# A yeast phenomic model for the influence of Warburg metabolism on genetic buffering of doxorubicin

**DOI:** 10.1101/517490

**Authors:** Sean M. Santos, John L. Hartman

**Affiliations:** University of Alabama at Birmingham, Department of Genetics, Birmingham, AL

**Keywords:** genetic buffering, yeast phenomics, quantitative high throughput cell array phenotyping (Q-HTCP), cell proliferation parameters (CPPs), doxorubicin, Warburg metabolism, differential gene interaction networks, recursive expectation-maximization clustering (REMc), pharmacogenomics, human-like / HL yeast media

## Abstract

**Background:** *Saccharomyces cerevisiae* represses respiration in the presence of adequate glucose, mimicking the Warburg effect, termed aerobic glycolysis. We conducted yeast phenomic experiments to characterize differential doxorubicin-gene interaction, in the context of respiration *vs.* glycolysis. The resulting systems level biology about doxorubicin cytotoxicity, including the influence of the Warburg effect, was integrated with cancer pharmacogenomics data to identify potentially causal correlations between differential gene expression and anti-cancer efficacy.

**Methods:** Quantitative high-throughput cell array phenotyping (Q-HTCP) was used to measure cell proliferation phenotypes (CPPs) of the yeast gene knockout/knockdown library, treated with escalating doxorubicin concentrations in fermentable and non-fermentable media. Doxorubicin-gene interaction was quantified by departure of the observed and expected phenotypes for the doxorubicin-treated mutant strain, with respect to phenotypes for the untreated mutant strain and both the treated and untreated reference strain. Recursive expectation-maximization clustering (REMc) and Gene Ontology-based analyses of interactions were used to identify functional biological modules that buffer doxorubicin cytotoxicity, and to characterize their Warburg-dependence. Yeast phenomic data was applied to cancer cell line pharmacogenomics data to predict differential gene expression that causally influences the anti-tumor efficacy, and potentially the anthracycline-associated host toxicity, of doxorubicin.

**Results:** Doxorubicin cytotoxicity was greater with respiration, suggesting the Warburg effect can influence therapeutic efficacy. Accordingly, doxorubicin drug-gene interaction was more extensive with respiration, including increased buffering by cellular processes related to chromatin organization, protein folding and modification, translation reinitiation, spermine metabolism, and fatty acid beta-oxidation. Pathway enrichment was less notable for glycolysis-specific buffering. Cellular processes exerting influence relatively independently, with respect to Warburg status, included homologous recombination, sphingolipid homeostasis, telomere tethering at nuclear periphery, and actin cortical patch localization. Causality for differential gene expression associated with doxorubicin cytotoxicity in tumor cells was predicted within the biological context of the phenomic model.

**Conclusions:** Warburg status influences the genetic requirements to buffer doxorubicin toxicity. Yeast phenomics provides an experimental platform to model the complexity of gene interaction networks that influence human disease phenotypes, as in this example of chemotherapy response. High-resolution, systems level yeast phenotyping is useful to predict the biological influence of functional variation on disease, offering the potential to fundamentally advance precision medicine.

## Background

Doxorubicin is used widely in oncology to treat both hematologic cancer and solid tumors [1]. Proposed mechanisms of doxorubicin cytotoxicity include topoisomerase II poisoning, DNA adduct formation, oxidative stress, and ceramide overproduction [1–6]. Topoisomerase II is an ATP-dependent enzyme that relieves the DNA torsional stress occurring with replication or transcription by catalyzing a double-stranded DNA (**dsDNA**) break, relaxing positive and negative DNA supercoiling, and finally re-ligating the DNA [7]. Inhibiting this activity can result in irreparable DNA damage and induction of apoptosis, selectively killing rapidly dividing proliferating cells [8–10]. Doxorubicin also causes histone eviction leading to chromatin trapping and damage [2, 11–13]. In addition to its potent anti-cancer therapeutic properties, doxorubicin is known for dose-limiting cardiomyocyte toxicity, causing cardiomyopathy and heart failure years post-treatment [14]. In this regard, topoisomerase IIB is highly expressed specifically in myocardiocytes, where tissue-specific deletion suppresses cardiac toxicity in mice [15]. Clinical guidelines recommend a maximum cumulative lifetime dose of 500 mg/m^2^; however, doxorubicin toxicity is variable and has a genetic basis [16]. Thus, a detailed understanding of drug-gene interaction could advance the rationale for more precisely prescribing doxorubicin (among other cytotoxic agents) and also predicting toxicity, based on the unique genetic context of each patient’s tumor genetic profile as well as germline functional variation.

This work establishes a yeast phenomic model to understand genetic pathways that buffer doxorubicin toxicity [17–23], and how the Warburg effect influences the doxorubicin-gene interaction network. Warburg won the Nobel Prize in 1931, yet there remains lack of consensus about how cancer cells undergo the Warburg transition and how aerobic glycolysis contributes to cancer [24–27]. In humans, aerobic glycolysis is considered a tumor-specific metabolic transition; however, yeast normally repress respiration in the presence of adequate glucose [28–30]. Thus, we wondered whether doxorubicin-gene interaction manifests differentially under glycolytic vs. respiratory conditions in yeast and if genetic insights from this model could lead to better understanding its variable anti-tumor efficacy between different patients [17]. We observed increased toxicity of doxorubicin in non-fermentable media, where yeast must respire to proliferate, suggesting the Warburg transition could confer resistance of tumor cells to doxorubicin, and perhaps help explain the dose-limiting toxicity observed in cardiomyocytes, which have respiratory rates among the highest of all cell types [31].

We conducted yeast phenomic analysis of doxorubicin-gene interaction, consisting of quantitative high throughput cell array phenotyping (**Q-HTCP**) of the yeast knockout and knockdown (**YKO/KD**) libraries, using multiple growth inhibitory concentrations of doxorubicin in either dextrose-(**HLD**) or ethanol/glycerol-based (**HLEG**) media. Q-HTCP provided cell proliferation parameters (**CPPs**) with which to quantify doxorubicin-gene interaction and determine its dependence on respiratory *vs.* glycolytic metabolism [32–34]. The yeast phenomic model was used to predict causality underlying correlations between doxorubicin sensitivity and increased or decreased expression of the homologous human gene in pharmacogenomics data from cancer cell lines. Thus, the work details genetic pathways for buffering doxorubicin toxicity in yeast and applies the information to predict interactions between doxorubicin and functional genetic variation manifest in cancers from different, individual patients.

## Methods

### Strains and media

The yeast gene knockout strain library (**YKO**) was obtained from Research Genetics (Huntsville, AL, USA). The knockdown (**KD**) collection, also known as the Decreased Abundance of mRNA Production (**DAmP**) library, was obtained from Open Biosystems (Huntsville, AL, USA). The genetic background for the YKO library was BY4741 (S288C MAT**a***ura3*-Δ*0 his3*-Δ*1 leu2*-Δ*0 met17*-Δ*0*). Additional information and lists of strains can be obtained at https://dharmacon.horizondiscovery.com/cdnas-and-orfs/non-mammalian-cdnas-and-orfs/yeast/#all. Some mutants appear multiple times in the library and they are treated independently in our analysis. HL yeast media, a modified synthetic complete media [20], was used with either 2% dextrose (HLD) or 3% ethanol and 3% glycerol (HLEG) as the carbon source.

### Quantitative high throughput cell array phenotyping (Q-HTCP)

Phenomic data was obtained by Q-HTCP, a custom, automated method of collecting growth curve phenotypes for the YKO/KD library arrayed onto agar media [34]. A Caliper Sciclone 3000 liquid handling robot was used for cell array printing, integrated with a custom imaging robot (Hartman laboratory) and Cytomat 6001 (Thermo Fisher Scientific, Asheville, NC, USA) incubator. 384-culture array images were obtained approximately every 2 hours and analyzed as previously described [21, 34]. To obtain CPPs, image data were fit to the logistic equation, G(t) = K/(1 + e^−r(t−l)^), assuming G(0) < K, where G(t) is the image intensity of a spotted culture vs. time, K is the carrying capacity, r is the maximum specific growth rate, and l is the moment of maximal absolute growth rate, occurring when G(t) = K/2 (the time to reach half of carrying capacity) [32]. The resulting CPPs were used as phenotypes to measure doxorubicin-gene interaction.

### Quantification of doxorubicin-gene interaction

Gene interaction was defined by departure of the corresponding YKO/KD strain from its expected phenotypic response to doxorubicin. The expected phenotype was determined by cell proliferation phenotypes of the mutant without doxorubicin, together with those of the reference strain with and without doxorubicin [17–19, 21]. The concentrations of doxorubicin (0, 2.5, 5, 7.5 and 15 ug/mL) were chosen based on phenotypic responses being functionally discriminating in the parental strain. We tested for effects of mating type or ploidy on doxorubicin growth inhibition (**Additional File 1, Fig. S1)**, and noted only small differences between the YKO/KD parental strain genotypes, BY4741 (MAT**a***ura3*-Δ*0 his3*-Δ*1 leu2*-Δ*0 met17*-Δ*0*), BY4742 (MATα *ura3*-Δ*0his3*-Δ*1 leu2*-Δ*0 lys2Δ0*), BY4741R (MAT**a***ura3*-Δ*0 his3*-Δ*1 leu2*-Δ*0 lys2*Δ*0*), BY4742R (MATα *ura3*-Δ*0 his3*-Δ*1 leu2*-Δ*0 met17*-Δ*0*), and diploid strains derived from these haploids. In this regard, haploid *MET17/lys2*-Δ*0* was associated with a lower carrying capacity in HLD media (**Additional File 1, Fig. S1**), but genome-wide experiments were not performed in this background.

Interaction scores were calculated as previously described [21], with slight modifications, as summarized below. Variables were defined as:

D_i_ = concentration (dose) of doxorubicin
R_i_ = observed mean growth parameter for parental Reference strain at D_i_
Y_i_ = observed growth parameter for the YKO/KD mutant strain at D_i_
K_i_ = Y_i_ – R_i_, the difference in growth parameter between the YKO/KD mutant (Y_i_) and Reference (R_i_) at D_i_
K_0_ = Y_0_ − R_0_, the effect of gene KO/KD on the observed phenotype in the absence of doxorubicin; this value is annotated as ‘shift’ and is subtracted from all K_i_ to obtain L_i_
L_i_ = K_i_ − K_0_, the interaction between (specific influence of) the KO/KD mutation on doxorubicin response, at D_i_

For cultures not generating a growth curve, Y_i_ = 0 for K and r, and the L parameter was assigned Y_i_ max, defined as the maximum observed Y_i_ among all cultures exhibiting a minimum carrying capacity (K) within 2 standard deviation (SD) of the parental reference strain mean at D_i_. Y_i_ max was also assigned to outlier values (*i.e.*, if Y_i_ > Y_i_ max).

Interaction was calculated by the following steps:

1. Compute the average value of the 768 reference cultures (R_i_) at each dose (D_i_):
2. Assign Y_i_ max (defined above) if growth curve is observed at D_0_, but not at D_i_, or if observed Y_i_ is greater than Y_i_ max.
3. Calculate K_i_ = Y_i_ − R_i_.
4. Calculate L_i_ = K_i_ − K_0_.
5. Fit data by linear regression (least squares): L_i_ = A + B*D_i_
6. Compute the interaction value ‘INT’ at the max dose: INT = L_i_-max = A + B*D_max_
7. Calculate the mean and standard deviation of interaction scores for reference strains, mean(REF_INT_) and SD(REF_INT_); mean(REF_INT_) is expected to be approximately zero, but SD(REF_INT_) is useful for standardizing against variance (**Additional Files 2-4**).
8. Calculate interaction z-scores (**Fig. 1D**):

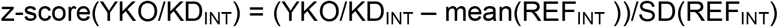

z-score(YKO/KD_INT_) > 2 for L or < −2 for K are referred to as gene deletion enhancers of doxorubicin cytotoxicity, and conversely, L interaction score < −2 or K interaction scores >2 are considered gene deletion suppressors (**Fig. 1E**).

**Figure 1.**
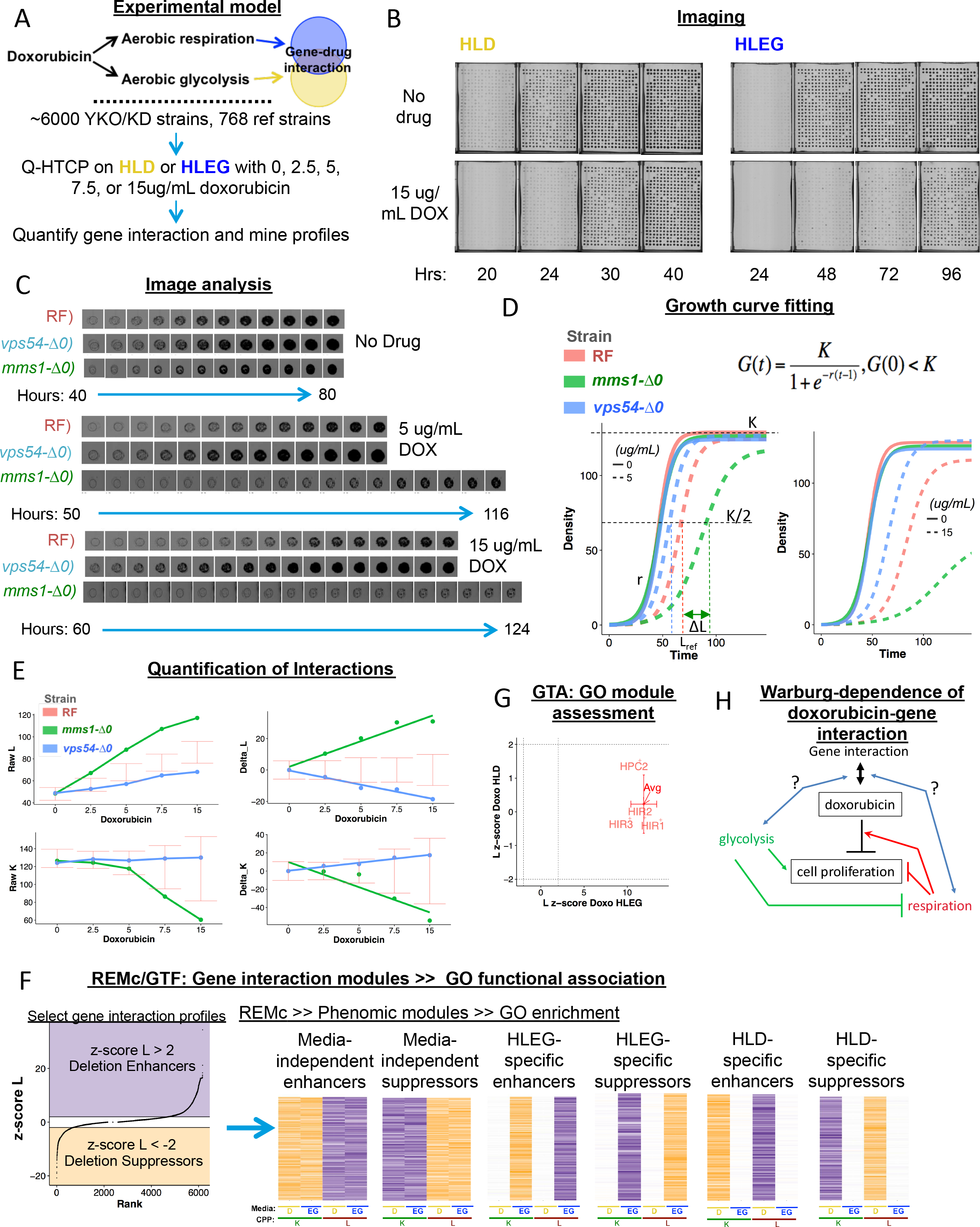
Experimental strategy to characterize differential doxorubicin-gene interaction, with respect to the Warburg metabolic transition. **(A)** The phenomic model incorporates treatment of individually grown cultures of the YKO/KD collection with increasing doxorubicin (0, 2.5, 5, 7.5, and 15 ug/mL) in “fermentable/glycolytic” (HLD) or “non-fermentable/respiratory” (HLEG) media. **(B)** Representative cell array images, treated and untreated with 15 ug/mL doxorubicin. **(C)** Time series of individual culture images, exemplifying gene deletion suppression (*vps54*-Δ*0*) and gene deletion enhancement (*mms1*-Δ*0*), relative to parental control (‘RF1’) in HLEG media with indicated concentrations (0, 5, and 15 ug/mL) of doxorubicin. **(D)** After image analysis, data time series are fit to a logistic growth function, G(*t*), to obtain the cell proliferation parameters (CPPs), *K* (carrying capacity), *L* (time at which K/2 is reached) and *r* (maximum specific rate) for each culture. ‘ΔL’ (left panel) indicates K_i_ (see methods). **(E)** Interaction is quantified by linear regression of Li (indicated ‘Delta_L’ and ‘Delta_K’ in right panels; see methods) across the entire dose range, which is converted to a z-score by dividing with the variance of the parental reference control (see methods). **(F)** Gene interaction profiles were grouped by recursive expectation-maximization clustering (REMc) to reveal deletion enhancing and deletion suppressing doxorubicin-gene interaction modules and the influence of the Warburg effect. Resulting clusters were analyzed with GOTermFinder (GTF) to identify enriched biological functions. **(G)** Gene Ontology Term Averaging (GTA) was used as a complement to REMc/GTF. **(H)** The model for genetic buffering of doxorubicin cytotoxicity incorporates primary and interaction effects involving glycolysis (green), and respiration (red), to explain the influence of Warburg context (blue) on doxorubicin-gene interaction (black).

### Recursive expectation-maximization clustering (REMc) and heatmap generation

REMc is a probability-based clustering method and was performed as previously described [35]. Clusters obtained by Weka 3.5, an EM-optimized Gaussian mixture-clustering module, were subjected to hierarchical clustering in R (http://www.r-project.org/) to further aid visualization with heatmaps. REMc was performed using L and K interaction z-scores (**Fig. 1F**). The effect of gene deletion on the CPP (in the absence of drug), termed ‘shift’ (K_0_), was not used for REMc, but was included for visualization in the final hierarchical clustering. **Additional File 5** contains REMc results in text files with associated data also displayed as heatmaps. In cases where a culture did not grow in the absence of drug, 0.0001 was assigned as the interaction score, and associated data were colored red (‘NA’) in the shift columns of the heatmaps.

### Gene ontology term finder (GTF)

A python script was used to format REMc clusters for analysis with the command line version of the GO Term Finder (GTF) tool downloaded from http://search.cpan.org/dist/GO-TermFinder/ [36]. GTF reports on enrichment of Gene Ontology (GO) terms by comparing the ratio of genes assigned to a term within a cluster to the respective ratio involving all genes tested. **Additional File 5** contains GTF analysis of all REMc clusters. GO-enriched terms from REMc were investigated with respect to genes representing the term and literature underlying their annotations [37].

### Gene ontology term averaging (GTA)

In addition to using GTF to survey functional enrichment in REMc clusters, we developed GTA as a complementary workflow, using the GO information on SGD at https://downloads.yeastgenome.org/curation/literature/ to perform the following analysis:

1. Calculate the average and SD for interaction values of all genes in a GO term.
2. Filter results to obtain terms having GTA value greater than 2 or less than −2.
3. Obtain GTA scores defined as |GTA value| - gtaSD; filter for GTA score > 2.

The GTA analysis is contained in **Additional File 6** as tables and interactive plots created using the R *plotly* package https://CRAN.R-project.org/package=plotly. GTA results were analyzed primarily using the L interaction scores, however GTA results with K interaction scores are included in **Additional File 6 (File D)**.

### Validation of doxorubicin-gene interaction

We retested 364 YKO/KD strains having human homologs in the P-POD database [38] and L interaction scores greater than 2 or less than −2 in at least one media type. Strains were struck to obtain four single colonies and arranged on replicate 384 well plates along with twenty reference strain controls and reanalyzed by Q-HTCP on HLD and HLEG, as in the genome-wide experiment. Results are summarized in **Fig 2S-T**, **Additional File 2 (Tables S5-S8), and Additional Files 3-4 (Files C-D)**.

**Figure 2.**
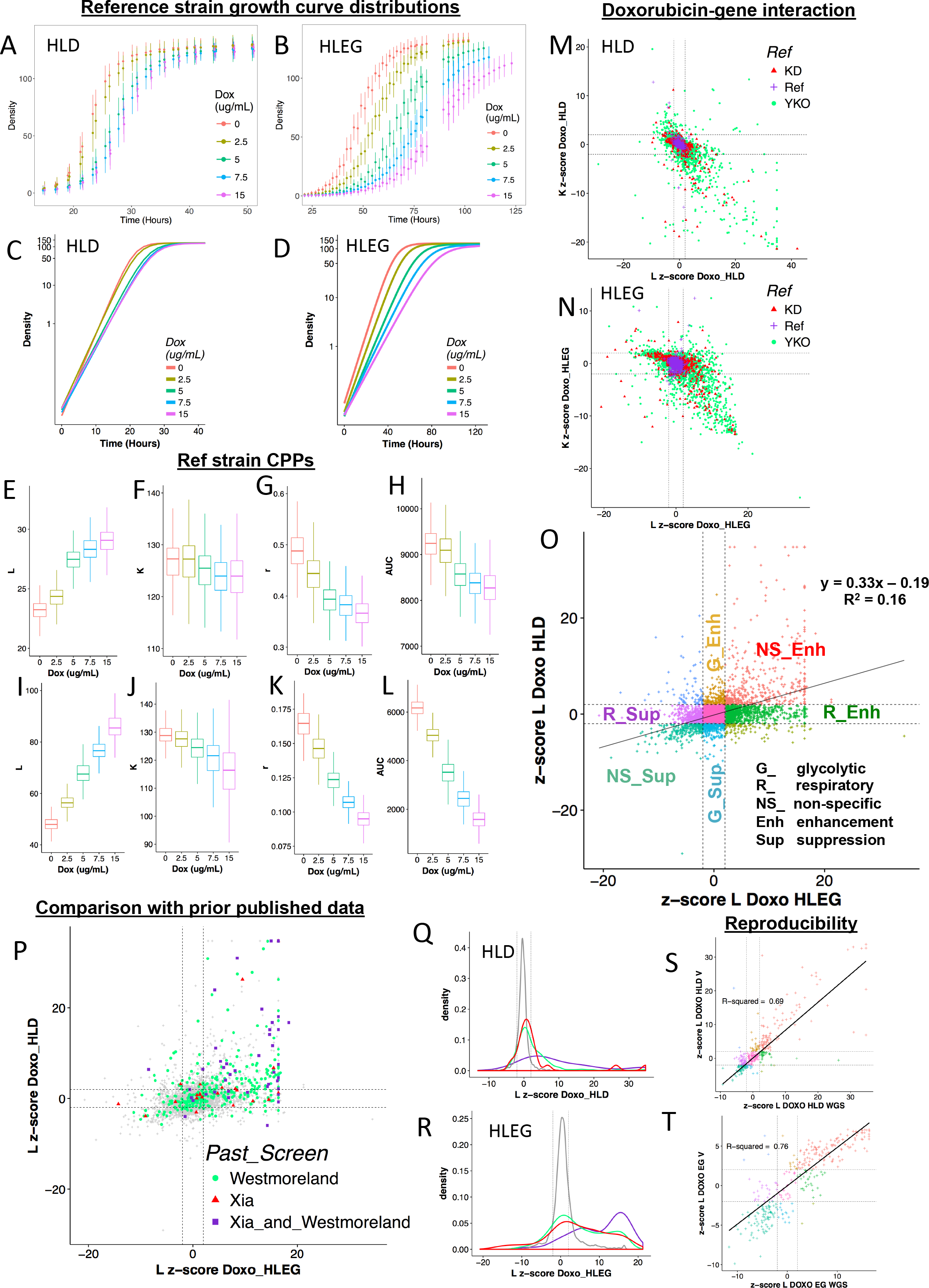
Q-HTCP provides cell proliferation parameters as phenotypes to quantify gene interaction. (**A, B**) Average pixel intensity and standard deviation for 768 reference strain cultures at indicated times after exposure to escalating doxorubicin concentrations in **(A)** HLD or **(B)** HLEG media. (**C-D**) Semi-log plots after fitting the data plotted above for **(C)** HLD or **(D)** HLEG to a logistic function (see Fig. 1D). (**E-L**) CPP distributions from data depicted in panels A-D for (**E-H**) HLD and (**I-J**) HLEG, including (**E, I**) L, (**F, J**) K, (**G, K**) r, and (**H, L**) AUC. **(M, N)** Comparison of doxorubicin-gene interaction scores using the L *vs.* K CPP in the context of either **(M)** HLD or **(N)** HLEG media. Score distributions of knockout (YKO, green), knock down / DAmP (YKD, Red), and non-mutant parental (Ref, Purple) strain cultures are indicated along with thresholds for deletion enhancement and suppression (dashed lines at +/− 2). **(O)** Differential doxorubicin-gene interaction (using L as the CPP) for HLD vs. HLEG, classified with respect to Warburg metabolism as non-specific (NS), respiratory-specific (R), or glycolysis-specific (G) deletion enhancement (Enh) or deletion suppression (Sup). **(P-R)** Comparisons between genome-wide studies of doxorubicin-gene interaction: **(P)** Genes reported from Westmoreland *et al.* (green), Xia *et al.* (Red) or both studies (purple) are plotted overlying L interaction scores (gray) in HLD *vs.* HLEG. (**Q-R**) L interaction scores (gray) for genes reported by Westmoreland *et al.* (green), Xia *et al.* (red), or both studies (purple) in (**Q**) HLD or (**R**) HLEG media. (**S-T**) Doxorubicin-gene interaction from genome wide (GWS) and validation (V) studies on **(S)** HLD or **(T)** HLEG media.

### Prediction of human homologs that influence tumor response to doxorubicin

PharmacoDB holds pharmacogenomics data from cancer cell lines, including transcriptomics and drug sensitivity [39]. The *PharmacoGx* R/Bioconductor package [40] was used to analyze the GDSC1000 (https://pharmacodb.pmgenomics.ca/datasets/5) and gCSI (https://pharmacodb.pmgenomics.ca/datasets/4) datasets, which contained transcriptomic and doxorubicin sensitivity results. A p-value < 0.05 was used for differential gene expression and doxorubicin sensitivity. For gene expression, the sign of the standardized coefficient denotes increased (+) or decreased (-) expression. The *biomaRt* R package [41, 42] was used with the Ensembl database [43] to match yeast and human homologs from the phenomic and transcriptomic data, classifying yeast-human homology as one to one, one to many, and many to many.

## Results

### Phenomic characterization of doxorubicin response genes

The workflow for analyzing doxorubicin-gene interaction and differential buffering of doxorubicin with respect to the Warburg effect is summarized in **Fig. 1**. Alternately, in a respiratory or glycolytic (HLEG or HLD media, respectively) context (**Fig. 1A**), Q-HTCP technology was used for high throughput kinetic imaging of 384-culture cell arrays plated on agar media (**Fig. 1B**), image analysis (**Fig. 1C**), and growth curve fitting (**Fig. 1D**) to obtain the CPPs, L (time to reach half carrying capacity), K (carrying capacity), and r (maximum specific rate) [21, 32, 34], which were used to measure doxorubicin-gene interaction across the entire YKO/KD library. Departure of the observed CPP from the expected doxorubicin response for each YKO/KD strain was derived using distributions from many replicate reference strain control cultures, and summarized across all doxorubicin concentrations by linear regression (**Fig. 1E**). Interaction scores with absolute value greater than two were considered as gene *deletion enhancement* (z-score_L ≥ 2 or z-score_K ≤ −2) or *deletion suppression* (z-score_L ≤ −2 or z-score_K ≥ 2) of doxorubicin cytotoxicity. Gene deletion enhancement (*e.g*., *mms1*-Δ*0*) and suppression (*e.g.*, *vps54*-Δ*0*) reveal functions that buffer or confer doxorubicin cytotoxicity, respectively. Doxorubicin-gene interaction profiles (selected if they contained L interaction scores with absolute value greater than two, in either HLD or HLEG media) were analyzed by REMc and assessed for GO Term enrichment (**Fig. 1F**). As a complement to clustering gene interaction profiles, functional enrichment was analyzed by GTA (see methods), systematically querying all GO processes, functions, and components (Fig. 1G **and methods**) with respect to CPPs and Warburg status. Taken together, REMc and GTA reveal genetic modules that buffer doxorubicin, and how they are influenced by Warburg metabolism (**Fig. 1H**).

Doxorubicin cytotoxicity was greater in HLEG than HLD media, evidenced by the reference strain being more growth inhibited (Fig. 2A-L, **Additional File 1, Fig. S1**). The ‘L’ parameter was the most sensitive CPP, while K reported larger phenotypic effects (**Fig. 2M-N) (Additional File 1, Fig. S2**). We noted positive correlation between doxorubicin-gene interaction in HLEG and HLD, however interaction was media-specific and more abundant in the context of respiration, i.e., with HLEG media (**Fig. 2O**).

We compared our results with two prior studies of doxorubicin cytotoxicity in the yeast knockout collections [44, 45]. One study was conducted in SC media with the haploid (BY4741) YKO library and identified 71 deletion enhancers of cytotoxicity [44]. A second study reported on the homozygous diploid (BY4743) YKO collection in YPD media, identifying 376 enhancers [45]. Overlap between these studies and ours is shown in **Fig. 2P-2R** and in **Additional File 7 (Table S9-10)**. While many genes overlapped between the studies, differing results were also observed, possibly attributable to strain background, media conditions, as well as methods for scoring interactions [20, 46]. To assess within-study reproducibility, we sub-cloned four colonies from glycerol stocks used in the first experiment and retested doxorubicin-gene interaction, revealing higher correlation and overall reproducibility within-study than between-study (**Fig. 2S-T**).

### Identification of functional gene interaction modules

Gene interaction profiles were analyzed by REMc (**Figs. 3, 4**), as described previously [35]. Briefly, REMc uses an expectation-maximization algorithm to define clusters probabilistically, and is applied recursively to resolve gene interaction profile clusters. REMc terminates when a round of clustering reveals no new clusters. The cluster naming convention is “A-B.C.D-X”, where ‘A’ = the round of clustering, ‘B’ = 0, and ‘C.D-X’ indicates the cluster pedigree. For example, 1-0-0 refers to the first cluster of the first round, 2-0.0-3 the fourth cluster derived from 1-0-0 (in round 2 of REMc), 3-0.0.3-1 indicates the second cluster derived from 2-0.0-3 (in round 3), and so on [35].

**Figure 3.**
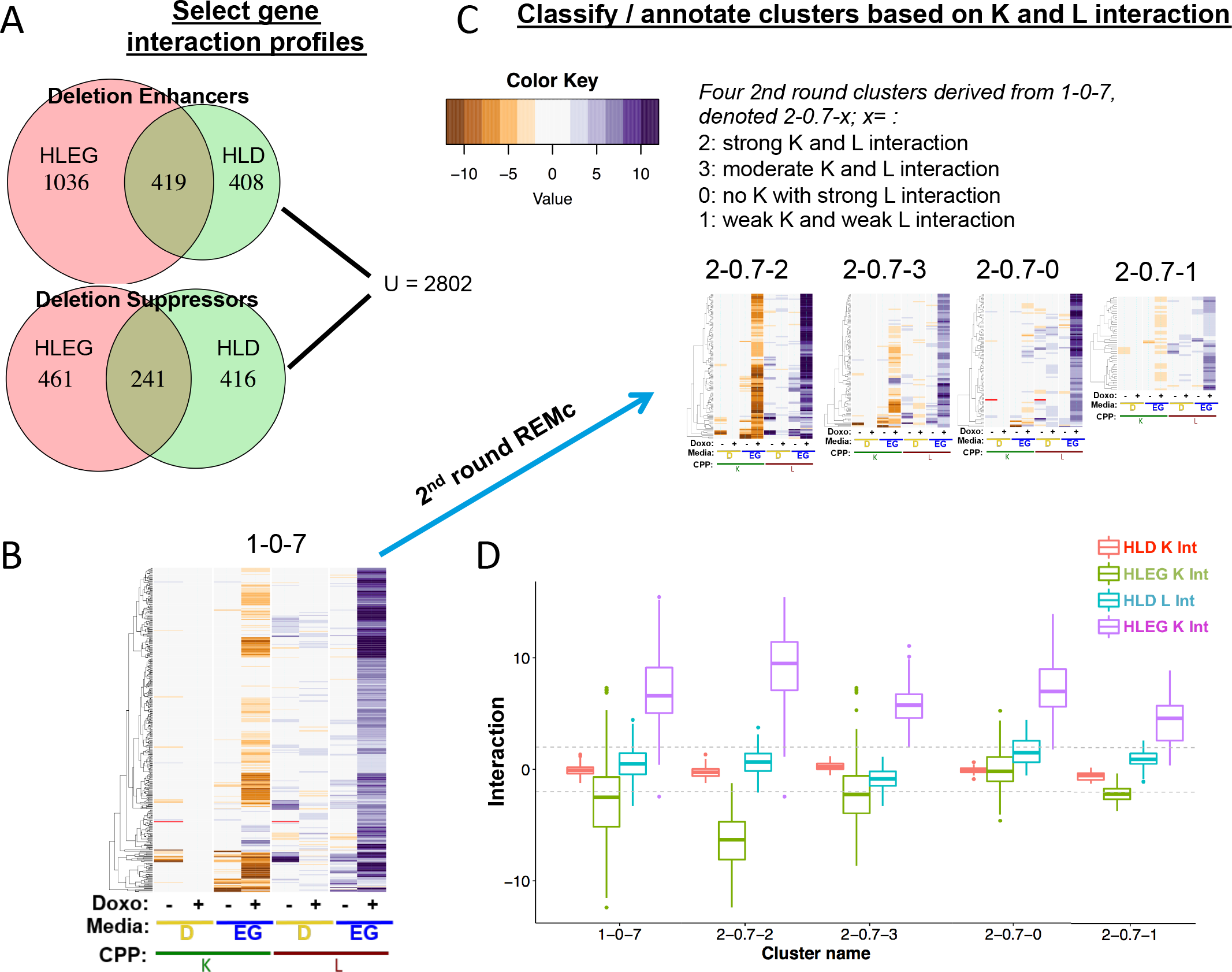
Characterization of Warburg-differential, doxorubicin-gene interaction profiles. **(A)** The union of enhancers (L z-score > 2) or suppressors (L z-score < −2) from the HLD and HLEG analyses totaled 2802 gene interaction profiles that were subjected to REMc (see methods). **(B-C)** The column order is the same for all heatmaps; ‘+’ indicates doxorubicin-gene interaction and ‘-‘ indicates ‘shift’ (K_0_; see methods). Interactions by K are negative (brown) if enhancing and positive (purple) if suppressing, while the signs of interaction are reversed for L (see methods). The heatmap color scale is incremented by twos; red indicates no growth curve in the absence of doxorubicin. **(B)** First round cluster 1-0-7 has a gene interaction profile indicative of HLEG-specific deletion enhancement. (**C**) Second round clusters (2-0.7-X) are ordered left to right by strength of influence. **(D)** The pattern of distributions for the different doxorubicin-gene interaction scores (‘+’ columns only from panel C) summarizes respective clusters from panel C. Deletion enhancement is considered to be qualitatively stronger if observed for K in addition to L.

**Figure 4.**
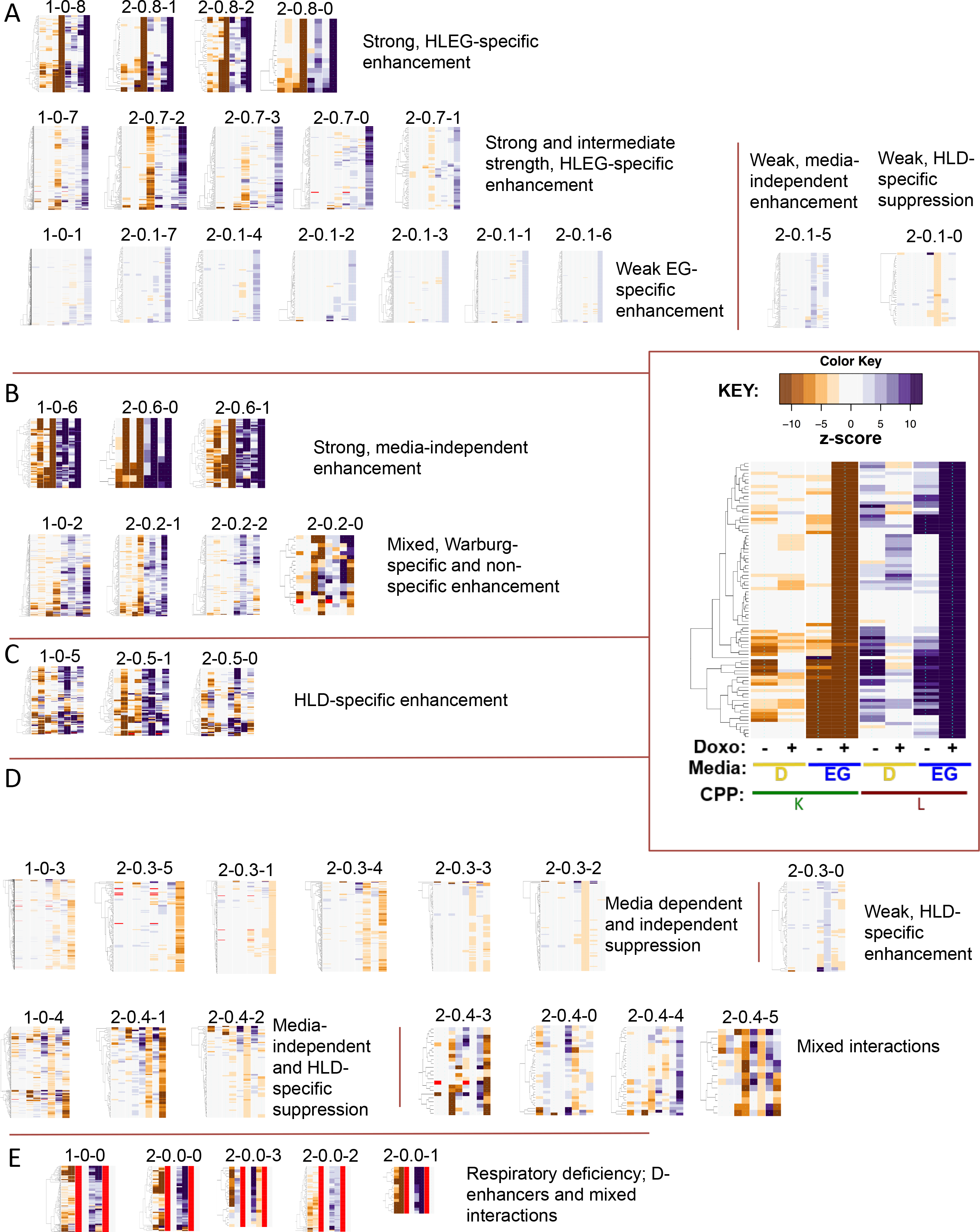
A summary of the first and second rounds of REMc. First round clusters are at the left end of each row of heatmap thumbnails; second round clusters derived from each first round cluster are ordered to the right by relative strength. Rows are grouped into panels by similarity in their gene interaction profiles. The columns in each heatmap have the same order from left to right (see inset panel), with K to the left and L to the right. Within the K and L groups, HLD is to the left and HLEG to the right. Within each of the CPP-media groupings, ‘shift’ (−) is left of the doxorubicin-gene interaction (+). **(A)** Respiration-specific enhancement. **(B)** Warburg-independent enhancement. **(C)** Glycolysis-specific enhancement. **(D)** HLD and HLEG suppression modules. **(E)** Respiratory deficiency.

The main effect of the gene KO or KD on cell proliferation, *i.e.*, K_i_ in the absence of doxorubicin (D_0_) is also referred to as ‘shift’ (see methods). ‘Shift’ was not subjected to REMc, but was included for hierarchical clustering and visualization by heatmaps after REMc (Fig. 3; **Additional File 5, File B)**. K_i_ is termed ‘shift’, because this value is subtracted from the data series for each YKO/KD to obtain L_i_ values, which are fit by linear regression for calculating drug-gene interaction (**Fig. 1E;** see methods).

GO TermFinder [36] was used to associate enrichment of biological functions with particular patterns of doxorubicin-gene interaction identified by REMc (Figs. 3–4; Table 1; **Additional File 5, File C**). In general, the first two rounds of REMc revealed distinctive profiles of gene interaction in respiratory vs. glycolytic media (**Fig. 4**). Later round clusters exhibited greater GO term enrichment in some cases; however, GO enrichment was other times reduced by further clustering (see **Additional File 8**).

**Table 1.**
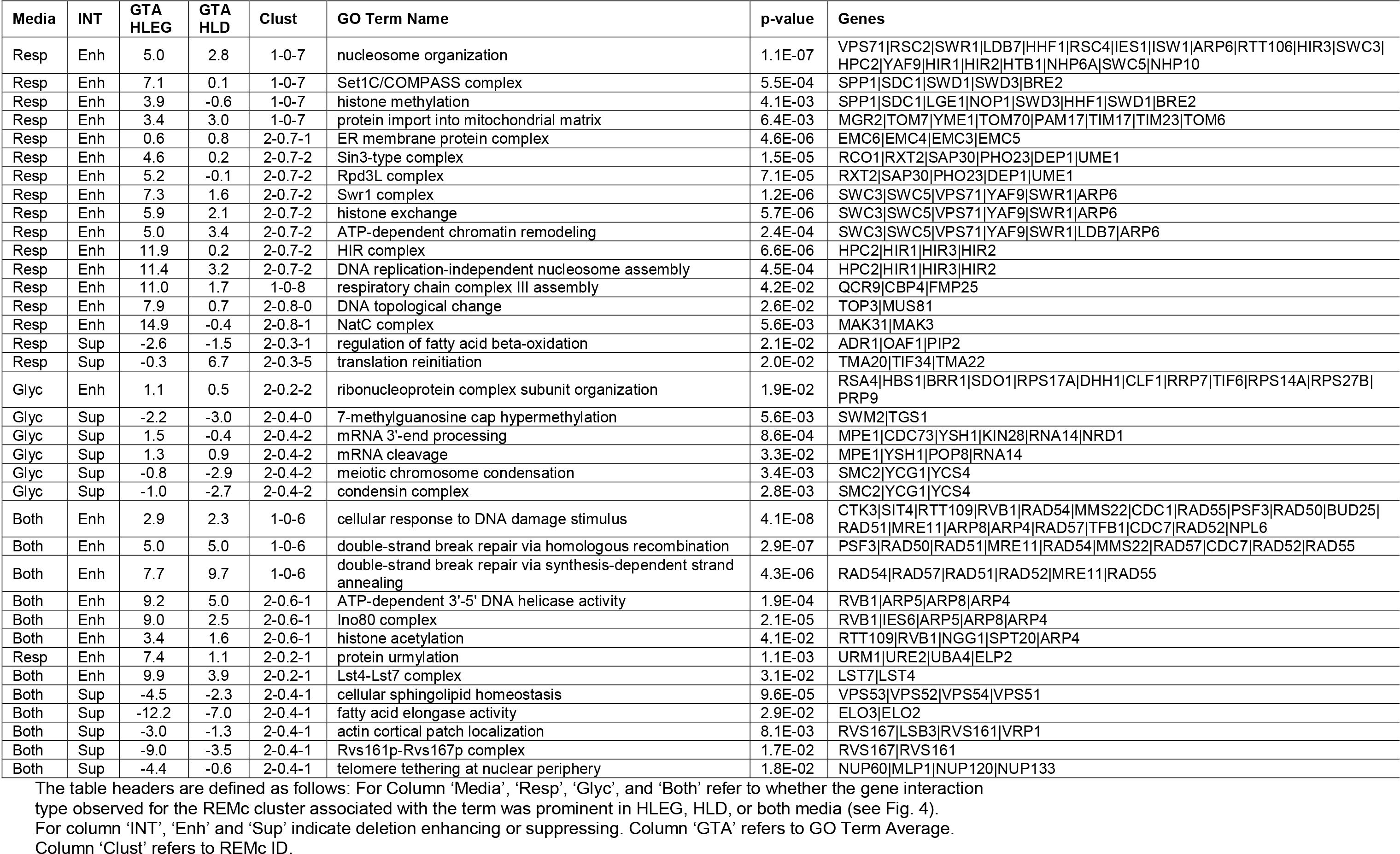
GO Terms enriched in REMc clusters.

GTA score revealed 129 GO terms, 39 of which were found by REMc/GTF (**Table 2** and **Additional File 6, Files A-C**). GTA identifies functions of smaller GO terms, *e.g.*, protein complexes. GTA with K interaction scores yielded only 35 GO terms (**Additional File 6, File D)**, with only 3 being unique from GTA with L interaction; thus, we focused on L interaction for GTA analysis. Interactive scatter plots (html files in which points contain embedded information) were used to visualize significant GO terms from both REMc and GTA (**Additional File 6, File B**). GO term-specific heatmaps further aided visualization of relationships between genes and the GO terms (see **Figs. 6-11** and **Additional File 9**) by systematically displaying, for all genes attributed to a parent term and its children, uniformity *vs.* pleiotropy of interaction effects across different conditions.

**Table 2.** GO terms identified by GTA.

| GO Term Name | Media | INT | HLEG<br>GTA | HLEG<br>gtaSD | HLD<br>GTA | HLD<br>gtaSD | Genes | REMc<br>related | p-value |
| --- | --- | --- | --- | --- | --- | --- | --- | --- | --- |
| HIR complex | Resp | Enh | 11.9 | 1.5 | 0.2 | 0.9 | HIR1 \| HIR2 \| HPC2 \| HIR3 | 2-0.7-2 | 6.6E-06 |
| histone monoubiquitination | Resp | Enh | 11.4 | 7.0 | 0.1 | 1.2 | RAD6 \| BRE1 | NA | NA |
| Ino80 complex | Resp | Enh | 9.0 | 6.8 | 2.5 | 7.7 | RVB1 \| IES6 \| ARP5 \| ARP8 \| ARP4 \| ARP7 \| IES5 \| IES3 \| NHP10 \| IES2 \| IES1 \| RVB2 \| IES4 \| TAF14 | 3-0.6.1-1 | 1.5E-06 |
| histone H4 acetylation | Resp | Enh | 8.0 | 4.8 | -0.8 | 2.1 | ESA1 \| NGG1 \| ELP4 \| EAF3 \| HAT1 | NA | NA |
| mitochondrial respiratory chain complex III assembly | Resp | Enh | 11.0 | 6.8 | 1.7 | 2.0 | QCR7 \| CBP6 \| CBP4 \| BCS1 \| QCR9 \| FMP25 \| FMP36 \| CBP3 | 1-0-8 | 4.2E-02 |
| mitochondrial respiratory chain supercomplex assembly | Resp | Enh | 15.9 | 0.6 | 0.8 | 0.1 | RCF1 \| COX13 | 1-0-8 | 7.0E-02 |
| mitochondrial outer membrane translocase complex | Resp | Enh | 9.1 | 6.4 | 0.7 | 3.1 | TOM22 \| TOM5 \| TOM6 \| TOM70 \| TOM7 \| TOM40 | NA | NA |
| protein urmylation | Resp | Enh | 7.4 | 2.6 | 1.1 | 0.9 | ELP2 \| URM1 \| NCS2 \| UBA4 \| ELP6 \| URE2 | 2-0.2-1 | 1.1E-03 |
| Elongator holoenzyme complex | Resp | Enh | 8.9 | 3.6 | 0.0 | 0.9 | TUP1 \| IKI3 \| ELP4 \| ELP2 \| ELP3 \| IKI1 \| ELP6 | 3-0.7.2-0 | 1.4E-04 |
| NatC complex | Resp | Enh | 14.9 | 1.7 | -0.4 | 0.6 | MAK31 \| MAK10 \| MAK3 | 2-0.8-1 | 5.6E-03 |
| DNA topological change | Resp | Enh | 7.9 | 5.7 | 0.7 | 2.6 | RFA2 \| TOP3 \| MUS81 \| RMI1 \| TOP1 \| SGS1 \| RFA1 \| RAD4 \| TOP2 | 2-0.8-0 | 2.6E-02 |
| tRNA (m1A) methyltransferase complex | Resp | Enh | 17.0 | 0.8 | 9.3 | 17.4 | GCD10 \| GCD14 | NA | NA |
| MUB1-RAD6-UBR2 ubiquitin ligase complex | Resp | Enh | 12.9 | 3.1 | 0.9 | 0.5 | RAD6 \| MUB1 \| UBR2 | NA | NA |
| malonyl-CoA biosynthetic process | Resp | Enh | 11.1 | 7.4 | 1.5 | 0.1 | HFA1 \| ACC1 | NA | NA |
| pyridoxal 5'-phosphate salvage | Resp | Enh | 11.1 | 8.7 | 1.5 | 5.3 | PDX3 \| BUD16 \| BUD17 | NA | NA |
| maintenance of transcriptional fidelity during DNA-templated transcription elongation from RNA polymerase II promoter | Resp | Enh | 11.1 | 7.5 | -0.4 | 4.2 | RPB9 \| DST1 | NA | NA |
| RNA polymerase II transcription corepressor activity | Resp | Enh | 11.0 | 7.6 | 2.2 | 1.7 | SIN3 \| MED8 \| SRB7 | NA | NA |
| pyruvate dehydrogenase activity | Resp | Enh | 10.6 | 6.4 | 2.8 | 0.9 | PDA1 \| LPD1 \| PDB1 | NA | NA |
| eukaryotic translation initiation factor 2 complex | Resp | Enh | 10.3 | 4.7 | 8.2 | 8.7 | SUI2 \| GCD11 | NA | NA |
| L-aspartate:2-oxoglutarate aminotransferase activity | Resp | Sup | -3.9 | 0.5 | -0.9 | 0.8 | AAT2 \| AAT1 | 2-0.4-3 | 5.9E-04 |
| nuclear pore outer ring | Resp | Sup | -6.3 | 3.7 | 1.4 | 7.6 | NUP145 \| SEH1 \| NUP84 \| NUP120 \| NUP133 | 3-0.4.1-0 | 9.7E-02 |
| positive regulation of fatty acid beta-oxidation | Resp | Sup | -2.6 | 0.5 | -1.5 | 0.2 | OAF1 \| ADR1 \| PIP2 | 2-0.3-1 | 2.1E-02 |
| EKC/KEOPS complex | Resp | Sup | -7.9 | 4.6 | -1.8 | 1.1 | KAE1 \| CGI121 \| GON7 \| BUD32 | NA | NA |
| spermine biosynthetic process | Resp | Sup | -2.6 | 0.3 | -0.3 | 0.7 | SPE4 \| SPE2 | NA | NA |
| Dom34-Hbs1 complex | Glyc | Enh | 0.3 | 2.1 | 2.7 | 0.3 | HBS1 \| DOM34 | NA | NA |
| Ubp3-Bre5 deubiquitination complex | Glyc | Enh | -1.1 | 3.2 | 8.8 | 2.2 | BRE5 \| UBP3 | NA | NA |
| Cul4-RING E3 ubiquitin ligase complex | Glyc | Enh | 1.8 | 4.1 | 4.6 | 2.3 | HRT1 \| PRP46 \| SOF1 | NA | NA |
| dTTP biosynthetic process | Glyc | Enh | -1.3 | 3.2 | 7.0 | 0.6 | CDC21 \| CDC8 | NA | NA |
| GDP-mannose transport | Glyc | Enh | 1.5 | 1.9 | 9.5 | 5.6 | VRG4 \| HVG1 | NA | NA |
| 7-methylguanosine cap hypermethylation | Glyc | Sup | -2.2 | 1.5 | -3.0 | 0.9 | SWM2 \| TGS1 | 2-0.4-0 | 5.6E-03 |
| meiotic chromosome condensation | Glyc | Sup | -0.8 | 0.9 | -2.9 | 0.9 | SMC2 \| YCG1 \| SMC4 \| YCS4 | 2-0.4-2 | 3.4E-03 |
| histone deubiquitination | Glyc | Sup | 1.8 | 1.9 | -3.4 | 1.1 | SEM1 \| UBP8 \| SGF73 \| SGF11 | NA | NA |
| HDA1 complex | Both | Enh | 8.9 | 0.3 | 4.0 | 1.1 | HDA2 \| HDA1 \| HDA3 | NA | NA |
| CTDK-1 complex | Both | Enh | 15.6 | 0.7 | 3.8 | 0.9 | CTK2 \| CTK3 \| CTK1 | 1-0-8 | 5.3E-02 |
| Cul8-RING ubiquitin ligase complex | Both | Enh | 9.1 | 4.5 | 6.1 | 1.1 | MMS22 \| MMS1 \| RTT101 \| HRT1 \| RTT107 | 1-0-2 | 1.0E-01 |
| Lst4-Lst7 complex | Both | Enh | 9.9 | 1.4 | 3.9 | 0.3 | LST7 \| LST4 | 2-0.2-1 | 3.1E-02 |
| MCM complex | Both | Enh | 4.2 | 1.4 | 4.9 | 2.6 | MCM7 \| MCM6 \| MCM5 \| MCM2 \| MCM3 | NA | NA |
| histone H3-K56 acetylation | Both | Enh | 10.3 | 7.0 | 8.6 | 4.3 | RTT109 \| SPT10 | NA | NA |
| fatty acid elongase activity | Both | Sup | -12.2 | 2.4 | -7.0 | 1.1 | SUR4 \| FEN1 | 2-0.4-1 | 2.9E-02 |
| GARP complex | Both | Sup | -6.8 | 0.9 | -3.5 | 0.8 | VPS53 \| VPS54 \| VPS52 \| VPS51 | 3-0.4.1-0 | 6.9E-07 |
| nuclear cap binding complex | Both | Sup | -4.7 | 0.5 | -3.4 | 0.5 | STO1 \| CBC2 | 3-0.4.1-0 | 9.9E-03 |
| Rvs161p-Rvs167p complex | Both | Sup | -9.0 | 0.4 | -3.5 | 0.2 | RVS167 \| RVS161 | 3-0.4.1-0 | 9.9E-03 |

In summary, we used REMc, GTA, and GO term-specific scatterplots and heatmaps to discover genetic modules that alternatively buffer (*i.e.*, deletion enhancing) or confer (*i.e.*, deletion suppressing) doxorubicin cytotoxicity, and to determine whether the Warburg-transition exerts influence on their effects (**Fig. 5)**.

**Figure 5.**
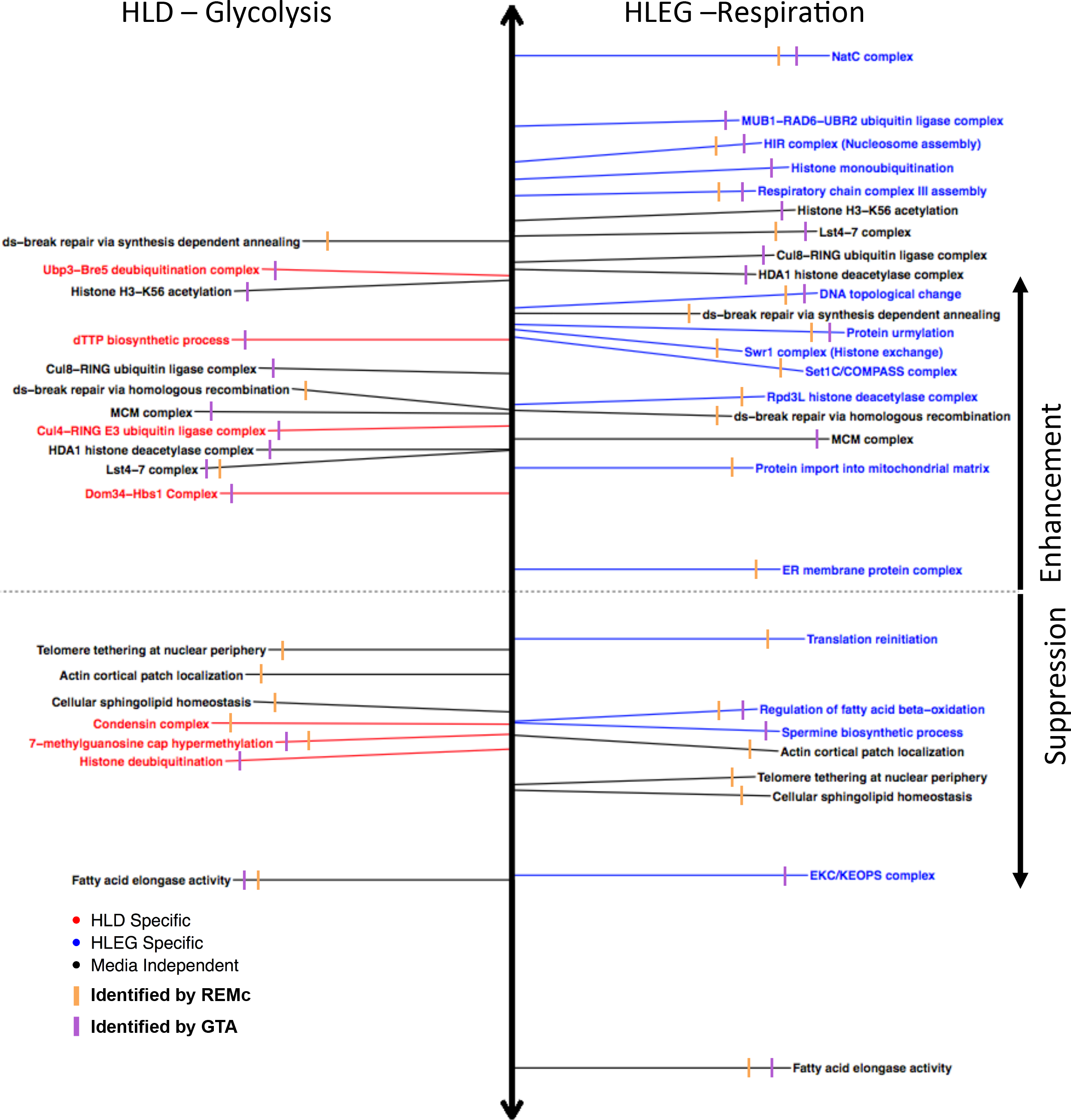
GO annotations associated with deletion enhancement or suppression of doxorubicin cytotoxicity, with respect to Warburg-dependence. Representative GO terms are listed, which were identified by REMc/GTF (orange), GTA (purple), or both methods, for HLD (left, red), HLEG (right, blue), or both media types (black), and for enhancement (above dashed line) or suppression (below dashed line) of doxorubicin cytotoxicity. Distance above or below the horizontal dashed line indicates the GTA value for terms identified by REMc or the GTA score if identified by GTA (see methods). See Additional Files 5 and 6, respectively, for all REMc and GTA results.

### Warburg transition-dependent doxorubicin gene interaction modules

#### Respiration-specific gene deletion enhancement

Respiration-specific deletion-enhancing clusters (see **Fig. 4**; 1-0-7 and 1-0-8) revealed GO Term enrichment for *histone modification and chromatin organization*, *respiratory chain complex III assembly*, *protein import into mitochondria, protein urmylation*, the *NatC complex*, *protein folding in endoplasmic reticulum*, *and DNA topological change* (Figs. 6–8; **Additional File 5, File C)**. Additional modules were identified using GTA (Fig. 8C **and Additional File 6, File A**).

**Figure 6.**
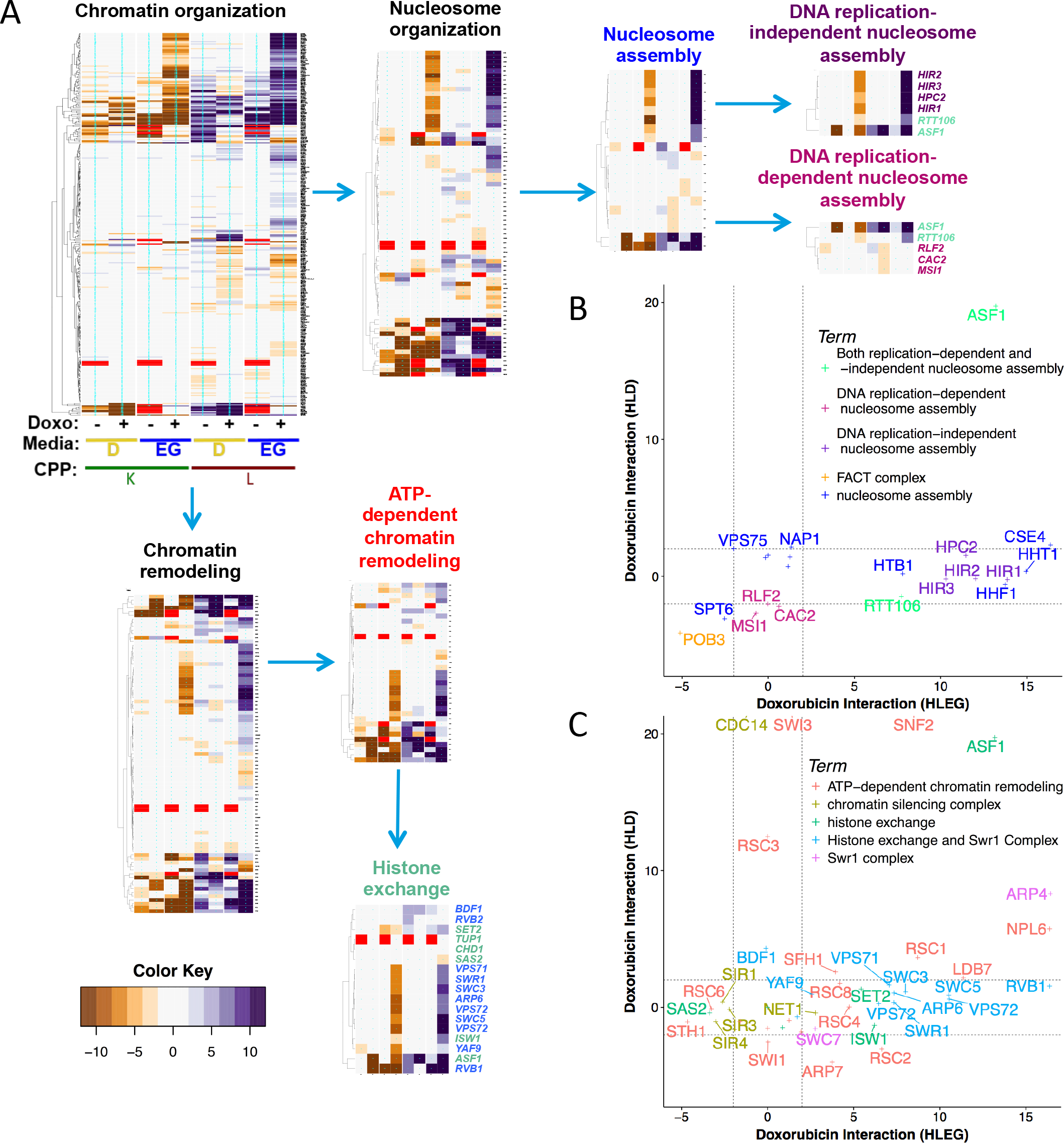
Respiration increases the role for chromatin organization in buffering doxorubicin toxicity. **(A)** GO term-specific heatmaps for chromatin organization and its child terms (indicated by arrows) clarify related but distinct biological functions that buffer doxorubicin, with respect to Warburg status. **(B-C)** L-based doxorubicin-gene interaction scores associated with GO terms that were enriched in cluster 2-0.7-2. Dashed lines indicate z-score thresholds for enhancers (>2) and suppressors (<-2). Sub-threshold gene interaction values are plotted, but not labeled.

#### Chromatin organization and histone modification

REMc/GTF and GTA identified several chromatin-related processes that buffer doxorubicin toxicity in a respiration-specific manner, including *DNA replication-independent nucleosome assembly*, *histone exchange*, *histone deacetylation*, and *histone methylation* (**Figs. 6 and 7)**.

**Figure 7.**
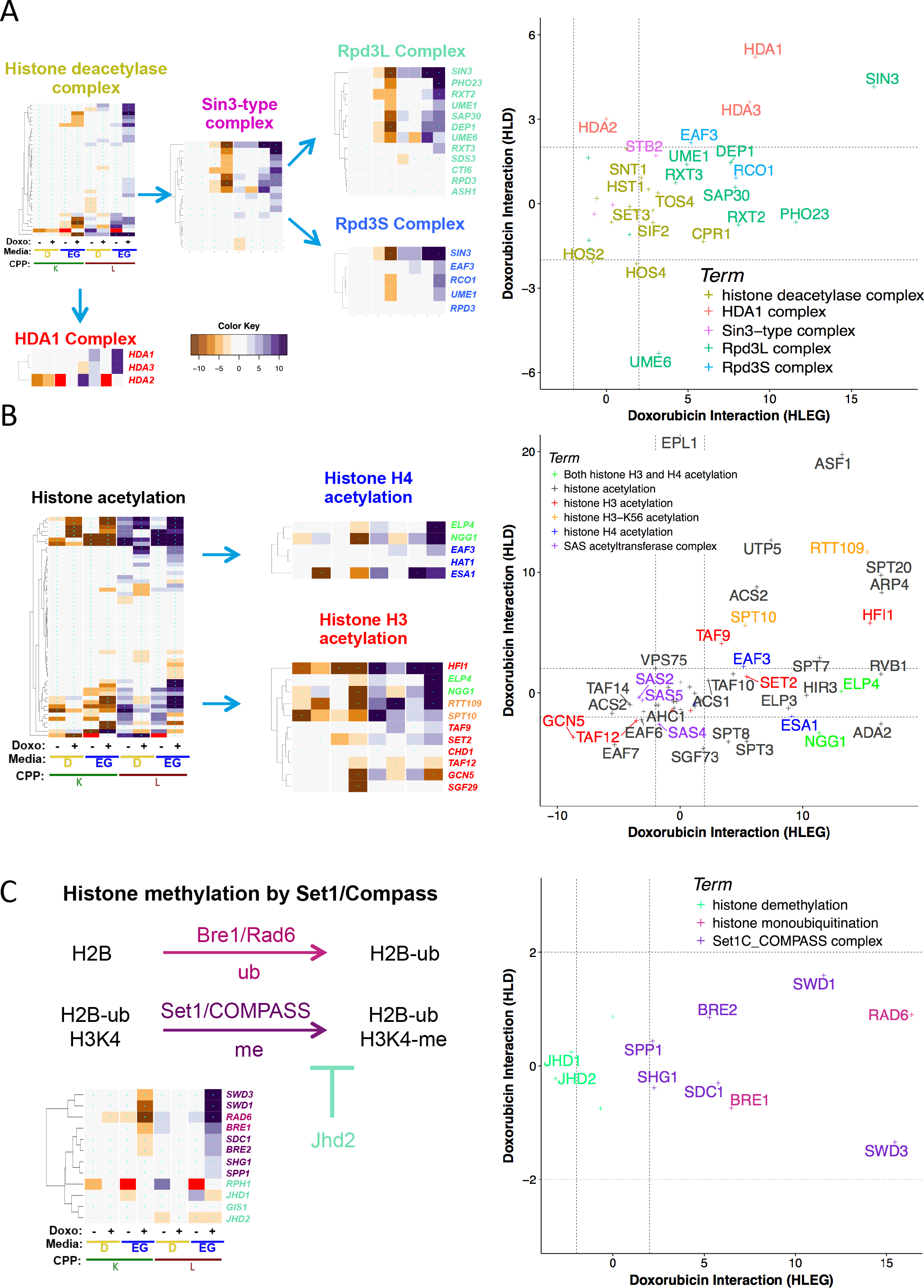
Distinct histone modifications differentially influence doxorubicin cytotoxicity. **(A)** Rpd3L and Rpd3S complexes exert strong HLEG-specific doxorubicin-enhancing influence relative to other Sin3-type histone deacetylases and the HDA1 complex. **(B)** In contrast to histone deacetylation (panel A), histone acetylation exhibits deletion enhancement that is Warburg-independent. **(C)** Histone H3K4 methylation by the Set1C/COMPASS complex, which requires histone mono-ubiquitination of H2B by the Bre1/Rad6 complex, is opposed by Jhd2, a histone H3K4 demethylase. The respiration-specific deletion enhancing interactions suggest the Warburg transition can protect tumors promoted by certain types of chromatin deregulation from doxorubicin.

##### (i) DNA replication-independent nucleosome assembly (HIR complex)

REMc/GTF identified the HIR complex (*HIR1-3* and *HPC2*), which functions as a histone chaperone in chromatin assembly and disassembly, in cluster 2-0.7-2 (**Fig. 4, Table 1**) [47]. Along with Asf1 and Rtt106, the HIR complex is involved in DNA replication-independent (*i.e.*, RNA transcriptional) histone deposition, and regulates transcription of three of the four histone genes [47–49]. Furthermore, genes encoding for *HTA1/HTB1, HHT1/2*, and *HHF1/2* were also respiratory-specific deletion enhancers. Asf1 and Rtt106 function in nucleosome assembly in both DNA replication and DNA replication-independent contexts. Asf1, which functions in the Rad53-dependent DNA damage response [50], enhanced doxorubicin toxicity in both respiratory and glycolytic media, like other DNA repair genes (see below). In further contrast, genes associated with *replication-dependent nucleosome assembly* (*RLF2, CAC2, MSI1*) by the chromatin assembly factor complex, CAF-1, [51] were HLD-specific suppressors (**Fig. 6A-B**).

Prior studies have reported enhanced doxorubicin cytotoxicity due to nucleosome disassembly and “chromatin trapping” by the FACT complex, referring to binding and resulting damage to disassembled chromatin in the context of doxorubicin exposure [13]. *POB3-DAmP*, the only member of the FACT complex represented in the YKO/KD library, resulted in suppression of doxorubicin cytotoxicity (**Fig. 6B**), presumably by suppressing its effect of trapping and damaging disassembled chromatin.

##### (ii) Histone exchange (Swr1 complex)

The Swr1 complex (enriched in cluster 2-0.7-2) uses ATP hydrolysis to replace the H2A nucleosome with the H2AZ variant [52]. Swr1 complex genes showing respiration-specific buffering of doxorubicin toxicity included *RVB1, SWC3, SWC5, ARP6, SWR1*, VPS71, and *VPS72* (**Fig. 6C**). Accordingly, the H2AZ variant, Htz1, which is enriched at most gene promoters in euchromatin [53–55], was also an HLEG-specific deletion enhancer. The Swr1 complex is recruited for repair of dsDNA breaks, where the H2AZ variant is incorporated [56]; however, the interaction profile of the Swr1 complex more closely resembles other respiratory specific enhancers involved in transcriptional regulation, whereas dsDNA-break repair by homologous recombination buffered doxorubicin toxicity independent of Warburg context (see cluster 1-0-6 from **Fig. 4**, **Table 1**, and descriptions below). The Swr1 complex can also inhibit subtelomeric spread of heterochromatin by impeding SIR-dependent silencing [57]. Consistent with knockout of Swr1 promoting silencing and having a deletion enhancing effect, deletion of *SIR1*, *SIR3* or *SIR4* (which disrupts chromatin silencing) also exerted respiratory-specific suppression of doxorubicin toxicity (**Fig. 6C**).

##### (iii) Histone deacetylation (Sin3-type and HDA1 complexes)

Deletion of genes functioning in the Rpd3L and Rpd3S histone deacetylase complexes (**HDAC**) was associated with strong respiratory enhancement of doxorubicin toxicity (cluster 2-0.7-2; **Fig. 7A**); however, genes constituting the Hda1 complex exerted weaker effects, but in both respiratory and glycolytic media (**Fig. 7A**, **Table 2**). The yeast Rpd3 deacetylase histone complexes are homologous to mammalian class I Rpd3-like proteins (Hdac1-3,8), while the yeast Hda1 complex is homologous to mammalian class II Hda1-like proteins (Hdac4-5,7,9) [58]. Hda1 and Rpd3 complexes both deacetylate histones H3 and H4; however, deletion of *RPD*3 *vs. HDA1* revealed different degrees of H4 lysine 5 and K12 hyperacetylation [59], implicating this functional distinction in Warburg-differential doxorubicin response.

Histone acetylation was GO-enriched in cluster 2-0.6-1, which displayed a Warburg-independent gene interaction profile (**Fig. 4, Table 1)**. GTA analysis confirmed H3K56 acetylation (*SPT10* and *RTT109*) and histone H3 acetylation (*TAF9* and *HFI1*) as media-independent, but also histone H4 acetylation (*EAF3, ESA1, NGG1*, and *ELP4*), which was relatively respiratory-specific in its deletion enhancement (**Fig. 7B**, **Table 2**). Rtt109 promotes H3K56 acetylation, which is associated with elongating RNA polymerase II [60], and can be persistent in the setting of DNA damage [61]. Warburg-independent deletion enhancement suggests its role in DNA repair becomes invoked.

The SAS acetyltransferase complex was deletion suppressing; *SAS2* and *SAS5* were HLEG-specific, and *SAS4* was HLD-specific (**Fig. 7B**). The Sas2 acetyltransferase complex creates a barrier against spread of heterochromatin at telomeres by opposing Sir protein deacetylation via effects on histone H4K16 [62]. The deacetylating SIR proteins (*SIR1, SIR3, SIR4*) were also HLEG-specific suppressors (**Fig. 6C**), suggesting dynamic regulation of telomeric histones (not simply acetylation or deacetylation), or perhaps a function of Sas2 acetyltransferase that is independent of SIR protein functions, confers doxorubicin cytotoxicity in respiring cells.

##### (iv) Histone methylation (Set1C/COMPASS complex)

Histone methylation differentially influences gene transcription, depending on the histone residues modified and the number of methyl groups added [63]. The Set1C/ COMPASS complex, which catalyzes mono-, di-, and tri-methylation of H3K4 [64–67], was enriched in cluster 1-0-7 (**Fig. 4; Table 1**). All genes tested from the Set1C/ COMPASS complex (*SPP1, SDS1, SWD1, SWD3, BRE2, SHG1; SET1* not in YKO/KD) were EG-specific deletion enhancers (**Fig. 7C**). The Set1C/COMPASS complex and H3K4 trimethylation localize at transcription start sites of actively transcribed genes [68, 69]. Furthermore, the Rad6-Bre1 complex, which mono-ubiquitinates histone H2B before Set1C/COMPASS methylates histone H3K4 [70–72], shared the same interaction profile, cross-implicating the Set1C/COMPASS and Rad6-Bre1 functions (**Fig. 7C**). The Rad6-Bre1 complex is additionally involved in the DNA damage response checkpoint to activate Rad53 [73], however, its HLEG-specific enhancing profile was more closely shared with transcriptional regulation modules, indicating its latter role is better related. *JHD1* and *JHD2* are JmjC domain family histone demethylases that act on H3-K36 and H3-K4 respectively, and their deletion suppression interactions are further evidence that histone methylation buffers doxorubicin cytotoxicity, especially in a respiratory context (**Fig. 7C**).

Taken together, the data suggest transcription-associated chromatin regulation buffers doxorubicin-mediated cellular toxicity, which is alleviated by the transition from respiratory to glycolytic metabolism. In contrast, Warburg-independent buffering by histone modifiers appears to be associated with functions related to DNA repair.

#### Mitochondrial functions

The abundance of deletion-enhancing doxorubicin-gene interactions in HLEG media (**Figure 2O**) caused us to closely examine genes annotated to mitochondrial function. Many mitochondrial gene deletion strains grew poorly on HLEG media, with petite-like proliferation defects on HLD media, as respiration is required to reach carrying capacity. Completely respiratory-deficient mutants clustered together in 1-0-0, however, many mitochondrial mutants maintained some or all respiratory capacity. For example, the *respiratory chain complex III assembly* and *protein import into mitochondrial matrix* terms were enriched in deletion enhancing clusters, 1-0-7 and 1-0-8 (Table 1, Figure 4, **Additional File 1**, **Fig. S3**). Some of these strains appeared respiratory sufficient yet the genes were required to buffer doxorubicin cytotoxicity under respiratory conditions. For example, evolutionarily conserved genes functioning in complex IV assembly (*RCF1/YML030W* and *COA6*) reached carrying capacity on HLEG media, yet exerted strong deletion enhancement of doxorubicin growth inhibition (**Additional File 1, Fig. S3A**). In contrast, other HLEG-specific deletion enhancing complex IV assembly components (*COA2*, *CMC1*, *RCF2*) and complex III assembly genes (*FMP25, FMP36, QCR9, CBP4*) were either not conserved in humans or exhibited strong respiratory defects (in absence of doxorubicin) (**Additional File 1**, **Fig. S3A-B**). Interactions specific to assembly of respiratory chain complexes may be informative for studies in cardiomyocytes regarding doxorubicin inhibition, depletion of cytochrome c and cardiolipin, reduced workload capacity, and accelerated aging [74, 75]. Functionally conserved (*TOM70*, *TIM10*, *TIM17, TIM23*, and *MGR2*) and yeast-specific (*TOM6* and *TOM7*) genes in *protein import into mitochondrial matrix* buffered doxorubicin cytotoxicity (**Additional File 1**, **Fig. S3C-E)**, possibly due to increased oxidative stress [76], which enhances doxorubicin toxicity [1, 4].

Systematic examination of the GO annotation *mitochondrion* (**Additional File 1**, **Fig. S4**) revealed several additional respiratory-competent gene-deletion strains exhibiting HLEG-specific enhancing interactions. *COX13* encodes subunit VIa of cytochrome c oxidase, which functions with Rcf1 in the formation of respirasomes (also called ‘supercomplexes’) [77, 78]. Others included *COX8*, encoding subunit VIII of cytochrome c oxidase [79]; *MPC1*, encoding a mitochondrial pyruvate carrier [80, 81]; *MME1*, encoding an inner mitochondrial membrane magnesium exporter [82]; *OMS1*, an inner membrane protein predicted to have methyltransferase activity [83]; *GUF1*, a matrix-localized GTPase that binds mitochondrial ribosomes and influences cytochrome oxidase assembly [84]; and *MIC10 (YCL057C-A)*, encoding a component of the MICOS complex, functioning in inner membrane organization and membrane contact site formation [85].

#### Protein folding, localization, and modification pathways

Protein biogenesis and modification pathways enriched in HLEG-specific enhancement clusters included the *endoplasmic reticulum membrane complex* (**EMC**) (2-0.7-1), *protein urmylation* (2-0.2-1), and N-terminal acetylation by the *NatC complex* (2-0.8-1) (**Figure 4, Table 1**).

##### (i) Protein folding in endoplasmic reticulum (ER membrane protein complex)

The ER membrane complex (*EMC1-6*, **Fig. 8A**) functions in protein folding in the ER [86] and together with the ER-mitochondria encounter structure (**ERMES**), the EMC enables ER-mitochondria phosphatidylserine transfer and tethering [87]. The EMC physically interacts with the mitochondrial translocase of the outer membrane (e.g., *TOM5, 6, 7, 22, 70;* described above) for the process of ER-mitochondria phosphatidylserine transfer [87]. The shared respiratory-specific, deletion-enhancing profiles suggest cooperative functions of the EMC and mitochondrial outer membrane translocase (**Additional File 1, Fig. S3D**) in buffering doxorubicin cytotoxicity. In contrast to the EMC, genes involved in the ERMES complex (1-0-0; **Additional File 5, File B-C**) were essential for respiration, and thus their influence on doxorubicin cytotoxicity could not be addressed with knockout mutants in HLEG media.

**Figure 8.**
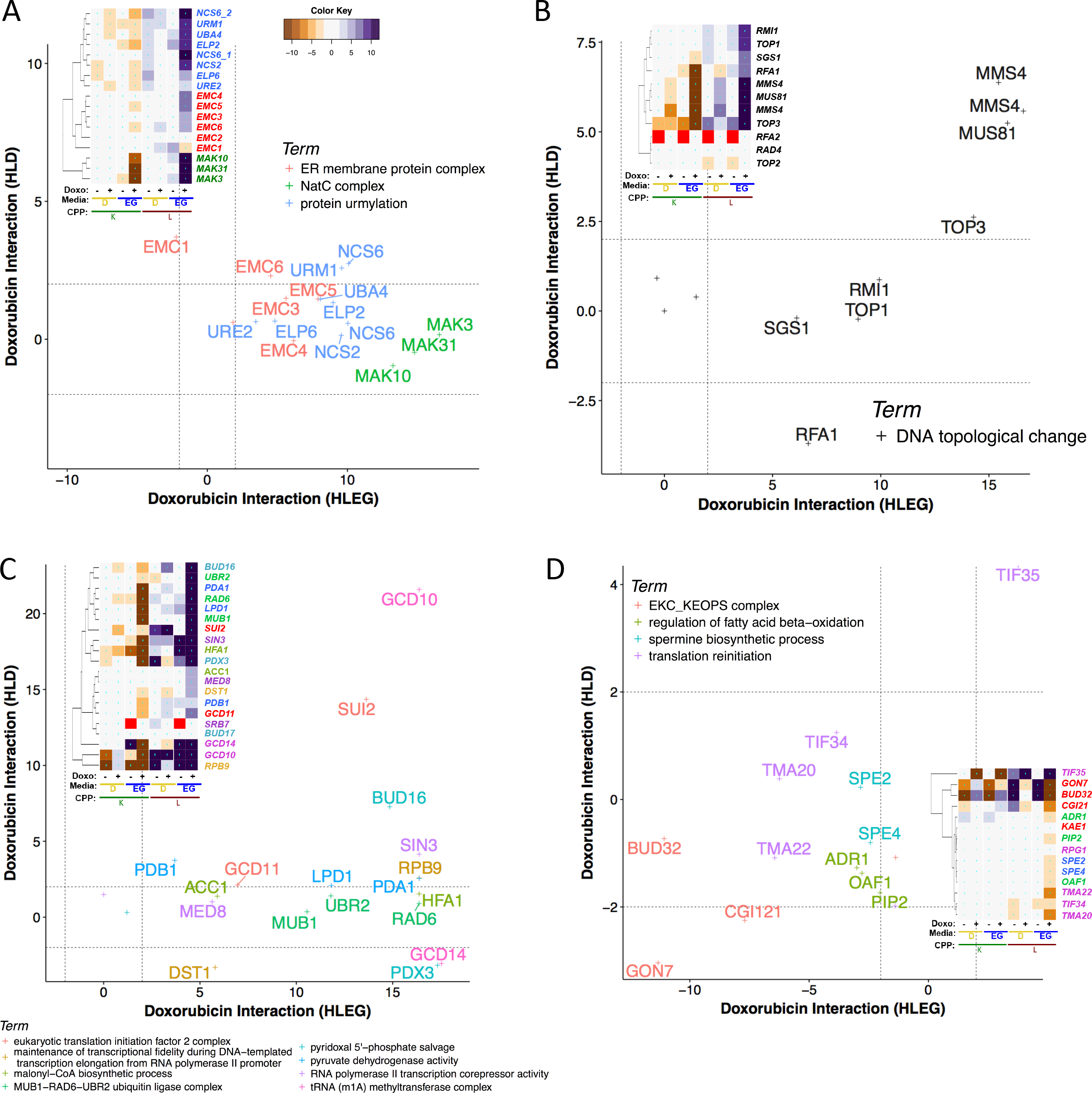
Additional respiration-specific deletion-enhancing and-suppressing functions that influence doxorubicin cytotoxicity. Heatmaps depicting complete phenotypic profiles are inset, corresponding to plots of L-based doxorubicin-gene interaction. (**A**) Protein folding in endoplasmic reticulum and the N-terminal protein-acetylating NatC complex are largely respiratory-dependent in their deletion-enhancing influence. **(B)** DNA topological change exerts deletion-enhancing interactions in both respiratory and glycolytic contexts. **(C)** GTA-identified terms tend to be smaller in number and display greater variability in the Warburg dependence among genes sharing the same functional annotation. **(D)** Functions implicated in respiratory-dependent deletion suppression of doxorubicin toxicity.

##### (ii) Protein urmylation, Elongator complex, and tRNA wobble uridine thiolation

*ELP2*, *UBA4*, *URM1*, and *URE2* clustered together in 2-0.2-1, constituting GO-enrichment in protein urmylation, the covalent modification of lysine residues with the ubiquitin-related modifier, Urm1 [88]. Other protein urmylation genes, *ELP6*, *NCS2*, and *NCS6/YGL211W*, displayed similar interaction profiles and clustered together in 1-0-7 (**Figure 8A**). *ELP2* and *ELP6* also function in the Elongator holoenzyme complex (*IKI1, IKI3, ELP2, ELP3, ELP4*, and *ELP6*) [89–91], associated with similar interaction profiles (**Additional File 1, Fig. S5**). *URM1*, *UBA4, NCS2*, and *NCS6* further function in tRNA wobble position uridine thiolation, where Urm1 functions as a sulfur carrier. Genes uniquely annotated to these terms (*IKI1, IKI3, ELP3, ELP4, TUM1, URE2*) also displayed related profiles (**Additional File 1, Fig. S5**). Thus, protein urmylation, Elongator complex function, and tRNA wobble thiolation appear to be distinct modules, comprised of shared genes, each buffering doxorubicin toxicity in a respiratory-specific way.

##### (iii) N-terminal acetylation by the NatC complex

The NatC complex (Mak3, Mak10, and Mak31) specifically acetylates methionine-starting hydrophobic N-terminal proteins (Met-Leu, Met-Phe, Met-Ile, Met-Tyr) [92], neutralizing positive charge on the alpha-amino group and impeding turnover by ubiquitination or other modifications [93]. N-acetylation occurs on around half of the soluble yeast proteome and over 80% in humans [94]. NatC-mediated N-terminal acetylation facilitates Golgi or inner nuclear membrane localization of some [95–98], but not most proteins [99]. The three genes encoding the NatC complex clustered together (**Fig. 8A**), however, NatC substrates were not enriched among doxorubicin-gene interactions (**Additional File 7, Table S11**). Perhaps a select few NatC targets or a novel function for NatC underlie its compensatory effects.

#### DNA topological change

DNA topological change, which refers to remodeling the turns of a double stranded DNA helix, was enriched in cluster 2-0.8-0 (**Figure 4, Table 1**). Representative genes were *SGS1*, *TOP1*, *RFA1*, *RMI1*, *TOP3, MMS4*, and *MUS81* (**Fig. 8B**). Types I and II topoisomerases resolve supercoiling during replication and transcription [100, 101]. Top1 is a type IB topoisomerase, which relaxes positive and negative supercoils [102, 103], compared to Top3, a type IA topoisomerase that specifically acts on negative supercoiling [104]. The Mms4-Mus81 endonuclease has overlapping functions with Top3 and Sgs1 in DNA repair [105]; however, their respective influences on doxorubicin toxicity were quantitatively distinct in both respiratory and glycolytic contexts, with a greater requirement for the *MMS4/MUS81* than *SGS1, TOP3*, *RFA1*, and *RMI1*; the latter four, functioning together for decatenation and unknotting of dsDNA [106].

#### GTA reveals additional biological functions that buffer doxorubicin toxicity

GTA scores revealed 71 respiratory-specific deletion enhancing GO terms, 24 of which were also found by REMc/GTF (see **Additional File 6, File A**). Strong enhancing terms (GTA value > 10) with functions relatively distinct from those identified above by REMc were *tRNA (m1A) methyltransferase complex*, *MUB1-RAD6-UBR2 ubiquitin ligase complex*, *malonyl-CoA biosynthetic process*, *pyridoxal 5'-phosphate salvage*, *maintenance of transcriptional fidelity during DNA-templated transcription elongation from RNA polymerase II promoter*, *RNA polymerase II transcription corepressor activity*, *pyruvate dehydrogenase activity*, and *eukaryotic translation initiation factor 2 complex* (**Fig. 8C**). Most terms identified by GTA consisted of 2-3 genes, and did not necessarily cluster together by REMc.

#### Respiration-specific gene deletion suppression of doxorubicin cytotoxicity

REMc clusters exhibiting respiration-dependent gene deletion suppression revealed GO Term enrichment for *regulation of fatty acid beta-oxidation*, (cluster 2-0.3-1) and *translation reinitiation* (cluster 2-0.3-5) **(Fig. 4, Table 1**). By GTA analysis, the *EKC/KEOPS complex* and *spermine biosynthetic process* were additionally found to confer HLEG-specific deletion suppression (**Fig. 8D, Table 2**).

##### Regulation of fatty acid beta-oxidation

*ADR1*, *OAF1*, and *PIP2* were grouped together in cluster 2-0.3-1 (**Fig. 4, Table 1**), displaying HLEG-specific gene deletion suppression (**Fig. 8D**). The Pip2-Oaf1 complex binds to oleate response elements, and along with *ADR1*, regulates transcription of peroxisomal genes [107, 108]. Doxorubicin inhibits beta-oxidation of long chain fatty acids in cardiac tissues, which is reversed by supplementing with propionyl-L-carnitine, and alleviates effects of doxorubicin cardiotoxicity [109]. Thus, the yeast model may be informative for investigating related gene networks in greater depth.

##### Translation reinitiation

In the respiratory-specific deletion suppressing cluster 2-0.3-5 (**Fig. 4)**, *TMA20*, *TMA22*, and *TIF34* represented enrichment for translation reinitiation, which is necessary after termination of short upstream open reading frames (**uORFs**) [110] (**Fig. 8D**). Some uORFs function in translational regulation of a downstream protein; for example *GCN4* expression is regulated in response to amino acid starvation [110]. However, using the Welsh two sample t-test we found no significant difference in means of interaction scores between the distribution of proteins regulated or not by uORFs [111] (p-value = 0.8357) (**Additional File 7, Table S12**).

##### Spermine biosynthetic process

Loss of spermine biosynthesis, specifically *SPE2* (S-adenosylmethionine decarboxylase) and *SPE4* (spermine synthase), suppressed doxorubicin toxicity in HLEG media (**Fig. 8D**). The pathways of polyamine metabolism and their physiologic effects on cancer are complex [112, 113], and although our data suggest spermine metabolism contributes to doxorubicin cytotoxicity, how this occurs mechanistically and specifically in respiring cells awaits further study [114].

##### EKC/KEOPS complex

GTA revealed the EKC/KEOPS complex (*CGI121, GON7*, and *BUD32*) as HLEG-specific deletion suppressing (**Fig. 8D**). The EKC/KEOPS complex is involved in threonyl carbamoyl adenosine (t6A) tRNA modification [115], which strengthens the A-U codon – anticodon interaction [116]. EKC/KEOPS has also been characterized with respect to telomere maintenance [117] and transcription [118]. Deletion of *GON7, BUD32*, or to a lesser extent, *CGI121*, inhibited cell proliferation in the absence of doxorubicin treatment, indicating that translational and/or transcriptional activity of the EKC/KEOPS complex function contributes to doxorubicin sensitivity.

#### Glycolysis-specific gene deletion enhancement of doxorubicin cytotoxicity

HLD-specific deletion enhancement of doxorubicin cytotoxicity could represent lethal vulnerabilities that emerge when a tumor undergoes the Warburg transition. In this regard, several genes, but few enriched GO terms were identified by REMc (**Fig. 4**, clusters 1-0-5, 2-0.3-0, and 2-0.2-2; **Additional File 5, File A**). *Ribonucleoprotein complex subunit organization* was suggested (**Table 1)**, however, the term-specific heatmap revealed doxorubicin-gene interaction within this cellular process to be pleiotropic (**Additional File 1, Fig. S6**).

##### Glycolysis-specific deletion enhancing terms identified by GTA

GTA analysis revealed HLD-specific deletion-enhancing genes encoding the Cul4-RING E3 ubiquitin ligase, the Dom34-Hbs1 complex, and the Ubp3-Bre5 deubiquitinase. *GDP-Mannose Transport* and *dTTP biosynthesis* were also revealed (Fig. 9A; **Supplemental File 6, File A**). *SOF1, HRT1*, and *PRP46* were computationally inferred to form the Cul4-RING E3 ubiquitin ligase complex [119]. Yeast Sof1 is an essential protein that is required for 40s ribosomal biogenesis, and overexpression of its human ortholog, *DCAF13/WDSOF1*, is associated with aggressive tumors and poorer survival in hepatocellular carcinoma [120]. *DOM34/PELO* and *HBS1/HBS1L* facilitate recycling of stalled ribosomes by promoting dissociation of large and small subunits through a process called no-go decay [121–123]. Knockdown by siRNA of either *WDSOF1* or *HBS1L* was synthetic lethal in a KRAS-driven tumor model [124]. The Ubp3-Bre5 deubiquitination complex regulates anterograde and retrograde transport between the ER and Golgi [125, 126]. Vrg4 and Hvg1 transport GDP-mannose into the Golgi lumen for protein glycosylation [127, 128]. Reduced dTTP pools, evidenced by *CDC8/DTYMK* and *CDC21/TYM*S, can increase doxorubicin cytotoxicity in cancer cell lines [129]. The human homologs of *UBP3*, *CDC8*, and *CDC21* were identified in genome-wide siRNA synthetic interaction studies in cancer cell line models [130–132].

**Figure 9.**
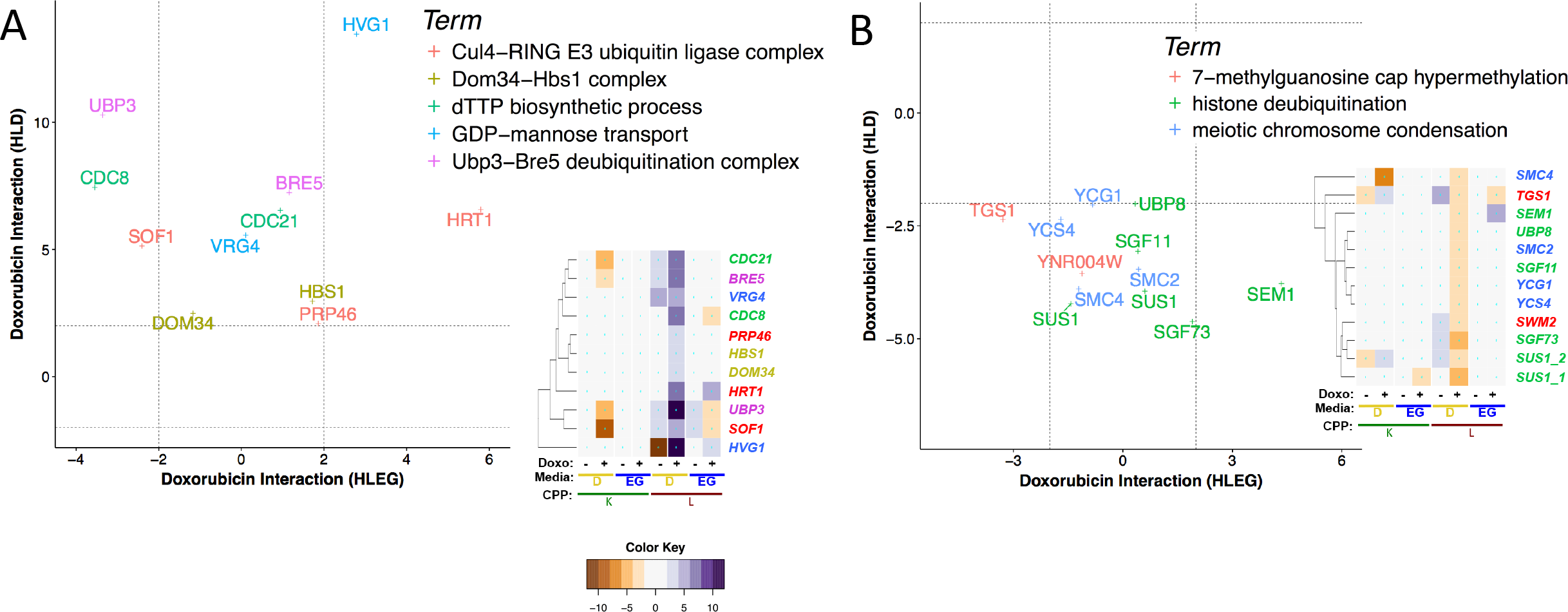
Glycolysis-specific enhancement and suppression of doxorubicin cytotoxicity. Doxorubicin-gene interaction profiles for HLD-specific GO terms identified by GTA are depicted for **(A)** deletion enhancement and **(B)** deletion suppression.

For several examples above, like *SOF1/DCAF13*, genes could be targeted as both a driver of the tumor and as a sensitizer to doxorubicin. To systematically identify all candidate vulnerabilities specific to glycolytic tumor cells (not constrained by GO enrichment), we filtered the overall data set, limiting the list to genes with human homologs and to YKO/KD strains that were growth sufficient (low shift on HLD) (**Additional File 1, Fig. S7)**. The human homologs, along with functional descriptions, are provided in **Additional File 10, Table S13**.

#### Glycolysis-specific gene deletion suppression of doxorubicin cytotoxicity

HLD-specific deletion suppression clusters (**Fig. 4**, clusters 2-0.1-0, 2-0.4-0, 2-0.4-2, and 3-0.3.3-1) had GO Term enrichment for terms related to mRNA processing and *meiotic chromosome condensation*. GTA also identified *histone deubiquitination* (**Table 2**). Deletion suppression points to genes that could potentially increase doxorubicin toxicity if overexpressed.

##### RNA processing

HLD-specific deletion suppression clusters (2-0.4-0, 2-0.4-2**; Fig. 4**) were enriched for mRNA processing-related terms including *mRNA 3’ end processing*, *mRNA cleavage*, and *7-methylguanosine cap hypermethylation* (**Table 1**), but the term-specific heatmaps revealed pleiotropic gene interaction profiles (**Additional File 1, Fig. S8**).

*SWM2/YNR004W* and *TGS1* function in 7-methylguanosine (m^7^G) cap trimethylation (cluster 2-0.4-0), however, the *tgs1*-Δ*0* allele also exerted deletion suppression in a respiratory context (**Fig. 9B**). m^7^G cap trimethylation protects small nuclear RNAs (**snRNAs**), and small nucleolar RNAs (**snoRNAs**) from degradation by exonucleases [133, 134], and promotes efficient pre-rRNA processing and ribosome biogenesis [135].

##### Meiotic chromosome condensation

*SMC2*, *SMC4*, *YCG1*, and *YCS4* constitute the nuclear condensin complex, which functions in chromosome condensation and segregation. The condensin complex associates with chromosomal sites bound by TFIIIC and the RNA Pol III transcription machinery [136], where it facilitates clustering of tRNA genes at the nucleolus [137] (**Fig. 9B**). The condensin complex has been suggested as a potential therapeutic target for cancer [138], and human homologs *YCG1/NCAPG2*, *YCS4/NCAPD2*, and *SMC4/SMC4* are synthetic lethal with the Ras oncogene [124].

##### Histone deubiquitination

Histone deubiquitination was identified by GTA and includes *SUS1, SGF11, SGF73, UBP8, and SEM1* (**Fig. 9B)**; all except *SEM1* are part of the DUBm complex, which mediates histone H2B deubiquitination and mRNA export [139]. Loss of histone H2B ubiquitination resulting in HLEG-specific enhancement (**Fig. 7C**) is consistent with loss of the DUBm deubiquitinase being suppressing. Together, they implicate regulation by histone ubiquitination as a mechanism of doxorubicin response. The human homologs of *UBP8, USP22* and *USP51*, were identified in an RNAi screen for resistance to ionizing radiation [140].

### Warburg transition-independent doxorubicin gene-interaction modules

Since many tumors have both respiratory and glycolytic cell populations, targeting Warburg-independent interactions could be especially efficacious, as described below.

#### Deletion enhancement

Cluster 1-0-6 (**Fig. 4**) had a strong deletion-enhancing profile in both metabolic contexts with GO Term enrichment for DNA repair (**Fig. 10**), as well as histone acetylation (discussed above, **Fig. 7B**). GTA analysis additionally revealed the Lst4-Lst7, the Cul8-RING ubiquitin ligase, and MCM complexes (**Fig. 10B**).

**Figure 10.**
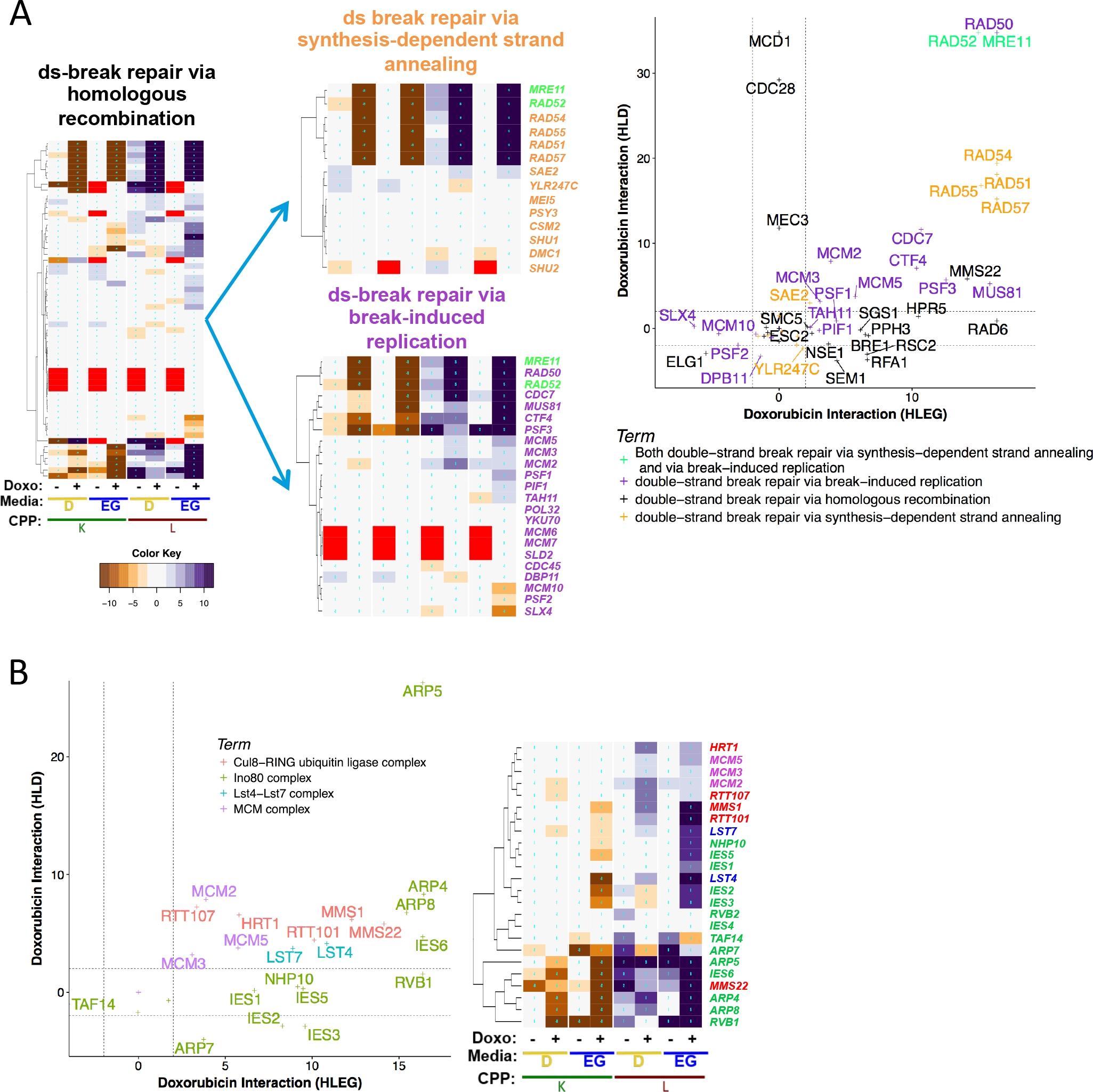
Warburg-independent deletion enhancement of doxorubicin cytotoxicity. Gene interaction profiles showing deletion enhancement in both respiratory and glycolytic context included: **(A)** *double-strand break repair via homologous recombination*, and its child terms (indicated by arrows), and **(B)** the *Cul8-RING ubiquitin ligase*, *Ino80 complex*, *Lst4-7 complex*, and *MCM complex*.

##### DNA repair

Warburg-independent, deletion-enhancing pathways included homologous recombination and break-induced replication repair (**Fig. 10A)**, along with the Ino80 complex (**Fig. 10B**), the latter explained by its role of histone acetylation in recruitment of DNA repair machinery to dsDNA break sites [52]. The Ino80 complex influences doxorubicin response in fission yeast [141, 142], further suggesting evolutionary conservation of this interaction, and thus potential relevance to mammalian systems [143]. DNA repair pathways, such as those involving *RAD52* and *INO80*, are evolutionarily conserved, involved in genome instability and tumorigenesis [144], and predictive of therapeutic response in some cancers [145], thus representing potential tumor-specific biomarkers for chemotherapeutic efficacy.

##### Complexes identified by GTA

Warburg-independent deletion enhancing modules identified by GTA were weaker, in many cases, than the dsDNA break repair pathways found by REMc, some of which had strong K parameter interactions (Fig. 10, **Additional File 9**). GTA-identified terms included: (1) The Cul8-RING ubiquitin ligase complex, which is encoded by *RTT101*, *RTT107*, *MMS1*, *MMS22*, and *HRT1*, and functions in replication-associated DNA repair [146]. Cul8/Rtt101, in fact, contributes to multiple complexes that regulate DNA damage responses, including Rtt101‐Mms1‐Mms22, which is required for Eco1-catalyzed Smc3 acetylation for normal sister chromatid cohesion establishment during S phase [147]; (2) The Lst4-Lst7 complex, which functions in general amino acid permease (*GAP1*) trafficking [148], threonine uptake, and maintenance of deoxyribonucleotide (**dNTP**) pools [19], clustered with *thr1*-Δ*0* (threonine biosynthesis) in 2-0.2-1 (**Additional File 5, File B**); and (3) the mini-chromosome maintenance (**MCM**) complex, which licenses and initiates DNA replication [149], was evidenced by the *mcm2-DAmP*, *mcm3-DAmP*, and *mcm5-DAmP* YKD strains (**Fig. 10B**). Work in pea plants showed that doxorubicin inhibits the *MCM6* DNA helicase activity [150]. Prior genome-wide experiments with doxorubicin did not analyze YKD mutants, thus the MCM complex highlights the utility of the DAmP collection in drug-gene interaction studies.

#### Media-independent deletion suppression

Genes functioning in processes that augment doxorubicin toxicity, when lost (*e.g.*, by deletion), can result in suppression. This was suggested in both respiratory and glycolytic contexts for *sphingolipid homeostasis*, *telomere tethering at nuclear periphery*, and *actin cortical patch localization* (**Fig. 4**, clusters 2-0.4-1 and 2-0.3-3). Conversely, their overexpression in cancer could potentiate toxicity and thereby therapeutic efficacy.

##### Sphingolipid homeostasis and metabolism

From cluster 2-0.4-1, *VPS51, VPS52, VPS53*, and *VPS54* (**Fig. 11A**) form the Golgi-associated retrograde protein (**GARP**) complex, which is required for endosome-to-Golgi retrograde vesicular transport. GARP deficiency results in accumulation of sphingolipid synthesis intermediates [151]. Also from this cluster came fatty acid elongase activity (*FEN1/ELO2* and *SUR4/ELO3*), which when deficient leads to reduced ceramide production and phytosphingosine accumulation [152, 153].

**Figure 11.**
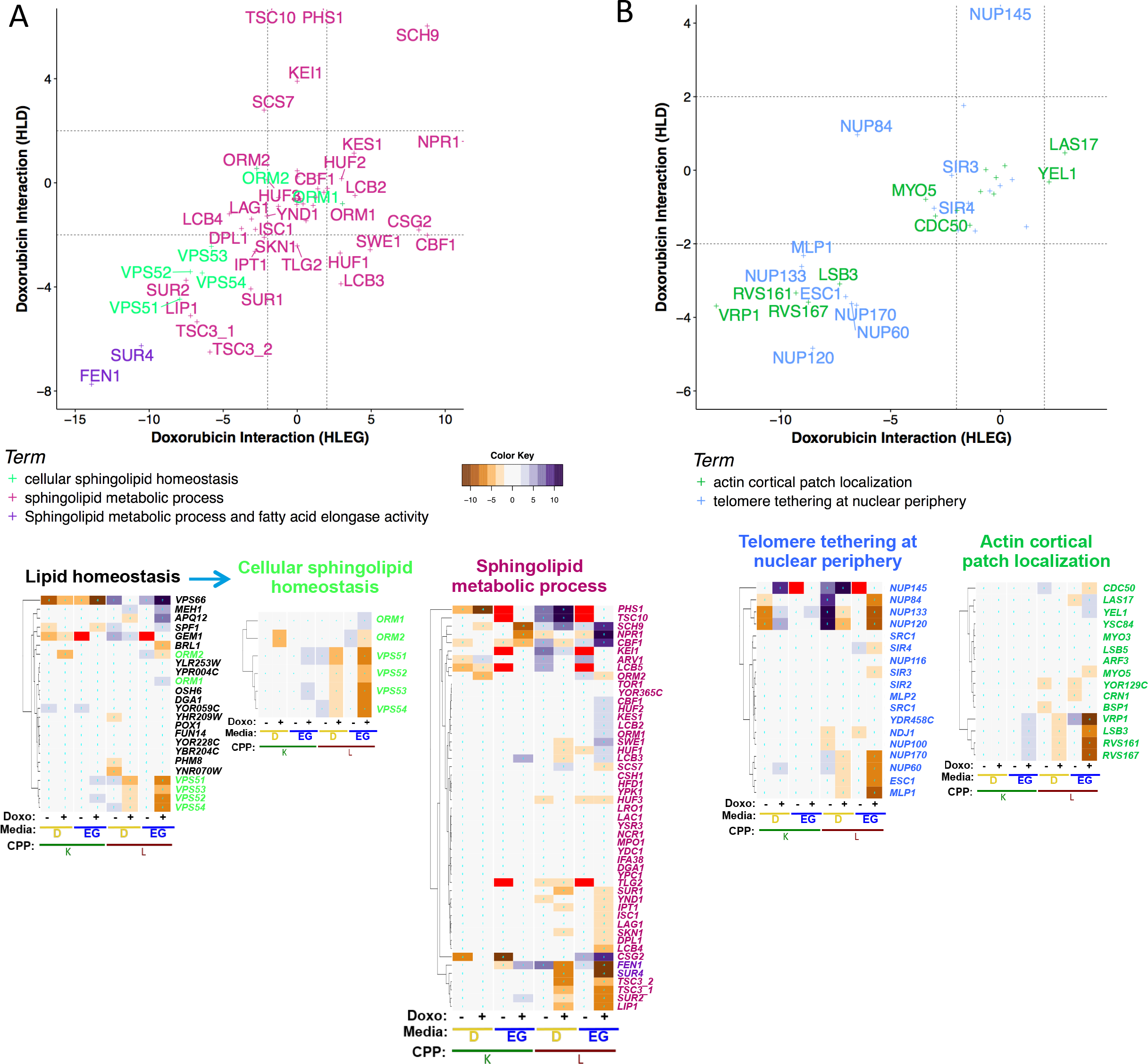
Warburg-independent deletion suppression of doxorubicin cytotoxicity. Doxorubicin-gene interaction profiles and L-interaction plots for genes associated with deletion suppression in HLEG or HLD media, including: (**A**) *Cellular sphingolipid homeostasis*, along with its parent term, *lipid homeostasis*, and related term *sphingolipid metabolism*; and **(B)** *actin cortical patch localization* and *telomere tethering at nuclear periphery*.

Since the GARP genes and fatty acid elongase activity genes function together in sphingolipid metabolism, we searched all genes annotated to this term and found other media-independent suppressors to include *TSC3*, *LIP1*, *SUR1*, *SUR2*, *IPT1*, and *SKN1* (**Fig. 11A**). Doxorubicin treatment induces accumulation of ceramide [5, 6], which mediates anti-proliferative responses and apoptosis in yeast and human and appears to mechanistically underlie the influence of this gene group [154] (**Additional File 1, Fig. S9**). These findings were further supported by the deletion enhancer, *SCH9*, which negatively regulates ceramide production by inducing ceramidases and negatively regulating *ISC1* (**Fig. 11A**) [155]. Multidrug-resistant HL-60/MX2 human promyelocytic leukemia cells are sensitized to doxorubicin by N,N-Dimethyl phytosphingosine [156].

Taken together, the model provides genetic detail regarding how disruption of sphingolipid metabolism increases resistance to doxorubicin, and that this occurs in a Warburg-independent manner, seemingly by reducing apoptosis associated with doxorubicin-induced ceramide overproduction [5, 157, 158].

##### Telomere tethering at nuclear periphery

Enrichment for *telomere tethering at nuclear periphery* in cluster 2-0.4-1 was comprised of *NUP60, NUP170, MLP1*, and *ESC1*. Paradoxically growth deficient on HLD media, *NUP84*, *NUP120*, and *NUP133* also exerted deletion suppression in HLEG (**Fig. 11B**). Nuclear pore functions include coordinating nuclear-cytoplasmic transport and localizing proteins and/or chromosomes at the nuclear periphery, which contributes to DNA repair, transcription, and chromatin silencing [159]. Thus, deletion of nuclear pore genes could influence doxorubicin resistance by multiple potential mechanisms involving altering chromatin states, transcriptional regulation, maintenance of telomeric regions, and DNA repair. Doxorubicin-gene interaction profiles for all nuclear pore-related genes are provided in **Additional File 1, Fig. S10 A**.

##### Actin cortical patch localization

Cluster 2-0.4-1 was enriched for *actin cortical patch localization*, including *RVS167*, *LSB3*, *RVS161*, and *VRP1* (**Fig. 11B**). Related terms (*Arp2/3 protein complex* and *actin cortical patch*) exhibited similar doxorubicin-gene interaction profiles, including *ARC15*, *ARC18*, *ARC35*, *INP52*, *INP53*, *ARP2*, *ARP3*, *GTS1*, *RSP5*, and *FKS1 (*see **Additional File 1, Fig. S10 B-C**). This result corroborates studies in mouse embryonic fibroblasts where deletion of *ROCK1* increased doxorubicin resistance by altering the actin cytoskeleton and protecting against apoptosis [160, 161]. Additional literature indicates an importance of actin-related processes for doxorubicin cytotoxicity [162–164], highlighting the utility of yeast phenomics to understand these effects in greater depth.

#### Respiratory-deficient doxorubicin-gene interaction modules

From cluster 1-0-0, we noted that respiratory deficient YKO/KD strains (those not generating a growth curve on HLEG) also had low K and/or increased L ‘shift’ values on HLD, as would be expected of petite strains [165]. Among strains in this category, those displaying doxorubicin-gene interaction tended to show deletion enhancement (**Fig. 4**).

Respiratory deficient deletion enhancers on HLD functioned primarily in mitochondrial processes (**Additional File 5, File C**; see GO enrichment for cluster 1-0-0 and derivative clusters), including *mitochondrial translation*, *mitochondrion-ER tethering*, *protein localization into mitochondria*, *mitochondrial genome maintenance*, *respiratory chain complex assembly*, and *proton transport*. Compromise of mitochondrial respiration leading to sensitization of cells to doxorubicin is of interest given recent findings that some glycolytic cancers are respiratory deficient [166, 167].

#### Phenomics-based predictions of doxorubicin-gene interaction in cancer cell lines

Differential gene expression, by itself, is a poor predictor of whether protein function affects proliferative response to a particular drug [168]. Thus, yeast phenomic data, which precisely measures enhancing and suppressing interactions with respect to growth phenotypes, could provide a systems model to prioritize candidate effectors of cancer cell line sensitivity and transcriptomic data [169, 170]. To investigate this possibility, yeast doxorubicin-gene interaction was matched by homology to differential gene expression in doxorubicin-sensitive cancer cell lines, using *PharmacoGx* [40] and *biomaRt* [41, 42]) in conjunction with the GDSC1000 [171, 172] or gCSI [173, 174] databases (**Fig. 12)**. Differential gene expression was analyzed for individual tissues and also aggregated across all tissues. Yeast gene deletion enhancers were matched to human homologs underexpressed in doxorubicin-sensitive cancer cell lines, termed ‘**UES**’. Conversely, yeast gene deletion suppressors were matched to human homologs overexpressed in doxorubicin sensitive cells, termed ‘**OES**’ (**Additional File 11)**.

**Figure 12.**
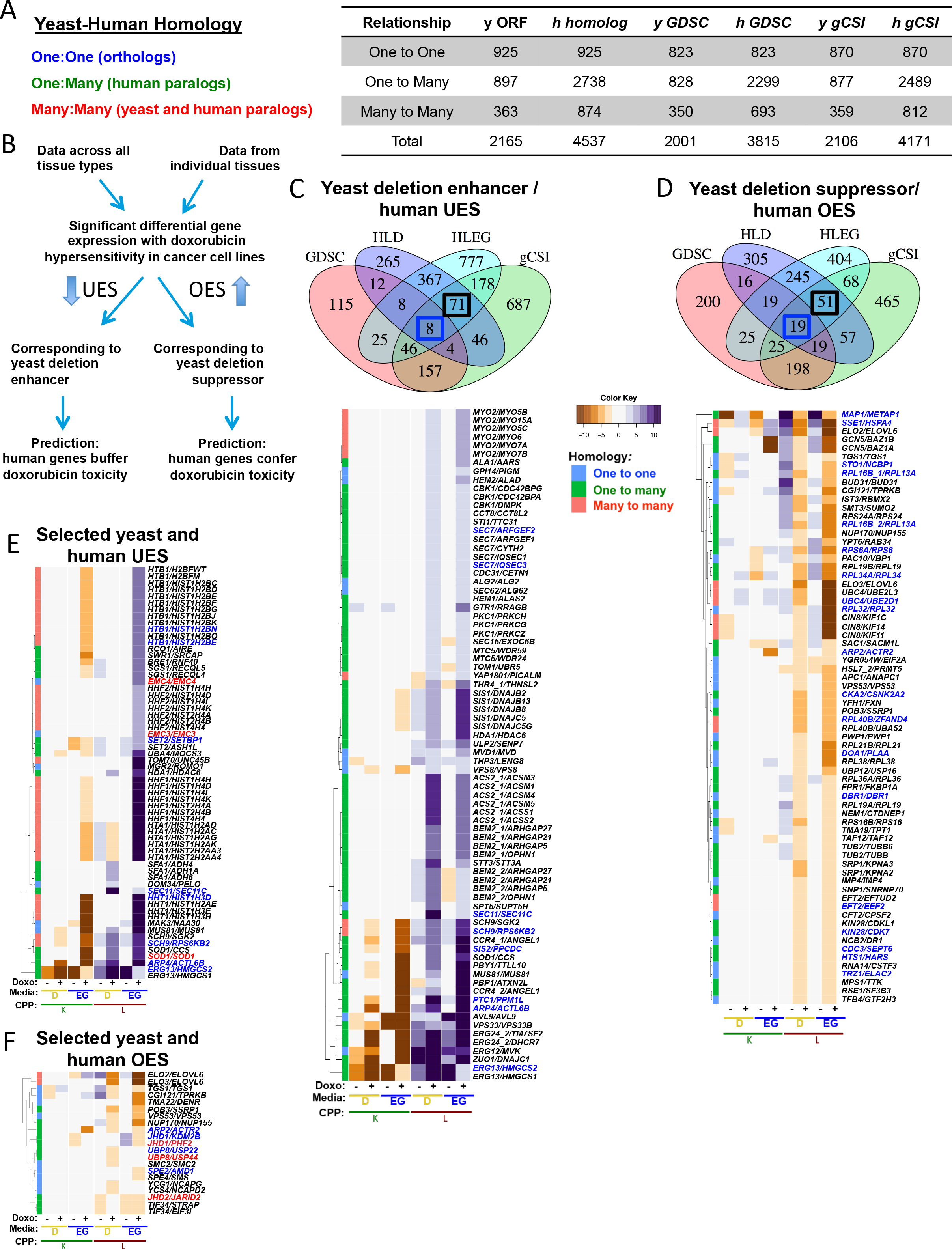
Use of the yeast phenomic model to predict doxorubicin-gene interaction in cancer cells. **(A)** *BiomaRt* was used to assign yeast-human gene homology for the GDSC and gCSI datasets. (**B**) *PharmacoGx* was used to retrieve differential gene expression for doxorubicin sensitive cell lines from the gCSI and GDSC databases, searching data from individual tissues or across data aggregated from all tissues. Human genes that are underexpressed in doxorubicin sensitive cell lines (UES) with yeast homologs that are deletion enhancers are predicted to be causal in their phenotypic association. Similarly, human genes that are overexpressed in doxorubicin sensitive cancer cell lines (OES) would be predicted to be causal if the yeast homolog was a deletion suppressor in the phenomic dataset. (**C-D**) Boxes inside of Venn diagrams indicate the genes for which gene interaction profiles are shown in the heatmaps below. Gene names are to the right of heatmaps, with **blue** labels indicating genes identified in both the GDSC and gCSI databases and **black** labels indicating genes found only in the gCSI dataset. The category of homology (see panel A) is indicated in the left column of each heatmap. **(C)** Deletion enhancement by yeast genes predicts human functions that buffer doxorubicin cytotoxicity, and thus, reduced expression of homologs in cancer cell lines is predicted to increase doxorubicin sensitivity. (**D**) Deletion suppression by yeast genes predicts functions that mediate cytotoxicity and is shown for human homologs having significant association of overexpression in cancer cell lines with increased doxorubicin sensitivity. (**E-F**) Genes representing enhancing or suppressing modules from REMc or GTA that are **(E)** UES or **(F)** OES in at least one of the two databases. **Red** labels indicate genes found only in the GDSC database. **Additional File 13** reports all results from the analysis described above, including assessment of individual tissues.

There was greater overlap in differential gene expression between the gCSI and GDSC databases for aggregated data (compared to data for individual tissues), of which agreement was greater for OES than UES. Among individual tissues, there was highest agreement between hematopoietic/lymphoid and lung. Differences could be partially explained by the two studies using different platforms for measuring gene expression and cell cytotoxicity (https://pharmacodb.pmgenomics.ca/drugs/273). The gCSI database also reported more UES and OES genes than GDSC (**Additional File 11, Files B and C**). Differentially expressed genes were mined with respect to p-value and matched to homologous yeast gene interactions (**Additional File 11, Files D-I**). Warburg status was not available for the cancer cell lines, so we first matched Warburg-independent yeast gene interactions to differentially expressed genes from aggregated data in both the gCSI and GDSC datasets, predicting eight UES (*ARP4/ACTL6B, ERG13/HMGCS2, PTC1/PPM1L, SCH9/RPS6KB2, SEC11/SEC11C, SEC7/ARFGEF2, SEC7/IQSEC3*, and *SIS2/PPCDC*) and 18 OES genes (*ARP2/ACTR2*, *CDC3/SEPT6*, *CKA2/CSNK2A2*, *DBR1/DBR1*, *DOA1/PLAA*, *EFT2/EEF2*, *HTS1/HARS*, *KIN28/CDK7*, *MAP1/METAP1*, *RPL16B/RPL13A*, *RPL32/RPL32, RPL34A/RPL34, RPL40B/ZFAND4*, RPS6A/RPS6, *SSE1/HSPA4*, *STO1/NCBP1*, *TRZ1/ELAC2*, and *UBC4/UBE2D1*) to have causal influences on the doxorubicin sensitivity phenotype (**Fig. 12C-D)**.

As detailed in **Tables 3–4** and described below, we expanded the analysis to genes representative of GO Term enrichments revealed by the yeast phenomic model, restricting to human genes differentially expressed across all cancer tissues, but without restricting by Warburg-independence or gCSI/GDSC co-evidence. Results for individual tissues are also provided in **Additional File 11, File A**.

**Table 3.**
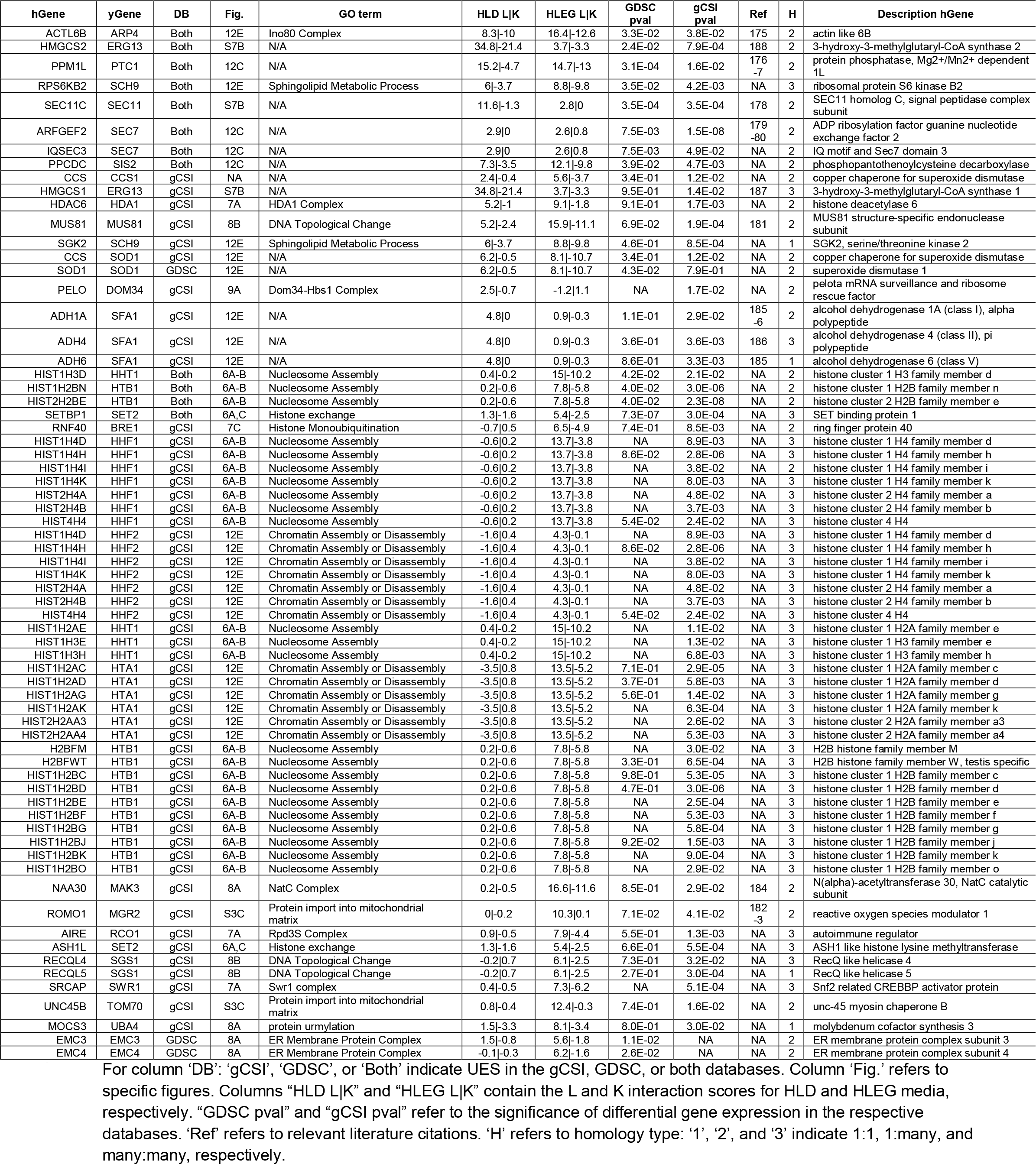
Yeast-human homologs with deletion enhancement and UES across all tissues.

**Table 4.**
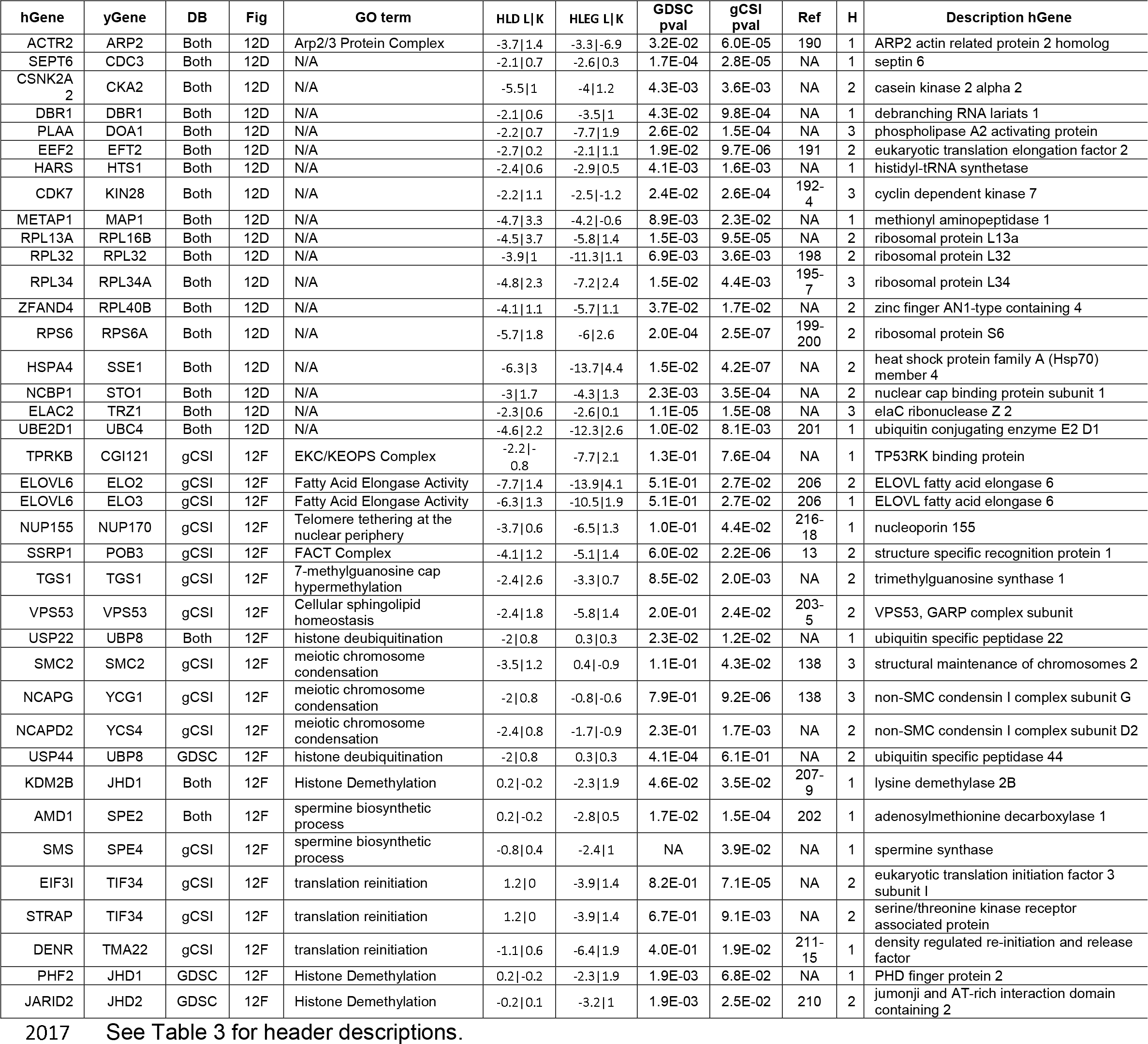
Yeast-human homologs with deletion suppression and OES across all tissues.

##### Deletion enhancers with UES homologs

Concordance between deletion-enhancing doxorubicin-gene interaction in yeast and UES observed for the corresponding human homologs in cancer cells suggests synergistic targets and biomarkers to increase therapeutic efficacy for doxorubicin, as summarized in **Table 3 and Fig. 12C**, and briefly discussed below.

Doxorubicin-enhancing interactions that were UES in both gCSI and GDSC included: *ACTL6B*, identified as a candidate tumor suppressor gene in primary hepatocellular carcinoma tissue [175]; *PPM1L*, which regulates ceramide trafficking at ER-Golgi membrane contact sites [176], and exhibits reduced expression in familial adenomatous polyposis [177]; *RPS6KB2*, which was UES in breast, ovarian and bone in gCSI, while *RPS6KA1*, *A2*, *A5* and *A6* were UES in select tissues in both databases (**Additional File 11, File A);***SEC11/SEC11C*, which is upregulated in response to hypoxia in non-small cell lung cancer tissue [178], and for which deletion enhancement was stronger in HLD media (**Additional File 1, Fig. S7**); *SEC7/ARFGEF2* (alias *BIG2*) exhibits increased gene and protein expression in pancreatic cancer [179], and shRNA knockdown of *ARFGEF2* can reduce Burkitt’s lymphoma cell survival [180].

We expanded the analysis above by matching yeast gene deletion enhancers to human UES genes in either database, i.e., not requiring that genes be significant in both datasets (**Figs 12E-F**). The result highlighted chromatin-related buffering processes, including nucleosome assembly (*HTA1, HTB1, HHF1, HHF2, HHT1, HHF1*), histone exchange (*SET2/SETBP1* and *SWR1/SRCAP*), and histone modifiers (*BRE1, HDA1, RCO1*) (**Fig. 12E, Table 3**). Other functions predicted by the yeast model to buffer doxorubicin toxicity in cancer cells included DNA topological change (*MUS81, SGS1*), mitochondrial maintenance (*MGR2, TOM70*), protein acetylation (*MAK3*), and metabolism (*SFA1, ERG13, SOD1*).

*MUS81* knockdown enhances sensitivity of colon cancer lines to cisplatin and other chemotherapy agents by activating the *CHK1* pathway [181]. *MGR2/ROMO1* is involved in protein import into the mitochondrial matrix and overexpression of *ROMO1* has been associated with poor prognosis in colorectal [182] and non-small cell lung cancer patients [183]. *MAK3/NAA30*, a component of the NatC complex (**Fig 8A**), induces p53-dependent apoptosis when knocked down in cancer cell lines [184]. The HLD-specific deletion enhancer, *SFA1*, has seven human homologs, of which three (*ADH4, ADH1A*, and *ADH6*) were UES in gCSI data (**Additional File 1, Figure S7)**‥ High expression of *ADH1A* or *ADH6* was predictive of improved prognosis for pancreatic adenocarcinoma [185] and high expression of *ADH1A* or *ADH4* had improved prognosis for non-small cell lung cancer [186]. The *ERG13* homolog, *GCS1*, has been suggested as a synthetic lethal target for BRAF^V600E^-positive human cancers [187], and *HMGCS2* plays a role in invasion and metastasis in colorectal and oral cancer [188]. Thus, doxorubicin treatment may have anti-tumor efficacy specifically in glycolytic tumors with reduced expression of *SFA1* and *ERG13* homologs.

##### Deletion suppressors with OES homologs

Choosing chemotherapeutic agents for patients based on their tumors exhibiting high expression of genes known to increase sensitivity represents a targeted strategy to increase therapeutic efficacy and could be particularly effective if the sensitizing overexpressed genes happen to also be drivers [189]. Human genes that are OES, homologous to yeast genes that are deletion suppressors, are highlighted in **Table 4** and **Fig. 12D**. *ARP2/ACTR2* is a member of the Arp2/3 protein complex (see **Additional File 1, Figure S10C)**, and silencing of the Arp2/3 protein complex reduces migration of pancreatic cancer cell lines [190]. *EEF2* protein is overexpressed in multiple cancer types, where shRNA knockdown inhibits growth [191]. *CDK7* overexpression in breast [192, 193] and gastric [194] cancer is predictive of poor prognosis. *RPL34* overexpression promotes proliferation, invasion, and metastasis in pancreatic [195], non-small cell lung [196], and squamous cell carcinoma [197], while *RPL32* was also overexpressed in a prostate cell cancer model [198]. In contrast to Rps6k family members being UES/deletion enhancing, Rps6 was OES/deletion suppressing in ovarian tissue. *RPS6* overexpression portends reduced survival for patients with renal carcinoma [199] and hyperphosphosphorylation of Rps6 confers poor prognosis in non-small cell lung cancer [200]. Overexpression of *UBE2D1* is associated with decreased survival in lung squamous cell carcinoma tissue [201], and numerous additional ubiquitin-conjugating enzyme family members were OES in analysis of individual tissues (**Additional File 11, File A**).

We expanded the analysis, similar to the way described above for the deletion enhancers, by relaxing the matching criteria in order to identify additional deletion suppressing pathways revealed by the yeast model (**Additional File 11**). The extended analysis identified yeast-human conserved functions in metabolism (*SPE2, SPE4, VPS53, ELO2, ELO4*), histone demethylation (*JHD1, JHD2*), translation reinitiation (*TMA22, TIF32*), the condensin complex (*YCG1*, *YCS4*, *SMC2*), and telomere tethering at the nuclear periphery (*NUP170*) (**Table 4, Fig. 12F)**. *SPE2/AMD1* is required for spermidine and spermine biosynthesis, and up-regulation of *AMD1* by mTORC1 rewires polyamine metabolism in prostate cancer cell lines and mouse models [202]. *VPS53*, a component of the GARP complex involved in sphingolipid homeostasis, is a tumor suppressor in hepatocellular carcinoma [203–205]. Inhibition of *ELOVL6* (homologous to yeast *ELO2* and *ELO3*) in mice reduces tumor growth and increases survival [206]. The histone demethylase, *JHD1/KDM2B*, is overexpressed in pancreatic cancer [207] and is associated with poor prognosis in glioma [208] and triple negative breast cancer [209]. A second homolog, *JHD2/JARID2*, is required for tumor initiation in bladder cancer [210]. The yeast model also predicts causality underlying OES associated with genes involved in translation reinitiation, *TMA22/DENR* (translation machinery associated) and *TIF32/EIF31. DENR-MCT-1* regulates a class of mRNAs encoding oncogenic kinases [211–213], and its overexpression in hepatocellular carcinoma is associated with metastasis [214]. *TMA22/DENR* also exerts evolutionarily conserved influence on telomeric function and cell proliferation [215]. *YCG1/NCAPG* and *SMC2/SMC2* are components of the condensin complex, which are overexpressed in cancer [138]. *NUP170/NUP155*, which functions in telomere tethering at the nuclear periphery (**Fig. 11B**), is hyper-methylated in association with breast cancer [216, 217], where its reduced expression contributes to a signature for bone metastasis [218].

## Discussion

Many genes are implicated in tumorigenesis and in chemotherapeutic response, with varying degrees of tissue-specific influence and yeast-human homology. The ability to assess mutation, differential gene expression, and other molecular correlates of cancer and chemotherapeutic efficacy is growing, but the direct assessment of drug-gene interaction (i.e., phenotypic/cell proliferative responses) remains a challenge due to the complex genetics and tissue-specific aspects of cancer. In stark contrast, yeast is a single-cell eukaryotic organism that is uniquely amenable to precise and genome-wide measures of drug-gene interaction, for which fundamental contributions to our understanding of human disease are well established [219–223]. Thus, we wondered whether phenomic analysis, using the yeast YKO/KD resource, might be informative about the potential of the Warburg effect to influence the anti-cancer efficacy of doxorubicin, and potentially other chemotherapeutic agents [25, 224]. From this unbiased systems perspective, we observed that a less extensive genetic network is required to buffer doxorubicin in glycolytic vs. respiring cells. The HLEG-specific doxorubicin-gene interaction network points to genetic vulnerabilities that respiratory tumors have, but that can be relieved of by the Warburg transition to glycolytic metabolism. Thus, the yeast phenomic model could be applied in the context of Warburg status and analysis of somatic mutations in an individual patient’s cancer, to aid in predicting doxorubicin therapeutic efficacy (**Fig. 13, Tables 3–4**). Cells can buffer the cytotoxic effects of doxorubicin in a glycolytic context with less reliance on pathways that can go awry in cancer and that influence doxorubicin cytotoxicity more in a respiratory context; examples include pathways of chromatin regulation, protein folding and modification, mitochondrial function, and DNA topological change. The yeast model also predicts that respiring tumors can better survive doxorubicin if functions for fatty acid beta-oxidation, spermine metabolism, and translation reinitiation are compromised by mutation (**Fig. 13, Table 3**). On the other hand, cells that transition to glycolytic metabolism need dTTP biosynthesis and protein complexes including the Cul4-RING E3 ubiquitin ligase, and the Ubp3-Bre5 deubuiquitinase, as well as Dom34-Hbs1, which functions in ‘no-go’ mRNA decay, in order to buffer doxorubicin. However, glycolytic cells become even more resistant if losing histone deubiquitination or the nuclear condensin complex (**Fig. 13, Table 3**). The yeast model also highlighted Warburg-independent pathways for which loss of function enhanced doxorubicin cytotoxicity, such as DNA repair and histone H3-K56 acetylation, along with deletion suppressing pathways, including sphingolipid homeostasis, actin cortical patch localization, and telomere tethering at the nuclear periphery (**Fig. 13, Table 3**).

**Figure 13.**
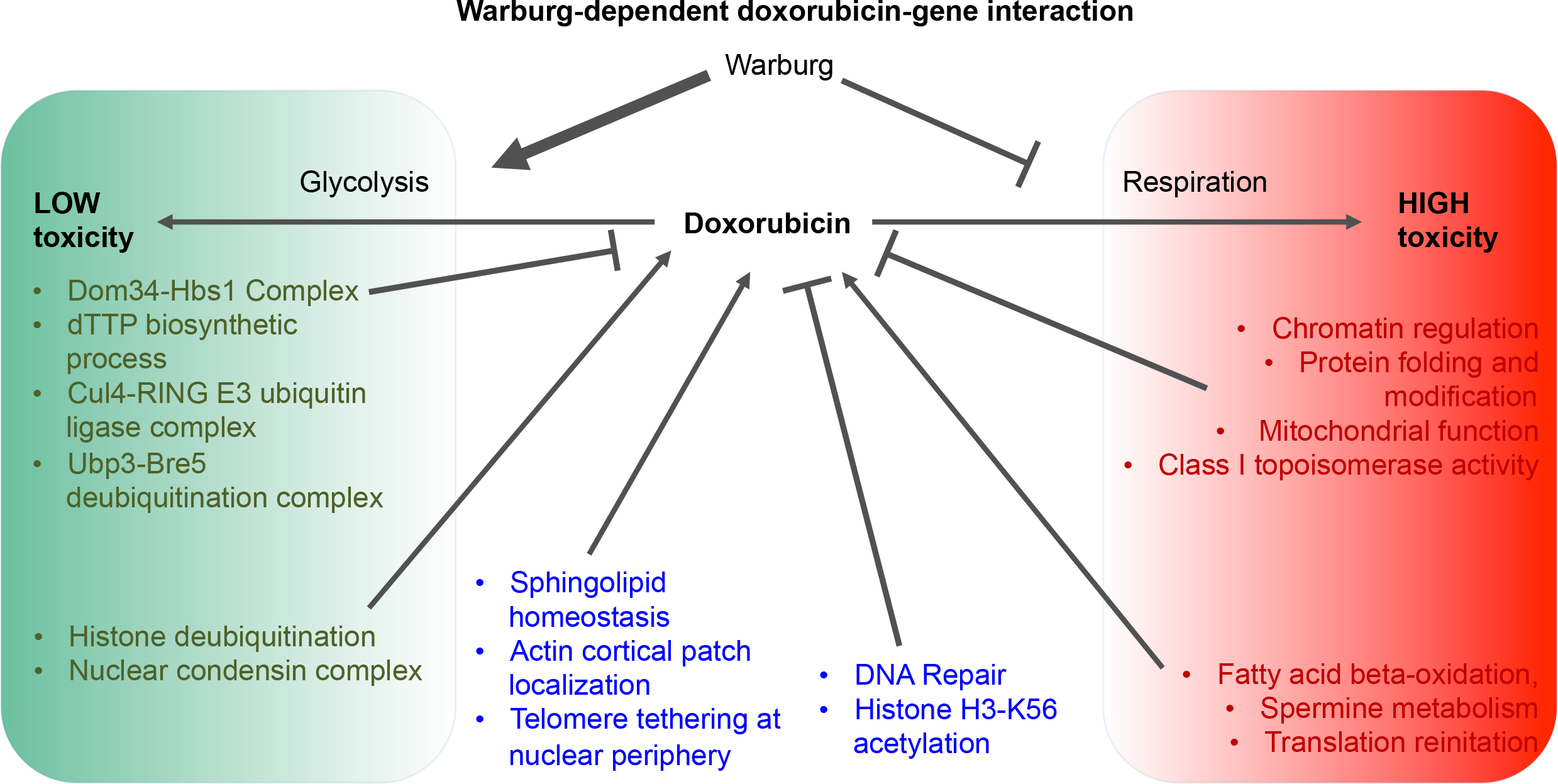
Yeast phenomic model for the influence of Warburg metabolism on doxorubicin-gene interaction. Shaded areas indicate influences that are relatively Warburg-dependent, being red or green if their effects are relatively specific to a respiratory or glycolytic context, respectively. Processes that influence doxorubicin cytotoxicity in a more Warburg-independent manner are unshaded. Arrowheads indicate processes for which genes predominantly transduce doxorubicin toxicity, based on their loss of function suppressing its growth inhibitory effects. Conversely, a perpendicular bar at the line head indicates a process that buffers doxorubicin toxicity, as genetic compromise of its function enhances the growth inhibitory effects of doxorubicin.

Studies in cancer cell lines, mice, and acute myeloid leukemia blast cells from patients were highlighted by the yeast phenomic model, suggesting histone eviction, increased mutation rates at active promoter sites [11, 12, 225], and accumulation of damage from chromatin trapping by the FACT complex as mechanisms of doxorubicin toxicity [13]. Further support of the importance of chromatin regulation was suggested by transcriptional control and assembly of histones, as well as histone modifications, which are all particularly important in a respiratory context. From a precision medicine perspective, tumors that are promoted by genetic compromise in chromatin regulation [226, 227] would be potentially more susceptible to treatment, but only if they have not undergone the Warburg transition to glycolysis. Analogously, patients with germline variation resulting in functional compromise of chromatin regulation may have normal tissue (e.g., cardiac muscle) that is susceptible to doxorubicin and thus may suffer greater toxic side effects of cancer treatment.

The examples of integrating yeast phenomic data with cancer cell line pharmacogenomics data to predict therapeutic efficacy are not limited to doxorubicin and/or the Warburg phenomenon. Analogous phenomic models could be generated for many cytotoxic agents and/or metabolic states. We found the global correlation of human UES and OES with yeast deletion suppressors and enhancers to be very low, consistent with yeast studies examining this expectation [168]. However, there were many examples of differential expression of individual genes, which were potentially explained biologically by the yeast phenomic model. These observations suggest that yeast phenomic models can be helpful and may even be necessary to associate differential gene expression and sensitivity of cancer cells to chemotherapy. We hope and anticipate that future integrative studies and ultimately clinical trials can further demonstrate whether yeast phenomic studies contribute useful information for personalizing clinical guidance and increasing therapeutic efficacy for patients.

The HDAC inhibitor, Abexinostat, enhanced doxorubicin cytotoxicity in cancer cell lines [228, 229], and a phase I clinical trial combining the agents in metastatic sarcomas showed some tumor responses [230]. Enhanced doxorubicin cardiotoxicity was observed with co-administration of HDAC inhibitors in mice [231]. The yeast phenomic model suggests it could be informative to monitor the Warburg status of cancer cells in such studies (**Fig. 7A**). The Sin3-type histone deacetylase complexes (Class I) exhibit respiration-specific deletion enhancement, however the influence of *HDA1/HDAC6* (Class II) is Warburg-independent, with *HDAC6* being UES (**Fig. 12E, Table 3**). *HDAC6* has a unique structure among histone deacetylases, increasing the ability to target it pharmacologically [232]. A clinical trial using Vorinostat in combination with paclitaxel and doxorubicin-cyclophosphamide to treat advanced breast cancer showed a positive response and reduced expression of *HDAC6* in the primary tumor [233]. Ricolinostat is a clinically safe *HDAC6*-specific inhibitor [234] that could enhance doxorubicin toxicity to cancer driven by epigenetic plasticity [226, 227], if the cancer undergoes the Warburg transition, as HDA1 complex mutants are less protected by glycolysis. Yet in a respiratory context, Sin3-type complexes exhibit stronger interaction (**Fig. 7A**). While speculative, these examples are intended to illustrate the potential power of yeast phenomic models to generate novel, testable hypotheses through integration of existing knowledge and new, unbiased experimental results.

In summary, we envision yeast phenomic drug-gene interaction models as a complement to existing cancer pharmacogenomics, providing an experimental platform to quantitatively derive drug-gene interaction network knowledge that can be integrated with DNA, RNA, protein, epigenetic, metabolite profiling, and/or cell proliferation data collected from tumors. Such predictions, applied to individual patients’ tumors, could be further used in conjunction with evaluation of tumor drug response; for example, before and after treatment to understand how recurrent cancer buffers the drug’s toxicities.

Analyses of patient-derived tumor organoids, for example, could include predictive modeling and experimental validation for development of treatment strategies, both initially and with recurrence [235–237]. The influence of Warburg status could also be integrated into such personalized models if monitoring its influence on responsiveness of cancer to chemotherapy proves useful for selectively killing tumors [238]. Moreover, yeast phenomic models could be tailored to individual patients to examine more complex interactions: for example, in the background of homologous recombination deficiency [145]. Yeast phenomics provides the experimental capabilities and genetic tractability to model genetic buffering networks relevant to human disease at high precision and resolution, and the biological relevance of yeast genetics to human disease is established; however, the extent to which yeast phenomics is predictive of human disease biology and complexity remains to be determined.

A major premise of precision medicine should be to systematically account for the contribution of genetic variance to phenotypes as well as influential interacting factors such as cell energy metabolism, age, drugs, or other environmental factors. However, functional genetic variation in human populations, and particularly for cancer, is essentially too abundant to resolve at a systems level with respect to drug-gene interaction. Thus, yeast phenomics, which can define gene interaction networks and genetic buffering in a highly tractable way [21, 239, 240], offers the potential to help resolve disease complexity [17, 241]. Although, the example of doxorubicin is a small sliver of biology, it exemplifies the potential of yeast phenomic modeling of human disease complexity. Lastly, doxorubicin and other cytotoxic agents are typically used in combination cocktails, and a future direction should be to develop yeast phenomic drug-gene interaction network models for buffering combination therapies.

## Conclusions

A yeast phenomic model for the influence of Warburg metabolism on doxorubicin cytotoxicity revealed that glycolysis reduces the cellular reliance on genetic buffering networks. The model reports gene deletion-enhancing and deletion-suppression pathways, and leverages yeast phenomic results to predict differentially expressed human genes that are causal in their association with doxorubicin killing from cancer cell line pharmacogenomics data. As such, this yeast model provides systems level information about gene networks that buffer doxorubicin, serving as example of how the YKO/KD enables experimental designs to quantify gene interaction globally at high resolution. In the case of doxorubicin, gene networks buffer cytotoxicity differentially with respect to Warburg metabolic status. Understanding cytotoxicity in terms of differential gene interaction networks has the potential to inform systems medicine by increasing the precision and rationale for personalizing the choice of cytotoxic agents, improving anti-tumor efficacy and thereby reducing host toxicity. Yeast phenomics is a scalable experimental platform that can, in principle, be expanded to other cytotoxic chemotherapeutic agents, singly or in combination, thus providing versatile, tractable models to map drug-gene interaction networks and understand their complex influence on cell proliferation.

## List of abbreviations

CPP: Cell proliferation parameter
DAmP: Decreased Abundance of mRNA Production
DE: Deletion enhancer
dNTP: deoxyribonucleotide triphosphate
DS: Deletion suppressor
dsDNA: double-stranded DNA
EMC: Endoplasmic reticulum membrane complex
ER: Endoplasmic reticulum
ERMES: ER-mitochondria encounter structure
GARP: Golgi-associated retrograde protein
GO: Gene ontology
GTF: Gene ontology term finder
GTA: Gene ontology term averaging
GTA value: Gene ontology term average value
gtaSD: standard deviation of GTA value
GTA score: (GTA value-gtaSD)
HDAC: Histone deacetylase complex
HLD: Human-like media with dextrose
HLEG: Human-like media with ethanol and glycerol
INT: Interaction score
m7G: 7-methylguanosine
MCM: Mini-chromosome maintenance
OES: Overexpressed in doxorubicin sensitive cells
Q-HTCP: Quantitative high throughput cell array phenotyping
Ref: Reference
REMc: Recursive expectation maximization clustering
ROS: Reactive oxygen species
RPA: Replication Protein A
SD: Standard deviation
SGD: Saccharomyces cerevisiae genome database
snoRNAs: Small nucleolar RNA
snRNA: Small nuclear RNA
t6A: Threonyl carbamoyl adenosine
UES: Underexpressed in doxorubicin sensitive cells
uORF: Upstream open reading frames
YKO: Yeast knockout
YKD: Yeast knockdown

## Declarations

### Ethics approval and consent to participate

-Not Applicable

### Consent for Publication

-Not Applicable

### Availability of data and materials

All data generated or analyzed during this study are either included in this published article and supplementary files or will be freely supplied upon request.

## Author’s contributions

SMS and JLH designed and conducted the experiments and analysis techniques, and wrote the manuscript.

## Acknowledgements

The authors thank Jingyu Guo and Brett McKinney for development of REMc tools, John Rodgers for help with Q-HTCP analysis, and Mary-Ann Bjornsti and Alex Stepanov for helpful discussions.

## Endnotes

Not Applicable

## Additional Files

**Additional File 1. Supplemental figures. Figure S1**. Doxorubicin dose responses of the YKO/KD parental strains, BY4741a, BY4742alpha, and BY4743a/alpha diploid. **Figure S2**. Correlation between interaction scores based on L *vs.* other CPPs (K, r, and AUC), for both HLD and HLEG media. **Figure S3**. Doxorubicin-gene interaction profiles for selected mitochondrial GO terms. **Figure S4**. Deletion of mitochondrial genes tends to influence doxorubicin-gene interaction in a respiratory (HLEG media) more so than a glycolytic (HLD media) context. **Figure S5**. Heatmaps for GO terms comprised of overlapping gene sets. **Figure S6**. Pleiotropic phenotypic influences from genetic perturbation of ribonucleoprotein complex subunit organization. **Figure S7**. HLD-specific deletion enhancement of doxorubicin toxicity by evolutionarily conserved genes. *See also Additional File 10 (Table S13).* **Figure S8**. GO term-specific heatmaps for *mRNA 3’ end processing* and *mRNA cleavage* gene interaction profiles. **Figure S9**. Suppression of doxorubicin cytotoxicity by perturbation of sphingolipid and ceramide metabolism. **Figure S10**. Deletion suppressing doxorubicin-gene interaction for nuclear pore and actin cortical patch functions is relatively Warburg-independent.

**Additional File 2. Doxorubicin-gene interaction data; Tables S1-S8. Tables S1-S4** are the genome-wide experiment: **Table S1**. YKO/KD strains in HLEG. **Table S2**. Reference cultures in HLEG. **Table S3**. YKO/KD strains in HLD. **Table S4**. Reference cultures in HLD. **Tables S5-S8** are the validation study: **Table S5**. YKO/KD strains in HLEG. **Table S6**. Reference cultures in HLEG. **Table S7**. YKO/KD strains in HLD. **Table S8**. Reference cultures in HLD.

**Additional File 3. Interaction plots for HLEG**. **(A, B)** Genome-wide and **(C, D)** validation analyses for **(A, C)** YKO/KD and **(B, D)** reference strains in HLEG. See also methods and Additional File 2.

**Additional File 4. Interaction plots for HLD**. **(A, B)** Genome-wide and **(C, D)** validation analyses. **(A, C)** YKO/KD and **(B, D)** reference strains in HLD media. See also methods and Additional File 2.

**Additional File 5. REMc results with doxorubicin-gene interaction profile heatmaps and Gene Ontology enrichment (GO Term Finder; GTF) results. File A** contains REMc results and associated gene interaction and shift data. **File B** is the heatmap representation of each REMc cluster after incorporating shift values and hierarchical clustering. **File C** contains the GTF results obtained for REMc clusters for the three ontologies – process, function, and component.

**Additional File 6. Gene Ontology Term Averaging (GTA) results and interactive plots. File A** contains all GTA values, cross-referenced with REMc-enriched terms. **File B** displays GTA values associated with above-threshold GTA scores (see note below) plotted for HLD *vs.* HLEG. GTA values for REMc-enriched terms are also included (regardless of whether |GTA score| >2). **File C** displays a subset of File B, containing only GO Terms with above-threshold GTA scores and that were enriched by REMc/GTF. **File D** reports GTA value using the K parameter. **Files B-D** should be opened in an Internet web browser so that embedded information from **Additional File 6A** can be viewed by scrolling over points on the graphs. Subsets in each of the plots can be toggled off and on by clicking on the respective legend label. In the embedded information, X1 represents HLEG and X2 represents HLD information. <u>Note</u>: The GTA score threshold (for L) indicates that GTA-gtaSD > 2 for enhancers or GTA+gtaSD < −2 for suppressors, in at least one media.

**Additional File 7. Systematic comparisons involving genome-wide studies of doxorubicin-gene interaction. Table S9**. Genes with deletion-enhancing doxorubicin-gene interaction from Xia et al. 2007 and Westmoreland et al. 2009. **Table S10**. Summary of experimental details associated with Table S9. **Table S11**. Test of enrichment for doxorubicin-gene interaction among genes encoding proteins predicted as substrates of the NatC complex. **Table S12**. Test of enrichment for doxorubicin-gene interaction among genes predicted to be regulated by conserved uORFs (*Cvijovic et al. 2007*).

**Additional File 8. Quantitative summaries of REMc clusters. File A** depicts REMc results, in terms of cluster distributions of L and K interaction (‘shift’ is not used for REMc and thus is not displayed), as a way to visualize cluster differences quantitatively. **File B** is organized by first round clusters and plots the change in p-value for significant terms with respect to round of clustering. Clusters derived from one another and sharing enrichment of the same GO term are connected by a line. Only GO terms with a background size of 500 or smaller are included. Scroll over a symbol to see embedded detail about each GO term. The square root of the p-value is used on the y-axis to evenly distribute data.

**Additional File 9. GO term-specific heatmaps for REMc/GTF-enriched clusters**. GO term-specific heatmaps for significant GO process terms were generated as described in methods and Figures 3 and 4. Any related child terms are presented in subsequent pages of the parent file name. GO terms with more than 100 children, with 2 or fewer genes annotated to the term, or a file size over 300KB are not shown. All heatmaps are generated with the same layout (see Figures 3 and 4).

**Additional File 10 (Table S13). HLD-specific gene deletion enhancement, not associated with ‘shift’ / growth deficiency**. Data were selected for yeast-human homologs if the respective YKO/KD strains generated growth curves in both HLD and HLEG media (in the absence doxorubicin), and either of the following two sets of criteria were met: (1) HLD L interaction > 2 and HLEG L interaction < 2; these data were further filtered for *HLD L Interaction* -*HLD L Shift* > 4, and are presented in Fig. S7A.; or (2) *HLD L Interaction* – *HLEG L interaction* > 4 and HLEG K interaction > - 10; these data were further filtered for *HLD L Interaction* - *HLD L Shift* > 4, and are presented in Fig. S7B. Data included in Fig. S7 are indicated in the last column.

**Additional File 11. Integration of yeast phenomic and cancer cell line pharmacogenomic data to predict human genes that modify doxorubicin toxicity in cancer cells**. **(A)** Tables of UES and OES human genes and whether their yeast homologs were found to be deletion enhancing or deletion suppressing, respectively. **(B-C)** Overlap between the gCSI and GDSC1000 databases with regard to UES and OES human genes (**B**) across all tissues or (**C**) for individual tissues. <u>Note</u>: the intersection of UES with OES between gCSI and GDSC was used as a negative control for assessing UES and OES overlap. **(D-E)** Yeast phenomic doxorubicin-gene interaction profiles for homologs of human UES or OES genes, sub-classified according to interaction type (deletion enhancing or suppressing) and Warburg-dependence of the interaction, for the **(D)** gCSI or **(E)** GDSC1000 databases. Similar to Figure 12, yeast-human homology relationships are shown to the left of heatmaps (blue - one to one; green - one to many; red - many to many). **(F-I)** Interactive plots for yeast-human homologs, comparing the p-value of human genes to L interaction scores for yeast counterparts in **(F, G)** HLD or **(H, I)** HLEG from **(F, H)** gCSI or **(G, I)** GDSC1000. For the standardized coefficient (‘estimate’ color gradient), a negative value (purple) indicates UES, while a positive value (orange) indicates OES. Thus, the model would predict causality for a human gene if its yeast homolog has a positive L interaction (deletion enhancing) and is colored purple (UES), or a negative L interaction (deletion suppressing) and colored orange (OES). Genes are only plotted if the human homolog was significant (p-value < 0.05).

