## Additional File 1 for "A yeast phenomic model for the influence of Warburg metabolism on genetic buffering of doxorubicin"

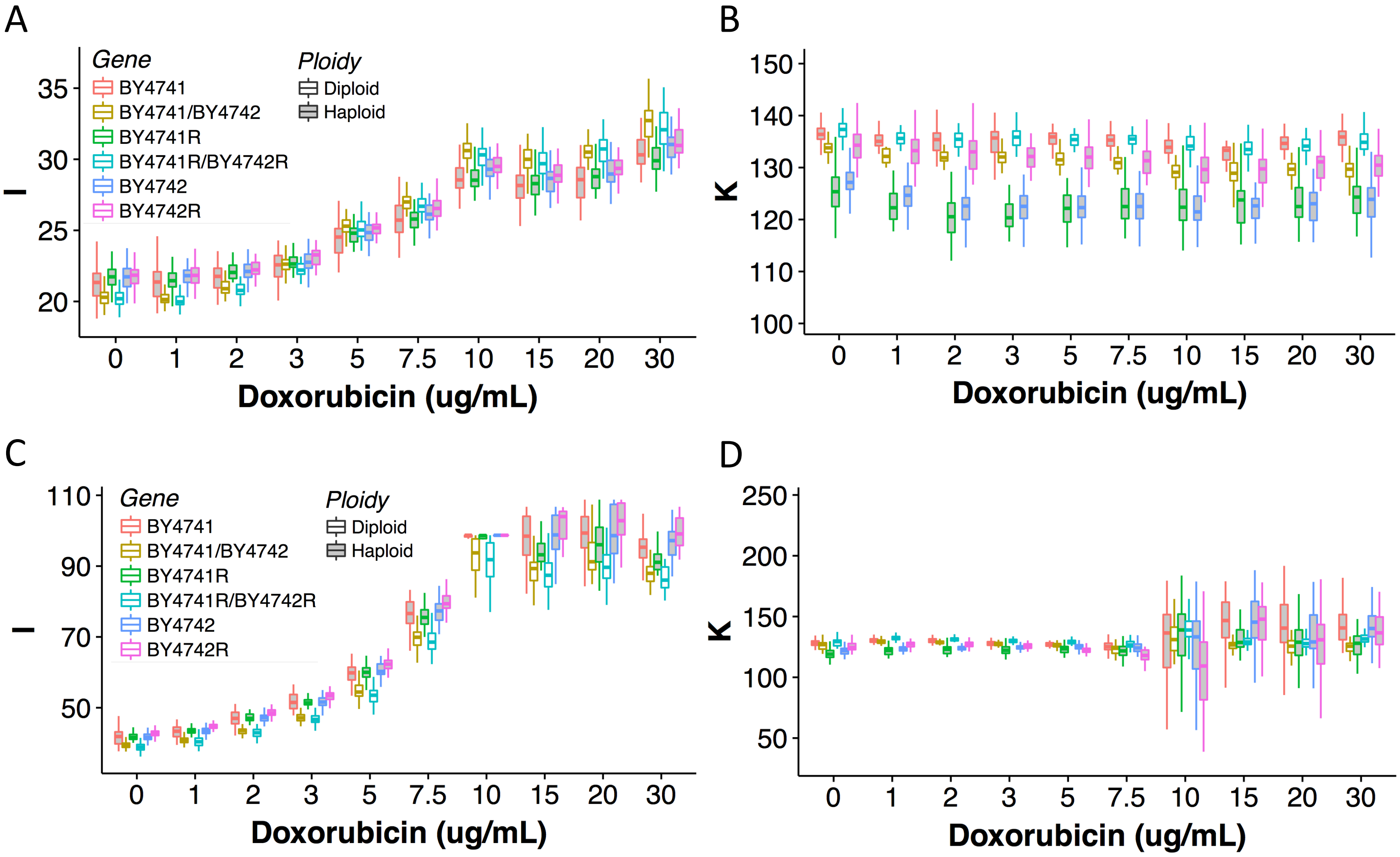


E

| Strain | MAT | LEU2 | URA3 | HIS3 | LYS2 | MET17 |
|---|---|---|---|---|---|---|
| BY4741 | a | *leu2-Δ0* | *ura3-Δ0* | *his3-Δ1* | *LYS2* | *met17-Δ0* |
| BY4742 | α | *leu2-Δ0* | *ura3-Δ0* | *his3-Δ1* | *lys2-Δ0* | *MET17* |
| BY4741R | a | *leu2-Δ0* | *ura3-Δ0* | *his3-Δ1* | *LYS2* | *met17-Δ0* |
| BY4742R | α | *leu2-Δ0* | *ura3-Δ0* | *his3-Δ1* | *lys2-Δ0* | *MET17* |
| BY4741/BY4742 | a/α | *leu2-Δ0/leu2-Δ0* | *ura3-Δ0/ura3-Δ0* | *his3-Δ1/his3-Δ1* | *LYS2/lys2-Δ0* | *MET17/met17-Δ0* |
| BY4741R/BY4742R | a/α | *leu2-Δ0/leu2-Δ0* | *ura3-Δ0/ura3-Δ0* | *his3-Δ1/his3-Δ1* | *LYS2/lys2-Δ0* | *MET17/met17-Δ0* |

**Figure S1. Doxorubicin dose response in the YKO/KD parental strains, BY4741a, BY4742alpha, and BY4743a/alpha diploid.** Reference strain responses to increasing doxorubicin concentrations in **(A, B)** HLD and **(C, D)** HLEG for the **(A, C)** L and **(B, D)** K cell proliferation parameters. **(E)** Genotypes of strains in panels A-D. BY4741 was the YKO background for this study. YKD (DAmP) strains have the common auxotrophic alleles (*leu2, ura3,* and *his3*) and the *LYS2* allele, but are clonally derived from *MET17/met17* heterozygous diploid cells, and thus the haploid DAmP library is mixed with respect to the *MET17* vs. *met17* background.

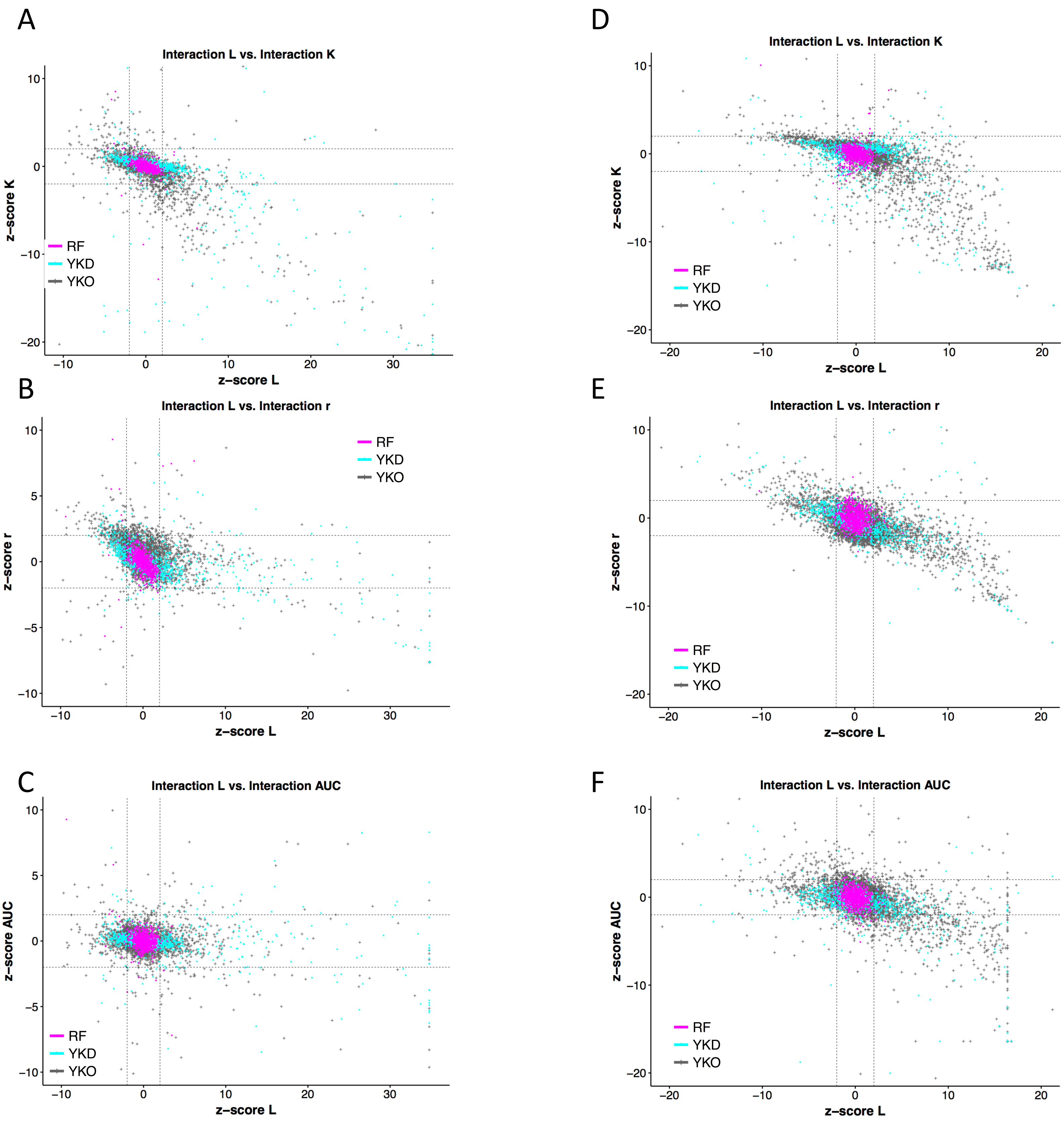


**Figure S2. Correlation between interaction scores based on L *vs.* other CPPs. (A-C)** Interaction scores in HLD for the L cell proliferation parameter vs. **(A)** K, **(B)** r, and **(C)** AUC. **(D-F)** Interaction scores in HLD for the L CPP vs. **(D)** K, **(E)** r, and **(F)** AUC. Dashed lines are drawn at +2 and -2 on the x and y axes. Reference (RF) strain data are overlain on yeast gene knockdown (YKD/DAmP) strain data, which are overlain on yeast gene knockout (YKO) data.

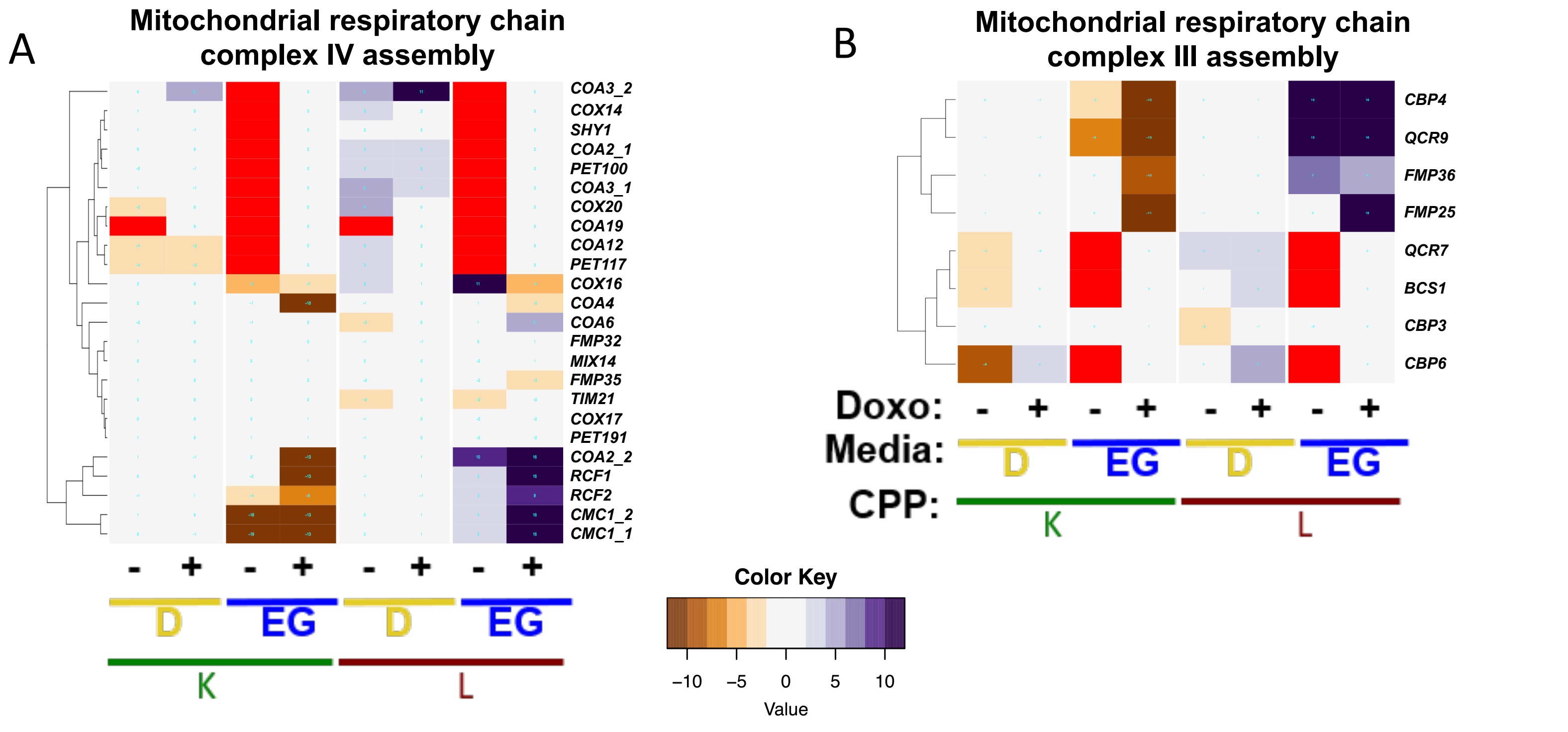


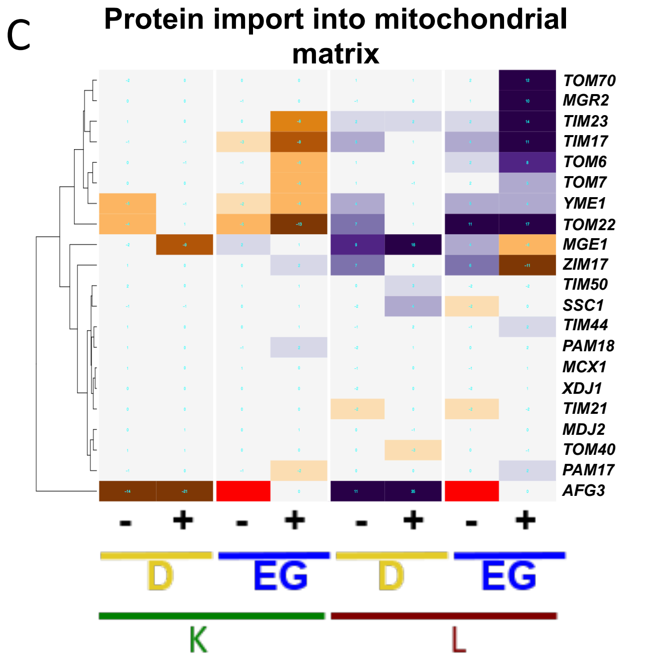


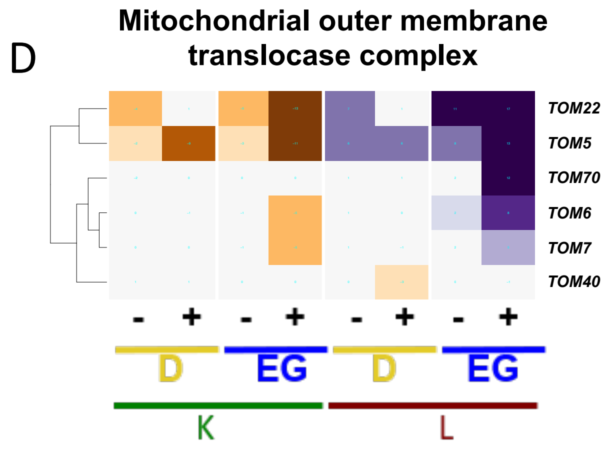


**Figure S3. Doxorubicin-gene interaction profiles for selected mitochondrial GO terms.** GO term-specific heatmaps for mitochondrial respiratory chain complex assembly for **(A)** complex IV and **(B)** complex III, and **(C-E)** terms related to mitochondrial protein import. For all terms, the respective gene deletions exhibit a range of respiratory insufficiency, quantified by the 'EG-' columns. HLEG-specific enhancing interactions are most prominent among these genes overall, indicated by the 'EG+' columns. However, all clusters are associated with pleiotropic interaction profiles.

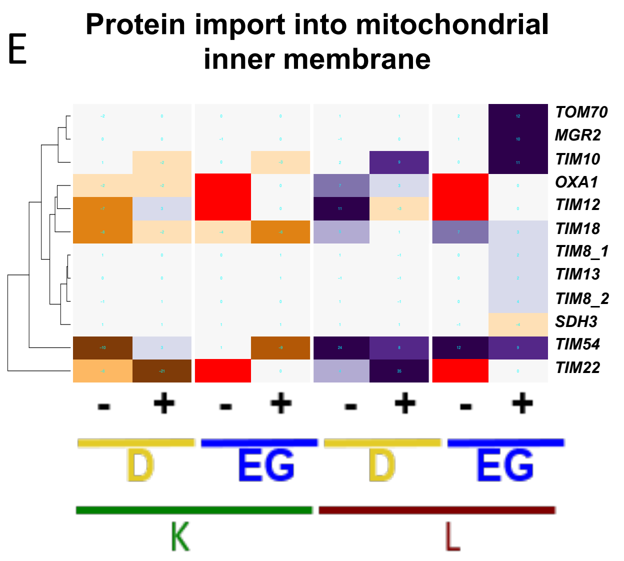

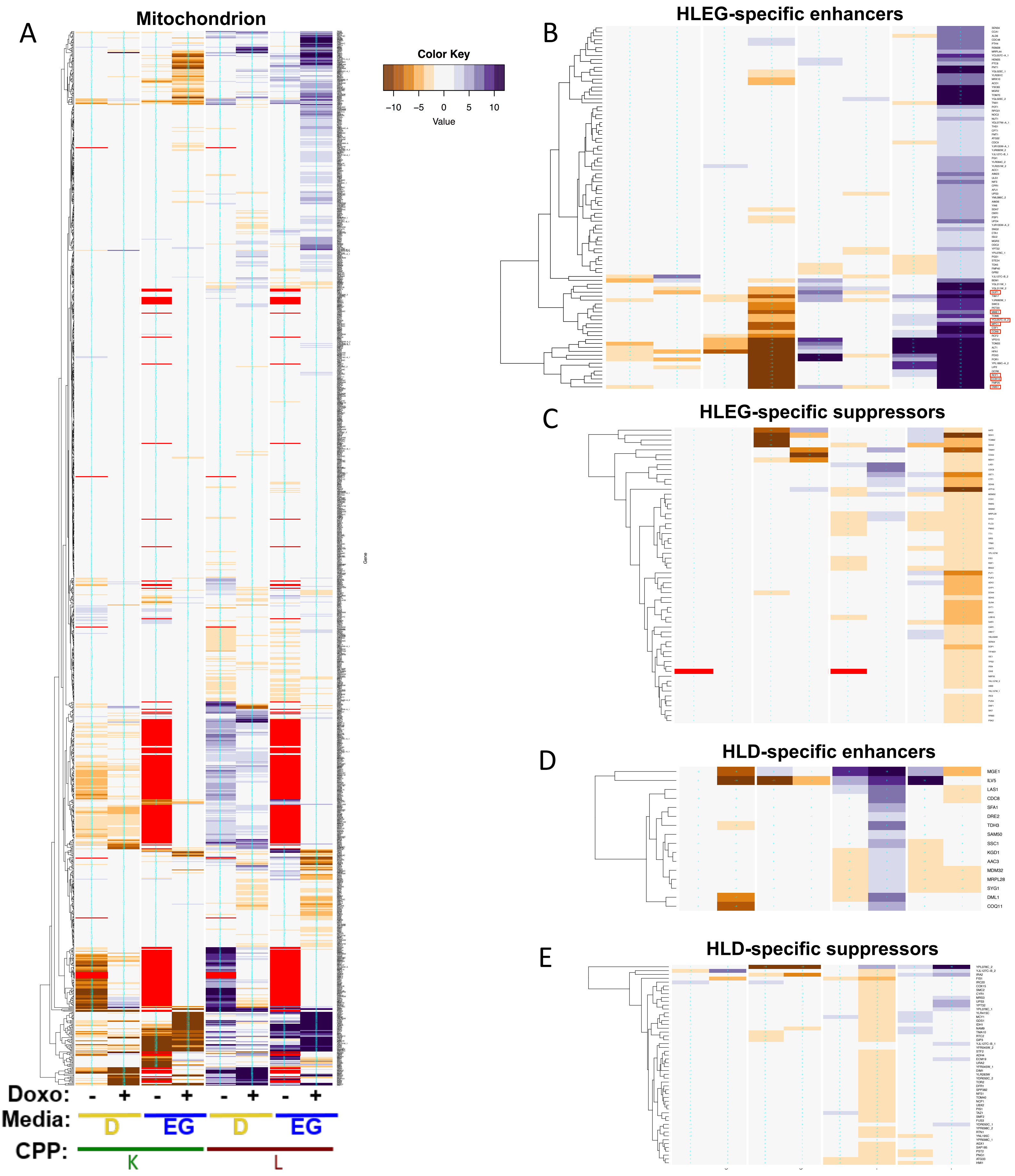


**Figure S4. Deletion of mitochondrial genes tends to result in HLEG-specific doxorubicin-gene interaction. (A)** Gene interaction profiles in the GO cellular component, mitochondrion. **(B-E)** Genes from Panel A were selected by the following criteria on **(B, C)** HLEG or **(D, E)** HLD, respectively: **(B, D)** deletion enhancers were selected if K shift was greater than negative 10 and if the L interaction was 4 greater than the L shift. **(C, E)** Deletion suppressors were selected if the L shift was less than 4. Genes indicated by red boxes in panel B are discussed in the main manuscript results.

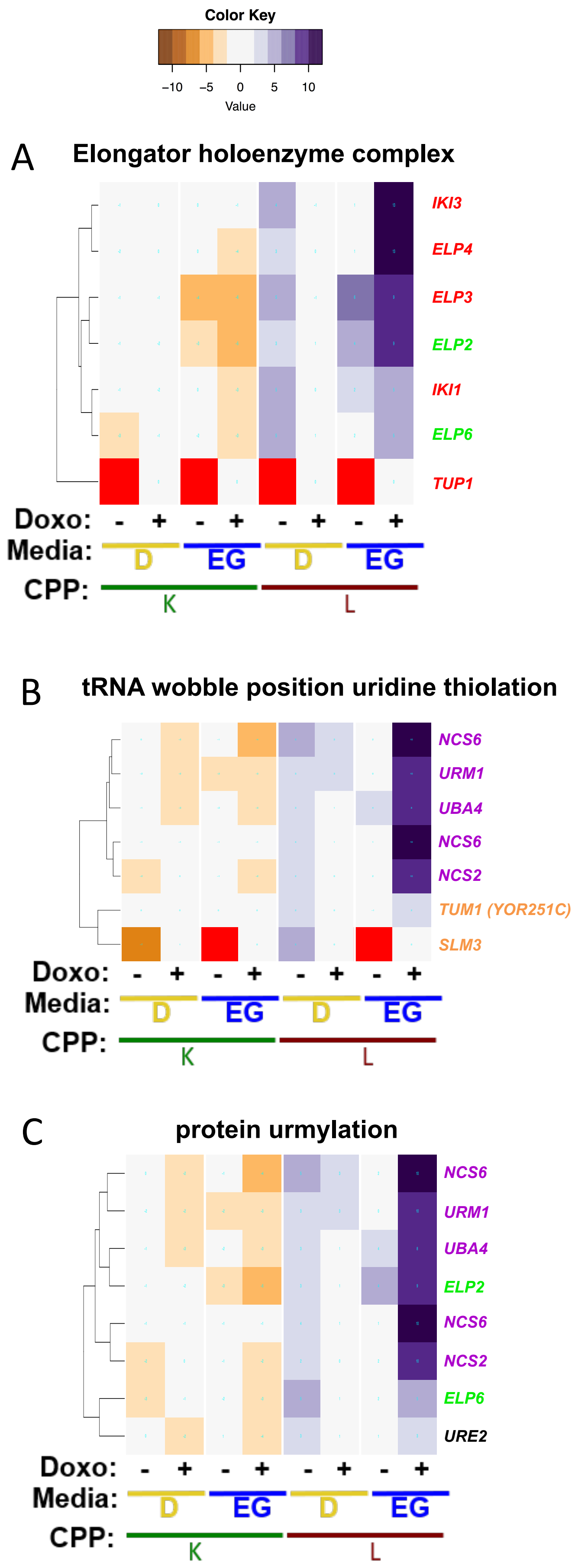


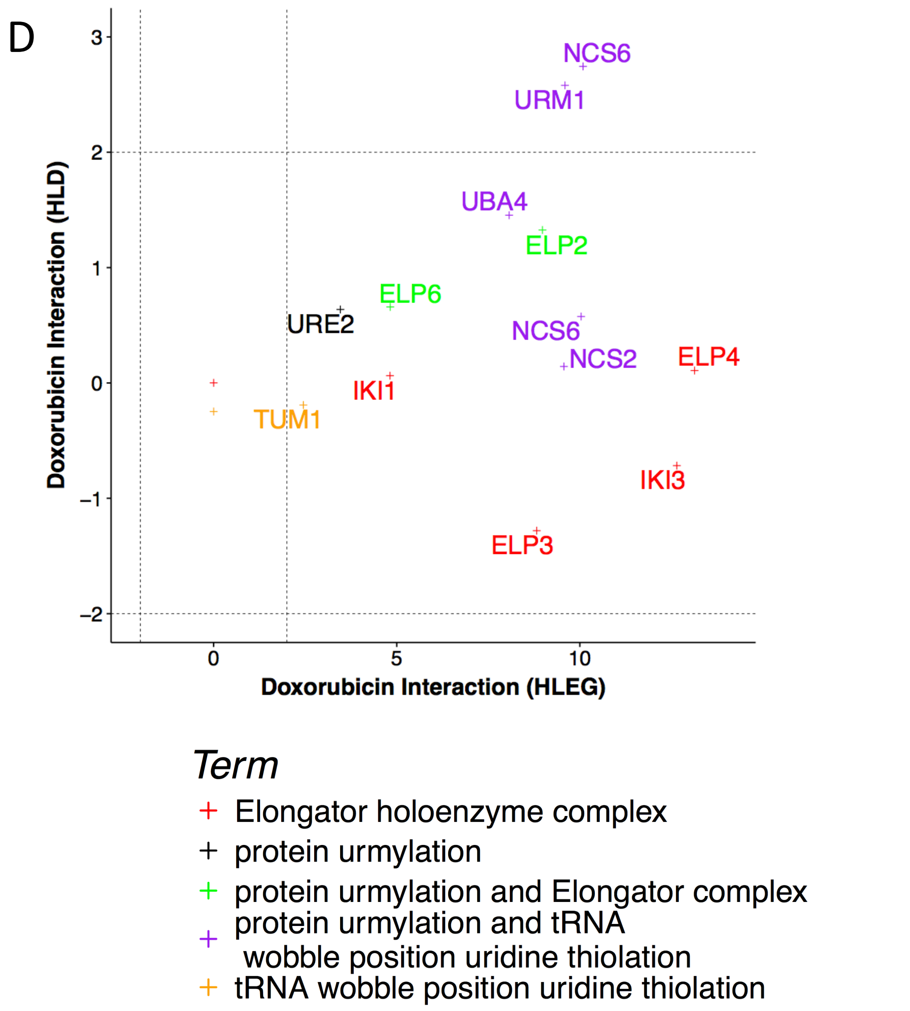


**Figure S5. Distinct GO terms comprised of overlapping gene sets.** Respiration-dependent, deletion-enhancing doxorubicin-gene interaction was enriched for **(A)** Elongator holoenzyme complex, **(B)** tRNA wobble position uridine thiolation, and **(C)** protein urmylation (see **Fig. 8A**, and related discussion).

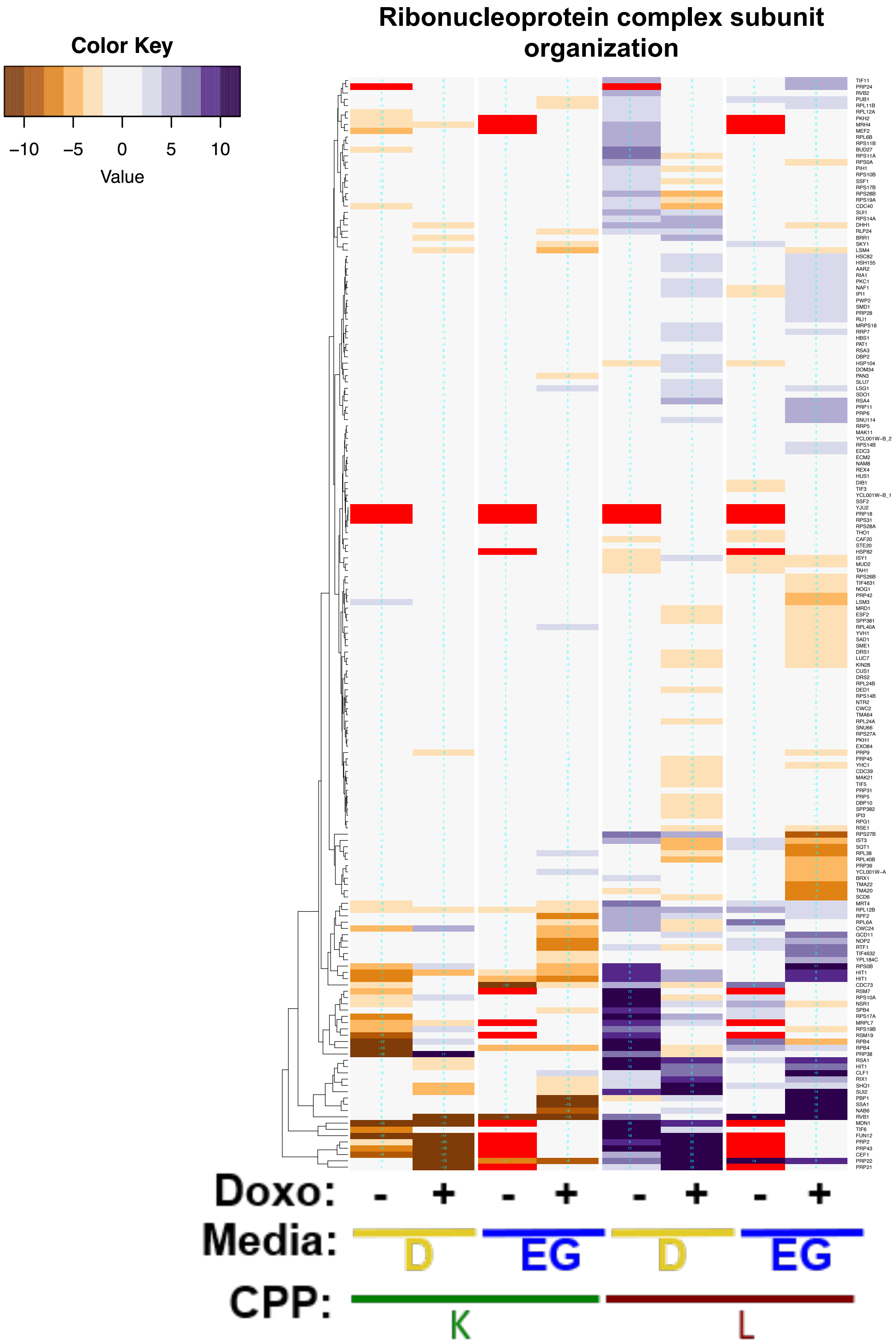


**Figure S6. Genetic perturbation of ribonucleoprotein complex subunit organization results in pleiotropic influences on cell proliferation and doxorubicin toxicity.** Cluster 2-0.2-2, which had an overall pattern of glycolysis-specific enhancement, was enriched for ribonucleoprotein complex subunit organization (**Fig. 4, Table 1**). However, the term-specific heatmap, i.e., all gene interaction profiles associated with the term, revealed a pleiotropic pattern of growth deficiency (no drug; '-' columns) and doxorubicin-gene interaction ('+' columns). Term-specific heatmaps thus provide a directed means to systematically examine and verify gene interaction profile patterns associated with GO terms discovered from the global, unbiased REMc and GTA analyses, and at the same time visualize the extent of growth deficiency attributable to the gene deletion alone.

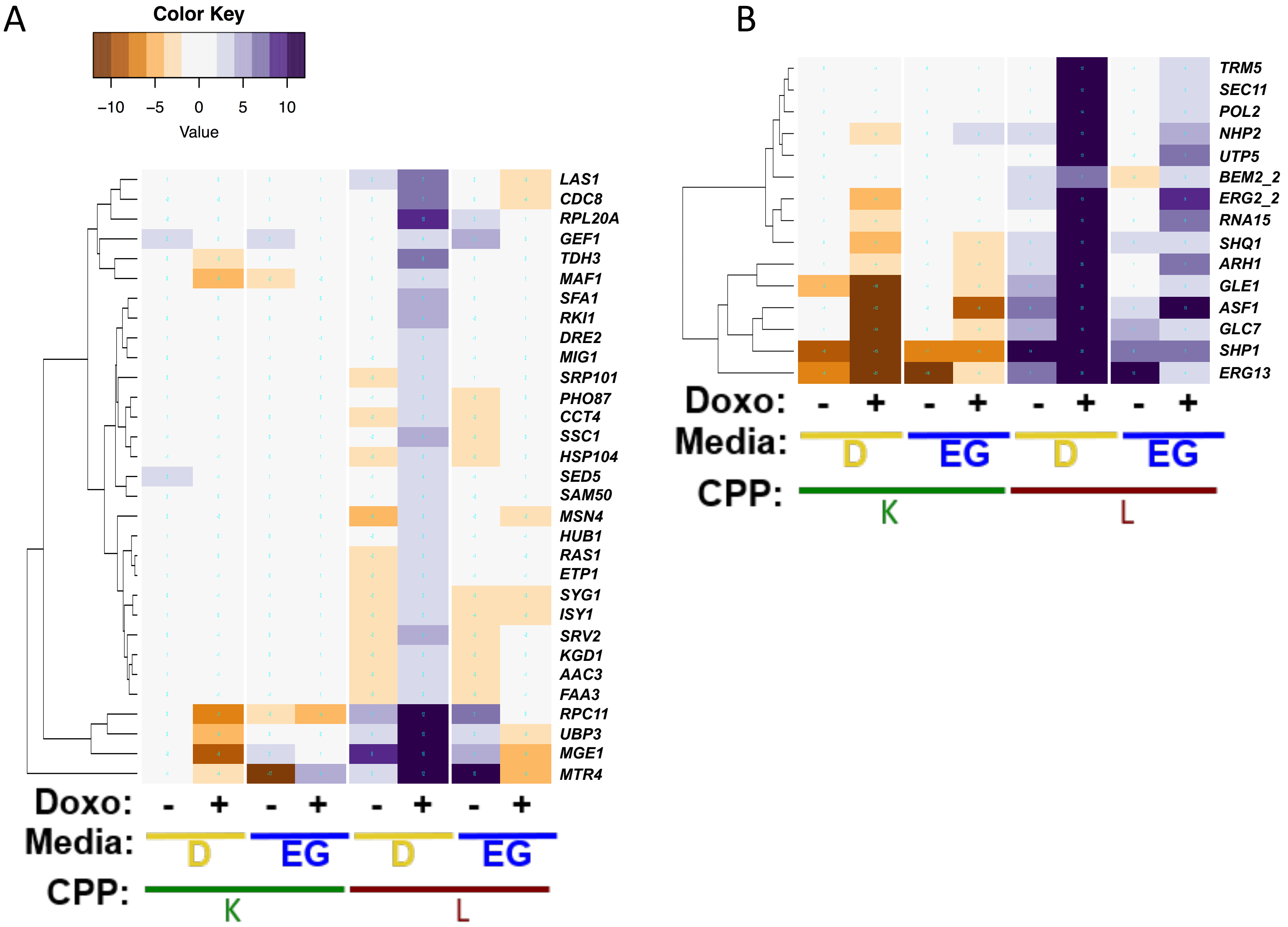


**Figure S7. HLD-specific deletion enhancement of doxorubicin toxicity by evolutionarily conserved genes.** To search systematically for HLD-specific deletion enhancers that could represent candidate vulnerabilities specific to glycolytic tumor cells, we focused on deletion of conserved genes that did not induce growth deficiency. Genes displayed in **panel A** were excluded from **panel B** if identified again. Additional details, including associated human homologs and descriptions of molecular functions are provided **Additional File 10, Table S13.**

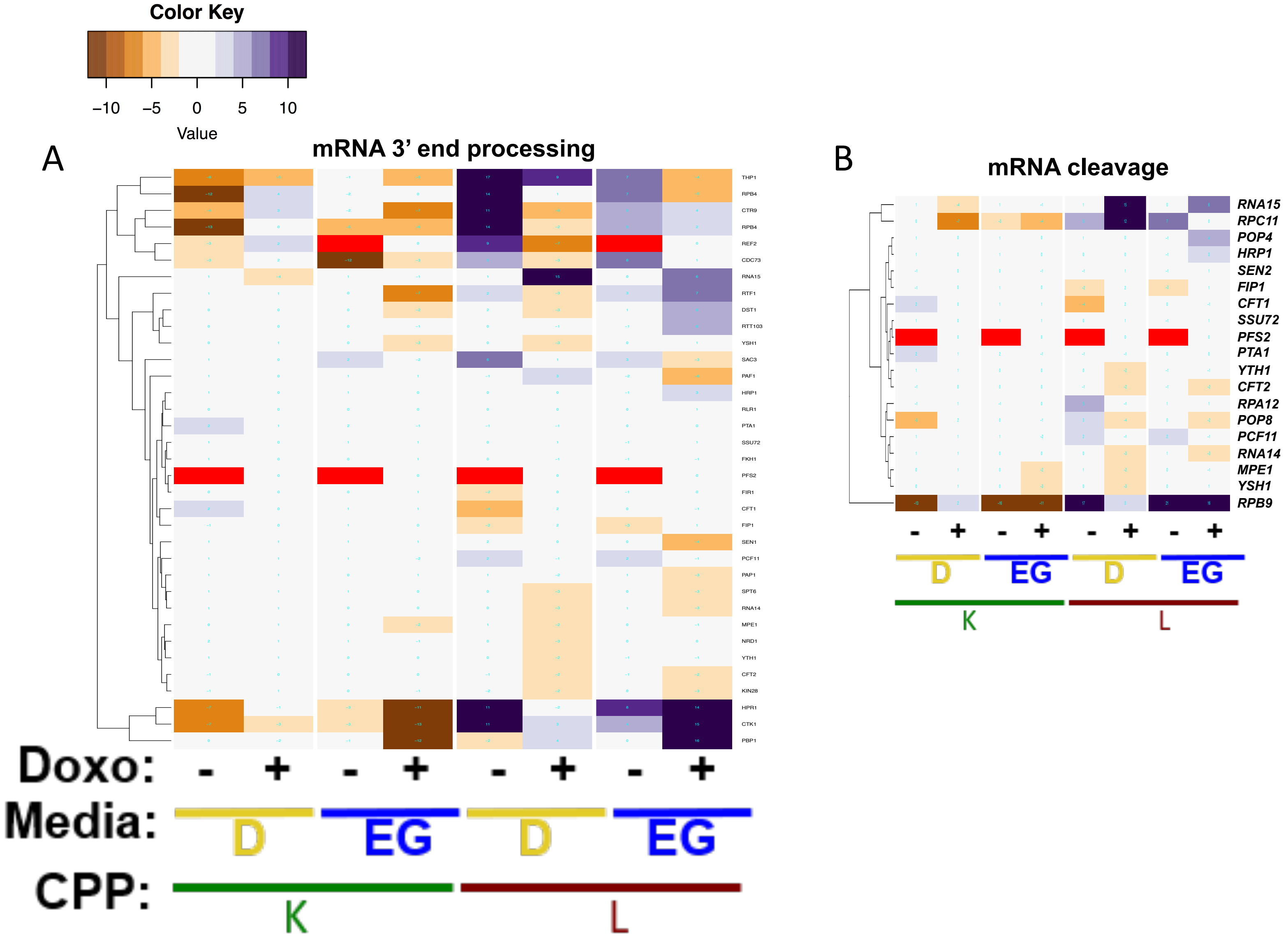


**Figure S8. The GO terms, *mRNA 3' end processing* and *mRNA cleavage,* are comprised of pleiotropic doxorubicin-gene interaction profiles**. Term-specific heat maps with hierarchical clustering for GO terms enriched in 2-0.4-2 including **(A)** mRNA processing and **(B)** mRNA cleavage.

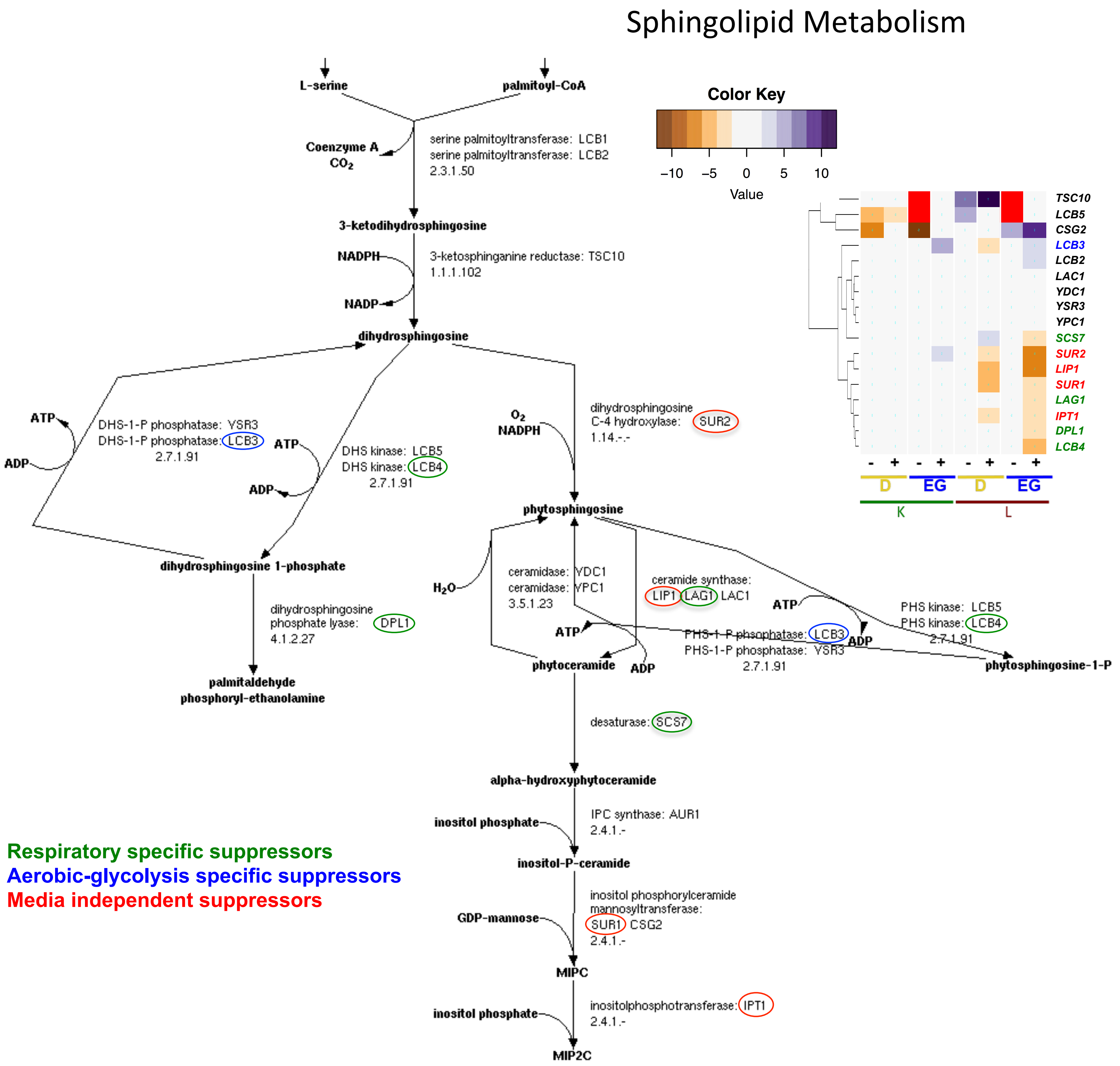


**Figure S9. Suppression of doxorubicin cytotoxicity by perturbation of sphingolipid and ceramide metabolism.** The diagram was modified from the SGD YeastPathways database at [https://pathway.yeastgenome.org/YEAST/NEW-IMAGE?type=PATHWAY&object=SPHINGOLIPID-SYN-PWY&detail-level=2&detail-level=1](https://pathway.yeastgenome.org/YEAST/NEW-IMAGE?type=PATHWAY&object=SPHINGOLIPID-SYN-PWY&detail-level=2&detail-level=1). Circled genes are color-coded by their Warburg-dependence. Interaction profiles for pathway genes are assembled in the heatmap inset.

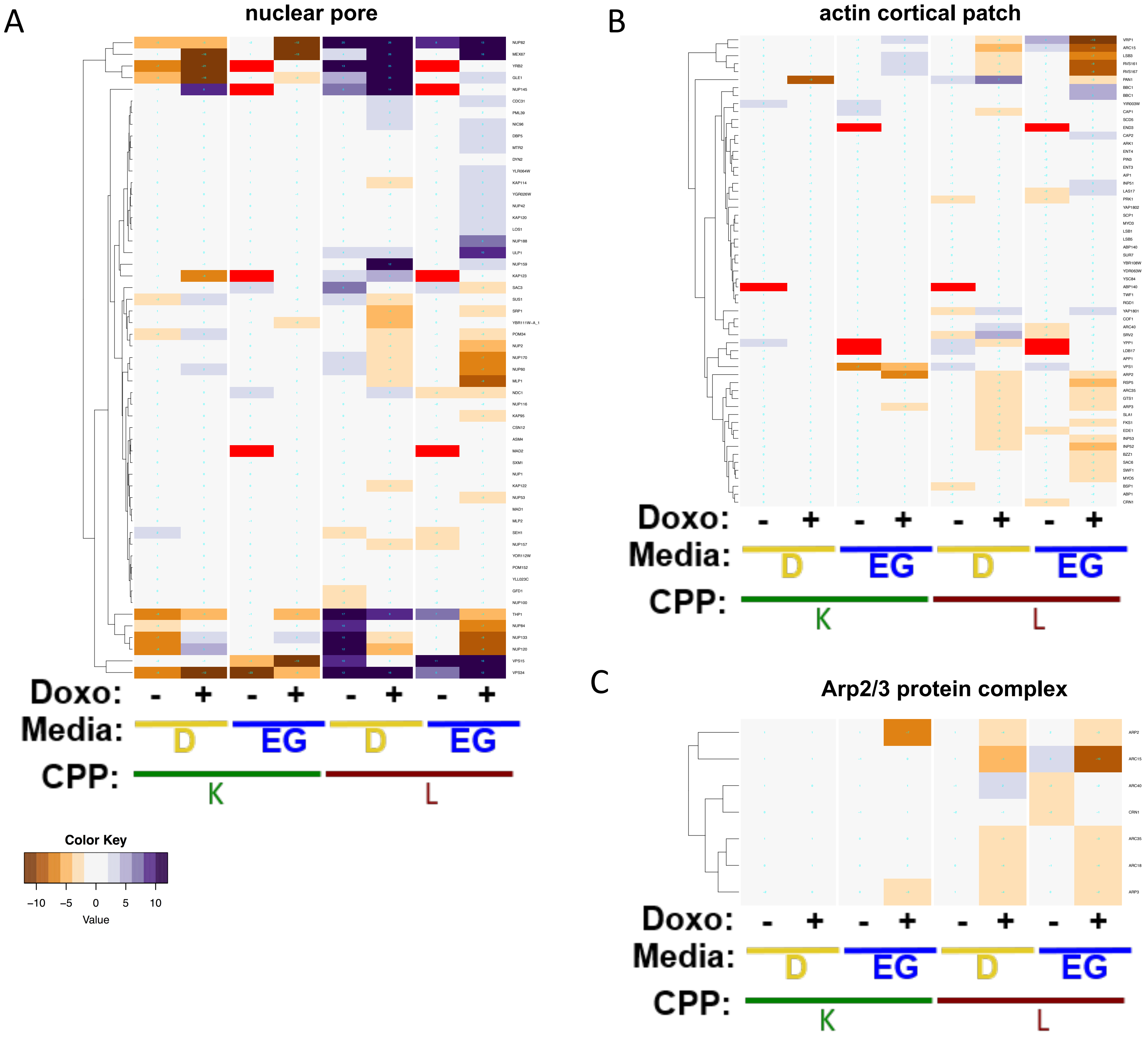


**Figure S10. Warburg-independence of deletion suppressing doxorubicin-gene interaction for nuclear pore and actin cortical patch functions.** Term specific heat maps for **(A)** nuclear pore, **(B)** actin cortical patch, and **(C)** Arp2/3 protein complex.
