## Additional File 3 for "A yeast phenomic model for the influence of Warburg metabolism on genetic buffering of doxorubicin": A - InteractionPlots_Doxo_HLEG.compressed.pdf

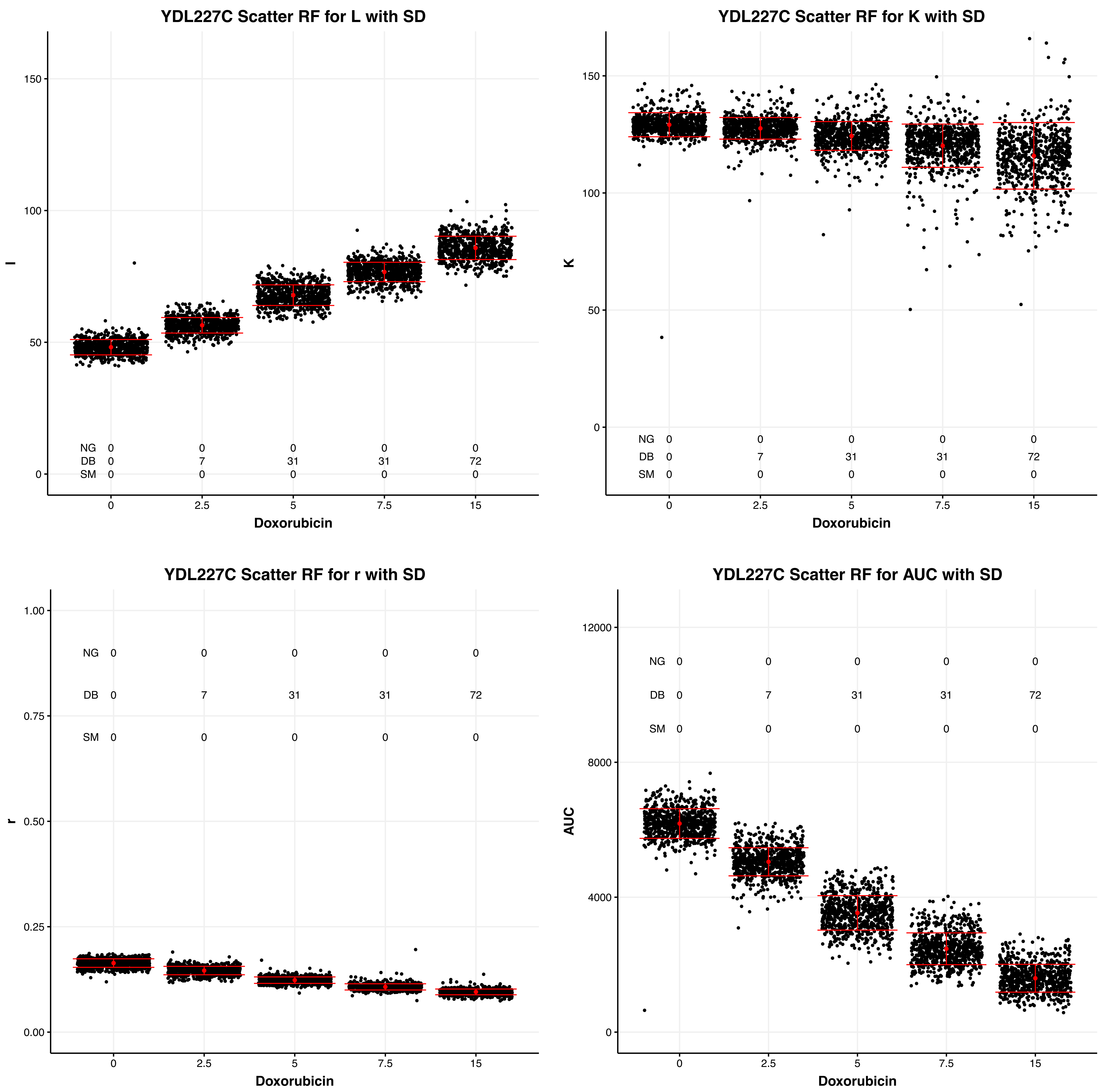

YDL227C Scatter RF for L with SD
l
Doxorubicin
NG 0 0 0 0 0
DB 0 7 31 31 72
SM 0 0 0 0 0
YDL227C Scatter RF for K with SD
K
Doxorubicin
NG 0 0 0 0 0
DB 0 7 31 31 72
SM 0 0 0 0 0
YDL227C Scatter RF for r with SD
r
Doxorubicin
NG 0 0 0 0 0
DB 0 7 31 31 72
SM 0 0 0 0 0
YDL227C Scatter RF for AUC with SD
AUC
Doxorubicin
NG 0 0 0 0 0
DB 0 7 31 31 72
SM 0 0 0 0 0

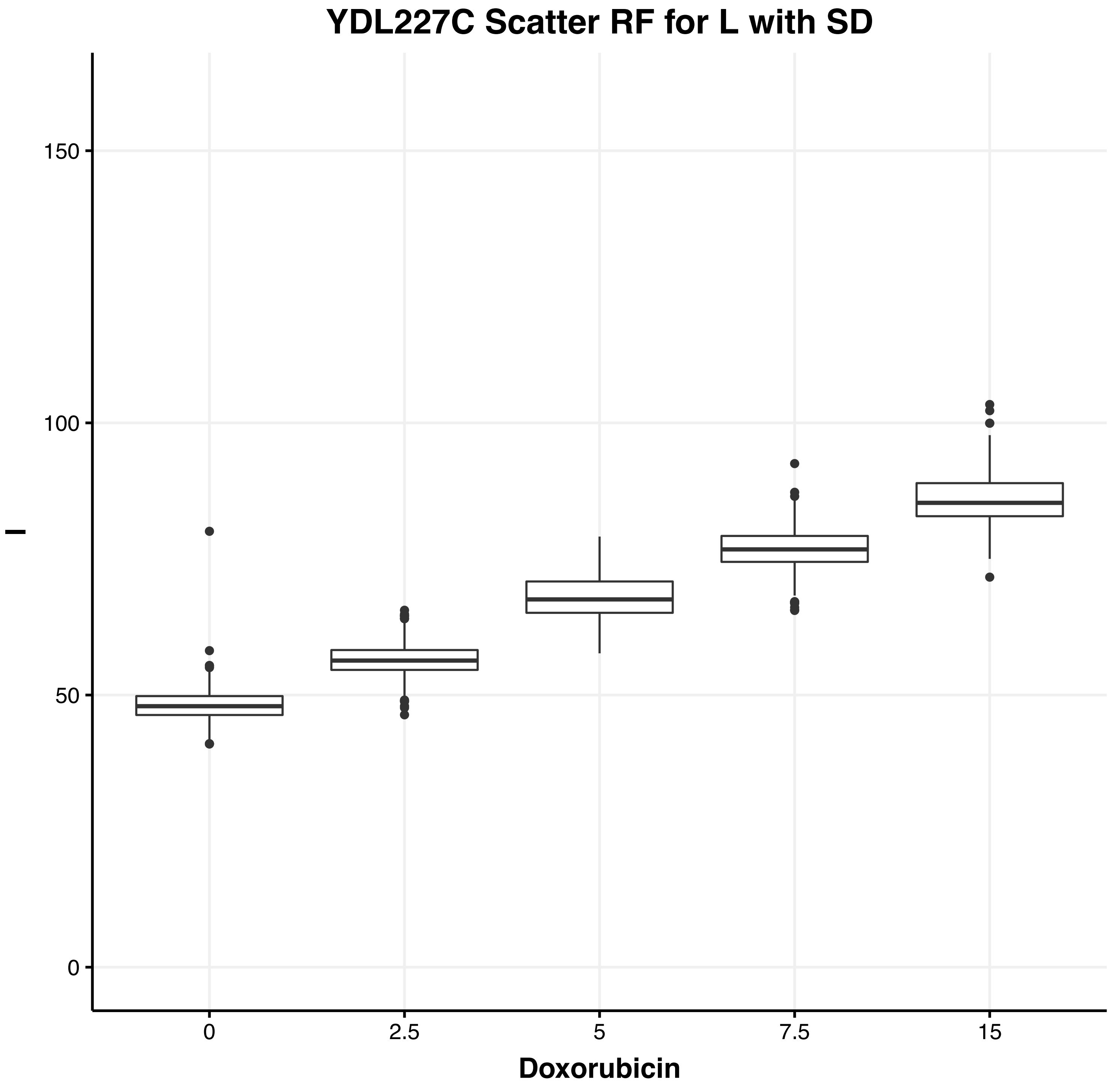
YDL227C Scatter RF for L with SD
l
0
50
100
150
0
2.5
5
7.5
15
Doxorubicin

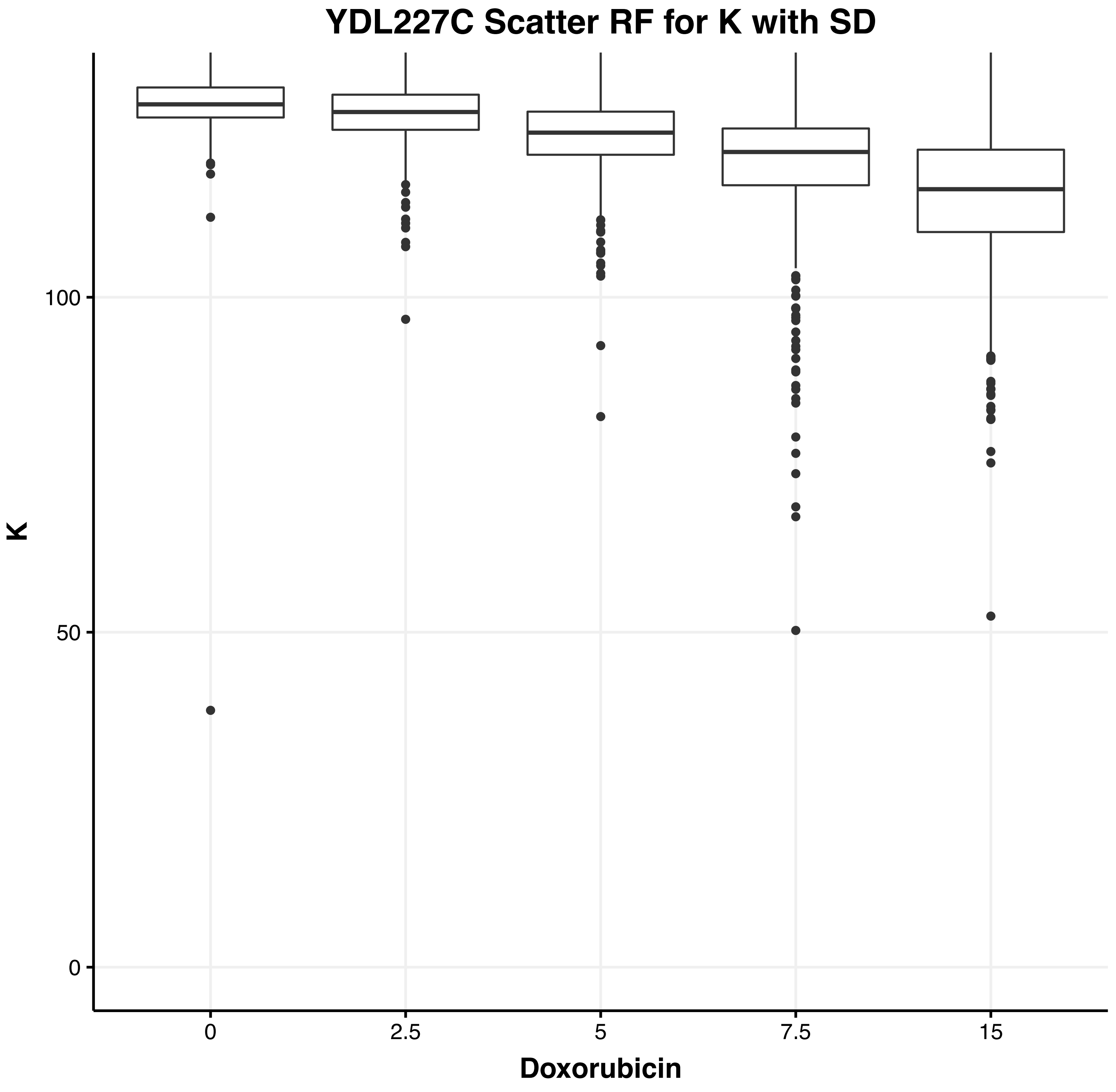
YDL227C Scatter RF for K with SD
K
0
50
100
0
2.5
5
7.5
15
Doxorubicin

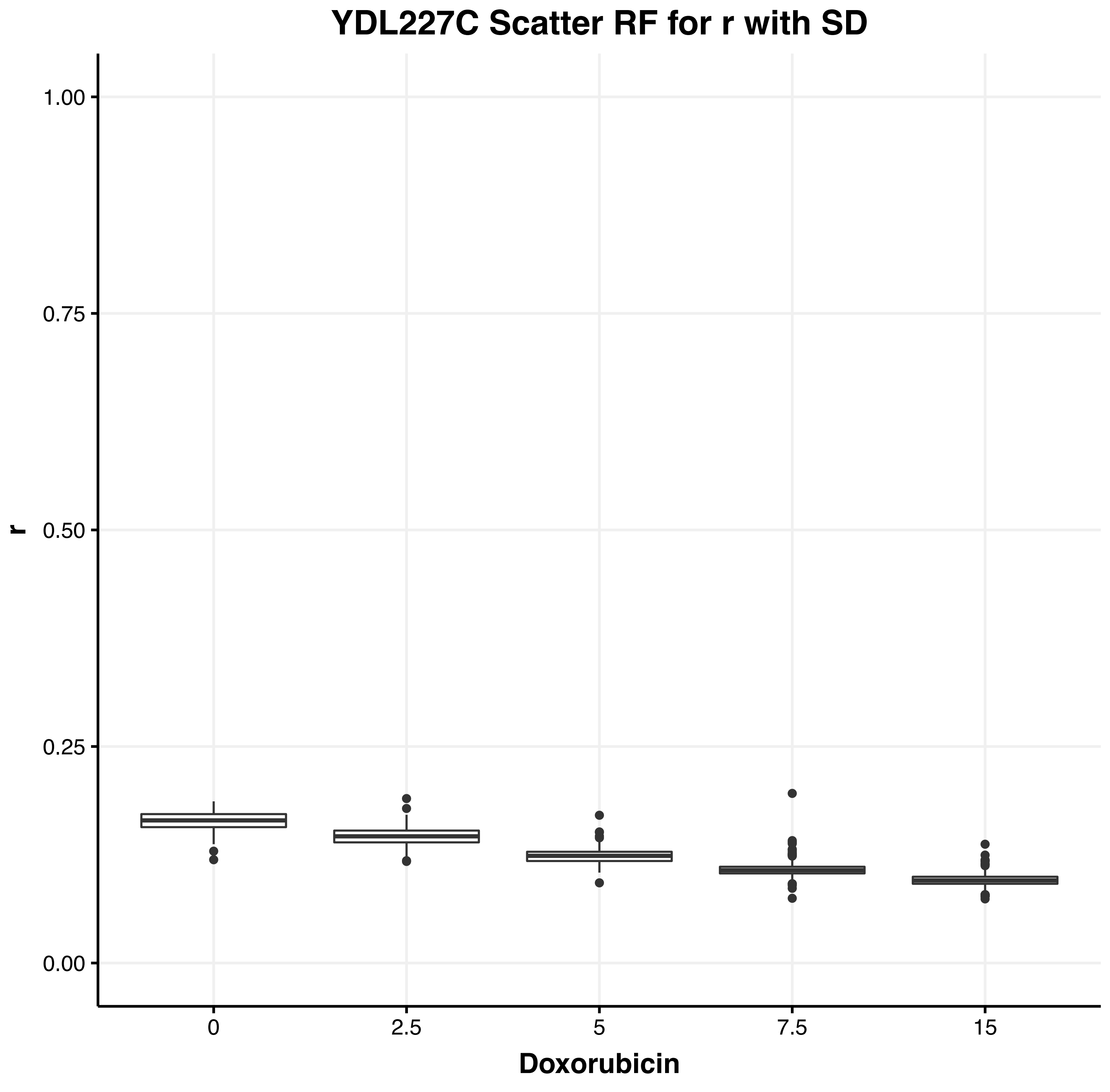
YDL227C Scatter RF for r with SD
r
0.00
0.25
0.50
0.75
1.00
0
2.5
5
7.5
15
Doxorubicin

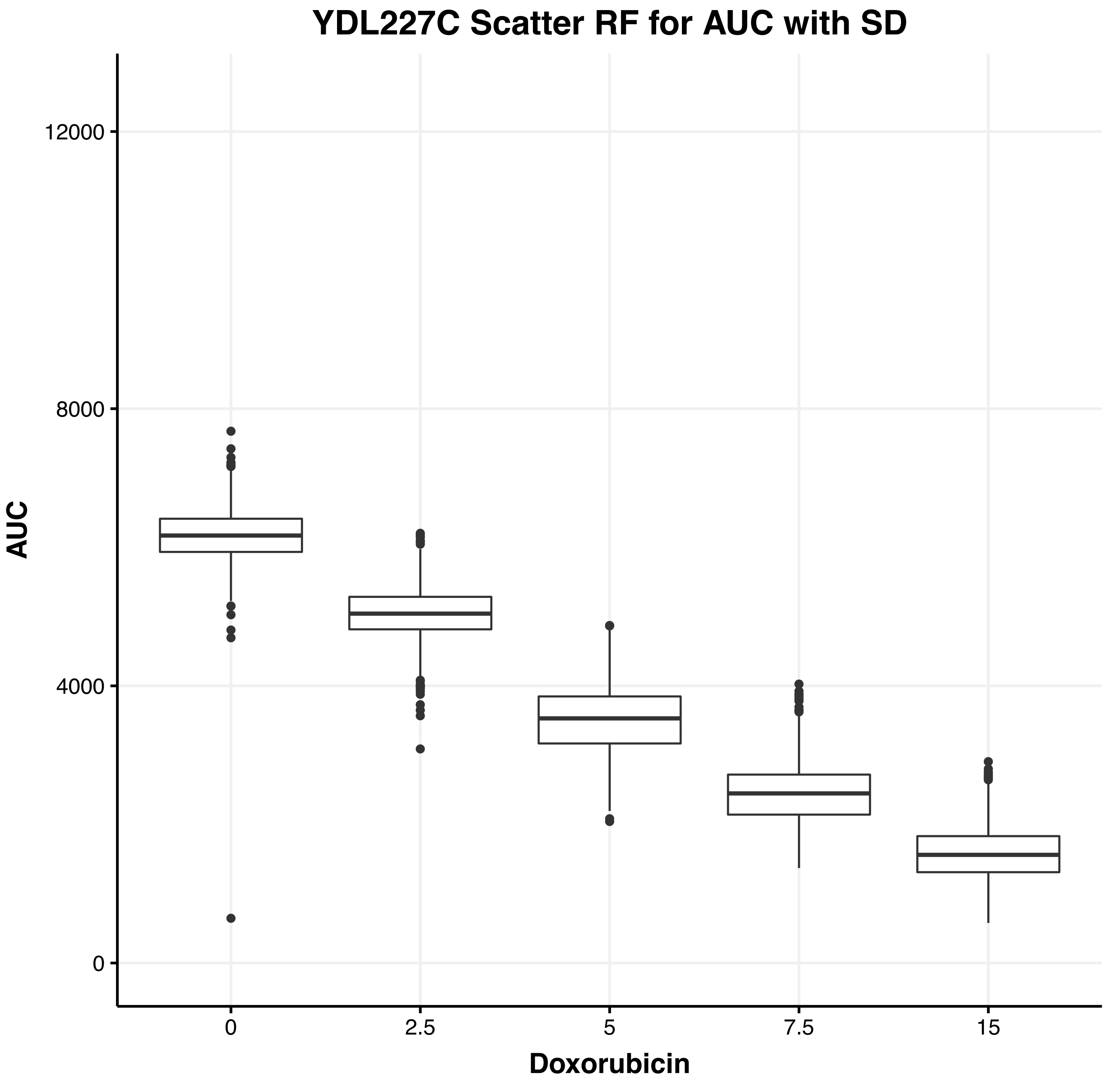
YDL227C Scatter RF for AUC with SD
AUC
0
4000
8000
12000
0
2.5
5
7.5
15
Doxorubicin

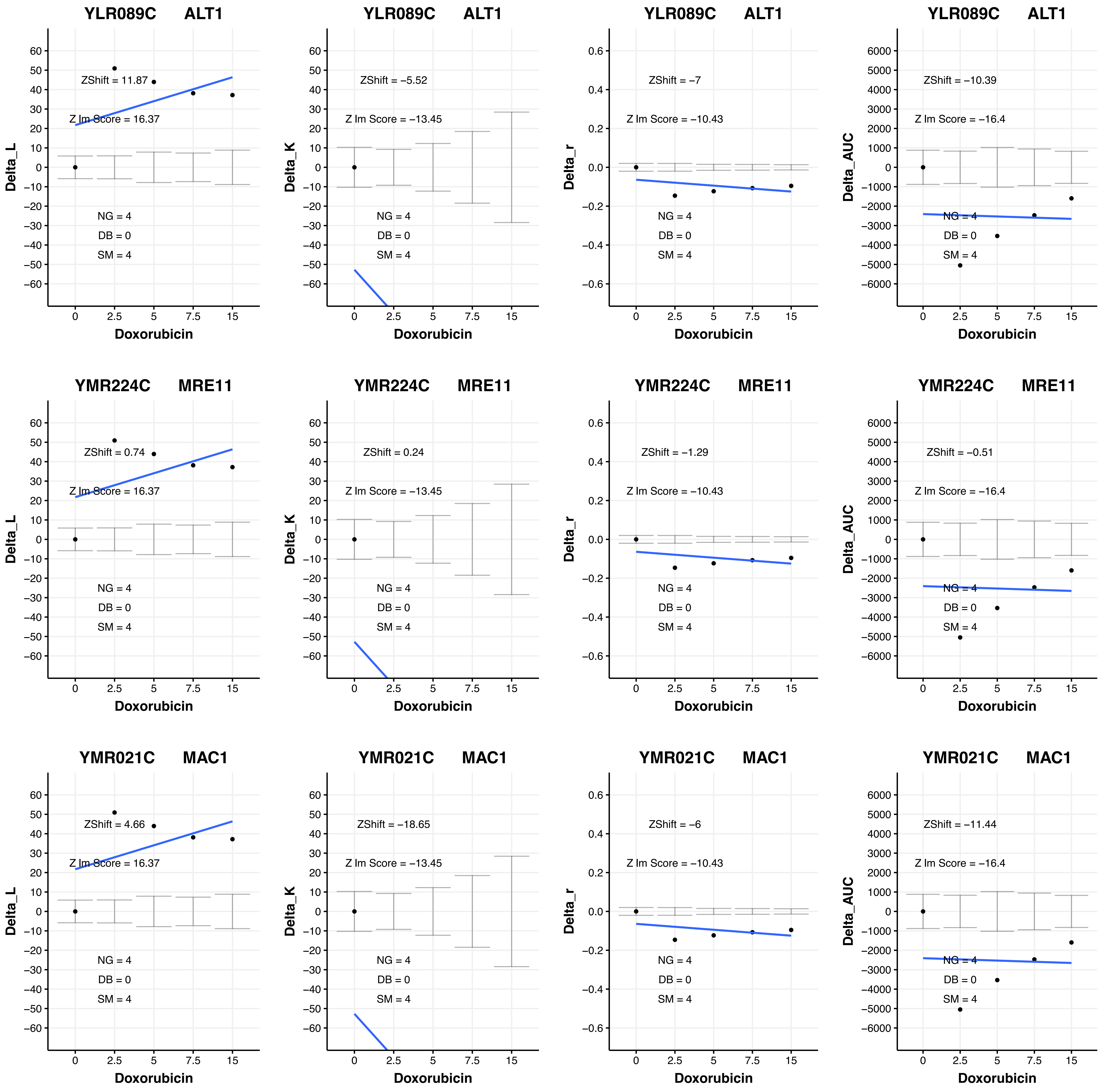

YLR089C ALT1
ZShift = 11.87
Z lm Score = 16.37
NG = 4
DB = 0
SM = 4
Delta_L
Doxorubicin
YLR089C ALT1
ZShift = −5.52
Z lm Score = −13.45
NG = 4
DB = 0
SM = 4
Delta_K
Doxorubicin
YLR089C ALT1
ZShift = −7
Z lm Score = −10.43
NG = 4
DB = 0
SM = 4
Delta_r
Doxorubicin
YLR089C ALT1
ZShift = −10.39
Z lm Score = −16.4
NG = 4
DB = 0
SM = 4
Delta_AUC
Doxorubicin
YMR224C MRE11
ZShift = 0.74
Z lm Score = 16.37
NG = 4
DB = 0
SM = 4
Delta_L
Doxorubicin
YMR224C MRE11
ZShift = 0.24
Z lm Score = −13.45
NG = 4
DB = 0
SM = 4
Delta_K
Doxorubicin
YMR224C MRE11
ZShift = −1.29
Z lm Score = −10.43
NG = 4
DB = 0
SM = 4
Delta_r
Doxorubicin
YMR224C MRE11
ZShift = −0.51
Z lm Score = −16.4
NG = 4
DB = 0
SM = 4
Delta_AUC
Doxorubicin
YMR021C MAC1
ZShift = 4.66
Z lm Score = 16.37
NG = 4
DB = 0
SM = 4
Delta_L
Doxorubicin
YMR021C MAC1
ZShift = −18.65
Z lm Score = −13.45
NG = 4
DB = 0
SM = 4
Delta_K
Doxorubicin
YMR021C MAC1
ZShift = −6
Z lm Score = −10.43
NG = 4
DB = 0
SM = 4
Delta_r
Doxorubicin
YMR021C MAC1
ZShift = −11.44
Z lm Score = −16.4
NG = 4
DB = 0
SM = 4
Delta_AUC
Doxorubicin

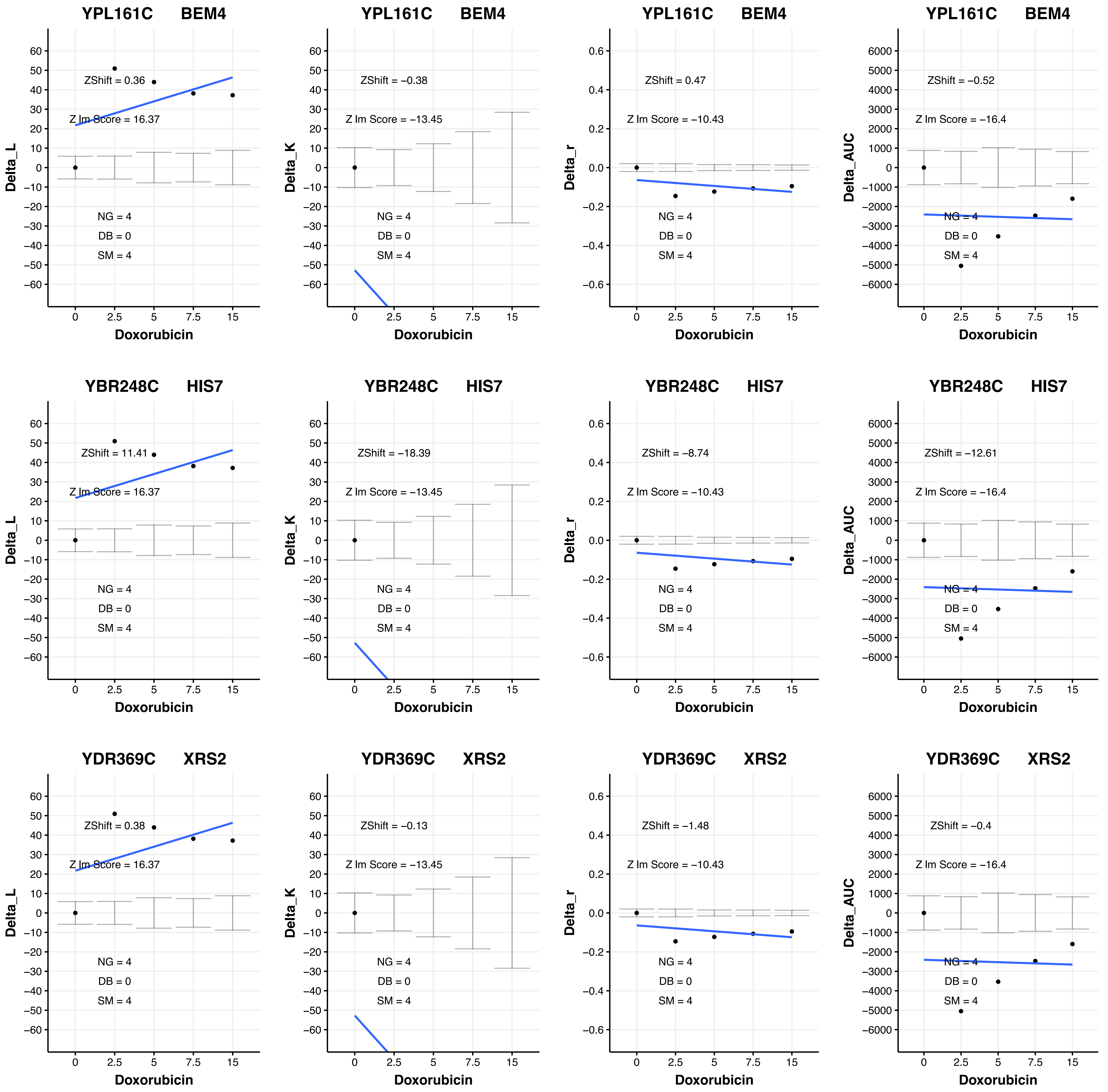
YPL161C BEM4
ZShift = 0.36
Z lm Score = 16.37
NG = 4
DB = 0
SM = 4
Delta_L
Doxorubicin
YPL161C BEM4
ZShift = −0.38
Z lm Score = −13.45
NG = 4
DB = 0
SM = 4
Delta_K
Doxorubicin
YPL161C BEM4
ZShift = 0.47
Z lm Score = −10.43
NG = 4
DB = 0
SM = 4
Delta_r
Doxorubicin
YPL161C BEM4
ZShift = −0.52
Z lm Score = −16.4
NG = 4
DB = 0
SM = 4
Delta_AUC
Doxorubicin
YBR248C HIS7
ZShift = 11.41
Z lm Score = 16.37
NG = 4
DB = 0
SM = 4
Delta_L
Doxorubicin
YBR248C HIS7
ZShift = −18.39
Z lm Score = −13.45
NG = 4
DB = 0
SM = 4
Delta_K
Doxorubicin
YBR248C HIS7
ZShift = −8.74
Z lm Score = −10.43
NG = 4
DB = 0
SM = 4
Delta_r
Doxorubicin
YBR248C HIS7
ZShift = −12.61
Z lm Score = −16.4
NG = 4
DB = 0
SM = 4
Delta_AUC
Doxorubicin
YDR369C XRS2
ZShift = 0.38
Z lm Score = 16.37
NG = 4
DB = 0
SM = 4
Delta_L
Doxorubicin
YDR369C XRS2
ZShift = −0.13
Z lm Score = −13.45
NG = 4
DB = 0
SM = 4
Delta_K
Doxorubicin
YDR369C XRS2
ZShift = −1.48
Z lm Score = −10.43
NG = 4
DB = 0
SM = 4
Delta_r
Doxorubicin
YDR369C XRS2
ZShift = −0.4
Z lm Score = −16.4
NG = 4
DB = 0
SM = 4
Delta_AUC
Doxorubicin

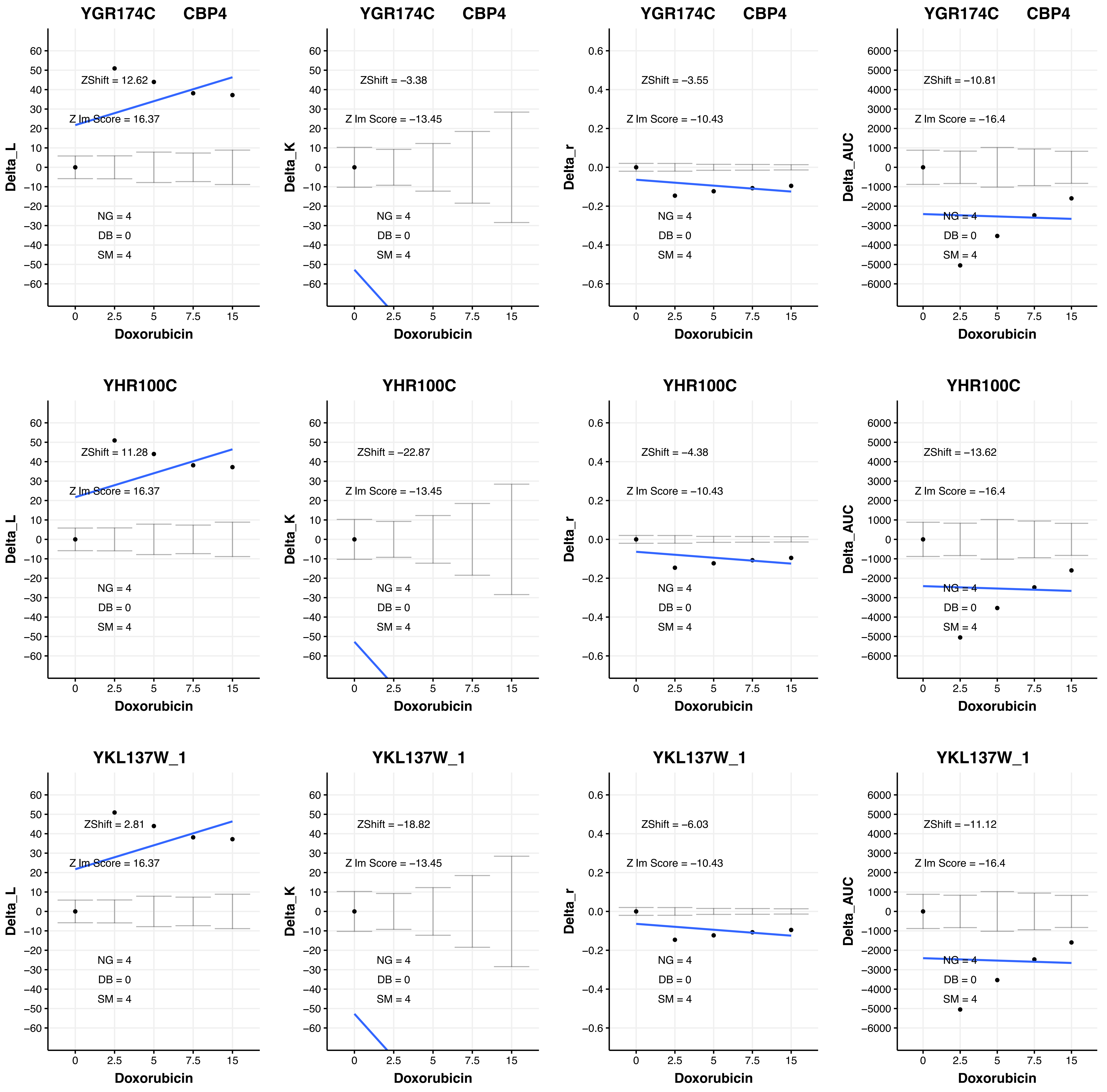
YGR174C CBP4
ZShift = 12.62
Z lm Score = 16.37
NG = 4
DB = 0
SM = 4
Delta_L
Doxorubicin
YGR174C CBP4
ZShift = −3.38
Z lm Score = −13.45
NG = 4
DB = 0
SM = 4
Delta_K
Doxorubicin
YGR174C CBP4
ZShift = −3.55
Z lm Score = −10.43
NG = 4
DB = 0
SM = 4
Delta_r
Doxorubicin
YGR174C CBP4
ZShift = −10.81
Z lm Score = −16.4
NG = 4
DB = 0
SM = 4
Delta_AUC
Doxorubicin
YHR100C
ZShift = 11.28
Z lm Score = 16.37
NG = 4
DB = 0
SM = 4
Delta_L
Doxorubicin
YHR100C
ZShift = −22.87
Z lm Score = −13.45
NG = 4
DB = 0
SM = 4
Delta_K
Doxorubicin
YHR100C
ZShift = −4.38
Z lm Score = −10.43
NG = 4
DB = 0
SM = 4
Delta_r
Doxorubicin
YHR100C
ZShift = −13.62
Z lm Score = −16.4
NG = 4
DB = 0
SM = 4
Delta_AUC
Doxorubicin
YKL137W_1
ZShift = 2.81
Z lm Score = 16.37
NG = 4
DB = 0
SM = 4
Delta_L
Doxorubicin
YKL137W_1
ZShift = −18.82
Z lm Score = −13.45
NG = 4
DB = 0
SM = 4
Delta_K
Doxorubicin
YKL137W_1
ZShift = −6.03
Z lm Score = −10.43
NG = 4
DB = 0
SM = 4
Delta_r
Doxorubicin
YKL137W_1
ZShift = −11.12
Z lm Score = −16.4
NG = 4
DB = 0
SM = 4
Delta_AUC
Doxorubicin

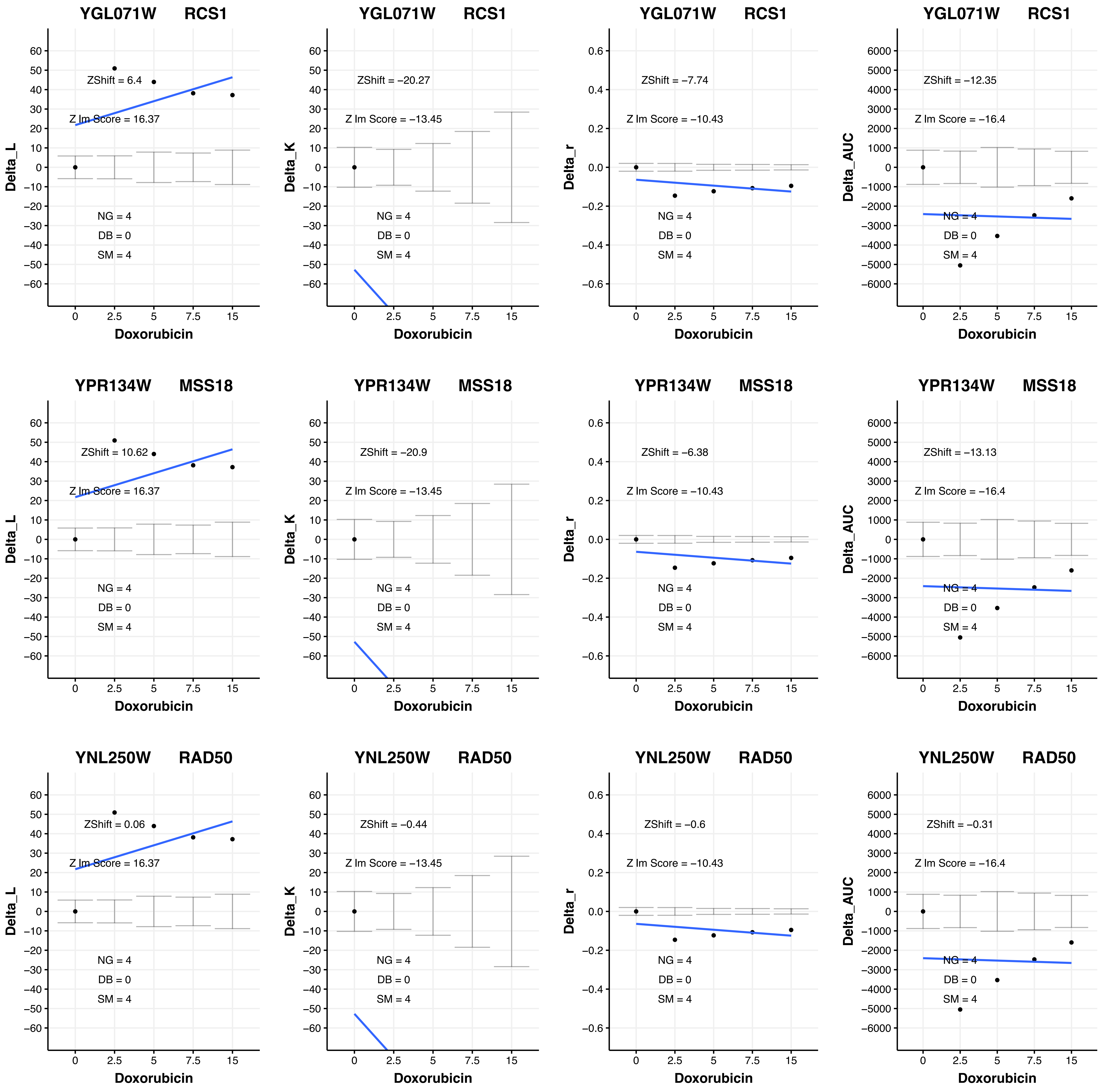
YGL071W RCS1
ZShift = 6.4
Z lm Score = 16.37
NG = 4
DB = 0
SM = 4
Delta_L
Doxorubicin
YGL071W RCS1
ZShift = −20.27
Z lm Score = −13.45
NG = 4
DB = 0
SM = 4
Delta_K
Doxorubicin
YGL071W RCS1
ZShift = −7.74
Z lm Score = −10.43
NG = 4
DB = 0
SM = 4
Delta_r
Doxorubicin
YGL071W RCS1
ZShift = −12.35
Z lm Score = −16.4
NG = 4
DB = 0
SM = 4
Delta_AUC
Doxorubicin
YPR134W MSS18
ZShift = 10.62
Z lm Score = 16.37
NG = 4
DB = 0
SM = 4
Delta_L
Doxorubicin
YPR134W MSS18
ZShift = −20.9
Z lm Score = −13.45
NG = 4
DB = 0
SM = 4
Delta_K
Doxorubicin
YPR134W MSS18
ZShift = −6.38
Z lm Score = −10.43
NG = 4
DB = 0
SM = 4
Delta_r
Doxorubicin
YPR134W MSS18
ZShift = −13.13
Z lm Score = −16.4
NG = 4
DB = 0
SM = 4
Delta_AUC
Doxorubicin
YNL250W RAD50
ZShift = 0.06
Z lm Score = 16.37
NG = 4
DB = 0
SM = 4
Delta_L
Doxorubicin
YNL250W RAD50
ZShift = −0.44
Z lm Score = −13.45
NG = 4
DB = 0
SM = 4
Delta_K
Doxorubicin
YNL250W RAD50
ZShift = −0.6
Z lm Score = −10.43
NG = 4
DB = 0
SM = 4
Delta_r
Doxorubicin
YNL250W RAD50
ZShift = −0.31
Z lm Score = −16.4
NG = 4
DB = 0
SM = 4
Delta_AUC
Doxorubicin

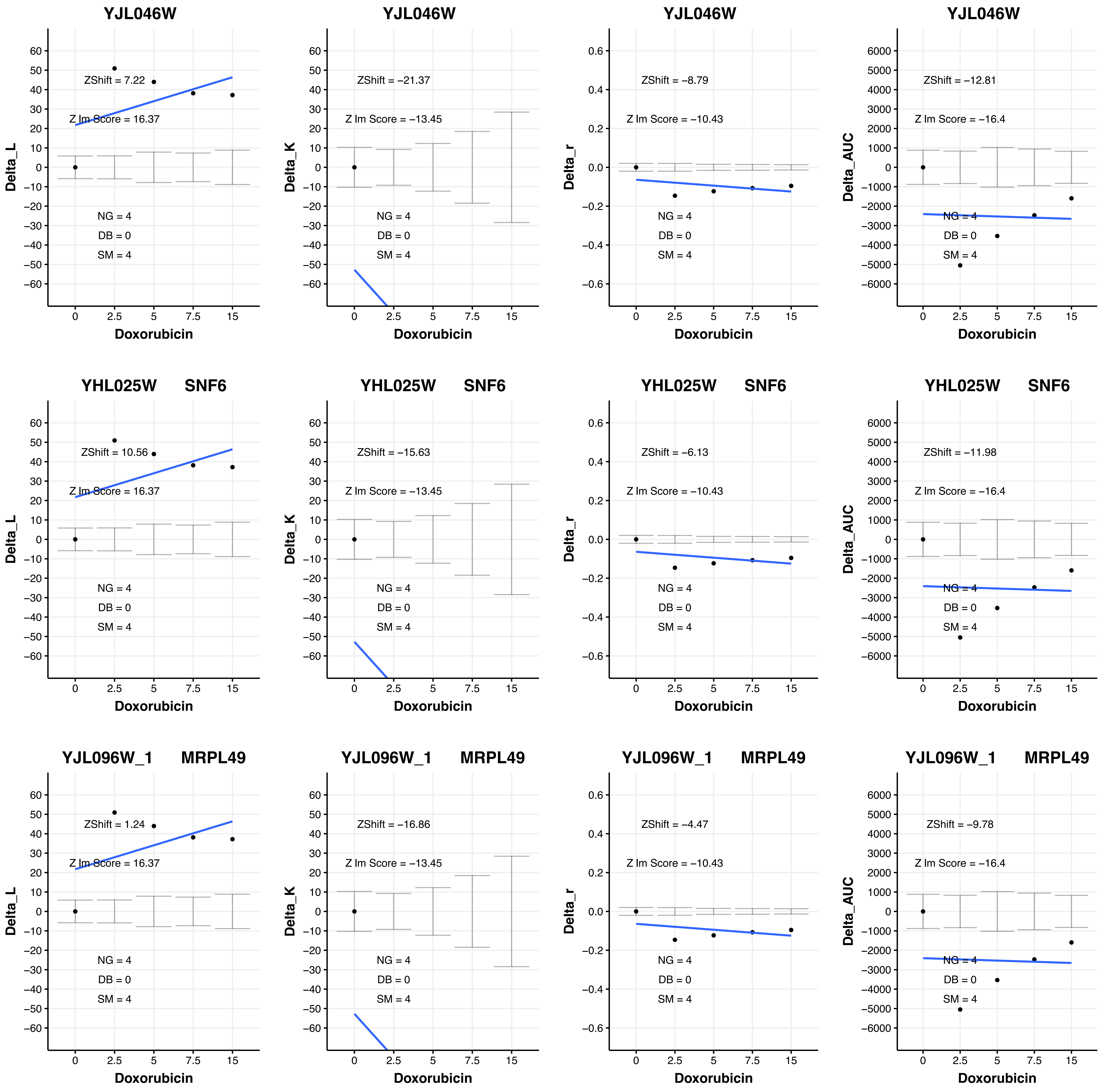
YJL046W
ZShift = 7.22
Z lm Score = 16.37
NG = 4
DB = 0
SM = 4
Delta_L
Doxorubicin
YJL046W
ZShift = −21.37
Z lm Score = −13.45
NG = 4
DB = 0
SM = 4
Delta_K
Doxorubicin
YJL046W
ZShift = −8.79
Z lm Score = −10.43
NG = 4
DB = 0
SM = 4
Delta_r
Doxorubicin
YJL046W
ZShift = −12.81
Z lm Score = −16.4
NG = 4
DB = 0
SM = 4
Delta_AUC
Doxorubicin
YHL025W SNF6
ZShift = 10.56
Z lm Score = 16.37
NG = 4
DB = 0
SM = 4
Delta_L
Doxorubicin
YHL025W SNF6
ZShift = −15.63
Z lm Score = −13.45
NG = 4
DB = 0
SM = 4
Delta_K
Doxorubicin
YHL025W SNF6
ZShift = −6.13
Z lm Score = −10.43
NG = 4
DB = 0
SM = 4
Delta_r
Doxorubicin
YHL025W SNF6
ZShift = −11.98
Z lm Score = −16.4
NG = 4
DB = 0
SM = 4
Delta_AUC
Doxorubicin
YJL096W_1 MRPL49
ZShift = 1.24
Z lm Score = 16.37
NG = 4
DB = 0
SM = 4
Delta_L
Doxorubicin
YJL096W_1 MRPL49
ZShift = −16.86
Z lm Score = −13.45
NG = 4
DB = 0
SM = 4
Delta_K
Doxorubicin
YJL096W_1 MRPL49
ZShift = −4.47
Z lm Score = −10.43
NG = 4
DB = 0
SM = 4
Delta_r
Doxorubicin
YJL096W_1 MRPL49
ZShift = −9.78
Z lm Score = −16.4
NG = 4
DB = 0
SM = 4
Delta_AUC
Doxorubicin

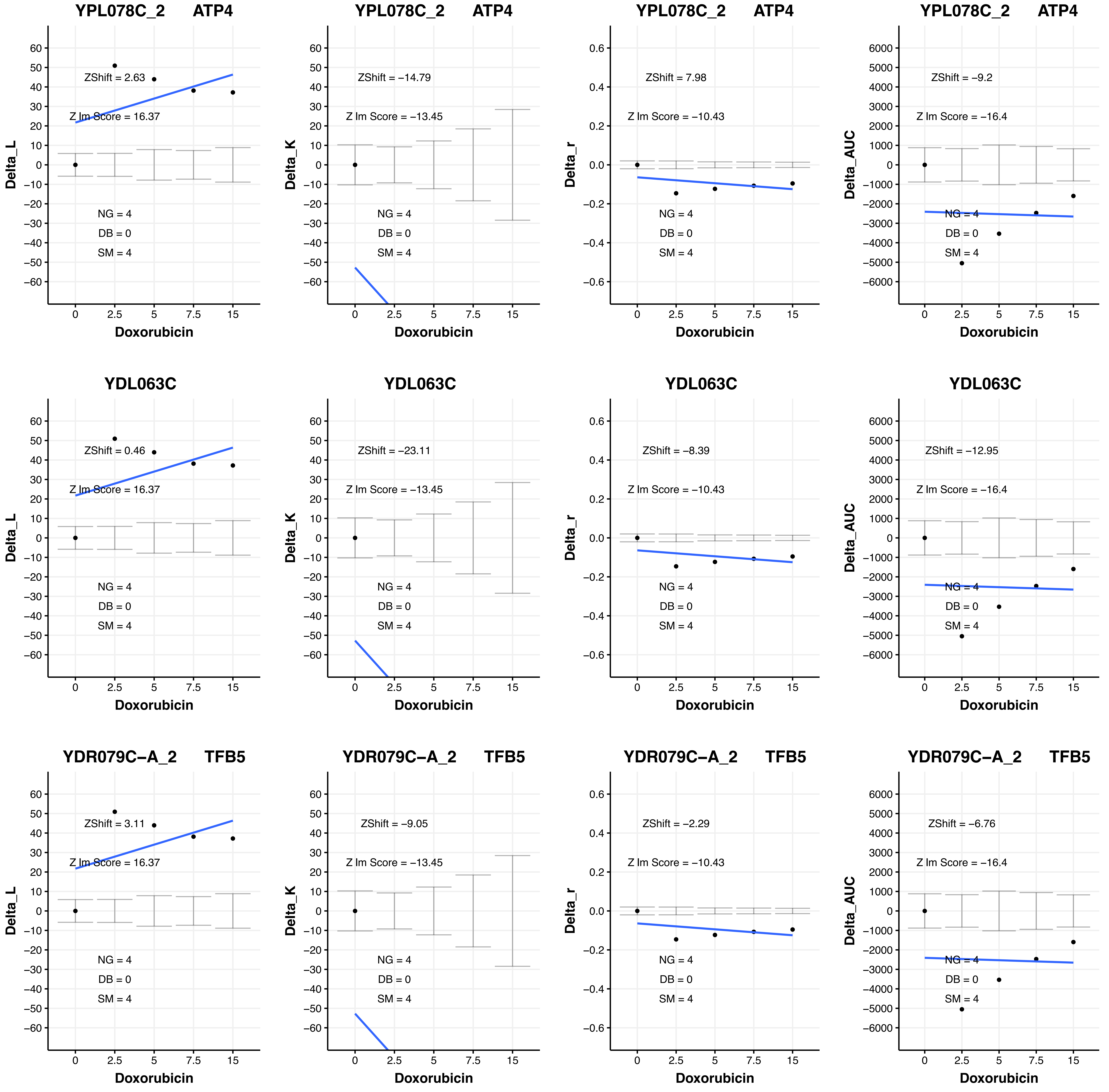
YPL078C_2 ATP4
ZShift = 2.63
Z lm Score = 16.37
NG = 4
DB = 0
SM = 4
Delta_L
Doxorubicin
YPL078C_2 ATP4
ZShift = −14.79
Z lm Score = −13.45
NG = 4
DB = 0
SM = 4
Delta_K
Doxorubicin
YPL078C_2 ATP4
ZShift = 7.98
Z lm Score = −10.43
NG = 4
DB = 0
SM = 4
Delta_r
Doxorubicin
YPL078C_2 ATP4
ZShift = −9.2
Z lm Score = −16.4
NG = 4
DB = 0
SM = 4
Delta_AUC
Doxorubicin
YDL063C
ZShift = 0.46
Z lm Score = 16.37
NG = 4
DB = 0
SM = 4
Delta_L
Doxorubicin
YDL063C
ZShift = −23.11
Z lm Score = −13.45
NG = 4
DB = 0
SM = 4
Delta_K
Doxorubicin
YDL063C
ZShift = −8.39
Z lm Score = −10.43
NG = 4
DB = 0
SM = 4
Delta_r
Doxorubicin
YDL063C
ZShift = −12.95
Z lm Score = −16.4
NG = 4
DB = 0
SM = 4
Delta_AUC
Doxorubicin
YDR079C−A_2 TFB5
ZShift = 3.11
Z lm Score = 16.37
NG = 4
DB = 0
SM = 4
Delta_L
Doxorubicin
YDR079C−A_2 TFB5
ZShift = −9.05
Z lm Score = −13.45
NG = 4
DB = 0
SM = 4
Delta_K
Doxorubicin
YDR079C−A_2 TFB5
ZShift = −2.29
Z lm Score = −10.43
NG = 4
DB = 0
SM = 4
Delta_r
Doxorubicin
YDR079C−A_2 TFB5
ZShift = −6.76
Z lm Score = −16.4
NG = 4
DB = 0
SM = 4
Delta_AUC
Doxorubicin

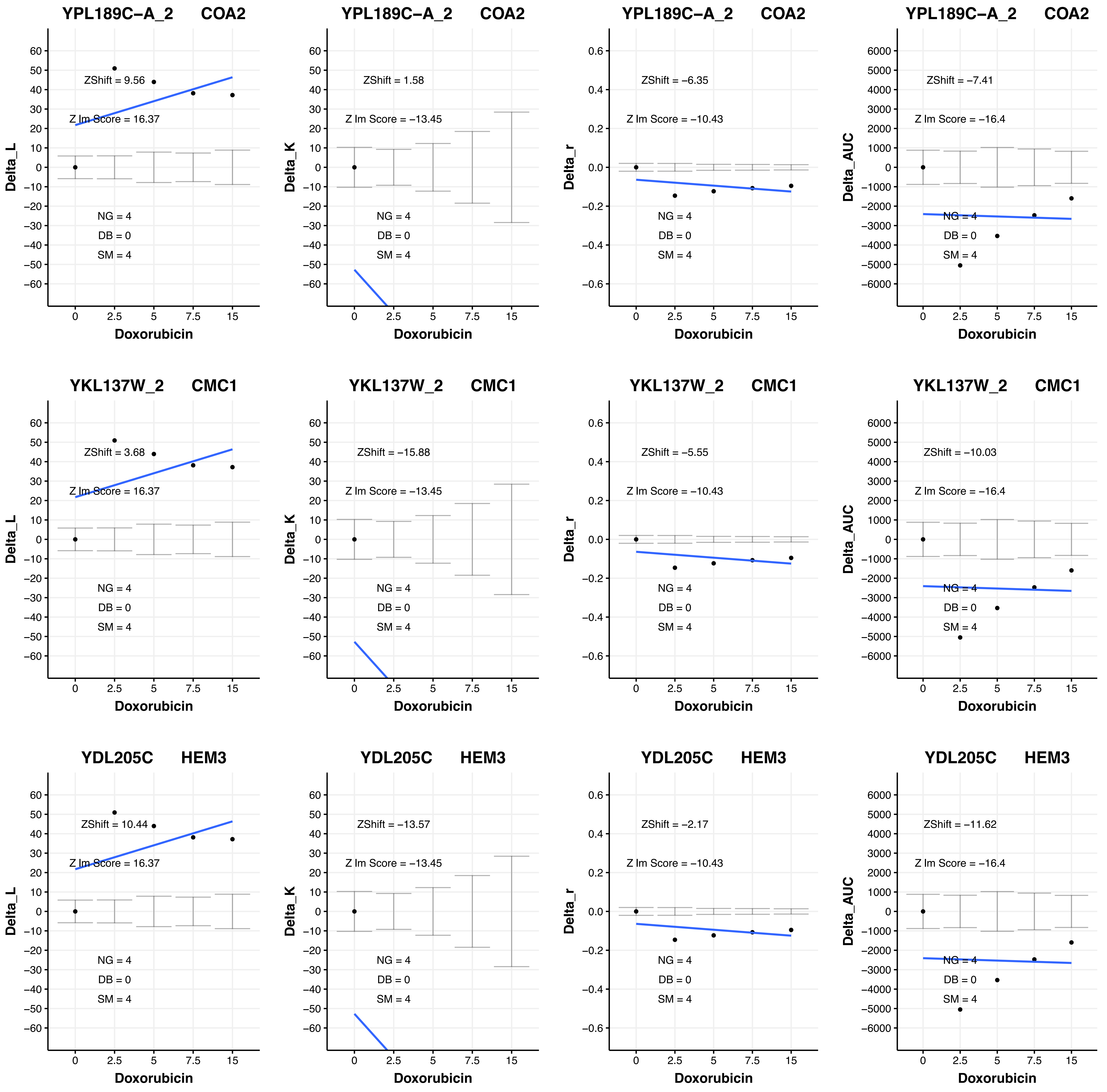
YPL189C−A_2 COA2
ZShift = 9.56
Z lm Score = 16.37
NG = 4
DB = 0
SM = 4
Delta_L
Doxorubicin
YPL189C−A_2 COA2
ZShift = 1.58
Z lm Score = −13.45
NG = 4
DB = 0
SM = 4
Delta_K
Doxorubicin
YPL189C−A_2 COA2
ZShift = −6.35
Z lm Score = −10.43
NG = 4
DB = 0
SM = 4
Delta_r
Doxorubicin
YPL189C−A_2 COA2
ZShift = −7.41
Z lm Score = −16.4
NG = 4
DB = 0
SM = 4
Delta_AUC
Doxorubicin
YKL137W_2 CMC1
ZShift = 3.68
Z lm Score = 16.37
NG = 4
DB = 0
SM = 4
Delta_L
Doxorubicin
YKL137W_2 CMC1
ZShift = −15.88
Z lm Score = −13.45
NG = 4
DB = 0
SM = 4
Delta_K
Doxorubicin
YKL137W_2 CMC1
ZShift = −5.55
Z lm Score = −10.43
NG = 4
DB = 0
SM = 4
Delta_r
Doxorubicin
YKL137W_2 CMC1
ZShift = −10.03
Z lm Score = −16.4
NG = 4
DB = 0
SM = 4
Delta_AUC
Doxorubicin
YDL205C HEM3
ZShift = 10.44
Z lm Score = 16.37
NG = 4
DB = 0
SM = 4
Delta_L
Doxorubicin
YDL205C HEM3
ZShift = −13.57
Z lm Score = −13.45
NG = 4
DB = 0
SM = 4
Delta_K
Doxorubicin
YDL205C HEM3
ZShift = −2.17
Z lm Score = −10.43
NG = 4
DB = 0
SM = 4
Delta_r
Doxorubicin
YDL205C HEM3
ZShift = −11.62
Z lm Score = −16.4
NG = 4
DB = 0
SM = 4
Delta_AUC
Doxorubicin

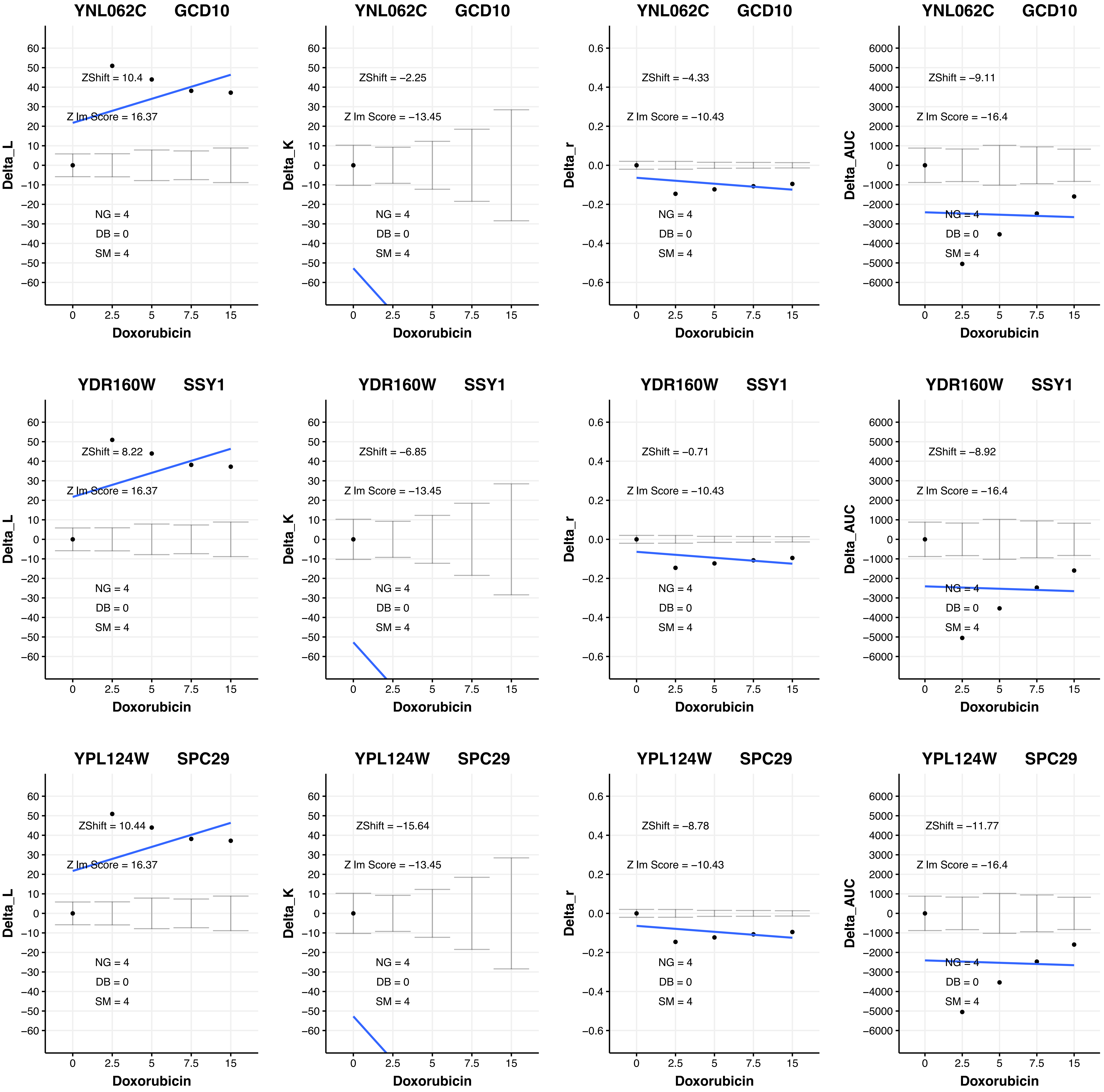

YNL062C GCD10
ZShift = 10.4
Z lm Score = 16.37
NG = 4
DB = 0
SM = 4
Delta_L
Doxorubicin
YNL062C GCD10
ZShift = −2.25
Z lm Score = −13.45
NG = 4
DB = 0
SM = 4
Delta_K
Doxorubicin
YNL062C GCD10
ZShift = −4.33
Z lm Score = −10.43
NG = 4
DB = 0
SM = 4
Delta_r
Doxorubicin
YNL062C GCD10
ZShift = −9.11
Z lm Score = −16.4
NG = 4
DB = 0
SM = 4
Delta_AUC
Doxorubicin
YDR160W SSY1
ZShift = 8.22
Z lm Score = 16.37
NG = 4
DB = 0
SM = 4
Delta_L
Doxorubicin
YDR160W SSY1
ZShift = −6.85
Z lm Score = −13.45
NG = 4
DB = 0
SM = 4
Delta_K
Doxorubicin
YDR160W SSY1
ZShift = −0.71
Z lm Score = −10.43
NG = 4
DB = 0
SM = 4
Delta_r
Doxorubicin
YDR160W SSY1
ZShift = −8.92
Z lm Score = −16.4
NG = 4
DB = 0
SM = 4
Delta_AUC
Doxorubicin
YPL124W SPC29
ZShift = 10.44
Z lm Score = 16.37
NG = 4
DB = 0
SM = 4
Delta_L
Doxorubicin
YPL124W SPC29
ZShift = −15.64
Z lm Score = −13.45
NG = 4
DB = 0
SM = 4
Delta_K
Doxorubicin
YPL124W SPC29
ZShift = −8.78
Z lm Score = −10.43
NG = 4
DB = 0
SM = 4
Delta_r
Doxorubicin
YPL124W SPC29
ZShift = −11.77
Z lm Score = −16.4
NG = 4
DB = 0
SM = 4
Delta_AUC
Doxorubicin

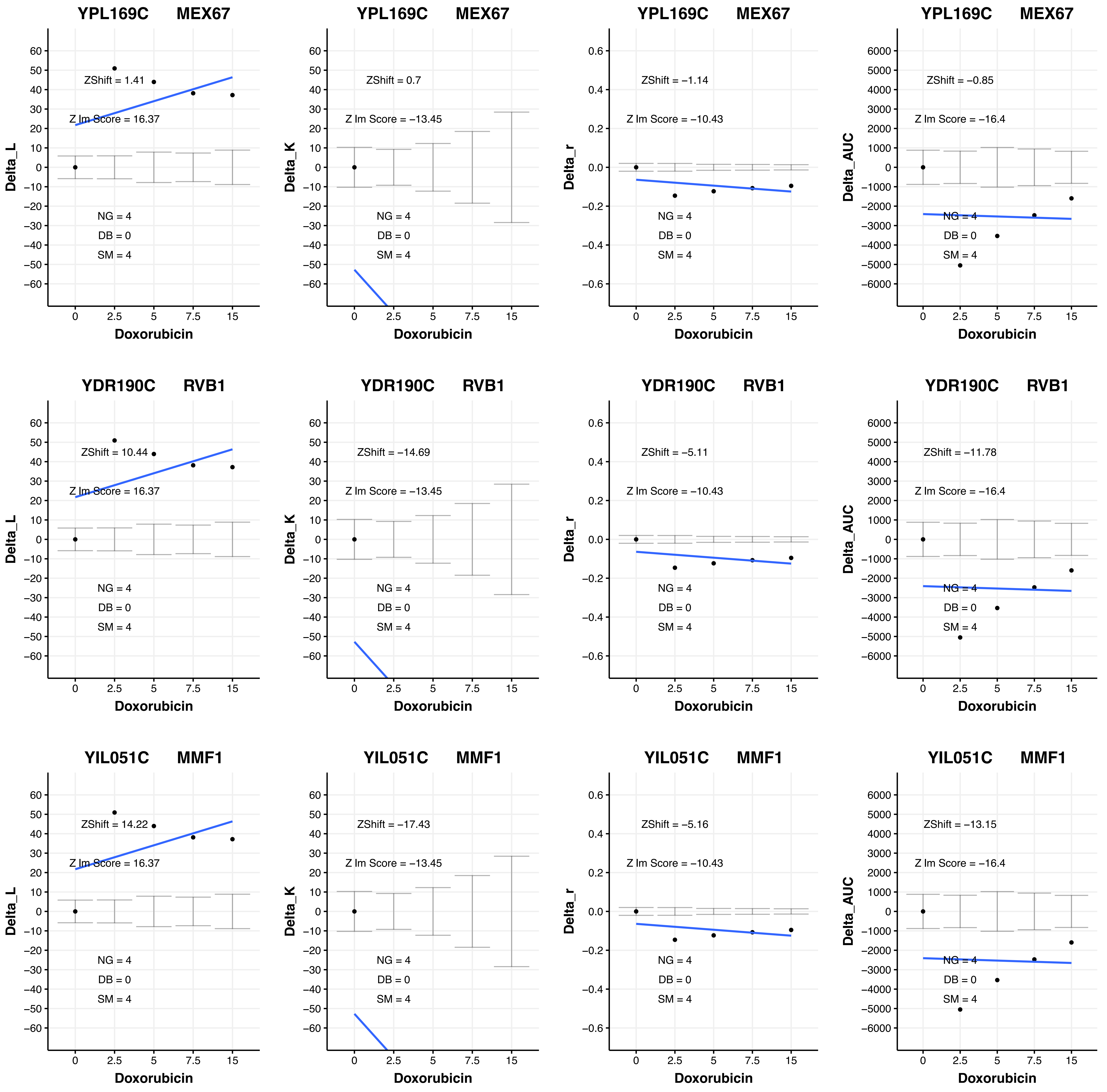
YPL169C MEX67
ZShift = 1.41
Z lm Score = 16.37
NG = 4
DB = 0
SM = 4
Delta_L
Doxorubicin
YPL169C MEX67
ZShift = 0.7
Z lm Score = −13.45
NG = 4
DB = 0
SM = 4
Delta_K
Doxorubicin
YPL169C MEX67
ZShift = −1.14
Z lm Score = −10.43
NG = 4
DB = 0
SM = 4
Delta_r
Doxorubicin
YPL169C MEX67
ZShift = −0.85
Z lm Score = −16.4
NG = 4
DB = 0
SM = 4
Delta_AUC
Doxorubicin
YDR190C RVB1
ZShift = 10.44
Z lm Score = 16.37
NG = 4
DB = 0
SM = 4
Delta_L
Doxorubicin
YDR190C RVB1
ZShift = −14.69
Z lm Score = −13.45
NG = 4
DB = 0
SM = 4
Delta_K
Doxorubicin
YDR190C RVB1
ZShift = −5.11
Z lm Score = −10.43
NG = 4
DB = 0
SM = 4
Delta_r
Doxorubicin
YDR190C RVB1
ZShift = −11.78
Z lm Score = −16.4
NG = 4
DB = 0
SM = 4
Delta_AUC
Doxorubicin
YIL051C MMF1
ZShift = 14.22
Z lm Score = 16.37
NG = 4
DB = 0
SM = 4
Delta_L
Doxorubicin
YIL051C MMF1
ZShift = −17.43
Z lm Score = −13.45
NG = 4
DB = 0
SM = 4
Delta_K
Doxorubicin
YIL051C MMF1
ZShift = −5.16
Z lm Score = −10.43
NG = 4
DB = 0
SM = 4
Delta_r
Doxorubicin
YIL051C MMF1
ZShift = −13.15
Z lm Score = −16.4
NG = 4
DB = 0
SM = 4
Delta_AUC
Doxorubicin

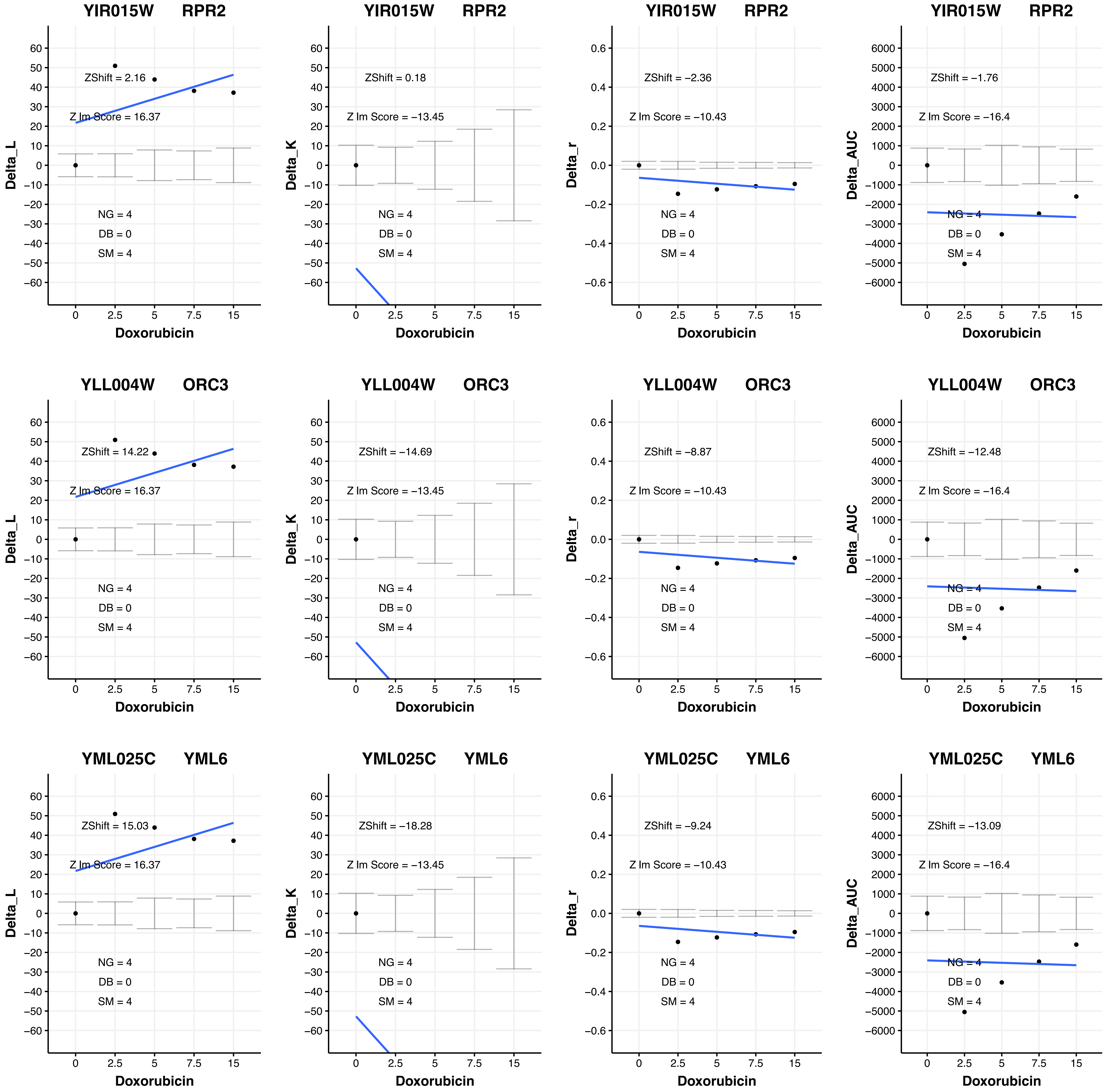
YIR015W RPR2
ZShift = 2.16
Z lm Score = 16.37
NG = 4
DB = 0
SM = 4
Delta_L
Doxorubicin
YIR015W RPR2
ZShift = 0.18
Z lm Score = −13.45
NG = 4
DB = 0
SM = 4
Delta_K
Doxorubicin
YIR015W RPR2
ZShift = −2.36
Z lm Score = −10.43
NG = 4
DB = 0
SM = 4
Delta_r
Doxorubicin
YIR015W RPR2
ZShift = −1.76
Z lm Score = −16.4
NG = 4
DB = 0
SM = 4
Delta_AUC
Doxorubicin
YLL004W ORC3
ZShift = 14.22
Z lm Score = 16.37
NG = 4
DB = 0
SM = 4
Delta_L
Doxorubicin
YLL004W ORC3
ZShift = −14.69
Z lm Score = −13.45
NG = 4
DB = 0
SM = 4
Delta_K
Doxorubicin
YLL004W ORC3
ZShift = −8.87
Z lm Score = −10.43
NG = 4
DB = 0
SM = 4
Delta_r
Doxorubicin
YLL004W ORC3
ZShift = −12.48
Z lm Score = −16.4
NG = 4
DB = 0
SM = 4
Delta_AUC
Doxorubicin
YML025C YML6
ZShift = 15.03
Z lm Score = 16.37
NG = 4
DB = 0
SM = 4
Delta_L
Doxorubicin
YML025C YML6
ZShift = −18.28
Z lm Score = −13.45
NG = 4
DB = 0
SM = 4
Delta_K
Doxorubicin
YML025C YML6
ZShift = −9.24
Z lm Score = −10.43
NG = 4
DB = 0
SM = 4
Delta_r
Doxorubicin
YML025C YML6
ZShift = −13.09
Z lm Score = −16.4
NG = 4
DB = 0
SM = 4
Delta_AUC
Doxorubicin

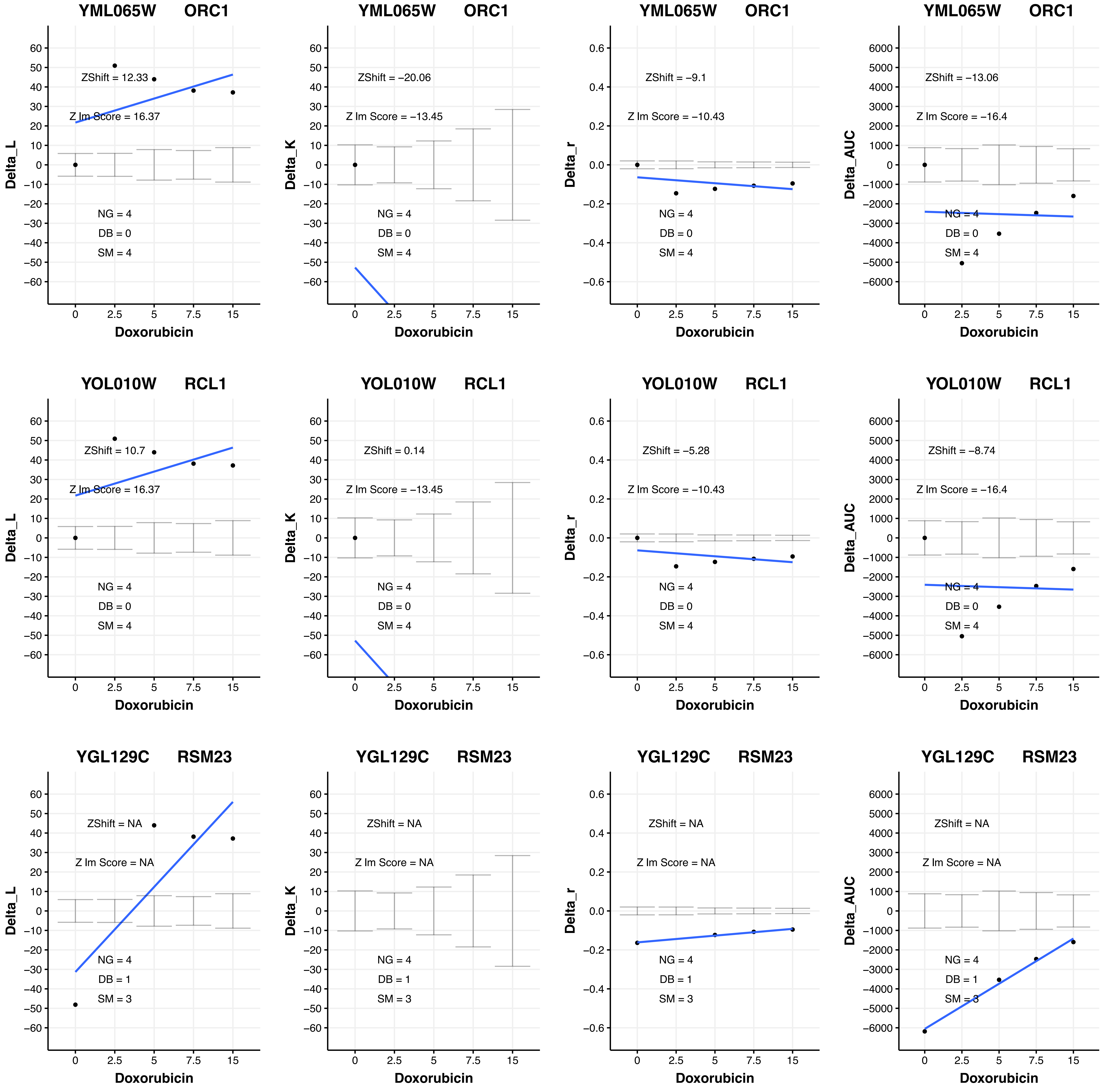

YML065W ORC1
ZShift = 12.33
Z lm Score = 16.37
NG = 4
DB = 0
SM = 4
Delta_L
Doxorubicin
YML065W ORC1
ZShift = −20.06
Z lm Score = −13.45
NG = 4
DB = 0
SM = 4
Delta_K
Doxorubicin
YML065W ORC1
ZShift = −9.1
Z lm Score = −10.43
NG = 4
DB = 0
SM = 4
Delta_r
Doxorubicin
YML065W ORC1
ZShift = −13.06
Z lm Score = −16.4
NG = 4
DB = 0
SM = 4
Delta_AUC
Doxorubicin
YOL010W RCL1
ZShift = 10.7
Z lm Score = 16.37
NG = 4
DB = 0
SM = 4
Delta_L
Doxorubicin
YOL010W RCL1
ZShift = 0.14
Z lm Score = −13.45
NG = 4
DB = 0
SM = 4
Delta_K
Doxorubicin
YOL010W RCL1
ZShift = −5.28
Z lm Score = −10.43
NG = 4
DB = 0
SM = 4
Delta_r
Doxorubicin
YOL010W RCL1
ZShift = −8.74
Z lm Score = −16.4
NG = 4
DB = 0
SM = 4
Delta_AUC
Doxorubicin
YGL129C RSM23
ZShift = NA
Z lm Score = NA
NG = 4
DB = 1
SM = 3
Delta_L
Doxorubicin
YGL129C RSM23
ZShift = NA
Z lm Score = NA
NG = 4
DB = 1
SM = 3
Delta_K
Doxorubicin
YGL129C RSM23
ZShift = NA
Z lm Score = NA
NG = 4
DB = 1
SM = 3
Delta_r
Doxorubicin
YGL129C RSM23
ZShift = NA
Z lm Score = NA
NG = 4
DB = 1
SM = 3
Delta_AUC
Doxorubicin

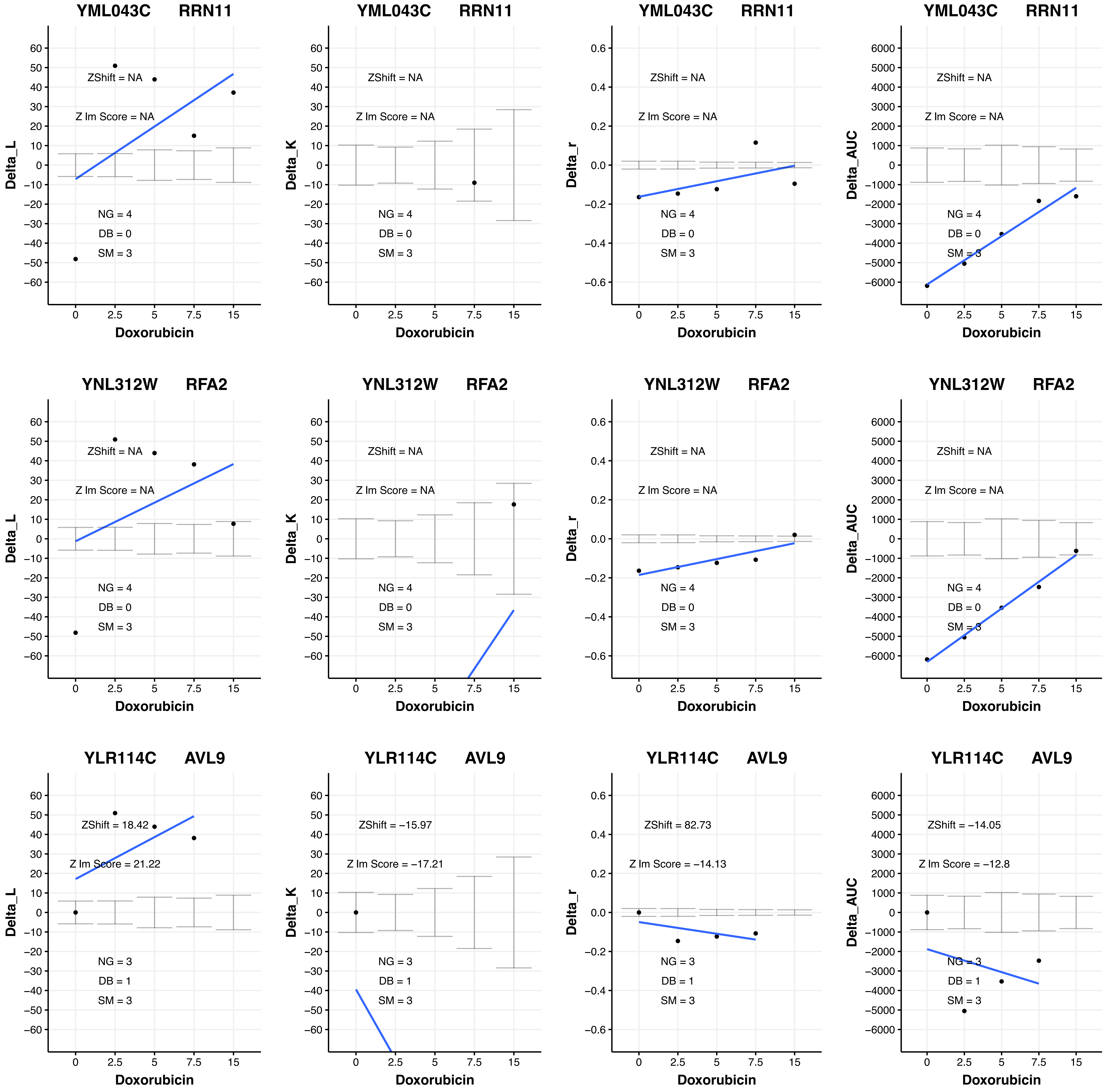
YML043C RRN11
ZShift = NA
Z lm Score = NA
NG = 4
DB = 0
SM = 3
Delta_L
Delta_K
Delta_r
Delta_AUC
Doxorubicin
YNL312W RFA2
YLR114C AVL9
ZShift = 18.42
Z lm Score = 21.22
ZShift = −15.97
Z lm Score = −17.21
ZShift = 82.73
Z lm Score = −14.13
ZShift = −14.05
Z lm Score = −12.8
NG = 3
DB = 1

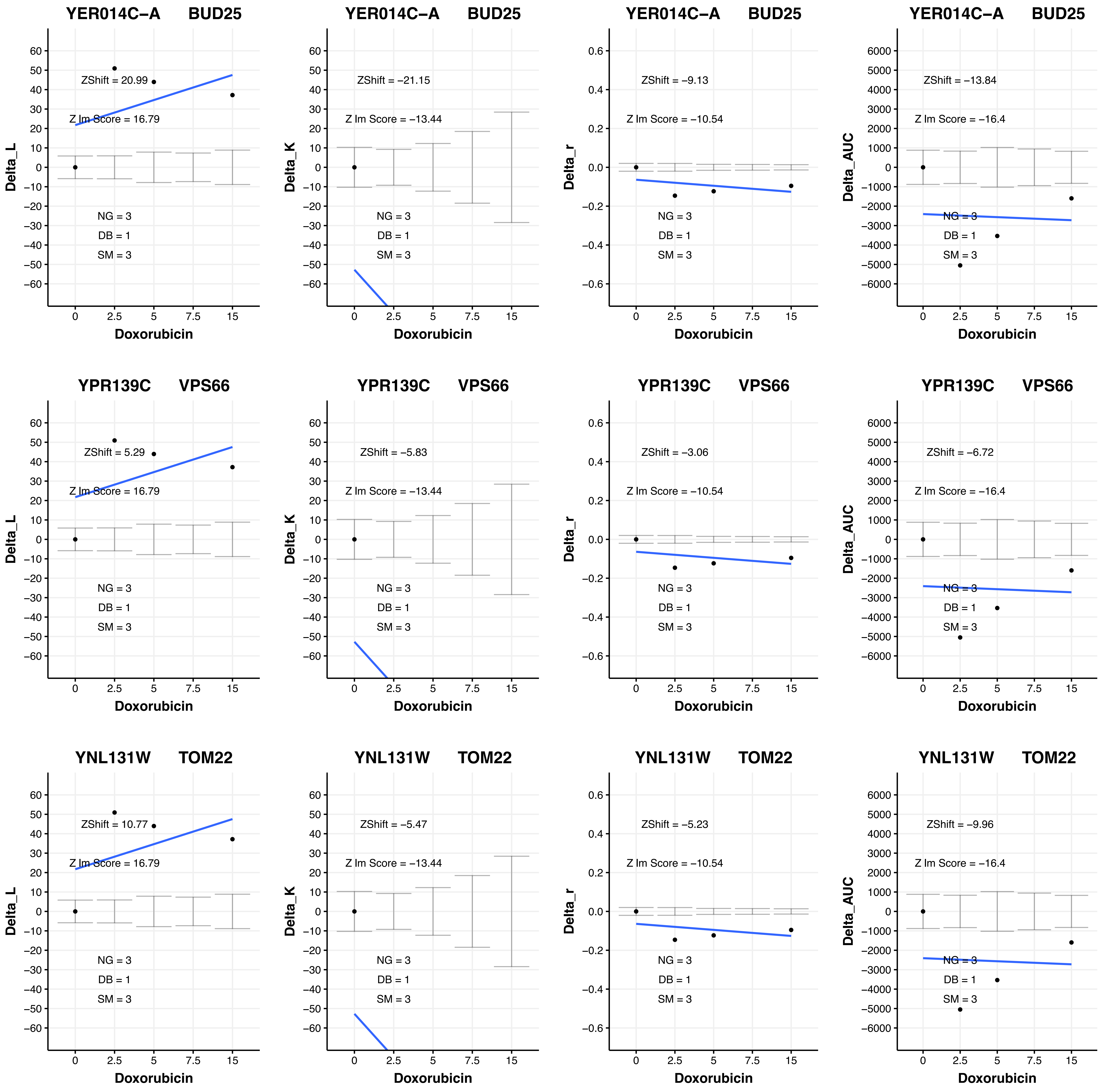
YER014C-A BUD25
ZShift = 20.99
Z lm Score = 16.79
NG = 3
DB = 1
SM = 3
Delta_L
Doxorubicin
0 2.5 5 7.5 15
YER014C-A BUD25
ZShift = −21.15
Z lm Score = −13.44
NG = 3
DB = 1
SM = 3
Delta_K
Doxorubicin
YER014C-A BUD25
ZShift = −9.13
Z lm Score = −10.54
NG = 3
DB = 1
SM = 3
Delta_r
Doxorubicin
YER014C-A BUD25
ZShift = −13.84
Z lm Score = −16.4
NG = 3
DB = 1
SM = 3
Delta_AUC
Doxorubicin
YPR139C VPS66
ZShift = 5.29
Z lm Score = 16.79
NG = 3
DB = 1
SM = 3
Delta_L
Doxorubicin
YPR139C VPS66
ZShift = −5.83
Z lm Score = −13.44
NG = 3
DB = 1
SM = 3
Delta_K
Doxorubicin
YPR139C VPS66
ZShift = −3.06
Z lm Score = −10.54
NG = 3
DB = 1
SM = 3
Delta_r
Doxorubicin
YPR139C VPS66
ZShift = −6.72
Z lm Score = −16.4
NG = 3
DB = 1
SM = 3
Delta_AUC
Doxorubicin
YNL131W TOM22
ZShift = 10.77
Z lm Score = 16.79
NG = 3
DB = 1
SM = 3
Delta_L
Doxorubicin
YNL131W TOM22
ZShift = −5.47
Z lm Score = −13.44
NG = 3
DB = 1
SM = 3
Delta_K
Doxorubicin
YNL131W TOM22
ZShift = −5.23
Z lm Score = −10.54
NG = 3
DB = 1
SM = 3
Delta_r
Doxorubicin
YNL131W TOM22
ZShift = −9.96
Z lm Score = −16.4
NG = 3
DB = 1
SM = 3
Delta_AUC
Doxorubicin

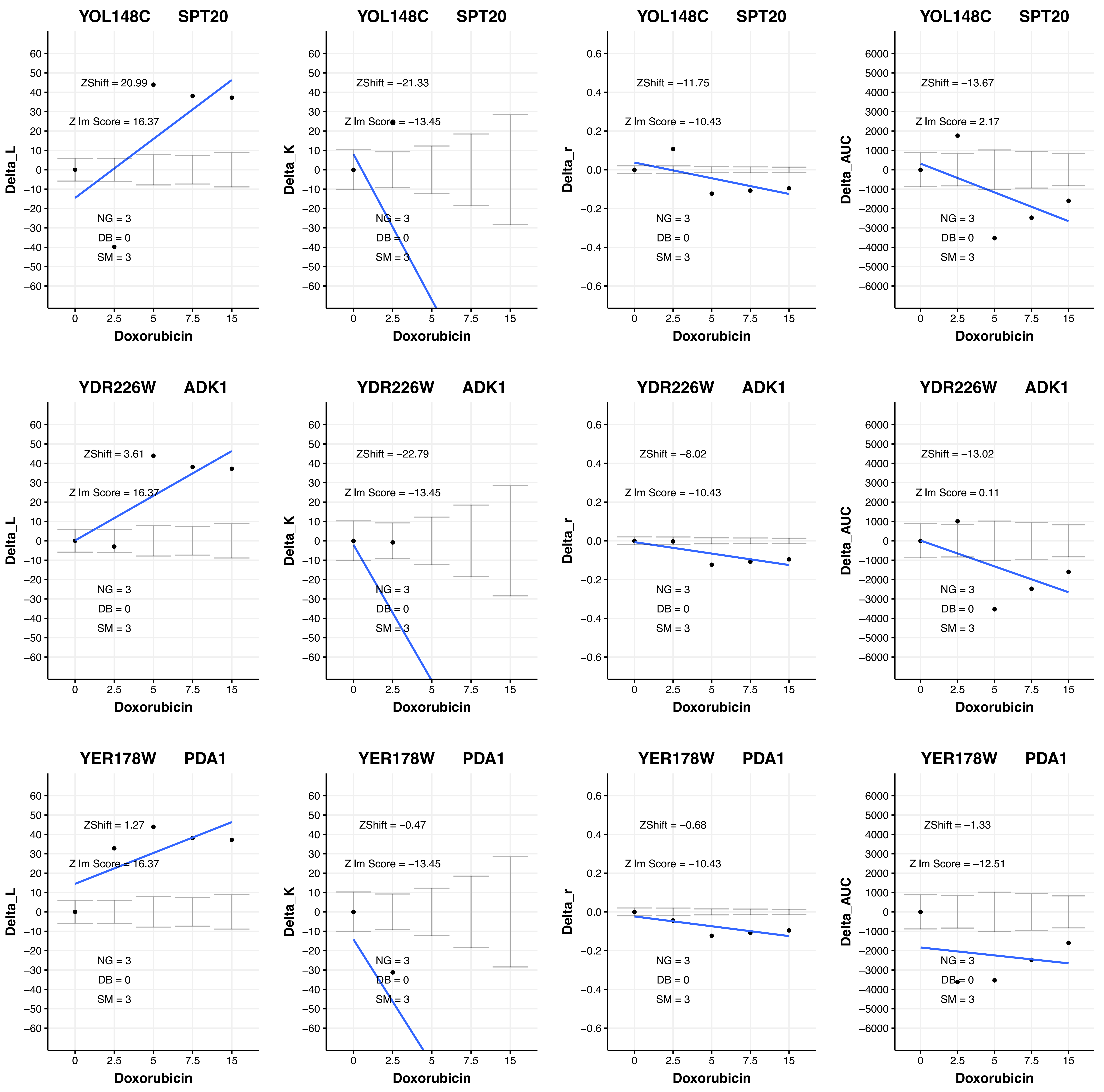

YOL148C SPT20
ZShift = 20.99
Z lm Score = 16.37
NG = 3
DB = 0
SM = 3
Delta_L
Doxorubicin
ZShift = −21.33
Z lm Score = −13.45
Delta_K
ZShift = −11.75
Z lm Score = −10.43
Delta_r
ZShift = −13.67
Z lm Score = 2.17
Delta_AUC
YDR226W ADK1
ZShift = 3.61
Z lm Score = 16.37
ZShift = −22.79
Z lm Score = −13.45
ZShift = −8.02
Z lm Score = −10.43
ZShift = −13.02
Z lm Score = 0.11
YER178W PDA1
ZShift = 1.27
Z lm Score = 16.37
ZShift = −0.47
Z lm Score = −13.45
ZShift = −0.68
Z lm Score = −10.43
ZShift = −1.33
Z lm Score = −12.51

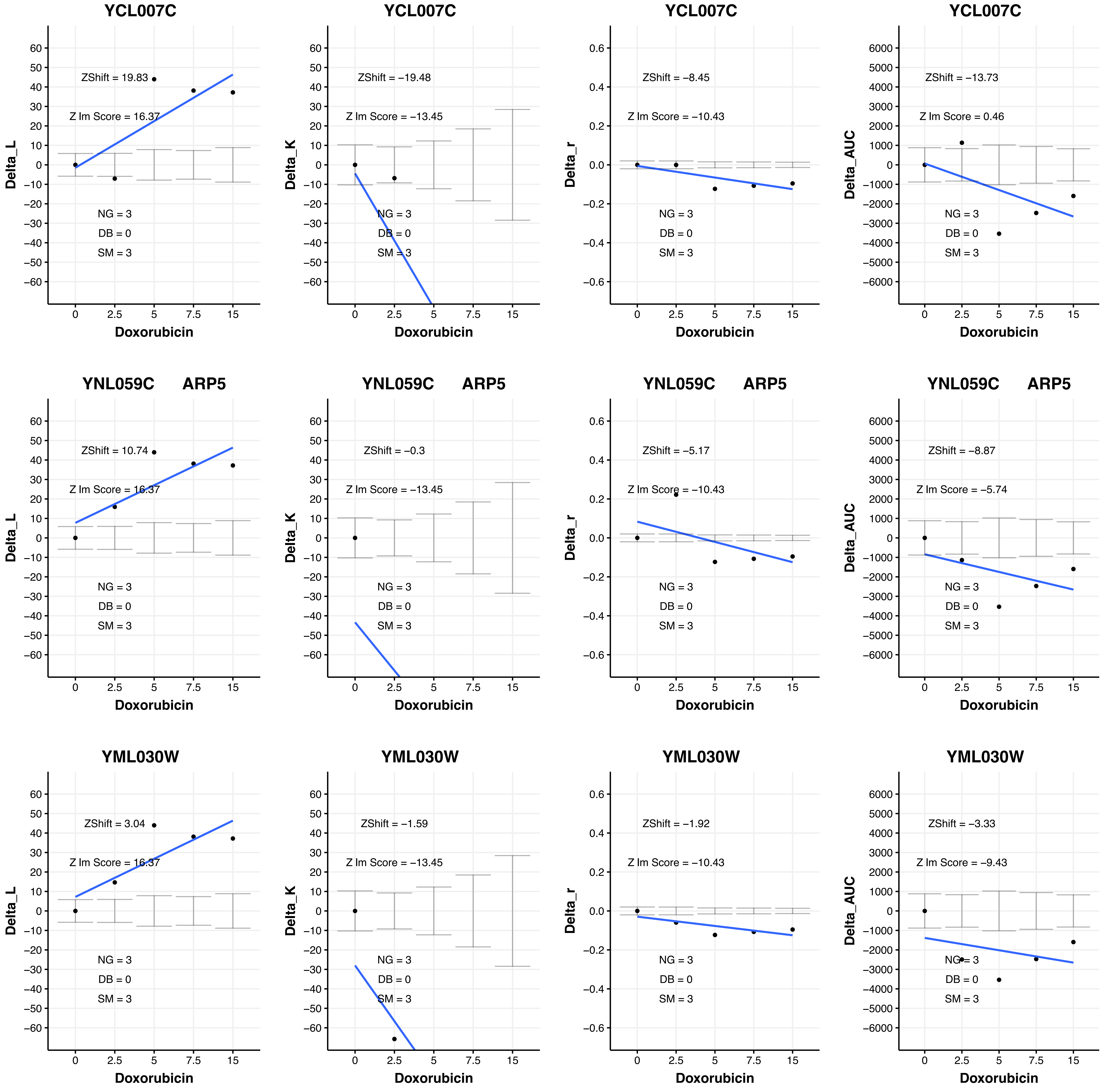
YCL007C
ZShift = 19.83
Z lm Score = 16.37
NG = 3
DB = 0
SM = 3
Delta_L
Doxorubicin
YCL007C
ZShift = −19.48
Z lm Score = −13.45
NG = 3
DB = 0
SM = 3
Delta_K
Doxorubicin
YCL007C
ZShift = −8.45
Z lm Score = −10.43
NG = 3
DB = 0
SM = 3
Delta_r
Doxorubicin
YCL007C
ZShift = −13.73
Z lm Score = 0.46
NG = 3
DB = 0
SM = 3
Delta_AUC
Doxorubicin
YNL059C ARP5
ZShift = 10.74
Z lm Score = 16.37
NG = 3
DB = 0
SM = 3
Delta_L
Doxorubicin
YNL059C ARP5
ZShift = −0.3
Z lm Score = −13.45
NG = 3
DB = 0
SM = 3
Delta_K
Doxorubicin
YNL059C ARP5
ZShift = −5.17
Z lm Score = −10.43
NG = 3
DB = 0
SM = 3
Delta_r
Doxorubicin
YNL059C ARP5
ZShift = −8.87
Z lm Score = −5.74
NG = 3
DB = 0
SM = 3
Delta_AUC
Doxorubicin
YML030W
ZShift = 3.04
Z lm Score = 16.37
NG = 3
DB = 0
SM = 3
Delta_L
Doxorubicin
YML030W
ZShift = −1.59
Z lm Score = −13.45
NG = 3
DB = 0
SM = 3
Delta_K
Doxorubicin
YML030W
ZShift = −1.92
Z lm Score = −10.43
NG = 3
DB = 0
SM = 3
Delta_r
Doxorubicin
YML030W
ZShift = −3.33
Z lm Score = −9.43
NG = 3
DB = 0
SM = 3
Delta_AUC
Doxorubicin

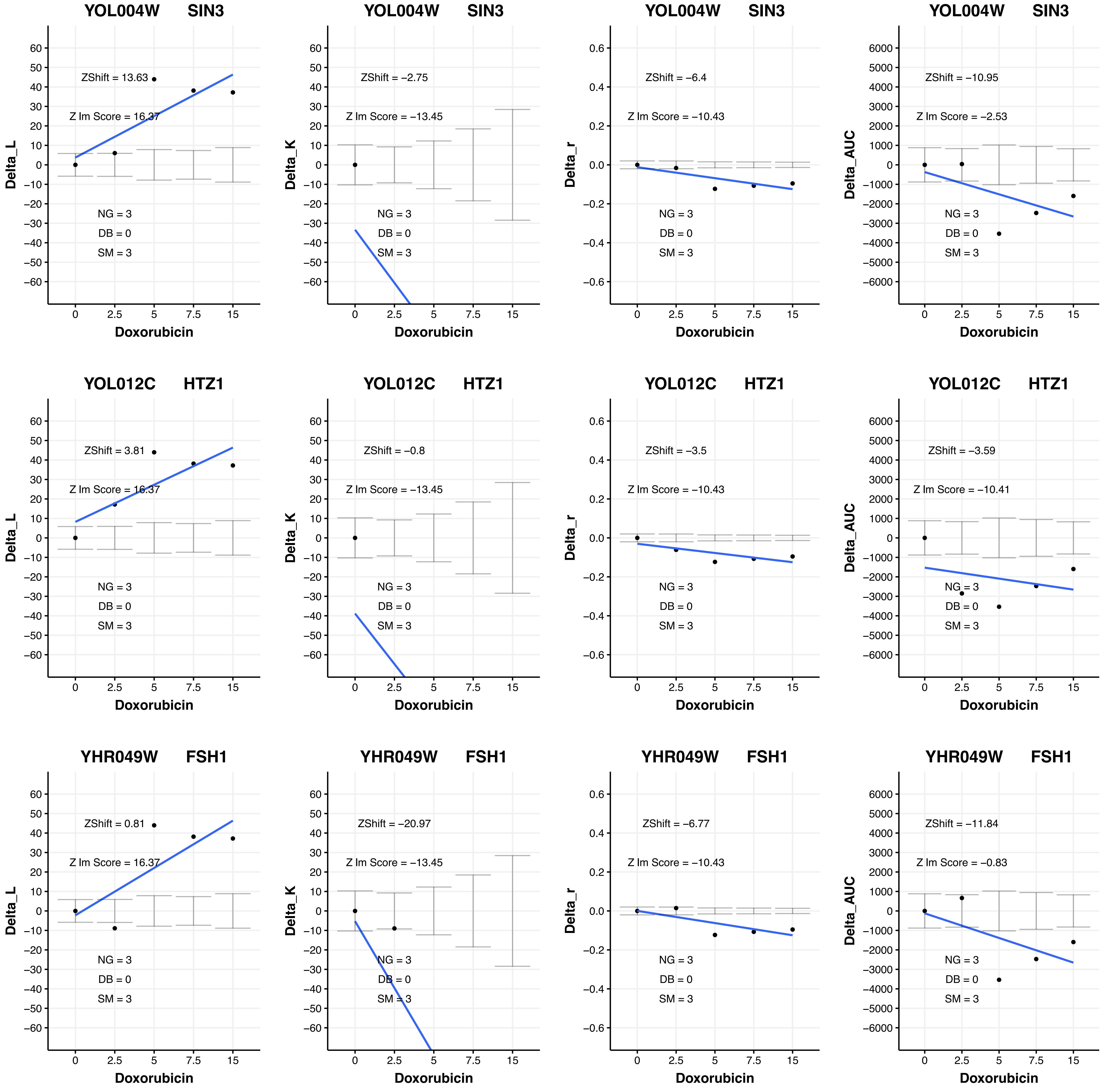

YOL004W SIN3
ZShift = 13.63
Z lm Score = 16.37
NG = 3
DB = 0
SM = 3
Delta_L
Doxorubicin
YOL004W SIN3
ZShift = −2.75
Z lm Score = −13.45
NG = 3
DB = 0
SM = 3
Delta_K
Doxorubicin
YOL004W SIN3
ZShift = −6.4
Z lm Score = −10.43
NG = 3
DB = 0
SM = 3
Delta_r
Doxorubicin
YOL004W SIN3
ZShift = −10.95
Z lm Score = −2.53
NG = 3
DB = 0
SM = 3
Delta_AUC
Doxorubicin
YOL012C HTZ1
ZShift = 3.81
Z lm Score = 16.37
NG = 3
DB = 0
SM = 3
Delta_L
Doxorubicin
YOL012C HTZ1
ZShift = −0.8
Z lm Score = −13.45
NG = 3
DB = 0
SM = 3
Delta_K
Doxorubicin
YOL012C HTZ1
ZShift = −3.5
Z lm Score = −10.43
NG = 3
DB = 0
SM = 3
Delta_r
Doxorubicin
YOL012C HTZ1
ZShift = −3.59
Z lm Score = −10.41
NG = 3
DB = 0
SM = 3
Delta_AUC
Doxorubicin
YHR049W FSH1
ZShift = 0.81
Z lm Score = 16.37
NG = 3
DB = 0
SM = 3
Delta_L
Doxorubicin
YHR049W FSH1
ZShift = −20.97
Z lm Score = −13.45
NG = 3
DB = 0
SM = 3
Delta_K
Doxorubicin
YHR049W FSH1
ZShift = −6.77
Z lm Score = −10.43
NG = 3
DB = 0
SM = 3
Delta_r
Doxorubicin
YHR049W FSH1
ZShift = −11.84
Z lm Score = −0.83
NG = 3
DB = 0
SM = 3
Delta_AUC
Doxorubicin

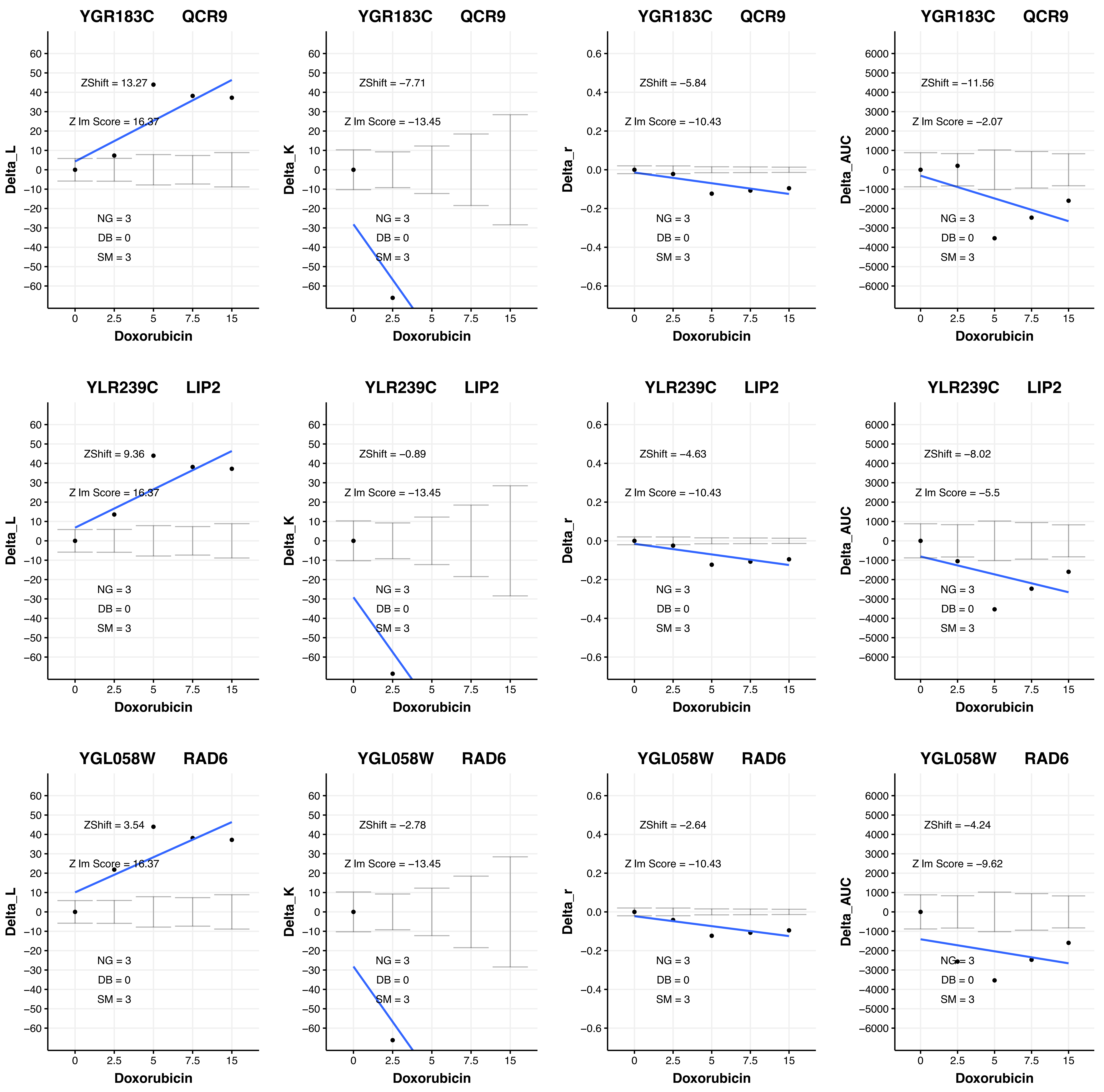

YGR183C QCR9
ZShift = 13.27
Z Im Score = 16.37
NG = 3
DB = 0
SM = 3
Delta_L
Doxorubicin
ZShift = −7.71
Z Im Score = −13.45
Delta_K
ZShift = −5.84
Z Im Score = −10.43
Delta_r
ZShift = −11.56
Z Im Score = −2.07
Delta_AUC
YLR239C LIP2
ZShift = 9.36
Z Im Score = 16.37
ZShift = −0.89
Z Im Score = −13.45
ZShift = −4.63
Z Im Score = −10.43
ZShift = −8.02
Z Im Score = −5.5
YGL058W RAD6
ZShift = 3.54
Z Im Score = 16.37
ZShift = −2.78
Z Im Score = −13.45
ZShift = −2.64
Z Im Score = −10.43
ZShift = −4.24
Z Im Score = −9.62

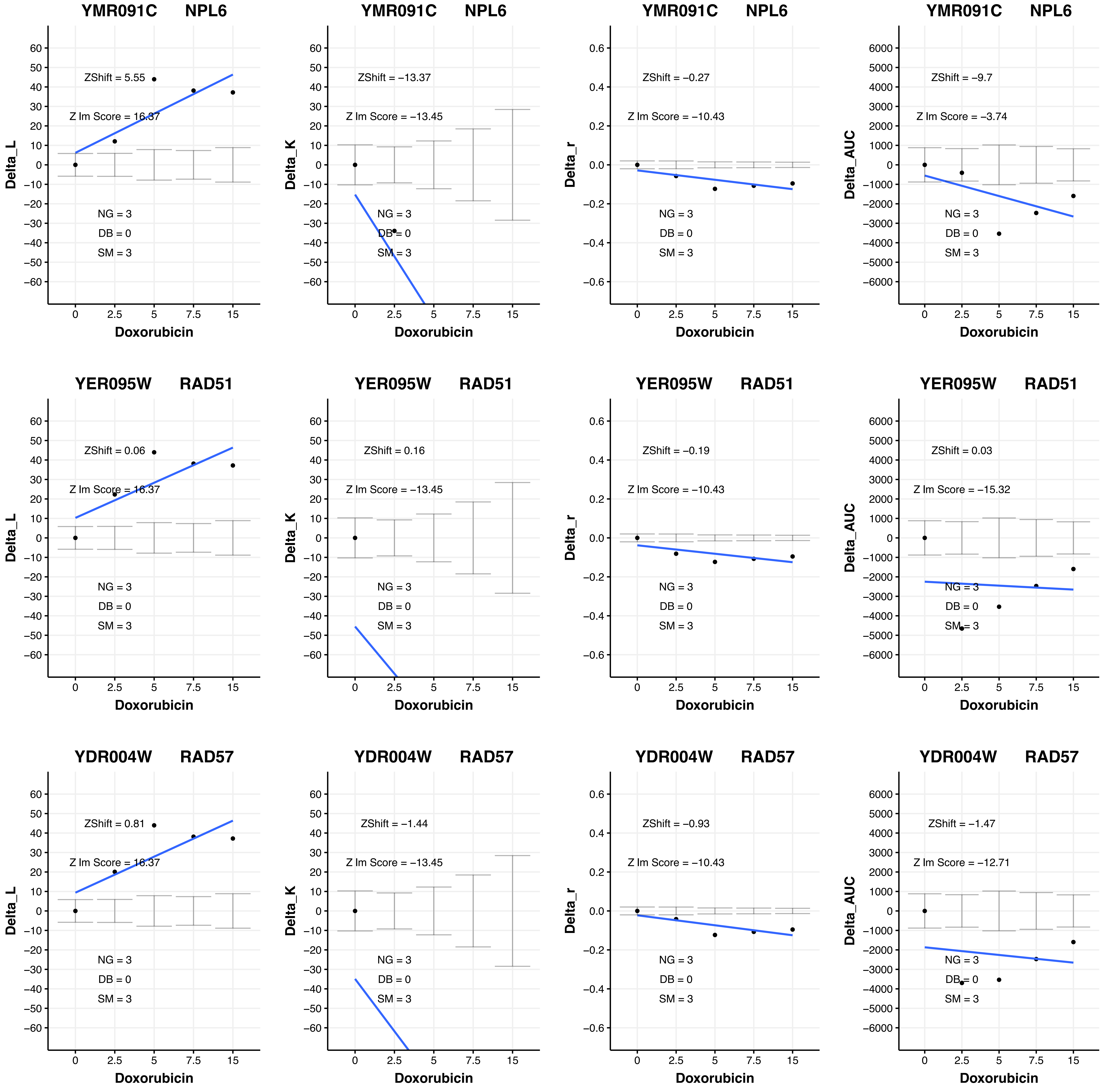
YMR091C NPL6
ZShift = 5.55
Z lm Score = 16.37
NG = 3
DB = 0
SM = 3
Delta_L
Doxorubicin
YMR091C NPL6
ZShift = −13.37
Z lm Score = −13.45
NG = 3
DB = 0
SM = 3
Delta_K
Doxorubicin
YMR091C NPL6
ZShift = −0.27
Z lm Score = −10.43
NG = 3
DB = 0
SM = 3
Delta_r
Doxorubicin
YMR091C NPL6
ZShift = −9.7
Z lm Score = −3.74
NG = 3
DB = 0
SM = 3
Delta_AUC
Doxorubicin
YER095W RAD51
ZShift = 0.06
Z lm Score = 16.37
NG = 3
DB = 0
SM = 3
Delta_L
Doxorubicin
YER095W RAD51
ZShift = 0.16
Z lm Score = −13.45
NG = 3
DB = 0
SM = 3
Delta_K
Doxorubicin
YER095W RAD51
ZShift = −0.19
Z lm Score = −10.43
NG = 3
DB = 0
SM = 3
Delta_r
Doxorubicin
YER095W RAD51
ZShift = 0.03
Z lm Score = −15.32
NG = 3
DB = 0
SM = 3
Delta_AUC
Doxorubicin
YDR004W RAD57
ZShift = 0.81
Z lm Score = 16.37
NG = 3
DB = 0
SM = 3
Delta_L
Doxorubicin
YDR004W RAD57
ZShift = −1.44
Z lm Score = −13.45
NG = 3
DB = 0
SM = 3
Delta_K
Doxorubicin
YDR004W RAD57
ZShift = −0.93
Z lm Score = −10.43
NG = 3
DB = 0
SM = 3
Delta_r
Doxorubicin
YDR004W RAD57
ZShift = −1.47
Z lm Score = −12.71
NG = 3
DB = 0
SM = 3
Delta_AUC
Doxorubicin

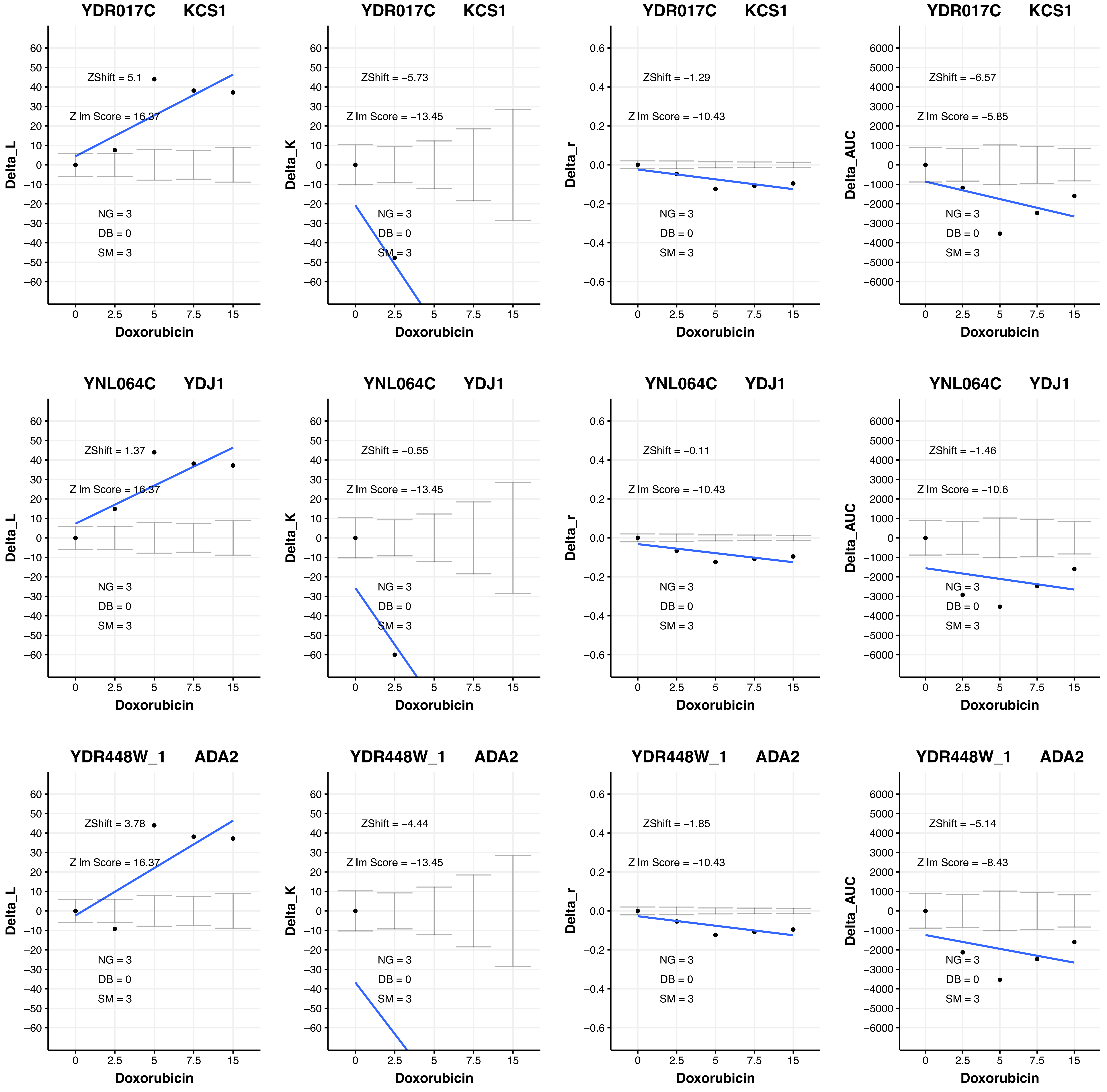

YDR017C KCS1
ZShift = 5.1
Z lm Score = 16.37
NG = 3
DB = 0
SM = 3
Delta_L
Doxorubicin
YDR017C KCS1
ZShift = −5.73
Z lm Score = −13.45
NG = 3
DB = 0
SM = 3
Delta_K
Doxorubicin
YDR017C KCS1
ZShift = −1.29
Z lm Score = −10.43
NG = 3
DB = 0
SM = 3
Delta_r
Doxorubicin
YDR017C KCS1
ZShift = −6.57
Z lm Score = −5.85
NG = 3
DB = 0
SM = 3
Delta_AUC
Doxorubicin
YNL064C YDJ1
ZShift = 1.37
Z lm Score = 16.37
NG = 3
DB = 0
SM = 3
Delta_L
Doxorubicin
YNL064C YDJ1
ZShift = −0.55
Z lm Score = −13.45
NG = 3
DB = 0
SM = 3
Delta_K
Doxorubicin
YNL064C YDJ1
ZShift = −0.11
Z lm Score = −10.43
NG = 3
DB = 0
SM = 3
Delta_r
Doxorubicin
YNL064C YDJ1
ZShift = −1.46
Z lm Score = −10.6
NG = 3
DB = 0
SM = 3
Delta_AUC
Doxorubicin
YDR448W_1 ADA2
ZShift = 3.78
Z lm Score = 16.37
NG = 3
DB = 0
SM = 3
Delta_L
Doxorubicin
YDR448W_1 ADA2
ZShift = −4.44
Z lm Score = −13.45
NG = 3
DB = 0
SM = 3
Delta_K
Doxorubicin
YDR448W_1 ADA2
ZShift = −1.85
Z lm Score = −10.43
NG = 3
DB = 0
SM = 3
Delta_r
Doxorubicin
YDR448W_1 ADA2
ZShift = −5.14
Z lm Score = −8.43
NG = 3
DB = 0
SM = 3
Delta_AUC
Doxorubicin

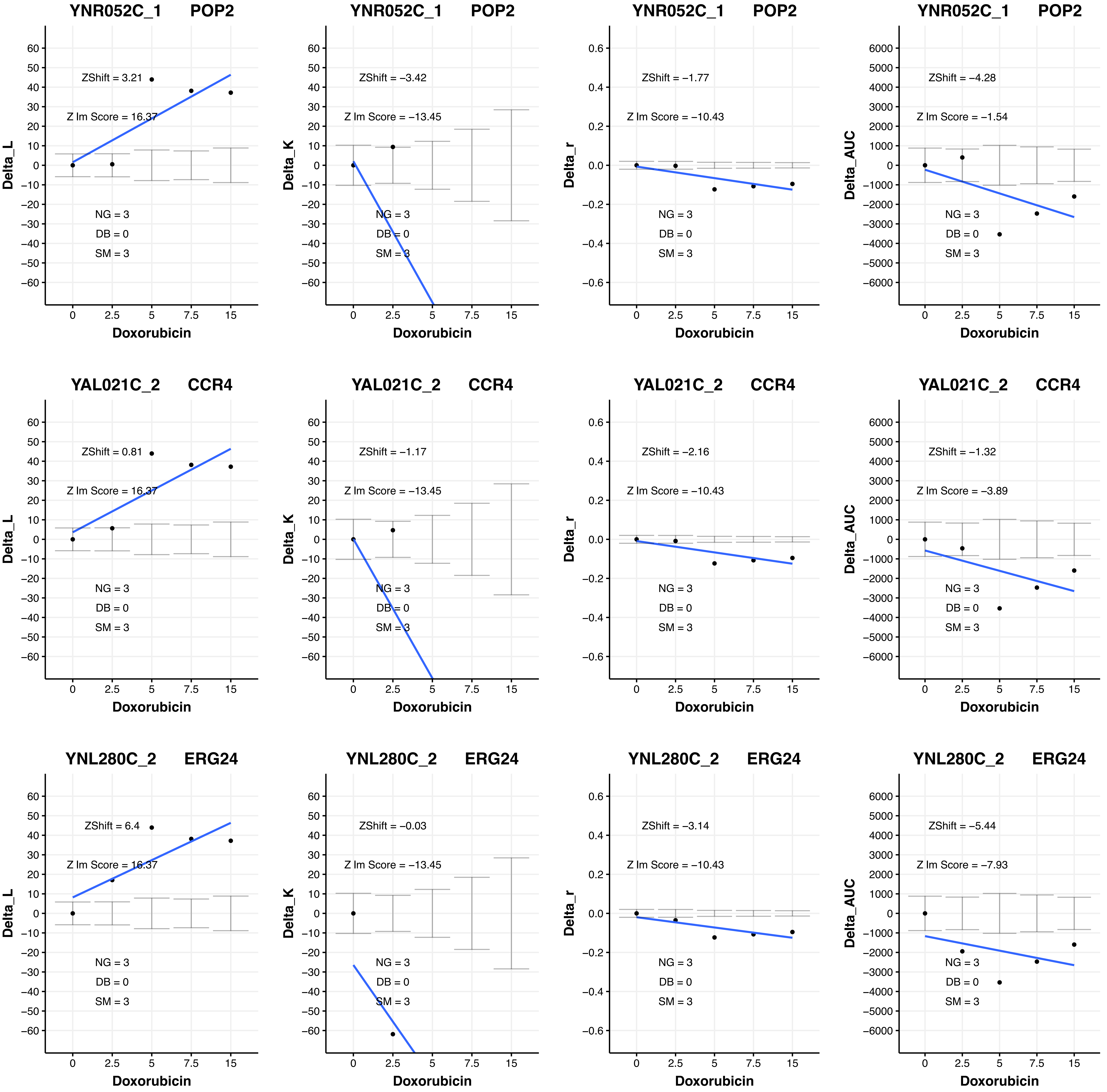

YNR052C_1 POP2
ZShift = 3.21
Z lm Score = 16.37
NG = 3
DB = 0
SM = 3
Delta_L
Doxorubicin
YNR052C_1 POP2
ZShift = −3.42
Z lm Score = −13.45
NG = 3
DB = 0
SM = 3
Delta_K
Doxorubicin
YNR052C_1 POP2
ZShift = −1.77
Z lm Score = −10.43
NG = 3
DB = 0
SM = 3
Delta_r
Doxorubicin
YNR052C_1 POP2
ZShift = −4.28
Z lm Score = −1.54
NG = 3
DB = 0
SM = 3
Delta_AUC
Doxorubicin
YAL021C_2 CCR4
ZShift = 0.81
Z lm Score = 16.37
NG = 3
DB = 0
SM = 3
Delta_L
Doxorubicin
YAL021C_2 CCR4
ZShift = −1.17
Z lm Score = −13.45
NG = 3
DB = 0
SM = 3
Delta_K
Doxorubicin
YAL021C_2 CCR4
ZShift = −2.16
Z lm Score = −10.43
NG = 3
DB = 0
SM = 3
Delta_r
Doxorubicin
YAL021C_2 CCR4
ZShift = −1.32
Z lm Score = −3.89
NG = 3
DB = 0
SM = 3
Delta_AUC
Doxorubicin
YNL280C_2 ERG24
ZShift = 6.4
Z lm Score = 16.37
NG = 3
DB = 0
SM = 3
Delta_L
Doxorubicin
YNL280C_2 ERG24
ZShift = −0.03
Z lm Score = −13.45
NG = 3
DB = 0
SM = 3
Delta_K
Doxorubicin
YNL280C_2 ERG24
ZShift = −3.14
Z lm Score = −10.43
NG = 3
DB = 0
SM = 3
Delta_r
Doxorubicin
YNL280C_2 ERG24
ZShift = −5.44
Z lm Score = −7.93
NG = 3
DB = 0
SM = 3
Delta_AUC
Doxorubicin

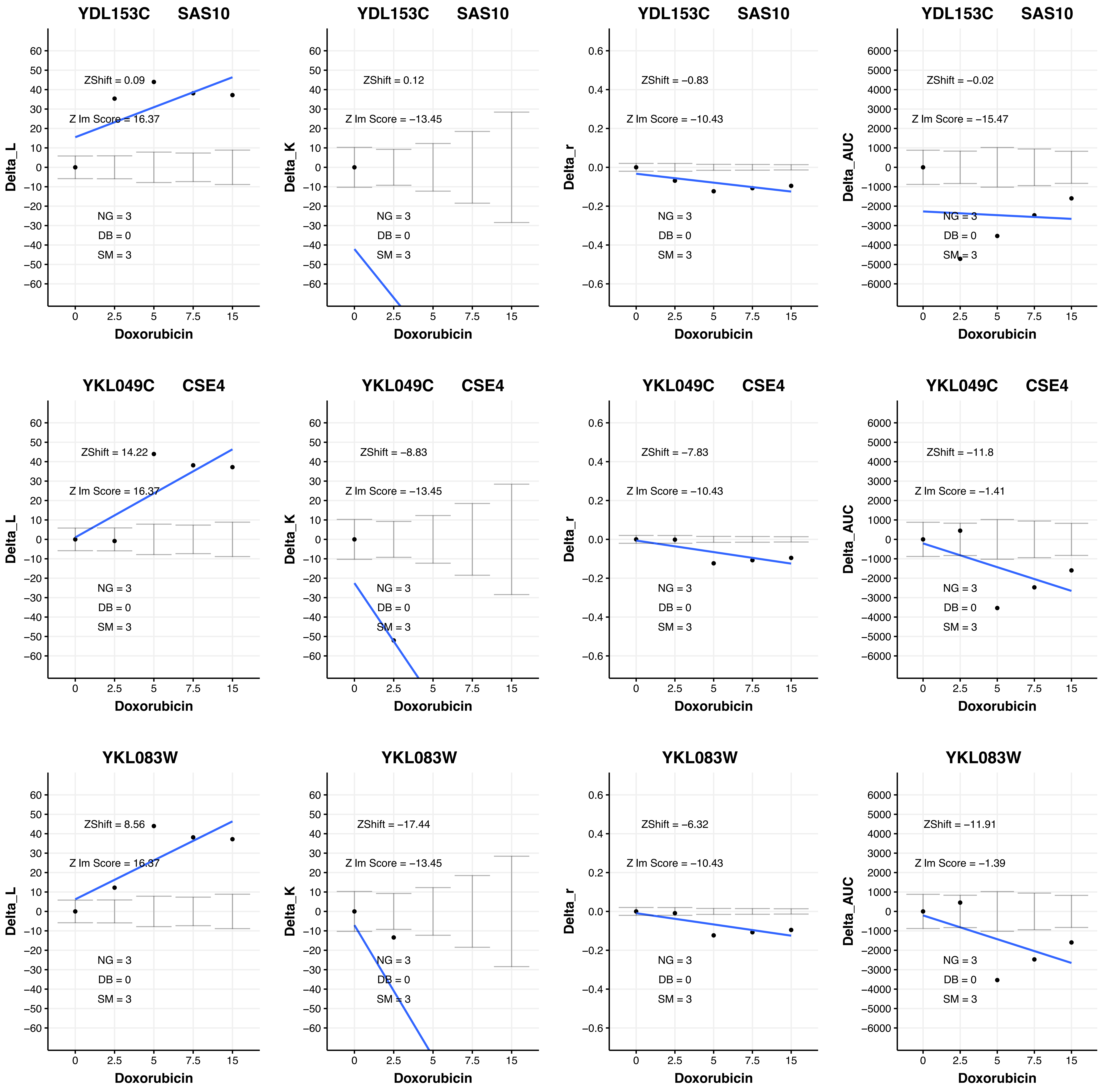
YDL153C SAS10
ZShift = 0.09
Z lm Score = 16.37
NG = 3
DB = 0
SM = 3
Delta_L
Doxorubicin
YDL153C SAS10
ZShift = 0.12
Z lm Score = −13.45
NG = 3
DB = 0
SM = 3
Delta_K
Doxorubicin
YDL153C SAS10
ZShift = −0.83
Z lm Score = −10.43
NG = 3
DB = 0
SM = 3
Delta_r
Doxorubicin
YDL153C SAS10
ZShift = −0.02
Z lm Score = −15.47
NG = 3
DB = 0
SM = 3
Delta_AUC
Doxorubicin
YKL049C CSE4
ZShift = 14.22
Z lm Score = 16.37
NG = 3
DB = 0
SM = 3
Delta_L
Doxorubicin
YKL049C CSE4
ZShift = −8.83
Z lm Score = −13.45
NG = 3
DB = 0
SM = 3
Delta_K
Doxorubicin
YKL049C CSE4
ZShift = −7.83
Z lm Score = −10.43
NG = 3
DB = 0
SM = 3
Delta_r
Doxorubicin
YKL049C CSE4
ZShift = −11.8
Z lm Score = −1.41
NG = 3
DB = 0
SM = 3
Delta_AUC
Doxorubicin
YKL083W
ZShift = 8.56
Z lm Score = 16.37
NG = 3
DB = 0
SM = 3
Delta_L
Doxorubicin
YKL083W
ZShift = −17.44
Z lm Score = −13.45
NG = 3
DB = 0
SM = 3
Delta_K
Doxorubicin
YKL083W
ZShift = −6.32
Z lm Score = −10.43
NG = 3
DB = 0
SM = 3
Delta_r
Doxorubicin
YKL083W
ZShift = −11.91
Z lm Score = −1.39
NG = 3
DB = 0
SM = 3
Delta_AUC
Doxorubicin

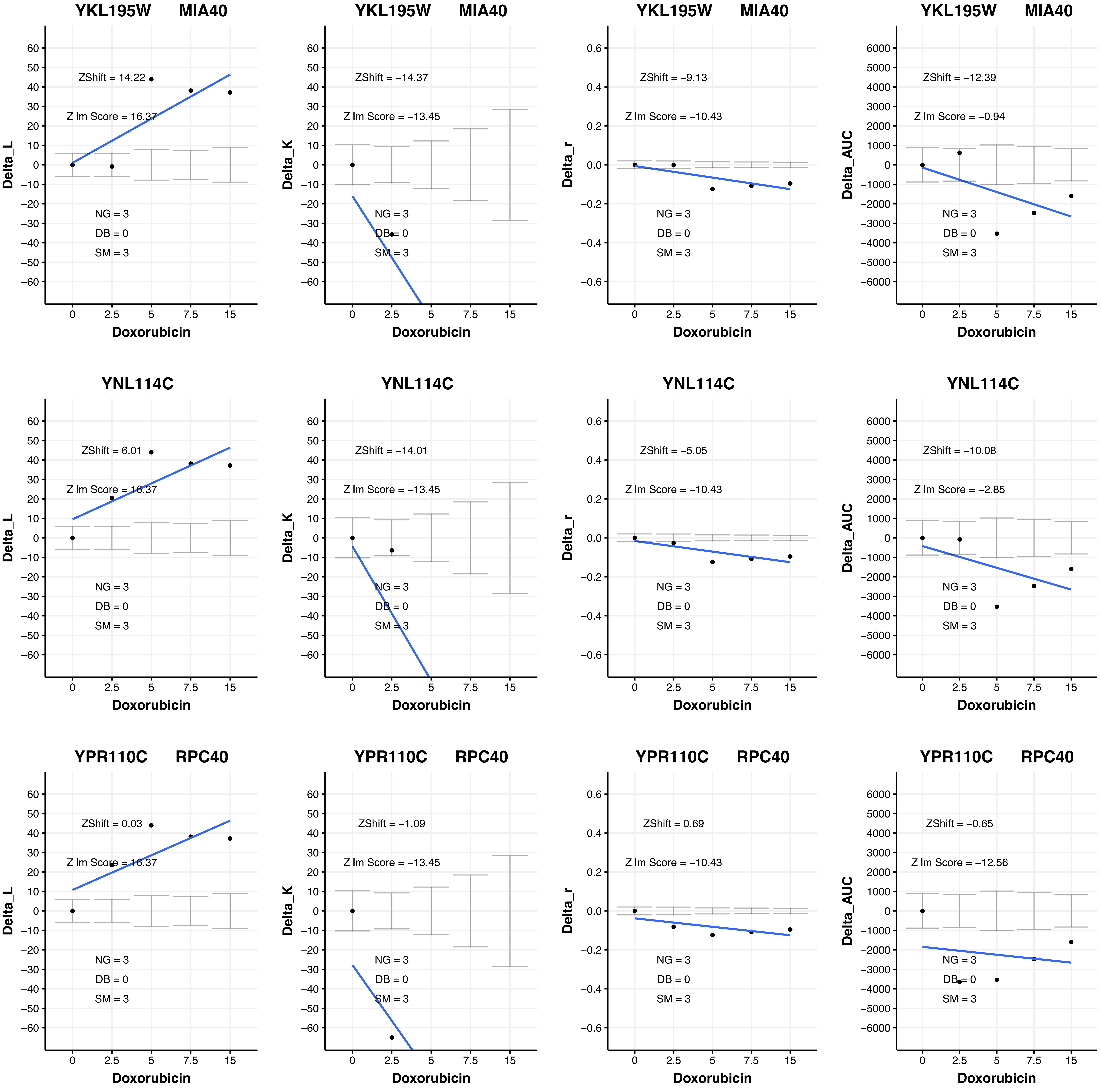

YKL195W MIA40
ZShift = 14.22
Z lm Score = 16.37
NG = 3
DB = 0
SM = 3
Delta_L
Doxorubicin
YKL195W MIA40
ZShift = −14.37
Z lm Score = −13.45
NG = 3
DB = 0
SM = 3
Delta_K
Doxorubicin
YKL195W MIA40
ZShift = −9.13
Z lm Score = −10.43
NG = 3
DB = 0
SM = 3
Delta_r
Doxorubicin
YKL195W MIA40
ZShift = −12.39
Z lm Score = −0.94
NG = 3
DB = 0
SM = 3
Delta_AUC
Doxorubicin
YNL114C
ZShift = 6.01
Z lm Score = 16.37
NG = 3
DB = 0
SM = 3
Delta_L
Doxorubicin
YNL114C
ZShift = −14.01
Z lm Score = −13.45
NG = 3
DB = 0
SM = 3
Delta_K
Doxorubicin
YNL114C
ZShift = −5.05
Z lm Score = −10.43
NG = 3
DB = 0
SM = 3
Delta_r
Doxorubicin
YNL114C
ZShift = −10.08
Z lm Score = −2.85
NG = 3
DB = 0
SM = 3
Delta_AUC
Doxorubicin
YPR110C RPC40
ZShift = 0.03
Z lm Score = 16.37
NG = 3
DB = 0
SM = 3
Delta_L
Doxorubicin
YPR110C RPC40
ZShift = −1.09
Z lm Score = −13.45
NG = 3
DB = 0
SM = 3
Delta_K
Doxorubicin
YPR110C RPC40
ZShift = 0.69
Z lm Score = −10.43
NG = 3
DB = 0
SM = 3
Delta_r
Doxorubicin
YPR110C RPC40
ZShift = −0.65
Z lm Score = −12.56
NG = 3
DB = 0
SM = 3
Delta_AUC
Doxorubicin

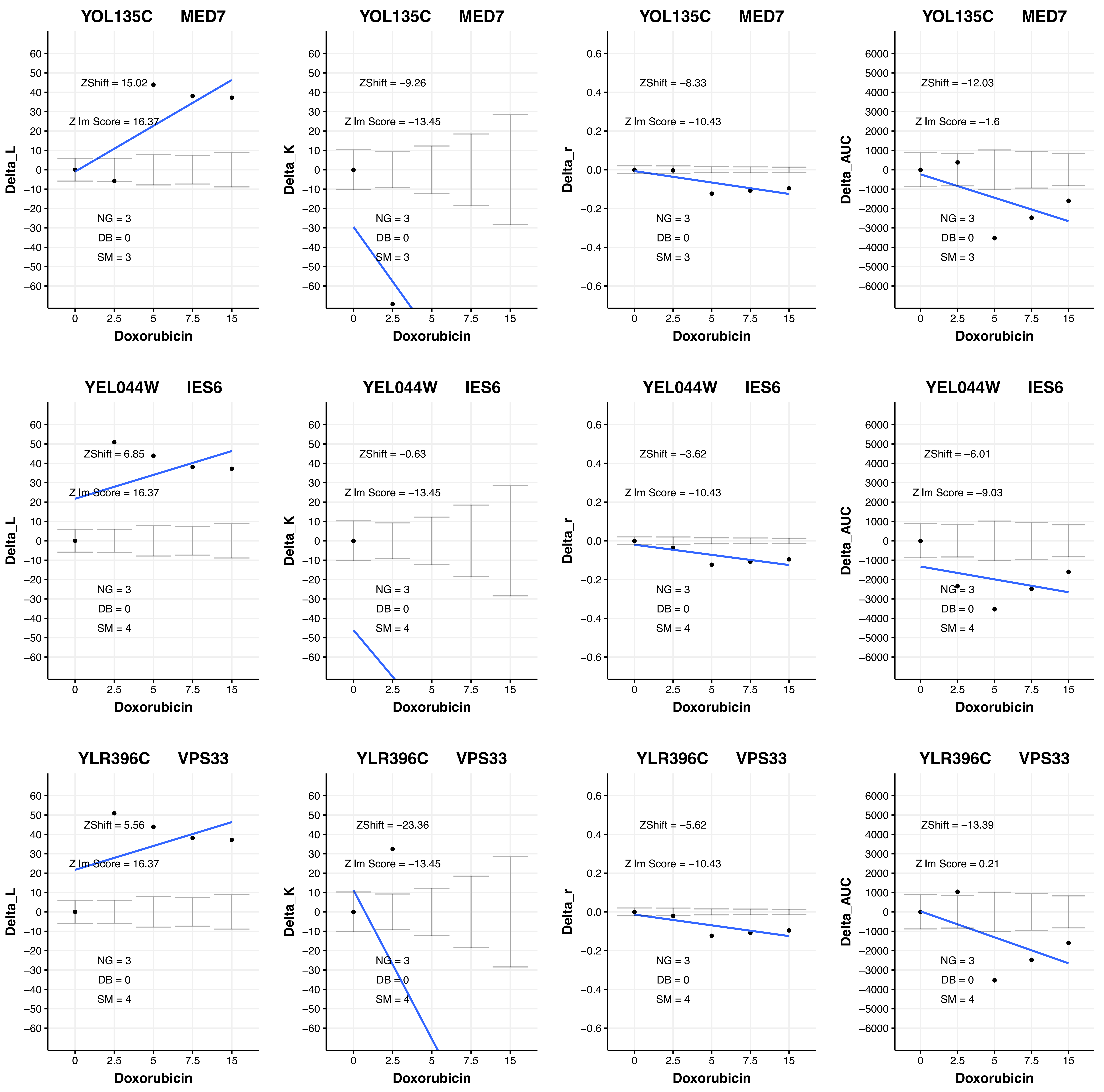
YOL135C MED7
ZShift = 15.02
Z lm Score = 16.37
NG = 3
DB = 0
SM = 3
Delta_L
Doxorubicin
YOL135C MED7
ZShift = −9.26
Z lm Score = −13.45
NG = 3
DB = 0
SM = 3
Delta_K
Doxorubicin
YOL135C MED7
ZShift = −8.33
Z lm Score = −10.43
NG = 3
DB = 0
SM = 3
Delta_r
Doxorubicin
YOL135C MED7
ZShift = −12.03
Z lm Score = −1.6
NG = 3
DB = 0
SM = 3
Delta_AUC
Doxorubicin
YEL044W IES6
ZShift = 6.85
Z lm Score = 16.37
NG = 3
DB = 0
SM = 4
Delta_L
Doxorubicin
YEL044W IES6
ZShift = −0.63
Z lm Score = −13.45
NG = 3
DB = 0
SM = 4
Delta_K
Doxorubicin
YEL044W IES6
ZShift = −3.62
Z lm Score = −10.43
NG = 3
DB = 0
SM = 4
Delta_r
Doxorubicin
YEL044W IES6
ZShift = −6.01
Z lm Score = −9.03
NG = 3
DB = 0
SM = 4
Delta_AUC
Doxorubicin
YLR396C VPS33
ZShift = 5.56
Z lm Score = 16.37
NG = 3
DB = 0
SM = 4
Delta_L
Doxorubicin
YLR396C VPS33
ZShift = −23.36
Z lm Score = −13.45
NG = 3
DB = 0
SM = 4
Delta_K
Doxorubicin
YLR396C VPS33
ZShift = −5.62
Z lm Score = −10.43
NG = 3
DB = 0
SM = 4
Delta_r
Doxorubicin
YLR396C VPS33
ZShift = −13.39
Z lm Score = 0.21
NG = 3
DB = 0
SM = 4
Delta_AUC
Doxorubicin

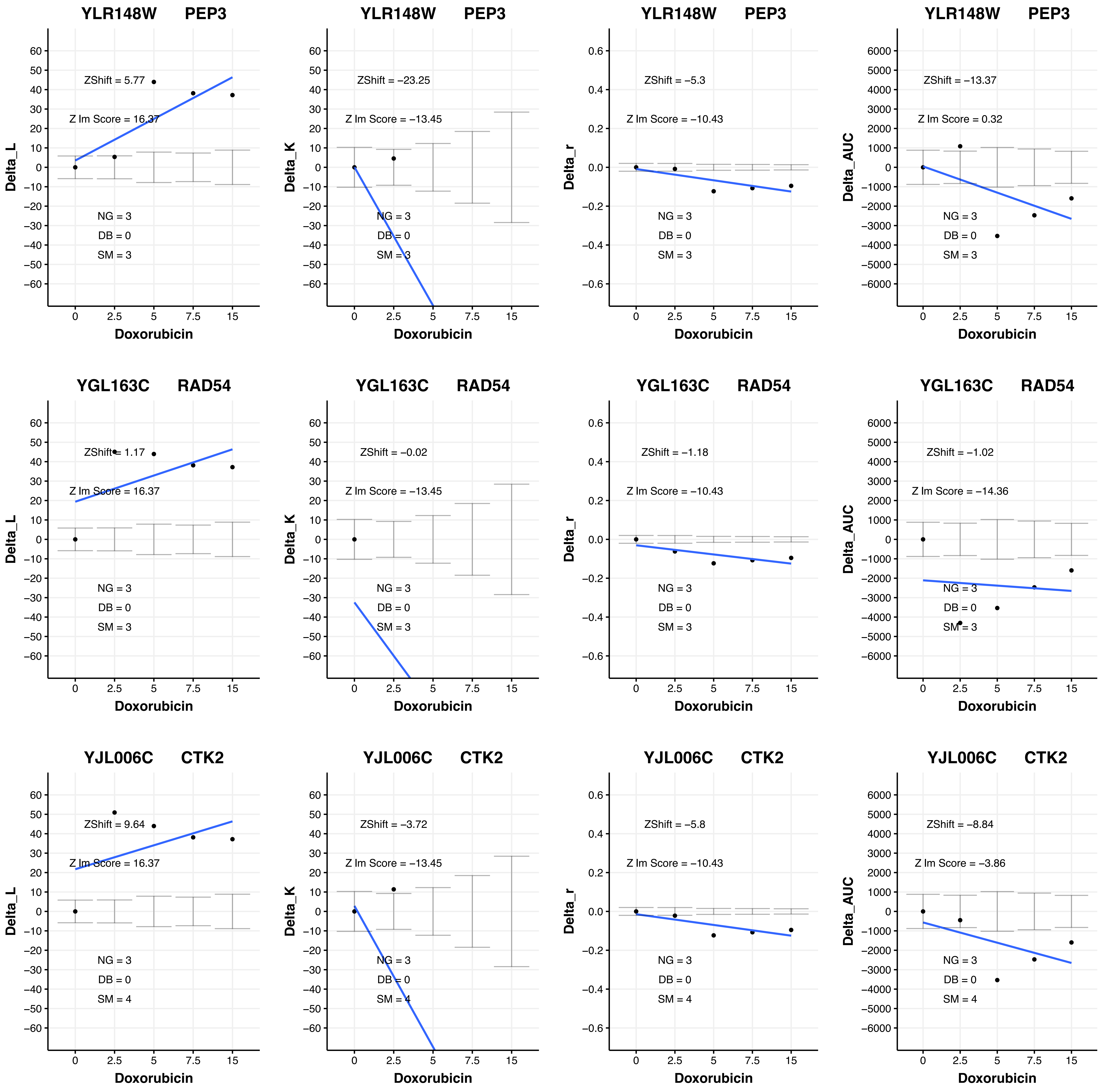
YLR148W PEP3
ZShift = 5.77
Z lm Score = 16.37
NG = 3
DB = 0
SM = 3
Delta_L
Doxorubicin
YLR148W PEP3
ZShift = −23.25
Z lm Score = −13.45
NG = 3
DB = 0
SM = 3
Delta_K
Doxorubicin
YLR148W PEP3
ZShift = −5.3
Z lm Score = −10.43
NG = 3
DB = 0
SM = 3
Delta_r
Doxorubicin
YLR148W PEP3
ZShift = −13.37
Z lm Score = 0.32
NG = 3
DB = 0
SM = 3
Delta_AUC
Doxorubicin
YGL163C RAD54
ZShift = 1.17
Z lm Score = 16.37
NG = 3
DB = 0
SM = 3
Delta_L
Doxorubicin
YGL163C RAD54
ZShift = −0.02
Z lm Score = −13.45
NG = 3
DB = 0
SM = 3
Delta_K
Doxorubicin
YGL163C RAD54
ZShift = −1.18
Z lm Score = −10.43
NG = 3
DB = 0
SM = 3
Delta_r
Doxorubicin
YGL163C RAD54
ZShift = −1.02
Z lm Score = −14.36
NG = 3
DB = 0
SM = 3
Delta_AUC
Doxorubicin
YJL006C CTK2
ZShift = 9.64
Z lm Score = 16.37
NG = 3
DB = 0
SM = 4
Delta_L
Doxorubicin
YJL006C CTK2
ZShift = −3.72
Z lm Score = −13.45
NG = 3
DB = 0
SM = 4
Delta_K
Doxorubicin
YJL006C CTK2
ZShift = −5.8
Z lm Score = −10.43
NG = 3
DB = 0
SM = 4
Delta_r
Doxorubicin
YJL006C CTK2
ZShift = −8.84
Z lm Score = −3.86
NG = 3
DB = 0
SM = 4
Delta_AUC
Doxorubicin

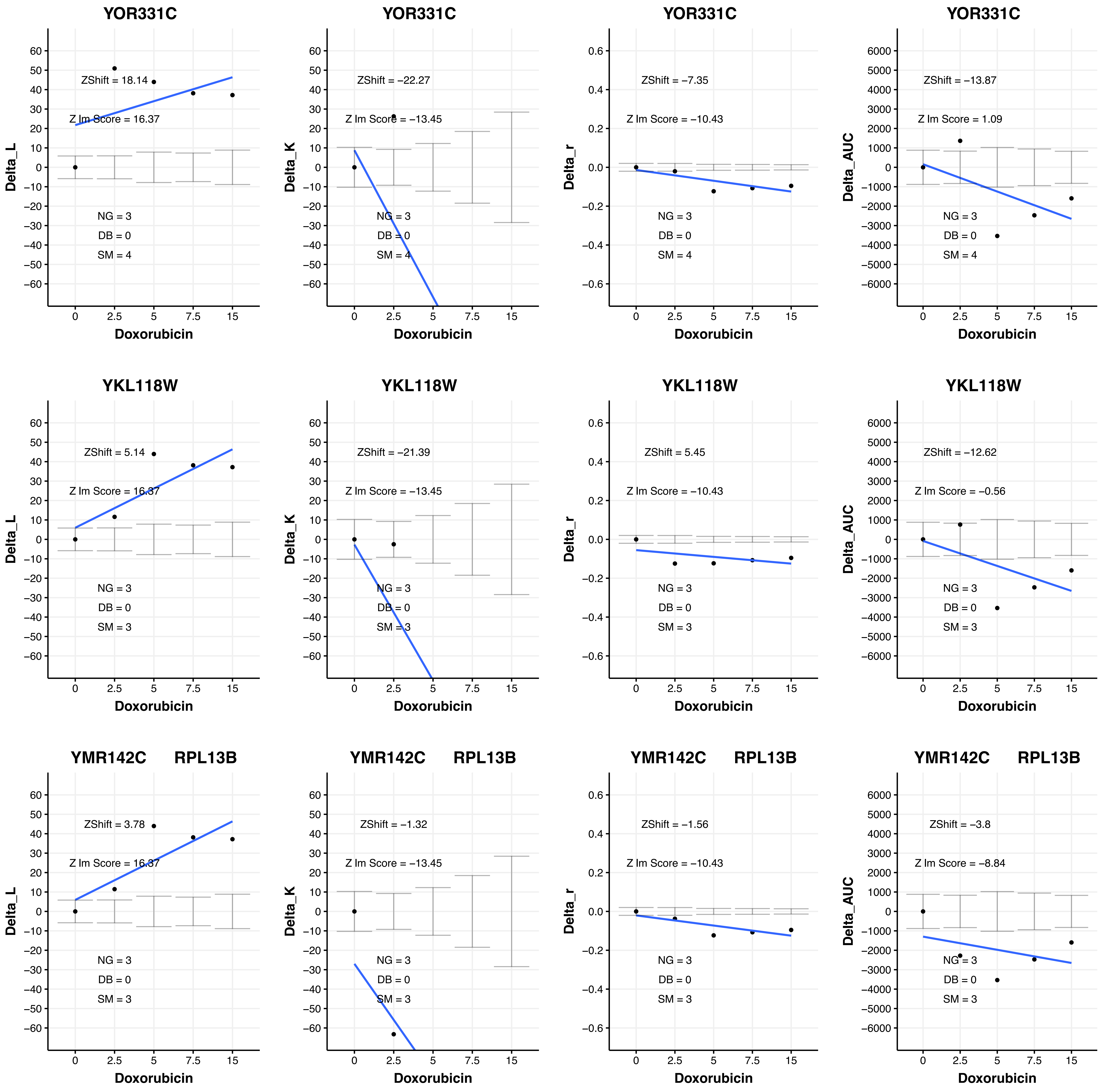
YOR331C
ZShift = 18.14
Z lm Score = 16.37
NG = 3
DB = 0
SM = 4
Delta_L
Doxorubicin
YOR331C
ZShift = −22.27
Z lm Score = −13.45
NG = 3
DB = 0
SM = 4
Delta_K
Doxorubicin
YOR331C
ZShift = −7.35
Z lm Score = −10.43
NG = 3
DB = 0
SM = 4
Delta_r
Doxorubicin
YOR331C
ZShift = −13.87
Z lm Score = 1.09
NG = 3
DB = 0
SM = 4
Delta_AUC
Doxorubicin
YKL118W
ZShift = 5.14
Z lm Score = 16.37
NG = 3
DB = 0
SM = 3
Delta_L
Doxorubicin
YKL118W
ZShift = −21.39
Z lm Score = −13.45
NG = 3
DB = 0
SM = 3
Delta_K
Doxorubicin
YKL118W
ZShift = 5.45
Z lm Score = −10.43
NG = 3
DB = 0
SM = 3
Delta_r
Doxorubicin
YKL118W
ZShift = −12.62
Z lm Score = −0.56
NG = 3
DB = 0
SM = 3
Delta_AUC
Doxorubicin
YMR142C RPL13B
ZShift = 3.78
Z lm Score = 16.37
NG = 3
DB = 0
SM = 3
Delta_L
Doxorubicin
YMR142C RPL13B
ZShift = −1.32
Z lm Score = −13.45
NG = 3
DB = 0
SM = 3
Delta_K
Doxorubicin
YMR142C RPL13B
ZShift = −1.56
Z lm Score = −10.43
NG = 3
DB = 0
SM = 3
Delta_r
Doxorubicin
YMR142C RPL13B
ZShift = −3.8
Z lm Score = −8.84
NG = 3
DB = 0
SM = 3
Delta_AUC
Doxorubicin

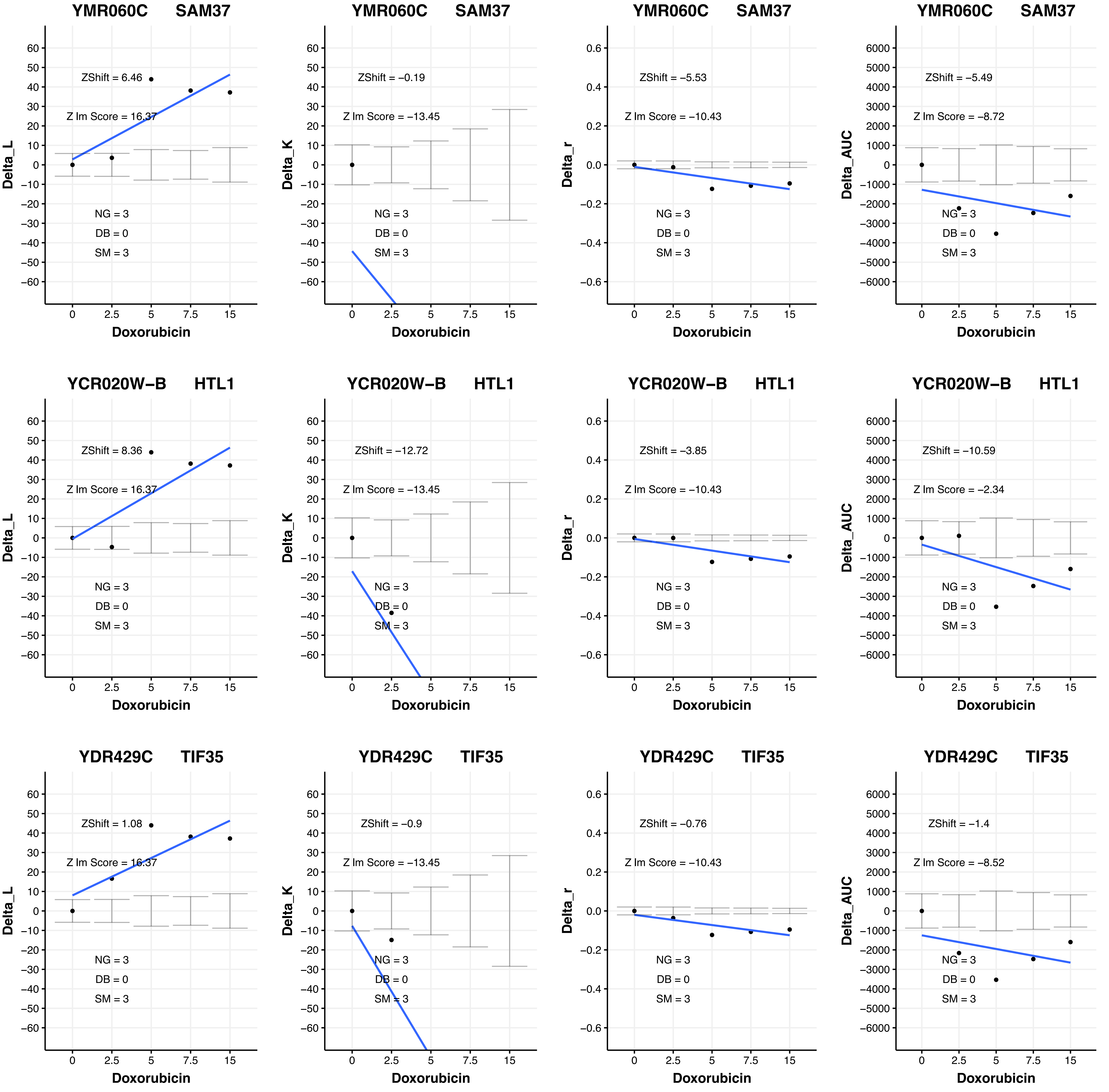
YMR060C SAM37
ZShift = 6.46
Z lm Score = 16.37
NG = 3
DB = 0
SM = 3
Delta_L
Doxorubicin
YMR060C SAM37
ZShift = −0.19
Z lm Score = −13.45
NG = 3
DB = 0
SM = 3
Delta_K
Doxorubicin
YMR060C SAM37
ZShift = −5.53
Z lm Score = −10.43
NG = 3
DB = 0
SM = 3
Delta_r
Doxorubicin
YMR060C SAM37
ZShift = −5.49
Z lm Score = −8.72
NG = 3
DB = 0
SM = 3
Delta_AUC
Doxorubicin
YCR020W−B HTL1
ZShift = 8.36
Z lm Score = 16.37
NG = 3
DB = 0
SM = 3
Delta_L
Doxorubicin
YCR020W−B HTL1
ZShift = −12.72
Z lm Score = −13.45
NG = 3
DB = 0
SM = 3
Delta_K
Doxorubicin
YCR020W−B HTL1
ZShift = −3.85
Z lm Score = −10.43
NG = 3
DB = 0
SM = 3
Delta_r
Doxorubicin
YCR020W−B HTL1
ZShift = −10.59
Z lm Score = −2.34
NG = 3
DB = 0
SM = 3
Delta_AUC
Doxorubicin
YDR429C TIF35
ZShift = 1.08
Z lm Score = 16.37
NG = 3
DB = 0
SM = 3
Delta_L
Doxorubicin
YDR429C TIF35
ZShift = −0.9
Z lm Score = −13.45
NG = 3
DB = 0
SM = 3
Delta_K
Doxorubicin
YDR429C TIF35
ZShift = −0.76
Z lm Score = −10.43
NG = 3
DB = 0
SM = 3
Delta_r
Doxorubicin
YDR429C TIF35
ZShift = −1.4
Z lm Score = −8.52
NG = 3
DB = 0
SM = 3
Delta_AUC
Doxorubicin

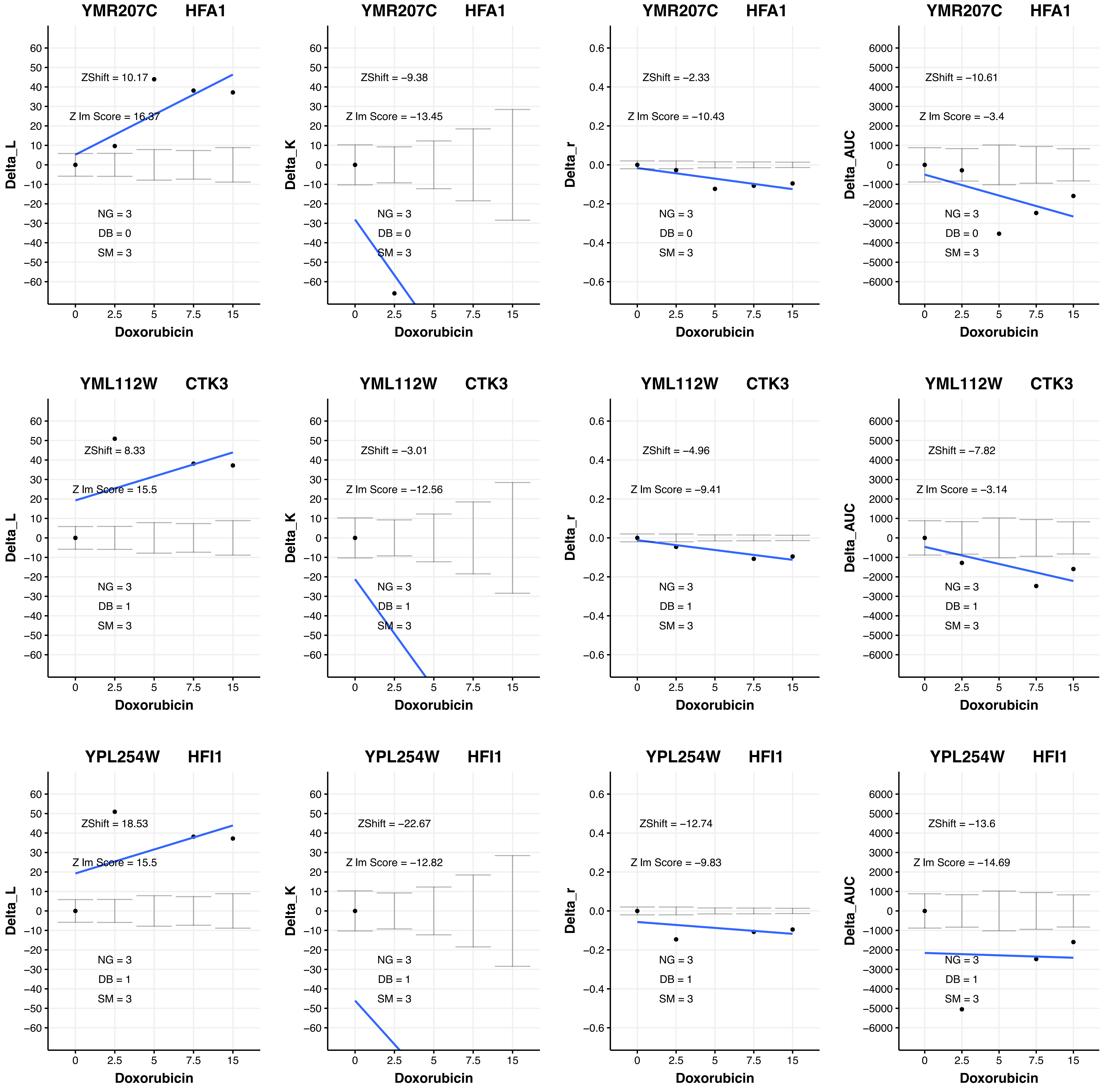
YMR207C HFA1
ZShift = 10.17
Z lm Score = 16.37
NG = 3
DB = 0
SM = 3
Delta_L
Doxorubicin
YMR207C HFA1
ZShift = −9.38
Z lm Score = −13.45
NG = 3
DB = 0
SM = 3
Delta_K
Doxorubicin
YMR207C HFA1
ZShift = −2.33
Z lm Score = −10.43
NG = 3
DB = 0
SM = 3
Delta_r
Doxorubicin
YMR207C HFA1
ZShift = −10.61
Z lm Score = −3.4
NG = 3
DB = 0
SM = 3
Delta_AUC
Doxorubicin
YML112W CTK3
ZShift = 8.33
Z lm Score = 15.5
NG = 3
DB = 1
SM = 3
Delta_L
Doxorubicin
YML112W CTK3
ZShift = −3.01
Z lm Score = −12.56
NG = 3
DB = 1
SM = 3
Delta_K
Doxorubicin
YML112W CTK3
ZShift = −4.96
Z lm Score = −9.41
NG = 3
DB = 1
SM = 3
Delta_r
Doxorubicin
YML112W CTK3
ZShift = −7.82
Z lm Score = −3.14
NG = 3
DB = 1
SM = 3
Delta_AUC
Doxorubicin
YPL254W HFI1
ZShift = 18.53
Z lm Score = 15.5
NG = 3
DB = 1
SM = 3
Delta_L
Doxorubicin
YPL254W HFI1
ZShift = −22.67
Z lm Score = −12.82
NG = 3
DB = 1
SM = 3
Delta_K
Doxorubicin
YPL254W HFI1
ZShift = −12.74
Z lm Score = −9.83
NG = 3
DB = 1
SM = 3
Delta_r
Doxorubicin
YPL254W HFI1
ZShift = −13.6
Z lm Score = −14.69
NG = 3
DB = 1
SM = 3
Delta_AUC
Doxorubicin

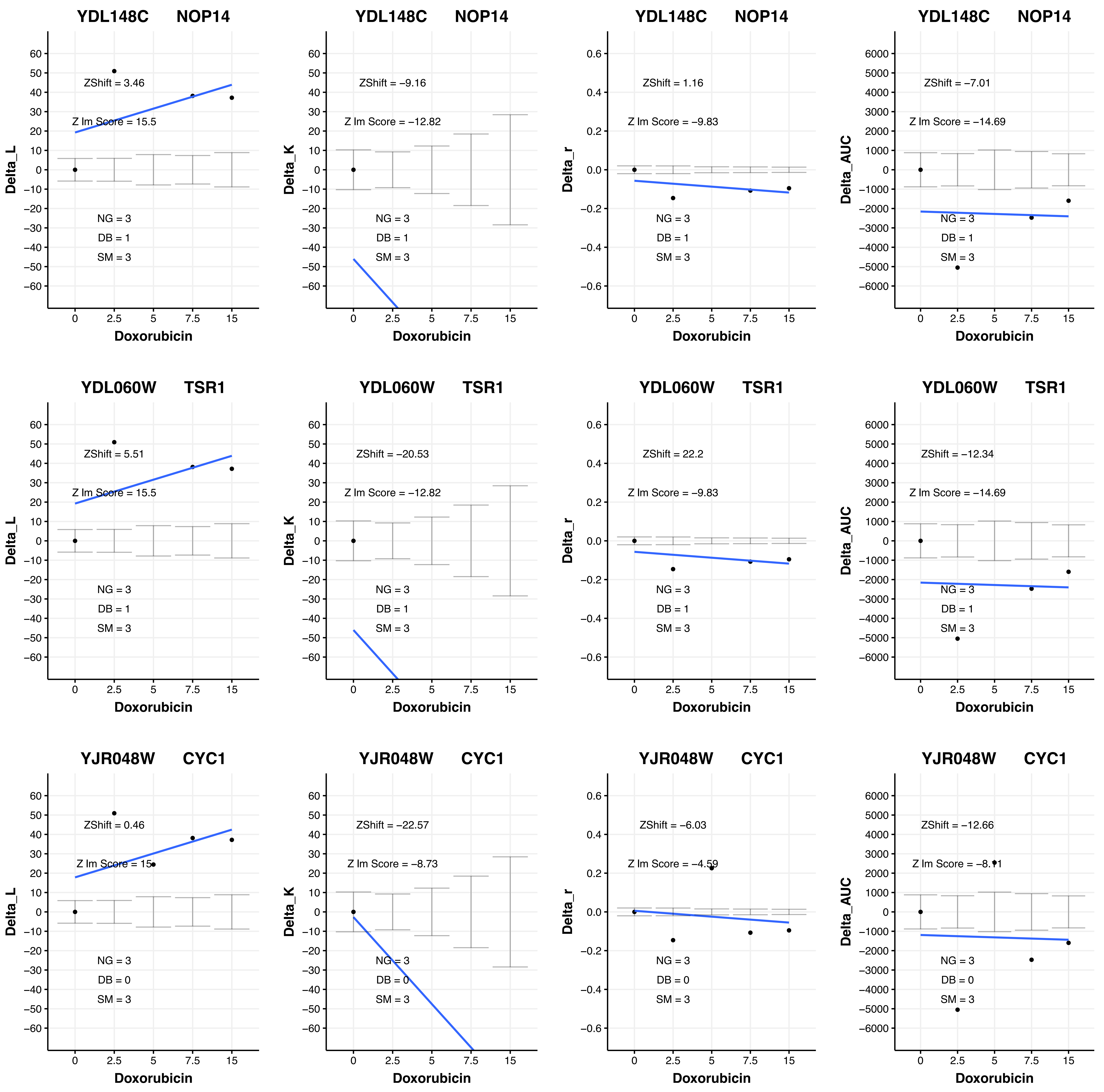
YDL148C NOP14
ZShift = 3.46
Z lm Score = 15.5
NG = 3
DB = 1
SM = 3
Delta_L
Doxorubicin
YDL148C NOP14
ZShift = −9.16
Z lm Score = −12.82
NG = 3
DB = 1
SM = 3
Delta_K
Doxorubicin
YDL148C NOP14
ZShift = 1.16
Z lm Score = −9.83
NG = 3
DB = 1
SM = 3
Delta_r
Doxorubicin
YDL148C NOP14
ZShift = −7.01
Z lm Score = −14.69
NG = 3
DB = 1
SM = 3
Delta_AUC
Doxorubicin
YDL060W TSR1
ZShift = 5.51
Z lm Score = 15.5
NG = 3
DB = 1
SM = 3
Delta_L
Doxorubicin
YDL060W TSR1
ZShift = −20.53
Z lm Score = −12.82
NG = 3
DB = 1
SM = 3
Delta_K
Doxorubicin
YDL060W TSR1
ZShift = 22.2
Z lm Score = −9.83
NG = 3
DB = 1
SM = 3
Delta_r
Doxorubicin
YDL060W TSR1
ZShift = −12.34
Z lm Score = −14.69
NG = 3
DB = 1
SM = 3
Delta_AUC
Doxorubicin
YJR048W CYC1
ZShift = 0.46
Z lm Score = 15
NG = 3
DB = 0
SM = 3
Delta_L
Doxorubicin
YJR048W CYC1
ZShift = −22.57
Z lm Score = −8.73
NG = 3
DB = 0
SM = 3
Delta_K
Doxorubicin
YJR048W CYC1
ZShift = −6.03
Z lm Score = −4.59
NG = 3
DB = 0
SM = 3
Delta_r
Doxorubicin
YJR048W CYC1
ZShift = −12.66
NG = 3
DB = 0
SM = 3
Delta_AUC
Doxorubicin

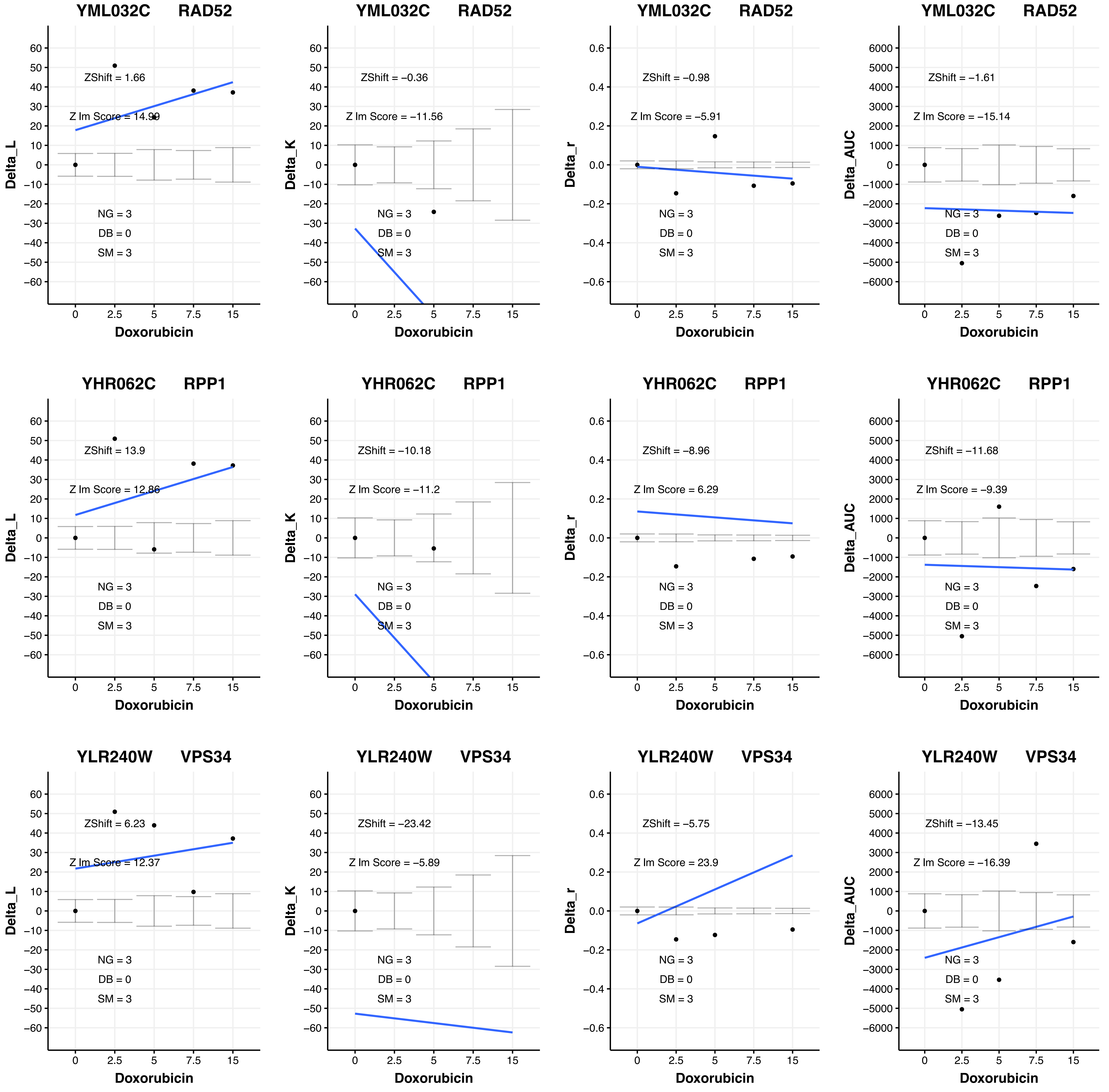
YML032C RAD52
ZShift = 1.66
Z lm Score = 14.99
NG = 3
DB = 0
SM = 3
Delta_L
Doxorubicin
YML032C RAD52
ZShift = −0.36
Z lm Score = −11.56
NG = 3
DB = 0
SM = 3
Delta_K
Doxorubicin
YML032C RAD52
ZShift = −0.98
Z lm Score = −5.91
NG = 3
DB = 0
SM = 3
Delta_r
Doxorubicin
YML032C RAD52
ZShift = −1.61
Z lm Score = −15.14
NG = 3
DB = 0
SM = 3
Delta_AUC
Doxorubicin
YHR062C RPP1
ZShift = 13.9
Z lm Score = 12.86
NG = 3
DB = 0
SM = 3
Delta_L
Doxorubicin
YHR062C RPP1
ZShift = −10.18
Z lm Score = −11.2
NG = 3
DB = 0
SM = 3
Delta_K
Doxorubicin
YHR062C RPP1
ZShift = −8.96
Z lm Score = 6.29
NG = 3
DB = 0
SM = 3
Delta_r
Doxorubicin
YHR062C RPP1
ZShift = −11.68
Z lm Score = −9.39
NG = 3
DB = 0
SM = 3
Delta_AUC
Doxorubicin
YLR240W VPS34
ZShift = 6.23
Z lm Score = 12.37
NG = 3
DB = 0
SM = 3
Delta_L
Doxorubicin
YLR240W VPS34
ZShift = −23.42
Z lm Score = −5.89
NG = 3
DB = 0
SM = 3
Delta_K
Doxorubicin
YLR240W VPS34
ZShift = −5.75
Z lm Score = 23.9
NG = 3
DB = 0
SM = 3
Delta_r
Doxorubicin
YLR240W VPS34
ZShift = −13.45
Z lm Score = −16.39
NG = 3
DB = 0
SM = 3
Delta_AUC
Doxorubicin

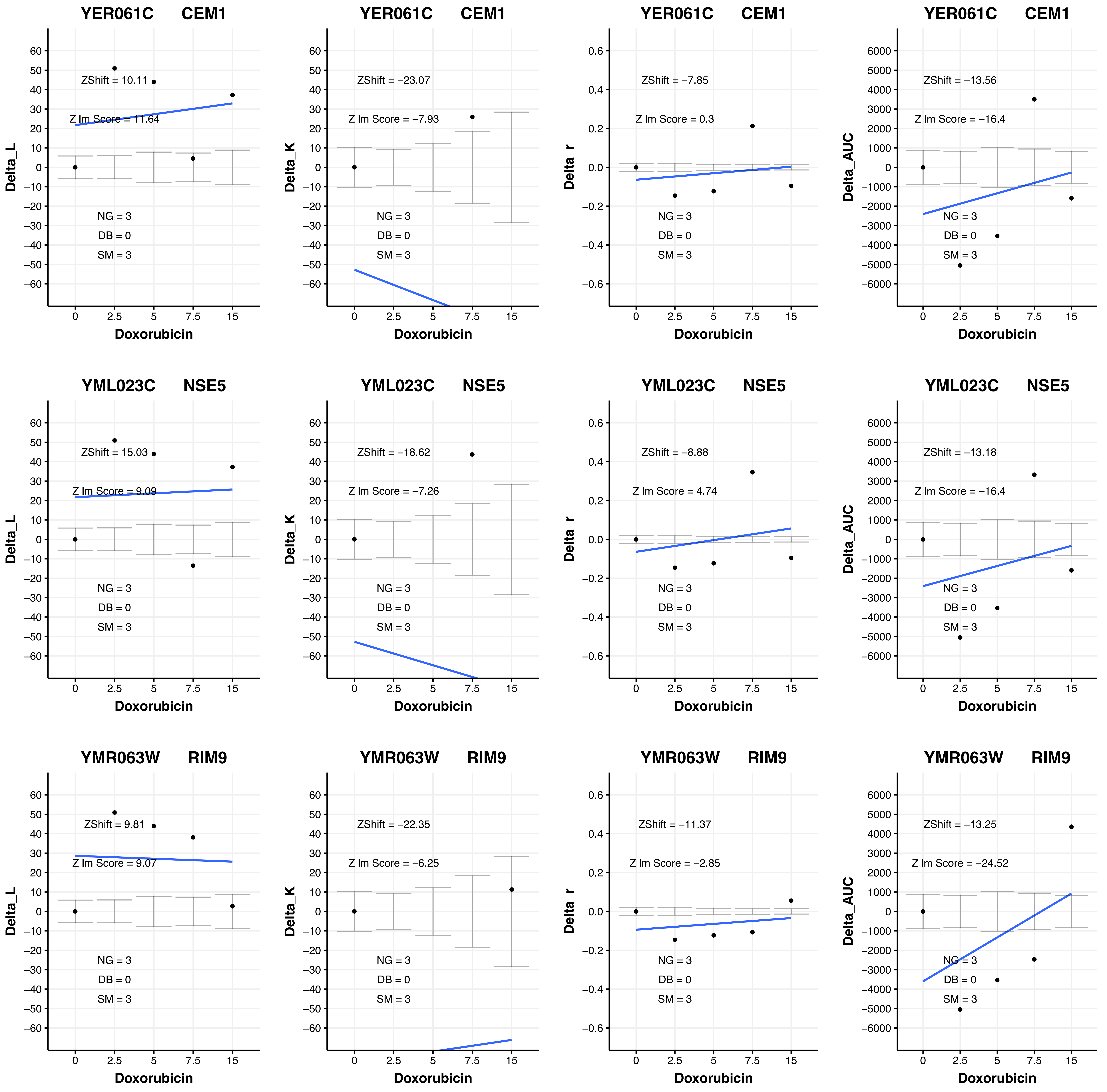

YER061C CEM1
ZShift = 10.11
Z lm Score = 11.64
NG = 3
DB = 0
SM = 3
Delta_L
Doxorubicin
YER061C CEM1
ZShift = −23.07
Z lm Score = −7.93
NG = 3
DB = 0
SM = 3
Delta_K
Doxorubicin
YER061C CEM1
ZShift = −7.85
Z lm Score = 0.3
NG = 3
DB = 0
SM = 3
Delta_r
Doxorubicin
YER061C CEM1
ZShift = −13.56
Z lm Score = −16.4
NG = 3
DB = 0
SM = 3
Delta_AUC
Doxorubicin
YML023C NSE5
ZShift = 15.03
Z lm Score = 9.09
NG = 3
DB = 0
SM = 3
Delta_L
Doxorubicin
YML023C NSE5
ZShift = −18.62
Z lm Score = −7.26
NG = 3
DB = 0
SM = 3
Delta_K
Doxorubicin
YML023C NSE5
ZShift = −8.88
Z lm Score = 4.74
NG = 3
DB = 0
SM = 3
Delta_r
Doxorubicin
YML023C NSE5
ZShift = −13.18
Z lm Score = −16.4
NG = 3
DB = 0
SM = 3
Delta_AUC
Doxorubicin
YMR063W RIM9
ZShift = 9.81
Z lm Score = 9.07
NG = 3
DB = 0
SM = 3
Delta_L
Doxorubicin
YMR063W RIM9
ZShift = −22.35
Z lm Score = −6.25
NG = 3
DB = 0
SM = 3
Delta_K
Doxorubicin
YMR063W RIM9
ZShift = −11.37
Z lm Score = −2.85
NG = 3
DB = 0
SM = 3
Delta_r
Doxorubicin
YMR063W RIM9
ZShift = −13.25
Z lm Score = −24.52
NG = 3
DB = 0
SM = 3
Delta_AUC
Doxorubicin

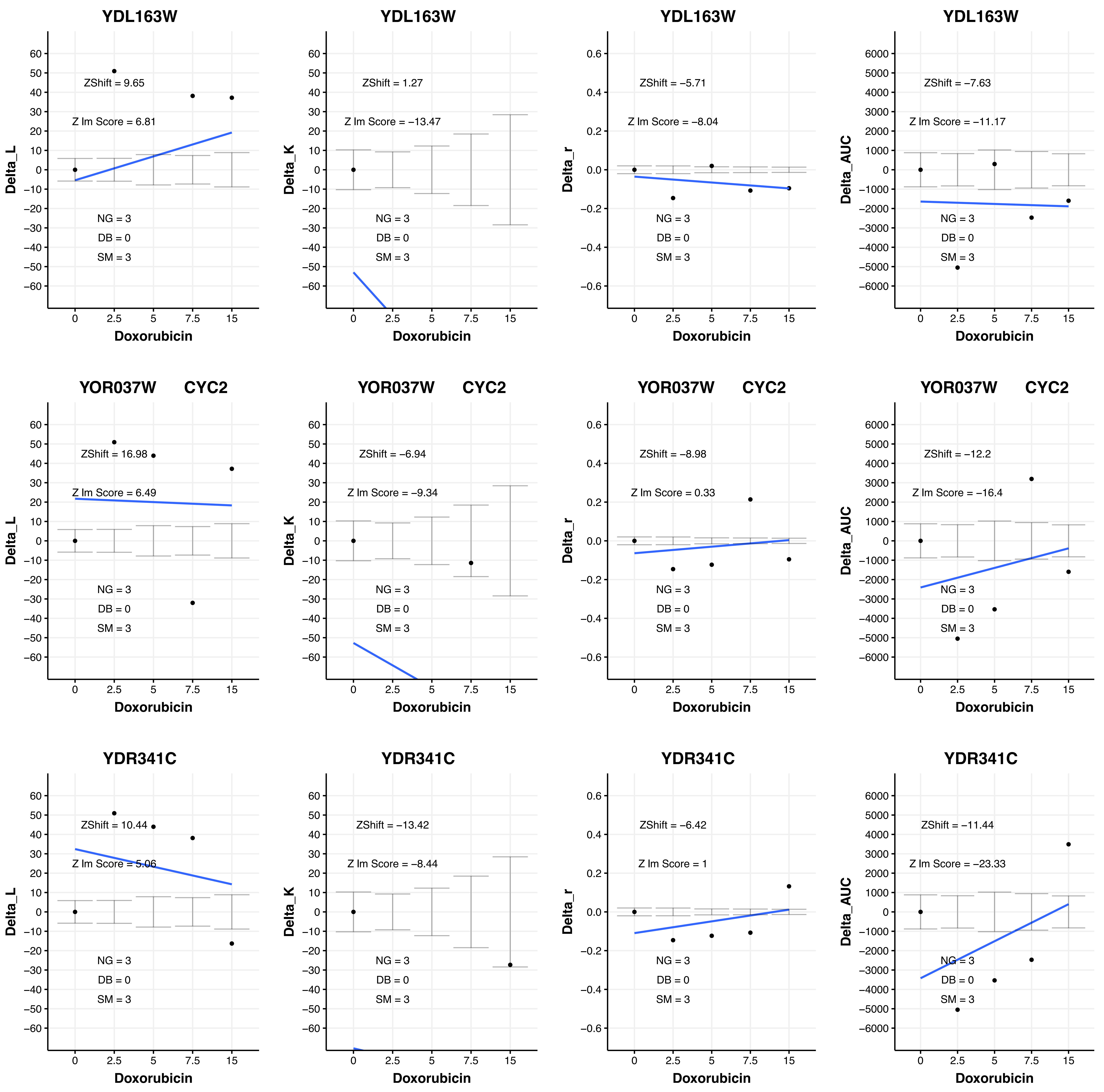
YDL163W
ZShift = 9.65
Z lm Score = 6.81
NG = 3
DB = 0
SM = 3
Delta_L
Doxorubicin
YDL163W
ZShift = 1.27
Z lm Score = −13.47
NG = 3
DB = 0
SM = 3
Delta_K
Doxorubicin
YDL163W
ZShift = −5.71
Z lm Score = −8.04
NG = 3
DB = 0
SM = 3
Delta_r
Doxorubicin
YDL163W
ZShift = −7.63
Z lm Score = −11.17
NG = 3
DB = 0
SM = 3
Delta_AUC
Doxorubicin
YOR037W CYC2
ZShift = 16.98
Z lm Score = 6.49
NG = 3
DB = 0
SM = 3
Delta_L
Doxorubicin
YOR037W CYC2
ZShift = −6.94
Z lm Score = −9.34
NG = 3
DB = 0
SM = 3
Delta_K
Doxorubicin
YOR037W CYC2
ZShift = −8.98
Z lm Score = 0.33
NG = 3
DB = 0
SM = 3
Delta_r
Doxorubicin
YOR037W CYC2
ZShift = −12.2
Z lm Score = −16.4
NG = 3
DB = 0
SM = 3
Delta_AUC
Doxorubicin
YDR341C
ZShift = 10.44
Z lm Score = 5.06
NG = 3
DB = 0
SM = 3
Delta_L
Doxorubicin
YDR341C
ZShift = −13.42
Z lm Score = −8.44
NG = 3
DB = 0
SM = 3
Delta_K
Doxorubicin
YDR341C
ZShift = −6.42
Z lm Score = 1
NG = 3
DB = 0
SM = 3
Delta_r
Doxorubicin
YDR341C
ZShift = −11.44
Z lm Score = −23.33
NG = 3
DB = 0
SM = 3
Delta_AUC
Doxorubicin

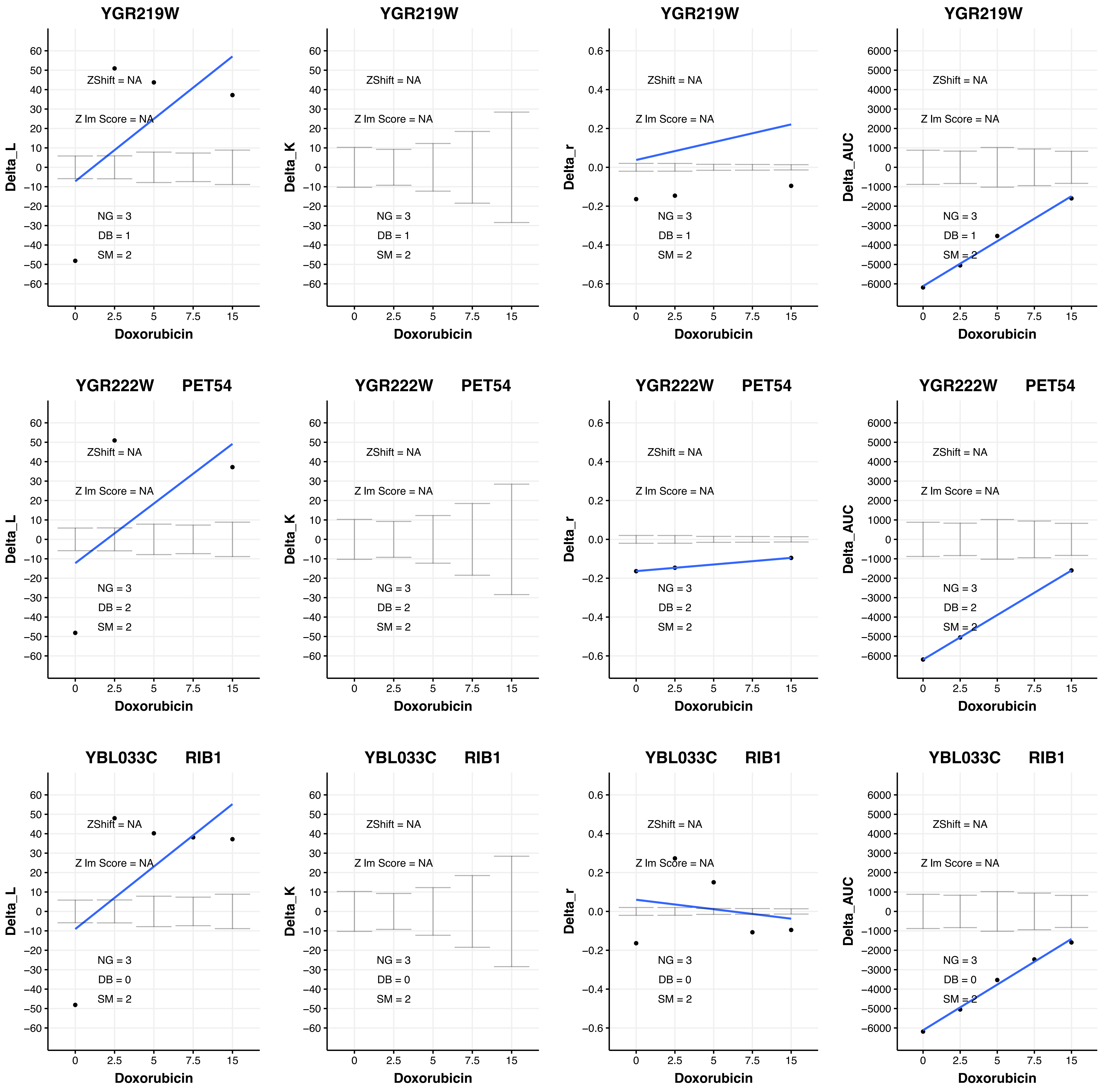
YGR219W
ZShift = NA
Z lm Score = NA
NG = 3
DB = 1
SM = 2
Delta_L
Delta_K
Delta_r
Delta_AUC
Doxorubicin
0
2.5
5
7.5
15
YGR222W PET54
ZShift = NA
Z lm Score = NA
NG = 3
DB = 2
SM = 2
Delta_L
Delta_K
Delta_r
Delta_AUC
Doxorubicin
YBL033C RIB1
ZShift = NA
Z lm Score = NA
NG = 3
DB = 0
SM = 2
Delta_L
Delta_K
Delta_r
Delta_AUC
Doxorubicin

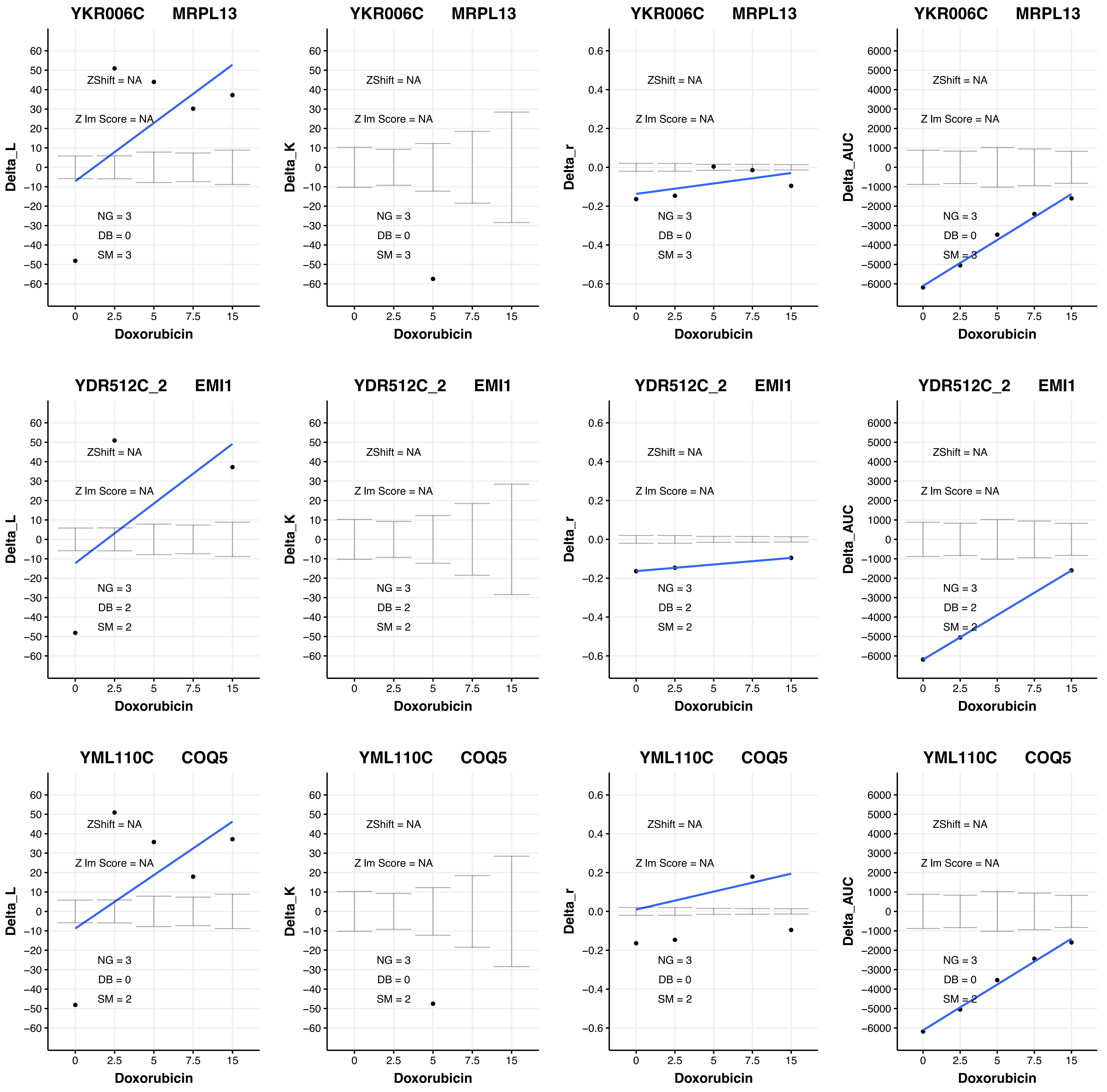

YKR006C MRPL13
ZShift = NA
Z lm Score = NA
NG = 3
DB = 0
SM = 3
Delta_L
Delta_K
Delta_r
Delta_AUC
Doxorubicin
0
2.5
5
7.5
15
YDR512C_2 EMI1
ZShift = NA
Z lm Score = NA
NG = 3
DB = 2
SM = 2
YML110C COQ5
ZShift = NA
Z lm Score = NA
NG = 3
DB = 0
SM = 2

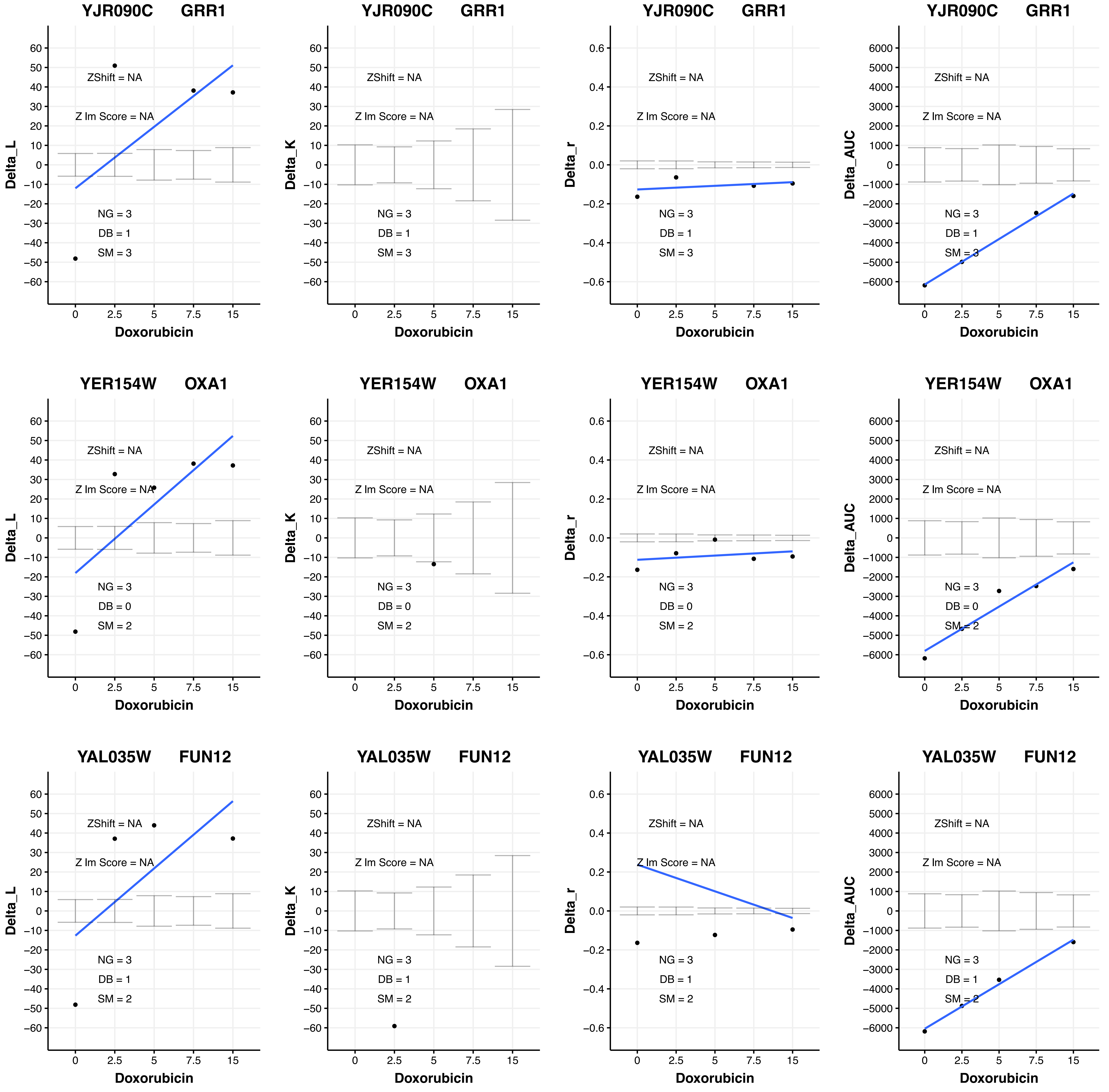
YJR090C GRR1
ZShift = NA
Z lm Score = NA
NG = 3
DB = 1
SM = 3
Delta_L
Delta_K
Delta_r
Delta_AUC
Doxorubicin
YER154W OXA1
ZShift = NA
Z lm Score = NA
NG = 3
DB = 0
SM = 2
YAL035W FUN12
ZShift = NA
Z lm Score = NA
NG = 3
DB = 1
SM = 2
0
2.5
5
7.5
15

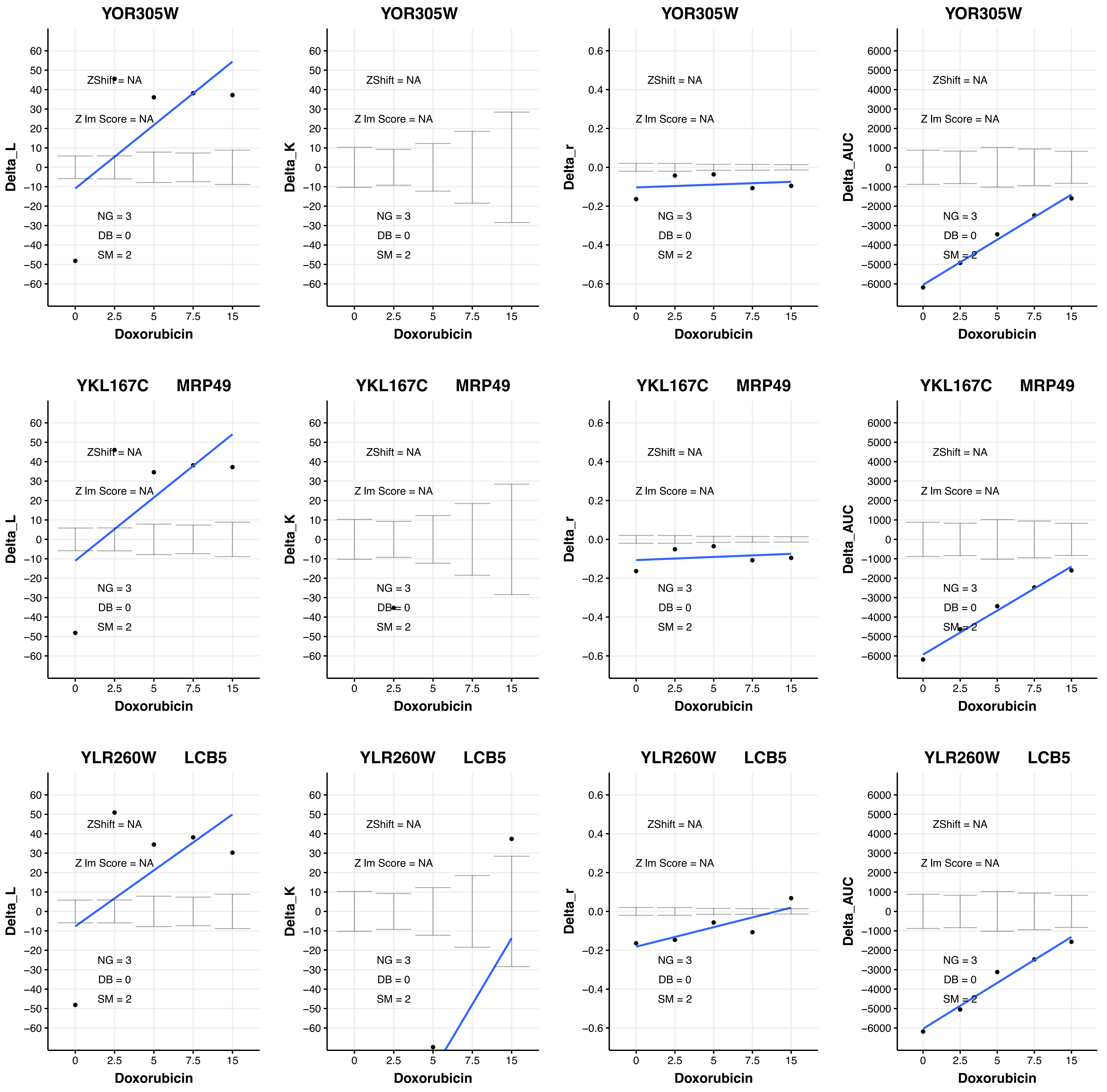

YOR305W
ZShift = NA
Z lm Score = NA
NG = 3
DB = 0
SM = 2
Delta_L
Doxorubicin
YOR305W
ZShift = NA
Z lm Score = NA
NG = 3
DB = 0
SM = 2
Delta_K
Doxorubicin
YOR305W
ZShift = NA
Z lm Score = NA
NG = 3
DB = 0
SM = 2
Delta_r
Doxorubicin
YOR305W
ZShift = NA
Z lm Score = NA
NG = 3
DB = 0
SM = 2
Delta_AUC
Doxorubicin
YKL167C MRP49
ZShift = NA
Z lm Score = NA
NG = 3
DB = 0
SM = 2
Delta_L
Doxorubicin
YKL167C MRP49
ZShift = NA
Z lm Score = NA
NG = 3
DB = 0
SM = 2
Delta_K
Doxorubicin
YKL167C MRP49
ZShift = NA
Z lm Score = NA
NG = 3
DB = 0
SM = 2
Delta_r
Doxorubicin
YKL167C MRP49
ZShift = NA
Z lm Score = NA
NG = 3
DB = 0
SM = 2
Delta_AUC
Doxorubicin
YLR260W LCB5
ZShift = NA
Z lm Score = NA
NG = 3
DB = 0
SM = 2
Delta_L
Doxorubicin
YLR260W LCB5
ZShift = NA
Z lm Score = NA
NG = 3
DB = 0
SM = 2
Delta_K
Doxorubicin
YLR260W LCB5
ZShift = NA
Z lm Score = NA
NG = 3
DB = 0
SM = 2
Delta_r
Doxorubicin
YLR260W LCB5
ZShift = NA
Z lm Score = NA
NG = 3
DB = 0
SM = 2
Delta_AUC
Doxorubicin

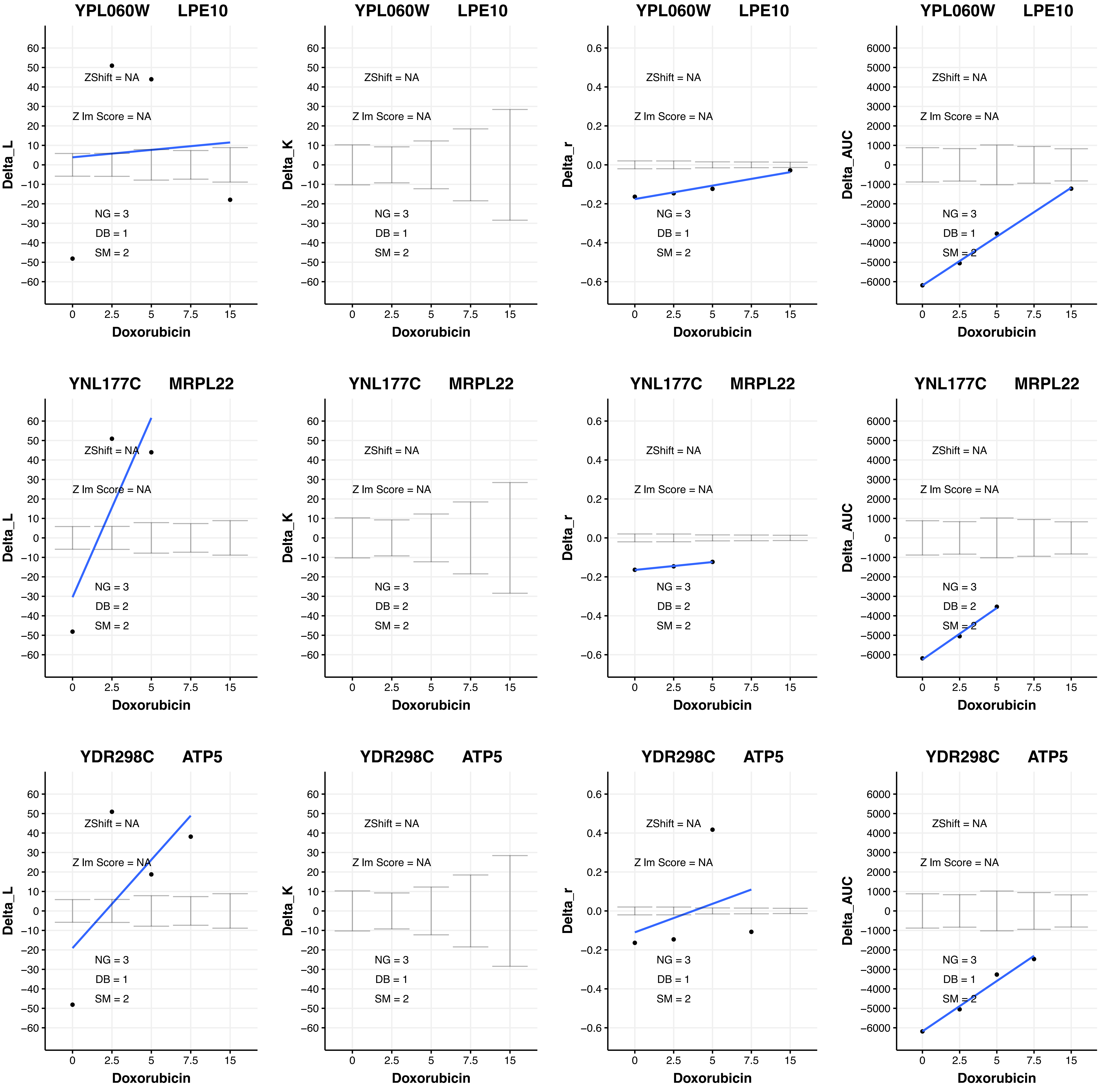

YPL060W LPE10
ZShift = NA
Z lm Score = NA
NG = 3
DB = 1
SM = 2
Delta_L
Delta_K
Delta_r
Delta_AUC
Doxorubicin
YNL177C MRPL22
ZShift = NA
Z lm Score = NA
NG = 3
DB = 2
SM = 2
YDR298C ATP5
ZShift = NA
Z lm Score = NA
NG = 3
DB = 1
SM = 2

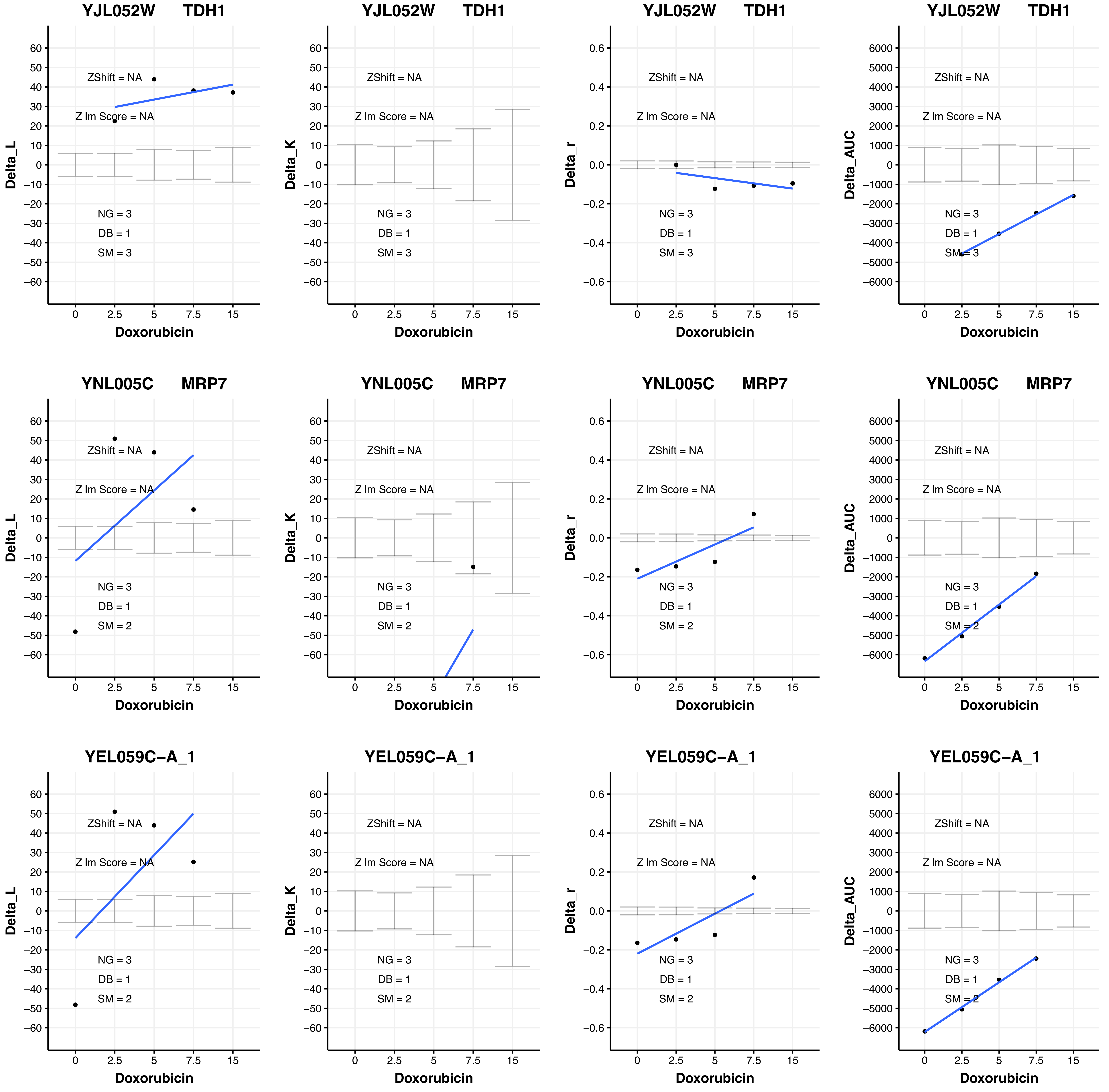
YJL052W TDH1
ZShift = NA
Z lm Score = NA
NG = 3
DB = 1
SM = 3
Delta_L
Delta_K
Delta_r
Delta_AUC
Doxorubicin
0
2.5
5
7.5
15
YNL005C MRP7
ZShift = NA
Z lm Score = NA
NG = 3
DB = 1
SM = 2
YEL059C−A_1
ZShift = NA
Z lm Score = NA
NG = 3
DB = 1
SM = 2

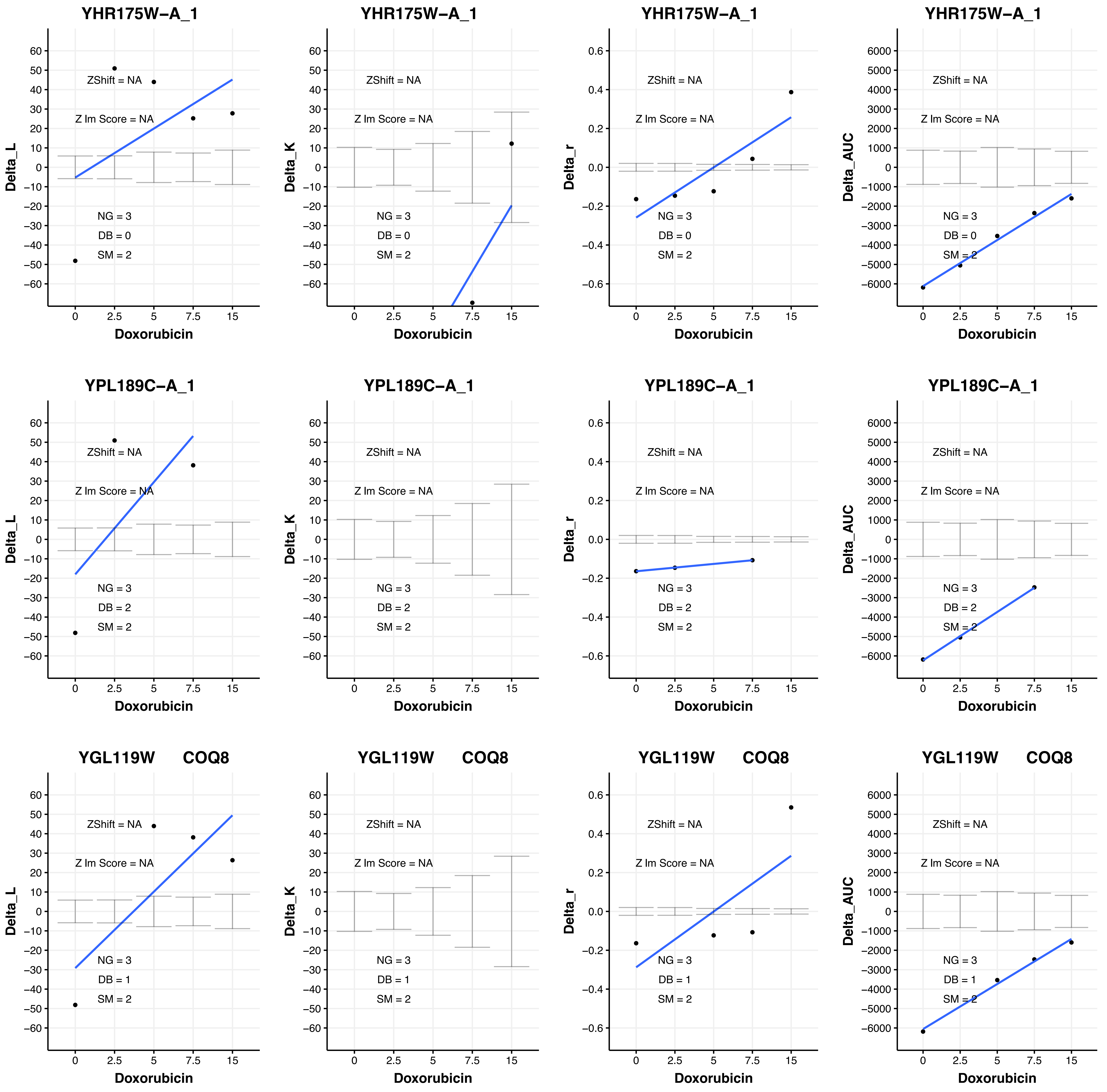

YHR175W−A_1
ZShift = NA
Z lm Score = NA
NG = 3
DB = 0
SM = 2
Delta_L
Delta_K
Delta_r
Delta_AUC
Doxorubicin
0
2.5
5
7.5
15
YPL189C−A_1
ZShift = NA
Z lm Score = NA
NG = 3
DB = 2
SM = 2
YGL119W COQ8
ZShift = NA
Z lm Score = NA
NG = 3
DB = 1
SM = 2

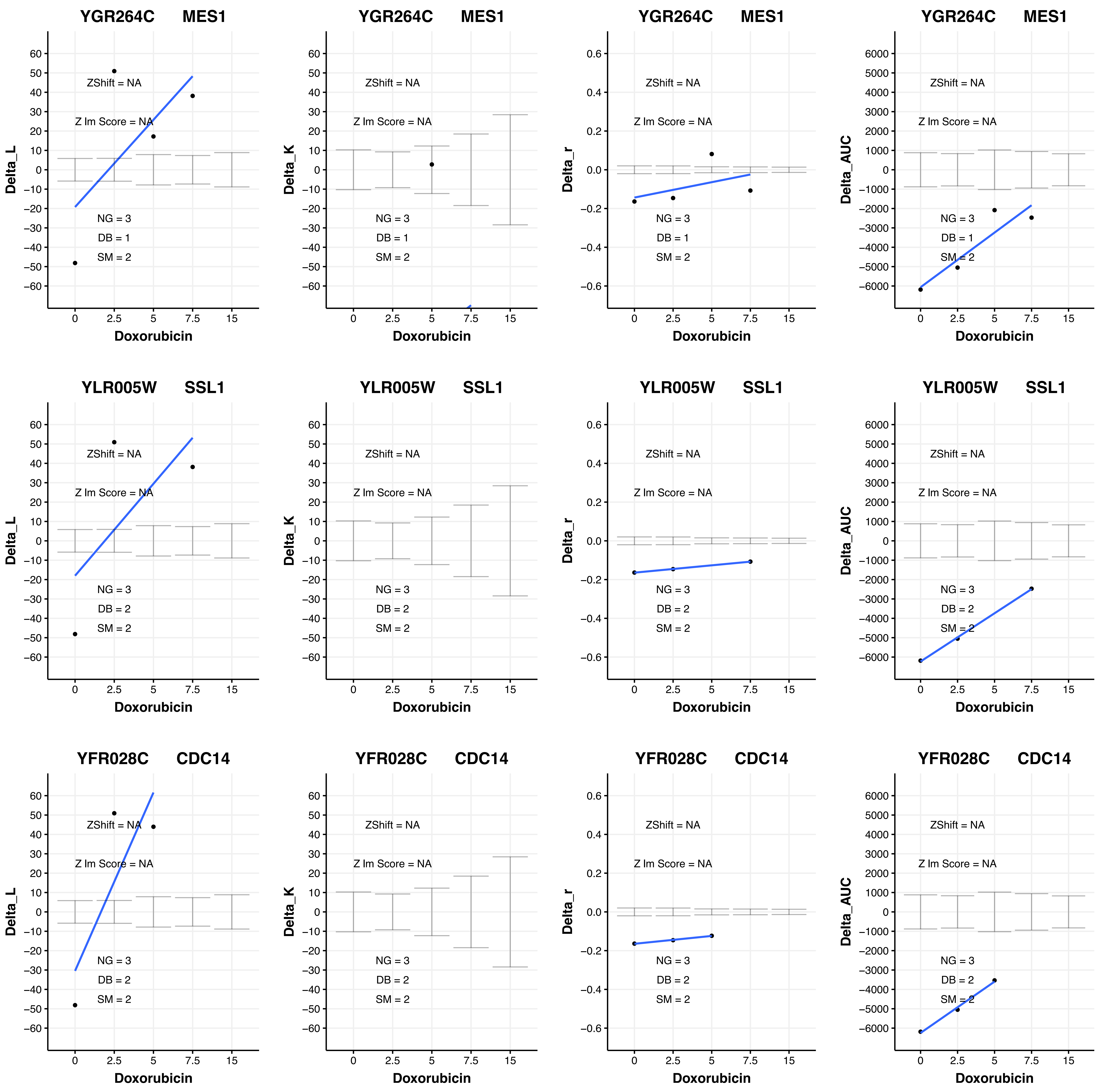

YGR264C MES1
ZShift = NA
Z lm Score = NA
NG = 3
DB = 1
SM = 2
Delta_L
Doxorubicin
YGR264C MES1
ZShift = NA
Z lm Score = NA
NG = 3
DB = 1
SM = 2
Delta_K
Doxorubicin
YGR264C MES1
ZShift = NA
Z lm Score = NA
NG = 3
DB = 1
SM = 2
Delta_r
Doxorubicin
YGR264C MES1
ZShift = NA
Z lm Score = NA
NG = 3
DB = 1
SM = 2
Delta_AUC
Doxorubicin
YLR005W SSL1
ZShift = NA
Z lm Score = NA
NG = 3
DB = 2
SM = 2
Delta_L
Doxorubicin
YLR005W SSL1
ZShift = NA
Z lm Score = NA
NG = 3
DB = 2
SM = 2
Delta_K
Doxorubicin
YLR005W SSL1
ZShift = NA
Z lm Score = NA
NG = 3
DB = 2
SM = 2
Delta_r
Doxorubicin
YLR005W SSL1
ZShift = NA
Z lm Score = NA
NG = 3
DB = 2
SM = 2
Delta_AUC
Doxorubicin
YFR028C CDC14
ZShift = NA
Z lm Score = NA
NG = 3
DB = 2
SM = 2
Delta_L
Doxorubicin
YFR028C CDC14
ZShift = NA
Z lm Score = NA
NG = 3
DB = 2
SM = 2
Delta_K
Doxorubicin
YFR028C CDC14
ZShift = NA
Z lm Score = NA
NG = 3
DB = 2
SM = 2
Delta_r
Doxorubicin
YFR028C CDC14
ZShift = NA
Z lm Score = NA
NG = 3
DB = 2
SM = 2
Delta_AUC
Doxorubicin

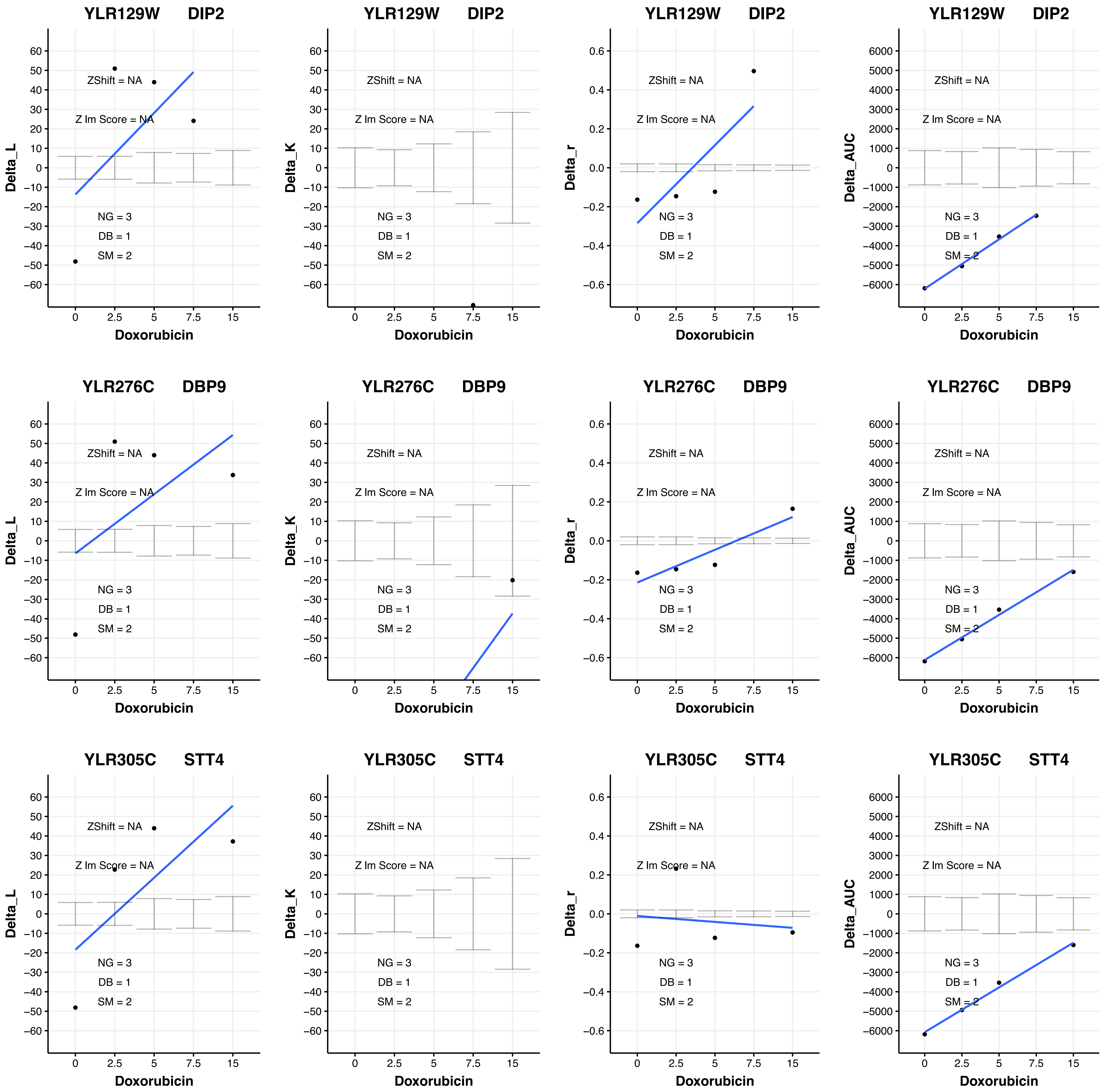

YLR129W DIP2
ZShift = NA
Z lm Score = NA
NG = 3
DB = 1
SM = 2
Delta_L
Delta_K
Delta_r
Delta_AUC
Doxorubicin
YLR276C DBP9
ZShift = NA
Z lm Score = NA
NG = 3
DB = 1
SM = 2
Delta_L
Delta_K
Delta_r
Delta_AUC
Doxorubicin
YLR305C STT4
ZShift = NA
Z lm Score = NA
NG = 3
DB = 1
SM = 2
Delta_L
Delta_K
Delta_r
Delta_AUC
Doxorubicin

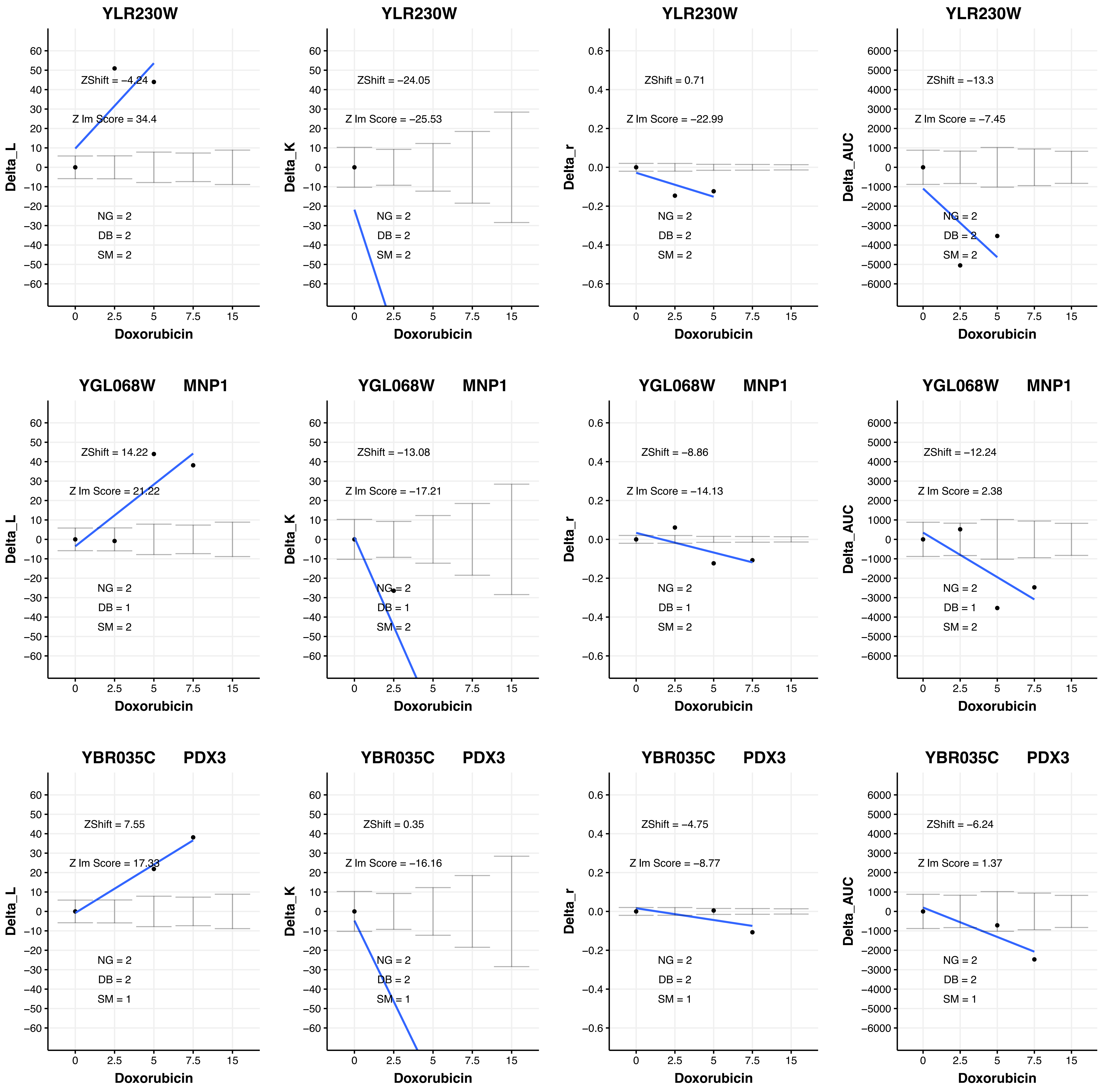
YLR230W
ZShift = −4.24
Z lm Score = 34.4
NG = 2
DB = 2
SM = 2
Delta_L
Doxorubicin
YLR230W
ZShift = −24.05
Z lm Score = −25.53
NG = 2
DB = 2
SM = 2
Delta_K
Doxorubicin
YLR230W
ZShift = 0.71
Z lm Score = −22.99
NG = 2
DB = 2
SM = 2
Delta_r
Doxorubicin
YLR230W
ZShift = −13.3
Z lm Score = −7.45
NG = 2
DB = 2
SM = 2
Delta_AUC
Doxorubicin
YGL068W MNP1
ZShift = 14.22
Z lm Score = 21.22
NG = 2
DB = 1
SM = 2
Delta_L
Doxorubicin
YGL068W MNP1
ZShift = −13.08
Z lm Score = −17.21
NG = 2
DB = 1
SM = 2
Delta_K
Doxorubicin
YGL068W MNP1
ZShift = −8.86
Z lm Score = −14.13
NG = 2
DB = 1
SM = 2
Delta_r
Doxorubicin
YGL068W MNP1
ZShift = −12.24
Z lm Score = 2.38
NG = 2
DB = 1
SM = 2
Delta_AUC
Doxorubicin
YBR035C PDX3
ZShift = 7.55
Z lm Score = 17.33
NG = 2
DB = 2
SM = 1
Delta_L
Doxorubicin
YBR035C PDX3
ZShift = 0.35
Z lm Score = −16.16
NG = 2
DB = 2
SM = 1
Delta_K
Doxorubicin
YBR035C PDX3
ZShift = −4.75
Z lm Score = −8.77
NG = 2
DB = 2
SM = 1
Delta_r
Doxorubicin
YBR035C PDX3
ZShift = −6.24
Z lm Score = 1.37
NG = 2
DB = 2
SM = 1
Delta_AUC
Doxorubicin

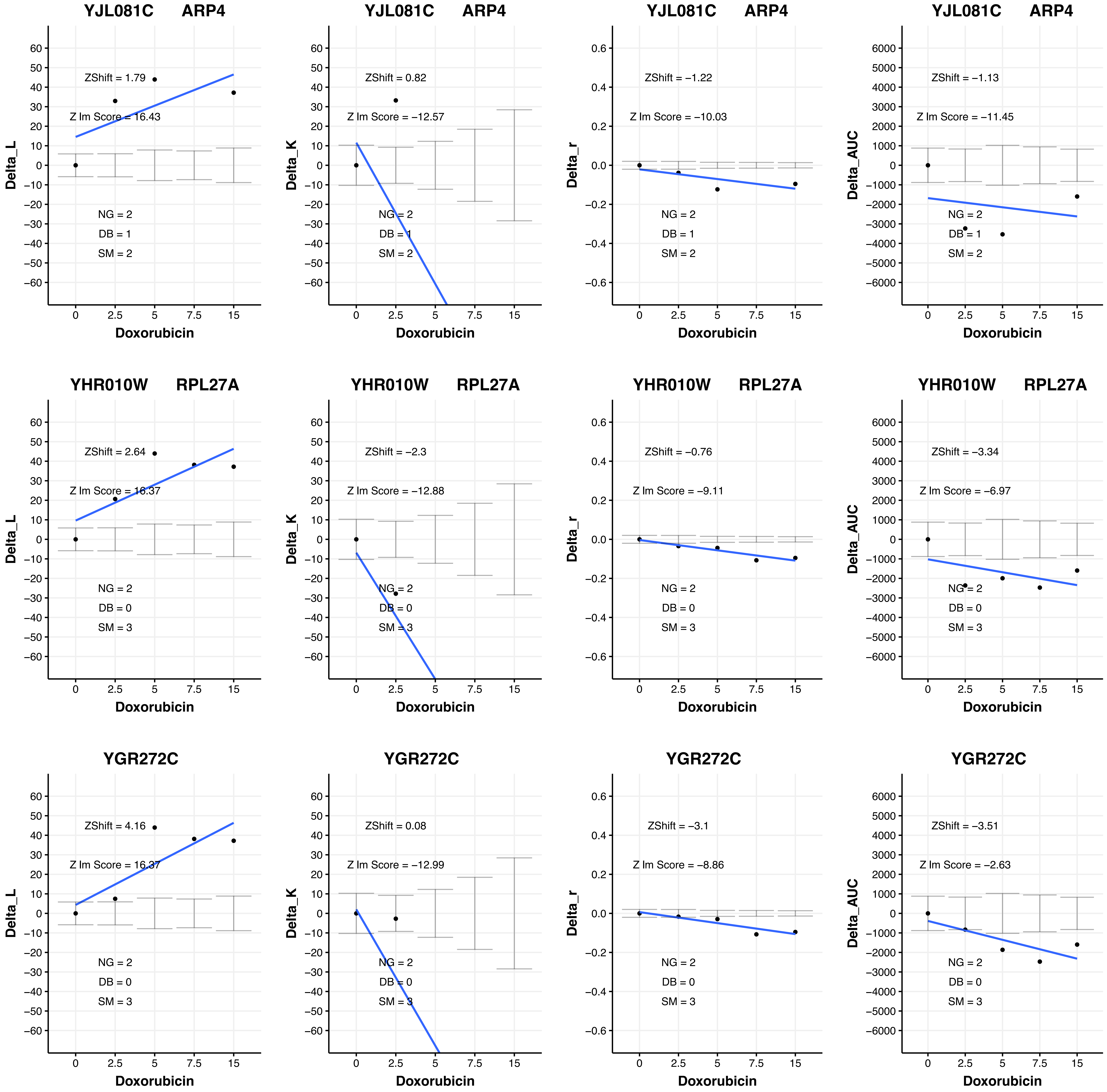
YJL081C ARP4
ZShift = 1.79
Z lm Score = 16.43
NG = 2
DB = 1
SM = 2
Delta_L
60
50
40
30
20
10
0
−10
−20
−30
−40
−50
−60
0
2.5
5
7.5
15
Doxorubicin
YJL081C ARP4
ZShift = 0.82
Z lm Score = −12.57
NG = 2
DB = 1
SM = 2
Delta_K
60
50
40
30
20
10
0
−10
−20
−30
−40
−50
−60
0
2.5
5
7.5
15
Doxorubicin
YJL081C ARP4
ZShift = −1.22
Z lm Score = −10.03
NG = 2
DB = 1
SM = 2
Delta_r
0.6
0.4
0.2
0.0
−0.2
−0.4
−0.6
0
2.5
5
7.5
15
Doxorubicin
YJL081C ARP4
ZShift = −1.13
Z lm Score = −11.45
NG = 2
DB = 1
SM = 2
Delta_AUC
6000
5000
4000
3000
2000
1000
0
−1000
−2000
−3000
−4000
−5000
−6000
0
2.5
5
7.5
15
Doxorubicin
YHR010W RPL27A
ZShift = 2.64
Z lm Score = 16.37
NG = 2
DB = 0
SM = 3
Delta_L
60
50
40
30
20
10
0
−10
−20
−30
−40
−50
−60
0
2.5
5
7.5
15
Doxorubicin
YHR010W RPL27A
ZShift = −2.3
Z lm Score = −12.88
NG = 2
DB = 0
SM = 3
Delta_K
60
50
40
30
20
10
0
−10
−20
−30
−40
−50
−60
0
2.5
5
7.5
15
Doxorubicin
YHR010W RPL27A
ZShift = −0.76
Z lm Score = −9.11
NG = 2
DB = 0
SM = 3
Delta_r
0.6
0.4
0.2
0.0
−0.2
−0.4
−0.6
0
2.5
5
7.5
15
Doxorubicin
YHR010W RPL27A
ZShift = −3.34
Z lm Score = −6.97
NG = 2
DB = 0
SM = 3
Delta_AUC
6000
5000
4000
3000
2000
1000
0
−1000
−2000
−3000
−4000
−5000
−6000
0
2.5
5
7.5
15
Doxorubicin
YGR272C
ZShift = 4.16
Z lm Score = 16.37
NG = 2
DB = 0
SM = 3
Delta_L
60
50
40
30
20
10
0
−10
−20
−30
−40
−50
−60
0
2.5
5
7.5
15
Doxorubicin
YGR272C
ZShift = 0.08
Z lm Score = −12.99
NG = 2
DB = 0
SM = 3
Delta_K
60
50
40
30
20
10
0
−10
−20
−30
−40
−50
−60
0
2.5
5
7.5
15
Doxorubicin
YGR272C
ZShift = −3.1
Z lm Score = −8.86
NG = 2
DB = 0
SM = 3
Delta_r
0.6
0.4
0.2
0.0
−0.2
−0.4
−0.6
0
2.5
5
7.5
15
Doxorubicin
YGR272C
ZShift = −3.51
Z lm Score = −2.63
NG = 2
DB = 0
SM = 3
Delta_AUC
6000
5000
4000
3000
2000
1000
0
−1000
−2000
−3000
−4000
−5000
−6000
0
2.5
5
7.5
15
Doxorubicin

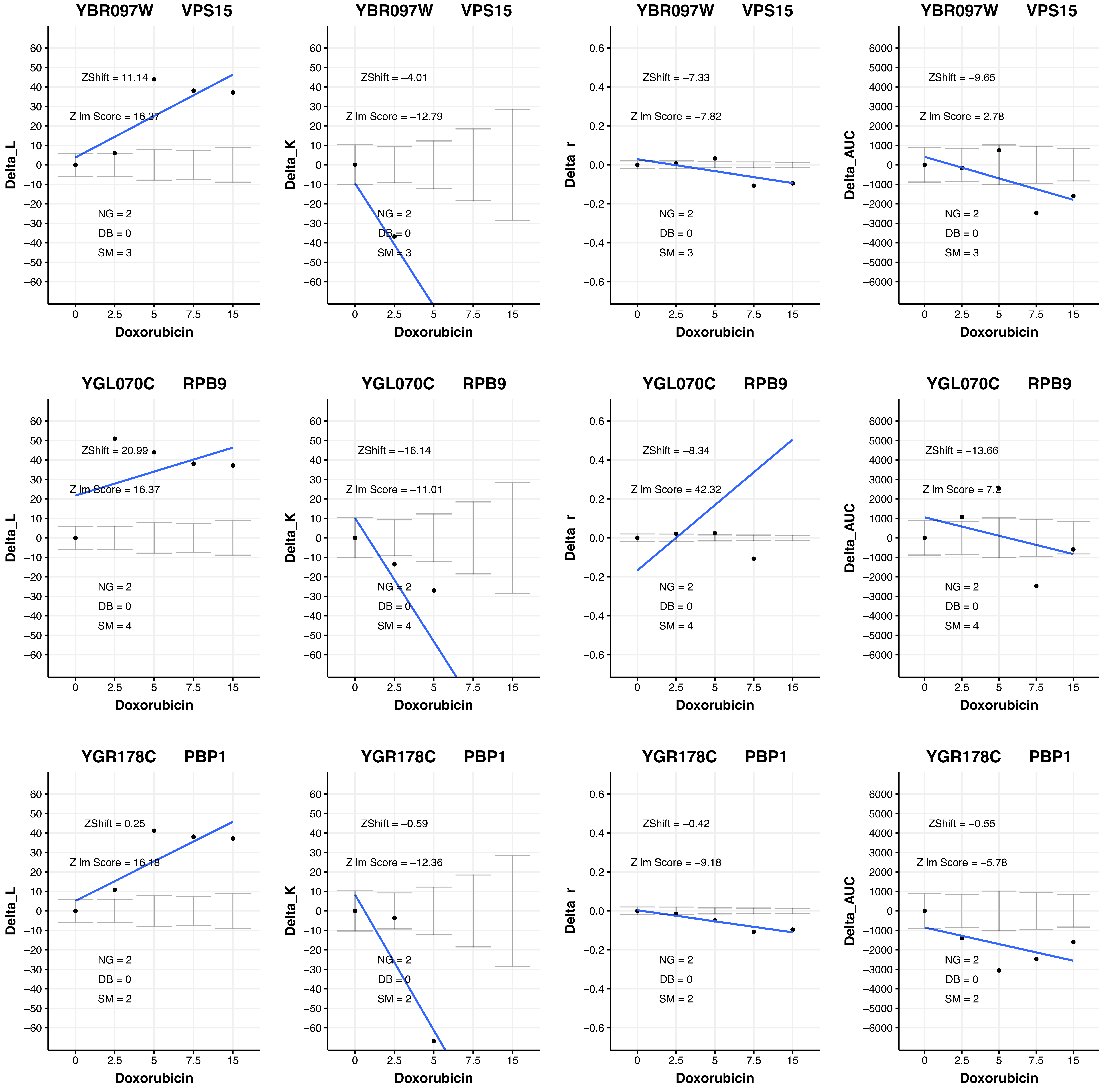
YBR097W VPS15
Delta_L
ZShift = 11.14
Z lm Score = 16.37
NG = 2
DB = 0
SM = 3
Doxorubicin
YBR097W VPS15
Delta_K
ZShift = −4.01
Z lm Score = −12.79
NG = 2
DB = 0
SM = 3
Doxorubicin
YBR097W VPS15
Delta_r
ZShift = −7.33
Z lm Score = −7.82
NG = 2
DB = 0
SM = 3
Doxorubicin
YBR097W VPS15
Delta_AUC
ZShift = −9.65
Z lm Score = 2.78
NG = 2
DB = 0
SM = 3
Doxorubicin
YGL070C RPB9
Delta_L
ZShift = 20.99
Z lm Score = 16.37
NG = 2
DB = 0
SM = 4
Doxorubicin
YGL070C RPB9
Delta_K
ZShift = −16.14
Z lm Score = −11.01
NG = 2
DB = 0
SM = 4
Doxorubicin
YGL070C RPB9
Delta_r
ZShift = −8.34
Z lm Score = 42.32
NG = 2
DB = 0
SM = 4
Doxorubicin
YGL070C RPB9
Delta_AUC
ZShift = −13.66
Z lm Score = 7.2
NG = 2
DB = 0
SM = 4
Doxorubicin
YGR178C PBP1
Delta_L
ZShift = 0.25
Z lm Score = 16.18
NG = 2
DB = 0
SM = 2
Doxorubicin
YGR178C PBP1
Delta_K
ZShift = −0.59
Z lm Score = −12.36
NG = 2
DB = 0
SM = 2
Doxorubicin
YGR178C PBP1
Delta_r
ZShift = −0.42
Z lm Score = −9.18
NG = 2
DB = 0
SM = 2
Doxorubicin
YGR178C PBP1
Delta_AUC
ZShift = −0.55
Z lm Score = −5.78
NG = 2
DB = 0
SM = 2
Doxorubicin

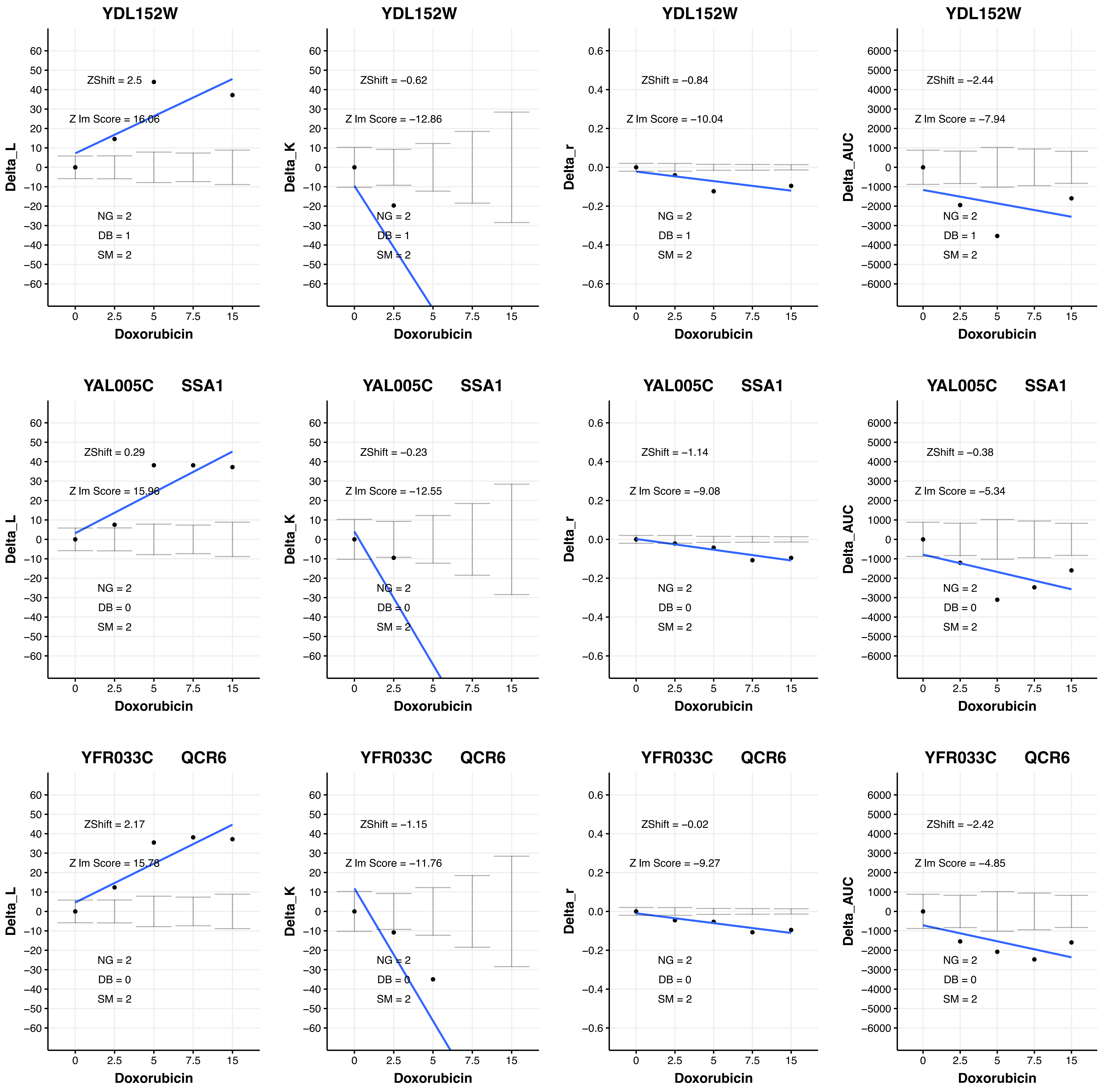

YDL152W
ZShift = 2.5
Z lm Score = 16.06
NG = 2
DB = 1
SM = 2
Delta_L
Doxorubicin
YDL152W
ZShift = −0.62
Z lm Score = −12.86
NG = 2
DB = 1
SM = 2
Delta_K
Doxorubicin
YDL152W
ZShift = −0.84
Z lm Score = −10.04
NG = 2
DB = 1
SM = 2
Delta_r
Doxorubicin
YDL152W
ZShift = −2.44
Z lm Score = −7.94
NG = 2
DB = 1
SM = 2
Delta_AUC
Doxorubicin
YAL005C SSA1
ZShift = 0.29
Z lm Score = 15.96
NG = 2
DB = 0
SM = 2
Delta_L
Doxorubicin
YAL005C SSA1
ZShift = −0.23
Z lm Score = −12.55
NG = 2
DB = 0
SM = 2
Delta_K
Doxorubicin
YAL005C SSA1
ZShift = −1.14
Z lm Score = −9.08
NG = 2
DB = 0
SM = 2
Delta_r
Doxorubicin
YAL005C SSA1
ZShift = −0.38
Z lm Score = −5.34
NG = 2
DB = 0
SM = 2
Delta_AUC
Doxorubicin
YFR033C QCR6
ZShift = 2.17
Z lm Score = 15.78
NG = 2
DB = 0
SM = 2
Delta_L
Doxorubicin
YFR033C QCR6
ZShift = −1.15
Z lm Score = −11.76
NG = 2
DB = 0
SM = 2
Delta_K
Doxorubicin
YFR033C QCR6
ZShift = −0.02
Z lm Score = −9.27
NG = 2
DB = 0
SM = 2
Delta_r
Doxorubicin
YFR033C QCR6
ZShift = −2.42
Z lm Score = −4.85
NG = 2
DB = 0
SM = 2
Delta_AUC
Doxorubicin

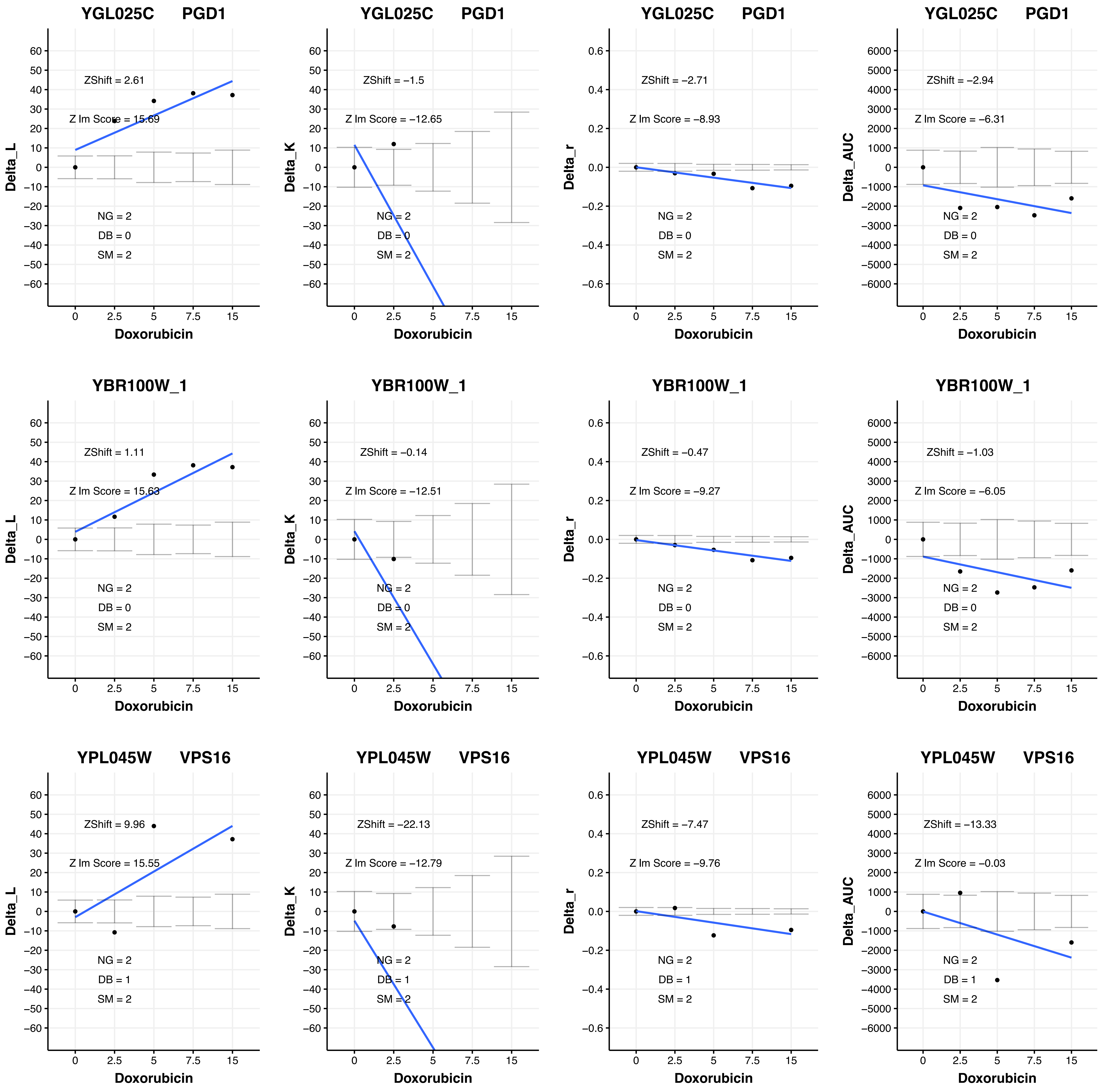
YGL025C PGD1
ZShift = 2.61
Z lm Score = 15.69
NG = 2
DB = 0
SM = 2
Delta_L
Doxorubicin
YGL025C PGD1
ZShift = −1.5
Z lm Score = −12.65
NG = 2
DB = 0
SM = 2
Delta_K
Doxorubicin
YGL025C PGD1
ZShift = −2.71
Z lm Score = −8.93
NG = 2
DB = 0
SM = 2
Delta_r
Doxorubicin
YGL025C PGD1
ZShift = −2.94
Z lm Score = −6.31
NG = 2
DB = 0
SM = 2
Delta_AUC
Doxorubicin
YBR100W_1
ZShift = 1.11
Z lm Score = 15.63
NG = 2
DB = 0
SM = 2
Delta_L
Doxorubicin
YBR100W_1
ZShift = −0.14
Z lm Score = −12.51
NG = 2
DB = 0
SM = 2
Delta_K
Doxorubicin
YBR100W_1
ZShift = −0.47
Z lm Score = −9.27
NG = 2
DB = 0
SM = 2
Delta_r
Doxorubicin
YBR100W_1
ZShift = −1.03
Z lm Score = −6.05
NG = 2
DB = 0
SM = 2
Delta_AUC
Doxorubicin
YPL045W VPS16
ZShift = 9.96
Z lm Score = 15.55
NG = 2
DB = 1
SM = 2
Delta_L
Doxorubicin
YPL045W VPS16
ZShift = −22.13
Z lm Score = −12.79
NG = 2
DB = 1
SM = 2
Delta_K
Doxorubicin
YPL045W VPS16
ZShift = −7.47
Z lm Score = −9.76
NG = 2
DB = 1
SM = 2
Delta_r
Doxorubicin
YPL045W VPS16
ZShift = −13.33
Z lm Score = −0.03
NG = 2
DB = 1
SM = 2
Delta_AUC
Doxorubicin

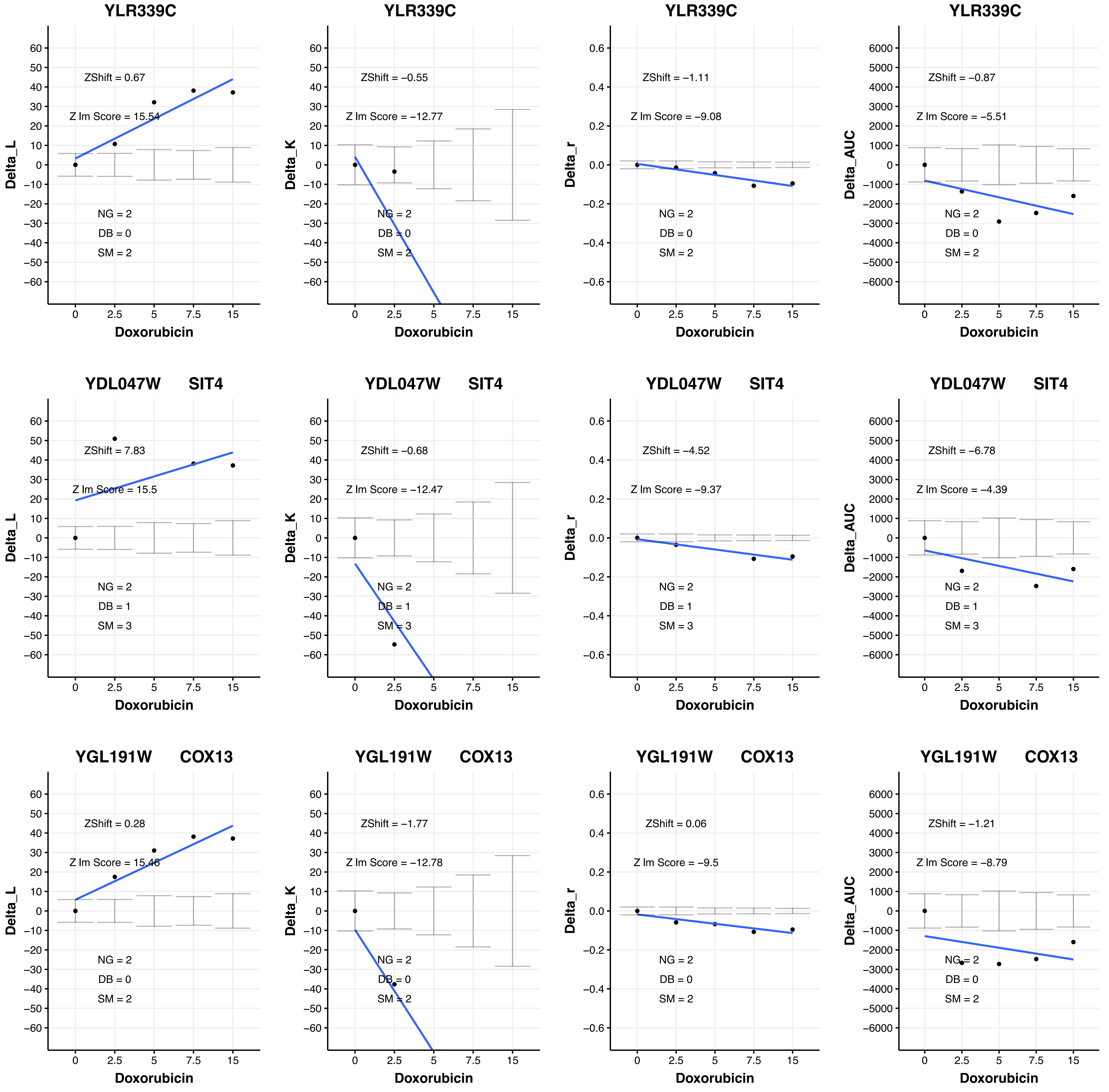
YLR339C
ZShift = 0.67
Z lm Score = 15.54
NG = 2
DB = 0
SM = 2
Delta_L
Doxorubicin
YLR339C
ZShift = −0.55
Z lm Score = −12.77
NG = 2
DB = 0
SM = 2
Delta_K
Doxorubicin
YLR339C
ZShift = −1.11
Z lm Score = −9.08
NG = 2
DB = 0
SM = 2
Delta_r
Doxorubicin
YLR339C
ZShift = −0.87
Z lm Score = −5.51
NG = 2
DB = 0
SM = 2
Delta_AUC
Doxorubicin
YDL047W SIT4
ZShift = 7.83
Z lm Score = 15.5
NG = 2
DB = 1
SM = 3
Delta_L
Doxorubicin
YDL047W SIT4
ZShift = −0.68
Z lm Score = −12.47
NG = 2
DB = 1
SM = 3
Delta_K
Doxorubicin
YDL047W SIT4
ZShift = −4.52
Z lm Score = −9.37
NG = 2
DB = 1
SM = 3
Delta_r
Doxorubicin
YDL047W SIT4
ZShift = −6.78
Z lm Score = −4.39
NG = 2
DB = 1
SM = 3
Delta_AUC
Doxorubicin
YGL191W COX13
ZShift = 0.28
Z lm Score = 15.46
NG = 2
DB = 0
SM = 2
Delta_L
Doxorubicin
YGL191W COX13
ZShift = −1.77
Z lm Score = −12.78
NG = 2
DB = 0
SM = 2
Delta_K
Doxorubicin
YGL191W COX13
ZShift = 0.06
Z lm Score = −9.5
NG = 2
DB = 0
SM = 2
Delta_r
Doxorubicin
YGL191W COX13
ZShift = −1.21
Z lm Score = −8.79
NG = 2
DB = 0
SM = 2
Delta_AUC
Doxorubicin

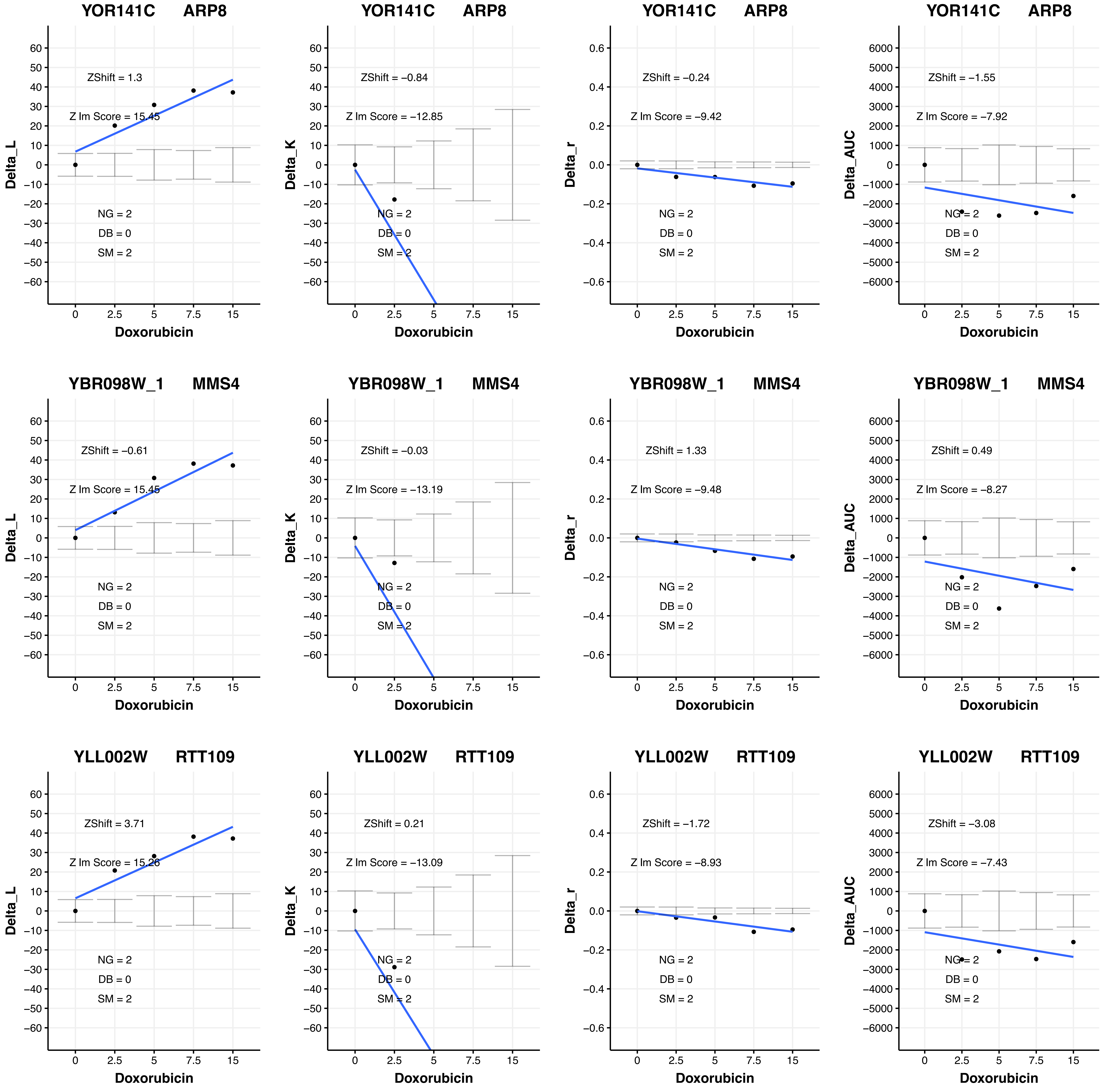

YOR141C ARP8
ZShift = 1.3
Z lm Score = 15.45
NG = 2
DB = 0
SM = 2
Delta_L
Doxorubicin
YOR141C ARP8
ZShift = −0.84
Z lm Score = −12.85
NG = 2
DB = 0
SM = 2
Delta_K
Doxorubicin
YOR141C ARP8
ZShift = −0.24
Z lm Score = −9.42
NG = 2
DB = 0
SM = 2
Delta_r
Doxorubicin
YOR141C ARP8
ZShift = −1.55
Z lm Score = −7.92
NG = 2
DB = 0
SM = 2
Delta_AUC
Doxorubicin
YBR098W_1 MMS4
ZShift = −0.61
Z lm Score = 15.45
NG = 2
DB = 0
SM = 2
Delta_L
Doxorubicin
YBR098W_1 MMS4
ZShift = −0.03
Z lm Score = −13.19
NG = 2
DB = 0
SM = 2
Delta_K
Doxorubicin
YBR098W_1 MMS4
ZShift = 1.33
Z lm Score = −9.48
NG = 2
DB = 0
SM = 2
Delta_r
Doxorubicin
YBR098W_1 MMS4
ZShift = 0.49
Z lm Score = −8.27
NG = 2
DB = 0
SM = 2
Delta_AUC
Doxorubicin
YLL002W RTT109
ZShift = 3.71
Z lm Score = 15.26
NG = 2
DB = 0
SM = 2
Delta_L
Doxorubicin
YLL002W RTT109
ZShift = 0.21
Z lm Score = −13.09
NG = 2
DB = 0
SM = 2
Delta_K
Doxorubicin
YLL002W RTT109
ZShift = −1.72
Z lm Score = −8.93
NG = 2
DB = 0
SM = 2
Delta_r
Doxorubicin
YLL002W RTT109
ZShift = −3.08
Z lm Score = −7.43
NG = 2
DB = 0
SM = 2
Delta_AUC
Doxorubicin

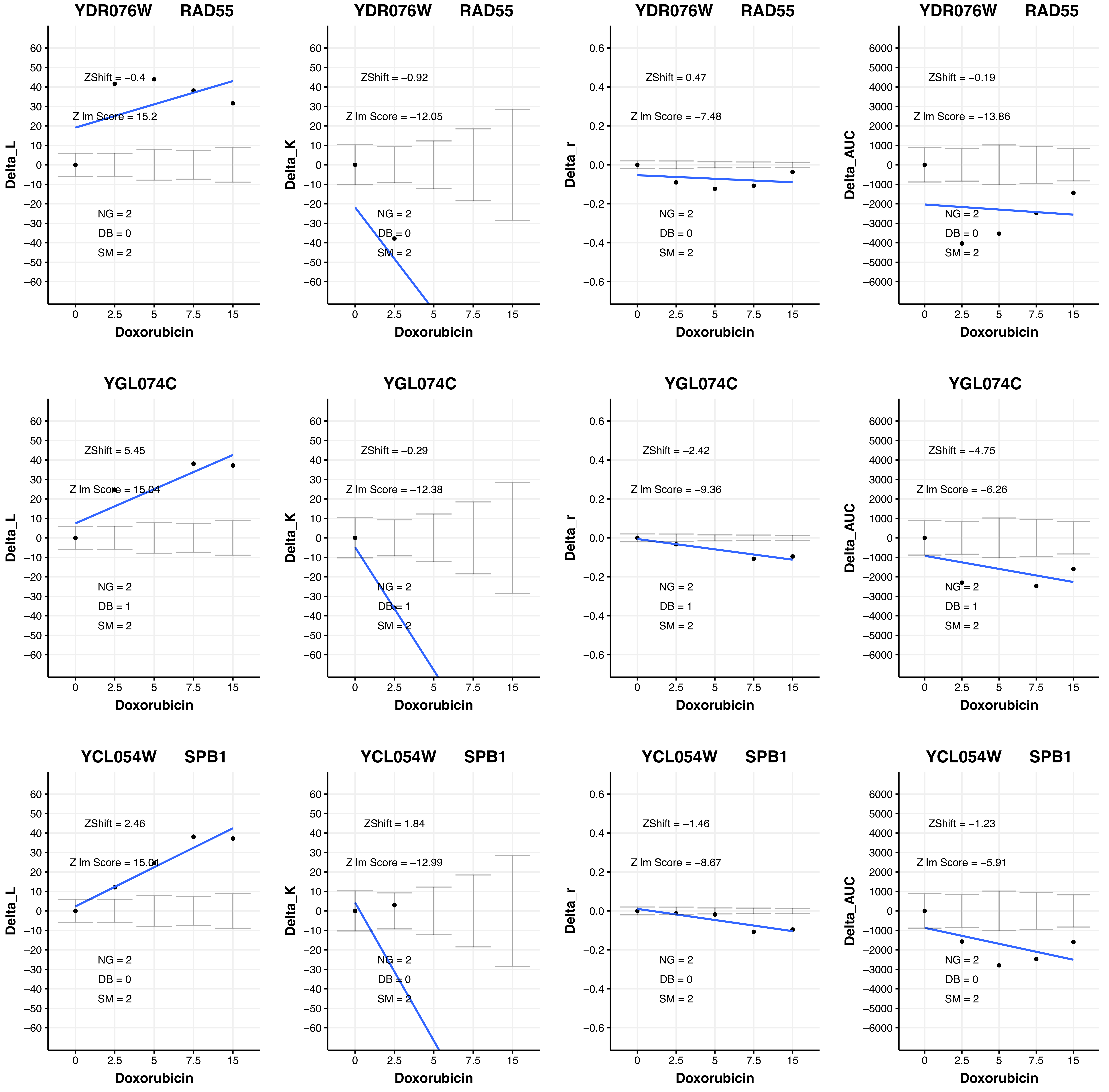
YDR076W RAD55
ZShift = −0.4
Z lm Score = 15.2
NG = 2
DB = 0
SM = 2
Delta_L
Doxorubicin
YDR076W RAD55
ZShift = −0.92
Z lm Score = −12.05
NG = 2
DB = 0
SM = 2
Delta_K
Doxorubicin
YDR076W RAD55
ZShift = 0.47
Z lm Score = −7.48
NG = 2
DB = 0
SM = 2
Delta_r
Doxorubicin
YDR076W RAD55
ZShift = −0.19
Z lm Score = −13.86
NG = 2
DB = 0
SM = 2
Delta_AUC
Doxorubicin
YGL074C
ZShift = 5.45
Z lm Score = 15.04
NG = 2
DB = 1
SM = 2
Delta_L
Doxorubicin
YGL074C
ZShift = −0.29
Z lm Score = −12.38
NG = 2
DB = 1
SM = 2
Delta_K
Doxorubicin
YGL074C
ZShift = −2.42
Z lm Score = −9.36
NG = 2
DB = 1
SM = 2
Delta_r
Doxorubicin
YGL074C
ZShift = −4.75
Z lm Score = −6.26
NG = 2
DB = 1
SM = 2
Delta_AUC
Doxorubicin
YCL054W SPB1
ZShift = 2.46
Z lm Score = 15.01
NG = 2
DB = 0
SM = 2
Delta_L
Doxorubicin
YCL054W SPB1
ZShift = 1.84
Z lm Score = −12.99
NG = 2
DB = 0
SM = 2
Delta_K
Doxorubicin
YCL054W SPB1
ZShift = −1.46
Z lm Score = −8.67
NG = 2
DB = 0
SM = 2
Delta_r
Doxorubicin
YCL054W SPB1
ZShift = −1.23
Z lm Score = −5.91
NG = 2
DB = 0
SM = 2
Delta_AUC
Doxorubicin

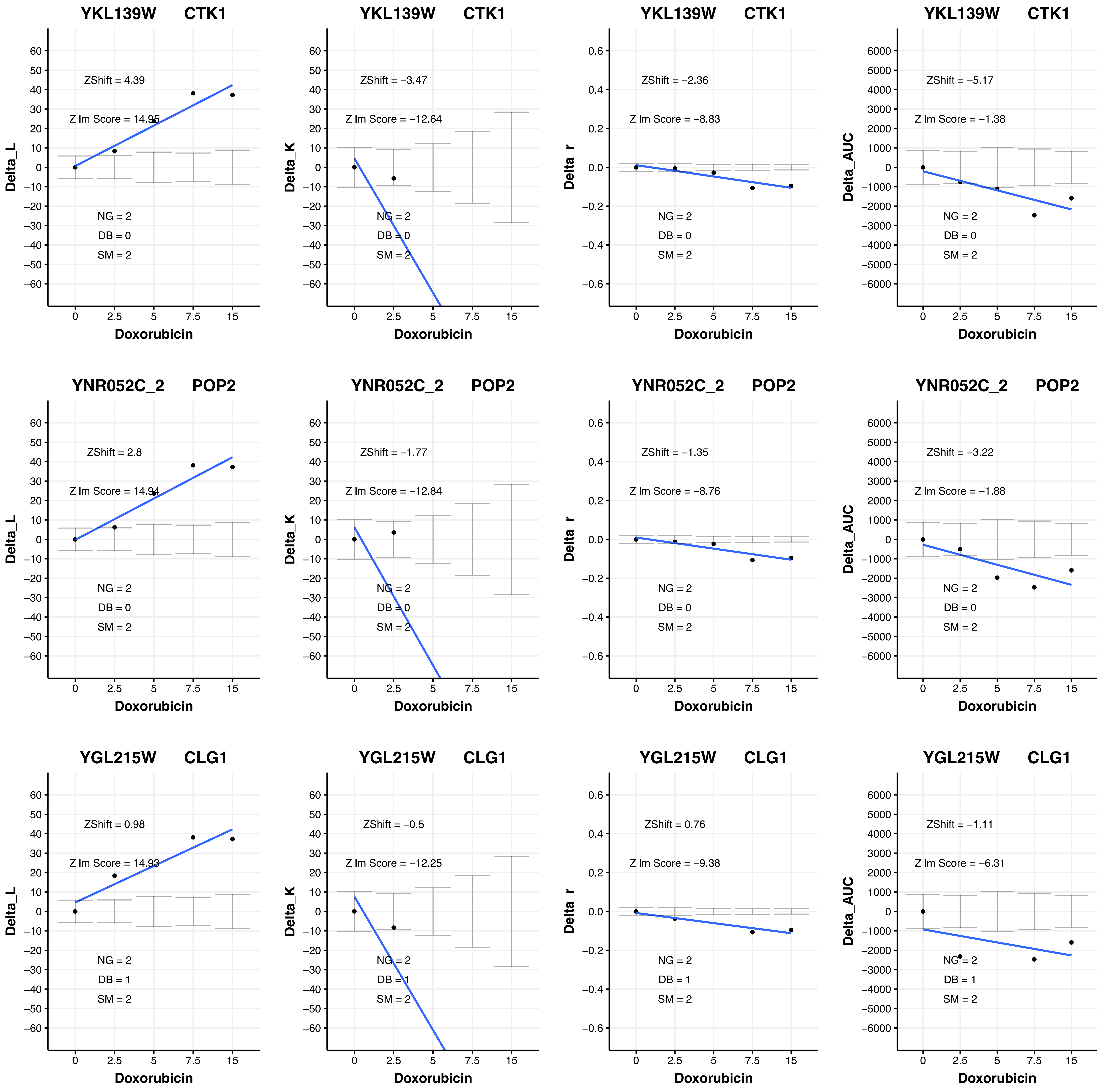
YKL139W CTK1
ZShift = 4.39
Z lm Score = 14.95
NG = 2
DB = 0
SM = 2
Delta_L
Doxorubicin
YKL139W CTK1
ZShift = −3.47
Z lm Score = −12.64
NG = 2
DB = 0
SM = 2
Delta_K
Doxorubicin
YKL139W CTK1
ZShift = −2.36
Z lm Score = −8.83
NG = 2
DB = 0
SM = 2
Delta_r
Doxorubicin
YKL139W CTK1
ZShift = −5.17
Z lm Score = −1.38
NG = 2
DB = 0
SM = 2
Delta_AUC
Doxorubicin
YNR052C_2 POP2
ZShift = 2.8
Z lm Score = 14.94
NG = 2
DB = 0
SM = 2
Delta_L
Doxorubicin
YNR052C_2 POP2
ZShift = −1.77
Z lm Score = −12.84
NG = 2
DB = 0
SM = 2
Delta_K
Doxorubicin
YNR052C_2 POP2
ZShift = −1.35
Z lm Score = −8.76
NG = 2
DB = 0
SM = 2
Delta_r
Doxorubicin
YNR052C_2 POP2
ZShift = −3.22
Z lm Score = −1.88
NG = 2
DB = 0
SM = 2
Delta_AUC
Doxorubicin
YGL215W CLG1
ZShift = 0.98
Z lm Score = 14.93
NG = 2
DB = 1
SM = 2
Delta_L
Doxorubicin
YGL215W CLG1
ZShift = −0.5
Z lm Score = −12.25
NG = 2
DB = 1
SM = 2
Delta_K
Doxorubicin
YGL215W CLG1
ZShift = 0.76
Z lm Score = −9.38
NG = 2
DB = 1
SM = 2
Delta_r
Doxorubicin
YGL215W CLG1
ZShift = −1.11
Z lm Score = −6.31
NG = 2
DB = 1
SM = 2
Delta_AUC
Doxorubicin

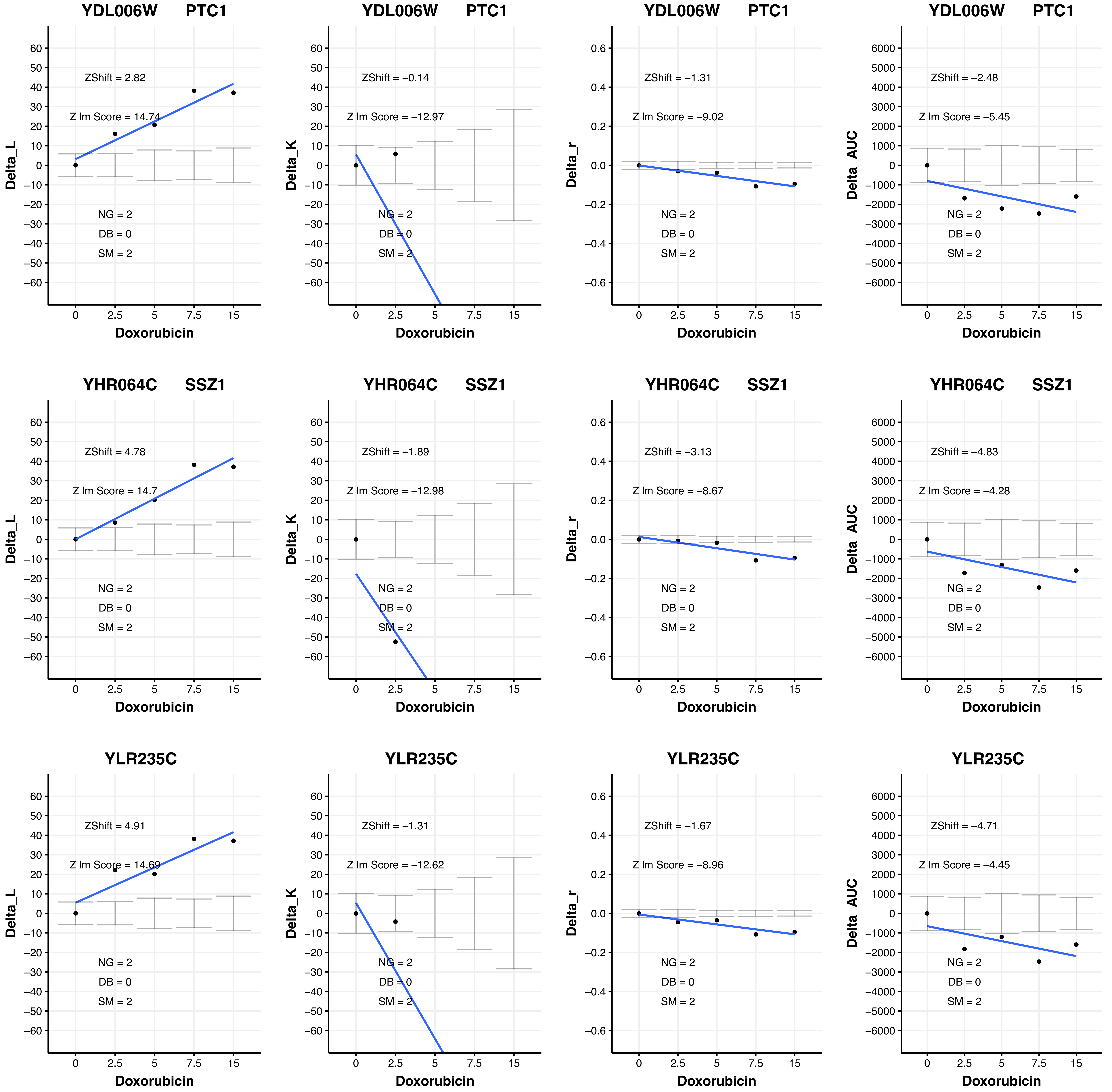

YDL006W PTC1
ZShift = 2.82
Z lm Score = 14.74
NG = 2
DB = 0
SM = 2
Delta_L
Doxorubicin
YDL006W PTC1
ZShift = −0.14
Z lm Score = −12.97
NG = 2
DB = 0
SM = 2
Delta_K
Doxorubicin
YDL006W PTC1
ZShift = −1.31
Z lm Score = −9.02
NG = 2
DB = 0
SM = 2
Delta_r
Doxorubicin
YDL006W PTC1
ZShift = −2.48
Z lm Score = −5.45
NG = 2
DB = 0
SM = 2
Delta_AUC
Doxorubicin
YHR064C SSZ1
ZShift = 4.78
Z lm Score = 14.7
NG = 2
DB = 0
SM = 2
Delta_L
Doxorubicin
YHR064C SSZ1
ZShift = −1.89
Z lm Score = −12.98
NG = 2
DB = 0
SM = 2
Delta_K
Doxorubicin
YHR064C SSZ1
ZShift = −3.13
Z lm Score = −8.67
NG = 2
DB = 0
SM = 2
Delta_r
Doxorubicin
YHR064C SSZ1
ZShift = −4.83
Z lm Score = −4.28
NG = 2
DB = 0
SM = 2
Delta_AUC
Doxorubicin
YLR235C
ZShift = 4.91
Z lm Score = 14.69
NG = 2
DB = 0
SM = 2
Delta_L
Doxorubicin
YLR235C
ZShift = −1.31
Z lm Score = −12.62
NG = 2
DB = 0
SM = 2
Delta_K
Doxorubicin
YLR235C
ZShift = −1.67
Z lm Score = −8.96
NG = 2
DB = 0
SM = 2
Delta_r
Doxorubicin
YLR235C
ZShift = −4.71
Z lm Score = −4.45
NG = 2
DB = 0
SM = 2
Delta_AUC
Doxorubicin

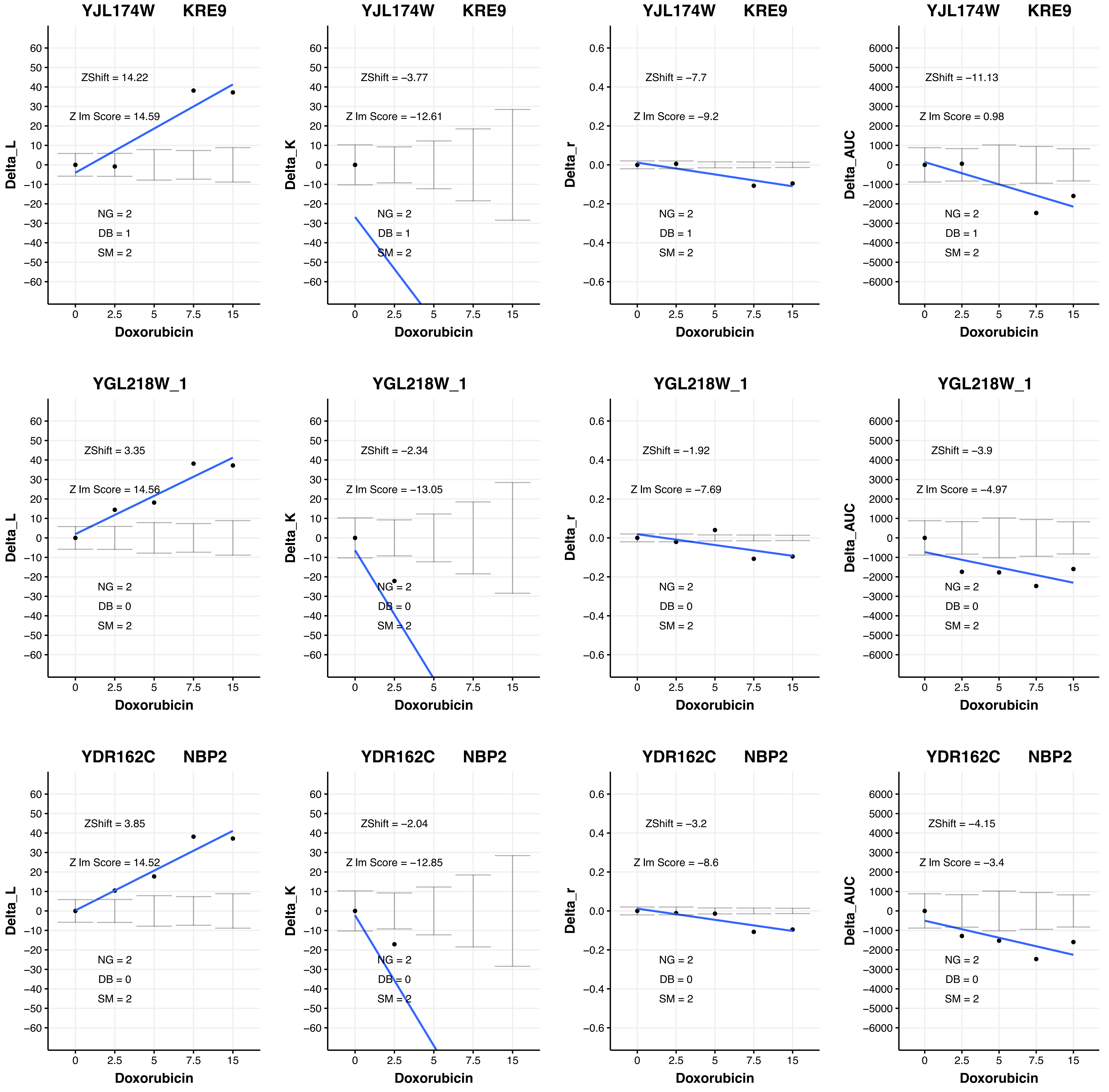

YJL174W KRE9
ZShift = 14.22
Z lm Score = 14.59
NG = 2
DB = 1
SM = 2
Delta_L
Doxorubicin
YJL174W KRE9
ZShift = −3.77
Z lm Score = −12.61
NG = 2
DB = 1
SM = 2
Delta_K
Doxorubicin
YJL174W KRE9
ZShift = −7.7
Z lm Score = −9.2
NG = 2
DB = 1
SM = 2
Delta_r
Doxorubicin
YJL174W KRE9
ZShift = −11.13
Z lm Score = 0.98
NG = 2
DB = 1
SM = 2
Delta_AUC
Doxorubicin
YGL218W_1
ZShift = 3.35
Z lm Score = 14.56
NG = 2
DB = 0
SM = 2
Delta_L
Doxorubicin
YGL218W_1
ZShift = −2.34
Z lm Score = −13.05
NG = 2
DB = 0
SM = 2
Delta_K
Doxorubicin
YGL218W_1
ZShift = −1.92
Z lm Score = −7.69
NG = 2
DB = 0
SM = 2
Delta_r
Doxorubicin
YGL218W_1
ZShift = −3.9
Z lm Score = −4.97
NG = 2
DB = 0
SM = 2
Delta_AUC
Doxorubicin
YDR162C NBP2
ZShift = 3.85
Z lm Score = 14.52
NG = 2
DB = 0
SM = 2
Delta_L
Doxorubicin
YDR162C NBP2
ZShift = −2.04
Z lm Score = −12.85
NG = 2
DB = 0
SM = 2
Delta_K
Doxorubicin
YDR162C NBP2
ZShift = −3.2
Z lm Score = −8.6
NG = 2
DB = 0
SM = 2
Delta_r
Doxorubicin
YDR162C NBP2
ZShift = −4.15
Z lm Score = −3.4
NG = 2
DB = 0
SM = 2
Delta_AUC
Doxorubicin

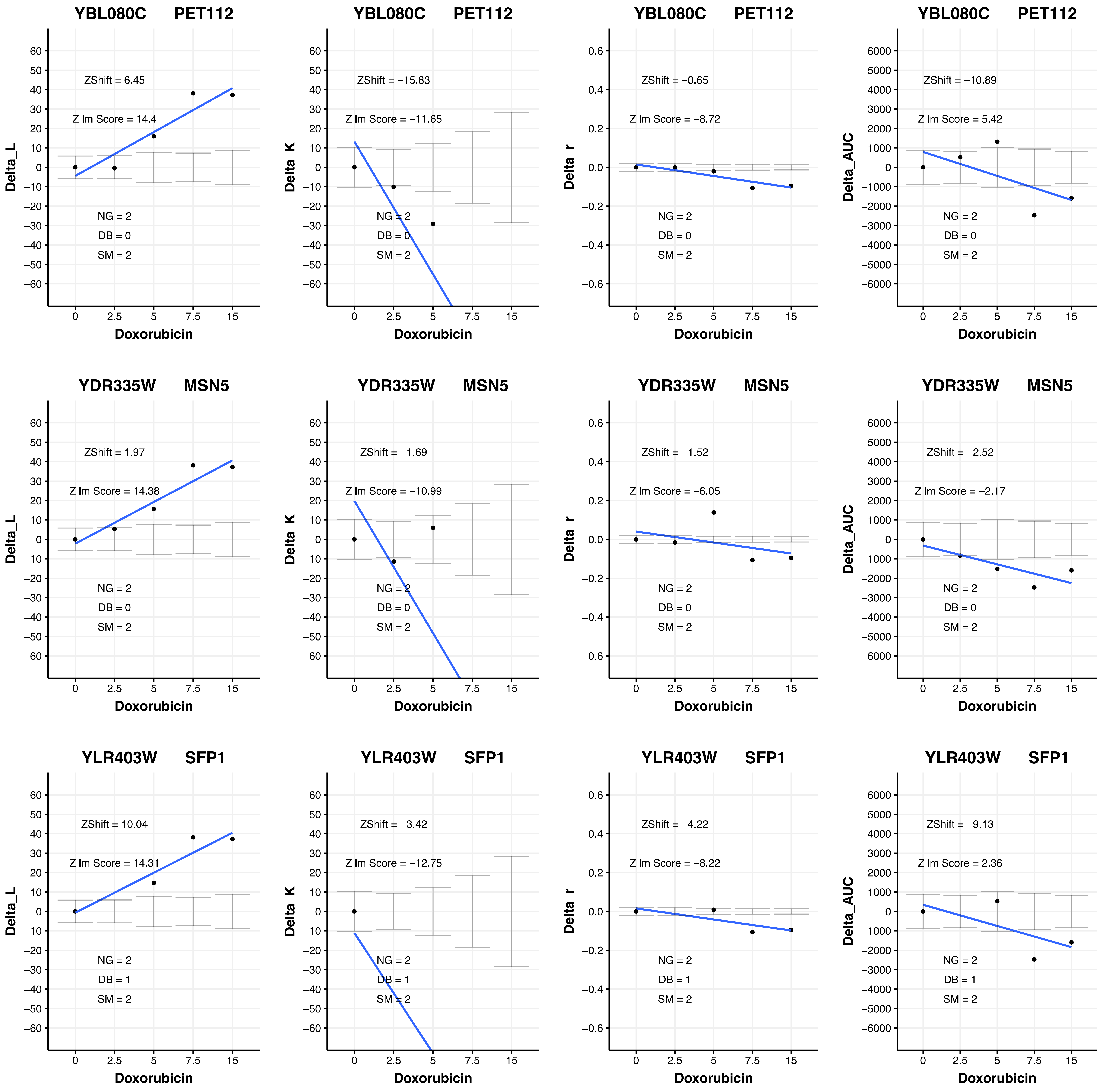
YBL080C PET112
ZShift = 6.45
Z lm Score = 14.4
NG = 2
DB = 0
SM = 2
Delta_L
Doxorubicin
YBL080C PET112
ZShift = −15.83
Z lm Score = −11.65
NG = 2
DB = 0
SM = 2
Delta_K
Doxorubicin
YBL080C PET112
ZShift = −0.65
Z lm Score = −8.72
NG = 2
DB = 0
SM = 2
Delta_r
Doxorubicin
YBL080C PET112
ZShift = −10.89
Z lm Score = 5.42
NG = 2
DB = 0
SM = 2
Delta_AUC
Doxorubicin
YDR335W MSN5
ZShift = 1.97
Z lm Score = 14.38
NG = 2
DB = 0
SM = 2
Delta_L
Doxorubicin
YDR335W MSN5
ZShift = −1.69
Z lm Score = −10.99
NG = 2
DB = 0
SM = 2
Delta_K
Doxorubicin
YDR335W MSN5
ZShift = −1.52
Z lm Score = −6.05
NG = 2
DB = 0
SM = 2
Delta_r
Doxorubicin
YDR335W MSN5
ZShift = −2.52
Z lm Score = −2.17
NG = 2
DB = 0
SM = 2
Delta_AUC
Doxorubicin
YLR403W SFP1
ZShift = 10.04
Z lm Score = 14.31
NG = 2
DB = 1
SM = 2
Delta_L
Doxorubicin
YLR403W SFP1
ZShift = −3.42
Z lm Score = −12.75
NG = 2
DB = 1
SM = 2
Delta_K
Doxorubicin
YLR403W SFP1
ZShift = −4.22
Z lm Score = −8.22
NG = 2
DB = 1
SM = 2
Delta_r
Doxorubicin
YLR403W SFP1
ZShift = −9.13
Z lm Score = 2.36
NG = 2
DB = 1
SM = 2
Delta_AUC
Doxorubicin

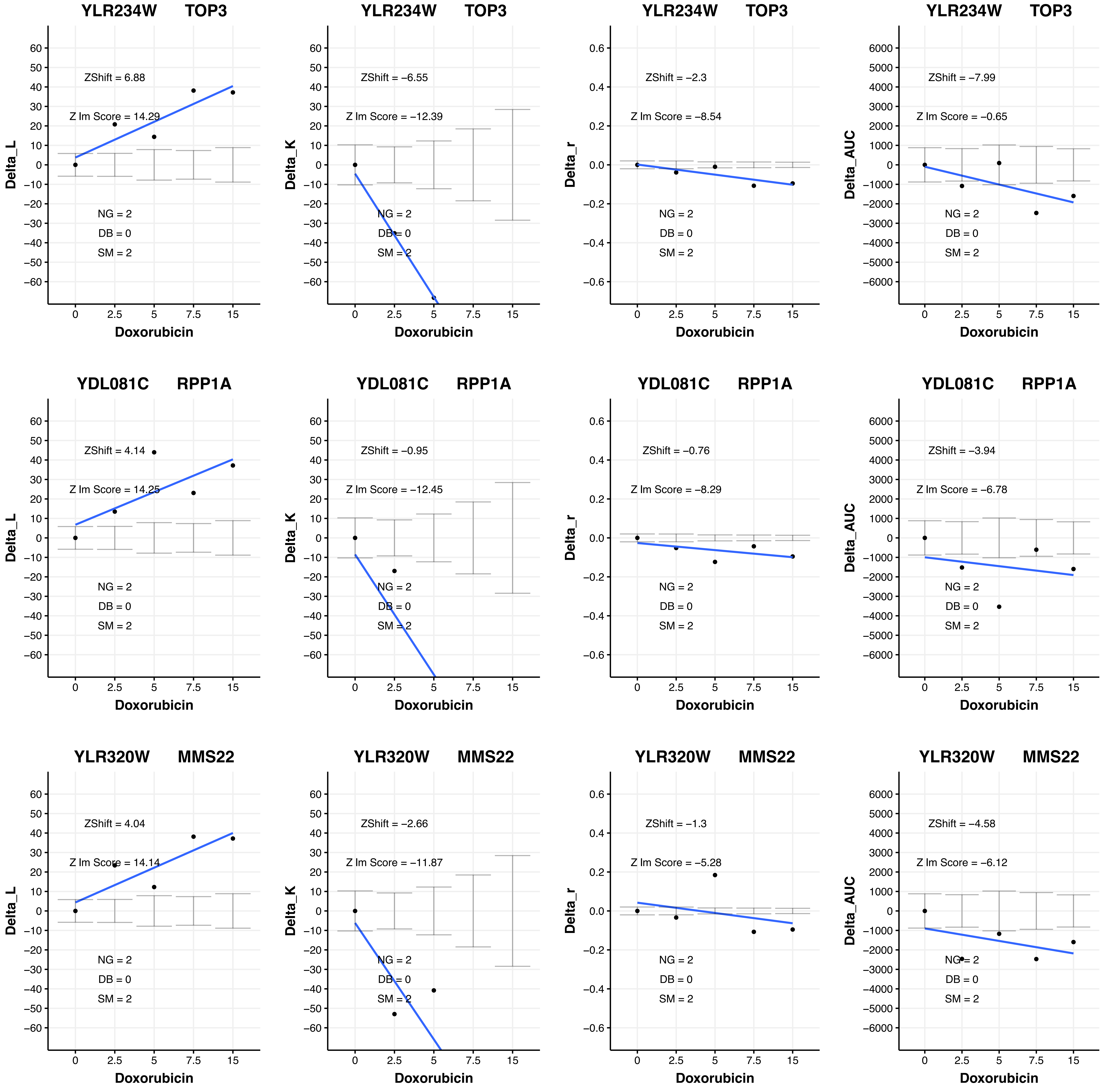
YLR234W TOP3
ZShift = 6.88
Z lm Score = 14.29
NG = 2
DB = 0
SM = 2
Delta_L
Doxorubicin
YLR234W TOP3
ZShift = −6.55
Z lm Score = −12.39
NG = 2
DB = 0
SM = 2
Delta_K
Doxorubicin
YLR234W TOP3
ZShift = −2.3
Z lm Score = −8.54
NG = 2
DB = 0
SM = 2
Delta_r
Doxorubicin
YLR234W TOP3
ZShift = −7.99
Z lm Score = −0.65
NG = 2
DB = 0
SM = 2
Delta_AUC
Doxorubicin
YDL081C RPP1A
ZShift = 4.14
Z lm Score = 14.25
NG = 2
DB = 0
SM = 2
Delta_L
Doxorubicin
YDL081C RPP1A
ZShift = −0.95
Z lm Score = −12.45
NG = 2
DB = 0
SM = 2
Delta_K
Doxorubicin
YDL081C RPP1A
ZShift = −0.76
Z lm Score = −8.29
NG = 2
DB = 0
SM = 2
Delta_r
Doxorubicin
YDL081C RPP1A
ZShift = −3.94
Z lm Score = −6.78
NG = 2
DB = 0
SM = 2
Delta_AUC
Doxorubicin
YLR320W MMS22
ZShift = 4.04
Z lm Score = 14.14
NG = 2
DB = 0
SM = 2
Delta_L
Doxorubicin
YLR320W MMS22
ZShift = −2.66
Z lm Score = −11.87
NG = 2
DB = 0
SM = 2
Delta_K
Doxorubicin
YLR320W MMS22
ZShift = −1.3
Z lm Score = −5.28
NG = 2
DB = 0
SM = 2
Delta_r
Doxorubicin
YLR320W MMS22
ZShift = −4.58
Z lm Score = −6.12
NG = 2
DB = 0
SM = 2
Delta_AUC
Doxorubicin

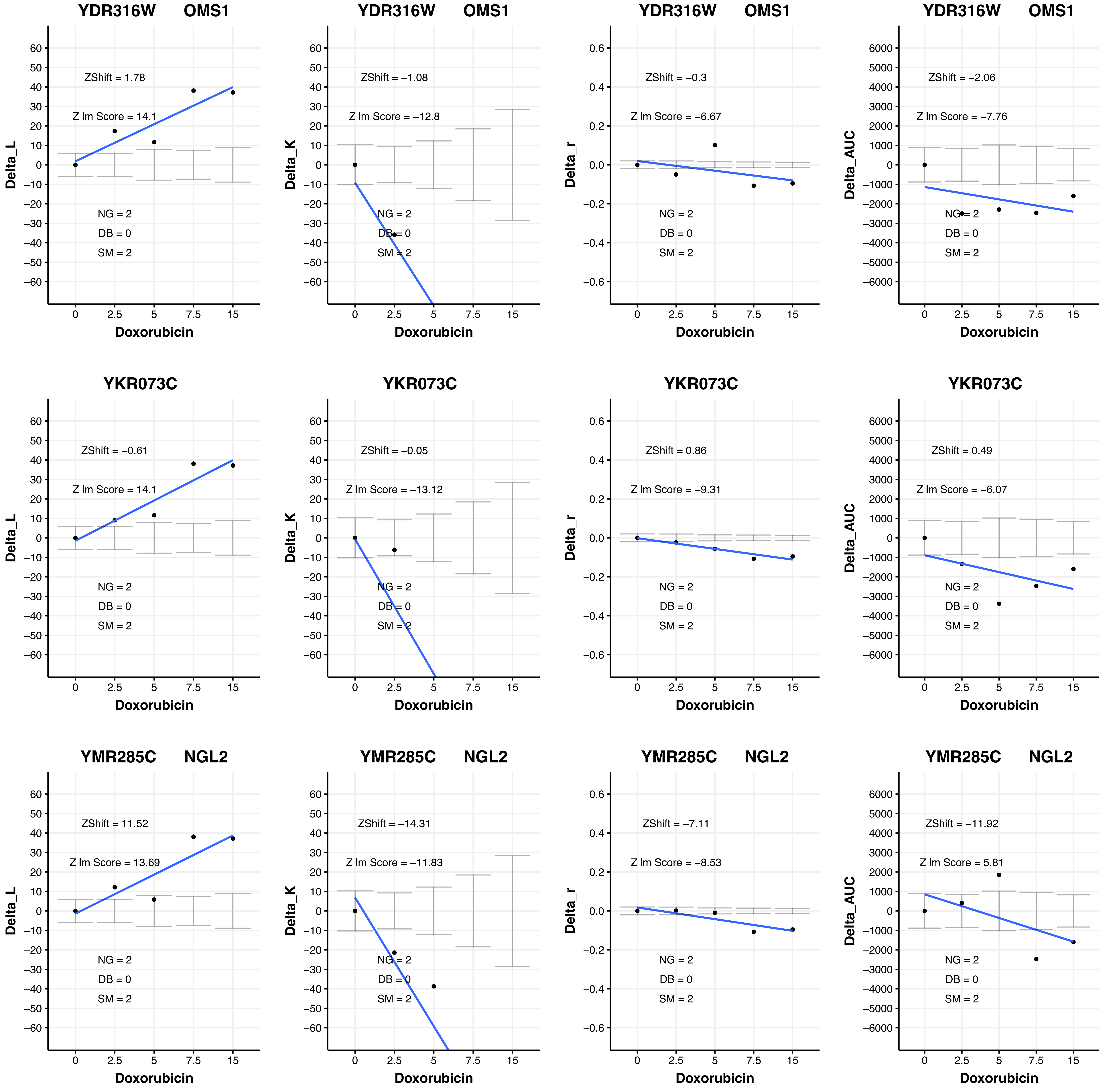
YDR316W OMS1
ZShift = 1.78
Z lm Score = 14.1
NG = 2
DB = 0
SM = 2
Delta_L
Doxorubicin
YDR316W OMS1
ZShift = −1.08
Z lm Score = −12.8
NG = 2
DB = 0
SM = 2
Delta_K
Doxorubicin
YDR316W OMS1
ZShift = −0.3
Z lm Score = −6.67
NG = 2
DB = 0
SM = 2
Delta_r
Doxorubicin
YDR316W OMS1
ZShift = −2.06
Z lm Score = −7.76
NG = 2
DB = 0
SM = 2
Delta_AUC
Doxorubicin
YKR073C
ZShift = −0.61
Z lm Score = 14.1
NG = 2
DB = 0
SM = 2
Delta_L
Doxorubicin
YKR073C
ZShift = −0.05
Z lm Score = −13.12
NG = 2
DB = 0
SM = 2
Delta_K
Doxorubicin
YKR073C
ZShift = 0.86
Z lm Score = −9.31
NG = 2
DB = 0
SM = 2
Delta_r
Doxorubicin
YKR073C
ZShift = 0.49
Z lm Score = −6.07
NG = 2
DB = 0
SM = 2
Delta_AUC
Doxorubicin
YMR285C NGL2
ZShift = 11.52
Z lm Score = 13.69
NG = 2
DB = 0
SM = 2
Delta_L
Doxorubicin
YMR285C NGL2
ZShift = −14.31
Z lm Score = −11.83
NG = 2
DB = 0
SM = 2
Delta_K
Doxorubicin
YMR285C NGL2
ZShift = −7.11
Z lm Score = −8.53
NG = 2
DB = 0
SM = 2
Delta_r
Doxorubicin
YMR285C NGL2
ZShift = −11.92
Z lm Score = 5.81
NG = 2
DB = 0
SM = 2
Delta_AUC
Doxorubicin

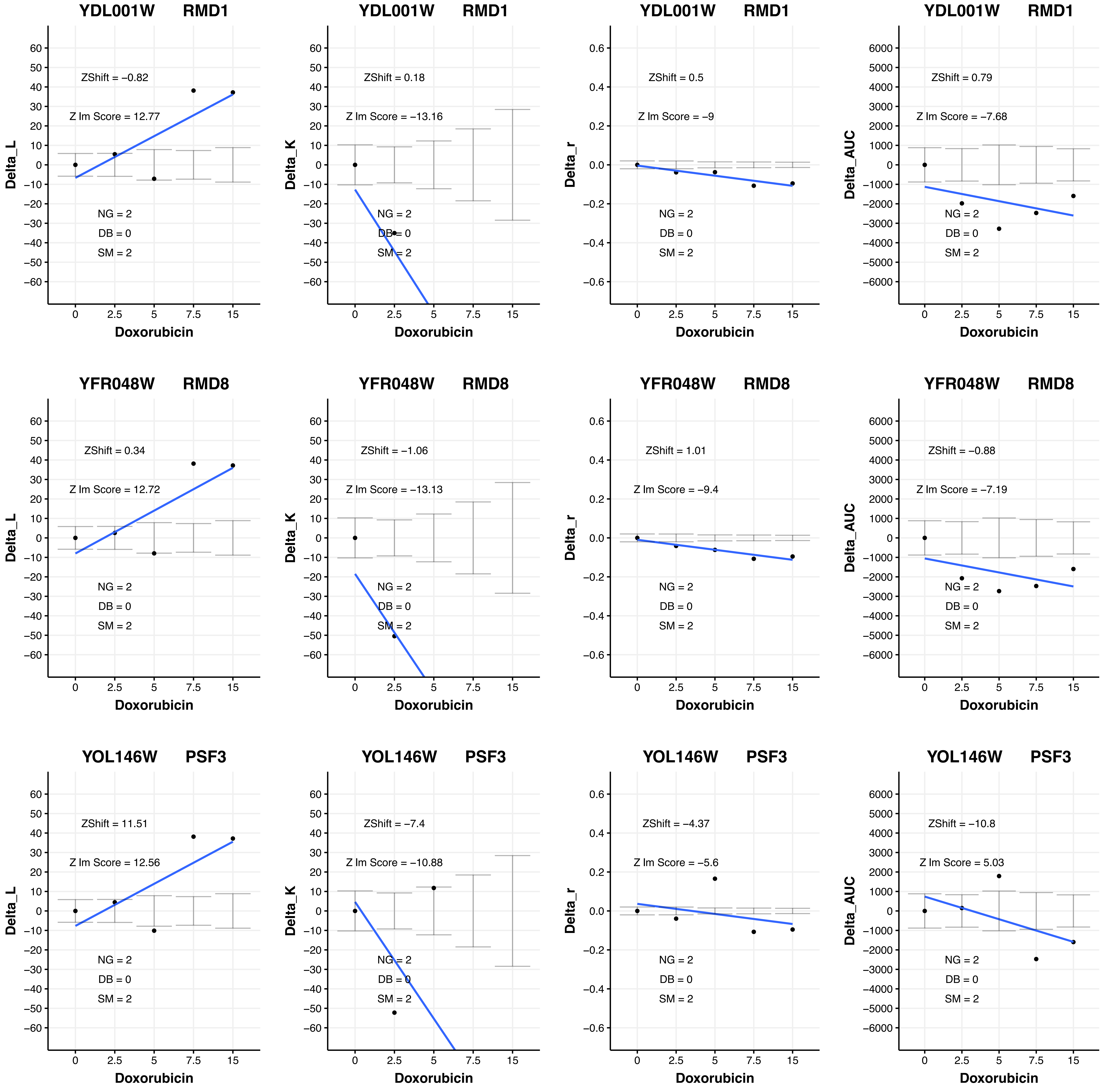
YDL001W RMD1
ZShift = −0.82
Z lm Score = 12.77
NG = 2
DB = 0
SM = 2
Delta_L
Doxorubicin
YDL001W RMD1
ZShift = 0.18
Z lm Score = −13.16
NG = 2
DB = 0
SM = 2
Delta_K
Doxorubicin
YDL001W RMD1
ZShift = 0.5
Z lm Score = −9
NG = 2
DB = 0
SM = 2
Delta_r
Doxorubicin
YDL001W RMD1
ZShift = 0.79
Z lm Score = −7.68
NG = 2
DB = 0
SM = 2
Delta_AUC
Doxorubicin
YFR048W RMD8
ZShift = 0.34
Z lm Score = 12.72
NG = 2
DB = 0
SM = 2
Delta_L
Doxorubicin
YFR048W RMD8
ZShift = −1.06
Z lm Score = −13.13
NG = 2
DB = 0
SM = 2
Delta_K
Doxorubicin
YFR048W RMD8
ZShift = 1.01
Z lm Score = −9.4
NG = 2
DB = 0
SM = 2
Delta_r
Doxorubicin
YFR048W RMD8
ZShift = −0.88
Z lm Score = −7.19
NG = 2
DB = 0
SM = 2
Delta_AUC
Doxorubicin
YOL146W PSF3
ZShift = 11.51
Z lm Score = 12.56
NG = 2
DB = 0
SM = 2
Delta_L
Doxorubicin
YOL146W PSF3
ZShift = −7.4
Z lm Score = −10.88
NG = 2
DB = 0
SM = 2
Delta_K
Doxorubicin
YOL146W PSF3
ZShift = −4.37
Z lm Score = −5.6
NG = 2
DB = 0
SM = 2
Delta_r
Doxorubicin
YOL146W PSF3
ZShift = −10.8
Z lm Score = 5.03
NG = 2
DB = 0
SM = 2
Delta_AUC
Doxorubicin

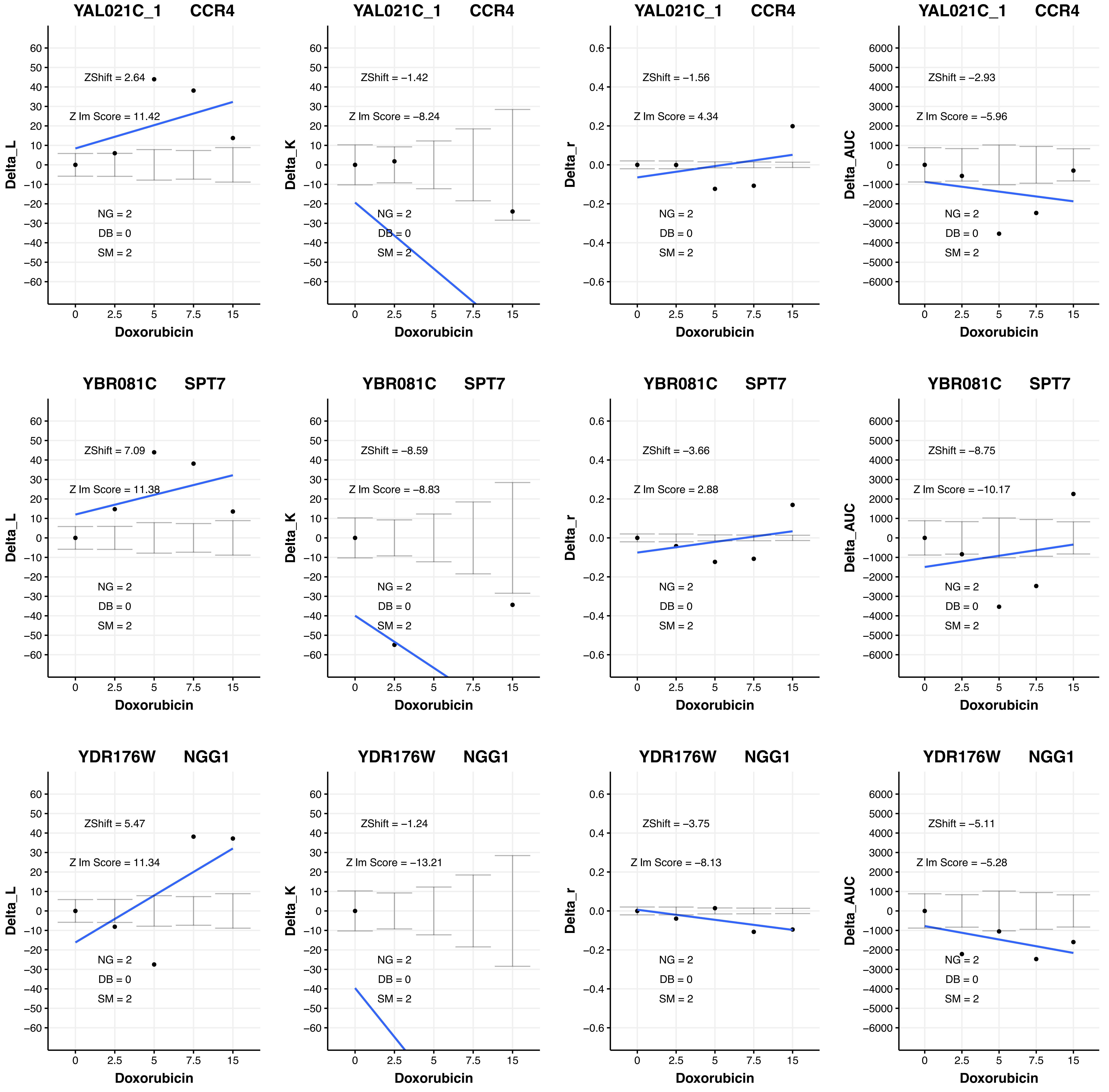

YAL021C_1 CCR4
ZShift = 2.64
Z lm Score = 11.42
NG = 2
DB = 0
SM = 2
Delta_L
Doxorubicin
YAL021C_1 CCR4
ZShift = −1.42
Z lm Score = −8.24
NG = 2
DB = 0
SM = 2
Delta_K
Doxorubicin
YAL021C_1 CCR4
ZShift = −1.56
Z lm Score = 4.34
NG = 2
DB = 0
SM = 2
Delta_r
Doxorubicin
YAL021C_1 CCR4
ZShift = −2.93
Z lm Score = −5.96
NG = 2
DB = 0
SM = 2
Delta_AUC
Doxorubicin
YBR081C SPT7
ZShift = 7.09
Z lm Score = 11.38
NG = 2
DB = 0
SM = 2
Delta_L
Doxorubicin
YBR081C SPT7
ZShift = −8.59
Z lm Score = −8.83
NG = 2
DB = 0
SM = 2
Delta_K
Doxorubicin
YBR081C SPT7
ZShift = −3.66
Z lm Score = 2.88
NG = 2
DB = 0
SM = 2
Delta_r
Doxorubicin
YBR081C SPT7
ZShift = −8.75
Z lm Score = −10.17
NG = 2
DB = 0
SM = 2
Delta_AUC
Doxorubicin
YDR176W NGG1
ZShift = 5.47
Z lm Score = 11.34
NG = 2
DB = 0
SM = 2
Delta_L
Doxorubicin
YDR176W NGG1
ZShift = −1.24
Z lm Score = −13.21
NG = 2
DB = 0
SM = 2
Delta_K
Doxorubicin
YDR176W NGG1
ZShift = −3.75
Z lm Score = −8.13
NG = 2
DB = 0
SM = 2
Delta_r
Doxorubicin
YDR176W NGG1
ZShift = −5.11
Z lm Score = −5.28
NG = 2
DB = 0
SM = 2
Delta_AUC
Doxorubicin

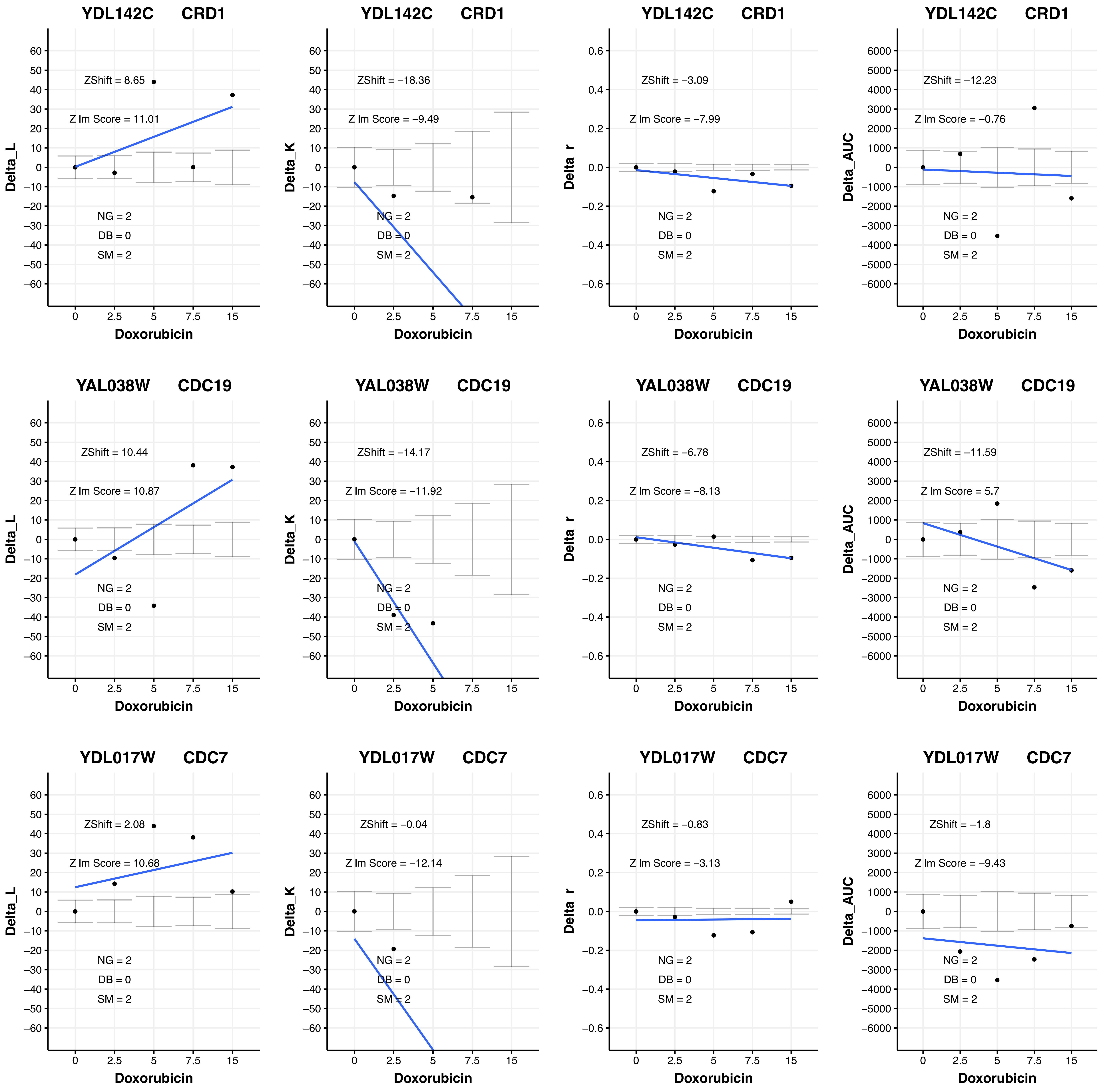

YDL142C CRD1
ZShift = 8.65
Z lm Score = 11.01
NG = 2
DB = 0
SM = 2
Delta_L
Doxorubicin
YDL142C CRD1
ZShift = −18.36
Z lm Score = −9.49
NG = 2
DB = 0
SM = 2
Delta_K
Doxorubicin
YDL142C CRD1
ZShift = −3.09
Z lm Score = −7.99
NG = 2
DB = 0
SM = 2
Delta_r
Doxorubicin
YDL142C CRD1
ZShift = −12.23
Z lm Score = −0.76
NG = 2
DB = 0
SM = 2
Delta_AUC
Doxorubicin
YAL038W CDC19
ZShift = 10.44
Z lm Score = 10.87
NG = 2
DB = 0
SM = 2
Delta_L
Doxorubicin
YAL038W CDC19
ZShift = −14.17
Z lm Score = −11.92
NG = 2
DB = 0
SM = 2
Delta_K
Doxorubicin
YAL038W CDC19
ZShift = −6.78
Z lm Score = −8.13
NG = 2
DB = 0
SM = 2
Delta_r
Doxorubicin
YAL038W CDC19
ZShift = −11.59
Z lm Score = 5.7
NG = 2
DB = 0
SM = 2
Delta_AUC
Doxorubicin
YDL017W CDC7
ZShift = 2.08
Z lm Score = 10.68
NG = 2
DB = 0
SM = 2
Delta_L
Doxorubicin
YDL017W CDC7
ZShift = −0.04
Z lm Score = −12.14
NG = 2
DB = 0
SM = 2
Delta_K
Doxorubicin
YDL017W CDC7
ZShift = −0.83
Z lm Score = −3.13
NG = 2
DB = 0
SM = 2
Delta_r
Doxorubicin
YDL017W CDC7
ZShift = −1.8
Z lm Score = −9.43
NG = 2
DB = 0
SM = 2
Delta_AUC
Doxorubicin

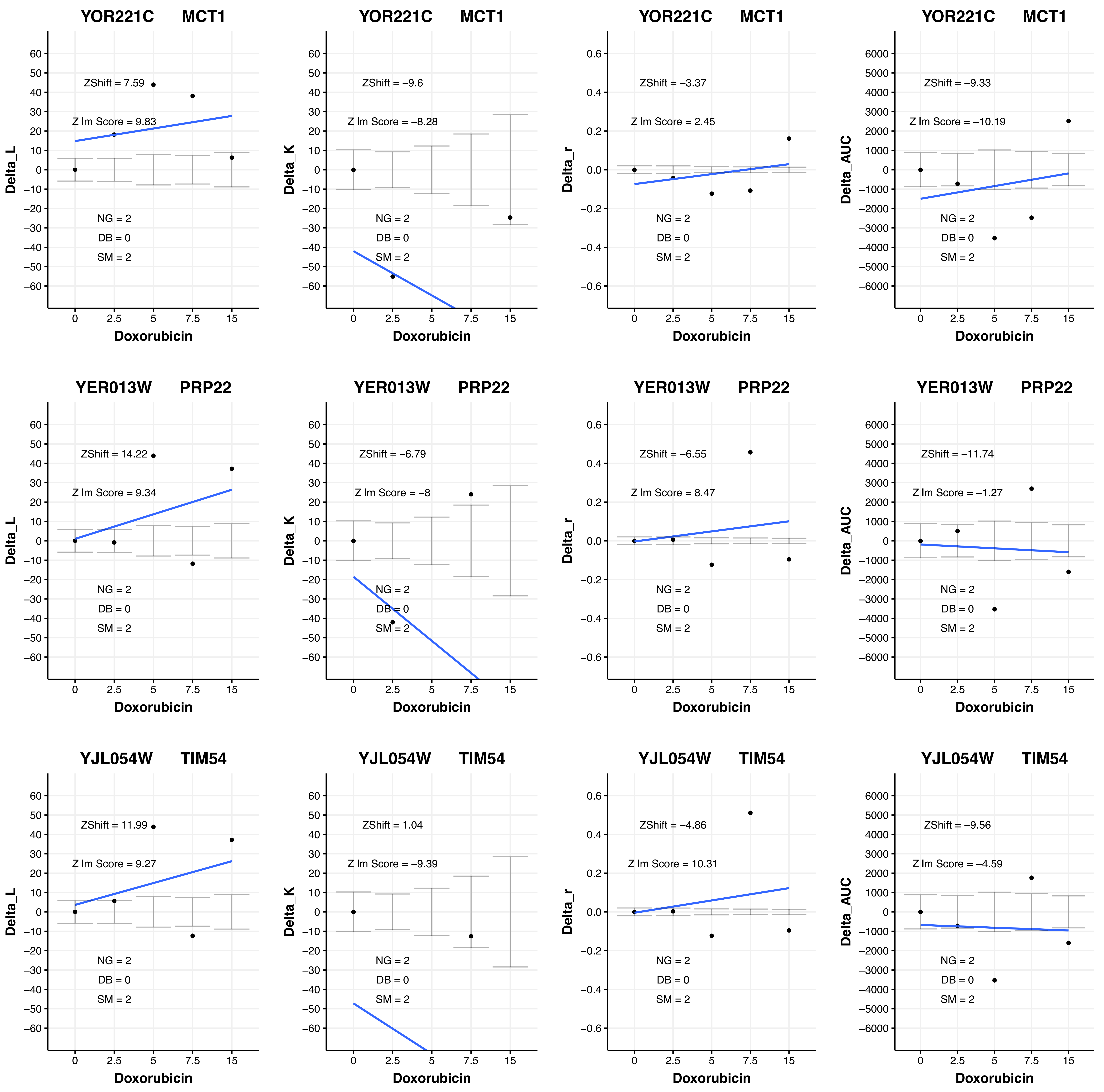

YOR221C MCT1
ZShift = 7.59
Z lm Score = 9.83
NG = 2
DB = 0
SM = 2
Delta_L
Doxorubicin
YOR221C MCT1
ZShift = −9.6
Z lm Score = −8.28
NG = 2
DB = 0
SM = 2
Delta_K
Doxorubicin
YOR221C MCT1
ZShift = −3.37
Z lm Score = 2.45
NG = 2
DB = 0
SM = 2
Delta_r
Doxorubicin
YOR221C MCT1
ZShift = −9.33
Z lm Score = −10.19
NG = 2
DB = 0
SM = 2
Delta_AUC
Doxorubicin
YER013W PRP22
ZShift = 14.22
Z lm Score = 9.34
NG = 2
DB = 0
SM = 2
Delta_L
Doxorubicin
YER013W PRP22
ZShift = −6.79
Z lm Score = −8
NG = 2
DB = 0
SM = 2
Delta_K
Doxorubicin
YER013W PRP22
ZShift = −6.55
Z lm Score = 8.47
NG = 2
DB = 0
SM = 2
Delta_r
Doxorubicin
YER013W PRP22
ZShift = −11.74
Z lm Score = −1.27
NG = 2
DB = 0
SM = 2
Delta_AUC
Doxorubicin
YJL054W TIM54
ZShift = 11.99
Z lm Score = 9.27
NG = 2
DB = 0
SM = 2
Delta_L
Doxorubicin
YJL054W TIM54
ZShift = 1.04
Z lm Score = −9.39
NG = 2
DB = 0
SM = 2
Delta_K
Doxorubicin
YJL054W TIM54
ZShift = −4.86
Z lm Score = 10.31
NG = 2
DB = 0
SM = 2
Delta_r
Doxorubicin
YJL054W TIM54
ZShift = −9.56
Z lm Score = −4.59
NG = 2
DB = 0
SM = 2
Delta_AUC
Doxorubicin

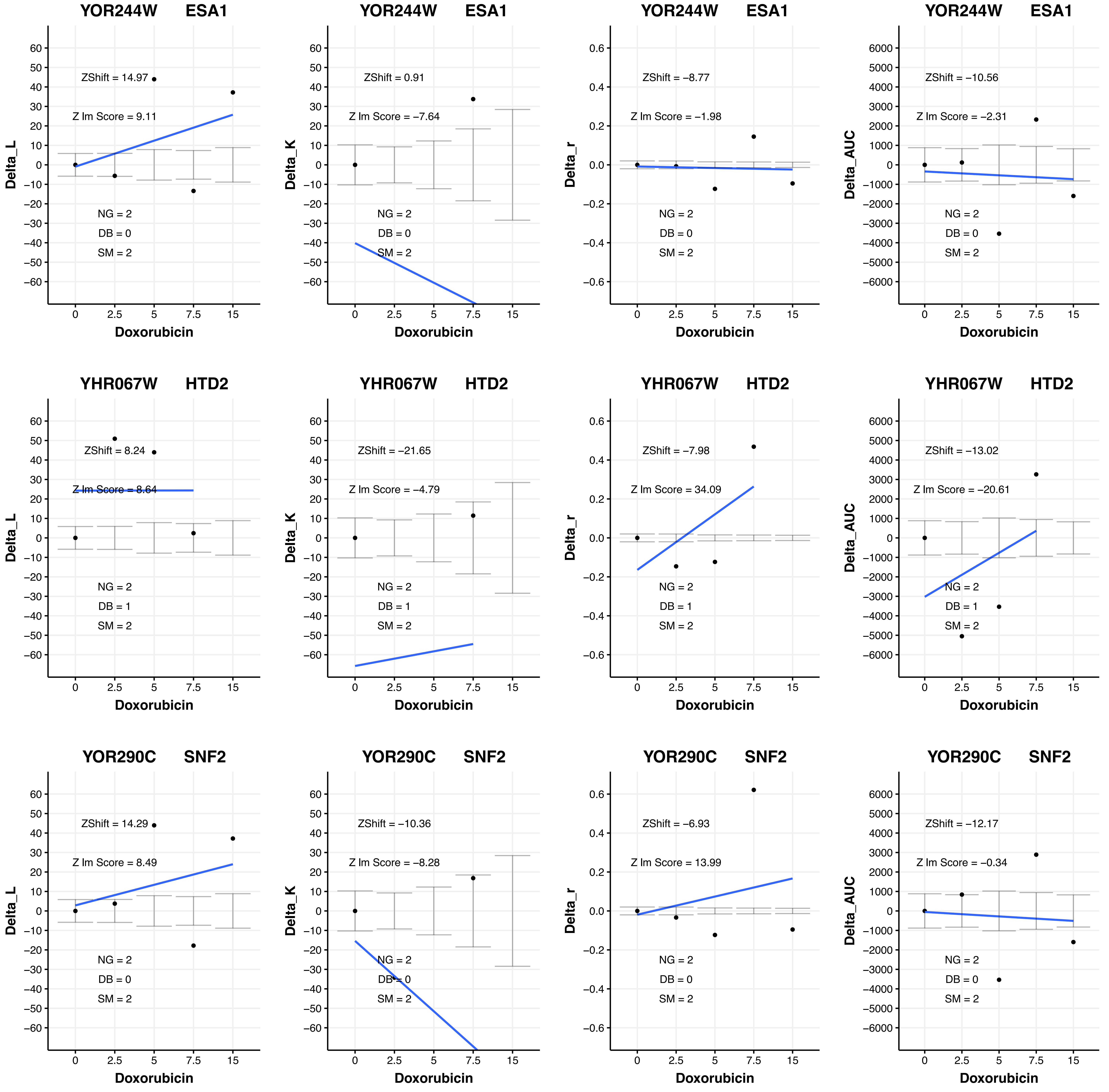
YOR244W ESA1
ZShift = 14.97
Z lm Score = 9.11
NG = 2
DB = 0
SM = 2
Delta_L
Doxorubicin
YOR244W ESA1
ZShift = 0.91
Z lm Score = −7.64
NG = 2
DB = 0
SM = 2
Delta_K
Doxorubicin
YOR244W ESA1
ZShift = −8.77
Z lm Score = −1.98
NG = 2
DB = 0
SM = 2
Delta_r
Doxorubicin
YOR244W ESA1
ZShift = −10.56
Z lm Score = −2.31
NG = 2
DB = 0
SM = 2
Delta_AUC
Doxorubicin
YHR067W HTD2
ZShift = 8.24
Z lm Score = 8.64
NG = 2
DB = 1
SM = 2
Delta_L
Doxorubicin
YHR067W HTD2
ZShift = −21.65
Z lm Score = −4.79
NG = 2
DB = 1
SM = 2
Delta_K
Doxorubicin
YHR067W HTD2
ZShift = −7.98
Z lm Score = 34.09
NG = 2
DB = 1
SM = 2
Delta_r
Doxorubicin
YHR067W HTD2
ZShift = −13.02
Z lm Score = −20.61
NG = 2
DB = 1
SM = 2
Delta_AUC
Doxorubicin
YOR290C SNF2
ZShift = 14.29
Z lm Score = 8.49
NG = 2
DB = 0
SM = 2
Delta_L
Doxorubicin
YOR290C SNF2
ZShift = −10.36
Z lm Score = −8.28
NG = 2
DB = 0
SM = 2
Delta_K
Doxorubicin
YOR290C SNF2
ZShift = −6.93
Z lm Score = 13.99
NG = 2
DB = 0
SM = 2
Delta_r
Doxorubicin
YOR290C SNF2
ZShift = −12.17
Z lm Score = −0.34
NG = 2
DB = 0
SM = 2
Delta_AUC
Doxorubicin

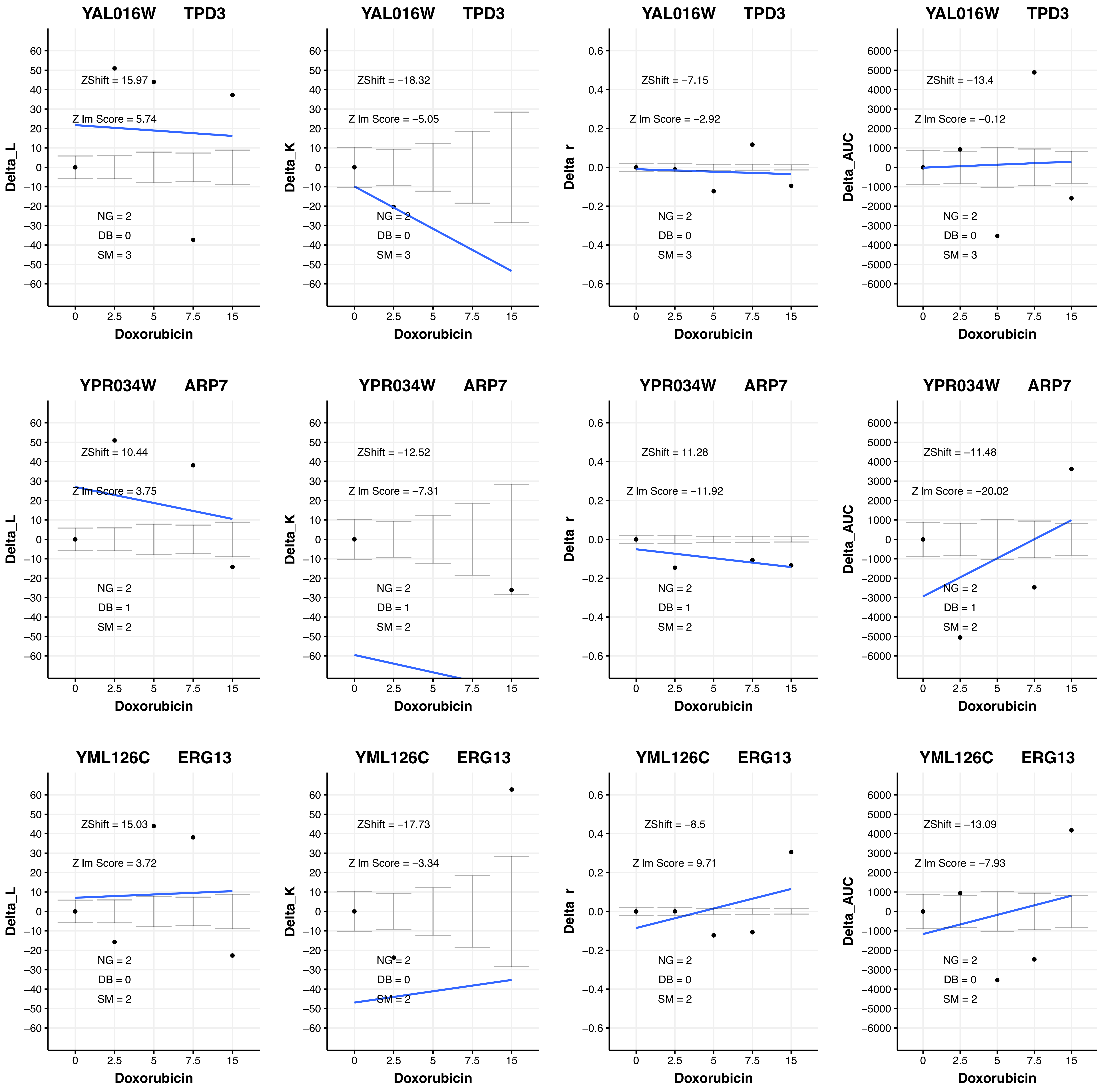
YAL016W TPD3
ZShift = 15.97
Z lm Score = 5.74
NG = 2
DB = 0
SM = 3
Delta_L
Doxorubicin
YAL016W TPD3
ZShift = −18.32
Z lm Score = −5.05
NG = 2
DB = 0
SM = 3
Delta_K
Doxorubicin
YAL016W TPD3
ZShift = −7.15
Z lm Score = −2.92
NG = 2
DB = 0
SM = 3
Delta_r
Doxorubicin
YAL016W TPD3
ZShift = −13.4
Z lm Score = −0.12
NG = 2
DB = 0
SM = 3
Delta_AUC
Doxorubicin
YPR034W ARP7
ZShift = 10.44
Z lm Score = 3.75
NG = 2
DB = 1
SM = 2
Delta_L
Doxorubicin
YPR034W ARP7
ZShift = −12.52
Z lm Score = −7.31
NG = 2
DB = 1
SM = 2
Delta_K
Doxorubicin
YPR034W ARP7
ZShift = 11.28
Z lm Score = −11.92
NG = 2
DB = 1
SM = 2
Delta_r
Doxorubicin
YPR034W ARP7
ZShift = −11.48
Z lm Score = −20.02
NG = 2
DB = 1
SM = 2
Delta_AUC
Doxorubicin
YML126C ERG13
ZShift = 15.03
Z lm Score = 3.72
NG = 2
DB = 0
SM = 2
Delta_L
Doxorubicin
YML126C ERG13
ZShift = −17.73
Z lm Score = −3.34
NG = 2
DB = 0
SM = 2
Delta_K
Doxorubicin
YML126C ERG13
ZShift = −8.5
Z lm Score = 9.71
NG = 2
DB = 0
SM = 2
Delta_r
Doxorubicin
YML126C ERG13
ZShift = −13.09
Z lm Score = −7.93
NG = 2
DB = 0
SM = 2
Delta_AUC
Doxorubicin

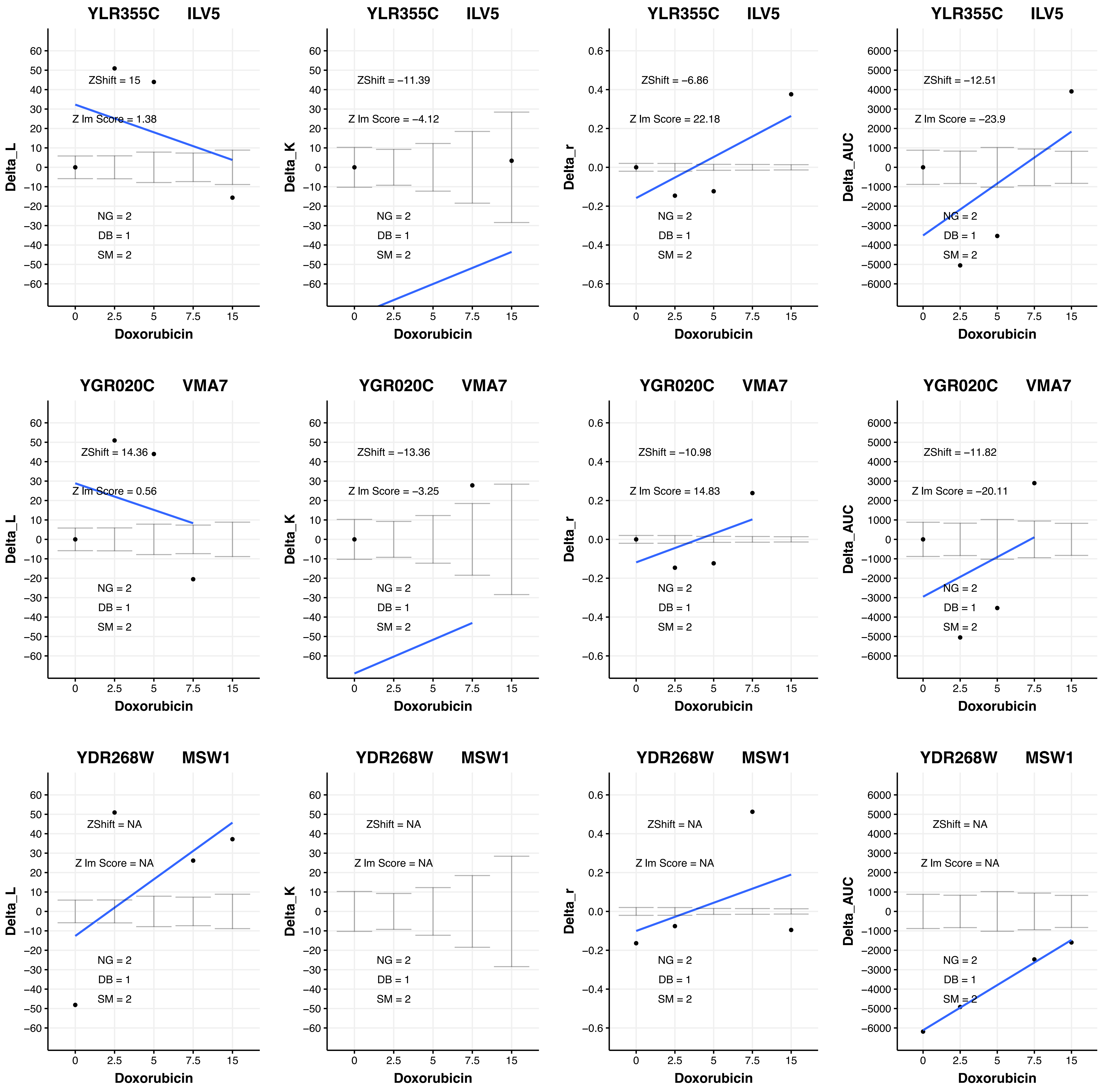
YLR355C ILV5
ZShift = 15
Z lm Score = 1.38
NG = 2
DB = 1
SM = 2
Delta_L
Doxorubicin
YLR355C ILV5
ZShift = −11.39
Z lm Score = −4.12
NG = 2
DB = 1
SM = 2
Delta_K
Doxorubicin
YLR355C ILV5
ZShift = −6.86
Z lm Score = 22.18
NG = 2
DB = 1
SM = 2
Delta_r
Doxorubicin
YLR355C ILV5
ZShift = −12.51
Z lm Score = −23.9
NG = 2
DB = 1
SM = 2
Delta_AUC
Doxorubicin
YGR020C VMA7
ZShift = 14.36
Z lm Score = 0.56
NG = 2
DB = 1
SM = 2
Delta_L
Doxorubicin
YGR020C VMA7
ZShift = −13.36
Z lm Score = −3.25
NG = 2
DB = 1
SM = 2
Delta_K
Doxorubicin
YGR020C VMA7
ZShift = −10.98
Z lm Score = 14.83
NG = 2
DB = 1
SM = 2
Delta_r
Doxorubicin
YGR020C VMA7
ZShift = −11.82
Z lm Score = −20.11
NG = 2
DB = 1
SM = 2
Delta_AUC
Doxorubicin
YDR268W MSW1
ZShift = NA
Z lm Score = NA
NG = 2
DB = 1
SM = 2
Delta_L
Doxorubicin
YDR268W MSW1
ZShift = NA
Z lm Score = NA
NG = 2
DB = 1
SM = 2
Delta_K
Doxorubicin
YDR268W MSW1
ZShift = NA
Z lm Score = NA
NG = 2
DB = 1
SM = 2
Delta_r
Doxorubicin
YDR268W MSW1
ZShift = NA
Z lm Score = NA
NG = 2
DB = 1
SM = 2
Delta_AUC
Doxorubicin

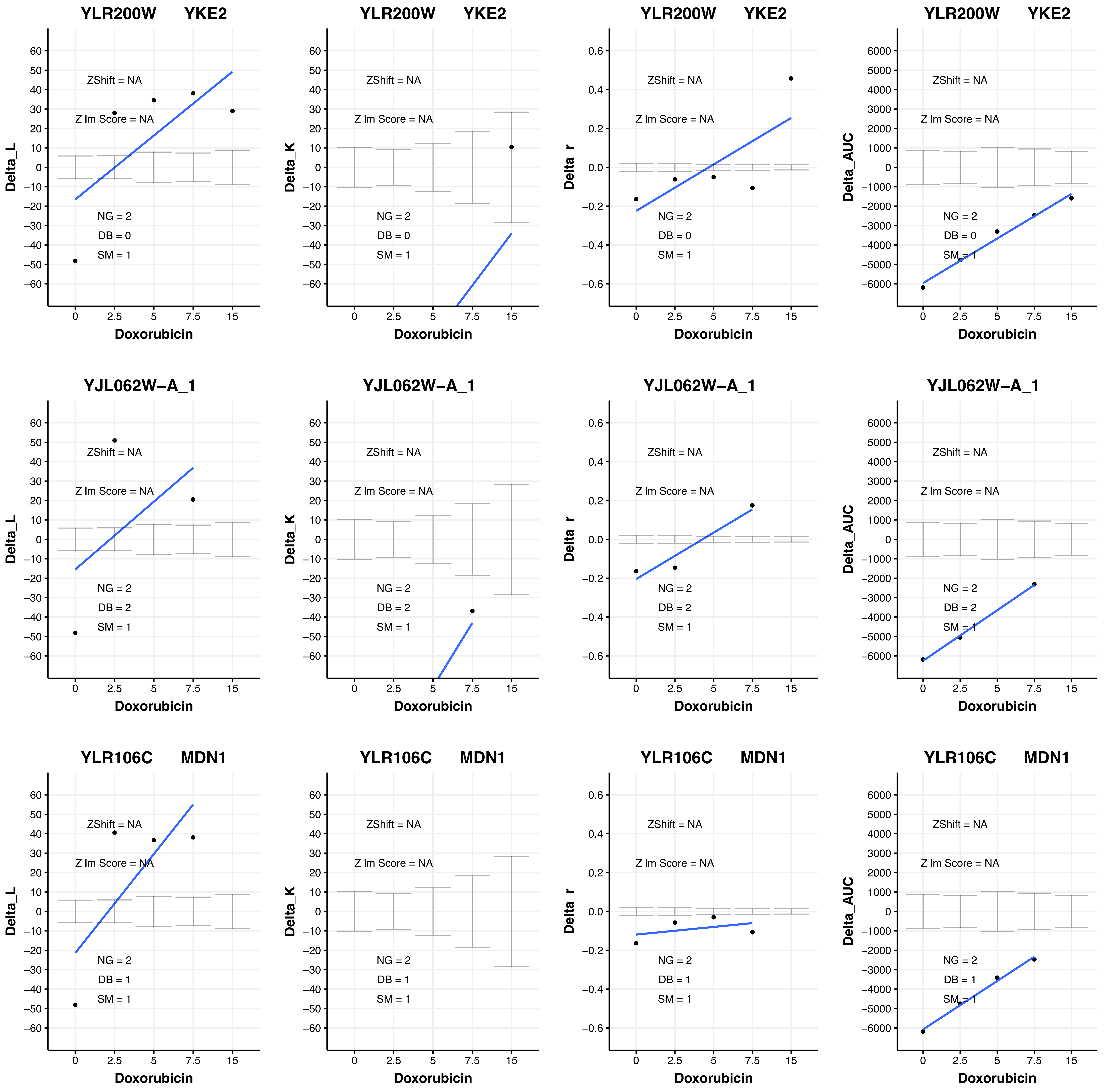
YLR200W YKE2
Delta_L
ZShift = NA
Z lm Score = NA
NG = 2
DB = 0
SM = 1
Doxorubicin
YLR200W YKE2
Delta_K
ZShift = NA
Z lm Score = NA
NG = 2
DB = 0
SM = 1
Doxorubicin
YLR200W YKE2
Delta_r
ZShift = NA
Z lm Score = NA
NG = 2
DB = 0
SM = 1
Doxorubicin
YLR200W YKE2
Delta_AUC
ZShift = NA
Z lm Score = NA
NG = 2
DB = 0
SM = 1
Doxorubicin
YJL062W−A_1
Delta_L
ZShift = NA
Z lm Score = NA
NG = 2
DB = 2
SM = 1
Doxorubicin
YJL062W−A_1
Delta_K
ZShift = NA
Z lm Score = NA
NG = 2
DB = 2
SM = 1
Doxorubicin
YJL062W−A_1
Delta_r
ZShift = NA
Z lm Score = NA
NG = 2
DB = 2
SM = 1
Doxorubicin
YJL062W−A_1
Delta_AUC
ZShift = NA
Z lm Score = NA
NG = 2
DB = 2
SM = 1
Doxorubicin
YLR106C MDN1
Delta_L
ZShift = NA
Z lm Score = NA
NG = 2
DB = 1
SM = 1
Doxorubicin
YLR106C MDN1
Delta_K
ZShift = NA
Z lm Score = NA
NG = 2
DB = 1
SM = 1
Doxorubicin
YLR106C MDN1
Delta_r
ZShift = NA
Z lm Score = NA
NG = 2
DB = 1
SM = 1
Doxorubicin
YLR106C MDN1
Delta_AUC
ZShift = NA
Z lm Score = NA
NG = 2
DB = 1
SM = 1
Doxorubicin

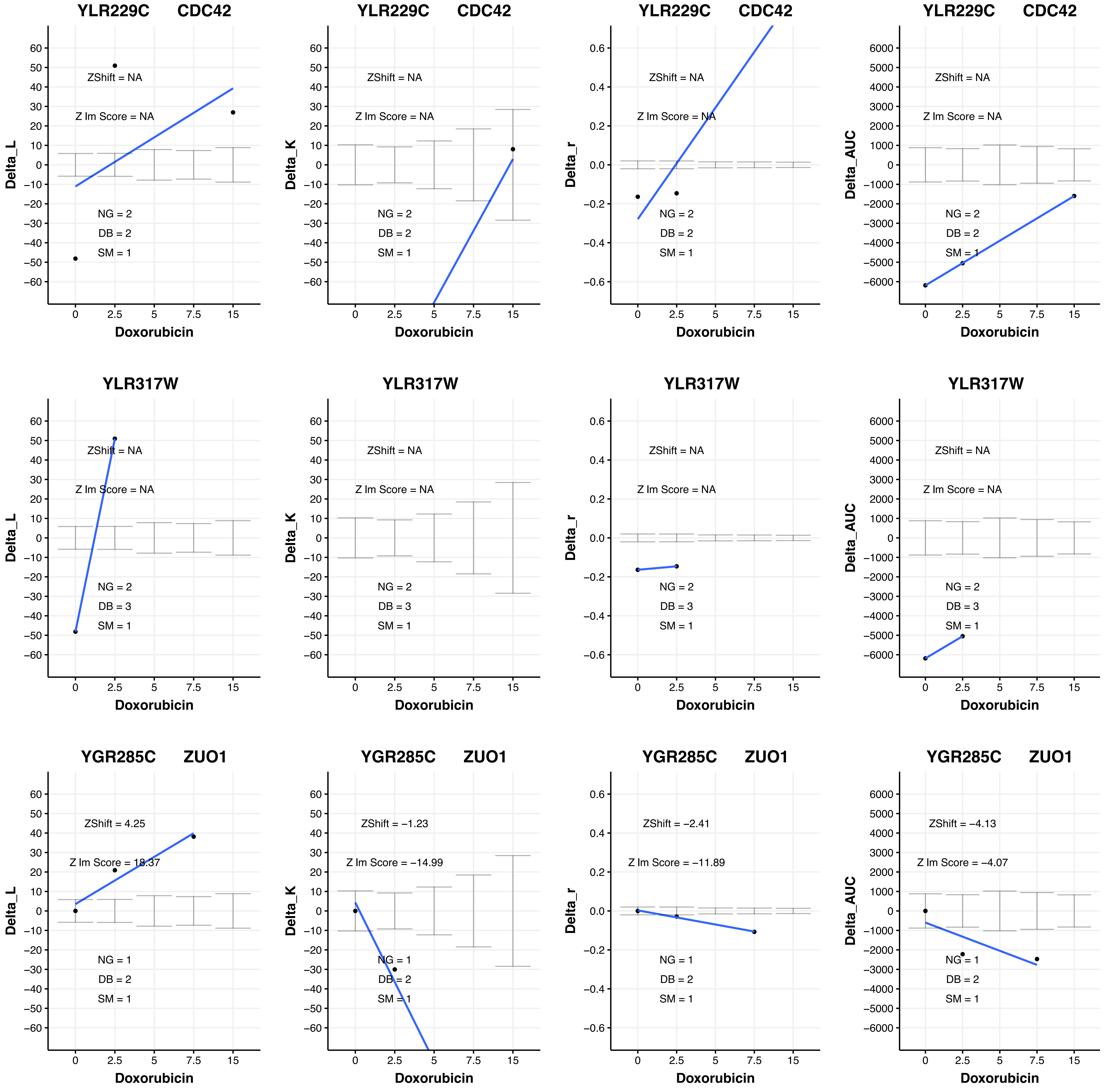
YLR229C CDC42
ZShift = NA
Z lm Score = NA
NG = 2
DB = 2
SM = 1
Delta_L
Doxorubicin
YLR229C CDC42
ZShift = NA
Z lm Score = NA
NG = 2
DB = 2
SM = 1
Delta_K
Doxorubicin
YLR229C CDC42
ZShift = NA
Z lm Score = NA
NG = 2
DB = 2
SM = 1
Delta_r
Doxorubicin
YLR229C CDC42
ZShift = NA
Z lm Score = NA
NG = 2
DB = 2
SM = 1
Delta_AUC
Doxorubicin
YLR317W
ZShift = NA
Z lm Score = NA
NG = 2
DB = 3
SM = 1
Delta_L
Doxorubicin
YLR317W
ZShift = NA
Z lm Score = NA
NG = 2
DB = 3
SM = 1
Delta_K
Doxorubicin
YLR317W
ZShift = NA
Z lm Score = NA
NG = 2
DB = 3
SM = 1
Delta_r
Doxorubicin
YLR317W
ZShift = NA
Z lm Score = NA
NG = 2
DB = 3
SM = 1
Delta_AUC
Doxorubicin
YGR285C ZUO1
ZShift = 4.25
Z lm Score = 18.37
NG = 1
DB = 2
SM = 1
Delta_L
Doxorubicin
YGR285C ZUO1
ZShift = −1.23
Z lm Score = −14.99
NG = 1
DB = 2
SM = 1
Delta_K
Doxorubicin
YGR285C ZUO1
ZShift = −2.41
Z lm Score = −11.89
NG = 1
DB = 2
SM = 1
Delta_r
Doxorubicin
YGR285C ZUO1
ZShift = −4.13
Z lm Score = −4.07
NG = 1
DB = 2
SM = 1
Delta_AUC
Doxorubicin

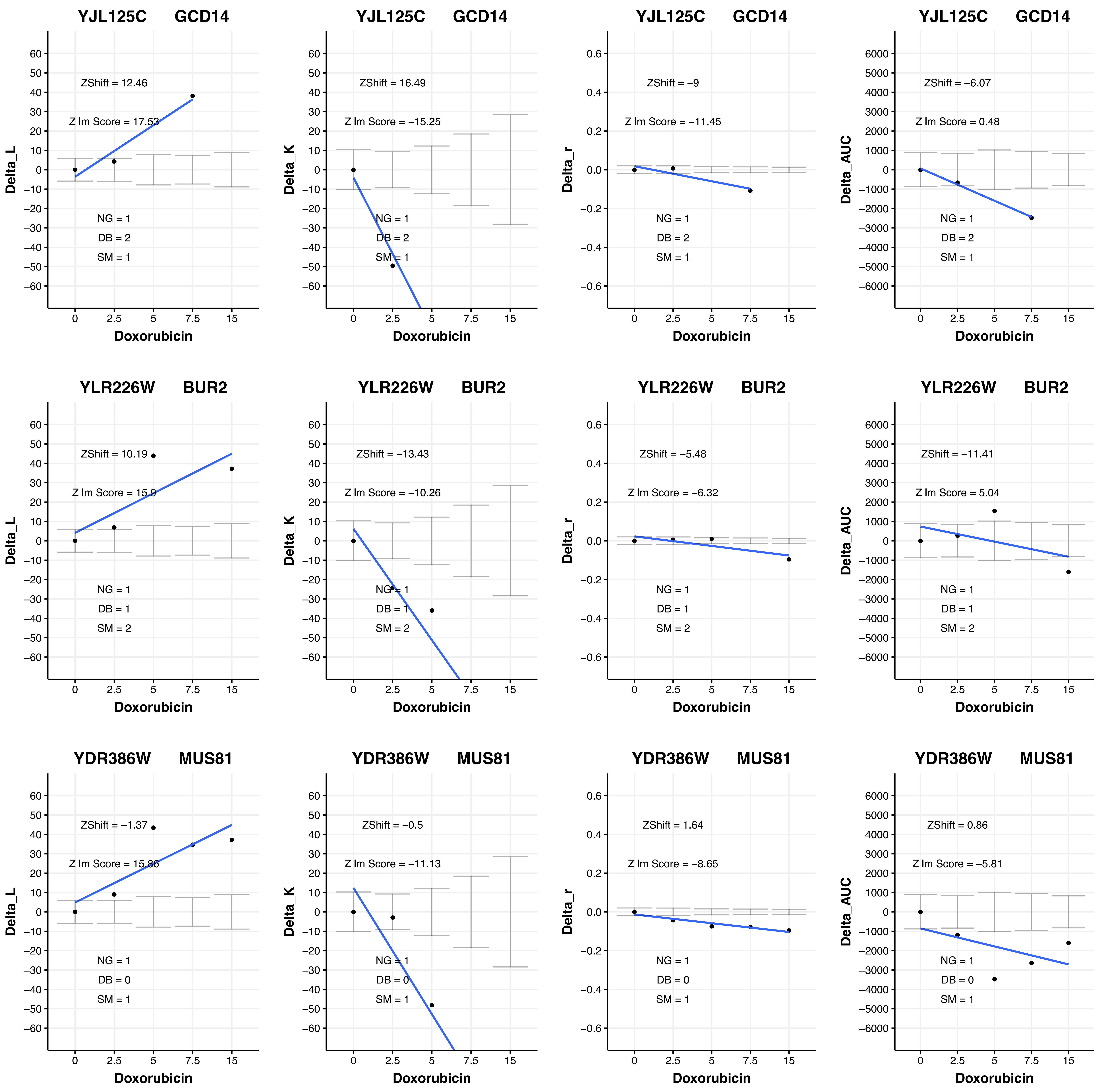
YJL125C GCD14
ZShift = 12.46
Z lm Score = 17.53
NG = 1
DB = 2
SM = 1
Delta_L
Doxorubicin
YJL125C GCD14
ZShift = 16.49
Z lm Score = −15.25
NG = 1
DB = 2
SM = 1
Delta_K
Doxorubicin
YJL125C GCD14
ZShift = −9
Z lm Score = −11.45
NG = 1
DB = 2
SM = 1
Delta_r
Doxorubicin
YJL125C GCD14
ZShift = −6.07
Z lm Score = 0.48
NG = 1
DB = 2
SM = 1
Delta_AUC
Doxorubicin
YLR226W BUR2
ZShift = 10.19
Z lm Score = 15.9
NG = 1
DB = 1
SM = 2
Delta_L
Doxorubicin
YLR226W BUR2
ZShift = −13.43
Z lm Score = −10.26
NG = 1
DB = 1
SM = 2
Delta_K
Doxorubicin
YLR226W BUR2
ZShift = −5.48
Z lm Score = −6.32
NG = 1
DB = 1
SM = 2
Delta_r
Doxorubicin
YLR226W BUR2
ZShift = −11.41
Z lm Score = 5.04
NG = 1
DB = 1
SM = 2
Delta_AUC
Doxorubicin
YDR386W MUS81
ZShift = −1.37
Z lm Score = 15.86
NG = 1
DB = 0
SM = 1
Delta_L
Doxorubicin
YDR386W MUS81
ZShift = −0.5
Z lm Score = −11.13
NG = 1
DB = 0
SM = 1
Delta_K
Doxorubicin
YDR386W MUS81
ZShift = 1.64
Z lm Score = −8.65
NG = 1
DB = 0
SM = 1
Delta_r
Doxorubicin
YDR386W MUS81
ZShift = 0.86
Z lm Score = −5.81
NG = 1
DB = 0
SM = 1
Delta_AUC
Doxorubicin

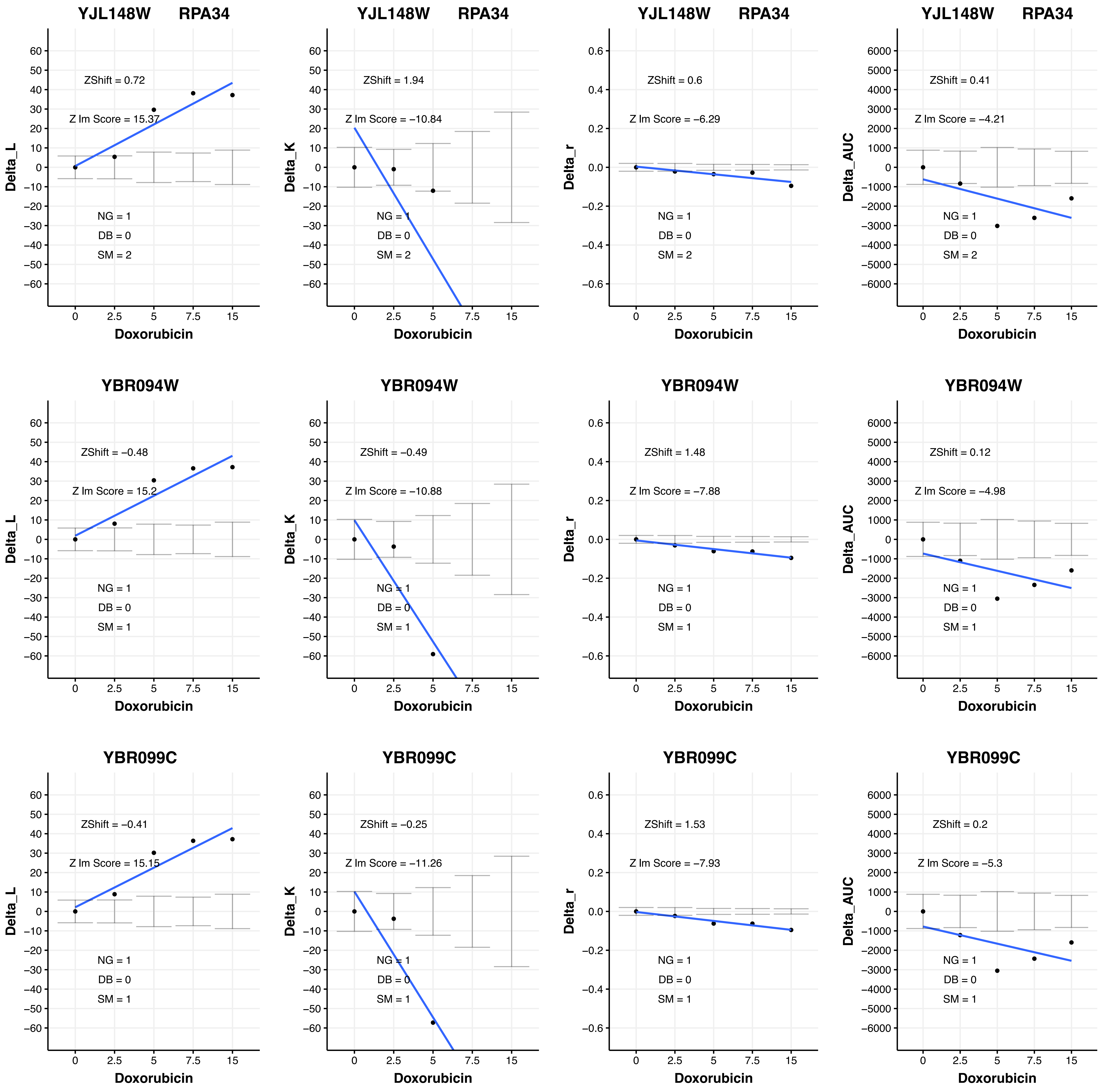
YJL148W RPA34
ZShift = 0.72
Z lm Score = 15.37
NG = 1
DB = 0
SM = 2
Delta_L
Doxorubicin
YJL148W RPA34
ZShift = 1.94
Z lm Score = −10.84
NG = 1
DB = 0
SM = 2
Delta_K
Doxorubicin
YJL148W RPA34
ZShift = 0.6
Z lm Score = −6.29
NG = 1
DB = 0
SM = 2
Delta_r
Doxorubicin
YJL148W RPA34
ZShift = 0.41
Z lm Score = −4.21
NG = 1
DB = 0
SM = 2
Delta_AUC
Doxorubicin
YBR094W
ZShift = −0.48
Z lm Score = 15.2
NG = 1
DB = 0
SM = 1
Delta_L
Doxorubicin
YBR094W
ZShift = −0.49
Z lm Score = −10.88
NG = 1
DB = 0
SM = 1
Delta_K
Doxorubicin
YBR094W
ZShift = 1.48
Z lm Score = −7.88
NG = 1
DB = 0
SM = 1
Delta_r
Doxorubicin
YBR094W
ZShift = 0.12
Z lm Score = −4.98
NG = 1
DB = 0
SM = 1
Delta_AUC
Doxorubicin
YBR099C
ZShift = −0.41
Z lm Score = 15.15
NG = 1
DB = 0
SM = 1
Delta_L
Doxorubicin
YBR099C
ZShift = −0.25
Z lm Score = −11.26
NG = 1
DB = 0
SM = 1
Delta_K
Doxorubicin
YBR099C
ZShift = 1.53
Z lm Score = −7.93
NG = 1
DB = 0
SM = 1
Delta_r
Doxorubicin
YBR099C
ZShift = 0.2
Z lm Score = −5.3
NG = 1
DB = 0
SM = 1
Delta_AUC
Doxorubicin

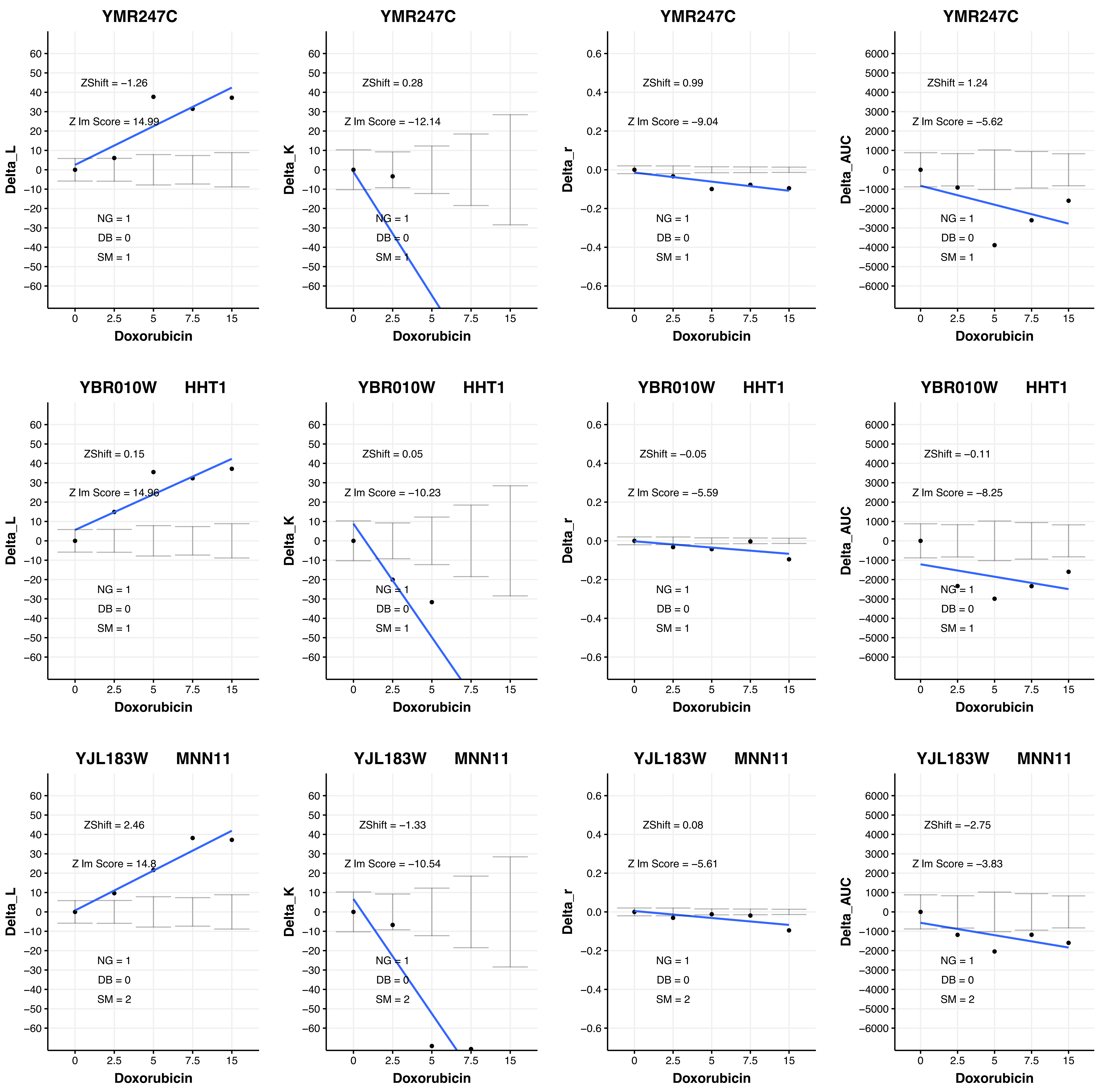

YMR247C
ZShift = −1.26
Z lm Score = 14.99
NG = 1
DB = 0
SM = 1
Delta_L
Doxorubicin
YMR247C
ZShift = 0.28
Z lm Score = −12.14
NG = 1
DB = 0
SM = 1
Delta_K
Doxorubicin
YMR247C
ZShift = 0.99
Z lm Score = −9.04
NG = 1
DB = 0
SM = 1
Delta_r
Doxorubicin
YMR247C
ZShift = 1.24
Z lm Score = −5.62
NG = 1
DB = 0
SM = 1
Delta_AUC
Doxorubicin
YBR010W HHT1
ZShift = 0.15
Z lm Score = 14.96
NG = 1
DB = 0
SM = 1
Delta_L
Doxorubicin
YBR010W HHT1
ZShift = 0.05
Z lm Score = −10.23
NG = 1
DB = 0
SM = 1
Delta_K
Doxorubicin
YBR010W HHT1
ZShift = −0.05
Z lm Score = −5.59
NG = 1
DB = 0
SM = 1
Delta_r
Doxorubicin
YBR010W HHT1
ZShift = −0.11
Z lm Score = −8.25
NG = 1
DB = 0
SM = 1
Delta_AUC
Doxorubicin
YJL183W MNN11
ZShift = 2.46
Z lm Score = 14.8
NG = 1
DB = 0
SM = 2
Delta_L
Doxorubicin
YJL183W MNN11
ZShift = −1.33
Z lm Score = −10.54
NG = 1
DB = 0
SM = 2
Delta_K
Doxorubicin
YJL183W MNN11
ZShift = 0.08
Z lm Score = −5.61
NG = 1
DB = 0
SM = 2
Delta_r
Doxorubicin
YJL183W MNN11
ZShift = −2.75
Z lm Score = −3.83
NG = 1
DB = 0
SM = 2
Delta_AUC
Doxorubicin

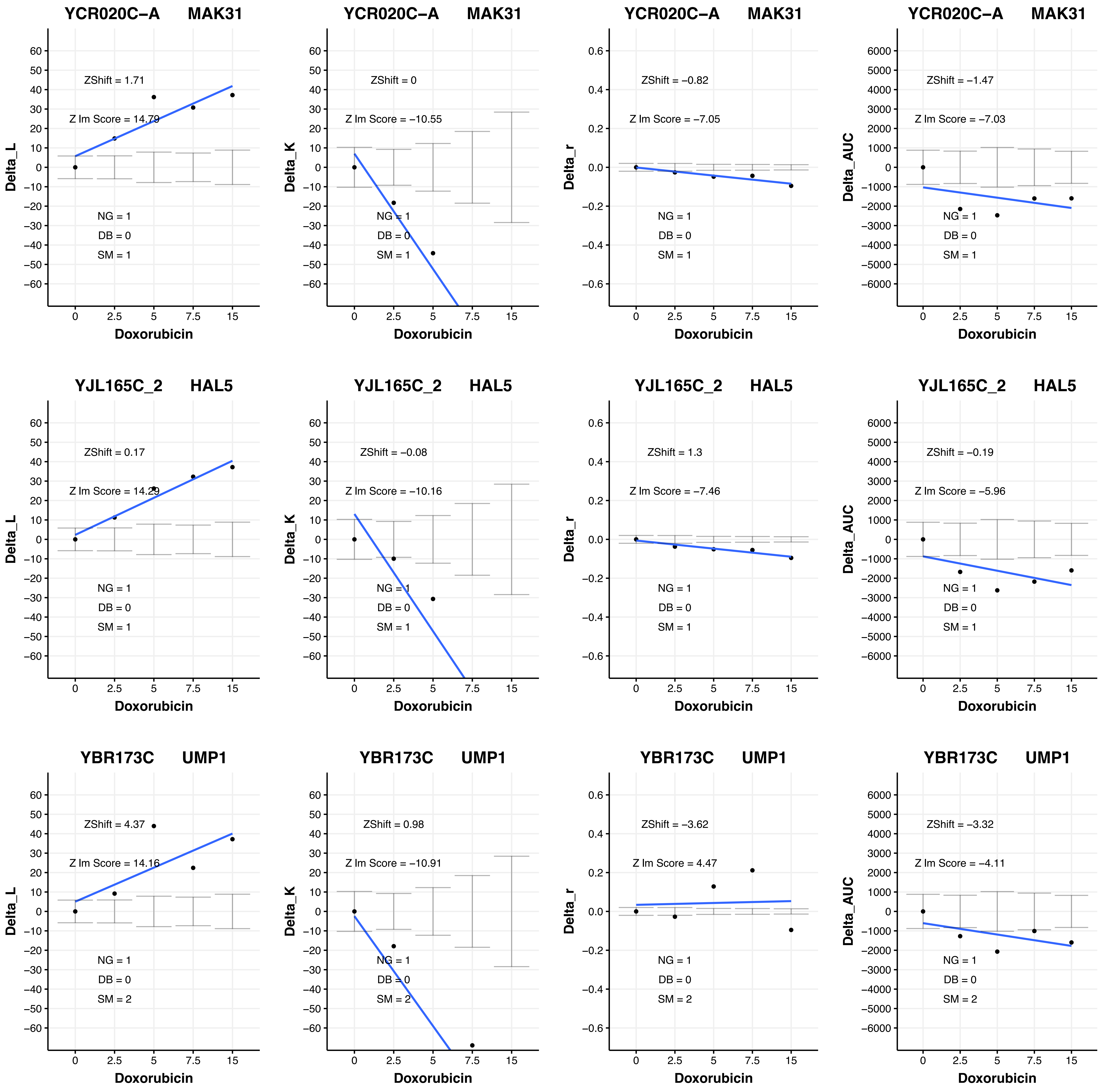

YCR020C-A MAK31
ZShift = 1.71
Z lm Score = 14.79
NG = 1
DB = 0
SM = 1
Delta_L
Doxorubicin
YCR020C-A MAK31
ZShift = 0
Z lm Score = −10.55
NG = 1
DB = 0
SM = 1
Delta_K
Doxorubicin
YCR020C-A MAK31
ZShift = −0.82
Z lm Score = −7.05
NG = 1
DB = 0
SM = 1
Delta_r
Doxorubicin
YCR020C-A MAK31
ZShift = −1.47
Z lm Score = −7.03
NG = 1
DB = 0
SM = 1
Delta_AUC
Doxorubicin
YJL165C_2 HAL5
ZShift = 0.17
Z lm Score = 14.29
NG = 1
DB = 0
SM = 1
Delta_L
Doxorubicin
YJL165C_2 HAL5
ZShift = −0.08
Z lm Score = −10.16
NG = 1
DB = 0
SM = 1
Delta_K
Doxorubicin
YJL165C_2 HAL5
ZShift = 1.3
Z lm Score = −7.46
NG = 1
DB = 0
SM = 1
Delta_r
Doxorubicin
YJL165C_2 HAL5
ZShift = −0.19
Z lm Score = −5.96
NG = 1
DB = 0
SM = 1
Delta_AUC
Doxorubicin
YBR173C UMP1
ZShift = 4.37
Z lm Score = 14.16
NG = 1
DB = 0
SM = 2
Delta_L
Doxorubicin
YBR173C UMP1
ZShift = 0.98
Z lm Score = −10.91
NG = 1
DB = 0
SM = 2
Delta_K
Doxorubicin
YBR173C UMP1
ZShift = −3.62
Z lm Score = 4.47
NG = 1
DB = 0
SM = 2
Delta_r
Doxorubicin
YBR173C UMP1
ZShift = −3.32
Z lm Score = −4.11
NG = 1
DB = 0
SM = 2
Delta_AUC
Doxorubicin

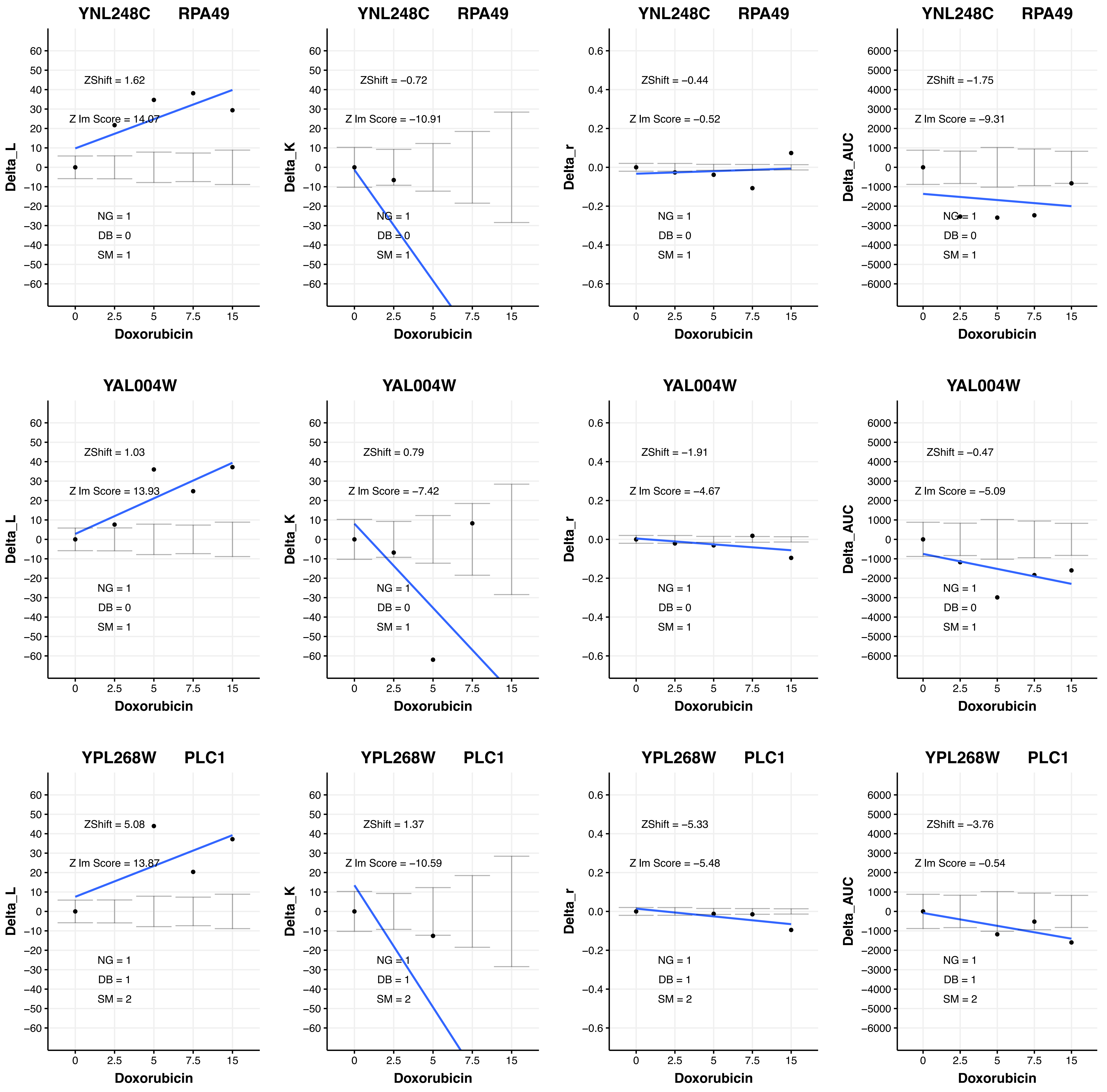

YNL248C RPA49
ZShift = 1.62
Z lm Score = 14.07
NG = 1
DB = 0
SM = 1
Delta_L
Doxorubicin
YNL248C RPA49
ZShift = −0.72
Z lm Score = −10.91
NG = 1
DB = 0
SM = 1
Delta_K
Doxorubicin
YNL248C RPA49
ZShift = −0.44
Z lm Score = −0.52
NG = 1
DB = 0
SM = 1
Delta_r
Doxorubicin
YNL248C RPA49
ZShift = −1.75
Z lm Score = −9.31
NG = 1
DB = 0
SM = 1
Delta_AUC
Doxorubicin
YAL004W
ZShift = 1.03
Z lm Score = 13.93
NG = 1
DB = 0
SM = 1
Delta_L
Doxorubicin
YAL004W
ZShift = 0.79
Z lm Score = −7.42
NG = 1
DB = 0
SM = 1
Delta_K
Doxorubicin
YAL004W
ZShift = −1.91
Z lm Score = −4.67
NG = 1
DB = 0
SM = 1
Delta_r
Doxorubicin
YAL004W
ZShift = −0.47
Z lm Score = −5.09
NG = 1
DB = 0
SM = 1
Delta_AUC
Doxorubicin
YPL268W PLC1
ZShift = 5.08
Z lm Score = 13.87
NG = 1
DB = 1
SM = 2
Delta_L
Doxorubicin
YPL268W PLC1
ZShift = 1.37
Z lm Score = −10.59
NG = 1
DB = 1
SM = 2
Delta_K
Doxorubicin
YPL268W PLC1
ZShift = −5.33
Z lm Score = −5.48
NG = 1
DB = 1
SM = 2
Delta_r
Doxorubicin
YPL268W PLC1
ZShift = −3.76
Z lm Score = −0.54
NG = 1
DB = 1
SM = 2
Delta_AUC
Doxorubicin

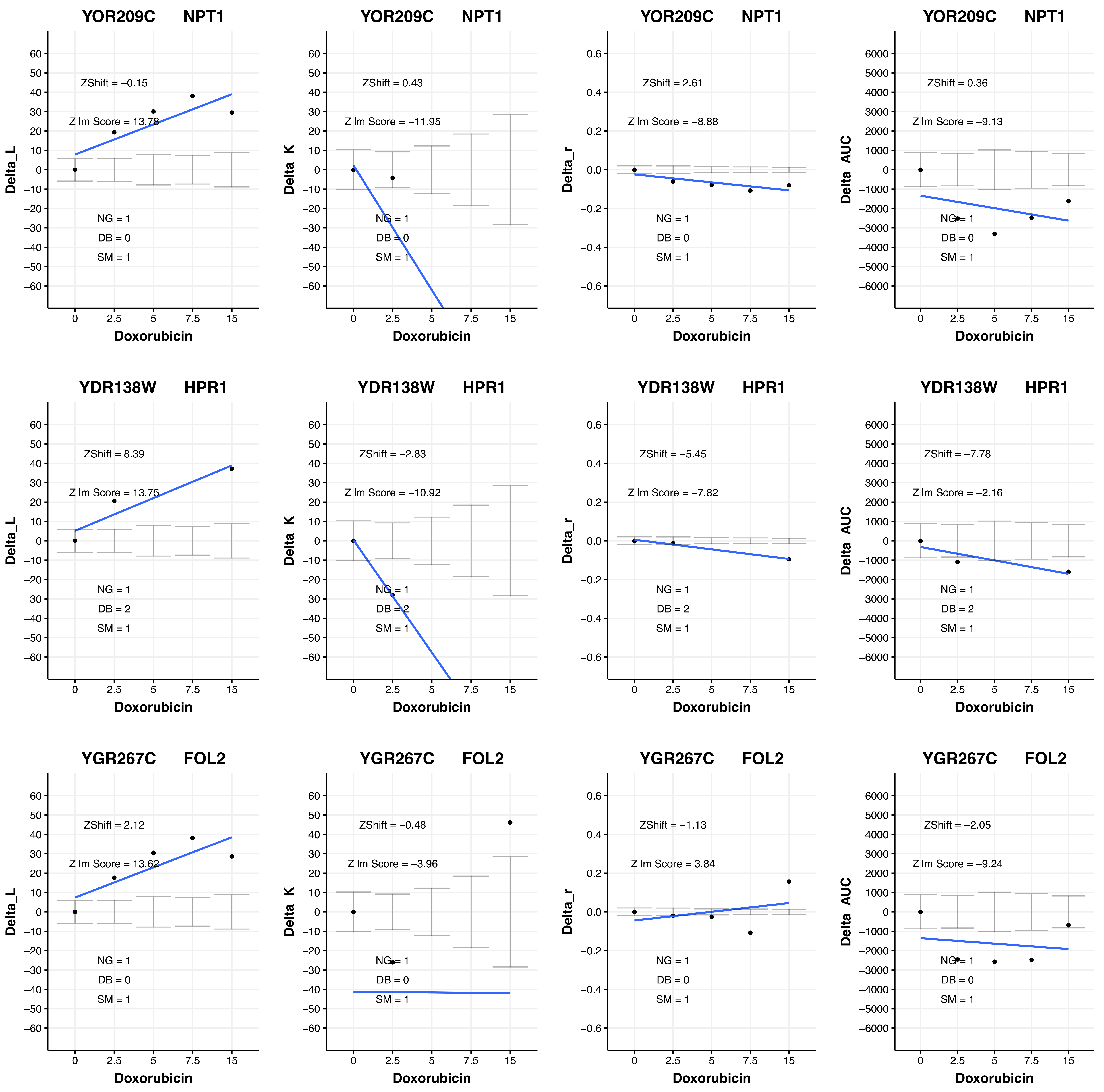
YOR209C NPT1
ZShift = −0.15
Z lm Score = 13.78
NG = 1
DB = 0
SM = 1
Delta_L
Doxorubicin
YOR209C NPT1
ZShift = 0.43
Z lm Score = −11.95
NG = 1
DB = 0
SM = 1
Delta_K
Doxorubicin
YOR209C NPT1
ZShift = 2.61
Z lm Score = −8.88
NG = 1
DB = 0
SM = 1
Delta_r
Doxorubicin
YOR209C NPT1
ZShift = 0.36
Z lm Score = −9.13
NG = 1
DB = 0
SM = 1
Delta_AUC
Doxorubicin
YDR138W HPR1
ZShift = 8.39
Z lm Score = 13.75
NG = 1
DB = 2
SM = 1
Delta_L
Doxorubicin
YDR138W HPR1
ZShift = −2.83
Z lm Score = −10.92
NG = 1
DB = 2
SM = 1
Delta_K
Doxorubicin
YDR138W HPR1
ZShift = −5.45
Z lm Score = −7.82
NG = 1
DB = 2
SM = 1
Delta_r
Doxorubicin
YDR138W HPR1
ZShift = −7.78
Z lm Score = −2.16
NG = 1
DB = 2
SM = 1
Delta_AUC
Doxorubicin
YGR267C FOL2
ZShift = 2.12
Z lm Score = 13.62
NG = 1
DB = 0
SM = 1
Delta_L
Doxorubicin
YGR267C FOL2
ZShift = −0.48
Z lm Score = −3.96
NG = 1
DB = 0
SM = 1
Delta_K
Doxorubicin
YGR267C FOL2
ZShift = −1.13
Z lm Score = 3.84
NG = 1
DB = 0
SM = 1
Delta_r
Doxorubicin
YGR267C FOL2
ZShift = −2.05
Z lm Score = −9.24
NG = 1
DB = 0
SM = 1
Delta_AUC
Doxorubicin

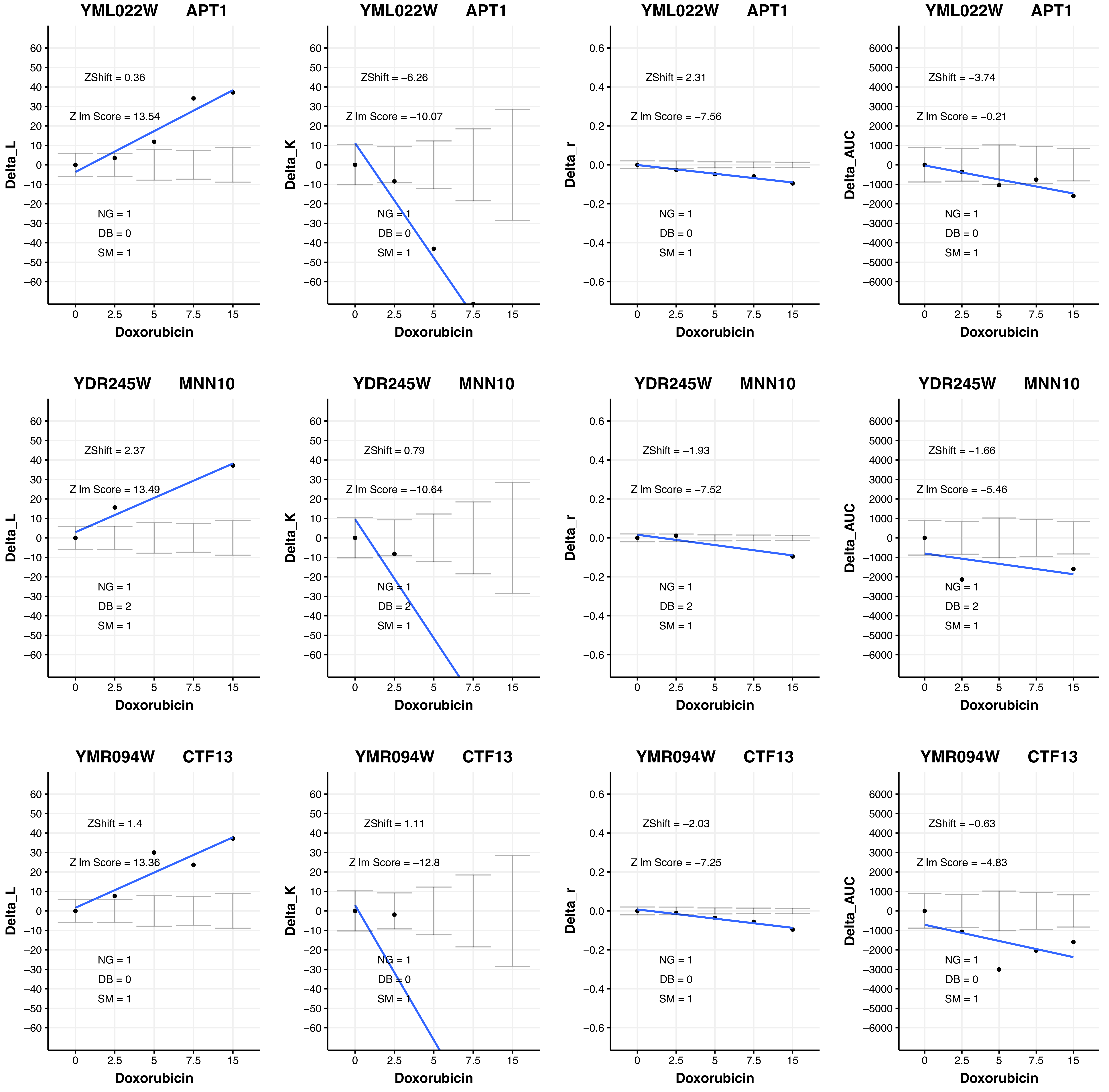
YML022W APT1
ZShift = 0.36
Z lm Score = 13.54
NG = 1
DB = 0
SM = 1
Delta_L
Doxorubicin
YML022W APT1
ZShift = −6.26
Z lm Score = −10.07
NG = 1
DB = 0
SM = 1
Delta_K
Doxorubicin
YML022W APT1
ZShift = 2.31
Z lm Score = −7.56
NG = 1
DB = 0
SM = 1
Delta_r
Doxorubicin
YML022W APT1
ZShift = −3.74
Z lm Score = −0.21
NG = 1
DB = 0
SM = 1
Delta_AUC
Doxorubicin
YDR245W MNN10
ZShift = 2.37
Z lm Score = 13.49
NG = 1
DB = 2
SM = 1
Delta_L
Doxorubicin
YDR245W MNN10
ZShift = 0.79
Z lm Score = −10.64
NG = 1
DB = 2
SM = 1
Delta_K
Doxorubicin
YDR245W MNN10
ZShift = −1.93
Z lm Score = −7.52
NG = 1
DB = 2
SM = 1
Delta_r
Doxorubicin
YDR245W MNN10
ZShift = −1.66
Z lm Score = −5.46
NG = 1
DB = 2
SM = 1
Delta_AUC
Doxorubicin
YMR094W CTF13
ZShift = 1.4
Z lm Score = 13.36
NG = 1
DB = 0
SM = 1
Delta_L
Doxorubicin
YMR094W CTF13
ZShift = 1.11
Z lm Score = −12.8
NG = 1
DB = 0
SM = 1
Delta_K
Doxorubicin
YMR094W CTF13
ZShift = −2.03
Z lm Score = −7.25
NG = 1
DB = 0
SM = 1
Delta_r
Doxorubicin
YMR094W CTF13
ZShift = −0.63
Z lm Score = −4.83
NG = 1
DB = 0
SM = 1
Delta_AUC
Doxorubicin

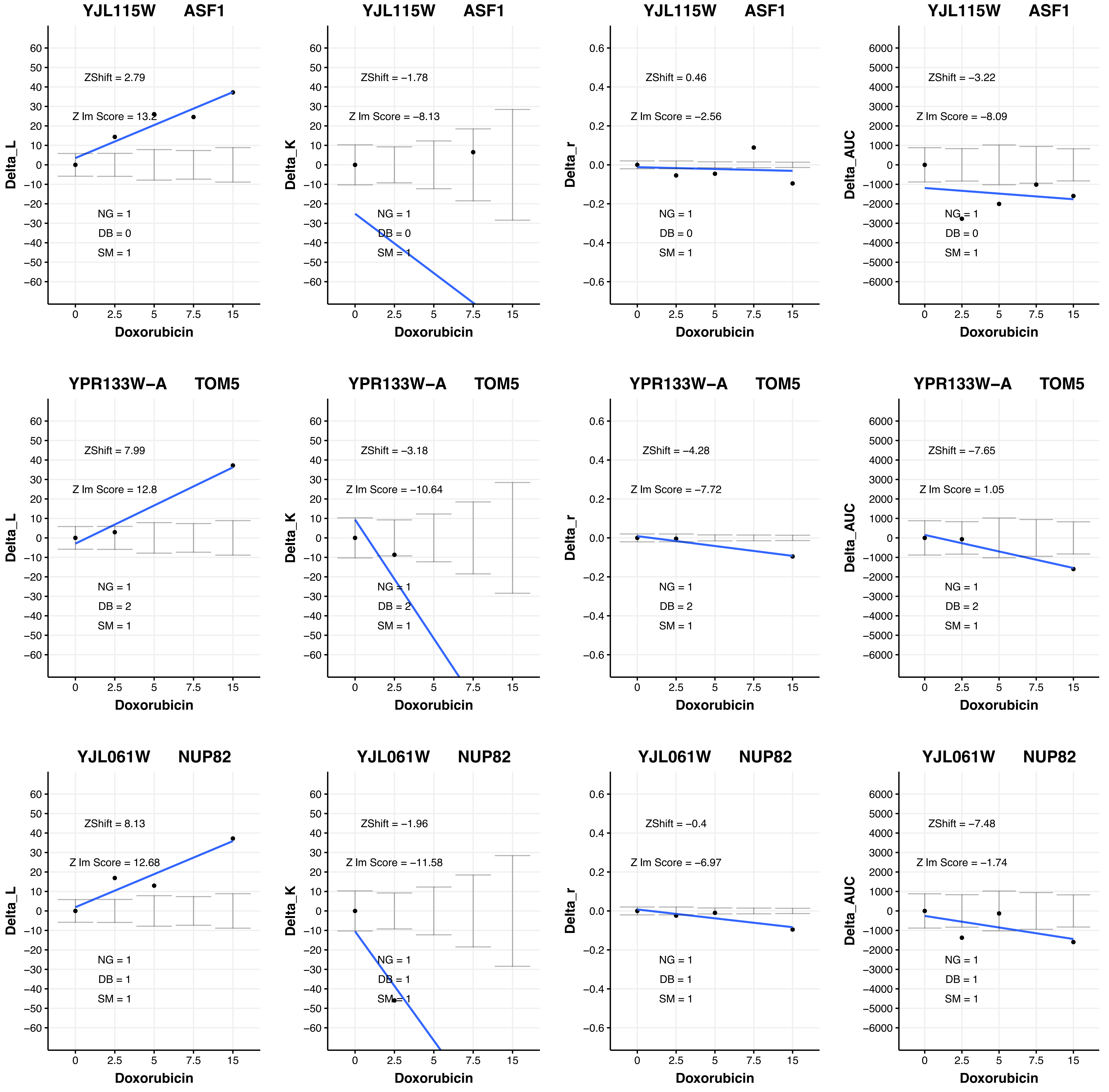
YJL115W ASF1
ZShift = 2.79
Z lm Score = 13.2
NG = 1
DB = 0
SM = 1
Delta_L
Doxorubicin
YJL115W ASF1
ZShift = −1.78
Z lm Score = −8.13
NG = 1
DB = 0
SM = 1
Delta_K
Doxorubicin
YJL115W ASF1
ZShift = 0.46
Z lm Score = −2.56
NG = 1
DB = 0
SM = 1
Delta_r
Doxorubicin
YJL115W ASF1
ZShift = −3.22
Z lm Score = −8.09
NG = 1
DB = 0
SM = 1
Delta_AUC
Doxorubicin
YPR133W−A TOM5
ZShift = 7.99
Z lm Score = 12.8
NG = 1
DB = 2
SM = 1
Delta_L
Doxorubicin
YPR133W−A TOM5
ZShift = −3.18
Z lm Score = −10.64
NG = 1
DB = 2
SM = 1
Delta_K
Doxorubicin
YPR133W−A TOM5
ZShift = −4.28
Z lm Score = −7.72
NG = 1
DB = 2
SM = 1
Delta_r
Doxorubicin
YPR133W−A TOM5
ZShift = −7.65
Z lm Score = 1.05
NG = 1
DB = 2
SM = 1
Delta_AUC
Doxorubicin
YJL061W NUP82
ZShift = 8.13
Z lm Score = 12.68
NG = 1
DB = 1
SM = 1
Delta_L
Doxorubicin
YJL061W NUP82
ZShift = −1.96
Z lm Score = −11.58
NG = 1
DB = 1
SM = 1
Delta_K
Doxorubicin
YJL061W NUP82
ZShift = −0.4
Z lm Score = −6.97
NG = 1
DB = 1
SM = 1
Delta_r
Doxorubicin
YJL061W NUP82
ZShift = −7.48
Z lm Score = −1.74
NG = 1
DB = 1
SM = 1
Delta_AUC
Doxorubicin

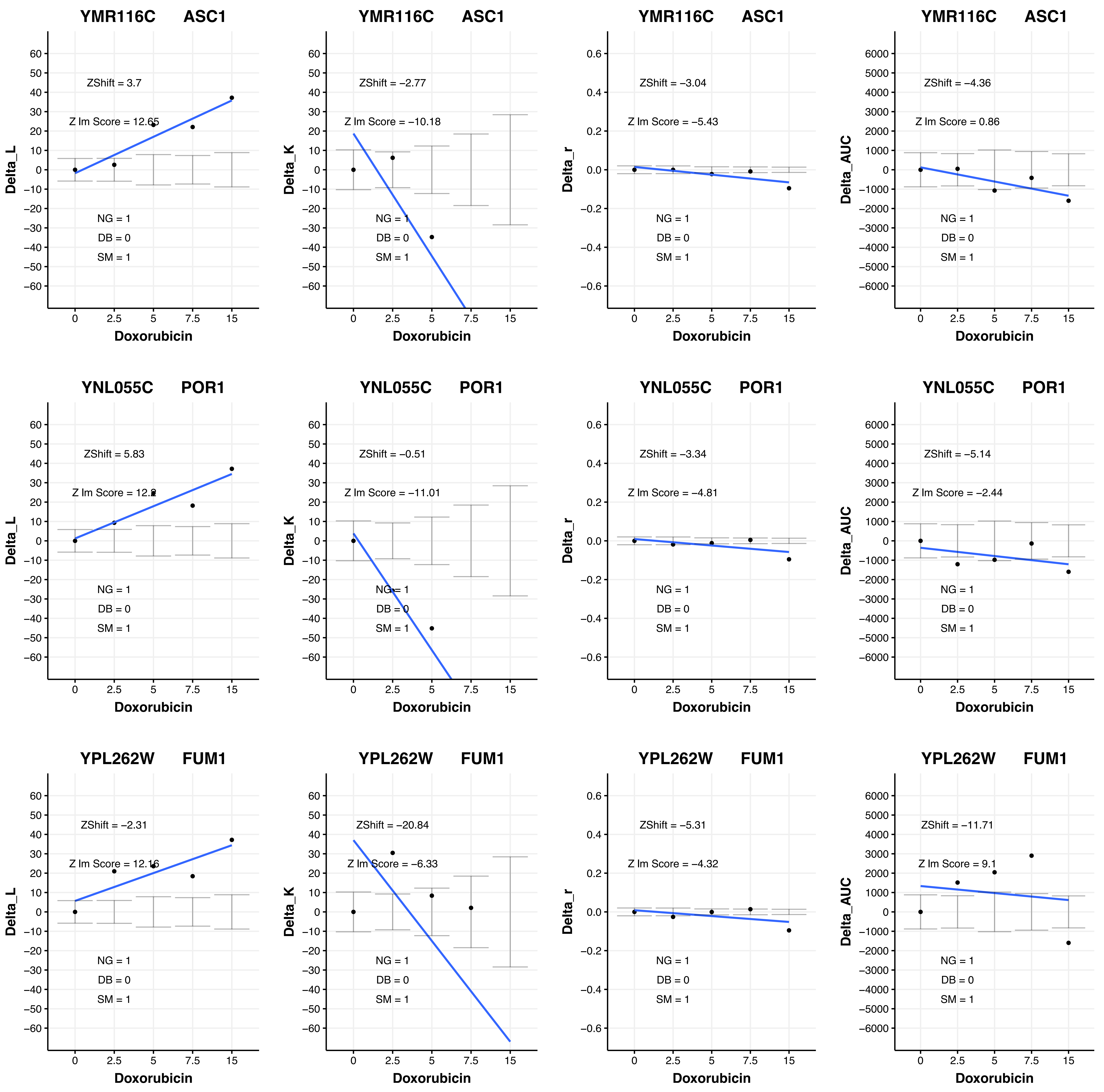
YMR116C ASC1
ZShift = 3.7
Z lm Score = 12.65
NG = 1
DB = 0
SM = 1
Delta_L
Doxorubicin
YMR116C ASC1
ZShift = −2.77
Z lm Score = −10.18
NG = 1
DB = 0
SM = 1
Delta_K
Doxorubicin
YMR116C ASC1
ZShift = −3.04
Z lm Score = −5.43
NG = 1
DB = 0
SM = 1
Delta_r
Doxorubicin
YMR116C ASC1
ZShift = −4.36
Z lm Score = 0.86
NG = 1
DB = 0
SM = 1
Delta_AUC
Doxorubicin
YNL055C POR1
ZShift = 5.83
Z lm Score = 12.2
NG = 1
DB = 0
SM = 1
Delta_L
Doxorubicin
YNL055C POR1
ZShift = −0.51
Z lm Score = −11.01
NG = 1
DB = 0
SM = 1
Delta_K
Doxorubicin
YNL055C POR1
ZShift = −3.34
Z lm Score = −4.81
NG = 1
DB = 0
SM = 1
Delta_r
Doxorubicin
YNL055C POR1
ZShift = −5.14
Z lm Score = −2.44
NG = 1
DB = 0
SM = 1
Delta_AUC
Doxorubicin
YPL262W FUM1
ZShift = −2.31
Z lm Score = 12.16
NG = 1
DB = 0
SM = 1
Delta_L
Doxorubicin
YPL262W FUM1
ZShift = −20.84
Z lm Score = −6.33
NG = 1
DB = 0
SM = 1
Delta_K
Doxorubicin
YPL262W FUM1
ZShift = −5.31
Z lm Score = −4.32
NG = 1
DB = 0
SM = 1
Delta_r
Doxorubicin
YPL262W FUM1
ZShift = −11.71
Z lm Score = 9.1
NG = 1
DB = 0
SM = 1
Delta_AUC
Doxorubicin

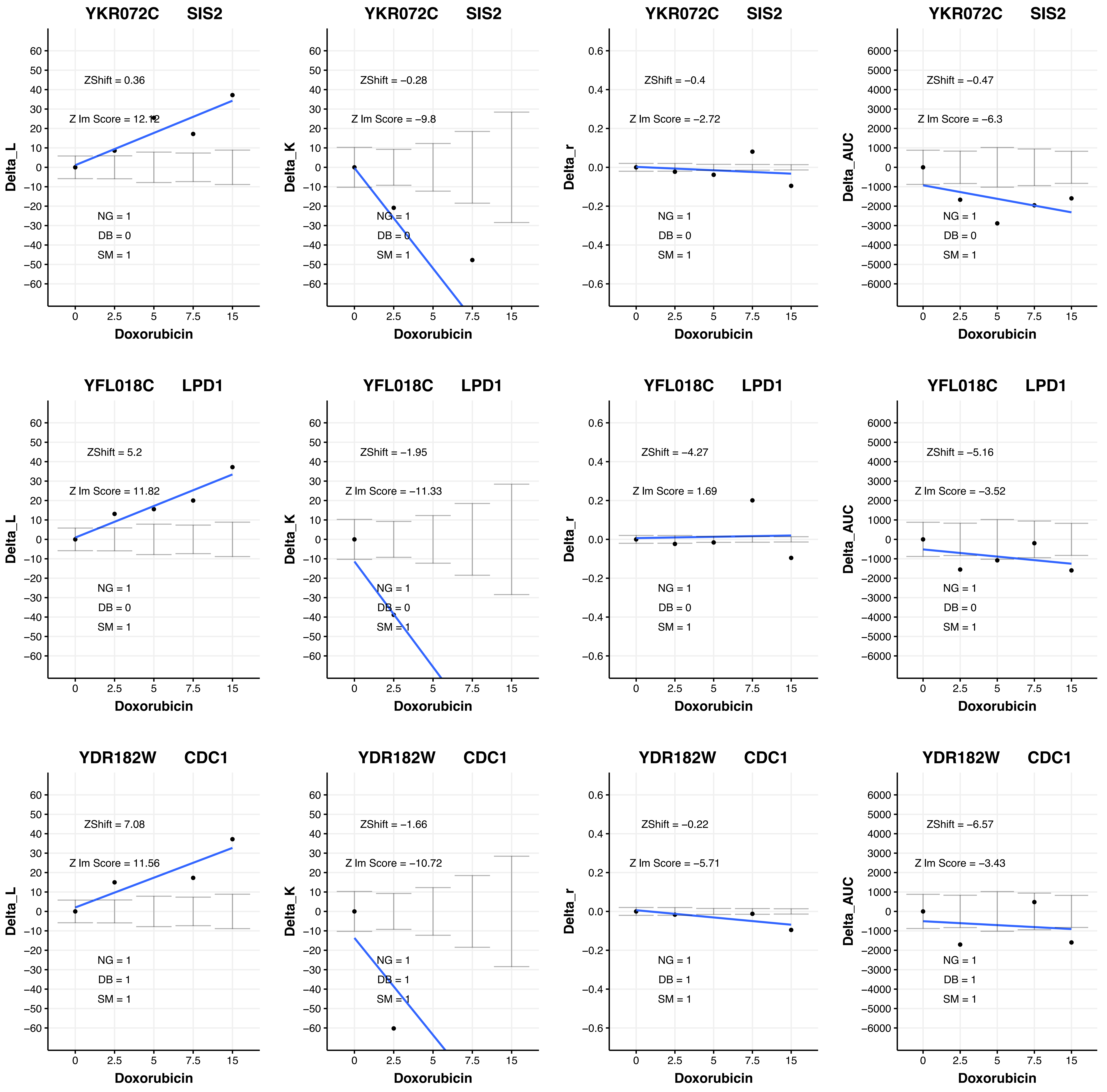

YKR072C SIS2
ZShift = 0.36
Z lm Score = 12.12
NG = 1
DB = 0
SM = 1
Delta_L
Doxorubicin
YKR072C SIS2
ZShift = −0.28
Z lm Score = −9.8
NG = 1
DB = 0
SM = 1
Delta_K
Doxorubicin
YKR072C SIS2
ZShift = −0.4
Z lm Score = −2.72
NG = 1
DB = 0
SM = 1
Delta_r
Doxorubicin
YKR072C SIS2
ZShift = −0.47
Z lm Score = −6.3
NG = 1
DB = 0
SM = 1
Delta_AUC
Doxorubicin
YFL018C LPD1
ZShift = 5.2
Z lm Score = 11.82
NG = 1
DB = 0
SM = 1
Delta_L
Doxorubicin
YFL018C LPD1
ZShift = −1.95
Z lm Score = −11.33
NG = 1
DB = 0
SM = 1
Delta_K
Doxorubicin
YFL018C LPD1
ZShift = −4.27
Z lm Score = 1.69
NG = 1
DB = 0
SM = 1
Delta_r
Doxorubicin
YFL018C LPD1
ZShift = −5.16
Z lm Score = −3.52
NG = 1
DB = 0
SM = 1
Delta_AUC
Doxorubicin
YDR182W CDC1
ZShift = 7.08
Z lm Score = 11.56
NG = 1
DB = 1
SM = 1
Delta_L
Doxorubicin
YDR182W CDC1
ZShift = −1.66
Z lm Score = −10.72
NG = 1
DB = 1
SM = 1
Delta_K
Doxorubicin
YDR182W CDC1
ZShift = −0.22
Z lm Score = −5.71
NG = 1
DB = 1
SM = 1
Delta_r
Doxorubicin
YDR182W CDC1
ZShift = −6.57
Z lm Score = −3.43
NG = 1
DB = 1
SM = 1
Delta_AUC
Doxorubicin

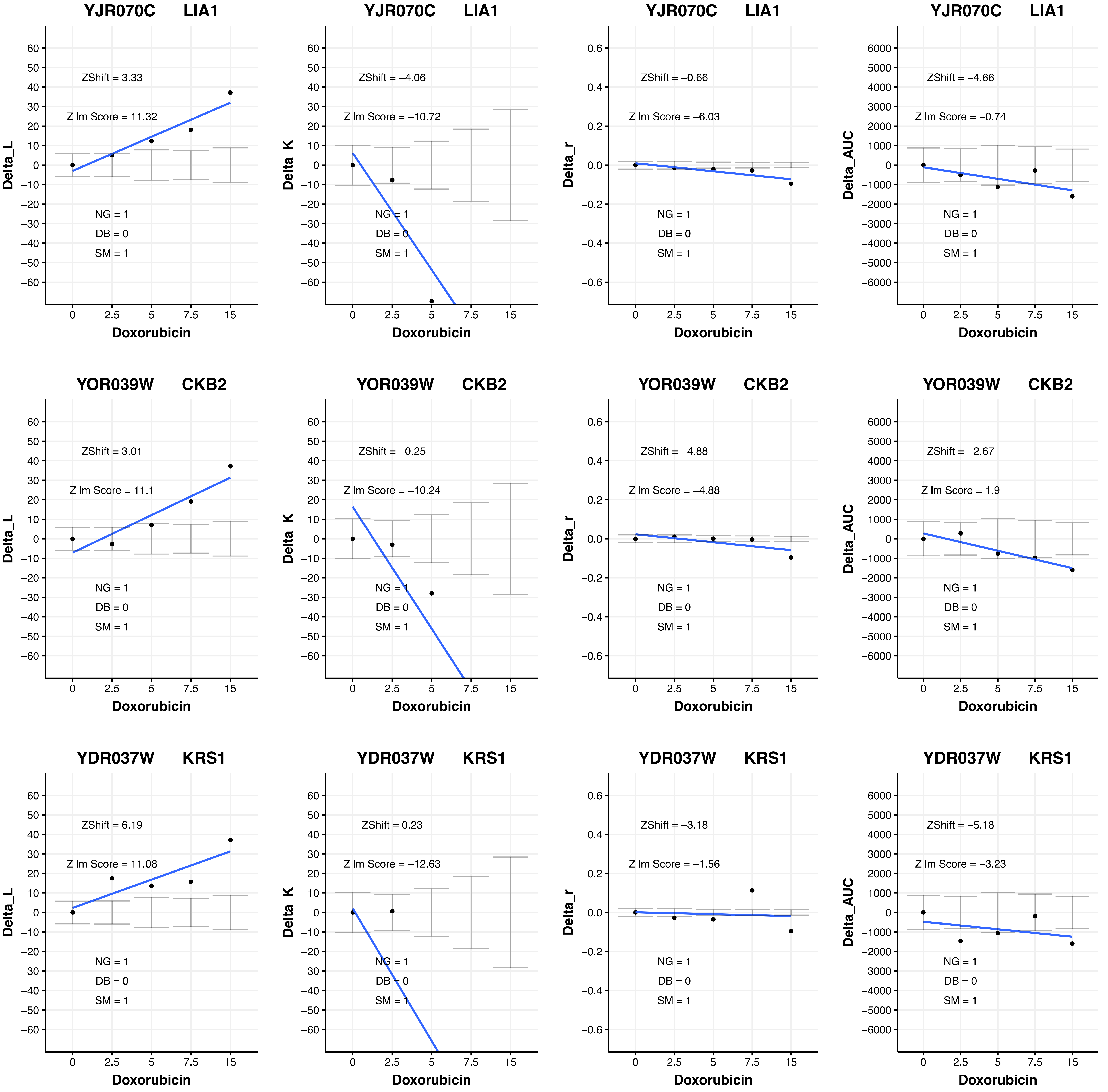

YJR070C LIA1
ZShift = 3.33
Z lm Score = 11.32
NG = 1
DB = 0
SM = 1
Delta_L
Doxorubicin
YJR070C LIA1
ZShift = −4.06
Z lm Score = −10.72
NG = 1
DB = 0
SM = 1
Delta_K
Doxorubicin
YJR070C LIA1
ZShift = −0.66
Z lm Score = −6.03
NG = 1
DB = 0
SM = 1
Delta_r
Doxorubicin
YJR070C LIA1
ZShift = −4.66
Z lm Score = −0.74
NG = 1
DB = 0
SM = 1
Delta_AUC
Doxorubicin
YOR039W CKB2
ZShift = 3.01
Z lm Score = 11.1
NG = 1
DB = 0
SM = 1
Delta_L
Doxorubicin
YOR039W CKB2
ZShift = −0.25
Z lm Score = −10.24
NG = 1
DB = 0
SM = 1
Delta_K
Doxorubicin
YOR039W CKB2
ZShift = −4.88
Z lm Score = −4.88
NG = 1
DB = 0
SM = 1
Delta_r
Doxorubicin
YOR039W CKB2
ZShift = −2.67
Z lm Score = 1.9
NG = 1
DB = 0
SM = 1
Delta_AUC
Doxorubicin
YDR037W KRS1
ZShift = 6.19
Z lm Score = 11.08
NG = 1
DB = 0
SM = 1
Delta_L
Doxorubicin
YDR037W KRS1
ZShift = 0.23
Z lm Score = −12.63
NG = 1
DB = 0
SM = 1
Delta_K
Doxorubicin
YDR037W KRS1
ZShift = −3.18
Z lm Score = −1.56
NG = 1
DB = 0
SM = 1
Delta_r
Doxorubicin
YDR037W KRS1
ZShift = −5.18
Z lm Score = −3.23
NG = 1
DB = 0
SM = 1
Delta_AUC
Doxorubicin

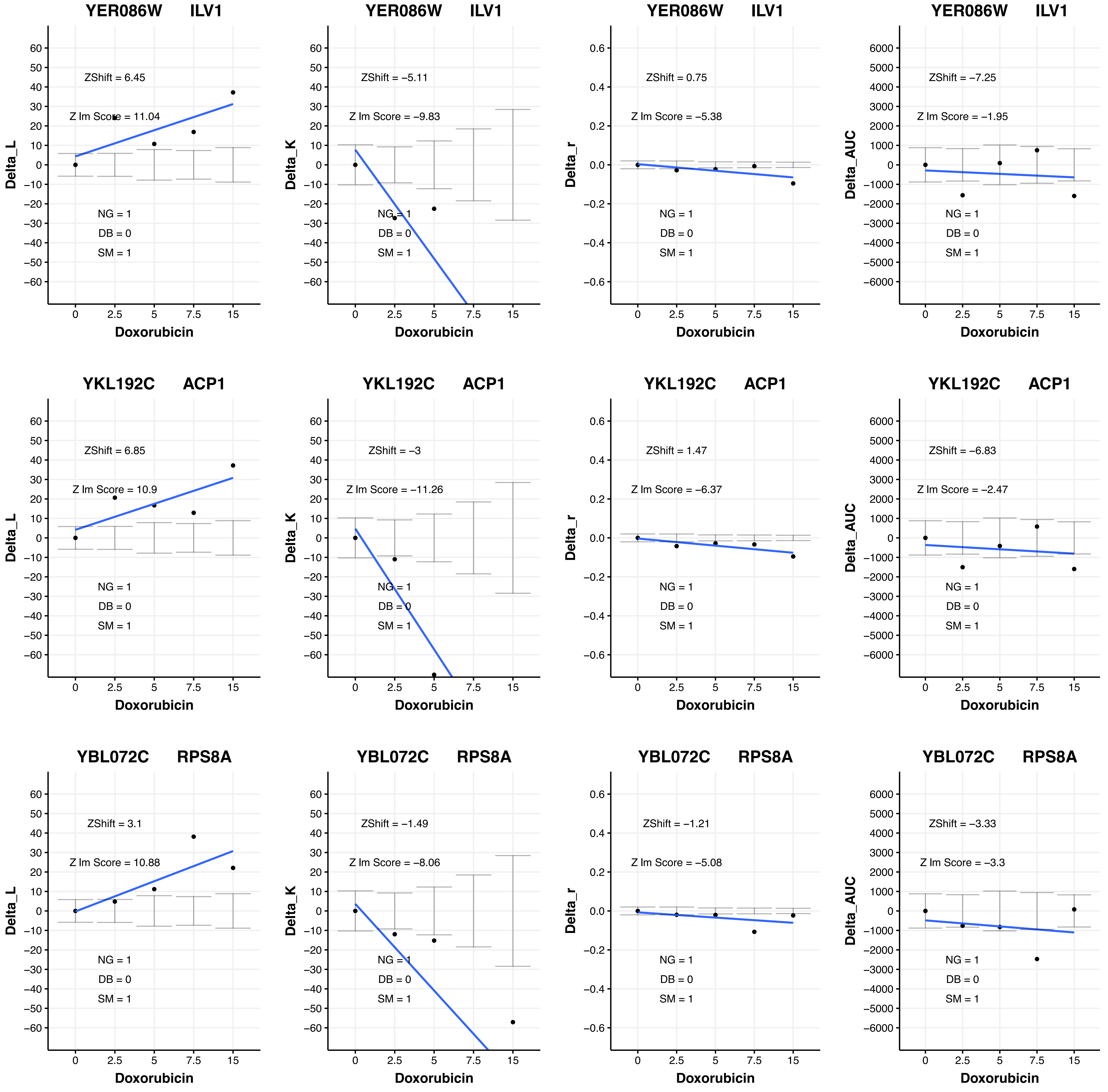

YER086W ILV1
ZShift = 6.45
Z lm Score = 11.04
NG = 1
DB = 0
SM = 1
Delta_L
Doxorubicin
YER086W ILV1
ZShift = −5.11
Z lm Score = −9.83
NG = 1
DB = 0
SM = 1
Delta_K
Doxorubicin
YER086W ILV1
ZShift = 0.75
Z lm Score = −5.38
NG = 1
DB = 0
SM = 1
Delta_r
Doxorubicin
YER086W ILV1
ZShift = −7.25
Z lm Score = −1.95
NG = 1
DB = 0
SM = 1
Delta_AUC
Doxorubicin
YKL192C ACP1
ZShift = 6.85
Z lm Score = 10.9
NG = 1
DB = 0
SM = 1
Delta_L
Doxorubicin
YKL192C ACP1
ZShift = −3
Z lm Score = −11.26
NG = 1
DB = 0
SM = 1
Delta_K
Doxorubicin
YKL192C ACP1
ZShift = 1.47
Z lm Score = −6.37
NG = 1
DB = 0
SM = 1
Delta_r
Doxorubicin
YKL192C ACP1
ZShift = −6.83
Z lm Score = −2.47
NG = 1
DB = 0
SM = 1
Delta_AUC
Doxorubicin
YBL072C RPS8A
ZShift = 3.1
Z lm Score = 10.88
NG = 1
DB = 0
SM = 1
Delta_L
Doxorubicin
YBL072C RPS8A
ZShift = −1.49
Z lm Score = −8.06
NG = 1
DB = 0
SM = 1
Delta_K
Doxorubicin
YBL072C RPS8A
ZShift = −1.21
Z lm Score = −5.08
NG = 1
DB = 0
SM = 1
Delta_r
Doxorubicin
YBL072C RPS8A
ZShift = −3.33
Z lm Score = −3.3
NG = 1
DB = 0
SM = 1
Delta_AUC
Doxorubicin

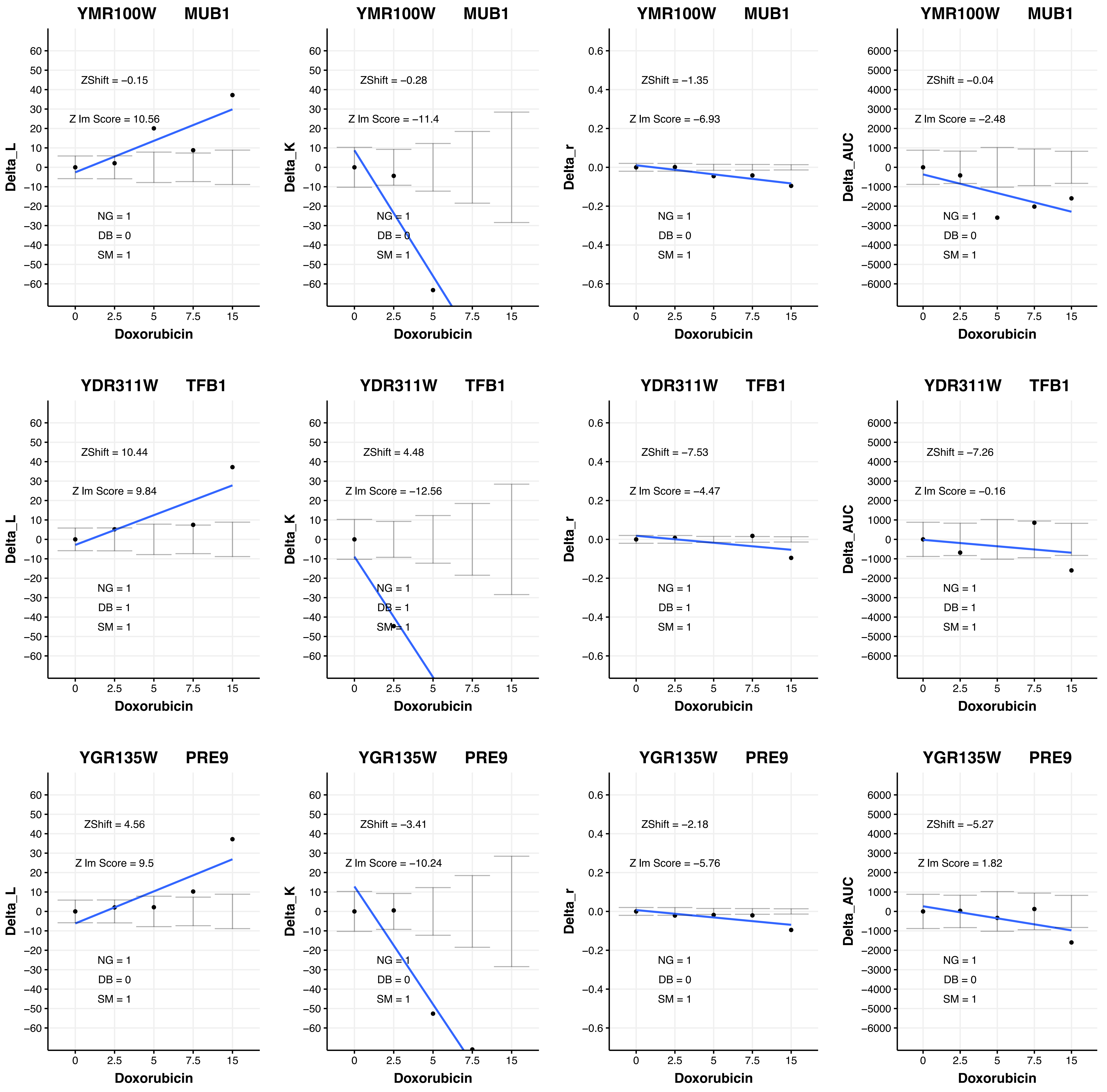
YMR100W MUB1
ZShift = −0.15
Z lm Score = 10.56
NG = 1
DB = 0
SM = 1
Delta_L
Doxorubicin
YMR100W MUB1
ZShift = −0.28
Z lm Score = −11.4
NG = 1
DB = 0
SM = 1
Delta_K
Doxorubicin
YMR100W MUB1
ZShift = −1.35
Z lm Score = −6.93
NG = 1
DB = 0
SM = 1
Delta_r
Doxorubicin
YMR100W MUB1
ZShift = −0.04
Z lm Score = −2.48
NG = 1
DB = 0
SM = 1
Delta_AUC
Doxorubicin
YDR311W TFB1
ZShift = 10.44
Z lm Score = 9.84
NG = 1
DB = 1
SM = 1
Delta_L
Doxorubicin
YDR311W TFB1
ZShift = 4.48
Z lm Score = −12.56
NG = 1
DB = 1
SM = 1
Delta_K
Doxorubicin
YDR311W TFB1
ZShift = −7.53
Z lm Score = −4.47
NG = 1
DB = 1
SM = 1
Delta_r
Doxorubicin
YDR311W TFB1
ZShift = −7.26
Z lm Score = −0.16
NG = 1
DB = 1
SM = 1
Delta_AUC
Doxorubicin
YGR135W PRE9
ZShift = 4.56
Z lm Score = 9.5
NG = 1
DB = 0
SM = 1
Delta_L
Doxorubicin
YGR135W PRE9
ZShift = −3.41
Z lm Score = −10.24
NG = 1
DB = 0
SM = 1
Delta_K
Doxorubicin
YGR135W PRE9
ZShift = −2.18
Z lm Score = −5.76
NG = 1
DB = 0
SM = 1
Delta_r
Doxorubicin
YGR135W PRE9
ZShift = −5.27
Z lm Score = 1.82
NG = 1
DB = 0
SM = 1
Delta_AUC
Doxorubicin

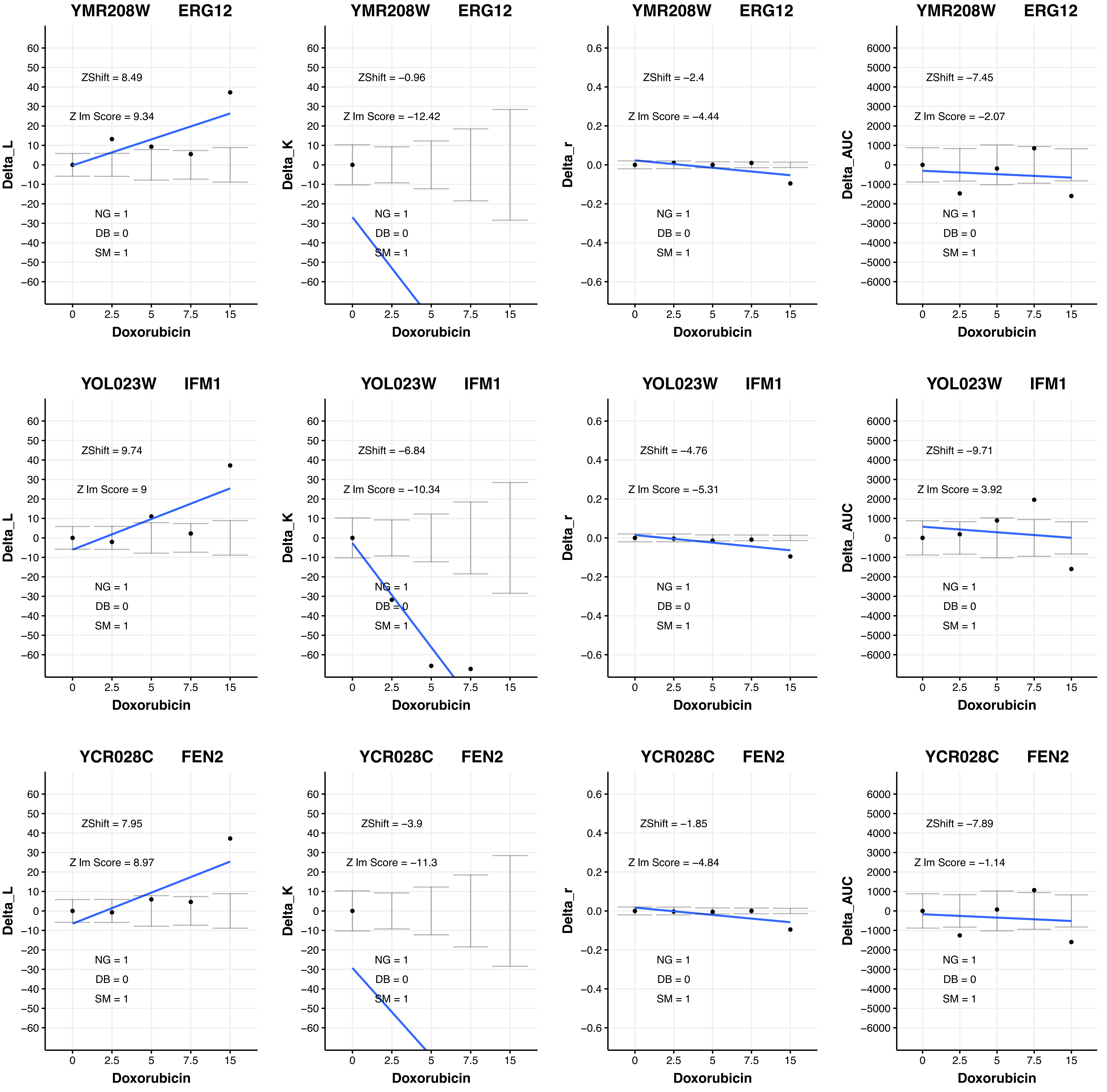
YMR208W ERG12
ZShift = 8.49
Z lm Score = 9.34
NG = 1
DB = 0
SM = 1
Delta_L
Doxorubicin
YMR208W ERG12
ZShift = −0.96
Z lm Score = −12.42
NG = 1
DB = 0
SM = 1
Delta_K
Doxorubicin
YMR208W ERG12
ZShift = −2.4
Z lm Score = −4.44
NG = 1
DB = 0
SM = 1
Delta_r
Doxorubicin
YMR208W ERG12
ZShift = −7.45
Z lm Score = −2.07
NG = 1
DB = 0
SM = 1
Delta_AUC
Doxorubicin
YOL023W IFM1
ZShift = 9.74
Z lm Score = 9
NG = 1
DB = 0
SM = 1
Delta_L
Doxorubicin
YOL023W IFM1
ZShift = −6.84
Z lm Score = −10.34
NG = 1
DB = 0
SM = 1
Delta_K
Doxorubicin
YOL023W IFM1
ZShift = −4.76
Z lm Score = −5.31
NG = 1
DB = 0
SM = 1
Delta_r
Doxorubicin
YOL023W IFM1
ZShift = −9.71
Z lm Score = 3.92
NG = 1
DB = 0
SM = 1
Delta_AUC
Doxorubicin
YCR028C FEN2
ZShift = 7.95
Z lm Score = 8.97
NG = 1
DB = 0
SM = 1
Delta_L
Doxorubicin
YCR028C FEN2
ZShift = −3.9
Z lm Score = −11.3
NG = 1
DB = 0
SM = 1
Delta_K
Doxorubicin
YCR028C FEN2
ZShift = −1.85
Z lm Score = −4.84
NG = 1
DB = 0
SM = 1
Delta_r
Doxorubicin
YCR028C FEN2
ZShift = −7.89
Z lm Score = −1.14
NG = 1
DB = 0
SM = 1
Delta_AUC
Doxorubicin

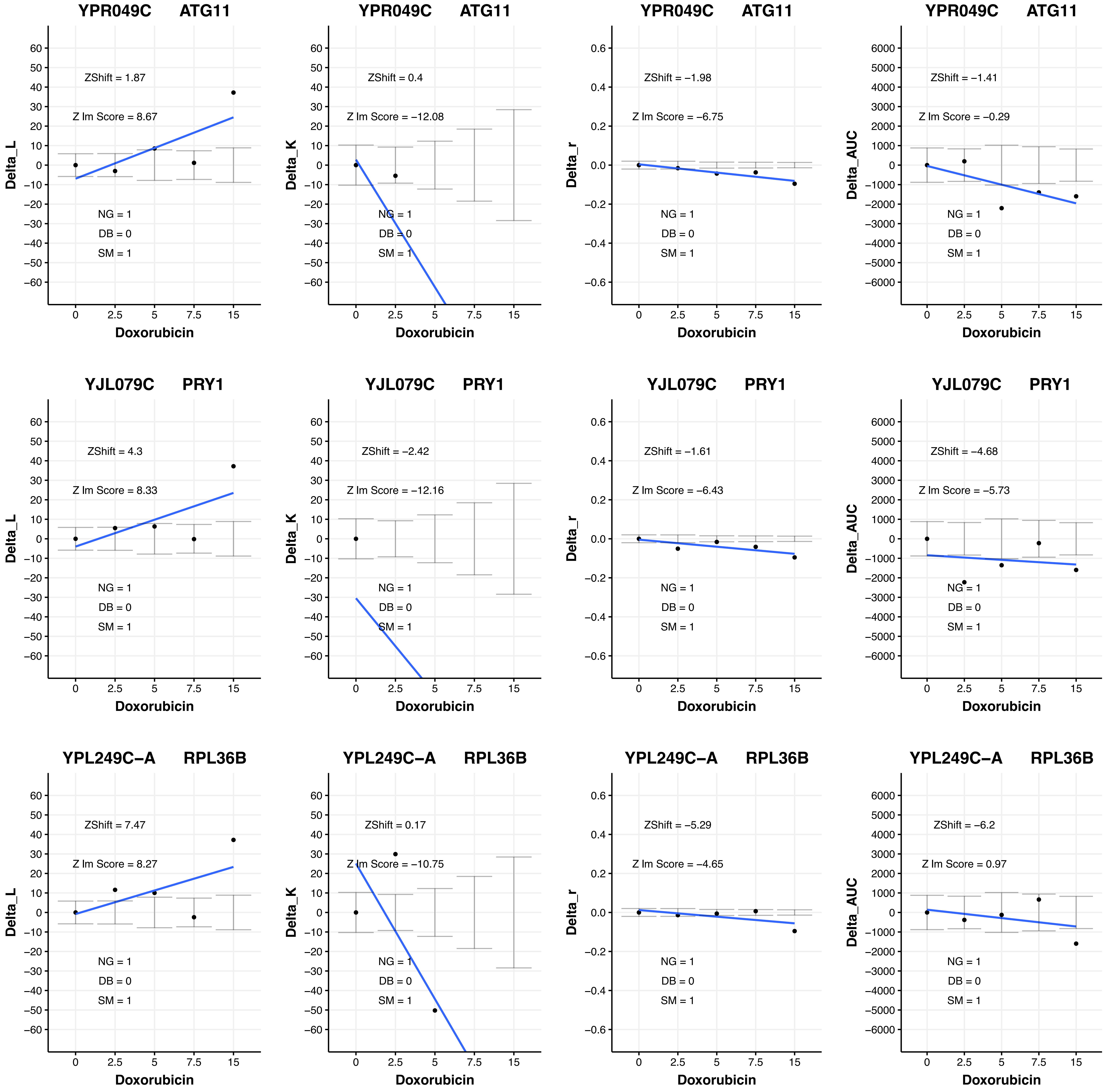

YPR049C ATG11
ZShift = 1.87
Z lm Score = 8.67
NG = 1
DB = 0
SM = 1
Delta_L
Doxorubicin
YPR049C ATG11
ZShift = 0.4
Z lm Score = −12.08
NG = 1
DB = 0
SM = 1
Delta_K
Doxorubicin
YPR049C ATG11
ZShift = −1.98
Z lm Score = −6.75
NG = 1
DB = 0
SM = 1
Delta_r
Doxorubicin
YPR049C ATG11
ZShift = −1.41
Z lm Score = −0.29
NG = 1
DB = 0
SM = 1
Delta_AUC
Doxorubicin
YJL079C PRY1
ZShift = 4.3
Z lm Score = 8.33
NG = 1
DB = 0
SM = 1
Delta_L
Doxorubicin
YJL079C PRY1
ZShift = −2.42
Z lm Score = −12.16
NG = 1
DB = 0
SM = 1
Delta_K
Doxorubicin
YJL079C PRY1
ZShift = −1.61
Z lm Score = −6.43
NG = 1
DB = 0
SM = 1
Delta_r
Doxorubicin
YJL079C PRY1
ZShift = −4.68
Z lm Score = −5.73
NG = 1
DB = 0
SM = 1
Delta_AUC
Doxorubicin
YPL249C−A RPL36B
ZShift = 7.47
Z lm Score = 8.27
NG = 1
DB = 0
SM = 1
Delta_L
Doxorubicin
YPL249C−A RPL36B
ZShift = 0.17
Z lm Score = −10.75
NG = 1
DB = 0
SM = 1
Delta_K
Doxorubicin
YPL249C−A RPL36B
ZShift = −5.29
Z lm Score = −4.65
NG = 1
DB = 0
SM = 1
Delta_r
Doxorubicin
YPL249C−A RPL36B
ZShift = −6.2
Z lm Score = 0.97
NG = 1
DB = 0
SM = 1
Delta_AUC
Doxorubicin

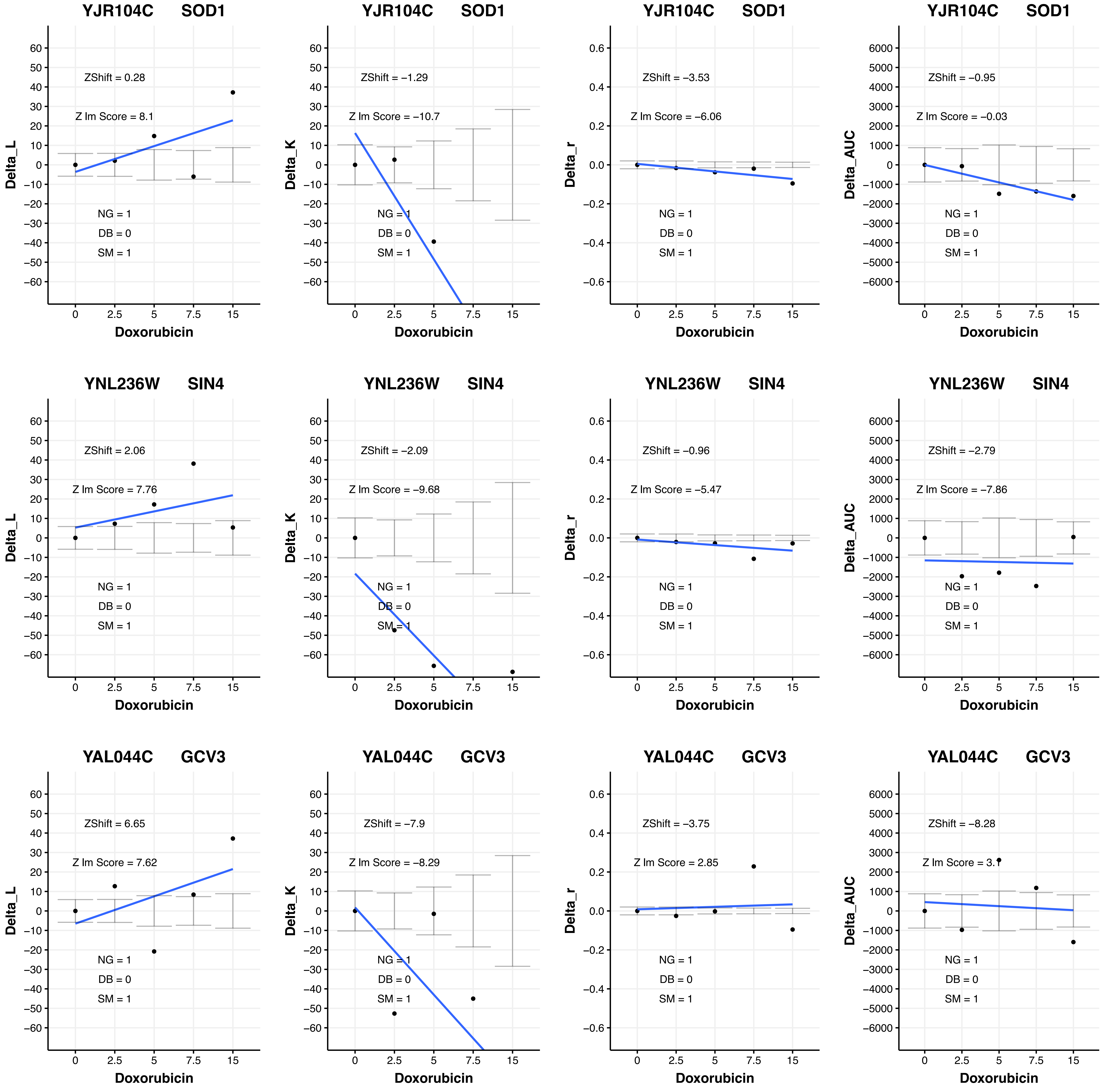

YJR104C SOD1
ZShift = 0.28
Z lm Score = 8.1
NG = 1
DB = 0
SM = 1
Delta_L
Doxorubicin
YJR104C SOD1
ZShift = −1.29
Z lm Score = −10.7
NG = 1
DB = 0
SM = 1
Delta_K
Doxorubicin
YJR104C SOD1
ZShift = −3.53
Z lm Score = −6.06
NG = 1
DB = 0
SM = 1
Delta_r
Doxorubicin
YJR104C SOD1
ZShift = −0.95
Z lm Score = −0.03
NG = 1
DB = 0
SM = 1
Delta_AUC
Doxorubicin
YNL236W SIN4
ZShift = 2.06
Z lm Score = 7.76
NG = 1
DB = 0
SM = 1
Delta_L
Doxorubicin
YNL236W SIN4
ZShift = −2.09
Z lm Score = −9.68
NG = 1
DB = 0
SM = 1
Delta_K
Doxorubicin
YNL236W SIN4
ZShift = −0.96
Z lm Score = −5.47
NG = 1
DB = 0
SM = 1
Delta_r
Doxorubicin
YNL236W SIN4
ZShift = −2.79
Z lm Score = −7.86
NG = 1
DB = 0
SM = 1
Delta_AUC
Doxorubicin
YAL044C GCV3
ZShift = 6.65
Z lm Score = 7.62
NG = 1
DB = 0
SM = 1
Delta_L
Doxorubicin
YAL044C GCV3
ZShift = −7.9
Z lm Score = −8.29
NG = 1
DB = 0
SM = 1
Delta_K
Doxorubicin
YAL044C GCV3
ZShift = −3.75
Z lm Score = 2.85
NG = 1
DB = 0
SM = 1
Delta_r
Doxorubicin
YAL044C GCV3
ZShift = −8.28
Z lm Score = 3.1
NG = 1
DB = 0
SM = 1
Delta_AUC
Doxorubicin

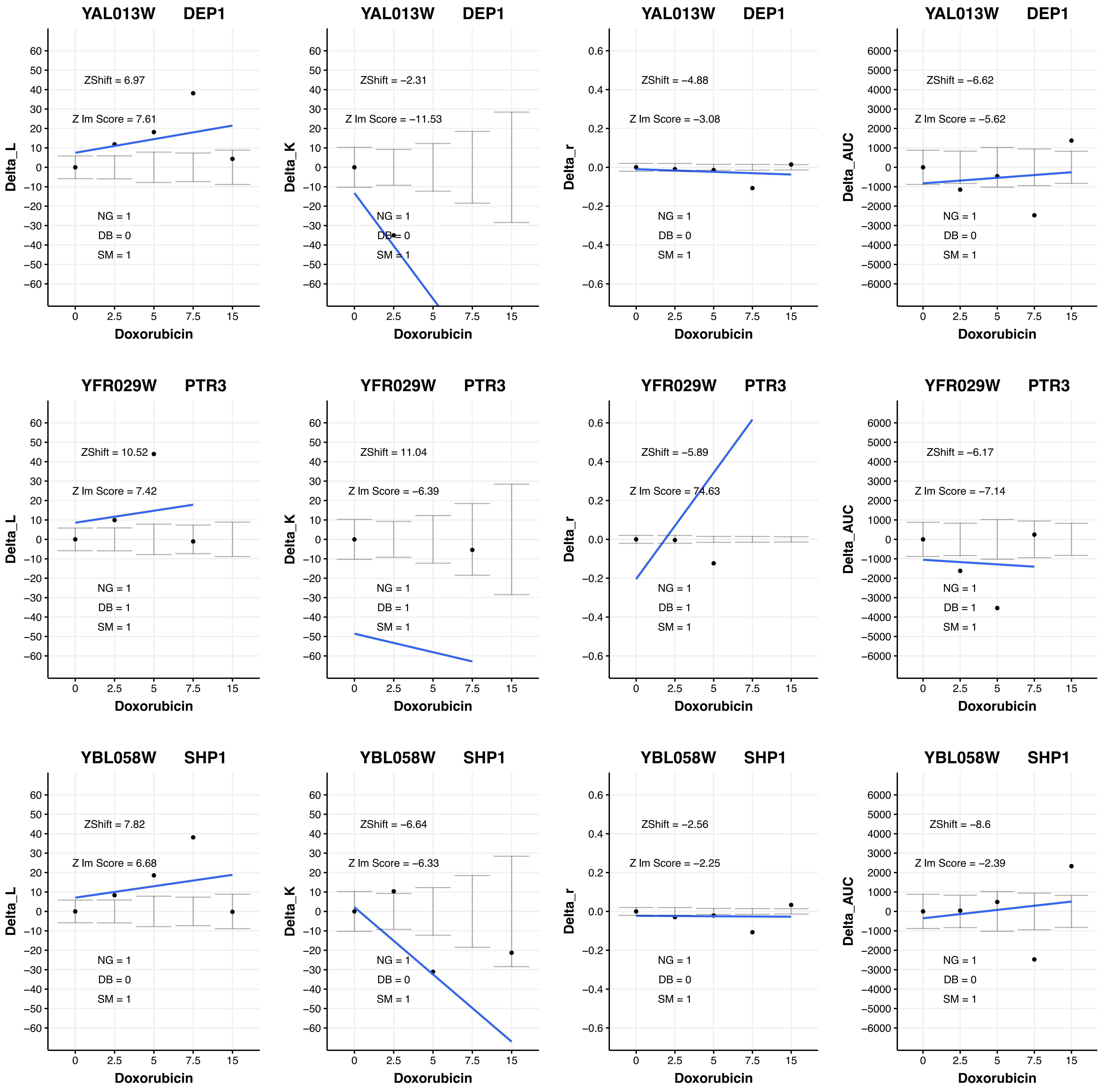
YAL013W DEP1
Delta_L
ZShift = 6.97
Z lm Score = 7.61
NG = 1
DB = 0
SM = 1
Doxorubicin
YAL013W DEP1
Delta_K
ZShift = −2.31
Z lm Score = −11.53
NG = 1
DB = 0
SM = 1
Doxorubicin
YAL013W DEP1
Delta_r
ZShift = −4.88
Z lm Score = −3.08
NG = 1
DB = 0
SM = 1
Doxorubicin
YAL013W DEP1
Delta_AUC
ZShift = −6.62
Z lm Score = −5.62
NG = 1
DB = 0
SM = 1
Doxorubicin
YFR029W PTR3
Delta_L
ZShift = 10.52
Z lm Score = 7.42
NG = 1
DB = 1
SM = 1
Doxorubicin
YFR029W PTR3
Delta_K
ZShift = 11.04
Z lm Score = −6.39
NG = 1
DB = 1
SM = 1
Doxorubicin
YFR029W PTR3
Delta_r
ZShift = −5.89
Z lm Score = 74.63
NG = 1
DB = 1
SM = 1
Doxorubicin
YFR029W PTR3
Delta_AUC
ZShift = −6.17
Z lm Score = −7.14
NG = 1
DB = 1
SM = 1
Doxorubicin
YBL058W SHP1
Delta_L
ZShift = 7.82
Z lm Score = 6.68
NG = 1
DB = 0
SM = 1
Doxorubicin
YBL058W SHP1
Delta_K
ZShift = −6.64
Z lm Score = −6.33
NG = 1
DB = 0
SM = 1
Doxorubicin
YBL058W SHP1
Delta_r
ZShift = −2.56
Z lm Score = −2.25
NG = 1
DB = 0
SM = 1
Doxorubicin
YBL058W SHP1
Delta_AUC
ZShift = −8.6
Z lm Score = −2.39
NG = 1
DB = 0
SM = 1
Doxorubicin

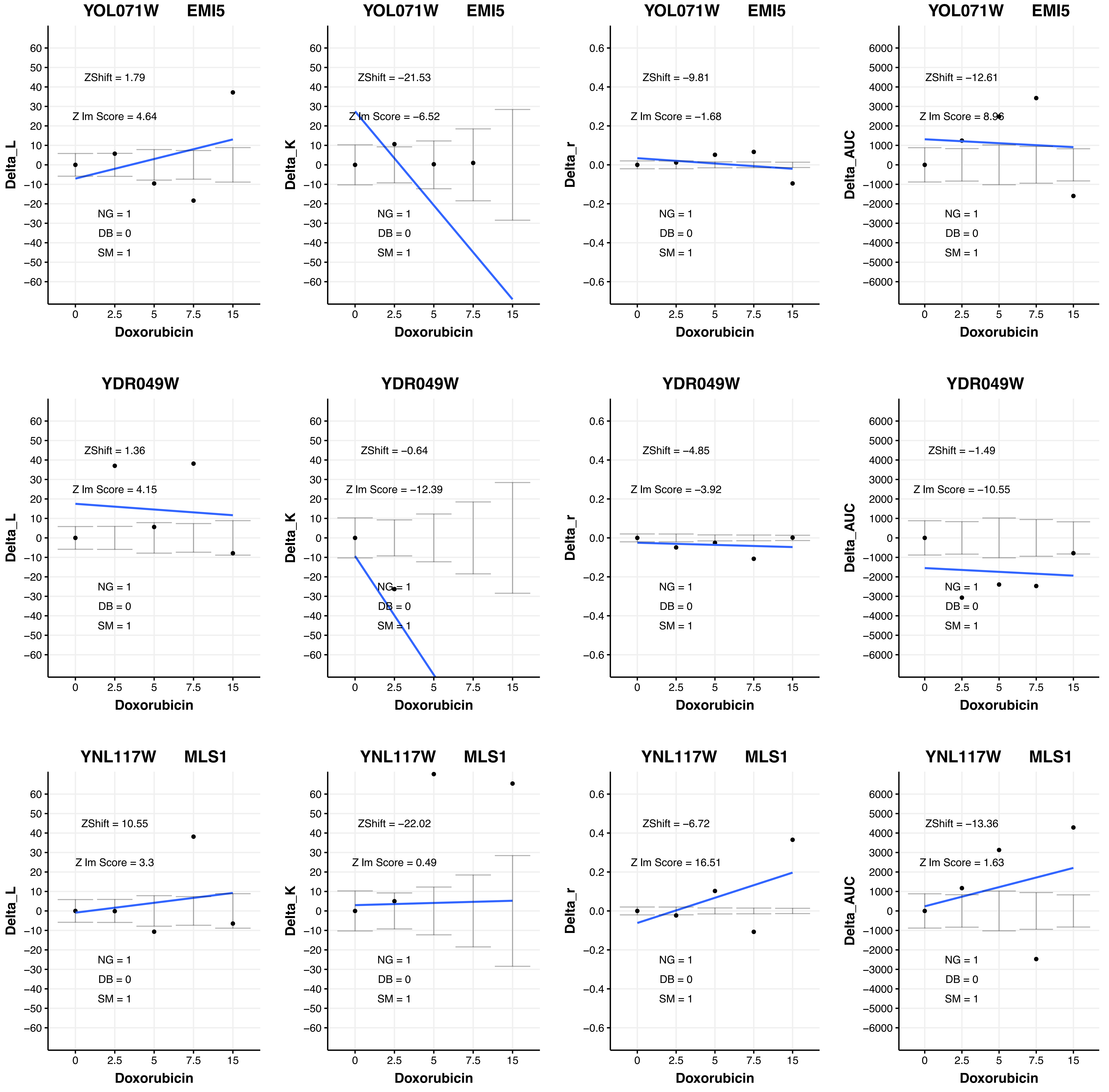
YOL071W EMI5
ZShift = 1.79
Z lm Score = 4.64
NG = 1
DB = 0
SM = 1
Delta_L
Doxorubicin
YOL071W EMI5
ZShift = −21.53
Z lm Score = −6.52
NG = 1
DB = 0
SM = 1
Delta_K
Doxorubicin
YOL071W EMI5
ZShift = −9.81
Z lm Score = −1.68
NG = 1
DB = 0
SM = 1
Delta_r
Doxorubicin
YOL071W EMI5
ZShift = −12.61
Z lm Score = 8.96
NG = 1
DB = 0
SM = 1
Delta_AUC
Doxorubicin
YDR049W
ZShift = 1.36
Z lm Score = 4.15
NG = 1
DB = 0
SM = 1
Delta_L
Doxorubicin
YDR049W
ZShift = −0.64
Z lm Score = −12.39
NG = 1
DB = 0
SM = 1
Delta_K
Doxorubicin
YDR049W
ZShift = −4.85
Z lm Score = −3.92
NG = 1
DB = 0
SM = 1
Delta_r
Doxorubicin
YDR049W
ZShift = −1.49
Z lm Score = −10.55
NG = 1
DB = 0
SM = 1
Delta_AUC
Doxorubicin
YNL117W MLS1
ZShift = 10.55
Z lm Score = 3.3
NG = 1
DB = 0
SM = 1
Delta_L
Doxorubicin
YNL117W MLS1
ZShift = −22.02
Z lm Score = 0.49
NG = 1
DB = 0
SM = 1
Delta_K
Doxorubicin
YNL117W MLS1
ZShift = −6.72
Z lm Score = 16.51
NG = 1
DB = 0
SM = 1
Delta_r
Doxorubicin
YNL117W MLS1
ZShift = −13.36
Z lm Score = 1.63
NG = 1
DB = 0
SM = 1
Delta_AUC
Doxorubicin

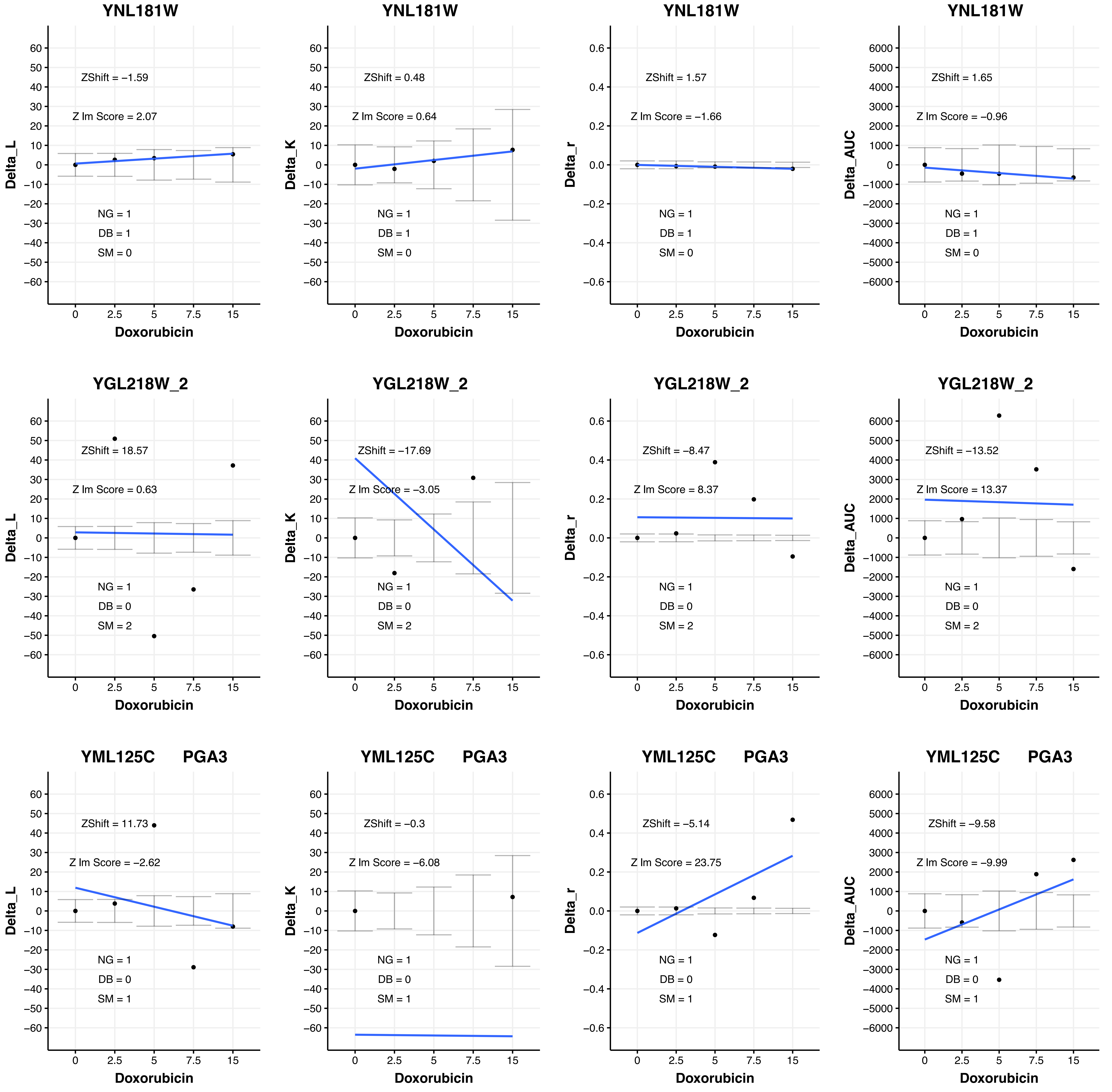
YNL181W
ZShift = −1.59
Z lm Score = 2.07
NG = 1
DB = 1
SM = 0
Delta_L
Doxorubicin
YNL181W
ZShift = 0.48
Z lm Score = 0.64
NG = 1
DB = 1
SM = 0
Delta_K
Doxorubicin
YNL181W
ZShift = 1.57
Z lm Score = −1.66
NG = 1
DB = 1
SM = 0
Delta_r
Doxorubicin
YNL181W
ZShift = 1.65
Z lm Score = −0.96
NG = 1
DB = 1
SM = 0
Delta_AUC
Doxorubicin
YGL218W_2
ZShift = 18.57
Z lm Score = 0.63
NG = 1
DB = 0
SM = 2
Delta_L
Doxorubicin
YGL218W_2
ZShift = −17.69
Z lm Score = −3.05
NG = 1
DB = 0
SM = 2
Delta_K
Doxorubicin
YGL218W_2
ZShift = −8.47
Z lm Score = 8.37
NG = 1
DB = 0
SM = 2
Delta_r
Doxorubicin
YGL218W_2
ZShift = −13.52
Z lm Score = 13.37
NG = 1
DB = 0
SM = 2
Delta_AUC
Doxorubicin
YML125C PGA3
ZShift = 11.73
Z lm Score = −2.62
NG = 1
DB = 0
SM = 1
Delta_L
Doxorubicin
YML125C PGA3
ZShift = −0.3
Z lm Score = −6.08
NG = 1
DB = 0
SM = 1
Delta_K
Doxorubicin
YML125C PGA3
ZShift = −5.14
Z lm Score = 23.75
NG = 1
DB = 0
SM = 1
Delta_r
Doxorubicin
YML125C PGA3
ZShift = −9.58
Z lm Score = −9.99
NG = 1
DB = 0
SM = 1
Delta_AUC
Doxorubicin

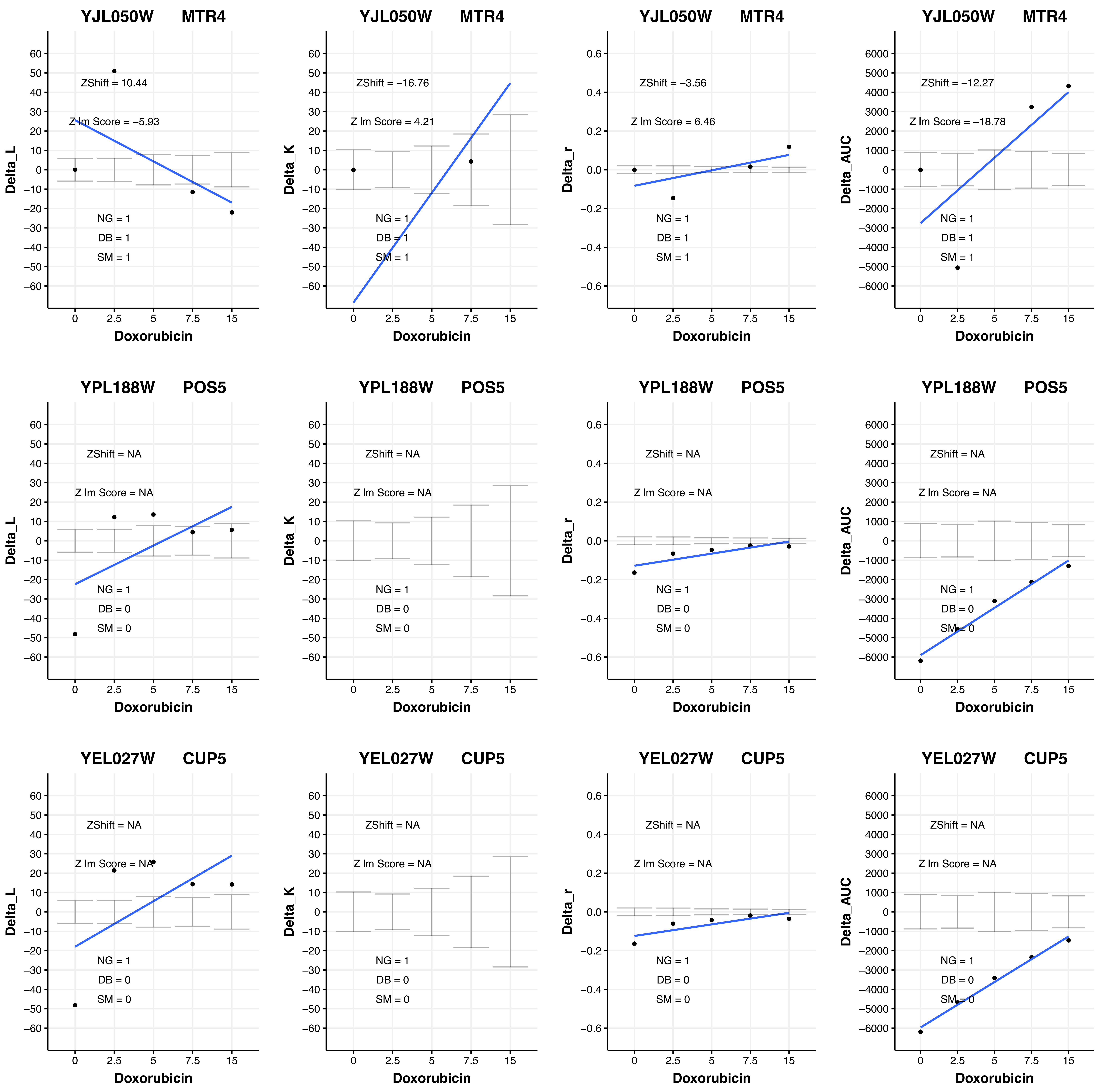
YJL050W MTR4
ZShift = 10.44
Z lm Score = −5.93
NG = 1
DB = 1
SM = 1
Delta_L
Doxorubicin
YJL050W MTR4
ZShift = −16.76
Z lm Score = 4.21
NG = 1
DB = 1
SM = 1
Delta_K
Doxorubicin
YJL050W MTR4
ZShift = −3.56
Z lm Score = 6.46
NG = 1
DB = 1
SM = 1
Delta_r
Doxorubicin
YJL050W MTR4
ZShift = −12.27
Z lm Score = −18.78
NG = 1
DB = 1
SM = 1
Delta_AUC
Doxorubicin
YPL188W POS5
ZShift = NA
Z lm Score = NA
NG = 1
DB = 0
SM = 0
Delta_L
Doxorubicin
YPL188W POS5
ZShift = NA
Z lm Score = NA
NG = 1
DB = 0
SM = 0
Delta_K
Doxorubicin
YPL188W POS5
ZShift = NA
Z lm Score = NA
NG = 1
DB = 0
SM = 0
Delta_r
Doxorubicin
YPL188W POS5
ZShift = NA
Z lm Score = NA
NG = 1
DB = 0
SM = 0
Delta_AUC
Doxorubicin
YEL027W CUP5
ZShift = NA
Z lm Score = NA
NG = 1
DB = 0
SM = 0
Delta_L
Doxorubicin
YEL027W CUP5
ZShift = NA
Z lm Score = NA
NG = 1
DB = 0
SM = 0
Delta_K
Doxorubicin
YEL027W CUP5
ZShift = NA
Z lm Score = NA
NG = 1
DB = 0
SM = 0
Delta_r
Doxorubicin
YEL027W CUP5
ZShift = NA
Z lm Score = NA
NG = 1
DB = 0
SM = 0
Delta_AUC
Doxorubicin

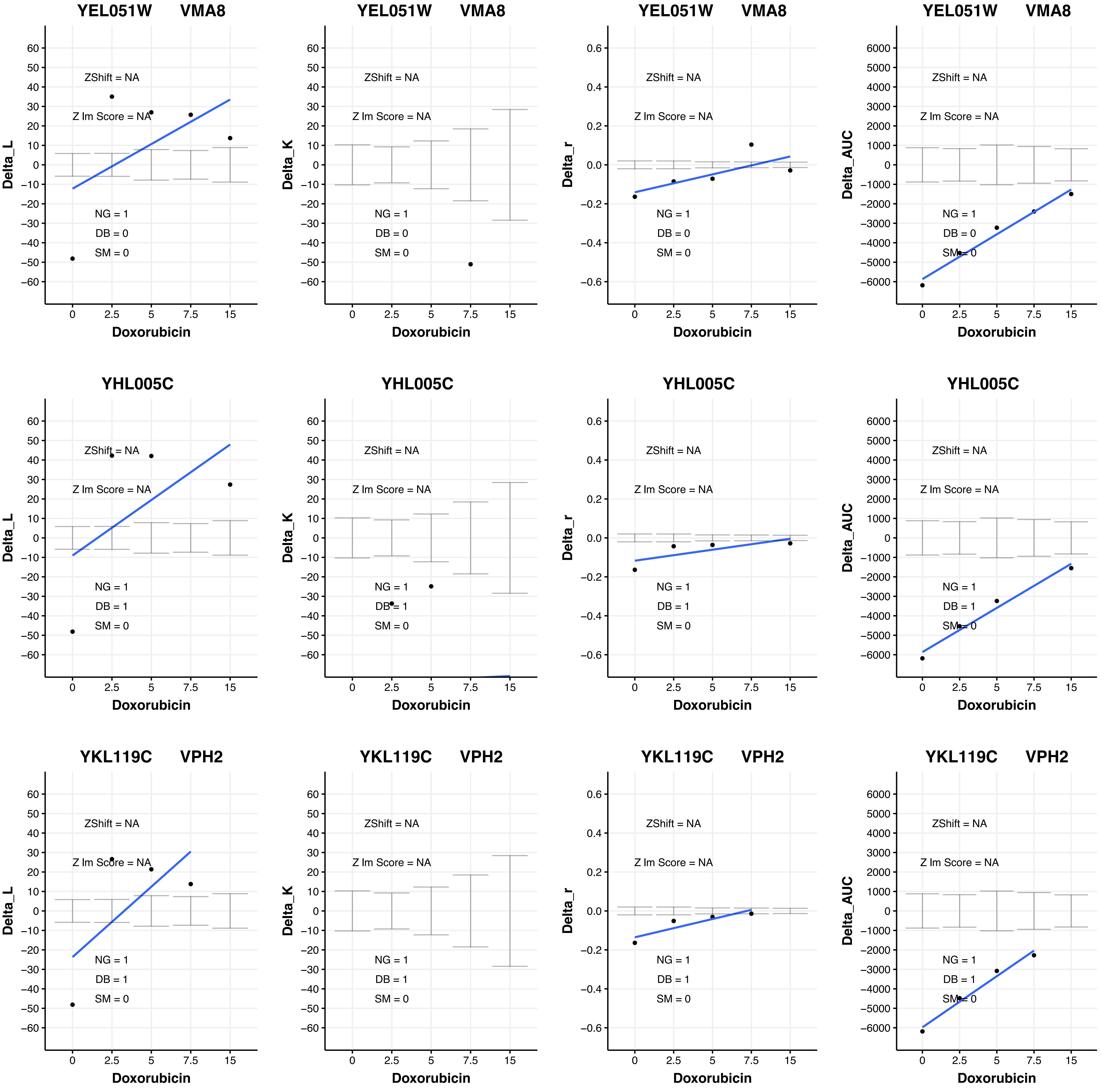
YEL051W VMA8
ZShift = NA
Z lm Score = NA
NG = 1
DB = 0
SM = 0
Delta_L
Doxorubicin
YEL051W VMA8
ZShift = NA
Z lm Score = NA
NG = 1
DB = 0
SM = 0
Delta_K
Doxorubicin
YEL051W VMA8
ZShift = NA
Z lm Score = NA
NG = 1
DB = 0
SM = 0
Delta_r
Doxorubicin
YEL051W VMA8
ZShift = NA
Z lm Score = NA
NG = 1
DB = 0
SM = 0
Delta_AUC
Doxorubicin
YHL005C
ZShift = NA
Z lm Score = NA
NG = 1
DB = 1
SM = 0
Delta_L
Doxorubicin
YHL005C
ZShift = NA
Z lm Score = NA
NG = 1
DB = 1
SM = 0
Delta_K
Doxorubicin
YHL005C
ZShift = NA
Z lm Score = NA
NG = 1
DB = 1
SM = 0
Delta_r
Doxorubicin
YHL005C
ZShift = NA
Z lm Score = NA
NG = 1
DB = 1
SM = 0
Delta_AUC
Doxorubicin
YKL119C VPH2
ZShift = NA
Z lm Score = NA
NG = 1
DB = 1
SM = 0
Delta_L
Doxorubicin
YKL119C VPH2
ZShift = NA
Z lm Score = NA
NG = 1
DB = 1
SM = 0
Delta_K
Doxorubicin
YKL119C VPH2
ZShift = NA
Z lm Score = NA
NG = 1
DB = 1
SM = 0
Delta_r
Doxorubicin
YKL119C VPH2
ZShift = NA
Z lm Score = NA
NG = 1
DB = 1
SM = 0
Delta_AUC
Doxorubicin

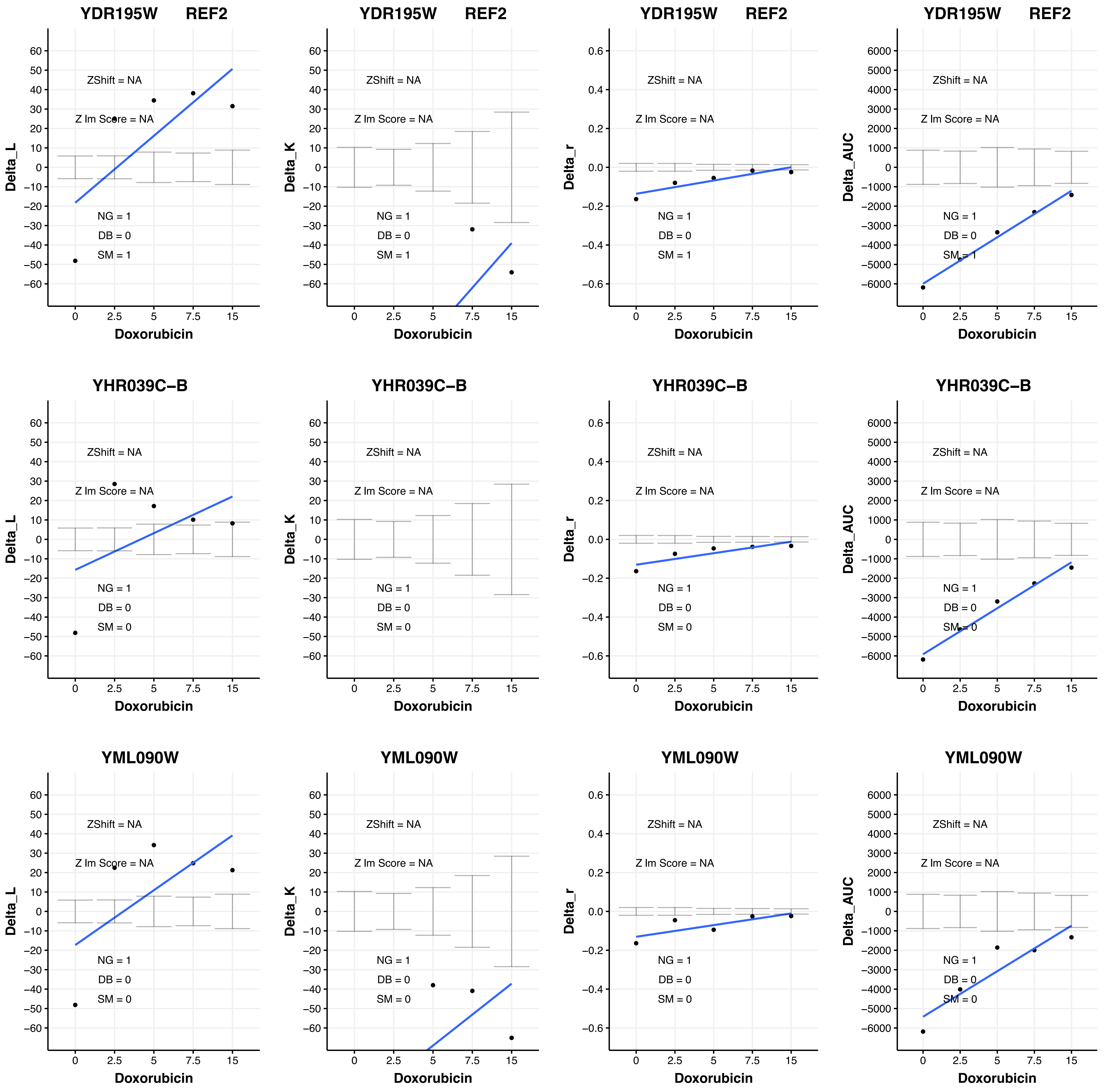

YDR195W REF2
ZShift = NA
Z lm Score = NA
NG = 1
DB = 0
SM = 1
Delta_L
Doxorubicin
YDR195W REF2
ZShift = NA
Z lm Score = NA
NG = 1
DB = 0
SM = 1
Delta_K
Doxorubicin
YDR195W REF2
ZShift = NA
Z lm Score = NA
NG = 1
DB = 0
SM = 1
Delta_r
Doxorubicin
YDR195W REF2
ZShift = NA
Z lm Score = NA
NG = 1
DB = 0
SM = 1
Delta_AUC
Doxorubicin
YHR039C−B
ZShift = NA
Z lm Score = NA
NG = 1
DB = 0
SM = 0
Delta_L
Doxorubicin
YHR039C−B
ZShift = NA
Z lm Score = NA
NG = 1
DB = 0
SM = 0
Delta_K
Doxorubicin
YHR039C−B
ZShift = NA
Z lm Score = NA
NG = 1
DB = 0
SM = 0
Delta_r
Doxorubicin
YHR039C−B
ZShift = NA
Z lm Score = NA
NG = 1
DB = 0
SM = 0
Delta_AUC
Doxorubicin
YML090W
ZShift = NA
Z lm Score = NA
NG = 1
DB = 0
SM = 0
Delta_L
Doxorubicin
YML090W
ZShift = NA
Z lm Score = NA
NG = 1
DB = 0
SM = 0
Delta_K
Doxorubicin
YML090W
ZShift = NA
Z lm Score = NA
NG = 1
DB = 0
SM = 0
Delta_r
Doxorubicin
YML090W
ZShift = NA
Z lm Score = NA
NG = 1
DB = 0
SM = 0
Delta_AUC
Doxorubicin

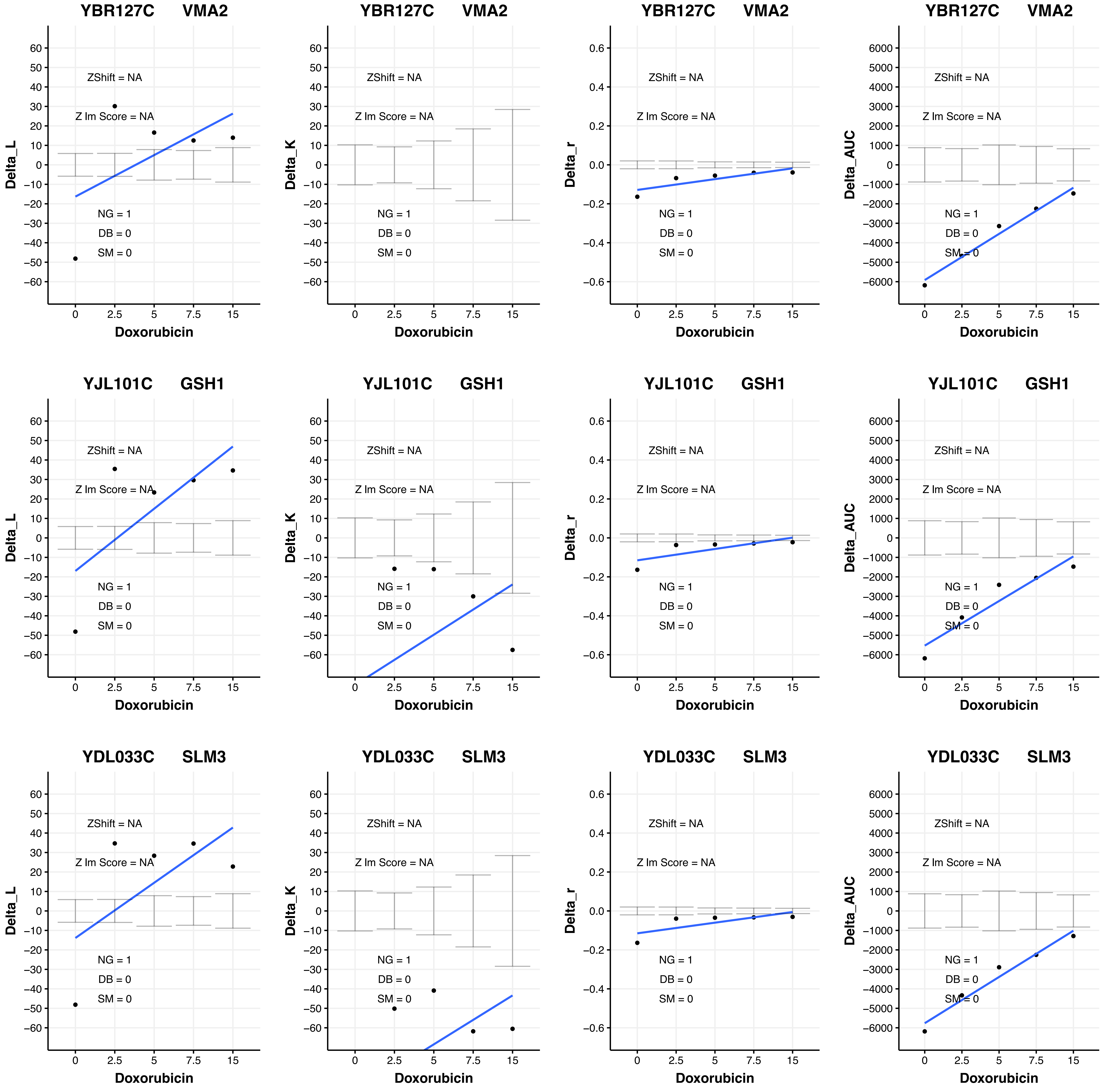

YBR127C VMA2
ZShift = NA
Z lm Score = NA
NG = 1
DB = 0
SM = 0
Delta_L
Delta_K
Delta_r
Delta_AUC
Doxorubicin
YJL101C GSH1
ZShift = NA
Z lm Score = NA
NG = 1
DB = 0
SM = 0
YDL033C SLM3
ZShift = NA
Z lm Score = NA
NG = 1
DB = 0
SM = 0

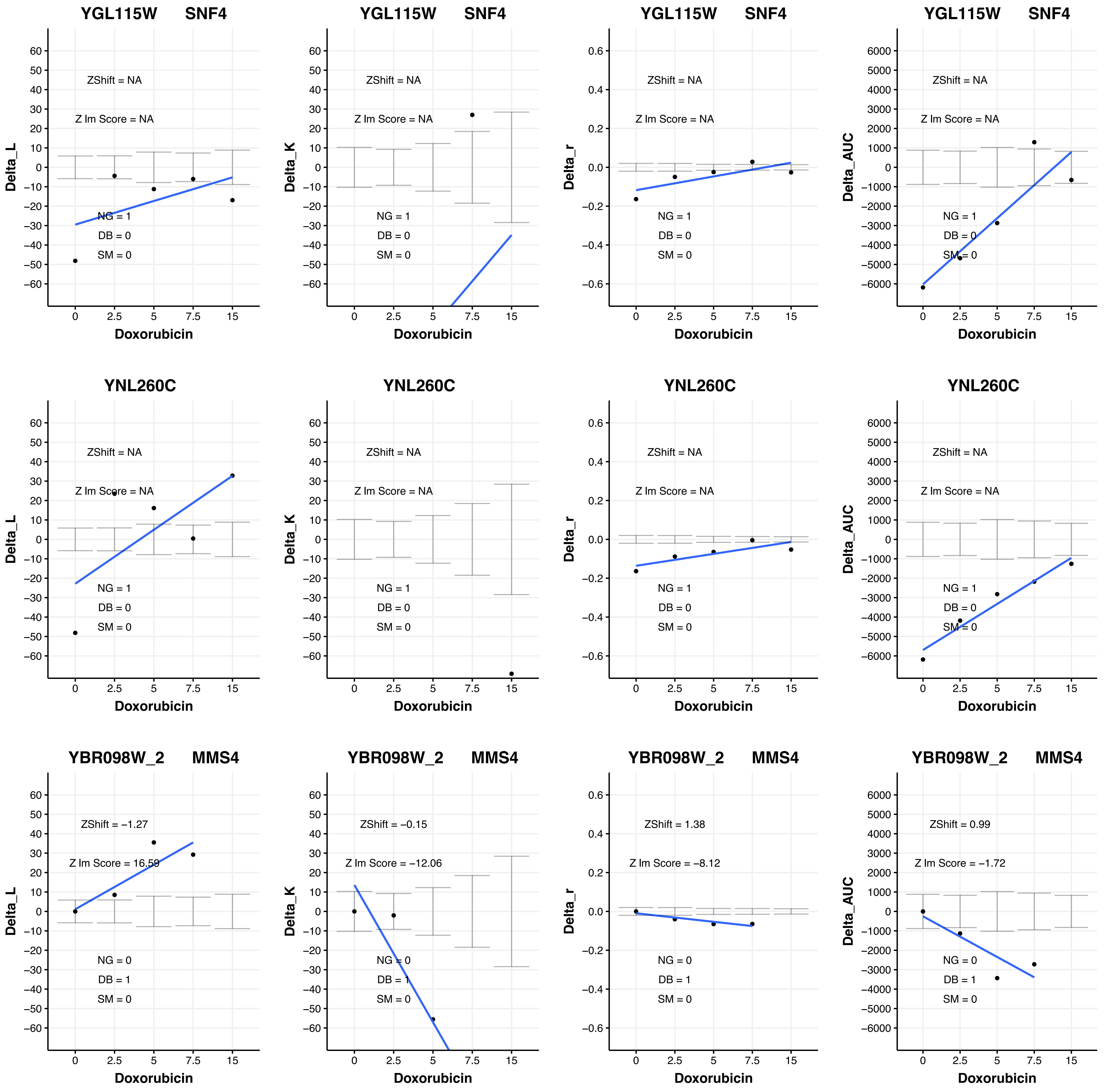

YGL115W SNF4
ZShift = NA
Z lm Score = NA
NG = 1
DB = 0
SM = 0
Delta_L
Delta_K
Delta_r
Delta_AUC
Doxorubicin
YNL260C
ZShift = NA
Z lm Score = NA
NG = 1
DB = 0
SM = 0
YBR098W_2 MMS4
ZShift = −1.27
Z lm Score = 16.59
ZShift = −0.15
Z lm Score = −12.06
ZShift = 1.38
Z lm Score = −8.12
ZShift = 0.99
Z lm Score = −1.72
NG = 0
DB = 1
SM = 0

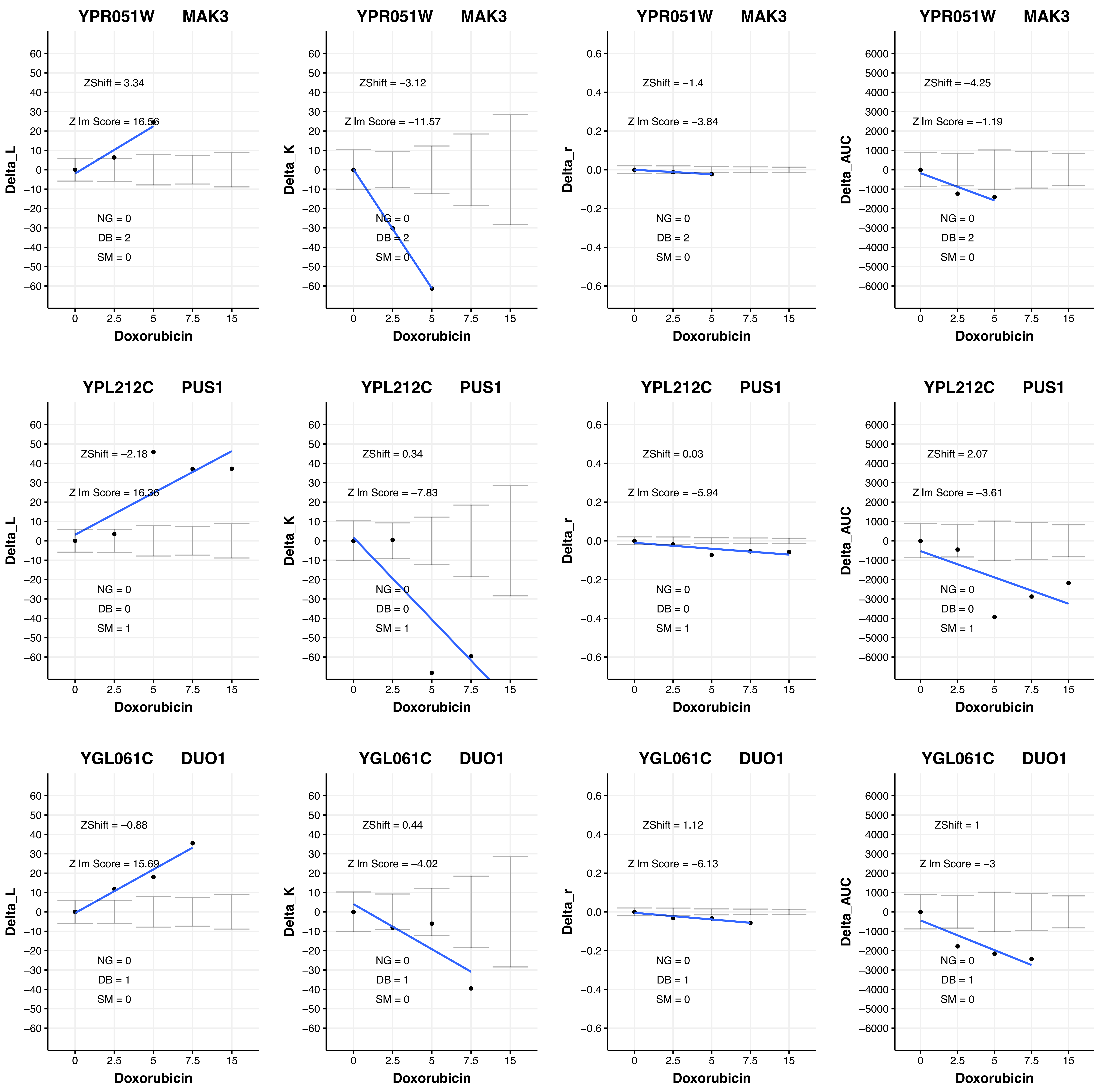
YPR051W MAK3
ZShift = 3.34
Z lm Score = 16.56
NG = 0
DB = 2
SM = 0
Delta_L
Doxorubicin
YPR051W MAK3
ZShift = −3.12
Z lm Score = −11.57
NG = 0
DB = 2
SM = 0
Delta_K
Doxorubicin
YPR051W MAK3
ZShift = −1.4
Z lm Score = −3.84
NG = 0
DB = 2
SM = 0
Delta_r
Doxorubicin
YPR051W MAK3
ZShift = −4.25
Z lm Score = −1.19
NG = 0
DB = 2
SM = 0
Delta_AUC
Doxorubicin
YPL212C PUS1
ZShift = −2.18
Z lm Score = 16.36
NG = 0
DB = 0
SM = 1
Delta_L
Doxorubicin
YPL212C PUS1
ZShift = 0.34
Z lm Score = −7.83
NG = 0
DB = 0
SM = 1
Delta_K
Doxorubicin
YPL212C PUS1
ZShift = 0.03
Z lm Score = −5.94
NG = 0
DB = 0
SM = 1
Delta_r
Doxorubicin
YPL212C PUS1
ZShift = 2.07
Z lm Score = −3.61
NG = 0
DB = 0
SM = 1
Delta_AUC
Doxorubicin
YGL061C DUO1
ZShift = −0.88
Z lm Score = 15.69
NG = 0
DB = 1
SM = 0
Delta_L
Doxorubicin
YGL061C DUO1
ZShift = 0.44
Z lm Score = −4.02
NG = 0
DB = 1
SM = 0
Delta_K
Doxorubicin
YGL061C DUO1
ZShift = 1.12
Z lm Score = −6.13
NG = 0
DB = 1
SM = 0
Delta_r
Doxorubicin
YGL061C DUO1
ZShift = 1
Z lm Score = −3
NG = 0
DB = 1
SM = 0
Delta_AUC
Doxorubicin

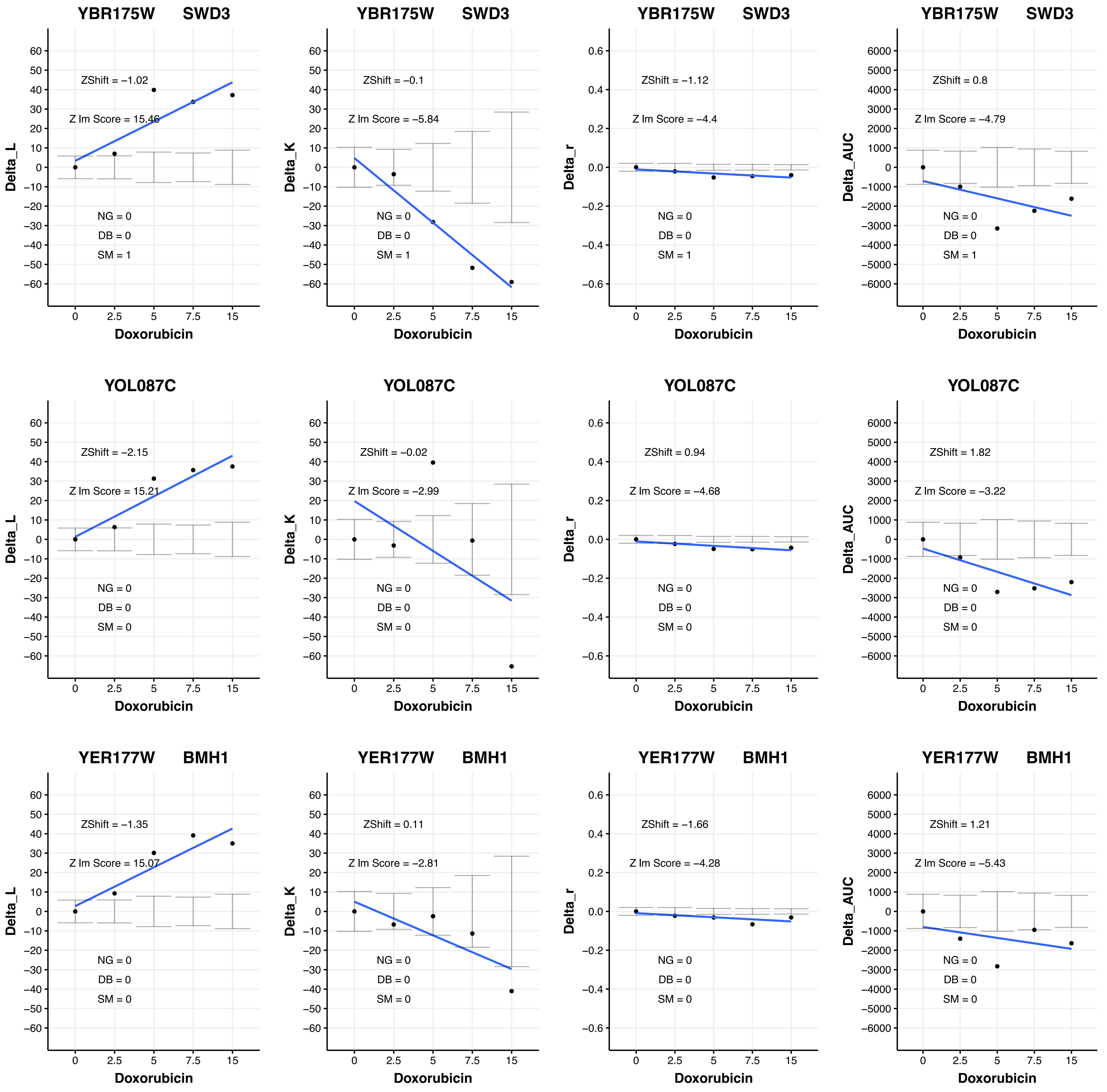

YBR175W SWD3
ZShift = −1.02
Z lm Score = 15.46
NG = 0
DB = 0
SM = 1
Delta_L
Doxorubicin
YBR175W SWD3
ZShift = −0.1
Z lm Score = −5.84
NG = 0
DB = 0
SM = 1
Delta_K
Doxorubicin
YBR175W SWD3
ZShift = −1.12
Z lm Score = −4.4
NG = 0
DB = 0
SM = 1
Delta_r
Doxorubicin
YBR175W SWD3
ZShift = 0.8
Z lm Score = −4.79
NG = 0
DB = 0
SM = 1
Delta_AUC
Doxorubicin
YOL087C
ZShift = −2.15
Z lm Score = 15.21
NG = 0
DB = 0
SM = 0
Delta_L
Doxorubicin
YOL087C
ZShift = −0.02
Z lm Score = −2.99
NG = 0
DB = 0
SM = 0
Delta_K
Doxorubicin
YOL087C
ZShift = 0.94
Z lm Score = −4.68
NG = 0
DB = 0
SM = 0
Delta_r
Doxorubicin
YOL087C
ZShift = 1.82
Z lm Score = −3.22
NG = 0
DB = 0
SM = 0
Delta_AUC
Doxorubicin
YER177W BMH1
ZShift = −1.35
Z lm Score = 15.07
NG = 0
DB = 0
SM = 0
Delta_L
Doxorubicin
YER177W BMH1
ZShift = 0.11
Z lm Score = −2.81
NG = 0
DB = 0
SM = 0
Delta_K
Doxorubicin
YER177W BMH1
ZShift = −1.66
Z lm Score = −4.28
NG = 0
DB = 0
SM = 0
Delta_r
Doxorubicin
YER177W BMH1
ZShift = 1.21
Z lm Score = −5.43
NG = 0
DB = 0
SM = 0
Delta_AUC
Doxorubicin

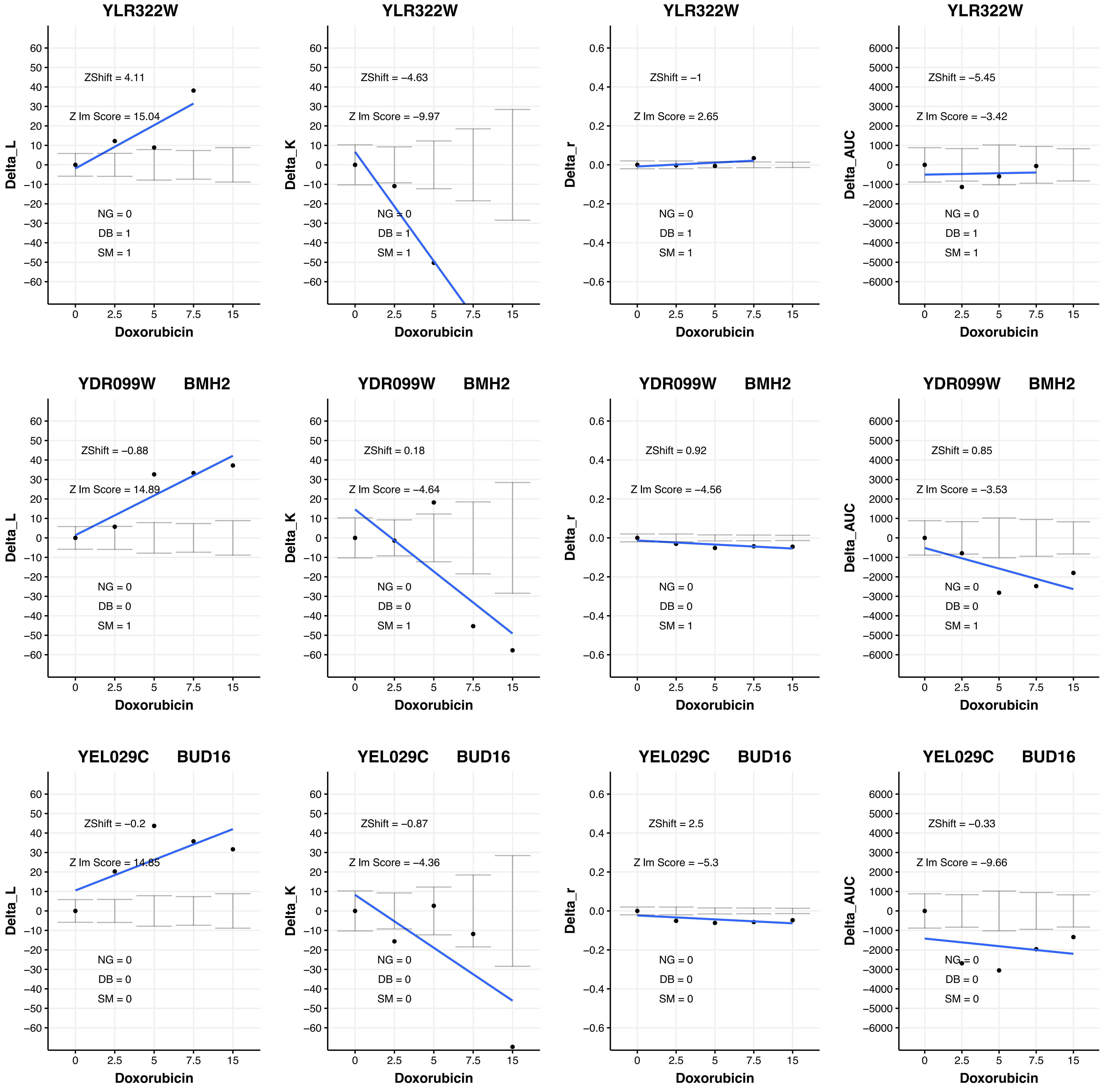

YLR322W
ZShift = 4.11
Z lm Score = 15.04
NG = 0
DB = 1
SM = 1
Delta_L
Doxorubicin
YLR322W
ZShift = −4.63
Z lm Score = −9.97
NG = 0
DB = 1
SM = 1
Delta_K
Doxorubicin
YLR322W
ZShift = −1
Z lm Score = 2.65
NG = 0
DB = 1
SM = 1
Delta_r
Doxorubicin
YLR322W
ZShift = −5.45
Z lm Score = −3.42
NG = 0
DB = 1
SM = 1
Delta_AUC
Doxorubicin
YDR099W BMH2
ZShift = −0.88
Z lm Score = 14.89
NG = 0
DB = 0
SM = 1
Delta_L
Doxorubicin
YDR099W BMH2
ZShift = 0.18
Z lm Score = −4.64
NG = 0
DB = 0
SM = 1
Delta_K
Doxorubicin
YDR099W BMH2
ZShift = 0.92
Z lm Score = −4.56
NG = 0
DB = 0
SM = 1
Delta_r
Doxorubicin
YDR099W BMH2
ZShift = 0.85
Z lm Score = −3.53
NG = 0
DB = 0
SM = 1
Delta_AUC
Doxorubicin
YEL029C BUD16
ZShift = −0.2
Z lm Score = 14.85
NG = 0
DB = 0
SM = 0
Delta_L
Doxorubicin
YEL029C BUD16
ZShift = −0.87
Z lm Score = −4.36
NG = 0
DB = 0
SM = 0
Delta_K
Doxorubicin
YEL029C BUD16
ZShift = 2.5
Z lm Score = −5.3
NG = 0
DB = 0
SM = 0
Delta_r
Doxorubicin
YEL029C BUD16
ZShift = −0.33
Z lm Score = −9.66
NG = 0
DB = 0
SM = 0
Delta_AUC
Doxorubicin

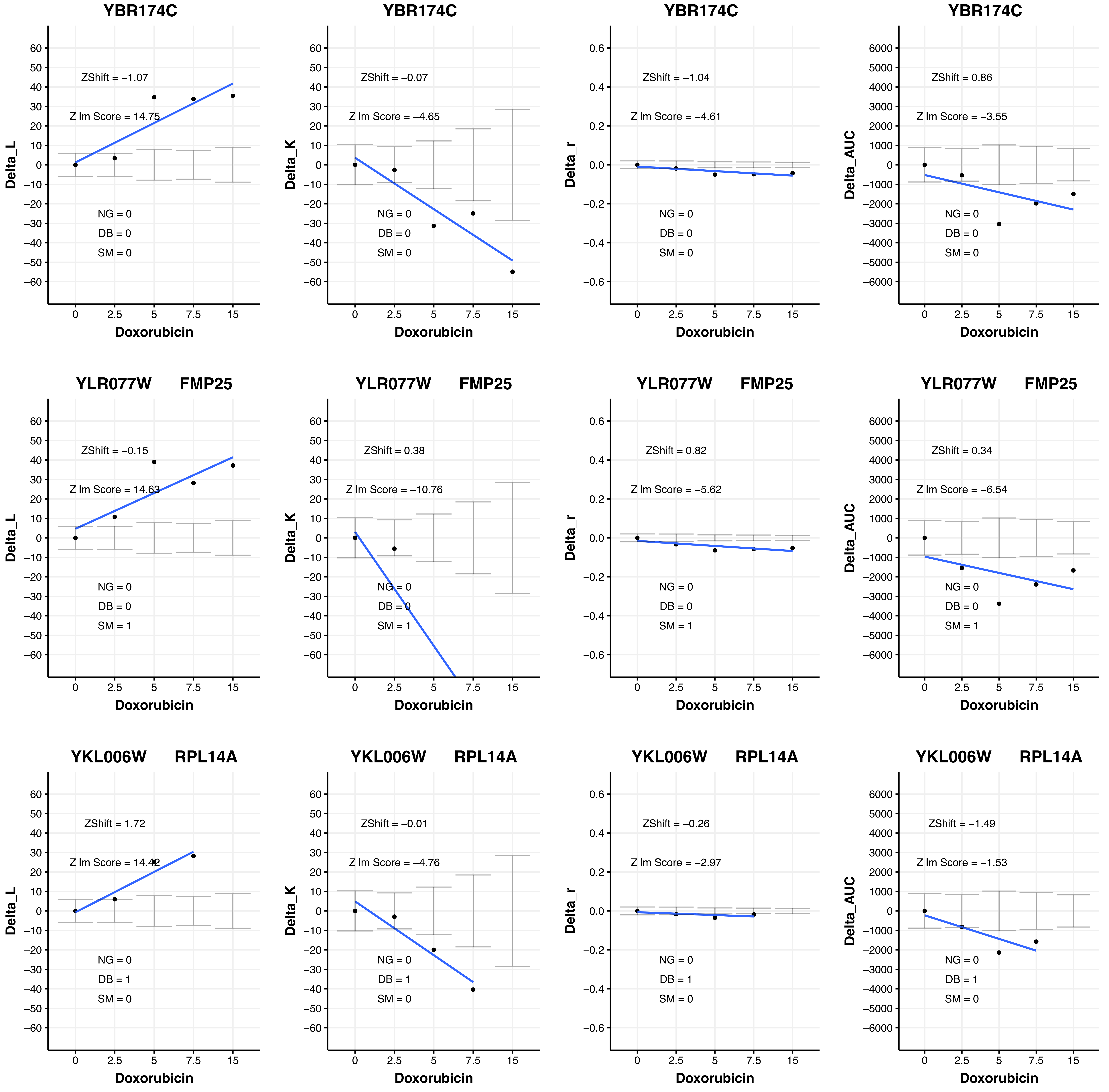
YBR174C
ZShift = −1.07
Z lm Score = 14.75
NG = 0
DB = 0
SM = 0
Delta_L
Doxorubicin
YBR174C
ZShift = −0.07
Z lm Score = −4.65
NG = 0
DB = 0
SM = 0
Delta_K
Doxorubicin
YBR174C
ZShift = −1.04
Z lm Score = −4.61
NG = 0
DB = 0
SM = 0
Delta_r
Doxorubicin
YBR174C
ZShift = 0.86
Z lm Score = −3.55
NG = 0
DB = 0
SM = 0
Delta_AUC
Doxorubicin
YLR077W FMP25
ZShift = −0.15
Z lm Score = 14.63
NG = 0
DB = 0
SM = 1
Delta_L
Doxorubicin
YLR077W FMP25
ZShift = 0.38
Z lm Score = −10.76
NG = 0
DB = 0
SM = 1
Delta_K
Doxorubicin
YLR077W FMP25
ZShift = 0.82
Z lm Score = −5.62
NG = 0
DB = 0
SM = 1
Delta_r
Doxorubicin
YLR077W FMP25
ZShift = 0.34
Z lm Score = −6.54
NG = 0
DB = 0
SM = 1
Delta_AUC
Doxorubicin
YKL006W RPL14A
ZShift = 1.72
Z lm Score = 14.42
NG = 0
DB = 1
SM = 0
Delta_L
Doxorubicin
YKL006W RPL14A
ZShift = −0.01
Z lm Score = −4.76
NG = 0
DB = 1
SM = 0
Delta_K
Doxorubicin
YKL006W RPL14A
ZShift = −0.26
Z lm Score = −2.97
NG = 0
DB = 1
SM = 0
Delta_r
Doxorubicin
YKL006W RPL14A
ZShift = −1.49
Z lm Score = −1.53
NG = 0
DB = 1
SM = 0
Delta_AUC
Doxorubicin

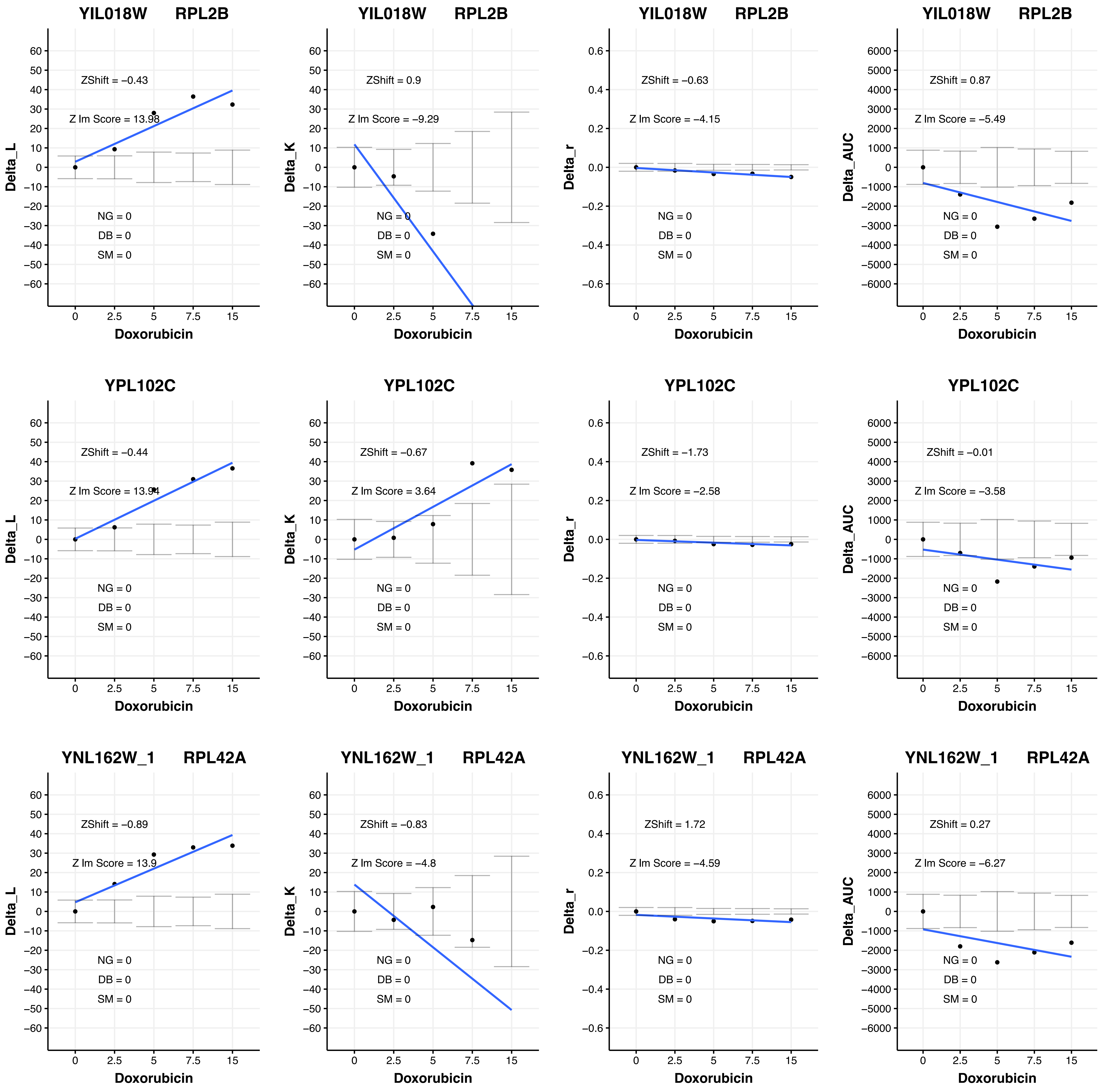

YIL018W RPL2B
ZShift = −0.43
Z lm Score = 13.98
NG = 0
DB = 0
SM = 0
Delta_L
Doxorubicin
YIL018W RPL2B
ZShift = 0.9
Z lm Score = −9.29
NG = 0
DB = 0
SM = 0
Delta_K
Doxorubicin
YIL018W RPL2B
ZShift = −0.63
Z lm Score = −4.15
NG = 0
DB = 0
SM = 0
Delta_r
Doxorubicin
YIL018W RPL2B
ZShift = 0.87
Z lm Score = −5.49
NG = 0
DB = 0
SM = 0
Delta_AUC
Doxorubicin
YPL102C
ZShift = −0.44
Z lm Score = 13.94
NG = 0
DB = 0
SM = 0
Delta_L
Doxorubicin
YPL102C
ZShift = −0.67
Z lm Score = 3.64
NG = 0
DB = 0
SM = 0
Delta_K
Doxorubicin
YPL102C
ZShift = −1.73
Z lm Score = −2.58
NG = 0
DB = 0
SM = 0
Delta_r
Doxorubicin
YPL102C
ZShift = −0.01
Z lm Score = −3.58
NG = 0
DB = 0
SM = 0
Delta_AUC
Doxorubicin
YNL162W_1 RPL42A
ZShift = −0.89
Z lm Score = 13.9
NG = 0
DB = 0
SM = 0
Delta_L
Doxorubicin
YNL162W_1 RPL42A
ZShift = −0.83
Z lm Score = −4.8
NG = 0
DB = 0
SM = 0
Delta_K
Doxorubicin
YNL162W_1 RPL42A
ZShift = 1.72
Z lm Score = −4.59
NG = 0
DB = 0
SM = 0
Delta_r
Doxorubicin
YNL162W_1 RPL42A
ZShift = 0.27
Z lm Score = −6.27
NG = 0
DB = 0
SM = 0
Delta_AUC
Doxorubicin

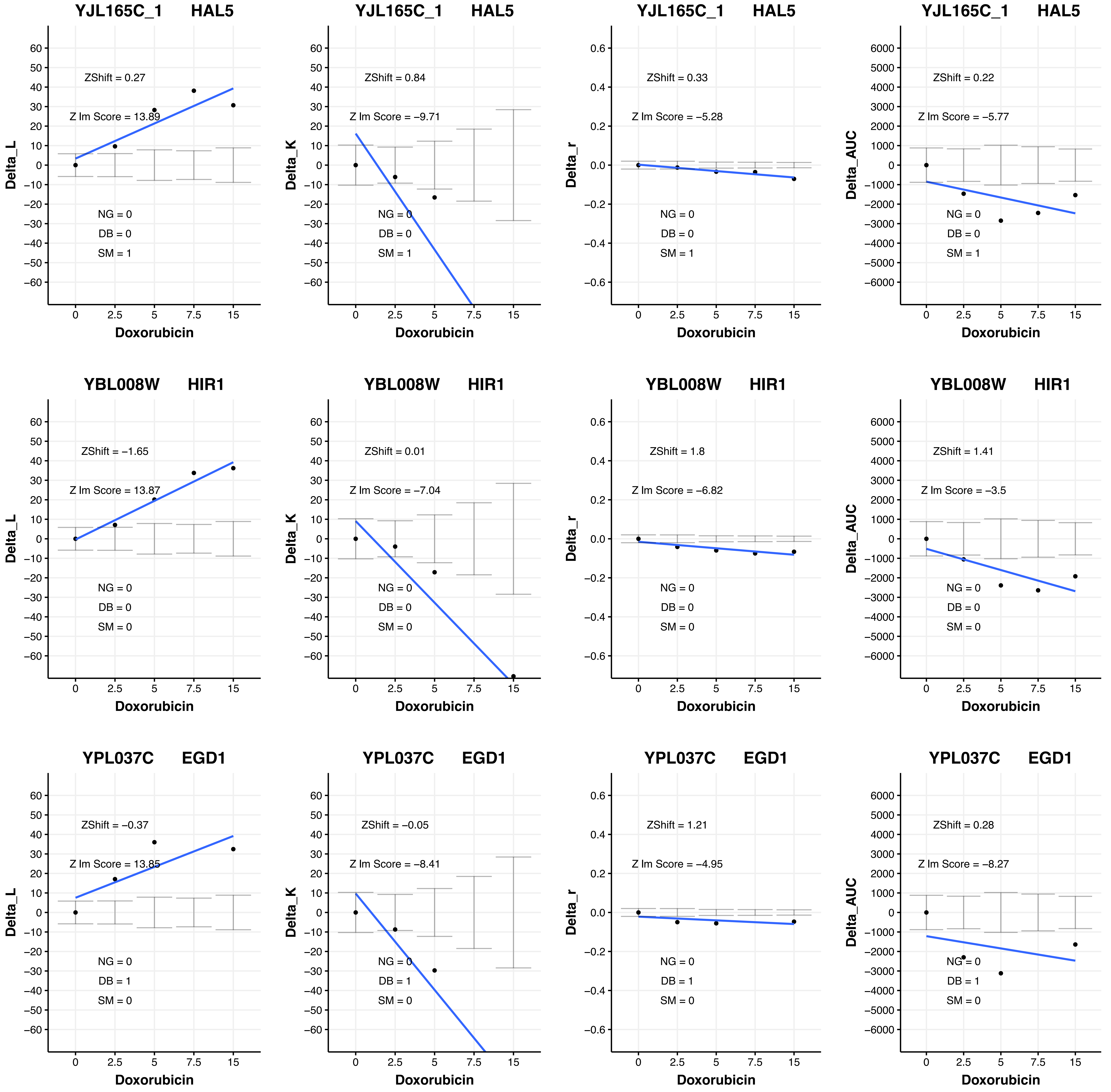
YJL165C_1 HAL5
ZShift = 0.27
Z lm Score = 13.89
NG = 0
DB = 0
SM = 1
Delta_L
Doxorubicin
YJL165C_1 HAL5
ZShift = 0.84
Z lm Score = −9.71
NG = 0
DB = 0
SM = 1
Delta_K
Doxorubicin
YJL165C_1 HAL5
ZShift = 0.33
Z lm Score = −5.28
NG = 0
DB = 0
SM = 1
Delta_r
Doxorubicin
YJL165C_1 HAL5
ZShift = 0.22
Z lm Score = −5.77
NG = 0
DB = 0
SM = 1
Delta_AUC
Doxorubicin
YBL008W HIR1
ZShift = −1.65
Z lm Score = 13.87
NG = 0
DB = 0
SM = 0
Delta_L
Doxorubicin
YBL008W HIR1
ZShift = 0.01
Z lm Score = −7.04
NG = 0
DB = 0
SM = 0
Delta_K
Doxorubicin
YBL008W HIR1
ZShift = 1.8
Z lm Score = −6.82
NG = 0
DB = 0
SM = 0
Delta_r
Doxorubicin
YBL008W HIR1
ZShift = 1.41
Z lm Score = −3.5
NG = 0
DB = 0
SM = 0
Delta_AUC
Doxorubicin
YPL037C EGD1
ZShift = −0.37
Z lm Score = 13.85
NG = 0
DB = 1
SM = 0
Delta_L
Doxorubicin
YPL037C EGD1
ZShift = −0.05
Z lm Score = −8.41
NG = 0
DB = 1
SM = 0
Delta_K
Doxorubicin
YPL037C EGD1
ZShift = 1.21
Z lm Score = −4.95
NG = 0
DB = 1
SM = 0
Delta_r
Doxorubicin
YPL037C EGD1
ZShift = 0.28
Z lm Score = −8.27
NG = 0
DB = 1
SM = 0
Delta_AUC
Doxorubicin

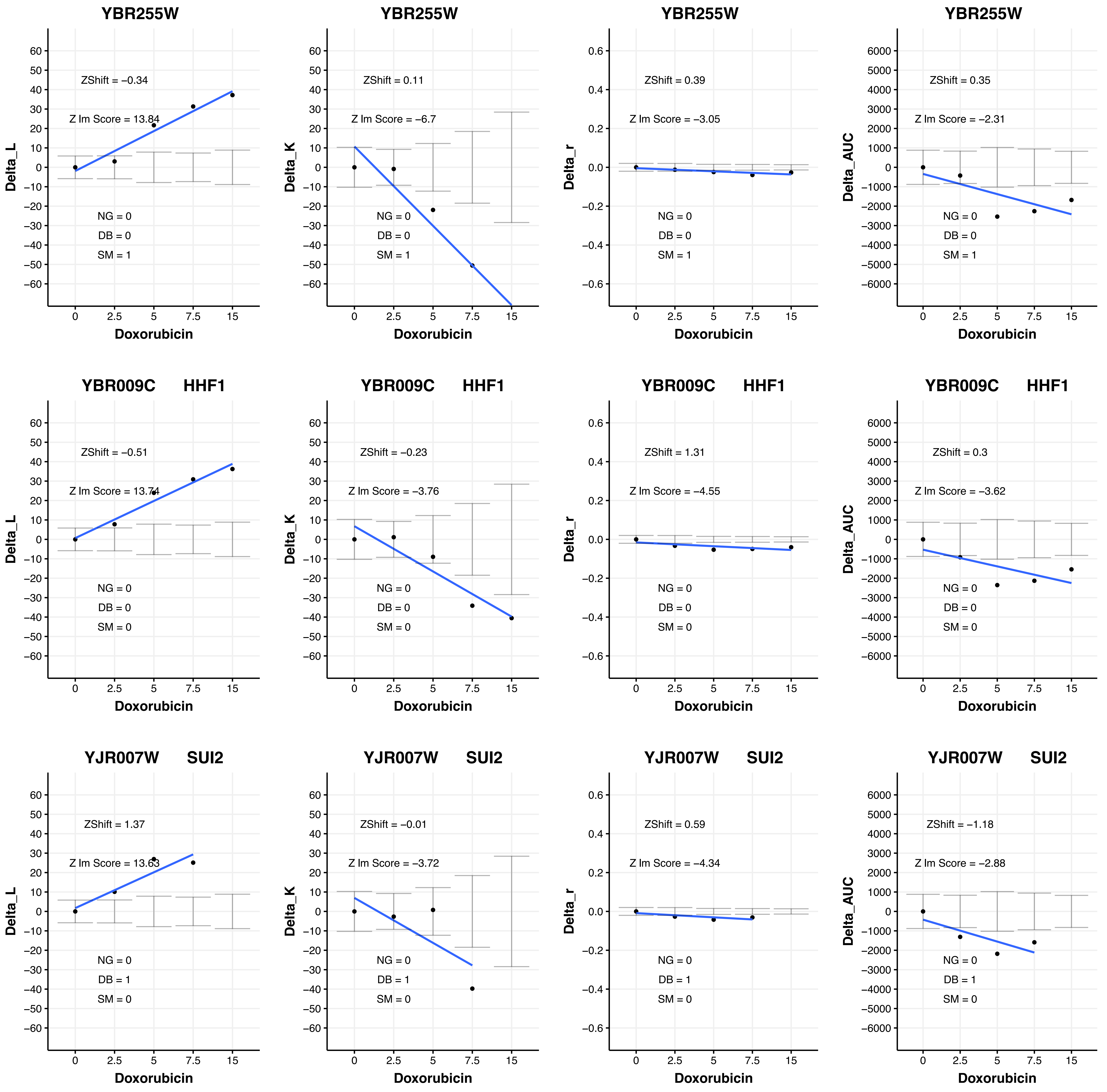
YBR255W
ZShift = −0.34
Z lm Score = 13.84
NG = 0
DB = 0
SM = 1
Delta_L
Doxorubicin
YBR255W
ZShift = 0.11
Z lm Score = −6.7
NG = 0
DB = 0
SM = 1
Delta_K
Doxorubicin
YBR255W
ZShift = 0.39
Z lm Score = −3.05
NG = 0
DB = 0
SM = 1
Delta_r
Doxorubicin
YBR255W
ZShift = 0.35
Z lm Score = −2.31
NG = 0
DB = 0
SM = 1
Delta_AUC
Doxorubicin
YBR009C HHF1
ZShift = −0.51
Z lm Score = 13.74
NG = 0
DB = 0
SM = 0
Delta_L
Doxorubicin
YBR009C HHF1
ZShift = −0.23
Z lm Score = −3.76
NG = 0
DB = 0
SM = 0
Delta_K
Doxorubicin
YBR009C HHF1
ZShift = 1.31
Z lm Score = −4.55
NG = 0
DB = 0
SM = 0
Delta_r
Doxorubicin
YBR009C HHF1
ZShift = 0.3
Z lm Score = −3.62
NG = 0
DB = 0
SM = 0
Delta_AUC
Doxorubicin
YJR007W SUI2
ZShift = 1.37
Z lm Score = 13.63
NG = 0
DB = 1
SM = 0
Delta_L
Doxorubicin
YJR007W SUI2
ZShift = −0.01
Z lm Score = −3.72
NG = 0
DB = 1
SM = 0
Delta_K
Doxorubicin
YJR007W SUI2
ZShift = 0.59
Z lm Score = −4.34
NG = 0
DB = 1
SM = 0
Delta_r
Doxorubicin
YJR007W SUI2
ZShift = −1.18
Z lm Score = −2.88
NG = 0
DB = 1
SM = 0
Delta_AUC
Doxorubicin

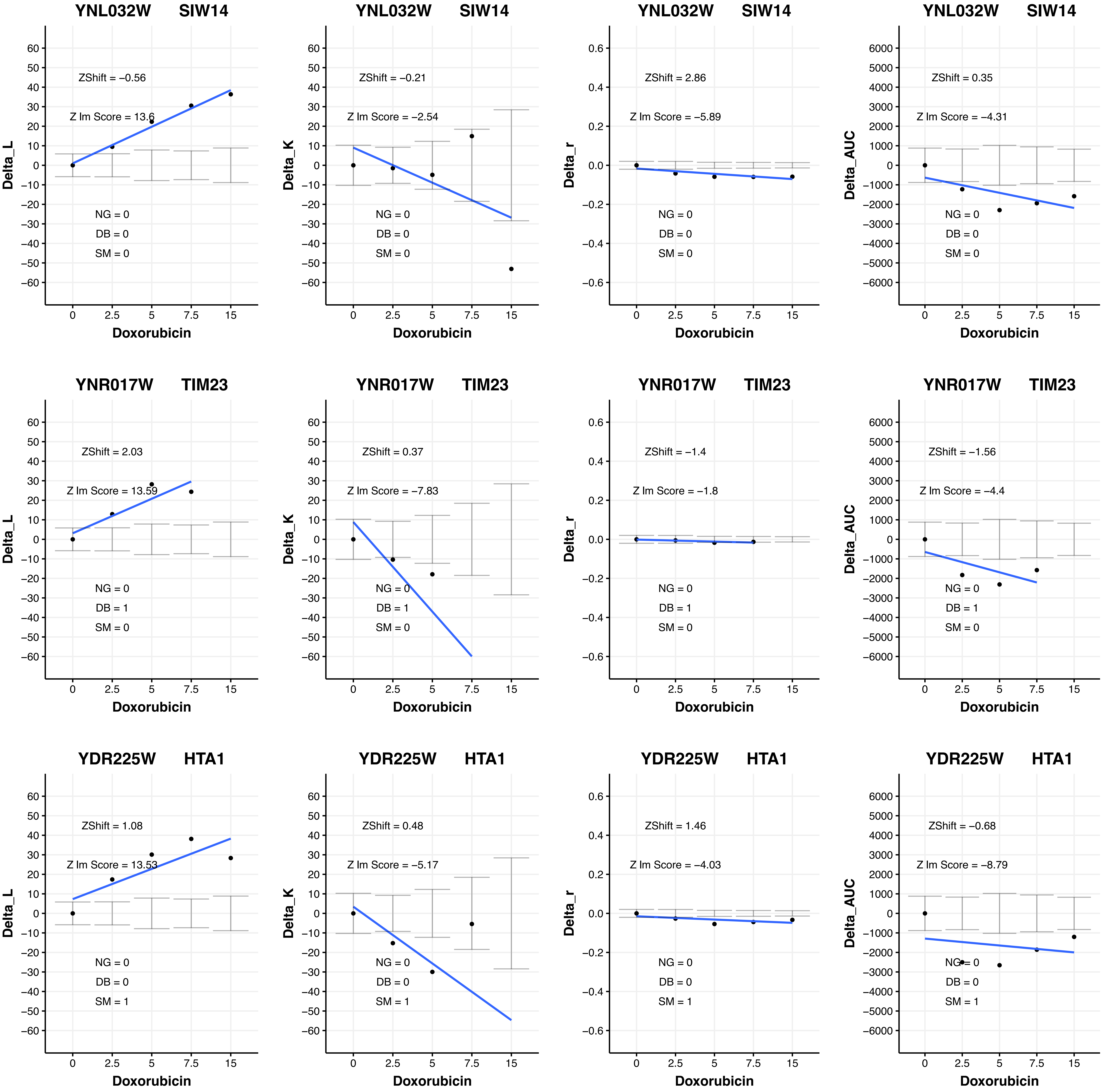

YNL032W SIW14
ZShift = −0.56
Z lm Score = 13.6
NG = 0
DB = 0
SM = 0
Delta_L
Doxorubicin
YNL032W SIW14
ZShift = −0.21
Z lm Score = −2.54
NG = 0
DB = 0
SM = 0
Delta_K
Doxorubicin
YNL032W SIW14
ZShift = 2.86
Z lm Score = −5.89
NG = 0
DB = 0
SM = 0
Delta_r
Doxorubicin
YNL032W SIW14
ZShift = 0.35
Z lm Score = −4.31
NG = 0
DB = 0
SM = 0
Delta_AUC
Doxorubicin
YNR017W TIM23
ZShift = 2.03
Z lm Score = 13.59
NG = 0
DB = 1
SM = 0
Delta_L
Doxorubicin
YNR017W TIM23
ZShift = 0.37
Z lm Score = −7.83
NG = 0
DB = 1
SM = 0
Delta_K
Doxorubicin
YNR017W TIM23
ZShift = −1.4
Z lm Score = −1.8
NG = 0
DB = 1
SM = 0
Delta_r
Doxorubicin
YNR017W TIM23
ZShift = −1.56
Z lm Score = −4.4
NG = 0
DB = 1
SM = 0
Delta_AUC
Doxorubicin
YDR225W HTA1
ZShift = 1.08
Z lm Score = 13.53
NG = 0
DB = 0
SM = 1
Delta_L
Doxorubicin
YDR225W HTA1
ZShift = 0.48
Z lm Score = −5.17
NG = 0
DB = 0
SM = 1
Delta_K
Doxorubicin
YDR225W HTA1
ZShift = 1.46
Z lm Score = −4.03
NG = 0
DB = 0
SM = 1
Delta_r
Doxorubicin
YDR225W HTA1
ZShift = −0.68
Z lm Score = −8.79
NG = 0
DB = 0
SM = 1
Delta_AUC
Doxorubicin

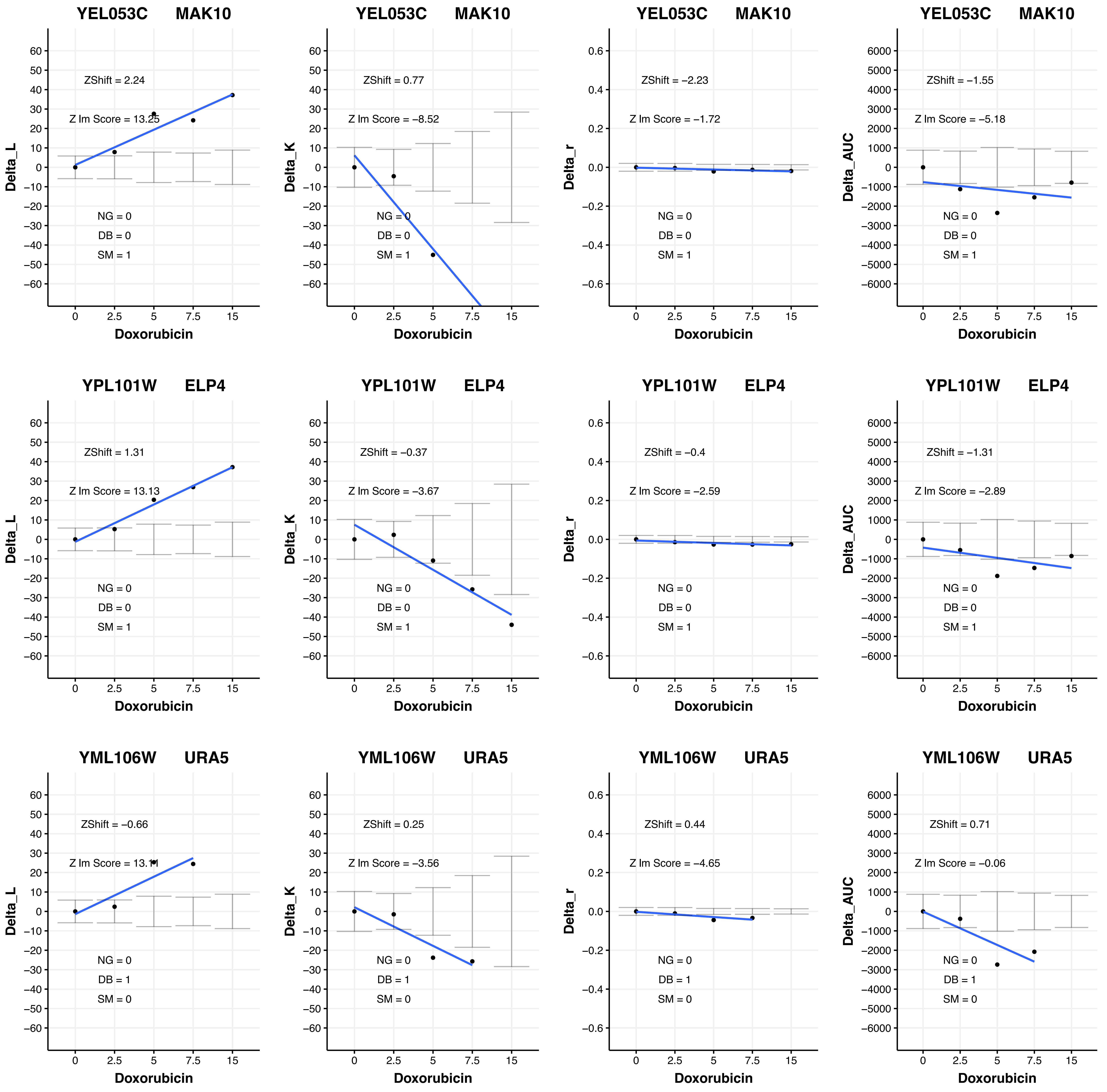
YEL053C MAK10
ZShift = 2.24
Z lm Score = 13.25
NG = 0
DB = 0
SM = 1
Delta_L
Doxorubicin
YEL053C MAK10
ZShift = 0.77
Z lm Score = −8.52
NG = 0
DB = 0
SM = 1
Delta_K
Doxorubicin
YEL053C MAK10
ZShift = −2.23
Z lm Score = −1.72
NG = 0
DB = 0
SM = 1
Delta_r
Doxorubicin
YEL053C MAK10
ZShift = −1.55
Z lm Score = −5.18
NG = 0
DB = 0
SM = 1
Delta_AUC
Doxorubicin
YPL101W ELP4
ZShift = 1.31
Z lm Score = 13.13
NG = 0
DB = 0
SM = 1
Delta_L
Doxorubicin
YPL101W ELP4
ZShift = −0.37
Z lm Score = −3.67
NG = 0
DB = 0
SM = 1
Delta_K
Doxorubicin
YPL101W ELP4
ZShift = −0.4
Z lm Score = −2.59
NG = 0
DB = 0
SM = 1
Delta_r
Doxorubicin
YPL101W ELP4
ZShift = −1.31
Z lm Score = −2.89
NG = 0
DB = 0
SM = 1
Delta_AUC
Doxorubicin
YML106W URA5
ZShift = −0.66
Z lm Score = 13.11
NG = 0
DB = 1
SM = 0
Delta_L
Doxorubicin
YML106W URA5
ZShift = 0.25
Z lm Score = −3.56
NG = 0
DB = 1
SM = 0
Delta_K
Doxorubicin
YML106W URA5
ZShift = 0.44
Z lm Score = −4.65
NG = 0
DB = 1
SM = 0
Delta_r
Doxorubicin
YML106W URA5
ZShift = 0.71
Z lm Score = −0.06
NG = 0
DB = 1
SM = 0
Delta_AUC
Doxorubicin

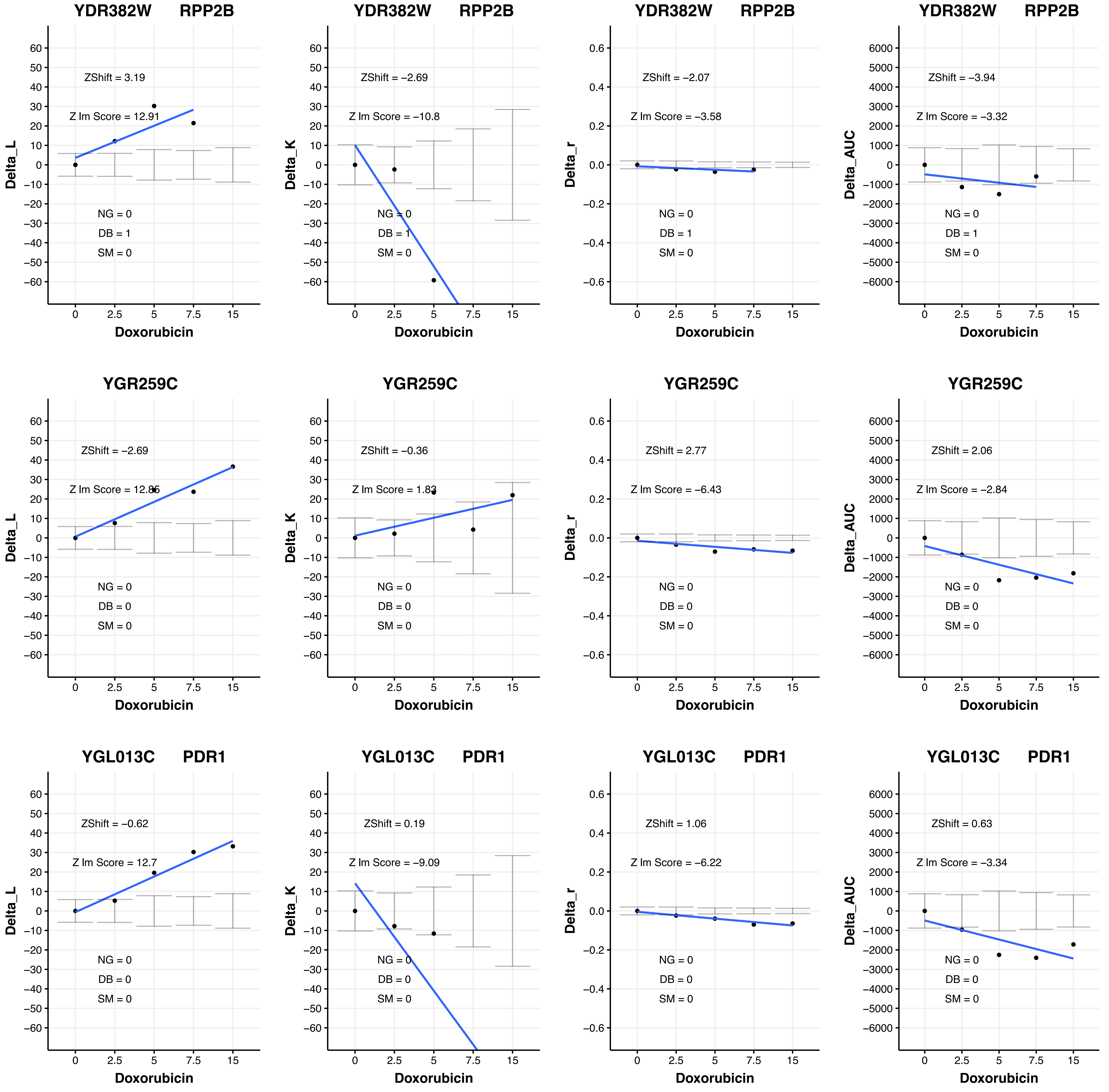

YDR382W RPP2B
ZShift = 3.19
Z lm Score = 12.91
NG = 0
DB = 1
SM = 0
Delta_L
Doxorubicin
YDR382W RPP2B
ZShift = −2.69
Z lm Score = −10.8
NG = 0
DB = 1
SM = 0
Delta_K
Doxorubicin
YDR382W RPP2B
ZShift = −2.07
Z lm Score = −3.58
NG = 0
DB = 1
SM = 0
Delta_r
Doxorubicin
YDR382W RPP2B
ZShift = −3.94
Z lm Score = −3.32
NG = 0
DB = 1
SM = 0
Delta_AUC
Doxorubicin
YGR259C
ZShift = −2.69
Z lm Score = 12.85
NG = 0
DB = 0
SM = 0
Delta_L
Doxorubicin
YGR259C
ZShift = −0.36
Z lm Score = 1.83
NG = 0
DB = 0
SM = 0
Delta_K
Doxorubicin
YGR259C
ZShift = 2.77
Z lm Score = −6.43
NG = 0
DB = 0
SM = 0
Delta_r
Doxorubicin
YGR259C
ZShift = 2.06
Z lm Score = −2.84
NG = 0
DB = 0
SM = 0
Delta_AUC
Doxorubicin
YGL013C PDR1
ZShift = −0.62
Z lm Score = 12.7
NG = 0
DB = 0
SM = 0
Delta_L
Doxorubicin
YGL013C PDR1
ZShift = 0.19
Z lm Score = −9.09
NG = 0
DB = 0
SM = 0
Delta_K
Doxorubicin
YGL013C PDR1
ZShift = 1.06
Z lm Score = −6.22
NG = 0
DB = 0
SM = 0
Delta_r
Doxorubicin
YGL013C PDR1
ZShift = 0.63
Z lm Score = −3.34
NG = 0
DB = 0
SM = 0
Delta_AUC
Doxorubicin

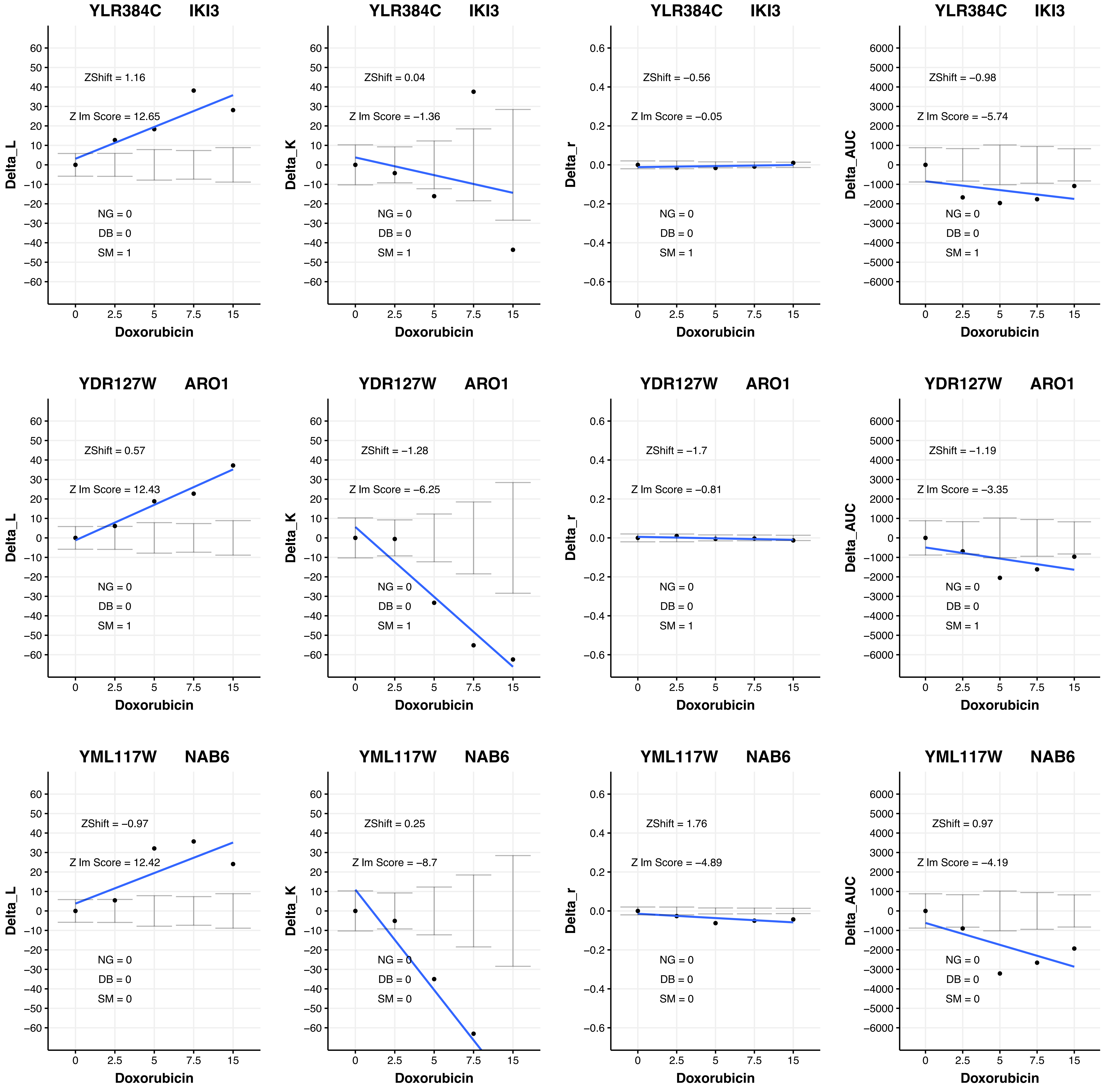

YLR384C IKI3
ZShift = 1.16
Z lm Score = 12.65
NG = 0
DB = 0
SM = 1
Delta_L
Doxorubicin
YLR384C IKI3
ZShift = 0.04
Z lm Score = −1.36
NG = 0
DB = 0
SM = 1
Delta_K
Doxorubicin
YLR384C IKI3
ZShift = −0.56
Z lm Score = −0.05
NG = 0
DB = 0
SM = 1
Delta_r
Doxorubicin
YLR384C IKI3
ZShift = −0.98
Z lm Score = −5.74
NG = 0
DB = 0
SM = 1
Delta_AUC
Doxorubicin
YDR127W ARO1
ZShift = 0.57
Z lm Score = 12.43
NG = 0
DB = 0
SM = 1
Delta_L
Doxorubicin
YDR127W ARO1
ZShift = −1.28
Z lm Score = −6.25
NG = 0
DB = 0
SM = 1
Delta_K
Doxorubicin
YDR127W ARO1
ZShift = −1.7
Z lm Score = −0.81
NG = 0
DB = 0
SM = 1
Delta_r
Doxorubicin
YDR127W ARO1
ZShift = −1.19
Z lm Score = −3.35
NG = 0
DB = 0
SM = 1
Delta_AUC
Doxorubicin
YML117W NAB6
ZShift = −0.97
Z lm Score = 12.42
NG = 0
DB = 0
SM = 0
Delta_L
Doxorubicin
YML117W NAB6
ZShift = 0.25
Z lm Score = −8.7
NG = 0
DB = 0
SM = 0
Delta_K
Doxorubicin
YML117W NAB6
ZShift = 1.76
Z lm Score = −4.89
NG = 0
DB = 0
SM = 0
Delta_r
Doxorubicin
YML117W NAB6
ZShift = 0.97
Z lm Score = −4.19
NG = 0
DB = 0
SM = 0
Delta_AUC
Doxorubicin

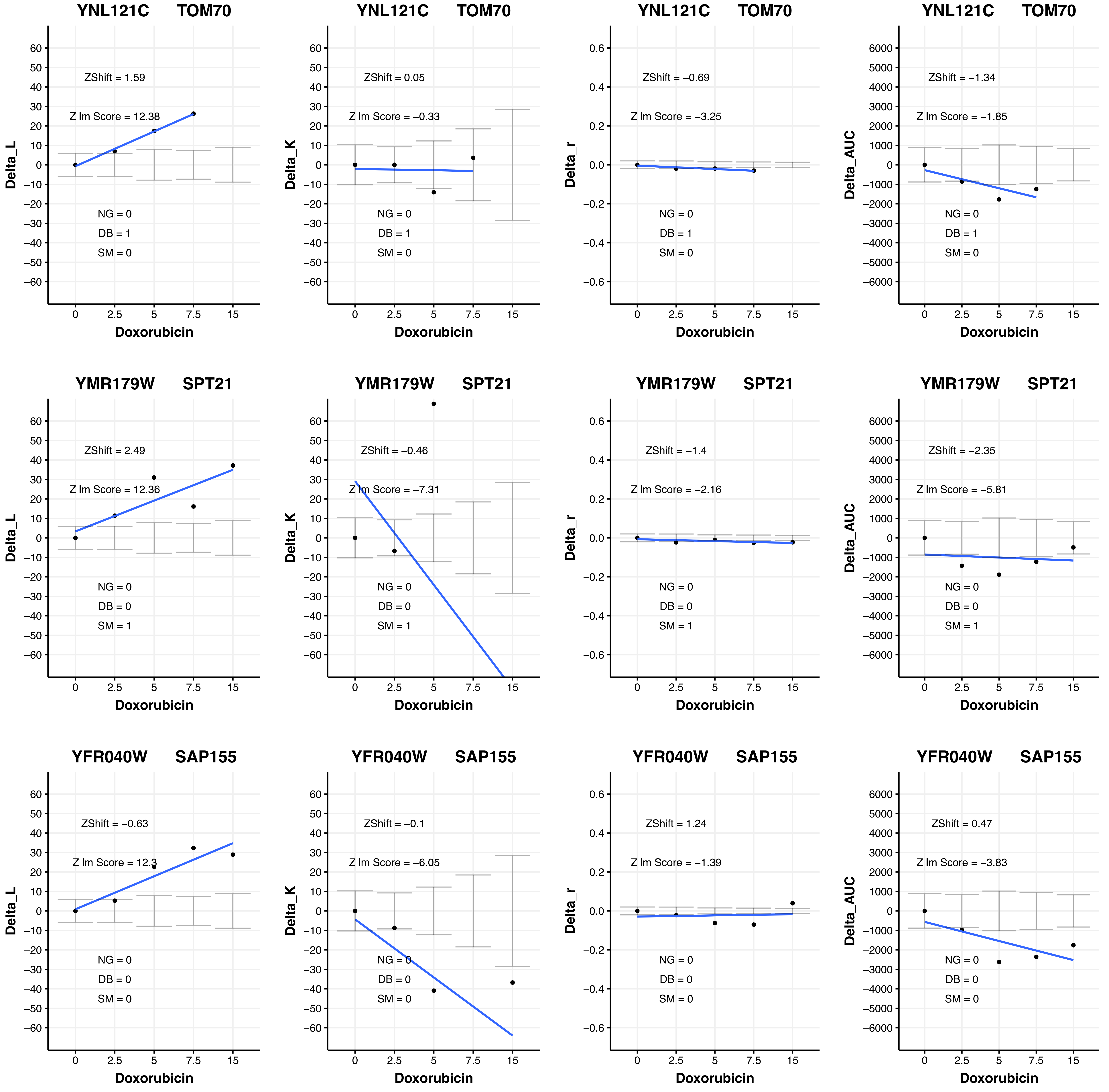

YNL121C TOM70
ZShift = 1.59
Z lm Score = 12.38
NG = 0
DB = 1
SM = 0
Delta_L
Doxorubicin
YNL121C TOM70
ZShift = 0.05
Z lm Score = −0.33
NG = 0
DB = 1
SM = 0
Delta_K
Doxorubicin
YNL121C TOM70
ZShift = −0.69
Z lm Score = −3.25
NG = 0
DB = 1
SM = 0
Delta_r
Doxorubicin
YNL121C TOM70
ZShift = −1.34
Z lm Score = −1.85
NG = 0
DB = 1
SM = 0
Delta_AUC
Doxorubicin
YMR179W SPT21
ZShift = 2.49
Z lm Score = 12.36
NG = 0
DB = 0
SM = 1
Delta_L
Doxorubicin
YMR179W SPT21
ZShift = −0.46
Z lm Score = −7.31
NG = 0
DB = 0
SM = 1
Delta_K
Doxorubicin
YMR179W SPT21
ZShift = −1.4
Z lm Score = −2.16
NG = 0
DB = 0
SM = 1
Delta_r
Doxorubicin
YMR179W SPT21
ZShift = −2.35
Z lm Score = −5.81
NG = 0
DB = 0
SM = 1
Delta_AUC
Doxorubicin
YFR040W SAP155
ZShift = −0.63
Z lm Score = 12.3
NG = 0
DB = 0
SM = 0
Delta_L
Doxorubicin
YFR040W SAP155
ZShift = −0.1
Z lm Score = −6.05
NG = 0
DB = 0
SM = 0
Delta_K
Doxorubicin
YFR040W SAP155
ZShift = 1.24
Z lm Score = −1.39
NG = 0
DB = 0
SM = 0
Delta_r
Doxorubicin
YFR040W SAP155
ZShift = 0.47
Z lm Score = −3.83
NG = 0
DB = 0
SM = 0
Delta_AUC
Doxorubicin

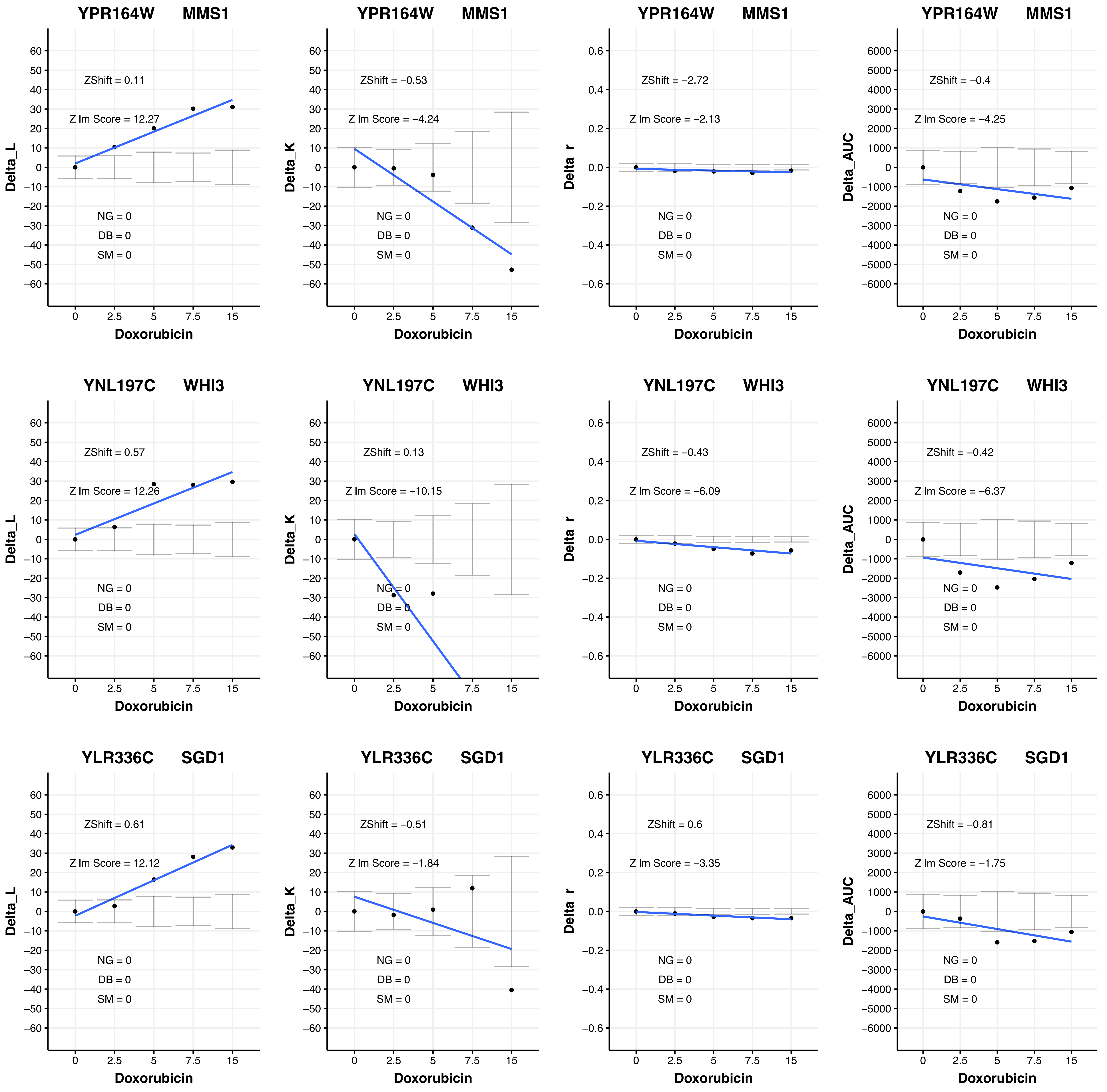

YPR164W MMS1
ZShift = 0.11
Z lm Score = 12.27
NG = 0
DB = 0
SM = 0
Delta_L
Doxorubicin
YPR164W MMS1
ZShift = −0.53
Z lm Score = −4.24
NG = 0
DB = 0
SM = 0
Delta_K
Doxorubicin
YPR164W MMS1
ZShift = −2.72
Z lm Score = −2.13
NG = 0
DB = 0
SM = 0
Delta_r
Doxorubicin
YPR164W MMS1
ZShift = −0.4
Z lm Score = −4.25
NG = 0
DB = 0
SM = 0
Delta_AUC
Doxorubicin
YNL197C WHI3
ZShift = 0.57
Z lm Score = 12.26
NG = 0
DB = 0
SM = 0
Delta_L
Doxorubicin
YNL197C WHI3
ZShift = 0.13
Z lm Score = −10.15
NG = 0
DB = 0
SM = 0
Delta_K
Doxorubicin
YNL197C WHI3
ZShift = −0.43
Z lm Score = −6.09
NG = 0
DB = 0
SM = 0
Delta_r
Doxorubicin
YNL197C WHI3
ZShift = −0.42
Z lm Score = −6.37
NG = 0
DB = 0
SM = 0
Delta_AUC
Doxorubicin
YLR336C SGD1
ZShift = 0.61
Z lm Score = 12.12
NG = 0
DB = 0
SM = 0
Delta_L
Doxorubicin
YLR336C SGD1
ZShift = −0.51
Z lm Score = −1.84
NG = 0
DB = 0
SM = 0
Delta_K
Doxorubicin
YLR336C SGD1
ZShift = 0.6
Z lm Score = −3.35
NG = 0
DB = 0
SM = 0
Delta_r
Doxorubicin
YLR336C SGD1
ZShift = −0.81
Z lm Score = −1.75
NG = 0
DB = 0
SM = 0
Delta_AUC
Doxorubicin

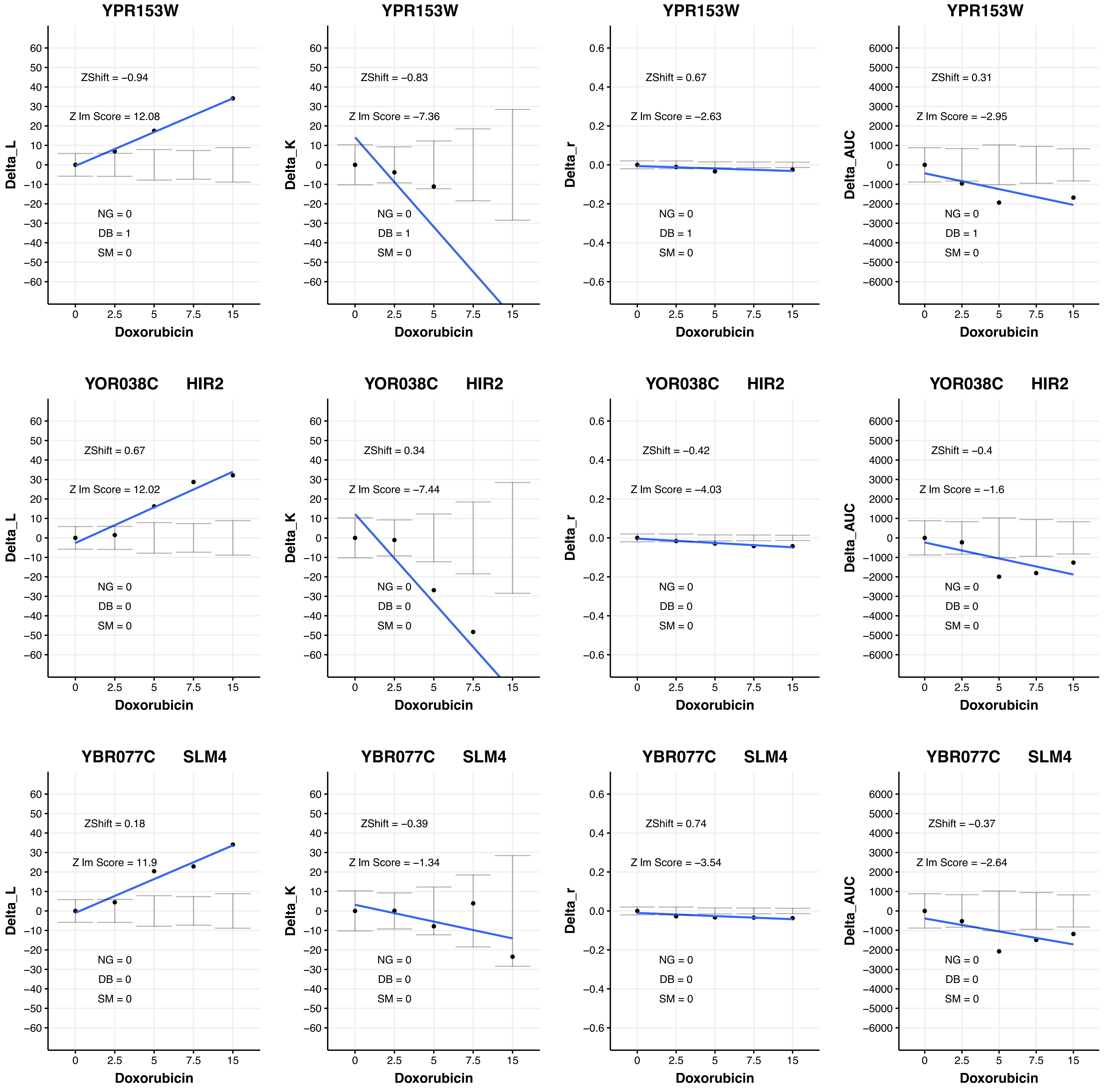
YPR153W
Delta_L
Doxorubicin
ZShift = −0.94
Z lm Score = 12.08
NG = 0
DB = 1
SM = 0
YPR153W
Delta_K
Doxorubicin
ZShift = −0.83
Z lm Score = −7.36
NG = 0
DB = 1
SM = 0
YPR153W
Delta_r
Doxorubicin
ZShift = 0.67
Z lm Score = −2.63
NG = 0
DB = 1
SM = 0
YPR153W
Delta_AUC
Doxorubicin
ZShift = 0.31
Z lm Score = −2.95
NG = 0
DB = 1
SM = 0
YOR038C HIR2
Delta_L
Doxorubicin
ZShift = 0.67
Z lm Score = 12.02
NG = 0
DB = 0
SM = 0
YOR038C HIR2
Delta_K
Doxorubicin
ZShift = 0.34
Z lm Score = −7.44
NG = 0
DB = 0
SM = 0
YOR038C HIR2
Delta_r
Doxorubicin
ZShift = −0.42
Z lm Score = −4.03
NG = 0
DB = 0
SM = 0
YOR038C HIR2
Delta_AUC
Doxorubicin
ZShift = −0.4
Z lm Score = −1.6
NG = 0
DB = 0
SM = 0
YBR077C SLM4
Delta_L
Doxorubicin
ZShift = 0.18
Z lm Score = 11.9
NG = 0
DB = 0
SM = 0
YBR077C SLM4
Delta_K
Doxorubicin
ZShift = −0.39
Z lm Score = −1.34
NG = 0
DB = 0
SM = 0
YBR077C SLM4
Delta_r
Doxorubicin
ZShift = 0.74
Z lm Score = −3.54
NG = 0
DB = 0
SM = 0
YBR077C SLM4
Delta_AUC
Doxorubicin
ZShift = −0.37
Z lm Score = −2.64
NG = 0
DB = 0
SM = 0

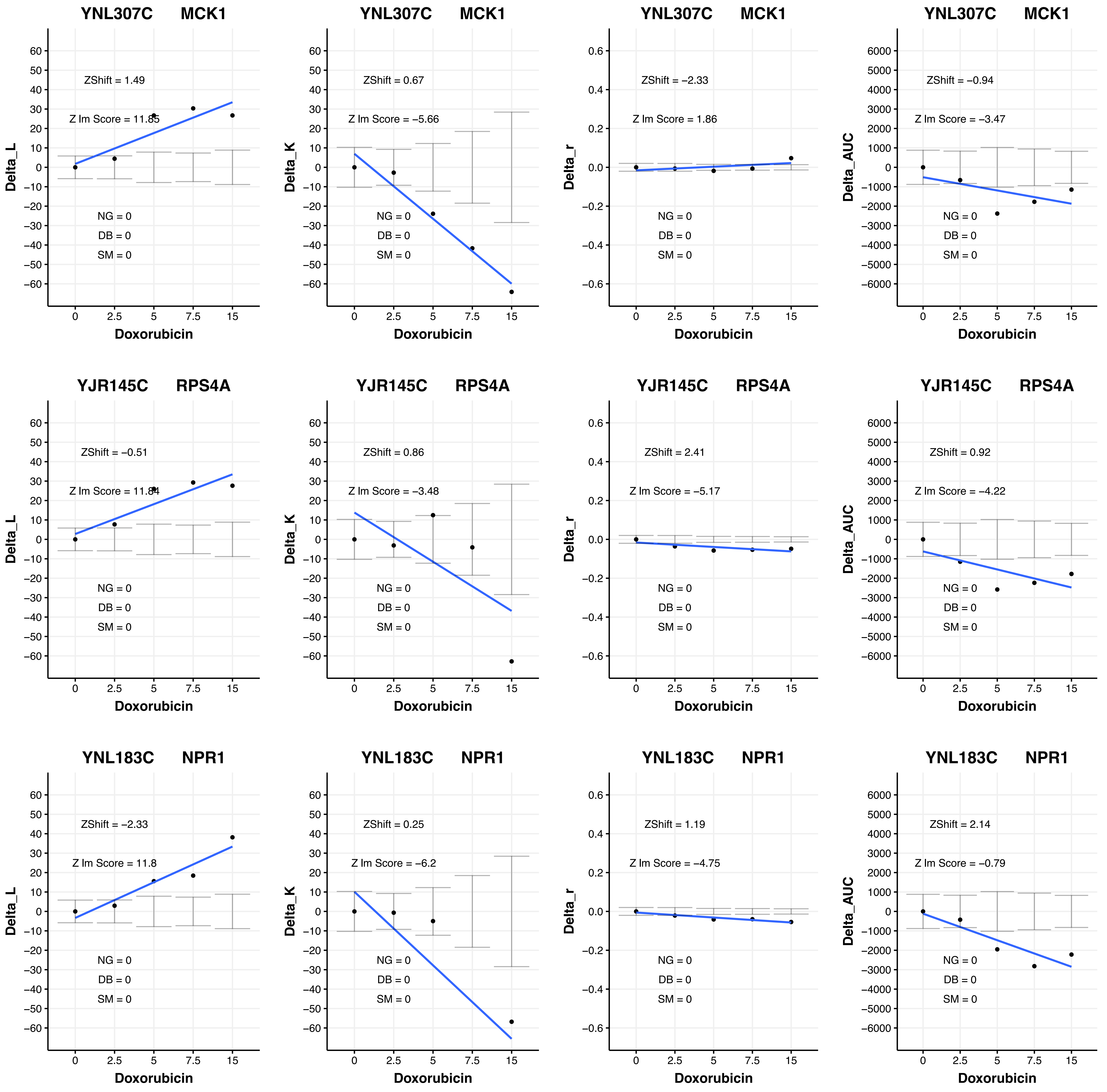

YNL307C MCK1
ZShift = 1.49
Z lm Score = 11.85
NG = 0
DB = 0
SM = 0
Delta_L
Doxorubicin
YNL307C MCK1
ZShift = 0.67
Z lm Score = −5.66
NG = 0
DB = 0
SM = 0
Delta_K
Doxorubicin
YNL307C MCK1
ZShift = −2.33
Z lm Score = 1.86
NG = 0
DB = 0
SM = 0
Delta_r
Doxorubicin
YNL307C MCK1
ZShift = −0.94
Z lm Score = −3.47
NG = 0
DB = 0
SM = 0
Delta_AUC
Doxorubicin
YJR145C RPS4A
ZShift = −0.51
Z lm Score = 11.84
NG = 0
DB = 0
SM = 0
Delta_L
Doxorubicin
YJR145C RPS4A
ZShift = 0.86
Z lm Score = −3.48
NG = 0
DB = 0
SM = 0
Delta_K
Doxorubicin
YJR145C RPS4A
ZShift = 2.41
Z lm Score = −5.17
NG = 0
DB = 0
SM = 0
Delta_r
Doxorubicin
YJR145C RPS4A
ZShift = 0.92
Z lm Score = −4.22
NG = 0
DB = 0
SM = 0
Delta_AUC
Doxorubicin
YNL183C NPR1
ZShift = −2.33
Z lm Score = 11.8
NG = 0
DB = 0
SM = 0
Delta_L
Doxorubicin
YNL183C NPR1
ZShift = 0.25
Z lm Score = −6.2
NG = 0
DB = 0
SM = 0
Delta_K
Doxorubicin
YNL183C NPR1
ZShift = 1.19
Z lm Score = −4.75
NG = 0
DB = 0
SM = 0
Delta_r
Doxorubicin
YNL183C NPR1
ZShift = 2.14
Z lm Score = −0.79
NG = 0
DB = 0
SM = 0
Delta_AUC
Doxorubicin

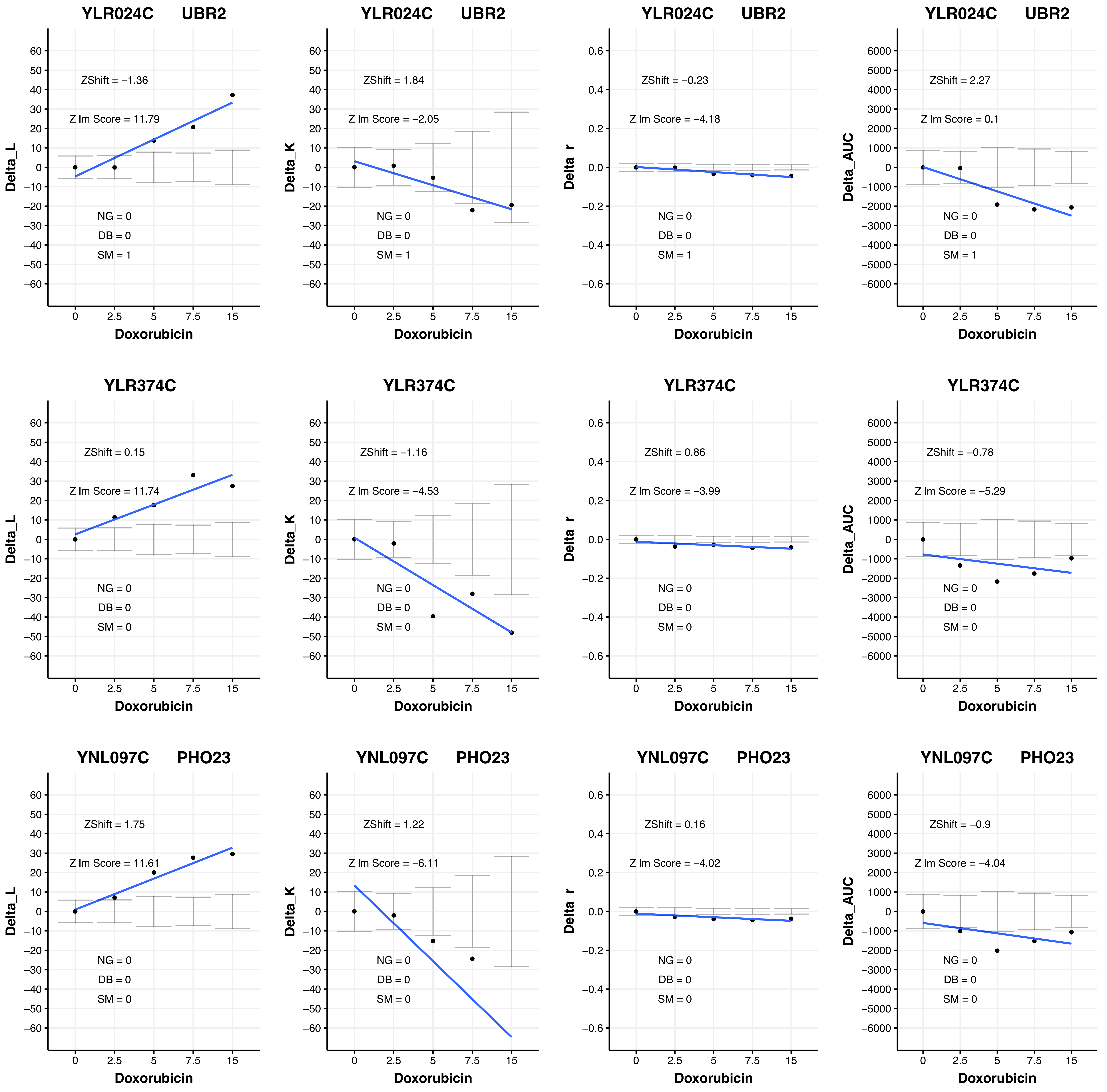
YLR024C UBR2
ZShift = −1.36
Z lm Score = 11.79
NG = 0
DB = 0
SM = 1
Delta_L
Doxorubicin
YLR024C UBR2
ZShift = 1.84
Z lm Score = −2.05
NG = 0
DB = 0
SM = 1
Delta_K
Doxorubicin
YLR024C UBR2
ZShift = −0.23
Z lm Score = −4.18
NG = 0
DB = 0
SM = 1
Delta_r
Doxorubicin
YLR024C UBR2
ZShift = 2.27
Z lm Score = 0.1
NG = 0
DB = 0
SM = 1
Delta_AUC
Doxorubicin
YLR374C
ZShift = 0.15
Z lm Score = 11.74
NG = 0
DB = 0
SM = 0
Delta_L
Doxorubicin
YLR374C
ZShift = −1.16
Z lm Score = −4.53
NG = 0
DB = 0
SM = 0
Delta_K
Doxorubicin
YLR374C
ZShift = 0.86
Z lm Score = −3.99
NG = 0
DB = 0
SM = 0
Delta_r
Doxorubicin
YLR374C
ZShift = −0.78
Z lm Score = −5.29
NG = 0
DB = 0
SM = 0
Delta_AUC
Doxorubicin
YNL097C PHO23
ZShift = 1.75
Z lm Score = 11.61
NG = 0
DB = 0
SM = 0
Delta_L
Doxorubicin
YNL097C PHO23
ZShift = 1.22
Z lm Score = −6.11
NG = 0
DB = 0
SM = 0
Delta_K
Doxorubicin
YNL097C PHO23
ZShift = 0.16
Z lm Score = −4.02
NG = 0
DB = 0
SM = 0
Delta_r
Doxorubicin
YNL097C PHO23
ZShift = −0.9
Z lm Score = −4.04
NG = 0
DB = 0
SM = 0
Delta_AUC
Doxorubicin

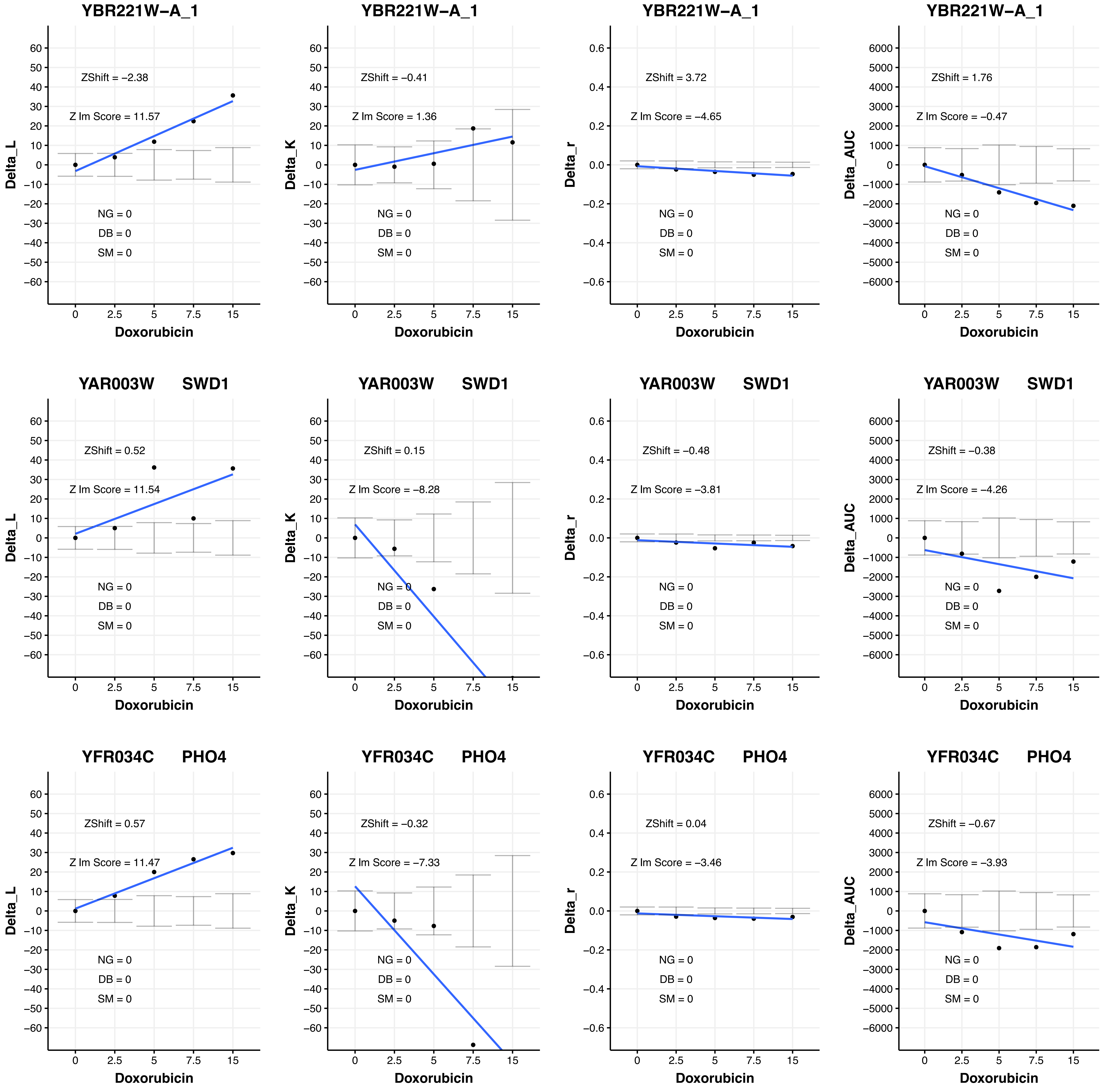

YBR221W−A_1
ZShift = −2.38
Z lm Score = 11.57
NG = 0
DB = 0
SM = 0
Delta_L
Doxorubicin
YBR221W−A_1
ZShift = −0.41
Z lm Score = 1.36
NG = 0
DB = 0
SM = 0
Delta_K
Doxorubicin
YBR221W−A_1
ZShift = 3.72
Z lm Score = −4.65
NG = 0
DB = 0
SM = 0
Delta_r
Doxorubicin
YBR221W−A_1
ZShift = 1.76
Z lm Score = −0.47
NG = 0
DB = 0
SM = 0
Delta_AUC
Doxorubicin
YAR003W SWD1
ZShift = 0.52
Z lm Score = 11.54
NG = 0
DB = 0
SM = 0
Delta_L
Doxorubicin
YAR003W SWD1
ZShift = 0.15
Z lm Score = −8.28
NG = 0
DB = 0
SM = 0
Delta_K
Doxorubicin
YAR003W SWD1
ZShift = −0.48
Z lm Score = −3.81
NG = 0
DB = 0
SM = 0
Delta_r
Doxorubicin
YAR003W SWD1
ZShift = −0.38
Z lm Score = −4.26
NG = 0
DB = 0
SM = 0
Delta_AUC
Doxorubicin
YFR034C PHO4
ZShift = 0.57
Z lm Score = 11.47
NG = 0
DB = 0
SM = 0
Delta_L
Doxorubicin
YFR034C PHO4
ZShift = −0.32
Z lm Score = −7.33
NG = 0
DB = 0
SM = 0
Delta_K
Doxorubicin
YFR034C PHO4
ZShift = 0.04
Z lm Score = −3.46
NG = 0
DB = 0
SM = 0
Delta_r
Doxorubicin
YFR034C PHO4
ZShift = −0.67
Z lm Score = −3.93
NG = 0
DB = 0
SM = 0
Delta_AUC
Doxorubicin

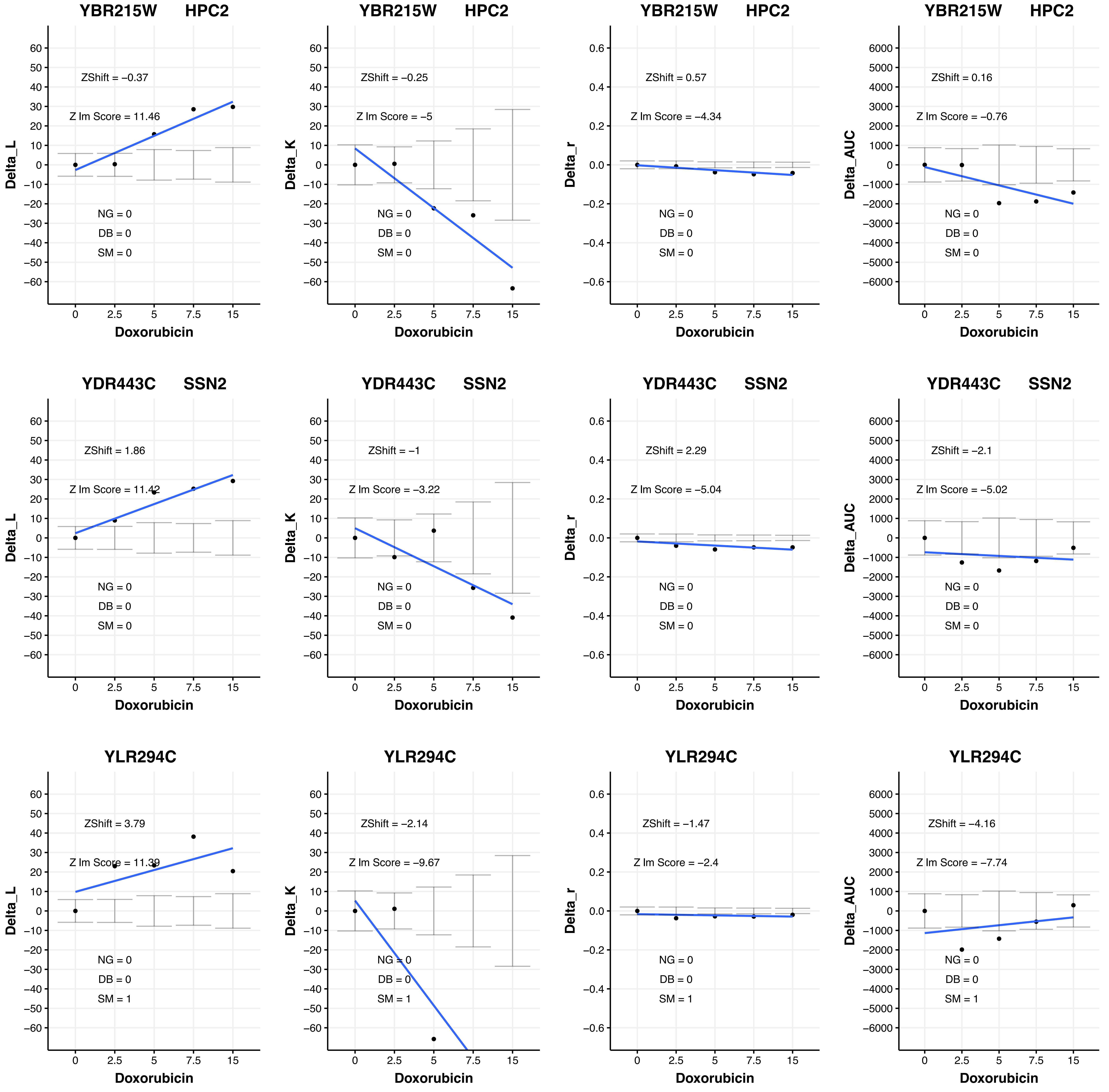

YBR215W HPC2
ZShift = −0.37
Z lm Score = 11.46
NG = 0
DB = 0
SM = 0
Delta_L
Doxorubicin
ZShift = −0.25
Z lm Score = −5
Delta_K
ZShift = 0.57
Z lm Score = −4.34
Delta_r
ZShift = 0.16
Z lm Score = −0.76
Delta_AUC
YDR443C SSN2
ZShift = 1.86
Z lm Score = 11.42
ZShift = −1
Z lm Score = −3.22
ZShift = 2.29
Z lm Score = −5.04
ZShift = −2.1
Z lm Score = −5.02
YLR294C
ZShift = 3.79
Z lm Score = 11.39
SM = 1
ZShift = −2.14
Z lm Score = −9.67
ZShift = −1.47
Z lm Score = −2.4
ZShift = −4.16
Z lm Score = −7.74

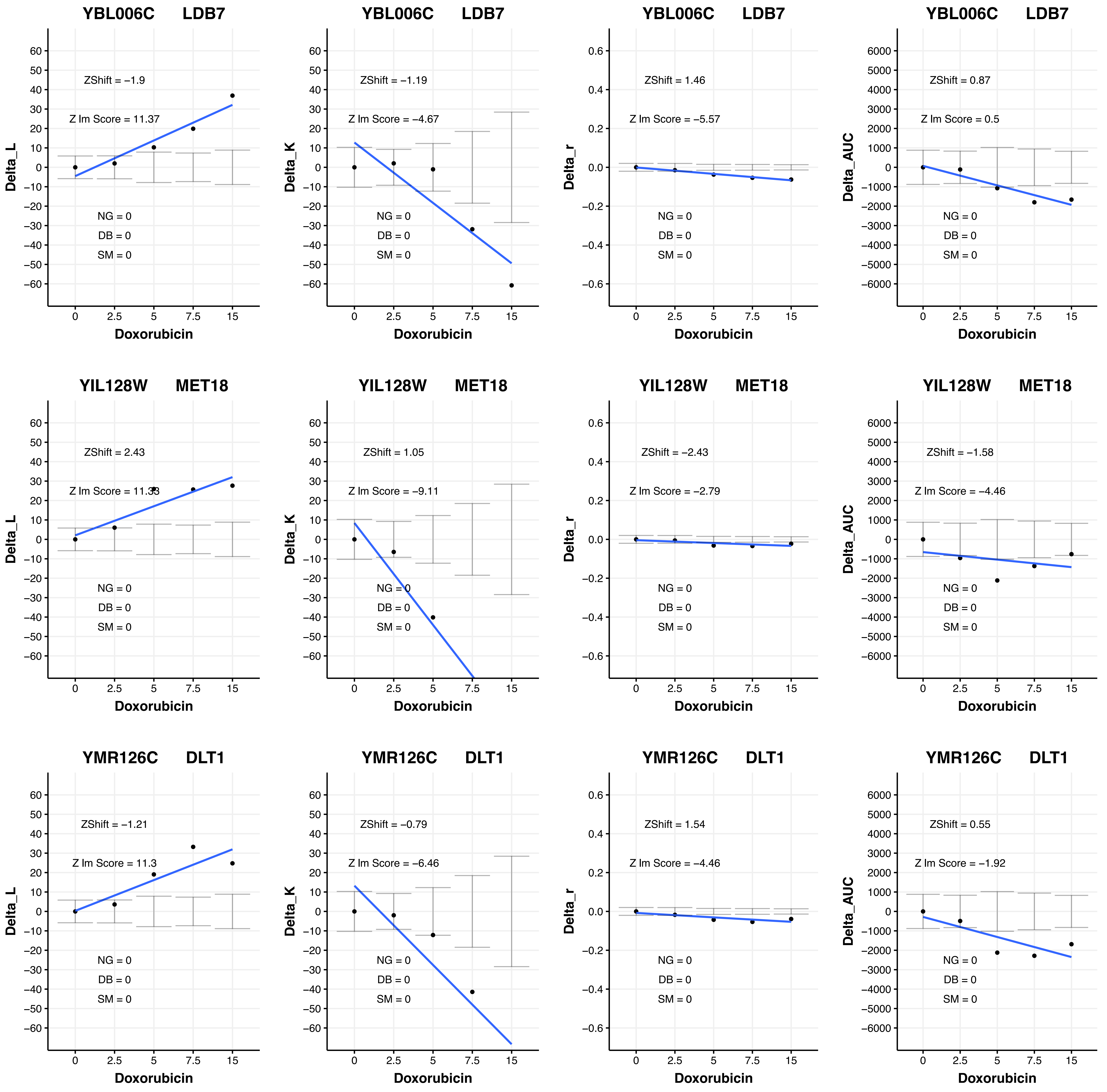

YBL006C LDB7
ZShift = −1.9
Z lm Score = 11.37
NG = 0
DB = 0
SM = 0
Delta_L
Doxorubicin
YBL006C LDB7
ZShift = −1.19
Z lm Score = −4.67
NG = 0
DB = 0
SM = 0
Delta_K
Doxorubicin
YBL006C LDB7
ZShift = 1.46
Z lm Score = −5.57
NG = 0
DB = 0
SM = 0
Delta_r
Doxorubicin
YBL006C LDB7
ZShift = 0.87
Z lm Score = 0.5
NG = 0
DB = 0
SM = 0
Delta_AUC
Doxorubicin
YIL128W MET18
ZShift = 2.43
Z lm Score = 11.33
NG = 0
DB = 0
SM = 0
Delta_L
Doxorubicin
YIL128W MET18
ZShift = 1.05
Z lm Score = −9.11
NG = 0
DB = 0
SM = 0
Delta_K
Doxorubicin
YIL128W MET18
ZShift = −2.43
Z lm Score = −2.79
NG = 0
DB = 0
SM = 0
Delta_r
Doxorubicin
YIL128W MET18
ZShift = −1.58
Z lm Score = −4.46
NG = 0
DB = 0
SM = 0
Delta_AUC
Doxorubicin
YMR126C DLT1
ZShift = −1.21
Z lm Score = 11.3
NG = 0
DB = 0
SM = 0
Delta_L
Doxorubicin
YMR126C DLT1
ZShift = −0.79
Z lm Score = −6.46
NG = 0
DB = 0
SM = 0
Delta_K
Doxorubicin
YMR126C DLT1
ZShift = 1.54
Z lm Score = −4.46
NG = 0
DB = 0
SM = 0
Delta_r
Doxorubicin
YMR126C DLT1
ZShift = 0.55
Z lm Score = −1.92
NG = 0
DB = 0
SM = 0
Delta_AUC
Doxorubicin

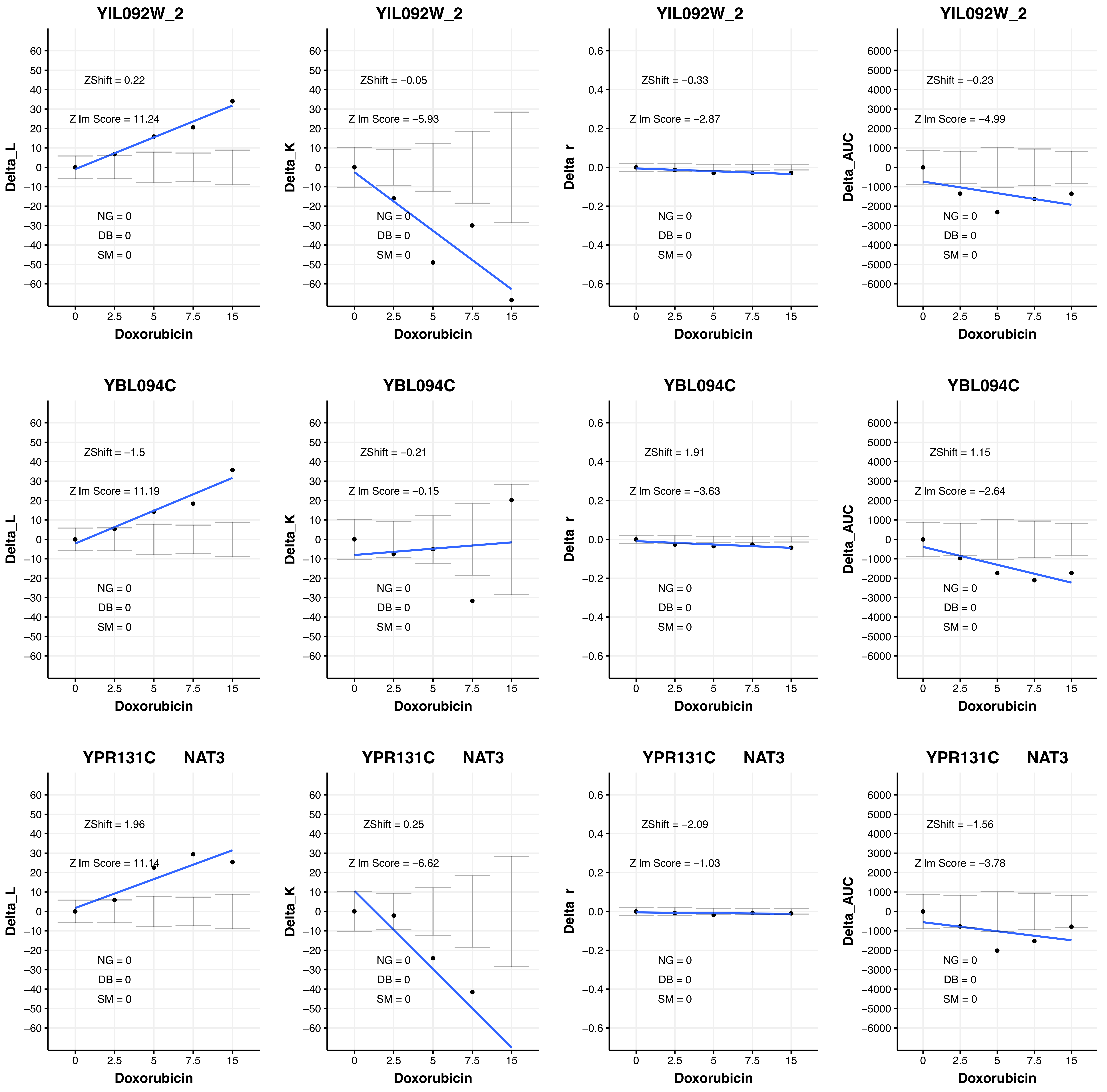

YIL092W_2
ZShift = 0.22
Z lm Score = 11.24
NG = 0
DB = 0
SM = 0
Delta_L
Doxorubicin
YIL092W_2
ZShift = −0.05
Z lm Score = −5.93
NG = 0
DB = 0
SM = 0
Delta_K
Doxorubicin
YIL092W_2
ZShift = −0.33
Z lm Score = −2.87
NG = 0
DB = 0
SM = 0
Delta_r
Doxorubicin
YIL092W_2
ZShift = −0.23
Z lm Score = −4.99
NG = 0
DB = 0
SM = 0
Delta_AUC
Doxorubicin
YBL094C
ZShift = −1.5
Z lm Score = 11.19
NG = 0
DB = 0
SM = 0
Delta_L
Doxorubicin
YBL094C
ZShift = −0.21
Z lm Score = −0.15
NG = 0
DB = 0
SM = 0
Delta_K
Doxorubicin
YBL094C
ZShift = 1.91
Z lm Score = −3.63
NG = 0
DB = 0
SM = 0
Delta_r
Doxorubicin
YBL094C
ZShift = 1.15
Z lm Score = −2.64
NG = 0
DB = 0
SM = 0
Delta_AUC
Doxorubicin
YPR131C NAT3
ZShift = 1.96
Z lm Score = 11.14
NG = 0
DB = 0
SM = 0
Delta_L
Doxorubicin
YPR131C NAT3
ZShift = 0.25
Z lm Score = −6.62
NG = 0
DB = 0
SM = 0
Delta_K
Doxorubicin
YPR131C NAT3
ZShift = −2.09
Z lm Score = −1.03
NG = 0
DB = 0
SM = 0
Delta_r
Doxorubicin
YPR131C NAT3
ZShift = −1.56
Z lm Score = −3.78
NG = 0
DB = 0
SM = 0
Delta_AUC
Doxorubicin

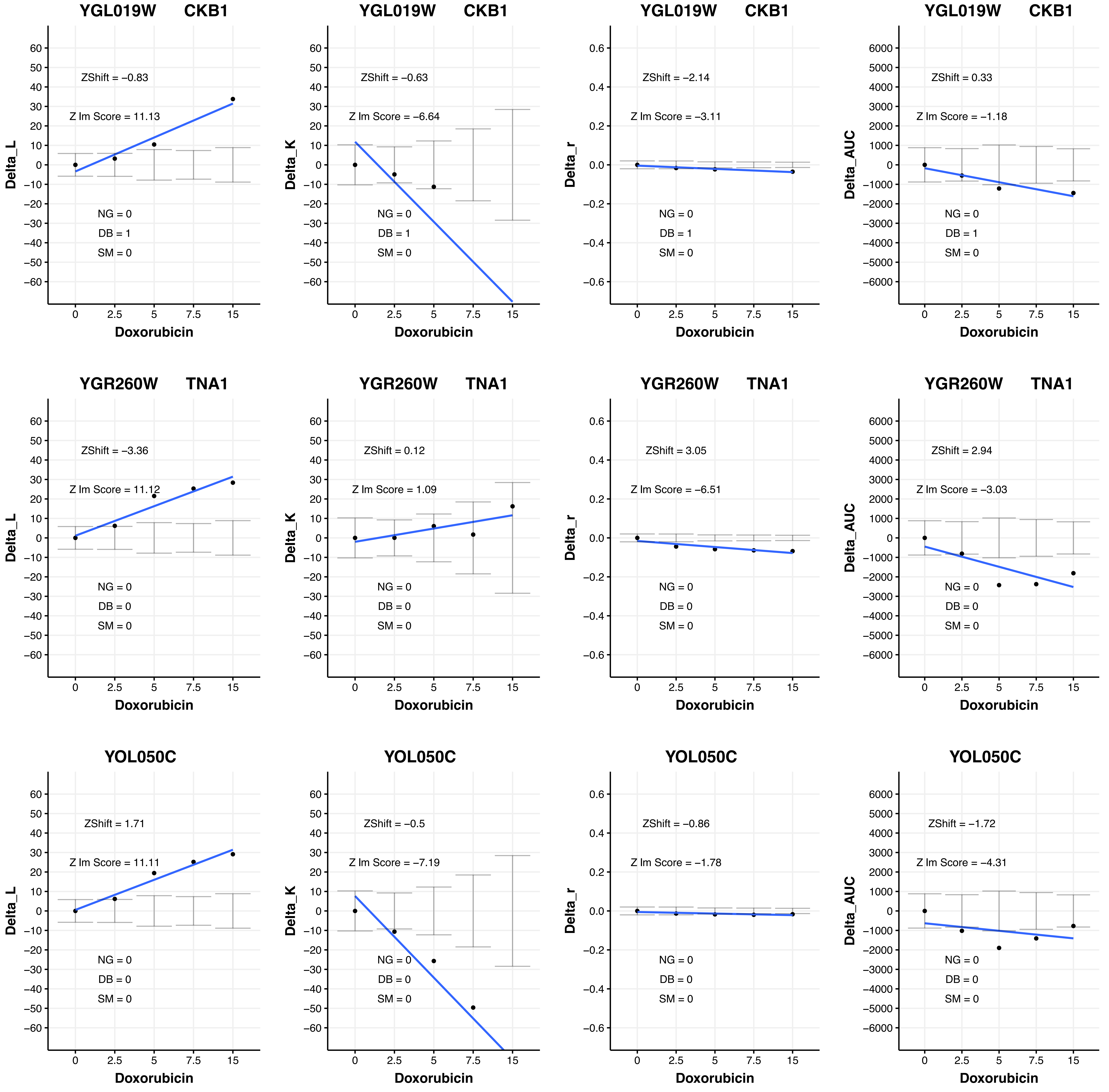
YGL019W CKB1
ZShift = −0.83
Z lm Score = 11.13
NG = 0
DB = 1
SM = 0
Delta_L
Doxorubicin
YGL019W CKB1
ZShift = −0.63
Z lm Score = −6.64
NG = 0
DB = 1
SM = 0
Delta_K
Doxorubicin
YGL019W CKB1
ZShift = −2.14
Z lm Score = −3.11
NG = 0
DB = 1
SM = 0
Delta_r
Doxorubicin
YGL019W CKB1
ZShift = 0.33
Z lm Score = −1.18
NG = 0
DB = 1
SM = 0
Delta_AUC
Doxorubicin
YGR260W TNA1
ZShift = −3.36
Z lm Score = 11.12
NG = 0
DB = 0
SM = 0
Delta_L
Doxorubicin
YGR260W TNA1
ZShift = 0.12
Z lm Score = 1.09
NG = 0
DB = 0
SM = 0
Delta_K
Doxorubicin
YGR260W TNA1
ZShift = 3.05
Z lm Score = −6.51
NG = 0
DB = 0
SM = 0
Delta_r
Doxorubicin
YGR260W TNA1
ZShift = 2.94
Z lm Score = −3.03
NG = 0
DB = 0
SM = 0
Delta_AUC
Doxorubicin
YOL050C
ZShift = 1.71
Z lm Score = 11.11
NG = 0
DB = 0
SM = 0
Delta_L
Doxorubicin
YOL050C
ZShift = −0.5
Z lm Score = −7.19
NG = 0
DB = 0
SM = 0
Delta_K
Doxorubicin
YOL050C
ZShift = −0.86
Z lm Score = −1.78
NG = 0
DB = 0
SM = 0
Delta_r
Doxorubicin
YOL050C
ZShift = −1.72
Z lm Score = −4.31
NG = 0
DB = 0
SM = 0
Delta_AUC
Doxorubicin

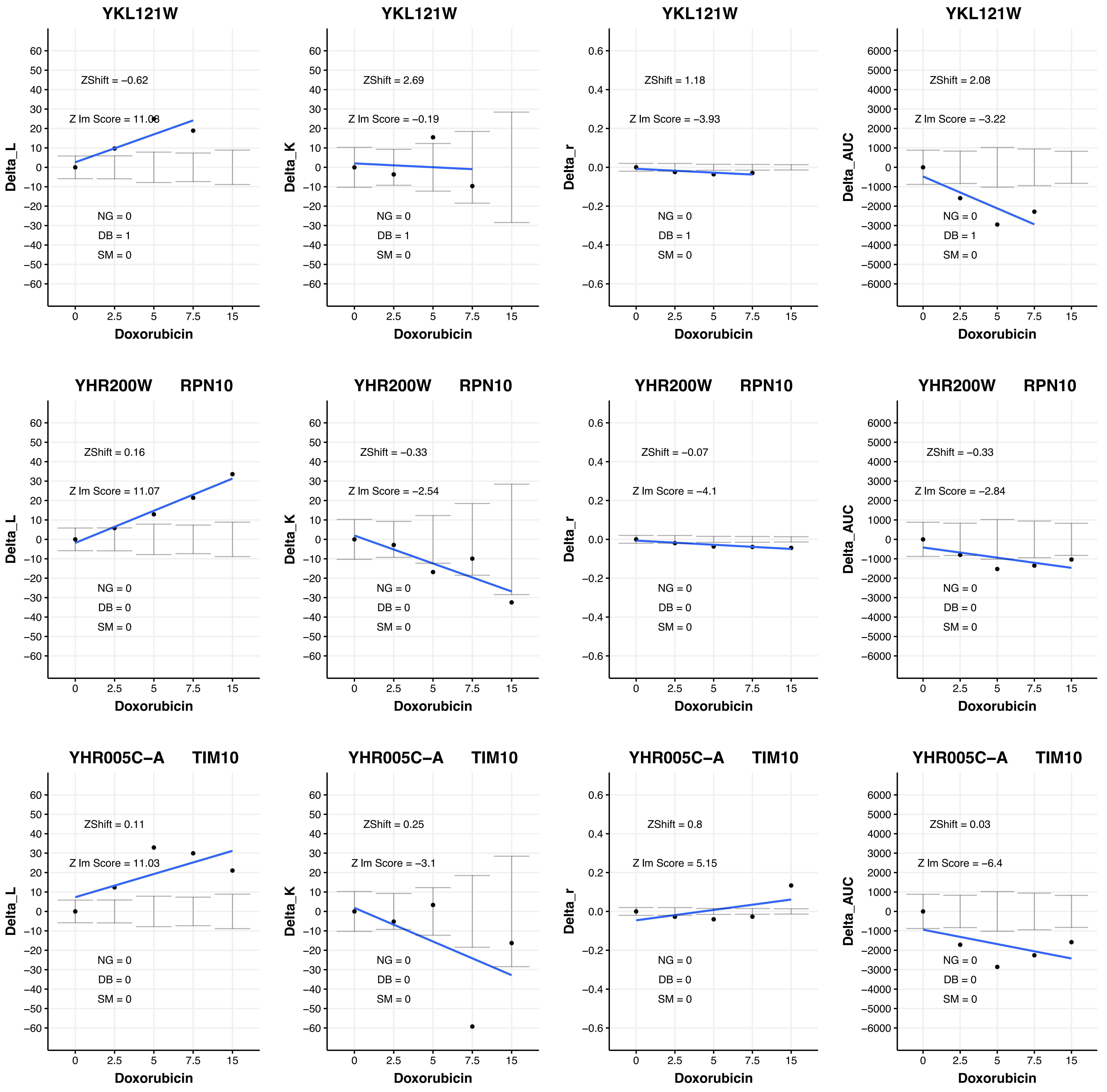
YKL121W
ZShift = −0.62
Z lm Score = 11.08
NG = 0
DB = 1
SM = 0
Delta_L
Doxorubicin
YKL121W
ZShift = 2.69
Z lm Score = −0.19
NG = 0
DB = 1
SM = 0
Delta_K
Doxorubicin
YKL121W
ZShift = 1.18
Z lm Score = −3.93
NG = 0
DB = 1
SM = 0
Delta_r
Doxorubicin
YKL121W
ZShift = 2.08
Z lm Score = −3.22
NG = 0
DB = 1
SM = 0
Delta_AUC
Doxorubicin
YHR200W RPN10
ZShift = 0.16
Z lm Score = 11.07
NG = 0
DB = 0
SM = 0
Delta_L
Doxorubicin
YHR200W RPN10
ZShift = −0.33
Z lm Score = −2.54
NG = 0
DB = 0
SM = 0
Delta_K
Doxorubicin
YHR200W RPN10
ZShift = −0.07
Z lm Score = −4.1
NG = 0
DB = 0
SM = 0
Delta_r
Doxorubicin
YHR200W RPN10
ZShift = −0.33
Z lm Score = −2.84
NG = 0
DB = 0
SM = 0
Delta_AUC
Doxorubicin
YHR005C−A TIM10
ZShift = 0.11
Z lm Score = 11.03
NG = 0
DB = 0
SM = 0
Delta_L
Doxorubicin
YHR005C−A TIM10
ZShift = 0.25
Z lm Score = −3.1
NG = 0
DB = 0
SM = 0
Delta_K
Doxorubicin
YHR005C−A TIM10
ZShift = 0.8
Z lm Score = 5.15
NG = 0
DB = 0
SM = 0
Delta_r
Doxorubicin
YHR005C−A TIM10
ZShift = 0.03
Z lm Score = −6.4
NG = 0
DB = 0
SM = 0
Delta_AUC
Doxorubicin

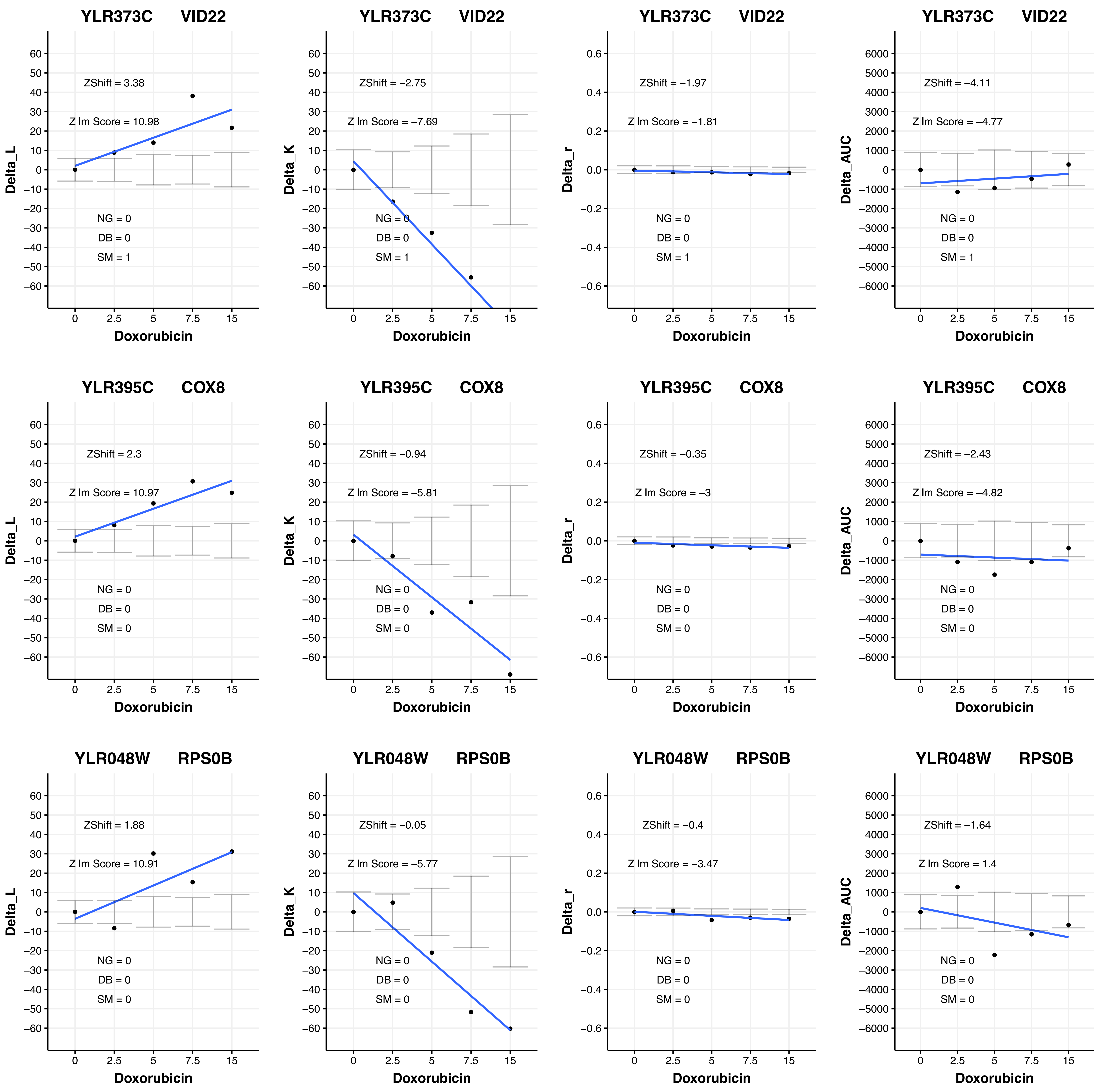

YLR373C VID22
ZShift = 3.38
Z lm Score = 10.98
NG = 0
DB = 0
SM = 1
Delta_L
Doxorubicin
ZShift = −2.75
Z lm Score = −7.69
NG = 0
DB = 0
SM = 1
Delta_K
ZShift = −1.97
Z lm Score = −1.81
NG = 0
DB = 0
SM = 1
Delta_r
ZShift = −4.11
Z lm Score = −4.77
NG = 0
DB = 0
SM = 1
Delta_AUC
YLR395C COX8
ZShift = 2.3
Z lm Score = 10.97
NG = 0
DB = 0
SM = 0
ZShift = −0.94
Z lm Score = −5.81
ZShift = −0.35
Z lm Score = −3
ZShift = −2.43
Z lm Score = −4.82
YLR048W RPS0B
ZShift = 1.88
Z lm Score = 10.91
ZShift = −0.05
Z lm Score = −5.77
ZShift = −0.4
Z lm Score = −3.47
ZShift = −1.64
Z lm Score = 1.4

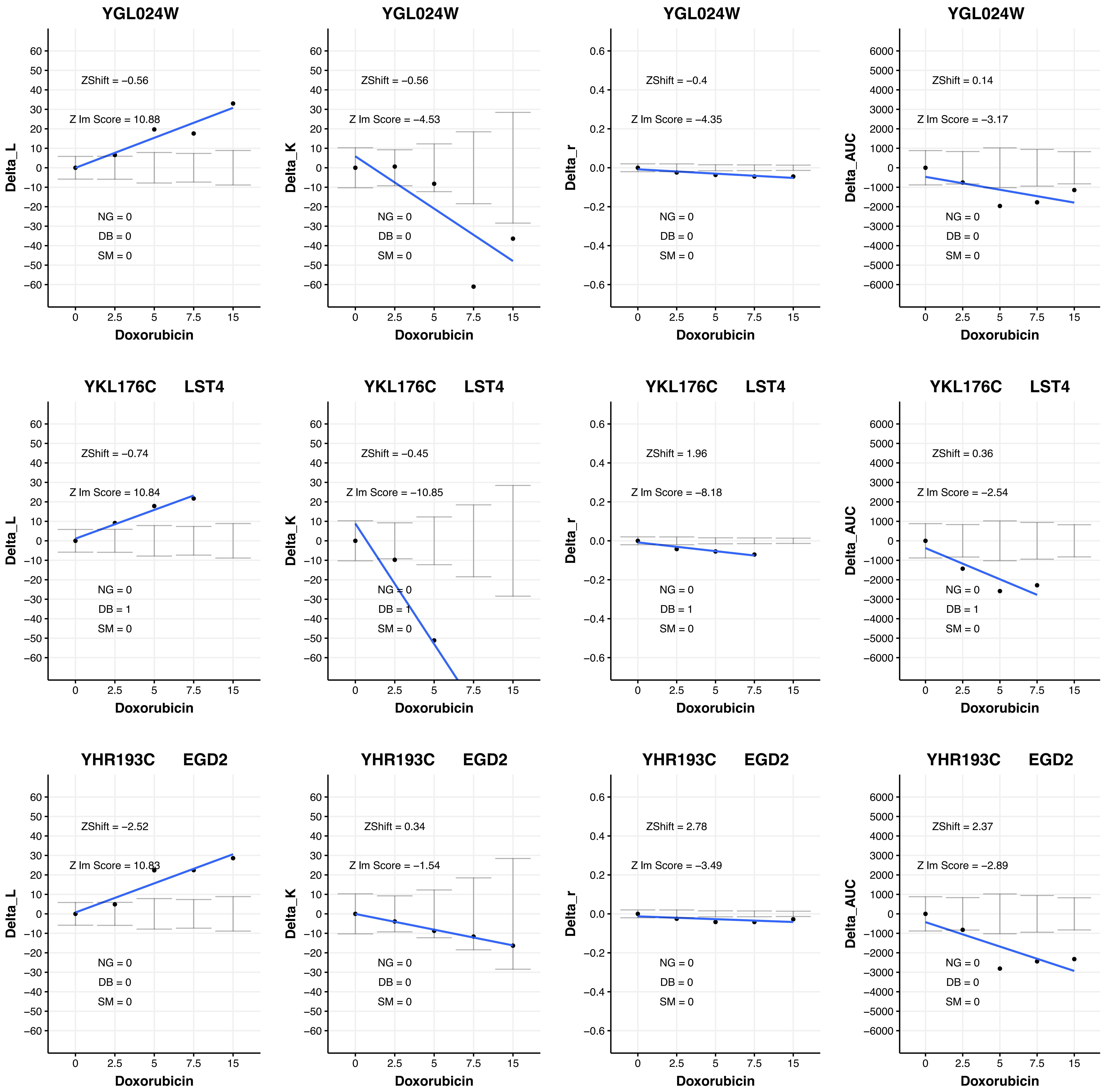
YGL024W
ZShift = −0.56
Z lm Score = 10.88
NG = 0
DB = 0
SM = 0
Delta_L
Doxorubicin
YGL024W
ZShift = −0.56
Z lm Score = −4.53
NG = 0
DB = 0
SM = 0
Delta_K
Doxorubicin
YGL024W
ZShift = −0.4
Z lm Score = −4.35
NG = 0
DB = 0
SM = 0
Delta_r
Doxorubicin
YGL024W
ZShift = 0.14
Z lm Score = −3.17
NG = 0
DB = 0
SM = 0
Delta_AUC
Doxorubicin
YKL176C LST4
ZShift = −0.74
Z lm Score = 10.84
NG = 0
DB = 1
SM = 0
Delta_L
Doxorubicin
YKL176C LST4
ZShift = −0.45
Z lm Score = −10.85
NG = 0
DB = 1
SM = 0
Delta_K
Doxorubicin
YKL176C LST4
ZShift = 1.96
Z lm Score = −8.18
NG = 0
DB = 1
SM = 0
Delta_r
Doxorubicin
YKL176C LST4
ZShift = 0.36
Z lm Score = −2.54
NG = 0
DB = 1
SM = 0
Delta_AUC
Doxorubicin
YHR193C EGD2
ZShift = −2.52
Z lm Score = 10.83
NG = 0
DB = 0
SM = 0
Delta_L
Doxorubicin
YHR193C EGD2
ZShift = 0.34
Z lm Score = −1.54
NG = 0
DB = 0
SM = 0
Delta_K
Doxorubicin
YHR193C EGD2
ZShift = 2.78
Z lm Score = −3.49
NG = 0
DB = 0
SM = 0
Delta_r
Doxorubicin
YHR193C EGD2
ZShift = 2.37
Z lm Score = −2.89
NG = 0
DB = 0
SM = 0
Delta_AUC
Doxorubicin

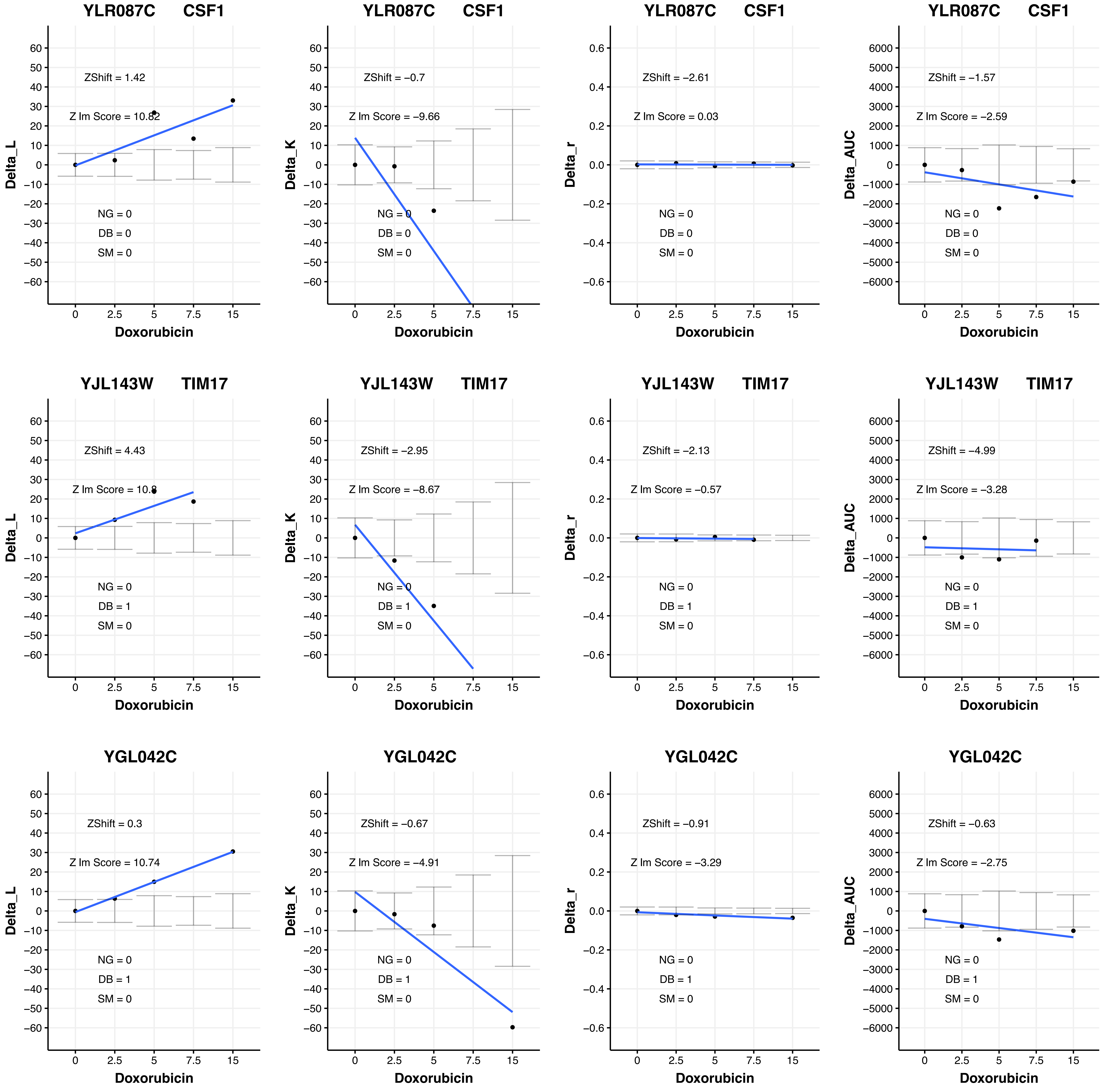
YLR087C CSF1
ZShift = 1.42
Z lm Score = 10.82
NG = 0
DB = 0
SM = 0
Delta_L
Doxorubicin
YLR087C CSF1
ZShift = −0.7
Z lm Score = −9.66
NG = 0
DB = 0
SM = 0
Delta_K
Doxorubicin
YLR087C CSF1
ZShift = −2.61
Z lm Score = 0.03
NG = 0
DB = 0
SM = 0
Delta_r
Doxorubicin
YLR087C CSF1
ZShift = −1.57
Z lm Score = −2.59
NG = 0
DB = 0
SM = 0
Delta_AUC
Doxorubicin
YJL143W TIM17
ZShift = 4.43
Z lm Score = 10.8
NG = 0
DB = 1
SM = 0
Delta_L
Doxorubicin
YJL143W TIM17
ZShift = −2.95
Z lm Score = −8.67
NG = 0
DB = 1
SM = 0
Delta_K
Doxorubicin
YJL143W TIM17
ZShift = −2.13
Z lm Score = −0.57
NG = 0
DB = 1
SM = 0
Delta_r
Doxorubicin
YJL143W TIM17
ZShift = −4.99
Z lm Score = −3.28
NG = 0
DB = 1
SM = 0
Delta_AUC
Doxorubicin
YGL042C
ZShift = 0.3
Z lm Score = 10.74
NG = 0
DB = 1
SM = 0
Delta_L
Doxorubicin
YGL042C
ZShift = −0.67
Z lm Score = −4.91
NG = 0
DB = 1
SM = 0
Delta_K
Doxorubicin
YGL042C
ZShift = −0.91
Z lm Score = −3.29
NG = 0
DB = 1
SM = 0
Delta_r
Doxorubicin
YGL042C
ZShift = −0.63
Z lm Score = −2.75
NG = 0
DB = 1
SM = 0
Delta_AUC
Doxorubicin

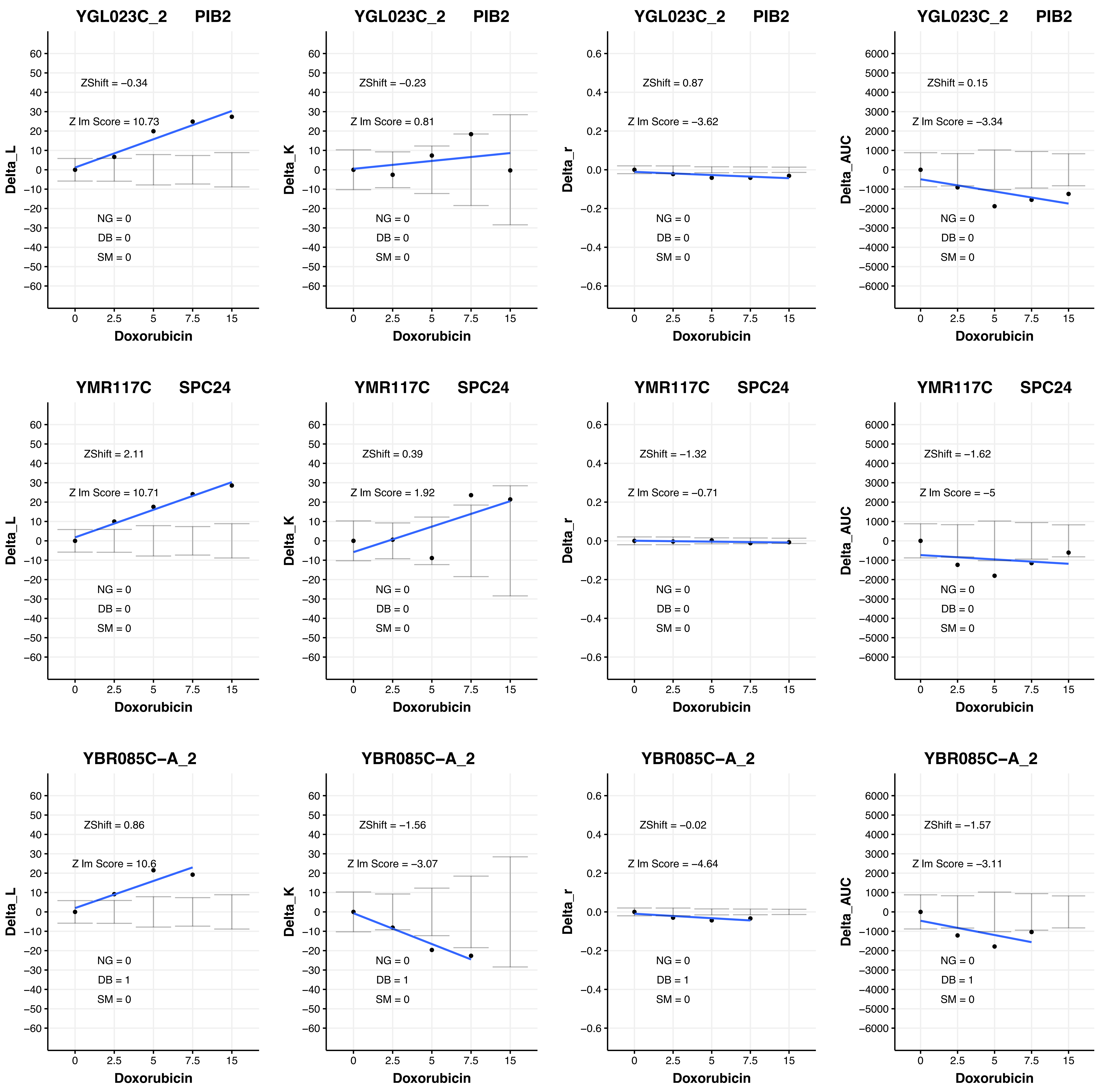
YGL023C_2 PIB2
ZShift = −0.34
Z lm Score = 10.73
NG = 0
DB = 0
SM = 0
Delta_L
Doxorubicin
YGL023C_2 PIB2
ZShift = −0.23
Z lm Score = 0.81
NG = 0
DB = 0
SM = 0
Delta_K
Doxorubicin
YGL023C_2 PIB2
ZShift = 0.87
Z lm Score = −3.62
NG = 0
DB = 0
SM = 0
Delta_r
Doxorubicin
YGL023C_2 PIB2
ZShift = 0.15
Z lm Score = −3.34
NG = 0
DB = 0
SM = 0
Delta_AUC
Doxorubicin
YMR117C SPC24
ZShift = 2.11
Z lm Score = 10.71
NG = 0
DB = 0
SM = 0
Delta_L
Doxorubicin
YMR117C SPC24
ZShift = 0.39
Z lm Score = 1.92
NG = 0
DB = 0
SM = 0
Delta_K
Doxorubicin
YMR117C SPC24
ZShift = −1.32
Z lm Score = −0.71
NG = 0
DB = 0
SM = 0
Delta_r
Doxorubicin
YMR117C SPC24
ZShift = −1.62
Z lm Score = −5
NG = 0
DB = 0
SM = 0
Delta_AUC
Doxorubicin
YBR085C−A_2
ZShift = 0.86
Z lm Score = 10.6
NG = 0
DB = 1
SM = 0
Delta_L
Doxorubicin
YBR085C−A_2
ZShift = −1.56
Z lm Score = −3.07
NG = 0
DB = 1
SM = 0
Delta_K
Doxorubicin
YBR085C−A_2
ZShift = −0.02
Z lm Score = −4.64
NG = 0
DB = 1
SM = 0
Delta_r
Doxorubicin
YBR085C−A_2
ZShift = −1.57
Z lm Score = −3.11
NG = 0
DB = 1
SM = 0
Delta_AUC
Doxorubicin

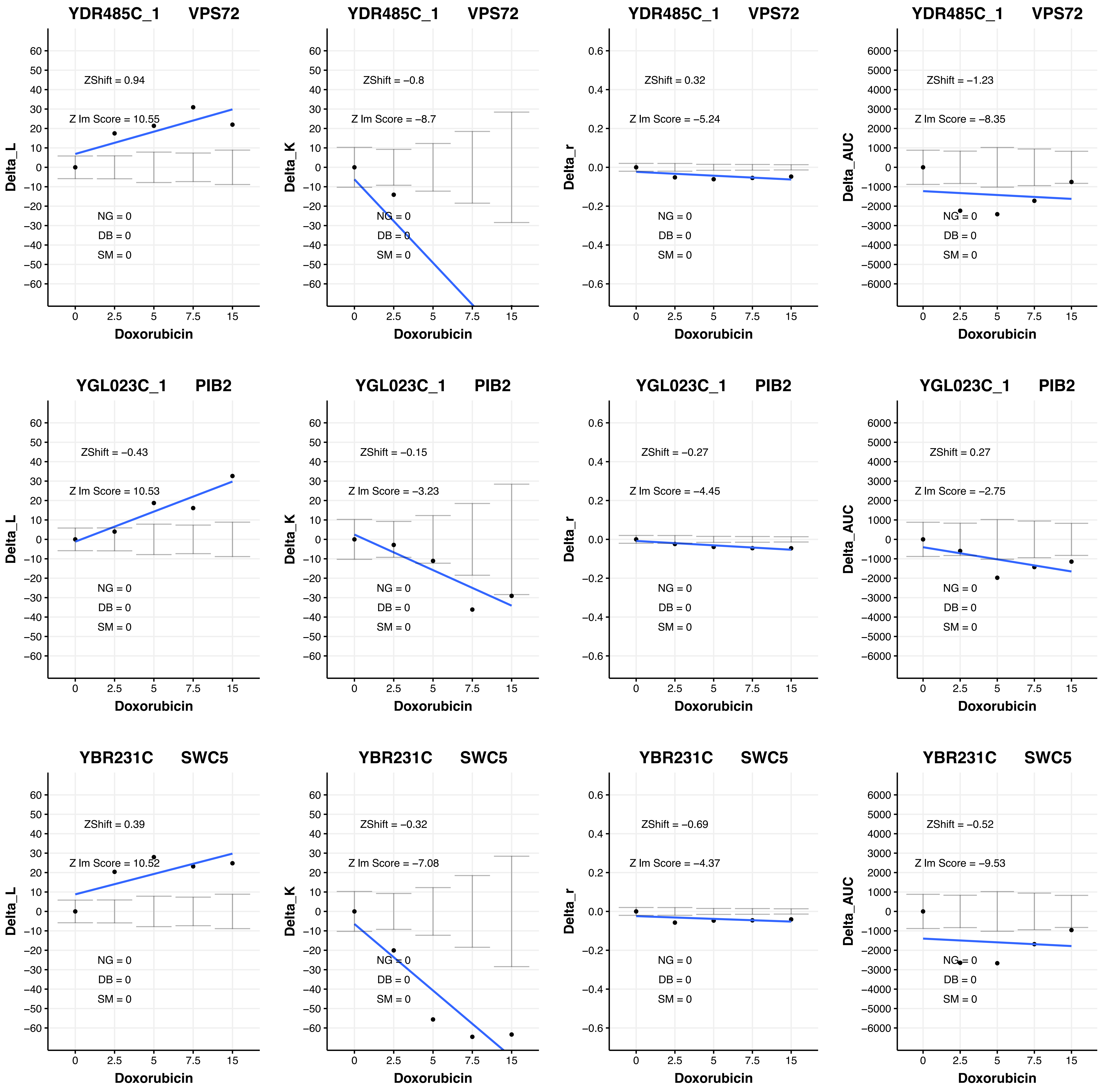
YDR485C_1 VPS72
Delta_L
ZShift = 0.94
Z lm Score = 10.55
NG = 0
DB = 0
SM = 0
Doxorubicin
YDR485C_1 VPS72
Delta_K
ZShift = −0.8
Z lm Score = −8.7
NG = 0
DB = 0
SM = 0
Doxorubicin
YDR485C_1 VPS72
Delta_r
ZShift = 0.32
Z lm Score = −5.24
NG = 0
DB = 0
SM = 0
Doxorubicin
YDR485C_1 VPS72
Delta_AUC
ZShift = −1.23
Z lm Score = −8.35
NG = 0
DB = 0
SM = 0
Doxorubicin
YGL023C_1 PIB2
Delta_L
ZShift = −0.43
Z lm Score = 10.53
NG = 0
DB = 0
SM = 0
Doxorubicin
YGL023C_1 PIB2
Delta_K
ZShift = −0.15
Z lm Score = −3.23
NG = 0
DB = 0
SM = 0
Doxorubicin
YGL023C_1 PIB2
Delta_r
ZShift = −0.27
Z lm Score = −4.45
NG = 0
DB = 0
SM = 0
Doxorubicin
YGL023C_1 PIB2
Delta_AUC
ZShift = 0.27
Z lm Score = −2.75
NG = 0
DB = 0
SM = 0
Doxorubicin
YBR231C SWC5
Delta_L
ZShift = 0.39
Z lm Score = 10.52
NG = 0
DB = 0
SM = 0
Doxorubicin
YBR231C SWC5
Delta_K
ZShift = −0.32
Z lm Score = −7.08
NG = 0
DB = 0
SM = 0
Doxorubicin
YBR231C SWC5
Delta_r
ZShift = −0.69
Z lm Score = −4.37
NG = 0
DB = 0
SM = 0
Doxorubicin
YBR231C SWC5
Delta_AUC
ZShift = −0.52
Z lm Score = −9.53
NG = 0
DB = 0
SM = 0
Doxorubicin

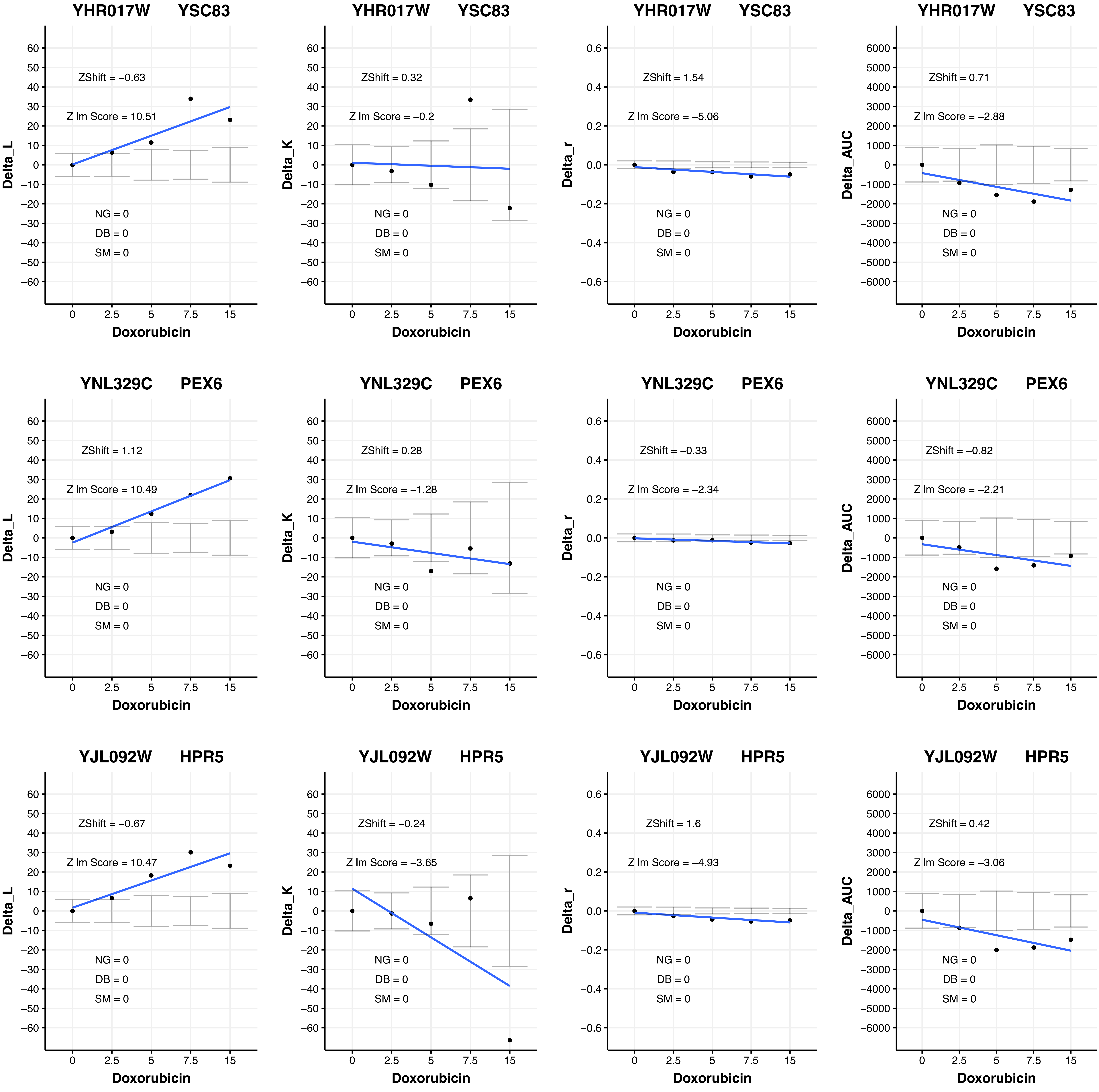
YHR017W YSC83
ZShift = −0.63
Z lm Score = 10.51
NG = 0
DB = 0
SM = 0
Delta_L
Doxorubicin
YHR017W YSC83
ZShift = 0.32
Z lm Score = −0.2
NG = 0
DB = 0
SM = 0
Delta_K
Doxorubicin
YHR017W YSC83
ZShift = 1.54
Z lm Score = −5.06
NG = 0
DB = 0
SM = 0
Delta_r
Doxorubicin
YHR017W YSC83
ZShift = 0.71
Z lm Score = −2.88
NG = 0
DB = 0
SM = 0
Delta_AUC
Doxorubicin
YNL329C PEX6
ZShift = 1.12
Z lm Score = 10.49
NG = 0
DB = 0
SM = 0
Delta_L
Doxorubicin
YNL329C PEX6
ZShift = 0.28
Z lm Score = −1.28
NG = 0
DB = 0
SM = 0
Delta_K
Doxorubicin
YNL329C PEX6
ZShift = −0.33
Z lm Score = −2.34
NG = 0
DB = 0
SM = 0
Delta_r
Doxorubicin
YNL329C PEX6
ZShift = −0.82
Z lm Score = −2.21
NG = 0
DB = 0
SM = 0
Delta_AUC
Doxorubicin
YJL092W HPR5
ZShift = −0.67
Z lm Score = 10.47
NG = 0
DB = 0
SM = 0
Delta_L
Doxorubicin
YJL092W HPR5
ZShift = −0.24
Z lm Score = −3.65
NG = 0
DB = 0
SM = 0
Delta_K
Doxorubicin
YJL092W HPR5
ZShift = 1.6
Z lm Score = −4.93
NG = 0
DB = 0
SM = 0
Delta_r
Doxorubicin
YJL092W HPR5
ZShift = 0.42
Z lm Score = −3.06
NG = 0
DB = 0
SM = 0
Delta_AUC
Doxorubicin

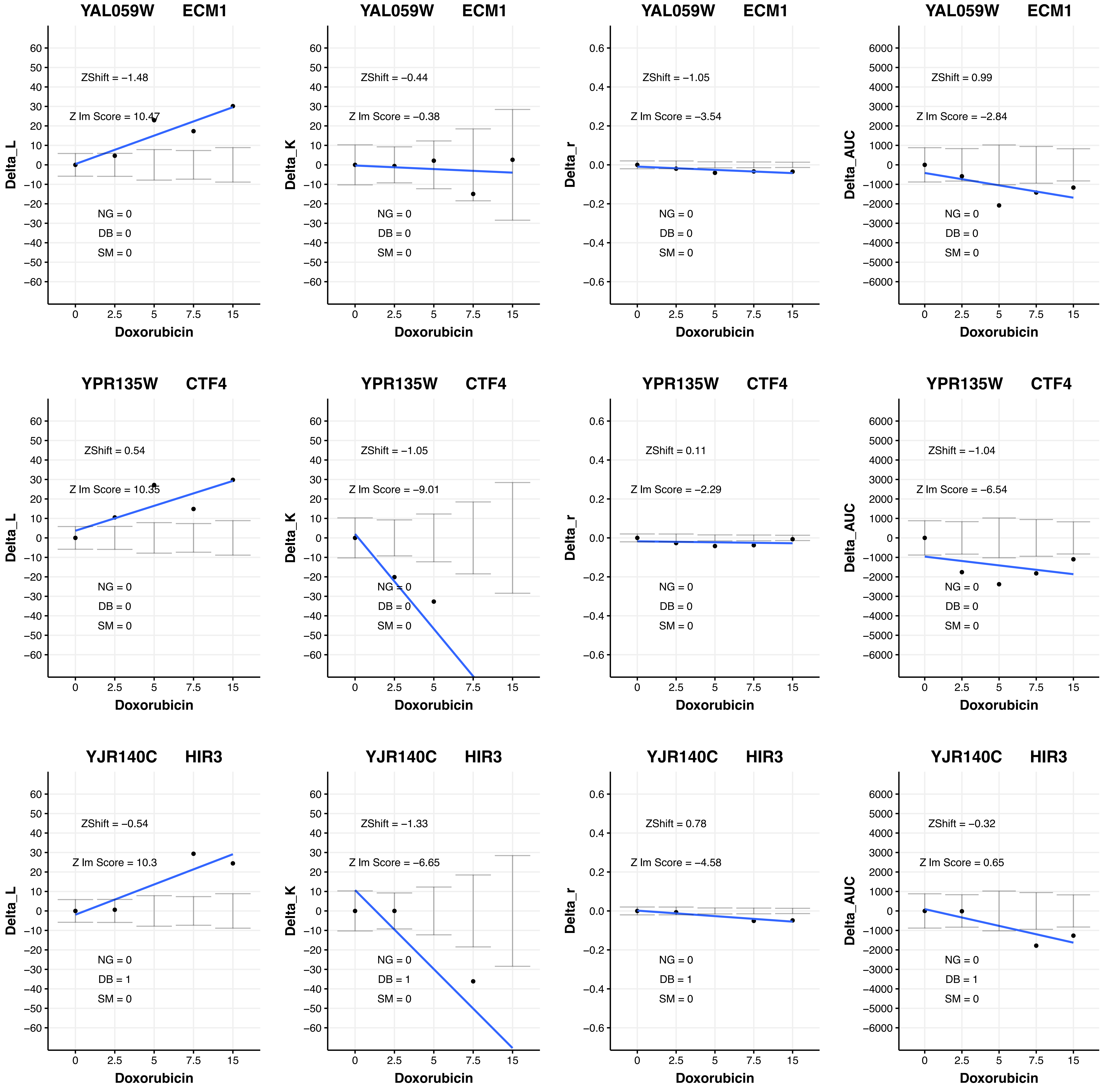

YAL059W ECM1
ZShift = −1.48
Z lm Score = 10.47
NG = 0
DB = 0
SM = 0
Delta_L
Doxorubicin
YAL059W ECM1
ZShift = −0.44
Z lm Score = −0.38
NG = 0
DB = 0
SM = 0
Delta_K
Doxorubicin
YAL059W ECM1
ZShift = −1.05
Z lm Score = −3.54
NG = 0
DB = 0
SM = 0
Delta_r
Doxorubicin
YAL059W ECM1
ZShift = 0.99
Z lm Score = −2.84
NG = 0
DB = 0
SM = 0
Delta_AUC
Doxorubicin
YPR135W CTF4
ZShift = 0.54
Z lm Score = 10.35
NG = 0
DB = 0
SM = 0
Delta_L
Doxorubicin
YPR135W CTF4
ZShift = −1.05
Z lm Score = −9.01
NG = 0
DB = 0
SM = 0
Delta_K
Doxorubicin
YPR135W CTF4
ZShift = 0.11
Z lm Score = −2.29
NG = 0
DB = 0
SM = 0
Delta_r
Doxorubicin
YPR135W CTF4
ZShift = −1.04
Z lm Score = −6.54
NG = 0
DB = 0
SM = 0
Delta_AUC
Doxorubicin
YJR140C HIR3
ZShift = −0.54
Z lm Score = 10.3
NG = 0
DB = 1
SM = 0
Delta_L
Doxorubicin
YJR140C HIR3
ZShift = −1.33
Z lm Score = −6.65
NG = 0
DB = 1
SM = 0
Delta_K
Doxorubicin
YJR140C HIR3
ZShift = 0.78
Z lm Score = −4.58
NG = 0
DB = 1
SM = 0
Delta_r
Doxorubicin
YJR140C HIR3
ZShift = −0.32
Z lm Score = 0.65
NG = 0
DB = 1
SM = 0
Delta_AUC
Doxorubicin

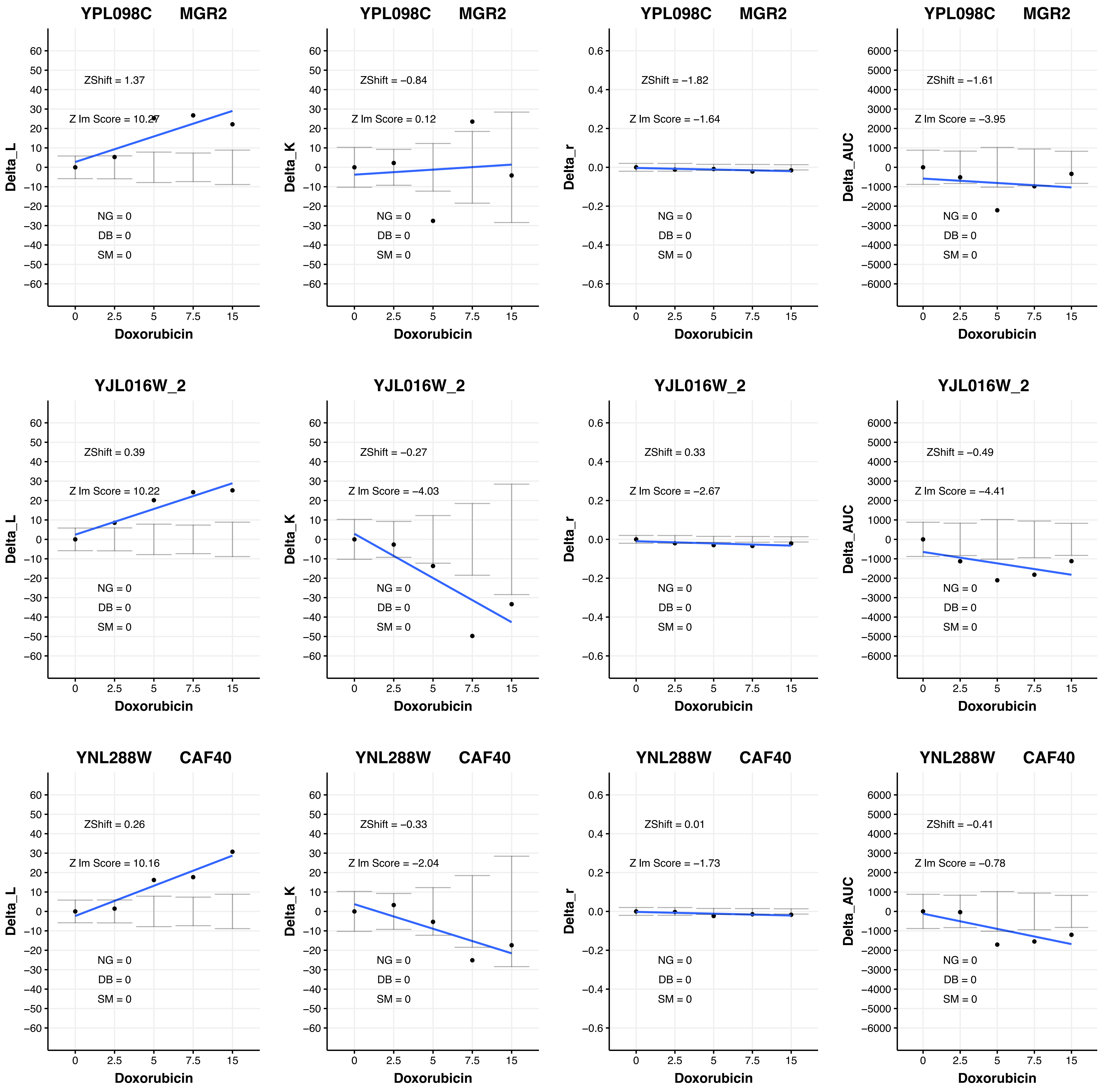

YPL098C MGR2
ZShift = 1.37
Z lm Score = 10.27
NG = 0
DB = 0
SM = 0
Delta_L
Doxorubicin
YPL098C MGR2
ZShift = −0.84
Z lm Score = 0.12
NG = 0
DB = 0
SM = 0
Delta_K
Doxorubicin
YPL098C MGR2
ZShift = −1.82
Z lm Score = −1.64
NG = 0
DB = 0
SM = 0
Delta_r
Doxorubicin
YPL098C MGR2
ZShift = −1.61
Z lm Score = −3.95
NG = 0
DB = 0
SM = 0
Delta_AUC
Doxorubicin
YJL016W_2
ZShift = 0.39
Z lm Score = 10.22
NG = 0
DB = 0
SM = 0
Delta_L
Doxorubicin
YJL016W_2
ZShift = −0.27
Z lm Score = −4.03
NG = 0
DB = 0
SM = 0
Delta_K
Doxorubicin
YJL016W_2
ZShift = 0.33
Z lm Score = −2.67
NG = 0
DB = 0
SM = 0
Delta_r
Doxorubicin
YJL016W_2
ZShift = −0.49
Z lm Score = −4.41
NG = 0
DB = 0
SM = 0
Delta_AUC
Doxorubicin
YNL288W CAF40
ZShift = 0.26
Z lm Score = 10.16
NG = 0
DB = 0
SM = 0
Delta_L
Doxorubicin
YNL288W CAF40
ZShift = −0.33
Z lm Score = −2.04
NG = 0
DB = 0
SM = 0
Delta_K
Doxorubicin
YNL288W CAF40
ZShift = 0.01
Z lm Score = −1.73
NG = 0
DB = 0
SM = 0
Delta_r
Doxorubicin
YNL288W CAF40
ZShift = −0.41
Z lm Score = −0.78
NG = 0
DB = 0
SM = 0
Delta_AUC
Doxorubicin

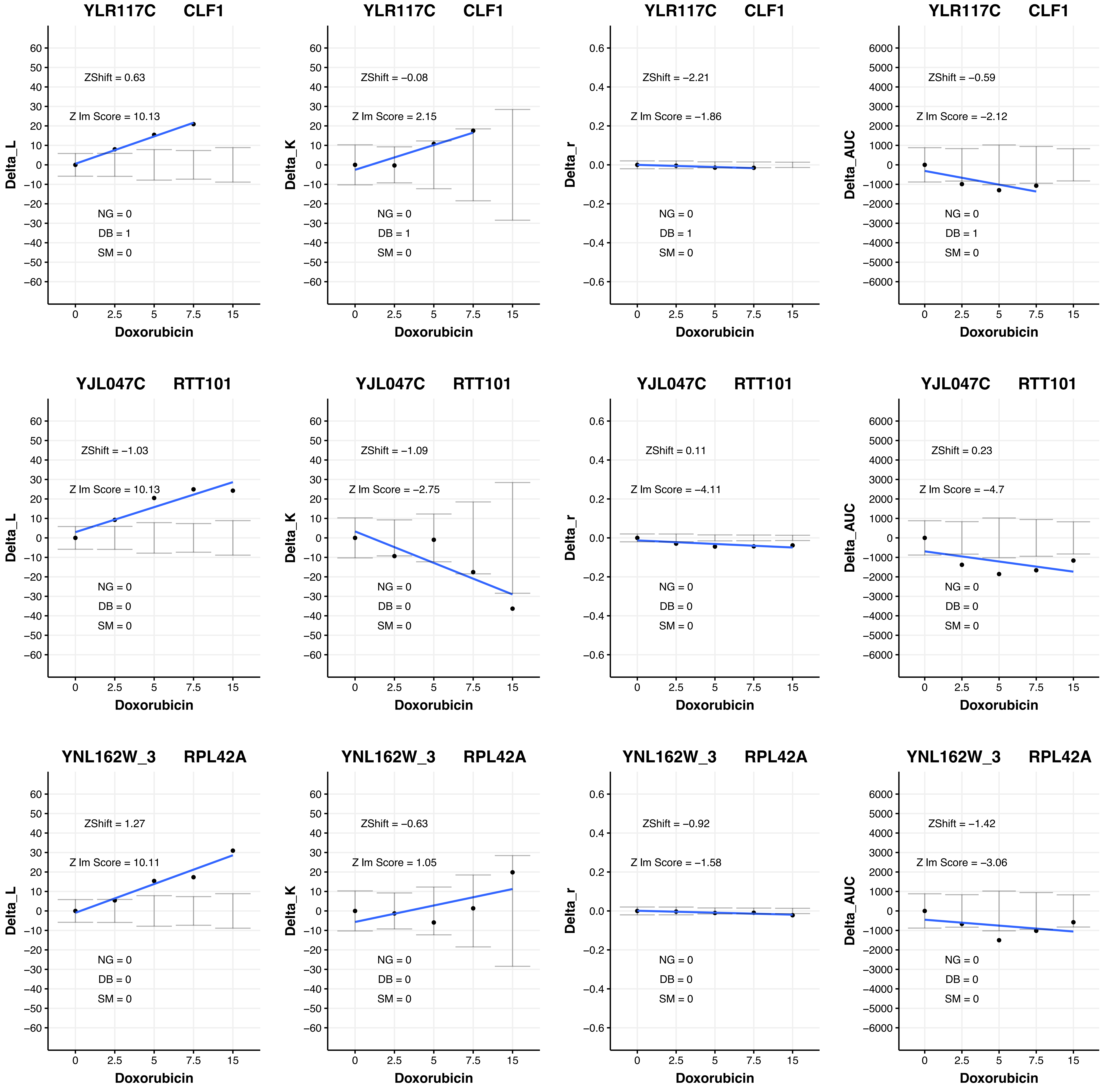
YLR117C CLF1
ZShift = 0.63
Z lm Score = 10.13
NG = 0
DB = 1
SM = 0
Delta_L
Doxorubicin
YLR117C CLF1
ZShift = −0.08
Z lm Score = 2.15
NG = 0
DB = 1
SM = 0
Delta_K
Doxorubicin
YLR117C CLF1
ZShift = −2.21
Z lm Score = −1.86
NG = 0
DB = 1
SM = 0
Delta_r
Doxorubicin
YLR117C CLF1
ZShift = −0.59
Z lm Score = −2.12
NG = 0
DB = 1
SM = 0
Delta_AUC
Doxorubicin
YJL047C RTT101
ZShift = −1.03
Z lm Score = 10.13
NG = 0
DB = 0
SM = 0
Delta_L
Doxorubicin
YJL047C RTT101
ZShift = −1.09
Z lm Score = −2.75
NG = 0
DB = 0
SM = 0
Delta_K
Doxorubicin
YJL047C RTT101
ZShift = 0.11
Z lm Score = −4.11
NG = 0
DB = 0
SM = 0
Delta_r
Doxorubicin
YJL047C RTT101
ZShift = 0.23
Z lm Score = −4.7
NG = 0
DB = 0
SM = 0
Delta_AUC
Doxorubicin
YNL162W_3 RPL42A
ZShift = 1.27
Z lm Score = 10.11
NG = 0
DB = 0
SM = 0
Delta_L
Doxorubicin
YNL162W_3 RPL42A
ZShift = −0.63
Z lm Score = 1.05
NG = 0
DB = 0
SM = 0
Delta_K
Doxorubicin
YNL162W_3 RPL42A
ZShift = −0.92
Z lm Score = −1.58
NG = 0
DB = 0
SM = 0
Delta_r
Doxorubicin
YNL162W_3 RPL42A
ZShift = −1.42
Z lm Score = −3.06
NG = 0
DB = 0
SM = 0
Delta_AUC
Doxorubicin

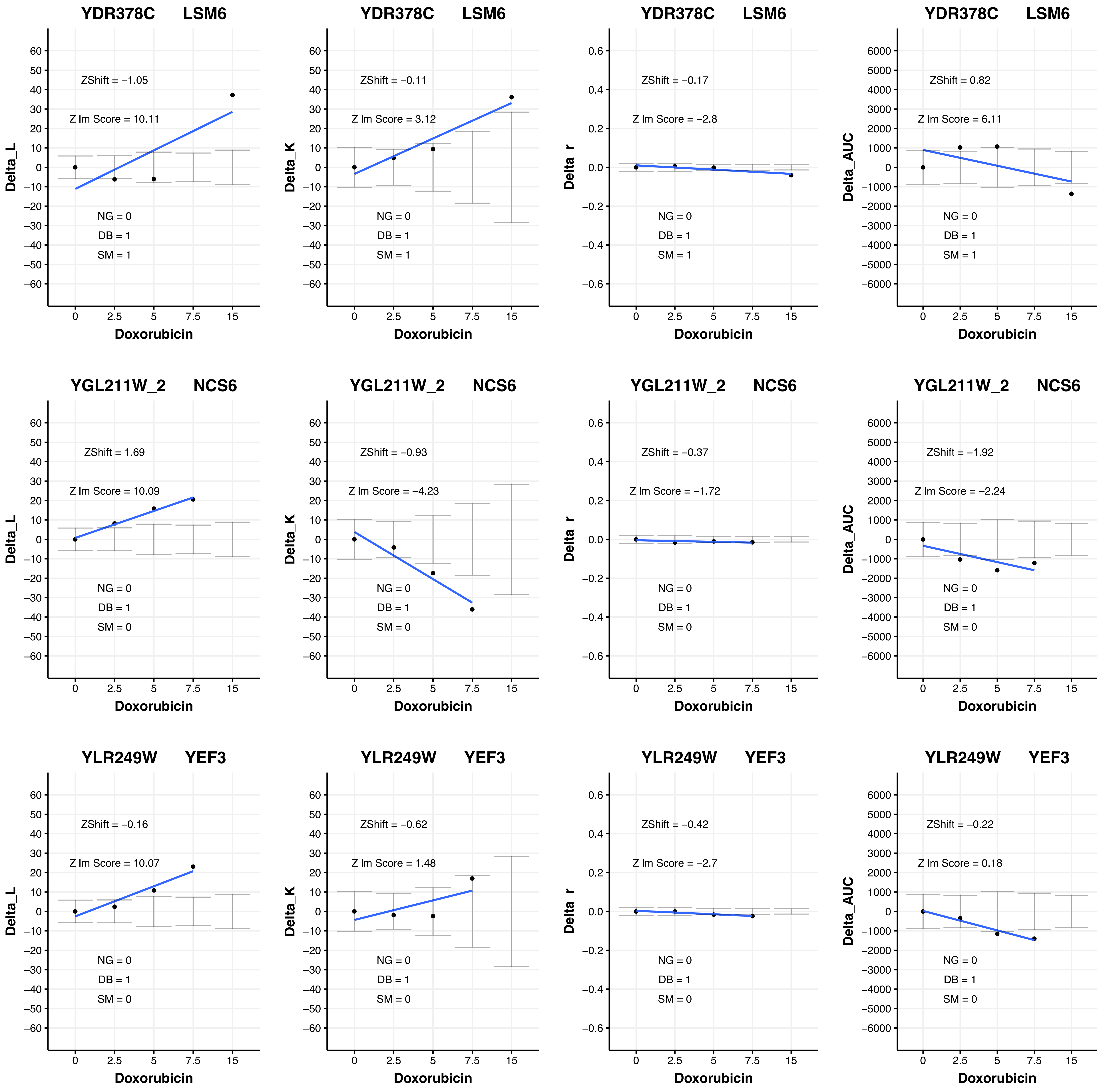

YDR378C LSM6
ZShift = −1.05
Z lm Score = 10.11
NG = 0
DB = 1
SM = 1
Delta_L
Doxorubicin
YDR378C LSM6
ZShift = −0.11
Z lm Score = 3.12
NG = 0
DB = 1
SM = 1
Delta_K
Doxorubicin
YDR378C LSM6
ZShift = −0.17
Z lm Score = −2.8
NG = 0
DB = 1
SM = 1
Delta_r
Doxorubicin
YDR378C LSM6
ZShift = 0.82
Z lm Score = 6.11
NG = 0
DB = 1
SM = 1
Delta_AUC
Doxorubicin
YGL211W_2 NCS6
ZShift = 1.69
Z lm Score = 10.09
NG = 0
DB = 1
SM = 0
Delta_L
Doxorubicin
YGL211W_2 NCS6
ZShift = −0.93
Z lm Score = −4.23
NG = 0
DB = 1
SM = 0
Delta_K
Doxorubicin
YGL211W_2 NCS6
ZShift = −0.37
Z lm Score = −1.72
NG = 0
DB = 1
SM = 0
Delta_r
Doxorubicin
YGL211W_2 NCS6
ZShift = −1.92
Z lm Score = −2.24
NG = 0
DB = 1
SM = 0
Delta_AUC
Doxorubicin
YLR249W YEF3
ZShift = −0.16
Z lm Score = 10.07
NG = 0
DB = 1
SM = 0
Delta_L
Doxorubicin
YLR249W YEF3
ZShift = −0.62
Z lm Score = 1.48
NG = 0
DB = 1
SM = 0
Delta_K
Doxorubicin
YLR249W YEF3
ZShift = −0.42
Z lm Score = −2.7
NG = 0
DB = 1
SM = 0
Delta_r
Doxorubicin
YLR249W YEF3
ZShift = −0.22
Z lm Score = 0.18
NG = 0
DB = 1
SM = 0
Delta_AUC
Doxorubicin

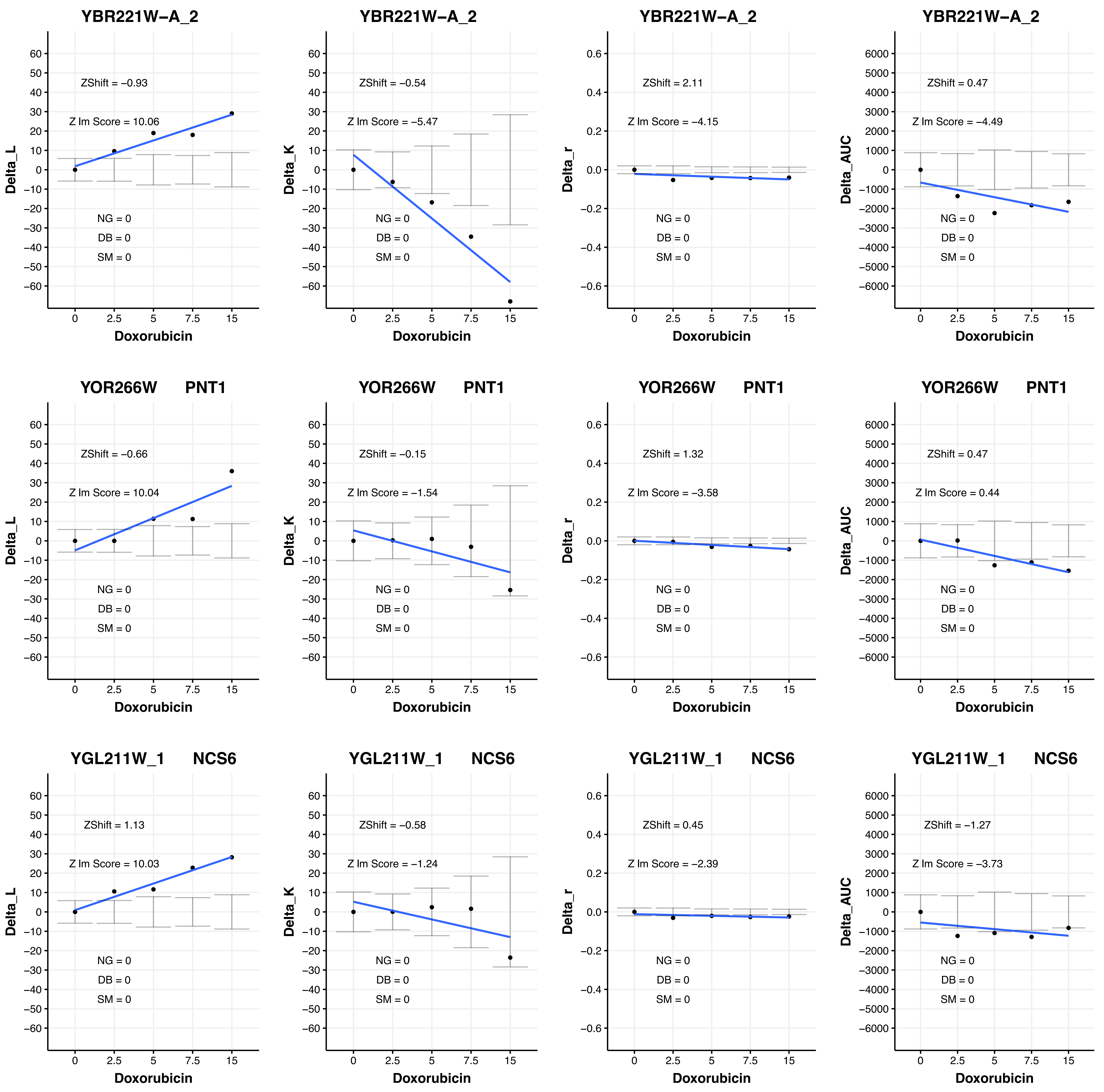

YBR221W−A_2
ZShift = −0.93
Z lm Score = 10.06
NG = 0
DB = 0
SM = 0
Delta_L
Doxorubicin
YBR221W−A_2
ZShift = −0.54
Z lm Score = −5.47
NG = 0
DB = 0
SM = 0
Delta_K
Doxorubicin
YBR221W−A_2
ZShift = 2.11
Z lm Score = −4.15
NG = 0
DB = 0
SM = 0
Delta_r
Doxorubicin
YBR221W−A_2
ZShift = 0.47
Z lm Score = −4.49
NG = 0
DB = 0
SM = 0
Delta_AUC
Doxorubicin
YOR266W PNT1
ZShift = −0.66
Z lm Score = 10.04
NG = 0
DB = 0
SM = 0
Delta_L
Doxorubicin
YOR266W PNT1
ZShift = −0.15
Z lm Score = −1.54
NG = 0
DB = 0
SM = 0
Delta_K
Doxorubicin
YOR266W PNT1
ZShift = 1.32
Z lm Score = −3.58
NG = 0
DB = 0
SM = 0
Delta_r
Doxorubicin
YOR266W PNT1
ZShift = 0.47
Z lm Score = 0.44
NG = 0
DB = 0
SM = 0
Delta_AUC
Doxorubicin
YGL211W_1 NCS6
ZShift = 1.13
Z lm Score = 10.03
NG = 0
DB = 0
SM = 0
Delta_L
Doxorubicin
YGL211W_1 NCS6
ZShift = −0.58
Z lm Score = −1.24
NG = 0
DB = 0
SM = 0
Delta_K
Doxorubicin
YGL211W_1 NCS6
ZShift = 0.45
Z lm Score = −2.39
NG = 0
DB = 0
SM = 0
Delta_r
Doxorubicin
YGL211W_1 NCS6
ZShift = −1.27
Z lm Score = −3.73
NG = 0
DB = 0
SM = 0
Delta_AUC
Doxorubicin

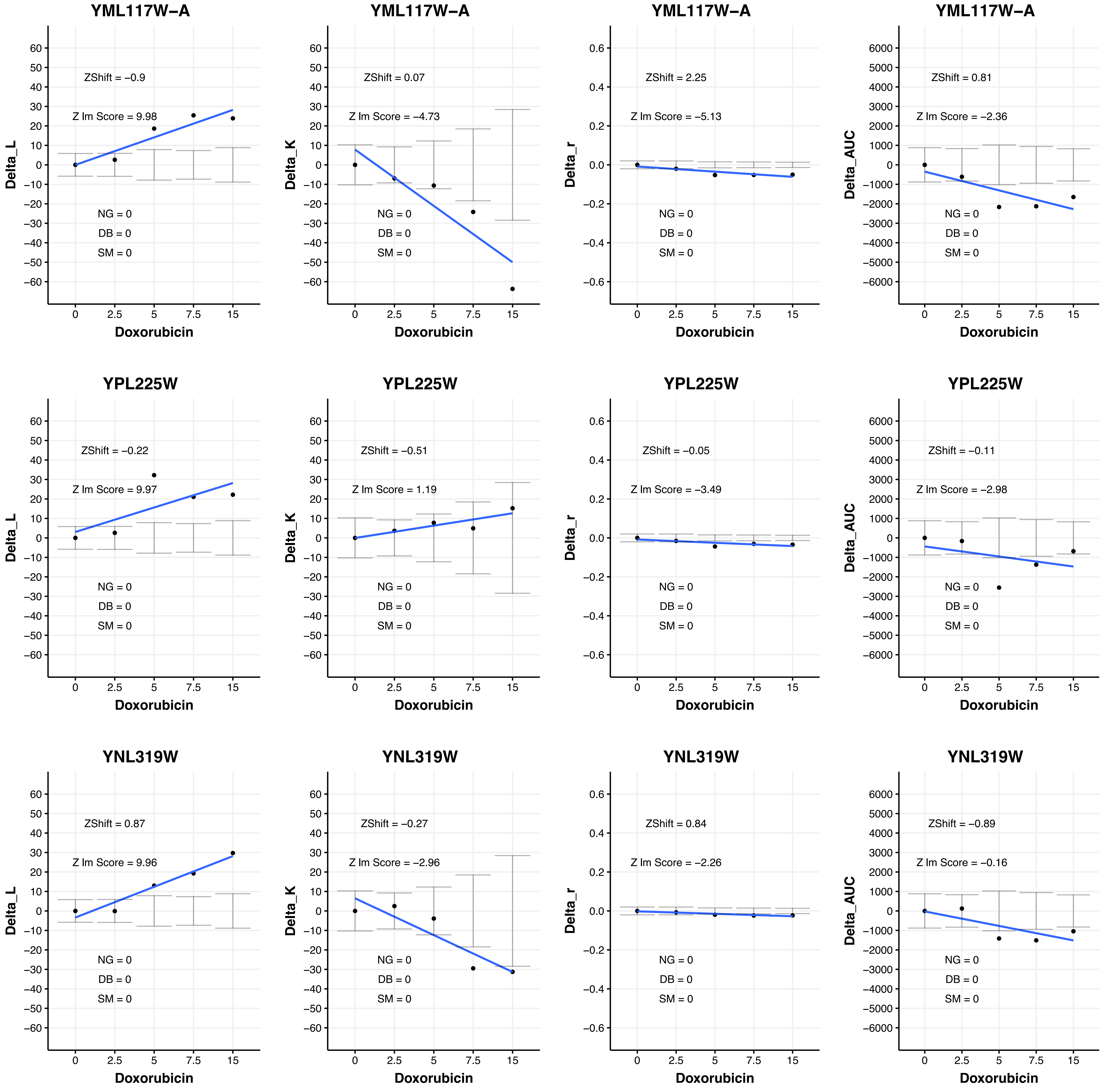
YML117W−A
ZShift = −0.9
Z lm Score = 9.98
NG = 0
DB = 0
SM = 0
Delta_L
Doxorubicin
YML117W−A
ZShift = 0.07
Z lm Score = −4.73
NG = 0
DB = 0
SM = 0
Delta_K
Doxorubicin
YML117W−A
ZShift = 2.25
Z lm Score = −5.13
NG = 0
DB = 0
SM = 0
Delta_r
Doxorubicin
YML117W−A
ZShift = 0.81
Z lm Score = −2.36
NG = 0
DB = 0
SM = 0
Delta_AUC
Doxorubicin
YPL225W
ZShift = −0.22
Z lm Score = 9.97
NG = 0
DB = 0
SM = 0
Delta_L
Doxorubicin
YPL225W
ZShift = −0.51
Z lm Score = 1.19
NG = 0
DB = 0
SM = 0
Delta_K
Doxorubicin
YPL225W
ZShift = −0.05
Z lm Score = −3.49
NG = 0
DB = 0
SM = 0
Delta_r
Doxorubicin
YPL225W
ZShift = −0.11
Z lm Score = −2.98
NG = 0
DB = 0
SM = 0
Delta_AUC
Doxorubicin
YNL319W
ZShift = 0.87
Z lm Score = 9.96
NG = 0
DB = 0
SM = 0
Delta_L
Doxorubicin
YNL319W
ZShift = −0.27
Z lm Score = −2.96
NG = 0
DB = 0
SM = 0
Delta_K
Doxorubicin
YNL319W
ZShift = 0.84
Z lm Score = −2.26
NG = 0
DB = 0
SM = 0
Delta_r
Doxorubicin
YNL319W
ZShift = −0.89
Z lm Score = −0.16
NG = 0
DB = 0
SM = 0
Delta_AUC
Doxorubicin

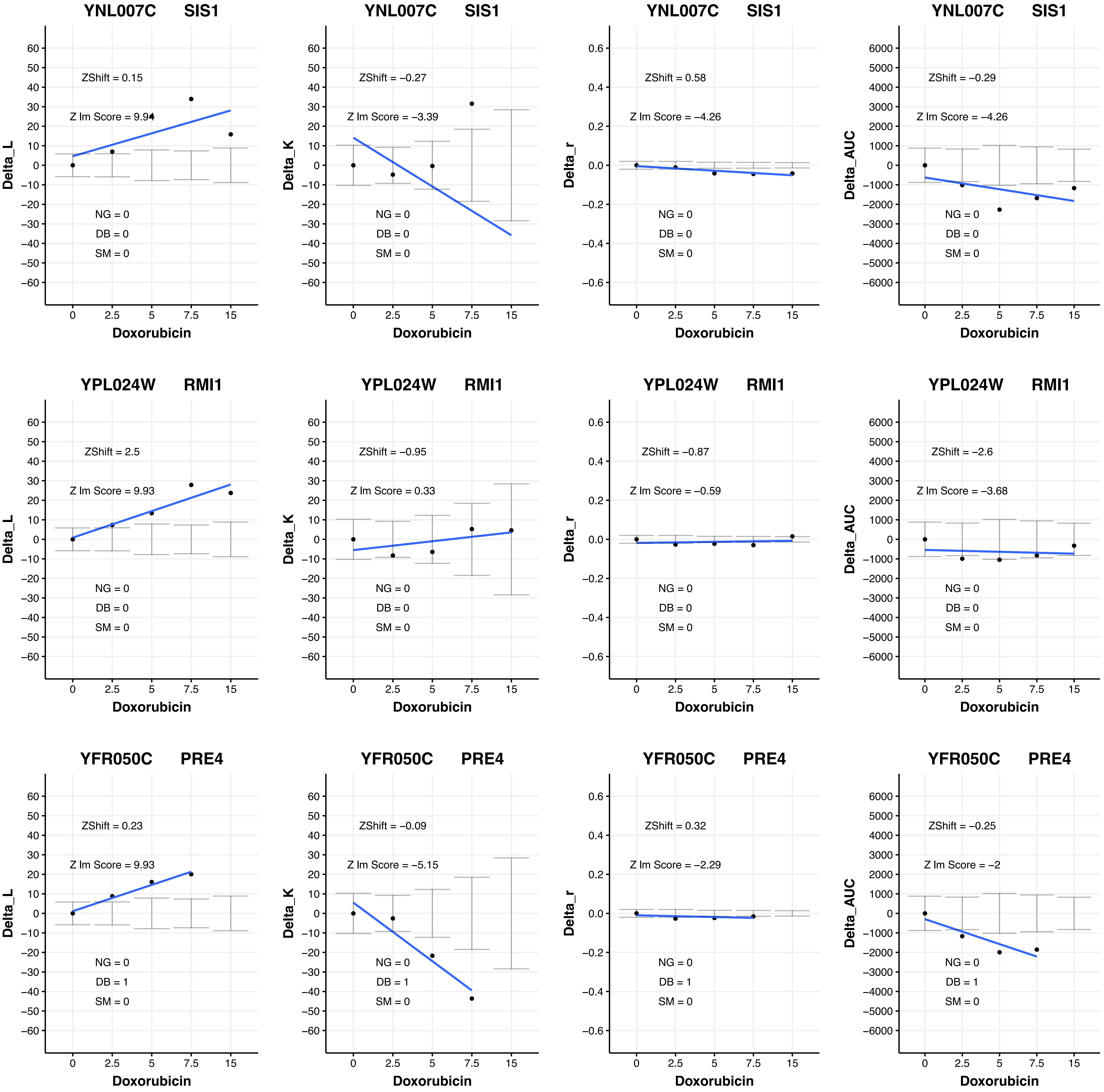

YNL007C SIS1
ZShift = 0.15
Z lm Score = 9.94
NG = 0
DB = 0
SM = 0
Delta_L
Doxorubicin
YNL007C SIS1
ZShift = −0.27
Z lm Score = −3.39
NG = 0
DB = 0
SM = 0
Delta_K
Doxorubicin
YNL007C SIS1
ZShift = 0.58
Z lm Score = −4.26
NG = 0
DB = 0
SM = 0
Delta_r
Doxorubicin
YNL007C SIS1
ZShift = −0.29
Z lm Score = −4.26
NG = 0
DB = 0
SM = 0
Delta_AUC
Doxorubicin
YPL024W RMI1
ZShift = 2.5
Z lm Score = 9.93
NG = 0
DB = 0
SM = 0
Delta_L
Doxorubicin
YPL024W RMI1
ZShift = −0.95
Z lm Score = 0.33
NG = 0
DB = 0
SM = 0
Delta_K
Doxorubicin
YPL024W RMI1
ZShift = −0.87
Z lm Score = −0.59
NG = 0
DB = 0
SM = 0
Delta_r
Doxorubicin
YPL024W RMI1
ZShift = −2.6
Z lm Score = −3.68
NG = 0
DB = 0
SM = 0
Delta_AUC
Doxorubicin
YFR050C PRE4
ZShift = 0.23
Z lm Score = 9.93
NG = 0
DB = 1
SM = 0
Delta_L
Doxorubicin
YFR050C PRE4
ZShift = −0.09
Z lm Score = −5.15
NG = 0
DB = 1
SM = 0
Delta_K
Doxorubicin
YFR050C PRE4
ZShift = 0.32
Z lm Score = −2.29
NG = 0
DB = 1
SM = 0
Delta_r
Doxorubicin
YFR050C PRE4
ZShift = −0.25
Z lm Score = −2
NG = 0
DB = 1
SM = 0
Delta_AUC
Doxorubicin

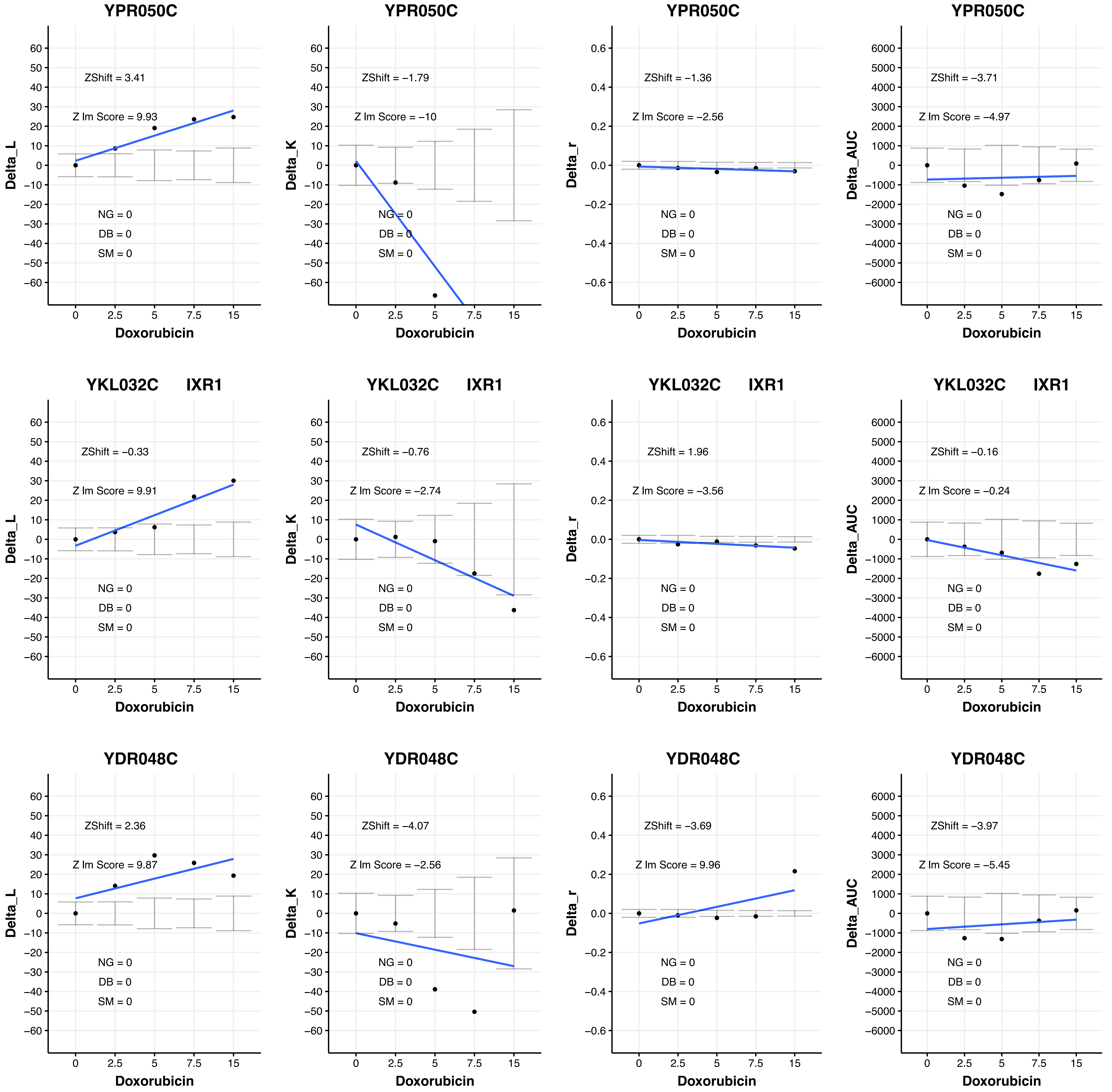

YPR050C
ZShift = 3.41
Z lm Score = 9.93
NG = 0
DB = 0
SM = 0
Delta_L
Doxorubicin
YPR050C
ZShift = −1.79
Z lm Score = −10
NG = 0
DB = 0
SM = 0
Delta_K
Doxorubicin
YPR050C
ZShift = −1.36
Z lm Score = −2.56
NG = 0
DB = 0
SM = 0
Delta_r
Doxorubicin
YPR050C
ZShift = −3.71
Z lm Score = −4.97
NG = 0
DB = 0
SM = 0
Delta_AUC
Doxorubicin
YKL032C IXR1
ZShift = −0.33
Z lm Score = 9.91
NG = 0
DB = 0
SM = 0
Delta_L
Doxorubicin
YKL032C IXR1
ZShift = −0.76
Z lm Score = −2.74
NG = 0
DB = 0
SM = 0
Delta_K
Doxorubicin
YKL032C IXR1
ZShift = 1.96
Z lm Score = −3.56
NG = 0
DB = 0
SM = 0
Delta_r
Doxorubicin
YKL032C IXR1
ZShift = −0.16
Z lm Score = −0.24
NG = 0
DB = 0
SM = 0
Delta_AUC
Doxorubicin
YDR048C
ZShift = 2.36
Z lm Score = 9.87
NG = 0
DB = 0
SM = 0
Delta_L
Doxorubicin
YDR048C
ZShift = −4.07
Z lm Score = −2.56
NG = 0
DB = 0
SM = 0
Delta_K
Doxorubicin
YDR048C
ZShift = −3.69
Z lm Score = 9.96
NG = 0
DB = 0
SM = 0
Delta_r
Doxorubicin
YDR048C
ZShift = −3.97
Z lm Score = −5.45
NG = 0
DB = 0
SM = 0
Delta_AUC
Doxorubicin

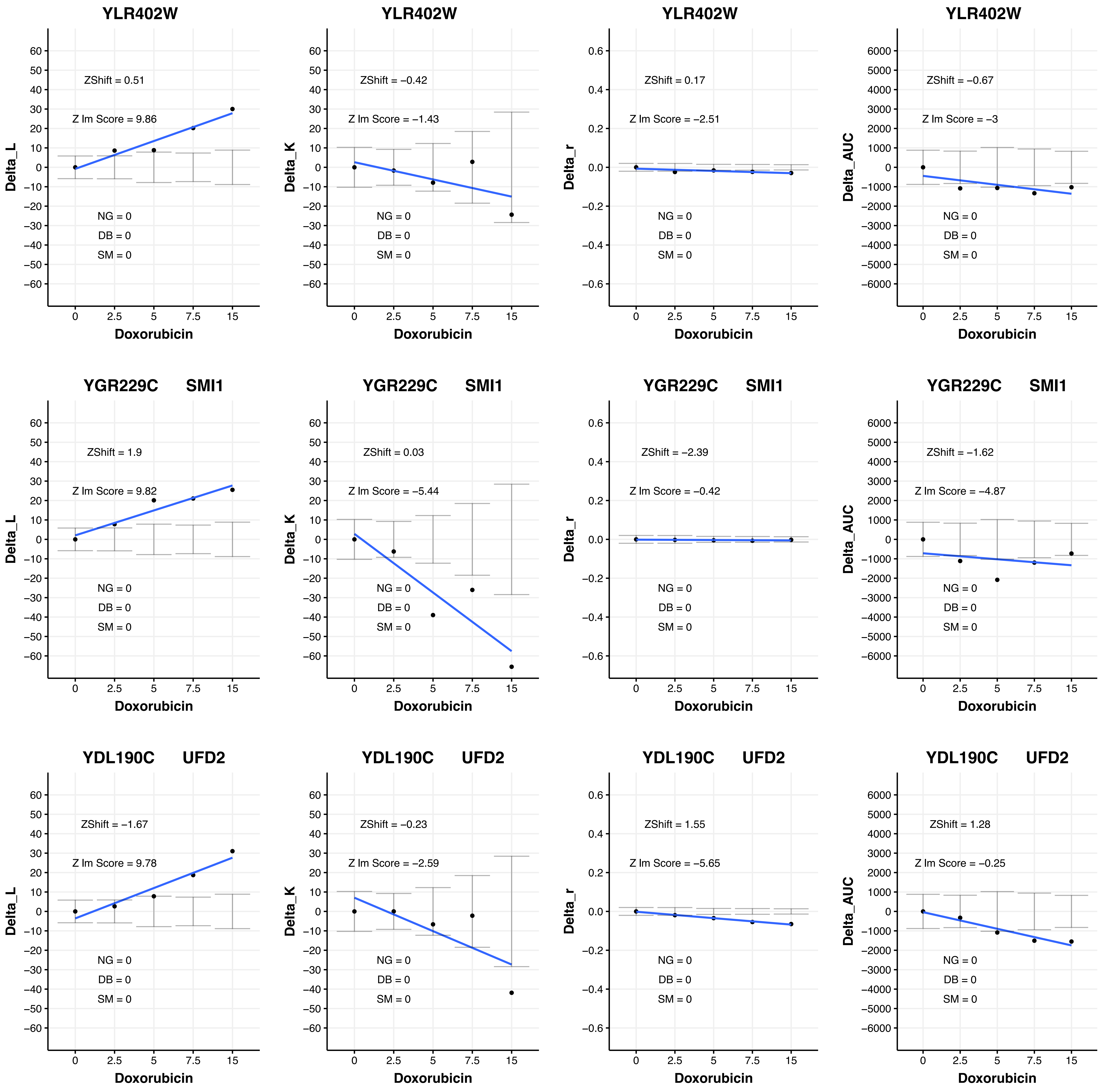
YLR402W
ZShift = 0.51
Z lm Score = 9.86
NG = 0
DB = 0
SM = 0
Delta_L
Doxorubicin
YLR402W
ZShift = −0.42
Z lm Score = −1.43
NG = 0
DB = 0
SM = 0
Delta_K
Doxorubicin
YLR402W
ZShift = 0.17
Z lm Score = −2.51
NG = 0
DB = 0
SM = 0
Delta_r
Doxorubicin
YLR402W
ZShift = −0.67
Z lm Score = −3
NG = 0
DB = 0
SM = 0
Delta_AUC
Doxorubicin
YGR229C SMI1
ZShift = 1.9
Z lm Score = 9.82
NG = 0
DB = 0
SM = 0
Delta_L
Doxorubicin
YGR229C SMI1
ZShift = 0.03
Z lm Score = −5.44
NG = 0
DB = 0
SM = 0
Delta_K
Doxorubicin
YGR229C SMI1
ZShift = −2.39
Z lm Score = −0.42
NG = 0
DB = 0
SM = 0
Delta_r
Doxorubicin
YGR229C SMI1
ZShift = −1.62
Z lm Score = −4.87
NG = 0
DB = 0
SM = 0
Delta_AUC
Doxorubicin
YDL190C UFD2
ZShift = −1.67
Z lm Score = 9.78
NG = 0
DB = 0
SM = 0
Delta_L
Doxorubicin
YDL190C UFD2
ZShift = −0.23
Z lm Score = −2.59
NG = 0
DB = 0
SM = 0
Delta_K
Doxorubicin
YDL190C UFD2
ZShift = 1.55
Z lm Score = −5.65
NG = 0
DB = 0
SM = 0
Delta_r
Doxorubicin
YDL190C UFD2
ZShift = 1.28
Z lm Score = −0.25
NG = 0
DB = 0
SM = 0
Delta_AUC
Doxorubicin

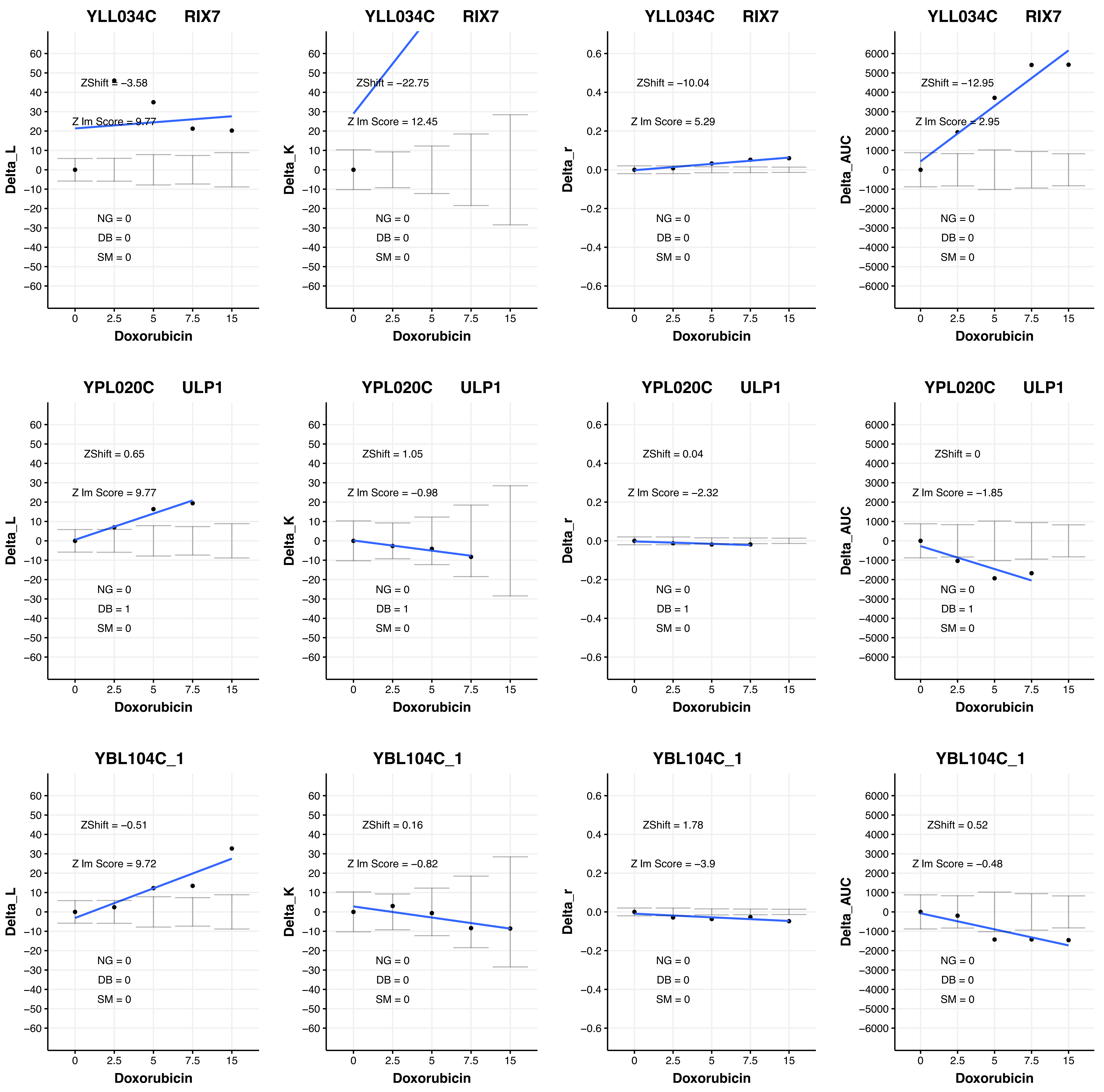
YLL034C RIX7
ZShift = −3.58
Z lm Score = 9.77
NG = 0
DB = 0
SM = 0
Delta_L
Doxorubicin
YLL034C RIX7
ZShift = −22.75
Z lm Score = 12.45
NG = 0
DB = 0
SM = 0
Delta_K
Doxorubicin
YLL034C RIX7
ZShift = −10.04
Z lm Score = 5.29
NG = 0
DB = 0
SM = 0
Delta_r
Doxorubicin
YLL034C RIX7
ZShift = −12.95
Z lm Score = 2.95
NG = 0
DB = 0
SM = 0
Delta_AUC
Doxorubicin
YPL020C ULP1
ZShift = 0.65
Z lm Score = 9.77
NG = 0
DB = 1
SM = 0
Delta_L
Doxorubicin
YPL020C ULP1
ZShift = 1.05
Z lm Score = −0.98
NG = 0
DB = 1
SM = 0
Delta_K
Doxorubicin
YPL020C ULP1
ZShift = 0.04
Z lm Score = −2.32
NG = 0
DB = 1
SM = 0
Delta_r
Doxorubicin
YPL020C ULP1
ZShift = 0
Z lm Score = −1.85
NG = 0
DB = 1
SM = 0
Delta_AUC
Doxorubicin
YBL104C_1
ZShift = −0.51
Z lm Score = 9.72
NG = 0
DB = 0
SM = 0
Delta_L
Doxorubicin
YBL104C_1
ZShift = 0.16
Z lm Score = −0.82
NG = 0
DB = 0
SM = 0
Delta_K
Doxorubicin
YBL104C_1
ZShift = 1.78
Z lm Score = −3.9
NG = 0
DB = 0
SM = 0
Delta_r
Doxorubicin
YBL104C_1
ZShift = 0.52
Z lm Score = −0.48
NG = 0
DB = 0
SM = 0
Delta_AUC
Doxorubicin

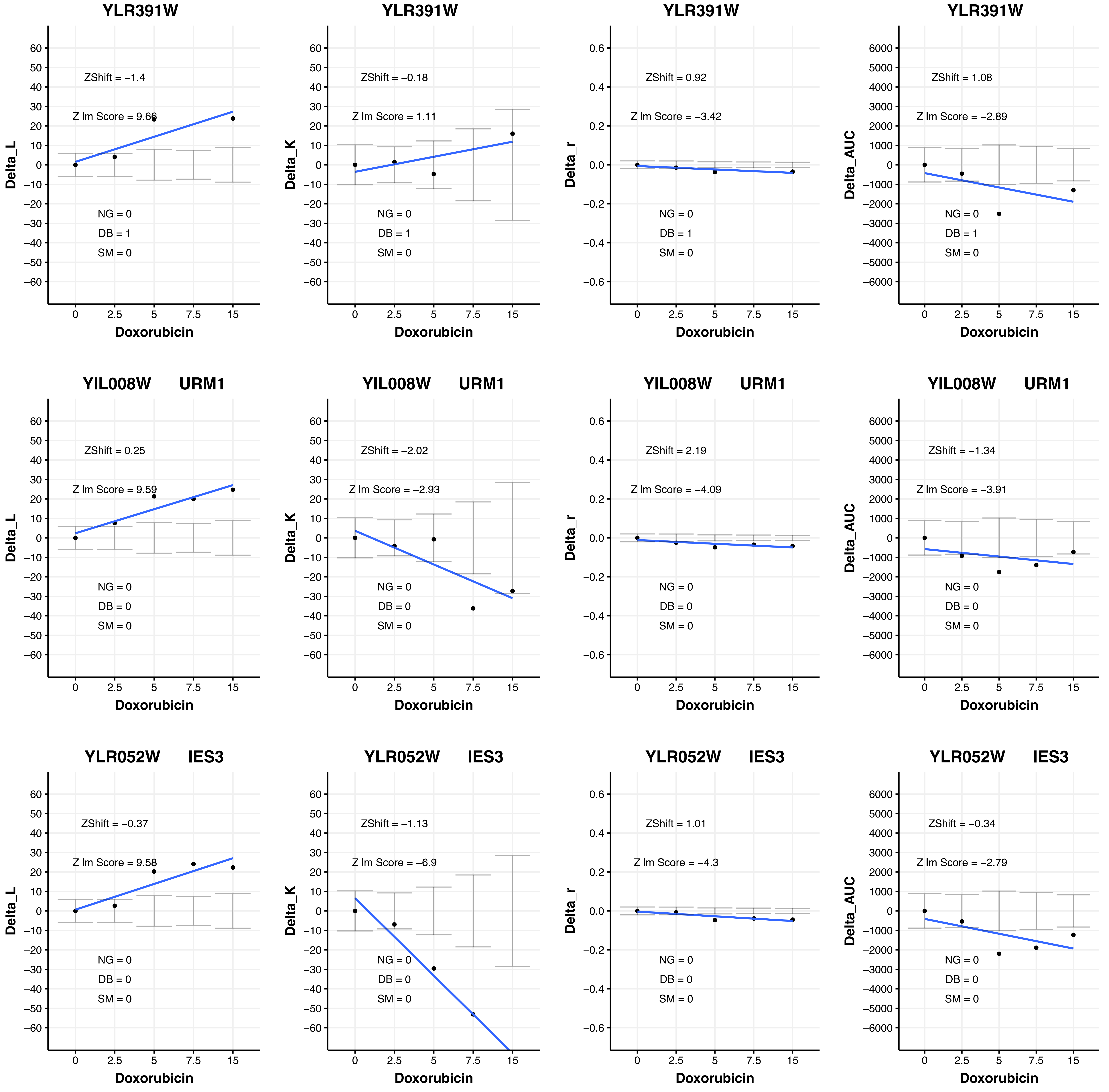
YLR391W
ZShift = −1.4
Z lm Score = 9.66
NG = 0
DB = 1
SM = 0
Delta_L
Doxorubicin
YLR391W
ZShift = −0.18
Z lm Score = 1.11
NG = 0
DB = 1
SM = 0
Delta_K
Doxorubicin
YLR391W
ZShift = 0.92
Z lm Score = −3.42
NG = 0
DB = 1
SM = 0
Delta_r
Doxorubicin
YLR391W
ZShift = 1.08
Z lm Score = −2.89
NG = 0
DB = 1
SM = 0
Delta_AUC
Doxorubicin
YIL008W URM1
ZShift = 0.25
Z lm Score = 9.59
NG = 0
DB = 0
SM = 0
Delta_L
Doxorubicin
YIL008W URM1
ZShift = −2.02
Z lm Score = −2.93
NG = 0
DB = 0
SM = 0
Delta_K
Doxorubicin
YIL008W URM1
ZShift = 2.19
Z lm Score = −4.09
NG = 0
DB = 0
SM = 0
Delta_r
Doxorubicin
YIL008W URM1
ZShift = −1.34
Z lm Score = −3.91
NG = 0
DB = 0
SM = 0
Delta_AUC
Doxorubicin
YLR052W IES3
ZShift = −0.37
Z lm Score = 9.58
NG = 0
DB = 0
SM = 0
Delta_L
Doxorubicin
YLR052W IES3
ZShift = −1.13
Z lm Score = −6.9
NG = 0
DB = 0
SM = 0
Delta_K
Doxorubicin
YLR052W IES3
ZShift = 1.01
Z lm Score = −4.3
NG = 0
DB = 0
SM = 0
Delta_r
Doxorubicin
YLR052W IES3
ZShift = −0.34
Z lm Score = −2.79
NG = 0
DB = 0
SM = 0
Delta_AUC
Doxorubicin

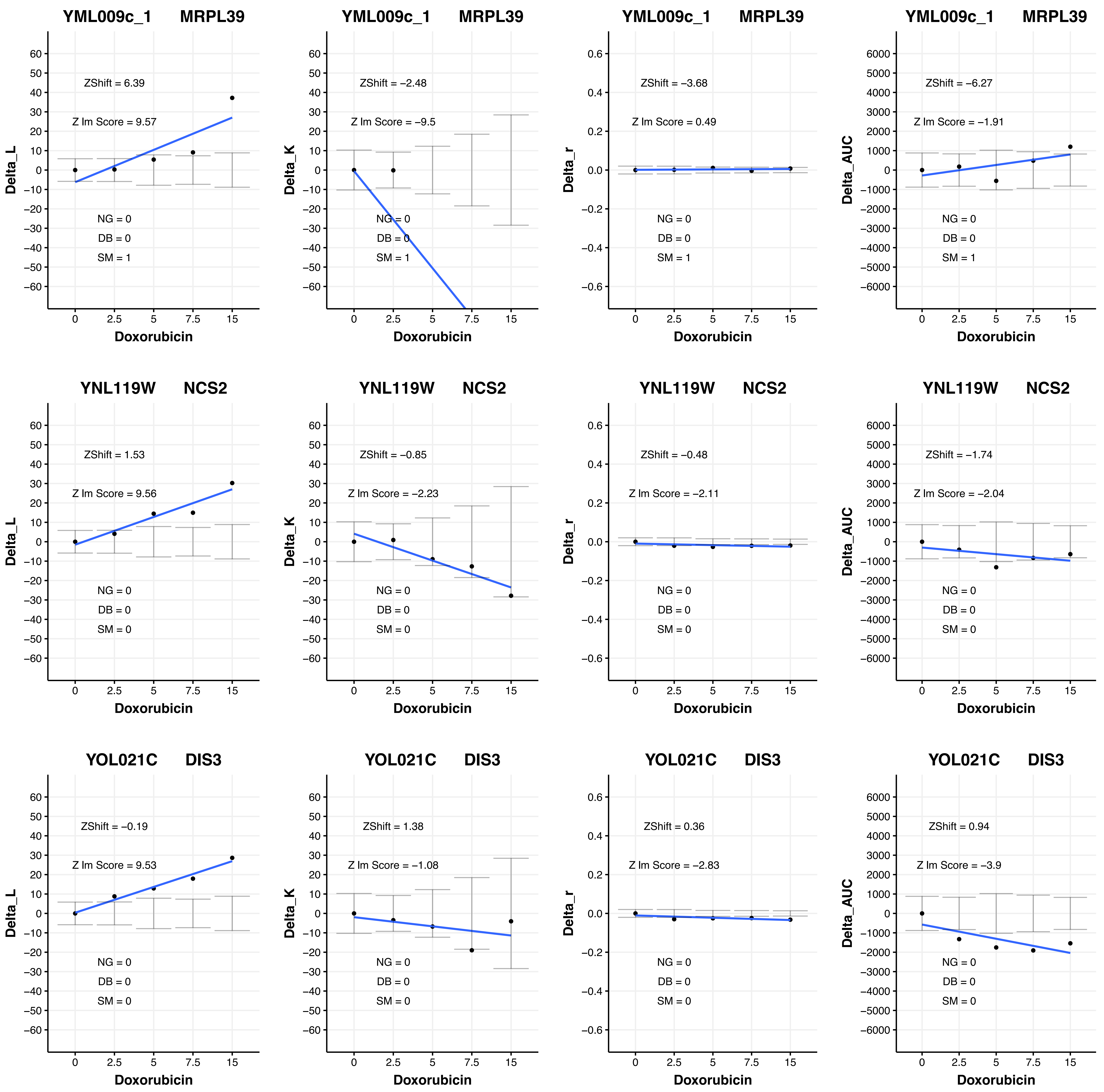
YML009c_1 MRPL39
ZShift = 6.39
Z lm Score = 9.57
NG = 0
DB = 0
SM = 1
Delta_L
Doxorubicin
YML009c_1 MRPL39
ZShift = −2.48
Z lm Score = −9.5
NG = 0
DB = 0
SM = 1
Delta_K
Doxorubicin
YML009c_1 MRPL39
ZShift = −3.68
Z lm Score = 0.49
NG = 0
DB = 0
SM = 1
Delta_r
Doxorubicin
YML009c_1 MRPL39
ZShift = −6.27
Z lm Score = −1.91
NG = 0
DB = 0
SM = 1
Delta_AUC
Doxorubicin
YNL119W NCS2
ZShift = 1.53
Z lm Score = 9.56
NG = 0
DB = 0
SM = 0
Delta_L
Doxorubicin
YNL119W NCS2
ZShift = −0.85
Z lm Score = −2.23
NG = 0
DB = 0
SM = 0
Delta_K
Doxorubicin
YNL119W NCS2
ZShift = −0.48
Z lm Score = −2.11
NG = 0
DB = 0
SM = 0
Delta_r
Doxorubicin
YNL119W NCS2
ZShift = −1.74
Z lm Score = −2.04
NG = 0
DB = 0
SM = 0
Delta_AUC
Doxorubicin
YOL021C DIS3
ZShift = −0.19
Z lm Score = 9.53
NG = 0
DB = 0
SM = 0
Delta_L
Doxorubicin
YOL021C DIS3
ZShift = 1.38
Z lm Score = −1.08
NG = 0
DB = 0
SM = 0
Delta_K
Doxorubicin
YOL021C DIS3
ZShift = 0.36
Z lm Score = −2.83
NG = 0
DB = 0
SM = 0
Delta_r
Doxorubicin
YOL021C DIS3
ZShift = 0.94
Z lm Score = −3.9
NG = 0
DB = 0
SM = 0
Delta_AUC
Doxorubicin

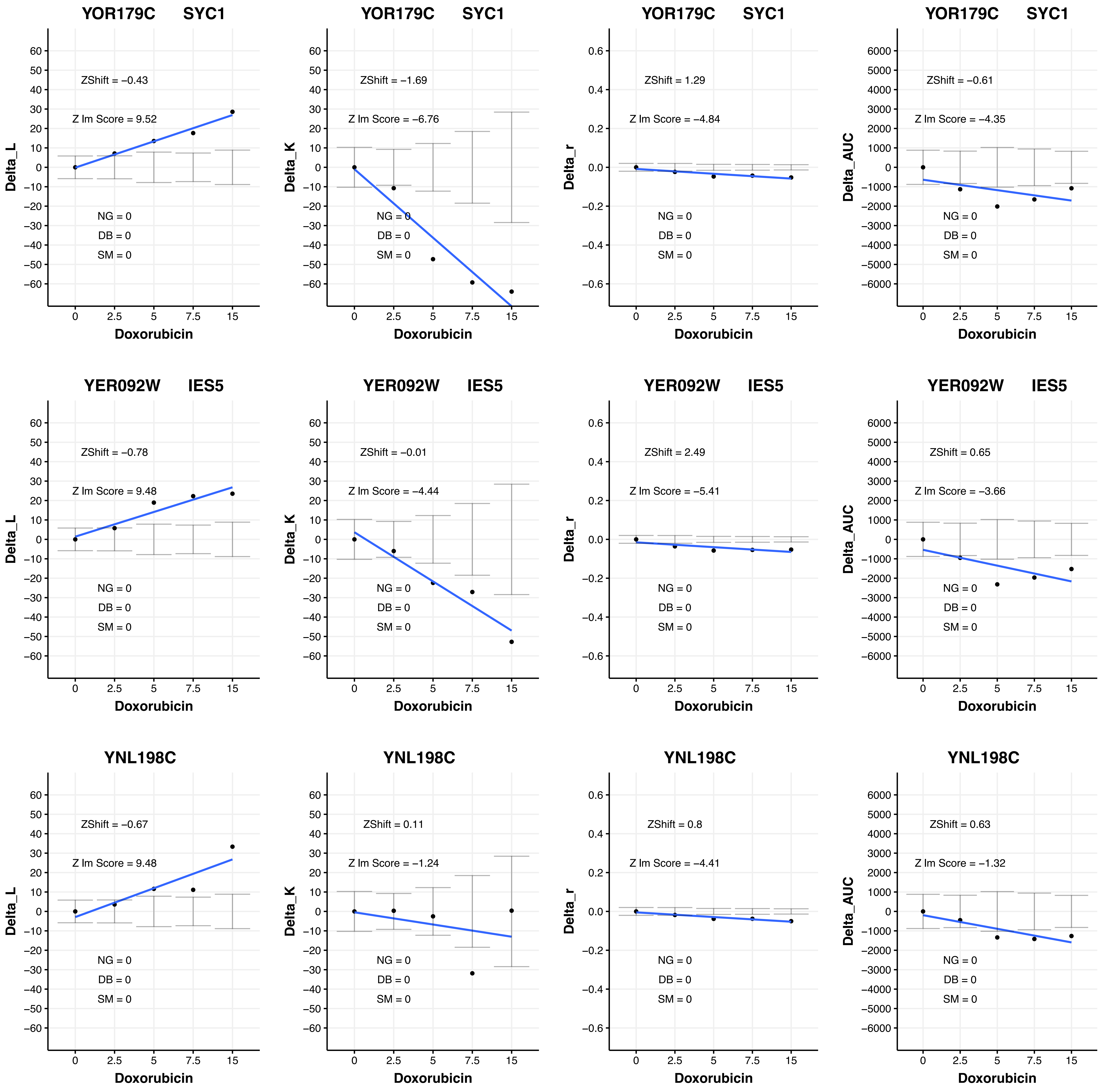

YOR179C SYC1
ZShift = −0.43
Z lm Score = 9.52
NG = 0
DB = 0
SM = 0
Delta_L
Doxorubicin
YOR179C SYC1
ZShift = −1.69
Z lm Score = −6.76
NG = 0
DB = 0
SM = 0
Delta_K
Doxorubicin
YOR179C SYC1
ZShift = 1.29
Z lm Score = −4.84
NG = 0
DB = 0
SM = 0
Delta_r
Doxorubicin
YOR179C SYC1
ZShift = −0.61
Z lm Score = −4.35
NG = 0
DB = 0
SM = 0
Delta_AUC
Doxorubicin
YER092W IES5
ZShift = −0.78
Z lm Score = 9.48
NG = 0
DB = 0
SM = 0
Delta_L
Doxorubicin
YER092W IES5
ZShift = −0.01
Z lm Score = −4.44
NG = 0
DB = 0
SM = 0
Delta_K
Doxorubicin
YER092W IES5
ZShift = 2.49
Z lm Score = −5.41
NG = 0
DB = 0
SM = 0
Delta_r
Doxorubicin
YER092W IES5
ZShift = 0.65
Z lm Score = −3.66
NG = 0
DB = 0
SM = 0
Delta_AUC
Doxorubicin
YNL198C
ZShift = −0.67
Z lm Score = 9.48
NG = 0
DB = 0
SM = 0
Delta_L
Doxorubicin
YNL198C
ZShift = 0.11
Z lm Score = −1.24
NG = 0
DB = 0
SM = 0
Delta_K
Doxorubicin
YNL198C
ZShift = 0.8
Z lm Score = −4.41
NG = 0
DB = 0
SM = 0
Delta_r
Doxorubicin
YNL198C
ZShift = 0.63
Z lm Score = −1.32
NG = 0
DB = 0
SM = 0
Delta_AUC
Doxorubicin

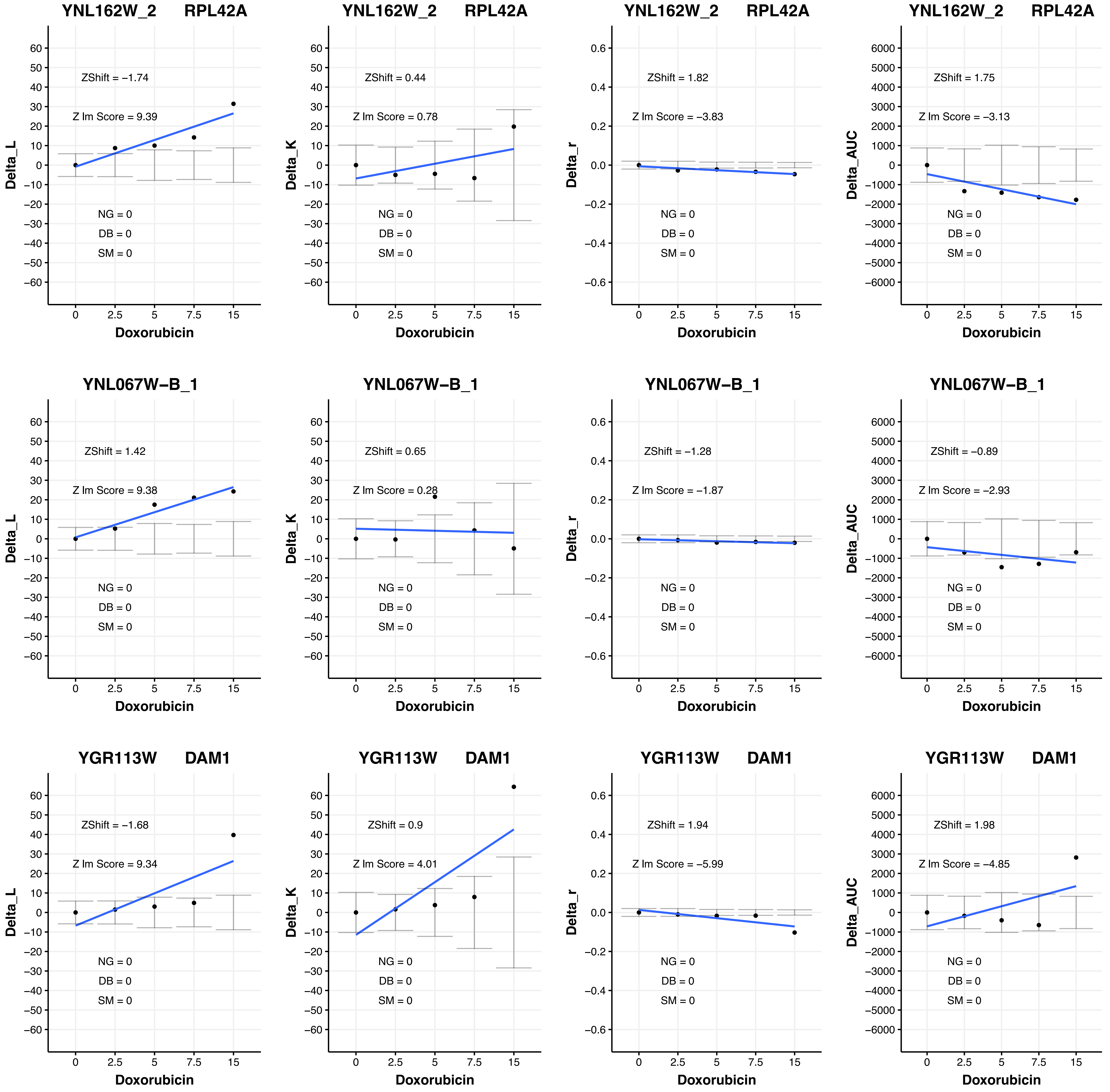
YNL162W_2 RPL42A
ZShift = −1.74
Z lm Score = 9.39
NG = 0
DB = 0
SM = 0
Delta_L
Doxorubicin
YNL162W_2 RPL42A
ZShift = 0.44
Z lm Score = 0.78
NG = 0
DB = 0
SM = 0
Delta_K
Doxorubicin
YNL162W_2 RPL42A
ZShift = 1.82
Z lm Score = −3.83
NG = 0
DB = 0
SM = 0
Delta_r
Doxorubicin
YNL162W_2 RPL42A
ZShift = 1.75
Z lm Score = −3.13
NG = 0
DB = 0
SM = 0
Delta_AUC
Doxorubicin
YNL067W−B_1
ZShift = 1.42
Z lm Score = 9.38
NG = 0
DB = 0
SM = 0
Delta_L
Doxorubicin
YNL067W−B_1
ZShift = 0.65
Z lm Score = 0.28
NG = 0
DB = 0
SM = 0
Delta_K
Doxorubicin
YNL067W−B_1
ZShift = −1.28
Z lm Score = −1.87
NG = 0
DB = 0
SM = 0
Delta_r
Doxorubicin
YNL067W−B_1
ZShift = −0.89
Z lm Score = −2.93
NG = 0
DB = 0
SM = 0
Delta_AUC
Doxorubicin
YGR113W DAM1
ZShift = −1.68
Z lm Score = 9.34
NG = 0
DB = 0
SM = 0
Delta_L
Doxorubicin
YGR113W DAM1
ZShift = 0.9
Z lm Score = 4.01
NG = 0
DB = 0
SM = 0
Delta_K
Doxorubicin
YGR113W DAM1
ZShift = 1.94
Z lm Score = −5.99
NG = 0
DB = 0
SM = 0
Delta_r
Doxorubicin
YGR113W DAM1
ZShift = 1.98
Z lm Score = −4.85
NG = 0
DB = 0
SM = 0
Delta_AUC
Doxorubicin

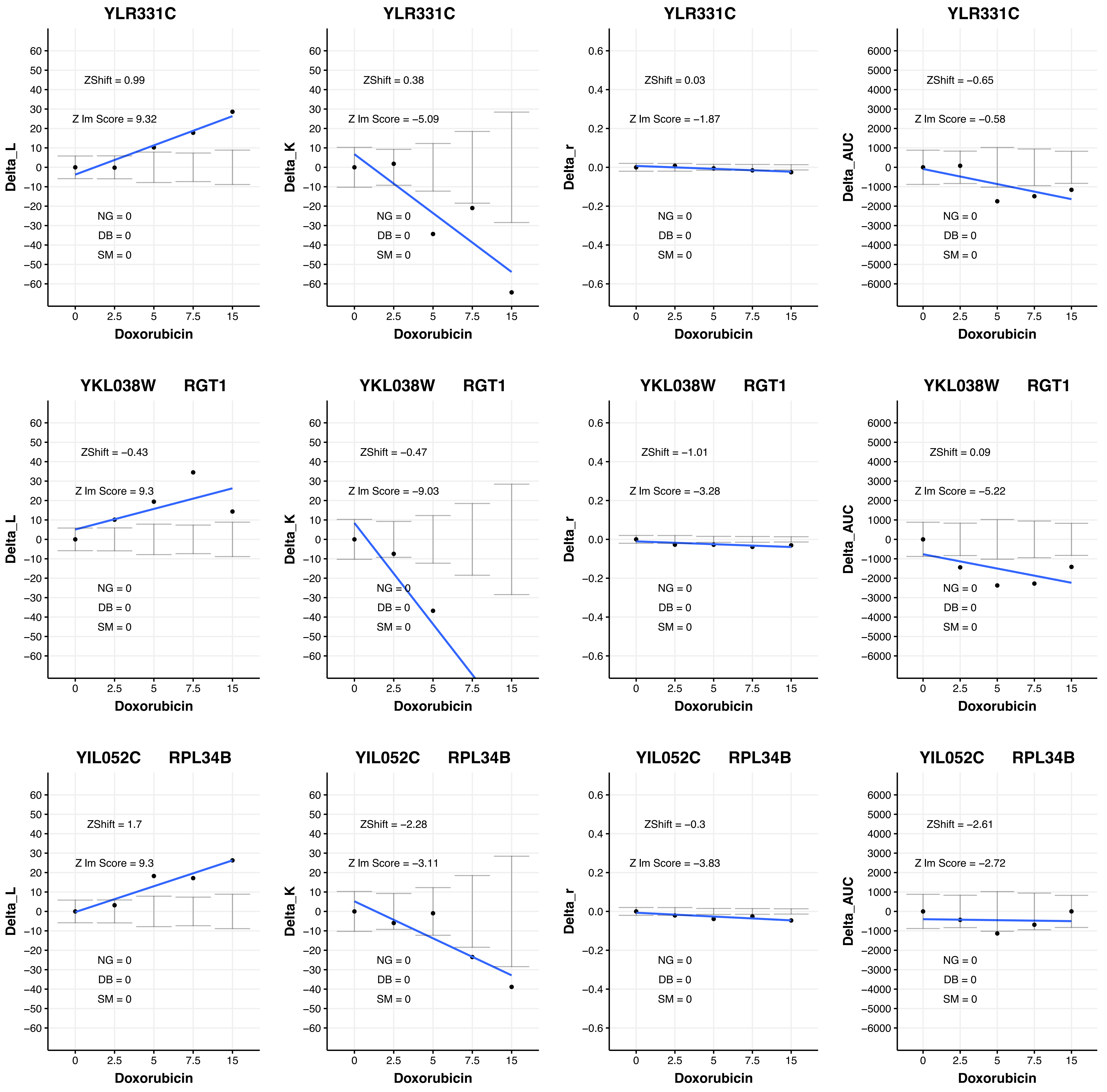
YLR331C
ZShift = 0.99
Z lm Score = 9.32
NG = 0
DB = 0
SM = 0
Delta_L
Doxorubicin
YLR331C
ZShift = 0.38
Z lm Score = −5.09
NG = 0
DB = 0
SM = 0
Delta_K
Doxorubicin
YLR331C
ZShift = 0.03
Z lm Score = −1.87
NG = 0
DB = 0
SM = 0
Delta_r
Doxorubicin
YLR331C
ZShift = −0.65
Z lm Score = −0.58
NG = 0
DB = 0
SM = 0
Delta_AUC
Doxorubicin
YKL038W RGT1
ZShift = −0.43
Z lm Score = 9.3
NG = 0
DB = 0
SM = 0
Delta_L
Doxorubicin
YKL038W RGT1
ZShift = −0.47
Z lm Score = −9.03
NG = 0
DB = 0
SM = 0
Delta_K
Doxorubicin
YKL038W RGT1
ZShift = −1.01
Z lm Score = −3.28
NG = 0
DB = 0
SM = 0
Delta_r
Doxorubicin
YKL038W RGT1
ZShift = 0.09
Z lm Score = −5.22
NG = 0
DB = 0
SM = 0
Delta_AUC
Doxorubicin
YIL052C RPL34B
ZShift = 1.7
Z lm Score = 9.3
NG = 0
DB = 0
SM = 0
Delta_L
Doxorubicin
YIL052C RPL34B
ZShift = −2.28
Z lm Score = −3.11
NG = 0
DB = 0
SM = 0
Delta_K
Doxorubicin
YIL052C RPL34B
ZShift = −0.3
Z lm Score = −3.83
NG = 0
DB = 0
SM = 0
Delta_r
Doxorubicin
YIL052C RPL34B
ZShift = −2.61
Z lm Score = −2.72
NG = 0
DB = 0
SM = 0
Delta_AUC
Doxorubicin

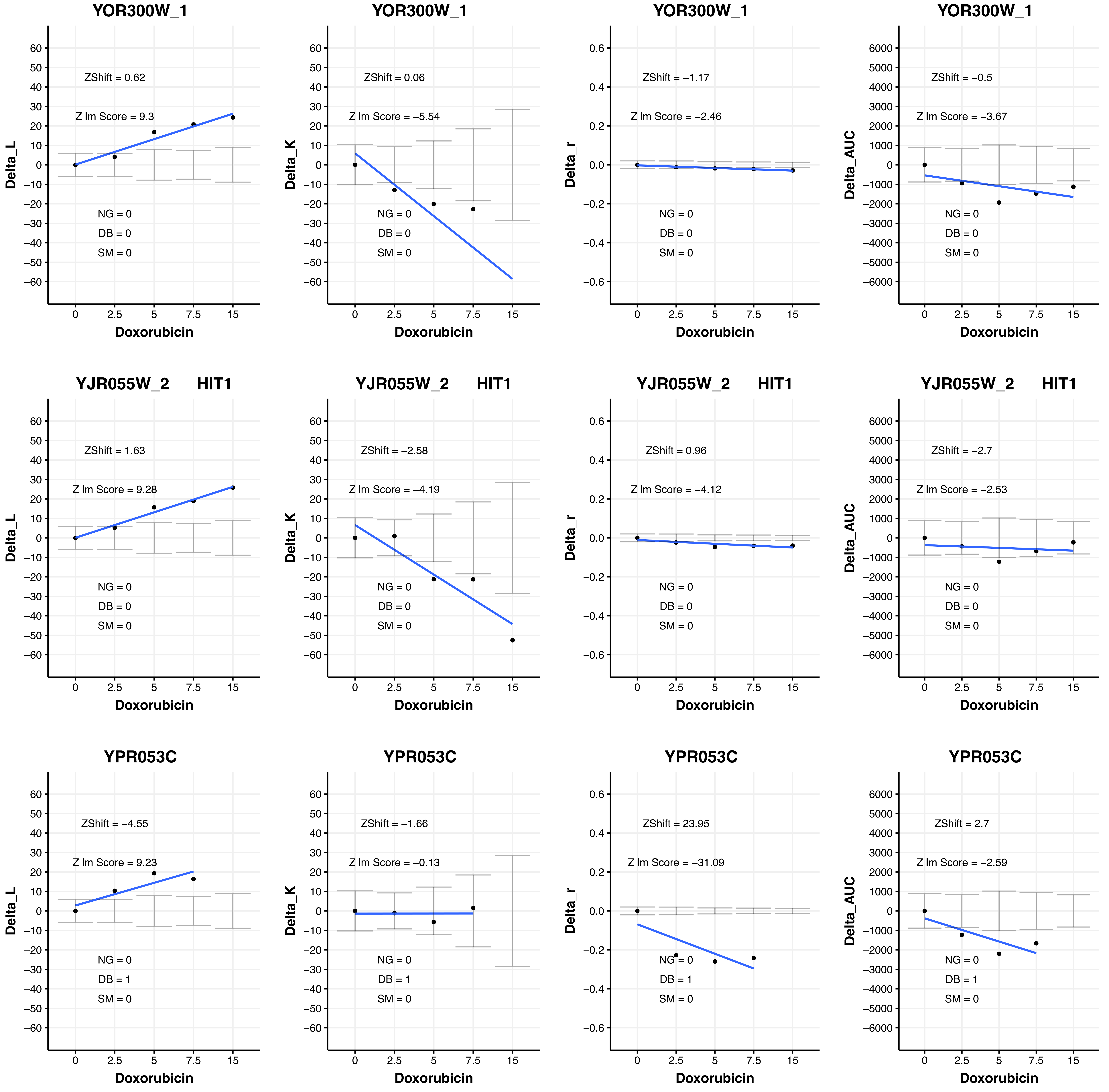
YOR300W_1
ZShift = 0.62
Z lm Score = 9.3
NG = 0
DB = 0
SM = 0
Delta_L
Doxorubicin
YOR300W_1
ZShift = 0.06
Z lm Score = −5.54
NG = 0
DB = 0
SM = 0
Delta_K
Doxorubicin
YOR300W_1
ZShift = −1.17
Z lm Score = −2.46
NG = 0
DB = 0
SM = 0
Delta_r
Doxorubicin
YOR300W_1
ZShift = −0.5
Z lm Score = −3.67
NG = 0
DB = 0
SM = 0
Delta_AUC
Doxorubicin
YJR055W_2 HIT1
ZShift = 1.63
Z lm Score = 9.28
NG = 0
DB = 0
SM = 0
Delta_L
Doxorubicin
YJR055W_2 HIT1
ZShift = −2.58
Z lm Score = −4.19
NG = 0
DB = 0
SM = 0
Delta_K
Doxorubicin
YJR055W_2 HIT1
ZShift = 0.96
Z lm Score = −4.12
NG = 0
DB = 0
SM = 0
Delta_r
Doxorubicin
YJR055W_2 HIT1
ZShift = −2.7
Z lm Score = −2.53
NG = 0
DB = 0
SM = 0
Delta_AUC
Doxorubicin
YPR053C
ZShift = −4.55
Z lm Score = 9.23
NG = 0
DB = 1
SM = 0
Delta_L
Doxorubicin
YPR053C
ZShift = −1.66
Z lm Score = −0.13
NG = 0
DB = 1
SM = 0
Delta_K
Doxorubicin
YPR053C
ZShift = 23.95
Z lm Score = −31.09
NG = 0
DB = 1
SM = 0
Delta_r
Doxorubicin
YPR053C
ZShift = 2.7
Z lm Score = −2.59
NG = 0
DB = 1
SM = 0
Delta_AUC
Doxorubicin

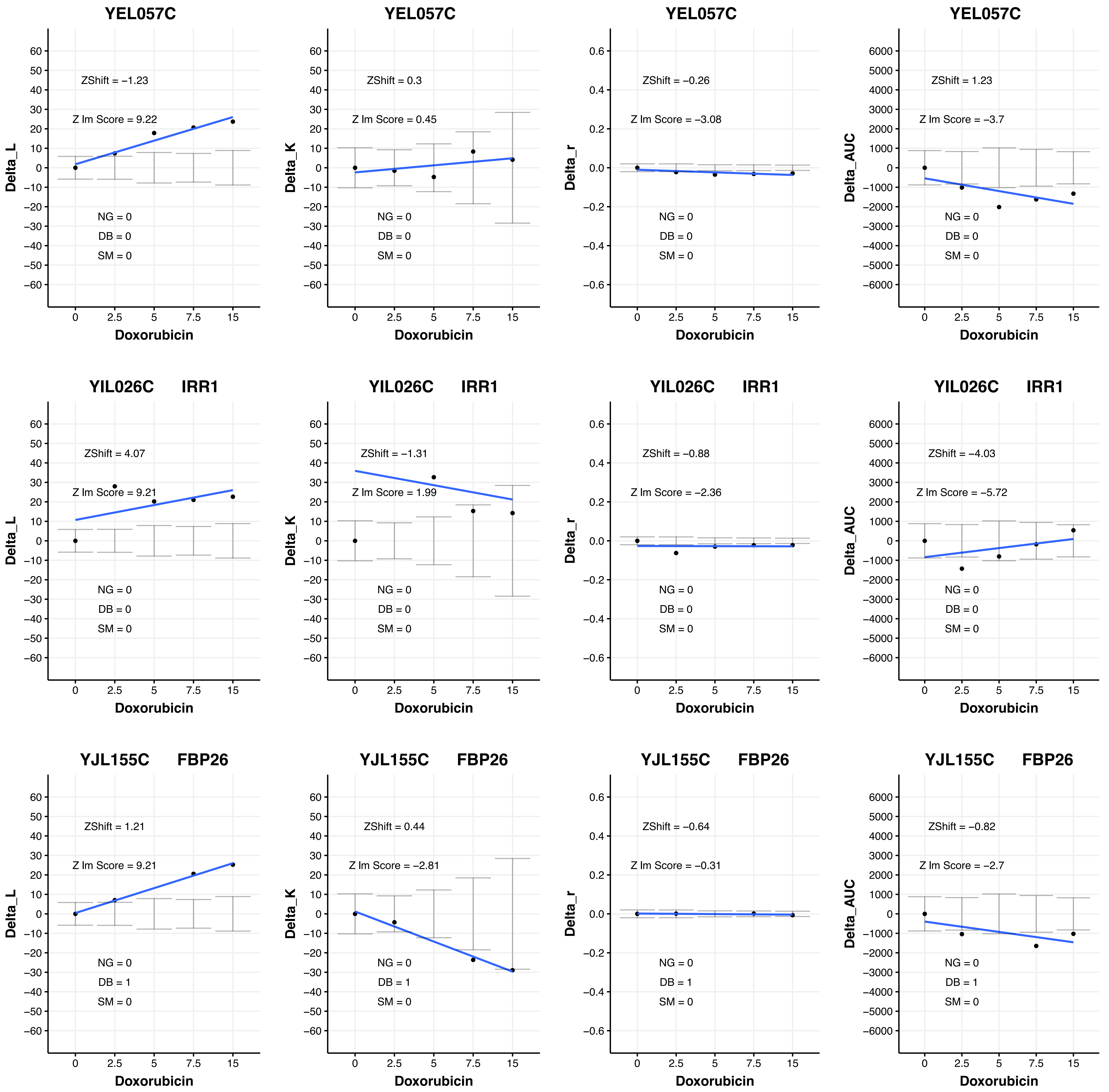
YEL057C
ZShift = −1.23
Z lm Score = 9.22
NG = 0
DB = 0
SM = 0
Delta_L
Doxorubicin
YEL057C
ZShift = 0.3
Z lm Score = 0.45
NG = 0
DB = 0
SM = 0
Delta_K
Doxorubicin
YEL057C
ZShift = −0.26
Z lm Score = −3.08
NG = 0
DB = 0
SM = 0
Delta_r
Doxorubicin
YEL057C
ZShift = 1.23
Z lm Score = −3.7
NG = 0
DB = 0
SM = 0
Delta_AUC
Doxorubicin
YIL026C IRR1
ZShift = 4.07
Z lm Score = 9.21
NG = 0
DB = 0
SM = 0
Delta_L
Doxorubicin
YIL026C IRR1
ZShift = −1.31
Z lm Score = 1.99
NG = 0
DB = 0
SM = 0
Delta_K
Doxorubicin
YIL026C IRR1
ZShift = −0.88
Z lm Score = −2.36
NG = 0
DB = 0
SM = 0
Delta_r
Doxorubicin
YIL026C IRR1
ZShift = −4.03
Z lm Score = −5.72
NG = 0
DB = 0
SM = 0
Delta_AUC
Doxorubicin
YJL155C FBP26
ZShift = 1.21
Z lm Score = 9.21
NG = 0
DB = 1
SM = 0
Delta_L
Doxorubicin
YJL155C FBP26
ZShift = 0.44
Z lm Score = −2.81
NG = 0
DB = 1
SM = 0
Delta_K
Doxorubicin
YJL155C FBP26
ZShift = −0.64
Z lm Score = −0.31
NG = 0
DB = 1
SM = 0
Delta_r
Doxorubicin
YJL155C FBP26
ZShift = −0.82
Z lm Score = −2.7
NG = 0
DB = 1
SM = 0
Delta_AUC
Doxorubicin

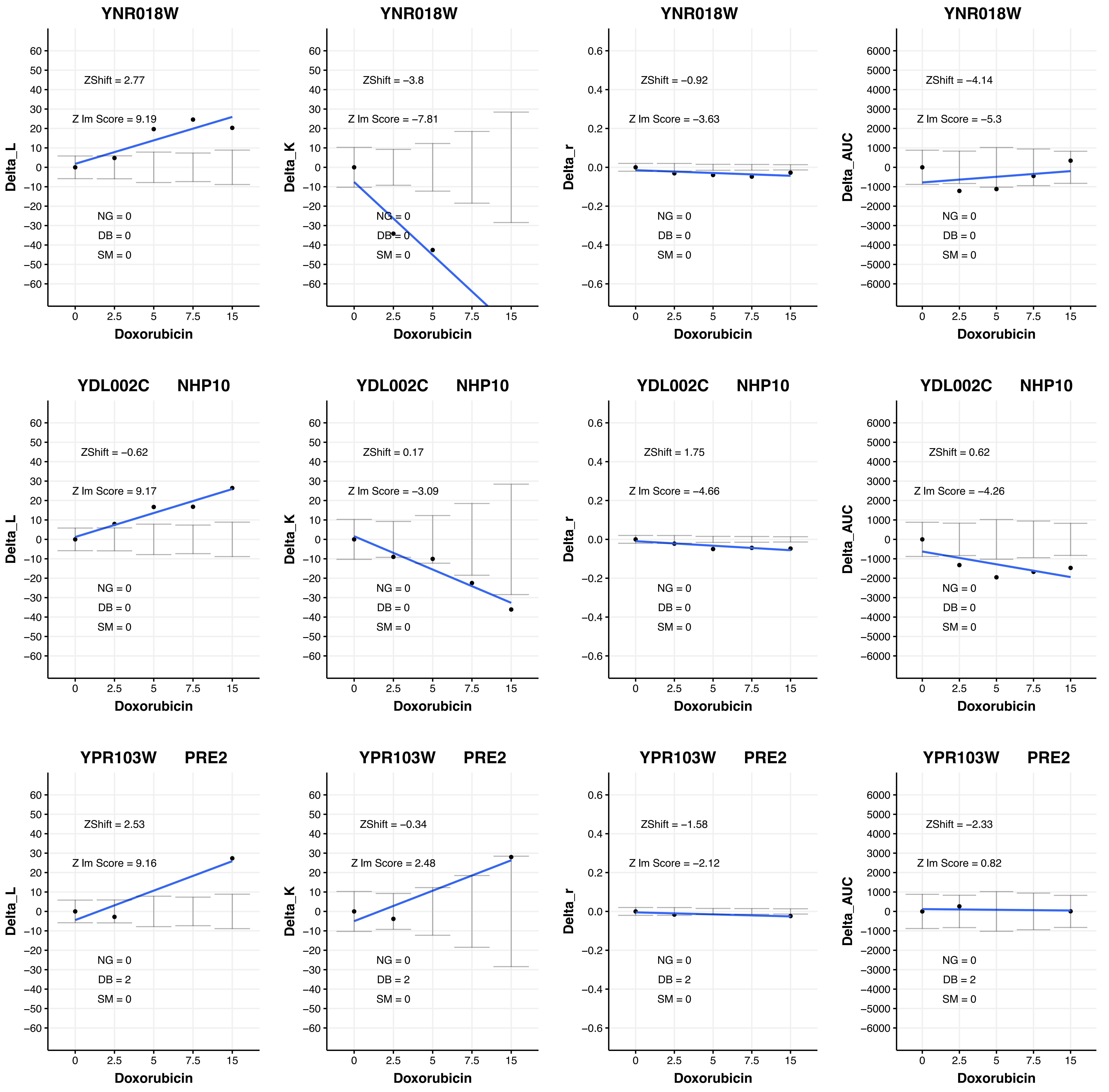
YNR018W
ZShift = 2.77
Z lm Score = 9.19
NG = 0
DB = 0
SM = 0
Delta_L
Doxorubicin
YNR018W
ZShift = −3.8
Z lm Score = −7.81
NG = 0
DB = 0
SM = 0
Delta_K
Doxorubicin
YNR018W
ZShift = −0.92
Z lm Score = −3.63
NG = 0
DB = 0
SM = 0
Delta_r
Doxorubicin
YNR018W
ZShift = −4.14
Z lm Score = −5.3
NG = 0
DB = 0
SM = 0
Delta_AUC
Doxorubicin
YDL002C NHP10
ZShift = −0.62
Z lm Score = 9.17
NG = 0
DB = 0
SM = 0
Delta_L
Doxorubicin
YDL002C NHP10
ZShift = 0.17
Z lm Score = −3.09
NG = 0
DB = 0
SM = 0
Delta_K
Doxorubicin
YDL002C NHP10
ZShift = 1.75
Z lm Score = −4.66
NG = 0
DB = 0
SM = 0
Delta_r
Doxorubicin
YDL002C NHP10
ZShift = 0.62
Z lm Score = −4.26
NG = 0
DB = 0
SM = 0
Delta_AUC
Doxorubicin
YPR103W PRE2
ZShift = 2.53
Z lm Score = 9.16
NG = 0
DB = 2
SM = 0
Delta_L
Doxorubicin
YPR103W PRE2
ZShift = −0.34
Z lm Score = 2.48
NG = 0
DB = 2
SM = 0
Delta_K
Doxorubicin
YPR103W PRE2
ZShift = −1.58
Z lm Score = −2.12
NG = 0
DB = 2
SM = 0
Delta_r
Doxorubicin
YPR103W PRE2
ZShift = −2.33
Z lm Score = 0.82
NG = 0
DB = 2
SM = 0
Delta_AUC
Doxorubicin

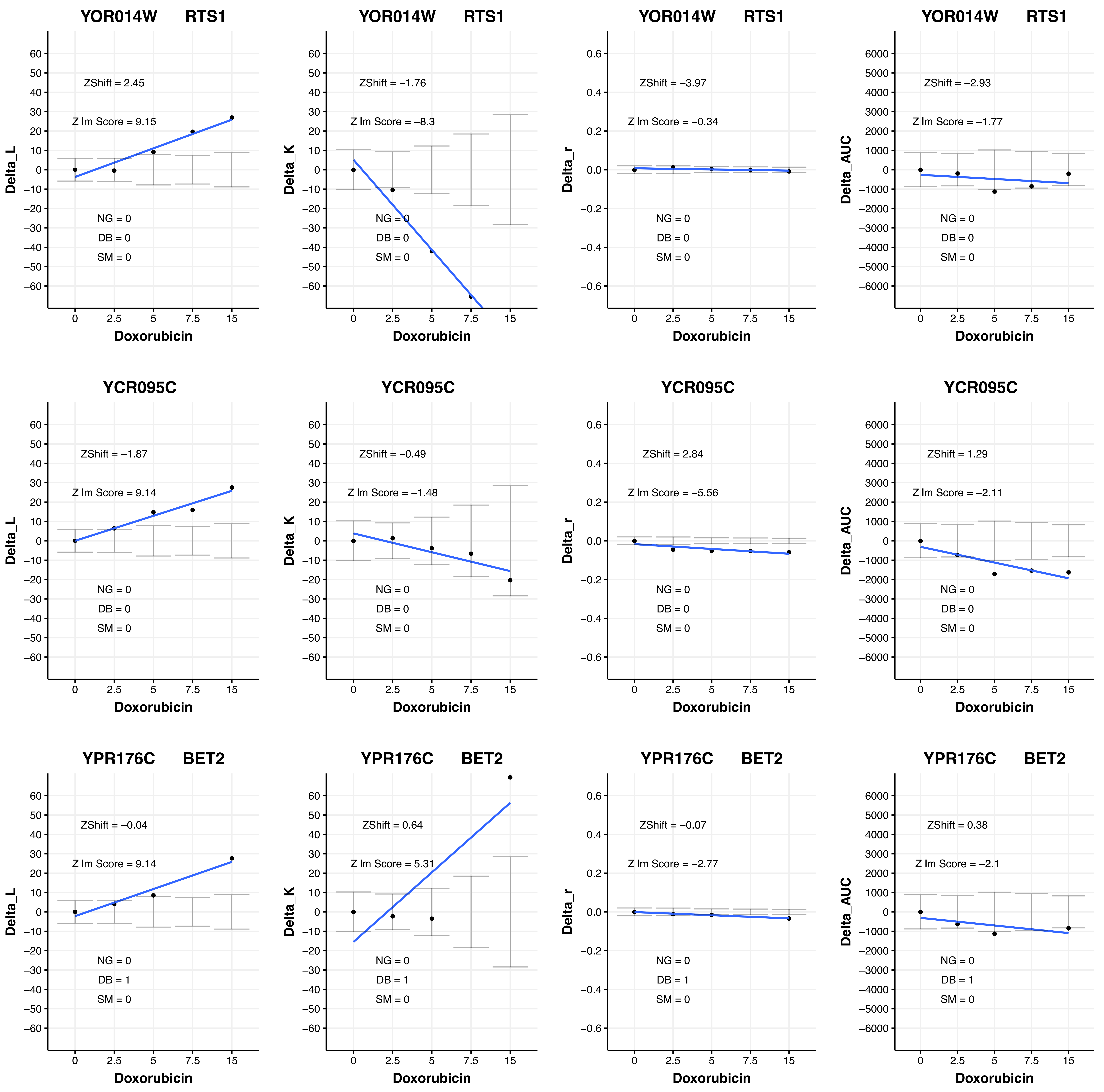
YOR014W RTS1
ZShift = 2.45
Z lm Score = 9.15
NG = 0
DB = 0
SM = 0
Delta_L
Doxorubicin
YOR014W RTS1
ZShift = −1.76
Z lm Score = −8.3
NG = 0
DB = 0
SM = 0
Delta_K
Doxorubicin
YOR014W RTS1
ZShift = −3.97
Z lm Score = −0.34
NG = 0
DB = 0
SM = 0
Delta_r
Doxorubicin
YOR014W RTS1
ZShift = −2.93
Z lm Score = −1.77
NG = 0
DB = 0
SM = 0
Delta_AUC
Doxorubicin
YCR095C
ZShift = −1.87
Z lm Score = 9.14
NG = 0
DB = 0
SM = 0
Delta_L
Doxorubicin
YCR095C
ZShift = −0.49
Z lm Score = −1.48
NG = 0
DB = 0
SM = 0
Delta_K
Doxorubicin
YCR095C
ZShift = 2.84
Z lm Score = −5.56
NG = 0
DB = 0
SM = 0
Delta_r
Doxorubicin
YCR095C
ZShift = 1.29
Z lm Score = −2.11
NG = 0
DB = 0
SM = 0
Delta_AUC
Doxorubicin
YPR176C BET2
ZShift = −0.04
Z lm Score = 9.14
NG = 0
DB = 1
SM = 0
Delta_L
Doxorubicin
YPR176C BET2
ZShift = 0.64
Z lm Score = 5.31
NG = 0
DB = 1
SM = 0
Delta_K
Doxorubicin
YPR176C BET2
ZShift = −0.07
Z lm Score = −2.77
NG = 0
DB = 1
SM = 0
Delta_r
Doxorubicin
YPR176C BET2
ZShift = 0.38
Z lm Score = −2.1
NG = 0
DB = 1
SM = 0
Delta_AUC
Doxorubicin

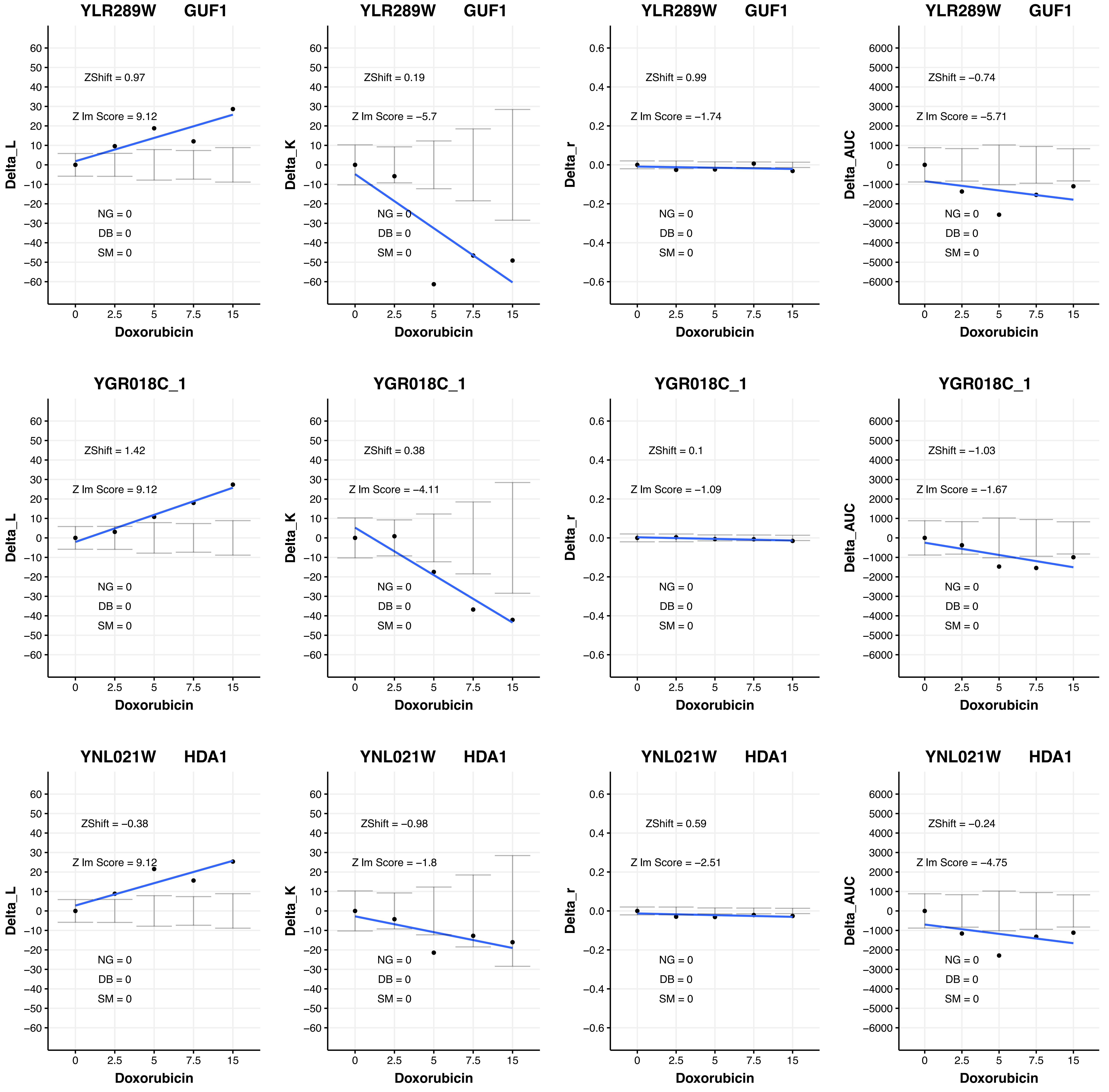
YLR289W GUF1
ZShift = 0.97
Z lm Score = 9.12
NG = 0
DB = 0
SM = 0
Delta_L
Doxorubicin
YLR289W GUF1
ZShift = 0.19
Z lm Score = −5.7
NG = 0
DB = 0
SM = 0
Delta_K
Doxorubicin
YLR289W GUF1
ZShift = 0.99
Z lm Score = −1.74
NG = 0
DB = 0
SM = 0
Delta_r
Doxorubicin
YLR289W GUF1
ZShift = −0.74
Z lm Score = −5.71
NG = 0
DB = 0
SM = 0
Delta_AUC
Doxorubicin
YGR018C_1
ZShift = 1.42
Z lm Score = 9.12
NG = 0
DB = 0
SM = 0
Delta_L
Doxorubicin
YGR018C_1
ZShift = 0.38
Z lm Score = −4.11
NG = 0
DB = 0
SM = 0
Delta_K
Doxorubicin
YGR018C_1
ZShift = 0.1
Z lm Score = −1.09
NG = 0
DB = 0
SM = 0
Delta_r
Doxorubicin
YGR018C_1
ZShift = −1.03
Z lm Score = −1.67
NG = 0
DB = 0
SM = 0
Delta_AUC
Doxorubicin
YNL021W HDA1
ZShift = −0.38
Z lm Score = 9.12
NG = 0
DB = 0
SM = 0
Delta_L
Doxorubicin
YNL021W HDA1
ZShift = −0.98
Z lm Score = −1.8
NG = 0
DB = 0
SM = 0
Delta_K
Doxorubicin
YNL021W HDA1
ZShift = 0.59
Z lm Score = −2.51
NG = 0
DB = 0
SM = 0
Delta_r
Doxorubicin
YNL021W HDA1
ZShift = −0.24
Z lm Score = −4.75
NG = 0
DB = 0
SM = 0
Delta_AUC
Doxorubicin

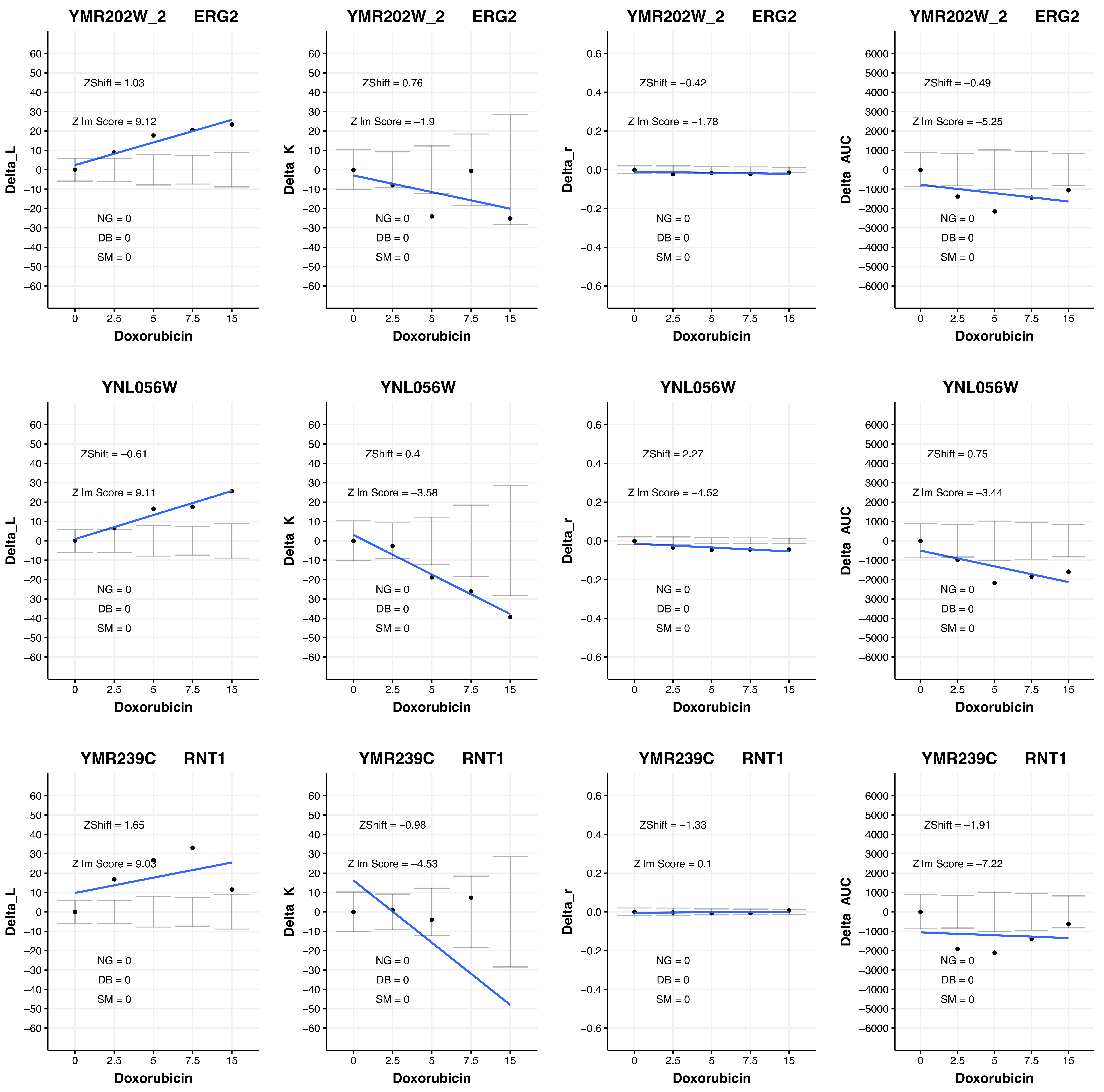
YMR202W_2 ERG2
ZShift = 1.03
Z lm Score = 9.12
NG = 0
DB = 0
SM = 0
Delta_L
Doxorubicin
YMR202W_2 ERG2
ZShift = 0.76
Z lm Score = −1.9
NG = 0
DB = 0
SM = 0
Delta_K
Doxorubicin
YMR202W_2 ERG2
ZShift = −0.42
Z lm Score = −1.78
NG = 0
DB = 0
SM = 0
Delta_r
Doxorubicin
YMR202W_2 ERG2
ZShift = −0.49
Z lm Score = −5.25
NG = 0
DB = 0
SM = 0
Delta_AUC
Doxorubicin
YNL056W
ZShift = −0.61
Z lm Score = 9.11
NG = 0
DB = 0
SM = 0
Delta_L
Doxorubicin
YNL056W
ZShift = 0.4
Z lm Score = −3.58
NG = 0
DB = 0
SM = 0
Delta_K
Doxorubicin
YNL056W
ZShift = 2.27
Z lm Score = −4.52
NG = 0
DB = 0
SM = 0
Delta_r
Doxorubicin
YNL056W
ZShift = 0.75
Z lm Score = −3.44
NG = 0
DB = 0
SM = 0
Delta_AUC
Doxorubicin
YMR239C RNT1
ZShift = 1.65
Z lm Score = 9.03
NG = 0
DB = 0
SM = 0
Delta_L
Doxorubicin
YMR239C RNT1
ZShift = −0.98
Z lm Score = −4.53
NG = 0
DB = 0
SM = 0
Delta_K
Doxorubicin
YMR239C RNT1
ZShift = −1.33
Z lm Score = 0.1
NG = 0
DB = 0
SM = 0
Delta_r
Doxorubicin
YMR239C RNT1
ZShift = −1.91
Z lm Score = −7.22
NG = 0
DB = 0
SM = 0
Delta_AUC
Doxorubicin

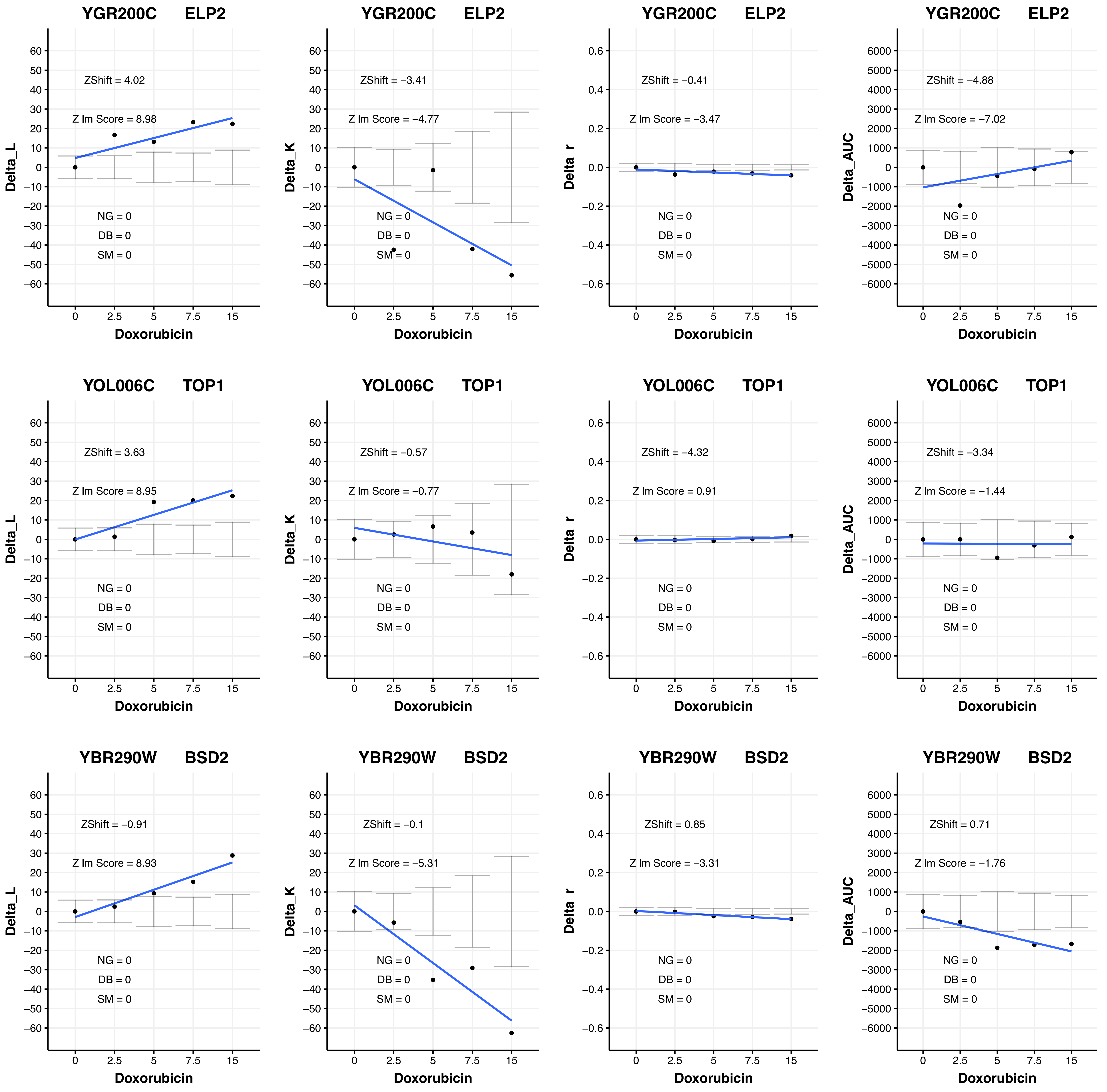

YGR200C ELP2
ZShift = 4.02
Z lm Score = 8.98
NG = 0
DB = 0
SM = 0
Delta_L
Doxorubicin
YGR200C ELP2
ZShift = −3.41
Z lm Score = −4.77
NG = 0
DB = 0
SM = 0
Delta_K
Doxorubicin
YGR200C ELP2
ZShift = −0.41
Z lm Score = −3.47
NG = 0
DB = 0
SM = 0
Delta_r
Doxorubicin
YGR200C ELP2
ZShift = −4.88
Z lm Score = −7.02
NG = 0
DB = 0
SM = 0
Delta_AUC
Doxorubicin
YOL006C TOP1
ZShift = 3.63
Z lm Score = 8.95
NG = 0
DB = 0
SM = 0
Delta_L
Doxorubicin
YOL006C TOP1
ZShift = −0.57
Z lm Score = −0.77
NG = 0
DB = 0
SM = 0
Delta_K
Doxorubicin
YOL006C TOP1
ZShift = −4.32
Z lm Score = 0.91
NG = 0
DB = 0
SM = 0
Delta_r
Doxorubicin
YOL006C TOP1
ZShift = −3.34
Z lm Score = −1.44
NG = 0
DB = 0
SM = 0
Delta_AUC
Doxorubicin
YBR290W BSD2
ZShift = −0.91
Z lm Score = 8.93
NG = 0
DB = 0
SM = 0
Delta_L
Doxorubicin
YBR290W BSD2
ZShift = −0.1
Z lm Score = −5.31
NG = 0
DB = 0
SM = 0
Delta_K
Doxorubicin
YBR290W BSD2
ZShift = 0.85
Z lm Score = −3.31
NG = 0
DB = 0
SM = 0
Delta_r
Doxorubicin
YBR290W BSD2
ZShift = 0.71
Z lm Score = −1.76
NG = 0
DB = 0
SM = 0
Delta_AUC
Doxorubicin

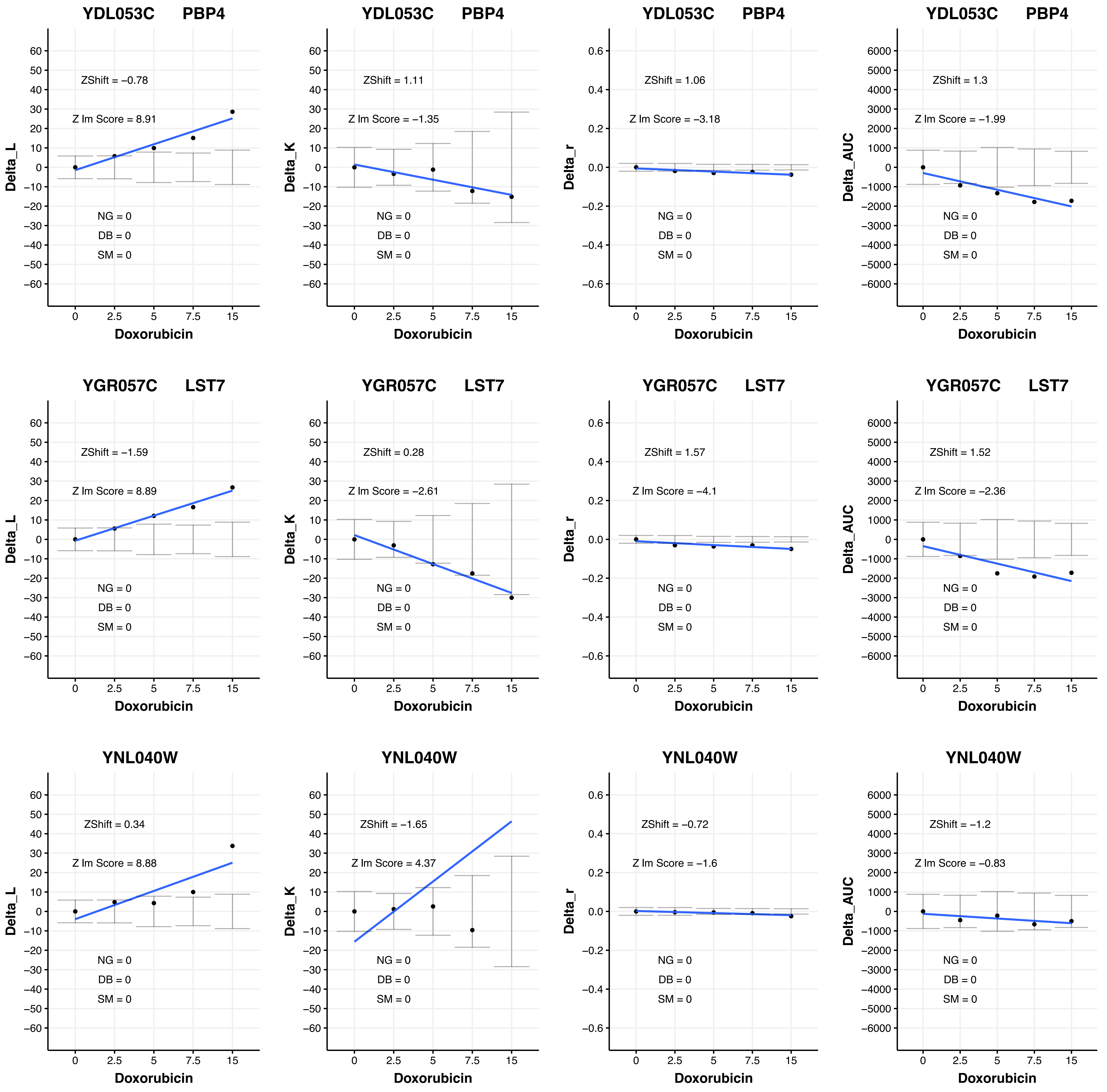
YDL053C PBP4
ZShift = −0.78
Z lm Score = 8.91
NG = 0
DB = 0
SM = 0
Delta_L
Doxorubicin
YDL053C PBP4
ZShift = 1.11
Z lm Score = −1.35
NG = 0
DB = 0
SM = 0
Delta_K
Doxorubicin
YDL053C PBP4
ZShift = 1.06
Z lm Score = −3.18
NG = 0
DB = 0
SM = 0
Delta_r
Doxorubicin
YDL053C PBP4
ZShift = 1.3
Z lm Score = −1.99
NG = 0
DB = 0
SM = 0
Delta_AUC
Doxorubicin
YGR057C LST7
ZShift = −1.59
Z lm Score = 8.89
NG = 0
DB = 0
SM = 0
Delta_L
Doxorubicin
YGR057C LST7
ZShift = 0.28
Z lm Score = −2.61
NG = 0
DB = 0
SM = 0
Delta_K
Doxorubicin
YGR057C LST7
ZShift = 1.57
Z lm Score = −4.1
NG = 0
DB = 0
SM = 0
Delta_r
Doxorubicin
YGR057C LST7
ZShift = 1.52
Z lm Score = −2.36
NG = 0
DB = 0
SM = 0
Delta_AUC
Doxorubicin
YNL040W
ZShift = 0.34
Z lm Score = 8.88
NG = 0
DB = 0
SM = 0
Delta_L
Doxorubicin
YNL040W
ZShift = −1.65
Z lm Score = 4.37
NG = 0
DB = 0
SM = 0
Delta_K
Doxorubicin
YNL040W
ZShift = −0.72
Z lm Score = −1.6
NG = 0
DB = 0
SM = 0
Delta_r
Doxorubicin
YNL040W
ZShift = −1.2
Z lm Score = −0.83
NG = 0
DB = 0
SM = 0
Delta_AUC
Doxorubicin

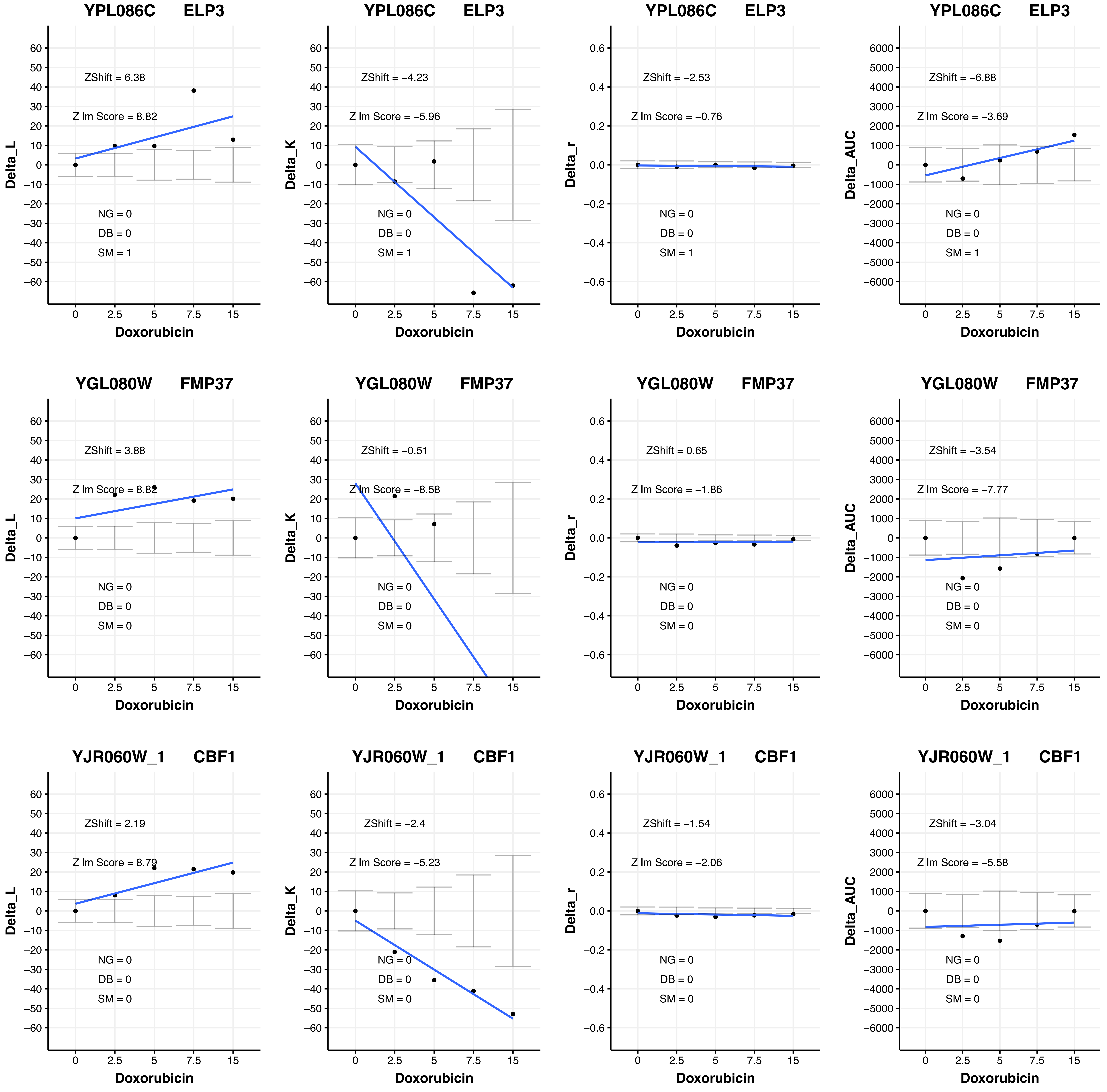
YPL086C ELP3
ZShift = 6.38
Z lm Score = 8.82
NG = 0
DB = 0
SM = 1
Delta_L
Doxorubicin
YPL086C ELP3
ZShift = −4.23
Z lm Score = −5.96
NG = 0
DB = 0
SM = 1
Delta_K
Doxorubicin
YPL086C ELP3
ZShift = −2.53
Z lm Score = −0.76
NG = 0
DB = 0
SM = 1
Delta_r
Doxorubicin
YPL086C ELP3
ZShift = −6.88
Z lm Score = −3.69
NG = 0
DB = 0
SM = 1
Delta_AUC
Doxorubicin
YGL080W FMP37
ZShift = 3.88
Z lm Score = 8.82
NG = 0
DB = 0
SM = 0
Delta_L
Doxorubicin
YGL080W FMP37
ZShift = −0.51
Z lm Score = −8.58
NG = 0
DB = 0
SM = 0
Delta_K
Doxorubicin
YGL080W FMP37
ZShift = 0.65
Z lm Score = −1.86
NG = 0
DB = 0
SM = 0
Delta_r
Doxorubicin
YGL080W FMP37
ZShift = −3.54
Z lm Score = −7.77
NG = 0
DB = 0
SM = 0
Delta_AUC
Doxorubicin
YJR060W_1 CBF1
ZShift = 2.19
Z lm Score = 8.79
NG = 0
DB = 0
SM = 0
Delta_L
Doxorubicin
YJR060W_1 CBF1
ZShift = −2.4
Z lm Score = −5.23
NG = 0
DB = 0
SM = 0
Delta_K
Doxorubicin
YJR060W_1 CBF1
ZShift = −1.54
Z lm Score = −2.06
NG = 0
DB = 0
SM = 0
Delta_r
Doxorubicin
YJR060W_1 CBF1
ZShift = −3.04
Z lm Score = −5.58
NG = 0
DB = 0
SM = 0
Delta_AUC
Doxorubicin

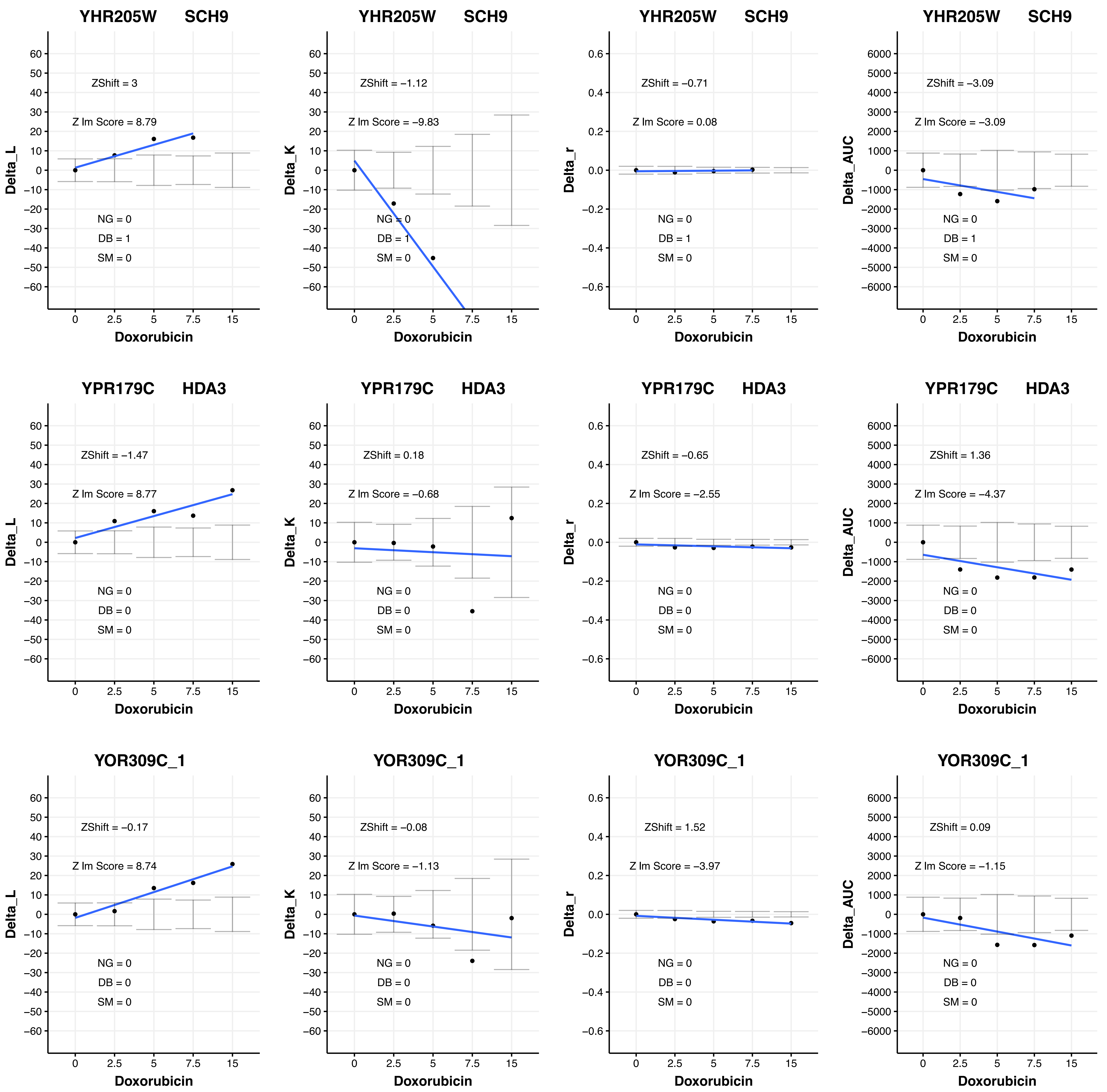

YHR205W SCH9
ZShift = 3
Z lm Score = 8.79
NG = 0
DB = 1
SM = 0
Delta_L
Doxorubicin
YHR205W SCH9
ZShift = −1.12
Z lm Score = −9.83
NG = 0
DB = 1
SM = 0
Delta_K
Doxorubicin
YHR205W SCH9
ZShift = −0.71
Z lm Score = 0.08
NG = 0
DB = 1
SM = 0
Delta_r
Doxorubicin
YHR205W SCH9
ZShift = −3.09
Z lm Score = −3.09
NG = 0
DB = 1
SM = 0
Delta_AUC
Doxorubicin
YPR179C HDA3
ZShift = −1.47
Z lm Score = 8.77
NG = 0
DB = 0
SM = 0
Delta_L
Doxorubicin
YPR179C HDA3
ZShift = 0.18
Z lm Score = −0.68
NG = 0
DB = 0
SM = 0
Delta_K
Doxorubicin
YPR179C HDA3
ZShift = −0.65
Z lm Score = −2.55
NG = 0
DB = 0
SM = 0
Delta_r
Doxorubicin
YPR179C HDA3
ZShift = 1.36
Z lm Score = −4.37
NG = 0
DB = 0
SM = 0
Delta_AUC
Doxorubicin
YOR309C_1
ZShift = −0.17
Z lm Score = 8.74
NG = 0
DB = 0
SM = 0
Delta_L
Doxorubicin
YOR309C_1
ZShift = −0.08
Z lm Score = −1.13
NG = 0
DB = 0
SM = 0
Delta_K
Doxorubicin
YOR309C_1
ZShift = 1.52
Z lm Score = −3.97
NG = 0
DB = 0
SM = 0
Delta_r
Doxorubicin
YOR309C_1
ZShift = 0.09
Z lm Score = −1.15
NG = 0
DB = 0
SM = 0
Delta_AUC
Doxorubicin
0 2.5 5 7.5 15

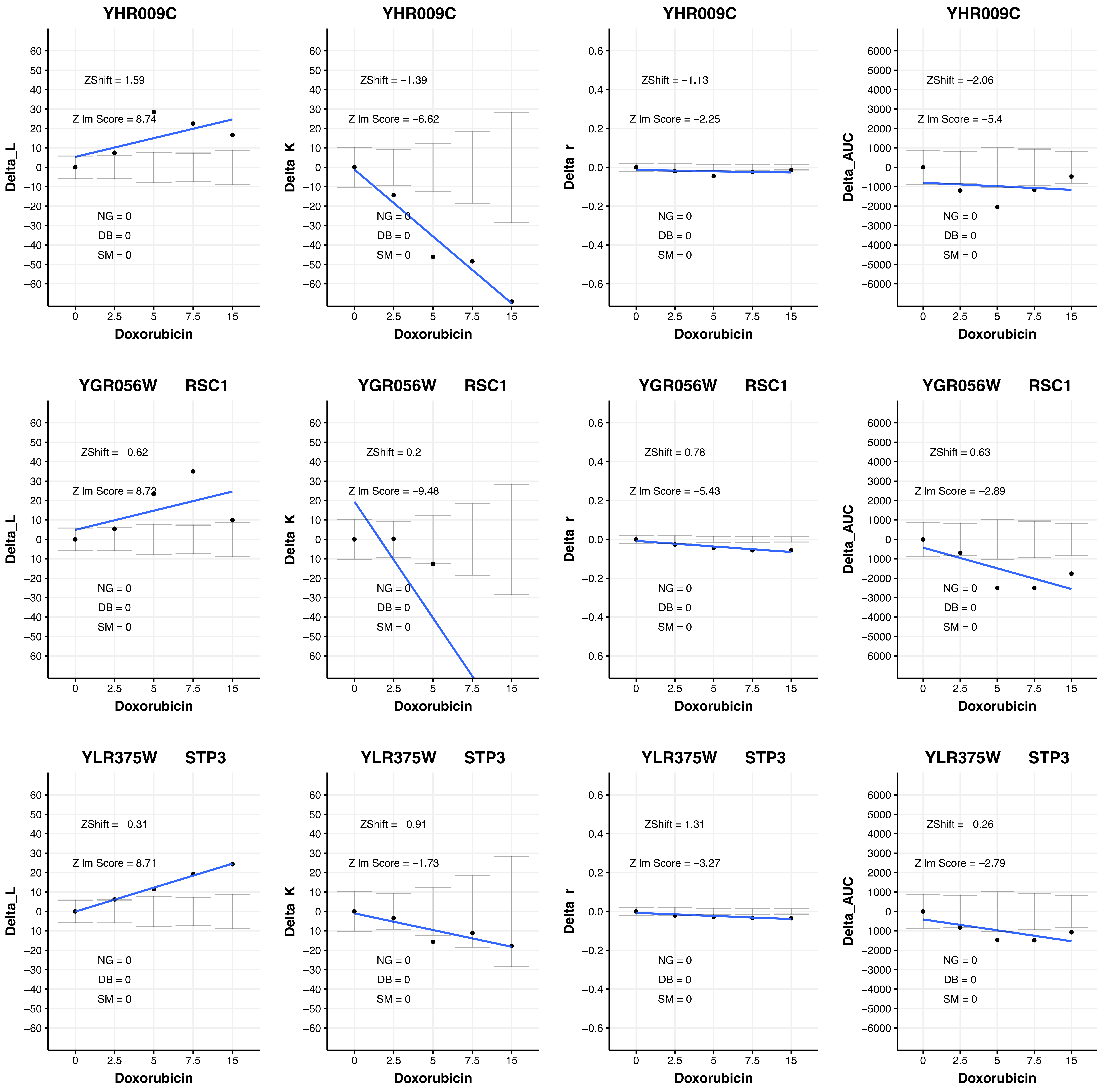
YHR009C
ZShift = 1.59
Z lm Score = 8.74
NG = 0
DB = 0
SM = 0
Delta_L
Doxorubicin
YHR009C
ZShift = −1.39
Z lm Score = −6.62
NG = 0
DB = 0
SM = 0
Delta_K
Doxorubicin
YHR009C
ZShift = −1.13
Z lm Score = −2.25
NG = 0
DB = 0
SM = 0
Delta_r
Doxorubicin
YHR009C
ZShift = −2.06
Z lm Score = −5.4
NG = 0
DB = 0
SM = 0
Delta_AUC
Doxorubicin
YGR056W RSC1
ZShift = −0.62
Z lm Score = 8.72
NG = 0
DB = 0
SM = 0
Delta_L
Doxorubicin
YGR056W RSC1
ZShift = 0.2
Z lm Score = −9.48
NG = 0
DB = 0
SM = 0
Delta_K
Doxorubicin
YGR056W RSC1
ZShift = 0.78
Z lm Score = −5.43
NG = 0
DB = 0
SM = 0
Delta_r
Doxorubicin
YGR056W RSC1
ZShift = 0.63
Z lm Score = −2.89
NG = 0
DB = 0
SM = 0
Delta_AUC
Doxorubicin
YLR375W STP3
ZShift = −0.31
Z lm Score = 8.71
NG = 0
DB = 0
SM = 0
Delta_L
Doxorubicin
YLR375W STP3
ZShift = −0.91
Z lm Score = −1.73
NG = 0
DB = 0
SM = 0
Delta_K
Doxorubicin
YLR375W STP3
ZShift = 1.31
Z lm Score = −3.27
NG = 0
DB = 0
SM = 0
Delta_r
Doxorubicin
YLR375W STP3
ZShift = −0.26
Z lm Score = −2.79
NG = 0
DB = 0
SM = 0
Delta_AUC
Doxorubicin

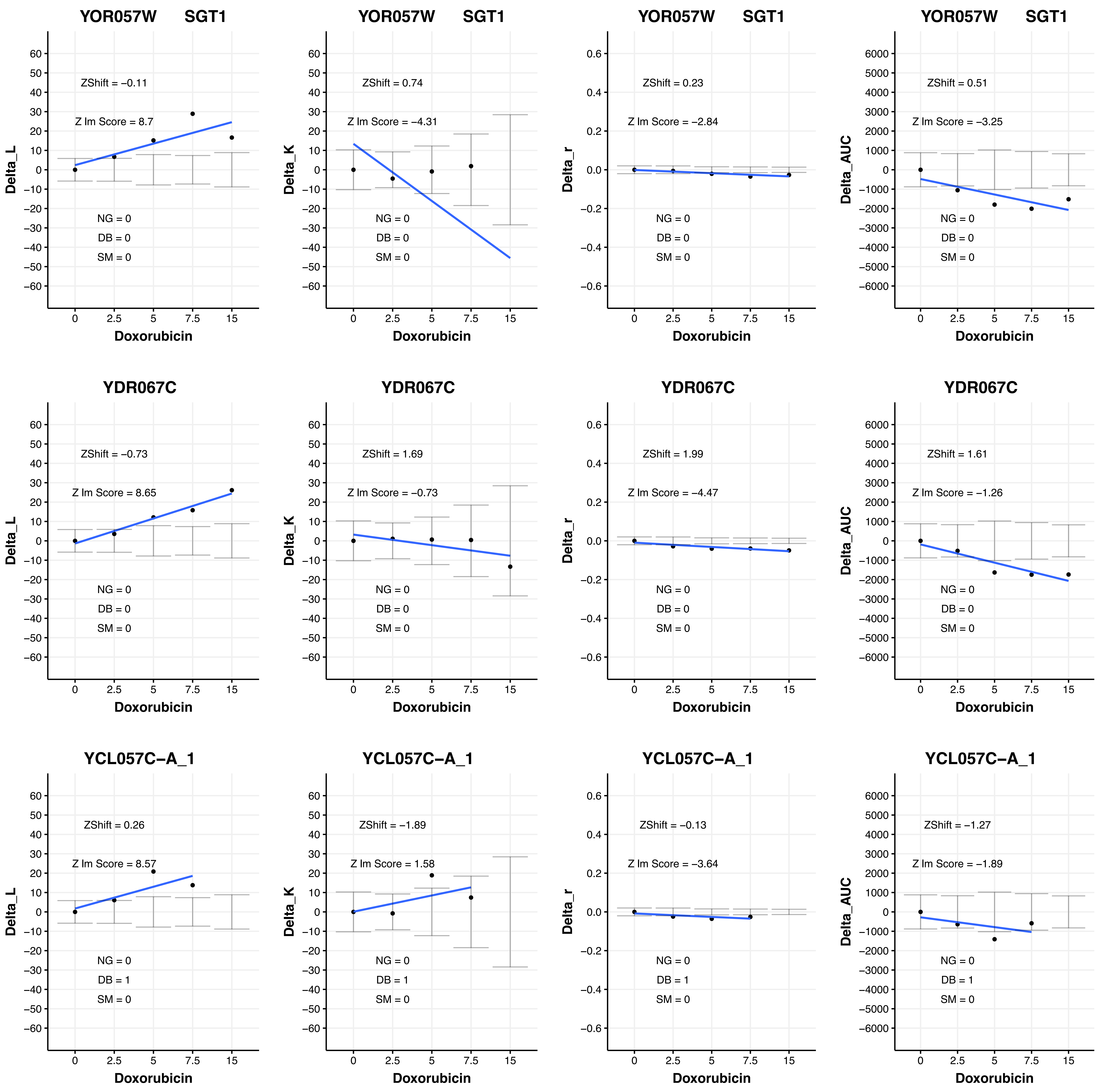

YOR057W SGT1
ZShift = −0.11
Z lm Score = 8.7
NG = 0
DB = 0
SM = 0
Delta_L
Doxorubicin
YOR057W SGT1
ZShift = 0.74
Z lm Score = −4.31
NG = 0
DB = 0
SM = 0
Delta_K
Doxorubicin
YOR057W SGT1
ZShift = 0.23
Z lm Score = −2.84
NG = 0
DB = 0
SM = 0
Delta_r
Doxorubicin
YOR057W SGT1
ZShift = 0.51
Z lm Score = −3.25
NG = 0
DB = 0
SM = 0
Delta_AUC
Doxorubicin
YDR067C
ZShift = −0.73
Z lm Score = 8.65
NG = 0
DB = 0
SM = 0
Delta_L
Doxorubicin
YDR067C
ZShift = 1.69
Z lm Score = −0.73
NG = 0
DB = 0
SM = 0
Delta_K
Doxorubicin
YDR067C
ZShift = 1.99
Z lm Score = −4.47
NG = 0
DB = 0
SM = 0
Delta_r
Doxorubicin
YDR067C
ZShift = 1.61
Z lm Score = −1.26
NG = 0
DB = 0
SM = 0
Delta_AUC
Doxorubicin
YCL057C−A_1
ZShift = 0.26
Z lm Score = 8.57
NG = 0
DB = 1
SM = 0
Delta_L
Doxorubicin
YCL057C−A_1
ZShift = −1.89
Z lm Score = 1.58
NG = 0
DB = 1
SM = 0
Delta_K
Doxorubicin
YCL057C−A_1
ZShift = −0.13
Z lm Score = −3.64
NG = 0
DB = 1
SM = 0
Delta_r
Doxorubicin
YCL057C−A_1
ZShift = −1.27
Z lm Score = −1.89
NG = 0
DB = 1
SM = 0
Delta_AUC
Doxorubicin

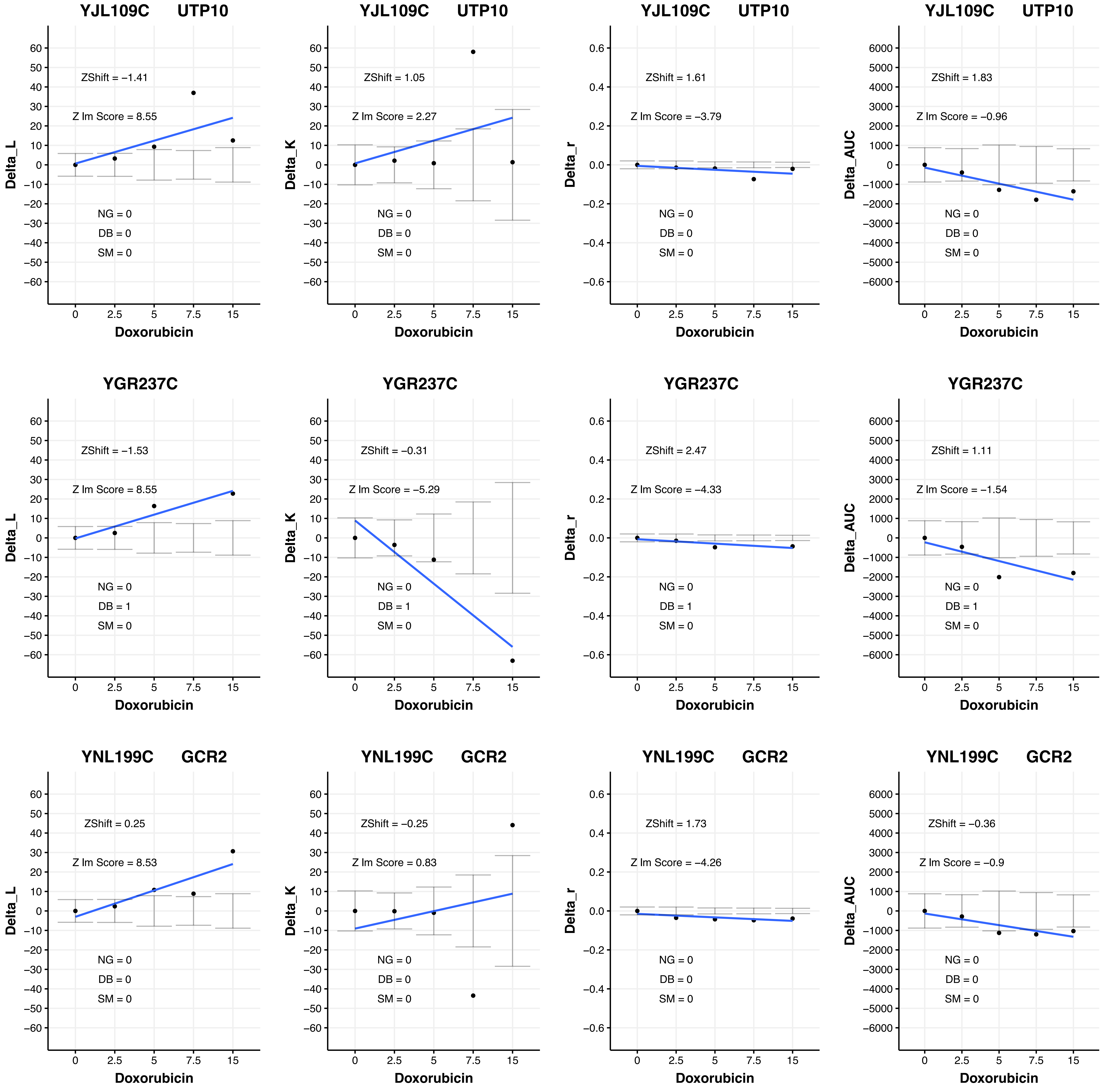
YJL109C UTP10
ZShift = −1.41
Z lm Score = 8.55
NG = 0
DB = 0
SM = 0
Delta_L
Doxorubicin
YJL109C UTP10
ZShift = 1.05
Z lm Score = 2.27
NG = 0
DB = 0
SM = 0
Delta_K
Doxorubicin
YJL109C UTP10
ZShift = 1.61
Z lm Score = −3.79
NG = 0
DB = 0
SM = 0
Delta_r
Doxorubicin
YJL109C UTP10
ZShift = 1.83
Z lm Score = −0.96
NG = 0
DB = 0
SM = 0
Delta_AUC
Doxorubicin
YGR237C
ZShift = −1.53
Z lm Score = 8.55
NG = 0
DB = 1
SM = 0
Delta_L
Doxorubicin
YGR237C
ZShift = −0.31
Z lm Score = −5.29
NG = 0
DB = 1
SM = 0
Delta_K
Doxorubicin
YGR237C
ZShift = 2.47
Z lm Score = −4.33
NG = 0
DB = 1
SM = 0
Delta_r
Doxorubicin
YGR237C
ZShift = 1.11
Z lm Score = −1.54
NG = 0
DB = 1
SM = 0
Delta_AUC
Doxorubicin
YNL199C GCR2
ZShift = 0.25
Z lm Score = 8.53
NG = 0
DB = 0
SM = 0
Delta_L
Doxorubicin
YNL199C GCR2
ZShift = −0.25
Z lm Score = 0.83
NG = 0
DB = 0
SM = 0
Delta_K
Doxorubicin
YNL199C GCR2
ZShift = 1.73
Z lm Score = −4.26
NG = 0
DB = 0
SM = 0
Delta_r
Doxorubicin
YNL199C GCR2
ZShift = −0.36
Z lm Score = −0.9
NG = 0
DB = 0
SM = 0
Delta_AUC
Doxorubicin

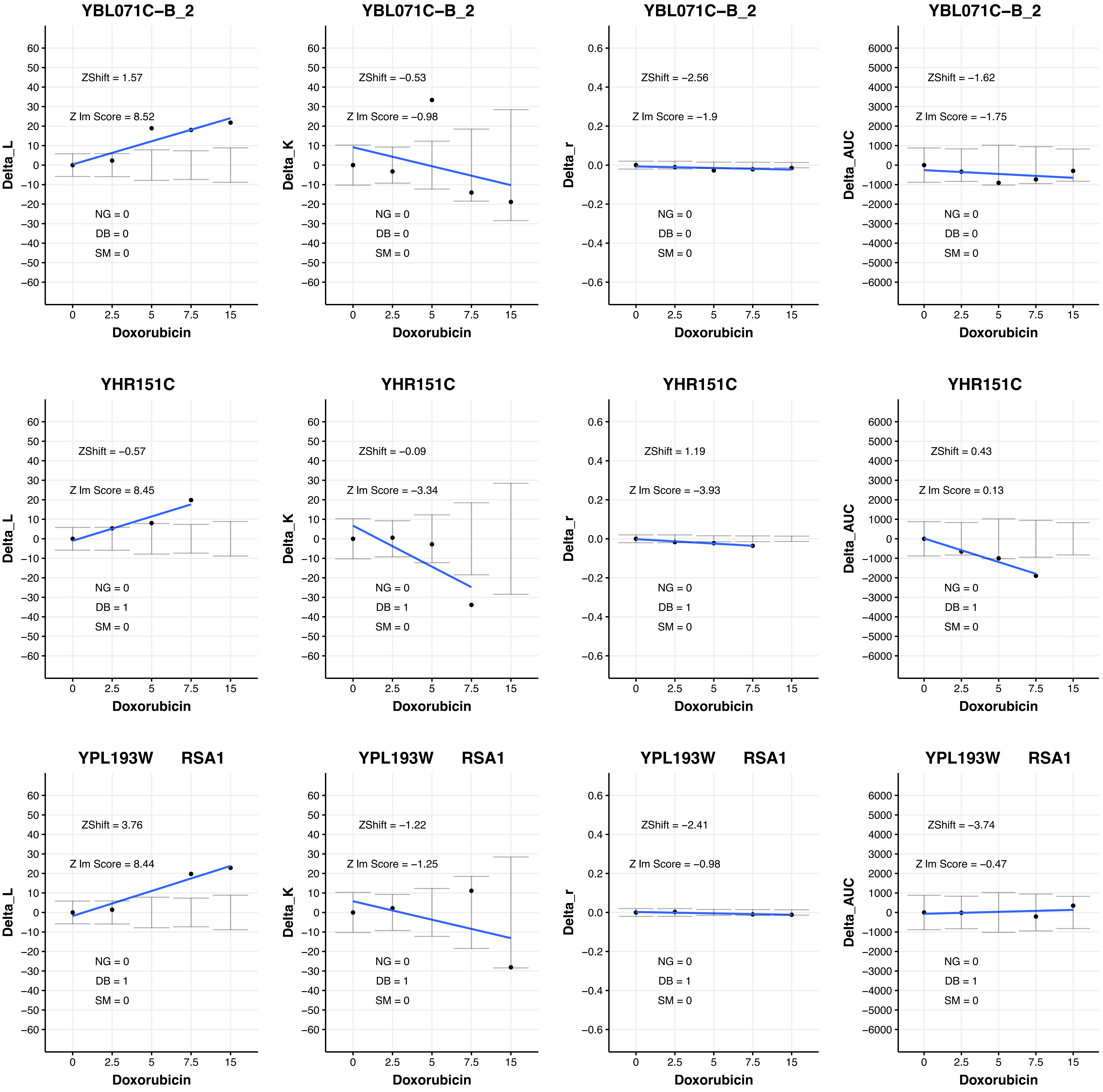

YBL071C−B_2
ZShift = 1.57
Z lm Score = 8.52
NG = 0
DB = 0
SM = 0
Delta_L
Doxorubicin
YBL071C−B_2
ZShift = −0.53
Z lm Score = −0.98
NG = 0
DB = 0
SM = 0
Delta_K
Doxorubicin
YBL071C−B_2
ZShift = −2.56
Z lm Score = −1.9
NG = 0
DB = 0
SM = 0
Delta_r
Doxorubicin
YBL071C−B_2
ZShift = −1.62
Z lm Score = −1.75
NG = 0
DB = 0
SM = 0
Delta_AUC
Doxorubicin
YHR151C
ZShift = −0.57
Z lm Score = 8.45
NG = 0
DB = 1
SM = 0
Delta_L
Doxorubicin
YHR151C
ZShift = −0.09
Z lm Score = −3.34
NG = 0
DB = 1
SM = 0
Delta_K
Doxorubicin
YHR151C
ZShift = 1.19
Z lm Score = −3.93
NG = 0
DB = 1
SM = 0
Delta_r
Doxorubicin
YHR151C
ZShift = 0.43
Z lm Score = 0.13
NG = 0
DB = 1
SM = 0
Delta_AUC
Doxorubicin
YPL193W RSA1
ZShift = 3.76
Z lm Score = 8.44
NG = 0
DB = 1
SM = 0
Delta_L
Doxorubicin
YPL193W RSA1
ZShift = −1.22
Z lm Score = −1.25
NG = 0
DB = 1
SM = 0
Delta_K
Doxorubicin
YPL193W RSA1
ZShift = −2.41
Z lm Score = −0.98
NG = 0
DB = 1
SM = 0
Delta_r
Doxorubicin
YPL193W RSA1
ZShift = −3.74
Z lm Score = −0.47
NG = 0
DB = 1
SM = 0
Delta_AUC
Doxorubicin

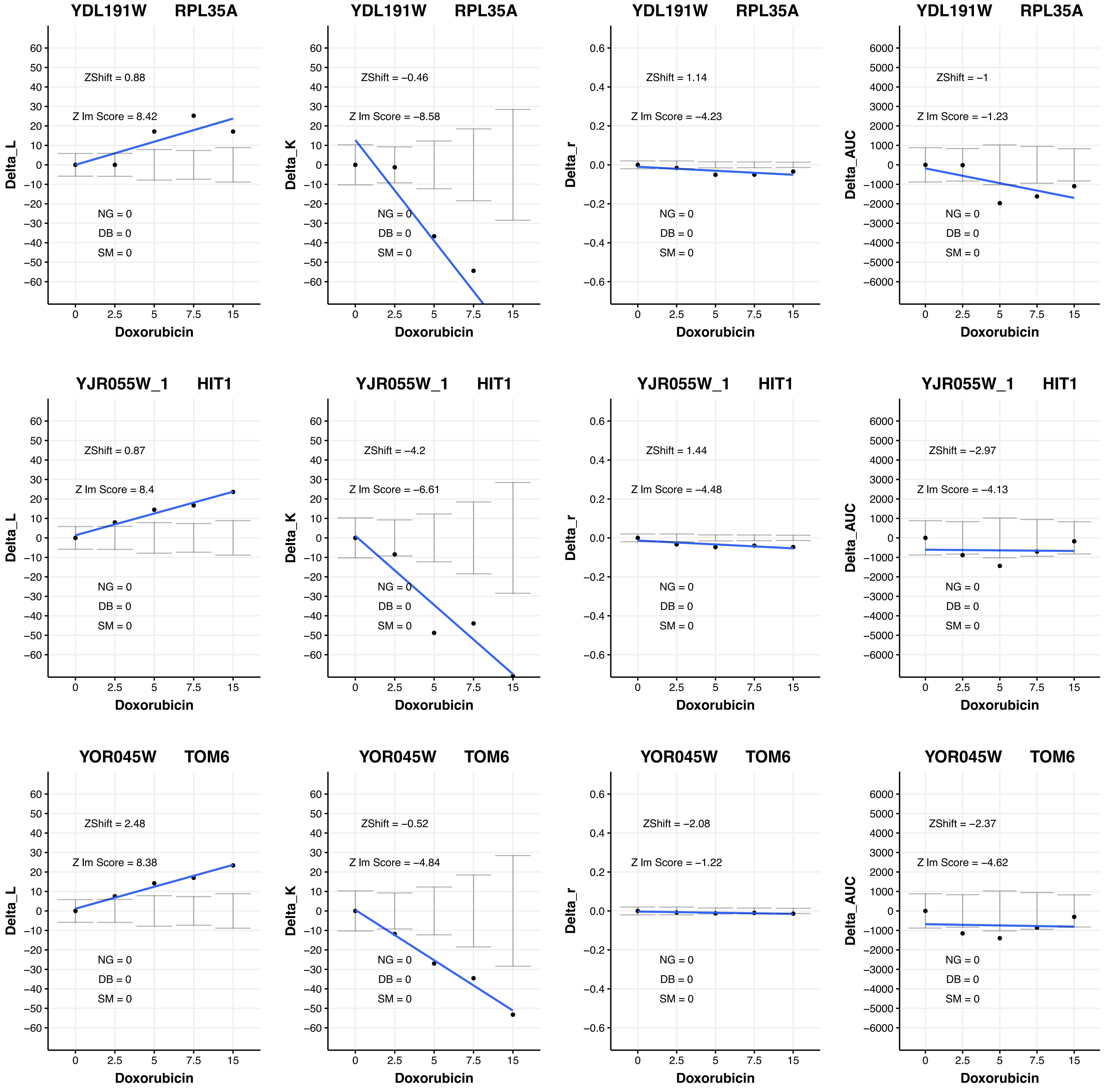
YDL191W RPL35A
ZShift = 0.88
Z lm Score = 8.42
NG = 0
DB = 0
SM = 0
Delta_L
Doxorubicin
YDL191W RPL35A
ZShift = −0.46
Z lm Score = −8.58
NG = 0
DB = 0
SM = 0
Delta_K
Doxorubicin
YDL191W RPL35A
ZShift = 1.14
Z lm Score = −4.23
NG = 0
DB = 0
SM = 0
Delta_r
Doxorubicin
YDL191W RPL35A
ZShift = −1
Z lm Score = −1.23
NG = 0
DB = 0
SM = 0
Delta_AUC
Doxorubicin
YJR055W_1 HIT1
ZShift = 0.87
Z lm Score = 8.4
NG = 0
DB = 0
SM = 0
Delta_L
Doxorubicin
YJR055W_1 HIT1
ZShift = −4.2
Z lm Score = −6.61
NG = 0
DB = 0
SM = 0
Delta_K
Doxorubicin
YJR055W_1 HIT1
ZShift = 1.44
Z lm Score = −4.48
NG = 0
DB = 0
SM = 0
Delta_r
Doxorubicin
YJR055W_1 HIT1
ZShift = −2.97
Z lm Score = −4.13
NG = 0
DB = 0
SM = 0
Delta_AUC
Doxorubicin
YOR045W TOM6
ZShift = 2.48
Z lm Score = 8.38
NG = 0
DB = 0
SM = 0
Delta_L
Doxorubicin
YOR045W TOM6
ZShift = −0.52
Z lm Score = −4.84
NG = 0
DB = 0
SM = 0
Delta_K
Doxorubicin
YOR045W TOM6
ZShift = −2.08
Z lm Score = −1.22
NG = 0
DB = 0
SM = 0
Delta_r
Doxorubicin
YOR045W TOM6
ZShift = −2.37
Z lm Score = −4.62
NG = 0
DB = 0
SM = 0
Delta_AUC
Doxorubicin

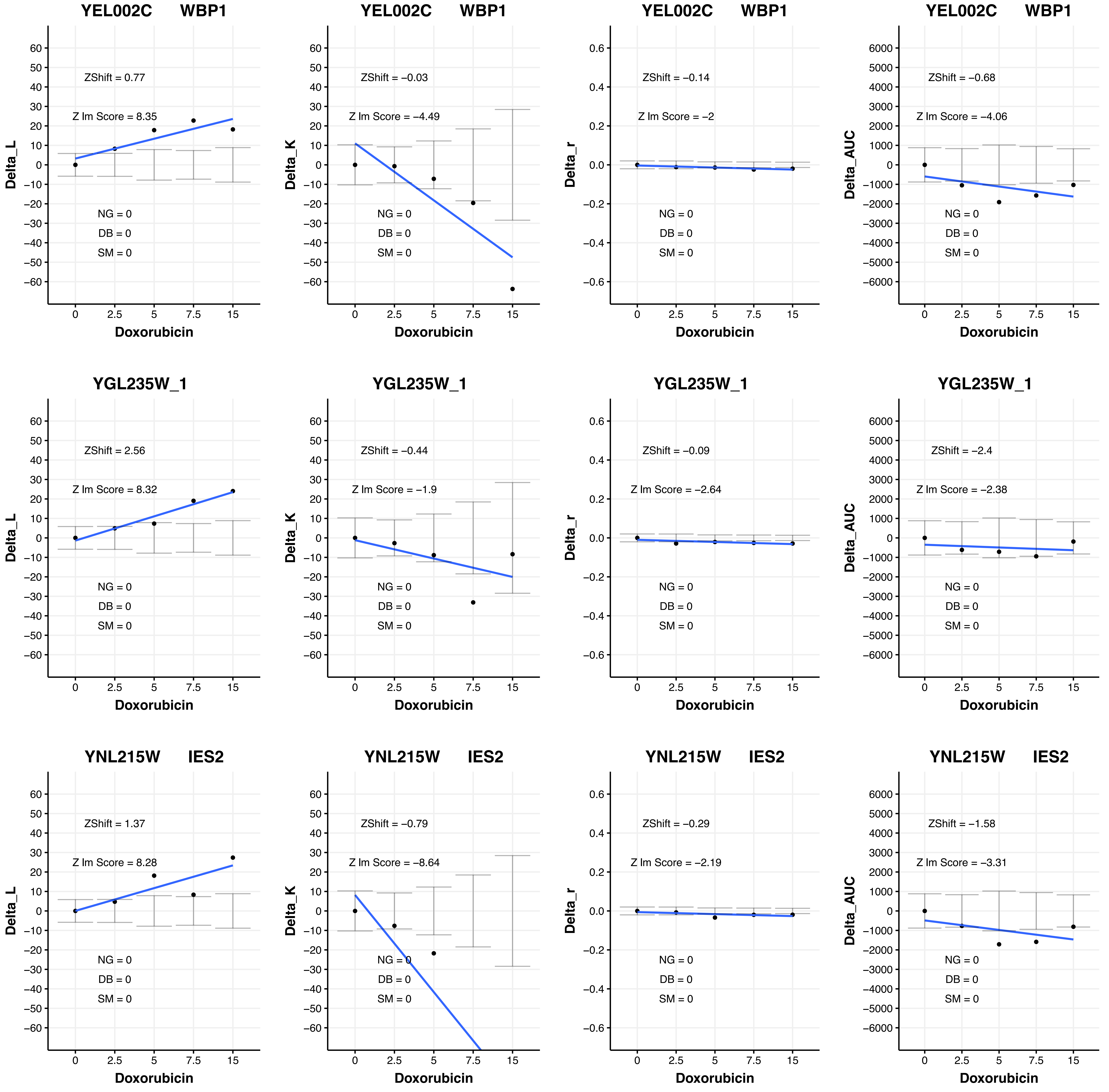
YEL002C WBP1
ZShift = 0.77
Z lm Score = 8.35
NG = 0
DB = 0
SM = 0
Delta_L
Doxorubicin
YEL002C WBP1
ZShift = −0.03
Z lm Score = −4.49
NG = 0
DB = 0
SM = 0
Delta_K
Doxorubicin
YEL002C WBP1
ZShift = −0.14
Z lm Score = −2
NG = 0
DB = 0
SM = 0
Delta_r
Doxorubicin
YEL002C WBP1
ZShift = −0.68
Z lm Score = −4.06
NG = 0
DB = 0
SM = 0
Delta_AUC
Doxorubicin
YGL235W_1
ZShift = 2.56
Z lm Score = 8.32
NG = 0
DB = 0
SM = 0
Delta_L
Doxorubicin
YGL235W_1
ZShift = −0.44
Z lm Score = −1.9
NG = 0
DB = 0
SM = 0
Delta_K
Doxorubicin
YGL235W_1
ZShift = −0.09
Z lm Score = −2.64
NG = 0
DB = 0
SM = 0
Delta_r
Doxorubicin
YGL235W_1
ZShift = −2.4
Z lm Score = −2.38
NG = 0
DB = 0
SM = 0
Delta_AUC
Doxorubicin
YNL215W IES2
ZShift = 1.37
Z lm Score = 8.28
NG = 0
DB = 0
SM = 0
Delta_L
Doxorubicin
YNL215W IES2
ZShift = −0.79
Z lm Score = −8.64
NG = 0
DB = 0
SM = 0
Delta_K
Doxorubicin
YNL215W IES2
ZShift = −0.29
Z lm Score = −2.19
NG = 0
DB = 0
SM = 0
Delta_r
Doxorubicin
YNL215W IES2
ZShift = −1.58
Z lm Score = −3.31
NG = 0
DB = 0
SM = 0
Delta_AUC
Doxorubicin

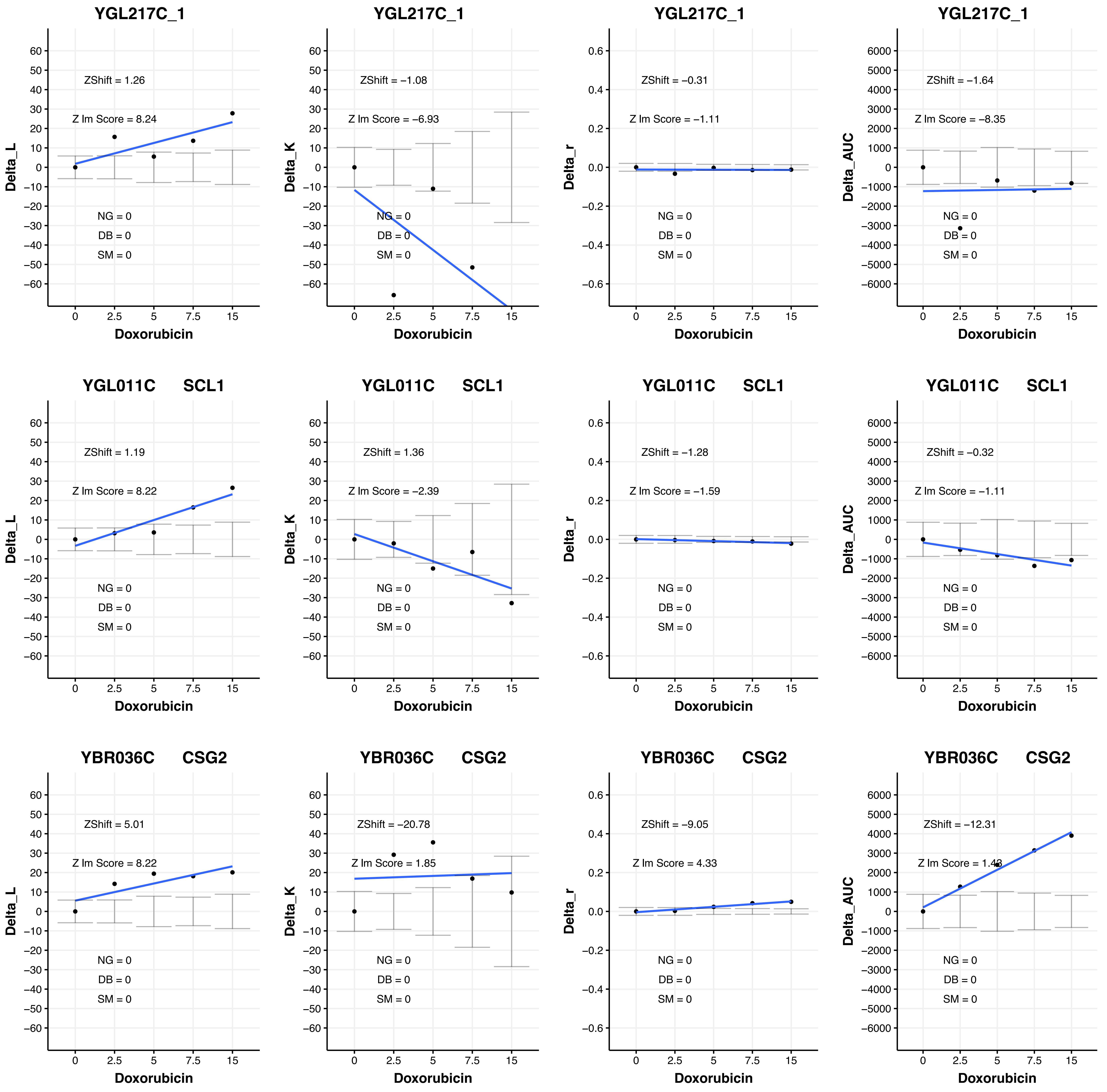

YGL217C_1
ZShift = 1.26
Z lm Score = 8.24
NG = 0
DB = 0
SM = 0
Delta_L
Doxorubicin
YGL217C_1
ZShift = −1.08
Z lm Score = −6.93
NG = 0
DB = 0
SM = 0
Delta_K
Doxorubicin
YGL217C_1
ZShift = −0.31
Z lm Score = −1.11
NG = 0
DB = 0
SM = 0
Delta_r
Doxorubicin
YGL217C_1
ZShift = −1.64
Z lm Score = −8.35
NG = 0
DB = 0
SM = 0
Delta_AUC
Doxorubicin
YGL011C SCL1
ZShift = 1.19
Z lm Score = 8.22
NG = 0
DB = 0
SM = 0
Delta_L
Doxorubicin
YGL011C SCL1
ZShift = 1.36
Z lm Score = −2.39
NG = 0
DB = 0
SM = 0
Delta_K
Doxorubicin
YGL011C SCL1
ZShift = −1.28
Z lm Score = −1.59
NG = 0
DB = 0
SM = 0
Delta_r
Doxorubicin
YGL011C SCL1
ZShift = −0.32
Z lm Score = −1.11
NG = 0
DB = 0
SM = 0
Delta_AUC
Doxorubicin
YBR036C CSG2
ZShift = 5.01
Z lm Score = 8.22
NG = 0
DB = 0
SM = 0
Delta_L
Doxorubicin
YBR036C CSG2
ZShift = −20.78
Z lm Score = 1.85
NG = 0
DB = 0
SM = 0
Delta_K
Doxorubicin
YBR036C CSG2
ZShift = −9.05
Z lm Score = 4.33
NG = 0
DB = 0
SM = 0
Delta_r
Doxorubicin
YBR036C CSG2
ZShift = −12.31
Z lm Score = 1.43
NG = 0
DB = 0
SM = 0
Delta_AUC
Doxorubicin

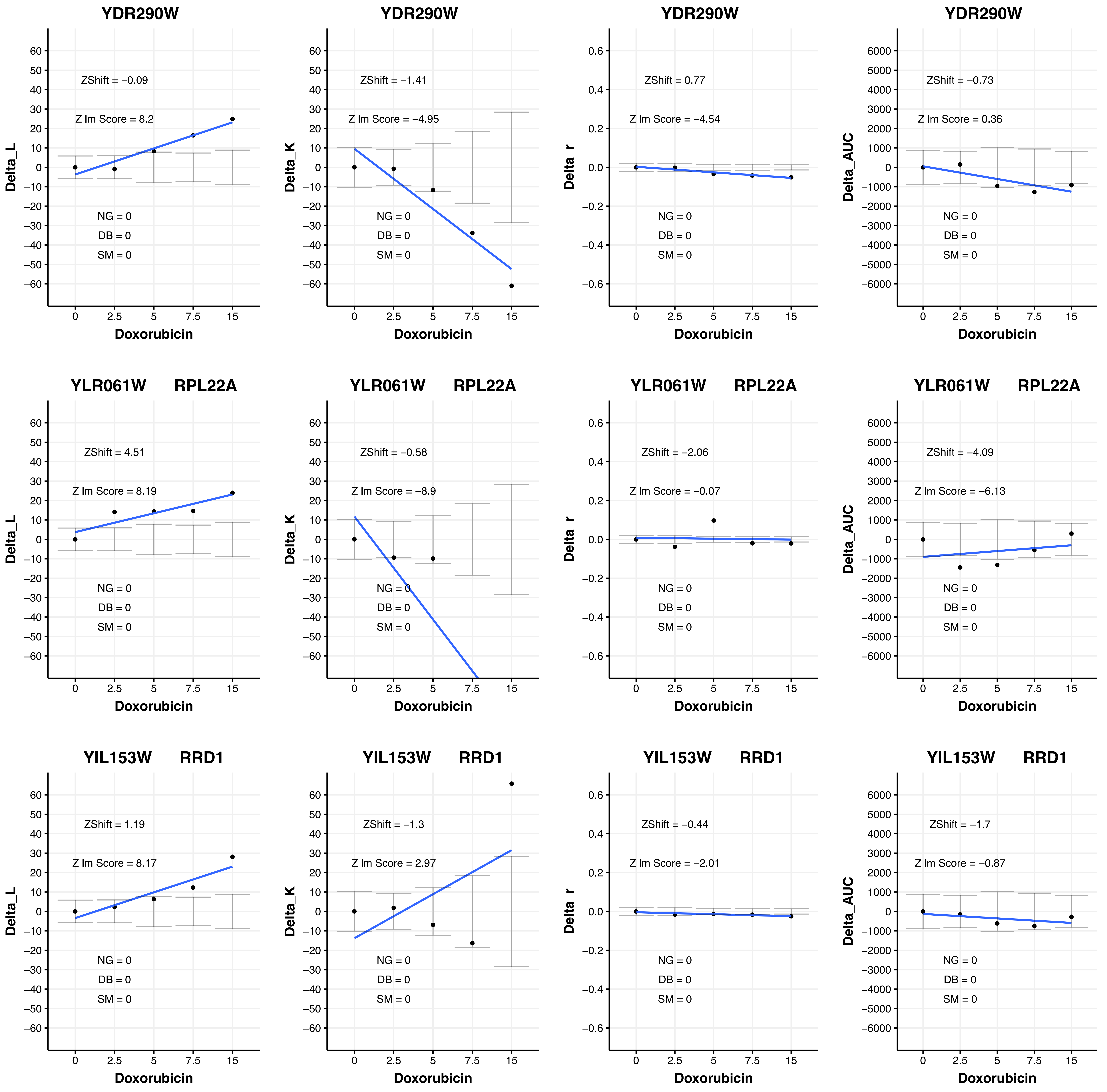
YDR290W
ZShift = −0.09
Z lm Score = 8.2
NG = 0
DB = 0
SM = 0
Delta_L
Doxorubicin
YDR290W
ZShift = −1.41
Z lm Score = −4.95
NG = 0
DB = 0
SM = 0
Delta_K
Doxorubicin
YDR290W
ZShift = 0.77
Z lm Score = −4.54
NG = 0
DB = 0
SM = 0
Delta_r
Doxorubicin
YDR290W
ZShift = −0.73
Z lm Score = 0.36
NG = 0
DB = 0
SM = 0
Delta_AUC
Doxorubicin
YLR061W RPL22A
ZShift = 4.51
Z lm Score = 8.19
NG = 0
DB = 0
SM = 0
Delta_L
Doxorubicin
YLR061W RPL22A
ZShift = −0.58
Z lm Score = −8.9
NG = 0
DB = 0
SM = 0
Delta_K
Doxorubicin
YLR061W RPL22A
ZShift = −2.06
Z lm Score = −0.07
NG = 0
DB = 0
SM = 0
Delta_r
Doxorubicin
YLR061W RPL22A
ZShift = −4.09
Z lm Score = −6.13
NG = 0
DB = 0
SM = 0
Delta_AUC
Doxorubicin
YIL153W RRD1
ZShift = 1.19
Z lm Score = 8.17
NG = 0
DB = 0
SM = 0
Delta_L
Doxorubicin
YIL153W RRD1
ZShift = −1.3
Z lm Score = 2.97
NG = 0
DB = 0
SM = 0
Delta_K
Doxorubicin
YIL153W RRD1
ZShift = −0.44
Z lm Score = −2.01
NG = 0
DB = 0
SM = 0
Delta_r
Doxorubicin
YIL153W RRD1
ZShift = −1.7
Z lm Score = −0.87
NG = 0
DB = 0
SM = 0
Delta_AUC
Doxorubicin

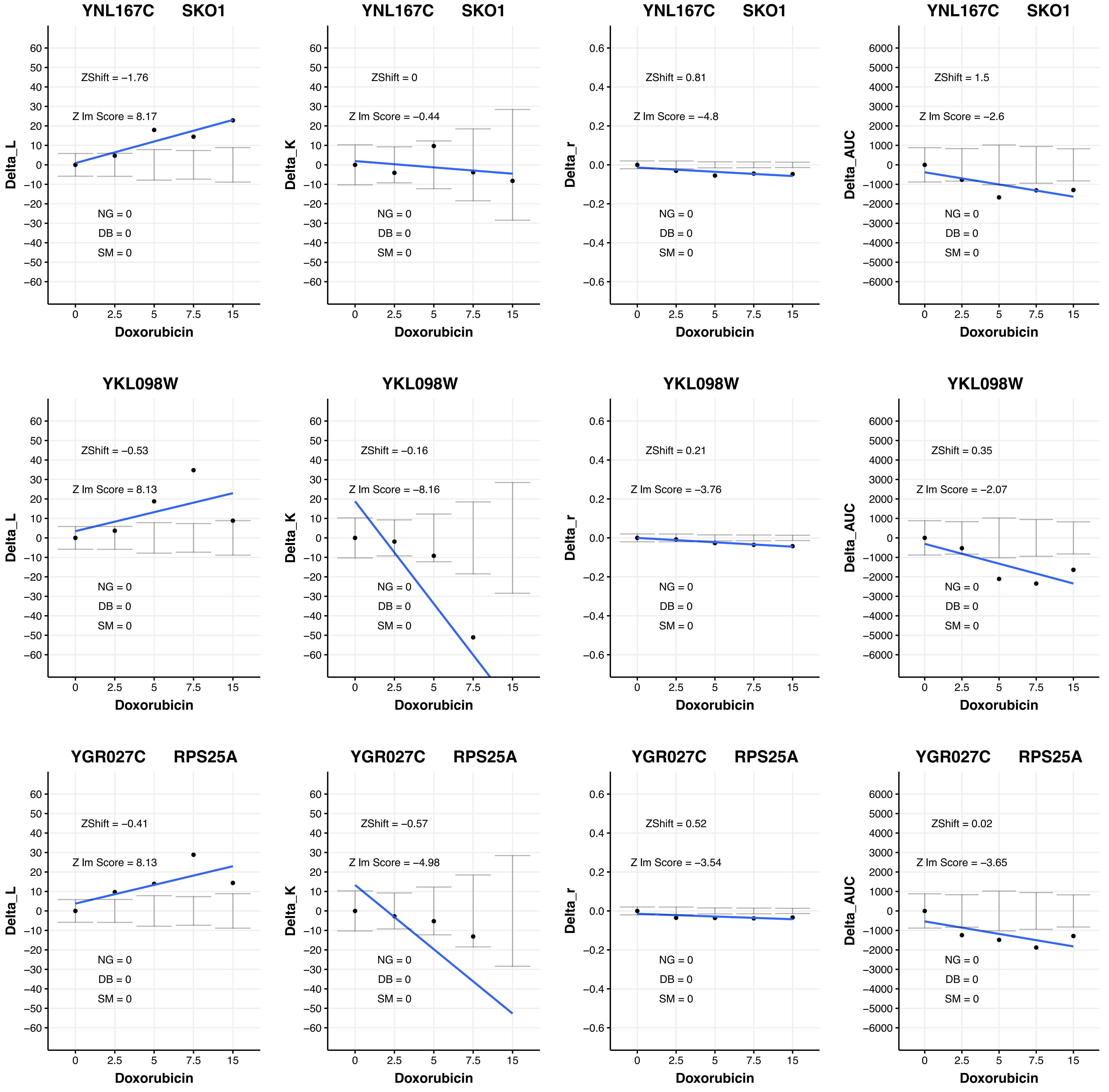
YNL167C SKO1
ZShift = −1.76
Z lm Score = 8.17
NG = 0
DB = 0
SM = 0
Delta_L
Doxorubicin
YNL167C SKO1
ZShift = 0
Z lm Score = −0.44
NG = 0
DB = 0
SM = 0
Delta_K
Doxorubicin
YNL167C SKO1
ZShift = 0.81
Z lm Score = −4.8
NG = 0
DB = 0
SM = 0
Delta_r
Doxorubicin
YNL167C SKO1
ZShift = 1.5
Z lm Score = −2.6
NG = 0
DB = 0
SM = 0
Delta_AUC
Doxorubicin
YKL098W
ZShift = −0.53
Z lm Score = 8.13
NG = 0
DB = 0
SM = 0
Delta_L
Doxorubicin
YKL098W
ZShift = −0.16
Z lm Score = −8.16
NG = 0
DB = 0
SM = 0
Delta_K
Doxorubicin
YKL098W
ZShift = 0.21
Z lm Score = −3.76
NG = 0
DB = 0
SM = 0
Delta_r
Doxorubicin
YKL098W
ZShift = 0.35
Z lm Score = −2.07
NG = 0
DB = 0
SM = 0
Delta_AUC
Doxorubicin
YGR027C RPS25A
ZShift = −0.41
Z lm Score = 8.13
NG = 0
DB = 0
SM = 0
Delta_L
Doxorubicin
YGR027C RPS25A
ZShift = −0.57
Z lm Score = −4.98
NG = 0
DB = 0
SM = 0
Delta_K
Doxorubicin
YGR027C RPS25A
ZShift = 0.52
Z lm Score = −3.54
NG = 0
DB = 0
SM = 0
Delta_r
Doxorubicin
YGR027C RPS25A
ZShift = 0.02
Z lm Score = −3.65
NG = 0
DB = 0
SM = 0
Delta_AUC
Doxorubicin

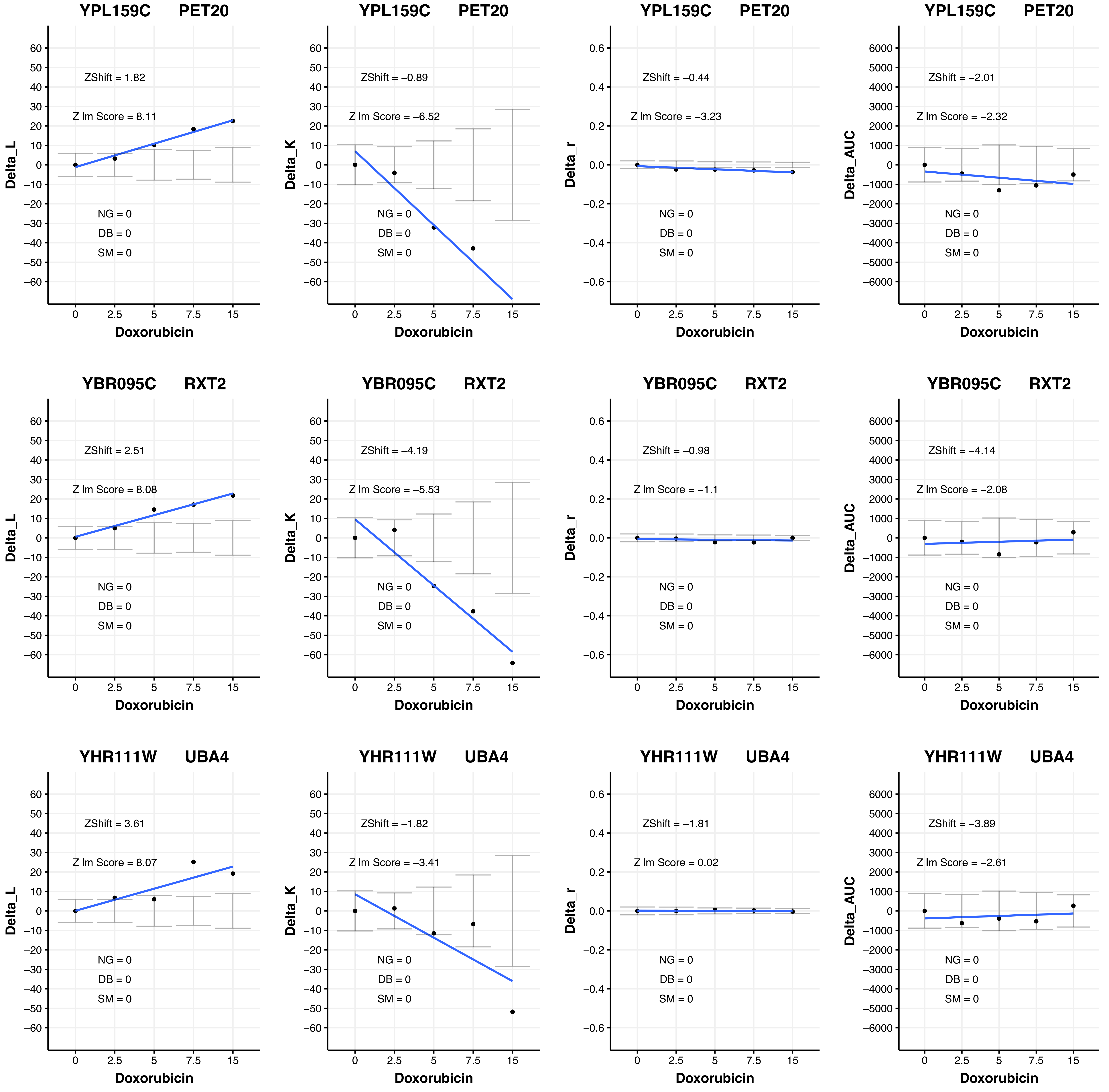
YPL159C PET20
ZShift = 1.82
Z lm Score = 8.11
NG = 0
DB = 0
SM = 0
Delta_L
Doxorubicin
YPL159C PET20
ZShift = −0.89
Z lm Score = −6.52
NG = 0
DB = 0
SM = 0
Delta_K
Doxorubicin
YPL159C PET20
ZShift = −0.44
Z lm Score = −3.23
NG = 0
DB = 0
SM = 0
Delta_r
Doxorubicin
YPL159C PET20
ZShift = −2.01
Z lm Score = −2.32
NG = 0
DB = 0
SM = 0
Delta_AUC
Doxorubicin
YBR095C RXT2
ZShift = 2.51
Z lm Score = 8.08
NG = 0
DB = 0
SM = 0
Delta_L
Doxorubicin
YBR095C RXT2
ZShift = −4.19
Z lm Score = −5.53
NG = 0
DB = 0
SM = 0
Delta_K
Doxorubicin
YBR095C RXT2
ZShift = −0.98
Z lm Score = −1.1
NG = 0
DB = 0
SM = 0
Delta_r
Doxorubicin
YBR095C RXT2
ZShift = −4.14
Z lm Score = −2.08
NG = 0
DB = 0
SM = 0
Delta_AUC
Doxorubicin
YHR111W UBA4
ZShift = 3.61
Z lm Score = 8.07
NG = 0
DB = 0
SM = 0
Delta_L
Doxorubicin
YHR111W UBA4
ZShift = −1.82
Z lm Score = −3.41
NG = 0
DB = 0
SM = 0
Delta_K
Doxorubicin
YHR111W UBA4
ZShift = −1.81
Z lm Score = 0.02
NG = 0
DB = 0
SM = 0
Delta_r
Doxorubicin
YHR111W UBA4
ZShift = −3.89
Z lm Score = −2.61
NG = 0
DB = 0
SM = 0
Delta_AUC
Doxorubicin

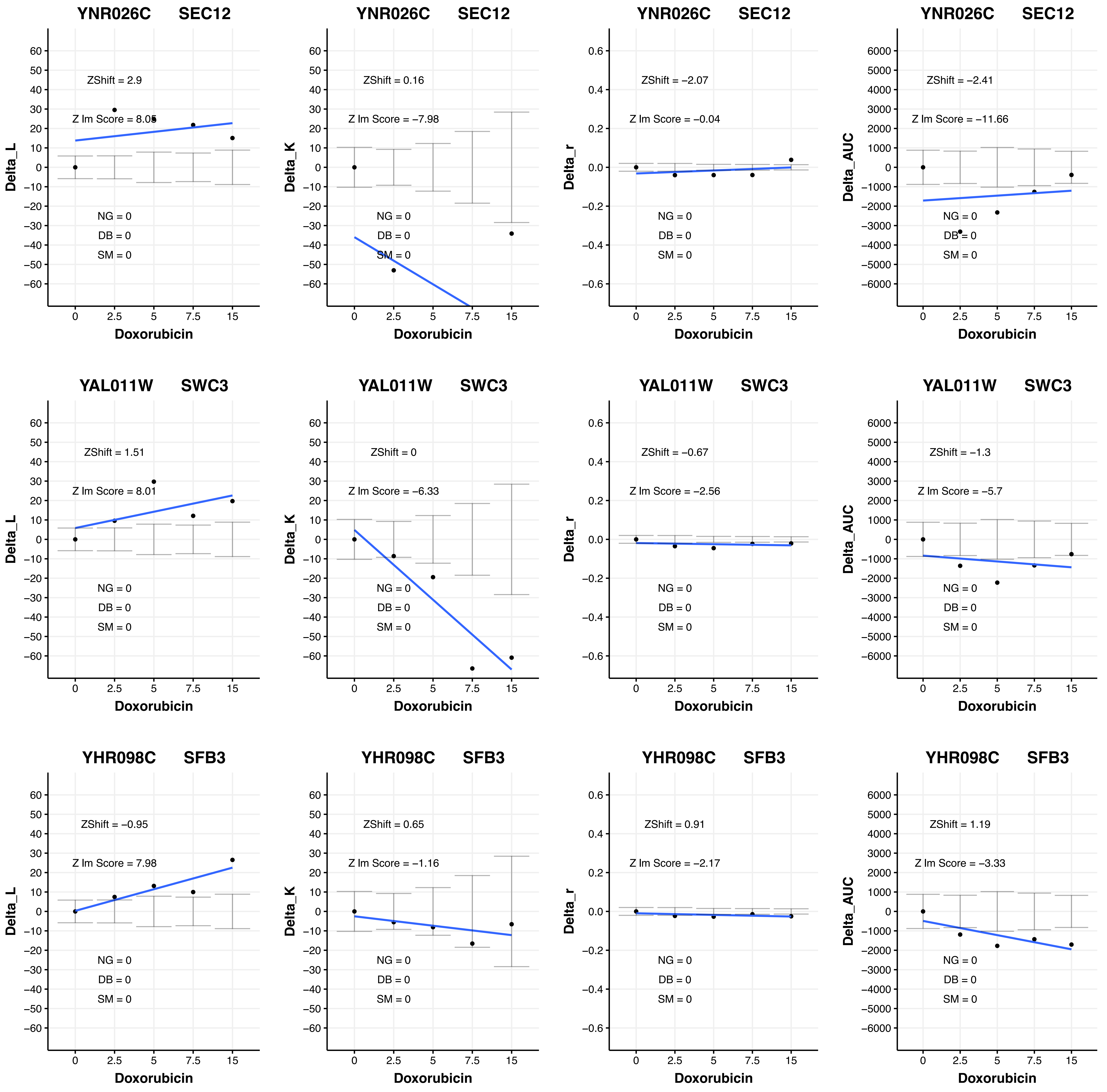
YNR026C SEC12
ZShift = 2.9
Z lm Score = 8.05
NG = 0
DB = 0
SM = 0
Delta_L
Doxorubicin
YNR026C SEC12
ZShift = 0.16
Z lm Score = −7.98
NG = 0
DB = 0
SM = 0
Delta_K
Doxorubicin
YNR026C SEC12
ZShift = −2.07
Z lm Score = −0.04
NG = 0
DB = 0
SM = 0
Delta_r
Doxorubicin
YNR026C SEC12
ZShift = −2.41
Z lm Score = −11.66
NG = 0
DB = 0
SM = 0
Delta_AUC
Doxorubicin
YAL011W SWC3
ZShift = 1.51
Z lm Score = 8.01
NG = 0
DB = 0
SM = 0
Delta_L
Doxorubicin
YAL011W SWC3
ZShift = 0
Z lm Score = −6.33
NG = 0
DB = 0
SM = 0
Delta_K
Doxorubicin
YAL011W SWC3
ZShift = −0.67
Z lm Score = −2.56
NG = 0
DB = 0
SM = 0
Delta_r
Doxorubicin
YAL011W SWC3
ZShift = −1.3
Z lm Score = −5.7
NG = 0
DB = 0
SM = 0
Delta_AUC
Doxorubicin
YHR098C SFB3
ZShift = −0.95
Z lm Score = 7.98
NG = 0
DB = 0
SM = 0
Delta_L
Doxorubicin
YHR098C SFB3
ZShift = 0.65
Z lm Score = −1.16
NG = 0
DB = 0
SM = 0
Delta_K
Doxorubicin
YHR098C SFB3
ZShift = 0.91
Z lm Score = −2.17
NG = 0
DB = 0
SM = 0
Delta_r
Doxorubicin
YHR098C SFB3
ZShift = 1.19
Z lm Score = −3.33
NG = 0
DB = 0
SM = 0
Delta_AUC
Doxorubicin

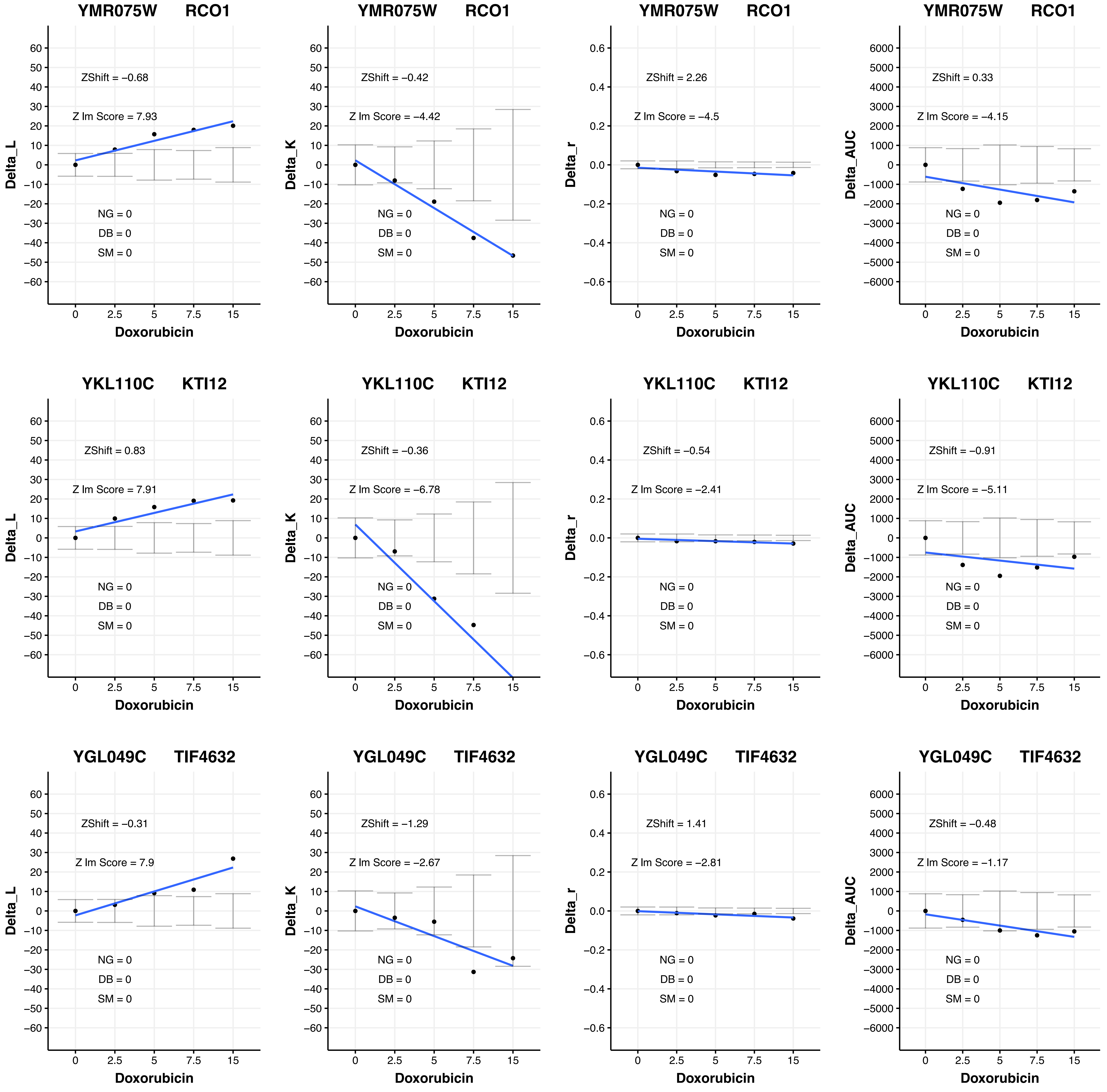
YMR075W RCO1
ZShift = −0.68
Z lm Score = 7.93
NG = 0
DB = 0
SM = 0
Delta_L
Doxorubicin
YMR075W RCO1
ZShift = −0.42
Z lm Score = −4.42
NG = 0
DB = 0
SM = 0
Delta_K
Doxorubicin
YMR075W RCO1
ZShift = 2.26
Z lm Score = −4.5
NG = 0
DB = 0
SM = 0
Delta_r
Doxorubicin
YMR075W RCO1
ZShift = 0.33
Z lm Score = −4.15
NG = 0
DB = 0
SM = 0
Delta_AUC
Doxorubicin
YKL110C KTI12
ZShift = 0.83
Z lm Score = 7.91
NG = 0
DB = 0
SM = 0
Delta_L
Doxorubicin
YKL110C KTI12
ZShift = −0.36
Z lm Score = −6.78
NG = 0
DB = 0
SM = 0
Delta_K
Doxorubicin
YKL110C KTI12
ZShift = −0.54
Z lm Score = −2.41
NG = 0
DB = 0
SM = 0
Delta_r
Doxorubicin
YKL110C KTI12
ZShift = −0.91
Z lm Score = −5.11
NG = 0
DB = 0
SM = 0
Delta_AUC
Doxorubicin
YGL049C TIF4632
ZShift = −0.31
Z lm Score = 7.9
NG = 0
DB = 0
SM = 0
Delta_L
Doxorubicin
YGL049C TIF4632
ZShift = −1.29
Z lm Score = −2.67
NG = 0
DB = 0
SM = 0
Delta_K
Doxorubicin
YGL049C TIF4632
ZShift = 1.41
Z lm Score = −2.81
NG = 0
DB = 0
SM = 0
Delta_r
Doxorubicin
YGL049C TIF4632
ZShift = −0.48
Z lm Score = −1.17
NG = 0
DB = 0
SM = 0
Delta_AUC
Doxorubicin

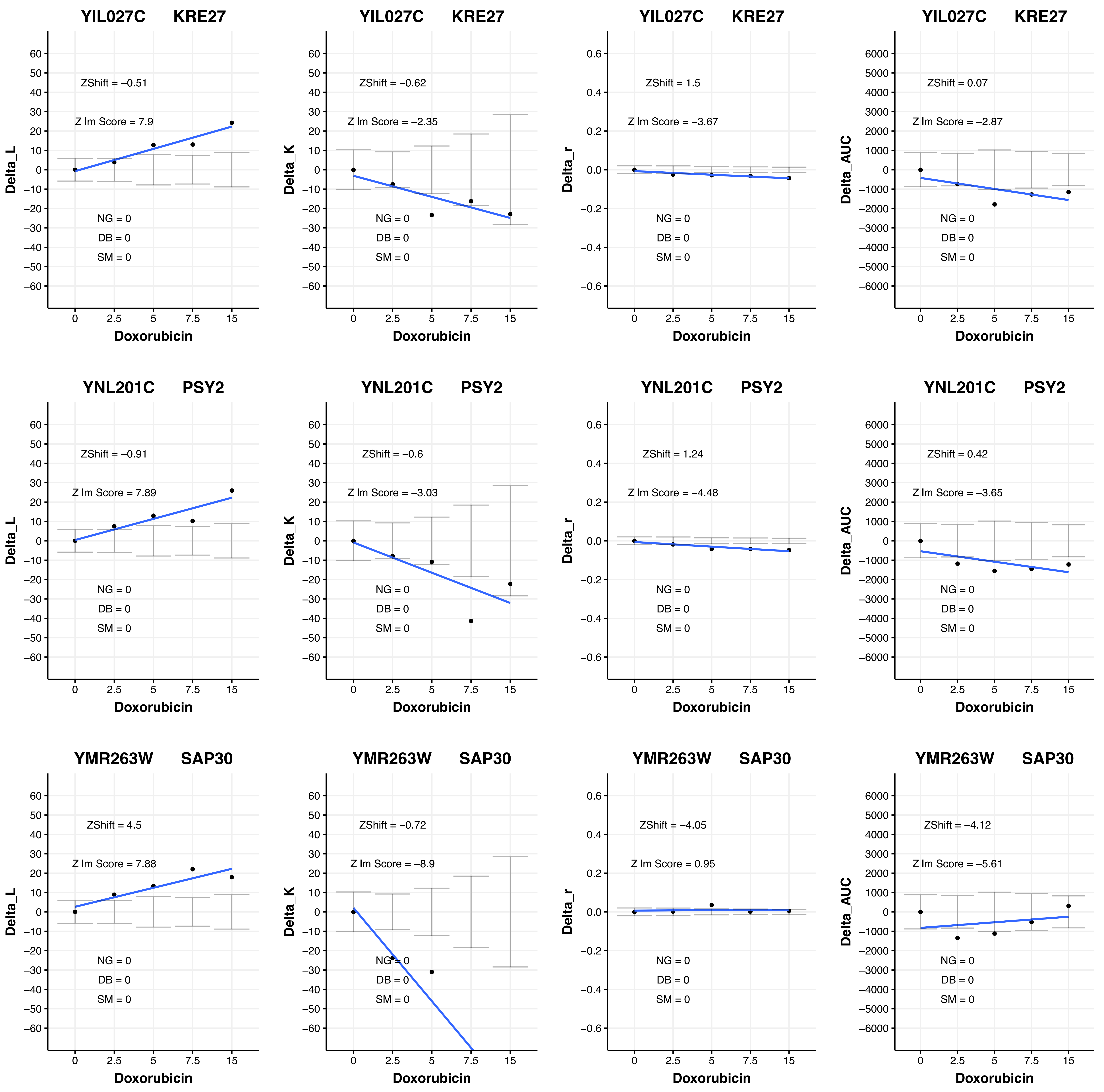

YIL027C KRE27
ZShift = −0.51
Z lm Score = 7.9
NG = 0
DB = 0
SM = 0
Delta_L
Doxorubicin
YIL027C KRE27
ZShift = −0.62
Z lm Score = −2.35
NG = 0
DB = 0
SM = 0
Delta_K
Doxorubicin
YIL027C KRE27
ZShift = 1.5
Z lm Score = −3.67
NG = 0
DB = 0
SM = 0
Delta_r
Doxorubicin
YIL027C KRE27
ZShift = 0.07
Z lm Score = −2.87
NG = 0
DB = 0
SM = 0
Delta_AUC
Doxorubicin
YNL201C PSY2
ZShift = −0.91
Z lm Score = 7.89
NG = 0
DB = 0
SM = 0
Delta_L
Doxorubicin
YNL201C PSY2
ZShift = −0.6
Z lm Score = −3.03
NG = 0
DB = 0
SM = 0
Delta_K
Doxorubicin
YNL201C PSY2
ZShift = 1.24
Z lm Score = −4.48
NG = 0
DB = 0
SM = 0
Delta_r
Doxorubicin
YNL201C PSY2
ZShift = 0.42
Z lm Score = −3.65
NG = 0
DB = 0
SM = 0
Delta_AUC
Doxorubicin
YMR263W SAP30
ZShift = 4.5
Z lm Score = 7.88
NG = 0
DB = 0
SM = 0
Delta_L
Doxorubicin
YMR263W SAP30
ZShift = −0.72
Z lm Score = −8.9
NG = 0
DB = 0
SM = 0
Delta_K
Doxorubicin
YMR263W SAP30
ZShift = −4.05
Z lm Score = 0.95
NG = 0
DB = 0
SM = 0
Delta_r
Doxorubicin
YMR263W SAP30
ZShift = −4.12
Z lm Score = −5.61
NG = 0
DB = 0
SM = 0
Delta_AUC
Doxorubicin

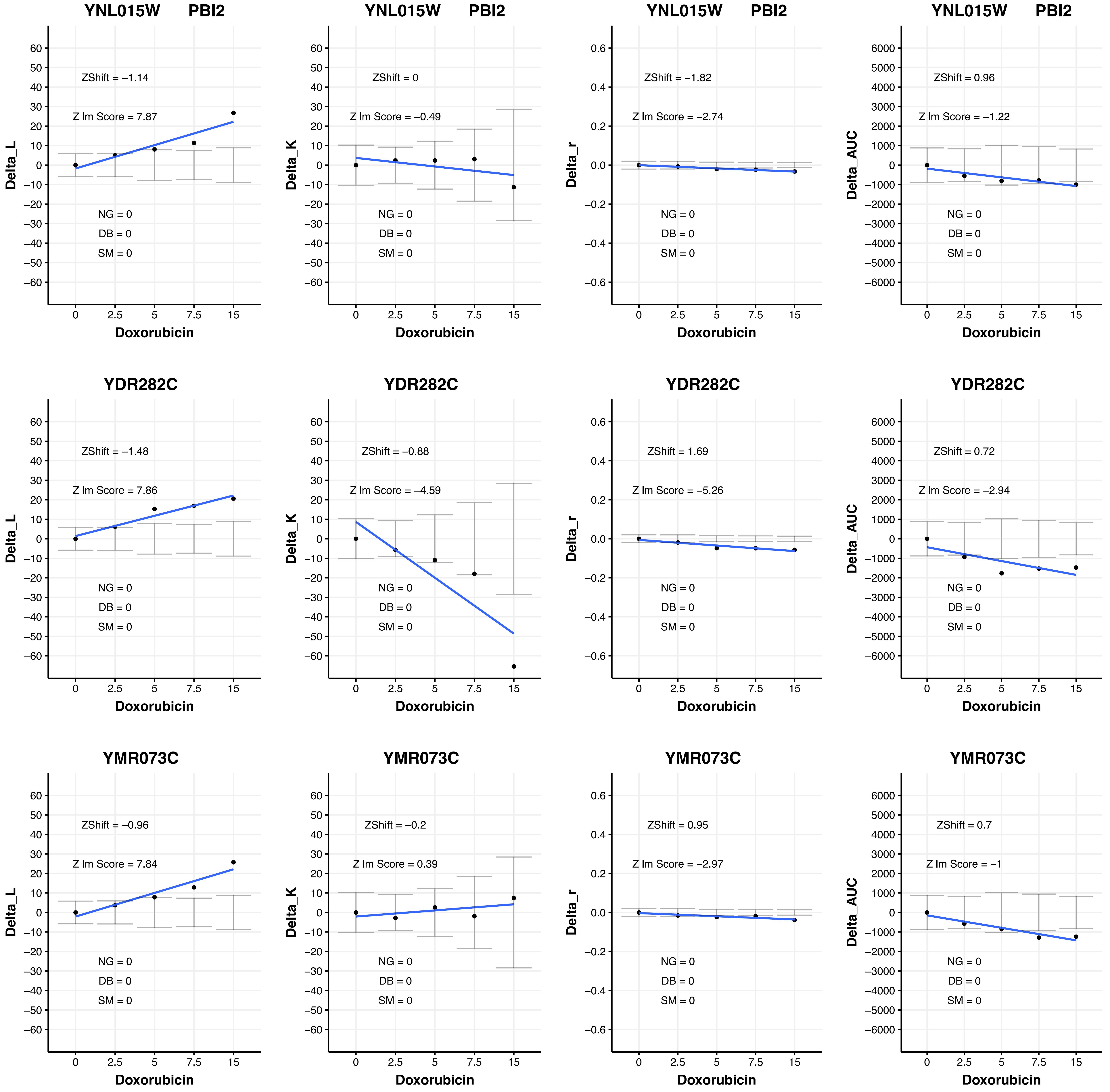
YNL015W PBI2
ZShift = −1.14
Z lm Score = 7.87
NG = 0
DB = 0
SM = 0
Delta_L
Doxorubicin
YNL015W PBI2
ZShift = 0
Z lm Score = −0.49
NG = 0
DB = 0
SM = 0
Delta_K
Doxorubicin
YNL015W PBI2
ZShift = −1.82
Z lm Score = −2.74
NG = 0
DB = 0
SM = 0
Delta_r
Doxorubicin
YNL015W PBI2
ZShift = 0.96
Z lm Score = −1.22
NG = 0
DB = 0
SM = 0
Delta_AUC
Doxorubicin
YDR282C
ZShift = −1.48
Z lm Score = 7.86
NG = 0
DB = 0
SM = 0
Delta_L
Doxorubicin
YDR282C
ZShift = −0.88
Z lm Score = −4.59
NG = 0
DB = 0
SM = 0
Delta_K
Doxorubicin
YDR282C
ZShift = 1.69
Z lm Score = −5.26
NG = 0
DB = 0
SM = 0
Delta_r
Doxorubicin
YDR282C
ZShift = 0.72
Z lm Score = −2.94
NG = 0
DB = 0
SM = 0
Delta_AUC
Doxorubicin
YMR073C
ZShift = −0.96
Z lm Score = 7.84
NG = 0
DB = 0
SM = 0
Delta_L
Doxorubicin
YMR073C
ZShift = −0.2
Z lm Score = 0.39
NG = 0
DB = 0
SM = 0
Delta_K
Doxorubicin
YMR073C
ZShift = 0.95
Z lm Score = −2.97
NG = 0
DB = 0
SM = 0
Delta_r
Doxorubicin
YMR073C
ZShift = 0.7
Z lm Score = −1
NG = 0
DB = 0
SM = 0
Delta_AUC
Doxorubicin

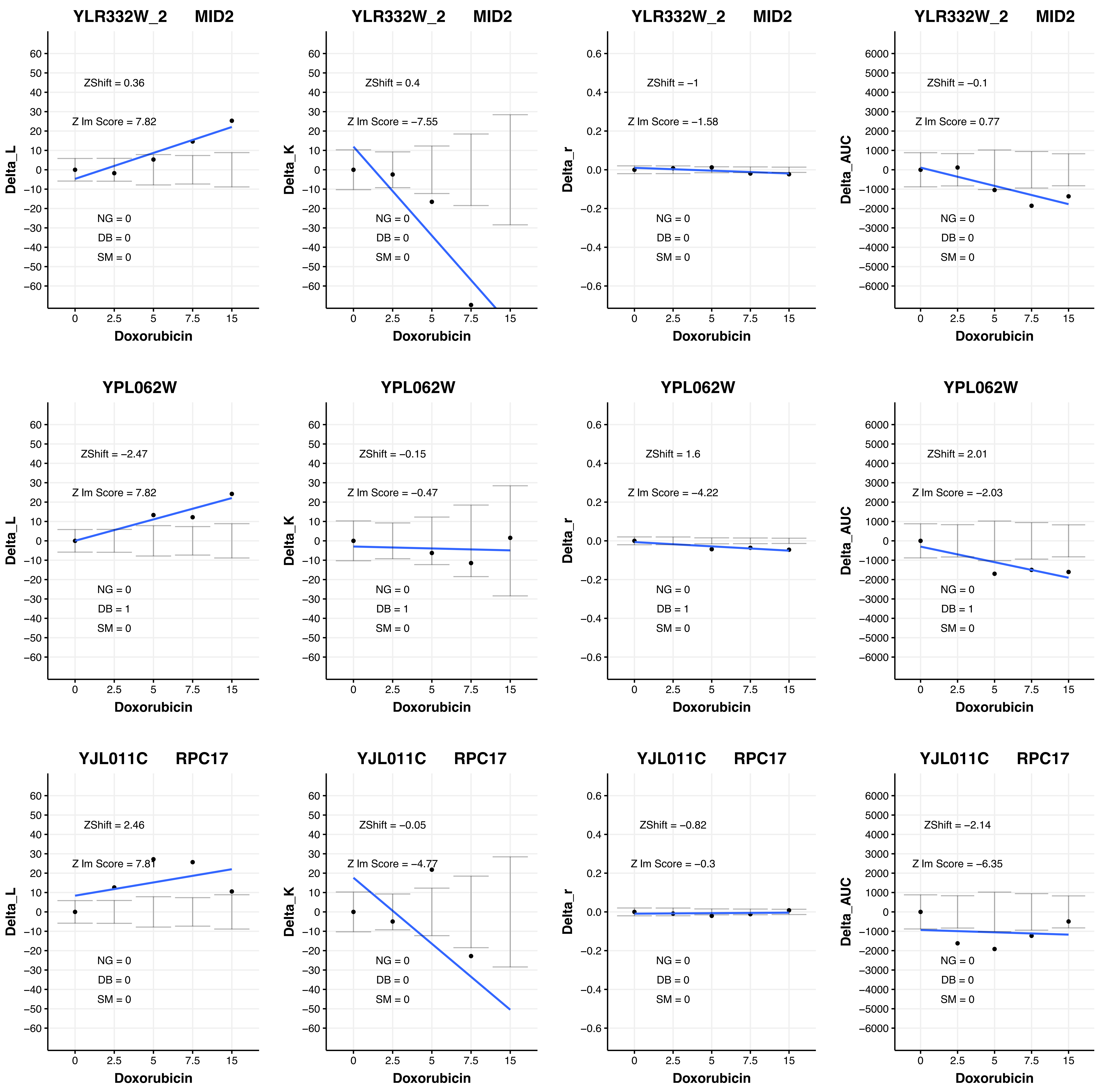

YLR332W_2 MID2
ZShift = 0.36
Z lm Score = 7.82
NG = 0
DB = 0
SM = 0
Delta_L
Doxorubicin
YLR332W_2 MID2
ZShift = 0.4
Z lm Score = −7.55
NG = 0
DB = 0
SM = 0
Delta_K
Doxorubicin
YLR332W_2 MID2
ZShift = −1
Z lm Score = −1.58
NG = 0
DB = 0
SM = 0
Delta_r
Doxorubicin
YLR332W_2 MID2
ZShift = −0.1
Z lm Score = 0.77
NG = 0
DB = 0
SM = 0
Delta_AUC
Doxorubicin
YPL062W
ZShift = −2.47
Z lm Score = 7.82
NG = 0
DB = 1
SM = 0
Delta_L
Doxorubicin
YPL062W
ZShift = −0.15
Z lm Score = −0.47
NG = 0
DB = 1
SM = 0
Delta_K
Doxorubicin
YPL062W
ZShift = 1.6
Z lm Score = −4.22
NG = 0
DB = 1
SM = 0
Delta_r
Doxorubicin
YPL062W
ZShift = 2.01
Z lm Score = −2.03
NG = 0
DB = 1
SM = 0
Delta_AUC
Doxorubicin
YJL011C RPC17
ZShift = 2.46
Z lm Score = 7.81
NG = 0
DB = 0
SM = 0
Delta_L
Doxorubicin
YJL011C RPC17
ZShift = −0.05
Z lm Score = −4.77
NG = 0
DB = 0
SM = 0
Delta_K
Doxorubicin
YJL011C RPC17
ZShift = −0.82
Z lm Score = −0.3
NG = 0
DB = 0
SM = 0
Delta_r
Doxorubicin
YJL011C RPC17
ZShift = −2.14
Z lm Score = −6.35
NG = 0
DB = 0
SM = 0
Delta_AUC
Doxorubicin

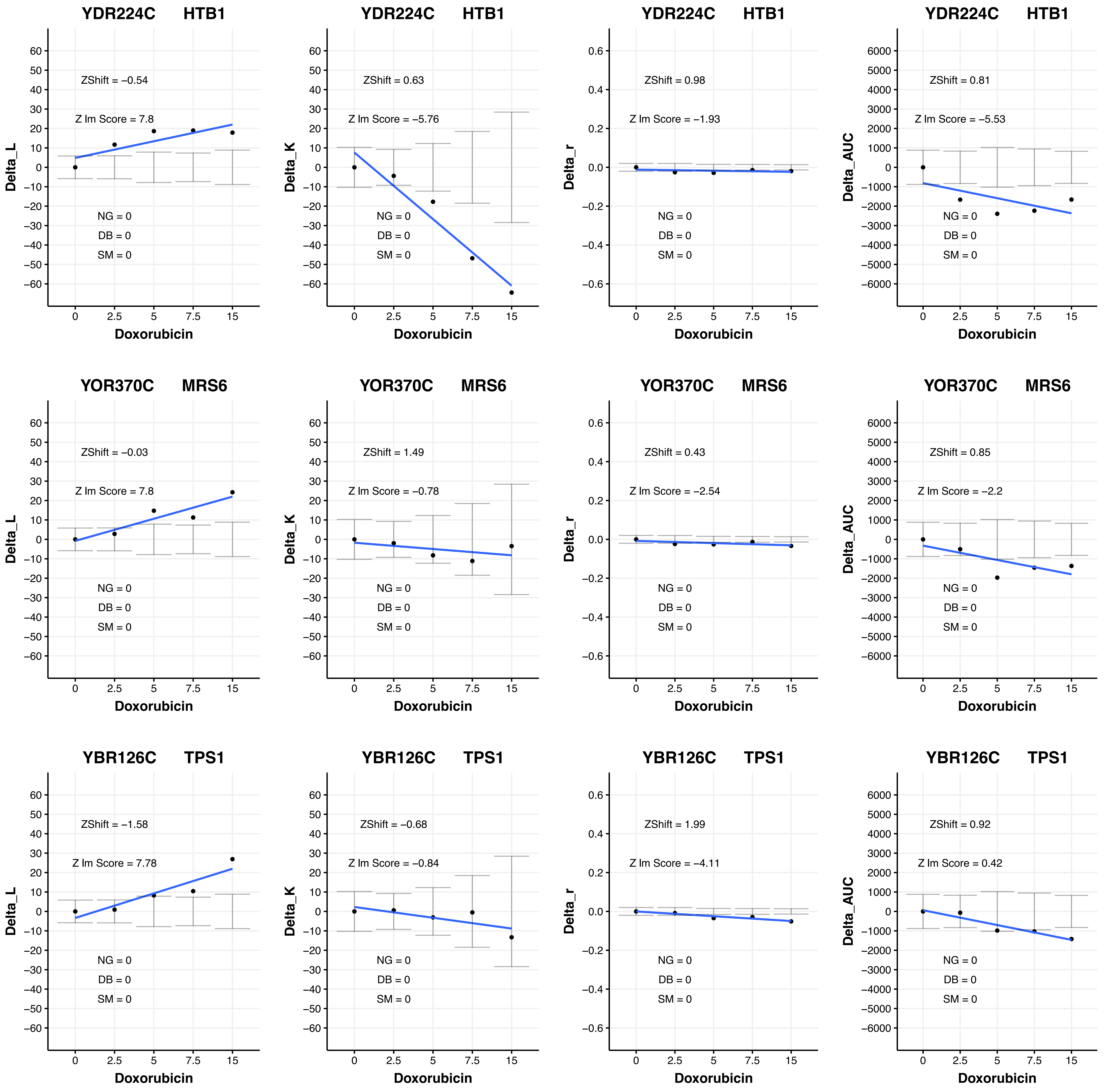

YDR224C HTB1
ZShift = −0.54
Z lm Score = 7.8
NG = 0
DB = 0
SM = 0
Delta_L
Doxorubicin
YDR224C HTB1
ZShift = 0.63
Z lm Score = −5.76
NG = 0
DB = 0
SM = 0
Delta_K
Doxorubicin
YDR224C HTB1
ZShift = 0.98
Z lm Score = −1.93
NG = 0
DB = 0
SM = 0
Delta_r
Doxorubicin
YDR224C HTB1
ZShift = 0.81
Z lm Score = −5.53
NG = 0
DB = 0
SM = 0
Delta_AUC
Doxorubicin
YOR370C MRS6
ZShift = −0.03
Z lm Score = 7.8
NG = 0
DB = 0
SM = 0
Delta_L
Doxorubicin
YOR370C MRS6
ZShift = 1.49
Z lm Score = −0.78
NG = 0
DB = 0
SM = 0
Delta_K
Doxorubicin
YOR370C MRS6
ZShift = 0.43
Z lm Score = −2.54
NG = 0
DB = 0
SM = 0
Delta_r
Doxorubicin
YOR370C MRS6
ZShift = 0.85
Z lm Score = −2.2
NG = 0
DB = 0
SM = 0
Delta_AUC
Doxorubicin
YBR126C TPS1
ZShift = −1.58
Z lm Score = 7.78
NG = 0
DB = 0
SM = 0
Delta_L
Doxorubicin
YBR126C TPS1
ZShift = −0.68
Z lm Score = −0.84
NG = 0
DB = 0
SM = 0
Delta_K
Doxorubicin
YBR126C TPS1
ZShift = 1.99
Z lm Score = −4.11
NG = 0
DB = 0
SM = 0
Delta_r
Doxorubicin
YBR126C TPS1
ZShift = 0.92
Z lm Score = 0.42
NG = 0
DB = 0
SM = 0
Delta_AUC
Doxorubicin

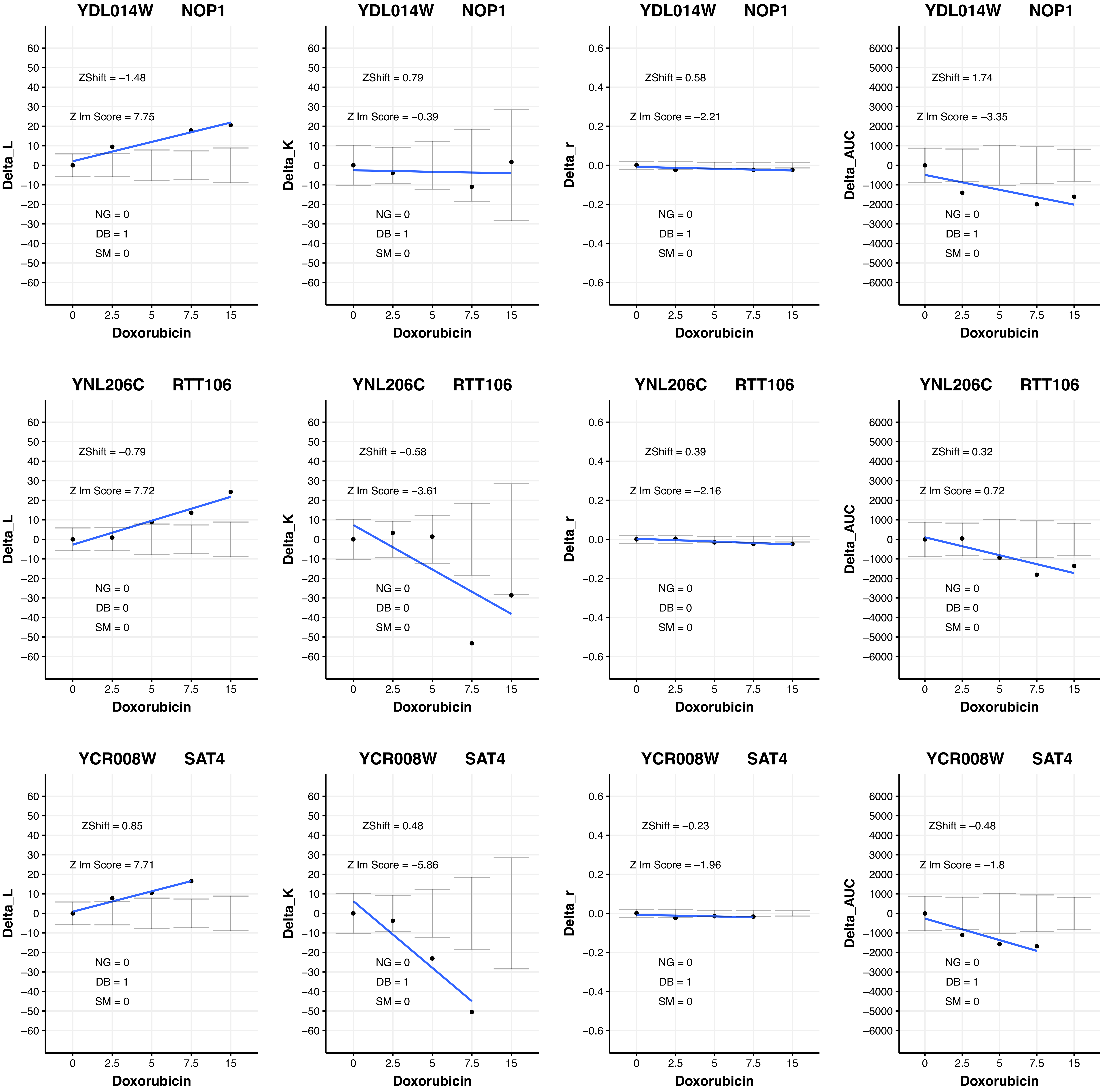
YDL014W NOP1
ZShift = −1.48
Z lm Score = 7.75
NG = 0
DB = 1
SM = 0
Delta_L
Doxorubicin
YDL014W NOP1
ZShift = 0.79
Z lm Score = −0.39
NG = 0
DB = 1
SM = 0
Delta_K
Doxorubicin
YDL014W NOP1
ZShift = 0.58
Z lm Score = −2.21
NG = 0
DB = 1
SM = 0
Delta_r
Doxorubicin
YDL014W NOP1
ZShift = 1.74
Z lm Score = −3.35
NG = 0
DB = 1
SM = 0
Delta_AUC
Doxorubicin
YNL206C RTT106
ZShift = −0.79
Z lm Score = 7.72
NG = 0
DB = 0
SM = 0
Delta_L
Doxorubicin
YNL206C RTT106
ZShift = −0.58
Z lm Score = −3.61
NG = 0
DB = 0
SM = 0
Delta_K
Doxorubicin
YNL206C RTT106
ZShift = 0.39
Z lm Score = −2.16
NG = 0
DB = 0
SM = 0
Delta_r
Doxorubicin
YNL206C RTT106
ZShift = 0.32
Z lm Score = 0.72
NG = 0
DB = 0
SM = 0
Delta_AUC
Doxorubicin
YCR008W SAT4
ZShift = 0.85
Z lm Score = 7.71
NG = 0
DB = 1
SM = 0
Delta_L
Doxorubicin
YCR008W SAT4
ZShift = 0.48
Z lm Score = −5.86
NG = 0
DB = 1
SM = 0
Delta_K
Doxorubicin
YCR008W SAT4
ZShift = −0.23
Z lm Score = −1.96
NG = 0
DB = 1
SM = 0
Delta_r
Doxorubicin
YCR008W SAT4
ZShift = −0.48
Z lm Score = −1.8
NG = 0
DB = 1
SM = 0
Delta_AUC
Doxorubicin

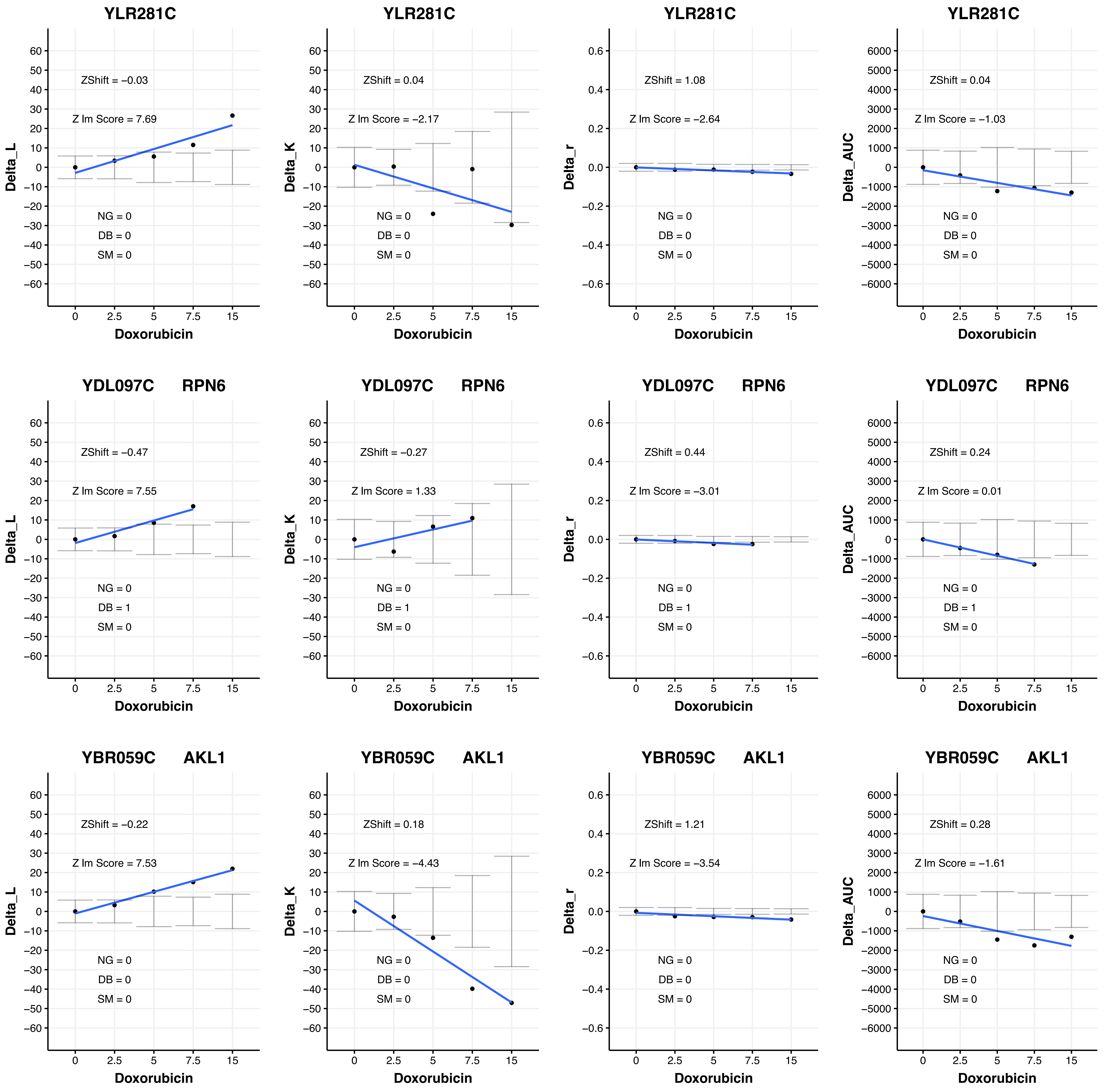
YLR281C
ZShift = −0.03
Z lm Score = 7.69
NG = 0
DB = 0
SM = 0
Delta_L
Doxorubicin
YLR281C
ZShift = 0.04
Z lm Score = −2.17
NG = 0
DB = 0
SM = 0
Delta_K
Doxorubicin
YLR281C
ZShift = 1.08
Z lm Score = −2.64
NG = 0
DB = 0
SM = 0
Delta_r
Doxorubicin
YLR281C
ZShift = 0.04
Z lm Score = −1.03
NG = 0
DB = 0
SM = 0
Delta_AUC
Doxorubicin
YDL097C RPN6
ZShift = −0.47
Z lm Score = 7.55
NG = 0
DB = 1
SM = 0
Delta_L
Doxorubicin
YDL097C RPN6
ZShift = −0.27
Z lm Score = 1.33
NG = 0
DB = 1
SM = 0
Delta_K
Doxorubicin
YDL097C RPN6
ZShift = 0.44
Z lm Score = −3.01
NG = 0
DB = 1
SM = 0
Delta_r
Doxorubicin
YDL097C RPN6
ZShift = 0.24
Z lm Score = 0.01
NG = 0
DB = 1
SM = 0
Delta_AUC
Doxorubicin
YBR059C AKL1
ZShift = −0.22
Z lm Score = 7.53
NG = 0
DB = 0
SM = 0
Delta_L
Doxorubicin
YBR059C AKL1
ZShift = 0.18
Z lm Score = −4.43
NG = 0
DB = 0
SM = 0
Delta_K
Doxorubicin
YBR059C AKL1
ZShift = 1.21
Z lm Score = −3.54
NG = 0
DB = 0
SM = 0
Delta_r
Doxorubicin
YBR059C AKL1
ZShift = 0.28
Z lm Score = −1.61
NG = 0
DB = 0
SM = 0
Delta_AUC
Doxorubicin

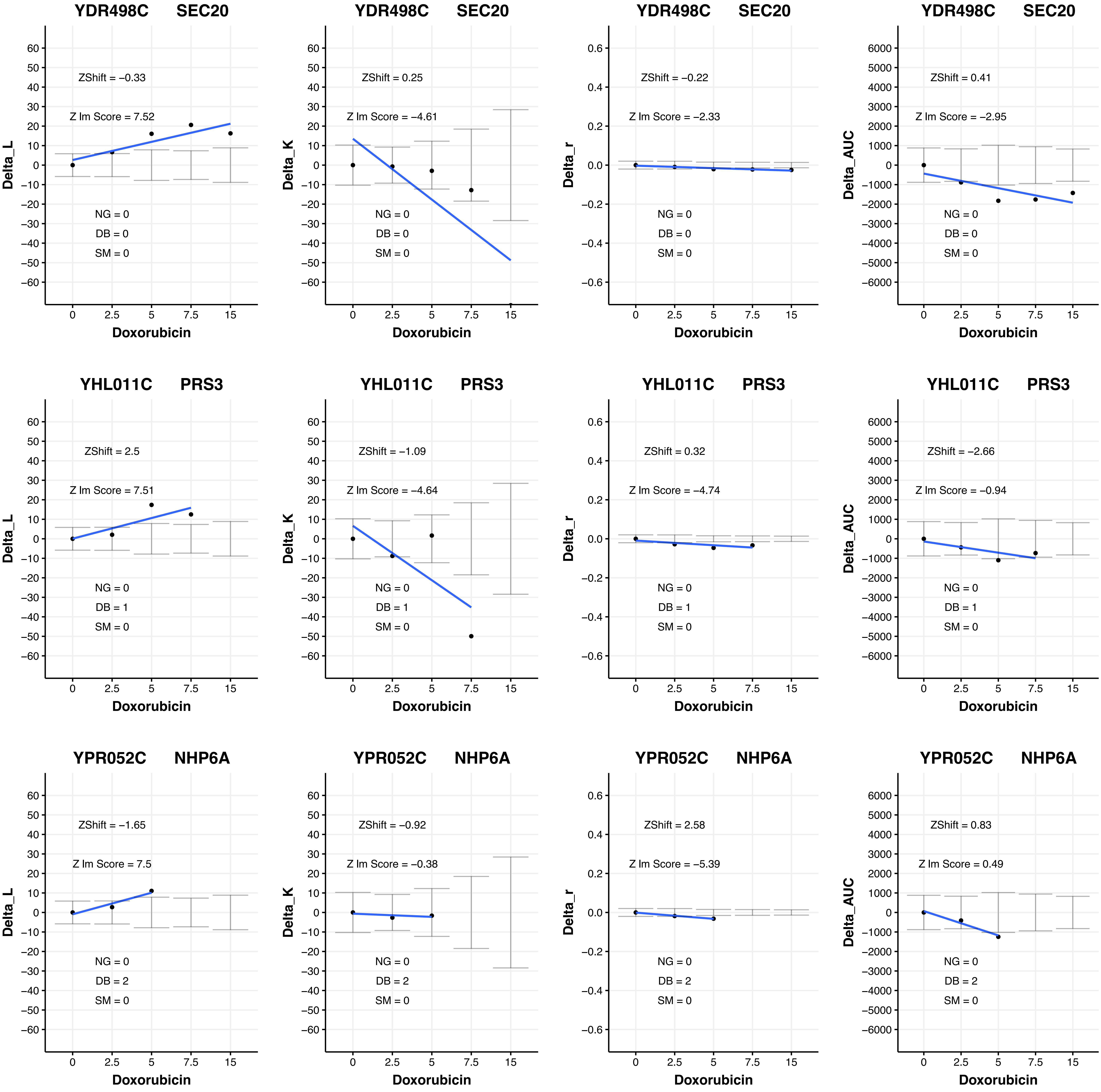

YDR498C SEC20
ZShift = −0.33
Z lm Score = 7.52
NG = 0
DB = 0
SM = 0
Delta_L
Doxorubicin
YDR498C SEC20
ZShift = 0.25
Z lm Score = −4.61
NG = 0
DB = 0
SM = 0
Delta_K
Doxorubicin
YDR498C SEC20
ZShift = −0.22
Z lm Score = −2.33
NG = 0
DB = 0
SM = 0
Delta_r
Doxorubicin
YDR498C SEC20
ZShift = 0.41
Z lm Score = −2.95
NG = 0
DB = 0
SM = 0
Delta_AUC
Doxorubicin
YHL011C PRS3
ZShift = 2.5
Z lm Score = 7.51
NG = 0
DB = 1
SM = 0
Delta_L
Doxorubicin
YHL011C PRS3
ZShift = −1.09
Z lm Score = −4.64
NG = 0
DB = 1
SM = 0
Delta_K
Doxorubicin
YHL011C PRS3
ZShift = 0.32
Z lm Score = −4.74
NG = 0
DB = 1
SM = 0
Delta_r
Doxorubicin
YHL011C PRS3
ZShift = −2.66
Z lm Score = −0.94
NG = 0
DB = 1
SM = 0
Delta_AUC
Doxorubicin
YPR052C NHP6A
ZShift = −1.65
Z lm Score = 7.5
NG = 0
DB = 2
SM = 0
Delta_L
Doxorubicin
YPR052C NHP6A
ZShift = −0.92
Z lm Score = −0.38
NG = 0
DB = 2
SM = 0
Delta_K
Doxorubicin
YPR052C NHP6A
ZShift = 2.58
Z lm Score = −5.39
NG = 0
DB = 2
SM = 0
Delta_r
Doxorubicin
YPR052C NHP6A
ZShift = 0.83
Z lm Score = 0.49
NG = 0
DB = 2
SM = 0
Delta_AUC
Doxorubicin

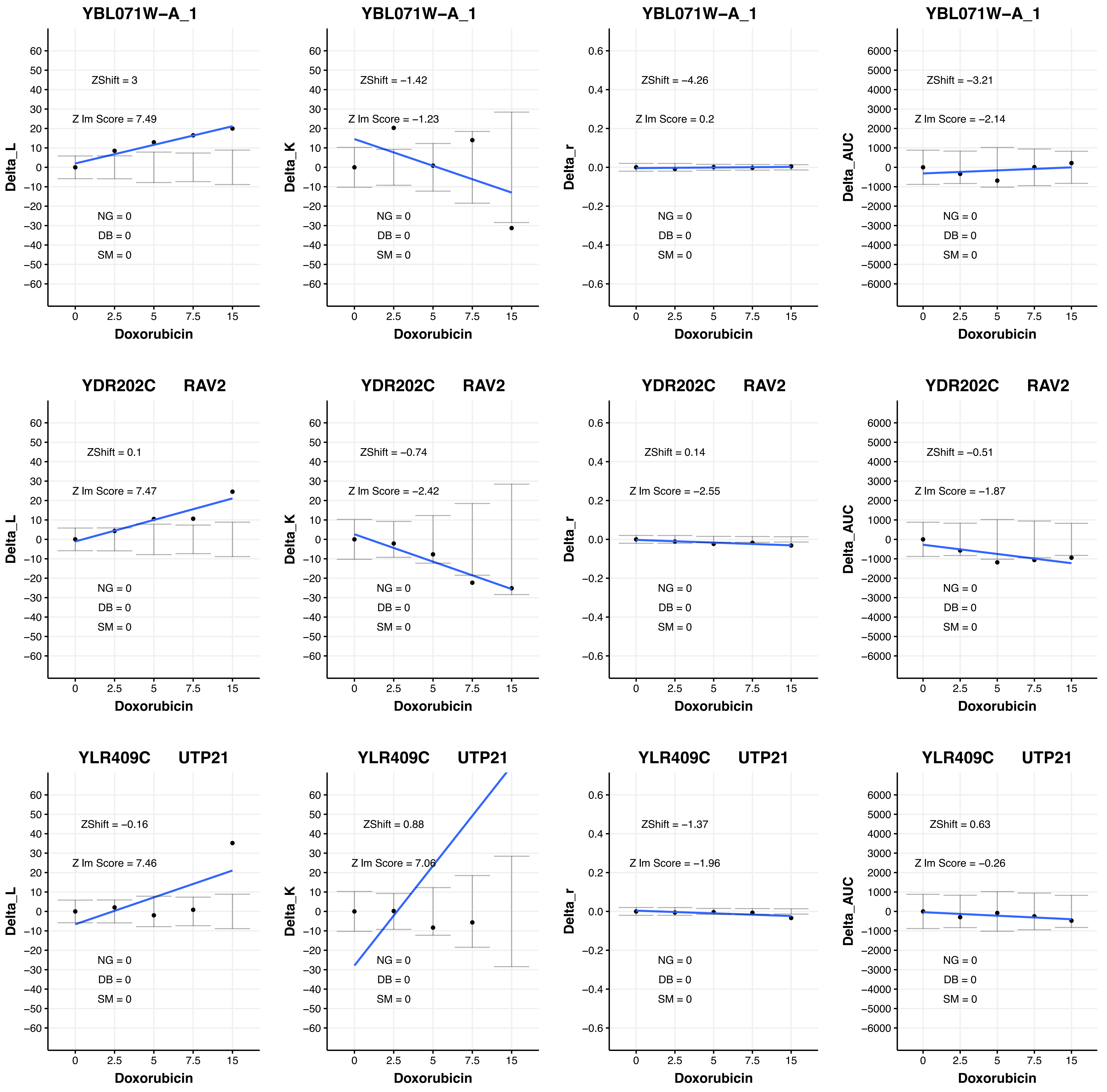
YBL071W−A_1
ZShift = 3
Z lm Score = 7.49
NG = 0
DB = 0
SM = 0
Delta_L
Doxorubicin
YBL071W−A_1
ZShift = −1.42
Z lm Score = −1.23
NG = 0
DB = 0
SM = 0
Delta_K
Doxorubicin
YBL071W−A_1
ZShift = −4.26
Z lm Score = 0.2
NG = 0
DB = 0
SM = 0
Delta_r
Doxorubicin
YBL071W−A_1
ZShift = −3.21
Z lm Score = −2.14
NG = 0
DB = 0
SM = 0
Delta_AUC
Doxorubicin
YDR202C RAV2
ZShift = 0.1
Z lm Score = 7.47
NG = 0
DB = 0
SM = 0
Delta_L
Doxorubicin
YDR202C RAV2
ZShift = −0.74
Z lm Score = −2.42
NG = 0
DB = 0
SM = 0
Delta_K
Doxorubicin
YDR202C RAV2
ZShift = 0.14
Z lm Score = −2.55
NG = 0
DB = 0
SM = 0
Delta_r
Doxorubicin
YDR202C RAV2
ZShift = −0.51
Z lm Score = −1.87
NG = 0
DB = 0
SM = 0
Delta_AUC
Doxorubicin
YLR409C UTP21
ZShift = −0.16
Z lm Score = 7.46
NG = 0
DB = 0
SM = 0
Delta_L
Doxorubicin
YLR409C UTP21
ZShift = 0.88
Z lm Score = 7.06
NG = 0
DB = 0
SM = 0
Delta_K
Doxorubicin
YLR409C UTP21
ZShift = −1.37
Z lm Score = −1.96
NG = 0
DB = 0
SM = 0
Delta_r
Doxorubicin
YLR409C UTP21
ZShift = 0.63
Z lm Score = −0.26
NG = 0
DB = 0
SM = 0
Delta_AUC
Doxorubicin

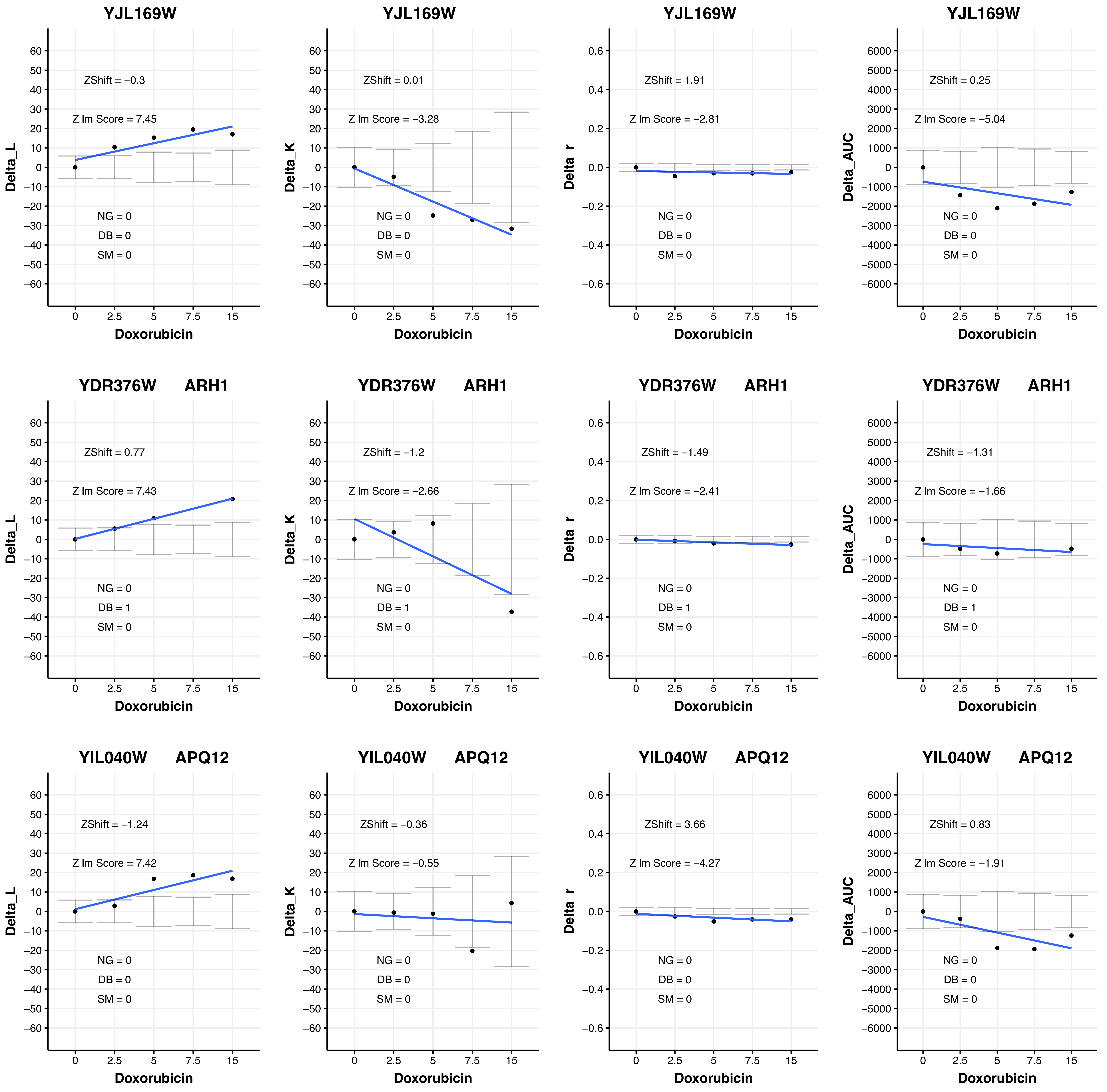
YJL169W
ZShift = −0.3
Z lm Score = 7.45
NG = 0
DB = 0
SM = 0
Delta_L
Doxorubicin
YJL169W
ZShift = 0.01
Z lm Score = −3.28
NG = 0
DB = 0
SM = 0
Delta_K
Doxorubicin
YJL169W
ZShift = 1.91
Z lm Score = −2.81
NG = 0
DB = 0
SM = 0
Delta_r
Doxorubicin
YJL169W
ZShift = 0.25
Z lm Score = −5.04
NG = 0
DB = 0
SM = 0
Delta_AUC
Doxorubicin
YDR376W ARH1
ZShift = 0.77
Z lm Score = 7.43
NG = 0
DB = 1
SM = 0
Delta_L
Doxorubicin
YDR376W ARH1
ZShift = −1.2
Z lm Score = −2.66
NG = 0
DB = 1
SM = 0
Delta_K
Doxorubicin
YDR376W ARH1
ZShift = −1.49
Z lm Score = −2.41
NG = 0
DB = 1
SM = 0
Delta_r
Doxorubicin
YDR376W ARH1
ZShift = −1.31
Z lm Score = −1.66
NG = 0
DB = 1
SM = 0
Delta_AUC
Doxorubicin
YIL040W APQ12
ZShift = −1.24
Z lm Score = 7.42
NG = 0
DB = 0
SM = 0
Delta_L
Doxorubicin
YIL040W APQ12
ZShift = −0.36
Z lm Score = −0.55
NG = 0
DB = 0
SM = 0
Delta_K
Doxorubicin
YIL040W APQ12
ZShift = 3.66
Z lm Score = −4.27
NG = 0
DB = 0
SM = 0
Delta_r
Doxorubicin
YIL040W APQ12
ZShift = 0.83
Z lm Score = −1.91
NG = 0
DB = 0
SM = 0
Delta_AUC
Doxorubicin

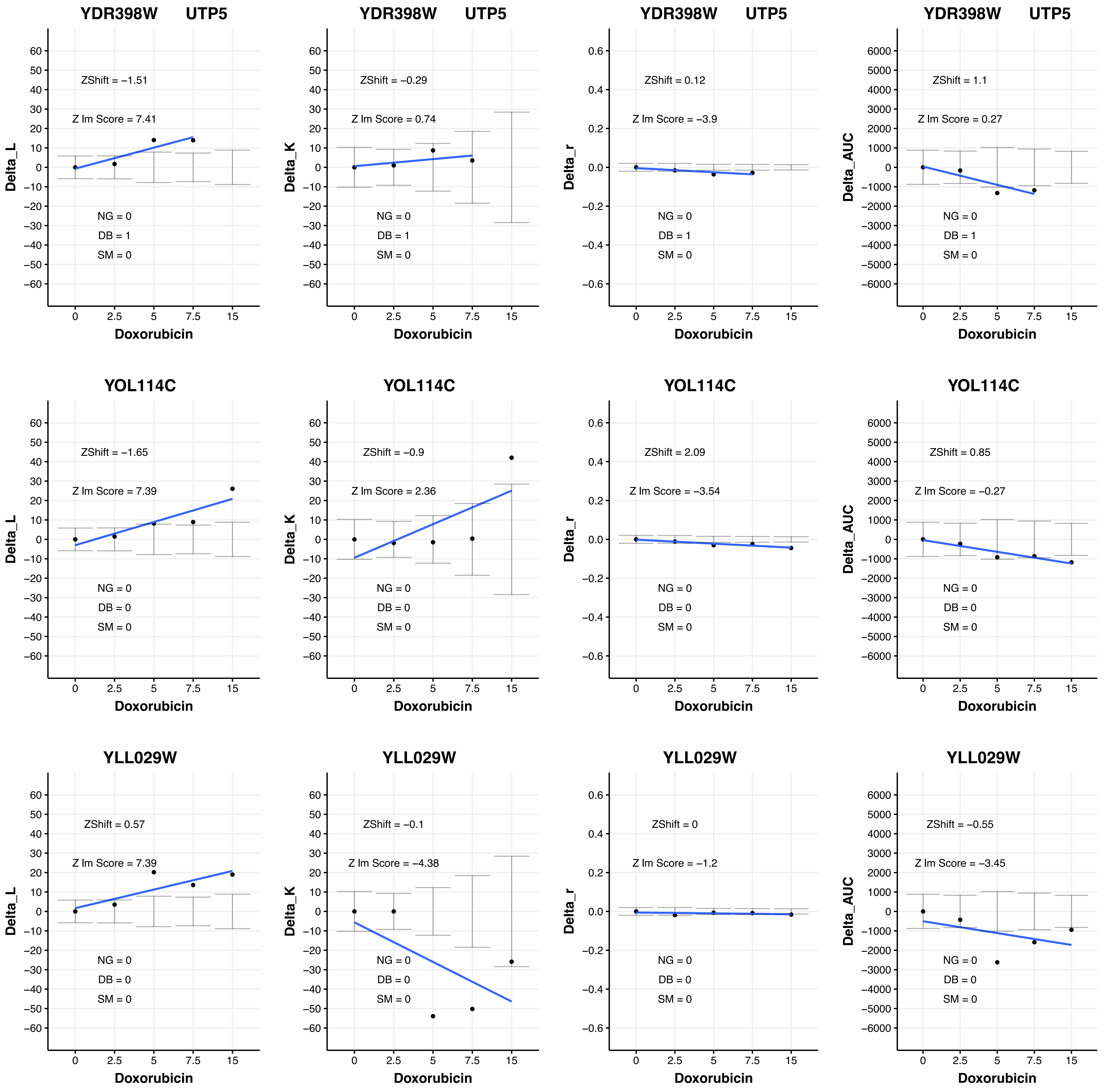

YDR398W UTP5
ZShift = −1.51
Z lm Score = 7.41
NG = 0
DB = 1
SM = 0
Delta_L
Doxorubicin
YDR398W UTP5
ZShift = −0.29
Z lm Score = 0.74
NG = 0
DB = 1
SM = 0
Delta_K
Doxorubicin
YDR398W UTP5
ZShift = 0.12
Z lm Score = −3.9
NG = 0
DB = 1
SM = 0
Delta_r
Doxorubicin
YDR398W UTP5
ZShift = 1.1
Z lm Score = 0.27
NG = 0
DB = 1
SM = 0
Delta_AUC
Doxorubicin
YOL114C
ZShift = −1.65
Z lm Score = 7.39
NG = 0
DB = 0
SM = 0
Delta_L
Doxorubicin
YOL114C
ZShift = −0.9
Z lm Score = 2.36
NG = 0
DB = 0
SM = 0
Delta_K
Doxorubicin
YOL114C
ZShift = 2.09
Z lm Score = −3.54
NG = 0
DB = 0
SM = 0
Delta_r
Doxorubicin
YOL114C
ZShift = 0.85
Z lm Score = −0.27
NG = 0
DB = 0
SM = 0
Delta_AUC
Doxorubicin
YLL029W
ZShift = 0.57
Z lm Score = 7.39
NG = 0
DB = 0
SM = 0
Delta_L
Doxorubicin
YLL029W
ZShift = −0.1
Z lm Score = −4.38
NG = 0
DB = 0
SM = 0
Delta_K
Doxorubicin
YLL029W
ZShift = 0
Z lm Score = −1.2
NG = 0
DB = 0
SM = 0
Delta_r
Doxorubicin
YLL029W
ZShift = −0.55
Z lm Score = −3.45
NG = 0
DB = 0
SM = 0
Delta_AUC
Doxorubicin

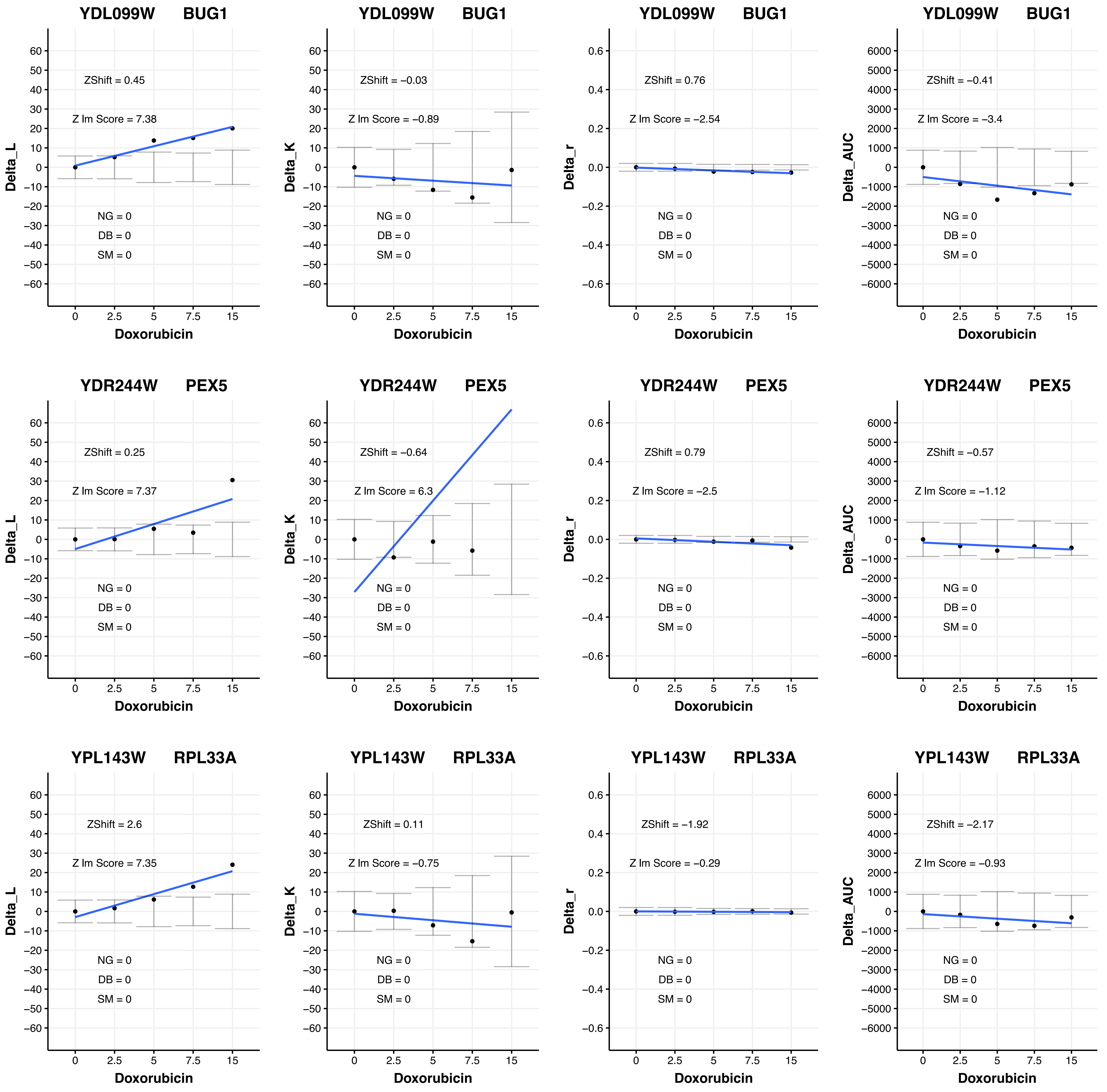
YDL099W BUG1
ZShift = 0.45
Z lm Score = 7.38
NG = 0
DB = 0
SM = 0
Delta_L
Doxorubicin
YDL099W BUG1
ZShift = −0.03
Z lm Score = −0.89
NG = 0
DB = 0
SM = 0
Delta_K
Doxorubicin
YDL099W BUG1
ZShift = 0.76
Z lm Score = −2.54
NG = 0
DB = 0
SM = 0
Delta_r
Doxorubicin
YDL099W BUG1
ZShift = −0.41
Z lm Score = −3.4
NG = 0
DB = 0
SM = 0
Delta_AUC
Doxorubicin
YDR244W PEX5
ZShift = 0.25
Z lm Score = 7.37
NG = 0
DB = 0
SM = 0
Delta_L
Doxorubicin
YDR244W PEX5
ZShift = −0.64
Z lm Score = 6.3
NG = 0
DB = 0
SM = 0
Delta_K
Doxorubicin
YDR244W PEX5
ZShift = 0.79
Z lm Score = −2.5
NG = 0
DB = 0
SM = 0
Delta_r
Doxorubicin
YDR244W PEX5
ZShift = −0.57
Z lm Score = −1.12
NG = 0
DB = 0
SM = 0
Delta_AUC
Doxorubicin
YPL143W RPL33A
ZShift = 2.6
Z lm Score = 7.35
NG = 0
DB = 0
SM = 0
Delta_L
Doxorubicin
YPL143W RPL33A
ZShift = 0.11
Z lm Score = −0.75
NG = 0
DB = 0
SM = 0
Delta_K
Doxorubicin
YPL143W RPL33A
ZShift = −1.92
Z lm Score = −0.29
NG = 0
DB = 0
SM = 0
Delta_r
Doxorubicin
YPL143W RPL33A
ZShift = −2.17
Z lm Score = −0.93
NG = 0
DB = 0
SM = 0
Delta_AUC
Doxorubicin

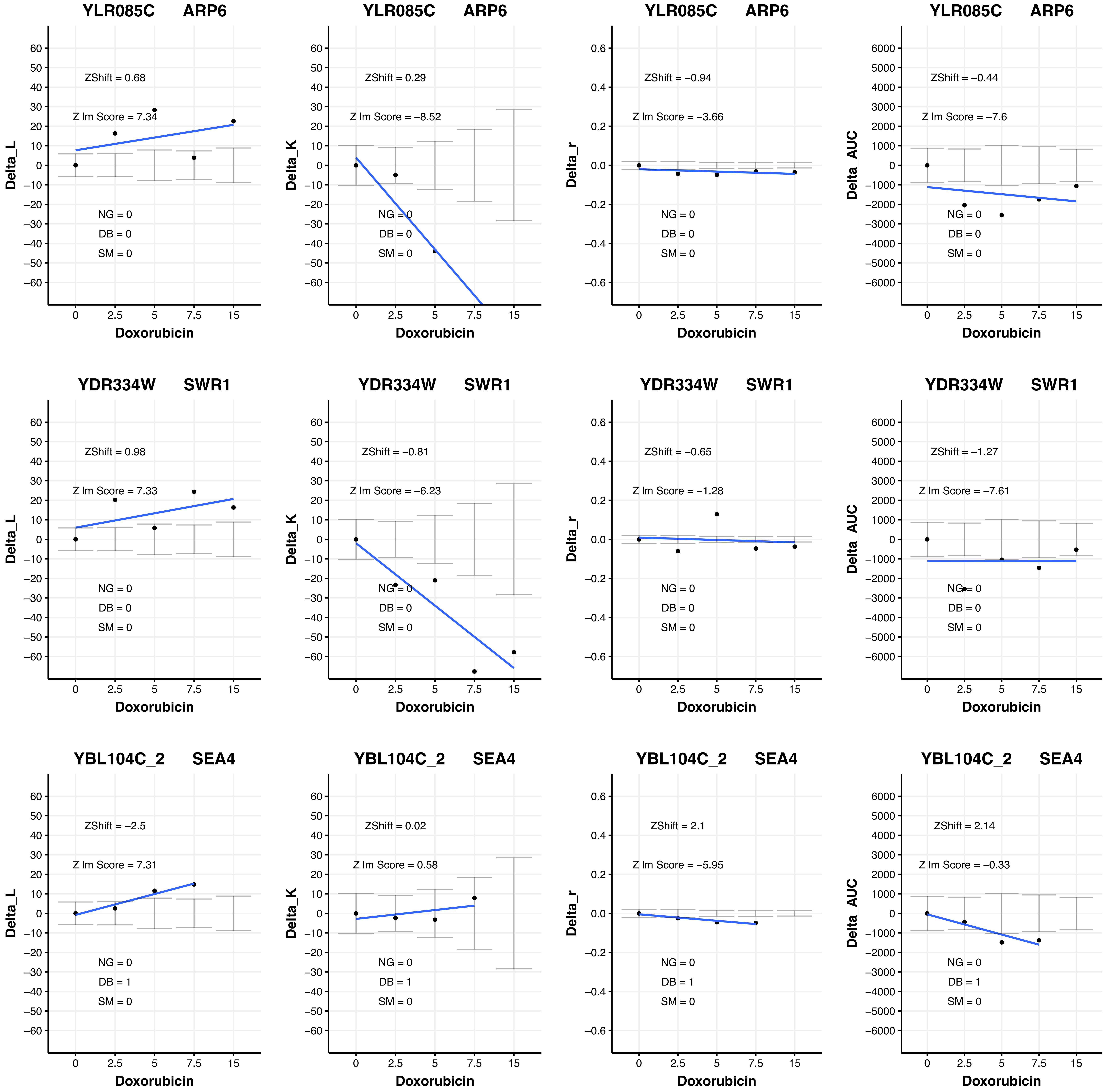
YLR085C ARP6
ZShift = 0.68
Z lm Score = 7.34
NG = 0
DB = 0
SM = 0
Delta_L
Doxorubicin
YLR085C ARP6
ZShift = 0.29
Z lm Score = −8.52
NG = 0
DB = 0
SM = 0
Delta_K
Doxorubicin
YLR085C ARP6
ZShift = −0.94
Z lm Score = −3.66
NG = 0
DB = 0
SM = 0
Delta_r
Doxorubicin
YLR085C ARP6
ZShift = −0.44
Z lm Score = −7.6
NG = 0
DB = 0
SM = 0
Delta_AUC
Doxorubicin
YDR334W SWR1
ZShift = 0.98
Z lm Score = 7.33
NG = 0
DB = 0
SM = 0
Delta_L
Doxorubicin
YDR334W SWR1
ZShift = −0.81
Z lm Score = −6.23
NG = 0
DB = 0
SM = 0
Delta_K
Doxorubicin
YDR334W SWR1
ZShift = −0.65
Z lm Score = −1.28
NG = 0
DB = 0
SM = 0
Delta_r
Doxorubicin
YDR334W SWR1
ZShift = −1.27
Z lm Score = −7.61
NG = 0
DB = 0
SM = 0
Delta_AUC
Doxorubicin
YBL104C_2 SEA4
ZShift = −2.5
Z lm Score = 7.31
NG = 0
DB = 1
SM = 0
Delta_L
Doxorubicin
YBL104C_2 SEA4
ZShift = 0.02
Z lm Score = 0.58
NG = 0
DB = 1
SM = 0
Delta_K
Doxorubicin
YBL104C_2 SEA4
ZShift = 2.1
Z lm Score = −5.95
NG = 0
DB = 1
SM = 0
Delta_r
Doxorubicin
YBL104C_2 SEA4
ZShift = 2.14
Z lm Score = −0.33
NG = 0
DB = 1
SM = 0
Delta_AUC
Doxorubicin

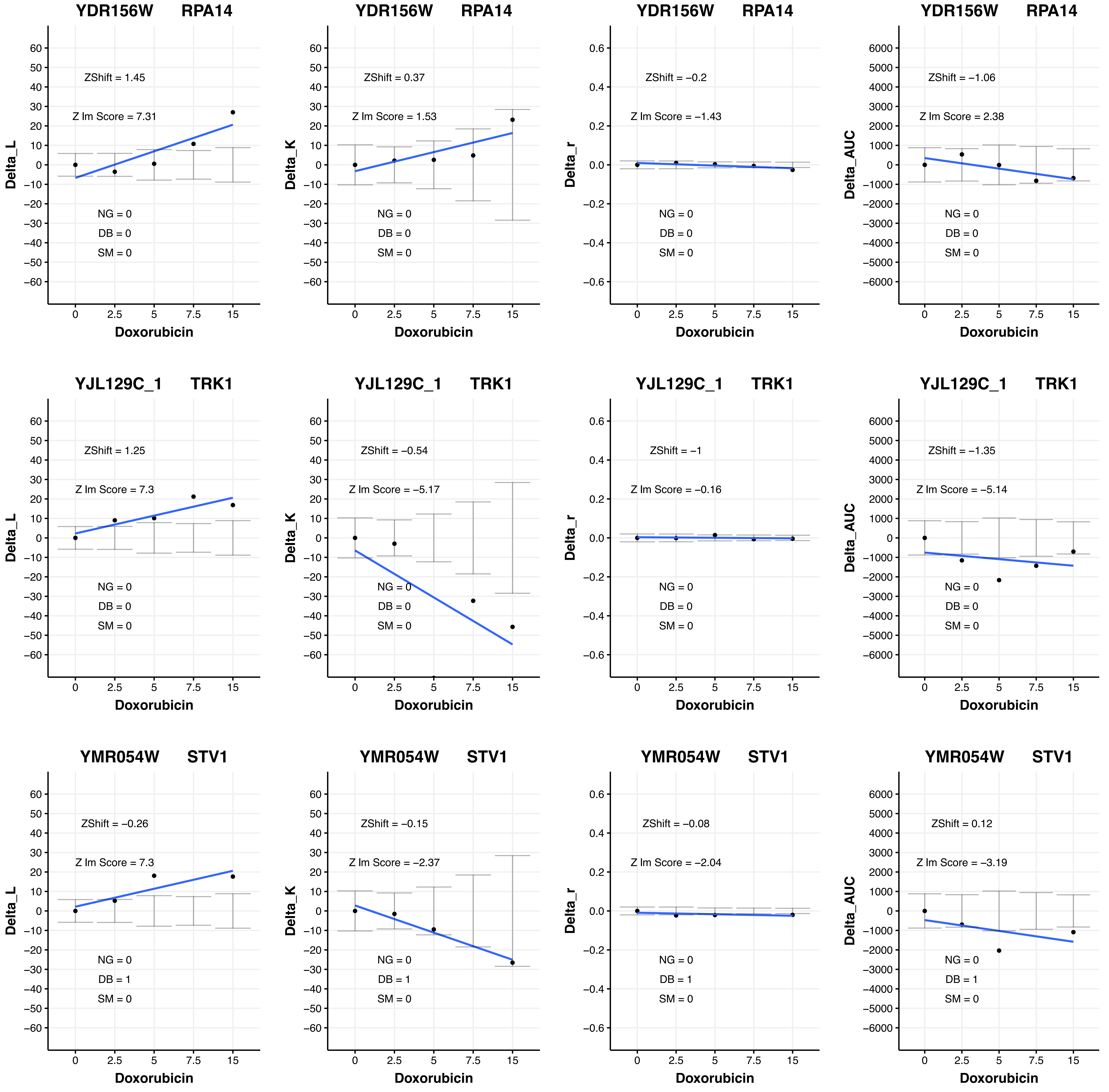
YDR156W RPA14
ZShift = 1.45
Z lm Score = 7.31
NG = 0
DB = 0
SM = 0
Delta_L
Doxorubicin
YDR156W RPA14
ZShift = 0.37
Z lm Score = 1.53
NG = 0
DB = 0
SM = 0
Delta_K
Doxorubicin
YDR156W RPA14
ZShift = −0.2
Z lm Score = −1.43
NG = 0
DB = 0
SM = 0
Delta_r
Doxorubicin
YDR156W RPA14
ZShift = −1.06
Z lm Score = 2.38
NG = 0
DB = 0
SM = 0
Delta_AUC
Doxorubicin
YJL129C_1 TRK1
ZShift = 1.25
Z lm Score = 7.3
NG = 0
DB = 0
SM = 0
Delta_L
Doxorubicin
YJL129C_1 TRK1
ZShift = −0.54
Z lm Score = −5.17
NG = 0
DB = 0
SM = 0
Delta_K
Doxorubicin
YJL129C_1 TRK1
ZShift = −1
Z lm Score = −0.16
NG = 0
DB = 0
SM = 0
Delta_r
Doxorubicin
YJL129C_1 TRK1
ZShift = −1.35
Z lm Score = −5.14
NG = 0
DB = 0
SM = 0
Delta_AUC
Doxorubicin
YMR054W STV1
ZShift = −0.26
Z lm Score = 7.3
NG = 0
DB = 1
SM = 0
Delta_L
Doxorubicin
YMR054W STV1
ZShift = −0.15
Z lm Score = −2.37
NG = 0
DB = 1
SM = 0
Delta_K
Doxorubicin
YMR054W STV1
ZShift = −0.08
Z lm Score = −2.04
NG = 0
DB = 1
SM = 0
Delta_r
Doxorubicin
YMR054W STV1
ZShift = 0.12
Z lm Score = −3.19
NG = 0
DB = 1
SM = 0
Delta_AUC
Doxorubicin

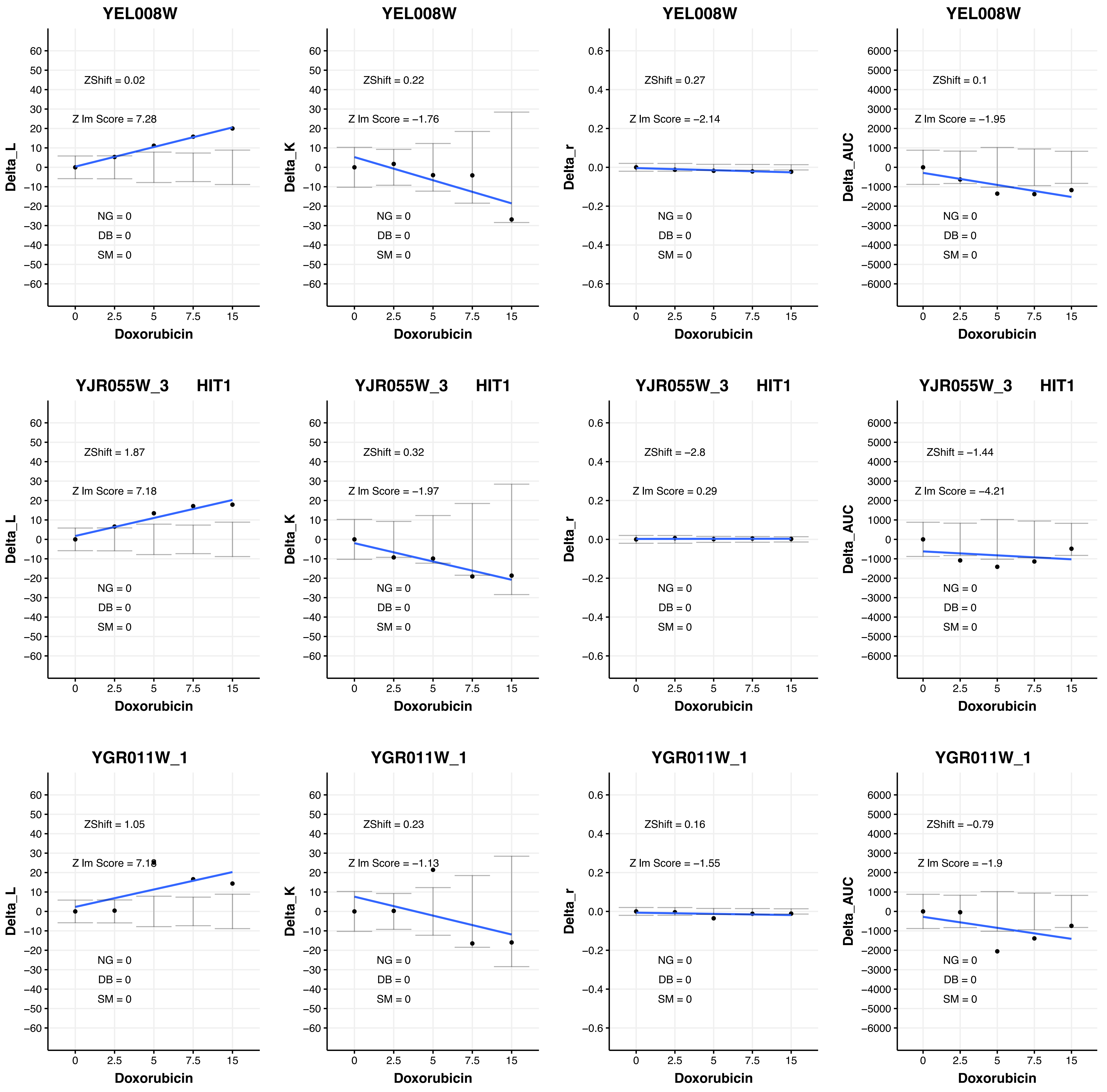

YEL008W
ZShift = 0.02
Z lm Score = 7.28
NG = 0
DB = 0
SM = 0
Delta_L
Doxorubicin
YEL008W
ZShift = 0.22
Z lm Score = −1.76
NG = 0
DB = 0
SM = 0
Delta_K
Doxorubicin
YEL008W
ZShift = 0.27
Z lm Score = −2.14
NG = 0
DB = 0
SM = 0
Delta_r
Doxorubicin
YEL008W
ZShift = 0.1
Z lm Score = −1.95
NG = 0
DB = 0
SM = 0
Delta_AUC
Doxorubicin
YJR055W_3 HIT1
ZShift = 1.87
Z lm Score = 7.18
NG = 0
DB = 0
SM = 0
Delta_L
Doxorubicin
YJR055W_3 HIT1
ZShift = 0.32
Z lm Score = −1.97
NG = 0
DB = 0
SM = 0
Delta_K
Doxorubicin
YJR055W_3 HIT1
ZShift = −2.8
Z lm Score = 0.29
NG = 0
DB = 0
SM = 0
Delta_r
Doxorubicin
YJR055W_3 HIT1
ZShift = −1.44
Z lm Score = −4.21
NG = 0
DB = 0
SM = 0
Delta_AUC
Doxorubicin
YGR011W_1
ZShift = 1.05
Z lm Score = 7.18
NG = 0
DB = 0
SM = 0
Delta_L
Doxorubicin
YGR011W_1
ZShift = 0.23
Z lm Score = −1.13
NG = 0
DB = 0
SM = 0
Delta_K
Doxorubicin
YGR011W_1
ZShift = 0.16
Z lm Score = −1.55
NG = 0
DB = 0
SM = 0
Delta_r
Doxorubicin
YGR011W_1
ZShift = −0.79
Z lm Score = −1.9
NG = 0
DB = 0
SM = 0
Delta_AUC
Doxorubicin

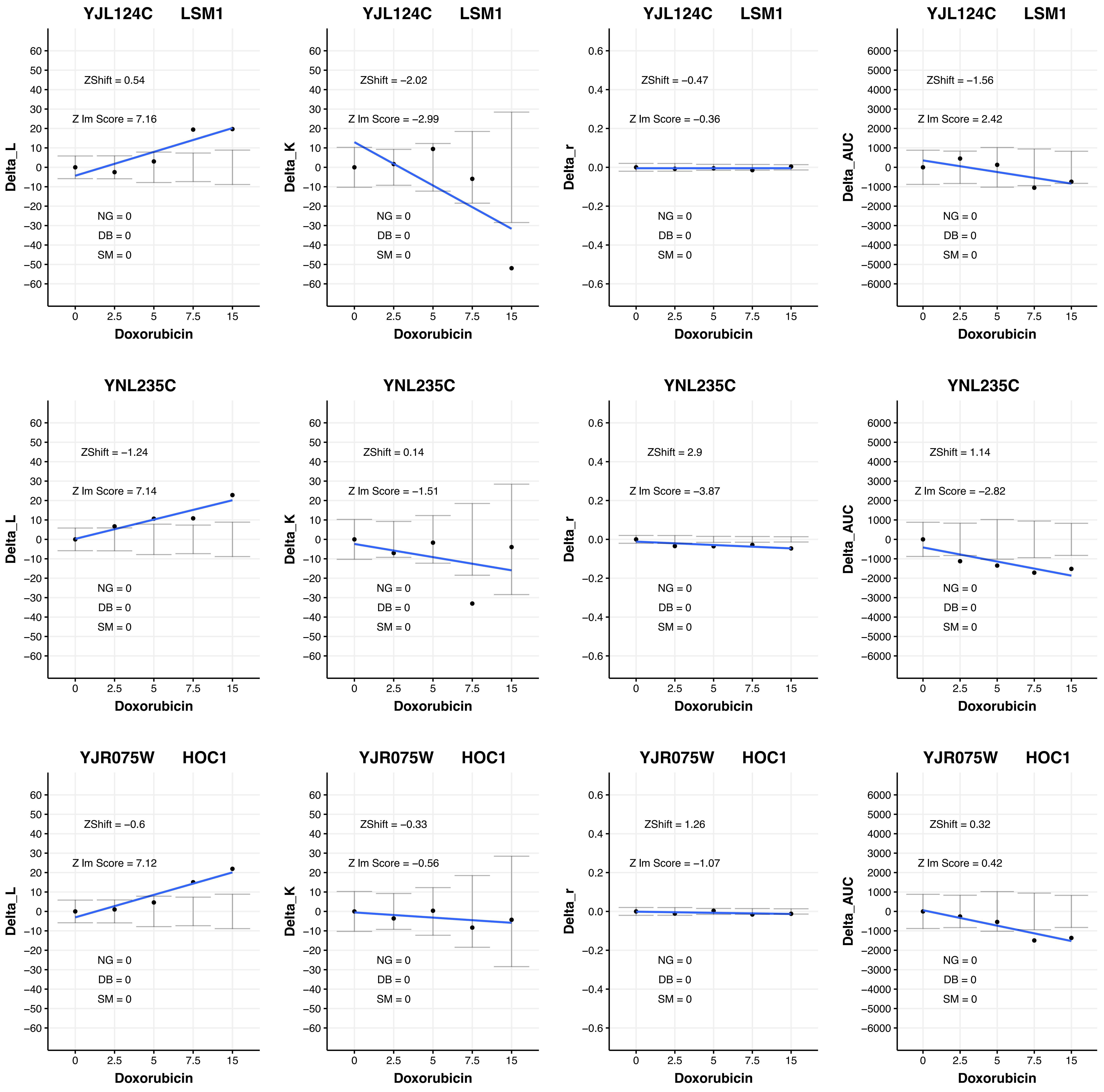

YJL124C LSM1
ZShift = 0.54
Z lm Score = 7.16
NG = 0
DB = 0
SM = 0
Delta_L
Doxorubicin
YJL124C LSM1
ZShift = −2.02
Z lm Score = −2.99
NG = 0
DB = 0
SM = 0
Delta_K
Doxorubicin
YJL124C LSM1
ZShift = −0.47
Z lm Score = −0.36
NG = 0
DB = 0
SM = 0
Delta_r
Doxorubicin
YJL124C LSM1
ZShift = −1.56
Z lm Score = 2.42
NG = 0
DB = 0
SM = 0
Delta_AUC
Doxorubicin
YNL235C
ZShift = −1.24
Z lm Score = 7.14
NG = 0
DB = 0
SM = 0
Delta_L
Doxorubicin
YNL235C
ZShift = 0.14
Z lm Score = −1.51
NG = 0
DB = 0
SM = 0
Delta_K
Doxorubicin
YNL235C
ZShift = 2.9
Z lm Score = −3.87
NG = 0
DB = 0
SM = 0
Delta_r
Doxorubicin
YNL235C
ZShift = 1.14
Z lm Score = −2.82
NG = 0
DB = 0
SM = 0
Delta_AUC
Doxorubicin
YJR075W HOC1
ZShift = −0.6
Z lm Score = 7.12
NG = 0
DB = 0
SM = 0
Delta_L
Doxorubicin
YJR075W HOC1
ZShift = −0.33
Z lm Score = −0.56
NG = 0
DB = 0
SM = 0
Delta_K
Doxorubicin
YJR075W HOC1
ZShift = 1.26
Z lm Score = −1.07
NG = 0
DB = 0
SM = 0
Delta_r
Doxorubicin
YJR075W HOC1
ZShift = 0.32
Z lm Score = 0.42
NG = 0
DB = 0
SM = 0
Delta_AUC
Doxorubicin

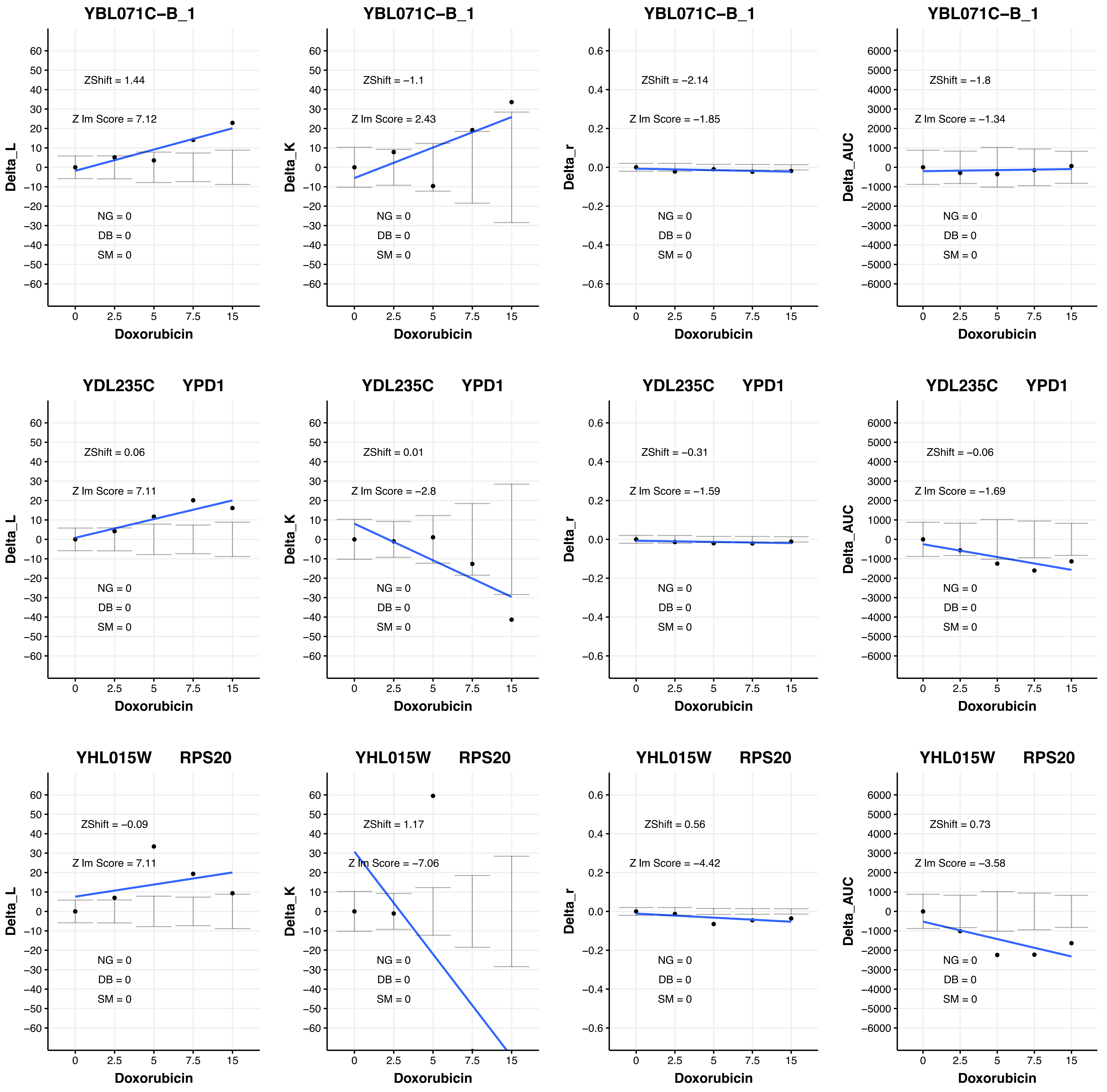
YBL071C−B_1
ZShift = 1.44
Z lm Score = 7.12
NG = 0
DB = 0
SM = 0
Delta_L
Doxorubicin
YBL071C−B_1
ZShift = −1.1
Z lm Score = 2.43
NG = 0
DB = 0
SM = 0
Delta_K
Doxorubicin
YBL071C−B_1
ZShift = −2.14
Z lm Score = −1.85
NG = 0
DB = 0
SM = 0
Delta_r
Doxorubicin
YBL071C−B_1
ZShift = −1.8
Z lm Score = −1.34
NG = 0
DB = 0
SM = 0
Delta_AUC
Doxorubicin
YDL235C YPD1
ZShift = 0.06
Z lm Score = 7.11
NG = 0
DB = 0
SM = 0
Delta_L
Doxorubicin
YDL235C YPD1
ZShift = 0.01
Z lm Score = −2.8
NG = 0
DB = 0
SM = 0
Delta_K
Doxorubicin
YDL235C YPD1
ZShift = −0.31
Z lm Score = −1.59
NG = 0
DB = 0
SM = 0
Delta_r
Doxorubicin
YDL235C YPD1
ZShift = −0.06
Z lm Score = −1.69
NG = 0
DB = 0
SM = 0
Delta_AUC
Doxorubicin
YHL015W RPS20
ZShift = −0.09
Z lm Score = 7.11
NG = 0
DB = 0
SM = 0
Delta_L
Doxorubicin
YHL015W RPS20
ZShift = 1.17
Z lm Score = −7.06
NG = 0
DB = 0
SM = 0
Delta_K
Doxorubicin
YHL015W RPS20
ZShift = 0.56
Z lm Score = −4.42
NG = 0
DB = 0
SM = 0
Delta_r
Doxorubicin
YHL015W RPS20
ZShift = 0.73
Z lm Score = −3.58
NG = 0
DB = 0
SM = 0
Delta_AUC
Doxorubicin

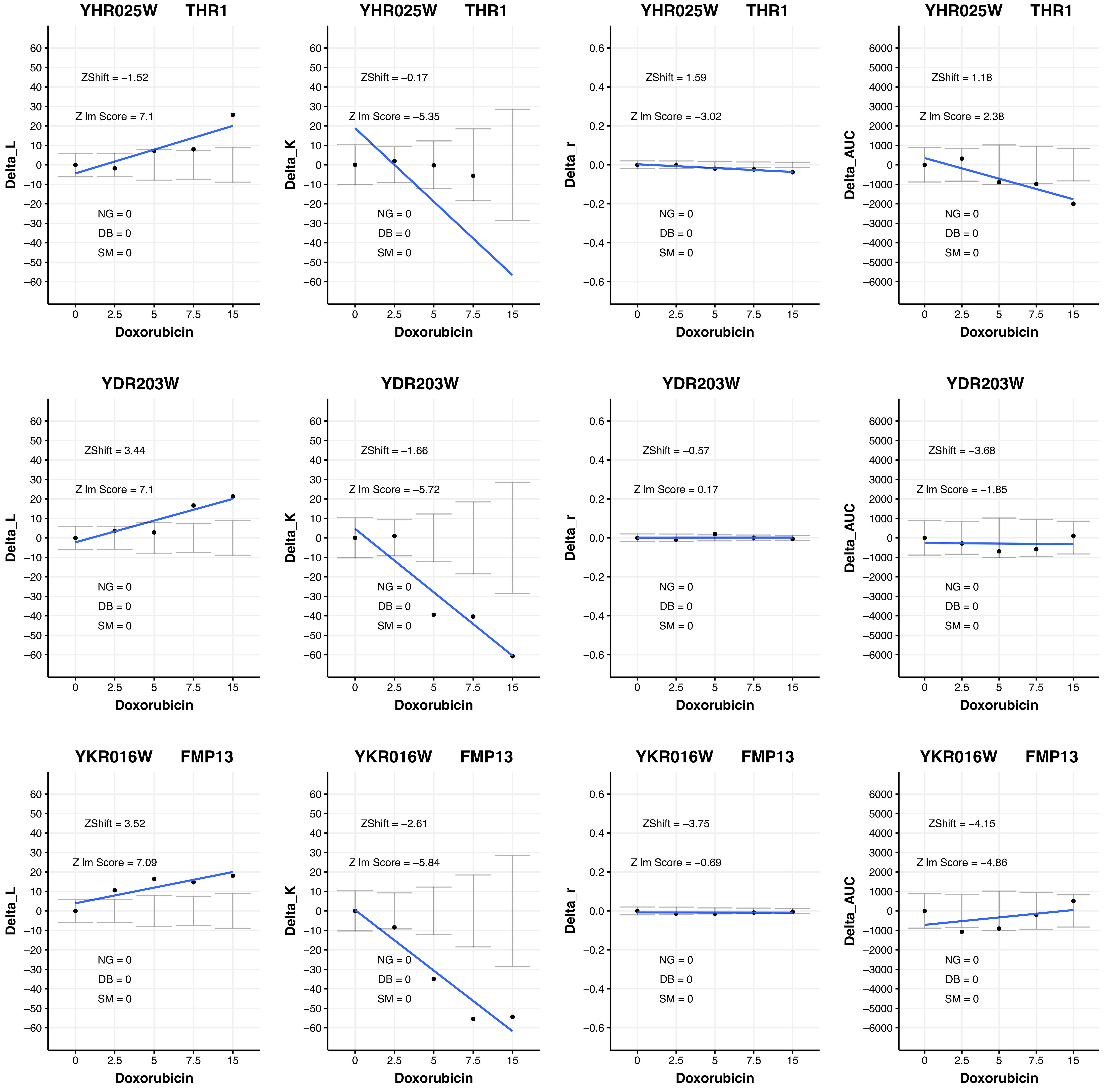

YHR025W THR1
ZShift = −1.52
Z lm Score = 7.1
NG = 0
DB = 0
SM = 0
Delta_L
Doxorubicin
YHR025W THR1
ZShift = −0.17
Z lm Score = −5.35
NG = 0
DB = 0
SM = 0
Delta_K
Doxorubicin
YHR025W THR1
ZShift = 1.59
Z lm Score = −3.02
NG = 0
DB = 0
SM = 0
Delta_r
Doxorubicin
YHR025W THR1
ZShift = 1.18
Z lm Score = 2.38
NG = 0
DB = 0
SM = 0
Delta_AUC
Doxorubicin
YDR203W
ZShift = 3.44
Z lm Score = 7.1
NG = 0
DB = 0
SM = 0
Delta_L
Doxorubicin
YDR203W
ZShift = −1.66
Z lm Score = −5.72
NG = 0
DB = 0
SM = 0
Delta_K
Doxorubicin
YDR203W
ZShift = −0.57
Z lm Score = 0.17
NG = 0
DB = 0
SM = 0
Delta_r
Doxorubicin
YDR203W
ZShift = −3.68
Z lm Score = −1.85
NG = 0
DB = 0
SM = 0
Delta_AUC
Doxorubicin
YKR016W FMP13
ZShift = 3.52
Z lm Score = 7.09
NG = 0
DB = 0
SM = 0
Delta_L
Doxorubicin
YKR016W FMP13
ZShift = −2.61
Z lm Score = −5.84
NG = 0
DB = 0
SM = 0
Delta_K
Doxorubicin
YKR016W FMP13
ZShift = −3.75
Z lm Score = −0.69
NG = 0
DB = 0
SM = 0
Delta_r
Doxorubicin
YKR016W FMP13
ZShift = −4.15
Z lm Score = −4.86
NG = 0
DB = 0
SM = 0
Delta_AUC
Doxorubicin

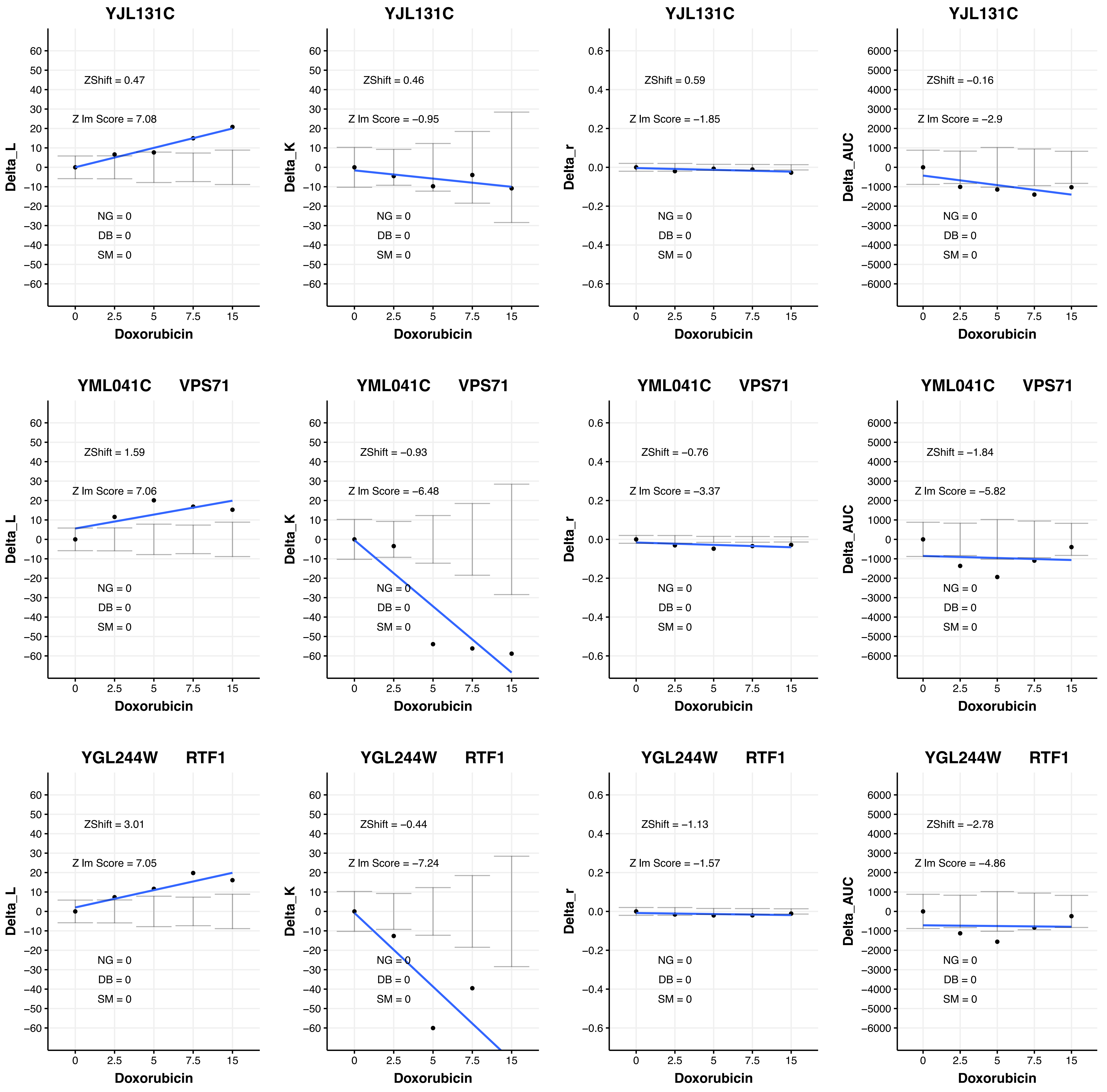

YJL131C
ZShift = 0.47
Z lm Score = 7.08
NG = 0
DB = 0
SM = 0
Delta_L
Doxorubicin
YJL131C
ZShift = 0.46
Z lm Score = −0.95
NG = 0
DB = 0
SM = 0
Delta_K
Doxorubicin
YJL131C
ZShift = 0.59
Z lm Score = −1.85
NG = 0
DB = 0
SM = 0
Delta_r
Doxorubicin
YJL131C
ZShift = −0.16
Z lm Score = −2.9
NG = 0
DB = 0
SM = 0
Delta_AUC
Doxorubicin
YML041C VPS71
ZShift = 1.59
Z lm Score = 7.06
NG = 0
DB = 0
SM = 0
Delta_L
Doxorubicin
YML041C VPS71
ZShift = −0.93
Z lm Score = −6.48
NG = 0
DB = 0
SM = 0
Delta_K
Doxorubicin
YML041C VPS71
ZShift = −0.76
Z lm Score = −3.37
NG = 0
DB = 0
SM = 0
Delta_r
Doxorubicin
YML041C VPS71
ZShift = −1.84
Z lm Score = −5.82
NG = 0
DB = 0
SM = 0
Delta_AUC
Doxorubicin
YGL244W RTF1
ZShift = 3.01
Z lm Score = 7.05
NG = 0
DB = 0
SM = 0
Delta_L
Doxorubicin
YGL244W RTF1
ZShift = −0.44
Z lm Score = −7.24
NG = 0
DB = 0
SM = 0
Delta_K
Doxorubicin
YGL244W RTF1
ZShift = −1.13
Z lm Score = −1.57
NG = 0
DB = 0
SM = 0
Delta_r
Doxorubicin
YGL244W RTF1
ZShift = −2.78
Z lm Score = −4.86
NG = 0
DB = 0
SM = 0
Delta_AUC
Doxorubicin

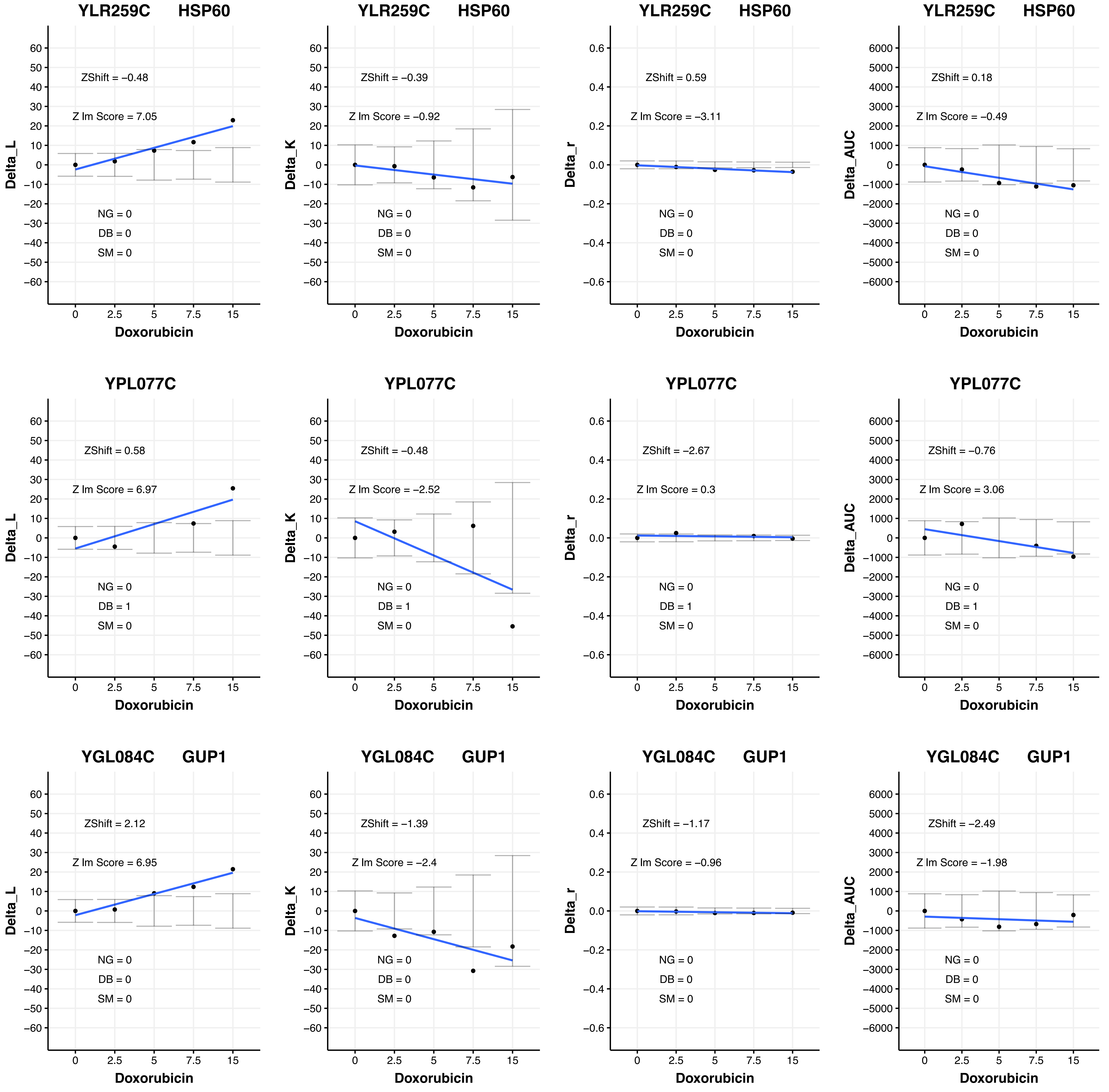

YLR259C HSP60
ZShift = −0.48
Z lm Score = 7.05
NG = 0
DB = 0
SM = 0
Delta_L
Doxorubicin
YLR259C HSP60
ZShift = −0.39
Z lm Score = −0.92
NG = 0
DB = 0
SM = 0
Delta_K
Doxorubicin
YLR259C HSP60
ZShift = 0.59
Z lm Score = −3.11
NG = 0
DB = 0
SM = 0
Delta_r
Doxorubicin
YLR259C HSP60
ZShift = 0.18
Z lm Score = −0.49
NG = 0
DB = 0
SM = 0
Delta_AUC
Doxorubicin
YPL077C
ZShift = 0.58
Z lm Score = 6.97
NG = 0
DB = 1
SM = 0
Delta_L
Doxorubicin
YPL077C
ZShift = −0.48
Z lm Score = −2.52
NG = 0
DB = 1
SM = 0
Delta_K
Doxorubicin
YPL077C
ZShift = −2.67
Z lm Score = 0.3
NG = 0
DB = 1
SM = 0
Delta_r
Doxorubicin
YPL077C
ZShift = −0.76
Z lm Score = 3.06
NG = 0
DB = 1
SM = 0
Delta_AUC
Doxorubicin
YGL084C GUP1
ZShift = 2.12
Z lm Score = 6.95
NG = 0
DB = 0
SM = 0
Delta_L
Doxorubicin
YGL084C GUP1
ZShift = −1.39
Z lm Score = −2.4
NG = 0
DB = 0
SM = 0
Delta_K
Doxorubicin
YGL084C GUP1
ZShift = −1.17
Z lm Score = −0.96
NG = 0
DB = 0
SM = 0
Delta_r
Doxorubicin
YGL084C GUP1
ZShift = −2.49
Z lm Score = −1.98
NG = 0
DB = 0
SM = 0
Delta_AUC
Doxorubicin

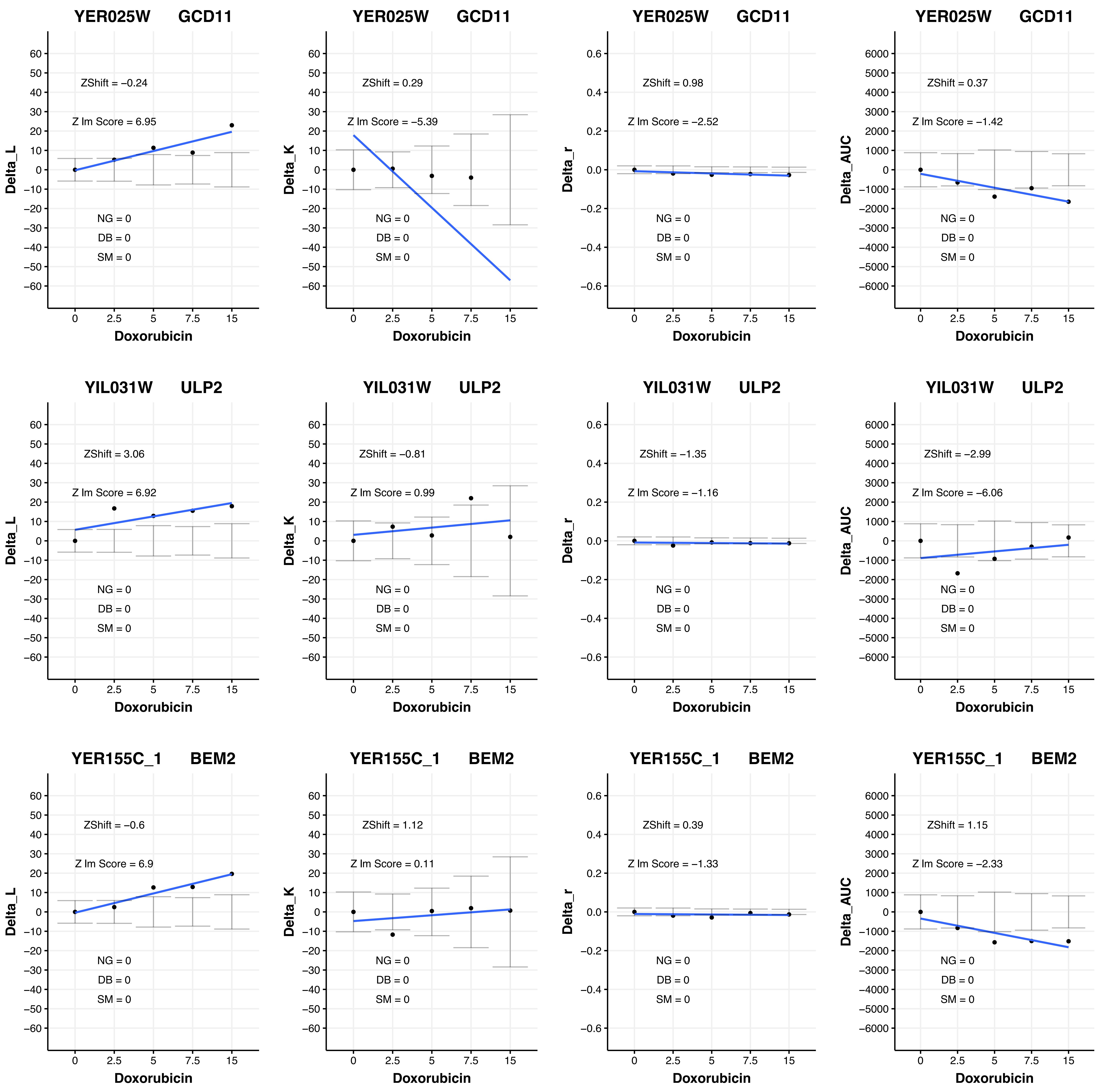

YER025W GCD11
ZShift = −0.24
Z lm Score = 6.95
NG = 0
DB = 0
SM = 0
Delta_L
Doxorubicin
YER025W GCD11
ZShift = 0.29
Z lm Score = −5.39
NG = 0
DB = 0
SM = 0
Delta_K
Doxorubicin
YER025W GCD11
ZShift = 0.98
Z lm Score = −2.52
NG = 0
DB = 0
SM = 0
Delta_r
Doxorubicin
YER025W GCD11
ZShift = 0.37
Z lm Score = −1.42
NG = 0
DB = 0
SM = 0
Delta_AUC
Doxorubicin
YIL031W ULP2
ZShift = 3.06
Z lm Score = 6.92
NG = 0
DB = 0
SM = 0
Delta_L
Doxorubicin
YIL031W ULP2
ZShift = −0.81
Z lm Score = 0.99
NG = 0
DB = 0
SM = 0
Delta_K
Doxorubicin
YIL031W ULP2
ZShift = −1.35
Z lm Score = −1.16
NG = 0
DB = 0
SM = 0
Delta_r
Doxorubicin
YIL031W ULP2
ZShift = −2.99
Z lm Score = −6.06
NG = 0
DB = 0
SM = 0
Delta_AUC
Doxorubicin
YER155C_1 BEM2
ZShift = −0.6
Z lm Score = 6.9
NG = 0
DB = 0
SM = 0
Delta_L
Doxorubicin
YER155C_1 BEM2
ZShift = 1.12
Z lm Score = 0.11
NG = 0
DB = 0
SM = 0
Delta_K
Doxorubicin
YER155C_1 BEM2
ZShift = 0.39
Z lm Score = −1.33
NG = 0
DB = 0
SM = 0
Delta_r
Doxorubicin
YER155C_1 BEM2
ZShift = 1.15
Z lm Score = −2.33
NG = 0
DB = 0
SM = 0
Delta_AUC
Doxorubicin

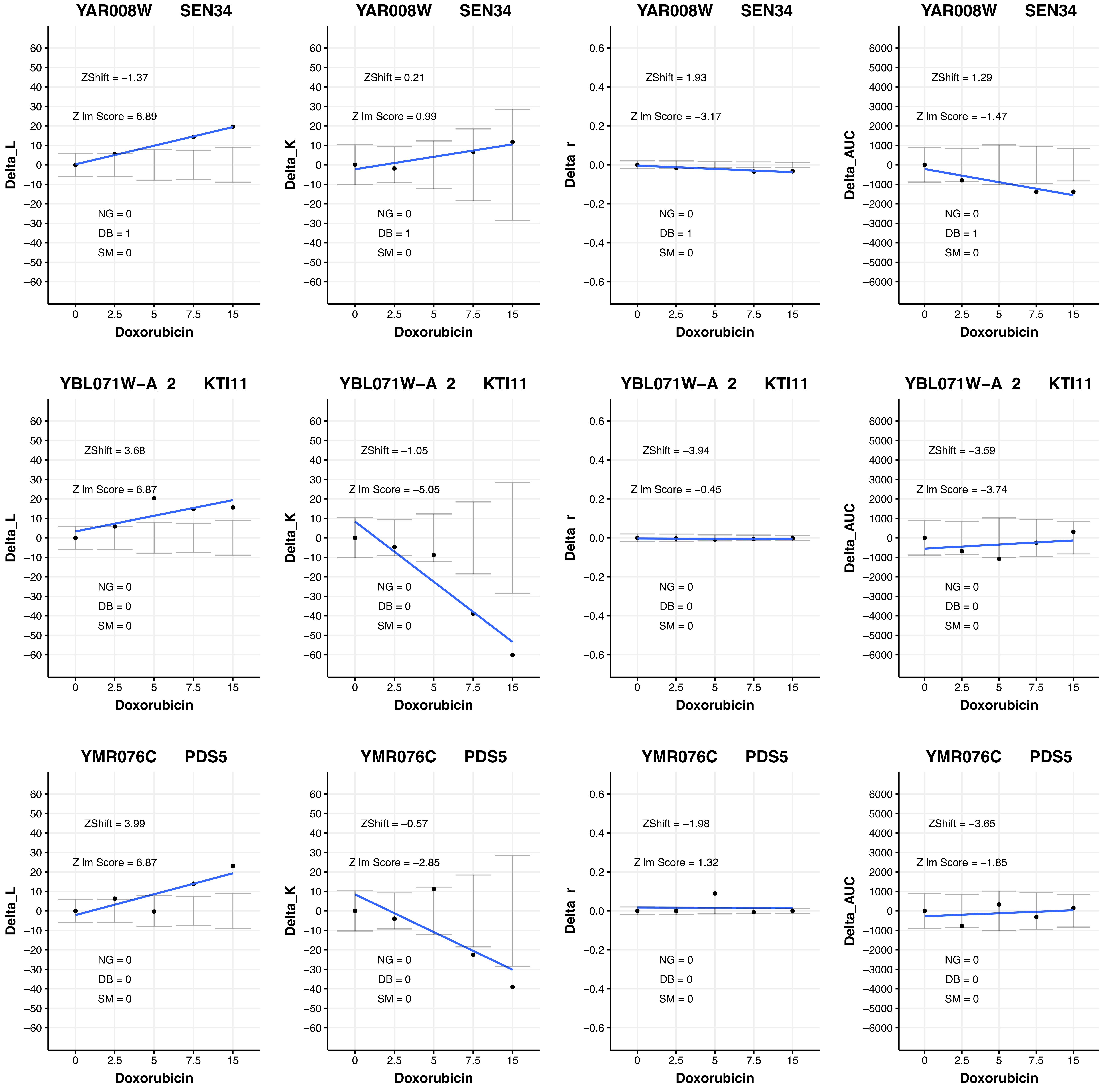

YAR008W SEN34
ZShift = −1.37
Z lm Score = 6.89
NG = 0
DB = 1
SM = 0
Delta_L
Doxorubicin
YAR008W SEN34
ZShift = 0.21
Z lm Score = 0.99
NG = 0
DB = 1
SM = 0
Delta_K
Doxorubicin
YAR008W SEN34
ZShift = 1.93
Z lm Score = −3.17
NG = 0
DB = 1
SM = 0
Delta_r
Doxorubicin
YAR008W SEN34
ZShift = 1.29
Z lm Score = −1.47
NG = 0
DB = 1
SM = 0
Delta_AUC
Doxorubicin
YBL071W−A_2 KTI11
ZShift = 3.68
Z lm Score = 6.87
NG = 0
DB = 0
SM = 0
Delta_L
Doxorubicin
YBL071W−A_2 KTI11
ZShift = −1.05
Z lm Score = −5.05
NG = 0
DB = 0
SM = 0
Delta_K
Doxorubicin
YBL071W−A_2 KTI11
ZShift = −3.94
Z lm Score = −0.45
NG = 0
DB = 0
SM = 0
Delta_r
Doxorubicin
YBL071W−A_2 KTI11
ZShift = −3.59
Z lm Score = −3.74
NG = 0
DB = 0
SM = 0
Delta_AUC
Doxorubicin
YMR076C PDS5
ZShift = 3.99
Z lm Score = 6.87
NG = 0
DB = 0
SM = 0
Delta_L
Doxorubicin
YMR076C PDS5
ZShift = −0.57
Z lm Score = −2.85
NG = 0
DB = 0
SM = 0
Delta_K
Doxorubicin
YMR076C PDS5
ZShift = −1.98
Z lm Score = 1.32
NG = 0
DB = 0
SM = 0
Delta_r
Doxorubicin
YMR076C PDS5
ZShift = −3.65
Z lm Score = −1.85
NG = 0
DB = 0
SM = 0
Delta_AUC
Doxorubicin

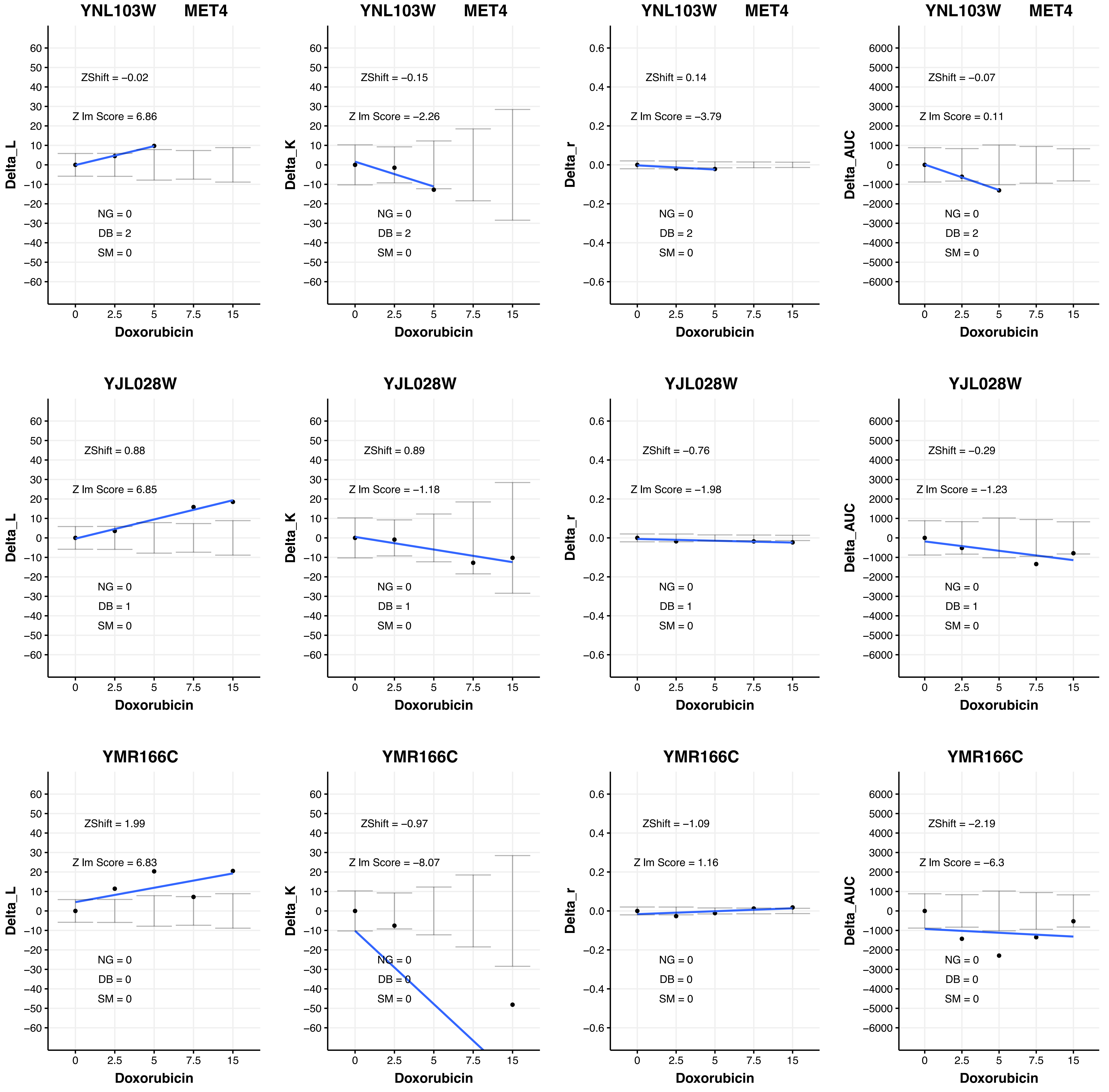
YNL103W MET4
ZShift = −0.02
Z lm Score = 6.86
NG = 0
DB = 2
SM = 0
Delta_L
Doxorubicin
YNL103W MET4
ZShift = −0.15
Z lm Score = −2.26
NG = 0
DB = 2
SM = 0
Delta_K
Doxorubicin
YNL103W MET4
ZShift = 0.14
Z lm Score = −3.79
NG = 0
DB = 2
SM = 0
Delta_r
Doxorubicin
YNL103W MET4
ZShift = −0.07
Z lm Score = 0.11
NG = 0
DB = 2
SM = 0
Delta_AUC
Doxorubicin
YJL028W
ZShift = 0.88
Z lm Score = 6.85
NG = 0
DB = 1
SM = 0
Delta_L
Doxorubicin
YJL028W
ZShift = 0.89
Z lm Score = −1.18
NG = 0
DB = 1
SM = 0
Delta_K
Doxorubicin
YJL028W
ZShift = −0.76
Z lm Score = −1.98
NG = 0
DB = 1
SM = 0
Delta_r
Doxorubicin
YJL028W
ZShift = −0.29
Z lm Score = −1.23
NG = 0
DB = 1
SM = 0
Delta_AUC
Doxorubicin
YMR166C
ZShift = 1.99
Z lm Score = 6.83
NG = 0
DB = 0
SM = 0
Delta_L
Doxorubicin
YMR166C
ZShift = −0.97
Z lm Score = −8.07
NG = 0
DB = 0
SM = 0
Delta_K
Doxorubicin
YMR166C
ZShift = −1.09
Z lm Score = 1.16
NG = 0
DB = 0
SM = 0
Delta_r
Doxorubicin
YMR166C
ZShift = −2.19
Z lm Score = −6.3
NG = 0
DB = 0
SM = 0
Delta_AUC
Doxorubicin

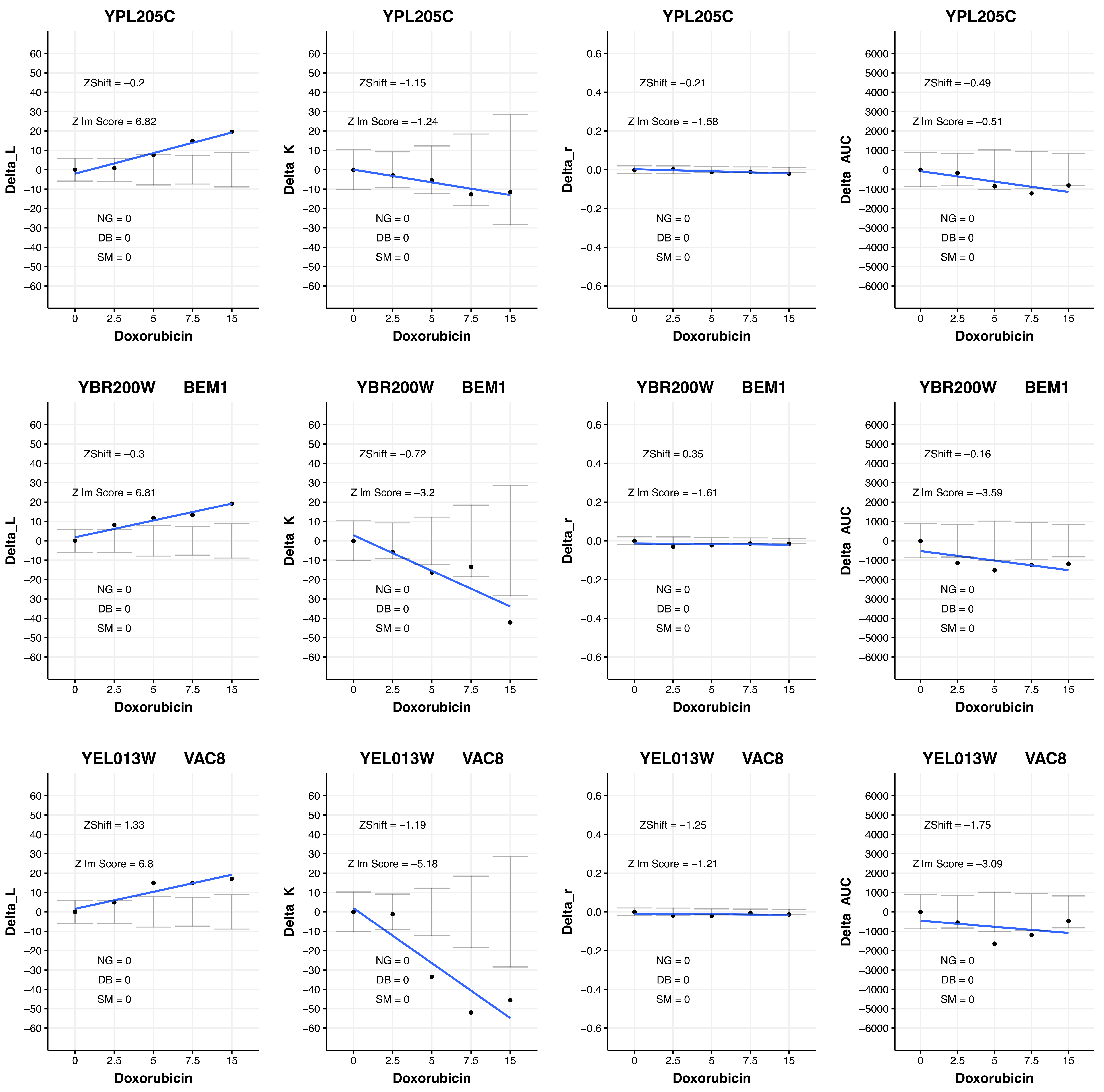

YPL205C
ZShift = −0.2
Z lm Score = 6.82
NG = 0
DB = 0
SM = 0
Delta_L
Doxorubicin
YPL205C
ZShift = −1.15
Z lm Score = −1.24
NG = 0
DB = 0
SM = 0
Delta_K
Doxorubicin
YPL205C
ZShift = −0.21
Z lm Score = −1.58
NG = 0
DB = 0
SM = 0
Delta_r
Doxorubicin
YPL205C
ZShift = −0.49
Z lm Score = −0.51
NG = 0
DB = 0
SM = 0
Delta_AUC
Doxorubicin
YBR200W BEM1
ZShift = −0.3
Z lm Score = 6.81
NG = 0
DB = 0
SM = 0
Delta_L
Doxorubicin
YBR200W BEM1
ZShift = −0.72
Z lm Score = −3.2
NG = 0
DB = 0
SM = 0
Delta_K
Doxorubicin
YBR200W BEM1
ZShift = 0.35
Z lm Score = −1.61
NG = 0
DB = 0
SM = 0
Delta_r
Doxorubicin
YBR200W BEM1
ZShift = −0.16
Z lm Score = −3.59
NG = 0
DB = 0
SM = 0
Delta_AUC
Doxorubicin
YEL013W VAC8
ZShift = 1.33
Z lm Score = 6.8
NG = 0
DB = 0
SM = 0
Delta_L
Doxorubicin
YEL013W VAC8
ZShift = −1.19
Z lm Score = −5.18
NG = 0
DB = 0
SM = 0
Delta_K
Doxorubicin
YEL013W VAC8
ZShift = −1.25
Z lm Score = −1.21
NG = 0
DB = 0
SM = 0
Delta_r
Doxorubicin
YEL013W VAC8
ZShift = −1.75
Z lm Score = −3.09
NG = 0
DB = 0
SM = 0
Delta_AUC
Doxorubicin

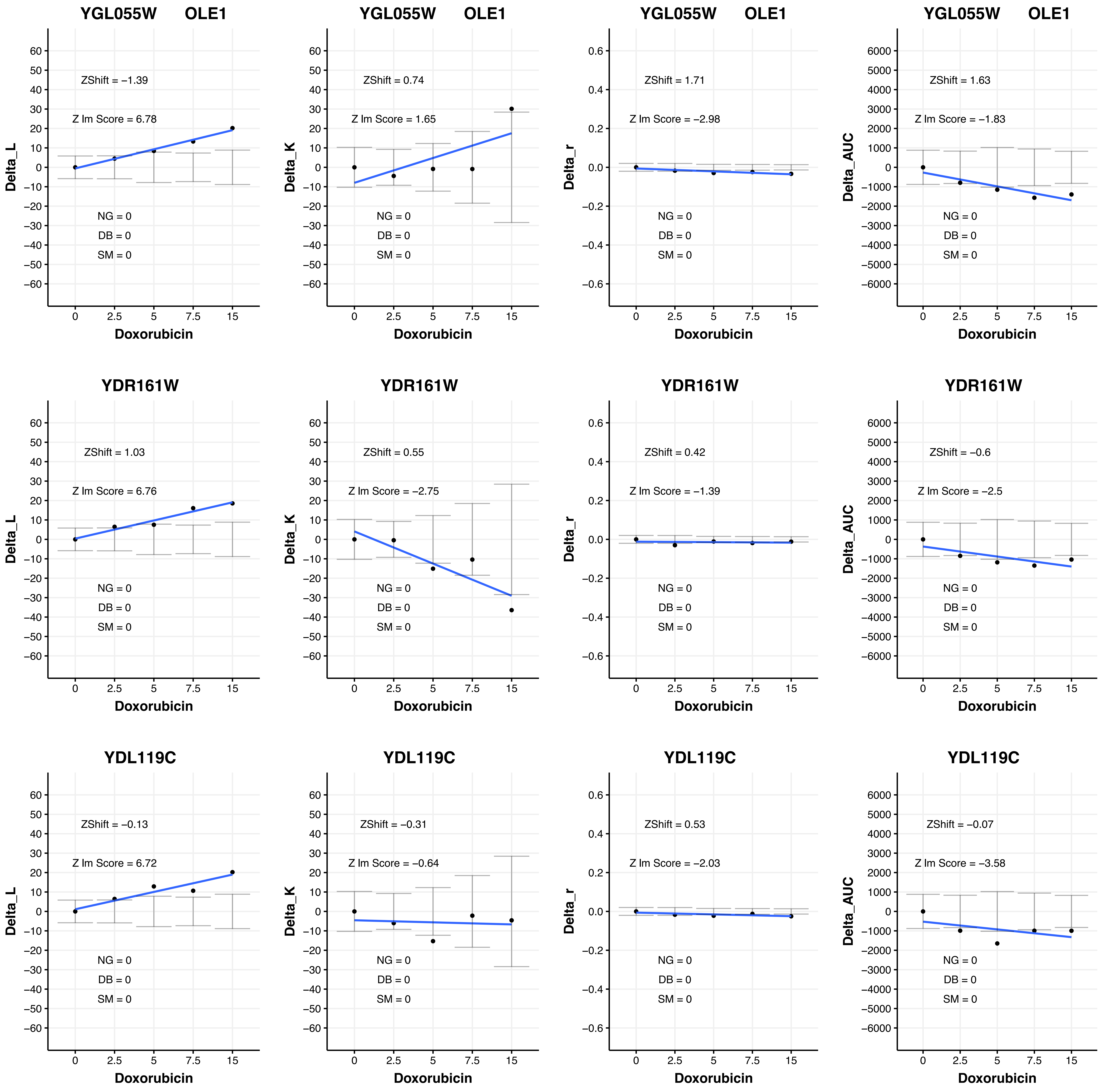
YGL055W OLE1
ZShift = −1.39
Z lm Score = 6.78
NG = 0
DB = 0
SM = 0
Delta_L
Doxorubicin
YGL055W OLE1
ZShift = 0.74
Z lm Score = 1.65
NG = 0
DB = 0
SM = 0
Delta_K
Doxorubicin
YGL055W OLE1
ZShift = 1.71
Z lm Score = −2.98
NG = 0
DB = 0
SM = 0
Delta_r
Doxorubicin
YGL055W OLE1
ZShift = 1.63
Z lm Score = −1.83
NG = 0
DB = 0
SM = 0
Delta_AUC
Doxorubicin
YDR161W
ZShift = 1.03
Z lm Score = 6.76
NG = 0
DB = 0
SM = 0
Delta_L
Doxorubicin
YDR161W
ZShift = 0.55
Z lm Score = −2.75
NG = 0
DB = 0
SM = 0
Delta_K
Doxorubicin
YDR161W
ZShift = 0.42
Z lm Score = −1.39
NG = 0
DB = 0
SM = 0
Delta_r
Doxorubicin
YDR161W
ZShift = −0.6
Z lm Score = −2.5
NG = 0
DB = 0
SM = 0
Delta_AUC
Doxorubicin
YDL119C
ZShift = −0.13
Z lm Score = 6.72
NG = 0
DB = 0
SM = 0
Delta_L
Doxorubicin
YDL119C
ZShift = −0.31
Z lm Score = −0.64
NG = 0
DB = 0
SM = 0
Delta_K
Doxorubicin
YDL119C
ZShift = 0.53
Z lm Score = −2.03
NG = 0
DB = 0
SM = 0
Delta_r
Doxorubicin
YDL119C
ZShift = −0.07
Z lm Score = −3.58
NG = 0
DB = 0
SM = 0
Delta_AUC
Doxorubicin

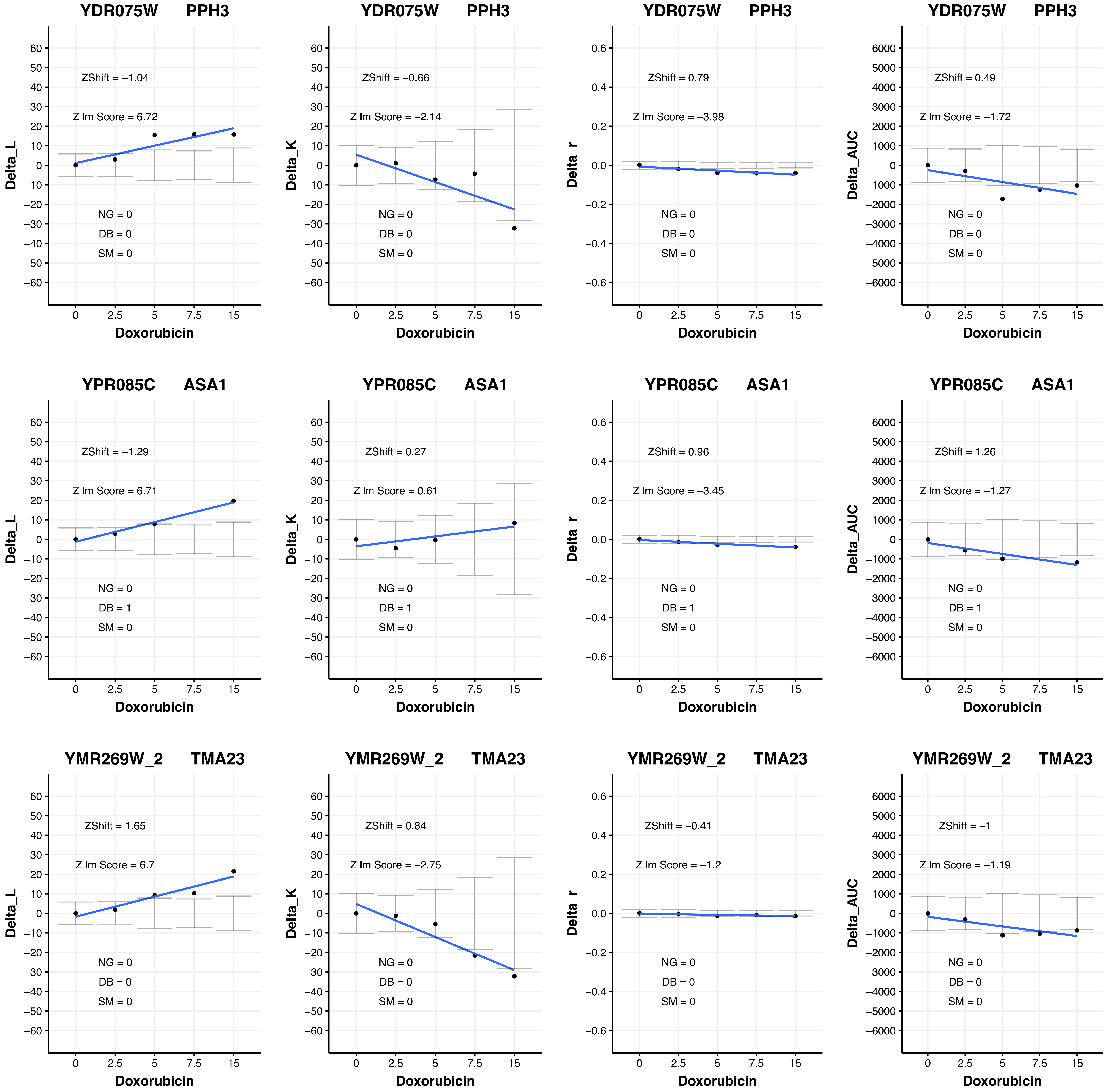
YDR075W PPH3
ZShift = −1.04
Z lm Score = 6.72
NG = 0
DB = 0
SM = 0
Delta_L
Doxorubicin
YDR075W PPH3
ZShift = −0.66
Z lm Score = −2.14
NG = 0
DB = 0
SM = 0
Delta_K
Doxorubicin
YDR075W PPH3
ZShift = 0.79
Z lm Score = −3.98
NG = 0
DB = 0
SM = 0
Delta_r
Doxorubicin
YDR075W PPH3
ZShift = 0.49
Z lm Score = −1.72
NG = 0
DB = 0
SM = 0
Delta_AUC
Doxorubicin
YPR085C ASA1
ZShift = −1.29
Z lm Score = 6.71
NG = 0
DB = 1
SM = 0
Delta_L
Doxorubicin
YPR085C ASA1
ZShift = 0.27
Z lm Score = 0.61
NG = 0
DB = 1
SM = 0
Delta_K
Doxorubicin
YPR085C ASA1
ZShift = 0.96
Z lm Score = −3.45
NG = 0
DB = 1
SM = 0
Delta_r
Doxorubicin
YPR085C ASA1
ZShift = 1.26
Z lm Score = −1.27
NG = 0
DB = 1
SM = 0
Delta_AUC
Doxorubicin
YMR269W_2 TMA23
ZShift = 1.65
Z lm Score = 6.7
NG = 0
DB = 0
SM = 0
Delta_L
Doxorubicin
YMR269W_2 TMA23
ZShift = 0.84
Z lm Score = −2.75
NG = 0
DB = 0
SM = 0
Delta_K
Doxorubicin
YMR269W_2 TMA23
ZShift = −0.41
Z lm Score = −1.2
NG = 0
DB = 0
SM = 0
Delta_r
Doxorubicin
YMR269W_2 TMA23
ZShift = −1
Z lm Score = −1.19
NG = 0
DB = 0
SM = 0
Delta_AUC
Doxorubicin

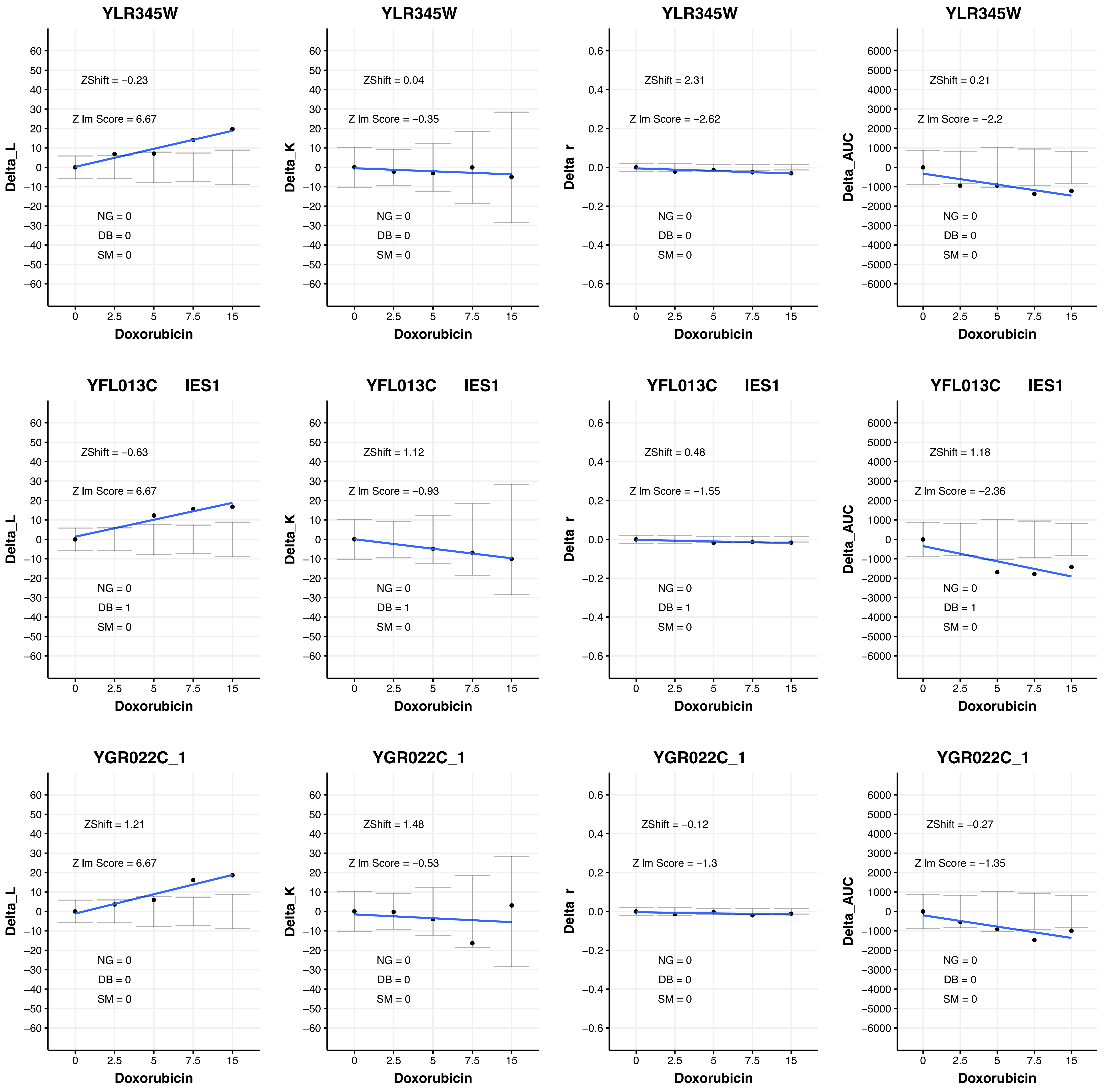
YLR345W
ZShift = −0.23
Z lm Score = 6.67
NG = 0
DB = 0
SM = 0
Delta_L
Doxorubicin
YLR345W
ZShift = 0.04
Z lm Score = −0.35
NG = 0
DB = 0
SM = 0
Delta_K
Doxorubicin
YLR345W
ZShift = 2.31
Z lm Score = −2.62
NG = 0
DB = 0
SM = 0
Delta_r
Doxorubicin
YLR345W
ZShift = 0.21
Z lm Score = −2.2
NG = 0
DB = 0
SM = 0
Delta_AUC
Doxorubicin
YFL013C IES1
ZShift = −0.63
Z lm Score = 6.67
NG = 0
DB = 1
SM = 0
Delta_L
Doxorubicin
YFL013C IES1
ZShift = 1.12
Z lm Score = −0.93
NG = 0
DB = 1
SM = 0
Delta_K
Doxorubicin
YFL013C IES1
ZShift = 0.48
Z lm Score = −1.55
NG = 0
DB = 1
SM = 0
Delta_r
Doxorubicin
YFL013C IES1
ZShift = 1.18
Z lm Score = −2.36
NG = 0
DB = 1
SM = 0
Delta_AUC
Doxorubicin
YGR022C_1
ZShift = 1.21
Z lm Score = 6.67
NG = 0
DB = 0
SM = 0
Delta_L
Doxorubicin
YGR022C_1
ZShift = 1.48
Z lm Score = −0.53
NG = 0
DB = 0
SM = 0
Delta_K
Doxorubicin
YGR022C_1
ZShift = −0.12
Z lm Score = −1.3
NG = 0
DB = 0
SM = 0
Delta_r
Doxorubicin
YGR022C_1
ZShift = −0.27
Z lm Score = −1.35
NG = 0
DB = 0
SM = 0
Delta_AUC
Doxorubicin

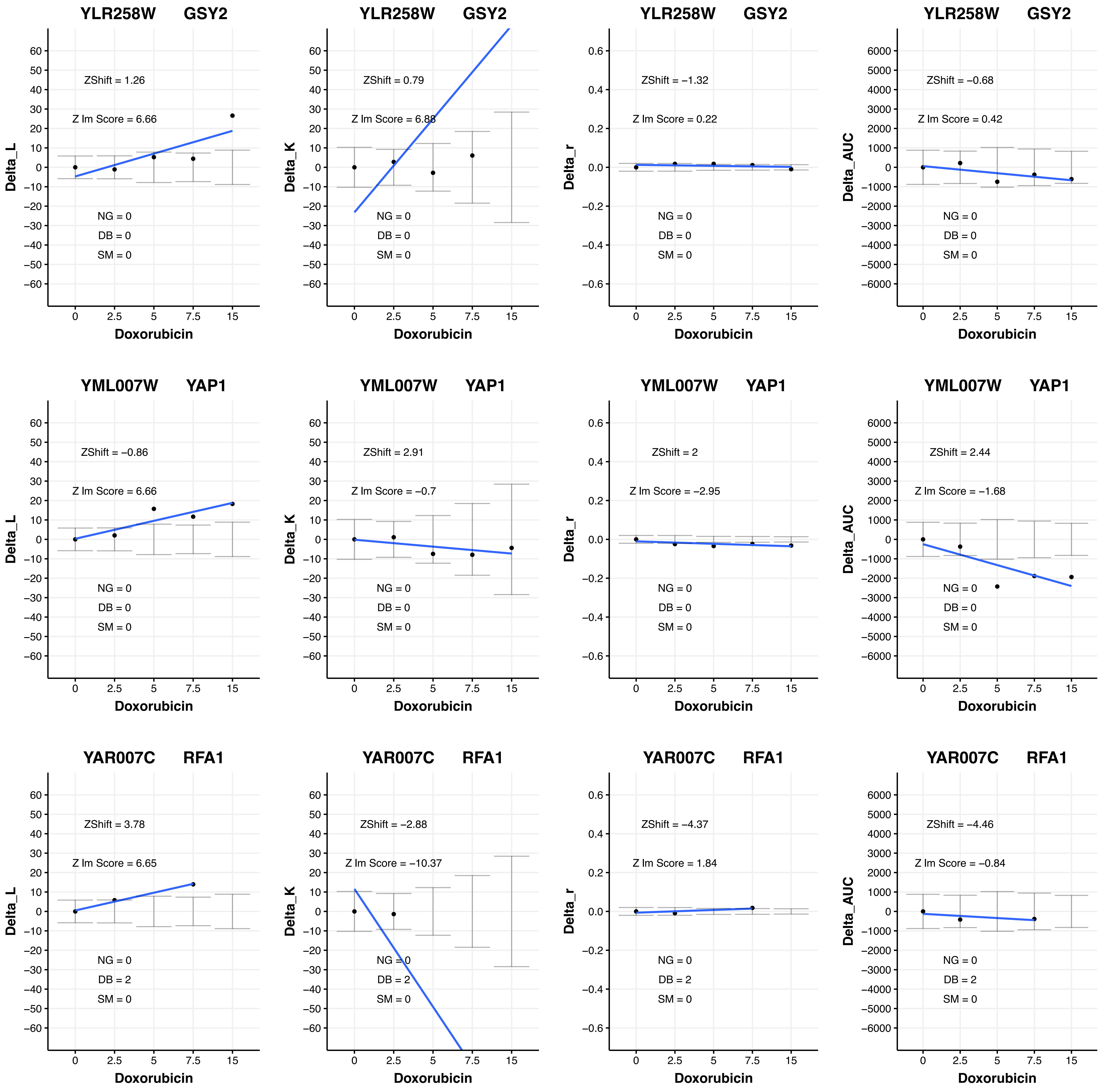

YLR258W GSY2
ZShift = 1.26
Z lm Score = 6.66
NG = 0
DB = 0
SM = 0
Delta_L
Doxorubicin
YLR258W GSY2
ZShift = 0.79
Z lm Score = 6.88
NG = 0
DB = 0
SM = 0
Delta_K
Doxorubicin
YLR258W GSY2
ZShift = −1.32
Z lm Score = 0.22
NG = 0
DB = 0
SM = 0
Delta_r
Doxorubicin
YLR258W GSY2
ZShift = −0.68
Z lm Score = 0.42
NG = 0
DB = 0
SM = 0
Delta_AUC
Doxorubicin
YML007W YAP1
ZShift = −0.86
Z lm Score = 6.66
NG = 0
DB = 0
SM = 0
Delta_L
Doxorubicin
YML007W YAP1
ZShift = 2.91
Z lm Score = −0.7
NG = 0
DB = 0
SM = 0
Delta_K
Doxorubicin
YML007W YAP1
ZShift = 2
Z lm Score = −2.95
NG = 0
DB = 0
SM = 0
Delta_r
Doxorubicin
YML007W YAP1
ZShift = 2.44
Z lm Score = −1.68
NG = 0
DB = 0
SM = 0
Delta_AUC
Doxorubicin
YAR007C RFA1
ZShift = 3.78
Z lm Score = 6.65
NG = 0
DB = 2
SM = 0
Delta_L
Doxorubicin
YAR007C RFA1
ZShift = −2.88
Z lm Score = −10.37
NG = 0
DB = 2
SM = 0
Delta_K
Doxorubicin
YAR007C RFA1
ZShift = −4.37
Z lm Score = 1.84
NG = 0
DB = 2
SM = 0
Delta_r
Doxorubicin
YAR007C RFA1
ZShift = −4.46
Z lm Score = −0.84
NG = 0
DB = 2
SM = 0
Delta_AUC
Doxorubicin

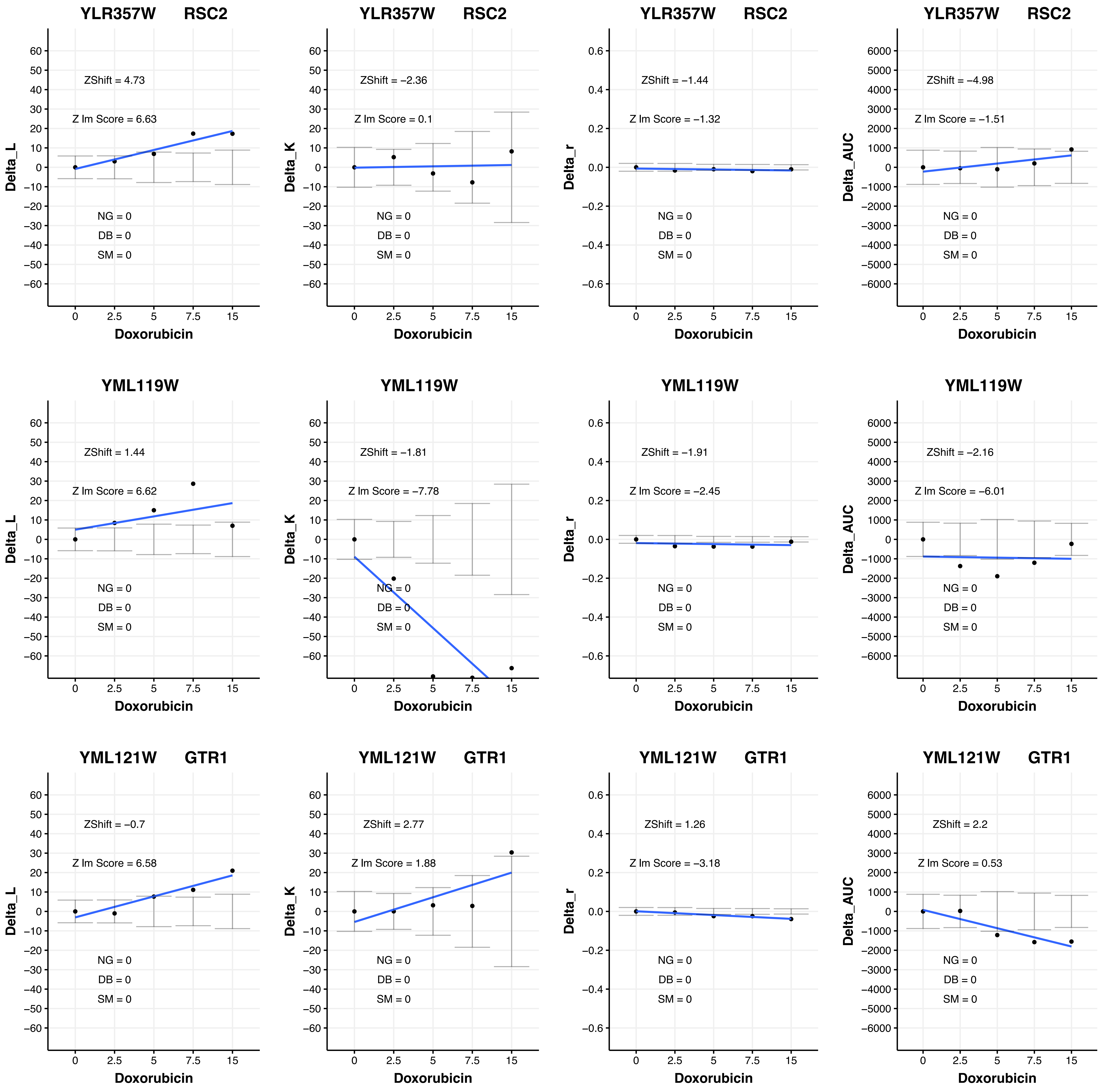
YLR357W RSC2
ZShift = 4.73
Z lm Score = 6.63
NG = 0
DB = 0
SM = 0
Delta_L
Doxorubicin
YLR357W RSC2
ZShift = −2.36
Z lm Score = 0.1
NG = 0
DB = 0
SM = 0
Delta_K
Doxorubicin
YLR357W RSC2
ZShift = −1.44
Z lm Score = −1.32
NG = 0
DB = 0
SM = 0
Delta_r
Doxorubicin
YLR357W RSC2
ZShift = −4.98
Z lm Score = −1.51
NG = 0
DB = 0
SM = 0
Delta_AUC
Doxorubicin
YML119W
ZShift = 1.44
Z lm Score = 6.62
NG = 0
DB = 0
SM = 0
Delta_L
Doxorubicin
YML119W
ZShift = −1.81
Z lm Score = −7.78
NG = 0
DB = 0
SM = 0
Delta_K
Doxorubicin
YML119W
ZShift = −1.91
Z lm Score = −2.45
NG = 0
DB = 0
SM = 0
Delta_r
Doxorubicin
YML119W
ZShift = −2.16
Z lm Score = −6.01
NG = 0
DB = 0
SM = 0
Delta_AUC
Doxorubicin
YML121W GTR1
ZShift = −0.7
Z lm Score = 6.58
NG = 0
DB = 0
SM = 0
Delta_L
Doxorubicin
YML121W GTR1
ZShift = 2.77
Z lm Score = 1.88
NG = 0
DB = 0
SM = 0
Delta_K
Doxorubicin
YML121W GTR1
ZShift = 1.26
Z lm Score = −3.18
NG = 0
DB = 0
SM = 0
Delta_r
Doxorubicin
YML121W GTR1
ZShift = 2.2
Z lm Score = 0.53
NG = 0
DB = 0
SM = 0
Delta_AUC
Doxorubicin

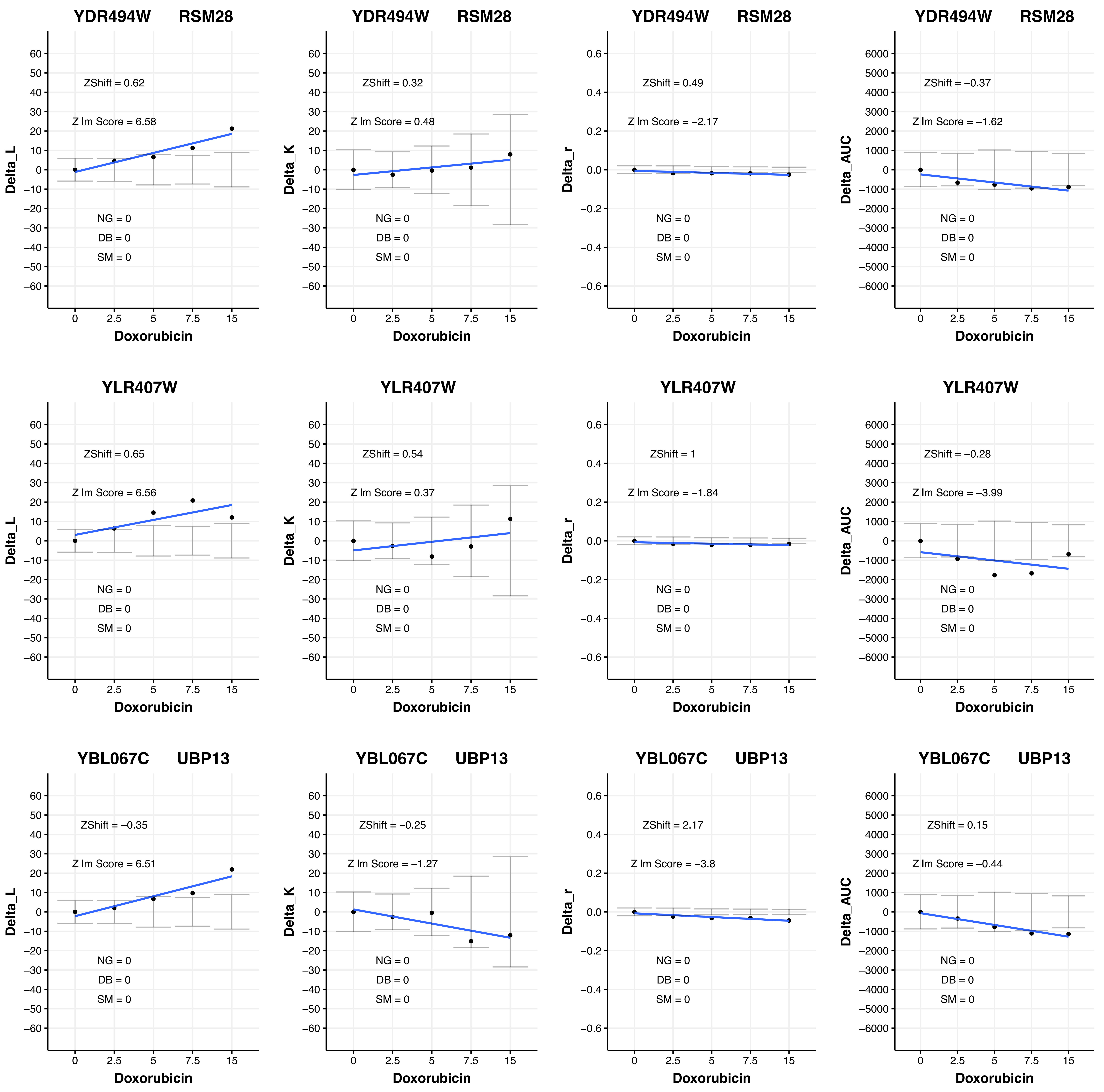
YDR494W RSM28
ZShift = 0.62
Z lm Score = 6.58
NG = 0
DB = 0
SM = 0
Delta_L
Doxorubicin
YDR494W RSM28
ZShift = 0.32
Z lm Score = 0.48
NG = 0
DB = 0
SM = 0
Delta_K
Doxorubicin
YDR494W RSM28
ZShift = 0.49
Z lm Score = −2.17
NG = 0
DB = 0
SM = 0
Delta_r
Doxorubicin
YDR494W RSM28
ZShift = −0.37
Z lm Score = −1.62
NG = 0
DB = 0
SM = 0
Delta_AUC
Doxorubicin
YLR407W
ZShift = 0.65
Z lm Score = 6.56
NG = 0
DB = 0
SM = 0
Delta_L
Doxorubicin
YLR407W
ZShift = 0.54
Z lm Score = 0.37
NG = 0
DB = 0
SM = 0
Delta_K
Doxorubicin
YLR407W
ZShift = 1
Z lm Score = −1.84
NG = 0
DB = 0
SM = 0
Delta_r
Doxorubicin
YLR407W
ZShift = −0.28
Z lm Score = −3.99
NG = 0
DB = 0
SM = 0
Delta_AUC
Doxorubicin
YBL067C UBP13
ZShift = −0.35
Z lm Score = 6.51
NG = 0
DB = 0
SM = 0
Delta_L
Doxorubicin
YBL067C UBP13
ZShift = −0.25
Z lm Score = −1.27
NG = 0
DB = 0
SM = 0
Delta_K
Doxorubicin
YBL067C UBP13
ZShift = 2.17
Z lm Score = −3.8
NG = 0
DB = 0
SM = 0
Delta_r
Doxorubicin
YBL067C UBP13
ZShift = 0.15
Z lm Score = −0.44
NG = 0
DB = 0
SM = 0
Delta_AUC
Doxorubicin

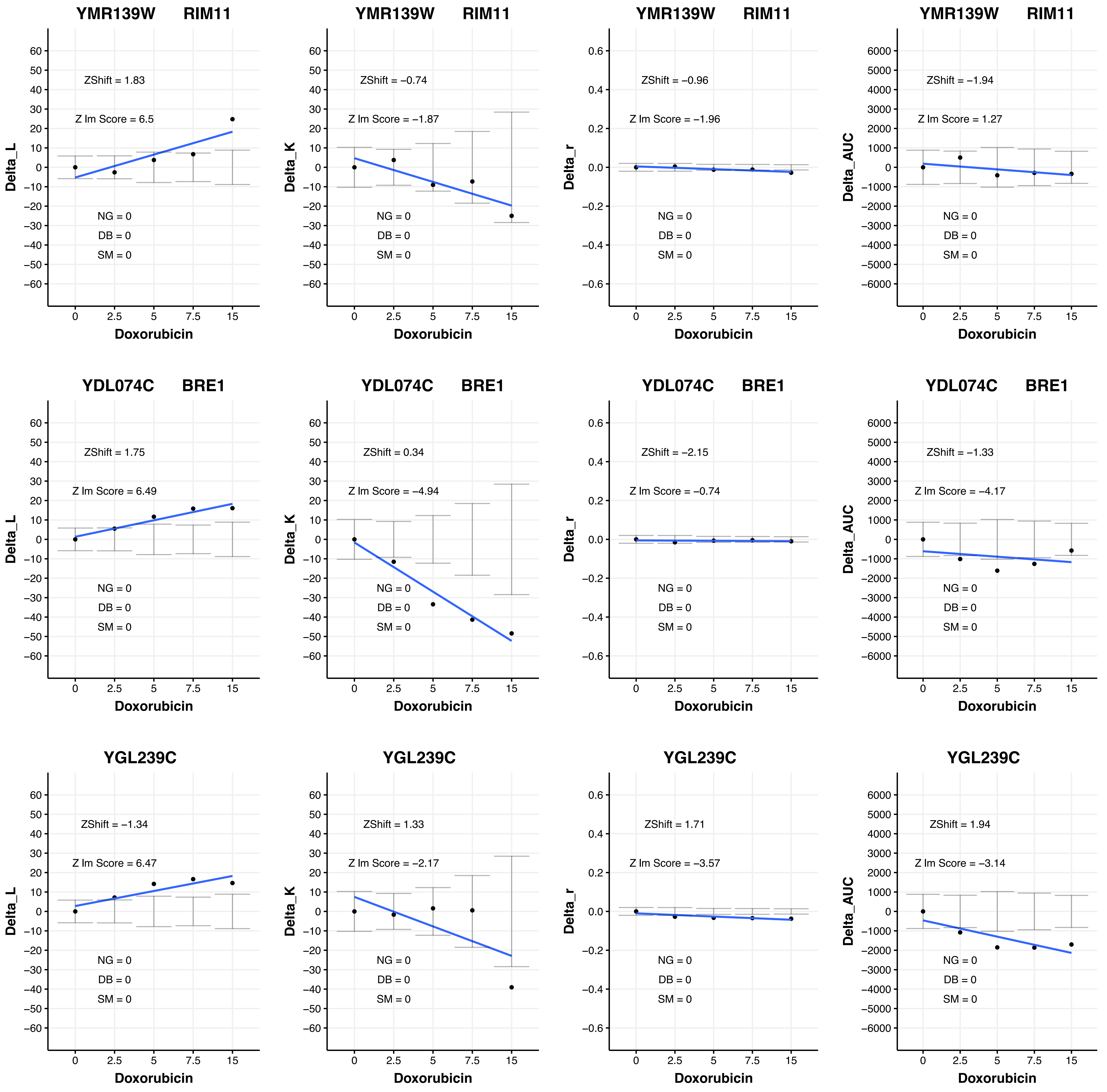
YMR139W RIM11
ZShift = 1.83
Z lm Score = 6.5
NG = 0
DB = 0
SM = 0
Delta_L
Doxorubicin
YMR139W RIM11
ZShift = −0.74
Z lm Score = −1.87
NG = 0
DB = 0
SM = 0
Delta_K
Doxorubicin
YMR139W RIM11
ZShift = −0.96
Z lm Score = −1.96
NG = 0
DB = 0
SM = 0
Delta_r
Doxorubicin
YMR139W RIM11
ZShift = −1.94
Z lm Score = 1.27
NG = 0
DB = 0
SM = 0
Delta_AUC
Doxorubicin
YDL074C BRE1
ZShift = 1.75
Z lm Score = 6.49
NG = 0
DB = 0
SM = 0
Delta_L
Doxorubicin
YDL074C BRE1
ZShift = 0.34
Z lm Score = −4.94
NG = 0
DB = 0
SM = 0
Delta_K
Doxorubicin
YDL074C BRE1
ZShift = −2.15
Z lm Score = −0.74
NG = 0
DB = 0
SM = 0
Delta_r
Doxorubicin
YDL074C BRE1
ZShift = −1.33
Z lm Score = −4.17
NG = 0
DB = 0
SM = 0
Delta_AUC
Doxorubicin
YGL239C
ZShift = −1.34
Z lm Score = 6.47
NG = 0
DB = 0
SM = 0
Delta_L
Doxorubicin
YGL239C
ZShift = 1.33
Z lm Score = −2.17
NG = 0
DB = 0
SM = 0
Delta_K
Doxorubicin
YGL239C
ZShift = 1.71
Z lm Score = −3.57
NG = 0
DB = 0
SM = 0
Delta_r
Doxorubicin
YGL239C
ZShift = 1.94
Z lm Score = −3.14
NG = 0
DB = 0
SM = 0
Delta_AUC
Doxorubicin

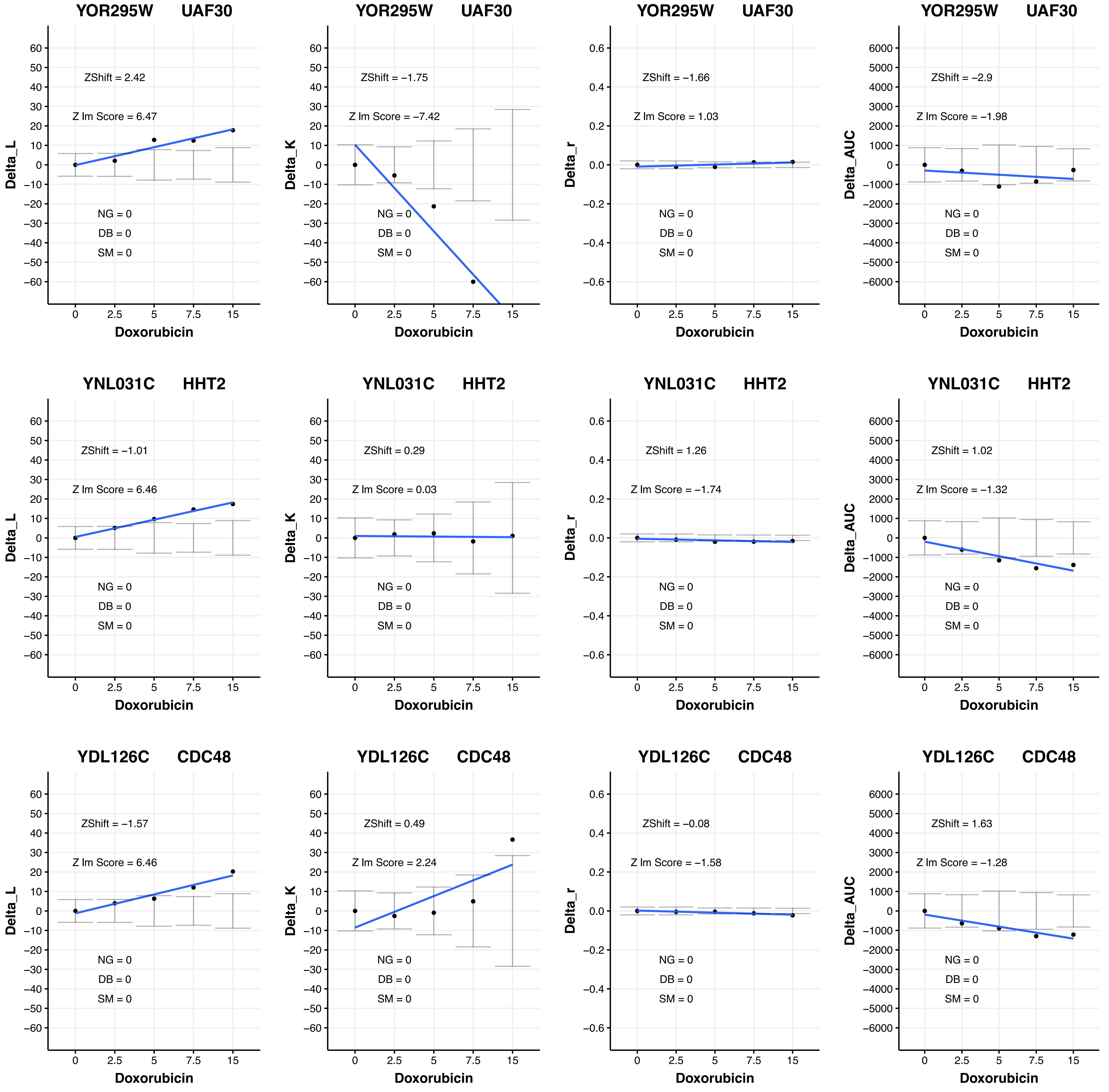

YOR295W UAF30
ZShift = 2.42
Z lm Score = 6.47
NG = 0
DB = 0
SM = 0
Delta_L
Doxorubicin
YOR295W UAF30
ZShift = −1.75
Z lm Score = −7.42
NG = 0
DB = 0
SM = 0
Delta_K
Doxorubicin
YOR295W UAF30
ZShift = −1.66
Z lm Score = 1.03
NG = 0
DB = 0
SM = 0
Delta_r
Doxorubicin
YOR295W UAF30
ZShift = −2.9
Z lm Score = −1.98
NG = 0
DB = 0
SM = 0
Delta_AUC
Doxorubicin
YNL031C HHT2
ZShift = −1.01
Z lm Score = 6.46
NG = 0
DB = 0
SM = 0
Delta_L
Doxorubicin
YNL031C HHT2
ZShift = 0.29
Z lm Score = 0.03
NG = 0
DB = 0
SM = 0
Delta_K
Doxorubicin
YNL031C HHT2
ZShift = 1.26
Z lm Score = −1.74
NG = 0
DB = 0
SM = 0
Delta_r
Doxorubicin
YNL031C HHT2
ZShift = 1.02
Z lm Score = −1.32
NG = 0
DB = 0
SM = 0
Delta_AUC
Doxorubicin
YDL126C CDC48
ZShift = −1.57
Z lm Score = 6.46
NG = 0
DB = 0
SM = 0
Delta_L
Doxorubicin
YDL126C CDC48
ZShift = 0.49
Z lm Score = 2.24
NG = 0
DB = 0
SM = 0
Delta_K
Doxorubicin
YDL126C CDC48
ZShift = −0.08
Z lm Score = −1.58
NG = 0
DB = 0
SM = 0
Delta_r
Doxorubicin
YDL126C CDC48
ZShift = 1.63
Z lm Score = −1.28
NG = 0
DB = 0
SM = 0
Delta_AUC
Doxorubicin

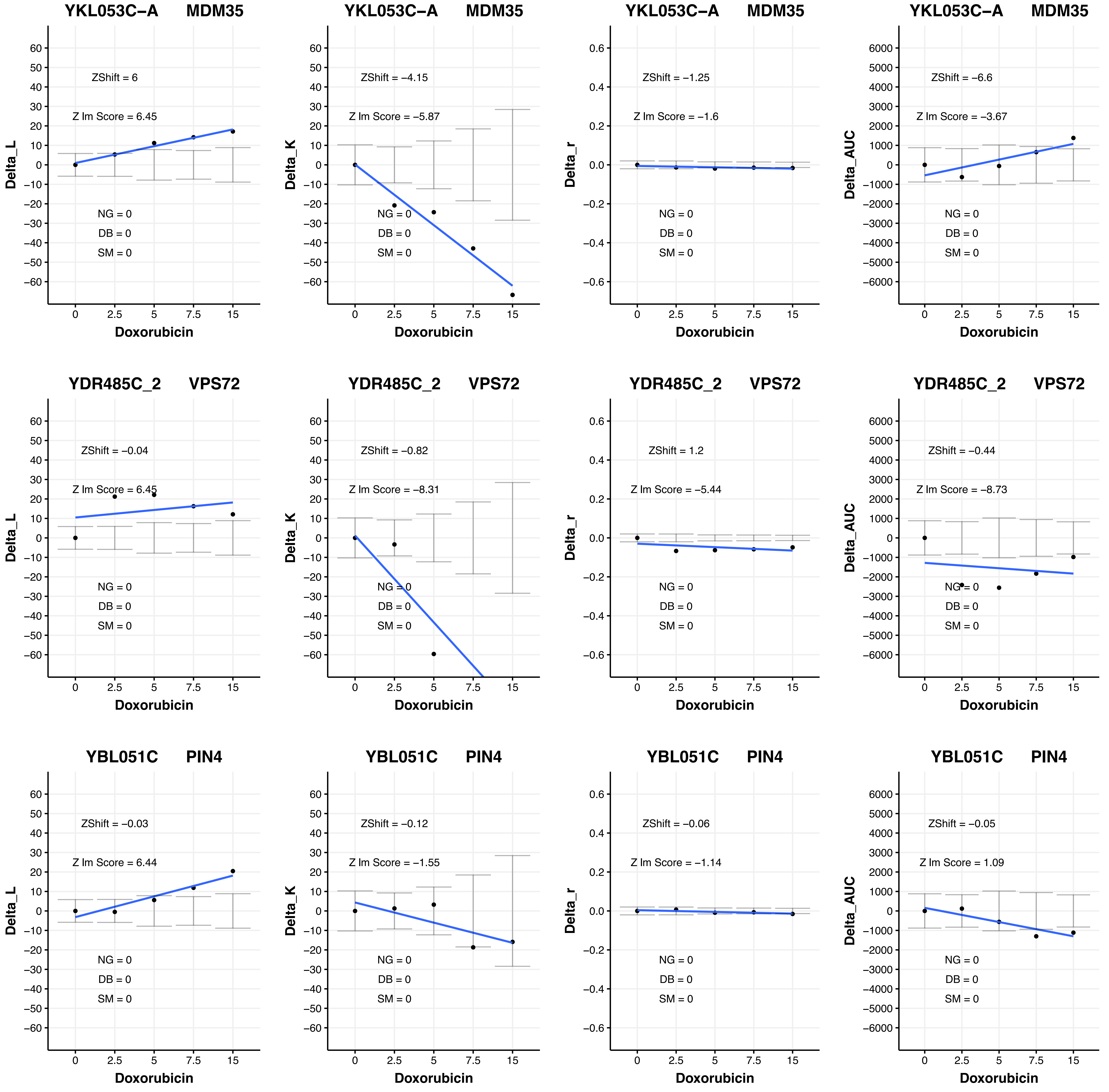

YKL053C–A MDM35
ZShift = 6
Z lm Score = 6.45
NG = 0
DB = 0
SM = 0
Delta_L
Doxorubicin
YKL053C–A MDM35
ZShift = –4.15
Z lm Score = –5.87
NG = 0
DB = 0
SM = 0
Delta_K
Doxorubicin
YKL053C–A MDM35
ZShift = –1.25
Z lm Score = –1.6
NG = 0
DB = 0
SM = 0
Delta_r
Doxorubicin
YKL053C–A MDM35
ZShift = –6.6
Z lm Score = –3.67
NG = 0
DB = 0
SM = 0
Delta_AUC
Doxorubicin
YDR485C_2 VPS72
ZShift = –0.04
Z lm Score = 6.45
NG = 0
DB = 0
SM = 0
Delta_L
Doxorubicin
YDR485C_2 VPS72
ZShift = –0.82
Z lm Score = –8.31
NG = 0
DB = 0
SM = 0
Delta_K
Doxorubicin
YDR485C_2 VPS72
ZShift = 1.2
Z lm Score = –5.44
NG = 0
DB = 0
SM = 0
Delta_r
Doxorubicin
YDR485C_2 VPS72
ZShift = –0.44
Z lm Score = –8.73
NG = 0
DB = 0
SM = 0
Delta_AUC
Doxorubicin
YBL051C PIN4
ZShift = –0.03
Z lm Score = 6.44
NG = 0
DB = 0
SM = 0
Delta_L
Doxorubicin
YBL051C PIN4
ZShift = –0.12
Z lm Score = –1.55
NG = 0
DB = 0
SM = 0
Delta_K
Doxorubicin
YBL051C PIN4
ZShift = –0.06
Z lm Score = –1.14
NG = 0
DB = 0
SM = 0
Delta_r
Doxorubicin
YBL051C PIN4
ZShift = –0.05
Z lm Score = 1.09
NG = 0
DB = 0
SM = 0
Delta_AUC
Doxorubicin

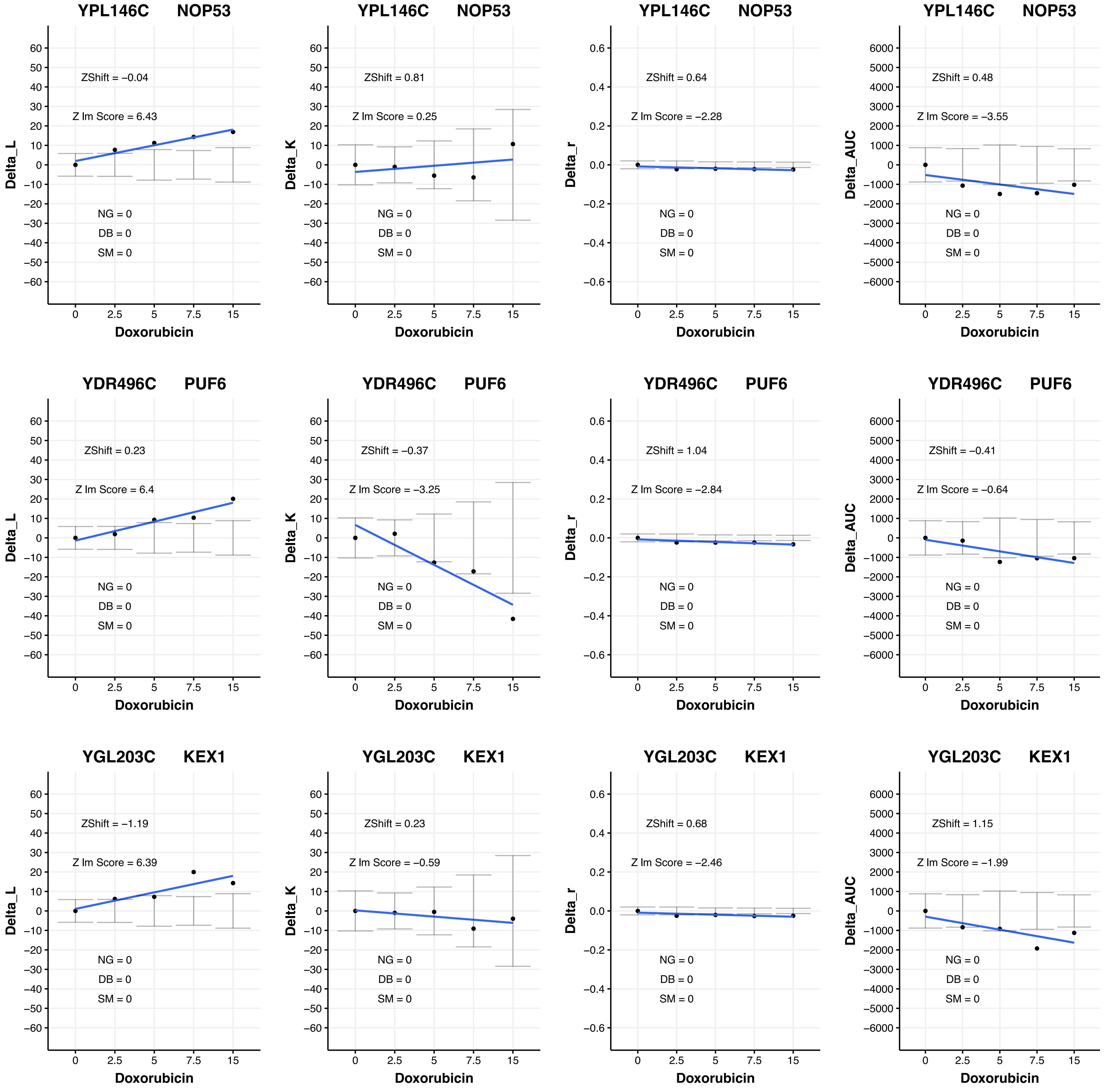
YPL146C NOP53
ZShift = −0.04
Z lm Score = 6.43
NG = 0
DB = 0
SM = 0
Delta_L
Doxorubicin
YPL146C NOP53
ZShift = 0.81
Z lm Score = 0.25
NG = 0
DB = 0
SM = 0
Delta_K
Doxorubicin
YPL146C NOP53
ZShift = 0.64
Z lm Score = −2.28
NG = 0
DB = 0
SM = 0
Delta_r
Doxorubicin
YPL146C NOP53
ZShift = 0.48
Z lm Score = −3.55
NG = 0
DB = 0
SM = 0
Delta_AUC
Doxorubicin
YDR496C PUF6
ZShift = 0.23
Z lm Score = 6.4
NG = 0
DB = 0
SM = 0
Delta_L
Doxorubicin
YDR496C PUF6
ZShift = −0.37
Z lm Score = −3.25
NG = 0
DB = 0
SM = 0
Delta_K
Doxorubicin
YDR496C PUF6
ZShift = 1.04
Z lm Score = −2.84
NG = 0
DB = 0
SM = 0
Delta_r
Doxorubicin
YDR496C PUF6
ZShift = −0.41
Z lm Score = −0.64
NG = 0
DB = 0
SM = 0
Delta_AUC
Doxorubicin
YGL203C KEX1
ZShift = −1.19
Z lm Score = 6.39
NG = 0
DB = 0
SM = 0
Delta_L
Doxorubicin
YGL203C KEX1
ZShift = 0.23
Z lm Score = −0.59
NG = 0
DB = 0
SM = 0
Delta_K
Doxorubicin
YGL203C KEX1
ZShift = 0.68
Z lm Score = −2.46
NG = 0
DB = 0
SM = 0
Delta_r
Doxorubicin
YGL203C KEX1
ZShift = 1.15
Z lm Score = −1.99
NG = 0
DB = 0
SM = 0
Delta_AUC
Doxorubicin

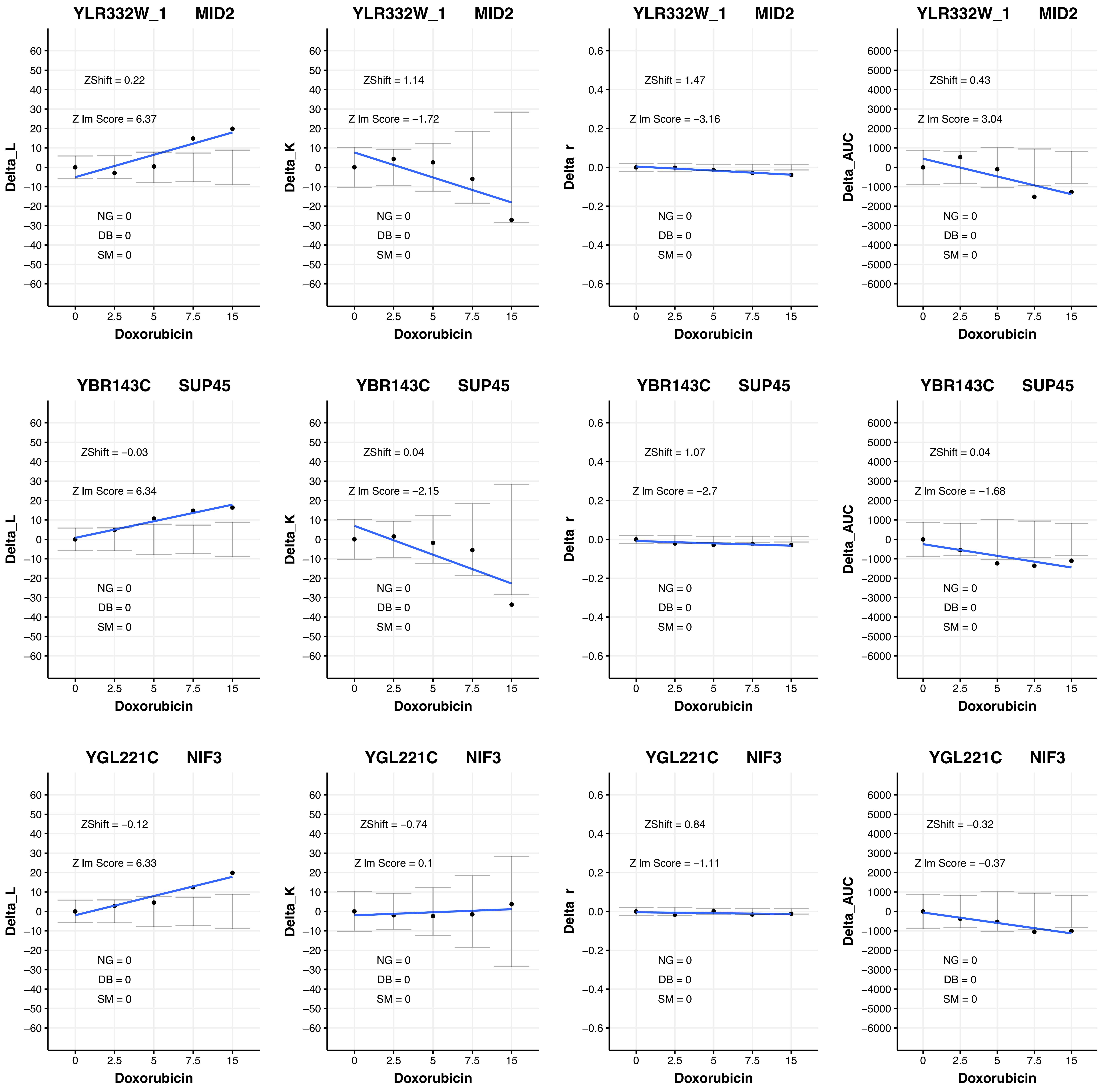
YLR332W_1 MID2
ZShift = 0.22
Z lm Score = 6.37
NG = 0
DB = 0
SM = 0
Delta_L
Doxorubicin
YLR332W_1 MID2
ZShift = 1.14
Z lm Score = −1.72
NG = 0
DB = 0
SM = 0
Delta_K
Doxorubicin
YLR332W_1 MID2
ZShift = 1.47
Z lm Score = −3.16
NG = 0
DB = 0
SM = 0
Delta_r
Doxorubicin
YLR332W_1 MID2
ZShift = 0.43
Z lm Score = 3.04
NG = 0
DB = 0
SM = 0
Delta_AUC
Doxorubicin
YBR143C SUP45
ZShift = −0.03
Z lm Score = 6.34
NG = 0
DB = 0
SM = 0
Delta_L
Doxorubicin
YBR143C SUP45
ZShift = 0.04
Z lm Score = −2.15
NG = 0
DB = 0
SM = 0
Delta_K
Doxorubicin
YBR143C SUP45
ZShift = 1.07
Z lm Score = −2.7
NG = 0
DB = 0
SM = 0
Delta_r
Doxorubicin
YBR143C SUP45
ZShift = 0.04
Z lm Score = −1.68
NG = 0
DB = 0
SM = 0
Delta_AUC
Doxorubicin
YGL221C NIF3
ZShift = −0.12
Z lm Score = 6.33
NG = 0
DB = 0
SM = 0
Delta_L
Doxorubicin
YGL221C NIF3
ZShift = −0.74
Z lm Score = 0.1
NG = 0
DB = 0
SM = 0
Delta_K
Doxorubicin
YGL221C NIF3
ZShift = 0.84
Z lm Score = −1.11
NG = 0
DB = 0
SM = 0
Delta_r
Doxorubicin
YGL221C NIF3
ZShift = −0.32
Z lm Score = −0.37
NG = 0
DB = 0
SM = 0
Delta_AUC
Doxorubicin

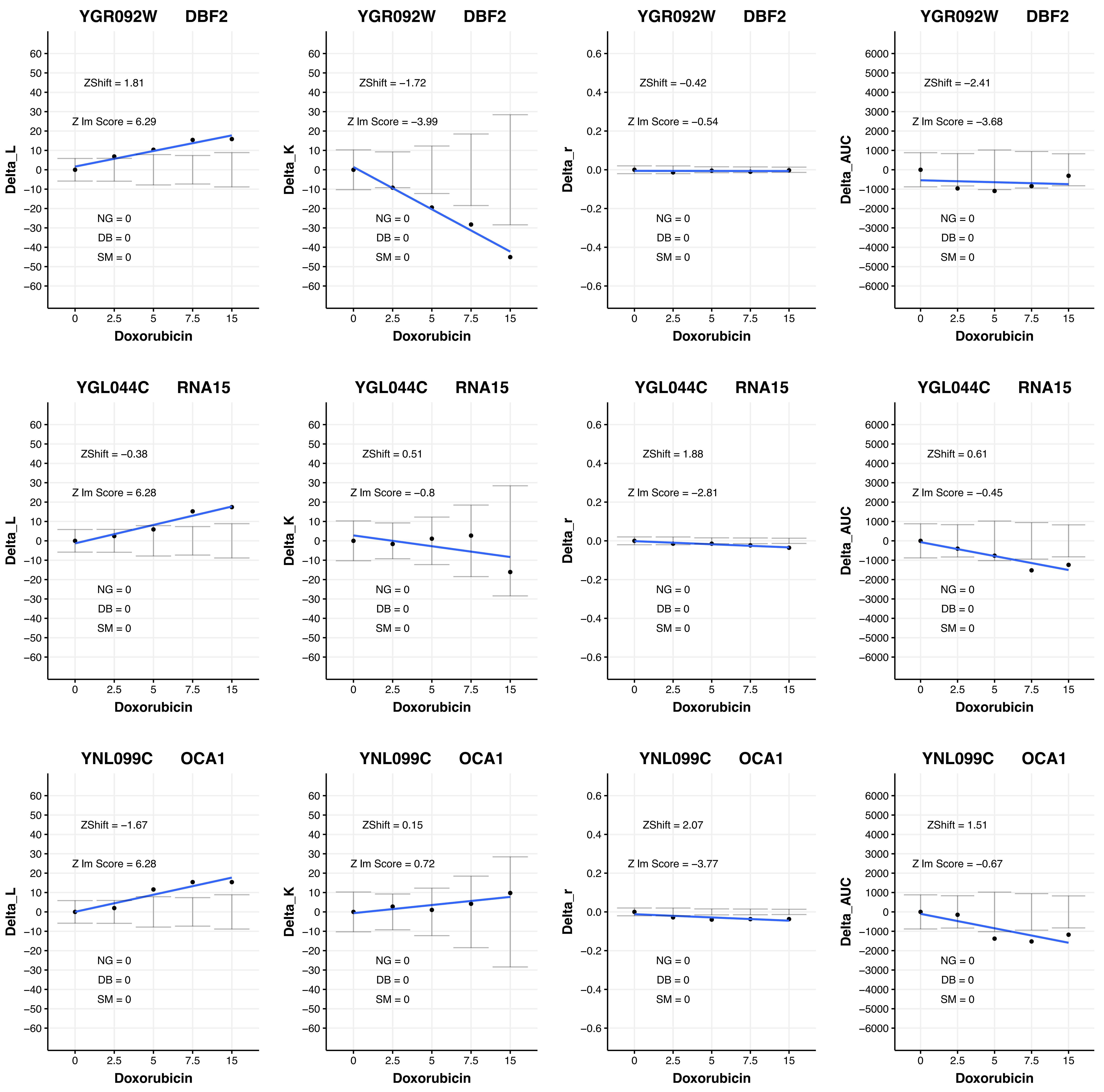
YGR092W DBF2
ZShift = 1.81
Z lm Score = 6.29
NG = 0
DB = 0
SM = 0
Delta_L
Doxorubicin
YGR092W DBF2
ZShift = −1.72
Z lm Score = −3.99
NG = 0
DB = 0
SM = 0
Delta_K
Doxorubicin
YGR092W DBF2
ZShift = −0.42
Z lm Score = −0.54
NG = 0
DB = 0
SM = 0
Delta_r
Doxorubicin
YGR092W DBF2
ZShift = −2.41
Z lm Score = −3.68
NG = 0
DB = 0
SM = 0
Delta_AUC
Doxorubicin
YGL044C RNA15
ZShift = −0.38
Z lm Score = 6.28
NG = 0
DB = 0
SM = 0
Delta_L
Doxorubicin
YGL044C RNA15
ZShift = 0.51
Z lm Score = −0.8
NG = 0
DB = 0
SM = 0
Delta_K
Doxorubicin
YGL044C RNA15
ZShift = 1.88
Z lm Score = −2.81
NG = 0
DB = 0
SM = 0
Delta_r
Doxorubicin
YGL044C RNA15
ZShift = 0.61
Z lm Score = −0.45
NG = 0
DB = 0
SM = 0
Delta_AUC
Doxorubicin
YNL099C OCA1
ZShift = −1.67
Z lm Score = 6.28
NG = 0
DB = 0
SM = 0
Delta_L
Doxorubicin
YNL099C OCA1
ZShift = 0.15
Z lm Score = 0.72
NG = 0
DB = 0
SM = 0
Delta_K
Doxorubicin
YNL099C OCA1
ZShift = 2.07
Z lm Score = −3.77
NG = 0
DB = 0
SM = 0
Delta_r
Doxorubicin
YNL099C OCA1
ZShift = 1.51
Z lm Score = −0.67
NG = 0
DB = 0
SM = 0
Delta_AUC
Doxorubicin

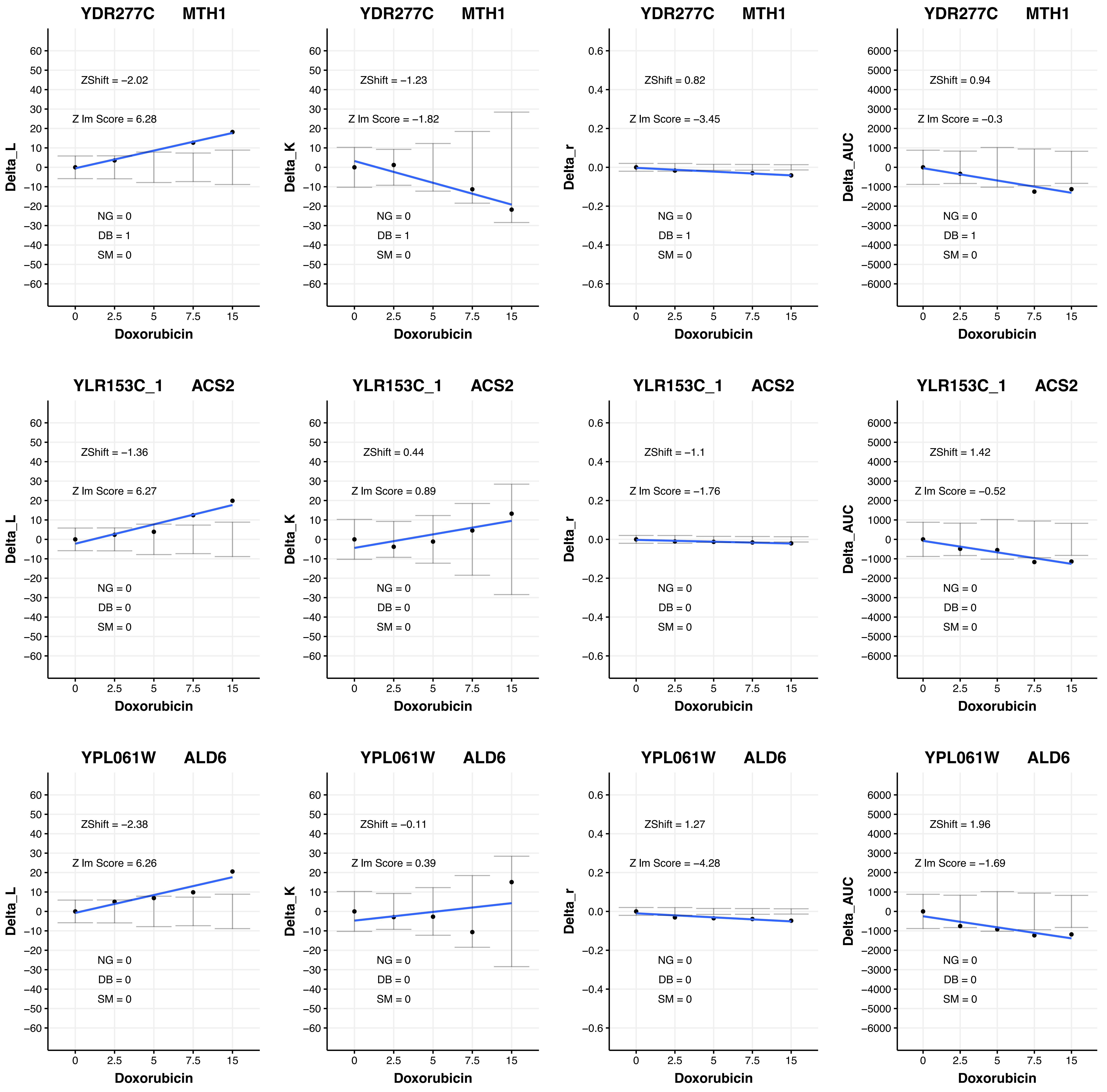
YDR277C MTH1
ZShift = −2.02
Z lm Score = 6.28
NG = 0
DB = 1
SM = 0
Delta_L
Doxorubicin
YDR277C MTH1
ZShift = −1.23
Z lm Score = −1.82
NG = 0
DB = 1
SM = 0
Delta_K
Doxorubicin
YDR277C MTH1
ZShift = 0.82
Z lm Score = −3.45
NG = 0
DB = 1
SM = 0
Delta_r
Doxorubicin
YDR277C MTH1
ZShift = 0.94
Z lm Score = −0.3
NG = 0
DB = 1
SM = 0
Delta_AUC
Doxorubicin
YLR153C_1 ACS2
ZShift = −1.36
Z lm Score = 6.27
NG = 0
DB = 0
SM = 0
Delta_L
Doxorubicin
YLR153C_1 ACS2
ZShift = 0.44
Z lm Score = 0.89
NG = 0
DB = 0
SM = 0
Delta_K
Doxorubicin
YLR153C_1 ACS2
ZShift = −1.1
Z lm Score = −1.76
NG = 0
DB = 0
SM = 0
Delta_r
Doxorubicin
YLR153C_1 ACS2
ZShift = 1.42
Z lm Score = −0.52
NG = 0
DB = 0
SM = 0
Delta_AUC
Doxorubicin
YPL061W ALD6
ZShift = −2.38
Z lm Score = 6.26
NG = 0
DB = 0
SM = 0
Delta_L
Doxorubicin
YPL061W ALD6
ZShift = −0.11
Z lm Score = 0.39
NG = 0
DB = 0
SM = 0
Delta_K
Doxorubicin
YPL061W ALD6
ZShift = 1.27
Z lm Score = −4.28
NG = 0
DB = 0
SM = 0
Delta_r
Doxorubicin
YPL061W ALD6
ZShift = 1.96
Z lm Score = −1.69
NG = 0
DB = 0
SM = 0
Delta_AUC
Doxorubicin

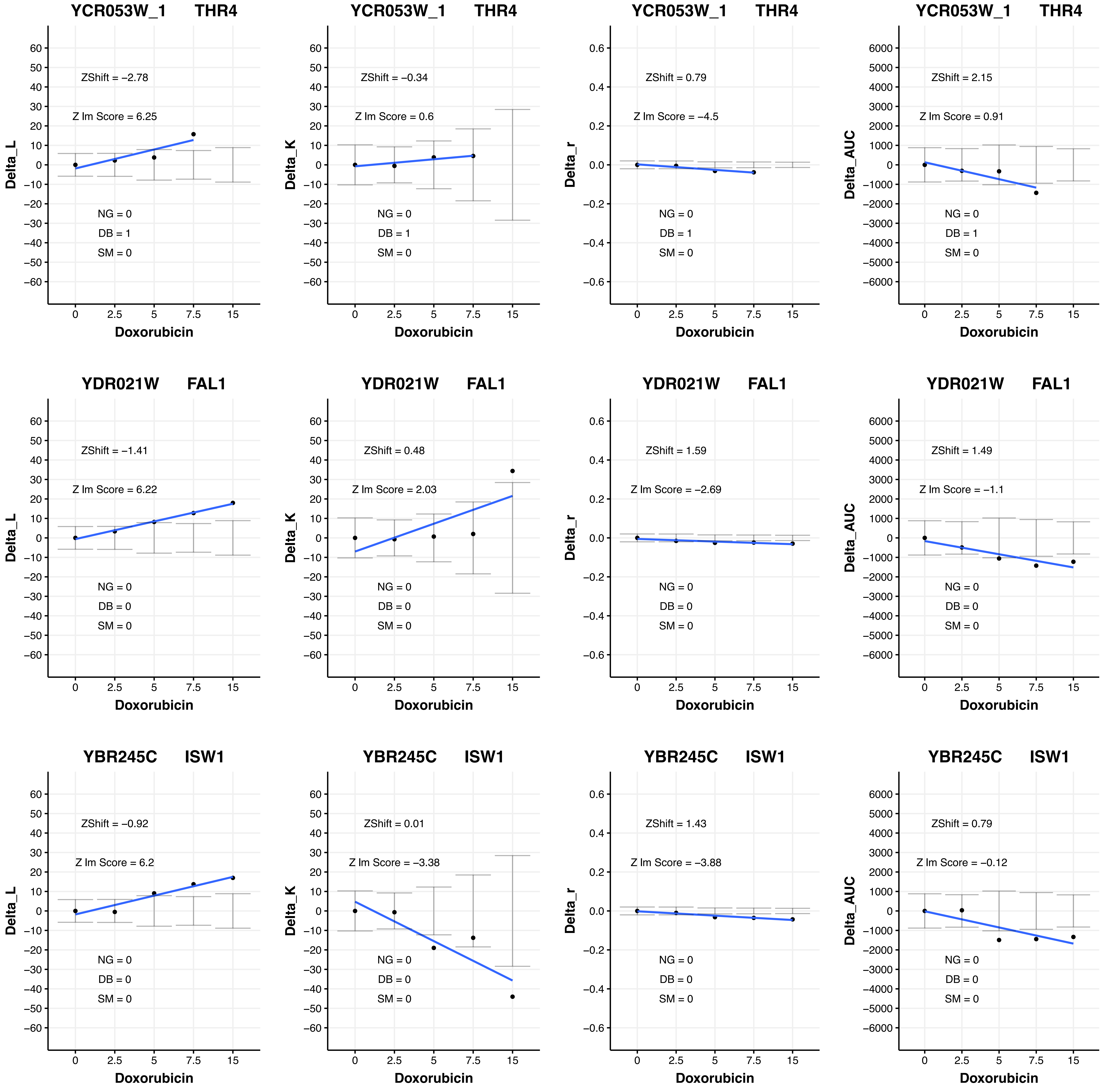
YCR053W_1 THR4
Delta_L
Delta_K
Delta_r
Delta_AUC
Doxorubicin
ZShift = −2.78
Z lm Score = 6.25
ZShift = −0.34
Z lm Score = 0.6
ZShift = 0.79
Z lm Score = −4.5
ZShift = 2.15
Z lm Score = 0.91
NG = 0
DB = 1
SM = 0
YDR021W FAL1
ZShift = −1.41
Z lm Score = 6.22
ZShift = 0.48
Z lm Score = 2.03
ZShift = 1.59
Z lm Score = −2.69
ZShift = 1.49
Z lm Score = −1.1
NG = 0
DB = 0
SM = 0
YBR245C ISW1
ZShift = −0.92
Z lm Score = 6.2
ZShift = 0.01
Z lm Score = −3.38
ZShift = 1.43
Z lm Score = −3.88
ZShift = 0.79
Z lm Score = −0.12
NG = 0
DB = 0
SM = 0

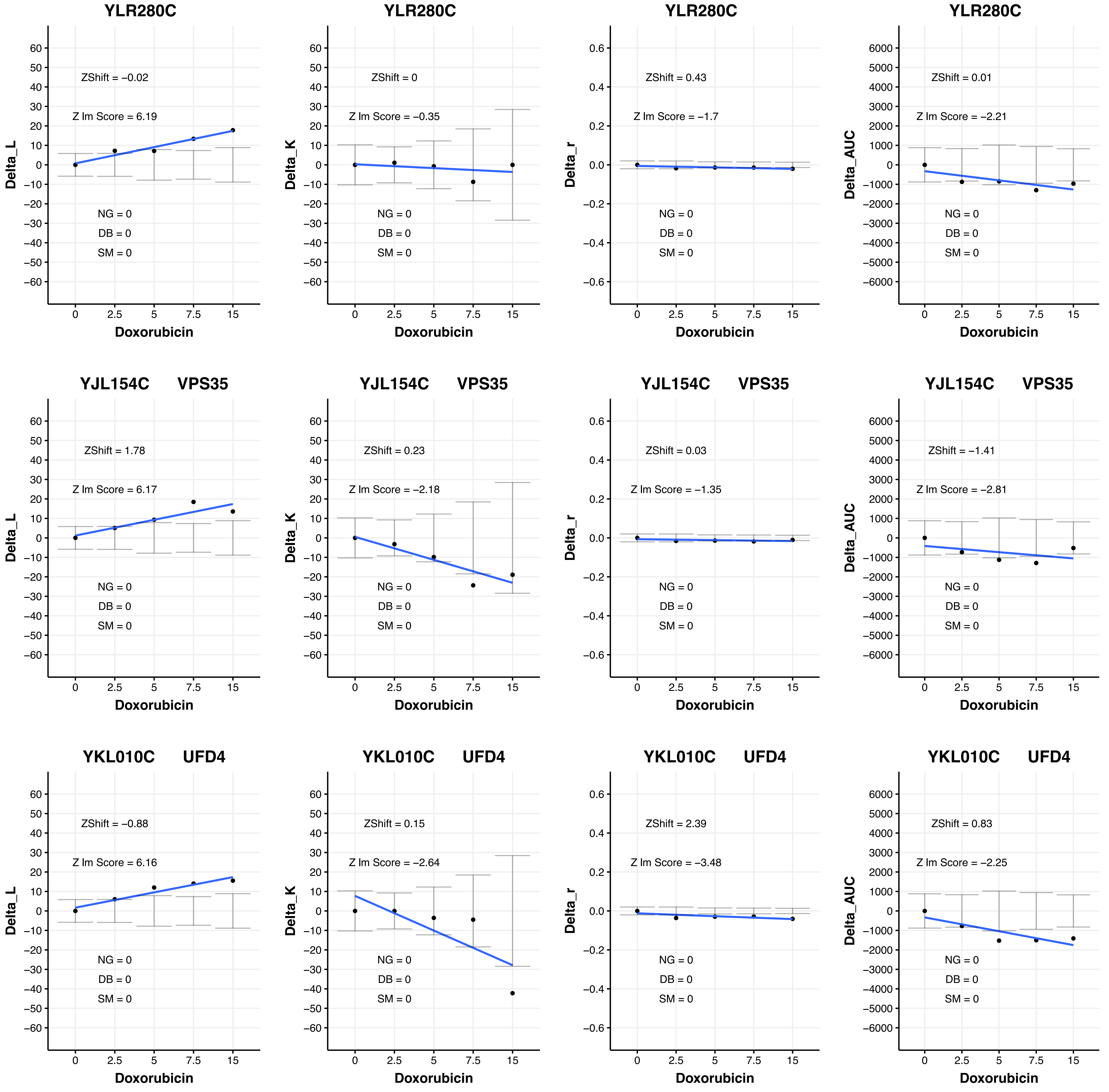
YLR280C
ZShift = −0.02
Z lm Score = 6.19
NG = 0
DB = 0
SM = 0
Delta_L
Doxorubicin
YLR280C
ZShift = 0
Z lm Score = −0.35
NG = 0
DB = 0
SM = 0
Delta_K
Doxorubicin
YLR280C
ZShift = 0.43
Z lm Score = −1.7
NG = 0
DB = 0
SM = 0
Delta_r
Doxorubicin
YLR280C
ZShift = 0.01
Z lm Score = −2.21
NG = 0
DB = 0
SM = 0
Delta_AUC
Doxorubicin
YJL154C VPS35
ZShift = 1.78
Z lm Score = 6.17
NG = 0
DB = 0
SM = 0
Delta_L
Doxorubicin
YJL154C VPS35
ZShift = 0.23
Z lm Score = −2.18
NG = 0
DB = 0
SM = 0
Delta_K
Doxorubicin
YJL154C VPS35
ZShift = 0.03
Z lm Score = −1.35
NG = 0
DB = 0
SM = 0
Delta_r
Doxorubicin
YJL154C VPS35
ZShift = −1.41
Z lm Score = −2.81
NG = 0
DB = 0
SM = 0
Delta_AUC
Doxorubicin
YKL010C UFD4
ZShift = −0.88
Z lm Score = 6.16
NG = 0
DB = 0
SM = 0
Delta_L
Doxorubicin
YKL010C UFD4
ZShift = 0.15
Z lm Score = −2.64
NG = 0
DB = 0
SM = 0
Delta_K
Doxorubicin
YKL010C UFD4
ZShift = 2.39
Z lm Score = −3.48
NG = 0
DB = 0
SM = 0
Delta_r
Doxorubicin
YKL010C UFD4
ZShift = 0.83
Z lm Score = −2.25
NG = 0
DB = 0
SM = 0
Delta_AUC
Doxorubicin

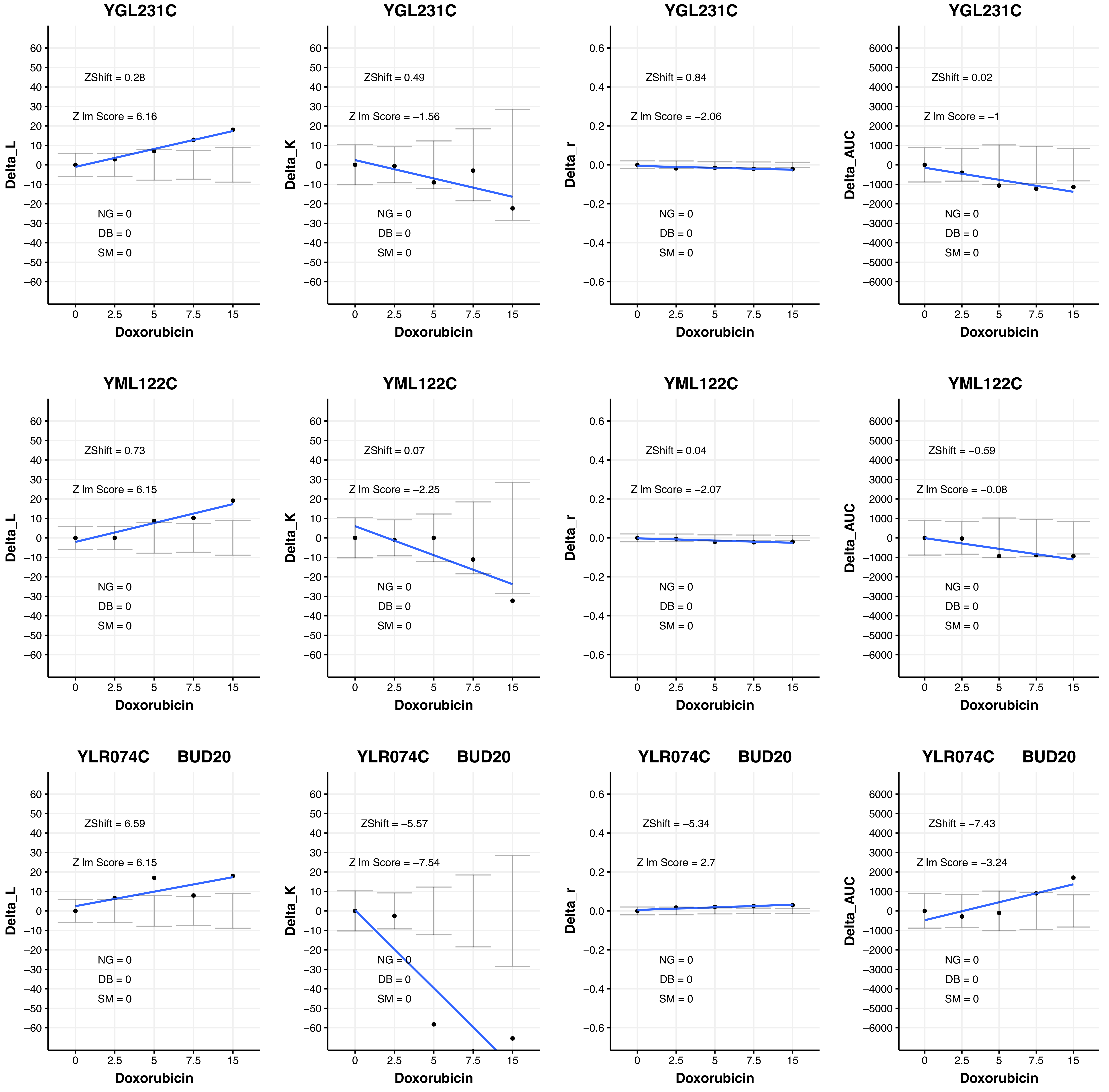

YGL231C
ZShift = 0.28
Z lm Score = 6.16
NG = 0
DB = 0
SM = 0
Delta_L
Doxorubicin
YGL231C
ZShift = 0.49
Z lm Score = −1.56
NG = 0
DB = 0
SM = 0
Delta_K
Doxorubicin
YGL231C
ZShift = 0.84
Z lm Score = −2.06
NG = 0
DB = 0
SM = 0
Delta_r
Doxorubicin
YGL231C
ZShift = 0.02
Z lm Score = −1
NG = 0
DB = 0
SM = 0
Delta_AUC
Doxorubicin
YML122C
ZShift = 0.73
Z lm Score = 6.15
NG = 0
DB = 0
SM = 0
Delta_L
Doxorubicin
YML122C
ZShift = 0.07
Z lm Score = −2.25
NG = 0
DB = 0
SM = 0
Delta_K
Doxorubicin
YML122C
ZShift = 0.04
Z lm Score = −2.07
NG = 0
DB = 0
SM = 0
Delta_r
Doxorubicin
YML122C
ZShift = −0.59
Z lm Score = −0.08
NG = 0
DB = 0
SM = 0
Delta_AUC
Doxorubicin
YLR074C BUD20
ZShift = 6.59
Z lm Score = 6.15
NG = 0
DB = 0
SM = 0
Delta_L
Doxorubicin
YLR074C BUD20
ZShift = −5.57
Z lm Score = −7.54
NG = 0
DB = 0
SM = 0
Delta_K
Doxorubicin
YLR074C BUD20
ZShift = −5.34
Z lm Score = 2.7
NG = 0
DB = 0
SM = 0
Delta_r
Doxorubicin
YLR074C BUD20
ZShift = −7.43
Z lm Score = −3.24
NG = 0
DB = 0
SM = 0
Delta_AUC
Doxorubicin

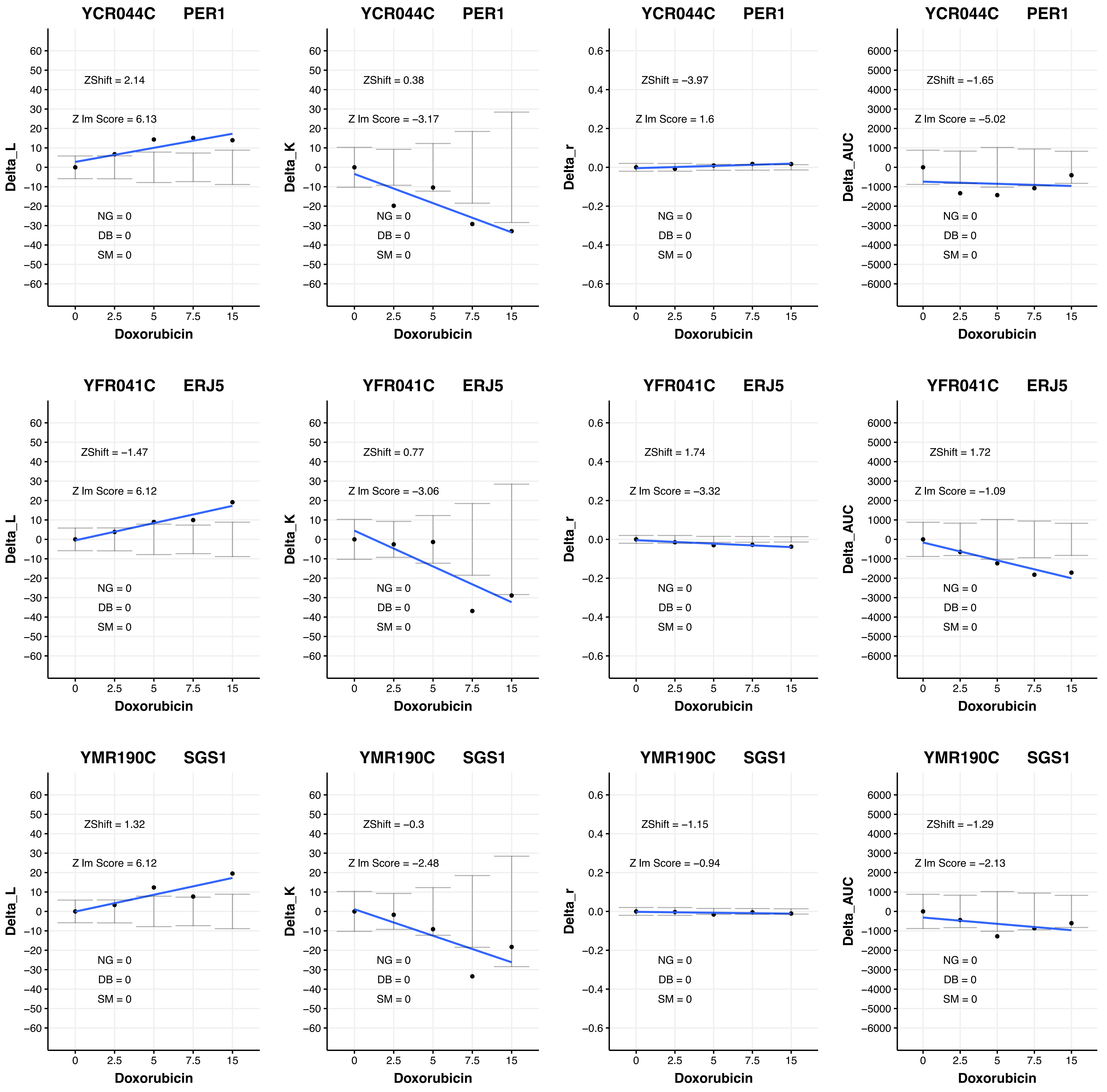
YCR044C PER1
ZShift = 2.14
Z lm Score = 6.13
NG = 0
DB = 0
SM = 0
Delta_L
Doxorubicin
YCR044C PER1
ZShift = 0.38
Z lm Score = −3.17
NG = 0
DB = 0
SM = 0
Delta_K
Doxorubicin
YCR044C PER1
ZShift = −3.97
Z lm Score = 1.6
NG = 0
DB = 0
SM = 0
Delta_r
Doxorubicin
YCR044C PER1
ZShift = −1.65
Z lm Score = −5.02
NG = 0
DB = 0
SM = 0
Delta_AUC
Doxorubicin
YFR041C ERJ5
ZShift = −1.47
Z lm Score = 6.12
NG = 0
DB = 0
SM = 0
Delta_L
Doxorubicin
YFR041C ERJ5
ZShift = 0.77
Z lm Score = −3.06
NG = 0
DB = 0
SM = 0
Delta_K
Doxorubicin
YFR041C ERJ5
ZShift = 1.74
Z lm Score = −3.32
NG = 0
DB = 0
SM = 0
Delta_r
Doxorubicin
YFR041C ERJ5
ZShift = 1.72
Z lm Score = −1.09
NG = 0
DB = 0
SM = 0
Delta_AUC
Doxorubicin
YMR190C SGS1
ZShift = 1.32
Z lm Score = 6.12
NG = 0
DB = 0
SM = 0
Delta_L
Doxorubicin
YMR190C SGS1
ZShift = −0.3
Z lm Score = −2.48
NG = 0
DB = 0
SM = 0
Delta_K
Doxorubicin
YMR190C SGS1
ZShift = −1.15
Z lm Score = −0.94
NG = 0
DB = 0
SM = 0
Delta_r
Doxorubicin
YMR190C SGS1
ZShift = −1.29
Z lm Score = −2.13
NG = 0
DB = 0
SM = 0
Delta_AUC
Doxorubicin

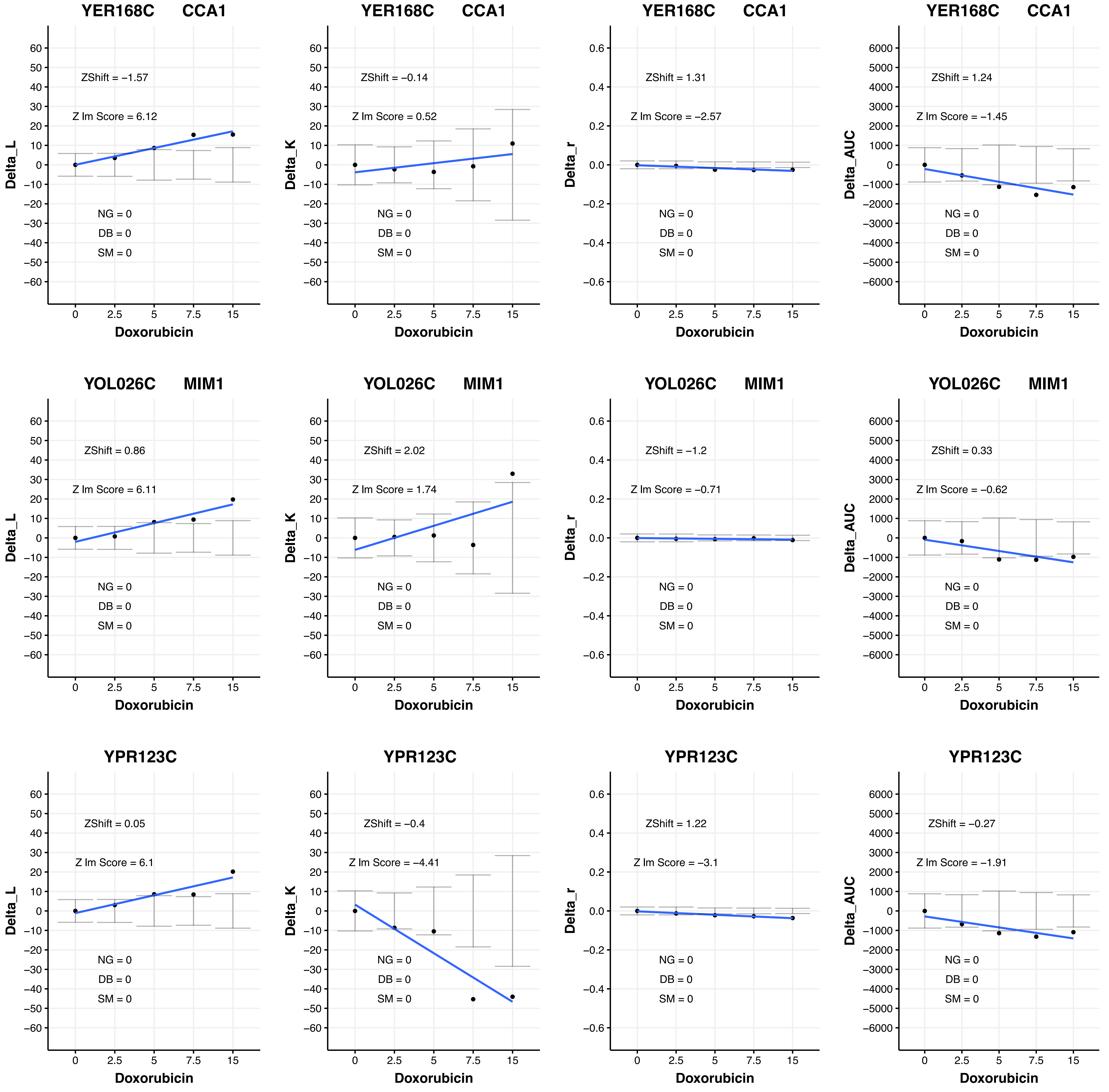
YER168C CCA1
ZShift = −1.57
Z lm Score = 6.12
NG = 0
DB = 0
SM = 0
Delta_L
Doxorubicin
YER168C CCA1
ZShift = −0.14
Z lm Score = 0.52
Delta_K
YER168C CCA1
ZShift = 1.31
Z lm Score = −2.57
Delta_r
YER168C CCA1
ZShift = 1.24
Z lm Score = −1.45
Delta_AUC
YOL026C MIM1
ZShift = 0.86
Z lm Score = 6.11
YOL026C MIM1
ZShift = 2.02
Z lm Score = 1.74
YOL026C MIM1
ZShift = −1.2
Z lm Score = −0.71
YOL026C MIM1
ZShift = 0.33
Z lm Score = −0.62
YPR123C
ZShift = 0.05
Z lm Score = 6.1
YPR123C
ZShift = −0.4
Z lm Score = −4.41
YPR123C
ZShift = 1.22
Z lm Score = −3.1
YPR123C
ZShift = −0.27
Z lm Score = −1.91

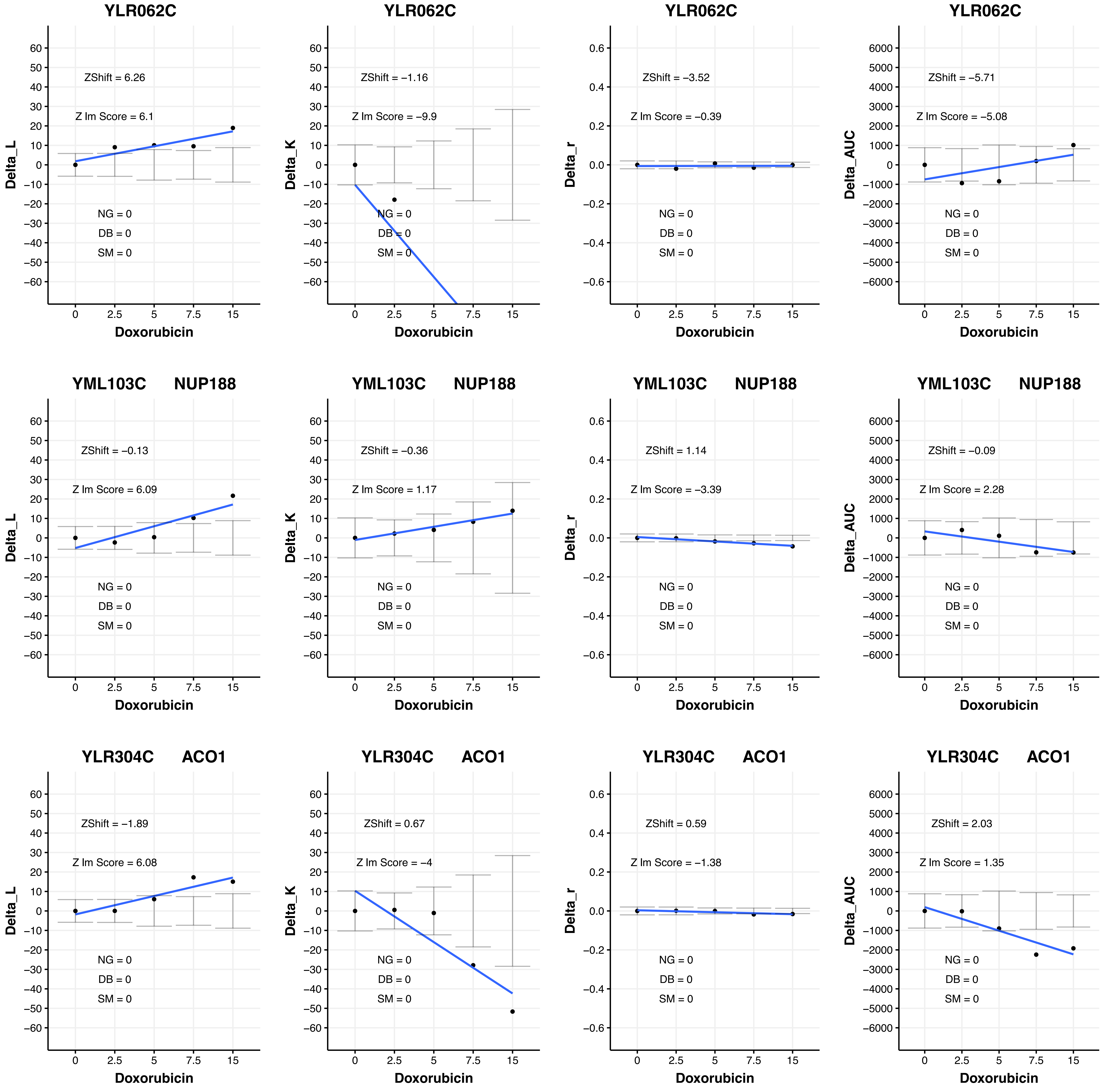

YLR062C
ZShift = 6.26
Z lm Score = 6.1
NG = 0
DB = 0
SM = 0
Delta_L
Doxorubicin
YLR062C
ZShift = −1.16
Z lm Score = −9.9
NG = 0
DB = 0
SM = 0
Delta_K
Doxorubicin
YLR062C
ZShift = −3.52
Z lm Score = −0.39
NG = 0
DB = 0
SM = 0
Delta_r
Doxorubicin
YLR062C
ZShift = −5.71
Z lm Score = −5.08
NG = 0
DB = 0
SM = 0
Delta_AUC
Doxorubicin
YML103C NUP188
ZShift = −0.13
Z lm Score = 6.09
NG = 0
DB = 0
SM = 0
Delta_L
Doxorubicin
YML103C NUP188
ZShift = −0.36
Z lm Score = 1.17
NG = 0
DB = 0
SM = 0
Delta_K
Doxorubicin
YML103C NUP188
ZShift = 1.14
Z lm Score = −3.39
NG = 0
DB = 0
SM = 0
Delta_r
Doxorubicin
YML103C NUP188
ZShift = −0.09
Z lm Score = 2.28
NG = 0
DB = 0
SM = 0
Delta_AUC
Doxorubicin
YLR304C ACO1
ZShift = −1.89
Z lm Score = 6.08
NG = 0
DB = 0
SM = 0
Delta_L
Doxorubicin
YLR304C ACO1
ZShift = 0.67
Z lm Score = −4
NG = 0
DB = 0
SM = 0
Delta_K
Doxorubicin
YLR304C ACO1
ZShift = 0.59
Z lm Score = −1.38
NG = 0
DB = 0
SM = 0
Delta_r
Doxorubicin
YLR304C ACO1
ZShift = 2.03
Z lm Score = 1.35
NG = 0
DB = 0
SM = 0
Delta_AUC
Doxorubicin

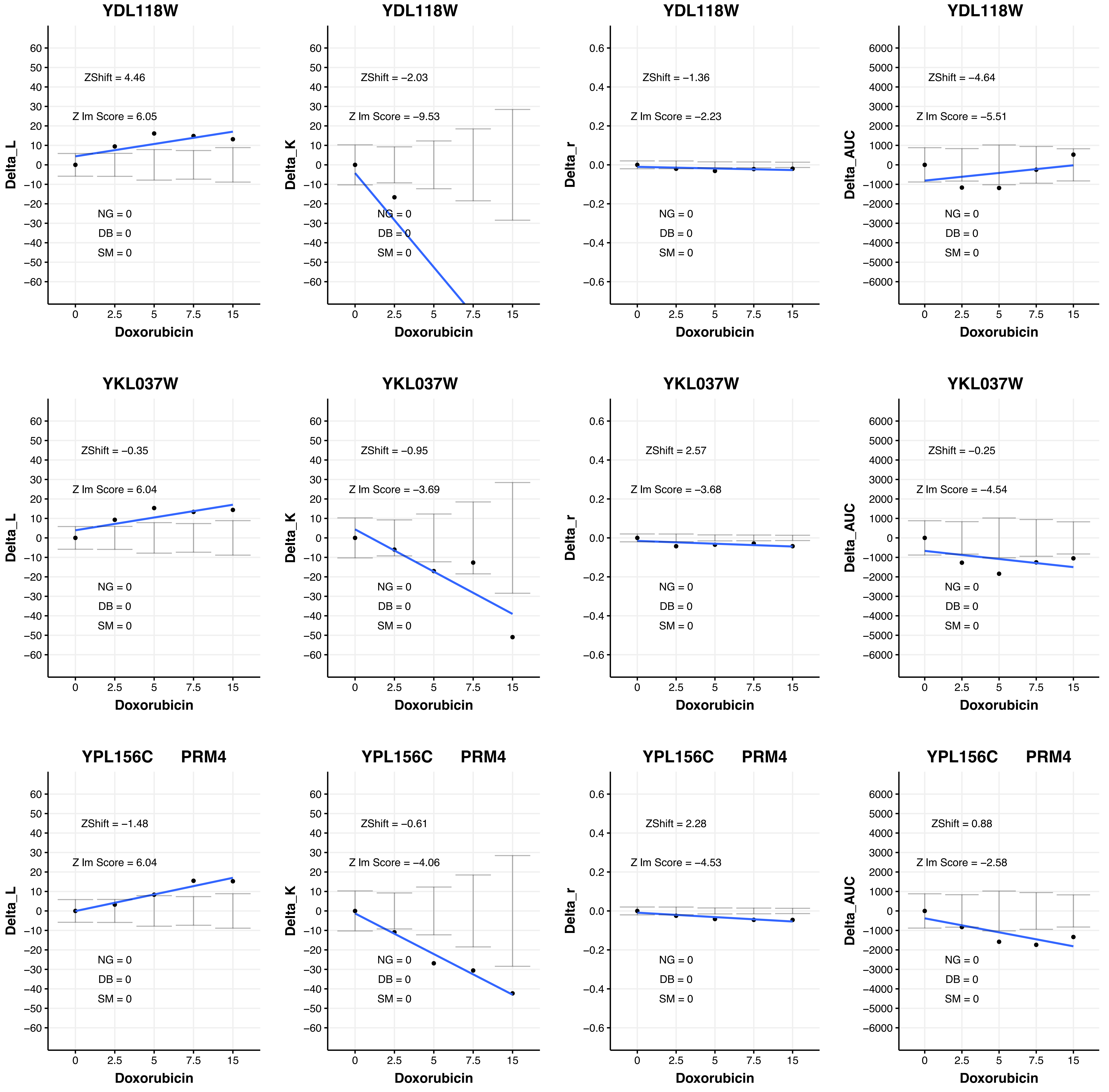
YDL118W
ZShift = 4.46
Z lm Score = 6.05
NG = 0
DB = 0
SM = 0
Delta_L
Doxorubicin
YDL118W
ZShift = −2.03
Z lm Score = −9.53
NG = 0
DB = 0
SM = 0
Delta_K
Doxorubicin
YDL118W
ZShift = −1.36
Z lm Score = −2.23
NG = 0
DB = 0
SM = 0
Delta_r
Doxorubicin
YDL118W
ZShift = −4.64
Z lm Score = −5.51
NG = 0
DB = 0
SM = 0
Delta_AUC
Doxorubicin
YKL037W
ZShift = −0.35
Z lm Score = 6.04
NG = 0
DB = 0
SM = 0
Delta_L
Doxorubicin
YKL037W
ZShift = −0.95
Z lm Score = −3.69
NG = 0
DB = 0
SM = 0
Delta_K
Doxorubicin
YKL037W
ZShift = 2.57
Z lm Score = −3.68
NG = 0
DB = 0
SM = 0
Delta_r
Doxorubicin
YKL037W
ZShift = −0.25
Z lm Score = −4.54
NG = 0
DB = 0
SM = 0
Delta_AUC
Doxorubicin
YPL156C PRM4
ZShift = −1.48
Z lm Score = 6.04
NG = 0
DB = 0
SM = 0
Delta_L
Doxorubicin
YPL156C PRM4
ZShift = −0.61
Z lm Score = −4.06
NG = 0
DB = 0
SM = 0
Delta_K
Doxorubicin
YPL156C PRM4
ZShift = 2.28
Z lm Score = −4.53
NG = 0
DB = 0
SM = 0
Delta_r
Doxorubicin
YPL156C PRM4
ZShift = 0.88
Z lm Score = −2.58
NG = 0
DB = 0
SM = 0
Delta_AUC
Doxorubicin

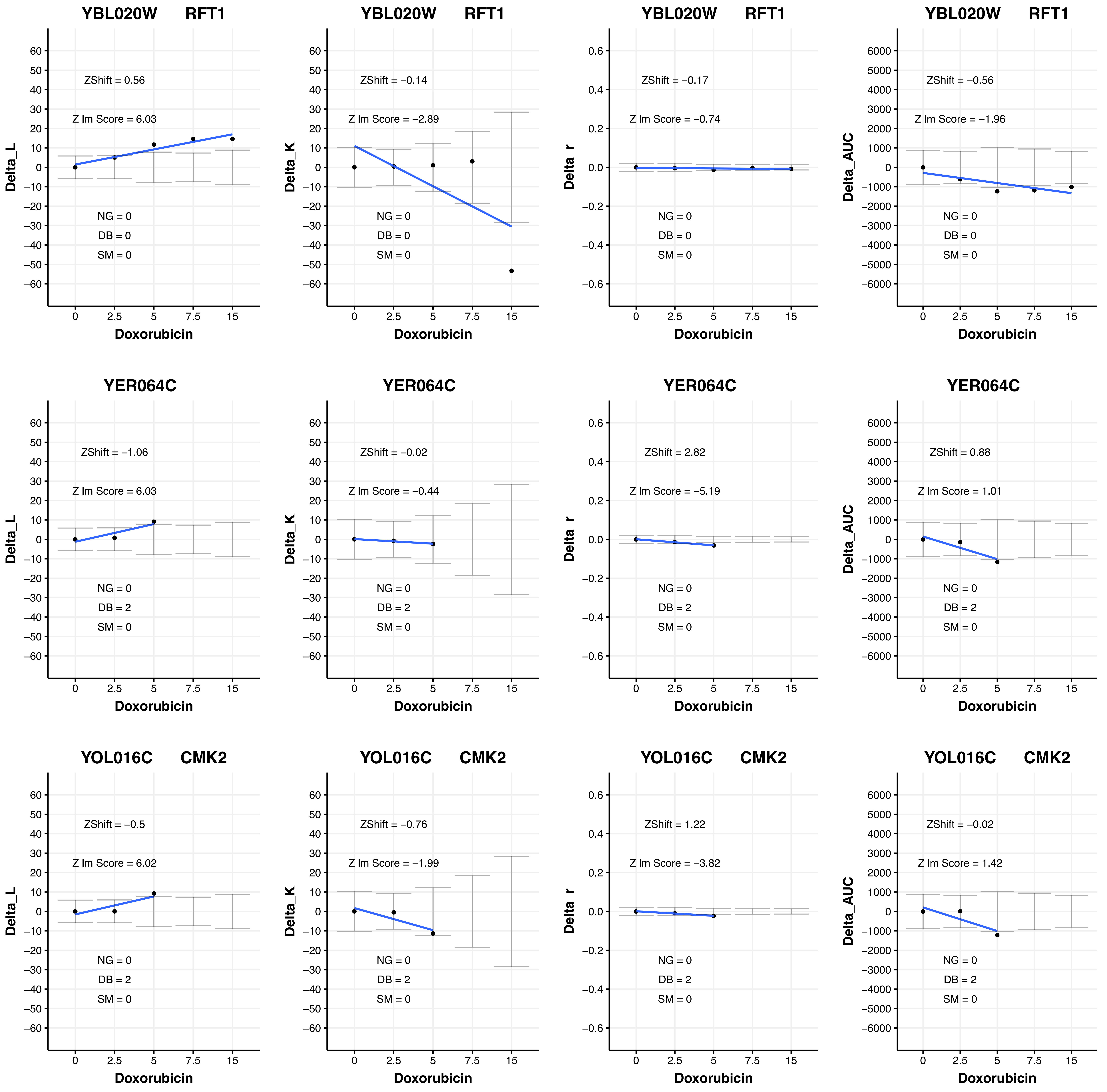
YBL020W RFT1
ZShift = 0.56
Z lm Score = 6.03
NG = 0
DB = 0
SM = 0
Delta_L
Doxorubicin
YBL020W RFT1
ZShift = −0.14
Z lm Score = −2.89
NG = 0
DB = 0
SM = 0
Delta_K
Doxorubicin
YBL020W RFT1
ZShift = −0.17
Z lm Score = −0.74
NG = 0
DB = 0
SM = 0
Delta_r
Doxorubicin
YBL020W RFT1
ZShift = −0.56
Z lm Score = −1.96
NG = 0
DB = 0
SM = 0
Delta_AUC
Doxorubicin
YER064C
ZShift = −1.06
Z lm Score = 6.03
NG = 0
DB = 2
SM = 0
Delta_L
Doxorubicin
YER064C
ZShift = −0.02
Z lm Score = −0.44
NG = 0
DB = 2
SM = 0
Delta_K
Doxorubicin
YER064C
ZShift = 2.82
Z lm Score = −5.19
NG = 0
DB = 2
SM = 0
Delta_r
Doxorubicin
YER064C
ZShift = 0.88
Z lm Score = 1.01
NG = 0
DB = 2
SM = 0
Delta_AUC
Doxorubicin
YOL016C CMK2
ZShift = −0.5
Z lm Score = 6.02
NG = 0
DB = 2
SM = 0
Delta_L
Doxorubicin
YOL016C CMK2
ZShift = −0.76
Z lm Score = −1.99
NG = 0
DB = 2
SM = 0
Delta_K
Doxorubicin
YOL016C CMK2
ZShift = 1.22
Z lm Score = −3.82
NG = 0
DB = 2
SM = 0
Delta_r
Doxorubicin
YOL016C CMK2
ZShift = −0.02
Z lm Score = 1.42
NG = 0
DB = 2
SM = 0
Delta_AUC
Doxorubicin

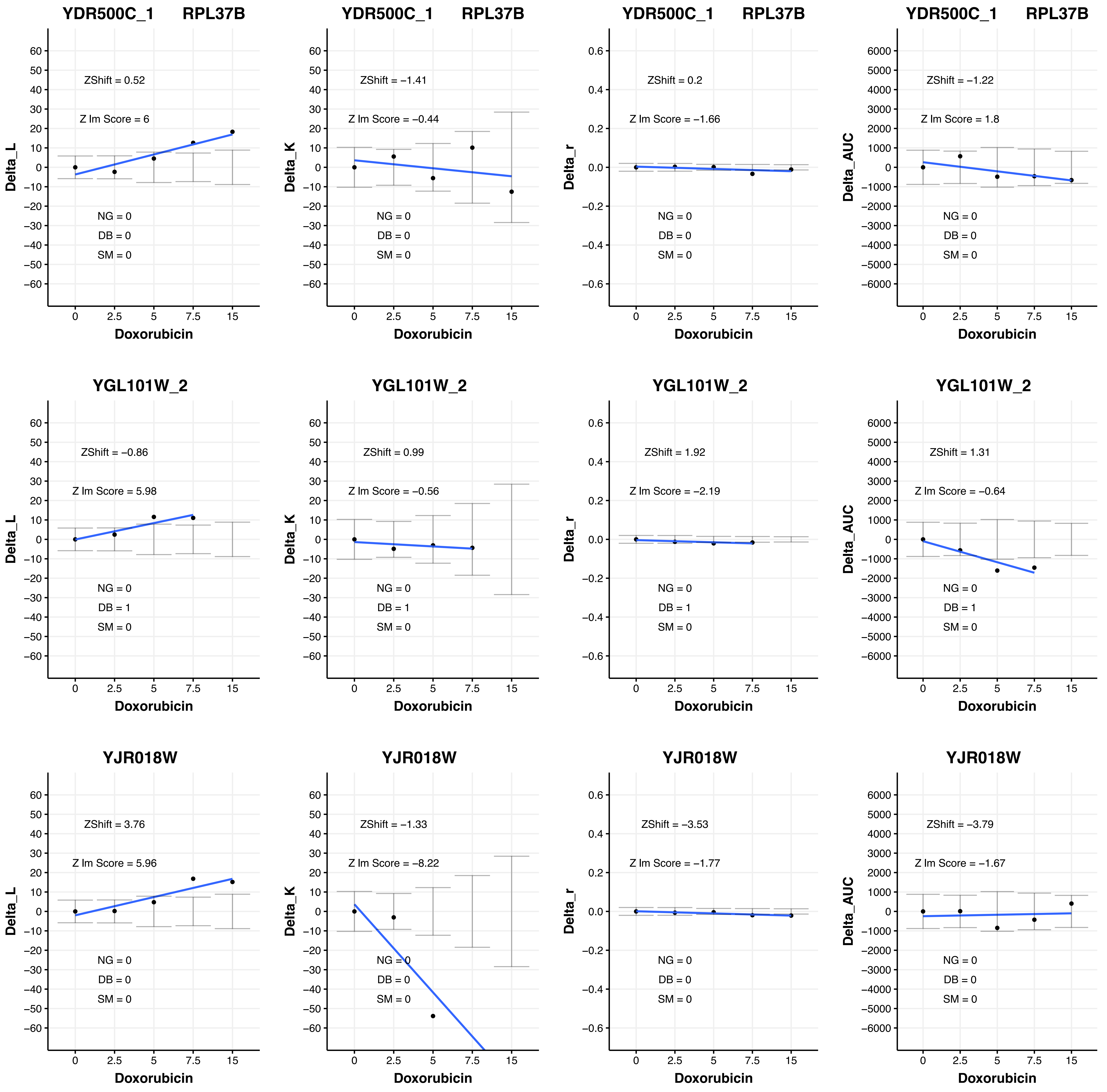
YDR500C_1 RPL37B
ZShift = 0.52
Z lm Score = 6
NG = 0
DB = 0
SM = 0
Delta_L
Doxorubicin
YDR500C_1 RPL37B
ZShift = −1.41
Z lm Score = −0.44
NG = 0
DB = 0
SM = 0
Delta_K
Doxorubicin
YDR500C_1 RPL37B
ZShift = 0.2
Z lm Score = −1.66
NG = 0
DB = 0
SM = 0
Delta_r
Doxorubicin
YDR500C_1 RPL37B
ZShift = −1.22
Z lm Score = 1.8
NG = 0
DB = 0
SM = 0
Delta_AUC
Doxorubicin
YGL101W_2
ZShift = −0.86
Z lm Score = 5.98
NG = 0
DB = 1
SM = 0
Delta_L
Doxorubicin
YGL101W_2
ZShift = 0.99
Z lm Score = −0.56
NG = 0
DB = 1
SM = 0
Delta_K
Doxorubicin
YGL101W_2
ZShift = 1.92
Z lm Score = −2.19
NG = 0
DB = 1
SM = 0
Delta_r
Doxorubicin
YGL101W_2
ZShift = 1.31
Z lm Score = −0.64
NG = 0
DB = 1
SM = 0
Delta_AUC
Doxorubicin
YJR018W
ZShift = 3.76
Z lm Score = 5.96
NG = 0
DB = 0
SM = 0
Delta_L
Doxorubicin
YJR018W
ZShift = −1.33
Z lm Score = −8.22
NG = 0
DB = 0
SM = 0
Delta_K
Doxorubicin
YJR018W
ZShift = −3.53
Z lm Score = −1.77
NG = 0
DB = 0
SM = 0
Delta_r
Doxorubicin
YJR018W
ZShift = −3.79
Z lm Score = −1.67
NG = 0
DB = 0
SM = 0
Delta_AUC
Doxorubicin

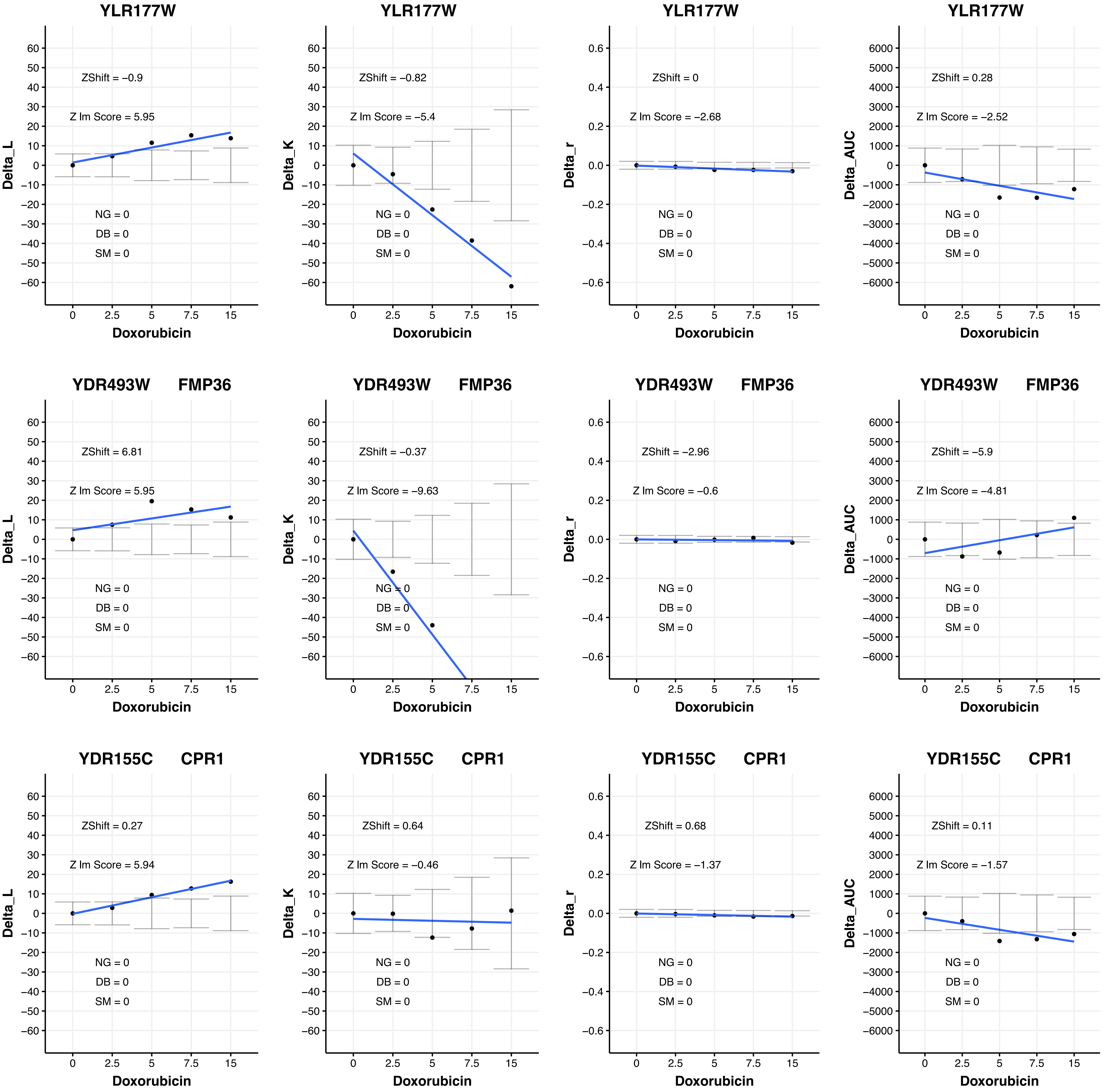
YLR177W
ZShift = −0.9
Z lm Score = 5.95
NG = 0
DB = 0
SM = 0
Delta_L
Doxorubicin
YLR177W
ZShift = −0.82
Z lm Score = −5.4
NG = 0
DB = 0
SM = 0
Delta_K
Doxorubicin
YLR177W
ZShift = 0
Z lm Score = −2.68
NG = 0
DB = 0
SM = 0
Delta_r
Doxorubicin
YLR177W
ZShift = 0.28
Z lm Score = −2.52
NG = 0
DB = 0
SM = 0
Delta_AUC
Doxorubicin
YDR493W FMP36
ZShift = 6.81
Z lm Score = 5.95
NG = 0
DB = 0
SM = 0
Delta_L
Doxorubicin
YDR493W FMP36
ZShift = −0.37
Z lm Score = −9.63
NG = 0
DB = 0
SM = 0
Delta_K
Doxorubicin
YDR493W FMP36
ZShift = −2.96
Z lm Score = −0.6
NG = 0
DB = 0
SM = 0
Delta_r
Doxorubicin
YDR493W FMP36
ZShift = −5.9
Z lm Score = −4.81
NG = 0
DB = 0
SM = 0
Delta_AUC
Doxorubicin
YDR155C CPR1
ZShift = 0.27
Z lm Score = 5.94
NG = 0
DB = 0
SM = 0
Delta_L
Doxorubicin
YDR155C CPR1
ZShift = 0.64
Z lm Score = −0.46
NG = 0
DB = 0
SM = 0
Delta_K
Doxorubicin
YDR155C CPR1
ZShift = 0.68
Z lm Score = −1.37
NG = 0
DB = 0
SM = 0
Delta_r
Doxorubicin
YDR155C CPR1
ZShift = 0.11
Z lm Score = −1.57
NG = 0
DB = 0
SM = 0
Delta_AUC
Doxorubicin

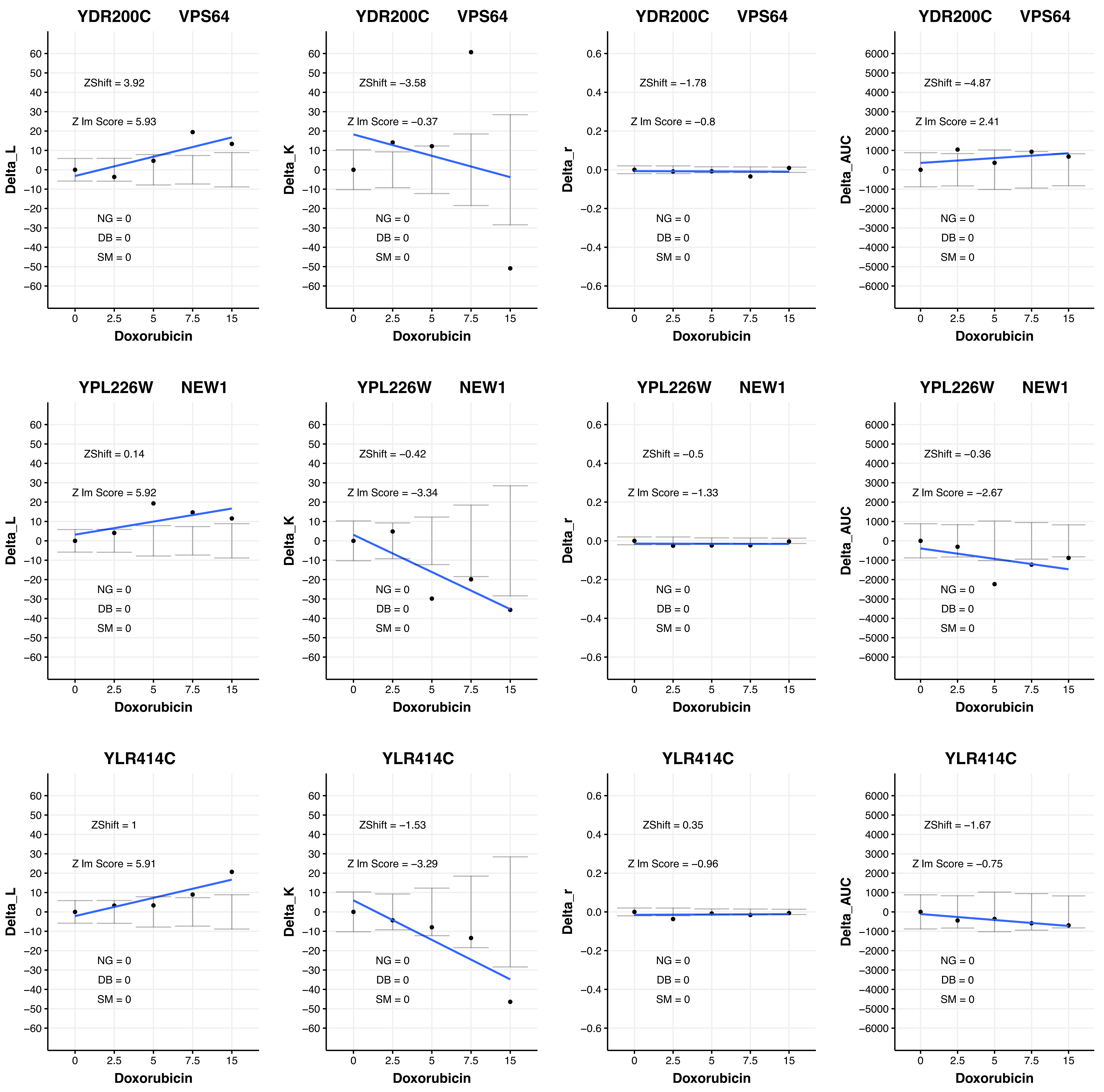

YDR200C VPS64
ZShift = 3.92
Z lm Score = 5.93
NG = 0
DB = 0
SM = 0
Delta_L
Doxorubicin
YDR200C VPS64
ZShift = −3.58
Z lm Score = −0.37
NG = 0
DB = 0
SM = 0
Delta_K
Doxorubicin
YDR200C VPS64
ZShift = −1.78
Z lm Score = −0.8
NG = 0
DB = 0
SM = 0
Delta_r
Doxorubicin
YDR200C VPS64
ZShift = −4.87
Z lm Score = 2.41
NG = 0
DB = 0
SM = 0
Delta_AUC
Doxorubicin
YPL226W NEW1
ZShift = 0.14
Z lm Score = 5.92
NG = 0
DB = 0
SM = 0
Delta_L
Doxorubicin
YPL226W NEW1
ZShift = −0.42
Z lm Score = −3.34
NG = 0
DB = 0
SM = 0
Delta_K
Doxorubicin
YPL226W NEW1
ZShift = −0.5
Z lm Score = −1.33
NG = 0
DB = 0
SM = 0
Delta_r
Doxorubicin
YPL226W NEW1
ZShift = −0.36
Z lm Score = −2.67
NG = 0
DB = 0
SM = 0
Delta_AUC
Doxorubicin
YLR414C
ZShift = 1
Z lm Score = 5.91
NG = 0
DB = 0
SM = 0
Delta_L
Doxorubicin
YLR414C
ZShift = −1.53
Z lm Score = −3.29
NG = 0
DB = 0
SM = 0
Delta_K
Doxorubicin
YLR414C
ZShift = 0.35
Z lm Score = −0.96
NG = 0
DB = 0
SM = 0
Delta_r
Doxorubicin
YLR414C
ZShift = −1.67
Z lm Score = −0.75
NG = 0
DB = 0
SM = 0
Delta_AUC
Doxorubicin

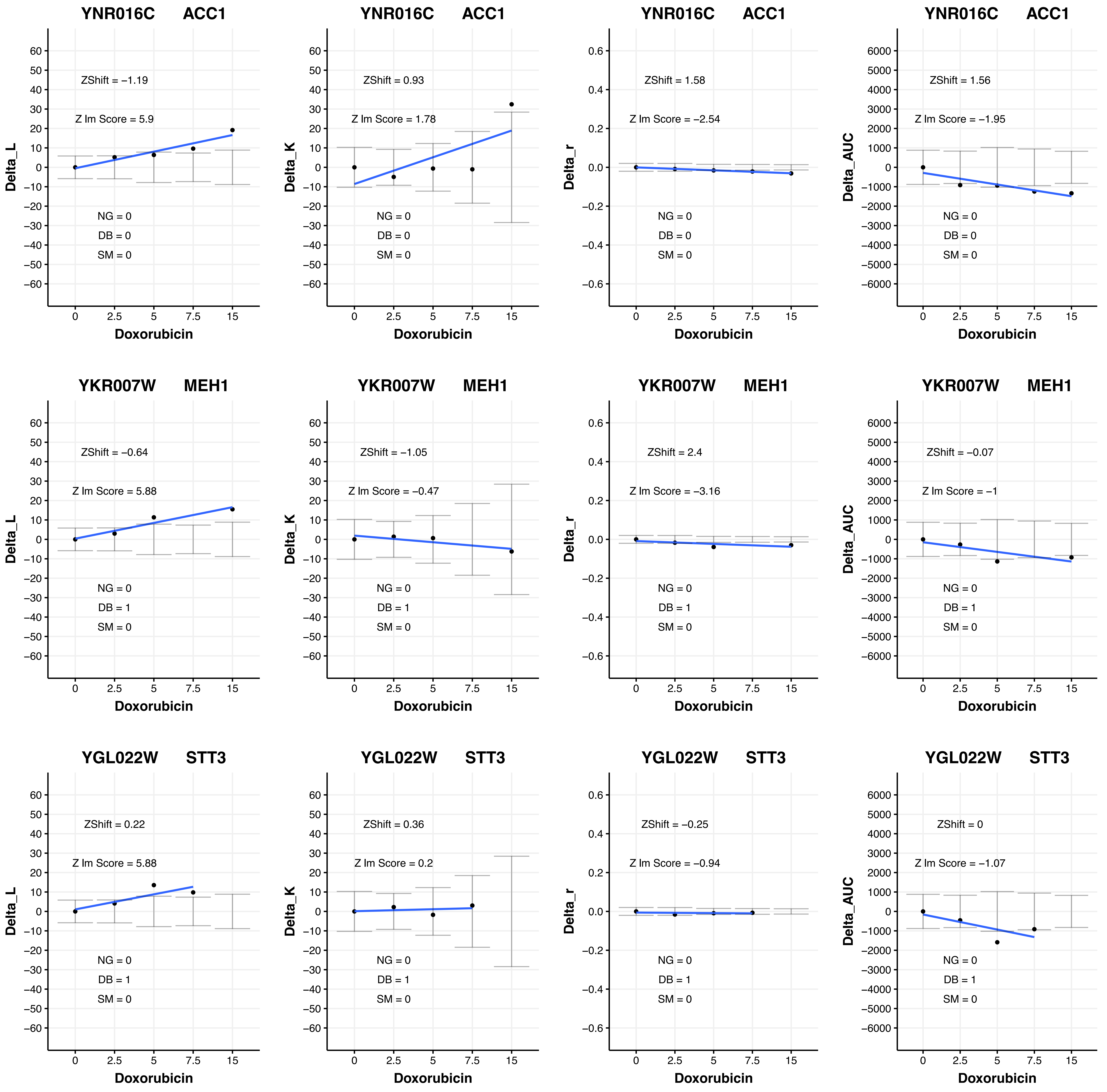

YNR016C ACC1
ZShift = −1.19
Z lm Score = 5.9
NG = 0
DB = 0
SM = 0
Delta_L
Doxorubicin
YNR016C ACC1
ZShift = 0.93
Z lm Score = 1.78
NG = 0
DB = 0
SM = 0
Delta_K
Doxorubicin
YNR016C ACC1
ZShift = 1.58
Z lm Score = −2.54
NG = 0
DB = 0
SM = 0
Delta_r
Doxorubicin
YNR016C ACC1
ZShift = 1.56
Z lm Score = −1.95
NG = 0
DB = 0
SM = 0
Delta_AUC
Doxorubicin
YKR007W MEH1
ZShift = −0.64
Z lm Score = 5.88
NG = 0
DB = 1
SM = 0
Delta_L
Doxorubicin
YKR007W MEH1
ZShift = −1.05
Z lm Score = −0.47
NG = 0
DB = 1
SM = 0
Delta_K
Doxorubicin
YKR007W MEH1
ZShift = 2.4
Z lm Score = −3.16
NG = 0
DB = 1
SM = 0
Delta_r
Doxorubicin
YKR007W MEH1
ZShift = −0.07
Z lm Score = −1
NG = 0
DB = 1
SM = 0
Delta_AUC
Doxorubicin
YGL022W STT3
ZShift = 0.22
Z lm Score = 5.88
NG = 0
DB = 1
SM = 0
Delta_L
Doxorubicin
YGL022W STT3
ZShift = 0.36
Z lm Score = 0.2
NG = 0
DB = 1
SM = 0
Delta_K
Doxorubicin
YGL022W STT3
ZShift = −0.25
Z lm Score = −0.94
NG = 0
DB = 1
SM = 0
Delta_r
Doxorubicin
YGL022W STT3
ZShift = 0
Z lm Score = −1.07
NG = 0
DB = 1
SM = 0
Delta_AUC
Doxorubicin

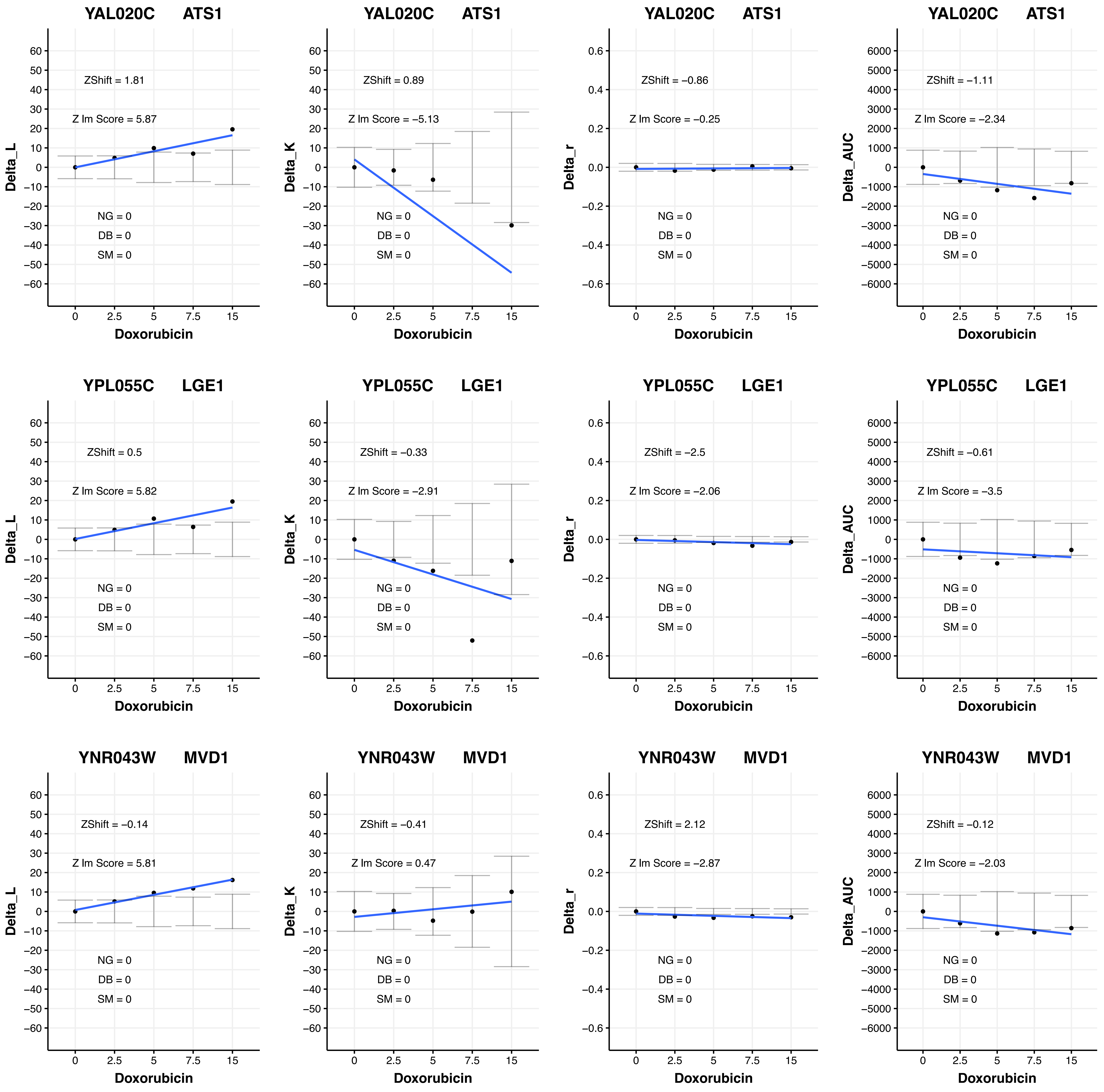
YAL020C ATS1
ZShift = 1.81
Z lm Score = 5.87
NG = 0
DB = 0
SM = 0
Delta_L
Doxorubicin
YAL020C ATS1
ZShift = 0.89
Z lm Score = −5.13
NG = 0
DB = 0
SM = 0
Delta_K
Doxorubicin
YAL020C ATS1
ZShift = −0.86
Z lm Score = −0.25
NG = 0
DB = 0
SM = 0
Delta_r
Doxorubicin
YAL020C ATS1
ZShift = −1.11
Z lm Score = −2.34
NG = 0
DB = 0
SM = 0
Delta_AUC
Doxorubicin
YPL055C LGE1
ZShift = 0.5
Z lm Score = 5.82
NG = 0
DB = 0
SM = 0
Delta_L
Doxorubicin
YPL055C LGE1
ZShift = −0.33
Z lm Score = −2.91
NG = 0
DB = 0
SM = 0
Delta_K
Doxorubicin
YPL055C LGE1
ZShift = −2.5
Z lm Score = −2.06
NG = 0
DB = 0
SM = 0
Delta_r
Doxorubicin
YPL055C LGE1
ZShift = −0.61
Z lm Score = −3.5
NG = 0
DB = 0
SM = 0
Delta_AUC
Doxorubicin
YNR043W MVD1
ZShift = −0.14
Z lm Score = 5.81
NG = 0
DB = 0
SM = 0
Delta_L
Doxorubicin
YNR043W MVD1
ZShift = −0.41
Z lm Score = 0.47
NG = 0
DB = 0
SM = 0
Delta_K
Doxorubicin
YNR043W MVD1
ZShift = 2.12
Z lm Score = −2.87
NG = 0
DB = 0
SM = 0
Delta_r
Doxorubicin
YNR043W MVD1
ZShift = −0.12
Z lm Score = −2.03
NG = 0
DB = 0
SM = 0
Delta_AUC
Doxorubicin

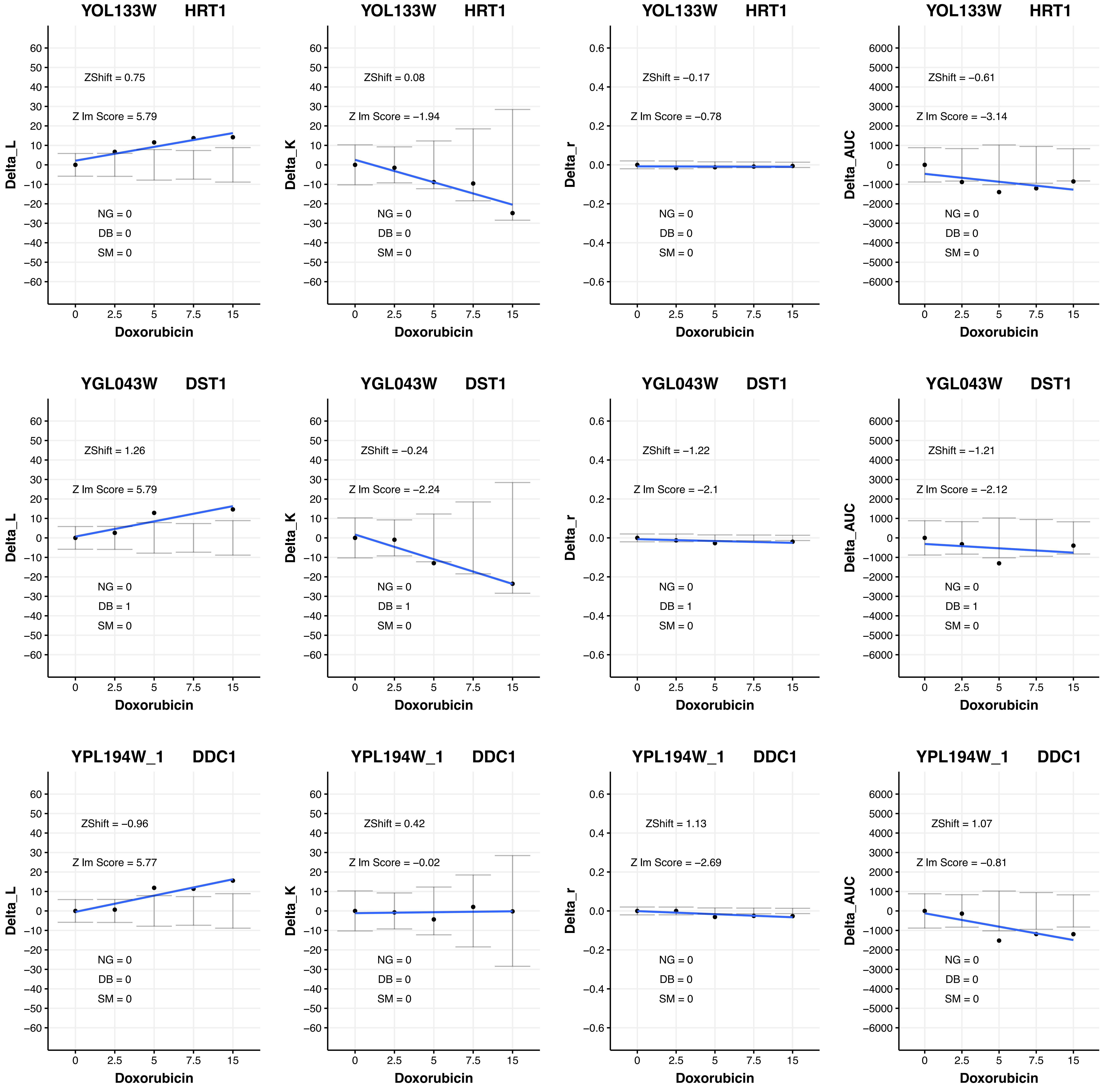
YOL133W HRT1
ZShift = 0.75
Z lm Score = 5.79
NG = 0
DB = 0
SM = 0
Delta_L
Doxorubicin
YOL133W HRT1
ZShift = 0.08
Z lm Score = −1.94
NG = 0
DB = 0
SM = 0
Delta_K
Doxorubicin
YOL133W HRT1
ZShift = −0.17
Z lm Score = −0.78
NG = 0
DB = 0
SM = 0
Delta_r
Doxorubicin
YOL133W HRT1
ZShift = −0.61
Z lm Score = −3.14
NG = 0
DB = 0
SM = 0
Delta_AUC
Doxorubicin
YGL043W DST1
ZShift = 1.26
Z lm Score = 5.79
NG = 0
DB = 1
SM = 0
Delta_L
Doxorubicin
YGL043W DST1
ZShift = −0.24
Z lm Score = −2.24
NG = 0
DB = 1
SM = 0
Delta_K
Doxorubicin
YGL043W DST1
ZShift = −1.22
Z lm Score = −2.1
NG = 0
DB = 1
SM = 0
Delta_r
Doxorubicin
YGL043W DST1
ZShift = −1.21
Z lm Score = −2.12
NG = 0
DB = 1
SM = 0
Delta_AUC
Doxorubicin
YPL194W_1 DDC1
ZShift = −0.96
Z lm Score = 5.77
NG = 0
DB = 0
SM = 0
Delta_L
Doxorubicin
YPL194W_1 DDC1
ZShift = 0.42
Z lm Score = −0.02
NG = 0
DB = 0
SM = 0
Delta_K
Doxorubicin
YPL194W_1 DDC1
ZShift = 1.13
Z lm Score = −2.69
NG = 0
DB = 0
SM = 0
Delta_r
Doxorubicin
YPL194W_1 DDC1
ZShift = 1.07
Z lm Score = −0.81
NG = 0
DB = 0
SM = 0
Delta_AUC
Doxorubicin

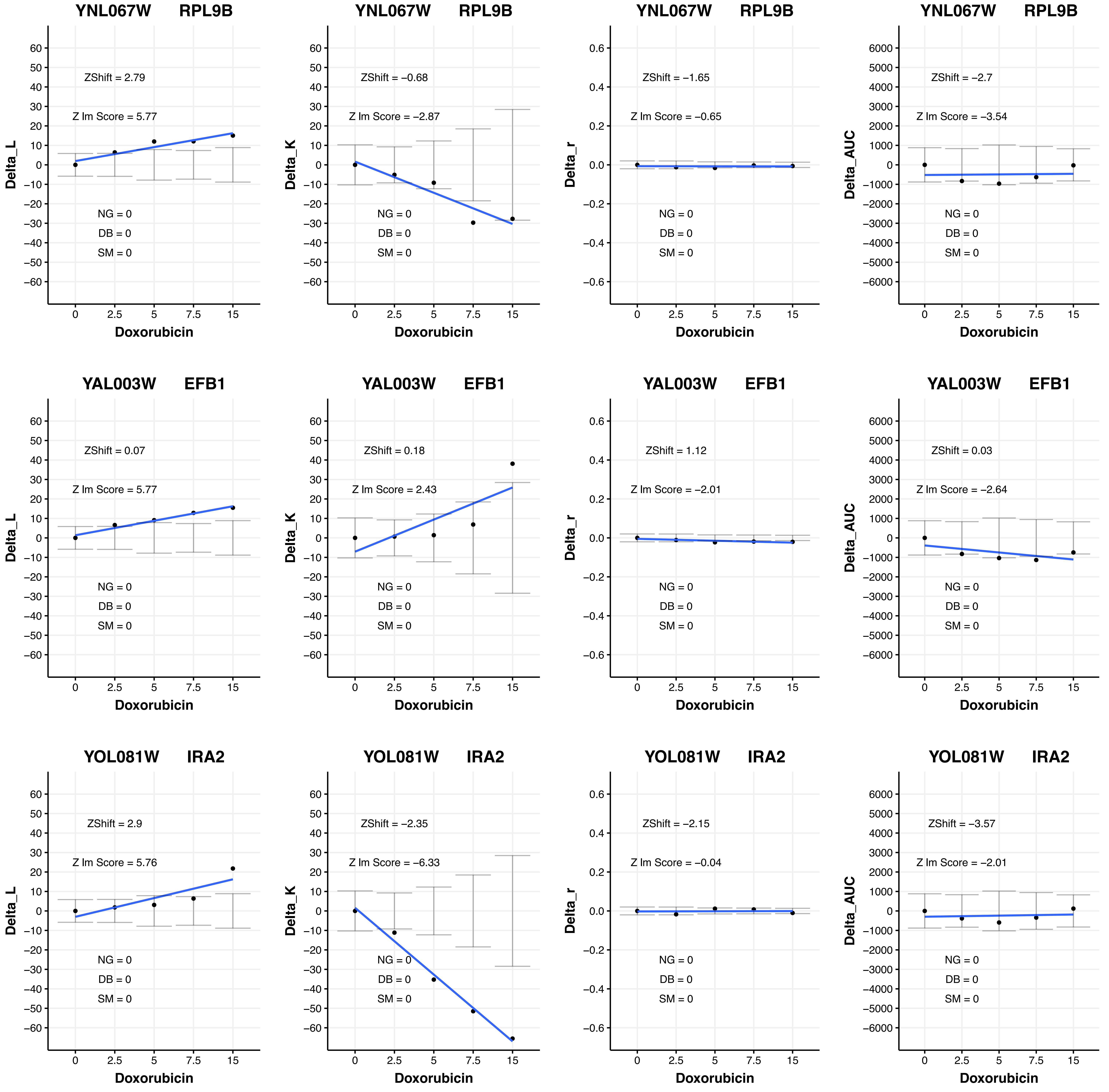

YNL067W RPL9B
ZShift = 2.79
Z lm Score = 5.77
NG = 0
DB = 0
SM = 0
Delta_L
Doxorubicin
YNL067W RPL9B
ZShift = −0.68
Z lm Score = −2.87
NG = 0
DB = 0
SM = 0
Delta_K
Doxorubicin
YNL067W RPL9B
ZShift = −1.65
Z lm Score = −0.65
NG = 0
DB = 0
SM = 0
Delta_r
Doxorubicin
YNL067W RPL9B
ZShift = −2.7
Z lm Score = −3.54
NG = 0
DB = 0
SM = 0
Delta_AUC
Doxorubicin
YAL003W EFB1
ZShift = 0.07
Z lm Score = 5.77
NG = 0
DB = 0
SM = 0
Delta_L
Doxorubicin
YAL003W EFB1
ZShift = 0.18
Z lm Score = 2.43
NG = 0
DB = 0
SM = 0
Delta_K
Doxorubicin
YAL003W EFB1
ZShift = 1.12
Z lm Score = −2.01
NG = 0
DB = 0
SM = 0
Delta_r
Doxorubicin
YAL003W EFB1
ZShift = 0.03
Z lm Score = −2.64
NG = 0
DB = 0
SM = 0
Delta_AUC
Doxorubicin
YOL081W IRA2
ZShift = 2.9
Z lm Score = 5.76
NG = 0
DB = 0
SM = 0
Delta_L
Doxorubicin
YOL081W IRA2
ZShift = −2.35
Z lm Score = −6.33
NG = 0
DB = 0
SM = 0
Delta_K
Doxorubicin
YOL081W IRA2
ZShift = −2.15
Z lm Score = −0.04
NG = 0
DB = 0
SM = 0
Delta_r
Doxorubicin
YOL081W IRA2
ZShift = −3.57
Z lm Score = −2.01
NG = 0
DB = 0
SM = 0
Delta_AUC
Doxorubicin

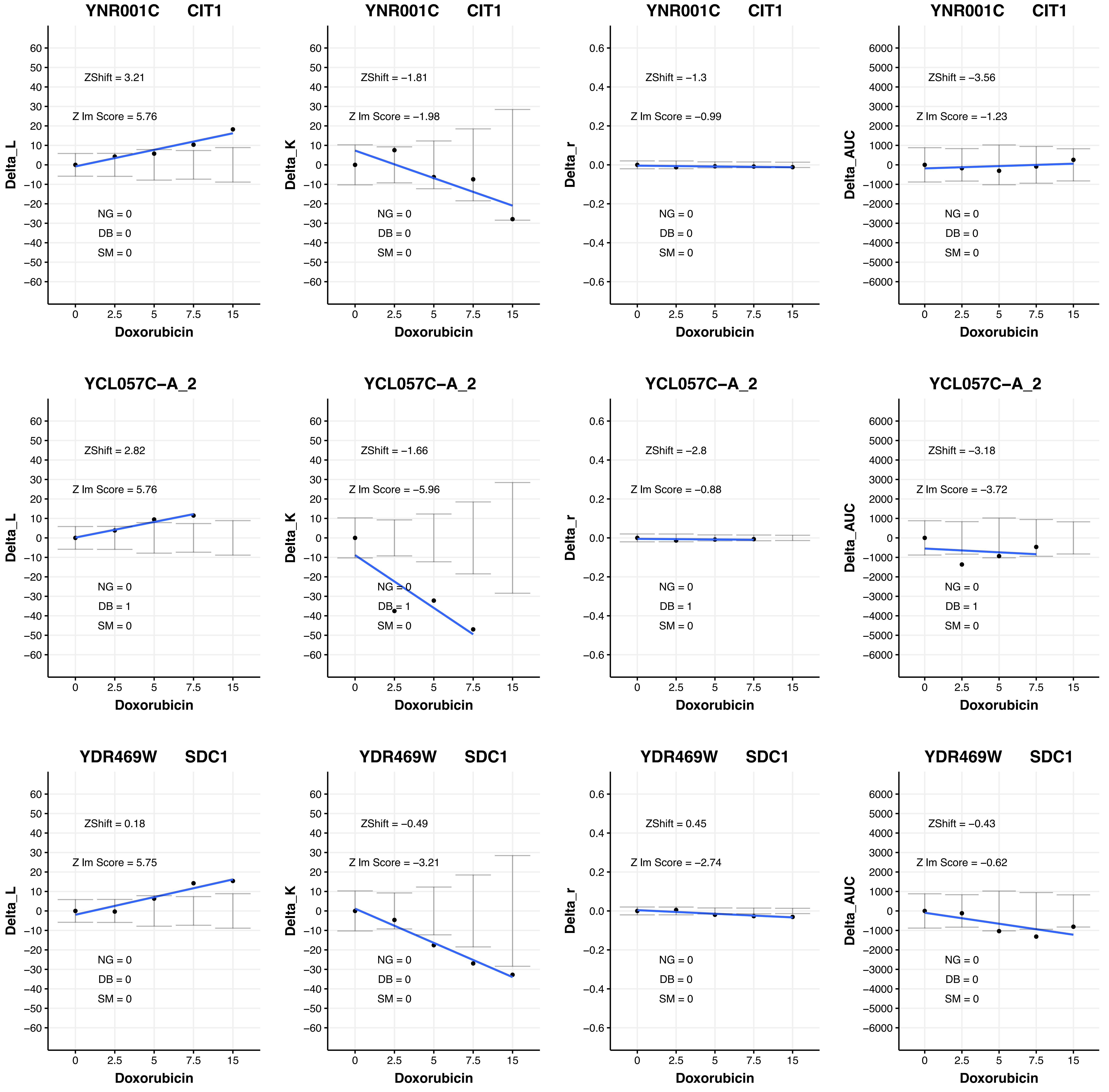

YNR001C CIT1
ZShift = 3.21
Z lm Score = 5.76
NG = 0
DB = 0
SM = 0
Delta_L
Doxorubicin
YNR001C CIT1
ZShift = −1.81
Z lm Score = −1.98
NG = 0
DB = 0
SM = 0
Delta_K
Doxorubicin
YNR001C CIT1
ZShift = −1.3
Z lm Score = −0.99
NG = 0
DB = 0
SM = 0
Delta_r
Doxorubicin
YNR001C CIT1
ZShift = −3.56
Z lm Score = −1.23
NG = 0
DB = 0
SM = 0
Delta_AUC
Doxorubicin
YCL057C−A_2
ZShift = 2.82
Z lm Score = 5.76
NG = 0
DB = 1
SM = 0
Delta_L
Doxorubicin
YCL057C−A_2
ZShift = −1.66
Z lm Score = −5.96
NG = 0
DB = 1
SM = 0
Delta_K
Doxorubicin
YCL057C−A_2
ZShift = −2.8
Z lm Score = −0.88
NG = 0
DB = 1
SM = 0
Delta_r
Doxorubicin
YCL057C−A_2
ZShift = −3.18
Z lm Score = −3.72
NG = 0
DB = 1
SM = 0
Delta_AUC
Doxorubicin
YDR469W SDC1
ZShift = 0.18
Z lm Score = 5.75
NG = 0
DB = 0
SM = 0
Delta_L
Doxorubicin
YDR469W SDC1
ZShift = −0.49
Z lm Score = −3.21
NG = 0
DB = 0
SM = 0
Delta_K
Doxorubicin
YDR469W SDC1
ZShift = 0.45
Z lm Score = −2.74
NG = 0
DB = 0
SM = 0
Delta_r
Doxorubicin
YDR469W SDC1
ZShift = −0.43
Z lm Score = −0.62
NG = 0
DB = 0
SM = 0
Delta_AUC
Doxorubicin

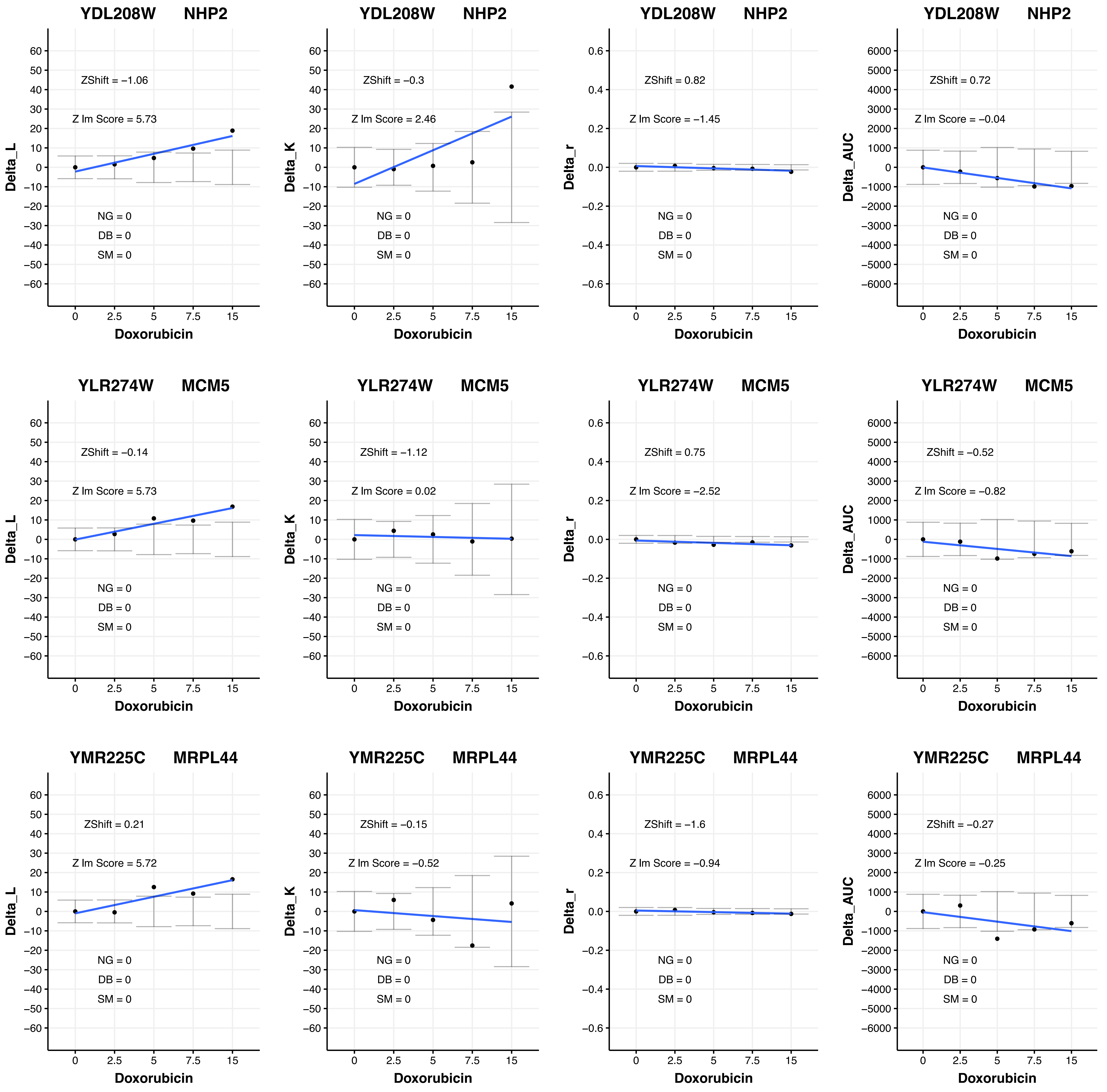
YDL208W NHP2
ZShift = −1.06
Z lm Score = 5.73
NG = 0
DB = 0
SM = 0
Delta_L
Doxorubicin
YDL208W NHP2
ZShift = −0.3
Z lm Score = 2.46
NG = 0
DB = 0
SM = 0
Delta_K
Doxorubicin
YDL208W NHP2
ZShift = 0.82
Z lm Score = −1.45
NG = 0
DB = 0
SM = 0
Delta_r
Doxorubicin
YDL208W NHP2
ZShift = 0.72
Z lm Score = −0.04
NG = 0
DB = 0
SM = 0
Delta_AUC
Doxorubicin
YLR274W MCM5
ZShift = −0.14
Z lm Score = 5.73
NG = 0
DB = 0
SM = 0
Delta_L
Doxorubicin
YLR274W MCM5
ZShift = −1.12
Z lm Score = 0.02
NG = 0
DB = 0
SM = 0
Delta_K
Doxorubicin
YLR274W MCM5
ZShift = 0.75
Z lm Score = −2.52
NG = 0
DB = 0
SM = 0
Delta_r
Doxorubicin
YLR274W MCM5
ZShift = −0.52
Z lm Score = −0.82
NG = 0
DB = 0
SM = 0
Delta_AUC
Doxorubicin
YMR225C MRPL44
ZShift = 0.21
Z lm Score = 5.72
NG = 0
DB = 0
SM = 0
Delta_L
Doxorubicin
YMR225C MRPL44
ZShift = −0.15
Z lm Score = −0.52
NG = 0
DB = 0
SM = 0
Delta_K
Doxorubicin
YMR225C MRPL44
ZShift = −1.6
Z lm Score = −0.94
NG = 0
DB = 0
SM = 0
Delta_r
Doxorubicin
YMR225C MRPL44
ZShift = −0.27
Z lm Score = −0.25
NG = 0
DB = 0
SM = 0
Delta_AUC
Doxorubicin

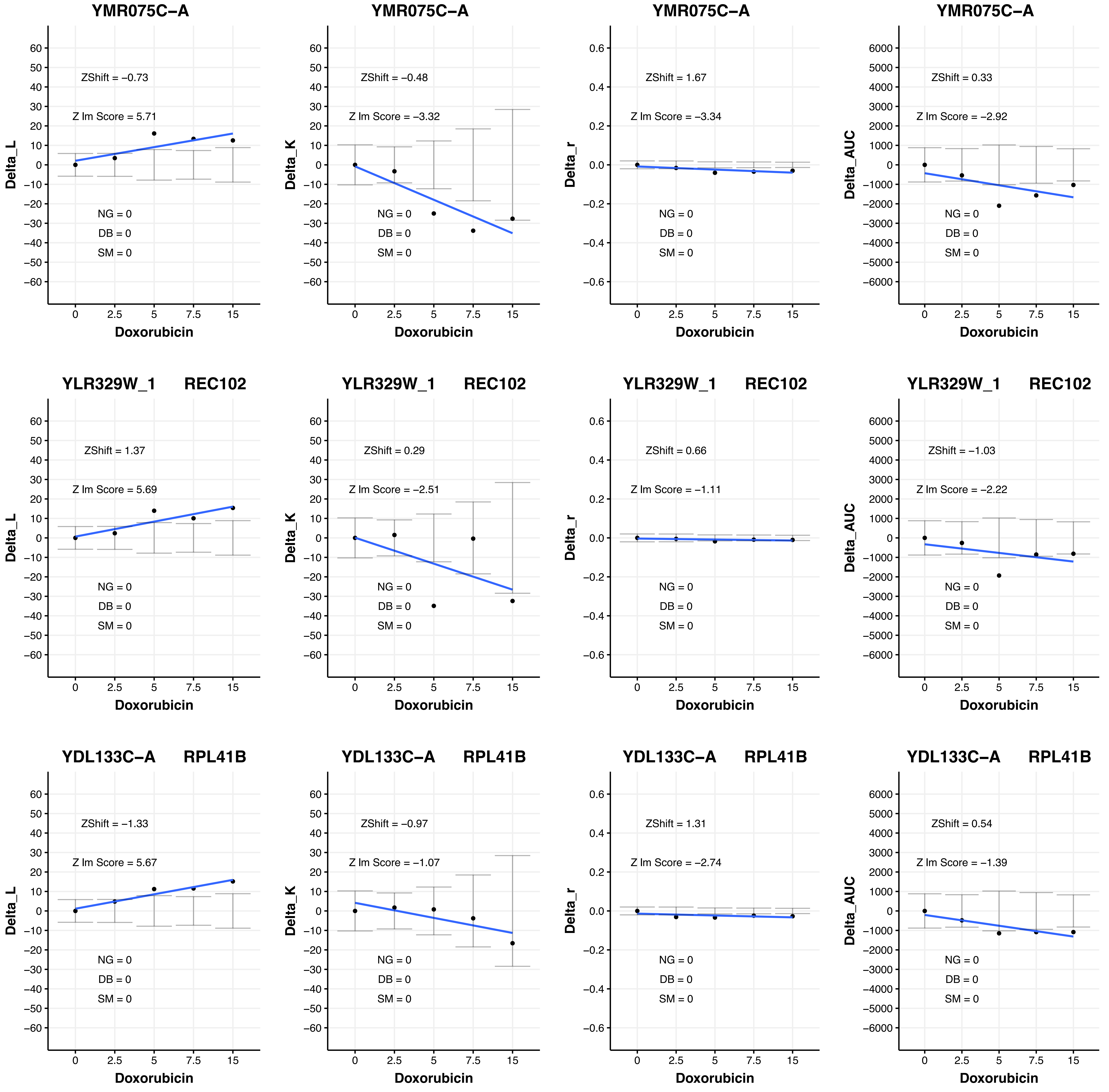
YMR075C−A
ZShift = −0.73
Z lm Score = 5.71
NG = 0
DB = 0
SM = 0
Delta_L
Doxorubicin
YMR075C−A
ZShift = −0.48
Z lm Score = −3.32
NG = 0
DB = 0
SM = 0
Delta_K
Doxorubicin
YMR075C−A
ZShift = 1.67
Z lm Score = −3.34
NG = 0
DB = 0
SM = 0
Delta_r
Doxorubicin
YMR075C−A
ZShift = 0.33
Z lm Score = −2.92
NG = 0
DB = 0
SM = 0
Delta_AUC
Doxorubicin
YLR329W_1 REC102
ZShift = 1.37
Z lm Score = 5.69
NG = 0
DB = 0
SM = 0
Delta_L
Doxorubicin
YLR329W_1 REC102
ZShift = 0.29
Z lm Score = −2.51
NG = 0
DB = 0
SM = 0
Delta_K
Doxorubicin
YLR329W_1 REC102
ZShift = 0.66
Z lm Score = −1.11
NG = 0
DB = 0
SM = 0
Delta_r
Doxorubicin
YLR329W_1 REC102
ZShift = −1.03
Z lm Score = −2.22
NG = 0
DB = 0
SM = 0
Delta_AUC
Doxorubicin
YDL133C−A RPL41B
ZShift = −1.33
Z lm Score = 5.67
NG = 0
DB = 0
SM = 0
Delta_L
Doxorubicin
YDL133C−A RPL41B
ZShift = −0.97
Z lm Score = −1.07
NG = 0
DB = 0
SM = 0
Delta_K
Doxorubicin
YDL133C−A RPL41B
ZShift = 1.31
Z lm Score = −2.74
NG = 0
DB = 0
SM = 0
Delta_r
Doxorubicin
YDL133C−A RPL41B
ZShift = 0.54
Z lm Score = −1.39
NG = 0
DB = 0
SM = 0
Delta_AUC
Doxorubicin

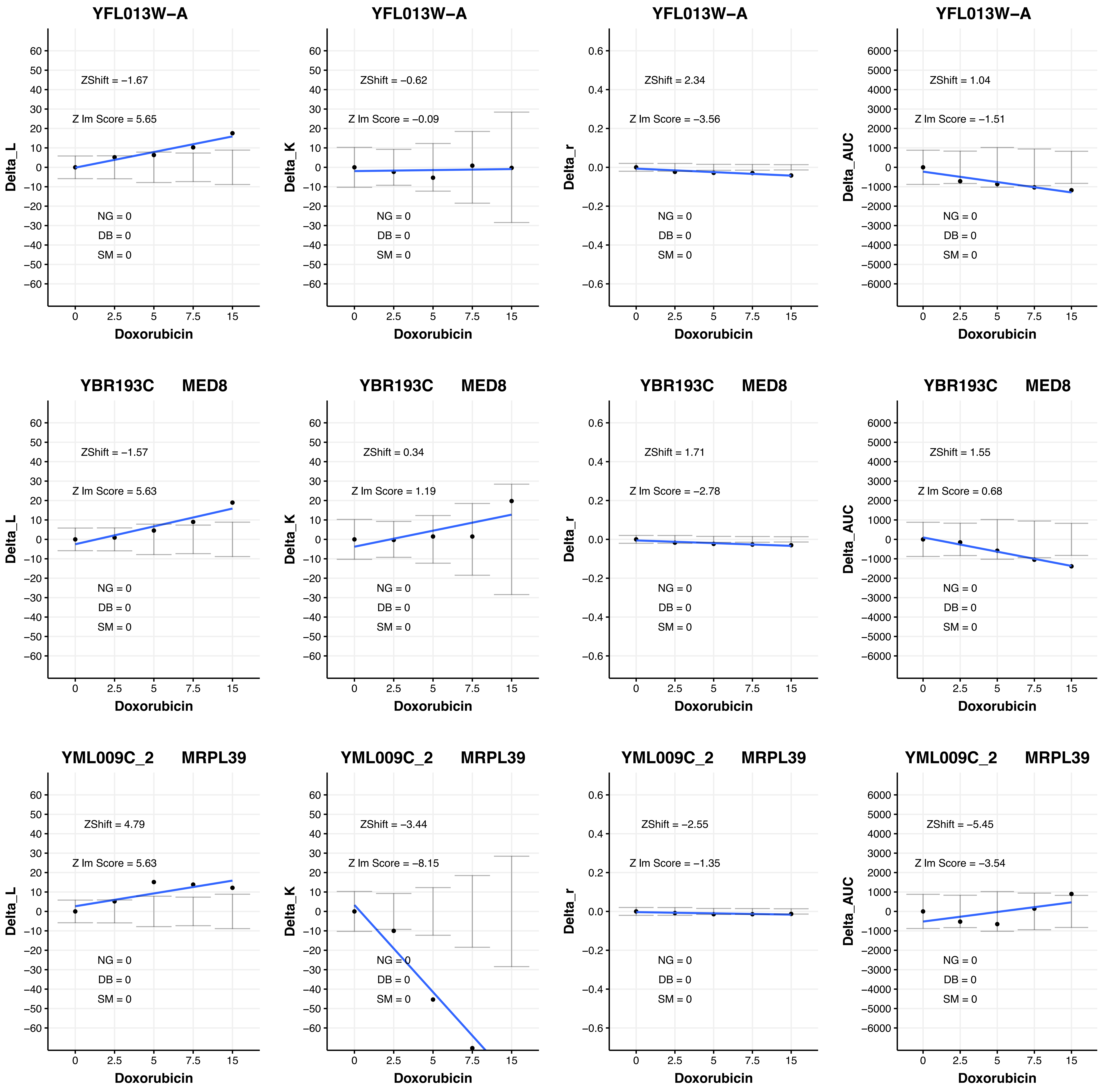
YFL013W−A
ZShift = −1.67
Z lm Score = 5.65
NG = 0
DB = 0
SM = 0
Delta_L
Doxorubicin
YFL013W−A
ZShift = −0.62
Z lm Score = −0.09
NG = 0
DB = 0
SM = 0
Delta_K
Doxorubicin
YFL013W−A
ZShift = 2.34
Z lm Score = −3.56
NG = 0
DB = 0
SM = 0
Delta_r
Doxorubicin
YFL013W−A
ZShift = 1.04
Z lm Score = −1.51
NG = 0
DB = 0
SM = 0
Delta_AUC
Doxorubicin
YBR193C MED8
ZShift = −1.57
Z lm Score = 5.63
NG = 0
DB = 0
SM = 0
Delta_L
Doxorubicin
YBR193C MED8
ZShift = 0.34
Z lm Score = 1.19
NG = 0
DB = 0
SM = 0
Delta_K
Doxorubicin
YBR193C MED8
ZShift = 1.71
Z lm Score = −2.78
NG = 0
DB = 0
SM = 0
Delta_r
Doxorubicin
YBR193C MED8
ZShift = 1.55
Z lm Score = 0.68
NG = 0
DB = 0
SM = 0
Delta_AUC
Doxorubicin
YML009C_2 MRPL39
ZShift = 4.79
Z lm Score = 5.63
NG = 0
DB = 0
SM = 0
Delta_L
Doxorubicin
YML009C_2 MRPL39
ZShift = −3.44
Z lm Score = −8.15
NG = 0
DB = 0
SM = 0
Delta_K
Doxorubicin
YML009C_2 MRPL39
ZShift = −2.55
Z lm Score = −1.35
NG = 0
DB = 0
SM = 0
Delta_r
Doxorubicin
YML009C_2 MRPL39
ZShift = −5.45
Z lm Score = −3.54
NG = 0
DB = 0
SM = 0
Delta_AUC
Doxorubicin

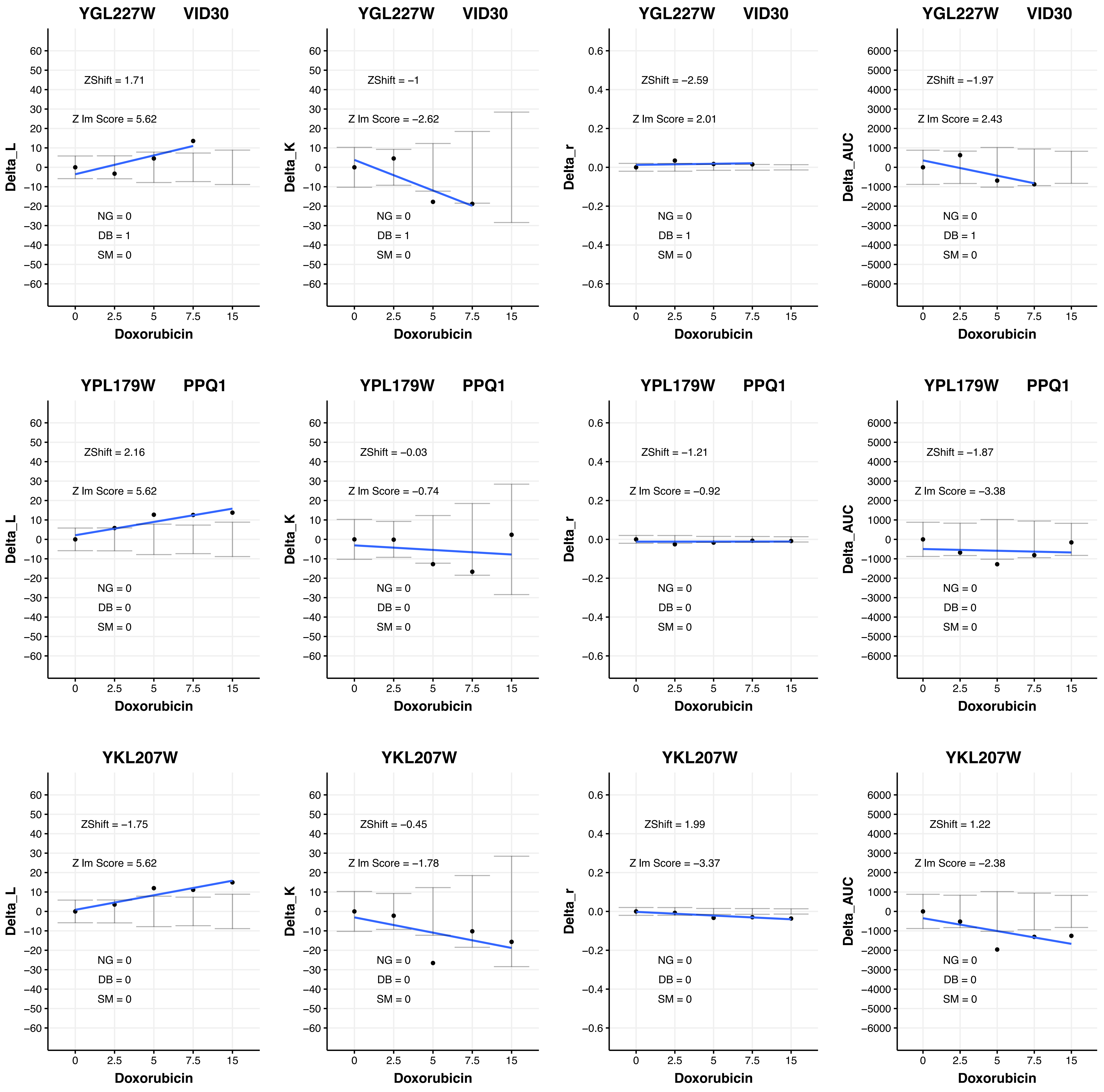

YGL227W VID30
ZShift = 1.71
Z lm Score = 5.62
NG = 0
DB = 1
SM = 0
Delta_L
Doxorubicin
YGL227W VID30
ZShift = −1
Z lm Score = −2.62
NG = 0
DB = 1
SM = 0
Delta_K
Doxorubicin
YGL227W VID30
ZShift = −2.59
Z lm Score = 2.01
NG = 0
DB = 1
SM = 0
Delta_r
Doxorubicin
YGL227W VID30
ZShift = −1.97
Z lm Score = 2.43
NG = 0
DB = 1
SM = 0
Delta_AUC
Doxorubicin
YPL179W PPQ1
ZShift = 2.16
Z lm Score = 5.62
NG = 0
DB = 0
SM = 0
Delta_L
Doxorubicin
YPL179W PPQ1
ZShift = −0.03
Z lm Score = −0.74
NG = 0
DB = 0
SM = 0
Delta_K
Doxorubicin
YPL179W PPQ1
ZShift = −1.21
Z lm Score = −0.92
NG = 0
DB = 0
SM = 0
Delta_r
Doxorubicin
YPL179W PPQ1
ZShift = −1.87
Z lm Score = −3.38
NG = 0
DB = 0
SM = 0
Delta_AUC
Doxorubicin
YKL207W
ZShift = −1.75
Z lm Score = 5.62
NG = 0
DB = 0
SM = 0
Delta_L
Doxorubicin
YKL207W
ZShift = −0.45
Z lm Score = −1.78
NG = 0
DB = 0
SM = 0
Delta_K
Doxorubicin
YKL207W
ZShift = 1.99
Z lm Score = −3.37
NG = 0
DB = 0
SM = 0
Delta_r
Doxorubicin
YKL207W
ZShift = 1.22
Z lm Score = −2.38
NG = 0
DB = 0
SM = 0
Delta_AUC
Doxorubicin

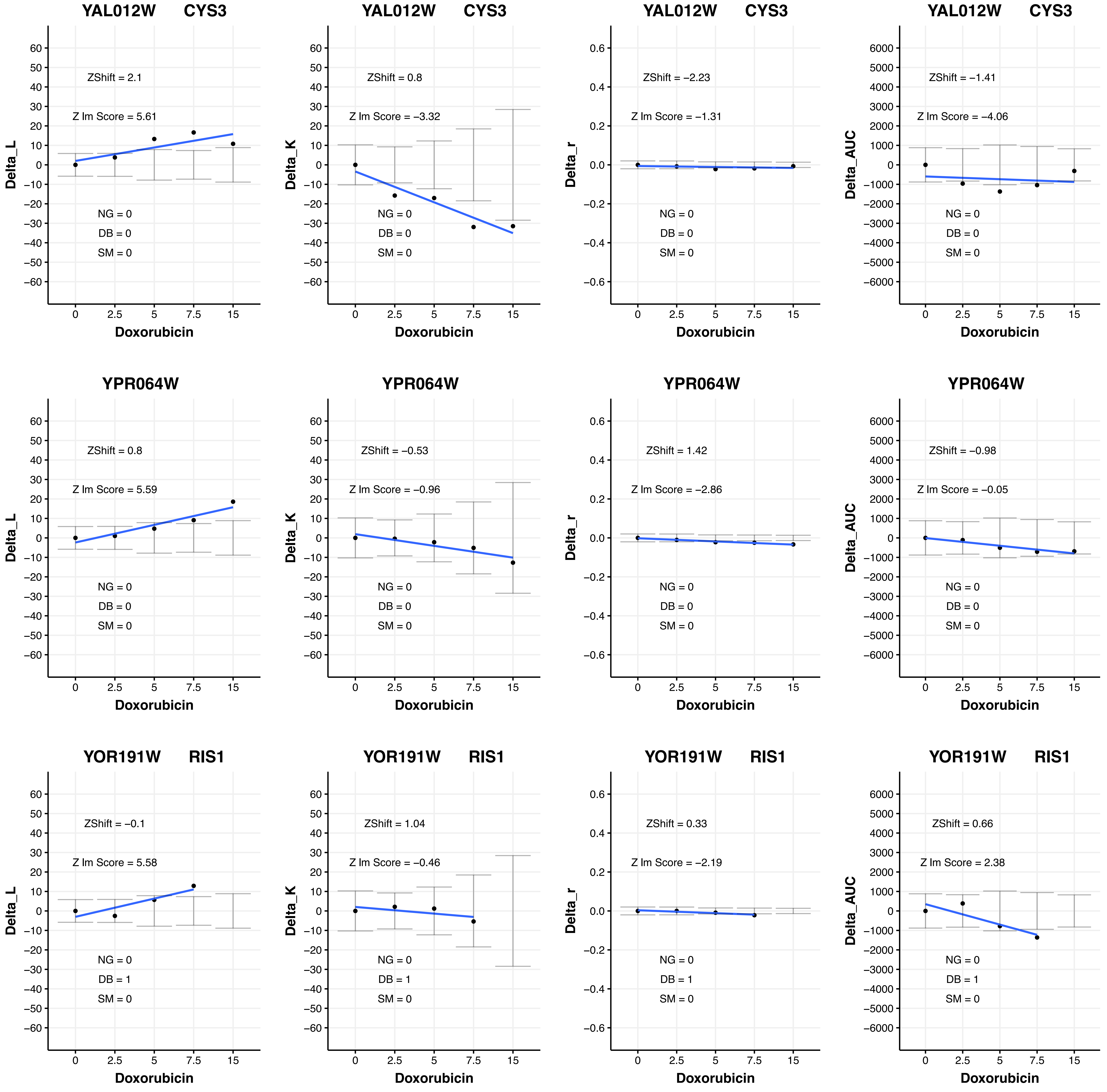
YAL012W CYS3
ZShift = 2.1
Z lm Score = 5.61
NG = 0
DB = 0
SM = 0
Delta_L
Doxorubicin
YAL012W CYS3
ZShift = 0.8
Z lm Score = −3.32
NG = 0
DB = 0
SM = 0
Delta_K
Doxorubicin
YAL012W CYS3
ZShift = −2.23
Z lm Score = −1.31
NG = 0
DB = 0
SM = 0
Delta_r
Doxorubicin
YAL012W CYS3
ZShift = −1.41
Z lm Score = −4.06
NG = 0
DB = 0
SM = 0
Delta_AUC
Doxorubicin
YPR064W
ZShift = 0.8
Z lm Score = 5.59
NG = 0
DB = 0
SM = 0
Delta_L
Doxorubicin
YPR064W
ZShift = −0.53
Z lm Score = −0.96
NG = 0
DB = 0
SM = 0
Delta_K
Doxorubicin
YPR064W
ZShift = 1.42
Z lm Score = −2.86
NG = 0
DB = 0
SM = 0
Delta_r
Doxorubicin
YPR064W
ZShift = −0.98
Z lm Score = −0.05
NG = 0
DB = 0
SM = 0
Delta_AUC
Doxorubicin
YOR191W RIS1
ZShift = −0.1
Z lm Score = 5.58
NG = 0
DB = 1
SM = 0
Delta_L
Doxorubicin
YOR191W RIS1
ZShift = 1.04
Z lm Score = −0.46
NG = 0
DB = 1
SM = 0
Delta_K
Doxorubicin
YOR191W RIS1
ZShift = 0.33
Z lm Score = −2.19
NG = 0
DB = 1
SM = 0
Delta_r
Doxorubicin
YOR191W RIS1
ZShift = 0.66
Z lm Score = 2.38
NG = 0
DB = 1
SM = 0
Delta_AUC
Doxorubicin

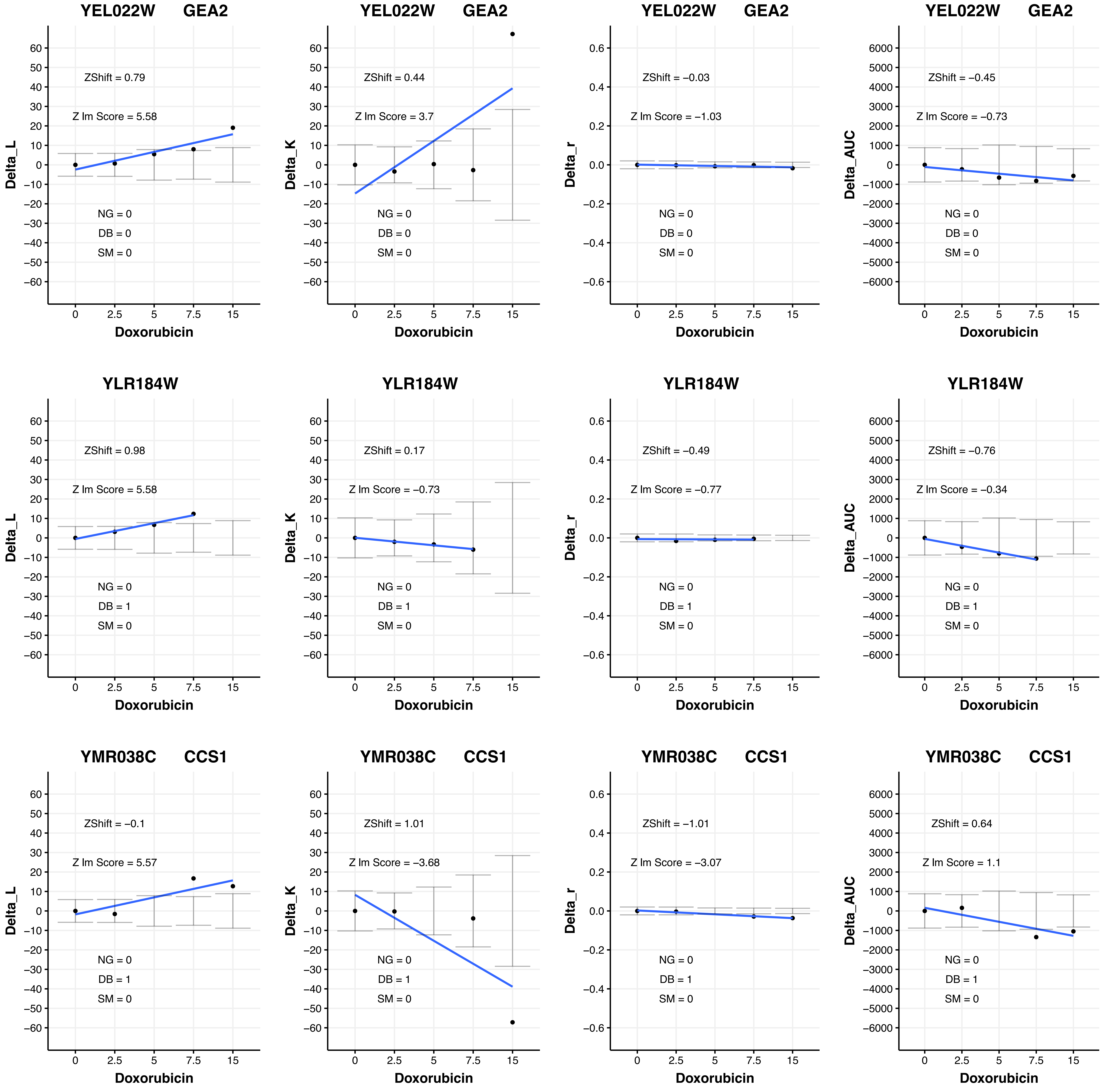
YEL022W GEA2
ZShift = 0.79
Z lm Score = 5.58
NG = 0
DB = 0
SM = 0
Delta_L
Doxorubicin
YEL022W GEA2
ZShift = 0.44
Z lm Score = 3.7
NG = 0
DB = 0
SM = 0
Delta_K
Doxorubicin
YEL022W GEA2
ZShift = −0.03
Z lm Score = −1.03
NG = 0
DB = 0
SM = 0
Delta_r
Doxorubicin
YEL022W GEA2
ZShift = −0.45
Z lm Score = −0.73
NG = 0
DB = 0
SM = 0
Delta_AUC
Doxorubicin
YLR184W
ZShift = 0.98
Z lm Score = 5.58
NG = 0
DB = 1
SM = 0
Delta_L
Doxorubicin
YLR184W
ZShift = 0.17
Z lm Score = −0.73
NG = 0
DB = 1
SM = 0
Delta_K
Doxorubicin
YLR184W
ZShift = −0.49
Z lm Score = −0.77
NG = 0
DB = 1
SM = 0
Delta_r
Doxorubicin
YLR184W
ZShift = −0.76
Z lm Score = −0.34
NG = 0
DB = 1
SM = 0
Delta_AUC
Doxorubicin
YMR038C CCS1
ZShift = −0.1
Z lm Score = 5.57
NG = 0
DB = 1
SM = 0
Delta_L
Doxorubicin
YMR038C CCS1
ZShift = 1.01
Z lm Score = −3.68
NG = 0
DB = 1
SM = 0
Delta_K
Doxorubicin
YMR038C CCS1
ZShift = −1.01
Z lm Score = −3.07
NG = 0
DB = 1
SM = 0
Delta_r
Doxorubicin
YMR038C CCS1
ZShift = 0.64
Z lm Score = 1.1
NG = 0
DB = 1
SM = 0
Delta_AUC
Doxorubicin

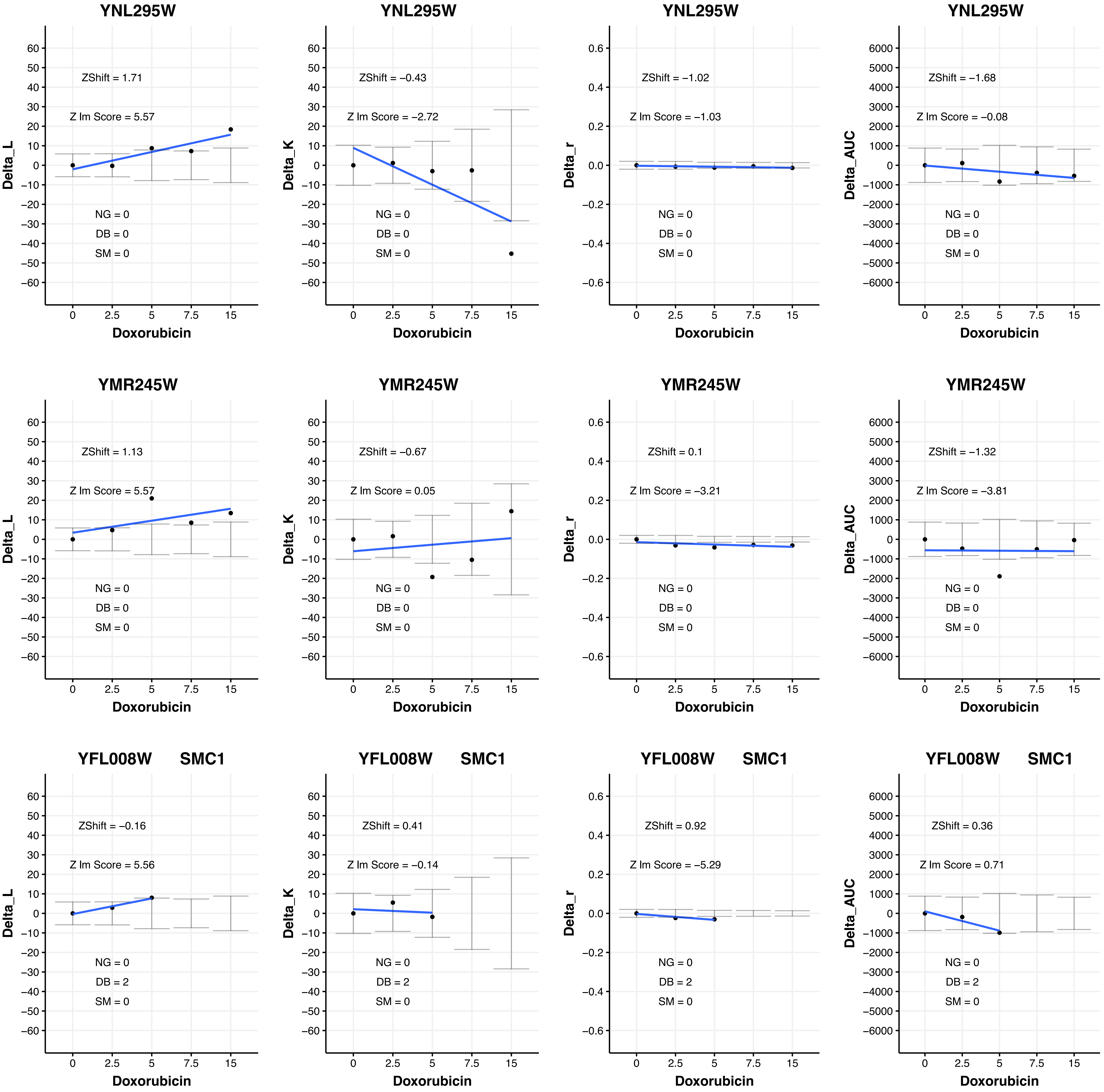
YNL295W
ZShift = 1.71
Z lm Score = 5.57
NG = 0
DB = 0
SM = 0
Delta_L
Doxorubicin
YNL295W
ZShift = −0.43
Z lm Score = −2.72
NG = 0
DB = 0
SM = 0
Delta_K
Doxorubicin
YNL295W
ZShift = −1.02
Z lm Score = −1.03
NG = 0
DB = 0
SM = 0
Delta_r
Doxorubicin
YNL295W
ZShift = −1.68
Z lm Score = −0.08
NG = 0
DB = 0
SM = 0
Delta_AUC
Doxorubicin
YMR245W
ZShift = 1.13
Z lm Score = 5.57
NG = 0
DB = 0
SM = 0
Delta_L
Doxorubicin
YMR245W
ZShift = −0.67
Z lm Score = 0.05
NG = 0
DB = 0
SM = 0
Delta_K
Doxorubicin
YMR245W
ZShift = 0.1
Z lm Score = −3.21
NG = 0
DB = 0
SM = 0
Delta_r
Doxorubicin
YMR245W
ZShift = −1.32
Z lm Score = −3.81
NG = 0
DB = 0
SM = 0
Delta_AUC
Doxorubicin
YFL008W SMC1
ZShift = −0.16
Z lm Score = 5.56
NG = 0
DB = 2
SM = 0
Delta_L
Doxorubicin
YFL008W SMC1
ZShift = 0.41
Z lm Score = −0.14
NG = 0
DB = 2
SM = 0
Delta_K
Doxorubicin
YFL008W SMC1
ZShift = 0.92
Z lm Score = −5.29
NG = 0
DB = 2
SM = 0
Delta_r
Doxorubicin
YFL008W SMC1
ZShift = 0.36
Z lm Score = 0.71
NG = 0
DB = 2
SM = 0
Delta_AUC
Doxorubicin

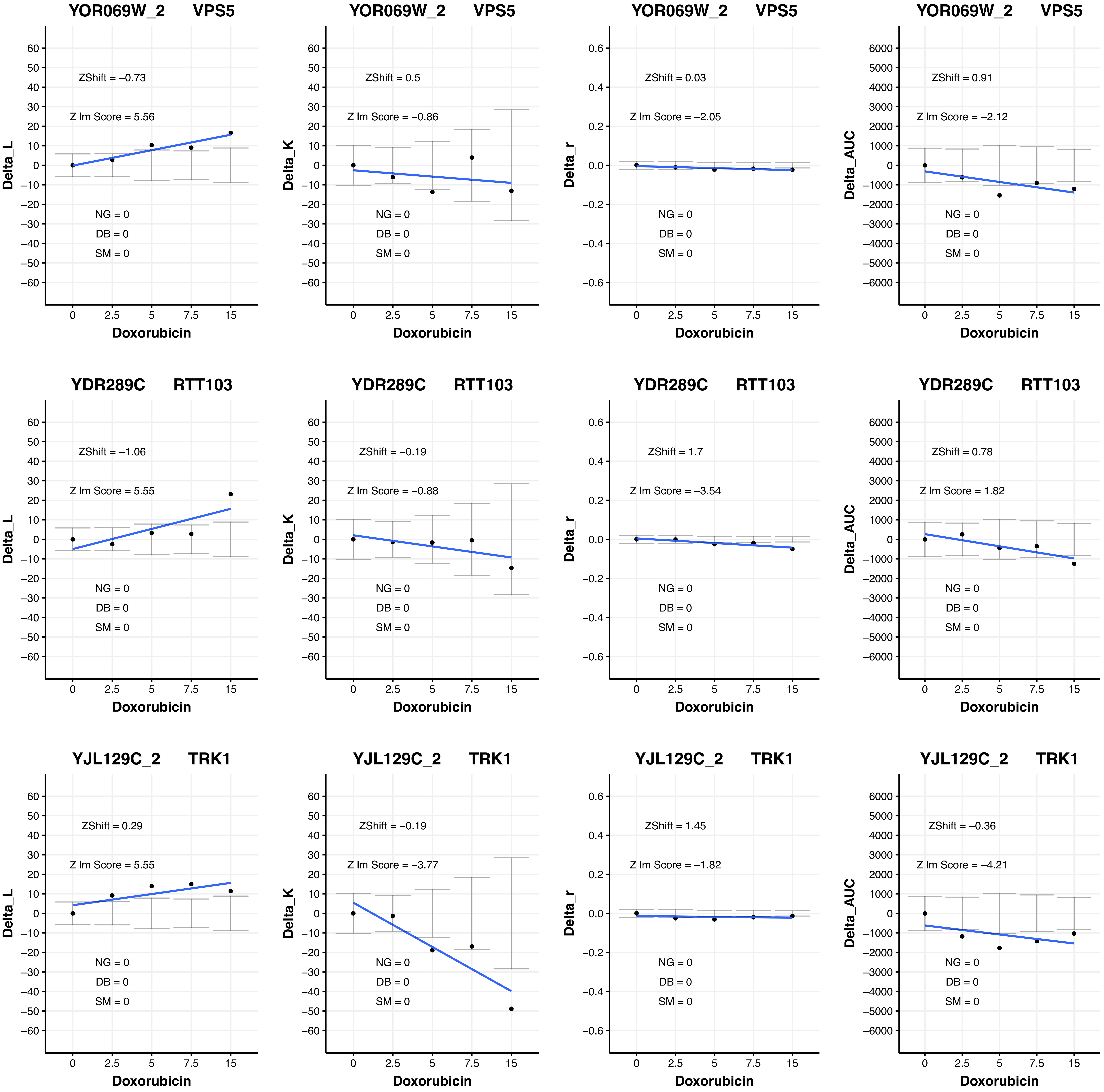
YOR069W_2 VPS5
ZShift = −0.73
Z lm Score = 5.56
NG = 0
DB = 0
SM = 0
Delta_L
Doxorubicin
YOR069W_2 VPS5
ZShift = 0.5
Z lm Score = −0.86
NG = 0
DB = 0
SM = 0
Delta_K
Doxorubicin
YOR069W_2 VPS5
ZShift = 0.03
Z lm Score = −2.05
NG = 0
DB = 0
SM = 0
Delta_r
Doxorubicin
YOR069W_2 VPS5
ZShift = 0.91
Z lm Score = −2.12
NG = 0
DB = 0
SM = 0
Delta_AUC
Doxorubicin
YDR289C RTT103
ZShift = −1.06
Z lm Score = 5.55
NG = 0
DB = 0
SM = 0
Delta_L
Doxorubicin
YDR289C RTT103
ZShift = −0.19
Z lm Score = −0.88
NG = 0
DB = 0
SM = 0
Delta_K
Doxorubicin
YDR289C RTT103
ZShift = 1.7
Z lm Score = −3.54
NG = 0
DB = 0
SM = 0
Delta_r
Doxorubicin
YDR289C RTT103
ZShift = 0.78
Z lm Score = 1.82
NG = 0
DB = 0
SM = 0
Delta_AUC
Doxorubicin
YJL129C_2 TRK1
ZShift = 0.29
Z lm Score = 5.55
NG = 0
DB = 0
SM = 0
Delta_L
Doxorubicin
YJL129C_2 TRK1
ZShift = −0.19
Z lm Score = −3.77
NG = 0
DB = 0
SM = 0
Delta_K
Doxorubicin
YJL129C_2 TRK1
ZShift = 1.45
Z lm Score = −1.82
NG = 0
DB = 0
SM = 0
Delta_r
Doxorubicin
YJL129C_2 TRK1
ZShift = −0.36
Z lm Score = −4.21
NG = 0
DB = 0
SM = 0
Delta_AUC
Doxorubicin

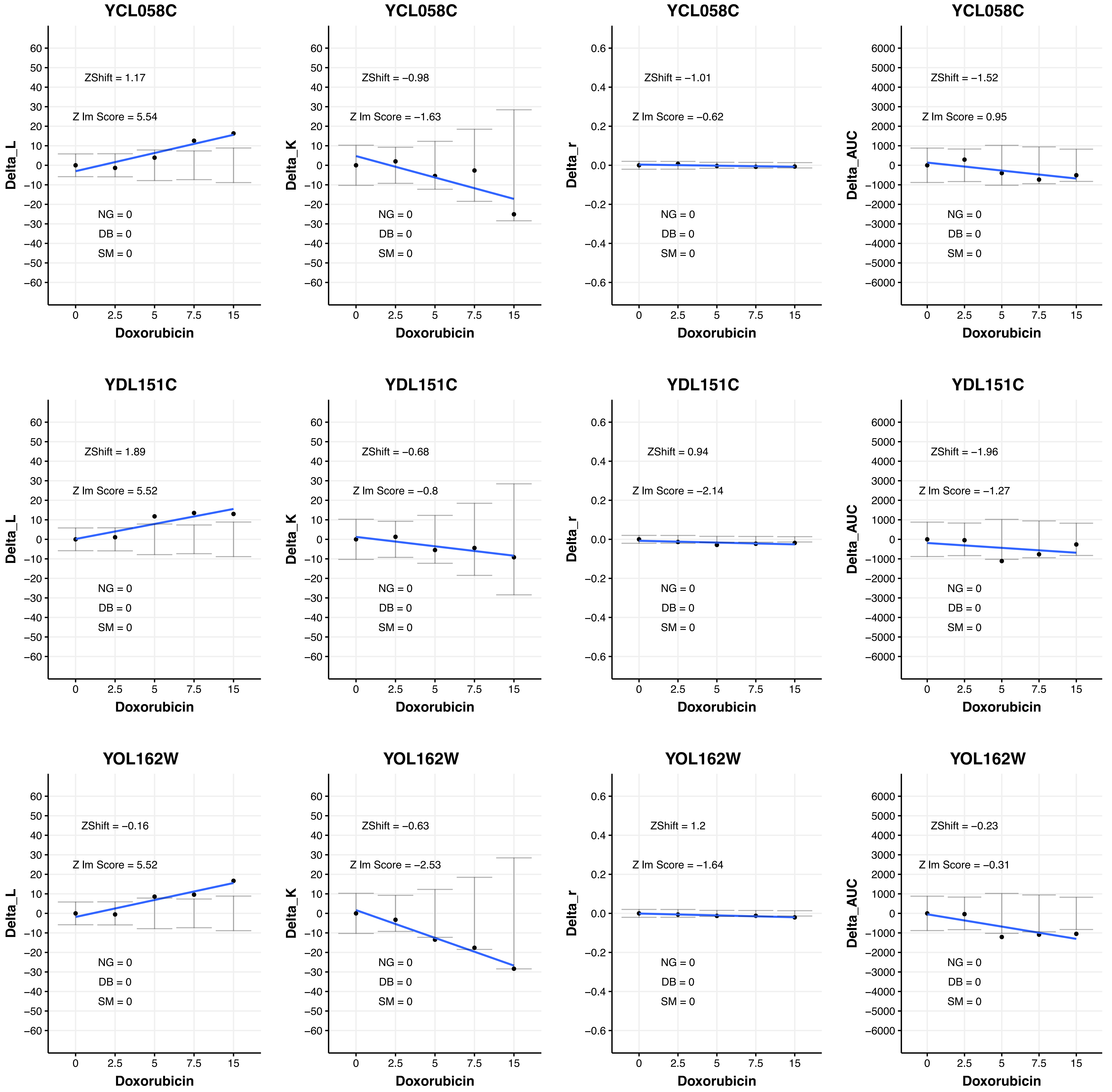
YCL058C
ZShift = 1.17
Z lm Score = 5.54
NG = 0
DB = 0
SM = 0
Delta_L
Doxorubicin
YCL058C
ZShift = −0.98
Z lm Score = −1.63
NG = 0
DB = 0
SM = 0
Delta_K
Doxorubicin
YCL058C
ZShift = −1.01
Z lm Score = −0.62
NG = 0
DB = 0
SM = 0
Delta_r
Doxorubicin
YCL058C
ZShift = −1.52
Z lm Score = 0.95
NG = 0
DB = 0
SM = 0
Delta_AUC
Doxorubicin
YDL151C
ZShift = 1.89
Z lm Score = 5.52
NG = 0
DB = 0
SM = 0
Delta_L
Doxorubicin
YDL151C
ZShift = −0.68
Z lm Score = −0.8
NG = 0
DB = 0
SM = 0
Delta_K
Doxorubicin
YDL151C
ZShift = 0.94
Z lm Score = −2.14
NG = 0
DB = 0
SM = 0
Delta_r
Doxorubicin
YDL151C
ZShift = −1.96
Z lm Score = −1.27
NG = 0
DB = 0
SM = 0
Delta_AUC
Doxorubicin
YOL162W
ZShift = −0.16
Z lm Score = 5.52
NG = 0
DB = 0
SM = 0
Delta_L
Doxorubicin
YOL162W
ZShift = −0.63
Z lm Score = −2.53
NG = 0
DB = 0
SM = 0
Delta_K
Doxorubicin
YOL162W
ZShift = 1.2
Z lm Score = −1.64
NG = 0
DB = 0
SM = 0
Delta_r
Doxorubicin
YOL162W
ZShift = −0.23
Z lm Score = −0.31
NG = 0
DB = 0
SM = 0
Delta_AUC
Doxorubicin

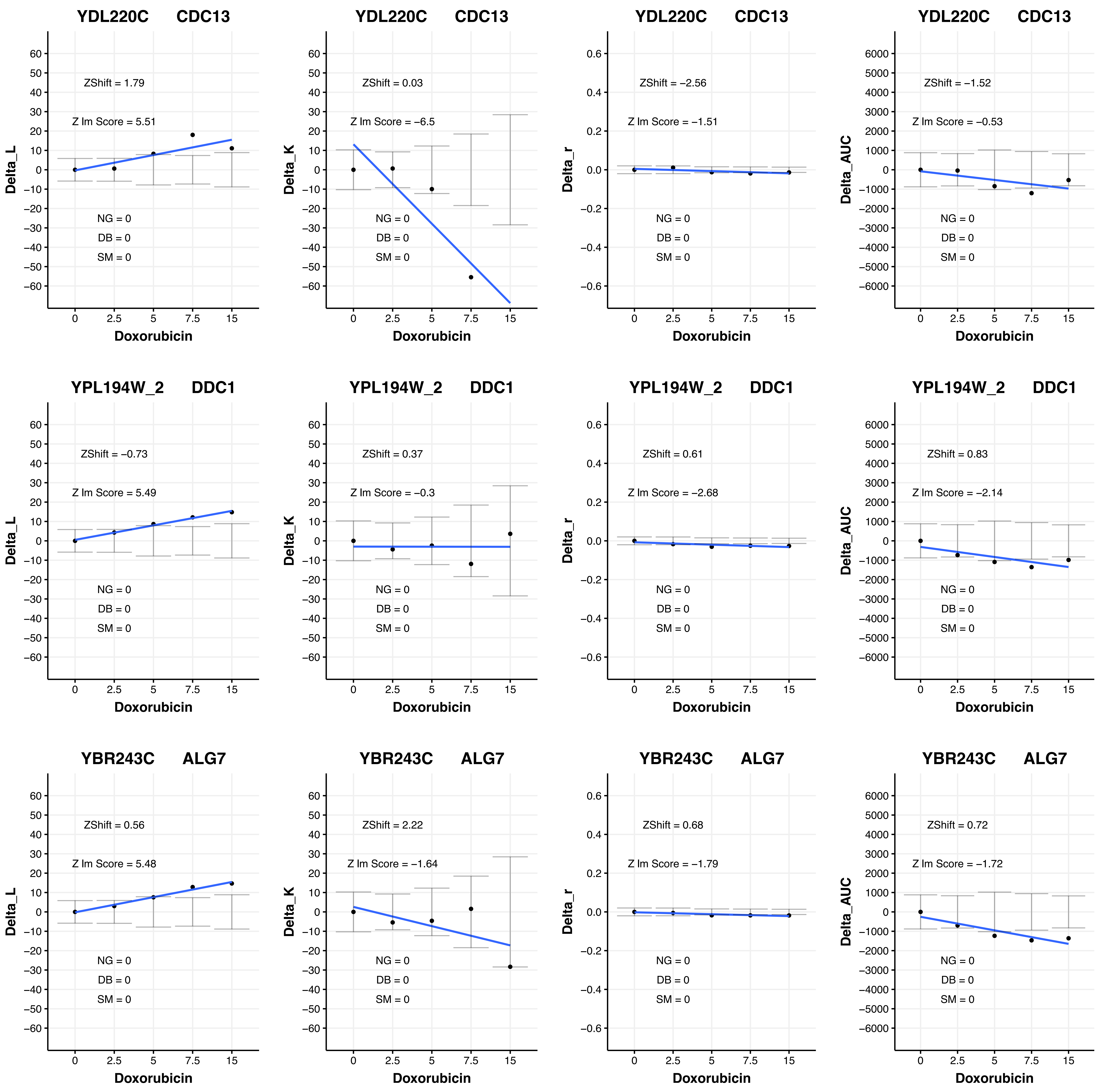
YDL220C CDC13
ZShift = 1.79
Z lm Score = 5.51
NG = 0
DB = 0
SM = 0
Delta_L
Doxorubicin
YDL220C CDC13
ZShift = 0.03
Z lm Score = −6.5
NG = 0
DB = 0
SM = 0
Delta_K
Doxorubicin
YDL220C CDC13
ZShift = −2.56
Z lm Score = −1.51
NG = 0
DB = 0
SM = 0
Delta_r
Doxorubicin
YDL220C CDC13
ZShift = −1.52
Z lm Score = −0.53
NG = 0
DB = 0
SM = 0
Delta_AUC
Doxorubicin
YPL194W_2 DDC1
ZShift = −0.73
Z lm Score = 5.49
NG = 0
DB = 0
SM = 0
Delta_L
Doxorubicin
YPL194W_2 DDC1
ZShift = 0.37
Z lm Score = −0.3
NG = 0
DB = 0
SM = 0
Delta_K
Doxorubicin
YPL194W_2 DDC1
ZShift = 0.61
Z lm Score = −2.68
NG = 0
DB = 0
SM = 0
Delta_r
Doxorubicin
YPL194W_2 DDC1
ZShift = 0.83
Z lm Score = −2.14
NG = 0
DB = 0
SM = 0
Delta_AUC
Doxorubicin
YBR243C ALG7
ZShift = 0.56
Z lm Score = 5.48
NG = 0
DB = 0
SM = 0
Delta_L
Doxorubicin
YBR243C ALG7
ZShift = 2.22
Z lm Score = −1.64
NG = 0
DB = 0
SM = 0
Delta_K
Doxorubicin
YBR243C ALG7
ZShift = 0.68
Z lm Score = −1.79
NG = 0
DB = 0
SM = 0
Delta_r
Doxorubicin
YBR243C ALG7
ZShift = 0.72
Z lm Score = −1.72
NG = 0
DB = 0
SM = 0
Delta_AUC
Doxorubicin

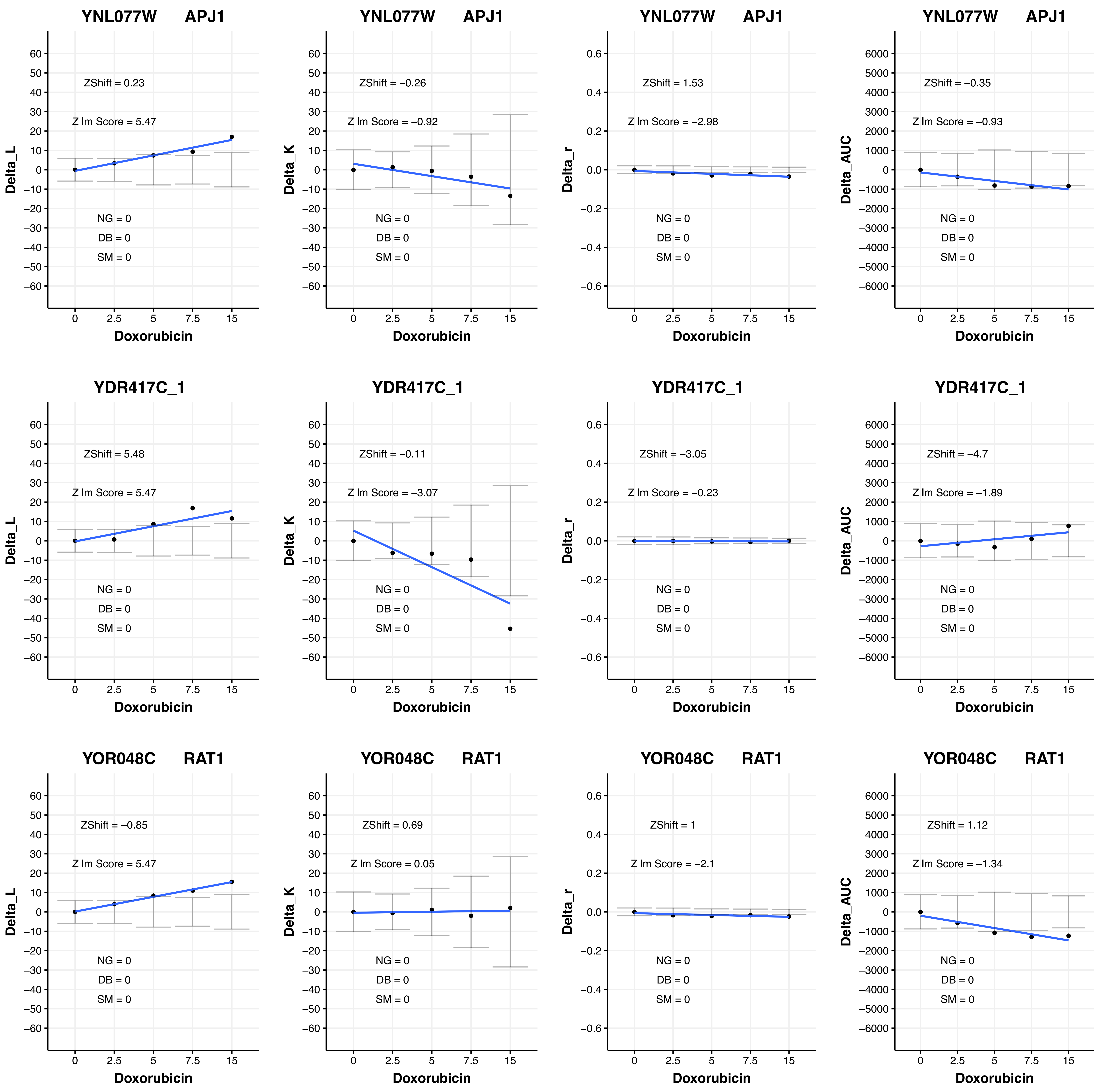
YNL077W APJ1
ZShift = 0.23
Z lm Score = 5.47
NG = 0
DB = 0
SM = 0
Delta_L
Doxorubicin
YNL077W APJ1
ZShift = −0.26
Z lm Score = −0.92
NG = 0
DB = 0
SM = 0
Delta_K
Doxorubicin
YNL077W APJ1
ZShift = 1.53
Z lm Score = −2.98
NG = 0
DB = 0
SM = 0
Delta_r
Doxorubicin
YNL077W APJ1
ZShift = −0.35
Z lm Score = −0.93
NG = 0
DB = 0
SM = 0
Delta_AUC
Doxorubicin
YDR417C_1
ZShift = 5.48
Z lm Score = 5.47
NG = 0
DB = 0
SM = 0
Delta_L
Doxorubicin
YDR417C_1
ZShift = −0.11
Z lm Score = −3.07
NG = 0
DB = 0
SM = 0
Delta_K
Doxorubicin
YDR417C_1
ZShift = −3.05
Z lm Score = −0.23
NG = 0
DB = 0
SM = 0
Delta_r
Doxorubicin
YDR417C_1
ZShift = −4.7
Z lm Score = −1.89
NG = 0
DB = 0
SM = 0
Delta_AUC
Doxorubicin
YOR048C RAT1
ZShift = −0.85
Z lm Score = 5.47
NG = 0
DB = 0
SM = 0
Delta_L
Doxorubicin
YOR048C RAT1
ZShift = 0.69
Z lm Score = 0.05
NG = 0
DB = 0
SM = 0
Delta_K
Doxorubicin
YOR048C RAT1
ZShift = 1
Z lm Score = −2.1
NG = 0
DB = 0
SM = 0
Delta_r
Doxorubicin
YOR048C RAT1
ZShift = 1.12
Z lm Score = −1.34
NG = 0
DB = 0
SM = 0
Delta_AUC
Doxorubicin

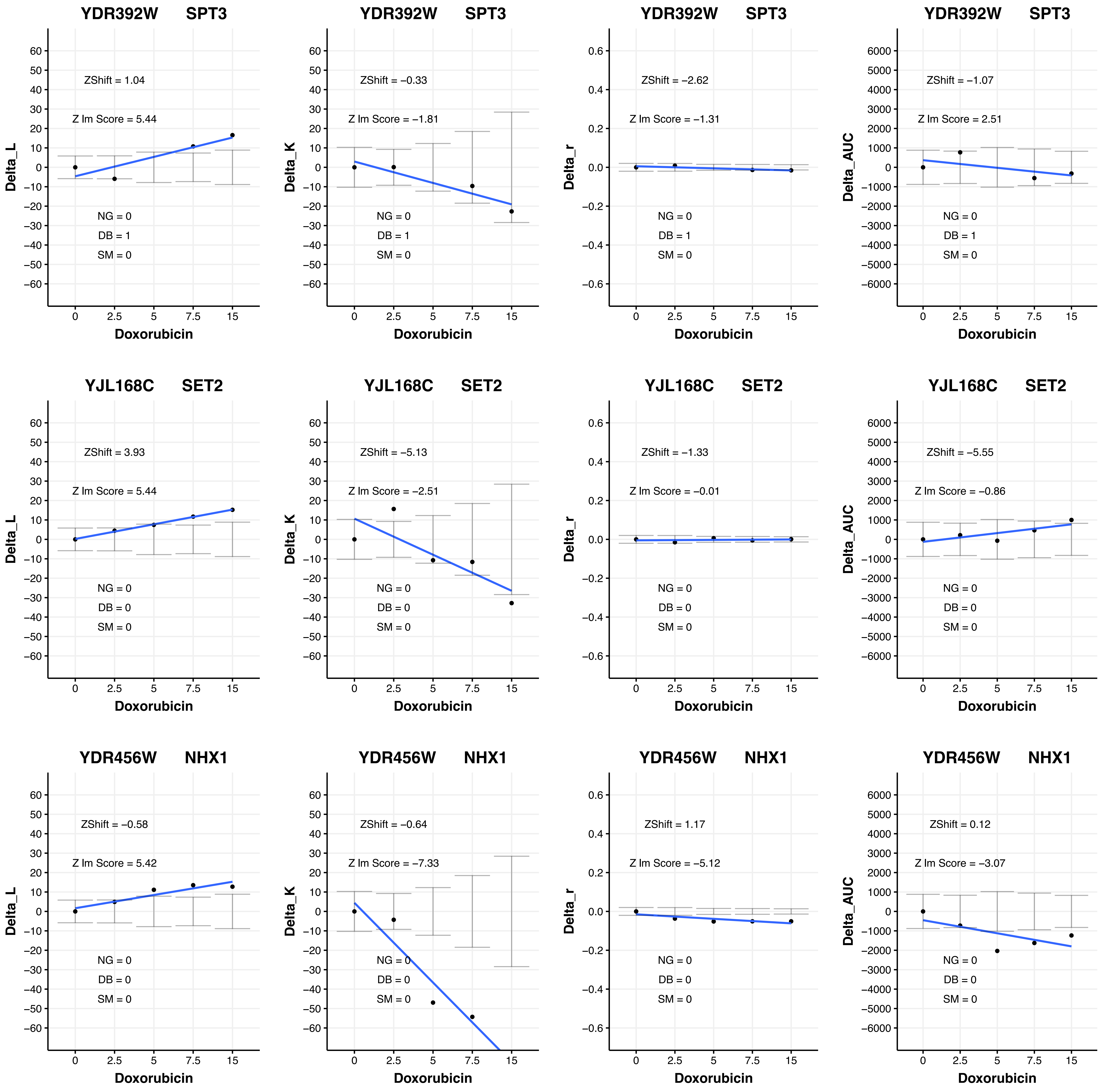
YDR392W SPT3
ZShift = 1.04
Z lm Score = 5.44
NG = 0
DB = 1
SM = 0
Delta_L
Doxorubicin
YDR392W SPT3
ZShift = −0.33
Z lm Score = −1.81
NG = 0
DB = 1
SM = 0
Delta_K
Doxorubicin
YDR392W SPT3
ZShift = −2.62
Z lm Score = −1.31
NG = 0
DB = 1
SM = 0
Delta_r
Doxorubicin
YDR392W SPT3
ZShift = −1.07
Z lm Score = 2.51
NG = 0
DB = 1
SM = 0
Delta_AUC
Doxorubicin
YJL168C SET2
ZShift = 3.93
Z lm Score = 5.44
NG = 0
DB = 0
SM = 0
Delta_L
Doxorubicin
YJL168C SET2
ZShift = −5.13
Z lm Score = −2.51
NG = 0
DB = 0
SM = 0
Delta_K
Doxorubicin
YJL168C SET2
ZShift = −1.33
Z lm Score = −0.01
NG = 0
DB = 0
SM = 0
Delta_r
Doxorubicin
YJL168C SET2
ZShift = −5.55
Z lm Score = −0.86
NG = 0
DB = 0
SM = 0
Delta_AUC
Doxorubicin
YDR456W NHX1
ZShift = −0.58
Z lm Score = 5.42
NG = 0
DB = 0
SM = 0
Delta_L
Doxorubicin
YDR456W NHX1
ZShift = −0.64
Z lm Score = −7.33
NG = 0
DB = 0
SM = 0
Delta_K
Doxorubicin
YDR456W NHX1
ZShift = 1.17
Z lm Score = −5.12
NG = 0
DB = 0
SM = 0
Delta_r
Doxorubicin
YDR456W NHX1
ZShift = 0.12
Z lm Score = −3.07
NG = 0
DB = 0
SM = 0
Delta_AUC
Doxorubicin

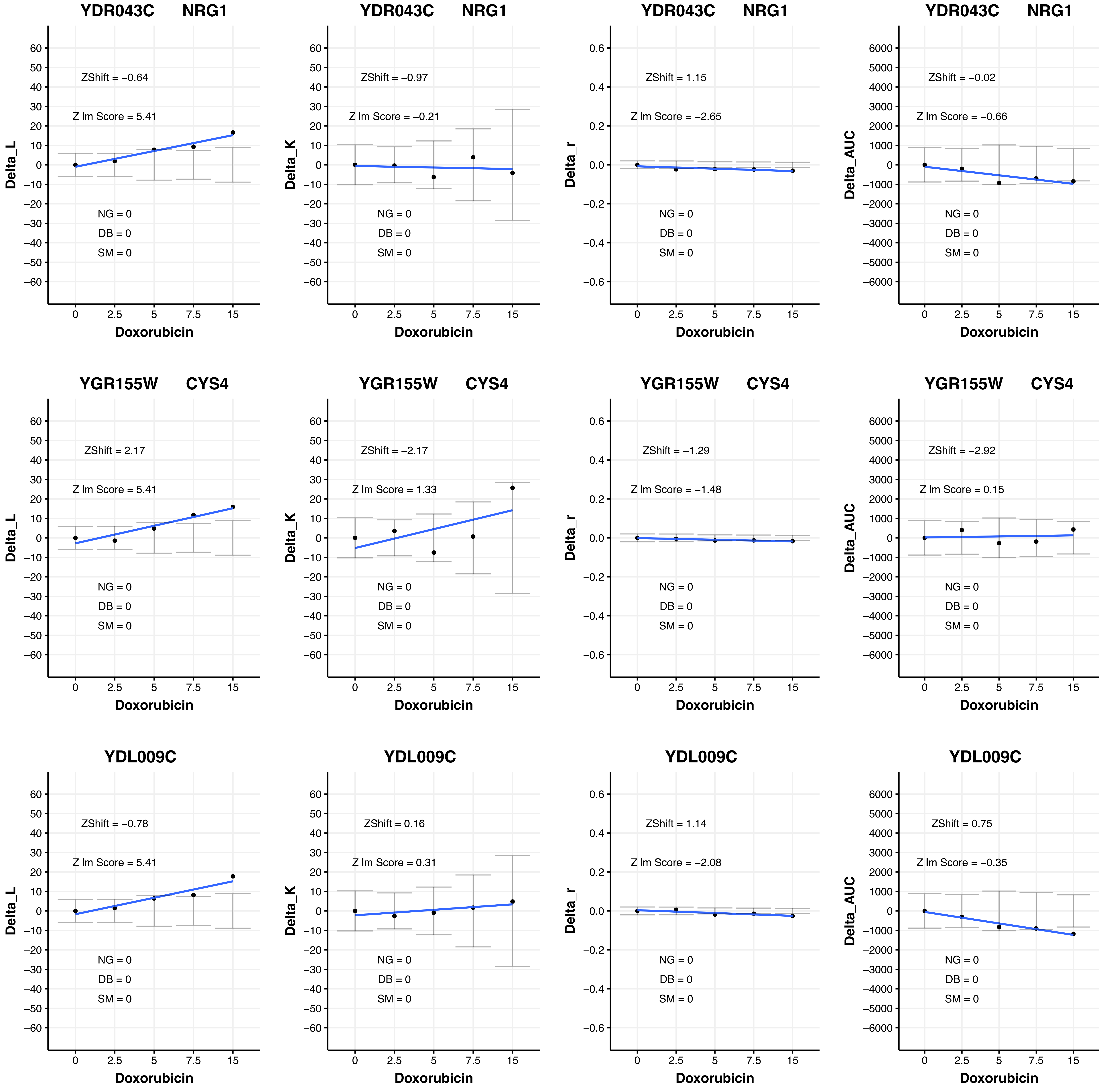
YDR043C NRG1
ZShift = −0.64
Z lm Score = 5.41
NG = 0
DB = 0
SM = 0
Delta_L
Doxorubicin
YDR043C NRG1
ZShift = −0.97
Z lm Score = −0.21
NG = 0
DB = 0
SM = 0
Delta_K
Doxorubicin
YDR043C NRG1
ZShift = 1.15
Z lm Score = −2.65
NG = 0
DB = 0
SM = 0
Delta_r
Doxorubicin
YDR043C NRG1
ZShift = −0.02
Z lm Score = −0.66
NG = 0
DB = 0
SM = 0
Delta_AUC
Doxorubicin
YGR155W CYS4
ZShift = 2.17
Z lm Score = 5.41
NG = 0
DB = 0
SM = 0
Delta_L
Doxorubicin
YGR155W CYS4
ZShift = −2.17
Z lm Score = 1.33
NG = 0
DB = 0
SM = 0
Delta_K
Doxorubicin
YGR155W CYS4
ZShift = −1.29
Z lm Score = −1.48
NG = 0
DB = 0
SM = 0
Delta_r
Doxorubicin
YGR155W CYS4
ZShift = −2.92
Z lm Score = 0.15
NG = 0
DB = 0
SM = 0
Delta_AUC
Doxorubicin
YDL009C
ZShift = −0.78
Z lm Score = 5.41
NG = 0
DB = 0
SM = 0
Delta_L
Doxorubicin
YDL009C
ZShift = 0.16
Z lm Score = 0.31
NG = 0
DB = 0
SM = 0
Delta_K
Doxorubicin
YDL009C
ZShift = 1.14
Z lm Score = −2.08
NG = 0
DB = 0
SM = 0
Delta_r
Doxorubicin
YDL009C
ZShift = 0.75
Z lm Score = −0.35
NG = 0
DB = 0
SM = 0
Delta_AUC
Doxorubicin

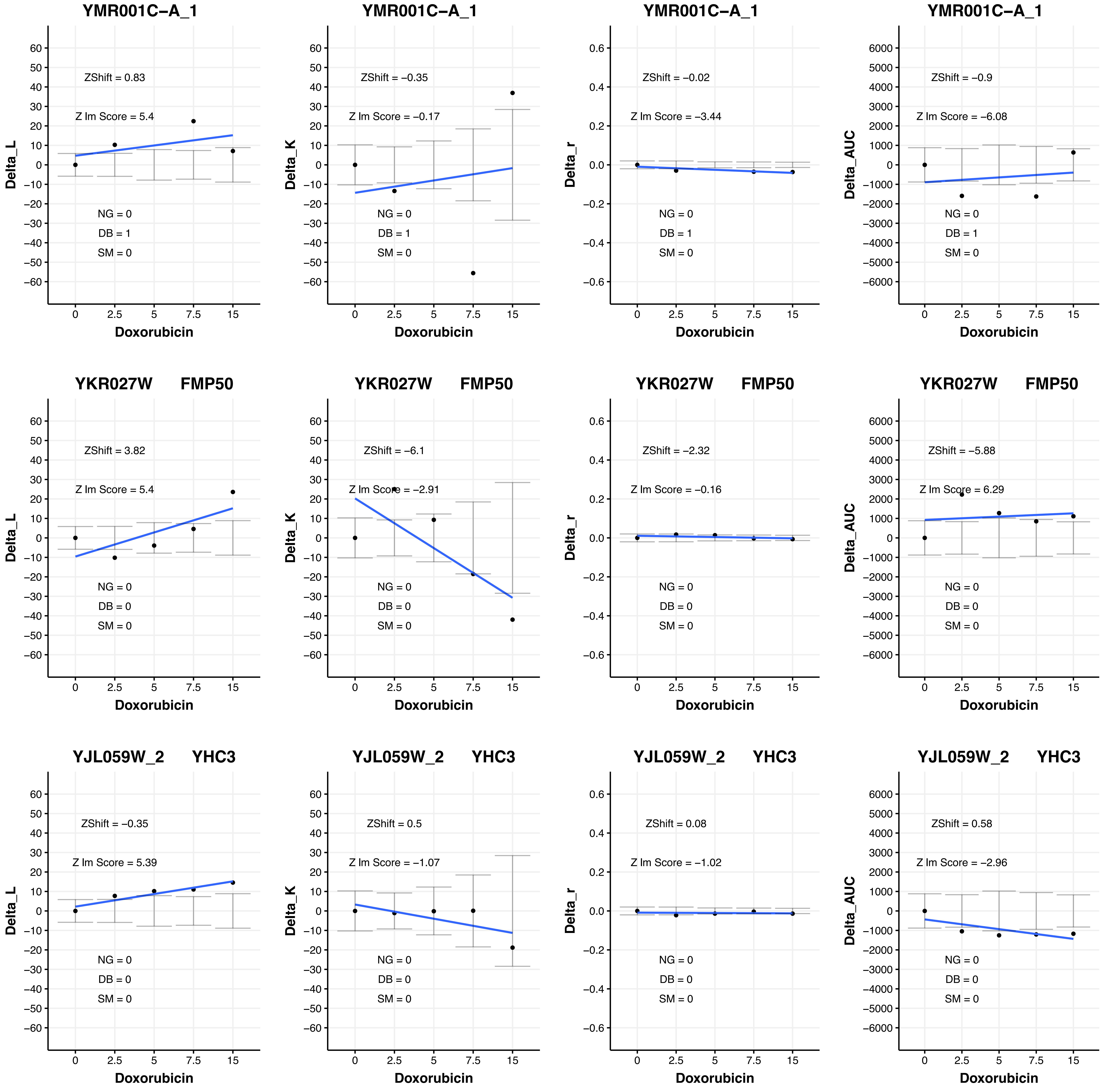
YMR001C−A_1
ZShift = 0.83
Z lm Score = 5.4
NG = 0
DB = 1
SM = 0
Delta_L
Doxorubicin
YMR001C−A_1
ZShift = −0.35
Z lm Score = −0.17
NG = 0
DB = 1
SM = 0
Delta_K
Doxorubicin
YMR001C−A_1
ZShift = −0.02
Z lm Score = −3.44
NG = 0
DB = 1
SM = 0
Delta_r
Doxorubicin
YMR001C−A_1
ZShift = −0.9
Z lm Score = −6.08
NG = 0
DB = 1
SM = 0
Delta_AUC
Doxorubicin
YKR027W FMP50
ZShift = 3.82
Z lm Score = 5.4
NG = 0
DB = 0
SM = 0
Delta_L
Doxorubicin
YKR027W FMP50
ZShift = −6.1
Z lm Score = −2.91
NG = 0
DB = 0
SM = 0
Delta_K
Doxorubicin
YKR027W FMP50
ZShift = −2.32
Z lm Score = −0.16
NG = 0
DB = 0
SM = 0
Delta_r
Doxorubicin
YKR027W FMP50
ZShift = −5.88
Z lm Score = 6.29
NG = 0
DB = 0
SM = 0
Delta_AUC
Doxorubicin
YJL059W_2 YHC3
ZShift = −0.35
Z lm Score = 5.39
NG = 0
DB = 0
SM = 0
Delta_L
Doxorubicin
YJL059W_2 YHC3
ZShift = 0.5
Z lm Score = −1.07
NG = 0
DB = 0
SM = 0
Delta_K
Doxorubicin
YJL059W_2 YHC3
ZShift = 0.08
Z lm Score = −1.02
NG = 0
DB = 0
SM = 0
Delta_r
Doxorubicin
YJL059W_2 YHC3
ZShift = 0.58
Z lm Score = −2.96
NG = 0
DB = 0
SM = 0
Delta_AUC
Doxorubicin

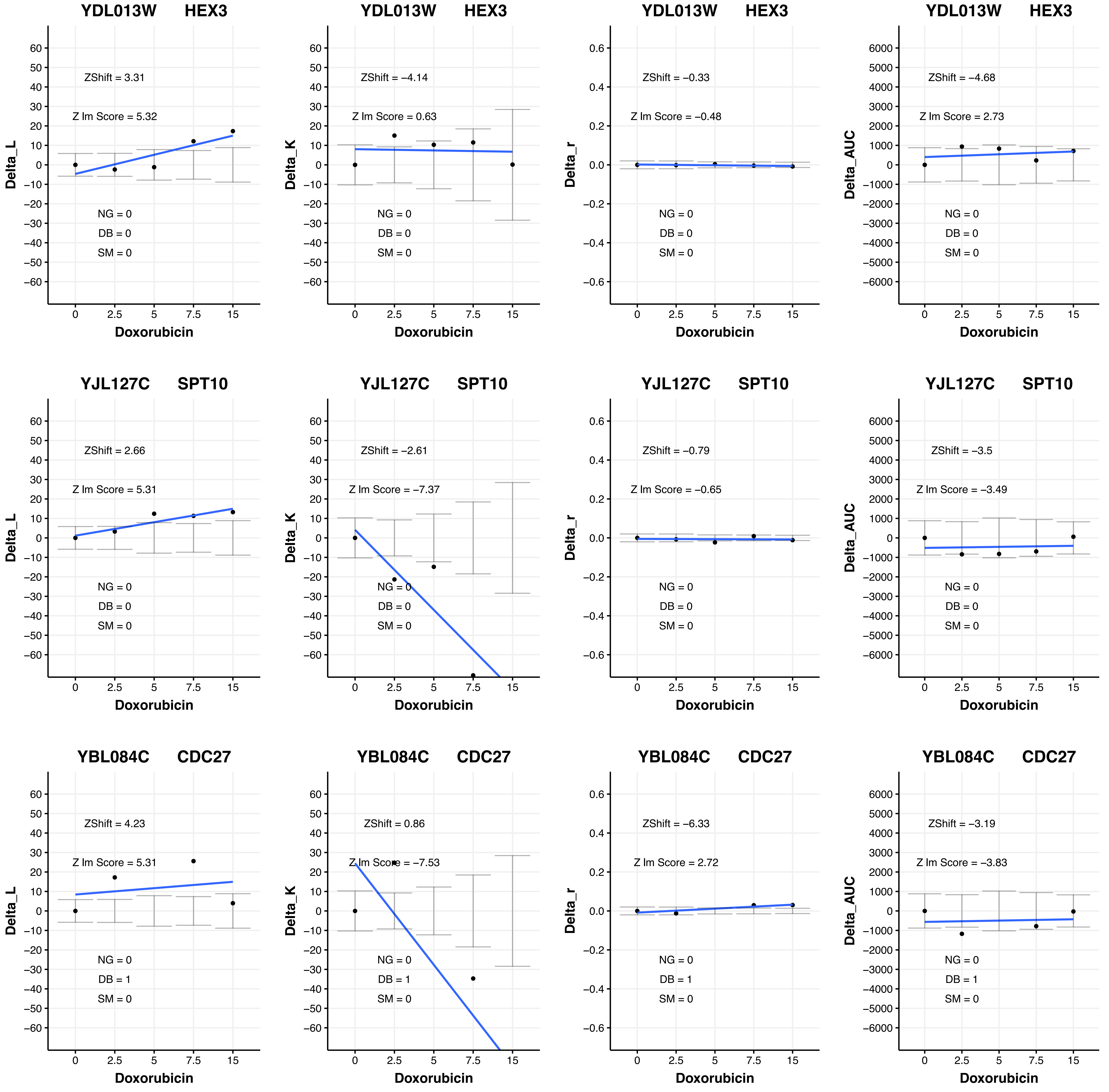
YDL013W HEX3
ZShift = 3.31
Z lm Score = 5.32
NG = 0
DB = 0
SM = 0
Delta_L
Doxorubicin
YDL013W HEX3
ZShift = −4.14
Z lm Score = 0.63
NG = 0
DB = 0
SM = 0
Delta_K
Doxorubicin
YDL013W HEX3
ZShift = −0.33
Z lm Score = −0.48
NG = 0
DB = 0
SM = 0
Delta_r
Doxorubicin
YDL013W HEX3
ZShift = −4.68
Z lm Score = 2.73
NG = 0
DB = 0
SM = 0
Delta_AUC
Doxorubicin
YJL127C SPT10
ZShift = 2.66
Z lm Score = 5.31
NG = 0
DB = 0
SM = 0
Delta_L
Doxorubicin
YJL127C SPT10
ZShift = −2.61
Z lm Score = −7.37
NG = 0
DB = 0
SM = 0
Delta_K
Doxorubicin
YJL127C SPT10
ZShift = −0.79
Z lm Score = −0.65
NG = 0
DB = 0
SM = 0
Delta_r
Doxorubicin
YJL127C SPT10
ZShift = −3.5
Z lm Score = −3.49
NG = 0
DB = 0
SM = 0
Delta_AUC
Doxorubicin
YBL084C CDC27
ZShift = 4.23
Z lm Score = 5.31
NG = 0
DB = 1
SM = 0
Delta_L
Doxorubicin
YBL084C CDC27
ZShift = 0.86
Z lm Score = −7.53
NG = 0
DB = 1
SM = 0
Delta_K
Doxorubicin
YBL084C CDC27
ZShift = −6.33
Z lm Score = 2.72
NG = 0
DB = 1
SM = 0
Delta_r
Doxorubicin
YBL084C CDC27
ZShift = −3.19
Z lm Score = −3.83
NG = 0
DB = 1
SM = 0
Delta_AUC
Doxorubicin

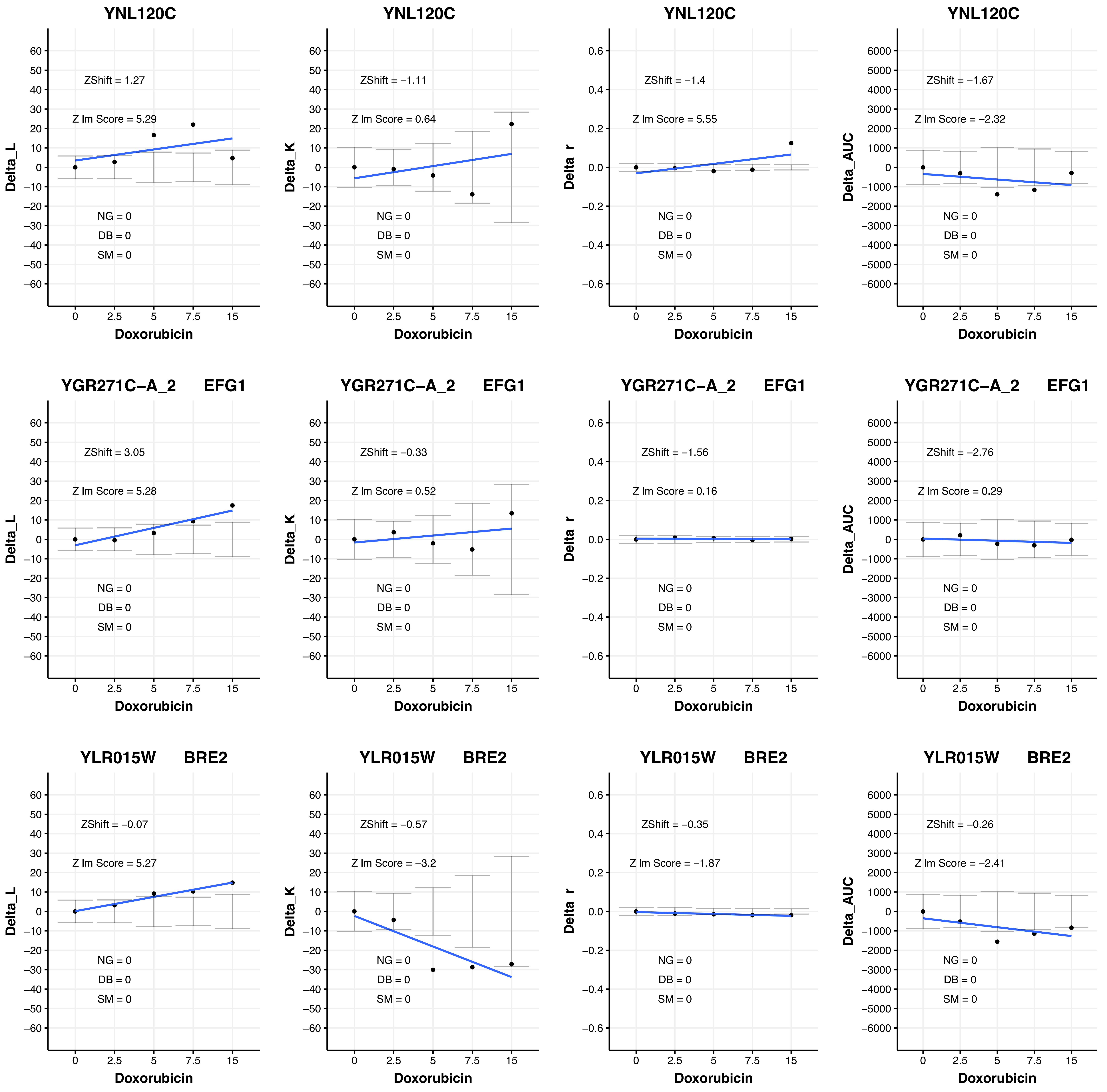
YNL120C
ZShift = 1.27
Z lm Score = 5.29
NG = 0
DB = 0
SM = 0
Delta_L
Doxorubicin
YNL120C
ZShift = −1.11
Z lm Score = 0.64
NG = 0
DB = 0
SM = 0
Delta_K
Doxorubicin
YNL120C
ZShift = −1.4
Z lm Score = 5.55
NG = 0
DB = 0
SM = 0
Delta_r
Doxorubicin
YNL120C
ZShift = −1.67
Z lm Score = −2.32
NG = 0
DB = 0
SM = 0
Delta_AUC
Doxorubicin
YGR271C−A_2 EFG1
ZShift = 3.05
Z lm Score = 5.28
NG = 0
DB = 0
SM = 0
Delta_L
Doxorubicin
YGR271C−A_2 EFG1
ZShift = −0.33
Z lm Score = 0.52
NG = 0
DB = 0
SM = 0
Delta_K
Doxorubicin
YGR271C−A_2 EFG1
ZShift = −1.56
Z lm Score = 0.16
NG = 0
DB = 0
SM = 0
Delta_r
Doxorubicin
YGR271C−A_2 EFG1
ZShift = −2.76
Z lm Score = 0.29
NG = 0
DB = 0
SM = 0
Delta_AUC
Doxorubicin
YLR015W BRE2
ZShift = −0.07
Z lm Score = 5.27
NG = 0
DB = 0
SM = 0
Delta_L
Doxorubicin
YLR015W BRE2
ZShift = −0.57
Z lm Score = −3.2
NG = 0
DB = 0
SM = 0
Delta_K
Doxorubicin
YLR015W BRE2
ZShift = −0.35
Z lm Score = −1.87
NG = 0
DB = 0
SM = 0
Delta_r
Doxorubicin
YLR015W BRE2
ZShift = −0.26
Z lm Score = −2.41
NG = 0
DB = 0
SM = 0
Delta_AUC
Doxorubicin

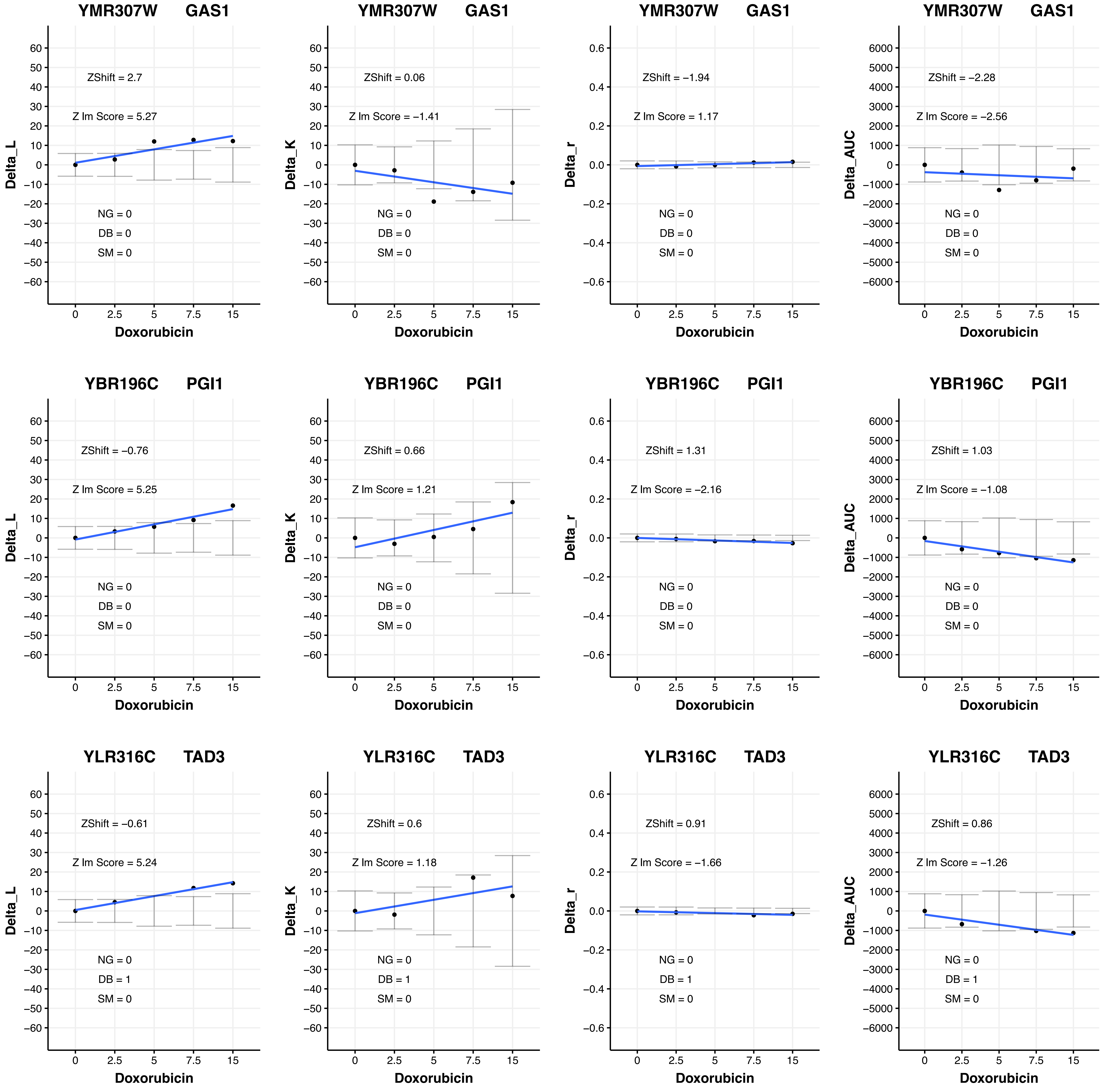

YMR307W GAS1
ZShift = 2.7
Z lm Score = 5.27
NG = 0
DB = 0
SM = 0
Delta_L
Doxorubicin
YMR307W GAS1
ZShift = 0.06
Z lm Score = −1.41
NG = 0
DB = 0
SM = 0
Delta_K
Doxorubicin
YMR307W GAS1
ZShift = −1.94
Z lm Score = 1.17
NG = 0
DB = 0
SM = 0
Delta_r
Doxorubicin
YMR307W GAS1
ZShift = −2.28
Z lm Score = −2.56
NG = 0
DB = 0
SM = 0
Delta_AUC
Doxorubicin
YBR196C PGI1
ZShift = −0.76
Z lm Score = 5.25
NG = 0
DB = 0
SM = 0
Delta_L
Doxorubicin
YBR196C PGI1
ZShift = 0.66
Z lm Score = 1.21
NG = 0
DB = 0
SM = 0
Delta_K
Doxorubicin
YBR196C PGI1
ZShift = 1.31
Z lm Score = −2.16
NG = 0
DB = 0
SM = 0
Delta_r
Doxorubicin
YBR196C PGI1
ZShift = 1.03
Z lm Score = −1.08
NG = 0
DB = 0
SM = 0
Delta_AUC
Doxorubicin
YLR316C TAD3
ZShift = −0.61
Z lm Score = 5.24
NG = 0
DB = 1
SM = 0
Delta_L
Doxorubicin
YLR316C TAD3
ZShift = 0.6
Z lm Score = 1.18
NG = 0
DB = 1
SM = 0
Delta_K
Doxorubicin
YLR316C TAD3
ZShift = 0.91
Z lm Score = −1.66
NG = 0
DB = 1
SM = 0
Delta_r
Doxorubicin
YLR316C TAD3
ZShift = 0.86
Z lm Score = −1.26
NG = 0
DB = 1
SM = 0
Delta_AUC
Doxorubicin

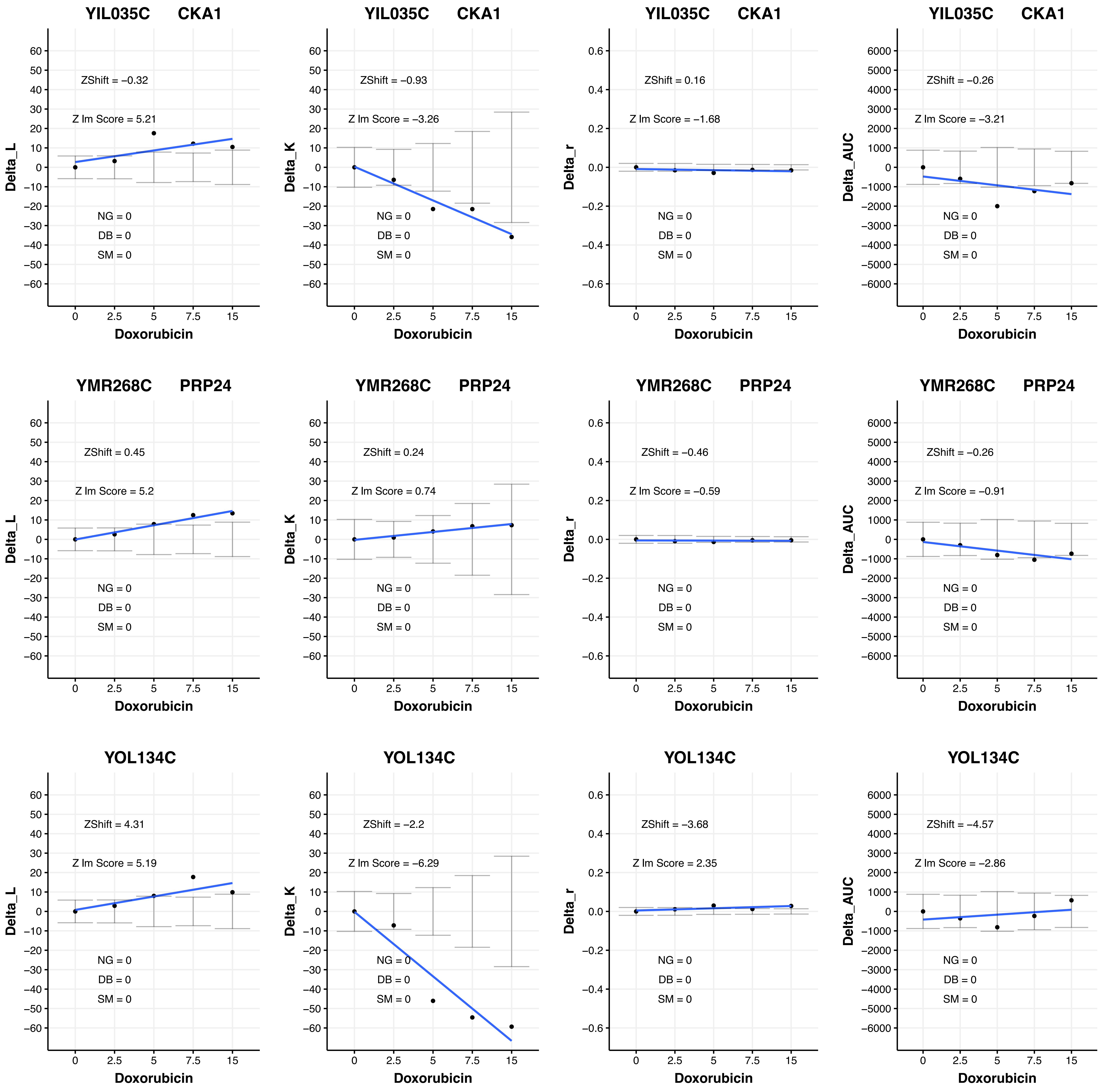
YIL035C CKA1
ZShift = −0.32
Z lm Score = 5.21
NG = 0
DB = 0
SM = 0
Delta_L
Doxorubicin
YIL035C CKA1
ZShift = −0.93
Z lm Score = −3.26
NG = 0
DB = 0
SM = 0
Delta_K
Doxorubicin
YIL035C CKA1
ZShift = 0.16
Z lm Score = −1.68
NG = 0
DB = 0
SM = 0
Delta_r
Doxorubicin
YIL035C CKA1
ZShift = −0.26
Z lm Score = −3.21
NG = 0
DB = 0
SM = 0
Delta_AUC
Doxorubicin
YMR268C PRP24
ZShift = 0.45
Z lm Score = 5.2
NG = 0
DB = 0
SM = 0
Delta_L
Doxorubicin
YMR268C PRP24
ZShift = 0.24
Z lm Score = 0.74
NG = 0
DB = 0
SM = 0
Delta_K
Doxorubicin
YMR268C PRP24
ZShift = −0.46
Z lm Score = −0.59
NG = 0
DB = 0
SM = 0
Delta_r
Doxorubicin
YMR268C PRP24
ZShift = −0.26
Z lm Score = −0.91
NG = 0
DB = 0
SM = 0
Delta_AUC
Doxorubicin
YOL134C
ZShift = 4.31
Z lm Score = 5.19
NG = 0
DB = 0
SM = 0
Delta_L
Doxorubicin
YOL134C
ZShift = −2.2
Z lm Score = −6.29
NG = 0
DB = 0
SM = 0
Delta_K
Doxorubicin
YOL134C
ZShift = −3.68
Z lm Score = 2.35
NG = 0
DB = 0
SM = 0
Delta_r
Doxorubicin
YOL134C
ZShift = −4.57
Z lm Score = −2.86
NG = 0
DB = 0
SM = 0
Delta_AUC
Doxorubicin

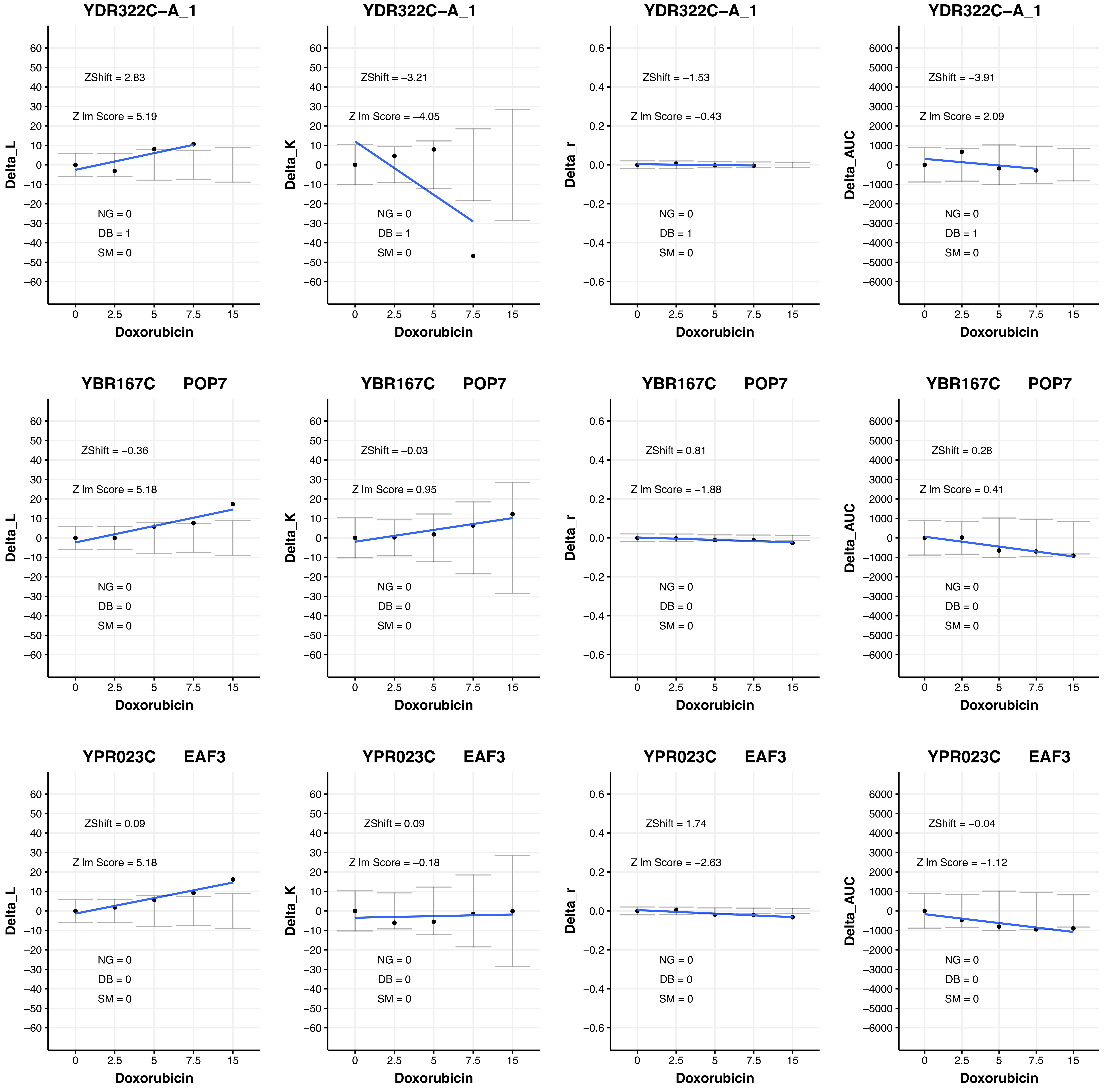
YDR322C−A_1
ZShift = 2.83
Z lm Score = 5.19
NG = 0
DB = 1
SM = 0
Delta_L
Doxorubicin
YDR322C−A_1
ZShift = −3.21
Z lm Score = −4.05
NG = 0
DB = 1
SM = 0
Delta_K
Doxorubicin
YDR322C−A_1
ZShift = −1.53
Z lm Score = −0.43
NG = 0
DB = 1
SM = 0
Delta_r
Doxorubicin
YDR322C−A_1
ZShift = −3.91
Z lm Score = 2.09
NG = 0
DB = 1
SM = 0
Delta_AUC
Doxorubicin
YBR167C POP7
ZShift = −0.36
Z lm Score = 5.18
NG = 0
DB = 0
SM = 0
Delta_L
Doxorubicin
YBR167C POP7
ZShift = −0.03
Z lm Score = 0.95
NG = 0
DB = 0
SM = 0
Delta_K
Doxorubicin
YBR167C POP7
ZShift = 0.81
Z lm Score = −1.88
NG = 0
DB = 0
SM = 0
Delta_r
Doxorubicin
YBR167C POP7
ZShift = 0.28
Z lm Score = 0.41
NG = 0
DB = 0
SM = 0
Delta_AUC
Doxorubicin
YPR023C EAF3
ZShift = 0.09
Z lm Score = 5.18
NG = 0
DB = 0
SM = 0
Delta_L
Doxorubicin
YPR023C EAF3
ZShift = 0.09
Z lm Score = −0.18
NG = 0
DB = 0
SM = 0
Delta_K
Doxorubicin
YPR023C EAF3
ZShift = 1.74
Z lm Score = −2.63
NG = 0
DB = 0
SM = 0
Delta_r
Doxorubicin
YPR023C EAF3
ZShift = −0.04
Z lm Score = −1.12
NG = 0
DB = 0
SM = 0
Delta_AUC
Doxorubicin

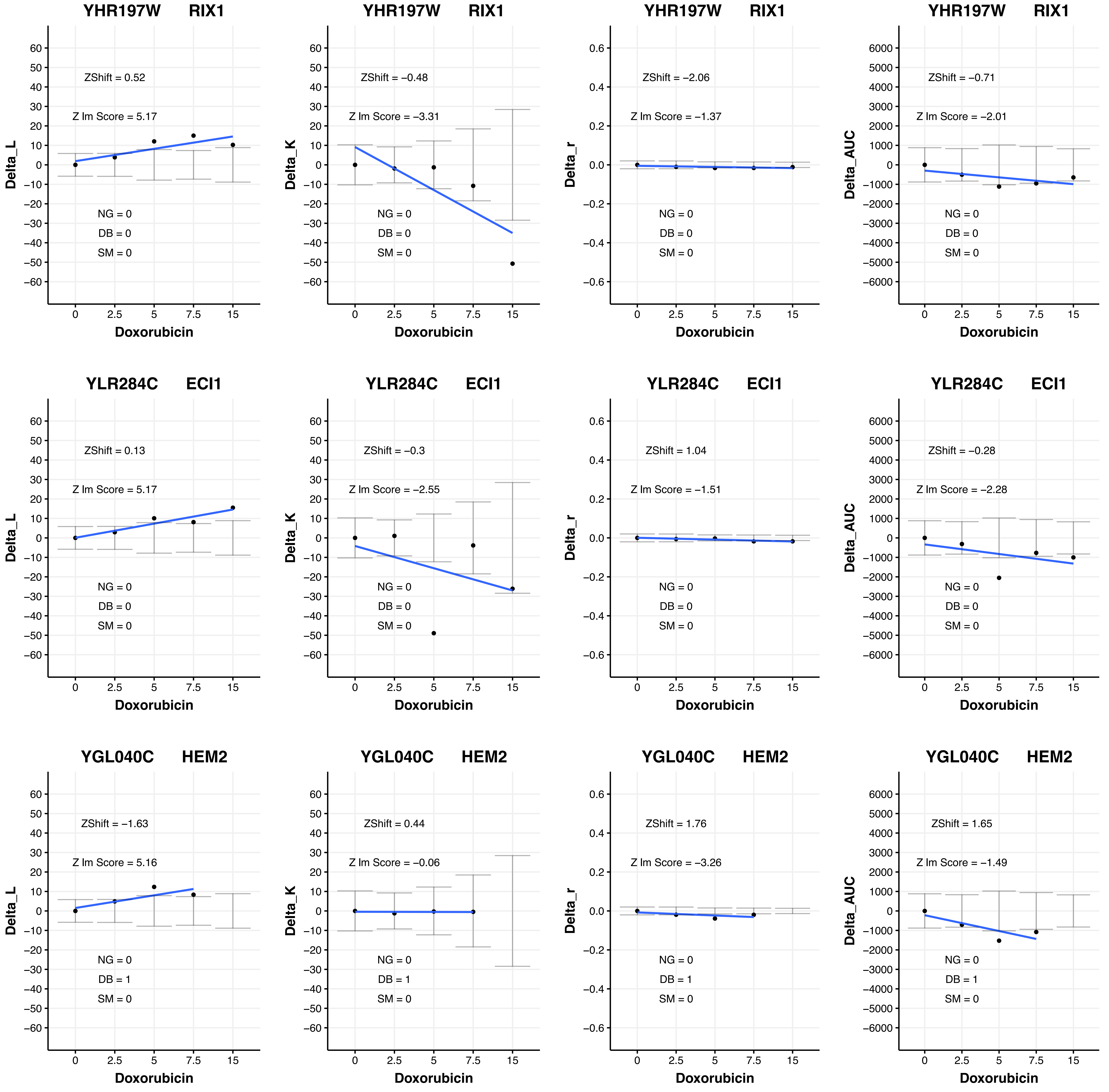
YHR197W RIX1
ZShift = 0.52
Z lm Score = 5.17
NG = 0
DB = 0
SM = 0
Delta_L
Doxorubicin
YHR197W RIX1
ZShift = −0.48
Z lm Score = −3.31
NG = 0
DB = 0
SM = 0
Delta_K
Doxorubicin
YHR197W RIX1
ZShift = −2.06
Z lm Score = −1.37
NG = 0
DB = 0
SM = 0
Delta_r
Doxorubicin
YHR197W RIX1
ZShift = −0.71
Z lm Score = −2.01
NG = 0
DB = 0
SM = 0
Delta_AUC
Doxorubicin
YLR284C ECI1
ZShift = 0.13
Z lm Score = 5.17
NG = 0
DB = 0
SM = 0
Delta_L
Doxorubicin
YLR284C ECI1
ZShift = −0.3
Z lm Score = −2.55
NG = 0
DB = 0
SM = 0
Delta_K
Doxorubicin
YLR284C ECI1
ZShift = 1.04
Z lm Score = −1.51
NG = 0
DB = 0
SM = 0
Delta_r
Doxorubicin
YLR284C ECI1
ZShift = −0.28
Z lm Score = −2.28
NG = 0
DB = 0
SM = 0
Delta_AUC
Doxorubicin
YGL040C HEM2
ZShift = −1.63
Z lm Score = 5.16
NG = 0
DB = 1
SM = 0
Delta_L
Doxorubicin
YGL040C HEM2
ZShift = 0.44
Z lm Score = −0.06
NG = 0
DB = 1
SM = 0
Delta_K
Doxorubicin
YGL040C HEM2
ZShift = 1.76
Z lm Score = −3.26
NG = 0
DB = 1
SM = 0
Delta_r
Doxorubicin
YGL040C HEM2
ZShift = 1.65
Z lm Score = −1.49
NG = 0
DB = 1
SM = 0
Delta_AUC
Doxorubicin

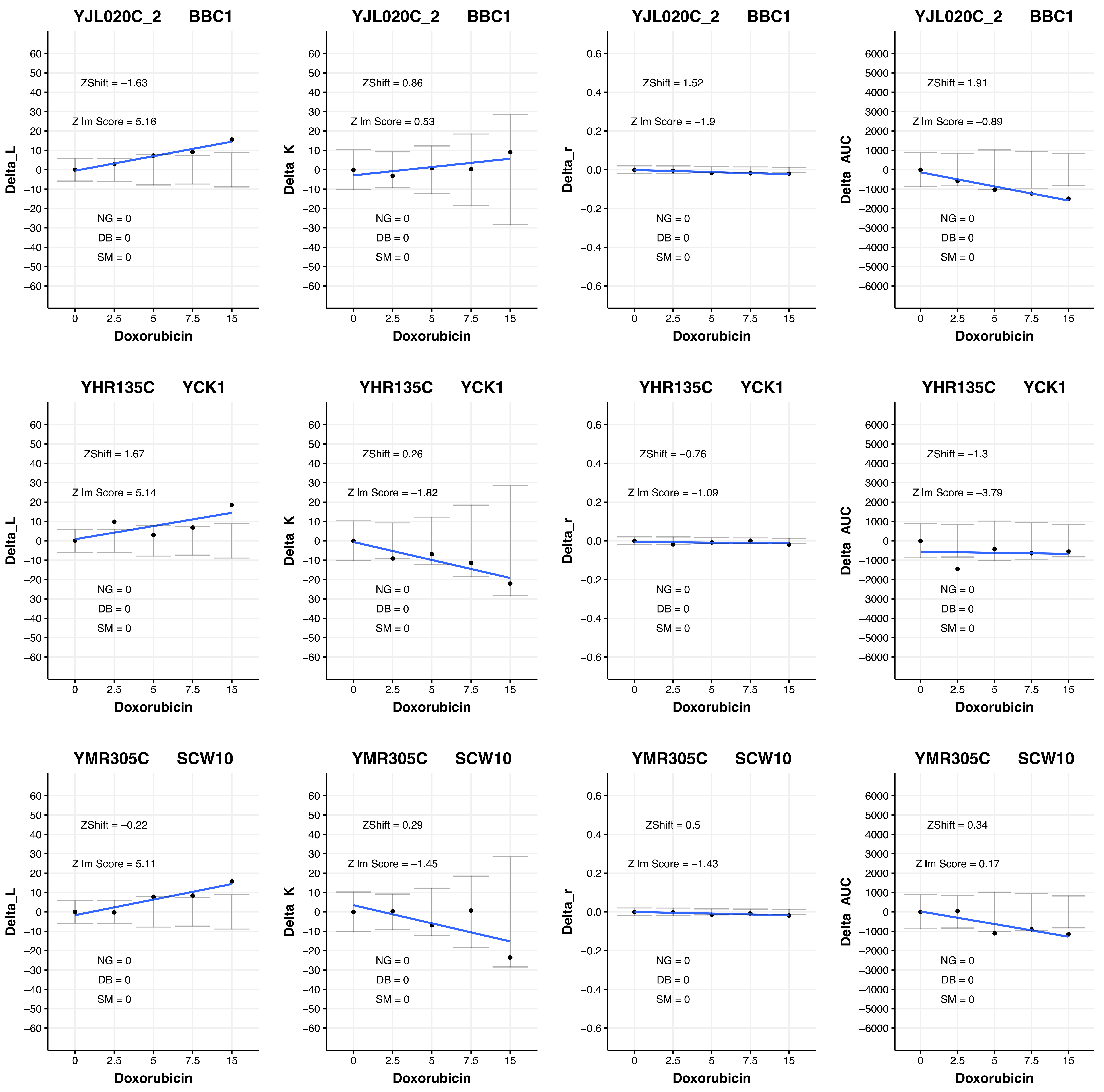

YJL020C_2 BBC1
ZShift = −1.63
Z lm Score = 5.16
Delta_L
NG = 0
DB = 0
SM = 0
Doxorubicin
YJL020C_2 BBC1
ZShift = 0.86
Z lm Score = 0.53
Delta_K
NG = 0
DB = 0
SM = 0
Doxorubicin
YJL020C_2 BBC1
ZShift = 1.52
Z lm Score = −1.9
Delta_r
NG = 0
DB = 0
SM = 0
Doxorubicin
YJL020C_2 BBC1
ZShift = 1.91
Z lm Score = −0.89
Delta_AUC
NG = 0
DB = 0
SM = 0
Doxorubicin
YHR135C YCK1
ZShift = 1.67
Z lm Score = 5.14
Delta_L
NG = 0
DB = 0
SM = 0
Doxorubicin
YHR135C YCK1
ZShift = 0.26
Z lm Score = −1.82
Delta_K
NG = 0
DB = 0
SM = 0
Doxorubicin
YHR135C YCK1
ZShift = −0.76
Z lm Score = −1.09
Delta_r
NG = 0
DB = 0
SM = 0
Doxorubicin
YHR135C YCK1
ZShift = −1.3
Z lm Score = −3.79
Delta_AUC
NG = 0
DB = 0
SM = 0
Doxorubicin
YMR305C SCW10
ZShift = −0.22
Z lm Score = 5.11
Delta_L
NG = 0
DB = 0
SM = 0
Doxorubicin
YMR305C SCW10
ZShift = 0.29
Z lm Score = −1.45
Delta_K
NG = 0
DB = 0
SM = 0
Doxorubicin
YMR305C SCW10
ZShift = 0.5
Z lm Score = −1.43
Delta_r
NG = 0
DB = 0
SM = 0
Doxorubicin
YMR305C SCW10
ZShift = 0.34
Z lm Score = 0.17
Delta_AUC
NG = 0
DB = 0
SM = 0
Doxorubicin

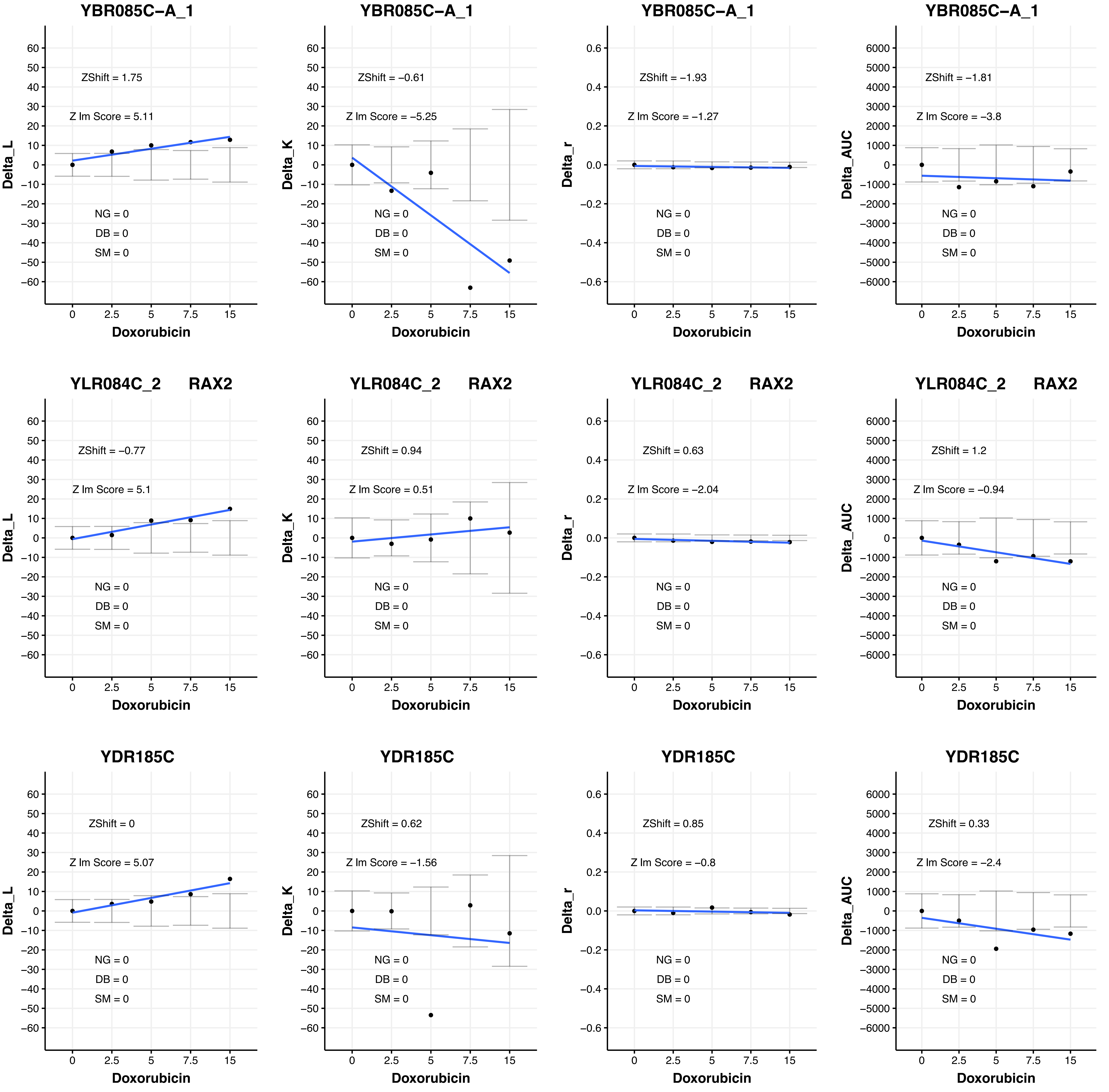
YBR085C−A_1
ZShift = 1.75
Z lm Score = 5.11
NG = 0
DB = 0
SM = 0
Delta_L
Doxorubicin
YBR085C−A_1
ZShift = −0.61
Z lm Score = −5.25
NG = 0
DB = 0
SM = 0
Delta_K
Doxorubicin
YBR085C−A_1
ZShift = −1.93
Z lm Score = −1.27
NG = 0
DB = 0
SM = 0
Delta_r
Doxorubicin
YBR085C−A_1
ZShift = −1.81
Z lm Score = −3.8
NG = 0
DB = 0
SM = 0
Delta_AUC
Doxorubicin
YLR084C_2 RAX2
ZShift = −0.77
Z lm Score = 5.1
NG = 0
DB = 0
SM = 0
Delta_L
Doxorubicin
YLR084C_2 RAX2
ZShift = 0.94
Z lm Score = 0.51
NG = 0
DB = 0
SM = 0
Delta_K
Doxorubicin
YLR084C_2 RAX2
ZShift = 0.63
Z lm Score = −2.04
NG = 0
DB = 0
SM = 0
Delta_r
Doxorubicin
YLR084C_2 RAX2
ZShift = 1.2
Z lm Score = −0.94
NG = 0
DB = 0
SM = 0
Delta_AUC
Doxorubicin
YDR185C
ZShift = 0
Z lm Score = 5.07
NG = 0
DB = 0
SM = 0
Delta_L
Doxorubicin
YDR185C
ZShift = 0.62
Z lm Score = −1.56
NG = 0
DB = 0
SM = 0
Delta_K
Doxorubicin
YDR185C
ZShift = 0.85
Z lm Score = −0.8
NG = 0
DB = 0
SM = 0
Delta_r
Doxorubicin
YDR185C
ZShift = 0.33
Z lm Score = −2.4
NG = 0
DB = 0
SM = 0
Delta_AUC
Doxorubicin

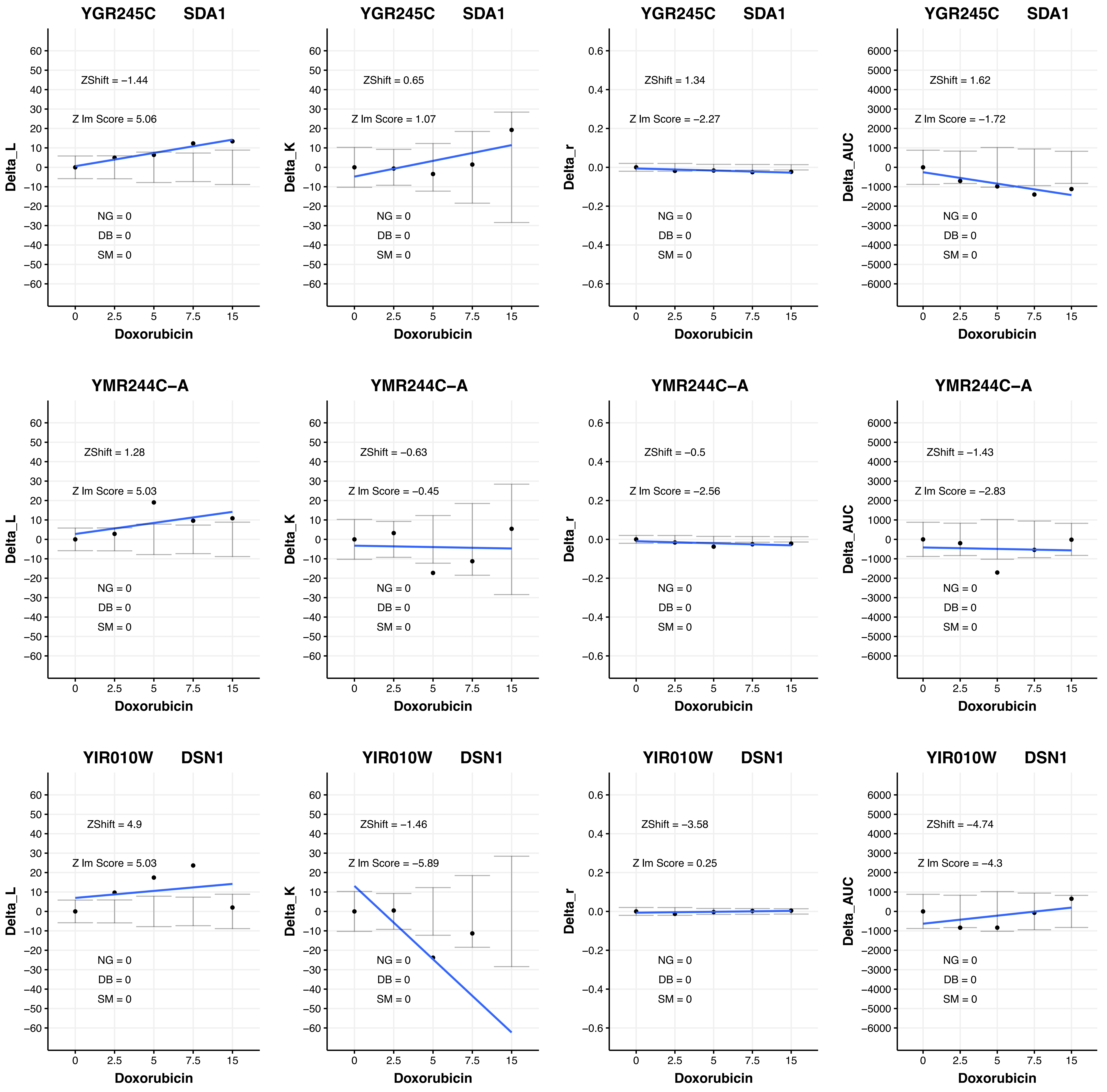
YGR245C SDA1
ZShift = −1.44
Z lm Score = 5.06
NG = 0
DB = 0
SM = 0
Delta_L
Doxorubicin
YGR245C SDA1
ZShift = 0.65
Z lm Score = 1.07
NG = 0
DB = 0
SM = 0
Delta_K
Doxorubicin
YGR245C SDA1
ZShift = 1.34
Z lm Score = −2.27
NG = 0
DB = 0
SM = 0
Delta_r
Doxorubicin
YGR245C SDA1
ZShift = 1.62
Z lm Score = −1.72
NG = 0
DB = 0
SM = 0
Delta_AUC
Doxorubicin
YMR244C−A
ZShift = 1.28
Z lm Score = 5.03
NG = 0
DB = 0
SM = 0
Delta_L
Doxorubicin
YMR244C−A
ZShift = −0.63
Z lm Score = −0.45
NG = 0
DB = 0
SM = 0
Delta_K
Doxorubicin
YMR244C−A
ZShift = −0.5
Z lm Score = −2.56
NG = 0
DB = 0
SM = 0
Delta_r
Doxorubicin
YMR244C−A
ZShift = −1.43
Z lm Score = −2.83
NG = 0
DB = 0
SM = 0
Delta_AUC
Doxorubicin
YIR010W DSN1
ZShift = 4.9
Z lm Score = 5.03
NG = 0
DB = 0
SM = 0
Delta_L
Doxorubicin
YIR010W DSN1
ZShift = −1.46
Z lm Score = −5.89
NG = 0
DB = 0
SM = 0
Delta_K
Doxorubicin
YIR010W DSN1
ZShift = −3.58
Z lm Score = 0.25
NG = 0
DB = 0
SM = 0
Delta_r
Doxorubicin
YIR010W DSN1
ZShift = −4.74
Z lm Score = −4.3
NG = 0
DB = 0
SM = 0
Delta_AUC
Doxorubicin

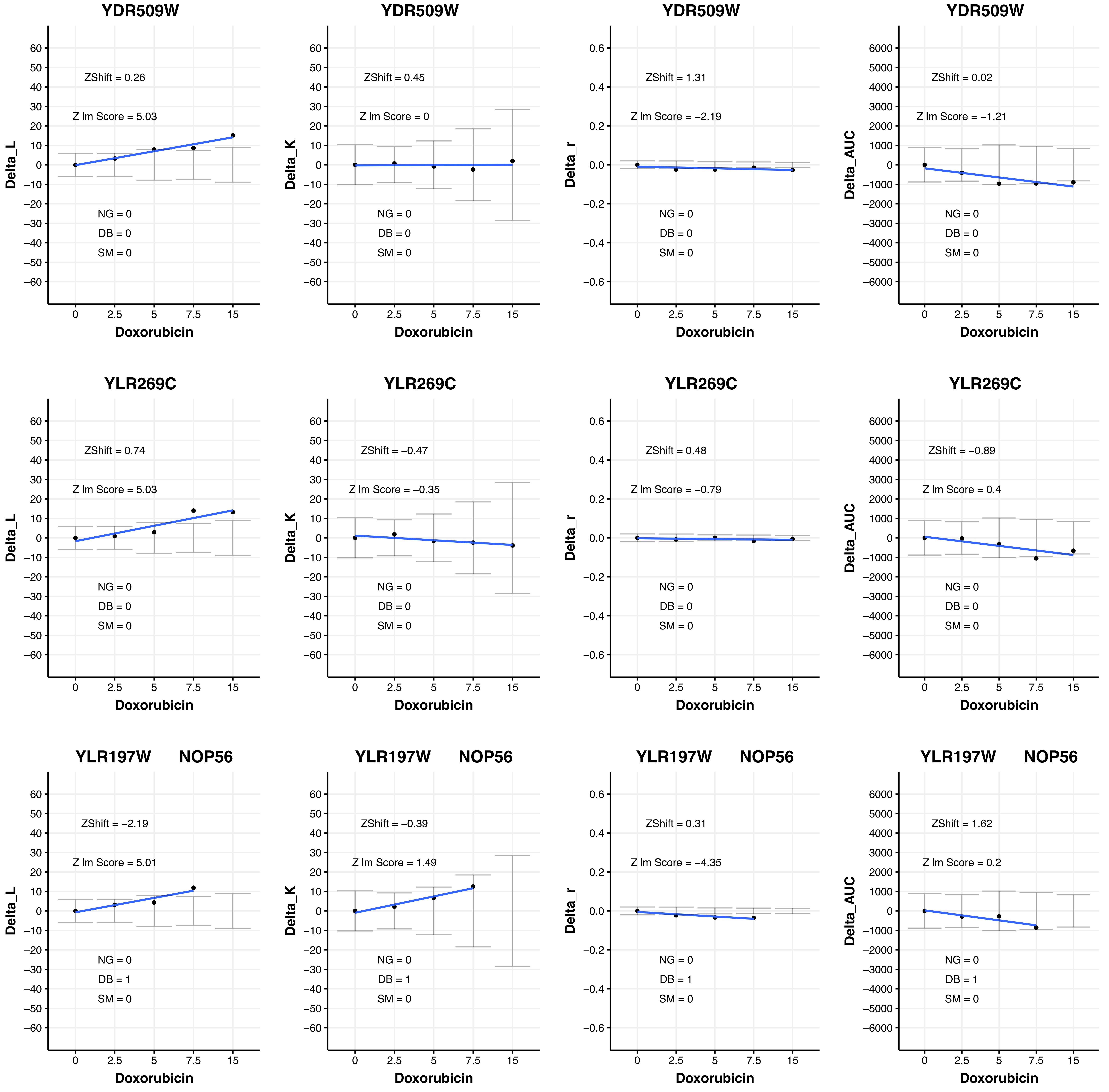

YDR509W
ZShift = 0.26
Z lm Score = 5.03
NG = 0
DB = 0
SM = 0
Delta_L
Doxorubicin
YDR509W
ZShift = 0.45
Z lm Score = 0
NG = 0
DB = 0
SM = 0
Delta_K
Doxorubicin
YDR509W
ZShift = 1.31
Z lm Score = −2.19
NG = 0
DB = 0
SM = 0
Delta_r
Doxorubicin
YDR509W
ZShift = 0.02
Z lm Score = −1.21
NG = 0
DB = 0
SM = 0
Delta_AUC
Doxorubicin
YLR269C
ZShift = 0.74
Z lm Score = 5.03
NG = 0
DB = 0
SM = 0
Delta_L
Doxorubicin
YLR269C
ZShift = −0.47
Z lm Score = −0.35
NG = 0
DB = 0
SM = 0
Delta_K
Doxorubicin
YLR269C
ZShift = 0.48
Z lm Score = −0.79
NG = 0
DB = 0
SM = 0
Delta_r
Doxorubicin
YLR269C
ZShift = −0.89
Z lm Score = 0.4
NG = 0
DB = 0
SM = 0
Delta_AUC
Doxorubicin
YLR197W NOP56
ZShift = −2.19
Z lm Score = 5.01
NG = 0
DB = 1
SM = 0
Delta_L
Doxorubicin
YLR197W NOP56
ZShift = −0.39
Z lm Score = 1.49
NG = 0
DB = 1
SM = 0
Delta_K
Doxorubicin
YLR197W NOP56
ZShift = 0.31
Z lm Score = −4.35
NG = 0
DB = 1
SM = 0
Delta_r
Doxorubicin
YLR197W NOP56
ZShift = 1.62
Z lm Score = 0.2
NG = 0
DB = 1
SM = 0
Delta_AUC
Doxorubicin

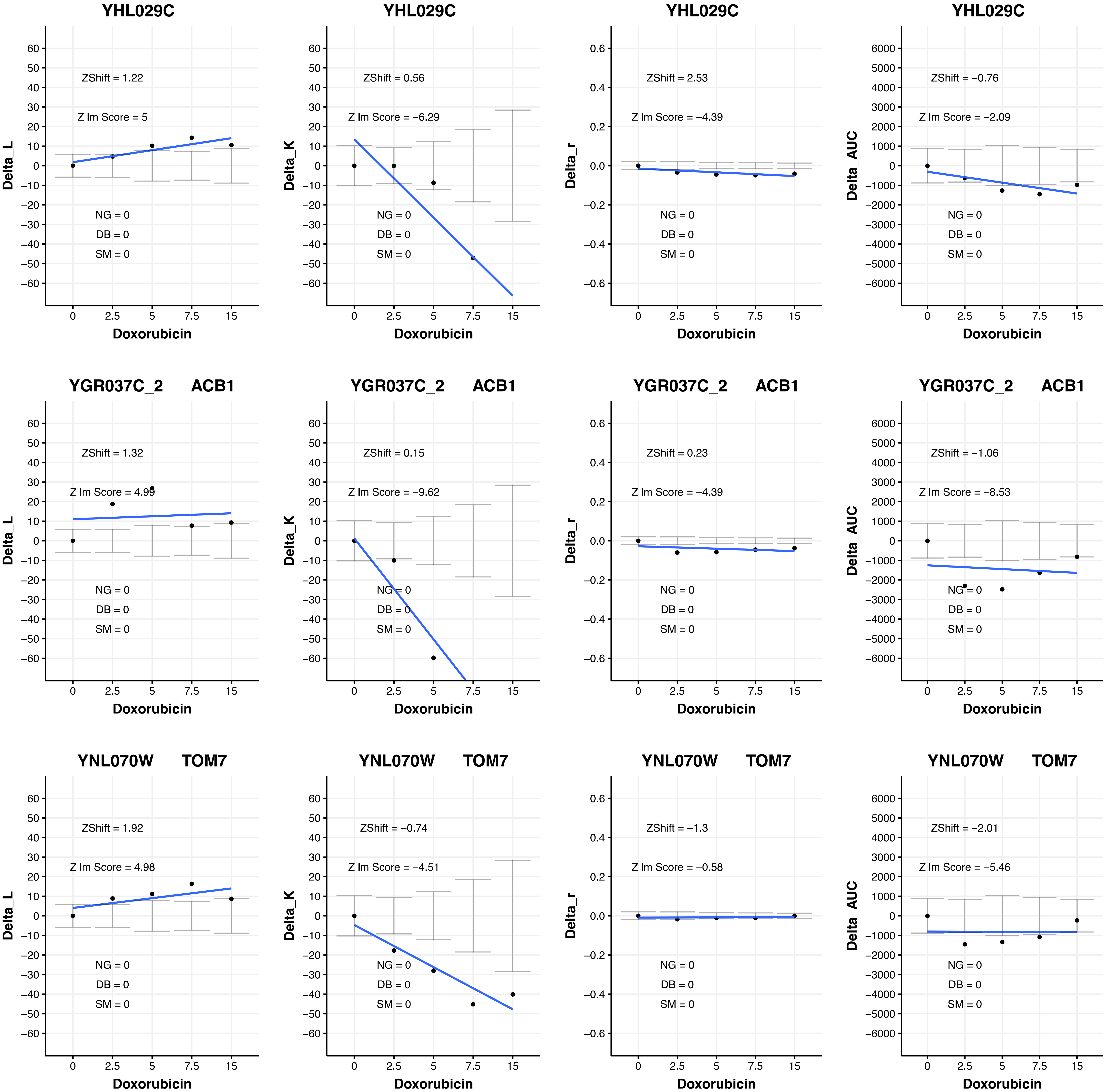

YHL029C
ZShift = 1.22
Z lm Score = 5
NG = 0
DB = 0
SM = 0
Delta_L
60 50 40 30 20 10 0 −10 −20 −30 −40 −50 −60
0 2.5 5 7.5 15
Doxorubicin
YHL029C
ZShift = 0.56
Z lm Score = −6.29
NG = 0
DB = 0
SM = 0
Delta_K
60 50 40 30 20 10 0 −10 −20 −30 −40 −50 −60
0 2.5 5 7.5 15
Doxorubicin
YHL029C
ZShift = 2.53
Z lm Score = −4.39
NG = 0
DB = 0
SM = 0
Delta_r
0.6 0.4 0.2 0.0 −0.2 −0.4 −0.6
0 2.5 5 7.5 15
Doxorubicin
YHL029C
ZShift = −0.76
Z lm Score = −2.09
NG = 0
DB = 0
SM = 0
Delta_AUC
6000 5000 4000 3000 2000 1000 0 −1000 −2000 −3000 −4000 −5000 −6000
0 2.5 5 7.5 15
Doxorubicin
YGR037C_2 ACB1
ZShift = 1.32
Z lm Score = 4.99
NG = 0
DB = 0
SM = 0
Delta_L
60 50 40 30 20 10 0 −10 −20 −30 −40 −50 −60
0 2.5 5 7.5 15
Doxorubicin
YGR037C_2 ACB1
ZShift = 0.15
Z lm Score = −9.62
NG = 0
DB = 0
SM = 0
Delta_K
60 50 40 30 20 10 0 −10 −20 −30 −40 −50 −60
0 2.5 5 7.5 15
Doxorubicin
YGR037C_2 ACB1
ZShift = 0.23
Z lm Score = −4.39
NG = 0
DB = 0
SM = 0
Delta_r
0.6 0.4 0.2 0.0 −0.2 −0.4 −0.6
0 2.5 5 7.5 15
Doxorubicin
YGR037C_2 ACB1
ZShift = −1.06
Z lm Score = −8.53
NG = 0
DB = 0
SM = 0
Delta_AUC
6000 5000 4000 3000 2000 1000 0 −1000 −2000 −3000 −4000 −5000 −6000
0 2.5 5 7.5 15
Doxorubicin
YNL070W TOM7
ZShift = 1.92
Z lm Score = 4.98
NG = 0
DB = 0
SM = 0
Delta_L
60 50 40 30 20 10 0 −10 −20 −30 −40 −50 −60
0 2.5 5 7.5 15
Doxorubicin
YNL070W TOM7
ZShift = −0.74
Z lm Score = −4.51
NG = 0
DB = 0
SM = 0
Delta_K
60 50 40 30 20 10 0 −10 −20 −30 −40 −50 −60
0 2.5 5 7.5 15
Doxorubicin
YNL070W TOM7
ZShift = −1.3
Z lm Score = −0.58
NG = 0
DB = 0
SM = 0
Delta_r
0.6 0.4 0.2 0.0 −0.2 −0.4 −0.6
0 2.5 5 7.5 15
Doxorubicin
YNL070W TOM7
ZShift = −2.01
Z lm Score = −5.46
NG = 0
DB = 0
SM = 0
Delta_AUC
6000 5000 4000 3000 2000 1000 0 −1000 −2000 −3000 −4000 −5000 −6000
0 2.5 5 7.5 15
Doxorubicin

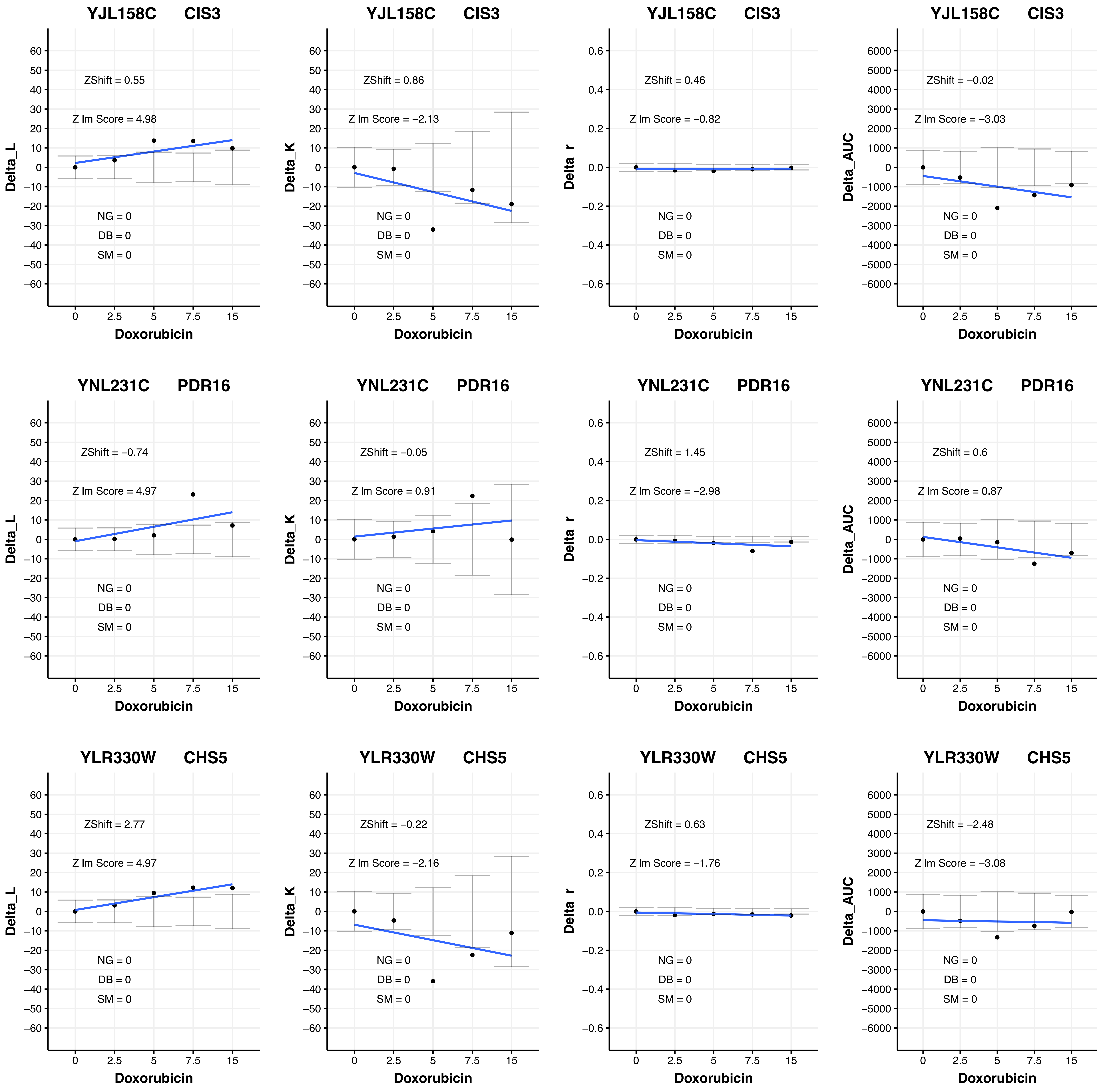

YJL158C CIS3
ZShift = 0.55
Z lm Score = 4.98
NG = 0
DB = 0
SM = 0
Delta_L
Doxorubicin
YJL158C CIS3
ZShift = 0.86
Z lm Score = −2.13
NG = 0
DB = 0
SM = 0
Delta_K
Doxorubicin
YJL158C CIS3
ZShift = 0.46
Z lm Score = −0.82
NG = 0
DB = 0
SM = 0
Delta_r
Doxorubicin
YJL158C CIS3
ZShift = −0.02
Z lm Score = −3.03
NG = 0
DB = 0
SM = 0
Delta_AUC
Doxorubicin
YNL231C PDR16
ZShift = −0.74
Z lm Score = 4.97
NG = 0
DB = 0
SM = 0
Delta_L
Doxorubicin
YNL231C PDR16
ZShift = −0.05
Z lm Score = 0.91
NG = 0
DB = 0
SM = 0
Delta_K
Doxorubicin
YNL231C PDR16
ZShift = 1.45
Z lm Score = −2.98
NG = 0
DB = 0
SM = 0
Delta_r
Doxorubicin
YNL231C PDR16
ZShift = 0.6
Z lm Score = 0.87
NG = 0
DB = 0
SM = 0
Delta_AUC
Doxorubicin
YLR330W CHS5
ZShift = 2.77
Z lm Score = 4.97
NG = 0
DB = 0
SM = 0
Delta_L
Doxorubicin
YLR330W CHS5
ZShift = −0.22
Z lm Score = −2.16
NG = 0
DB = 0
SM = 0
Delta_K
Doxorubicin
YLR330W CHS5
ZShift = 0.63
Z lm Score = −1.76
NG = 0
DB = 0
SM = 0
Delta_r
Doxorubicin
YLR330W CHS5
ZShift = −2.48
Z lm Score = −3.08
NG = 0
DB = 0
SM = 0
Delta_AUC
Doxorubicin

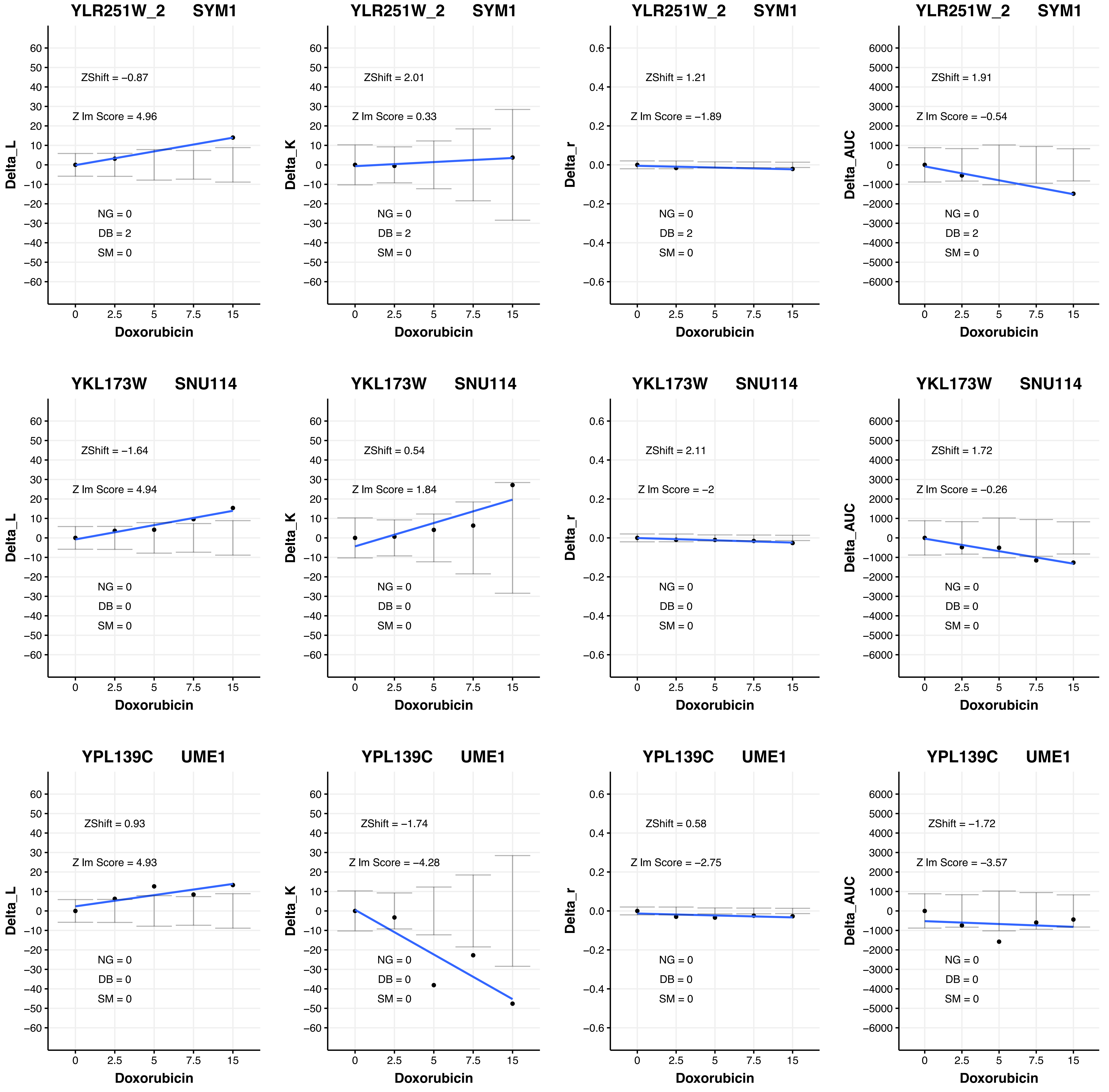

YLR251W_2 SYM1
ZShift = −0.87
Z lm Score = 4.96
NG = 0
DB = 2
SM = 0
Delta_L
Doxorubicin
YLR251W_2 SYM1
ZShift = 2.01
Z lm Score = 0.33
NG = 0
DB = 2
SM = 0
Delta_K
Doxorubicin
YLR251W_2 SYM1
ZShift = 1.21
Z lm Score = −1.89
NG = 0
DB = 2
SM = 0
Delta_r
Doxorubicin
YLR251W_2 SYM1
ZShift = 1.91
Z lm Score = −0.54
NG = 0
DB = 2
SM = 0
Delta_AUC
Doxorubicin
YKL173W SNU114
ZShift = −1.64
Z lm Score = 4.94
NG = 0
DB = 0
SM = 0
Delta_L
Doxorubicin
YKL173W SNU114
ZShift = 0.54
Z lm Score = 1.84
NG = 0
DB = 0
SM = 0
Delta_K
Doxorubicin
YKL173W SNU114
ZShift = 2.11
Z lm Score = −2
NG = 0
DB = 0
SM = 0
Delta_r
Doxorubicin
YKL173W SNU114
ZShift = 1.72
Z lm Score = −0.26
NG = 0
DB = 0
SM = 0
Delta_AUC
Doxorubicin
YPL139C UME1
ZShift = 0.93
Z lm Score = 4.93
NG = 0
DB = 0
SM = 0
Delta_L
Doxorubicin
YPL139C UME1
ZShift = −1.74
Z lm Score = −4.28
NG = 0
DB = 0
SM = 0
Delta_K
Doxorubicin
YPL139C UME1
ZShift = 0.58
Z lm Score = −2.75
NG = 0
DB = 0
SM = 0
Delta_r
Doxorubicin
YPL139C UME1
ZShift = −1.72
Z lm Score = −3.57
NG = 0
DB = 0
SM = 0
Delta_AUC
Doxorubicin

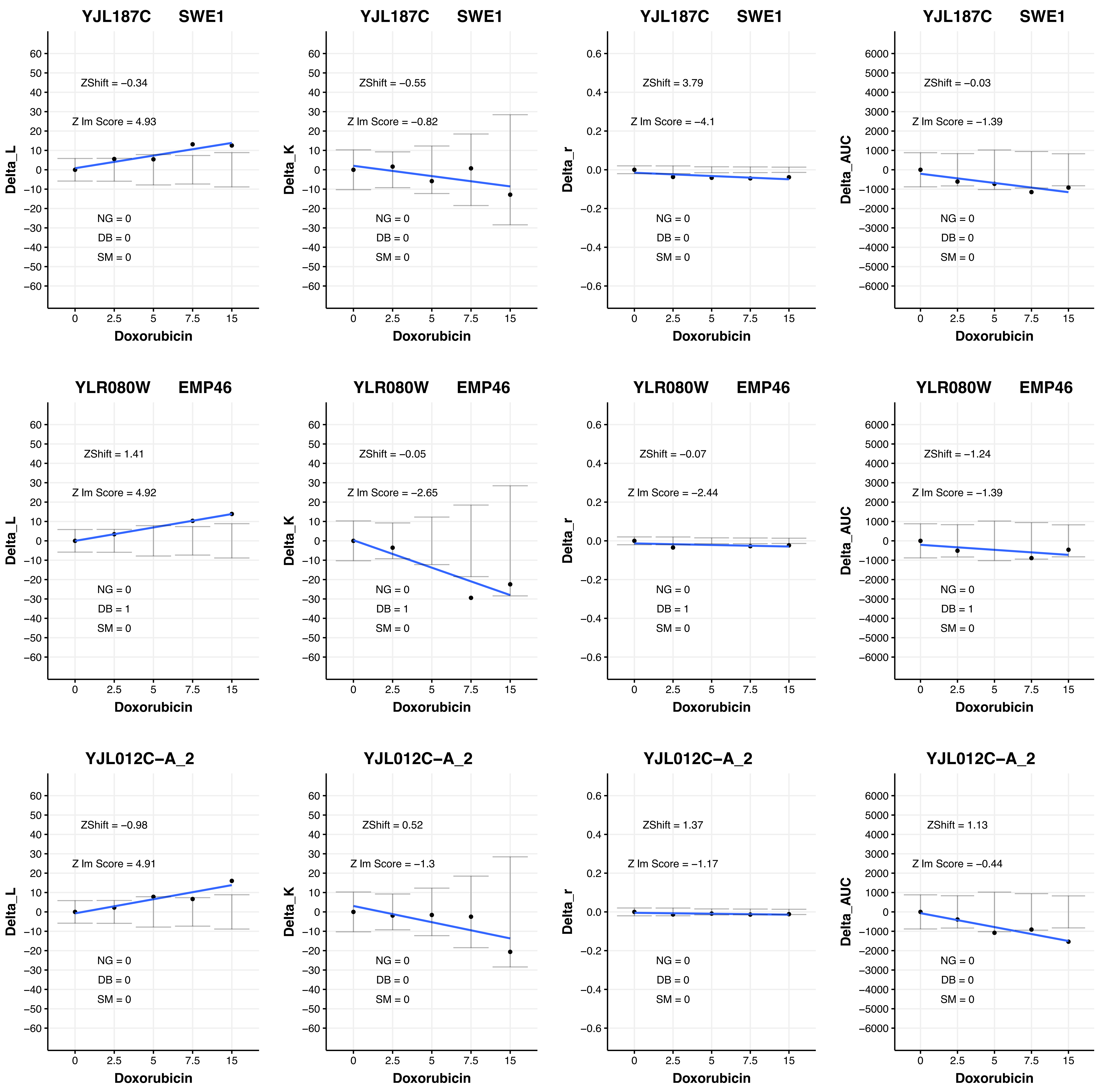
YJL187C SWE1
ZShift = −0.34
Z lm Score = 4.93
NG = 0
DB = 0
SM = 0
Delta_L
Doxorubicin
YJL187C SWE1
ZShift = −0.55
Z lm Score = −0.82
NG = 0
DB = 0
SM = 0
Delta_K
Doxorubicin
YJL187C SWE1
ZShift = 3.79
Z lm Score = −4.1
NG = 0
DB = 0
SM = 0
Delta_r
Doxorubicin
YJL187C SWE1
ZShift = −0.03
Z lm Score = −1.39
NG = 0
DB = 0
SM = 0
Delta_AUC
Doxorubicin
YLR080W EMP46
ZShift = 1.41
Z lm Score = 4.92
NG = 0
DB = 1
SM = 0
Delta_L
Doxorubicin
YLR080W EMP46
ZShift = −0.05
Z lm Score = −2.65
NG = 0
DB = 1
SM = 0
Delta_K
Doxorubicin
YLR080W EMP46
ZShift = −0.07
Z lm Score = −2.44
NG = 0
DB = 1
SM = 0
Delta_r
Doxorubicin
YLR080W EMP46
ZShift = −1.24
Z lm Score = −1.39
NG = 0
DB = 1
SM = 0
Delta_AUC
Doxorubicin
YJL012C−A_2
ZShift = −0.98
Z lm Score = 4.91
NG = 0
DB = 0
SM = 0
Delta_L
Doxorubicin
YJL012C−A_2
ZShift = 0.52
Z lm Score = −1.3
NG = 0
DB = 0
SM = 0
Delta_K
Doxorubicin
YJL012C−A_2
ZShift = 1.37
Z lm Score = −1.17
NG = 0
DB = 0
SM = 0
Delta_r
Doxorubicin
YJL012C−A_2
ZShift = 1.13
Z lm Score = −0.44
NG = 0
DB = 0
SM = 0
Delta_AUC
Doxorubicin

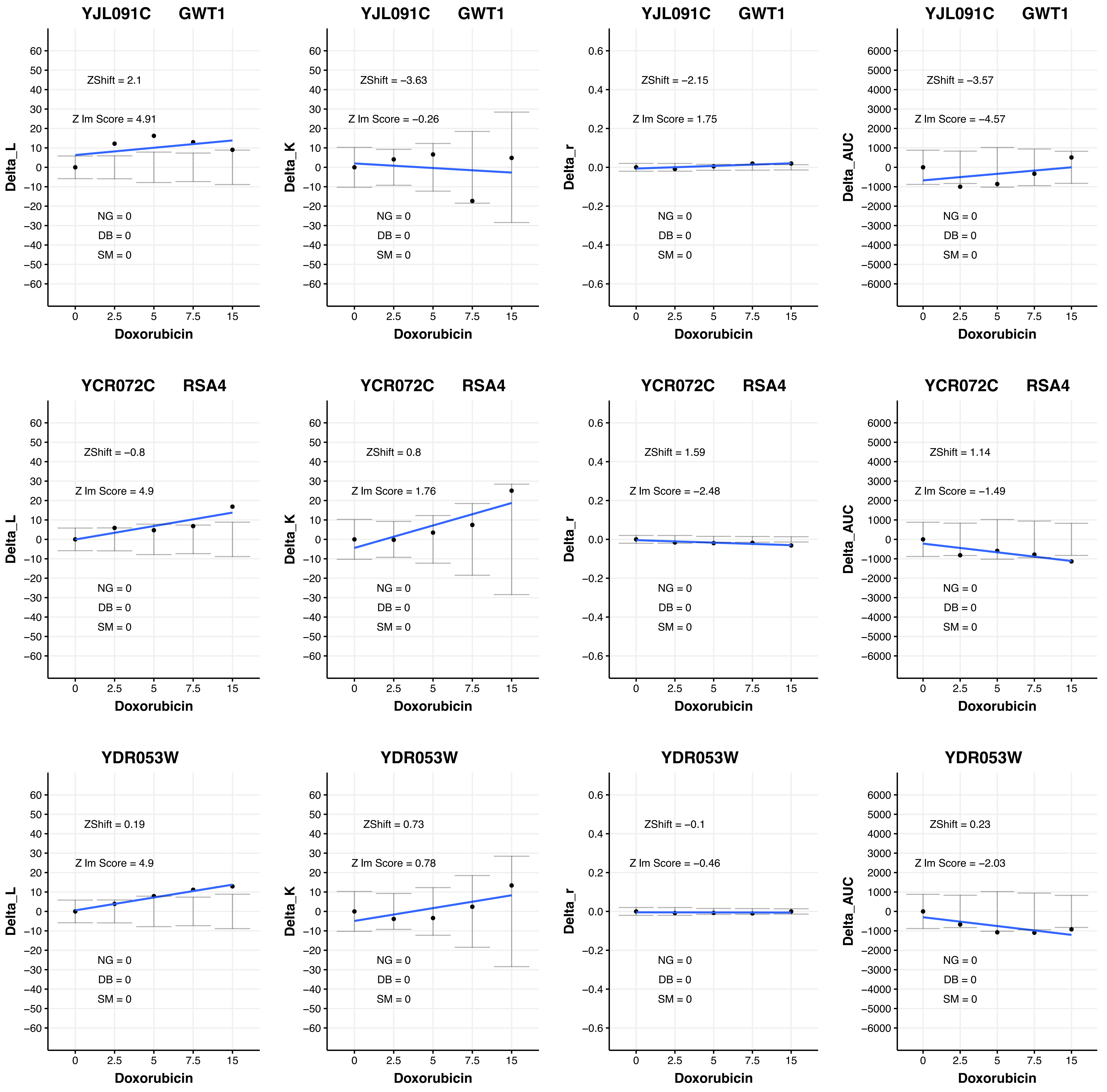
YJL091C GWT1
ZShift = 2.1
Z lm Score = 4.91
NG = 0
DB = 0
SM = 0
Delta_L
Doxorubicin
YJL091C GWT1
ZShift = −3.63
Z lm Score = −0.26
NG = 0
DB = 0
SM = 0
Delta_K
Doxorubicin
YJL091C GWT1
ZShift = −2.15
Z lm Score = 1.75
NG = 0
DB = 0
SM = 0
Delta_r
Doxorubicin
YJL091C GWT1
ZShift = −3.57
Z lm Score = −4.57
NG = 0
DB = 0
SM = 0
Delta_AUC
Doxorubicin
YCR072C RSA4
ZShift = −0.8
Z lm Score = 4.9
NG = 0
DB = 0
SM = 0
Delta_L
Doxorubicin
YCR072C RSA4
ZShift = 0.8
Z lm Score = 1.76
NG = 0
DB = 0
SM = 0
Delta_K
Doxorubicin
YCR072C RSA4
ZShift = 1.59
Z lm Score = −2.48
NG = 0
DB = 0
SM = 0
Delta_r
Doxorubicin
YCR072C RSA4
ZShift = 1.14
Z lm Score = −1.49
NG = 0
DB = 0
SM = 0
Delta_AUC
Doxorubicin
YDR053W
ZShift = 0.19
Z lm Score = 4.9
NG = 0
DB = 0
SM = 0
Delta_L
Doxorubicin
YDR053W
ZShift = 0.73
Z lm Score = 0.78
NG = 0
DB = 0
SM = 0
Delta_K
Doxorubicin
YDR053W
ZShift = −0.1
Z lm Score = −0.46
NG = 0
DB = 0
SM = 0
Delta_r
Doxorubicin
YDR053W
ZShift = 0.23
Z lm Score = −2.03
NG = 0
DB = 0
SM = 0
Delta_AUC
Doxorubicin

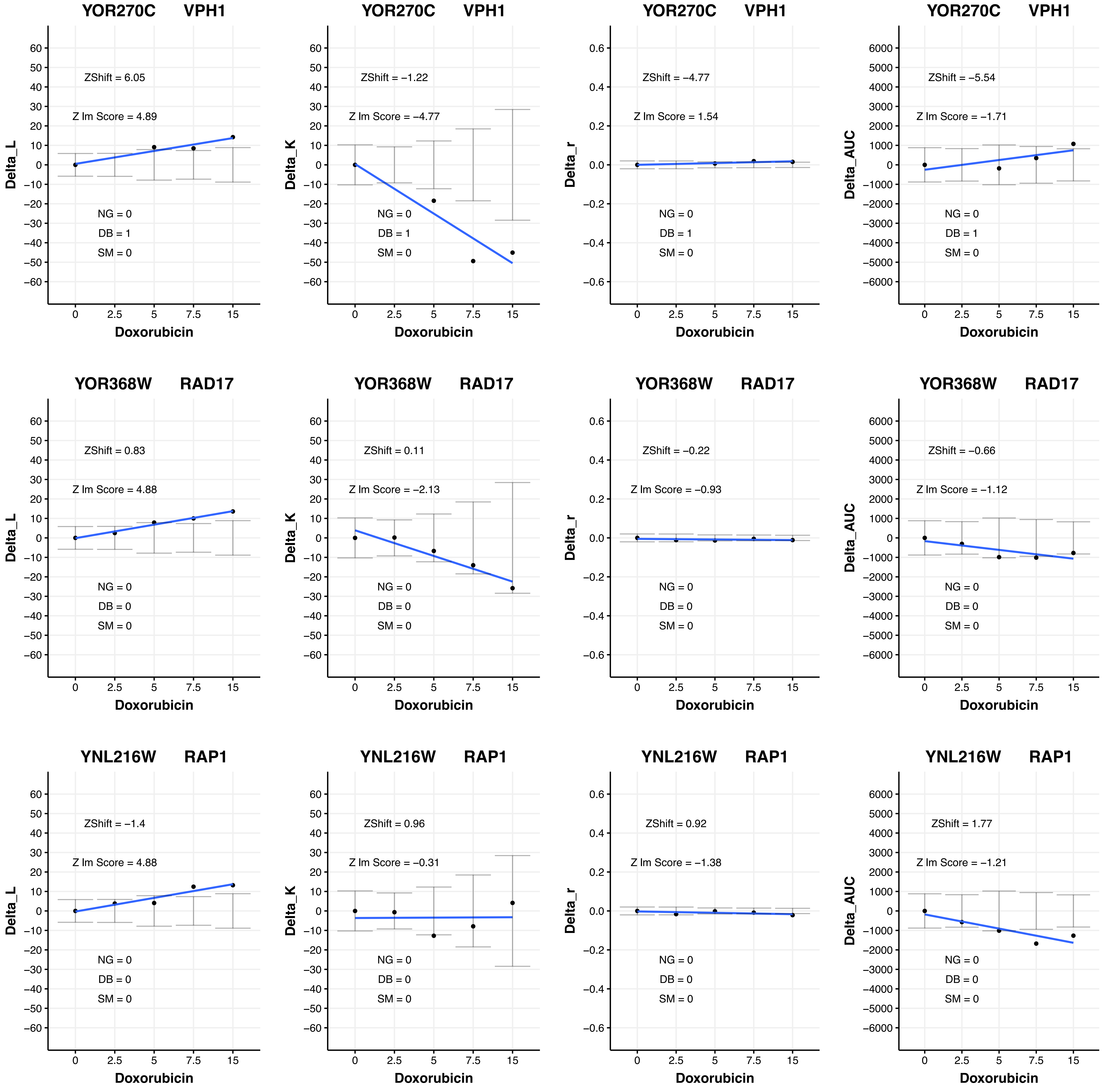
YOR270C VPH1
ZShift = 6.05
Z lm Score = 4.89
NG = 0
DB = 1
SM = 0
Delta_L
Doxorubicin
YOR270C VPH1
ZShift = −1.22
Z lm Score = −4.77
NG = 0
DB = 1
SM = 0
Delta_K
Doxorubicin
YOR270C VPH1
ZShift = −4.77
Z lm Score = 1.54
NG = 0
DB = 1
SM = 0
Delta_r
Doxorubicin
YOR270C VPH1
ZShift = −5.54
Z lm Score = −1.71
NG = 0
DB = 1
SM = 0
Delta_AUC
Doxorubicin
YOR368W RAD17
ZShift = 0.83
Z lm Score = 4.88
NG = 0
DB = 0
SM = 0
Delta_L
Doxorubicin
YOR368W RAD17
ZShift = 0.11
Z lm Score = −2.13
NG = 0
DB = 0
SM = 0
Delta_K
Doxorubicin
YOR368W RAD17
ZShift = −0.22
Z lm Score = −0.93
NG = 0
DB = 0
SM = 0
Delta_r
Doxorubicin
YOR368W RAD17
ZShift = −0.66
Z lm Score = −1.12
NG = 0
DB = 0
SM = 0
Delta_AUC
Doxorubicin
YNL216W RAP1
ZShift = −1.4
Z lm Score = 4.88
NG = 0
DB = 0
SM = 0
Delta_L
Doxorubicin
YNL216W RAP1
ZShift = 0.96
Z lm Score = −0.31
NG = 0
DB = 0
SM = 0
Delta_K
Doxorubicin
YNL216W RAP1
ZShift = 0.92
Z lm Score = −1.38
NG = 0
DB = 0
SM = 0
Delta_r
Doxorubicin
YNL216W RAP1
ZShift = 1.77
Z lm Score = −1.21
NG = 0
DB = 0
SM = 0
Delta_AUC
Doxorubicin

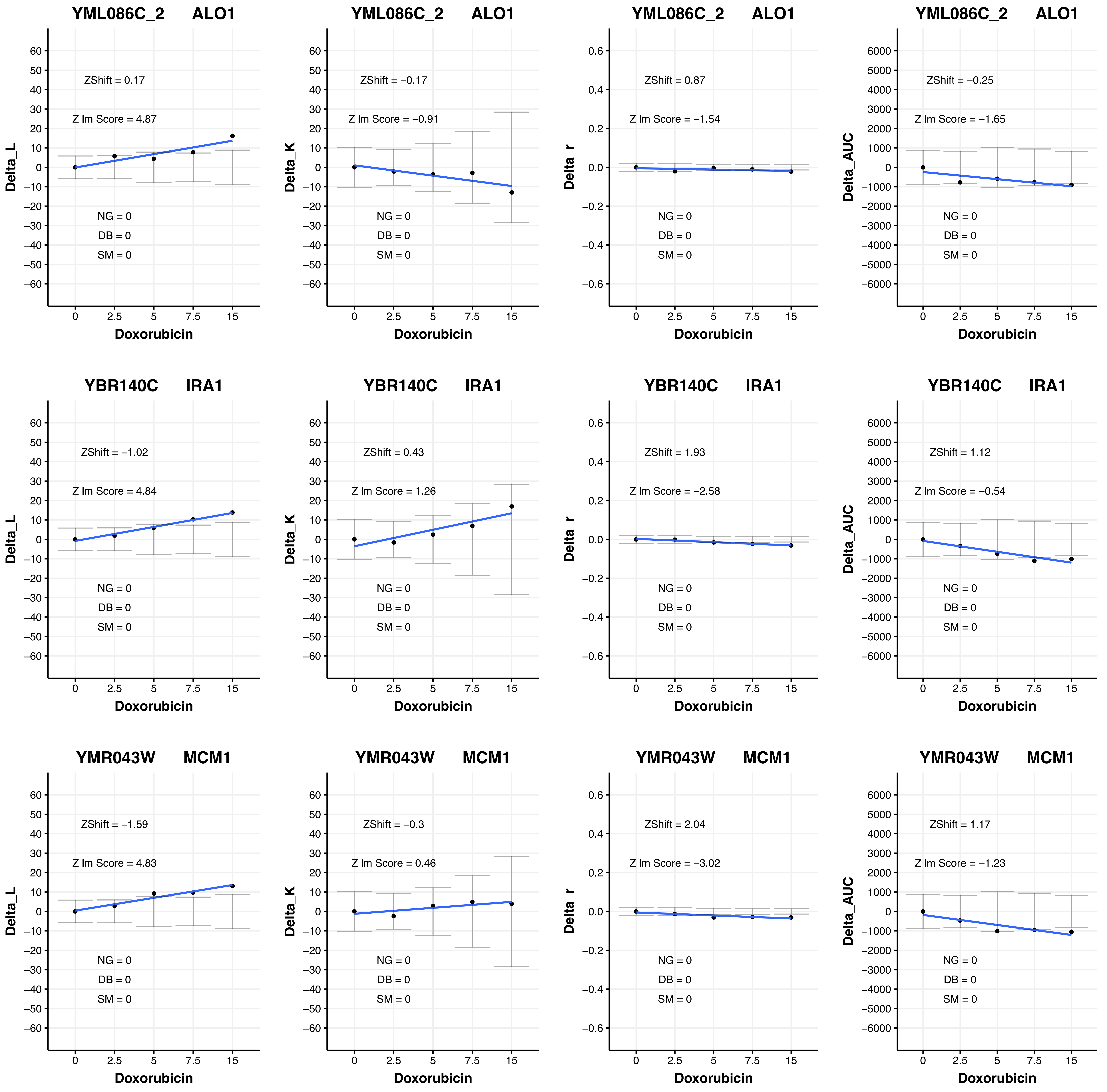

YML086C_2 ALO1
ZShift = 0.17
Z lm Score = 4.87
NG = 0
DB = 0
SM = 0
Delta_L
Doxorubicin
YML086C_2 ALO1
ZShift = −0.17
Z lm Score = −0.91
NG = 0
DB = 0
SM = 0
Delta_K
Doxorubicin
YML086C_2 ALO1
ZShift = 0.87
Z lm Score = −1.54
NG = 0
DB = 0
SM = 0
Delta_r
Doxorubicin
YML086C_2 ALO1
ZShift = −0.25
Z lm Score = −1.65
NG = 0
DB = 0
SM = 0
Delta_AUC
Doxorubicin
YBR140C IRA1
ZShift = −1.02
Z lm Score = 4.84
NG = 0
DB = 0
SM = 0
Delta_L
Doxorubicin
YBR140C IRA1
ZShift = 0.43
Z lm Score = 1.26
NG = 0
DB = 0
SM = 0
Delta_K
Doxorubicin
YBR140C IRA1
ZShift = 1.93
Z lm Score = −2.58
NG = 0
DB = 0
SM = 0
Delta_r
Doxorubicin
YBR140C IRA1
ZShift = 1.12
Z lm Score = −0.54
NG = 0
DB = 0
SM = 0
Delta_AUC
Doxorubicin
YMR043W MCM1
ZShift = −1.59
Z lm Score = 4.83
NG = 0
DB = 0
SM = 0
Delta_L
Doxorubicin
YMR043W MCM1
ZShift = −0.3
Z lm Score = 0.46
NG = 0
DB = 0
SM = 0
Delta_K
Doxorubicin
YMR043W MCM1
ZShift = 2.04
Z lm Score = −3.02
NG = 0
DB = 0
SM = 0
Delta_r
Doxorubicin
YMR043W MCM1
ZShift = 1.17
Z lm Score = −1.23
NG = 0
DB = 0
SM = 0
Delta_AUC
Doxorubicin

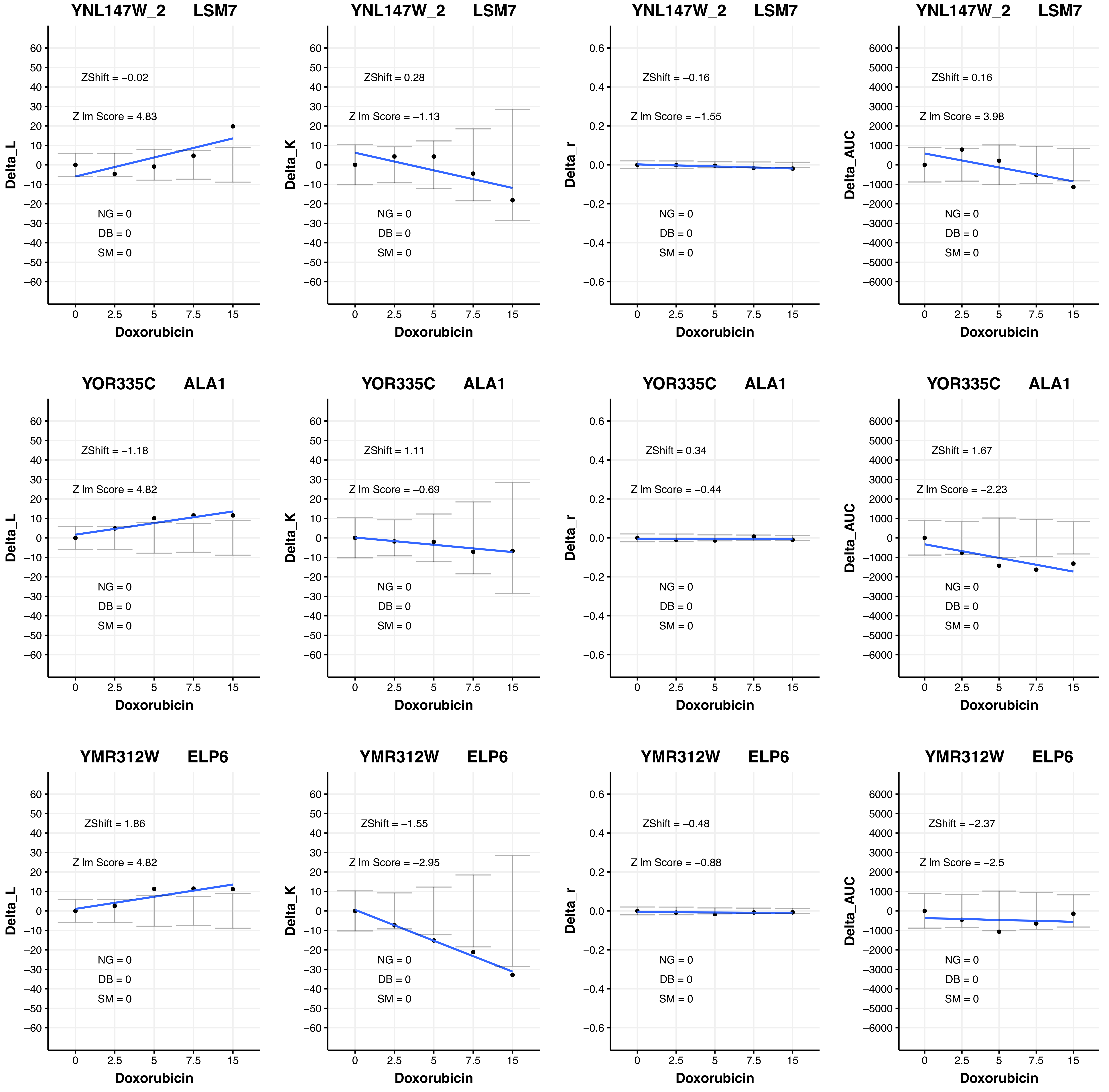
YNL147W_2 LSM7
ZShift = −0.02
Z lm Score = 4.83
NG = 0
DB = 0
SM = 0
Delta_L
Doxorubicin
YNL147W_2 LSM7
ZShift = 0.28
Z lm Score = −1.13
NG = 0
DB = 0
SM = 0
Delta_K
Doxorubicin
YNL147W_2 LSM7
ZShift = −0.16
Z lm Score = −1.55
NG = 0
DB = 0
SM = 0
Delta_r
Doxorubicin
YNL147W_2 LSM7
ZShift = 0.16
Z lm Score = 3.98
NG = 0
DB = 0
SM = 0
Delta_AUC
Doxorubicin
YOR335C ALA1
ZShift = −1.18
Z lm Score = 4.82
NG = 0
DB = 0
SM = 0
Delta_L
Doxorubicin
YOR335C ALA1
ZShift = 1.11
Z lm Score = −0.69
NG = 0
DB = 0
SM = 0
Delta_K
Doxorubicin
YOR335C ALA1
ZShift = 0.34
Z lm Score = −0.44
NG = 0
DB = 0
SM = 0
Delta_r
Doxorubicin
YOR335C ALA1
ZShift = 1.67
Z lm Score = −2.23
NG = 0
DB = 0
SM = 0
Delta_AUC
Doxorubicin
YMR312W ELP6
ZShift = 1.86
Z lm Score = 4.82
NG = 0
DB = 0
SM = 0
Delta_L
Doxorubicin
YMR312W ELP6
ZShift = −1.55
Z lm Score = −2.95
NG = 0
DB = 0
SM = 0
Delta_K
Doxorubicin
YMR312W ELP6
ZShift = −0.48
Z lm Score = −0.88
NG = 0
DB = 0
SM = 0
Delta_r
Doxorubicin
YMR312W ELP6
ZShift = −2.37
Z lm Score = −2.5
NG = 0
DB = 0
SM = 0
Delta_AUC
Doxorubicin

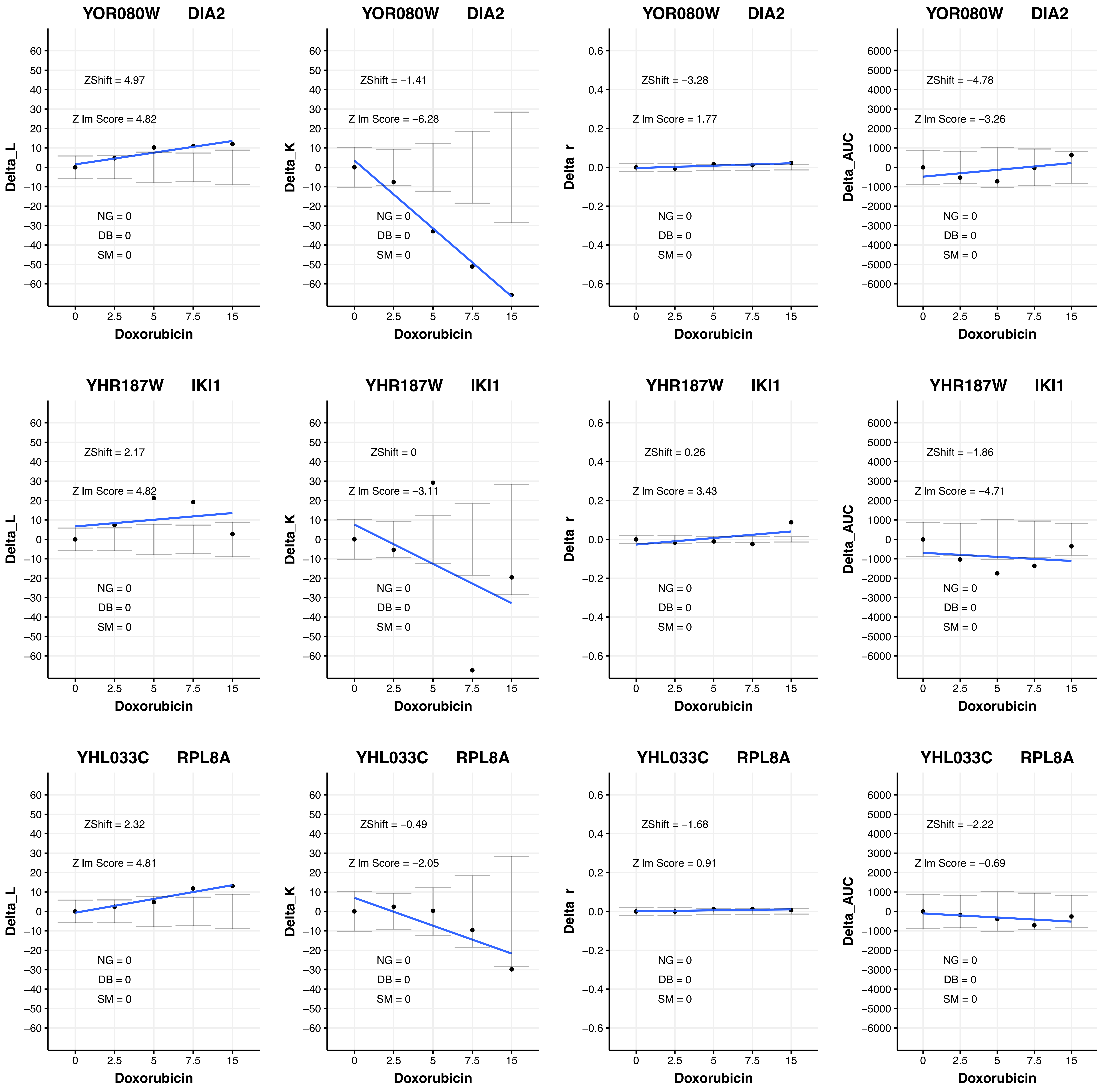
YOR080W DIA2
ZShift = 4.97
Z lm Score = 4.82
NG = 0
DB = 0
SM = 0
Delta_L
Doxorubicin
YOR080W DIA2
ZShift = −1.41
Z lm Score = −6.28
NG = 0
DB = 0
SM = 0
Delta_K
Doxorubicin
YOR080W DIA2
ZShift = −3.28
Z lm Score = 1.77
NG = 0
DB = 0
SM = 0
Delta_r
Doxorubicin
YOR080W DIA2
ZShift = −4.78
Z lm Score = −3.26
NG = 0
DB = 0
SM = 0
Delta_AUC
Doxorubicin
YHR187W IKI1
ZShift = 2.17
Z lm Score = 4.82
NG = 0
DB = 0
SM = 0
Delta_L
Doxorubicin
YHR187W IKI1
ZShift = 0
Z lm Score = −3.11
NG = 0
DB = 0
SM = 0
Delta_K
Doxorubicin
YHR187W IKI1
ZShift = 0.26
Z lm Score = 3.43
NG = 0
DB = 0
SM = 0
Delta_r
Doxorubicin
YHR187W IKI1
ZShift = −1.86
Z lm Score = −4.71
NG = 0
DB = 0
SM = 0
Delta_AUC
Doxorubicin
YHL033C RPL8A
ZShift = 2.32
Z lm Score = 4.81
NG = 0
DB = 0
SM = 0
Delta_L
Doxorubicin
YHL033C RPL8A
ZShift = −0.49
Z lm Score = −2.05
NG = 0
DB = 0
SM = 0
Delta_K
Doxorubicin
YHL033C RPL8A
ZShift = −1.68
Z lm Score = 0.91
NG = 0
DB = 0
SM = 0
Delta_r
Doxorubicin
YHL033C RPL8A
ZShift = −2.22
Z lm Score = −0.69
NG = 0
DB = 0
SM = 0
Delta_AUC
Doxorubicin

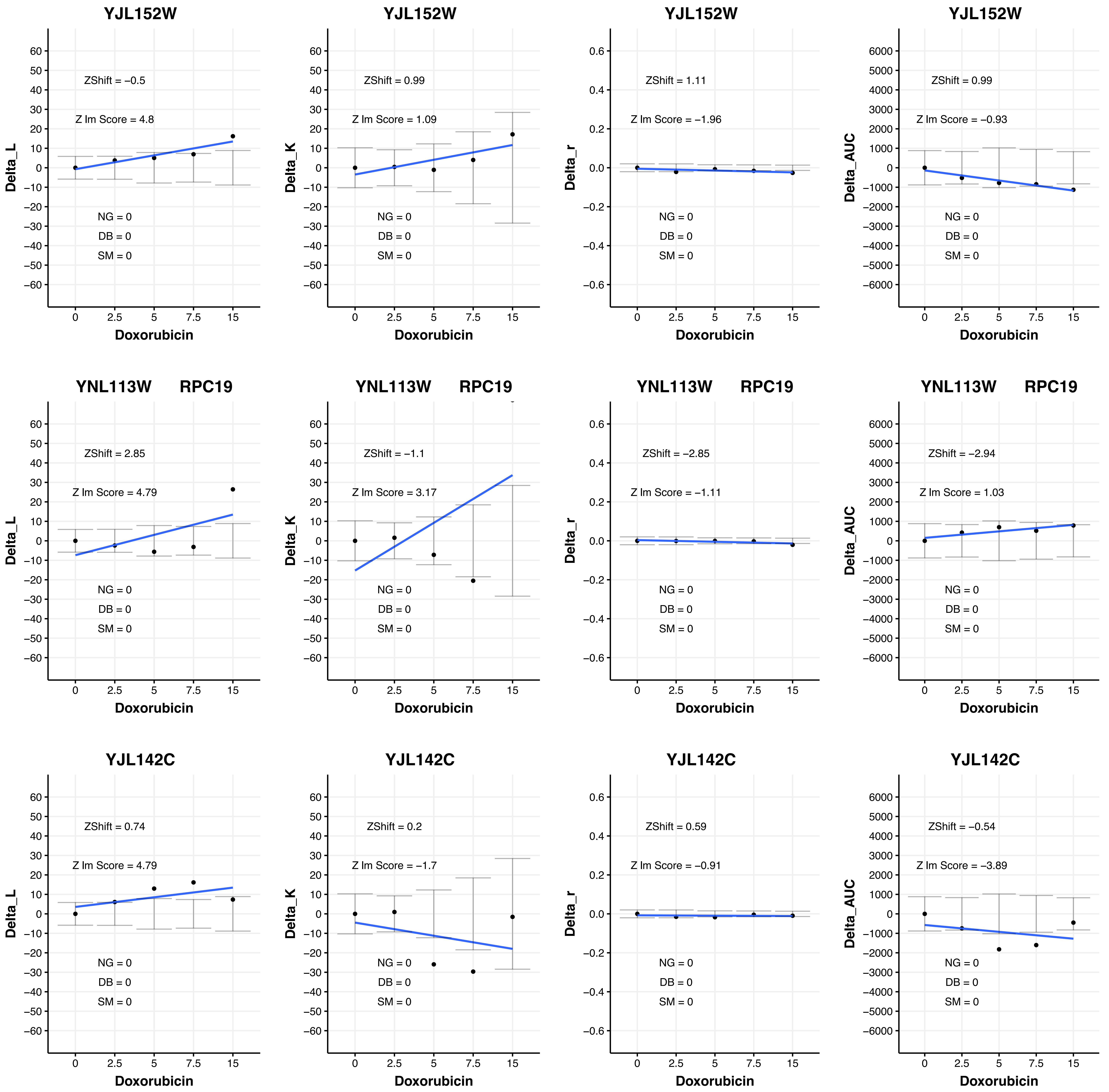

YJL152W
ZShift = −0.5
Z lm Score = 4.8
NG = 0
DB = 0
SM = 0
Delta_L
Doxorubicin
YJL152W
ZShift = 0.99
Z lm Score = 1.09
Delta_K
YJL152W
ZShift = 1.11
Z lm Score = −1.96
Delta_r
YJL152W
ZShift = 0.99
Z lm Score = −0.93
Delta_AUC
YNL113W RPC19
ZShift = 2.85
Z lm Score = 4.79
ZShift = −1.1
Z lm Score = 3.17
ZShift = −2.85
Z lm Score = −1.11
ZShift = −2.94
Z lm Score = 1.03
YJL142C
ZShift = 0.74
Z lm Score = 4.79
ZShift = 0.2
Z lm Score = −1.7
ZShift = 0.59
Z lm Score = −0.91
ZShift = −0.54
Z lm Score = −3.89

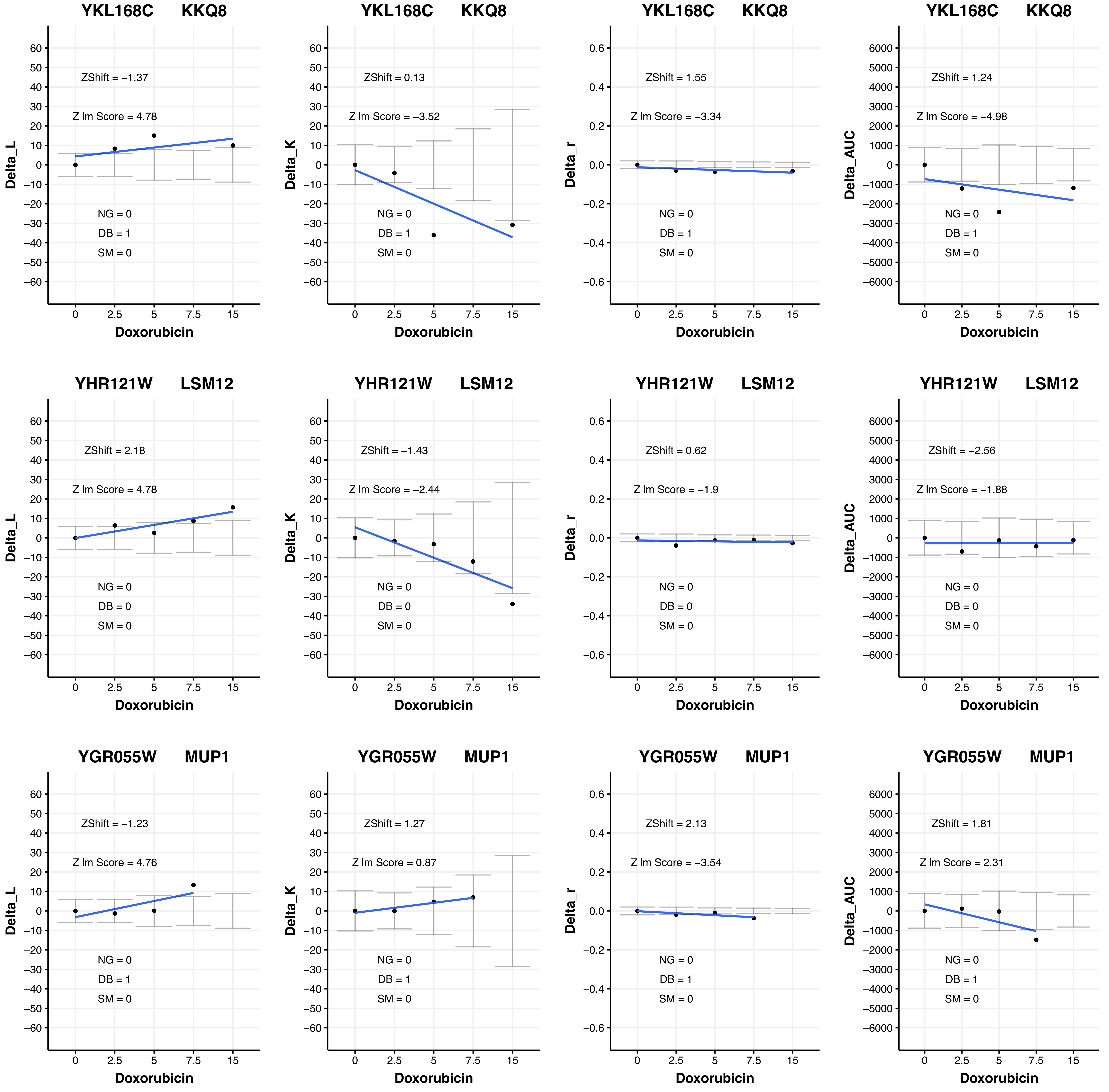
YKL168C KKQ8
ZShift = −1.37
Z lm Score = 4.78
NG = 0
DB = 1
SM = 0
Delta_L
Doxorubicin
YKL168C KKQ8
ZShift = 0.13
Z lm Score = −3.52
NG = 0
DB = 1
SM = 0
Delta_K
Doxorubicin
YKL168C KKQ8
ZShift = 1.55
Z lm Score = −3.34
NG = 0
DB = 1
SM = 0
Delta_r
Doxorubicin
YKL168C KKQ8
ZShift = 1.24
Z lm Score = −4.98
NG = 0
DB = 1
SM = 0
Delta_AUC
Doxorubicin
YHR121W LSM12
ZShift = 2.18
Z lm Score = 4.78
NG = 0
DB = 0
SM = 0
Delta_L
Doxorubicin
YHR121W LSM12
ZShift = −1.43
Z lm Score = −2.44
NG = 0
DB = 0
SM = 0
Delta_K
Doxorubicin
YHR121W LSM12
ZShift = 0.62
Z lm Score = −1.9
NG = 0
DB = 0
SM = 0
Delta_r
Doxorubicin
YHR121W LSM12
ZShift = −2.56
Z lm Score = −1.88
NG = 0
DB = 0
SM = 0
Delta_AUC
Doxorubicin
YGR055W MUP1
ZShift = −1.23
Z lm Score = 4.76
NG = 0
DB = 1
SM = 0
Delta_L
Doxorubicin
YGR055W MUP1
ZShift = 1.27
Z lm Score = 0.87
NG = 0
DB = 1
SM = 0
Delta_K
Doxorubicin
YGR055W MUP1
ZShift = 2.13
Z lm Score = −3.54
NG = 0
DB = 1
SM = 0
Delta_r
Doxorubicin
YGR055W MUP1
ZShift = 1.81
Z lm Score = 2.31
NG = 0
DB = 1
SM = 0
Delta_AUC
Doxorubicin

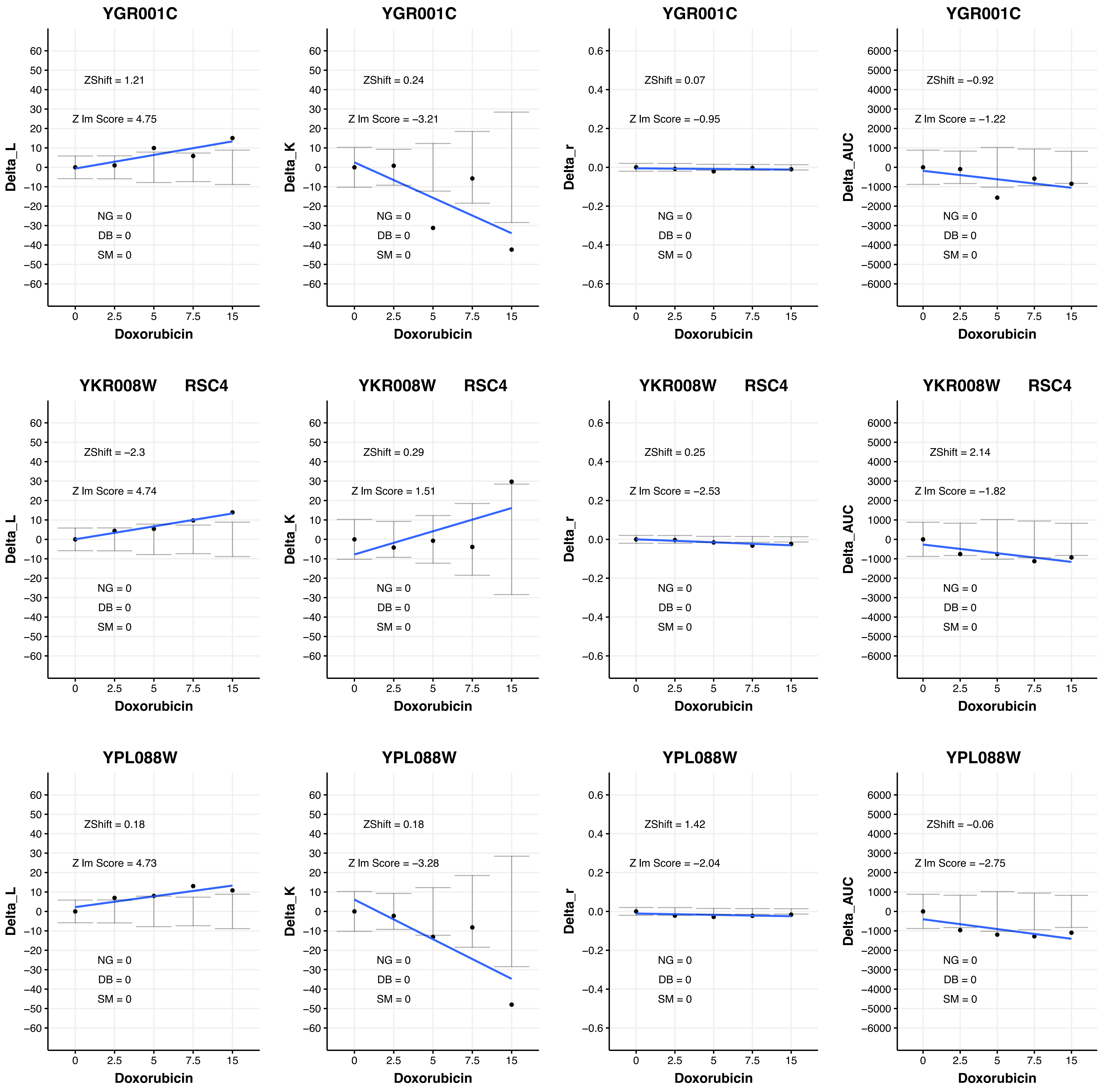

YGR001C
ZShift = 1.21
Z lm Score = 4.75
NG = 0
DB = 0
SM = 0
Delta_L
Doxorubicin
YGR001C
ZShift = 0.24
Z lm Score = −3.21
NG = 0
DB = 0
SM = 0
Delta_K
Doxorubicin
YGR001C
ZShift = 0.07
Z lm Score = −0.95
NG = 0
DB = 0
SM = 0
Delta_r
Doxorubicin
YGR001C
ZShift = −0.92
Z lm Score = −1.22
NG = 0
DB = 0
SM = 0
Delta_AUC
Doxorubicin
YKR008W RSC4
ZShift = −2.3
Z lm Score = 4.74
NG = 0
DB = 0
SM = 0
Delta_L
Doxorubicin
YKR008W RSC4
ZShift = 0.29
Z lm Score = 1.51
NG = 0
DB = 0
SM = 0
Delta_K
Doxorubicin
YKR008W RSC4
ZShift = 0.25
Z lm Score = −2.53
NG = 0
DB = 0
SM = 0
Delta_r
Doxorubicin
YKR008W RSC4
ZShift = 2.14
Z lm Score = −1.82
NG = 0
DB = 0
SM = 0
Delta_AUC
Doxorubicin
YPL088W
ZShift = 0.18
Z lm Score = 4.73
NG = 0
DB = 0
SM = 0
Delta_L
Doxorubicin
YPL088W
ZShift = 0.18
Z lm Score = −3.28
NG = 0
DB = 0
SM = 0
Delta_K
Doxorubicin
YPL088W
ZShift = 1.42
Z lm Score = −2.04
NG = 0
DB = 0
SM = 0
Delta_r
Doxorubicin
YPL088W
ZShift = −0.06
Z lm Score = −2.75
NG = 0
DB = 0
SM = 0
Delta_AUC
Doxorubicin

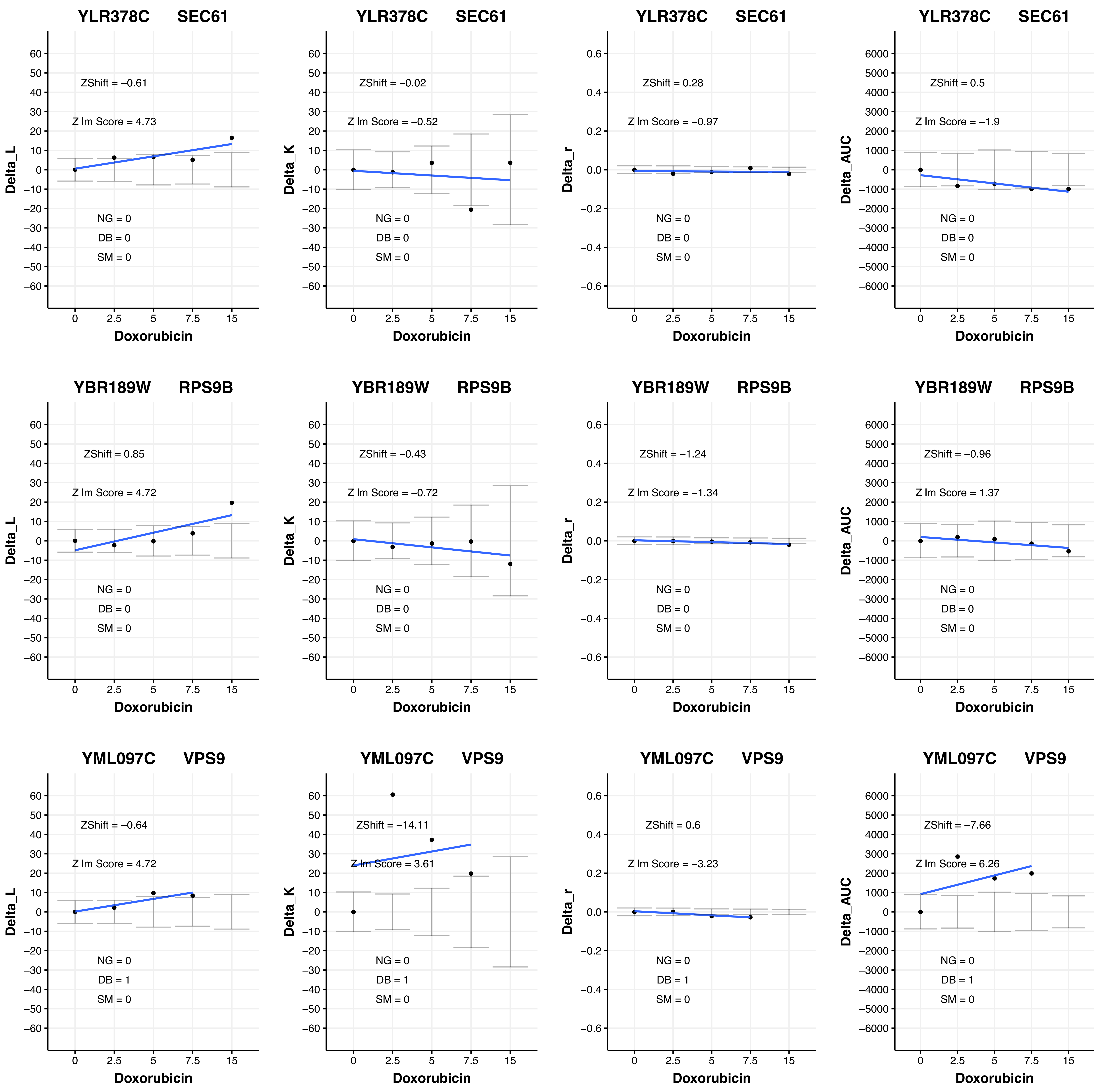
YLR378C SEC61
ZShift = −0.61
Z lm Score = 4.73
NG = 0
DB = 0
SM = 0
Delta_L
Doxorubicin
YLR378C SEC61
ZShift = −0.02
Z lm Score = −0.52
NG = 0
DB = 0
SM = 0
Delta_K
Doxorubicin
YLR378C SEC61
ZShift = 0.28
Z lm Score = −0.97
NG = 0
DB = 0
SM = 0
Delta_r
Doxorubicin
YLR378C SEC61
ZShift = 0.5
Z lm Score = −1.9
NG = 0
DB = 0
SM = 0
Delta_AUC
Doxorubicin
YBR189W RPS9B
ZShift = 0.85
Z lm Score = 4.72
NG = 0
DB = 0
SM = 0
Delta_L
Doxorubicin
YBR189W RPS9B
ZShift = −0.43
Z lm Score = −0.72
NG = 0
DB = 0
SM = 0
Delta_K
Doxorubicin
YBR189W RPS9B
ZShift = −1.24
Z lm Score = −1.34
NG = 0
DB = 0
SM = 0
Delta_r
Doxorubicin
YBR189W RPS9B
ZShift = −0.96
Z lm Score = 1.37
NG = 0
DB = 0
SM = 0
Delta_AUC
Doxorubicin
YML097C VPS9
ZShift = −0.64
Z lm Score = 4.72
NG = 0
DB = 1
SM = 0
Delta_L
Doxorubicin
YML097C VPS9
ZShift = −14.11
Z lm Score = 3.61
NG = 0
DB = 1
SM = 0
Delta_K
Doxorubicin
YML097C VPS9
ZShift = 0.6
Z lm Score = −3.23
NG = 0
DB = 1
SM = 0
Delta_r
Doxorubicin
YML097C VPS9
ZShift = −7.66
Z lm Score = 6.26
NG = 0
DB = 1
SM = 0
Delta_AUC
Doxorubicin

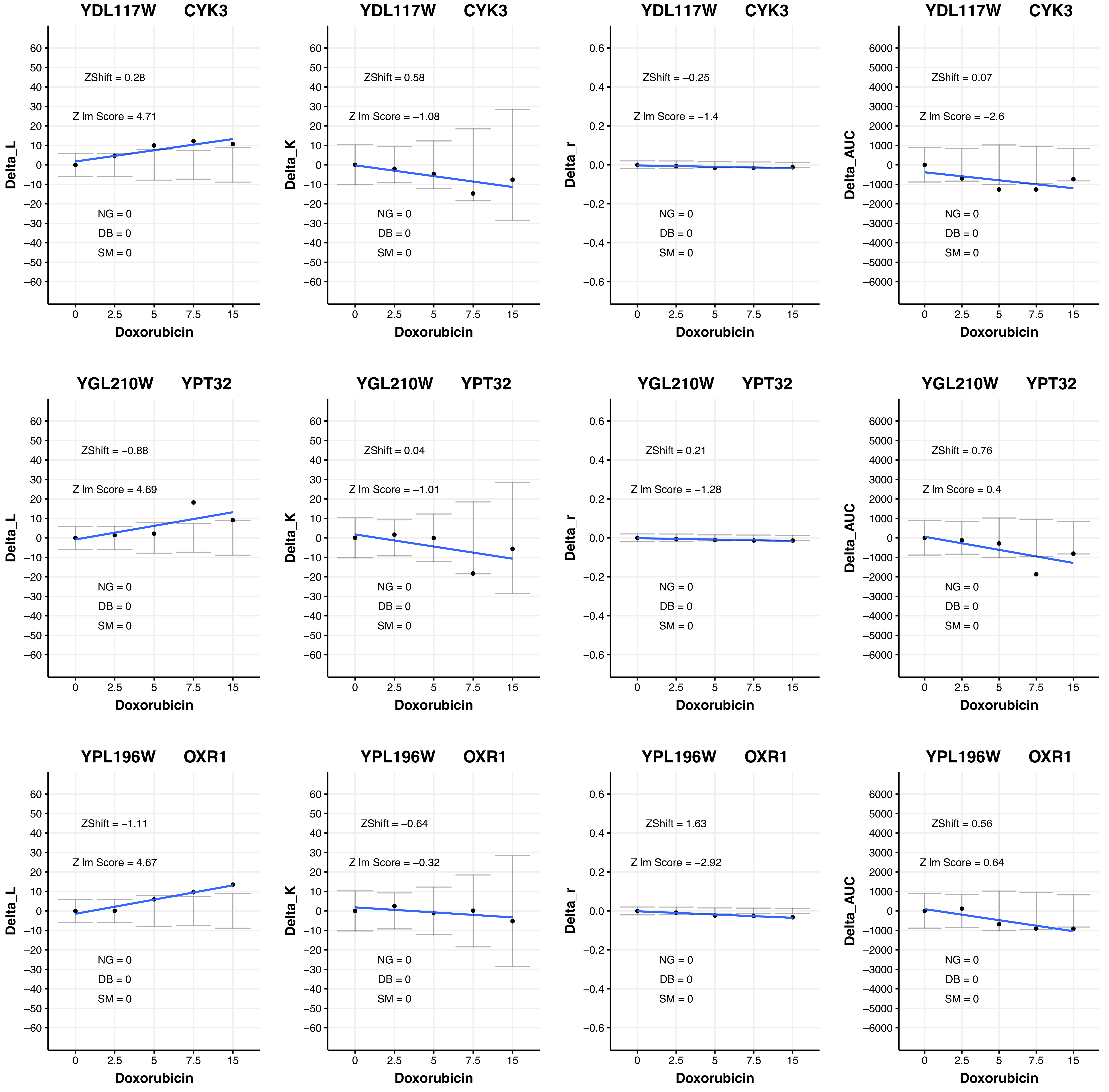

YDL117W CYK3
ZShift = 0.28
Z lm Score = 4.71
NG = 0
DB = 0
SM = 0
Delta_L
Doxorubicin
YDL117W CYK3
ZShift = 0.58
Z lm Score = −1.08
NG = 0
DB = 0
SM = 0
Delta_K
Doxorubicin
YDL117W CYK3
ZShift = −0.25
Z lm Score = −1.4
NG = 0
DB = 0
SM = 0
Delta_r
Doxorubicin
YDL117W CYK3
ZShift = 0.07
Z lm Score = −2.6
NG = 0
DB = 0
SM = 0
Delta_AUC
Doxorubicin
YGL210W YPT32
ZShift = −0.88
Z lm Score = 4.69
NG = 0
DB = 0
SM = 0
Delta_L
Doxorubicin
YGL210W YPT32
ZShift = 0.04
Z lm Score = −1.01
NG = 0
DB = 0
SM = 0
Delta_K
Doxorubicin
YGL210W YPT32
ZShift = 0.21
Z lm Score = −1.28
NG = 0
DB = 0
SM = 0
Delta_r
Doxorubicin
YGL210W YPT32
ZShift = 0.76
Z lm Score = 0.4
NG = 0
DB = 0
SM = 0
Delta_AUC
Doxorubicin
YPL196W OXR1
ZShift = −1.11
Z lm Score = 4.67
NG = 0
DB = 0
SM = 0
Delta_L
Doxorubicin
YPL196W OXR1
ZShift = −0.64
Z lm Score = −0.32
NG = 0
DB = 0
SM = 0
Delta_K
Doxorubicin
YPL196W OXR1
ZShift = 1.63
Z lm Score = −2.92
NG = 0
DB = 0
SM = 0
Delta_r
Doxorubicin
YPL196W OXR1
ZShift = 0.56
Z lm Score = 0.64
NG = 0
DB = 0
SM = 0
Delta_AUC
Doxorubicin

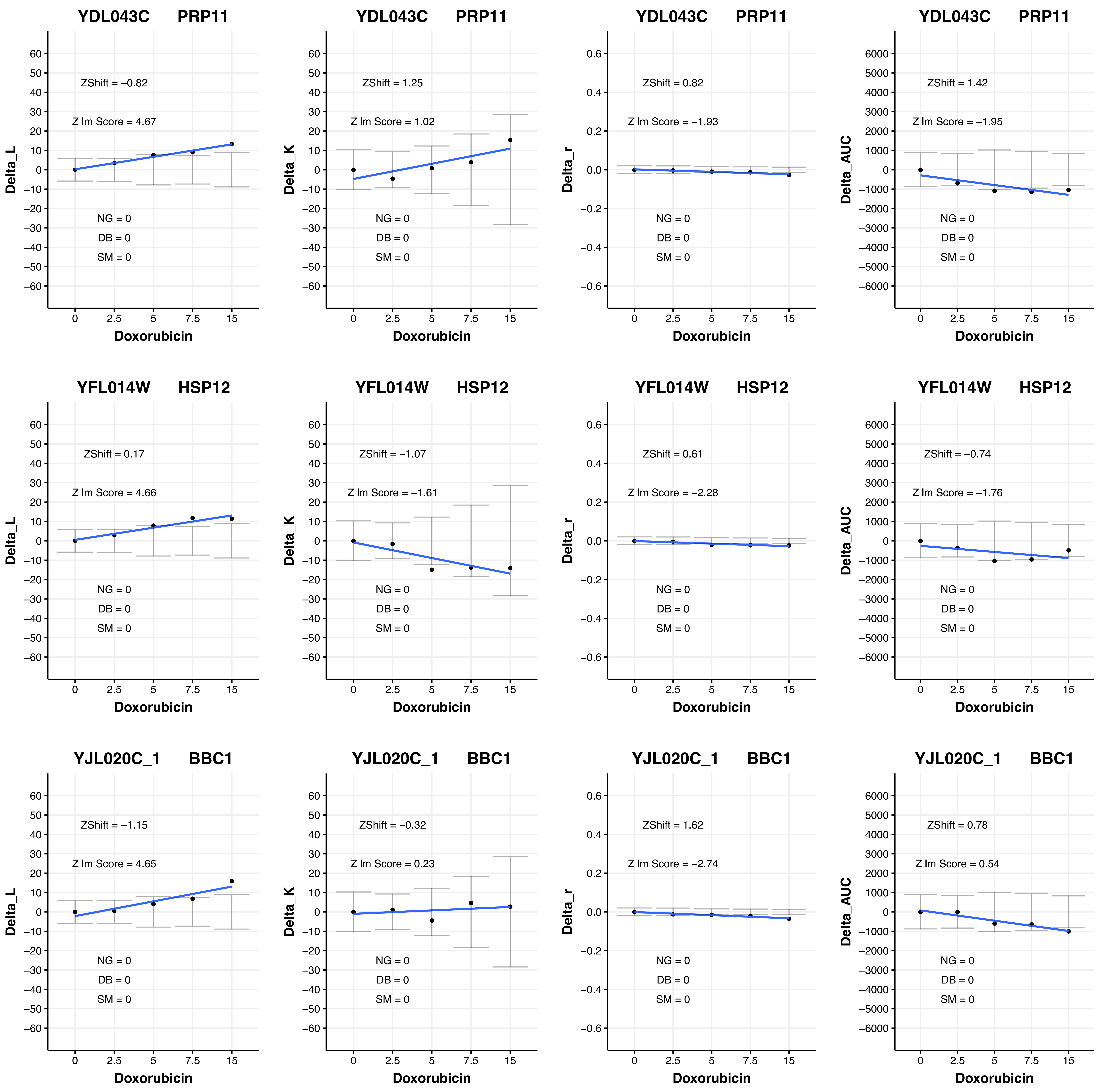

YDL043C PRP11
ZShift = −0.82
Z lm Score = 4.67
NG = 0
DB = 0
SM = 0
Delta_L
Doxorubicin
YDL043C PRP11
ZShift = 1.25
Z lm Score = 1.02
NG = 0
DB = 0
SM = 0
Delta_K
Doxorubicin
YDL043C PRP11
ZShift = 0.82
Z lm Score = −1.93
NG = 0
DB = 0
SM = 0
Delta_r
Doxorubicin
YDL043C PRP11
ZShift = 1.42
Z lm Score = −1.95
NG = 0
DB = 0
SM = 0
Delta_AUC
Doxorubicin
YFL014W HSP12
ZShift = 0.17
Z lm Score = 4.66
NG = 0
DB = 0
SM = 0
Delta_L
Doxorubicin
YFL014W HSP12
ZShift = −1.07
Z lm Score = −1.61
NG = 0
DB = 0
SM = 0
Delta_K
Doxorubicin
YFL014W HSP12
ZShift = 0.61
Z lm Score = −2.28
NG = 0
DB = 0
SM = 0
Delta_r
Doxorubicin
YFL014W HSP12
ZShift = −0.74
Z lm Score = −1.76
NG = 0
DB = 0
SM = 0
Delta_AUC
Doxorubicin
YJL020C_1 BBC1
ZShift = −1.15
Z lm Score = 4.65
NG = 0
DB = 0
SM = 0
Delta_L
Doxorubicin
YJL020C_1 BBC1
ZShift = −0.32
Z lm Score = 0.23
NG = 0
DB = 0
SM = 0
Delta_K
Doxorubicin
YJL020C_1 BBC1
ZShift = 1.62
Z lm Score = −2.74
NG = 0
DB = 0
SM = 0
Delta_r
Doxorubicin
YJL020C_1 BBC1
ZShift = 0.78
Z lm Score = 0.54
NG = 0
DB = 0
SM = 0
Delta_AUC
Doxorubicin

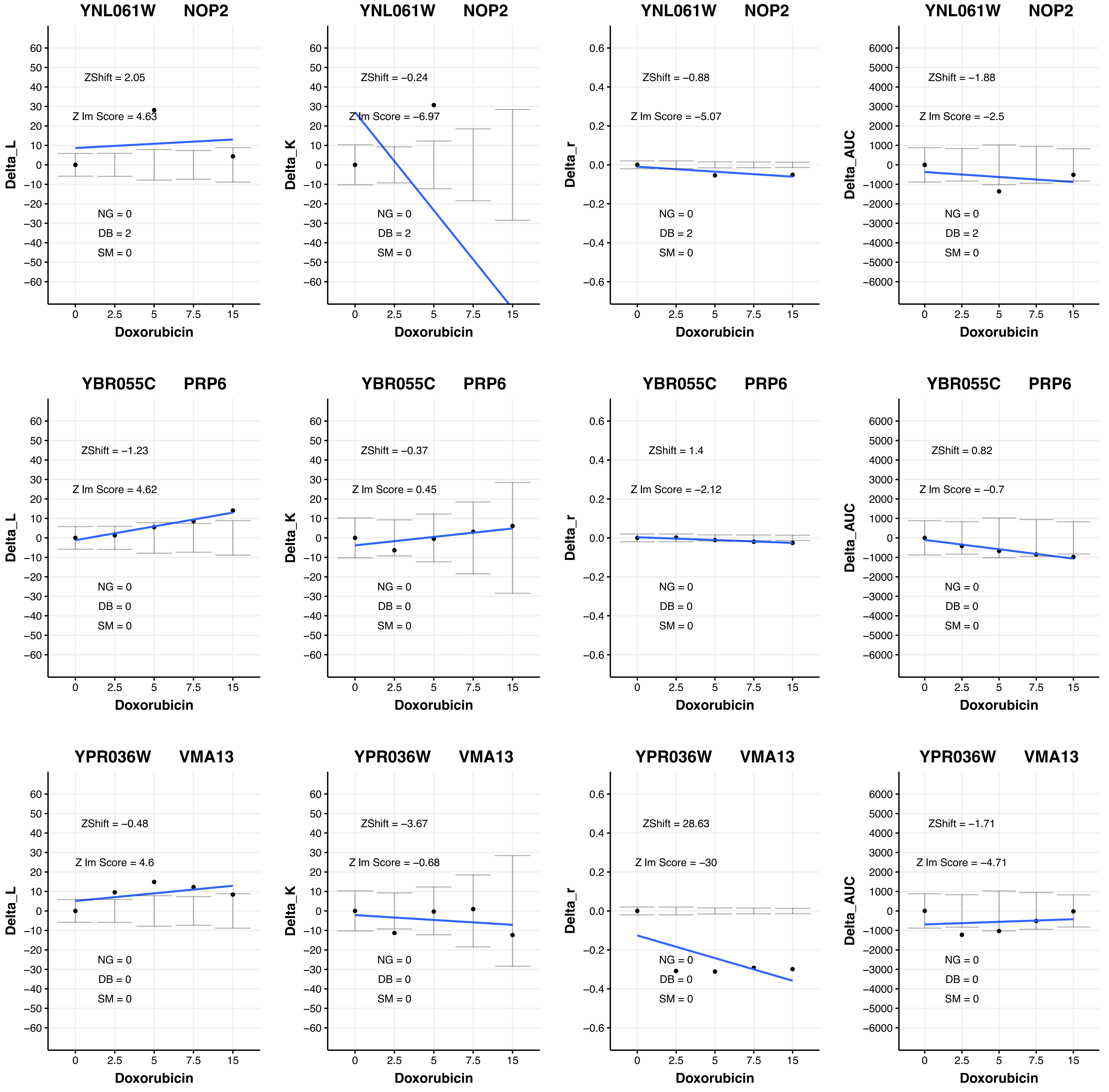
YNL061W NOP2
ZShift = 2.05
Z lm Score = 4.63
NG = 0
DB = 2
SM = 0
Delta_L
Doxorubicin
YNL061W NOP2
ZShift = −0.24
Z lm Score = −6.97
NG = 0
DB = 2
SM = 0
Delta_K
Doxorubicin
YNL061W NOP2
ZShift = −0.88
Z lm Score = −5.07
NG = 0
DB = 2
SM = 0
Delta_r
Doxorubicin
YNL061W NOP2
ZShift = −1.88
Z lm Score = −2.5
NG = 0
DB = 2
SM = 0
Delta_AUC
Doxorubicin
YBR055C PRP6
ZShift = −1.23
Z lm Score = 4.62
NG = 0
DB = 0
SM = 0
Delta_L
Doxorubicin
YBR055C PRP6
ZShift = −0.37
Z lm Score = 0.45
NG = 0
DB = 0
SM = 0
Delta_K
Doxorubicin
YBR055C PRP6
ZShift = 1.4
Z lm Score = −2.12
NG = 0
DB = 0
SM = 0
Delta_r
Doxorubicin
YBR055C PRP6
ZShift = 0.82
Z lm Score = −0.7
NG = 0
DB = 0
SM = 0
Delta_AUC
Doxorubicin
YPR036W VMA13
ZShift = −0.48
Z lm Score = 4.6
NG = 0
DB = 0
SM = 0
Delta_L
Doxorubicin
YPR036W VMA13
ZShift = −3.67
Z lm Score = −0.68
NG = 0
DB = 0
SM = 0
Delta_K
Doxorubicin
YPR036W VMA13
ZShift = 28.63
Z lm Score = −30
NG = 0
DB = 0
SM = 0
Delta_r
Doxorubicin
YPR036W VMA13
ZShift = −1.71
Z lm Score = −4.71
NG = 0
DB = 0
SM = 0
Delta_AUC
Doxorubicin

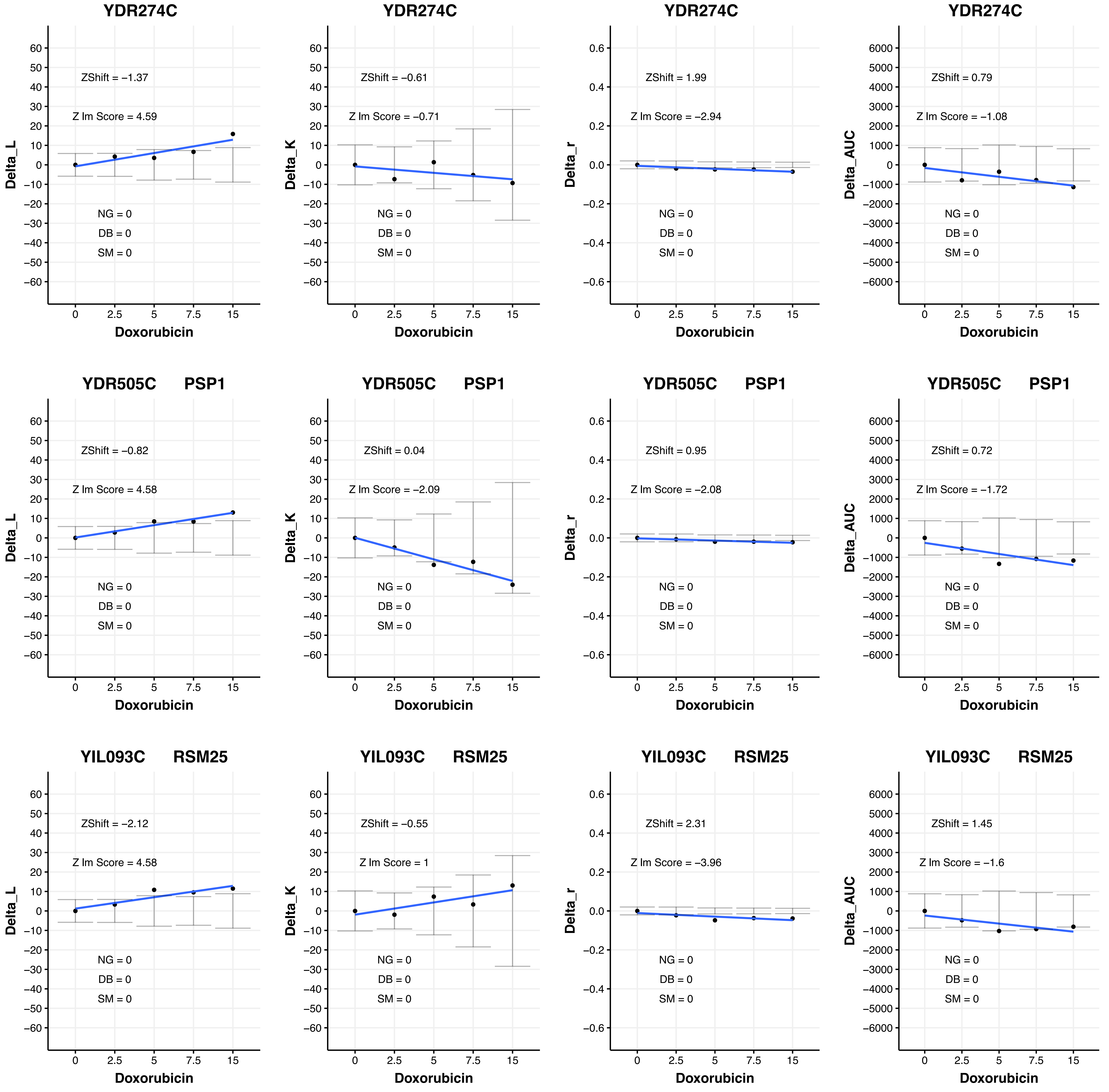

YDR274C
ZShift = −1.37
Z lm Score = 4.59
NG = 0
DB = 0
SM = 0
Delta_L
Doxorubicin
YDR274C
ZShift = −0.61
Z lm Score = −0.71
NG = 0
DB = 0
SM = 0
Delta_K
Doxorubicin
YDR274C
ZShift = 1.99
Z lm Score = −2.94
NG = 0
DB = 0
SM = 0
Delta_r
Doxorubicin
YDR274C
ZShift = 0.79
Z lm Score = −1.08
NG = 0
DB = 0
SM = 0
Delta_AUC
Doxorubicin
YDR505C PSP1
ZShift = −0.82
Z lm Score = 4.58
NG = 0
DB = 0
SM = 0
Delta_L
Doxorubicin
YDR505C PSP1
ZShift = 0.04
Z lm Score = −2.09
NG = 0
DB = 0
SM = 0
Delta_K
Doxorubicin
YDR505C PSP1
ZShift = 0.95
Z lm Score = −2.08
NG = 0
DB = 0
SM = 0
Delta_r
Doxorubicin
YDR505C PSP1
ZShift = 0.72
Z lm Score = −1.72
NG = 0
DB = 0
SM = 0
Delta_AUC
Doxorubicin
YIL093C RSM25
ZShift = −2.12
Z lm Score = 4.58
NG = 0
DB = 0
SM = 0
Delta_L
Doxorubicin
YIL093C RSM25
ZShift = −0.55
Z lm Score = 1
NG = 0
DB = 0
SM = 0
Delta_K
Doxorubicin
YIL093C RSM25
ZShift = 2.31
Z lm Score = −3.96
NG = 0
DB = 0
SM = 0
Delta_r
Doxorubicin
YIL093C RSM25
ZShift = 1.45
Z lm Score = −1.6
NG = 0
DB = 0
SM = 0
Delta_AUC
Doxorubicin

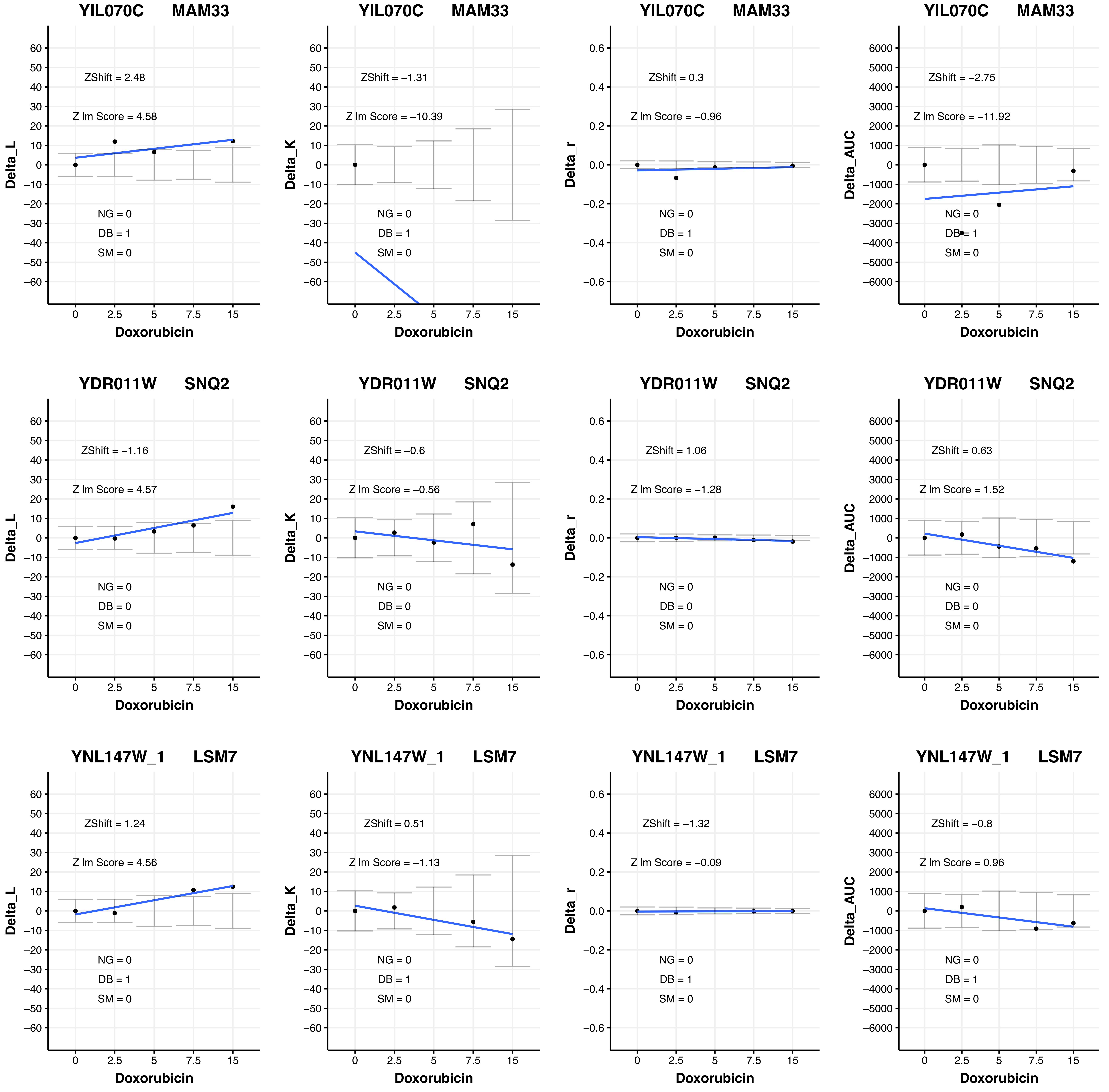
YIL070C MAM33
ZShift = 2.48
Z lm Score = 4.58
NG = 0
DB = 1
SM = 0
Delta_L
Doxorubicin
YIL070C MAM33
ZShift = −1.31
Z lm Score = −10.39
NG = 0
DB = 1
SM = 0
Delta_K
Doxorubicin
YIL070C MAM33
ZShift = 0.3
Z lm Score = −0.96
NG = 0
DB = 1
SM = 0
Delta_r
Doxorubicin
YIL070C MAM33
ZShift = −2.75
Z lm Score = −11.92
NG = 0
DB = 1
SM = 0
Delta_AUC
Doxorubicin
YDR011W SNQ2
ZShift = −1.16
Z lm Score = 4.57
NG = 0
DB = 0
SM = 0
Delta_L
Doxorubicin
YDR011W SNQ2
ZShift = −0.6
Z lm Score = −0.56
NG = 0
DB = 0
SM = 0
Delta_K
Doxorubicin
YDR011W SNQ2
ZShift = 1.06
Z lm Score = −1.28
NG = 0
DB = 0
SM = 0
Delta_r
Doxorubicin
YDR011W SNQ2
ZShift = 0.63
Z lm Score = 1.52
NG = 0
DB = 0
SM = 0
Delta_AUC
Doxorubicin
YNL147W_1 LSM7
ZShift = 1.24
Z lm Score = 4.56
NG = 0
DB = 1
SM = 0
Delta_L
Doxorubicin
YNL147W_1 LSM7
ZShift = 0.51
Z lm Score = −1.13
NG = 0
DB = 1
SM = 0
Delta_K
Doxorubicin
YNL147W_1 LSM7
ZShift = −1.32
Z lm Score = −0.09
NG = 0
DB = 1
SM = 0
Delta_r
Doxorubicin
YNL147W_1 LSM7
ZShift = −0.8
Z lm Score = 0.96
NG = 0
DB = 1
SM = 0
Delta_AUC
Doxorubicin

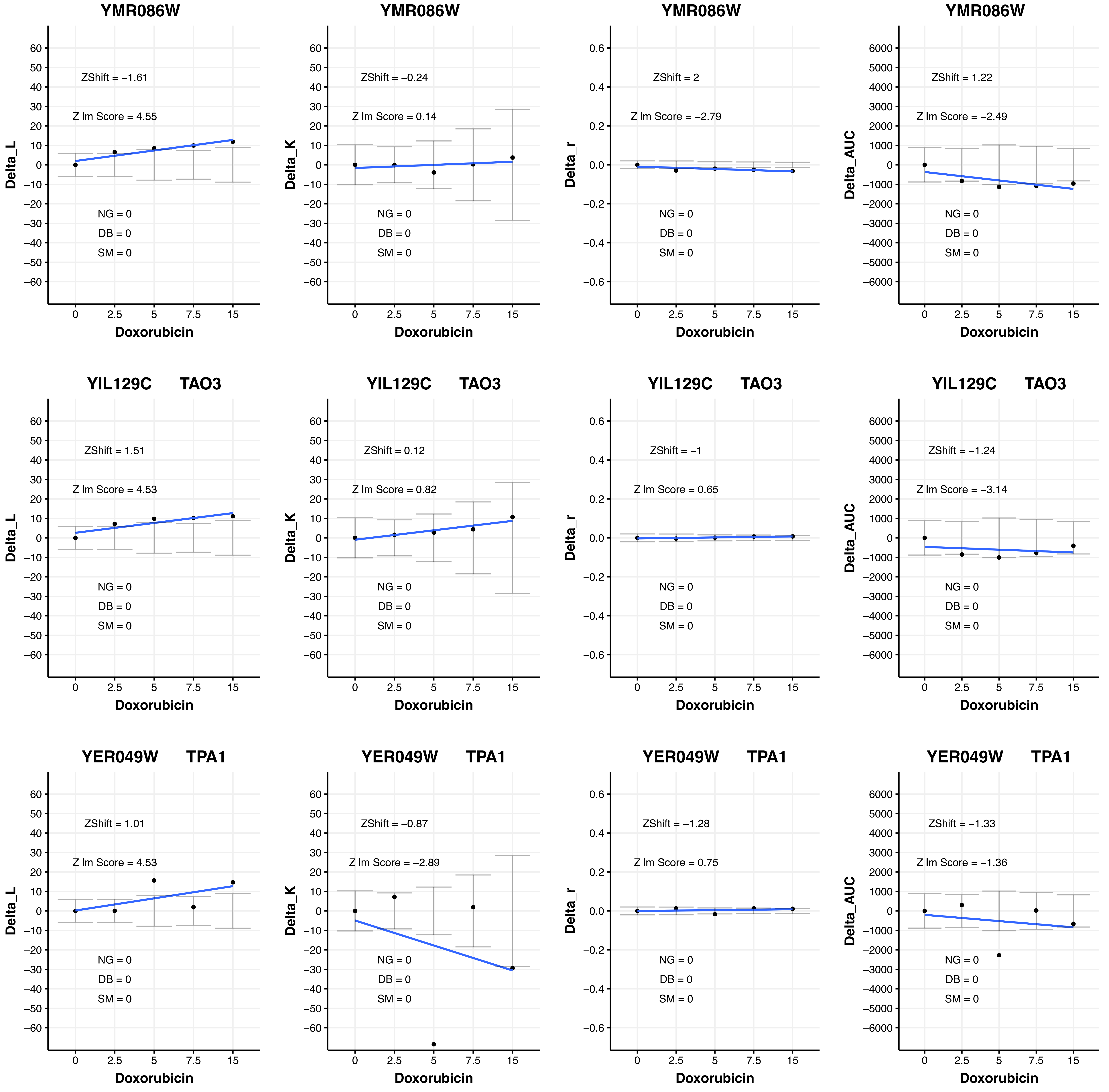
YMR086W
ZShift = −1.61
Z lm Score = 4.55
NG = 0
DB = 0
SM = 0
Delta_L
Doxorubicin
YMR086W
ZShift = −0.24
Z lm Score = 0.14
NG = 0
DB = 0
SM = 0
Delta_K
Doxorubicin
YMR086W
ZShift = 2
Z lm Score = −2.79
NG = 0
DB = 0
SM = 0
Delta_r
Doxorubicin
YMR086W
ZShift = 1.22
Z lm Score = −2.49
NG = 0
DB = 0
SM = 0
Delta_AUC
Doxorubicin
YIL129C TAO3
ZShift = 1.51
Z lm Score = 4.53
NG = 0
DB = 0
SM = 0
Delta_L
Doxorubicin
YIL129C TAO3
ZShift = 0.12
Z lm Score = 0.82
NG = 0
DB = 0
SM = 0
Delta_K
Doxorubicin
YIL129C TAO3
ZShift = −1
Z lm Score = 0.65
NG = 0
DB = 0
SM = 0
Delta_r
Doxorubicin
YIL129C TAO3
ZShift = −1.24
Z lm Score = −3.14
NG = 0
DB = 0
SM = 0
Delta_AUC
Doxorubicin
YER049W TPA1
ZShift = 1.01
Z lm Score = 4.53
NG = 0
DB = 0
SM = 0
Delta_L
Doxorubicin
YER049W TPA1
ZShift = −0.87
Z lm Score = −2.89
NG = 0
DB = 0
SM = 0
Delta_K
Doxorubicin
YER049W TPA1
ZShift = −1.28
Z lm Score = 0.75
NG = 0
DB = 0
SM = 0
Delta_r
Doxorubicin
YER049W TPA1
ZShift = −1.33
Z lm Score = −1.36
NG = 0
DB = 0
SM = 0
Delta_AUC
Doxorubicin

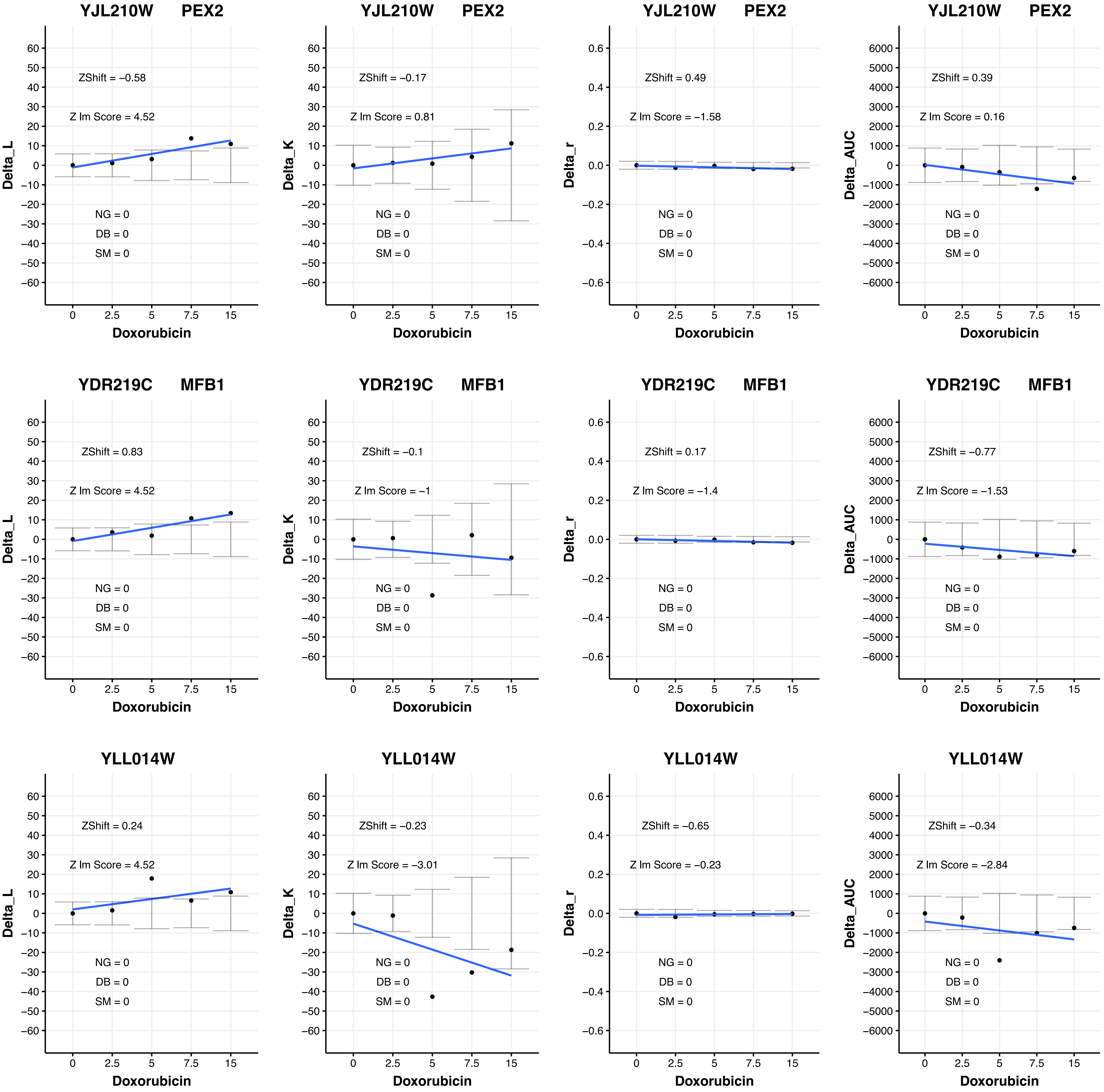

YJL210W PEX2
ZShift = −0.58
Z lm Score = 4.52
NG = 0
DB = 0
SM = 0
Delta_L
Doxorubicin
YJL210W PEX2
ZShift = −0.17
Z lm Score = 0.81
NG = 0
DB = 0
SM = 0
Delta_K
Doxorubicin
YJL210W PEX2
ZShift = 0.49
Z lm Score = −1.58
NG = 0
DB = 0
SM = 0
Delta_r
Doxorubicin
YJL210W PEX2
ZShift = 0.39
Z lm Score = 0.16
NG = 0
DB = 0
SM = 0
Delta_AUC
Doxorubicin
YDR219C MFB1
ZShift = 0.83
Z lm Score = 4.52
NG = 0
DB = 0
SM = 0
Delta_L
Doxorubicin
YDR219C MFB1
ZShift = −0.1
Z lm Score = −1
NG = 0
DB = 0
SM = 0
Delta_K
Doxorubicin
YDR219C MFB1
ZShift = 0.17
Z lm Score = −1.4
NG = 0
DB = 0
SM = 0
Delta_r
Doxorubicin
YDR219C MFB1
ZShift = −0.77
Z lm Score = −1.53
NG = 0
DB = 0
SM = 0
Delta_AUC
Doxorubicin
YLL014W
ZShift = 0.24
Z lm Score = 4.52
NG = 0
DB = 0
SM = 0
Delta_L
Doxorubicin
YLL014W
ZShift = −0.23
Z lm Score = −3.01
NG = 0
DB = 0
SM = 0
Delta_K
Doxorubicin
YLL014W
ZShift = −0.65
Z lm Score = −0.23
NG = 0
DB = 0
SM = 0
Delta_r
Doxorubicin
YLL014W
ZShift = −0.34
Z lm Score = −2.84
NG = 0
DB = 0
SM = 0
Delta_AUC
Doxorubicin

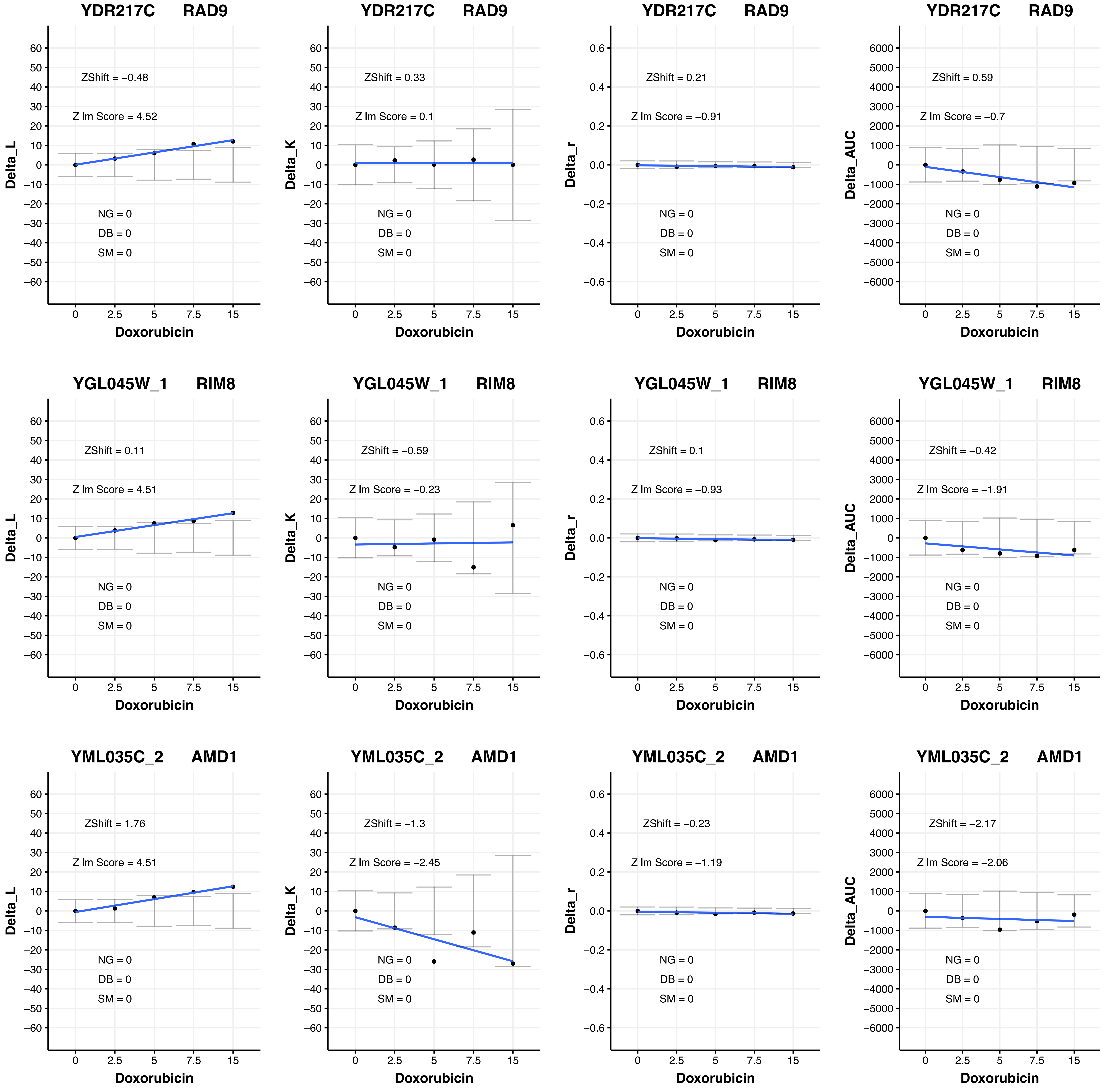

YDR217C RAD9
ZShift = −0.48
Z lm Score = 4.52
NG = 0
DB = 0
SM = 0
Delta_L
Doxorubicin
YDR217C RAD9
ZShift = 0.33
Z lm Score = 0.1
NG = 0
DB = 0
SM = 0
Delta_K
Doxorubicin
YDR217C RAD9
ZShift = 0.21
Z lm Score = −0.91
NG = 0
DB = 0
SM = 0
Delta_r
Doxorubicin
YDR217C RAD9
ZShift = 0.59
Z lm Score = −0.7
NG = 0
DB = 0
SM = 0
Delta_AUC
Doxorubicin
YGL045W_1 RIM8
ZShift = 0.11
Z lm Score = 4.51
NG = 0
DB = 0
SM = 0
Delta_L
Doxorubicin
YGL045W_1 RIM8
ZShift = −0.59
Z lm Score = −0.23
NG = 0
DB = 0
SM = 0
Delta_K
Doxorubicin
YGL045W_1 RIM8
ZShift = 0.1
Z lm Score = −0.93
NG = 0
DB = 0
SM = 0
Delta_r
Doxorubicin
YGL045W_1 RIM8
ZShift = −0.42
Z lm Score = −1.91
NG = 0
DB = 0
SM = 0
Delta_AUC
Doxorubicin
YML035C_2 AMD1
ZShift = 1.76
Z lm Score = 4.51
NG = 0
DB = 0
SM = 0
Delta_L
Doxorubicin
YML035C_2 AMD1
ZShift = −1.3
Z lm Score = −2.45
NG = 0
DB = 0
SM = 0
Delta_K
Doxorubicin
YML035C_2 AMD1
ZShift = −0.23
Z lm Score = −1.19
NG = 0
DB = 0
SM = 0
Delta_r
Doxorubicin
YML035C_2 AMD1
ZShift = −2.17
Z lm Score = −2.06
NG = 0
DB = 0
SM = 0
Delta_AUC
Doxorubicin

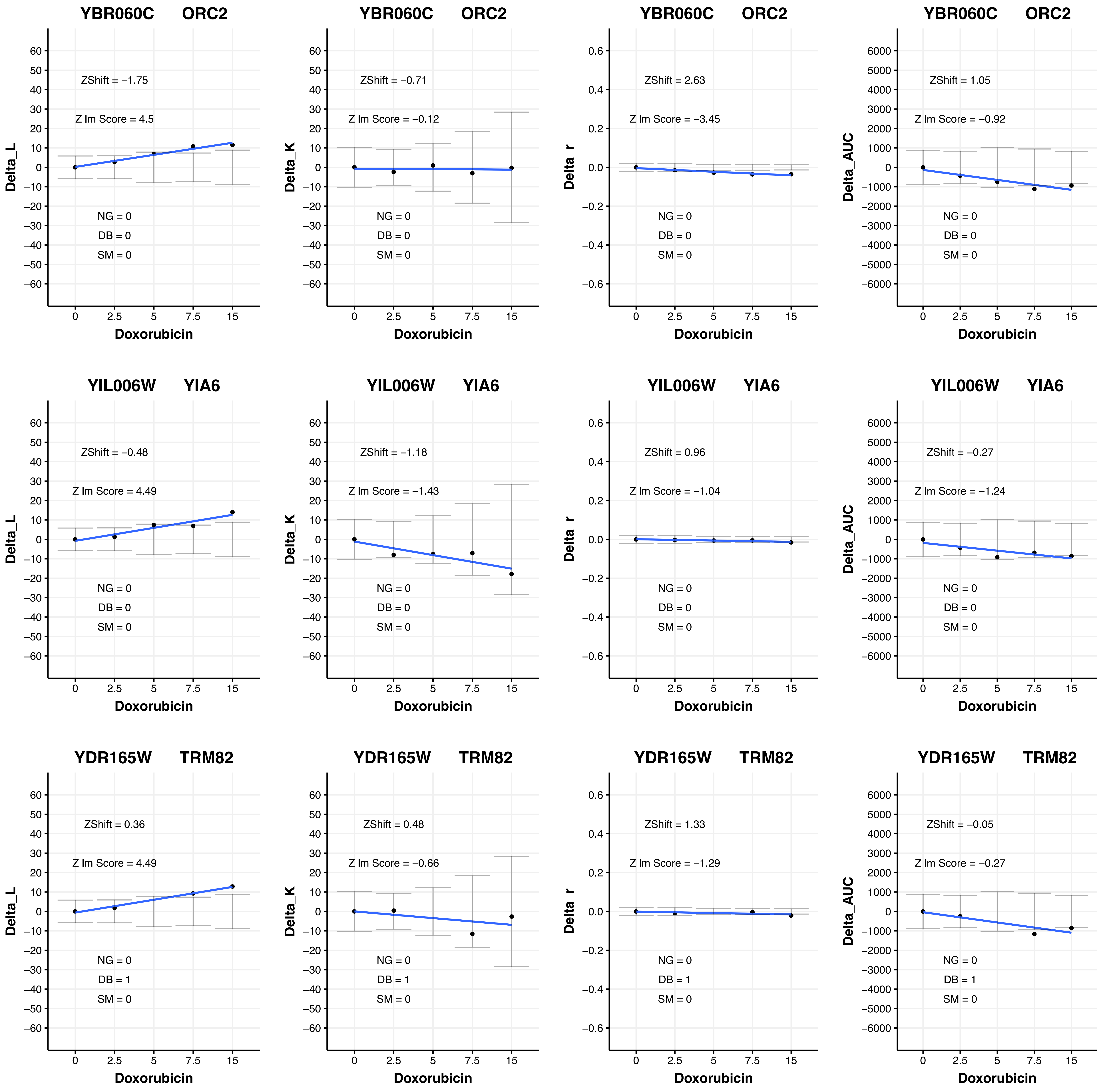

YBR060C ORC2
ZShift = −1.75
Z lm Score = 4.5
NG = 0
DB = 0
SM = 0
Delta_L
Doxorubicin
YBR060C ORC2
ZShift = −0.71
Z lm Score = −0.12
NG = 0
DB = 0
SM = 0
Delta_K
Doxorubicin
YBR060C ORC2
ZShift = 2.63
Z lm Score = −3.45
NG = 0
DB = 0
SM = 0
Delta_r
Doxorubicin
YBR060C ORC2
ZShift = 1.05
Z lm Score = −0.92
NG = 0
DB = 0
SM = 0
Delta_AUC
Doxorubicin
YIL006W YIA6
ZShift = −0.48
Z lm Score = 4.49
NG = 0
DB = 0
SM = 0
Delta_L
Doxorubicin
YIL006W YIA6
ZShift = −1.18
Z lm Score = −1.43
NG = 0
DB = 0
SM = 0
Delta_K
Doxorubicin
YIL006W YIA6
ZShift = 0.96
Z lm Score = −1.04
NG = 0
DB = 0
SM = 0
Delta_r
Doxorubicin
YIL006W YIA6
ZShift = −0.27
Z lm Score = −1.24
NG = 0
DB = 0
SM = 0
Delta_AUC
Doxorubicin
YDR165W TRM82
ZShift = 0.36
Z lm Score = 4.49
NG = 0
DB = 1
SM = 0
Delta_L
Doxorubicin
YDR165W TRM82
ZShift = 0.48
Z lm Score = −0.66
NG = 0
DB = 1
SM = 0
Delta_K
Doxorubicin
YDR165W TRM82
ZShift = 1.33
Z lm Score = −1.29
NG = 0
DB = 1
SM = 0
Delta_r
Doxorubicin
YDR165W TRM82
ZShift = −0.05
Z lm Score = −0.27
NG = 0
DB = 1
SM = 0
Delta_AUC
Doxorubicin

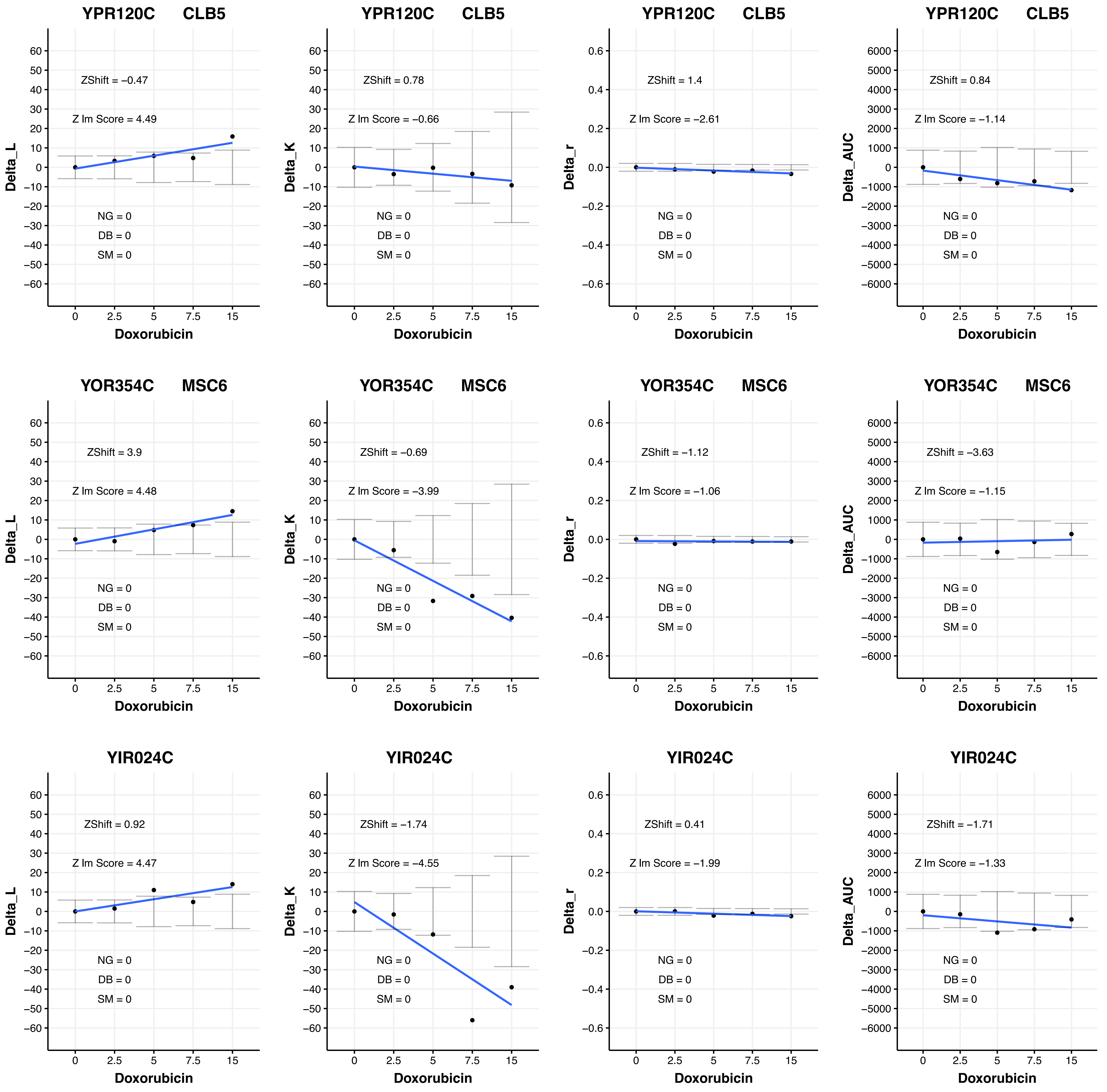
YPR120C CLB5
ZShift = −0.47
Z lm Score = 4.49
NG = 0
DB = 0
SM = 0
Delta_L
Doxorubicin
YPR120C CLB5
ZShift = 0.78
Z lm Score = −0.66
NG = 0
DB = 0
SM = 0
Delta_K
Doxorubicin
YPR120C CLB5
ZShift = 1.4
Z lm Score = −2.61
NG = 0
DB = 0
SM = 0
Delta_r
Doxorubicin
YPR120C CLB5
ZShift = 0.84
Z lm Score = −1.14
NG = 0
DB = 0
SM = 0
Delta_AUC
Doxorubicin
YOR354C MSC6
ZShift = 3.9
Z lm Score = 4.48
NG = 0
DB = 0
SM = 0
Delta_L
Doxorubicin
YOR354C MSC6
ZShift = −0.69
Z lm Score = −3.99
NG = 0
DB = 0
SM = 0
Delta_K
Doxorubicin
YOR354C MSC6
ZShift = −1.12
Z lm Score = −1.06
NG = 0
DB = 0
SM = 0
Delta_r
Doxorubicin
YOR354C MSC6
ZShift = −3.63
Z lm Score = −1.15
NG = 0
DB = 0
SM = 0
Delta_AUC
Doxorubicin
YIR024C
ZShift = 0.92
Z lm Score = 4.47
NG = 0
DB = 0
SM = 0
Delta_L
Doxorubicin
YIR024C
ZShift = −1.74
Z lm Score = −4.55
NG = 0
DB = 0
SM = 0
Delta_K
Doxorubicin
YIR024C
ZShift = 0.41
Z lm Score = −1.99
NG = 0
DB = 0
SM = 0
Delta_r
Doxorubicin
YIR024C
ZShift = −1.71
Z lm Score = −1.33
NG = 0
DB = 0
SM = 0
Delta_AUC
Doxorubicin

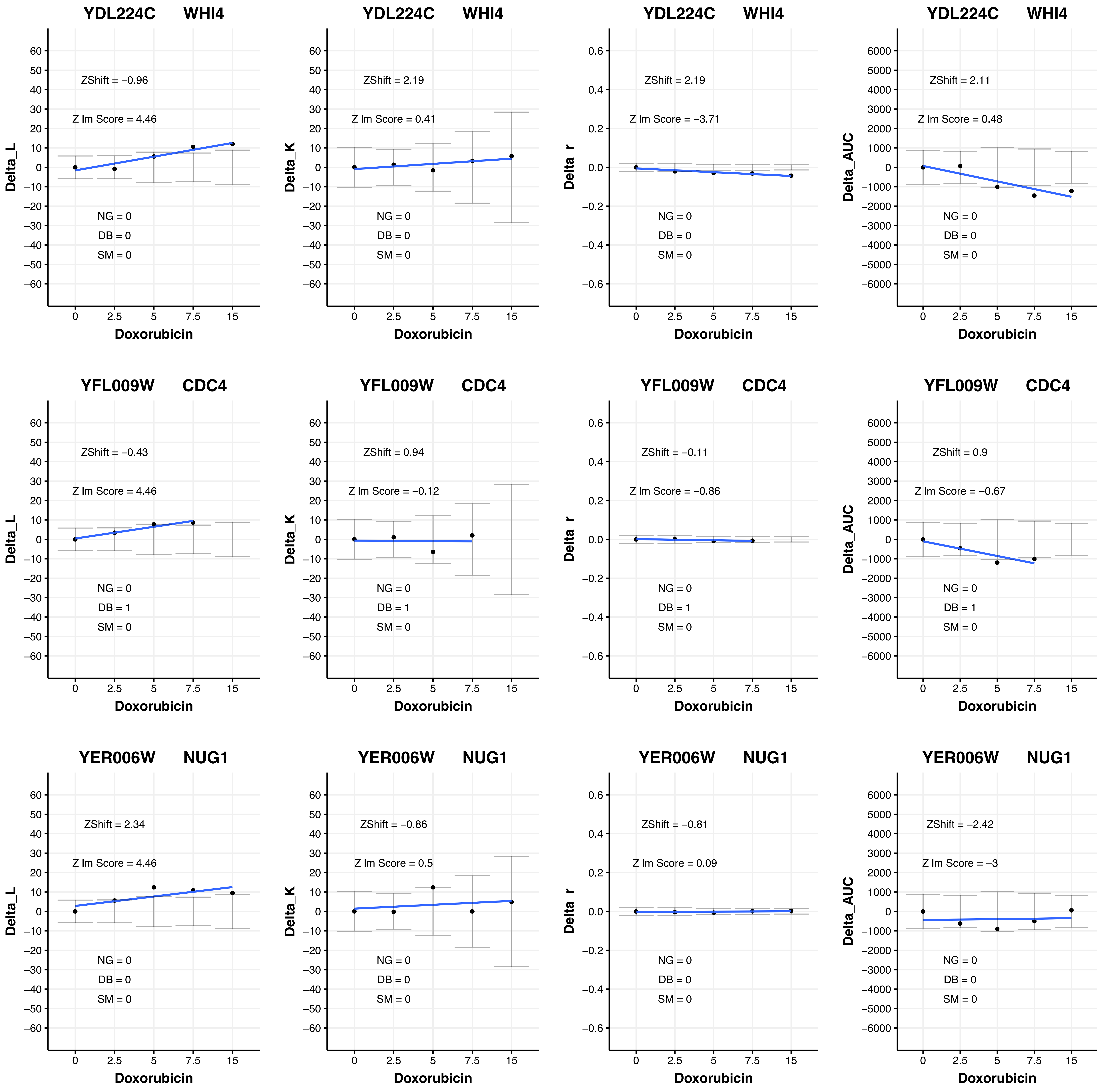

YDL224C WHI4
ZShift = −0.96
Z lm Score = 4.46
NG = 0
DB = 0
SM = 0
Delta_L
Doxorubicin
YDL224C WHI4
ZShift = 2.19
Z lm Score = 0.41
NG = 0
DB = 0
SM = 0
Delta_K
Doxorubicin
YDL224C WHI4
ZShift = 2.19
Z lm Score = −3.71
NG = 0
DB = 0
SM = 0
Delta_r
Doxorubicin
YDL224C WHI4
ZShift = 2.11
Z lm Score = 0.48
NG = 0
DB = 0
SM = 0
Delta_AUC
Doxorubicin
YFL009W CDC4
ZShift = −0.43
Z lm Score = 4.46
NG = 0
DB = 1
SM = 0
Delta_L
Doxorubicin
YFL009W CDC4
ZShift = 0.94
Z lm Score = −0.12
NG = 0
DB = 1
SM = 0
Delta_K
Doxorubicin
YFL009W CDC4
ZShift = −0.11
Z lm Score = −0.86
NG = 0
DB = 1
SM = 0
Delta_r
Doxorubicin
YFL009W CDC4
ZShift = 0.9
Z lm Score = −0.67
NG = 0
DB = 1
SM = 0
Delta_AUC
Doxorubicin
YER006W NUG1
ZShift = 2.34
Z lm Score = 4.46
NG = 0
DB = 0
SM = 0
Delta_L
Doxorubicin
YER006W NUG1
ZShift = −0.86
Z lm Score = 0.5
NG = 0
DB = 0
SM = 0
Delta_K
Doxorubicin
YER006W NUG1
ZShift = −0.81
Z lm Score = 0.09
NG = 0
DB = 0
SM = 0
Delta_r
Doxorubicin
YER006W NUG1
ZShift = −2.42
Z lm Score = −3
NG = 0
DB = 0
SM = 0
Delta_AUC
Doxorubicin

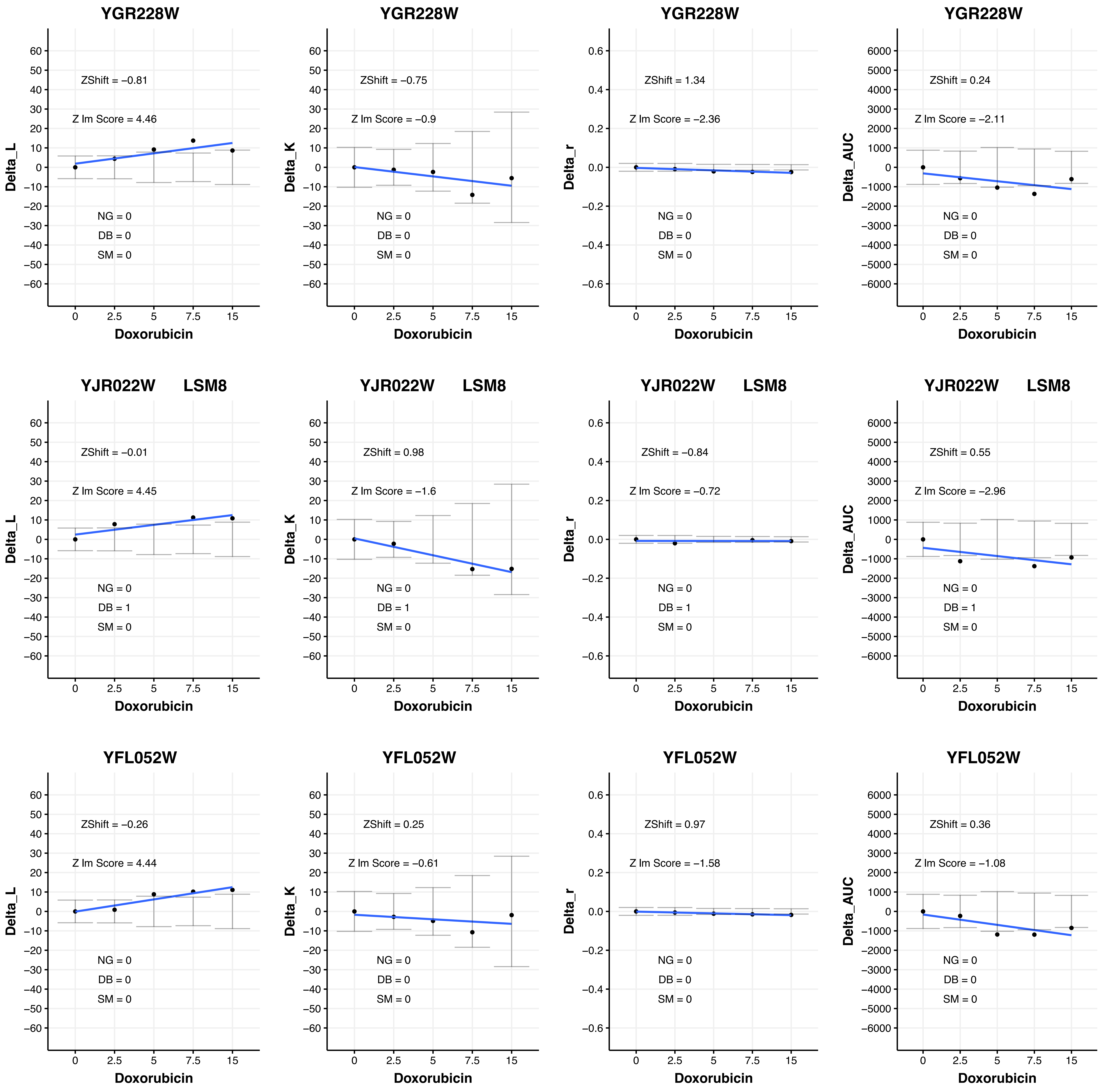

YGR228W
ZShift = −0.81
Z lm Score = 4.46
NG = 0
DB = 0
SM = 0
Delta_L
Doxorubicin
YGR228W
ZShift = −0.75
Z lm Score = −0.9
NG = 0
DB = 0
SM = 0
Delta_K
Doxorubicin
YGR228W
ZShift = 1.34
Z lm Score = −2.36
NG = 0
DB = 0
SM = 0
Delta_r
Doxorubicin
YGR228W
ZShift = 0.24
Z lm Score = −2.11
NG = 0
DB = 0
SM = 0
Delta_AUC
Doxorubicin
YJR022W LSM8
ZShift = −0.01
Z lm Score = 4.45
NG = 0
DB = 1
SM = 0
Delta_L
Doxorubicin
YJR022W LSM8
ZShift = 0.98
Z lm Score = −1.6
NG = 0
DB = 1
SM = 0
Delta_K
Doxorubicin
YJR022W LSM8
ZShift = −0.84
Z lm Score = −0.72
NG = 0
DB = 1
SM = 0
Delta_r
Doxorubicin
YJR022W LSM8
ZShift = 0.55
Z lm Score = −2.96
NG = 0
DB = 1
SM = 0
Delta_AUC
Doxorubicin
YFL052W
ZShift = −0.26
Z lm Score = 4.44
NG = 0
DB = 0
SM = 0
Delta_L
Doxorubicin
YFL052W
ZShift = 0.25
Z lm Score = −0.61
NG = 0
DB = 0
SM = 0
Delta_K
Doxorubicin
YFL052W
ZShift = 0.97
Z lm Score = −1.58
NG = 0
DB = 0
SM = 0
Delta_r
Doxorubicin
YFL052W
ZShift = 0.36
Z lm Score = −1.08
NG = 0
DB = 0
SM = 0
Delta_AUC
Doxorubicin

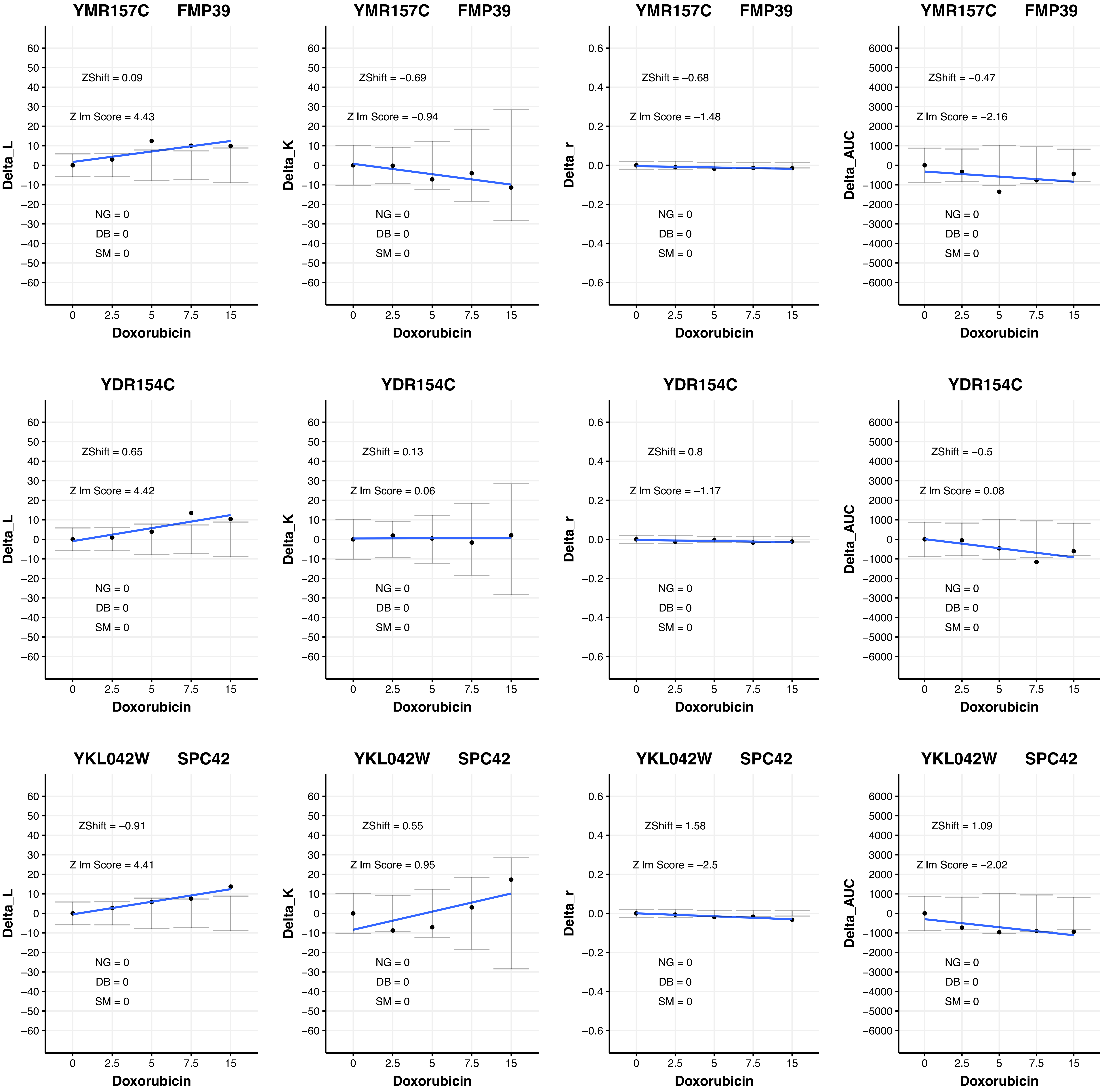
YMR157C FMP39
ZShift = 0.09
Z lm Score = 4.43
NG = 0
DB = 0
SM = 0
Delta_L
Doxorubicin
YMR157C FMP39
ZShift = −0.69
Z lm Score = −0.94
NG = 0
DB = 0
SM = 0
Delta_K
Doxorubicin
YMR157C FMP39
ZShift = −0.68
Z lm Score = −1.48
NG = 0
DB = 0
SM = 0
Delta_r
Doxorubicin
YMR157C FMP39
ZShift = −0.47
Z lm Score = −2.16
NG = 0
DB = 0
SM = 0
Delta_AUC
Doxorubicin
YDR154C
ZShift = 0.65
Z lm Score = 4.42
NG = 0
DB = 0
SM = 0
Delta_L
Doxorubicin
YDR154C
ZShift = 0.13
Z lm Score = 0.06
NG = 0
DB = 0
SM = 0
Delta_K
Doxorubicin
YDR154C
ZShift = 0.8
Z lm Score = −1.17
NG = 0
DB = 0
SM = 0
Delta_r
Doxorubicin
YDR154C
ZShift = −0.5
Z lm Score = 0.08
NG = 0
DB = 0
SM = 0
Delta_AUC
Doxorubicin
YKL042W SPC42
ZShift = −0.91
Z lm Score = 4.41
NG = 0
DB = 0
SM = 0
Delta_L
Doxorubicin
YKL042W SPC42
ZShift = 0.55
Z lm Score = 0.95
NG = 0
DB = 0
SM = 0
Delta_K
Doxorubicin
YKL042W SPC42
ZShift = 1.58
Z lm Score = −2.5
NG = 0
DB = 0
SM = 0
Delta_r
Doxorubicin
YKL042W SPC42
ZShift = 1.09
Z lm Score = −2.02
NG = 0
DB = 0
SM = 0
Delta_AUC
Doxorubicin

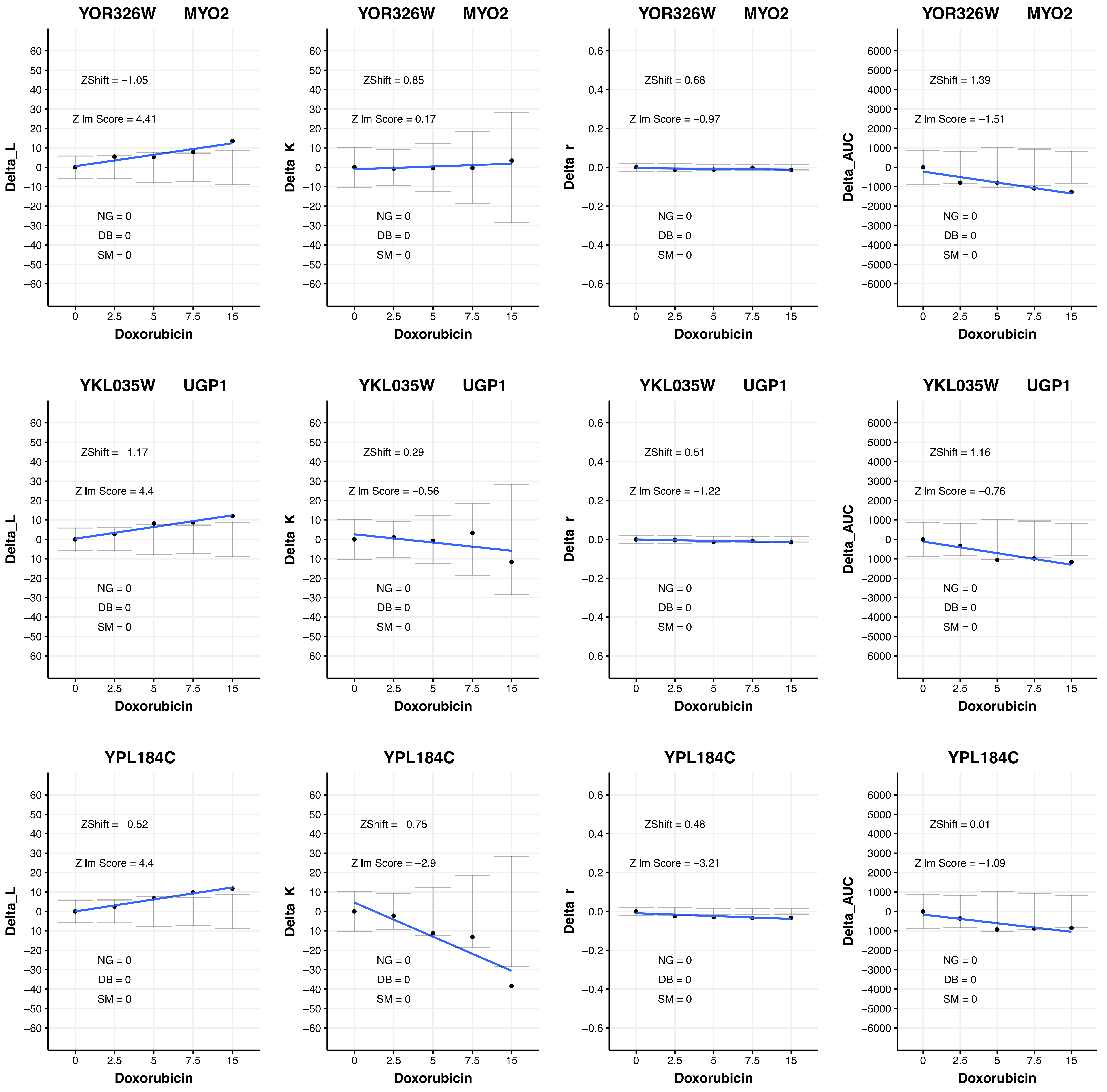
YOR326W MYO2
ZShift = −1.05
Z lm Score = 4.41
NG = 0
DB = 0
SM = 0
Delta_L
Doxorubicin
YOR326W MYO2
ZShift = 0.85
Z lm Score = 0.17
NG = 0
DB = 0
SM = 0
Delta_K
Doxorubicin
YOR326W MYO2
ZShift = 0.68
Z lm Score = −0.97
NG = 0
DB = 0
SM = 0
Delta_r
Doxorubicin
YOR326W MYO2
ZShift = 1.39
Z lm Score = −1.51
NG = 0
DB = 0
SM = 0
Delta_AUC
Doxorubicin
YKL035W UGP1
ZShift = −1.17
Z lm Score = 4.4
NG = 0
DB = 0
SM = 0
Delta_L
Doxorubicin
YKL035W UGP1
ZShift = 0.29
Z lm Score = −0.56
NG = 0
DB = 0
SM = 0
Delta_K
Doxorubicin
YKL035W UGP1
ZShift = 0.51
Z lm Score = −1.22
NG = 0
DB = 0
SM = 0
Delta_r
Doxorubicin
YKL035W UGP1
ZShift = 1.16
Z lm Score = −0.76
NG = 0
DB = 0
SM = 0
Delta_AUC
Doxorubicin
YPL184C
ZShift = −0.52
Z lm Score = 4.4
NG = 0
DB = 0
SM = 0
Delta_L
Doxorubicin
YPL184C
ZShift = −0.75
Z lm Score = −2.9
NG = 0
DB = 0
SM = 0
Delta_K
Doxorubicin
YPL184C
ZShift = 0.48
Z lm Score = −3.21
NG = 0
DB = 0
SM = 0
Delta_r
Doxorubicin
YPL184C
ZShift = 0.01
Z lm Score = −1.09
NG = 0
DB = 0
SM = 0
Delta_AUC
Doxorubicin

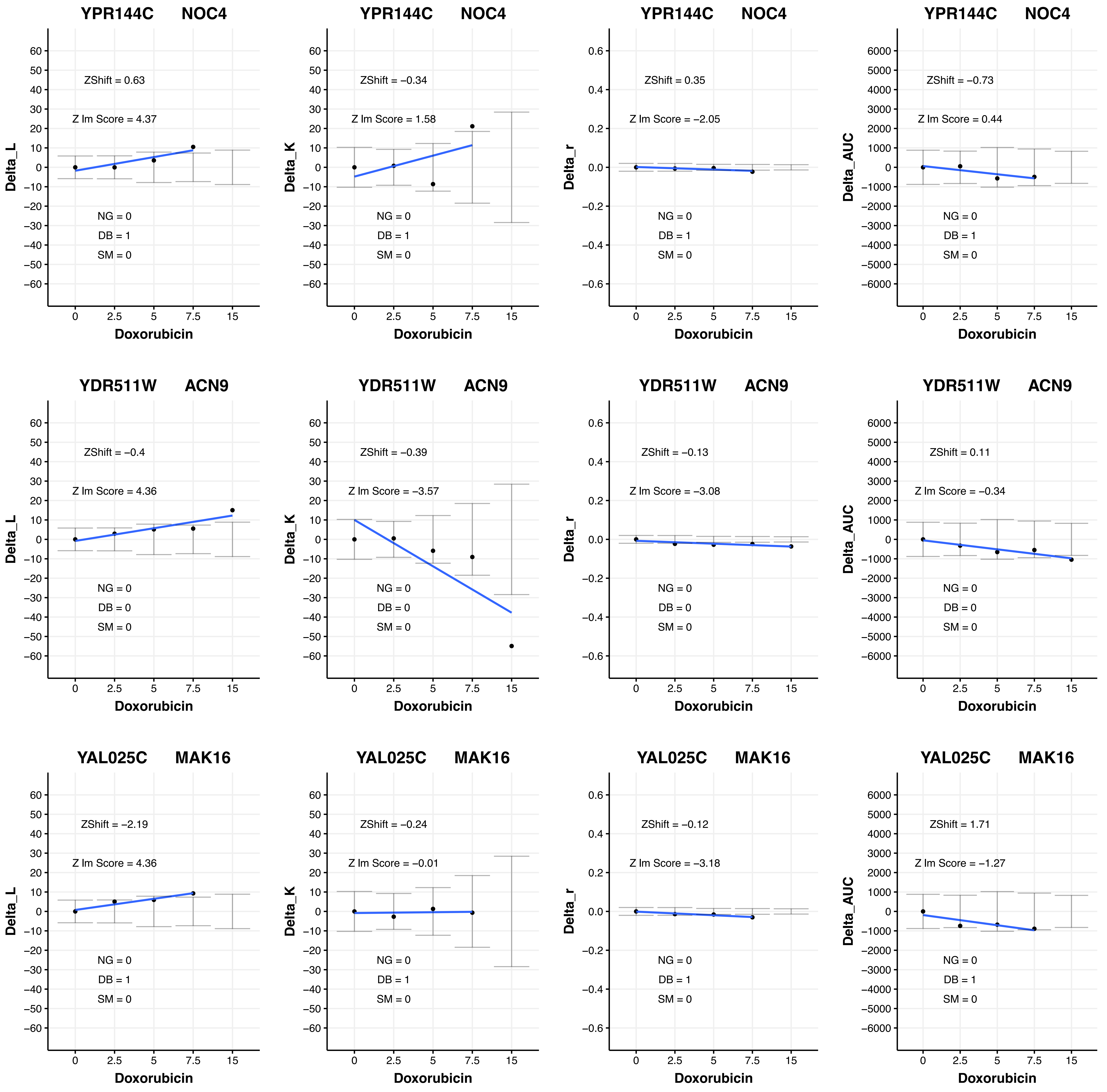
YPR144C NOC4
ZShift = 0.63
Z lm Score = 4.37
NG = 0
DB = 1
SM = 0
Delta_L
Doxorubicin
YPR144C NOC4
ZShift = −0.34
Z lm Score = 1.58
NG = 0
DB = 1
SM = 0
Delta_K
Doxorubicin
YPR144C NOC4
ZShift = 0.35
Z lm Score = −2.05
NG = 0
DB = 1
SM = 0
Delta_r
Doxorubicin
YPR144C NOC4
ZShift = −0.73
Z lm Score = 0.44
NG = 0
DB = 1
SM = 0
Delta_AUC
Doxorubicin
YDR511W ACN9
ZShift = −0.4
Z lm Score = 4.36
NG = 0
DB = 0
SM = 0
Delta_L
Doxorubicin
YDR511W ACN9
ZShift = −0.39
Z lm Score = −3.57
NG = 0
DB = 0
SM = 0
Delta_K
Doxorubicin
YDR511W ACN9
ZShift = −0.13
Z lm Score = −3.08
NG = 0
DB = 0
SM = 0
Delta_r
Doxorubicin
YDR511W ACN9
ZShift = 0.11
Z lm Score = −0.34
NG = 0
DB = 0
SM = 0
Delta_AUC
Doxorubicin
YAL025C MAK16
ZShift = −2.19
Z lm Score = 4.36
NG = 0
DB = 1
SM = 0
Delta_L
Doxorubicin
YAL025C MAK16
ZShift = −0.24
Z lm Score = −0.01
NG = 0
DB = 1
SM = 0
Delta_K
Doxorubicin
YAL025C MAK16
ZShift = −0.12
Z lm Score = −3.18
NG = 0
DB = 1
SM = 0
Delta_r
Doxorubicin
YAL025C MAK16
ZShift = 1.71
Z lm Score = −1.27
NG = 0
DB = 1
SM = 0
Delta_AUC
Doxorubicin

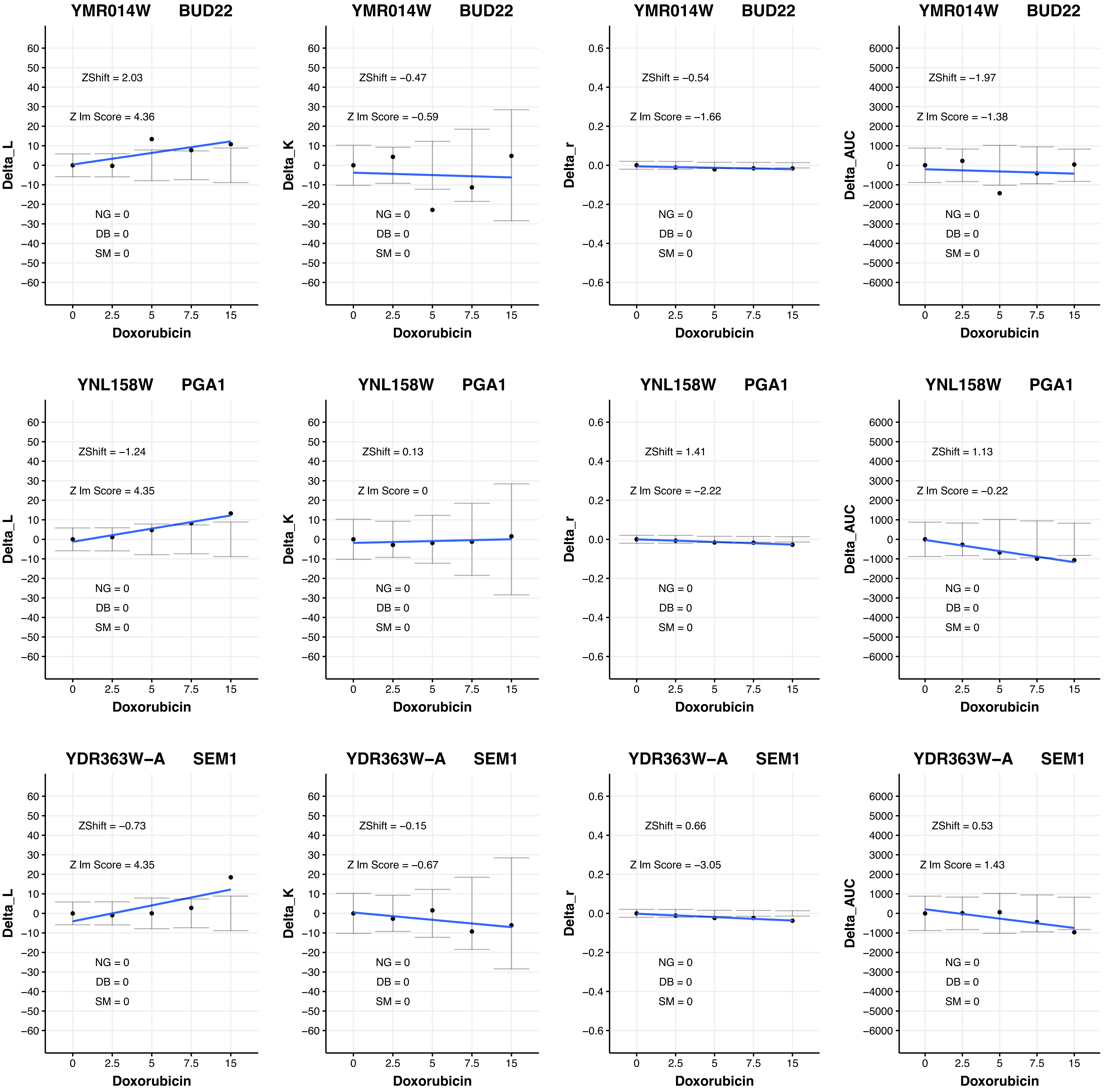

YMR014W BUD22
ZShift = 2.03
Z lm Score = 4.36
NG = 0
DB = 0
SM = 0
Delta_L
Doxorubicin
YMR014W BUD22
ZShift = −0.47
Z lm Score = −0.59
NG = 0
DB = 0
SM = 0
Delta_K
Doxorubicin
YMR014W BUD22
ZShift = −0.54
Z lm Score = −1.66
NG = 0
DB = 0
SM = 0
Delta_r
Doxorubicin
YMR014W BUD22
ZShift = −1.97
Z lm Score = −1.38
NG = 0
DB = 0
SM = 0
Delta_AUC
Doxorubicin
YNL158W PGA1
ZShift = −1.24
Z lm Score = 4.35
NG = 0
DB = 0
SM = 0
Delta_L
Doxorubicin
YNL158W PGA1
ZShift = 0.13
Z lm Score = 0
NG = 0
DB = 0
SM = 0
Delta_K
Doxorubicin
YNL158W PGA1
ZShift = 1.41
Z lm Score = −2.22
NG = 0
DB = 0
SM = 0
Delta_r
Doxorubicin
YNL158W PGA1
ZShift = 1.13
Z lm Score = −0.22
NG = 0
DB = 0
SM = 0
Delta_AUC
Doxorubicin
YDR363W−A SEM1
ZShift = −0.73
Z lm Score = 4.35
NG = 0
DB = 0
SM = 0
Delta_L
Doxorubicin
YDR363W−A SEM1
ZShift = −0.15
Z lm Score = −0.67
NG = 0
DB = 0
SM = 0
Delta_K
Doxorubicin
YDR363W−A SEM1
ZShift = 0.66
Z lm Score = −3.05
NG = 0
DB = 0
SM = 0
Delta_r
Doxorubicin
YDR363W−A SEM1
ZShift = 0.53
Z lm Score = 1.43
NG = 0
DB = 0
SM = 0
Delta_AUC
Doxorubicin

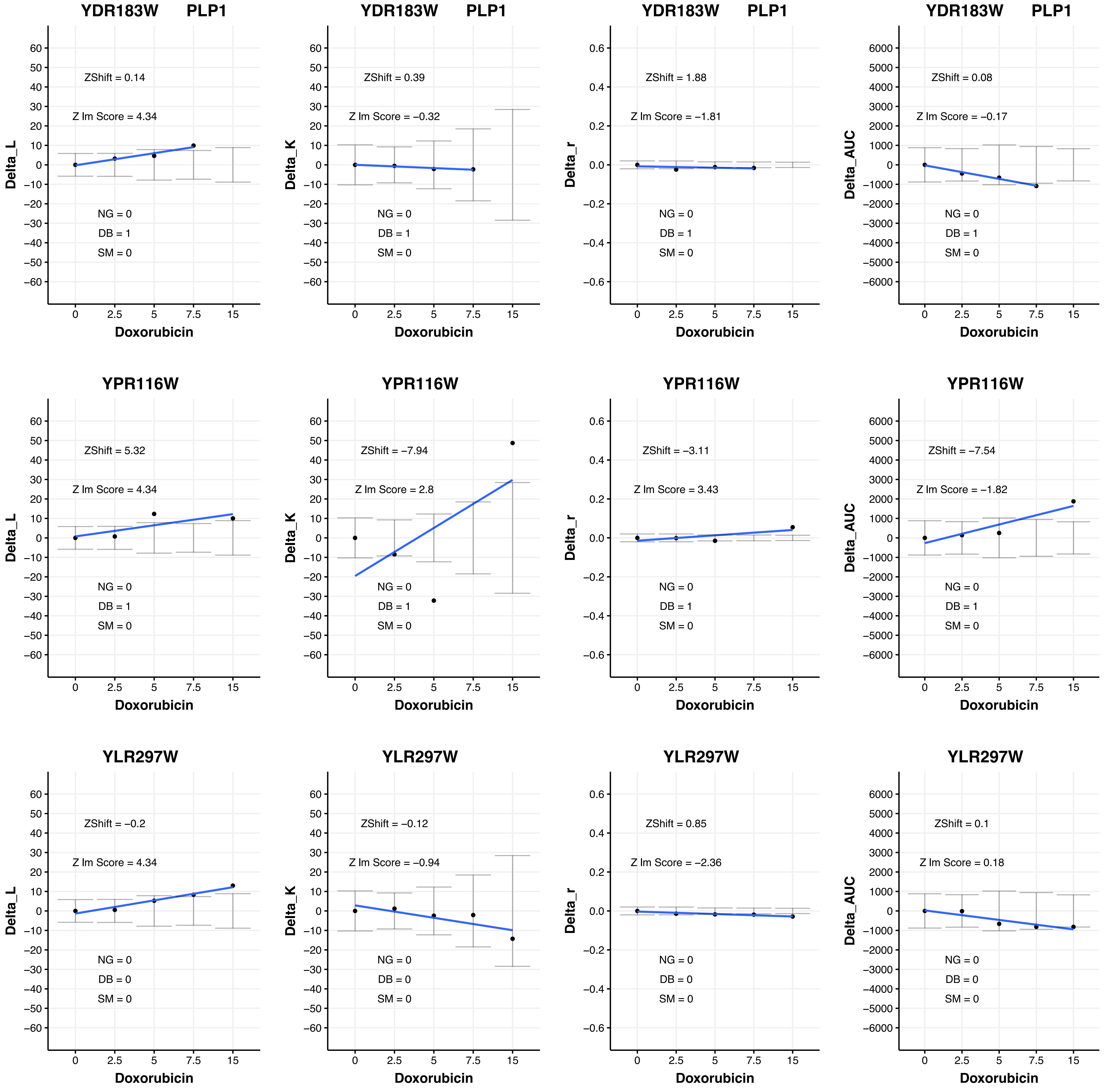
YDR183W PLP1
ZShift = 0.14
Z lm Score = 4.34
NG = 0
DB = 1
SM = 0
Delta_L
Doxorubicin
YDR183W PLP1
ZShift = 0.39
Z lm Score = −0.32
NG = 0
DB = 1
SM = 0
Delta_K
Doxorubicin
YDR183W PLP1
ZShift = 1.88
Z lm Score = −1.81
NG = 0
DB = 1
SM = 0
Delta_r
Doxorubicin
YDR183W PLP1
ZShift = 0.08
Z lm Score = −0.17
NG = 0
DB = 1
SM = 0
Delta_AUC
Doxorubicin
YPR116W
ZShift = 5.32
Z lm Score = 4.34
NG = 0
DB = 1
SM = 0
Delta_L
Doxorubicin
YPR116W
ZShift = −7.94
Z lm Score = 2.8
NG = 0
DB = 1
SM = 0
Delta_K
Doxorubicin
YPR116W
ZShift = −3.11
Z lm Score = 3.43
NG = 0
DB = 1
SM = 0
Delta_r
Doxorubicin
YPR116W
ZShift = −7.54
Z lm Score = −1.82
NG = 0
DB = 1
SM = 0
Delta_AUC
Doxorubicin
YLR297W
ZShift = −0.2
Z lm Score = 4.34
NG = 0
DB = 0
SM = 0
Delta_L
Doxorubicin
YLR297W
ZShift = −0.12
Z lm Score = −0.94
NG = 0
DB = 0
SM = 0
Delta_K
Doxorubicin
YLR297W
ZShift = 0.85
Z lm Score = −2.36
NG = 0
DB = 0
SM = 0
Delta_r
Doxorubicin
YLR297W
ZShift = 0.1
Z lm Score = 0.18
NG = 0
DB = 0
SM = 0
Delta_AUC
Doxorubicin

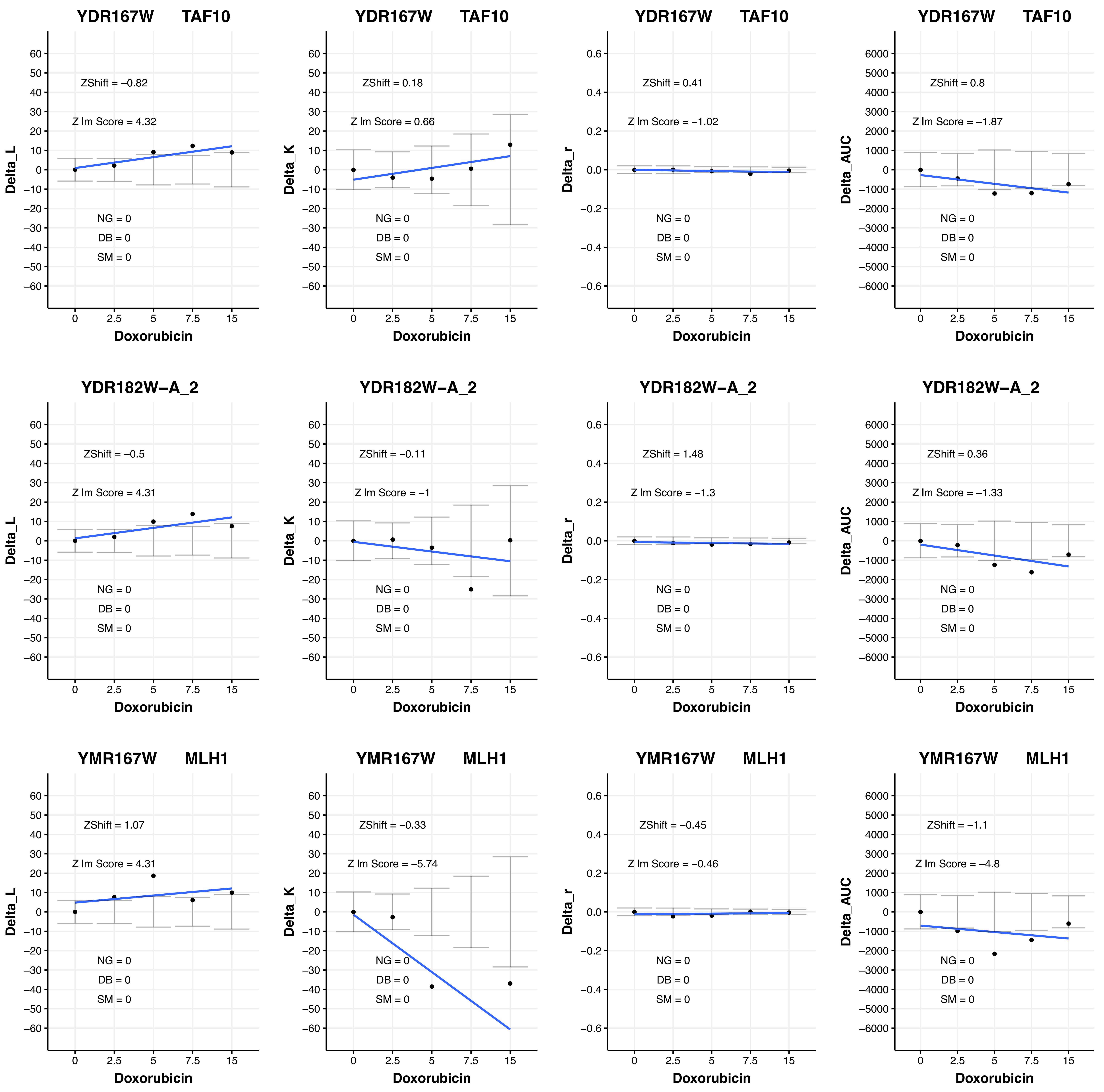

YDR167W TAF10
ZShift = −0.82
Z lm Score = 4.32
NG = 0
DB = 0
SM = 0
Delta_L
Doxorubicin
YDR167W TAF10
ZShift = 0.18
Z lm Score = 0.66
NG = 0
DB = 0
SM = 0
Delta_K
Doxorubicin
YDR167W TAF10
ZShift = 0.41
Z lm Score = −1.02
NG = 0
DB = 0
SM = 0
Delta_r
Doxorubicin
YDR167W TAF10
ZShift = 0.8
Z lm Score = −1.87
NG = 0
DB = 0
SM = 0
Delta_AUC
Doxorubicin
YDR182W−A_2
ZShift = −0.5
Z lm Score = 4.31
NG = 0
DB = 0
SM = 0
Delta_L
Doxorubicin
YDR182W−A_2
ZShift = −0.11
Z lm Score = −1
NG = 0
DB = 0
SM = 0
Delta_K
Doxorubicin
YDR182W−A_2
ZShift = 1.48
Z lm Score = −1.3
NG = 0
DB = 0
SM = 0
Delta_r
Doxorubicin
YDR182W−A_2
ZShift = 0.36
Z lm Score = −1.33
NG = 0
DB = 0
SM = 0
Delta_AUC
Doxorubicin
YMR167W MLH1
ZShift = 1.07
Z lm Score = 4.31
NG = 0
DB = 0
SM = 0
Delta_L
Doxorubicin
YMR167W MLH1
ZShift = −0.33
Z lm Score = −5.74
NG = 0
DB = 0
SM = 0
Delta_K
Doxorubicin
YMR167W MLH1
ZShift = −0.45
Z lm Score = −0.46
NG = 0
DB = 0
SM = 0
Delta_r
Doxorubicin
YMR167W MLH1
ZShift = −1.1
Z lm Score = −4.8
NG = 0
DB = 0
SM = 0
Delta_AUC
Doxorubicin

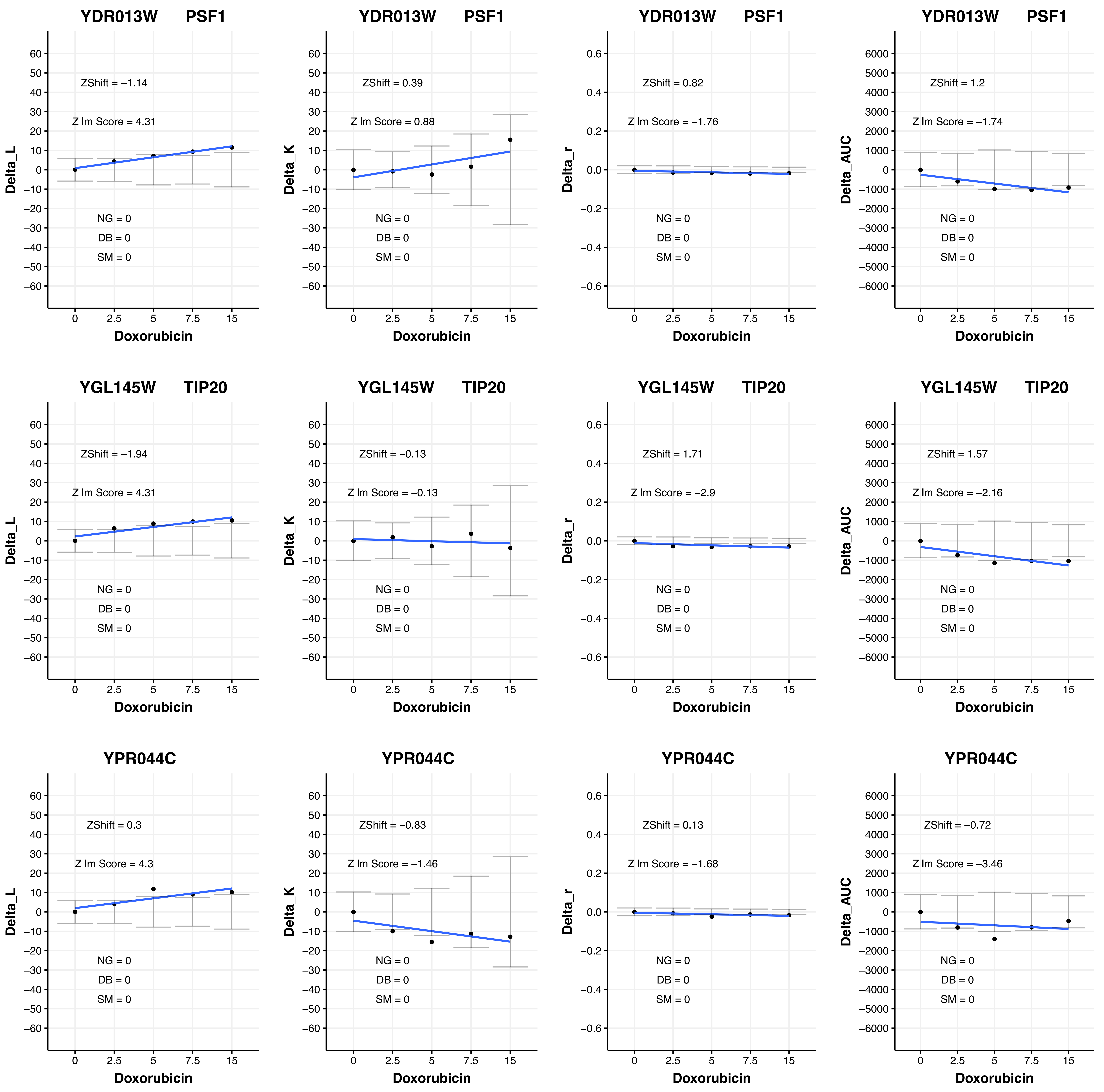
YDR013W PSF1
ZShift = −1.14
Z lm Score = 4.31
NG = 0
DB = 0
SM = 0
Delta_L
Doxorubicin
YDR013W PSF1
ZShift = 0.39
Z lm Score = 0.88
NG = 0
DB = 0
SM = 0
Delta_K
Doxorubicin
YDR013W PSF1
ZShift = 0.82
Z lm Score = −1.76
NG = 0
DB = 0
SM = 0
Delta_r
Doxorubicin
YDR013W PSF1
ZShift = 1.2
Z lm Score = −1.74
NG = 0
DB = 0
SM = 0
Delta_AUC
Doxorubicin
YGL145W TIP20
ZShift = −1.94
Z lm Score = 4.31
NG = 0
DB = 0
SM = 0
Delta_L
Doxorubicin
YGL145W TIP20
ZShift = −0.13
Z lm Score = −0.13
NG = 0
DB = 0
SM = 0
Delta_K
Doxorubicin
YGL145W TIP20
ZShift = 1.71
Z lm Score = −2.9
NG = 0
DB = 0
SM = 0
Delta_r
Doxorubicin
YGL145W TIP20
ZShift = 1.57
Z lm Score = −2.16
NG = 0
DB = 0
SM = 0
Delta_AUC
Doxorubicin
YPR044C
ZShift = 0.3
Z lm Score = 4.3
NG = 0
DB = 0
SM = 0
Delta_L
Doxorubicin
YPR044C
ZShift = −0.83
Z lm Score = −1.46
NG = 0
DB = 0
SM = 0
Delta_K
Doxorubicin
YPR044C
ZShift = 0.13
Z lm Score = −1.68
NG = 0
DB = 0
SM = 0
Delta_r
Doxorubicin
YPR044C
ZShift = −0.72
Z lm Score = −3.46
NG = 0
DB = 0
SM = 0
Delta_AUC
Doxorubicin

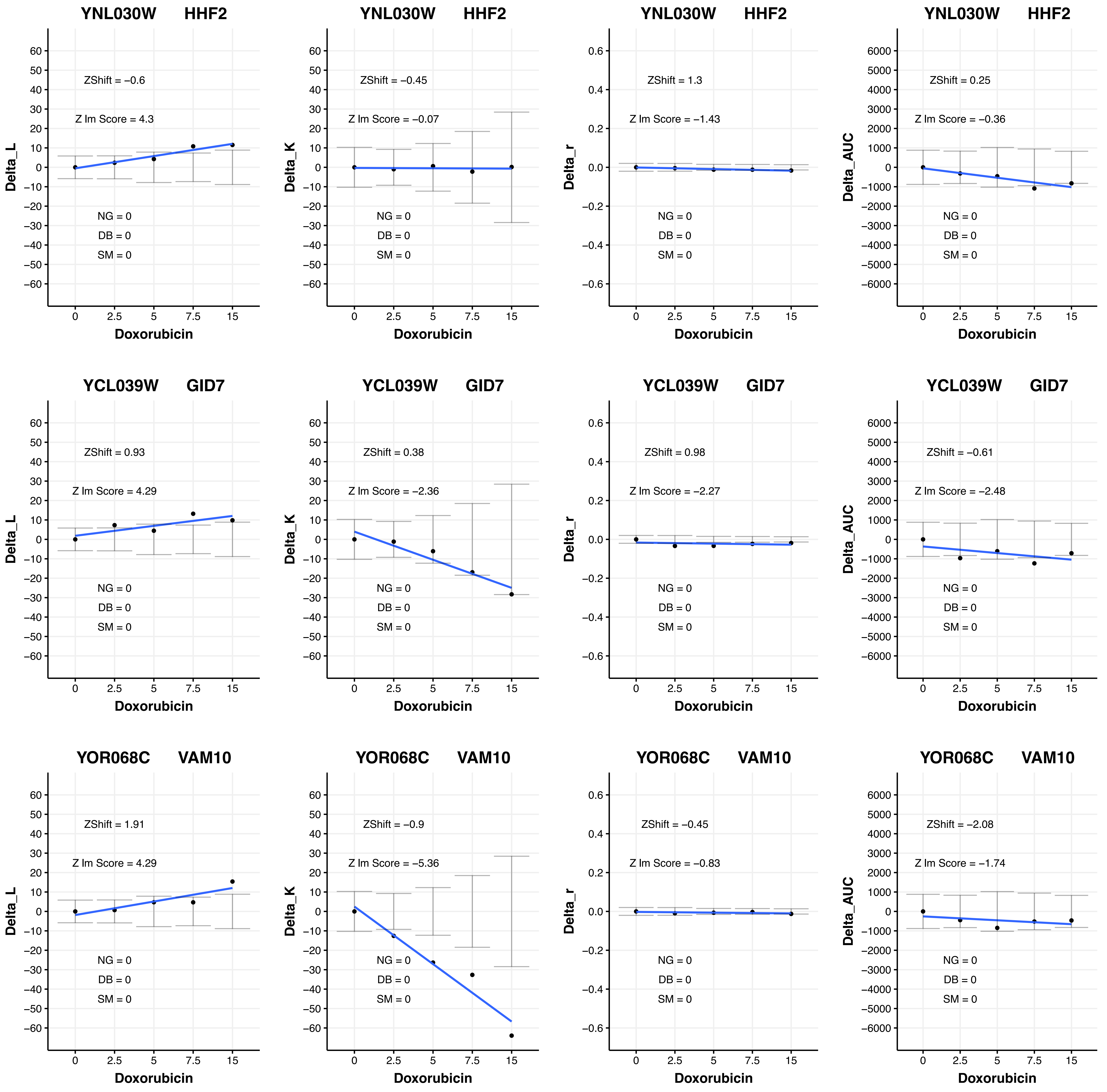

YNL030W HHF2
ZShift = −0.6
Z lm Score = 4.3
NG = 0
DB = 0
SM = 0
Delta_L
Doxorubicin
YNL030W HHF2
ZShift = −0.45
Z lm Score = −0.07
NG = 0
DB = 0
SM = 0
Delta_K
Doxorubicin
YNL030W HHF2
ZShift = 1.3
Z lm Score = −1.43
NG = 0
DB = 0
SM = 0
Delta_r
Doxorubicin
YNL030W HHF2
ZShift = 0.25
Z lm Score = −0.36
NG = 0
DB = 0
SM = 0
Delta_AUC
Doxorubicin
YCL039W GID7
ZShift = 0.93
Z lm Score = 4.29
NG = 0
DB = 0
SM = 0
Delta_L
Doxorubicin
YCL039W GID7
ZShift = 0.38
Z lm Score = −2.36
NG = 0
DB = 0
SM = 0
Delta_K
Doxorubicin
YCL039W GID7
ZShift = 0.98
Z lm Score = −2.27
NG = 0
DB = 0
SM = 0
Delta_r
Doxorubicin
YCL039W GID7
ZShift = −0.61
Z lm Score = −2.48
NG = 0
DB = 0
SM = 0
Delta_AUC
Doxorubicin
YOR068C VAM10
ZShift = 1.91
Z lm Score = 4.29
NG = 0
DB = 0
SM = 0
Delta_L
Doxorubicin
YOR068C VAM10
ZShift = −0.9
Z lm Score = −5.36
NG = 0
DB = 0
SM = 0
Delta_K
Doxorubicin
YOR068C VAM10
ZShift = −0.45
Z lm Score = −0.83
NG = 0
DB = 0
SM = 0
Delta_r
Doxorubicin
YOR068C VAM10
ZShift = −2.08
Z lm Score = −1.74
NG = 0
DB = 0
SM = 0
Delta_AUC
Doxorubicin

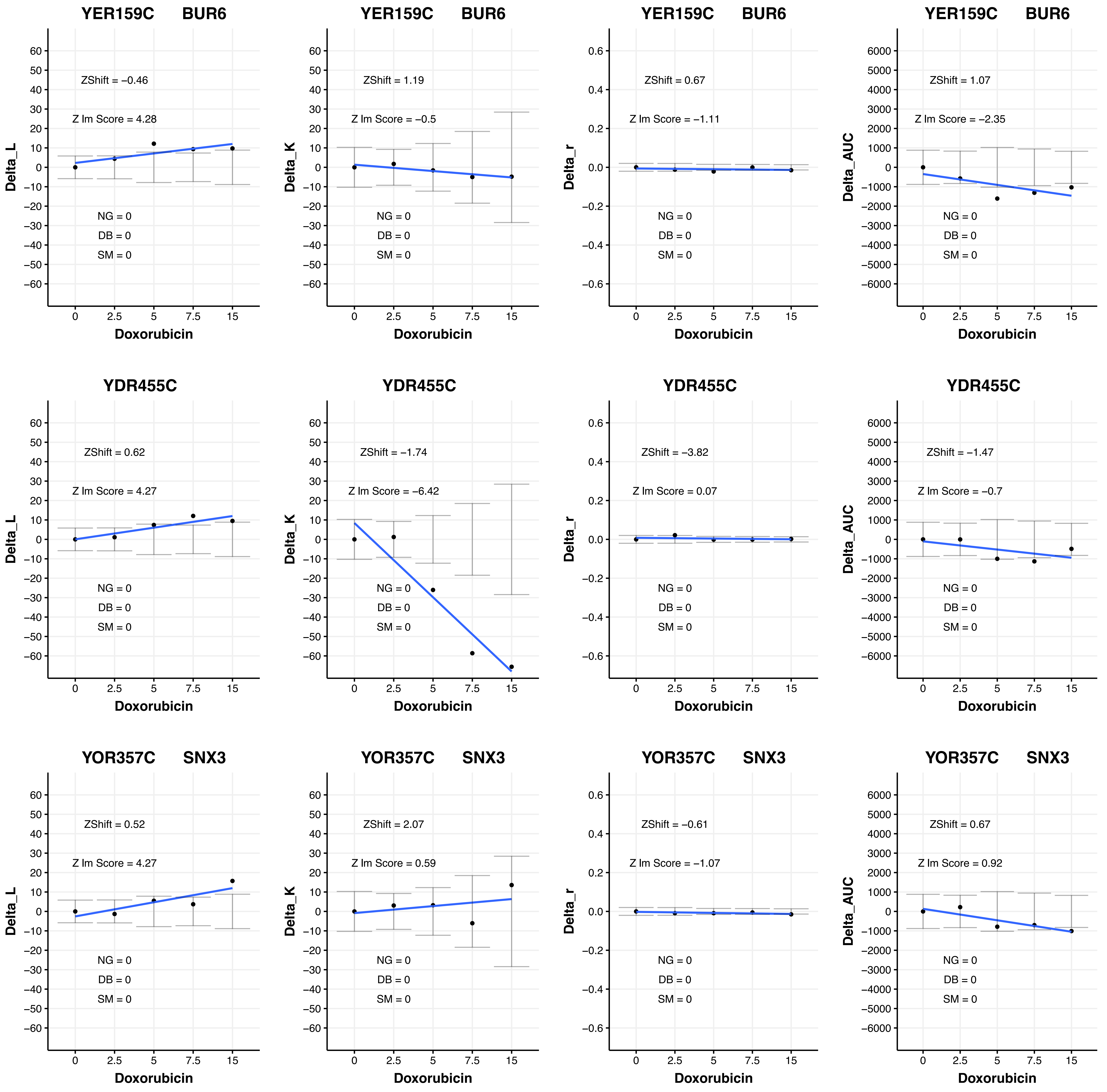
YER159C BUR6
ZShift = −0.46
Z lm Score = 4.28
NG = 0
DB = 0
SM = 0
Delta_L
Doxorubicin
YER159C BUR6
ZShift = 1.19
Z lm Score = −0.5
NG = 0
DB = 0
SM = 0
Delta_K
Doxorubicin
YER159C BUR6
ZShift = 0.67
Z lm Score = −1.11
NG = 0
DB = 0
SM = 0
Delta_r
Doxorubicin
YER159C BUR6
ZShift = 1.07
Z lm Score = −2.35
NG = 0
DB = 0
SM = 0
Delta_AUC
Doxorubicin
YDR455C
ZShift = 0.62
Z lm Score = 4.27
NG = 0
DB = 0
SM = 0
Delta_L
Doxorubicin
YDR455C
ZShift = −1.74
Z lm Score = −6.42
NG = 0
DB = 0
SM = 0
Delta_K
Doxorubicin
YDR455C
ZShift = −3.82
Z lm Score = 0.07
NG = 0
DB = 0
SM = 0
Delta_r
Doxorubicin
YDR455C
ZShift = −1.47
Z lm Score = −0.7
NG = 0
DB = 0
SM = 0
Delta_AUC
Doxorubicin
YOR357C SNX3
ZShift = 0.52
Z lm Score = 4.27
NG = 0
DB = 0
SM = 0
Delta_L
Doxorubicin
YOR357C SNX3
ZShift = 2.07
Z lm Score = 0.59
NG = 0
DB = 0
SM = 0
Delta_K
Doxorubicin
YOR357C SNX3
ZShift = −0.61
Z lm Score = −1.07
NG = 0
DB = 0
SM = 0
Delta_r
Doxorubicin
YOR357C SNX3
ZShift = 0.67
Z lm Score = 0.92
NG = 0
DB = 0
SM = 0
Delta_AUC
Doxorubicin

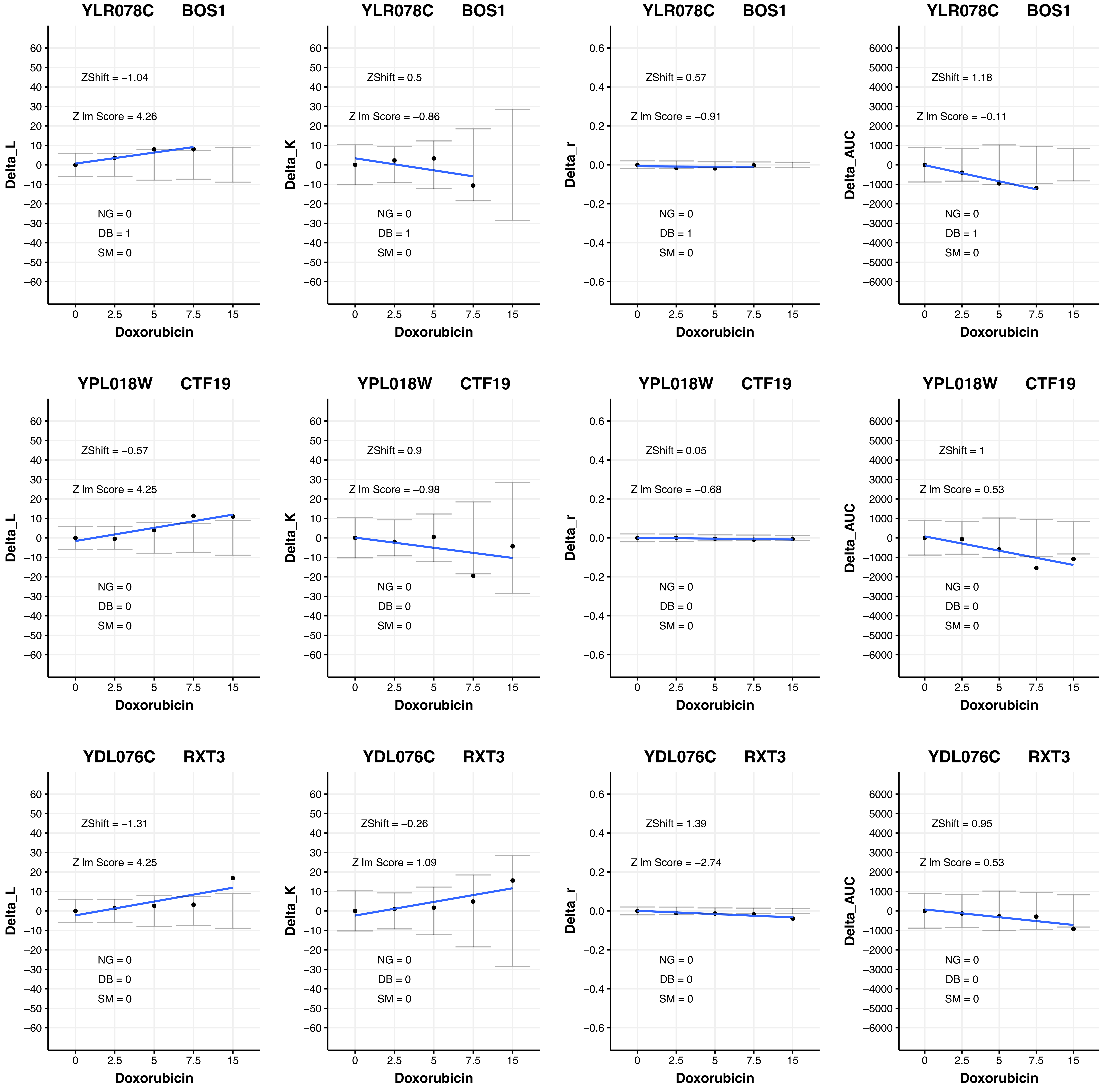
YLR078C BOS1
ZShift = −1.04
Z lm Score = 4.26
NG = 0
DB = 1
SM = 0
Delta_L
Doxorubicin
YLR078C BOS1
ZShift = 0.5
Z lm Score = −0.86
NG = 0
DB = 1
SM = 0
Delta_K
Doxorubicin
YLR078C BOS1
ZShift = 0.57
Z lm Score = −0.91
NG = 0
DB = 1
SM = 0
Delta_r
Doxorubicin
YLR078C BOS1
ZShift = 1.18
Z lm Score = −0.11
NG = 0
DB = 1
SM = 0
Delta_AUC
Doxorubicin
YPL018W CTF19
ZShift = −0.57
Z lm Score = 4.25
NG = 0
DB = 0
SM = 0
Delta_L
Doxorubicin
YPL018W CTF19
ZShift = 0.9
Z lm Score = −0.98
NG = 0
DB = 0
SM = 0
Delta_K
Doxorubicin
YPL018W CTF19
ZShift = 0.05
Z lm Score = −0.68
NG = 0
DB = 0
SM = 0
Delta_r
Doxorubicin
YPL018W CTF19
ZShift = 1
Z lm Score = 0.53
NG = 0
DB = 0
SM = 0
Delta_AUC
Doxorubicin
YDL076C RXT3
ZShift = −1.31
Z lm Score = 4.25
NG = 0
DB = 0
SM = 0
Delta_L
Doxorubicin
YDL076C RXT3
ZShift = −0.26
Z lm Score = 1.09
NG = 0
DB = 0
SM = 0
Delta_K
Doxorubicin
YDL076C RXT3
ZShift = 1.39
Z lm Score = −2.74
NG = 0
DB = 0
SM = 0
Delta_r
Doxorubicin
YDL076C RXT3
ZShift = 0.95
Z lm Score = 0.53
NG = 0
DB = 0
SM = 0
Delta_AUC
Doxorubicin

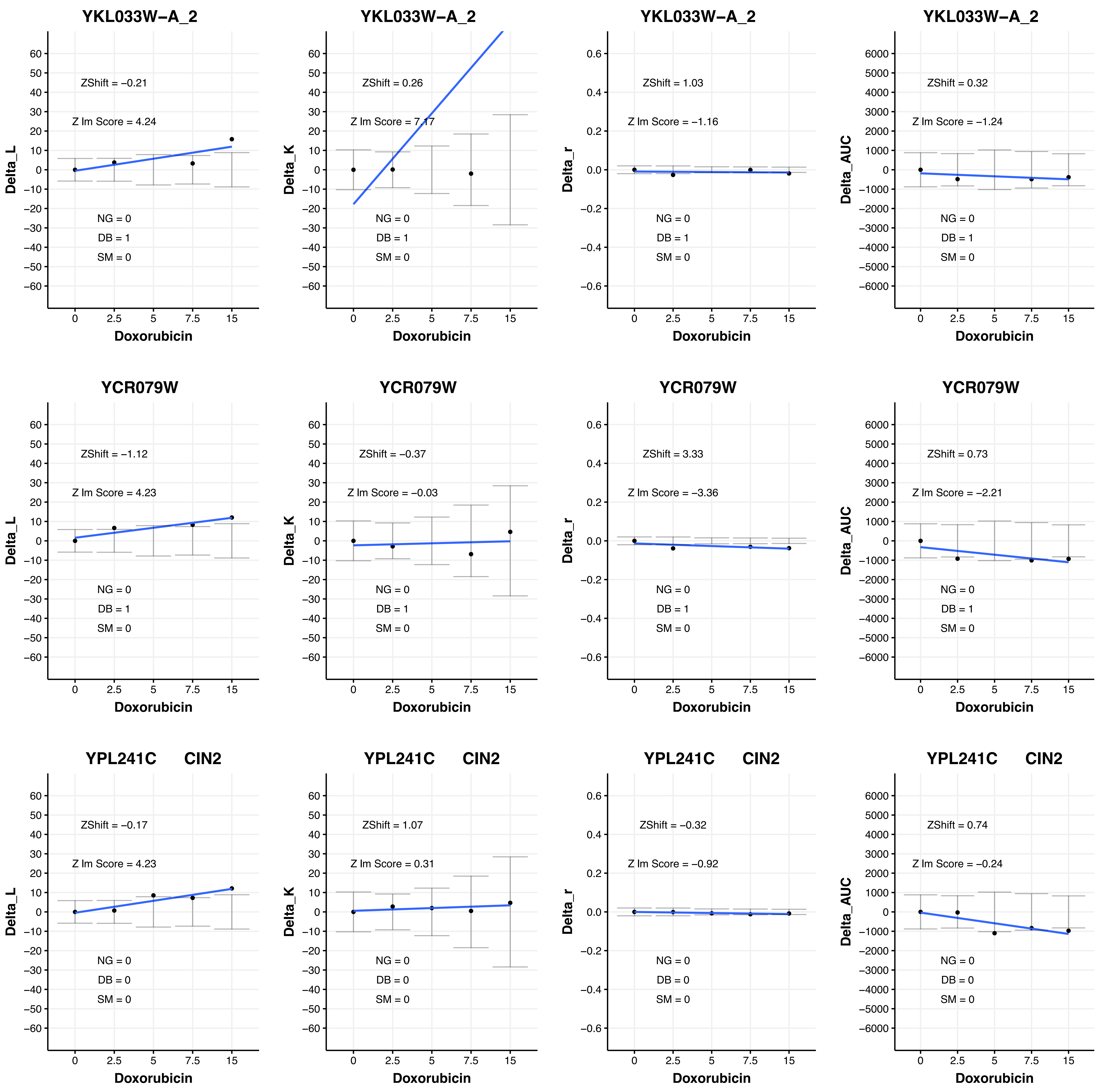
YKL033W−A_2
ZShift = −0.21
Z lm Score = 4.24
NG = 0
DB = 1
SM = 0
Delta_L
Doxorubicin
YKL033W−A_2
ZShift = 0.26
Z lm Score = 7.17
NG = 0
DB = 1
SM = 0
Delta_K
Doxorubicin
YKL033W−A_2
ZShift = 1.03
Z lm Score = −1.16
NG = 0
DB = 1
SM = 0
Delta_r
Doxorubicin
YKL033W−A_2
ZShift = 0.32
Z lm Score = −1.24
NG = 0
DB = 1
SM = 0
Delta_AUC
Doxorubicin
YCR079W
ZShift = −1.12
Z lm Score = 4.23
NG = 0
DB = 1
SM = 0
Delta_L
Doxorubicin
YCR079W
ZShift = −0.37
Z lm Score = −0.03
NG = 0
DB = 1
SM = 0
Delta_K
Doxorubicin
YCR079W
ZShift = 3.33
Z lm Score = −3.36
NG = 0
DB = 1
SM = 0
Delta_r
Doxorubicin
YCR079W
ZShift = 0.73
Z lm Score = −2.21
NG = 0
DB = 1
SM = 0
Delta_AUC
Doxorubicin
YPL241C CIN2
ZShift = −0.17
Z lm Score = 4.23
NG = 0
DB = 0
SM = 0
Delta_L
Doxorubicin
YPL241C CIN2
ZShift = 1.07
Z lm Score = 0.31
NG = 0
DB = 0
SM = 0
Delta_K
Doxorubicin
YPL241C CIN2
ZShift = −0.32
Z lm Score = −0.92
NG = 0
DB = 0
SM = 0
Delta_r
Doxorubicin
YPL241C CIN2
ZShift = 0.74
Z lm Score = −0.24
NG = 0
DB = 0
SM = 0
Delta_AUC
Doxorubicin

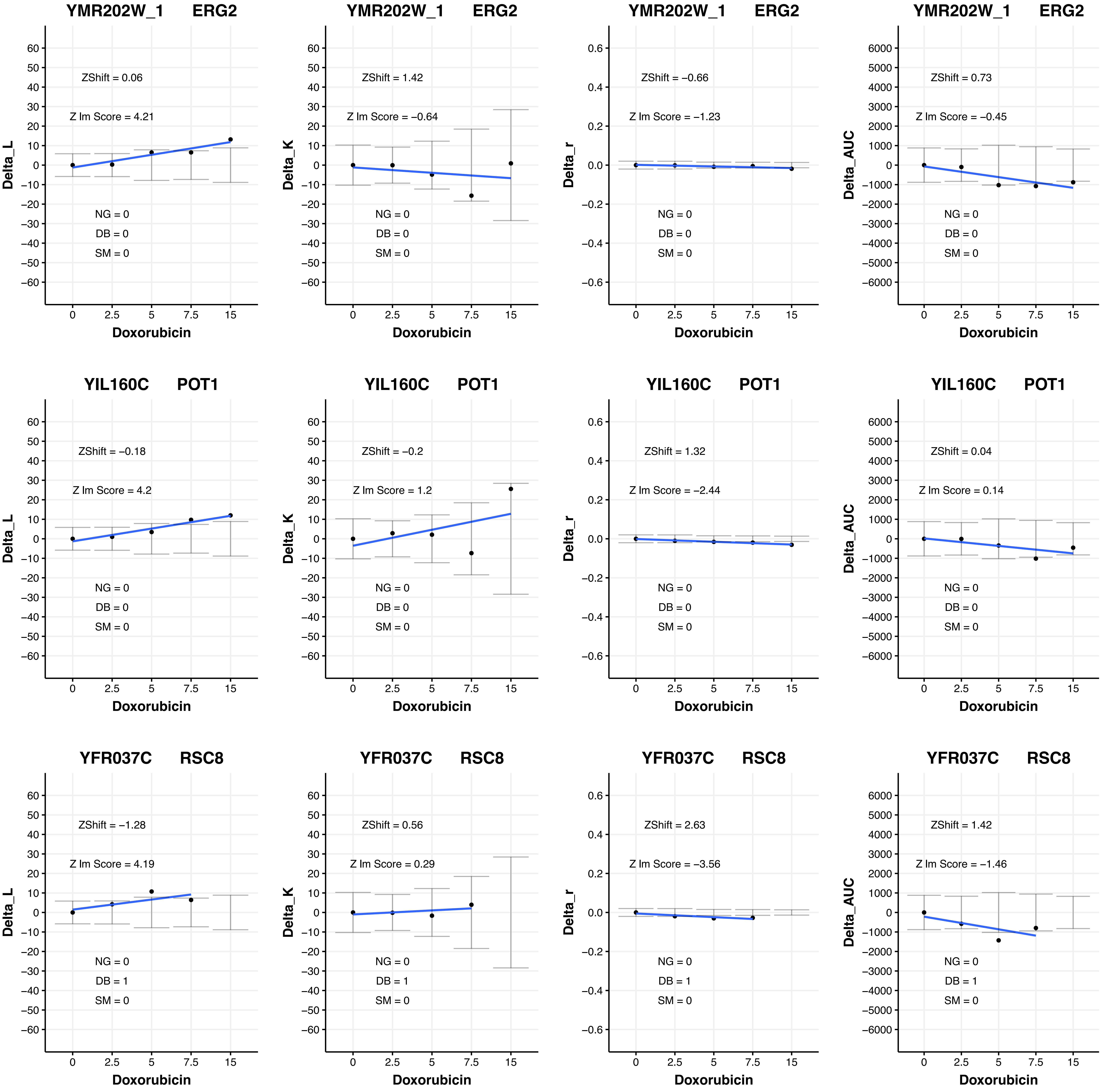
YMR202W_1 ERG2
ZShift = 0.06
Z lm Score = 4.21
NG = 0
DB = 0
SM = 0
Delta_L
Doxorubicin
YMR202W_1 ERG2
ZShift = 1.42
Z lm Score = −0.64
NG = 0
DB = 0
SM = 0
Delta_K
Doxorubicin
YMR202W_1 ERG2
ZShift = −0.66
Z lm Score = −1.23
NG = 0
DB = 0
SM = 0
Delta_r
Doxorubicin
YMR202W_1 ERG2
ZShift = 0.73
Z lm Score = −0.45
NG = 0
DB = 0
SM = 0
Delta_AUC
Doxorubicin
YIL160C POT1
ZShift = −0.18
Z lm Score = 4.2
NG = 0
DB = 0
SM = 0
Delta_L
Doxorubicin
YIL160C POT1
ZShift = −0.2
Z lm Score = 1.2
NG = 0
DB = 0
SM = 0
Delta_K
Doxorubicin
YIL160C POT1
ZShift = 1.32
Z lm Score = −2.44
NG = 0
DB = 0
SM = 0
Delta_r
Doxorubicin
YIL160C POT1
ZShift = 0.04
Z lm Score = 0.14
NG = 0
DB = 0
SM = 0
Delta_AUC
Doxorubicin
YFR037C RSC8
ZShift = −1.28
Z lm Score = 4.19
NG = 0
DB = 1
SM = 0
Delta_L
Doxorubicin
YFR037C RSC8
ZShift = 0.56
Z lm Score = 0.29
NG = 0
DB = 1
SM = 0
Delta_K
Doxorubicin
YFR037C RSC8
ZShift = 2.63
Z lm Score = −3.56
NG = 0
DB = 1
SM = 0
Delta_r
Doxorubicin
YFR037C RSC8
ZShift = 1.42
Z lm Score = −1.46
NG = 0
DB = 1
SM = 0
Delta_AUC
Doxorubicin

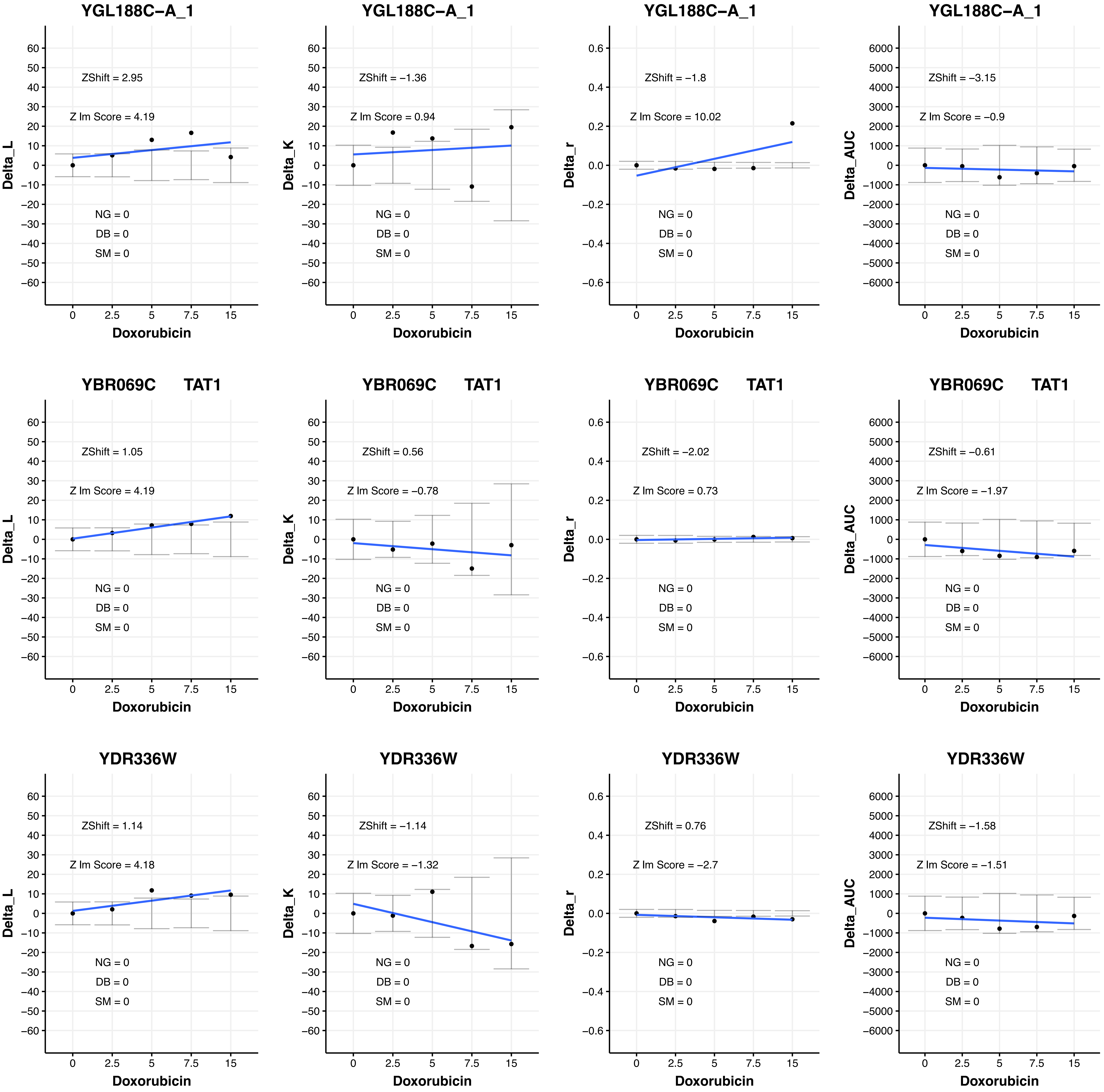

YGL188C−A_1
ZShift = 2.95
Z lm Score = 4.19
NG = 0
DB = 0
SM = 0
Delta_L
Doxorubicin
YGL188C−A_1
ZShift = −1.36
Z lm Score = 0.94
NG = 0
DB = 0
SM = 0
Delta_K
Doxorubicin
YGL188C−A_1
ZShift = −1.8
Z lm Score = 10.02
NG = 0
DB = 0
SM = 0
Delta_r
Doxorubicin
YGL188C−A_1
ZShift = −3.15
Z lm Score = −0.9
NG = 0
DB = 0
SM = 0
Delta_AUC
Doxorubicin
YBR069C TAT1
ZShift = 1.05
Z lm Score = 4.19
NG = 0
DB = 0
SM = 0
Delta_L
Doxorubicin
YBR069C TAT1
ZShift = 0.56
Z lm Score = −0.78
NG = 0
DB = 0
SM = 0
Delta_K
Doxorubicin
YBR069C TAT1
ZShift = −2.02
Z lm Score = 0.73
NG = 0
DB = 0
SM = 0
Delta_r
Doxorubicin
YBR069C TAT1
ZShift = −0.61
Z lm Score = −1.97
NG = 0
DB = 0
SM = 0
Delta_AUC
Doxorubicin
YDR336W
ZShift = 1.14
Z lm Score = 4.18
NG = 0
DB = 0
SM = 0
Delta_L
Doxorubicin
YDR336W
ZShift = −1.14
Z lm Score = −1.32
NG = 0
DB = 0
SM = 0
Delta_K
Doxorubicin
YDR336W
ZShift = 0.76
Z lm Score = −2.7
NG = 0
DB = 0
SM = 0
Delta_r
Doxorubicin
YDR336W
ZShift = −1.58
Z lm Score = −1.51
NG = 0
DB = 0
SM = 0
Delta_AUC
Doxorubicin

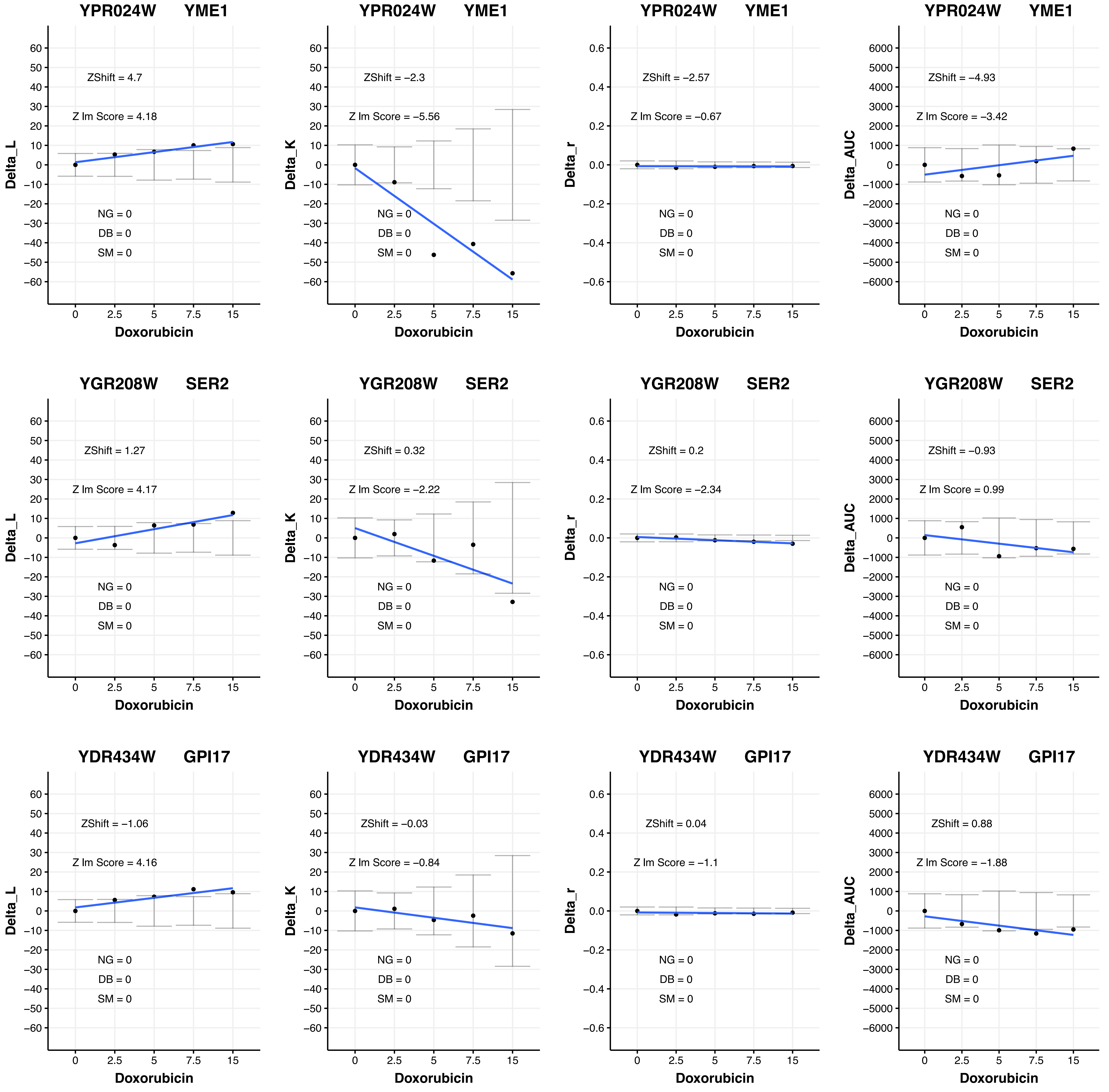

YPR024W YME1
ZShift = 4.7
Z lm Score = 4.18
NG = 0
DB = 0
SM = 0
Delta_L
Doxorubicin
YPR024W YME1
ZShift = −2.3
Z lm Score = −5.56
NG = 0
DB = 0
SM = 0
Delta_K
Doxorubicin
YPR024W YME1
ZShift = −2.57
Z lm Score = −0.67
NG = 0
DB = 0
SM = 0
Delta_r
Doxorubicin
YPR024W YME1
ZShift = −4.93
Z lm Score = −3.42
NG = 0
DB = 0
SM = 0
Delta_AUC
Doxorubicin
YGR208W SER2
ZShift = 1.27
Z lm Score = 4.17
NG = 0
DB = 0
SM = 0
Delta_L
Doxorubicin
YGR208W SER2
ZShift = 0.32
Z lm Score = −2.22
NG = 0
DB = 0
SM = 0
Delta_K
Doxorubicin
YGR208W SER2
ZShift = 0.2
Z lm Score = −2.34
NG = 0
DB = 0
SM = 0
Delta_r
Doxorubicin
YGR208W SER2
ZShift = −0.93
Z lm Score = 0.99
NG = 0
DB = 0
SM = 0
Delta_AUC
Doxorubicin
YDR434W GPI17
ZShift = −1.06
Z lm Score = 4.16
NG = 0
DB = 0
SM = 0
Delta_L
Doxorubicin
YDR434W GPI17
ZShift = −0.03
Z lm Score = −0.84
NG = 0
DB = 0
SM = 0
Delta_K
Doxorubicin
YDR434W GPI17
ZShift = 0.04
Z lm Score = −1.1
NG = 0
DB = 0
SM = 0
Delta_r
Doxorubicin
YDR434W GPI17
ZShift = 0.88
Z lm Score = −1.88
NG = 0
DB = 0
SM = 0
Delta_AUC
Doxorubicin

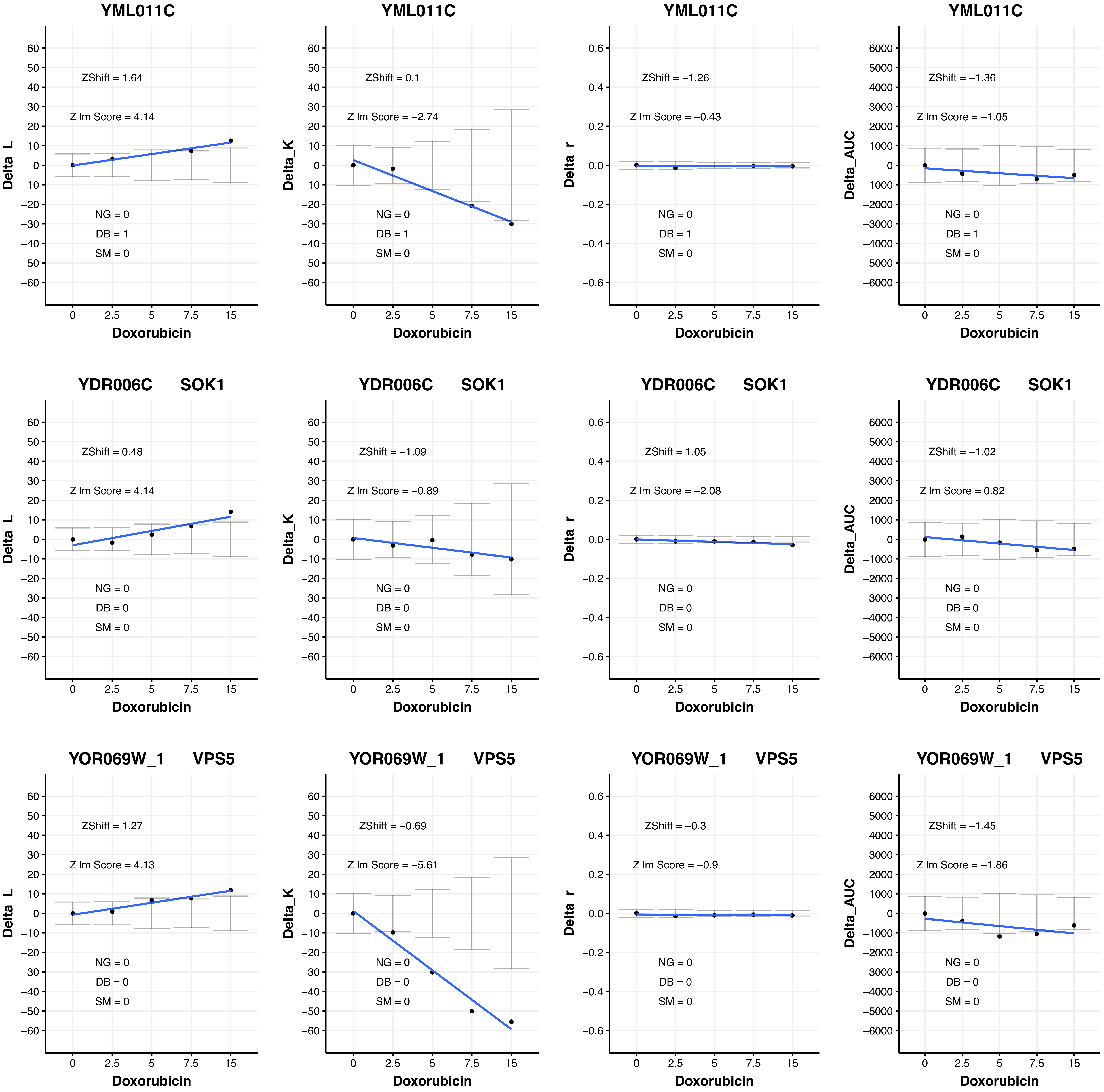
YML011C
ZShift = 1.64
Z lm Score = 4.14
NG = 0
DB = 1
SM = 0
Delta_L
Doxorubicin
YML011C
ZShift = 0.1
Z lm Score = −2.74
NG = 0
DB = 1
SM = 0
Delta_K
Doxorubicin
YML011C
ZShift = −1.26
Z lm Score = −0.43
NG = 0
DB = 1
SM = 0
Delta_r
Doxorubicin
YML011C
ZShift = −1.36
Z lm Score = −1.05
NG = 0
DB = 1
SM = 0
Delta_AUC
Doxorubicin
YDR006C SOK1
ZShift = 0.48
Z lm Score = 4.14
NG = 0
DB = 0
SM = 0
Delta_L
Doxorubicin
YDR006C SOK1
ZShift = −1.09
Z lm Score = −0.89
NG = 0
DB = 0
SM = 0
Delta_K
Doxorubicin
YDR006C SOK1
ZShift = 1.05
Z lm Score = −2.08
NG = 0
DB = 0
SM = 0
Delta_r
Doxorubicin
YDR006C SOK1
ZShift = −1.02
Z lm Score = 0.82
NG = 0
DB = 0
SM = 0
Delta_AUC
Doxorubicin
YOR069W_1 VPS5
ZShift = 1.27
Z lm Score = 4.13
NG = 0
DB = 0
SM = 0
Delta_L
Doxorubicin
YOR069W_1 VPS5
ZShift = −0.69
Z lm Score = −5.61
NG = 0
DB = 0
SM = 0
Delta_K
Doxorubicin
YOR069W_1 VPS5
ZShift = −0.3
Z lm Score = −0.9
NG = 0
DB = 0
SM = 0
Delta_r
Doxorubicin
YOR069W_1 VPS5
ZShift = −1.45
Z lm Score = −1.86
NG = 0
DB = 0
SM = 0
Delta_AUC
Doxorubicin

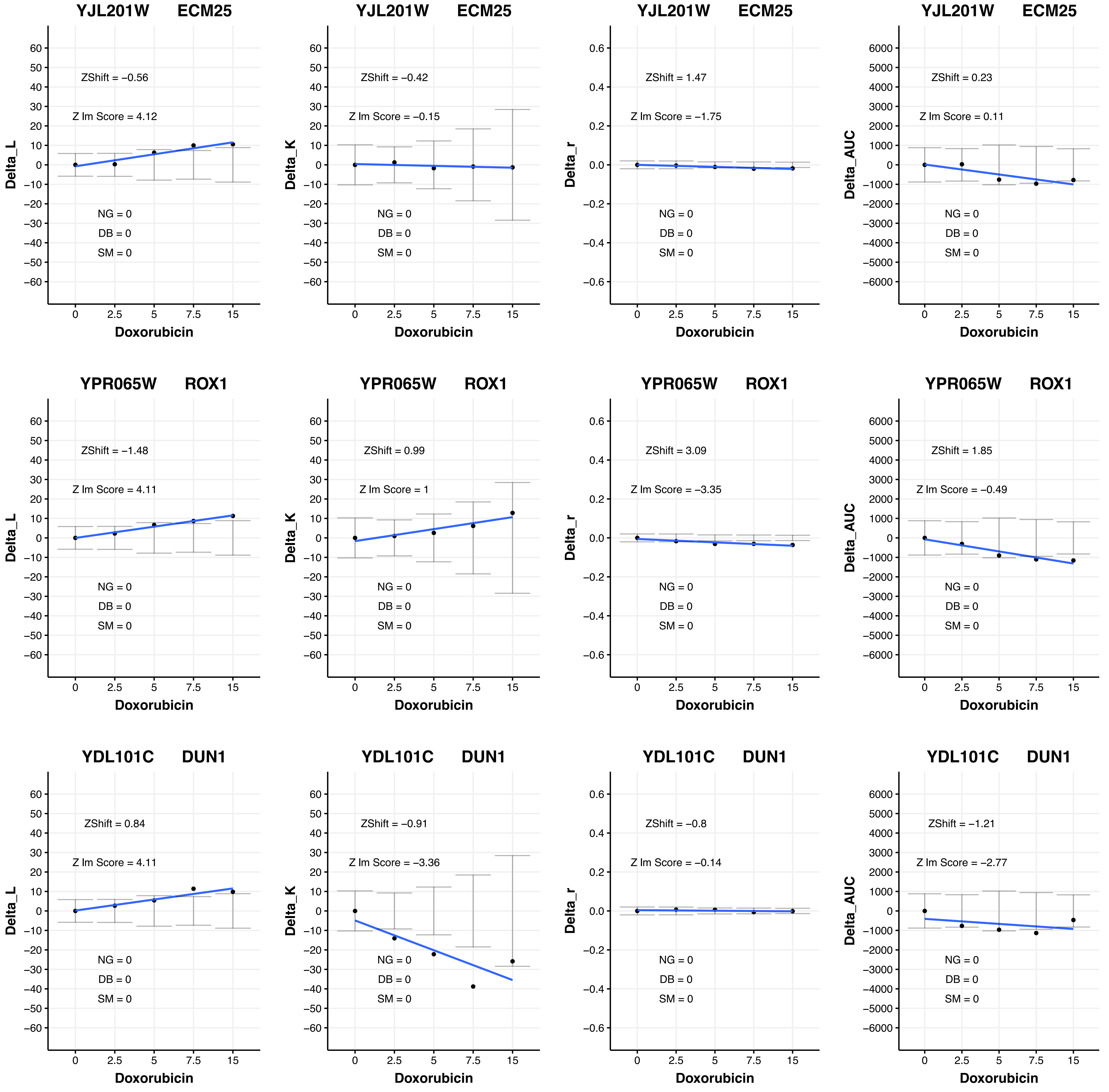
YJL201W ECM25
ZShift = −0.56
Z lm Score = 4.12
NG = 0
DB = 0
SM = 0
Delta_L
Doxorubicin
YJL201W ECM25
ZShift = −0.42
Z lm Score = −0.15
NG = 0
DB = 0
SM = 0
Delta_K
Doxorubicin
YJL201W ECM25
ZShift = 1.47
Z lm Score = −1.75
NG = 0
DB = 0
SM = 0
Delta_r
Doxorubicin
YJL201W ECM25
ZShift = 0.23
Z lm Score = 0.11
NG = 0
DB = 0
SM = 0
Delta_AUC
Doxorubicin
YPR065W ROX1
ZShift = −1.48
Z lm Score = 4.11
NG = 0
DB = 0
SM = 0
Delta_L
Doxorubicin
YPR065W ROX1
ZShift = 0.99
Z lm Score = 1
NG = 0
DB = 0
SM = 0
Delta_K
Doxorubicin
YPR065W ROX1
ZShift = 3.09
Z lm Score = −3.35
NG = 0
DB = 0
SM = 0
Delta_r
Doxorubicin
YPR065W ROX1
ZShift = 1.85
Z lm Score = −0.49
NG = 0
DB = 0
SM = 0
Delta_AUC
Doxorubicin
YDL101C DUN1
ZShift = 0.84
Z lm Score = 4.11
NG = 0
DB = 0
SM = 0
Delta_L
Doxorubicin
YDL101C DUN1
ZShift = −0.91
Z lm Score = −3.36
NG = 0
DB = 0
SM = 0
Delta_K
Doxorubicin
YDL101C DUN1
ZShift = −0.8
Z lm Score = −0.14
NG = 0
DB = 0
SM = 0
Delta_r
Doxorubicin
YDL101C DUN1
ZShift = −1.21
Z lm Score = −2.77
NG = 0
DB = 0
SM = 0
Delta_AUC
Doxorubicin

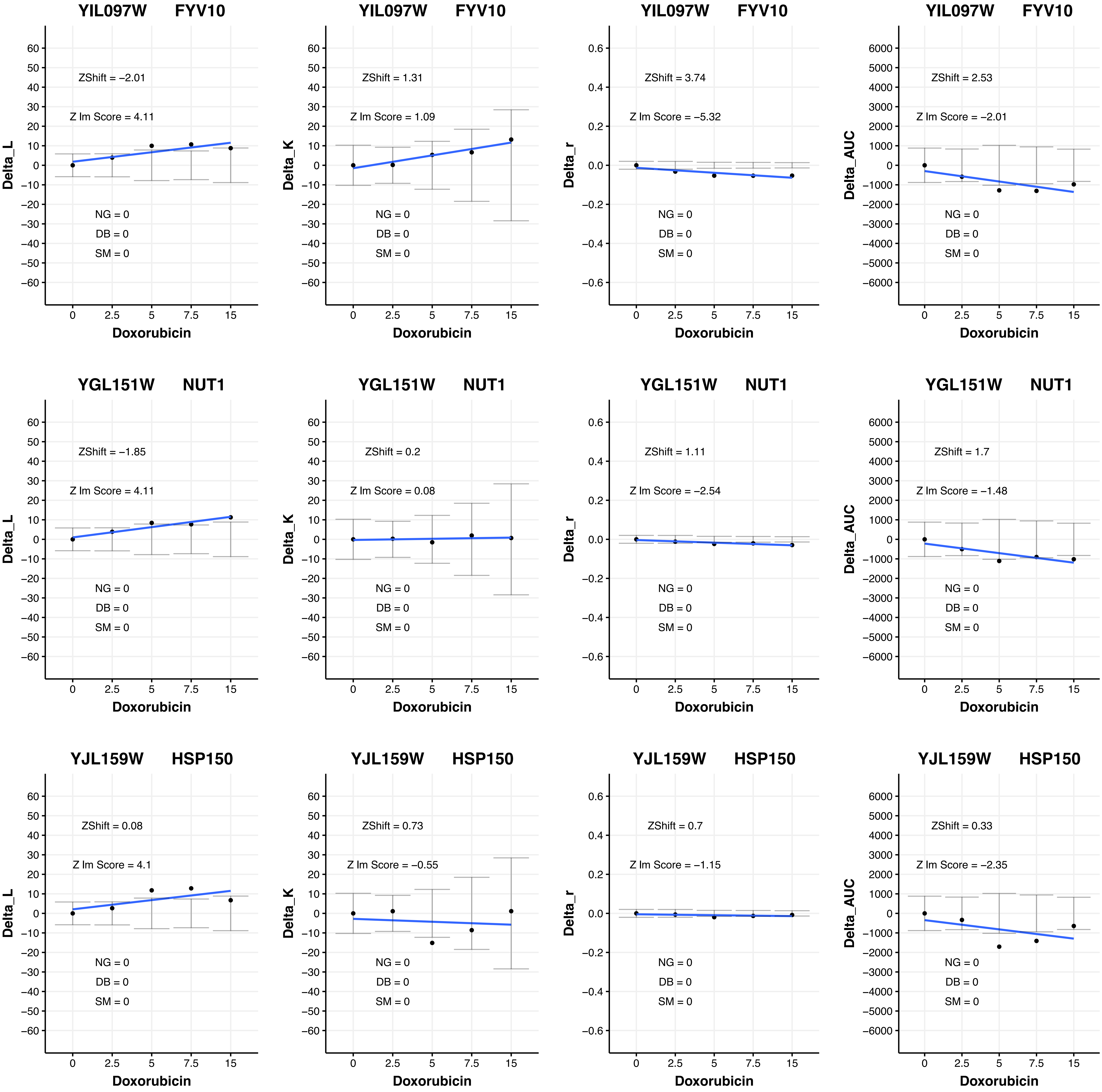
YIL097W FYV10
Delta_L
ZShift = −2.01
Z lm Score = 4.11
NG = 0
DB = 0
SM = 0
Doxorubicin
YIL097W FYV10
Delta_K
ZShift = 1.31
Z lm Score = 1.09
NG = 0
DB = 0
SM = 0
Doxorubicin
YIL097W FYV10
Delta_r
ZShift = 3.74
Z lm Score = −5.32
NG = 0
DB = 0
SM = 0
Doxorubicin
YIL097W FYV10
Delta_AUC
ZShift = 2.53
Z lm Score = −2.01
NG = 0
DB = 0
SM = 0
Doxorubicin
YGL151W NUT1
Delta_L
ZShift = −1.85
Z lm Score = 4.11
NG = 0
DB = 0
SM = 0
Doxorubicin
YGL151W NUT1
Delta_K
ZShift = 0.2
Z lm Score = 0.08
NG = 0
DB = 0
SM = 0
Doxorubicin
YGL151W NUT1
Delta_r
ZShift = 1.11
Z lm Score = −2.54
NG = 0
DB = 0
SM = 0
Doxorubicin
YGL151W NUT1
Delta_AUC
ZShift = 1.7
Z lm Score = −1.48
NG = 0
DB = 0
SM = 0
Doxorubicin
YJL159W HSP150
Delta_L
ZShift = 0.08
Z lm Score = 4.1
NG = 0
DB = 0
SM = 0
Doxorubicin
YJL159W HSP150
Delta_K
ZShift = 0.73
Z lm Score = −0.55
NG = 0
DB = 0
SM = 0
Doxorubicin
YJL159W HSP150
Delta_r
ZShift = 0.7
Z lm Score = −1.15
NG = 0
DB = 0
SM = 0
Doxorubicin
YJL159W HSP150
Delta_AUC
ZShift = 0.33
Z lm Score = −2.35
NG = 0
DB = 0
SM = 0
Doxorubicin

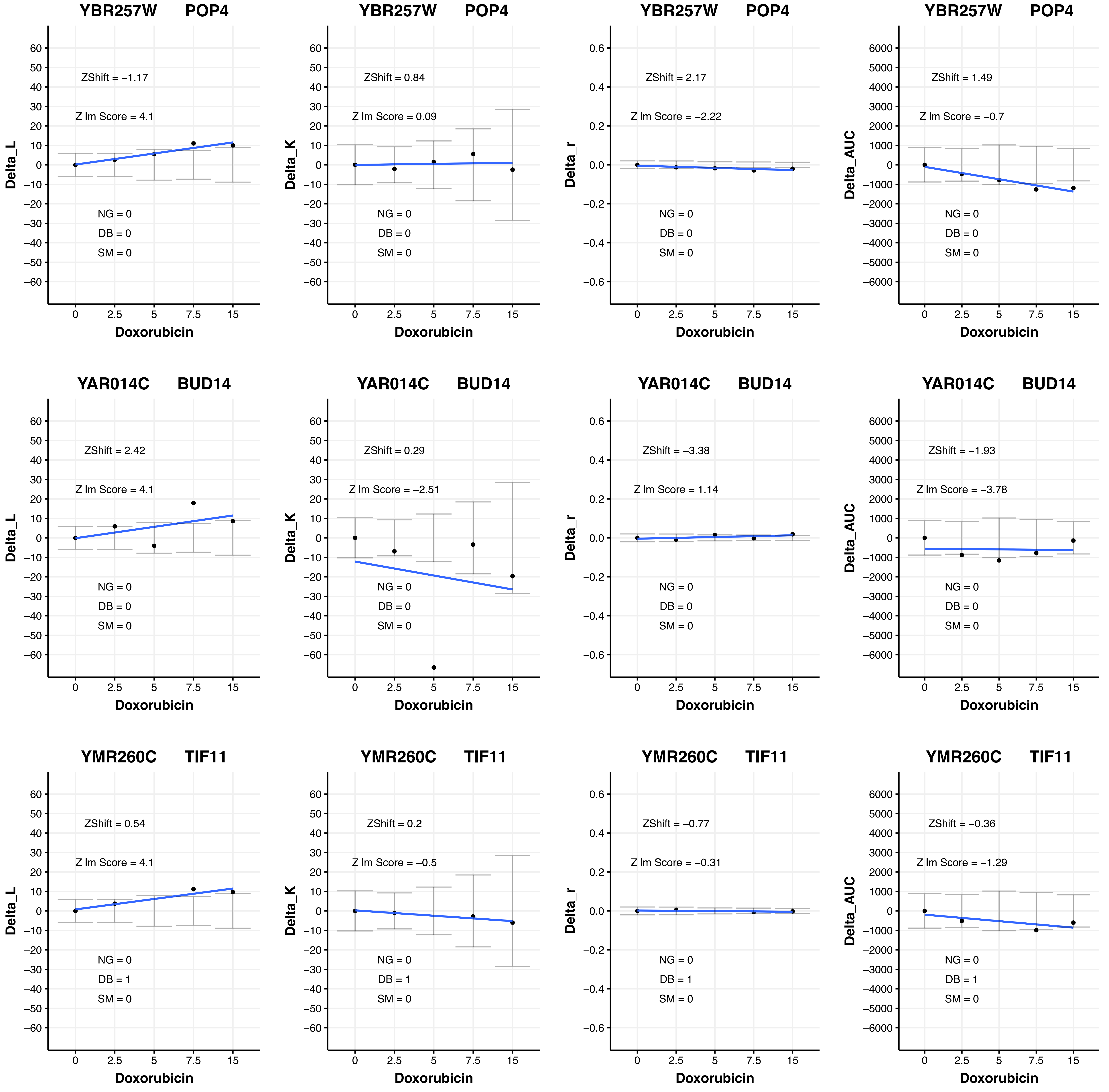
YBR257W POP4
ZShift = −1.17
Z lm Score = 4.1
NG = 0
DB = 0
SM = 0
Delta_L
Doxorubicin
YBR257W POP4
ZShift = 0.84
Z lm Score = 0.09
NG = 0
DB = 0
SM = 0
Delta_K
Doxorubicin
YBR257W POP4
ZShift = 2.17
Z lm Score = −2.22
NG = 0
DB = 0
SM = 0
Delta_r
Doxorubicin
YBR257W POP4
ZShift = 1.49
Z lm Score = −0.7
NG = 0
DB = 0
SM = 0
Delta_AUC
Doxorubicin
YAR014C BUD14
ZShift = 2.42
Z lm Score = 4.1
NG = 0
DB = 0
SM = 0
Delta_L
Doxorubicin
YAR014C BUD14
ZShift = 0.29
Z lm Score = −2.51
NG = 0
DB = 0
SM = 0
Delta_K
Doxorubicin
YAR014C BUD14
ZShift = −3.38
Z lm Score = 1.14
NG = 0
DB = 0
SM = 0
Delta_r
Doxorubicin
YAR014C BUD14
ZShift = −1.93
Z lm Score = −3.78
NG = 0
DB = 0
SM = 0
Delta_AUC
Doxorubicin
YMR260C TIF11
ZShift = 0.54
Z lm Score = 4.1
NG = 0
DB = 1
SM = 0
Delta_L
Doxorubicin
YMR260C TIF11
ZShift = 0.2
Z lm Score = −0.5
NG = 0
DB = 1
SM = 0
Delta_K
Doxorubicin
YMR260C TIF11
ZShift = −0.77
Z lm Score = −0.31
NG = 0
DB = 1
SM = 0
Delta_r
Doxorubicin
YMR260C TIF11
ZShift = −0.36
Z lm Score = −1.29
NG = 0
DB = 1
SM = 0
Delta_AUC
Doxorubicin

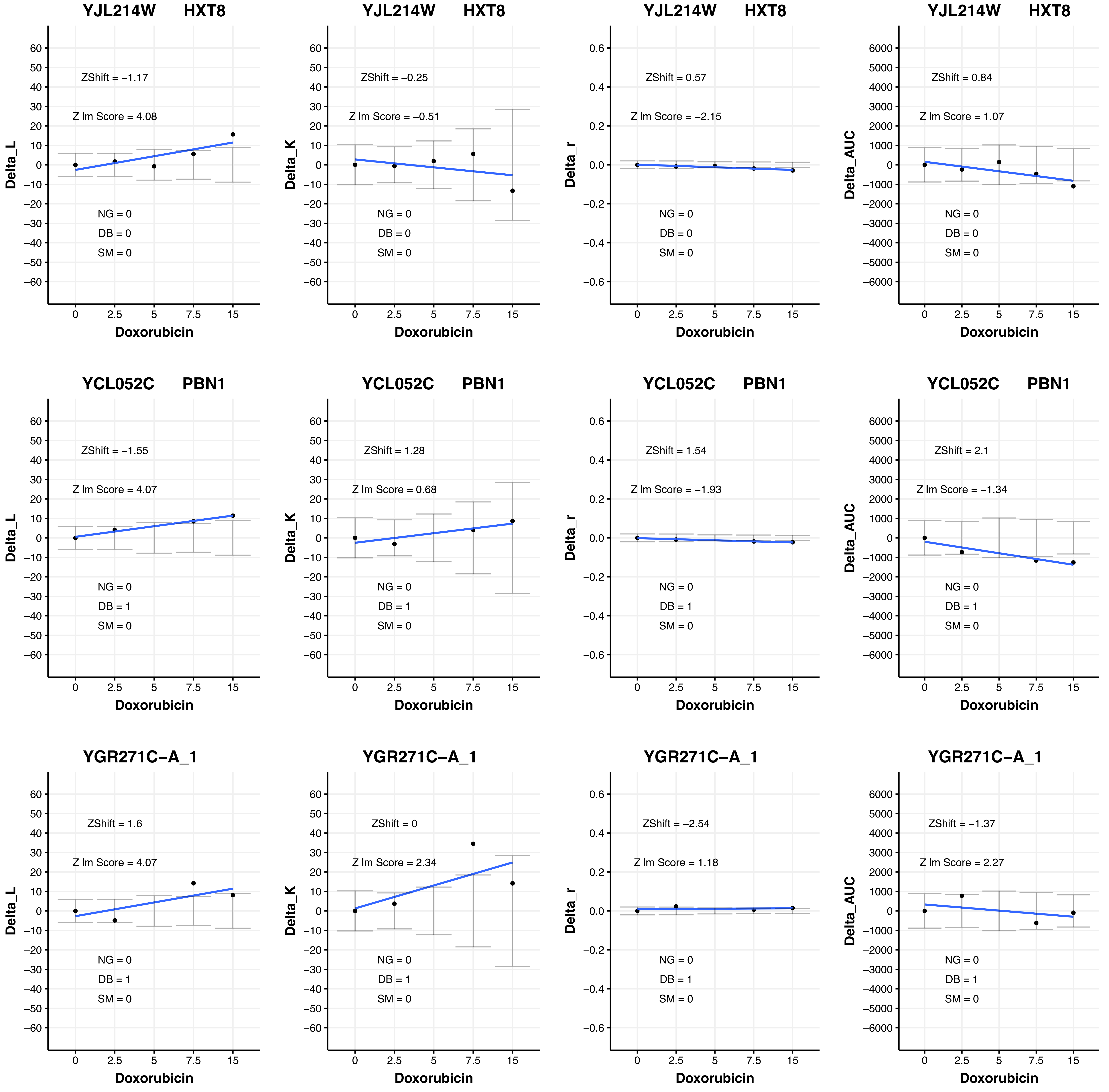
YJL214W HXT8
ZShift = −1.17
Z lm Score = 4.08
NG = 0
DB = 0
SM = 0
Delta_L
Doxorubicin
YJL214W HXT8
ZShift = −0.25
Z lm Score = −0.51
NG = 0
DB = 0
SM = 0
Delta_K
Doxorubicin
YJL214W HXT8
ZShift = 0.57
Z lm Score = −2.15
NG = 0
DB = 0
SM = 0
Delta_r
Doxorubicin
YJL214W HXT8
ZShift = 0.84
Z lm Score = 1.07
NG = 0
DB = 0
SM = 0
Delta_AUC
Doxorubicin
YCL052C PBN1
ZShift = −1.55
Z lm Score = 4.07
NG = 0
DB = 1
SM = 0
Delta_L
Doxorubicin
YCL052C PBN1
ZShift = 1.28
Z lm Score = 0.68
NG = 0
DB = 1
SM = 0
Delta_K
Doxorubicin
YCL052C PBN1
ZShift = 1.54
Z lm Score = −1.93
NG = 0
DB = 1
SM = 0
Delta_r
Doxorubicin
YCL052C PBN1
ZShift = 2.1
Z lm Score = −1.34
NG = 0
DB = 1
SM = 0
Delta_AUC
Doxorubicin
YGR271C−A_1
ZShift = 1.6
Z lm Score = 4.07
NG = 0
DB = 1
SM = 0
Delta_L
Doxorubicin
YGR271C−A_1
ZShift = 0
Z lm Score = 2.34
NG = 0
DB = 1
SM = 0
Delta_K
Doxorubicin
YGR271C−A_1
ZShift = −2.54
Z lm Score = 1.18
NG = 0
DB = 1
SM = 0
Delta_r
Doxorubicin
YGR271C−A_1
ZShift = −1.37
Z lm Score = 2.27
NG = 0
DB = 1
SM = 0
Delta_AUC
Doxorubicin

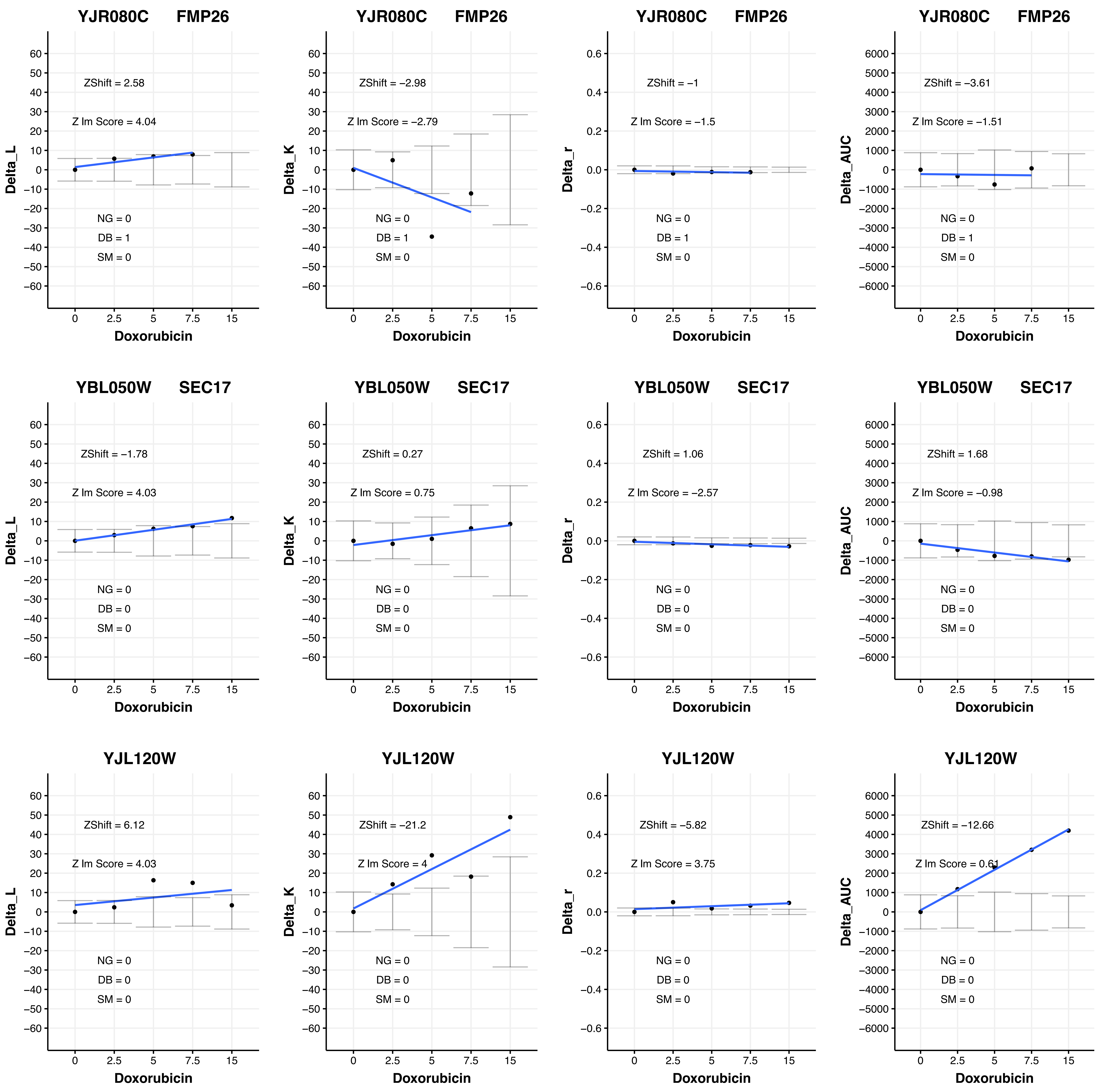
YJR080C FMP26
Delta_L
ZShift = 2.58
Z lm Score = 4.04
NG = 0
DB = 1
SM = 0
Doxorubicin
YJR080C FMP26
Delta_K
ZShift = −2.98
Z lm Score = −2.79
NG = 0
DB = 1
SM = 0
Doxorubicin
YJR080C FMP26
Delta_r
ZShift = −1
Z lm Score = −1.5
NG = 0
DB = 1
SM = 0
Doxorubicin
YJR080C FMP26
Delta_AUC
ZShift = −3.61
Z lm Score = −1.51
NG = 0
DB = 1
SM = 0
Doxorubicin
YBL050W SEC17
Delta_L
ZShift = −1.78
Z lm Score = 4.03
NG = 0
DB = 0
SM = 0
Doxorubicin
YBL050W SEC17
Delta_K
ZShift = 0.27
Z lm Score = 0.75
NG = 0
DB = 0
SM = 0
Doxorubicin
YBL050W SEC17
Delta_r
ZShift = 1.06
Z lm Score = −2.57
NG = 0
DB = 0
SM = 0
Doxorubicin
YBL050W SEC17
Delta_AUC
ZShift = 1.68
Z lm Score = −0.98
NG = 0
DB = 0
SM = 0
Doxorubicin
YJL120W
Delta_L
ZShift = 6.12
Z lm Score = 4.03
NG = 0
DB = 0
SM = 0
Doxorubicin
YJL120W
Delta_K
ZShift = −21.2
Z lm Score = 4
NG = 0
DB = 0
SM = 0
Doxorubicin
YJL120W
Delta_r
ZShift = −5.82
Z lm Score = 3.75
NG = 0
DB = 0
SM = 0
Doxorubicin
YJL120W
Delta_AUC
ZShift = −12.66
Z lm Score = 0.61
NG = 0
DB = 0
SM = 0
Doxorubicin

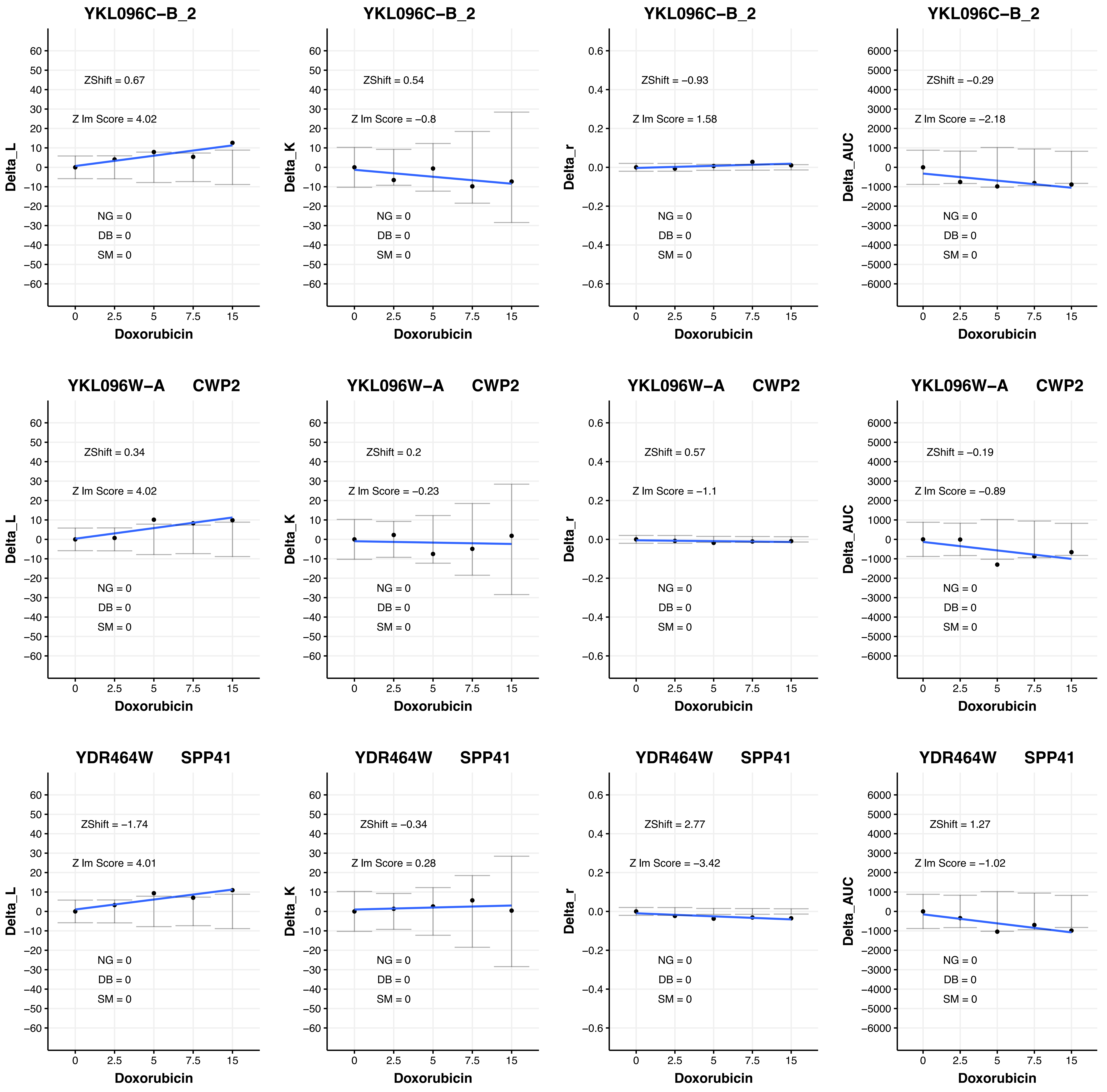

YKL096C−B_2
ZShift = 0.67
Z lm Score = 4.02
NG = 0
DB = 0
SM = 0
Delta_L
Doxorubicin
YKL096C−B_2
ZShift = 0.54
Z lm Score = −0.8
NG = 0
DB = 0
SM = 0
Delta_K
Doxorubicin
YKL096C−B_2
ZShift = −0.93
Z lm Score = 1.58
NG = 0
DB = 0
SM = 0
Delta_r
Doxorubicin
YKL096C−B_2
ZShift = −0.29
Z lm Score = −2.18
NG = 0
DB = 0
SM = 0
Delta_AUC
Doxorubicin
YKL096W−A CWP2
ZShift = 0.34
Z lm Score = 4.02
NG = 0
DB = 0
SM = 0
Delta_L
Doxorubicin
YKL096W−A CWP2
ZShift = 0.2
Z lm Score = −0.23
NG = 0
DB = 0
SM = 0
Delta_K
Doxorubicin
YKL096W−A CWP2
ZShift = 0.57
Z lm Score = −1.1
NG = 0
DB = 0
SM = 0
Delta_r
Doxorubicin
YKL096W−A CWP2
ZShift = −0.19
Z lm Score = −0.89
NG = 0
DB = 0
SM = 0
Delta_AUC
Doxorubicin
YDR464W SPP41
ZShift = −1.74
Z lm Score = 4.01
NG = 0
DB = 0
SM = 0
Delta_L
Doxorubicin
YDR464W SPP41
ZShift = −0.34
Z lm Score = 0.28
NG = 0
DB = 0
SM = 0
Delta_K
Doxorubicin
YDR464W SPP41
ZShift = 2.77
Z lm Score = −3.42
NG = 0
DB = 0
SM = 0
Delta_r
Doxorubicin
YDR464W SPP41
ZShift = 1.27
Z lm Score = −1.02
NG = 0
DB = 0
SM = 0
Delta_AUC
Doxorubicin

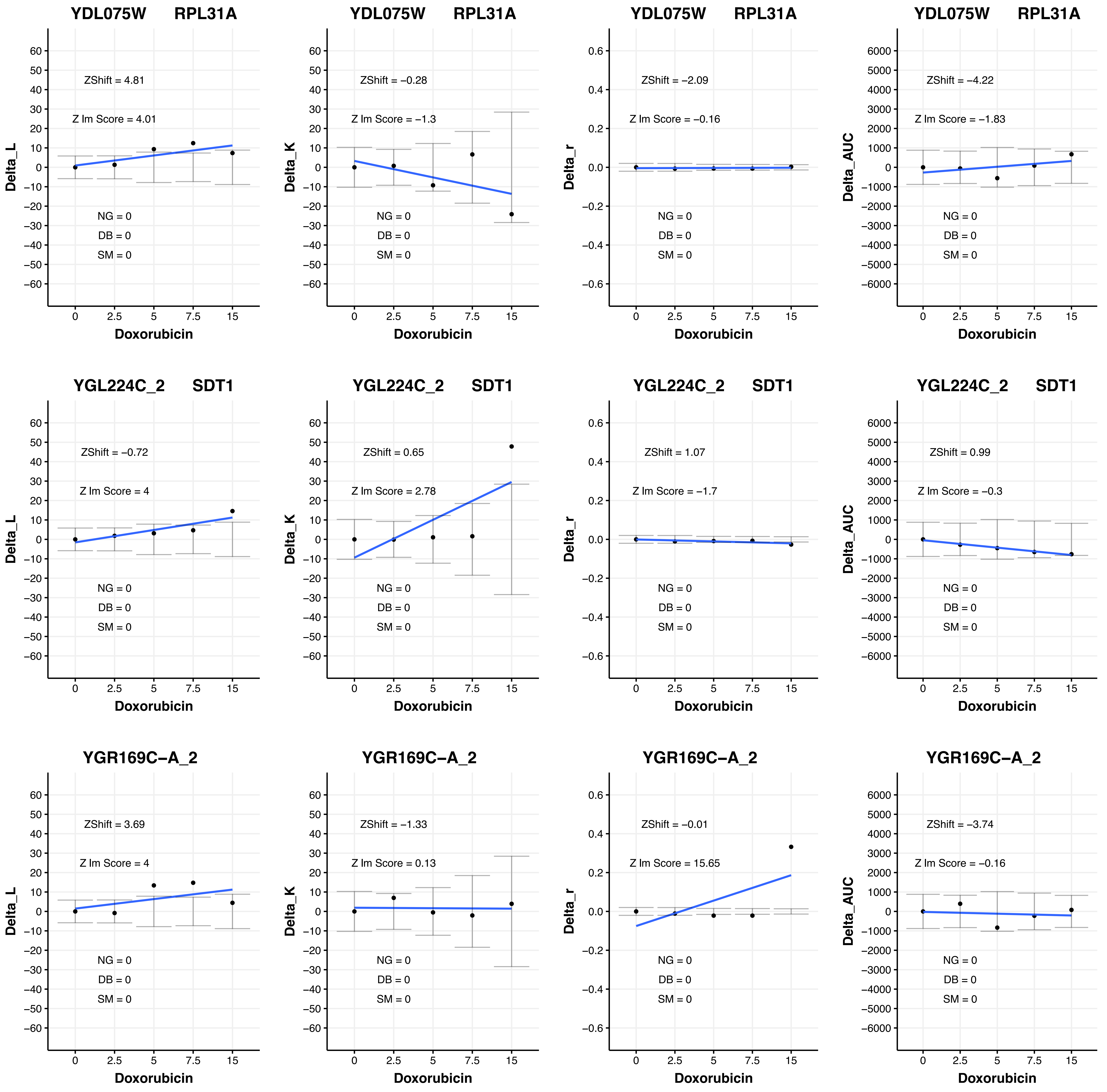

YDL075W RPL31A
ZShift = 4.81
Z lm Score = 4.01
NG = 0
DB = 0
SM = 0
Delta_L
Doxorubicin
YDL075W RPL31A
ZShift = −0.28
Z lm Score = −1.3
NG = 0
DB = 0
SM = 0
Delta_K
Doxorubicin
YDL075W RPL31A
ZShift = −2.09
Z lm Score = −0.16
NG = 0
DB = 0
SM = 0
Delta_r
Doxorubicin
YDL075W RPL31A
ZShift = −4.22
Z lm Score = −1.83
NG = 0
DB = 0
SM = 0
Delta_AUC
Doxorubicin
YGL224C_2 SDT1
ZShift = −0.72
Z lm Score = 4
NG = 0
DB = 0
SM = 0
Delta_L
Doxorubicin
YGL224C_2 SDT1
ZShift = 0.65
Z lm Score = 2.78
NG = 0
DB = 0
SM = 0
Delta_K
Doxorubicin
YGL224C_2 SDT1
ZShift = 1.07
Z lm Score = −1.7
NG = 0
DB = 0
SM = 0
Delta_r
Doxorubicin
YGL224C_2 SDT1
ZShift = 0.99
Z lm Score = −0.3
NG = 0
DB = 0
SM = 0
Delta_AUC
Doxorubicin
YGR169C−A_2
ZShift = 3.69
Z lm Score = 4
NG = 0
DB = 0
SM = 0
Delta_L
Doxorubicin
YGR169C−A_2
ZShift = −1.33
Z lm Score = 0.13
NG = 0
DB = 0
SM = 0
Delta_K
Doxorubicin
YGR169C−A_2
ZShift = −0.01
Z lm Score = 15.65
NG = 0
DB = 0
SM = 0
Delta_r
Doxorubicin
YGR169C−A_2
ZShift = −3.74
Z lm Score = −0.16
NG = 0
DB = 0
SM = 0
Delta_AUC
Doxorubicin

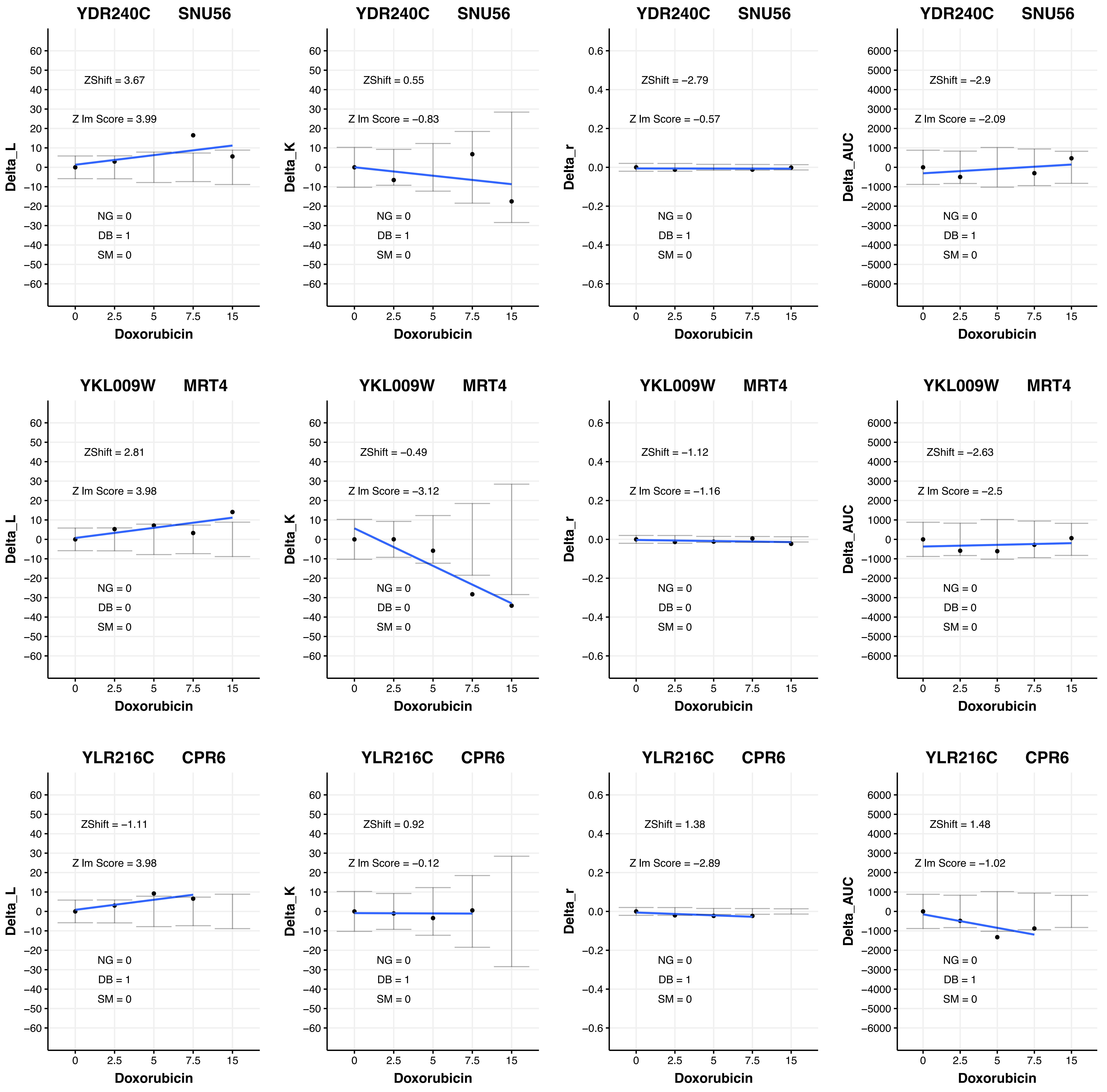
YDR240C SNU56
ZShift = 3.67
Z lm Score = 3.99
NG = 0
DB = 1
SM = 0
Delta_L
Doxorubicin
YDR240C SNU56
ZShift = 0.55
Z lm Score = −0.83
NG = 0
DB = 1
SM = 0
Delta_K
Doxorubicin
YDR240C SNU56
ZShift = −2.79
Z lm Score = −0.57
NG = 0
DB = 1
SM = 0
Delta_r
Doxorubicin
YDR240C SNU56
ZShift = −2.9
Z lm Score = −2.09
NG = 0
DB = 1
SM = 0
Delta_AUC
Doxorubicin
YKL009W MRT4
ZShift = 2.81
Z lm Score = 3.98
NG = 0
DB = 0
SM = 0
Delta_L
Doxorubicin
YKL009W MRT4
ZShift = −0.49
Z lm Score = −3.12
NG = 0
DB = 0
SM = 0
Delta_K
Doxorubicin
YKL009W MRT4
ZShift = −1.12
Z lm Score = −1.16
NG = 0
DB = 0
SM = 0
Delta_r
Doxorubicin
YKL009W MRT4
ZShift = −2.63
Z lm Score = −2.5
NG = 0
DB = 0
SM = 0
Delta_AUC
Doxorubicin
YLR216C CPR6
ZShift = −1.11
Z lm Score = 3.98
NG = 0
DB = 1
SM = 0
Delta_L
Doxorubicin
YLR216C CPR6
ZShift = 0.92
Z lm Score = −0.12
NG = 0
DB = 1
SM = 0
Delta_K
Doxorubicin
YLR216C CPR6
ZShift = 1.38
Z lm Score = −2.89
NG = 0
DB = 1
SM = 0
Delta_r
Doxorubicin
YLR216C CPR6
ZShift = 1.48
Z lm Score = −1.02
NG = 0
DB = 1
SM = 0
Delta_AUC
Doxorubicin

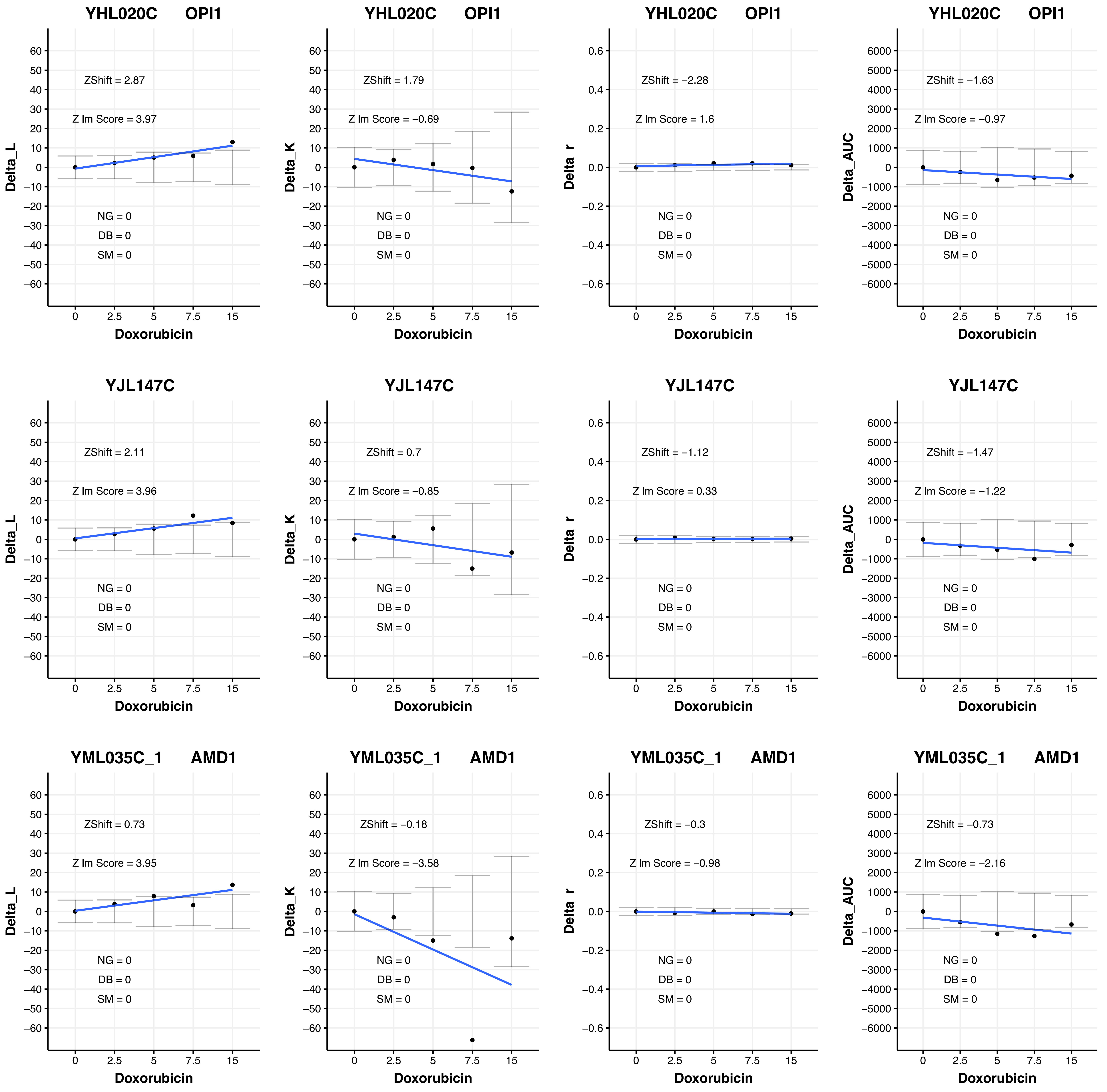
YHL020C OPI1
ZShift = 2.87
Z lm Score = 3.97
NG = 0
DB = 0
SM = 0
Delta_L
Doxorubicin
YHL020C OPI1
ZShift = 1.79
Z lm Score = −0.69
NG = 0
DB = 0
SM = 0
Delta_K
Doxorubicin
YHL020C OPI1
ZShift = −2.28
Z lm Score = 1.6
NG = 0
DB = 0
SM = 0
Delta_r
Doxorubicin
YHL020C OPI1
ZShift = −1.63
Z lm Score = −0.97
NG = 0
DB = 0
SM = 0
Delta_AUC
Doxorubicin
YJL147C
ZShift = 2.11
Z lm Score = 3.96
NG = 0
DB = 0
SM = 0
Delta_L
Doxorubicin
YJL147C
ZShift = 0.7
Z lm Score = −0.85
NG = 0
DB = 0
SM = 0
Delta_K
Doxorubicin
YJL147C
ZShift = −1.12
Z lm Score = 0.33
NG = 0
DB = 0
SM = 0
Delta_r
Doxorubicin
YJL147C
ZShift = −1.47
Z lm Score = −1.22
NG = 0
DB = 0
SM = 0
Delta_AUC
Doxorubicin
YML035C_1 AMD1
ZShift = 0.73
Z lm Score = 3.95
NG = 0
DB = 0
SM = 0
Delta_L
Doxorubicin
YML035C_1 AMD1
ZShift = −0.18
Z lm Score = −3.58
NG = 0
DB = 0
SM = 0
Delta_K
Doxorubicin
YML035C_1 AMD1
ZShift = −0.3
Z lm Score = −0.98
NG = 0
DB = 0
SM = 0
Delta_r
Doxorubicin
YML035C_1 AMD1
ZShift = −0.73
Z lm Score = −2.16
NG = 0
DB = 0
SM = 0
Delta_AUC
Doxorubicin

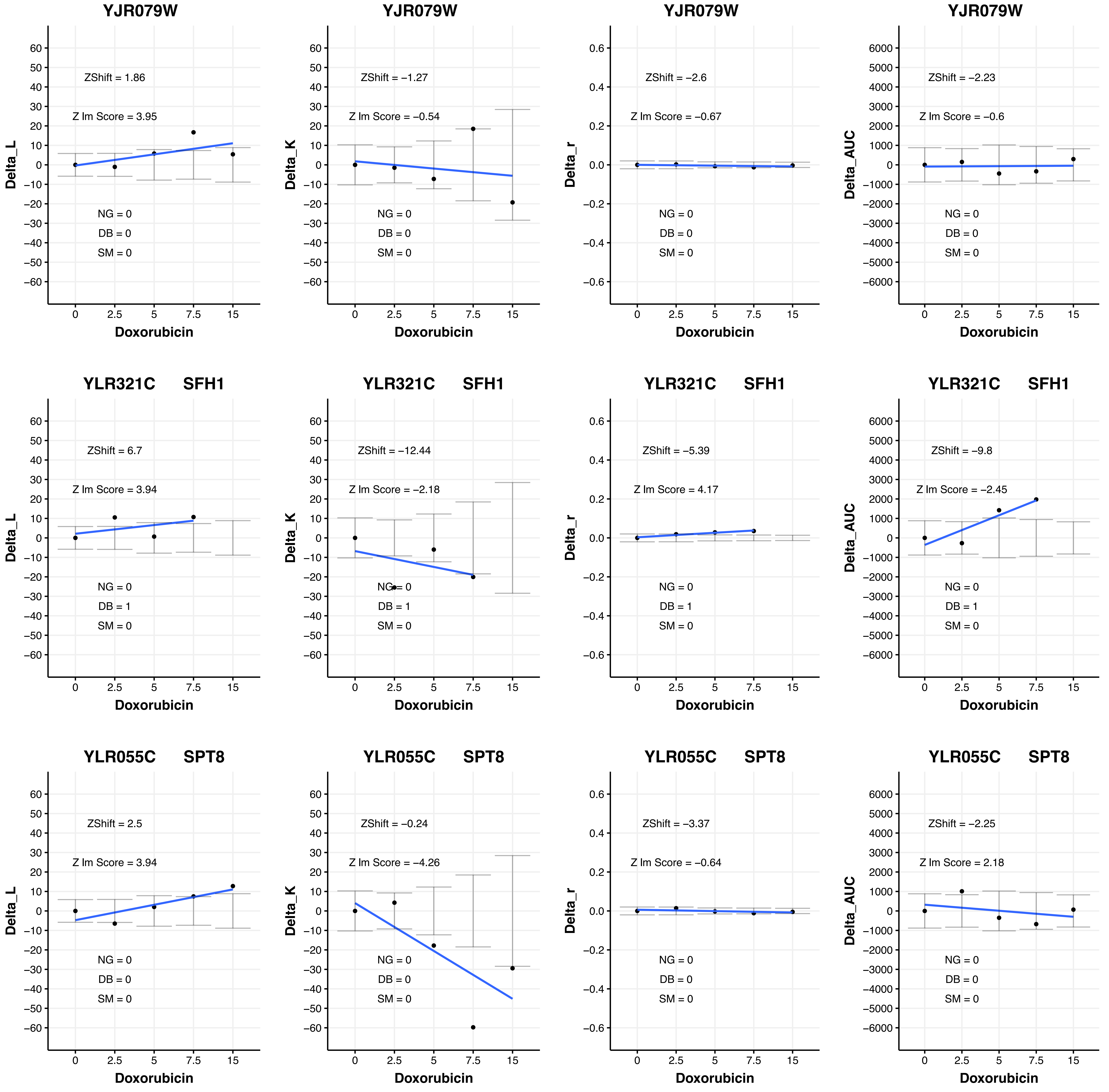

YJR079W
ZShift = 1.86
Z lm Score = 3.95
NG = 0
DB = 0
SM = 0
Delta_L
Doxorubicin
YJR079W
ZShift = −1.27
Z lm Score = −0.54
NG = 0
DB = 0
SM = 0
Delta_K
Doxorubicin
YJR079W
ZShift = −2.6
Z lm Score = −0.67
NG = 0
DB = 0
SM = 0
Delta_r
Doxorubicin
YJR079W
ZShift = −2.23
Z lm Score = −0.6
NG = 0
DB = 0
SM = 0
Delta_AUC
Doxorubicin
YLR321C SFH1
ZShift = 6.7
Z lm Score = 3.94
NG = 0
DB = 1
SM = 0
Delta_L
Doxorubicin
YLR321C SFH1
ZShift = −12.44
Z lm Score = −2.18
NG = 0
DB = 1
SM = 0
Delta_K
Doxorubicin
YLR321C SFH1
ZShift = −5.39
Z lm Score = 4.17
NG = 0
DB = 1
SM = 0
Delta_r
Doxorubicin
YLR321C SFH1
ZShift = −9.8
Z lm Score = −2.45
NG = 0
DB = 1
SM = 0
Delta_AUC
Doxorubicin
YLR055C SPT8
ZShift = 2.5
Z lm Score = 3.94
NG = 0
DB = 0
SM = 0
Delta_L
Doxorubicin
YLR055C SPT8
ZShift = −0.24
Z lm Score = −4.26
NG = 0
DB = 0
SM = 0
Delta_K
Doxorubicin
YLR055C SPT8
ZShift = −3.37
Z lm Score = −0.64
NG = 0
DB = 0
SM = 0
Delta_r
Doxorubicin
YLR055C SPT8
ZShift = −2.25
Z lm Score = 2.18
NG = 0
DB = 0
SM = 0
Delta_AUC
Doxorubicin

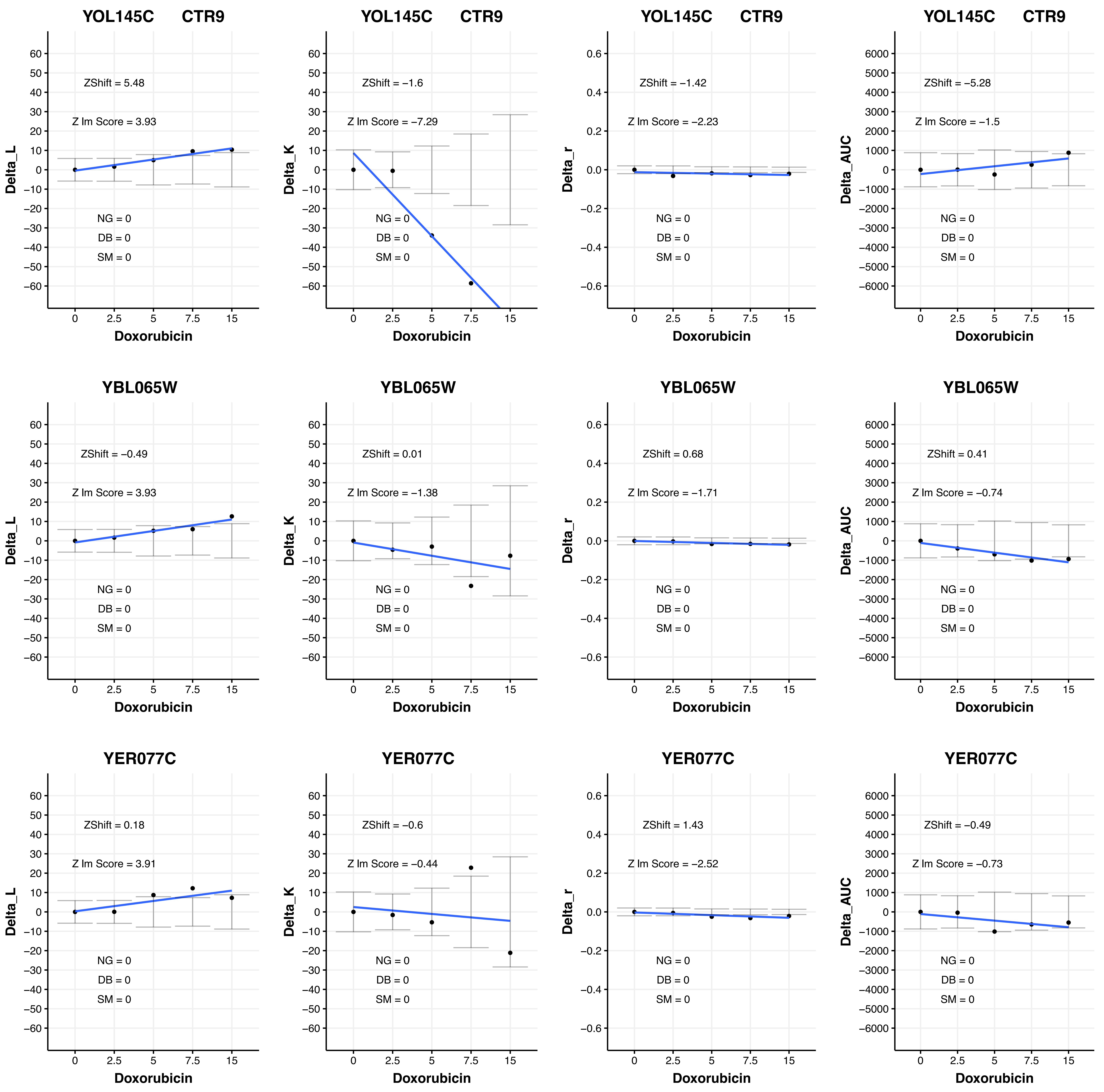
YOL145C CTR9
ZShift = 5.48
Z lm Score = 3.93
NG = 0
DB = 0
SM = 0
Delta_L
Doxorubicin
YOL145C CTR9
ZShift = −1.6
Z lm Score = −7.29
NG = 0
DB = 0
SM = 0
Delta_K
Doxorubicin
YOL145C CTR9
ZShift = −1.42
Z lm Score = −2.23
NG = 0
DB = 0
SM = 0
Delta_r
Doxorubicin
YOL145C CTR9
ZShift = −5.28
Z lm Score = −1.5
NG = 0
DB = 0
SM = 0
Delta_AUC
Doxorubicin
YBL065W
ZShift = −0.49
Z lm Score = 3.93
NG = 0
DB = 0
SM = 0
Delta_L
Doxorubicin
YBL065W
ZShift = 0.01
Z lm Score = −1.38
NG = 0
DB = 0
SM = 0
Delta_K
Doxorubicin
YBL065W
ZShift = 0.68
Z lm Score = −1.71
NG = 0
DB = 0
SM = 0
Delta_r
Doxorubicin
YBL065W
ZShift = 0.41
Z lm Score = −0.74
NG = 0
DB = 0
SM = 0
Delta_AUC
Doxorubicin
YER077C
ZShift = 0.18
Z lm Score = 3.91
NG = 0
DB = 0
SM = 0
Delta_L
Doxorubicin
YER077C
ZShift = −0.6
Z lm Score = −0.44
NG = 0
DB = 0
SM = 0
Delta_K
Doxorubicin
YER077C
ZShift = 1.43
Z lm Score = −2.52
NG = 0
DB = 0
SM = 0
Delta_r
Doxorubicin
YER077C
ZShift = −0.49
Z lm Score = −0.73
NG = 0
DB = 0
SM = 0
Delta_AUC
Doxorubicin

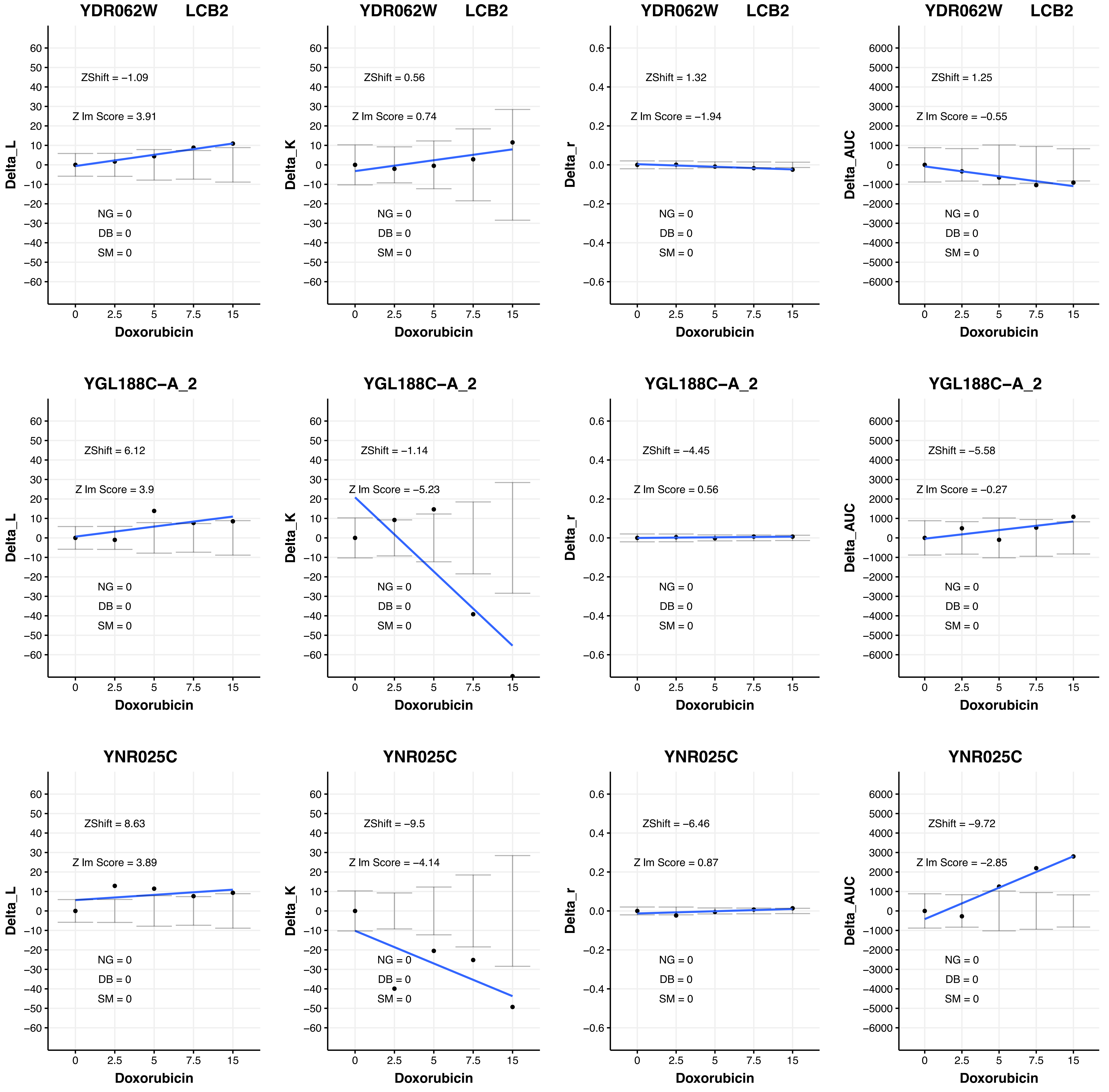
YDR062W LCB2
ZShift = −1.09
Z lm Score = 3.91
NG = 0
DB = 0
SM = 0
Delta_L
Doxorubicin
YDR062W LCB2
ZShift = 0.56
Z lm Score = 0.74
NG = 0
DB = 0
SM = 0
Delta_K
Doxorubicin
YDR062W LCB2
ZShift = 1.32
Z lm Score = −1.94
NG = 0
DB = 0
SM = 0
Delta_r
Doxorubicin
YDR062W LCB2
ZShift = 1.25
Z lm Score = −0.55
NG = 0
DB = 0
SM = 0
Delta_AUC
Doxorubicin
YGL188C−A_2
ZShift = 6.12
Z lm Score = 3.9
NG = 0
DB = 0
SM = 0
Delta_L
Doxorubicin
YGL188C−A_2
ZShift = −1.14
Z lm Score = −5.23
NG = 0
DB = 0
SM = 0
Delta_K
Doxorubicin
YGL188C−A_2
ZShift = −4.45
Z lm Score = 0.56
NG = 0
DB = 0
SM = 0
Delta_r
Doxorubicin
YGL188C−A_2
ZShift = −5.58
Z lm Score = −0.27
NG = 0
DB = 0
SM = 0
Delta_AUC
Doxorubicin
YNR025C
ZShift = 8.63
Z lm Score = 3.89
NG = 0
DB = 0
SM = 0
Delta_L
Doxorubicin
YNR025C
ZShift = −9.5
Z lm Score = −4.14
NG = 0
DB = 0
SM = 0
Delta_K
Doxorubicin
YNR025C
ZShift = −6.46
Z lm Score = 0.87
NG = 0
DB = 0
SM = 0
Delta_r
Doxorubicin
YNR025C
ZShift = −9.72
Z lm Score = −2.85
NG = 0
DB = 0
SM = 0
Delta_AUC
Doxorubicin

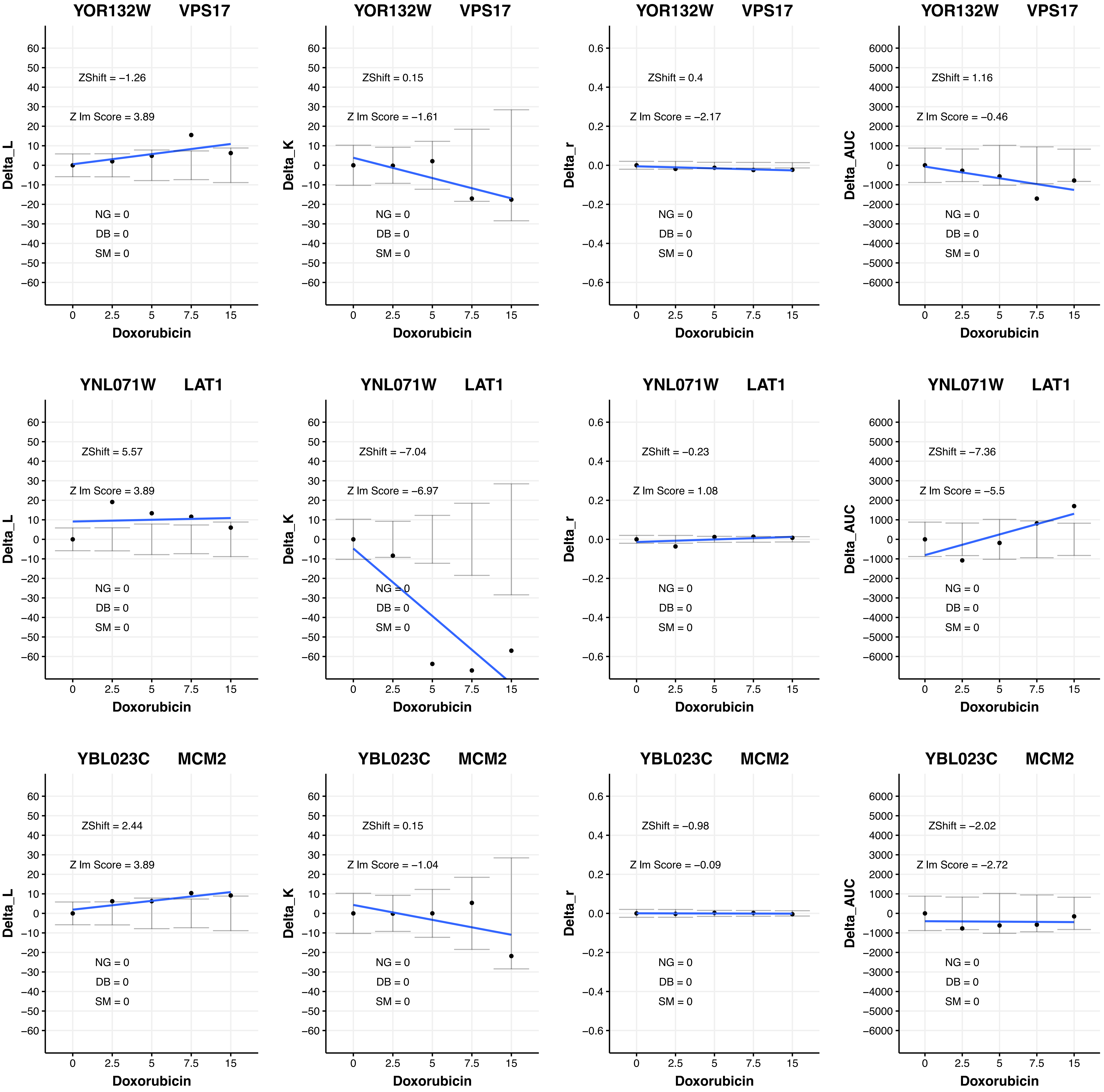

YOR132W VPS17
ZShift = −1.26
Z lm Score = 3.89
NG = 0
DB = 0
SM = 0
Delta_L
Doxorubicin
YOR132W VPS17
ZShift = 0.15
Z lm Score = −1.61
NG = 0
DB = 0
SM = 0
Delta_K
Doxorubicin
YOR132W VPS17
ZShift = 0.4
Z lm Score = −2.17
NG = 0
DB = 0
SM = 0
Delta_r
Doxorubicin
YOR132W VPS17
ZShift = 1.16
Z lm Score = −0.46
NG = 0
DB = 0
SM = 0
Delta_AUC
Doxorubicin
YNL071W LAT1
ZShift = 5.57
Z lm Score = 3.89
NG = 0
DB = 0
SM = 0
Delta_L
Doxorubicin
YNL071W LAT1
ZShift = −7.04
Z lm Score = −6.97
NG = 0
DB = 0
SM = 0
Delta_K
Doxorubicin
YNL071W LAT1
ZShift = −0.23
Z lm Score = 1.08
NG = 0
DB = 0
SM = 0
Delta_r
Doxorubicin
YNL071W LAT1
ZShift = −7.36
Z lm Score = −5.5
NG = 0
DB = 0
SM = 0
Delta_AUC
Doxorubicin
YBL023C MCM2
ZShift = 2.44
Z lm Score = 3.89
NG = 0
DB = 0
SM = 0
Delta_L
Doxorubicin
YBL023C MCM2
ZShift = 0.15
Z lm Score = −1.04
NG = 0
DB = 0
SM = 0
Delta_K
Doxorubicin
YBL023C MCM2
ZShift = −0.98
Z lm Score = −0.09
NG = 0
DB = 0
SM = 0
Delta_r
Doxorubicin
YBL023C MCM2
ZShift = −2.02
Z lm Score = −2.72
NG = 0
DB = 0
SM = 0
Delta_AUC
Doxorubicin

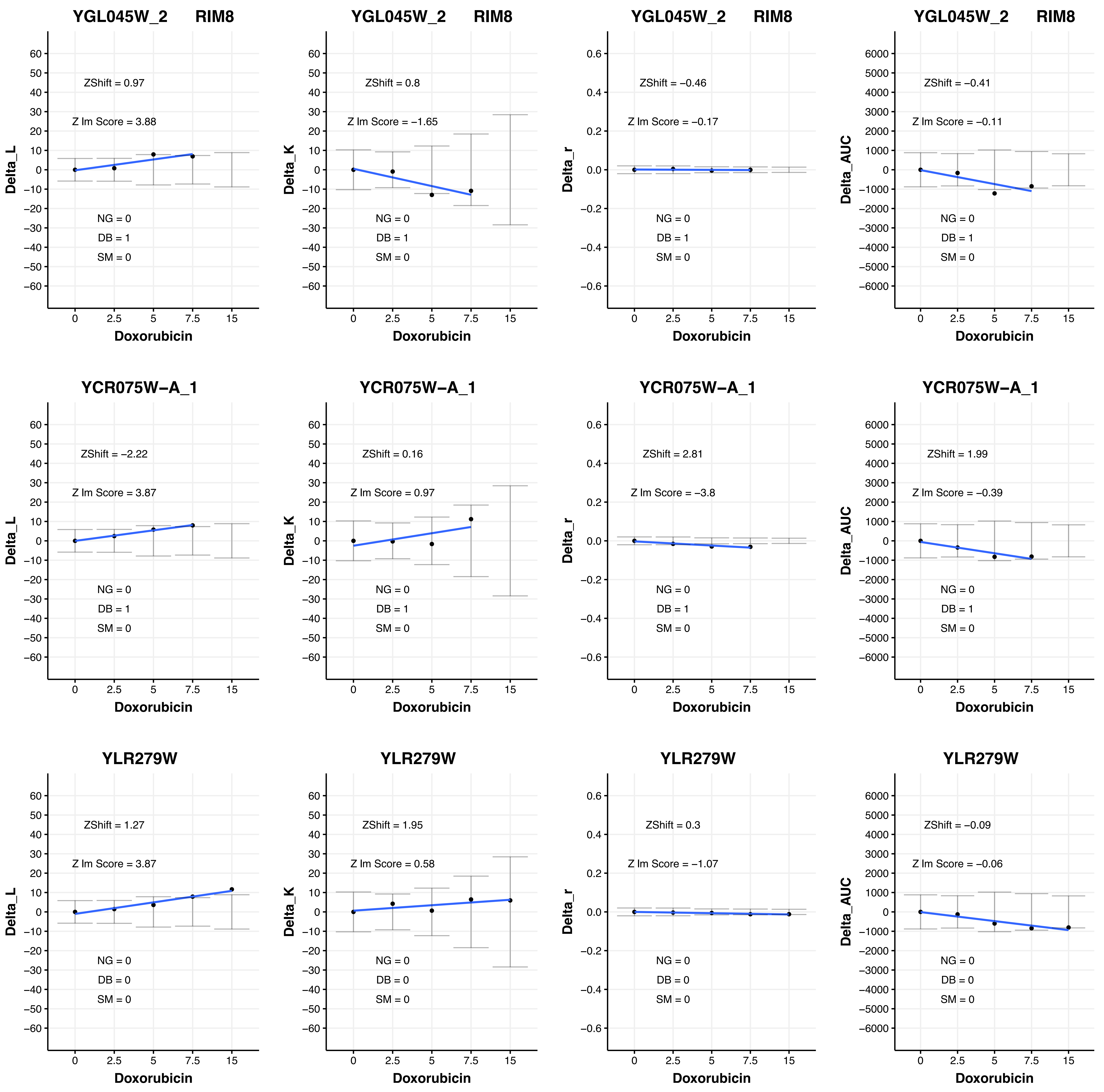
YGL045W_2 RIM8
ZShift = 0.97
Z lm Score = 3.88
NG = 0
DB = 1
SM = 0
Delta_L
Doxorubicin
YGL045W_2 RIM8
ZShift = 0.8
Z lm Score = −1.65
NG = 0
DB = 1
SM = 0
Delta_K
Doxorubicin
YGL045W_2 RIM8
ZShift = −0.46
Z lm Score = −0.17
NG = 0
DB = 1
SM = 0
Delta_r
Doxorubicin
YGL045W_2 RIM8
ZShift = −0.41
Z lm Score = −0.11
NG = 0
DB = 1
SM = 0
Delta_AUC
Doxorubicin
YCR075W−A_1
ZShift = −2.22
Z lm Score = 3.87
NG = 0
DB = 1
SM = 0
Delta_L
Doxorubicin
YCR075W−A_1
ZShift = 0.16
Z lm Score = 0.97
NG = 0
DB = 1
SM = 0
Delta_K
Doxorubicin
YCR075W−A_1
ZShift = 2.81
Z lm Score = −3.8
NG = 0
DB = 1
SM = 0
Delta_r
Doxorubicin
YCR075W−A_1
ZShift = 1.99
Z lm Score = −0.39
NG = 0
DB = 1
SM = 0
Delta_AUC
Doxorubicin
YLR279W
ZShift = 1.27
Z lm Score = 3.87
NG = 0
DB = 0
SM = 0
Delta_L
Doxorubicin
YLR279W
ZShift = 1.95
Z lm Score = 0.58
NG = 0
DB = 0
SM = 0
Delta_K
Doxorubicin
YLR279W
ZShift = 0.3
Z lm Score = −1.07
NG = 0
DB = 0
SM = 0
Delta_r
Doxorubicin
YLR279W
ZShift = −0.09
Z lm Score = −0.06
NG = 0
DB = 0
SM = 0
Delta_AUC
Doxorubicin

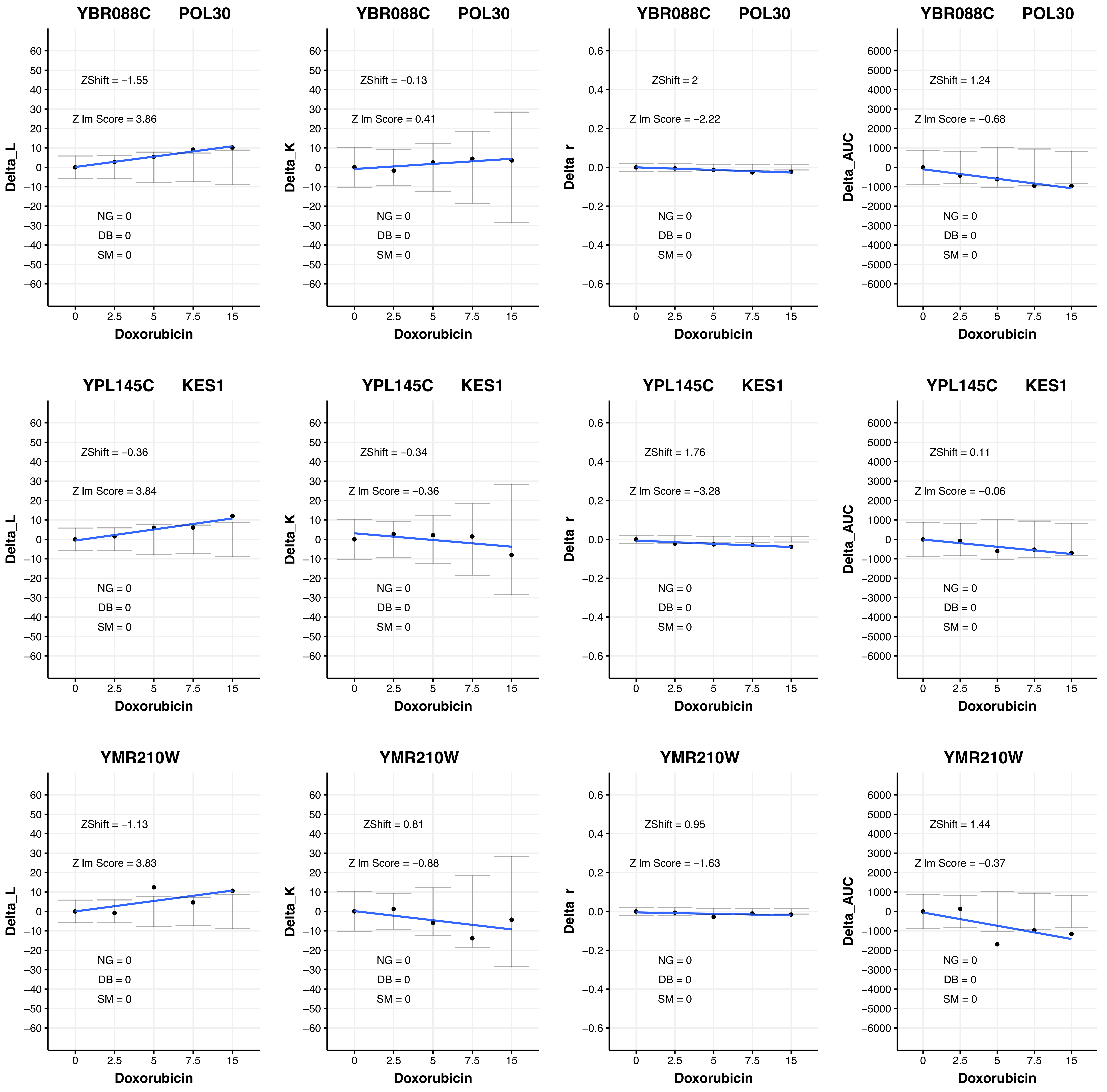

YBR088C POL30
ZShift = −1.55
Z lm Score = 3.86
NG = 0
DB = 0
SM = 0
Delta_L
Doxorubicin
YBR088C POL30
ZShift = −0.13
Z lm Score = 0.41
NG = 0
DB = 0
SM = 0
Delta_K
Doxorubicin
YBR088C POL30
ZShift = 2
Z lm Score = −2.22
NG = 0
DB = 0
SM = 0
Delta_r
Doxorubicin
YBR088C POL30
ZShift = 1.24
Z lm Score = −0.68
NG = 0
DB = 0
SM = 0
Delta_AUC
Doxorubicin
YPL145C KES1
ZShift = −0.36
Z lm Score = 3.84
NG = 0
DB = 0
SM = 0
Delta_L
Doxorubicin
YPL145C KES1
ZShift = −0.34
Z lm Score = −0.36
NG = 0
DB = 0
SM = 0
Delta_K
Doxorubicin
YPL145C KES1
ZShift = 1.76
Z lm Score = −3.28
NG = 0
DB = 0
SM = 0
Delta_r
Doxorubicin
YPL145C KES1
ZShift = 0.11
Z lm Score = −0.06
NG = 0
DB = 0
SM = 0
Delta_AUC
Doxorubicin
YMR210W
ZShift = −1.13
Z lm Score = 3.83
NG = 0
DB = 0
SM = 0
Delta_L
Doxorubicin
YMR210W
ZShift = 0.81
Z lm Score = −0.88
NG = 0
DB = 0
SM = 0
Delta_K
Doxorubicin
YMR210W
ZShift = 0.95
Z lm Score = −1.63
NG = 0
DB = 0
SM = 0
Delta_r
Doxorubicin
YMR210W
ZShift = 1.44
Z lm Score = −0.37
NG = 0
DB = 0
SM = 0
Delta_AUC
Doxorubicin

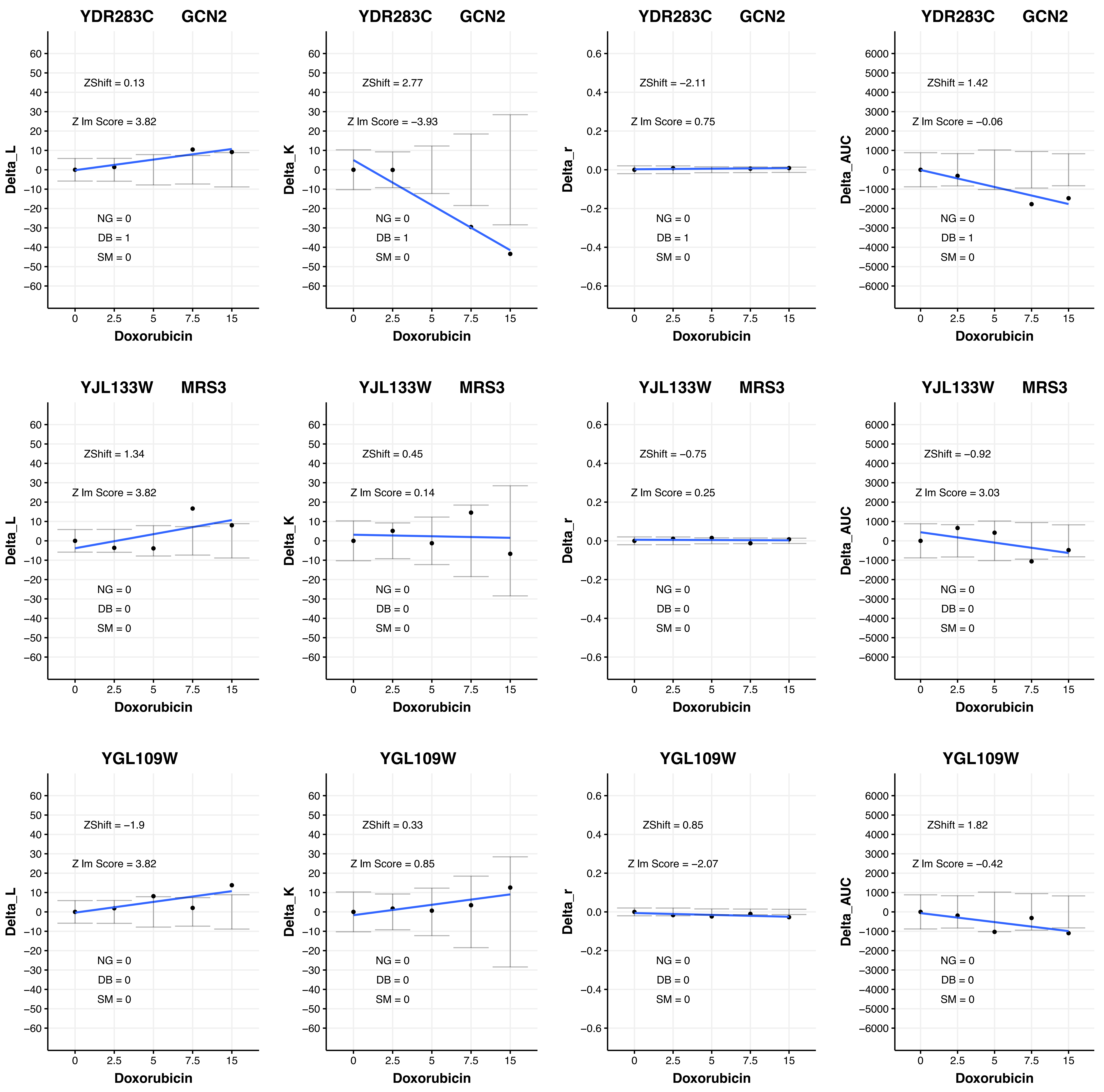
YDR283C GCN2
ZShift = 0.13
Z lm Score = 3.82
NG = 0
DB = 1
SM = 0
Delta_L
Doxorubicin
YDR283C GCN2
ZShift = 2.77
Z lm Score = −3.93
NG = 0
DB = 1
SM = 0
Delta_K
Doxorubicin
YDR283C GCN2
ZShift = −2.11
Z lm Score = 0.75
NG = 0
DB = 1
SM = 0
Delta_r
Doxorubicin
YDR283C GCN2
ZShift = 1.42
Z lm Score = −0.06
NG = 0
DB = 1
SM = 0
Delta_AUC
Doxorubicin
YJL133W MRS3
ZShift = 1.34
Z lm Score = 3.82
NG = 0
DB = 0
SM = 0
Delta_L
Doxorubicin
YJL133W MRS3
ZShift = 0.45
Z lm Score = 0.14
NG = 0
DB = 0
SM = 0
Delta_K
Doxorubicin
YJL133W MRS3
ZShift = −0.75
Z lm Score = 0.25
NG = 0
DB = 0
SM = 0
Delta_r
Doxorubicin
YJL133W MRS3
ZShift = −0.92
Z lm Score = 3.03
NG = 0
DB = 0
SM = 0
Delta_AUC
Doxorubicin
YGL109W
ZShift = −1.9
Z lm Score = 3.82
NG = 0
DB = 0
SM = 0
Delta_L
Doxorubicin
YGL109W
ZShift = 0.33
Z lm Score = 0.85
NG = 0
DB = 0
SM = 0
Delta_K
Doxorubicin
YGL109W
ZShift = 0.85
Z lm Score = −2.07
NG = 0
DB = 0
SM = 0
Delta_r
Doxorubicin
YGL109W
ZShift = 1.82
Z lm Score = −0.42
NG = 0
DB = 0
SM = 0
Delta_AUC
Doxorubicin

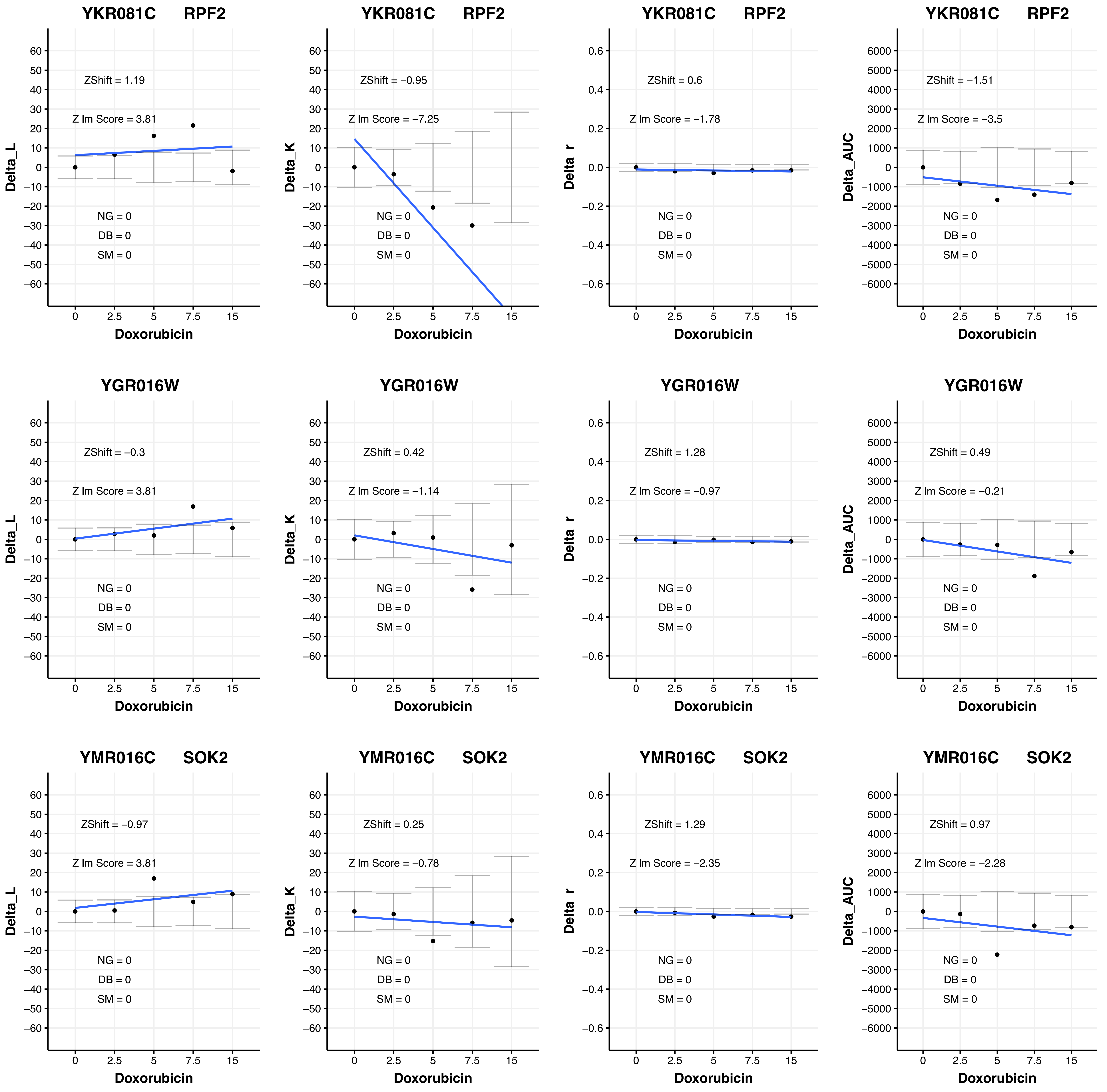
YKR081C RPF2
ZShift = 1.19
Z lm Score = 3.81
NG = 0
DB = 0
SM = 0
Delta_L
Doxorubicin
YKR081C RPF2
ZShift = −0.95
Z lm Score = −7.25
NG = 0
DB = 0
SM = 0
Delta_K
Doxorubicin
YKR081C RPF2
ZShift = 0.6
Z lm Score = −1.78
NG = 0
DB = 0
SM = 0
Delta_r
Doxorubicin
YKR081C RPF2
ZShift = −1.51
Z lm Score = −3.5
NG = 0
DB = 0
SM = 0
Delta_AUC
Doxorubicin
YGR016W
ZShift = −0.3
Z lm Score = 3.81
NG = 0
DB = 0
SM = 0
Delta_L
Doxorubicin
YGR016W
ZShift = 0.42
Z lm Score = −1.14
NG = 0
DB = 0
SM = 0
Delta_K
Doxorubicin
YGR016W
ZShift = 1.28
Z lm Score = −0.97
NG = 0
DB = 0
SM = 0
Delta_r
Doxorubicin
YGR016W
ZShift = 0.49
Z lm Score = −0.21
NG = 0
DB = 0
SM = 0
Delta_AUC
Doxorubicin
YMR016C SOK2
ZShift = −0.97
Z lm Score = 3.81
NG = 0
DB = 0
SM = 0
Delta_L
Doxorubicin
YMR016C SOK2
ZShift = 0.25
Z lm Score = −0.78
NG = 0
DB = 0
SM = 0
Delta_K
Doxorubicin
YMR016C SOK2
ZShift = 1.29
Z lm Score = −2.35
NG = 0
DB = 0
SM = 0
Delta_r
Doxorubicin
YMR016C SOK2
ZShift = 0.97
Z lm Score = −2.28
NG = 0
DB = 0
SM = 0
Delta_AUC
Doxorubicin

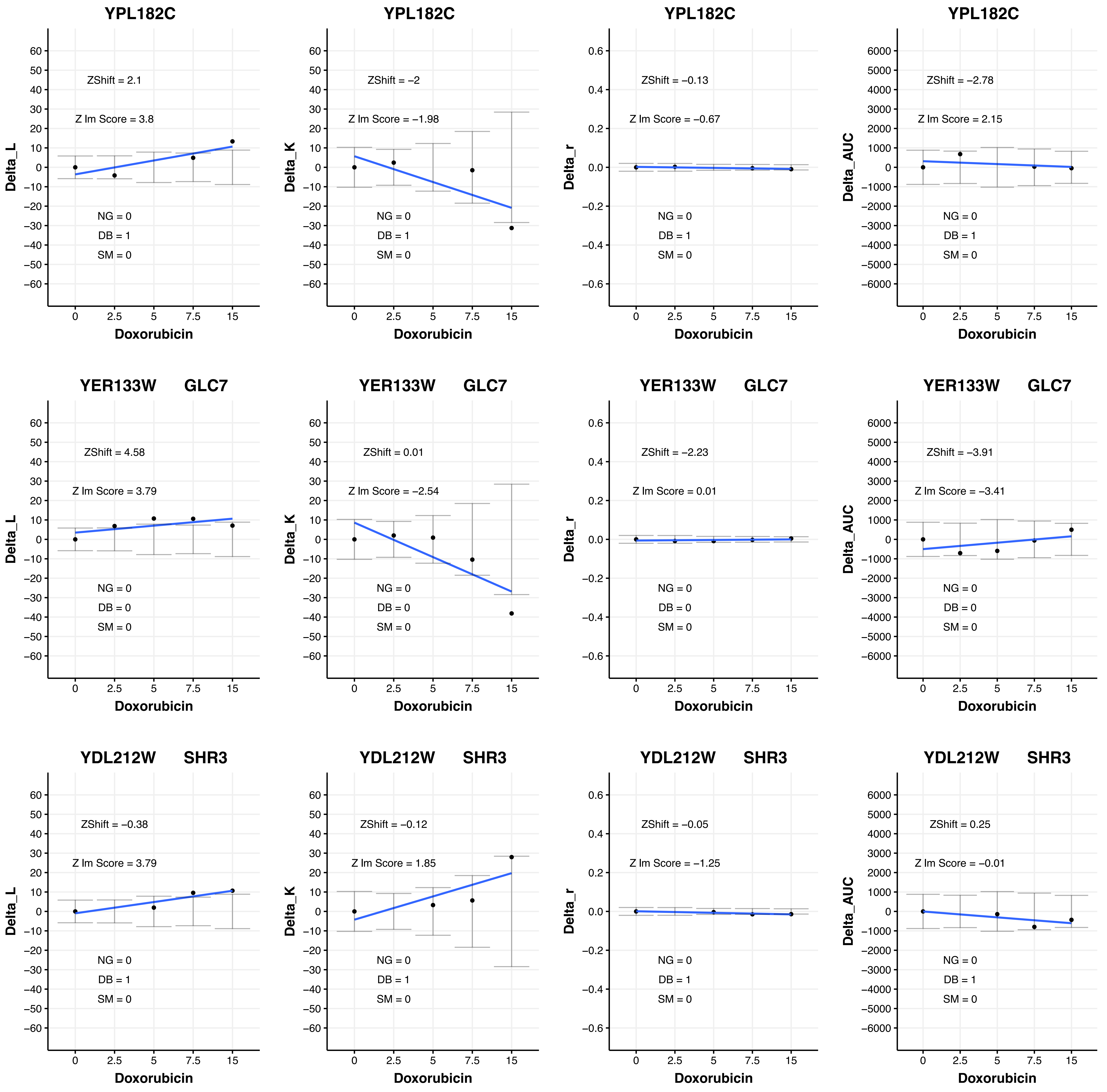
YPL182C
ZShift = 2.1
Z lm Score = 3.8
NG = 0
DB = 1
SM = 0
Delta_L
Doxorubicin
YPL182C
ZShift = −2
Z lm Score = −1.98
NG = 0
DB = 1
SM = 0
Delta_K
Doxorubicin
YPL182C
ZShift = −0.13
Z lm Score = −0.67
NG = 0
DB = 1
SM = 0
Delta_r
Doxorubicin
YPL182C
ZShift = −2.78
Z lm Score = 2.15
NG = 0
DB = 1
SM = 0
Delta_AUC
Doxorubicin
YER133W GLC7
ZShift = 4.58
Z lm Score = 3.79
NG = 0
DB = 0
SM = 0
Delta_L
Doxorubicin
YER133W GLC7
ZShift = 0.01
Z lm Score = −2.54
NG = 0
DB = 0
SM = 0
Delta_K
Doxorubicin
YER133W GLC7
ZShift = −2.23
Z lm Score = 0.01
NG = 0
DB = 0
SM = 0
Delta_r
Doxorubicin
YER133W GLC7
ZShift = −3.91
Z lm Score = −3.41
NG = 0
DB = 0
SM = 0
Delta_AUC
Doxorubicin
YDL212W SHR3
ZShift = −0.38
Z lm Score = 3.79
NG = 0
DB = 1
SM = 0
Delta_L
Doxorubicin
YDL212W SHR3
ZShift = −0.12
Z lm Score = 1.85
NG = 0
DB = 1
SM = 0
Delta_K
Doxorubicin
YDL212W SHR3
ZShift = −0.05
Z lm Score = −1.25
NG = 0
DB = 1
SM = 0
Delta_r
Doxorubicin
YDL212W SHR3
ZShift = 0.25
Z lm Score = −0.01
NG = 0
DB = 1
SM = 0
Delta_AUC
Doxorubicin

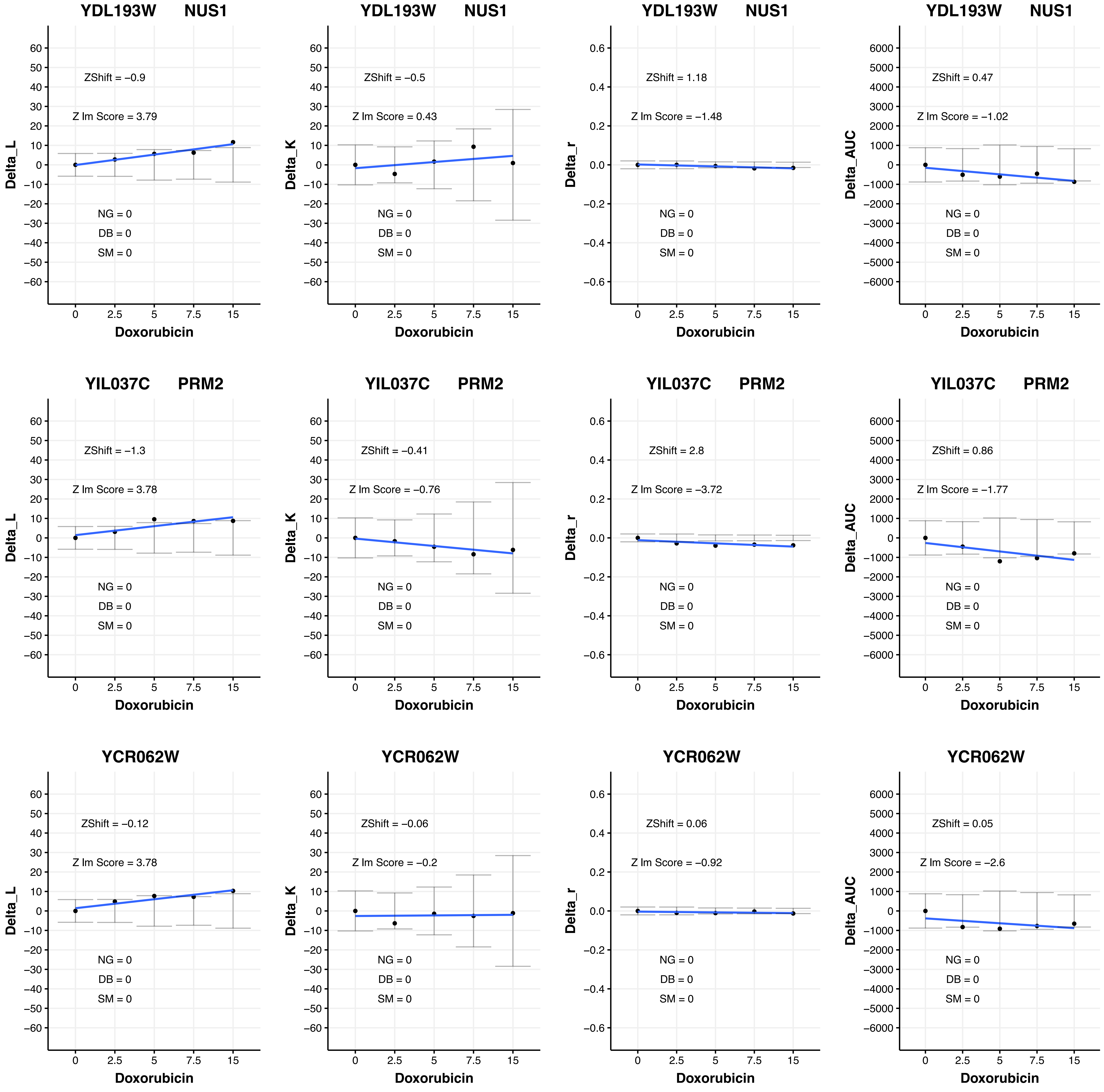
YDL193W NUS1
ZShift = −0.9
Z lm Score = 3.79
NG = 0
DB = 0
SM = 0
Delta_L
Doxorubicin
YDL193W NUS1
ZShift = −0.5
Z lm Score = 0.43
NG = 0
DB = 0
SM = 0
Delta_K
Doxorubicin
YDL193W NUS1
ZShift = 1.18
Z lm Score = −1.48
NG = 0
DB = 0
SM = 0
Delta_r
Doxorubicin
YDL193W NUS1
ZShift = 0.47
Z lm Score = −1.02
NG = 0
DB = 0
SM = 0
Delta_AUC
Doxorubicin
YIL037C PRM2
ZShift = −1.3
Z lm Score = 3.78
NG = 0
DB = 0
SM = 0
Delta_L
Doxorubicin
YIL037C PRM2
ZShift = −0.41
Z lm Score = −0.76
NG = 0
DB = 0
SM = 0
Delta_K
Doxorubicin
YIL037C PRM2
ZShift = 2.8
Z lm Score = −3.72
NG = 0
DB = 0
SM = 0
Delta_r
Doxorubicin
YIL037C PRM2
ZShift = 0.86
Z lm Score = −1.77
NG = 0
DB = 0
SM = 0
Delta_AUC
Doxorubicin
YCR062W
ZShift = −0.12
Z lm Score = 3.78
NG = 0
DB = 0
SM = 0
Delta_L
Doxorubicin
YCR062W
ZShift = −0.06
Z lm Score = −0.2
NG = 0
DB = 0
SM = 0
Delta_K
Doxorubicin
YCR062W
ZShift = 0.06
Z lm Score = −0.92
NG = 0
DB = 0
SM = 0
Delta_r
Doxorubicin
YCR062W
ZShift = 0.05
Z lm Score = −2.6
NG = 0
DB = 0
SM = 0
Delta_AUC
Doxorubicin

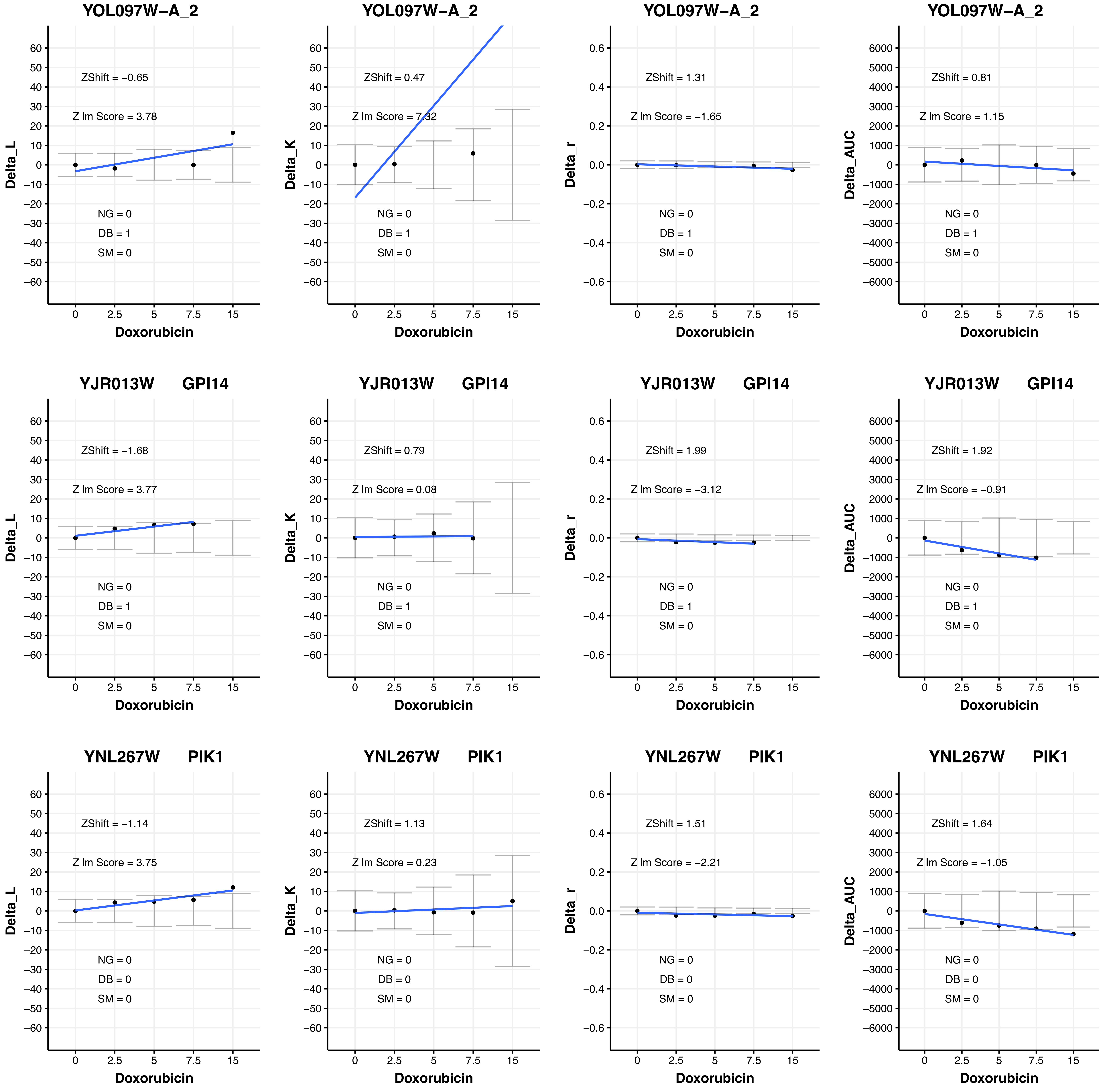
YOL097W-A_2
ZShift = −0.65
Z lm Score = 3.78
NG = 0
DB = 1
SM = 0
Delta_L
Doxorubicin
YOL097W-A_2
ZShift = 0.47
Z lm Score = 7.32
NG = 0
DB = 1
SM = 0
Delta_K
Doxorubicin
YOL097W-A_2
ZShift = 1.31
Z lm Score = −1.65
NG = 0
DB = 1
SM = 0
Delta_r
Doxorubicin
YOL097W-A_2
ZShift = 0.81
Z lm Score = 1.15
NG = 0
DB = 1
SM = 0
Delta_AUC
Doxorubicin
YJR013W GPI14
ZShift = −1.68
Z lm Score = 3.77
NG = 0
DB = 1
SM = 0
Delta_L
Doxorubicin
YJR013W GPI14
ZShift = 0.79
Z lm Score = 0.08
NG = 0
DB = 1
SM = 0
Delta_K
Doxorubicin
YJR013W GPI14
ZShift = 1.99
Z lm Score = −3.12
NG = 0
DB = 1
SM = 0
Delta_r
Doxorubicin
YJR013W GPI14
ZShift = 1.92
Z lm Score = −0.91
NG = 0
DB = 1
SM = 0
Delta_AUC
Doxorubicin
YNL267W PIK1
ZShift = −1.14
Z lm Score = 3.75
NG = 0
DB = 0
SM = 0
Delta_L
Doxorubicin
YNL267W PIK1
ZShift = 1.13
Z lm Score = 0.23
NG = 0
DB = 0
SM = 0
Delta_K
Doxorubicin
YNL267W PIK1
ZShift = 1.51
Z lm Score = −2.21
NG = 0
DB = 0
SM = 0
Delta_r
Doxorubicin
YNL267W PIK1
ZShift = 1.64
Z lm Score = −1.05
NG = 0
DB = 0
SM = 0
Delta_AUC
Doxorubicin

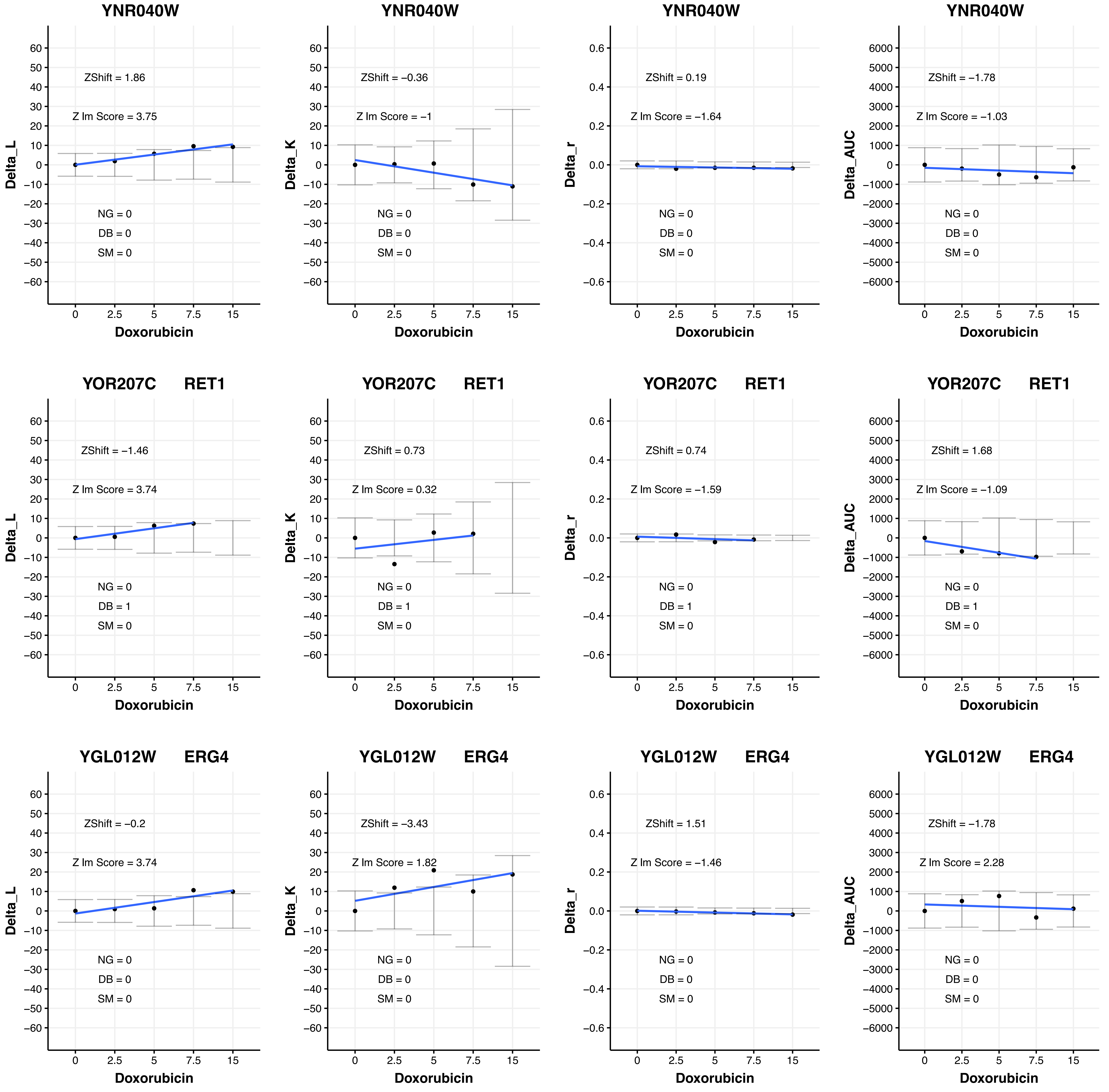
YNR040W
ZShift = 1.86
Z lm Score = 3.75
NG = 0
DB = 0
SM = 0
Delta_L
Doxorubicin
YNR040W
ZShift = −0.36
Z lm Score = −1
NG = 0
DB = 0
SM = 0
Delta_K
Doxorubicin
YNR040W
ZShift = 0.19
Z lm Score = −1.64
NG = 0
DB = 0
SM = 0
Delta_r
Doxorubicin
YNR040W
ZShift = −1.78
Z lm Score = −1.03
NG = 0
DB = 0
SM = 0
Delta_AUC
Doxorubicin
YOR207C RET1
ZShift = −1.46
Z lm Score = 3.74
NG = 0
DB = 1
SM = 0
Delta_L
Doxorubicin
YOR207C RET1
ZShift = 0.73
Z lm Score = 0.32
NG = 0
DB = 1
SM = 0
Delta_K
Doxorubicin
YOR207C RET1
ZShift = 0.74
Z lm Score = −1.59
NG = 0
DB = 1
SM = 0
Delta_r
Doxorubicin
YOR207C RET1
ZShift = 1.68
Z lm Score = −1.09
NG = 0
DB = 1
SM = 0
Delta_AUC
Doxorubicin
YGL012W ERG4
ZShift = −0.2
Z lm Score = 3.74
NG = 0
DB = 0
SM = 0
Delta_L
Doxorubicin
YGL012W ERG4
ZShift = −3.43
Z lm Score = 1.82
NG = 0
DB = 0
SM = 0
Delta_K
Doxorubicin
YGL012W ERG4
ZShift = 1.51
Z lm Score = −1.46
NG = 0
DB = 0
SM = 0
Delta_r
Doxorubicin
YGL012W ERG4
ZShift = −1.78
Z lm Score = 2.28
NG = 0
DB = 0
SM = 0
Delta_AUC
Doxorubicin

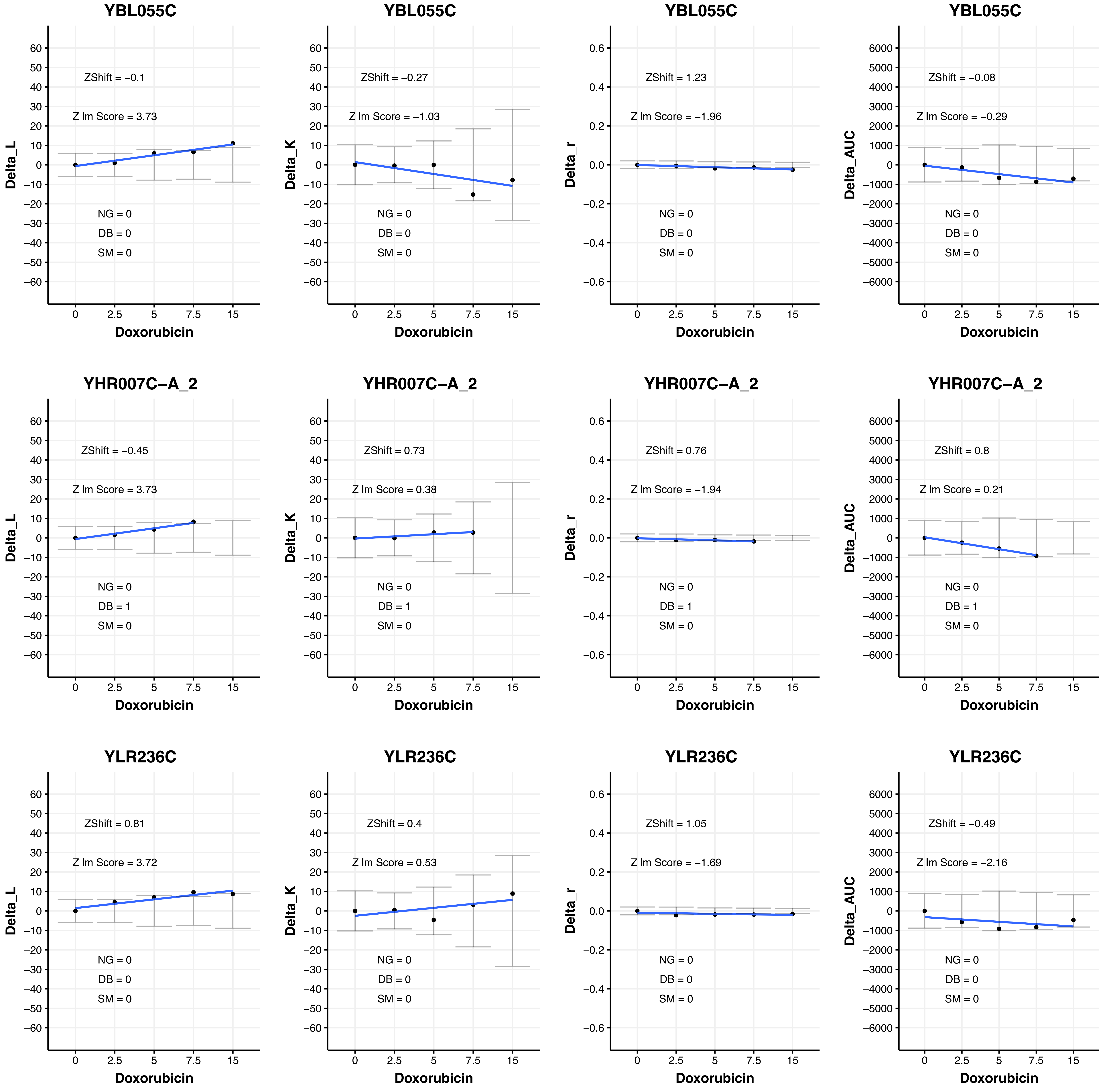
YBL055C
ZShift = −0.1
Z lm Score = 3.73
NG = 0
DB = 0
SM = 0
Delta_L
Doxorubicin
YBL055C
ZShift = −0.27
Z lm Score = −1.03
NG = 0
DB = 0
SM = 0
Delta_K
Doxorubicin
YBL055C
ZShift = 1.23
Z lm Score = −1.96
NG = 0
DB = 0
SM = 0
Delta_r
Doxorubicin
YBL055C
ZShift = −0.08
Z lm Score = −0.29
NG = 0
DB = 0
SM = 0
Delta_AUC
Doxorubicin
YHR007C−A_2
ZShift = −0.45
Z lm Score = 3.73
NG = 0
DB = 1
SM = 0
Delta_L
Doxorubicin
YHR007C−A_2
ZShift = 0.73
Z lm Score = 0.38
NG = 0
DB = 1
SM = 0
Delta_K
Doxorubicin
YHR007C−A_2
ZShift = 0.76
Z lm Score = −1.94
NG = 0
DB = 1
SM = 0
Delta_r
Doxorubicin
YHR007C−A_2
ZShift = 0.8
Z lm Score = 0.21
NG = 0
DB = 1
SM = 0
Delta_AUC
Doxorubicin
YLR236C
ZShift = 0.81
Z lm Score = 3.72
NG = 0
DB = 0
SM = 0
Delta_L
Doxorubicin
YLR236C
ZShift = 0.4
Z lm Score = 0.53
NG = 0
DB = 0
SM = 0
Delta_K
Doxorubicin
YLR236C
ZShift = 1.05
Z lm Score = −1.69
NG = 0
DB = 0
SM = 0
Delta_r
Doxorubicin
YLR236C
ZShift = −0.49
Z lm Score = −2.16
NG = 0
DB = 0
SM = 0
Delta_AUC
Doxorubicin

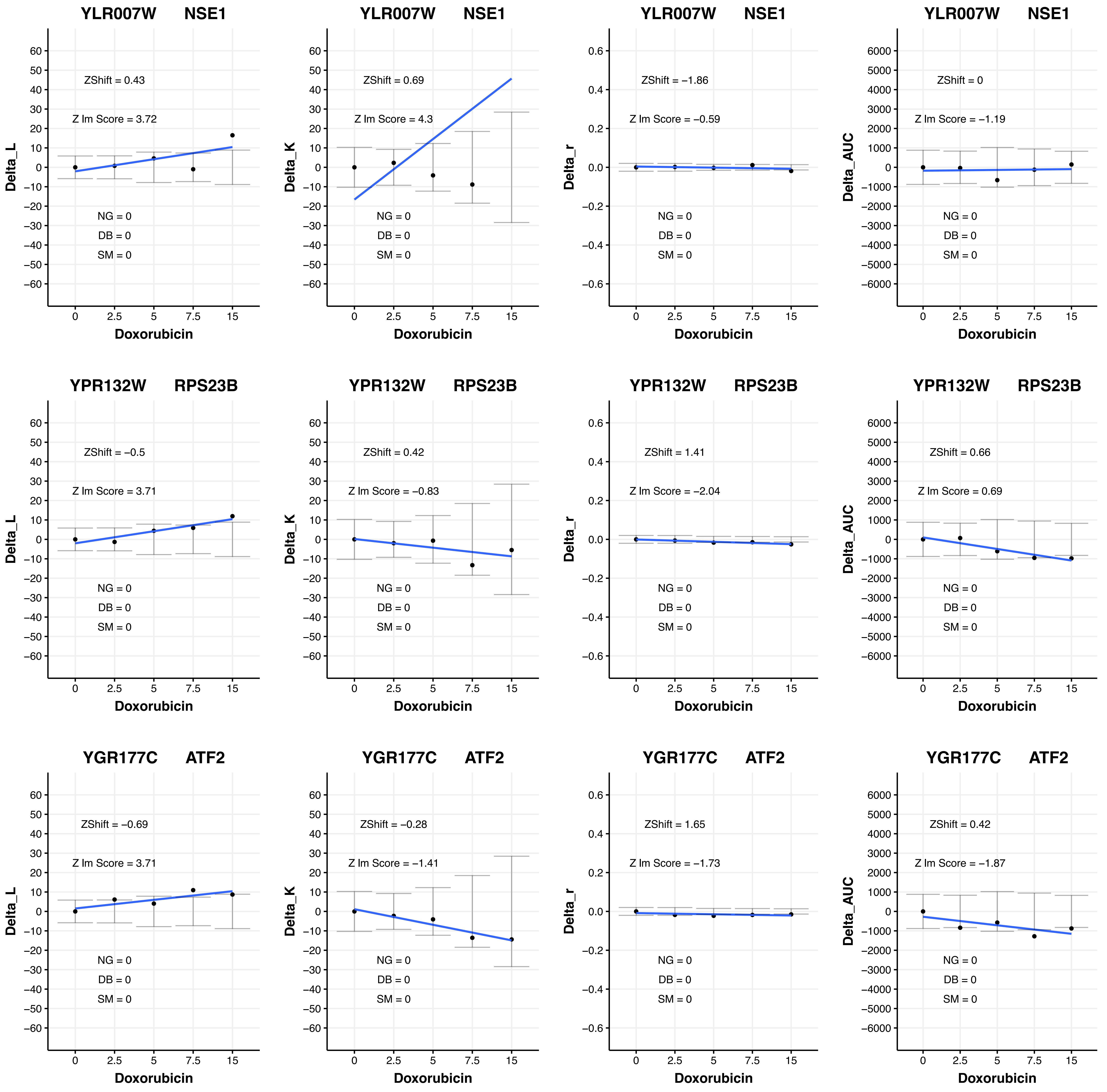

YLR007W NSE1
ZShift = 0.43
Z lm Score = 3.72
NG = 0
DB = 0
SM = 0
Delta_L
Doxorubicin
YLR007W NSE1
ZShift = 0.69
Z lm Score = 4.3
NG = 0
DB = 0
SM = 0
Delta_K
Doxorubicin
YLR007W NSE1
ZShift = −1.86
Z lm Score = −0.59
NG = 0
DB = 0
SM = 0
Delta_r
Doxorubicin
YLR007W NSE1
ZShift = 0
Z lm Score = −1.19
NG = 0
DB = 0
SM = 0
Delta_AUC
Doxorubicin
YPR132W RPS23B
ZShift = −0.5
Z lm Score = 3.71
NG = 0
DB = 0
SM = 0
Delta_L
Doxorubicin
YPR132W RPS23B
ZShift = 0.42
Z lm Score = −0.83
NG = 0
DB = 0
SM = 0
Delta_K
Doxorubicin
YPR132W RPS23B
ZShift = 1.41
Z lm Score = −2.04
NG = 0
DB = 0
SM = 0
Delta_r
Doxorubicin
YPR132W RPS23B
ZShift = 0.66
Z lm Score = 0.69
NG = 0
DB = 0
SM = 0
Delta_AUC
Doxorubicin
YGR177C ATF2
ZShift = −0.69
Z lm Score = 3.71
NG = 0
DB = 0
SM = 0
Delta_L
Doxorubicin
YGR177C ATF2
ZShift = −0.28
Z lm Score = −1.41
NG = 0
DB = 0
SM = 0
Delta_K
Doxorubicin
YGR177C ATF2
ZShift = 1.65
Z lm Score = −1.73
NG = 0
DB = 0
SM = 0
Delta_r
Doxorubicin
YGR177C ATF2
ZShift = 0.42
Z lm Score = −1.87
NG = 0
DB = 0
SM = 0
Delta_AUC
Doxorubicin

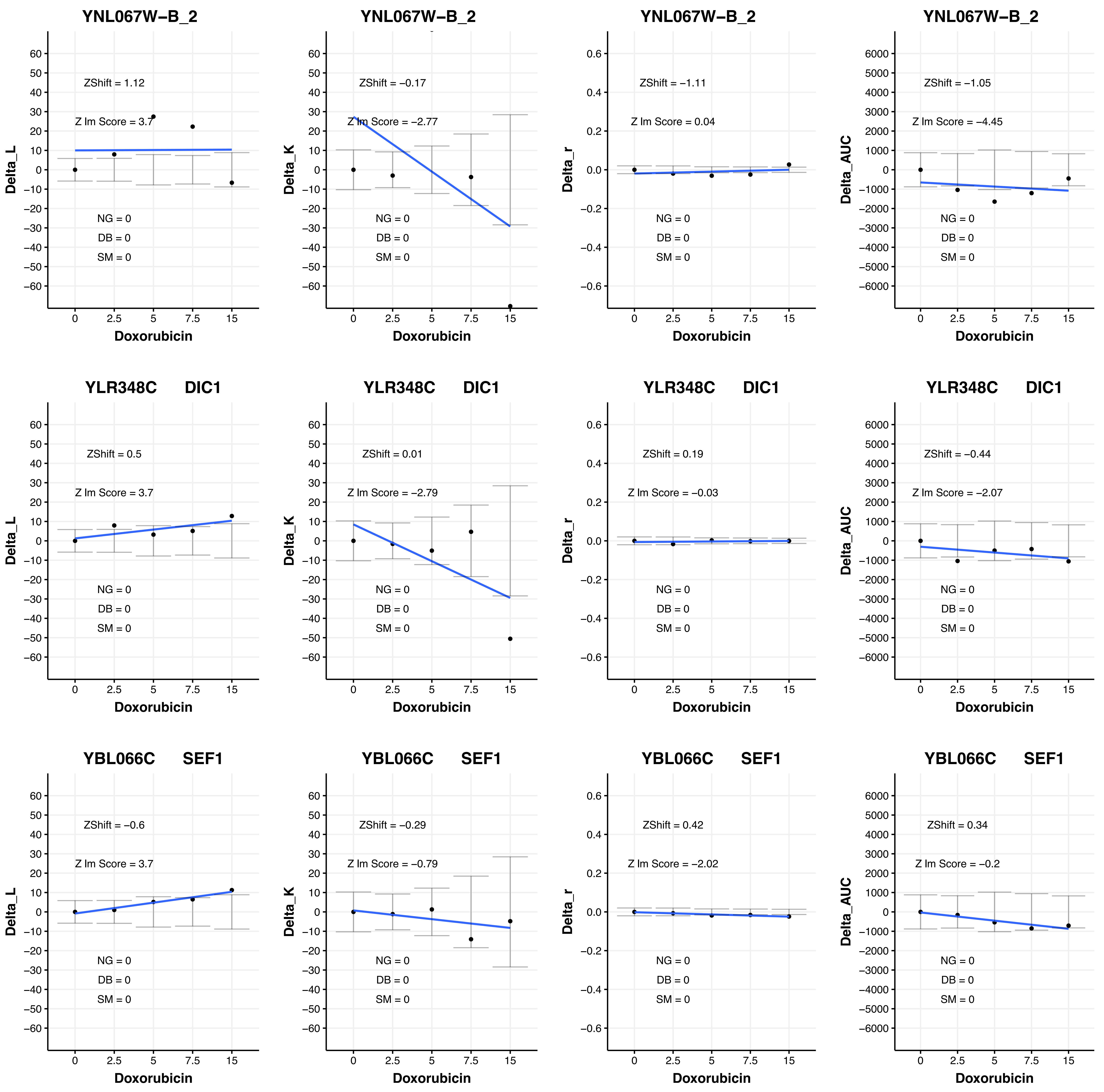
YNL067W−B_2
ZShift = 1.12
Z lm Score = 3.7
NG = 0
DB = 0
SM = 0
Delta_L
Doxorubicin
YNL067W−B_2
ZShift = −0.17
Z lm Score = −2.77
NG = 0
DB = 0
SM = 0
Delta_K
Doxorubicin
YNL067W−B_2
ZShift = −1.11
Z lm Score = 0.04
NG = 0
DB = 0
SM = 0
Delta_r
Doxorubicin
YNL067W−B_2
ZShift = −1.05
Z lm Score = −4.45
NG = 0
DB = 0
SM = 0
Delta_AUC
Doxorubicin
YLR348C DIC1
ZShift = 0.5
Z lm Score = 3.7
NG = 0
DB = 0
SM = 0
Delta_L
Doxorubicin
YLR348C DIC1
ZShift = 0.01
Z lm Score = −2.79
NG = 0
DB = 0
SM = 0
Delta_K
Doxorubicin
YLR348C DIC1
ZShift = 0.19
Z lm Score = −0.03
NG = 0
DB = 0
SM = 0
Delta_r
Doxorubicin
YLR348C DIC1
ZShift = −0.44
Z lm Score = −2.07
NG = 0
DB = 0
SM = 0
Delta_AUC
Doxorubicin
YBL066C SEF1
ZShift = −0.6
Z lm Score = 3.7
NG = 0
DB = 0
SM = 0
Delta_L
Doxorubicin
YBL066C SEF1
ZShift = −0.29
Z lm Score = −0.79
NG = 0
DB = 0
SM = 0
Delta_K
Doxorubicin
YBL066C SEF1
ZShift = 0.42
Z lm Score = −2.02
NG = 0
DB = 0
SM = 0
Delta_r
Doxorubicin
YBL066C SEF1
ZShift = 0.34
Z lm Score = −0.2
NG = 0
DB = 0
SM = 0
Delta_AUC
Doxorubicin

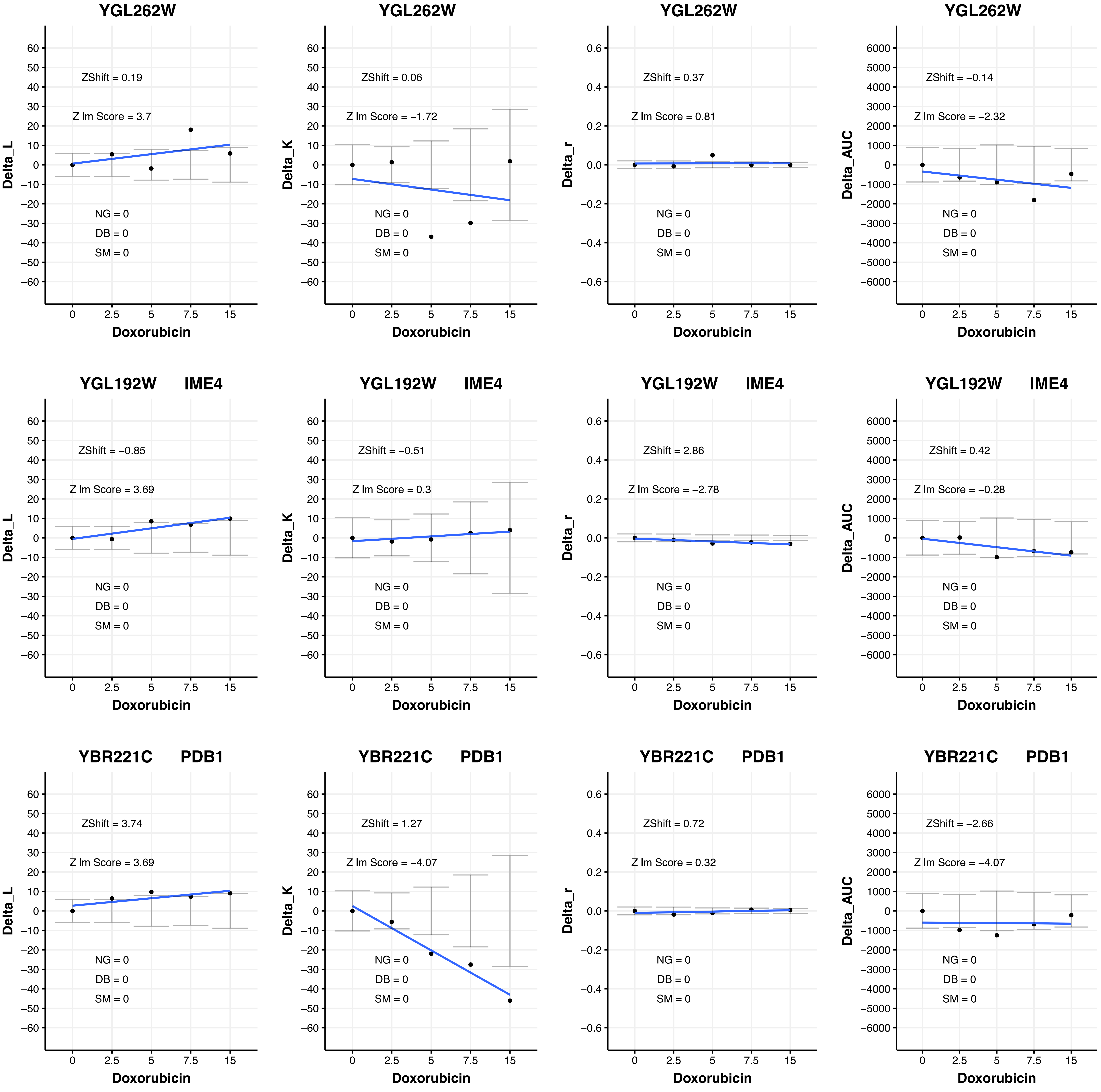

YGL262W
ZShift = 0.19
Z lm Score = 3.7
NG = 0
DB = 0
SM = 0
Delta_L
Doxorubicin
YGL262W
ZShift = 0.06
Z lm Score = −1.72
NG = 0
DB = 0
SM = 0
Delta_K
Doxorubicin
YGL262W
ZShift = 0.37
Z lm Score = 0.81
NG = 0
DB = 0
SM = 0
Delta_r
Doxorubicin
YGL262W
ZShift = −0.14
Z lm Score = −2.32
NG = 0
DB = 0
SM = 0
Delta_AUC
Doxorubicin
YGL192W IME4
ZShift = −0.85
Z lm Score = 3.69
NG = 0
DB = 0
SM = 0
Delta_L
Doxorubicin
YGL192W IME4
ZShift = −0.51
Z lm Score = 0.3
NG = 0
DB = 0
SM = 0
Delta_K
Doxorubicin
YGL192W IME4
ZShift = 2.86
Z lm Score = −2.78
NG = 0
DB = 0
SM = 0
Delta_r
Doxorubicin
YGL192W IME4
ZShift = 0.42
Z lm Score = −0.28
NG = 0
DB = 0
SM = 0
Delta_AUC
Doxorubicin
YBR221C PDB1
ZShift = 3.74
Z lm Score = 3.69
NG = 0
DB = 0
SM = 0
Delta_L
Doxorubicin
YBR221C PDB1
ZShift = 1.27
Z lm Score = −4.07
NG = 0
DB = 0
SM = 0
Delta_K
Doxorubicin
YBR221C PDB1
ZShift = 0.72
Z lm Score = 0.32
NG = 0
DB = 0
SM = 0
Delta_r
Doxorubicin
YBR221C PDB1
ZShift = −2.66
Z lm Score = −4.07
NG = 0
DB = 0
SM = 0
Delta_AUC
Doxorubicin

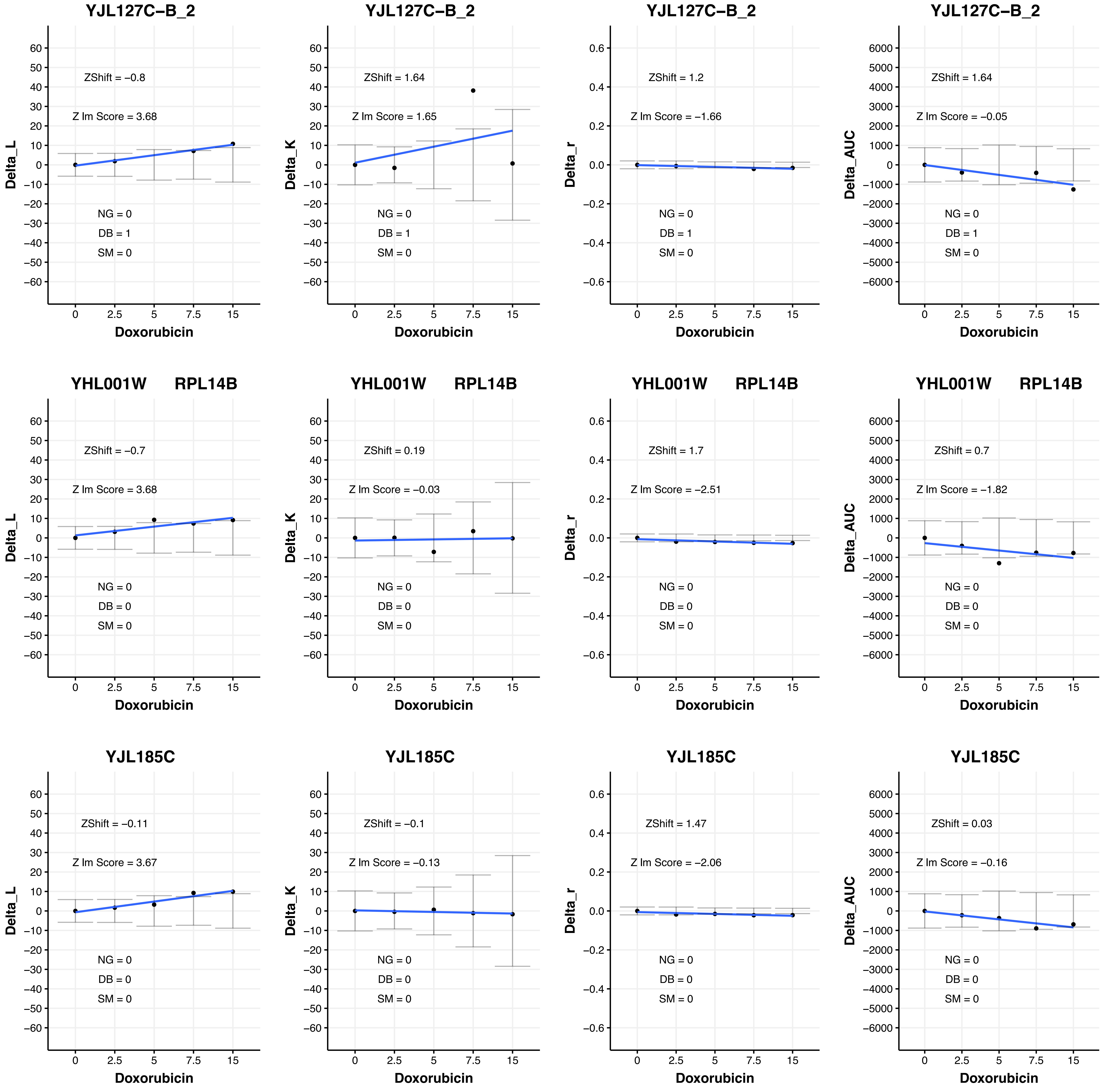
YJL127C–B_2
ZShift = –0.8
Z lm Score = 3.68
NG = 0
DB = 1
SM = 0
Delta_L
Doxorubicin
YJL127C–B_2
ZShift = 1.64
Z lm Score = 1.65
NG = 0
DB = 1
SM = 0
Delta_K
Doxorubicin
YJL127C–B_2
ZShift = 1.2
Z lm Score = –1.66
NG = 0
DB = 1
SM = 0
Delta_r
Doxorubicin
YJL127C–B_2
ZShift = 1.64
Z lm Score = –0.05
NG = 0
DB = 1
SM = 0
Delta_AUC
Doxorubicin
YHL001W RPL14B
ZShift = –0.7
Z lm Score = 3.68
NG = 0
DB = 0
SM = 0
Delta_L
Doxorubicin
YHL001W RPL14B
ZShift = 0.19
Z lm Score = –0.03
NG = 0
DB = 0
SM = 0
Delta_K
Doxorubicin
YHL001W RPL14B
ZShift = 1.7
Z lm Score = –2.51
NG = 0
DB = 0
SM = 0
Delta_r
Doxorubicin
YHL001W RPL14B
ZShift = 0.7
Z lm Score = –1.82
NG = 0
DB = 0
SM = 0
Delta_AUC
Doxorubicin
YJL185C
ZShift = –0.11
Z lm Score = 3.67
NG = 0
DB = 0
SM = 0
Delta_L
Doxorubicin
YJL185C
ZShift = –0.1
Z lm Score = –0.13
NG = 0
DB = 0
SM = 0
Delta_K
Doxorubicin
YJL185C
ZShift = 1.47
Z lm Score = –2.06
NG = 0
DB = 0
SM = 0
Delta_r
Doxorubicin
YJL185C
ZShift = 0.03
Z lm Score = –0.16
NG = 0
DB = 0
SM = 0
Delta_AUC
Doxorubicin

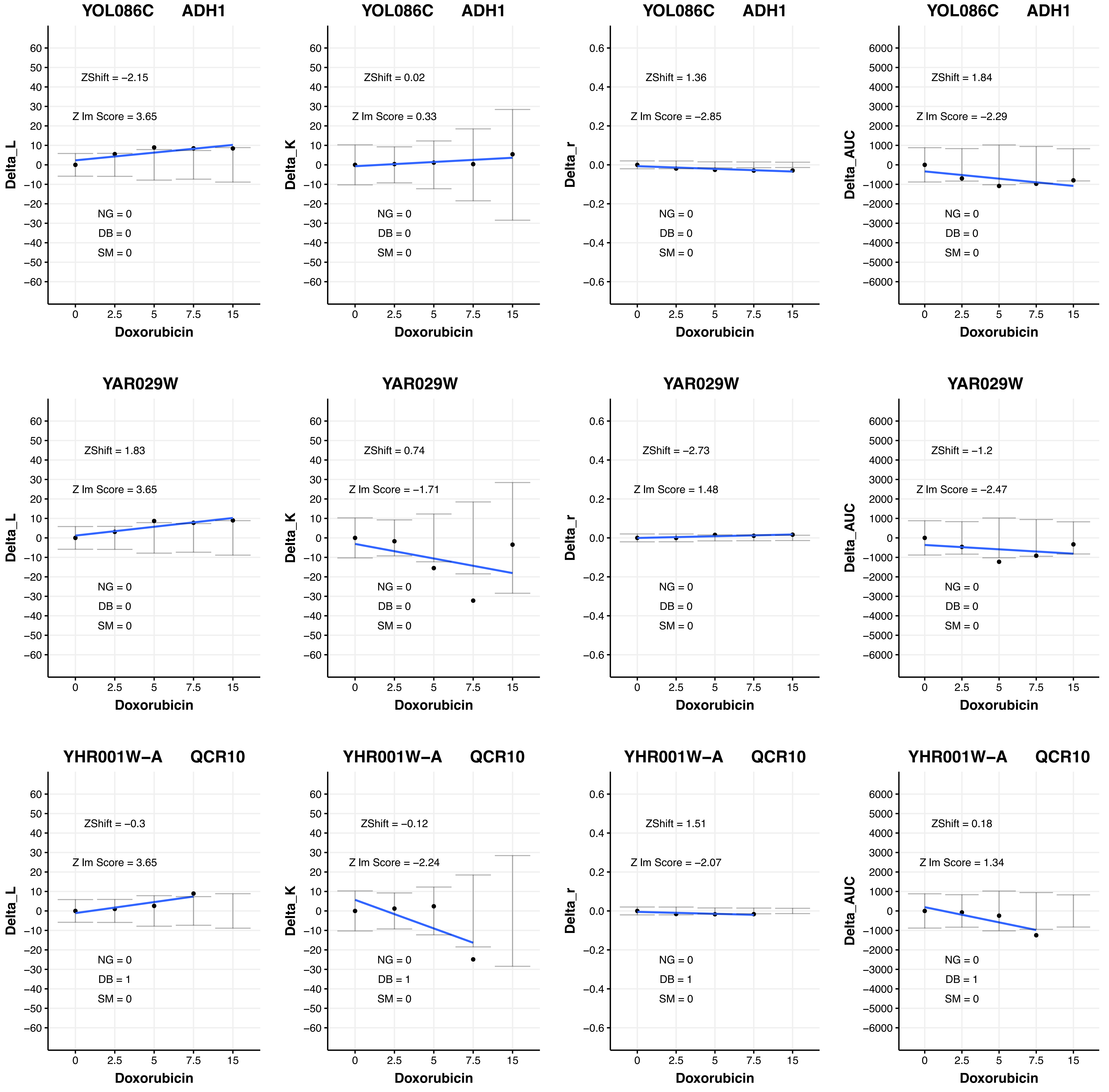
YOL086C ADH1
ZShift = −2.15
Z lm Score = 3.65
NG = 0
DB = 0
SM = 0
Delta_L
Doxorubicin
YOL086C ADH1
ZShift = 0.02
Z lm Score = 0.33
NG = 0
DB = 0
SM = 0
Delta_K
Doxorubicin
YOL086C ADH1
ZShift = 1.36
Z lm Score = −2.85
NG = 0
DB = 0
SM = 0
Delta_r
Doxorubicin
YOL086C ADH1
ZShift = 1.84
Z lm Score = −2.29
NG = 0
DB = 0
SM = 0
Delta_AUC
Doxorubicin
YAR029W
ZShift = 1.83
Z lm Score = 3.65
NG = 0
DB = 0
SM = 0
Delta_L
Doxorubicin
YAR029W
ZShift = 0.74
Z lm Score = −1.71
NG = 0
DB = 0
SM = 0
Delta_K
Doxorubicin
YAR029W
ZShift = −2.73
Z lm Score = 1.48
NG = 0
DB = 0
SM = 0
Delta_r
Doxorubicin
YAR029W
ZShift = −1.2
Z lm Score = −2.47
NG = 0
DB = 0
SM = 0
Delta_AUC
Doxorubicin
YHR001W−A QCR10
ZShift = −0.3
Z lm Score = 3.65
NG = 0
DB = 1
SM = 0
Delta_L
Doxorubicin
YHR001W−A QCR10
ZShift = −0.12
Z lm Score = −2.24
NG = 0
DB = 1
SM = 0
Delta_K
Doxorubicin
YHR001W−A QCR10
ZShift = 1.51
Z lm Score = −2.07
NG = 0
DB = 1
SM = 0
Delta_r
Doxorubicin
YHR001W−A QCR10
ZShift = 0.18
Z lm Score = 1.34
NG = 0
DB = 1
SM = 0
Delta_AUC
Doxorubicin

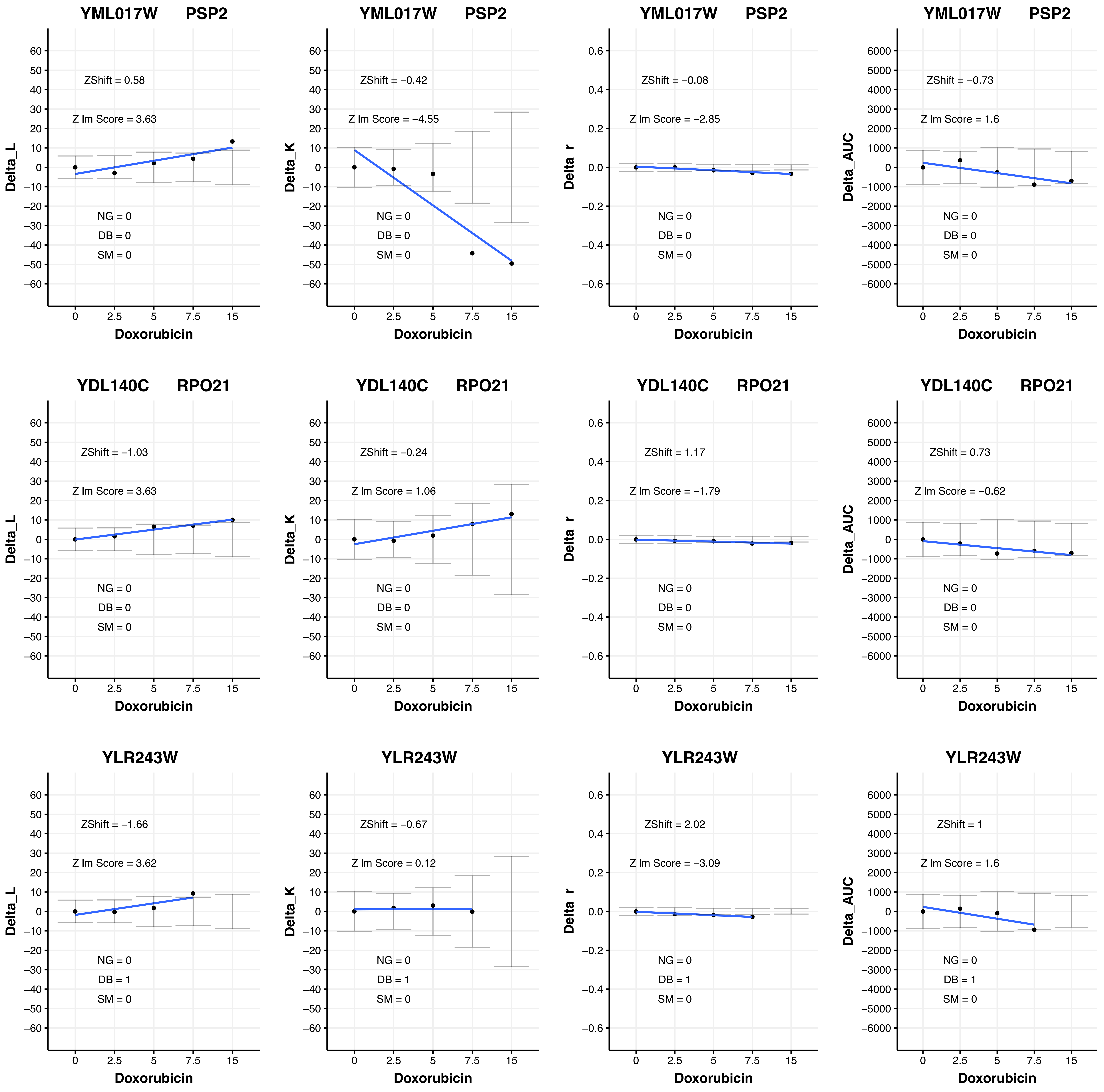
YML017W PSP2
ZShift = 0.58
Z lm Score = 3.63
NG = 0
DB = 0
SM = 0
Delta_L
Doxorubicin
YML017W PSP2
ZShift = −0.42
Z lm Score = −4.55
NG = 0
DB = 0
SM = 0
Delta_K
Doxorubicin
YML017W PSP2
ZShift = −0.08
Z lm Score = −2.85
NG = 0
DB = 0
SM = 0
Delta_r
Doxorubicin
YML017W PSP2
ZShift = −0.73
Z lm Score = 1.6
NG = 0
DB = 0
SM = 0
Delta_AUC
Doxorubicin
YDL140C RPO21
ZShift = −1.03
Z lm Score = 3.63
NG = 0
DB = 0
SM = 0
Delta_L
Doxorubicin
YDL140C RPO21
ZShift = −0.24
Z lm Score = 1.06
NG = 0
DB = 0
SM = 0
Delta_K
Doxorubicin
YDL140C RPO21
ZShift = 1.17
Z lm Score = −1.79
NG = 0
DB = 0
SM = 0
Delta_r
Doxorubicin
YDL140C RPO21
ZShift = 0.73
Z lm Score = −0.62
NG = 0
DB = 0
SM = 0
Delta_AUC
Doxorubicin
YLR243W
ZShift = −1.66
Z lm Score = 3.62
NG = 0
DB = 1
SM = 0
Delta_L
Doxorubicin
YLR243W
ZShift = −0.67
Z lm Score = 0.12
NG = 0
DB = 1
SM = 0
Delta_K
Doxorubicin
YLR243W
ZShift = 2.02
Z lm Score = −3.09
NG = 0
DB = 1
SM = 0
Delta_r
Doxorubicin
YLR243W
ZShift = 1
Z lm Score = 1.6
NG = 0
DB = 1
SM = 0
Delta_AUC
Doxorubicin

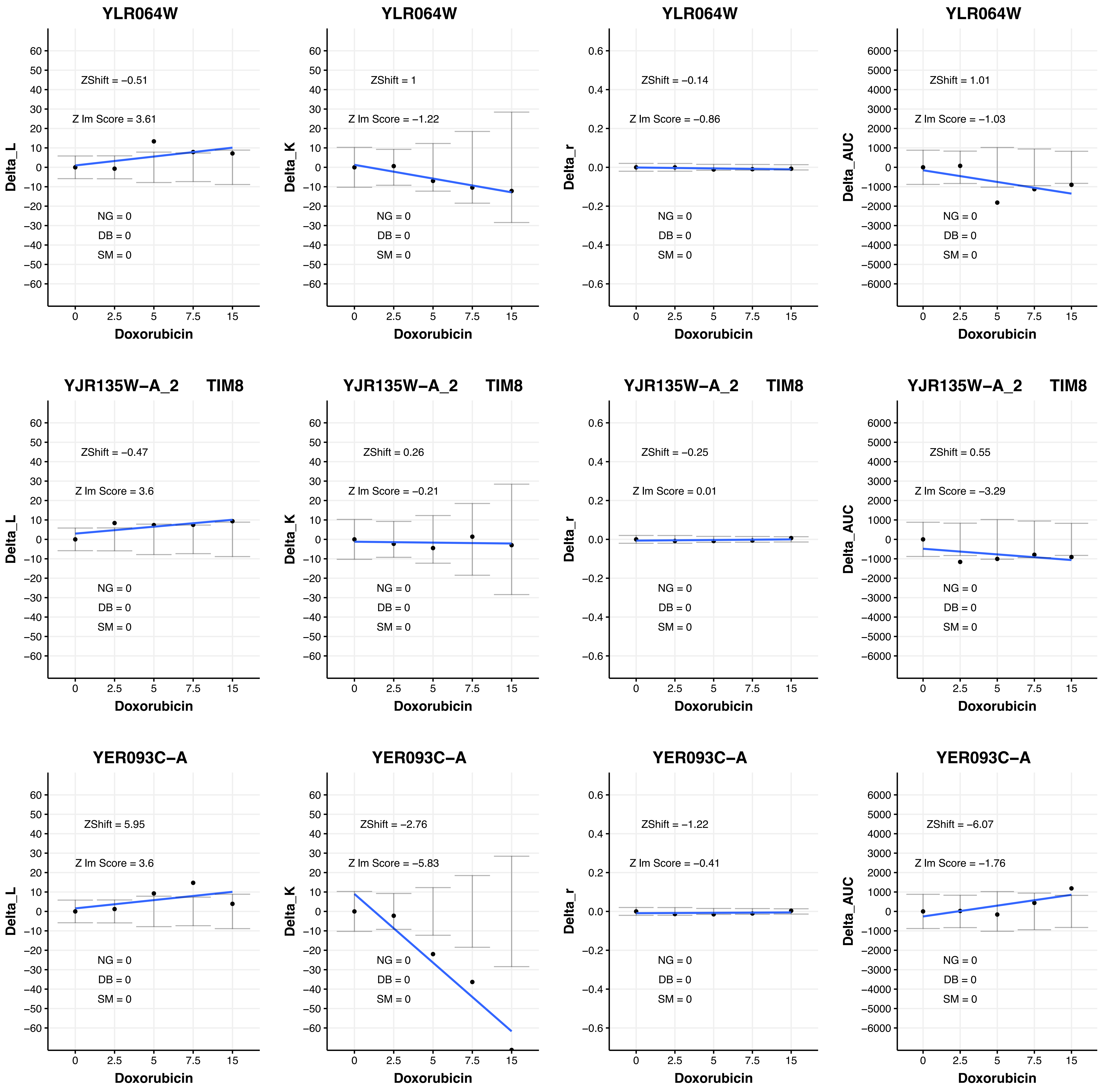
YLR064W
ZShift = −0.51
Z lm Score = 3.61
NG = 0
DB = 0
SM = 0
Delta_L
Doxorubicin
YLR064W
ZShift = 1
Z lm Score = −1.22
NG = 0
DB = 0
SM = 0
Delta_K
Doxorubicin
YLR064W
ZShift = −0.14
Z lm Score = −0.86
NG = 0
DB = 0
SM = 0
Delta_r
Doxorubicin
YLR064W
ZShift = 1.01
Z lm Score = −1.03
NG = 0
DB = 0
SM = 0
Delta_AUC
Doxorubicin
YJR135W−A_2 TIM8
ZShift = −0.47
Z lm Score = 3.6
NG = 0
DB = 0
SM = 0
Delta_L
Doxorubicin
YJR135W−A_2 TIM8
ZShift = 0.26
Z lm Score = −0.21
NG = 0
DB = 0
SM = 0
Delta_K
Doxorubicin
YJR135W−A_2 TIM8
ZShift = −0.25
Z lm Score = 0.01
NG = 0
DB = 0
SM = 0
Delta_r
Doxorubicin
YJR135W−A_2 TIM8
ZShift = 0.55
Z lm Score = −3.29
NG = 0
DB = 0
SM = 0
Delta_AUC
Doxorubicin
YER093C−A
ZShift = 5.95
Z lm Score = 3.6
NG = 0
DB = 0
SM = 0
Delta_L
Doxorubicin
YER093C−A
ZShift = −2.76
Z lm Score = −5.83
NG = 0
DB = 0
SM = 0
Delta_K
Doxorubicin
YER093C−A
ZShift = −1.22
Z lm Score = −0.41
NG = 0
DB = 0
SM = 0
Delta_r
Doxorubicin
YER093C−A
ZShift = −6.07
Z lm Score = −1.76
NG = 0
DB = 0
SM = 0
Delta_AUC
Doxorubicin

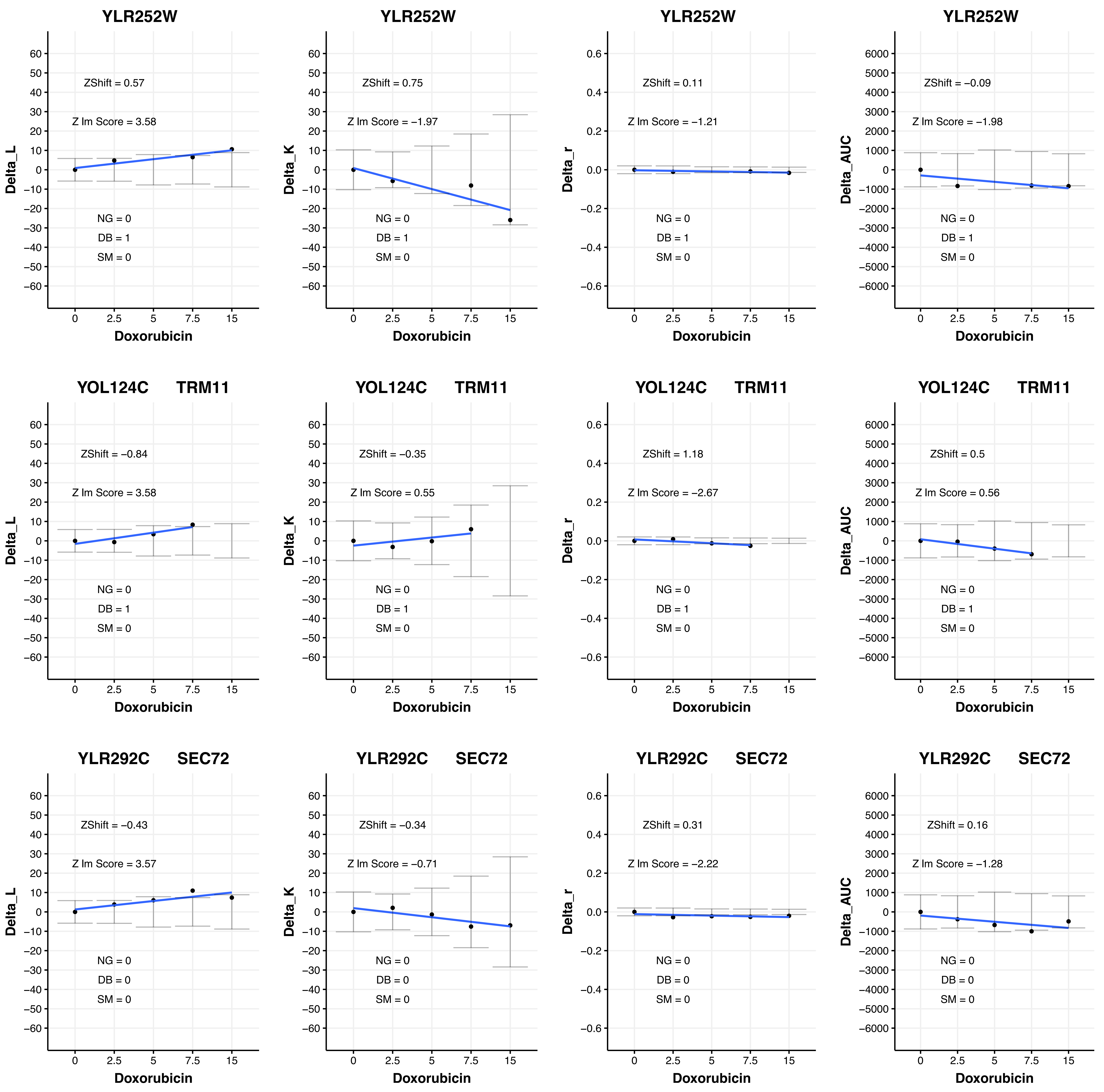

YLR252W
ZShift = 0.57
Z lm Score = 3.58
NG = 0
DB = 1
SM = 0
Delta_L
Doxorubicin
YLR252W
ZShift = 0.75
Z lm Score = −1.97
NG = 0
DB = 1
SM = 0
Delta_K
Doxorubicin
YLR252W
ZShift = 0.11
Z lm Score = −1.21
NG = 0
DB = 1
SM = 0
Delta_r
Doxorubicin
YLR252W
ZShift = −0.09
Z lm Score = −1.98
NG = 0
DB = 1
SM = 0
Delta_AUC
Doxorubicin
YOL124C TRM11
ZShift = −0.84
Z lm Score = 3.58
NG = 0
DB = 1
SM = 0
Delta_L
Doxorubicin
YOL124C TRM11
ZShift = −0.35
Z lm Score = 0.55
NG = 0
DB = 1
SM = 0
Delta_K
Doxorubicin
YOL124C TRM11
ZShift = 1.18
Z lm Score = −2.67
NG = 0
DB = 1
SM = 0
Delta_r
Doxorubicin
YOL124C TRM11
ZShift = 0.5
Z lm Score = 0.56
NG = 0
DB = 1
SM = 0
Delta_AUC
Doxorubicin
YLR292C SEC72
ZShift = −0.43
Z lm Score = 3.57
NG = 0
DB = 0
SM = 0
Delta_L
Doxorubicin
YLR292C SEC72
ZShift = −0.34
Z lm Score = −0.71
NG = 0
DB = 0
SM = 0
Delta_K
Doxorubicin
YLR292C SEC72
ZShift = 0.31
Z lm Score = −2.22
NG = 0
DB = 0
SM = 0
Delta_r
Doxorubicin
YLR292C SEC72
ZShift = 0.16
Z lm Score = −1.28
NG = 0
DB = 0
SM = 0
Delta_AUC
Doxorubicin

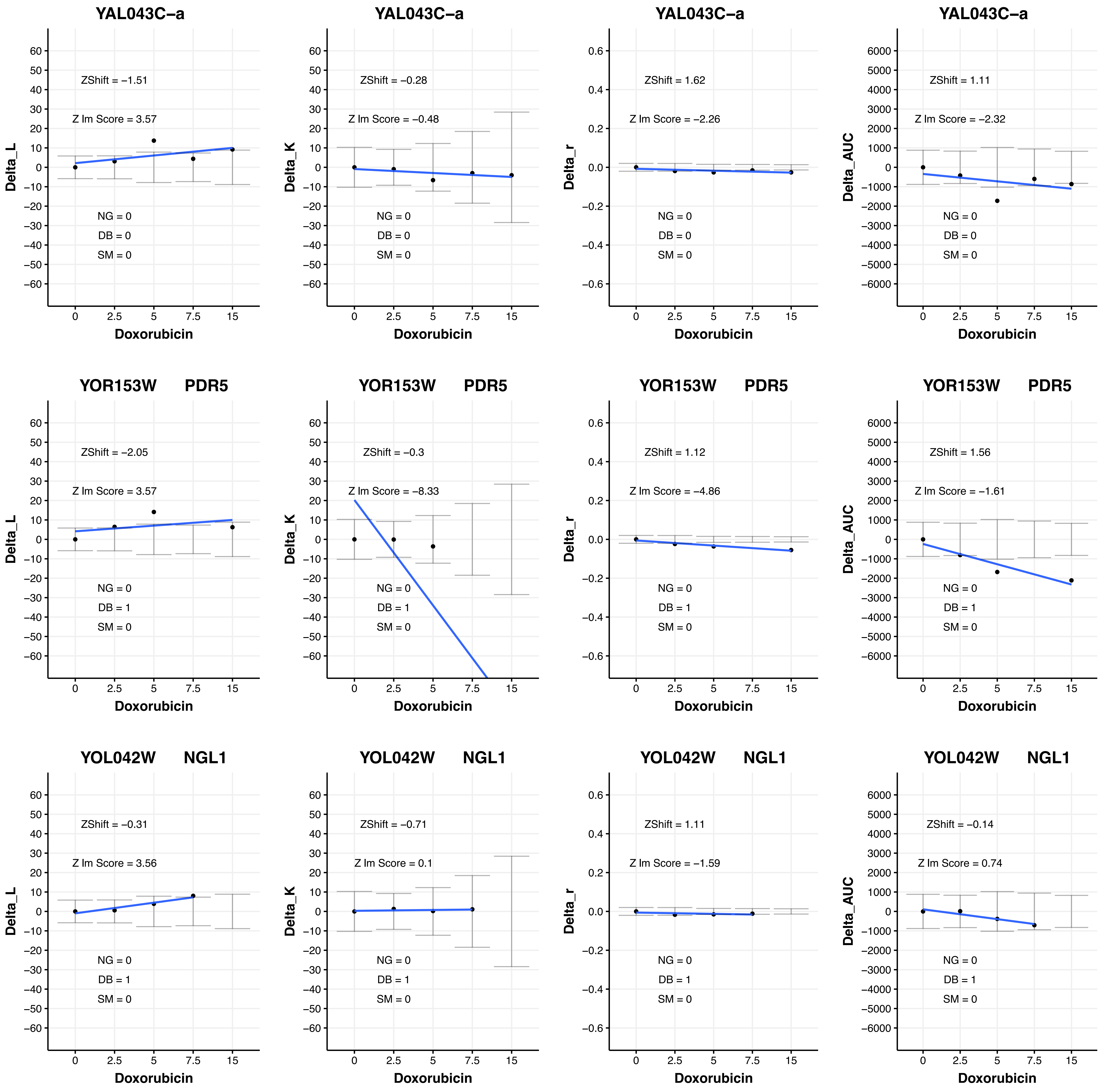
YAL043C–a
ZShift = –1.51
Z lm Score = 3.57
NG = 0
DB = 0
SM = 0
Delta_L
Doxorubicin
YAL043C–a
ZShift = –0.28
Z lm Score = –0.48
NG = 0
DB = 0
SM = 0
Delta_K
Doxorubicin
YAL043C–a
ZShift = 1.62
Z lm Score = –2.26
NG = 0
DB = 0
SM = 0
Delta_r
Doxorubicin
YAL043C–a
ZShift = 1.11
Z lm Score = –2.32
NG = 0
DB = 0
SM = 0
Delta_AUC
Doxorubicin
YOR153W PDR5
ZShift = –2.05
Z lm Score = 3.57
NG = 0
DB = 1
SM = 0
Delta_L
Doxorubicin
YOR153W PDR5
ZShift = –0.3
Z lm Score = –8.33
NG = 0
DB = 1
SM = 0
Delta_K
Doxorubicin
YOR153W PDR5
ZShift = 1.12
Z lm Score = –4.86
NG = 0
DB = 1
SM = 0
Delta_r
Doxorubicin
YOR153W PDR5
ZShift = 1.56
Z lm Score = –1.61
NG = 0
DB = 1
SM = 0
Delta_AUC
Doxorubicin
YOL042W NGL1
ZShift = –0.31
Z lm Score = 3.56
NG = 0
DB = 1
SM = 0
Delta_L
Doxorubicin
YOL042W NGL1
ZShift = –0.71
Z lm Score = 0.1
NG = 0
DB = 1
SM = 0
Delta_K
Doxorubicin
YOL042W NGL1
ZShift = 1.11
Z lm Score = –1.59
NG = 0
DB = 1
SM = 0
Delta_r
Doxorubicin
YOL042W NGL1
ZShift = –0.14
Z lm Score = 0.74
NG = 0
DB = 1
SM = 0
Delta_AUC
Doxorubicin

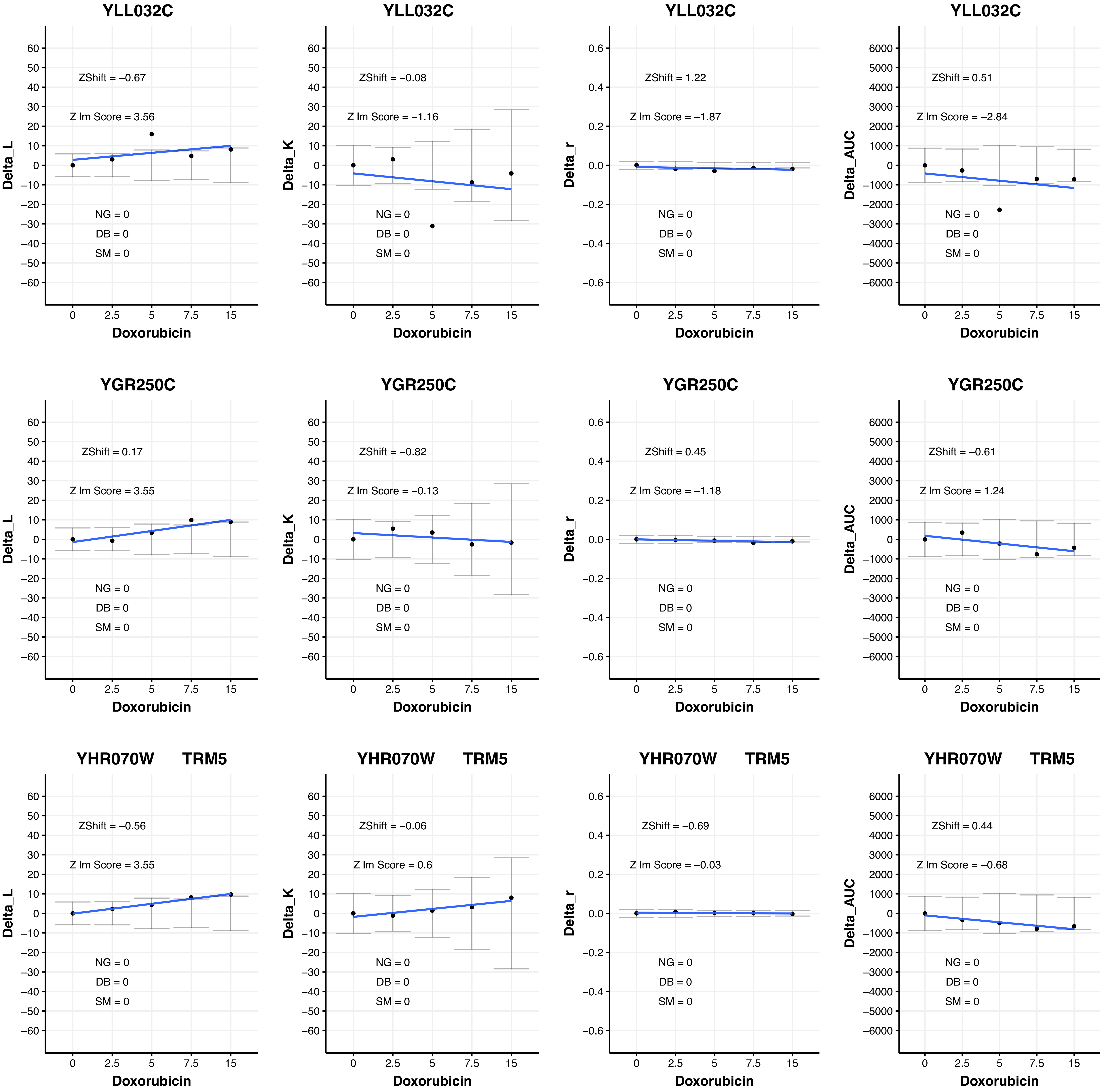

YLL032C
ZShift = −0.67
Z lm Score = 3.56
NG = 0
DB = 0
SM = 0
Delta_L
Doxorubicin
YLL032C
ZShift = −0.08
Z lm Score = −1.16
NG = 0
DB = 0
SM = 0
Delta_K
Doxorubicin
YLL032C
ZShift = 1.22
Z lm Score = −1.87
NG = 0
DB = 0
SM = 0
Delta_r
Doxorubicin
YLL032C
ZShift = 0.51
Z lm Score = −2.84
NG = 0
DB = 0
SM = 0
Delta_AUC
Doxorubicin
YGR250C
ZShift = 0.17
Z lm Score = 3.55
NG = 0
DB = 0
SM = 0
Delta_L
Doxorubicin
YGR250C
ZShift = −0.82
Z lm Score = −0.13
NG = 0
DB = 0
SM = 0
Delta_K
Doxorubicin
YGR250C
ZShift = 0.45
Z lm Score = −1.18
NG = 0
DB = 0
SM = 0
Delta_r
Doxorubicin
YGR250C
ZShift = −0.61
Z lm Score = 1.24
NG = 0
DB = 0
SM = 0
Delta_AUC
Doxorubicin
YHR070W TRM5
ZShift = −0.56
Z lm Score = 3.55
NG = 0
DB = 0
SM = 0
Delta_L
Doxorubicin
YHR070W TRM5
ZShift = −0.06
Z lm Score = 0.6
NG = 0
DB = 0
SM = 0
Delta_K
Doxorubicin
YHR070W TRM5
ZShift = −0.69
Z lm Score = −0.03
NG = 0
DB = 0
SM = 0
Delta_r
Doxorubicin
YHR070W TRM5
ZShift = 0.44
Z lm Score = −0.68
NG = 0
DB = 0
SM = 0
Delta_AUC
Doxorubicin

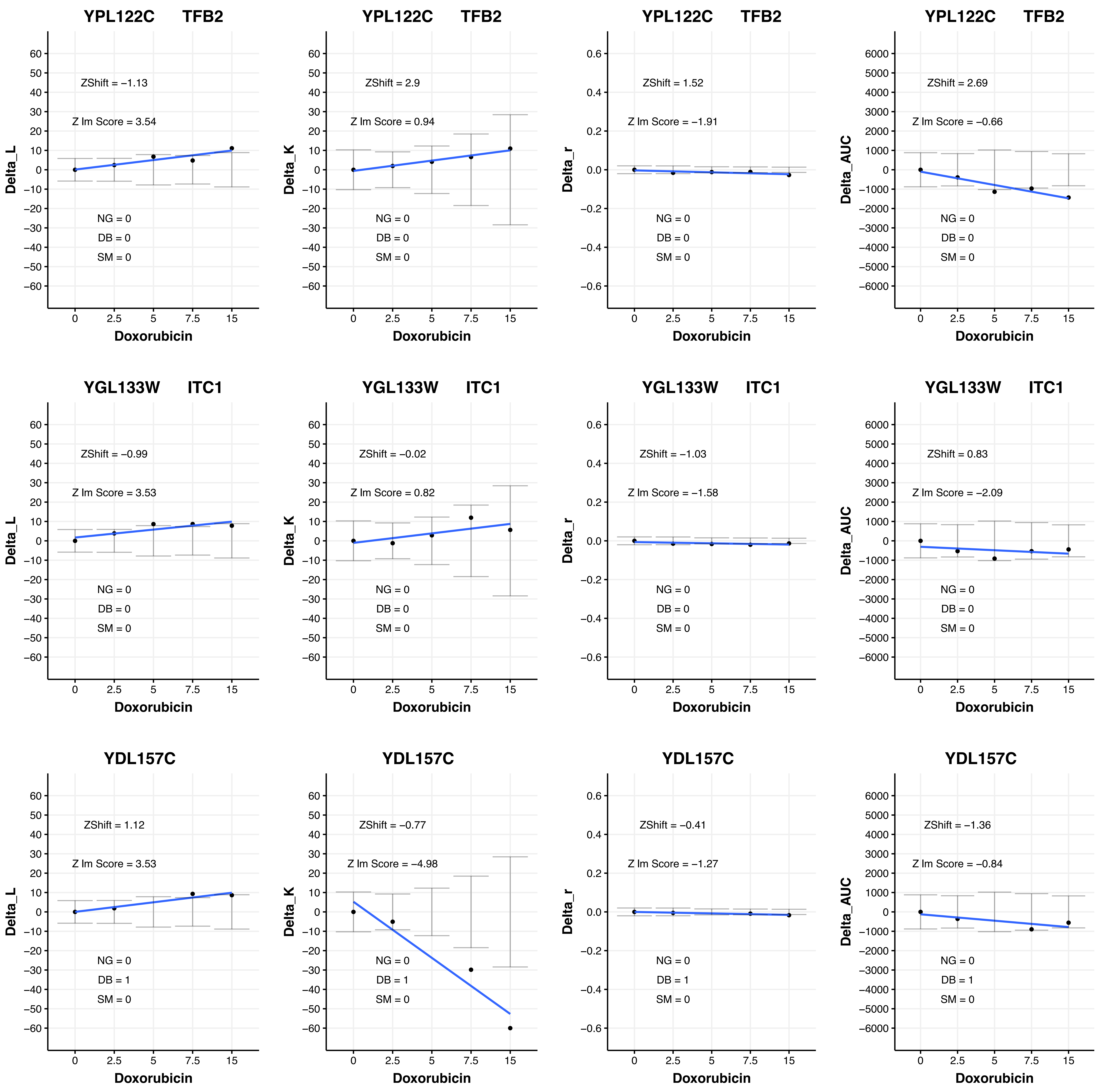

YPL122C TFB2
ZShift = −1.13
Z lm Score = 3.54
NG = 0
DB = 0
SM = 0
Delta_L
Doxorubicin
YPL122C TFB2
ZShift = 2.9
Z lm Score = 0.94
NG = 0
DB = 0
SM = 0
Delta_K
Doxorubicin
YPL122C TFB2
ZShift = 1.52
Z lm Score = −1.91
NG = 0
DB = 0
SM = 0
Delta_r
Doxorubicin
YPL122C TFB2
ZShift = 2.69
Z lm Score = −0.66
NG = 0
DB = 0
SM = 0
Delta_AUC
Doxorubicin
YGL133W ITC1
ZShift = −0.99
Z lm Score = 3.53
NG = 0
DB = 0
SM = 0
Delta_L
Doxorubicin
YGL133W ITC1
ZShift = −0.02
Z lm Score = 0.82
NG = 0
DB = 0
SM = 0
Delta_K
Doxorubicin
YGL133W ITC1
ZShift = −1.03
Z lm Score = −1.58
NG = 0
DB = 0
SM = 0
Delta_r
Doxorubicin
YGL133W ITC1
ZShift = 0.83
Z lm Score = −2.09
NG = 0
DB = 0
SM = 0
Delta_AUC
Doxorubicin
YDL157C
ZShift = 1.12
Z lm Score = 3.53
NG = 0
DB = 1
SM = 0
Delta_L
Doxorubicin
YDL157C
ZShift = −0.77
Z lm Score = −4.98
NG = 0
DB = 1
SM = 0
Delta_K
Doxorubicin
YDL157C
ZShift = −0.41
Z lm Score = −1.27
NG = 0
DB = 1
SM = 0
Delta_r
Doxorubicin
YDL157C
ZShift = −1.36
Z lm Score = −0.84
NG = 0
DB = 1
SM = 0
Delta_AUC
Doxorubicin

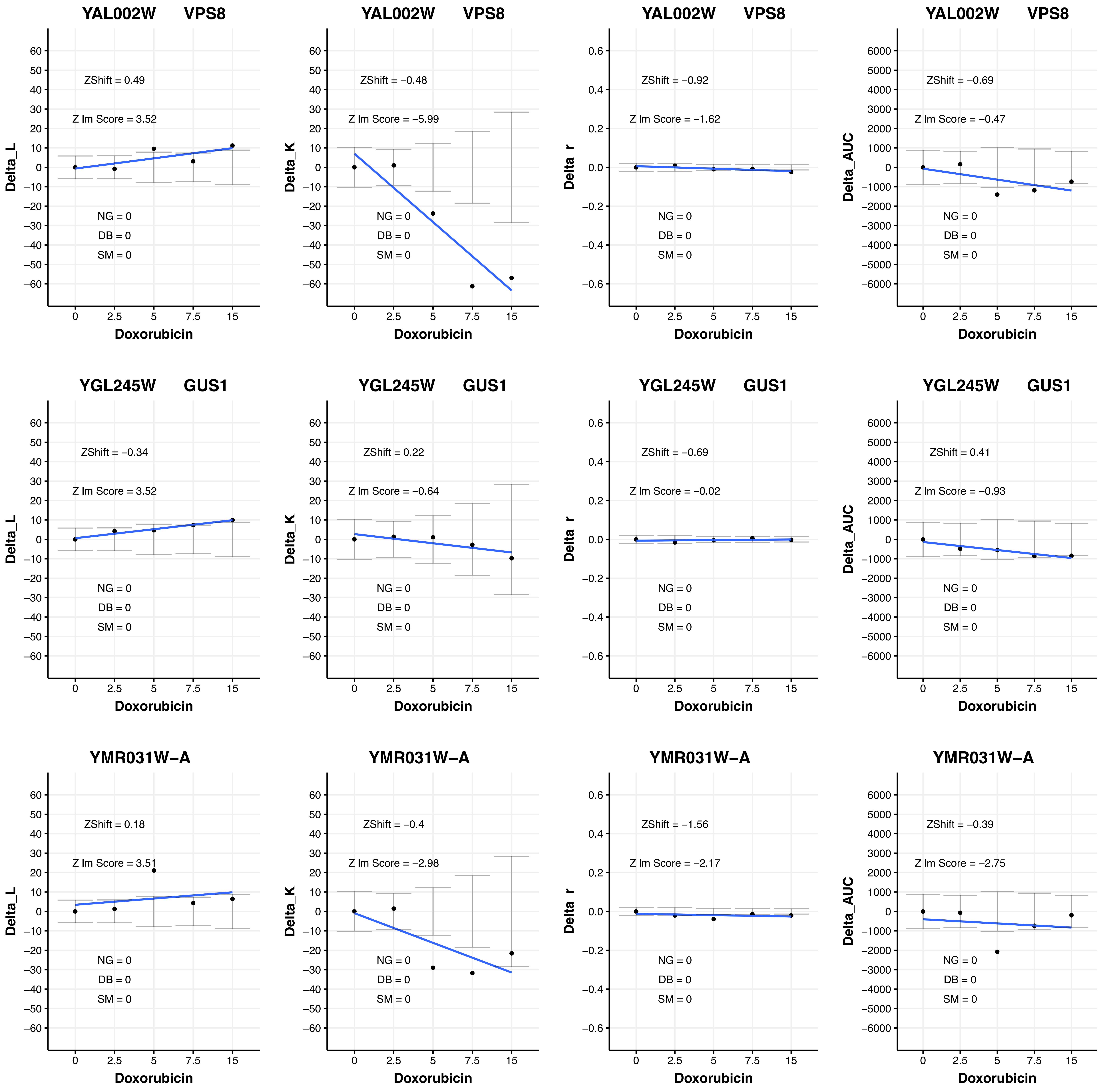
YAL002W VPS8
ZShift = 0.49
Z lm Score = 3.52
NG = 0
DB = 0
SM = 0
Delta_L
Doxorubicin
YAL002W VPS8
ZShift = −0.48
Z lm Score = −5.99
NG = 0
DB = 0
SM = 0
Delta_K
Doxorubicin
YAL002W VPS8
ZShift = −0.92
Z lm Score = −1.62
NG = 0
DB = 0
SM = 0
Delta_r
Doxorubicin
YAL002W VPS8
ZShift = −0.69
Z lm Score = −0.47
NG = 0
DB = 0
SM = 0
Delta_AUC
Doxorubicin
YGL245W GUS1
ZShift = −0.34
Z lm Score = 3.52
NG = 0
DB = 0
SM = 0
Delta_L
Doxorubicin
YGL245W GUS1
ZShift = 0.22
Z lm Score = −0.64
NG = 0
DB = 0
SM = 0
Delta_K
Doxorubicin
YGL245W GUS1
ZShift = −0.69
Z lm Score = −0.02
NG = 0
DB = 0
SM = 0
Delta_r
Doxorubicin
YGL245W GUS1
ZShift = 0.41
Z lm Score = −0.93
NG = 0
DB = 0
SM = 0
Delta_AUC
Doxorubicin
YMR031W−A
ZShift = 0.18
Z lm Score = 3.51
NG = 0
DB = 0
SM = 0
Delta_L
Doxorubicin
YMR031W−A
ZShift = −0.4
Z lm Score = −2.98
NG = 0
DB = 0
SM = 0
Delta_K
Doxorubicin
YMR031W−A
ZShift = −1.56
Z lm Score = −2.17
NG = 0
DB = 0
SM = 0
Delta_r
Doxorubicin
YMR031W−A
ZShift = −0.39
Z lm Score = −2.75
NG = 0
DB = 0
SM = 0
Delta_AUC
Doxorubicin

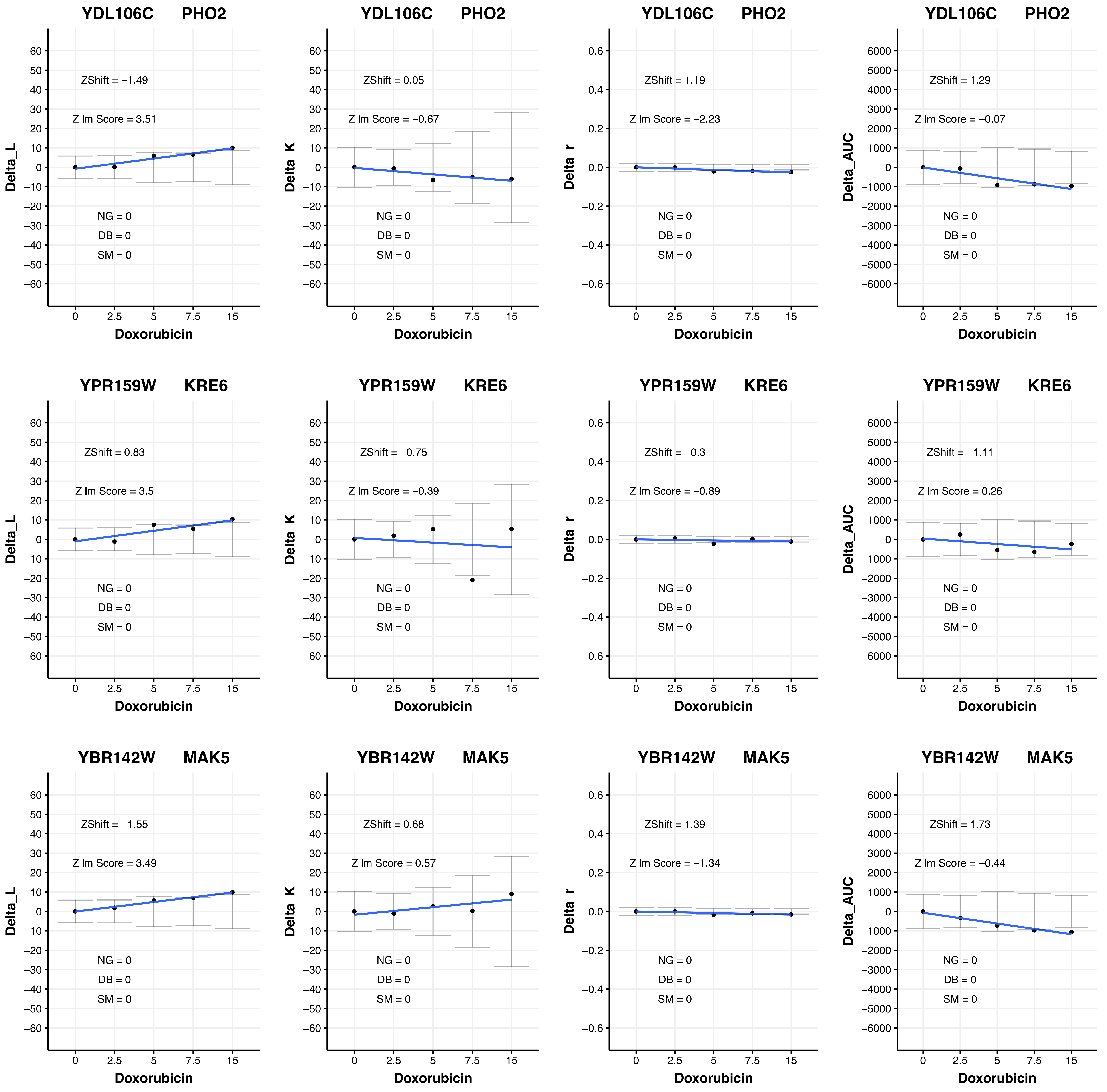
YDL106C PHO2
Delta_L
ZShift = −1.49
Z lm Score = 3.51
NG = 0
DB = 0
SM = 0
Doxorubicin
YDL106C PHO2
Delta_K
ZShift = 0.05
Z lm Score = −0.67
NG = 0
DB = 0
SM = 0
Doxorubicin
YDL106C PHO2
Delta_r
ZShift = 1.19
Z lm Score = −2.23
NG = 0
DB = 0
SM = 0
Doxorubicin
YDL106C PHO2
Delta_AUC
ZShift = 1.29
Z lm Score = −0.07
NG = 0
DB = 0
SM = 0
Doxorubicin
YPR159W KRE6
Delta_L
ZShift = 0.83
Z lm Score = 3.5
NG = 0
DB = 0
SM = 0
Doxorubicin
YPR159W KRE6
Delta_K
ZShift = −0.75
Z lm Score = −0.39
NG = 0
DB = 0
SM = 0
Doxorubicin
YPR159W KRE6
Delta_r
ZShift = −0.3
Z lm Score = −0.89
NG = 0
DB = 0
SM = 0
Doxorubicin
YPR159W KRE6
Delta_AUC
ZShift = −1.11
Z lm Score = 0.26
NG = 0
DB = 0
SM = 0
Doxorubicin
YBR142W MAK5
Delta_L
ZShift = −1.55
Z lm Score = 3.49
NG = 0
DB = 0
SM = 0
Doxorubicin
YBR142W MAK5
Delta_K
ZShift = 0.68
Z lm Score = 0.57
NG = 0
DB = 0
SM = 0
Doxorubicin
YBR142W MAK5
Delta_r
ZShift = 1.39
Z lm Score = −1.34
NG = 0
DB = 0
SM = 0
Doxorubicin
YBR142W MAK5
Delta_AUC
ZShift = 1.73
Z lm Score = −0.44
NG = 0
DB = 0
SM = 0
Doxorubicin

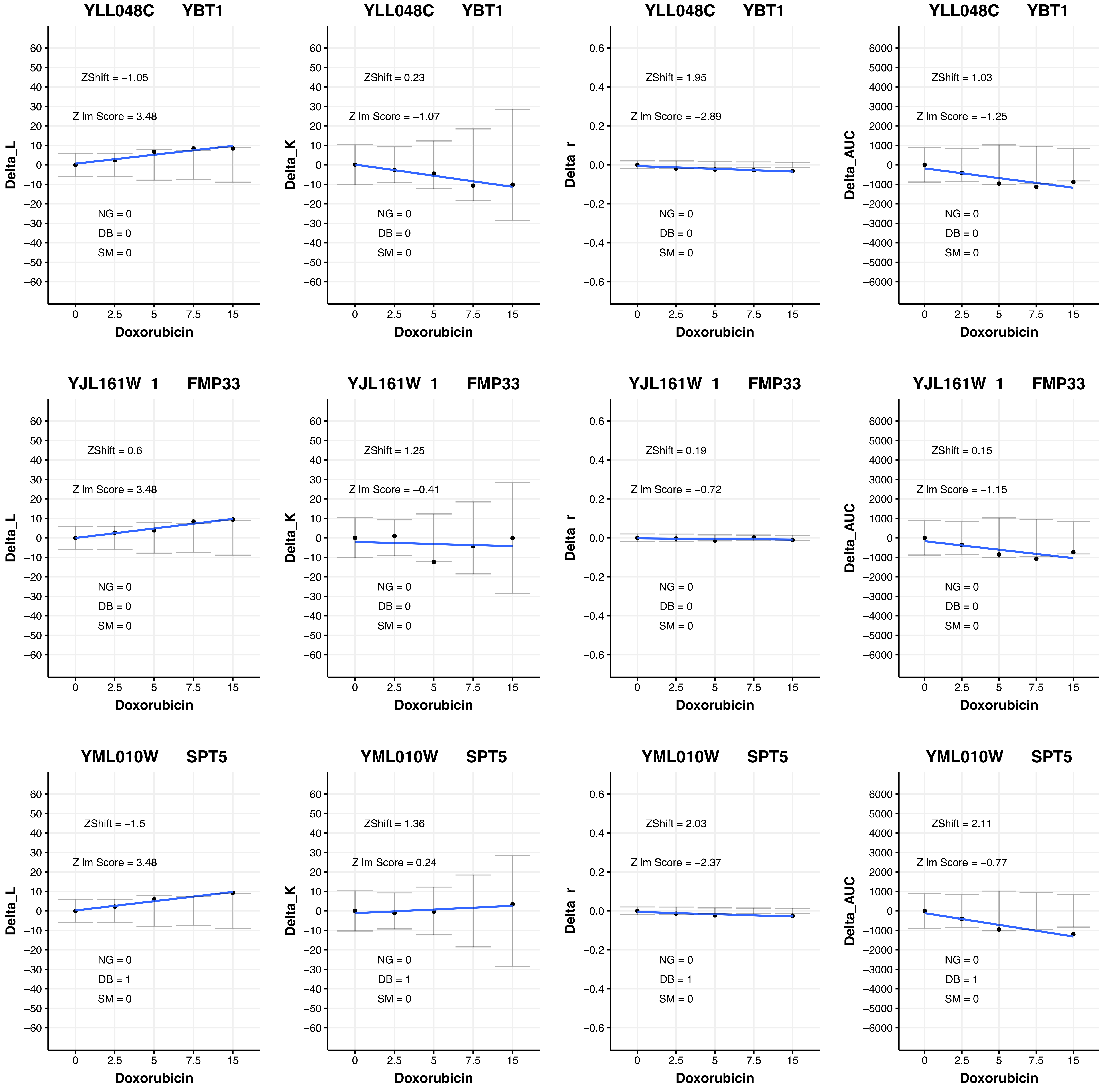
YLL048C YBT1
ZShift = −1.05
Z lm Score = 3.48
NG = 0
DB = 0
SM = 0
Delta_L
Doxorubicin
YLL048C YBT1
ZShift = 0.23
Z lm Score = −1.07
NG = 0
DB = 0
SM = 0
Delta_K
Doxorubicin
YLL048C YBT1
ZShift = 1.95
Z lm Score = −2.89
NG = 0
DB = 0
SM = 0
Delta_r
Doxorubicin
YLL048C YBT1
ZShift = 1.03
Z lm Score = −1.25
NG = 0
DB = 0
SM = 0
Delta_AUC
Doxorubicin
YJL161W_1 FMP33
ZShift = 0.6
Z lm Score = 3.48
NG = 0
DB = 0
SM = 0
Delta_L
Doxorubicin
YJL161W_1 FMP33
ZShift = 1.25
Z lm Score = −0.41
NG = 0
DB = 0
SM = 0
Delta_K
Doxorubicin
YJL161W_1 FMP33
ZShift = 0.19
Z lm Score = −0.72
NG = 0
DB = 0
SM = 0
Delta_r
Doxorubicin
YJL161W_1 FMP33
ZShift = 0.15
Z lm Score = −1.15
NG = 0
DB = 0
SM = 0
Delta_AUC
Doxorubicin
YML010W SPT5
ZShift = −1.5
Z lm Score = 3.48
NG = 0
DB = 1
SM = 0
Delta_L
Doxorubicin
YML010W SPT5
ZShift = 1.36
Z lm Score = 0.24
NG = 0
DB = 1
SM = 0
Delta_K
Doxorubicin
YML010W SPT5
ZShift = 2.03
Z lm Score = −2.37
NG = 0
DB = 1
SM = 0
Delta_r
Doxorubicin
YML010W SPT5
ZShift = 2.11
Z lm Score = −0.77
NG = 0
DB = 1
SM = 0
Delta_AUC
Doxorubicin

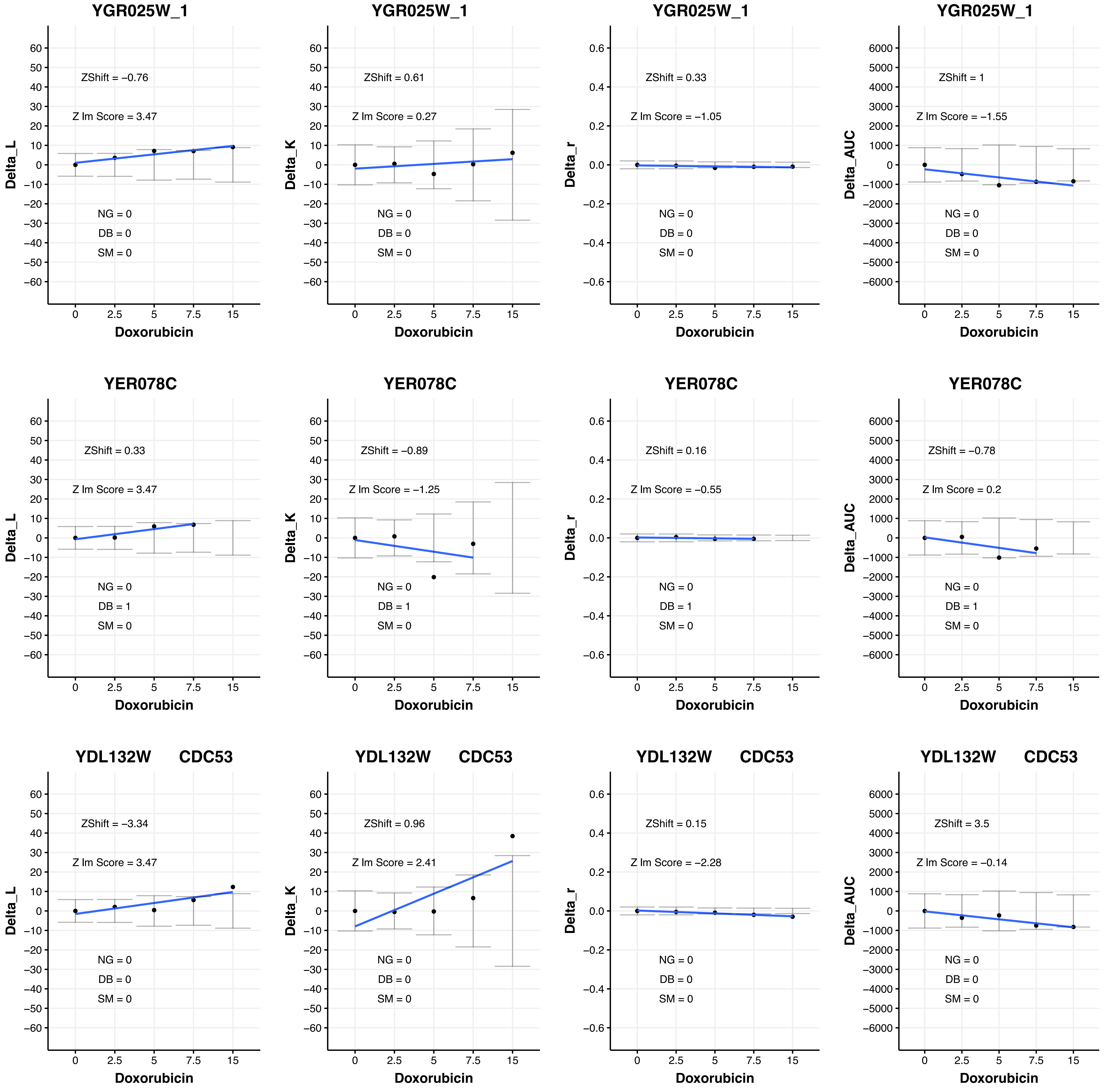
YGR025W_1
ZShift = −0.76
Z lm Score = 3.47
NG = 0
DB = 0
SM = 0
Delta_L
Doxorubicin
YGR025W_1
ZShift = 0.61
Z lm Score = 0.27
NG = 0
DB = 0
SM = 0
Delta_K
Doxorubicin
YGR025W_1
ZShift = 0.33
Z lm Score = −1.05
NG = 0
DB = 0
SM = 0
Delta_r
Doxorubicin
YGR025W_1
ZShift = 1
Z lm Score = −1.55
NG = 0
DB = 0
SM = 0
Delta_AUC
Doxorubicin
YER078C
ZShift = 0.33
Z lm Score = 3.47
NG = 0
DB = 1
SM = 0
Delta_L
Doxorubicin
YER078C
ZShift = −0.89
Z lm Score = −1.25
NG = 0
DB = 1
SM = 0
Delta_K
Doxorubicin
YER078C
ZShift = 0.16
Z lm Score = −0.55
NG = 0
DB = 1
SM = 0
Delta_r
Doxorubicin
YER078C
ZShift = −0.78
Z lm Score = 0.2
NG = 0
DB = 1
SM = 0
Delta_AUC
Doxorubicin
YDL132W CDC53
ZShift = −3.34
Z lm Score = 3.47
NG = 0
DB = 0
SM = 0
Delta_L
Doxorubicin
YDL132W CDC53
ZShift = 0.96
Z lm Score = 2.41
NG = 0
DB = 0
SM = 0
Delta_K
Doxorubicin
YDL132W CDC53
ZShift = 0.15
Z lm Score = −2.28
NG = 0
DB = 0
SM = 0
Delta_r
Doxorubicin
YDL132W CDC53
ZShift = 3.5
Z lm Score = −0.14
NG = 0
DB = 0
SM = 0
Delta_AUC
Doxorubicin

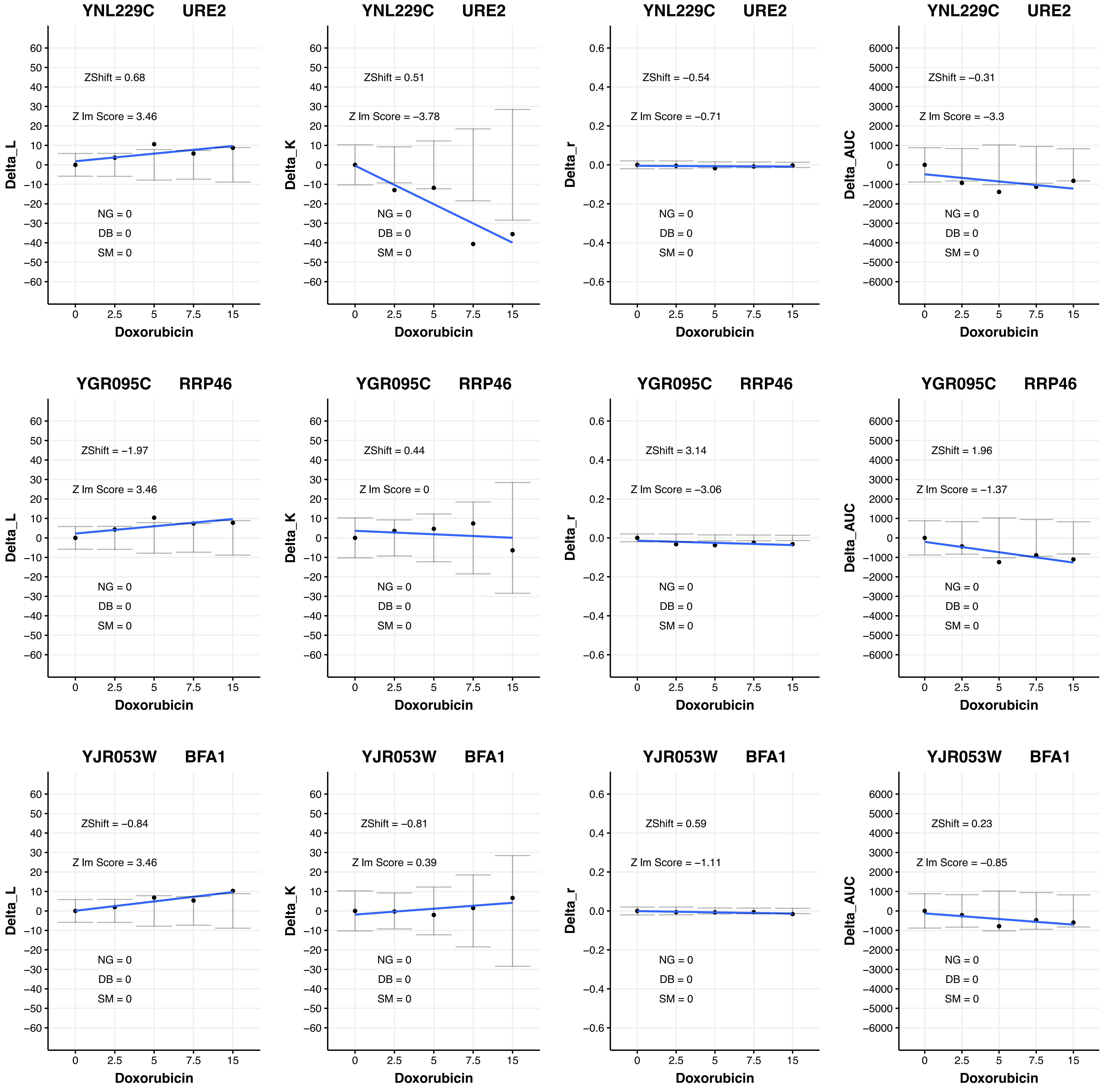
YNL229C URE2
ZShift = 0.68
Z lm Score = 3.46
NG = 0
DB = 0
SM = 0
Delta_L
Doxorubicin
YNL229C URE2
ZShift = 0.51
Z lm Score = −3.78
NG = 0
DB = 0
SM = 0
Delta_K
Doxorubicin
YNL229C URE2
ZShift = −0.54
Z lm Score = −0.71
NG = 0
DB = 0
SM = 0
Delta_r
Doxorubicin
YNL229C URE2
ZShift = −0.31
Z lm Score = −3.3
NG = 0
DB = 0
SM = 0
Delta_AUC
Doxorubicin
YGR095C RRP46
ZShift = −1.97
Z lm Score = 3.46
NG = 0
DB = 0
SM = 0
Delta_L
Doxorubicin
YGR095C RRP46
ZShift = 0.44
Z lm Score = 0
NG = 0
DB = 0
SM = 0
Delta_K
Doxorubicin
YGR095C RRP46
ZShift = 3.14
Z lm Score = −3.06
NG = 0
DB = 0
SM = 0
Delta_r
Doxorubicin
YGR095C RRP46
ZShift = 1.96
Z lm Score = −1.37
NG = 0
DB = 0
SM = 0
Delta_AUC
Doxorubicin
YJR053W BFA1
ZShift = −0.84
Z lm Score = 3.46
NG = 0
DB = 0
SM = 0
Delta_L
Doxorubicin
YJR053W BFA1
ZShift = −0.81
Z lm Score = 0.39
NG = 0
DB = 0
SM = 0
Delta_K
Doxorubicin
YJR053W BFA1
ZShift = 0.59
Z lm Score = −1.11
NG = 0
DB = 0
SM = 0
Delta_r
Doxorubicin
YJR053W BFA1
ZShift = 0.23
Z lm Score = −0.85
NG = 0
DB = 0
SM = 0
Delta_AUC
Doxorubicin

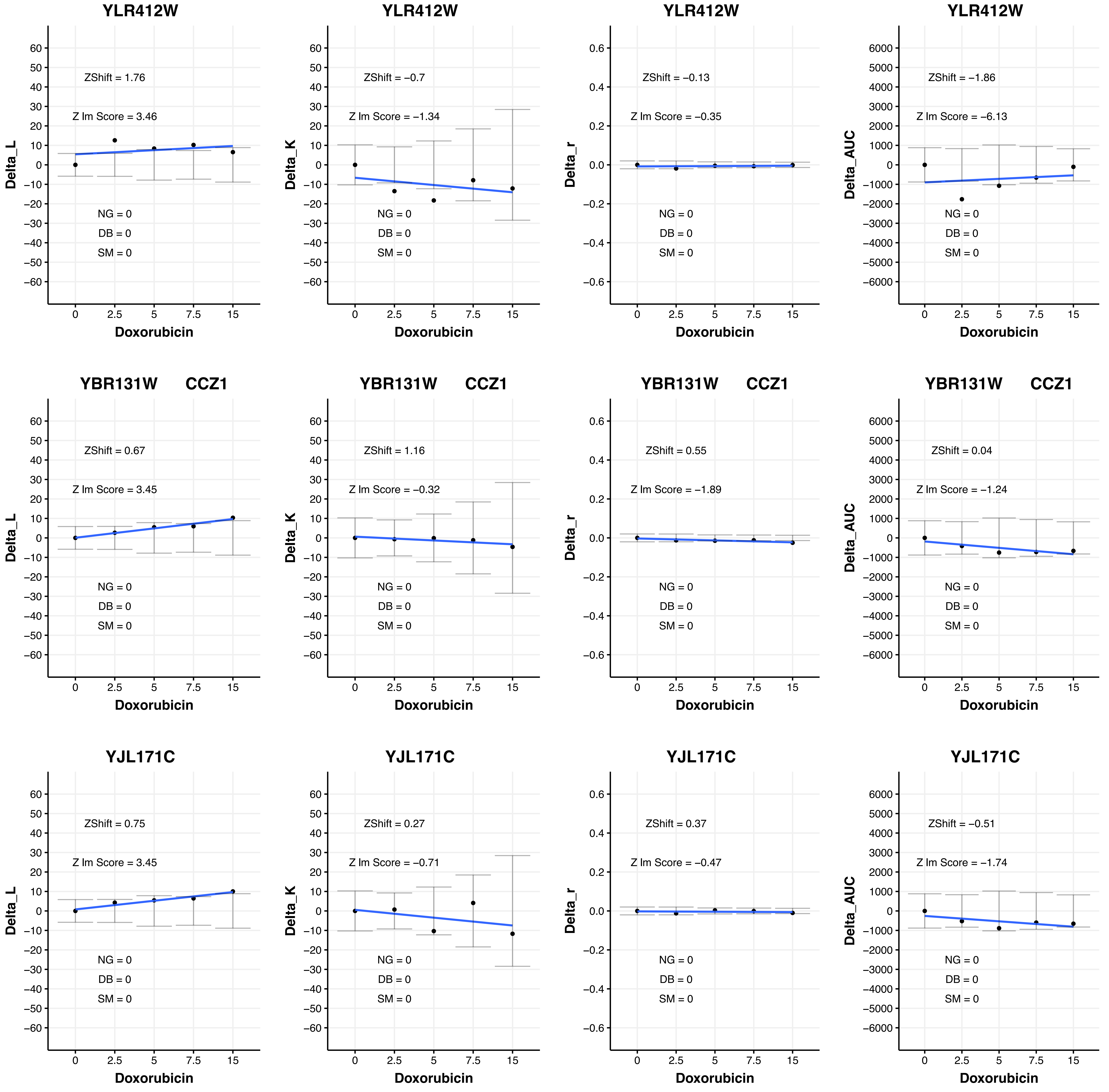
YLR412W
ZShift = 1.76
Z lm Score = 3.46
NG = 0
DB = 0
SM = 0
Delta_L
Doxorubicin
YLR412W
ZShift = −0.7
Z lm Score = −1.34
Delta_K
YLR412W
ZShift = −0.13
Z lm Score = −0.35
Delta_r
YLR412W
ZShift = −1.86
Z lm Score = −6.13
Delta_AUC
YBR131W CCZ1
ZShift = 0.67
Z lm Score = 3.45
ZShift = 1.16
Z lm Score = −0.32
ZShift = 0.55
Z lm Score = −1.89
ZShift = 0.04
Z lm Score = −1.24
YJL171C
ZShift = 0.75
Z lm Score = 3.45
ZShift = 0.27
Z lm Score = −0.71
ZShift = 0.37
Z lm Score = −0.47
ZShift = −0.51
Z lm Score = −1.74

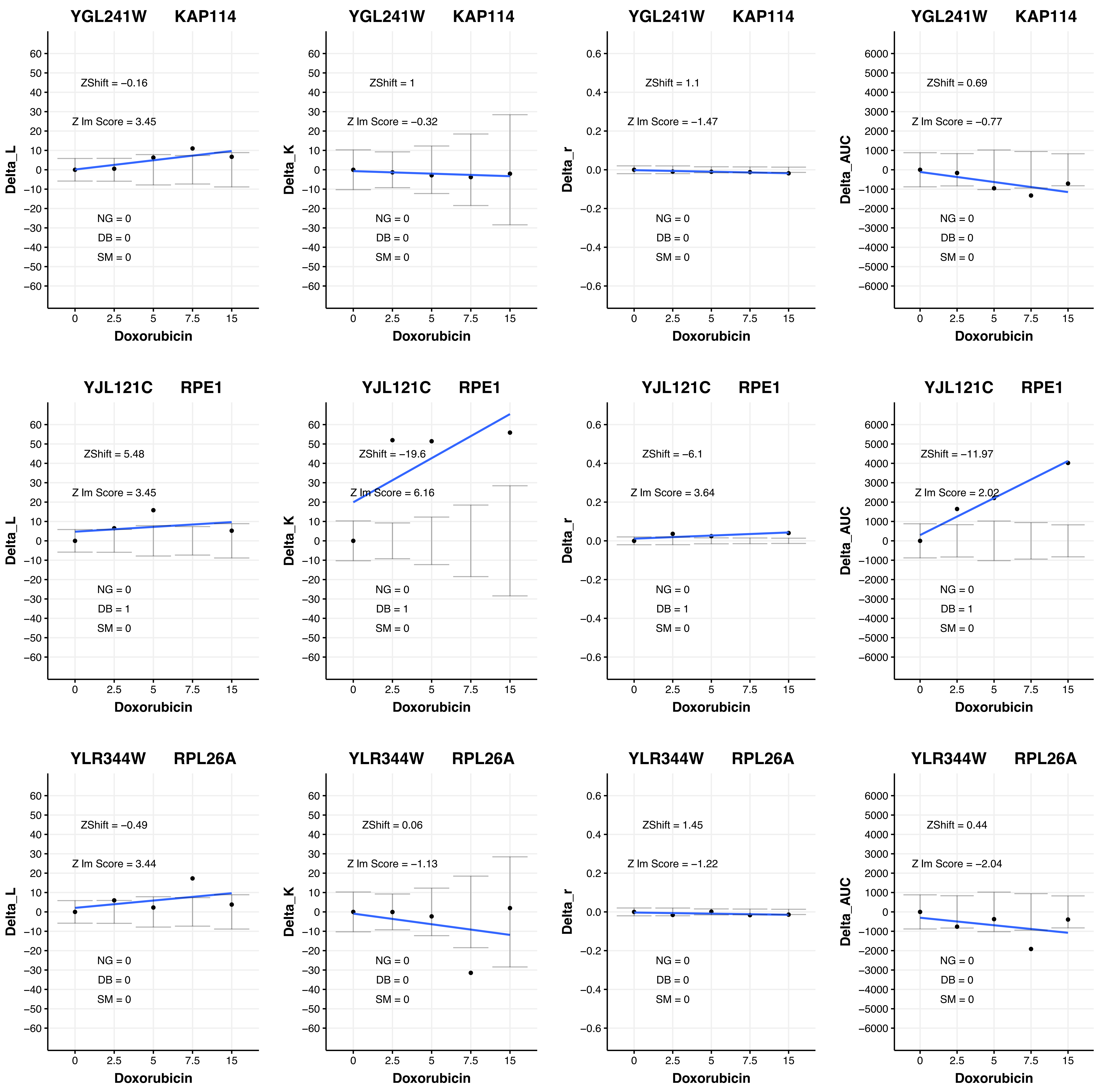
YGL241W KAP114
ZShift = −0.16
Z lm Score = 3.45
NG = 0
DB = 0
SM = 0
Delta_L
Doxorubicin
YGL241W KAP114
ZShift = 1
Z lm Score = −0.32
NG = 0
DB = 0
SM = 0
Delta_K
Doxorubicin
YGL241W KAP114
ZShift = 1.1
Z lm Score = −1.47
NG = 0
DB = 0
SM = 0
Delta_r
Doxorubicin
YGL241W KAP114
ZShift = 0.69
Z lm Score = −0.77
NG = 0
DB = 0
SM = 0
Delta_AUC
Doxorubicin
YJL121C RPE1
ZShift = 5.48
Z lm Score = 3.45
NG = 0
DB = 1
SM = 0
Delta_L
Doxorubicin
YJL121C RPE1
ZShift = −19.6
Z lm Score = 6.16
NG = 0
DB = 1
SM = 0
Delta_K
Doxorubicin
YJL121C RPE1
ZShift = −6.1
Z lm Score = 3.64
NG = 0
DB = 1
SM = 0
Delta_r
Doxorubicin
YJL121C RPE1
ZShift = −11.97
Z lm Score = 2.02
NG = 0
DB = 1
SM = 0
Delta_AUC
Doxorubicin
YLR344W RPL26A
ZShift = −0.49
Z lm Score = 3.44
NG = 0
DB = 0
SM = 0
Delta_L
Doxorubicin
YLR344W RPL26A
ZShift = 0.06
Z lm Score = −1.13
NG = 0
DB = 0
SM = 0
Delta_K
Doxorubicin
YLR344W RPL26A
ZShift = 1.45
Z lm Score = −1.22
NG = 0
DB = 0
SM = 0
Delta_r
Doxorubicin
YLR344W RPL26A
ZShift = 0.44
Z lm Score = −2.04
NG = 0
DB = 0
SM = 0
Delta_AUC
Doxorubicin

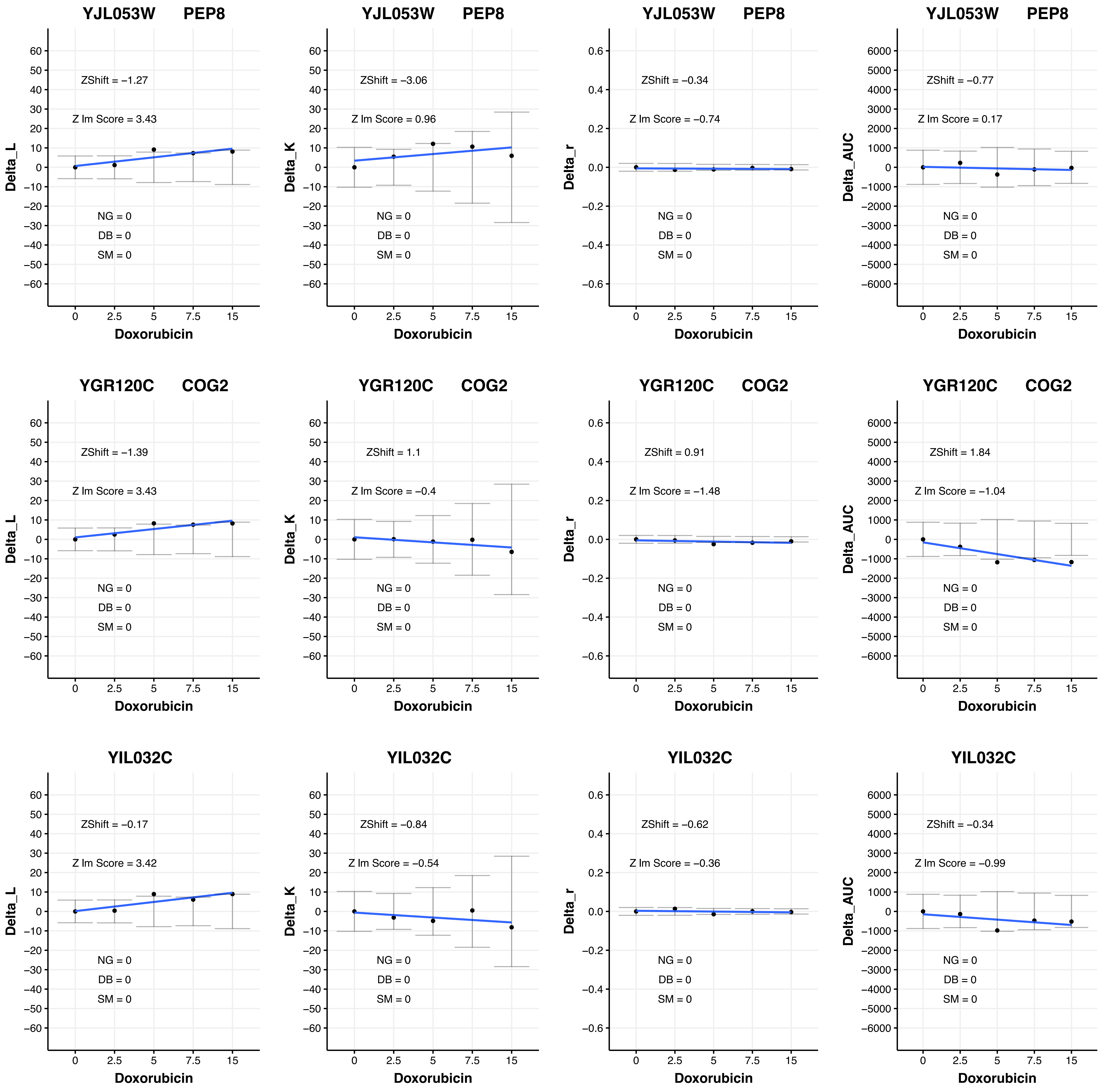
YJL053W PEP8
ZShift = −1.27
Z lm Score = 3.43
NG = 0
DB = 0
SM = 0
Delta_L
Doxorubicin
YJL053W PEP8
ZShift = −3.06
Z lm Score = 0.96
NG = 0
DB = 0
SM = 0
Delta_K
Doxorubicin
YJL053W PEP8
ZShift = −0.34
Z lm Score = −0.74
NG = 0
DB = 0
SM = 0
Delta_r
Doxorubicin
YJL053W PEP8
ZShift = −0.77
Z lm Score = 0.17
NG = 0
DB = 0
SM = 0
Delta_AUC
Doxorubicin
YGR120C COG2
ZShift = −1.39
Z lm Score = 3.43
NG = 0
DB = 0
SM = 0
Delta_L
Doxorubicin
YGR120C COG2
ZShift = 1.1
Z lm Score = −0.4
NG = 0
DB = 0
SM = 0
Delta_K
Doxorubicin
YGR120C COG2
ZShift = 0.91
Z lm Score = −1.48
NG = 0
DB = 0
SM = 0
Delta_r
Doxorubicin
YGR120C COG2
ZShift = 1.84
Z lm Score = −1.04
NG = 0
DB = 0
SM = 0
Delta_AUC
Doxorubicin
YIL032C
ZShift = −0.17
Z lm Score = 3.42
NG = 0
DB = 0
SM = 0
Delta_L
Doxorubicin
YIL032C
ZShift = −0.84
Z lm Score = −0.54
NG = 0
DB = 0
SM = 0
Delta_K
Doxorubicin
YIL032C
ZShift = −0.62
Z lm Score = −0.36
NG = 0
DB = 0
SM = 0
Delta_r
Doxorubicin
YIL032C
ZShift = −0.34
Z lm Score = −0.99
NG = 0
DB = 0
SM = 0
Delta_AUC
Doxorubicin

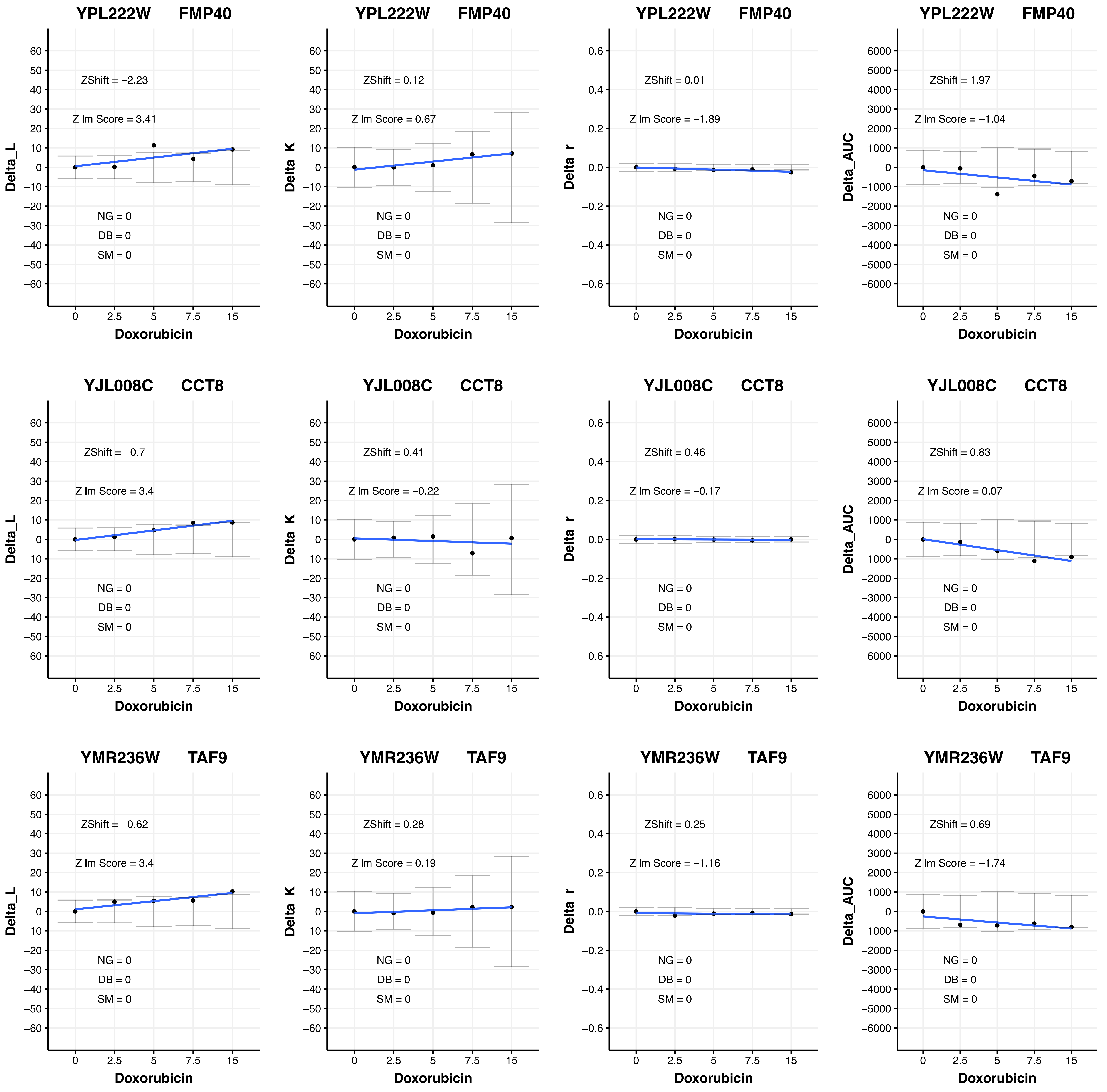
YPL222W FMP40
ZShift = −2.23
Z lm Score = 3.41
NG = 0
DB = 0
SM = 0
Delta_L
Doxorubicin
YPL222W FMP40
ZShift = 0.12
Z lm Score = 0.67
NG = 0
DB = 0
SM = 0
Delta_K
Doxorubicin
YPL222W FMP40
ZShift = 0.01
Z lm Score = −1.89
NG = 0
DB = 0
SM = 0
Delta_r
Doxorubicin
YPL222W FMP40
ZShift = 1.97
Z lm Score = −1.04
NG = 0
DB = 0
SM = 0
Delta_AUC
Doxorubicin
YJL008C CCT8
ZShift = −0.7
Z lm Score = 3.4
NG = 0
DB = 0
SM = 0
Delta_L
Doxorubicin
YJL008C CCT8
ZShift = 0.41
Z lm Score = −0.22
NG = 0
DB = 0
SM = 0
Delta_K
Doxorubicin
YJL008C CCT8
ZShift = 0.46
Z lm Score = −0.17
NG = 0
DB = 0
SM = 0
Delta_r
Doxorubicin
YJL008C CCT8
ZShift = 0.83
Z lm Score = 0.07
NG = 0
DB = 0
SM = 0
Delta_AUC
Doxorubicin
YMR236W TAF9
ZShift = −0.62
Z lm Score = 3.4
NG = 0
DB = 0
SM = 0
Delta_L
Doxorubicin
YMR236W TAF9
ZShift = 0.28
Z lm Score = 0.19
NG = 0
DB = 0
SM = 0
Delta_K
Doxorubicin
YMR236W TAF9
ZShift = 0.25
Z lm Score = −1.16
NG = 0
DB = 0
SM = 0
Delta_r
Doxorubicin
YMR236W TAF9
ZShift = 0.69
Z lm Score = −1.74
NG = 0
DB = 0
SM = 0
Delta_AUC
Doxorubicin

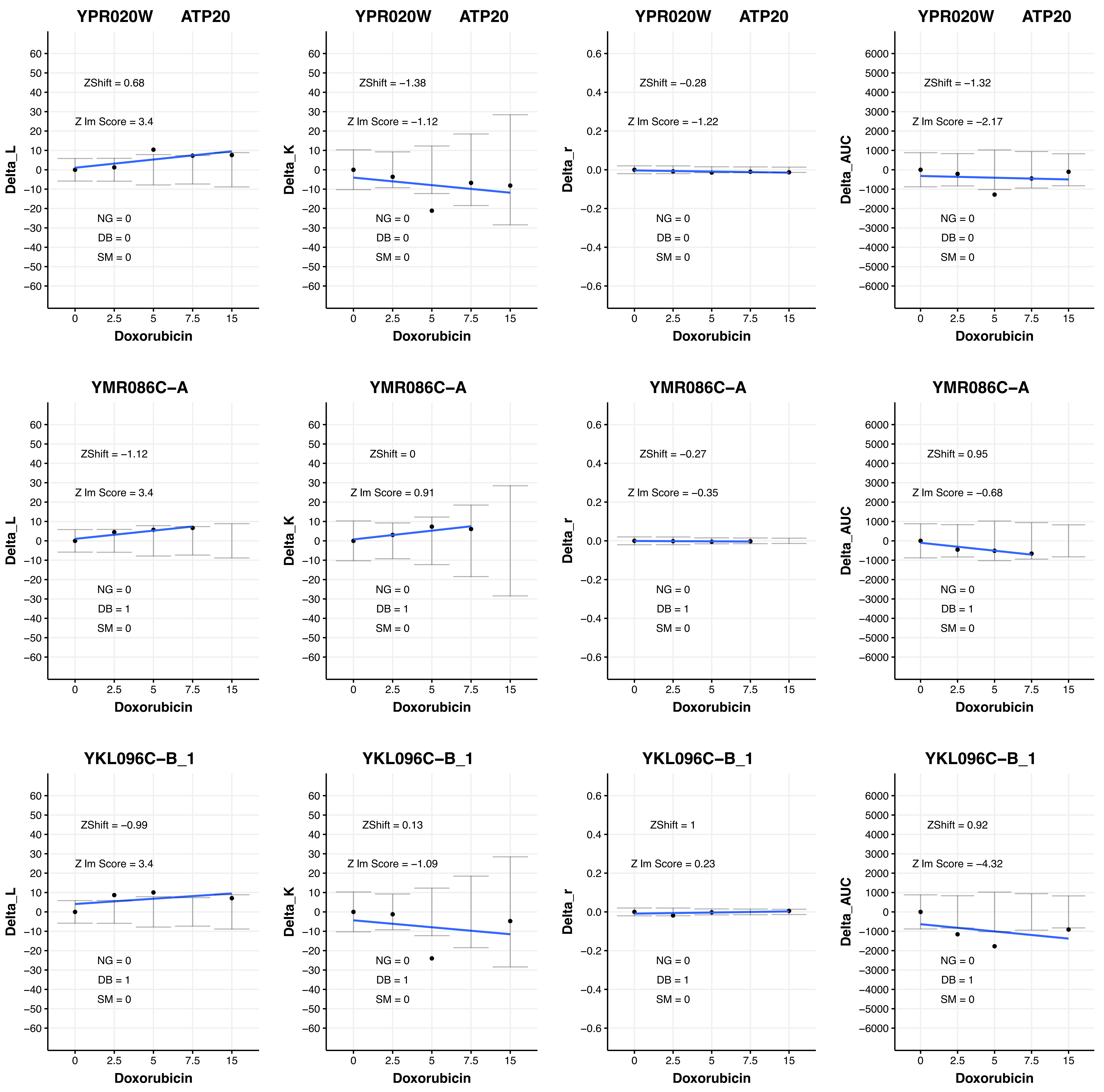
YPR020W ATP20
ZShift = 0.68
Z lm Score = 3.4
NG = 0
DB = 0
SM = 0
Delta_L
Doxorubicin
YPR020W ATP20
ZShift = −1.38
Z lm Score = −1.12
NG = 0
DB = 0
SM = 0
Delta_K
Doxorubicin
YPR020W ATP20
ZShift = −0.28
Z lm Score = −1.22
NG = 0
DB = 0
SM = 0
Delta_r
Doxorubicin
YPR020W ATP20
ZShift = −1.32
Z lm Score = −2.17
NG = 0
DB = 0
SM = 0
Delta_AUC
Doxorubicin
YMR086C−A
ZShift = −1.12
Z lm Score = 3.4
NG = 0
DB = 1
SM = 0
Delta_L
Doxorubicin
YMR086C−A
ZShift = 0
Z lm Score = 0.91
NG = 0
DB = 1
SM = 0
Delta_K
Doxorubicin
YMR086C−A
ZShift = −0.27
Z lm Score = −0.35
NG = 0
DB = 1
SM = 0
Delta_r
Doxorubicin
YMR086C−A
ZShift = 0.95
Z lm Score = −0.68
NG = 0
DB = 1
SM = 0
Delta_AUC
Doxorubicin
YKL096C−B_1
ZShift = −0.99
Z lm Score = 3.4
NG = 0
DB = 1
SM = 0
Delta_L
Doxorubicin
YKL096C−B_1
ZShift = 0.13
Z lm Score = −1.09
NG = 0
DB = 1
SM = 0
Delta_K
Doxorubicin
YKL096C−B_1
ZShift = 1
Z lm Score = 0.23
NG = 0
DB = 1
SM = 0
Delta_r
Doxorubicin
YKL096C−B_1
ZShift = 0.92
Z lm Score = −4.32
NG = 0
DB = 1
SM = 0
Delta_AUC
Doxorubicin

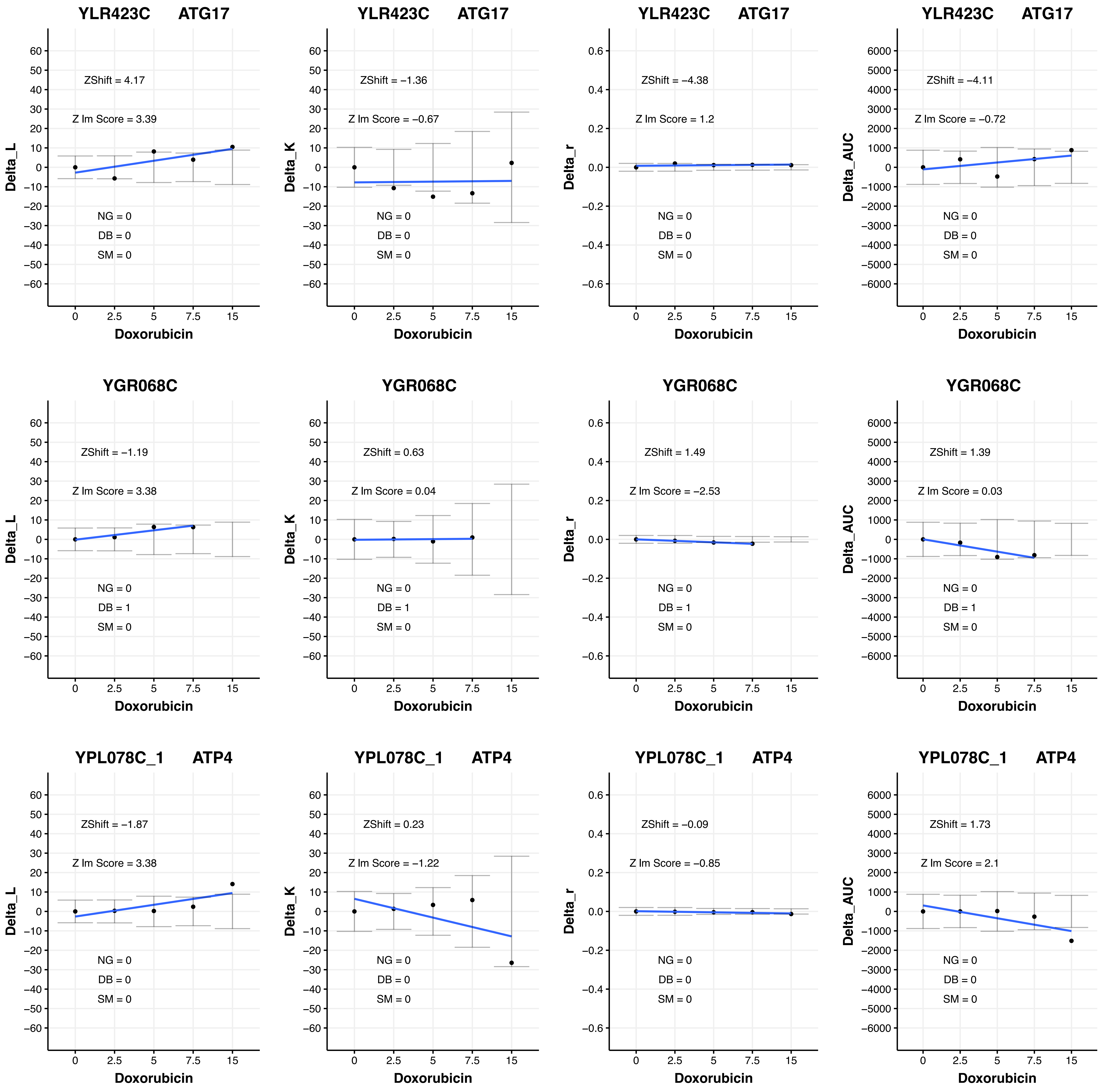
YLR423C ATG17
ZShift = 4.17
Z lm Score = 3.39
NG = 0
DB = 0
SM = 0
Delta_L
Doxorubicin
YLR423C ATG17
ZShift = −1.36
Z lm Score = −0.67
NG = 0
DB = 0
SM = 0
Delta_K
Doxorubicin
YLR423C ATG17
ZShift = −4.38
Z lm Score = 1.2
NG = 0
DB = 0
SM = 0
Delta_r
Doxorubicin
YLR423C ATG17
ZShift = −4.11
Z lm Score = −0.72
NG = 0
DB = 0
SM = 0
Delta_AUC
Doxorubicin
YGR068C
ZShift = −1.19
Z lm Score = 3.38
NG = 0
DB = 1
SM = 0
Delta_L
Doxorubicin
YGR068C
ZShift = 0.63
Z lm Score = 0.04
NG = 0
DB = 1
SM = 0
Delta_K
Doxorubicin
YGR068C
ZShift = 1.49
Z lm Score = −2.53
NG = 0
DB = 1
SM = 0
Delta_r
Doxorubicin
YGR068C
ZShift = 1.39
Z lm Score = 0.03
NG = 0
DB = 1
SM = 0
Delta_AUC
Doxorubicin
YPL078C_1 ATP4
ZShift = −1.87
Z lm Score = 3.38
NG = 0
DB = 0
SM = 0
Delta_L
Doxorubicin
YPL078C_1 ATP4
ZShift = 0.23
Z lm Score = −1.22
NG = 0
DB = 0
SM = 0
Delta_K
Doxorubicin
YPL078C_1 ATP4
ZShift = −0.09
Z lm Score = −0.85
NG = 0
DB = 0
SM = 0
Delta_r
Doxorubicin
YPL078C_1 ATP4
ZShift = 1.73
Z lm Score = 2.1
NG = 0
DB = 0
SM = 0
Delta_AUC
Doxorubicin

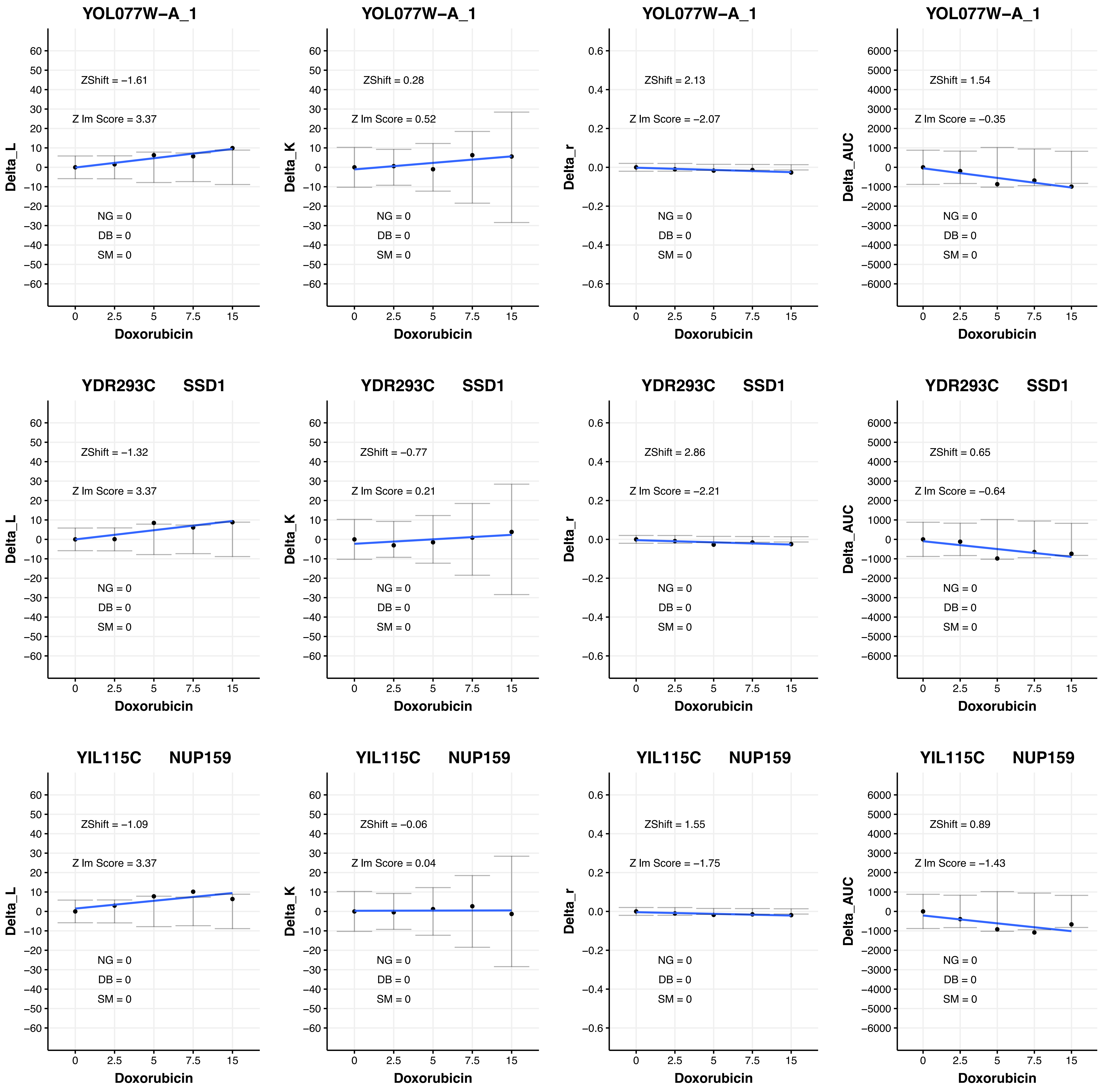

YOL077W−A_1
ZShift = −1.61
Z lm Score = 3.37
NG = 0
DB = 0
SM = 0
Delta_L
Doxorubicin
YOL077W−A_1
ZShift = 0.28
Z lm Score = 0.52
NG = 0
DB = 0
SM = 0
Delta_K
Doxorubicin
YOL077W−A_1
ZShift = 2.13
Z lm Score = −2.07
NG = 0
DB = 0
SM = 0
Delta_r
Doxorubicin
YOL077W−A_1
ZShift = 1.54
Z lm Score = −0.35
NG = 0
DB = 0
SM = 0
Delta_AUC
Doxorubicin
YDR293C SSD1
ZShift = −1.32
Z lm Score = 3.37
NG = 0
DB = 0
SM = 0
Delta_L
Doxorubicin
YDR293C SSD1
ZShift = −0.77
Z lm Score = 0.21
NG = 0
DB = 0
SM = 0
Delta_K
Doxorubicin
YDR293C SSD1
ZShift = 2.86
Z lm Score = −2.21
NG = 0
DB = 0
SM = 0
Delta_r
Doxorubicin
YDR293C SSD1
ZShift = 0.65
Z lm Score = −0.64
NG = 0
DB = 0
SM = 0
Delta_AUC
Doxorubicin
YIL115C NUP159
ZShift = −1.09
Z lm Score = 3.37
NG = 0
DB = 0
SM = 0
Delta_L
Doxorubicin
YIL115C NUP159
ZShift = −0.06
Z lm Score = 0.04
NG = 0
DB = 0
SM = 0
Delta_K
Doxorubicin
YIL115C NUP159
ZShift = 1.55
Z lm Score = −1.75
NG = 0
DB = 0
SM = 0
Delta_r
Doxorubicin
YIL115C NUP159
ZShift = 0.89
Z lm Score = −1.43
NG = 0
DB = 0
SM = 0
Delta_AUC
Doxorubicin

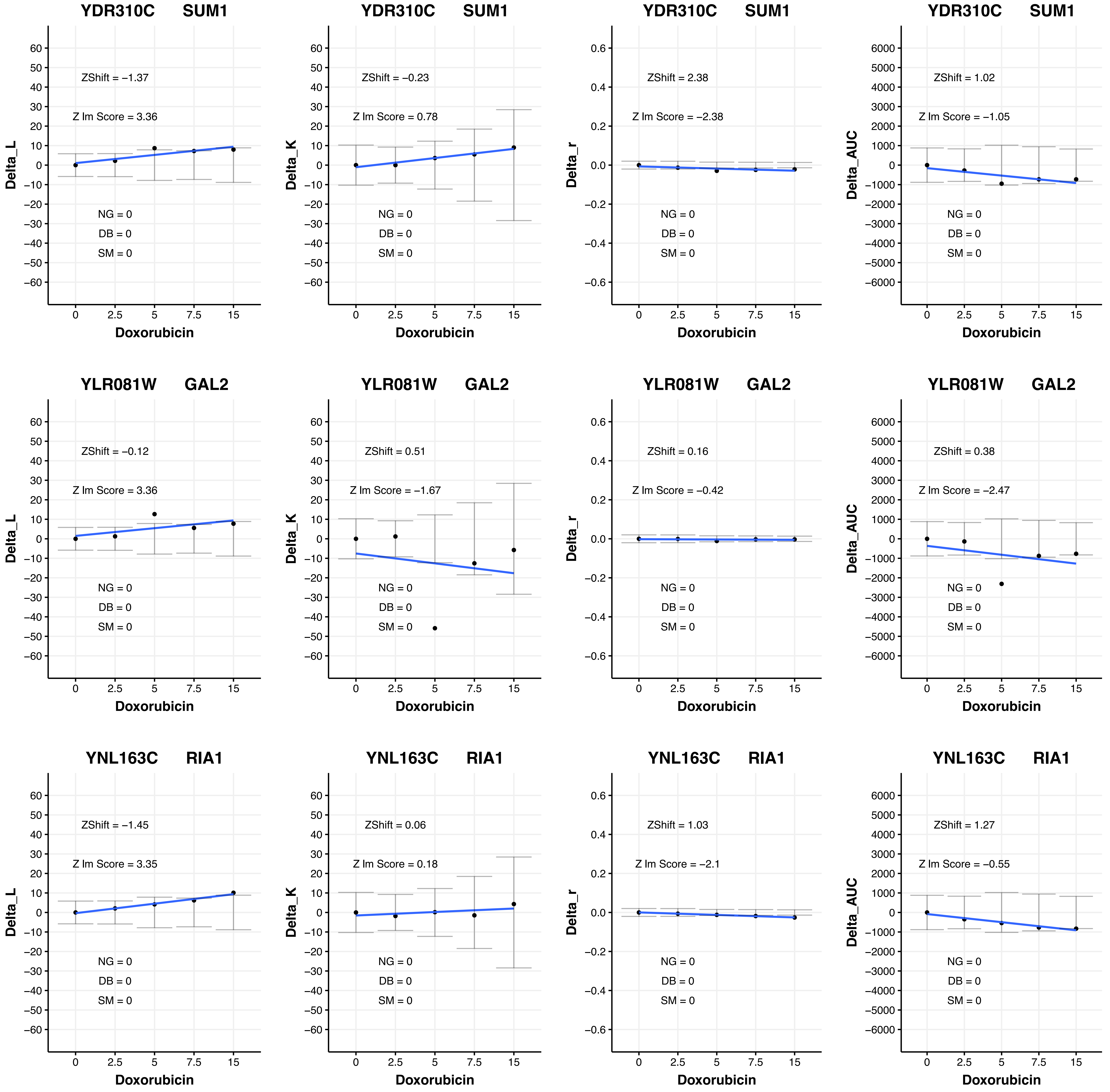
YDR310C SUM1
ZShift = –1.37
Z lm Score = 3.36
NG = 0
DB = 0
SM = 0
Delta_L
Doxorubicin
YDR310C SUM1
ZShift = –0.23
Z lm Score = 0.78
NG = 0
DB = 0
SM = 0
Delta_K
Doxorubicin
YDR310C SUM1
ZShift = 2.38
Z lm Score = –2.38
NG = 0
DB = 0
SM = 0
Delta_r
Doxorubicin
YDR310C SUM1
ZShift = 1.02
Z lm Score = –1.05
NG = 0
DB = 0
SM = 0
Delta_AUC
Doxorubicin
YLR081W GAL2
ZShift = –0.12
Z lm Score = 3.36
NG = 0
DB = 0
SM = 0
Delta_L
Doxorubicin
YLR081W GAL2
ZShift = 0.51
Z lm Score = –1.67
NG = 0
DB = 0
SM = 0
Delta_K
Doxorubicin
YLR081W GAL2
ZShift = 0.16
Z lm Score = –0.42
NG = 0
DB = 0
SM = 0
Delta_r
Doxorubicin
YLR081W GAL2
ZShift = 0.38
Z lm Score = –2.47
NG = 0
DB = 0
SM = 0
Delta_AUC
Doxorubicin
YNL163C RIA1
ZShift = –1.45
Z lm Score = 3.35
NG = 0
DB = 0
SM = 0
Delta_L
Doxorubicin
YNL163C RIA1
ZShift = 0.06
Z lm Score = 0.18
NG = 0
DB = 0
SM = 0
Delta_K
Doxorubicin
YNL163C RIA1
ZShift = 1.03
Z lm Score = –2.1
NG = 0
DB = 0
SM = 0
Delta_r
Doxorubicin
YNL163C RIA1
ZShift = 1.27
Z lm Score = –0.55
NG = 0
DB = 0
SM = 0
Delta_AUC
Doxorubicin

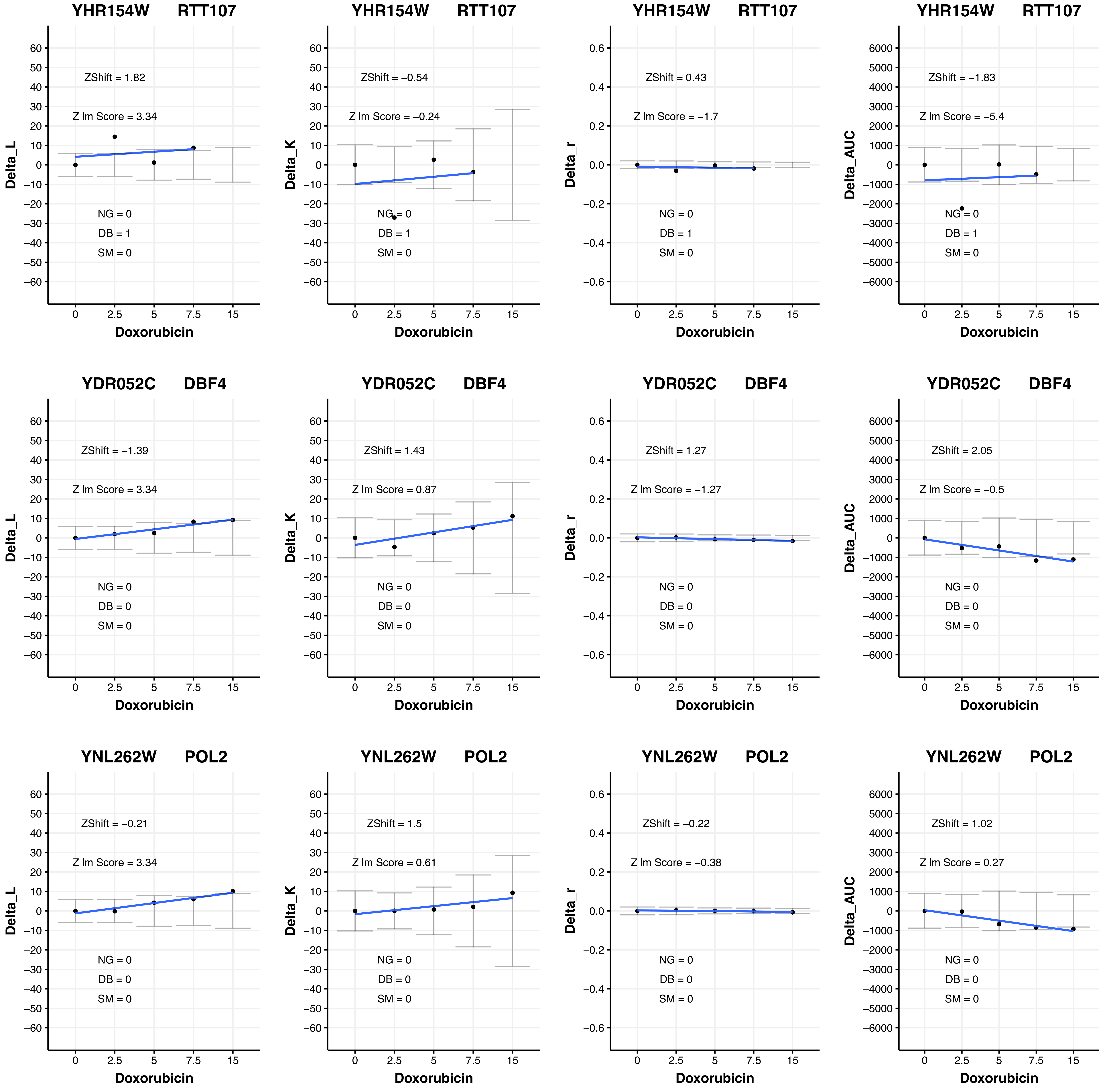

YHR154W RTT107
ZShift = 1.82
Z lm Score = 3.34
NG = 0
DB = 1
SM = 0
Delta_L
Doxorubicin
YHR154W RTT107
ZShift = −0.54
Z lm Score = −0.24
NG = 0
DB = 1
SM = 0
Delta_K
Doxorubicin
YHR154W RTT107
ZShift = 0.43
Z lm Score = −1.7
NG = 0
DB = 1
SM = 0
Delta_r
Doxorubicin
YHR154W RTT107
ZShift = −1.83
Z lm Score = −5.4
NG = 0
DB = 1
SM = 0
Delta_AUC
Doxorubicin
YDR052C DBF4
ZShift = −1.39
Z lm Score = 3.34
NG = 0
DB = 0
SM = 0
Delta_L
Doxorubicin
YDR052C DBF4
ZShift = 1.43
Z lm Score = 0.87
NG = 0
DB = 0
SM = 0
Delta_K
Doxorubicin
YDR052C DBF4
ZShift = 1.27
Z lm Score = −1.27
NG = 0
DB = 0
SM = 0
Delta_r
Doxorubicin
YDR052C DBF4
ZShift = 2.05
Z lm Score = −0.5
NG = 0
DB = 0
SM = 0
Delta_AUC
Doxorubicin
YNL262W POL2
ZShift = −0.21
Z lm Score = 3.34
NG = 0
DB = 0
SM = 0
Delta_L
Doxorubicin
YNL262W POL2
ZShift = 1.5
Z lm Score = 0.61
NG = 0
DB = 0
SM = 0
Delta_K
Doxorubicin
YNL262W POL2
ZShift = −0.22
Z lm Score = −0.38
NG = 0
DB = 0
SM = 0
Delta_r
Doxorubicin
YNL262W POL2
ZShift = 1.02
Z lm Score = 0.27
NG = 0
DB = 0
SM = 0
Delta_AUC
Doxorubicin

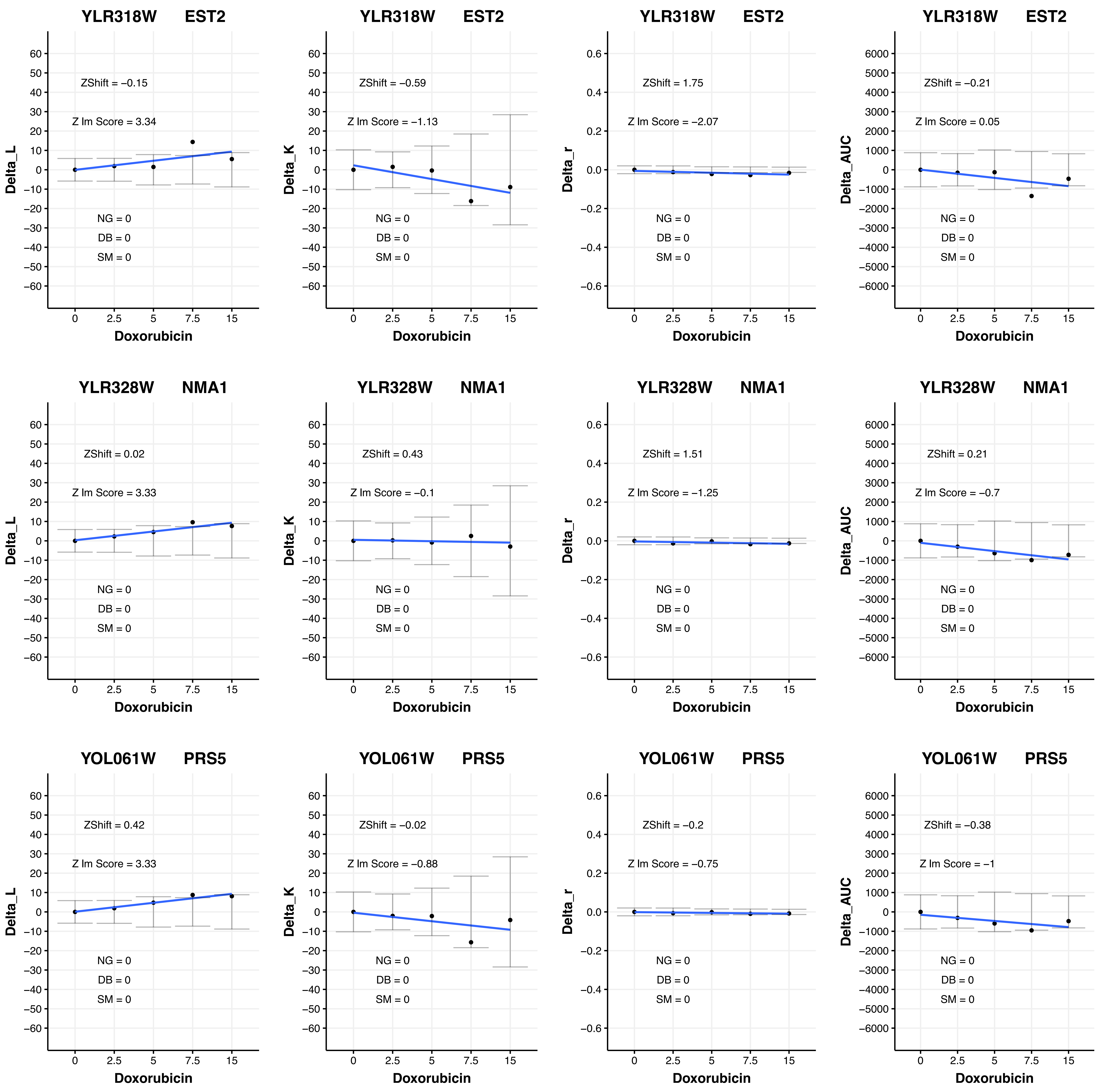
YLR318W EST2
ZShift = −0.15
Z lm Score = 3.34
NG = 0
DB = 0
SM = 0
Delta_L
Doxorubicin
YLR318W EST2
ZShift = −0.59
Z lm Score = −1.13
NG = 0
DB = 0
SM = 0
Delta_K
Doxorubicin
YLR318W EST2
ZShift = 1.75
Z lm Score = −2.07
NG = 0
DB = 0
SM = 0
Delta_r
Doxorubicin
YLR318W EST2
ZShift = −0.21
Z lm Score = 0.05
NG = 0
DB = 0
SM = 0
Delta_AUC
Doxorubicin
YLR328W NMA1
ZShift = 0.02
Z lm Score = 3.33
NG = 0
DB = 0
SM = 0
Delta_L
Doxorubicin
YLR328W NMA1
ZShift = 0.43
Z lm Score = −0.1
NG = 0
DB = 0
SM = 0
Delta_K
Doxorubicin
YLR328W NMA1
ZShift = 1.51
Z lm Score = −1.25
NG = 0
DB = 0
SM = 0
Delta_r
Doxorubicin
YLR328W NMA1
ZShift = 0.21
Z lm Score = −0.7
NG = 0
DB = 0
SM = 0
Delta_AUC
Doxorubicin
YOL061W PRS5
ZShift = 0.42
Z lm Score = 3.33
NG = 0
DB = 0
SM = 0
Delta_L
Doxorubicin
YOL061W PRS5
ZShift = −0.02
Z lm Score = −0.88
NG = 0
DB = 0
SM = 0
Delta_K
Doxorubicin
YOL061W PRS5
ZShift = −0.2
Z lm Score = −0.75
NG = 0
DB = 0
SM = 0
Delta_r
Doxorubicin
YOL061W PRS5
ZShift = −0.38
Z lm Score = −1
NG = 0
DB = 0
SM = 0
Delta_AUC
Doxorubicin

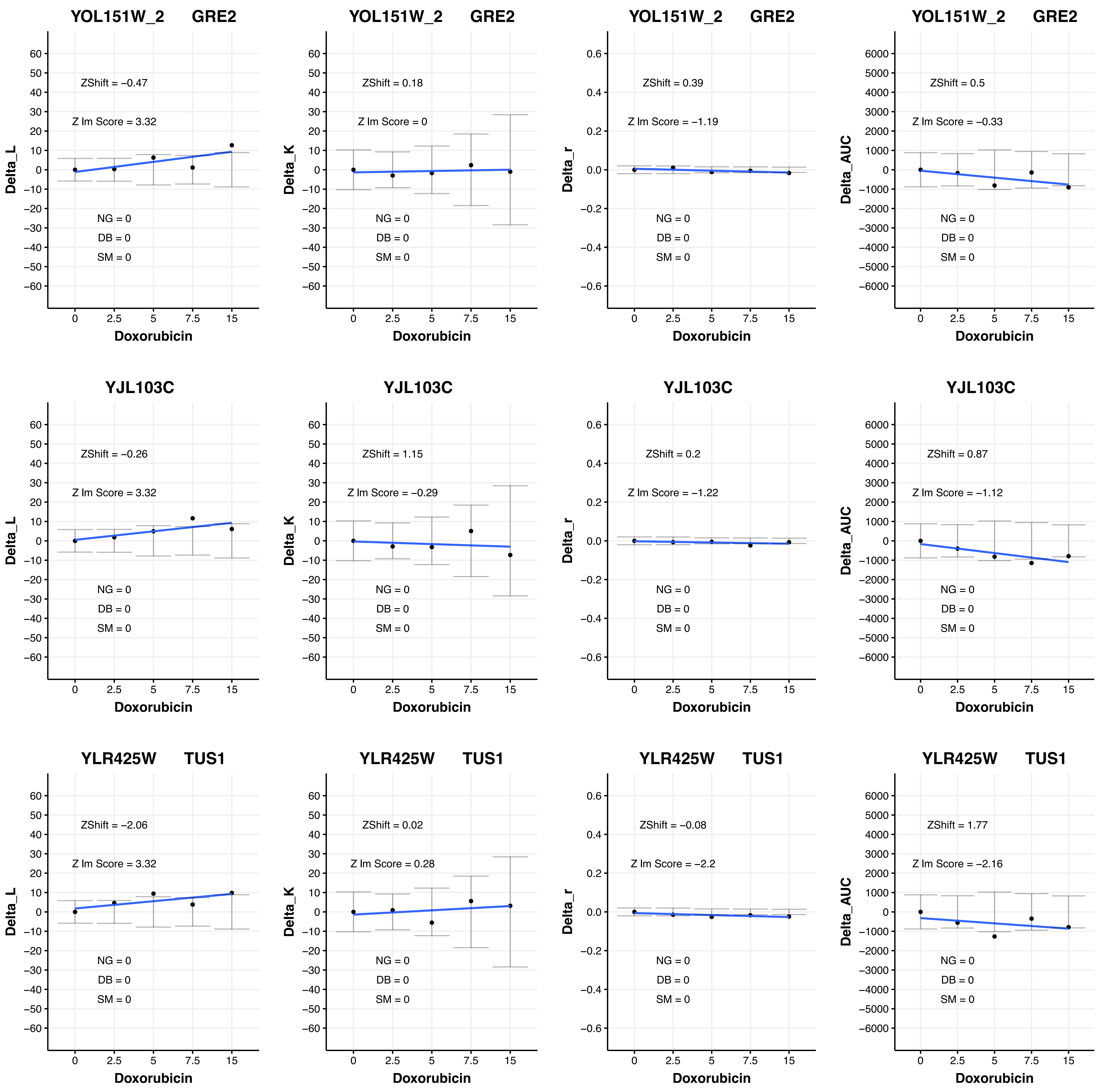

YOL151W_2 GRE2
ZShift = −0.47
Z lm Score = 3.32
NG = 0
DB = 0
SM = 0
Delta_L
Doxorubicin
YOL151W_2 GRE2
ZShift = 0.18
Z lm Score = 0
NG = 0
DB = 0
SM = 0
Delta_K
Doxorubicin
YOL151W_2 GRE2
ZShift = 0.39
Z lm Score = −1.19
NG = 0
DB = 0
SM = 0
Delta_r
Doxorubicin
YOL151W_2 GRE2
ZShift = 0.5
Z lm Score = −0.33
NG = 0
DB = 0
SM = 0
Delta_AUC
Doxorubicin
YJL103C
ZShift = −0.26
Z lm Score = 3.32
NG = 0
DB = 0
SM = 0
Delta_L
Doxorubicin
YJL103C
ZShift = 1.15
Z lm Score = −0.29
NG = 0
DB = 0
SM = 0
Delta_K
Doxorubicin
YJL103C
ZShift = 0.2
Z lm Score = −1.22
NG = 0
DB = 0
SM = 0
Delta_r
Doxorubicin
YJL103C
ZShift = 0.87
Z lm Score = −1.12
NG = 0
DB = 0
SM = 0
Delta_AUC
Doxorubicin
YLR425W TUS1
ZShift = −2.06
Z lm Score = 3.32
NG = 0
DB = 0
SM = 0
Delta_L
Doxorubicin
YLR425W TUS1
ZShift = 0.02
Z lm Score = 0.28
NG = 0
DB = 0
SM = 0
Delta_K
Doxorubicin
YLR425W TUS1
ZShift = −0.08
Z lm Score = −2.2
NG = 0
DB = 0
SM = 0
Delta_r
Doxorubicin
YLR425W TUS1
ZShift = 1.77
Z lm Score = −2.16
NG = 0
DB = 0
SM = 0
Delta_AUC
Doxorubicin

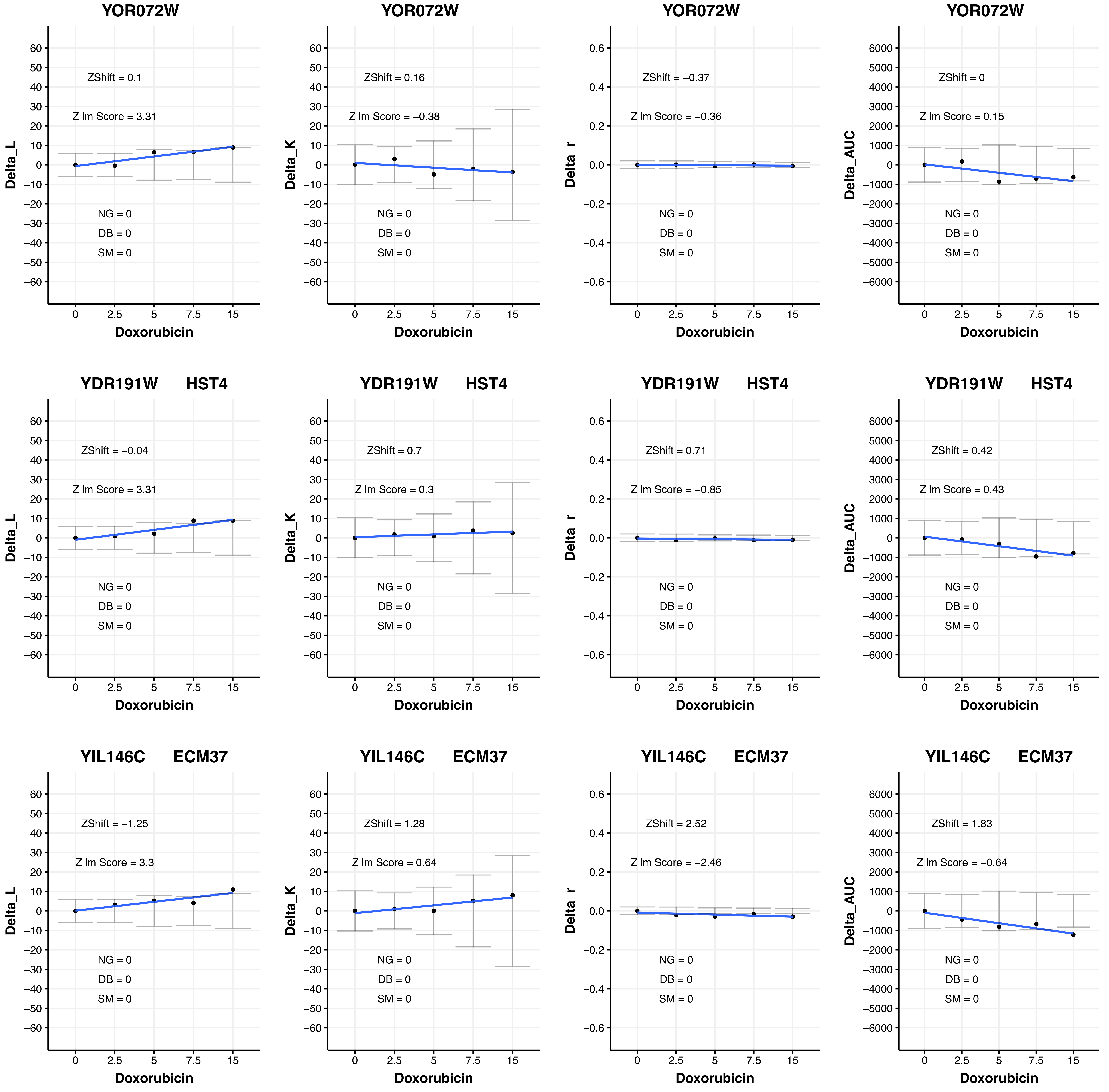
YOR072W
ZShift = 0.1
Z lm Score = 3.31
NG = 0
DB = 0
SM = 0
Delta_L
Doxorubicin
YOR072W
ZShift = 0.16
Z lm Score = −0.38
NG = 0
DB = 0
SM = 0
Delta_K
Doxorubicin
YOR072W
ZShift = −0.37
Z lm Score = −0.36
NG = 0
DB = 0
SM = 0
Delta_r
Doxorubicin
YOR072W
ZShift = 0
Z lm Score = 0.15
NG = 0
DB = 0
SM = 0
Delta_AUC
Doxorubicin
YDR191W HST4
ZShift = −0.04
Z lm Score = 3.31
NG = 0
DB = 0
SM = 0
Delta_L
Doxorubicin
YDR191W HST4
ZShift = 0.7
Z lm Score = 0.3
NG = 0
DB = 0
SM = 0
Delta_K
Doxorubicin
YDR191W HST4
ZShift = 0.71
Z lm Score = −0.85
NG = 0
DB = 0
SM = 0
Delta_r
Doxorubicin
YDR191W HST4
ZShift = 0.42
Z lm Score = 0.43
NG = 0
DB = 0
SM = 0
Delta_AUC
Doxorubicin
YIL146C ECM37
ZShift = −1.25
Z lm Score = 3.3
NG = 0
DB = 0
SM = 0
Delta_L
Doxorubicin
YIL146C ECM37
ZShift = 1.28
Z lm Score = 0.64
NG = 0
DB = 0
SM = 0
Delta_K
Doxorubicin
YIL146C ECM37
ZShift = 2.52
Z lm Score = −2.46
NG = 0
DB = 0
SM = 0
Delta_r
Doxorubicin
YIL146C ECM37
ZShift = 1.83
Z lm Score = −0.64
NG = 0
DB = 0
SM = 0
Delta_AUC
Doxorubicin

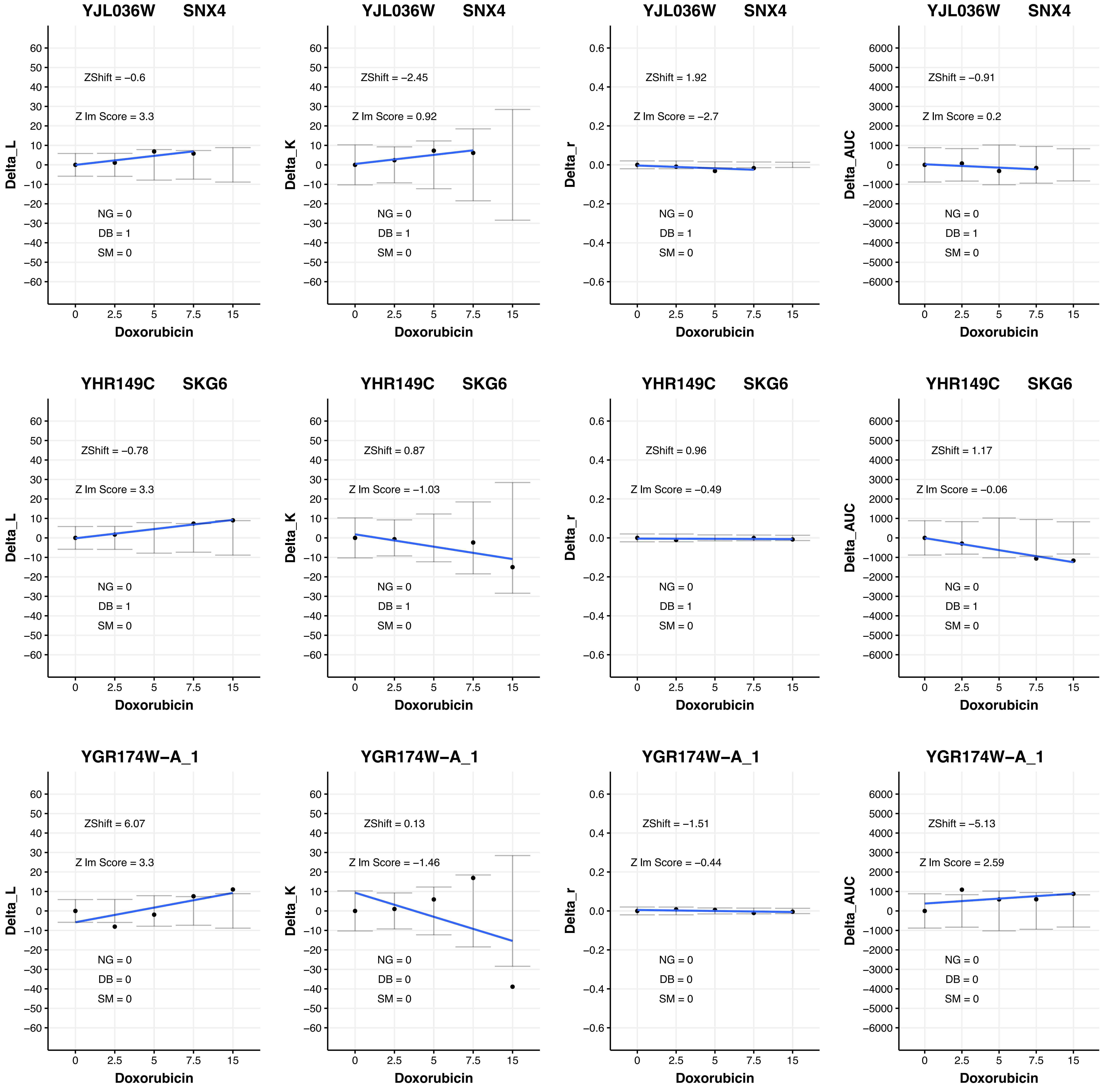
YJL036W SNX4
ZShift = −0.6
Z lm Score = 3.3
NG = 0
DB = 1
SM = 0
Delta_L
Doxorubicin
YJL036W SNX4
ZShift = −2.45
Z lm Score = 0.92
NG = 0
DB = 1
SM = 0
Delta_K
Doxorubicin
YJL036W SNX4
ZShift = 1.92
Z lm Score = −2.7
NG = 0
DB = 1
SM = 0
Delta_r
Doxorubicin
YJL036W SNX4
ZShift = −0.91
Z lm Score = 0.2
NG = 0
DB = 1
SM = 0
Delta_AUC
Doxorubicin
YHR149C SKG6
ZShift = −0.78
Z lm Score = 3.3
NG = 0
DB = 1
SM = 0
Delta_L
Doxorubicin
YHR149C SKG6
ZShift = 0.87
Z lm Score = −1.03
NG = 0
DB = 1
SM = 0
Delta_K
Doxorubicin
YHR149C SKG6
ZShift = 0.96
Z lm Score = −0.49
NG = 0
DB = 1
SM = 0
Delta_r
Doxorubicin
YHR149C SKG6
ZShift = 1.17
Z lm Score = −0.06
NG = 0
DB = 1
SM = 0
Delta_AUC
Doxorubicin
YGR174W−A_1
ZShift = 6.07
Z lm Score = 3.3
NG = 0
DB = 0
SM = 0
Delta_L
Doxorubicin
YGR174W−A_1
ZShift = 0.13
Z lm Score = −1.46
NG = 0
DB = 0
SM = 0
Delta_K
Doxorubicin
YGR174W−A_1
ZShift = −1.51
Z lm Score = −0.44
NG = 0
DB = 0
SM = 0
Delta_r
Doxorubicin
YGR174W−A_1
ZShift = −5.13
Z lm Score = 2.59
NG = 0
DB = 0
SM = 0
Delta_AUC
Doxorubicin

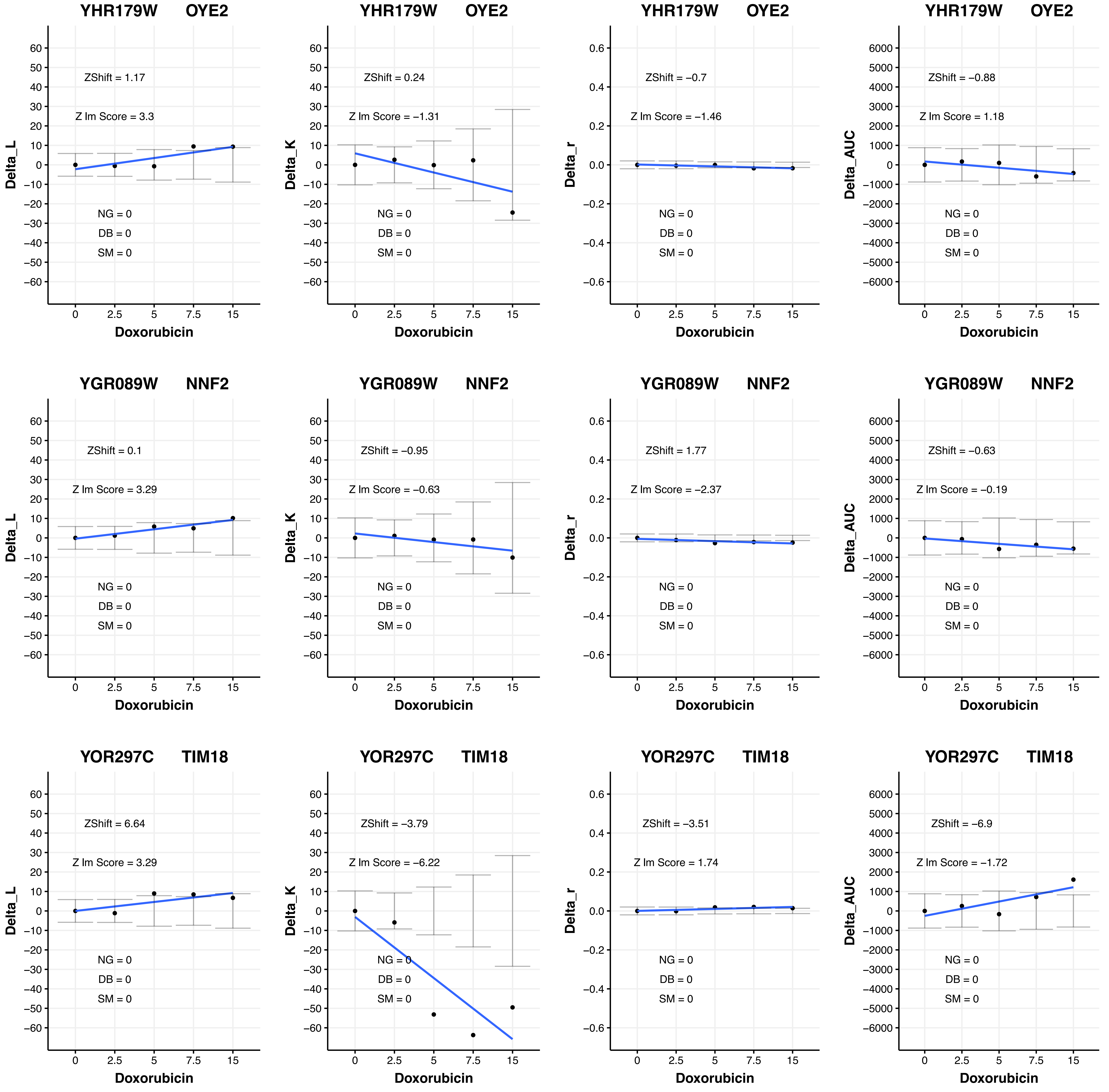
YHR179W OYE2
ZShift = 1.17
Z lm Score = 3.3
NG = 0
DB = 0
SM = 0
Delta_L
Doxorubicin
YHR179W OYE2
ZShift = 0.24
Z lm Score = −1.31
NG = 0
DB = 0
SM = 0
Delta_K
Doxorubicin
YHR179W OYE2
ZShift = −0.7
Z lm Score = −1.46
NG = 0
DB = 0
SM = 0
Delta_r
Doxorubicin
YHR179W OYE2
ZShift = −0.88
Z lm Score = 1.18
NG = 0
DB = 0
SM = 0
Delta_AUC
Doxorubicin
YGR089W NNF2
ZShift = 0.1
Z lm Score = 3.29
NG = 0
DB = 0
SM = 0
Delta_L
Doxorubicin
YGR089W NNF2
ZShift = −0.95
Z lm Score = −0.63
NG = 0
DB = 0
SM = 0
Delta_K
Doxorubicin
YGR089W NNF2
ZShift = 1.77
Z lm Score = −2.37
NG = 0
DB = 0
SM = 0
Delta_r
Doxorubicin
YGR089W NNF2
ZShift = −0.63
Z lm Score = −0.19
NG = 0
DB = 0
SM = 0
Delta_AUC
Doxorubicin
YOR297C TIM18
ZShift = 6.64
Z lm Score = 3.29
NG = 0
DB = 0
SM = 0
Delta_L
Doxorubicin
YOR297C TIM18
ZShift = −3.79
Z lm Score = −6.22
NG = 0
DB = 0
SM = 0
Delta_K
Doxorubicin
YOR297C TIM18
ZShift = −3.51
Z lm Score = 1.74
NG = 0
DB = 0
SM = 0
Delta_r
Doxorubicin
YOR297C TIM18
ZShift = −6.9
Z lm Score = −1.72
NG = 0
DB = 0
SM = 0
Delta_AUC
Doxorubicin

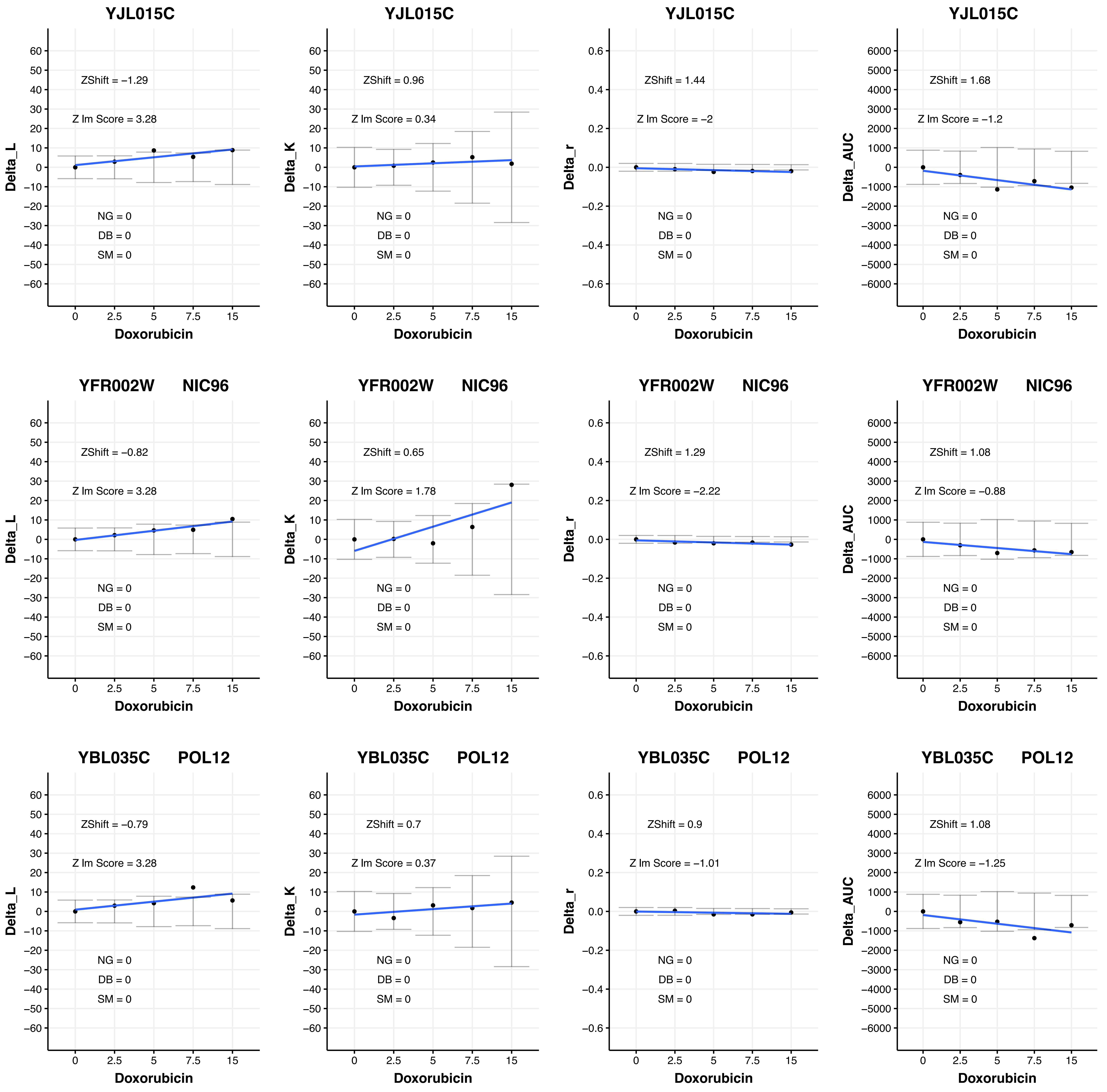

YJL015C
Delta_L
ZShift = −1.29
Z lm Score = 3.28
NG = 0
DB = 0
SM = 0
Doxorubicin
YJL015C
Delta_K
ZShift = 0.96
Z lm Score = 0.34
NG = 0
DB = 0
SM = 0
Doxorubicin
YJL015C
Delta_r
ZShift = 1.44
Z lm Score = −2
NG = 0
DB = 0
SM = 0
Doxorubicin
YJL015C
Delta_AUC
ZShift = 1.68
Z lm Score = −1.2
NG = 0
DB = 0
SM = 0
Doxorubicin
YFR002W NIC96
Delta_L
ZShift = −0.82
Z lm Score = 3.28
NG = 0
DB = 0
SM = 0
Doxorubicin
YFR002W NIC96
Delta_K
ZShift = 0.65
Z lm Score = 1.78
NG = 0
DB = 0
SM = 0
Doxorubicin
YFR002W NIC96
Delta_r
ZShift = 1.29
Z lm Score = −2.22
NG = 0
DB = 0
SM = 0
Doxorubicin
YFR002W NIC96
Delta_AUC
ZShift = 1.08
Z lm Score = −0.88
NG = 0
DB = 0
SM = 0
Doxorubicin
YBL035C POL12
Delta_L
ZShift = −0.79
Z lm Score = 3.28
NG = 0
DB = 0
SM = 0
Doxorubicin
YBL035C POL12
Delta_K
ZShift = 0.7
Z lm Score = 0.37
NG = 0
DB = 0
SM = 0
Doxorubicin
YBL035C POL12
Delta_r
ZShift = 0.9
Z lm Score = −1.01
NG = 0
DB = 0
SM = 0
Doxorubicin
YBL035C POL12
Delta_AUC
ZShift = 1.08
Z lm Score = −1.25
NG = 0
DB = 0
SM = 0
Doxorubicin

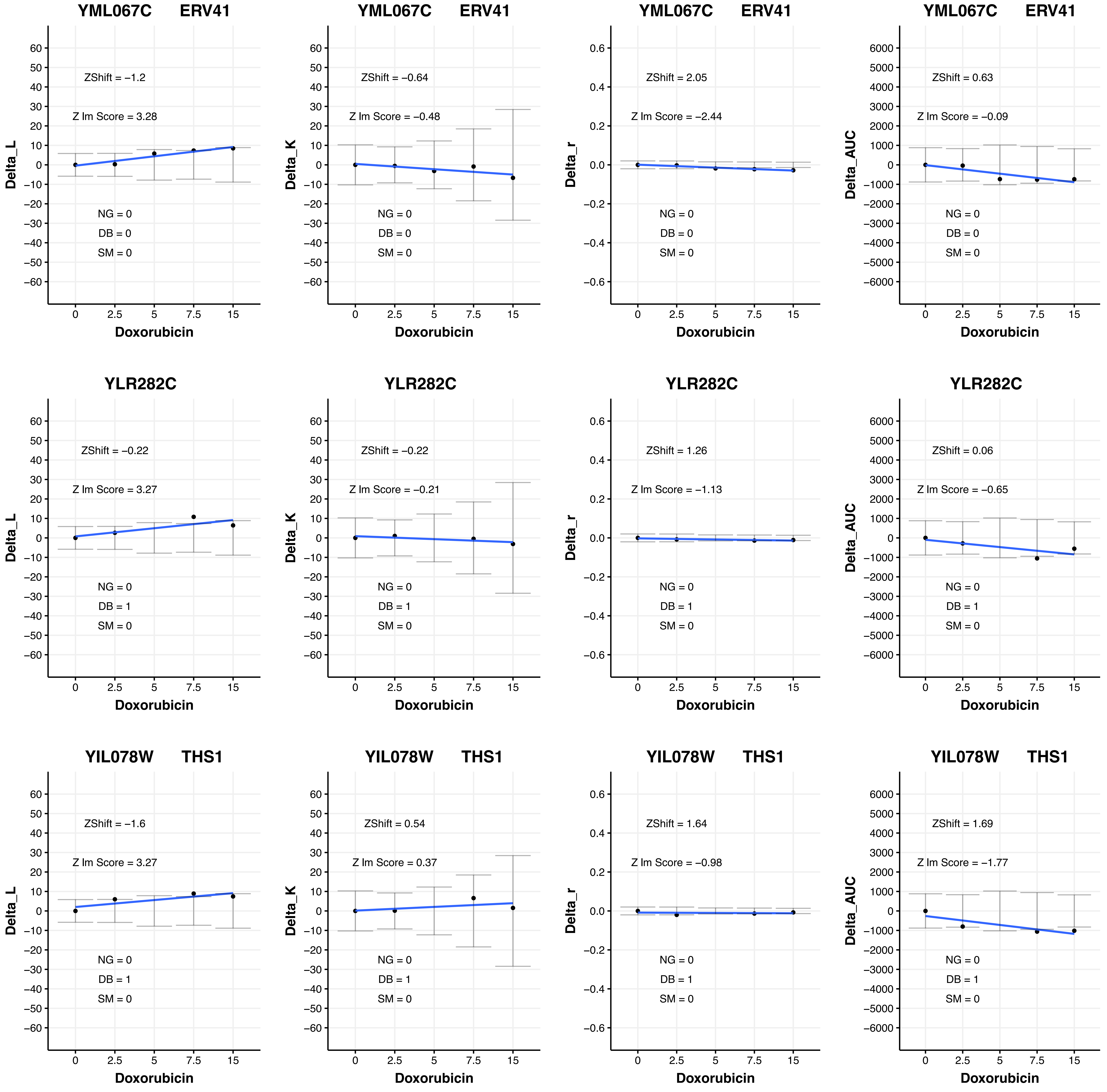
YML067C ERV41
ZShift = −1.2
Z lm Score = 3.28
NG = 0
DB = 0
SM = 0
Delta_L
Doxorubicin
YML067C ERV41
ZShift = −0.64
Z lm Score = −0.48
NG = 0
DB = 0
SM = 0
Delta_K
Doxorubicin
YML067C ERV41
ZShift = 2.05
Z lm Score = −2.44
NG = 0
DB = 0
SM = 0
Delta_r
Doxorubicin
YML067C ERV41
ZShift = 0.63
Z lm Score = −0.09
NG = 0
DB = 0
SM = 0
Delta_AUC
Doxorubicin
YLR282C
ZShift = −0.22
Z lm Score = 3.27
NG = 0
DB = 1
SM = 0
Delta_L
Doxorubicin
YLR282C
ZShift = −0.22
Z lm Score = −0.21
NG = 0
DB = 1
SM = 0
Delta_K
Doxorubicin
YLR282C
ZShift = 1.26
Z lm Score = −1.13
NG = 0
DB = 1
SM = 0
Delta_r
Doxorubicin
YLR282C
ZShift = 0.06
Z lm Score = −0.65
NG = 0
DB = 1
SM = 0
Delta_AUC
Doxorubicin
YIL078W THS1
ZShift = −1.6
Z lm Score = 3.27
NG = 0
DB = 1
SM = 0
Delta_L
Doxorubicin
YIL078W THS1
ZShift = 0.54
Z lm Score = 0.37
NG = 0
DB = 1
SM = 0
Delta_K
Doxorubicin
YIL078W THS1
ZShift = 1.64
Z lm Score = −0.98
NG = 0
DB = 1
SM = 0
Delta_r
Doxorubicin
YIL078W THS1
ZShift = 1.69
Z lm Score = −1.77
NG = 0
DB = 1
SM = 0
Delta_AUC
Doxorubicin

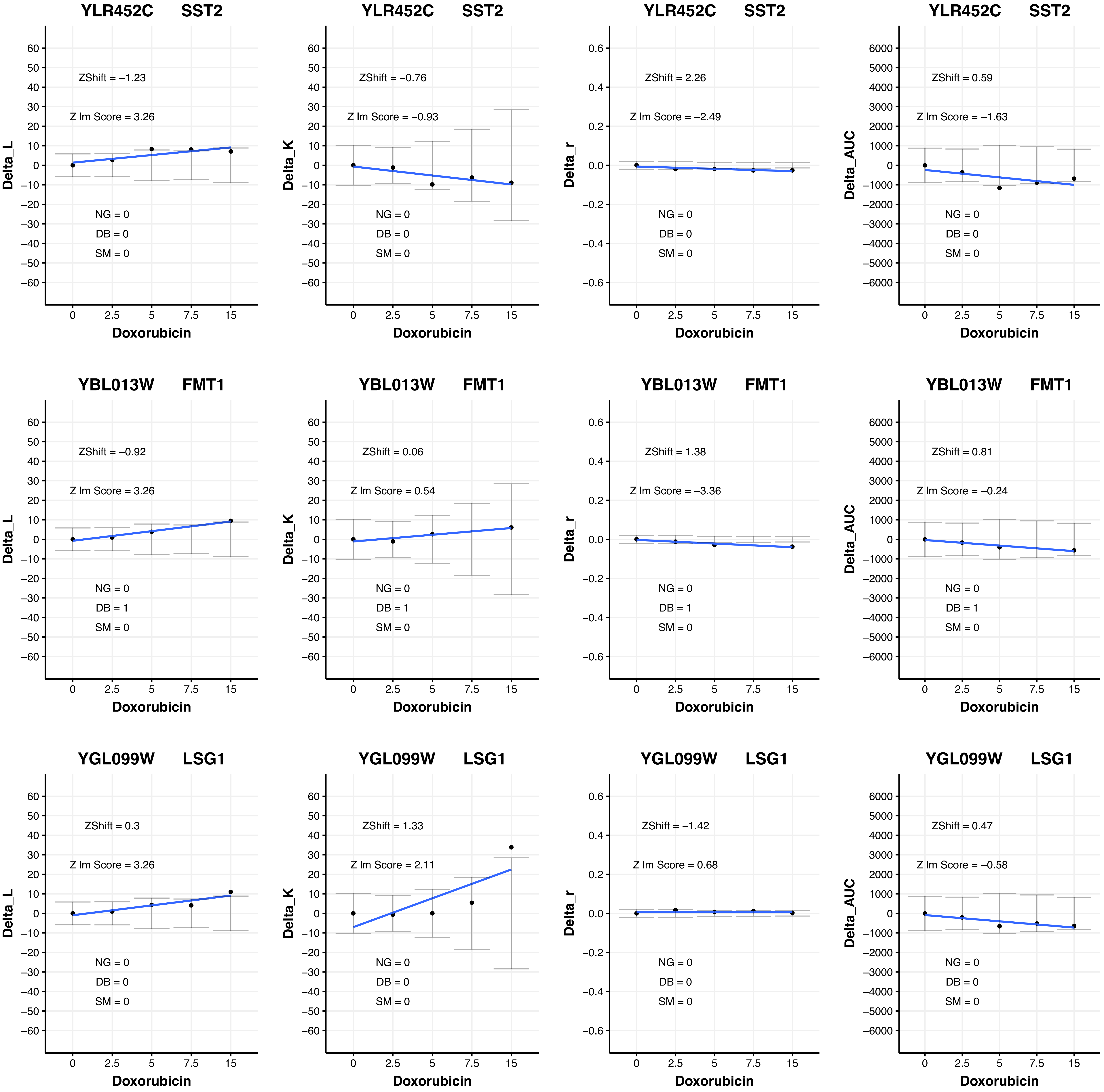
YLR452C SST2
ZShift = −1.23
Z lm Score = 3.26
NG = 0
DB = 0
SM = 0
Delta_L
Doxorubicin
YLR452C SST2
ZShift = −0.76
Z lm Score = −0.93
NG = 0
DB = 0
SM = 0
Delta_K
Doxorubicin
YLR452C SST2
ZShift = 2.26
Z lm Score = −2.49
NG = 0
DB = 0
SM = 0
Delta_r
Doxorubicin
YLR452C SST2
ZShift = 0.59
Z lm Score = −1.63
NG = 0
DB = 0
SM = 0
Delta_AUC
Doxorubicin
YBL013W FMT1
ZShift = −0.92
Z lm Score = 3.26
NG = 0
DB = 1
SM = 0
Delta_L
Doxorubicin
YBL013W FMT1
ZShift = 0.06
Z lm Score = 0.54
NG = 0
DB = 1
SM = 0
Delta_K
Doxorubicin
YBL013W FMT1
ZShift = 1.38
Z lm Score = −3.36
NG = 0
DB = 1
SM = 0
Delta_r
Doxorubicin
YBL013W FMT1
ZShift = 0.81
Z lm Score = −0.24
NG = 0
DB = 1
SM = 0
Delta_AUC
Doxorubicin
YGL099W LSG1
ZShift = 0.3
Z lm Score = 3.26
NG = 0
DB = 0
SM = 0
Delta_L
Doxorubicin
YGL099W LSG1
ZShift = 1.33
Z lm Score = 2.11
NG = 0
DB = 0
SM = 0
Delta_K
Doxorubicin
YGL099W LSG1
ZShift = −1.42
Z lm Score = 0.68
NG = 0
DB = 0
SM = 0
Delta_r
Doxorubicin
YGL099W LSG1
ZShift = 0.47
Z lm Score = −0.58
NG = 0
DB = 0
SM = 0
Delta_AUC
Doxorubicin

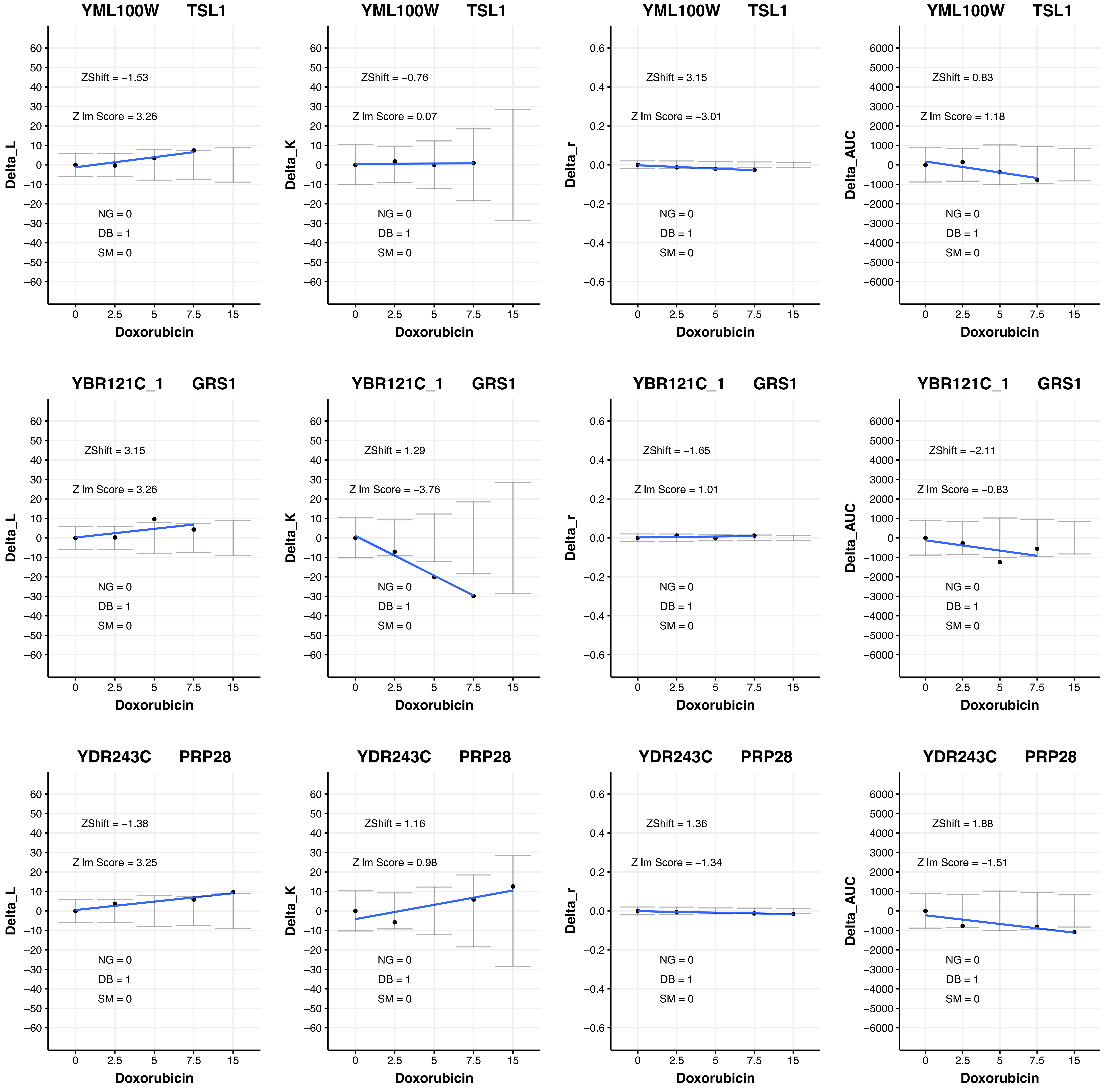
YML100W TSL1
ZShift = −1.53
Z lm Score = 3.26
NG = 0
DB = 1
SM = 0
Delta_L
Doxorubicin
YML100W TSL1
ZShift = −0.76
Z lm Score = 0.07
NG = 0
DB = 1
SM = 0
Delta_K
Doxorubicin
YML100W TSL1
ZShift = 3.15
Z lm Score = −3.01
NG = 0
DB = 1
SM = 0
Delta_r
Doxorubicin
YML100W TSL1
ZShift = 0.83
Z lm Score = 1.18
NG = 0
DB = 1
SM = 0
Delta_AUC
Doxorubicin
YBR121C_1 GRS1
ZShift = 3.15
Z lm Score = 3.26
NG = 0
DB = 1
SM = 0
Delta_L
Doxorubicin
YBR121C_1 GRS1
ZShift = 1.29
Z lm Score = −3.76
NG = 0
DB = 1
SM = 0
Delta_K
Doxorubicin
YBR121C_1 GRS1
ZShift = −1.65
Z lm Score = 1.01
NG = 0
DB = 1
SM = 0
Delta_r
Doxorubicin
YBR121C_1 GRS1
ZShift = −2.11
Z lm Score = −0.83
NG = 0
DB = 1
SM = 0
Delta_AUC
Doxorubicin
YDR243C PRP28
ZShift = −1.38
Z lm Score = 3.25
NG = 0
DB = 1
SM = 0
Delta_L
Doxorubicin
YDR243C PRP28
ZShift = 1.16
Z lm Score = 0.98
NG = 0
DB = 1
SM = 0
Delta_K
Doxorubicin
YDR243C PRP28
ZShift = 1.36
Z lm Score = −1.34
NG = 0
DB = 1
SM = 0
Delta_r
Doxorubicin
YDR243C PRP28
ZShift = 1.88
Z lm Score = −1.51
NG = 0
DB = 1
SM = 0
Delta_AUC
Doxorubicin
0
2.5
5
7.5
15

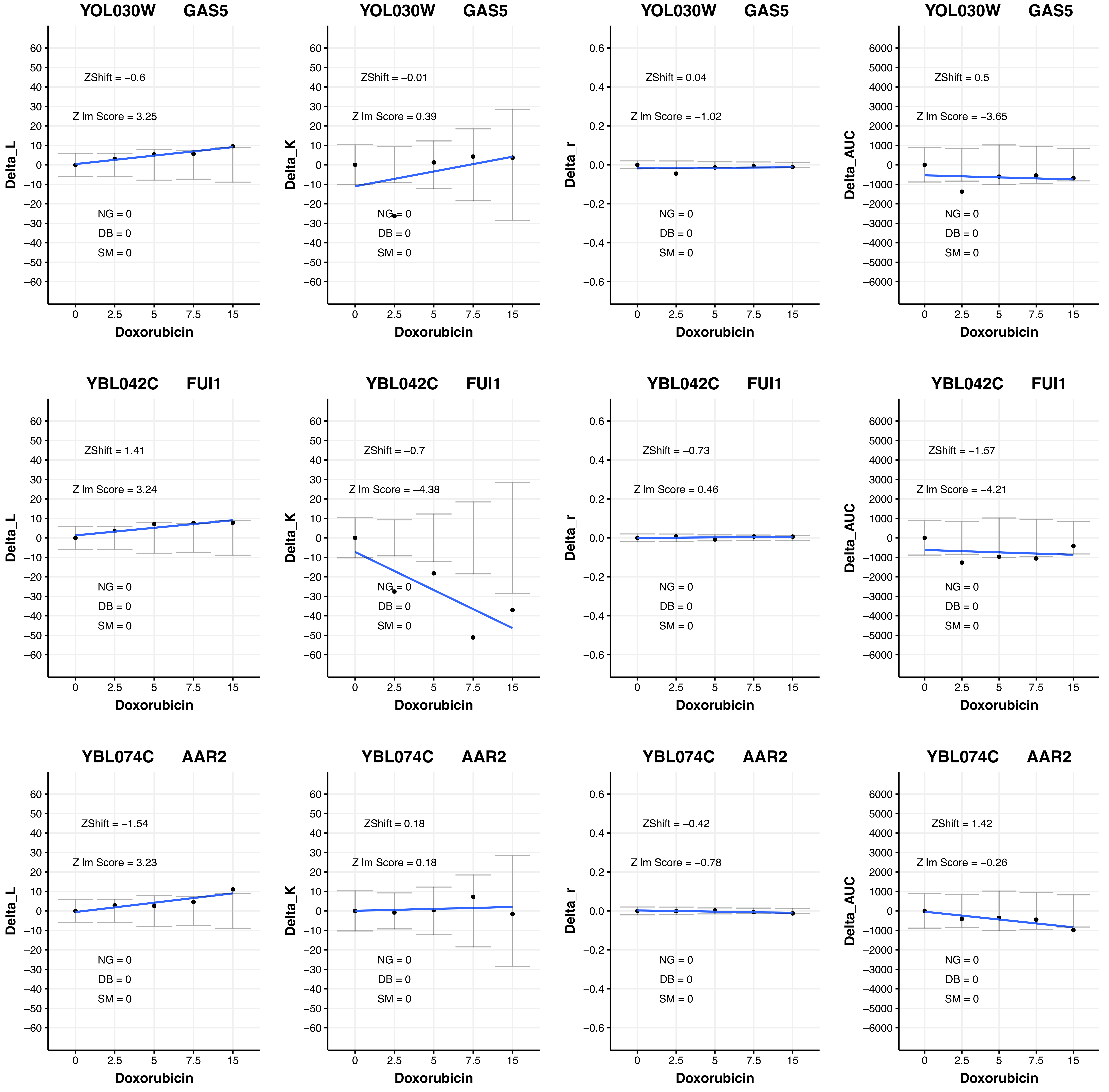
YOL030W GAS5
Delta_L
Doxorubicin
ZShift = −0.6
Z lm Score = 3.25
NG = 0
DB = 0
SM = 0
YOL030W GAS5
Delta_K
Doxorubicin
ZShift = −0.01
Z lm Score = 0.39
NG = 0
DB = 0
SM = 0
YOL030W GAS5
Delta_r
Doxorubicin
ZShift = 0.04
Z lm Score = −1.02
NG = 0
DB = 0
SM = 0
YOL030W GAS5
Delta_AUC
Doxorubicin
ZShift = 0.5
Z lm Score = −3.65
NG = 0
DB = 0
SM = 0
YBL042C FUI1
Delta_L
Doxorubicin
ZShift = 1.41
Z lm Score = 3.24
NG = 0
DB = 0
SM = 0
YBL042C FUI1
Delta_K
Doxorubicin
ZShift = −0.7
Z lm Score = −4.38
NG = 0
DB = 0
SM = 0
YBL042C FUI1
Delta_r
Doxorubicin
ZShift = −0.73
Z lm Score = 0.46
NG = 0
DB = 0
SM = 0
YBL042C FUI1
Delta_AUC
Doxorubicin
ZShift = −1.57
Z lm Score = −4.21
NG = 0
DB = 0
SM = 0
YBL074C AAR2
Delta_L
Doxorubicin
ZShift = −1.54
Z lm Score = 3.23
NG = 0
DB = 0
SM = 0
YBL074C AAR2
Delta_K
Doxorubicin
ZShift = 0.18
Z lm Score = 0.18
NG = 0
DB = 0
SM = 0
YBL074C AAR2
Delta_r
Doxorubicin
ZShift = −0.42
Z lm Score = −0.78
NG = 0
DB = 0
SM = 0
YBL074C AAR2
Delta_AUC
Doxorubicin
ZShift = 1.42
Z lm Score = −0.26
NG = 0
DB = 0
SM = 0

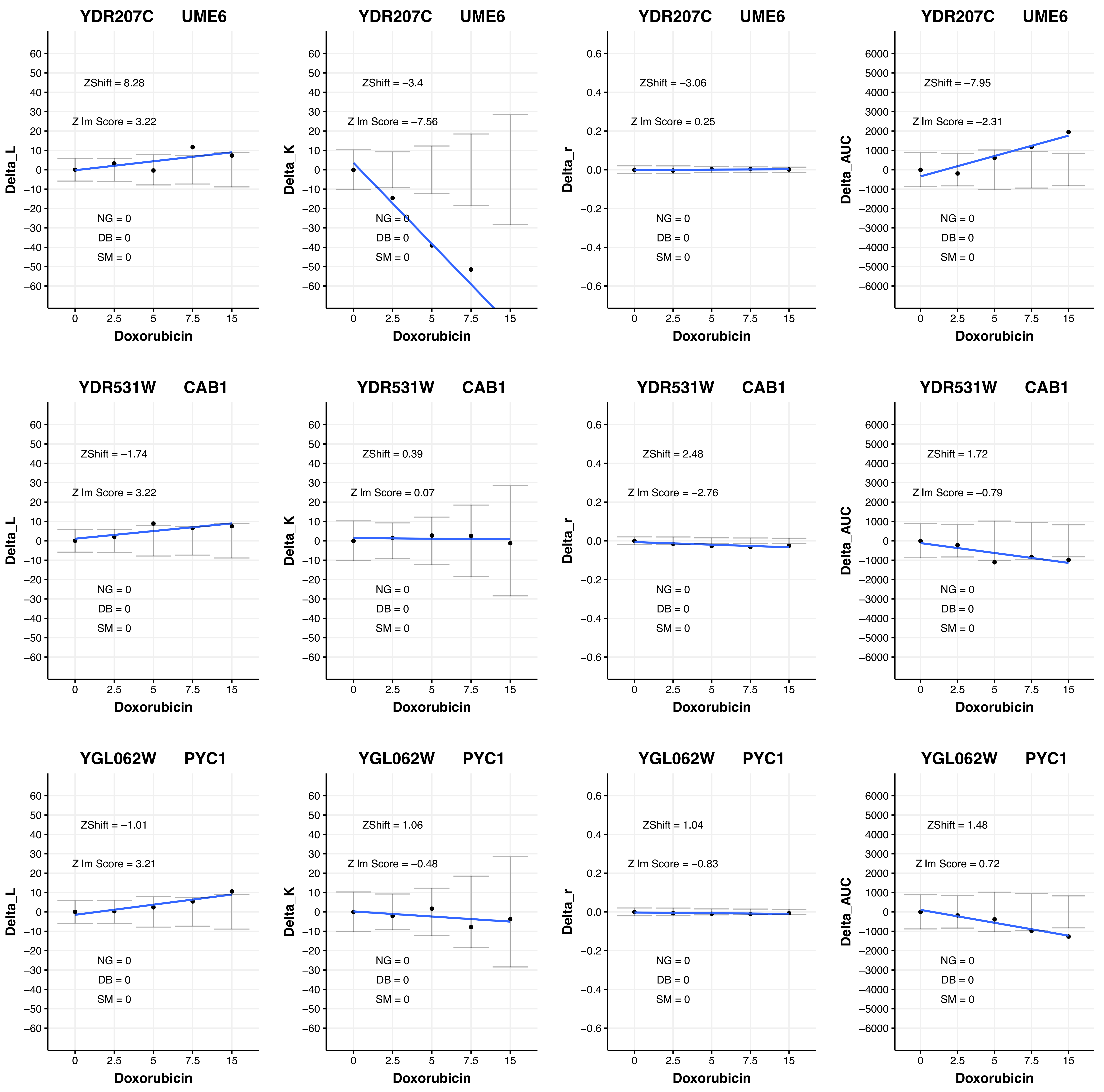
YDR207C UME6
ZShift = 8.28
Z lm Score = 3.22
NG = 0
DB = 0
SM = 0
Delta_L
Doxorubicin
YDR207C UME6
ZShift = −3.4
Z lm Score = −7.56
NG = 0
DB = 0
SM = 0
Delta_K
Doxorubicin
YDR207C UME6
ZShift = −3.06
Z lm Score = 0.25
NG = 0
DB = 0
SM = 0
Delta_r
Doxorubicin
YDR207C UME6
ZShift = −7.95
Z lm Score = −2.31
NG = 0
DB = 0
SM = 0
Delta_AUC
Doxorubicin
YDR531W CAB1
ZShift = −1.74
Z lm Score = 3.22
NG = 0
DB = 0
SM = 0
Delta_L
Doxorubicin
YDR531W CAB1
ZShift = 0.39
Z lm Score = 0.07
NG = 0
DB = 0
SM = 0
Delta_K
Doxorubicin
YDR531W CAB1
ZShift = 2.48
Z lm Score = −2.76
NG = 0
DB = 0
SM = 0
Delta_r
Doxorubicin
YDR531W CAB1
ZShift = 1.72
Z lm Score = −0.79
NG = 0
DB = 0
SM = 0
Delta_AUC
Doxorubicin
YGL062W PYC1
ZShift = −1.01
Z lm Score = 3.21
NG = 0
DB = 0
SM = 0
Delta_L
Doxorubicin
YGL062W PYC1
ZShift = 1.06
Z lm Score = −0.48
NG = 0
DB = 0
SM = 0
Delta_K
Doxorubicin
YGL062W PYC1
ZShift = 1.04
Z lm Score = −0.83
NG = 0
DB = 0
SM = 0
Delta_r
Doxorubicin
YGL062W PYC1
ZShift = 1.48
Z lm Score = 0.72
NG = 0
DB = 0
SM = 0
Delta_AUC
Doxorubicin

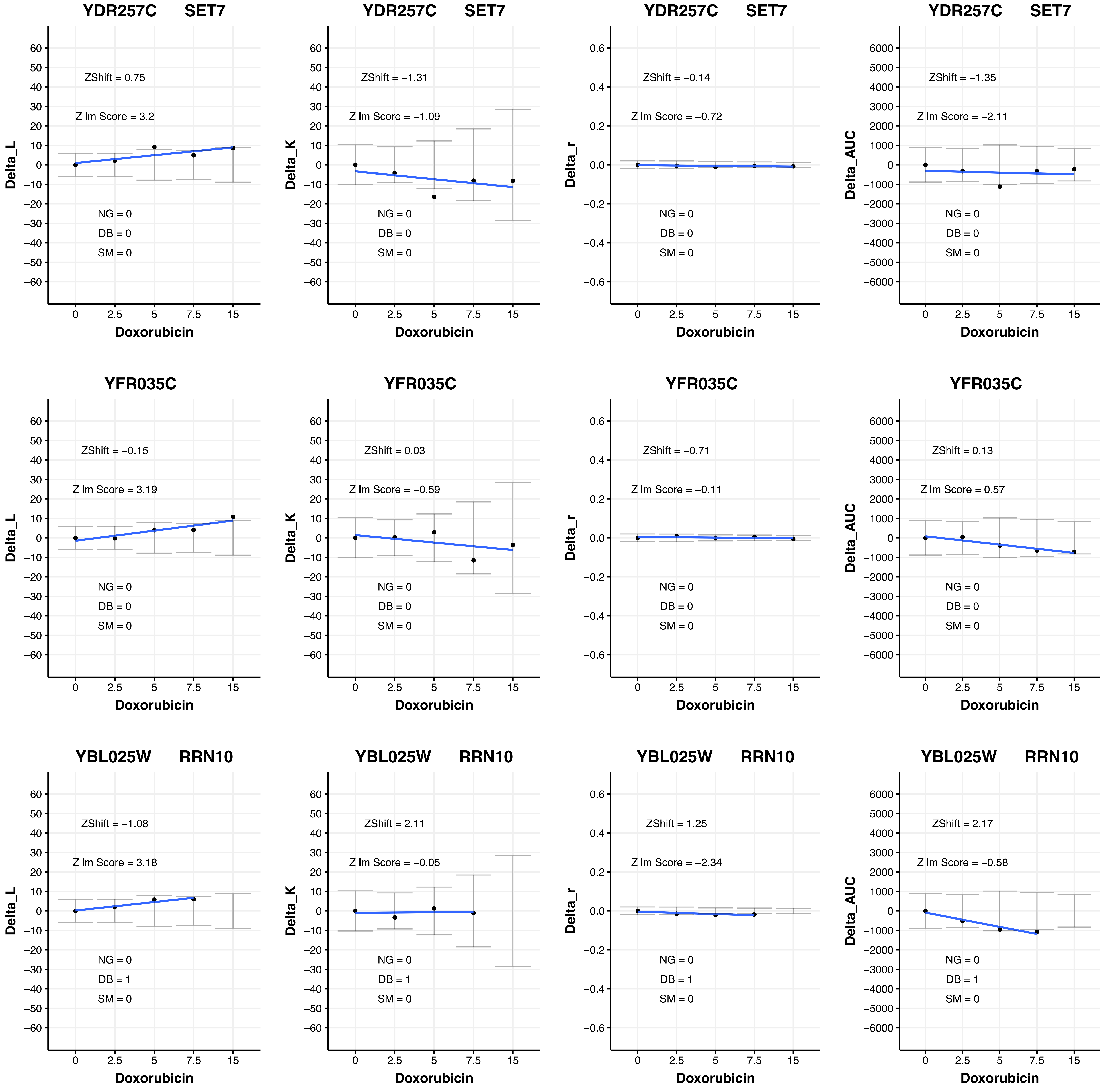
YDR257C SET7
ZShift = 0.75
Z lm Score = 3.2
NG = 0
DB = 0
SM = 0
Delta_L
Doxorubicin
YDR257C SET7
ZShift = −1.31
Z lm Score = −1.09
NG = 0
DB = 0
SM = 0
Delta_K
Doxorubicin
YDR257C SET7
ZShift = −0.14
Z lm Score = −0.72
NG = 0
DB = 0
SM = 0
Delta_r
Doxorubicin
YDR257C SET7
ZShift = −1.35
Z lm Score = −2.11
NG = 0
DB = 0
SM = 0
Delta_AUC
Doxorubicin
YFR035C
ZShift = −0.15
Z lm Score = 3.19
NG = 0
DB = 0
SM = 0
Delta_L
Doxorubicin
YFR035C
ZShift = 0.03
Z lm Score = −0.59
NG = 0
DB = 0
SM = 0
Delta_K
Doxorubicin
YFR035C
ZShift = −0.71
Z lm Score = −0.11
NG = 0
DB = 0
SM = 0
Delta_r
Doxorubicin
YFR035C
ZShift = 0.13
Z lm Score = 0.57
NG = 0
DB = 0
SM = 0
Delta_AUC
Doxorubicin
YBL025W RRN10
ZShift = −1.08
Z lm Score = 3.18
NG = 0
DB = 1
SM = 0
Delta_L
Doxorubicin
YBL025W RRN10
ZShift = 2.11
Z lm Score = −0.05
NG = 0
DB = 1
SM = 0
Delta_K
Doxorubicin
YBL025W RRN10
ZShift = 1.25
Z lm Score = −2.34
NG = 0
DB = 1
SM = 0
Delta_r
Doxorubicin
YBL025W RRN10
ZShift = 2.17
Z lm Score = −0.58
NG = 0
DB = 1
SM = 0
Delta_AUC
Doxorubicin

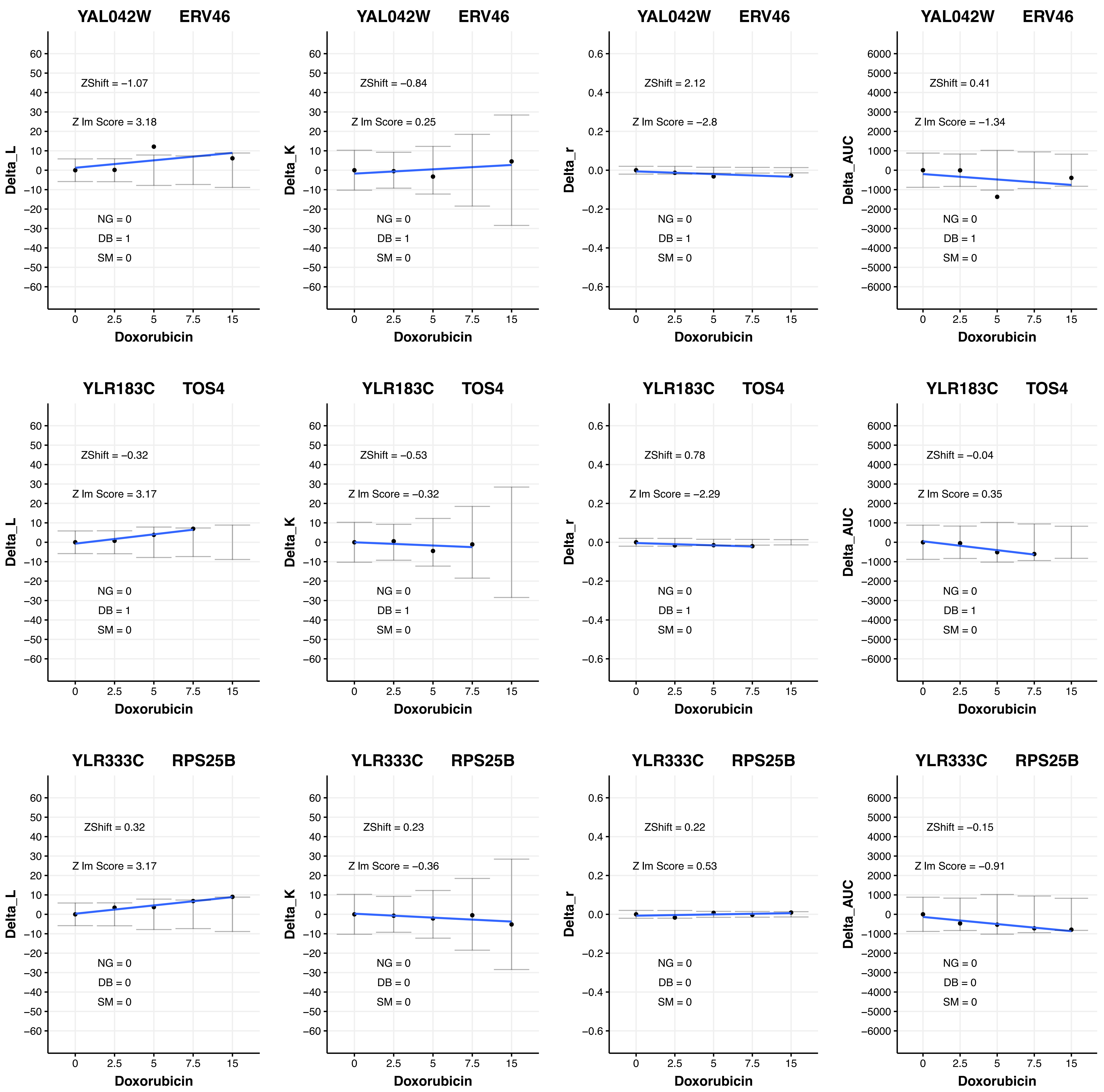
YAL042W ERV46
ZShift = −1.07
Z lm Score = 3.18
NG = 0
DB = 1
SM = 0
Delta_L
Doxorubicin
YAL042W ERV46
ZShift = −0.84
Z lm Score = 0.25
NG = 0
DB = 1
SM = 0
Delta_K
Doxorubicin
YAL042W ERV46
ZShift = 2.12
Z lm Score = −2.8
NG = 0
DB = 1
SM = 0
Delta_r
Doxorubicin
YAL042W ERV46
ZShift = 0.41
Z lm Score = −1.34
NG = 0
DB = 1
SM = 0
Delta_AUC
Doxorubicin
YLR183C TOS4
ZShift = −0.32
Z lm Score = 3.17
NG = 0
DB = 1
SM = 0
Delta_L
Doxorubicin
YLR183C TOS4
ZShift = −0.53
Z lm Score = −0.32
NG = 0
DB = 1
SM = 0
Delta_K
Doxorubicin
YLR183C TOS4
ZShift = 0.78
Z lm Score = −2.29
NG = 0
DB = 1
SM = 0
Delta_r
Doxorubicin
YLR183C TOS4
ZShift = −0.04
Z lm Score = 0.35
NG = 0
DB = 1
SM = 0
Delta_AUC
Doxorubicin
YLR333C RPS25B
ZShift = 0.32
Z lm Score = 3.17
NG = 0
DB = 0
SM = 0
Delta_L
Doxorubicin
YLR333C RPS25B
ZShift = 0.23
Z lm Score = −0.36
NG = 0
DB = 0
SM = 0
Delta_K
Doxorubicin
YLR333C RPS25B
ZShift = 0.22
Z lm Score = 0.53
NG = 0
DB = 0
SM = 0
Delta_r
Doxorubicin
YLR333C RPS25B
ZShift = −0.15
Z lm Score = −0.91
NG = 0
DB = 0
SM = 0
Delta_AUC
Doxorubicin

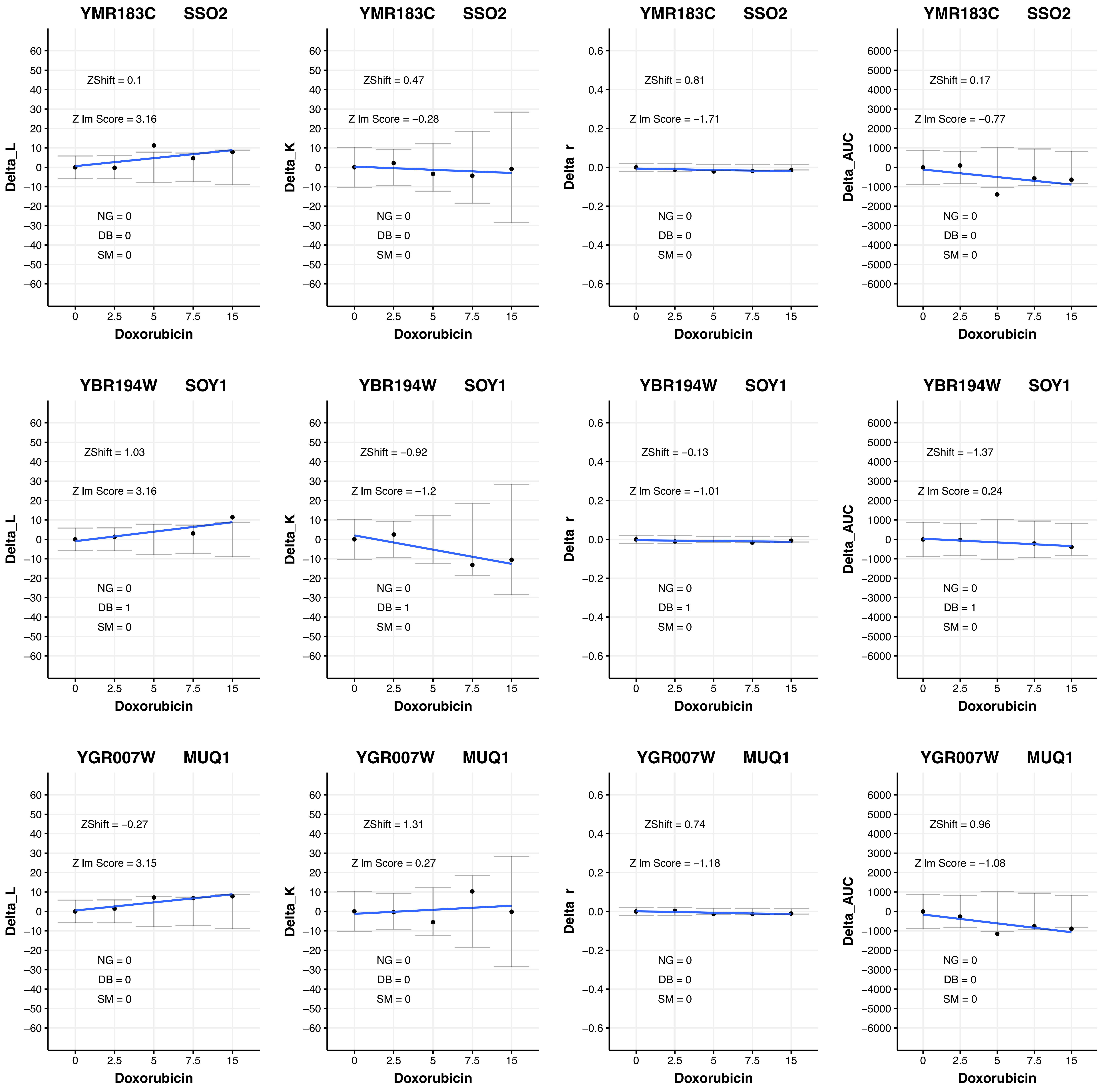
YMR183C SSO2
ZShift = 0.1
Z lm Score = 3.16
NG = 0
DB = 0
SM = 0
Delta_L
Doxorubicin
YMR183C SSO2
ZShift = 0.47
Z lm Score = −0.28
NG = 0
DB = 0
SM = 0
Delta_K
Doxorubicin
YMR183C SSO2
ZShift = 0.81
Z lm Score = −1.71
NG = 0
DB = 0
SM = 0
Delta_r
Doxorubicin
YMR183C SSO2
ZShift = 0.17
Z lm Score = −0.77
NG = 0
DB = 0
SM = 0
Delta_AUC
Doxorubicin
YBR194W SOY1
ZShift = 1.03
Z lm Score = 3.16
NG = 0
DB = 1
SM = 0
Delta_L
Doxorubicin
YBR194W SOY1
ZShift = −0.92
Z lm Score = −1.2
NG = 0
DB = 1
SM = 0
Delta_K
Doxorubicin
YBR194W SOY1
ZShift = −0.13
Z lm Score = −1.01
NG = 0
DB = 1
SM = 0
Delta_r
Doxorubicin
YBR194W SOY1
ZShift = −1.37
Z lm Score = 0.24
NG = 0
DB = 1
SM = 0
Delta_AUC
Doxorubicin
YGR007W MUQ1
ZShift = −0.27
Z lm Score = 3.15
NG = 0
DB = 0
SM = 0
Delta_L
Doxorubicin
YGR007W MUQ1
ZShift = 1.31
Z lm Score = 0.27
NG = 0
DB = 0
SM = 0
Delta_K
Doxorubicin
YGR007W MUQ1
ZShift = 0.74
Z lm Score = −1.18
NG = 0
DB = 0
SM = 0
Delta_r
Doxorubicin
YGR007W MUQ1
ZShift = 0.96
Z lm Score = −1.08
NG = 0
DB = 0
SM = 0
Delta_AUC
Doxorubicin

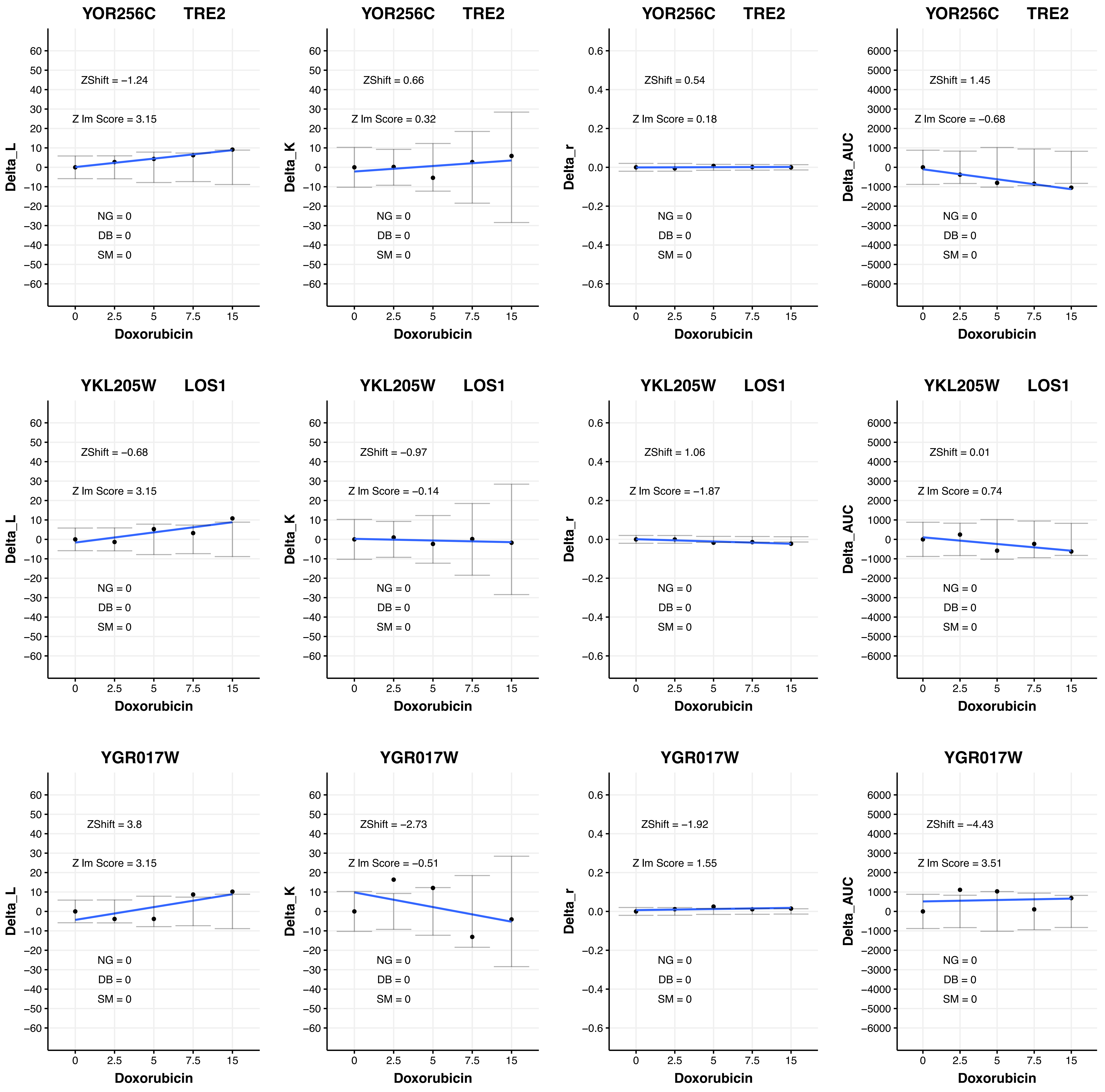
YOR256C TRE2
ZShift = −1.24
Z lm Score = 3.15
NG = 0
DB = 0
SM = 0
Delta_L
Doxorubicin
YOR256C TRE2
ZShift = 0.66
Z lm Score = 0.32
NG = 0
DB = 0
SM = 0
Delta_K
Doxorubicin
YOR256C TRE2
ZShift = 0.54
Z lm Score = 0.18
NG = 0
DB = 0
SM = 0
Delta_r
Doxorubicin
YOR256C TRE2
ZShift = 1.45
Z lm Score = −0.68
NG = 0
DB = 0
SM = 0
Delta_AUC
Doxorubicin
YKL205W LOS1
ZShift = −0.68
Z lm Score = 3.15
NG = 0
DB = 0
SM = 0
Delta_L
Doxorubicin
YKL205W LOS1
ZShift = −0.97
Z lm Score = −0.14
NG = 0
DB = 0
SM = 0
Delta_K
Doxorubicin
YKL205W LOS1
ZShift = 1.06
Z lm Score = −1.87
NG = 0
DB = 0
SM = 0
Delta_r
Doxorubicin
YKL205W LOS1
ZShift = 0.01
Z lm Score = 0.74
NG = 0
DB = 0
SM = 0
Delta_AUC
Doxorubicin
YGR017W
ZShift = 3.8
Z lm Score = 3.15
NG = 0
DB = 0
SM = 0
Delta_L
Doxorubicin
YGR017W
ZShift = −2.73
Z lm Score = −0.51
NG = 0
DB = 0
SM = 0
Delta_K
Doxorubicin
YGR017W
ZShift = −1.92
Z lm Score = 1.55
NG = 0
DB = 0
SM = 0
Delta_r
Doxorubicin
YGR017W
ZShift = −4.43
Z lm Score = 3.51
NG = 0
DB = 0
SM = 0
Delta_AUC
Doxorubicin

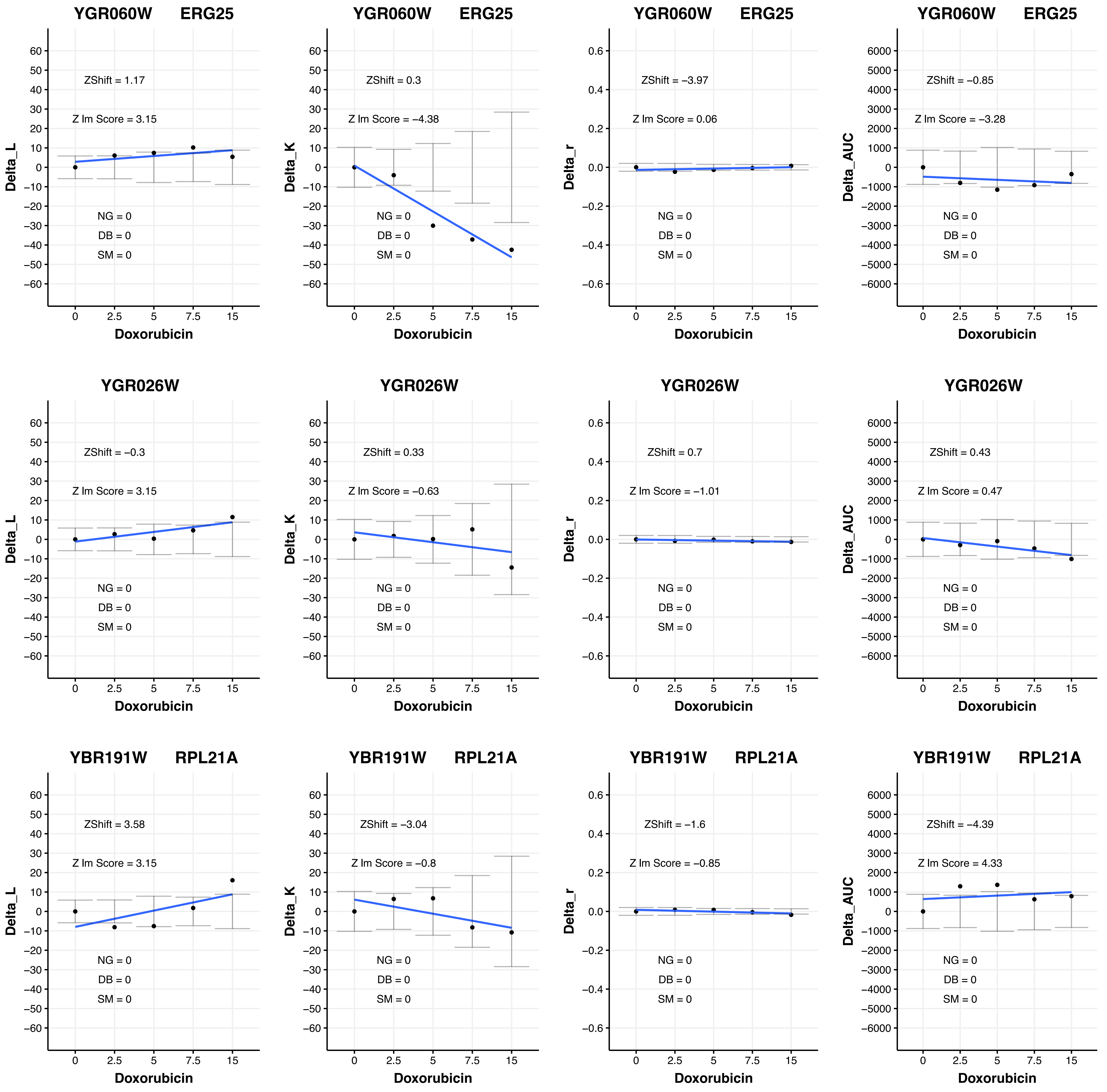
YGR060W ERG25
ZShift = 1.17
Z lm Score = 3.15
NG = 0
DB = 0
SM = 0
Delta_L
Doxorubicin
YGR060W ERG25
ZShift = 0.3
Z lm Score = −4.38
NG = 0
DB = 0
SM = 0
Delta_K
Doxorubicin
YGR060W ERG25
ZShift = −3.97
Z lm Score = 0.06
NG = 0
DB = 0
SM = 0
Delta_r
Doxorubicin
YGR060W ERG25
ZShift = −0.85
Z lm Score = −3.28
NG = 0
DB = 0
SM = 0
Delta_AUC
Doxorubicin
YGR026W
ZShift = −0.3
Z lm Score = 3.15
NG = 0
DB = 0
SM = 0
Delta_L
Doxorubicin
YGR026W
ZShift = 0.33
Z lm Score = −0.63
NG = 0
DB = 0
SM = 0
Delta_K
Doxorubicin
YGR026W
ZShift = 0.7
Z lm Score = −1.01
NG = 0
DB = 0
SM = 0
Delta_r
Doxorubicin
YGR026W
ZShift = 0.43
Z lm Score = 0.47
NG = 0
DB = 0
SM = 0
Delta_AUC
Doxorubicin
YBR191W RPL21A
ZShift = 3.58
Z lm Score = 3.15
NG = 0
DB = 0
SM = 0
Delta_L
Doxorubicin
YBR191W RPL21A
ZShift = −3.04
Z lm Score = −0.8
NG = 0
DB = 0
SM = 0
Delta_K
Doxorubicin
YBR191W RPL21A
ZShift = −1.6
Z lm Score = −0.85
NG = 0
DB = 0
SM = 0
Delta_r
Doxorubicin
YBR191W RPL21A
ZShift = −4.39
Z lm Score = 4.33
NG = 0
DB = 0
SM = 0
Delta_AUC
Doxorubicin

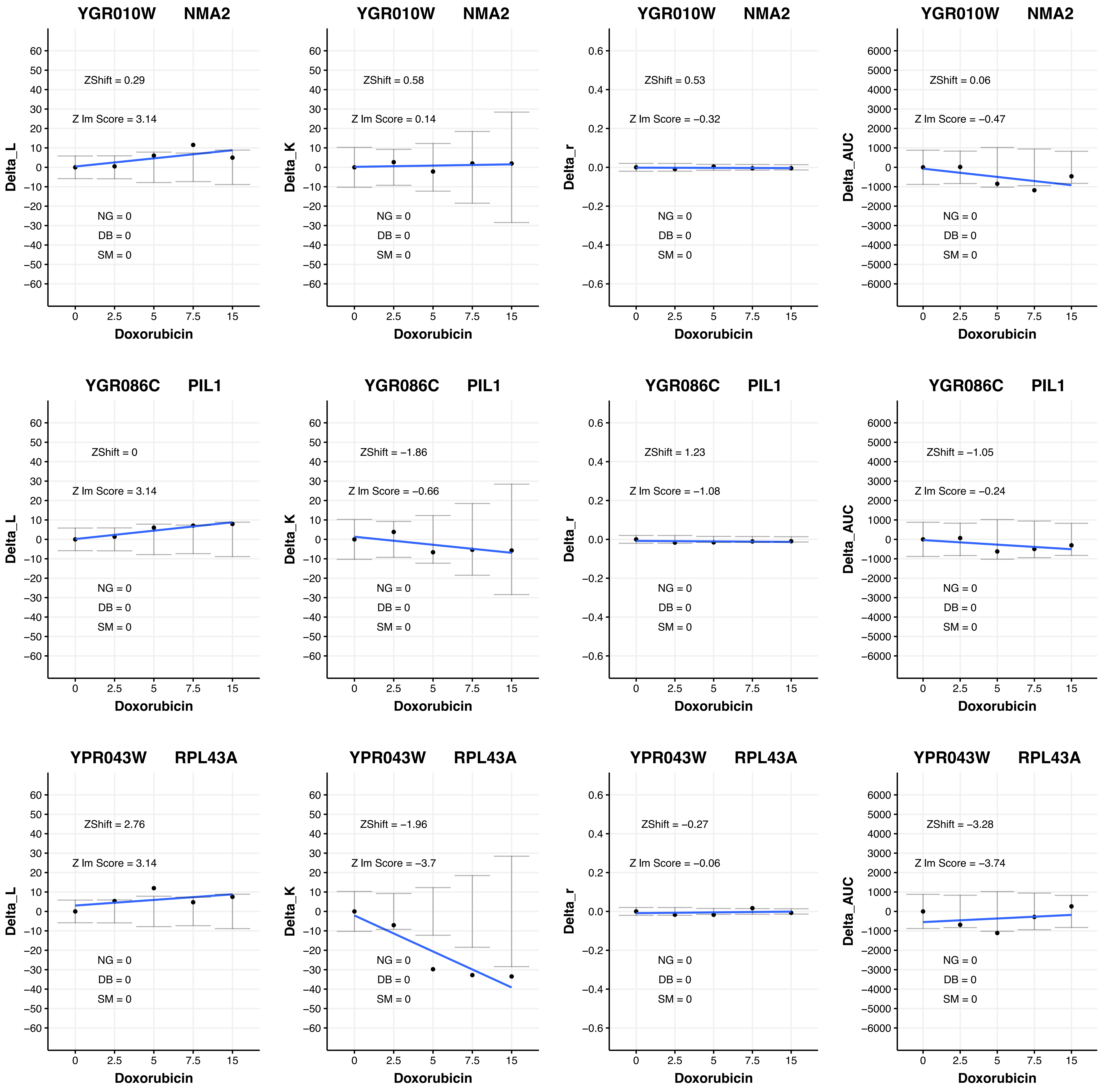
YGR010W NMA2
ZShift = 0.29
Z lm Score = 3.14
NG = 0
DB = 0
SM = 0
Delta_L
Doxorubicin
YGR010W NMA2
ZShift = 0.58
Z lm Score = 0.14
NG = 0
DB = 0
SM = 0
Delta_K
Doxorubicin
YGR010W NMA2
ZShift = 0.53
Z lm Score = −0.32
NG = 0
DB = 0
SM = 0
Delta_r
Doxorubicin
YGR010W NMA2
ZShift = 0.06
Z lm Score = −0.47
NG = 0
DB = 0
SM = 0
Delta_AUC
Doxorubicin
YGR086C PIL1
ZShift = 0
Z lm Score = 3.14
NG = 0
DB = 0
SM = 0
Delta_L
Doxorubicin
YGR086C PIL1
ZShift = −1.86
Z lm Score = −0.66
NG = 0
DB = 0
SM = 0
Delta_K
Doxorubicin
YGR086C PIL1
ZShift = 1.23
Z lm Score = −1.08
NG = 0
DB = 0
SM = 0
Delta_r
Doxorubicin
YGR086C PIL1
ZShift = −1.05
Z lm Score = −0.24
NG = 0
DB = 0
SM = 0
Delta_AUC
Doxorubicin
YPR043W RPL43A
ZShift = 2.76
Z lm Score = 3.14
NG = 0
DB = 0
SM = 0
Delta_L
Doxorubicin
YPR043W RPL43A
ZShift = −1.96
Z lm Score = −3.7
NG = 0
DB = 0
SM = 0
Delta_K
Doxorubicin
YPR043W RPL43A
ZShift = −0.27
Z lm Score = −0.06
NG = 0
DB = 0
SM = 0
Delta_r
Doxorubicin
YPR043W RPL43A
ZShift = −3.28
Z lm Score = −3.74
NG = 0
DB = 0
SM = 0
Delta_AUC
Doxorubicin
0
2.5
5
7.5
15

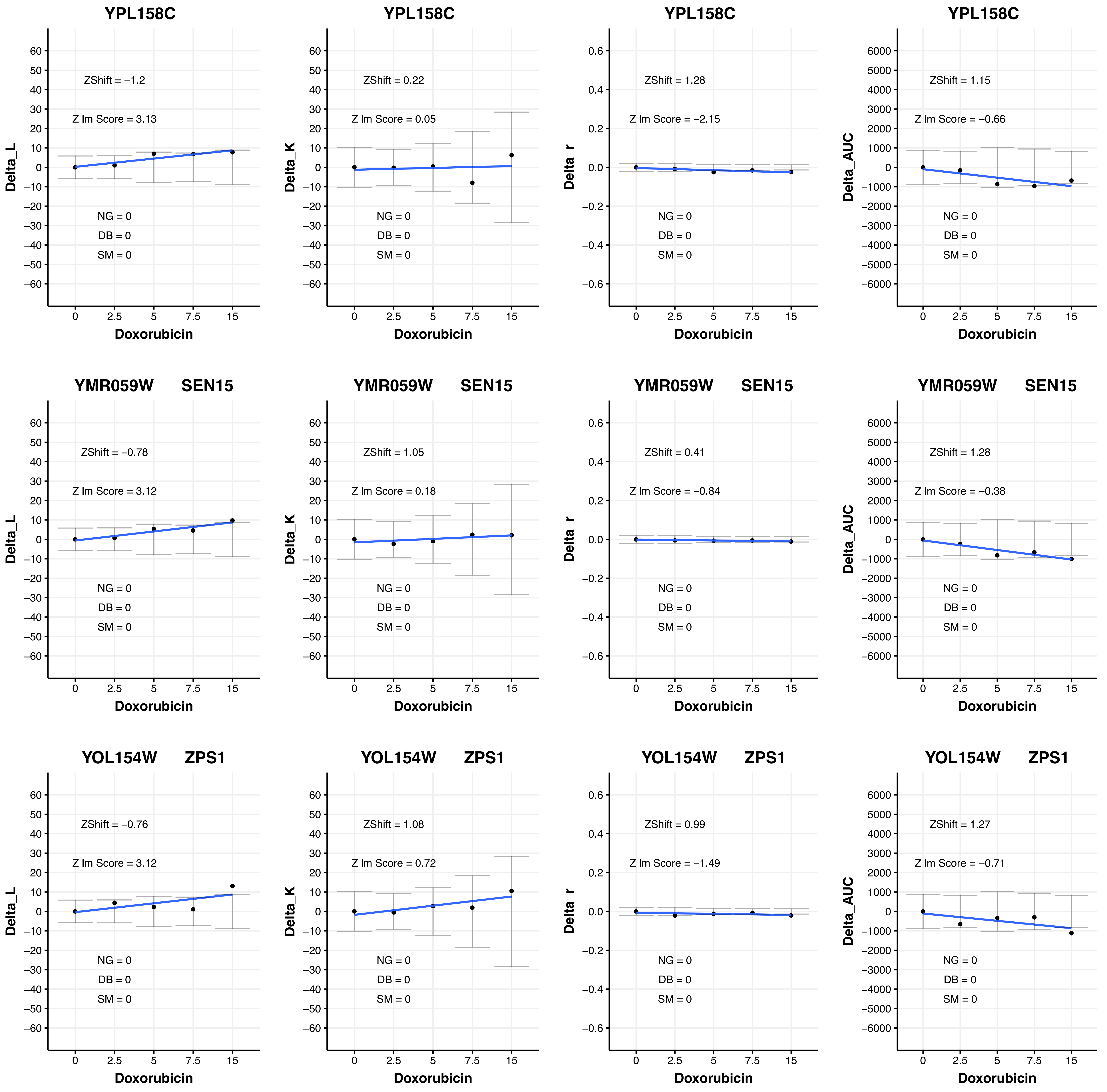
YPL158C
ZShift = −1.2
Z lm Score = 3.13
NG = 0
DB = 0
SM = 0
Delta_L
Doxorubicin
YPL158C
ZShift = 0.22
Z lm Score = 0.05
NG = 0
DB = 0
SM = 0
Delta_K
Doxorubicin
YPL158C
ZShift = 1.28
Z lm Score = −2.15
NG = 0
DB = 0
SM = 0
Delta_r
Doxorubicin
YPL158C
ZShift = 1.15
Z lm Score = −0.66
NG = 0
DB = 0
SM = 0
Delta_AUC
Doxorubicin
YMR059W SEN15
ZShift = −0.78
Z lm Score = 3.12
NG = 0
DB = 0
SM = 0
Delta_L
Doxorubicin
YMR059W SEN15
ZShift = 1.05
Z lm Score = 0.18
NG = 0
DB = 0
SM = 0
Delta_K
Doxorubicin
YMR059W SEN15
ZShift = 0.41
Z lm Score = −0.84
NG = 0
DB = 0
SM = 0
Delta_r
Doxorubicin
YMR059W SEN15
ZShift = 1.28
Z lm Score = −0.38
NG = 0
DB = 0
SM = 0
Delta_AUC
Doxorubicin
YOL154W ZPS1
ZShift = −0.76
Z lm Score = 3.12
NG = 0
DB = 0
SM = 0
Delta_L
Doxorubicin
YOL154W ZPS1
ZShift = 1.08
Z lm Score = 0.72
NG = 0
DB = 0
SM = 0
Delta_K
Doxorubicin
YOL154W ZPS1
ZShift = 0.99
Z lm Score = −1.49
NG = 0
DB = 0
SM = 0
Delta_r
Doxorubicin
YOL154W ZPS1
ZShift = 1.27
Z lm Score = −0.71
NG = 0
DB = 0
SM = 0
Delta_AUC
Doxorubicin

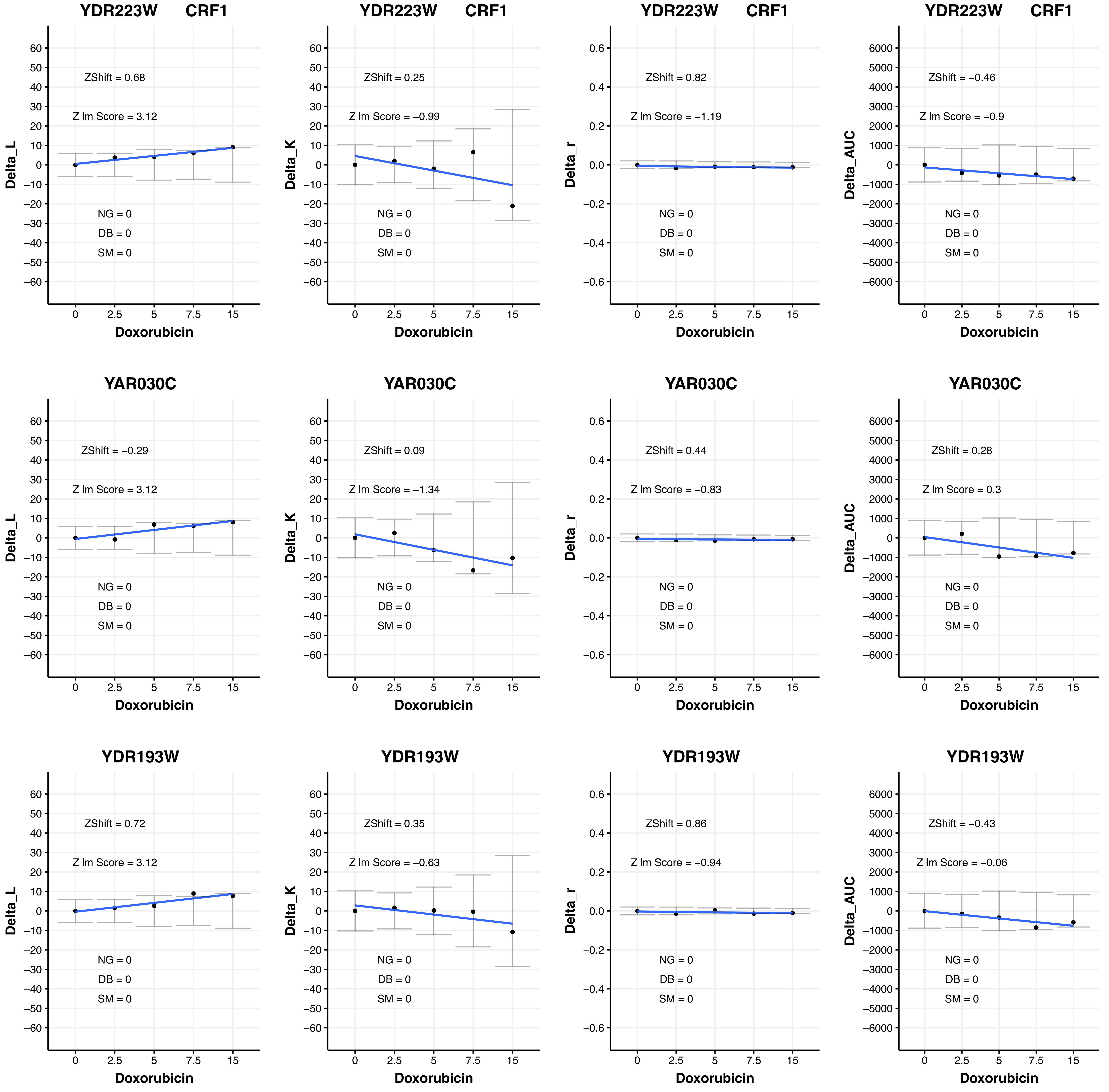

YDR223W CRF1
ZShift = 0.68
Z lm Score = 3.12
NG = 0
DB = 0
SM = 0
Delta_L
Doxorubicin
YDR223W CRF1
ZShift = 0.25
Z lm Score = −0.99
NG = 0
DB = 0
SM = 0
Delta_K
Doxorubicin
YDR223W CRF1
ZShift = 0.82
Z lm Score = −1.19
NG = 0
DB = 0
SM = 0
Delta_r
Doxorubicin
YDR223W CRF1
ZShift = −0.46
Z lm Score = −0.9
NG = 0
DB = 0
SM = 0
Delta_AUC
Doxorubicin
YAR030C
ZShift = −0.29
Z lm Score = 3.12
NG = 0
DB = 0
SM = 0
Delta_L
Doxorubicin
YAR030C
ZShift = 0.09
Z lm Score = −1.34
NG = 0
DB = 0
SM = 0
Delta_K
Doxorubicin
YAR030C
ZShift = 0.44
Z lm Score = −0.83
NG = 0
DB = 0
SM = 0
Delta_r
Doxorubicin
YAR030C
ZShift = 0.28
Z lm Score = 0.3
NG = 0
DB = 0
SM = 0
Delta_AUC
Doxorubicin
YDR193W
ZShift = 0.72
Z lm Score = 3.12
NG = 0
DB = 0
SM = 0
Delta_L
Doxorubicin
YDR193W
ZShift = 0.35
Z lm Score = −0.63
NG = 0
DB = 0
SM = 0
Delta_K
Doxorubicin
YDR193W
ZShift = 0.86
Z lm Score = −0.94
NG = 0
DB = 0
SM = 0
Delta_r
Doxorubicin
YDR193W
ZShift = −0.43
Z lm Score = −0.06
NG = 0
DB = 0
SM = 0
Delta_AUC
Doxorubicin

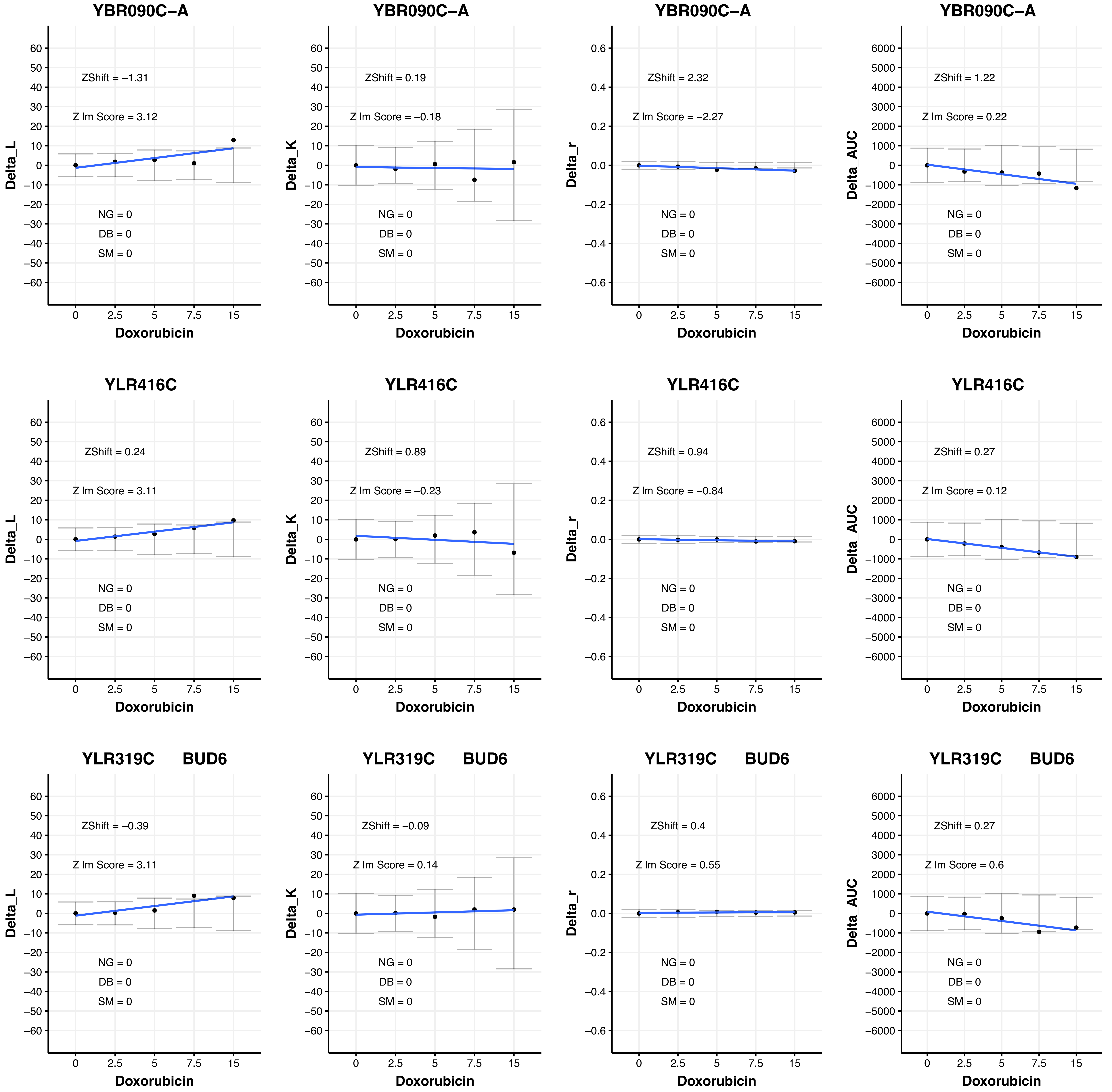
YBR090C−A
ZShift = −1.31
Z lm Score = 3.12
NG = 0
DB = 0
SM = 0
Delta_L
Doxorubicin
YBR090C−A
ZShift = 0.19
Z lm Score = −0.18
NG = 0
DB = 0
SM = 0
Delta_K
Doxorubicin
YBR090C−A
ZShift = 2.32
Z lm Score = −2.27
NG = 0
DB = 0
SM = 0
Delta_r
Doxorubicin
YBR090C−A
ZShift = 1.22
Z lm Score = 0.22
NG = 0
DB = 0
SM = 0
Delta_AUC
Doxorubicin
YLR416C
ZShift = 0.24
Z lm Score = 3.11
NG = 0
DB = 0
SM = 0
Delta_L
Doxorubicin
YLR416C
ZShift = 0.89
Z lm Score = −0.23
NG = 0
DB = 0
SM = 0
Delta_K
Doxorubicin
YLR416C
ZShift = 0.94
Z lm Score = −0.84
NG = 0
DB = 0
SM = 0
Delta_r
Doxorubicin
YLR416C
ZShift = 0.27
Z lm Score = 0.12
NG = 0
DB = 0
SM = 0
Delta_AUC
Doxorubicin
YLR319C BUD6
ZShift = −0.39
Z lm Score = 3.11
NG = 0
DB = 0
SM = 0
Delta_L
Doxorubicin
YLR319C BUD6
ZShift = −0.09
Z lm Score = 0.14
NG = 0
DB = 0
SM = 0
Delta_K
Doxorubicin
YLR319C BUD6
ZShift = 0.4
Z lm Score = 0.55
NG = 0
DB = 0
SM = 0
Delta_r
Doxorubicin
YLR319C BUD6
ZShift = 0.27
Z lm Score = 0.6
NG = 0
DB = 0
SM = 0
Delta_AUC
Doxorubicin

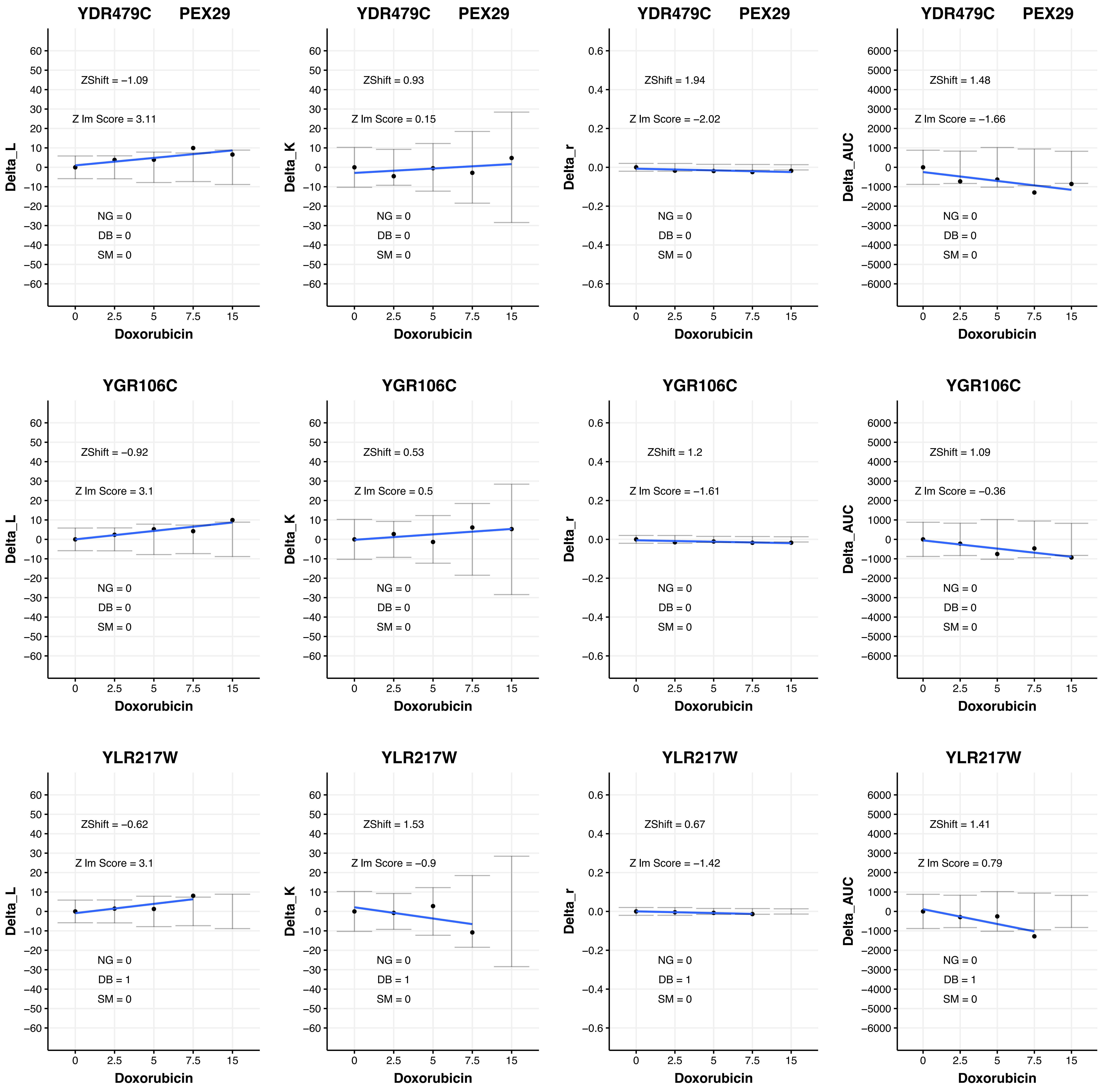
YDR479C PEX29
ZShift = −1.09
Z lm Score = 3.11
NG = 0
DB = 0
SM = 0
Delta_L
Doxorubicin
YDR479C PEX29
ZShift = 0.93
Z lm Score = 0.15
NG = 0
DB = 0
SM = 0
Delta_K
Doxorubicin
YDR479C PEX29
ZShift = 1.94
Z lm Score = −2.02
NG = 0
DB = 0
SM = 0
Delta_r
Doxorubicin
YDR479C PEX29
ZShift = 1.48
Z lm Score = −1.66
NG = 0
DB = 0
SM = 0
Delta_AUC
Doxorubicin
YGR106C
ZShift = −0.92
Z lm Score = 3.1
NG = 0
DB = 0
SM = 0
Delta_L
Doxorubicin
YGR106C
ZShift = 0.53
Z lm Score = 0.5
NG = 0
DB = 0
SM = 0
Delta_K
Doxorubicin
YGR106C
ZShift = 1.2
Z lm Score = −1.61
NG = 0
DB = 0
SM = 0
Delta_r
Doxorubicin
YGR106C
ZShift = 1.09
Z lm Score = −0.36
NG = 0
DB = 0
SM = 0
Delta_AUC
Doxorubicin
YLR217W
ZShift = −0.62
Z lm Score = 3.1
NG = 0
DB = 1
SM = 0
Delta_L
Doxorubicin
YLR217W
ZShift = 1.53
Z lm Score = −0.9
NG = 0
DB = 1
SM = 0
Delta_K
Doxorubicin
YLR217W
ZShift = 0.67
Z lm Score = −1.42
NG = 0
DB = 1
SM = 0
Delta_r
Doxorubicin
YLR217W
ZShift = 1.41
Z lm Score = 0.79
NG = 0
DB = 1
SM = 0
Delta_AUC
Doxorubicin

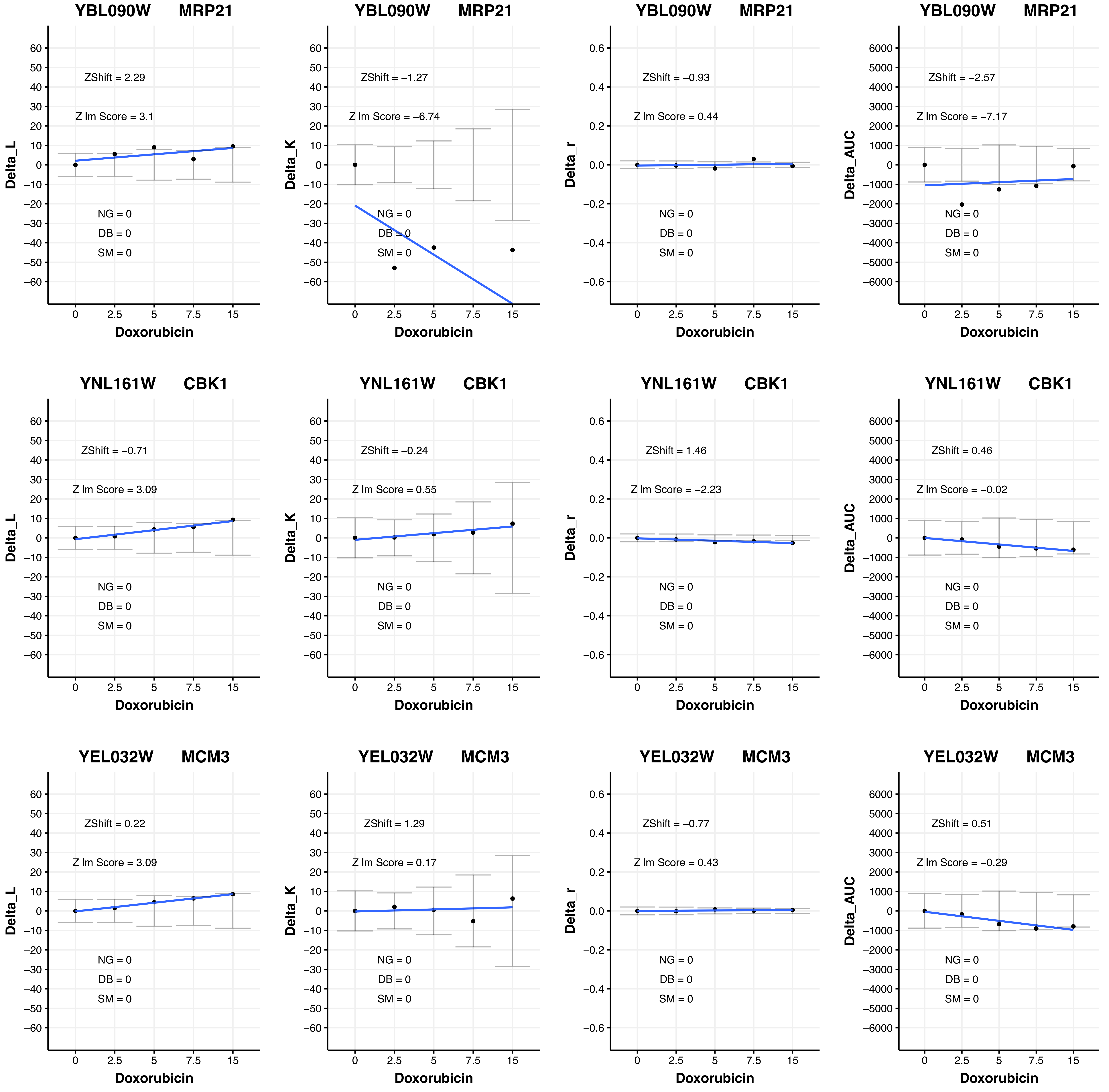
YBL090W MRP21
ZShift = 2.29
Z lm Score = 3.1
NG = 0
DB = 0
SM = 0
Delta_L
Doxorubicin
YBL090W MRP21
ZShift = −1.27
Z lm Score = −6.74
NG = 0
DB = 0
SM = 0
Delta_K
Doxorubicin
YBL090W MRP21
ZShift = −0.93
Z lm Score = 0.44
NG = 0
DB = 0
SM = 0
Delta_r
Doxorubicin
YBL090W MRP21
ZShift = −2.57
Z lm Score = −7.17
NG = 0
DB = 0
SM = 0
Delta_AUC
Doxorubicin
YNL161W CBK1
ZShift = −0.71
Z lm Score = 3.09
NG = 0
DB = 0
SM = 0
Delta_L
Doxorubicin
YNL161W CBK1
ZShift = −0.24
Z lm Score = 0.55
NG = 0
DB = 0
SM = 0
Delta_K
Doxorubicin
YNL161W CBK1
ZShift = 1.46
Z lm Score = −2.23
NG = 0
DB = 0
SM = 0
Delta_r
Doxorubicin
YNL161W CBK1
ZShift = 0.46
Z lm Score = −0.02
NG = 0
DB = 0
SM = 0
Delta_AUC
Doxorubicin
YEL032W MCM3
ZShift = 0.22
Z lm Score = 3.09
NG = 0
DB = 0
SM = 0
Delta_L
Doxorubicin
YEL032W MCM3
ZShift = 1.29
Z lm Score = 0.17
NG = 0
DB = 0
SM = 0
Delta_K
Doxorubicin
YEL032W MCM3
ZShift = −0.77
Z lm Score = 0.43
NG = 0
DB = 0
SM = 0
Delta_r
Doxorubicin
YEL032W MCM3
ZShift = 0.51
Z lm Score = −0.29
NG = 0
DB = 0
SM = 0
Delta_AUC
Doxorubicin

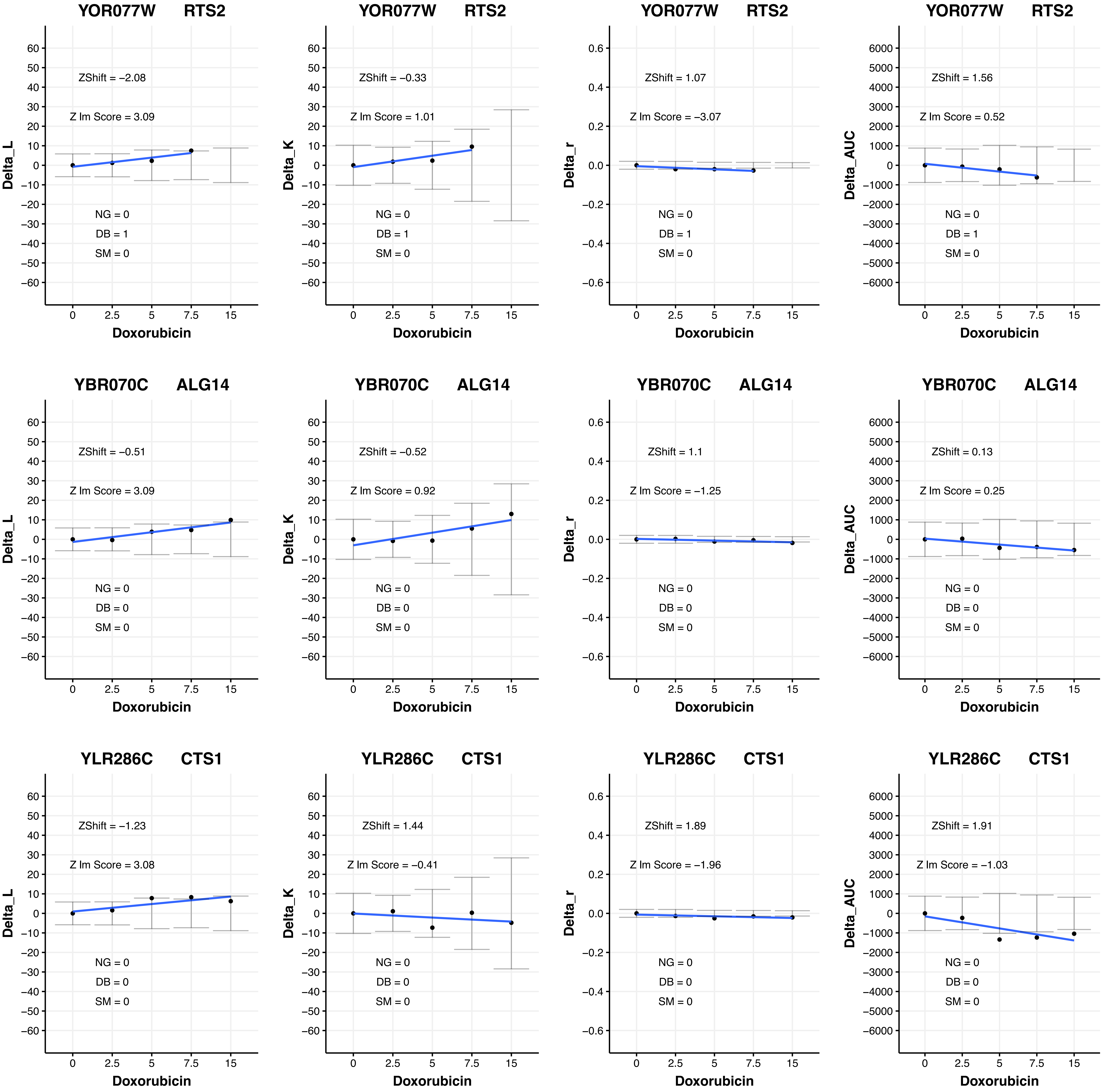
YOR077W RTS2
ZShift = −2.08
Z lm Score = 3.09
NG = 0
DB = 1
SM = 0
Delta_L
Doxorubicin
YOR077W RTS2
ZShift = −0.33
Z lm Score = 1.01
NG = 0
DB = 1
SM = 0
Delta_K
Doxorubicin
YOR077W RTS2
ZShift = 1.07
Z lm Score = −3.07
NG = 0
DB = 1
SM = 0
Delta_r
Doxorubicin
YOR077W RTS2
ZShift = 1.56
Z lm Score = 0.52
NG = 0
DB = 1
SM = 0
Delta_AUC
Doxorubicin
YBR070C ALG14
ZShift = −0.51
Z lm Score = 3.09
NG = 0
DB = 0
SM = 0
Delta_L
Doxorubicin
YBR070C ALG14
ZShift = −0.52
Z lm Score = 0.92
NG = 0
DB = 0
SM = 0
Delta_K
Doxorubicin
YBR070C ALG14
ZShift = 1.1
Z lm Score = −1.25
NG = 0
DB = 0
SM = 0
Delta_r
Doxorubicin
YBR070C ALG14
ZShift = 0.13
Z lm Score = 0.25
NG = 0
DB = 0
SM = 0
Delta_AUC
Doxorubicin
YLR286C CTS1
ZShift = −1.23
Z lm Score = 3.08
NG = 0
DB = 0
SM = 0
Delta_L
Doxorubicin
YLR286C CTS1
ZShift = 1.44
Z lm Score = −0.41
NG = 0
DB = 0
SM = 0
Delta_K
Doxorubicin
YLR286C CTS1
ZShift = 1.89
Z lm Score = −1.96
NG = 0
DB = 0
SM = 0
Delta_r
Doxorubicin
YLR286C CTS1
ZShift = 1.91
Z lm Score = −1.03
NG = 0
DB = 0
SM = 0
Delta_AUC
Doxorubicin

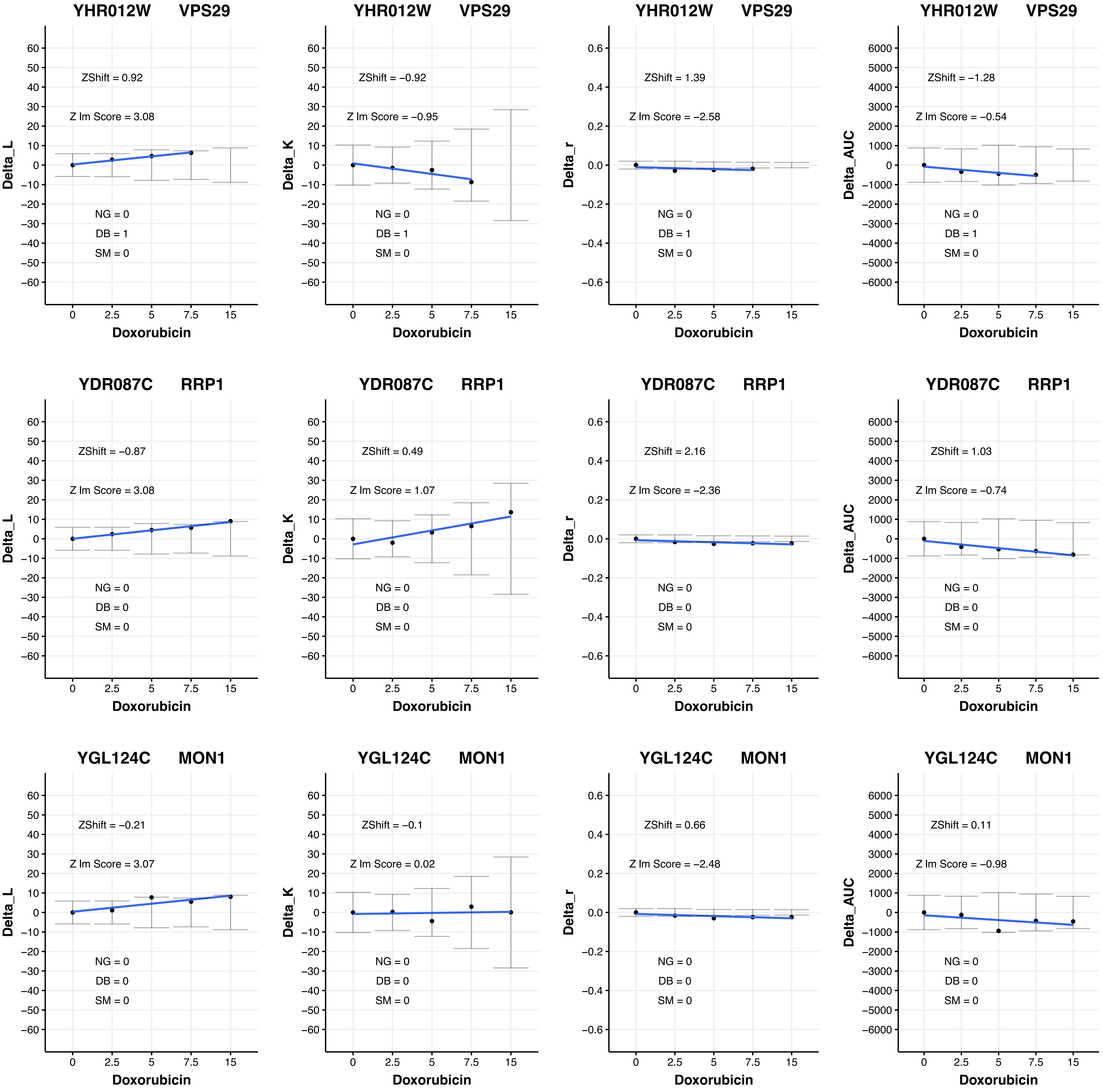
YHR012W VPS29
ZShift = 0.92
Z lm Score = 3.08
NG = 0
DB = 1
SM = 0
Delta_L
Doxorubicin
YHR012W VPS29
ZShift = −0.92
Z lm Score = −0.95
NG = 0
DB = 1
SM = 0
Delta_K
Doxorubicin
YHR012W VPS29
ZShift = 1.39
Z lm Score = −2.58
NG = 0
DB = 1
SM = 0
Delta_r
Doxorubicin
YHR012W VPS29
ZShift = −1.28
Z lm Score = −0.54
NG = 0
DB = 1
SM = 0
Delta_AUC
Doxorubicin
YDR087C RRP1
ZShift = −0.87
Z lm Score = 3.08
NG = 0
DB = 0
SM = 0
Delta_L
Doxorubicin
YDR087C RRP1
ZShift = 0.49
Z lm Score = 1.07
NG = 0
DB = 0
SM = 0
Delta_K
Doxorubicin
YDR087C RRP1
ZShift = 2.16
Z lm Score = −2.36
NG = 0
DB = 0
SM = 0
Delta_r
Doxorubicin
YDR087C RRP1
ZShift = 1.03
Z lm Score = −0.74
NG = 0
DB = 0
SM = 0
Delta_AUC
Doxorubicin
YGL124C MON1
ZShift = −0.21
Z lm Score = 3.07
NG = 0
DB = 0
SM = 0
Delta_L
Doxorubicin
YGL124C MON1
ZShift = −0.1
Z lm Score = 0.02
NG = 0
DB = 0
SM = 0
Delta_K
Doxorubicin
YGL124C MON1
ZShift = 0.66
Z lm Score = −2.48
NG = 0
DB = 0
SM = 0
Delta_r
Doxorubicin
YGL124C MON1
ZShift = 0.11
Z lm Score = −0.98
NG = 0
DB = 0
SM = 0
Delta_AUC
Doxorubicin

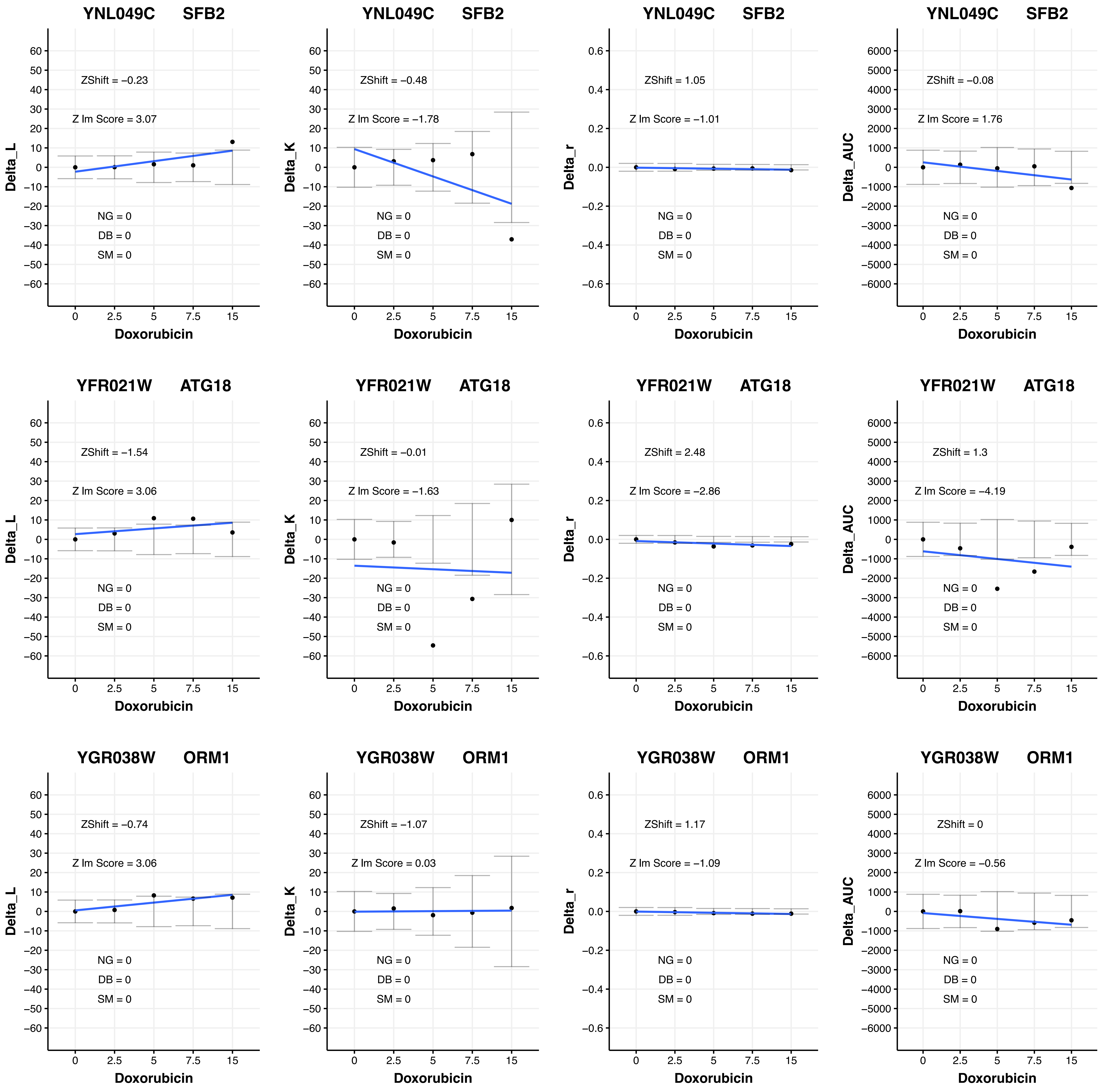
YNL049C SFB2
ZShift = −0.23
Z lm Score = 3.07
NG = 0
DB = 0
SM = 0
Delta_L
Doxorubicin
YNL049C SFB2
ZShift = −0.48
Z lm Score = −1.78
NG = 0
DB = 0
SM = 0
Delta_K
Doxorubicin
YNL049C SFB2
ZShift = 1.05
Z lm Score = −1.01
NG = 0
DB = 0
SM = 0
Delta_r
Doxorubicin
YNL049C SFB2
ZShift = −0.08
Z lm Score = 1.76
NG = 0
DB = 0
SM = 0
Delta_AUC
Doxorubicin
YFR021W ATG18
ZShift = −1.54
Z lm Score = 3.06
NG = 0
DB = 0
SM = 0
Delta_L
Doxorubicin
YFR021W ATG18
ZShift = −0.01
Z lm Score = −1.63
NG = 0
DB = 0
SM = 0
Delta_K
Doxorubicin
YFR021W ATG18
ZShift = 2.48
Z lm Score = −2.86
NG = 0
DB = 0
SM = 0
Delta_r
Doxorubicin
YFR021W ATG18
ZShift = 1.3
Z lm Score = −4.19
NG = 0
DB = 0
SM = 0
Delta_AUC
Doxorubicin
YGR038W ORM1
ZShift = −0.74
Z lm Score = 3.06
NG = 0
DB = 0
SM = 0
Delta_L
Doxorubicin
YGR038W ORM1
ZShift = −1.07
Z lm Score = 0.03
NG = 0
DB = 0
SM = 0
Delta_K
Doxorubicin
YGR038W ORM1
ZShift = 1.17
Z lm Score = −1.09
NG = 0
DB = 0
SM = 0
Delta_r
Doxorubicin
YGR038W ORM1
ZShift = 0
Z lm Score = −0.56
NG = 0
DB = 0
SM = 0
Delta_AUC
Doxorubicin

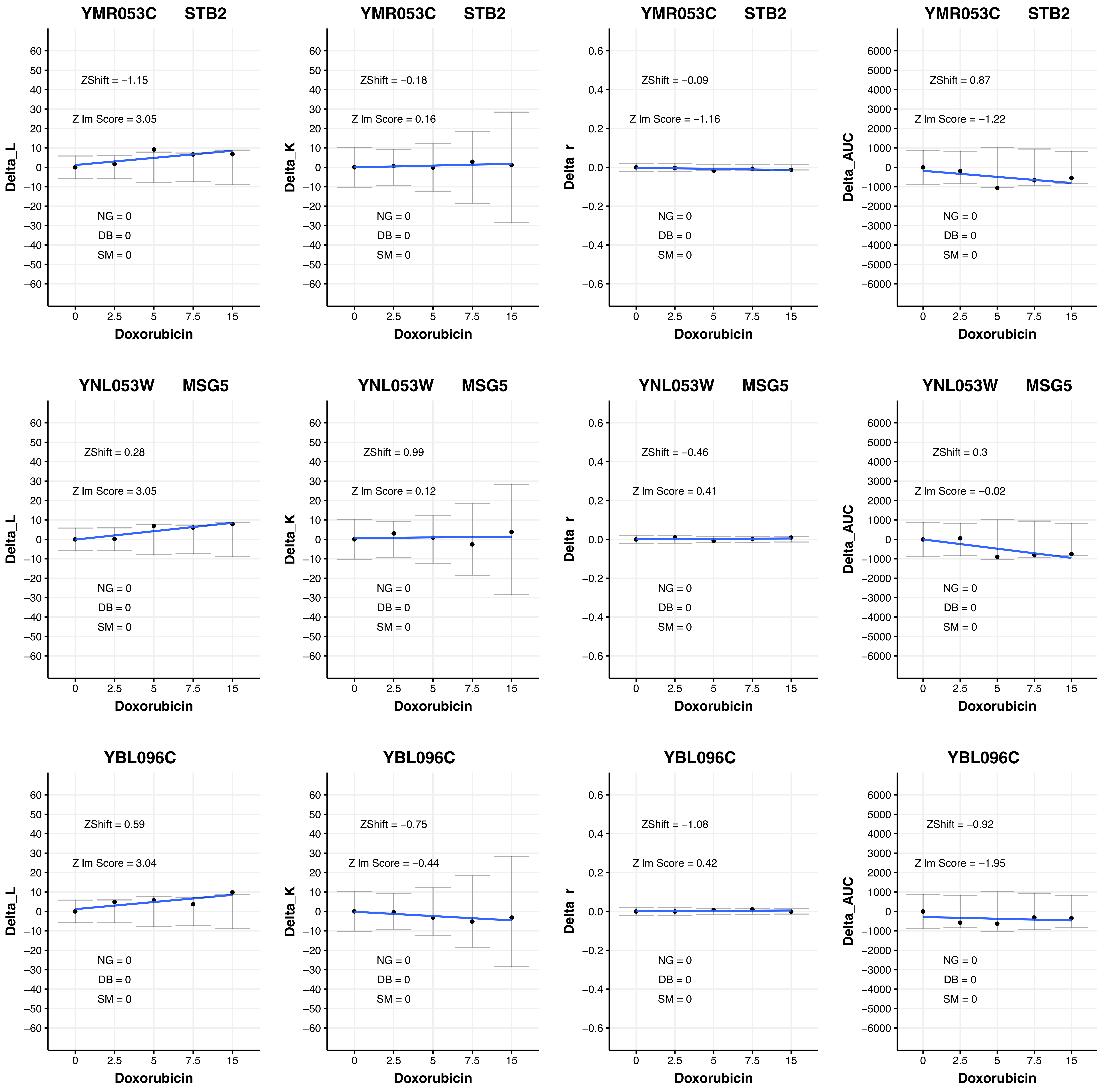

YMR053C STB2
ZShift = −1.15
Z lm Score = 3.05
NG = 0
DB = 0
SM = 0
Delta_L
Doxorubicin
YMR053C STB2
ZShift = −0.18
Z lm Score = 0.16
NG = 0
DB = 0
SM = 0
Delta_K
Doxorubicin
YMR053C STB2
ZShift = −0.09
Z lm Score = −1.16
NG = 0
DB = 0
SM = 0
Delta_r
Doxorubicin
YMR053C STB2
ZShift = 0.87
Z lm Score = −1.22
NG = 0
DB = 0
SM = 0
Delta_AUC
Doxorubicin
YNL053W MSG5
ZShift = 0.28
Z lm Score = 3.05
NG = 0
DB = 0
SM = 0
Delta_L
Doxorubicin
YNL053W MSG5
ZShift = 0.99
Z lm Score = 0.12
NG = 0
DB = 0
SM = 0
Delta_K
Doxorubicin
YNL053W MSG5
ZShift = −0.46
Z lm Score = 0.41
NG = 0
DB = 0
SM = 0
Delta_r
Doxorubicin
YNL053W MSG5
ZShift = 0.3
Z lm Score = −0.02
NG = 0
DB = 0
SM = 0
Delta_AUC
Doxorubicin
YBL096C
ZShift = 0.59
Z lm Score = 3.04
NG = 0
DB = 0
SM = 0
Delta_L
Doxorubicin
YBL096C
ZShift = −0.75
Z lm Score = −0.44
NG = 0
DB = 0
SM = 0
Delta_K
Doxorubicin
YBL096C
ZShift = −1.08
Z lm Score = 0.42
NG = 0
DB = 0
SM = 0
Delta_r
Doxorubicin
YBL096C
ZShift = −0.92
Z lm Score = −1.95
NG = 0
DB = 0
SM = 0
Delta_AUC
Doxorubicin

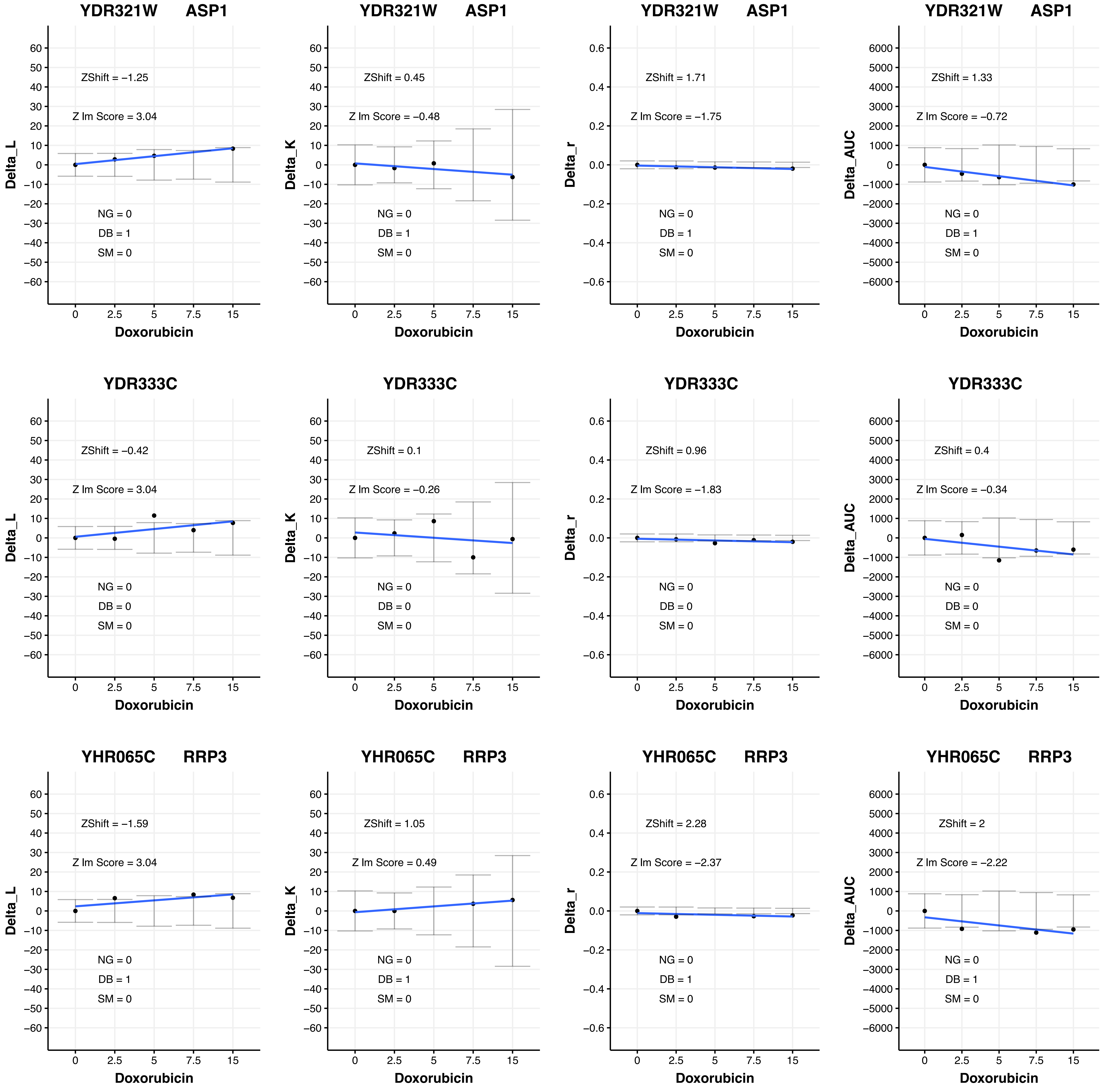
YDR321W ASP1
ZShift = −1.25
Z lm Score = 3.04
NG = 0
DB = 1
SM = 0
Delta_L
Doxorubicin
YDR321W ASP1
ZShift = 0.45
Z lm Score = −0.48
NG = 0
DB = 1
SM = 0
Delta_K
Doxorubicin
YDR321W ASP1
ZShift = 1.71
Z lm Score = −1.75
NG = 0
DB = 1
SM = 0
Delta_r
Doxorubicin
YDR321W ASP1
ZShift = 1.33
Z lm Score = −0.72
NG = 0
DB = 1
SM = 0
Delta_AUC
Doxorubicin
YDR333C
ZShift = −0.42
Z lm Score = 3.04
NG = 0
DB = 0
SM = 0
Delta_L
Doxorubicin
YDR333C
ZShift = 0.1
Z lm Score = −0.26
NG = 0
DB = 0
SM = 0
Delta_K
Doxorubicin
YDR333C
ZShift = 0.96
Z lm Score = −1.83
NG = 0
DB = 0
SM = 0
Delta_r
Doxorubicin
YDR333C
ZShift = 0.4
Z lm Score = −0.34
NG = 0
DB = 0
SM = 0
Delta_AUC
Doxorubicin
YHR065C RRP3
ZShift = −1.59
Z lm Score = 3.04
NG = 0
DB = 1
SM = 0
Delta_L
Doxorubicin
YHR065C RRP3
ZShift = 1.05
Z lm Score = 0.49
NG = 0
DB = 1
SM = 0
Delta_K
Doxorubicin
YHR065C RRP3
ZShift = 2.28
Z lm Score = −2.37
NG = 0
DB = 1
SM = 0
Delta_r
Doxorubicin
YHR065C RRP3
ZShift = 2
Z lm Score = −2.22
NG = 0
DB = 1
SM = 0
Delta_AUC
Doxorubicin

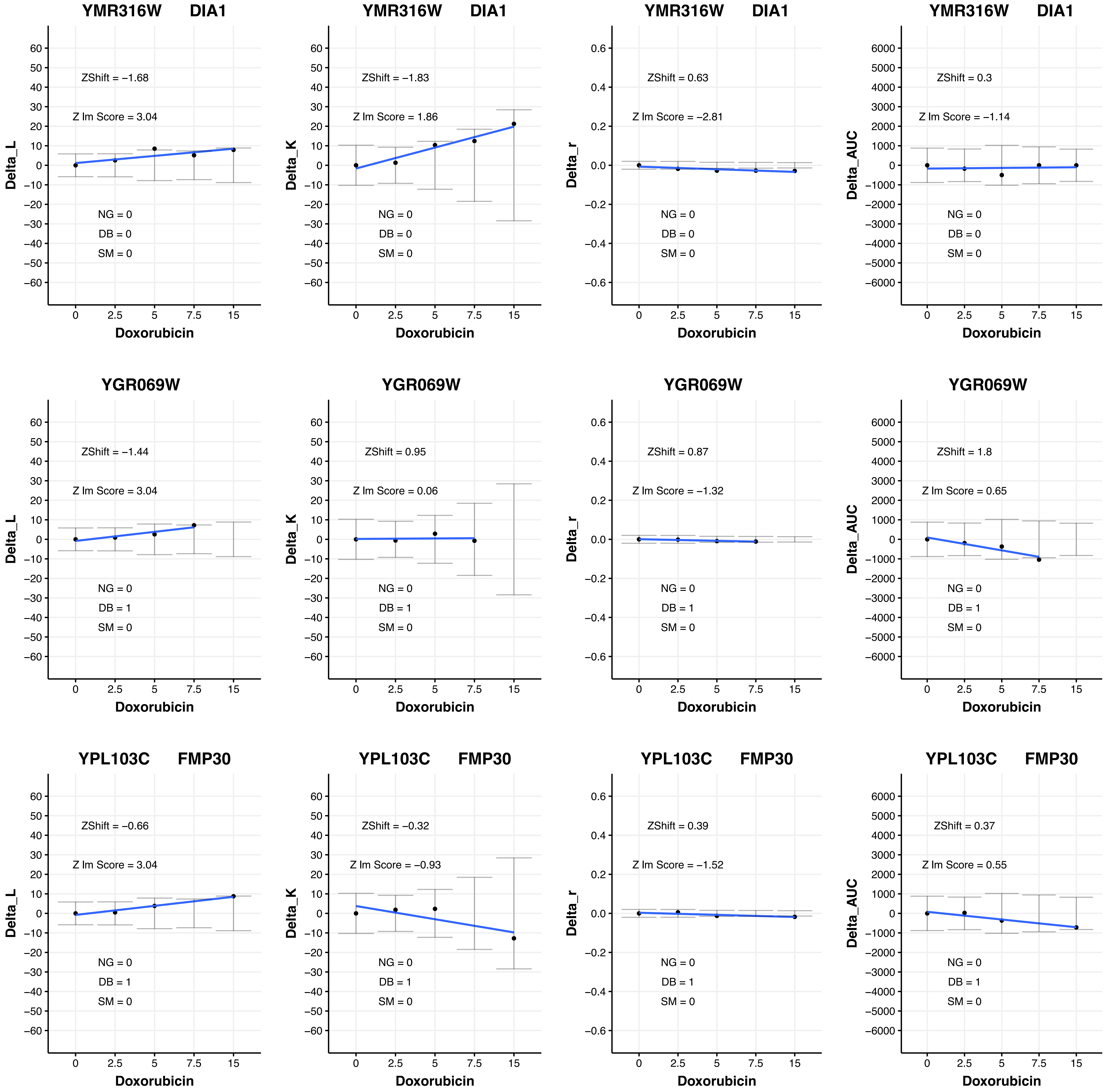
YMR316W DIA1
ZShift = −1.68
Z lm Score = 3.04
NG = 0
DB = 0
SM = 0
Delta_L
Doxorubicin
YMR316W DIA1
ZShift = −1.83
Z lm Score = 1.86
NG = 0
DB = 0
SM = 0
Delta_K
Doxorubicin
YMR316W DIA1
ZShift = 0.63
Z lm Score = −2.81
NG = 0
DB = 0
SM = 0
Delta_r
Doxorubicin
YMR316W DIA1
ZShift = 0.3
Z lm Score = −1.14
NG = 0
DB = 0
SM = 0
Delta_AUC
Doxorubicin
YGR069W
ZShift = −1.44
Z lm Score = 3.04
NG = 0
DB = 1
SM = 0
Delta_L
Doxorubicin
YGR069W
ZShift = 0.95
Z lm Score = 0.06
NG = 0
DB = 1
SM = 0
Delta_K
Doxorubicin
YGR069W
ZShift = 0.87
Z lm Score = −1.32
NG = 0
DB = 1
SM = 0
Delta_r
Doxorubicin
YGR069W
ZShift = 1.8
Z lm Score = 0.65
NG = 0
DB = 1
SM = 0
Delta_AUC
Doxorubicin
YPL103C FMP30
ZShift = −0.66
Z lm Score = 3.04
NG = 0
DB = 1
SM = 0
Delta_L
Doxorubicin
YPL103C FMP30
ZShift = −0.32
Z lm Score = −0.93
NG = 0
DB = 1
SM = 0
Delta_K
Doxorubicin
YPL103C FMP30
ZShift = 0.39
Z lm Score = −1.52
NG = 0
DB = 1
SM = 0
Delta_r
Doxorubicin
YPL103C FMP30
ZShift = 0.37
Z lm Score = 0.55
NG = 0
DB = 1
SM = 0
Delta_AUC
Doxorubicin

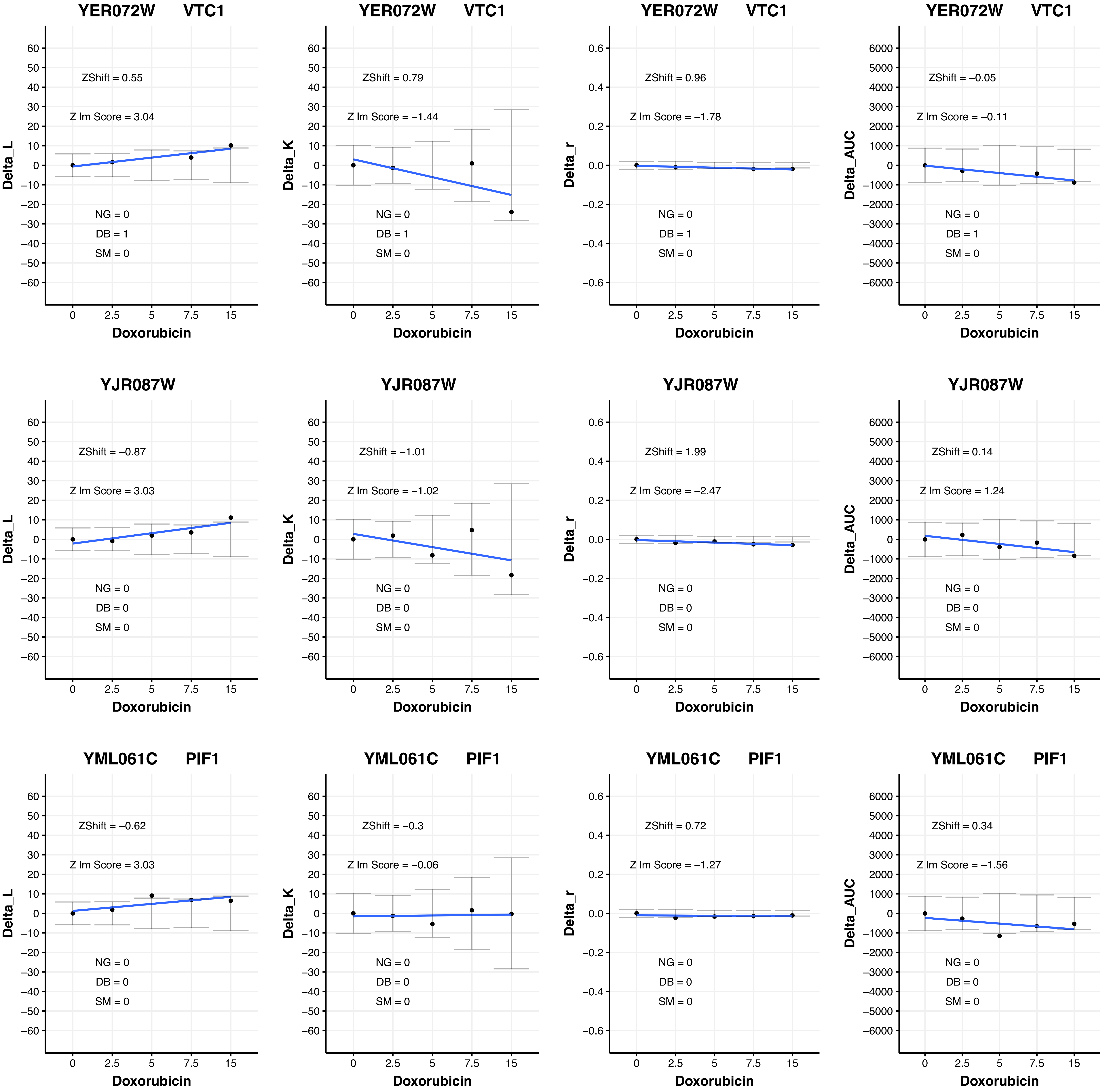
YER072W VTC1
ZShift = 0.55
Z lm Score = 3.04
NG = 0
DB = 1
SM = 0
Delta_L
Doxorubicin
YER072W VTC1
ZShift = 0.79
Z lm Score = −1.44
NG = 0
DB = 1
SM = 0
Delta_K
Doxorubicin
YER072W VTC1
ZShift = 0.96
Z lm Score = −1.78
NG = 0
DB = 1
SM = 0
Delta_r
Doxorubicin
YER072W VTC1
ZShift = −0.05
Z lm Score = −0.11
NG = 0
DB = 1
SM = 0
Delta_AUC
Doxorubicin
YJR087W
ZShift = −0.87
Z lm Score = 3.03
NG = 0
DB = 0
SM = 0
Delta_L
Doxorubicin
YJR087W
ZShift = −1.01
Z lm Score = −1.02
NG = 0
DB = 0
SM = 0
Delta_K
Doxorubicin
YJR087W
ZShift = 1.99
Z lm Score = −2.47
NG = 0
DB = 0
SM = 0
Delta_r
Doxorubicin
YJR087W
ZShift = 0.14
Z lm Score = 1.24
NG = 0
DB = 0
SM = 0
Delta_AUC
Doxorubicin
YML061C PIF1
ZShift = −0.62
Z lm Score = 3.03
NG = 0
DB = 0
SM = 0
Delta_L
Doxorubicin
YML061C PIF1
ZShift = −0.3
Z lm Score = −0.06
NG = 0
DB = 0
SM = 0
Delta_K
Doxorubicin
YML061C PIF1
ZShift = 0.72
Z lm Score = −1.27
NG = 0
DB = 0
SM = 0
Delta_r
Doxorubicin
YML061C PIF1
ZShift = 0.34
Z lm Score = −1.56
NG = 0
DB = 0
SM = 0
Delta_AUC
Doxorubicin

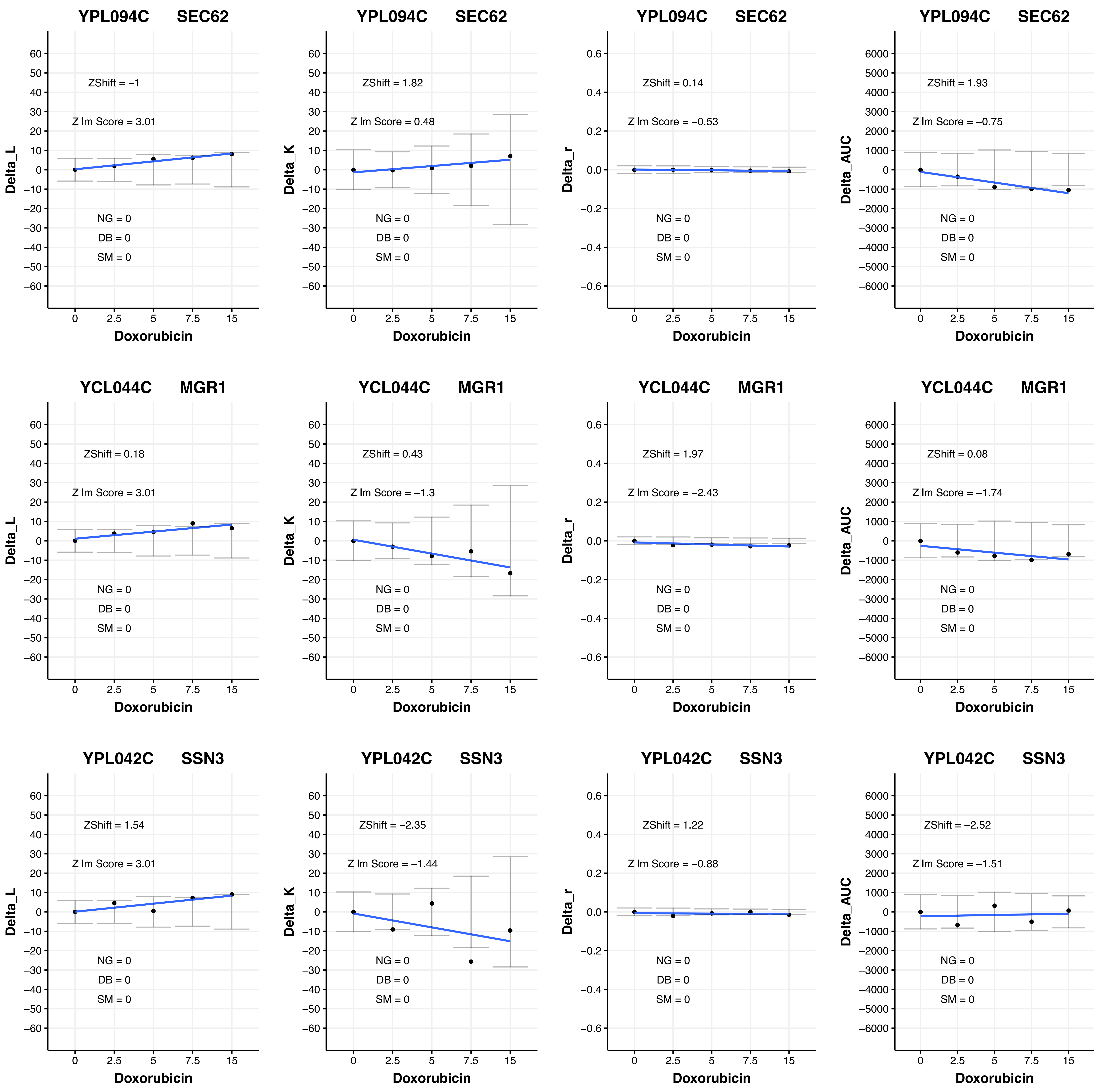
YPL094C SEC62
ZShift = −1
Z lm Score = 3.01
NG = 0
DB = 0
SM = 0
Delta_L
Doxorubicin
YPL094C SEC62
ZShift = 1.82
Z lm Score = 0.48
NG = 0
DB = 0
SM = 0
Delta_K
Doxorubicin
YPL094C SEC62
ZShift = 0.14
Z lm Score = −0.53
NG = 0
DB = 0
SM = 0
Delta_r
Doxorubicin
YPL094C SEC62
ZShift = 1.93
Z lm Score = −0.75
NG = 0
DB = 0
SM = 0
Delta_AUC
Doxorubicin
YCL044C MGR1
ZShift = 0.18
Z lm Score = 3.01
NG = 0
DB = 0
SM = 0
Delta_L
Doxorubicin
YCL044C MGR1
ZShift = 0.43
Z lm Score = −1.3
NG = 0
DB = 0
SM = 0
Delta_K
Doxorubicin
YCL044C MGR1
ZShift = 1.97
Z lm Score = −2.43
NG = 0
DB = 0
SM = 0
Delta_r
Doxorubicin
YCL044C MGR1
ZShift = 0.08
Z lm Score = −1.74
NG = 0
DB = 0
SM = 0
Delta_AUC
Doxorubicin
YPL042C SSN3
ZShift = 1.54
Z lm Score = 3.01
NG = 0
DB = 0
SM = 0
Delta_L
Doxorubicin
YPL042C SSN3
ZShift = −2.35
Z lm Score = −1.44
NG = 0
DB = 0
SM = 0
Delta_K
Doxorubicin
YPL042C SSN3
ZShift = 1.22
Z lm Score = −0.88
NG = 0
DB = 0
SM = 0
Delta_r
Doxorubicin
YPL042C SSN3
ZShift = −2.52
Z lm Score = −1.51
NG = 0
DB = 0
SM = 0
Delta_AUC
Doxorubicin

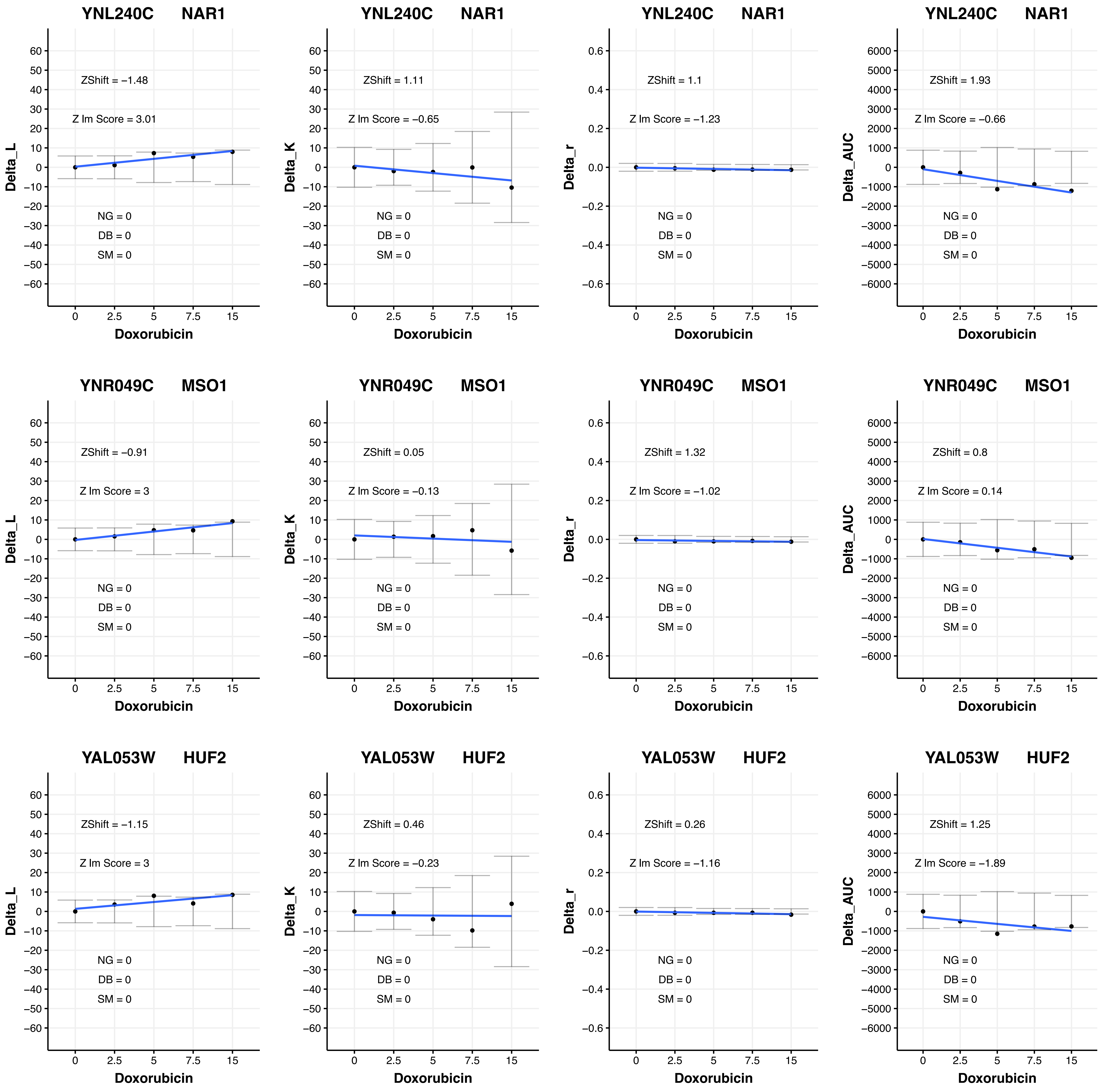
YNL240C NAR1
ZShift = −1.48
Z lm Score = 3.01
NG = 0
DB = 0
SM = 0
Delta_L
Doxorubicin
YNL240C NAR1
ZShift = 1.11
Z lm Score = −0.65
NG = 0
DB = 0
SM = 0
Delta_K
Doxorubicin
YNL240C NAR1
ZShift = 1.1
Z lm Score = −1.23
NG = 0
DB = 0
SM = 0
Delta_r
Doxorubicin
YNL240C NAR1
ZShift = 1.93
Z lm Score = −0.66
NG = 0
DB = 0
SM = 0
Delta_AUC
Doxorubicin
YNR049C MSO1
ZShift = −0.91
Z lm Score = 3
NG = 0
DB = 0
SM = 0
Delta_L
Doxorubicin
YNR049C MSO1
ZShift = 0.05
Z lm Score = −0.13
NG = 0
DB = 0
SM = 0
Delta_K
Doxorubicin
YNR049C MSO1
ZShift = 1.32
Z lm Score = −1.02
NG = 0
DB = 0
SM = 0
Delta_r
Doxorubicin
YNR049C MSO1
ZShift = 0.8
Z lm Score = 0.14
NG = 0
DB = 0
SM = 0
Delta_AUC
Doxorubicin
YAL053W HUF2
ZShift = −1.15
Z lm Score = 3
NG = 0
DB = 0
SM = 0
Delta_L
Doxorubicin
YAL053W HUF2
ZShift = 0.46
Z lm Score = −0.23
NG = 0
DB = 0
SM = 0
Delta_K
Doxorubicin
YAL053W HUF2
ZShift = 0.26
Z lm Score = −1.16
NG = 0
DB = 0
SM = 0
Delta_r
Doxorubicin
YAL053W HUF2
ZShift = 1.25
Z lm Score = −1.89
NG = 0
DB = 0
SM = 0
Delta_AUC
Doxorubicin

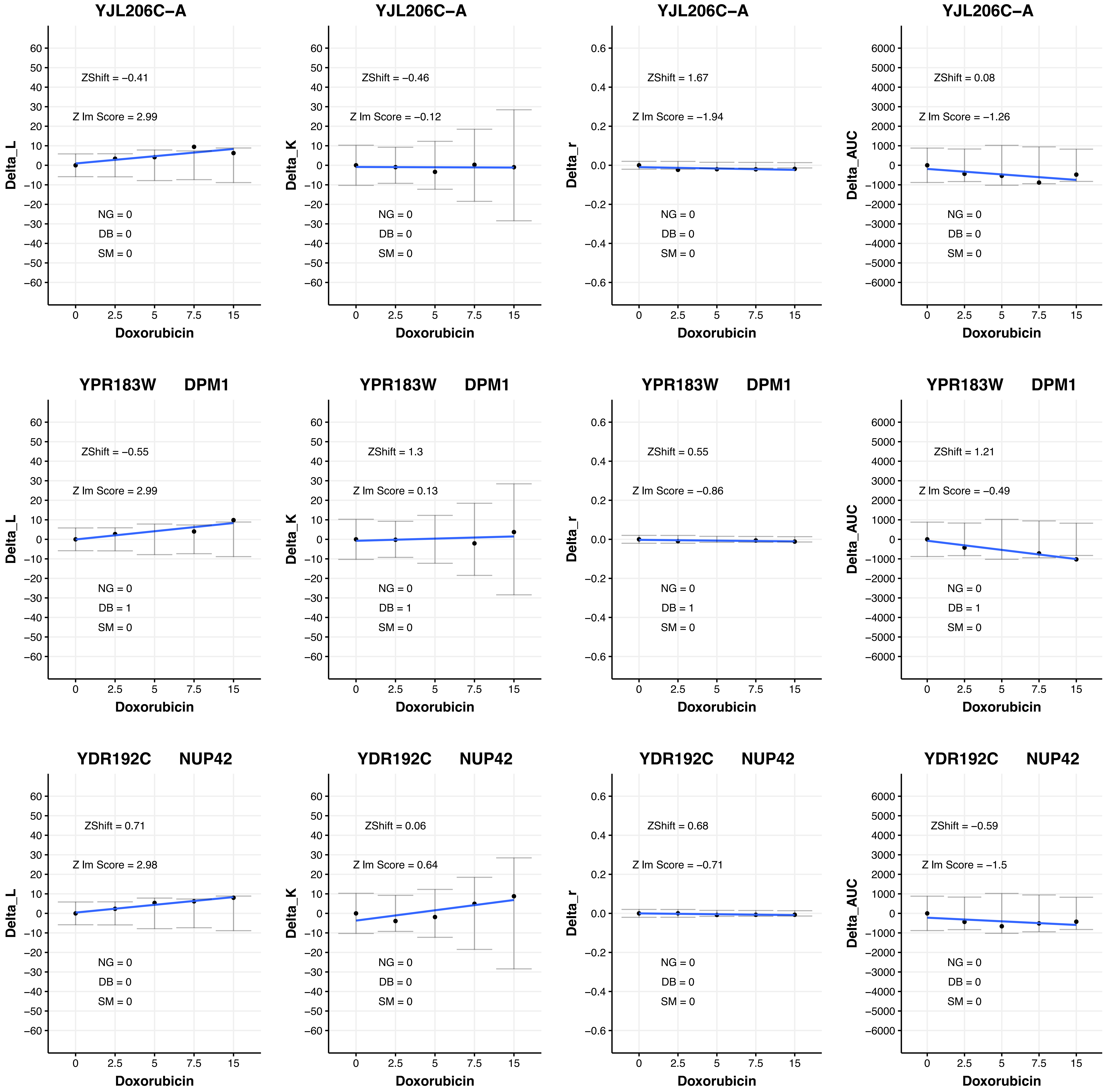

YJL206C−A
ZShift = −0.41
Z lm Score = 2.99
NG = 0
DB = 0
SM = 0
Delta_L
Doxorubicin
YJL206C−A
ZShift = −0.46
Z lm Score = −0.12
NG = 0
DB = 0
SM = 0
Delta_K
Doxorubicin
YJL206C−A
ZShift = 1.67
Z lm Score = −1.94
NG = 0
DB = 0
SM = 0
Delta_r
Doxorubicin
YJL206C−A
ZShift = 0.08
Z lm Score = −1.26
NG = 0
DB = 0
SM = 0
Delta_AUC
Doxorubicin
YPR183W DPM1
ZShift = −0.55
Z lm Score = 2.99
NG = 0
DB = 1
SM = 0
Delta_L
Doxorubicin
YPR183W DPM1
ZShift = 1.3
Z lm Score = 0.13
NG = 0
DB = 1
SM = 0
Delta_K
Doxorubicin
YPR183W DPM1
ZShift = 0.55
Z lm Score = −0.86
NG = 0
DB = 1
SM = 0
Delta_r
Doxorubicin
YPR183W DPM1
ZShift = 1.21
Z lm Score = −0.49
NG = 0
DB = 1
SM = 0
Delta_AUC
Doxorubicin
YDR192C NUP42
ZShift = 0.71
Z lm Score = 2.98
NG = 0
DB = 0
SM = 0
Delta_L
Doxorubicin
YDR192C NUP42
ZShift = 0.06
Z lm Score = 0.64
NG = 0
DB = 0
SM = 0
Delta_K
Doxorubicin
YDR192C NUP42
ZShift = 0.68
Z lm Score = −0.71
NG = 0
DB = 0
SM = 0
Delta_r
Doxorubicin
YDR192C NUP42
ZShift = −0.59
Z lm Score = −1.5
NG = 0
DB = 0
SM = 0
Delta_AUC
Doxorubicin

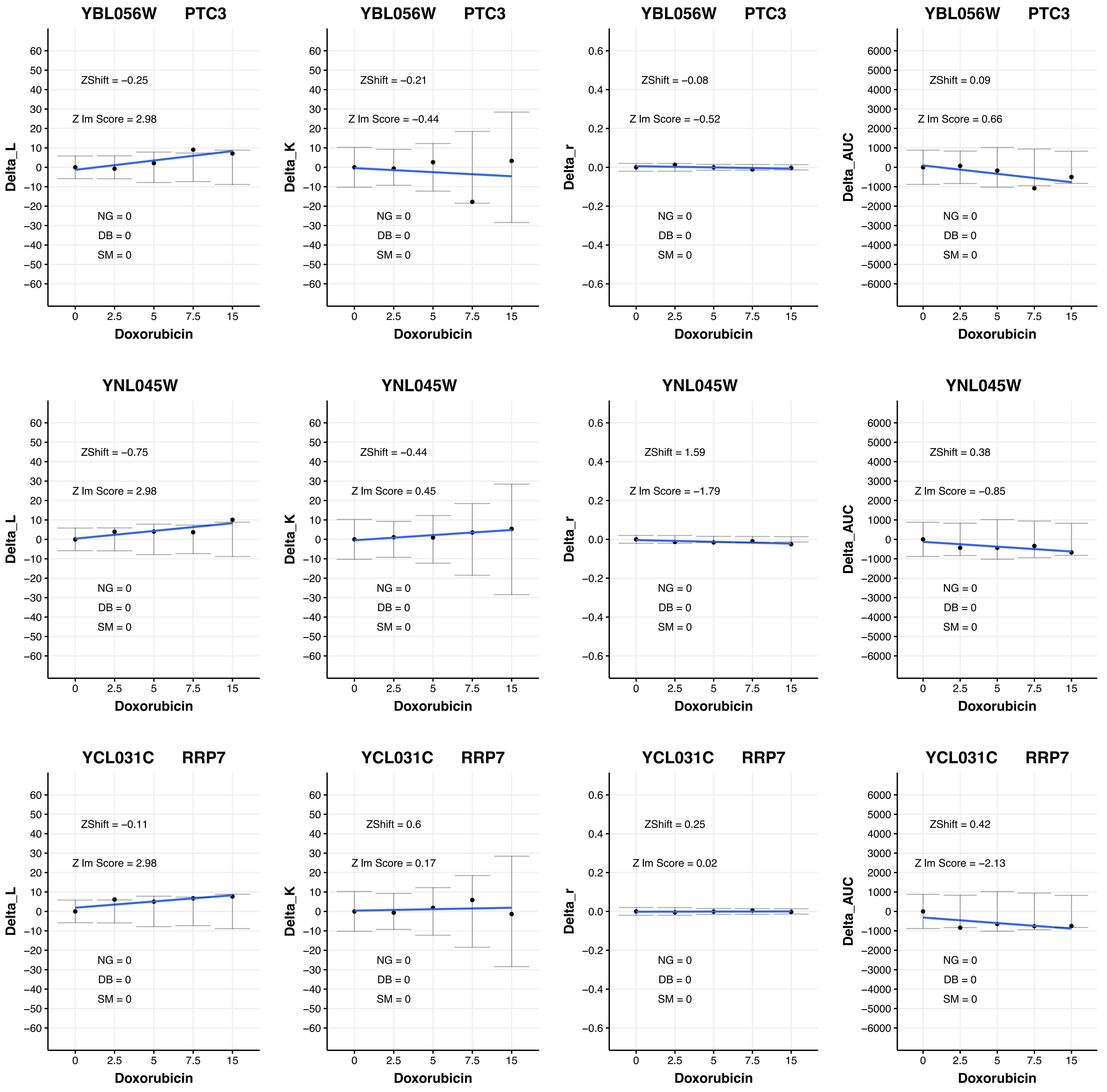
YBL056W PTC3
ZShift = −0.25
Z lm Score = 2.98
NG = 0
DB = 0
SM = 0
Delta_L
Doxorubicin
YBL056W PTC3
ZShift = −0.21
Z lm Score = −0.44
NG = 0
DB = 0
SM = 0
Delta_K
Doxorubicin
YBL056W PTC3
ZShift = −0.08
Z lm Score = −0.52
NG = 0
DB = 0
SM = 0
Delta_r
Doxorubicin
YBL056W PTC3
ZShift = 0.09
Z lm Score = 0.66
NG = 0
DB = 0
SM = 0
Delta_AUC
Doxorubicin
YNL045W
ZShift = −0.75
Z lm Score = 2.98
NG = 0
DB = 0
SM = 0
Delta_L
Doxorubicin
YNL045W
ZShift = −0.44
Z lm Score = 0.45
NG = 0
DB = 0
SM = 0
Delta_K
Doxorubicin
YNL045W
ZShift = 1.59
Z lm Score = −1.79
NG = 0
DB = 0
SM = 0
Delta_r
Doxorubicin
YNL045W
ZShift = 0.38
Z lm Score = −0.85
NG = 0
DB = 0
SM = 0
Delta_AUC
Doxorubicin
YCL031C RRP7
ZShift = −0.11
Z lm Score = 2.98
NG = 0
DB = 0
SM = 0
Delta_L
Doxorubicin
YCL031C RRP7
ZShift = 0.6
Z lm Score = 0.17
NG = 0
DB = 0
SM = 0
Delta_K
Doxorubicin
YCL031C RRP7
ZShift = 0.25
Z lm Score = 0.02
NG = 0
DB = 0
SM = 0
Delta_r
Doxorubicin
YCL031C RRP7
ZShift = 0.42
Z lm Score = −2.13
NG = 0
DB = 0
SM = 0
Delta_AUC
Doxorubicin

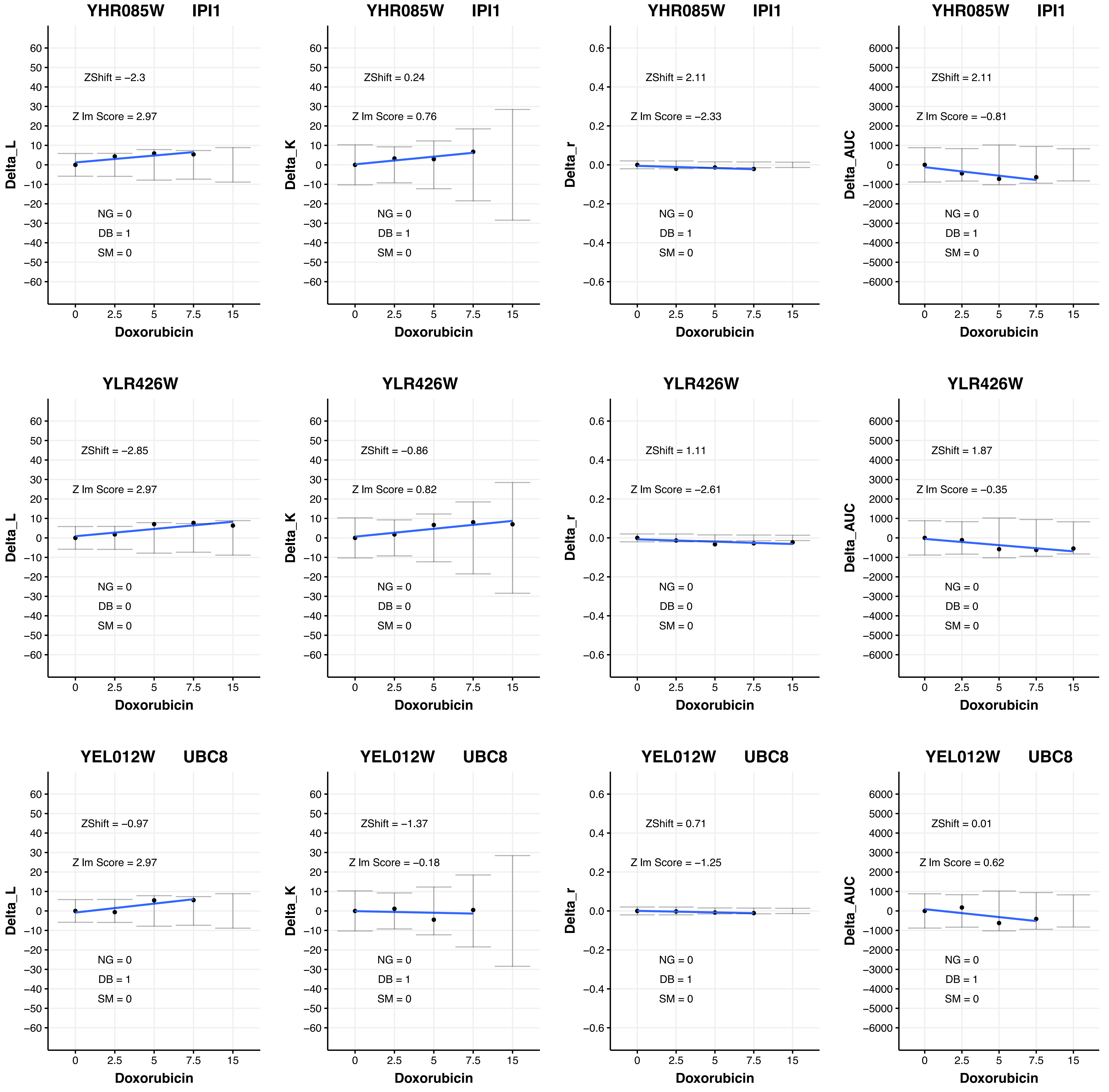
YHR085W IPI1
ZShift = −2.3
Z lm Score = 2.97
NG = 0
DB = 1
SM = 0
Delta_L
Doxorubicin
YHR085W IPI1
ZShift = 0.24
Z lm Score = 0.76
NG = 0
DB = 1
SM = 0
Delta_K
Doxorubicin
YHR085W IPI1
ZShift = 2.11
Z lm Score = −2.33
NG = 0
DB = 1
SM = 0
Delta_r
Doxorubicin
YHR085W IPI1
ZShift = 2.11
Z lm Score = −0.81
NG = 0
DB = 1
SM = 0
Delta_AUC
Doxorubicin
YLR426W
ZShift = −2.85
Z lm Score = 2.97
NG = 0
DB = 0
SM = 0
Delta_L
Doxorubicin
YLR426W
ZShift = −0.86
Z lm Score = 0.82
NG = 0
DB = 0
SM = 0
Delta_K
Doxorubicin
YLR426W
ZShift = 1.11
Z lm Score = −2.61
NG = 0
DB = 0
SM = 0
Delta_r
Doxorubicin
YLR426W
ZShift = 1.87
Z lm Score = −0.35
NG = 0
DB = 0
SM = 0
Delta_AUC
Doxorubicin
YEL012W UBC8
ZShift = −0.97
Z lm Score = 2.97
NG = 0
DB = 1
SM = 0
Delta_L
Doxorubicin
YEL012W UBC8
ZShift = −1.37
Z lm Score = −0.18
NG = 0
DB = 1
SM = 0
Delta_K
Doxorubicin
YEL012W UBC8
ZShift = 0.71
Z lm Score = −1.25
NG = 0
DB = 1
SM = 0
Delta_r
Doxorubicin
YEL012W UBC8
ZShift = 0.01
Z lm Score = 0.62
NG = 0
DB = 1
SM = 0
Delta_AUC
Doxorubicin

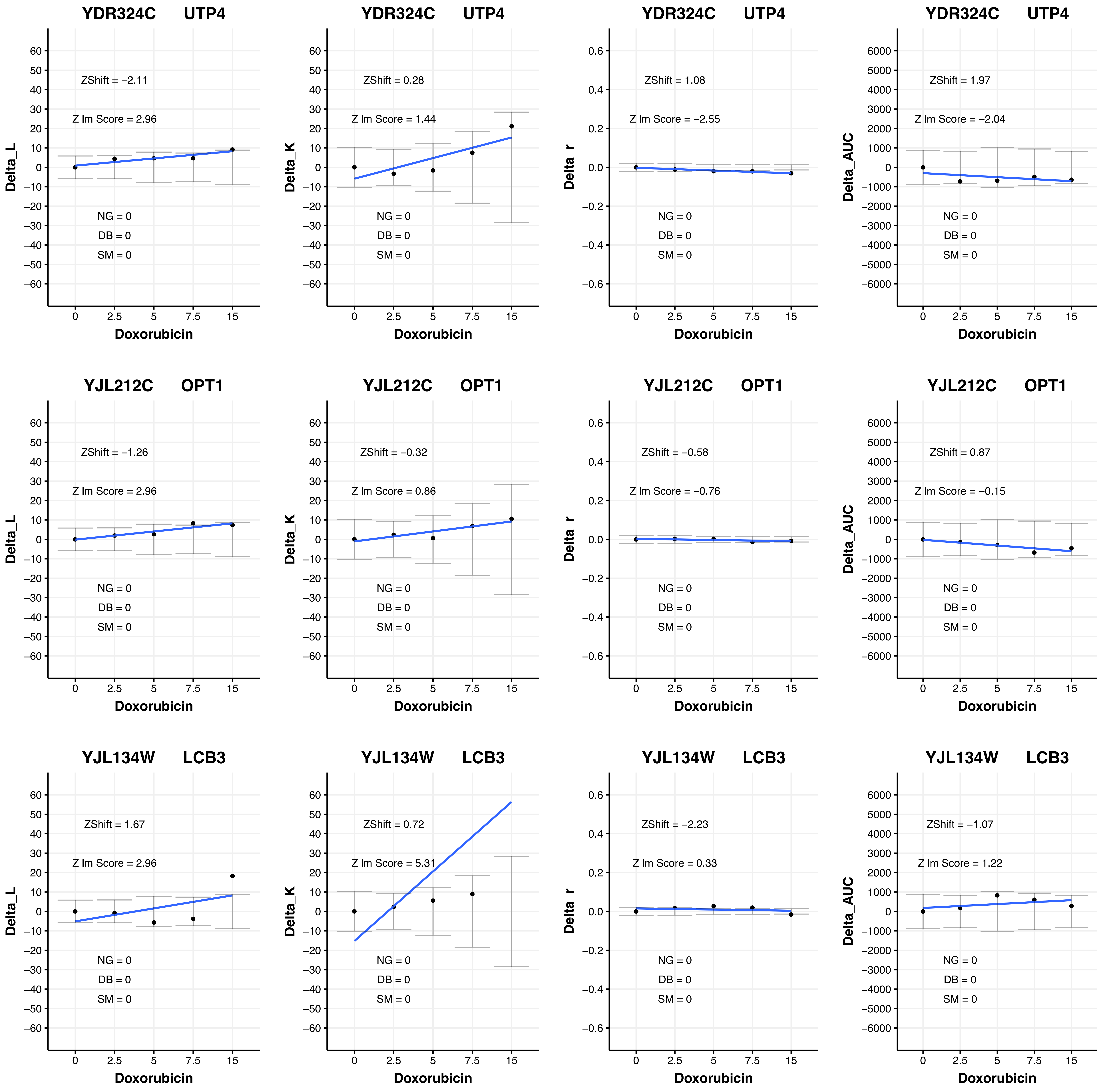
YDR324C UTP4
ZShift = −2.11
Z lm Score = 2.96
NG = 0
DB = 0
SM = 0
Delta_L
Doxorubicin
YDR324C UTP4
ZShift = 0.28
Z lm Score = 1.44
NG = 0
DB = 0
SM = 0
Delta_K
Doxorubicin
YDR324C UTP4
ZShift = 1.08
Z lm Score = −2.55
NG = 0
DB = 0
SM = 0
Delta_r
Doxorubicin
YDR324C UTP4
ZShift = 1.97
Z lm Score = −2.04
NG = 0
DB = 0
SM = 0
Delta_AUC
Doxorubicin
YJL212C OPT1
ZShift = −1.26
Z lm Score = 2.96
NG = 0
DB = 0
SM = 0
Delta_L
Doxorubicin
YJL212C OPT1
ZShift = −0.32
Z lm Score = 0.86
NG = 0
DB = 0
SM = 0
Delta_K
Doxorubicin
YJL212C OPT1
ZShift = −0.58
Z lm Score = −0.76
NG = 0
DB = 0
SM = 0
Delta_r
Doxorubicin
YJL212C OPT1
ZShift = 0.87
Z lm Score = −0.15
NG = 0
DB = 0
SM = 0
Delta_AUC
Doxorubicin
YJL134W LCB3
ZShift = 1.67
Z lm Score = 2.96
NG = 0
DB = 0
SM = 0
Delta_L
Doxorubicin
YJL134W LCB3
ZShift = 0.72
Z lm Score = 5.31
NG = 0
DB = 0
SM = 0
Delta_K
Doxorubicin
YJL134W LCB3
ZShift = −2.23
Z lm Score = 0.33
NG = 0
DB = 0
SM = 0
Delta_r
Doxorubicin
YJL134W LCB3
ZShift = −1.07
Z lm Score = 1.22
NG = 0
DB = 0
SM = 0
Delta_AUC
Doxorubicin

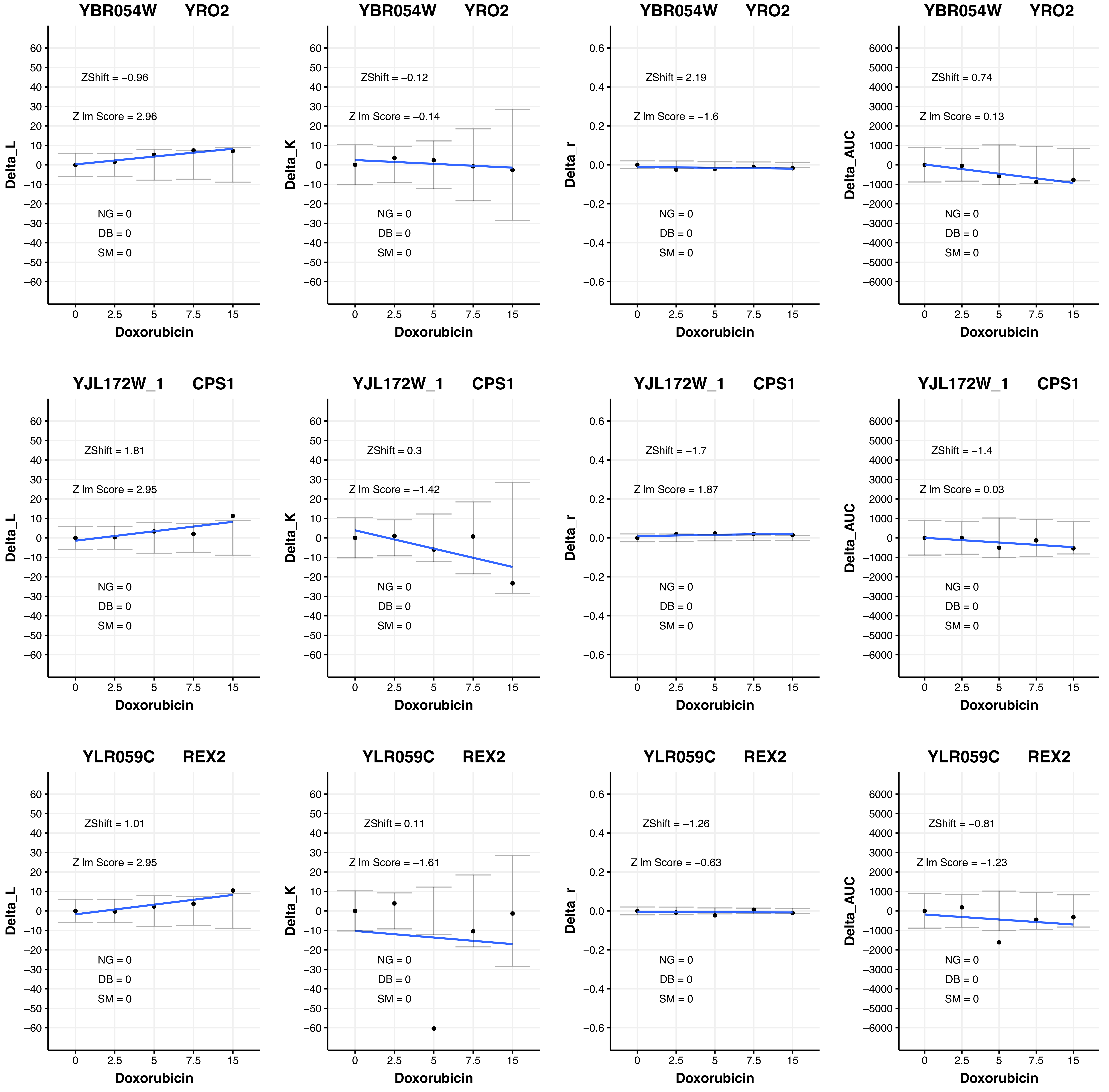
YBR054W YRO2
ZShift = −0.96
Z lm Score = 2.96
NG = 0
DB = 0
SM = 0
Delta_L
Doxorubicin
YBR054W YRO2
ZShift = −0.12
Z lm Score = −0.14
NG = 0
DB = 0
SM = 0
Delta_K
Doxorubicin
YBR054W YRO2
ZShift = 2.19
Z lm Score = −1.6
NG = 0
DB = 0
SM = 0
Delta_r
Doxorubicin
YBR054W YRO2
ZShift = 0.74
Z lm Score = 0.13
NG = 0
DB = 0
SM = 0
Delta_AUC
Doxorubicin
YJL172W_1 CPS1
ZShift = 1.81
Z lm Score = 2.95
NG = 0
DB = 0
SM = 0
Delta_L
Doxorubicin
YJL172W_1 CPS1
ZShift = 0.3
Z lm Score = −1.42
NG = 0
DB = 0
SM = 0
Delta_K
Doxorubicin
YJL172W_1 CPS1
ZShift = −1.7
Z lm Score = 1.87
NG = 0
DB = 0
SM = 0
Delta_r
Doxorubicin
YJL172W_1 CPS1
ZShift = −1.4
Z lm Score = 0.03
NG = 0
DB = 0
SM = 0
Delta_AUC
Doxorubicin
YLR059C REX2
ZShift = 1.01
Z lm Score = 2.95
NG = 0
DB = 0
SM = 0
Delta_L
Doxorubicin
YLR059C REX2
ZShift = 0.11
Z lm Score = −1.61
NG = 0
DB = 0
SM = 0
Delta_K
Doxorubicin
YLR059C REX2
ZShift = −1.26
Z lm Score = −0.63
NG = 0
DB = 0
SM = 0
Delta_r
Doxorubicin
YLR059C REX2
ZShift = −0.81
Z lm Score = −1.23
NG = 0
DB = 0
SM = 0
Delta_AUC
Doxorubicin

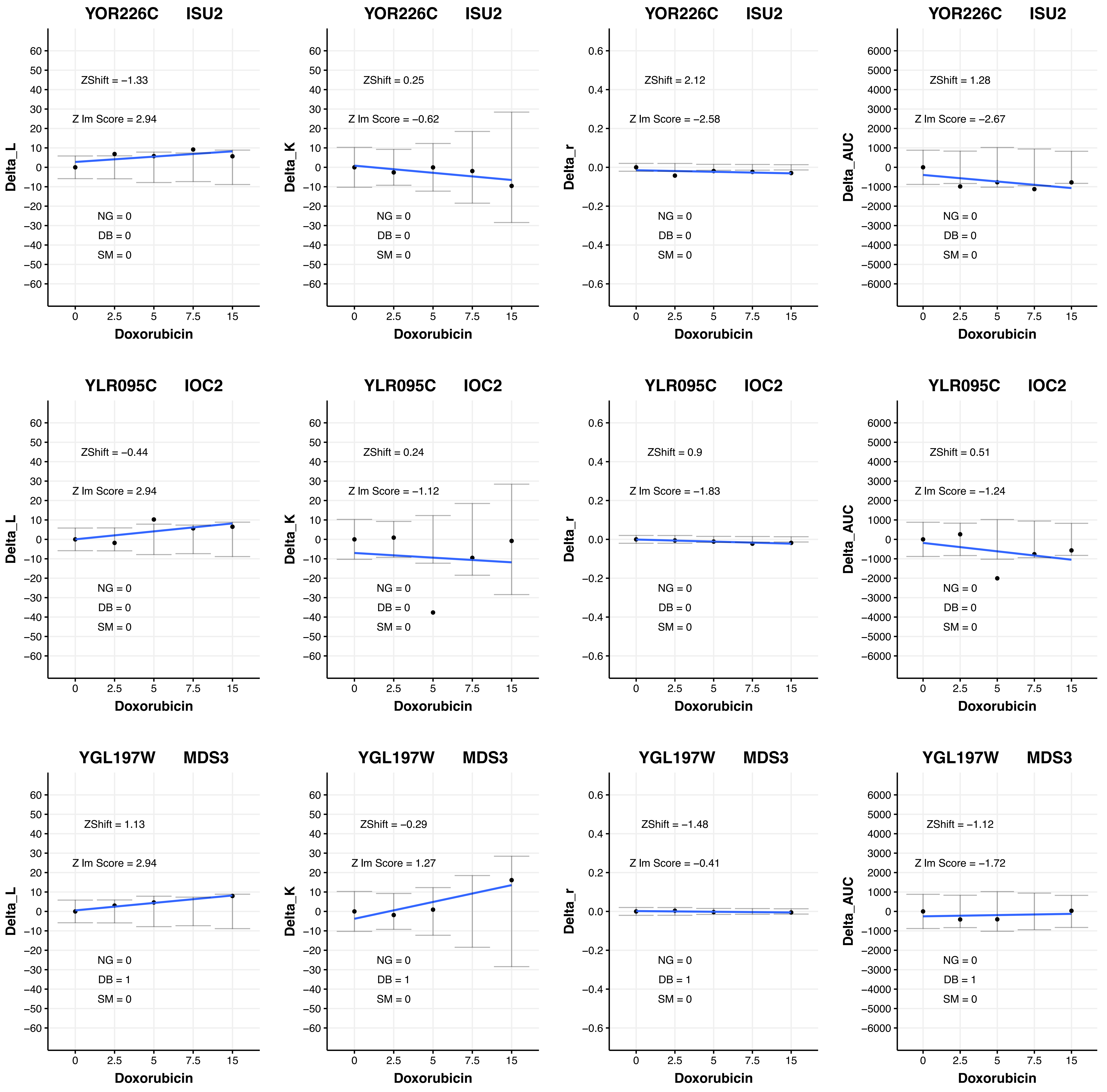

YOR226C ISU2
ZShift = −1.33
Z lm Score = 2.94
NG = 0
DB = 0
SM = 0
Delta_L
Doxorubicin
YOR226C ISU2
ZShift = 0.25
Z lm Score = −0.62
NG = 0
DB = 0
SM = 0
Delta_K
Doxorubicin
YOR226C ISU2
ZShift = 2.12
Z lm Score = −2.58
NG = 0
DB = 0
SM = 0
Delta_r
Doxorubicin
YOR226C ISU2
ZShift = 1.28
Z lm Score = −2.67
NG = 0
DB = 0
SM = 0
Delta_AUC
Doxorubicin
YLR095C IOC2
ZShift = −0.44
Z lm Score = 2.94
NG = 0
DB = 0
SM = 0
Delta_L
Doxorubicin
YLR095C IOC2
ZShift = 0.24
Z lm Score = −1.12
NG = 0
DB = 0
SM = 0
Delta_K
Doxorubicin
YLR095C IOC2
ZShift = 0.9
Z lm Score = −1.83
NG = 0
DB = 0
SM = 0
Delta_r
Doxorubicin
YLR095C IOC2
ZShift = 0.51
Z lm Score = −1.24
NG = 0
DB = 0
SM = 0
Delta_AUC
Doxorubicin
YGL197W MDS3
ZShift = 1.13
Z lm Score = 2.94
NG = 0
DB = 1
SM = 0
Delta_L
Doxorubicin
YGL197W MDS3
ZShift = −0.29
Z lm Score = 1.27
NG = 0
DB = 1
SM = 0
Delta_K
Doxorubicin
YGL197W MDS3
ZShift = −1.48
Z lm Score = −0.41
NG = 0
DB = 1
SM = 0
Delta_r
Doxorubicin
YGL197W MDS3
ZShift = −1.12
Z lm Score = −1.72
NG = 0
DB = 1
SM = 0
Delta_AUC
Doxorubicin

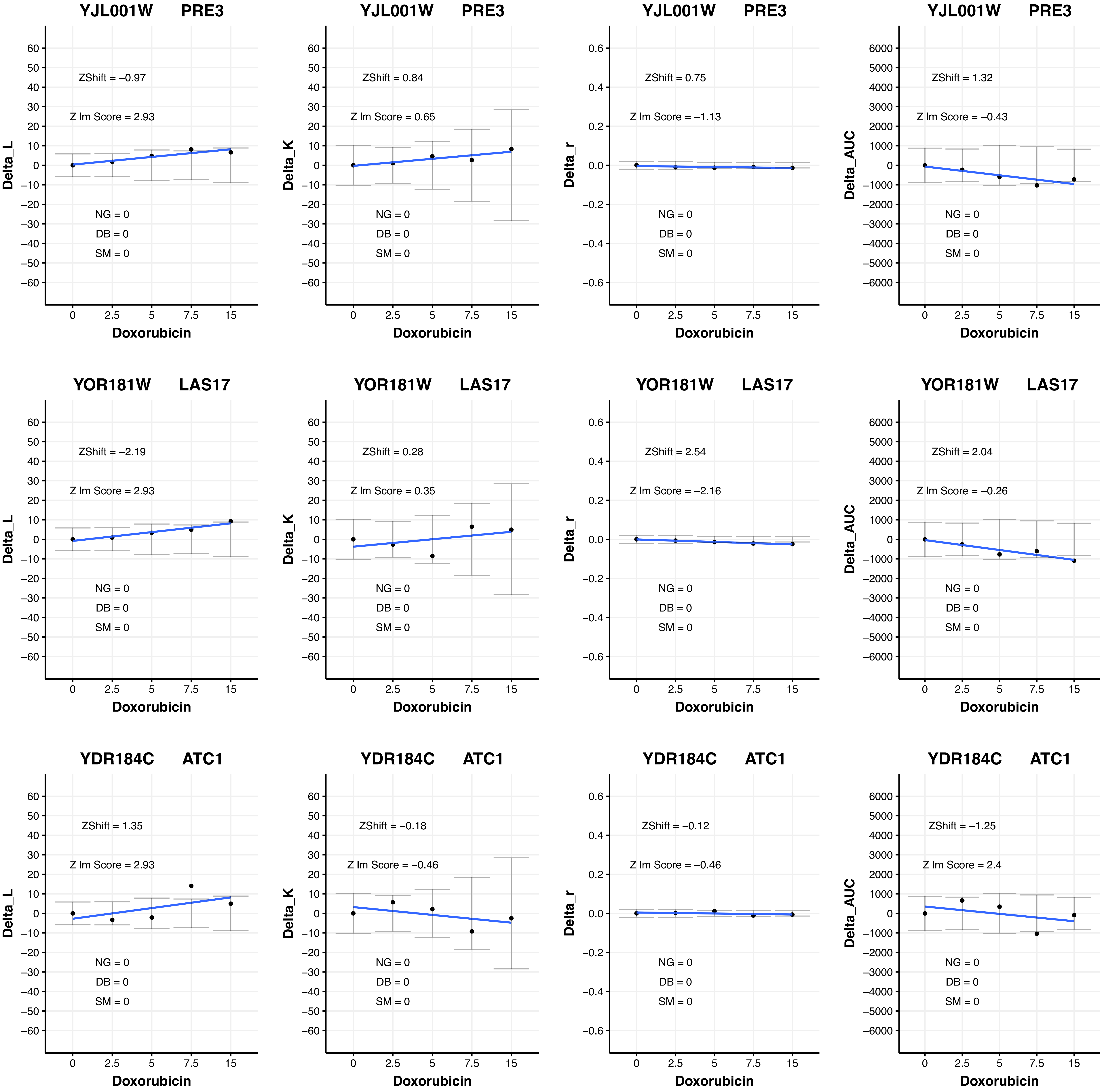
YJL001W PRE3
ZShift = −0.97
Z lm Score = 2.93
NG = 0
DB = 0
SM = 0
Delta_L
Doxorubicin
YJL001W PRE3
ZShift = 0.84
Z lm Score = 0.65
NG = 0
DB = 0
SM = 0
Delta_K
Doxorubicin
YJL001W PRE3
ZShift = 0.75
Z lm Score = −1.13
NG = 0
DB = 0
SM = 0
Delta_r
Doxorubicin
YJL001W PRE3
ZShift = 1.32
Z lm Score = −0.43
NG = 0
DB = 0
SM = 0
Delta_AUC
Doxorubicin
YOR181W LAS17
ZShift = −2.19
Z lm Score = 2.93
NG = 0
DB = 0
SM = 0
Delta_L
Doxorubicin
YOR181W LAS17
ZShift = 0.28
Z lm Score = 0.35
NG = 0
DB = 0
SM = 0
Delta_K
Doxorubicin
YOR181W LAS17
ZShift = 2.54
Z lm Score = −2.16
NG = 0
DB = 0
SM = 0
Delta_r
Doxorubicin
YOR181W LAS17
ZShift = 2.04
Z lm Score = −0.26
NG = 0
DB = 0
SM = 0
Delta_AUC
Doxorubicin
YDR184C ATC1
ZShift = 1.35
Z lm Score = 2.93
NG = 0
DB = 0
SM = 0
Delta_L
Doxorubicin
YDR184C ATC1
ZShift = −0.18
Z lm Score = −0.46
NG = 0
DB = 0
SM = 0
Delta_K
Doxorubicin
YDR184C ATC1
ZShift = −0.12
Z lm Score = −0.46
NG = 0
DB = 0
SM = 0
Delta_r
Doxorubicin
YDR184C ATC1
ZShift = −1.25
Z lm Score = 2.4
NG = 0
DB = 0
SM = 0
Delta_AUC
Doxorubicin

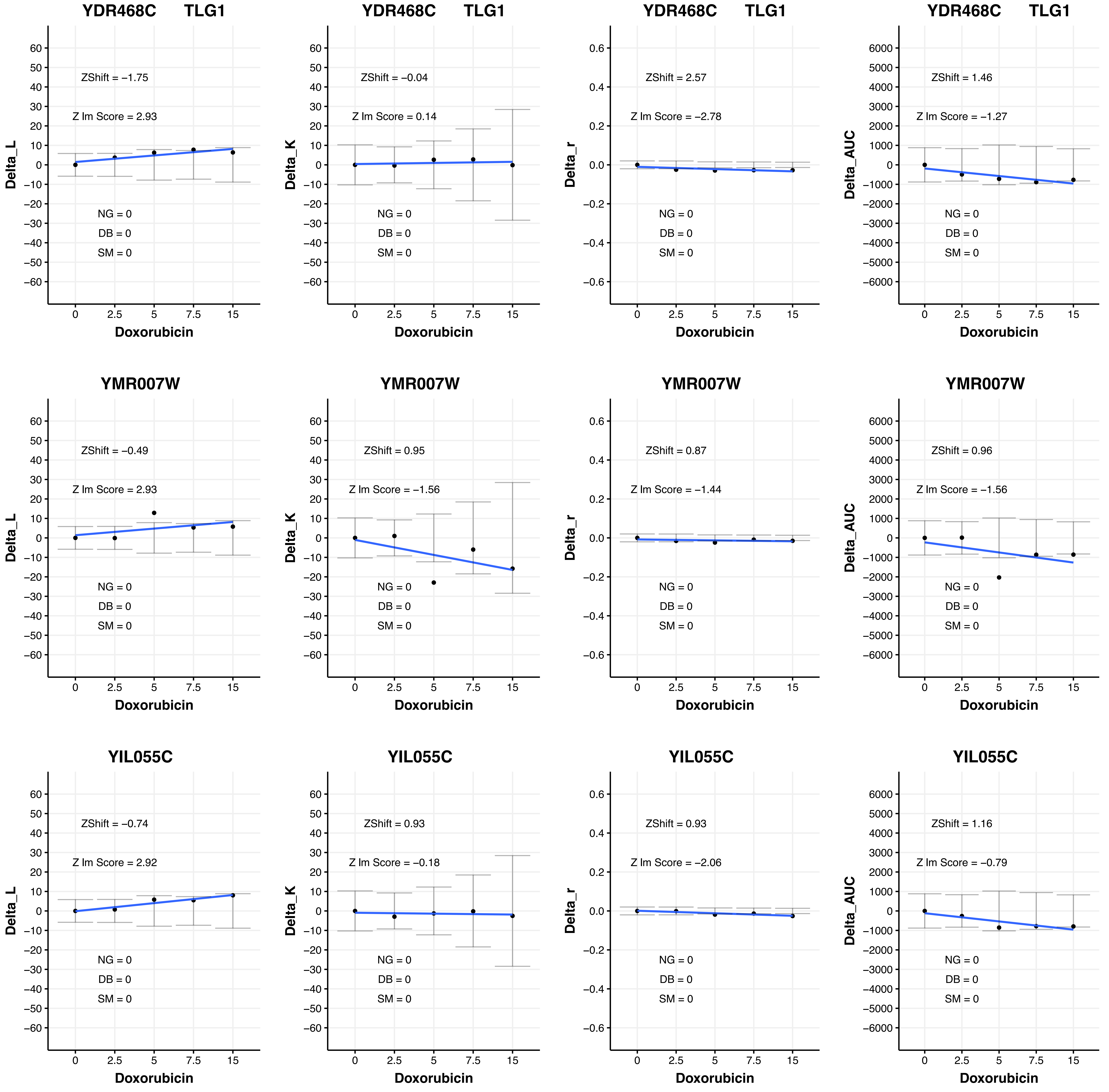

YDR468C TLG1
ZShift = −1.75
Z lm Score = 2.93
NG = 0
DB = 0
SM = 0
Delta_L
Doxorubicin
YDR468C TLG1
ZShift = −0.04
Z lm Score = 0.14
NG = 0
DB = 0
SM = 0
Delta_K
Doxorubicin
YDR468C TLG1
ZShift = 2.57
Z lm Score = −2.78
NG = 0
DB = 0
SM = 0
Delta_r
Doxorubicin
YDR468C TLG1
ZShift = 1.46
Z lm Score = −1.27
NG = 0
DB = 0
SM = 0
Delta_AUC
Doxorubicin
YMR007W
ZShift = −0.49
Z lm Score = 2.93
NG = 0
DB = 0
SM = 0
Delta_L
Doxorubicin
YMR007W
ZShift = 0.95
Z lm Score = −1.56
NG = 0
DB = 0
SM = 0
Delta_K
Doxorubicin
YMR007W
ZShift = 0.87
Z lm Score = −1.44
NG = 0
DB = 0
SM = 0
Delta_r
Doxorubicin
YMR007W
ZShift = 0.96
Z lm Score = −1.56
NG = 0
DB = 0
SM = 0
Delta_AUC
Doxorubicin
YIL055C
ZShift = −0.74
Z lm Score = 2.92
NG = 0
DB = 0
SM = 0
Delta_L
Doxorubicin
YIL055C
ZShift = 0.93
Z lm Score = −0.18
NG = 0
DB = 0
SM = 0
Delta_K
Doxorubicin
YIL055C
ZShift = 0.93
Z lm Score = −2.06
NG = 0
DB = 0
SM = 0
Delta_r
Doxorubicin
YIL055C
ZShift = 1.16
Z lm Score = −0.79
NG = 0
DB = 0
SM = 0
Delta_AUC
Doxorubicin

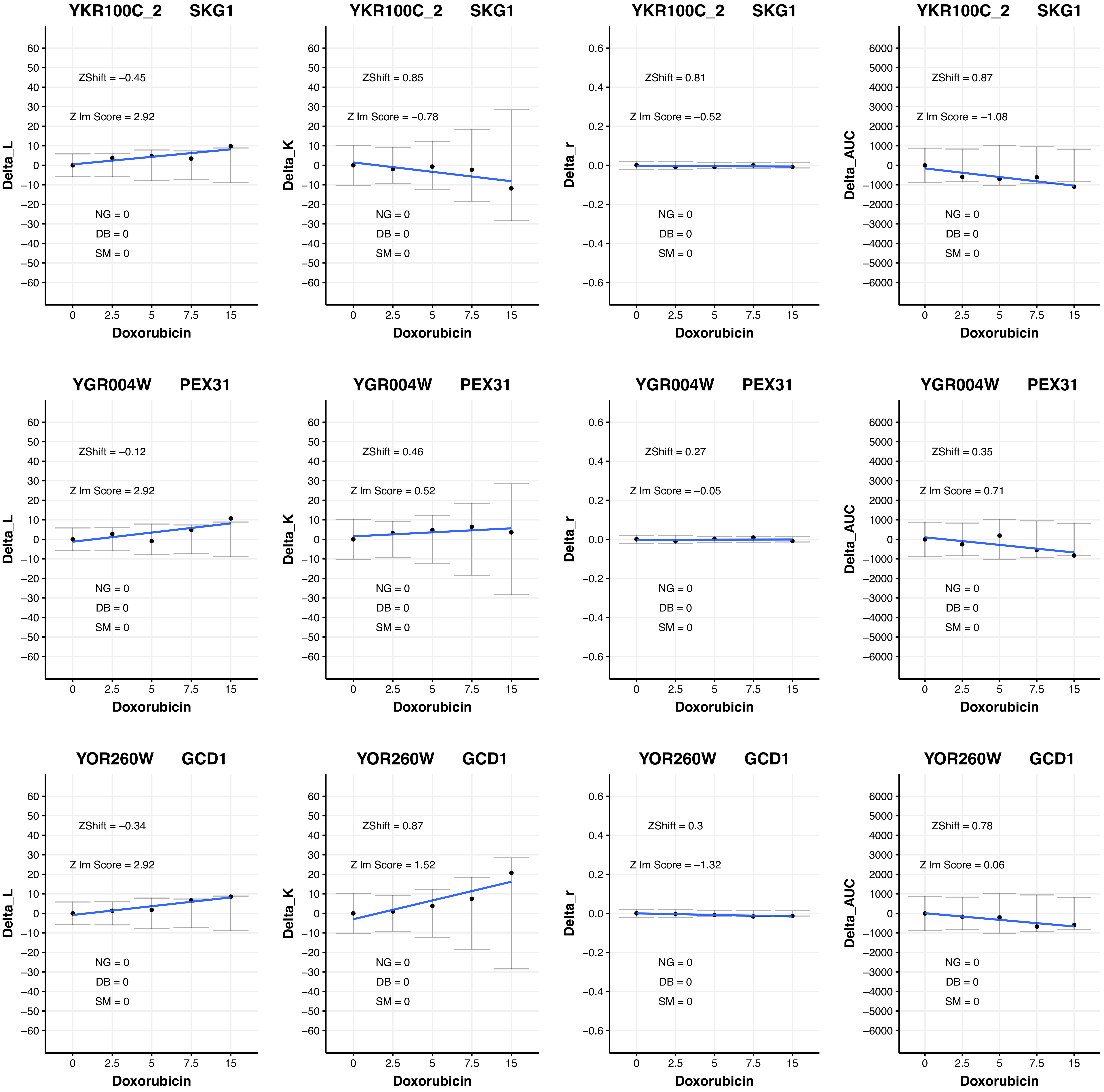
YKR100C_2 SKG1
ZShift = −0.45
Z lm Score = 2.92
NG = 0
DB = 0
SM = 0
Delta_L
Doxorubicin
YKR100C_2 SKG1
ZShift = 0.85
Z lm Score = −0.78
NG = 0
DB = 0
SM = 0
Delta_K
Doxorubicin
YKR100C_2 SKG1
ZShift = 0.81
Z lm Score = −0.52
NG = 0
DB = 0
SM = 0
Delta_r
Doxorubicin
YKR100C_2 SKG1
ZShift = 0.87
Z lm Score = −1.08
NG = 0
DB = 0
SM = 0
Delta_AUC
Doxorubicin
YGR004W PEX31
ZShift = −0.12
Z lm Score = 2.92
NG = 0
DB = 0
SM = 0
Delta_L
Doxorubicin
YGR004W PEX31
ZShift = 0.46
Z lm Score = 0.52
NG = 0
DB = 0
SM = 0
Delta_K
Doxorubicin
YGR004W PEX31
ZShift = 0.27
Z lm Score = −0.05
NG = 0
DB = 0
SM = 0
Delta_r
Doxorubicin
YGR004W PEX31
ZShift = 0.35
Z lm Score = 0.71
NG = 0
DB = 0
SM = 0
Delta_AUC
Doxorubicin
YOR260W GCD1
ZShift = −0.34
Z lm Score = 2.92
NG = 0
DB = 0
SM = 0
Delta_L
Doxorubicin
YOR260W GCD1
ZShift = 0.87
Z lm Score = 1.52
NG = 0
DB = 0
SM = 0
Delta_K
Doxorubicin
YOR260W GCD1
ZShift = 0.3
Z lm Score = −1.32
NG = 0
DB = 0
SM = 0
Delta_r
Doxorubicin
YOR260W GCD1
ZShift = 0.78
Z lm Score = 0.06
NG = 0
DB = 0
SM = 0
Delta_AUC
Doxorubicin

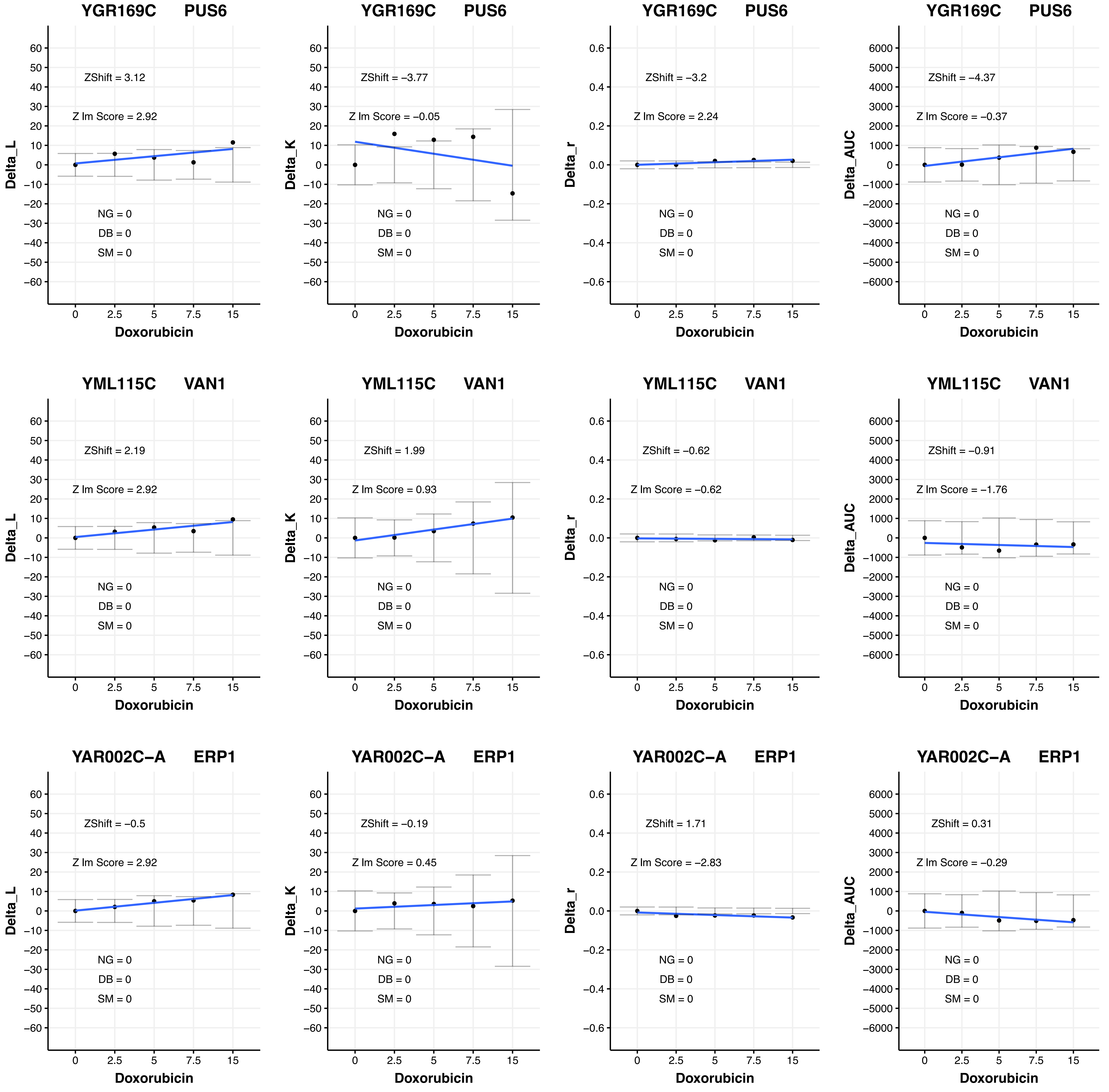
YGR169C PUS6
ZShift = 3.12
Z lm Score = 2.92
NG = 0
DB = 0
SM = 0
Delta_L
Doxorubicin
YGR169C PUS6
ZShift = −3.77
Z lm Score = −0.05
NG = 0
DB = 0
SM = 0
Delta_K
Doxorubicin
YGR169C PUS6
ZShift = −3.2
Z lm Score = 2.24
NG = 0
DB = 0
SM = 0
Delta_r
Doxorubicin
YGR169C PUS6
ZShift = −4.37
Z lm Score = −0.37
NG = 0
DB = 0
SM = 0
Delta_AUC
Doxorubicin
YML115C VAN1
ZShift = 2.19
Z lm Score = 2.92
NG = 0
DB = 0
SM = 0
Delta_L
Doxorubicin
YML115C VAN1
ZShift = 1.99
Z lm Score = 0.93
NG = 0
DB = 0
SM = 0
Delta_K
Doxorubicin
YML115C VAN1
ZShift = −0.62
Z lm Score = −0.62
NG = 0
DB = 0
SM = 0
Delta_r
Doxorubicin
YML115C VAN1
ZShift = −0.91
Z lm Score = −1.76
NG = 0
DB = 0
SM = 0
Delta_AUC
Doxorubicin
YAR002C−A ERP1
ZShift = −0.5
Z lm Score = 2.92
NG = 0
DB = 0
SM = 0
Delta_L
Doxorubicin
YAR002C−A ERP1
ZShift = −0.19
Z lm Score = 0.45
NG = 0
DB = 0
SM = 0
Delta_K
Doxorubicin
YAR002C−A ERP1
ZShift = 1.71
Z lm Score = −2.83
NG = 0
DB = 0
SM = 0
Delta_r
Doxorubicin
YAR002C−A ERP1
ZShift = 0.31
Z lm Score = −0.29
NG = 0
DB = 0
SM = 0
Delta_AUC
Doxorubicin

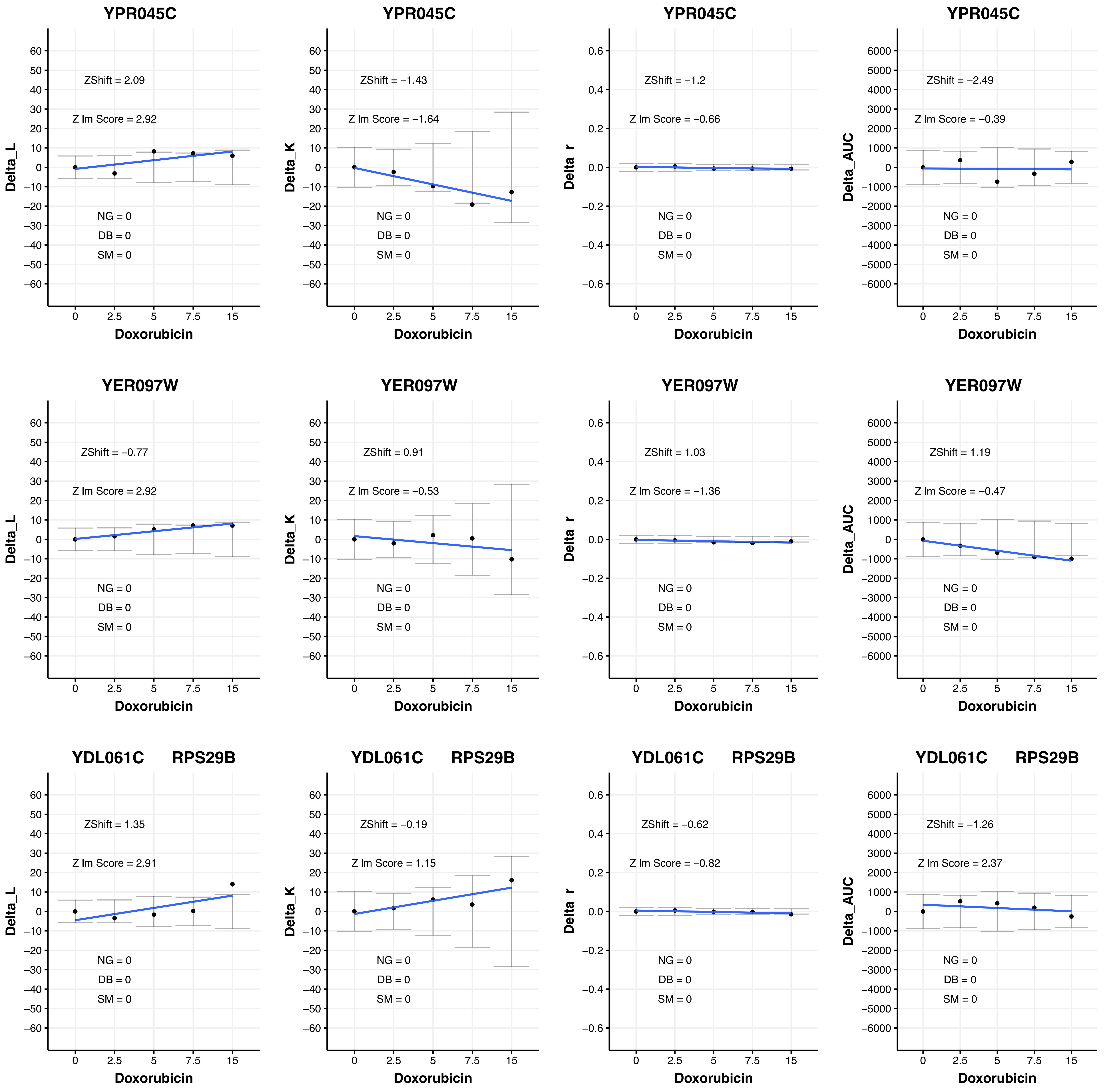
YPR045C
ZShift = 2.09
Z lm Score = 2.92
NG = 0
DB = 0
SM = 0
Delta_L
Doxorubicin
YPR045C
ZShift = −1.43
Z lm Score = −1.64
NG = 0
DB = 0
SM = 0
Delta_K
Doxorubicin
YPR045C
ZShift = −1.2
Z lm Score = −0.66
NG = 0
DB = 0
SM = 0
Delta_r
Doxorubicin
YPR045C
ZShift = −2.49
Z lm Score = −0.39
NG = 0
DB = 0
SM = 0
Delta_AUC
Doxorubicin
YER097W
ZShift = −0.77
Z lm Score = 2.92
NG = 0
DB = 0
SM = 0
Delta_L
Doxorubicin
YER097W
ZShift = 0.91
Z lm Score = −0.53
NG = 0
DB = 0
SM = 0
Delta_K
Doxorubicin
YER097W
ZShift = 1.03
Z lm Score = −1.36
NG = 0
DB = 0
SM = 0
Delta_r
Doxorubicin
YER097W
ZShift = 1.19
Z lm Score = −0.47
NG = 0
DB = 0
SM = 0
Delta_AUC
Doxorubicin
YDL061C RPS29B
ZShift = 1.35
Z lm Score = 2.91
NG = 0
DB = 0
SM = 0
Delta_L
Doxorubicin
YDL061C RPS29B
ZShift = −0.19
Z lm Score = 1.15
NG = 0
DB = 0
SM = 0
Delta_K
Doxorubicin
YDL061C RPS29B
ZShift = −0.62
Z lm Score = −0.82
NG = 0
DB = 0
SM = 0
Delta_r
Doxorubicin
YDL061C RPS29B
ZShift = −1.26
Z lm Score = 2.37
NG = 0
DB = 0
SM = 0
Delta_AUC
Doxorubicin

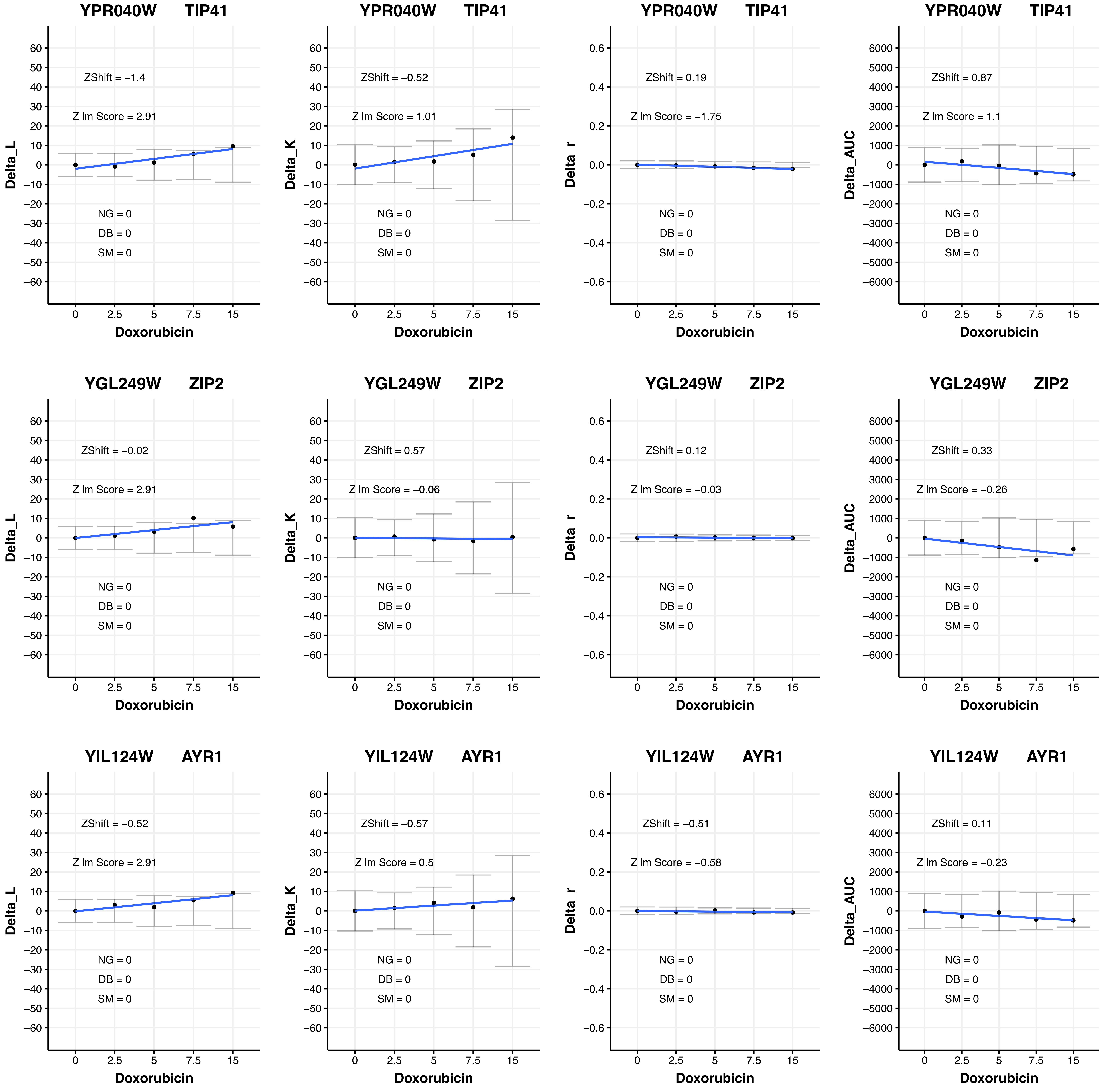

YPR040W TIP41
ZShift = −1.4
Z lm Score = 2.91
NG = 0
DB = 0
SM = 0
Delta_L
Doxorubicin
YPR040W TIP41
ZShift = −0.52
Z lm Score = 1.01
NG = 0
DB = 0
SM = 0
Delta_K
Doxorubicin
YPR040W TIP41
ZShift = 0.19
Z lm Score = −1.75
NG = 0
DB = 0
SM = 0
Delta_r
Doxorubicin
YPR040W TIP41
ZShift = 0.87
Z lm Score = 1.1
NG = 0
DB = 0
SM = 0
Delta_AUC
Doxorubicin
YGL249W ZIP2
ZShift = −0.02
Z lm Score = 2.91
NG = 0
DB = 0
SM = 0
Delta_L
Doxorubicin
YGL249W ZIP2
ZShift = 0.57
Z lm Score = −0.06
NG = 0
DB = 0
SM = 0
Delta_K
Doxorubicin
YGL249W ZIP2
ZShift = 0.12
Z lm Score = −0.03
NG = 0
DB = 0
SM = 0
Delta_r
Doxorubicin
YGL249W ZIP2
ZShift = 0.33
Z lm Score = −0.26
NG = 0
DB = 0
SM = 0
Delta_AUC
Doxorubicin
YIL124W AYR1
ZShift = −0.52
Z lm Score = 2.91
NG = 0
DB = 0
SM = 0
Delta_L
Doxorubicin
YIL124W AYR1
ZShift = −0.57
Z lm Score = 0.5
NG = 0
DB = 0
SM = 0
Delta_K
Doxorubicin
YIL124W AYR1
ZShift = −0.51
Z lm Score = −0.58
NG = 0
DB = 0
SM = 0
Delta_r
Doxorubicin
YIL124W AYR1
ZShift = 0.11
Z lm Score = −0.23
NG = 0
DB = 0
SM = 0
Delta_AUC
Doxorubicin

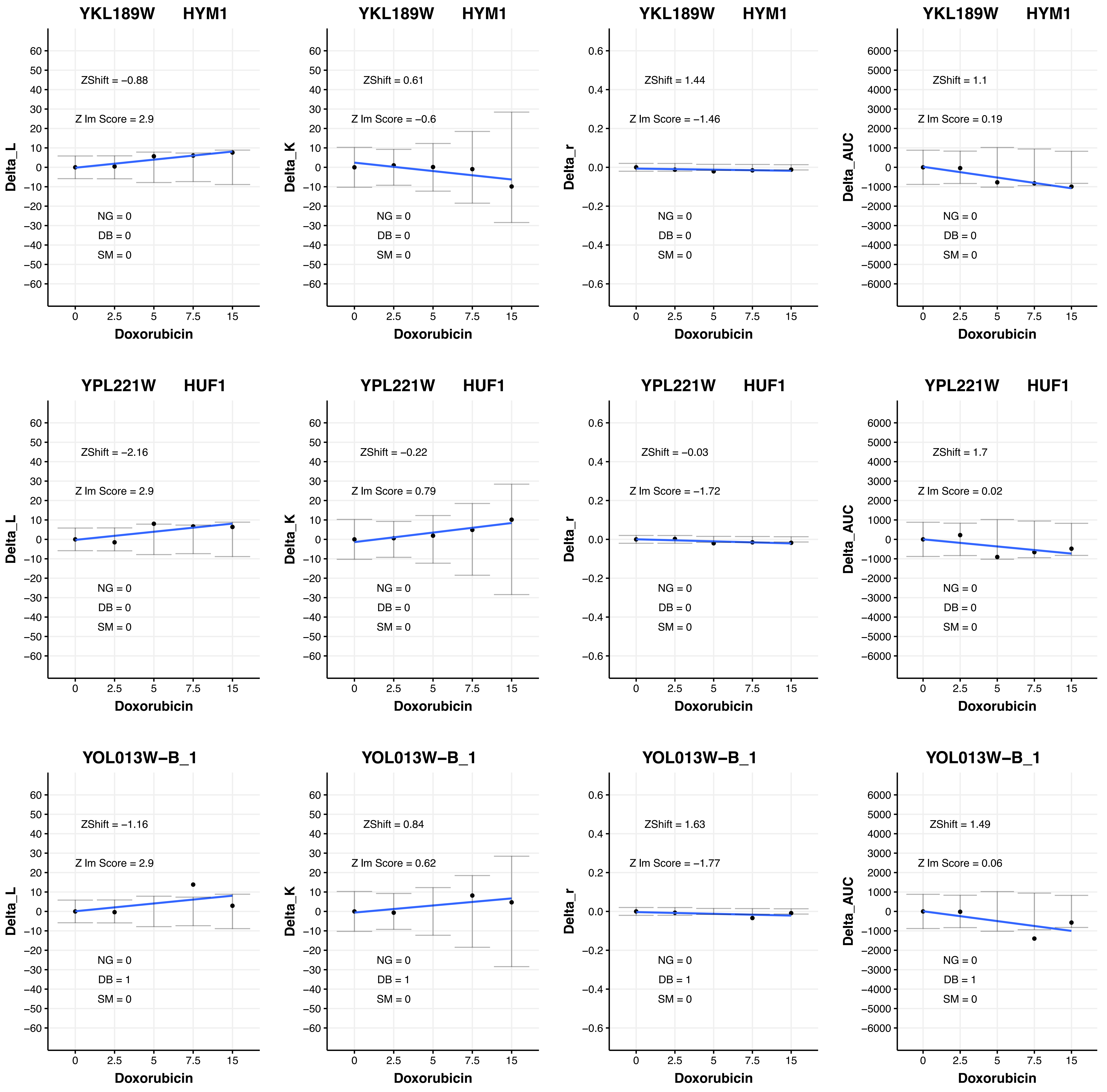
YKL189W HYM1
ZShift = −0.88
Z lm Score = 2.9
NG = 0
DB = 0
SM = 0
Delta_L
Doxorubicin
YKL189W HYM1
ZShift = 0.61
Z lm Score = −0.6
NG = 0
DB = 0
SM = 0
Delta_K
Doxorubicin
YKL189W HYM1
ZShift = 1.44
Z lm Score = −1.46
NG = 0
DB = 0
SM = 0
Delta_r
Doxorubicin
YKL189W HYM1
ZShift = 1.1
Z lm Score = 0.19
NG = 0
DB = 0
SM = 0
Delta_AUC
Doxorubicin
YPL221W HUF1
ZShift = −2.16
Z lm Score = 2.9
NG = 0
DB = 0
SM = 0
Delta_L
Doxorubicin
YPL221W HUF1
ZShift = −0.22
Z lm Score = 0.79
NG = 0
DB = 0
SM = 0
Delta_K
Doxorubicin
YPL221W HUF1
ZShift = −0.03
Z lm Score = −1.72
NG = 0
DB = 0
SM = 0
Delta_r
Doxorubicin
YPL221W HUF1
ZShift = 1.7
Z lm Score = 0.02
NG = 0
DB = 0
SM = 0
Delta_AUC
Doxorubicin
YOL013W−B_1
ZShift = −1.16
Z lm Score = 2.9
NG = 0
DB = 1
SM = 0
Delta_L
Doxorubicin
YOL013W−B_1
ZShift = 0.84
Z lm Score = 0.62
NG = 0
DB = 1
SM = 0
Delta_K
Doxorubicin
YOL013W−B_1
ZShift = 1.63
Z lm Score = −1.77
NG = 0
DB = 1
SM = 0
Delta_r
Doxorubicin
YOL013W−B_1
ZShift = 1.49
Z lm Score = 0.06
NG = 0
DB = 1
SM = 0
Delta_AUC
Doxorubicin

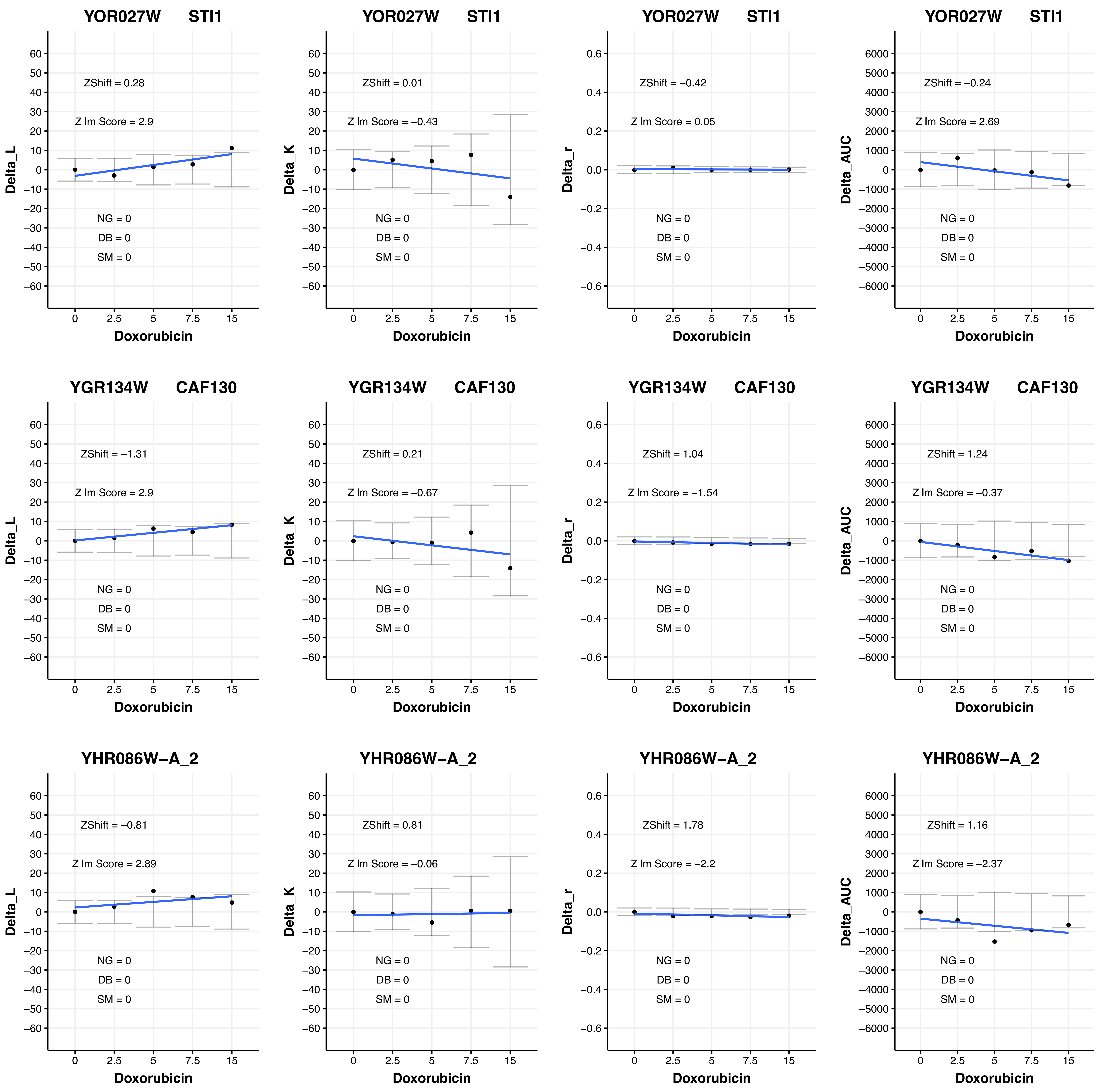

YOR027W STI1
ZShift = 0.28
Z lm Score = 2.9
NG = 0
DB = 0
SM = 0
Delta_L
Doxorubicin
YOR027W STI1
ZShift = 0.01
Z lm Score = −0.43
NG = 0
DB = 0
SM = 0
Delta_K
Doxorubicin
YOR027W STI1
ZShift = −0.42
Z lm Score = 0.05
NG = 0
DB = 0
SM = 0
Delta_r
Doxorubicin
YOR027W STI1
ZShift = −0.24
Z lm Score = 2.69
NG = 0
DB = 0
SM = 0
Delta_AUC
Doxorubicin
YGR134W CAF130
ZShift = −1.31
Z lm Score = 2.9
NG = 0
DB = 0
SM = 0
Delta_L
Doxorubicin
YGR134W CAF130
ZShift = 0.21
Z lm Score = −0.67
NG = 0
DB = 0
SM = 0
Delta_K
Doxorubicin
YGR134W CAF130
ZShift = 1.04
Z lm Score = −1.54
NG = 0
DB = 0
SM = 0
Delta_r
Doxorubicin
YGR134W CAF130
ZShift = 1.24
Z lm Score = −0.37
NG = 0
DB = 0
SM = 0
Delta_AUC
Doxorubicin
YHR086W−A_2
ZShift = −0.81
Z lm Score = 2.89
NG = 0
DB = 0
SM = 0
Delta_L
Doxorubicin
YHR086W−A_2
ZShift = 0.81
Z lm Score = −0.06
NG = 0
DB = 0
SM = 0
Delta_K
Doxorubicin
YHR086W−A_2
ZShift = 1.78
Z lm Score = −2.2
NG = 0
DB = 0
SM = 0
Delta_r
Doxorubicin
YHR086W−A_2
ZShift = 1.16
Z lm Score = −2.37
NG = 0
DB = 0
SM = 0
Delta_AUC
Doxorubicin

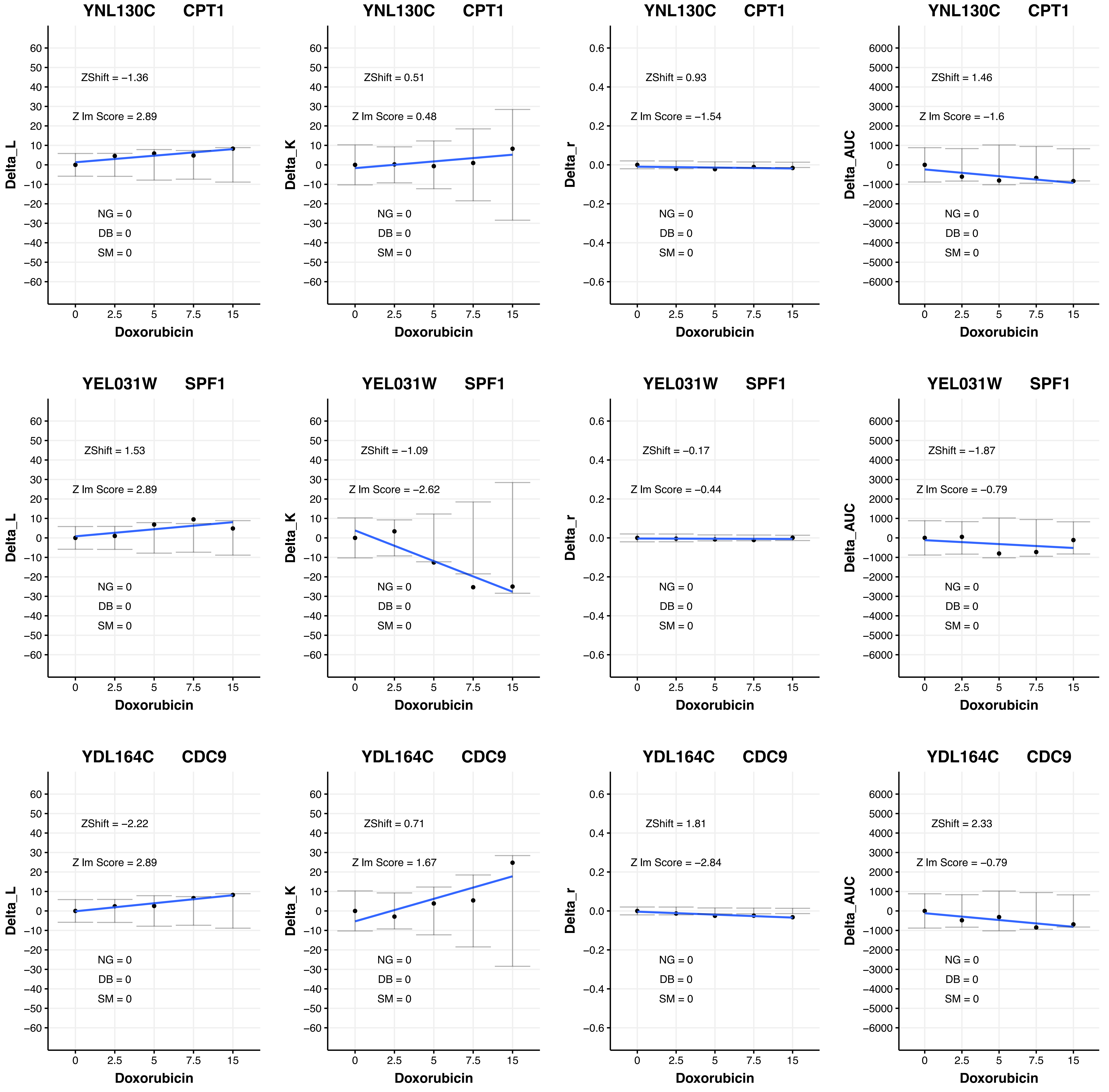

YNL130C CPT1
ZShift = −1.36
Z lm Score = 2.89
NG = 0
DB = 0
SM = 0
Delta_L
Doxorubicin
YNL130C CPT1
ZShift = 0.51
Z lm Score = 0.48
NG = 0
DB = 0
SM = 0
Delta_K
Doxorubicin
YNL130C CPT1
ZShift = 0.93
Z lm Score = −1.54
NG = 0
DB = 0
SM = 0
Delta_r
Doxorubicin
YNL130C CPT1
ZShift = 1.46
Z lm Score = −1.6
NG = 0
DB = 0
SM = 0
Delta_AUC
Doxorubicin
YEL031W SPF1
ZShift = 1.53
Z lm Score = 2.89
NG = 0
DB = 0
SM = 0
Delta_L
Doxorubicin
YEL031W SPF1
ZShift = −1.09
Z lm Score = −2.62
NG = 0
DB = 0
SM = 0
Delta_K
Doxorubicin
YEL031W SPF1
ZShift = −0.17
Z lm Score = −0.44
NG = 0
DB = 0
SM = 0
Delta_r
Doxorubicin
YEL031W SPF1
ZShift = −1.87
Z lm Score = −0.79
NG = 0
DB = 0
SM = 0
Delta_AUC
Doxorubicin
YDL164C CDC9
ZShift = −2.22
Z lm Score = 2.89
NG = 0
DB = 0
SM = 0
Delta_L
Doxorubicin
YDL164C CDC9
ZShift = 0.71
Z lm Score = 1.67
NG = 0
DB = 0
SM = 0
Delta_K
Doxorubicin
YDL164C CDC9
ZShift = 1.81
Z lm Score = −2.84
NG = 0
DB = 0
SM = 0
Delta_r
Doxorubicin
YDL164C CDC9
ZShift = 2.33
Z lm Score = −0.79
NG = 0
DB = 0
SM = 0
Delta_AUC
Doxorubicin

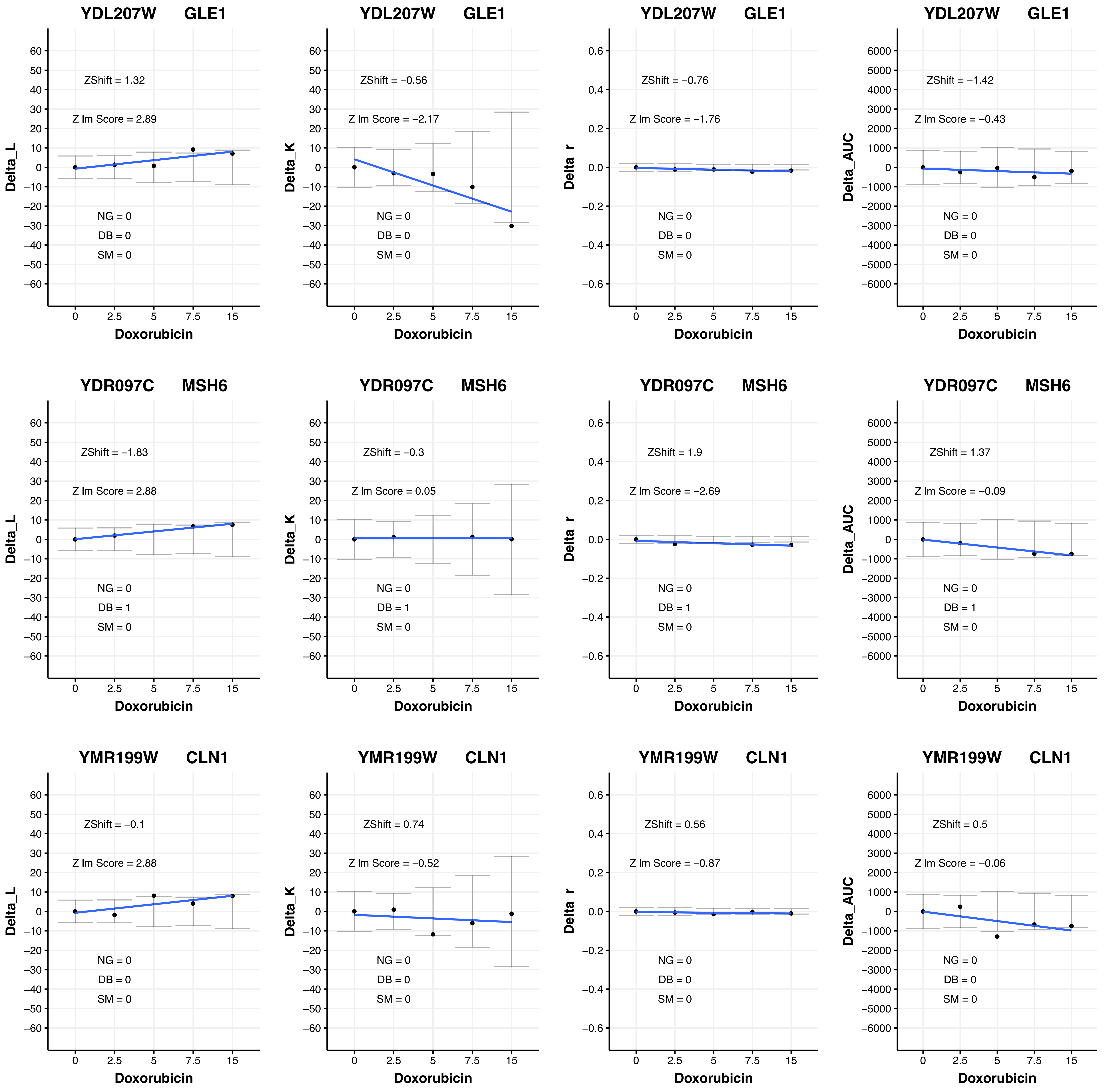

YDL207W GLE1
ZShift = 1.32
Z lm Score = 2.89
Delta_L
NG = 0
DB = 0
SM = 0
Doxorubicin
YDL207W GLE1
ZShift = −0.56
Z lm Score = −2.17
Delta_K
NG = 0
DB = 0
SM = 0
Doxorubicin
YDL207W GLE1
ZShift = −0.76
Z lm Score = −1.76
Delta_r
NG = 0
DB = 0
SM = 0
Doxorubicin
YDL207W GLE1
ZShift = −1.42
Z lm Score = −0.43
Delta_AUC
NG = 0
DB = 0
SM = 0
Doxorubicin
YDR097C MSH6
ZShift = −1.83
Z lm Score = 2.88
Delta_L
NG = 0
DB = 1
SM = 0
Doxorubicin
YDR097C MSH6
ZShift = −0.3
Z lm Score = 0.05
Delta_K
NG = 0
DB = 1
SM = 0
Doxorubicin
YDR097C MSH6
ZShift = 1.9
Z lm Score = −2.69
Delta_r
NG = 0
DB = 1
SM = 0
Doxorubicin
YDR097C MSH6
ZShift = 1.37
Z lm Score = −0.09
Delta_AUC
NG = 0
DB = 1
SM = 0
Doxorubicin
YMR199W CLN1
ZShift = −0.1
Z lm Score = 2.88
Delta_L
NG = 0
DB = 0
SM = 0
Doxorubicin
YMR199W CLN1
ZShift = 0.74
Z lm Score = −0.52
Delta_K
NG = 0
DB = 0
SM = 0
Doxorubicin
YMR199W CLN1
ZShift = 0.56
Z lm Score = −0.87
Delta_r
NG = 0
DB = 0
SM = 0
Doxorubicin
YMR199W CLN1
ZShift = 0.5
Z lm Score = −0.06
Delta_AUC
NG = 0
DB = 0
SM = 0
Doxorubicin

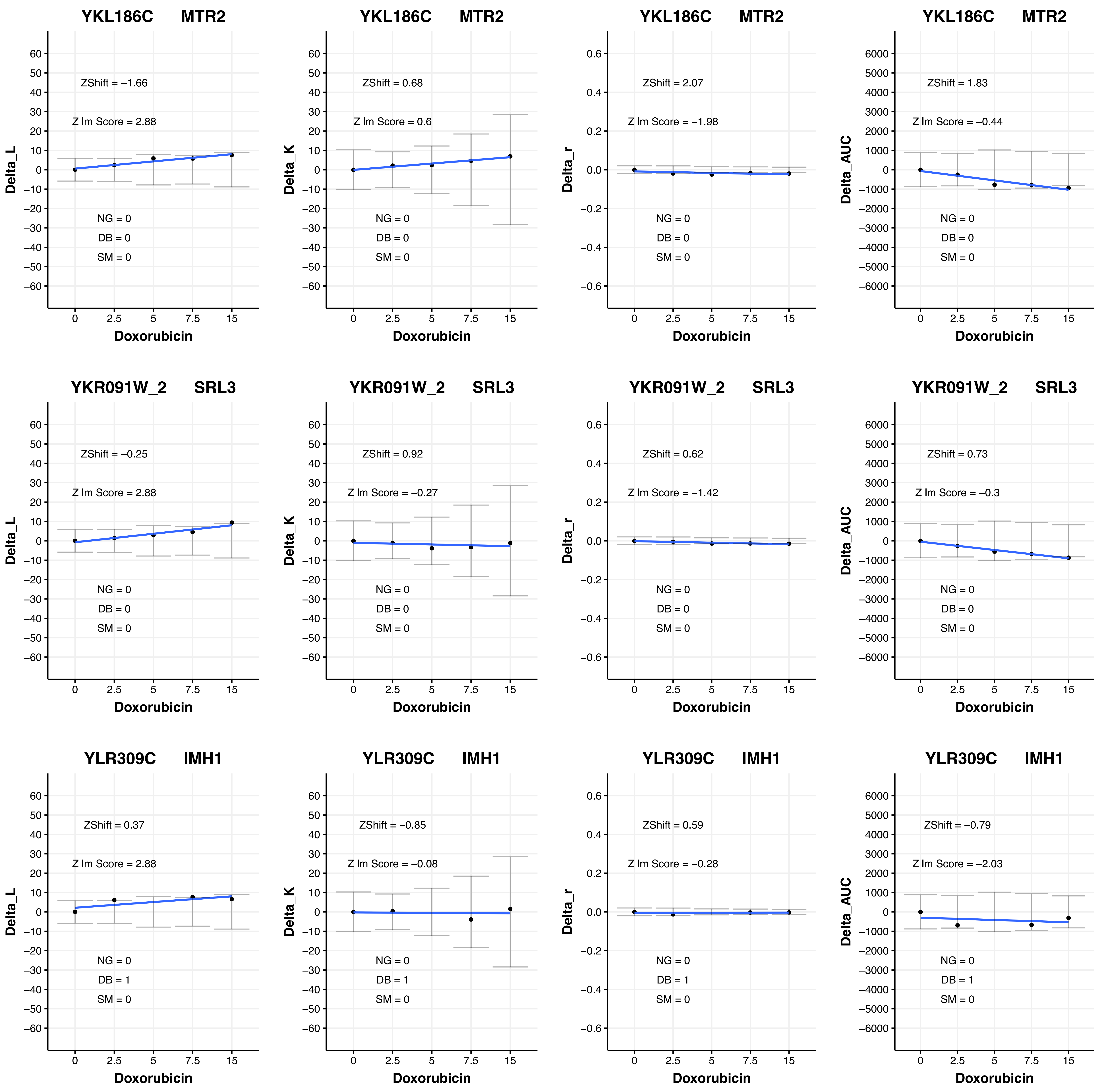

YKL186C MTR2
ZShift = −1.66
Z lm Score = 2.88
NG = 0
DB = 0
SM = 0
Delta_L
Doxorubicin
YKL186C MTR2
ZShift = 0.68
Z lm Score = 0.6
NG = 0
DB = 0
SM = 0
Delta_K
Doxorubicin
YKL186C MTR2
ZShift = 2.07
Z lm Score = −1.98
NG = 0
DB = 0
SM = 0
Delta_r
Doxorubicin
YKL186C MTR2
ZShift = 1.83
Z lm Score = −0.44
NG = 0
DB = 0
SM = 0
Delta_AUC
Doxorubicin
YKR091W_2 SRL3
ZShift = −0.25
Z lm Score = 2.88
NG = 0
DB = 0
SM = 0
Delta_L
Doxorubicin
YKR091W_2 SRL3
ZShift = 0.92
Z lm Score = −0.27
NG = 0
DB = 0
SM = 0
Delta_K
Doxorubicin
YKR091W_2 SRL3
ZShift = 0.62
Z lm Score = −1.42
NG = 0
DB = 0
SM = 0
Delta_r
Doxorubicin
YKR091W_2 SRL3
ZShift = 0.73
Z lm Score = −0.3
NG = 0
DB = 0
SM = 0
Delta_AUC
Doxorubicin
YLR309C IMH1
ZShift = 0.37
Z lm Score = 2.88
NG = 0
DB = 1
SM = 0
Delta_L
Doxorubicin
YLR309C IMH1
ZShift = −0.85
Z lm Score = −0.08
NG = 0
DB = 1
SM = 0
Delta_K
Doxorubicin
YLR309C IMH1
ZShift = 0.59
Z lm Score = −0.28
NG = 0
DB = 1
SM = 0
Delta_r
Doxorubicin
YLR309C IMH1
ZShift = −0.79
Z lm Score = −2.03
NG = 0
DB = 1
SM = 0
Delta_AUC
Doxorubicin

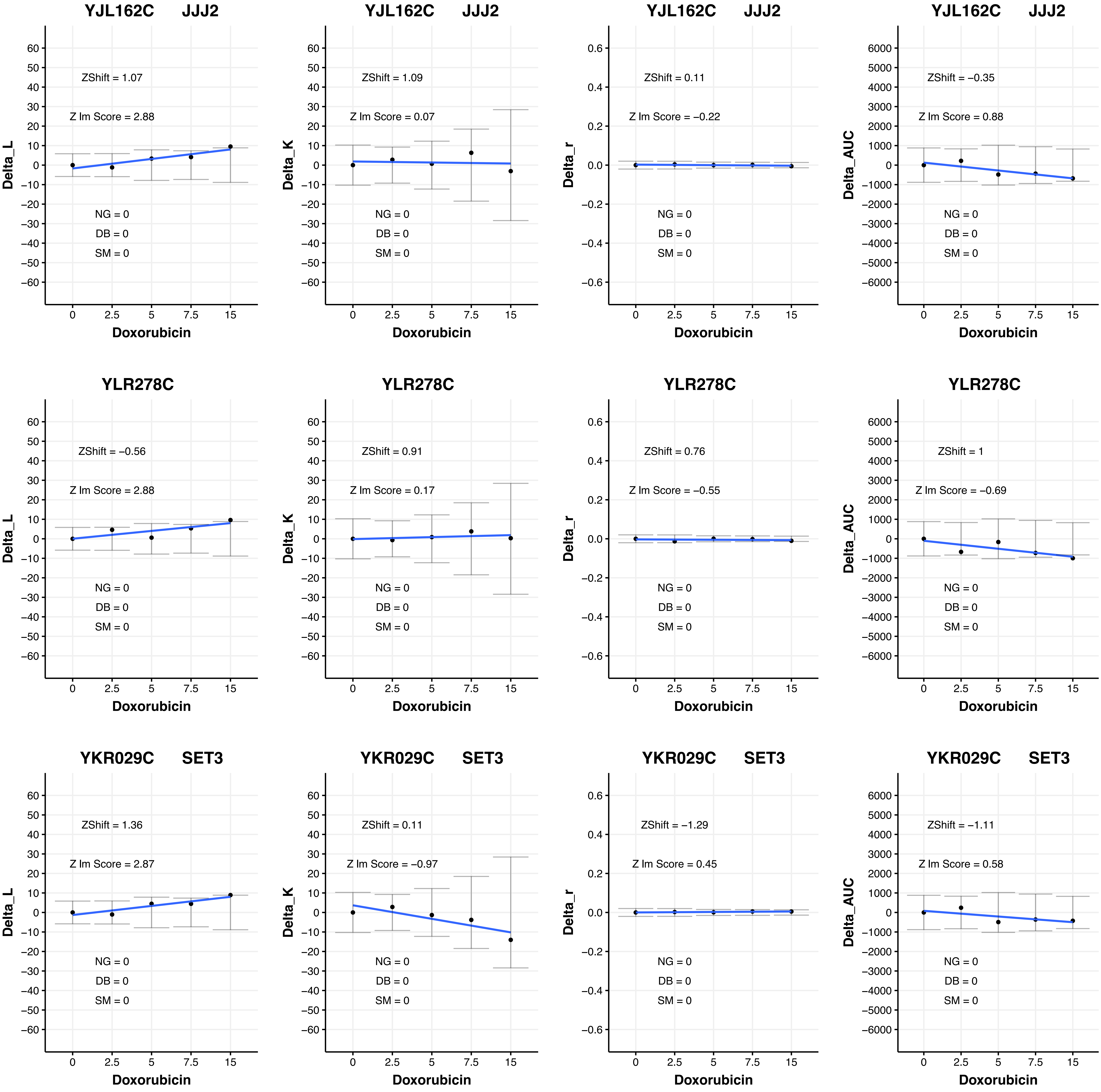
YJL162C JJJ2
ZShift = 1.07
Z lm Score = 2.88
NG = 0
DB = 0
SM = 0
Delta_L
Doxorubicin
YJL162C JJJ2
ZShift = 1.09
Z lm Score = 0.07
NG = 0
DB = 0
SM = 0
Delta_K
Doxorubicin
YJL162C JJJ2
ZShift = 0.11
Z lm Score = −0.22
NG = 0
DB = 0
SM = 0
Delta_r
Doxorubicin
YJL162C JJJ2
ZShift = −0.35
Z lm Score = 0.88
NG = 0
DB = 0
SM = 0
Delta_AUC
Doxorubicin
YLR278C
ZShift = −0.56
Z lm Score = 2.88
NG = 0
DB = 0
SM = 0
Delta_L
Doxorubicin
YLR278C
ZShift = 0.91
Z lm Score = 0.17
NG = 0
DB = 0
SM = 0
Delta_K
Doxorubicin
YLR278C
ZShift = 0.76
Z lm Score = −0.55
NG = 0
DB = 0
SM = 0
Delta_r
Doxorubicin
YLR278C
ZShift = 1
Z lm Score = −0.69
NG = 0
DB = 0
SM = 0
Delta_AUC
Doxorubicin
YKR029C SET3
ZShift = 1.36
Z lm Score = 2.87
NG = 0
DB = 0
SM = 0
Delta_L
Doxorubicin
YKR029C SET3
ZShift = 0.11
Z lm Score = −0.97
NG = 0
DB = 0
SM = 0
Delta_K
Doxorubicin
YKR029C SET3
ZShift = −1.29
Z lm Score = 0.45
NG = 0
DB = 0
SM = 0
Delta_r
Doxorubicin
YKR029C SET3
ZShift = −1.11
Z lm Score = 0.58
NG = 0
DB = 0
SM = 0
Delta_AUC
Doxorubicin

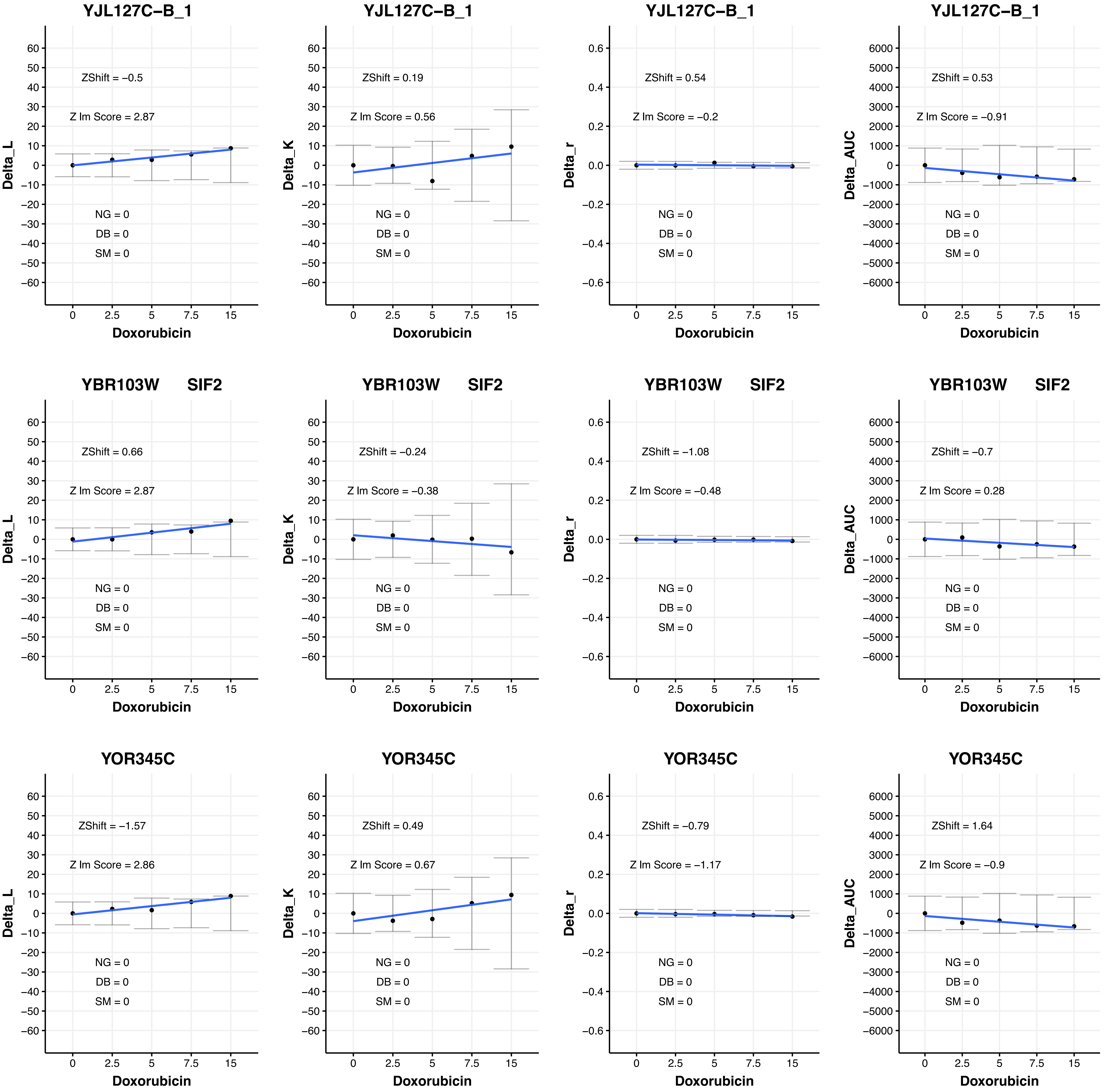

YJL127C−B_1
ZShift = −0.5
Z lm Score = 2.87
NG = 0
DB = 0
SM = 0
Delta_L
Doxorubicin
YJL127C−B_1
ZShift = 0.19
Z lm Score = 0.56
NG = 0
DB = 0
SM = 0
Delta_K
Doxorubicin
YJL127C−B_1
ZShift = 0.54
Z lm Score = −0.2
NG = 0
DB = 0
SM = 0
Delta_r
Doxorubicin
YJL127C−B_1
ZShift = 0.53
Z lm Score = −0.91
NG = 0
DB = 0
SM = 0
Delta_AUC
Doxorubicin
YBR103W SIF2
ZShift = 0.66
Z lm Score = 2.87
NG = 0
DB = 0
SM = 0
Delta_L
Doxorubicin
YBR103W SIF2
ZShift = −0.24
Z lm Score = −0.38
NG = 0
DB = 0
SM = 0
Delta_K
Doxorubicin
YBR103W SIF2
ZShift = −1.08
Z lm Score = −0.48
NG = 0
DB = 0
SM = 0
Delta_r
Doxorubicin
YBR103W SIF2
ZShift = −0.7
Z lm Score = 0.28
NG = 0
DB = 0
SM = 0
Delta_AUC
Doxorubicin
YOR345C
ZShift = −1.57
Z lm Score = 2.86
NG = 0
DB = 0
SM = 0
Delta_L
Doxorubicin
YOR345C
ZShift = 0.49
Z lm Score = 0.67
NG = 0
DB = 0
SM = 0
Delta_K
Doxorubicin
YOR345C
ZShift = −0.79
Z lm Score = −1.17
NG = 0
DB = 0
SM = 0
Delta_r
Doxorubicin
YOR345C
ZShift = 1.64
Z lm Score = −0.9
NG = 0
DB = 0
SM = 0
Delta_AUC
Doxorubicin

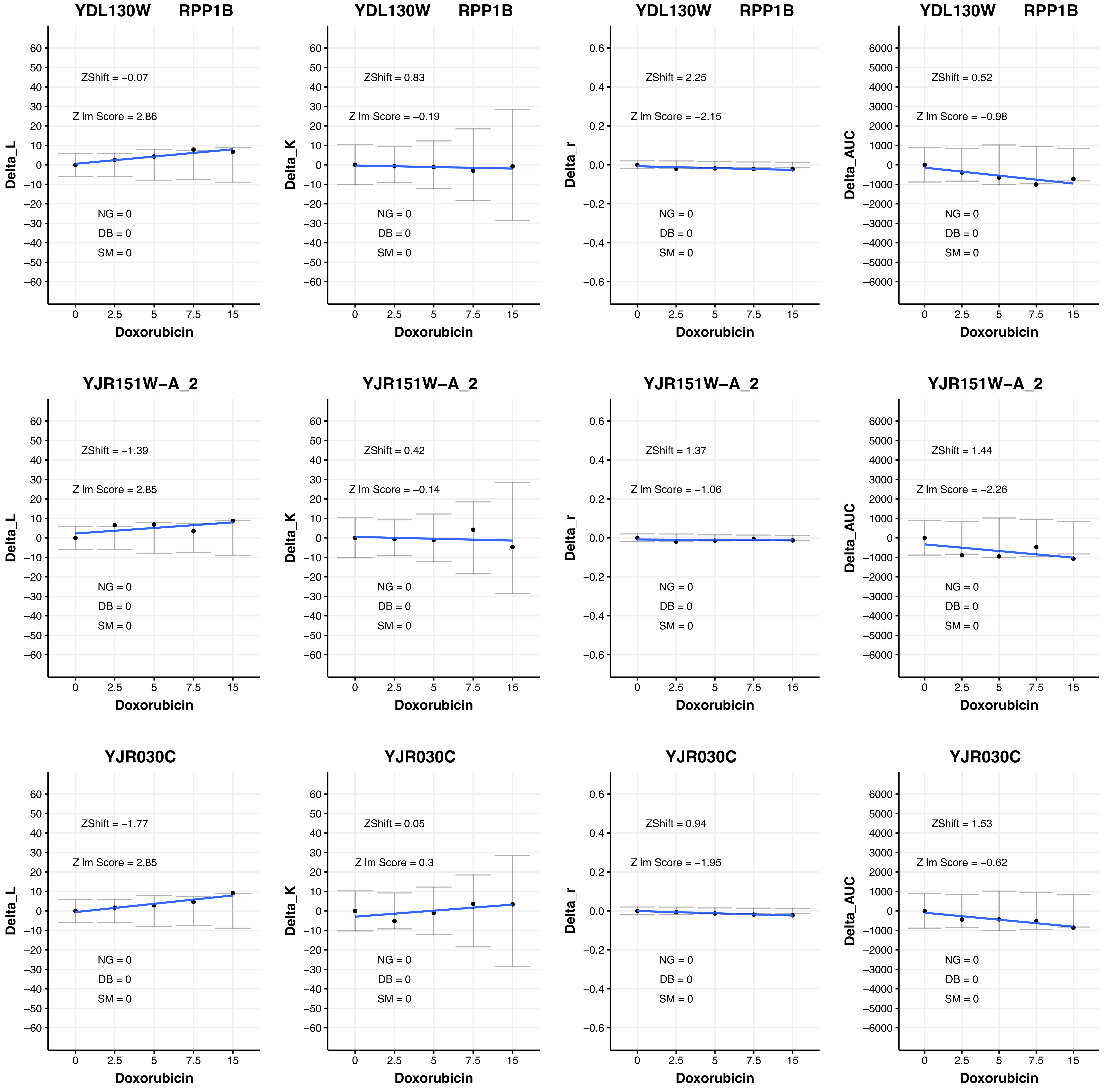

YDL130W RPP1B
ZShift = −0.07
Z lm Score = 2.86
NG = 0
DB = 0
SM = 0
Delta_L
Doxorubicin
YDL130W RPP1B
ZShift = 0.83
Z lm Score = −0.19
NG = 0
DB = 0
SM = 0
Delta_K
Doxorubicin
YDL130W RPP1B
ZShift = 2.25
Z lm Score = −2.15
NG = 0
DB = 0
SM = 0
Delta_r
Doxorubicin
YDL130W RPP1B
ZShift = 0.52
Z lm Score = −0.98
NG = 0
DB = 0
SM = 0
Delta_AUC
Doxorubicin
YJR151W−A_2
ZShift = −1.39
Z lm Score = 2.85
NG = 0
DB = 0
SM = 0
Delta_L
Doxorubicin
YJR151W−A_2
ZShift = 0.42
Z lm Score = −0.14
NG = 0
DB = 0
SM = 0
Delta_K
Doxorubicin
YJR151W−A_2
ZShift = 1.37
Z lm Score = −1.06
NG = 0
DB = 0
SM = 0
Delta_r
Doxorubicin
YJR151W−A_2
ZShift = 1.44
Z lm Score = −2.26
NG = 0
DB = 0
SM = 0
Delta_AUC
Doxorubicin
YJR030C
ZShift = −1.77
Z lm Score = 2.85
NG = 0
DB = 0
SM = 0
Delta_L
Doxorubicin
YJR030C
ZShift = 0.05
Z lm Score = 0.3
NG = 0
DB = 0
SM = 0
Delta_K
Doxorubicin
YJR030C
ZShift = 0.94
Z lm Score = −1.95
NG = 0
DB = 0
SM = 0
Delta_r
Doxorubicin
YJR030C
ZShift = 1.53
Z lm Score = −0.62
NG = 0
DB = 0
SM = 0
Delta_AUC
Doxorubicin

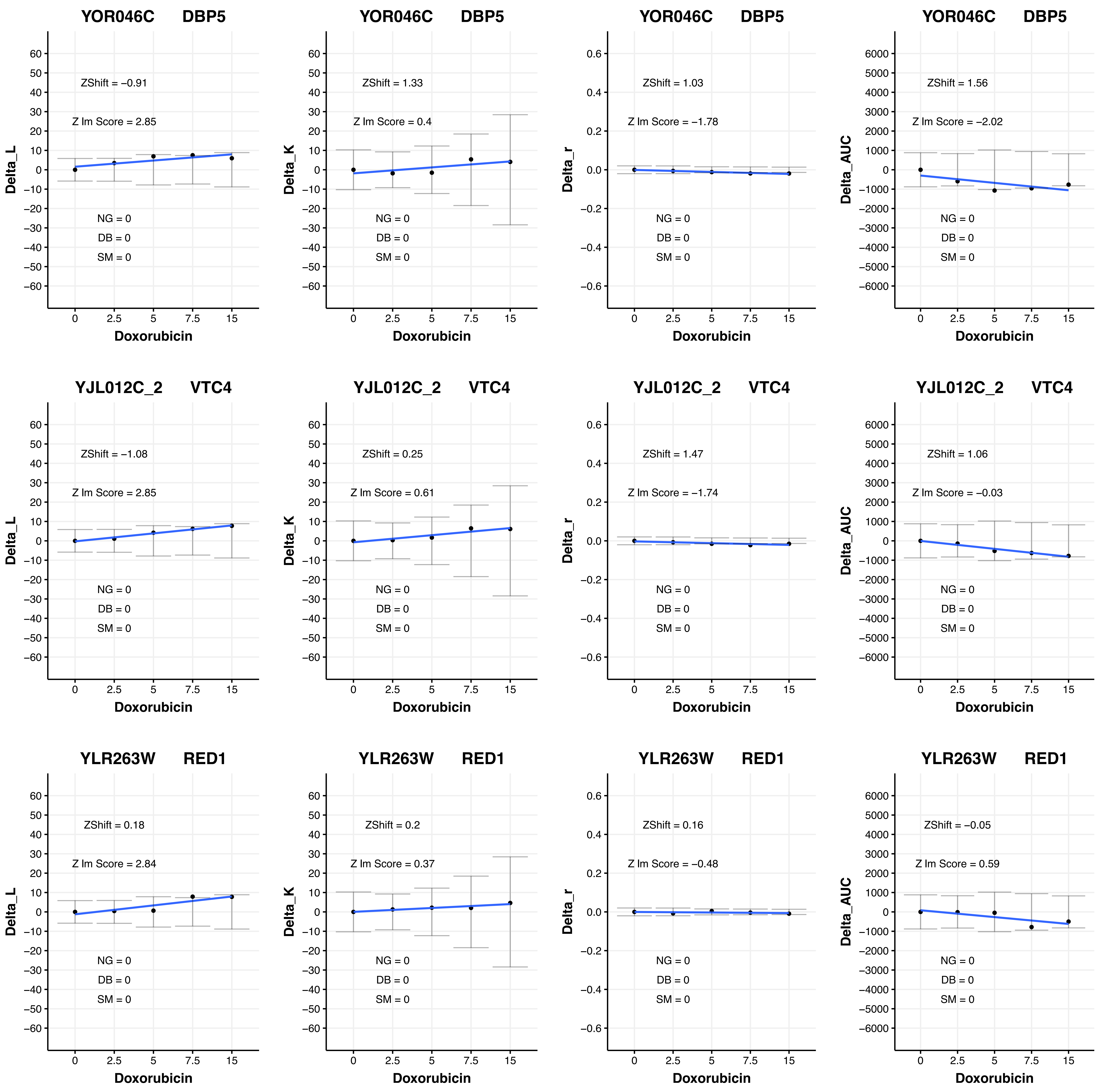

YOR046C DBP5
ZShift = −0.91
Z lm Score = 2.85
NG = 0
DB = 0
SM = 0
Delta_L
Doxorubicin
YOR046C DBP5
ZShift = 1.33
Z lm Score = 0.4
NG = 0
DB = 0
SM = 0
Delta_K
Doxorubicin
YOR046C DBP5
ZShift = 1.03
Z lm Score = −1.78
NG = 0
DB = 0
SM = 0
Delta_r
Doxorubicin
YOR046C DBP5
ZShift = 1.56
Z lm Score = −2.02
NG = 0
DB = 0
SM = 0
Delta_AUC
Doxorubicin
YJL012C_2 VTC4
ZShift = −1.08
Z lm Score = 2.85
NG = 0
DB = 0
SM = 0
Delta_L
Doxorubicin
YJL012C_2 VTC4
ZShift = 0.25
Z lm Score = 0.61
NG = 0
DB = 0
SM = 0
Delta_K
Doxorubicin
YJL012C_2 VTC4
ZShift = 1.47
Z lm Score = −1.74
NG = 0
DB = 0
SM = 0
Delta_r
Doxorubicin
YJL012C_2 VTC4
ZShift = 1.06
Z lm Score = −0.03
NG = 0
DB = 0
SM = 0
Delta_AUC
Doxorubicin
YLR263W RED1
ZShift = 0.18
Z lm Score = 2.84
NG = 0
DB = 0
SM = 0
Delta_L
Doxorubicin
YLR263W RED1
ZShift = 0.2
Z lm Score = 0.37
NG = 0
DB = 0
SM = 0
Delta_K
Doxorubicin
YLR263W RED1
ZShift = 0.16
Z lm Score = −0.48
NG = 0
DB = 0
SM = 0
Delta_r
Doxorubicin
YLR263W RED1
ZShift = −0.05
Z lm Score = 0.59
NG = 0
DB = 0
SM = 0
Delta_AUC
Doxorubicin

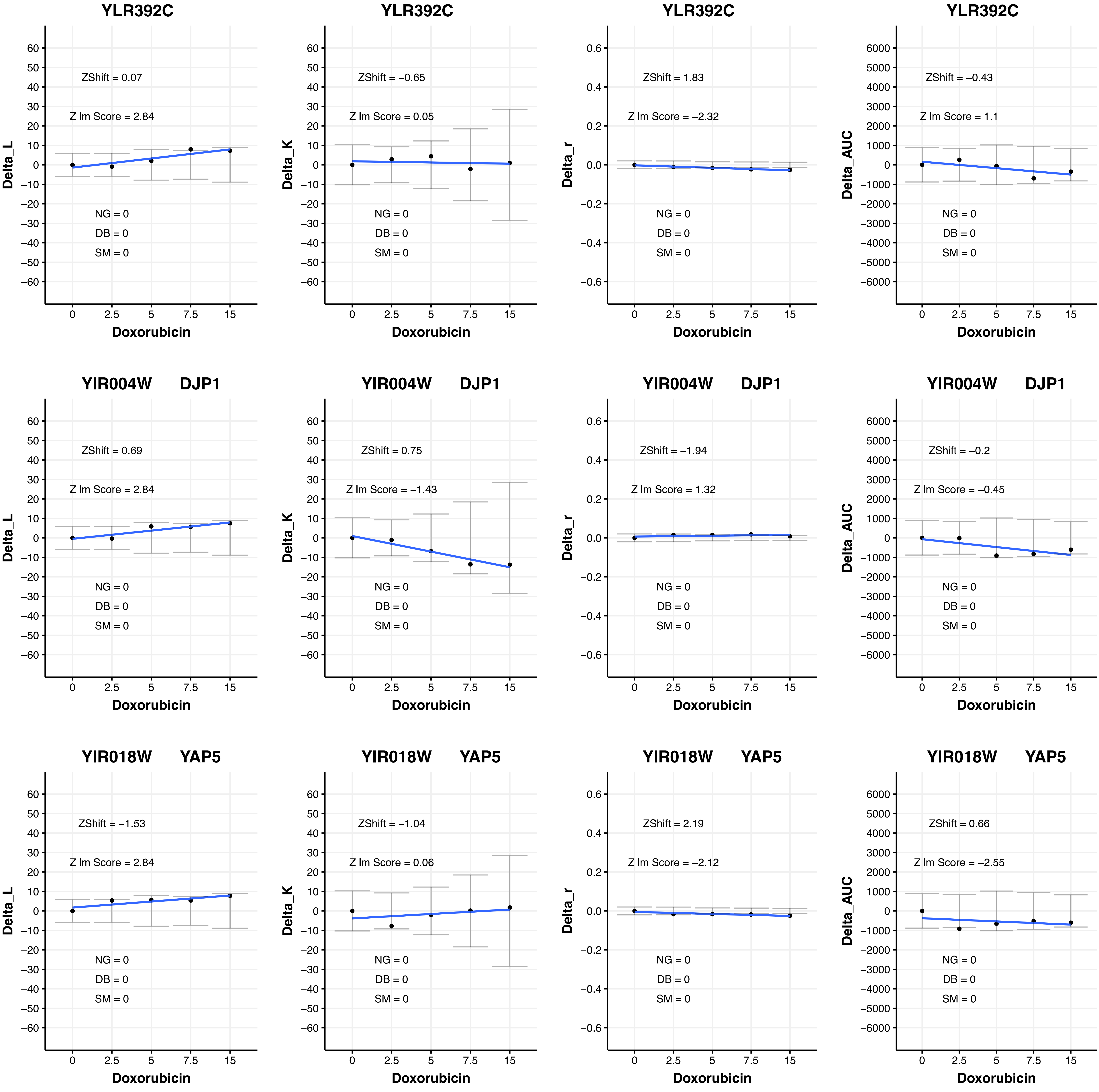

YLR392C
ZShift = 0.07
Z lm Score = 2.84
NG = 0
DB = 0
SM = 0
Delta_L
Doxorubicin
YLR392C
ZShift = −0.65
Z lm Score = 0.05
NG = 0
DB = 0
SM = 0
Delta_K
Doxorubicin
YLR392C
ZShift = 1.83
Z lm Score = −2.32
NG = 0
DB = 0
SM = 0
Delta_r
Doxorubicin
YLR392C
ZShift = −0.43
Z lm Score = 1.1
NG = 0
DB = 0
SM = 0
Delta_AUC
Doxorubicin
YIR004W DJP1
ZShift = 0.69
Z lm Score = 2.84
NG = 0
DB = 0
SM = 0
Delta_L
Doxorubicin
YIR004W DJP1
ZShift = 0.75
Z lm Score = −1.43
NG = 0
DB = 0
SM = 0
Delta_K
Doxorubicin
YIR004W DJP1
ZShift = −1.94
Z lm Score = 1.32
NG = 0
DB = 0
SM = 0
Delta_r
Doxorubicin
YIR004W DJP1
ZShift = −0.2
Z lm Score = −0.45
NG = 0
DB = 0
SM = 0
Delta_AUC
Doxorubicin
YIR018W YAP5
ZShift = −1.53
Z lm Score = 2.84
NG = 0
DB = 0
SM = 0
Delta_L
Doxorubicin
YIR018W YAP5
ZShift = −1.04
Z lm Score = 0.06
NG = 0
DB = 0
SM = 0
Delta_K
Doxorubicin
YIR018W YAP5
ZShift = 2.19
Z lm Score = −2.12
NG = 0
DB = 0
SM = 0
Delta_r
Doxorubicin
YIR018W YAP5
ZShift = 0.66
Z lm Score = −2.55
NG = 0
DB = 0
SM = 0
Delta_AUC
Doxorubicin

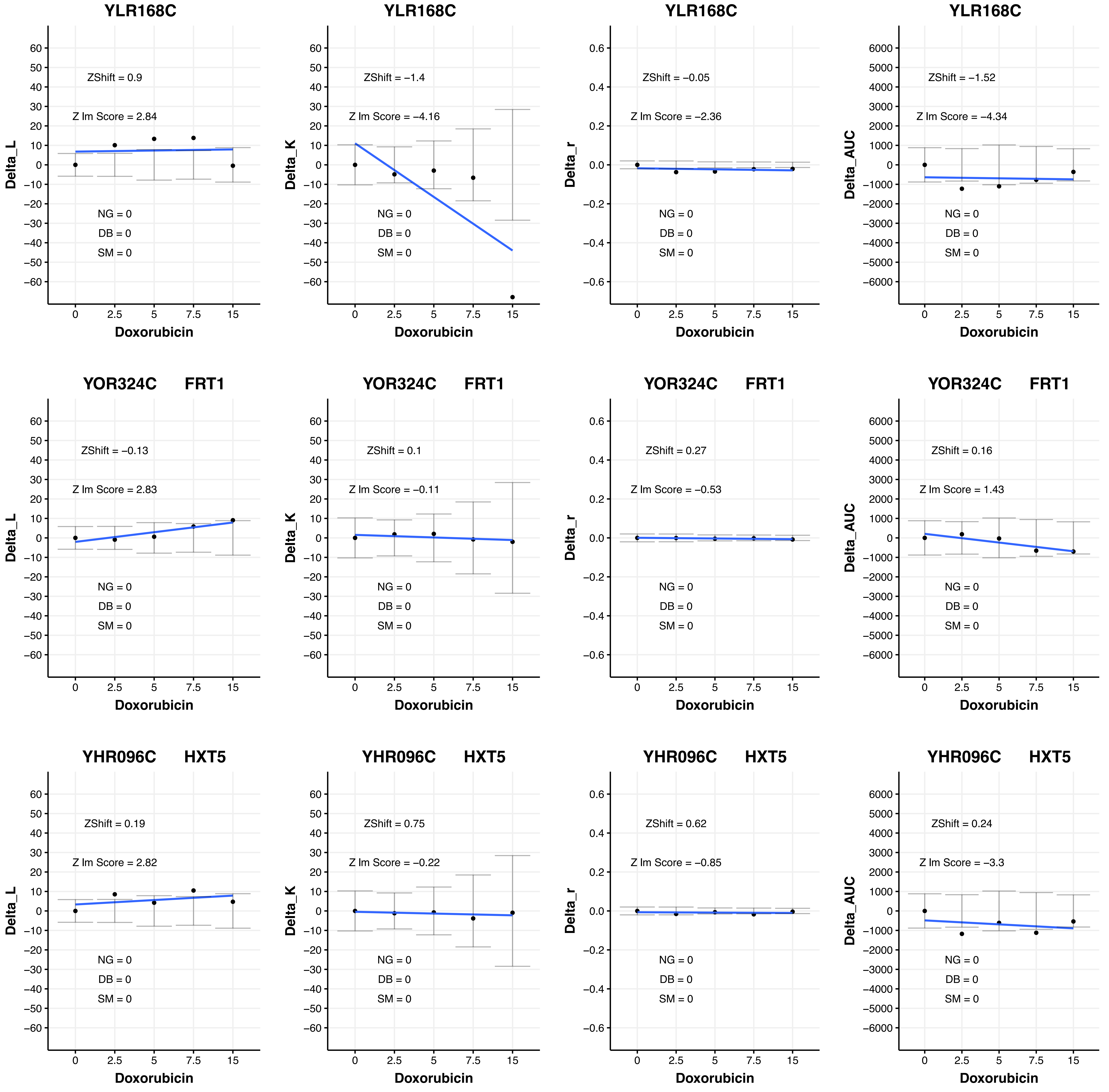

YLR168C
ZShift = 0.9
Z lm Score = 2.84
NG = 0
DB = 0
SM = 0
Delta_L
Doxorubicin
YLR168C
ZShift = −1.4
Z lm Score = −4.16
NG = 0
DB = 0
SM = 0
Delta_K
Doxorubicin
YLR168C
ZShift = −0.05
Z lm Score = −2.36
NG = 0
DB = 0
SM = 0
Delta_r
Doxorubicin
YLR168C
ZShift = −1.52
Z lm Score = −4.34
NG = 0
DB = 0
SM = 0
Delta_AUC
Doxorubicin
YOR324C FRT1
ZShift = −0.13
Z lm Score = 2.83
NG = 0
DB = 0
SM = 0
Delta_L
Doxorubicin
YOR324C FRT1
ZShift = 0.1
Z lm Score = −0.11
NG = 0
DB = 0
SM = 0
Delta_K
Doxorubicin
YOR324C FRT1
ZShift = 0.27
Z lm Score = −0.53
NG = 0
DB = 0
SM = 0
Delta_r
Doxorubicin
YOR324C FRT1
ZShift = 0.16
Z lm Score = 1.43
NG = 0
DB = 0
SM = 0
Delta_AUC
Doxorubicin
YHR096C HXT5
ZShift = 0.19
Z lm Score = 2.82
NG = 0
DB = 0
SM = 0
Delta_L
Doxorubicin
YHR096C HXT5
ZShift = 0.75
Z lm Score = −0.22
NG = 0
DB = 0
SM = 0
Delta_K
Doxorubicin
YHR096C HXT5
ZShift = 0.62
Z lm Score = −0.85
NG = 0
DB = 0
SM = 0
Delta_r
Doxorubicin
YHR096C HXT5
ZShift = 0.24
Z lm Score = −3.3
NG = 0
DB = 0
SM = 0
Delta_AUC
Doxorubicin

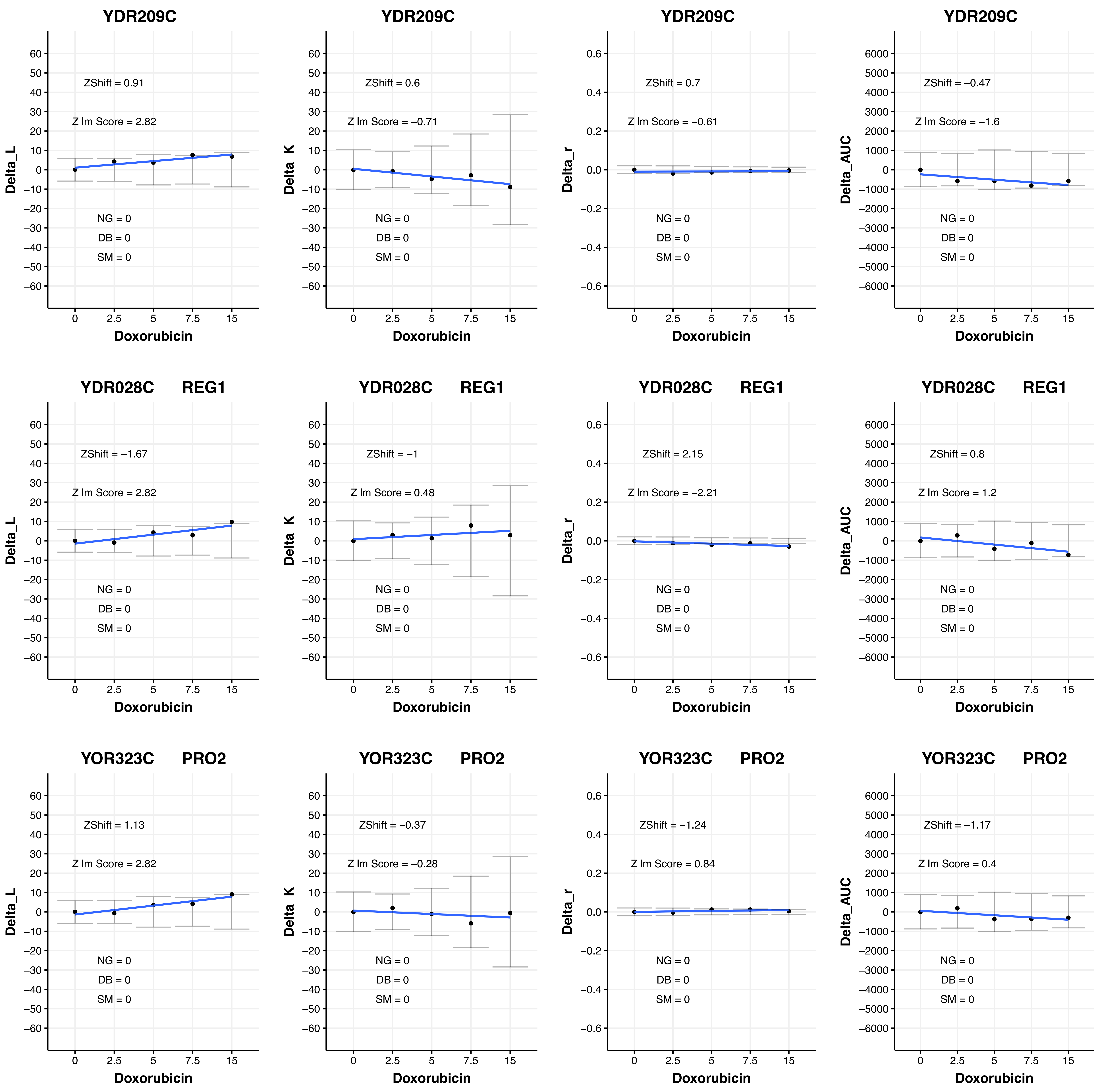
YDR209C
ZShift = 0.91
Z lm Score = 2.82
NG = 0
DB = 0
SM = 0
Delta_L
Doxorubicin
YDR209C
ZShift = 0.6
Z lm Score = −0.71
NG = 0
DB = 0
SM = 0
Delta_K
Doxorubicin
YDR209C
ZShift = 0.7
Z lm Score = −0.61
NG = 0
DB = 0
SM = 0
Delta_r
Doxorubicin
YDR209C
ZShift = −0.47
Z lm Score = −1.6
NG = 0
DB = 0
SM = 0
Delta_AUC
Doxorubicin
YDR028C REG1
ZShift = −1.67
Z lm Score = 2.82
NG = 0
DB = 0
SM = 0
Delta_L
Doxorubicin
YDR028C REG1
ZShift = −1
Z lm Score = 0.48
NG = 0
DB = 0
SM = 0
Delta_K
Doxorubicin
YDR028C REG1
ZShift = 2.15
Z lm Score = −2.21
NG = 0
DB = 0
SM = 0
Delta_r
Doxorubicin
YDR028C REG1
ZShift = 0.8
Z lm Score = 1.2
NG = 0
DB = 0
SM = 0
Delta_AUC
Doxorubicin
YOR323C PRO2
ZShift = 1.13
Z lm Score = 2.82
NG = 0
DB = 0
SM = 0
Delta_L
Doxorubicin
YOR323C PRO2
ZShift = −0.37
Z lm Score = −0.28
NG = 0
DB = 0
SM = 0
Delta_K
Doxorubicin
YOR323C PRO2
ZShift = −1.24
Z lm Score = 0.84
NG = 0
DB = 0
SM = 0
Delta_r
Doxorubicin
YOR323C PRO2
ZShift = −1.17
Z lm Score = 0.4
NG = 0
DB = 0
SM = 0
Delta_AUC
Doxorubicin

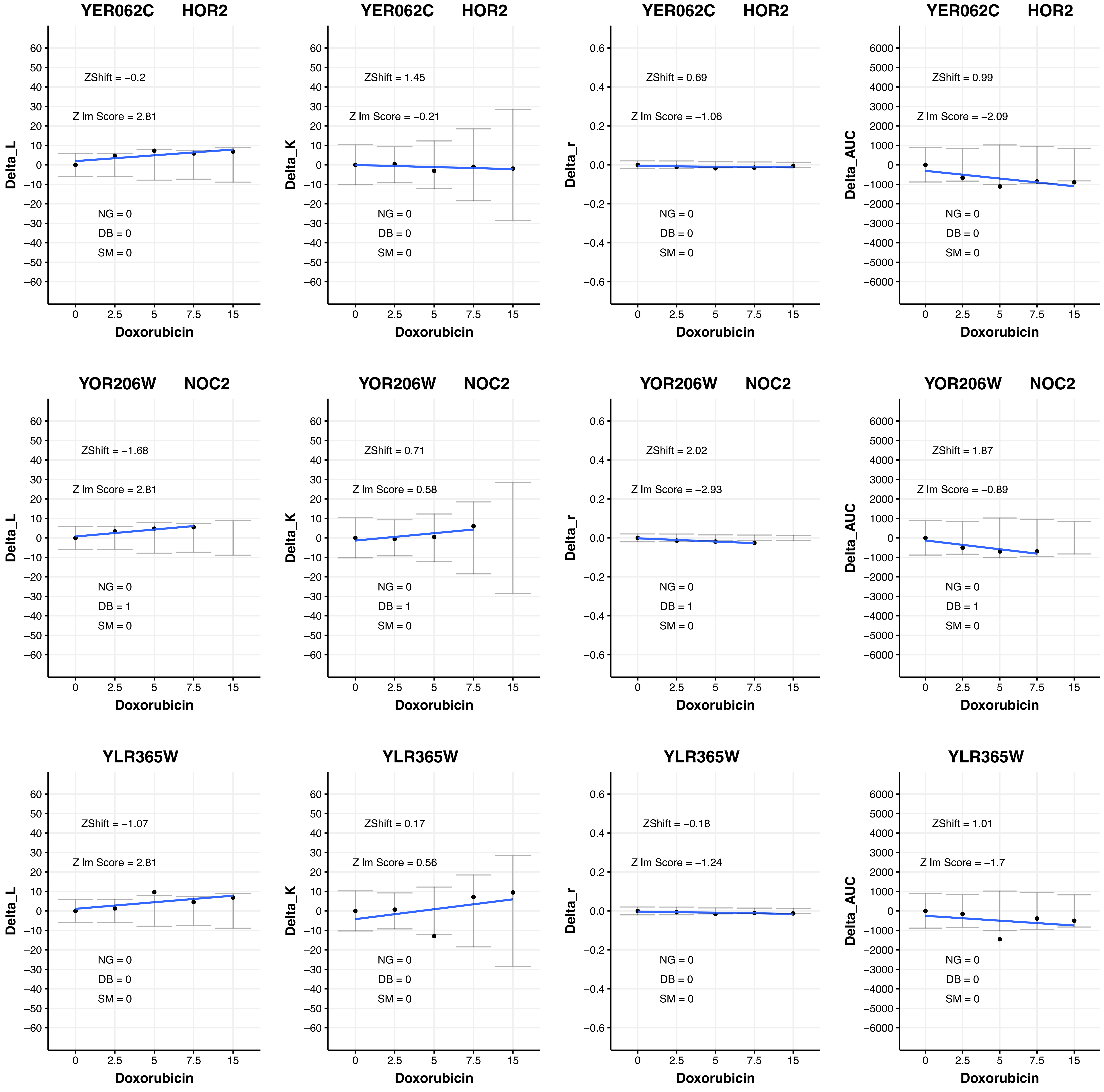
YER062C HOR2
ZShift = −0.2
Z lm Score = 2.81
NG = 0
DB = 0
SM = 0
Delta_L
Doxorubicin
YER062C HOR2
ZShift = 1.45
Z lm Score = −0.21
NG = 0
DB = 0
SM = 0
Delta_K
Doxorubicin
YER062C HOR2
ZShift = 0.69
Z lm Score = −1.06
NG = 0
DB = 0
SM = 0
Delta_r
Doxorubicin
YER062C HOR2
ZShift = 0.99
Z lm Score = −2.09
NG = 0
DB = 0
SM = 0
Delta_AUC
Doxorubicin
YOR206W NOC2
ZShift = −1.68
Z lm Score = 2.81
NG = 0
DB = 1
SM = 0
Delta_L
Doxorubicin
YOR206W NOC2
ZShift = 0.71
Z lm Score = 0.58
NG = 0
DB = 1
SM = 0
Delta_K
Doxorubicin
YOR206W NOC2
ZShift = 2.02
Z lm Score = −2.93
NG = 0
DB = 1
SM = 0
Delta_r
Doxorubicin
YOR206W NOC2
ZShift = 1.87
Z lm Score = −0.89
NG = 0
DB = 1
SM = 0
Delta_AUC
Doxorubicin
YLR365W
ZShift = −1.07
Z lm Score = 2.81
NG = 0
DB = 0
SM = 0
Delta_L
Doxorubicin
YLR365W
ZShift = 0.17
Z lm Score = 0.56
NG = 0
DB = 0
SM = 0
Delta_K
Doxorubicin
YLR365W
ZShift = −0.18
Z lm Score = −1.24
NG = 0
DB = 0
SM = 0
Delta_r
Doxorubicin
YLR365W
ZShift = 1.01
Z lm Score = −1.7
NG = 0
DB = 0
SM = 0
Delta_AUC
Doxorubicin

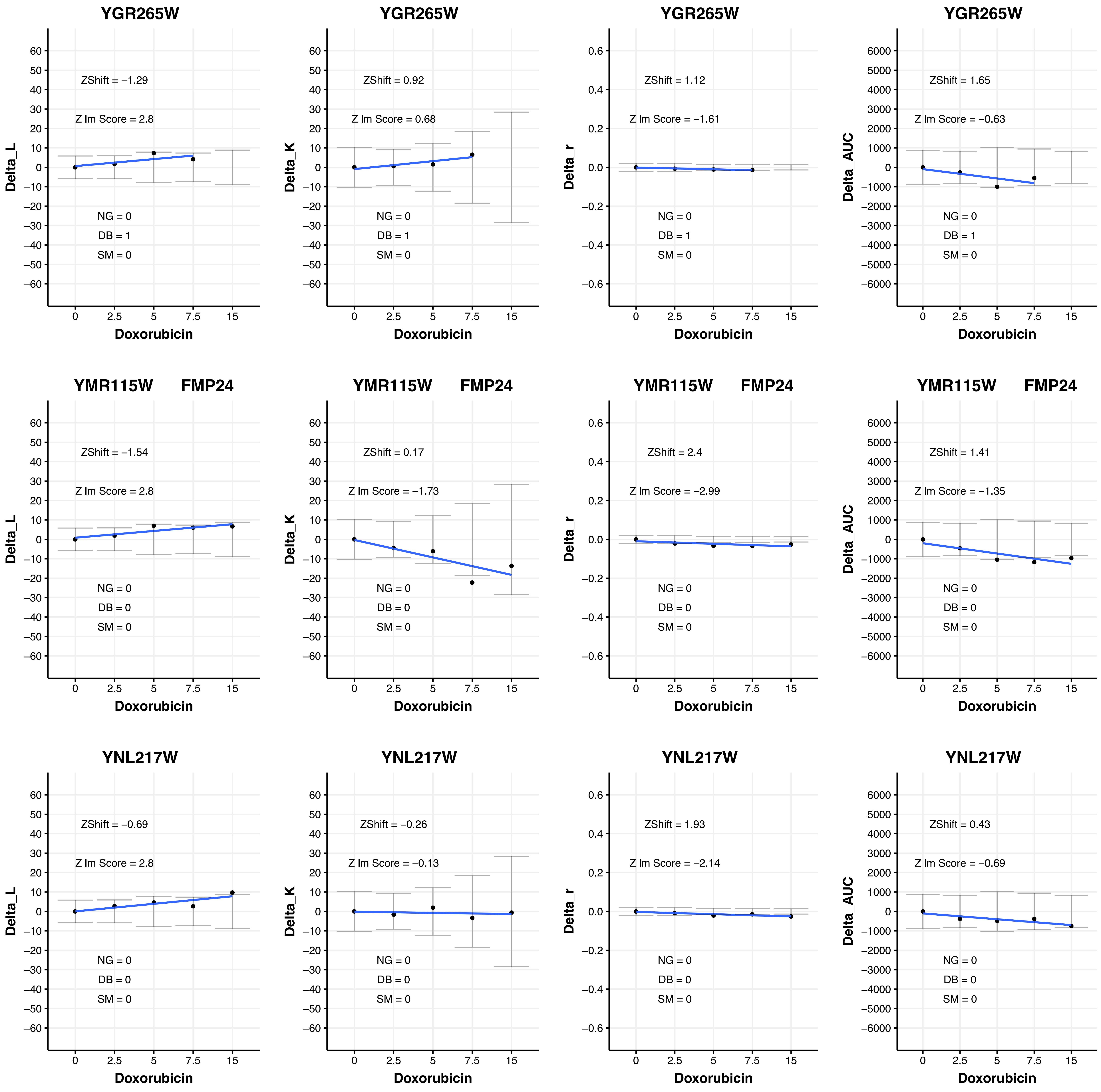
YGR265W
ZShift = −1.29
Z lm Score = 2.8
NG = 0
DB = 1
SM = 0
Delta_L
Doxorubicin
YGR265W
ZShift = 0.92
Z lm Score = 0.68
NG = 0
DB = 1
SM = 0
Delta_K
Doxorubicin
YGR265W
ZShift = 1.12
Z lm Score = −1.61
NG = 0
DB = 1
SM = 0
Delta_r
Doxorubicin
YGR265W
ZShift = 1.65
Z lm Score = −0.63
NG = 0
DB = 1
SM = 0
Delta_AUC
Doxorubicin
YMR115W FMP24
ZShift = −1.54
Z lm Score = 2.8
NG = 0
DB = 0
SM = 0
Delta_L
Doxorubicin
YMR115W FMP24
ZShift = 0.17
Z lm Score = −1.73
NG = 0
DB = 0
SM = 0
Delta_K
Doxorubicin
YMR115W FMP24
ZShift = 2.4
Z lm Score = −2.99
NG = 0
DB = 0
SM = 0
Delta_r
Doxorubicin
YMR115W FMP24
ZShift = 1.41
Z lm Score = −1.35
NG = 0
DB = 0
SM = 0
Delta_AUC
Doxorubicin
YNL217W
ZShift = −0.69
Z lm Score = 2.8
NG = 0
DB = 0
SM = 0
Delta_L
Doxorubicin
YNL217W
ZShift = −0.26
Z lm Score = −0.13
NG = 0
DB = 0
SM = 0
Delta_K
Doxorubicin
YNL217W
ZShift = 1.93
Z lm Score = −2.14
NG = 0
DB = 0
SM = 0
Delta_r
Doxorubicin
YNL217W
ZShift = 0.43
Z lm Score = −0.69
NG = 0
DB = 0
SM = 0
Delta_AUC
Doxorubicin
0 2.5 5 7.5 15

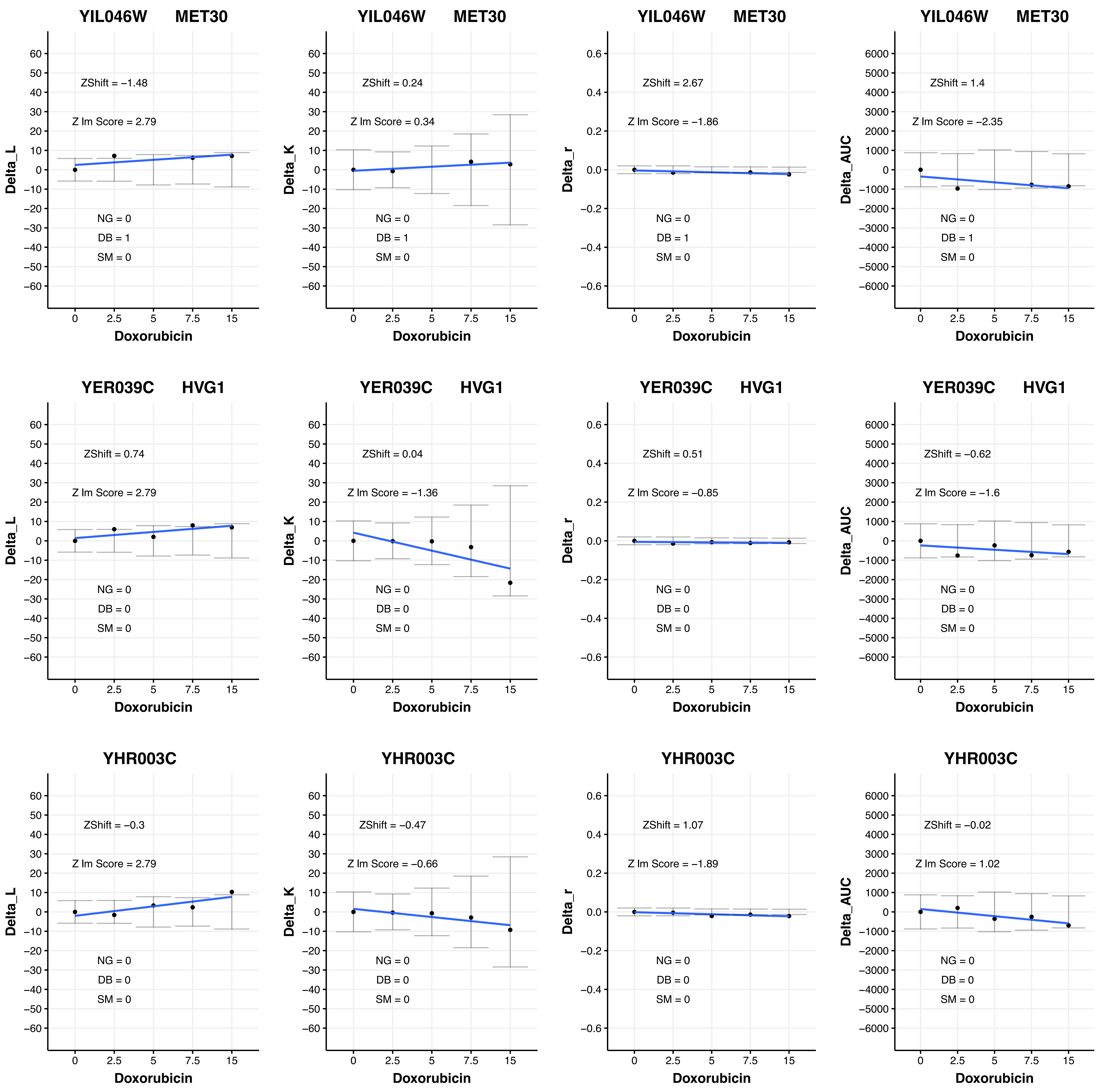
YIL046W MET30
ZShift = −1.48
Z lm Score = 2.79
NG = 0
DB = 1
SM = 0
Delta_L
Doxorubicin
YIL046W MET30
ZShift = 0.24
Z lm Score = 0.34
NG = 0
DB = 1
SM = 0
Delta_K
Doxorubicin
YIL046W MET30
ZShift = 2.67
Z lm Score = −1.86
NG = 0
DB = 1
SM = 0
Delta_r
Doxorubicin
YIL046W MET30
ZShift = 1.4
Z lm Score = −2.35
NG = 0
DB = 1
SM = 0
Delta_AUC
Doxorubicin
YER039C HVG1
ZShift = 0.74
Z lm Score = 2.79
NG = 0
DB = 0
SM = 0
Delta_L
Doxorubicin
YER039C HVG1
ZShift = 0.04
Z lm Score = −1.36
NG = 0
DB = 0
SM = 0
Delta_K
Doxorubicin
YER039C HVG1
ZShift = 0.51
Z lm Score = −0.85
NG = 0
DB = 0
SM = 0
Delta_r
Doxorubicin
YER039C HVG1
ZShift = −0.62
Z lm Score = −1.6
NG = 0
DB = 0
SM = 0
Delta_AUC
Doxorubicin
YHR003C
ZShift = −0.3
Z lm Score = 2.79
NG = 0
DB = 0
SM = 0
Delta_L
Doxorubicin
YHR003C
ZShift = −0.47
Z lm Score = −0.66
NG = 0
DB = 0
SM = 0
Delta_K
Doxorubicin
YHR003C
ZShift = 1.07
Z lm Score = −1.89
NG = 0
DB = 0
SM = 0
Delta_r
Doxorubicin
YHR003C
ZShift = −0.02
Z lm Score = 1.02
NG = 0
DB = 0
SM = 0
Delta_AUC
Doxorubicin

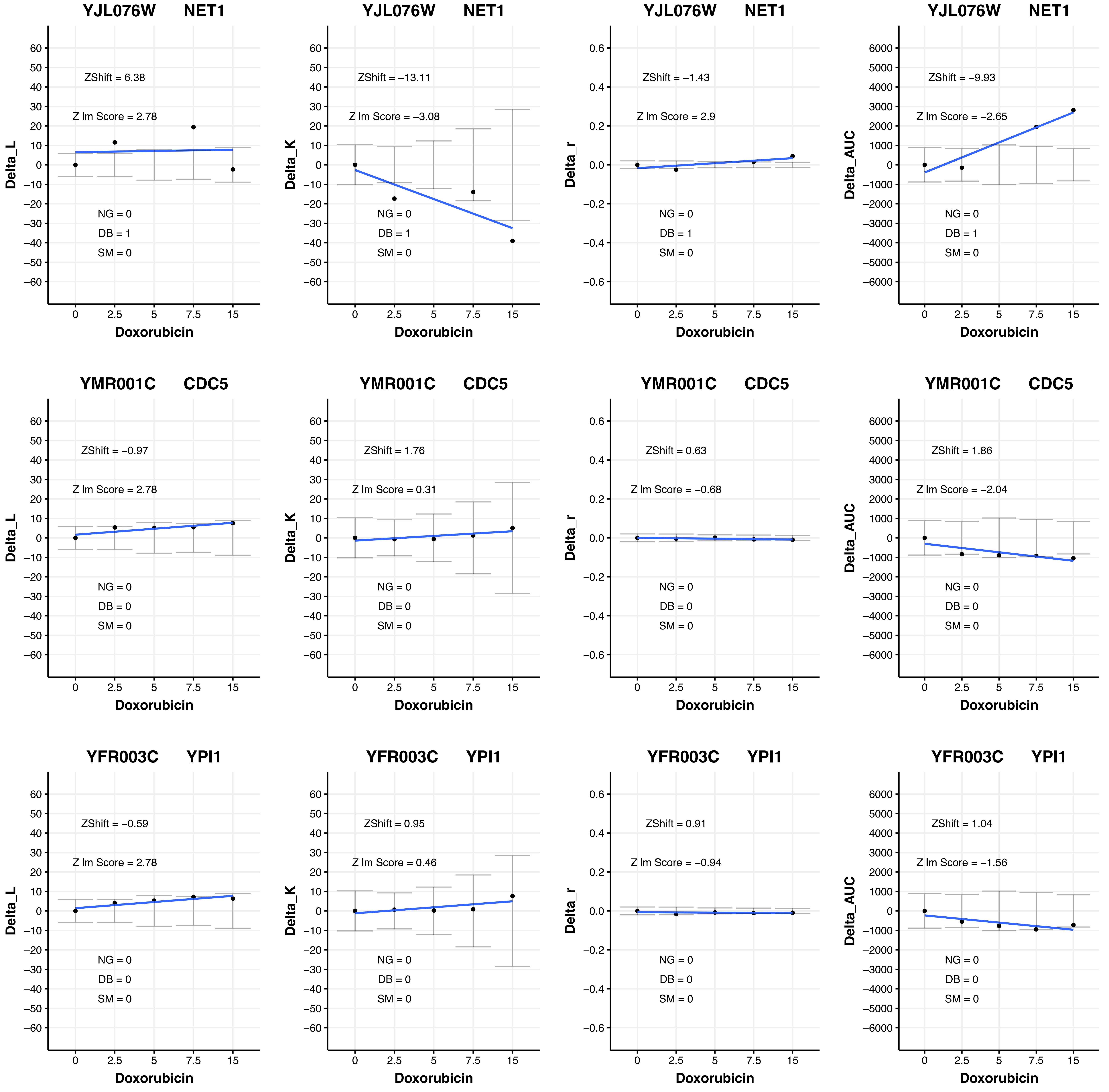
YJL076W NET1
ZShift = 6.38
Z lm Score = 2.78
NG = 0
DB = 1
SM = 0
Delta_L
Doxorubicin
YJL076W NET1
ZShift = −13.11
Z lm Score = −3.08
NG = 0
DB = 1
SM = 0
Delta_K
Doxorubicin
YJL076W NET1
ZShift = −1.43
Z lm Score = 2.9
NG = 0
DB = 1
SM = 0
Delta_r
Doxorubicin
YJL076W NET1
ZShift = −9.93
Z lm Score = −2.65
NG = 0
DB = 1
SM = 0
Delta_AUC
Doxorubicin
YMR001C CDC5
ZShift = −0.97
Z lm Score = 2.78
NG = 0
DB = 0
SM = 0
Delta_L
Doxorubicin
YMR001C CDC5
ZShift = 1.76
Z lm Score = 0.31
NG = 0
DB = 0
SM = 0
Delta_K
Doxorubicin
YMR001C CDC5
ZShift = 0.63
Z lm Score = −0.68
NG = 0
DB = 0
SM = 0
Delta_r
Doxorubicin
YMR001C CDC5
ZShift = 1.86
Z lm Score = −2.04
NG = 0
DB = 0
SM = 0
Delta_AUC
Doxorubicin
YFR003C YPI1
ZShift = −0.59
Z lm Score = 2.78
NG = 0
DB = 0
SM = 0
Delta_L
Doxorubicin
YFR003C YPI1
ZShift = 0.95
Z lm Score = 0.46
NG = 0
DB = 0
SM = 0
Delta_K
Doxorubicin
YFR003C YPI1
ZShift = 0.91
Z lm Score = −0.94
NG = 0
DB = 0
SM = 0
Delta_r
Doxorubicin
YFR003C YPI1
ZShift = 1.04
Z lm Score = −1.56
NG = 0
DB = 0
SM = 0
Delta_AUC
Doxorubicin

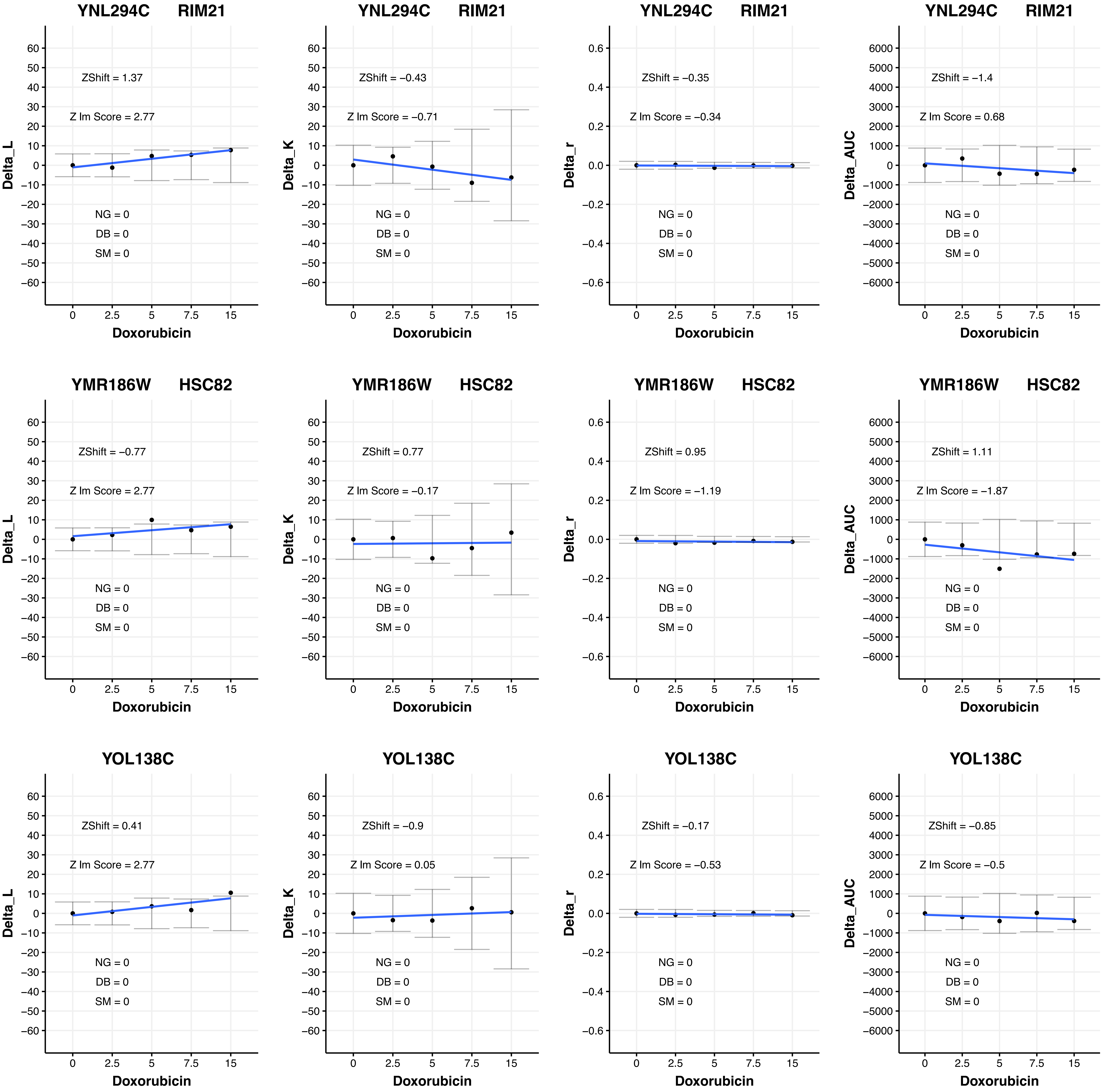

YNL294C RIM21
ZShift = 1.37
Z lm Score = 2.77
Delta_L
NG = 0
DB = 0
SM = 0
Doxorubicin
YNL294C RIM21
ZShift = −0.43
Z lm Score = −0.71
Delta_K
NG = 0
DB = 0
SM = 0
Doxorubicin
YNL294C RIM21
ZShift = −0.35
Z lm Score = −0.34
Delta_r
NG = 0
DB = 0
SM = 0
Doxorubicin
YNL294C RIM21
ZShift = −1.4
Z lm Score = 0.68
Delta_AUC
NG = 0
DB = 0
SM = 0
Doxorubicin
YMR186W HSC82
ZShift = −0.77
Z lm Score = 2.77
Delta_L
NG = 0
DB = 0
SM = 0
Doxorubicin
YMR186W HSC82
ZShift = 0.77
Z lm Score = −0.17
Delta_K
NG = 0
DB = 0
SM = 0
Doxorubicin
YMR186W HSC82
ZShift = 0.95
Z lm Score = −1.19
Delta_r
NG = 0
DB = 0
SM = 0
Doxorubicin
YMR186W HSC82
ZShift = 1.11
Z lm Score = −1.87
Delta_AUC
NG = 0
DB = 0
SM = 0
Doxorubicin
YOL138C
ZShift = 0.41
Z lm Score = 2.77
Delta_L
NG = 0
DB = 0
SM = 0
Doxorubicin
YOL138C
ZShift = −0.9
Z lm Score = 0.05
Delta_K
NG = 0
DB = 0
SM = 0
Doxorubicin
YOL138C
ZShift = −0.17
Z lm Score = −0.53
Delta_r
NG = 0
DB = 0
SM = 0
Doxorubicin
YOL138C
ZShift = −0.85
Z lm Score = −0.5
Delta_AUC
NG = 0
DB = 0
SM = 0
Doxorubicin

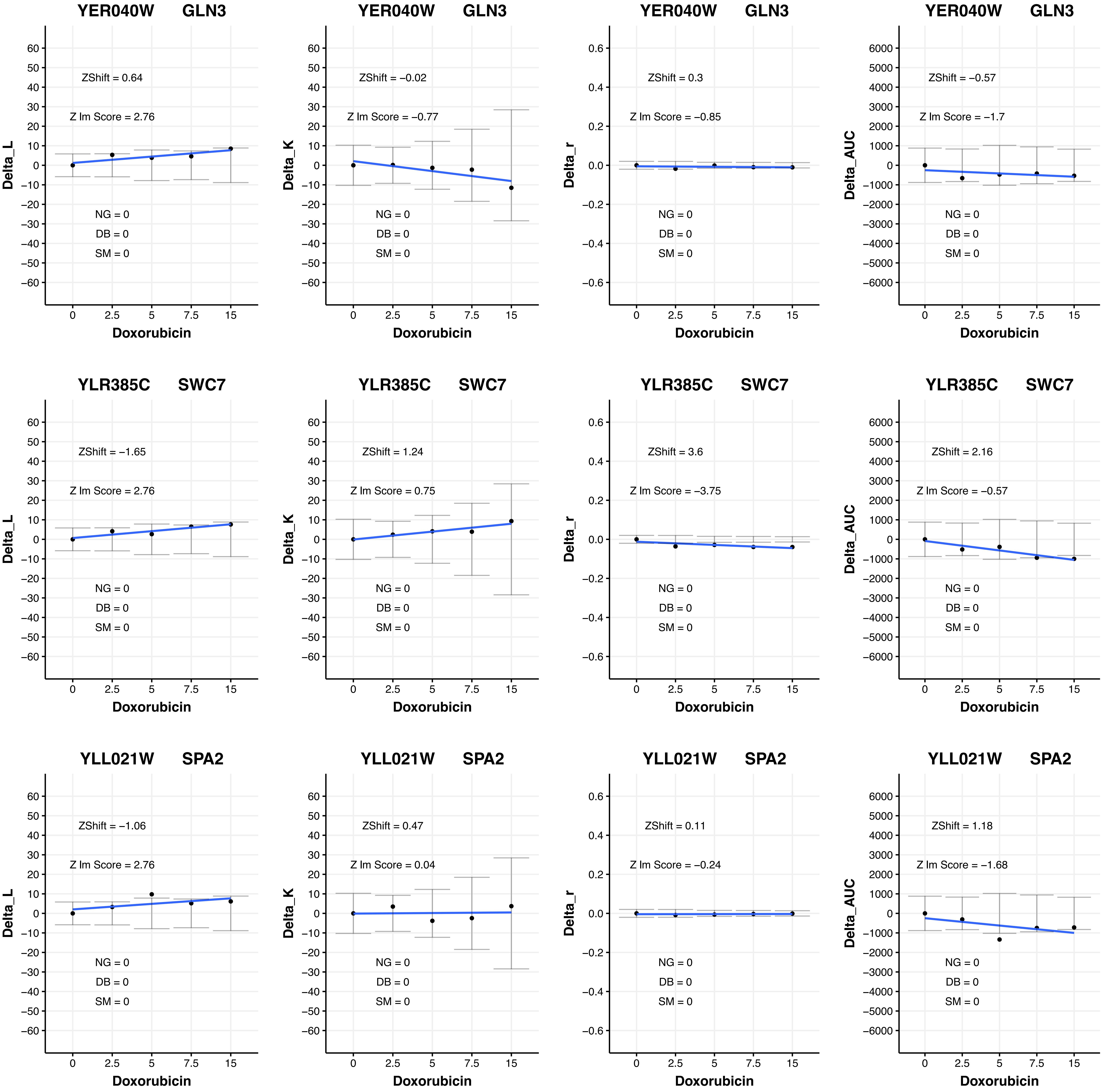
YER040W GLN3
ZShift = 0.64
Z lm Score = 2.76
NG = 0
DB = 0
SM = 0
Delta_L
Doxorubicin
YER040W GLN3
ZShift = −0.02
Z lm Score = −0.77
NG = 0
DB = 0
SM = 0
Delta_K
Doxorubicin
YER040W GLN3
ZShift = 0.3
Z lm Score = −0.85
NG = 0
DB = 0
SM = 0
Delta_r
Doxorubicin
YER040W GLN3
ZShift = −0.57
Z lm Score = −1.7
NG = 0
DB = 0
SM = 0
Delta_AUC
Doxorubicin
YLR385C SWC7
ZShift = −1.65
Z lm Score = 2.76
NG = 0
DB = 0
SM = 0
Delta_L
Doxorubicin
YLR385C SWC7
ZShift = 1.24
Z lm Score = 0.75
NG = 0
DB = 0
SM = 0
Delta_K
Doxorubicin
YLR385C SWC7
ZShift = 3.6
Z lm Score = −3.75
NG = 0
DB = 0
SM = 0
Delta_r
Doxorubicin
YLR385C SWC7
ZShift = 2.16
Z lm Score = −0.57
NG = 0
DB = 0
SM = 0
Delta_AUC
Doxorubicin
YLL021W SPA2
ZShift = −1.06
Z lm Score = 2.76
NG = 0
DB = 0
SM = 0
Delta_L
Doxorubicin
YLL021W SPA2
ZShift = 0.47
Z lm Score = 0.04
NG = 0
DB = 0
SM = 0
Delta_K
Doxorubicin
YLL021W SPA2
ZShift = 0.11
Z lm Score = −0.24
NG = 0
DB = 0
SM = 0
Delta_r
Doxorubicin
YLL021W SPA2
ZShift = 1.18
Z lm Score = −1.68
NG = 0
DB = 0
SM = 0
Delta_AUC
Doxorubicin

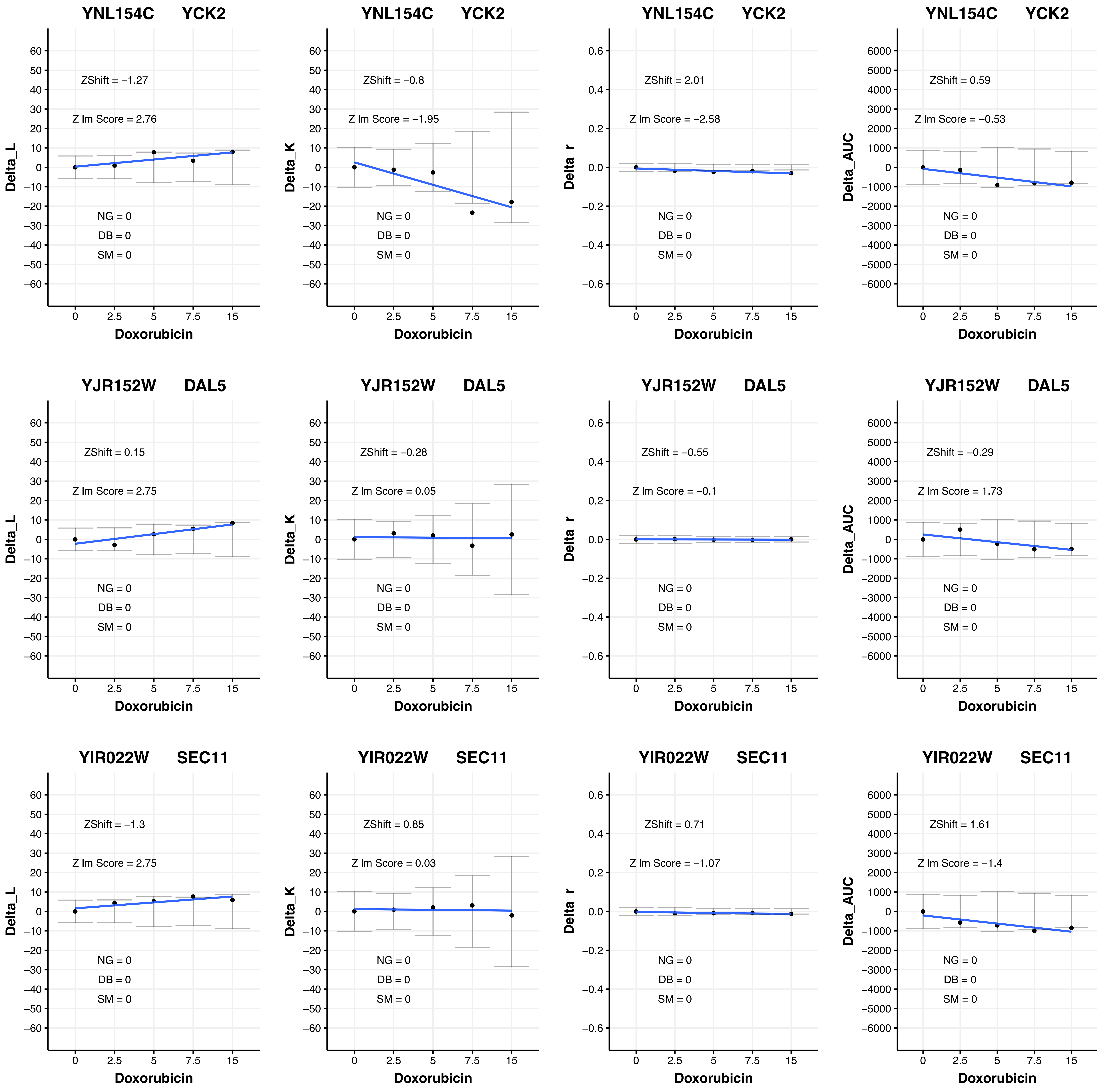
YNL154C YCK2
ZShift = −1.27
Z lm Score = 2.76
NG = 0
DB = 0
SM = 0
Delta_L
Doxorubicin
YNL154C YCK2
ZShift = −0.8
Z lm Score = −1.95
NG = 0
DB = 0
SM = 0
Delta_K
Doxorubicin
YNL154C YCK2
ZShift = 2.01
Z lm Score = −2.58
NG = 0
DB = 0
SM = 0
Delta_r
Doxorubicin
YNL154C YCK2
ZShift = 0.59
Z lm Score = −0.53
NG = 0
DB = 0
SM = 0
Delta_AUC
Doxorubicin
YJR152W DAL5
ZShift = 0.15
Z lm Score = 2.75
NG = 0
DB = 0
SM = 0
Delta_L
Doxorubicin
YJR152W DAL5
ZShift = −0.28
Z lm Score = 0.05
NG = 0
DB = 0
SM = 0
Delta_K
Doxorubicin
YJR152W DAL5
ZShift = −0.55
Z lm Score = −0.1
NG = 0
DB = 0
SM = 0
Delta_r
Doxorubicin
YJR152W DAL5
ZShift = −0.29
Z lm Score = 1.73
NG = 0
DB = 0
SM = 0
Delta_AUC
Doxorubicin
YIR022W SEC11
ZShift = −1.3
Z lm Score = 2.75
NG = 0
DB = 0
SM = 0
Delta_L
Doxorubicin
YIR022W SEC11
ZShift = 0.85
Z lm Score = 0.03
NG = 0
DB = 0
SM = 0
Delta_K
Doxorubicin
YIR022W SEC11
ZShift = 0.71
Z lm Score = −1.07
NG = 0
DB = 0
SM = 0
Delta_r
Doxorubicin
YIR022W SEC11
ZShift = 1.61
Z lm Score = −1.4
NG = 0
DB = 0
SM = 0
Delta_AUC
Doxorubicin

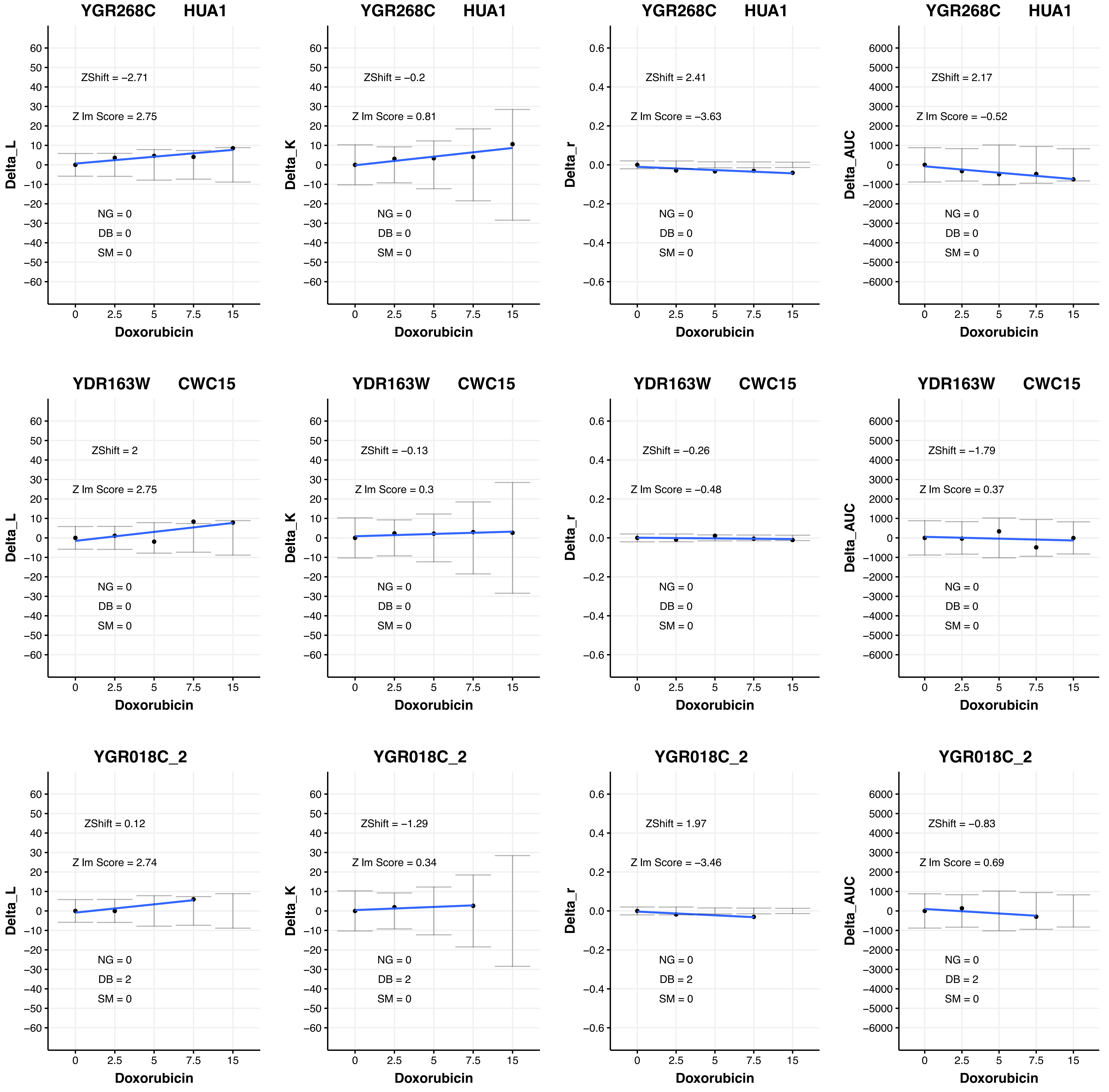

YGR268C HUA1
ZShift = −2.71
Z lm Score = 2.75
NG = 0
DB = 0
SM = 0
Delta_L
Doxorubicin
YGR268C HUA1
ZShift = −0.2
Z lm Score = 0.81
NG = 0
DB = 0
SM = 0
Delta_K
Doxorubicin
YGR268C HUA1
ZShift = 2.41
Z lm Score = −3.63
NG = 0
DB = 0
SM = 0
Delta_r
Doxorubicin
YGR268C HUA1
ZShift = 2.17
Z lm Score = −0.52
NG = 0
DB = 0
SM = 0
Delta_AUC
Doxorubicin
YDR163W CWC15
ZShift = 2
Z lm Score = 2.75
NG = 0
DB = 0
SM = 0
Delta_L
Doxorubicin
YDR163W CWC15
ZShift = −0.13
Z lm Score = 0.3
NG = 0
DB = 0
SM = 0
Delta_K
Doxorubicin
YDR163W CWC15
ZShift = −0.26
Z lm Score = −0.48
NG = 0
DB = 0
SM = 0
Delta_r
Doxorubicin
YDR163W CWC15
ZShift = −1.79
Z lm Score = 0.37
NG = 0
DB = 0
SM = 0
Delta_AUC
Doxorubicin
YGR018C_2
ZShift = 0.12
Z lm Score = 2.74
NG = 0
DB = 2
SM = 0
Delta_L
Doxorubicin
YGR018C_2
ZShift = −1.29
Z lm Score = 0.34
NG = 0
DB = 2
SM = 0
Delta_K
Doxorubicin
YGR018C_2
ZShift = 1.97
Z lm Score = −3.46
NG = 0
DB = 2
SM = 0
Delta_r
Doxorubicin
YGR018C_2
ZShift = −0.83
Z lm Score = 0.69
NG = 0
DB = 2
SM = 0
Delta_AUC
Doxorubicin

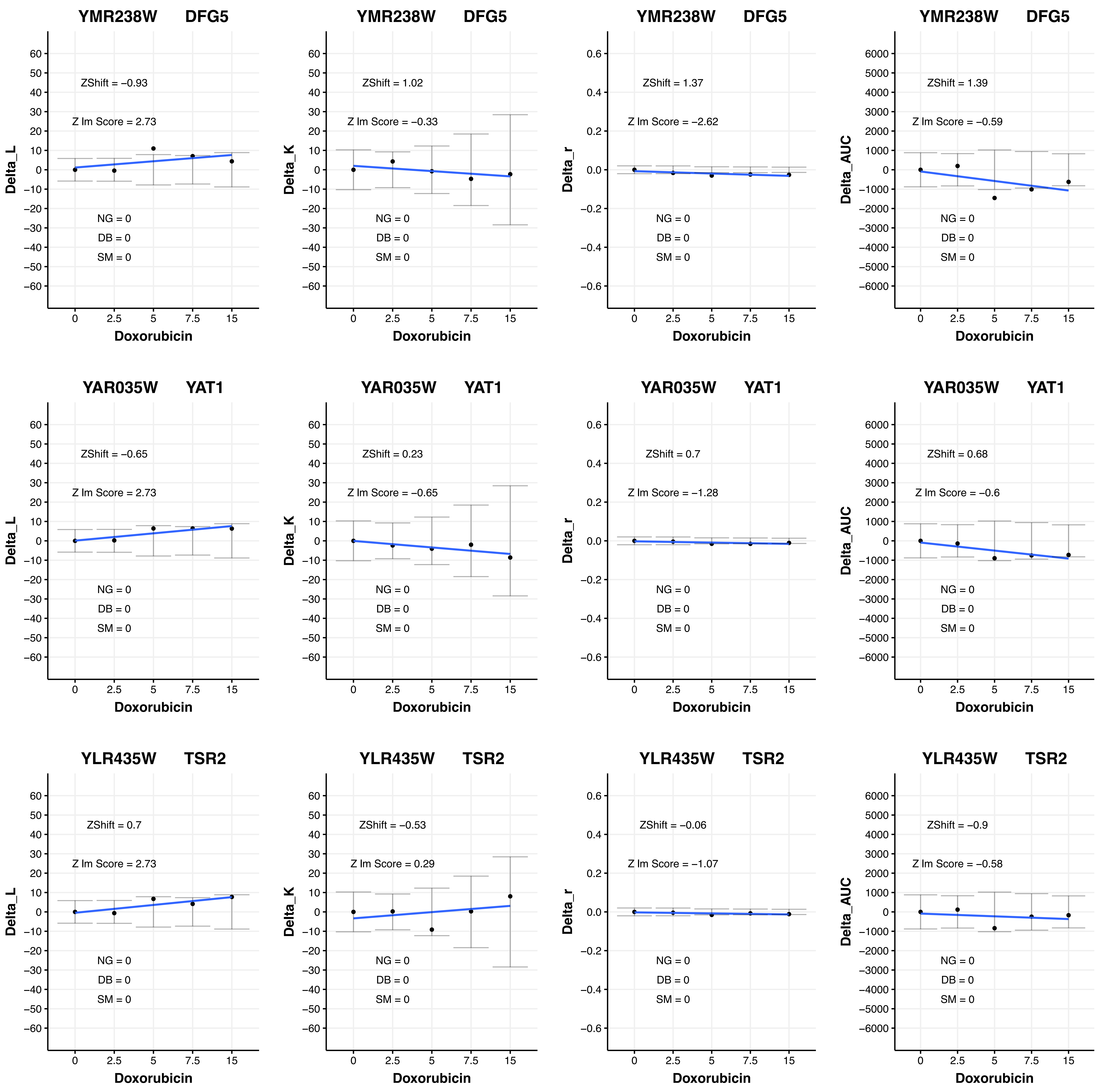
YMR238W DFG5
ZShift = −0.93
Z lm Score = 2.73
NG = 0
DB = 0
SM = 0
Delta_L
Doxorubicin
YMR238W DFG5
ZShift = 1.02
Z lm Score = −0.33
NG = 0
DB = 0
SM = 0
Delta_K
Doxorubicin
YMR238W DFG5
ZShift = 1.37
Z lm Score = −2.62
NG = 0
DB = 0
SM = 0
Delta_r
Doxorubicin
YMR238W DFG5
ZShift = 1.39
Z lm Score = −0.59
NG = 0
DB = 0
SM = 0
Delta_AUC
Doxorubicin
YAR035W YAT1
ZShift = −0.65
Z lm Score = 2.73
NG = 0
DB = 0
SM = 0
Delta_L
Doxorubicin
YAR035W YAT1
ZShift = 0.23
Z lm Score = −0.65
NG = 0
DB = 0
SM = 0
Delta_K
Doxorubicin
YAR035W YAT1
ZShift = 0.7
Z lm Score = −1.28
NG = 0
DB = 0
SM = 0
Delta_r
Doxorubicin
YAR035W YAT1
ZShift = 0.68
Z lm Score = −0.6
NG = 0
DB = 0
SM = 0
Delta_AUC
Doxorubicin
YLR435W TSR2
ZShift = 0.7
Z lm Score = 2.73
NG = 0
DB = 0
SM = 0
Delta_L
Doxorubicin
YLR435W TSR2
ZShift = −0.53
Z lm Score = 0.29
NG = 0
DB = 0
SM = 0
Delta_K
Doxorubicin
YLR435W TSR2
ZShift = −0.06
Z lm Score = −1.07
NG = 0
DB = 0
SM = 0
Delta_r
Doxorubicin
YLR435W TSR2
ZShift = −0.9
Z lm Score = −0.58
NG = 0
DB = 0
SM = 0
Delta_AUC
Doxorubicin

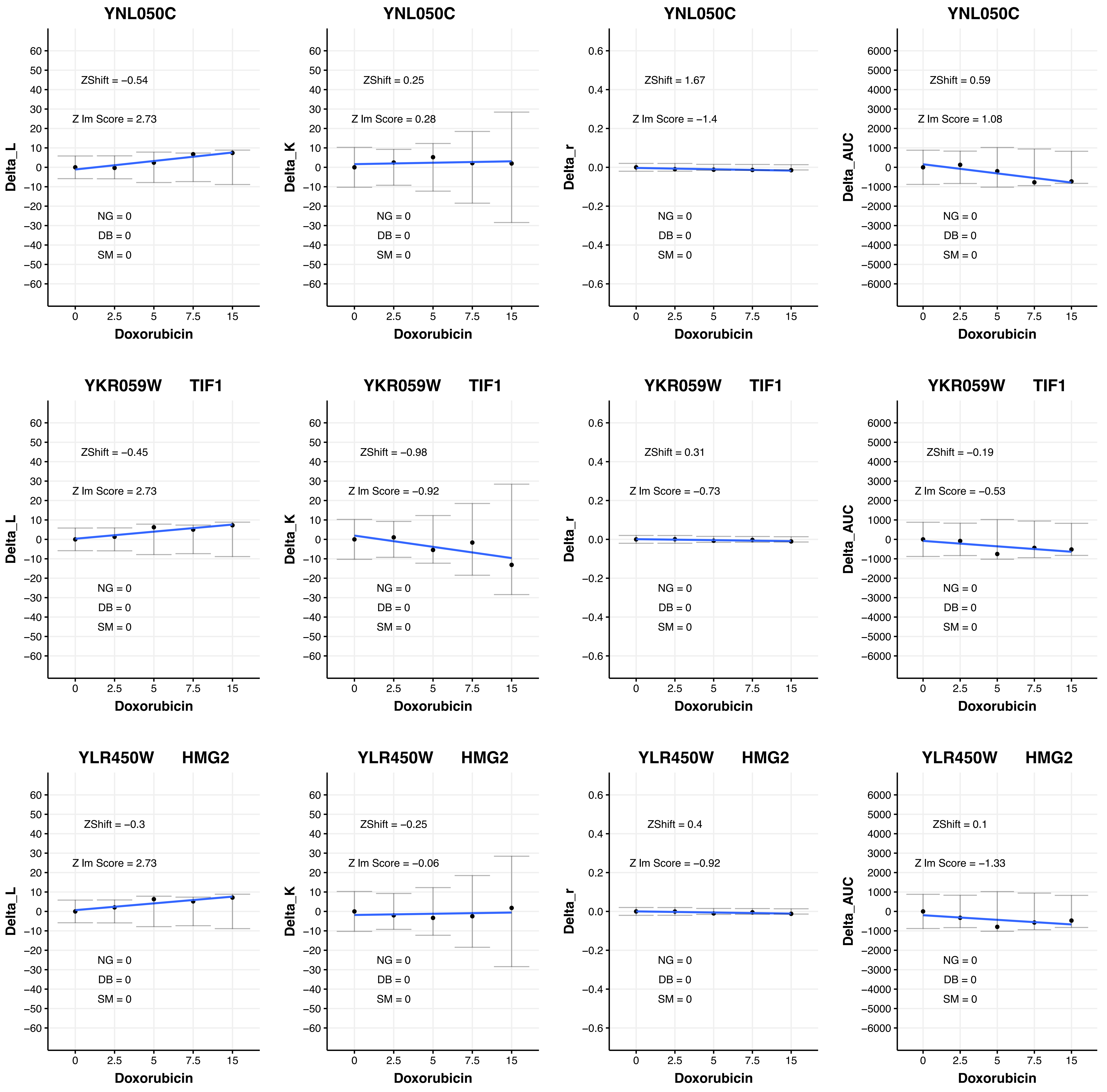
YNL050C
ZShift = −0.54
Z lm Score = 2.73
NG = 0
DB = 0
SM = 0
Delta_L
Doxorubicin
YNL050C
ZShift = 0.25
Z lm Score = 0.28
NG = 0
DB = 0
SM = 0
Delta_K
Doxorubicin
YNL050C
ZShift = 1.67
Z lm Score = −1.4
NG = 0
DB = 0
SM = 0
Delta_r
Doxorubicin
YNL050C
ZShift = 0.59
Z lm Score = 1.08
NG = 0
DB = 0
SM = 0
Delta_AUC
Doxorubicin
YKR059W TIF1
ZShift = −0.45
Z lm Score = 2.73
NG = 0
DB = 0
SM = 0
Delta_L
Doxorubicin
YKR059W TIF1
ZShift = −0.98
Z lm Score = −0.92
NG = 0
DB = 0
SM = 0
Delta_K
Doxorubicin
YKR059W TIF1
ZShift = 0.31
Z lm Score = −0.73
NG = 0
DB = 0
SM = 0
Delta_r
Doxorubicin
YKR059W TIF1
ZShift = −0.19
Z lm Score = −0.53
NG = 0
DB = 0
SM = 0
Delta_AUC
Doxorubicin
YLR450W HMG2
ZShift = −0.3
Z lm Score = 2.73
NG = 0
DB = 0
SM = 0
Delta_L
Doxorubicin
YLR450W HMG2
ZShift = −0.25
Z lm Score = −0.06
NG = 0
DB = 0
SM = 0
Delta_K
Doxorubicin
YLR450W HMG2
ZShift = 0.4
Z lm Score = −0.92
NG = 0
DB = 0
SM = 0
Delta_r
Doxorubicin
YLR450W HMG2
ZShift = 0.1
Z lm Score = −1.33
NG = 0
DB = 0
SM = 0
Delta_AUC
Doxorubicin

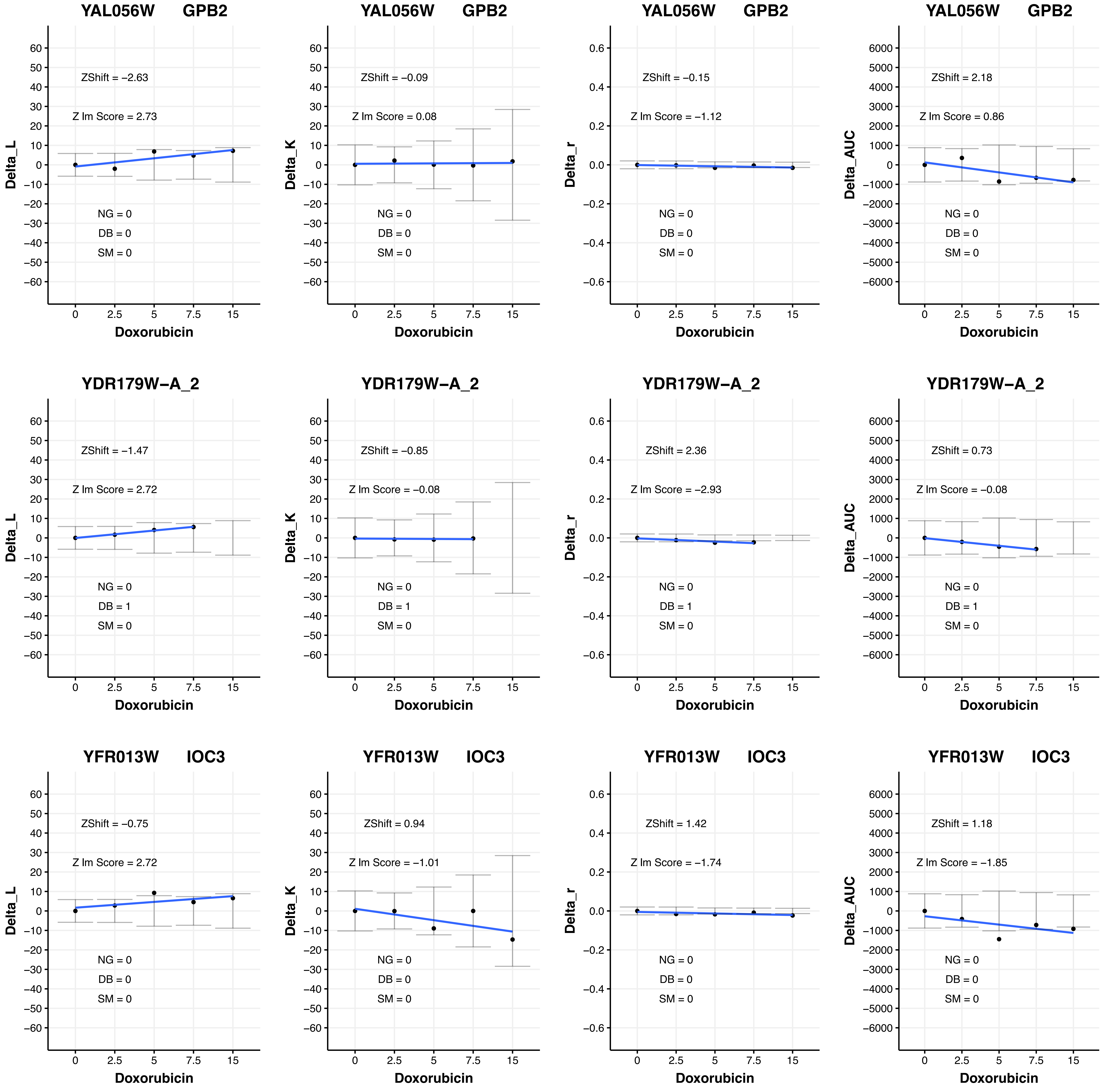
YAL056W GPB2
ZShift = −2.63
Z lm Score = 2.73
NG = 0
DB = 0
SM = 0
Delta_L
Doxorubicin
YAL056W GPB2
ZShift = −0.09
Z lm Score = 0.08
NG = 0
DB = 0
SM = 0
Delta_K
Doxorubicin
YAL056W GPB2
ZShift = −0.15
Z lm Score = −1.12
NG = 0
DB = 0
SM = 0
Delta_r
Doxorubicin
YAL056W GPB2
ZShift = 2.18
Z lm Score = 0.86
NG = 0
DB = 0
SM = 0
Delta_AUC
Doxorubicin
YDR179W−A_2
ZShift = −1.47
Z lm Score = 2.72
NG = 0
DB = 1
SM = 0
Delta_L
Doxorubicin
YDR179W−A_2
ZShift = −0.85
Z lm Score = −0.08
NG = 0
DB = 1
SM = 0
Delta_K
Doxorubicin
YDR179W−A_2
ZShift = 2.36
Z lm Score = −2.93
NG = 0
DB = 1
SM = 0
Delta_r
Doxorubicin
YDR179W−A_2
ZShift = 0.73
Z lm Score = −0.08
NG = 0
DB = 1
SM = 0
Delta_AUC
Doxorubicin
YFR013W IOC3
ZShift = −0.75
Z lm Score = 2.72
NG = 0
DB = 0
SM = 0
Delta_L
Doxorubicin
YFR013W IOC3
ZShift = 0.94
Z lm Score = −1.01
NG = 0
DB = 0
SM = 0
Delta_K
Doxorubicin
YFR013W IOC3
ZShift = 1.42
Z lm Score = −1.74
NG = 0
DB = 0
SM = 0
Delta_r
Doxorubicin
YFR013W IOC3
ZShift = 1.18
Z lm Score = −1.85
NG = 0
DB = 0
SM = 0
Delta_AUC
Doxorubicin

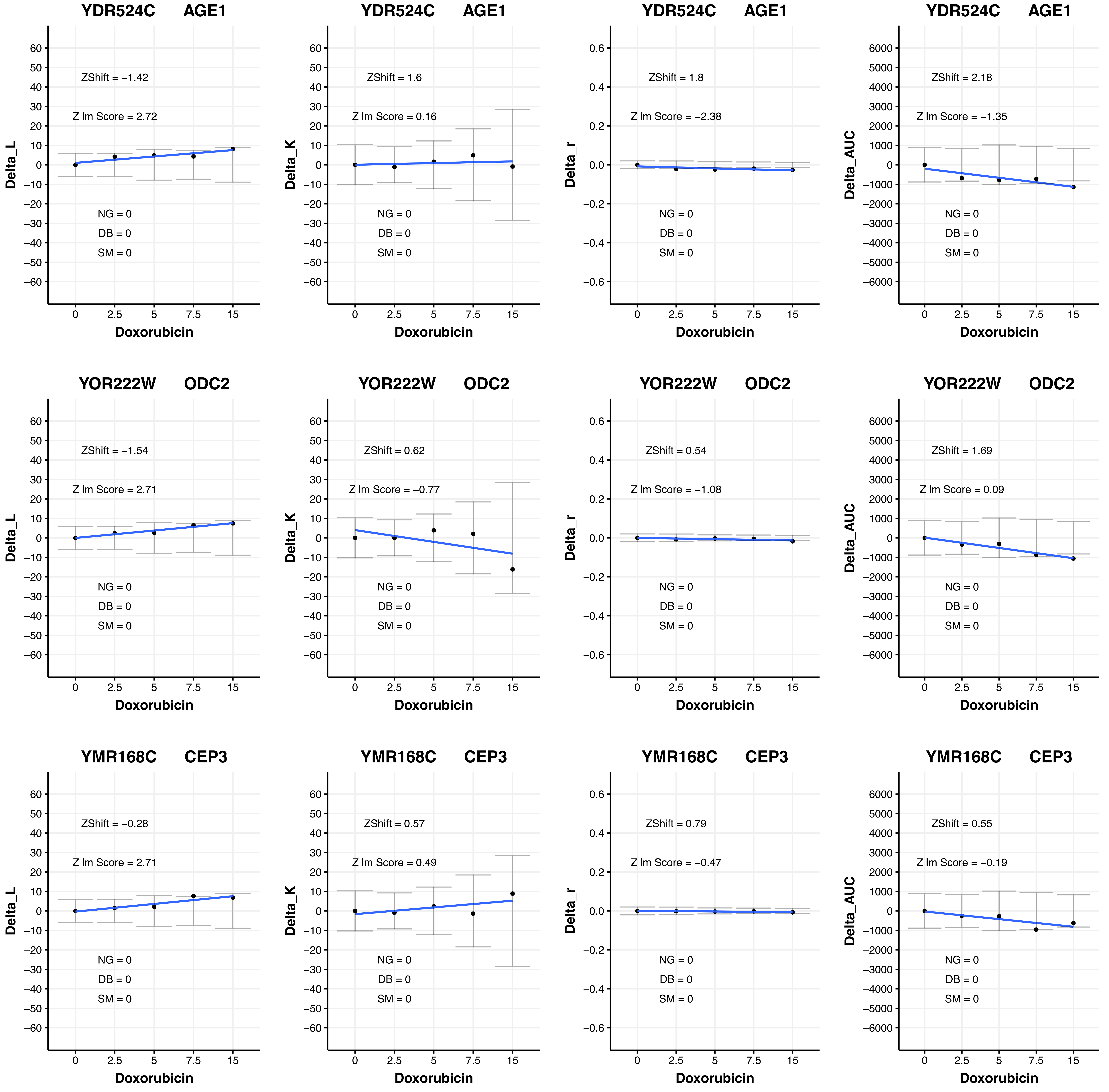

YDR524C AGE1
ZShift = −1.42
Z lm Score = 2.72
NG = 0
DB = 0
SM = 0
Delta_L
Doxorubicin
YDR524C AGE1
ZShift = 1.6
Z lm Score = 0.16
NG = 0
DB = 0
SM = 0
Delta_K
Doxorubicin
YDR524C AGE1
ZShift = 1.8
Z lm Score = −2.38
NG = 0
DB = 0
SM = 0
Delta_r
Doxorubicin
YDR524C AGE1
ZShift = 2.18
Z lm Score = −1.35
NG = 0
DB = 0
SM = 0
Delta_AUC
Doxorubicin
YOR222W ODC2
ZShift = −1.54
Z lm Score = 2.71
NG = 0
DB = 0
SM = 0
Delta_L
Doxorubicin
YOR222W ODC2
ZShift = 0.62
Z lm Score = −0.77
NG = 0
DB = 0
SM = 0
Delta_K
Doxorubicin
YOR222W ODC2
ZShift = 0.54
Z lm Score = −1.08
NG = 0
DB = 0
SM = 0
Delta_r
Doxorubicin
YOR222W ODC2
ZShift = 1.69
Z lm Score = 0.09
NG = 0
DB = 0
SM = 0
Delta_AUC
Doxorubicin
YMR168C CEP3
ZShift = −0.28
Z lm Score = 2.71
NG = 0
DB = 0
SM = 0
Delta_L
Doxorubicin
YMR168C CEP3
ZShift = 0.57
Z lm Score = 0.49
NG = 0
DB = 0
SM = 0
Delta_K
Doxorubicin
YMR168C CEP3
ZShift = 0.79
Z lm Score = −0.47
NG = 0
DB = 0
SM = 0
Delta_r
Doxorubicin
YMR168C CEP3
ZShift = 0.55
Z lm Score = −0.19
NG = 0
DB = 0
SM = 0
Delta_AUC
Doxorubicin

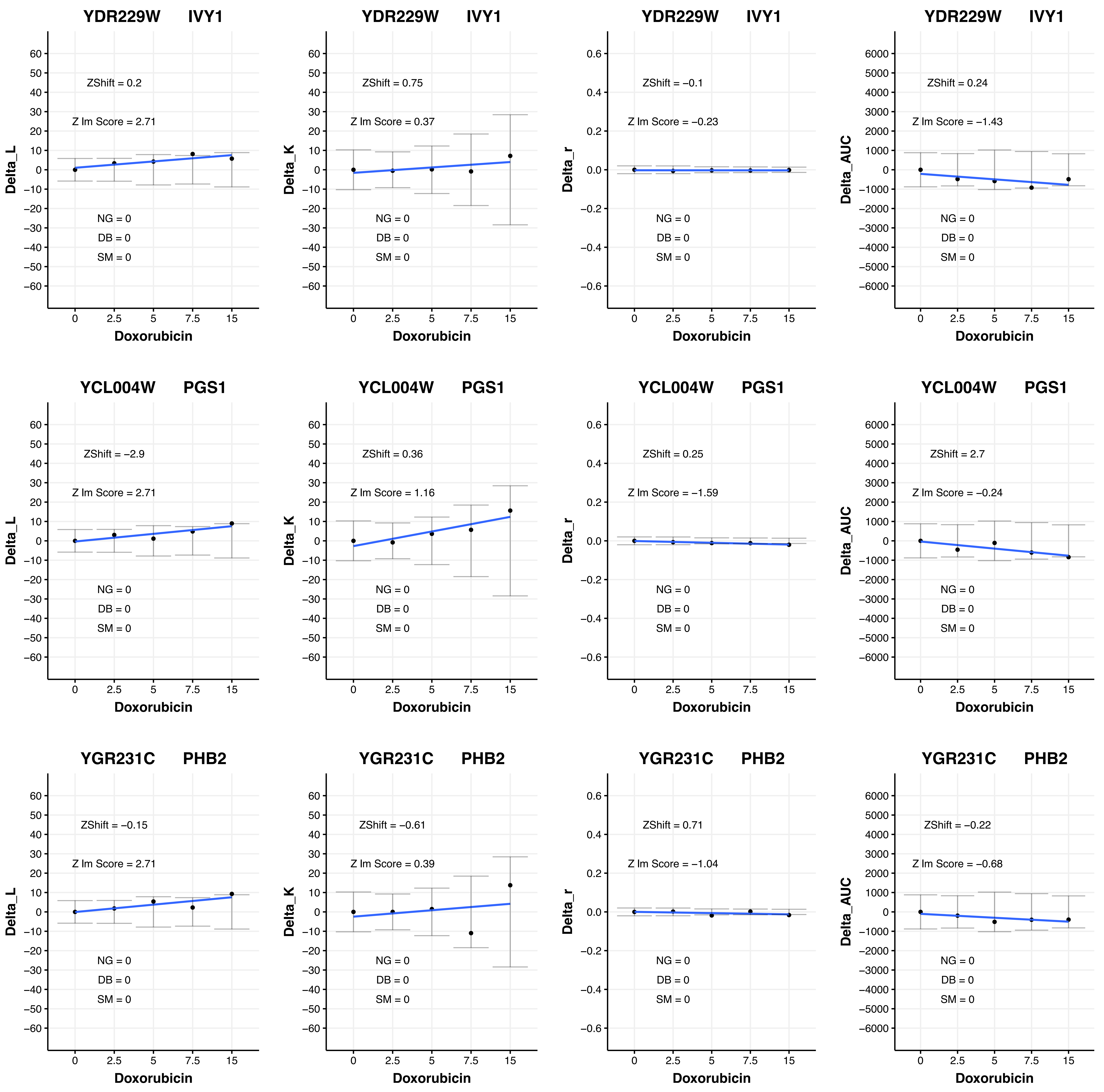

YDR229W IVY1
ZShift = 0.2
Z lm Score = 2.71
NG = 0
DB = 0
SM = 0
Delta_L
Doxorubicin
YDR229W IVY1
ZShift = 0.75
Z lm Score = 0.37
NG = 0
DB = 0
SM = 0
Delta_K
Doxorubicin
YDR229W IVY1
ZShift = −0.1
Z lm Score = −0.23
NG = 0
DB = 0
SM = 0
Delta_r
Doxorubicin
YDR229W IVY1
ZShift = 0.24
Z lm Score = −1.43
NG = 0
DB = 0
SM = 0
Delta_AUC
Doxorubicin
YCL004W PGS1
ZShift = −2.9
Z lm Score = 2.71
NG = 0
DB = 0
SM = 0
Delta_L
Doxorubicin
YCL004W PGS1
ZShift = 0.36
Z lm Score = 1.16
NG = 0
DB = 0
SM = 0
Delta_K
Doxorubicin
YCL004W PGS1
ZShift = 0.25
Z lm Score = −1.59
NG = 0
DB = 0
SM = 0
Delta_r
Doxorubicin
YCL004W PGS1
ZShift = 2.7
Z lm Score = −0.24
NG = 0
DB = 0
SM = 0
Delta_AUC
Doxorubicin
YGR231C PHB2
ZShift = −0.15
Z lm Score = 2.71
NG = 0
DB = 0
SM = 0
Delta_L
Doxorubicin
YGR231C PHB2
ZShift = −0.61
Z lm Score = 0.39
NG = 0
DB = 0
SM = 0
Delta_K
Doxorubicin
YGR231C PHB2
ZShift = 0.71
Z lm Score = −1.04
NG = 0
DB = 0
SM = 0
Delta_r
Doxorubicin
YGR231C PHB2
ZShift = −0.22
Z lm Score = −0.68
NG = 0
DB = 0
SM = 0
Delta_AUC
Doxorubicin

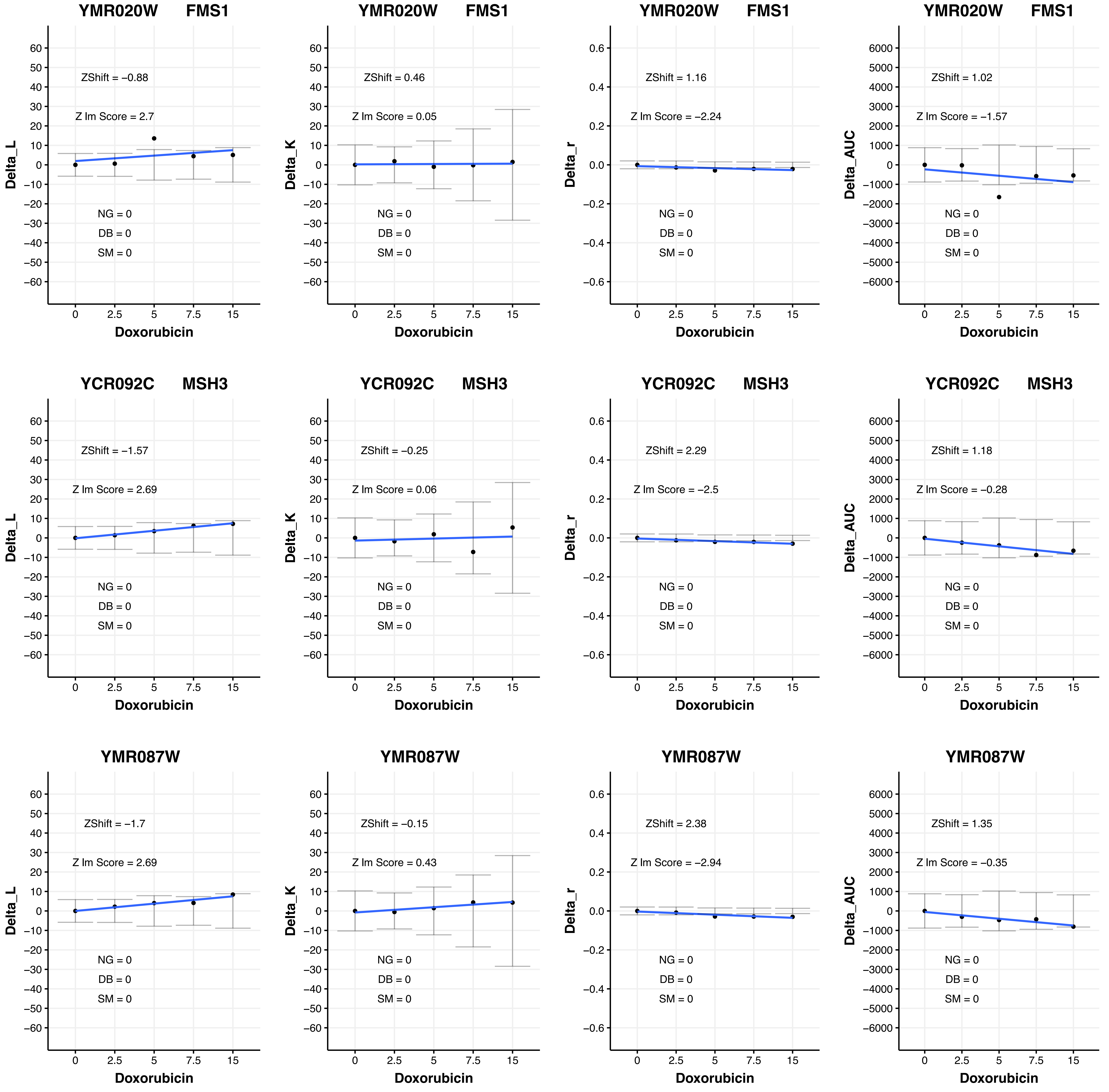

YMR020W FMS1
ZShift = −0.88
Z lm Score = 2.7
NG = 0
DB = 0
SM = 0
Delta_L
Doxorubicin
YMR020W FMS1
ZShift = 0.46
Z lm Score = 0.05
NG = 0
DB = 0
SM = 0
Delta_K
Doxorubicin
YMR020W FMS1
ZShift = 1.16
Z lm Score = −2.24
NG = 0
DB = 0
SM = 0
Delta_r
Doxorubicin
YMR020W FMS1
ZShift = 1.02
Z lm Score = −1.57
NG = 0
DB = 0
SM = 0
Delta_AUC
Doxorubicin
YCR092C MSH3
ZShift = −1.57
Z lm Score = 2.69
NG = 0
DB = 0
SM = 0
Delta_L
Doxorubicin
YCR092C MSH3
ZShift = −0.25
Z lm Score = 0.06
NG = 0
DB = 0
SM = 0
Delta_K
Doxorubicin
YCR092C MSH3
ZShift = 2.29
Z lm Score = −2.5
NG = 0
DB = 0
SM = 0
Delta_r
Doxorubicin
YCR092C MSH3
ZShift = 1.18
Z lm Score = −0.28
NG = 0
DB = 0
SM = 0
Delta_AUC
Doxorubicin
YMR087W
ZShift = −1.7
Z lm Score = 2.69
NG = 0
DB = 0
SM = 0
Delta_L
Doxorubicin
YMR087W
ZShift = −0.15
Z lm Score = 0.43
NG = 0
DB = 0
SM = 0
Delta_K
Doxorubicin
YMR087W
ZShift = 2.38
Z lm Score = −2.94
NG = 0
DB = 0
SM = 0
Delta_r
Doxorubicin
YMR087W
ZShift = 1.35
Z lm Score = −0.35
NG = 0
DB = 0
SM = 0
Delta_AUC
Doxorubicin

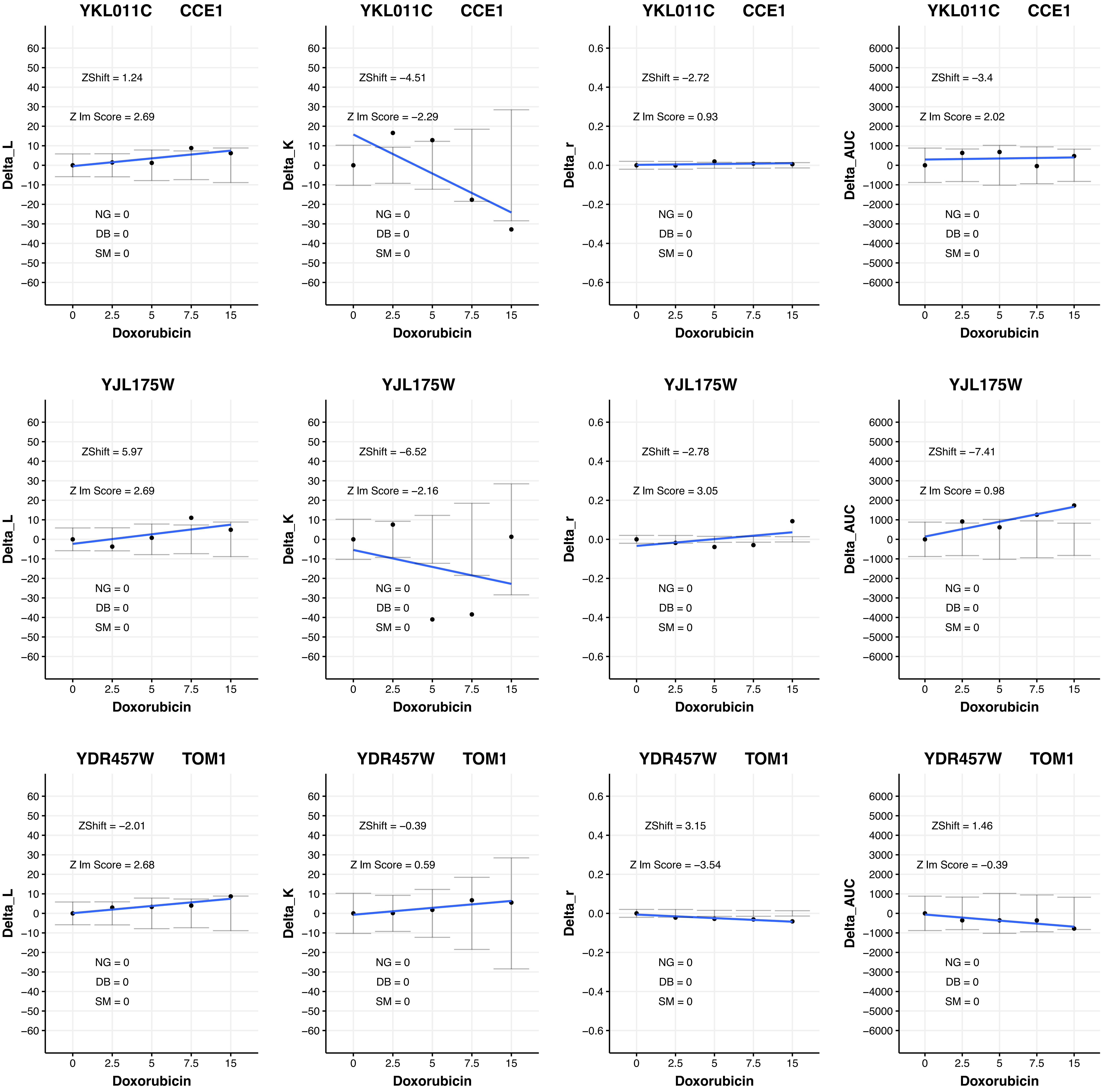
YKL011C CCE1
ZShift = 1.24
Z lm Score = 2.69
NG = 0
DB = 0
SM = 0
Delta_L
Doxorubicin
YKL011C CCE1
ZShift = −4.51
Z lm Score = −2.29
NG = 0
DB = 0
SM = 0
Delta_K
Doxorubicin
YKL011C CCE1
ZShift = −2.72
Z lm Score = 0.93
NG = 0
DB = 0
SM = 0
Delta_r
Doxorubicin
YKL011C CCE1
ZShift = −3.4
Z lm Score = 2.02
NG = 0
DB = 0
SM = 0
Delta_AUC
Doxorubicin
YJL175W
ZShift = 5.97
Z lm Score = 2.69
NG = 0
DB = 0
SM = 0
Delta_L
Doxorubicin
YJL175W
ZShift = −6.52
Z lm Score = −2.16
NG = 0
DB = 0
SM = 0
Delta_K
Doxorubicin
YJL175W
ZShift = −2.78
Z lm Score = 3.05
NG = 0
DB = 0
SM = 0
Delta_r
Doxorubicin
YJL175W
ZShift = −7.41
Z lm Score = 0.98
NG = 0
DB = 0
SM = 0
Delta_AUC
Doxorubicin
YDR457W TOM1
ZShift = −2.01
Z lm Score = 2.68
NG = 0
DB = 0
SM = 0
Delta_L
Doxorubicin
YDR457W TOM1
ZShift = −0.39
Z lm Score = 0.59
NG = 0
DB = 0
SM = 0
Delta_K
Doxorubicin
YDR457W TOM1
ZShift = 3.15
Z lm Score = −3.54
NG = 0
DB = 0
SM = 0
Delta_r
Doxorubicin
YDR457W TOM1
ZShift = 1.46
Z lm Score = −0.39
NG = 0
DB = 0
SM = 0
Delta_AUC
Doxorubicin

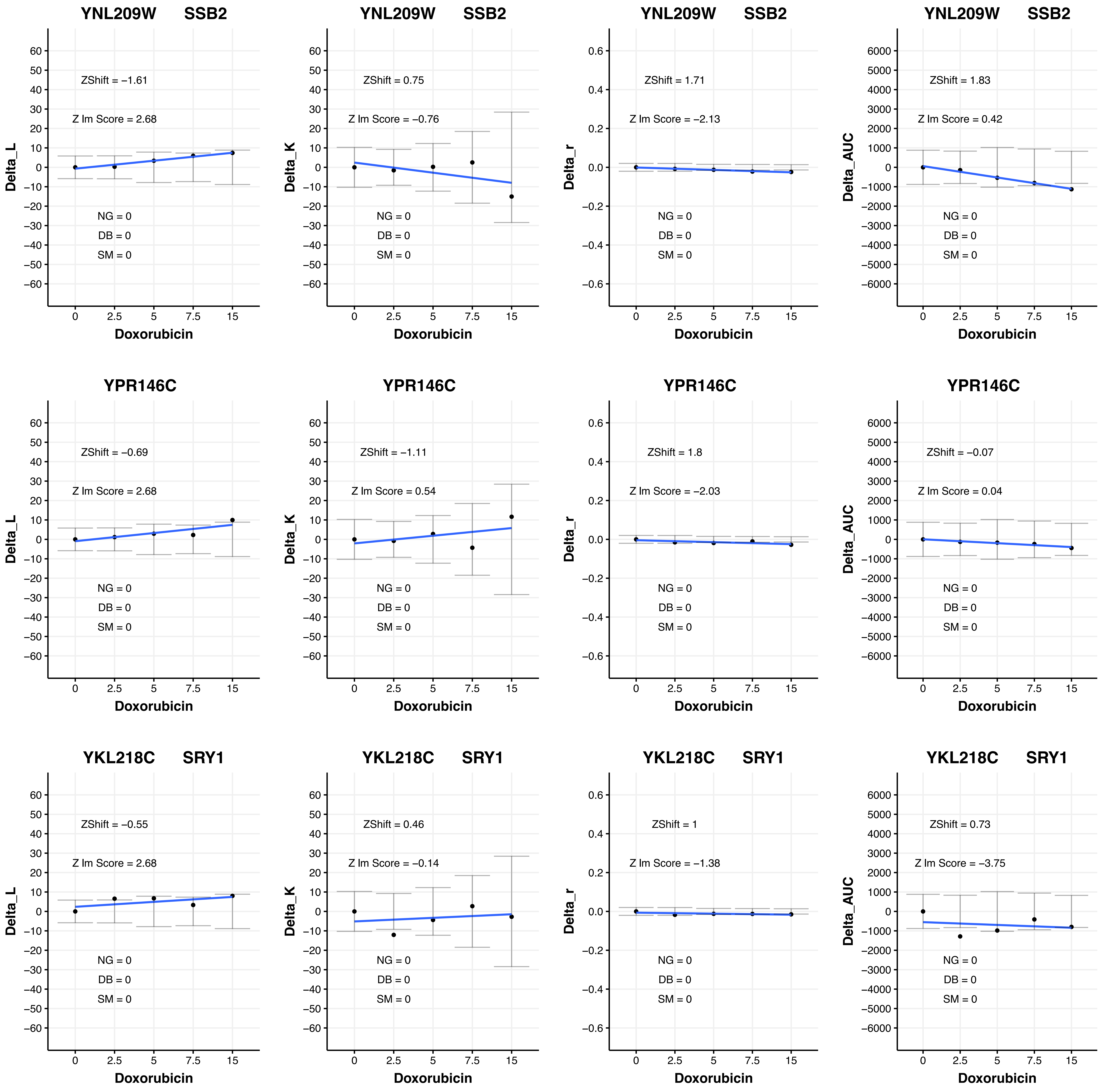

YNL209W SSB2
ZShift = −1.61
Z lm Score = 2.68
NG = 0
DB = 0
SM = 0
Delta_L
Doxorubicin
YNL209W SSB2
ZShift = 0.75
Z lm Score = −0.76
NG = 0
DB = 0
SM = 0
Delta_K
Doxorubicin
YNL209W SSB2
ZShift = 1.71
Z lm Score = −2.13
NG = 0
DB = 0
SM = 0
Delta_r
Doxorubicin
YNL209W SSB2
ZShift = 1.83
Z lm Score = 0.42
NG = 0
DB = 0
SM = 0
Delta_AUC
Doxorubicin
YPR146C
ZShift = −0.69
Z lm Score = 2.68
NG = 0
DB = 0
SM = 0
Delta_L
Doxorubicin
YPR146C
ZShift = −1.11
Z lm Score = 0.54
NG = 0
DB = 0
SM = 0
Delta_K
Doxorubicin
YPR146C
ZShift = 1.8
Z lm Score = −2.03
NG = 0
DB = 0
SM = 0
Delta_r
Doxorubicin
YPR146C
ZShift = −0.07
Z lm Score = 0.04
NG = 0
DB = 0
SM = 0
Delta_AUC
Doxorubicin
YKL218C SRY1
ZShift = −0.55
Z lm Score = 2.68
NG = 0
DB = 0
SM = 0
Delta_L
Doxorubicin
YKL218C SRY1
ZShift = 0.46
Z lm Score = −0.14
NG = 0
DB = 0
SM = 0
Delta_K
Doxorubicin
YKL218C SRY1
ZShift = 1
Z lm Score = −1.38
NG = 0
DB = 0
SM = 0
Delta_r
Doxorubicin
YKL218C SRY1
ZShift = 0.73
Z lm Score = −3.75
NG = 0
DB = 0
SM = 0
Delta_AUC
Doxorubicin

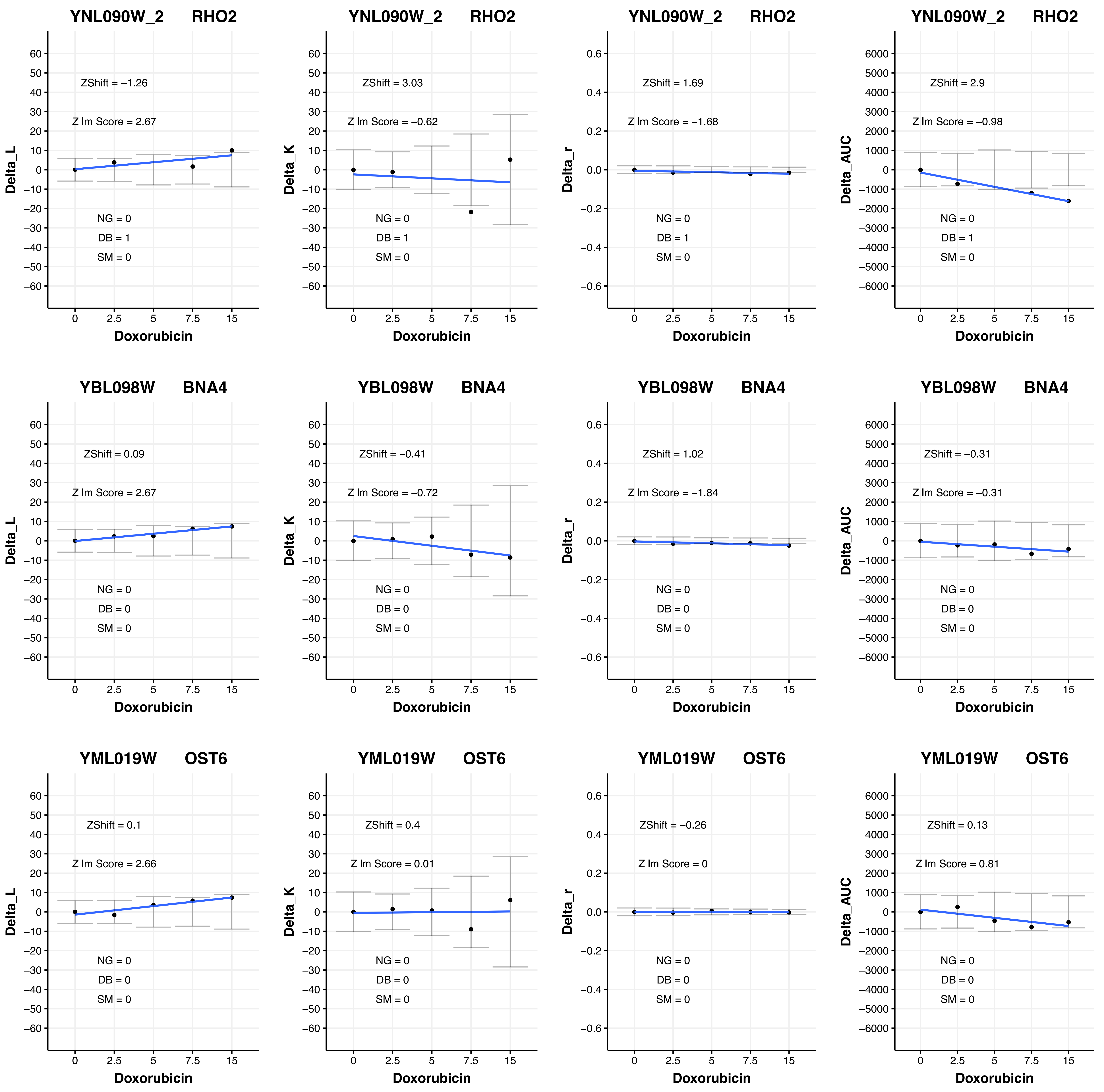

YNL090W_2 RHO2
ZShift = −1.26
Z lm Score = 2.67
NG = 0
DB = 1
SM = 0
Delta_L
Doxorubicin
YNL090W_2 RHO2
ZShift = 3.03
Z lm Score = −0.62
NG = 0
DB = 1
SM = 0
Delta_K
Doxorubicin
YNL090W_2 RHO2
ZShift = 1.69
Z lm Score = −1.68
NG = 0
DB = 1
SM = 0
Delta_r
Doxorubicin
YNL090W_2 RHO2
ZShift = 2.9
Z lm Score = −0.98
NG = 0
DB = 1
SM = 0
Delta_AUC
Doxorubicin
YBL098W BNA4
ZShift = 0.09
Z lm Score = 2.67
NG = 0
DB = 0
SM = 0
Delta_L
Doxorubicin
YBL098W BNA4
ZShift = −0.41
Z lm Score = −0.72
NG = 0
DB = 0
SM = 0
Delta_K
Doxorubicin
YBL098W BNA4
ZShift = 1.02
Z lm Score = −1.84
NG = 0
DB = 0
SM = 0
Delta_r
Doxorubicin
YBL098W BNA4
ZShift = −0.31
Z lm Score = −0.31
NG = 0
DB = 0
SM = 0
Delta_AUC
Doxorubicin
YML019W OST6
ZShift = 0.1
Z lm Score = 2.66
NG = 0
DB = 0
SM = 0
Delta_L
Doxorubicin
YML019W OST6
ZShift = 0.4
Z lm Score = 0.01
NG = 0
DB = 0
SM = 0
Delta_K
Doxorubicin
YML019W OST6
ZShift = −0.26
Z lm Score = 0
NG = 0
DB = 0
SM = 0
Delta_r
Doxorubicin
YML019W OST6
ZShift = 0.13
Z lm Score = 0.81
NG = 0
DB = 0
SM = 0
Delta_AUC
Doxorubicin

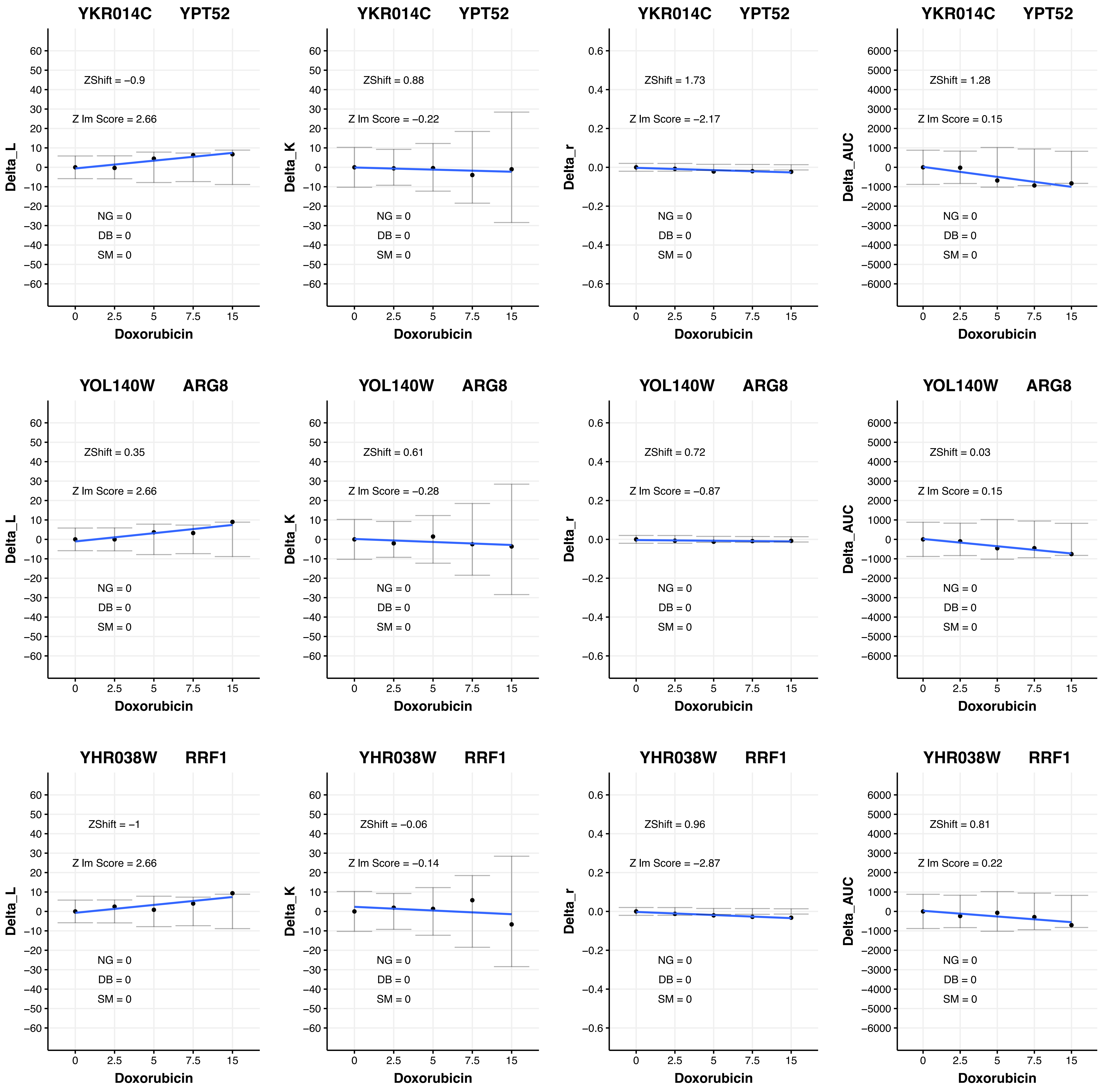
YKR014C YPT52
ZShift = −0.9
Z lm Score = 2.66
NG = 0
DB = 0
SM = 0
Delta_L
Doxorubicin
YKR014C YPT52
ZShift = 0.88
Z lm Score = −0.22
NG = 0
DB = 0
SM = 0
Delta_K
Doxorubicin
YKR014C YPT52
ZShift = 1.73
Z lm Score = −2.17
NG = 0
DB = 0
SM = 0
Delta_r
Doxorubicin
YKR014C YPT52
ZShift = 1.28
Z lm Score = 0.15
NG = 0
DB = 0
SM = 0
Delta_AUC
Doxorubicin
YOL140W ARG8
ZShift = 0.35
Z lm Score = 2.66
NG = 0
DB = 0
SM = 0
Delta_L
Doxorubicin
YOL140W ARG8
ZShift = 0.61
Z lm Score = −0.28
NG = 0
DB = 0
SM = 0
Delta_K
Doxorubicin
YOL140W ARG8
ZShift = 0.72
Z lm Score = −0.87
NG = 0
DB = 0
SM = 0
Delta_r
Doxorubicin
YOL140W ARG8
ZShift = 0.03
Z lm Score = 0.15
NG = 0
DB = 0
SM = 0
Delta_AUC
Doxorubicin
YHR038W RRF1
ZShift = −1
Z lm Score = 2.66
NG = 0
DB = 0
SM = 0
Delta_L
Doxorubicin
YHR038W RRF1
ZShift = −0.06
Z lm Score = −0.14
NG = 0
DB = 0
SM = 0
Delta_K
Doxorubicin
YHR038W RRF1
ZShift = 0.96
Z lm Score = −2.87
NG = 0
DB = 0
SM = 0
Delta_r
Doxorubicin
YHR038W RRF1
ZShift = 0.81
Z lm Score = 0.22
NG = 0
DB = 0
SM = 0
Delta_AUC
Doxorubicin

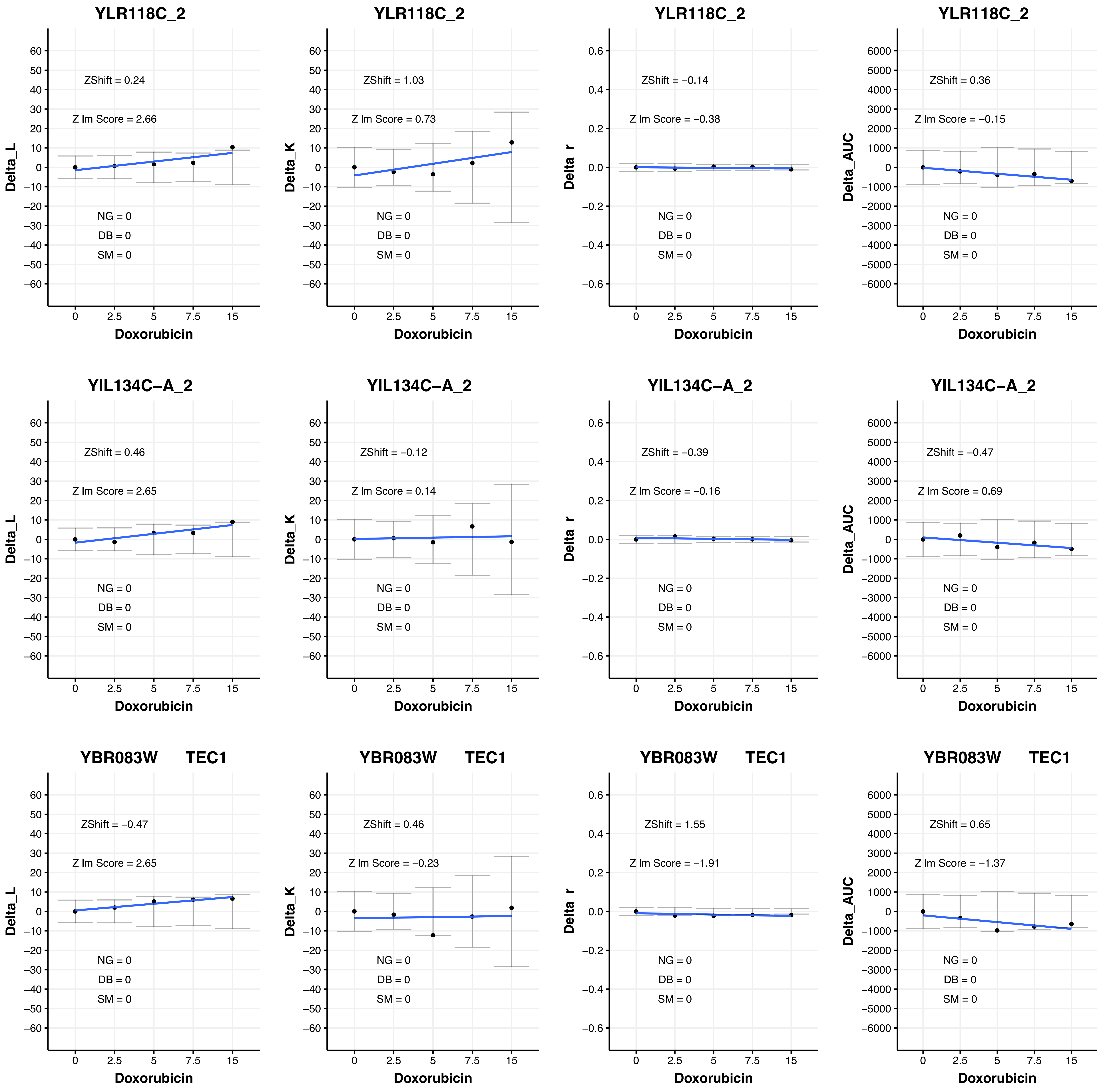
YLR118C_2
ZShift = 0.24
Z lm Score = 2.66
NG = 0
DB = 0
SM = 0
Delta_L
Doxorubicin
YLR118C_2
ZShift = 1.03
Z lm Score = 0.73
NG = 0
DB = 0
SM = 0
Delta_K
Doxorubicin
YLR118C_2
ZShift = −0.14
Z lm Score = −0.38
NG = 0
DB = 0
SM = 0
Delta_r
Doxorubicin
YLR118C_2
ZShift = 0.36
Z lm Score = −0.15
NG = 0
DB = 0
SM = 0
Delta_AUC
Doxorubicin
YIL134C−A_2
ZShift = 0.46
Z lm Score = 2.65
NG = 0
DB = 0
SM = 0
Delta_L
Doxorubicin
YIL134C−A_2
ZShift = −0.12
Z lm Score = 0.14
NG = 0
DB = 0
SM = 0
Delta_K
Doxorubicin
YIL134C−A_2
ZShift = −0.39
Z lm Score = −0.16
NG = 0
DB = 0
SM = 0
Delta_r
Doxorubicin
YIL134C−A_2
ZShift = −0.47
Z lm Score = 0.69
NG = 0
DB = 0
SM = 0
Delta_AUC
Doxorubicin
YBR083W TEC1
ZShift = −0.47
Z lm Score = 2.65
NG = 0
DB = 0
SM = 0
Delta_L
Doxorubicin
YBR083W TEC1
ZShift = 0.46
Z lm Score = −0.23
NG = 0
DB = 0
SM = 0
Delta_K
Doxorubicin
YBR083W TEC1
ZShift = 1.55
Z lm Score = −1.91
NG = 0
DB = 0
SM = 0
Delta_r
Doxorubicin
YBR083W TEC1
ZShift = 0.65
Z lm Score = −1.37
NG = 0
DB = 0
SM = 0
Delta_AUC
Doxorubicin

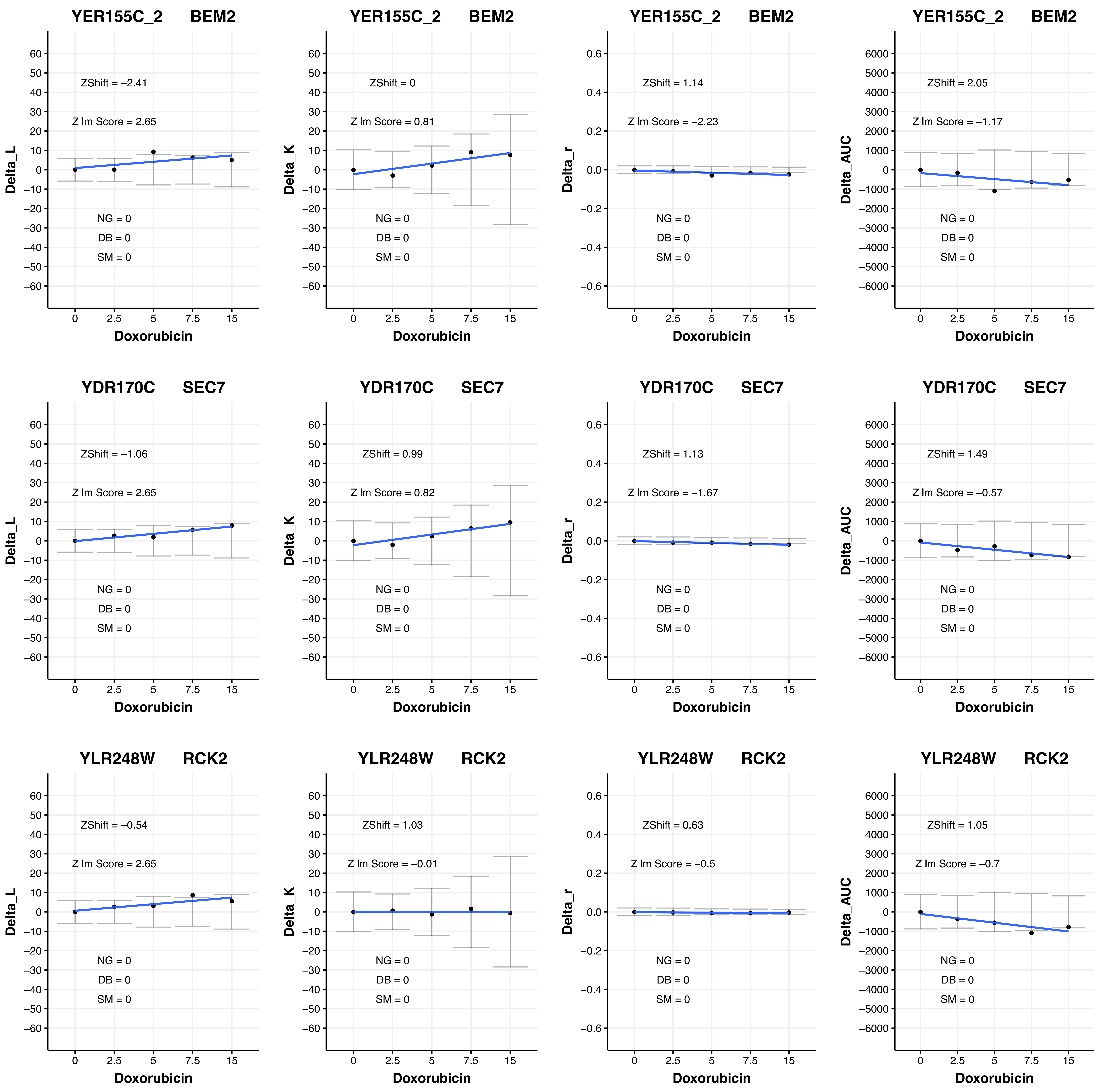
YER155C_2 BEM2
ZShift = −2.41
Z lm Score = 2.65
Delta_L
NG = 0
DB = 0
SM = 0
Doxorubicin
YER155C_2 BEM2
ZShift = 0
Z lm Score = 0.81
Delta_K
NG = 0
DB = 0
SM = 0
Doxorubicin
YER155C_2 BEM2
ZShift = 1.14
Z lm Score = −2.23
Delta_r
NG = 0
DB = 0
SM = 0
Doxorubicin
YER155C_2 BEM2
ZShift = 2.05
Z lm Score = −1.17
Delta_AUC
NG = 0
DB = 0
SM = 0
Doxorubicin
YDR170C SEC7
ZShift = −1.06
Z lm Score = 2.65
Delta_L
NG = 0
DB = 0
SM = 0
Doxorubicin
YDR170C SEC7
ZShift = 0.99
Z lm Score = 0.82
Delta_K
NG = 0
DB = 0
SM = 0
Doxorubicin
YDR170C SEC7
ZShift = 1.13
Z lm Score = −1.67
Delta_r
NG = 0
DB = 0
SM = 0
Doxorubicin
YDR170C SEC7
ZShift = 1.49
Z lm Score = −0.57
Delta_AUC
NG = 0
DB = 0
SM = 0
Doxorubicin
YLR248W RCK2
ZShift = −0.54
Z lm Score = 2.65
Delta_L
NG = 0
DB = 0
SM = 0
Doxorubicin
YLR248W RCK2
ZShift = 1.03
Z lm Score = −0.01
Delta_K
NG = 0
DB = 0
SM = 0
Doxorubicin
YLR248W RCK2
ZShift = 0.63
Z lm Score = −0.5
Delta_r
NG = 0
DB = 0
SM = 0
Doxorubicin
YLR248W RCK2
ZShift = 1.05
Z lm Score = −0.7
Delta_AUC
NG = 0
DB = 0
SM = 0
Doxorubicin

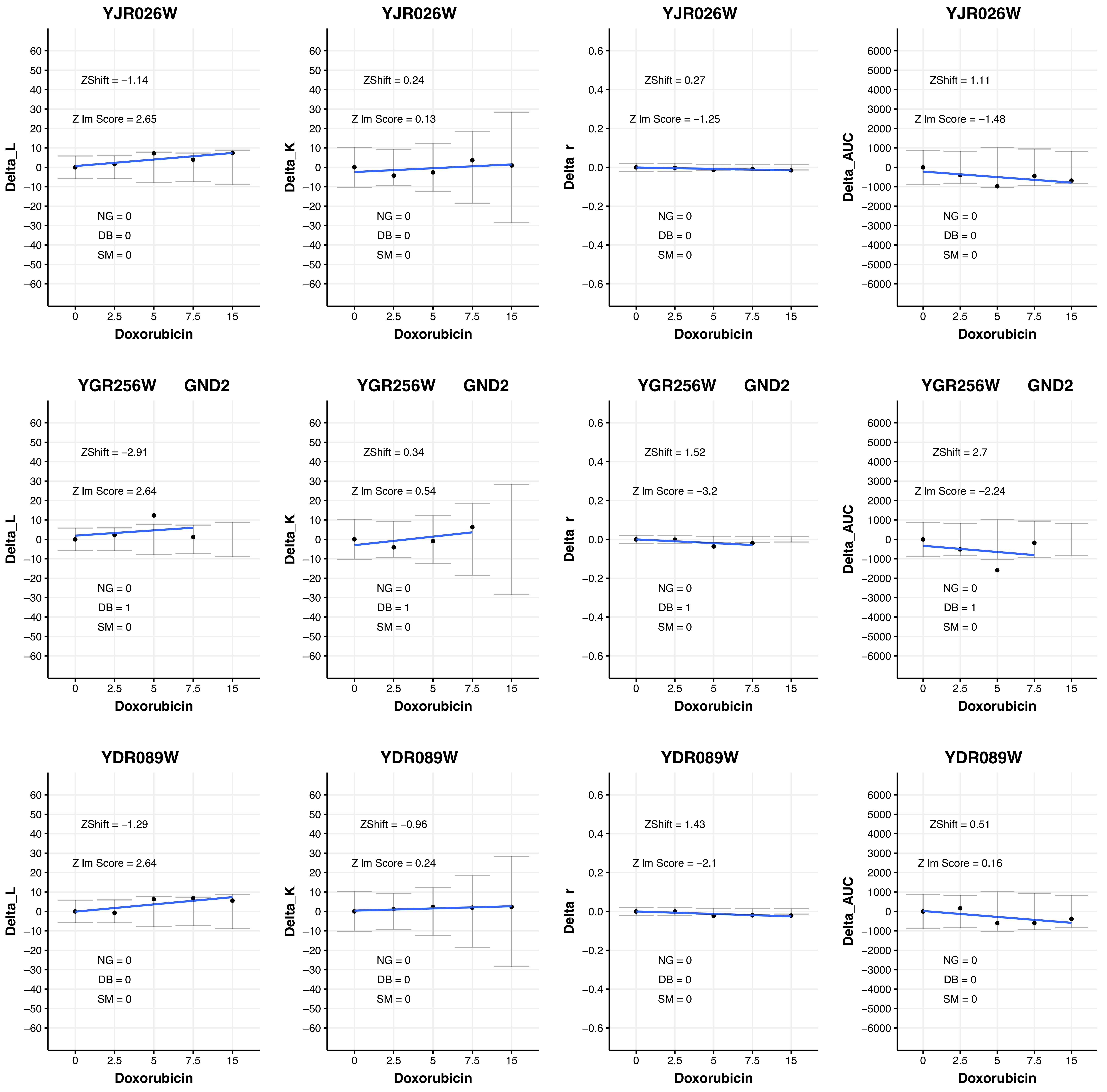
YJR026W
ZShift = −1.14
Z lm Score = 2.65
NG = 0
DB = 0
SM = 0
Delta_L
Doxorubicin
YJR026W
ZShift = 0.24
Z lm Score = 0.13
NG = 0
DB = 0
SM = 0
Delta_K
Doxorubicin
YJR026W
ZShift = 0.27
Z lm Score = −1.25
NG = 0
DB = 0
SM = 0
Delta_r
Doxorubicin
YJR026W
ZShift = 1.11
Z lm Score = −1.48
NG = 0
DB = 0
SM = 0
Delta_AUC
Doxorubicin
YGR256W GND2
ZShift = −2.91
Z lm Score = 2.64
NG = 0
DB = 1
SM = 0
Delta_L
Doxorubicin
YGR256W GND2
ZShift = 0.34
Z lm Score = 0.54
NG = 0
DB = 1
SM = 0
Delta_K
Doxorubicin
YGR256W GND2
ZShift = 1.52
Z lm Score = −3.2
NG = 0
DB = 1
SM = 0
Delta_r
Doxorubicin
YGR256W GND2
ZShift = 2.7
Z lm Score = −2.24
NG = 0
DB = 1
SM = 0
Delta_AUC
Doxorubicin
YDR089W
ZShift = −1.29
Z lm Score = 2.64
NG = 0
DB = 0
SM = 0
Delta_L
Doxorubicin
YDR089W
ZShift = −0.96
Z lm Score = 0.24
NG = 0
DB = 0
SM = 0
Delta_K
Doxorubicin
YDR089W
ZShift = 1.43
Z lm Score = −2.1
NG = 0
DB = 0
SM = 0
Delta_r
Doxorubicin
YDR089W
ZShift = 0.51
Z lm Score = 0.16
NG = 0
DB = 0
SM = 0
Delta_AUC
Doxorubicin

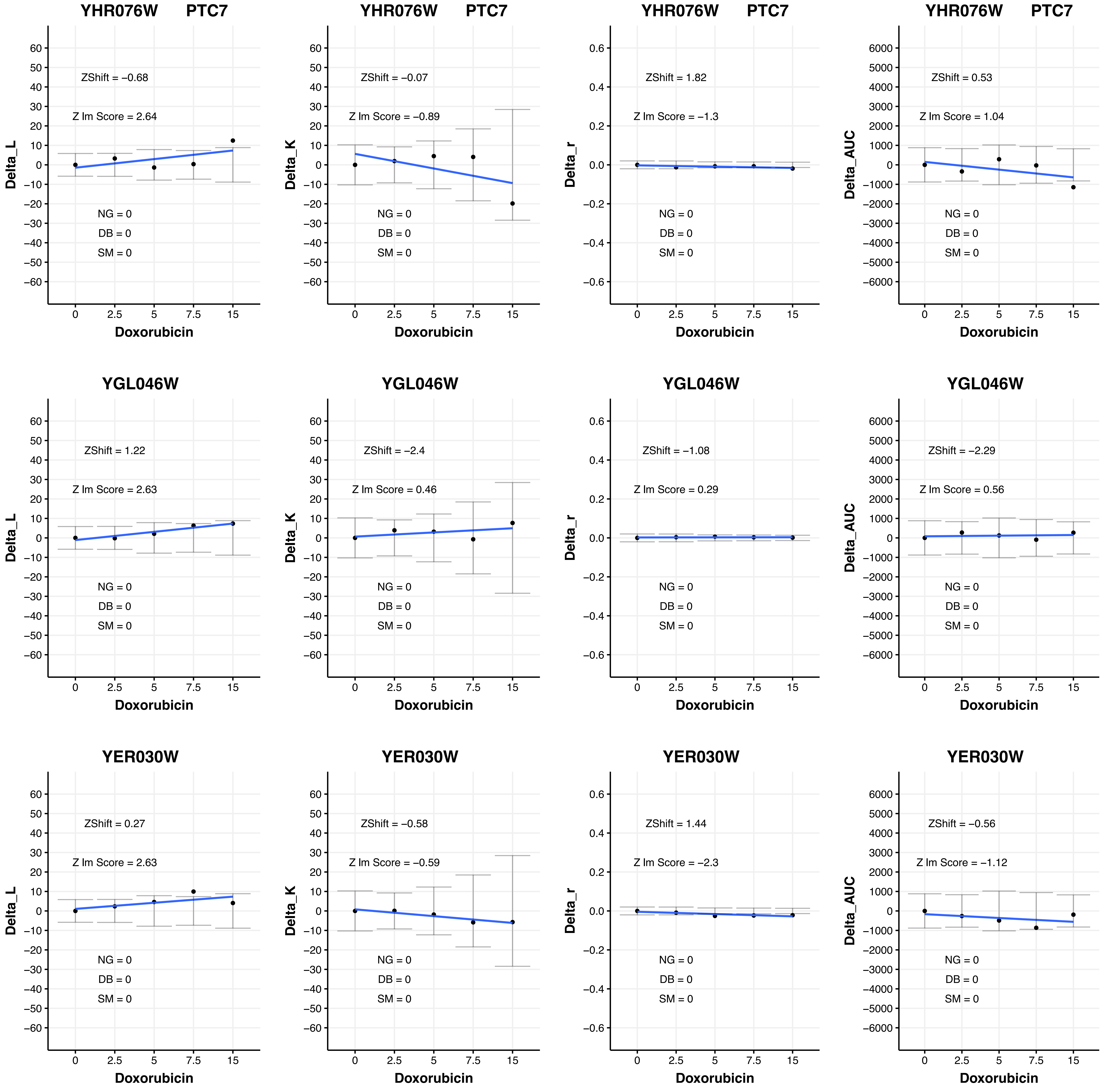

YHR076W PTC7
ZShift = −0.68
Z lm Score = 2.64
NG = 0
DB = 0
SM = 0
Delta_L
Doxorubicin
YHR076W PTC7
ZShift = −0.07
Z lm Score = −0.89
NG = 0
DB = 0
SM = 0
Delta_K
Doxorubicin
YHR076W PTC7
ZShift = 1.82
Z lm Score = −1.3
NG = 0
DB = 0
SM = 0
Delta_r
Doxorubicin
YHR076W PTC7
ZShift = 0.53
Z lm Score = 1.04
NG = 0
DB = 0
SM = 0
Delta_AUC
Doxorubicin
YGL046W
ZShift = 1.22
Z lm Score = 2.63
NG = 0
DB = 0
SM = 0
Delta_L
Doxorubicin
YGL046W
ZShift = −2.4
Z lm Score = 0.46
NG = 0
DB = 0
SM = 0
Delta_K
Doxorubicin
YGL046W
ZShift = −1.08
Z lm Score = 0.29
NG = 0
DB = 0
SM = 0
Delta_r
Doxorubicin
YGL046W
ZShift = −2.29
Z lm Score = 0.56
NG = 0
DB = 0
SM = 0
Delta_AUC
Doxorubicin
YER030W
ZShift = 0.27
Z lm Score = 2.63
NG = 0
DB = 0
SM = 0
Delta_L
Doxorubicin
YER030W
ZShift = −0.58
Z lm Score = −0.59
NG = 0
DB = 0
SM = 0
Delta_K
Doxorubicin
YER030W
ZShift = 1.44
Z lm Score = −2.3
NG = 0
DB = 0
SM = 0
Delta_r
Doxorubicin
YER030W
ZShift = −0.56
Z lm Score = −1.12
NG = 0
DB = 0
SM = 0
Delta_AUC
Doxorubicin

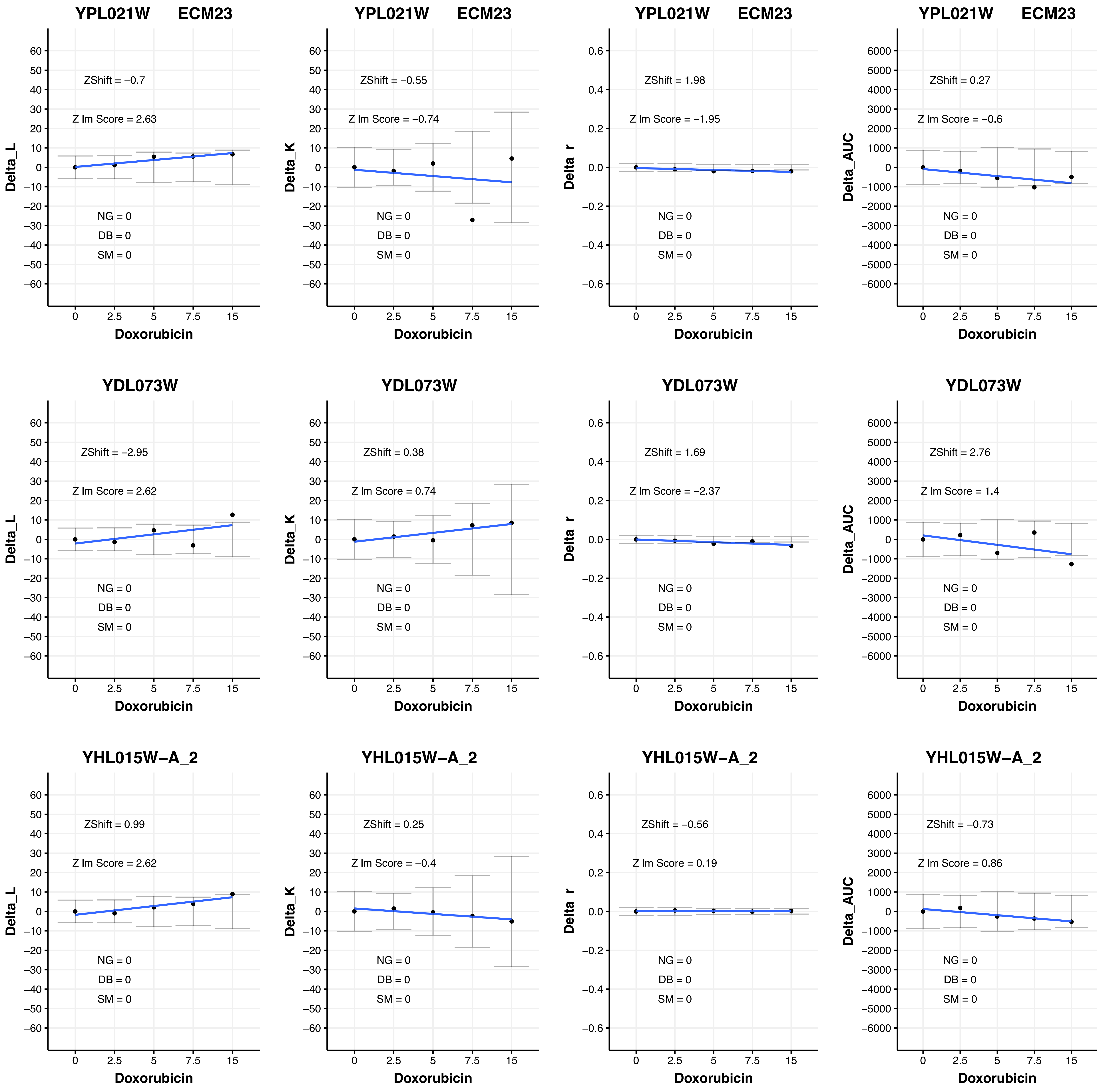
YPL021W ECM23
ZShift = −0.7
Z lm Score = 2.63
Delta_L
NG = 0
DB = 0
SM = 0
Doxorubicin
YPL021W ECM23
ZShift = −0.55
Z lm Score = −0.74
Delta_K
NG = 0
DB = 0
SM = 0
Doxorubicin
YPL021W ECM23
ZShift = 1.98
Z lm Score = −1.95
Delta_r
NG = 0
DB = 0
SM = 0
Doxorubicin
YPL021W ECM23
ZShift = 0.27
Z lm Score = −0.6
Delta_AUC
NG = 0
DB = 0
SM = 0
Doxorubicin
YDL073W
ZShift = −2.95
Z lm Score = 2.62
Delta_L
NG = 0
DB = 0
SM = 0
Doxorubicin
YDL073W
ZShift = 0.38
Z lm Score = 0.74
Delta_K
NG = 0
DB = 0
SM = 0
Doxorubicin
YDL073W
ZShift = 1.69
Z lm Score = −2.37
Delta_r
NG = 0
DB = 0
SM = 0
Doxorubicin
YDL073W
ZShift = 2.76
Z lm Score = 1.4
Delta_AUC
NG = 0
DB = 0
SM = 0
Doxorubicin
YHL015W−A_2
ZShift = 0.99
Z lm Score = 2.62
Delta_L
NG = 0
DB = 0
SM = 0
Doxorubicin
YHL015W−A_2
ZShift = 0.25
Z lm Score = −0.4
Delta_K
NG = 0
DB = 0
SM = 0
Doxorubicin
YHL015W−A_2
ZShift = −0.56
Z lm Score = 0.19
Delta_r
NG = 0
DB = 0
SM = 0
Doxorubicin
YHL015W−A_2
ZShift = −0.73
Z lm Score = 0.86
Delta_AUC
NG = 0
DB = 0
SM = 0
Doxorubicin

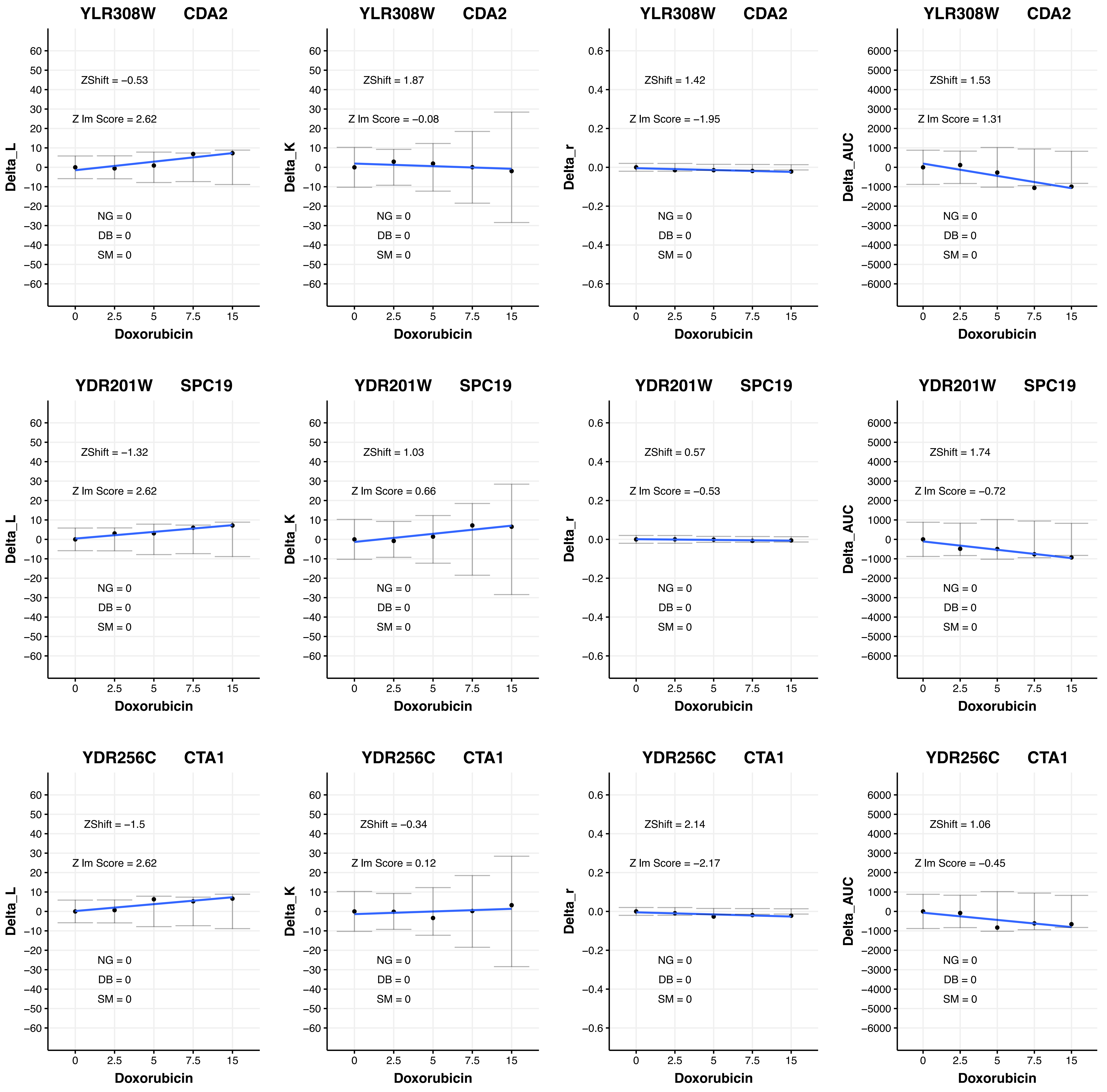

YLR308W CDA2
ZShift = −0.53
Z lm Score = 2.62
NG = 0
DB = 0
SM = 0
Delta_L
Doxorubicin
YLR308W CDA2
ZShift = 1.87
Z lm Score = −0.08
NG = 0
DB = 0
SM = 0
Delta_K
Doxorubicin
YLR308W CDA2
ZShift = 1.42
Z lm Score = −1.95
NG = 0
DB = 0
SM = 0
Delta_r
Doxorubicin
YLR308W CDA2
ZShift = 1.53
Z lm Score = 1.31
NG = 0
DB = 0
SM = 0
Delta_AUC
Doxorubicin
YDR201W SPC19
ZShift = −1.32
Z lm Score = 2.62
NG = 0
DB = 0
SM = 0
Delta_L
Doxorubicin
YDR201W SPC19
ZShift = 1.03
Z lm Score = 0.66
NG = 0
DB = 0
SM = 0
Delta_K
Doxorubicin
YDR201W SPC19
ZShift = 0.57
Z lm Score = −0.53
NG = 0
DB = 0
SM = 0
Delta_r
Doxorubicin
YDR201W SPC19
ZShift = 1.74
Z lm Score = −0.72
NG = 0
DB = 0
SM = 0
Delta_AUC
Doxorubicin
YDR256C CTA1
ZShift = −1.5
Z lm Score = 2.62
NG = 0
DB = 0
SM = 0
Delta_L
Doxorubicin
YDR256C CTA1
ZShift = −0.34
Z lm Score = 0.12
NG = 0
DB = 0
SM = 0
Delta_K
Doxorubicin
YDR256C CTA1
ZShift = 2.14
Z lm Score = −2.17
NG = 0
DB = 0
SM = 0
Delta_r
Doxorubicin
YDR256C CTA1
ZShift = 1.06
Z lm Score = −0.45
NG = 0
DB = 0
SM = 0
Delta_AUC
Doxorubicin

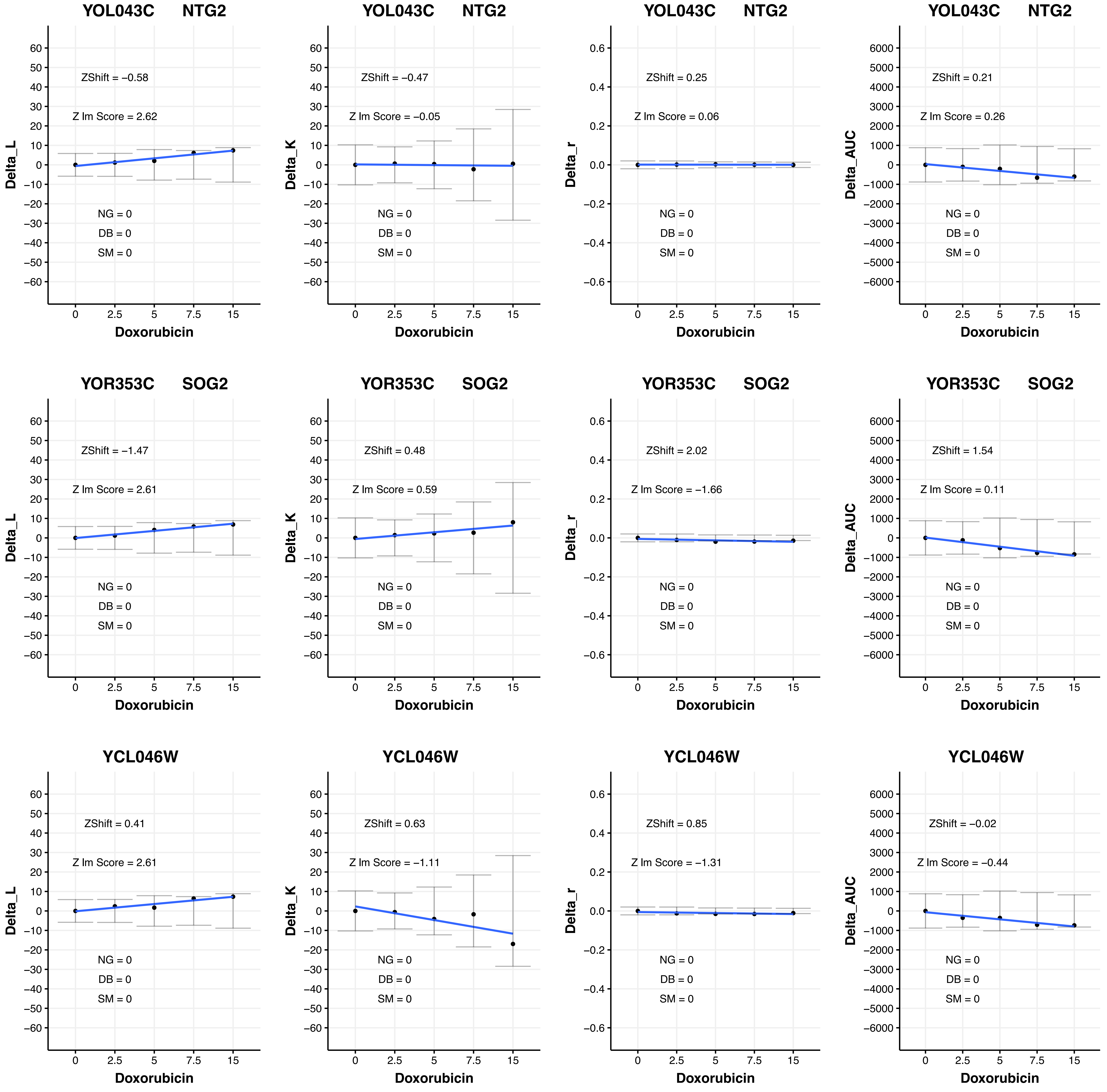
YOL043C NTG2
ZShift = −0.58
Z lm Score = 2.62
NG = 0
DB = 0
SM = 0
Delta_L
Doxorubicin
YOL043C NTG2
ZShift = −0.47
Z lm Score = −0.05
NG = 0
DB = 0
SM = 0
Delta_K
Doxorubicin
YOL043C NTG2
ZShift = 0.25
Z lm Score = 0.06
NG = 0
DB = 0
SM = 0
Delta_r
Doxorubicin
YOL043C NTG2
ZShift = 0.21
Z lm Score = 0.26
NG = 0
DB = 0
SM = 0
Delta_AUC
Doxorubicin
YOR353C SOG2
ZShift = −1.47
Z lm Score = 2.61
NG = 0
DB = 0
SM = 0
Delta_L
Doxorubicin
YOR353C SOG2
ZShift = 0.48
Z lm Score = 0.59
NG = 0
DB = 0
SM = 0
Delta_K
Doxorubicin
YOR353C SOG2
ZShift = 2.02
Z lm Score = −1.66
NG = 0
DB = 0
SM = 0
Delta_r
Doxorubicin
YOR353C SOG2
ZShift = 1.54
Z lm Score = 0.11
NG = 0
DB = 0
SM = 0
Delta_AUC
Doxorubicin
YCL046W
ZShift = 0.41
Z lm Score = 2.61
NG = 0
DB = 0
SM = 0
Delta_L
Doxorubicin
YCL046W
ZShift = 0.63
Z lm Score = −1.11
NG = 0
DB = 0
SM = 0
Delta_K
Doxorubicin
YCL046W
ZShift = 0.85
Z lm Score = −1.31
NG = 0
DB = 0
SM = 0
Delta_r
Doxorubicin
YCL046W
ZShift = −0.02
Z lm Score = −0.44
NG = 0
DB = 0
SM = 0
Delta_AUC
Doxorubicin

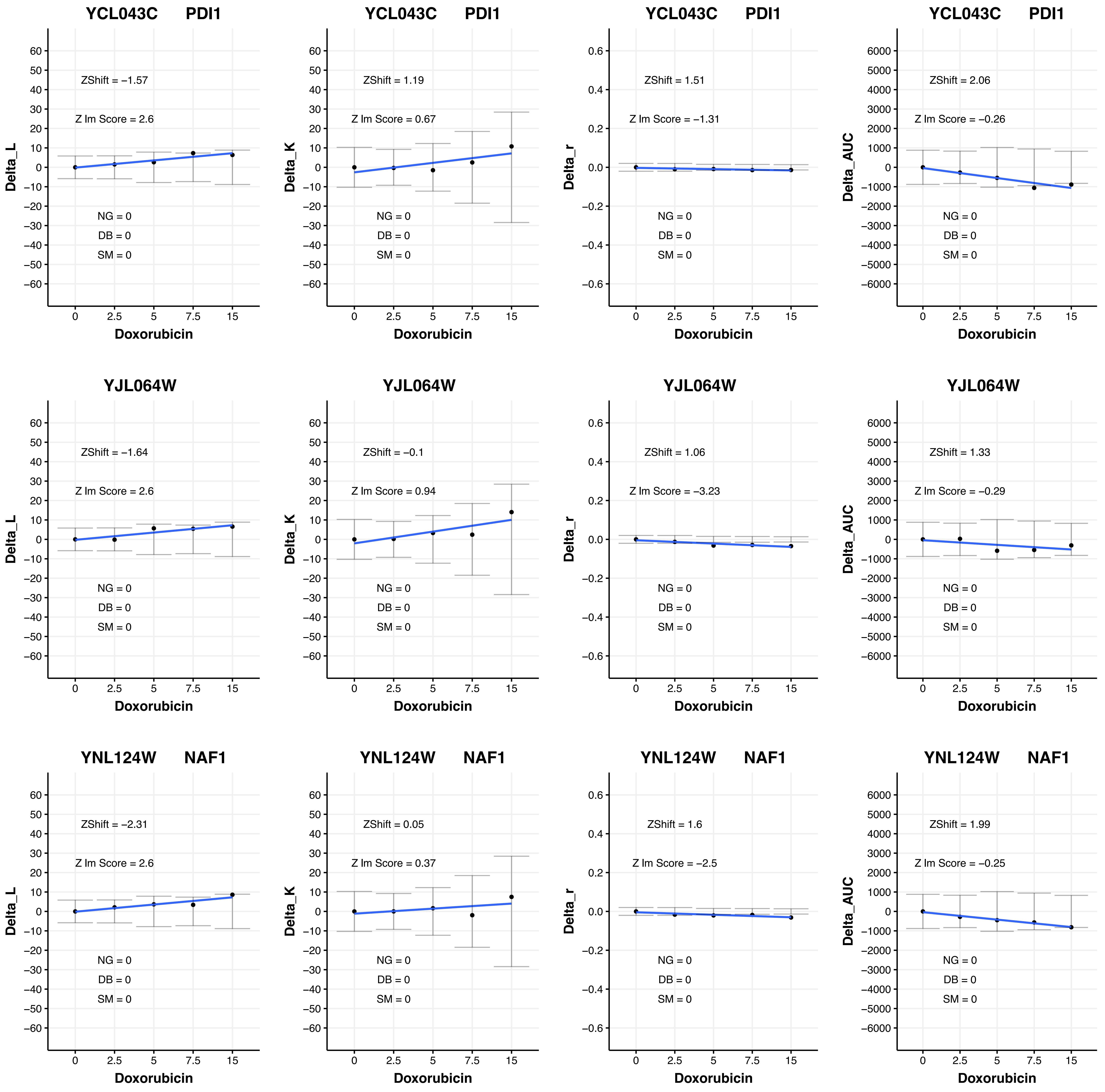
YCL043C PDI1
ZShift = −1.57
Z lm Score = 2.6
NG = 0
DB = 0
SM = 0
Delta_L
Doxorubicin
YCL043C PDI1
ZShift = 1.19
Z lm Score = 0.67
NG = 0
DB = 0
SM = 0
Delta_K
Doxorubicin
YCL043C PDI1
ZShift = 1.51
Z lm Score = −1.31
NG = 0
DB = 0
SM = 0
Delta_r
Doxorubicin
YCL043C PDI1
ZShift = 2.06
Z lm Score = −0.26
NG = 0
DB = 0
SM = 0
Delta_AUC
Doxorubicin
YJL064W
ZShift = −1.64
Z lm Score = 2.6
NG = 0
DB = 0
SM = 0
Delta_L
Doxorubicin
YJL064W
ZShift = −0.1
Z lm Score = 0.94
NG = 0
DB = 0
SM = 0
Delta_K
Doxorubicin
YJL064W
ZShift = 1.06
Z lm Score = −3.23
NG = 0
DB = 0
SM = 0
Delta_r
Doxorubicin
YJL064W
ZShift = 1.33
Z lm Score = −0.29
NG = 0
DB = 0
SM = 0
Delta_AUC
Doxorubicin
YNL124W NAF1
ZShift = −2.31
Z lm Score = 2.6
NG = 0
DB = 0
SM = 0
Delta_L
Doxorubicin
YNL124W NAF1
ZShift = 0.05
Z lm Score = 0.37
NG = 0
DB = 0
SM = 0
Delta_K
Doxorubicin
YNL124W NAF1
ZShift = 1.6
Z lm Score = −2.5
NG = 0
DB = 0
SM = 0
Delta_r
Doxorubicin
YNL124W NAF1
ZShift = 1.99
Z lm Score = −0.25
NG = 0
DB = 0
SM = 0
Delta_AUC
Doxorubicin

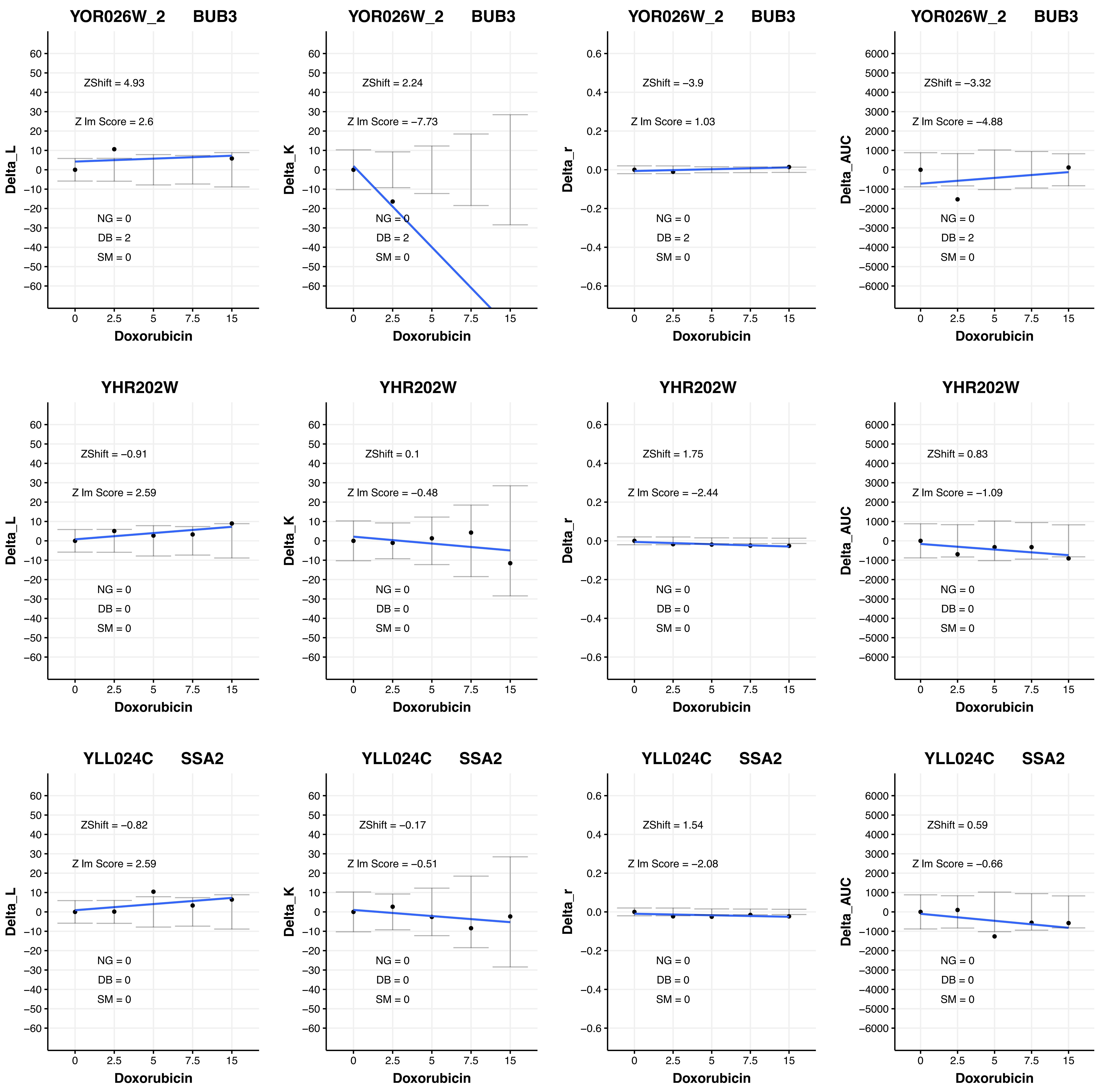
YOR026W_2 BUB3
ZShift = 4.93
Z lm Score = 2.6
NG = 0
DB = 2
SM = 0
Delta_L
Doxorubicin
YOR026W_2 BUB3
ZShift = 2.24
Z lm Score = −7.73
NG = 0
DB = 2
SM = 0
Delta_K
Doxorubicin
YOR026W_2 BUB3
ZShift = −3.9
Z lm Score = 1.03
NG = 0
DB = 2
SM = 0
Delta_r
Doxorubicin
YOR026W_2 BUB3
ZShift = −3.32
Z lm Score = −4.88
NG = 0
DB = 2
SM = 0
Delta_AUC
Doxorubicin
YHR202W
ZShift = −0.91
Z lm Score = 2.59
NG = 0
DB = 0
SM = 0
Delta_L
Doxorubicin
YHR202W
ZShift = 0.1
Z lm Score = −0.48
NG = 0
DB = 0
SM = 0
Delta_K
Doxorubicin
YHR202W
ZShift = 1.75
Z lm Score = −2.44
NG = 0
DB = 0
SM = 0
Delta_r
Doxorubicin
YHR202W
ZShift = 0.83
Z lm Score = −1.09
NG = 0
DB = 0
SM = 0
Delta_AUC
Doxorubicin
YLL024C SSA2
ZShift = −0.82
Z lm Score = 2.59
NG = 0
DB = 0
SM = 0
Delta_L
Doxorubicin
YLL024C SSA2
ZShift = −0.17
Z lm Score = −0.51
NG = 0
DB = 0
SM = 0
Delta_K
Doxorubicin
YLL024C SSA2
ZShift = 1.54
Z lm Score = −2.08
NG = 0
DB = 0
SM = 0
Delta_r
Doxorubicin
YLL024C SSA2
ZShift = 0.59
Z lm Score = −0.66
NG = 0
DB = 0
SM = 0
Delta_AUC
Doxorubicin

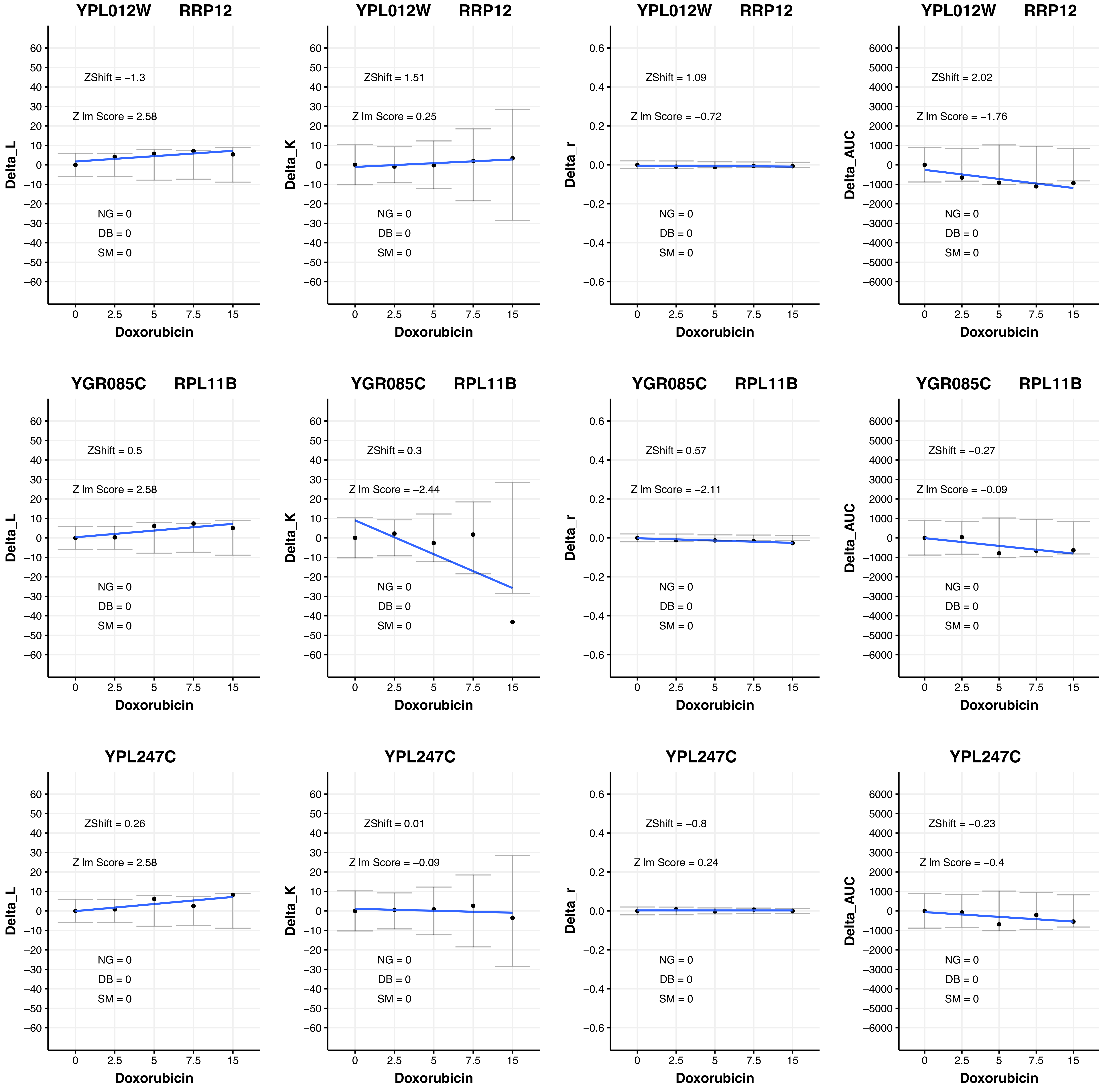

YPL012W RRP12
ZShift = −1.3
Z lm Score = 2.58
NG = 0
DB = 0
SM = 0
Delta_L
Doxorubicin
YPL012W RRP12
ZShift = 1.51
Z lm Score = 0.25
NG = 0
DB = 0
SM = 0
Delta_K
Doxorubicin
YPL012W RRP12
ZShift = 1.09
Z lm Score = −0.72
NG = 0
DB = 0
SM = 0
Delta_r
Doxorubicin
YPL012W RRP12
ZShift = 2.02
Z lm Score = −1.76
NG = 0
DB = 0
SM = 0
Delta_AUC
Doxorubicin
YGR085C RPL11B
ZShift = 0.5
Z lm Score = 2.58
NG = 0
DB = 0
SM = 0
Delta_L
Doxorubicin
YGR085C RPL11B
ZShift = 0.3
Z lm Score = −2.44
NG = 0
DB = 0
SM = 0
Delta_K
Doxorubicin
YGR085C RPL11B
ZShift = 0.57
Z lm Score = −2.11
NG = 0
DB = 0
SM = 0
Delta_r
Doxorubicin
YGR085C RPL11B
ZShift = −0.27
Z lm Score = −0.09
NG = 0
DB = 0
SM = 0
Delta_AUC
Doxorubicin
YPL247C
ZShift = 0.26
Z lm Score = 2.58
NG = 0
DB = 0
SM = 0
Delta_L
Doxorubicin
YPL247C
ZShift = 0.01
Z lm Score = −0.09
NG = 0
DB = 0
SM = 0
Delta_K
Doxorubicin
YPL247C
ZShift = −0.8
Z lm Score = 0.24
NG = 0
DB = 0
SM = 0
Delta_r
Doxorubicin
YPL247C
ZShift = −0.23
Z lm Score = −0.4
NG = 0
DB = 0
SM = 0
Delta_AUC
Doxorubicin

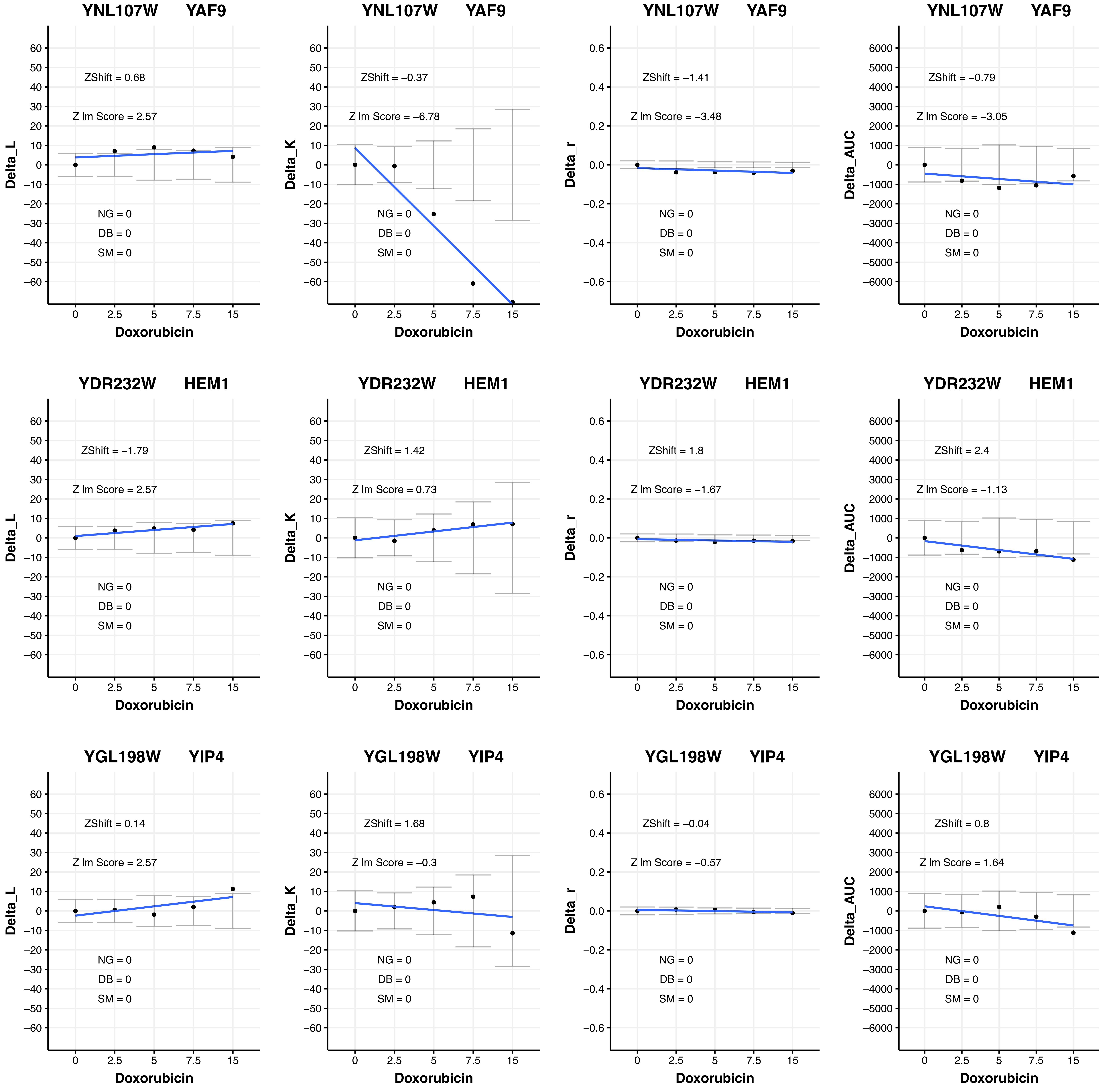

YNL107W YAF9
ZShift = 0.68
Z lm Score = 2.57
NG = 0
DB = 0
SM = 0
Delta_L
Doxorubicin
YNL107W YAF9
ZShift = −0.37
Z lm Score = −6.78
NG = 0
DB = 0
SM = 0
Delta_K
Doxorubicin
YNL107W YAF9
ZShift = −1.41
Z lm Score = −3.48
NG = 0
DB = 0
SM = 0
Delta_r
Doxorubicin
YNL107W YAF9
ZShift = −0.79
Z lm Score = −3.05
NG = 0
DB = 0
SM = 0
Delta_AUC
Doxorubicin
YDR232W HEM1
ZShift = −1.79
Z lm Score = 2.57
NG = 0
DB = 0
SM = 0
Delta_L
Doxorubicin
YDR232W HEM1
ZShift = 1.42
Z lm Score = 0.73
NG = 0
DB = 0
SM = 0
Delta_K
Doxorubicin
YDR232W HEM1
ZShift = 1.8
Z lm Score = −1.67
NG = 0
DB = 0
SM = 0
Delta_r
Doxorubicin
YDR232W HEM1
ZShift = 2.4
Z lm Score = −1.13
NG = 0
DB = 0
SM = 0
Delta_AUC
Doxorubicin
YGL198W YIP4
ZShift = 0.14
Z lm Score = 2.57
NG = 0
DB = 0
SM = 0
Delta_L
Doxorubicin
YGL198W YIP4
ZShift = 1.68
Z lm Score = −0.3
NG = 0
DB = 0
SM = 0
Delta_K
Doxorubicin
YGL198W YIP4
ZShift = −0.04
Z lm Score = −0.57
NG = 0
DB = 0
SM = 0
Delta_r
Doxorubicin
YGL198W YIP4
ZShift = 0.8
Z lm Score = 1.64
NG = 0
DB = 0
SM = 0
Delta_AUC
Doxorubicin

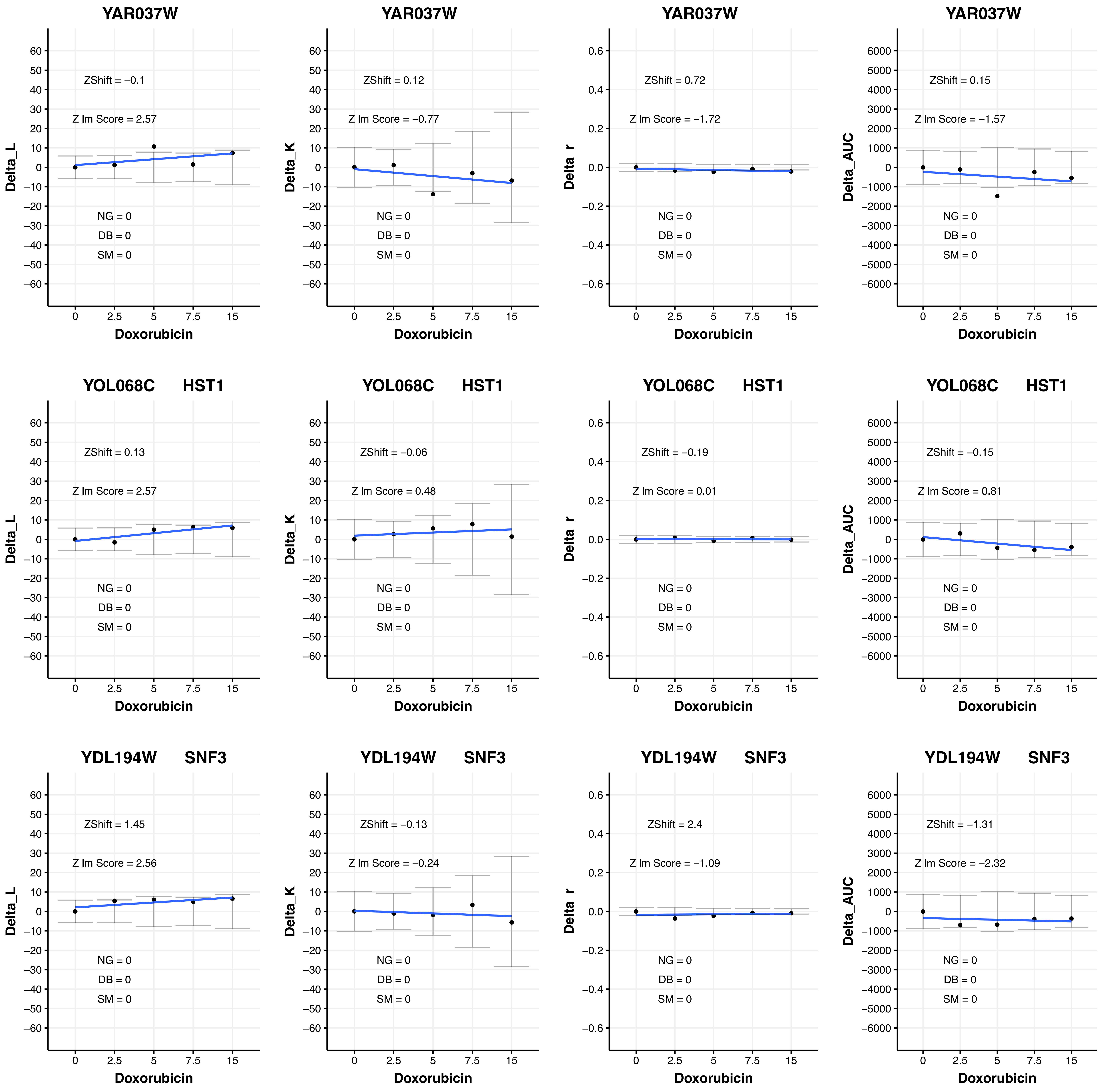
YAR037W
ZShift = −0.1
Z lm Score = 2.57
NG = 0
DB = 0
SM = 0
Delta_L
Doxorubicin
YAR037W
ZShift = 0.12
Z lm Score = −0.77
NG = 0
DB = 0
SM = 0
Delta_K
Doxorubicin
YAR037W
ZShift = 0.72
Z lm Score = −1.72
NG = 0
DB = 0
SM = 0
Delta_r
Doxorubicin
YAR037W
ZShift = 0.15
Z lm Score = −1.57
NG = 0
DB = 0
SM = 0
Delta_AUC
Doxorubicin
YOL068C HST1
ZShift = 0.13
Z lm Score = 2.57
NG = 0
DB = 0
SM = 0
Delta_L
Doxorubicin
YOL068C HST1
ZShift = −0.06
Z lm Score = 0.48
NG = 0
DB = 0
SM = 0
Delta_K
Doxorubicin
YOL068C HST1
ZShift = −0.19
Z lm Score = 0.01
NG = 0
DB = 0
SM = 0
Delta_r
Doxorubicin
YOL068C HST1
ZShift = −0.15
Z lm Score = 0.81
NG = 0
DB = 0
SM = 0
Delta_AUC
Doxorubicin
YDL194W SNF3
ZShift = 1.45
Z lm Score = 2.56
NG = 0
DB = 0
SM = 0
Delta_L
Doxorubicin
YDL194W SNF3
ZShift = −0.13
Z lm Score = −0.24
NG = 0
DB = 0
SM = 0
Delta_K
Doxorubicin
YDL194W SNF3
ZShift = 2.4
Z lm Score = −1.09
NG = 0
DB = 0
SM = 0
Delta_r
Doxorubicin
YDL194W SNF3
ZShift = −1.31
Z lm Score = −2.32
NG = 0
DB = 0
SM = 0
Delta_AUC
Doxorubicin

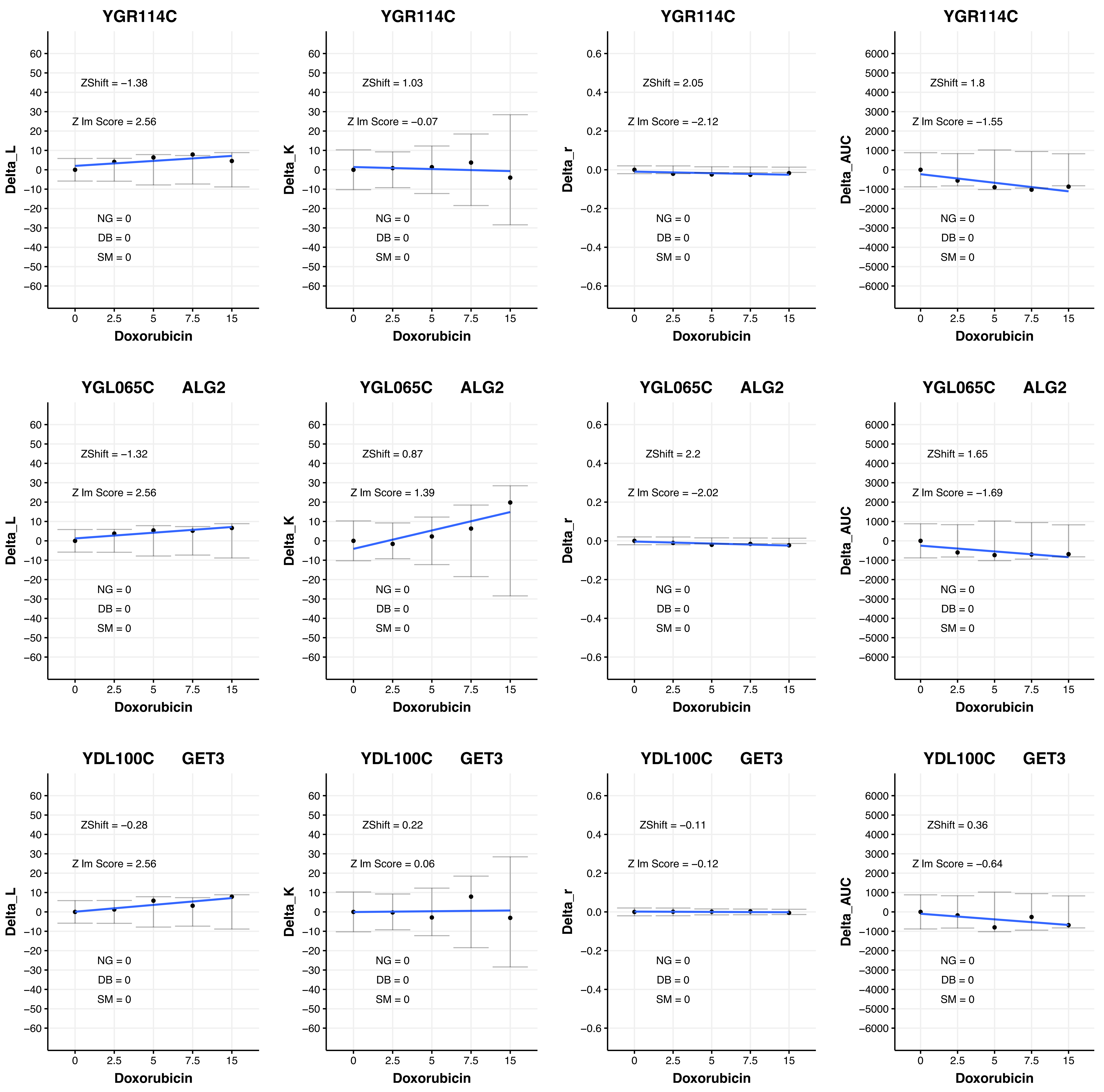
YGR114C
ZShift = −1.38
Z lm Score = 2.56
NG = 0
DB = 0
SM = 0
Delta_L
Doxorubicin
YGR114C
ZShift = 1.03
Z lm Score = −0.07
NG = 0
DB = 0
SM = 0
Delta_K
Doxorubicin
YGR114C
ZShift = 2.05
Z lm Score = −2.12
NG = 0
DB = 0
SM = 0
Delta_r
Doxorubicin
YGR114C
ZShift = 1.8
Z lm Score = −1.55
NG = 0
DB = 0
SM = 0
Delta_AUC
Doxorubicin
YGL065C ALG2
ZShift = −1.32
Z lm Score = 2.56
NG = 0
DB = 0
SM = 0
Delta_L
Doxorubicin
YGL065C ALG2
ZShift = 0.87
Z lm Score = 1.39
NG = 0
DB = 0
SM = 0
Delta_K
Doxorubicin
YGL065C ALG2
ZShift = 2.2
Z lm Score = −2.02
NG = 0
DB = 0
SM = 0
Delta_r
Doxorubicin
YGL065C ALG2
ZShift = 1.65
Z lm Score = −1.69
NG = 0
DB = 0
SM = 0
Delta_AUC
Doxorubicin
YDL100C GET3
ZShift = −0.28
Z lm Score = 2.56
NG = 0
DB = 0
SM = 0
Delta_L
Doxorubicin
YDL100C GET3
ZShift = 0.22
Z lm Score = 0.06
NG = 0
DB = 0
SM = 0
Delta_K
Doxorubicin
YDL100C GET3
ZShift = −0.11
Z lm Score = −0.12
NG = 0
DB = 0
SM = 0
Delta_r
Doxorubicin
YDL100C GET3
ZShift = 0.36
Z lm Score = −0.64
NG = 0
DB = 0
SM = 0
Delta_AUC
Doxorubicin

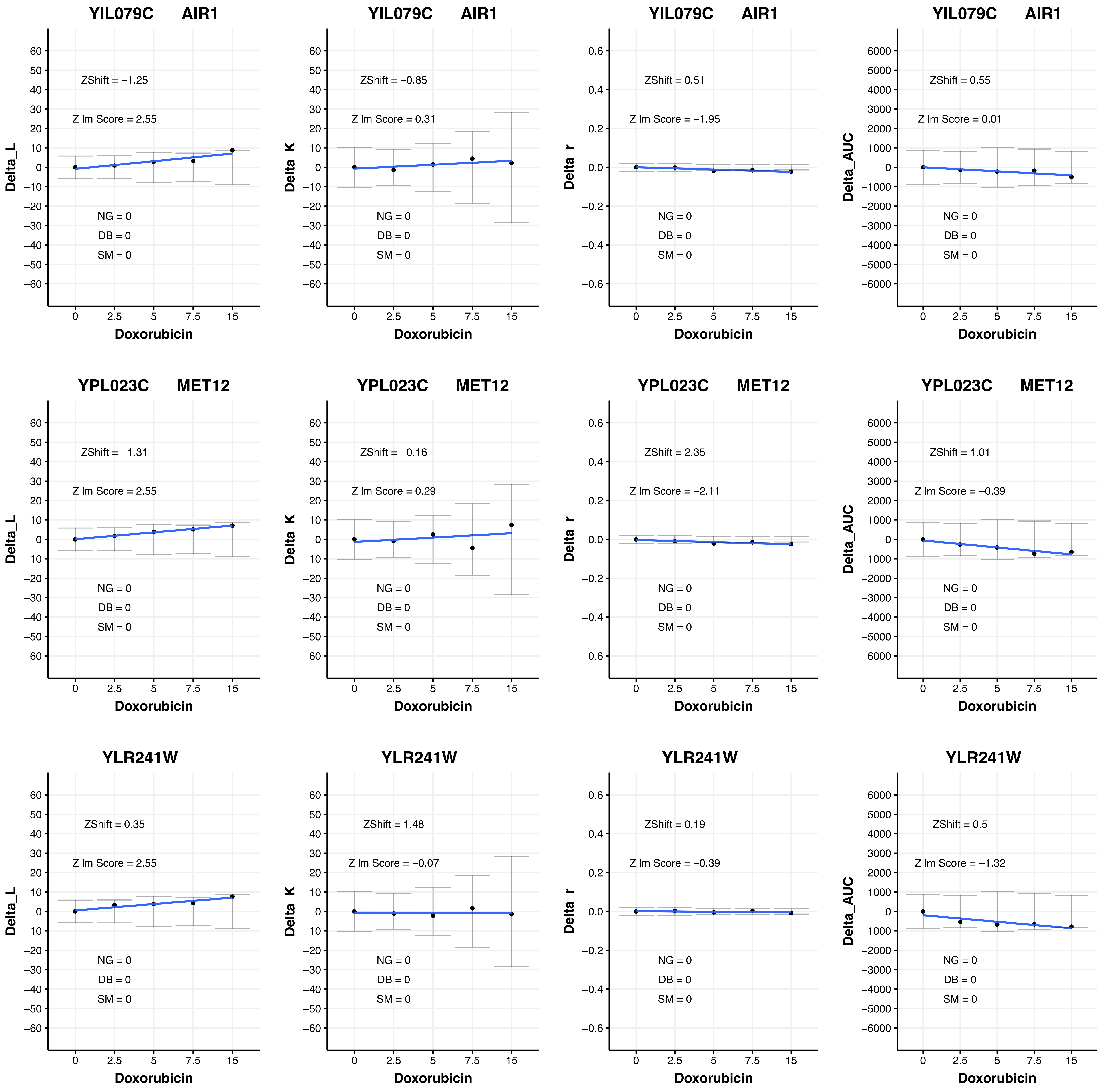

YIL079C AIR1
ZShift = −1.25
Z lm Score = 2.55
NG = 0
DB = 0
SM = 0
Delta_L
Doxorubicin
YIL079C AIR1
ZShift = −0.85
Z lm Score = 0.31
NG = 0
DB = 0
SM = 0
Delta_K
Doxorubicin
YIL079C AIR1
ZShift = 0.51
Z lm Score = −1.95
NG = 0
DB = 0
SM = 0
Delta_r
Doxorubicin
YIL079C AIR1
ZShift = 0.55
Z lm Score = 0.01
NG = 0
DB = 0
SM = 0
Delta_AUC
Doxorubicin
YPL023C MET12
ZShift = −1.31
Z lm Score = 2.55
NG = 0
DB = 0
SM = 0
Delta_L
Doxorubicin
YPL023C MET12
ZShift = −0.16
Z lm Score = 0.29
NG = 0
DB = 0
SM = 0
Delta_K
Doxorubicin
YPL023C MET12
ZShift = 2.35
Z lm Score = −2.11
NG = 0
DB = 0
SM = 0
Delta_r
Doxorubicin
YPL023C MET12
ZShift = 1.01
Z lm Score = −0.39
NG = 0
DB = 0
SM = 0
Delta_AUC
Doxorubicin
YLR241W
ZShift = 0.35
Z lm Score = 2.55
NG = 0
DB = 0
SM = 0
Delta_L
Doxorubicin
YLR241W
ZShift = 1.48
Z lm Score = −0.07
NG = 0
DB = 0
SM = 0
Delta_K
Doxorubicin
YLR241W
ZShift = 0.19
Z lm Score = −0.39
NG = 0
DB = 0
SM = 0
Delta_r
Doxorubicin
YLR241W
ZShift = 0.5
Z lm Score = −1.32
NG = 0
DB = 0
SM = 0
Delta_AUC
Doxorubicin

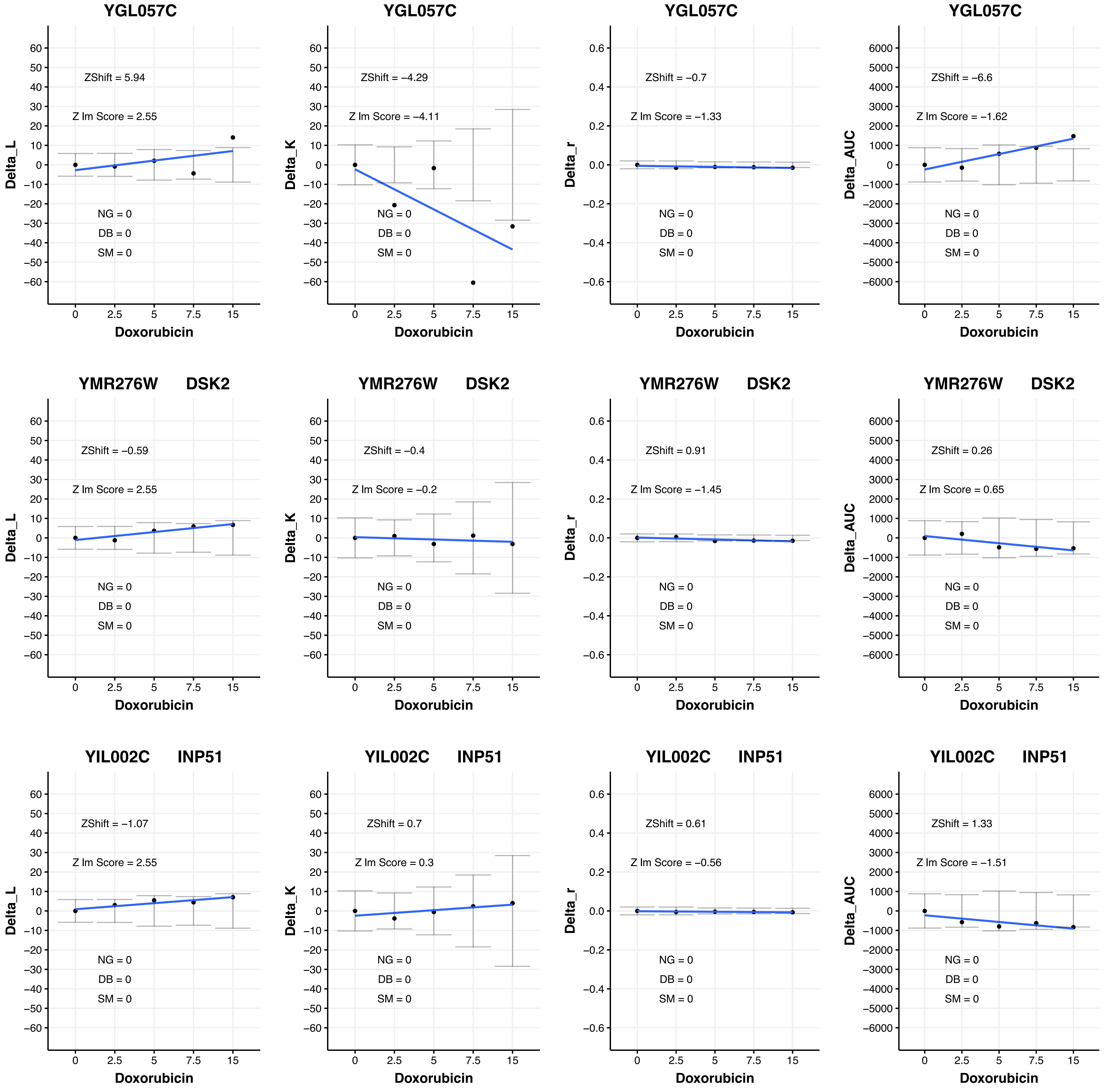

YGL057C
ZShift = 5.94
Z lm Score = 2.55
NG = 0
DB = 0
SM = 0
Delta_L
Doxorubicin
YGL057C
ZShift = −4.29
Z lm Score = −4.11
NG = 0
DB = 0
SM = 0
Delta_K
Doxorubicin
YGL057C
ZShift = −0.7
Z lm Score = −1.33
NG = 0
DB = 0
SM = 0
Delta_r
Doxorubicin
YGL057C
ZShift = −6.6
Z lm Score = −1.62
NG = 0
DB = 0
SM = 0
Delta_AUC
Doxorubicin
YMR276W DSK2
ZShift = −0.59
Z lm Score = 2.55
NG = 0
DB = 0
SM = 0
Delta_L
Doxorubicin
YMR276W DSK2
ZShift = −0.4
Z lm Score = −0.2
NG = 0
DB = 0
SM = 0
Delta_K
Doxorubicin
YMR276W DSK2
ZShift = 0.91
Z lm Score = −1.45
NG = 0
DB = 0
SM = 0
Delta_r
Doxorubicin
YMR276W DSK2
ZShift = 0.26
Z lm Score = 0.65
NG = 0
DB = 0
SM = 0
Delta_AUC
Doxorubicin
YIL002C INP51
ZShift = −1.07
Z lm Score = 2.55
NG = 0
DB = 0
SM = 0
Delta_L
Doxorubicin
YIL002C INP51
ZShift = 0.7
Z lm Score = 0.3
NG = 0
DB = 0
SM = 0
Delta_K
Doxorubicin
YIL002C INP51
ZShift = 0.61
Z lm Score = −0.56
NG = 0
DB = 0
SM = 0
Delta_r
Doxorubicin
YIL002C INP51
ZShift = 1.33
Z lm Score = −1.51
NG = 0
DB = 0
SM = 0
Delta_AUC
Doxorubicin

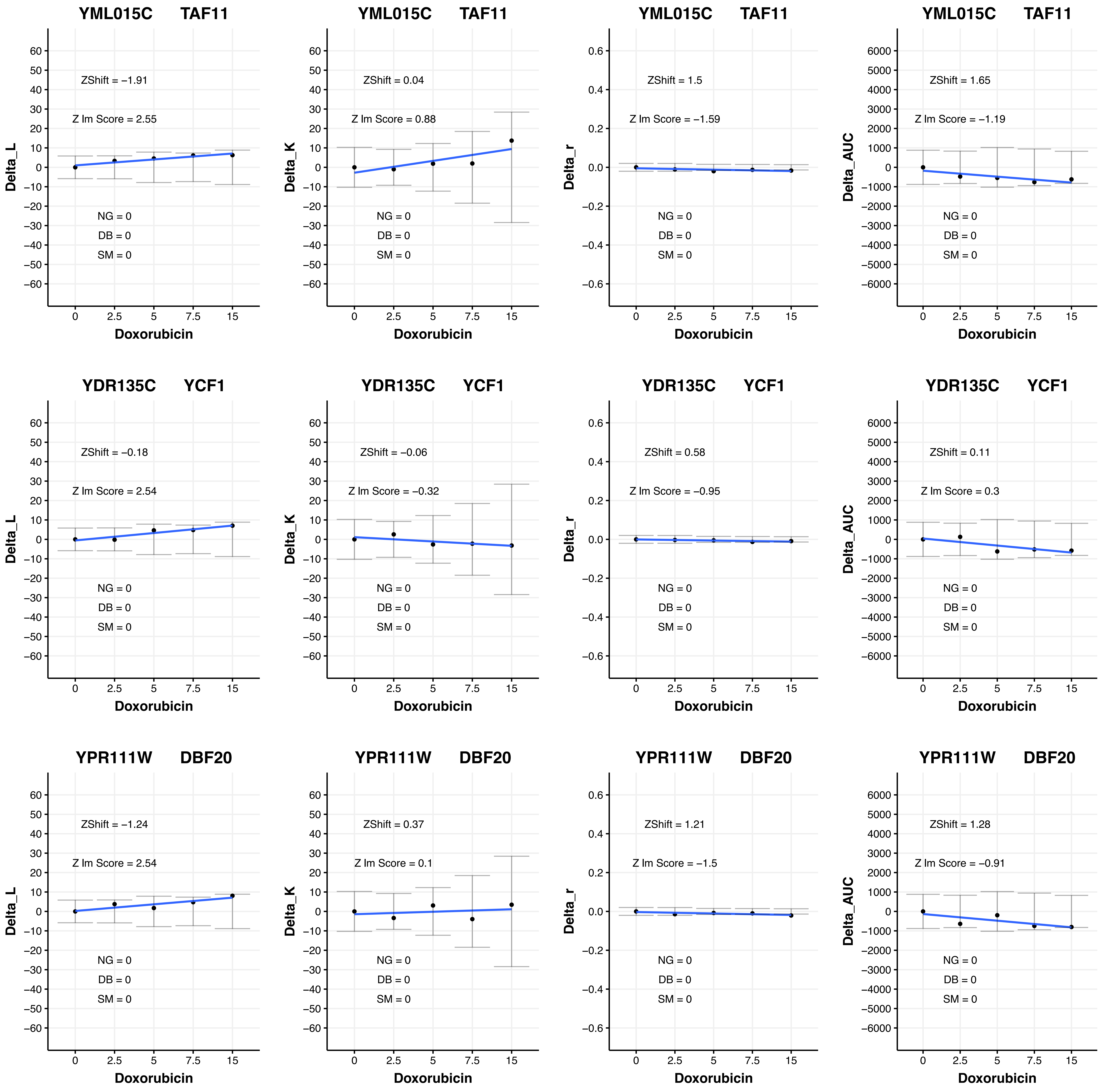
YML015C TAF11
ZShift = −1.91
Z lm Score = 2.55
NG = 0
DB = 0
SM = 0
Delta_L
Doxorubicin
YML015C TAF11
ZShift = 0.04
Z lm Score = 0.88
NG = 0
DB = 0
SM = 0
Delta_K
Doxorubicin
YML015C TAF11
ZShift = 1.5
Z lm Score = −1.59
NG = 0
DB = 0
SM = 0
Delta_r
Doxorubicin
YML015C TAF11
ZShift = 1.65
Z lm Score = −1.19
NG = 0
DB = 0
SM = 0
Delta_AUC
Doxorubicin
YDR135C YCF1
ZShift = −0.18
Z lm Score = 2.54
NG = 0
DB = 0
SM = 0
Delta_L
Doxorubicin
YDR135C YCF1
ZShift = −0.06
Z lm Score = −0.32
NG = 0
DB = 0
SM = 0
Delta_K
Doxorubicin
YDR135C YCF1
ZShift = 0.58
Z lm Score = −0.95
NG = 0
DB = 0
SM = 0
Delta_r
Doxorubicin
YDR135C YCF1
ZShift = 0.11
Z lm Score = 0.3
NG = 0
DB = 0
SM = 0
Delta_AUC
Doxorubicin
YPR111W DBF20
ZShift = −1.24
Z lm Score = 2.54
NG = 0
DB = 0
SM = 0
Delta_L
Doxorubicin
YPR111W DBF20
ZShift = 0.37
Z lm Score = 0.1
NG = 0
DB = 0
SM = 0
Delta_K
Doxorubicin
YPR111W DBF20
ZShift = 1.21
Z lm Score = −1.5
NG = 0
DB = 0
SM = 0
Delta_r
Doxorubicin
YPR111W DBF20
ZShift = 1.28
Z lm Score = −0.91
NG = 0
DB = 0
SM = 0
Delta_AUC
Doxorubicin

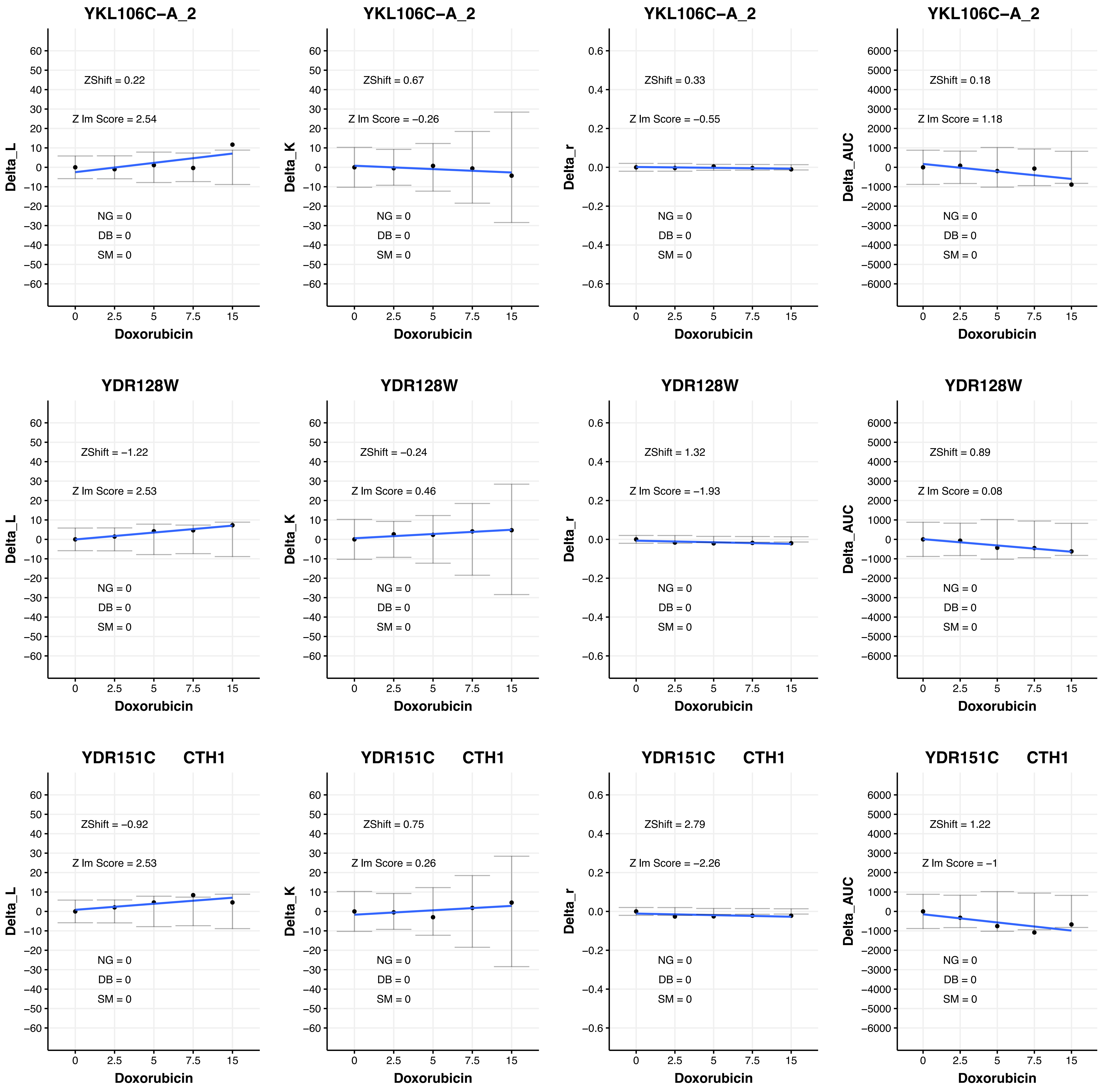
YKL106C−A_2
ZShift = 0.22
Z lm Score = 2.54
NG = 0
DB = 0
SM = 0
Delta_L
Doxorubicin
YKL106C−A_2
ZShift = 0.67
Z lm Score = −0.26
NG = 0
DB = 0
SM = 0
Delta_K
Doxorubicin
YKL106C−A_2
ZShift = 0.33
Z lm Score = −0.55
NG = 0
DB = 0
SM = 0
Delta_r
Doxorubicin
YKL106C−A_2
ZShift = 0.18
Z lm Score = 1.18
NG = 0
DB = 0
SM = 0
Delta_AUC
Doxorubicin
YDR128W
ZShift = −1.22
Z lm Score = 2.53
NG = 0
DB = 0
SM = 0
Delta_L
Doxorubicin
YDR128W
ZShift = −0.24
Z lm Score = 0.46
NG = 0
DB = 0
SM = 0
Delta_K
Doxorubicin
YDR128W
ZShift = 1.32
Z lm Score = −1.93
NG = 0
DB = 0
SM = 0
Delta_r
Doxorubicin
YDR128W
ZShift = 0.89
Z lm Score = 0.08
NG = 0
DB = 0
SM = 0
Delta_AUC
Doxorubicin
YDR151C CTH1
ZShift = −0.92
Z lm Score = 2.53
NG = 0
DB = 0
SM = 0
Delta_L
Doxorubicin
YDR151C CTH1
ZShift = 0.75
Z lm Score = 0.26
NG = 0
DB = 0
SM = 0
Delta_K
Doxorubicin
YDR151C CTH1
ZShift = 2.79
Z lm Score = −2.26
NG = 0
DB = 0
SM = 0
Delta_r
Doxorubicin
YDR151C CTH1
ZShift = 1.22
Z lm Score = −1
NG = 0
DB = 0
SM = 0
Delta_AUC
Doxorubicin

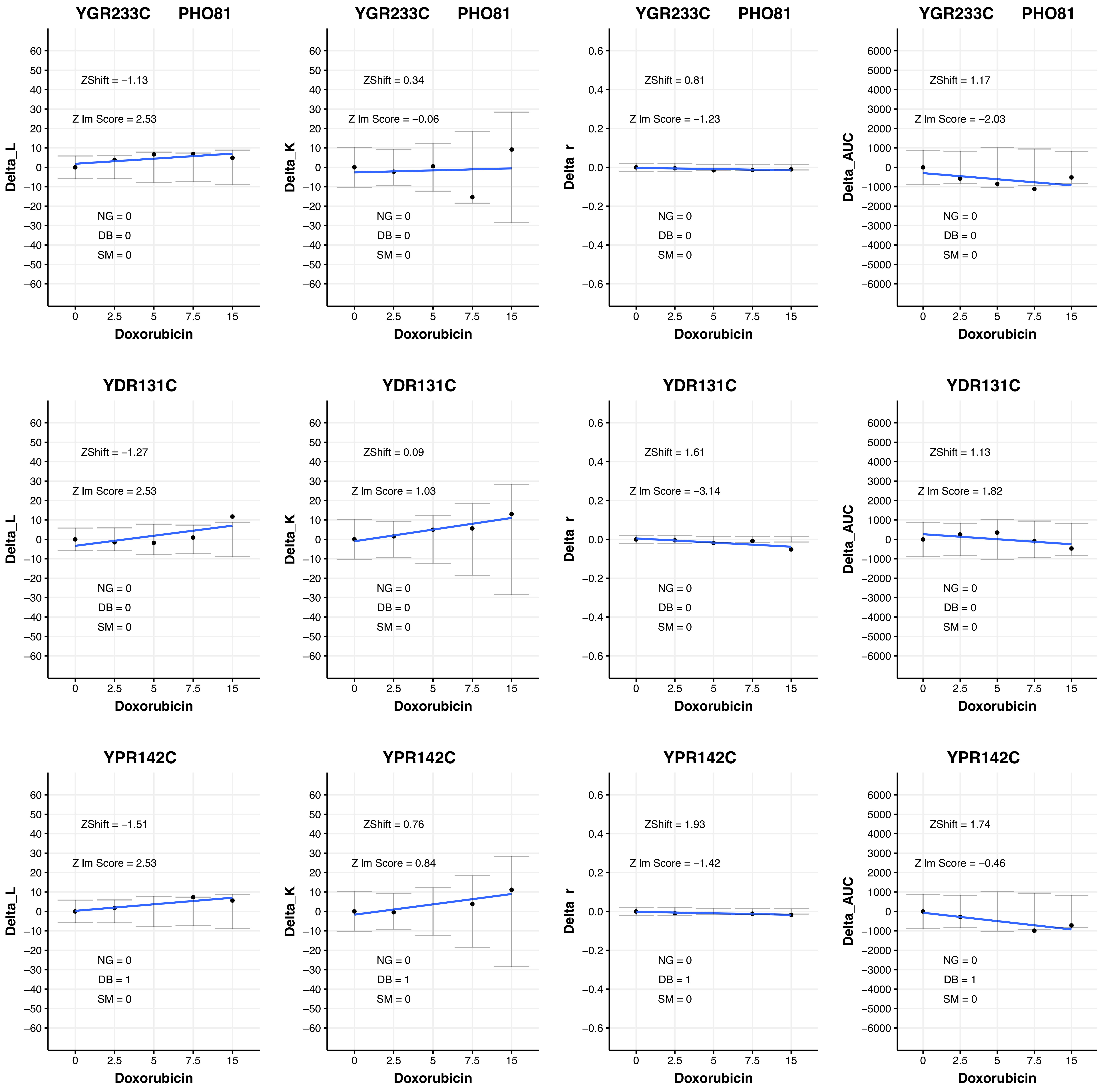
YGR233C PHO81
ZShift = −1.13
Z lm Score = 2.53
NG = 0
DB = 0
SM = 0
Delta_L
Doxorubicin
YGR233C PHO81
ZShift = 0.34
Z lm Score = −0.06
NG = 0
DB = 0
SM = 0
Delta_K
Doxorubicin
YGR233C PHO81
ZShift = 0.81
Z lm Score = −1.23
NG = 0
DB = 0
SM = 0
Delta_r
Doxorubicin
YGR233C PHO81
ZShift = 1.17
Z lm Score = −2.03
NG = 0
DB = 0
SM = 0
Delta_AUC
Doxorubicin
YDR131C
ZShift = −1.27
Z lm Score = 2.53
NG = 0
DB = 0
SM = 0
Delta_L
Doxorubicin
YDR131C
ZShift = 0.09
Z lm Score = 1.03
NG = 0
DB = 0
SM = 0
Delta_K
Doxorubicin
YDR131C
ZShift = 1.61
Z lm Score = −3.14
NG = 0
DB = 0
SM = 0
Delta_r
Doxorubicin
YDR131C
ZShift = 1.13
Z lm Score = 1.82
NG = 0
DB = 0
SM = 0
Delta_AUC
Doxorubicin
YPR142C
ZShift = −1.51
Z lm Score = 2.53
NG = 0
DB = 1
SM = 0
Delta_L
Doxorubicin
YPR142C
ZShift = 0.76
Z lm Score = 0.84
NG = 0
DB = 1
SM = 0
Delta_K
Doxorubicin
YPR142C
ZShift = 1.93
Z lm Score = −1.42
NG = 0
DB = 1
SM = 0
Delta_r
Doxorubicin
YPR142C
ZShift = 1.74
Z lm Score = −0.46
NG = 0
DB = 1
SM = 0
Delta_AUC
Doxorubicin

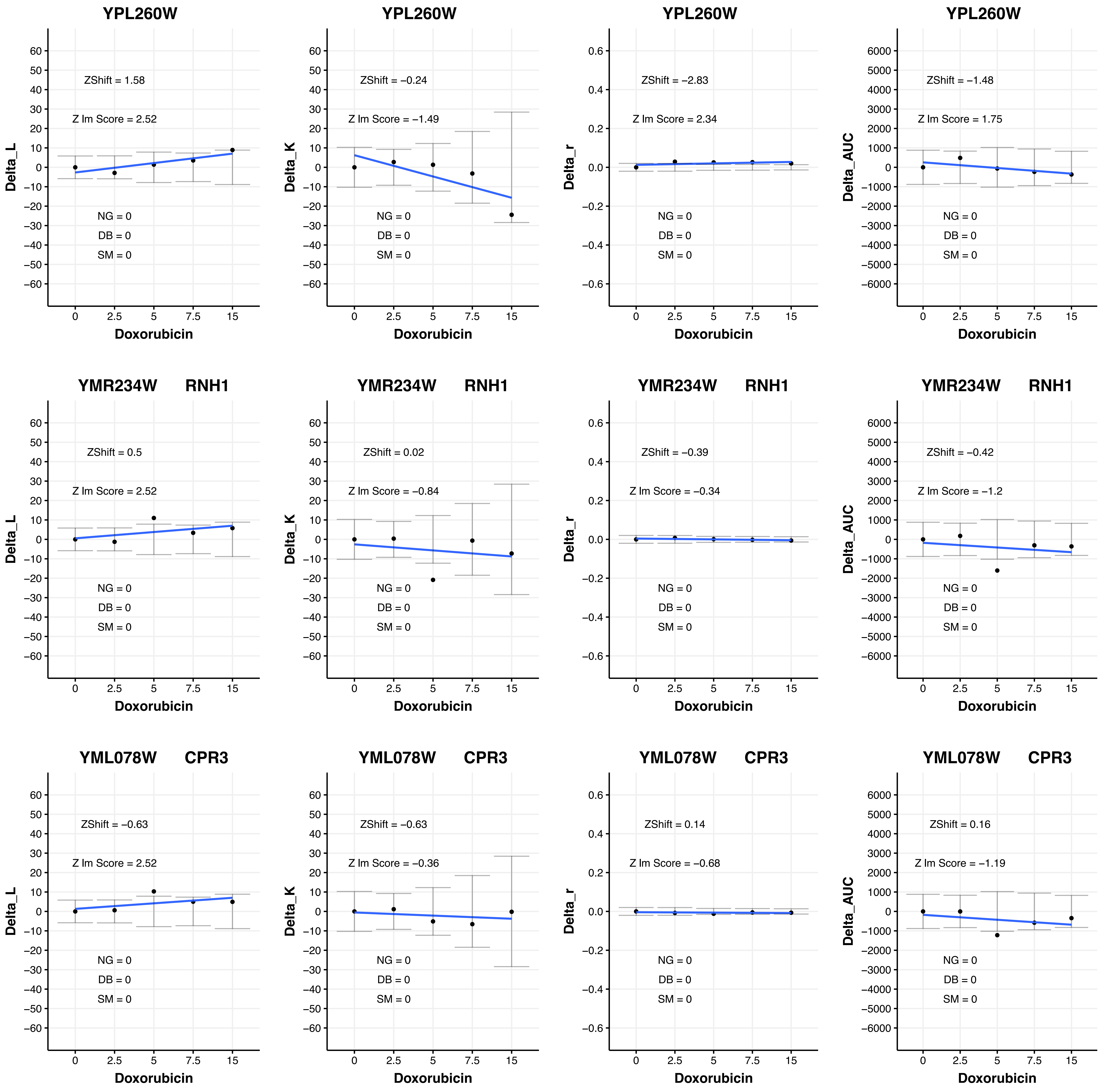
YPL260W
ZShift = 1.58
Z lm Score = 2.52
NG = 0
DB = 0
SM = 0
Delta_L
Doxorubicin
YPL260W
ZShift = −0.24
Z lm Score = −1.49
NG = 0
DB = 0
SM = 0
Delta_K
Doxorubicin
YPL260W
ZShift = −2.83
Z lm Score = 2.34
NG = 0
DB = 0
SM = 0
Delta_r
Doxorubicin
YPL260W
ZShift = −1.48
Z lm Score = 1.75
NG = 0
DB = 0
SM = 0
Delta_AUC
Doxorubicin
YMR234W RNH1
ZShift = 0.5
Z lm Score = 2.52
NG = 0
DB = 0
SM = 0
Delta_L
Doxorubicin
YMR234W RNH1
ZShift = 0.02
Z lm Score = −0.84
NG = 0
DB = 0
SM = 0
Delta_K
Doxorubicin
YMR234W RNH1
ZShift = −0.39
Z lm Score = −0.34
NG = 0
DB = 0
SM = 0
Delta_r
Doxorubicin
YMR234W RNH1
ZShift = −0.42
Z lm Score = −1.2
NG = 0
DB = 0
SM = 0
Delta_AUC
Doxorubicin
YML078W CPR3
ZShift = −0.63
Z lm Score = 2.52
NG = 0
DB = 0
SM = 0
Delta_L
Doxorubicin
YML078W CPR3
ZShift = −0.63
Z lm Score = −0.36
NG = 0
DB = 0
SM = 0
Delta_K
Doxorubicin
YML078W CPR3
ZShift = 0.14
Z lm Score = −0.68
NG = 0
DB = 0
SM = 0
Delta_r
Doxorubicin
YML078W CPR3
ZShift = 0.16
Z lm Score = −1.19
NG = 0
DB = 0
SM = 0
Delta_AUC
Doxorubicin

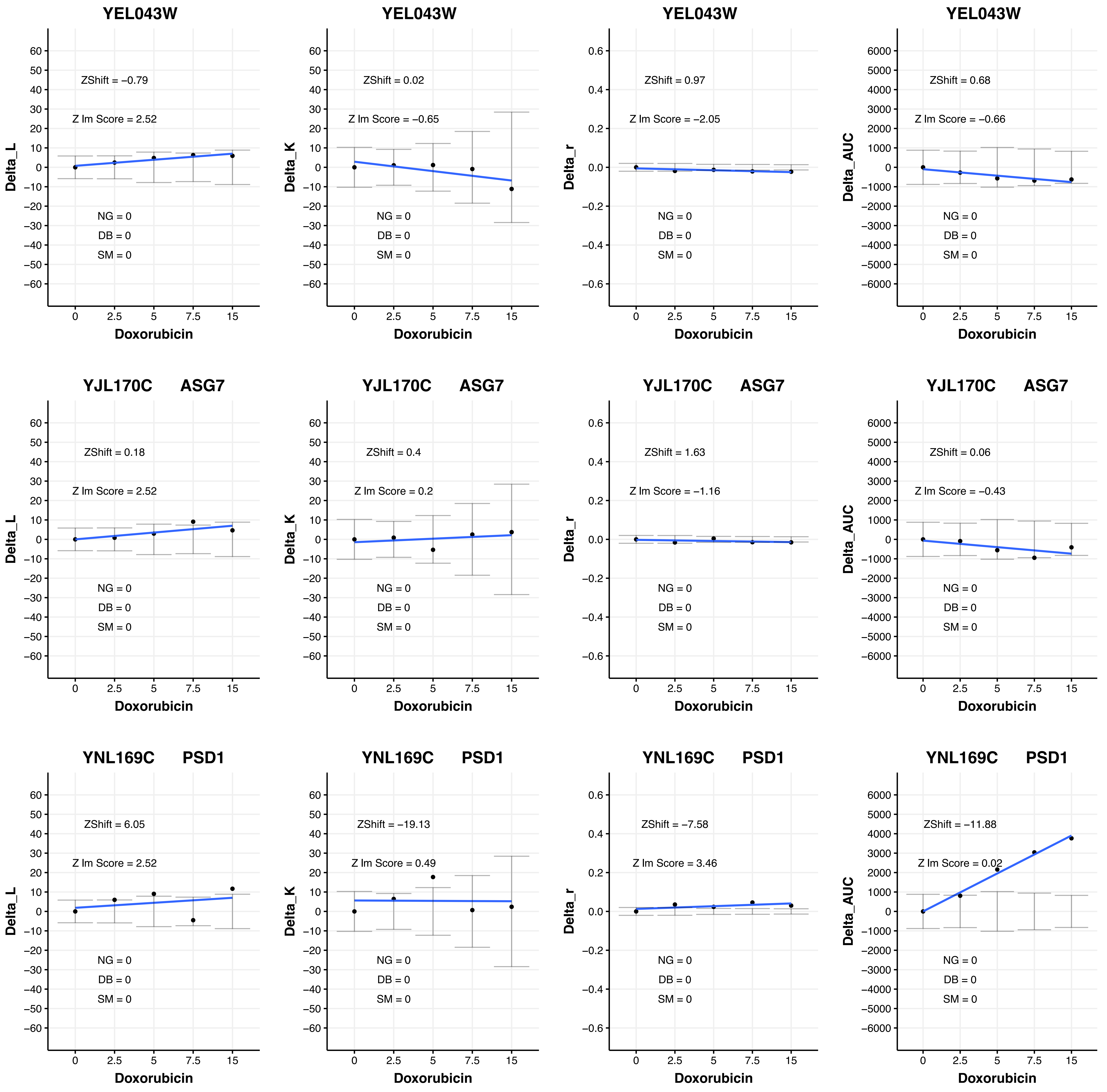
YEL043W
ZShift = −0.79
Z lm Score = 2.52
NG = 0
DB = 0
SM = 0
Delta_L
Doxorubicin
YEL043W
ZShift = 0.02
Z lm Score = −0.65
NG = 0
DB = 0
SM = 0
Delta_K
Doxorubicin
YEL043W
ZShift = 0.97
Z lm Score = −2.05
NG = 0
DB = 0
SM = 0
Delta_r
Doxorubicin
YEL043W
ZShift = 0.68
Z lm Score = −0.66
NG = 0
DB = 0
SM = 0
Delta_AUC
Doxorubicin
YJL170C ASG7
ZShift = 0.18
Z lm Score = 2.52
NG = 0
DB = 0
SM = 0
Delta_L
Doxorubicin
YJL170C ASG7
ZShift = 0.4
Z lm Score = 0.2
NG = 0
DB = 0
SM = 0
Delta_K
Doxorubicin
YJL170C ASG7
ZShift = 1.63
Z lm Score = −1.16
NG = 0
DB = 0
SM = 0
Delta_r
Doxorubicin
YJL170C ASG7
ZShift = 0.06
Z lm Score = −0.43
NG = 0
DB = 0
SM = 0
Delta_AUC
Doxorubicin
YNL169C PSD1
ZShift = 6.05
Z lm Score = 2.52
NG = 0
DB = 0
SM = 0
Delta_L
Doxorubicin
YNL169C PSD1
ZShift = −19.13
Z lm Score = 0.49
NG = 0
DB = 0
SM = 0
Delta_K
Doxorubicin
YNL169C PSD1
ZShift = −7.58
Z lm Score = 3.46
NG = 0
DB = 0
SM = 0
Delta_r
Doxorubicin
YNL169C PSD1
ZShift = −11.88
Z lm Score = 0.02
NG = 0
DB = 0
SM = 0
Delta_AUC
Doxorubicin

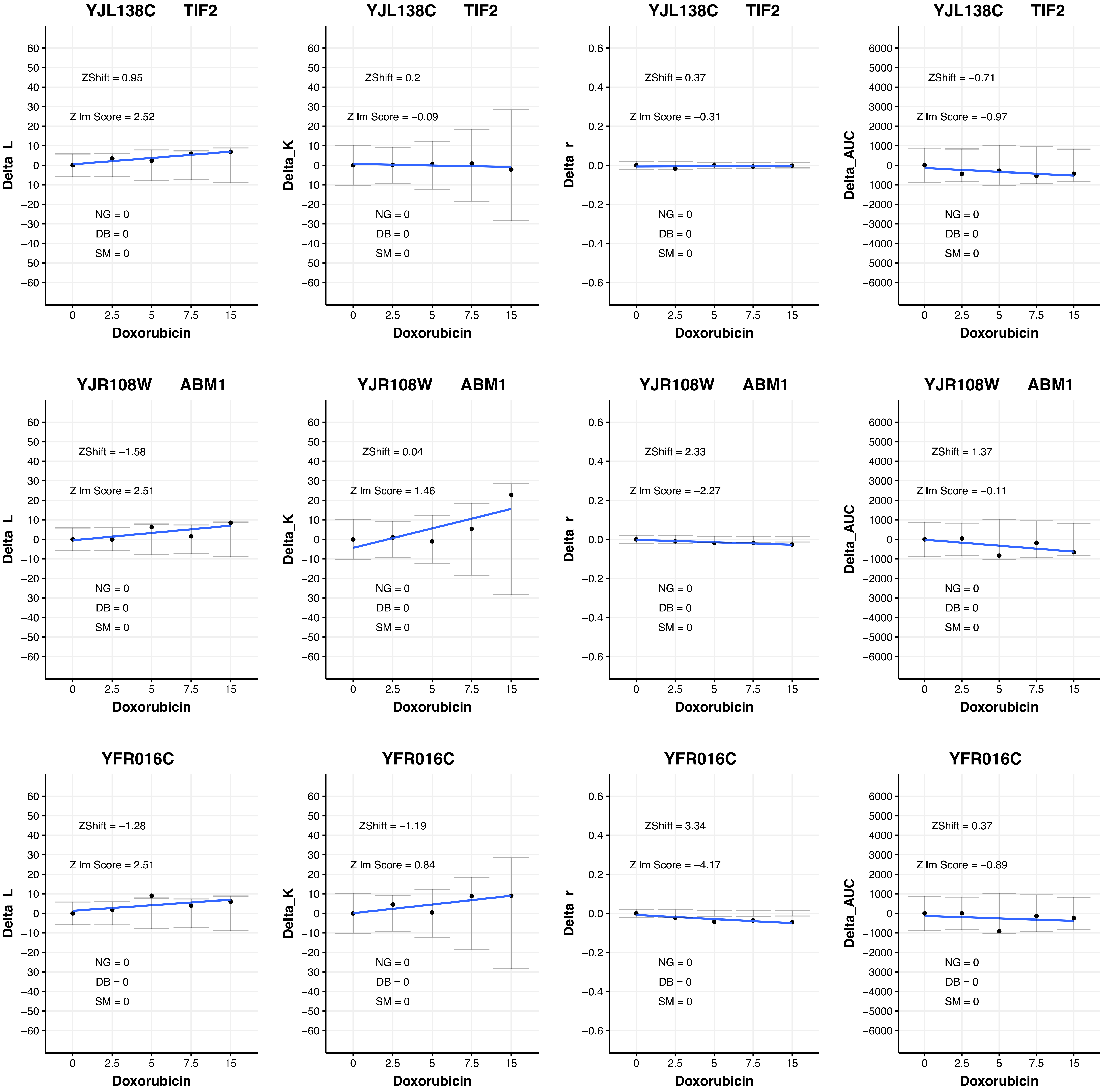
YJL138C TIF2
ZShift = 0.95
Z lm Score = 2.52
NG = 0
DB = 0
SM = 0
Delta_L
Doxorubicin
YJL138C TIF2
ZShift = 0.2
Z lm Score = −0.09
NG = 0
DB = 0
SM = 0
Delta_K
Doxorubicin
YJL138C TIF2
ZShift = 0.37
Z lm Score = −0.31
NG = 0
DB = 0
SM = 0
Delta_r
Doxorubicin
YJL138C TIF2
ZShift = −0.71
Z lm Score = −0.97
NG = 0
DB = 0
SM = 0
Delta_AUC
Doxorubicin
YJR108W ABM1
ZShift = −1.58
Z lm Score = 2.51
NG = 0
DB = 0
SM = 0
Delta_L
Doxorubicin
YJR108W ABM1
ZShift = 0.04
Z lm Score = 1.46
NG = 0
DB = 0
SM = 0
Delta_K
Doxorubicin
YJR108W ABM1
ZShift = 2.33
Z lm Score = −2.27
NG = 0
DB = 0
SM = 0
Delta_r
Doxorubicin
YJR108W ABM1
ZShift = 1.37
Z lm Score = −0.11
NG = 0
DB = 0
SM = 0
Delta_AUC
Doxorubicin
YFR016C
ZShift = −1.28
Z lm Score = 2.51
NG = 0
DB = 0
SM = 0
Delta_L
Doxorubicin
YFR016C
ZShift = −1.19
Z lm Score = 0.84
NG = 0
DB = 0
SM = 0
Delta_K
Doxorubicin
YFR016C
ZShift = 3.34
Z lm Score = −4.17
NG = 0
DB = 0
SM = 0
Delta_r
Doxorubicin
YFR016C
ZShift = 0.37
Z lm Score = −0.89
NG = 0
DB = 0
SM = 0
Delta_AUC
Doxorubicin

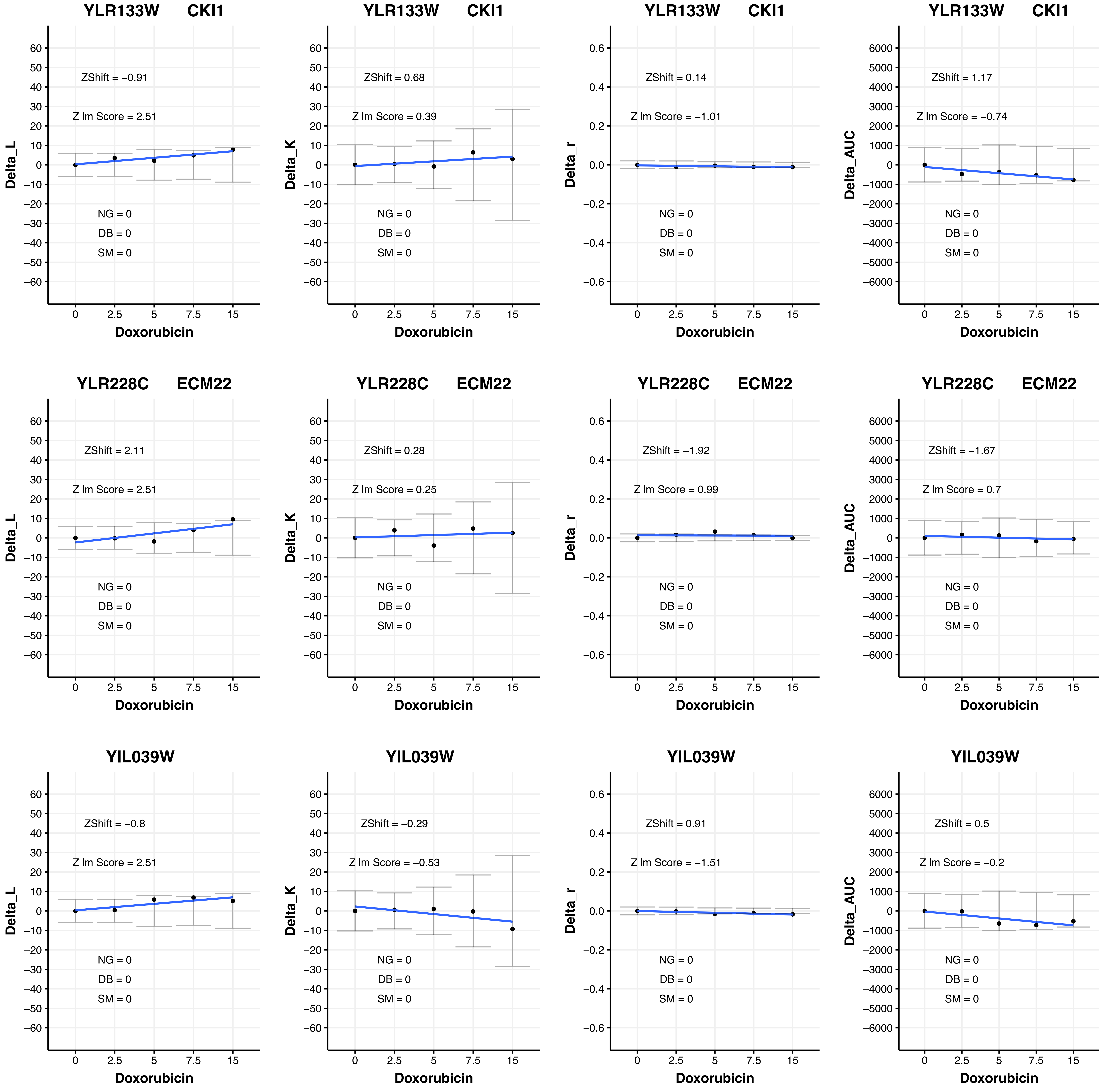

YLR133W CKI1
ZShift = −0.91
Z lm Score = 2.51
NG = 0
DB = 0
SM = 0
Delta_L
Doxorubicin
YLR133W CKI1
ZShift = 0.68
Z lm Score = 0.39
NG = 0
DB = 0
SM = 0
Delta_K
Doxorubicin
YLR133W CKI1
ZShift = 0.14
Z lm Score = −1.01
NG = 0
DB = 0
SM = 0
Delta_r
Doxorubicin
YLR133W CKI1
ZShift = 1.17
Z lm Score = −0.74
NG = 0
DB = 0
SM = 0
Delta_AUC
Doxorubicin
YLR228C ECM22
ZShift = 2.11
Z lm Score = 2.51
NG = 0
DB = 0
SM = 0
Delta_L
Doxorubicin
YLR228C ECM22
ZShift = 0.28
Z lm Score = 0.25
NG = 0
DB = 0
SM = 0
Delta_K
Doxorubicin
YLR228C ECM22
ZShift = −1.92
Z lm Score = 0.99
NG = 0
DB = 0
SM = 0
Delta_r
Doxorubicin
YLR228C ECM22
ZShift = −1.67
Z lm Score = 0.7
NG = 0
DB = 0
SM = 0
Delta_AUC
Doxorubicin
YIL039W
ZShift = −0.8
Z lm Score = 2.51
NG = 0
DB = 0
SM = 0
Delta_L
Doxorubicin
YIL039W
ZShift = −0.29
Z lm Score = −0.53
NG = 0
DB = 0
SM = 0
Delta_K
Doxorubicin
YIL039W
ZShift = 0.91
Z lm Score = −1.51
NG = 0
DB = 0
SM = 0
Delta_r
Doxorubicin
YIL039W
ZShift = 0.5
Z lm Score = −0.2
NG = 0
DB = 0
SM = 0
Delta_AUC
Doxorubicin

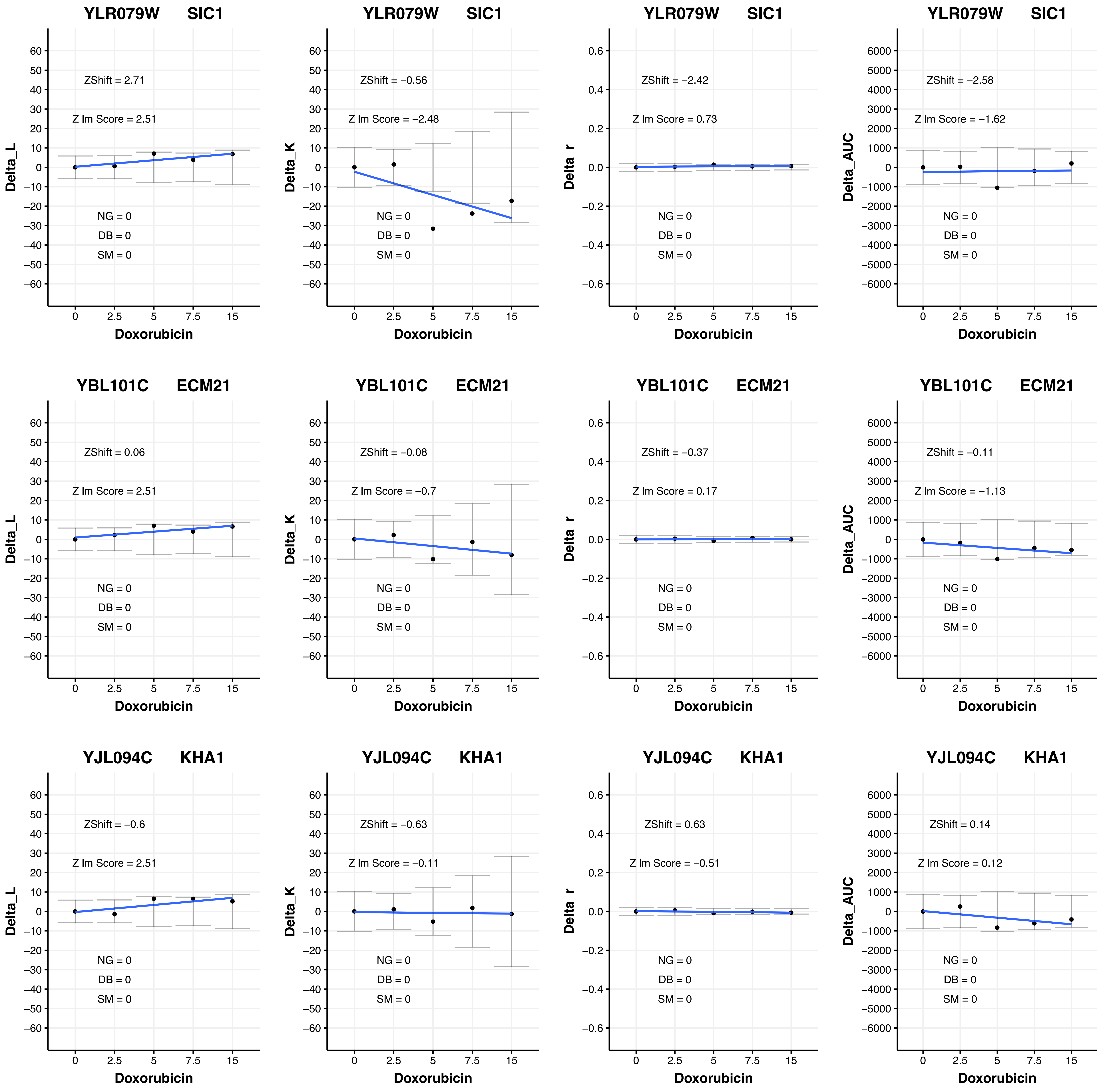
YLR079W SIC1
ZShift = 2.71
Z lm Score = 2.51
NG = 0
DB = 0
SM = 0
Delta_L
Doxorubicin
YLR079W SIC1
ZShift = −0.56
Z lm Score = −2.48
NG = 0
DB = 0
SM = 0
Delta_K
Doxorubicin
YLR079W SIC1
ZShift = −2.42
Z lm Score = 0.73
NG = 0
DB = 0
SM = 0
Delta_r
Doxorubicin
YLR079W SIC1
ZShift = −2.58
Z lm Score = −1.62
NG = 0
DB = 0
SM = 0
Delta_AUC
Doxorubicin
YBL101C ECM21
ZShift = 0.06
Z lm Score = 2.51
NG = 0
DB = 0
SM = 0
Delta_L
Doxorubicin
YBL101C ECM21
ZShift = −0.08
Z lm Score = −0.7
NG = 0
DB = 0
SM = 0
Delta_K
Doxorubicin
YBL101C ECM21
ZShift = −0.37
Z lm Score = 0.17
NG = 0
DB = 0
SM = 0
Delta_r
Doxorubicin
YBL101C ECM21
ZShift = −0.11
Z lm Score = −1.13
NG = 0
DB = 0
SM = 0
Delta_AUC
Doxorubicin
YJL094C KHA1
ZShift = −0.6
Z lm Score = 2.51
NG = 0
DB = 0
SM = 0
Delta_L
Doxorubicin
YJL094C KHA1
ZShift = −0.63
Z lm Score = −0.11
NG = 0
DB = 0
SM = 0
Delta_K
Doxorubicin
YJL094C KHA1
ZShift = 0.63
Z lm Score = −0.51
NG = 0
DB = 0
SM = 0
Delta_r
Doxorubicin
YJL094C KHA1
ZShift = 0.14
Z lm Score = 0.12
NG = 0
DB = 0
SM = 0
Delta_AUC
Doxorubicin

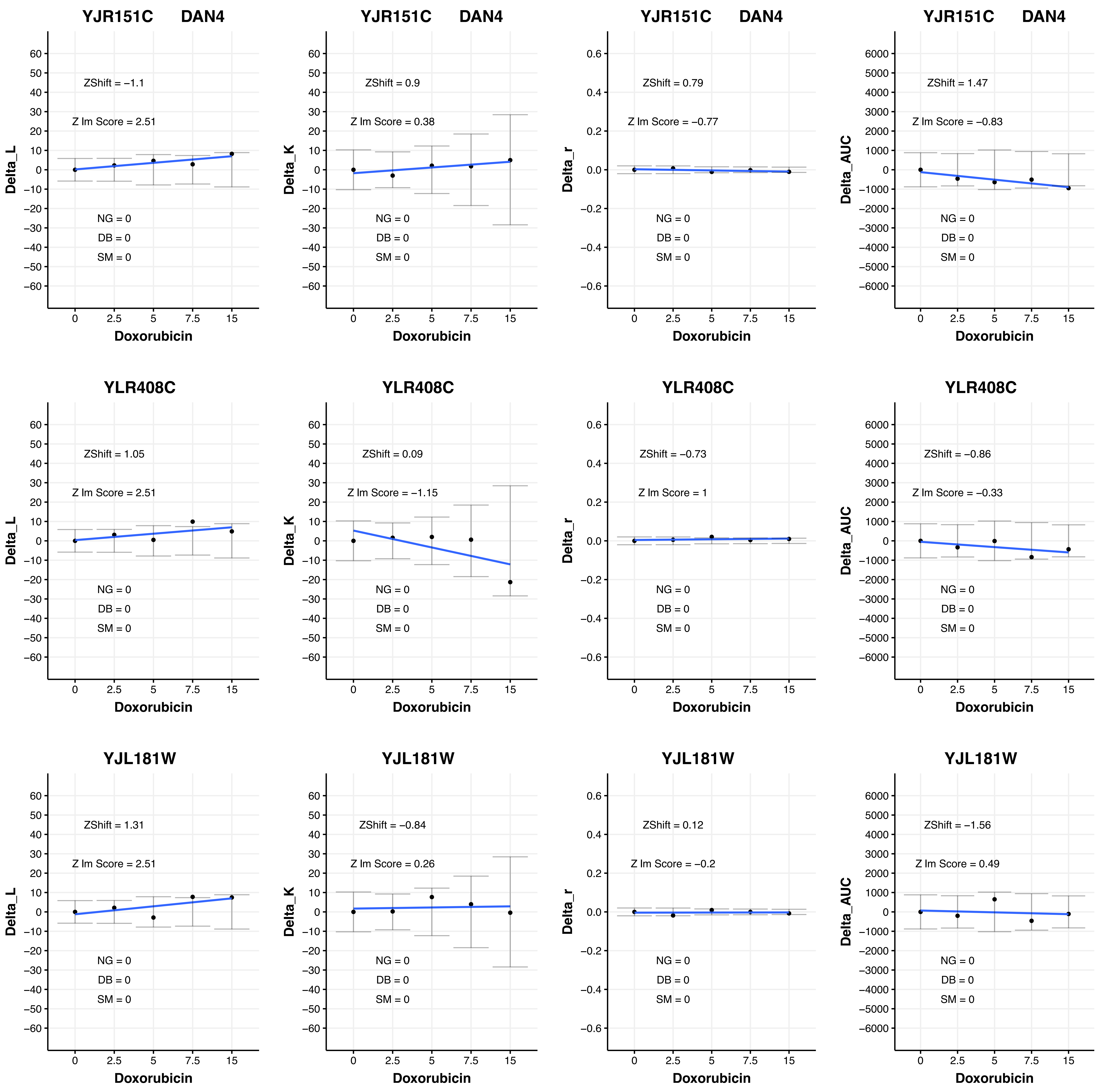

YJR151C DAN4
ZShift = −1.1
Z lm Score = 2.51
NG = 0
DB = 0
SM = 0
Delta_L
Doxorubicin
YJR151C DAN4
ZShift = 0.9
Z lm Score = 0.38
NG = 0
DB = 0
SM = 0
Delta_K
Doxorubicin
YJR151C DAN4
ZShift = 0.79
Z lm Score = −0.77
NG = 0
DB = 0
SM = 0
Delta_r
Doxorubicin
YJR151C DAN4
ZShift = 1.47
Z lm Score = −0.83
NG = 0
DB = 0
SM = 0
Delta_AUC
Doxorubicin
YLR408C
ZShift = 1.05
Z lm Score = 2.51
NG = 0
DB = 0
SM = 0
Delta_L
Doxorubicin
YLR408C
ZShift = 0.09
Z lm Score = −1.15
NG = 0
DB = 0
SM = 0
Delta_K
Doxorubicin
YLR408C
ZShift = −0.73
Z lm Score = 1
NG = 0
DB = 0
SM = 0
Delta_r
Doxorubicin
YLR408C
ZShift = −0.86
Z lm Score = −0.33
NG = 0
DB = 0
SM = 0
Delta_AUC
Doxorubicin
YJL181W
ZShift = 1.31
Z lm Score = 2.51
NG = 0
DB = 0
SM = 0
Delta_L
Doxorubicin
YJL181W
ZShift = −0.84
Z lm Score = 0.26
NG = 0
DB = 0
SM = 0
Delta_K
Doxorubicin
YJL181W
ZShift = 0.12
Z lm Score = −0.2
NG = 0
DB = 0
SM = 0
Delta_r
Doxorubicin
YJL181W
ZShift = −1.56
Z lm Score = 0.49
NG = 0
DB = 0
SM = 0
Delta_AUC
Doxorubicin

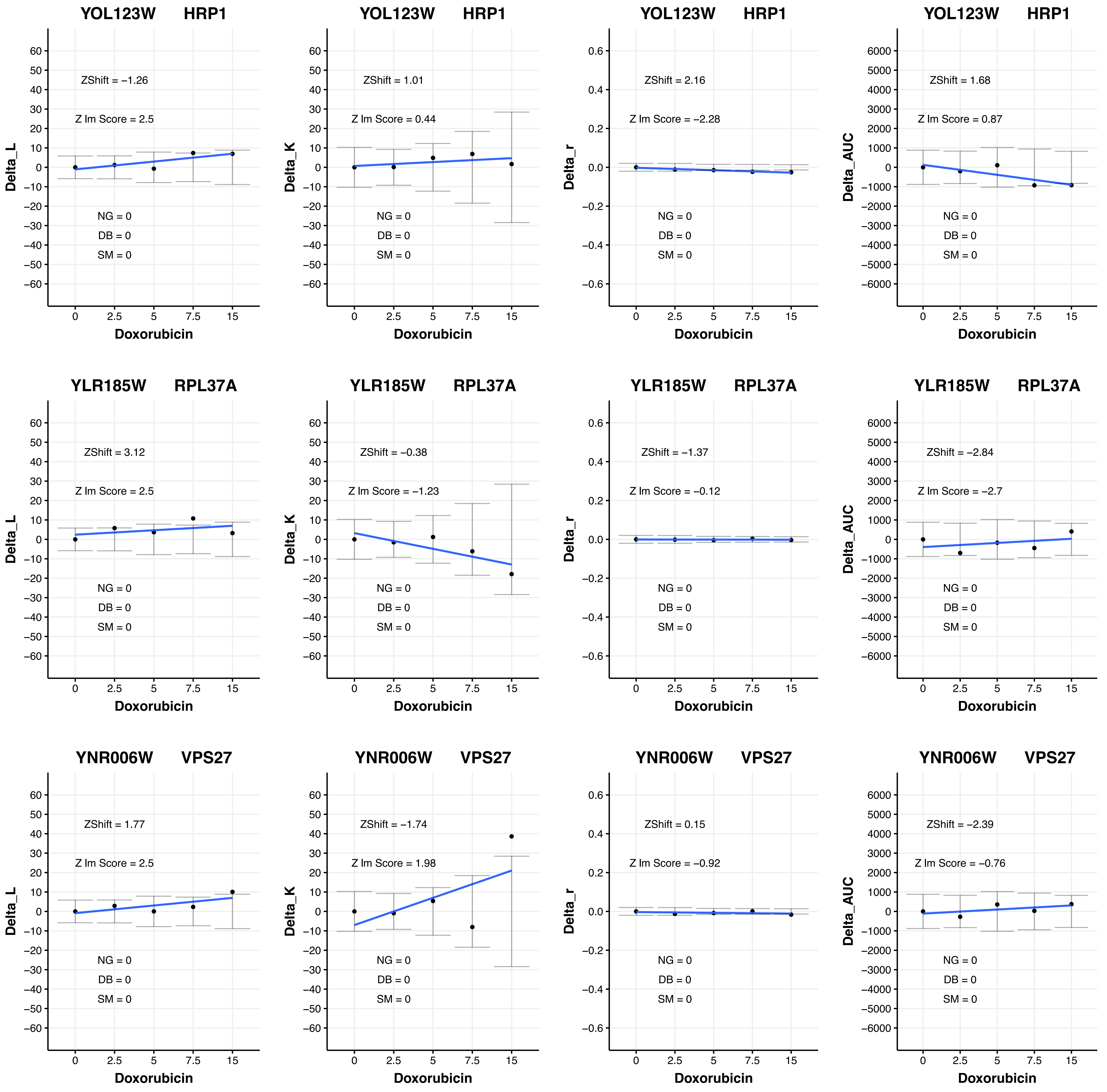

YOL123W HRP1
ZShift = −1.26
Z lm Score = 2.5
NG = 0
DB = 0
SM = 0
Delta_L
Doxorubicin
YOL123W HRP1
ZShift = 1.01
Z lm Score = 0.44
NG = 0
DB = 0
SM = 0
Delta_K
Doxorubicin
YOL123W HRP1
ZShift = 2.16
Z lm Score = −2.28
NG = 0
DB = 0
SM = 0
Delta_r
Doxorubicin
YOL123W HRP1
ZShift = 1.68
Z lm Score = 0.87
NG = 0
DB = 0
SM = 0
Delta_AUC
Doxorubicin
YLR185W RPL37A
ZShift = 3.12
Z lm Score = 2.5
NG = 0
DB = 0
SM = 0
Delta_L
Doxorubicin
YLR185W RPL37A
ZShift = −0.38
Z lm Score = −1.23
NG = 0
DB = 0
SM = 0
Delta_K
Doxorubicin
YLR185W RPL37A
ZShift = −1.37
Z lm Score = −0.12
NG = 0
DB = 0
SM = 0
Delta_r
Doxorubicin
YLR185W RPL37A
ZShift = −2.84
Z lm Score = −2.7
NG = 0
DB = 0
SM = 0
Delta_AUC
Doxorubicin
YNR006W VPS27
ZShift = 1.77
Z lm Score = 2.5
NG = 0
DB = 0
SM = 0
Delta_L
Doxorubicin
YNR006W VPS27
ZShift = −1.74
Z lm Score = 1.98
NG = 0
DB = 0
SM = 0
Delta_K
Doxorubicin
YNR006W VPS27
ZShift = 0.15
Z lm Score = −0.92
NG = 0
DB = 0
SM = 0
Delta_r
Doxorubicin
YNR006W VPS27
ZShift = −2.39
Z lm Score = −0.76
NG = 0
DB = 0
SM = 0
Delta_AUC
Doxorubicin

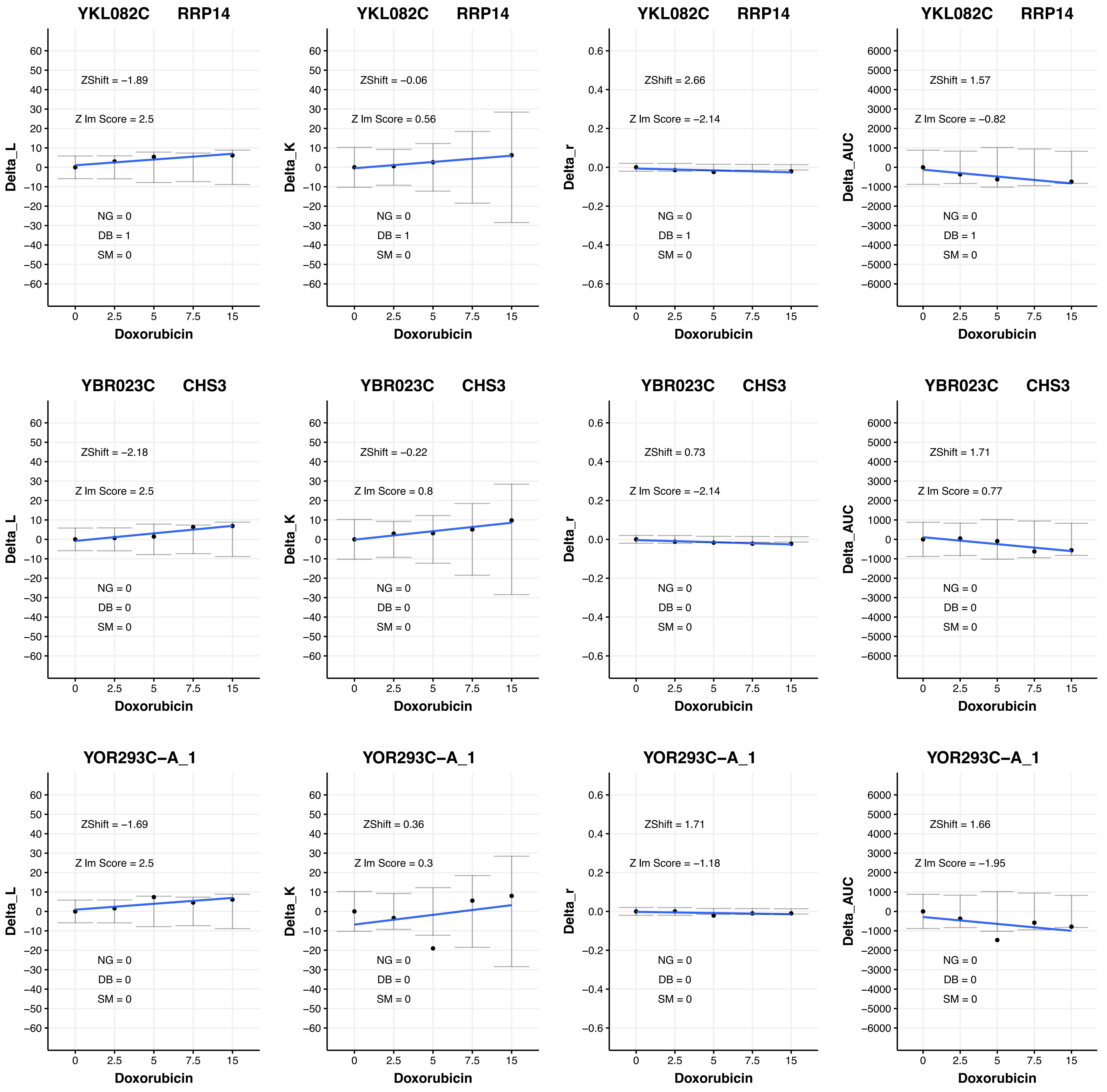

YKL082C RRP14
ZShift = −1.89
Z lm Score = 2.5
NG = 0
DB = 1
SM = 0
Delta_L
Doxorubicin
YKL082C RRP14
ZShift = −0.06
Z lm Score = 0.56
NG = 0
DB = 1
SM = 0
Delta_K
Doxorubicin
YKL082C RRP14
ZShift = 2.66
Z lm Score = −2.14
NG = 0
DB = 1
SM = 0
Delta_r
Doxorubicin
YKL082C RRP14
ZShift = 1.57
Z lm Score = −0.82
NG = 0
DB = 1
SM = 0
Delta_AUC
Doxorubicin
YBR023C CHS3
ZShift = −2.18
Z lm Score = 2.5
NG = 0
DB = 0
SM = 0
Delta_L
Doxorubicin
YBR023C CHS3
ZShift = −0.22
Z lm Score = 0.8
NG = 0
DB = 0
SM = 0
Delta_K
Doxorubicin
YBR023C CHS3
ZShift = 0.73
Z lm Score = −2.14
NG = 0
DB = 0
SM = 0
Delta_r
Doxorubicin
YBR023C CHS3
ZShift = 1.71
Z lm Score = 0.77
NG = 0
DB = 0
SM = 0
Delta_AUC
Doxorubicin
YOR293C−A_1
ZShift = −1.69
Z lm Score = 2.5
NG = 0
DB = 0
SM = 0
Delta_L
Doxorubicin
YOR293C−A_1
ZShift = 0.36
Z lm Score = 0.3
NG = 0
DB = 0
SM = 0
Delta_K
Doxorubicin
YOR293C−A_1
ZShift = 1.71
Z lm Score = −1.18
NG = 0
DB = 0
SM = 0
Delta_r
Doxorubicin
YOR293C−A_1
ZShift = 1.66
Z lm Score = −1.95
NG = 0
DB = 0
SM = 0
Delta_AUC
Doxorubicin

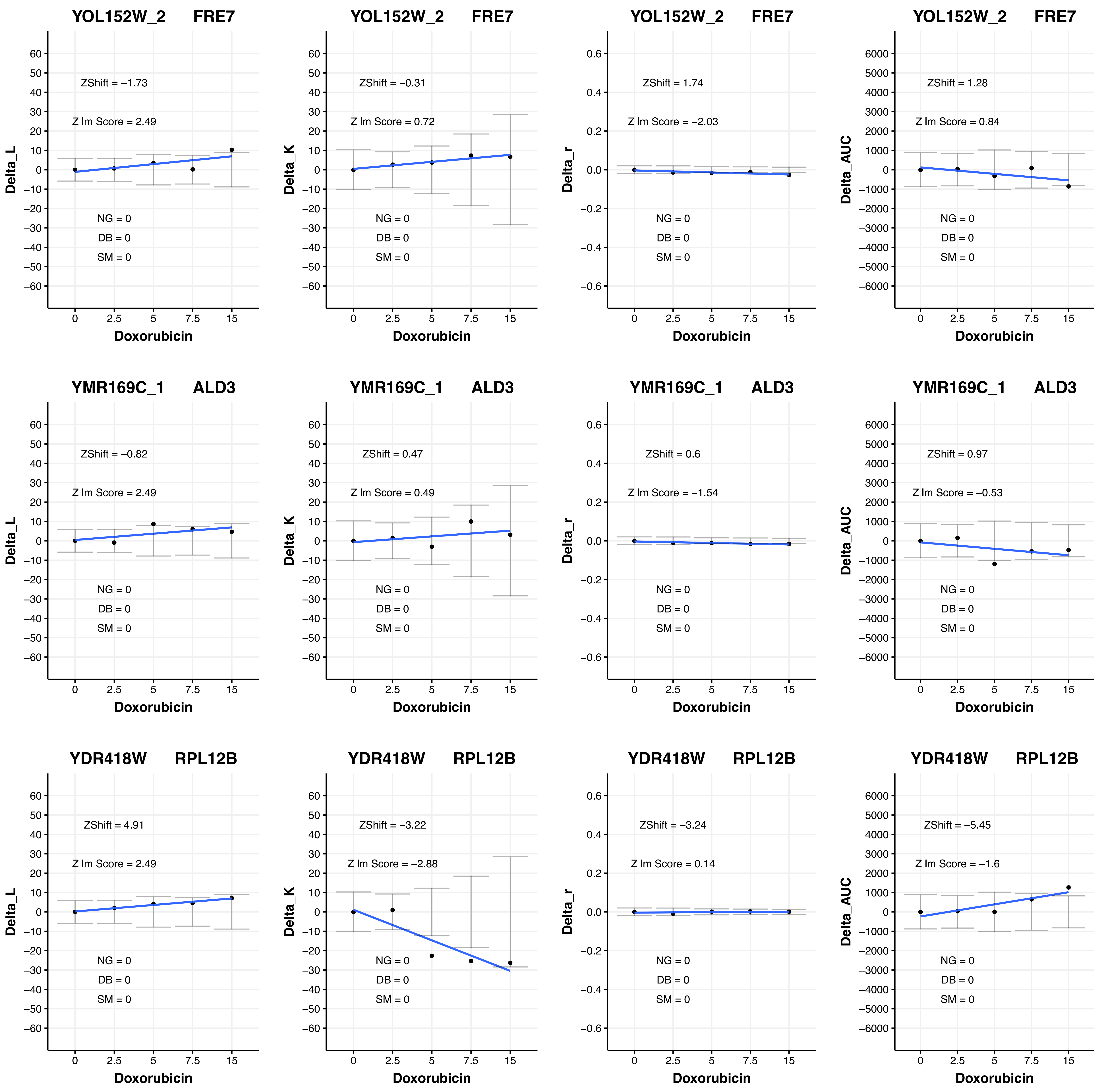
YOL152W_2 FRE7
ZShift = −1.73
Z lm Score = 2.49
NG = 0
DB = 0
SM = 0
Delta_L
Doxorubicin
YOL152W_2 FRE7
ZShift = −0.31
Z lm Score = 0.72
NG = 0
DB = 0
SM = 0
Delta_K
Doxorubicin
YOL152W_2 FRE7
ZShift = 1.74
Z lm Score = −2.03
NG = 0
DB = 0
SM = 0
Delta_r
Doxorubicin
YOL152W_2 FRE7
ZShift = 1.28
Z lm Score = 0.84
NG = 0
DB = 0
SM = 0
Delta_AUC
Doxorubicin
YMR169C_1 ALD3
ZShift = −0.82
Z lm Score = 2.49
NG = 0
DB = 0
SM = 0
Delta_L
Doxorubicin
YMR169C_1 ALD3
ZShift = 0.47
Z lm Score = 0.49
NG = 0
DB = 0
SM = 0
Delta_K
Doxorubicin
YMR169C_1 ALD3
ZShift = 0.6
Z lm Score = −1.54
NG = 0
DB = 0
SM = 0
Delta_r
Doxorubicin
YMR169C_1 ALD3
ZShift = 0.97
Z lm Score = −0.53
NG = 0
DB = 0
SM = 0
Delta_AUC
Doxorubicin
YDR418W RPL12B
ZShift = 4.91
Z lm Score = 2.49
NG = 0
DB = 0
SM = 0
Delta_L
Doxorubicin
YDR418W RPL12B
ZShift = −3.22
Z lm Score = −2.88
NG = 0
DB = 0
SM = 0
Delta_K
Doxorubicin
YDR418W RPL12B
ZShift = −3.24
Z lm Score = 0.14
NG = 0
DB = 0
SM = 0
Delta_r
Doxorubicin
YDR418W RPL12B
ZShift = −5.45
Z lm Score = −1.6
NG = 0
DB = 0
SM = 0
Delta_AUC
Doxorubicin

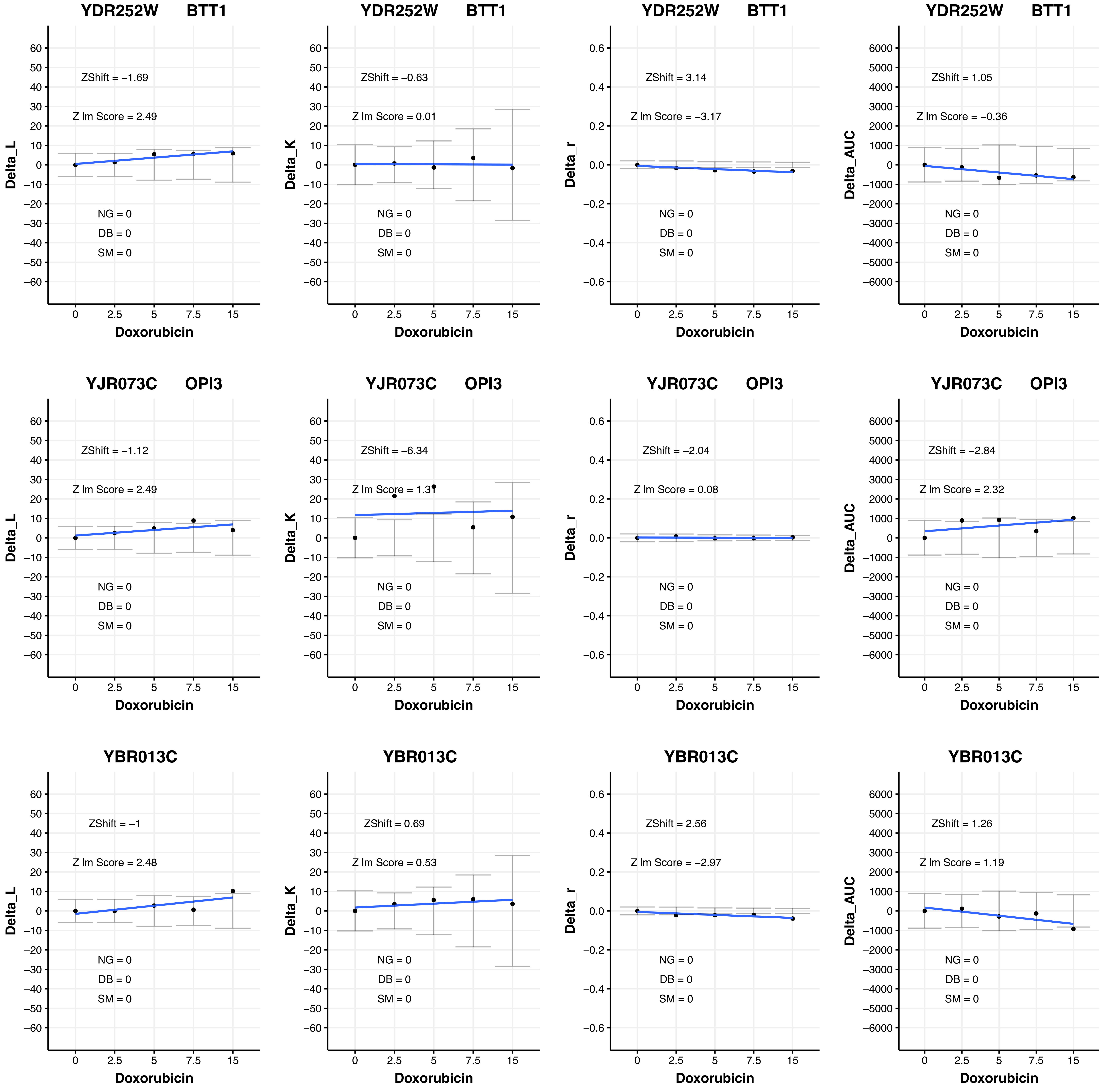

YDR252W BTT1
ZShift = −1.69
Z lm Score = 2.49
NG = 0
DB = 0
SM = 0
Delta_L
Doxorubicin
YDR252W BTT1
ZShift = −0.63
Z lm Score = 0.01
NG = 0
DB = 0
SM = 0
Delta_K
Doxorubicin
YDR252W BTT1
ZShift = 3.14
Z lm Score = −3.17
NG = 0
DB = 0
SM = 0
Delta_r
Doxorubicin
YDR252W BTT1
ZShift = 1.05
Z lm Score = −0.36
NG = 0
DB = 0
SM = 0
Delta_AUC
Doxorubicin
YJR073C OPI3
ZShift = −1.12
Z lm Score = 2.49
NG = 0
DB = 0
SM = 0
Delta_L
Doxorubicin
YJR073C OPI3
ZShift = −6.34
Z lm Score = 1.31
NG = 0
DB = 0
SM = 0
Delta_K
Doxorubicin
YJR073C OPI3
ZShift = −2.04
Z lm Score = 0.08
NG = 0
DB = 0
SM = 0
Delta_r
Doxorubicin
YJR073C OPI3
ZShift = −2.84
Z lm Score = 2.32
NG = 0
DB = 0
SM = 0
Delta_AUC
Doxorubicin
YBR013C
ZShift = −1
Z lm Score = 2.48
NG = 0
DB = 0
SM = 0
Delta_L
Doxorubicin
YBR013C
ZShift = 0.69
Z lm Score = 0.53
NG = 0
DB = 0
SM = 0
Delta_K
Doxorubicin
YBR013C
ZShift = 2.56
Z lm Score = −2.97
NG = 0
DB = 0
SM = 0
Delta_r
Doxorubicin
YBR013C
ZShift = 1.26
Z lm Score = 1.19
NG = 0
DB = 0
SM = 0
Delta_AUC
Doxorubicin

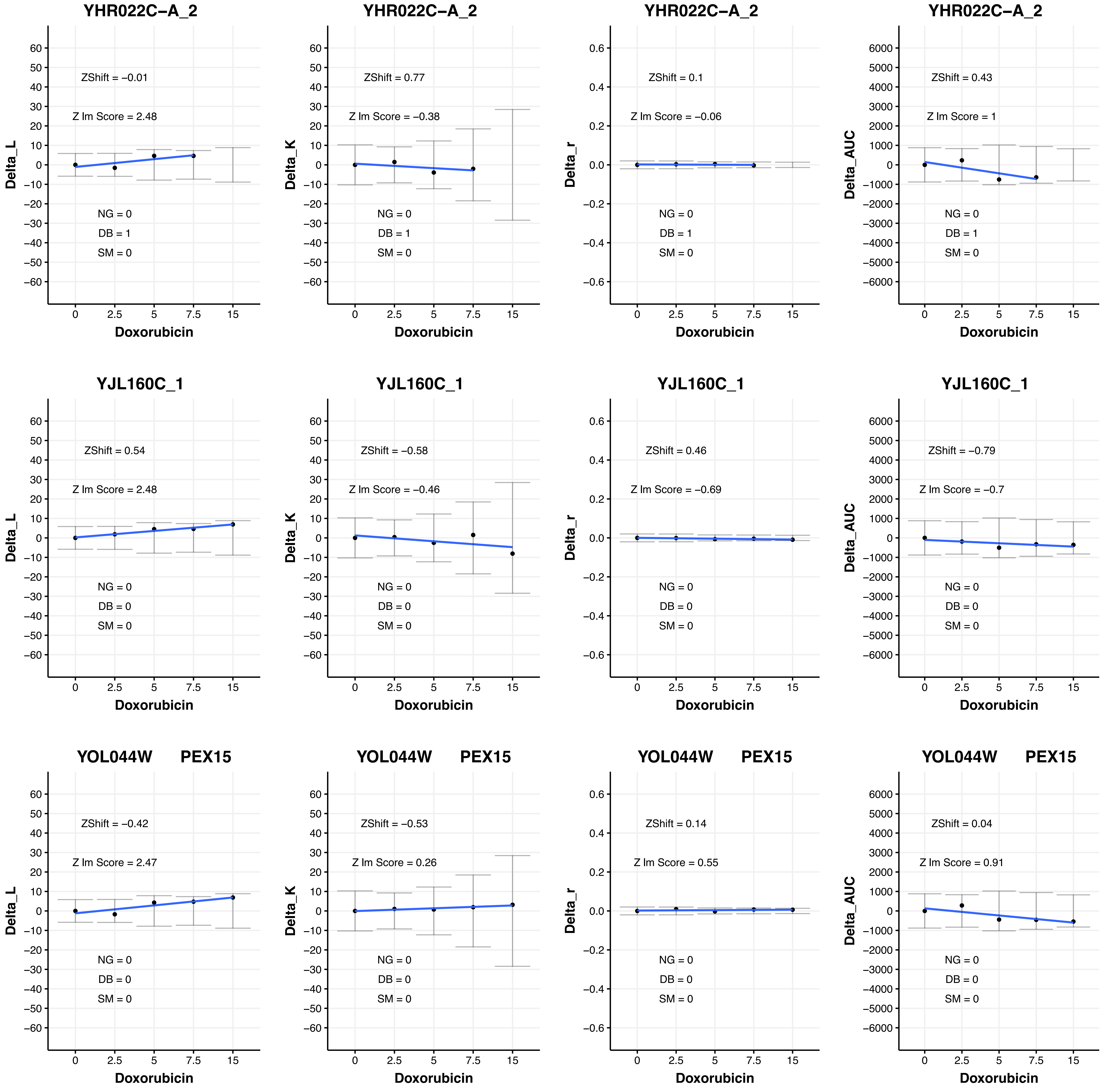
YHR022C−A_2
ZShift = −0.01
Z lm Score = 2.48
NG = 0
DB = 1
SM = 0
Delta_L
Doxorubicin
YHR022C−A_2
ZShift = 0.77
Z lm Score = −0.38
NG = 0
DB = 1
SM = 0
Delta_K
Doxorubicin
YHR022C−A_2
ZShift = 0.1
Z lm Score = −0.06
NG = 0
DB = 1
SM = 0
Delta_r
Doxorubicin
YHR022C−A_2
ZShift = 0.43
Z lm Score = 1
NG = 0
DB = 1
SM = 0
Delta_AUC
Doxorubicin
YJL160C_1
ZShift = 0.54
Z lm Score = 2.48
NG = 0
DB = 0
SM = 0
Delta_L
Doxorubicin
YJL160C_1
ZShift = −0.58
Z lm Score = −0.46
NG = 0
DB = 0
SM = 0
Delta_K
Doxorubicin
YJL160C_1
ZShift = 0.46
Z lm Score = −0.69
NG = 0
DB = 0
SM = 0
Delta_r
Doxorubicin
YJL160C_1
ZShift = −0.79
Z lm Score = −0.7
NG = 0
DB = 0
SM = 0
Delta_AUC
Doxorubicin
YOL044W PEX15
ZShift = −0.42
Z lm Score = 2.47
NG = 0
DB = 0
SM = 0
Delta_L
Doxorubicin
YOL044W PEX15
ZShift = −0.53
Z lm Score = 0.26
NG = 0
DB = 0
SM = 0
Delta_K
Doxorubicin
YOL044W PEX15
ZShift = 0.14
Z lm Score = 0.55
NG = 0
DB = 0
SM = 0
Delta_r
Doxorubicin
YOL044W PEX15
ZShift = 0.04
Z lm Score = 0.91
NG = 0
DB = 0
SM = 0
Delta_AUC
Doxorubicin

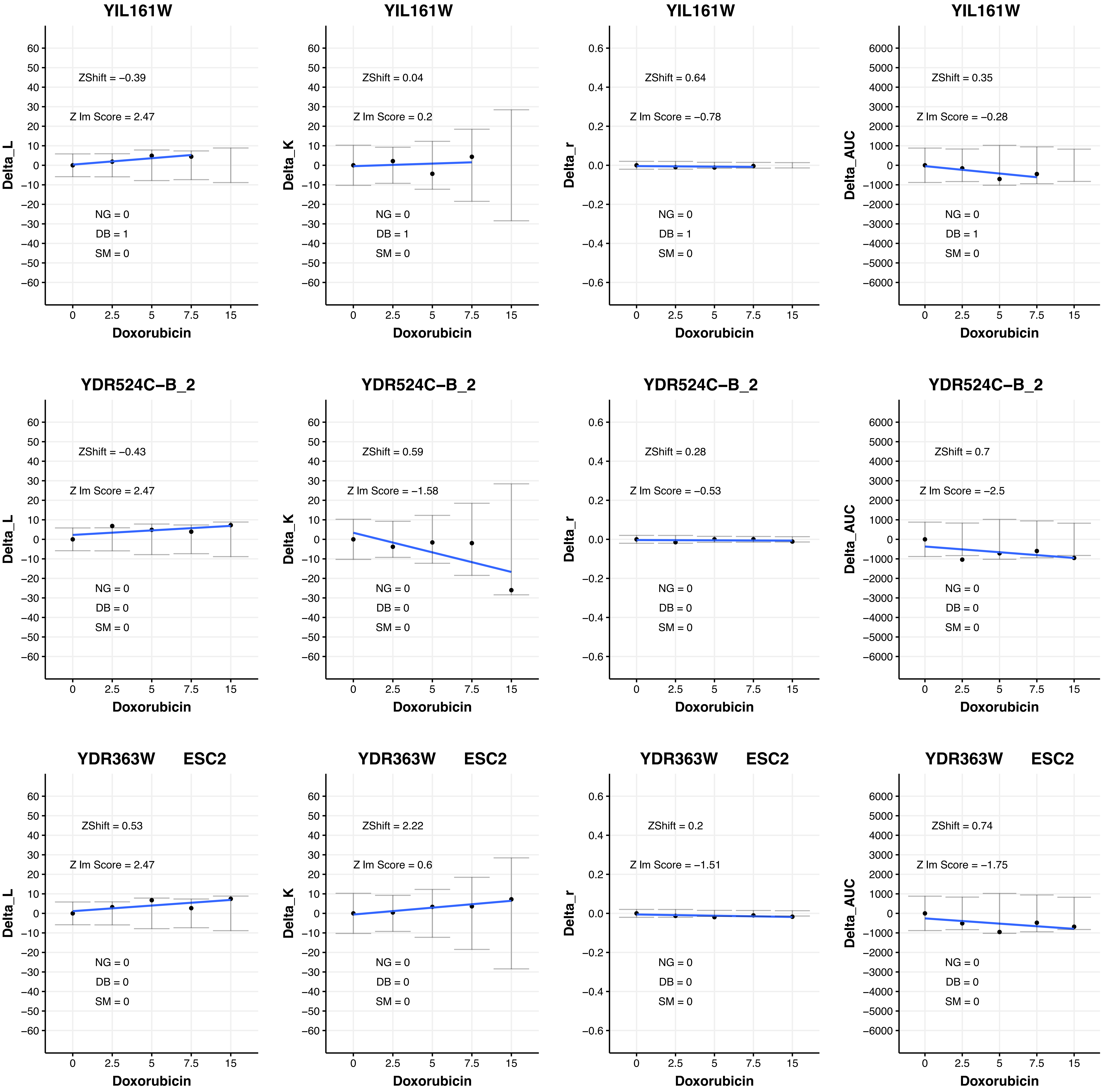

YIL161W
ZShift = −0.39
Z lm Score = 2.47
NG = 0
DB = 1
SM = 0
Delta_L
Doxorubicin
YIL161W
ZShift = 0.04
Z lm Score = 0.2
NG = 0
DB = 1
SM = 0
Delta_K
Doxorubicin
YIL161W
ZShift = 0.64
Z lm Score = −0.78
NG = 0
DB = 1
SM = 0
Delta_r
Doxorubicin
YIL161W
ZShift = 0.35
Z lm Score = −0.28
NG = 0
DB = 1
SM = 0
Delta_AUC
Doxorubicin
YDR524C−B_2
ZShift = −0.43
Z lm Score = 2.47
NG = 0
DB = 0
SM = 0
Delta_L
Doxorubicin
YDR524C−B_2
ZShift = 0.59
Z lm Score = −1.58
NG = 0
DB = 0
SM = 0
Delta_K
Doxorubicin
YDR524C−B_2
ZShift = 0.28
Z lm Score = −0.53
NG = 0
DB = 0
SM = 0
Delta_r
Doxorubicin
YDR524C−B_2
ZShift = 0.7
Z lm Score = −2.5
NG = 0
DB = 0
SM = 0
Delta_AUC
Doxorubicin
YDR363W ESC2
ZShift = 0.53
Z lm Score = 2.47
NG = 0
DB = 0
SM = 0
Delta_L
Doxorubicin
YDR363W ESC2
ZShift = 2.22
Z lm Score = 0.6
NG = 0
DB = 0
SM = 0
Delta_K
Doxorubicin
YDR363W ESC2
ZShift = 0.2
Z lm Score = −1.51
NG = 0
DB = 0
SM = 0
Delta_r
Doxorubicin
YDR363W ESC2
ZShift = 0.74
Z lm Score = −1.75
NG = 0
DB = 0
SM = 0
Delta_AUC
Doxorubicin

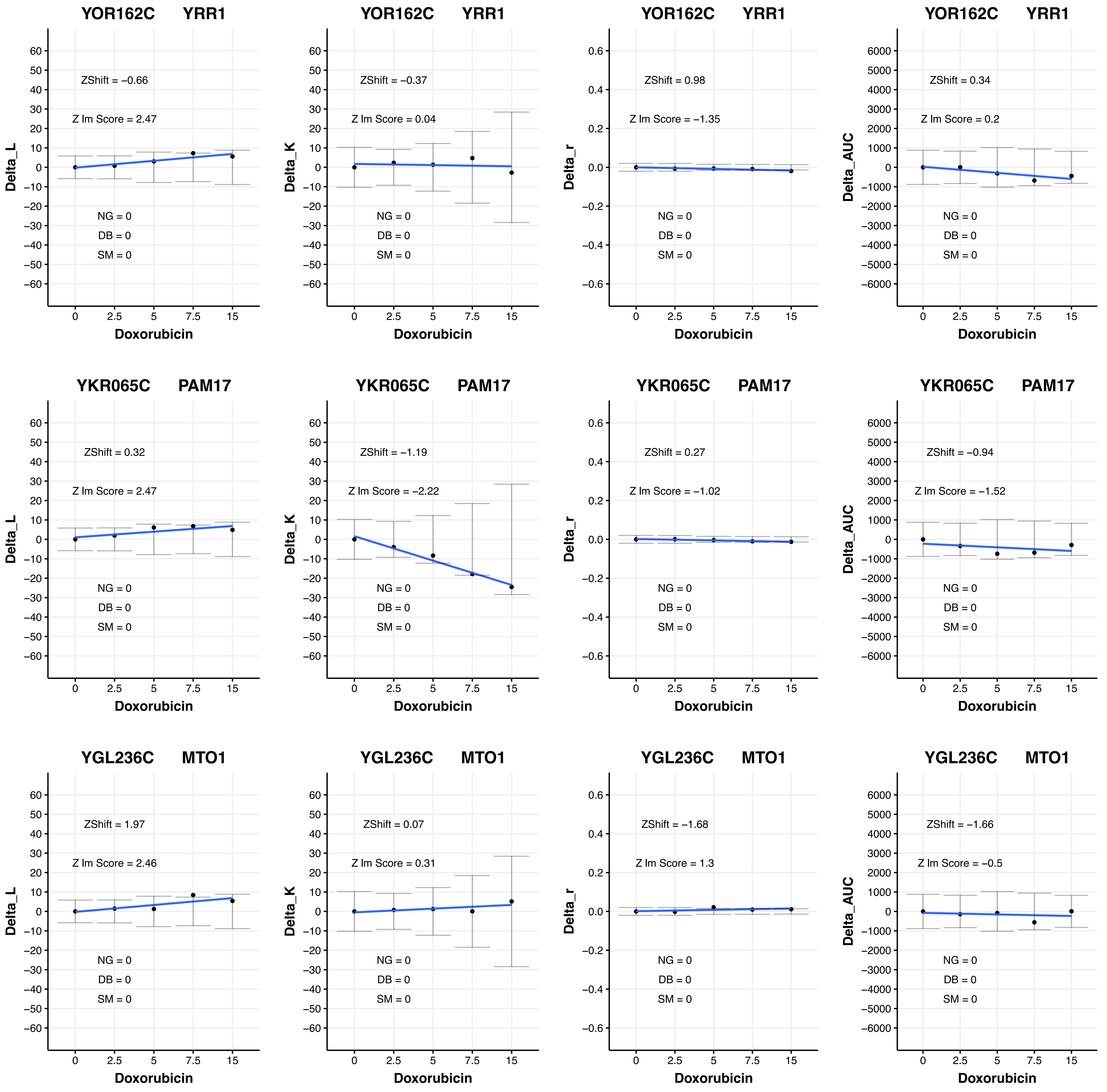

YOR162C YRR1
ZShift = −0.66
Z lm Score = 2.47
NG = 0
DB = 0
SM = 0
Delta_L
Doxorubicin
YOR162C YRR1
ZShift = −0.37
Z lm Score = 0.04
NG = 0
DB = 0
SM = 0
Delta_K
Doxorubicin
YOR162C YRR1
ZShift = 0.98
Z lm Score = −1.35
NG = 0
DB = 0
SM = 0
Delta_r
Doxorubicin
YOR162C YRR1
ZShift = 0.34
Z lm Score = 0.2
NG = 0
DB = 0
SM = 0
Delta_AUC
Doxorubicin
YKR065C PAM17
ZShift = 0.32
Z lm Score = 2.47
NG = 0
DB = 0
SM = 0
Delta_L
Doxorubicin
YKR065C PAM17
ZShift = −1.19
Z lm Score = −2.22
NG = 0
DB = 0
SM = 0
Delta_K
Doxorubicin
YKR065C PAM17
ZShift = 0.27
Z lm Score = −1.02
NG = 0
DB = 0
SM = 0
Delta_r
Doxorubicin
YKR065C PAM17
ZShift = −0.94
Z lm Score = −1.52
NG = 0
DB = 0
SM = 0
Delta_AUC
Doxorubicin
YGL236C MTO1
ZShift = 1.97
Z lm Score = 2.46
NG = 0
DB = 0
SM = 0
Delta_L
Doxorubicin
YGL236C MTO1
ZShift = 0.07
Z lm Score = 0.31
NG = 0
DB = 0
SM = 0
Delta_K
Doxorubicin
YGL236C MTO1
ZShift = −1.68
Z lm Score = 1.3
NG = 0
DB = 0
SM = 0
Delta_r
Doxorubicin
YGL236C MTO1
ZShift = −1.66
Z lm Score = −0.5
NG = 0
DB = 0
SM = 0
Delta_AUC
Doxorubicin

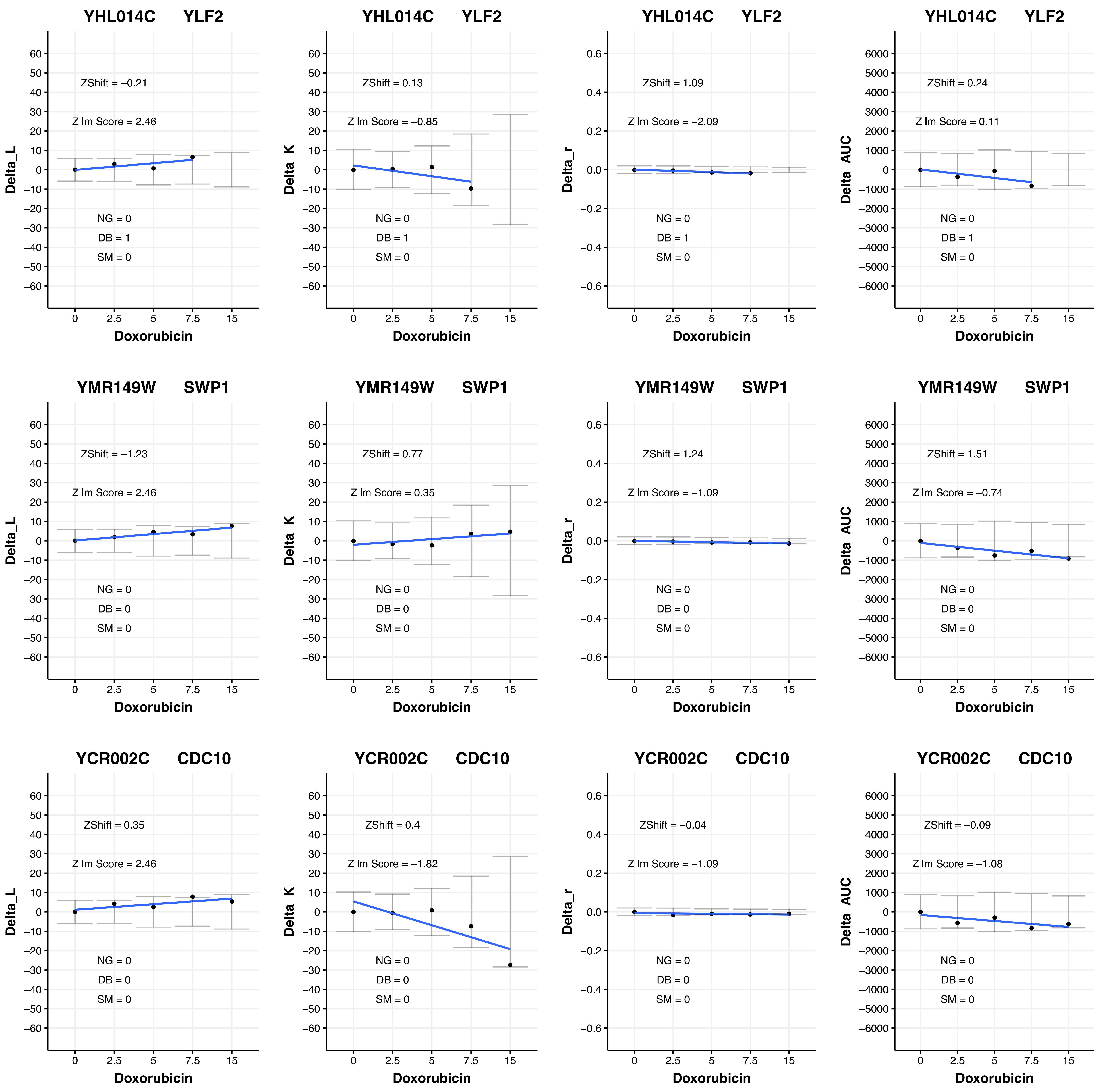

YHL014C YLF2
ZShift = −0.21
Z lm Score = 2.46
NG = 0
DB = 1
SM = 0
Delta_L
Doxorubicin
YHL014C YLF2
ZShift = 0.13
Z lm Score = −0.85
NG = 0
DB = 1
SM = 0
Delta_K
Doxorubicin
YHL014C YLF2
ZShift = 1.09
Z lm Score = −2.09
NG = 0
DB = 1
SM = 0
Delta_r
Doxorubicin
YHL014C YLF2
ZShift = 0.24
Z lm Score = 0.11
NG = 0
DB = 1
SM = 0
Delta_AUC
Doxorubicin
YMR149W SWP1
ZShift = −1.23
Z lm Score = 2.46
NG = 0
DB = 0
SM = 0
Delta_L
Doxorubicin
YMR149W SWP1
ZShift = 0.77
Z lm Score = 0.35
NG = 0
DB = 0
SM = 0
Delta_K
Doxorubicin
YMR149W SWP1
ZShift = 1.24
Z lm Score = −1.09
NG = 0
DB = 0
SM = 0
Delta_r
Doxorubicin
YMR149W SWP1
ZShift = 1.51
Z lm Score = −0.74
NG = 0
DB = 0
SM = 0
Delta_AUC
Doxorubicin
YCR002C CDC10
ZShift = 0.35
Z lm Score = 2.46
NG = 0
DB = 0
SM = 0
Delta_L
Doxorubicin
YCR002C CDC10
ZShift = 0.4
Z lm Score = −1.82
NG = 0
DB = 0
SM = 0
Delta_K
Doxorubicin
YCR002C CDC10
ZShift = −0.04
Z lm Score = −1.09
NG = 0
DB = 0
SM = 0
Delta_r
Doxorubicin
YCR002C CDC10
ZShift = −0.09
Z lm Score = −1.08
NG = 0
DB = 0
SM = 0
Delta_AUC
Doxorubicin

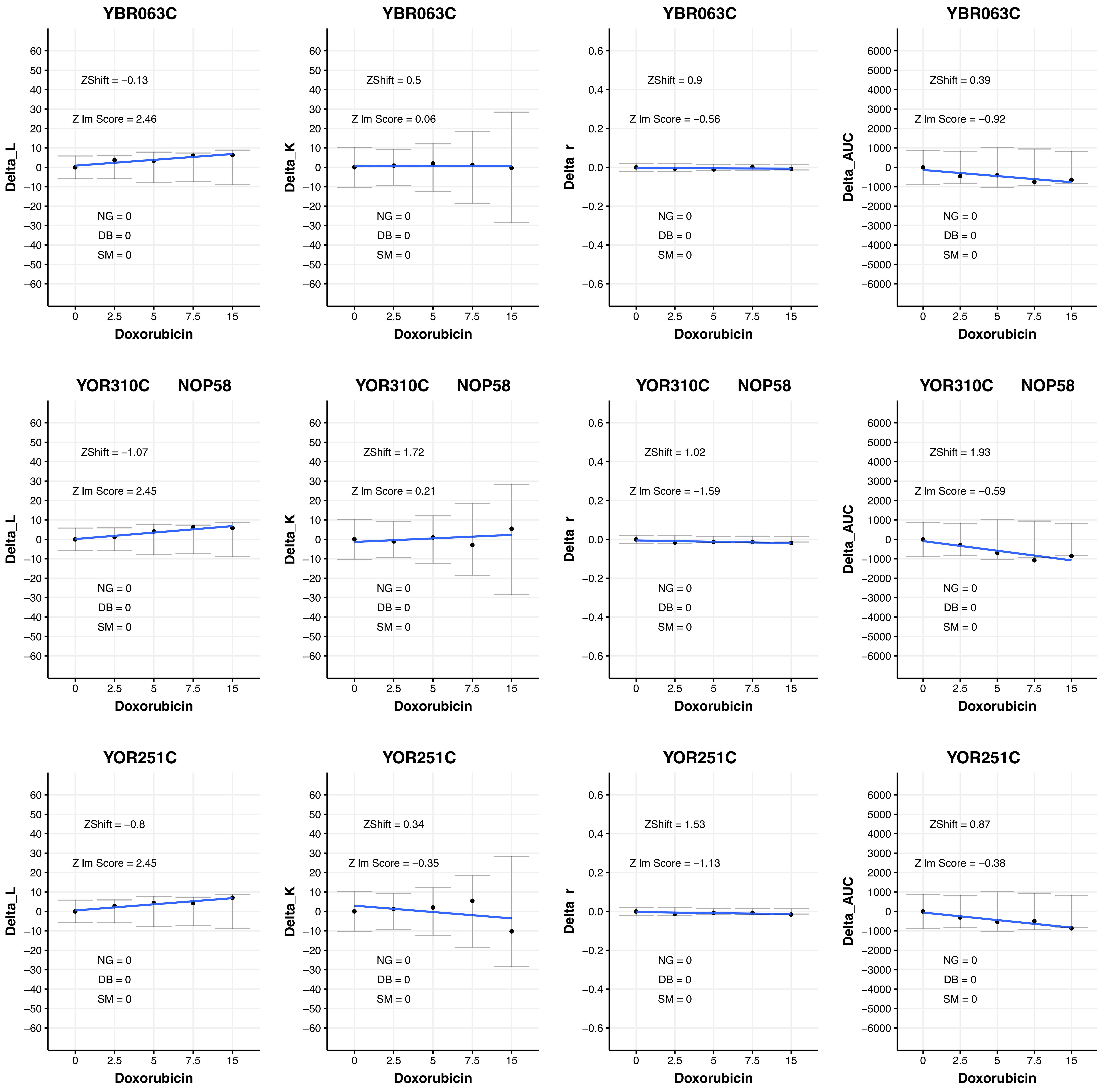
YBR063C
ZShift = −0.13
Z lm Score = 2.46
NG = 0
DB = 0
SM = 0
Delta_L
Doxorubicin
YBR063C
ZShift = 0.5
Z lm Score = 0.06
NG = 0
DB = 0
SM = 0
Delta_K
Doxorubicin
YBR063C
ZShift = 0.9
Z lm Score = −0.56
NG = 0
DB = 0
SM = 0
Delta_r
Doxorubicin
YBR063C
ZShift = 0.39
Z lm Score = −0.92
NG = 0
DB = 0
SM = 0
Delta_AUC
Doxorubicin
YOR310C NOP58
ZShift = −1.07
Z lm Score = 2.45
NG = 0
DB = 0
SM = 0
Delta_L
Doxorubicin
YOR310C NOP58
ZShift = 1.72
Z lm Score = 0.21
NG = 0
DB = 0
SM = 0
Delta_K
Doxorubicin
YOR310C NOP58
ZShift = 1.02
Z lm Score = −1.59
NG = 0
DB = 0
SM = 0
Delta_r
Doxorubicin
YOR310C NOP58
ZShift = 1.93
Z lm Score = −0.59
NG = 0
DB = 0
SM = 0
Delta_AUC
Doxorubicin
YOR251C
ZShift = −0.8
Z lm Score = 2.45
NG = 0
DB = 0
SM = 0
Delta_L
Doxorubicin
YOR251C
ZShift = 0.34
Z lm Score = −0.35
NG = 0
DB = 0
SM = 0
Delta_K
Doxorubicin
YOR251C
ZShift = 1.53
Z lm Score = −1.13
NG = 0
DB = 0
SM = 0
Delta_r
Doxorubicin
YOR251C
ZShift = 0.87
Z lm Score = −0.38
NG = 0
DB = 0
SM = 0
Delta_AUC
Doxorubicin

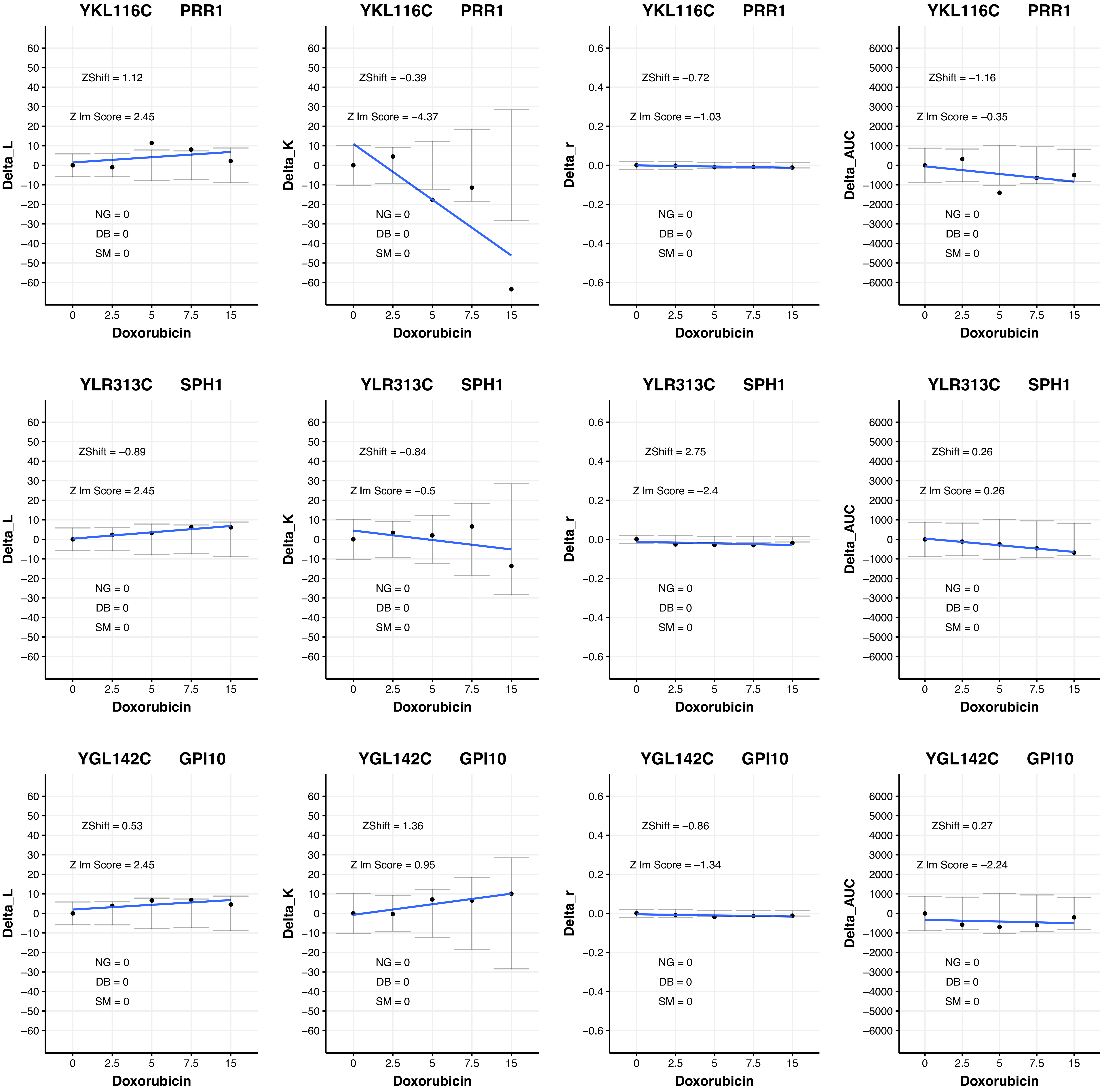

YKL116C PRR1
ZShift = 1.12
Z lm Score = 2.45
NG = 0
DB = 0
SM = 0
Delta_L
Doxorubicin
YKL116C PRR1
ZShift = −0.39
Z lm Score = −4.37
NG = 0
DB = 0
SM = 0
Delta_K
Doxorubicin
YKL116C PRR1
ZShift = −0.72
Z lm Score = −1.03
NG = 0
DB = 0
SM = 0
Delta_r
Doxorubicin
YKL116C PRR1
ZShift = −1.16
Z lm Score = −0.35
NG = 0
DB = 0
SM = 0
Delta_AUC
Doxorubicin
YLR313C SPH1
ZShift = −0.89
Z lm Score = 2.45
NG = 0
DB = 0
SM = 0
Delta_L
Doxorubicin
YLR313C SPH1
ZShift = −0.84
Z lm Score = −0.5
NG = 0
DB = 0
SM = 0
Delta_K
Doxorubicin
YLR313C SPH1
ZShift = 2.75
Z lm Score = −2.4
NG = 0
DB = 0
SM = 0
Delta_r
Doxorubicin
YLR313C SPH1
ZShift = 0.26
Z lm Score = 0.26
NG = 0
DB = 0
SM = 0
Delta_AUC
Doxorubicin
YGL142C GPI10
ZShift = 0.53
Z lm Score = 2.45
NG = 0
DB = 0
SM = 0
Delta_L
Doxorubicin
YGL142C GPI10
ZShift = 1.36
Z lm Score = 0.95
NG = 0
DB = 0
SM = 0
Delta_K
Doxorubicin
YGL142C GPI10
ZShift = −0.86
Z lm Score = −1.34
NG = 0
DB = 0
SM = 0
Delta_r
Doxorubicin
YGL142C GPI10
ZShift = 0.27
Z lm Score = −2.24
NG = 0
DB = 0
SM = 0
Delta_AUC
Doxorubicin

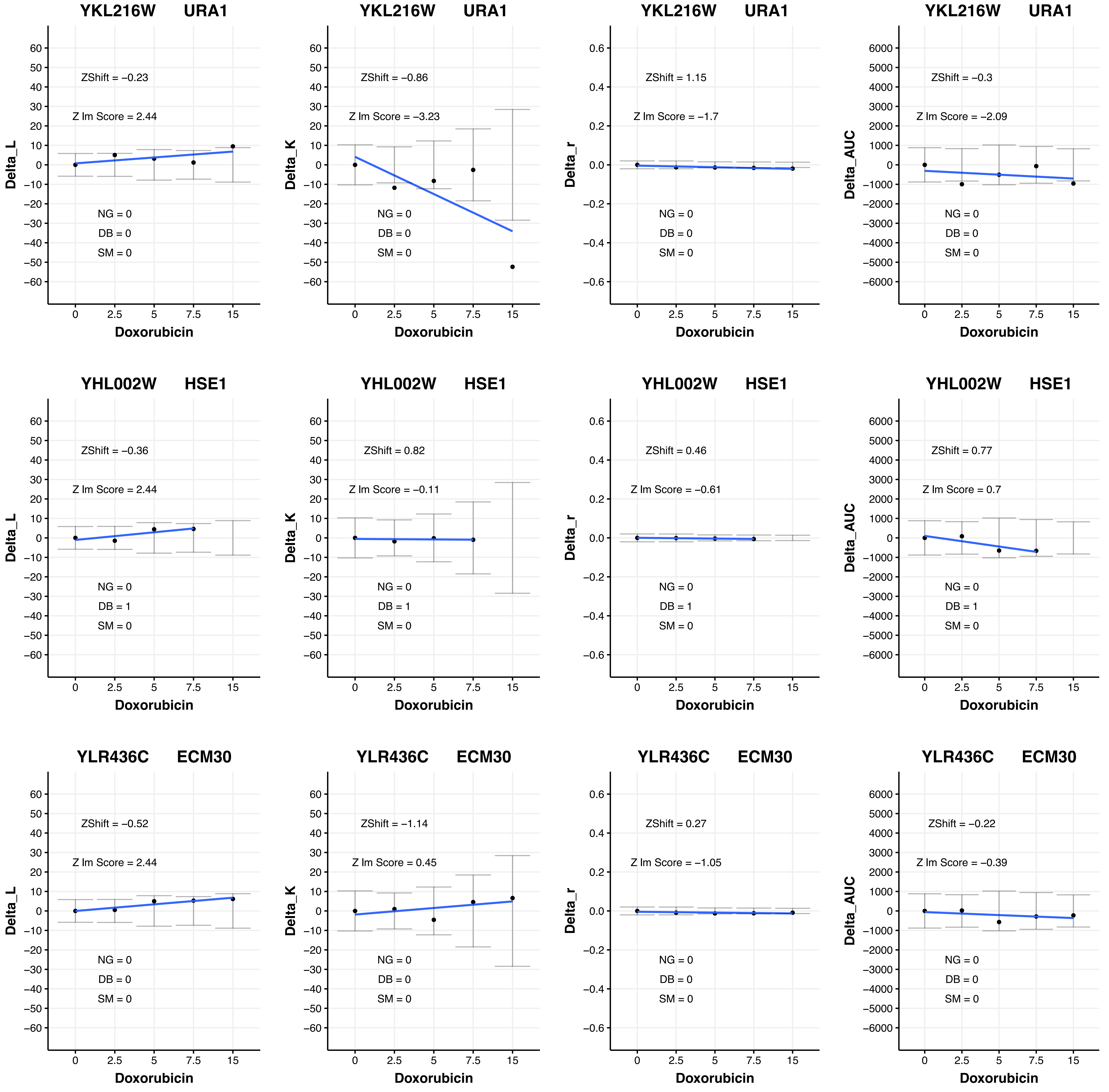
YKL216W URA1
ZShift = −0.23
Z lm Score = 2.44
NG = 0
DB = 0
SM = 0
Delta_L
Doxorubicin
YKL216W URA1
ZShift = −0.86
Z lm Score = −3.23
NG = 0
DB = 0
SM = 0
Delta_K
Doxorubicin
YKL216W URA1
ZShift = 1.15
Z lm Score = −1.7
NG = 0
DB = 0
SM = 0
Delta_r
Doxorubicin
YKL216W URA1
ZShift = −0.3
Z lm Score = −2.09
NG = 0
DB = 0
SM = 0
Delta_AUC
Doxorubicin
YHL002W HSE1
ZShift = −0.36
Z lm Score = 2.44
NG = 0
DB = 1
SM = 0
Delta_L
Doxorubicin
YHL002W HSE1
ZShift = 0.82
Z lm Score = −0.11
NG = 0
DB = 1
SM = 0
Delta_K
Doxorubicin
YHL002W HSE1
ZShift = 0.46
Z lm Score = −0.61
NG = 0
DB = 1
SM = 0
Delta_r
Doxorubicin
YHL002W HSE1
ZShift = 0.77
Z lm Score = 0.7
NG = 0
DB = 1
SM = 0
Delta_AUC
Doxorubicin
YLR436C ECM30
ZShift = −0.52
Z lm Score = 2.44
NG = 0
DB = 0
SM = 0
Delta_L
Doxorubicin
YLR436C ECM30
ZShift = −1.14
Z lm Score = 0.45
NG = 0
DB = 0
SM = 0
Delta_K
Doxorubicin
YLR436C ECM30
ZShift = 0.27
Z lm Score = −1.05
NG = 0
DB = 0
SM = 0
Delta_r
Doxorubicin
YLR436C ECM30
ZShift = −0.22
Z lm Score = −0.39
NG = 0
DB = 0
SM = 0
Delta_AUC
Doxorubicin

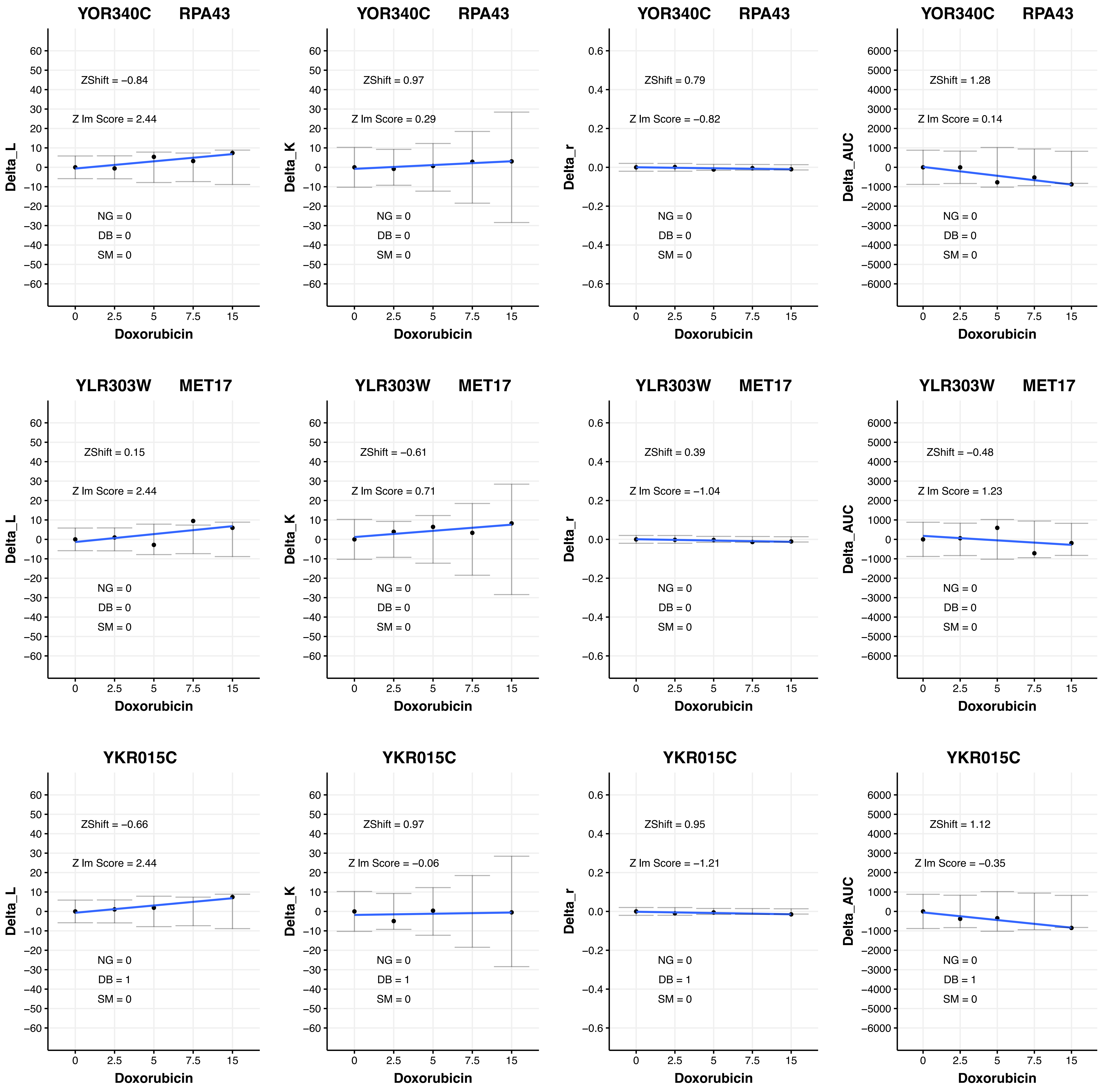
YOR340C RPA43
ZShift = −0.84
Z lm Score = 2.44
NG = 0
DB = 0
SM = 0
Delta_L
Doxorubicin
YOR340C RPA43
ZShift = 0.97
Z lm Score = 0.29
NG = 0
DB = 0
SM = 0
Delta_K
Doxorubicin
YOR340C RPA43
ZShift = 0.79
Z lm Score = −0.82
NG = 0
DB = 0
SM = 0
Delta_r
Doxorubicin
YOR340C RPA43
ZShift = 1.28
Z lm Score = 0.14
NG = 0
DB = 0
SM = 0
Delta_AUC
Doxorubicin
YLR303W MET17
ZShift = 0.15
Z lm Score = 2.44
NG = 0
DB = 0
SM = 0
Delta_L
Doxorubicin
YLR303W MET17
ZShift = −0.61
Z lm Score = 0.71
NG = 0
DB = 0
SM = 0
Delta_K
Doxorubicin
YLR303W MET17
ZShift = 0.39
Z lm Score = −1.04
NG = 0
DB = 0
SM = 0
Delta_r
Doxorubicin
YLR303W MET17
ZShift = −0.48
Z lm Score = 1.23
NG = 0
DB = 0
SM = 0
Delta_AUC
Doxorubicin
YKR015C
ZShift = −0.66
Z lm Score = 2.44
NG = 0
DB = 1
SM = 0
Delta_L
Doxorubicin
YKR015C
ZShift = 0.97
Z lm Score = −0.06
NG = 0
DB = 1
SM = 0
Delta_K
Doxorubicin
YKR015C
ZShift = 0.95
Z lm Score = −1.21
NG = 0
DB = 1
SM = 0
Delta_r
Doxorubicin
YKR015C
ZShift = 1.12
Z lm Score = −0.35
NG = 0
DB = 1
SM = 0
Delta_AUC
Doxorubicin

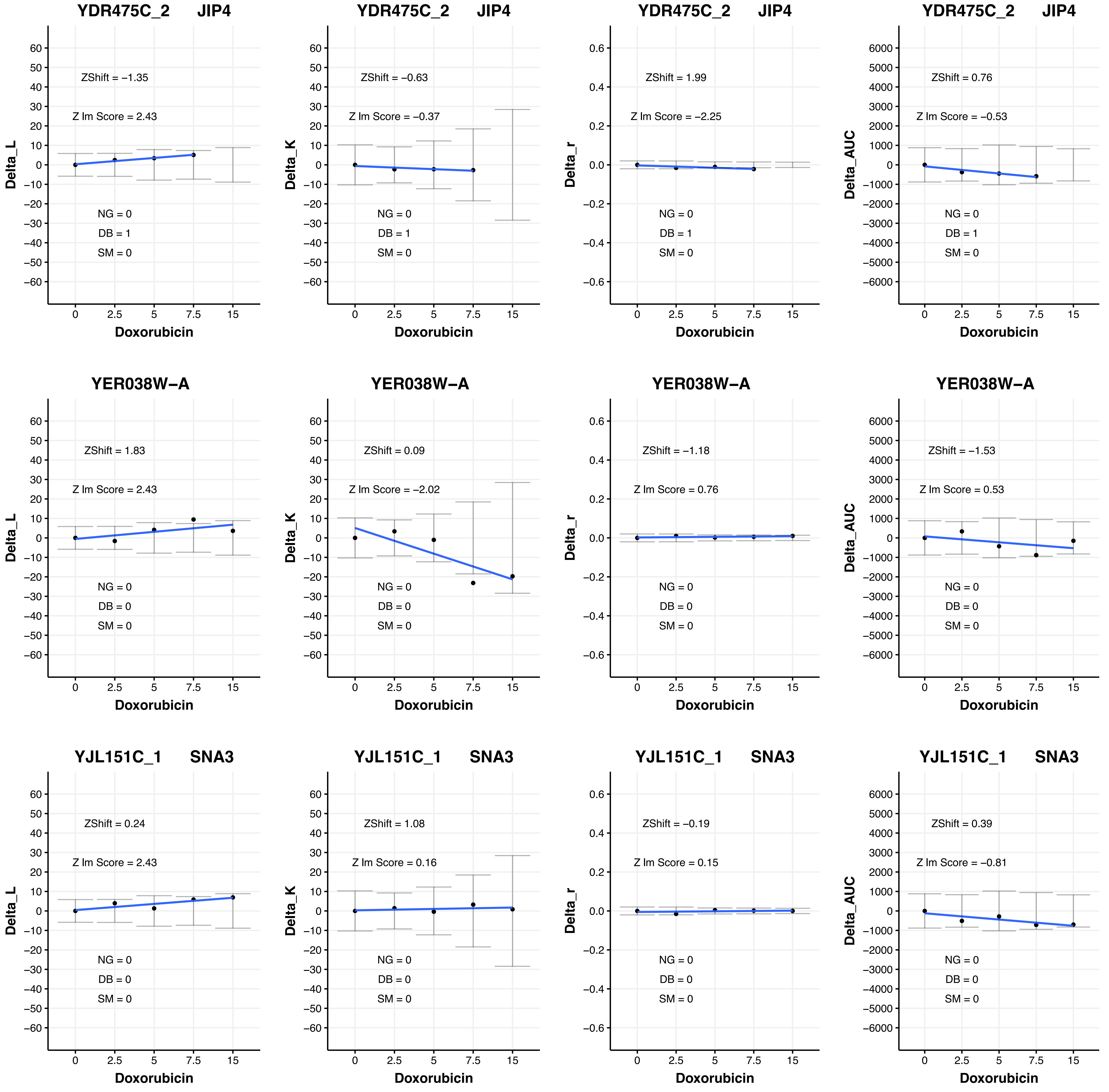

YDR475C_2 JIP4
ZShift = −1.35
Z lm Score = 2.43
NG = 0
DB = 1
SM = 0
Delta_L
Doxorubicin
YDR475C_2 JIP4
ZShift = −0.63
Z lm Score = −0.37
NG = 0
DB = 1
SM = 0
Delta_K
Doxorubicin
YDR475C_2 JIP4
ZShift = 1.99
Z lm Score = −2.25
NG = 0
DB = 1
SM = 0
Delta_r
Doxorubicin
YDR475C_2 JIP4
ZShift = 0.76
Z lm Score = −0.53
NG = 0
DB = 1
SM = 0
Delta_AUC
Doxorubicin
YER038W−A
ZShift = 1.83
Z lm Score = 2.43
NG = 0
DB = 0
SM = 0
Delta_L
Doxorubicin
YER038W−A
ZShift = 0.09
Z lm Score = −2.02
NG = 0
DB = 0
SM = 0
Delta_K
Doxorubicin
YER038W−A
ZShift = −1.18
Z lm Score = 0.76
NG = 0
DB = 0
SM = 0
Delta_r
Doxorubicin
YER038W−A
ZShift = −1.53
Z lm Score = 0.53
NG = 0
DB = 0
SM = 0
Delta_AUC
Doxorubicin
YJL151C_1 SNA3
ZShift = 0.24
Z lm Score = 2.43
NG = 0
DB = 0
SM = 0
Delta_L
Doxorubicin
YJL151C_1 SNA3
ZShift = 1.08
Z lm Score = 0.16
NG = 0
DB = 0
SM = 0
Delta_K
Doxorubicin
YJL151C_1 SNA3
ZShift = −0.19
Z lm Score = 0.15
NG = 0
DB = 0
SM = 0
Delta_r
Doxorubicin
YJL151C_1 SNA3
ZShift = 0.39
Z lm Score = −0.81
NG = 0
DB = 0
SM = 0
Delta_AUC
Doxorubicin

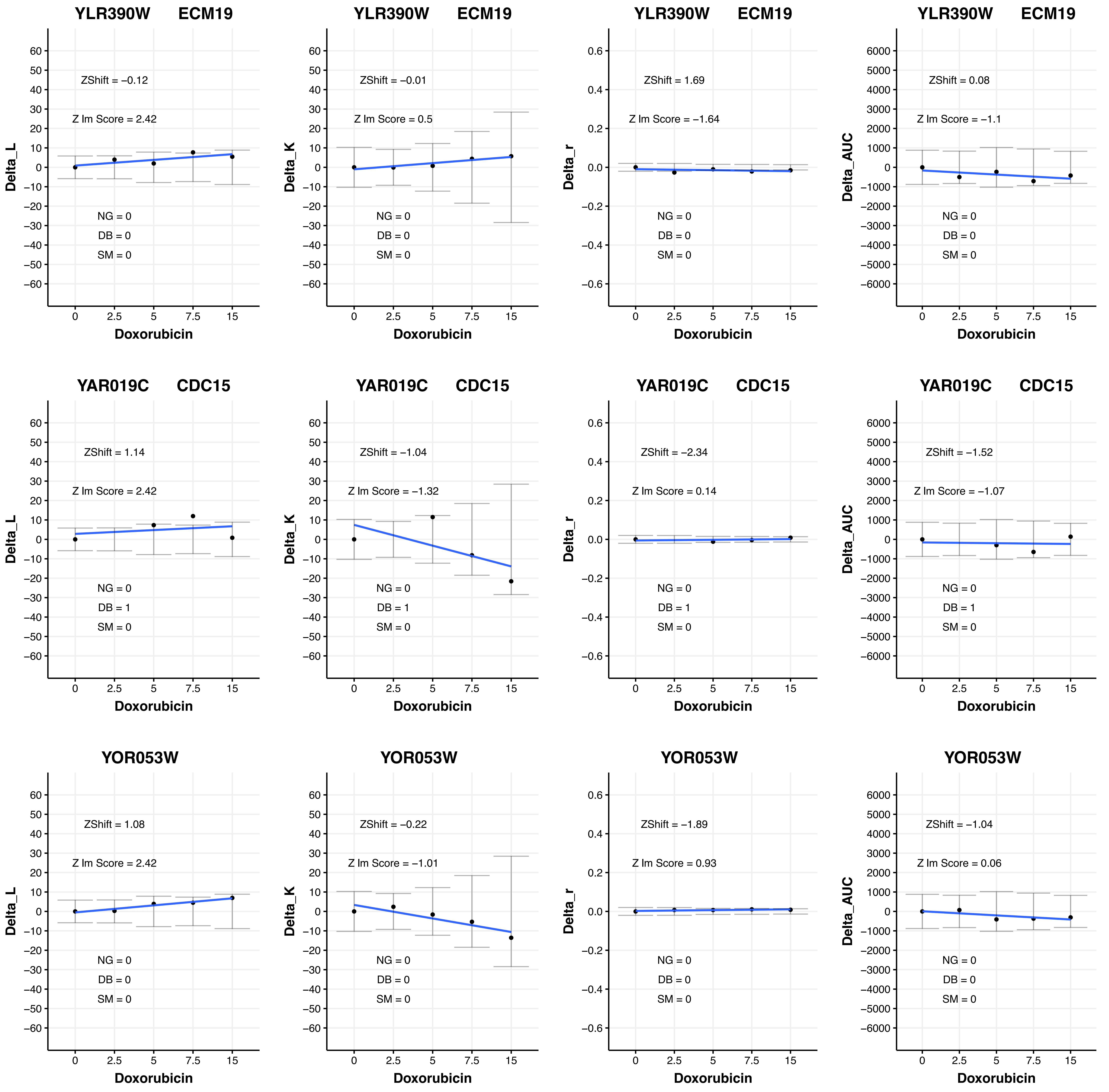

YLR390W ECM19
ZShift = −0.12
Z lm Score = 2.42
NG = 0
DB = 0
SM = 0
Delta_L
Doxorubicin
YLR390W ECM19
ZShift = −0.01
Z lm Score = 0.5
NG = 0
DB = 0
SM = 0
Delta_K
Doxorubicin
YLR390W ECM19
ZShift = 1.69
Z lm Score = −1.64
NG = 0
DB = 0
SM = 0
Delta_r
Doxorubicin
YLR390W ECM19
ZShift = 0.08
Z lm Score = −1.1
NG = 0
DB = 0
SM = 0
Delta_AUC
Doxorubicin
YAR019C CDC15
ZShift = 1.14
Z lm Score = 2.42
NG = 0
DB = 1
SM = 0
Delta_L
Doxorubicin
YAR019C CDC15
ZShift = −1.04
Z lm Score = −1.32
NG = 0
DB = 1
SM = 0
Delta_K
Doxorubicin
YAR019C CDC15
ZShift = −2.34
Z lm Score = 0.14
NG = 0
DB = 1
SM = 0
Delta_r
Doxorubicin
YAR019C CDC15
ZShift = −1.52
Z lm Score = −1.07
NG = 0
DB = 1
SM = 0
Delta_AUC
Doxorubicin
YOR053W
ZShift = 1.08
Z lm Score = 2.42
NG = 0
DB = 0
SM = 0
Delta_L
Doxorubicin
YOR053W
ZShift = −0.22
Z lm Score = −1.01
NG = 0
DB = 0
SM = 0
Delta_K
Doxorubicin
YOR053W
ZShift = −1.89
Z lm Score = 0.93
NG = 0
DB = 0
SM = 0
Delta_r
Doxorubicin
YOR053W
ZShift = −1.04
Z lm Score = 0.06
NG = 0
DB = 0
SM = 0
Delta_AUC
Doxorubicin

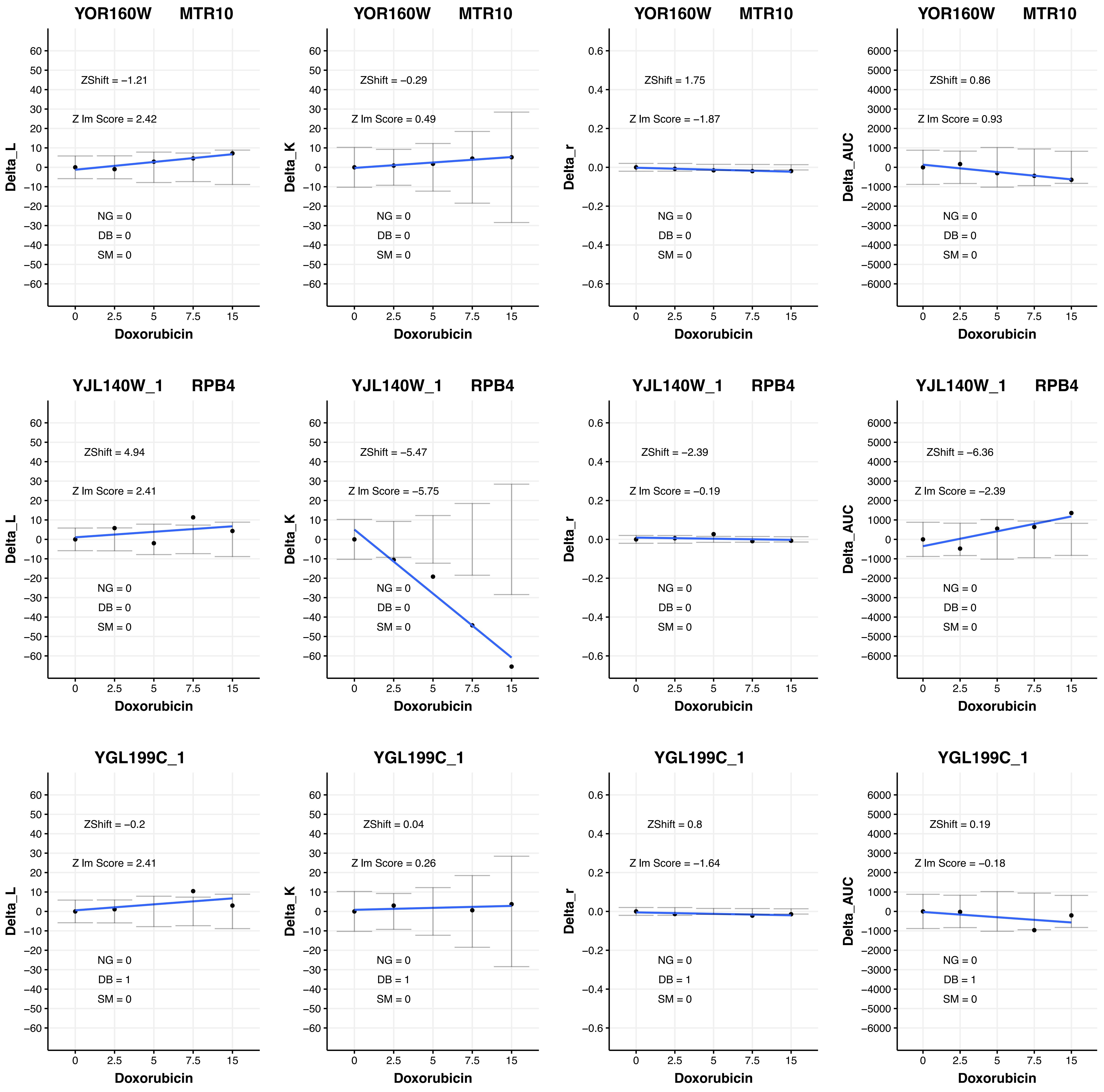
YOR160W MTR10
ZShift = −1.21
Z lm Score = 2.42
NG = 0
DB = 0
SM = 0
Delta_L
Doxorubicin
YOR160W MTR10
ZShift = −0.29
Z lm Score = 0.49
NG = 0
DB = 0
SM = 0
Delta_K
Doxorubicin
YOR160W MTR10
ZShift = 1.75
Z lm Score = −1.87
NG = 0
DB = 0
SM = 0
Delta_r
Doxorubicin
YOR160W MTR10
ZShift = 0.86
Z lm Score = 0.93
NG = 0
DB = 0
SM = 0
Delta_AUC
Doxorubicin
YJL140W_1 RPB4
ZShift = 4.94
Z lm Score = 2.41
NG = 0
DB = 0
SM = 0
Delta_L
Doxorubicin
YJL140W_1 RPB4
ZShift = −5.47
Z lm Score = −5.75
NG = 0
DB = 0
SM = 0
Delta_K
Doxorubicin
YJL140W_1 RPB4
ZShift = −2.39
Z lm Score = −0.19
NG = 0
DB = 0
SM = 0
Delta_r
Doxorubicin
YJL140W_1 RPB4
ZShift = −6.36
Z lm Score = −2.39
NG = 0
DB = 0
SM = 0
Delta_AUC
Doxorubicin
YGL199C_1
ZShift = −0.2
Z lm Score = 2.41
NG = 0
DB = 1
SM = 0
Delta_L
Doxorubicin
YGL199C_1
ZShift = 0.04
Z lm Score = 0.26
NG = 0
DB = 1
SM = 0
Delta_K
Doxorubicin
YGL199C_1
ZShift = 0.8
Z lm Score = −1.64
NG = 0
DB = 1
SM = 0
Delta_r
Doxorubicin
YGL199C_1
ZShift = 0.19
Z lm Score = −0.18
NG = 0
DB = 1
SM = 0
Delta_AUC
Doxorubicin

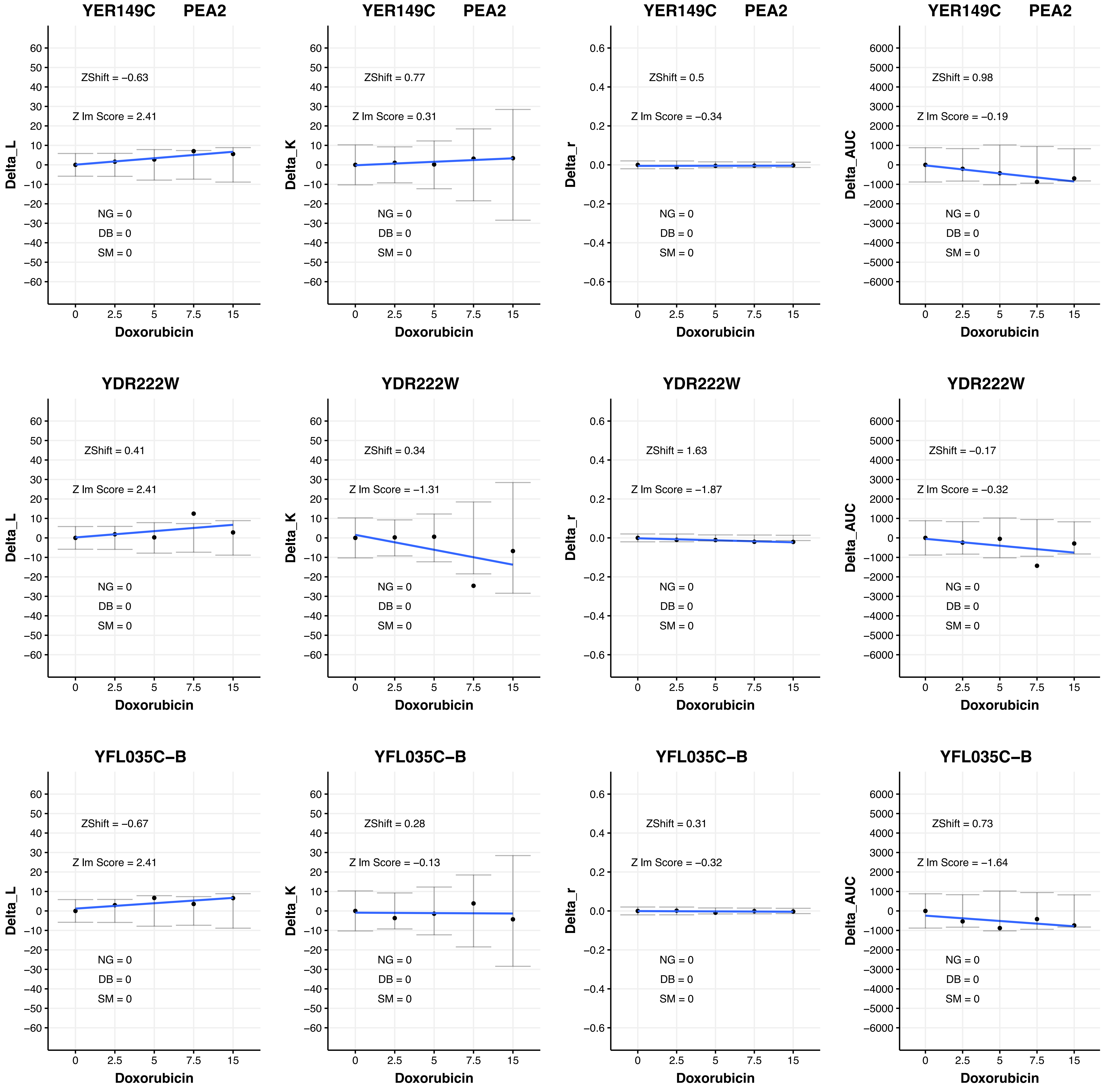
YER149C PEA2
ZShift = −0.63
Z lm Score = 2.41
NG = 0
DB = 0
SM = 0
Delta_L
Doxorubicin
YER149C PEA2
ZShift = 0.77
Z lm Score = 0.31
NG = 0
DB = 0
SM = 0
Delta_K
Doxorubicin
YER149C PEA2
ZShift = 0.5
Z lm Score = −0.34
NG = 0
DB = 0
SM = 0
Delta_r
Doxorubicin
YER149C PEA2
ZShift = 0.98
Z lm Score = −0.19
NG = 0
DB = 0
SM = 0
Delta_AUC
Doxorubicin
YDR222W
ZShift = 0.41
Z lm Score = 2.41
NG = 0
DB = 0
SM = 0
Delta_L
Doxorubicin
YDR222W
ZShift = 0.34
Z lm Score = −1.31
NG = 0
DB = 0
SM = 0
Delta_K
Doxorubicin
YDR222W
ZShift = 1.63
Z lm Score = −1.87
NG = 0
DB = 0
SM = 0
Delta_r
Doxorubicin
YDR222W
ZShift = −0.17
Z lm Score = −0.32
NG = 0
DB = 0
SM = 0
Delta_AUC
Doxorubicin
YFL035C−B
ZShift = −0.67
Z lm Score = 2.41
NG = 0
DB = 0
SM = 0
Delta_L
Doxorubicin
YFL035C−B
ZShift = 0.28
Z lm Score = −0.13
NG = 0
DB = 0
SM = 0
Delta_K
Doxorubicin
YFL035C−B
ZShift = 0.31
Z lm Score = −0.32
NG = 0
DB = 0
SM = 0
Delta_r
Doxorubicin
YFL035C−B
ZShift = 0.73
Z lm Score = −1.64
NG = 0
DB = 0
SM = 0
Delta_AUC
Doxorubicin

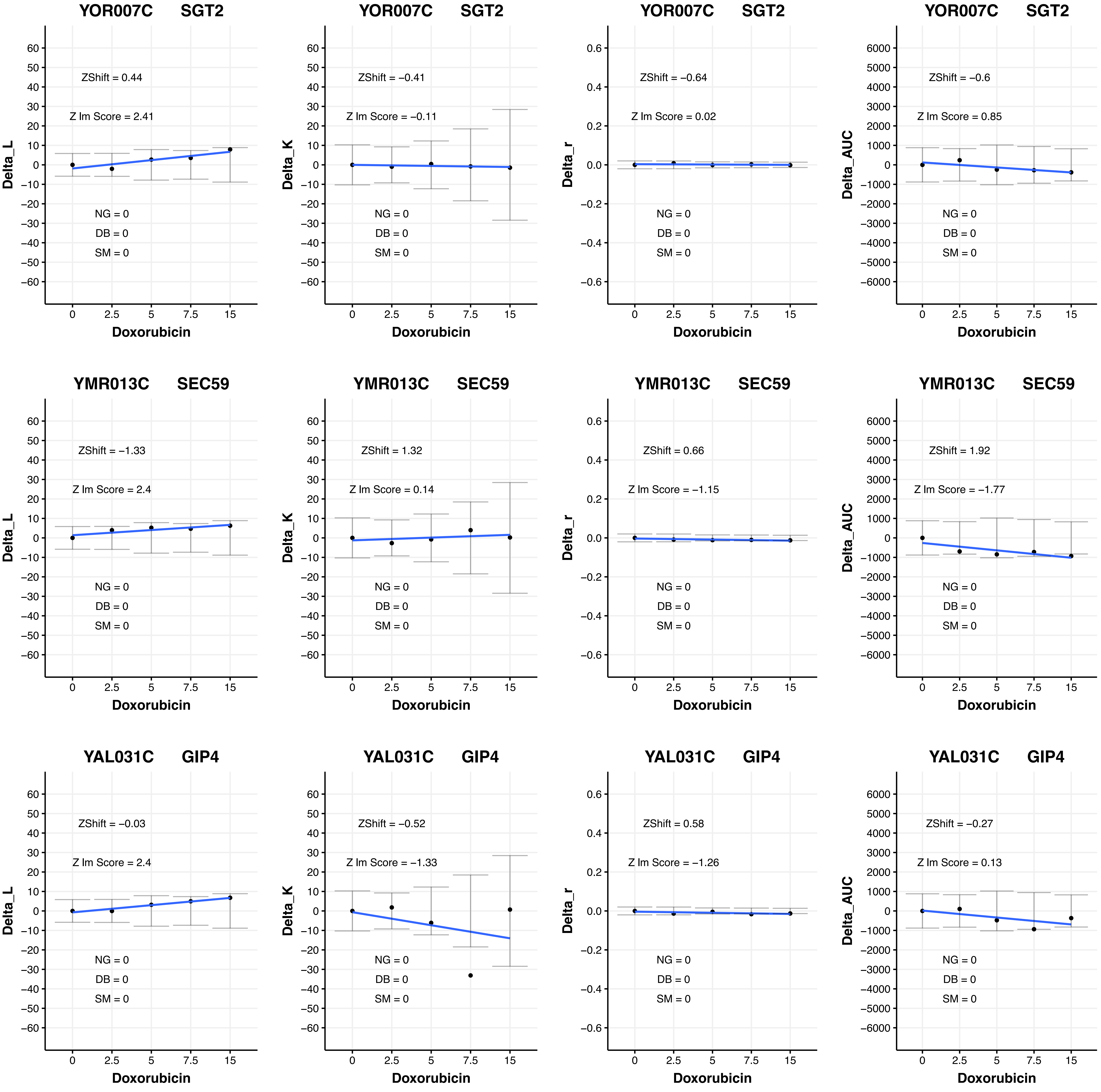
YOR007C SGT2
ZShift = 0.44
Z lm Score = 2.41
NG = 0
DB = 0
SM = 0
Delta_L
Doxorubicin
YOR007C SGT2
ZShift = −0.41
Z lm Score = −0.11
NG = 0
DB = 0
SM = 0
Delta_K
Doxorubicin
YOR007C SGT2
ZShift = −0.64
Z lm Score = 0.02
NG = 0
DB = 0
SM = 0
Delta_r
Doxorubicin
YOR007C SGT2
ZShift = −0.6
Z lm Score = 0.85
NG = 0
DB = 0
SM = 0
Delta_AUC
Doxorubicin
YMR013C SEC59
ZShift = −1.33
Z lm Score = 2.4
NG = 0
DB = 0
SM = 0
Delta_L
Doxorubicin
YMR013C SEC59
ZShift = 1.32
Z lm Score = 0.14
NG = 0
DB = 0
SM = 0
Delta_K
Doxorubicin
YMR013C SEC59
ZShift = 0.66
Z lm Score = −1.15
NG = 0
DB = 0
SM = 0
Delta_r
Doxorubicin
YMR013C SEC59
ZShift = 1.92
Z lm Score = −1.77
NG = 0
DB = 0
SM = 0
Delta_AUC
Doxorubicin
YAL031C GIP4
ZShift = −0.03
Z lm Score = 2.4
NG = 0
DB = 0
SM = 0
Delta_L
Doxorubicin
YAL031C GIP4
ZShift = −0.52
Z lm Score = −1.33
NG = 0
DB = 0
SM = 0
Delta_K
Doxorubicin
YAL031C GIP4
ZShift = 0.58
Z lm Score = −1.26
NG = 0
DB = 0
SM = 0
Delta_r
Doxorubicin
YAL031C GIP4
ZShift = −0.27
Z lm Score = 0.13
NG = 0
DB = 0
SM = 0
Delta_AUC
Doxorubicin

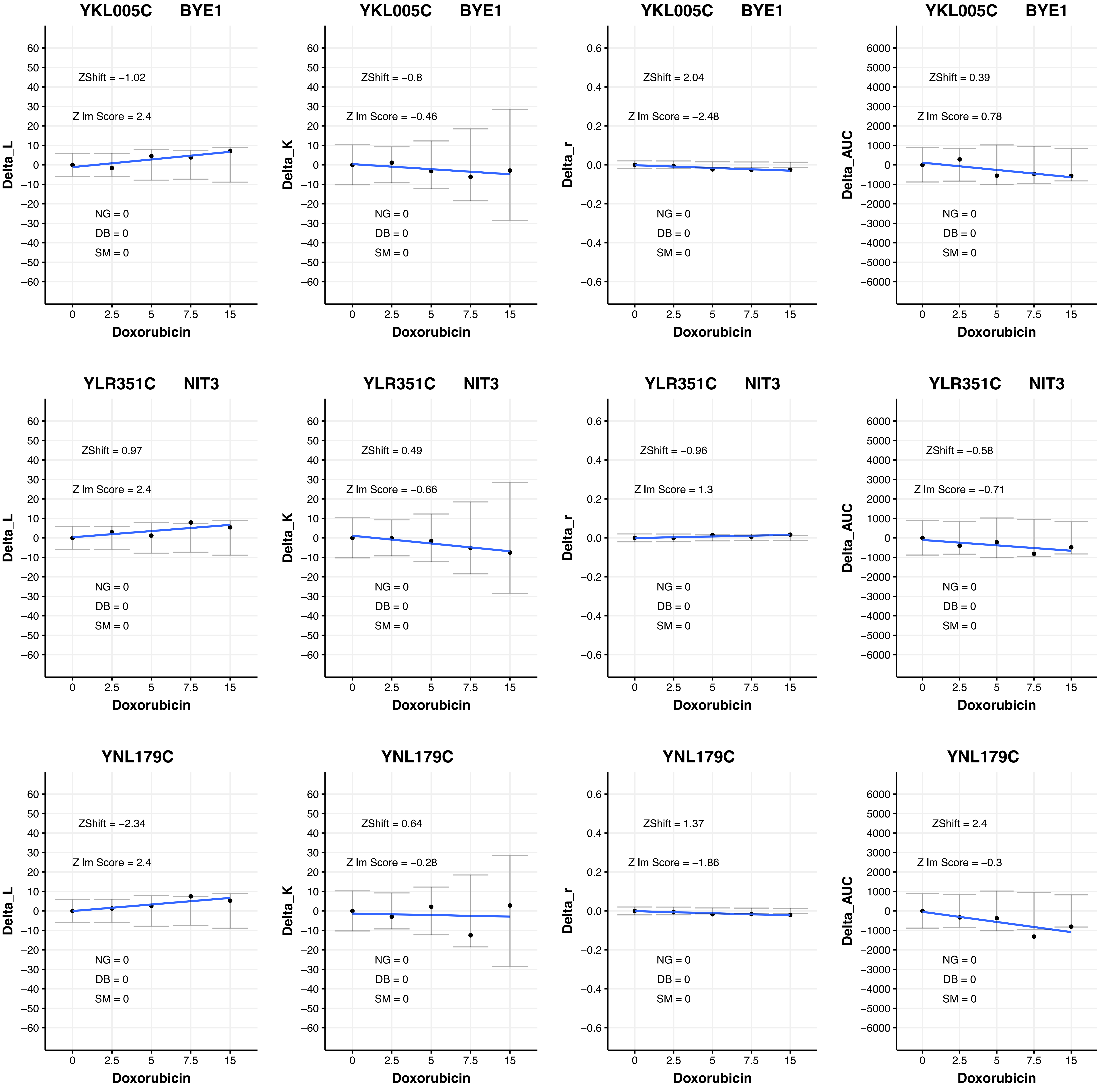
YKL005C BYE1
ZShift = −1.02
Z lm Score = 2.4
NG = 0
DB = 0
SM = 0
Delta_L
Doxorubicin
YKL005C BYE1
ZShift = −0.8
Z lm Score = −0.46
NG = 0
DB = 0
SM = 0
Delta_K
Doxorubicin
YKL005C BYE1
ZShift = 2.04
Z lm Score = −2.48
NG = 0
DB = 0
SM = 0
Delta_r
Doxorubicin
YKL005C BYE1
ZShift = 0.39
Z lm Score = 0.78
NG = 0
DB = 0
SM = 0
Delta_AUC
Doxorubicin
YLR351C NIT3
ZShift = 0.97
Z lm Score = 2.4
NG = 0
DB = 0
SM = 0
Delta_L
Doxorubicin
YLR351C NIT3
ZShift = 0.49
Z lm Score = −0.66
NG = 0
DB = 0
SM = 0
Delta_K
Doxorubicin
YLR351C NIT3
ZShift = −0.96
Z lm Score = 1.3
NG = 0
DB = 0
SM = 0
Delta_r
Doxorubicin
YLR351C NIT3
ZShift = −0.58
Z lm Score = −0.71
NG = 0
DB = 0
SM = 0
Delta_AUC
Doxorubicin
YNL179C
ZShift = −2.34
Z lm Score = 2.4
NG = 0
DB = 0
SM = 0
Delta_L
Doxorubicin
YNL179C
ZShift = 0.64
Z lm Score = −0.28
NG = 0
DB = 0
SM = 0
Delta_K
Doxorubicin
YNL179C
ZShift = 1.37
Z lm Score = −1.86
NG = 0
DB = 0
SM = 0
Delta_r
Doxorubicin
YNL179C
ZShift = 2.4
Z lm Score = −0.3
NG = 0
DB = 0
SM = 0
Delta_AUC
Doxorubicin

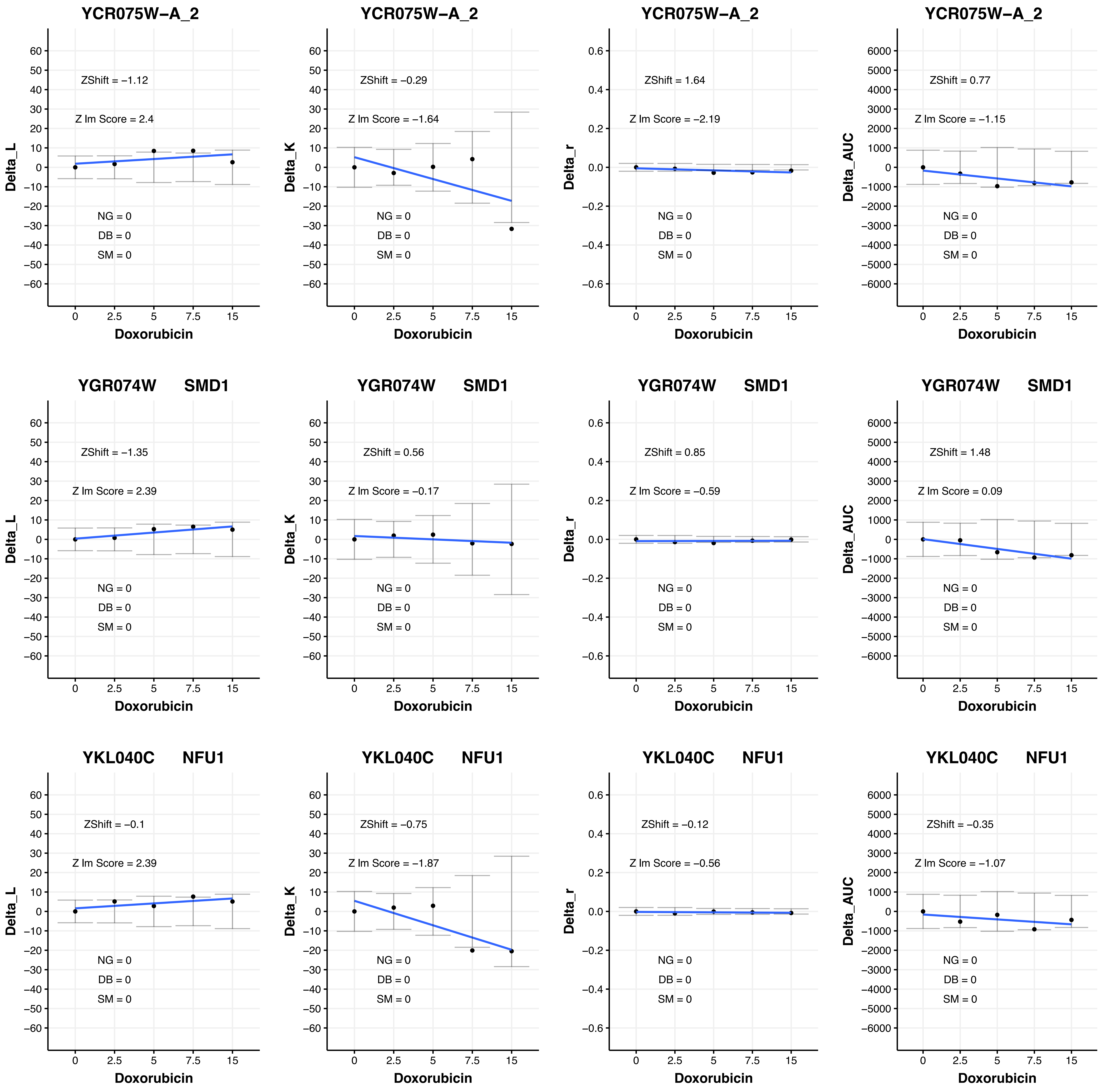

YCR075W−A_2
ZShift = −1.12
Z lm Score = 2.4
NG = 0
DB = 0
SM = 0
Delta_L
YCR075W−A_2
ZShift = −0.29
Z lm Score = −1.64
NG = 0
DB = 0
SM = 0
Delta_K
YCR075W−A_2
ZShift = 1.64
Z lm Score = −2.19
NG = 0
DB = 0
SM = 0
Delta_r
YCR075W−A_2
ZShift = 0.77
Z lm Score = −1.15
NG = 0
DB = 0
SM = 0
Delta_AUC
YGR074W SMD1
ZShift = −1.35
Z lm Score = 2.39
NG = 0
DB = 0
SM = 0
Delta_L
YGR074W SMD1
ZShift = 0.56
Z lm Score = −0.17
NG = 0
DB = 0
SM = 0
Delta_K
YGR074W SMD1
ZShift = 0.85
Z lm Score = −0.59
NG = 0
DB = 0
SM = 0
Delta_r
YGR074W SMD1
ZShift = 1.48
Z lm Score = 0.09
NG = 0
DB = 0
SM = 0
Delta_AUC
YKL040C NFU1
ZShift = −0.1
Z lm Score = 2.39
NG = 0
DB = 0
SM = 0
Delta_L
YKL040C NFU1
ZShift = −0.75
Z lm Score = −1.87
NG = 0
DB = 0
SM = 0
Delta_K
YKL040C NFU1
ZShift = −0.12
Z lm Score = −0.56
NG = 0
DB = 0
SM = 0
Delta_r
YKL040C NFU1
ZShift = −0.35
Z lm Score = −1.07
NG = 0
DB = 0
SM = 0
Delta_AUC
Doxorubicin
0
2.5
5
7.5
15

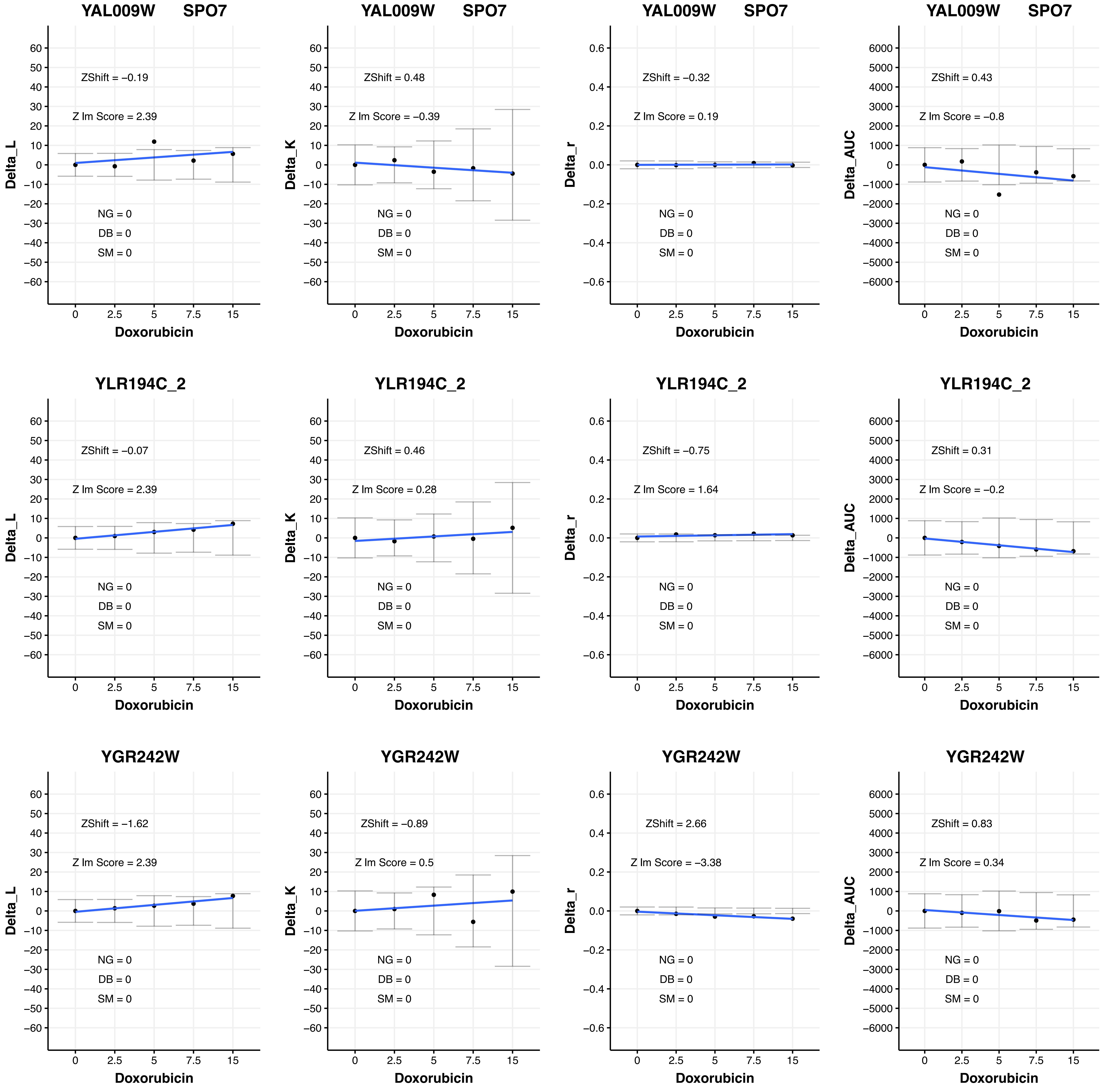
YAL009W SPO7
ZShift = −0.19
Z lm Score = 2.39
NG = 0
DB = 0
SM = 0
Delta_L
Doxorubicin
YAL009W SPO7
ZShift = 0.48
Z lm Score = −0.39
NG = 0
DB = 0
SM = 0
Delta_K
Doxorubicin
YAL009W SPO7
ZShift = −0.32
Z lm Score = 0.19
NG = 0
DB = 0
SM = 0
Delta_r
Doxorubicin
YAL009W SPO7
ZShift = 0.43
Z lm Score = −0.8
NG = 0
DB = 0
SM = 0
Delta_AUC
Doxorubicin
YLR194C_2
ZShift = −0.07
Z lm Score = 2.39
NG = 0
DB = 0
SM = 0
Delta_L
Doxorubicin
YLR194C_2
ZShift = 0.46
Z lm Score = 0.28
NG = 0
DB = 0
SM = 0
Delta_K
Doxorubicin
YLR194C_2
ZShift = −0.75
Z lm Score = 1.64
NG = 0
DB = 0
SM = 0
Delta_r
Doxorubicin
YLR194C_2
ZShift = 0.31
Z lm Score = −0.2
NG = 0
DB = 0
SM = 0
Delta_AUC
Doxorubicin
YGR242W
ZShift = −1.62
Z lm Score = 2.39
NG = 0
DB = 0
SM = 0
Delta_L
Doxorubicin
YGR242W
ZShift = −0.89
Z lm Score = 0.5
NG = 0
DB = 0
SM = 0
Delta_K
Doxorubicin
YGR242W
ZShift = 2.66
Z lm Score = −3.38
NG = 0
DB = 0
SM = 0
Delta_r
Doxorubicin
YGR242W
ZShift = 0.83
Z lm Score = 0.34
NG = 0
DB = 0
SM = 0
Delta_AUC
Doxorubicin

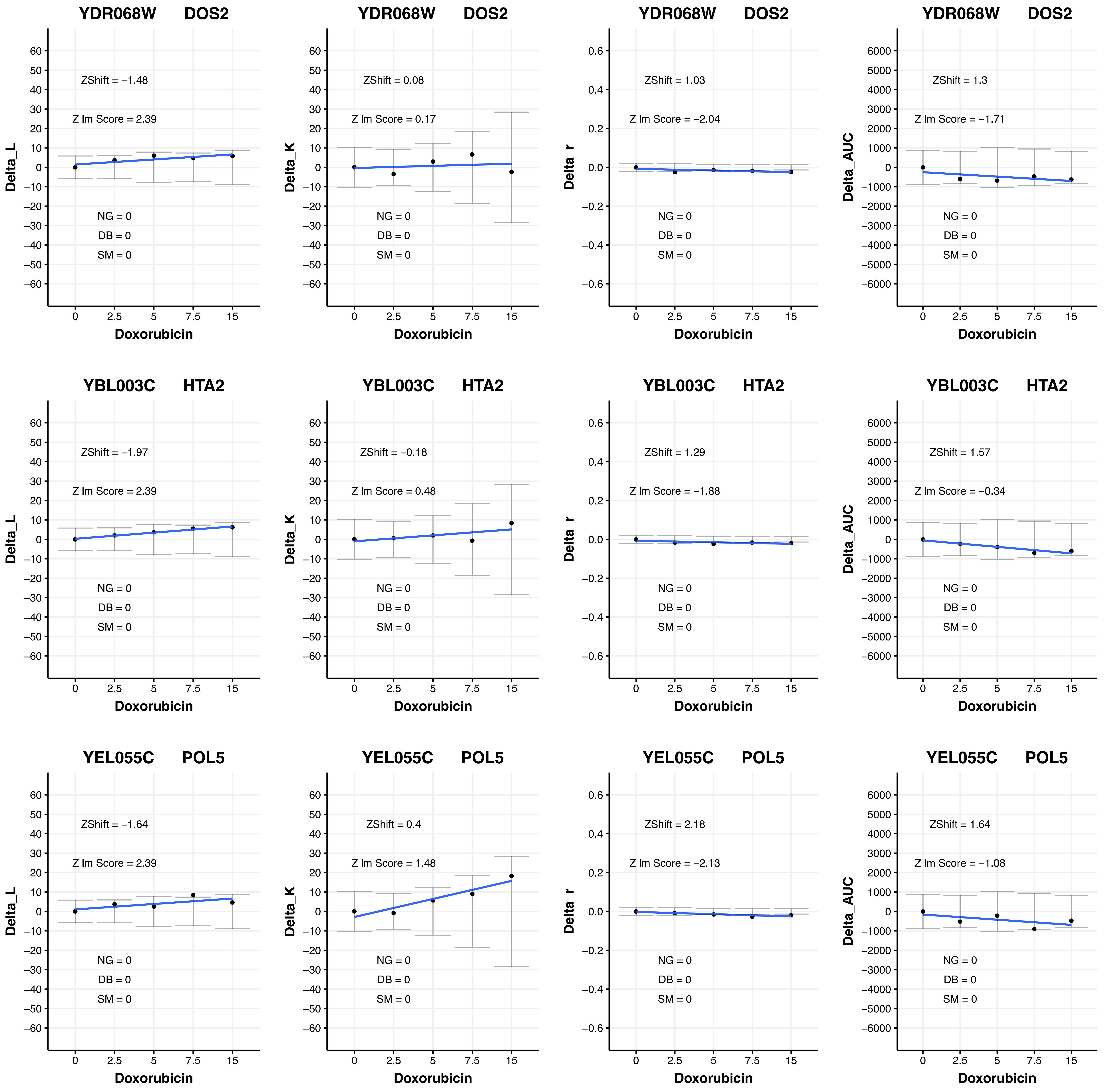

YDR068W DOS2
ZShift = −1.48
Z lm Score = 2.39
NG = 0
DB = 0
SM = 0
Delta_L
Doxorubicin
YDR068W DOS2
ZShift = 0.08
Z lm Score = 0.17
NG = 0
DB = 0
SM = 0
Delta_K
Doxorubicin
YDR068W DOS2
ZShift = 1.03
Z lm Score = −2.04
NG = 0
DB = 0
SM = 0
Delta_r
Doxorubicin
YDR068W DOS2
ZShift = 1.3
Z lm Score = −1.71
NG = 0
DB = 0
SM = 0
Delta_AUC
Doxorubicin
YBL003C HTA2
ZShift = −1.97
Z lm Score = 2.39
NG = 0
DB = 0
SM = 0
Delta_L
Doxorubicin
YBL003C HTA2
ZShift = −0.18
Z lm Score = 0.48
NG = 0
DB = 0
SM = 0
Delta_K
Doxorubicin
YBL003C HTA2
ZShift = 1.29
Z lm Score = −1.88
NG = 0
DB = 0
SM = 0
Delta_r
Doxorubicin
YBL003C HTA2
ZShift = 1.57
Z lm Score = −0.34
NG = 0
DB = 0
SM = 0
Delta_AUC
Doxorubicin
YEL055C POL5
ZShift = −1.64
Z lm Score = 2.39
NG = 0
DB = 0
SM = 0
Delta_L
Doxorubicin
YEL055C POL5
ZShift = 0.4
Z lm Score = 1.48
NG = 0
DB = 0
SM = 0
Delta_K
Doxorubicin
YEL055C POL5
ZShift = 2.18
Z lm Score = −2.13
NG = 0
DB = 0
SM = 0
Delta_r
Doxorubicin
YEL055C POL5
ZShift = 1.64
Z lm Score = −1.08
NG = 0
DB = 0
SM = 0
Delta_AUC
Doxorubicin

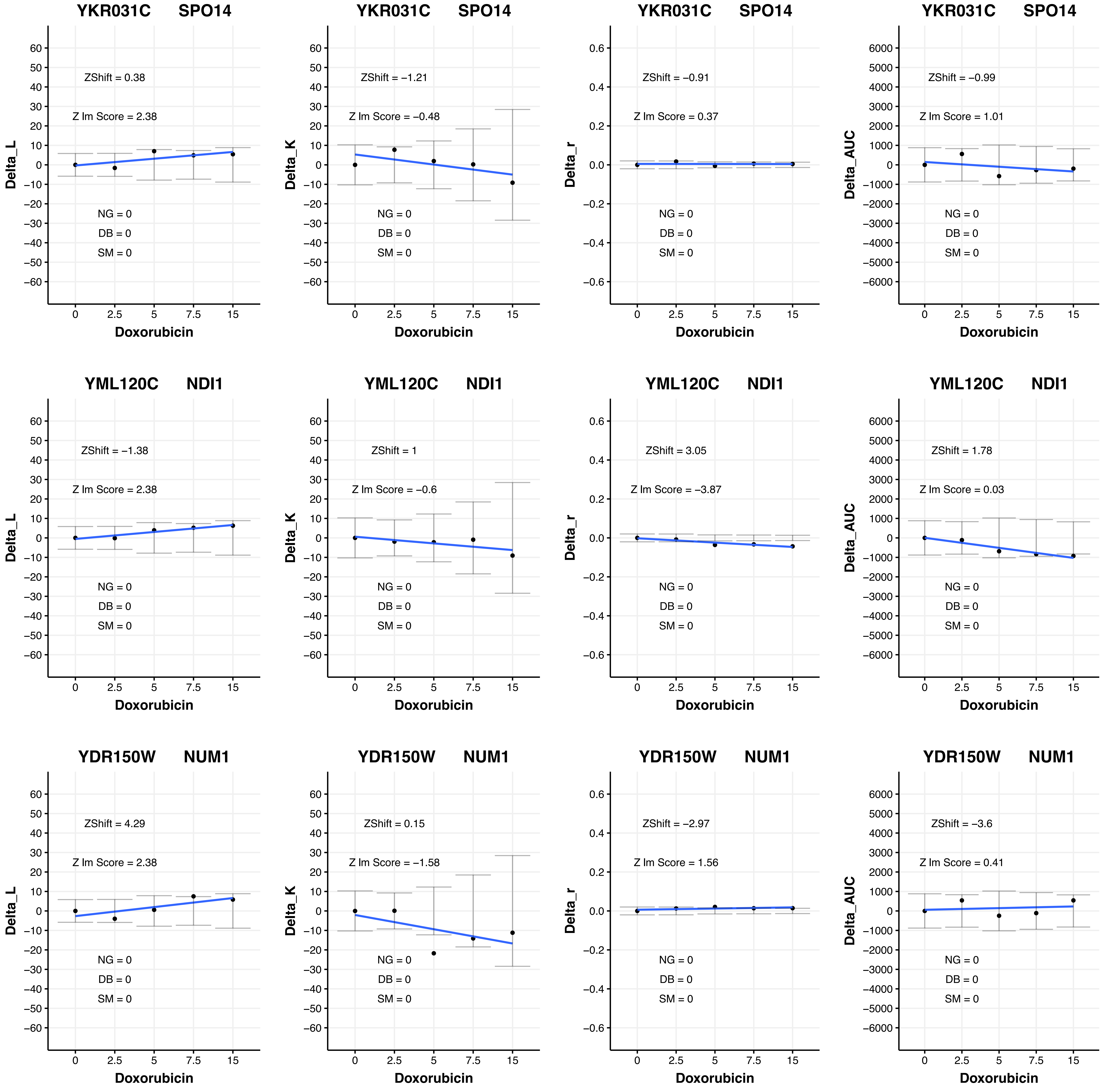

YKR031C SPO14
ZShift = 0.38
Z lm Score = 2.38
NG = 0
DB = 0
SM = 0
Delta_L
Doxorubicin
YKR031C SPO14
ZShift = −1.21
Z lm Score = −0.48
NG = 0
DB = 0
SM = 0
Delta_K
Doxorubicin
YKR031C SPO14
ZShift = −0.91
Z lm Score = 0.37
NG = 0
DB = 0
SM = 0
Delta_r
Doxorubicin
YKR031C SPO14
ZShift = −0.99
Z lm Score = 1.01
NG = 0
DB = 0
SM = 0
Delta_AUC
Doxorubicin
YML120C NDI1
ZShift = −1.38
Z lm Score = 2.38
NG = 0
DB = 0
SM = 0
Delta_L
Doxorubicin
YML120C NDI1
ZShift = 1
Z lm Score = −0.6
NG = 0
DB = 0
SM = 0
Delta_K
Doxorubicin
YML120C NDI1
ZShift = 3.05
Z lm Score = −3.87
NG = 0
DB = 0
SM = 0
Delta_r
Doxorubicin
YML120C NDI1
ZShift = 1.78
Z lm Score = 0.03
NG = 0
DB = 0
SM = 0
Delta_AUC
Doxorubicin
YDR150W NUM1
ZShift = 4.29
Z lm Score = 2.38
NG = 0
DB = 0
SM = 0
Delta_L
Doxorubicin
YDR150W NUM1
ZShift = 0.15
Z lm Score = −1.58
NG = 0
DB = 0
SM = 0
Delta_K
Doxorubicin
YDR150W NUM1
ZShift = −2.97
Z lm Score = 1.56
NG = 0
DB = 0
SM = 0
Delta_r
Doxorubicin
YDR150W NUM1
ZShift = −3.6
Z lm Score = 0.41
NG = 0
DB = 0
SM = 0
Delta_AUC
Doxorubicin

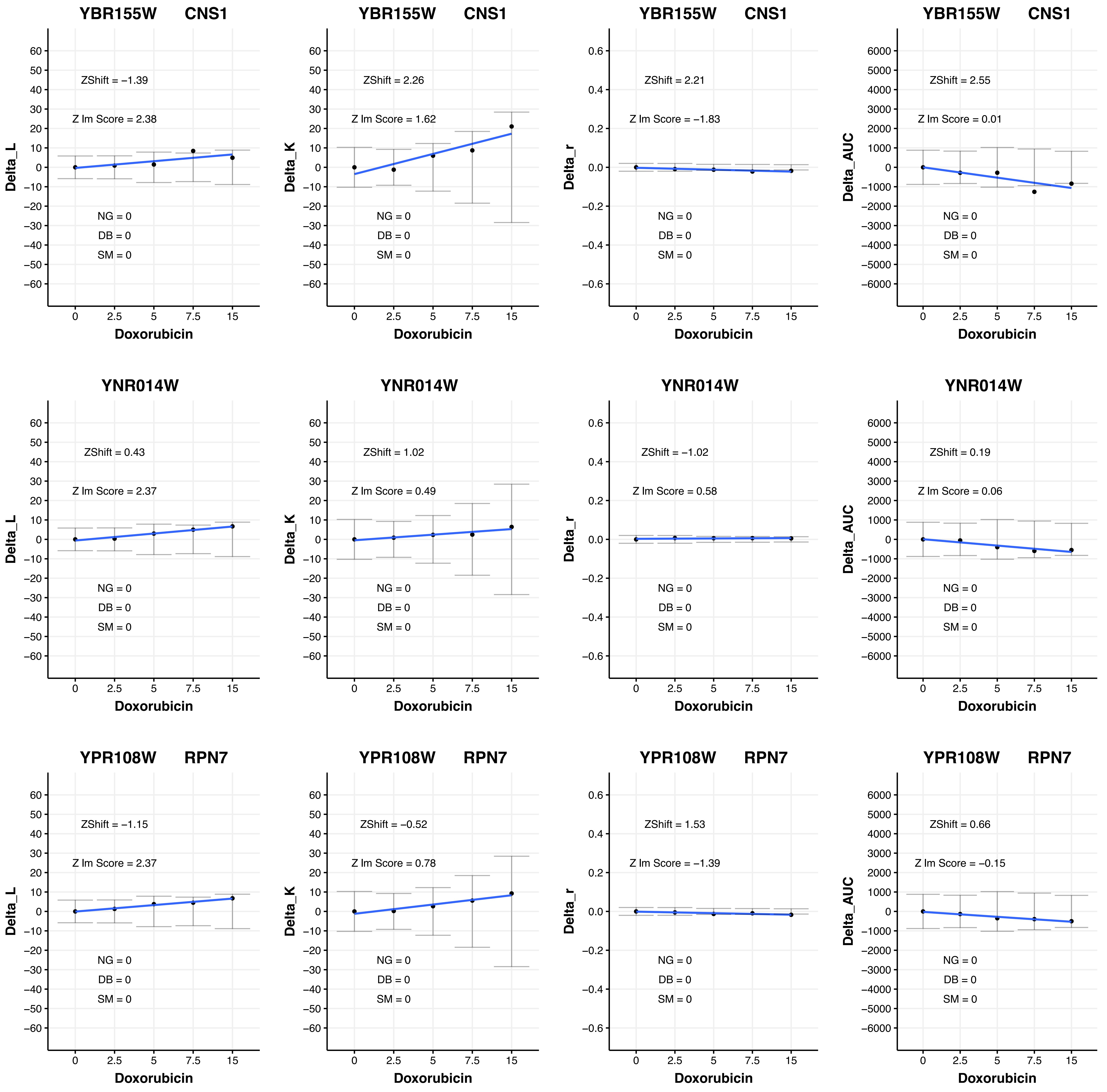
YBR155W CNS1
ZShift = −1.39
Z lm Score = 2.38
NG = 0
DB = 0
SM = 0
Delta_L
Doxorubicin
YBR155W CNS1
ZShift = 2.26
Z lm Score = 1.62
NG = 0
DB = 0
SM = 0
Delta_K
Doxorubicin
YBR155W CNS1
ZShift = 2.21
Z lm Score = −1.83
NG = 0
DB = 0
SM = 0
Delta_r
Doxorubicin
YBR155W CNS1
ZShift = 2.55
Z lm Score = 0.01
NG = 0
DB = 0
SM = 0
Delta_AUC
Doxorubicin
YNR014W
ZShift = 0.43
Z lm Score = 2.37
NG = 0
DB = 0
SM = 0
Delta_L
Doxorubicin
YNR014W
ZShift = 1.02
Z lm Score = 0.49
NG = 0
DB = 0
SM = 0
Delta_K
Doxorubicin
YNR014W
ZShift = −1.02
Z lm Score = 0.58
NG = 0
DB = 0
SM = 0
Delta_r
Doxorubicin
YNR014W
ZShift = 0.19
Z lm Score = 0.06
NG = 0
DB = 0
SM = 0
Delta_AUC
Doxorubicin
YPR108W RPN7
ZShift = −1.15
Z lm Score = 2.37
NG = 0
DB = 0
SM = 0
Delta_L
Doxorubicin
YPR108W RPN7
ZShift = −0.52
Z lm Score = 0.78
NG = 0
DB = 0
SM = 0
Delta_K
Doxorubicin
YPR108W RPN7
ZShift = 1.53
Z lm Score = −1.39
NG = 0
DB = 0
SM = 0
Delta_r
Doxorubicin
YPR108W RPN7
ZShift = 0.66
Z lm Score = −0.15
NG = 0
DB = 0
SM = 0
Delta_AUC
Doxorubicin

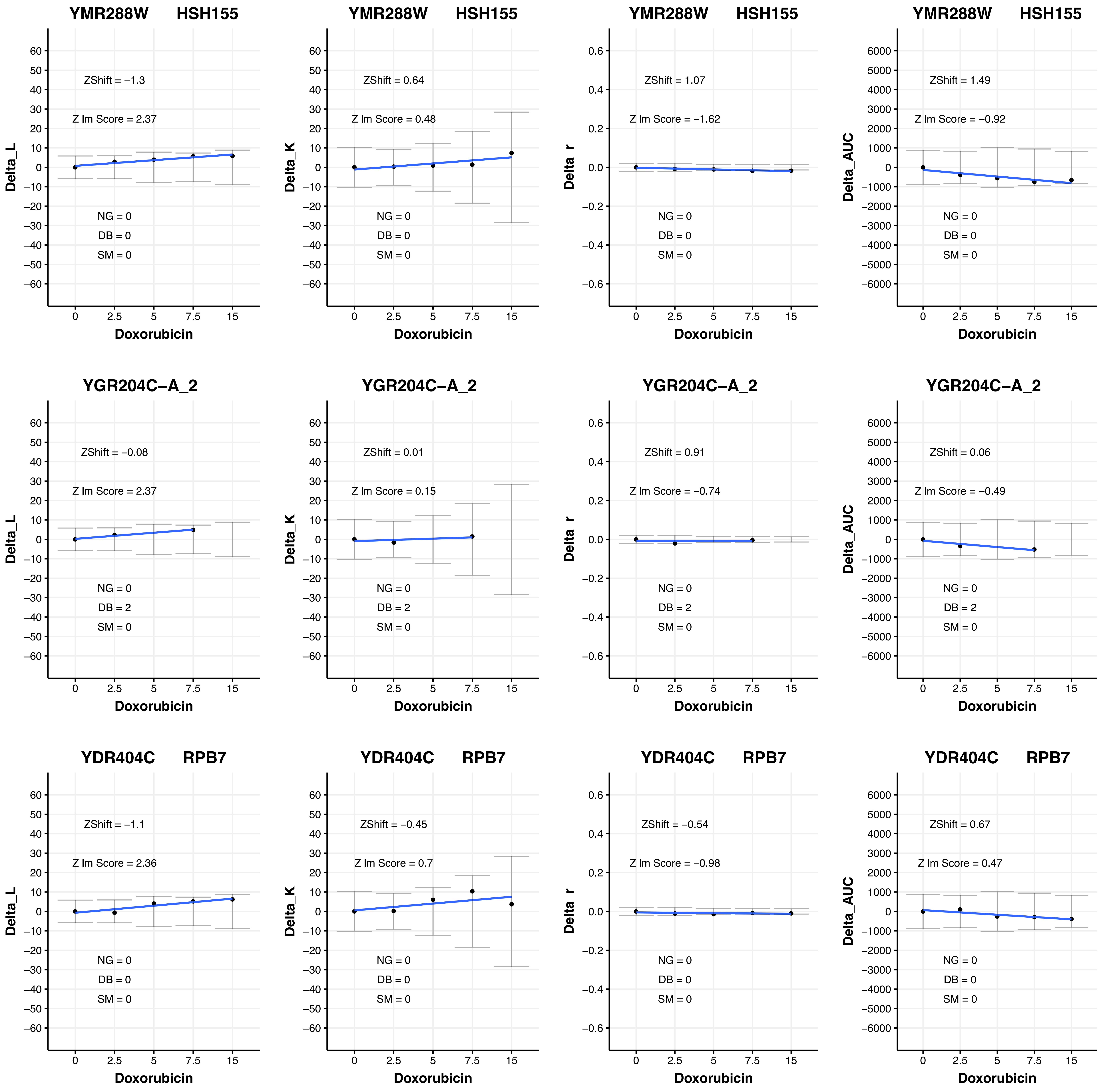

YMR288W HSH155
ZShift = −1.3
Z lm Score = 2.37
NG = 0
DB = 0
SM = 0
Delta_L
Doxorubicin
YMR288W HSH155
ZShift = 0.64
Z lm Score = 0.48
NG = 0
DB = 0
SM = 0
Delta_K
Doxorubicin
YMR288W HSH155
ZShift = 1.07
Z lm Score = −1.62
NG = 0
DB = 0
SM = 0
Delta_r
Doxorubicin
YMR288W HSH155
ZShift = 1.49
Z lm Score = −0.92
NG = 0
DB = 0
SM = 0
Delta_AUC
Doxorubicin
YGR204C−A_2
ZShift = −0.08
Z lm Score = 2.37
NG = 0
DB = 2
SM = 0
Delta_L
Doxorubicin
YGR204C−A_2
ZShift = 0.01
Z lm Score = 0.15
NG = 0
DB = 2
SM = 0
Delta_K
Doxorubicin
YGR204C−A_2
ZShift = 0.91
Z lm Score = −0.74
NG = 0
DB = 2
SM = 0
Delta_r
Doxorubicin
YGR204C−A_2
ZShift = 0.06
Z lm Score = −0.49
NG = 0
DB = 2
SM = 0
Delta_AUC
Doxorubicin
YDR404C RPB7
ZShift = −1.1
Z lm Score = 2.36
NG = 0
DB = 0
SM = 0
Delta_L
Doxorubicin
YDR404C RPB7
ZShift = −0.45
Z lm Score = 0.7
NG = 0
DB = 0
SM = 0
Delta_K
Doxorubicin
YDR404C RPB7
ZShift = −0.54
Z lm Score = −0.98
NG = 0
DB = 0
SM = 0
Delta_r
Doxorubicin
YDR404C RPB7
ZShift = 0.67
Z lm Score = 0.47
NG = 0
DB = 0
SM = 0
Delta_AUC
Doxorubicin

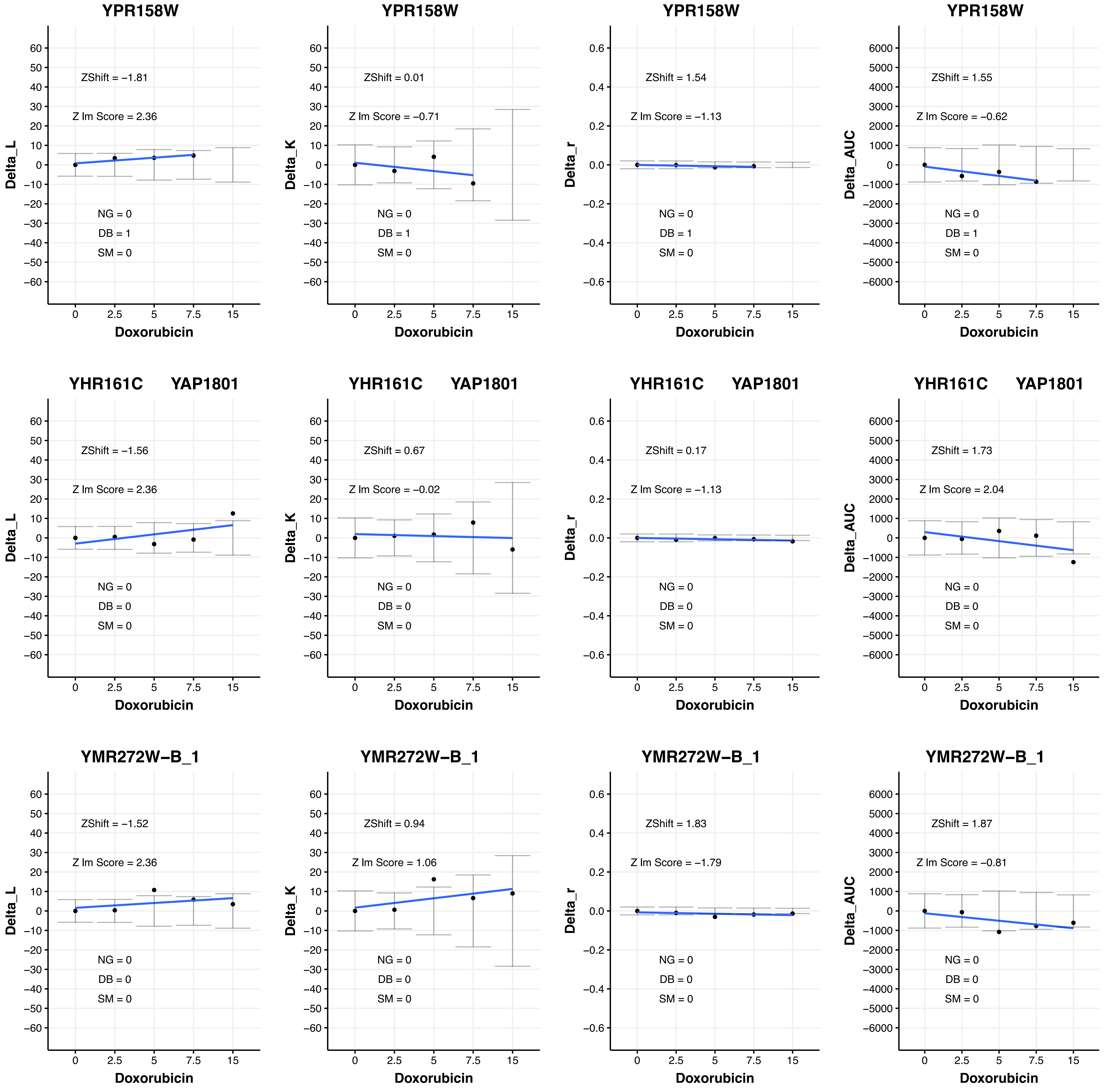

YPR158W
ZShift = −1.81
Z lm Score = 2.36
NG = 0
DB = 1
SM = 0
Delta_L
Doxorubicin
YPR158W
ZShift = 0.01
Z lm Score = −0.71
NG = 0
DB = 1
SM = 0
Delta_K
Doxorubicin
YPR158W
ZShift = 1.54
Z lm Score = −1.13
NG = 0
DB = 1
SM = 0
Delta_r
Doxorubicin
YPR158W
ZShift = 1.55
Z lm Score = −0.62
NG = 0
DB = 1
SM = 0
Delta_AUC
Doxorubicin
YHR161C YAP1801
ZShift = −1.56
Z lm Score = 2.36
NG = 0
DB = 0
SM = 0
Delta_L
Doxorubicin
YHR161C YAP1801
ZShift = 0.67
Z lm Score = −0.02
NG = 0
DB = 0
SM = 0
Delta_K
Doxorubicin
YHR161C YAP1801
ZShift = 0.17
Z lm Score = −1.13
NG = 0
DB = 0
SM = 0
Delta_r
Doxorubicin
YHR161C YAP1801
ZShift = 1.73
Z lm Score = 2.04
NG = 0
DB = 0
SM = 0
Delta_AUC
Doxorubicin
YMR272W−B_1
ZShift = −1.52
Z lm Score = 2.36
NG = 0
DB = 0
SM = 0
Delta_L
Doxorubicin
YMR272W−B_1
ZShift = 0.94
Z lm Score = 1.06
NG = 0
DB = 0
SM = 0
Delta_K
Doxorubicin
YMR272W−B_1
ZShift = 1.83
Z lm Score = −1.79
NG = 0
DB = 0
SM = 0
Delta_r
Doxorubicin
YMR272W−B_1
ZShift = 1.87
Z lm Score = −0.81
NG = 0
DB = 0
SM = 0
Delta_AUC
Doxorubicin

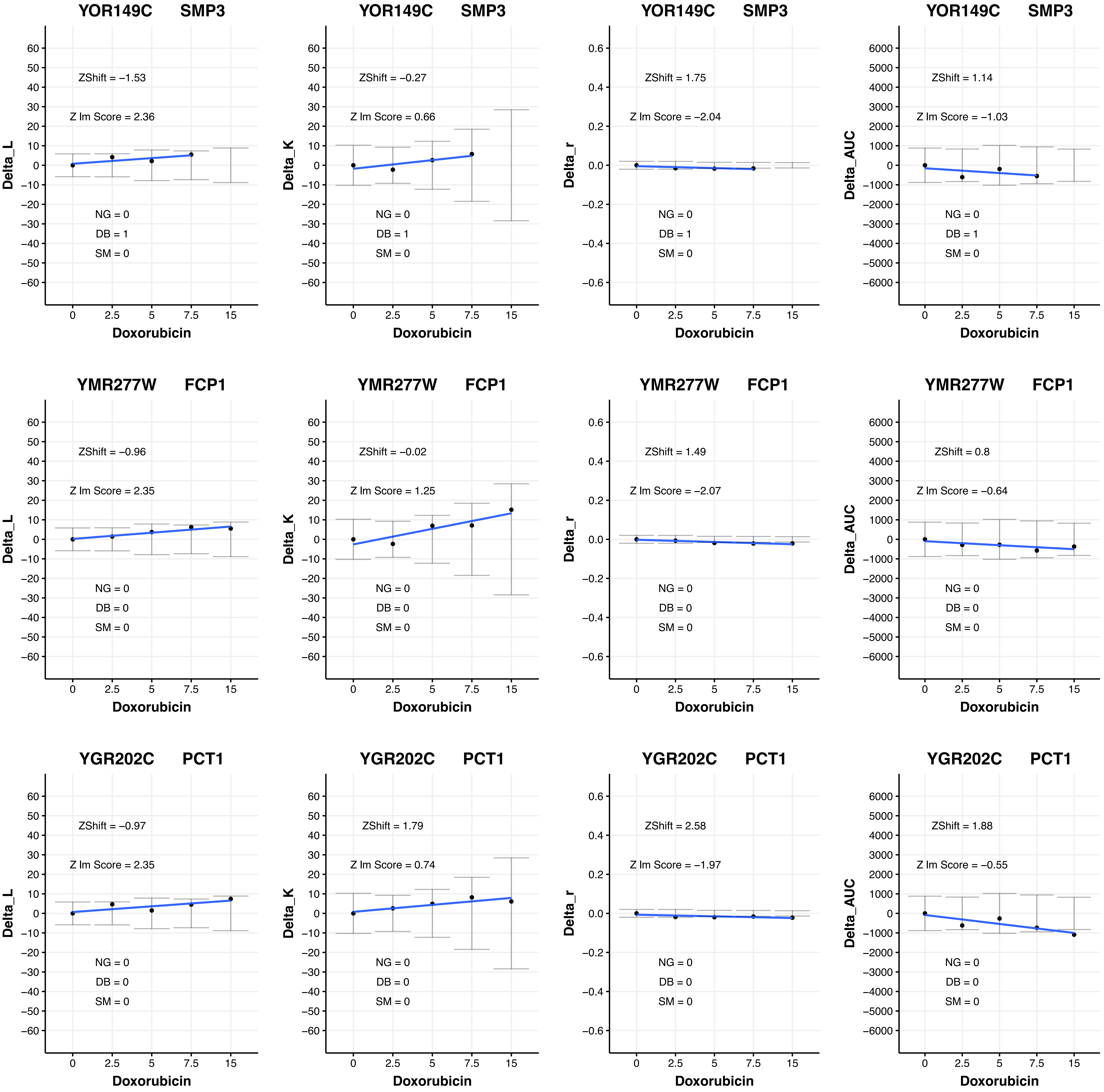
YOR149C SMP3
ZShift = –1.53
Z lm Score = 2.36
NG = 0
DB = 1
SM = 0
Delta_L
Doxorubicin
YOR149C SMP3
ZShift = –0.27
Z lm Score = 0.66
NG = 0
DB = 1
SM = 0
Delta_K
Doxorubicin
YOR149C SMP3
ZShift = 1.75
Z lm Score = –2.04
NG = 0
DB = 1
SM = 0
Delta_r
Doxorubicin
YOR149C SMP3
ZShift = 1.14
Z lm Score = –1.03
NG = 0
DB = 1
SM = 0
Delta_AUC
Doxorubicin
YMR277W FCP1
ZShift = –0.96
Z lm Score = 2.35
NG = 0
DB = 0
SM = 0
Delta_L
Doxorubicin
YMR277W FCP1
ZShift = –0.02
Z lm Score = 1.25
NG = 0
DB = 0
SM = 0
Delta_K
Doxorubicin
YMR277W FCP1
ZShift = 1.49
Z lm Score = –2.07
NG = 0
DB = 0
SM = 0
Delta_r
Doxorubicin
YMR277W FCP1
ZShift = 0.8
Z lm Score = –0.64
NG = 0
DB = 0
SM = 0
Delta_AUC
Doxorubicin
YGR202C PCT1
ZShift = –0.97
Z lm Score = 2.35
NG = 0
DB = 0
SM = 0
Delta_L
Doxorubicin
YGR202C PCT1
ZShift = 1.79
Z lm Score = 0.74
NG = 0
DB = 0
SM = 0
Delta_K
Doxorubicin
YGR202C PCT1
ZShift = 2.58
Z lm Score = –1.97
NG = 0
DB = 0
SM = 0
Delta_r
Doxorubicin
YGR202C PCT1
ZShift = 1.88
Z lm Score = –0.55
NG = 0
DB = 0
SM = 0
Delta_AUC
Doxorubicin

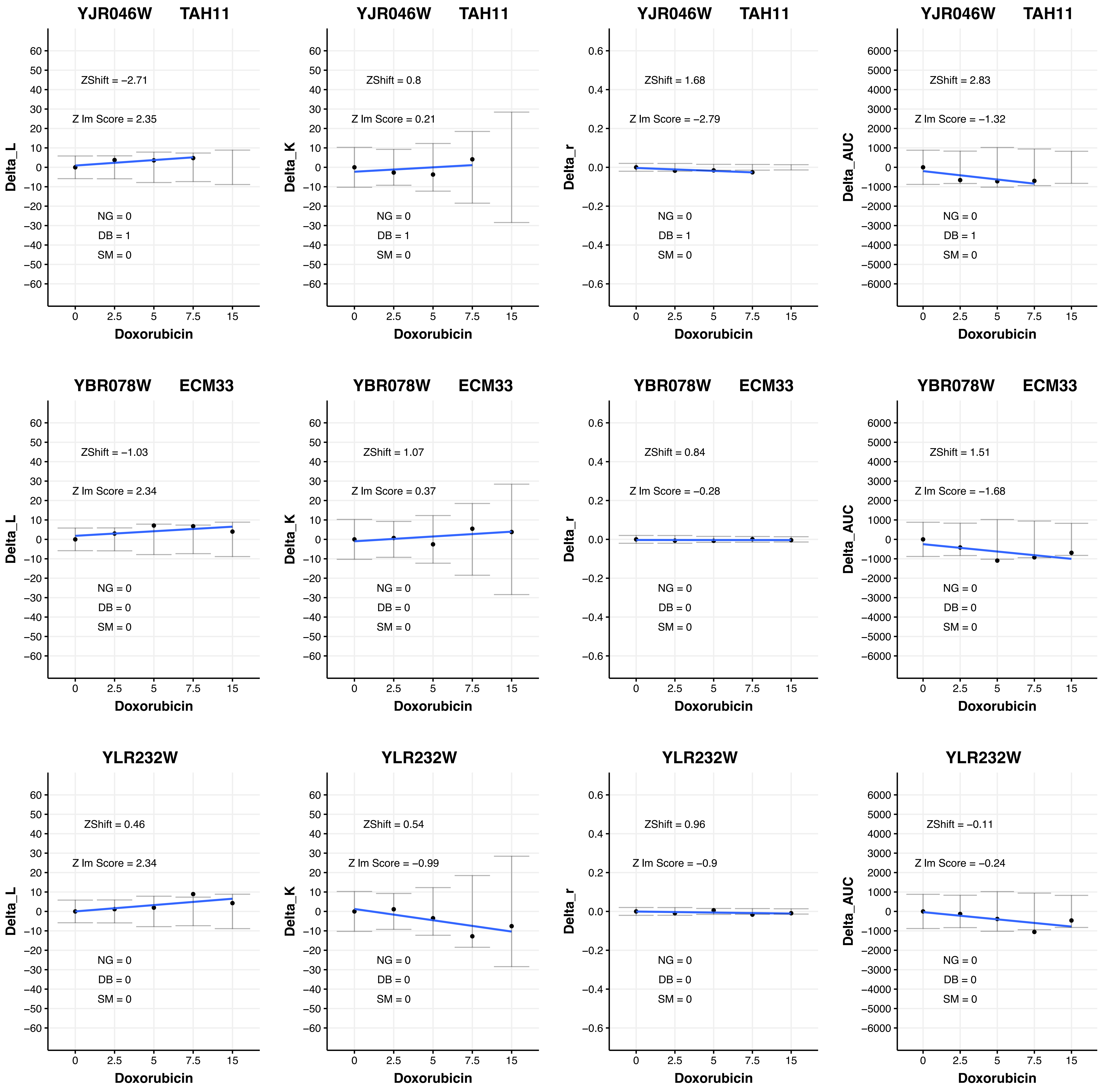
YJR046W TAH11
ZShift = −2.71
Z lm Score = 2.35
NG = 0
DB = 1
SM = 0
Delta_L
Doxorubicin
YJR046W TAH11
ZShift = 0.8
Z lm Score = 0.21
NG = 0
DB = 1
SM = 0
Delta_K
Doxorubicin
YJR046W TAH11
ZShift = 1.68
Z lm Score = −2.79
NG = 0
DB = 1
SM = 0
Delta_r
Doxorubicin
YJR046W TAH11
ZShift = 2.83
Z lm Score = −1.32
NG = 0
DB = 1
SM = 0
Delta_AUC
Doxorubicin
YBR078W ECM33
ZShift = −1.03
Z lm Score = 2.34
NG = 0
DB = 0
SM = 0
Delta_L
Doxorubicin
YBR078W ECM33
ZShift = 1.07
Z lm Score = 0.37
NG = 0
DB = 0
SM = 0
Delta_K
Doxorubicin
YBR078W ECM33
ZShift = 0.84
Z lm Score = −0.28
NG = 0
DB = 0
SM = 0
Delta_r
Doxorubicin
YBR078W ECM33
ZShift = 1.51
Z lm Score = −1.68
NG = 0
DB = 0
SM = 0
Delta_AUC
Doxorubicin
YLR232W
ZShift = 0.46
Z lm Score = 2.34
NG = 0
DB = 0
SM = 0
Delta_L
Doxorubicin
YLR232W
ZShift = 0.54
Z lm Score = −0.99
NG = 0
DB = 0
SM = 0
Delta_K
Doxorubicin
YLR232W
ZShift = 0.96
Z lm Score = −0.9
NG = 0
DB = 0
SM = 0
Delta_r
Doxorubicin
YLR232W
ZShift = −0.11
Z lm Score = −0.24
NG = 0
DB = 0
SM = 0
Delta_AUC
Doxorubicin

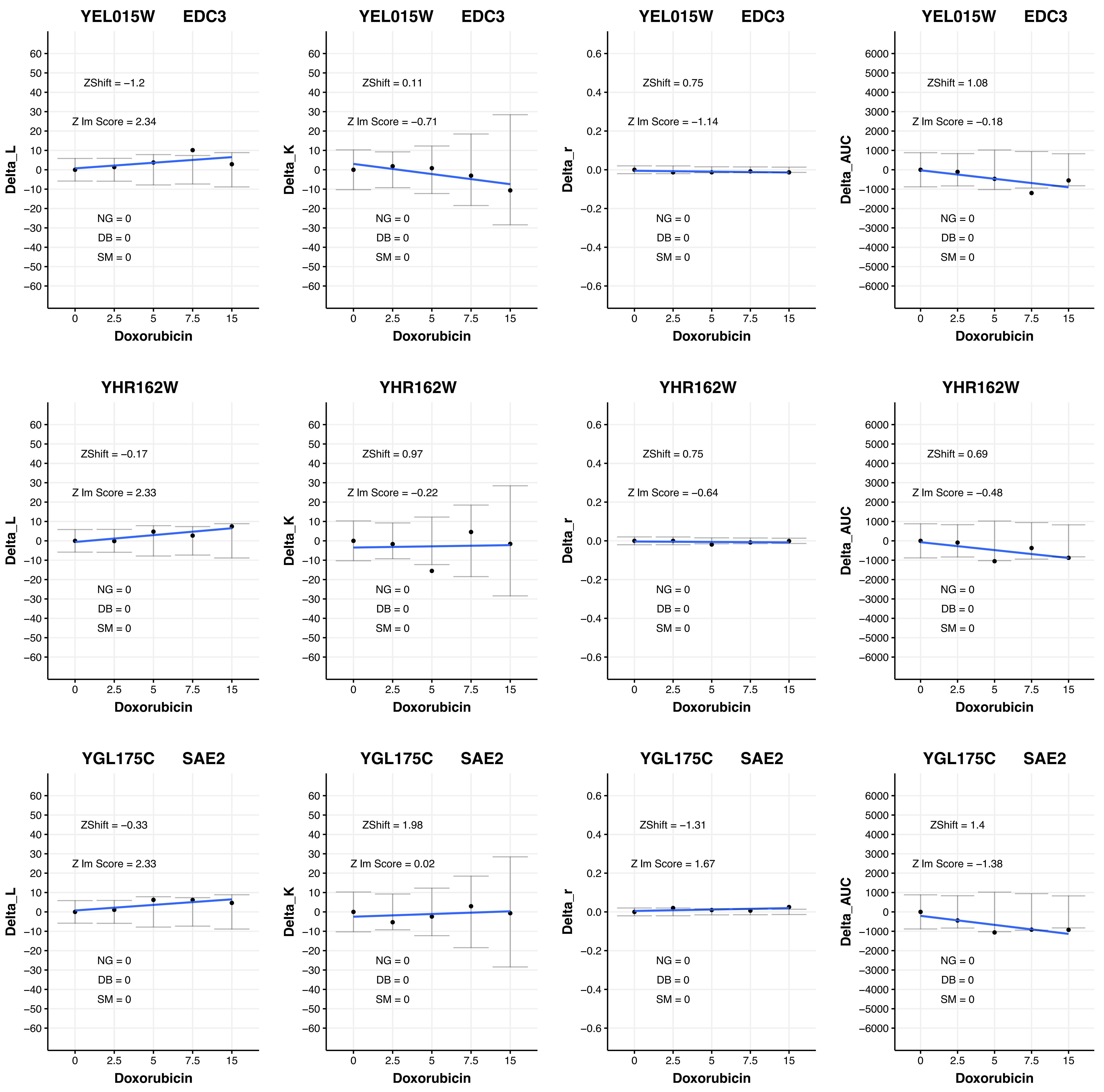

YEL015W EDC3
ZShift = −1.2
Z lm Score = 2.34
NG = 0
DB = 0
SM = 0
Delta_L
Doxorubicin
YEL015W EDC3
ZShift = 0.11
Z lm Score = −0.71
NG = 0
DB = 0
SM = 0
Delta_K
Doxorubicin
YEL015W EDC3
ZShift = 0.75
Z lm Score = −1.14
NG = 0
DB = 0
SM = 0
Delta_r
Doxorubicin
YEL015W EDC3
ZShift = 1.08
Z lm Score = −0.18
NG = 0
DB = 0
SM = 0
Delta_AUC
Doxorubicin
YHR162W
ZShift = −0.17
Z lm Score = 2.33
NG = 0
DB = 0
SM = 0
Delta_L
Doxorubicin
YHR162W
ZShift = 0.97
Z lm Score = −0.22
NG = 0
DB = 0
SM = 0
Delta_K
Doxorubicin
YHR162W
ZShift = 0.75
Z lm Score = −0.64
NG = 0
DB = 0
SM = 0
Delta_r
Doxorubicin
YHR162W
ZShift = 0.69
Z lm Score = −0.48
NG = 0
DB = 0
SM = 0
Delta_AUC
Doxorubicin
YGL175C SAE2
ZShift = −0.33
Z lm Score = 2.33
NG = 0
DB = 0
SM = 0
Delta_L
Doxorubicin
YGL175C SAE2
ZShift = 1.98
Z lm Score = 0.02
NG = 0
DB = 0
SM = 0
Delta_K
Doxorubicin
YGL175C SAE2
ZShift = −1.31
Z lm Score = 1.67
NG = 0
DB = 0
SM = 0
Delta_r
Doxorubicin
YGL175C SAE2
ZShift = 1.4
Z lm Score = −1.38
NG = 0
DB = 0
SM = 0
Delta_AUC
Doxorubicin

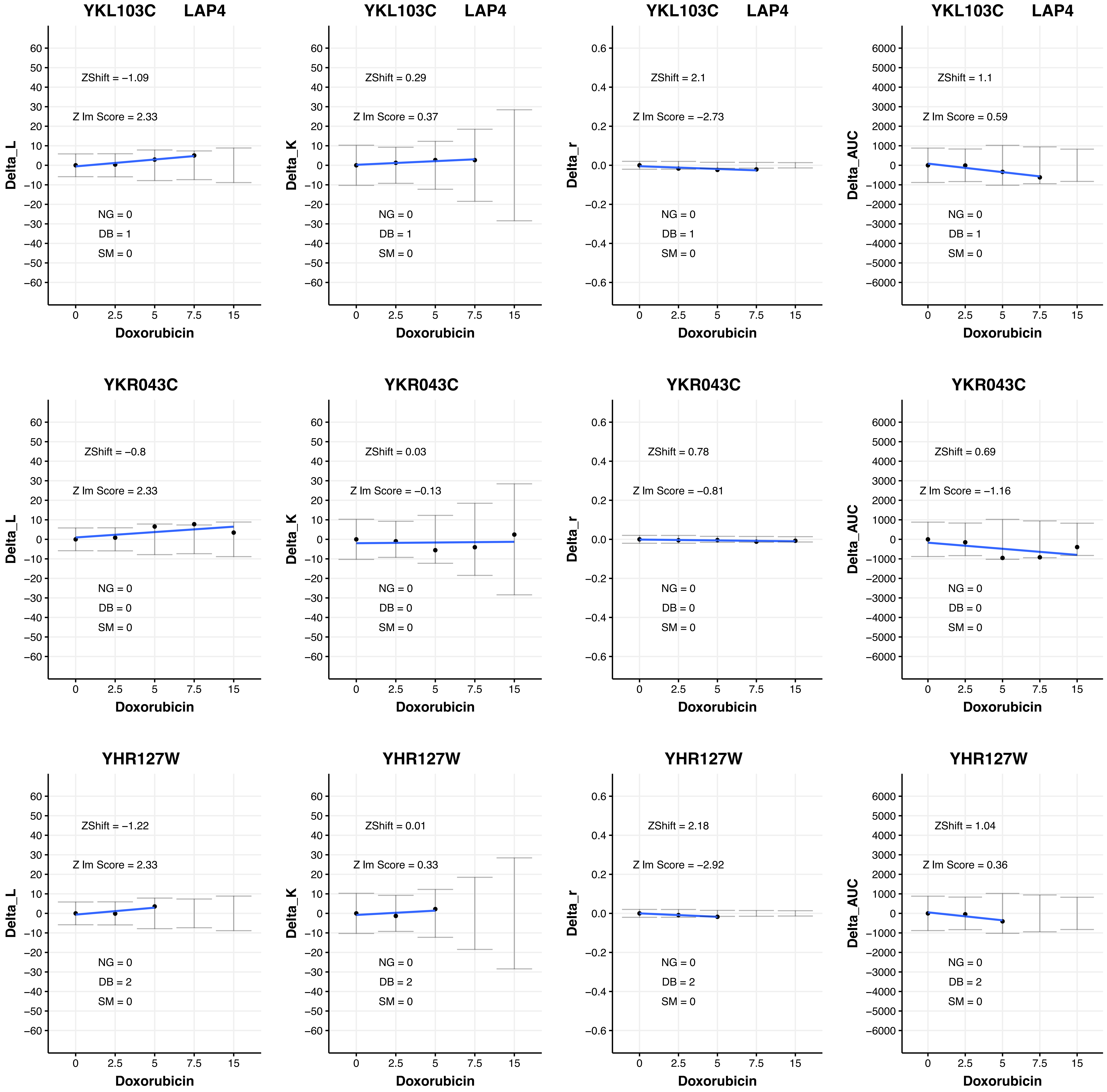

YKL103C LAP4
ZShift = −1.09
Z lm Score = 2.33
NG = 0
DB = 1
SM = 0
Delta_L
Doxorubicin
YKL103C LAP4
ZShift = 0.29
Z lm Score = 0.37
NG = 0
DB = 1
SM = 0
Delta_K
Doxorubicin
YKL103C LAP4
ZShift = 2.1
Z lm Score = −2.73
NG = 0
DB = 1
SM = 0
Delta_r
Doxorubicin
YKL103C LAP4
ZShift = 1.1
Z lm Score = 0.59
NG = 0
DB = 1
SM = 0
Delta_AUC
Doxorubicin
YKR043C
ZShift = −0.8
Z lm Score = 2.33
NG = 0
DB = 0
SM = 0
Delta_L
Doxorubicin
YKR043C
ZShift = 0.03
Z lm Score = −0.13
NG = 0
DB = 0
SM = 0
Delta_K
Doxorubicin
YKR043C
ZShift = 0.78
Z lm Score = −0.81
NG = 0
DB = 0
SM = 0
Delta_r
Doxorubicin
YKR043C
ZShift = 0.69
Z lm Score = −1.16
NG = 0
DB = 0
SM = 0
Delta_AUC
Doxorubicin
YHR127W
ZShift = −1.22
Z lm Score = 2.33
NG = 0
DB = 2
SM = 0
Delta_L
Doxorubicin
YHR127W
ZShift = 0.01
Z lm Score = 0.33
NG = 0
DB = 2
SM = 0
Delta_K
Doxorubicin
YHR127W
ZShift = 2.18
Z lm Score = −2.92
NG = 0
DB = 2
SM = 0
Delta_r
Doxorubicin
YHR127W
ZShift = 1.04
Z lm Score = 0.36
NG = 0
DB = 2
SM = 0
Delta_AUC
Doxorubicin

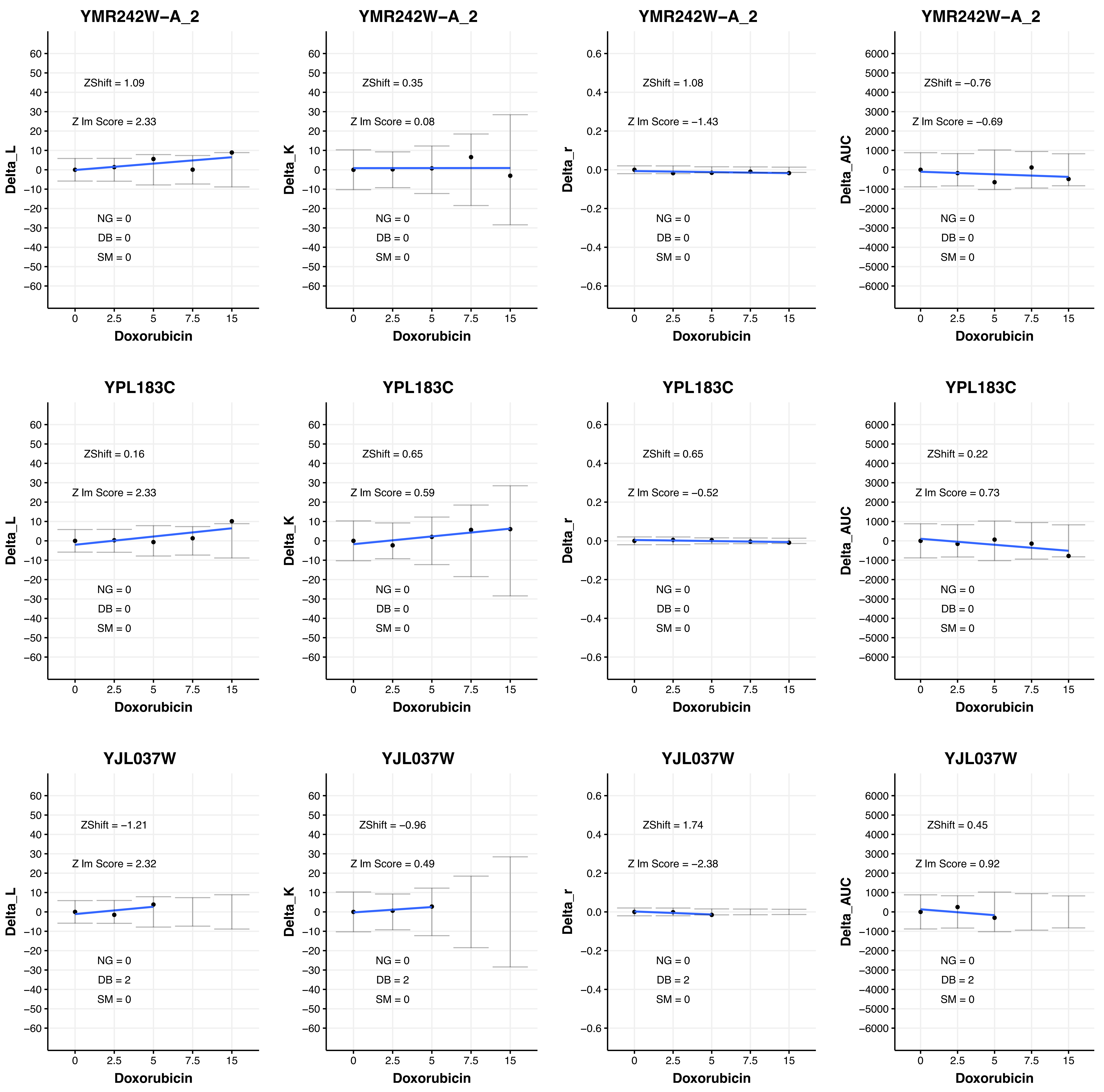
YMR242W−A_2
Delta_L
ZShift = 1.09
Z lm Score = 2.33
NG = 0
DB = 0
SM = 0
Doxorubicin
YMR242W−A_2
Delta_K
ZShift = 0.35
Z lm Score = 0.08
NG = 0
DB = 0
SM = 0
Doxorubicin
YMR242W−A_2
Delta_r
ZShift = 1.08
Z lm Score = −1.43
NG = 0
DB = 0
SM = 0
Doxorubicin
YMR242W−A_2
Delta_AUC
ZShift = −0.76
Z lm Score = −0.69
NG = 0
DB = 0
SM = 0
Doxorubicin
YPL183C
Delta_L
ZShift = 0.16
Z lm Score = 2.33
NG = 0
DB = 0
SM = 0
Doxorubicin
YPL183C
Delta_K
ZShift = 0.65
Z lm Score = 0.59
NG = 0
DB = 0
SM = 0
Doxorubicin
YPL183C
Delta_r
ZShift = 0.65
Z lm Score = −0.52
NG = 0
DB = 0
SM = 0
Doxorubicin
YPL183C
Delta_AUC
ZShift = 0.22
Z lm Score = 0.73
NG = 0
DB = 0
SM = 0
Doxorubicin
YJL037W
Delta_L
ZShift = −1.21
Z lm Score = 2.32
NG = 0
DB = 2
SM = 0
Doxorubicin
YJL037W
Delta_K
ZShift = −0.96
Z lm Score = 0.49
NG = 0
DB = 2
SM = 0
Doxorubicin
YJL037W
Delta_r
ZShift = 1.74
Z lm Score = −2.38
NG = 0
DB = 2
SM = 0
Doxorubicin
YJL037W
Delta_AUC
ZShift = 0.45
Z lm Score = 0.92
NG = 0
DB = 2
SM = 0
Doxorubicin

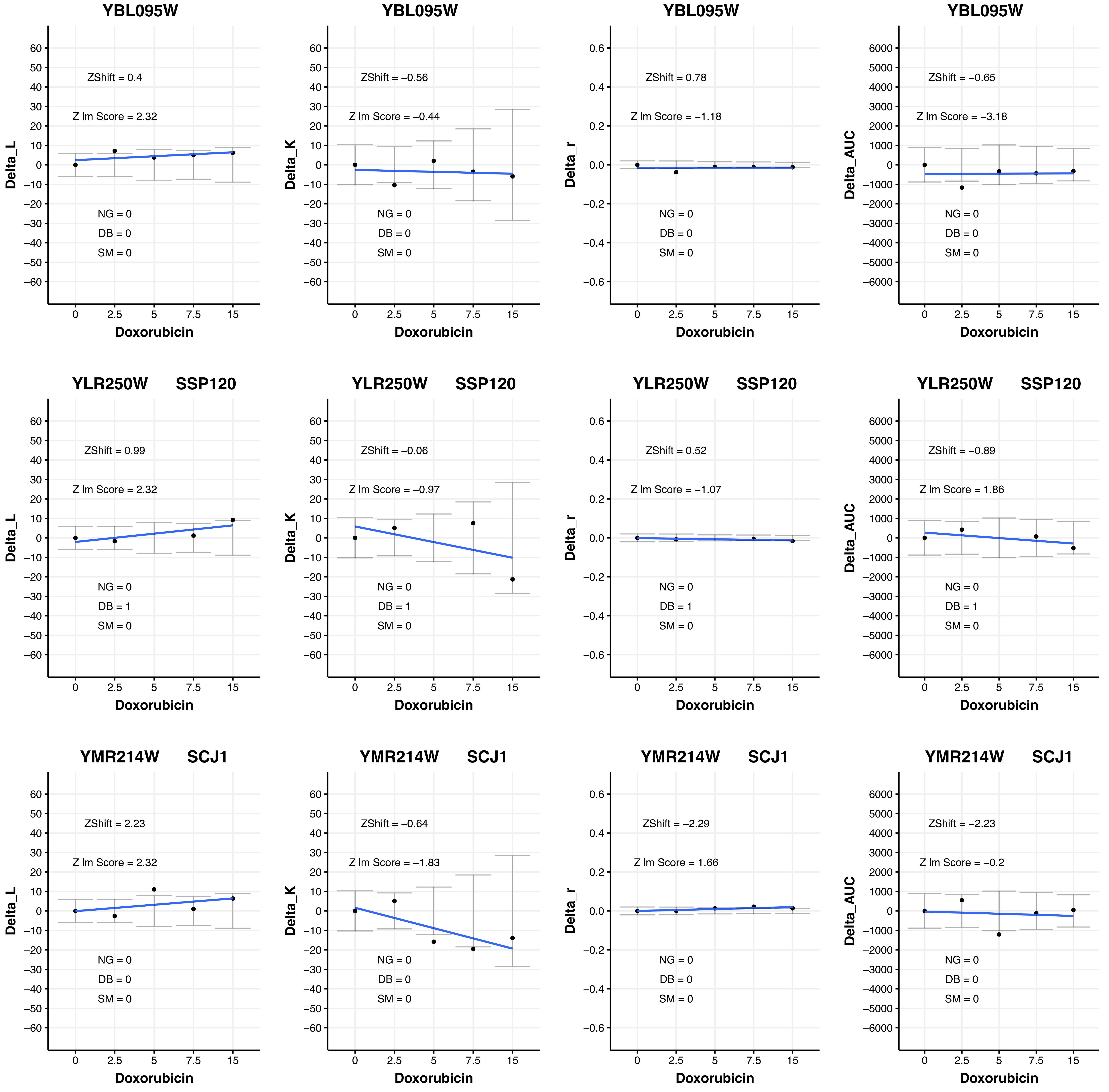

YBL095W
ZShift = 0.4
Z lm Score = 2.32
NG = 0
DB = 0
SM = 0
Delta_L
Doxorubicin
YBL095W
ZShift = −0.56
Z lm Score = −0.44
NG = 0
DB = 0
SM = 0
Delta_K
Doxorubicin
YBL095W
ZShift = 0.78
Z lm Score = −1.18
NG = 0
DB = 0
SM = 0
Delta_r
Doxorubicin
YBL095W
ZShift = −0.65
Z lm Score = −3.18
NG = 0
DB = 0
SM = 0
Delta_AUC
Doxorubicin
YLR250W SSP120
ZShift = 0.99
Z lm Score = 2.32
NG = 0
DB = 1
SM = 0
Delta_L
Doxorubicin
YLR250W SSP120
ZShift = −0.06
Z lm Score = −0.97
NG = 0
DB = 1
SM = 0
Delta_K
Doxorubicin
YLR250W SSP120
ZShift = 0.52
Z lm Score = −1.07
NG = 0
DB = 1
SM = 0
Delta_r
Doxorubicin
YLR250W SSP120
ZShift = −0.89
Z lm Score = 1.86
NG = 0
DB = 1
SM = 0
Delta_AUC
Doxorubicin
YMR214W SCJ1
ZShift = 2.23
Z lm Score = 2.32
NG = 0
DB = 0
SM = 0
Delta_L
Doxorubicin
YMR214W SCJ1
ZShift = −0.64
Z lm Score = −1.83
NG = 0
DB = 0
SM = 0
Delta_K
Doxorubicin
YMR214W SCJ1
ZShift = −2.29
Z lm Score = 1.66
NG = 0
DB = 0
SM = 0
Delta_r
Doxorubicin
YMR214W SCJ1
ZShift = −2.23
Z lm Score = −0.2
NG = 0
DB = 0
SM = 0
Delta_AUC
Doxorubicin

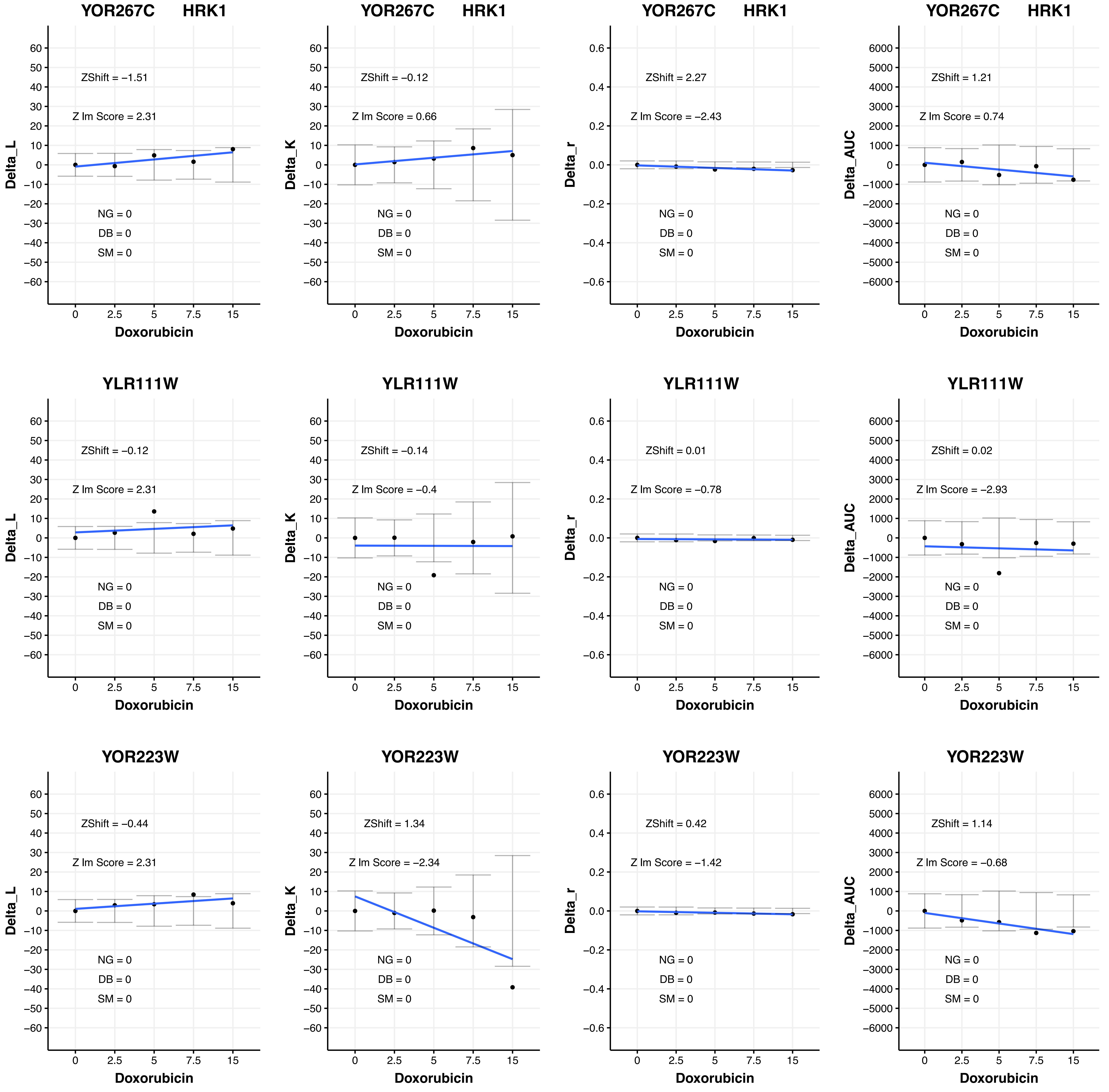

YOR267C HRK1
ZShift = −1.51
Z lm Score = 2.31
NG = 0
DB = 0
SM = 0
Delta_L
Doxorubicin
YOR267C HRK1
ZShift = −0.12
Z lm Score = 0.66
NG = 0
DB = 0
SM = 0
Delta_K
Doxorubicin
YOR267C HRK1
ZShift = 2.27
Z lm Score = −2.43
NG = 0
DB = 0
SM = 0
Delta_r
Doxorubicin
YOR267C HRK1
ZShift = 1.21
Z lm Score = 0.74
NG = 0
DB = 0
SM = 0
Delta_AUC
Doxorubicin
YLR111W
ZShift = −0.12
Z lm Score = 2.31
NG = 0
DB = 0
SM = 0
Delta_L
Doxorubicin
YLR111W
ZShift = −0.14
Z lm Score = −0.4
NG = 0
DB = 0
SM = 0
Delta_K
Doxorubicin
YLR111W
ZShift = 0.01
Z lm Score = −0.78
NG = 0
DB = 0
SM = 0
Delta_r
Doxorubicin
YLR111W
ZShift = 0.02
Z lm Score = −2.93
NG = 0
DB = 0
SM = 0
Delta_AUC
Doxorubicin
YOR223W
ZShift = −0.44
Z lm Score = 2.31
NG = 0
DB = 0
SM = 0
Delta_L
Doxorubicin
YOR223W
ZShift = 1.34
Z lm Score = −2.34
NG = 0
DB = 0
SM = 0
Delta_K
Doxorubicin
YOR223W
ZShift = 0.42
Z lm Score = −1.42
NG = 0
DB = 0
SM = 0
Delta_r
Doxorubicin
YOR223W
ZShift = 1.14
Z lm Score = −0.68
NG = 0
DB = 0
SM = 0
Delta_AUC
Doxorubicin

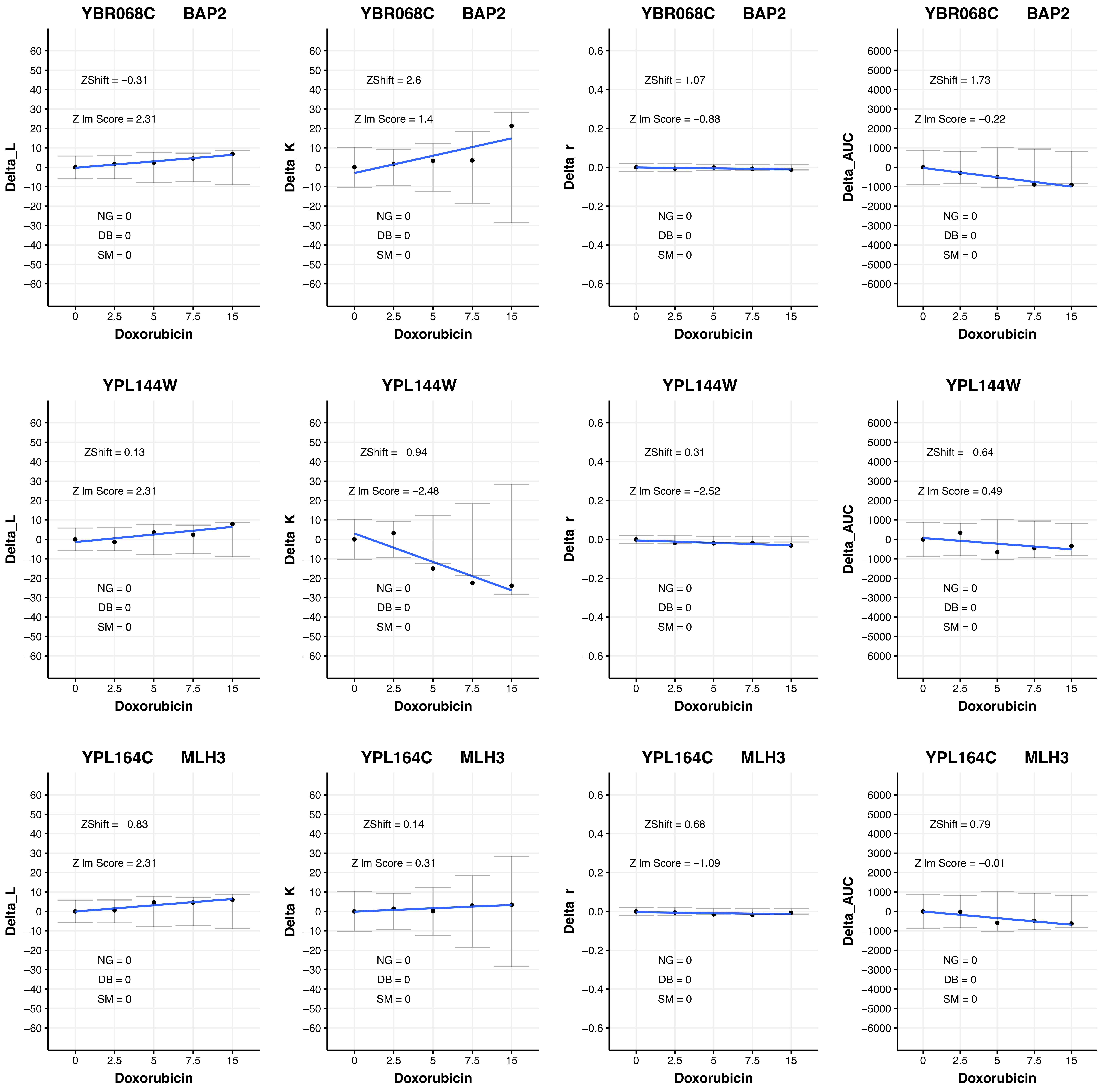
YBR068C BAP2
ZShift = −0.31
Z lm Score = 2.31
NG = 0
DB = 0
SM = 0
Delta_L
Doxorubicin
YBR068C BAP2
ZShift = 2.6
Z lm Score = 1.4
NG = 0
DB = 0
SM = 0
Delta_K
Doxorubicin
YBR068C BAP2
ZShift = 1.07
Z lm Score = −0.88
NG = 0
DB = 0
SM = 0
Delta_r
Doxorubicin
YBR068C BAP2
ZShift = 1.73
Z lm Score = −0.22
NG = 0
DB = 0
SM = 0
Delta_AUC
Doxorubicin
YPL144W
ZShift = 0.13
Z lm Score = 2.31
NG = 0
DB = 0
SM = 0
Delta_L
Doxorubicin
YPL144W
ZShift = −0.94
Z lm Score = −2.48
NG = 0
DB = 0
SM = 0
Delta_K
Doxorubicin
YPL144W
ZShift = 0.31
Z lm Score = −2.52
NG = 0
DB = 0
SM = 0
Delta_r
Doxorubicin
YPL144W
ZShift = −0.64
Z lm Score = 0.49
NG = 0
DB = 0
SM = 0
Delta_AUC
Doxorubicin
YPL164C MLH3
ZShift = −0.83
Z lm Score = 2.31
NG = 0
DB = 0
SM = 0
Delta_L
Doxorubicin
YPL164C MLH3
ZShift = 0.14
Z lm Score = 0.31
NG = 0
DB = 0
SM = 0
Delta_K
Doxorubicin
YPL164C MLH3
ZShift = 0.68
Z lm Score = −1.09
NG = 0
DB = 0
SM = 0
Delta_r
Doxorubicin
YPL164C MLH3
ZShift = 0.79
Z lm Score = −0.01
NG = 0
DB = 0
SM = 0
Delta_AUC
Doxorubicin

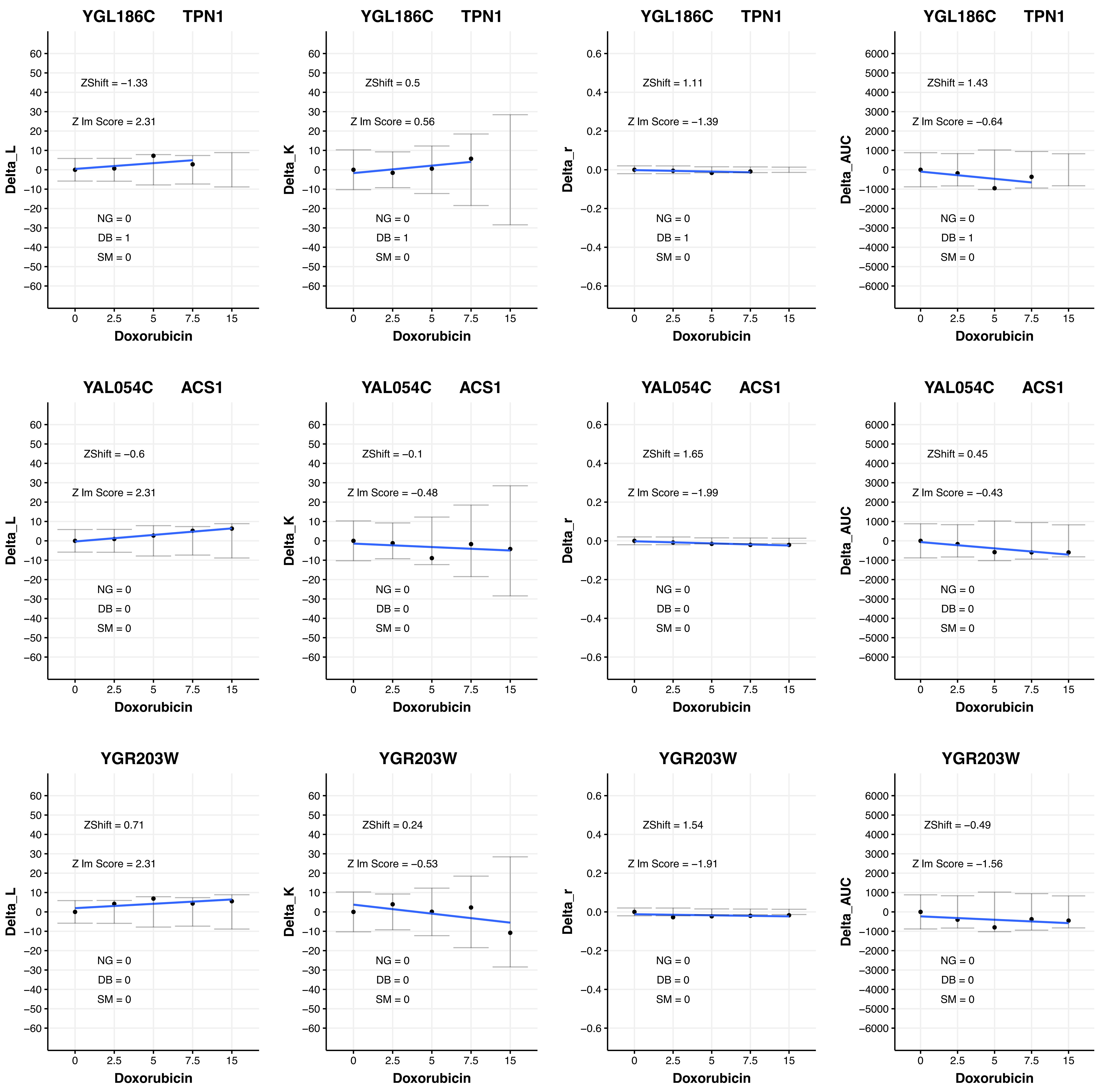
YGL186C TPN1
ZShift = −1.33
Z lm Score = 2.31
NG = 0
DB = 1
SM = 0
Delta_L
Doxorubicin
YGL186C TPN1
ZShift = 0.5
Z lm Score = 0.56
NG = 0
DB = 1
SM = 0
Delta_K
Doxorubicin
YGL186C TPN1
ZShift = 1.11
Z lm Score = −1.39
NG = 0
DB = 1
SM = 0
Delta_r
Doxorubicin
YGL186C TPN1
ZShift = 1.43
Z lm Score = −0.64
NG = 0
DB = 1
SM = 0
Delta_AUC
Doxorubicin
YAL054C ACS1
ZShift = −0.6
Z lm Score = 2.31
NG = 0
DB = 0
SM = 0
Delta_L
Doxorubicin
YAL054C ACS1
ZShift = −0.1
Z lm Score = −0.48
NG = 0
DB = 0
SM = 0
Delta_K
Doxorubicin
YAL054C ACS1
ZShift = 1.65
Z lm Score = −1.99
NG = 0
DB = 0
SM = 0
Delta_r
Doxorubicin
YAL054C ACS1
ZShift = 0.45
Z lm Score = −0.43
NG = 0
DB = 0
SM = 0
Delta_AUC
Doxorubicin
YGR203W
ZShift = 0.71
Z lm Score = 2.31
NG = 0
DB = 0
SM = 0
Delta_L
Doxorubicin
YGR203W
ZShift = 0.24
Z lm Score = −0.53
NG = 0
DB = 0
SM = 0
Delta_K
Doxorubicin
YGR203W
ZShift = 1.54
Z lm Score = −1.91
NG = 0
DB = 0
SM = 0
Delta_r
Doxorubicin
YGR203W
ZShift = −0.49
Z lm Score = −1.56
NG = 0
DB = 0
SM = 0
Delta_AUC
Doxorubicin

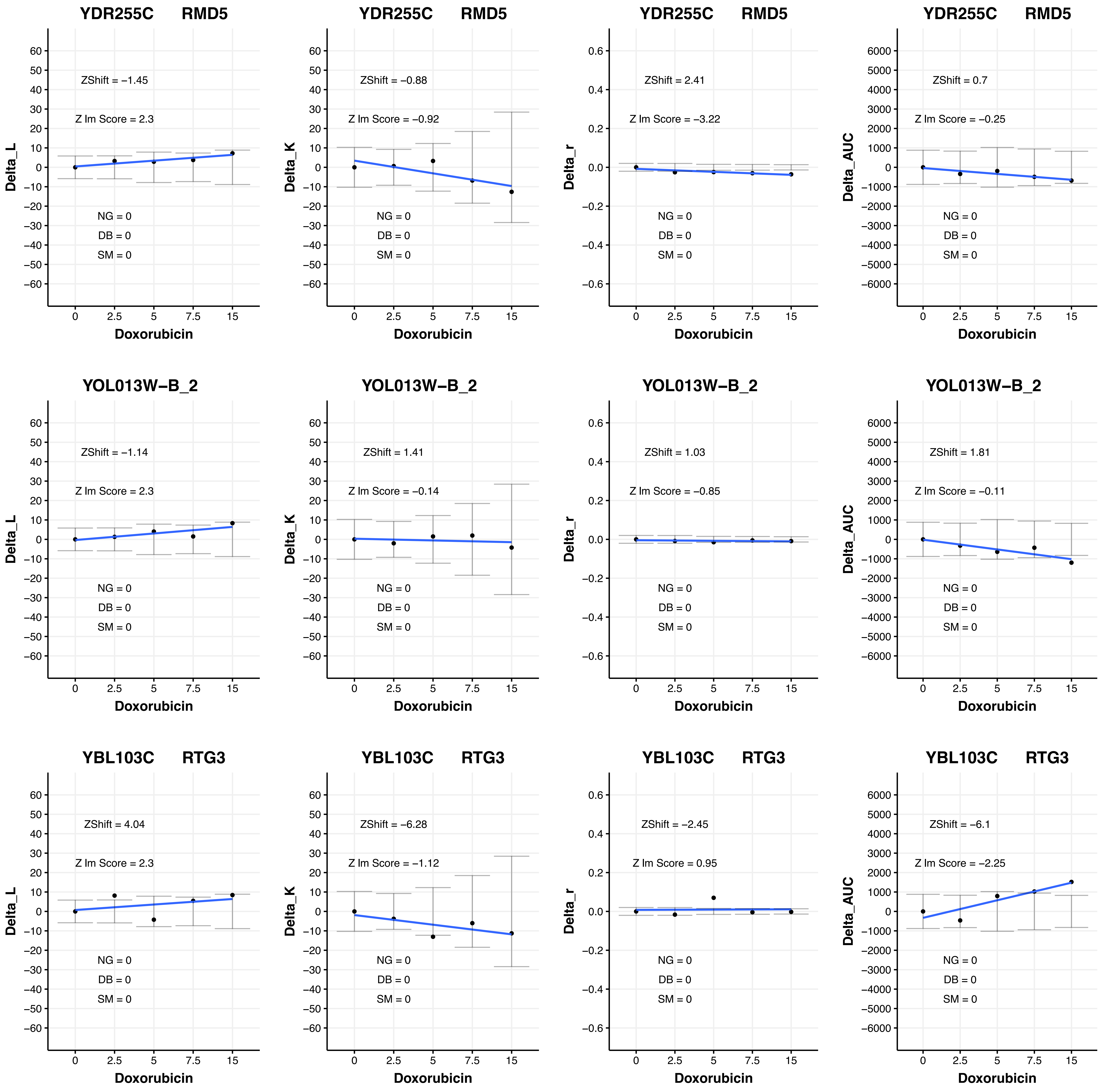

YDR255C RMD5
ZShift = −1.45
Z lm Score = 2.3
NG = 0
DB = 0
SM = 0
Delta_L
Doxorubicin
YDR255C RMD5
ZShift = −0.88
Z lm Score = −0.92
NG = 0
DB = 0
SM = 0
Delta_K
Doxorubicin
YDR255C RMD5
ZShift = 2.41
Z lm Score = −3.22
NG = 0
DB = 0
SM = 0
Delta_r
Doxorubicin
YDR255C RMD5
ZShift = 0.7
Z lm Score = −0.25
NG = 0
DB = 0
SM = 0
Delta_AUC
Doxorubicin
YOL013W−B_2
ZShift = −1.14
Z lm Score = 2.3
NG = 0
DB = 0
SM = 0
Delta_L
Doxorubicin
YOL013W−B_2
ZShift = 1.41
Z lm Score = −0.14
NG = 0
DB = 0
SM = 0
Delta_K
Doxorubicin
YOL013W−B_2
ZShift = 1.03
Z lm Score = −0.85
NG = 0
DB = 0
SM = 0
Delta_r
Doxorubicin
YOL013W−B_2
ZShift = 1.81
Z lm Score = −0.11
NG = 0
DB = 0
SM = 0
Delta_AUC
Doxorubicin
YBL103C RTG3
ZShift = 4.04
Z lm Score = 2.3
NG = 0
DB = 0
SM = 0
Delta_L
Doxorubicin
YBL103C RTG3
ZShift = −6.28
Z lm Score = −1.12
NG = 0
DB = 0
SM = 0
Delta_K
Doxorubicin
YBL103C RTG3
ZShift = −2.45
Z lm Score = 0.95
NG = 0
DB = 0
SM = 0
Delta_r
Doxorubicin
YBL103C RTG3
ZShift = −6.1
Z lm Score = −2.25
NG = 0
DB = 0
SM = 0
Delta_AUC
Doxorubicin

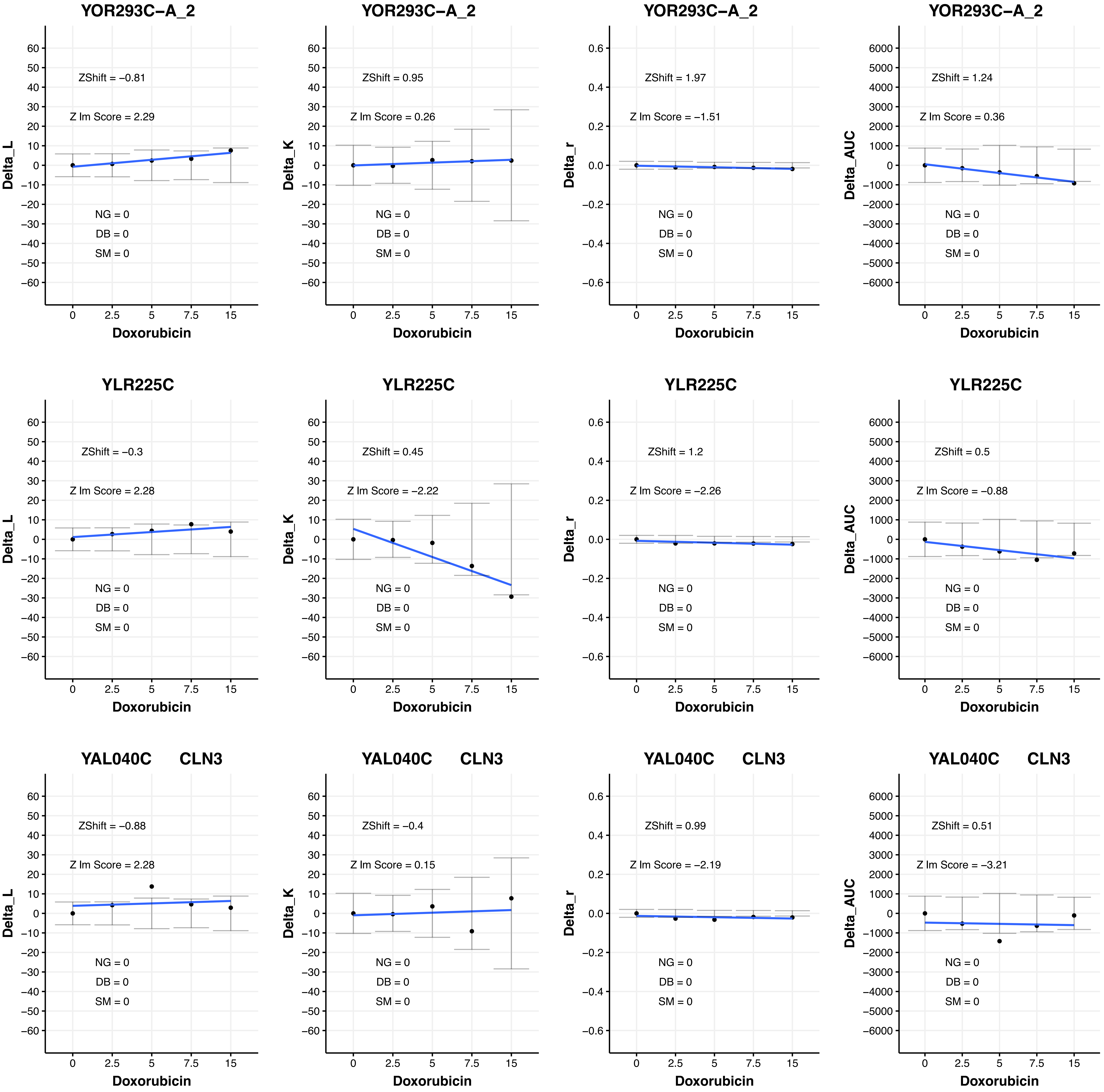
YOR293C−A_2
ZShift = −0.81
Z lm Score = 2.29
NG = 0
DB = 0
SM = 0
Delta_L
Doxorubicin
YOR293C−A_2
ZShift = 0.95
Z lm Score = 0.26
NG = 0
DB = 0
SM = 0
Delta_K
Doxorubicin
YOR293C−A_2
ZShift = 1.97
Z lm Score = −1.51
NG = 0
DB = 0
SM = 0
Delta_r
Doxorubicin
YOR293C−A_2
ZShift = 1.24
Z lm Score = 0.36
NG = 0
DB = 0
SM = 0
Delta_AUC
Doxorubicin
YLR225C
ZShift = −0.3
Z lm Score = 2.28
NG = 0
DB = 0
SM = 0
Delta_L
Doxorubicin
YLR225C
ZShift = 0.45
Z lm Score = −2.22
NG = 0
DB = 0
SM = 0
Delta_K
Doxorubicin
YLR225C
ZShift = 1.2
Z lm Score = −2.26
NG = 0
DB = 0
SM = 0
Delta_r
Doxorubicin
YLR225C
ZShift = 0.5
Z lm Score = −0.88
NG = 0
DB = 0
SM = 0
Delta_AUC
Doxorubicin
YAL040C CLN3
ZShift = −0.88
Z lm Score = 2.28
NG = 0
DB = 0
SM = 0
Delta_L
Doxorubicin
YAL040C CLN3
ZShift = −0.4
Z lm Score = 0.15
NG = 0
DB = 0
SM = 0
Delta_K
Doxorubicin
YAL040C CLN3
ZShift = 0.99
Z lm Score = −2.19
NG = 0
DB = 0
SM = 0
Delta_r
Doxorubicin
YAL040C CLN3
ZShift = 0.51
Z lm Score = −3.21
NG = 0
DB = 0
SM = 0
Delta_AUC
Doxorubicin

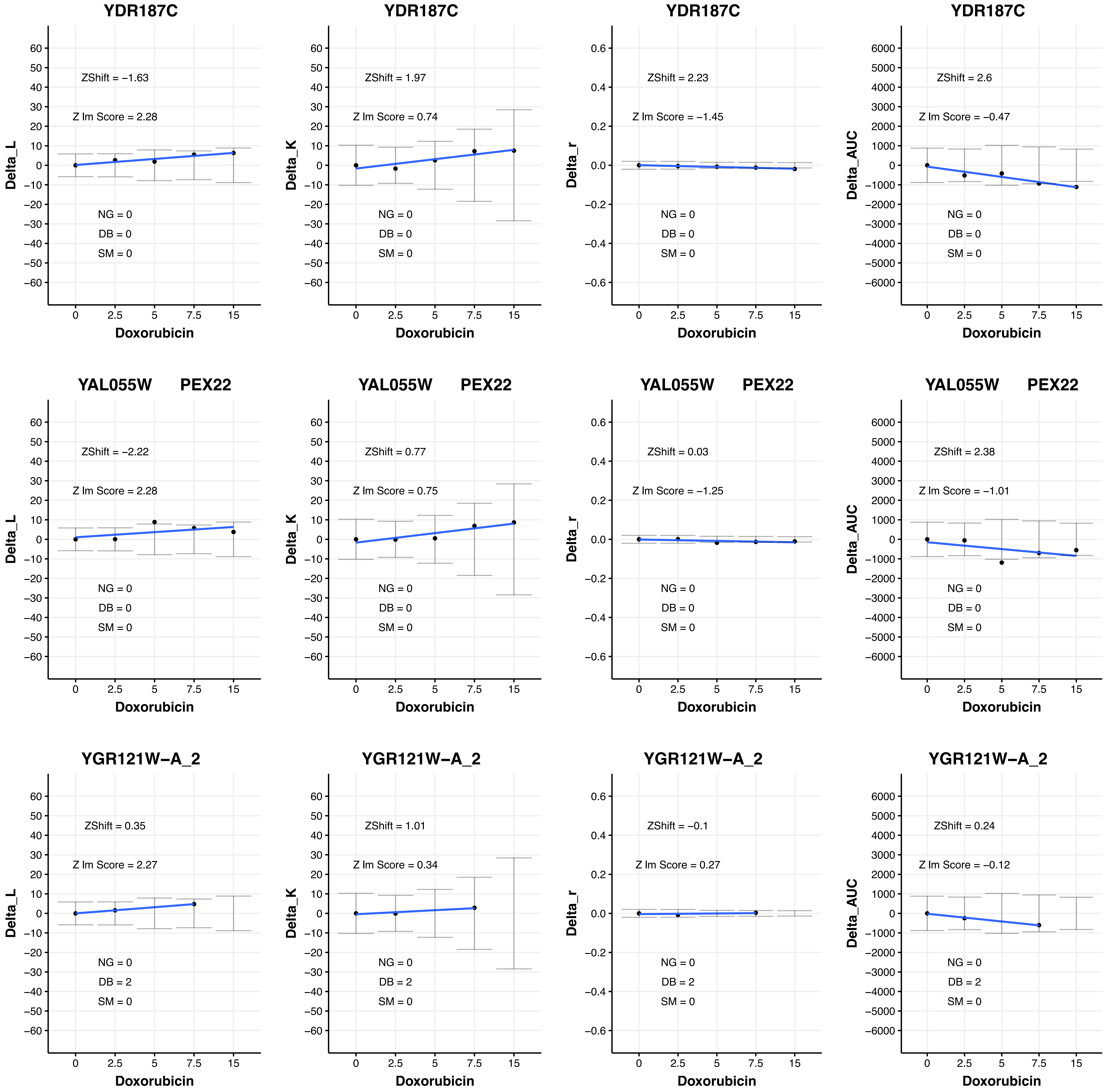

YDR187C
ZShift = −1.63
Z lm Score = 2.28
NG = 0
DB = 0
SM = 0
Delta_L
Doxorubicin
YDR187C
ZShift = 1.97
Z lm Score = 0.74
NG = 0
DB = 0
SM = 0
Delta_K
Doxorubicin
YDR187C
ZShift = 2.23
Z lm Score = −1.45
NG = 0
DB = 0
SM = 0
Delta_r
Doxorubicin
YDR187C
ZShift = 2.6
Z lm Score = −0.47
NG = 0
DB = 0
SM = 0
Delta_AUC
Doxorubicin
YAL055W PEX22
ZShift = −2.22
Z lm Score = 2.28
NG = 0
DB = 0
SM = 0
Delta_L
Doxorubicin
YAL055W PEX22
ZShift = 0.77
Z lm Score = 0.75
NG = 0
DB = 0
SM = 0
Delta_K
Doxorubicin
YAL055W PEX22
ZShift = 0.03
Z lm Score = −1.25
NG = 0
DB = 0
SM = 0
Delta_r
Doxorubicin
YAL055W PEX22
ZShift = 2.38
Z lm Score = −1.01
NG = 0
DB = 0
SM = 0
Delta_AUC
Doxorubicin
YGR121W−A_2
ZShift = 0.35
Z lm Score = 2.27
NG = 0
DB = 2
SM = 0
Delta_L
Doxorubicin
YGR121W−A_2
ZShift = 1.01
Z lm Score = 0.34
NG = 0
DB = 2
SM = 0
Delta_K
Doxorubicin
YGR121W−A_2
ZShift = −0.1
Z lm Score = 0.27
NG = 0
DB = 2
SM = 0
Delta_r
Doxorubicin
YGR121W−A_2
ZShift = 0.24
Z lm Score = −0.12
NG = 0
DB = 2
SM = 0
Delta_AUC
Doxorubicin

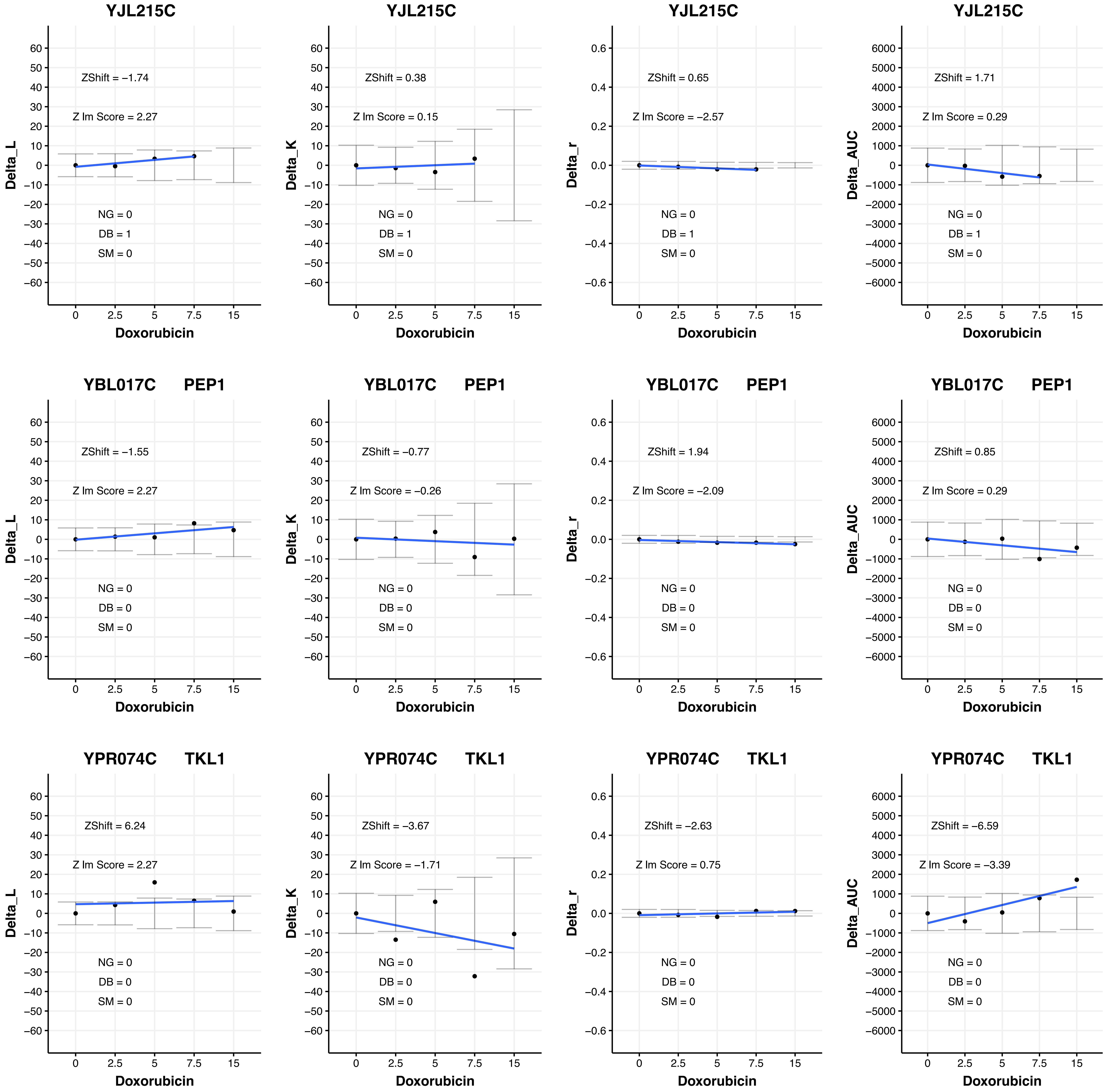

YJL215C
ZShift = −1.74
Z lm Score = 2.27
NG = 0
DB = 1
SM = 0
Delta_L
Doxorubicin
YJL215C
ZShift = 0.38
Z lm Score = 0.15
NG = 0
DB = 1
SM = 0
Delta_K
Doxorubicin
YJL215C
ZShift = 0.65
Z lm Score = −2.57
NG = 0
DB = 1
SM = 0
Delta_r
Doxorubicin
YJL215C
ZShift = 1.71
Z lm Score = 0.29
NG = 0
DB = 1
SM = 0
Delta_AUC
Doxorubicin
YBL017C PEP1
ZShift = −1.55
Z lm Score = 2.27
NG = 0
DB = 0
SM = 0
Delta_L
Doxorubicin
YBL017C PEP1
ZShift = −0.77
Z lm Score = −0.26
NG = 0
DB = 0
SM = 0
Delta_K
Doxorubicin
YBL017C PEP1
ZShift = 1.94
Z lm Score = −2.09
NG = 0
DB = 0
SM = 0
Delta_r
Doxorubicin
YBL017C PEP1
ZShift = 0.85
Z lm Score = 0.29
NG = 0
DB = 0
SM = 0
Delta_AUC
Doxorubicin
YPR074C TKL1
ZShift = 6.24
Z lm Score = 2.27
NG = 0
DB = 0
SM = 0
Delta_L
Doxorubicin
YPR074C TKL1
ZShift = −3.67
Z lm Score = −1.71
NG = 0
DB = 0
SM = 0
Delta_K
Doxorubicin
YPR074C TKL1
ZShift = −2.63
Z lm Score = 0.75
NG = 0
DB = 0
SM = 0
Delta_r
Doxorubicin
YPR074C TKL1
ZShift = −6.59
Z lm Score = −3.39
NG = 0
DB = 0
SM = 0
Delta_AUC
Doxorubicin

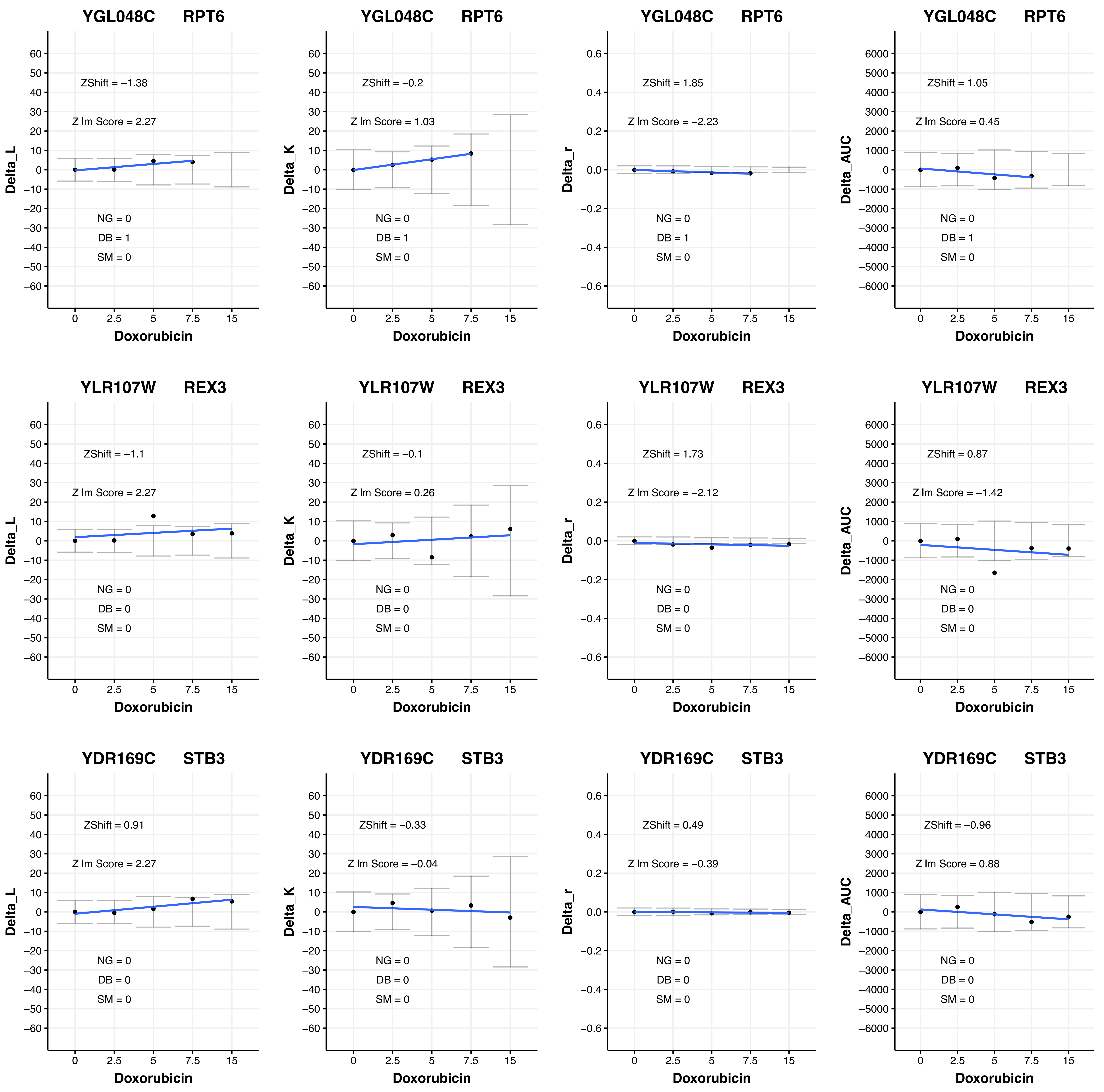

YGL048C RPT6
ZShift = −1.38
Z lm Score = 2.27
NG = 0
DB = 1
SM = 0
Delta_L
Doxorubicin
YGL048C RPT6
ZShift = −0.2
Z lm Score = 1.03
NG = 0
DB = 1
SM = 0
Delta_K
Doxorubicin
YGL048C RPT6
ZShift = 1.85
Z lm Score = −2.23
NG = 0
DB = 1
SM = 0
Delta_r
Doxorubicin
YGL048C RPT6
ZShift = 1.05
Z lm Score = 0.45
NG = 0
DB = 1
SM = 0
Delta_AUC
Doxorubicin
YLR107W REX3
ZShift = −1.1
Z lm Score = 2.27
NG = 0
DB = 0
SM = 0
Delta_L
Doxorubicin
YLR107W REX3
ZShift = −0.1
Z lm Score = 0.26
NG = 0
DB = 0
SM = 0
Delta_K
Doxorubicin
YLR107W REX3
ZShift = 1.73
Z lm Score = −2.12
NG = 0
DB = 0
SM = 0
Delta_r
Doxorubicin
YLR107W REX3
ZShift = 0.87
Z lm Score = −1.42
NG = 0
DB = 0
SM = 0
Delta_AUC
Doxorubicin
YDR169C STB3
ZShift = 0.91
Z lm Score = 2.27
NG = 0
DB = 0
SM = 0
Delta_L
Doxorubicin
YDR169C STB3
ZShift = −0.33
Z lm Score = −0.04
NG = 0
DB = 0
SM = 0
Delta_K
Doxorubicin
YDR169C STB3
ZShift = 0.49
Z lm Score = −0.39
NG = 0
DB = 0
SM = 0
Delta_r
Doxorubicin
YDR169C STB3
ZShift = −0.96
Z lm Score = 0.88
NG = 0
DB = 0
SM = 0
Delta_AUC
Doxorubicin

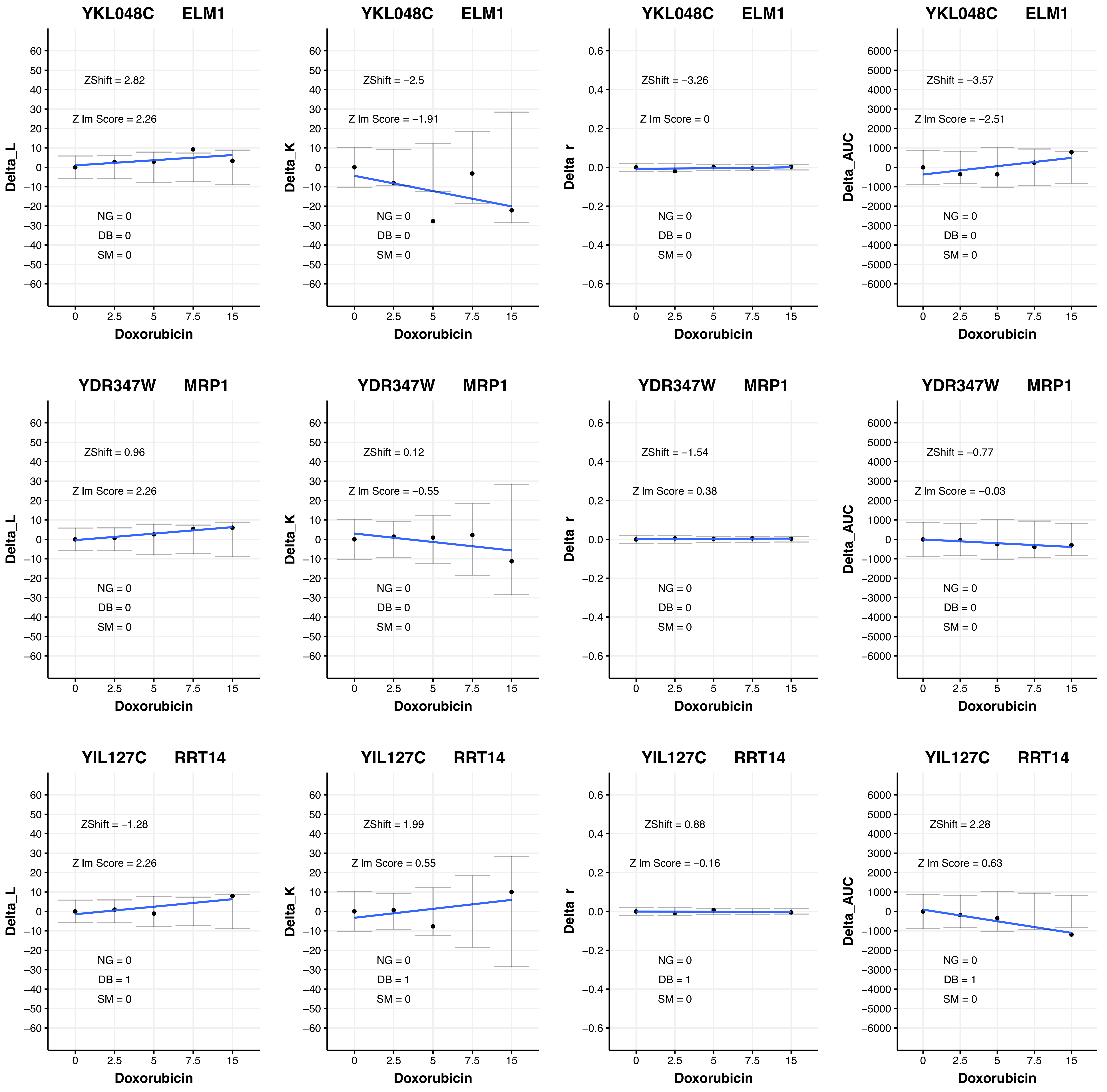
YKL048C ELM1
ZShift = 2.82
Z lm Score = 2.26
NG = 0
DB = 0
SM = 0
Delta_L
Doxorubicin
YKL048C ELM1
ZShift = −2.5
Z lm Score = −1.91
NG = 0
DB = 0
SM = 0
Delta_K
Doxorubicin
YKL048C ELM1
ZShift = −3.26
Z lm Score = 0
NG = 0
DB = 0
SM = 0
Delta_r
Doxorubicin
YKL048C ELM1
ZShift = −3.57
Z lm Score = −2.51
NG = 0
DB = 0
SM = 0
Delta_AUC
Doxorubicin
YDR347W MRP1
ZShift = 0.96
Z lm Score = 2.26
NG = 0
DB = 0
SM = 0
Delta_L
Doxorubicin
YDR347W MRP1
ZShift = 0.12
Z lm Score = −0.55
NG = 0
DB = 0
SM = 0
Delta_K
Doxorubicin
YDR347W MRP1
ZShift = −1.54
Z lm Score = 0.38
NG = 0
DB = 0
SM = 0
Delta_r
Doxorubicin
YDR347W MRP1
ZShift = −0.77
Z lm Score = −0.03
NG = 0
DB = 0
SM = 0
Delta_AUC
Doxorubicin
YIL127C RRT14
ZShift = −1.28
Z lm Score = 2.26
NG = 0
DB = 1
SM = 0
Delta_L
Doxorubicin
YIL127C RRT14
ZShift = 1.99
Z lm Score = 0.55
NG = 0
DB = 1
SM = 0
Delta_K
Doxorubicin
YIL127C RRT14
ZShift = 0.88
Z lm Score = −0.16
NG = 0
DB = 1
SM = 0
Delta_r
Doxorubicin
YIL127C RRT14
ZShift = 2.28
Z lm Score = 0.63
NG = 0
DB = 1
SM = 0
Delta_AUC
Doxorubicin

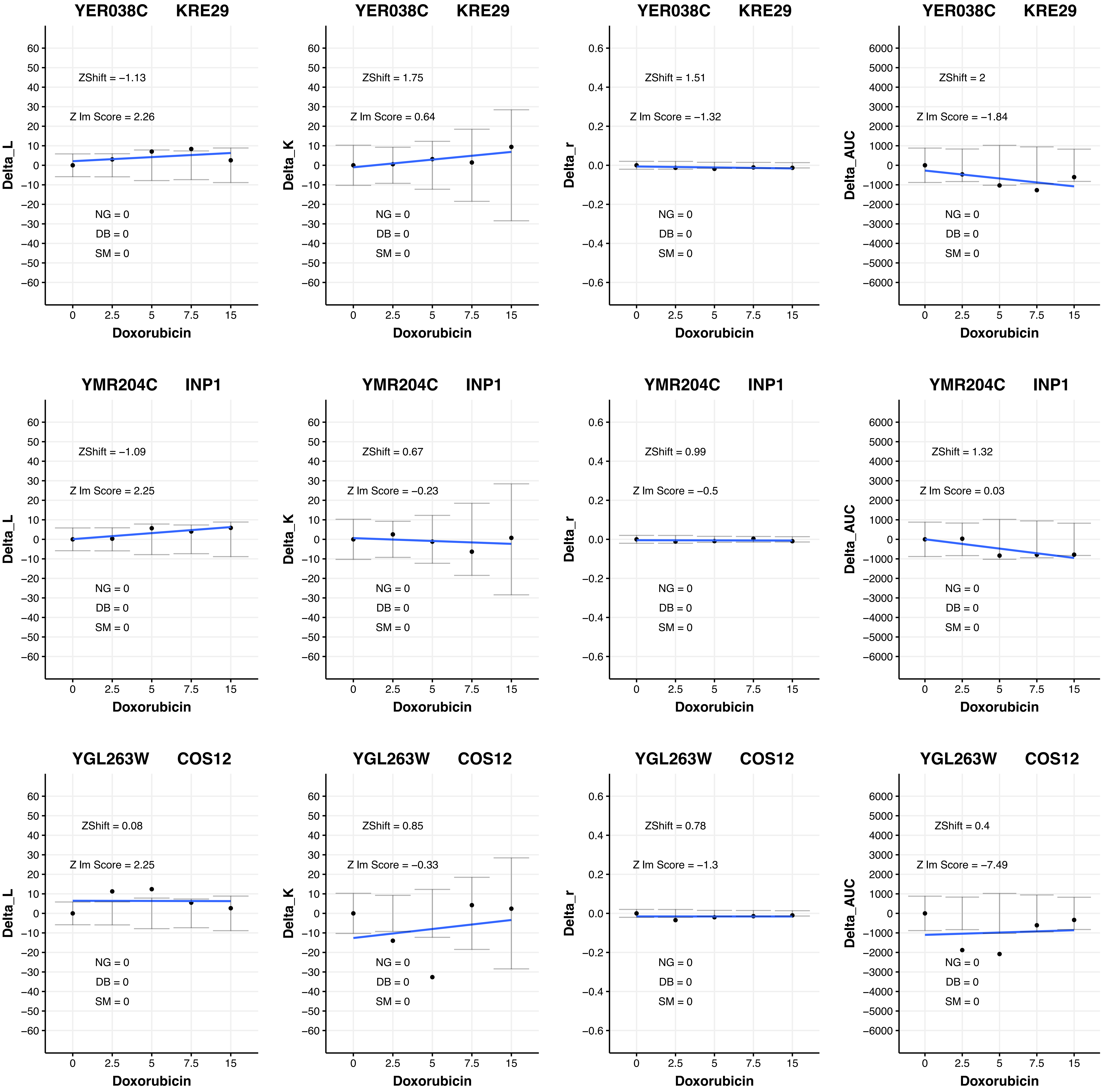
YER038C KRE29
ZShift = −1.13
Z lm Score = 2.26
NG = 0
DB = 0
SM = 0
Delta_L
Doxorubicin
YER038C KRE29
ZShift = 1.75
Z lm Score = 0.64
NG = 0
DB = 0
SM = 0
Delta_K
Doxorubicin
YER038C KRE29
ZShift = 1.51
Z lm Score = −1.32
NG = 0
DB = 0
SM = 0
Delta_r
Doxorubicin
YER038C KRE29
ZShift = 2
Z lm Score = −1.84
NG = 0
DB = 0
SM = 0
Delta_AUC
Doxorubicin
YMR204C INP1
ZShift = −1.09
Z lm Score = 2.25
NG = 0
DB = 0
SM = 0
Delta_L
Doxorubicin
YMR204C INP1
ZShift = 0.67
Z lm Score = −0.23
NG = 0
DB = 0
SM = 0
Delta_K
Doxorubicin
YMR204C INP1
ZShift = 0.99
Z lm Score = −0.5
NG = 0
DB = 0
SM = 0
Delta_r
Doxorubicin
YMR204C INP1
ZShift = 1.32
Z lm Score = 0.03
NG = 0
DB = 0
SM = 0
Delta_AUC
Doxorubicin
YGL263W COS12
ZShift = 0.08
Z lm Score = 2.25
NG = 0
DB = 0
SM = 0
Delta_L
Doxorubicin
YGL263W COS12
ZShift = 0.85
Z lm Score = −0.33
NG = 0
DB = 0
SM = 0
Delta_K
Doxorubicin
YGL263W COS12
ZShift = 0.78
Z lm Score = −1.3
NG = 0
DB = 0
SM = 0
Delta_r
Doxorubicin
YGL263W COS12
ZShift = 0.4
Z lm Score = −7.49
NG = 0
DB = 0
SM = 0
Delta_AUC
Doxorubicin

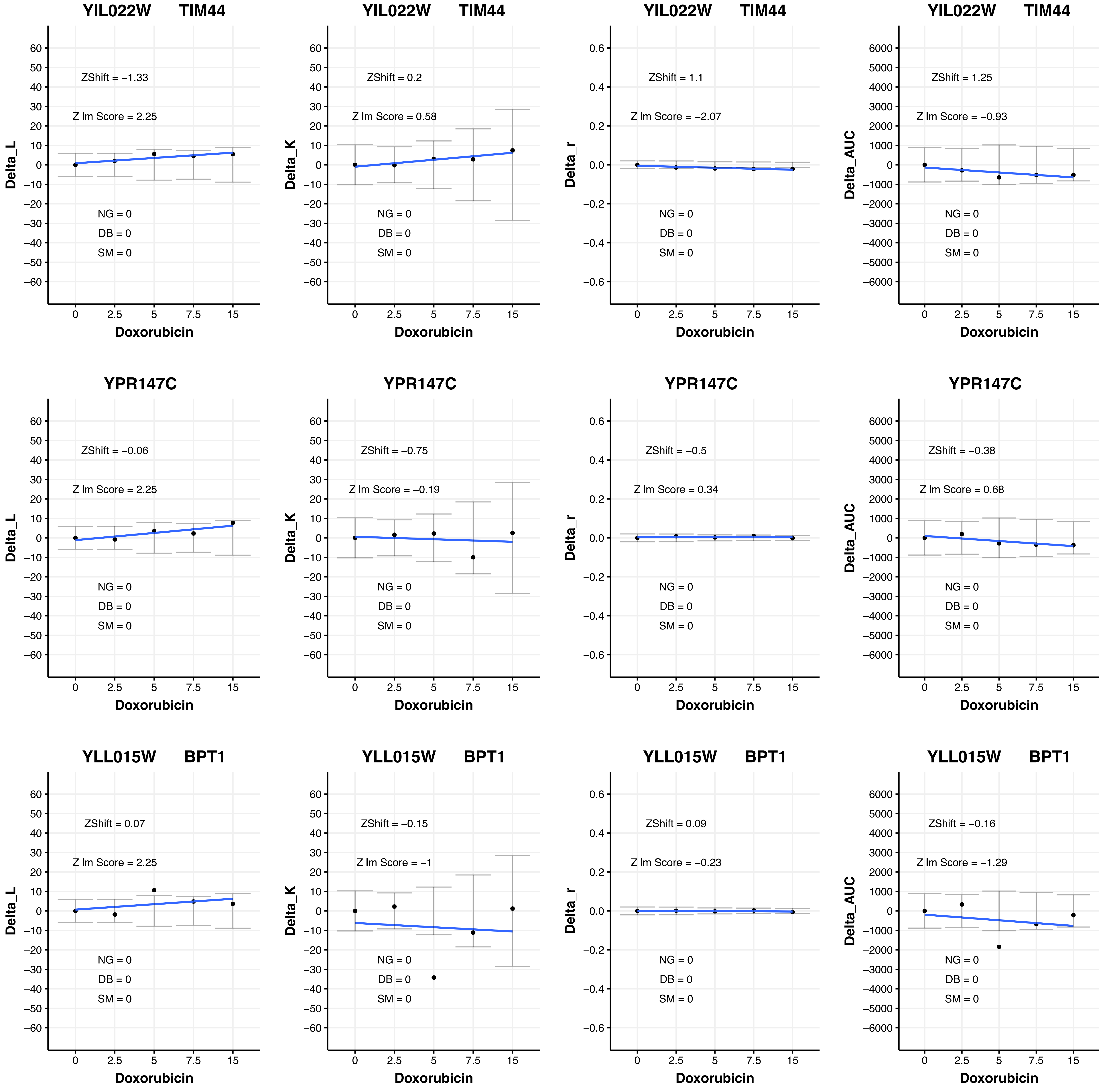

YIL022W TIM44
ZShift = −1.33
Z lm Score = 2.25
NG = 0
DB = 0
SM = 0
Delta_L
Doxorubicin
YIL022W TIM44
ZShift = 0.2
Z lm Score = 0.58
NG = 0
DB = 0
SM = 0
Delta_K
Doxorubicin
YIL022W TIM44
ZShift = 1.1
Z lm Score = −2.07
NG = 0
DB = 0
SM = 0
Delta_r
Doxorubicin
YIL022W TIM44
ZShift = 1.25
Z lm Score = −0.93
NG = 0
DB = 0
SM = 0
Delta_AUC
Doxorubicin
YPR147C
ZShift = −0.06
Z lm Score = 2.25
NG = 0
DB = 0
SM = 0
Delta_L
Doxorubicin
YPR147C
ZShift = −0.75
Z lm Score = −0.19
NG = 0
DB = 0
SM = 0
Delta_K
Doxorubicin
YPR147C
ZShift = −0.5
Z lm Score = 0.34
NG = 0
DB = 0
SM = 0
Delta_r
Doxorubicin
YPR147C
ZShift = −0.38
Z lm Score = 0.68
NG = 0
DB = 0
SM = 0
Delta_AUC
Doxorubicin
YLL015W BPT1
ZShift = 0.07
Z lm Score = 2.25
NG = 0
DB = 0
SM = 0
Delta_L
Doxorubicin
YLL015W BPT1
ZShift = −0.15
Z lm Score = −1
NG = 0
DB = 0
SM = 0
Delta_K
Doxorubicin
YLL015W BPT1
ZShift = 0.09
Z lm Score = −0.23
NG = 0
DB = 0
SM = 0
Delta_r
Doxorubicin
YLL015W BPT1
ZShift = −0.16
Z lm Score = −1.29
NG = 0
DB = 0
SM = 0
Delta_AUC
Doxorubicin

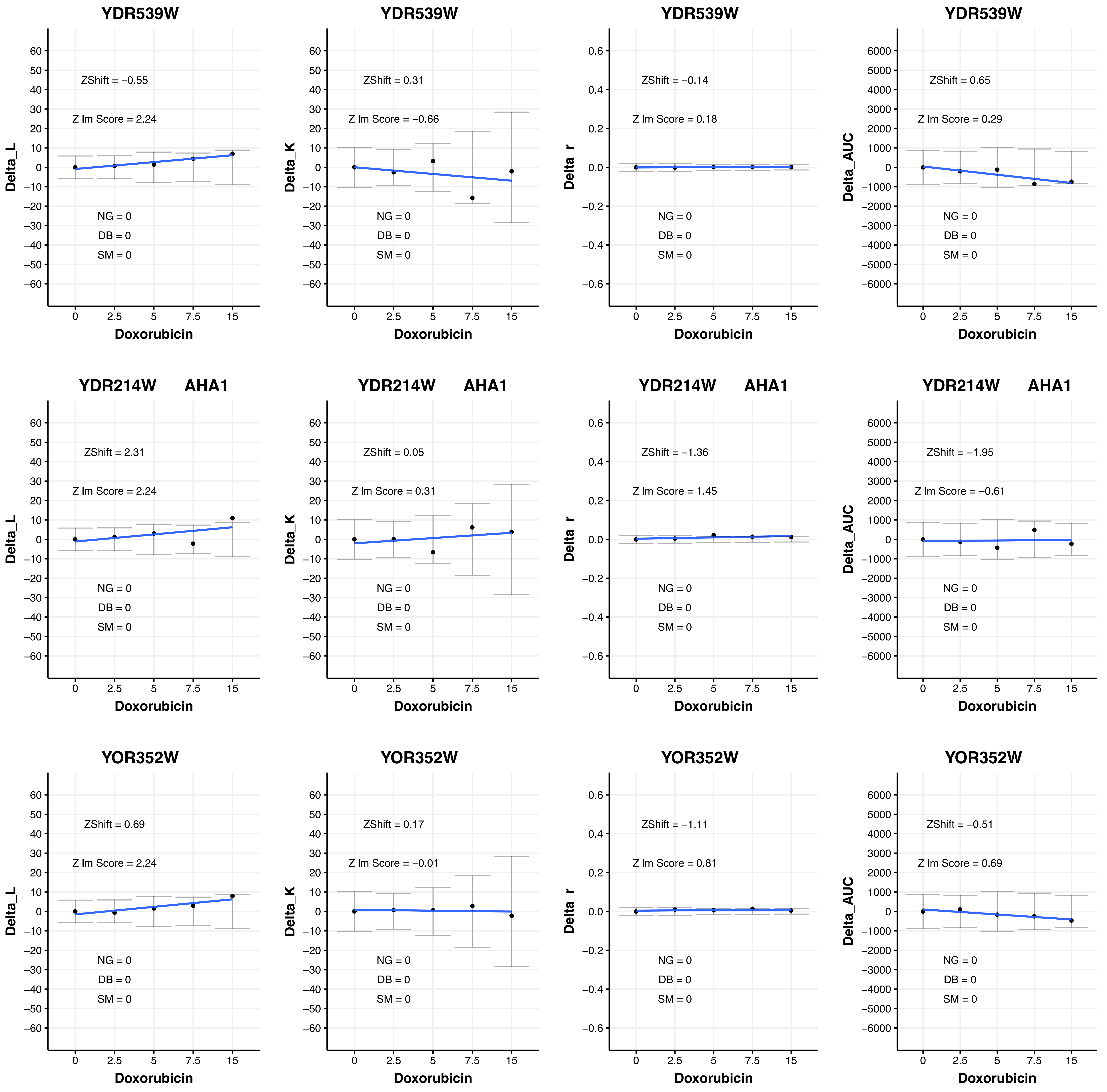
YDR539W
ZShift = −0.55
Z lm Score = 2.24
NG = 0
DB = 0
SM = 0
Delta_L
Doxorubicin
YDR539W
ZShift = 0.31
Z lm Score = −0.66
NG = 0
DB = 0
SM = 0
Delta_K
Doxorubicin
YDR539W
ZShift = −0.14
Z lm Score = 0.18
NG = 0
DB = 0
SM = 0
Delta_r
Doxorubicin
YDR539W
ZShift = 0.65
Z lm Score = 0.29
NG = 0
DB = 0
SM = 0
Delta_AUC
Doxorubicin
YDR214W AHA1
ZShift = 2.31
Z lm Score = 2.24
NG = 0
DB = 0
SM = 0
Delta_L
Doxorubicin
YDR214W AHA1
ZShift = 0.05
Z lm Score = 0.31
NG = 0
DB = 0
SM = 0
Delta_K
Doxorubicin
YDR214W AHA1
ZShift = −1.36
Z lm Score = 1.45
NG = 0
DB = 0
SM = 0
Delta_r
Doxorubicin
YDR214W AHA1
ZShift = −1.95
Z lm Score = −0.61
NG = 0
DB = 0
SM = 0
Delta_AUC
Doxorubicin
YOR352W
ZShift = 0.69
Z lm Score = 2.24
NG = 0
DB = 0
SM = 0
Delta_L
Doxorubicin
YOR352W
ZShift = 0.17
Z lm Score = −0.01
NG = 0
DB = 0
SM = 0
Delta_K
Doxorubicin
YOR352W
ZShift = −1.11
Z lm Score = 0.81
NG = 0
DB = 0
SM = 0
Delta_r
Doxorubicin
YOR352W
ZShift = −0.51
Z lm Score = 0.69
NG = 0
DB = 0
SM = 0
Delta_AUC
Doxorubicin

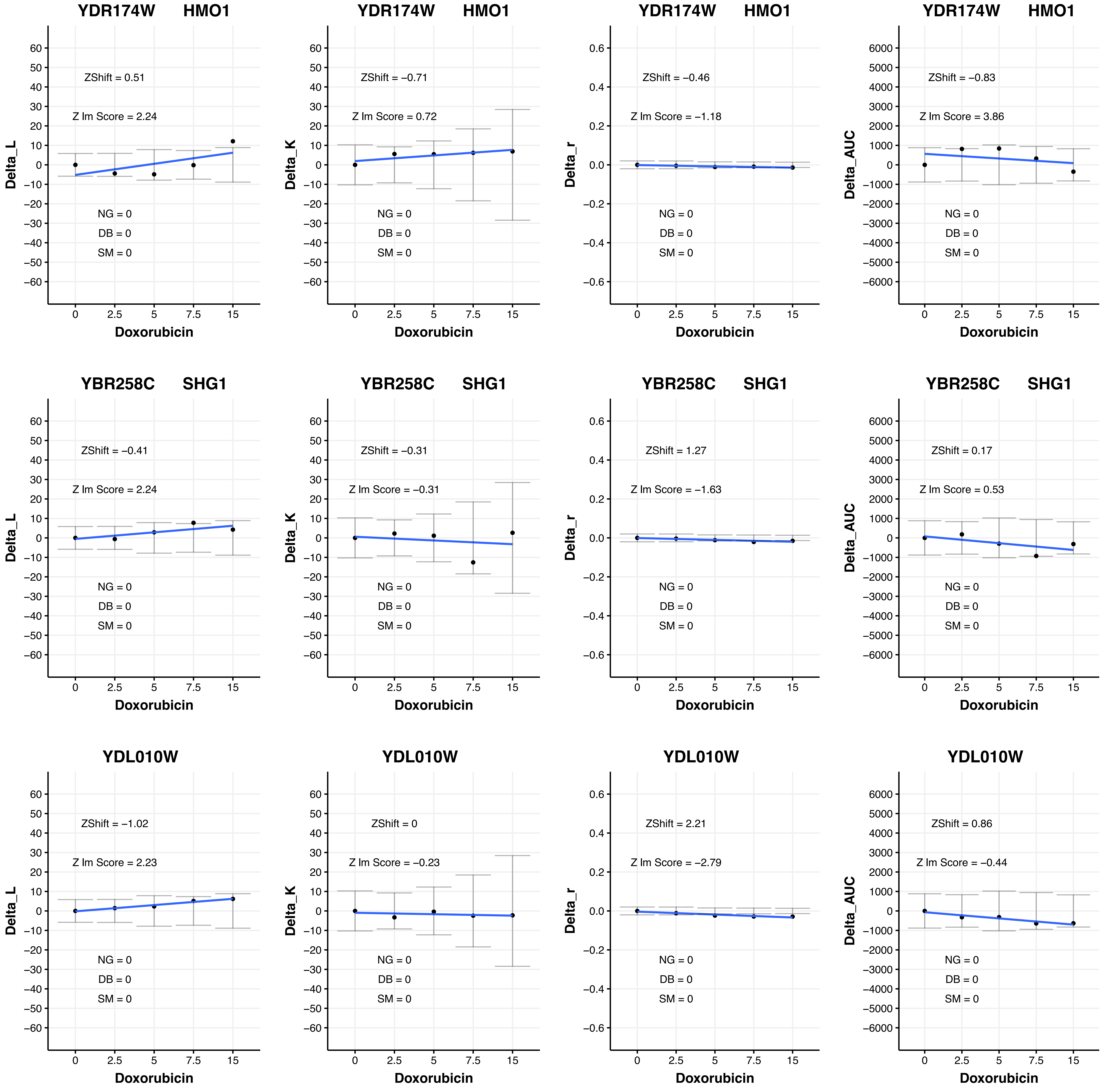
YDR174W HMO1
ZShift = 0.51
Z lm Score = 2.24
NG = 0
DB = 0
SM = 0
Delta_L
Doxorubicin
YDR174W HMO1
ZShift = −0.71
Z lm Score = 0.72
NG = 0
DB = 0
SM = 0
Delta_K
Doxorubicin
YDR174W HMO1
ZShift = −0.46
Z lm Score = −1.18
NG = 0
DB = 0
SM = 0
Delta_r
Doxorubicin
YDR174W HMO1
ZShift = −0.83
Z lm Score = 3.86
NG = 0
DB = 0
SM = 0
Delta_AUC
Doxorubicin
YBR258C SHG1
ZShift = −0.41
Z lm Score = 2.24
NG = 0
DB = 0
SM = 0
Delta_L
Doxorubicin
YBR258C SHG1
ZShift = −0.31
Z lm Score = −0.31
NG = 0
DB = 0
SM = 0
Delta_K
Doxorubicin
YBR258C SHG1
ZShift = 1.27
Z lm Score = −1.63
NG = 0
DB = 0
SM = 0
Delta_r
Doxorubicin
YBR258C SHG1
ZShift = 0.17
Z lm Score = 0.53
NG = 0
DB = 0
SM = 0
Delta_AUC
Doxorubicin
YDL010W
ZShift = −1.02
Z lm Score = 2.23
NG = 0
DB = 0
SM = 0
Delta_L
Doxorubicin
YDL010W
ZShift = 0
Z lm Score = −0.23
NG = 0
DB = 0
SM = 0
Delta_K
Doxorubicin
YDL010W
ZShift = 2.21
Z lm Score = −2.79
NG = 0
DB = 0
SM = 0
Delta_r
Doxorubicin
YDL010W
ZShift = 0.86
Z lm Score = −0.44
NG = 0
DB = 0
SM = 0
Delta_AUC
Doxorubicin

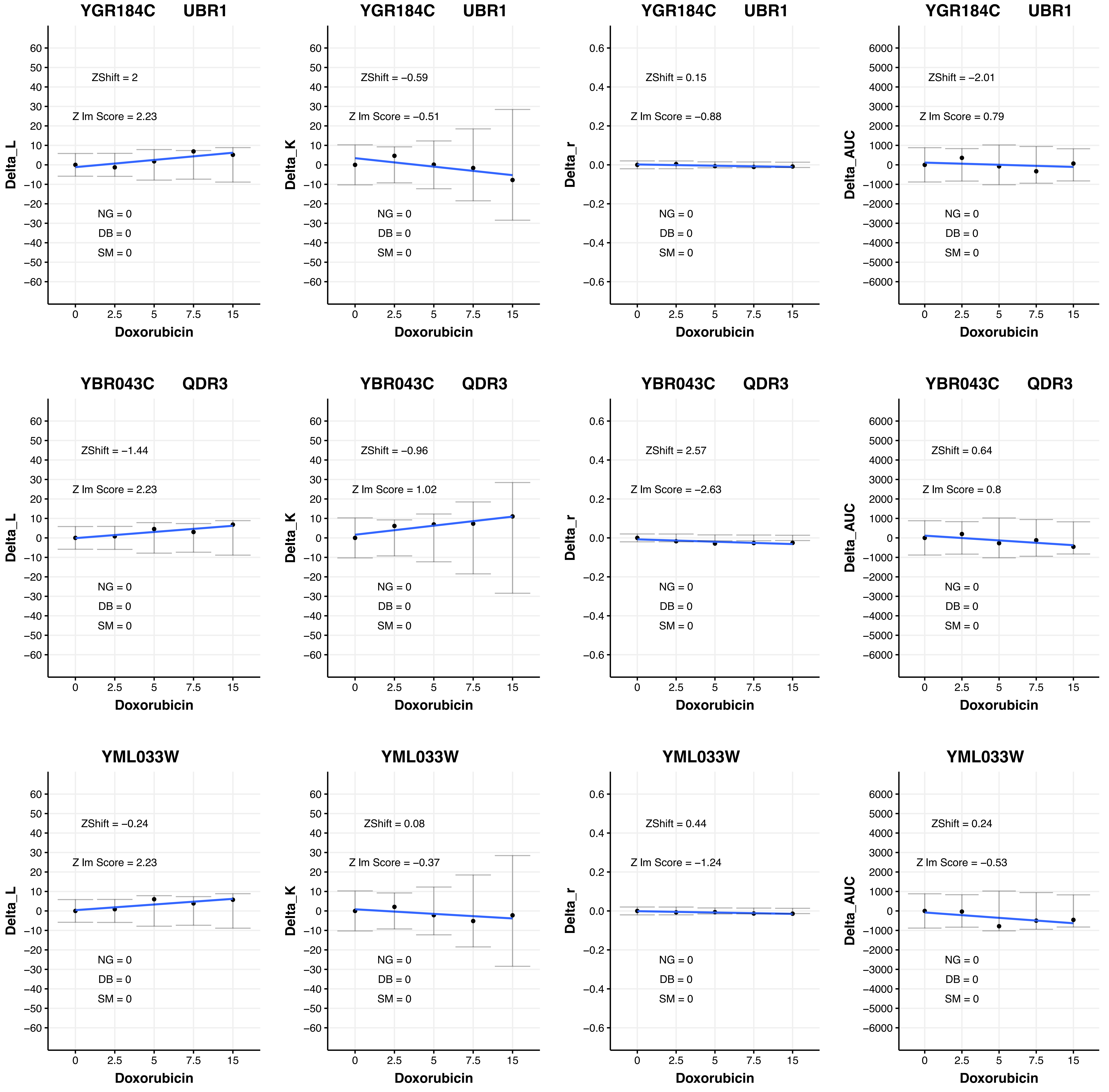
YGR184C UBR1
ZShift = 2
Z lm Score = 2.23
NG = 0
DB = 0
SM = 0
Delta_L
Doxorubicin
YGR184C UBR1
ZShift = −0.59
Z lm Score = −0.51
NG = 0
DB = 0
SM = 0
Delta_K
Doxorubicin
YGR184C UBR1
ZShift = 0.15
Z lm Score = −0.88
NG = 0
DB = 0
SM = 0
Delta_r
Doxorubicin
YGR184C UBR1
ZShift = −2.01
Z lm Score = 0.79
NG = 0
DB = 0
SM = 0
Delta_AUC
Doxorubicin
YBR043C QDR3
ZShift = −1.44
Z lm Score = 2.23
NG = 0
DB = 0
SM = 0
Delta_L
Doxorubicin
YBR043C QDR3
ZShift = −0.96
Z lm Score = 1.02
NG = 0
DB = 0
SM = 0
Delta_K
Doxorubicin
YBR043C QDR3
ZShift = 2.57
Z lm Score = −2.63
NG = 0
DB = 0
SM = 0
Delta_r
Doxorubicin
YBR043C QDR3
ZShift = 0.64
Z lm Score = 0.8
NG = 0
DB = 0
SM = 0
Delta_AUC
Doxorubicin
YML033W
ZShift = −0.24
Z lm Score = 2.23
NG = 0
DB = 0
SM = 0
Delta_L
Doxorubicin
YML033W
ZShift = 0.08
Z lm Score = −0.37
NG = 0
DB = 0
SM = 0
Delta_K
Doxorubicin
YML033W
ZShift = 0.44
Z lm Score = −1.24
NG = 0
DB = 0
SM = 0
Delta_r
Doxorubicin
YML033W
ZShift = 0.24
Z lm Score = −0.53
NG = 0
DB = 0
SM = 0
Delta_AUC
Doxorubicin

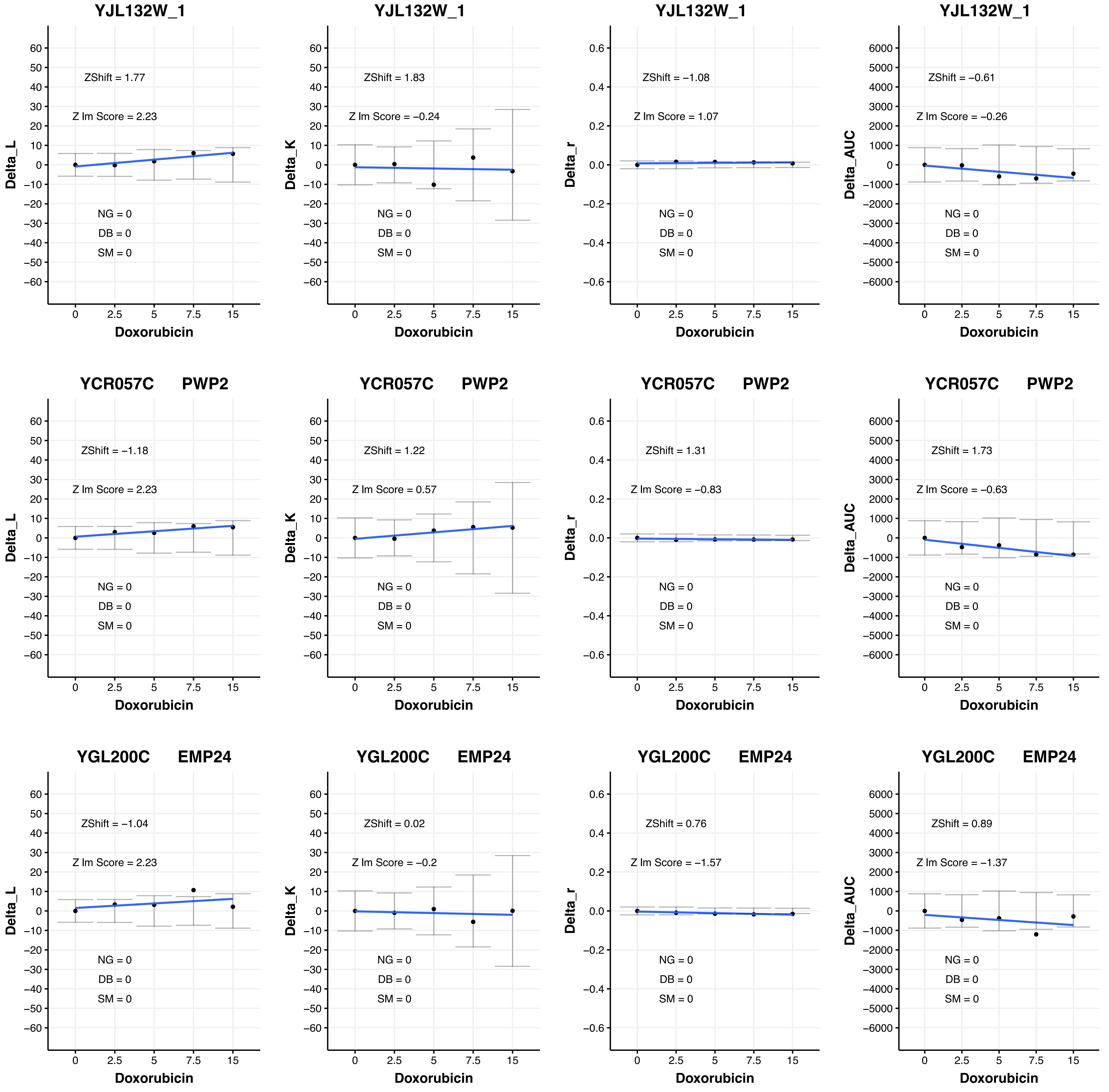

YJL132W_1
ZShift = 1.77
Z lm Score = 2.23
NG = 0
DB = 0
SM = 0
Delta_L
Doxorubicin
YJL132W_1
ZShift = 1.83
Z lm Score = −0.24
NG = 0
DB = 0
SM = 0
Delta_K
Doxorubicin
YJL132W_1
ZShift = −1.08
Z lm Score = 1.07
NG = 0
DB = 0
SM = 0
Delta_r
Doxorubicin
YJL132W_1
ZShift = −0.61
Z lm Score = −0.26
NG = 0
DB = 0
SM = 0
Delta_AUC
Doxorubicin
YCR057C PWP2
ZShift = −1.18
Z lm Score = 2.23
NG = 0
DB = 0
SM = 0
Delta_L
Doxorubicin
YCR057C PWP2
ZShift = 1.22
Z lm Score = 0.57
NG = 0
DB = 0
SM = 0
Delta_K
Doxorubicin
YCR057C PWP2
ZShift = 1.31
Z lm Score = −0.83
NG = 0
DB = 0
SM = 0
Delta_r
Doxorubicin
YCR057C PWP2
ZShift = 1.73
Z lm Score = −0.63
NG = 0
DB = 0
SM = 0
Delta_AUC
Doxorubicin
YGL200C EMP24
ZShift = −1.04
Z lm Score = 2.23
NG = 0
DB = 0
SM = 0
Delta_L
Doxorubicin
YGL200C EMP24
ZShift = 0.02
Z lm Score = −0.2
NG = 0
DB = 0
SM = 0
Delta_K
Doxorubicin
YGL200C EMP24
ZShift = 0.76
Z lm Score = −1.57
NG = 0
DB = 0
SM = 0
Delta_r
Doxorubicin
YGL200C EMP24
ZShift = 0.89
Z lm Score = −1.37
NG = 0
DB = 0
SM = 0
Delta_AUC
Doxorubicin

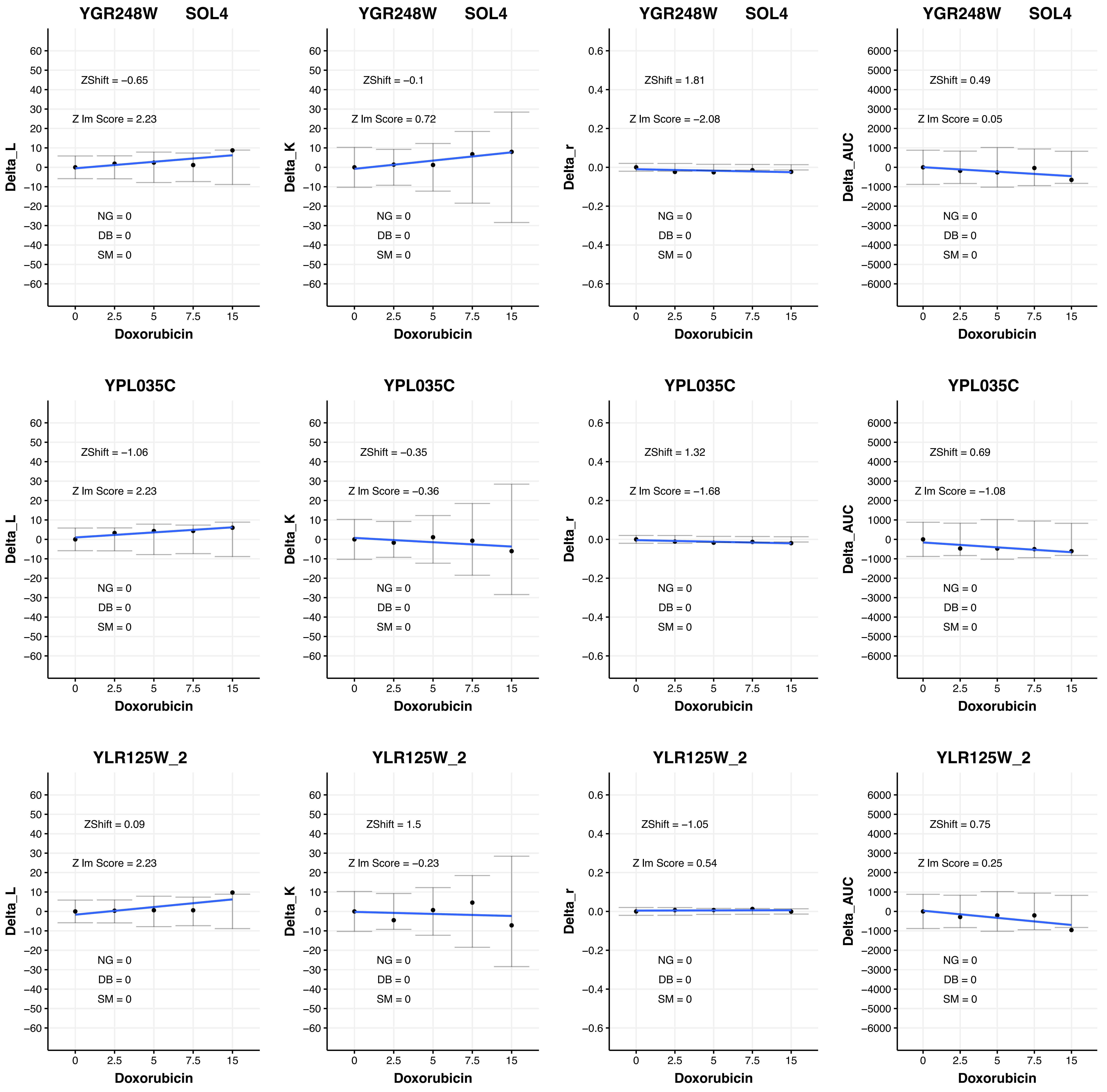
YGR248W SOL4
ZShift = −0.65
Z lm Score = 2.23
NG = 0
DB = 0
SM = 0
Delta_L
Doxorubicin
YGR248W SOL4
ZShift = −0.1
Z lm Score = 0.72
NG = 0
DB = 0
SM = 0
Delta_K
Doxorubicin
YGR248W SOL4
ZShift = 1.81
Z lm Score = −2.08
NG = 0
DB = 0
SM = 0
Delta_r
Doxorubicin
YGR248W SOL4
ZShift = 0.49
Z lm Score = 0.05
NG = 0
DB = 0
SM = 0
Delta_AUC
Doxorubicin
YPL035C
ZShift = −1.06
Z lm Score = 2.23
NG = 0
DB = 0
SM = 0
Delta_L
Doxorubicin
YPL035C
ZShift = −0.35
Z lm Score = −0.36
NG = 0
DB = 0
SM = 0
Delta_K
Doxorubicin
YPL035C
ZShift = 1.32
Z lm Score = −1.68
NG = 0
DB = 0
SM = 0
Delta_r
Doxorubicin
YPL035C
ZShift = 0.69
Z lm Score = −1.08
NG = 0
DB = 0
SM = 0
Delta_AUC
Doxorubicin
YLR125W_2
ZShift = 0.09
Z lm Score = 2.23
NG = 0
DB = 0
SM = 0
Delta_L
Doxorubicin
YLR125W_2
ZShift = 1.5
Z lm Score = −0.23
NG = 0
DB = 0
SM = 0
Delta_K
Doxorubicin
YLR125W_2
ZShift = −1.05
Z lm Score = 0.54
NG = 0
DB = 0
SM = 0
Delta_r
Doxorubicin
YLR125W_2
ZShift = 0.75
Z lm Score = 0.25
NG = 0
DB = 0
SM = 0
Delta_AUC
Doxorubicin

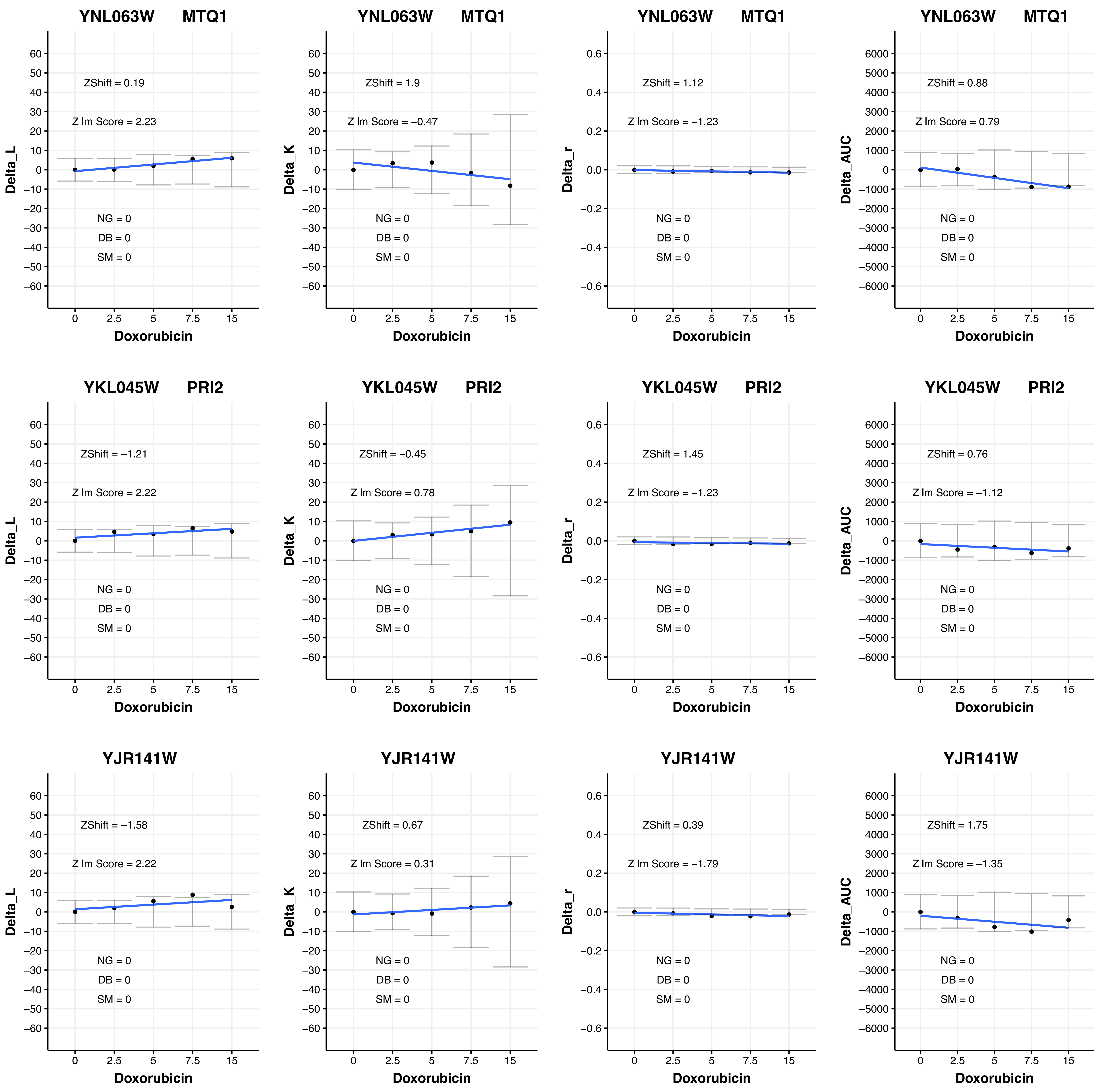

YNL063W MTQ1
ZShift = 0.19
Z lm Score = 2.23
NG = 0
DB = 0
SM = 0
Delta_L
Doxorubicin
YNL063W MTQ1
ZShift = 1.9
Z lm Score = −0.47
NG = 0
DB = 0
SM = 0
Delta_K
Doxorubicin
YNL063W MTQ1
ZShift = 1.12
Z lm Score = −1.23
NG = 0
DB = 0
SM = 0
Delta_r
Doxorubicin
YNL063W MTQ1
ZShift = 0.88
Z lm Score = 0.79
NG = 0
DB = 0
SM = 0
Delta_AUC
Doxorubicin
YKL045W PRI2
ZShift = −1.21
Z lm Score = 2.22
NG = 0
DB = 0
SM = 0
Delta_L
Doxorubicin
YKL045W PRI2
ZShift = −0.45
Z lm Score = 0.78
NG = 0
DB = 0
SM = 0
Delta_K
Doxorubicin
YKL045W PRI2
ZShift = 1.45
Z lm Score = −1.23
NG = 0
DB = 0
SM = 0
Delta_r
Doxorubicin
YKL045W PRI2
ZShift = 0.76
Z lm Score = −1.12
NG = 0
DB = 0
SM = 0
Delta_AUC
Doxorubicin
YJR141W
ZShift = −1.58
Z lm Score = 2.22
NG = 0
DB = 0
SM = 0
Delta_L
Doxorubicin
YJR141W
ZShift = 0.67
Z lm Score = 0.31
NG = 0
DB = 0
SM = 0
Delta_K
Doxorubicin
YJR141W
ZShift = 0.39
Z lm Score = −1.79
NG = 0
DB = 0
SM = 0
Delta_r
Doxorubicin
YJR141W
ZShift = 1.75
Z lm Score = −1.35
NG = 0
DB = 0
SM = 0
Delta_AUC
Doxorubicin

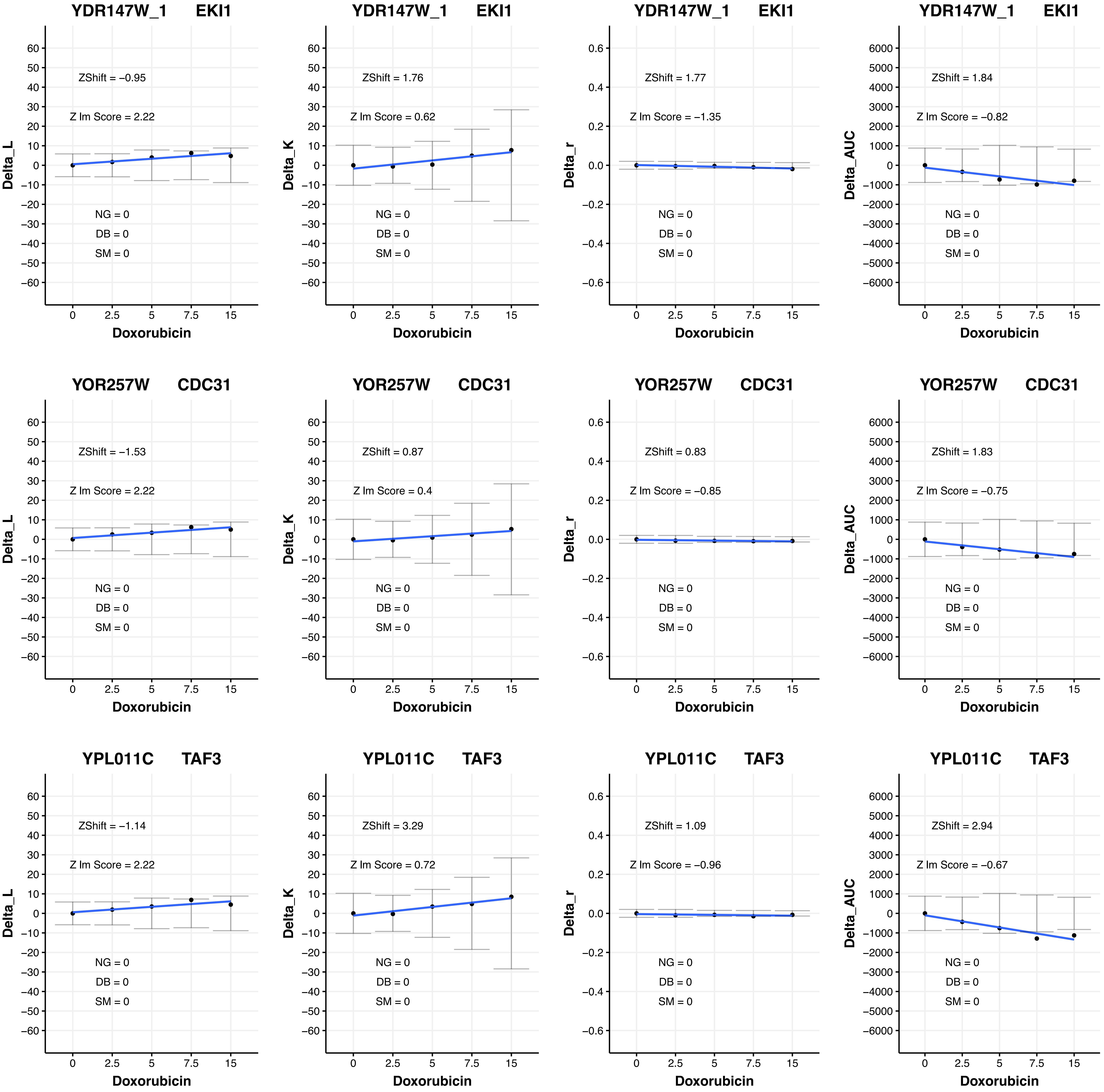

YDR147W_1 EKI1
ZShift = −0.95
Z lm Score = 2.22
NG = 0
DB = 0
SM = 0
Delta_L
Doxorubicin
YDR147W_1 EKI1
ZShift = 1.76
Z lm Score = 0.62
NG = 0
DB = 0
SM = 0
Delta_K
Doxorubicin
YDR147W_1 EKI1
ZShift = 1.77
Z lm Score = −1.35
NG = 0
DB = 0
SM = 0
Delta_r
Doxorubicin
YDR147W_1 EKI1
ZShift = 1.84
Z lm Score = −0.82
NG = 0
DB = 0
SM = 0
Delta_AUC
Doxorubicin
YOR257W CDC31
ZShift = −1.53
Z lm Score = 2.22
NG = 0
DB = 0
SM = 0
Delta_L
Doxorubicin
YOR257W CDC31
ZShift = 0.87
Z lm Score = 0.4
NG = 0
DB = 0
SM = 0
Delta_K
Doxorubicin
YOR257W CDC31
ZShift = 0.83
Z lm Score = −0.85
NG = 0
DB = 0
SM = 0
Delta_r
Doxorubicin
YOR257W CDC31
ZShift = 1.83
Z lm Score = −0.75
NG = 0
DB = 0
SM = 0
Delta_AUC
Doxorubicin
YPL011C TAF3
ZShift = −1.14
Z lm Score = 2.22
NG = 0
DB = 0
SM = 0
Delta_L
Doxorubicin
YPL011C TAF3
ZShift = 3.29
Z lm Score = 0.72
NG = 0
DB = 0
SM = 0
Delta_K
Doxorubicin
YPL011C TAF3
ZShift = 1.09
Z lm Score = −0.96
NG = 0
DB = 0
SM = 0
Delta_r
Doxorubicin
YPL011C TAF3
ZShift = 2.94
Z lm Score = −0.67
NG = 0
DB = 0
SM = 0
Delta_AUC
Doxorubicin

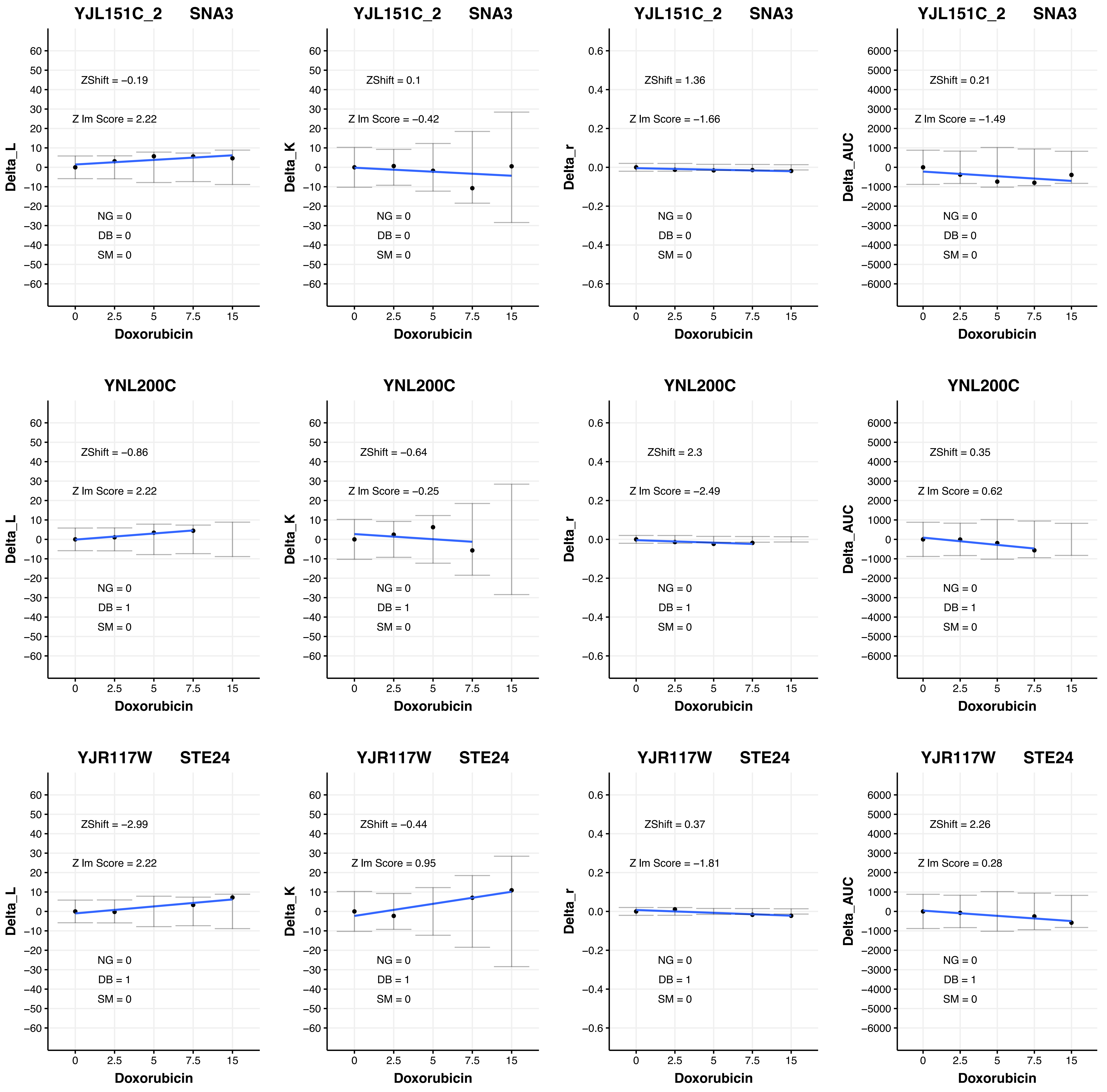
YJL151C_2 SNA3
ZShift = −0.19
Z lm Score = 2.22
NG = 0
DB = 0
SM = 0
Delta_L
Doxorubicin
YJL151C_2 SNA3
ZShift = 0.1
Z lm Score = −0.42
NG = 0
DB = 0
SM = 0
Delta_K
Doxorubicin
YJL151C_2 SNA3
ZShift = 1.36
Z lm Score = −1.66
NG = 0
DB = 0
SM = 0
Delta_r
Doxorubicin
YJL151C_2 SNA3
ZShift = 0.21
Z lm Score = −1.49
NG = 0
DB = 0
SM = 0
Delta_AUC
Doxorubicin
YNL200C
ZShift = −0.86
Z lm Score = 2.22
NG = 0
DB = 1
SM = 0
Delta_L
Doxorubicin
YNL200C
ZShift = −0.64
Z lm Score = −0.25
NG = 0
DB = 1
SM = 0
Delta_K
Doxorubicin
YNL200C
ZShift = 2.3
Z lm Score = −2.49
NG = 0
DB = 1
SM = 0
Delta_r
Doxorubicin
YNL200C
ZShift = 0.35
Z lm Score = 0.62
NG = 0
DB = 1
SM = 0
Delta_AUC
Doxorubicin
YJR117W STE24
ZShift = −2.99
Z lm Score = 2.22
NG = 0
DB = 1
SM = 0
Delta_L
Doxorubicin
YJR117W STE24
ZShift = −0.44
Z lm Score = 0.95
NG = 0
DB = 1
SM = 0
Delta_K
Doxorubicin
YJR117W STE24
ZShift = 0.37
Z lm Score = −1.81
NG = 0
DB = 1
SM = 0
Delta_r
Doxorubicin
YJR117W STE24
ZShift = 2.26
Z lm Score = 0.28
NG = 0
DB = 1
SM = 0
Delta_AUC
Doxorubicin

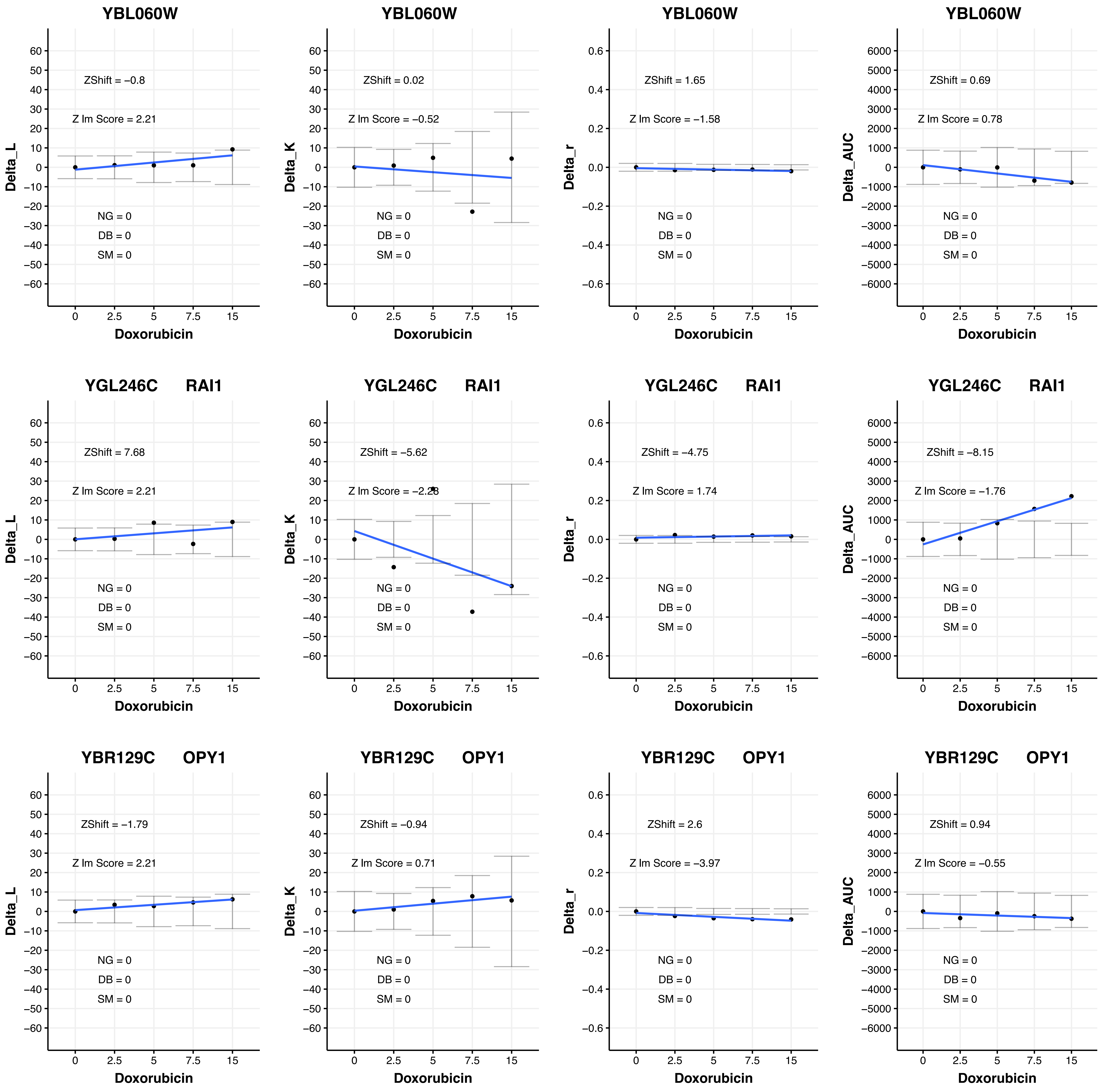
YBL060W
ZShift = −0.8
Z lm Score = 2.21
NG = 0
DB = 0
SM = 0
Delta_L
Doxorubicin
YBL060W
ZShift = 0.02
Z lm Score = −0.52
NG = 0
DB = 0
SM = 0
Delta_K
Doxorubicin
YBL060W
ZShift = 1.65
Z lm Score = −1.58
NG = 0
DB = 0
SM = 0
Delta_r
Doxorubicin
YBL060W
ZShift = 0.69
Z lm Score = 0.78
NG = 0
DB = 0
SM = 0
Delta_AUC
Doxorubicin
YGL246C RAI1
ZShift = 7.68
Z lm Score = 2.21
NG = 0
DB = 0
SM = 0
Delta_L
Doxorubicin
YGL246C RAI1
ZShift = −5.62
Z lm Score = −2.28
NG = 0
DB = 0
SM = 0
Delta_K
Doxorubicin
YGL246C RAI1
ZShift = −4.75
Z lm Score = 1.74
NG = 0
DB = 0
SM = 0
Delta_r
Doxorubicin
YGL246C RAI1
ZShift = −8.15
Z lm Score = −1.76
NG = 0
DB = 0
SM = 0
Delta_AUC
Doxorubicin
YBR129C OPY1
ZShift = −1.79
Z lm Score = 2.21
NG = 0
DB = 0
SM = 0
Delta_L
Doxorubicin
YBR129C OPY1
ZShift = −0.94
Z lm Score = 0.71
NG = 0
DB = 0
SM = 0
Delta_K
Doxorubicin
YBR129C OPY1
ZShift = 2.6
Z lm Score = −3.97
NG = 0
DB = 0
SM = 0
Delta_r
Doxorubicin
YBR129C OPY1
ZShift = 0.94
Z lm Score = −0.55
NG = 0
DB = 0
SM = 0
Delta_AUC
Doxorubicin

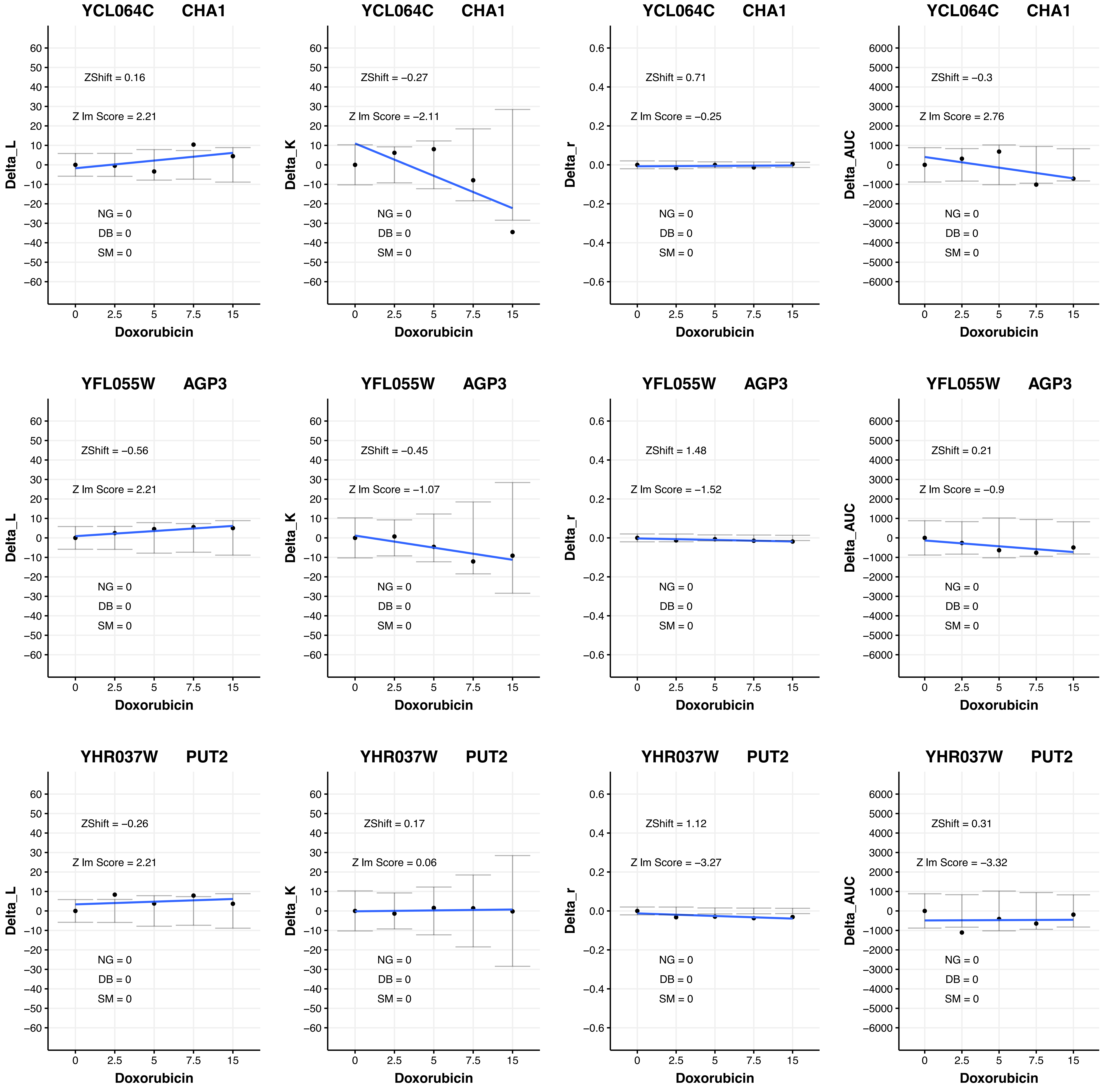

YCL064C CHA1
ZShift = 0.16
Z lm Score = 2.21
NG = 0
DB = 0
SM = 0
Delta_L
Doxorubicin
YCL064C CHA1
ZShift = −0.27
Z lm Score = −2.11
NG = 0
DB = 0
SM = 0
Delta_K
Doxorubicin
YCL064C CHA1
ZShift = 0.71
Z lm Score = −0.25
NG = 0
DB = 0
SM = 0
Delta_r
Doxorubicin
YCL064C CHA1
ZShift = −0.3
Z lm Score = 2.76
NG = 0
DB = 0
SM = 0
Delta_AUC
Doxorubicin
YFL055W AGP3
ZShift = −0.56
Z lm Score = 2.21
NG = 0
DB = 0
SM = 0
Delta_L
Doxorubicin
YFL055W AGP3
ZShift = −0.45
Z lm Score = −1.07
NG = 0
DB = 0
SM = 0
Delta_K
Doxorubicin
YFL055W AGP3
ZShift = 1.48
Z lm Score = −1.52
NG = 0
DB = 0
SM = 0
Delta_r
Doxorubicin
YFL055W AGP3
ZShift = 0.21
Z lm Score = −0.9
NG = 0
DB = 0
SM = 0
Delta_AUC
Doxorubicin
YHR037W PUT2
ZShift = −0.26
Z lm Score = 2.21
NG = 0
DB = 0
SM = 0
Delta_L
Doxorubicin
YHR037W PUT2
ZShift = 0.17
Z lm Score = 0.06
NG = 0
DB = 0
SM = 0
Delta_K
Doxorubicin
YHR037W PUT2
ZShift = 1.12
Z lm Score = −3.27
NG = 0
DB = 0
SM = 0
Delta_r
Doxorubicin
YHR037W PUT2
ZShift = 0.31
Z lm Score = −3.32
NG = 0
DB = 0
SM = 0
Delta_AUC
Doxorubicin

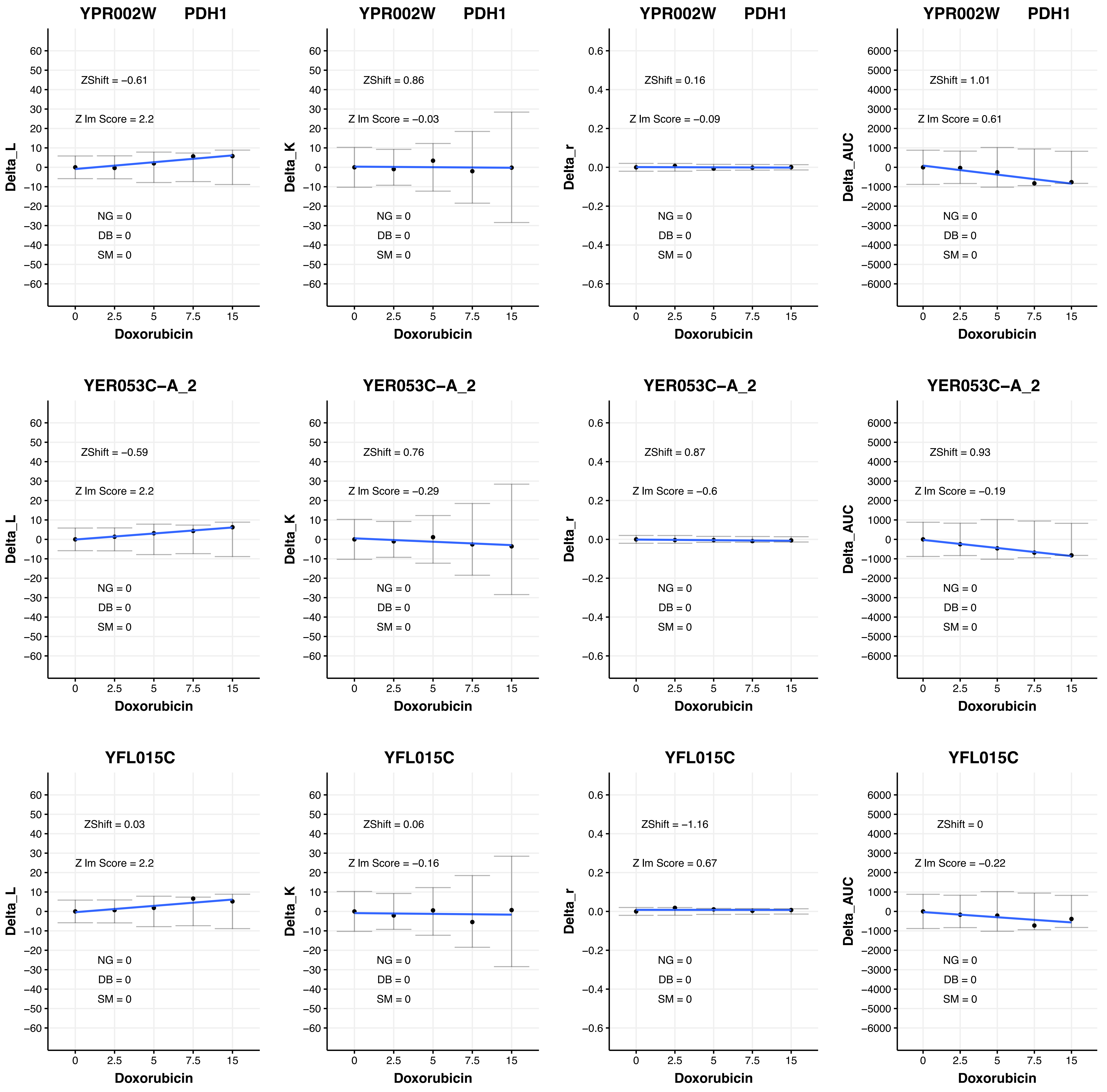
YPR002W PDH1
ZShift = −0.61
Z lm Score = 2.2
NG = 0
DB = 0
SM = 0
Delta_L
Doxorubicin
YPR002W PDH1
ZShift = 0.86
Z lm Score = −0.03
NG = 0
DB = 0
SM = 0
Delta_K
Doxorubicin
YPR002W PDH1
ZShift = 0.16
Z lm Score = −0.09
NG = 0
DB = 0
SM = 0
Delta_r
Doxorubicin
YPR002W PDH1
ZShift = 1.01
Z lm Score = 0.61
NG = 0
DB = 0
SM = 0
Delta_AUC
Doxorubicin
YER053C−A_2
ZShift = −0.59
Z lm Score = 2.2
NG = 0
DB = 0
SM = 0
Delta_L
Doxorubicin
YER053C−A_2
ZShift = 0.76
Z lm Score = −0.29
NG = 0
DB = 0
SM = 0
Delta_K
Doxorubicin
YER053C−A_2
ZShift = 0.87
Z lm Score = −0.6
NG = 0
DB = 0
SM = 0
Delta_r
Doxorubicin
YER053C−A_2
ZShift = 0.93
Z lm Score = −0.19
NG = 0
DB = 0
SM = 0
Delta_AUC
Doxorubicin
YFL015C
ZShift = 0.03
Z lm Score = 2.2
NG = 0
DB = 0
SM = 0
Delta_L
Doxorubicin
YFL015C
ZShift = 0.06
Z lm Score = −0.16
NG = 0
DB = 0
SM = 0
Delta_K
Doxorubicin
YFL015C
ZShift = −1.16
Z lm Score = 0.67
NG = 0
DB = 0
SM = 0
Delta_r
Doxorubicin
YFL015C
ZShift = 0
Z lm Score = −0.22
NG = 0
DB = 0
SM = 0
Delta_AUC
Doxorubicin

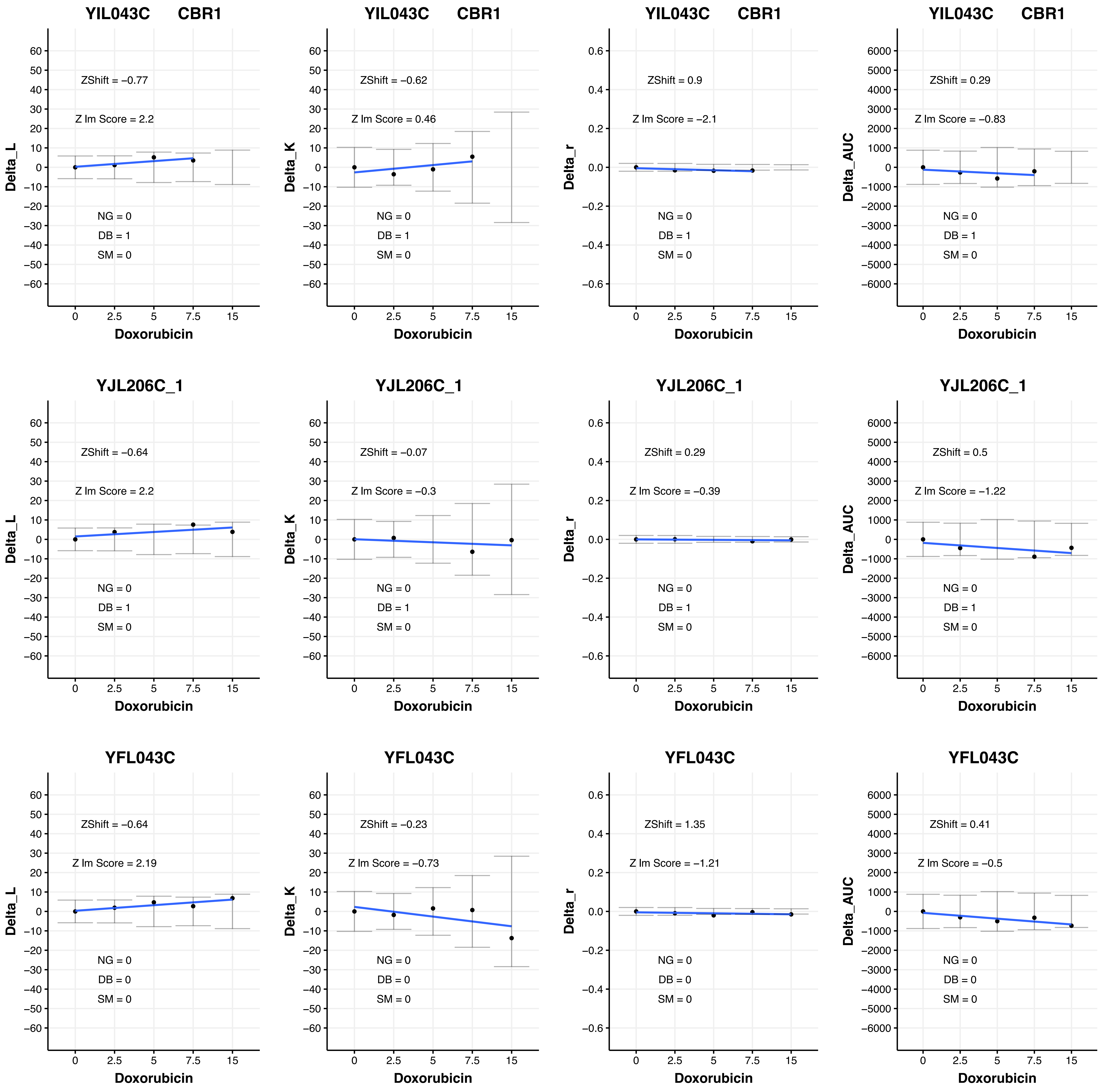

YIL043C CBR1
ZShift = −0.77
Z Im Score = 2.2
NG = 0
DB = 1
SM = 0
Delta_L
Doxorubicin
YIL043C CBR1
ZShift = −0.62
Z Im Score = 0.46
NG = 0
DB = 1
SM = 0
Delta_K
Doxorubicin
YIL043C CBR1
ZShift = 0.9
Z Im Score = −2.1
NG = 0
DB = 1
SM = 0
Delta_r
Doxorubicin
YIL043C CBR1
ZShift = 0.29
Z Im Score = −0.83
NG = 0
DB = 1
SM = 0
Delta_AUC
Doxorubicin
YJL206C_1
ZShift = −0.64
Z Im Score = 2.2
NG = 0
DB = 1
SM = 0
Delta_L
Doxorubicin
YJL206C_1
ZShift = −0.07
Z Im Score = −0.3
NG = 0
DB = 1
SM = 0
Delta_K
Doxorubicin
YJL206C_1
ZShift = 0.29
Z Im Score = −0.39
NG = 0
DB = 1
SM = 0
Delta_r
Doxorubicin
YJL206C_1
ZShift = 0.5
Z Im Score = −1.22
NG = 0
DB = 1
SM = 0
Delta_AUC
Doxorubicin
YFL043C
ZShift = −0.64
Z Im Score = 2.19
NG = 0
DB = 0
SM = 0
Delta_L
Doxorubicin
YFL043C
ZShift = −0.23
Z Im Score = −0.73
NG = 0
DB = 0
SM = 0
Delta_K
Doxorubicin
YFL043C
ZShift = 1.35
Z Im Score = −1.21
NG = 0
DB = 0
SM = 0
Delta_r
Doxorubicin
YFL043C
ZShift = 0.41
Z Im Score = −0.5
NG = 0
DB = 0
SM = 0
Delta_AUC
Doxorubicin

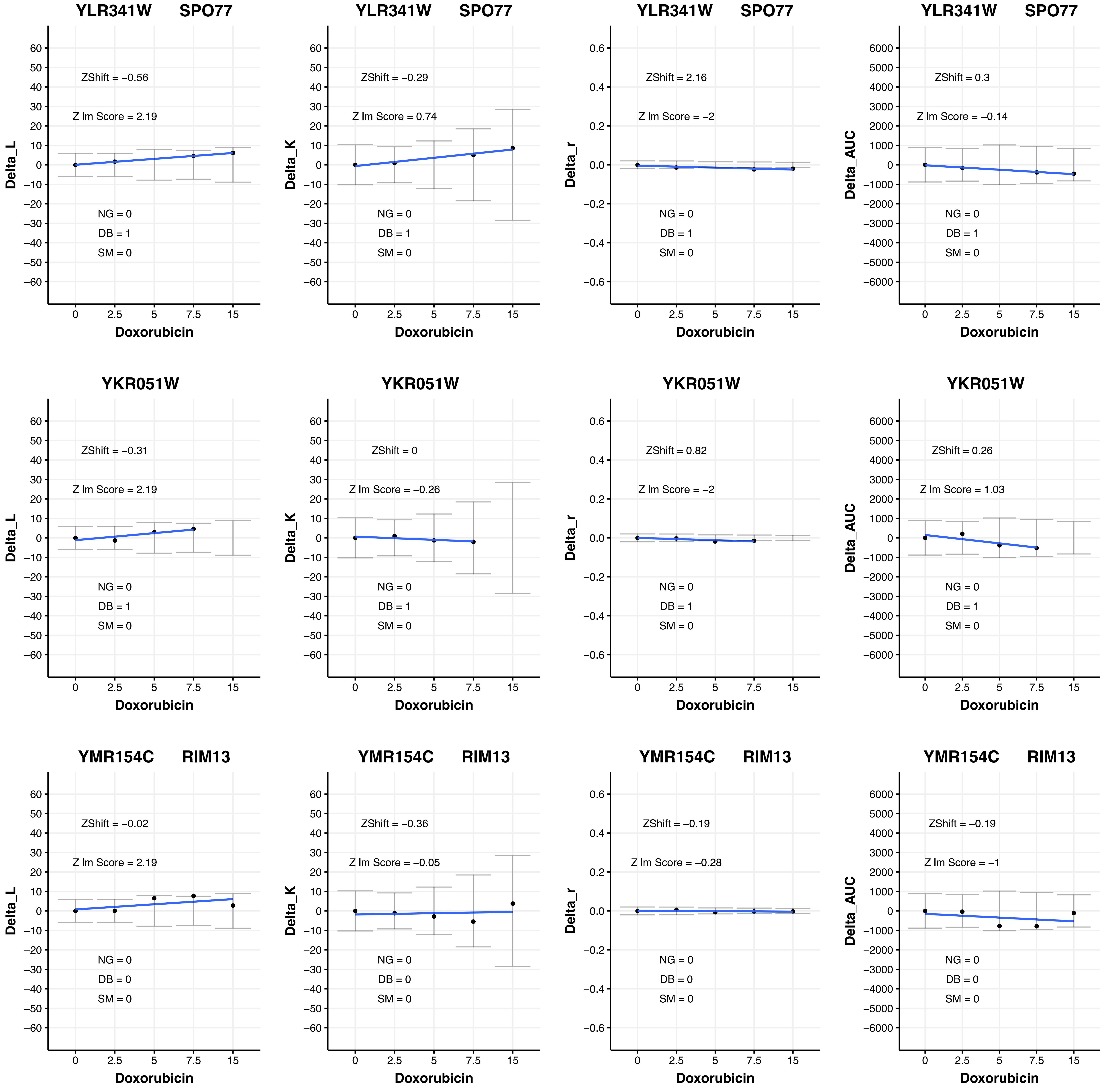
YLR341W SPO77
ZShift = −0.56
Z lm Score = 2.19
NG = 0
DB = 1
SM = 0
Delta_L
Doxorubicin
YLR341W SPO77
ZShift = −0.29
Z lm Score = 0.74
NG = 0
DB = 1
SM = 0
Delta_K
Doxorubicin
YLR341W SPO77
ZShift = 2.16
Z lm Score = −2
NG = 0
DB = 1
SM = 0
Delta_r
Doxorubicin
YLR341W SPO77
ZShift = 0.3
Z lm Score = −0.14
NG = 0
DB = 1
SM = 0
Delta_AUC
Doxorubicin
YKR051W
ZShift = −0.31
Z lm Score = 2.19
NG = 0
DB = 1
SM = 0
Delta_L
Doxorubicin
YKR051W
ZShift = 0
Z lm Score = −0.26
NG = 0
DB = 1
SM = 0
Delta_K
Doxorubicin
YKR051W
ZShift = 0.82
Z lm Score = −2
NG = 0
DB = 1
SM = 0
Delta_r
Doxorubicin
YKR051W
ZShift = 0.26
Z lm Score = 1.03
NG = 0
DB = 1
SM = 0
Delta_AUC
Doxorubicin
YMR154C RIM13
ZShift = −0.02
Z lm Score = 2.19
NG = 0
DB = 0
SM = 0
Delta_L
Doxorubicin
YMR154C RIM13
ZShift = −0.36
Z lm Score = −0.05
NG = 0
DB = 0
SM = 0
Delta_K
Doxorubicin
YMR154C RIM13
ZShift = −0.19
Z lm Score = −0.28
NG = 0
DB = 0
SM = 0
Delta_r
Doxorubicin
YMR154C RIM13
ZShift = −0.19
Z lm Score = −1
NG = 0
DB = 0
SM = 0
Delta_AUC
Doxorubicin

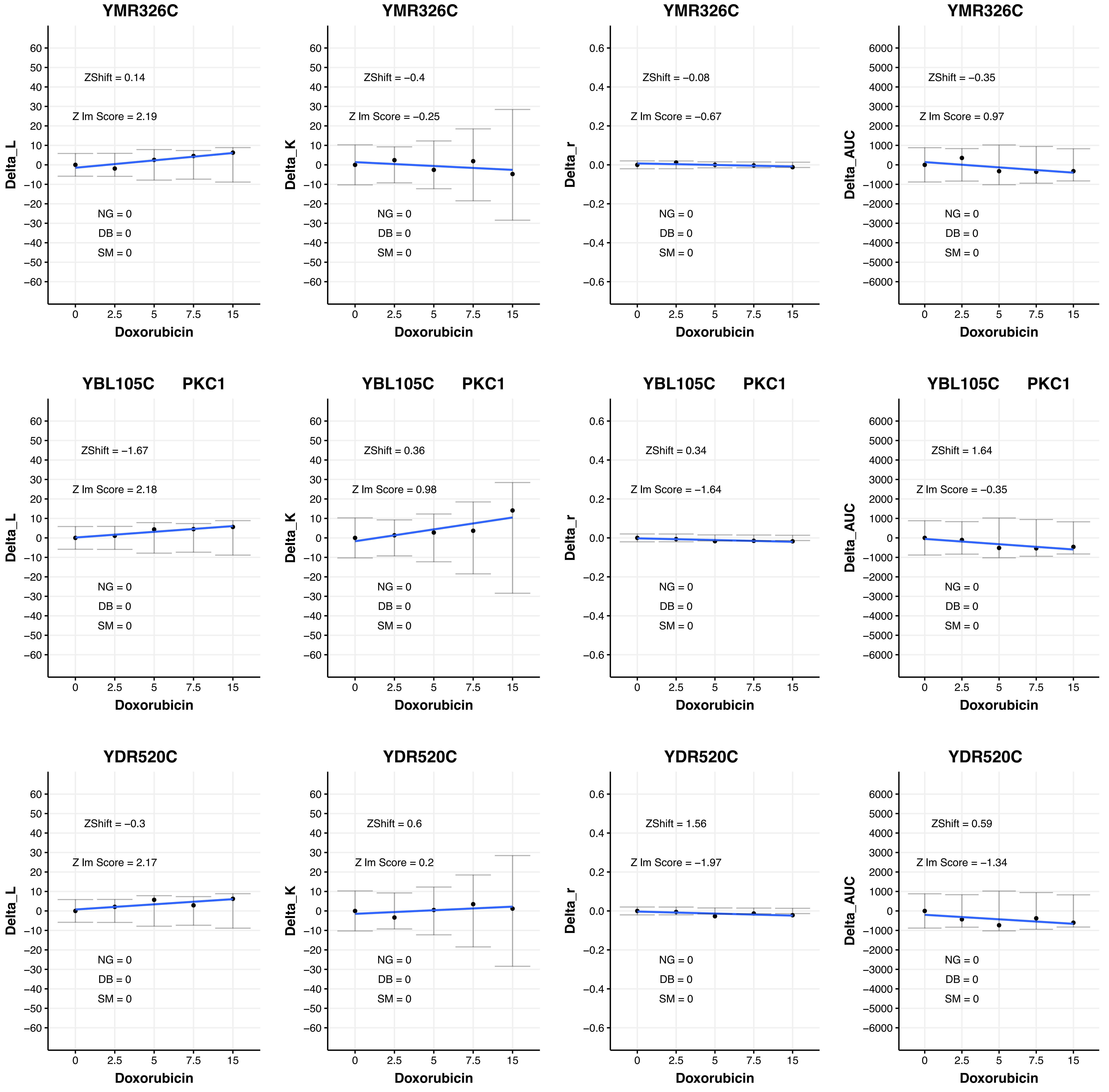

YMR326C
ZShift = 0.14
Z lm Score = 2.19
NG = 0
DB = 0
SM = 0
Delta_L
Doxorubicin
YMR326C
ZShift = −0.4
Z lm Score = −0.25
NG = 0
DB = 0
SM = 0
Delta_K
Doxorubicin
YMR326C
ZShift = −0.08
Z lm Score = −0.67
NG = 0
DB = 0
SM = 0
Delta_r
Doxorubicin
YMR326C
ZShift = −0.35
Z lm Score = 0.97
NG = 0
DB = 0
SM = 0
Delta_AUC
Doxorubicin
YBL105C PKC1
ZShift = −1.67
Z lm Score = 2.18
NG = 0
DB = 0
SM = 0
Delta_L
Doxorubicin
YBL105C PKC1
ZShift = 0.36
Z lm Score = 0.98
NG = 0
DB = 0
SM = 0
Delta_K
Doxorubicin
YBL105C PKC1
ZShift = 0.34
Z lm Score = −1.64
NG = 0
DB = 0
SM = 0
Delta_r
Doxorubicin
YBL105C PKC1
ZShift = 1.64
Z lm Score = −0.35
NG = 0
DB = 0
SM = 0
Delta_AUC
Doxorubicin
YDR520C
ZShift = −0.3
Z lm Score = 2.17
NG = 0
DB = 0
SM = 0
Delta_L
Doxorubicin
YDR520C
ZShift = 0.6
Z lm Score = 0.2
NG = 0
DB = 0
SM = 0
Delta_K
Doxorubicin
YDR520C
ZShift = 1.56
Z lm Score = −1.97
NG = 0
DB = 0
SM = 0
Delta_r
Doxorubicin
YDR520C
ZShift = 0.59
Z lm Score = −1.34
NG = 0
DB = 0
SM = 0
Delta_AUC
Doxorubicin

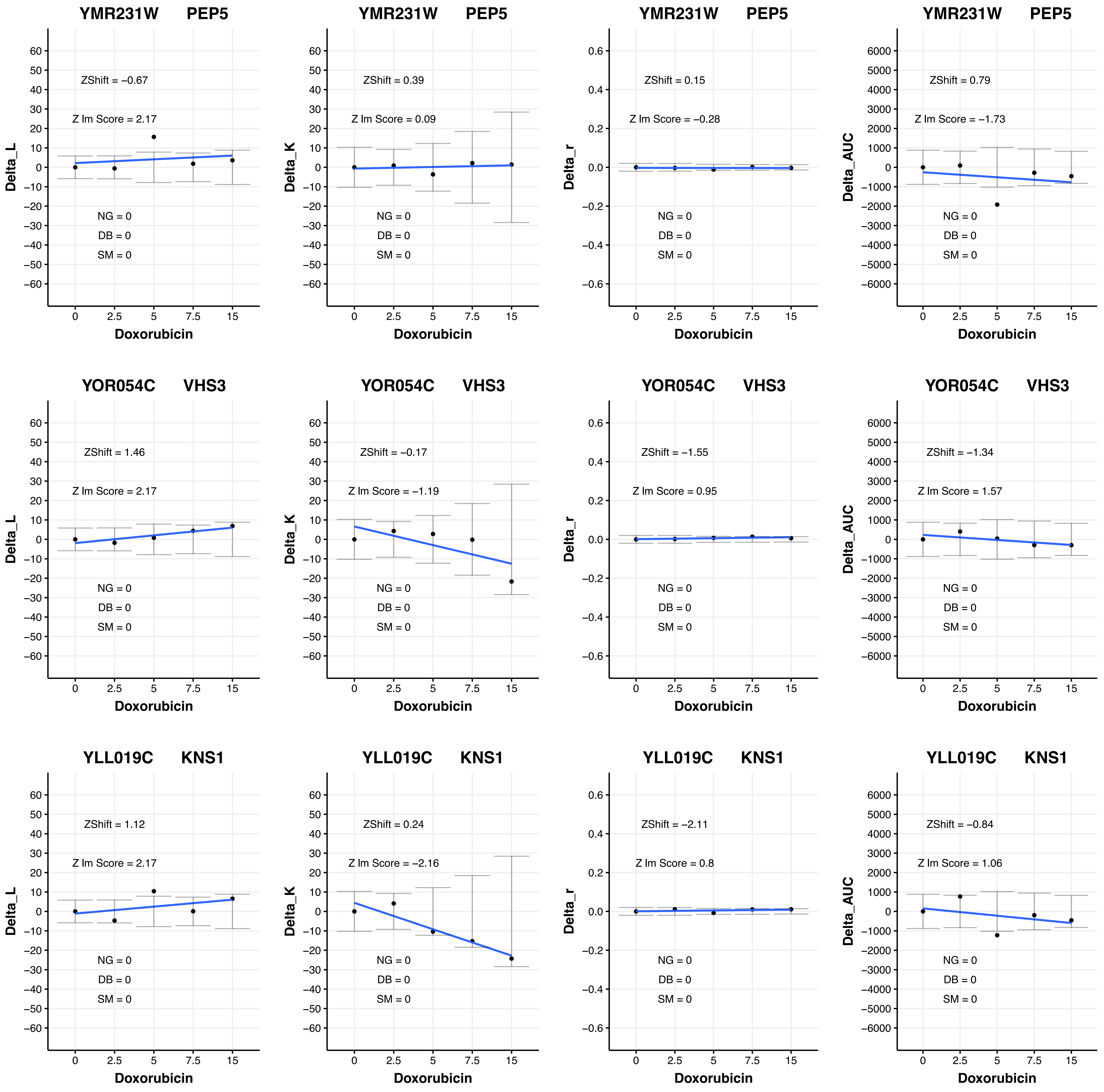
YMR231W PEP5
ZShift = −0.67
Z lm Score = 2.17
NG = 0
DB = 0
SM = 0
Delta_L
Doxorubicin
YMR231W PEP5
ZShift = 0.39
Z lm Score = 0.09
NG = 0
DB = 0
SM = 0
Delta_K
Doxorubicin
YMR231W PEP5
ZShift = 0.15
Z lm Score = −0.28
NG = 0
DB = 0
SM = 0
Delta_r
Doxorubicin
YMR231W PEP5
ZShift = 0.79
Z lm Score = −1.73
NG = 0
DB = 0
SM = 0
Delta_AUC
Doxorubicin
YOR054C VHS3
ZShift = 1.46
Z lm Score = 2.17
NG = 0
DB = 0
SM = 0
Delta_L
Doxorubicin
YOR054C VHS3
ZShift = −0.17
Z lm Score = −1.19
NG = 0
DB = 0
SM = 0
Delta_K
Doxorubicin
YOR054C VHS3
ZShift = −1.55
Z lm Score = 0.95
NG = 0
DB = 0
SM = 0
Delta_r
Doxorubicin
YOR054C VHS3
ZShift = −1.34
Z lm Score = 1.57
NG = 0
DB = 0
SM = 0
Delta_AUC
Doxorubicin
YLL019C KNS1
ZShift = 1.12
Z lm Score = 2.17
NG = 0
DB = 0
SM = 0
Delta_L
Doxorubicin
YLL019C KNS1
ZShift = 0.24
Z lm Score = −2.16
NG = 0
DB = 0
SM = 0
Delta_K
Doxorubicin
YLL019C KNS1
ZShift = −2.11
Z lm Score = 0.8
NG = 0
DB = 0
SM = 0
Delta_r
Doxorubicin
YLL019C KNS1
ZShift = −0.84
Z lm Score = 1.06
NG = 0
DB = 0
SM = 0
Delta_AUC
Doxorubicin

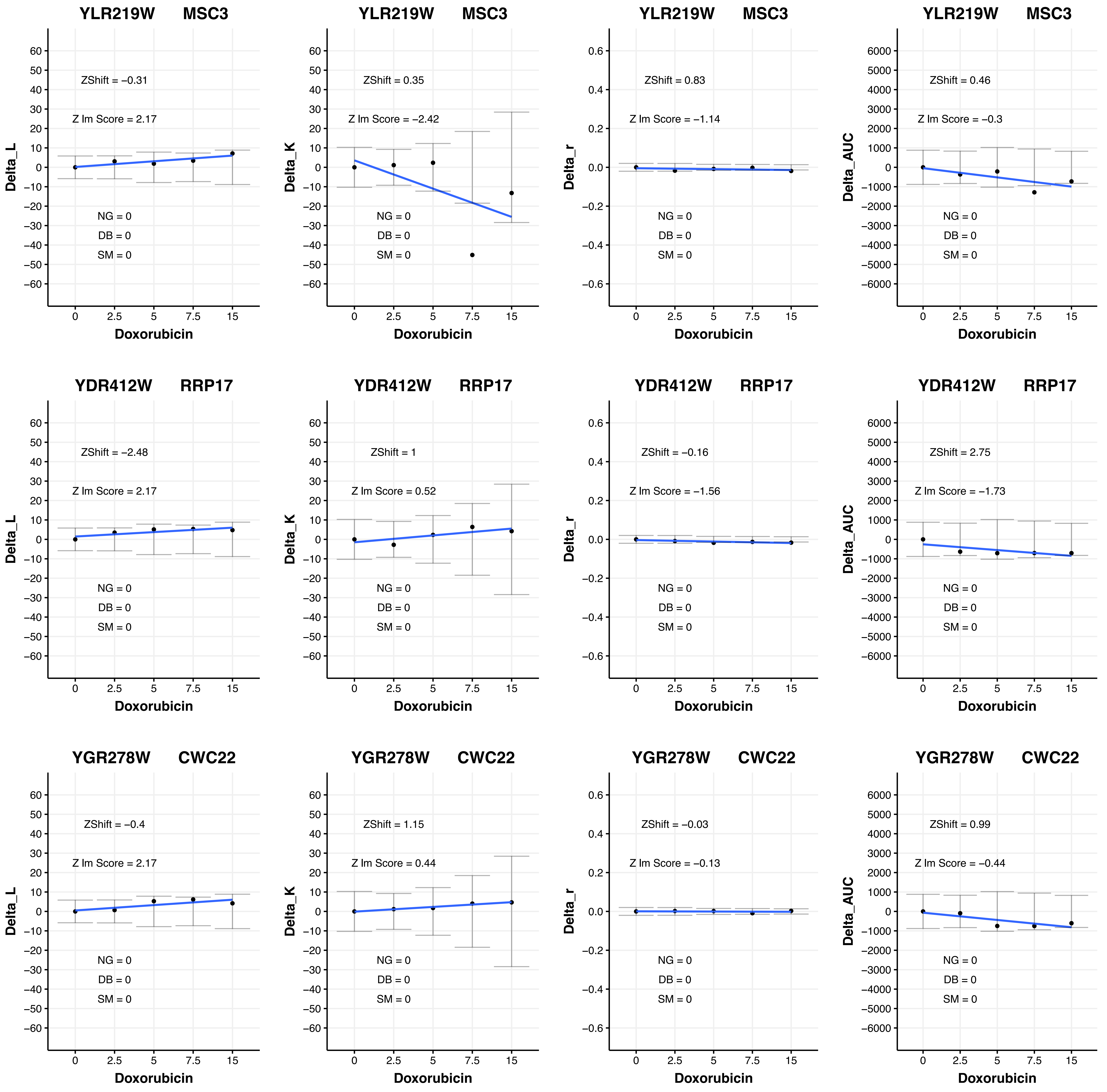
YLR219W MSC3
ZShift = −0.31
Z lm Score = 2.17
NG = 0
DB = 0
SM = 0
Delta_L
Doxorubicin
YLR219W MSC3
ZShift = 0.35
Z lm Score = −2.42
NG = 0
DB = 0
SM = 0
Delta_K
Doxorubicin
YLR219W MSC3
ZShift = 0.83
Z lm Score = −1.14
NG = 0
DB = 0
SM = 0
Delta_r
Doxorubicin
YLR219W MSC3
ZShift = 0.46
Z lm Score = −0.3
NG = 0
DB = 0
SM = 0
Delta_AUC
Doxorubicin
YDR412W RRP17
ZShift = −2.48
Z lm Score = 2.17
NG = 0
DB = 0
SM = 0
Delta_L
Doxorubicin
YDR412W RRP17
ZShift = 1
Z lm Score = 0.52
NG = 0
DB = 0
SM = 0
Delta_K
Doxorubicin
YDR412W RRP17
ZShift = −0.16
Z lm Score = −1.56
NG = 0
DB = 0
SM = 0
Delta_r
Doxorubicin
YDR412W RRP17
ZShift = 2.75
Z lm Score = −1.73
NG = 0
DB = 0
SM = 0
Delta_AUC
Doxorubicin
YGR278W CWC22
ZShift = −0.4
Z lm Score = 2.17
NG = 0
DB = 0
SM = 0
Delta_L
Doxorubicin
YGR278W CWC22
ZShift = 1.15
Z lm Score = 0.44
NG = 0
DB = 0
SM = 0
Delta_K
Doxorubicin
YGR278W CWC22
ZShift = −0.03
Z lm Score = −0.13
NG = 0
DB = 0
SM = 0
Delta_r
Doxorubicin
YGR278W CWC22
ZShift = 0.99
Z lm Score = −0.44
NG = 0
DB = 0
SM = 0
Delta_AUC
Doxorubicin

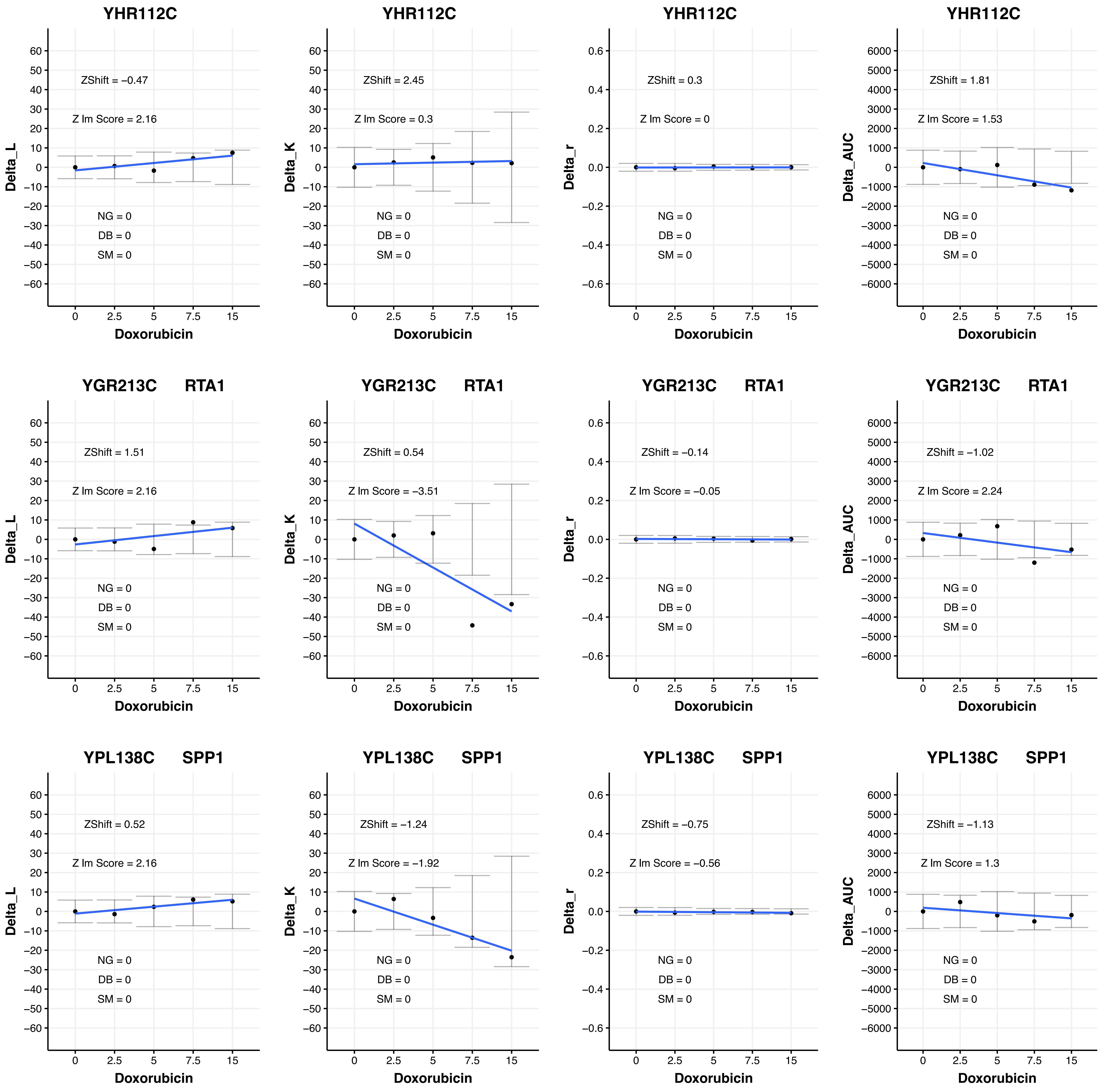
YHR112C
ZShift = −0.47
Z lm Score = 2.16
NG = 0
DB = 0
SM = 0
Delta_L
Doxorubicin
YHR112C
ZShift = 2.45
Z lm Score = 0.3
NG = 0
DB = 0
SM = 0
Delta_K
Doxorubicin
YHR112C
ZShift = 0.3
Z lm Score = 0
NG = 0
DB = 0
SM = 0
Delta_r
Doxorubicin
YHR112C
ZShift = 1.81
Z lm Score = 1.53
NG = 0
DB = 0
SM = 0
Delta_AUC
Doxorubicin
YGR213C RTA1
ZShift = 1.51
Z lm Score = 2.16
NG = 0
DB = 0
SM = 0
Delta_L
Doxorubicin
YGR213C RTA1
ZShift = 0.54
Z lm Score = −3.51
NG = 0
DB = 0
SM = 0
Delta_K
Doxorubicin
YGR213C RTA1
ZShift = −0.14
Z lm Score = −0.05
NG = 0
DB = 0
SM = 0
Delta_r
Doxorubicin
YGR213C RTA1
ZShift = −1.02
Z lm Score = 2.24
NG = 0
DB = 0
SM = 0
Delta_AUC
Doxorubicin
YPL138C SPP1
ZShift = 0.52
Z lm Score = 2.16
NG = 0
DB = 0
SM = 0
Delta_L
Doxorubicin
YPL138C SPP1
ZShift = −1.24
Z lm Score = −1.92
NG = 0
DB = 0
SM = 0
Delta_K
Doxorubicin
YPL138C SPP1
ZShift = −0.75
Z lm Score = −0.56
NG = 0
DB = 0
SM = 0
Delta_r
Doxorubicin
YPL138C SPP1
ZShift = −1.13
Z lm Score = 1.3
NG = 0
DB = 0
SM = 0
Delta_AUC
Doxorubicin

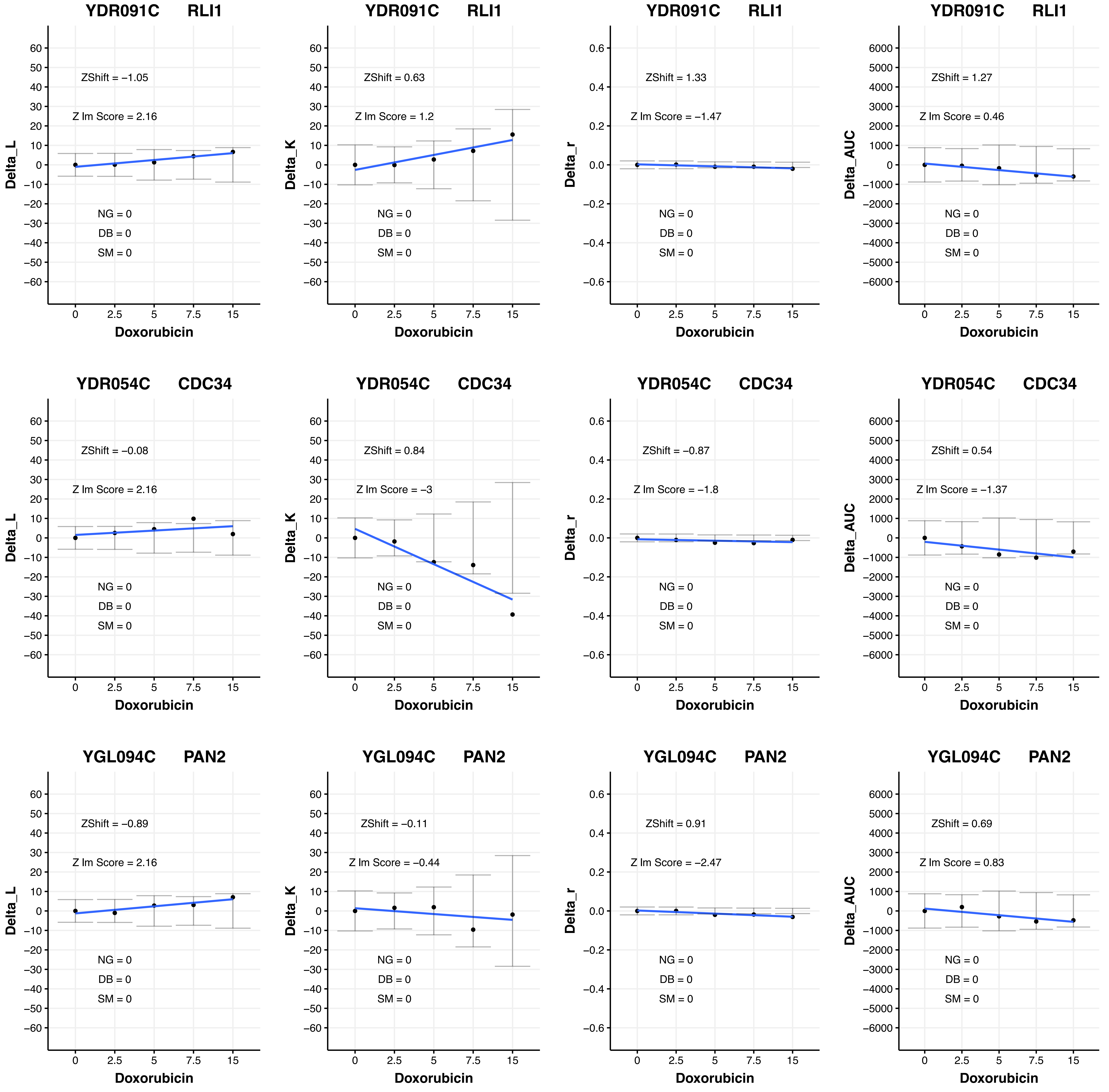

YDR091C RLI1
ZShift = −1.05
Z lm Score = 2.16
NG = 0
DB = 0
SM = 0
Delta_L
Doxorubicin
YDR091C RLI1
ZShift = 0.63
Z lm Score = 1.2
NG = 0
DB = 0
SM = 0
Delta_K
Doxorubicin
YDR091C RLI1
ZShift = 1.33
Z lm Score = −1.47
NG = 0
DB = 0
SM = 0
Delta_r
Doxorubicin
YDR091C RLI1
ZShift = 1.27
Z lm Score = 0.46
NG = 0
DB = 0
SM = 0
Delta_AUC
Doxorubicin
YDR054C CDC34
ZShift = −0.08
Z lm Score = 2.16
NG = 0
DB = 0
SM = 0
Delta_L
Doxorubicin
YDR054C CDC34
ZShift = 0.84
Z lm Score = −3
NG = 0
DB = 0
SM = 0
Delta_K
Doxorubicin
YDR054C CDC34
ZShift = −0.87
Z lm Score = −1.8
NG = 0
DB = 0
SM = 0
Delta_r
Doxorubicin
YDR054C CDC34
ZShift = 0.54
Z lm Score = −1.37
NG = 0
DB = 0
SM = 0
Delta_AUC
Doxorubicin
YGL094C PAN2
ZShift = −0.89
Z lm Score = 2.16
NG = 0
DB = 0
SM = 0
Delta_L
Doxorubicin
YGL094C PAN2
ZShift = −0.11
Z lm Score = −0.44
NG = 0
DB = 0
SM = 0
Delta_K
Doxorubicin
YGL094C PAN2
ZShift = 0.91
Z lm Score = −2.47
NG = 0
DB = 0
SM = 0
Delta_r
Doxorubicin
YGL094C PAN2
ZShift = 0.69
Z lm Score = 0.83
NG = 0
DB = 0
SM = 0
Delta_AUC
Doxorubicin

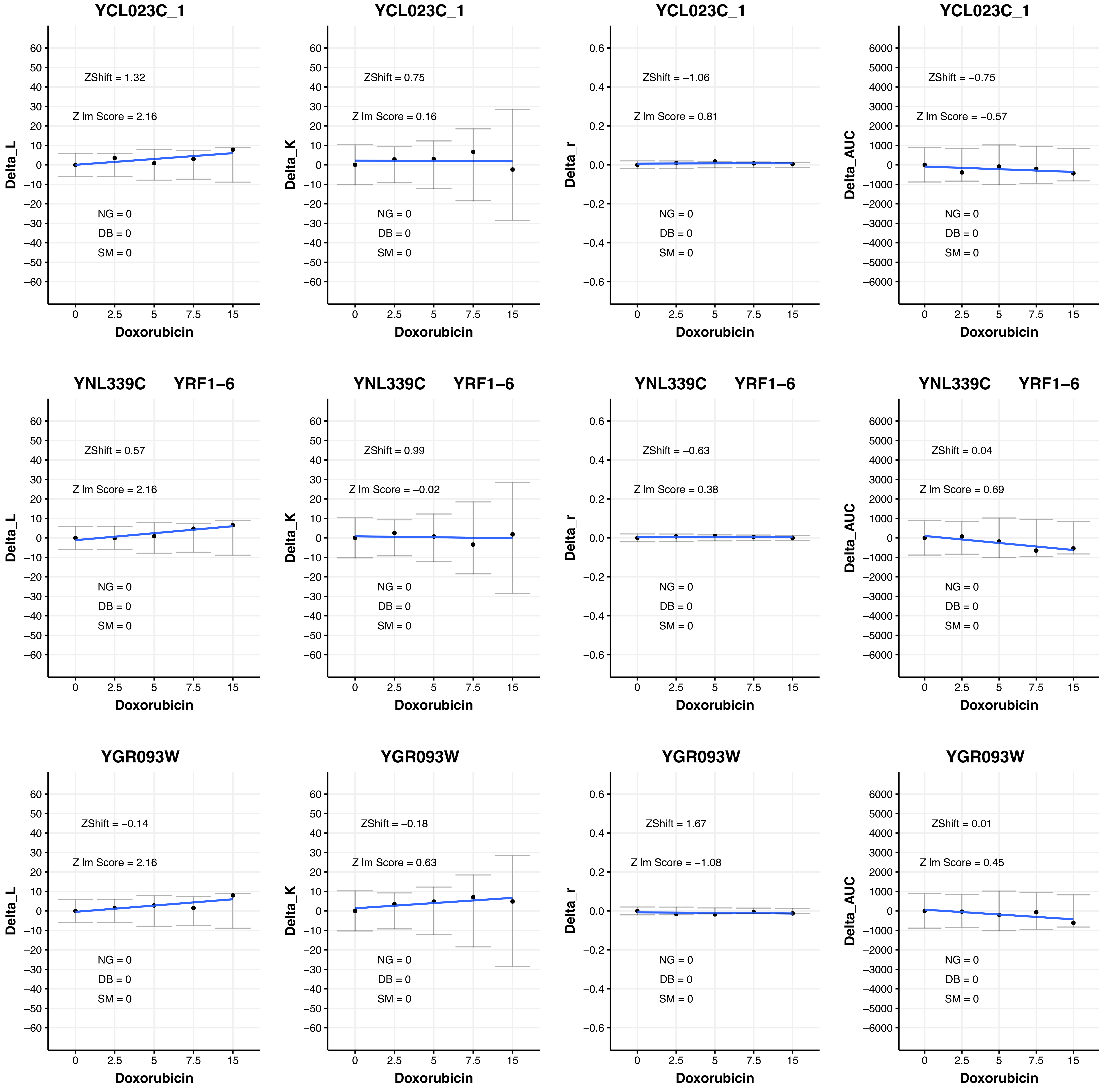
YCL023C_1
ZShift = 1.32
Z lm Score = 2.16
NG = 0
DB = 0
SM = 0
Delta_L
Doxorubicin
YCL023C_1
ZShift = 0.75
Z lm Score = 0.16
NG = 0
DB = 0
SM = 0
Delta_K
Doxorubicin
YCL023C_1
ZShift = −1.06
Z lm Score = 0.81
NG = 0
DB = 0
SM = 0
Delta_r
Doxorubicin
YCL023C_1
ZShift = −0.75
Z lm Score = −0.57
NG = 0
DB = 0
SM = 0
Delta_AUC
Doxorubicin
YNL339C YRF1−6
ZShift = 0.57
Z lm Score = 2.16
NG = 0
DB = 0
SM = 0
Delta_L
Doxorubicin
YNL339C YRF1−6
ZShift = 0.99
Z lm Score = −0.02
NG = 0
DB = 0
SM = 0
Delta_K
Doxorubicin
YNL339C YRF1−6
ZShift = −0.63
Z lm Score = 0.38
NG = 0
DB = 0
SM = 0
Delta_r
Doxorubicin
YNL339C YRF1−6
ZShift = 0.04
Z lm Score = 0.69
NG = 0
DB = 0
SM = 0
Delta_AUC
Doxorubicin
YGR093W
ZShift = −0.14
Z lm Score = 2.16
NG = 0
DB = 0
SM = 0
Delta_L
Doxorubicin
YGR093W
ZShift = −0.18
Z lm Score = 0.63
NG = 0
DB = 0
SM = 0
Delta_K
Doxorubicin
YGR093W
ZShift = 1.67
Z lm Score = −1.08
NG = 0
DB = 0
SM = 0
Delta_r
Doxorubicin
YGR093W
ZShift = 0.01
Z lm Score = 0.45
NG = 0
DB = 0
SM = 0
Delta_AUC
Doxorubicin

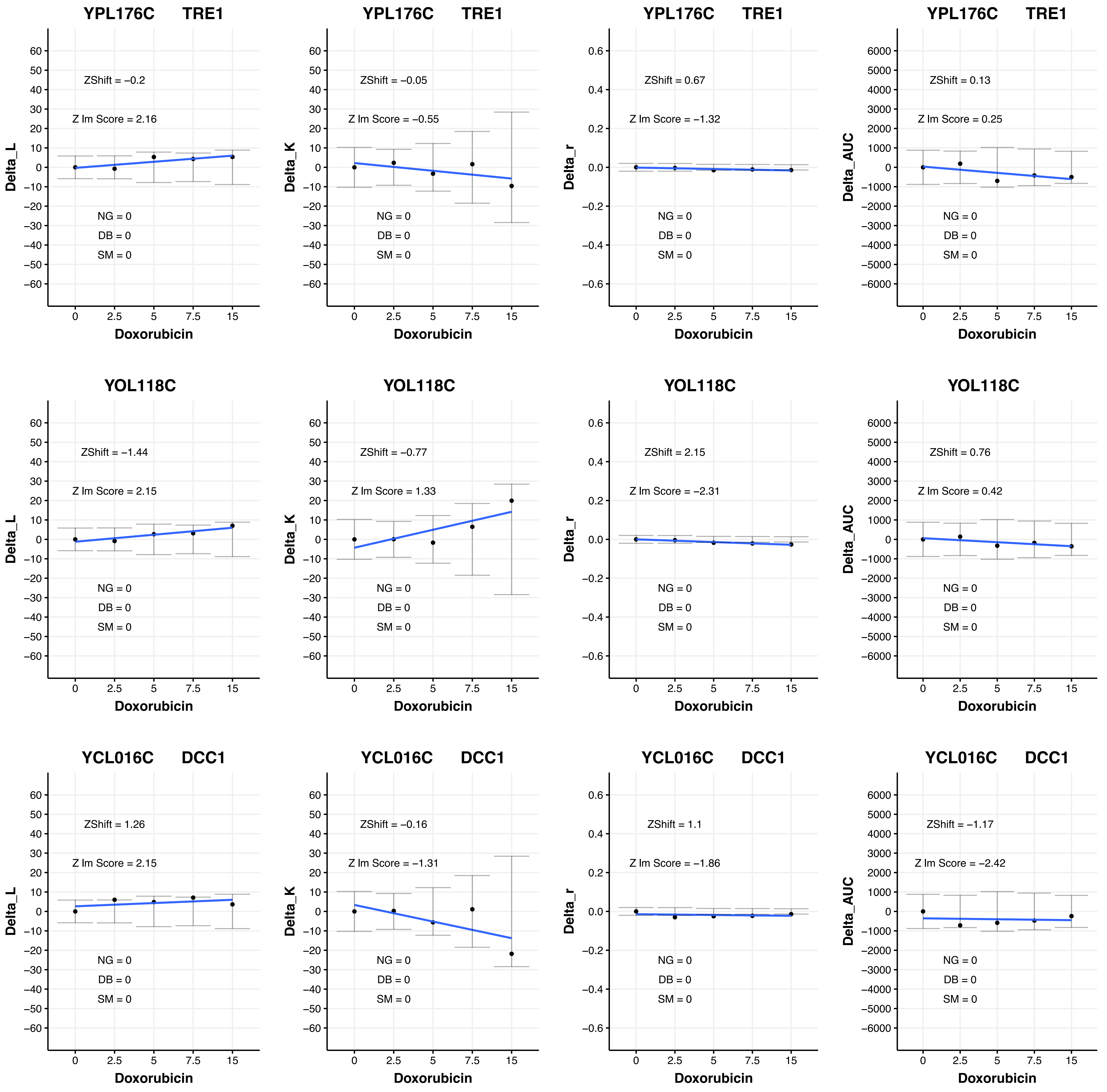

YPL176C TRE1
ZShift = −0.2
Z lm Score = 2.16
NG = 0
DB = 0
SM = 0
Delta_L
Doxorubicin
YPL176C TRE1
ZShift = −0.05
Z lm Score = −0.55
NG = 0
DB = 0
SM = 0
Delta_K
Doxorubicin
YPL176C TRE1
ZShift = 0.67
Z lm Score = −1.32
NG = 0
DB = 0
SM = 0
Delta_r
Doxorubicin
YPL176C TRE1
ZShift = 0.13
Z lm Score = 0.25
NG = 0
DB = 0
SM = 0
Delta_AUC
Doxorubicin
YOL118C
ZShift = −1.44
Z lm Score = 2.15
NG = 0
DB = 0
SM = 0
Delta_L
Doxorubicin
YOL118C
ZShift = −0.77
Z lm Score = 1.33
NG = 0
DB = 0
SM = 0
Delta_K
Doxorubicin
YOL118C
ZShift = 2.15
Z lm Score = −2.31
NG = 0
DB = 0
SM = 0
Delta_r
Doxorubicin
YOL118C
ZShift = 0.76
Z lm Score = 0.42
NG = 0
DB = 0
SM = 0
Delta_AUC
Doxorubicin
YCL016C DCC1
ZShift = 1.26
Z lm Score = 2.15
NG = 0
DB = 0
SM = 0
Delta_L
Doxorubicin
YCL016C DCC1
ZShift = −0.16
Z lm Score = −1.31
NG = 0
DB = 0
SM = 0
Delta_K
Doxorubicin
YCL016C DCC1
ZShift = 1.1
Z lm Score = −1.86
NG = 0
DB = 0
SM = 0
Delta_r
Doxorubicin
YCL016C DCC1
ZShift = −1.17
Z lm Score = −2.42
NG = 0
DB = 0
SM = 0
Delta_AUC
Doxorubicin

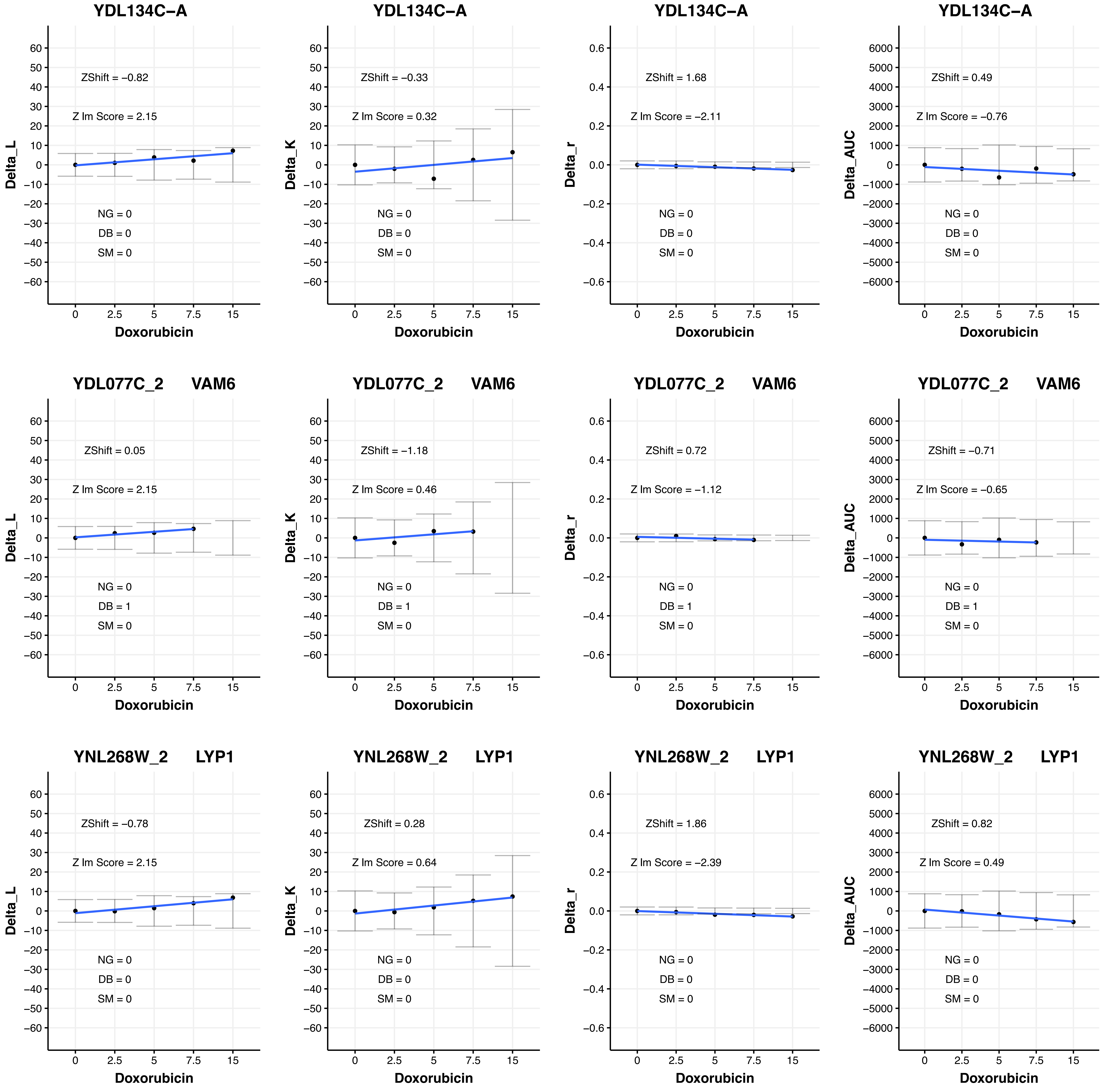
YDL134C−A
ZShift = −0.82
Z lm Score = 2.15
NG = 0
DB = 0
SM = 0
Delta_L
Doxorubicin
YDL134C−A
ZShift = −0.33
Z lm Score = 0.32
NG = 0
DB = 0
SM = 0
Delta_K
Doxorubicin
YDL134C−A
ZShift = 1.68
Z lm Score = −2.11
NG = 0
DB = 0
SM = 0
Delta_r
Doxorubicin
YDL134C−A
ZShift = 0.49
Z lm Score = −0.76
NG = 0
DB = 0
SM = 0
Delta_AUC
Doxorubicin
YDL077C_2 VAM6
ZShift = 0.05
Z lm Score = 2.15
NG = 0
DB = 1
SM = 0
Delta_L
Doxorubicin
YDL077C_2 VAM6
ZShift = −1.18
Z lm Score = 0.46
NG = 0
DB = 1
SM = 0
Delta_K
Doxorubicin
YDL077C_2 VAM6
ZShift = 0.72
Z lm Score = −1.12
NG = 0
DB = 1
SM = 0
Delta_r
Doxorubicin
YDL077C_2 VAM6
ZShift = −0.71
Z lm Score = −0.65
NG = 0
DB = 1
SM = 0
Delta_AUC
Doxorubicin
YNL268W_2 LYP1
ZShift = −0.78
Z lm Score = 2.15
NG = 0
DB = 0
SM = 0
Delta_L
Doxorubicin
YNL268W_2 LYP1
ZShift = 0.28
Z lm Score = 0.64
NG = 0
DB = 0
SM = 0
Delta_K
Doxorubicin
YNL268W_2 LYP1
ZShift = 1.86
Z lm Score = −2.39
NG = 0
DB = 0
SM = 0
Delta_r
Doxorubicin
YNL268W_2 LYP1
ZShift = 0.82
Z lm Score = 0.49
NG = 0
DB = 0
SM = 0
Delta_AUC
Doxorubicin

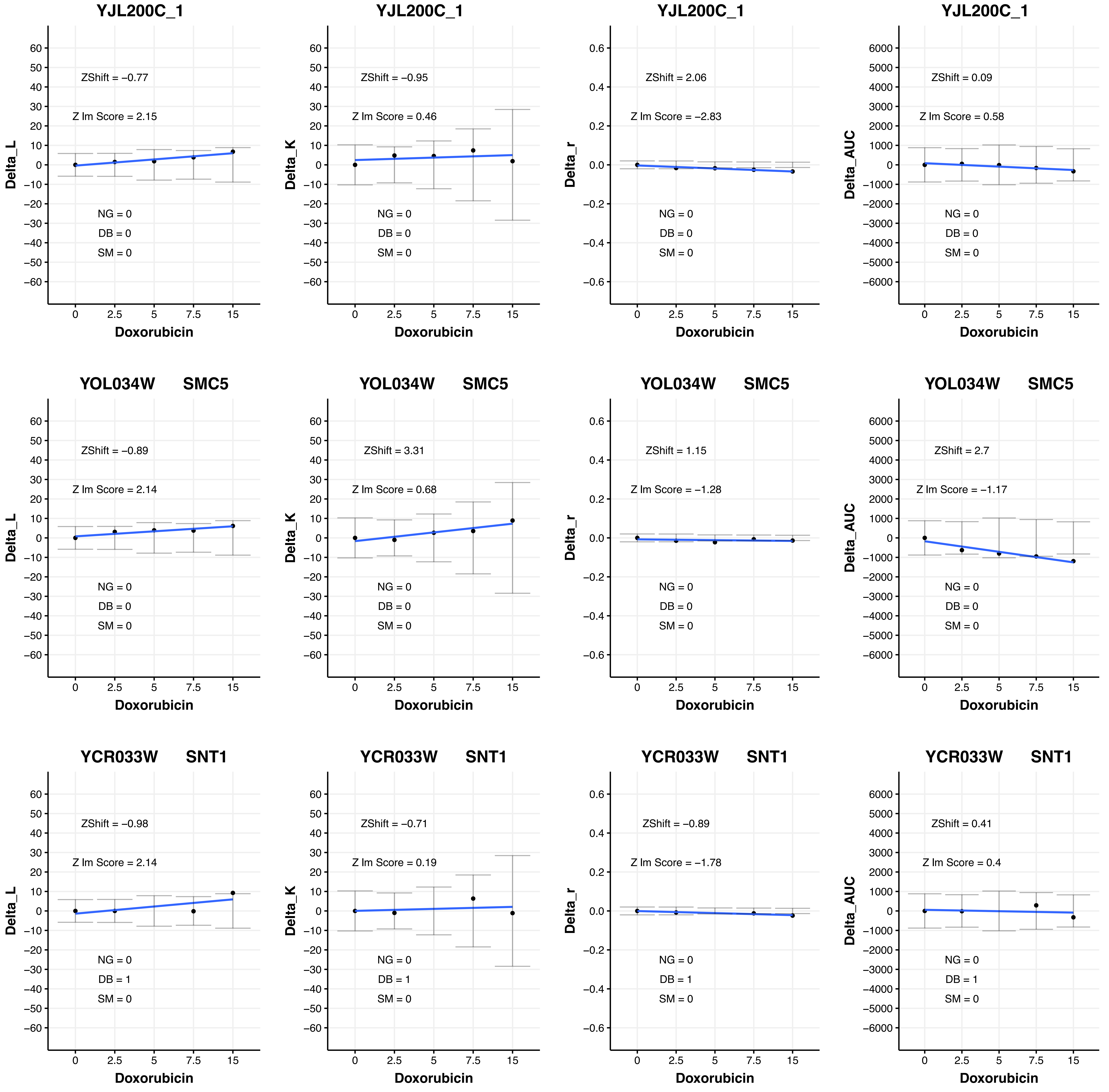
YJL200C_1
ZShift = −0.77
Z lm Score = 2.15
NG = 0
DB = 0
SM = 0
Delta_L
Doxorubicin
YJL200C_1
ZShift = −0.95
Z lm Score = 0.46
NG = 0
DB = 0
SM = 0
Delta_K
Doxorubicin
YJL200C_1
ZShift = 2.06
Z lm Score = −2.83
NG = 0
DB = 0
SM = 0
Delta_r
Doxorubicin
YJL200C_1
ZShift = 0.09
Z lm Score = 0.58
NG = 0
DB = 0
SM = 0
Delta_AUC
Doxorubicin
YOL034W SMC5
ZShift = −0.89
Z lm Score = 2.14
NG = 0
DB = 0
SM = 0
Delta_L
Doxorubicin
YOL034W SMC5
ZShift = 3.31
Z lm Score = 0.68
NG = 0
DB = 0
SM = 0
Delta_K
Doxorubicin
YOL034W SMC5
ZShift = 1.15
Z lm Score = −1.28
NG = 0
DB = 0
SM = 0
Delta_r
Doxorubicin
YOL034W SMC5
ZShift = 2.7
Z lm Score = −1.17
NG = 0
DB = 0
SM = 0
Delta_AUC
Doxorubicin
YCR033W SNT1
ZShift = −0.98
Z lm Score = 2.14
NG = 0
DB = 1
SM = 0
Delta_L
Doxorubicin
YCR033W SNT1
ZShift = −0.71
Z lm Score = 0.19
NG = 0
DB = 1
SM = 0
Delta_K
Doxorubicin
YCR033W SNT1
ZShift = −0.89
Z lm Score = −1.78
NG = 0
DB = 1
SM = 0
Delta_r
Doxorubicin
YCR033W SNT1
ZShift = 0.41
Z lm Score = 0.4
NG = 0
DB = 1
SM = 0
Delta_AUC
Doxorubicin

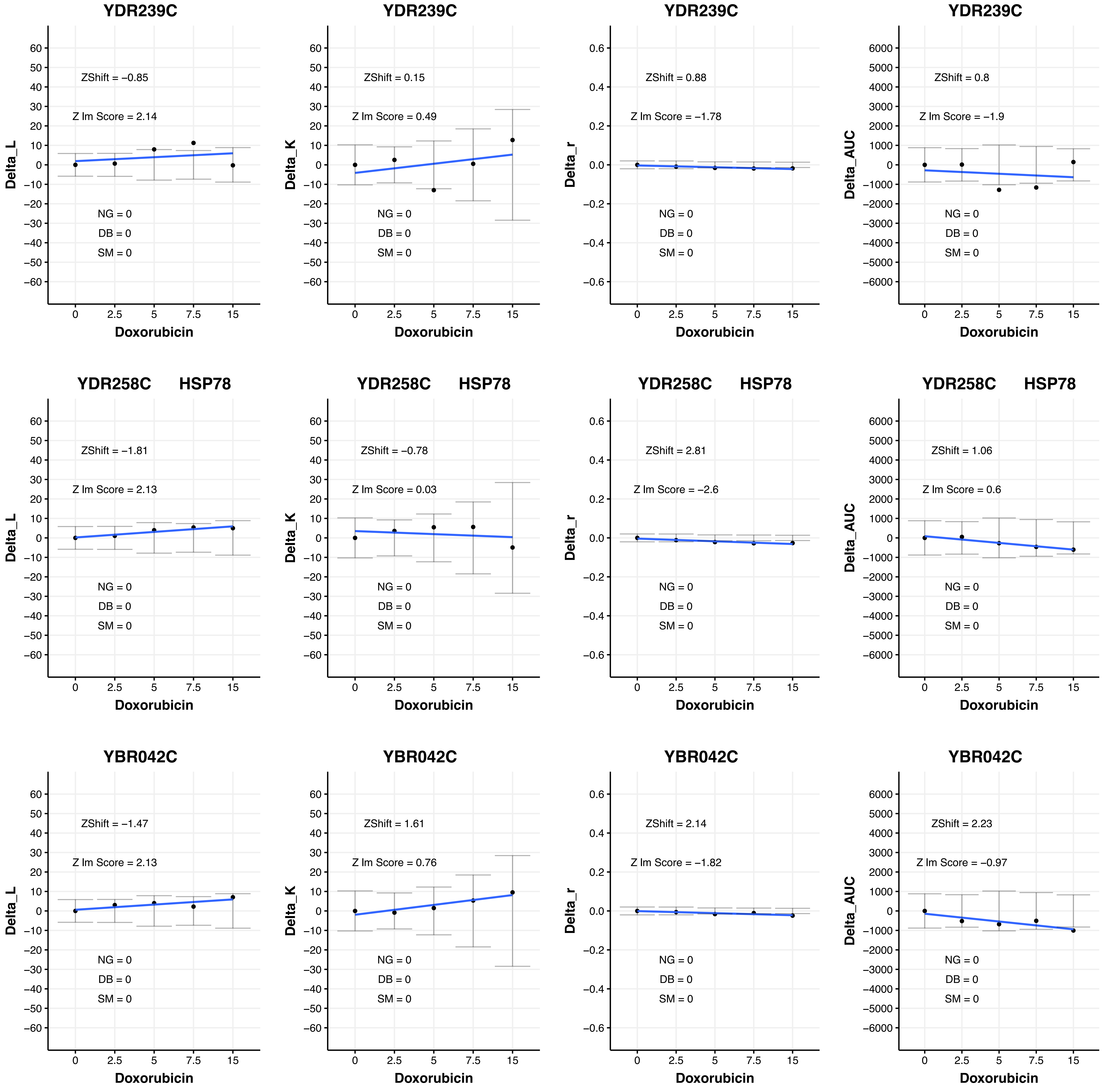

YDR239C
ZShift = −0.85
Z lm Score = 2.14
NG = 0
DB = 0
SM = 0
Delta_L
Doxorubicin
YDR239C
ZShift = 0.15
Z lm Score = 0.49
NG = 0
DB = 0
SM = 0
Delta_K
Doxorubicin
YDR239C
ZShift = 0.88
Z lm Score = −1.78
NG = 0
DB = 0
SM = 0
Delta_r
Doxorubicin
YDR239C
ZShift = 0.8
Z lm Score = −1.9
NG = 0
DB = 0
SM = 0
Delta_AUC
Doxorubicin
YDR258C HSP78
ZShift = −1.81
Z lm Score = 2.13
NG = 0
DB = 0
SM = 0
Delta_L
Doxorubicin
YDR258C HSP78
ZShift = −0.78
Z lm Score = 0.03
NG = 0
DB = 0
SM = 0
Delta_K
Doxorubicin
YDR258C HSP78
ZShift = 2.81
Z lm Score = −2.6
NG = 0
DB = 0
SM = 0
Delta_r
Doxorubicin
YDR258C HSP78
ZShift = 1.06
Z lm Score = 0.6
NG = 0
DB = 0
SM = 0
Delta_AUC
Doxorubicin
YBR042C
ZShift = −1.47
Z lm Score = 2.13
NG = 0
DB = 0
SM = 0
Delta_L
Doxorubicin
YBR042C
ZShift = 1.61
Z lm Score = 0.76
NG = 0
DB = 0
SM = 0
Delta_K
Doxorubicin
YBR042C
ZShift = 2.14
Z lm Score = −1.82
NG = 0
DB = 0
SM = 0
Delta_r
Doxorubicin
YBR042C
ZShift = 2.23
Z lm Score = −0.97
NG = 0
DB = 0
SM = 0
Delta_AUC
Doxorubicin

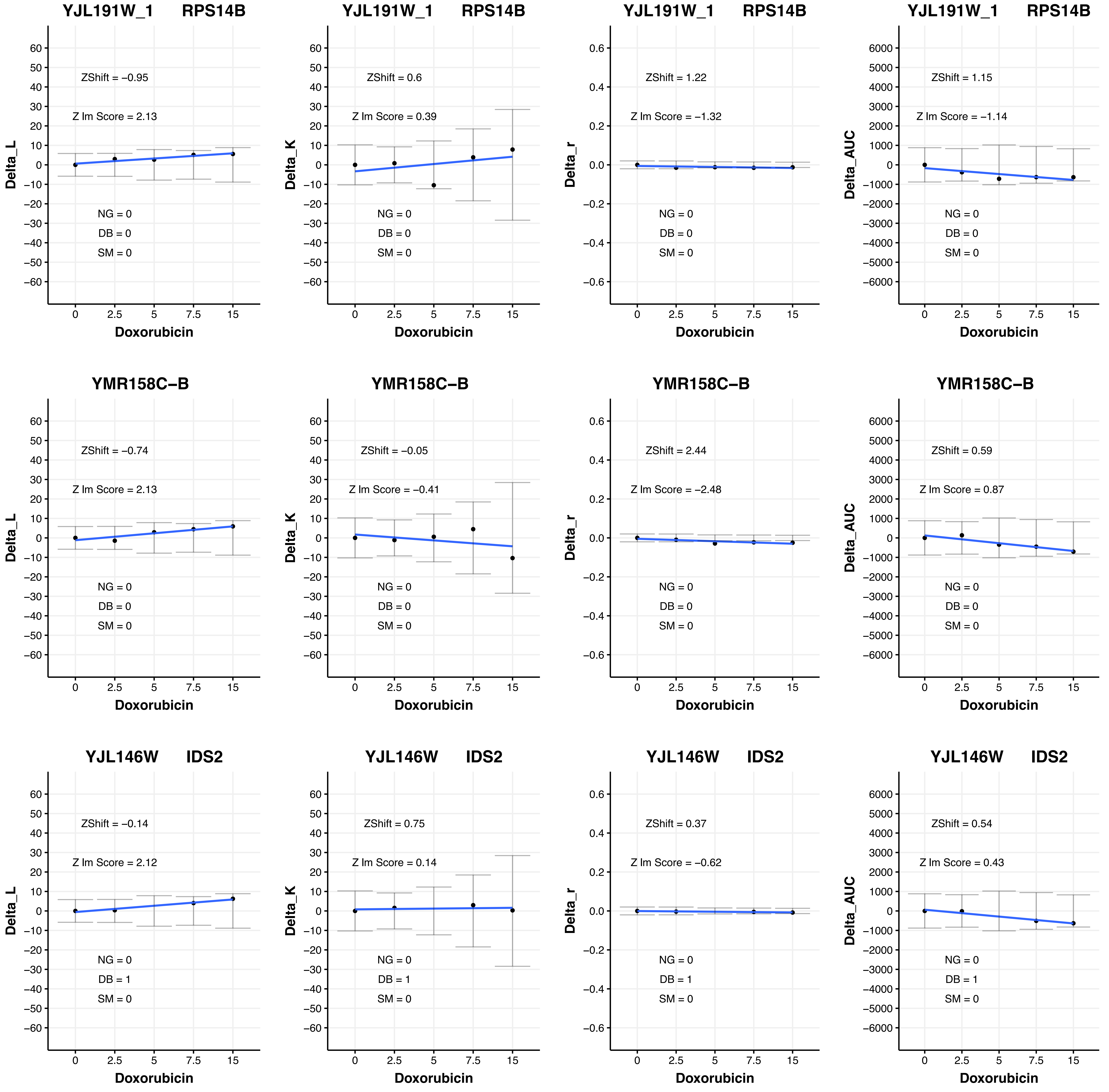

YJL191W_1 RPS14B
ZShift = −0.95
Z lm Score = 2.13
NG = 0
DB = 0
SM = 0
Delta_L
Doxorubicin
YJL191W_1 RPS14B
ZShift = 0.6
Z lm Score = 0.39
NG = 0
DB = 0
SM = 0
Delta_K
Doxorubicin
YJL191W_1 RPS14B
ZShift = 1.22
Z lm Score = −1.32
NG = 0
DB = 0
SM = 0
Delta_r
Doxorubicin
YJL191W_1 RPS14B
ZShift = 1.15
Z lm Score = −1.14
NG = 0
DB = 0
SM = 0
Delta_AUC
Doxorubicin
YMR158C−B
ZShift = −0.74
Z lm Score = 2.13
NG = 0
DB = 0
SM = 0
Delta_L
Doxorubicin
YMR158C−B
ZShift = −0.05
Z lm Score = −0.41
NG = 0
DB = 0
SM = 0
Delta_K
Doxorubicin
YMR158C−B
ZShift = 2.44
Z lm Score = −2.48
NG = 0
DB = 0
SM = 0
Delta_r
Doxorubicin
YMR158C−B
ZShift = 0.59
Z lm Score = 0.87
NG = 0
DB = 0
SM = 0
Delta_AUC
Doxorubicin
YJL146W IDS2
ZShift = −0.14
Z lm Score = 2.12
NG = 0
DB = 1
SM = 0
Delta_L
Doxorubicin
YJL146W IDS2
ZShift = 0.75
Z lm Score = 0.14
NG = 0
DB = 1
SM = 0
Delta_K
Doxorubicin
YJL146W IDS2
ZShift = 0.37
Z lm Score = −0.62
NG = 0
DB = 1
SM = 0
Delta_r
Doxorubicin
YJL146W IDS2
ZShift = 0.54
Z lm Score = 0.43
NG = 0
DB = 1
SM = 0
Delta_AUC
Doxorubicin

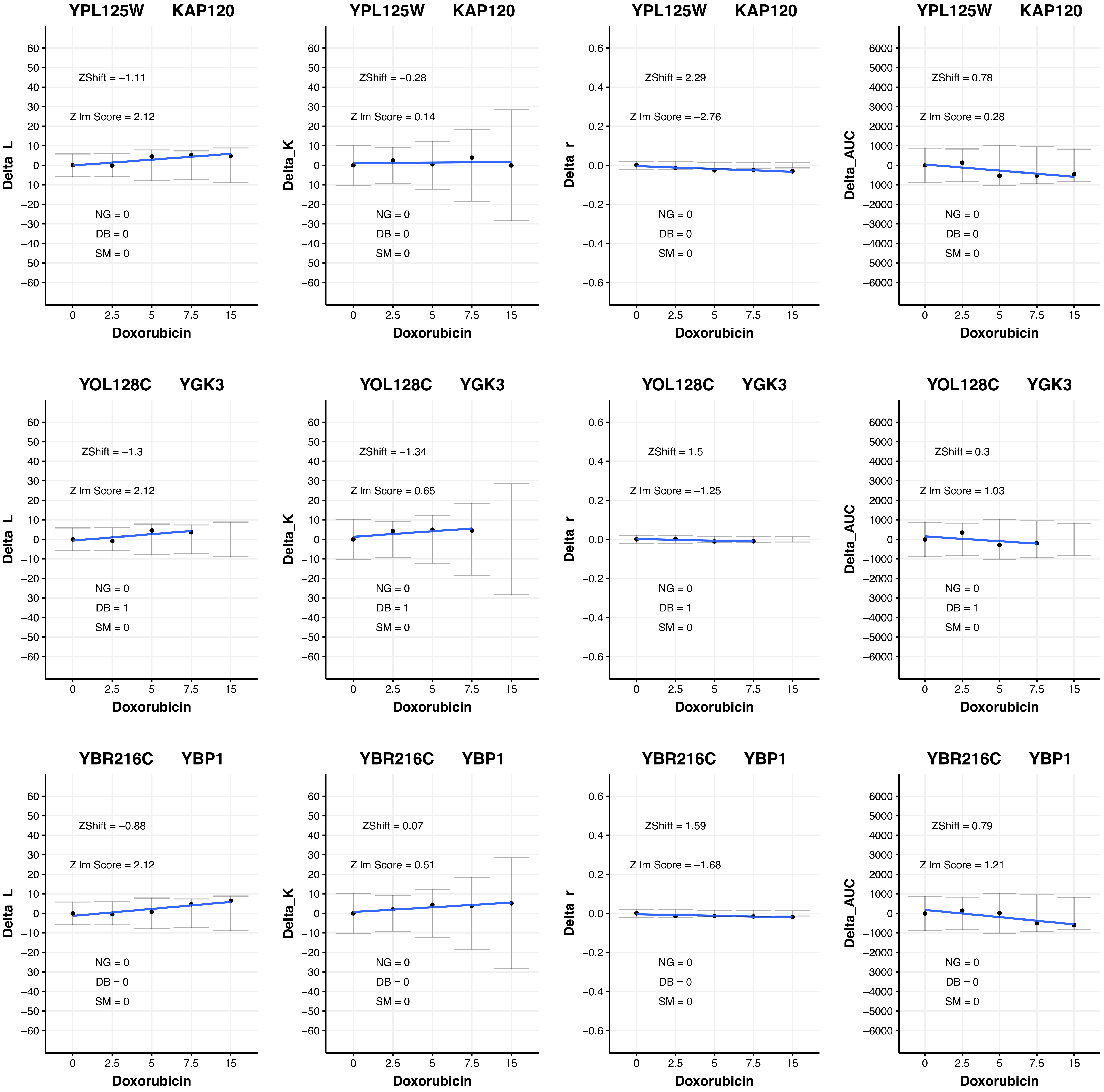
YPL125W KAP120
ZShift = −1.11
Z lm Score = 2.12
NG = 0
DB = 0
SM = 0
Delta_L
Doxorubicin
YPL125W KAP120
ZShift = −0.28
Z lm Score = 0.14
NG = 0
DB = 0
SM = 0
Delta_K
Doxorubicin
YPL125W KAP120
ZShift = 2.29
Z lm Score = −2.76
NG = 0
DB = 0
SM = 0
Delta_r
Doxorubicin
YPL125W KAP120
ZShift = 0.78
Z lm Score = 0.28
NG = 0
DB = 0
SM = 0
Delta_AUC
Doxorubicin
YOL128C YGK3
ZShift = −1.3
Z lm Score = 2.12
NG = 0
DB = 1
SM = 0
Delta_L
Doxorubicin
YOL128C YGK3
ZShift = −1.34
Z lm Score = 0.65
NG = 0
DB = 1
SM = 0
Delta_K
Doxorubicin
YOL128C YGK3
ZShift = 1.5
Z lm Score = −1.25
NG = 0
DB = 1
SM = 0
Delta_r
Doxorubicin
YOL128C YGK3
ZShift = 0.3
Z lm Score = 1.03
NG = 0
DB = 1
SM = 0
Delta_AUC
Doxorubicin
YBR216C YBP1
ZShift = −0.88
Z lm Score = 2.12
NG = 0
DB = 0
SM = 0
Delta_L
Doxorubicin
YBR216C YBP1
ZShift = 0.07
Z lm Score = 0.51
NG = 0
DB = 0
SM = 0
Delta_K
Doxorubicin
YBR216C YBP1
ZShift = 1.59
Z lm Score = −1.68
NG = 0
DB = 0
SM = 0
Delta_r
Doxorubicin
YBR216C YBP1
ZShift = 0.79
Z lm Score = 1.21
NG = 0
DB = 0
SM = 0
Delta_AUC
Doxorubicin

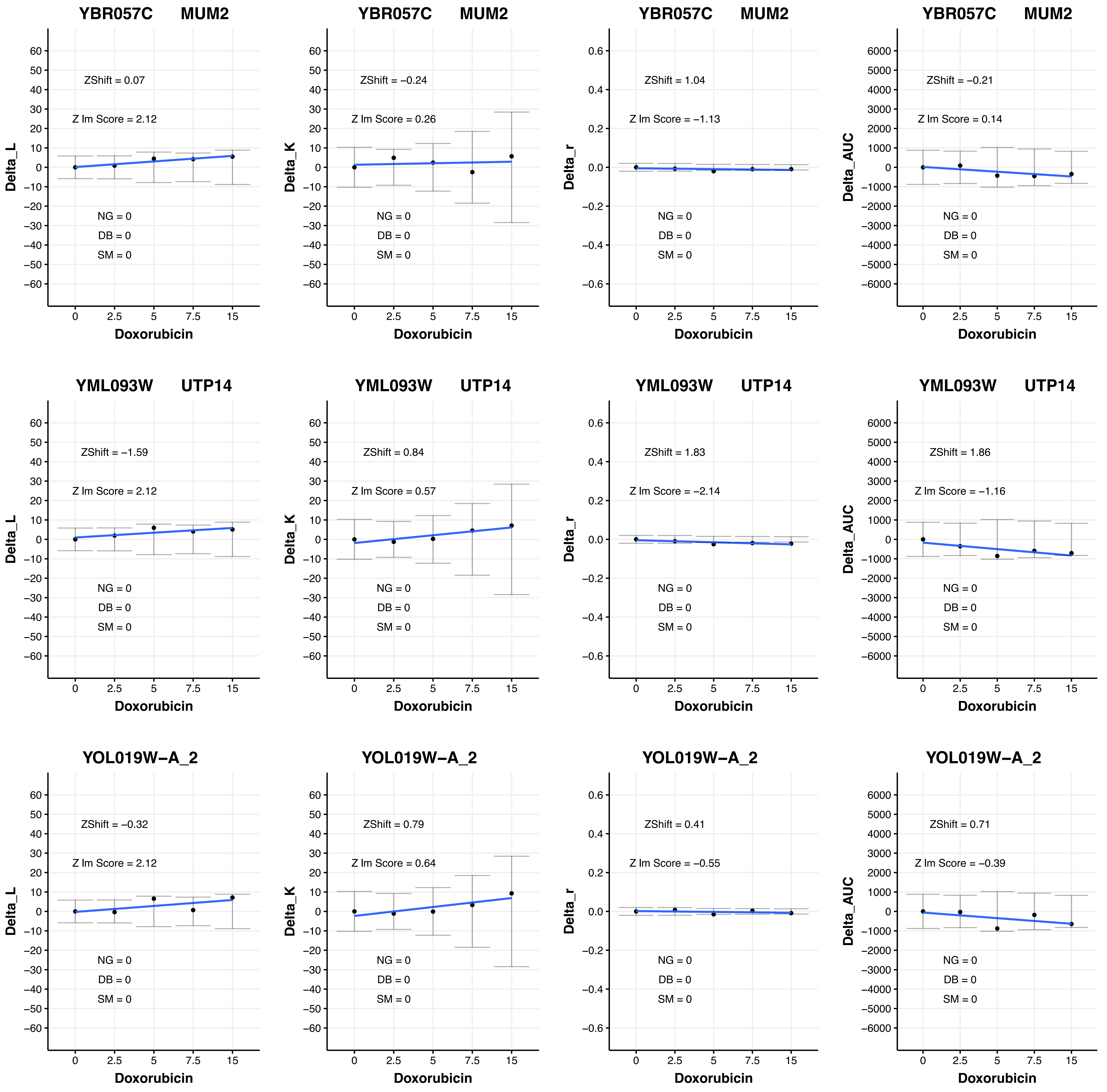
YBR057C MUM2
ZShift = 0.07
Z lm Score = 2.12
NG = 0
DB = 0
SM = 0
Delta_L
Doxorubicin
YBR057C MUM2
ZShift = −0.24
Z lm Score = 0.26
NG = 0
DB = 0
SM = 0
Delta_K
Doxorubicin
YBR057C MUM2
ZShift = 1.04
Z lm Score = −1.13
NG = 0
DB = 0
SM = 0
Delta_r
Doxorubicin
YBR057C MUM2
ZShift = −0.21
Z lm Score = 0.14
NG = 0
DB = 0
SM = 0
Delta_AUC
Doxorubicin
YML093W UTP14
ZShift = −1.59
Z lm Score = 2.12
NG = 0
DB = 0
SM = 0
Delta_L
Doxorubicin
YML093W UTP14
ZShift = 0.84
Z lm Score = 0.57
NG = 0
DB = 0
SM = 0
Delta_K
Doxorubicin
YML093W UTP14
ZShift = 1.83
Z lm Score = −2.14
NG = 0
DB = 0
SM = 0
Delta_r
Doxorubicin
YML093W UTP14
ZShift = 1.86
Z lm Score = −1.16
NG = 0
DB = 0
SM = 0
Delta_AUC
Doxorubicin
YOL019W−A_2
ZShift = −0.32
Z lm Score = 2.12
NG = 0
DB = 0
SM = 0
Delta_L
Doxorubicin
YOL019W−A_2
ZShift = 0.79
Z lm Score = 0.64
NG = 0
DB = 0
SM = 0
Delta_K
Doxorubicin
YOL019W−A_2
ZShift = 0.41
Z lm Score = −0.55
NG = 0
DB = 0
SM = 0
Delta_r
Doxorubicin
YOL019W−A_2
ZShift = 0.71
Z lm Score = −0.39
NG = 0
DB = 0
SM = 0
Delta_AUC
Doxorubicin

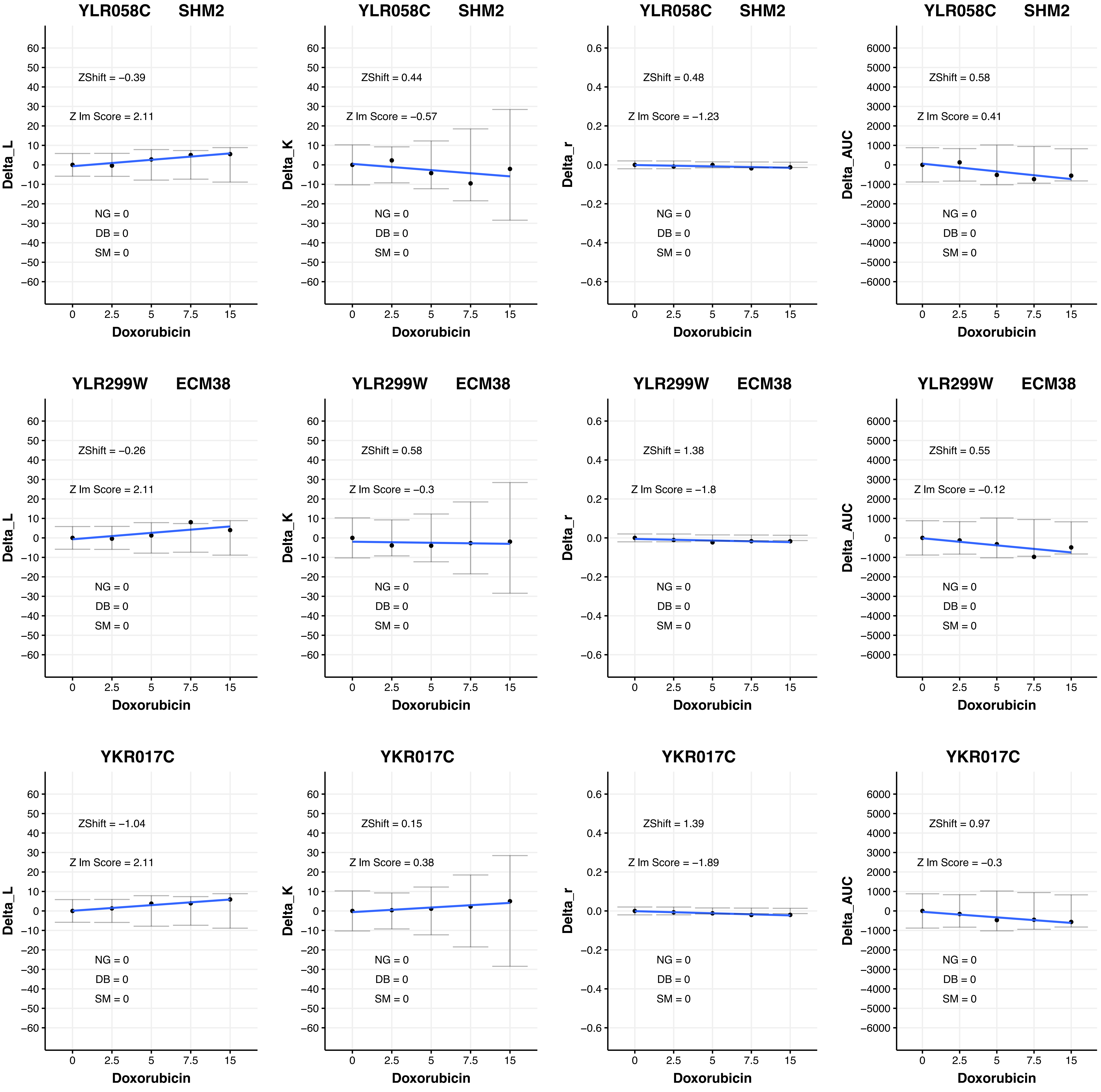

YLR058C SHM2
ZShift = −0.39
Z lm Score = 2.11
NG = 0
DB = 0
SM = 0
Delta_L
Doxorubicin
YLR058C SHM2
ZShift = 0.44
Z lm Score = −0.57
NG = 0
DB = 0
SM = 0
Delta_K
Doxorubicin
YLR058C SHM2
ZShift = 0.48
Z lm Score = −1.23
NG = 0
DB = 0
SM = 0
Delta_r
Doxorubicin
YLR058C SHM2
ZShift = 0.58
Z lm Score = 0.41
NG = 0
DB = 0
SM = 0
Delta_AUC
Doxorubicin
YLR299W ECM38
ZShift = −0.26
Z lm Score = 2.11
NG = 0
DB = 0
SM = 0
Delta_L
Doxorubicin
YLR299W ECM38
ZShift = 0.58
Z lm Score = −0.3
NG = 0
DB = 0
SM = 0
Delta_K
Doxorubicin
YLR299W ECM38
ZShift = 1.38
Z lm Score = −1.8
NG = 0
DB = 0
SM = 0
Delta_r
Doxorubicin
YLR299W ECM38
ZShift = 0.55
Z lm Score = −0.12
NG = 0
DB = 0
SM = 0
Delta_AUC
Doxorubicin
YKR017C
ZShift = −1.04
Z lm Score = 2.11
NG = 0
DB = 0
SM = 0
Delta_L
Doxorubicin
YKR017C
ZShift = 0.15
Z lm Score = 0.38
NG = 0
DB = 0
SM = 0
Delta_K
Doxorubicin
YKR017C
ZShift = 1.39
Z lm Score = −1.89
NG = 0
DB = 0
SM = 0
Delta_r
Doxorubicin
YKR017C
ZShift = 0.97
Z lm Score = −0.3
NG = 0
DB = 0
SM = 0
Delta_AUC
Doxorubicin

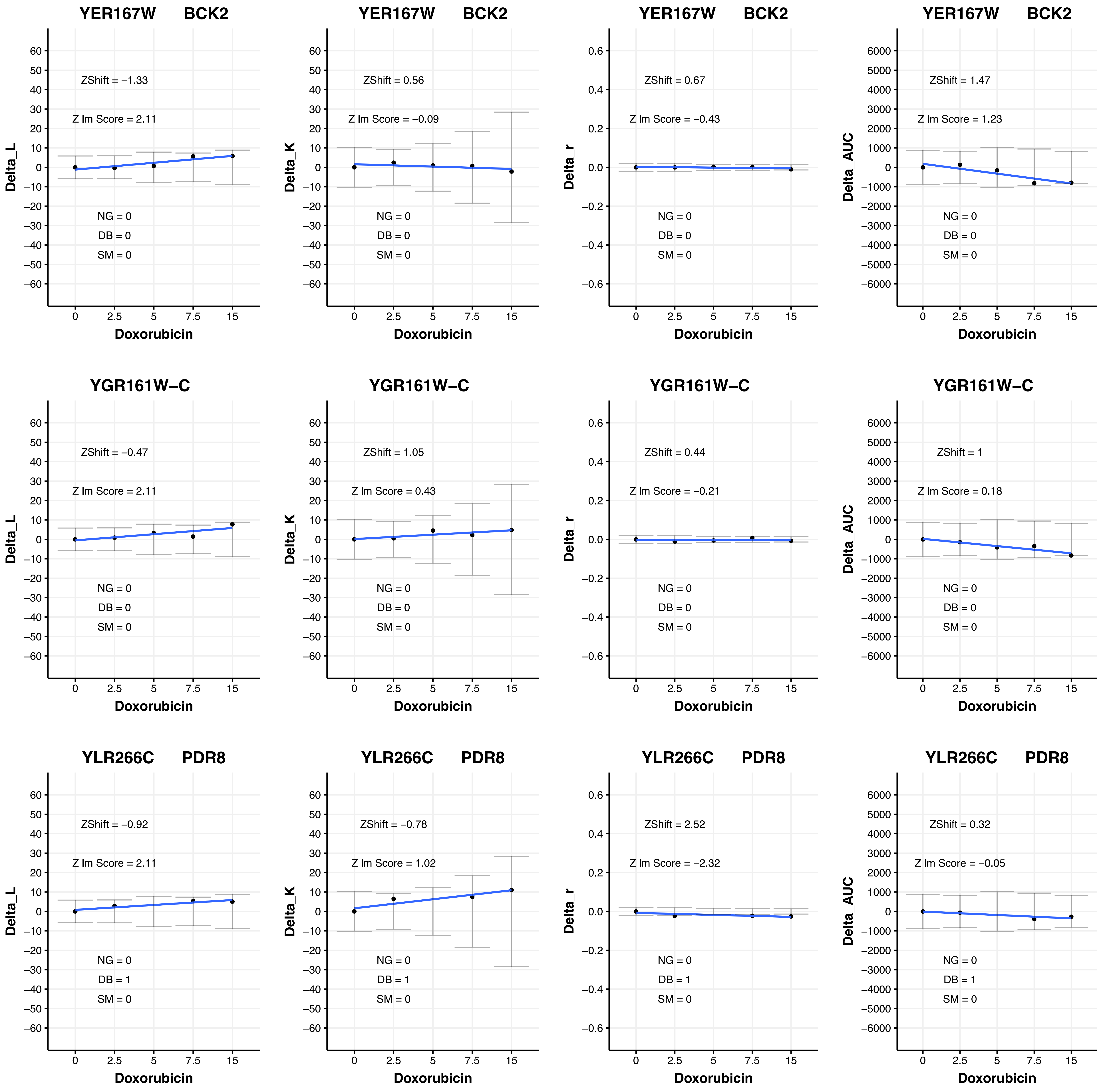
YER167W BCK2
ZShift = −1.33
Z lm Score = 2.11
NG = 0
DB = 0
SM = 0
Delta_L
Doxorubicin
YER167W BCK2
ZShift = 0.56
Z lm Score = −0.09
NG = 0
DB = 0
SM = 0
Delta_K
Doxorubicin
YER167W BCK2
ZShift = 0.67
Z lm Score = −0.43
NG = 0
DB = 0
SM = 0
Delta_r
Doxorubicin
YER167W BCK2
ZShift = 1.47
Z lm Score = 1.23
NG = 0
DB = 0
SM = 0
Delta_AUC
Doxorubicin
YGR161W−C
ZShift = −0.47
Z lm Score = 2.11
NG = 0
DB = 0
SM = 0
Delta_L
Doxorubicin
YGR161W−C
ZShift = 1.05
Z lm Score = 0.43
NG = 0
DB = 0
SM = 0
Delta_K
Doxorubicin
YGR161W−C
ZShift = 0.44
Z lm Score = −0.21
NG = 0
DB = 0
SM = 0
Delta_r
Doxorubicin
YGR161W−C
ZShift = 1
Z lm Score = 0.18
NG = 0
DB = 0
SM = 0
Delta_AUC
Doxorubicin
YLR266C PDR8
ZShift = −0.92
Z lm Score = 2.11
NG = 0
DB = 1
SM = 0
Delta_L
Doxorubicin
YLR266C PDR8
ZShift = −0.78
Z lm Score = 1.02
NG = 0
DB = 1
SM = 0
Delta_K
Doxorubicin
YLR266C PDR8
ZShift = 2.52
Z lm Score = −2.32
NG = 0
DB = 1
SM = 0
Delta_r
Doxorubicin
YLR266C PDR8
ZShift = 0.32
Z lm Score = −0.05
NG = 0
DB = 1
SM = 0
Delta_AUC
Doxorubicin

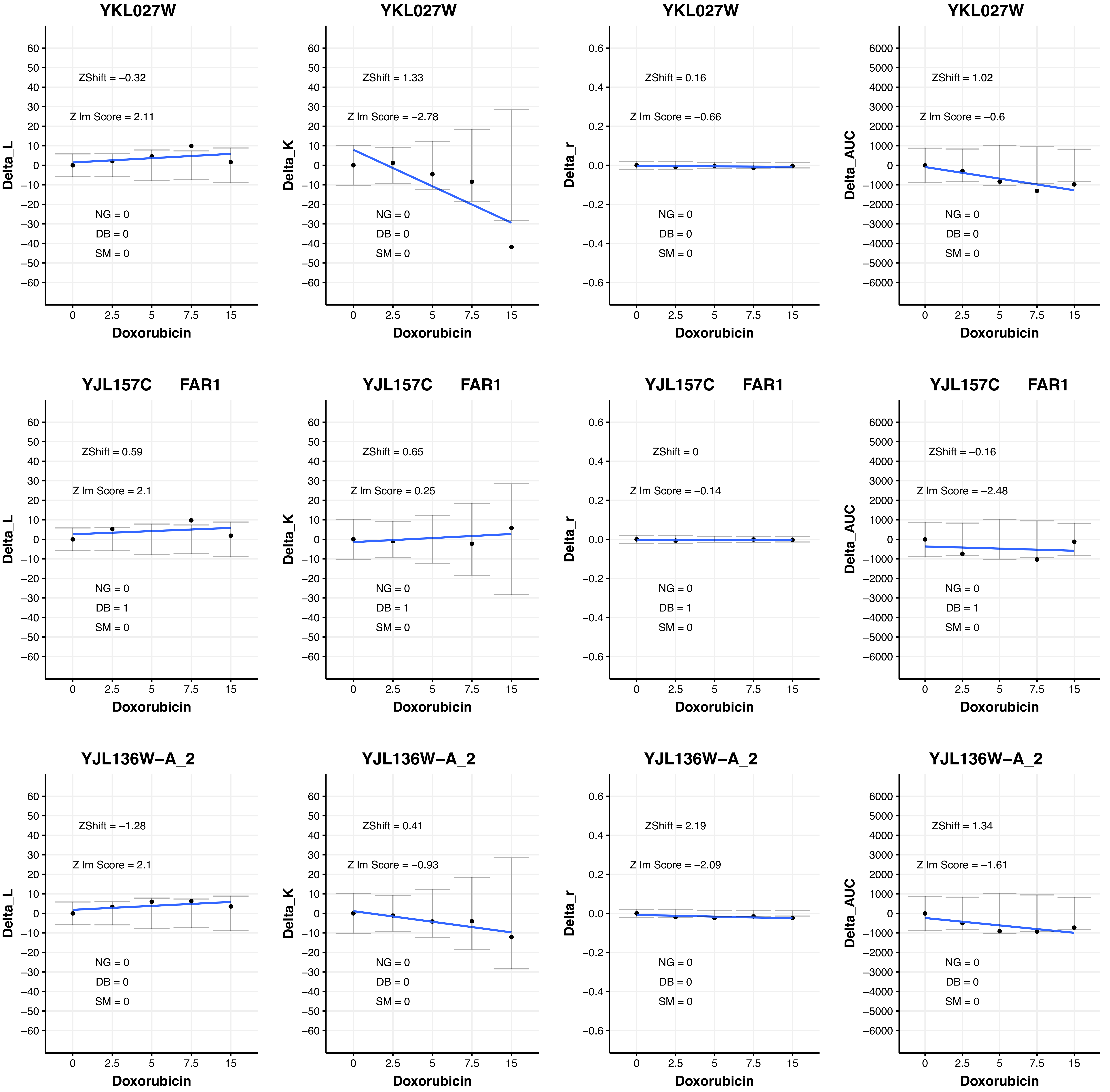
YKL027W
ZShift = −0.32
Z lm Score = 2.11
NG = 0
DB = 0
SM = 0
Delta_L
Doxorubicin
YKL027W
ZShift = 1.33
Z lm Score = −2.78
NG = 0
DB = 0
SM = 0
Delta_K
Doxorubicin
YKL027W
ZShift = 0.16
Z lm Score = −0.66
NG = 0
DB = 0
SM = 0
Delta_r
Doxorubicin
YKL027W
ZShift = 1.02
Z lm Score = −0.6
NG = 0
DB = 0
SM = 0
Delta_AUC
Doxorubicin
YJL157C FAR1
ZShift = 0.59
Z lm Score = 2.1
NG = 0
DB = 1
SM = 0
Delta_L
Doxorubicin
YJL157C FAR1
ZShift = 0.65
Z lm Score = 0.25
NG = 0
DB = 1
SM = 0
Delta_K
Doxorubicin
YJL157C FAR1
ZShift = 0
Z lm Score = −0.14
NG = 0
DB = 1
SM = 0
Delta_r
Doxorubicin
YJL157C FAR1
ZShift = −0.16
Z lm Score = −2.48
NG = 0
DB = 1
SM = 0
Delta_AUC
Doxorubicin
YJL136W−A_2
ZShift = −1.28
Z lm Score = 2.1
NG = 0
DB = 0
SM = 0
Delta_L
Doxorubicin
YJL136W−A_2
ZShift = 0.41
Z lm Score = −0.93
NG = 0
DB = 0
SM = 0
Delta_K
Doxorubicin
YJL136W−A_2
ZShift = 2.19
Z lm Score = −2.09
NG = 0
DB = 0
SM = 0
Delta_r
Doxorubicin
YJL136W−A_2
ZShift = 1.34
Z lm Score = −1.61
NG = 0
DB = 0
SM = 0
Delta_AUC
Doxorubicin

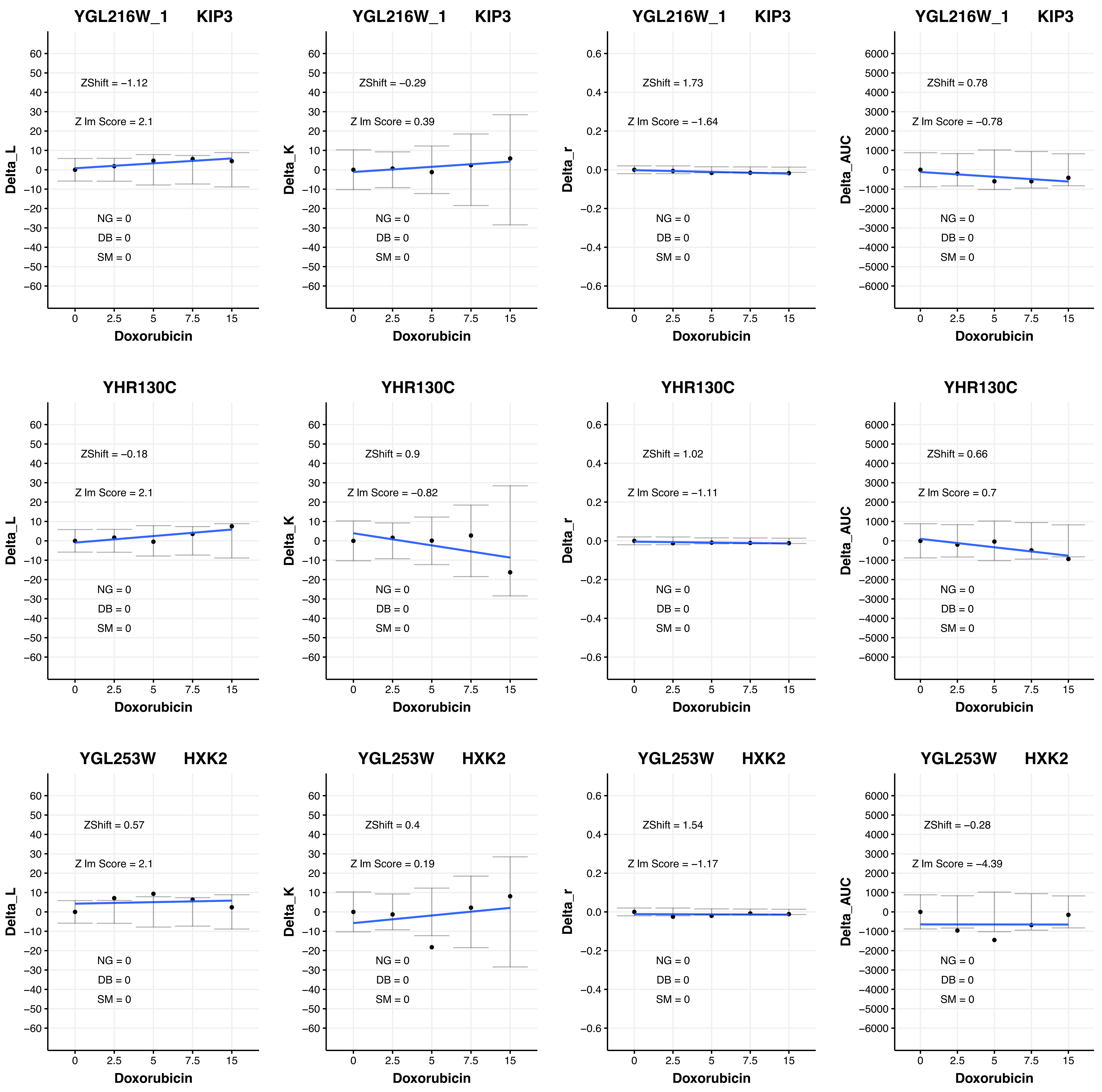
YGL216W_1 KIP3
Delta_L
ZShift = −1.12
Z lm Score = 2.1
NG = 0
DB = 0
SM = 0
Doxorubicin
YGL216W_1 KIP3
Delta_K
ZShift = −0.29
Z lm Score = 0.39
NG = 0
DB = 0
SM = 0
Doxorubicin
YGL216W_1 KIP3
Delta_r
ZShift = 1.73
Z lm Score = −1.64
NG = 0
DB = 0
SM = 0
Doxorubicin
YGL216W_1 KIP3
Delta_AUC
ZShift = 0.78
Z lm Score = −0.78
NG = 0
DB = 0
SM = 0
Doxorubicin
YHR130C
Delta_L
ZShift = −0.18
Z lm Score = 2.1
NG = 0
DB = 0
SM = 0
Doxorubicin
YHR130C
Delta_K
ZShift = 0.9
Z lm Score = −0.82
NG = 0
DB = 0
SM = 0
Doxorubicin
YHR130C
Delta_r
ZShift = 1.02
Z lm Score = −1.11
NG = 0
DB = 0
SM = 0
Doxorubicin
YHR130C
Delta_AUC
ZShift = 0.66
Z lm Score = 0.7
NG = 0
DB = 0
SM = 0
Doxorubicin
YGL253W HXK2
Delta_L
ZShift = 0.57
Z lm Score = 2.1
NG = 0
DB = 0
SM = 0
Doxorubicin
YGL253W HXK2
Delta_K
ZShift = 0.4
Z lm Score = 0.19
NG = 0
DB = 0
SM = 0
Doxorubicin
YGL253W HXK2
Delta_r
ZShift = 1.54
Z lm Score = −1.17
NG = 0
DB = 0
SM = 0
Doxorubicin
YGL253W HXK2
Delta_AUC
ZShift = −0.28
Z lm Score = −4.39
NG = 0
DB = 0
SM = 0
Doxorubicin

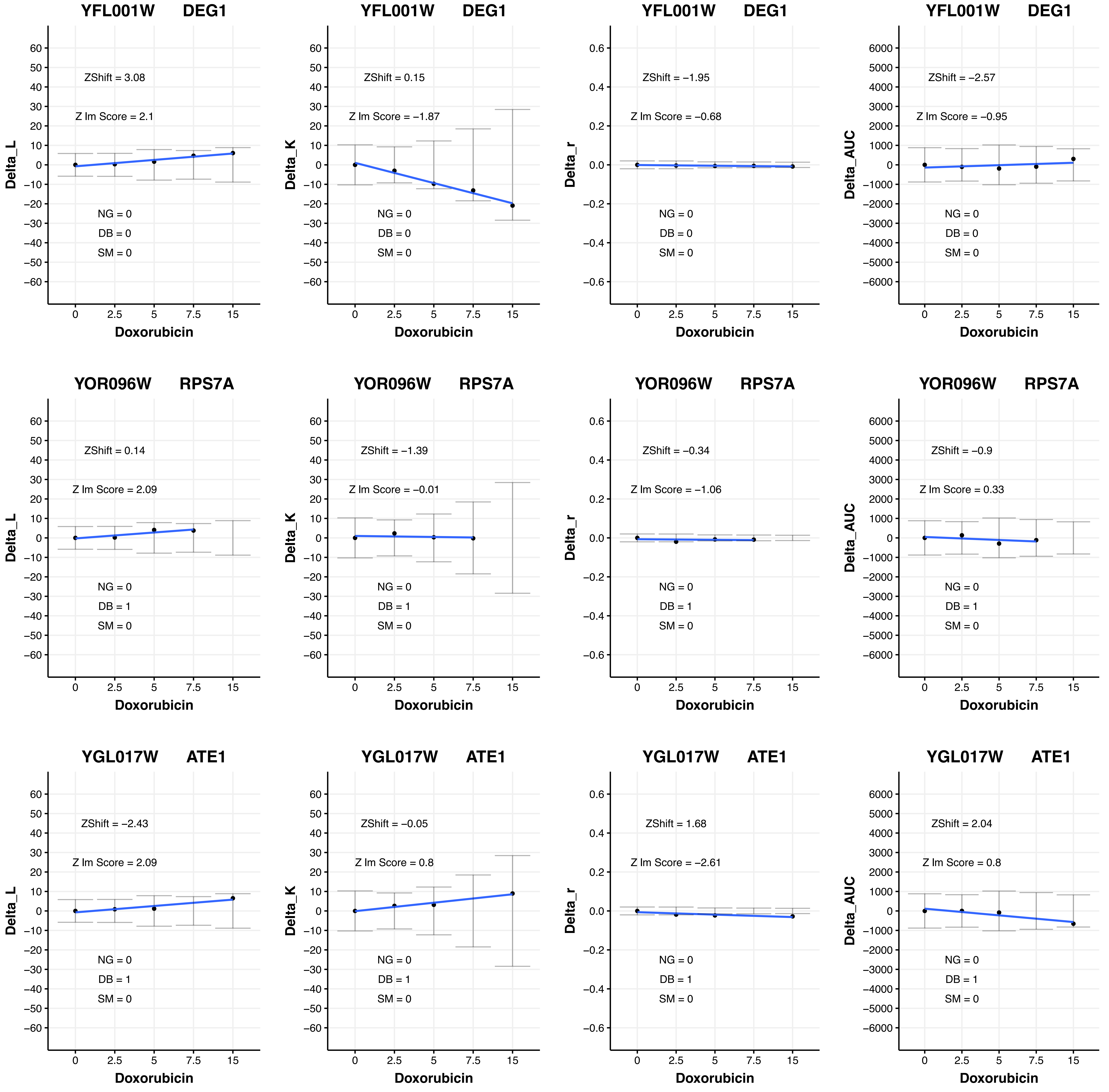

YFL001W DEG1
ZShift = 3.08
Z lm Score = 2.1
NG = 0
DB = 0
SM = 0
Delta_L
Doxorubicin
YFL001W DEG1
ZShift = 0.15
Z lm Score = −1.87
NG = 0
DB = 0
SM = 0
Delta_K
Doxorubicin
YFL001W DEG1
ZShift = −1.95
Z lm Score = −0.68
NG = 0
DB = 0
SM = 0
Delta_r
Doxorubicin
YFL001W DEG1
ZShift = −2.57
Z lm Score = −0.95
NG = 0
DB = 0
SM = 0
Delta_AUC
Doxorubicin
YOR096W RPS7A
ZShift = 0.14
Z lm Score = 2.09
NG = 0
DB = 1
SM = 0
Delta_L
Doxorubicin
YOR096W RPS7A
ZShift = −1.39
Z lm Score = −0.01
NG = 0
DB = 1
SM = 0
Delta_K
Doxorubicin
YOR096W RPS7A
ZShift = −0.34
Z lm Score = −1.06
NG = 0
DB = 1
SM = 0
Delta_r
Doxorubicin
YOR096W RPS7A
ZShift = −0.9
Z lm Score = 0.33
NG = 0
DB = 1
SM = 0
Delta_AUC
Doxorubicin
YGL017W ATE1
ZShift = −2.43
Z lm Score = 2.09
NG = 0
DB = 1
SM = 0
Delta_L
Doxorubicin
YGL017W ATE1
ZShift = −0.05
Z lm Score = 0.8
NG = 0
DB = 1
SM = 0
Delta_K
Doxorubicin
YGL017W ATE1
ZShift = 1.68
Z lm Score = −2.61
NG = 0
DB = 1
SM = 0
Delta_r
Doxorubicin
YGL017W ATE1
ZShift = 2.04
Z lm Score = 0.8
NG = 0
DB = 1
SM = 0
Delta_AUC
Doxorubicin

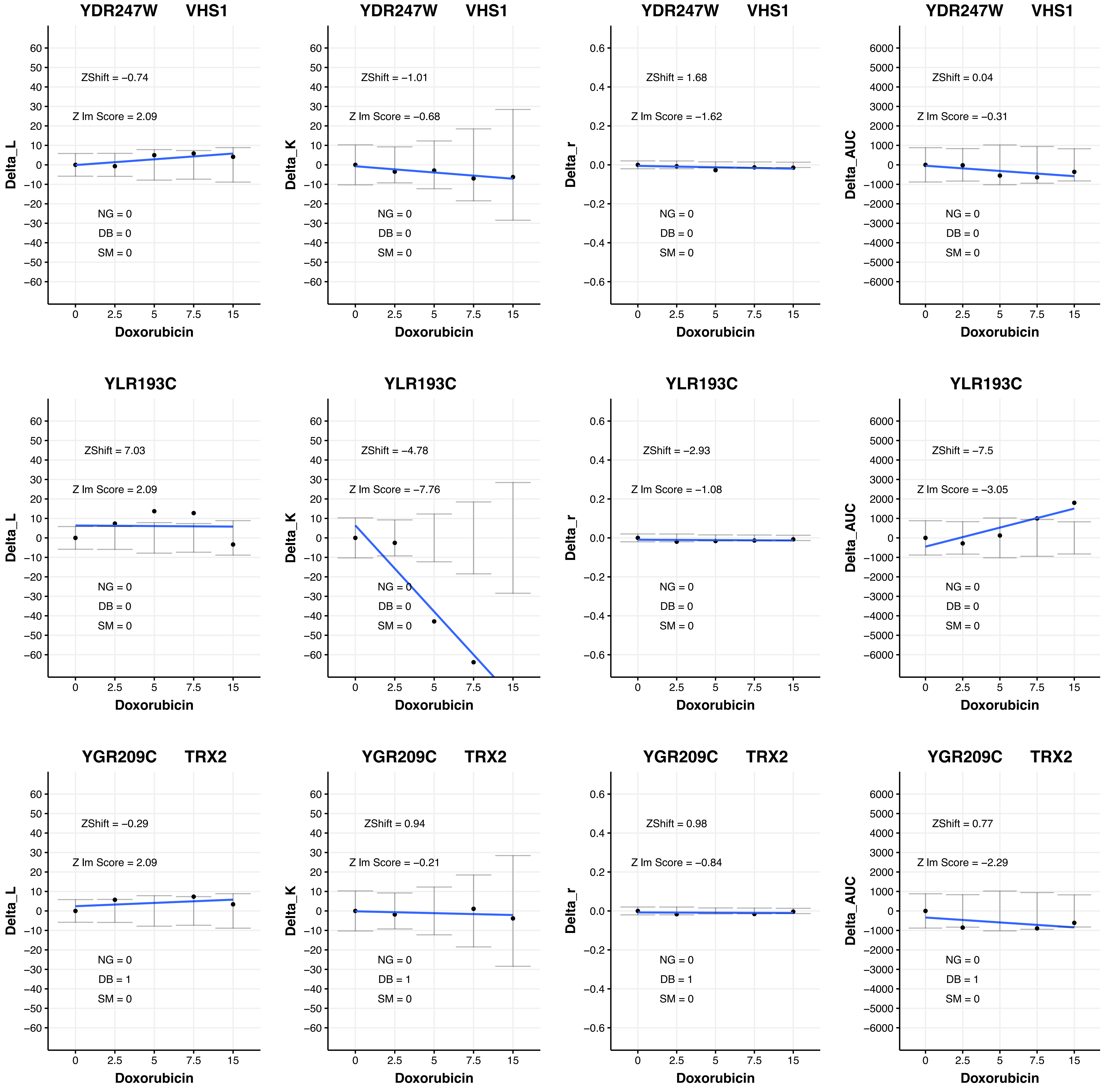

YDR247W VHS1
Delta_L
ZShift = −0.74
Z lm Score = 2.09
NG = 0
DB = 0
SM = 0
Doxorubicin
YDR247W VHS1
Delta_K
ZShift = −1.01
Z lm Score = −0.68
NG = 0
DB = 0
SM = 0
Doxorubicin
YDR247W VHS1
Delta_r
ZShift = 1.68
Z lm Score = −1.62
NG = 0
DB = 0
SM = 0
Doxorubicin
YDR247W VHS1
Delta_AUC
ZShift = 0.04
Z lm Score = −0.31
NG = 0
DB = 0
SM = 0
Doxorubicin
YLR193C
Delta_L
ZShift = 7.03
Z lm Score = 2.09
NG = 0
DB = 0
SM = 0
Doxorubicin
YLR193C
Delta_K
ZShift = −4.78
Z lm Score = −7.76
NG = 0
DB = 0
SM = 0
Doxorubicin
YLR193C
Delta_r
ZShift = −2.93
Z lm Score = −1.08
NG = 0
DB = 0
SM = 0
Doxorubicin
YLR193C
Delta_AUC
ZShift = −7.5
Z lm Score = −3.05
NG = 0
DB = 0
SM = 0
Doxorubicin
YGR209C TRX2
Delta_L
ZShift = −0.29
Z lm Score = 2.09
NG = 0
DB = 1
SM = 0
Doxorubicin
YGR209C TRX2
Delta_K
ZShift = 0.94
Z lm Score = −0.21
NG = 0
DB = 1
SM = 0
Doxorubicin
YGR209C TRX2
Delta_r
ZShift = 0.98
Z lm Score = −0.84
NG = 0
DB = 1
SM = 0
Doxorubicin
YGR209C TRX2
Delta_AUC
ZShift = 0.77
Z lm Score = −2.29
NG = 0
DB = 1
SM = 0
Doxorubicin

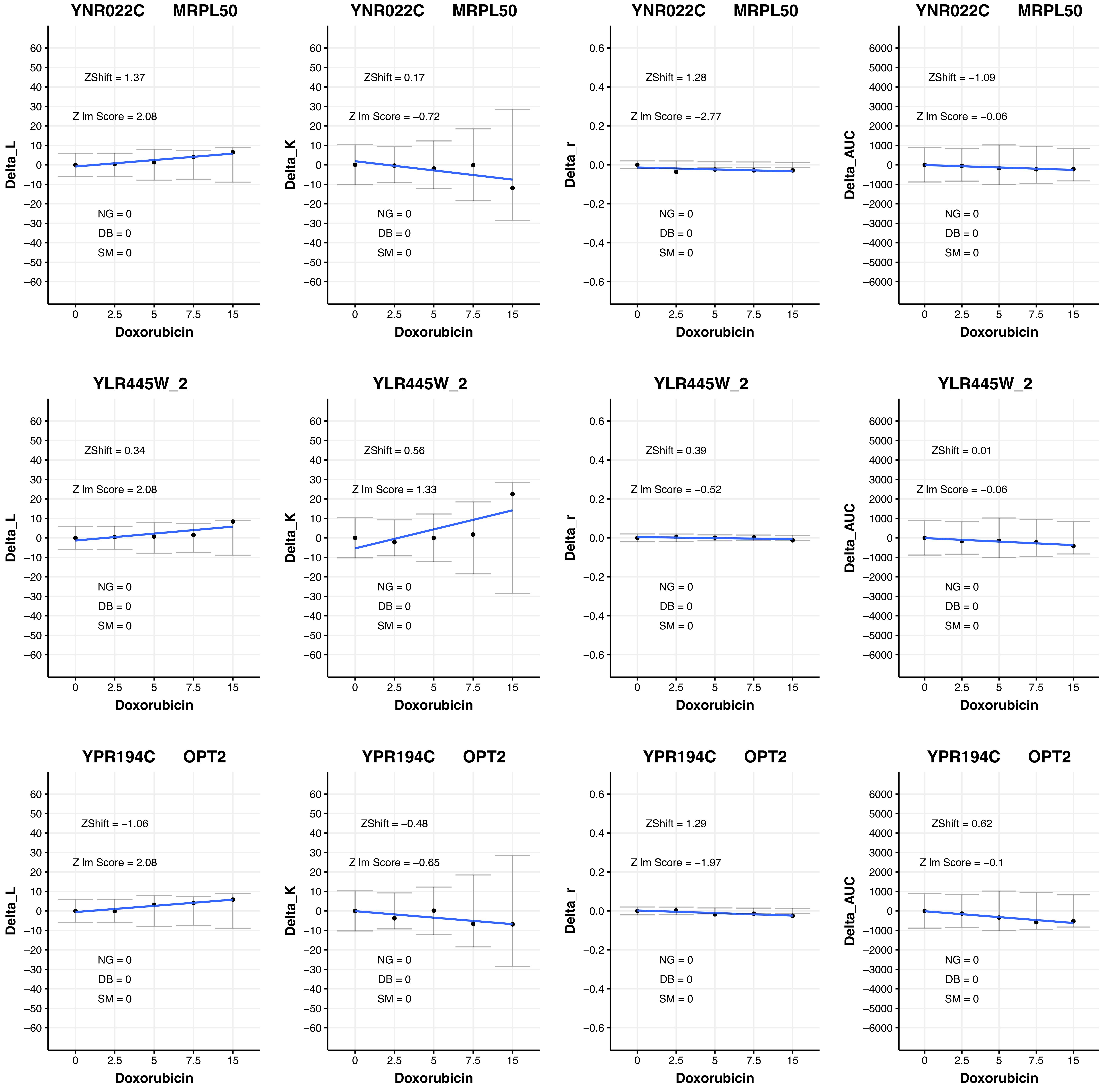

YNR022C MRPL50
ZShift = 1.37
Z lm Score = 2.08
NG = 0
DB = 0
SM = 0
Delta_L
Doxorubicin
YNR022C MRPL50
ZShift = 0.17
Z lm Score = −0.72
NG = 0
DB = 0
SM = 0
Delta_K
Doxorubicin
YNR022C MRPL50
ZShift = 1.28
Z lm Score = −2.77
NG = 0
DB = 0
SM = 0
Delta_r
Doxorubicin
YNR022C MRPL50
ZShift = −1.09
Z lm Score = −0.06
NG = 0
DB = 0
SM = 0
Delta_AUC
Doxorubicin
YLR445W_2
ZShift = 0.34
Z lm Score = 2.08
NG = 0
DB = 0
SM = 0
Delta_L
Doxorubicin
YLR445W_2
ZShift = 0.56
Z lm Score = 1.33
NG = 0
DB = 0
SM = 0
Delta_K
Doxorubicin
YLR445W_2
ZShift = 0.39
Z lm Score = −0.52
NG = 0
DB = 0
SM = 0
Delta_r
Doxorubicin
YLR445W_2
ZShift = 0.01
Z lm Score = −0.06
NG = 0
DB = 0
SM = 0
Delta_AUC
Doxorubicin
YPR194C OPT2
ZShift = −1.06
Z lm Score = 2.08
NG = 0
DB = 0
SM = 0
Delta_L
Doxorubicin
YPR194C OPT2
ZShift = −0.48
Z lm Score = −0.65
NG = 0
DB = 0
SM = 0
Delta_K
Doxorubicin
YPR194C OPT2
ZShift = 1.29
Z lm Score = −1.97
NG = 0
DB = 0
SM = 0
Delta_r
Doxorubicin
YPR194C OPT2
ZShift = 0.62
Z lm Score = −0.1
NG = 0
DB = 0
SM = 0
Delta_AUC
Doxorubicin

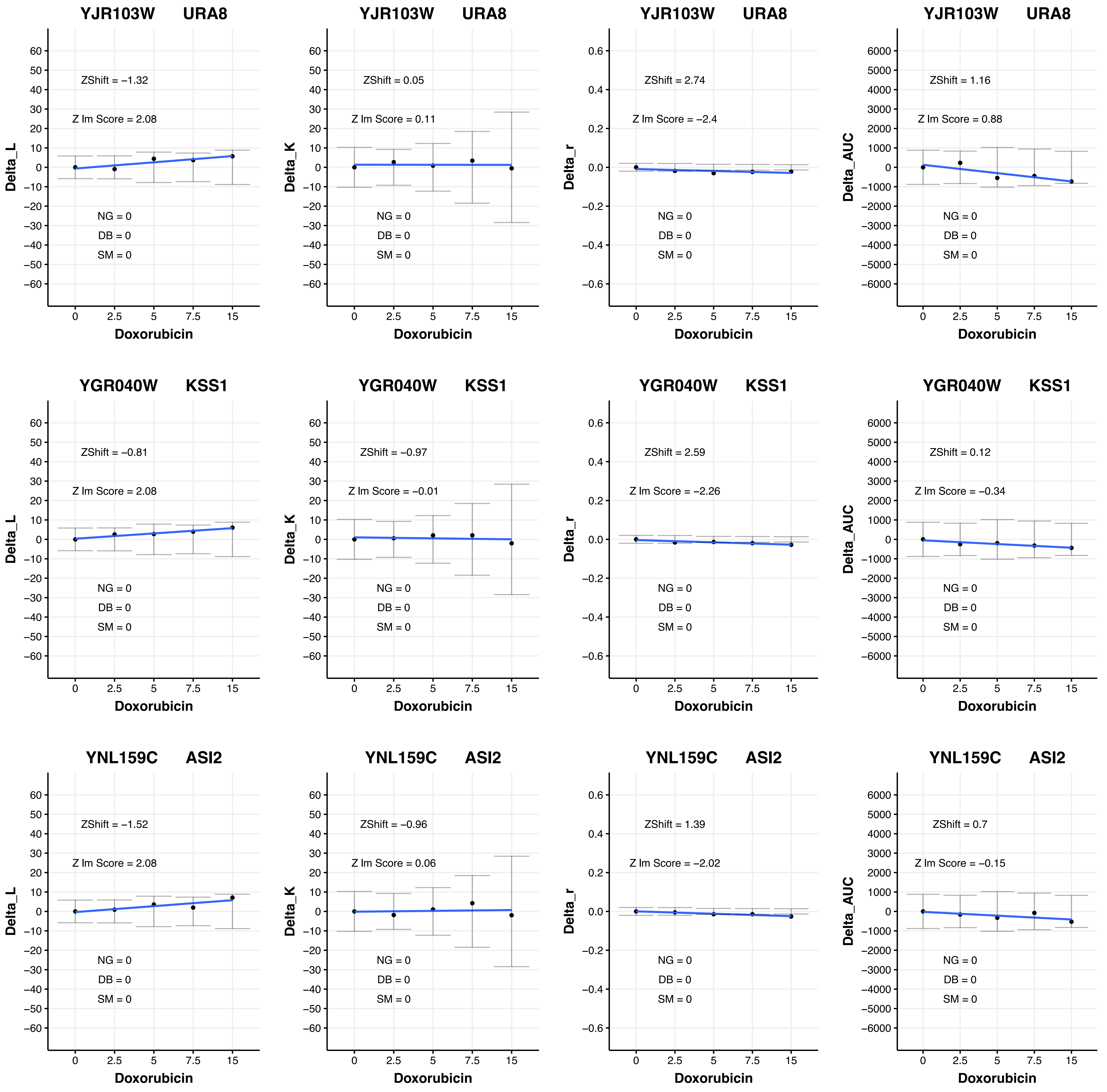
YJR103W URA8
ZShift = −1.32
Z lm Score = 2.08
NG = 0
DB = 0
SM = 0
Delta_L
Doxorubicin
YJR103W URA8
ZShift = 0.05
Z lm Score = 0.11
NG = 0
DB = 0
SM = 0
Delta_K
Doxorubicin
YJR103W URA8
ZShift = 2.74
Z lm Score = −2.4
NG = 0
DB = 0
SM = 0
Delta_r
Doxorubicin
YJR103W URA8
ZShift = 1.16
Z lm Score = 0.88
NG = 0
DB = 0
SM = 0
Delta_AUC
Doxorubicin
YGR040W KSS1
ZShift = −0.81
Z lm Score = 2.08
NG = 0
DB = 0
SM = 0
Delta_L
Doxorubicin
YGR040W KSS1
ZShift = −0.97
Z lm Score = −0.01
NG = 0
DB = 0
SM = 0
Delta_K
Doxorubicin
YGR040W KSS1
ZShift = 2.59
Z lm Score = −2.26
NG = 0
DB = 0
SM = 0
Delta_r
Doxorubicin
YGR040W KSS1
ZShift = 0.12
Z lm Score = −0.34
NG = 0
DB = 0
SM = 0
Delta_AUC
Doxorubicin
YNL159C ASI2
ZShift = −1.52
Z lm Score = 2.08
NG = 0
DB = 0
SM = 0
Delta_L
Doxorubicin
YNL159C ASI2
ZShift = −0.96
Z lm Score = 0.06
NG = 0
DB = 0
SM = 0
Delta_K
Doxorubicin
YNL159C ASI2
ZShift = 1.39
Z lm Score = −2.02
NG = 0
DB = 0
SM = 0
Delta_r
Doxorubicin
YNL159C ASI2
ZShift = 0.7
Z lm Score = −0.15
NG = 0
DB = 0
SM = 0
Delta_AUC
Doxorubicin

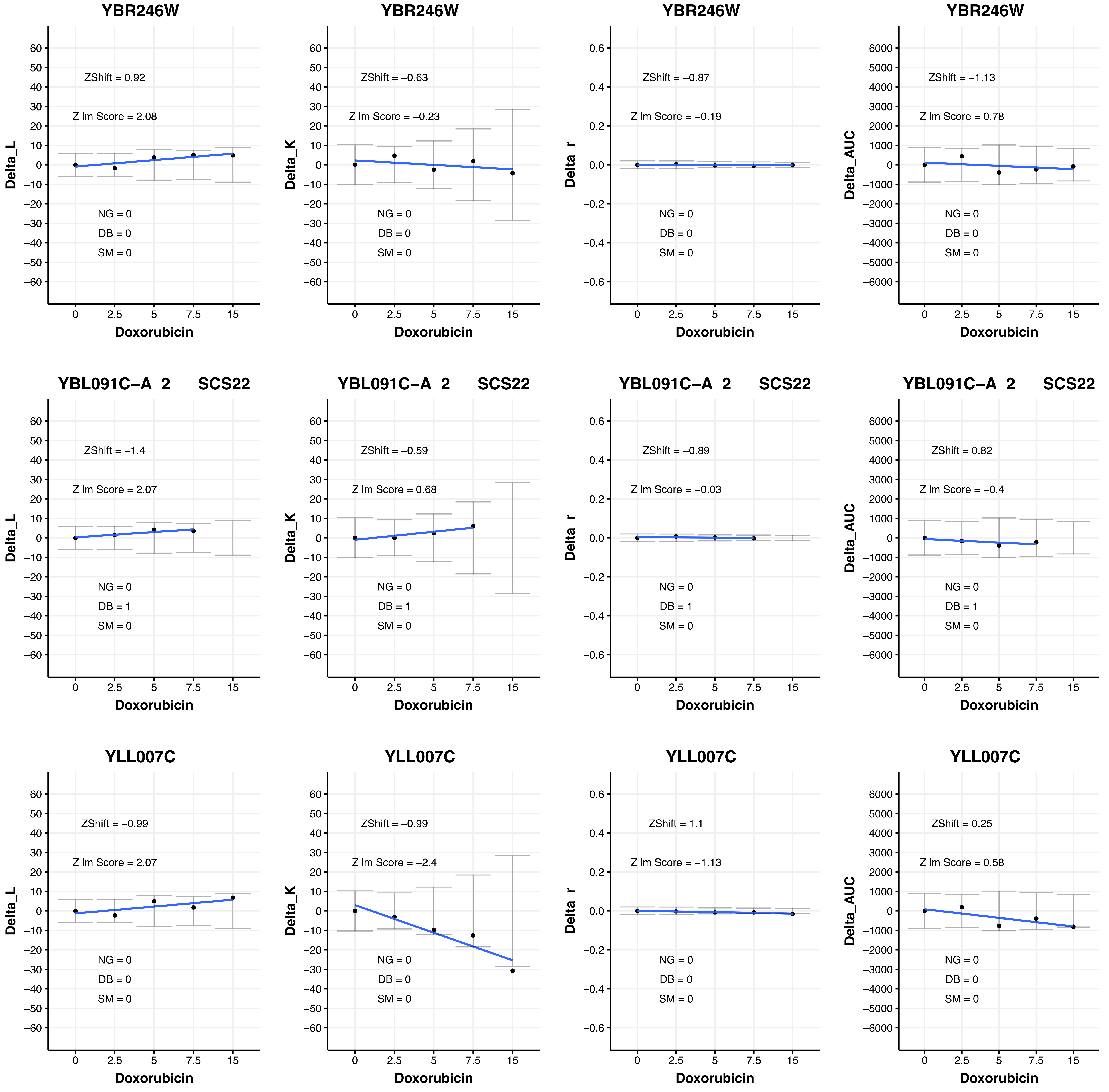
YBR246W
ZShift = 0.92
Z lm Score = 2.08
NG = 0
DB = 0
SM = 0
Delta_L
Doxorubicin
YBR246W
ZShift = −0.63
Z lm Score = −0.23
NG = 0
DB = 0
SM = 0
Delta_K
Doxorubicin
YBR246W
ZShift = −0.87
Z lm Score = −0.19
NG = 0
DB = 0
SM = 0
Delta_r
Doxorubicin
YBR246W
ZShift = −1.13
Z lm Score = 0.78
NG = 0
DB = 0
SM = 0
Delta_AUC
Doxorubicin
YBL091C−A_2 SCS22
ZShift = −1.4
Z lm Score = 2.07
NG = 0
DB = 1
SM = 0
Delta_L
Doxorubicin
YBL091C−A_2 SCS22
ZShift = −0.59
Z lm Score = 0.68
NG = 0
DB = 1
SM = 0
Delta_K
Doxorubicin
YBL091C−A_2 SCS22
ZShift = −0.89
Z lm Score = −0.03
NG = 0
DB = 1
SM = 0
Delta_r
Doxorubicin
YBL091C−A_2 SCS22
ZShift = 0.82
Z lm Score = −0.4
NG = 0
DB = 1
SM = 0
Delta_AUC
Doxorubicin
YLL007C
ZShift = −0.99
Z lm Score = 2.07
NG = 0
DB = 0
SM = 0
Delta_L
Doxorubicin
YLL007C
ZShift = −0.99
Z lm Score = −2.4
NG = 0
DB = 0
SM = 0
Delta_K
Doxorubicin
YLL007C
ZShift = 1.1
Z lm Score = −1.13
NG = 0
DB = 0
SM = 0
Delta_r
Doxorubicin
YLL007C
ZShift = 0.25
Z lm Score = 0.58
NG = 0
DB = 0
SM = 0
Delta_AUC
Doxorubicin

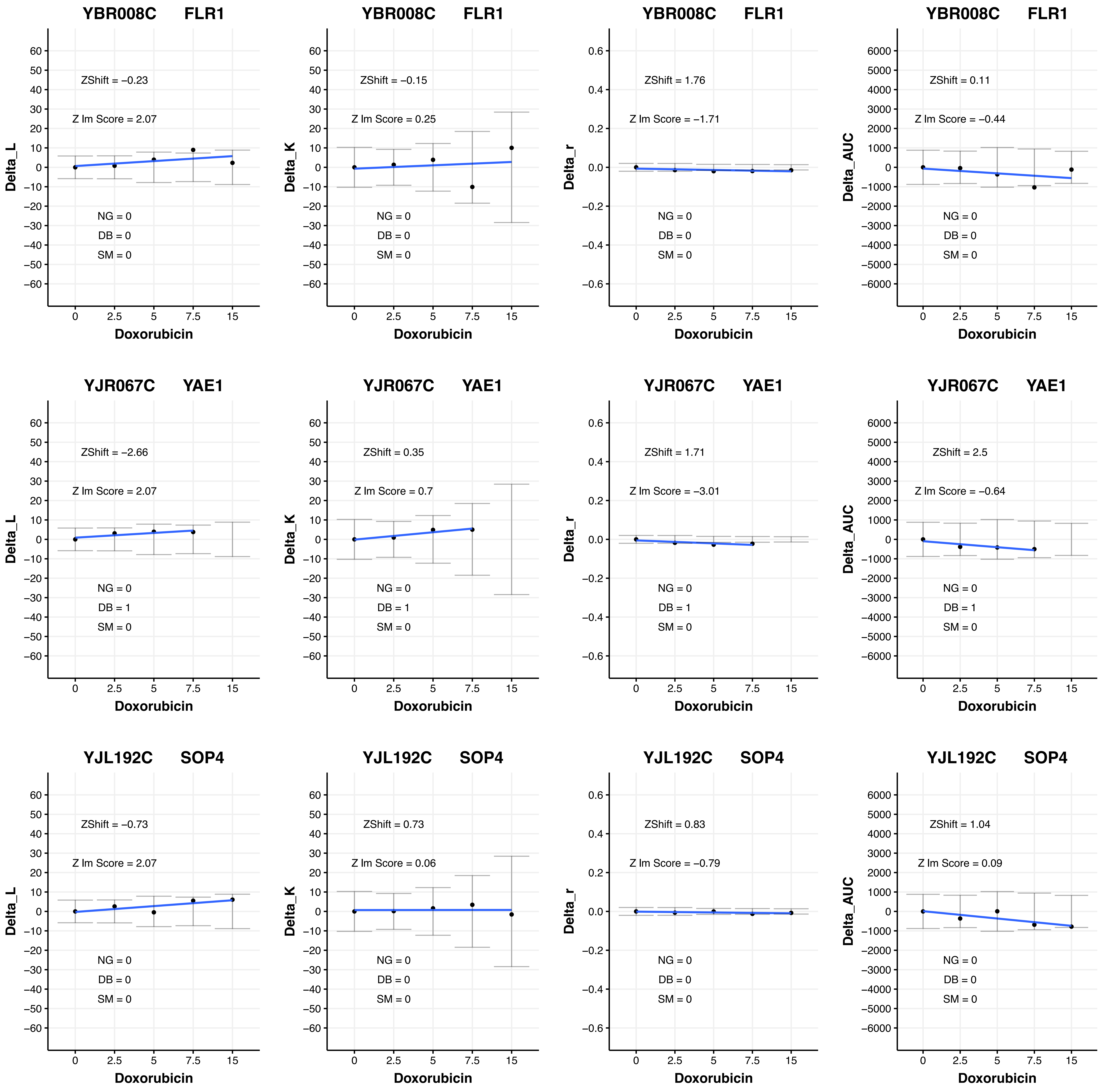
YBR008C FLR1
ZShift = −0.23
Z lm Score = 2.07
NG = 0
DB = 0
SM = 0
Delta_L
Doxorubicin
YBR008C FLR1
ZShift = −0.15
Z lm Score = 0.25
NG = 0
DB = 0
SM = 0
Delta_K
Doxorubicin
YBR008C FLR1
ZShift = 1.76
Z lm Score = −1.71
NG = 0
DB = 0
SM = 0
Delta_r
Doxorubicin
YBR008C FLR1
ZShift = 0.11
Z lm Score = −0.44
NG = 0
DB = 0
SM = 0
Delta_AUC
Doxorubicin
YJR067C YAE1
ZShift = −2.66
Z lm Score = 2.07
NG = 0
DB = 1
SM = 0
Delta_L
Doxorubicin
YJR067C YAE1
ZShift = 0.35
Z lm Score = 0.7
NG = 0
DB = 1
SM = 0
Delta_K
Doxorubicin
YJR067C YAE1
ZShift = 1.71
Z lm Score = −3.01
NG = 0
DB = 1
SM = 0
Delta_r
Doxorubicin
YJR067C YAE1
ZShift = 2.5
Z lm Score = −0.64
NG = 0
DB = 1
SM = 0
Delta_AUC
Doxorubicin
YJL192C SOP4
ZShift = −0.73
Z lm Score = 2.07
NG = 0
DB = 0
SM = 0
Delta_L
Doxorubicin
YJL192C SOP4
ZShift = 0.73
Z lm Score = 0.06
NG = 0
DB = 0
SM = 0
Delta_K
Doxorubicin
YJL192C SOP4
ZShift = 0.83
Z lm Score = −0.79
NG = 0
DB = 0
SM = 0
Delta_r
Doxorubicin
YJL192C SOP4
ZShift = 1.04
Z lm Score = 0.09
NG = 0
DB = 0
SM = 0
Delta_AUC
Doxorubicin

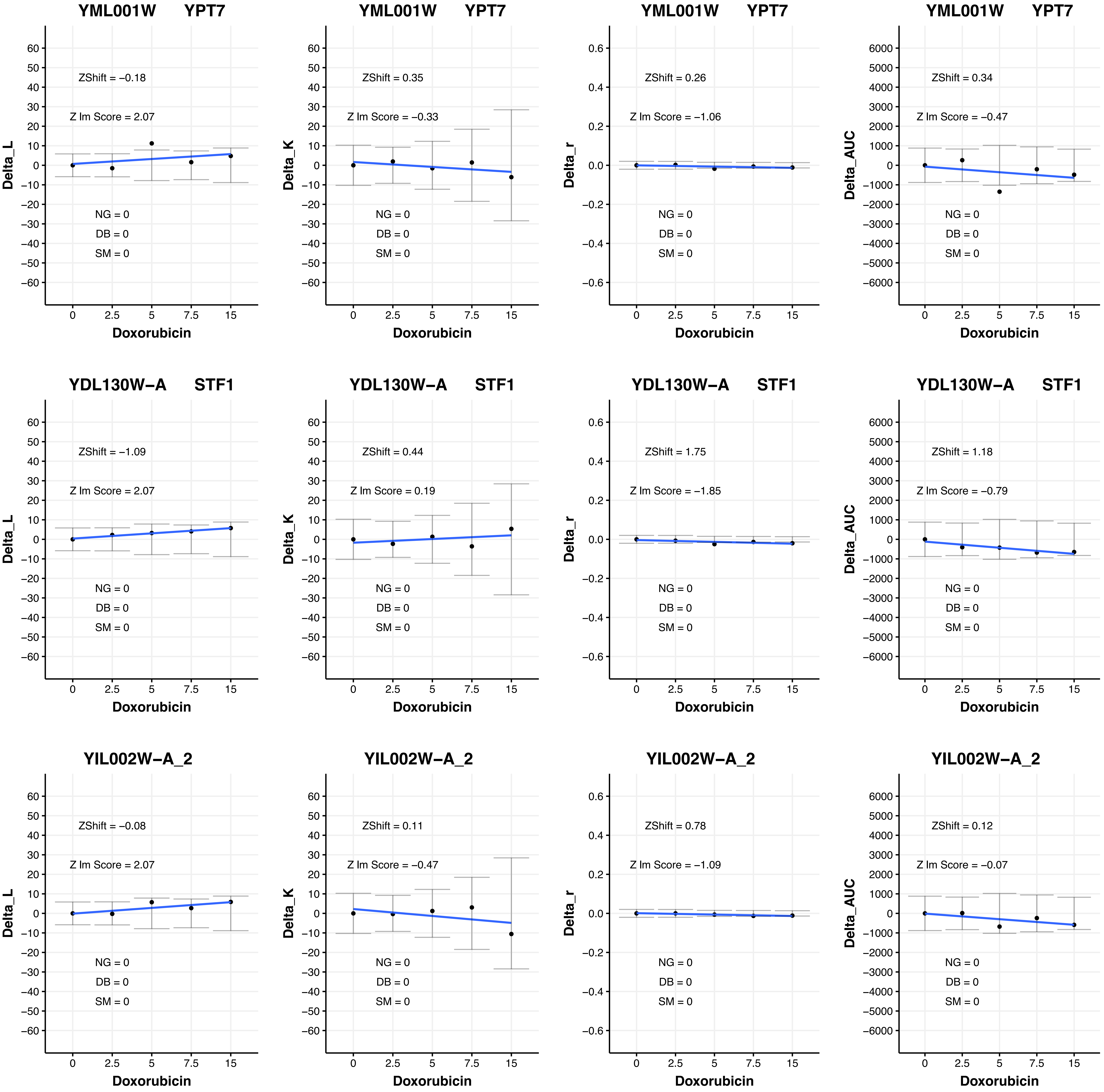

YML001W YPT7
Delta_L
ZShift = −0.18
Z lm Score = 2.07
NG = 0
DB = 0
SM = 0
Doxorubicin
YML001W YPT7
Delta_K
ZShift = 0.35
Z lm Score = −0.33
NG = 0
DB = 0
SM = 0
Doxorubicin
YML001W YPT7
Delta_r
ZShift = 0.26
Z lm Score = −1.06
NG = 0
DB = 0
SM = 0
Doxorubicin
YML001W YPT7
Delta_AUC
ZShift = 0.34
Z lm Score = −0.47
NG = 0
DB = 0
SM = 0
Doxorubicin
YDL130W−A STF1
Delta_L
ZShift = −1.09
Z lm Score = 2.07
NG = 0
DB = 0
SM = 0
Doxorubicin
YDL130W−A STF1
Delta_K
ZShift = 0.44
Z lm Score = 0.19
NG = 0
DB = 0
SM = 0
Doxorubicin
YDL130W−A STF1
Delta_r
ZShift = 1.75
Z lm Score = −1.85
NG = 0
DB = 0
SM = 0
Doxorubicin
YDL130W−A STF1
Delta_AUC
ZShift = 1.18
Z lm Score = −0.79
NG = 0
DB = 0
SM = 0
Doxorubicin
YIL002W−A_2
Delta_L
ZShift = −0.08
Z lm Score = 2.07
NG = 0
DB = 0
SM = 0
Doxorubicin
YIL002W−A_2
Delta_K
ZShift = 0.11
Z lm Score = −0.47
NG = 0
DB = 0
SM = 0
Doxorubicin
YIL002W−A_2
Delta_r
ZShift = 0.78
Z lm Score = −1.09
NG = 0
DB = 0
SM = 0
Doxorubicin
YIL002W−A_2
Delta_AUC
ZShift = 0.12
Z lm Score = −0.07
NG = 0
DB = 0
SM = 0
Doxorubicin

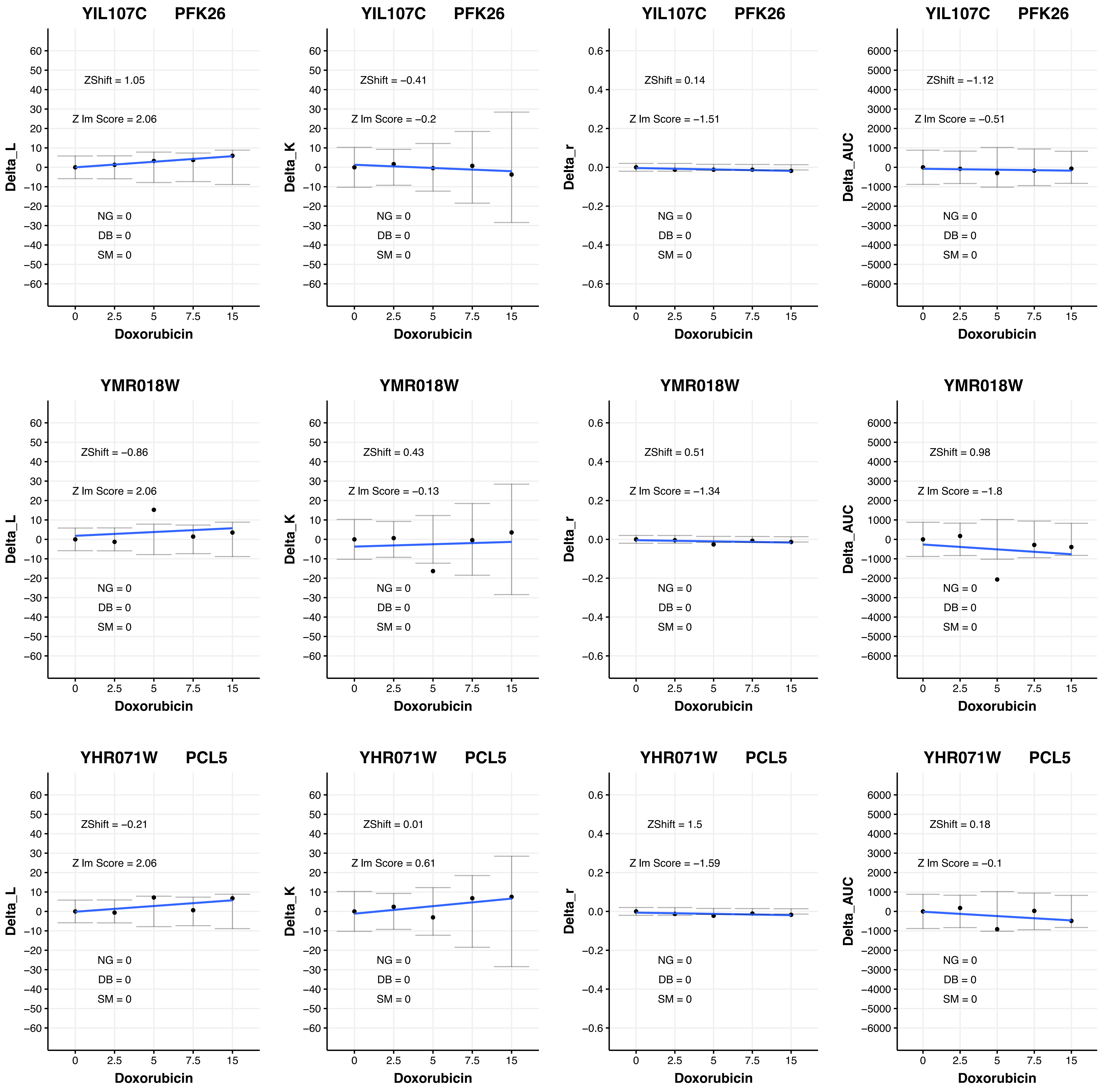
YIL107C PFK26
ZShift = 1.05
Z lm Score = 2.06
NG = 0
DB = 0
SM = 0
Delta_L
Doxorubicin
YIL107C PFK26
ZShift = −0.41
Z lm Score = −0.2
NG = 0
DB = 0
SM = 0
Delta_K
Doxorubicin
YIL107C PFK26
ZShift = 0.14
Z lm Score = −1.51
NG = 0
DB = 0
SM = 0
Delta_r
Doxorubicin
YIL107C PFK26
ZShift = −1.12
Z lm Score = −0.51
NG = 0
DB = 0
SM = 0
Delta_AUC
Doxorubicin
YMR018W
ZShift = −0.86
Z lm Score = 2.06
NG = 0
DB = 0
SM = 0
Delta_L
Doxorubicin
YMR018W
ZShift = 0.43
Z lm Score = −0.13
NG = 0
DB = 0
SM = 0
Delta_K
Doxorubicin
YMR018W
ZShift = 0.51
Z lm Score = −1.34
NG = 0
DB = 0
SM = 0
Delta_r
Doxorubicin
YMR018W
ZShift = 0.98
Z lm Score = −1.8
NG = 0
DB = 0
SM = 0
Delta_AUC
Doxorubicin
YHR071W PCL5
ZShift = −0.21
Z lm Score = 2.06
NG = 0
DB = 0
SM = 0
Delta_L
Doxorubicin
YHR071W PCL5
ZShift = 0.01
Z lm Score = 0.61
NG = 0
DB = 0
SM = 0
Delta_K
Doxorubicin
YHR071W PCL5
ZShift = 1.5
Z lm Score = −1.59
NG = 0
DB = 0
SM = 0
Delta_r
Doxorubicin
YHR071W PCL5
ZShift = 0.18
Z lm Score = −0.1
NG = 0
DB = 0
SM = 0
Delta_AUC
Doxorubicin

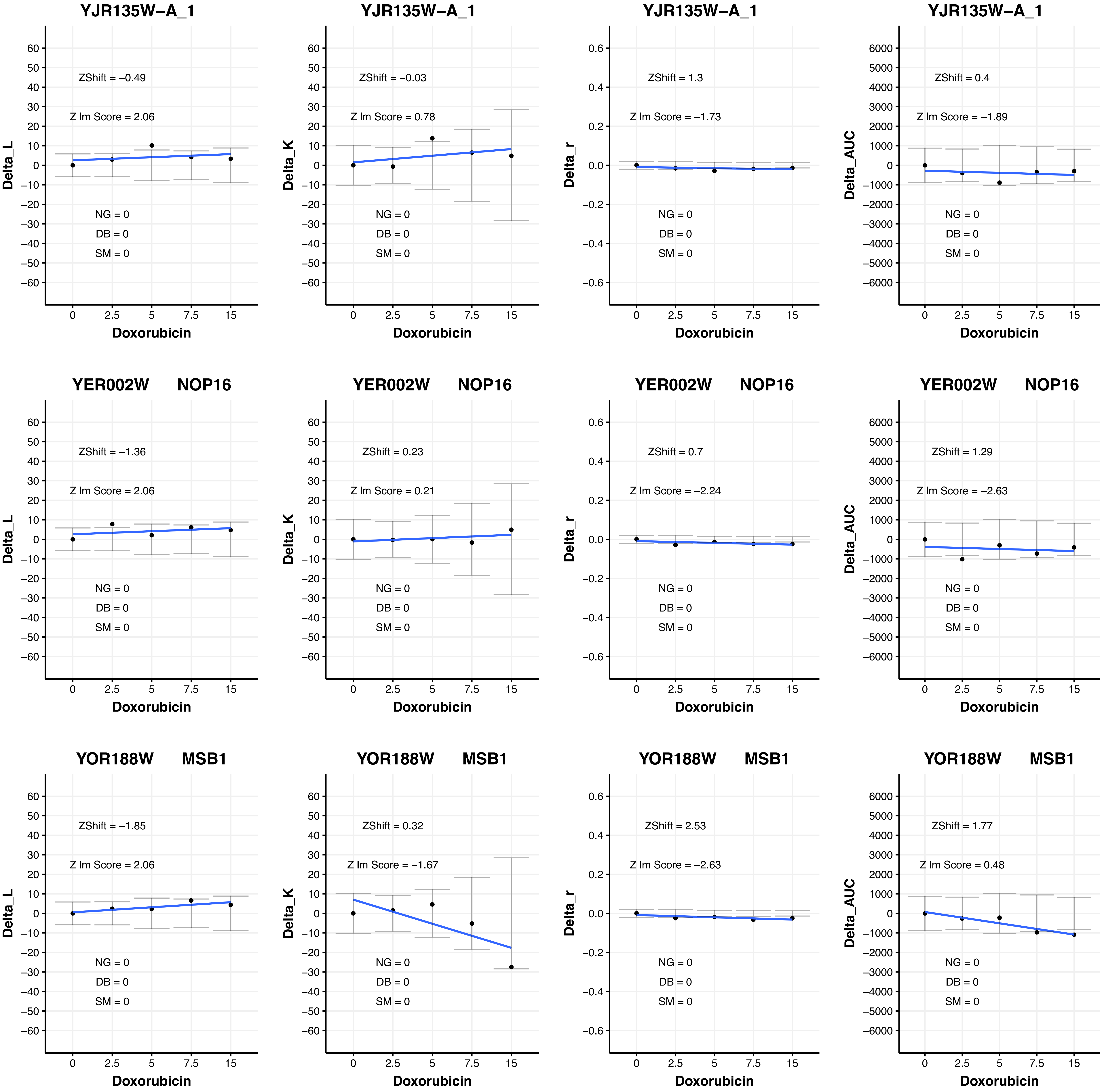

YJR135W−A_1
ZShift = −0.49
Z lm Score = 2.06
NG = 0
DB = 0
SM = 0
Delta_L
Doxorubicin
YJR135W−A_1
ZShift = −0.03
Z lm Score = 0.78
NG = 0
DB = 0
SM = 0
Delta_K
Doxorubicin
YJR135W−A_1
ZShift = 1.3
Z lm Score = −1.73
NG = 0
DB = 0
SM = 0
Delta_r
Doxorubicin
YJR135W−A_1
ZShift = 0.4
Z lm Score = −1.89
NG = 0
DB = 0
SM = 0
Delta_AUC
Doxorubicin
YER002W NOP16
ZShift = −1.36
Z lm Score = 2.06
NG = 0
DB = 0
SM = 0
Delta_L
Doxorubicin
YER002W NOP16
ZShift = 0.23
Z lm Score = 0.21
NG = 0
DB = 0
SM = 0
Delta_K
Doxorubicin
YER002W NOP16
ZShift = 0.7
Z lm Score = −2.24
NG = 0
DB = 0
SM = 0
Delta_r
Doxorubicin
YER002W NOP16
ZShift = 1.29
Z lm Score = −2.63
NG = 0
DB = 0
SM = 0
Delta_AUC
Doxorubicin
YOR188W MSB1
ZShift = −1.85
Z lm Score = 2.06
NG = 0
DB = 0
SM = 0
Delta_L
Doxorubicin
YOR188W MSB1
ZShift = 0.32
Z lm Score = −1.67
NG = 0
DB = 0
SM = 0
Delta_K
Doxorubicin
YOR188W MSB1
ZShift = 2.53
Z lm Score = −2.63
NG = 0
DB = 0
SM = 0
Delta_r
Doxorubicin
YOR188W MSB1
ZShift = 1.77
Z lm Score = 0.48
NG = 0
DB = 0
SM = 0
Delta_AUC
Doxorubicin

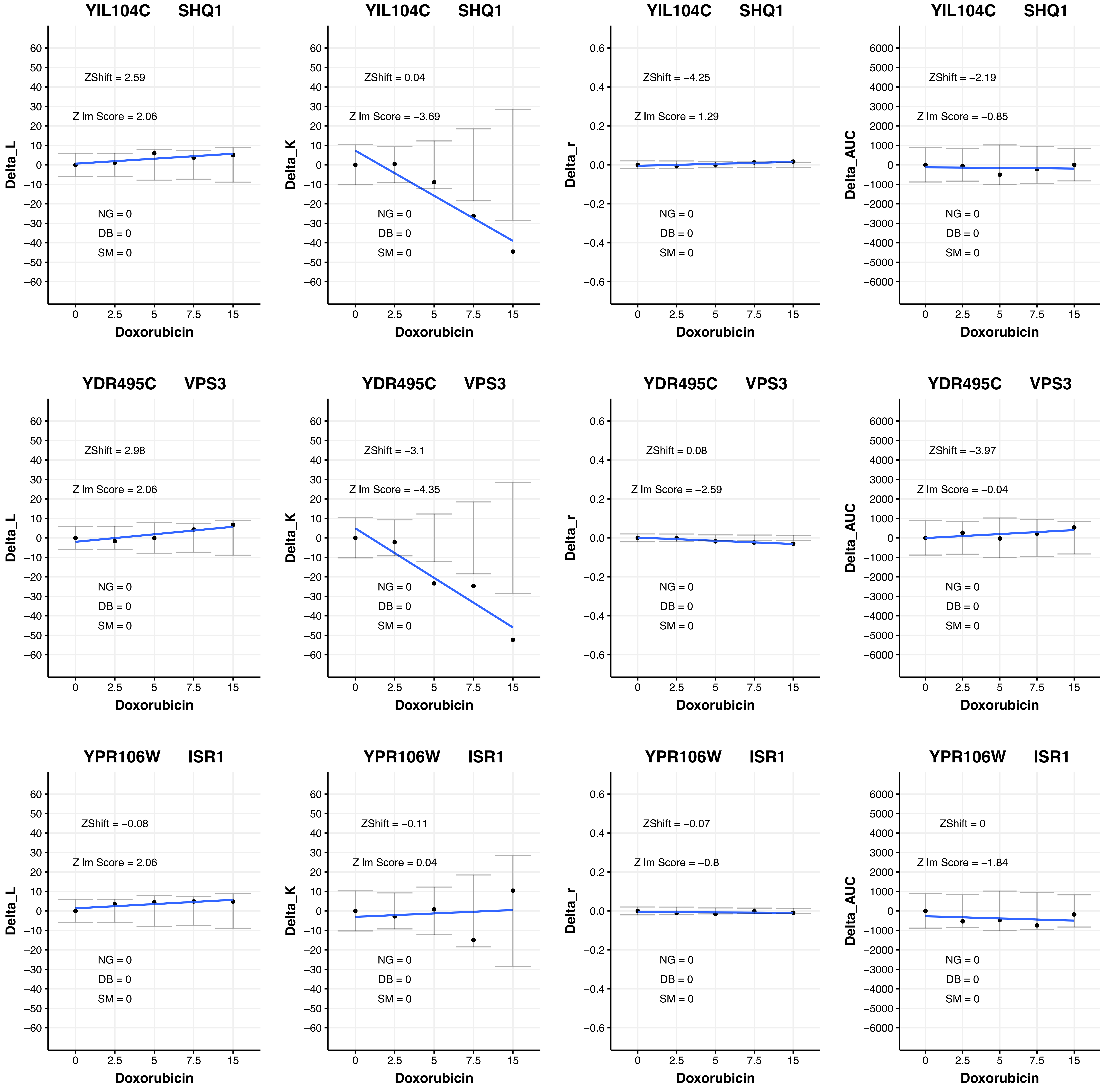
YIL104C SHQ1
ZShift = 2.59
Z lm Score = 2.06
NG = 0
DB = 0
SM = 0
Delta_L
Doxorubicin
YIL104C SHQ1
ZShift = 0.04
Z lm Score = −3.69
NG = 0
DB = 0
SM = 0
Delta_K
Doxorubicin
YIL104C SHQ1
ZShift = −4.25
Z lm Score = 1.29
NG = 0
DB = 0
SM = 0
Delta_r
Doxorubicin
YIL104C SHQ1
ZShift = −2.19
Z lm Score = −0.85
NG = 0
DB = 0
SM = 0
Delta_AUC
Doxorubicin
YDR495C VPS3
ZShift = 2.98
Z lm Score = 2.06
NG = 0
DB = 0
SM = 0
Delta_L
Doxorubicin
YDR495C VPS3
ZShift = −3.1
Z lm Score = −4.35
NG = 0
DB = 0
SM = 0
Delta_K
Doxorubicin
YDR495C VPS3
ZShift = 0.08
Z lm Score = −2.59
NG = 0
DB = 0
SM = 0
Delta_r
Doxorubicin
YDR495C VPS3
ZShift = −3.97
Z lm Score = −0.04
NG = 0
DB = 0
SM = 0
Delta_AUC
Doxorubicin
YPR106W ISR1
ZShift = −0.08
Z lm Score = 2.06
NG = 0
DB = 0
SM = 0
Delta_L
Doxorubicin
YPR106W ISR1
ZShift = −0.11
Z lm Score = 0.04
NG = 0
DB = 0
SM = 0
Delta_K
Doxorubicin
YPR106W ISR1
ZShift = −0.07
Z lm Score = −0.8
NG = 0
DB = 0
SM = 0
Delta_r
Doxorubicin
YPR106W ISR1
ZShift = 0
Z lm Score = −1.84
NG = 0
DB = 0
SM = 0
Delta_AUC
Doxorubicin

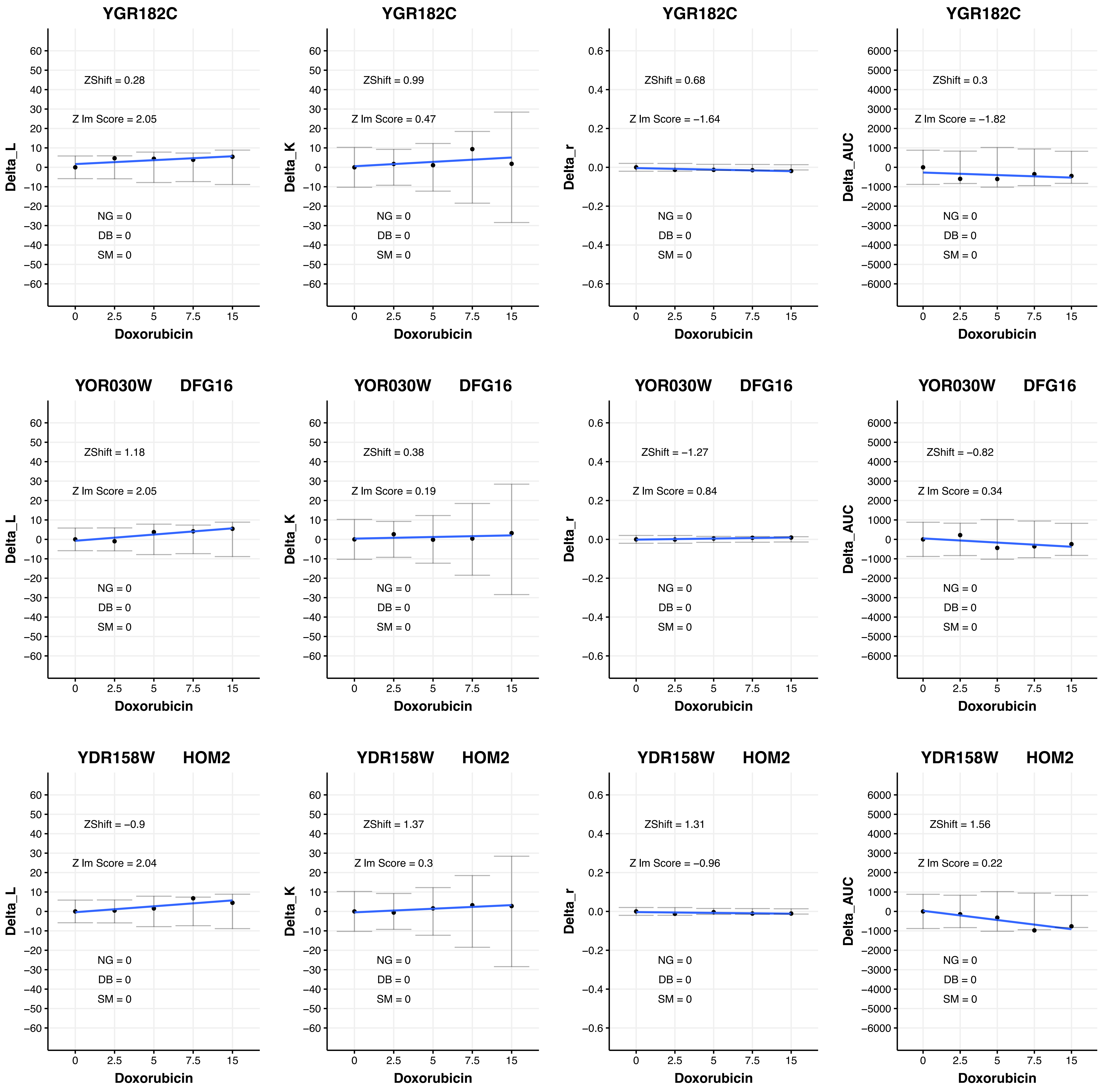
YGR182C
ZShift = 0.28
Z lm Score = 2.05
NG = 0
DB = 0
SM = 0
Delta_L
Doxorubicin
YGR182C
ZShift = 0.99
Z lm Score = 0.47
NG = 0
DB = 0
SM = 0
Delta_K
Doxorubicin
YGR182C
ZShift = 0.68
Z lm Score = −1.64
NG = 0
DB = 0
SM = 0
Delta_r
Doxorubicin
YGR182C
ZShift = 0.3
Z lm Score = −1.82
NG = 0
DB = 0
SM = 0
Delta_AUC
Doxorubicin
YOR030W DFG16
ZShift = 1.18
Z lm Score = 2.05
NG = 0
DB = 0
SM = 0
Delta_L
Doxorubicin
YOR030W DFG16
ZShift = 0.38
Z lm Score = 0.19
NG = 0
DB = 0
SM = 0
Delta_K
Doxorubicin
YOR030W DFG16
ZShift = −1.27
Z lm Score = 0.84
NG = 0
DB = 0
SM = 0
Delta_r
Doxorubicin
YOR030W DFG16
ZShift = −0.82
Z lm Score = 0.34
NG = 0
DB = 0
SM = 0
Delta_AUC
Doxorubicin
YDR158W HOM2
ZShift = −0.9
Z lm Score = 2.04
NG = 0
DB = 0
SM = 0
Delta_L
Doxorubicin
YDR158W HOM2
ZShift = 1.37
Z lm Score = 0.3
NG = 0
DB = 0
SM = 0
Delta_K
Doxorubicin
YDR158W HOM2
ZShift = 1.31
Z lm Score = −0.96
NG = 0
DB = 0
SM = 0
Delta_r
Doxorubicin
YDR158W HOM2
ZShift = 1.56
Z lm Score = 0.22
NG = 0
DB = 0
SM = 0
Delta_AUC
Doxorubicin

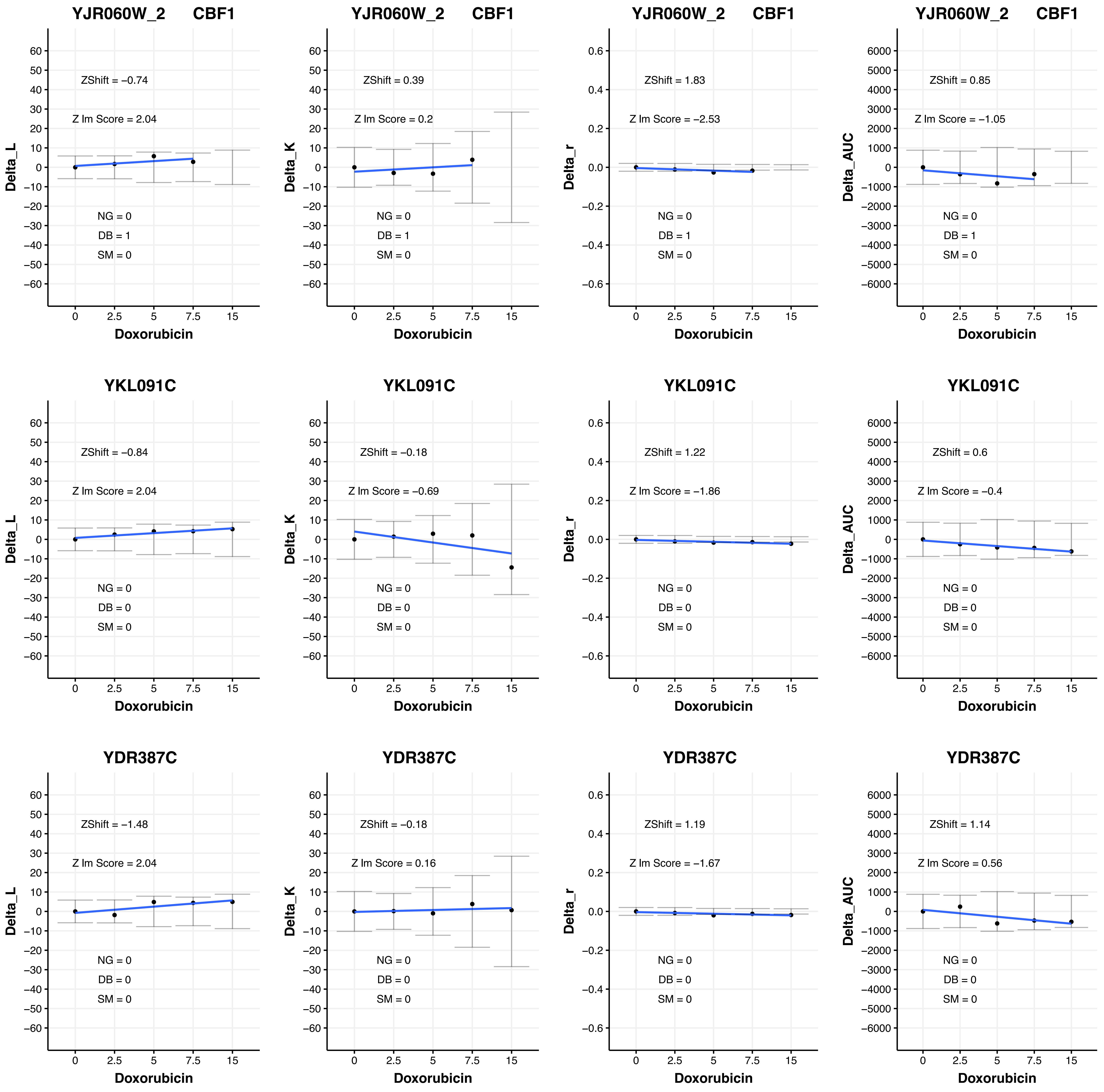
YJR060W_2 CBF1
ZShift = −0.74
Z lm Score = 2.04
NG = 0
DB = 1
SM = 0
Delta_L
Doxorubicin
YJR060W_2 CBF1
ZShift = 0.39
Z lm Score = 0.2
NG = 0
DB = 1
SM = 0
Delta_K
Doxorubicin
YJR060W_2 CBF1
ZShift = 1.83
Z lm Score = −2.53
NG = 0
DB = 1
SM = 0
Delta_r
Doxorubicin
YJR060W_2 CBF1
ZShift = 0.85
Z lm Score = −1.05
NG = 0
DB = 1
SM = 0
Delta_AUC
Doxorubicin
YKL091C
ZShift = −0.84
Z lm Score = 2.04
NG = 0
DB = 0
SM = 0
Delta_L
Doxorubicin
YKL091C
ZShift = −0.18
Z lm Score = −0.69
NG = 0
DB = 0
SM = 0
Delta_K
Doxorubicin
YKL091C
ZShift = 1.22
Z lm Score = −1.86
NG = 0
DB = 0
SM = 0
Delta_r
Doxorubicin
YKL091C
ZShift = 0.6
Z lm Score = −0.4
NG = 0
DB = 0
SM = 0
Delta_AUC
Doxorubicin
YDR387C
ZShift = −1.48
Z lm Score = 2.04
NG = 0
DB = 0
SM = 0
Delta_L
Doxorubicin
YDR387C
ZShift = −0.18
Z lm Score = 0.16
NG = 0
DB = 0
SM = 0
Delta_K
Doxorubicin
YDR387C
ZShift = 1.19
Z lm Score = −1.67
NG = 0
DB = 0
SM = 0
Delta_r
Doxorubicin
YDR387C
ZShift = 1.14
Z lm Score = 0.56
NG = 0
DB = 0
SM = 0
Delta_AUC
Doxorubicin

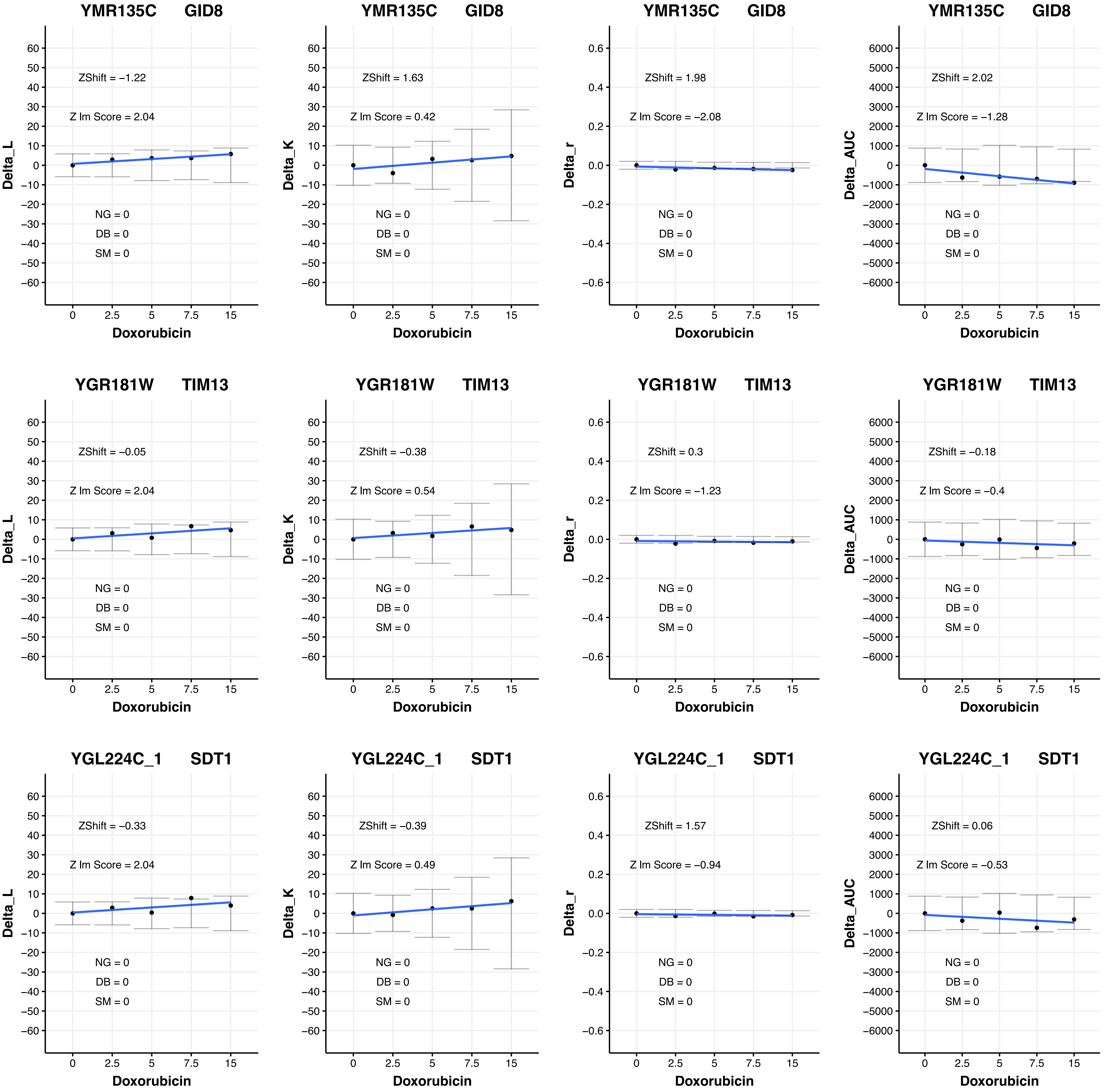
YMR135C GID8
ZShift = −1.22
Z lm Score = 2.04
NG = 0
DB = 0
SM = 0
Delta_L
Doxorubicin
YMR135C GID8
ZShift = 1.63
Z lm Score = 0.42
NG = 0
DB = 0
SM = 0
Delta_K
Doxorubicin
YMR135C GID8
ZShift = 1.98
Z lm Score = −2.08
NG = 0
DB = 0
SM = 0
Delta_r
Doxorubicin
YMR135C GID8
ZShift = 2.02
Z lm Score = −1.28
NG = 0
DB = 0
SM = 0
Delta_AUC
Doxorubicin
YGR181W TIM13
ZShift = −0.05
Z lm Score = 2.04
NG = 0
DB = 0
SM = 0
Delta_L
Doxorubicin
YGR181W TIM13
ZShift = −0.38
Z lm Score = 0.54
NG = 0
DB = 0
SM = 0
Delta_K
Doxorubicin
YGR181W TIM13
ZShift = 0.3
Z lm Score = −1.23
NG = 0
DB = 0
SM = 0
Delta_r
Doxorubicin
YGR181W TIM13
ZShift = −0.18
Z lm Score = −0.4
NG = 0
DB = 0
SM = 0
Delta_AUC
Doxorubicin
YGL224C_1 SDT1
ZShift = −0.33
Z lm Score = 2.04
NG = 0
DB = 0
SM = 0
Delta_L
Doxorubicin
YGL224C_1 SDT1
ZShift = −0.39
Z lm Score = 0.49
NG = 0
DB = 0
SM = 0
Delta_K
Doxorubicin
YGL224C_1 SDT1
ZShift = 1.57
Z lm Score = −0.94
NG = 0
DB = 0
SM = 0
Delta_r
Doxorubicin
YGL224C_1 SDT1
ZShift = 0.06
Z lm Score = −0.53
NG = 0
DB = 0
SM = 0
Delta_AUC
Doxorubicin

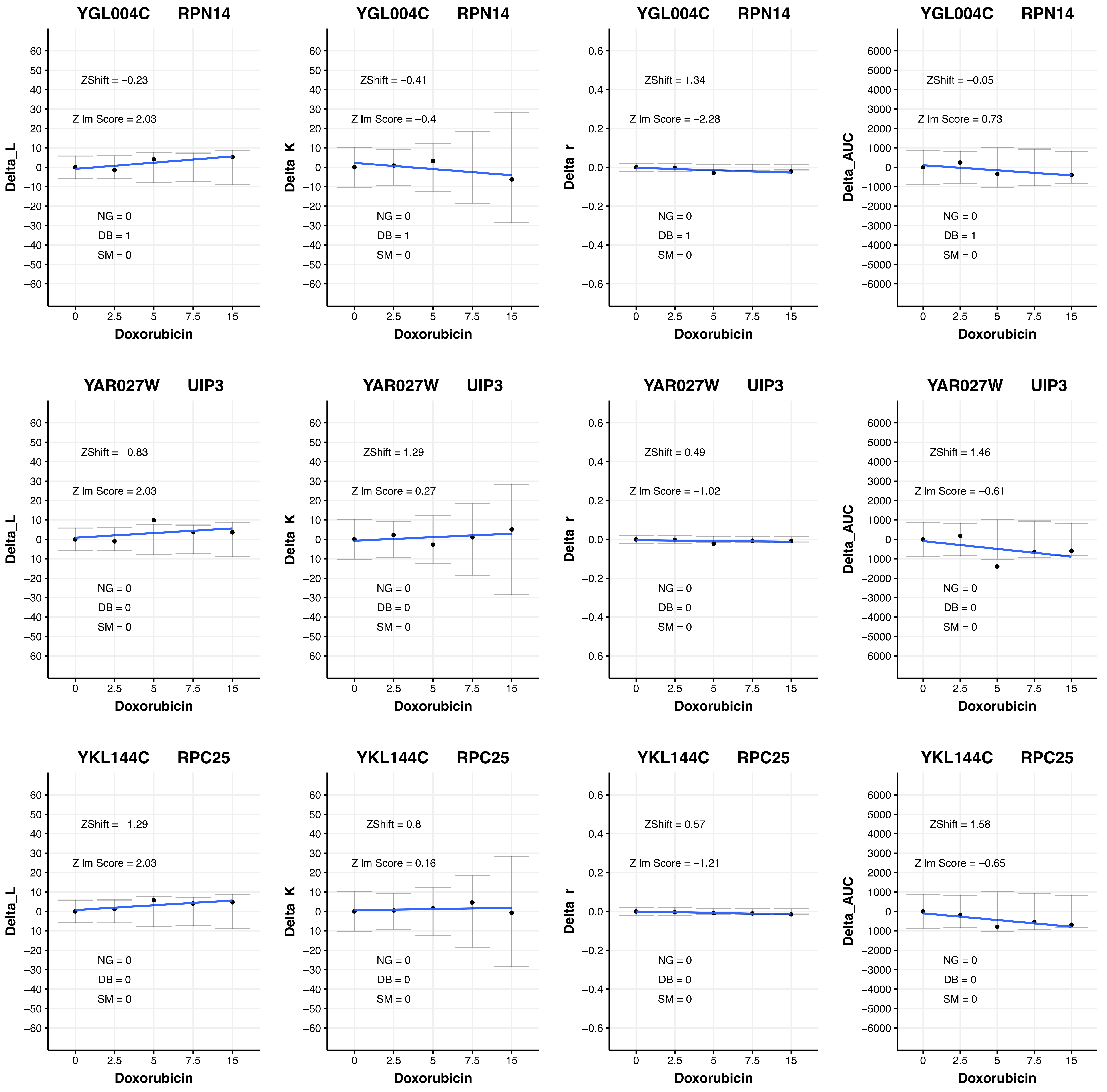
YGL004C RPN14
Delta_L
ZShift = −0.23
Z lm Score = 2.03
NG = 0
DB = 1
SM = 0
Doxorubicin
YGL004C RPN14
Delta_K
ZShift = −0.41
Z lm Score = −0.4
NG = 0
DB = 1
SM = 0
Doxorubicin
YGL004C RPN14
Delta_r
ZShift = 1.34
Z lm Score = −2.28
NG = 0
DB = 1
SM = 0
Doxorubicin
YGL004C RPN14
Delta_AUC
ZShift = −0.05
Z lm Score = 0.73
NG = 0
DB = 1
SM = 0
Doxorubicin
YAR027W UIP3
Delta_L
ZShift = −0.83
Z lm Score = 2.03
NG = 0
DB = 0
SM = 0
Doxorubicin
YAR027W UIP3
Delta_K
ZShift = 1.29
Z lm Score = 0.27
NG = 0
DB = 0
SM = 0
Doxorubicin
YAR027W UIP3
Delta_r
ZShift = 0.49
Z lm Score = −1.02
NG = 0
DB = 0
SM = 0
Doxorubicin
YAR027W UIP3
Delta_AUC
ZShift = 1.46
Z lm Score = −0.61
NG = 0
DB = 0
SM = 0
Doxorubicin
YKL144C RPC25
Delta_L
ZShift = −1.29
Z lm Score = 2.03
NG = 0
DB = 0
SM = 0
Doxorubicin
YKL144C RPC25
Delta_K
ZShift = 0.8
Z lm Score = 0.16
NG = 0
DB = 0
SM = 0
Doxorubicin
YKL144C RPC25
Delta_r
ZShift = 0.57
Z lm Score = −1.21
NG = 0
DB = 0
SM = 0
Doxorubicin
YKL144C RPC25
Delta_AUC
ZShift = 1.58
Z lm Score = −0.65
NG = 0
DB = 0
SM = 0
Doxorubicin

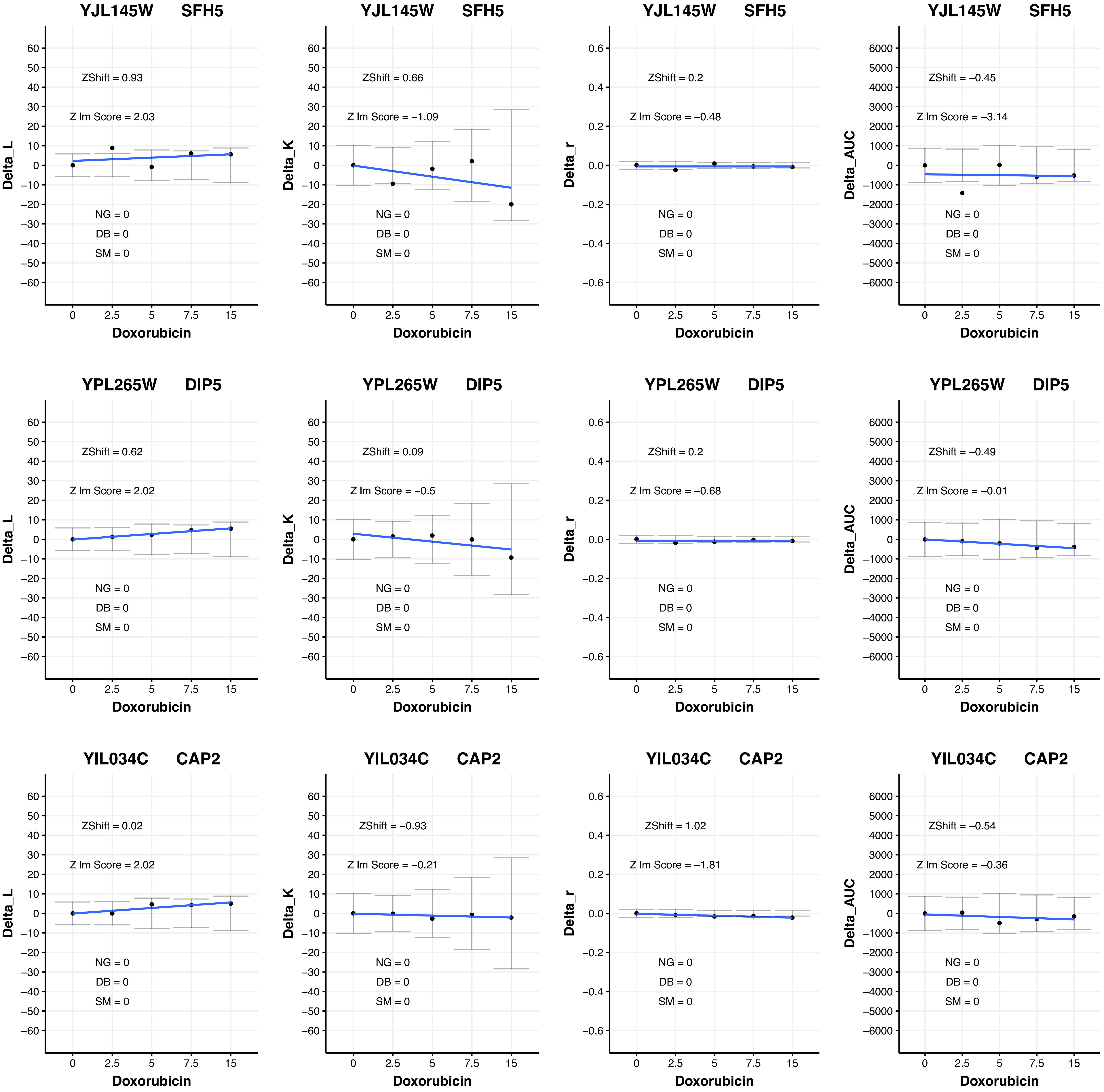

YJL145W SFH5
ZShift = 0.93
Z lm Score = 2.03
NG = 0
DB = 0
SM = 0
Delta_L
Doxorubicin
YJL145W SFH5
ZShift = 0.66
Z lm Score = −1.09
NG = 0
DB = 0
SM = 0
Delta_K
Doxorubicin
YJL145W SFH5
ZShift = 0.2
Z lm Score = −0.48
NG = 0
DB = 0
SM = 0
Delta_r
Doxorubicin
YJL145W SFH5
ZShift = −0.45
Z lm Score = −3.14
NG = 0
DB = 0
SM = 0
Delta_AUC
Doxorubicin
YPL265W DIP5
ZShift = 0.62
Z lm Score = 2.02
NG = 0
DB = 0
SM = 0
Delta_L
Doxorubicin
YPL265W DIP5
ZShift = 0.09
Z lm Score = −0.5
NG = 0
DB = 0
SM = 0
Delta_K
Doxorubicin
YPL265W DIP5
ZShift = 0.2
Z lm Score = −0.68
NG = 0
DB = 0
SM = 0
Delta_r
Doxorubicin
YPL265W DIP5
ZShift = −0.49
Z lm Score = −0.01
NG = 0
DB = 0
SM = 0
Delta_AUC
Doxorubicin
YIL034C CAP2
ZShift = 0.02
Z lm Score = 2.02
NG = 0
DB = 0
SM = 0
Delta_L
Doxorubicin
YIL034C CAP2
ZShift = −0.93
Z lm Score = −0.21
NG = 0
DB = 0
SM = 0
Delta_K
Doxorubicin
YIL034C CAP2
ZShift = 1.02
Z lm Score = −1.81
NG = 0
DB = 0
SM = 0
Delta_r
Doxorubicin
YIL034C CAP2
ZShift = −0.54
Z lm Score = −0.36
NG = 0
DB = 0
SM = 0
Delta_AUC
Doxorubicin

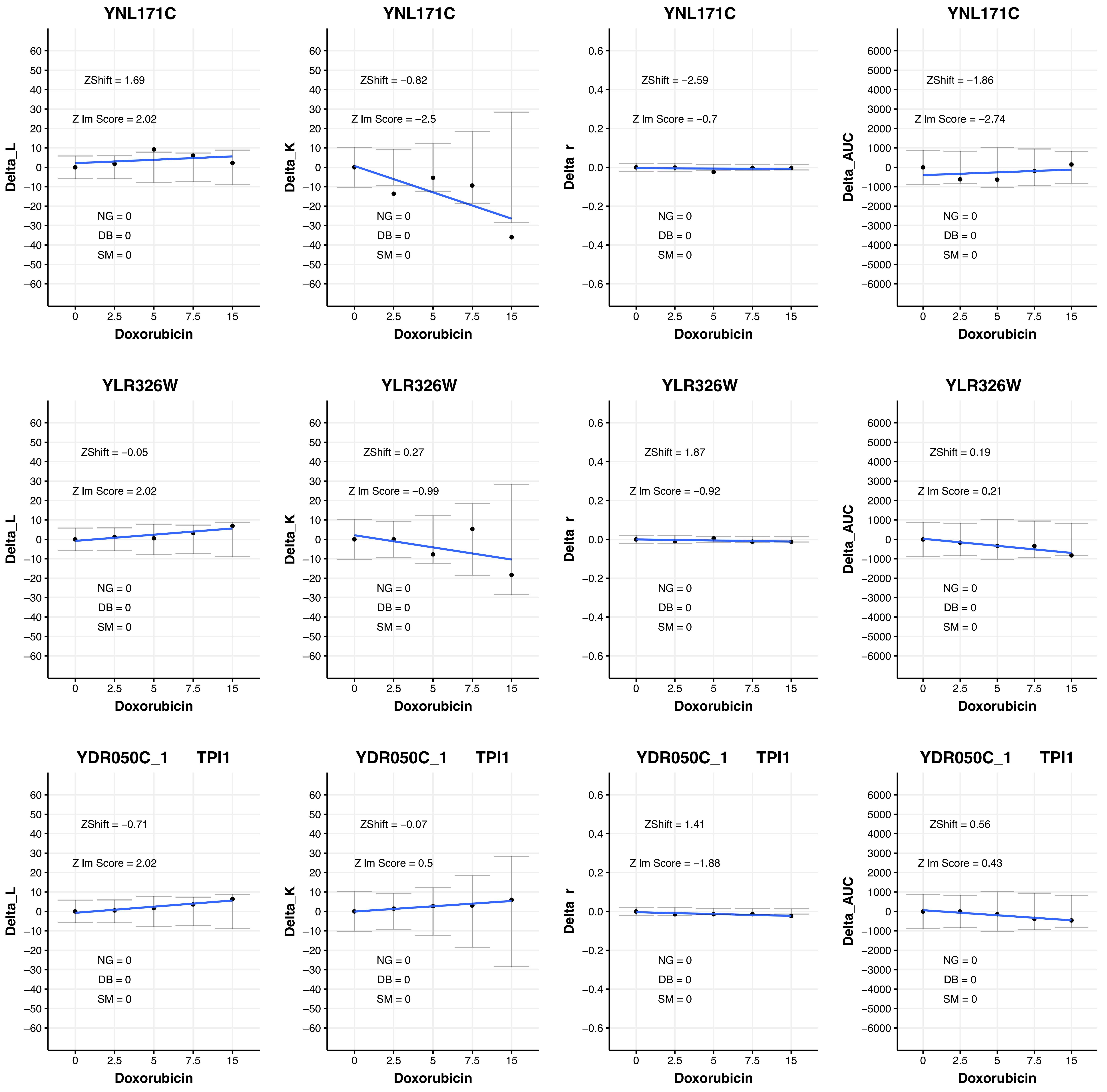

YNL171C
ZShift = 1.69
Z lm Score = 2.02
NG = 0
DB = 0
SM = 0
Delta_L
Doxorubicin
YNL171C
ZShift = −0.82
Z lm Score = −2.5
NG = 0
DB = 0
SM = 0
Delta_K
Doxorubicin
YNL171C
ZShift = −2.59
Z lm Score = −0.7
NG = 0
DB = 0
SM = 0
Delta_r
Doxorubicin
YNL171C
ZShift = −1.86
Z lm Score = −2.74
NG = 0
DB = 0
SM = 0
Delta_AUC
Doxorubicin
YLR326W
ZShift = −0.05
Z lm Score = 2.02
NG = 0
DB = 0
SM = 0
Delta_L
Doxorubicin
YLR326W
ZShift = 0.27
Z lm Score = −0.99
NG = 0
DB = 0
SM = 0
Delta_K
Doxorubicin
YLR326W
ZShift = 1.87
Z lm Score = −0.92
NG = 0
DB = 0
SM = 0
Delta_r
Doxorubicin
YLR326W
ZShift = 0.19
Z lm Score = 0.21
NG = 0
DB = 0
SM = 0
Delta_AUC
Doxorubicin
YDR050C_1 TPI1
ZShift = −0.71
Z lm Score = 2.02
NG = 0
DB = 0
SM = 0
Delta_L
Doxorubicin
YDR050C_1 TPI1
ZShift = −0.07
Z lm Score = 0.5
NG = 0
DB = 0
SM = 0
Delta_K
Doxorubicin
YDR050C_1 TPI1
ZShift = 1.41
Z lm Score = −1.88
NG = 0
DB = 0
SM = 0
Delta_r
Doxorubicin
YDR050C_1 TPI1
ZShift = 0.56
Z lm Score = 0.43
NG = 0
DB = 0
SM = 0
Delta_AUC
Doxorubicin

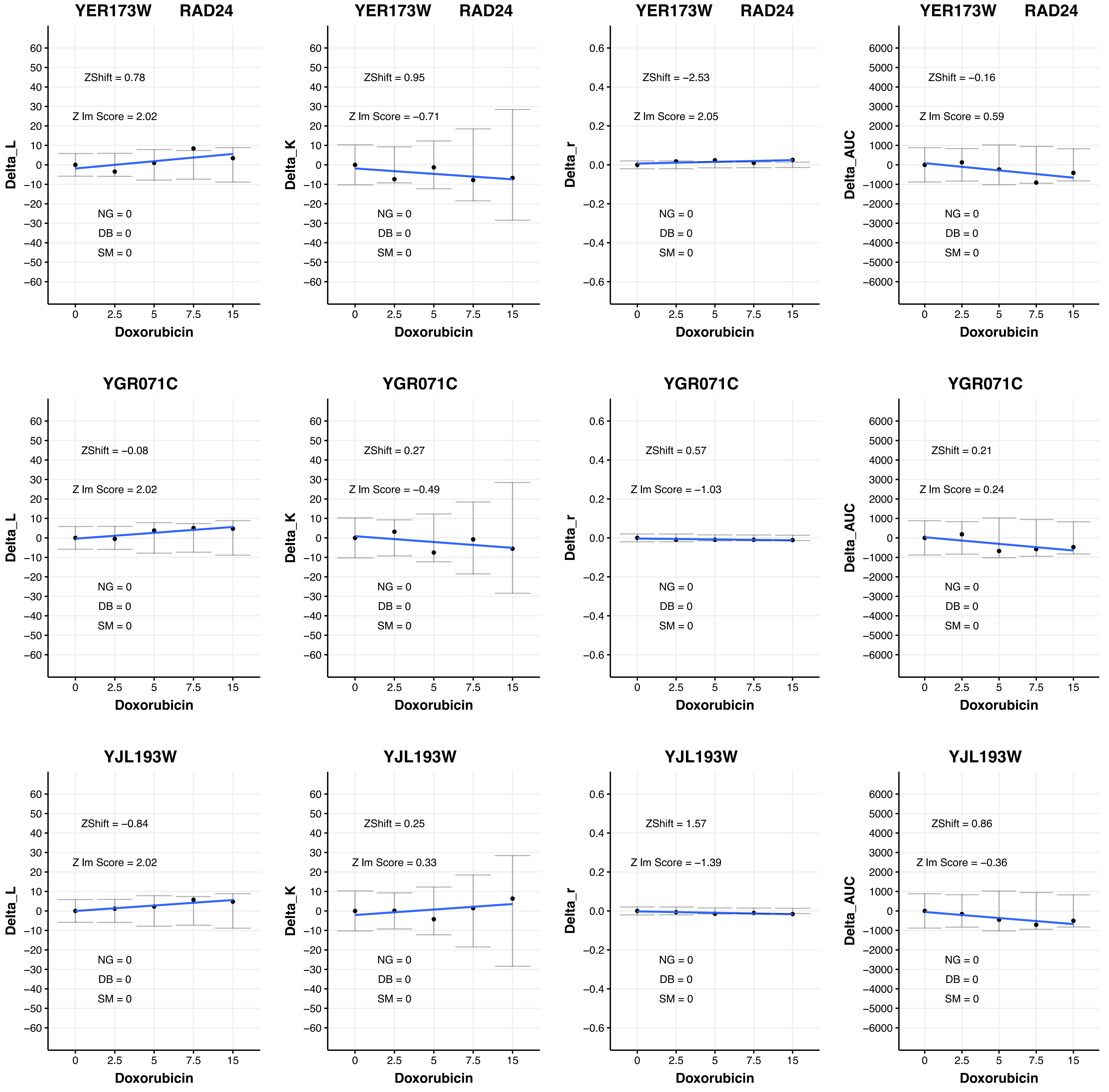

YER173W RAD24
ZShift = 0.78
Z lm Score = 2.02
NG = 0
DB = 0
SM = 0
Delta_L
Doxorubicin
YER173W RAD24
ZShift = 0.95
Z lm Score = −0.71
NG = 0
DB = 0
SM = 0
Delta_K
Doxorubicin
YER173W RAD24
ZShift = −2.53
Z lm Score = 2.05
NG = 0
DB = 0
SM = 0
Delta_r
Doxorubicin
YER173W RAD24
ZShift = −0.16
Z lm Score = 0.59
NG = 0
DB = 0
SM = 0
Delta_AUC
Doxorubicin
YGR071C
ZShift = −0.08
Z lm Score = 2.02
NG = 0
DB = 0
SM = 0
Delta_L
Doxorubicin
YGR071C
ZShift = 0.27
Z lm Score = −0.49
NG = 0
DB = 0
SM = 0
Delta_K
Doxorubicin
YGR071C
ZShift = 0.57
Z lm Score = −1.03
NG = 0
DB = 0
SM = 0
Delta_r
Doxorubicin
YGR071C
ZShift = 0.21
Z lm Score = 0.24
NG = 0
DB = 0
SM = 0
Delta_AUC
Doxorubicin
YJL193W
ZShift = −0.84
Z lm Score = 2.02
NG = 0
DB = 0
SM = 0
Delta_L
Doxorubicin
YJL193W
ZShift = 0.25
Z lm Score = 0.33
NG = 0
DB = 0
SM = 0
Delta_K
Doxorubicin
YJL193W
ZShift = 1.57
Z lm Score = −1.39
NG = 0
DB = 0
SM = 0
Delta_r
Doxorubicin
YJL193W
ZShift = 0.86
Z lm Score = −0.36
NG = 0
DB = 0
SM = 0
Delta_AUC
Doxorubicin

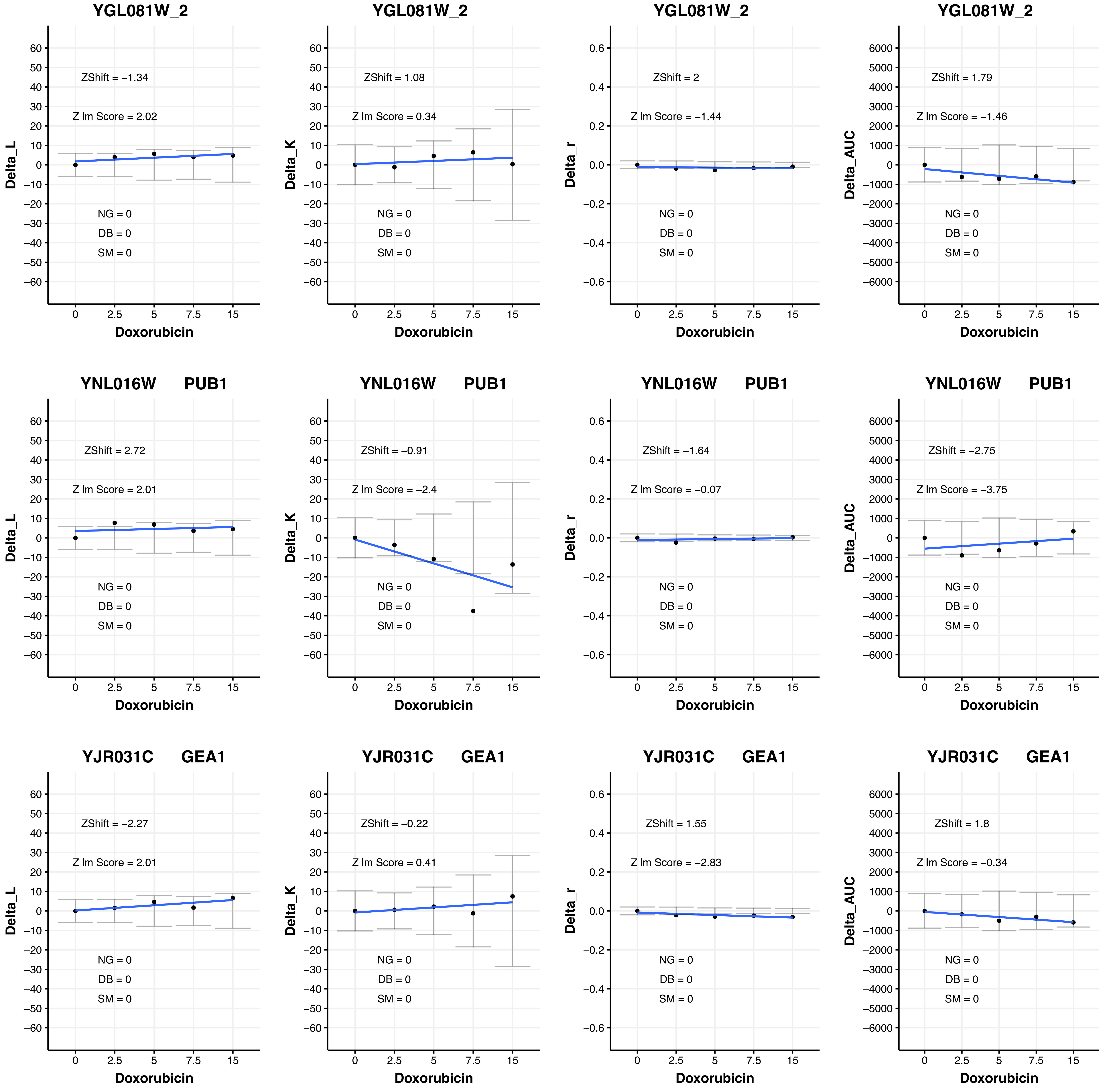

YGL081W_2
ZShift = −1.34
Z lm Score = 2.02
ZShift = 1.08
Z lm Score = 0.34
ZShift = 2
Z lm Score = −1.44
ZShift = 1.79
Z lm Score = −1.46
YNL016W PUB1
ZShift = 2.72
Z lm Score = 2.01
ZShift = −0.91
Z lm Score = −2.4
ZShift = −1.64
Z lm Score = −0.07
ZShift = −2.75
Z lm Score = −3.75
YJR031C GEA1
ZShift = −2.27
Z lm Score = 2.01
ZShift = −0.22
Z lm Score = 0.41
ZShift = 1.55
Z lm Score = −2.83
ZShift = 1.8
Z lm Score = −0.34
NG = 0
DB = 0
SM = 0
Delta_L
Delta_K
Delta_r
Delta_AUC
Doxorubicin

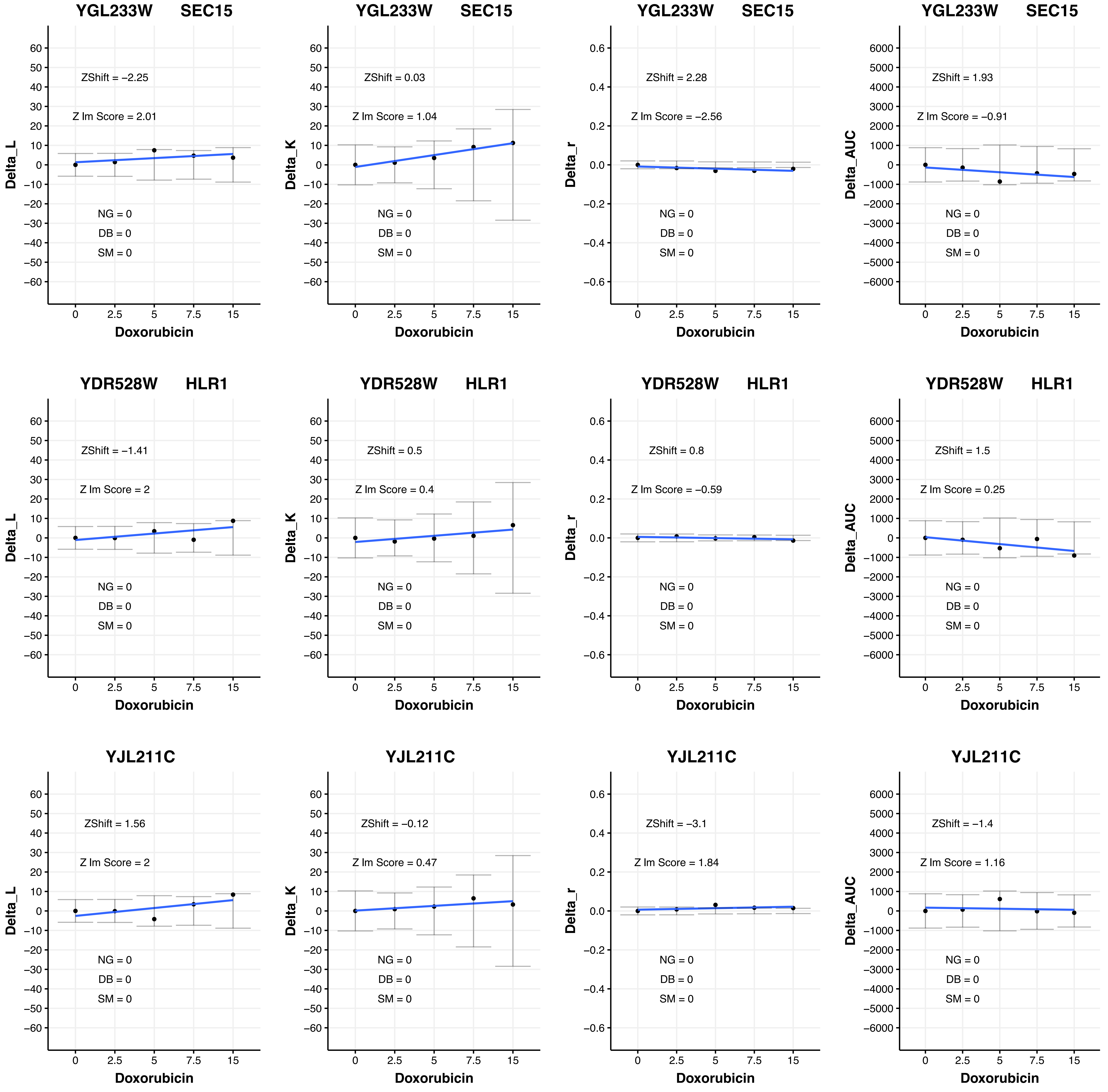

YGL233W SEC15
ZShift = −2.25
Z lm Score = 2.01
NG = 0
DB = 0
SM = 0
Delta_L
Doxorubicin
YGL233W SEC15
ZShift = 0.03
Z lm Score = 1.04
NG = 0
DB = 0
SM = 0
Delta_K
Doxorubicin
YGL233W SEC15
ZShift = 2.28
Z lm Score = −2.56
NG = 0
DB = 0
SM = 0
Delta_r
Doxorubicin
YGL233W SEC15
ZShift = 1.93
Z lm Score = −0.91
NG = 0
DB = 0
SM = 0
Delta_AUC
Doxorubicin
YDR528W HLR1
ZShift = −1.41
Z lm Score = 2
NG = 0
DB = 0
SM = 0
Delta_L
Doxorubicin
YDR528W HLR1
ZShift = 0.5
Z lm Score = 0.4
NG = 0
DB = 0
SM = 0
Delta_K
Doxorubicin
YDR528W HLR1
ZShift = 0.8
Z lm Score = −0.59
NG = 0
DB = 0
SM = 0
Delta_r
Doxorubicin
YDR528W HLR1
ZShift = 1.5
Z lm Score = 0.25
NG = 0
DB = 0
SM = 0
Delta_AUC
Doxorubicin
YJL211C
ZShift = 1.56
Z lm Score = 2
NG = 0
DB = 0
SM = 0
Delta_L
Doxorubicin
YJL211C
ZShift = −0.12
Z lm Score = 0.47
NG = 0
DB = 0
SM = 0
Delta_K
Doxorubicin
YJL211C
ZShift = −3.1
Z lm Score = 1.84
NG = 0
DB = 0
SM = 0
Delta_r
Doxorubicin
YJL211C
ZShift = −1.4
Z lm Score = 1.16
NG = 0
DB = 0
SM = 0
Delta_AUC
Doxorubicin

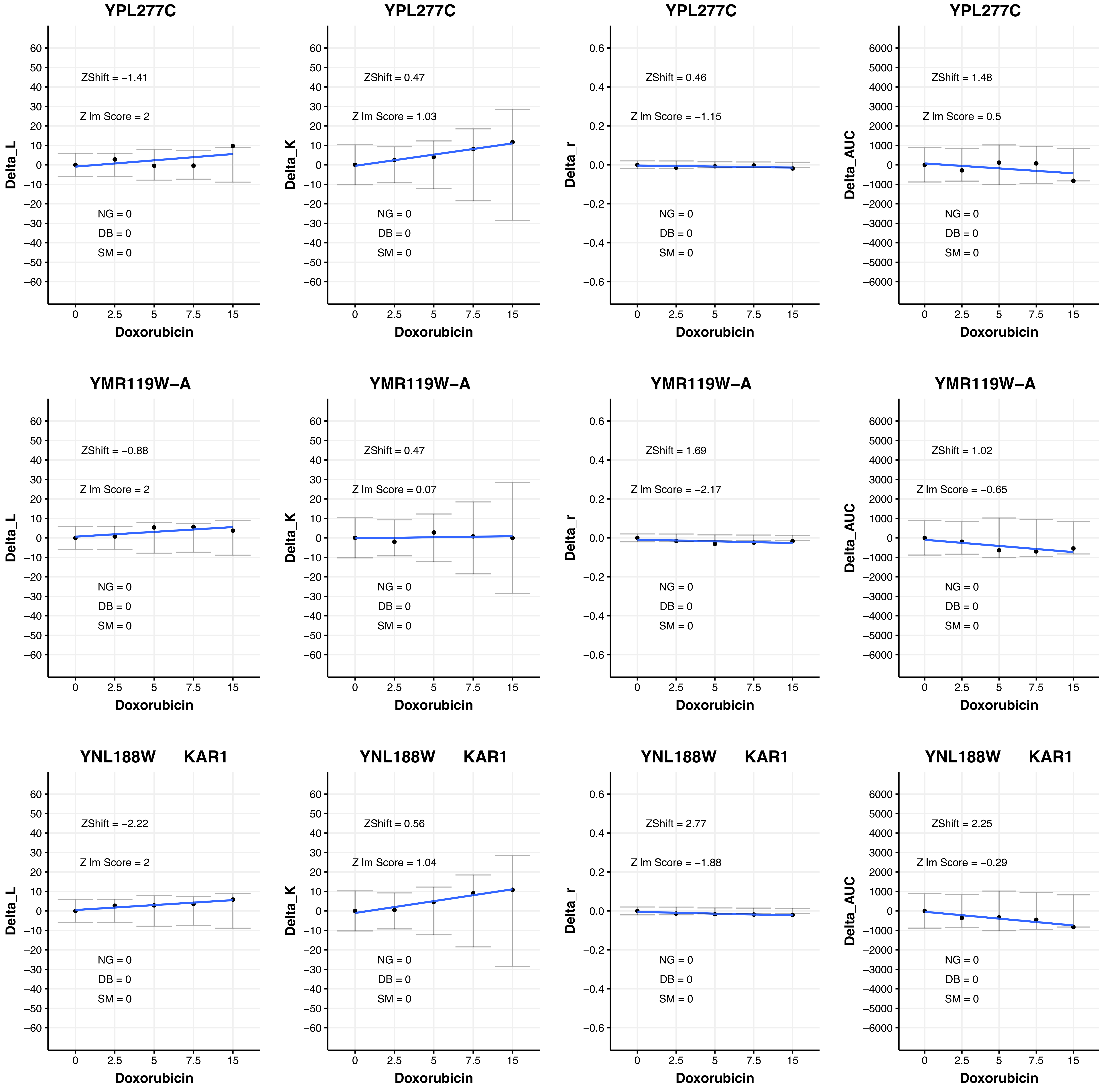

YPL277C
ZShift = −1.41
Z lm Score = 2
NG = 0
DB = 0
SM = 0
Delta_L
Doxorubicin
YPL277C
ZShift = 0.47
Z lm Score = 1.03
NG = 0
DB = 0
SM = 0
Delta_K
Doxorubicin
YPL277C
ZShift = 0.46
Z lm Score = −1.15
NG = 0
DB = 0
SM = 0
Delta_r
Doxorubicin
YPL277C
ZShift = 1.48
Z lm Score = 0.5
NG = 0
DB = 0
SM = 0
Delta_AUC
Doxorubicin
YMR119W−A
ZShift = −0.88
Z lm Score = 2
NG = 0
DB = 0
SM = 0
Delta_L
Doxorubicin
YMR119W−A
ZShift = 0.47
Z lm Score = 0.07
NG = 0
DB = 0
SM = 0
Delta_K
Doxorubicin
YMR119W−A
ZShift = 1.69
Z lm Score = −2.17
NG = 0
DB = 0
SM = 0
Delta_r
Doxorubicin
YMR119W−A
ZShift = 1.02
Z lm Score = −0.65
NG = 0
DB = 0
SM = 0
Delta_AUC
Doxorubicin
YNL188W KAR1
ZShift = −2.22
Z lm Score = 2
NG = 0
DB = 0
SM = 0
Delta_L
Doxorubicin
YNL188W KAR1
ZShift = 0.56
Z lm Score = 1.04
NG = 0
DB = 0
SM = 0
Delta_K
Doxorubicin
YNL188W KAR1
ZShift = 2.77
Z lm Score = −1.88
NG = 0
DB = 0
SM = 0
Delta_r
Doxorubicin
YNL188W KAR1
ZShift = 2.25
Z lm Score = −0.29
NG = 0
DB = 0
SM = 0
Delta_AUC
Doxorubicin

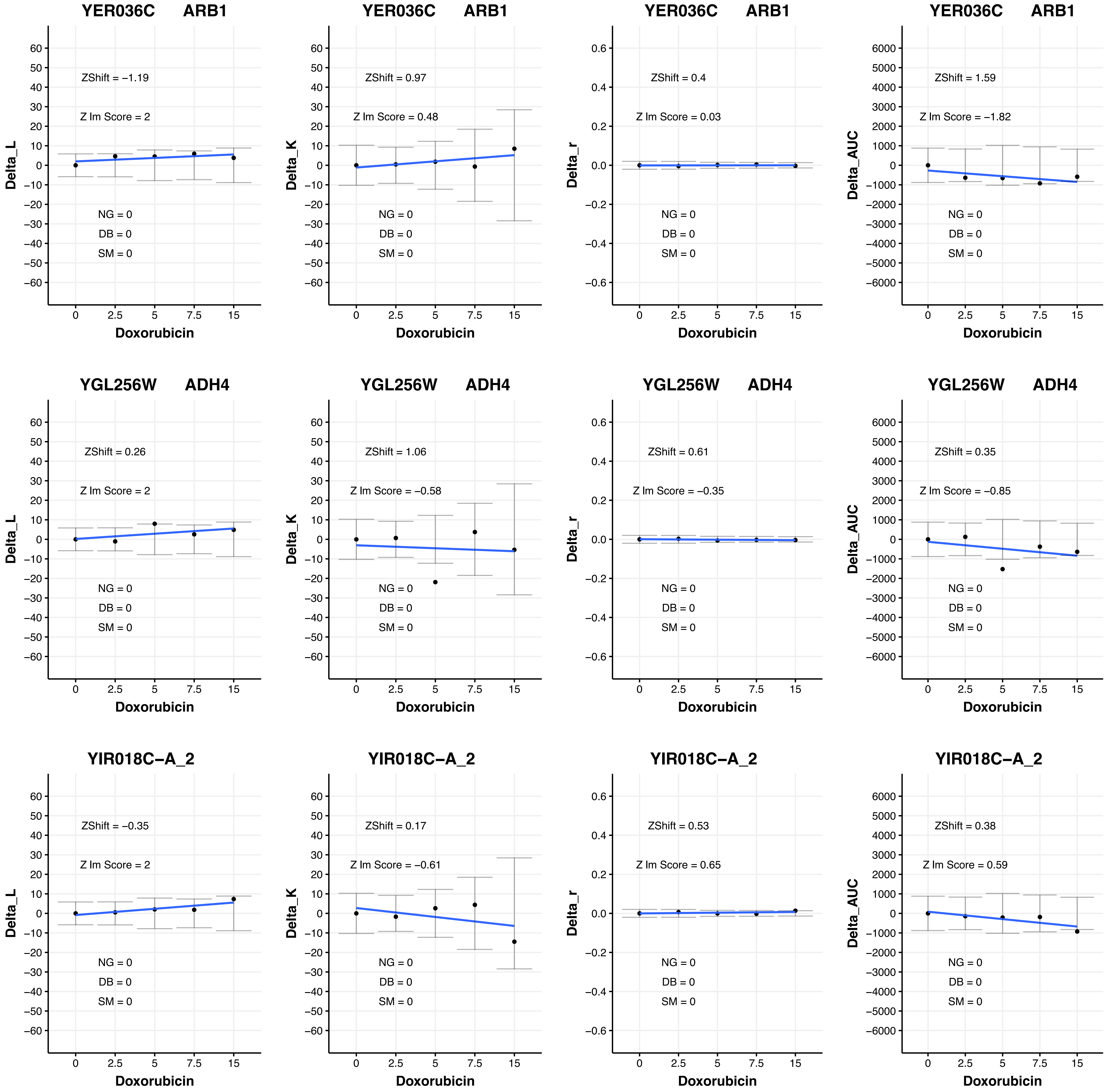
YER036C ARB1
ZShift = −1.19
Z lm Score = 2
NG = 0
DB = 0
SM = 0
Delta_L
Doxorubicin
YER036C ARB1
ZShift = 0.97
Z lm Score = 0.48
NG = 0
DB = 0
SM = 0
Delta_K
Doxorubicin
YER036C ARB1
ZShift = 0.4
Z lm Score = 0.03
NG = 0
DB = 0
SM = 0
Delta_r
Doxorubicin
YER036C ARB1
ZShift = 1.59
Z lm Score = −1.82
NG = 0
DB = 0
SM = 0
Delta_AUC
Doxorubicin
YGL256W ADH4
ZShift = 0.26
Z lm Score = 2
NG = 0
DB = 0
SM = 0
Delta_L
Doxorubicin
YGL256W ADH4
ZShift = 1.06
Z lm Score = −0.58
NG = 0
DB = 0
SM = 0
Delta_K
Doxorubicin
YGL256W ADH4
ZShift = 0.61
Z lm Score = −0.35
NG = 0
DB = 0
SM = 0
Delta_r
Doxorubicin
YGL256W ADH4
ZShift = 0.35
Z lm Score = −0.85
NG = 0
DB = 0
SM = 0
Delta_AUC
Doxorubicin
YIR018C−A_2
ZShift = −0.35
Z lm Score = 2
NG = 0
DB = 0
SM = 0
Delta_L
Doxorubicin
YIR018C−A_2
ZShift = 0.17
Z lm Score = −0.61
NG = 0
DB = 0
SM = 0
Delta_K
Doxorubicin
YIR018C−A_2
ZShift = 0.53
Z lm Score = 0.65
NG = 0
DB = 0
SM = 0
Delta_r
Doxorubicin
YIR018C−A_2
ZShift = 0.38
Z lm Score = 0.59
NG = 0
DB = 0
SM = 0
Delta_AUC
Doxorubicin

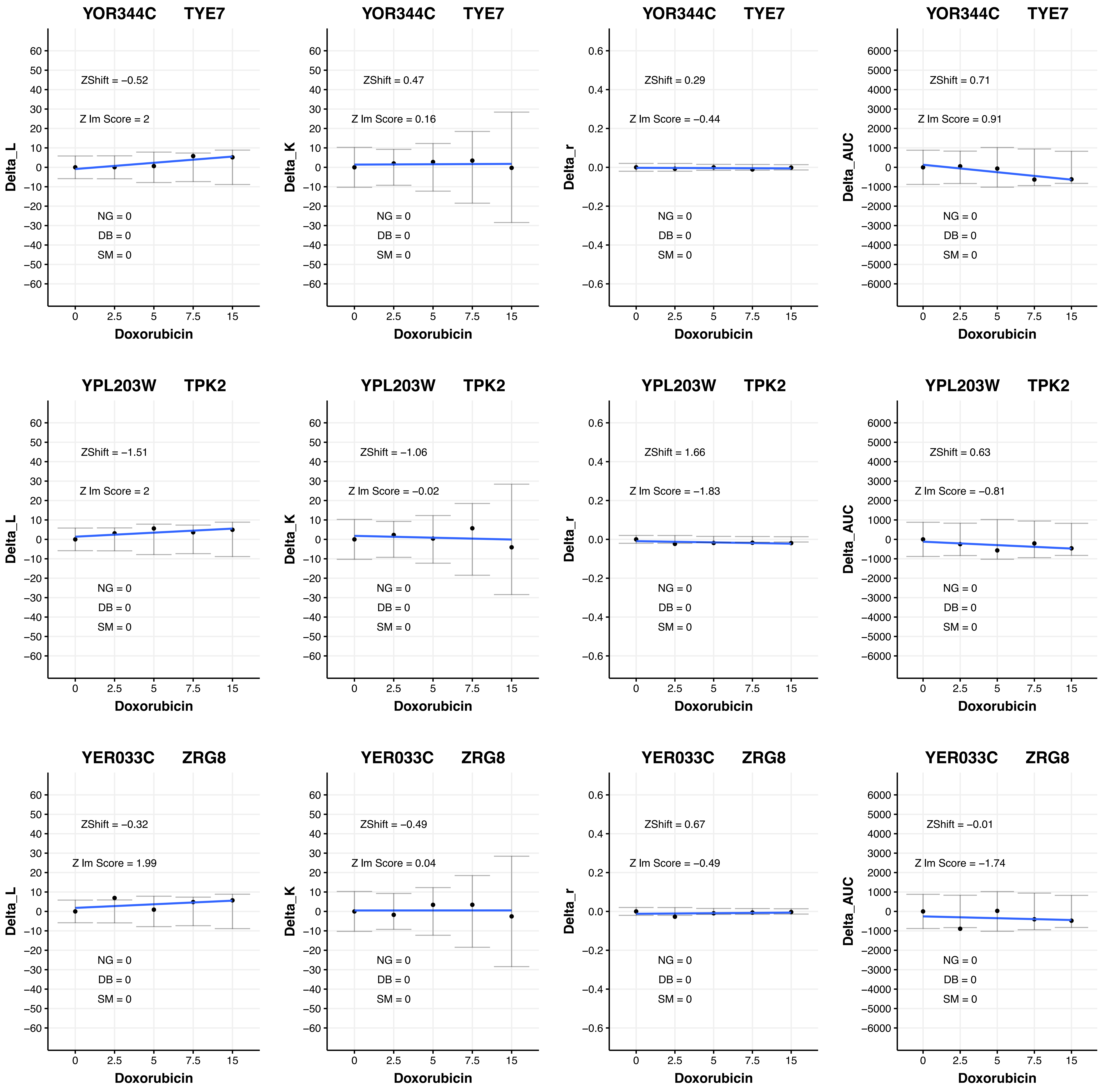
YOR344C TYE7
ZShift = −0.52
Z lm Score = 2
NG = 0
DB = 0
SM = 0
Delta_L
Doxorubicin
YOR344C TYE7
ZShift = 0.47
Z lm Score = 0.16
NG = 0
DB = 0
SM = 0
Delta_K
Doxorubicin
YOR344C TYE7
ZShift = 0.29
Z lm Score = −0.44
NG = 0
DB = 0
SM = 0
Delta_r
Doxorubicin
YOR344C TYE7
ZShift = 0.71
Z lm Score = 0.91
NG = 0
DB = 0
SM = 0
Delta_AUC
Doxorubicin
YPL203W TPK2
ZShift = −1.51
Z lm Score = 2
NG = 0
DB = 0
SM = 0
Delta_L
Doxorubicin
YPL203W TPK2
ZShift = −1.06
Z lm Score = −0.02
NG = 0
DB = 0
SM = 0
Delta_K
Doxorubicin
YPL203W TPK2
ZShift = 1.66
Z lm Score = −1.83
NG = 0
DB = 0
SM = 0
Delta_r
Doxorubicin
YPL203W TPK2
ZShift = 0.63
Z lm Score = −0.81
NG = 0
DB = 0
SM = 0
Delta_AUC
Doxorubicin
YER033C ZRG8
ZShift = −0.32
Z lm Score = 1.99
NG = 0
DB = 0
SM = 0
Delta_L
Doxorubicin
YER033C ZRG8
ZShift = −0.49
Z lm Score = 0.04
NG = 0
DB = 0
SM = 0
Delta_K
Doxorubicin
YER033C ZRG8
ZShift = 0.67
Z lm Score = −0.49
NG = 0
DB = 0
SM = 0
Delta_r
Doxorubicin
YER033C ZRG8
ZShift = −0.01
Z lm Score = −1.74
NG = 0
DB = 0
SM = 0
Delta_AUC
Doxorubicin

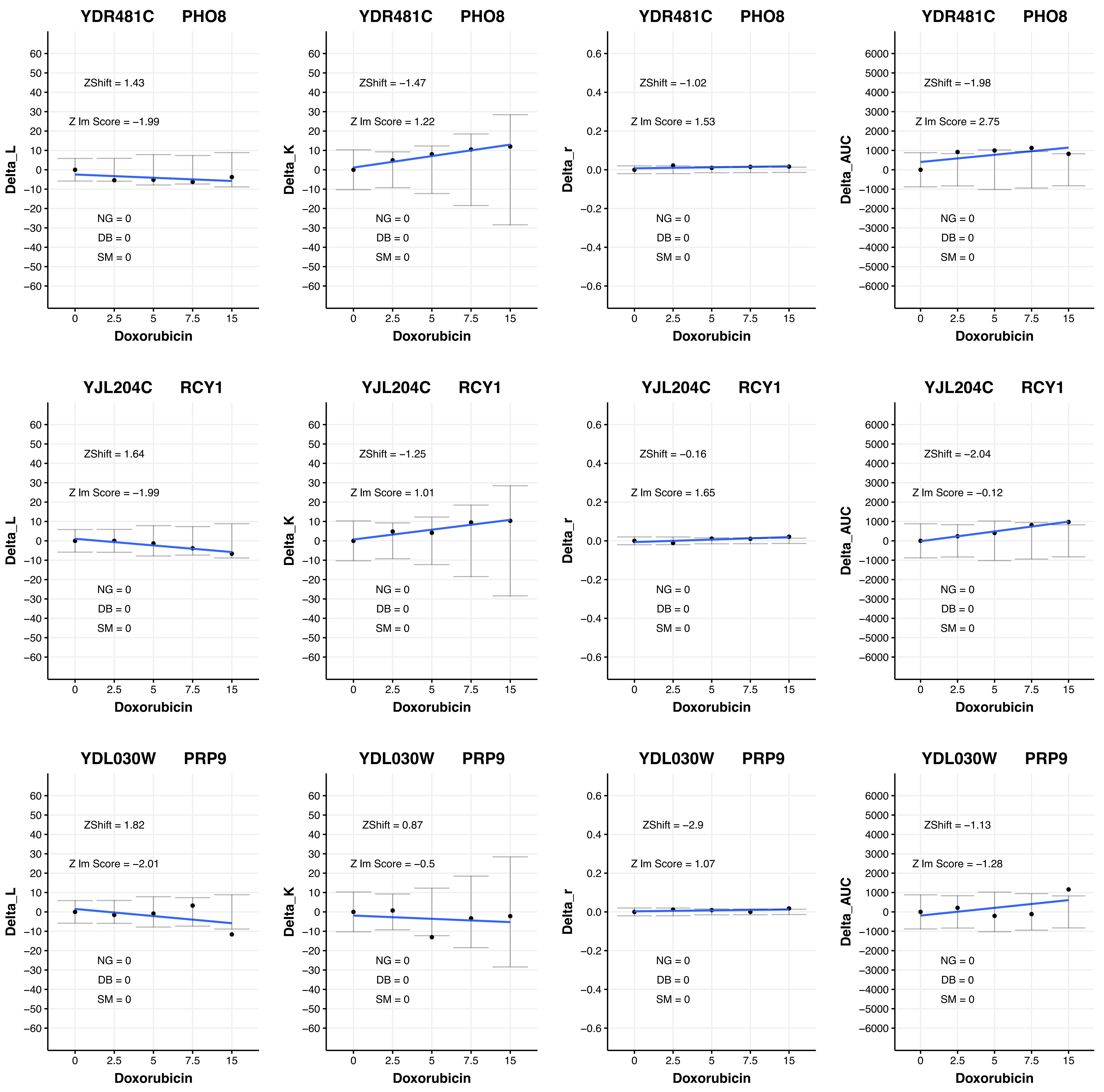
YDR481C PHO8
ZShift = 1.43
Z lm Score = −1.99
Delta_L
NG = 0
DB = 0
SM = 0
Doxorubicin
YDR481C PHO8
ZShift = −1.47
Z lm Score = 1.22
Delta_K
NG = 0
DB = 0
SM = 0
Doxorubicin
YDR481C PHO8
ZShift = −1.02
Z lm Score = 1.53
Delta_r
NG = 0
DB = 0
SM = 0
Doxorubicin
YDR481C PHO8
ZShift = −1.98
Z lm Score = 2.75
Delta_AUC
NG = 0
DB = 0
SM = 0
Doxorubicin
YJL204C RCY1
ZShift = 1.64
Z lm Score = −1.99
Delta_L
NG = 0
DB = 0
SM = 0
Doxorubicin
YJL204C RCY1
ZShift = −1.25
Z lm Score = 1.01
Delta_K
NG = 0
DB = 0
SM = 0
Doxorubicin
YJL204C RCY1
ZShift = −0.16
Z lm Score = 1.65
Delta_r
NG = 0
DB = 0
SM = 0
Doxorubicin
YJL204C RCY1
ZShift = −2.04
Z lm Score = −0.12
Delta_AUC
NG = 0
DB = 0
SM = 0
Doxorubicin
YDL030W PRP9
ZShift = 1.82
Z lm Score = −2.01
Delta_L
NG = 0
DB = 0
SM = 0
Doxorubicin
YDL030W PRP9
ZShift = 0.87
Z lm Score = −0.5
Delta_K
NG = 0
DB = 0
SM = 0
Doxorubicin
YDL030W PRP9
ZShift = −2.9
Z lm Score = 1.07
Delta_r
NG = 0
DB = 0
SM = 0
Doxorubicin
YDL030W PRP9
ZShift = −1.13
Z lm Score = −1.28
Delta_AUC
NG = 0
DB = 0
SM = 0
Doxorubicin

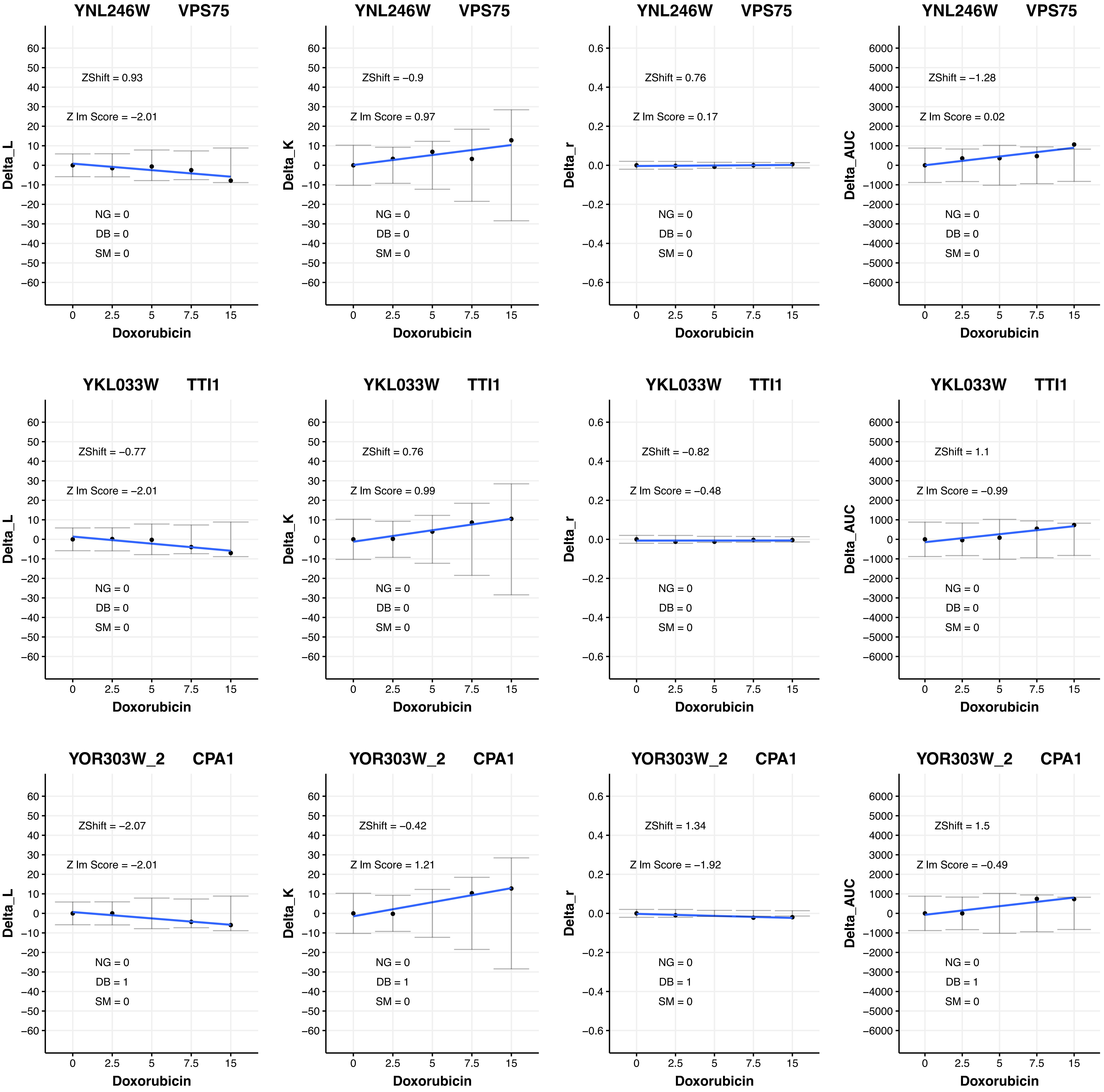
YNL246W VPS75
ZShift = 0.93
Z lm Score = −2.01
NG = 0
DB = 0
SM = 0
Delta_L
Doxorubicin
YNL246W VPS75
ZShift = −0.9
Z lm Score = 0.97
NG = 0
DB = 0
SM = 0
Delta_K
Doxorubicin
YNL246W VPS75
ZShift = 0.76
Z lm Score = 0.17
NG = 0
DB = 0
SM = 0
Delta_r
Doxorubicin
YNL246W VPS75
ZShift = −1.28
Z lm Score = 0.02
NG = 0
DB = 0
SM = 0
Delta_AUC
Doxorubicin
YKL033W TTI1
ZShift = −0.77
Z lm Score = −2.01
NG = 0
DB = 0
SM = 0
Delta_L
Doxorubicin
YKL033W TTI1
ZShift = 0.76
Z lm Score = 0.99
NG = 0
DB = 0
SM = 0
Delta_K
Doxorubicin
YKL033W TTI1
ZShift = −0.82
Z lm Score = −0.48
NG = 0
DB = 0
SM = 0
Delta_r
Doxorubicin
YKL033W TTI1
ZShift = 1.1
Z lm Score = −0.99
NG = 0
DB = 0
SM = 0
Delta_AUC
Doxorubicin
YOR303W_2 CPA1
ZShift = −2.07
Z lm Score = −2.01
NG = 0
DB = 1
SM = 0
Delta_L
Doxorubicin
YOR303W_2 CPA1
ZShift = −0.42
Z lm Score = 1.21
NG = 0
DB = 1
SM = 0
Delta_K
Doxorubicin
YOR303W_2 CPA1
ZShift = 1.34
Z lm Score = −1.92
NG = 0
DB = 1
SM = 0
Delta_r
Doxorubicin
YOR303W_2 CPA1
ZShift = 1.5
Z lm Score = −0.49
NG = 0
DB = 1
SM = 0
Delta_AUC
Doxorubicin

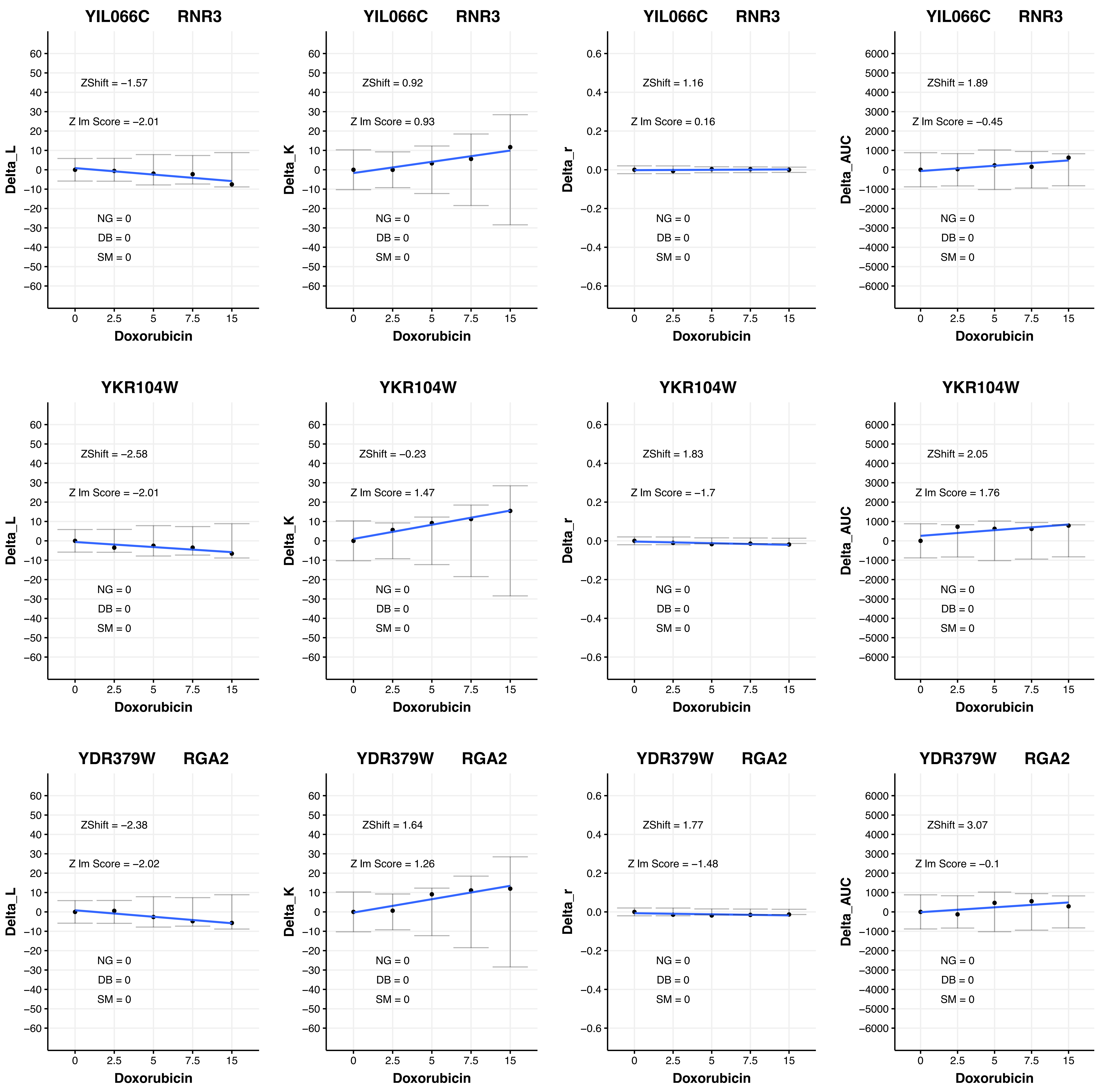
YIL066C RNR3
ZShift = −1.57
Z lm Score = −2.01
Delta_L
NG = 0
DB = 0
SM = 0
Doxorubicin
YIL066C RNR3
ZShift = 0.92
Z lm Score = 0.93
Delta_K
NG = 0
DB = 0
SM = 0
Doxorubicin
YIL066C RNR3
ZShift = 1.16
Z lm Score = 0.16
Delta_r
NG = 0
DB = 0
SM = 0
Doxorubicin
YIL066C RNR3
ZShift = 1.89
Z lm Score = −0.45
Delta_AUC
NG = 0
DB = 0
SM = 0
Doxorubicin
YKR104W
ZShift = −2.58
Z lm Score = −2.01
Delta_L
NG = 0
DB = 0
SM = 0
Doxorubicin
YKR104W
ZShift = −0.23
Z lm Score = 1.47
Delta_K
NG = 0
DB = 0
SM = 0
Doxorubicin
YKR104W
ZShift = 1.83
Z lm Score = −1.7
Delta_r
NG = 0
DB = 0
SM = 0
Doxorubicin
YKR104W
ZShift = 2.05
Z lm Score = 1.76
Delta_AUC
NG = 0
DB = 0
SM = 0
Doxorubicin
YDR379W RGA2
ZShift = −2.38
Z lm Score = −2.02
Delta_L
NG = 0
DB = 0
SM = 0
Doxorubicin
YDR379W RGA2
ZShift = 1.64
Z lm Score = 1.26
Delta_K
NG = 0
DB = 0
SM = 0
Doxorubicin
YDR379W RGA2
ZShift = 1.77
Z lm Score = −1.48
Delta_r
NG = 0
DB = 0
SM = 0
Doxorubicin
YDR379W RGA2
ZShift = 3.07
Z lm Score = −0.1
Delta_AUC
NG = 0
DB = 0
SM = 0
Doxorubicin

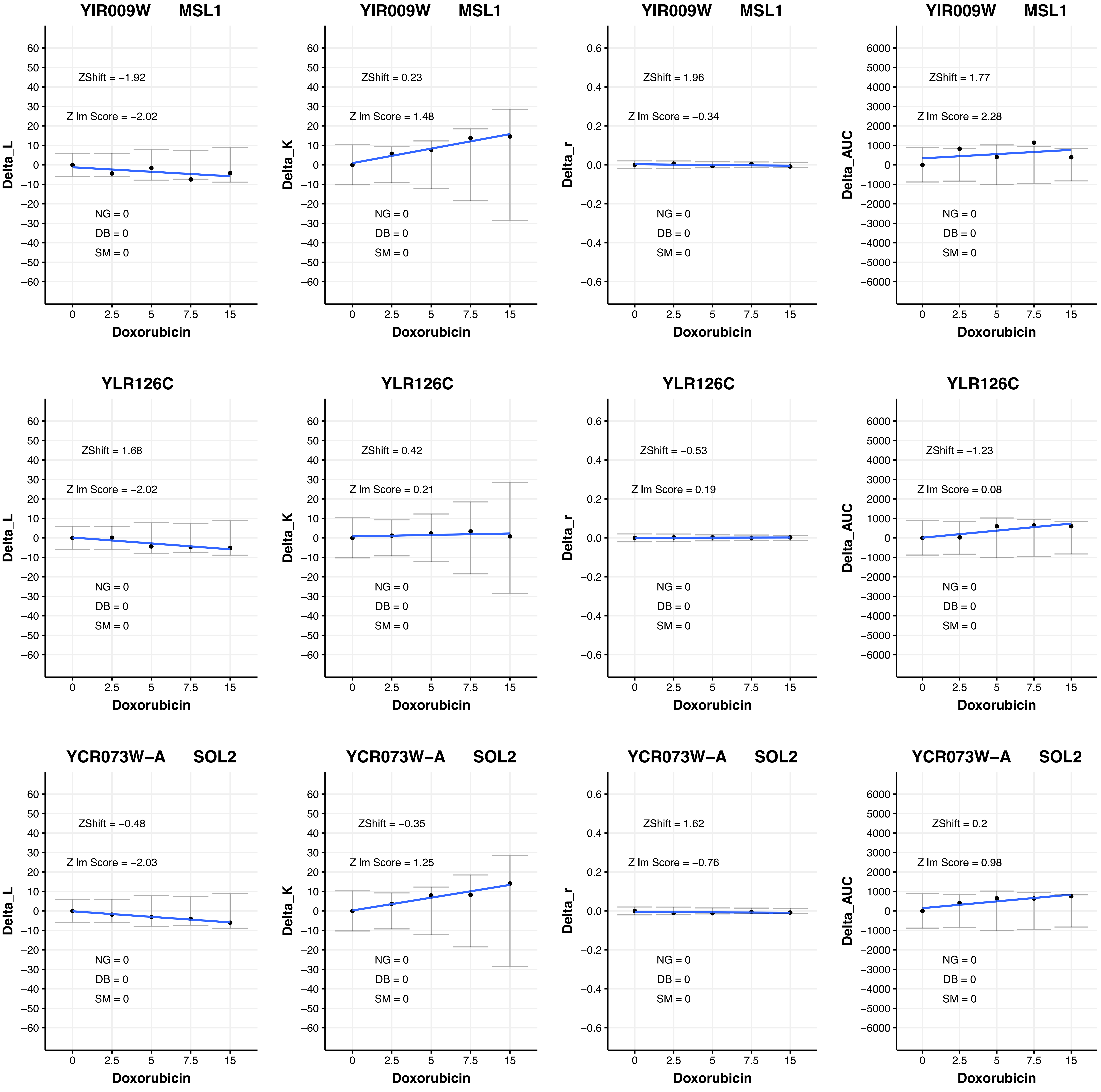

YIR009W MSL1
ZShift = −1.92
Z lm Score = −2.02
NG = 0
DB = 0
SM = 0
Delta_L
Doxorubicin
YIR009W MSL1
ZShift = 0.23
Z lm Score = 1.48
NG = 0
DB = 0
SM = 0
Delta_K
Doxorubicin
YIR009W MSL1
ZShift = 1.96
Z lm Score = −0.34
NG = 0
DB = 0
SM = 0
Delta_r
Doxorubicin
YIR009W MSL1
ZShift = 1.77
Z lm Score = 2.28
NG = 0
DB = 0
SM = 0
Delta_AUC
Doxorubicin
YLR126C
ZShift = 1.68
Z lm Score = −2.02
NG = 0
DB = 0
SM = 0
Delta_L
Doxorubicin
YLR126C
ZShift = 0.42
Z lm Score = 0.21
NG = 0
DB = 0
SM = 0
Delta_K
Doxorubicin
YLR126C
ZShift = −0.53
Z lm Score = 0.19
NG = 0
DB = 0
SM = 0
Delta_r
Doxorubicin
YLR126C
ZShift = −1.23
Z lm Score = 0.08
NG = 0
DB = 0
SM = 0
Delta_AUC
Doxorubicin
YCR073W−A SOL2
ZShift = −0.48
Z lm Score = −2.03
NG = 0
DB = 0
SM = 0
Delta_L
Doxorubicin
YCR073W−A SOL2
ZShift = −0.35
Z lm Score = 1.25
NG = 0
DB = 0
SM = 0
Delta_K
Doxorubicin
YCR073W−A SOL2
ZShift = 1.62
Z lm Score = −0.76
NG = 0
DB = 0
SM = 0
Delta_r
Doxorubicin
YCR073W−A SOL2
ZShift = 0.2
Z lm Score = 0.98
NG = 0
DB = 0
SM = 0
Delta_AUC
Doxorubicin

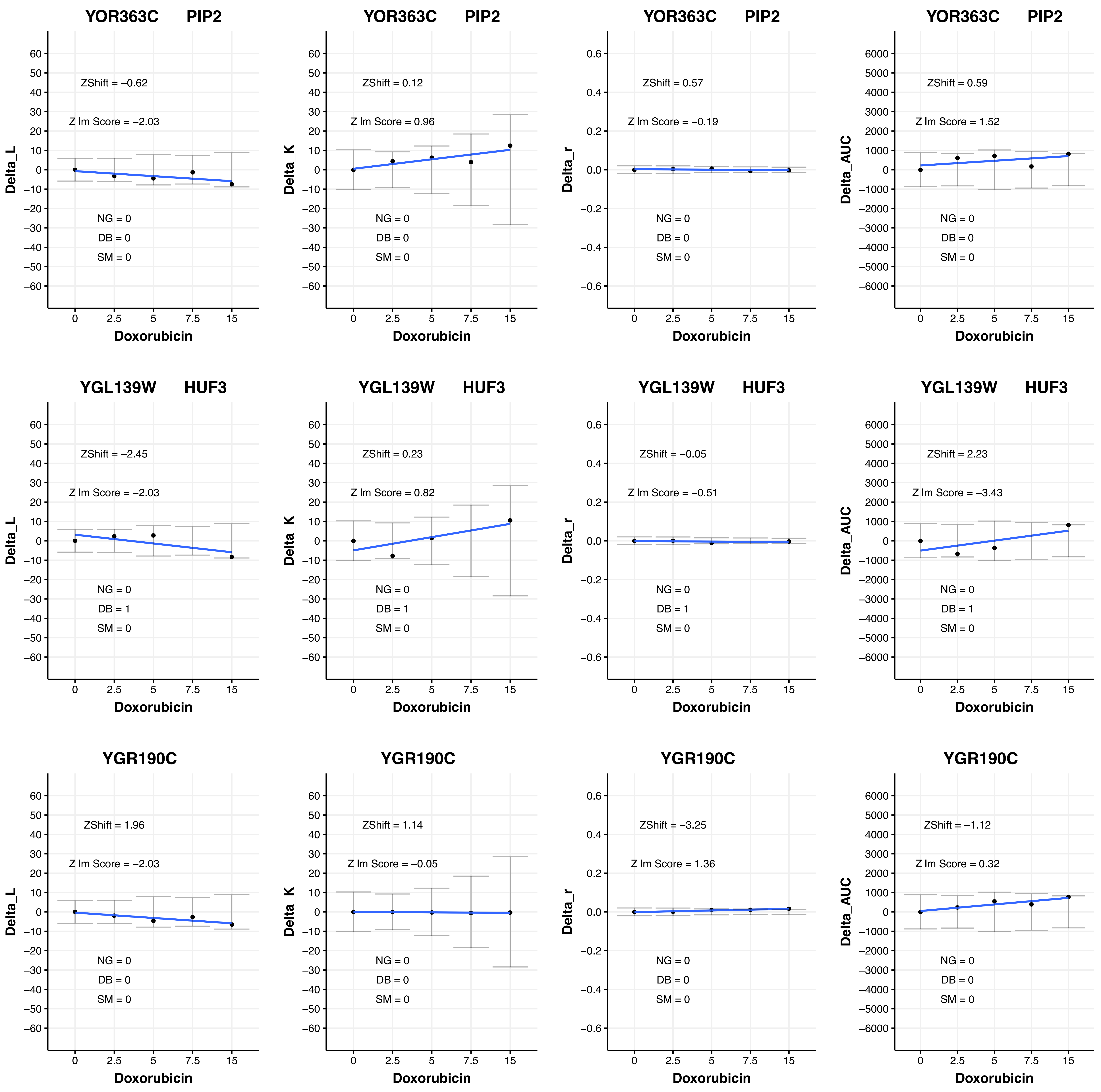
YOR363C PIP2
ZShift = −0.62
Z lm Score = −2.03
NG = 0
DB = 0
SM = 0
Delta_L
Doxorubicin
YOR363C PIP2
ZShift = 0.12
Z lm Score = 0.96
NG = 0
DB = 0
SM = 0
Delta_K
Doxorubicin
YOR363C PIP2
ZShift = 0.57
Z lm Score = −0.19
NG = 0
DB = 0
SM = 0
Delta_r
Doxorubicin
YOR363C PIP2
ZShift = 0.59
Z lm Score = 1.52
NG = 0
DB = 0
SM = 0
Delta_AUC
Doxorubicin
YGL139W HUF3
ZShift = −2.45
Z lm Score = −2.03
NG = 0
DB = 1
SM = 0
Delta_L
Doxorubicin
YGL139W HUF3
ZShift = 0.23
Z lm Score = 0.82
NG = 0
DB = 1
SM = 0
Delta_K
Doxorubicin
YGL139W HUF3
ZShift = −0.05
Z lm Score = −0.51
NG = 0
DB = 1
SM = 0
Delta_r
Doxorubicin
YGL139W HUF3
ZShift = 2.23
Z lm Score = −3.43
NG = 0
DB = 1
SM = 0
Delta_AUC
Doxorubicin
YGR190C
ZShift = 1.96
Z lm Score = −2.03
NG = 0
DB = 0
SM = 0
Delta_L
Doxorubicin
YGR190C
ZShift = 1.14
Z lm Score = −0.05
NG = 0
DB = 0
SM = 0
Delta_K
Doxorubicin
YGR190C
ZShift = −3.25
Z lm Score = 1.36
NG = 0
DB = 0
SM = 0
Delta_r
Doxorubicin
YGR190C
ZShift = −1.12
Z lm Score = 0.32
NG = 0
DB = 0
SM = 0
Delta_AUC
Doxorubicin

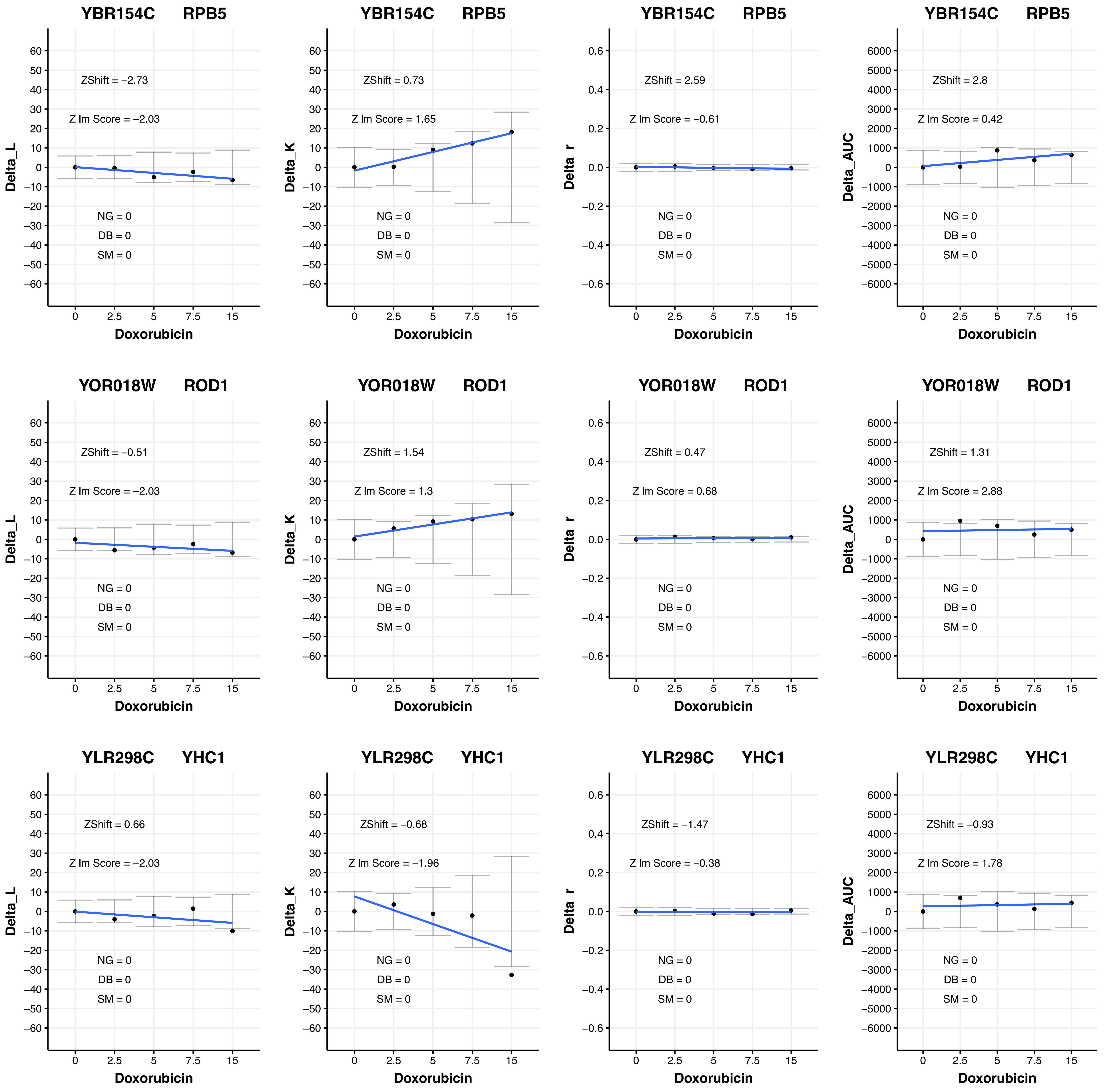
YBR154C RPB5
ZShift = −2.73
Z lm Score = −2.03
NG = 0
DB = 0
SM = 0
Delta_L
Doxorubicin
YBR154C RPB5
ZShift = 0.73
Z lm Score = 1.65
NG = 0
DB = 0
SM = 0
Delta_K
Doxorubicin
YBR154C RPB5
ZShift = 2.59
Z lm Score = −0.61
NG = 0
DB = 0
SM = 0
Delta_r
Doxorubicin
YBR154C RPB5
ZShift = 2.8
Z lm Score = 0.42
NG = 0
DB = 0
SM = 0
Delta_AUC
Doxorubicin
YOR018W ROD1
ZShift = −0.51
Z lm Score = −2.03
NG = 0
DB = 0
SM = 0
Delta_L
Doxorubicin
YOR018W ROD1
ZShift = 1.54
Z lm Score = 1.3
NG = 0
DB = 0
SM = 0
Delta_K
Doxorubicin
YOR018W ROD1
ZShift = 0.47
Z lm Score = 0.68
NG = 0
DB = 0
SM = 0
Delta_r
Doxorubicin
YOR018W ROD1
ZShift = 1.31
Z lm Score = 2.88
NG = 0
DB = 0
SM = 0
Delta_AUC
Doxorubicin
YLR298C YHC1
ZShift = 0.66
Z lm Score = −2.03
NG = 0
DB = 0
SM = 0
Delta_L
Doxorubicin
YLR298C YHC1
ZShift = −0.68
Z lm Score = −1.96
NG = 0
DB = 0
SM = 0
Delta_K
Doxorubicin
YLR298C YHC1
ZShift = −1.47
Z lm Score = −0.38
NG = 0
DB = 0
SM = 0
Delta_r
Doxorubicin
YLR298C YHC1
ZShift = −0.93
Z lm Score = 1.78
NG = 0
DB = 0
SM = 0
Delta_AUC
Doxorubicin

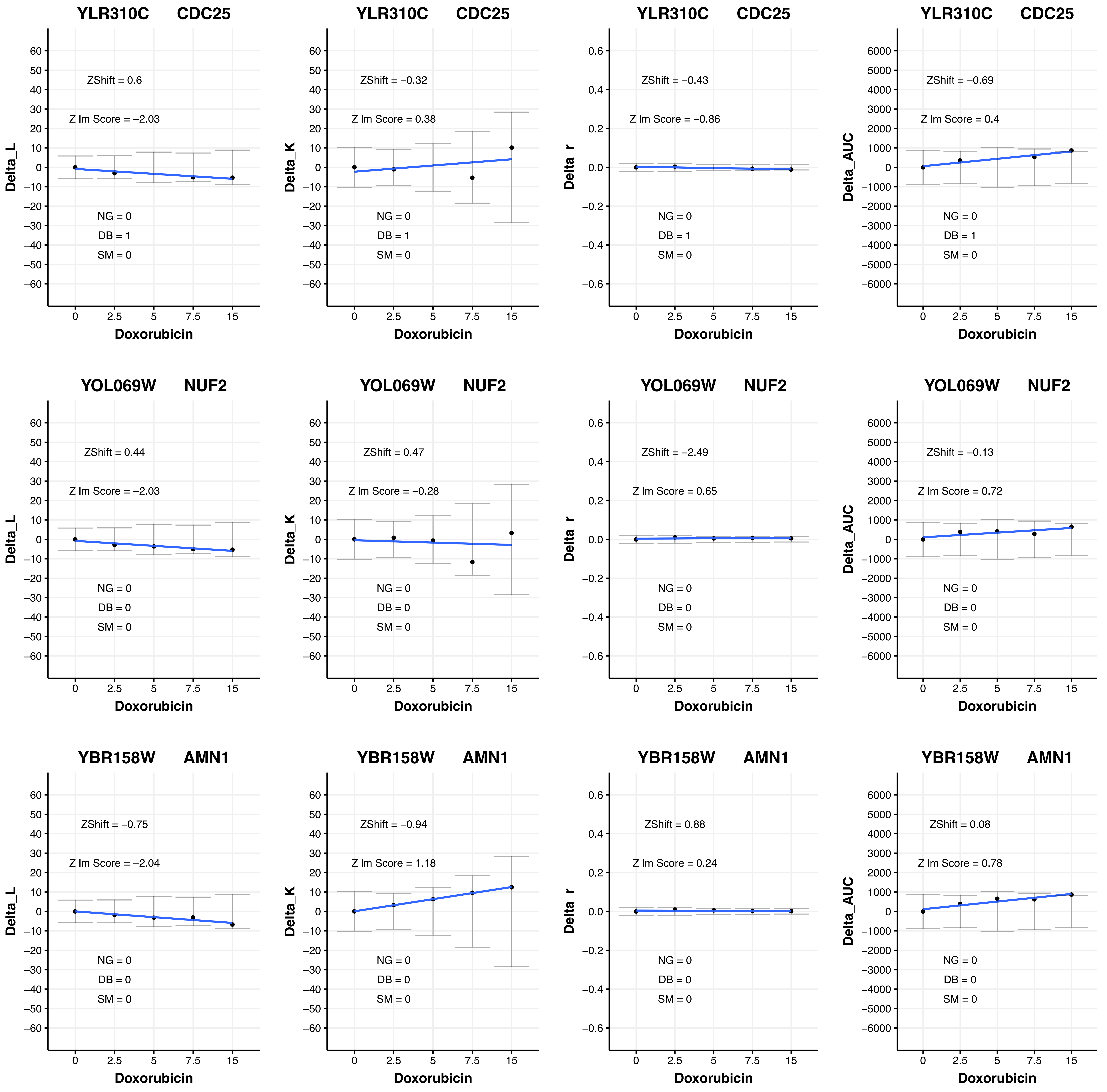
YLR310C CDC25
ZShift = 0.6
Z lm Score = −2.03
NG = 0
DB = 1
SM = 0
Delta_L
Doxorubicin
YLR310C CDC25
ZShift = −0.32
Z lm Score = 0.38
NG = 0
DB = 1
SM = 0
Delta_K
Doxorubicin
YLR310C CDC25
ZShift = −0.43
Z lm Score = −0.86
NG = 0
DB = 1
SM = 0
Delta_r
Doxorubicin
YLR310C CDC25
ZShift = −0.69
Z lm Score = 0.4
NG = 0
DB = 1
SM = 0
Delta_AUC
Doxorubicin
YOL069W NUF2
ZShift = 0.44
Z lm Score = −2.03
NG = 0
DB = 0
SM = 0
Delta_L
Doxorubicin
YOL069W NUF2
ZShift = 0.47
Z lm Score = −0.28
NG = 0
DB = 0
SM = 0
Delta_K
Doxorubicin
YOL069W NUF2
ZShift = −2.49
Z lm Score = 0.65
NG = 0
DB = 0
SM = 0
Delta_r
Doxorubicin
YOL069W NUF2
ZShift = −0.13
Z lm Score = 0.72
NG = 0
DB = 0
SM = 0
Delta_AUC
Doxorubicin
YBR158W AMN1
ZShift = −0.75
Z lm Score = −2.04
NG = 0
DB = 0
SM = 0
Delta_L
Doxorubicin
YBR158W AMN1
ZShift = −0.94
Z lm Score = 1.18
NG = 0
DB = 0
SM = 0
Delta_K
Doxorubicin
YBR158W AMN1
ZShift = 0.88
Z lm Score = 0.24
NG = 0
DB = 0
SM = 0
Delta_r
Doxorubicin
YBR158W AMN1
ZShift = 0.08
Z lm Score = 0.78
NG = 0
DB = 0
SM = 0
Delta_AUC
Doxorubicin

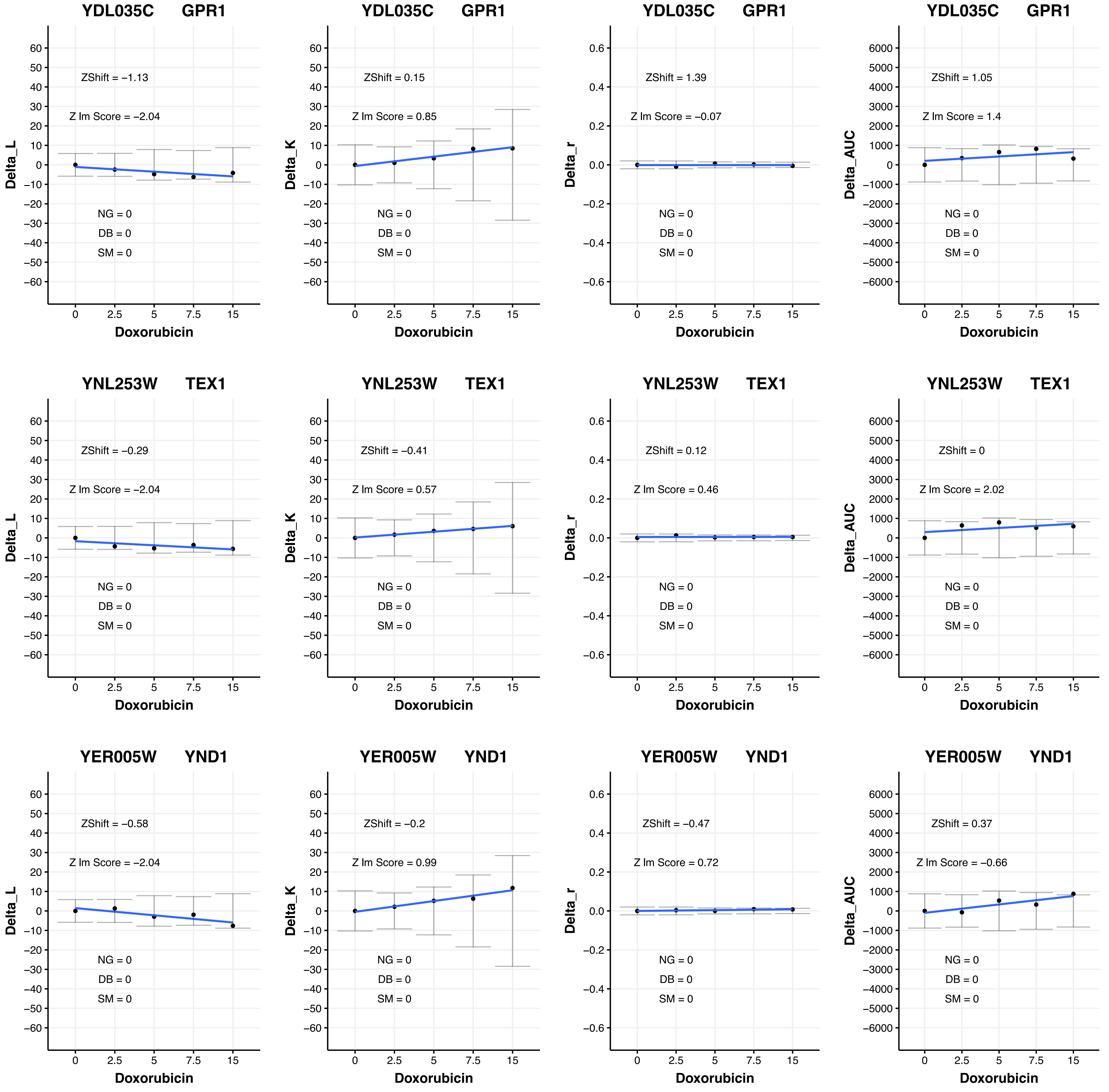

YDL035C GPR1
Delta_L
Doxorubicin
ZShift = −1.13
Z lm Score = −2.04
NG = 0
DB = 0
SM = 0
YDL035C GPR1
Delta_K
Doxorubicin
ZShift = 0.15
Z lm Score = 0.85
NG = 0
DB = 0
SM = 0
YDL035C GPR1
Delta_r
Doxorubicin
ZShift = 1.39
Z lm Score = −0.07
NG = 0
DB = 0
SM = 0
YDL035C GPR1
Delta_AUC
Doxorubicin
ZShift = 1.05
Z lm Score = 1.4
NG = 0
DB = 0
SM = 0
YNL253W TEX1
Delta_L
Doxorubicin
ZShift = −0.29
Z lm Score = −2.04
NG = 0
DB = 0
SM = 0
YNL253W TEX1
Delta_K
Doxorubicin
ZShift = −0.41
Z lm Score = 0.57
NG = 0
DB = 0
SM = 0
YNL253W TEX1
Delta_r
Doxorubicin
ZShift = 0.12
Z lm Score = 0.46
NG = 0
DB = 0
SM = 0
YNL253W TEX1
Delta_AUC
Doxorubicin
ZShift = 0
Z lm Score = 2.02
NG = 0
DB = 0
SM = 0
YER005W YND1
Delta_L
Doxorubicin
ZShift = −0.58
Z lm Score = −2.04
NG = 0
DB = 0
SM = 0
YER005W YND1
Delta_K
Doxorubicin
ZShift = −0.2
Z lm Score = 0.99
NG = 0
DB = 0
SM = 0
YER005W YND1
Delta_r
Doxorubicin
ZShift = −0.47
Z lm Score = 0.72
NG = 0
DB = 0
SM = 0
YER005W YND1
Delta_AUC
Doxorubicin
ZShift = 0.37
Z lm Score = −0.66
NG = 0
DB = 0
SM = 0

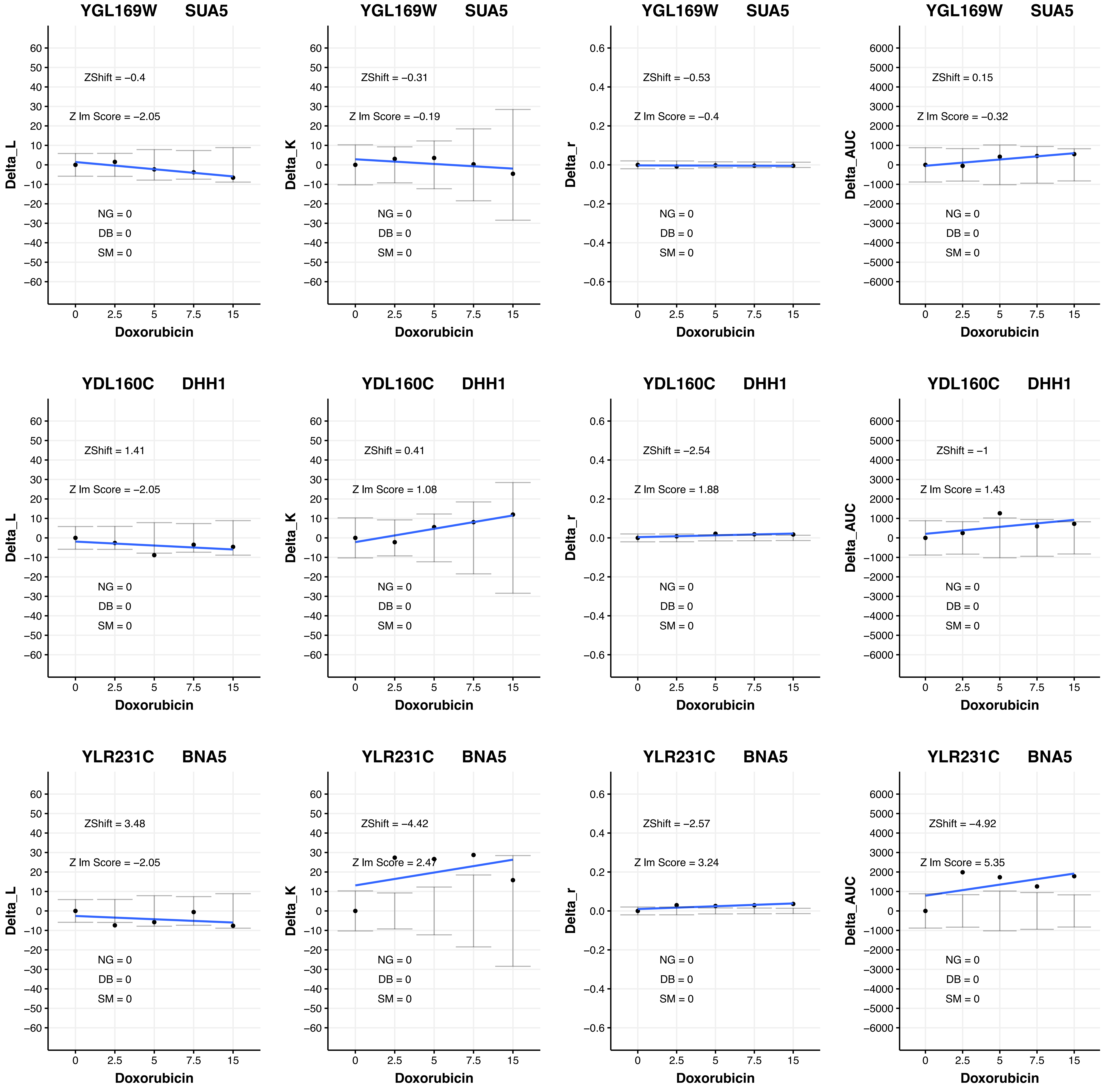
YGL169W SUA5
ZShift = −0.4
Z lm Score = −2.05
NG = 0
DB = 0
SM = 0
Delta_L
Doxorubicin
YGL169W SUA5
ZShift = −0.31
Z lm Score = −0.19
NG = 0
DB = 0
SM = 0
Delta_K
Doxorubicin
YGL169W SUA5
ZShift = −0.53
Z lm Score = −0.4
NG = 0
DB = 0
SM = 0
Delta_r
Doxorubicin
YGL169W SUA5
ZShift = 0.15
Z lm Score = −0.32
NG = 0
DB = 0
SM = 0
Delta_AUC
Doxorubicin
YDL160C DHH1
ZShift = 1.41
Z lm Score = −2.05
NG = 0
DB = 0
SM = 0
Delta_L
Doxorubicin
YDL160C DHH1
ZShift = 0.41
Z lm Score = 1.08
NG = 0
DB = 0
SM = 0
Delta_K
Doxorubicin
YDL160C DHH1
ZShift = −2.54
Z lm Score = 1.88
NG = 0
DB = 0
SM = 0
Delta_r
Doxorubicin
YDL160C DHH1
ZShift = −1
Z lm Score = 1.43
NG = 0
DB = 0
SM = 0
Delta_AUC
Doxorubicin
YLR231C BNA5
ZShift = 3.48
Z lm Score = −2.05
NG = 0
DB = 0
SM = 0
Delta_L
Doxorubicin
YLR231C BNA5
ZShift = −4.42
Z lm Score = 2.47
NG = 0
DB = 0
SM = 0
Delta_K
Doxorubicin
YLR231C BNA5
ZShift = −2.57
Z lm Score = 3.24
NG = 0
DB = 0
SM = 0
Delta_r
Doxorubicin
YLR231C BNA5
ZShift = −4.92
Z lm Score = 5.35
NG = 0
DB = 0
SM = 0
Delta_AUC
Doxorubicin

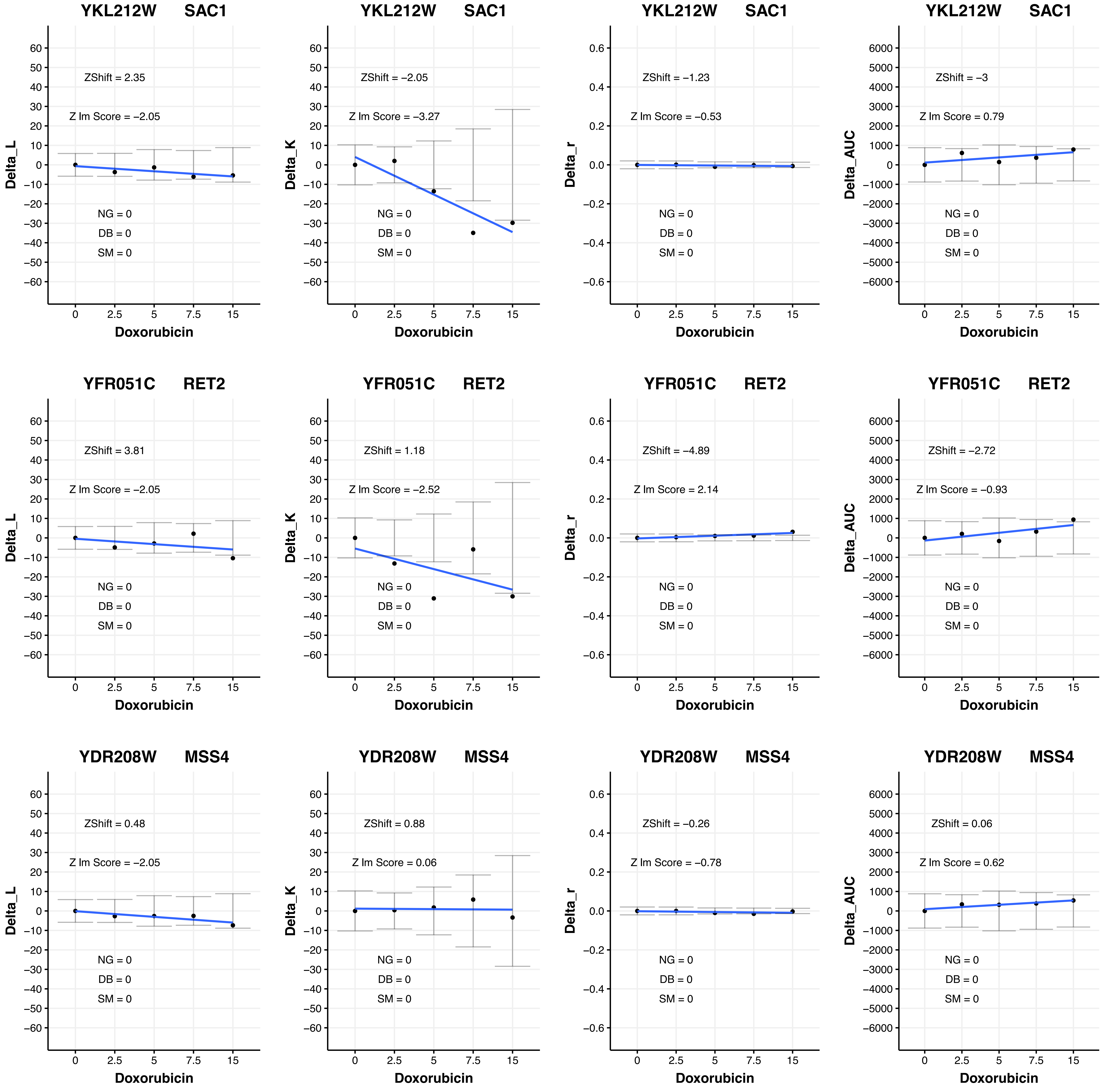
YKL212W SAC1
ZShift = 2.35
Z lm Score = −2.05
NG = 0
DB = 0
SM = 0
Delta_L
Doxorubicin
YKL212W SAC1
ZShift = −2.05
Z lm Score = −3.27
NG = 0
DB = 0
SM = 0
Delta_K
Doxorubicin
YKL212W SAC1
ZShift = −1.23
Z lm Score = −0.53
NG = 0
DB = 0
SM = 0
Delta_r
Doxorubicin
YKL212W SAC1
ZShift = −3
Z lm Score = 0.79
NG = 0
DB = 0
SM = 0
Delta_AUC
Doxorubicin
YFR051C RET2
ZShift = 3.81
Z lm Score = −2.05
NG = 0
DB = 0
SM = 0
Delta_L
Doxorubicin
YFR051C RET2
ZShift = 1.18
Z lm Score = −2.52
NG = 0
DB = 0
SM = 0
Delta_K
Doxorubicin
YFR051C RET2
ZShift = −4.89
Z lm Score = 2.14
NG = 0
DB = 0
SM = 0
Delta_r
Doxorubicin
YFR051C RET2
ZShift = −2.72
Z lm Score = −0.93
NG = 0
DB = 0
SM = 0
Delta_AUC
Doxorubicin
YDR208W MSS4
ZShift = 0.48
Z lm Score = −2.05
NG = 0
DB = 0
SM = 0
Delta_L
Doxorubicin
YDR208W MSS4
ZShift = 0.88
Z lm Score = 0.06
NG = 0
DB = 0
SM = 0
Delta_K
Doxorubicin
YDR208W MSS4
ZShift = −0.26
Z lm Score = −0.78
NG = 0
DB = 0
SM = 0
Delta_r
Doxorubicin
YDR208W MSS4
ZShift = 0.06
Z lm Score = 0.62
NG = 0
DB = 0
SM = 0
Delta_AUC
Doxorubicin

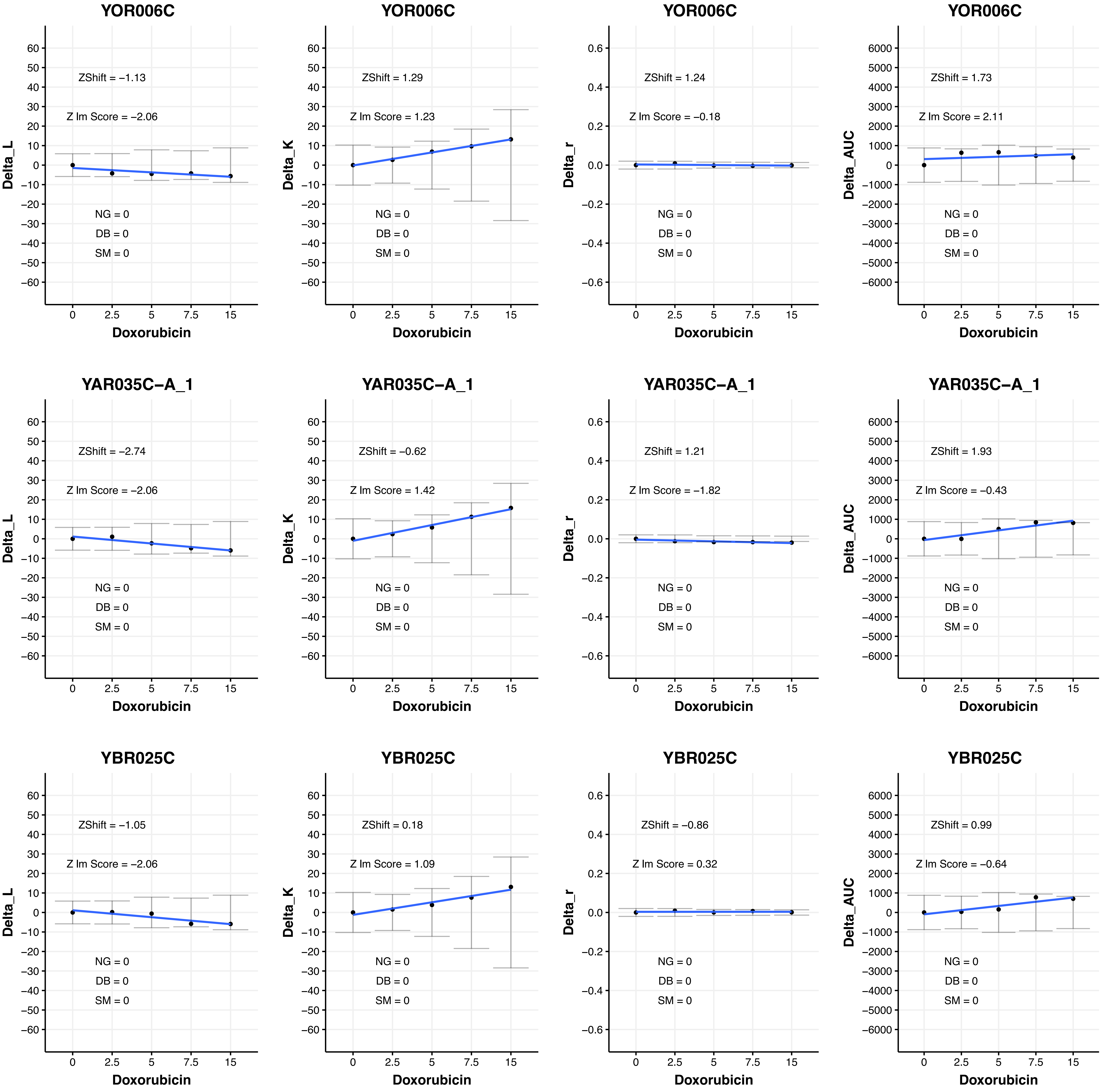
YOR006C
ZShift = −1.13
Z lm Score = −2.06
NG = 0
DB = 0
SM = 0
Delta_L
Doxorubicin
YOR006C
ZShift = 1.29
Z lm Score = 1.23
NG = 0
DB = 0
SM = 0
Delta_K
Doxorubicin
YOR006C
ZShift = 1.24
Z lm Score = −0.18
NG = 0
DB = 0
SM = 0
Delta_r
Doxorubicin
YOR006C
ZShift = 1.73
Z lm Score = 2.11
NG = 0
DB = 0
SM = 0
Delta_AUC
Doxorubicin
YAR035C−A_1
ZShift = −2.74
Z lm Score = −2.06
NG = 0
DB = 0
SM = 0
Delta_L
Doxorubicin
YAR035C−A_1
ZShift = −0.62
Z lm Score = 1.42
NG = 0
DB = 0
SM = 0
Delta_K
Doxorubicin
YAR035C−A_1
ZShift = 1.21
Z lm Score = −1.82
NG = 0
DB = 0
SM = 0
Delta_r
Doxorubicin
YAR035C−A_1
ZShift = 1.93
Z lm Score = −0.43
NG = 0
DB = 0
SM = 0
Delta_AUC
Doxorubicin
YBR025C
ZShift = −1.05
Z lm Score = −2.06
NG = 0
DB = 0
SM = 0
Delta_L
Doxorubicin
YBR025C
ZShift = 0.18
Z lm Score = 1.09
NG = 0
DB = 0
SM = 0
Delta_K
Doxorubicin
YBR025C
ZShift = −0.86
Z lm Score = 0.32
NG = 0
DB = 0
SM = 0
Delta_r
Doxorubicin
YBR025C
ZShift = 0.99
Z lm Score = −0.64
NG = 0
DB = 0
SM = 0
Delta_AUC
Doxorubicin

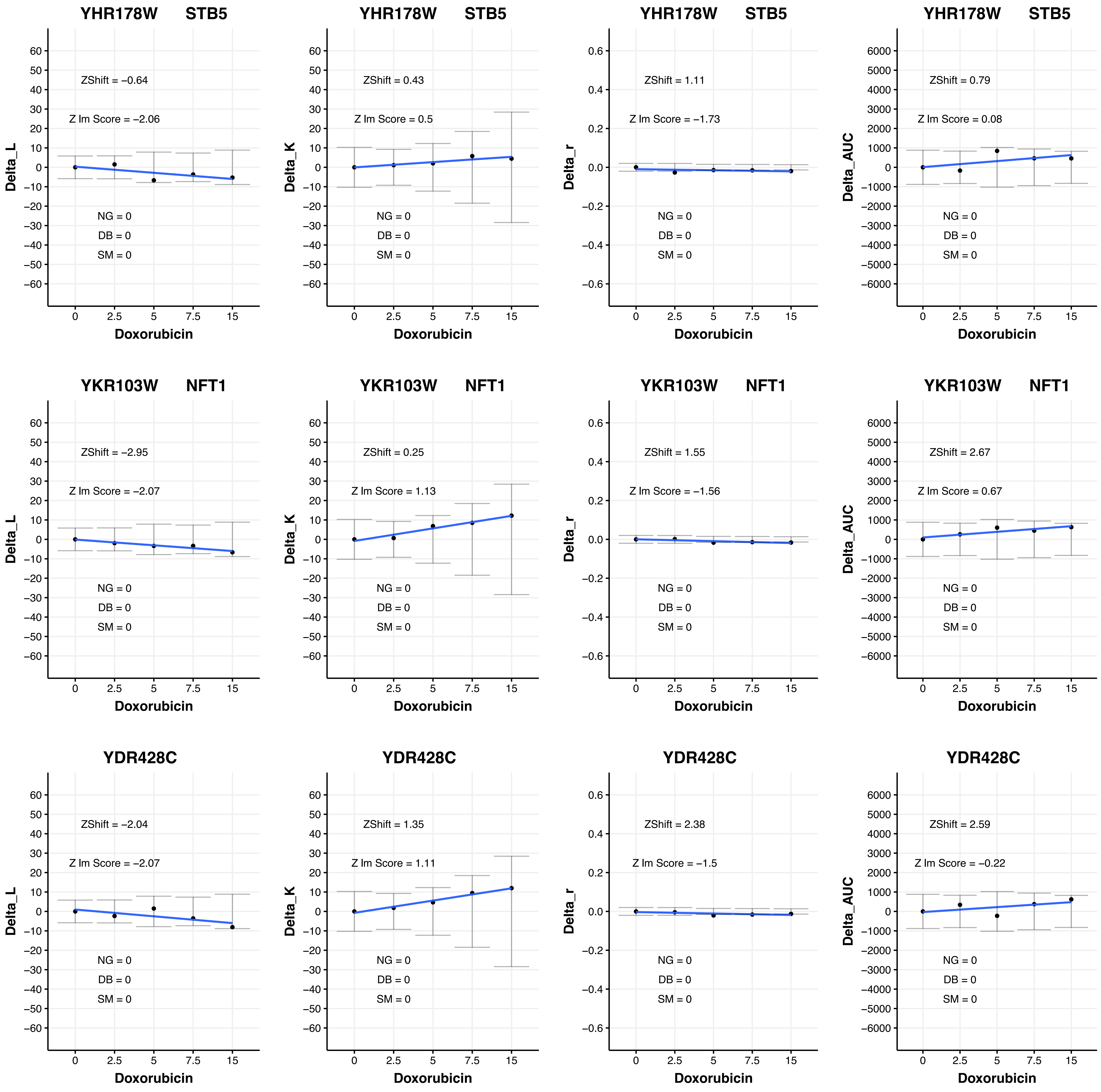

YHR178W STB5
ZShift = −0.64
Z lm Score = −2.06
Delta_L
NG = 0
DB = 0
SM = 0
Doxorubicin
YHR178W STB5
ZShift = 0.43
Z lm Score = 0.5
Delta_K
NG = 0
DB = 0
SM = 0
Doxorubicin
YHR178W STB5
ZShift = 1.11
Z lm Score = −1.73
Delta_r
NG = 0
DB = 0
SM = 0
Doxorubicin
YHR178W STB5
ZShift = 0.79
Z lm Score = 0.08
Delta_AUC
NG = 0
DB = 0
SM = 0
Doxorubicin
YKR103W NFT1
ZShift = −2.95
Z lm Score = −2.07
Delta_L
NG = 0
DB = 0
SM = 0
Doxorubicin
YKR103W NFT1
ZShift = 0.25
Z lm Score = 1.13
Delta_K
NG = 0
DB = 0
SM = 0
Doxorubicin
YKR103W NFT1
ZShift = 1.55
Z lm Score = −1.56
Delta_r
NG = 0
DB = 0
SM = 0
Doxorubicin
YKR103W NFT1
ZShift = 2.67
Z lm Score = 0.67
Delta_AUC
NG = 0
DB = 0
SM = 0
Doxorubicin
YDR428C
ZShift = −2.04
Z lm Score = −2.07
Delta_L
NG = 0
DB = 0
SM = 0
Doxorubicin
YDR428C
ZShift = 1.35
Z lm Score = 1.11
Delta_K
NG = 0
DB = 0
SM = 0
Doxorubicin
YDR428C
ZShift = 2.38
Z lm Score = −1.5
Delta_r
NG = 0
DB = 0
SM = 0
Doxorubicin
YDR428C
ZShift = 2.59
Z lm Score = −0.22
Delta_AUC
NG = 0
DB = 0
SM = 0
Doxorubicin

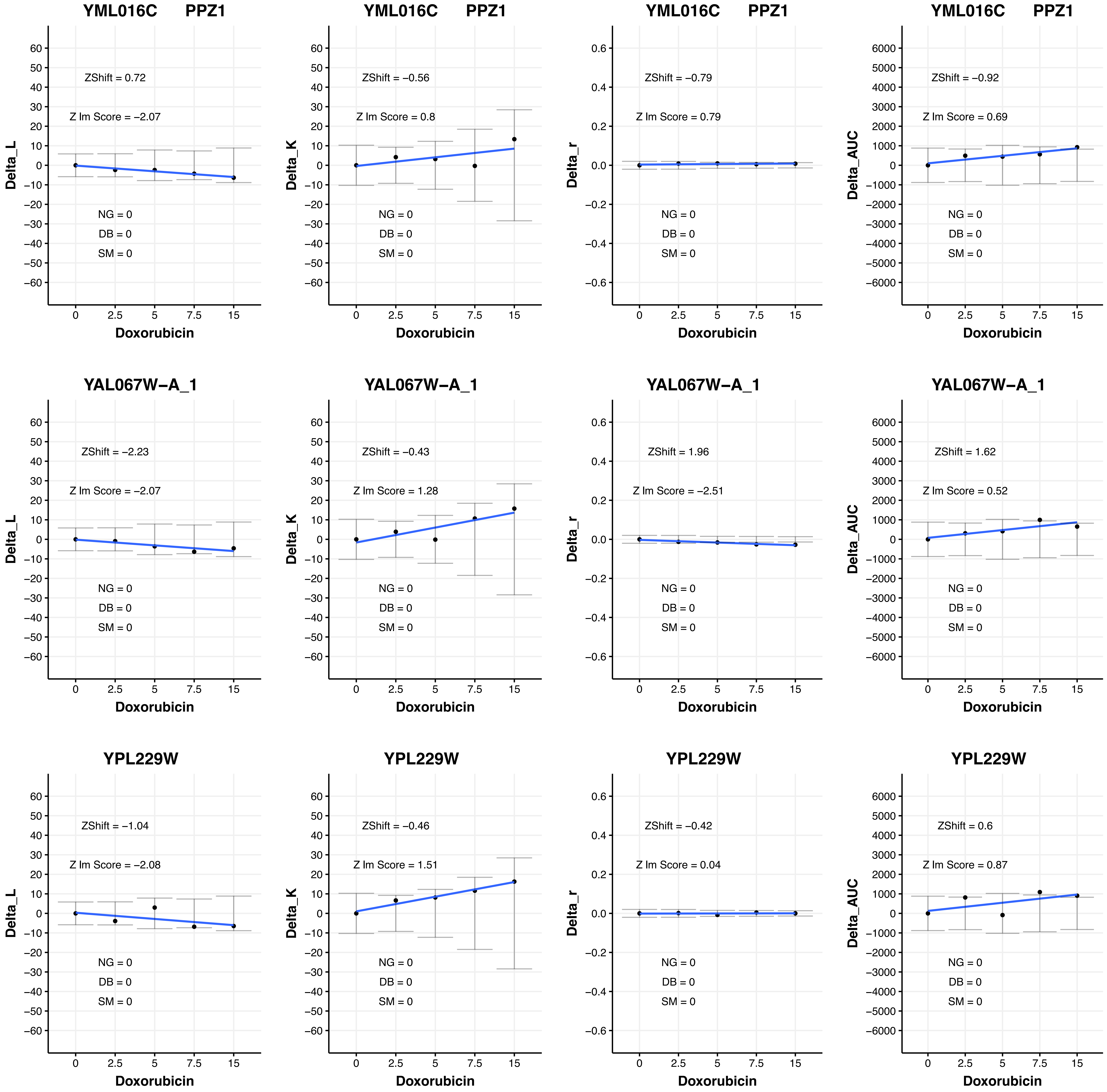

YML016C PPZ1
ZShift = 0.72
Z lm Score = −2.07
NG = 0
DB = 0
SM = 0
Delta_L
Doxorubicin
YML016C PPZ1
ZShift = −0.56
Z lm Score = 0.8
NG = 0
DB = 0
SM = 0
Delta_K
Doxorubicin
YML016C PPZ1
ZShift = −0.79
Z lm Score = 0.79
NG = 0
DB = 0
SM = 0
Delta_r
Doxorubicin
YML016C PPZ1
ZShift = −0.92
Z lm Score = 0.69
NG = 0
DB = 0
SM = 0
Delta_AUC
Doxorubicin
YAL067W−A_1
ZShift = −2.23
Z lm Score = −2.07
NG = 0
DB = 0
SM = 0
Delta_L
Doxorubicin
YAL067W−A_1
ZShift = −0.43
Z lm Score = 1.28
NG = 0
DB = 0
SM = 0
Delta_K
Doxorubicin
YAL067W−A_1
ZShift = 1.96
Z lm Score = −2.51
NG = 0
DB = 0
SM = 0
Delta_r
Doxorubicin
YAL067W−A_1
ZShift = 1.62
Z lm Score = 0.52
NG = 0
DB = 0
SM = 0
Delta_AUC
Doxorubicin
YPL229W
ZShift = −1.04
Z lm Score = −2.08
NG = 0
DB = 0
SM = 0
Delta_L
Doxorubicin
YPL229W
ZShift = −0.46
Z lm Score = 1.51
NG = 0
DB = 0
SM = 0
Delta_K
Doxorubicin
YPL229W
ZShift = −0.42
Z lm Score = 0.04
NG = 0
DB = 0
SM = 0
Delta_r
Doxorubicin
YPL229W
ZShift = 0.6
Z lm Score = 0.87
NG = 0
DB = 0
SM = 0
Delta_AUC
Doxorubicin

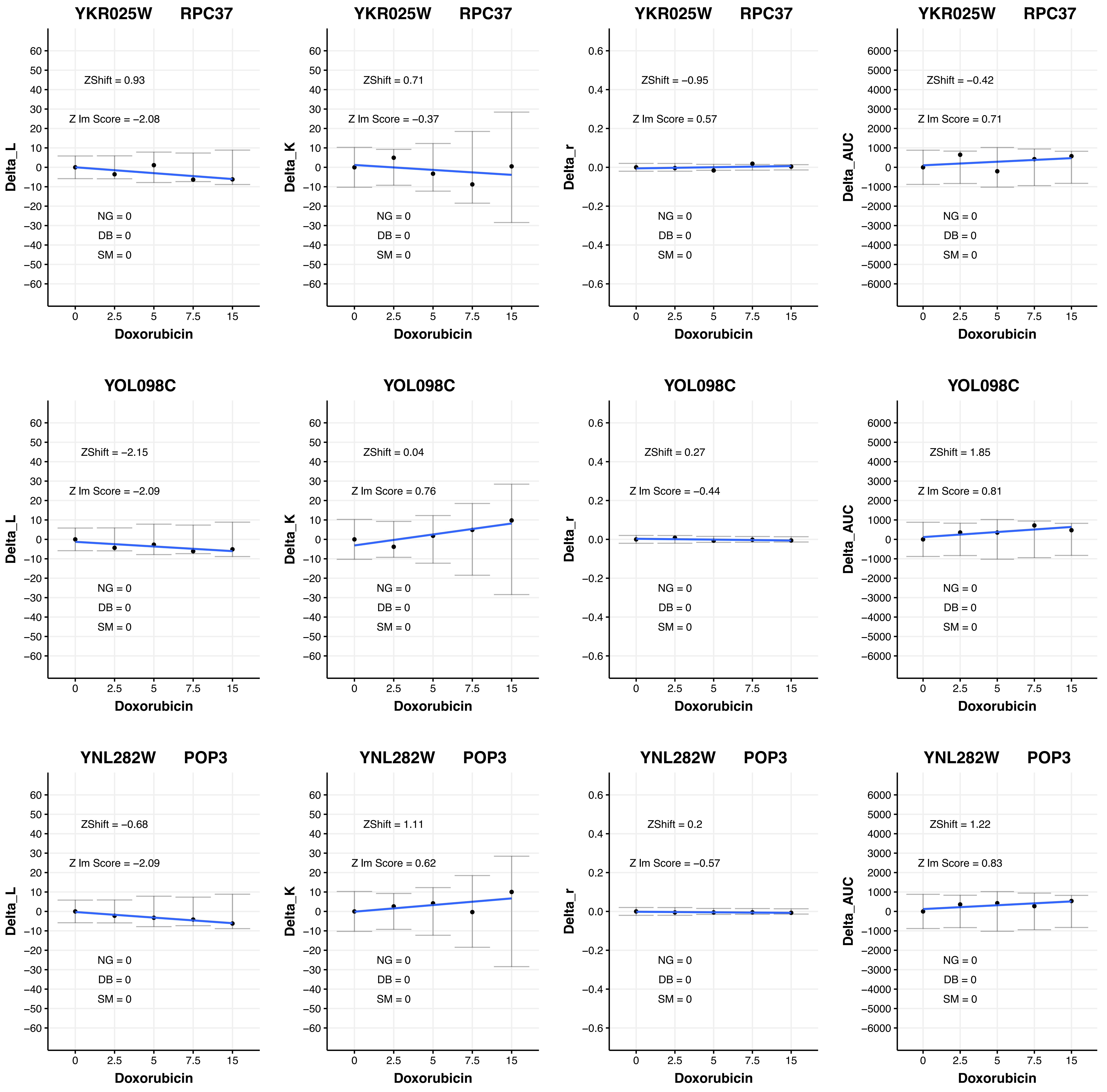
YKR025W RPC37
ZShift = 0.93
Z lm Score = −2.08
NG = 0
DB = 0
SM = 0
Delta_L
Doxorubicin
YKR025W RPC37
ZShift = 0.71
Z lm Score = −0.37
NG = 0
DB = 0
SM = 0
Delta_K
Doxorubicin
YKR025W RPC37
ZShift = −0.95
Z lm Score = 0.57
NG = 0
DB = 0
SM = 0
Delta_r
Doxorubicin
YKR025W RPC37
ZShift = −0.42
Z lm Score = 0.71
NG = 0
DB = 0
SM = 0
Delta_AUC
Doxorubicin
YOL098C
ZShift = −2.15
Z lm Score = −2.09
NG = 0
DB = 0
SM = 0
Delta_L
Doxorubicin
YOL098C
ZShift = 0.04
Z lm Score = 0.76
NG = 0
DB = 0
SM = 0
Delta_K
Doxorubicin
YOL098C
ZShift = 0.27
Z lm Score = −0.44
NG = 0
DB = 0
SM = 0
Delta_r
Doxorubicin
YOL098C
ZShift = 1.85
Z lm Score = 0.81
NG = 0
DB = 0
SM = 0
Delta_AUC
Doxorubicin
YNL282W POP3
ZShift = −0.68
Z lm Score = −2.09
NG = 0
DB = 0
SM = 0
Delta_L
Doxorubicin
YNL282W POP3
ZShift = 1.11
Z lm Score = 0.62
NG = 0
DB = 0
SM = 0
Delta_K
Doxorubicin
YNL282W POP3
ZShift = 0.2
Z lm Score = −0.57
NG = 0
DB = 0
SM = 0
Delta_r
Doxorubicin
YNL282W POP3
ZShift = 1.22
Z lm Score = 0.83
NG = 0
DB = 0
SM = 0
Delta_AUC
Doxorubicin

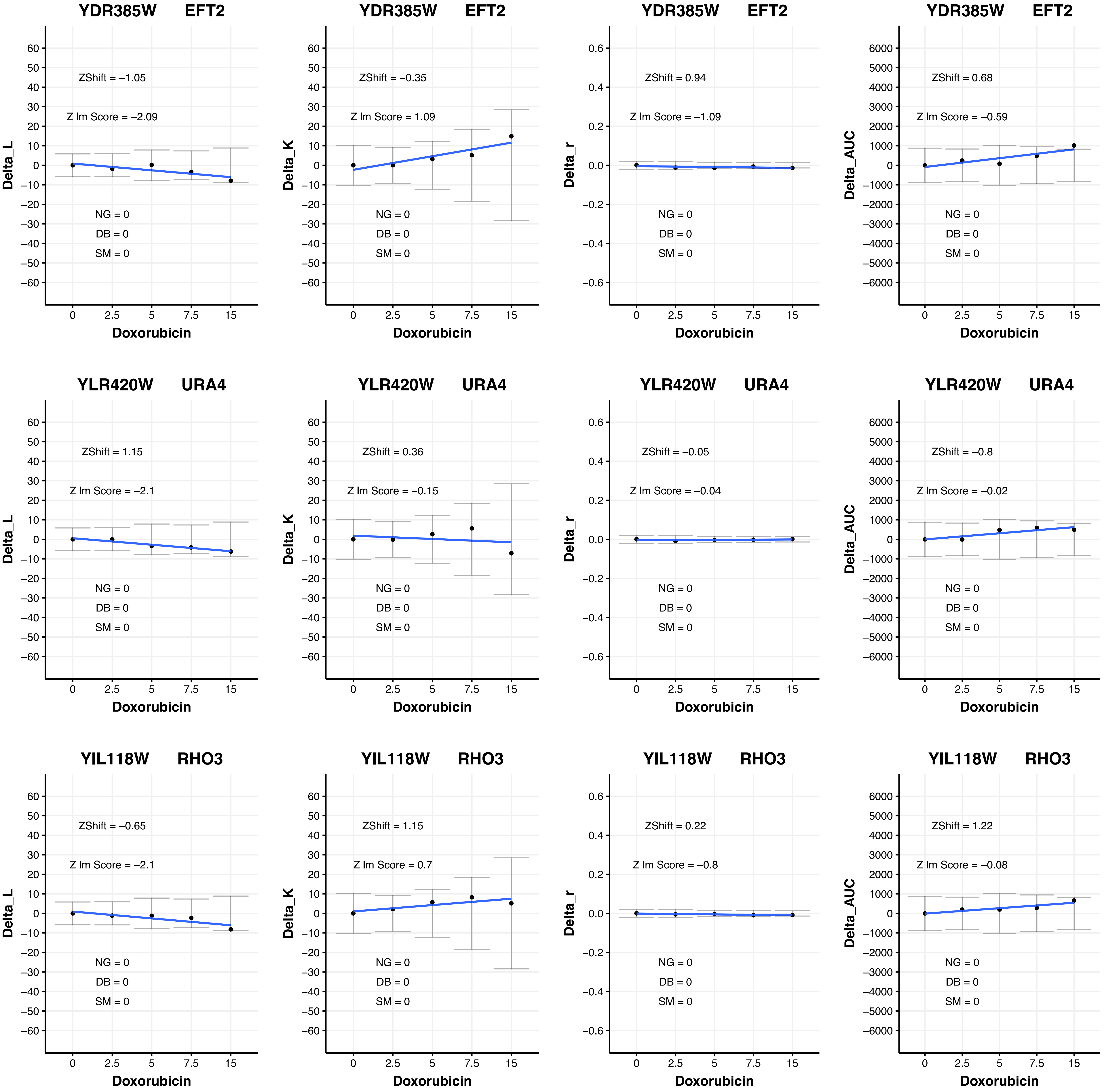

YDR385W EFT2
ZShift = −1.05
Z lm Score = −2.09
NG = 0
DB = 0
SM = 0
Delta_L
Doxorubicin
YDR385W EFT2
ZShift = −0.35
Z lm Score = 1.09
NG = 0
DB = 0
SM = 0
Delta_K
Doxorubicin
YDR385W EFT2
ZShift = 0.94
Z lm Score = −1.09
NG = 0
DB = 0
SM = 0
Delta_r
Doxorubicin
YDR385W EFT2
ZShift = 0.68
Z lm Score = −0.59
NG = 0
DB = 0
SM = 0
Delta_AUC
Doxorubicin
YLR420W URA4
ZShift = 1.15
Z lm Score = −2.1
NG = 0
DB = 0
SM = 0
Delta_L
Doxorubicin
YLR420W URA4
ZShift = 0.36
Z lm Score = −0.15
NG = 0
DB = 0
SM = 0
Delta_K
Doxorubicin
YLR420W URA4
ZShift = −0.05
Z lm Score = −0.04
NG = 0
DB = 0
SM = 0
Delta_r
Doxorubicin
YLR420W URA4
ZShift = −0.8
Z lm Score = −0.02
NG = 0
DB = 0
SM = 0
Delta_AUC
Doxorubicin
YIL118W RHO3
ZShift = −0.65
Z lm Score = −2.1
NG = 0
DB = 0
SM = 0
Delta_L
Doxorubicin
YIL118W RHO3
ZShift = 1.15
Z lm Score = 0.7
NG = 0
DB = 0
SM = 0
Delta_K
Doxorubicin
YIL118W RHO3
ZShift = 0.22
Z lm Score = −0.8
NG = 0
DB = 0
SM = 0
Delta_r
Doxorubicin
YIL118W RHO3
ZShift = 1.22
Z lm Score = −0.08
NG = 0
DB = 0
SM = 0
Delta_AUC
Doxorubicin

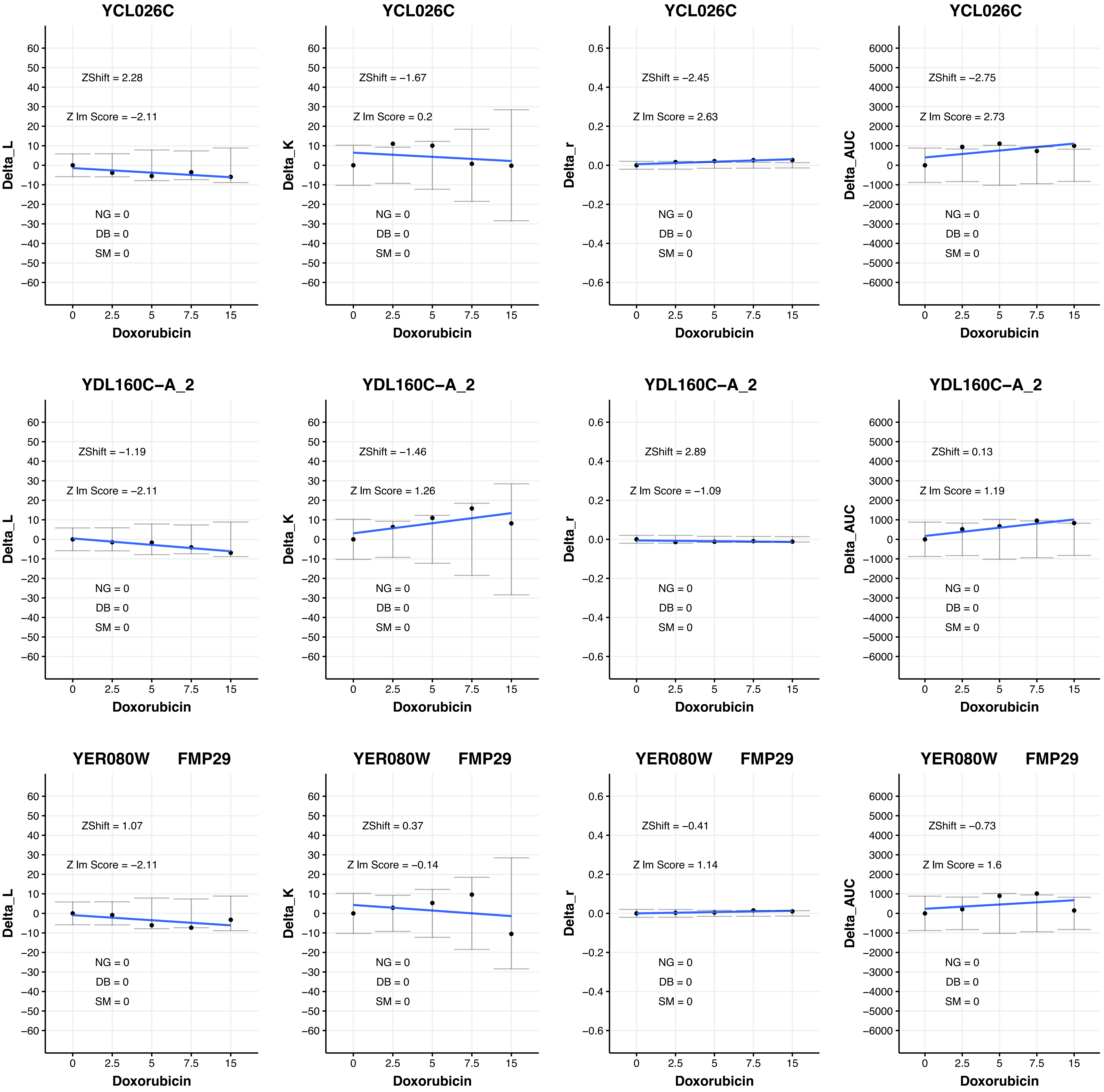

YCL026C
ZShift = 2.28
Z lm Score = −2.11
NG = 0
DB = 0
SM = 0
Delta_L
Doxorubicin
YCL026C
ZShift = −1.67
Z lm Score = 0.2
NG = 0
DB = 0
SM = 0
Delta_K
Doxorubicin
YCL026C
ZShift = −2.45
Z lm Score = 2.63
NG = 0
DB = 0
SM = 0
Delta_r
Doxorubicin
YCL026C
ZShift = −2.75
Z lm Score = 2.73
NG = 0
DB = 0
SM = 0
Delta_AUC
Doxorubicin
YDL160C−A_2
ZShift = −1.19
Z lm Score = −2.11
NG = 0
DB = 0
SM = 0
Delta_L
Doxorubicin
YDL160C−A_2
ZShift = −1.46
Z lm Score = 1.26
NG = 0
DB = 0
SM = 0
Delta_K
Doxorubicin
YDL160C−A_2
ZShift = 2.89
Z lm Score = −1.09
NG = 0
DB = 0
SM = 0
Delta_r
Doxorubicin
YDL160C−A_2
ZShift = 0.13
Z lm Score = 1.19
NG = 0
DB = 0
SM = 0
Delta_AUC
Doxorubicin
YER080W FMP29
ZShift = 1.07
Z lm Score = −2.11
NG = 0
DB = 0
SM = 0
Delta_L
Doxorubicin
YER080W FMP29
ZShift = 0.37
Z lm Score = −0.14
NG = 0
DB = 0
SM = 0
Delta_K
Doxorubicin
YER080W FMP29
ZShift = −0.41
Z lm Score = 1.14
NG = 0
DB = 0
SM = 0
Delta_r
Doxorubicin
YER080W FMP29
ZShift = −0.73
Z lm Score = 1.6
NG = 0
DB = 0
SM = 0
Delta_AUC
Doxorubicin

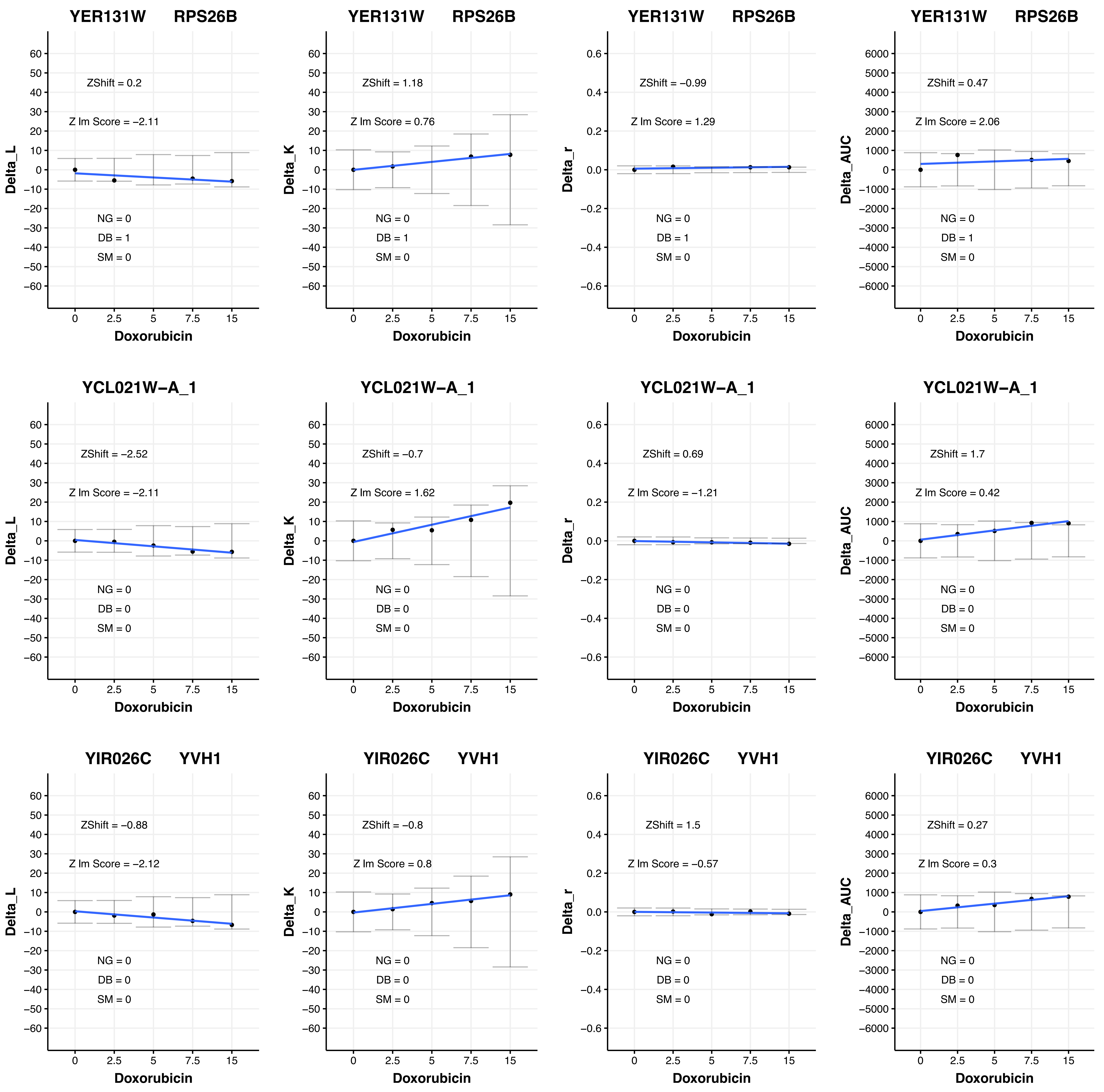
YER131W RPS26B
ZShift = 0.2
Z lm Score = −2.11
NG = 0
DB = 1
SM = 0
Delta_L
Doxorubicin
YER131W RPS26B
ZShift = 1.18
Z lm Score = 0.76
NG = 0
DB = 1
SM = 0
Delta_K
Doxorubicin
YER131W RPS26B
ZShift = −0.99
Z lm Score = 1.29
NG = 0
DB = 1
SM = 0
Delta_r
Doxorubicin
YER131W RPS26B
ZShift = 0.47
Z lm Score = 2.06
NG = 0
DB = 1
SM = 0
Delta_AUC
Doxorubicin
YCL021W−A_1
ZShift = −2.52
Z lm Score = −2.11
NG = 0
DB = 0
SM = 0
Delta_L
Doxorubicin
YCL021W−A_1
ZShift = −0.7
Z lm Score = 1.62
NG = 0
DB = 0
SM = 0
Delta_K
Doxorubicin
YCL021W−A_1
ZShift = 0.69
Z lm Score = −1.21
NG = 0
DB = 0
SM = 0
Delta_r
Doxorubicin
YCL021W−A_1
ZShift = 1.7
Z lm Score = 0.42
NG = 0
DB = 0
SM = 0
Delta_AUC
Doxorubicin
YIR026C YVH1
ZShift = −0.88
Z lm Score = −2.12
NG = 0
DB = 0
SM = 0
Delta_L
Doxorubicin
YIR026C YVH1
ZShift = −0.8
Z lm Score = 0.8
NG = 0
DB = 0
SM = 0
Delta_K
Doxorubicin
YIR026C YVH1
ZShift = 1.5
Z lm Score = −0.57
NG = 0
DB = 0
SM = 0
Delta_r
Doxorubicin
YIR026C YVH1
ZShift = 0.27
Z lm Score = 0.3
NG = 0
DB = 0
SM = 0
Delta_AUC
Doxorubicin

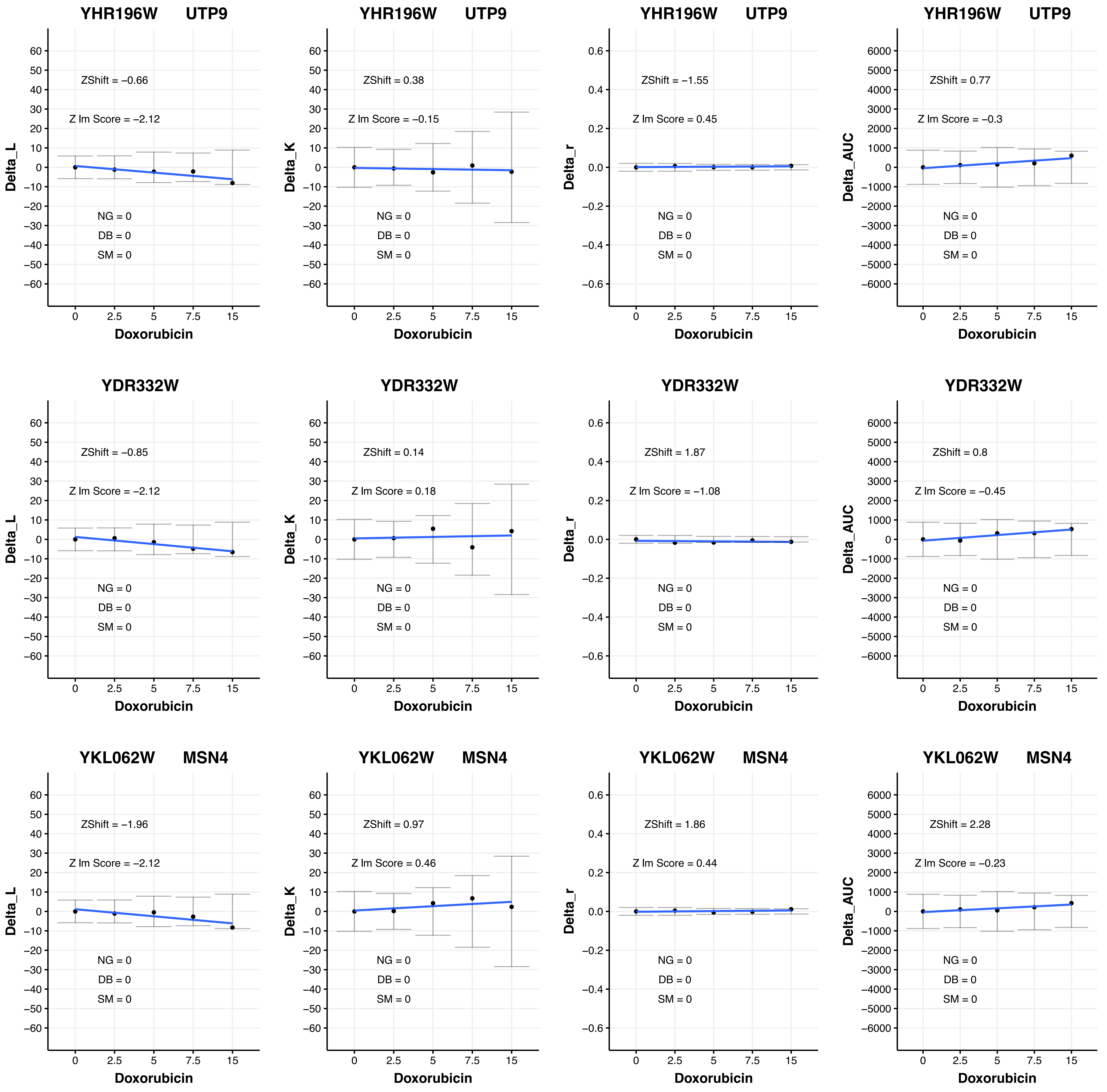
YHR196W UTP9
ZShift = −0.66
Z lm Score = −2.12
NG = 0
DB = 0
SM = 0
Delta_L
Doxorubicin
YHR196W UTP9
ZShift = 0.38
Z lm Score = −0.15
NG = 0
DB = 0
SM = 0
Delta_K
Doxorubicin
YHR196W UTP9
ZShift = −1.55
Z lm Score = 0.45
NG = 0
DB = 0
SM = 0
Delta_r
Doxorubicin
YHR196W UTP9
ZShift = 0.77
Z lm Score = −0.3
NG = 0
DB = 0
SM = 0
Delta_AUC
Doxorubicin
YDR332W
ZShift = −0.85
Z lm Score = −2.12
NG = 0
DB = 0
SM = 0
Delta_L
Doxorubicin
YDR332W
ZShift = 0.14
Z lm Score = 0.18
NG = 0
DB = 0
SM = 0
Delta_K
Doxorubicin
YDR332W
ZShift = 1.87
Z lm Score = −1.08
NG = 0
DB = 0
SM = 0
Delta_r
Doxorubicin
YDR332W
ZShift = 0.8
Z lm Score = −0.45
NG = 0
DB = 0
SM = 0
Delta_AUC
Doxorubicin
YKL062W MSN4
ZShift = −1.96
Z lm Score = −2.12
NG = 0
DB = 0
SM = 0
Delta_L
Doxorubicin
YKL062W MSN4
ZShift = 0.97
Z lm Score = 0.46
NG = 0
DB = 0
SM = 0
Delta_K
Doxorubicin
YKL062W MSN4
ZShift = 1.86
Z lm Score = 0.44
NG = 0
DB = 0
SM = 0
Delta_r
Doxorubicin
YKL062W MSN4
ZShift = 2.28
Z lm Score = −0.23
NG = 0
DB = 0
SM = 0
Delta_AUC
Doxorubicin

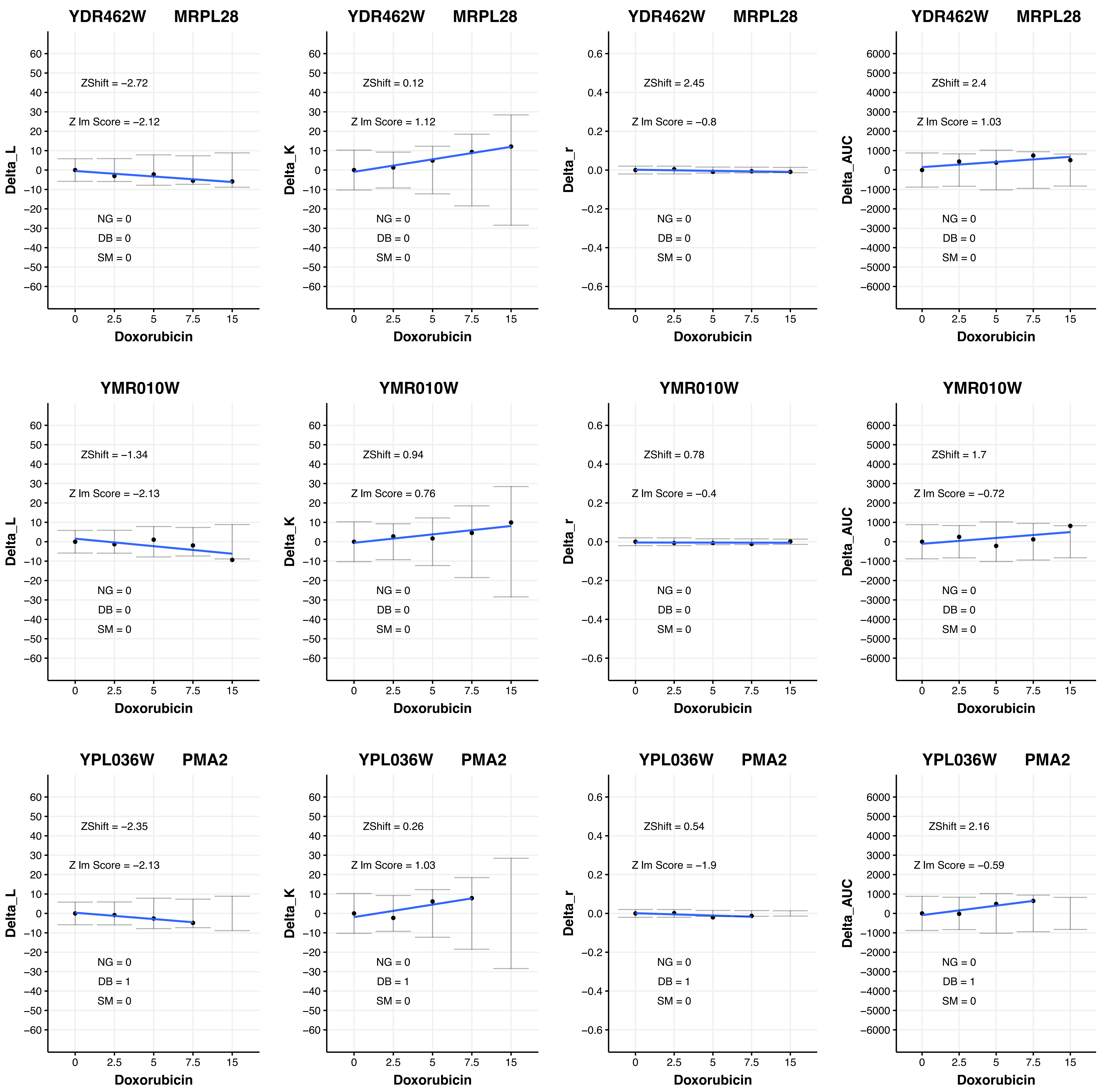

YDR462W MRPL28
ZShift = −2.72
Z lm Score = −2.12
NG = 0
DB = 0
SM = 0
Delta_L
Doxorubicin
YDR462W MRPL28
ZShift = 0.12
Z lm Score = 1.12
NG = 0
DB = 0
SM = 0
Delta_K
Doxorubicin
YDR462W MRPL28
ZShift = 2.45
Z lm Score = −0.8
NG = 0
DB = 0
SM = 0
Delta_r
Doxorubicin
YDR462W MRPL28
ZShift = 2.4
Z lm Score = 1.03
NG = 0
DB = 0
SM = 0
Delta_AUC
Doxorubicin
YMR010W
ZShift = −1.34
Z lm Score = −2.13
NG = 0
DB = 0
SM = 0
Delta_L
Doxorubicin
YMR010W
ZShift = 0.94
Z lm Score = 0.76
NG = 0
DB = 0
SM = 0
Delta_K
Doxorubicin
YMR010W
ZShift = 0.78
Z lm Score = −0.4
NG = 0
DB = 0
SM = 0
Delta_r
Doxorubicin
YMR010W
ZShift = 1.7
Z lm Score = −0.72
NG = 0
DB = 0
SM = 0
Delta_AUC
Doxorubicin
YPL036W PMA2
ZShift = −2.35
Z lm Score = −2.13
NG = 0
DB = 1
SM = 0
Delta_L
Doxorubicin
YPL036W PMA2
ZShift = 0.26
Z lm Score = 1.03
NG = 0
DB = 1
SM = 0
Delta_K
Doxorubicin
YPL036W PMA2
ZShift = 0.54
Z lm Score = −1.9
NG = 0
DB = 1
SM = 0
Delta_r
Doxorubicin
YPL036W PMA2
ZShift = 2.16
Z lm Score = −0.59
NG = 0
DB = 1
SM = 0
Delta_AUC
Doxorubicin

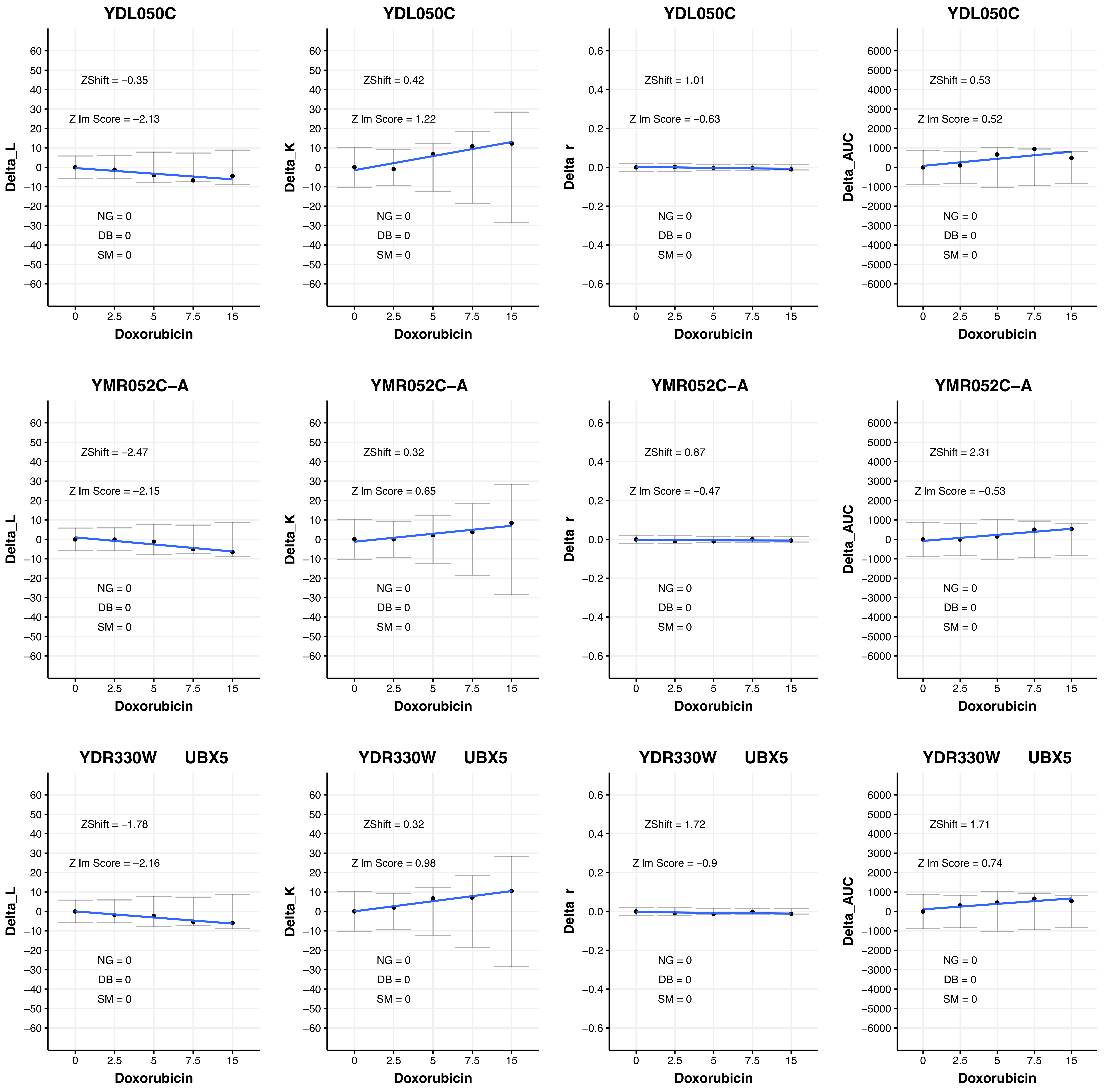
YDL050C
ZShift = −0.35
Z lm Score = −2.13
NG = 0
DB = 0
SM = 0
Delta_L
Doxorubicin
YDL050C
ZShift = 0.42
Z lm Score = 1.22
NG = 0
DB = 0
SM = 0
Delta_K
Doxorubicin
YDL050C
ZShift = 1.01
Z lm Score = −0.63
NG = 0
DB = 0
SM = 0
Delta_r
Doxorubicin
YDL050C
ZShift = 0.53
Z lm Score = 0.52
NG = 0
DB = 0
SM = 0
Delta_AUC
Doxorubicin
YMR052C−A
ZShift = −2.47
Z lm Score = −2.15
NG = 0
DB = 0
SM = 0
Delta_L
Doxorubicin
YMR052C−A
ZShift = 0.32
Z lm Score = 0.65
NG = 0
DB = 0
SM = 0
Delta_K
Doxorubicin
YMR052C−A
ZShift = 0.87
Z lm Score = −0.47
NG = 0
DB = 0
SM = 0
Delta_r
Doxorubicin
YMR052C−A
ZShift = 2.31
Z lm Score = −0.53
NG = 0
DB = 0
SM = 0
Delta_AUC
Doxorubicin
YDR330W UBX5
ZShift = −1.78
Z lm Score = −2.16
NG = 0
DB = 0
SM = 0
Delta_L
Doxorubicin
YDR330W UBX5
ZShift = 0.32
Z lm Score = 0.98
NG = 0
DB = 0
SM = 0
Delta_K
Doxorubicin
YDR330W UBX5
ZShift = 1.72
Z lm Score = −0.9
NG = 0
DB = 0
SM = 0
Delta_r
Doxorubicin
YDR330W UBX5
ZShift = 1.71
Z lm Score = 0.74
NG = 0
DB = 0
SM = 0
Delta_AUC
Doxorubicin

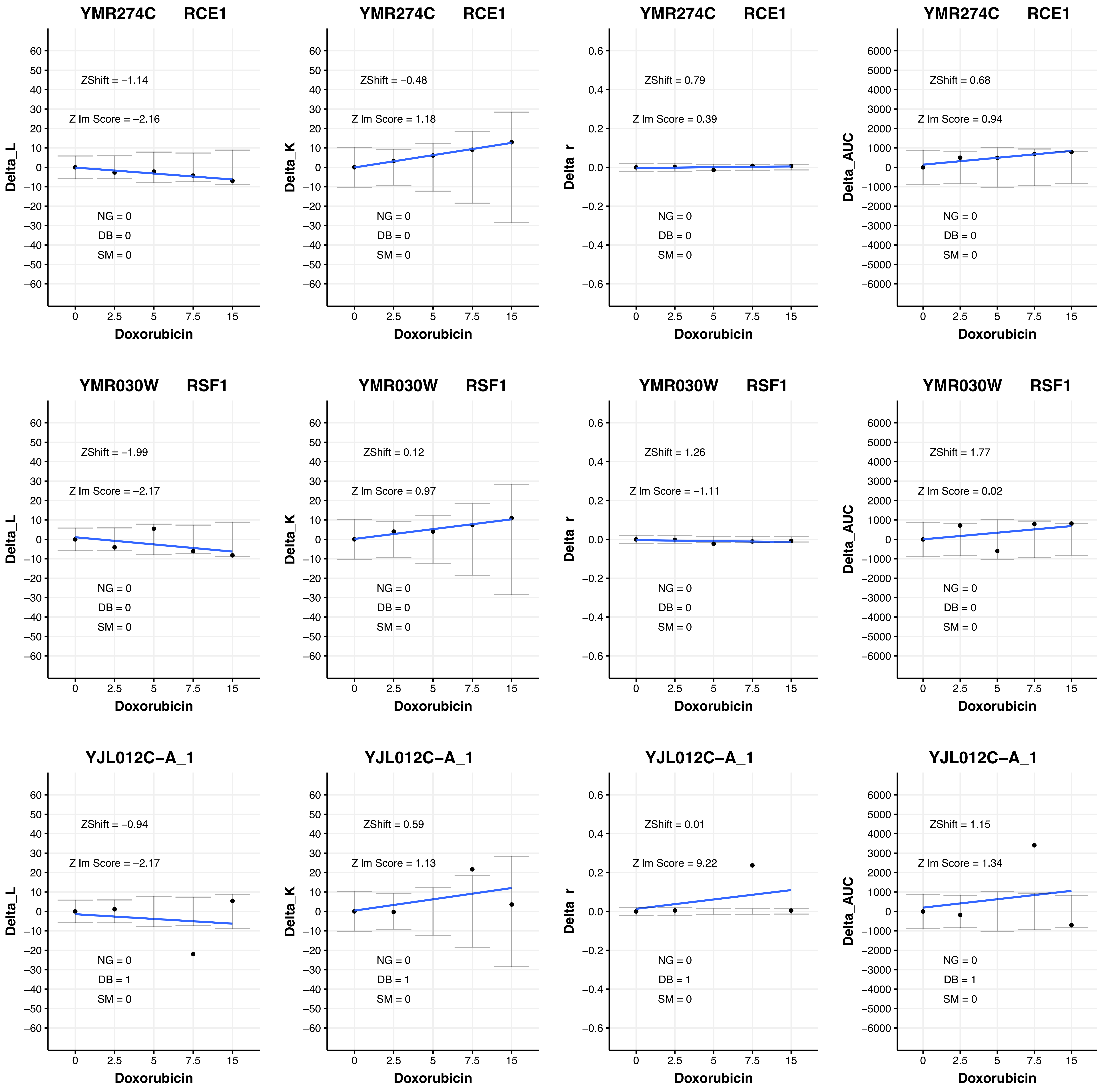
YMR274C RCE1
ZShift = −1.14
Z lm Score = −2.16
NG = 0
DB = 0
SM = 0
Delta_L
Doxorubicin
YMR274C RCE1
ZShift = −0.48
Z lm Score = 1.18
NG = 0
DB = 0
SM = 0
Delta_K
Doxorubicin
YMR274C RCE1
ZShift = 0.79
Z lm Score = 0.39
NG = 0
DB = 0
SM = 0
Delta_r
Doxorubicin
YMR274C RCE1
ZShift = 0.68
Z lm Score = 0.94
NG = 0
DB = 0
SM = 0
Delta_AUC
Doxorubicin
YMR030W RSF1
ZShift = −1.99
Z lm Score = −2.17
NG = 0
DB = 0
SM = 0
Delta_L
Doxorubicin
YMR030W RSF1
ZShift = 0.12
Z lm Score = 0.97
NG = 0
DB = 0
SM = 0
Delta_K
Doxorubicin
YMR030W RSF1
ZShift = 1.26
Z lm Score = −1.11
NG = 0
DB = 0
SM = 0
Delta_r
Doxorubicin
YMR030W RSF1
ZShift = 1.77
Z lm Score = 0.02
NG = 0
DB = 0
SM = 0
Delta_AUC
Doxorubicin
YJL012C−A_1
ZShift = −0.94
Z lm Score = −2.17
NG = 0
DB = 1
SM = 0
Delta_L
Doxorubicin
YJL012C−A_1
ZShift = 0.59
Z lm Score = 1.13
NG = 0
DB = 1
SM = 0
Delta_K
Doxorubicin
YJL012C−A_1
ZShift = 0.01
Z lm Score = 9.22
NG = 0
DB = 1
SM = 0
Delta_r
Doxorubicin
YJL012C−A_1
ZShift = 1.15
Z lm Score = 1.34
NG = 0
DB = 1
SM = 0
Delta_AUC
Doxorubicin

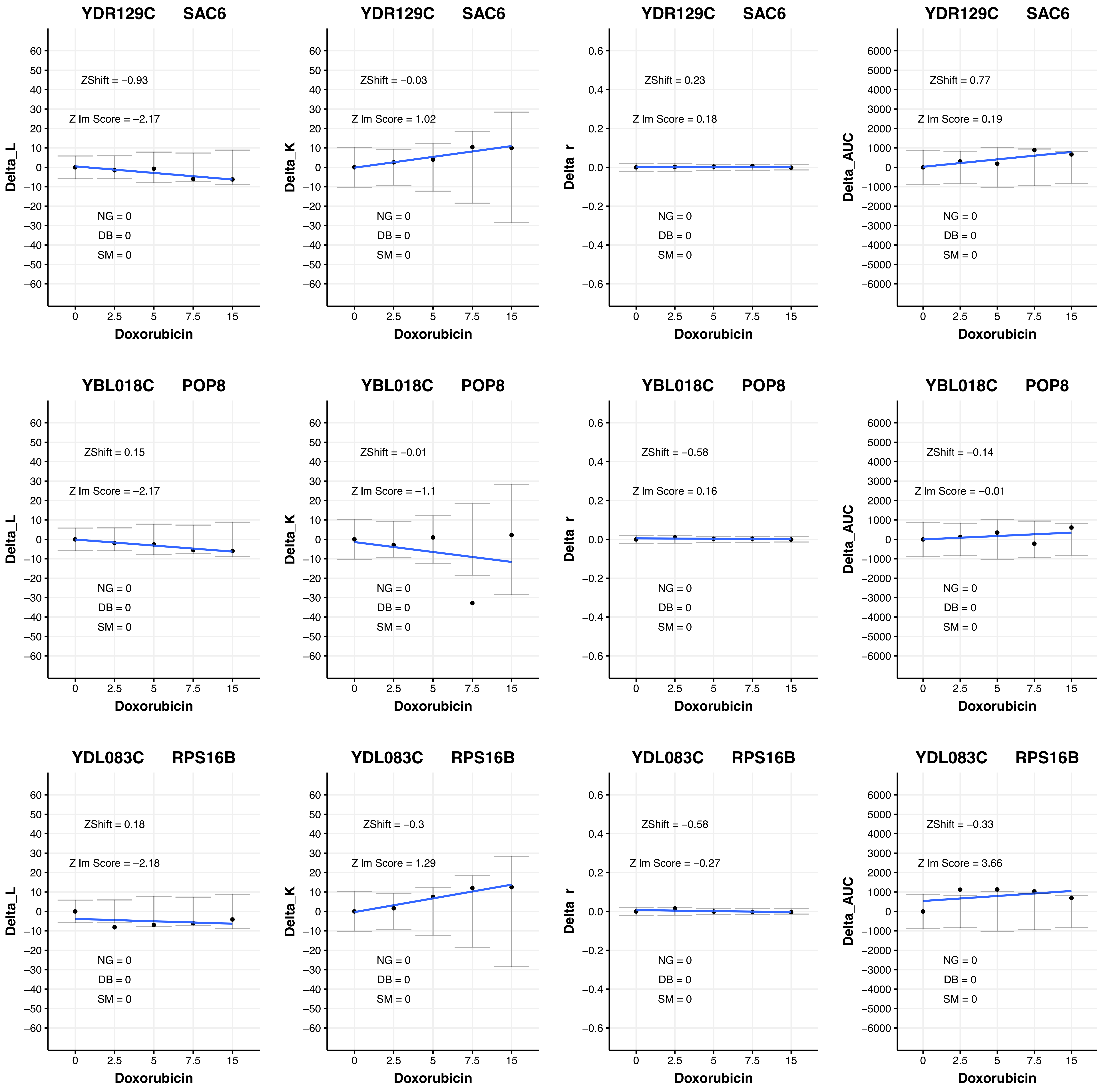
YDR129C SAC6
ZShift = −0.93
Z lm Score = −2.17
NG = 0
DB = 0
SM = 0
Delta_L
Doxorubicin
YDR129C SAC6
ZShift = −0.03
Z lm Score = 1.02
NG = 0
DB = 0
SM = 0
Delta_K
Doxorubicin
YDR129C SAC6
ZShift = 0.23
Z lm Score = 0.18
NG = 0
DB = 0
SM = 0
Delta_r
Doxorubicin
YDR129C SAC6
ZShift = 0.77
Z lm Score = 0.19
NG = 0
DB = 0
SM = 0
Delta_AUC
Doxorubicin
YBL018C POP8
ZShift = 0.15
Z lm Score = −2.17
NG = 0
DB = 0
SM = 0
Delta_L
Doxorubicin
YBL018C POP8
ZShift = −0.01
Z lm Score = −1.1
NG = 0
DB = 0
SM = 0
Delta_K
Doxorubicin
YBL018C POP8
ZShift = −0.58
Z lm Score = 0.16
NG = 0
DB = 0
SM = 0
Delta_r
Doxorubicin
YBL018C POP8
ZShift = −0.14
Z lm Score = −0.01
NG = 0
DB = 0
SM = 0
Delta_AUC
Doxorubicin
YDL083C RPS16B
ZShift = 0.18
Z lm Score = −2.18
NG = 0
DB = 0
SM = 0
Delta_L
Doxorubicin
YDL083C RPS16B
ZShift = −0.3
Z lm Score = 1.29
NG = 0
DB = 0
SM = 0
Delta_K
Doxorubicin
YDL083C RPS16B
ZShift = −0.58
Z lm Score = −0.27
NG = 0
DB = 0
SM = 0
Delta_r
Doxorubicin
YDL083C RPS16B
ZShift = −0.33
Z lm Score = 3.66
NG = 0
DB = 0
SM = 0
Delta_AUC
Doxorubicin

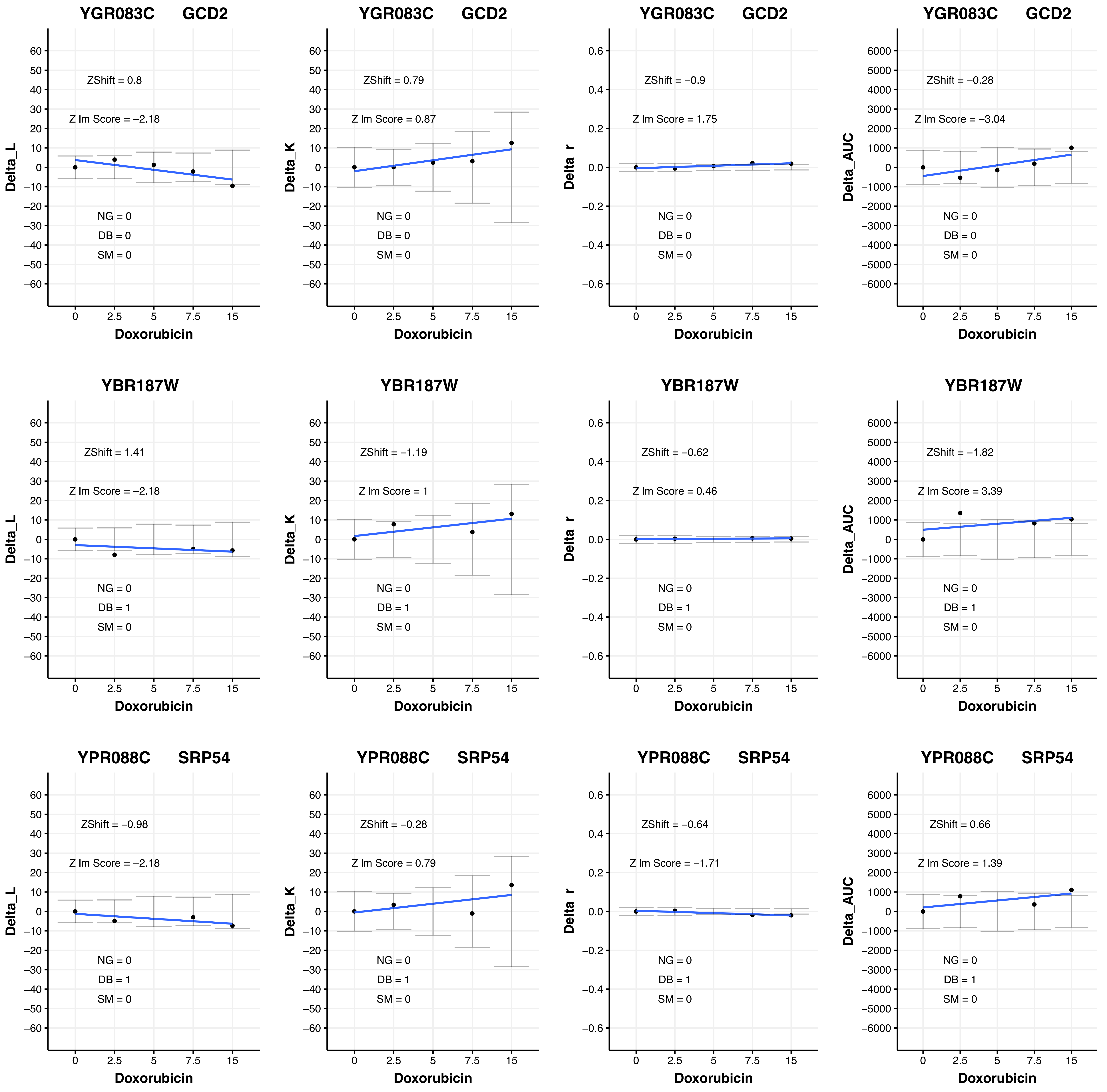

YGR083C GCD2
ZShift = 0.8
Z lm Score = −2.18
NG = 0
DB = 0
SM = 0
Delta_L
Doxorubicin
YGR083C GCD2
ZShift = 0.79
Z lm Score = 0.87
NG = 0
DB = 0
SM = 0
Delta_K
Doxorubicin
YGR083C GCD2
ZShift = −0.9
Z lm Score = 1.75
NG = 0
DB = 0
SM = 0
Delta_r
Doxorubicin
YGR083C GCD2
ZShift = −0.28
Z lm Score = −3.04
NG = 0
DB = 0
SM = 0
Delta_AUC
Doxorubicin
YBR187W
ZShift = 1.41
Z lm Score = −2.18
NG = 0
DB = 1
SM = 0
Delta_L
Doxorubicin
YBR187W
ZShift = −1.19
Z lm Score = 1
NG = 0
DB = 1
SM = 0
Delta_K
Doxorubicin
YBR187W
ZShift = −0.62
Z lm Score = 0.46
NG = 0
DB = 1
SM = 0
Delta_r
Doxorubicin
YBR187W
ZShift = −1.82
Z lm Score = 3.39
NG = 0
DB = 1
SM = 0
Delta_AUC
Doxorubicin
YPR088C SRP54
ZShift = −0.98
Z lm Score = −2.18
NG = 0
DB = 1
SM = 0
Delta_L
Doxorubicin
YPR088C SRP54
ZShift = −0.28
Z lm Score = 0.79
NG = 0
DB = 1
SM = 0
Delta_K
Doxorubicin
YPR088C SRP54
ZShift = −0.64
Z lm Score = −1.71
NG = 0
DB = 1
SM = 0
Delta_r
Doxorubicin
YPR088C SRP54
ZShift = 0.66
Z lm Score = 1.39
NG = 0
DB = 1
SM = 0
Delta_AUC
Doxorubicin

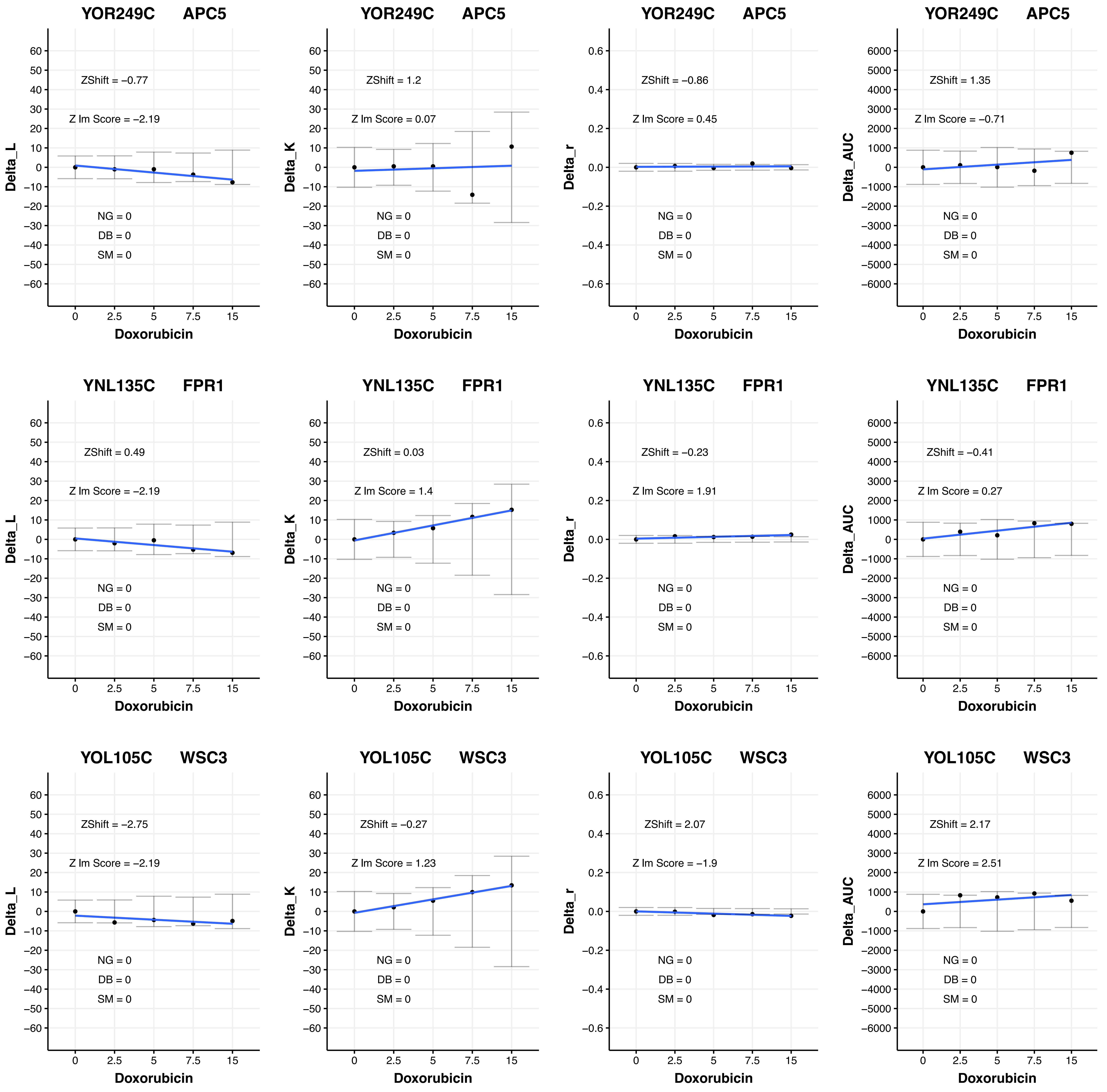
YOR249C APC5
ZShift = −0.77
Z lm Score = −2.19
NG = 0
DB = 0
SM = 0
Delta_L
Doxorubicin
YOR249C APC5
ZShift = 1.2
Z lm Score = 0.07
NG = 0
DB = 0
SM = 0
Delta_K
Doxorubicin
YOR249C APC5
ZShift = −0.86
Z lm Score = 0.45
NG = 0
DB = 0
SM = 0
Delta_r
Doxorubicin
YOR249C APC5
ZShift = 1.35
Z lm Score = −0.71
NG = 0
DB = 0
SM = 0
Delta_AUC
Doxorubicin
YNL135C FPR1
ZShift = 0.49
Z lm Score = −2.19
NG = 0
DB = 0
SM = 0
Delta_L
Doxorubicin
YNL135C FPR1
ZShift = 0.03
Z lm Score = 1.4
NG = 0
DB = 0
SM = 0
Delta_K
Doxorubicin
YNL135C FPR1
ZShift = −0.23
Z lm Score = 1.91
NG = 0
DB = 0
SM = 0
Delta_r
Doxorubicin
YNL135C FPR1
ZShift = −0.41
Z lm Score = 0.27
NG = 0
DB = 0
SM = 0
Delta_AUC
Doxorubicin
YOL105C WSC3
ZShift = −2.75
Z lm Score = −2.19
NG = 0
DB = 0
SM = 0
Delta_L
Doxorubicin
YOL105C WSC3
ZShift = −0.27
Z lm Score = 1.23
NG = 0
DB = 0
SM = 0
Delta_K
Doxorubicin
YOL105C WSC3
ZShift = 2.07
Z lm Score = −1.9
NG = 0
DB = 0
SM = 0
Delta_r
Doxorubicin
YOL105C WSC3
ZShift = 2.17
Z lm Score = 2.51
NG = 0
DB = 0
SM = 0
Delta_AUC
Doxorubicin

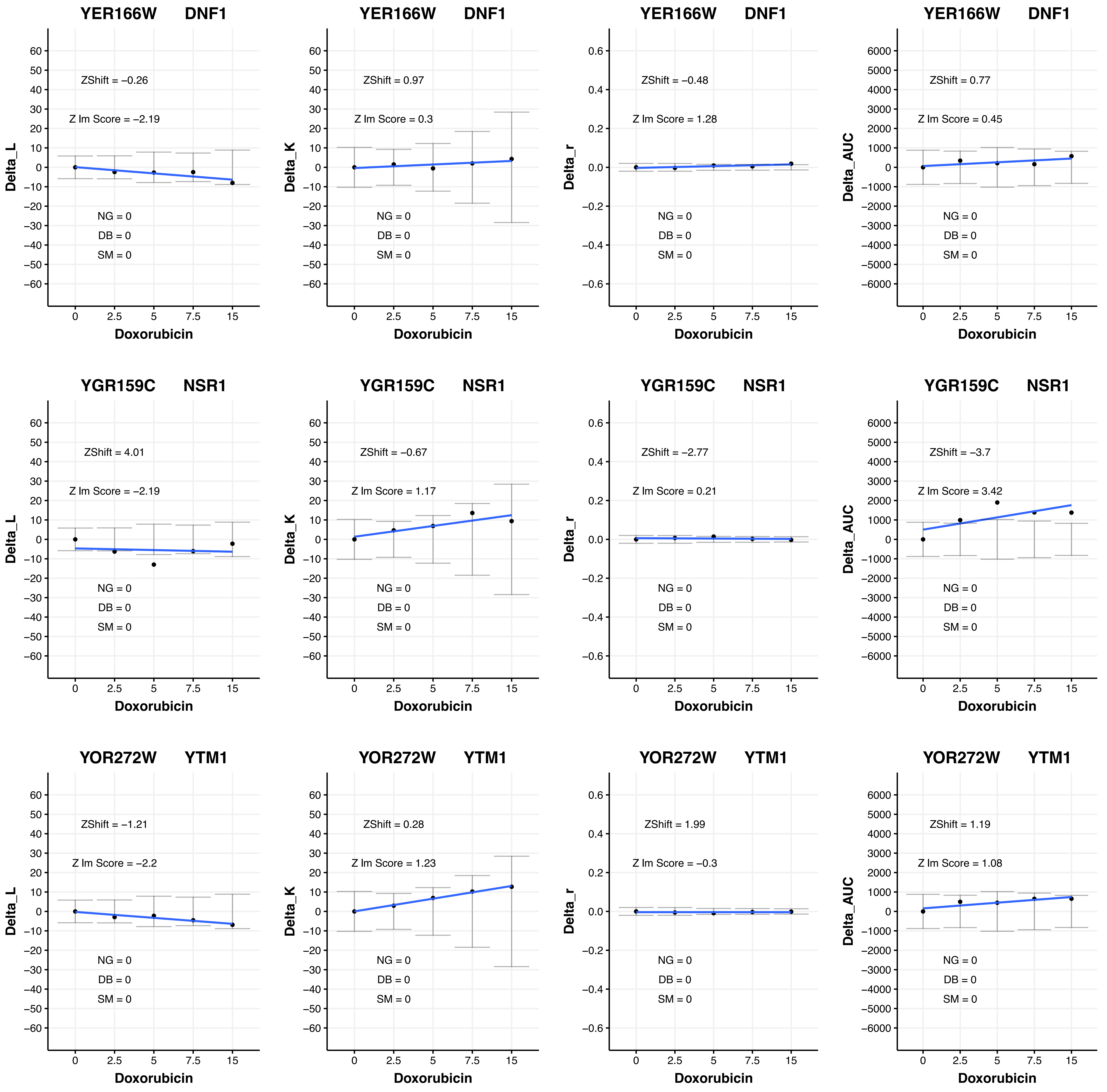

YER166W DNF1
ZShift = −0.26
Z lm Score = −2.19
NG = 0
DB = 0
SM = 0
Delta_L
Doxorubicin
YER166W DNF1
ZShift = 0.97
Z lm Score = 0.3
NG = 0
DB = 0
SM = 0
Delta_K
Doxorubicin
YER166W DNF1
ZShift = −0.48
Z lm Score = 1.28
NG = 0
DB = 0
SM = 0
Delta_r
Doxorubicin
YER166W DNF1
ZShift = 0.77
Z lm Score = 0.45
NG = 0
DB = 0
SM = 0
Delta_AUC
Doxorubicin
YGR159C NSR1
ZShift = 4.01
Z lm Score = −2.19
NG = 0
DB = 0
SM = 0
Delta_L
Doxorubicin
YGR159C NSR1
ZShift = −0.67
Z lm Score = 1.17
NG = 0
DB = 0
SM = 0
Delta_K
Doxorubicin
YGR159C NSR1
ZShift = −2.77
Z lm Score = 0.21
NG = 0
DB = 0
SM = 0
Delta_r
Doxorubicin
YGR159C NSR1
ZShift = −3.7
Z lm Score = 3.42
NG = 0
DB = 0
SM = 0
Delta_AUC
Doxorubicin
YOR272W YTM1
ZShift = −1.21
Z lm Score = −2.2
NG = 0
DB = 0
SM = 0
Delta_L
Doxorubicin
YOR272W YTM1
ZShift = 0.28
Z lm Score = 1.23
NG = 0
DB = 0
SM = 0
Delta_K
Doxorubicin
YOR272W YTM1
ZShift = 1.99
Z lm Score = −0.3
NG = 0
DB = 0
SM = 0
Delta_r
Doxorubicin
YOR272W YTM1
ZShift = 1.19
Z lm Score = 1.08
NG = 0
DB = 0
SM = 0
Delta_AUC
Doxorubicin

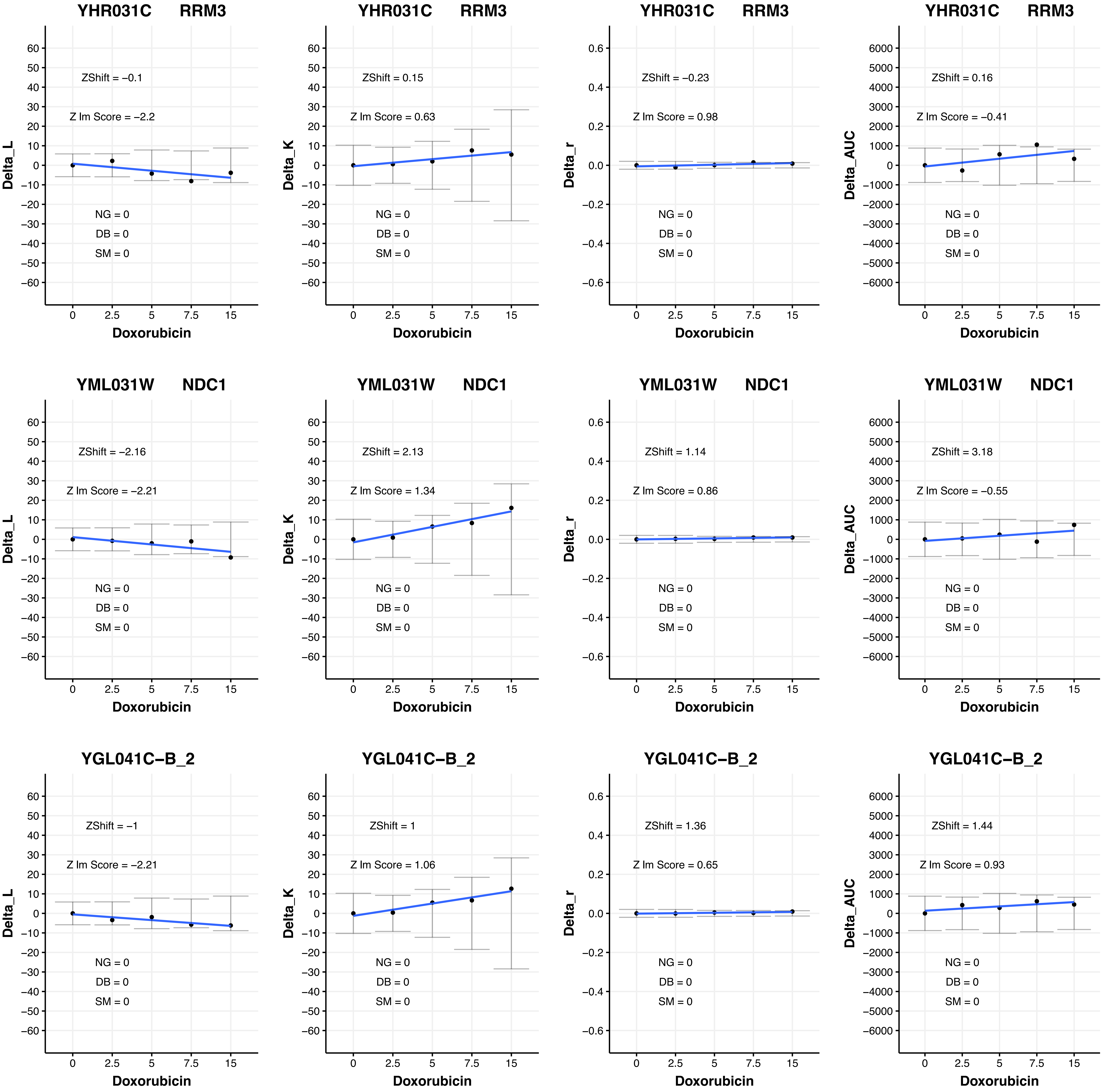

YHR031C RRM3
ZShift = −0.1
Z lm Score = −2.2
NG = 0
DB = 0
SM = 0
Delta_L
Doxorubicin
YHR031C RRM3
ZShift = 0.15
Z lm Score = 0.63
NG = 0
DB = 0
SM = 0
Delta_K
Doxorubicin
YHR031C RRM3
ZShift = −0.23
Z lm Score = 0.98
NG = 0
DB = 0
SM = 0
Delta_r
Doxorubicin
YHR031C RRM3
ZShift = 0.16
Z lm Score = −0.41
NG = 0
DB = 0
SM = 0
Delta_AUC
Doxorubicin
YML031W NDC1
ZShift = −2.16
Z lm Score = −2.21
NG = 0
DB = 0
SM = 0
Delta_L
Doxorubicin
YML031W NDC1
ZShift = 2.13
Z lm Score = 1.34
NG = 0
DB = 0
SM = 0
Delta_K
Doxorubicin
YML031W NDC1
ZShift = 1.14
Z lm Score = 0.86
NG = 0
DB = 0
SM = 0
Delta_r
Doxorubicin
YML031W NDC1
ZShift = 3.18
Z lm Score = −0.55
NG = 0
DB = 0
SM = 0
Delta_AUC
Doxorubicin
YGL041C−B_2
ZShift = −1
Z lm Score = −2.21
NG = 0
DB = 0
SM = 0
Delta_L
Doxorubicin
YGL041C−B_2
ZShift = 1
Z lm Score = 1.06
NG = 0
DB = 0
SM = 0
Delta_K
Doxorubicin
YGL041C−B_2
ZShift = 1.36
Z lm Score = 0.65
NG = 0
DB = 0
SM = 0
Delta_r
Doxorubicin
YGL041C−B_2
ZShift = 1.44
Z lm Score = 0.93
NG = 0
DB = 0
SM = 0
Delta_AUC
Doxorubicin

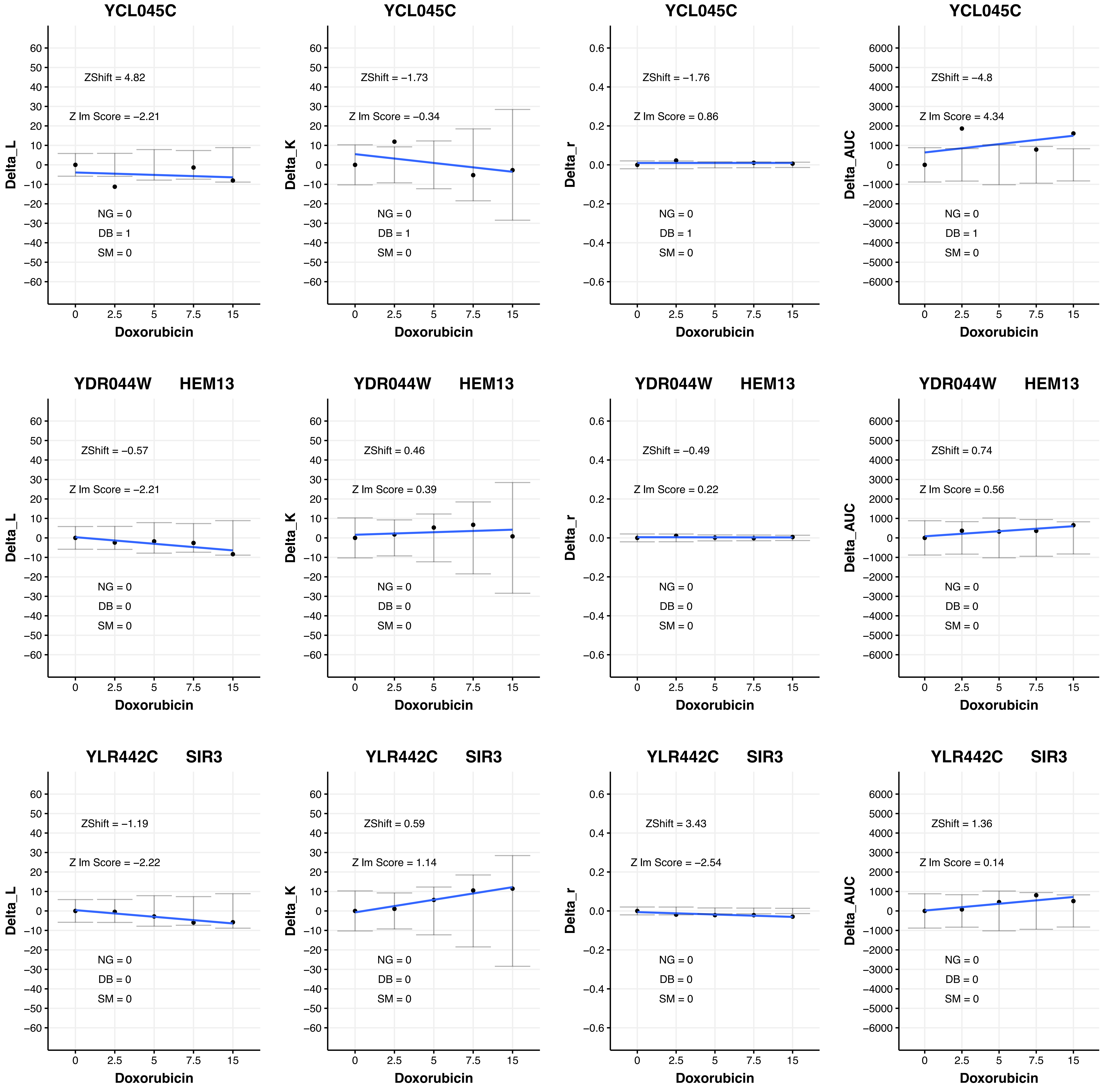

YCL045C
ZShift = 4.82
Z lm Score = −2.21
NG = 0
DB = 1
SM = 0
Delta_L
Doxorubicin
YCL045C
ZShift = −1.73
Z lm Score = −0.34
NG = 0
DB = 1
SM = 0
Delta_K
Doxorubicin
YCL045C
ZShift = −1.76
Z lm Score = 0.86
NG = 0
DB = 1
SM = 0
Delta_r
Doxorubicin
YCL045C
ZShift = −4.8
Z lm Score = 4.34
NG = 0
DB = 1
SM = 0
Delta_AUC
Doxorubicin
YDR044W HEM13
ZShift = −0.57
Z lm Score = −2.21
NG = 0
DB = 0
SM = 0
Delta_L
Doxorubicin
YDR044W HEM13
ZShift = 0.46
Z lm Score = 0.39
NG = 0
DB = 0
SM = 0
Delta_K
Doxorubicin
YDR044W HEM13
ZShift = −0.49
Z lm Score = 0.22
NG = 0
DB = 0
SM = 0
Delta_r
Doxorubicin
YDR044W HEM13
ZShift = 0.74
Z lm Score = 0.56
NG = 0
DB = 0
SM = 0
Delta_AUC
Doxorubicin
YLR442C SIR3
ZShift = −1.19
Z lm Score = −2.22
NG = 0
DB = 0
SM = 0
Delta_L
Doxorubicin
YLR442C SIR3
ZShift = 0.59
Z lm Score = 1.14
NG = 0
DB = 0
SM = 0
Delta_K
Doxorubicin
YLR442C SIR3
ZShift = 3.43
Z lm Score = −2.54
NG = 0
DB = 0
SM = 0
Delta_r
Doxorubicin
YLR442C SIR3
ZShift = 1.36
Z lm Score = 0.14
NG = 0
DB = 0
SM = 0
Delta_AUC
Doxorubicin

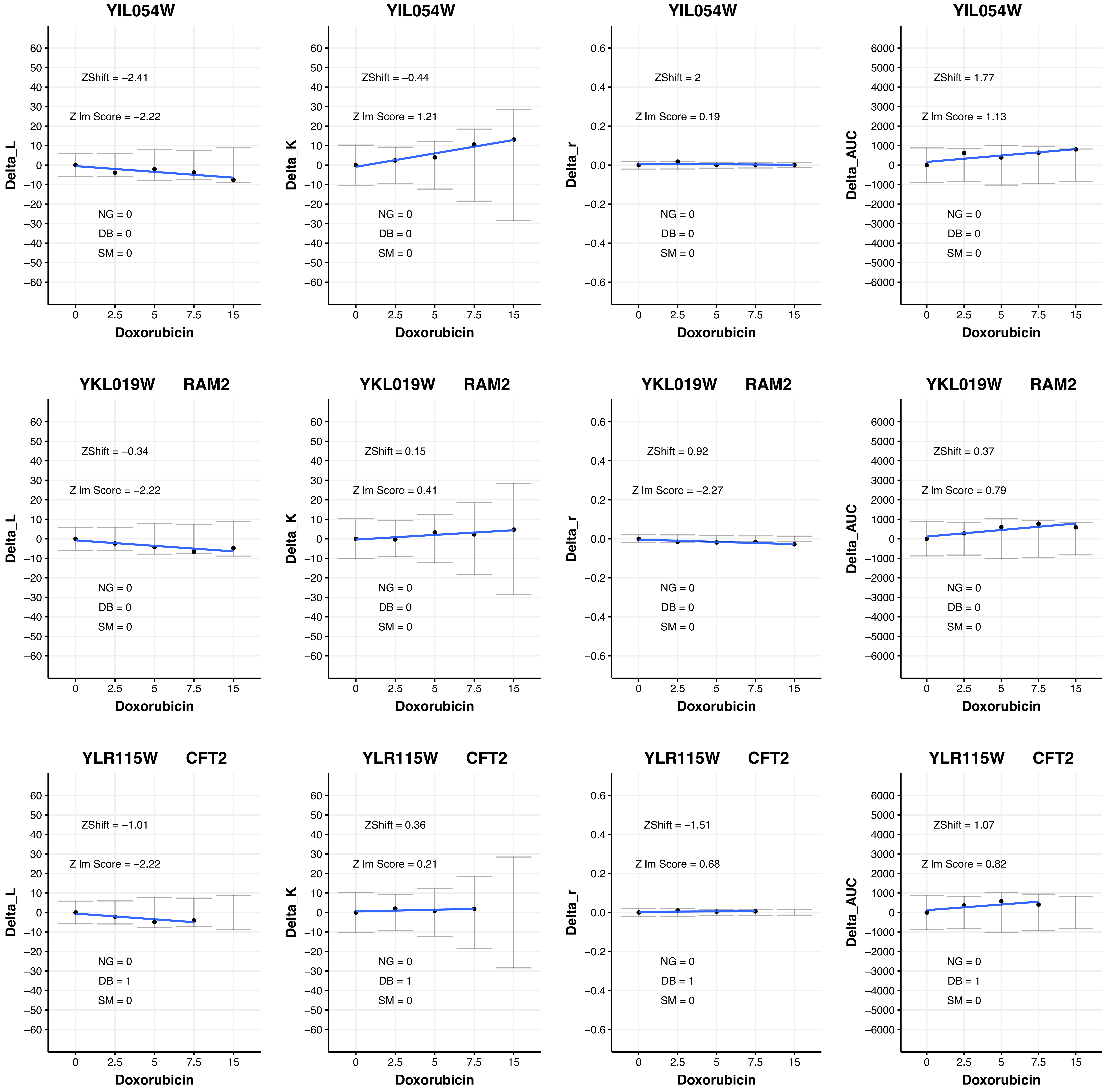
YIL054W
ZShift = −2.41
Z lm Score = −2.22
NG = 0
DB = 0
SM = 0
Delta_L
Doxorubicin
YIL054W
ZShift = −0.44
Z lm Score = 1.21
NG = 0
DB = 0
SM = 0
Delta_K
Doxorubicin
YIL054W
ZShift = 2
Z lm Score = 0.19
NG = 0
DB = 0
SM = 0
Delta_r
Doxorubicin
YIL054W
ZShift = 1.77
Z lm Score = 1.13
NG = 0
DB = 0
SM = 0
Delta_AUC
Doxorubicin
YKL019W RAM2
ZShift = −0.34
Z lm Score = −2.22
NG = 0
DB = 0
SM = 0
Delta_L
Doxorubicin
YKL019W RAM2
ZShift = 0.15
Z lm Score = 0.41
NG = 0
DB = 0
SM = 0
Delta_K
Doxorubicin
YKL019W RAM2
ZShift = 0.92
Z lm Score = −2.27
NG = 0
DB = 0
SM = 0
Delta_r
Doxorubicin
YKL019W RAM2
ZShift = 0.37
Z lm Score = 0.79
NG = 0
DB = 0
SM = 0
Delta_AUC
Doxorubicin
YLR115W CFT2
ZShift = −1.01
Z lm Score = −2.22
NG = 0
DB = 1
SM = 0
Delta_L
Doxorubicin
YLR115W CFT2
ZShift = 0.36
Z lm Score = 0.21
NG = 0
DB = 1
SM = 0
Delta_K
Doxorubicin
YLR115W CFT2
ZShift = −1.51
Z lm Score = 0.68
NG = 0
DB = 1
SM = 0
Delta_r
Doxorubicin
YLR115W CFT2
ZShift = 1.07
Z lm Score = 0.82
NG = 0
DB = 1
SM = 0
Delta_AUC
Doxorubicin

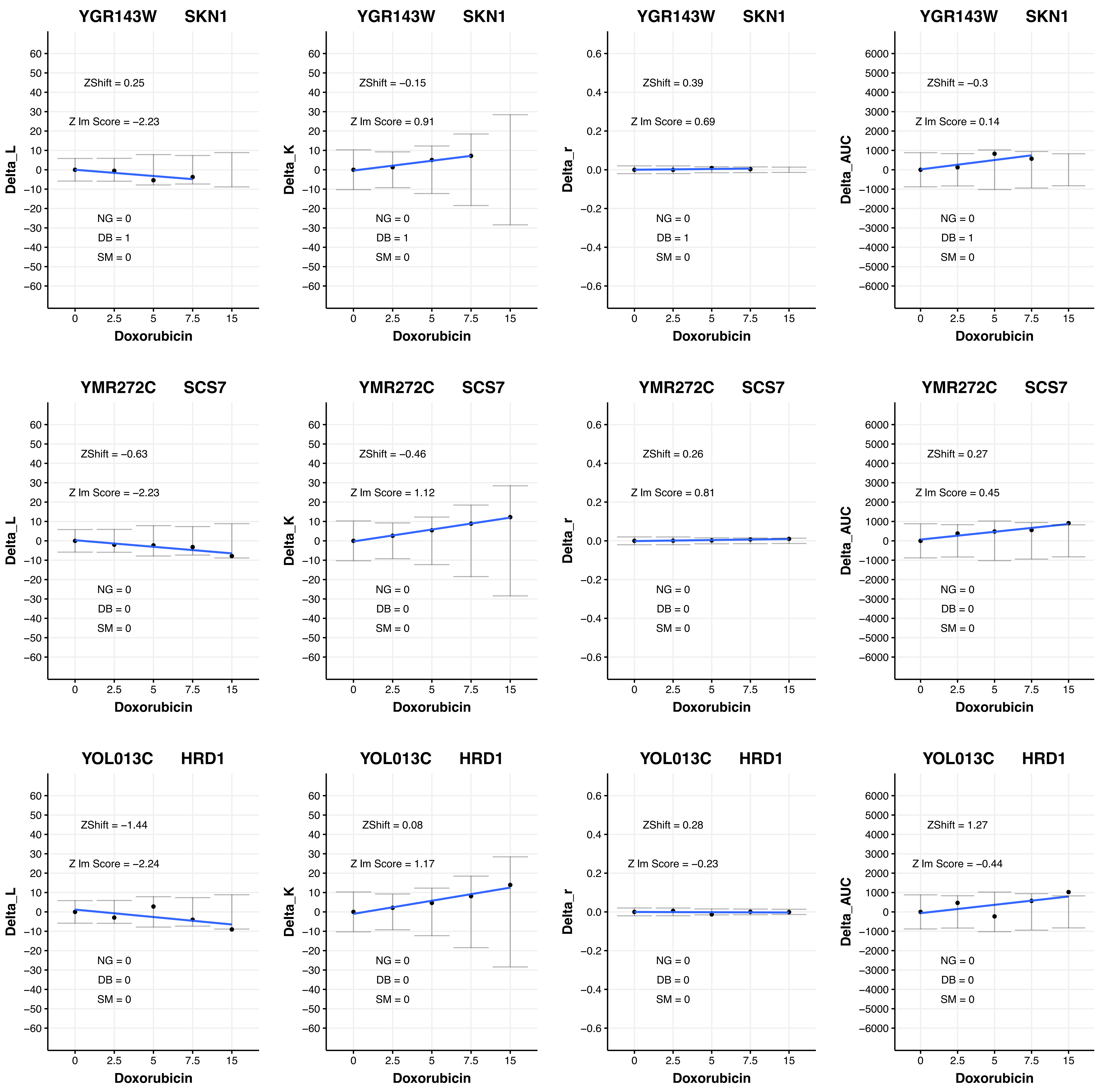
YGR143W SKN1
ZShift = 0.25
Z lm Score = −2.23
NG = 0
DB = 1
SM = 0
Delta_L
Doxorubicin
YGR143W SKN1
ZShift = −0.15
Z lm Score = 0.91
NG = 0
DB = 1
SM = 0
Delta_K
Doxorubicin
YGR143W SKN1
ZShift = 0.39
Z lm Score = 0.69
NG = 0
DB = 1
SM = 0
Delta_r
Doxorubicin
YGR143W SKN1
ZShift = −0.3
Z lm Score = 0.14
NG = 0
DB = 1
SM = 0
Delta_AUC
Doxorubicin
YMR272C SCS7
ZShift = −0.63
Z lm Score = −2.23
NG = 0
DB = 0
SM = 0
Delta_L
Doxorubicin
YMR272C SCS7
ZShift = −0.46
Z lm Score = 1.12
NG = 0
DB = 0
SM = 0
Delta_K
Doxorubicin
YMR272C SCS7
ZShift = 0.26
Z lm Score = 0.81
NG = 0
DB = 0
SM = 0
Delta_r
Doxorubicin
YMR272C SCS7
ZShift = 0.27
Z lm Score = 0.45
NG = 0
DB = 0
SM = 0
Delta_AUC
Doxorubicin
YOL013C HRD1
ZShift = −1.44
Z lm Score = −2.24
NG = 0
DB = 0
SM = 0
Delta_L
Doxorubicin
YOL013C HRD1
ZShift = 0.08
Z lm Score = 1.17
NG = 0
DB = 0
SM = 0
Delta_K
Doxorubicin
YOL013C HRD1
ZShift = 0.28
Z lm Score = −0.23
NG = 0
DB = 0
SM = 0
Delta_r
Doxorubicin
YOL013C HRD1
ZShift = 1.27
Z lm Score = −0.44
NG = 0
DB = 0
SM = 0
Delta_AUC
Doxorubicin

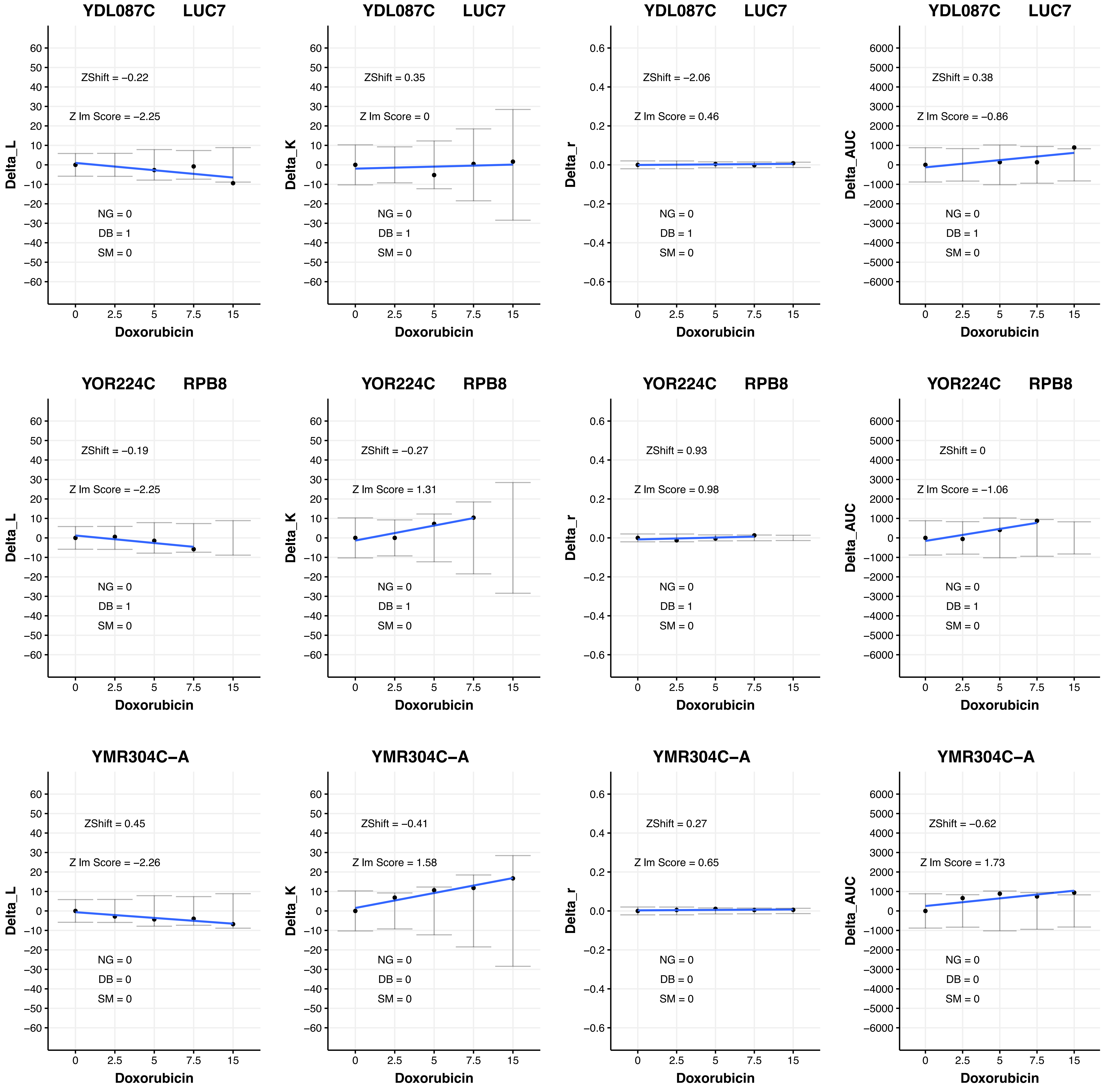

YDL087C LUC7
ZShift = −0.22
Z lm Score = −2.25
NG = 0
DB = 1
SM = 0
Delta_L
Doxorubicin
YDL087C LUC7
ZShift = 0.35
Z lm Score = 0
NG = 0
DB = 1
SM = 0
Delta_K
Doxorubicin
YDL087C LUC7
ZShift = −2.06
Z lm Score = 0.46
NG = 0
DB = 1
SM = 0
Delta_r
Doxorubicin
YDL087C LUC7
ZShift = 0.38
Z lm Score = −0.86
NG = 0
DB = 1
SM = 0
Delta_AUC
Doxorubicin
YOR224C RPB8
ZShift = −0.19
Z lm Score = −2.25
NG = 0
DB = 1
SM = 0
Delta_L
Doxorubicin
YOR224C RPB8
ZShift = −0.27
Z lm Score = 1.31
NG = 0
DB = 1
SM = 0
Delta_K
Doxorubicin
YOR224C RPB8
ZShift = 0.93
Z lm Score = 0.98
NG = 0
DB = 1
SM = 0
Delta_r
Doxorubicin
YOR224C RPB8
ZShift = 0
Z lm Score = −1.06
NG = 0
DB = 1
SM = 0
Delta_AUC
Doxorubicin
YMR304C−A
ZShift = 0.45
Z lm Score = −2.26
NG = 0
DB = 0
SM = 0
Delta_L
Doxorubicin
YMR304C−A
ZShift = −0.41
Z lm Score = 1.58
NG = 0
DB = 0
SM = 0
Delta_K
Doxorubicin
YMR304C−A
ZShift = 0.27
Z lm Score = 0.65
NG = 0
DB = 0
SM = 0
Delta_r
Doxorubicin
YMR304C−A
ZShift = −0.62
Z lm Score = 1.73
NG = 0
DB = 0
SM = 0
Delta_AUC
Doxorubicin

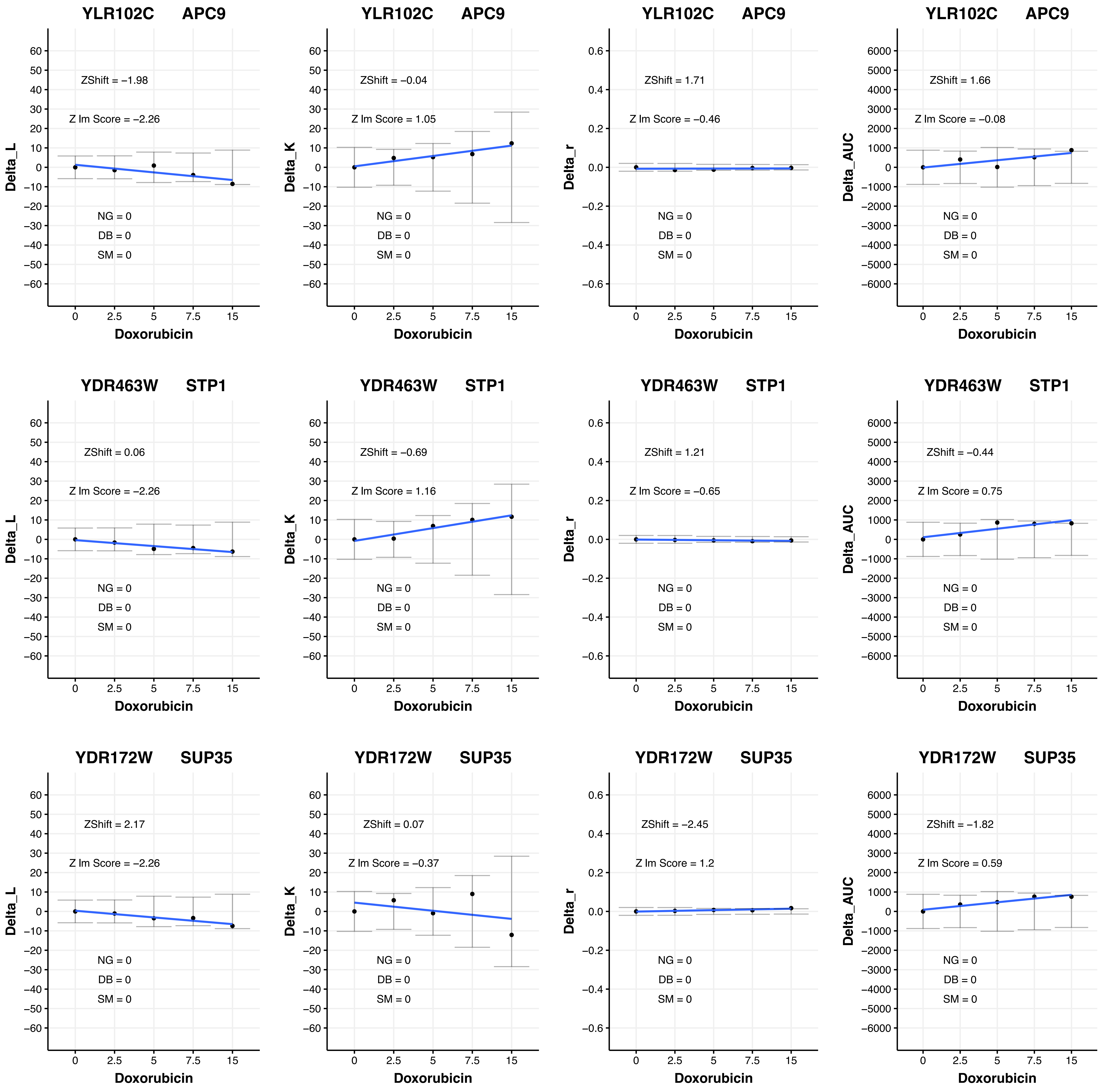
YLR102C APC9
ZShift = −1.98
Z lm Score = −2.26
NG = 0
DB = 0
SM = 0
Delta_L
Doxorubicin
YLR102C APC9
ZShift = −0.04
Z lm Score = 1.05
NG = 0
DB = 0
SM = 0
Delta_K
Doxorubicin
YLR102C APC9
ZShift = 1.71
Z lm Score = −0.46
NG = 0
DB = 0
SM = 0
Delta_r
Doxorubicin
YLR102C APC9
ZShift = 1.66
Z lm Score = −0.08
NG = 0
DB = 0
SM = 0
Delta_AUC
Doxorubicin
YDR463W STP1
ZShift = 0.06
Z lm Score = −2.26
NG = 0
DB = 0
SM = 0
Delta_L
Doxorubicin
YDR463W STP1
ZShift = −0.69
Z lm Score = 1.16
NG = 0
DB = 0
SM = 0
Delta_K
Doxorubicin
YDR463W STP1
ZShift = 1.21
Z lm Score = −0.65
NG = 0
DB = 0
SM = 0
Delta_r
Doxorubicin
YDR463W STP1
ZShift = −0.44
Z lm Score = 0.75
NG = 0
DB = 0
SM = 0
Delta_AUC
Doxorubicin
YDR172W SUP35
ZShift = 2.17
Z lm Score = −2.26
NG = 0
DB = 0
SM = 0
Delta_L
Doxorubicin
YDR172W SUP35
ZShift = 0.07
Z lm Score = −0.37
NG = 0
DB = 0
SM = 0
Delta_K
Doxorubicin
YDR172W SUP35
ZShift = −2.45
Z lm Score = 1.2
NG = 0
DB = 0
SM = 0
Delta_r
Doxorubicin
YDR172W SUP35
ZShift = −1.82
Z lm Score = 0.59
NG = 0
DB = 0
SM = 0
Delta_AUC
Doxorubicin

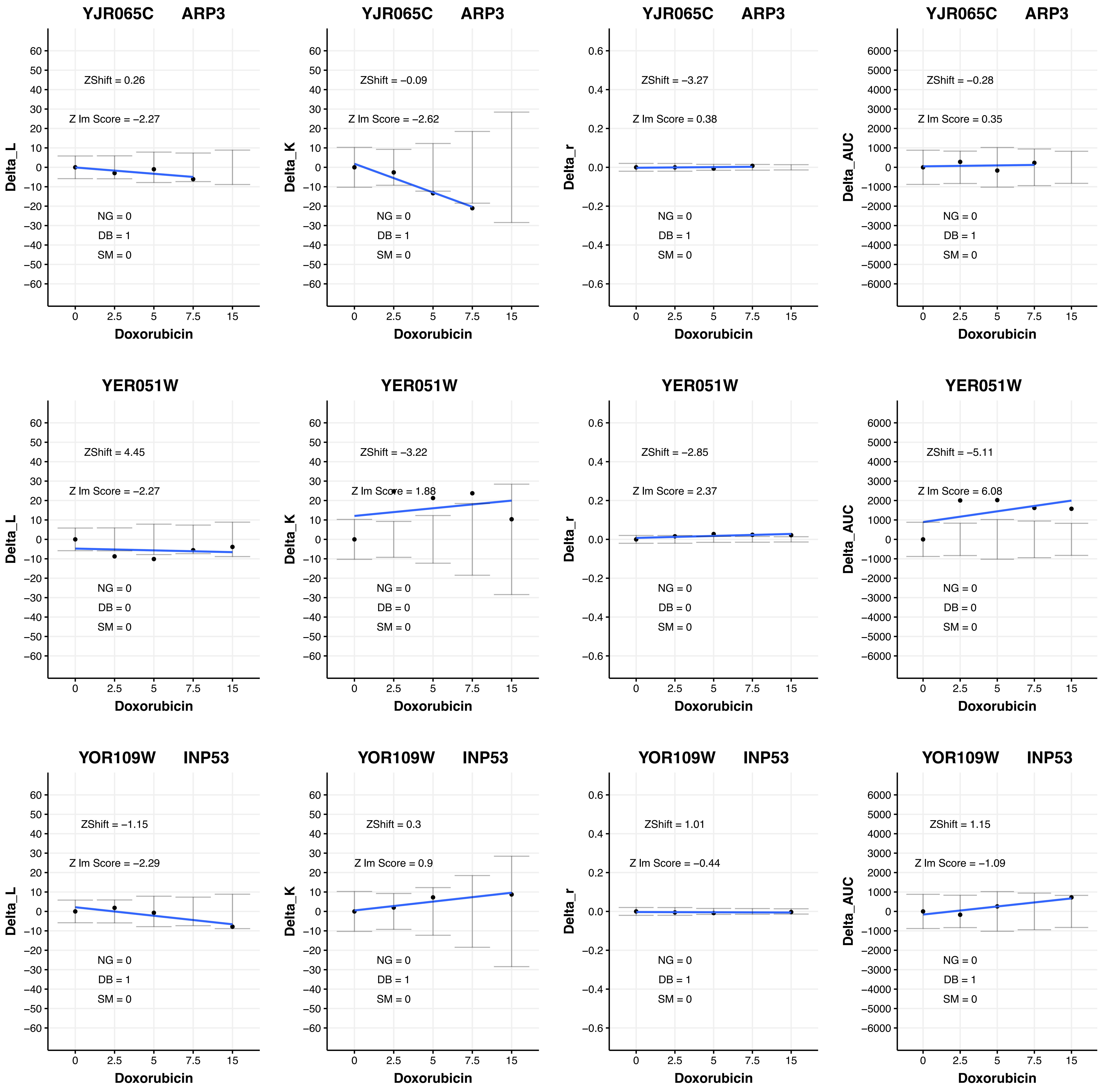

YJR065C ARP3
ZShift = 0.26
Z lm Score = −2.27
NG = 0
DB = 1
SM = 0
Delta_L
Doxorubicin
YJR065C ARP3
ZShift = −0.09
Z lm Score = −2.62
NG = 0
DB = 1
SM = 0
Delta_K
Doxorubicin
YJR065C ARP3
ZShift = −3.27
Z lm Score = 0.38
NG = 0
DB = 1
SM = 0
Delta_r
Doxorubicin
YJR065C ARP3
ZShift = −0.28
Z lm Score = 0.35
NG = 0
DB = 1
SM = 0
Delta_AUC
Doxorubicin
YER051W
ZShift = 4.45
Z lm Score = −2.27
NG = 0
DB = 0
SM = 0
Delta_L
Doxorubicin
YER051W
ZShift = −3.22
Z lm Score = 1.88
NG = 0
DB = 0
SM = 0
Delta_K
Doxorubicin
YER051W
ZShift = −2.85
Z lm Score = 2.37
NG = 0
DB = 0
SM = 0
Delta_r
Doxorubicin
YER051W
ZShift = −5.11
Z lm Score = 6.08
NG = 0
DB = 0
SM = 0
Delta_AUC
Doxorubicin
YOR109W INP53
ZShift = −1.15
Z lm Score = −2.29
NG = 0
DB = 1
SM = 0
Delta_L
Doxorubicin
YOR109W INP53
ZShift = 0.3
Z lm Score = 0.9
NG = 0
DB = 1
SM = 0
Delta_K
Doxorubicin
YOR109W INP53
ZShift = 1.01
Z lm Score = −0.44
NG = 0
DB = 1
SM = 0
Delta_r
Doxorubicin
YOR109W INP53
ZShift = 1.15
Z lm Score = −1.09
NG = 0
DB = 1
SM = 0
Delta_AUC
Doxorubicin

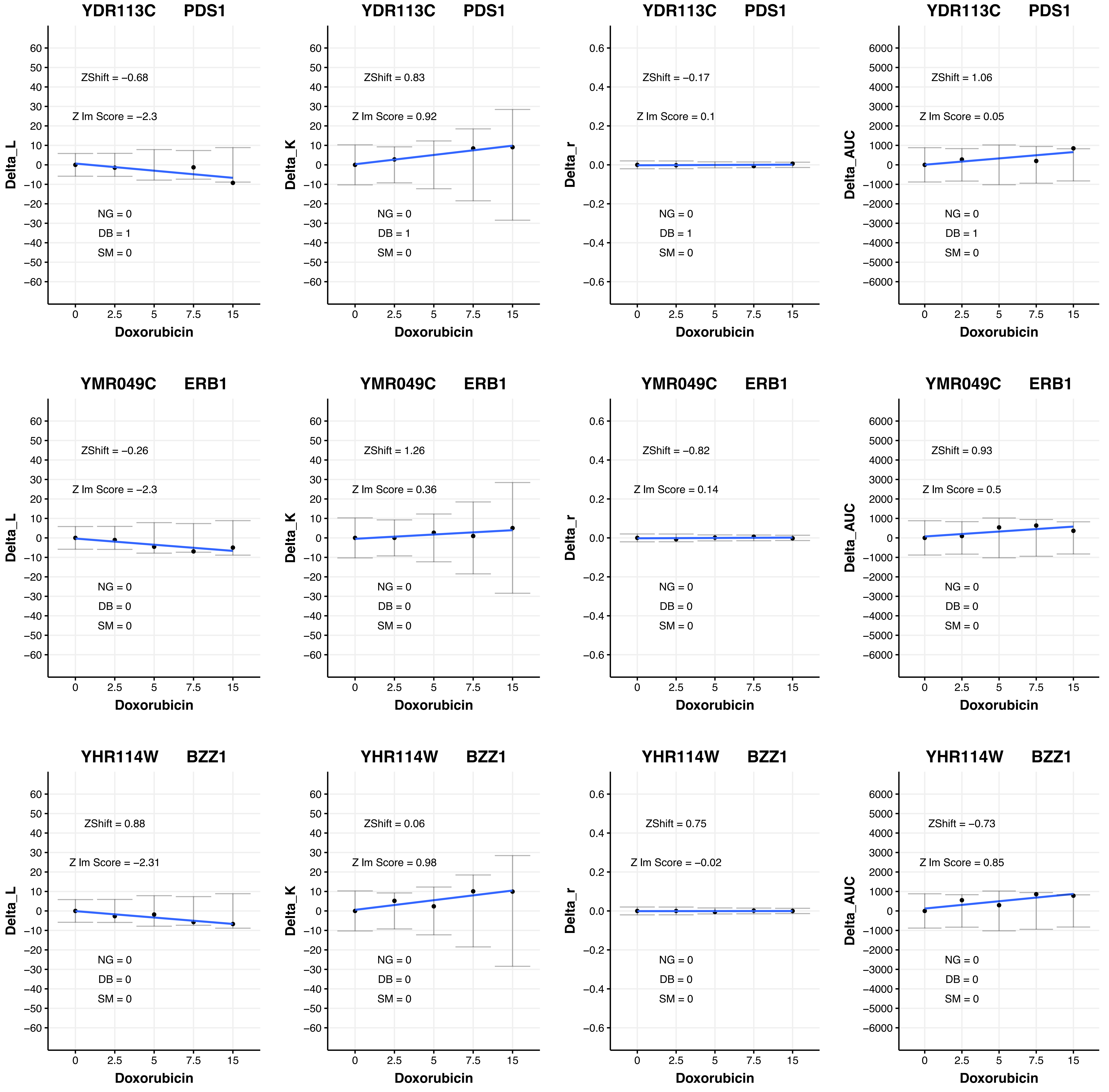
YDR113C PDS1
ZShift = −0.68
Z lm Score = −2.3
NG = 0
DB = 1
SM = 0
Delta_L
Doxorubicin
YDR113C PDS1
ZShift = 0.83
Z lm Score = 0.92
NG = 0
DB = 1
SM = 0
Delta_K
Doxorubicin
YDR113C PDS1
ZShift = −0.17
Z lm Score = 0.1
NG = 0
DB = 1
SM = 0
Delta_r
Doxorubicin
YDR113C PDS1
ZShift = 1.06
Z lm Score = 0.05
NG = 0
DB = 1
SM = 0
Delta_AUC
Doxorubicin
YMR049C ERB1
ZShift = −0.26
Z lm Score = −2.3
NG = 0
DB = 0
SM = 0
Delta_L
Doxorubicin
YMR049C ERB1
ZShift = 1.26
Z lm Score = 0.36
NG = 0
DB = 0
SM = 0
Delta_K
Doxorubicin
YMR049C ERB1
ZShift = −0.82
Z lm Score = 0.14
NG = 0
DB = 0
SM = 0
Delta_r
Doxorubicin
YMR049C ERB1
ZShift = 0.93
Z lm Score = 0.5
NG = 0
DB = 0
SM = 0
Delta_AUC
Doxorubicin
YHR114W BZZ1
ZShift = 0.88
Z lm Score = −2.31
NG = 0
DB = 0
SM = 0
Delta_L
Doxorubicin
YHR114W BZZ1
ZShift = 0.06
Z lm Score = 0.98
NG = 0
DB = 0
SM = 0
Delta_K
Doxorubicin
YHR114W BZZ1
ZShift = 0.75
Z lm Score = −0.02
NG = 0
DB = 0
SM = 0
Delta_r
Doxorubicin
YHR114W BZZ1
ZShift = −0.73
Z lm Score = 0.85
NG = 0
DB = 0
SM = 0
Delta_AUC
Doxorubicin

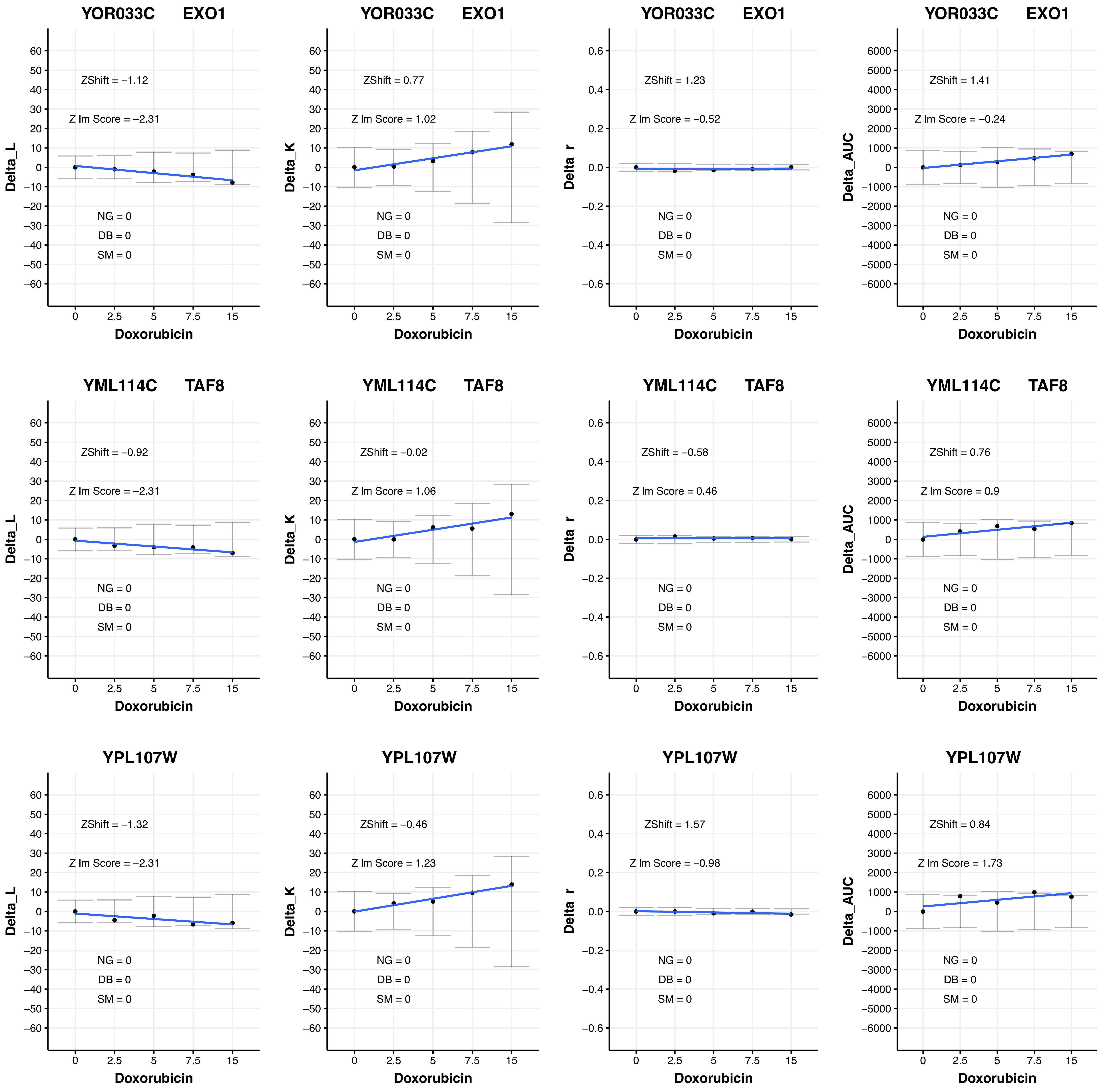
YOR033C EXO1
ZShift = −1.12
Z lm Score = −2.31
NG = 0
DB = 0
SM = 0
Delta_L
Doxorubicin
YOR033C EXO1
ZShift = 0.77
Z lm Score = 1.02
NG = 0
DB = 0
SM = 0
Delta_K
Doxorubicin
YOR033C EXO1
ZShift = 1.23
Z lm Score = −0.52
NG = 0
DB = 0
SM = 0
Delta_r
Doxorubicin
YOR033C EXO1
ZShift = 1.41
Z lm Score = −0.24
NG = 0
DB = 0
SM = 0
Delta_AUC
Doxorubicin
YML114C TAF8
ZShift = −0.92
Z lm Score = −2.31
NG = 0
DB = 0
SM = 0
Delta_L
Doxorubicin
YML114C TAF8
ZShift = −0.02
Z lm Score = 1.06
NG = 0
DB = 0
SM = 0
Delta_K
Doxorubicin
YML114C TAF8
ZShift = −0.58
Z lm Score = 0.46
NG = 0
DB = 0
SM = 0
Delta_r
Doxorubicin
YML114C TAF8
ZShift = 0.76
Z lm Score = 0.9
NG = 0
DB = 0
SM = 0
Delta_AUC
Doxorubicin
YPL107W
ZShift = −1.32
Z lm Score = −2.31
NG = 0
DB = 0
SM = 0
Delta_L
Doxorubicin
YPL107W
ZShift = −0.46
Z lm Score = 1.23
NG = 0
DB = 0
SM = 0
Delta_K
Doxorubicin
YPL107W
ZShift = 1.57
Z lm Score = −0.98
NG = 0
DB = 0
SM = 0
Delta_r
Doxorubicin
YPL107W
ZShift = 0.84
Z lm Score = 1.73
NG = 0
DB = 0
SM = 0
Delta_AUC
Doxorubicin

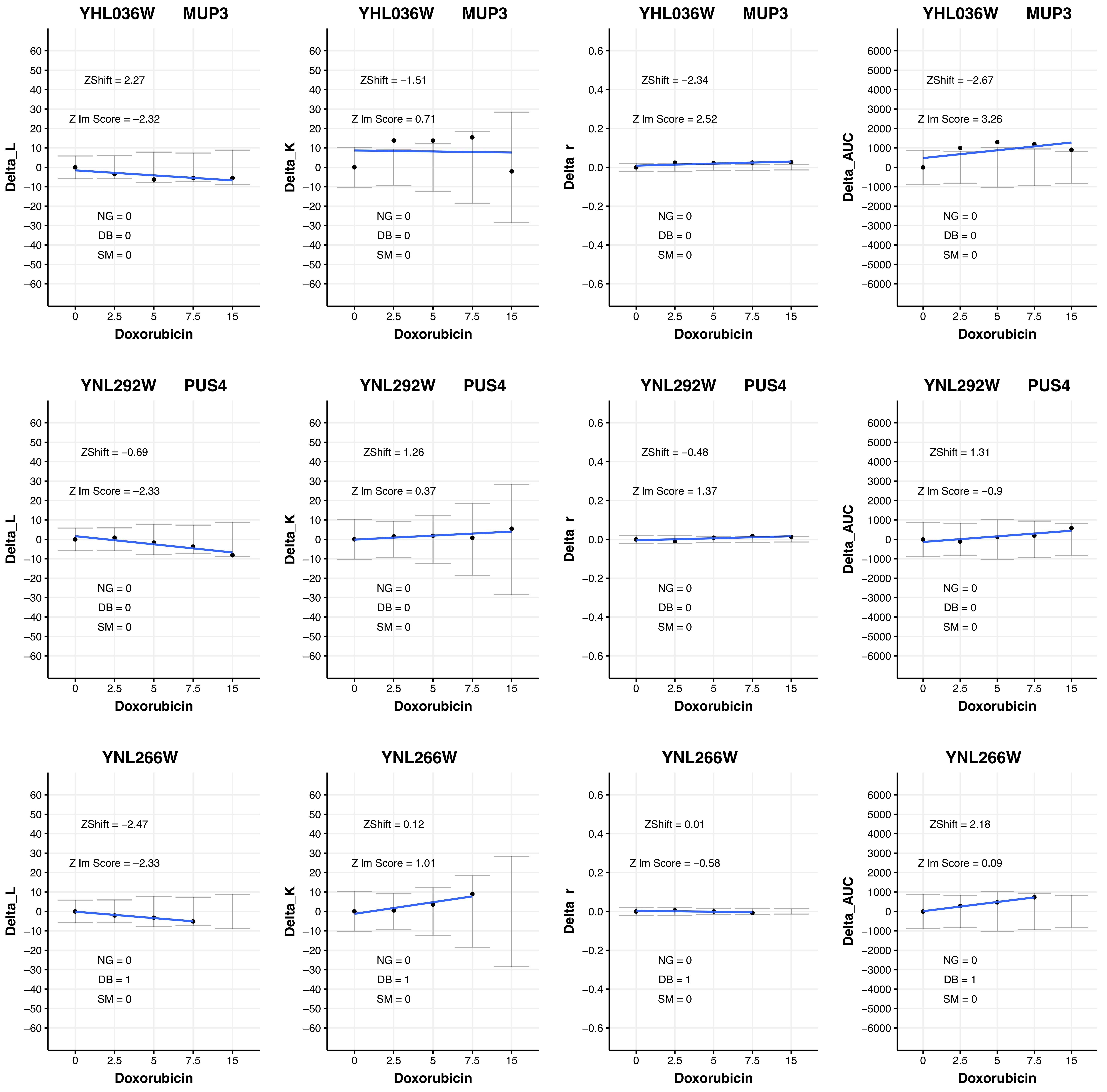
YHL036W MUP3
ZShift = 2.27
Z lm Score = −2.32
Delta_L
NG = 0
DB = 0
SM = 0
Doxorubicin
YHL036W MUP3
ZShift = −1.51
Z lm Score = 0.71
Delta_K
NG = 0
DB = 0
SM = 0
Doxorubicin
YHL036W MUP3
ZShift = −2.34
Z lm Score = 2.52
Delta_r
NG = 0
DB = 0
SM = 0
Doxorubicin
YHL036W MUP3
ZShift = −2.67
Z lm Score = 3.26
Delta_AUC
NG = 0
DB = 0
SM = 0
Doxorubicin
YNL292W PUS4
ZShift = −0.69
Z lm Score = −2.33
Delta_L
NG = 0
DB = 0
SM = 0
Doxorubicin
YNL292W PUS4
ZShift = 1.26
Z lm Score = 0.37
Delta_K
NG = 0
DB = 0
SM = 0
Doxorubicin
YNL292W PUS4
ZShift = −0.48
Z lm Score = 1.37
Delta_r
NG = 0
DB = 0
SM = 0
Doxorubicin
YNL292W PUS4
ZShift = 1.31
Z lm Score = −0.9
Delta_AUC
NG = 0
DB = 0
SM = 0
Doxorubicin
YNL266W
ZShift = −2.47
Z lm Score = −2.33
Delta_L
NG = 0
DB = 1
SM = 0
Doxorubicin
YNL266W
ZShift = 0.12
Z lm Score = 1.01
Delta_K
NG = 0
DB = 1
SM = 0
Doxorubicin
YNL266W
ZShift = 0.01
Z lm Score = −0.58
Delta_r
NG = 0
DB = 1
SM = 0
Doxorubicin
YNL266W
ZShift = 2.18
Z lm Score = 0.09
Delta_AUC
NG = 0
DB = 1
SM = 0
Doxorubicin

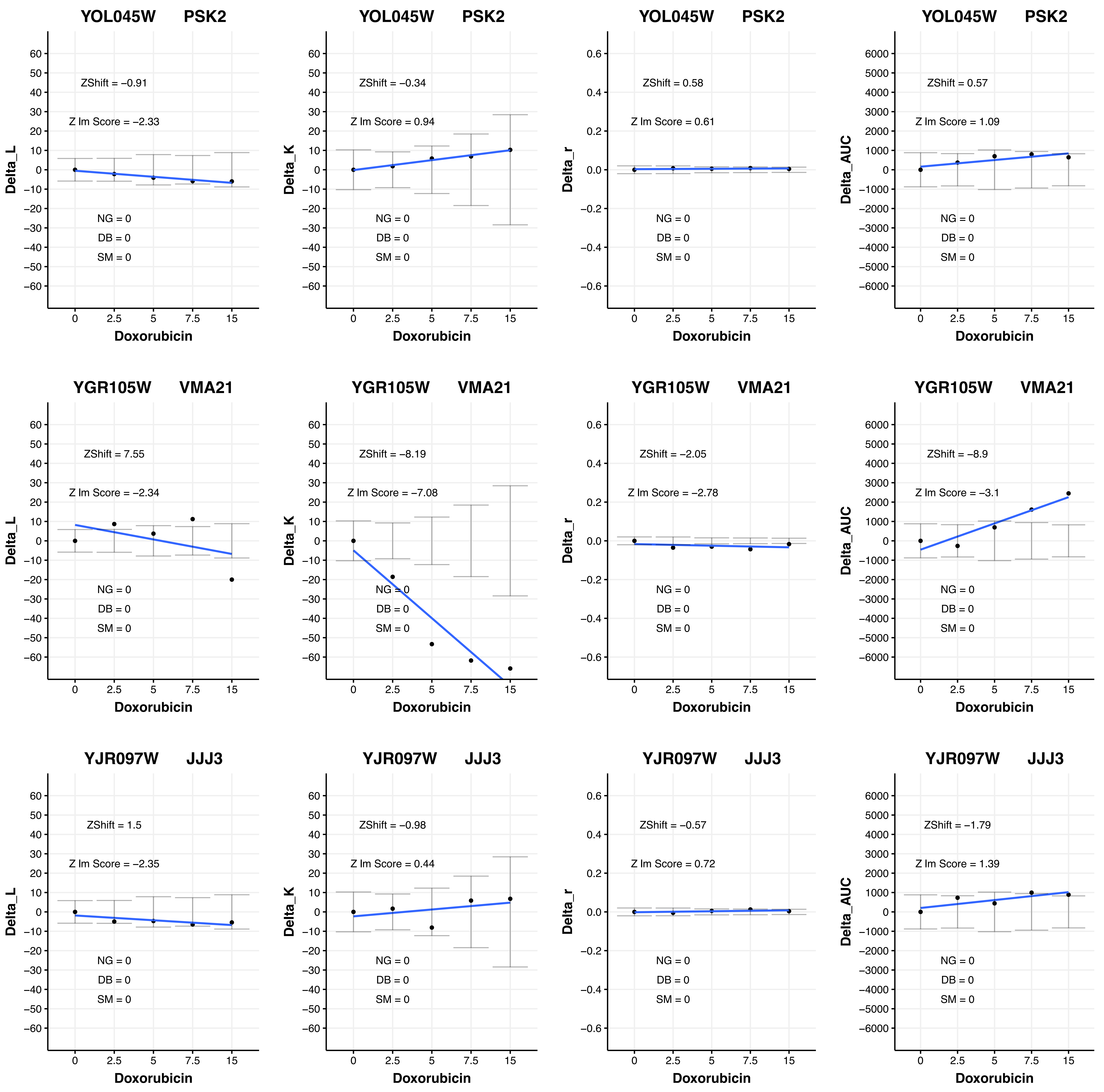

YOL045W PSK2
ZShift = −0.91
Z lm Score = −2.33
NG = 0
DB = 0
SM = 0
Delta_L
Doxorubicin
YOL045W PSK2
ZShift = −0.34
Z lm Score = 0.94
NG = 0
DB = 0
SM = 0
Delta_K
Doxorubicin
YOL045W PSK2
ZShift = 0.58
Z lm Score = 0.61
NG = 0
DB = 0
SM = 0
Delta_r
Doxorubicin
YOL045W PSK2
ZShift = 0.57
Z lm Score = 1.09
NG = 0
DB = 0
SM = 0
Delta_AUC
Doxorubicin
YGR105W VMA21
ZShift = 7.55
Z lm Score = −2.34
NG = 0
DB = 0
SM = 0
Delta_L
Doxorubicin
YGR105W VMA21
ZShift = −8.19
Z lm Score = −7.08
NG = 0
DB = 0
SM = 0
Delta_K
Doxorubicin
YGR105W VMA21
ZShift = −2.05
Z lm Score = −2.78
NG = 0
DB = 0
SM = 0
Delta_r
Doxorubicin
YGR105W VMA21
ZShift = −8.9
Z lm Score = −3.1
NG = 0
DB = 0
SM = 0
Delta_AUC
Doxorubicin
YJR097W JJJ3
ZShift = 1.5
Z lm Score = −2.35
NG = 0
DB = 0
SM = 0
Delta_L
Doxorubicin
YJR097W JJJ3
ZShift = −0.98
Z lm Score = 0.44
NG = 0
DB = 0
SM = 0
Delta_K
Doxorubicin
YJR097W JJJ3
ZShift = −0.57
Z lm Score = 0.72
NG = 0
DB = 0
SM = 0
Delta_r
Doxorubicin
YJR097W JJJ3
ZShift = −1.79
Z lm Score = 1.39
NG = 0
DB = 0
SM = 0
Delta_AUC
Doxorubicin

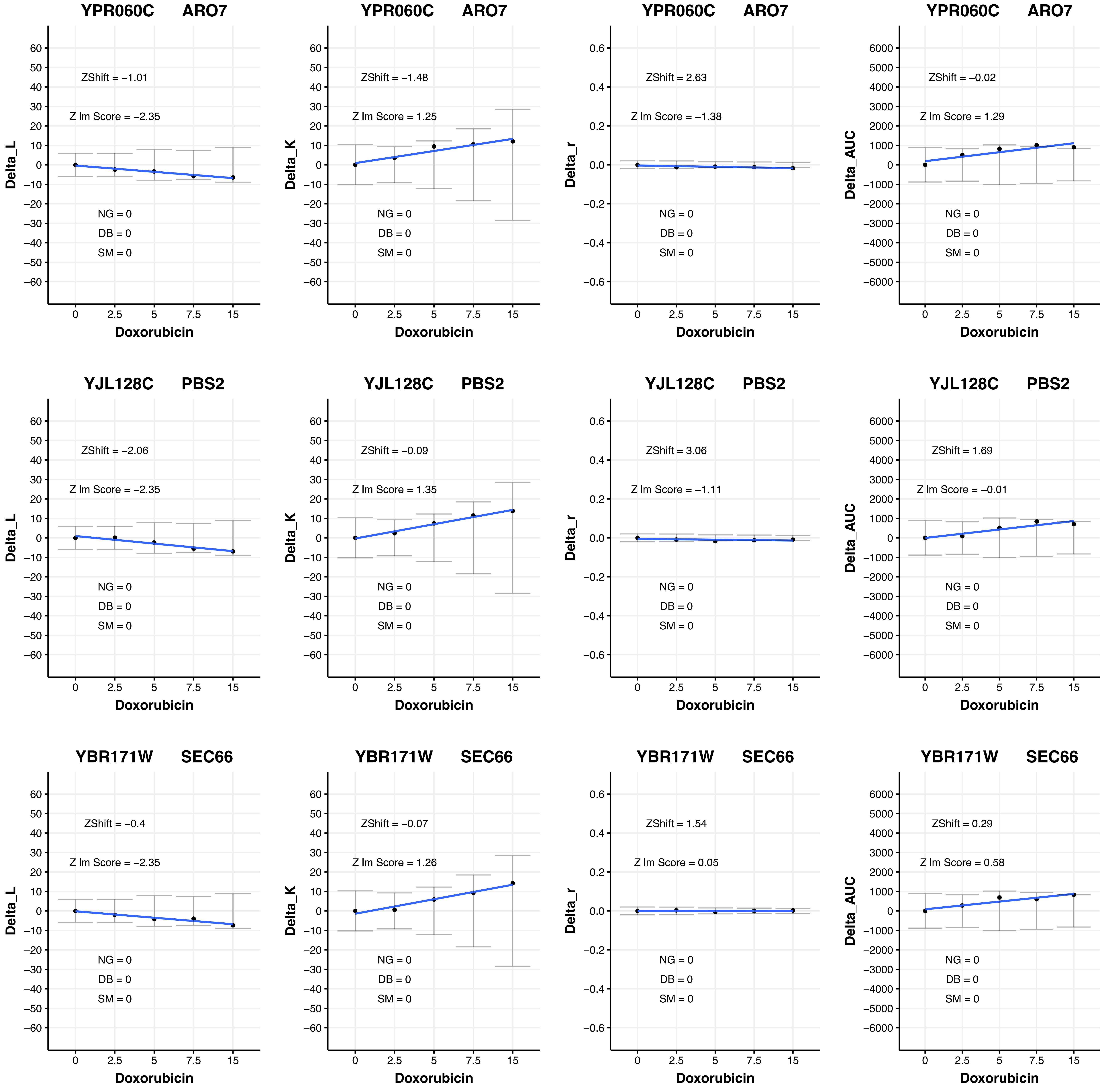
YPR060C ARO7
ZShift = −1.01
Z lm Score = −2.35
NG = 0
DB = 0
SM = 0
Delta_L
Doxorubicin
YPR060C ARO7
ZShift = −1.48
Z lm Score = 1.25
NG = 0
DB = 0
SM = 0
Delta_K
Doxorubicin
YPR060C ARO7
ZShift = 2.63
Z lm Score = −1.38
NG = 0
DB = 0
SM = 0
Delta_r
Doxorubicin
YPR060C ARO7
ZShift = −0.02
Z lm Score = 1.29
NG = 0
DB = 0
SM = 0
Delta_AUC
Doxorubicin
YJL128C PBS2
ZShift = −2.06
Z lm Score = −2.35
NG = 0
DB = 0
SM = 0
Delta_L
Doxorubicin
YJL128C PBS2
ZShift = −0.09
Z lm Score = 1.35
NG = 0
DB = 0
SM = 0
Delta_K
Doxorubicin
YJL128C PBS2
ZShift = 3.06
Z lm Score = −1.11
NG = 0
DB = 0
SM = 0
Delta_r
Doxorubicin
YJL128C PBS2
ZShift = 1.69
Z lm Score = −0.01
NG = 0
DB = 0
SM = 0
Delta_AUC
Doxorubicin
YBR171W SEC66
ZShift = −0.4
Z lm Score = −2.35
NG = 0
DB = 0
SM = 0
Delta_L
Doxorubicin
YBR171W SEC66
ZShift = −0.07
Z lm Score = 1.26
NG = 0
DB = 0
SM = 0
Delta_K
Doxorubicin
YBR171W SEC66
ZShift = 1.54
Z lm Score = 0.05
NG = 0
DB = 0
SM = 0
Delta_r
Doxorubicin
YBR171W SEC66
ZShift = 0.29
Z lm Score = 0.58
NG = 0
DB = 0
SM = 0
Delta_AUC
Doxorubicin

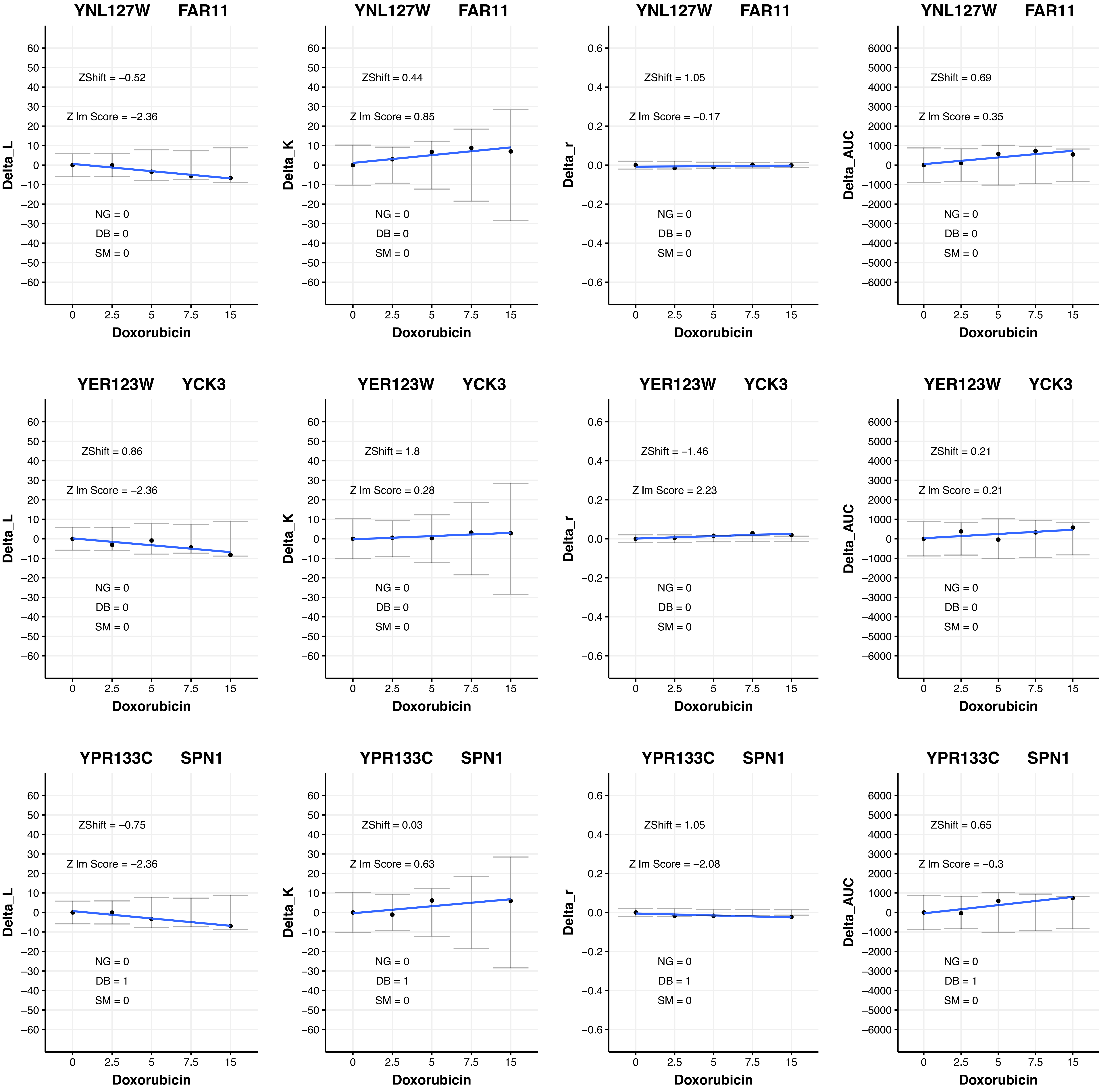
YNL127W FAR11
ZShift = −0.52
Z lm Score = −2.36
NG = 0
DB = 0
SM = 0
Delta_L
Doxorubicin
YNL127W FAR11
ZShift = 0.44
Z lm Score = 0.85
NG = 0
DB = 0
SM = 0
Delta_K
Doxorubicin
YNL127W FAR11
ZShift = 1.05
Z lm Score = −0.17
NG = 0
DB = 0
SM = 0
Delta_r
Doxorubicin
YNL127W FAR11
ZShift = 0.69
Z lm Score = 0.35
NG = 0
DB = 0
SM = 0
Delta_AUC
Doxorubicin
YER123W YCK3
ZShift = 0.86
Z lm Score = −2.36
NG = 0
DB = 0
SM = 0
Delta_L
Doxorubicin
YER123W YCK3
ZShift = 1.8
Z lm Score = 0.28
NG = 0
DB = 0
SM = 0
Delta_K
Doxorubicin
YER123W YCK3
ZShift = −1.46
Z lm Score = 2.23
NG = 0
DB = 0
SM = 0
Delta_r
Doxorubicin
YER123W YCK3
ZShift = 0.21
Z lm Score = 0.21
NG = 0
DB = 0
SM = 0
Delta_AUC
Doxorubicin
YPR133C SPN1
ZShift = −0.75
Z lm Score = −2.36
NG = 0
DB = 1
SM = 0
Delta_L
Doxorubicin
YPR133C SPN1
ZShift = 0.03
Z lm Score = 0.63
NG = 0
DB = 1
SM = 0
Delta_K
Doxorubicin
YPR133C SPN1
ZShift = 1.05
Z lm Score = −2.08
NG = 0
DB = 1
SM = 0
Delta_r
Doxorubicin
YPR133C SPN1
ZShift = 0.65
Z lm Score = −0.3
NG = 0
DB = 1
SM = 0
Delta_AUC
Doxorubicin

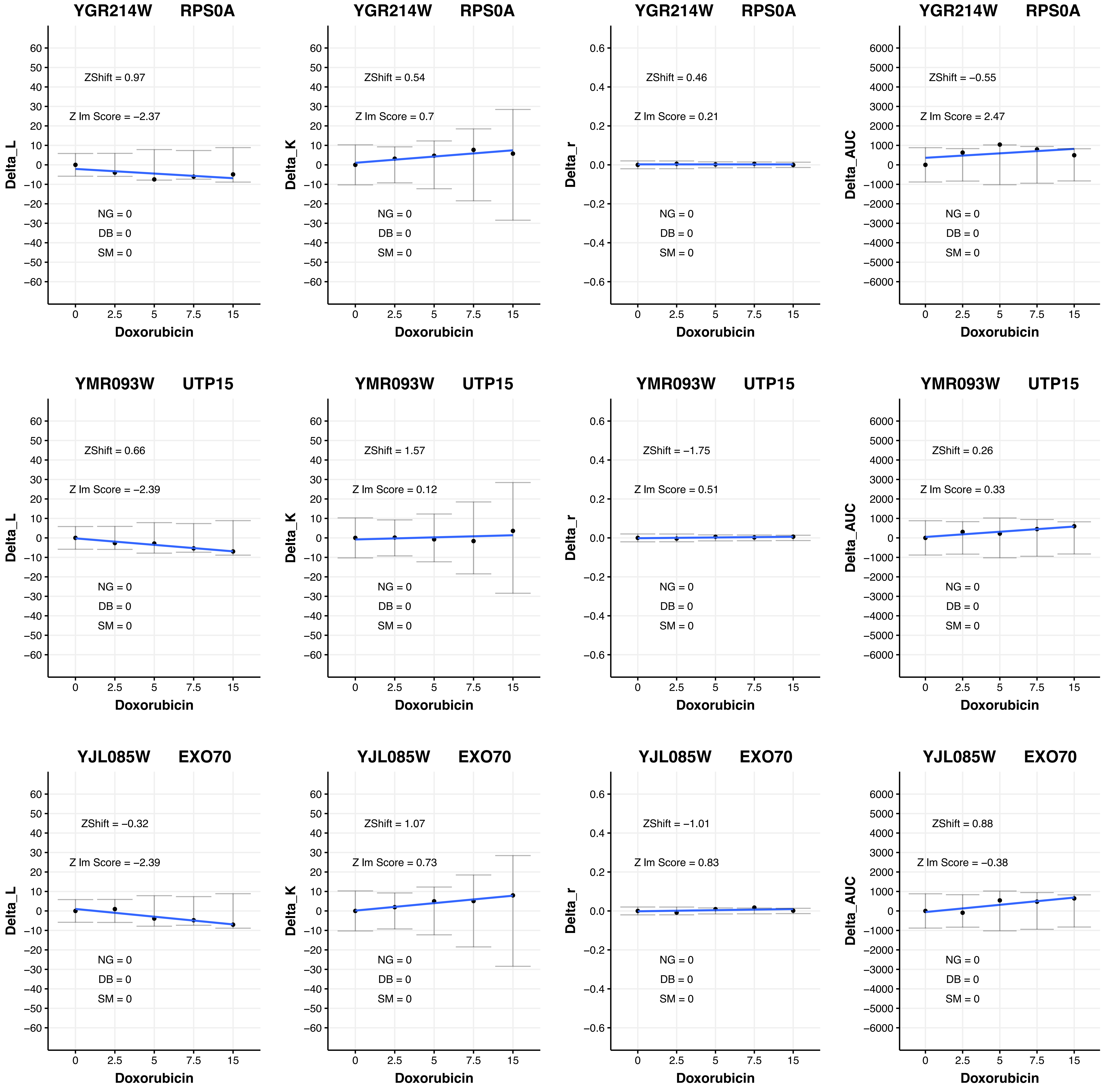
YGR214W RPS0A
ZShift = 0.97
Z lm Score = −2.37
NG = 0
DB = 0
SM = 0
Delta_L
Doxorubicin
YGR214W RPS0A
ZShift = 0.54
Z lm Score = 0.7
NG = 0
DB = 0
SM = 0
Delta_K
Doxorubicin
YGR214W RPS0A
ZShift = 0.46
Z lm Score = 0.21
NG = 0
DB = 0
SM = 0
Delta_r
Doxorubicin
YGR214W RPS0A
ZShift = −0.55
Z lm Score = 2.47
NG = 0
DB = 0
SM = 0
Delta_AUC
Doxorubicin
YMR093W UTP15
ZShift = 0.66
Z lm Score = −2.39
NG = 0
DB = 0
SM = 0
Delta_L
Doxorubicin
YMR093W UTP15
ZShift = 1.57
Z lm Score = 0.12
NG = 0
DB = 0
SM = 0
Delta_K
Doxorubicin
YMR093W UTP15
ZShift = −1.75
Z lm Score = 0.51
NG = 0
DB = 0
SM = 0
Delta_r
Doxorubicin
YMR093W UTP15
ZShift = 0.26
Z lm Score = 0.33
NG = 0
DB = 0
SM = 0
Delta_AUC
Doxorubicin
YJL085W EXO70
ZShift = −0.32
Z lm Score = −2.39
NG = 0
DB = 0
SM = 0
Delta_L
Doxorubicin
YJL085W EXO70
ZShift = 1.07
Z lm Score = 0.73
NG = 0
DB = 0
SM = 0
Delta_K
Doxorubicin
YJL085W EXO70
ZShift = −1.01
Z lm Score = 0.83
NG = 0
DB = 0
SM = 0
Delta_r
Doxorubicin
YJL085W EXO70
ZShift = 0.88
Z lm Score = −0.38
NG = 0
DB = 0
SM = 0
Delta_AUC
Doxorubicin

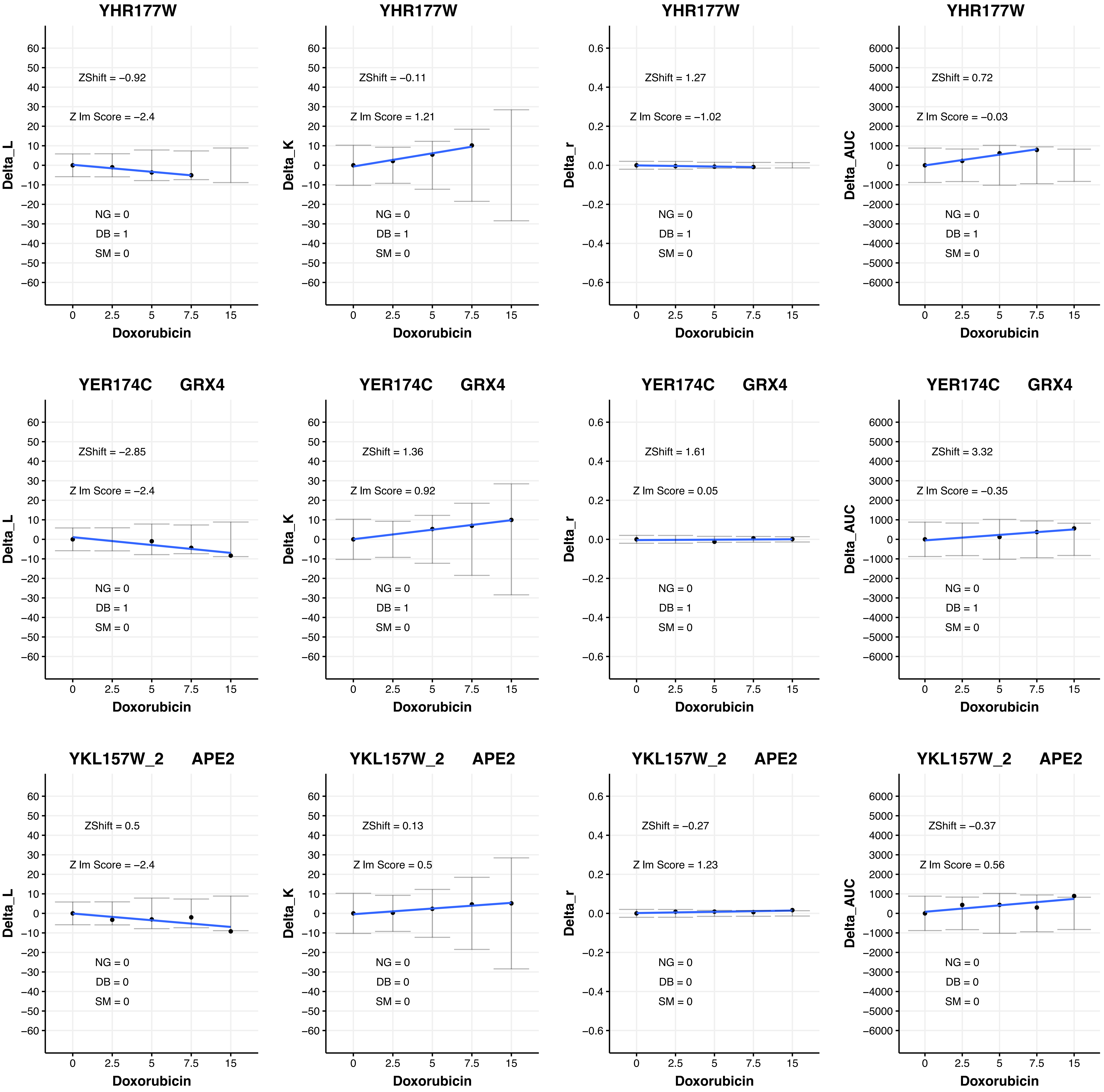
YHR177W
ZShift = −0.92
Z lm Score = −2.4
NG = 0
DB = 1
SM = 0
Delta_L
Doxorubicin
YHR177W
ZShift = −0.11
Z lm Score = 1.21
NG = 0
DB = 1
SM = 0
Delta_K
Doxorubicin
YHR177W
ZShift = 1.27
Z lm Score = −1.02
NG = 0
DB = 1
SM = 0
Delta_r
Doxorubicin
YHR177W
ZShift = 0.72
Z lm Score = −0.03
NG = 0
DB = 1
SM = 0
Delta_AUC
Doxorubicin
YER174C GRX4
ZShift = −2.85
Z lm Score = −2.4
NG = 0
DB = 1
SM = 0
Delta_L
Doxorubicin
YER174C GRX4
ZShift = 1.36
Z lm Score = 0.92
NG = 0
DB = 1
SM = 0
Delta_K
Doxorubicin
YER174C GRX4
ZShift = 1.61
Z lm Score = 0.05
NG = 0
DB = 1
SM = 0
Delta_r
Doxorubicin
YER174C GRX4
ZShift = 3.32
Z lm Score = −0.35
NG = 0
DB = 1
SM = 0
Delta_AUC
Doxorubicin
YKL157W_2 APE2
ZShift = 0.5
Z lm Score = −2.4
NG = 0
DB = 0
SM = 0
Delta_L
Doxorubicin
YKL157W_2 APE2
ZShift = 0.13
Z lm Score = 0.5
NG = 0
DB = 0
SM = 0
Delta_K
Doxorubicin
YKL157W_2 APE2
ZShift = −0.27
Z lm Score = 1.23
NG = 0
DB = 0
SM = 0
Delta_r
Doxorubicin
YKL157W_2 APE2
ZShift = −0.37
Z lm Score = 0.56
NG = 0
DB = 0
SM = 0
Delta_AUC
Doxorubicin

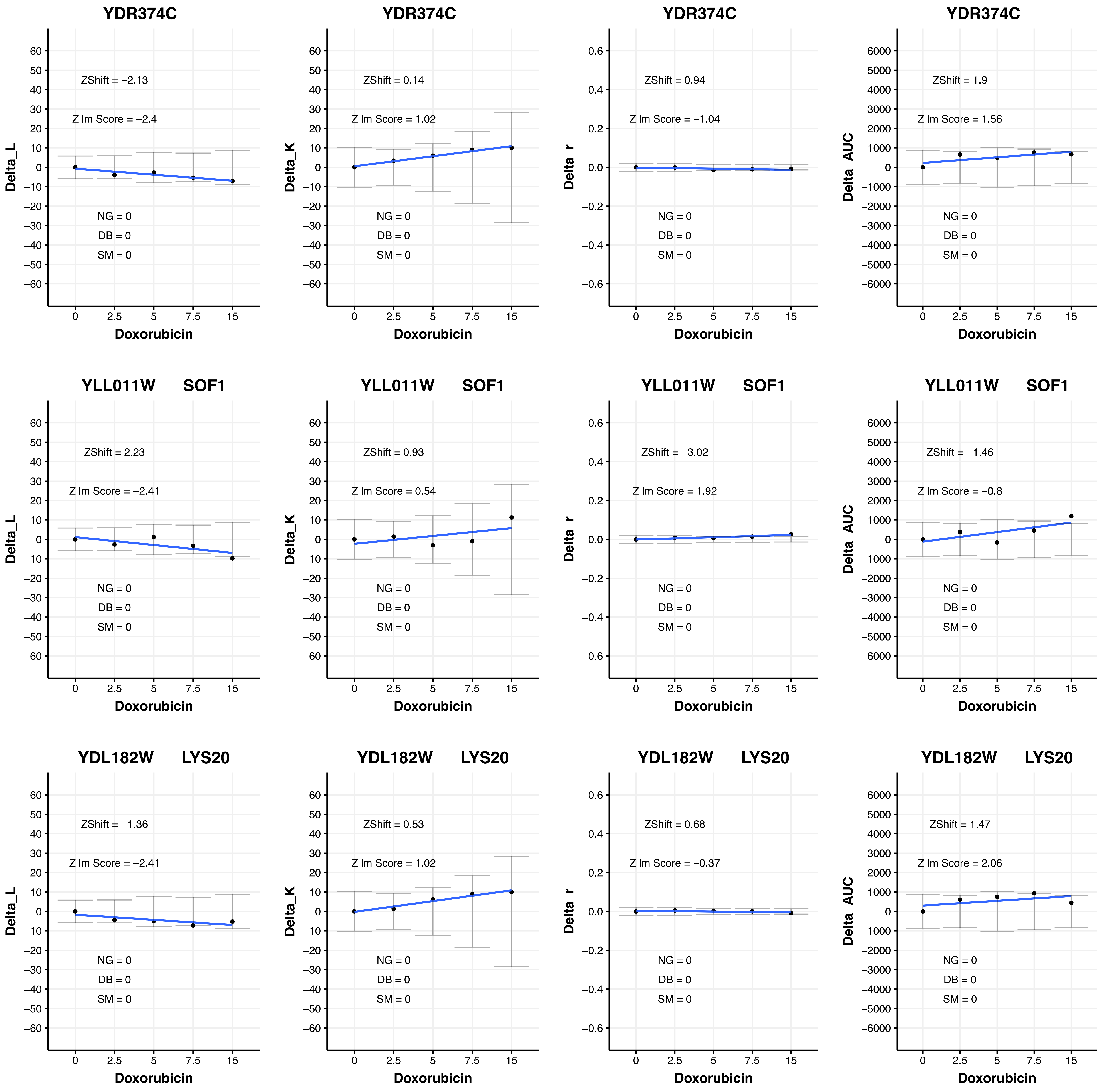
YDR374C
ZShift = −2.13
Z lm Score = −2.4
NG = 0
DB = 0
SM = 0
Delta_L
Doxorubicin
YDR374C
ZShift = 0.14
Z lm Score = 1.02
NG = 0
DB = 0
SM = 0
Delta_K
Doxorubicin
YDR374C
ZShift = 0.94
Z lm Score = −1.04
NG = 0
DB = 0
SM = 0
Delta_r
Doxorubicin
YDR374C
ZShift = 1.9
Z lm Score = 1.56
NG = 0
DB = 0
SM = 0
Delta_AUC
Doxorubicin
YLL011W SOF1
ZShift = 2.23
Z lm Score = −2.41
NG = 0
DB = 0
SM = 0
Delta_L
Doxorubicin
YLL011W SOF1
ZShift = 0.93
Z lm Score = 0.54
NG = 0
DB = 0
SM = 0
Delta_K
Doxorubicin
YLL011W SOF1
ZShift = −3.02
Z lm Score = 1.92
NG = 0
DB = 0
SM = 0
Delta_r
Doxorubicin
YLL011W SOF1
ZShift = −1.46
Z lm Score = −0.8
NG = 0
DB = 0
SM = 0
Delta_AUC
Doxorubicin
YDL182W LYS20
ZShift = −1.36
Z lm Score = −2.41
NG = 0
DB = 0
SM = 0
Delta_L
Doxorubicin
YDL182W LYS20
ZShift = 0.53
Z lm Score = 1.02
NG = 0
DB = 0
SM = 0
Delta_K
Doxorubicin
YDL182W LYS20
ZShift = 0.68
Z lm Score = −0.37
NG = 0
DB = 0
SM = 0
Delta_r
Doxorubicin
YDL182W LYS20
ZShift = 1.47
Z lm Score = 2.06
NG = 0
DB = 0
SM = 0
Delta_AUC
Doxorubicin

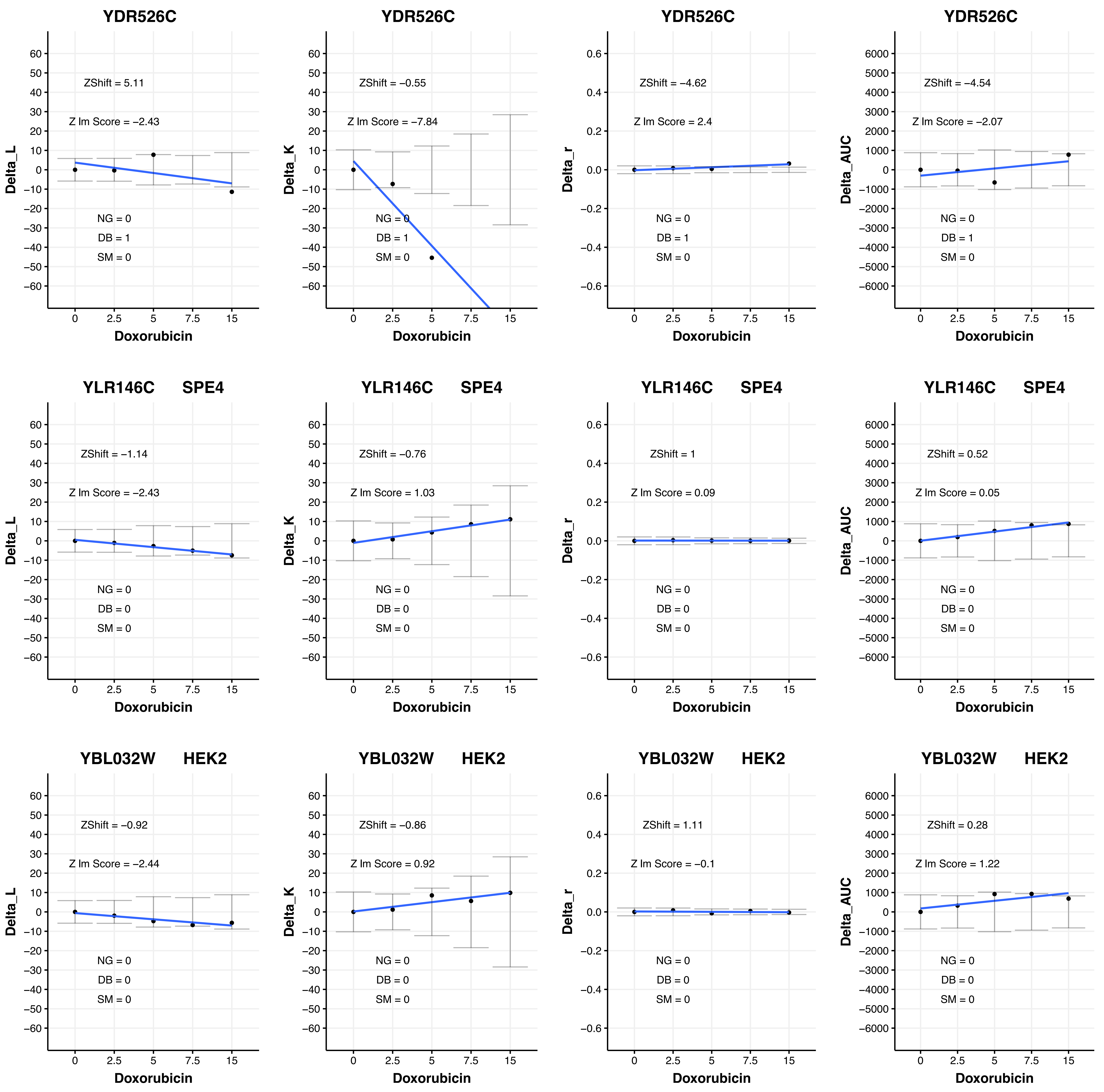
YDR526C
ZShift = 5.11
Z lm Score = −2.43
NG = 0
DB = 1
SM = 0
Delta_L
Doxorubicin
YDR526C
ZShift = −0.55
Z lm Score = −7.84
NG = 0
DB = 1
SM = 0
Delta_K
Doxorubicin
YDR526C
ZShift = −4.62
Z lm Score = 2.4
NG = 0
DB = 1
SM = 0
Delta_r
Doxorubicin
YDR526C
ZShift = −4.54
Z lm Score = −2.07
NG = 0
DB = 1
SM = 0
Delta_AUC
Doxorubicin
YLR146C SPE4
ZShift = −1.14
Z lm Score = −2.43
NG = 0
DB = 0
SM = 0
Delta_L
Doxorubicin
YLR146C SPE4
ZShift = −0.76
Z lm Score = 1.03
NG = 0
DB = 0
SM = 0
Delta_K
Doxorubicin
YLR146C SPE4
ZShift = 1
Z lm Score = 0.09
NG = 0
DB = 0
SM = 0
Delta_r
Doxorubicin
YLR146C SPE4
ZShift = 0.52
Z lm Score = 0.05
NG = 0
DB = 0
SM = 0
Delta_AUC
Doxorubicin
YBL032W HEK2
ZShift = −0.92
Z lm Score = −2.44
NG = 0
DB = 0
SM = 0
Delta_L
Doxorubicin
YBL032W HEK2
ZShift = −0.86
Z lm Score = 0.92
NG = 0
DB = 0
SM = 0
Delta_K
Doxorubicin
YBL032W HEK2
ZShift = 1.11
Z lm Score = −0.1
NG = 0
DB = 0
SM = 0
Delta_r
Doxorubicin
YBL032W HEK2
ZShift = 0.28
Z lm Score = 1.22
NG = 0
DB = 0
SM = 0
Delta_AUC
Doxorubicin

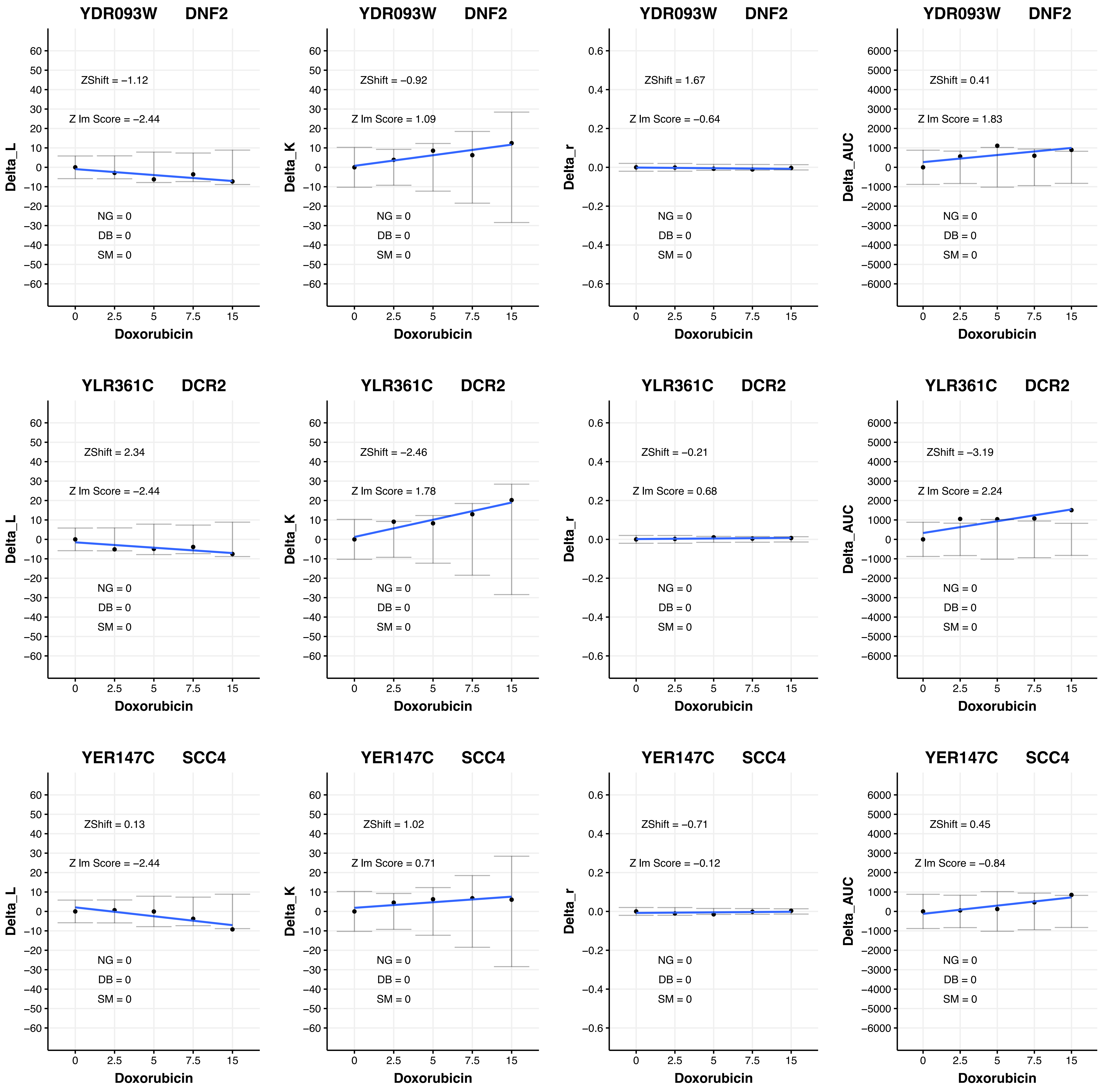
YDR093W DNF2
ZShift = −1.12
Z lm Score = −2.44
NG = 0
DB = 0
SM = 0
Delta_L
Doxorubicin
YDR093W DNF2
ZShift = −0.92
Z lm Score = 1.09
NG = 0
DB = 0
SM = 0
Delta_K
Doxorubicin
YDR093W DNF2
ZShift = 1.67
Z lm Score = −0.64
NG = 0
DB = 0
SM = 0
Delta_r
Doxorubicin
YDR093W DNF2
ZShift = 0.41
Z lm Score = 1.83
NG = 0
DB = 0
SM = 0
Delta_AUC
Doxorubicin
YLR361C DCR2
ZShift = 2.34
Z lm Score = −2.44
NG = 0
DB = 0
SM = 0
Delta_L
Doxorubicin
YLR361C DCR2
ZShift = −2.46
Z lm Score = 1.78
NG = 0
DB = 0
SM = 0
Delta_K
Doxorubicin
YLR361C DCR2
ZShift = −0.21
Z lm Score = 0.68
NG = 0
DB = 0
SM = 0
Delta_r
Doxorubicin
YLR361C DCR2
ZShift = −3.19
Z lm Score = 2.24
NG = 0
DB = 0
SM = 0
Delta_AUC
Doxorubicin
YER147C SCC4
ZShift = 0.13
Z lm Score = −2.44
NG = 0
DB = 0
SM = 0
Delta_L
Doxorubicin
YER147C SCC4
ZShift = 1.02
Z lm Score = 0.71
NG = 0
DB = 0
SM = 0
Delta_K
Doxorubicin
YER147C SCC4
ZShift = −0.71
Z lm Score = −0.12
NG = 0
DB = 0
SM = 0
Delta_r
Doxorubicin
YER147C SCC4
ZShift = 0.45
Z lm Score = −0.84
NG = 0
DB = 0
SM = 0
Delta_AUC
Doxorubicin

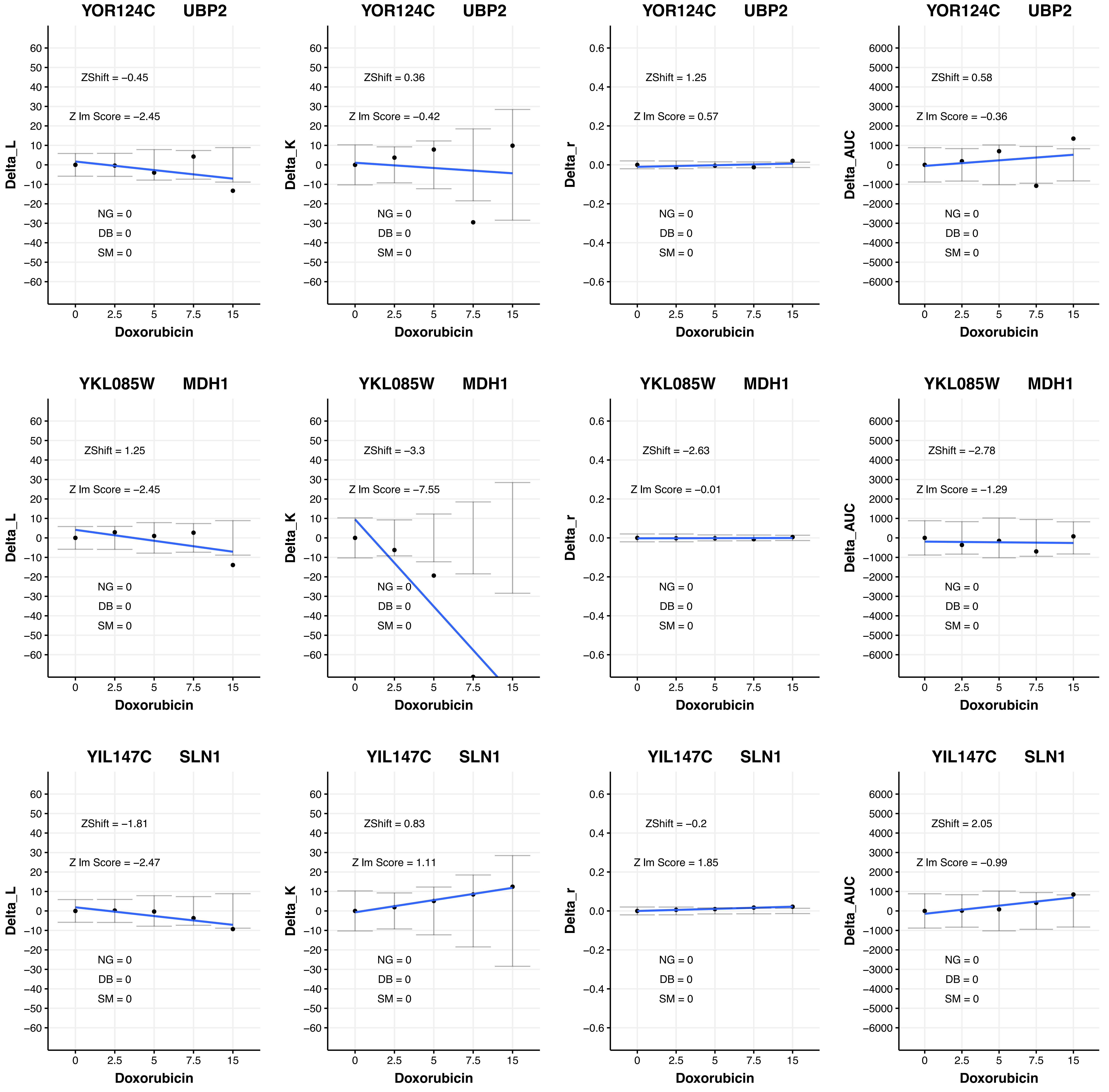
YOR124C UBP2
ZShift = −0.45
Z lm Score = −2.45
NG = 0
DB = 0
SM = 0
Delta_L
Doxorubicin
YOR124C UBP2
ZShift = 0.36
Z lm Score = −0.42
NG = 0
DB = 0
SM = 0
Delta_K
Doxorubicin
YOR124C UBP2
ZShift = 1.25
Z lm Score = 0.57
NG = 0
DB = 0
SM = 0
Delta_r
Doxorubicin
YOR124C UBP2
ZShift = 0.58
Z lm Score = −0.36
NG = 0
DB = 0
SM = 0
Delta_AUC
Doxorubicin
YKL085W MDH1
ZShift = 1.25
Z lm Score = −2.45
NG = 0
DB = 0
SM = 0
Delta_L
Doxorubicin
YKL085W MDH1
ZShift = −3.3
Z lm Score = −7.55
NG = 0
DB = 0
SM = 0
Delta_K
Doxorubicin
YKL085W MDH1
ZShift = −2.63
Z lm Score = −0.01
NG = 0
DB = 0
SM = 0
Delta_r
Doxorubicin
YKL085W MDH1
ZShift = −2.78
Z lm Score = −1.29
NG = 0
DB = 0
SM = 0
Delta_AUC
Doxorubicin
YIL147C SLN1
ZShift = −1.81
Z lm Score = −2.47
NG = 0
DB = 0
SM = 0
Delta_L
Doxorubicin
YIL147C SLN1
ZShift = 0.83
Z lm Score = 1.11
NG = 0
DB = 0
SM = 0
Delta_K
Doxorubicin
YIL147C SLN1
ZShift = −0.2
Z lm Score = 1.85
NG = 0
DB = 0
SM = 0
Delta_r
Doxorubicin
YIL147C SLN1
ZShift = 2.05
Z lm Score = −0.99
NG = 0
DB = 0
SM = 0
Delta_AUC
Doxorubicin

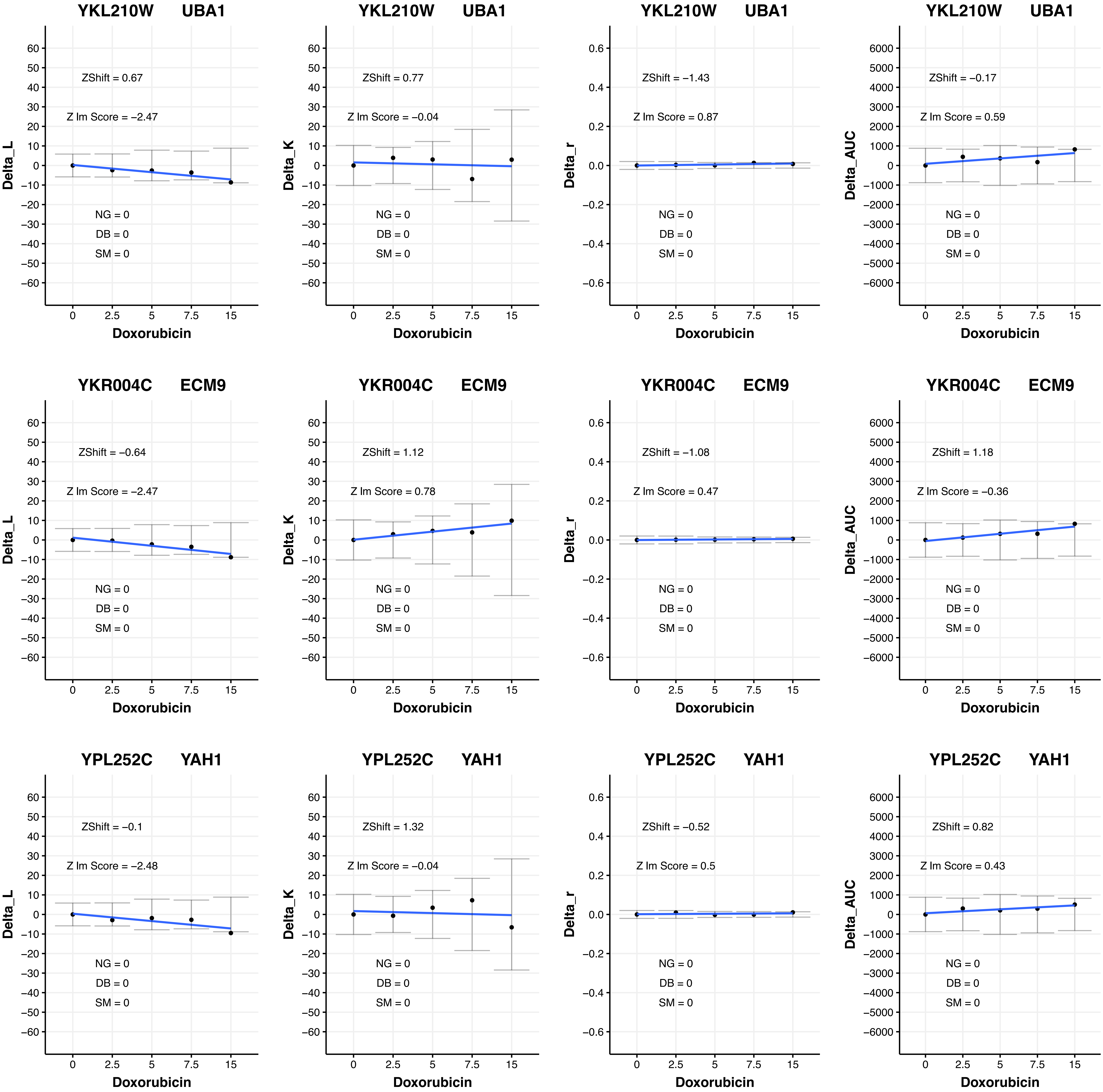

YKL210W UBA1
ZShift = 0.67
Z lm Score = −2.47
NG = 0
DB = 0
SM = 0
Delta_L
Doxorubicin
YKL210W UBA1
ZShift = 0.77
Z lm Score = −0.04
NG = 0
DB = 0
SM = 0
Delta_K
Doxorubicin
YKL210W UBA1
ZShift = −1.43
Z lm Score = 0.87
NG = 0
DB = 0
SM = 0
Delta_r
Doxorubicin
YKL210W UBA1
ZShift = −0.17
Z lm Score = 0.59
NG = 0
DB = 0
SM = 0
Delta_AUC
Doxorubicin
YKR004C ECM9
ZShift = −0.64
Z lm Score = −2.47
NG = 0
DB = 0
SM = 0
Delta_L
Doxorubicin
YKR004C ECM9
ZShift = 1.12
Z lm Score = 0.78
NG = 0
DB = 0
SM = 0
Delta_K
Doxorubicin
YKR004C ECM9
ZShift = −1.08
Z lm Score = 0.47
NG = 0
DB = 0
SM = 0
Delta_r
Doxorubicin
YKR004C ECM9
ZShift = 1.18
Z lm Score = −0.36
NG = 0
DB = 0
SM = 0
Delta_AUC
Doxorubicin
YPL252C YAH1
ZShift = −0.1
Z lm Score = −2.48
NG = 0
DB = 0
SM = 0
Delta_L
Doxorubicin
YPL252C YAH1
ZShift = 1.32
Z lm Score = −0.04
NG = 0
DB = 0
SM = 0
Delta_K
Doxorubicin
YPL252C YAH1
ZShift = −0.52
Z lm Score = 0.5
NG = 0
DB = 0
SM = 0
Delta_r
Doxorubicin
YPL252C YAH1
ZShift = 0.82
Z lm Score = 0.43
NG = 0
DB = 0
SM = 0
Delta_AUC
Doxorubicin

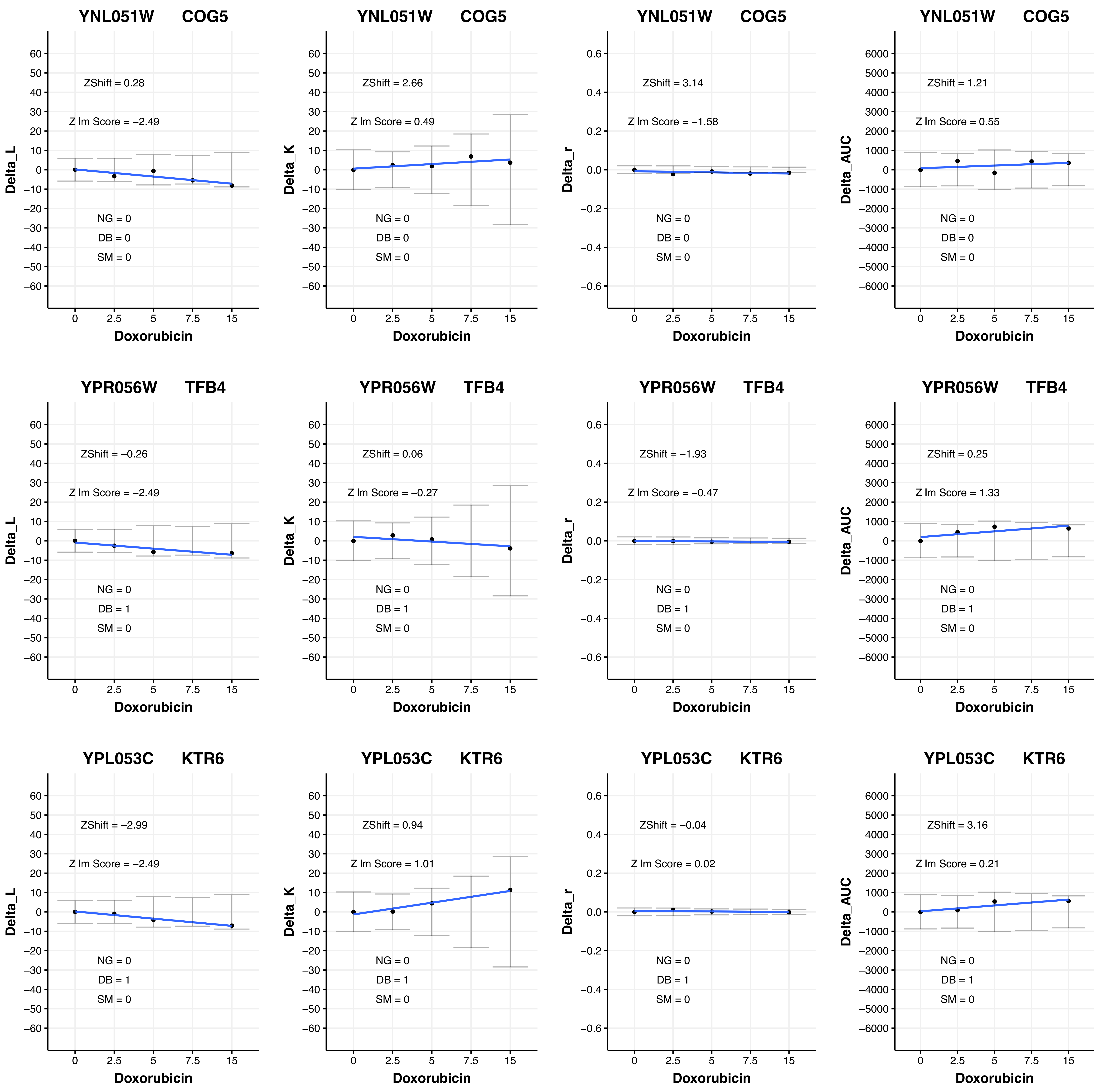
YNL051W COG5
ZShift = 0.28
Z lm Score = −2.49
NG = 0
DB = 0
SM = 0
Delta_L
Doxorubicin
YNL051W COG5
ZShift = 2.66
Z lm Score = 0.49
NG = 0
DB = 0
SM = 0
Delta_K
Doxorubicin
YNL051W COG5
ZShift = 3.14
Z lm Score = −1.58
NG = 0
DB = 0
SM = 0
Delta_r
Doxorubicin
YNL051W COG5
ZShift = 1.21
Z lm Score = 0.55
NG = 0
DB = 0
SM = 0
Delta_AUC
Doxorubicin
YPR056W TFB4
ZShift = −0.26
Z lm Score = −2.49
NG = 0
DB = 1
SM = 0
Delta_L
Doxorubicin
YPR056W TFB4
ZShift = 0.06
Z lm Score = −0.27
NG = 0
DB = 1
SM = 0
Delta_K
Doxorubicin
YPR056W TFB4
ZShift = −1.93
Z lm Score = −0.47
NG = 0
DB = 1
SM = 0
Delta_r
Doxorubicin
YPR056W TFB4
ZShift = 0.25
Z lm Score = 1.33
NG = 0
DB = 1
SM = 0
Delta_AUC
Doxorubicin
YPL053C KTR6
ZShift = −2.99
Z lm Score = −2.49
NG = 0
DB = 1
SM = 0
Delta_L
Doxorubicin
YPL053C KTR6
ZShift = 0.94
Z lm Score = 1.01
NG = 0
DB = 1
SM = 0
Delta_K
Doxorubicin
YPL053C KTR6
ZShift = −0.04
Z lm Score = 0.02
NG = 0
DB = 1
SM = 0
Delta_r
Doxorubicin
YPL053C KTR6
ZShift = 3.16
Z lm Score = 0.21
NG = 0
DB = 1
SM = 0
Delta_AUC
Doxorubicin

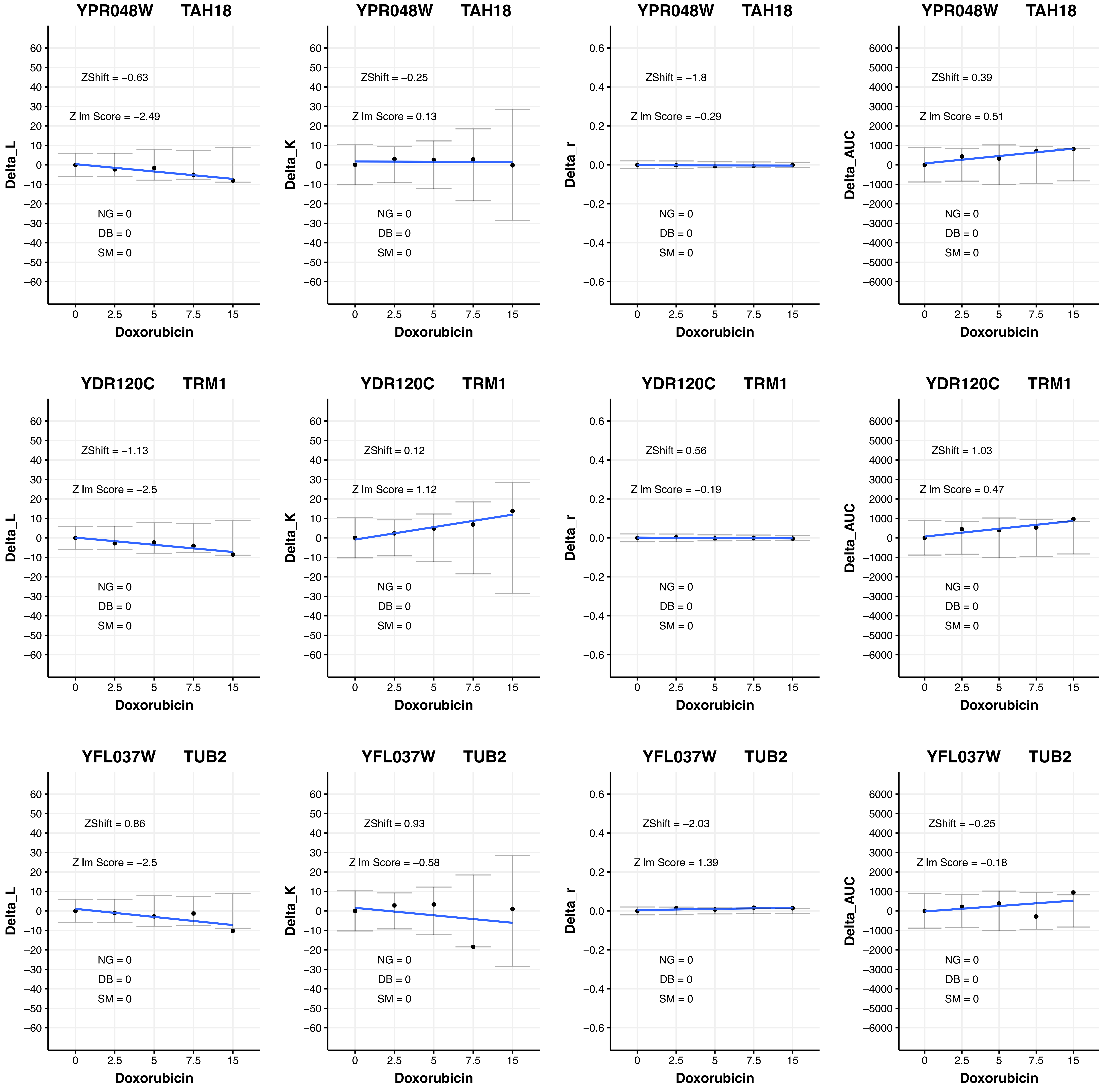
YPR048W TAH18
ZShift = −0.63
Z lm Score = −2.49
NG = 0
DB = 0
SM = 0
Delta_L
Doxorubicin
YPR048W TAH18
ZShift = −0.25
Z lm Score = 0.13
NG = 0
DB = 0
SM = 0
Delta_K
Doxorubicin
YPR048W TAH18
ZShift = −1.8
Z lm Score = −0.29
NG = 0
DB = 0
SM = 0
Delta_r
Doxorubicin
YPR048W TAH18
ZShift = 0.39
Z lm Score = 0.51
NG = 0
DB = 0
SM = 0
Delta_AUC
Doxorubicin
YDR120C TRM1
ZShift = −1.13
Z lm Score = −2.5
NG = 0
DB = 0
SM = 0
Delta_L
Doxorubicin
YDR120C TRM1
ZShift = 0.12
Z lm Score = 1.12
NG = 0
DB = 0
SM = 0
Delta_K
Doxorubicin
YDR120C TRM1
ZShift = 0.56
Z lm Score = −0.19
NG = 0
DB = 0
SM = 0
Delta_r
Doxorubicin
YDR120C TRM1
ZShift = 1.03
Z lm Score = 0.47
NG = 0
DB = 0
SM = 0
Delta_AUC
Doxorubicin
YFL037W TUB2
ZShift = 0.86
Z lm Score = −2.5
NG = 0
DB = 0
SM = 0
Delta_L
Doxorubicin
YFL037W TUB2
ZShift = 0.93
Z lm Score = −0.58
NG = 0
DB = 0
SM = 0
Delta_K
Doxorubicin
YFL037W TUB2
ZShift = −2.03
Z lm Score = 1.39
NG = 0
DB = 0
SM = 0
Delta_r
Doxorubicin
YFL037W TUB2
ZShift = −0.25
Z lm Score = −0.18
NG = 0
DB = 0
SM = 0
Delta_AUC
Doxorubicin

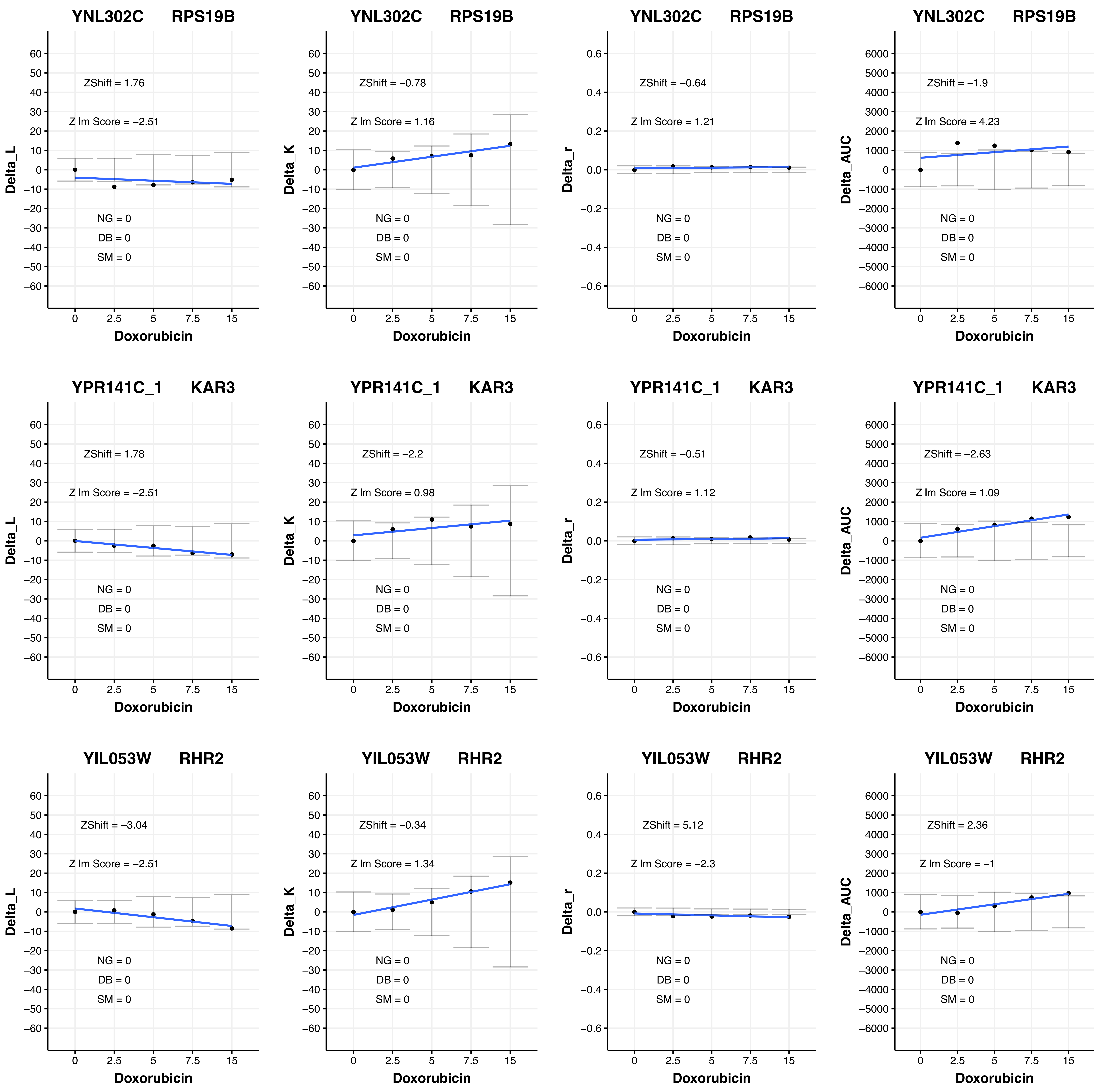

YNL302C RPS19B
ZShift = 1.76
Z lm Score = −2.51
NG = 0
DB = 0
SM = 0
Delta_L
Doxorubicin
YNL302C RPS19B
ZShift = −0.78
Z lm Score = 1.16
NG = 0
DB = 0
SM = 0
Delta_K
Doxorubicin
YNL302C RPS19B
ZShift = −0.64
Z lm Score = 1.21
NG = 0
DB = 0
SM = 0
Delta_r
Doxorubicin
YNL302C RPS19B
ZShift = −1.9
Z lm Score = 4.23
NG = 0
DB = 0
SM = 0
Delta_AUC
Doxorubicin
YPR141C_1 KAR3
ZShift = 1.78
Z lm Score = −2.51
NG = 0
DB = 0
SM = 0
Delta_L
Doxorubicin
YPR141C_1 KAR3
ZShift = −2.2
Z lm Score = 0.98
NG = 0
DB = 0
SM = 0
Delta_K
Doxorubicin
YPR141C_1 KAR3
ZShift = −0.51
Z lm Score = 1.12
NG = 0
DB = 0
SM = 0
Delta_r
Doxorubicin
YPR141C_1 KAR3
ZShift = −2.63
Z lm Score = 1.09
NG = 0
DB = 0
SM = 0
Delta_AUC
Doxorubicin
YIL053W RHR2
ZShift = −3.04
Z lm Score = −2.51
NG = 0
DB = 0
SM = 0
Delta_L
Doxorubicin
YIL053W RHR2
ZShift = −0.34
Z lm Score = 1.34
NG = 0
DB = 0
SM = 0
Delta_K
Doxorubicin
YIL053W RHR2
ZShift = 5.12
Z lm Score = −2.3
NG = 0
DB = 0
SM = 0
Delta_r
Doxorubicin
YIL053W RHR2
ZShift = 2.36
Z lm Score = −1
NG = 0
DB = 0
SM = 0
Delta_AUC
Doxorubicin

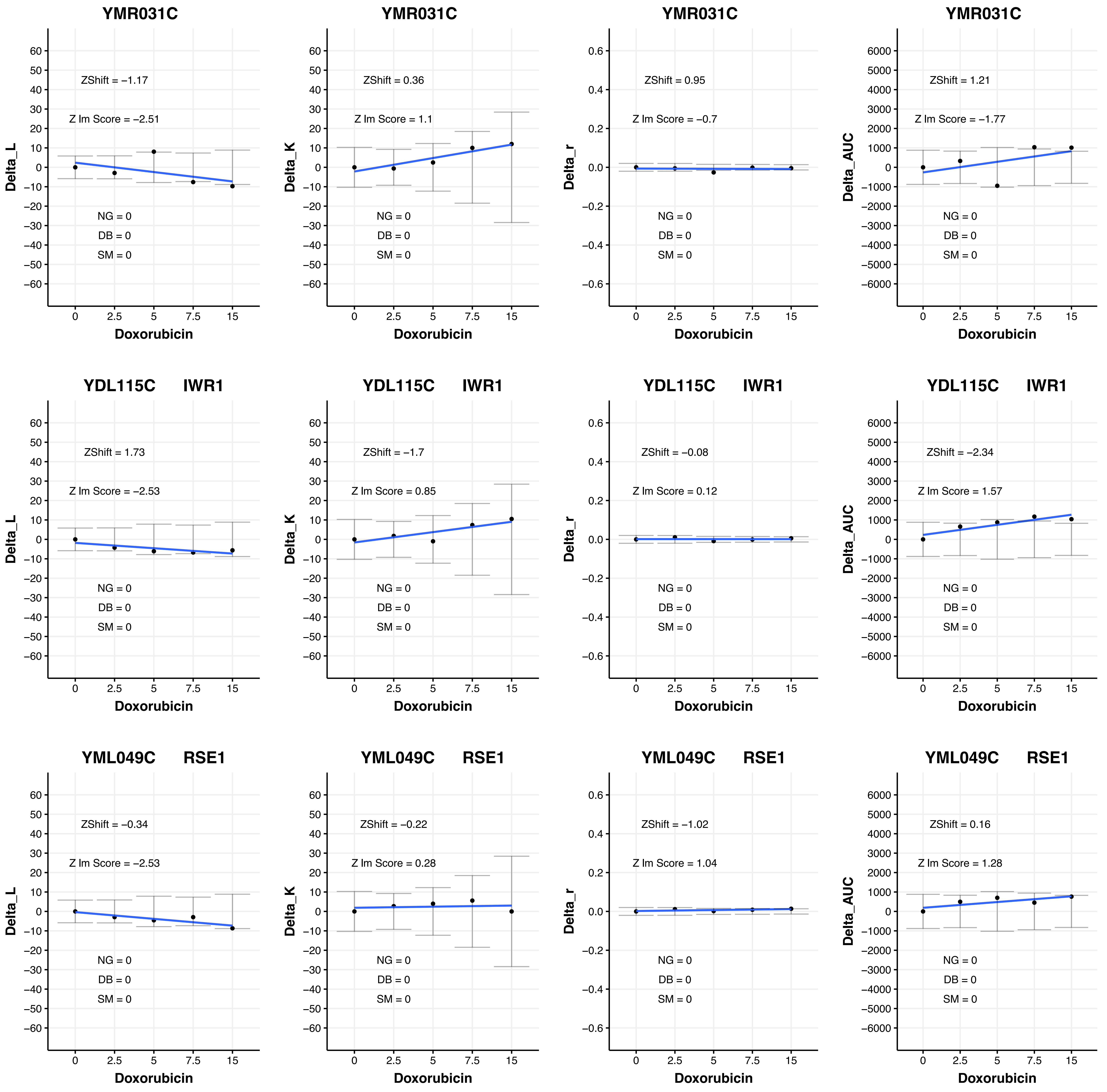
YMR031C
ZShift = −1.17
Z lm Score = −2.51
NG = 0
DB = 0
SM = 0
Delta_L
Doxorubicin
YMR031C
ZShift = 0.36
Z lm Score = 1.1
NG = 0
DB = 0
SM = 0
Delta_K
Doxorubicin
YMR031C
ZShift = 0.95
Z lm Score = −0.7
NG = 0
DB = 0
SM = 0
Delta_r
Doxorubicin
YMR031C
ZShift = 1.21
Z lm Score = −1.77
NG = 0
DB = 0
SM = 0
Delta_AUC
Doxorubicin
YDL115C IWR1
ZShift = 1.73
Z lm Score = −2.53
NG = 0
DB = 0
SM = 0
Delta_L
Doxorubicin
YDL115C IWR1
ZShift = −1.7
Z lm Score = 0.85
NG = 0
DB = 0
SM = 0
Delta_K
Doxorubicin
YDL115C IWR1
ZShift = −0.08
Z lm Score = 0.12
NG = 0
DB = 0
SM = 0
Delta_r
Doxorubicin
YDL115C IWR1
ZShift = −2.34
Z lm Score = 1.57
NG = 0
DB = 0
SM = 0
Delta_AUC
Doxorubicin
YML049C RSE1
ZShift = −0.34
Z lm Score = −2.53
NG = 0
DB = 0
SM = 0
Delta_L
Doxorubicin
YML049C RSE1
ZShift = −0.22
Z lm Score = 0.28
NG = 0
DB = 0
SM = 0
Delta_K
Doxorubicin
YML049C RSE1
ZShift = −1.02
Z lm Score = 1.04
NG = 0
DB = 0
SM = 0
Delta_r
Doxorubicin
YML049C RSE1
ZShift = 0.16
Z lm Score = 1.28
NG = 0
DB = 0
SM = 0
Delta_AUC
Doxorubicin

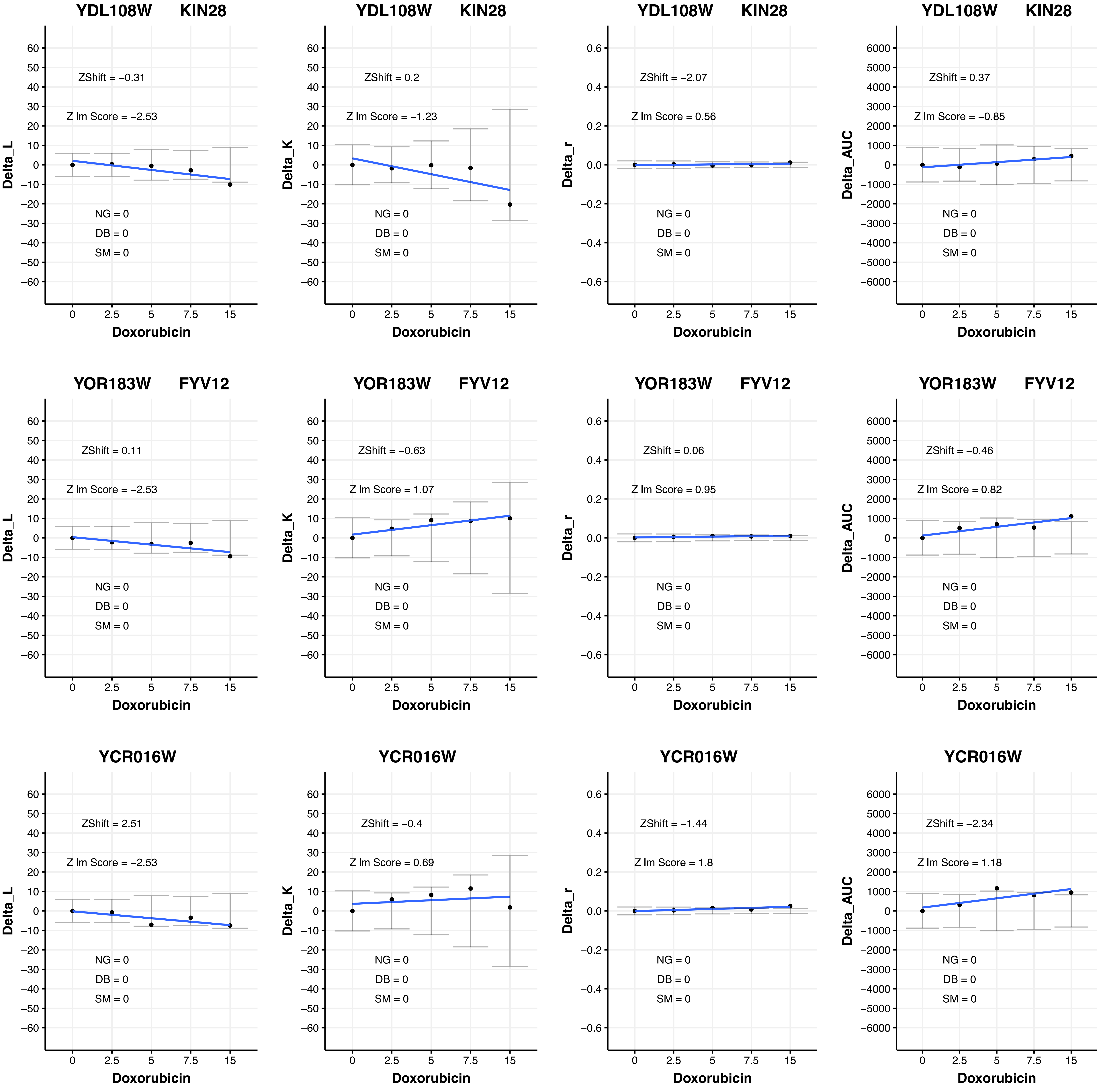
YDL108W KIN28
ZShift = −0.31
Z lm Score = −2.53
NG = 0
DB = 0
SM = 0
Delta_L
Doxorubicin
YDL108W KIN28
ZShift = 0.2
Z lm Score = −1.23
NG = 0
DB = 0
SM = 0
Delta_K
Doxorubicin
YDL108W KIN28
ZShift = −2.07
Z lm Score = 0.56
NG = 0
DB = 0
SM = 0
Delta_r
Doxorubicin
YDL108W KIN28
ZShift = 0.37
Z lm Score = −0.85
NG = 0
DB = 0
SM = 0
Delta_AUC
Doxorubicin
YOR183W FYV12
ZShift = 0.11
Z lm Score = −2.53
NG = 0
DB = 0
SM = 0
Delta_L
Doxorubicin
YOR183W FYV12
ZShift = −0.63
Z lm Score = 1.07
NG = 0
DB = 0
SM = 0
Delta_K
Doxorubicin
YOR183W FYV12
ZShift = 0.06
Z lm Score = 0.95
NG = 0
DB = 0
SM = 0
Delta_r
Doxorubicin
YOR183W FYV12
ZShift = −0.46
Z lm Score = 0.82
NG = 0
DB = 0
SM = 0
Delta_AUC
Doxorubicin
YCR016W
ZShift = 2.51
Z lm Score = −2.53
NG = 0
DB = 0
SM = 0
Delta_L
Doxorubicin
YCR016W
ZShift = −0.4
Z lm Score = 0.69
NG = 0
DB = 0
SM = 0
Delta_K
Doxorubicin
YCR016W
ZShift = −1.44
Z lm Score = 1.8
NG = 0
DB = 0
SM = 0
Delta_r
Doxorubicin
YCR016W
ZShift = −2.34
Z lm Score = 1.18
NG = 0
DB = 0
SM = 0
Delta_AUC
Doxorubicin

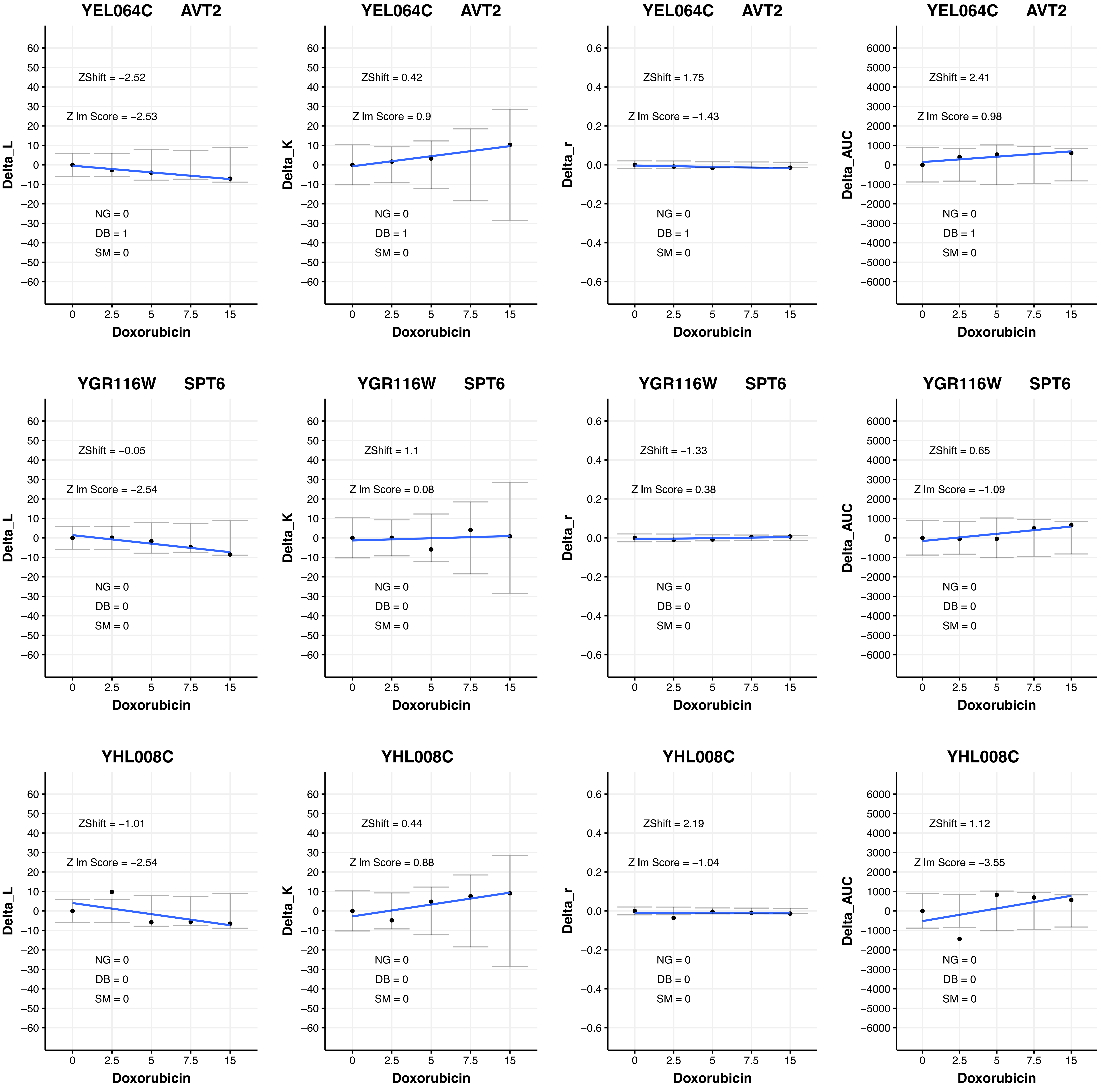

YEL064C AVT2
ZShift = −2.52
Z lm Score = −2.53
NG = 0
DB = 1
SM = 0
Delta_L
Doxorubicin
YEL064C AVT2
ZShift = 0.42
Z lm Score = 0.9
NG = 0
DB = 1
SM = 0
Delta_K
Doxorubicin
YEL064C AVT2
ZShift = 1.75
Z lm Score = −1.43
NG = 0
DB = 1
SM = 0
Delta_r
Doxorubicin
YEL064C AVT2
ZShift = 2.41
Z lm Score = 0.98
NG = 0
DB = 1
SM = 0
Delta_AUC
Doxorubicin
YGR116W SPT6
ZShift = −0.05
Z lm Score = −2.54
NG = 0
DB = 0
SM = 0
Delta_L
Doxorubicin
YGR116W SPT6
ZShift = 1.1
Z lm Score = 0.08
NG = 0
DB = 0
SM = 0
Delta_K
Doxorubicin
YGR116W SPT6
ZShift = −1.33
Z lm Score = 0.38
NG = 0
DB = 0
SM = 0
Delta_r
Doxorubicin
YGR116W SPT6
ZShift = 0.65
Z lm Score = −1.09
NG = 0
DB = 0
SM = 0
Delta_AUC
Doxorubicin
YHL008C
ZShift = −1.01
Z lm Score = −2.54
NG = 0
DB = 0
SM = 0
Delta_L
Doxorubicin
YHL008C
ZShift = 0.44
Z lm Score = 0.88
NG = 0
DB = 0
SM = 0
Delta_K
Doxorubicin
YHL008C
ZShift = 2.19
Z lm Score = −1.04
NG = 0
DB = 0
SM = 0
Delta_r
Doxorubicin
YHL008C
ZShift = 1.12
Z lm Score = −3.55
NG = 0
DB = 0
SM = 0
Delta_AUC
Doxorubicin

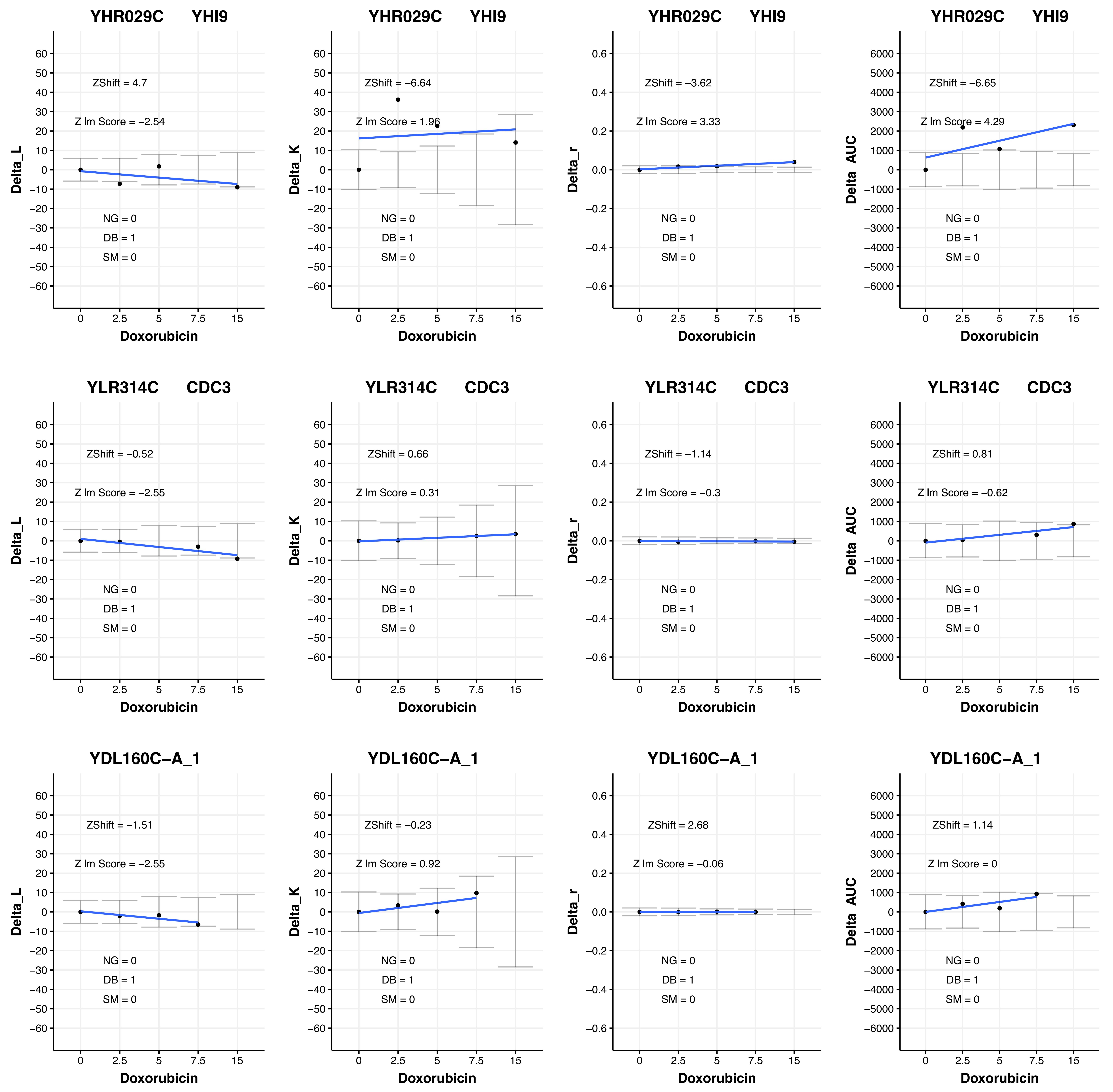
YHR029C YHI9
ZShift = 4.7
Z lm Score = −2.54
NG = 0
DB = 1
SM = 0
Delta_L
Doxorubicin
YHR029C YHI9
ZShift = −6.64
Z lm Score = 1.96
NG = 0
DB = 1
SM = 0
Delta_K
Doxorubicin
YHR029C YHI9
ZShift = −3.62
Z lm Score = 3.33
NG = 0
DB = 1
SM = 0
Delta_r
Doxorubicin
YHR029C YHI9
ZShift = −6.65
Z lm Score = 4.29
NG = 0
DB = 1
SM = 0
Delta_AUC
Doxorubicin
YLR314C CDC3
ZShift = −0.52
Z lm Score = −2.55
NG = 0
DB = 1
SM = 0
Delta_L
Doxorubicin
YLR314C CDC3
ZShift = 0.66
Z lm Score = 0.31
NG = 0
DB = 1
SM = 0
Delta_K
Doxorubicin
YLR314C CDC3
ZShift = −1.14
Z lm Score = −0.3
NG = 0
DB = 1
SM = 0
Delta_r
Doxorubicin
YLR314C CDC3
ZShift = 0.81
Z lm Score = −0.62
NG = 0
DB = 1
SM = 0
Delta_AUC
Doxorubicin
YDL160C−A_1
ZShift = −1.51
Z lm Score = −2.55
NG = 0
DB = 1
SM = 0
Delta_L
Doxorubicin
YDL160C−A_1
ZShift = −0.23
Z lm Score = 0.92
NG = 0
DB = 1
SM = 0
Delta_K
Doxorubicin
YDL160C−A_1
ZShift = 2.68
Z lm Score = −0.06
NG = 0
DB = 1
SM = 0
Delta_r
Doxorubicin
YDL160C−A_1
ZShift = 1.14
Z lm Score = 0
NG = 0
DB = 1
SM = 0
Delta_AUC
Doxorubicin

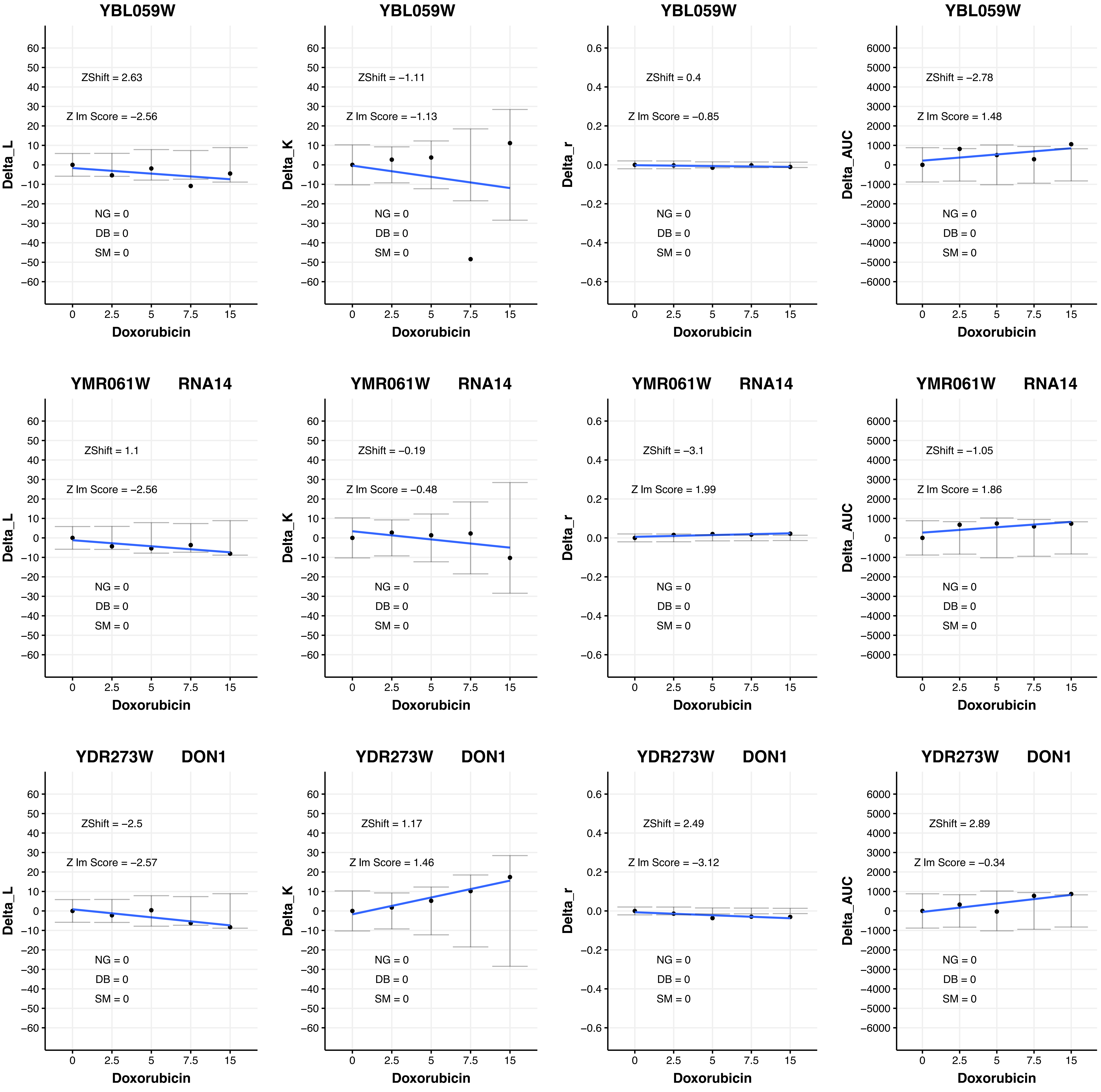
YBL059W
ZShift = 2.63
Z lm Score = −2.56
NG = 0
DB = 0
SM = 0
Delta_L
Doxorubicin
YBL059W
ZShift = −1.11
Z lm Score = −1.13
NG = 0
DB = 0
SM = 0
Delta_K
Doxorubicin
YBL059W
ZShift = 0.4
Z lm Score = −0.85
NG = 0
DB = 0
SM = 0
Delta_r
Doxorubicin
YBL059W
ZShift = −2.78
Z lm Score = 1.48
NG = 0
DB = 0
SM = 0
Delta_AUC
Doxorubicin
YMR061W RNA14
ZShift = 1.1
Z lm Score = −2.56
NG = 0
DB = 0
SM = 0
Delta_L
Doxorubicin
YMR061W RNA14
ZShift = −0.19
Z lm Score = −0.48
NG = 0
DB = 0
SM = 0
Delta_K
Doxorubicin
YMR061W RNA14
ZShift = −3.1
Z lm Score = 1.99
NG = 0
DB = 0
SM = 0
Delta_r
Doxorubicin
YMR061W RNA14
ZShift = −1.05
Z lm Score = 1.86
NG = 0
DB = 0
SM = 0
Delta_AUC
Doxorubicin
YDR273W DON1
ZShift = −2.5
Z lm Score = −2.57
NG = 0
DB = 0
SM = 0
Delta_L
Doxorubicin
YDR273W DON1
ZShift = 1.17
Z lm Score = 1.46
NG = 0
DB = 0
SM = 0
Delta_K
Doxorubicin
YDR273W DON1
ZShift = 2.49
Z lm Score = −3.12
NG = 0
DB = 0
SM = 0
Delta_r
Doxorubicin
YDR273W DON1
ZShift = 2.89
Z lm Score = −0.34
NG = 0
DB = 0
SM = 0
Delta_AUC
Doxorubicin

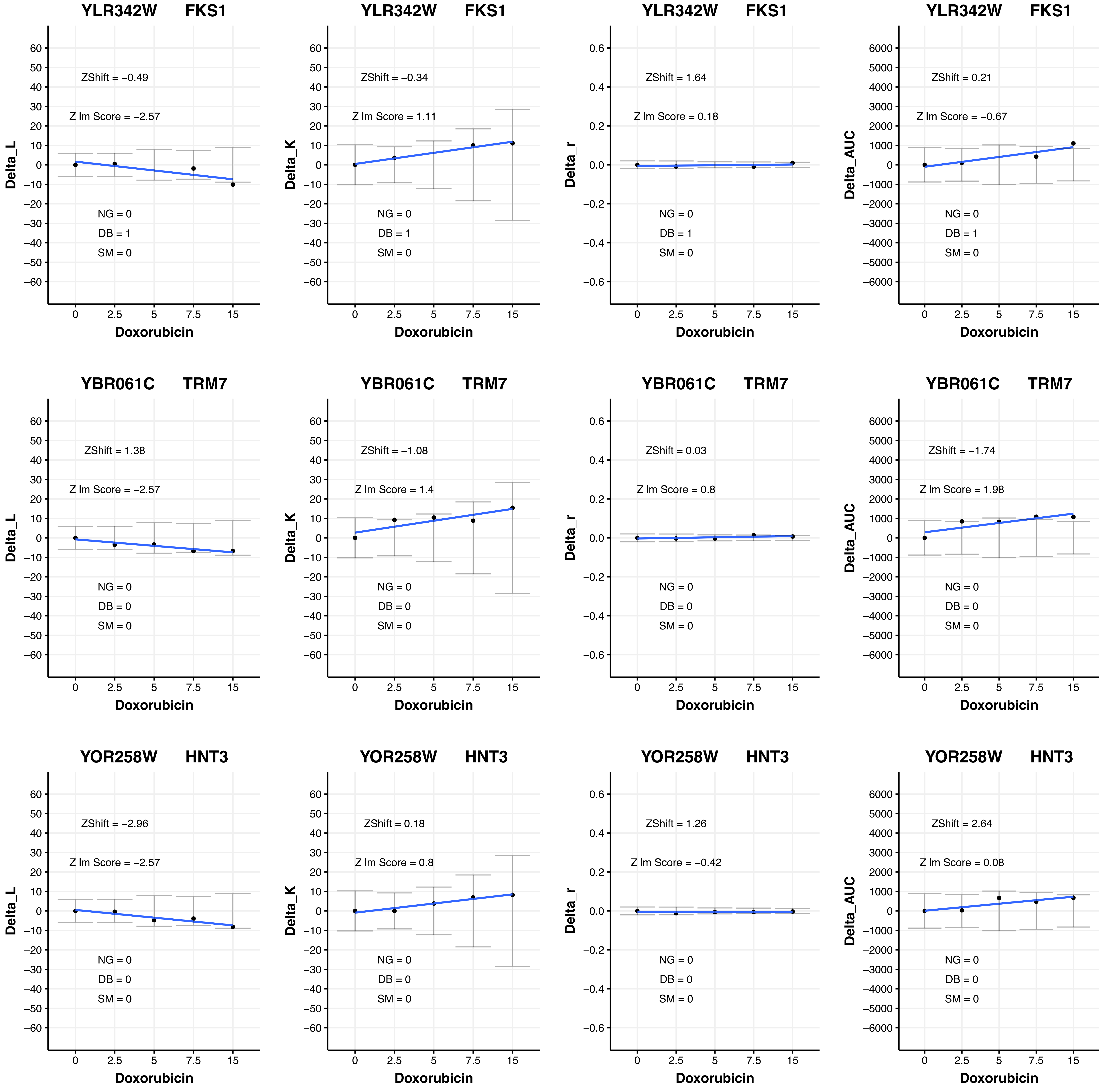

YLR342W FKS1
ZShift = −0.49
Z lm Score = −2.57
NG = 0
DB = 1
SM = 0
Delta_L
Doxorubicin
YLR342W FKS1
ZShift = −0.34
Z lm Score = 1.11
NG = 0
DB = 1
SM = 0
Delta_K
Doxorubicin
YLR342W FKS1
ZShift = 1.64
Z lm Score = 0.18
NG = 0
DB = 1
SM = 0
Delta_r
Doxorubicin
YLR342W FKS1
ZShift = 0.21
Z lm Score = −0.67
NG = 0
DB = 1
SM = 0
Delta_AUC
Doxorubicin
YBR061C TRM7
ZShift = 1.38
Z lm Score = −2.57
NG = 0
DB = 0
SM = 0
Delta_L
Doxorubicin
YBR061C TRM7
ZShift = −1.08
Z lm Score = 1.4
NG = 0
DB = 0
SM = 0
Delta_K
Doxorubicin
YBR061C TRM7
ZShift = 0.03
Z lm Score = 0.8
NG = 0
DB = 0
SM = 0
Delta_r
Doxorubicin
YBR061C TRM7
ZShift = −1.74
Z lm Score = 1.98
NG = 0
DB = 0
SM = 0
Delta_AUC
Doxorubicin
YOR258W HNT3
ZShift = −2.96
Z lm Score = −2.57
NG = 0
DB = 0
SM = 0
Delta_L
Doxorubicin
YOR258W HNT3
ZShift = 0.18
Z lm Score = 0.8
NG = 0
DB = 0
SM = 0
Delta_K
Doxorubicin
YOR258W HNT3
ZShift = 1.26
Z lm Score = −0.42
NG = 0
DB = 0
SM = 0
Delta_r
Doxorubicin
YOR258W HNT3
ZShift = 2.64
Z lm Score = 0.08
NG = 0
DB = 0
SM = 0
Delta_AUC
Doxorubicin

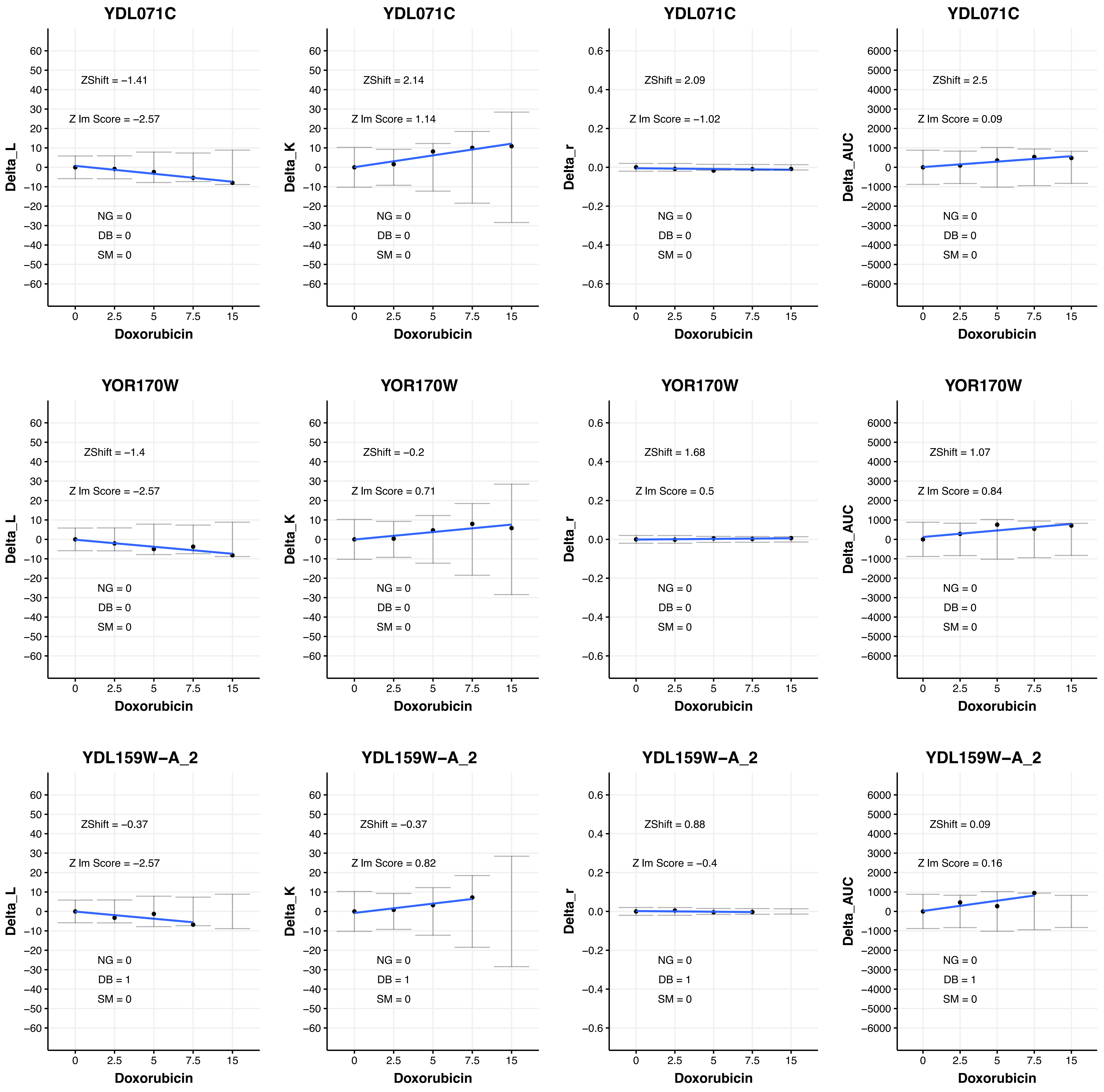

YDL071C
ZShift = −1.41
Z lm Score = −2.57
NG = 0
DB = 0
SM = 0
Delta_L
Doxorubicin
YDL071C
ZShift = 2.14
Z lm Score = 1.14
NG = 0
DB = 0
SM = 0
Delta_K
Doxorubicin
YDL071C
ZShift = 2.09
Z lm Score = −1.02
NG = 0
DB = 0
SM = 0
Delta_r
Doxorubicin
YDL071C
ZShift = 2.5
Z lm Score = 0.09
NG = 0
DB = 0
SM = 0
Delta_AUC
Doxorubicin
YOR170W
ZShift = −1.4
Z lm Score = −2.57
NG = 0
DB = 0
SM = 0
Delta_L
Doxorubicin
YOR170W
ZShift = −0.2
Z lm Score = 0.71
NG = 0
DB = 0
SM = 0
Delta_K
Doxorubicin
YOR170W
ZShift = 1.68
Z lm Score = 0.5
NG = 0
DB = 0
SM = 0
Delta_r
Doxorubicin
YOR170W
ZShift = 1.07
Z lm Score = 0.84
NG = 0
DB = 0
SM = 0
Delta_AUC
Doxorubicin
YDL159W−A_2
ZShift = −0.37
Z lm Score = −2.57
NG = 0
DB = 1
SM = 0
Delta_L
Doxorubicin
YDL159W−A_2
ZShift = −0.37
Z lm Score = 0.82
NG = 0
DB = 1
SM = 0
Delta_K
Doxorubicin
YDL159W−A_2
ZShift = 0.88
Z lm Score = −0.4
NG = 0
DB = 1
SM = 0
Delta_r
Doxorubicin
YDL159W−A_2
ZShift = 0.09
Z lm Score = 0.16
NG = 0
DB = 1
SM = 0
Delta_AUC
Doxorubicin

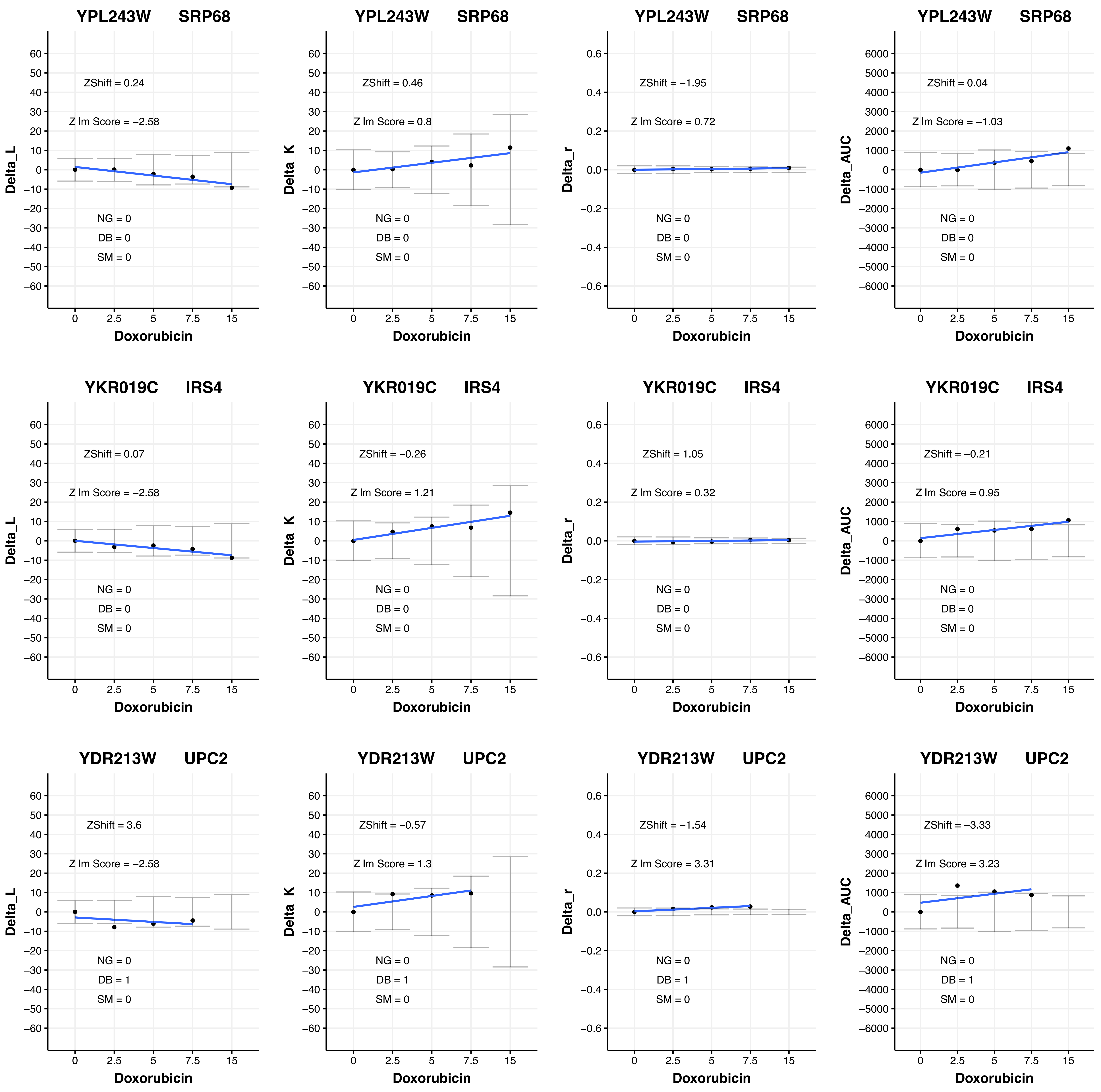

YPL243W SRP68
ZShift = 0.24
Z lm Score = −2.58
NG = 0
DB = 0
SM = 0
Delta_L
Doxorubicin
YPL243W SRP68
ZShift = 0.46
Z lm Score = 0.8
NG = 0
DB = 0
SM = 0
Delta_K
Doxorubicin
YPL243W SRP68
ZShift = −1.95
Z lm Score = 0.72
NG = 0
DB = 0
SM = 0
Delta_r
Doxorubicin
YPL243W SRP68
ZShift = 0.04
Z lm Score = −1.03
NG = 0
DB = 0
SM = 0
Delta_AUC
Doxorubicin
YKR019C IRS4
ZShift = 0.07
Z lm Score = −2.58
NG = 0
DB = 0
SM = 0
Delta_L
Doxorubicin
YKR019C IRS4
ZShift = −0.26
Z lm Score = 1.21
NG = 0
DB = 0
SM = 0
Delta_K
Doxorubicin
YKR019C IRS4
ZShift = 1.05
Z lm Score = 0.32
NG = 0
DB = 0
SM = 0
Delta_r
Doxorubicin
YKR019C IRS4
ZShift = −0.21
Z lm Score = 0.95
NG = 0
DB = 0
SM = 0
Delta_AUC
Doxorubicin
YDR213W UPC2
ZShift = 3.6
Z lm Score = −2.58
NG = 0
DB = 1
SM = 0
Delta_L
Doxorubicin
YDR213W UPC2
ZShift = −0.57
Z lm Score = 1.3
NG = 0
DB = 1
SM = 0
Delta_K
Doxorubicin
YDR213W UPC2
ZShift = −1.54
Z lm Score = 3.31
NG = 0
DB = 1
SM = 0
Delta_r
Doxorubicin
YDR213W UPC2
ZShift = −3.33
Z lm Score = 3.23
NG = 0
DB = 1
SM = 0
Delta_AUC
Doxorubicin

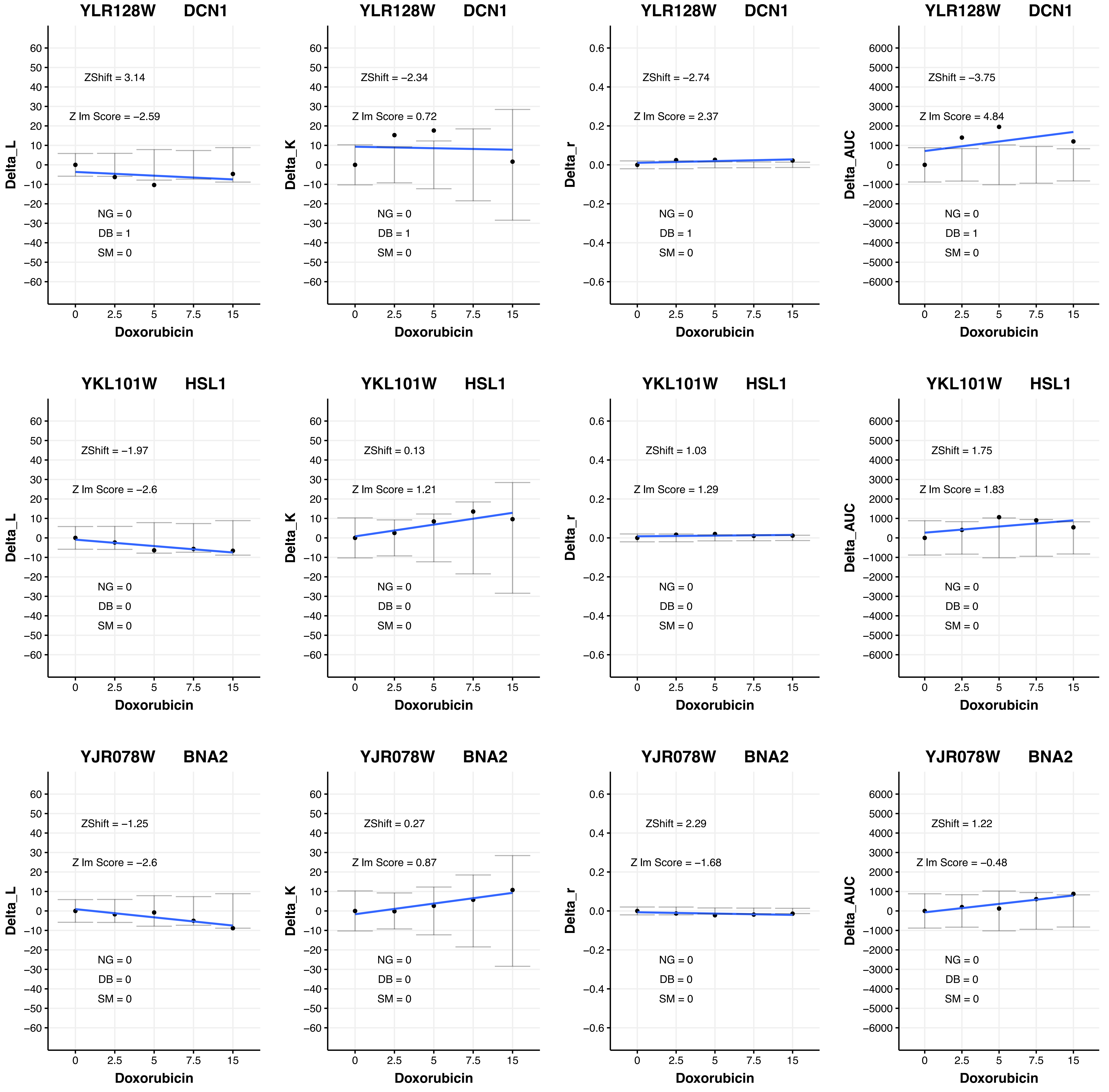

YLR128W DCN1
ZShift = 3.14
Z lm Score = −2.59
NG = 0
DB = 1
SM = 0
Delta_L
Doxorubicin
YLR128W DCN1
ZShift = −2.34
Z lm Score = 0.72
NG = 0
DB = 1
SM = 0
Delta_K
Doxorubicin
YLR128W DCN1
ZShift = −2.74
Z lm Score = 2.37
NG = 0
DB = 1
SM = 0
Delta_r
Doxorubicin
YLR128W DCN1
ZShift = −3.75
Z lm Score = 4.84
NG = 0
DB = 1
SM = 0
Delta_AUC
Doxorubicin
YKL101W HSL1
ZShift = −1.97
Z lm Score = −2.6
NG = 0
DB = 0
SM = 0
Delta_L
Doxorubicin
YKL101W HSL1
ZShift = 0.13
Z lm Score = 1.21
NG = 0
DB = 0
SM = 0
Delta_K
Doxorubicin
YKL101W HSL1
ZShift = 1.03
Z lm Score = 1.29
NG = 0
DB = 0
SM = 0
Delta_r
Doxorubicin
YKL101W HSL1
ZShift = 1.75
Z lm Score = 1.83
NG = 0
DB = 0
SM = 0
Delta_AUC
Doxorubicin
YJR078W BNA2
ZShift = −1.25
Z lm Score = −2.6
NG = 0
DB = 0
SM = 0
Delta_L
Doxorubicin
YJR078W BNA2
ZShift = 0.27
Z lm Score = 0.87
NG = 0
DB = 0
SM = 0
Delta_K
Doxorubicin
YJR078W BNA2
ZShift = 2.29
Z lm Score = −1.68
NG = 0
DB = 0
SM = 0
Delta_r
Doxorubicin
YJR078W BNA2
ZShift = 1.22
Z lm Score = −0.48
NG = 0
DB = 0
SM = 0
Delta_AUC
Doxorubicin

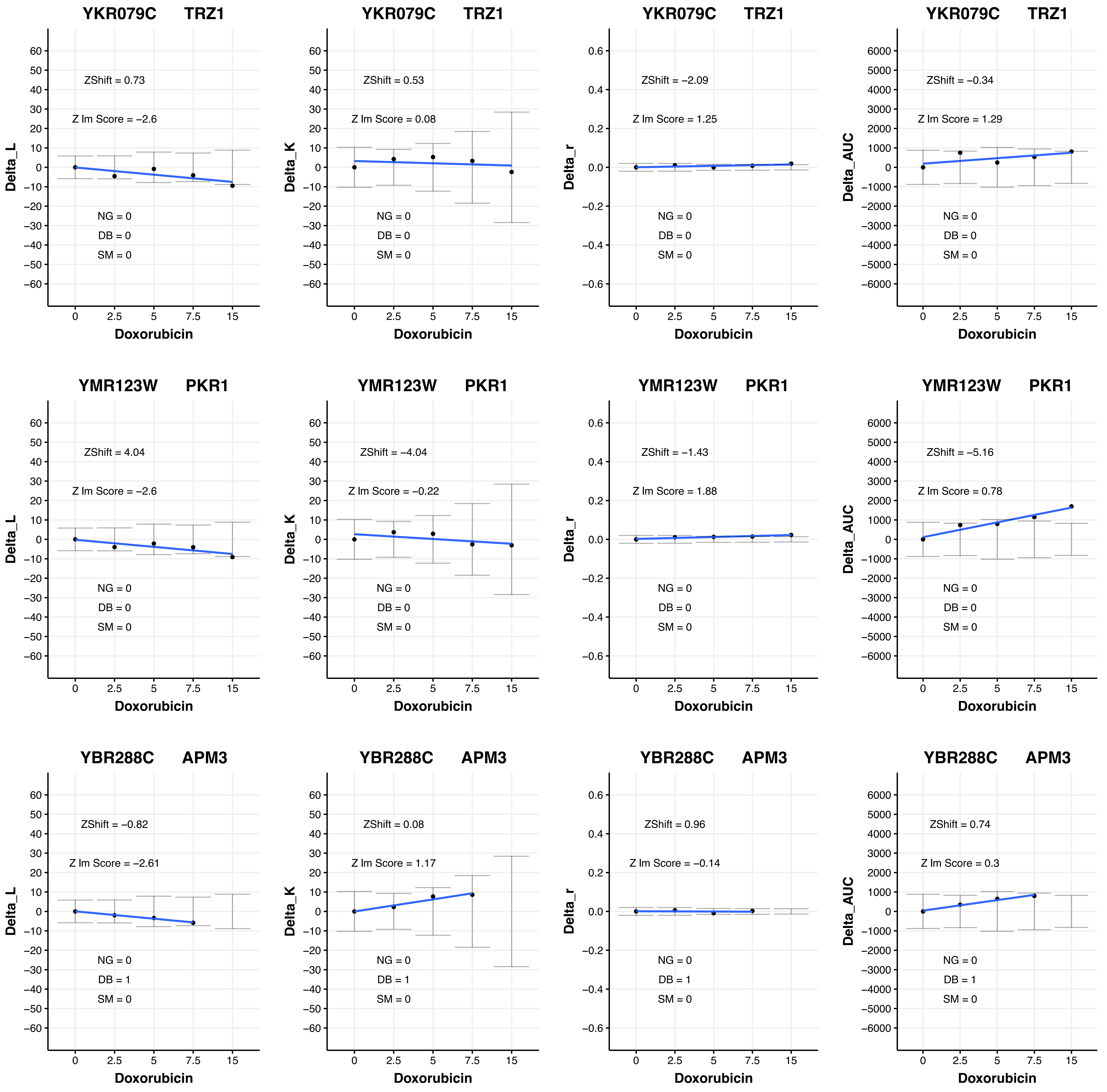

YKR079C TRZ1
ZShift = 0.73
Z lm Score = −2.6
NG = 0
DB = 0
SM = 0
Delta_L
Doxorubicin
YKR079C TRZ1
ZShift = 0.53
Z lm Score = 0.08
NG = 0
DB = 0
SM = 0
Delta_K
Doxorubicin
YKR079C TRZ1
ZShift = −2.09
Z lm Score = 1.25
NG = 0
DB = 0
SM = 0
Delta_r
Doxorubicin
YKR079C TRZ1
ZShift = −0.34
Z lm Score = 1.29
NG = 0
DB = 0
SM = 0
Delta_AUC
Doxorubicin
YMR123W PKR1
ZShift = 4.04
Z lm Score = −2.6
NG = 0
DB = 0
SM = 0
Delta_L
Doxorubicin
YMR123W PKR1
ZShift = −4.04
Z lm Score = −0.22
NG = 0
DB = 0
SM = 0
Delta_K
Doxorubicin
YMR123W PKR1
ZShift = −1.43
Z lm Score = 1.88
NG = 0
DB = 0
SM = 0
Delta_r
Doxorubicin
YMR123W PKR1
ZShift = −5.16
Z lm Score = 0.78
NG = 0
DB = 0
SM = 0
Delta_AUC
Doxorubicin
YBR288C APM3
ZShift = −0.82
Z lm Score = −2.61
NG = 0
DB = 1
SM = 0
Delta_L
Doxorubicin
YBR288C APM3
ZShift = 0.08
Z lm Score = 1.17
NG = 0
DB = 1
SM = 0
Delta_K
Doxorubicin
YBR288C APM3
ZShift = 0.96
Z lm Score = −0.14
NG = 0
DB = 1
SM = 0
Delta_r
Doxorubicin
YBR288C APM3
ZShift = 0.74
Z lm Score = 0.3
NG = 0
DB = 1
SM = 0
Delta_AUC
Doxorubicin

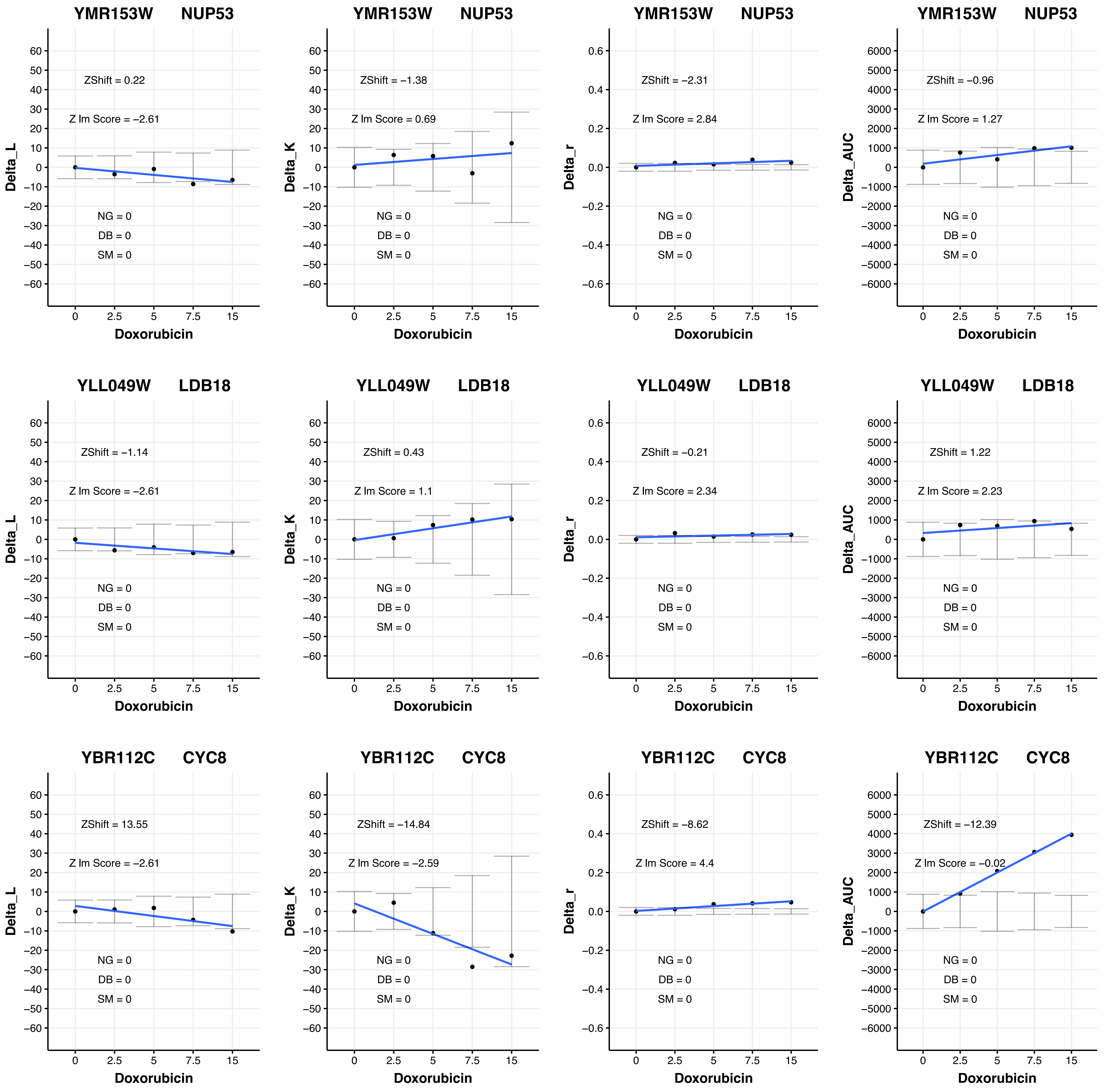
YMR153W NUP53
ZShift = 0.22
Z lm Score = −2.61
NG = 0
DB = 0
SM = 0
Delta_L
Doxorubicin
YMR153W NUP53
ZShift = −1.38
Z lm Score = 0.69
NG = 0
DB = 0
SM = 0
Delta_K
Doxorubicin
YMR153W NUP53
ZShift = −2.31
Z lm Score = 2.84
NG = 0
DB = 0
SM = 0
Delta_r
Doxorubicin
YMR153W NUP53
ZShift = −0.96
Z lm Score = 1.27
NG = 0
DB = 0
SM = 0
Delta_AUC
Doxorubicin
YLL049W LDB18
ZShift = −1.14
Z lm Score = −2.61
NG = 0
DB = 0
SM = 0
Delta_L
Doxorubicin
YLL049W LDB18
ZShift = 0.43
Z lm Score = 1.1
NG = 0
DB = 0
SM = 0
Delta_K
Doxorubicin
YLL049W LDB18
ZShift = −0.21
Z lm Score = 2.34
NG = 0
DB = 0
SM = 0
Delta_r
Doxorubicin
YLL049W LDB18
ZShift = 1.22
Z lm Score = 2.23
NG = 0
DB = 0
SM = 0
Delta_AUC
Doxorubicin
YBR112C CYC8
ZShift = 13.55
Z lm Score = −2.61
NG = 0
DB = 0
SM = 0
Delta_L
Doxorubicin
YBR112C CYC8
ZShift = −14.84
Z lm Score = −2.59
NG = 0
DB = 0
SM = 0
Delta_K
Doxorubicin
YBR112C CYC8
ZShift = −8.62
Z lm Score = 4.4
NG = 0
DB = 0
SM = 0
Delta_r
Doxorubicin
YBR112C CYC8
ZShift = −12.39
Z lm Score = −0.02
NG = 0
DB = 0
SM = 0
Delta_AUC
Doxorubicin

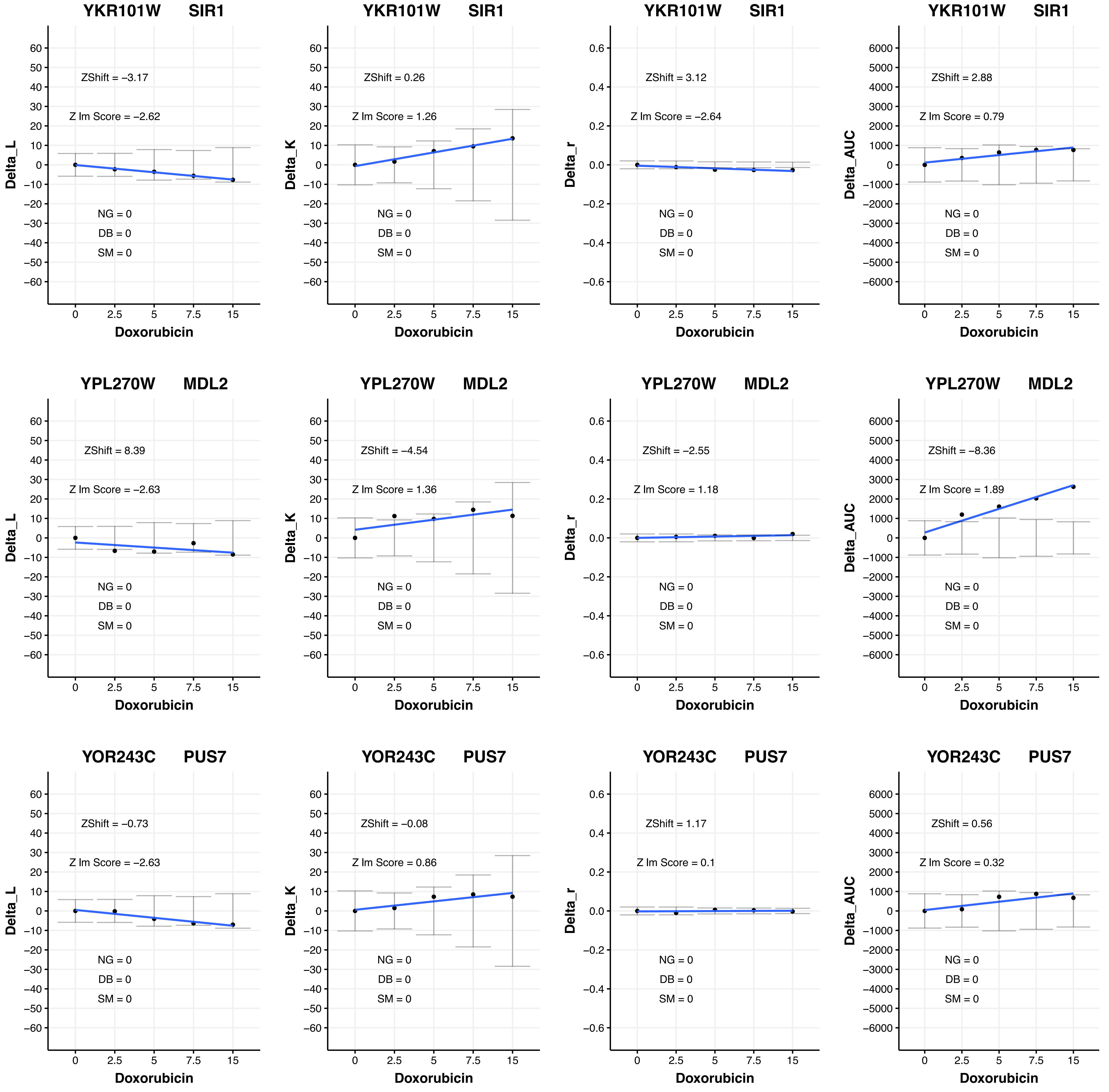
YKR101W SIR1
ZShift = −3.17
Z lm Score = −2.62
NG = 0
DB = 0
SM = 0
Delta_L
Doxorubicin
YKR101W SIR1
ZShift = 0.26
Z lm Score = 1.26
NG = 0
DB = 0
SM = 0
Delta_K
Doxorubicin
YKR101W SIR1
ZShift = 3.12
Z lm Score = −2.64
NG = 0
DB = 0
SM = 0
Delta_r
Doxorubicin
YKR101W SIR1
ZShift = 2.88
Z lm Score = 0.79
NG = 0
DB = 0
SM = 0
Delta_AUC
Doxorubicin
YPL270W MDL2
ZShift = 8.39
Z lm Score = −2.63
NG = 0
DB = 0
SM = 0
Delta_L
Doxorubicin
YPL270W MDL2
ZShift = −4.54
Z lm Score = 1.36
NG = 0
DB = 0
SM = 0
Delta_K
Doxorubicin
YPL270W MDL2
ZShift = −2.55
Z lm Score = 1.18
NG = 0
DB = 0
SM = 0
Delta_r
Doxorubicin
YPL270W MDL2
ZShift = −8.36
Z lm Score = 1.89
NG = 0
DB = 0
SM = 0
Delta_AUC
Doxorubicin
YOR243C PUS7
ZShift = −0.73
Z lm Score = −2.63
NG = 0
DB = 0
SM = 0
Delta_L
Doxorubicin
YOR243C PUS7
ZShift = −0.08
Z lm Score = 0.86
NG = 0
DB = 0
SM = 0
Delta_K
Doxorubicin
YOR243C PUS7
ZShift = 1.17
Z lm Score = 0.1
NG = 0
DB = 0
SM = 0
Delta_r
Doxorubicin
YOR243C PUS7
ZShift = 0.56
Z lm Score = 0.32
NG = 0
DB = 0
SM = 0
Delta_AUC
Doxorubicin

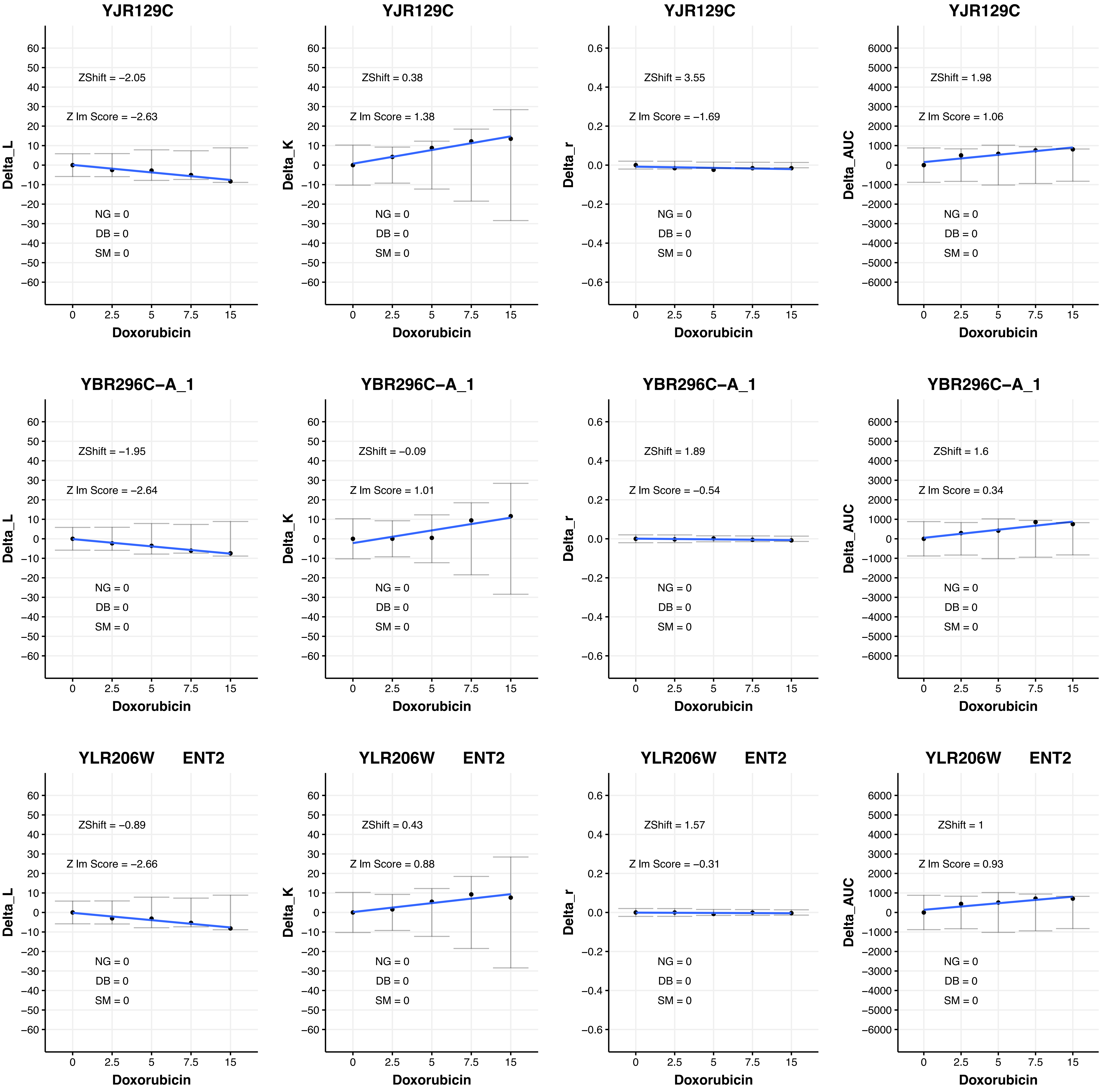

YJR129C
ZShift = −2.05
Z lm Score = −2.63
NG = 0
DB = 0
SM = 0
Delta_L
Doxorubicin
YJR129C
ZShift = 0.38
Z lm Score = 1.38
NG = 0
DB = 0
SM = 0
Delta_K
Doxorubicin
YJR129C
ZShift = 3.55
Z lm Score = −1.69
NG = 0
DB = 0
SM = 0
Delta_r
Doxorubicin
YJR129C
ZShift = 1.98
Z lm Score = 1.06
NG = 0
DB = 0
SM = 0
Delta_AUC
Doxorubicin
YBR296C−A_1
ZShift = −1.95
Z lm Score = −2.64
NG = 0
DB = 0
SM = 0
Delta_L
Doxorubicin
YBR296C−A_1
ZShift = −0.09
Z lm Score = 1.01
NG = 0
DB = 0
SM = 0
Delta_K
Doxorubicin
YBR296C−A_1
ZShift = 1.89
Z lm Score = −0.54
NG = 0
DB = 0
SM = 0
Delta_r
Doxorubicin
YBR296C−A_1
ZShift = 1.6
Z lm Score = 0.34
NG = 0
DB = 0
SM = 0
Delta_AUC
Doxorubicin
YLR206W ENT2
ZShift = −0.89
Z lm Score = −2.66
NG = 0
DB = 0
SM = 0
Delta_L
Doxorubicin
YLR206W ENT2
ZShift = 0.43
Z lm Score = 0.88
NG = 0
DB = 0
SM = 0
Delta_K
Doxorubicin
YLR206W ENT2
ZShift = 1.57
Z lm Score = −0.31
NG = 0
DB = 0
SM = 0
Delta_r
Doxorubicin
YLR206W ENT2
ZShift = 1
Z lm Score = 0.93
NG = 0
DB = 0
SM = 0
Delta_AUC
Doxorubicin

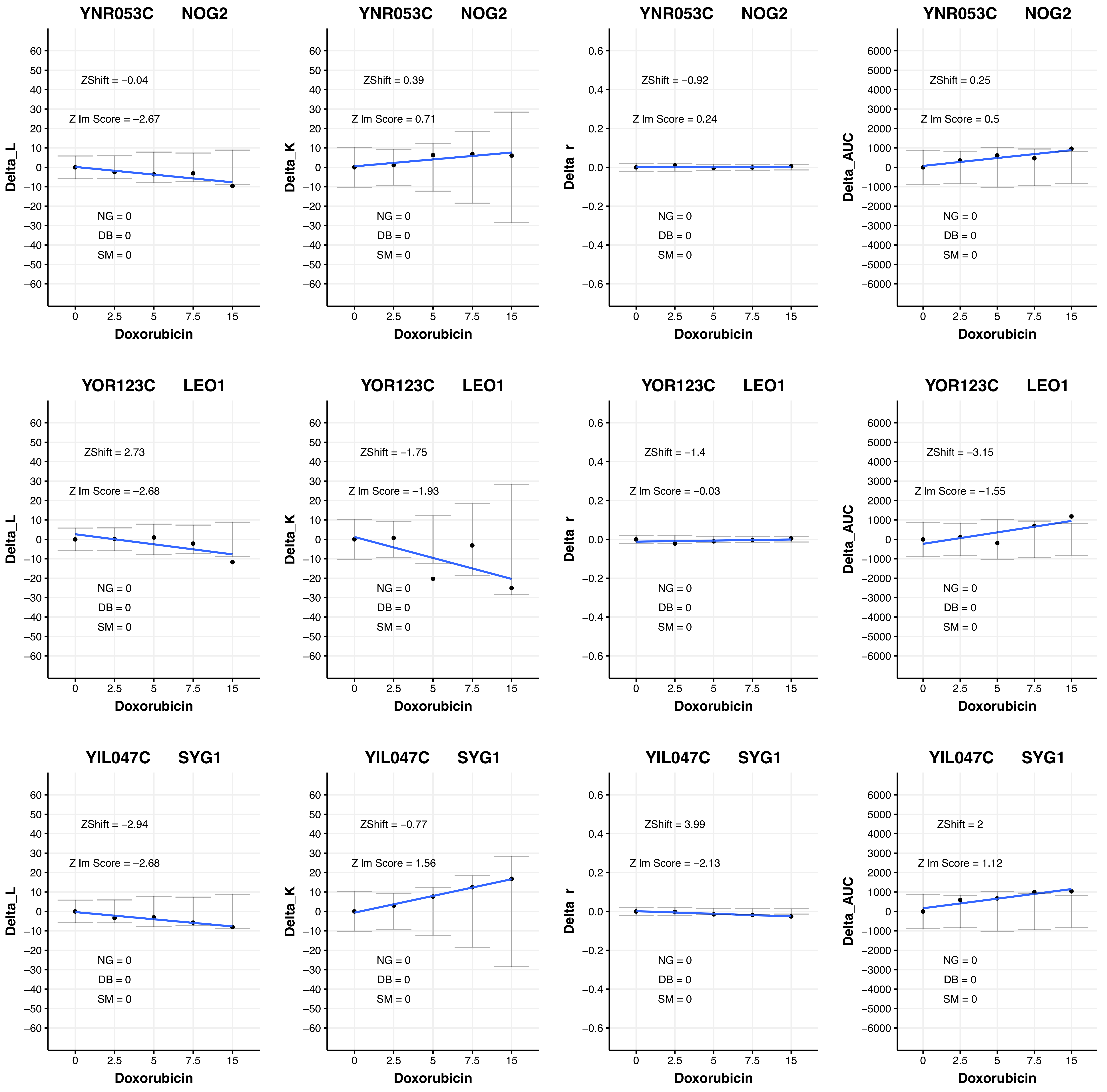
YNR053C NOG2
ZShift = −0.04
Z lm Score = −2.67
NG = 0
DB = 0
SM = 0
Delta_L
Doxorubicin
YNR053C NOG2
ZShift = 0.39
Z lm Score = 0.71
NG = 0
DB = 0
SM = 0
Delta_K
Doxorubicin
YNR053C NOG2
ZShift = −0.92
Z lm Score = 0.24
NG = 0
DB = 0
SM = 0
Delta_r
Doxorubicin
YNR053C NOG2
ZShift = 0.25
Z lm Score = 0.5
NG = 0
DB = 0
SM = 0
Delta_AUC
Doxorubicin
YOR123C LEO1
ZShift = 2.73
Z lm Score = −2.68
NG = 0
DB = 0
SM = 0
Delta_L
Doxorubicin
YOR123C LEO1
ZShift = −1.75
Z lm Score = −1.93
NG = 0
DB = 0
SM = 0
Delta_K
Doxorubicin
YOR123C LEO1
ZShift = −1.4
Z lm Score = −0.03
NG = 0
DB = 0
SM = 0
Delta_r
Doxorubicin
YOR123C LEO1
ZShift = −3.15
Z lm Score = −1.55
NG = 0
DB = 0
SM = 0
Delta_AUC
Doxorubicin
YIL047C SYG1
ZShift = −2.94
Z lm Score = −2.68
NG = 0
DB = 0
SM = 0
Delta_L
Doxorubicin
YIL047C SYG1
ZShift = −0.77
Z lm Score = 1.56
NG = 0
DB = 0
SM = 0
Delta_K
Doxorubicin
YIL047C SYG1
ZShift = 3.99
Z lm Score = −2.13
NG = 0
DB = 0
SM = 0
Delta_r
Doxorubicin
YIL047C SYG1
ZShift = 2
Z lm Score = 1.12
NG = 0
DB = 0
SM = 0
Delta_AUC
Doxorubicin

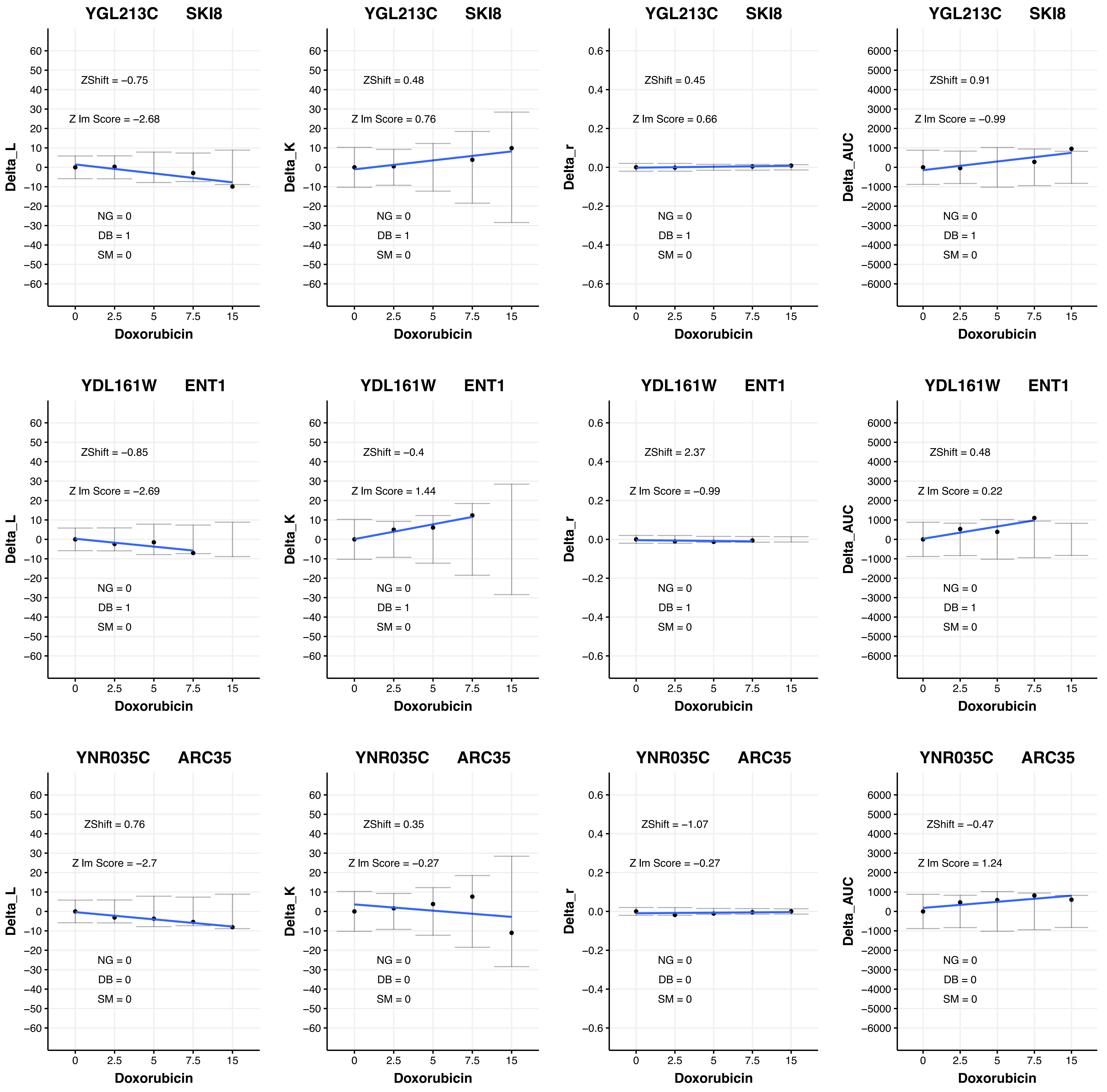

YGL213C SKI8
ZShift = −0.75
Z lm Score = −2.68
NG = 0
DB = 1
SM = 0
Delta_L
Doxorubicin
YGL213C SKI8
ZShift = 0.48
Z lm Score = 0.76
NG = 0
DB = 1
SM = 0
Delta_K
Doxorubicin
YGL213C SKI8
ZShift = 0.45
Z lm Score = 0.66
NG = 0
DB = 1
SM = 0
Delta_r
Doxorubicin
YGL213C SKI8
ZShift = 0.91
Z lm Score = −0.99
NG = 0
DB = 1
SM = 0
Delta_AUC
Doxorubicin
YDL161W ENT1
ZShift = −0.85
Z lm Score = −2.69
NG = 0
DB = 1
SM = 0
Delta_L
Doxorubicin
YDL161W ENT1
ZShift = −0.4
Z lm Score = 1.44
NG = 0
DB = 1
SM = 0
Delta_K
Doxorubicin
YDL161W ENT1
ZShift = 2.37
Z lm Score = −0.99
NG = 0
DB = 1
SM = 0
Delta_r
Doxorubicin
YDL161W ENT1
ZShift = 0.48
Z lm Score = 0.22
NG = 0
DB = 1
SM = 0
Delta_AUC
Doxorubicin
YNR035C ARC35
ZShift = 0.76
Z lm Score = −2.7
NG = 0
DB = 0
SM = 0
Delta_L
Doxorubicin
YNR035C ARC35
ZShift = 0.35
Z lm Score = −0.27
NG = 0
DB = 0
SM = 0
Delta_K
Doxorubicin
YNR035C ARC35
ZShift = −1.07
Z lm Score = −0.27
NG = 0
DB = 0
SM = 0
Delta_r
Doxorubicin
YNR035C ARC35
ZShift = −0.47
Z lm Score = 1.24
NG = 0
DB = 0
SM = 0
Delta_AUC
Doxorubicin

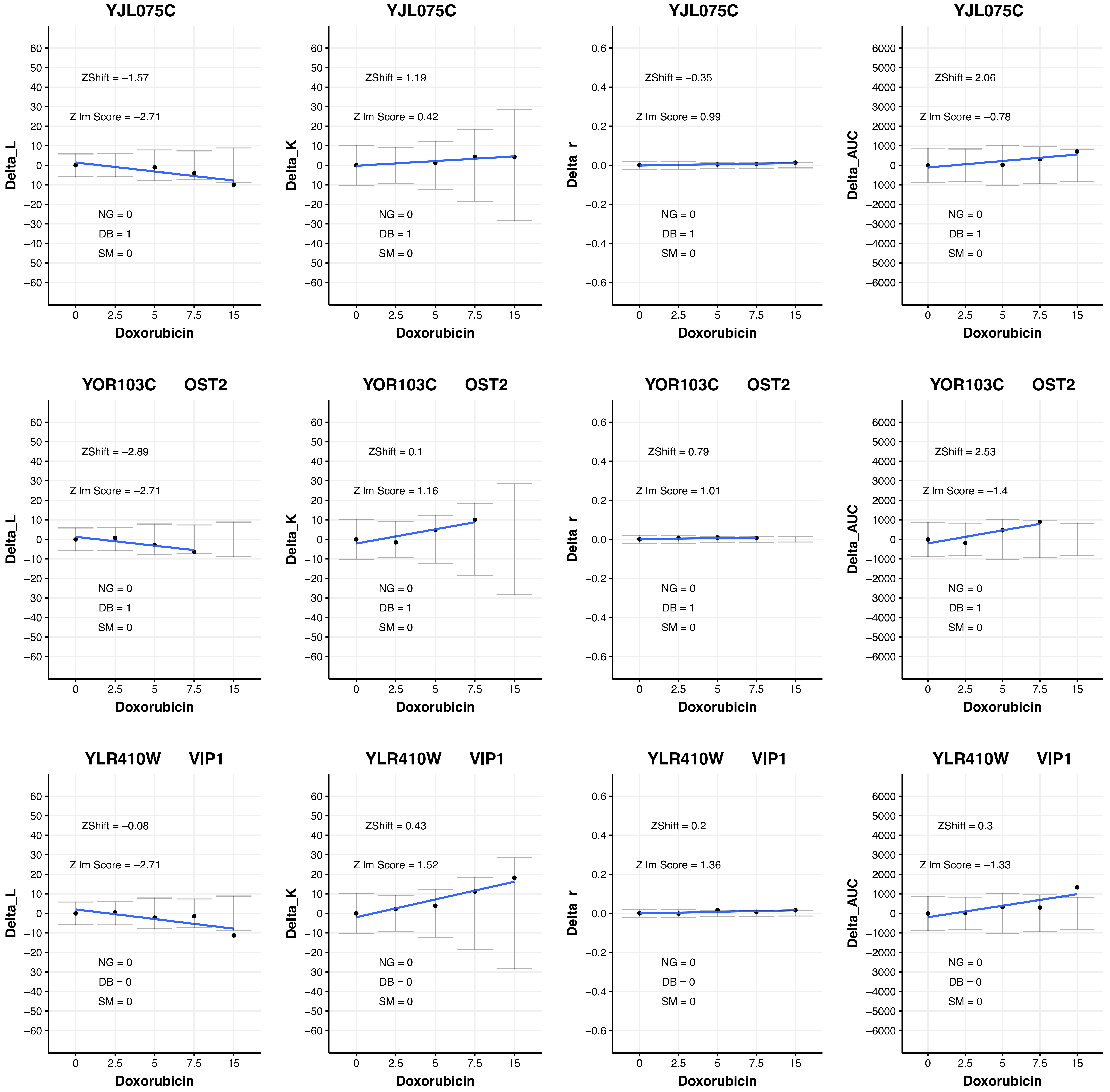
YJL075C
ZShift = −1.57
Z lm Score = −2.71
NG = 0
DB = 1
SM = 0
Delta_L
Doxorubicin
YJL075C
ZShift = 1.19
Z lm Score = 0.42
NG = 0
DB = 1
SM = 0
Delta_K
Doxorubicin
YJL075C
ZShift = −0.35
Z lm Score = 0.99
NG = 0
DB = 1
SM = 0
Delta_r
Doxorubicin
YJL075C
ZShift = 2.06
Z lm Score = −0.78
NG = 0
DB = 1
SM = 0
Delta_AUC
Doxorubicin
YOR103C OST2
ZShift = −2.89
Z lm Score = −2.71
NG = 0
DB = 1
SM = 0
Delta_L
Doxorubicin
YOR103C OST2
ZShift = 0.1
Z lm Score = 1.16
NG = 0
DB = 1
SM = 0
Delta_K
Doxorubicin
YOR103C OST2
ZShift = 0.79
Z lm Score = 1.01
NG = 0
DB = 1
SM = 0
Delta_r
Doxorubicin
YOR103C OST2
ZShift = 2.53
Z lm Score = −1.4
NG = 0
DB = 1
SM = 0
Delta_AUC
Doxorubicin
YLR410W VIP1
ZShift = −0.08
Z lm Score = −2.71
NG = 0
DB = 0
SM = 0
Delta_L
Doxorubicin
YLR410W VIP1
ZShift = 0.43
Z lm Score = 1.52
NG = 0
DB = 0
SM = 0
Delta_K
Doxorubicin
YLR410W VIP1
ZShift = 0.2
Z lm Score = 1.36
NG = 0
DB = 0
SM = 0
Delta_r
Doxorubicin
YLR410W VIP1
ZShift = 0.3
Z lm Score = −1.33
NG = 0
DB = 0
SM = 0
Delta_AUC
Doxorubicin

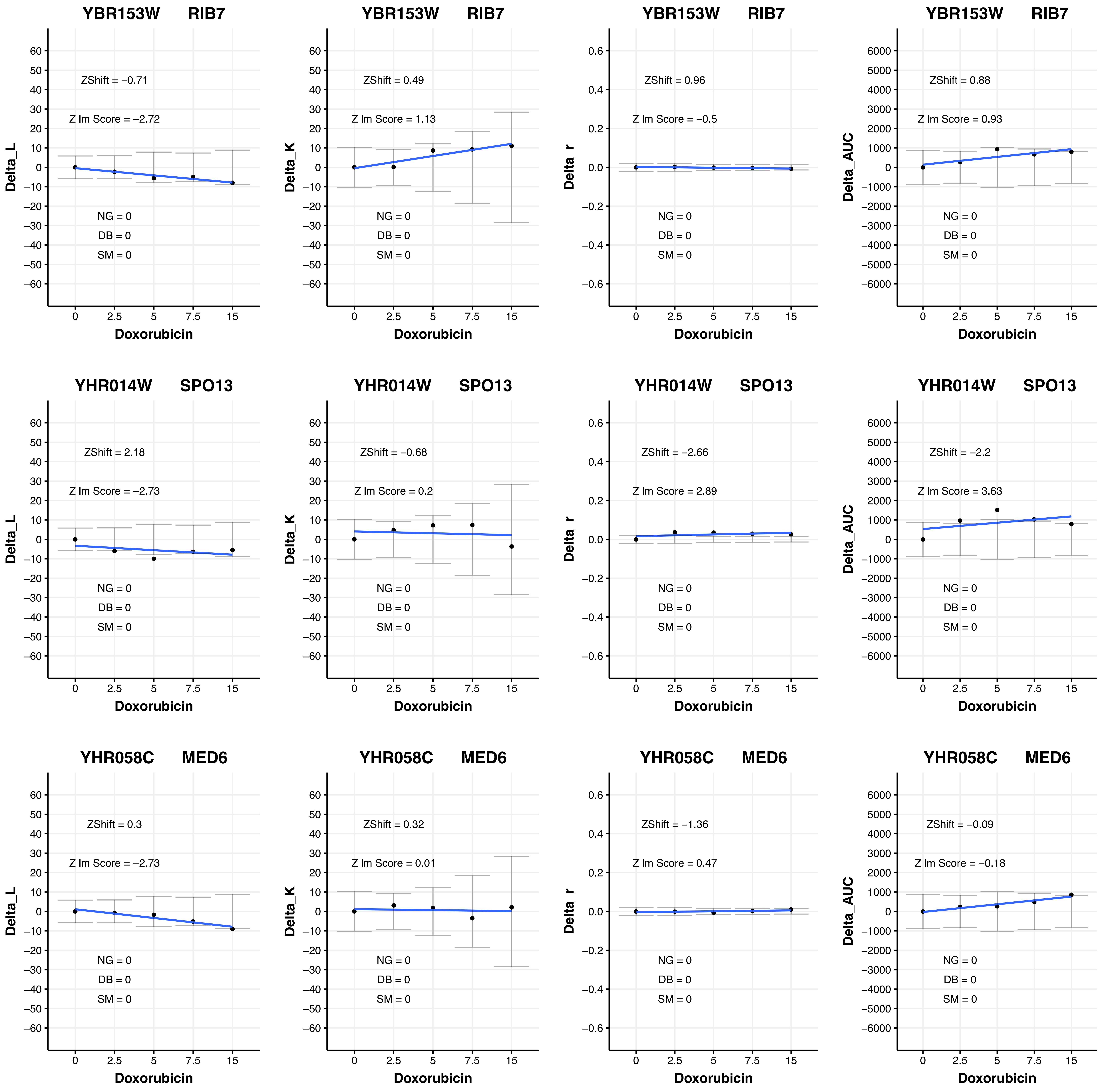

YBR153W RIB7
ZShift = −0.71
Z lm Score = −2.72
NG = 0
DB = 0
SM = 0
Delta_L
Doxorubicin
YBR153W RIB7
ZShift = 0.49
Z lm Score = 1.13
NG = 0
DB = 0
SM = 0
Delta_K
Doxorubicin
YBR153W RIB7
ZShift = 0.96
Z lm Score = −0.5
NG = 0
DB = 0
SM = 0
Delta_r
Doxorubicin
YBR153W RIB7
ZShift = 0.88
Z lm Score = 0.93
NG = 0
DB = 0
SM = 0
Delta_AUC
Doxorubicin
YHR014W SPO13
ZShift = 2.18
Z lm Score = −2.73
NG = 0
DB = 0
SM = 0
Delta_L
Doxorubicin
YHR014W SPO13
ZShift = −0.68
Z lm Score = 0.2
NG = 0
DB = 0
SM = 0
Delta_K
Doxorubicin
YHR014W SPO13
ZShift = −2.66
Z lm Score = 2.89
NG = 0
DB = 0
SM = 0
Delta_r
Doxorubicin
YHR014W SPO13
ZShift = −2.2
Z lm Score = 3.63
NG = 0
DB = 0
SM = 0
Delta_AUC
Doxorubicin
YHR058C MED6
ZShift = 0.3
Z lm Score = −2.73
NG = 0
DB = 0
SM = 0
Delta_L
Doxorubicin
YHR058C MED6
ZShift = 0.32
Z lm Score = 0.01
NG = 0
DB = 0
SM = 0
Delta_K
Doxorubicin
YHR058C MED6
ZShift = −1.36
Z lm Score = 0.47
NG = 0
DB = 0
SM = 0
Delta_r
Doxorubicin
YHR058C MED6
ZShift = −0.09
Z lm Score = −0.18
NG = 0
DB = 0
SM = 0
Delta_AUC
Doxorubicin

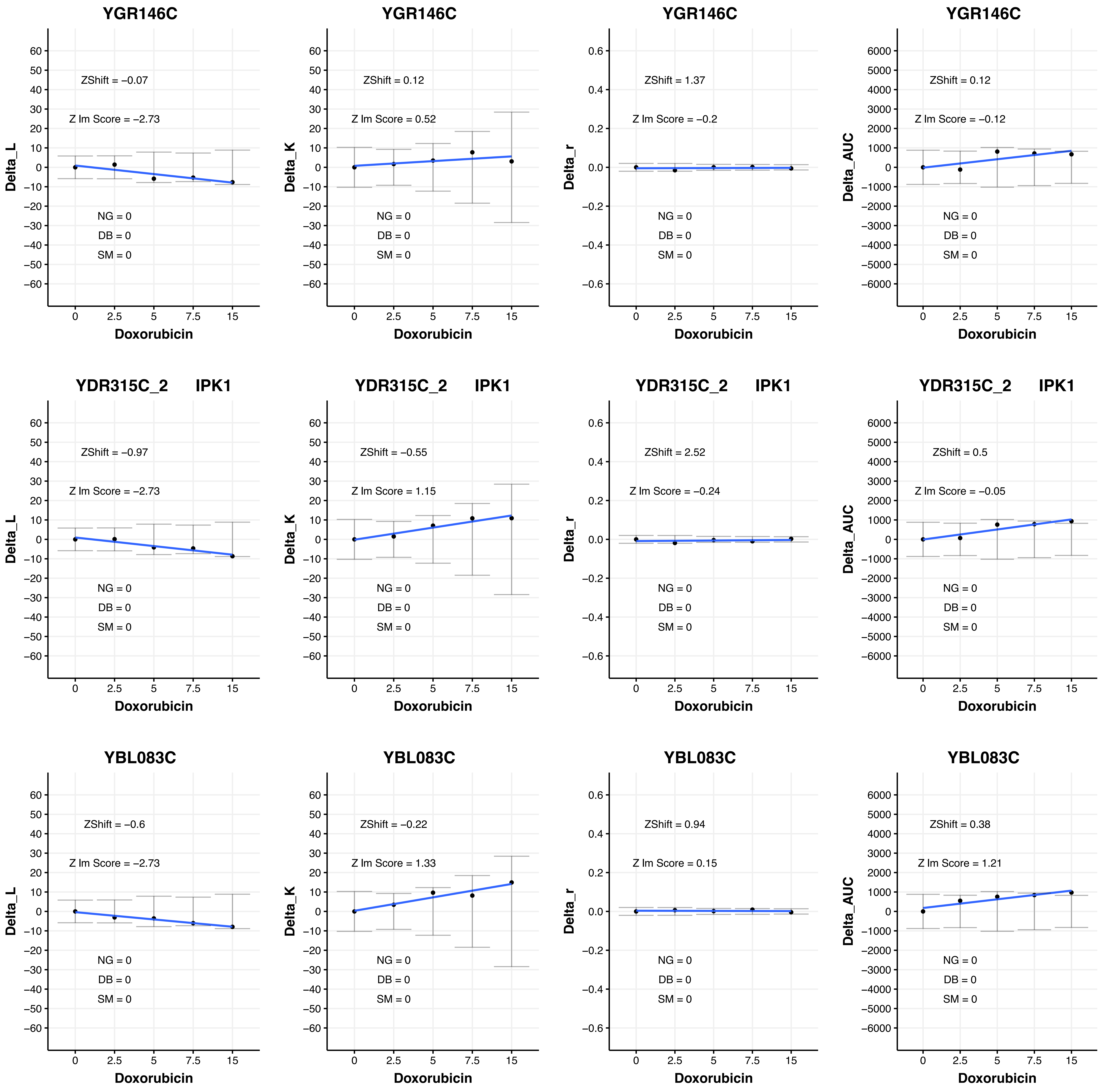
YGR146C
Delta_L
ZShift = −0.07
Z lm Score = −2.73
NG = 0
DB = 0
SM = 0
Doxorubicin
YGR146C
Delta_K
ZShift = 0.12
Z lm Score = 0.52
NG = 0
DB = 0
SM = 0
Doxorubicin
YGR146C
Delta_r
ZShift = 1.37
Z lm Score = −0.2
NG = 0
DB = 0
SM = 0
Doxorubicin
YGR146C
Delta_AUC
ZShift = 0.12
Z lm Score = −0.12
NG = 0
DB = 0
SM = 0
Doxorubicin
YDR315C_2 IPK1
Delta_L
ZShift = −0.97
Z lm Score = −2.73
NG = 0
DB = 0
SM = 0
Doxorubicin
YDR315C_2 IPK1
Delta_K
ZShift = −0.55
Z lm Score = 1.15
NG = 0
DB = 0
SM = 0
Doxorubicin
YDR315C_2 IPK1
Delta_r
ZShift = 2.52
Z lm Score = −0.24
NG = 0
DB = 0
SM = 0
Doxorubicin
YDR315C_2 IPK1
Delta_AUC
ZShift = 0.5
Z lm Score = −0.05
NG = 0
DB = 0
SM = 0
Doxorubicin
YBL083C
Delta_L
ZShift = −0.6
Z lm Score = −2.73
NG = 0
DB = 0
SM = 0
Doxorubicin
YBL083C
Delta_K
ZShift = −0.22
Z lm Score = 1.33
NG = 0
DB = 0
SM = 0
Doxorubicin
YBL083C
Delta_r
ZShift = 0.94
Z lm Score = 0.15
NG = 0
DB = 0
SM = 0
Doxorubicin
YBL083C
Delta_AUC
ZShift = 0.38
Z lm Score = 1.21
NG = 0
DB = 0
SM = 0
Doxorubicin

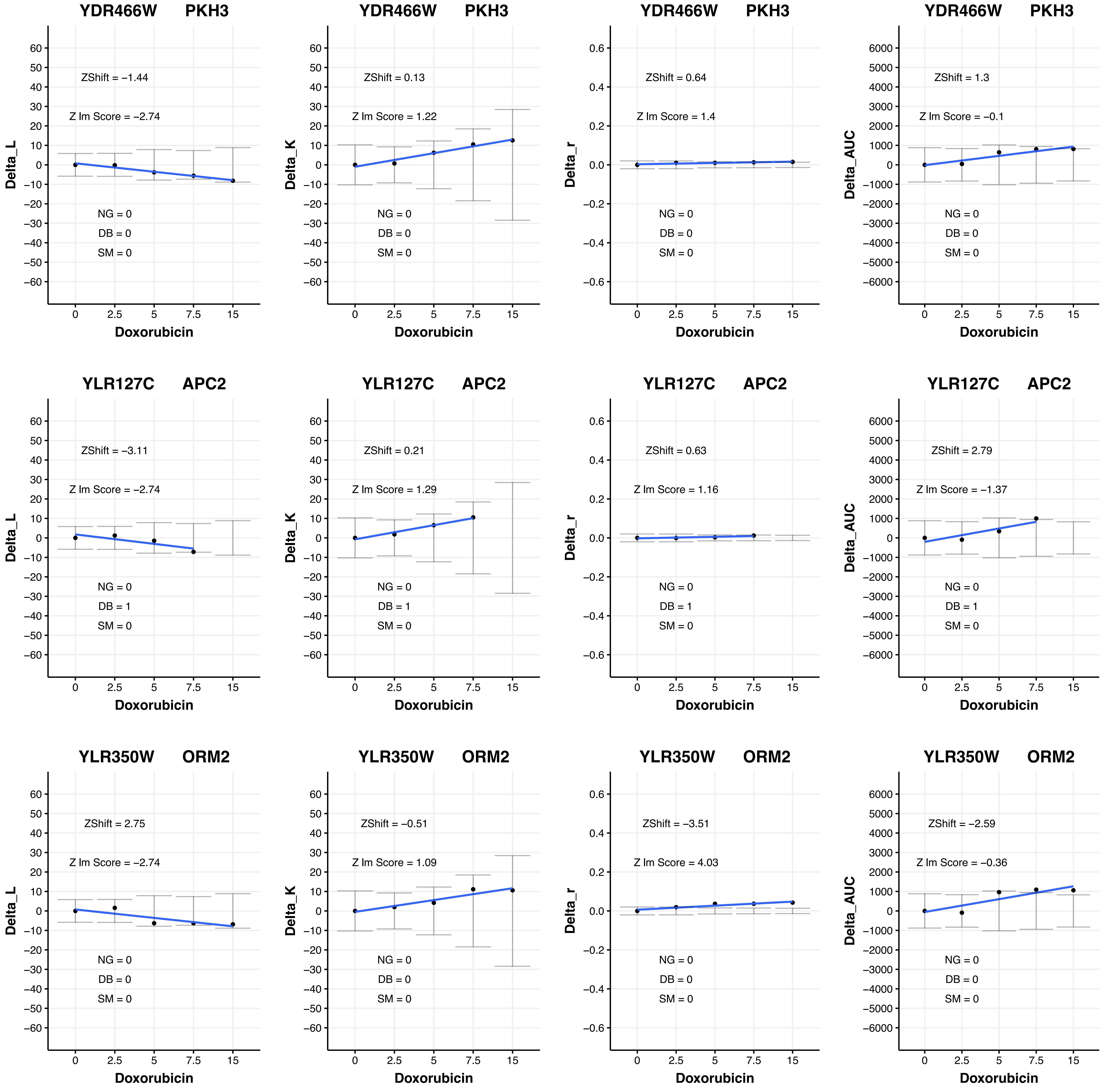
YDR466W PKH3
ZShift = −1.44
Z lm Score = −2.74
NG = 0
DB = 0
SM = 0
Delta_L
Doxorubicin
YDR466W PKH3
ZShift = 0.13
Z lm Score = 1.22
NG = 0
DB = 0
SM = 0
Delta_K
Doxorubicin
YDR466W PKH3
ZShift = 0.64
Z lm Score = 1.4
NG = 0
DB = 0
SM = 0
Delta_r
Doxorubicin
YDR466W PKH3
ZShift = 1.3
Z lm Score = −0.1
NG = 0
DB = 0
SM = 0
Delta_AUC
Doxorubicin
YLR127C APC2
ZShift = −3.11
Z lm Score = −2.74
NG = 0
DB = 1
SM = 0
Delta_L
Doxorubicin
YLR127C APC2
ZShift = 0.21
Z lm Score = 1.29
NG = 0
DB = 1
SM = 0
Delta_K
Doxorubicin
YLR127C APC2
ZShift = 0.63
Z lm Score = 1.16
NG = 0
DB = 1
SM = 0
Delta_r
Doxorubicin
YLR127C APC2
ZShift = 2.79
Z lm Score = −1.37
NG = 0
DB = 1
SM = 0
Delta_AUC
Doxorubicin
YLR350W ORM2
ZShift = 2.75
Z lm Score = −2.74
NG = 0
DB = 0
SM = 0
Delta_L
Doxorubicin
YLR350W ORM2
ZShift = −0.51
Z lm Score = 1.09
NG = 0
DB = 0
SM = 0
Delta_K
Doxorubicin
YLR350W ORM2
ZShift = −3.51
Z lm Score = 4.03
NG = 0
DB = 0
SM = 0
Delta_r
Doxorubicin
YLR350W ORM2
ZShift = −2.59
Z lm Score = −0.36
NG = 0
DB = 0
SM = 0
Delta_AUC
Doxorubicin

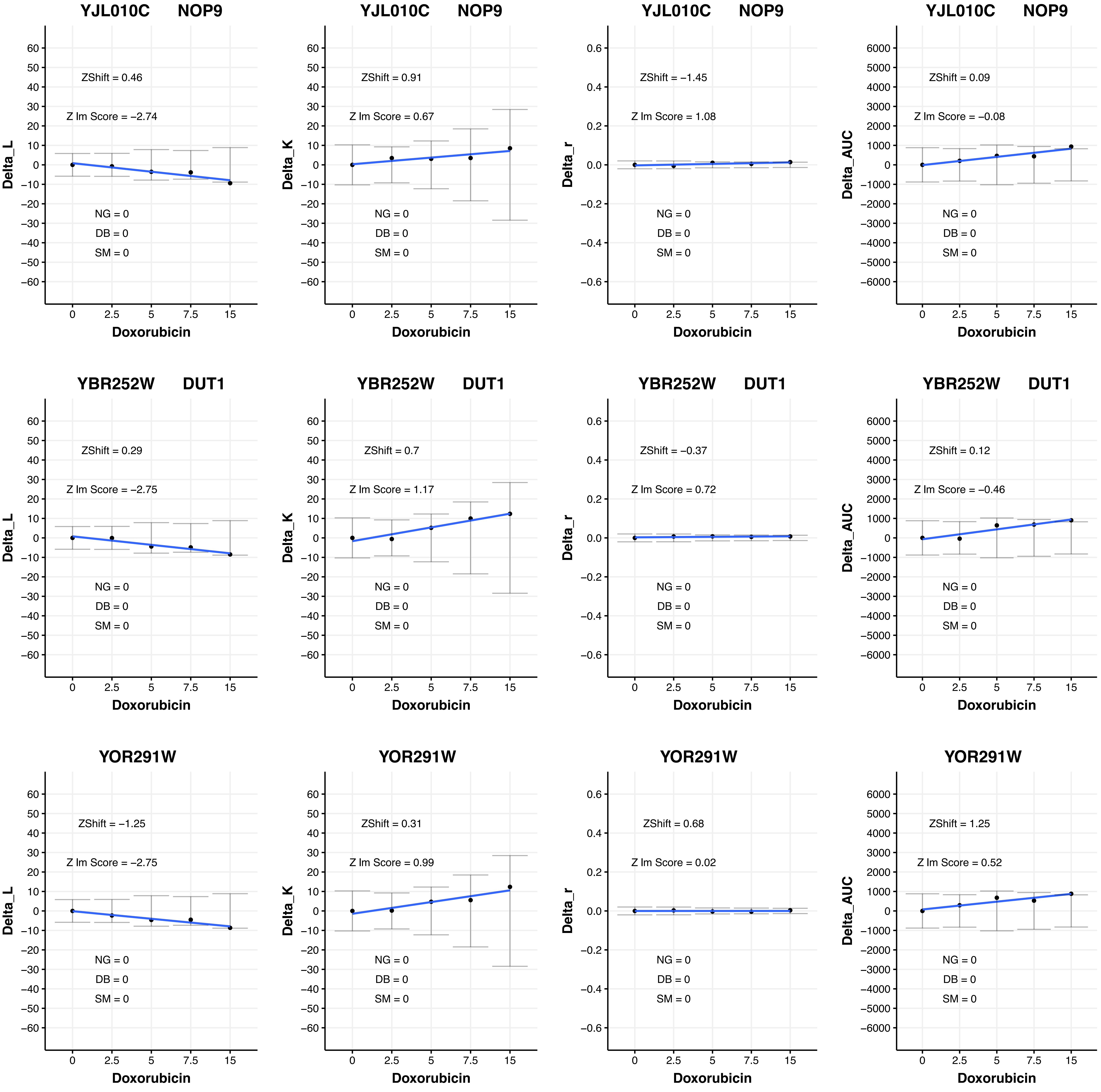
YJL010C NOP9
ZShift = 0.46
Z lm Score = −2.74
NG = 0
DB = 0
SM = 0
Delta_L
Doxorubicin
YJL010C NOP9
ZShift = 0.91
Z lm Score = 0.67
NG = 0
DB = 0
SM = 0
Delta_K
Doxorubicin
YJL010C NOP9
ZShift = −1.45
Z lm Score = 1.08
NG = 0
DB = 0
SM = 0
Delta_r
Doxorubicin
YJL010C NOP9
ZShift = 0.09
Z lm Score = −0.08
NG = 0
DB = 0
SM = 0
Delta_AUC
Doxorubicin
YBR252W DUT1
ZShift = 0.29
Z lm Score = −2.75
NG = 0
DB = 0
SM = 0
Delta_L
Doxorubicin
YBR252W DUT1
ZShift = 0.7
Z lm Score = 1.17
NG = 0
DB = 0
SM = 0
Delta_K
Doxorubicin
YBR252W DUT1
ZShift = −0.37
Z lm Score = 0.72
NG = 0
DB = 0
SM = 0
Delta_r
Doxorubicin
YBR252W DUT1
ZShift = 0.12
Z lm Score = −0.46
NG = 0
DB = 0
SM = 0
Delta_AUC
Doxorubicin
YOR291W
ZShift = −1.25
Z lm Score = −2.75
NG = 0
DB = 0
SM = 0
Delta_L
Doxorubicin
YOR291W
ZShift = 0.31
Z lm Score = 0.99
NG = 0
DB = 0
SM = 0
Delta_K
Doxorubicin
YOR291W
ZShift = 0.68
Z lm Score = 0.02
NG = 0
DB = 0
SM = 0
Delta_r
Doxorubicin
YOR291W
ZShift = 1.25
Z lm Score = 0.52
NG = 0
DB = 0
SM = 0
Delta_AUC
Doxorubicin

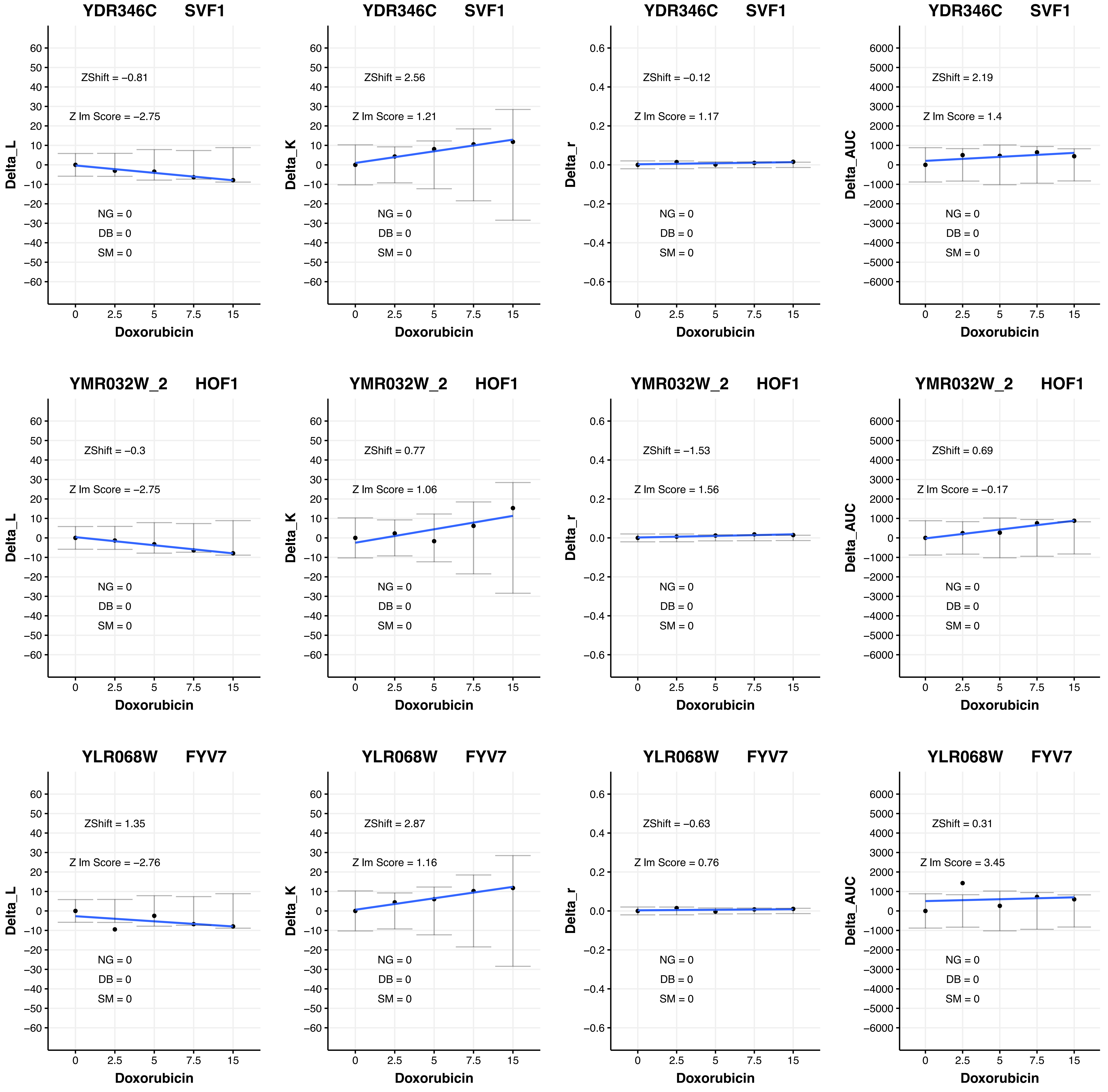
YDR346C SVF1
ZShift = −0.81
Z lm Score = −2.75
NG = 0
DB = 0
SM = 0
Delta_L
Doxorubicin
YDR346C SVF1
ZShift = 2.56
Z lm Score = 1.21
NG = 0
DB = 0
SM = 0
Delta_K
Doxorubicin
YDR346C SVF1
ZShift = −0.12
Z lm Score = 1.17
NG = 0
DB = 0
SM = 0
Delta_r
Doxorubicin
YDR346C SVF1
ZShift = 2.19
Z lm Score = 1.4
NG = 0
DB = 0
SM = 0
Delta_AUC
Doxorubicin
YMR032W_2 HOF1
ZShift = −0.3
Z lm Score = −2.75
NG = 0
DB = 0
SM = 0
Delta_L
Doxorubicin
YMR032W_2 HOF1
ZShift = 0.77
Z lm Score = 1.06
NG = 0
DB = 0
SM = 0
Delta_K
Doxorubicin
YMR032W_2 HOF1
ZShift = −1.53
Z lm Score = 1.56
NG = 0
DB = 0
SM = 0
Delta_r
Doxorubicin
YMR032W_2 HOF1
ZShift = 0.69
Z lm Score = −0.17
NG = 0
DB = 0
SM = 0
Delta_AUC
Doxorubicin
YLR068W FYV7
ZShift = 1.35
Z lm Score = −2.76
NG = 0
DB = 0
SM = 0
Delta_L
Doxorubicin
YLR068W FYV7
ZShift = 2.87
Z lm Score = 1.16
NG = 0
DB = 0
SM = 0
Delta_K
Doxorubicin
YLR068W FYV7
ZShift = −0.63
Z lm Score = 0.76
NG = 0
DB = 0
SM = 0
Delta_r
Doxorubicin
YLR068W FYV7
ZShift = 0.31
Z lm Score = 3.45
NG = 0
DB = 0
SM = 0
Delta_AUC
Doxorubicin

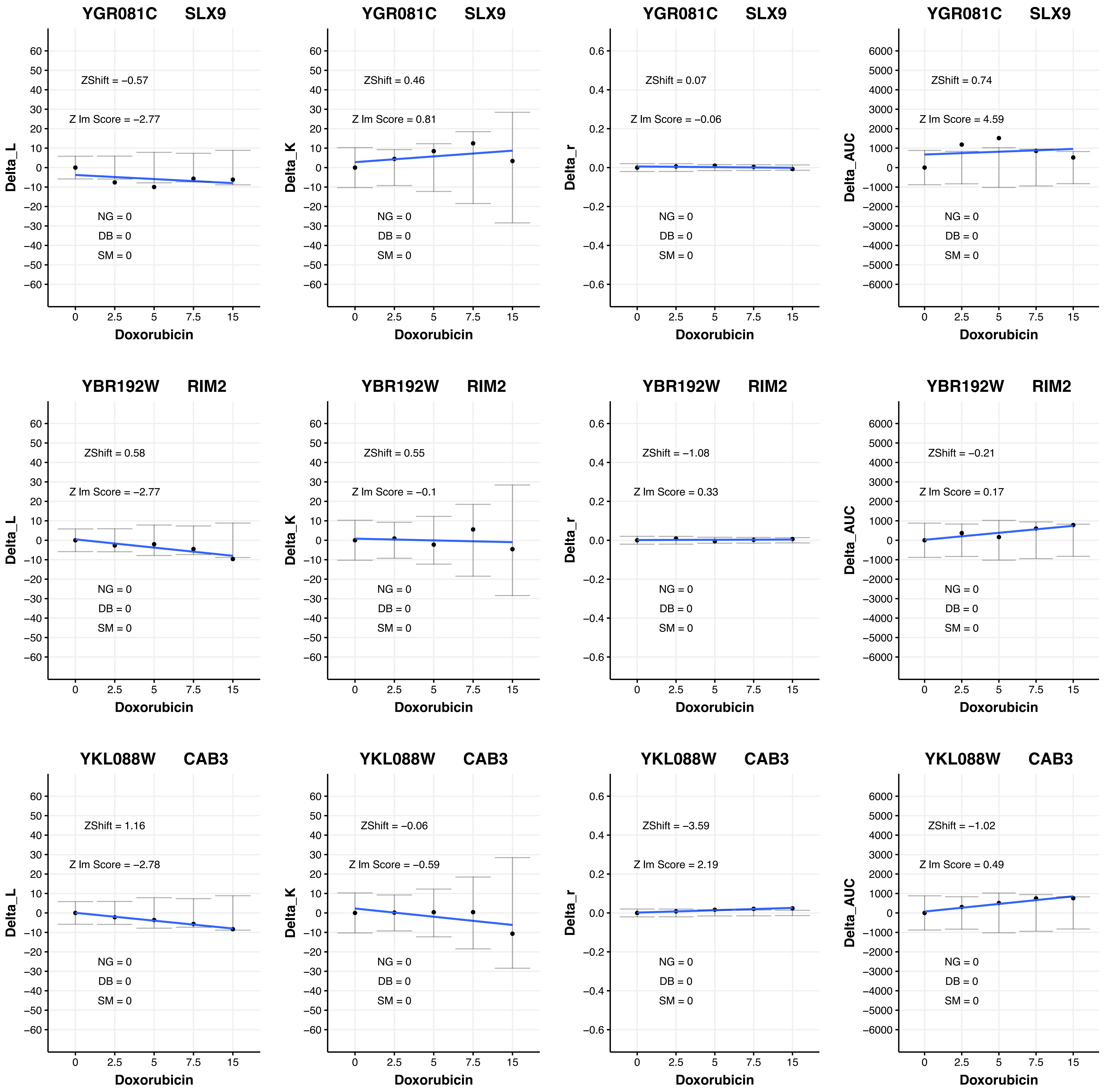

YGR081C SLX9
ZShift = −0.57
Z lm Score = −2.77
NG = 0
DB = 0
SM = 0
Delta_L
Doxorubicin
YGR081C SLX9
ZShift = 0.46
Z lm Score = 0.81
NG = 0
DB = 0
SM = 0
Delta_K
Doxorubicin
YGR081C SLX9
ZShift = 0.07
Z lm Score = −0.06
NG = 0
DB = 0
SM = 0
Delta_r
Doxorubicin
YGR081C SLX9
ZShift = 0.74
Z lm Score = 4.59
NG = 0
DB = 0
SM = 0
Delta_AUC
Doxorubicin
YBR192W RIM2
ZShift = 0.58
Z lm Score = −2.77
NG = 0
DB = 0
SM = 0
Delta_L
Doxorubicin
YBR192W RIM2
ZShift = 0.55
Z lm Score = −0.1
NG = 0
DB = 0
SM = 0
Delta_K
Doxorubicin
YBR192W RIM2
ZShift = −1.08
Z lm Score = 0.33
NG = 0
DB = 0
SM = 0
Delta_r
Doxorubicin
YBR192W RIM2
ZShift = −0.21
Z lm Score = 0.17
NG = 0
DB = 0
SM = 0
Delta_AUC
Doxorubicin
YKL088W CAB3
ZShift = 1.16
Z lm Score = −2.78
NG = 0
DB = 0
SM = 0
Delta_L
Doxorubicin
YKL088W CAB3
ZShift = −0.06
Z lm Score = −0.59
NG = 0
DB = 0
SM = 0
Delta_K
Doxorubicin
YKL088W CAB3
ZShift = −3.59
Z lm Score = 2.19
NG = 0
DB = 0
SM = 0
Delta_r
Doxorubicin
YKL088W CAB3
ZShift = −1.02
Z lm Score = 0.49
NG = 0
DB = 0
SM = 0
Delta_AUC
Doxorubicin

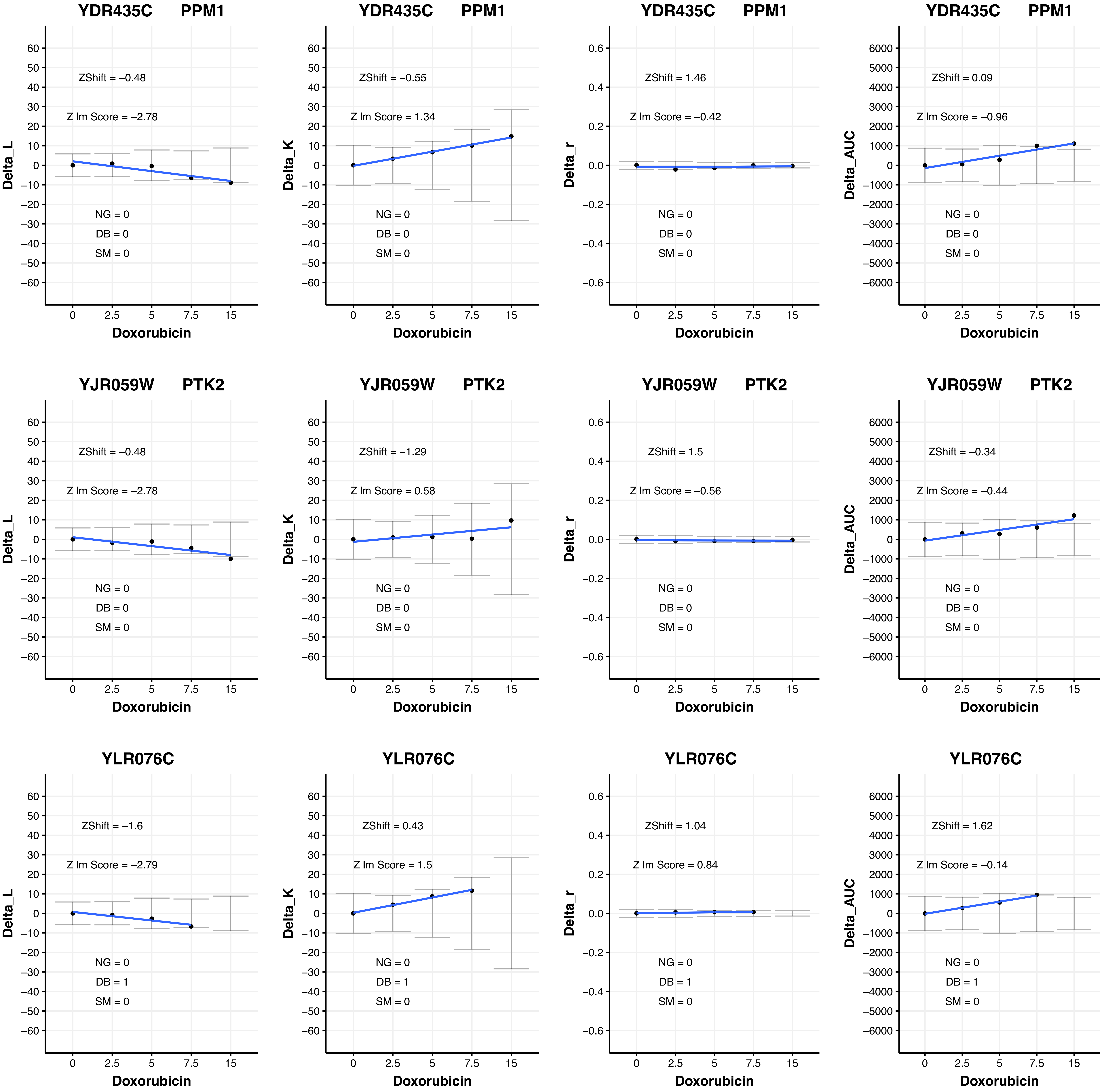

YDR435C PPM1
ZShift = −0.48
Z lm Score = −2.78
NG = 0
DB = 0
SM = 0
Delta_L
Doxorubicin
YDR435C PPM1
ZShift = −0.55
Z lm Score = 1.34
NG = 0
DB = 0
SM = 0
Delta_K
Doxorubicin
YDR435C PPM1
ZShift = 1.46
Z lm Score = −0.42
NG = 0
DB = 0
SM = 0
Delta_r
Doxorubicin
YDR435C PPM1
ZShift = 0.09
Z lm Score = −0.96
NG = 0
DB = 0
SM = 0
Delta_AUC
Doxorubicin
YJR059W PTK2
ZShift = −0.48
Z lm Score = −2.78
NG = 0
DB = 0
SM = 0
Delta_L
Doxorubicin
YJR059W PTK2
ZShift = −1.29
Z lm Score = 0.58
NG = 0
DB = 0
SM = 0
Delta_K
Doxorubicin
YJR059W PTK2
ZShift = 1.5
Z lm Score = −0.56
NG = 0
DB = 0
SM = 0
Delta_r
Doxorubicin
YJR059W PTK2
ZShift = −0.34
Z lm Score = −0.44
NG = 0
DB = 0
SM = 0
Delta_AUC
Doxorubicin
YLR076C
ZShift = −1.6
Z lm Score = −2.79
NG = 0
DB = 1
SM = 0
Delta_L
Doxorubicin
YLR076C
ZShift = 0.43
Z lm Score = 1.5
NG = 0
DB = 1
SM = 0
Delta_K
Doxorubicin
YLR076C
ZShift = 1.04
Z lm Score = 0.84
NG = 0
DB = 1
SM = 0
Delta_r
Doxorubicin
YLR076C
ZShift = 1.62
Z lm Score = −0.14
NG = 0
DB = 1
SM = 0
Delta_AUC
Doxorubicin

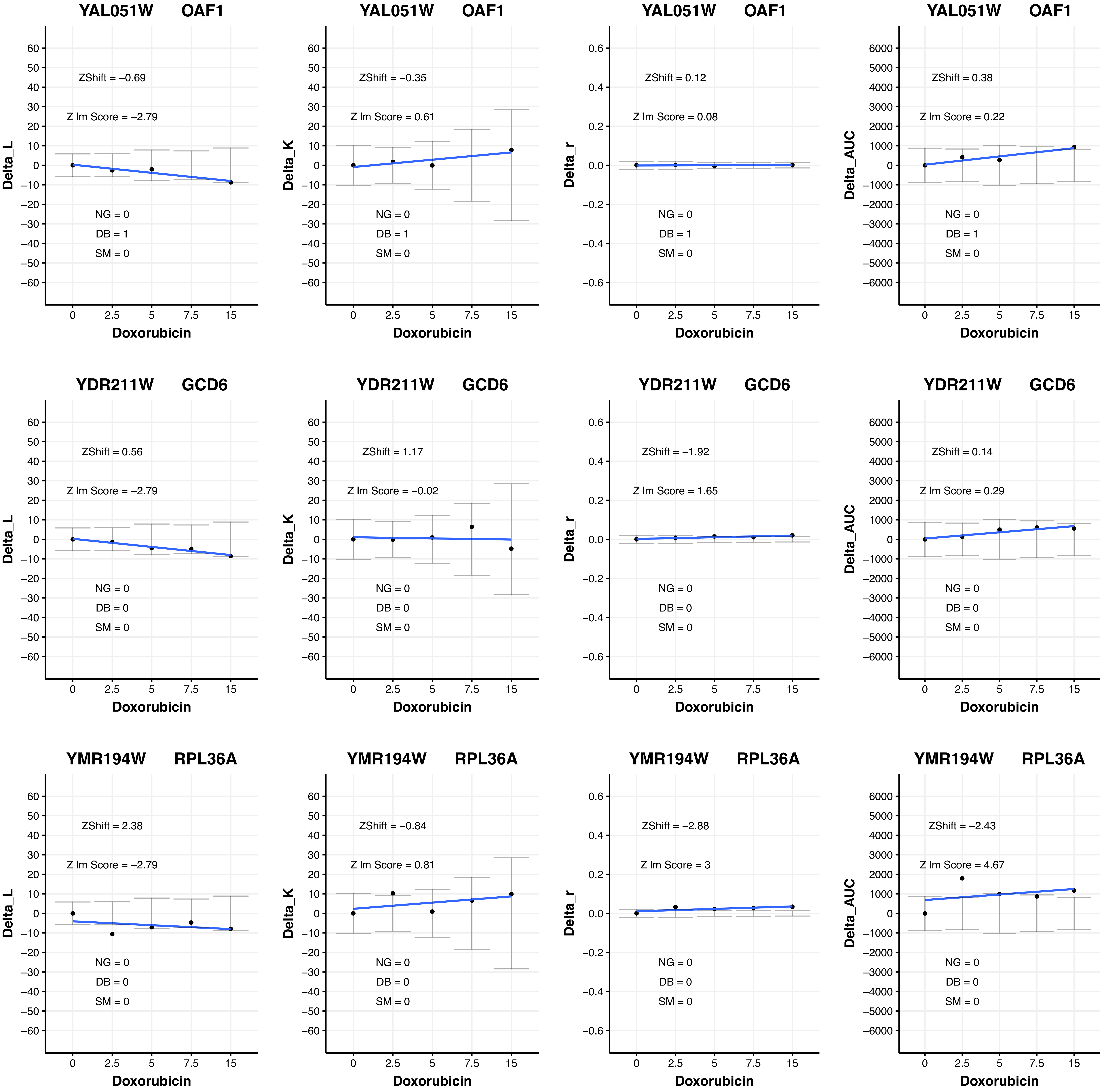
YAL051W OAF1
ZShift = −0.69
Z Im Score = −2.79
NG = 0
DB = 1
SM = 0
Delta_L
Doxorubicin
YAL051W OAF1
ZShift = −0.35
Z Im Score = 0.61
NG = 0
DB = 1
SM = 0
Delta_K
Doxorubicin
YAL051W OAF1
ZShift = 0.12
Z Im Score = 0.08
NG = 0
DB = 1
SM = 0
Delta_r
Doxorubicin
YAL051W OAF1
ZShift = 0.38
Z Im Score = 0.22
NG = 0
DB = 1
SM = 0
Delta_AUC
Doxorubicin
YDR211W GCD6
ZShift = 0.56
Z Im Score = −2.79
NG = 0
DB = 0
SM = 0
Delta_L
Doxorubicin
YDR211W GCD6
ZShift = 1.17
Z Im Score = −0.02
NG = 0
DB = 0
SM = 0
Delta_K
Doxorubicin
YDR211W GCD6
ZShift = −1.92
Z Im Score = 1.65
NG = 0
DB = 0
SM = 0
Delta_r
Doxorubicin
YDR211W GCD6
ZShift = 0.14
Z Im Score = 0.29
NG = 0
DB = 0
SM = 0
Delta_AUC
Doxorubicin
YMR194W RPL36A
ZShift = 2.38
Z Im Score = −2.79
NG = 0
DB = 0
SM = 0
Delta_L
Doxorubicin
YMR194W RPL36A
ZShift = −0.84
Z Im Score = 0.81
NG = 0
DB = 0
SM = 0
Delta_K
Doxorubicin
YMR194W RPL36A
ZShift = −2.88
Z Im Score = 3
NG = 0
DB = 0
SM = 0
Delta_r
Doxorubicin
YMR194W RPL36A
ZShift = −2.43
Z Im Score = 4.67
NG = 0
DB = 0
SM = 0
Delta_AUC
Doxorubicin

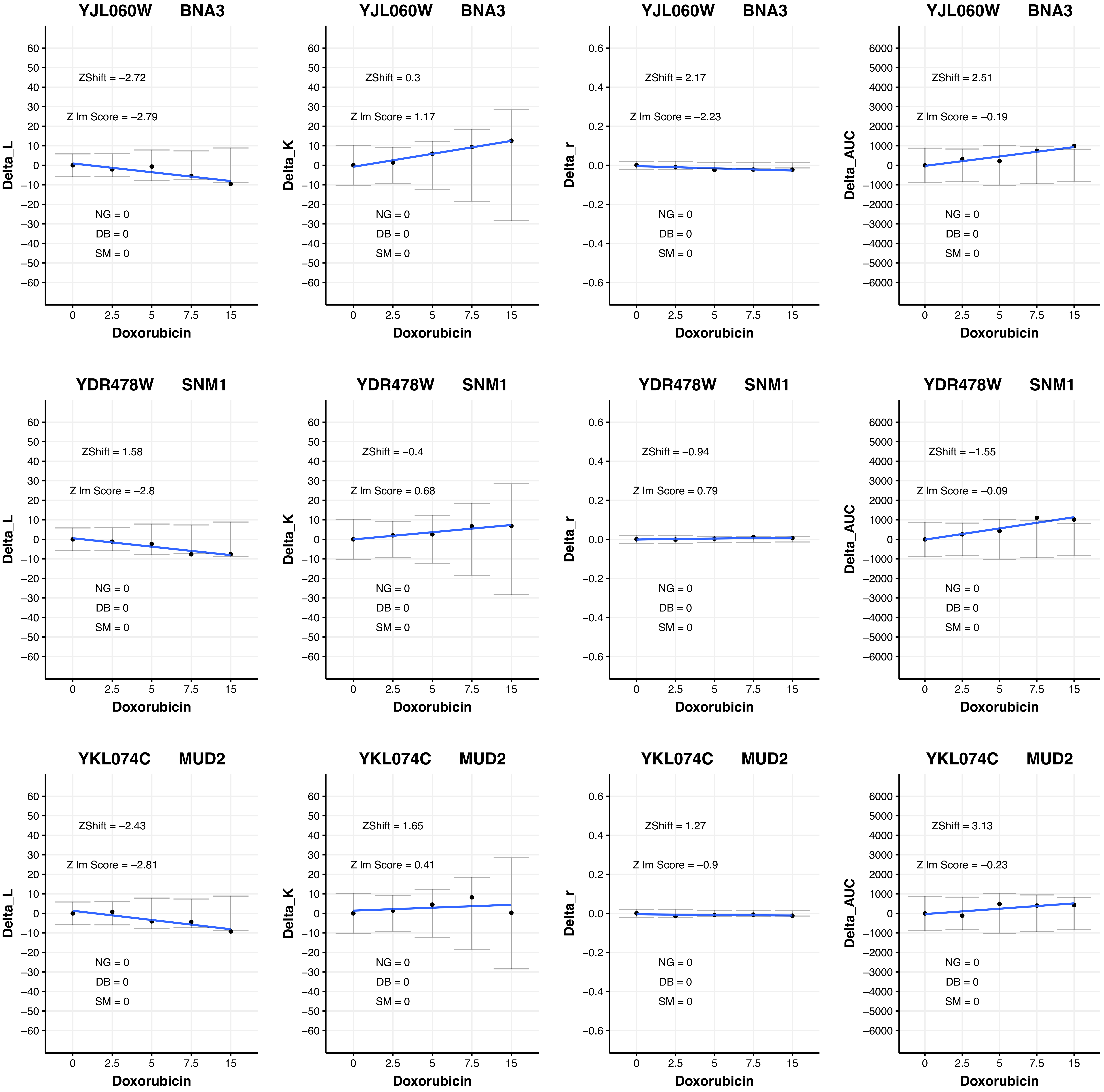

YJL060W BNA3
ZShift = −2.72
Z lm Score = −2.79
NG = 0
DB = 0
SM = 0
Delta_L
Doxorubicin
YJL060W BNA3
ZShift = 0.3
Z lm Score = 1.17
NG = 0
DB = 0
SM = 0
Delta_K
Doxorubicin
YJL060W BNA3
ZShift = 2.17
Z lm Score = −2.23
NG = 0
DB = 0
SM = 0
Delta_r
Doxorubicin
YJL060W BNA3
ZShift = 2.51
Z lm Score = −0.19
NG = 0
DB = 0
SM = 0
Delta_AUC
Doxorubicin
YDR478W SNM1
ZShift = 1.58
Z lm Score = −2.8
NG = 0
DB = 0
SM = 0
Delta_L
Doxorubicin
YDR478W SNM1
ZShift = −0.4
Z lm Score = 0.68
NG = 0
DB = 0
SM = 0
Delta_K
Doxorubicin
YDR478W SNM1
ZShift = −0.94
Z lm Score = 0.79
NG = 0
DB = 0
SM = 0
Delta_r
Doxorubicin
YDR478W SNM1
ZShift = −1.55
Z lm Score = −0.09
NG = 0
DB = 0
SM = 0
Delta_AUC
Doxorubicin
YKL074C MUD2
ZShift = −2.43
Z lm Score = −2.81
NG = 0
DB = 0
SM = 0
Delta_L
Doxorubicin
YKL074C MUD2
ZShift = 1.65
Z lm Score = 0.41
NG = 0
DB = 0
SM = 0
Delta_K
Doxorubicin
YKL074C MUD2
ZShift = 1.27
Z lm Score = −0.9
NG = 0
DB = 0
SM = 0
Delta_r
Doxorubicin
YKL074C MUD2
ZShift = 3.13
Z lm Score = −0.23
NG = 0
DB = 0
SM = 0
Delta_AUC
Doxorubicin

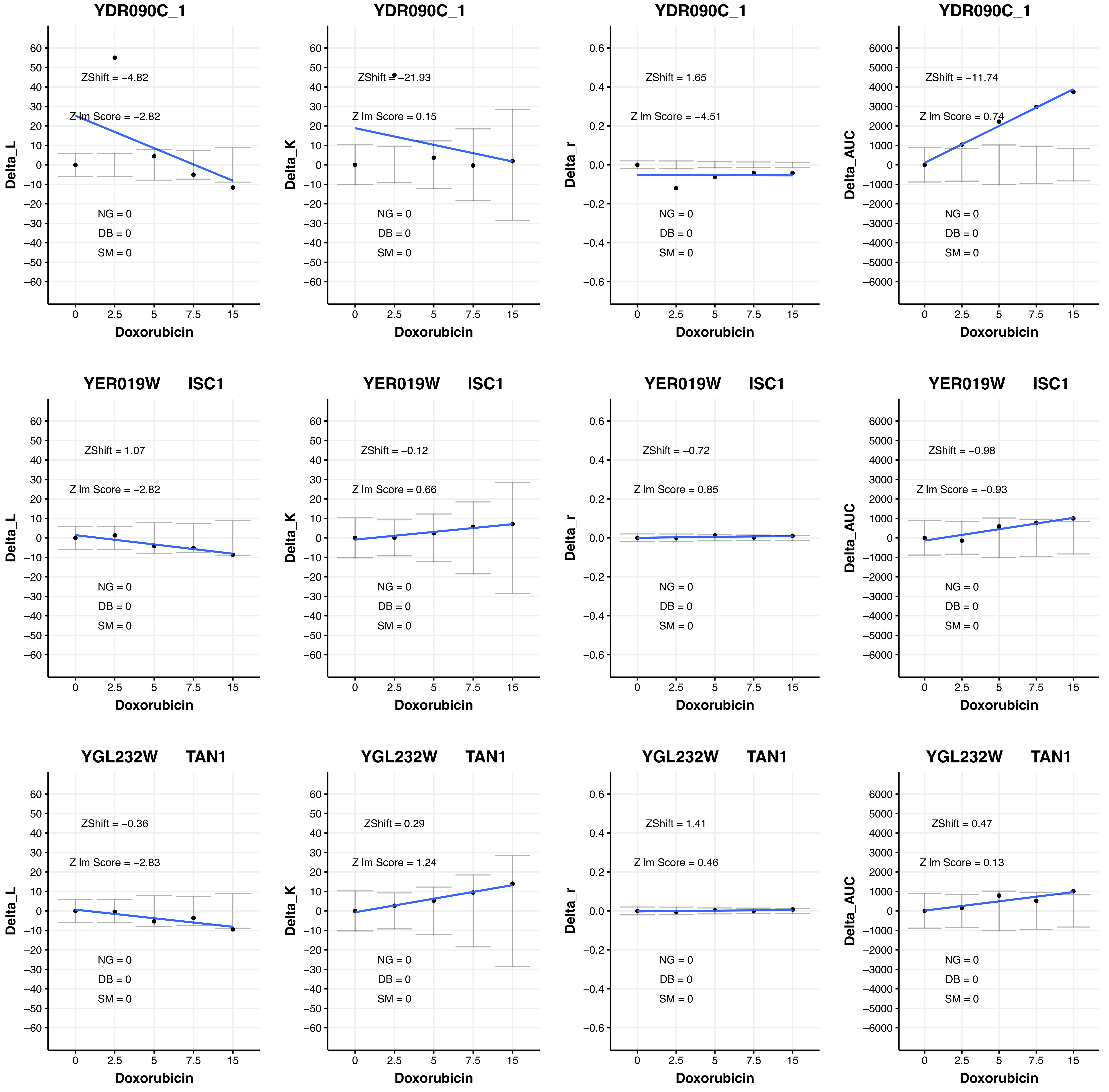
YDR090C_1
ZShift = −4.82
Z lm Score = −2.82
NG = 0
DB = 0
SM = 0
Delta_L
Doxorubicin
YDR090C_1
ZShift = −21.93
Z lm Score = 0.15
Delta_K
YDR090C_1
ZShift = 1.65
Z lm Score = −4.51
Delta_r
YDR090C_1
ZShift = −11.74
Z lm Score = 0.74
Delta_AUC
YER019W ISC1
ZShift = 1.07
Z lm Score = −2.82
ZShift = −0.12
Z lm Score = 0.66
ZShift = −0.72
Z lm Score = 0.85
ZShift = −0.98
Z lm Score = −0.93
YGL232W TAN1
ZShift = −0.36
Z lm Score = −2.83
ZShift = 0.29
Z lm Score = 1.24
ZShift = 1.41
Z lm Score = 0.46
ZShift = 0.47
Z lm Score = 0.13

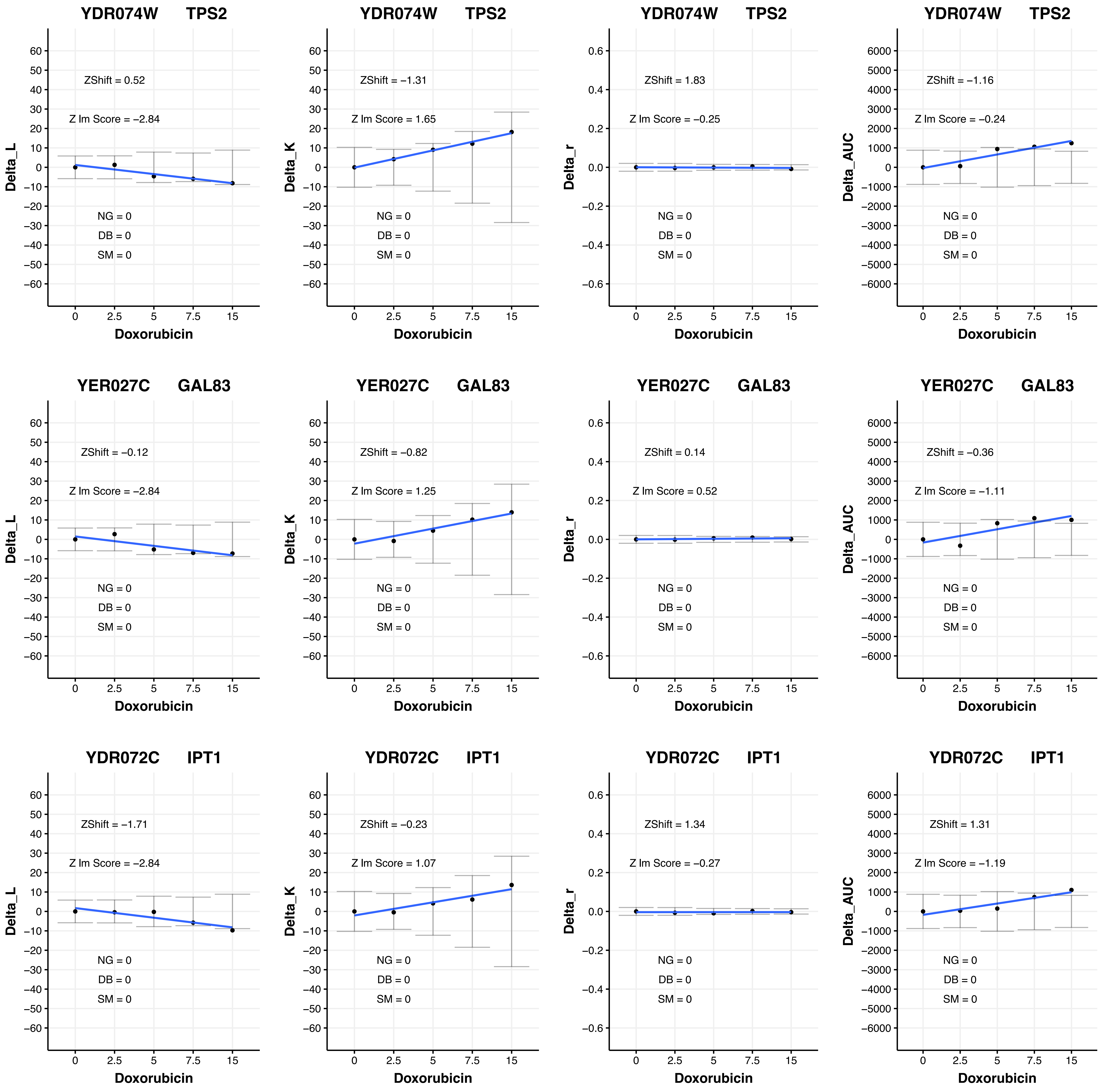
YDR074W TPS2
Delta_L
ZShift = 0.52
Z lm Score = −2.84
NG = 0
DB = 0
SM = 0
Doxorubicin
YDR074W TPS2
Delta_K
ZShift = −1.31
Z lm Score = 1.65
NG = 0
DB = 0
SM = 0
Doxorubicin
YDR074W TPS2
Delta_r
ZShift = 1.83
Z lm Score = −0.25
NG = 0
DB = 0
SM = 0
Doxorubicin
YDR074W TPS2
Delta_AUC
ZShift = −1.16
Z lm Score = −0.24
NG = 0
DB = 0
SM = 0
Doxorubicin
YER027C GAL83
Delta_L
ZShift = −0.12
Z lm Score = −2.84
NG = 0
DB = 0
SM = 0
Doxorubicin
YER027C GAL83
Delta_K
ZShift = −0.82
Z lm Score = 1.25
NG = 0
DB = 0
SM = 0
Doxorubicin
YER027C GAL83
Delta_r
ZShift = 0.14
Z lm Score = 0.52
NG = 0
DB = 0
SM = 0
Doxorubicin
YER027C GAL83
Delta_AUC
ZShift = −0.36
Z lm Score = −1.11
NG = 0
DB = 0
SM = 0
Doxorubicin
YDR072C IPT1
Delta_L
ZShift = −1.71
Z lm Score = −2.84
NG = 0
DB = 0
SM = 0
Doxorubicin
YDR072C IPT1
Delta_K
ZShift = −0.23
Z lm Score = 1.07
NG = 0
DB = 0
SM = 0
Doxorubicin
YDR072C IPT1
Delta_r
ZShift = 1.34
Z lm Score = −0.27
NG = 0
DB = 0
SM = 0
Doxorubicin
YDR072C IPT1
Delta_AUC
ZShift = 1.31
Z lm Score = −1.19
NG = 0
DB = 0
SM = 0
Doxorubicin

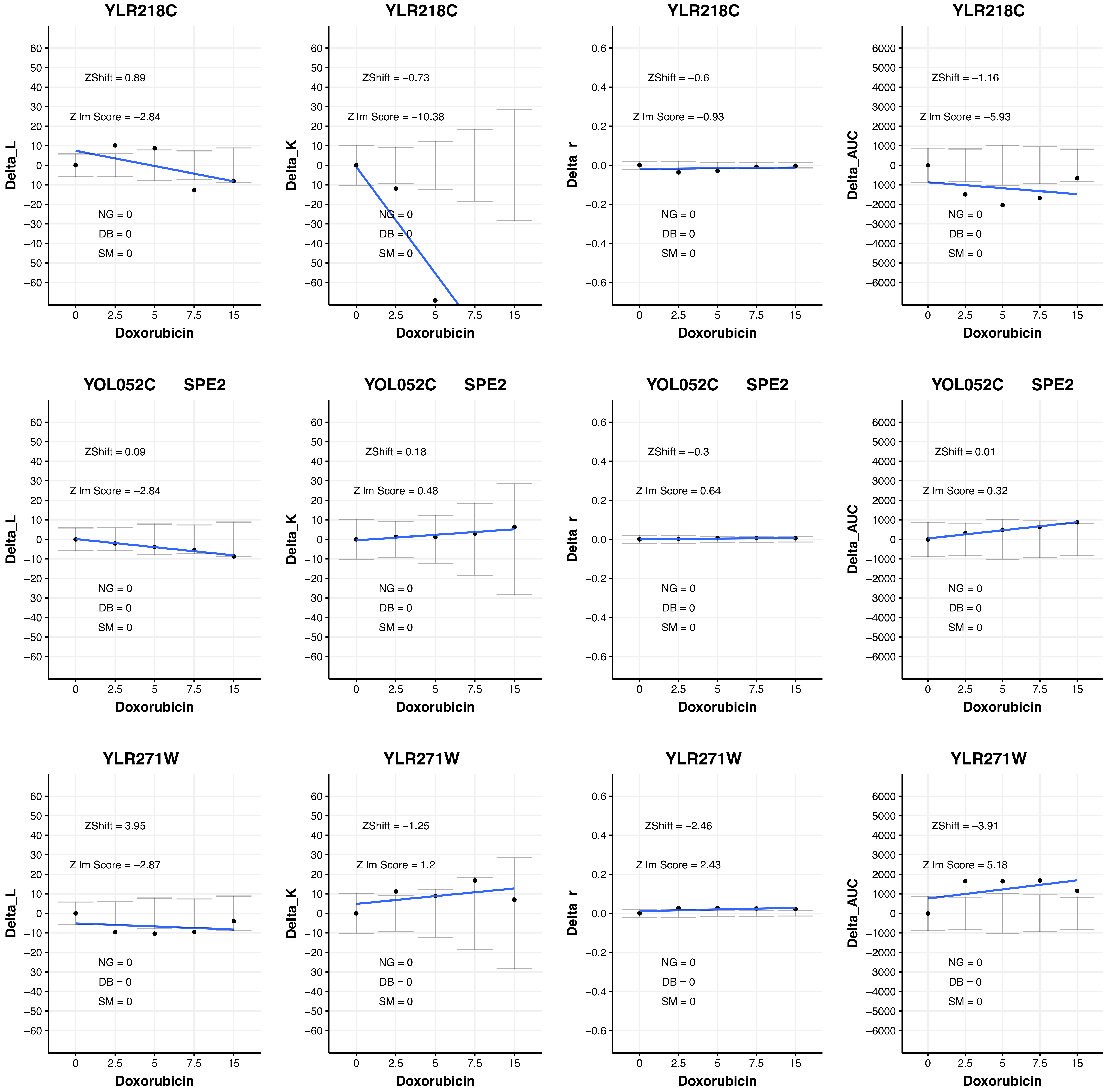
YLR218C
ZShift = 0.89
Z lm Score = −2.84
NG = 0
DB = 0
SM = 0
Delta_L
Doxorubicin
YLR218C
ZShift = −0.73
Z lm Score = −10.38
NG = 0
DB = 0
SM = 0
Delta_K
Doxorubicin
YLR218C
ZShift = −0.6
Z lm Score = −0.93
NG = 0
DB = 0
SM = 0
Delta_r
Doxorubicin
YLR218C
ZShift = −1.16
Z lm Score = −5.93
NG = 0
DB = 0
SM = 0
Delta_AUC
Doxorubicin
YOL052C SPE2
ZShift = 0.09
Z lm Score = −2.84
NG = 0
DB = 0
SM = 0
Delta_L
Doxorubicin
YOL052C SPE2
ZShift = 0.18
Z lm Score = 0.48
NG = 0
DB = 0
SM = 0
Delta_K
Doxorubicin
YOL052C SPE2
ZShift = −0.3
Z lm Score = 0.64
NG = 0
DB = 0
SM = 0
Delta_r
Doxorubicin
YOL052C SPE2
ZShift = 0.01
Z lm Score = 0.32
NG = 0
DB = 0
SM = 0
Delta_AUC
Doxorubicin
YLR271W
ZShift = 3.95
Z lm Score = −2.87
NG = 0
DB = 0
SM = 0
Delta_L
Doxorubicin
YLR271W
ZShift = −1.25
Z lm Score = 1.2
NG = 0
DB = 0
SM = 0
Delta_K
Doxorubicin
YLR271W
ZShift = −2.46
Z lm Score = 2.43
NG = 0
DB = 0
SM = 0
Delta_r
Doxorubicin
YLR271W
ZShift = −3.91
Z lm Score = 5.18
NG = 0
DB = 0
SM = 0
Delta_AUC
Doxorubicin

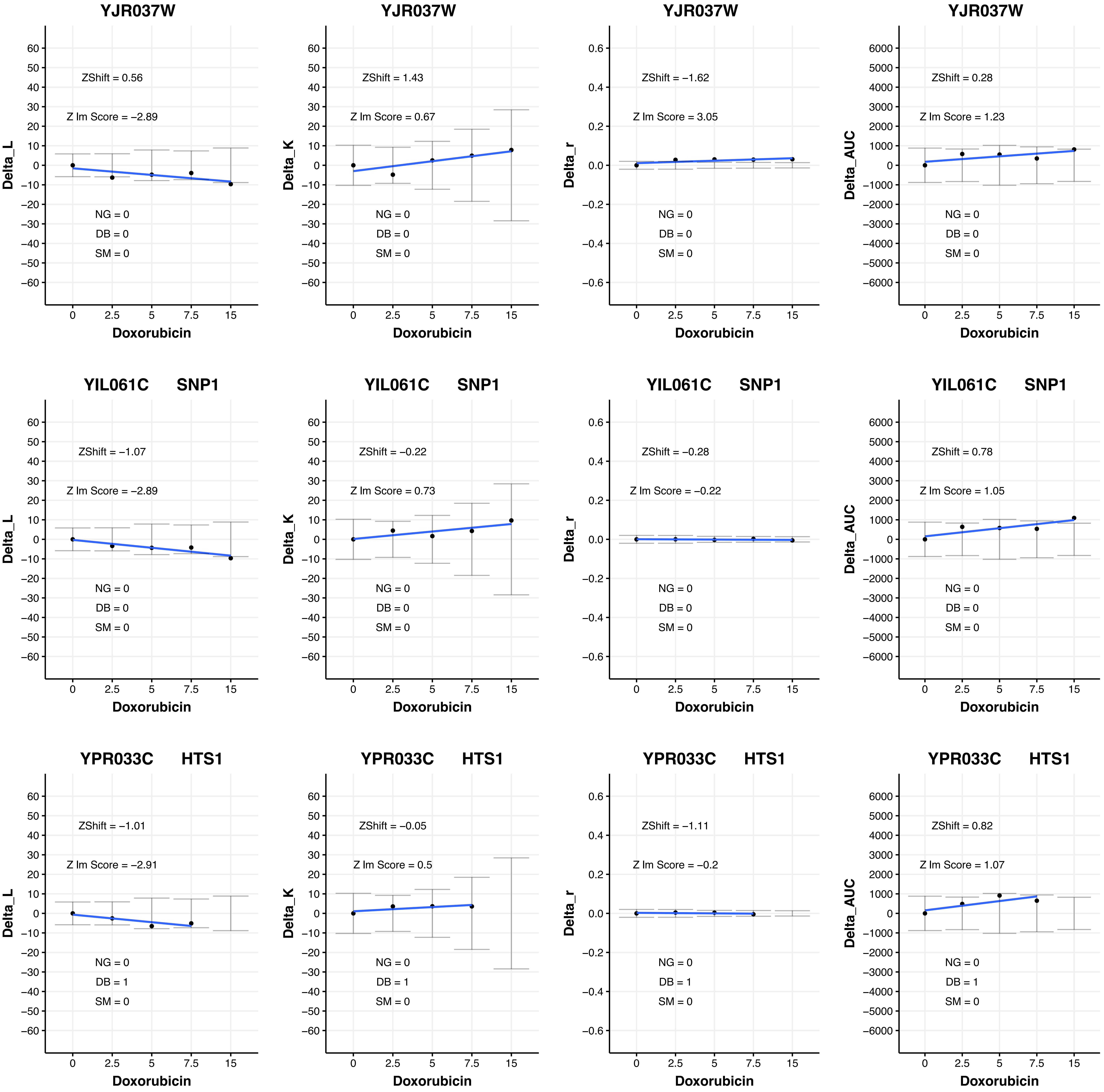
YJR037W
ZShift = 0.56
Z lm Score = −2.89
NG = 0
DB = 0
SM = 0
Delta_L
Doxorubicin
YJR037W
ZShift = 1.43
Z lm Score = 0.67
NG = 0
DB = 0
SM = 0
Delta_K
Doxorubicin
YJR037W
ZShift = −1.62
Z lm Score = 3.05
NG = 0
DB = 0
SM = 0
Delta_r
Doxorubicin
YJR037W
ZShift = 0.28
Z lm Score = 1.23
NG = 0
DB = 0
SM = 0
Delta_AUC
Doxorubicin
YIL061C SNP1
ZShift = −1.07
Z lm Score = −2.89
NG = 0
DB = 0
SM = 0
Delta_L
Doxorubicin
YIL061C SNP1
ZShift = −0.22
Z lm Score = 0.73
NG = 0
DB = 0
SM = 0
Delta_K
Doxorubicin
YIL061C SNP1
ZShift = −0.28
Z lm Score = −0.22
NG = 0
DB = 0
SM = 0
Delta_r
Doxorubicin
YIL061C SNP1
ZShift = 0.78
Z lm Score = 1.05
NG = 0
DB = 0
SM = 0
Delta_AUC
Doxorubicin
YPR033C HTS1
ZShift = −1.01
Z lm Score = −2.91
NG = 0
DB = 1
SM = 0
Delta_L
Doxorubicin
YPR033C HTS1
ZShift = −0.05
Z lm Score = 0.5
NG = 0
DB = 1
SM = 0
Delta_K
Doxorubicin
YPR033C HTS1
ZShift = −1.11
Z lm Score = −0.2
NG = 0
DB = 1
SM = 0
Delta_r
Doxorubicin
YPR033C HTS1
ZShift = 0.82
Z lm Score = 1.07
NG = 0
DB = 1
SM = 0
Delta_AUC
Doxorubicin

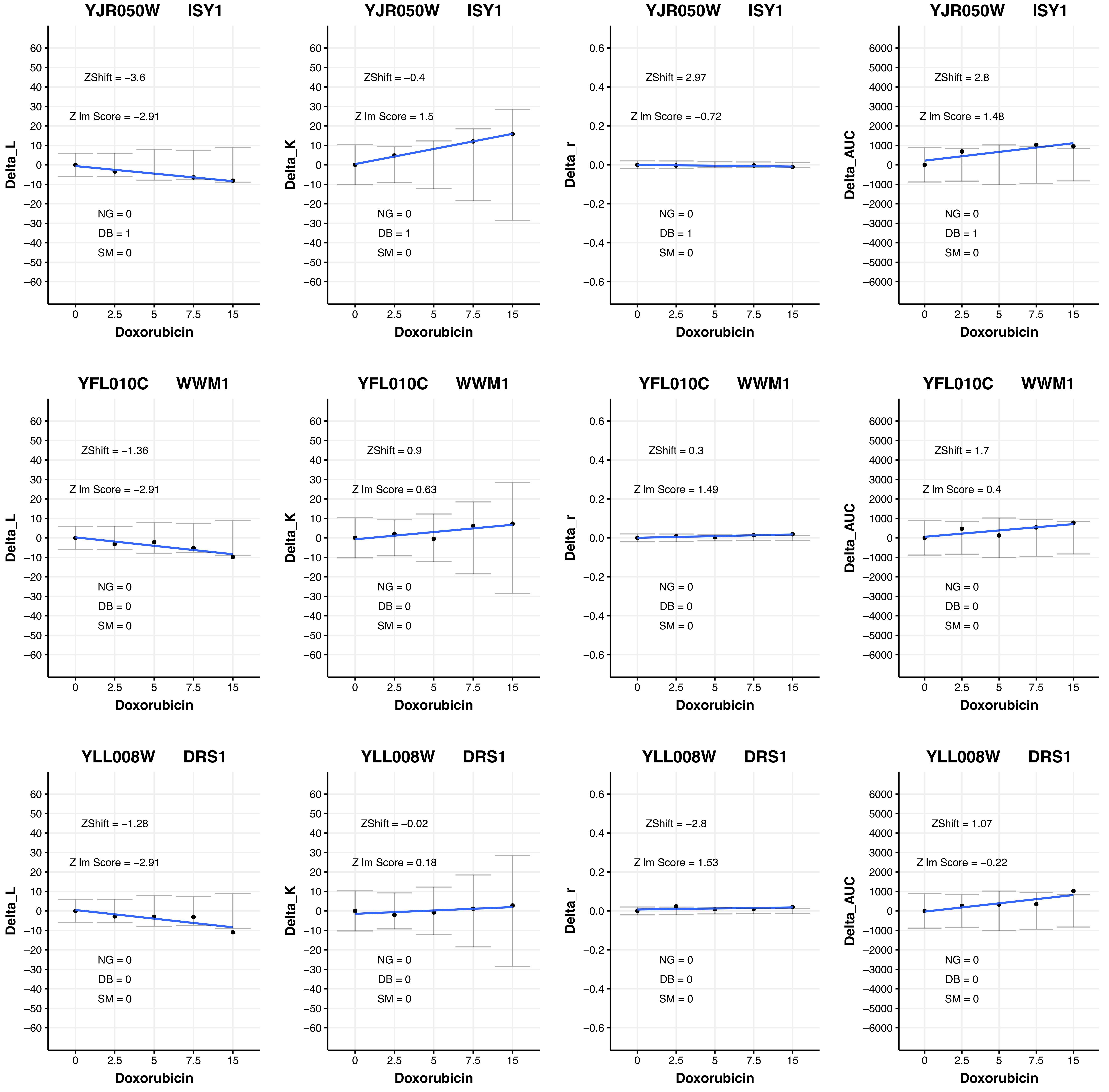
YJR050W ISY1
ZShift = −3.6
Z lm Score = −2.91
NG = 0
DB = 1
SM = 0
Delta_L
Doxorubicin
YJR050W ISY1
ZShift = −0.4
Z lm Score = 1.5
NG = 0
DB = 1
SM = 0
Delta_K
Doxorubicin
YJR050W ISY1
ZShift = 2.97
Z lm Score = −0.72
NG = 0
DB = 1
SM = 0
Delta_r
Doxorubicin
YJR050W ISY1
ZShift = 2.8
Z lm Score = 1.48
NG = 0
DB = 1
SM = 0
Delta_AUC
Doxorubicin
YFL010C WWM1
ZShift = −1.36
Z lm Score = −2.91
NG = 0
DB = 0
SM = 0
Delta_L
Doxorubicin
YFL010C WWM1
ZShift = 0.9
Z lm Score = 0.63
NG = 0
DB = 0
SM = 0
Delta_K
Doxorubicin
YFL010C WWM1
ZShift = 0.3
Z lm Score = 1.49
NG = 0
DB = 0
SM = 0
Delta_r
Doxorubicin
YFL010C WWM1
ZShift = 1.7
Z lm Score = 0.4
NG = 0
DB = 0
SM = 0
Delta_AUC
Doxorubicin
YLL008W DRS1
ZShift = −1.28
Z lm Score = −2.91
NG = 0
DB = 0
SM = 0
Delta_L
Doxorubicin
YLL008W DRS1
ZShift = −0.02
Z lm Score = 0.18
NG = 0
DB = 0
SM = 0
Delta_K
Doxorubicin
YLL008W DRS1
ZShift = −2.8
Z lm Score = 1.53
NG = 0
DB = 0
SM = 0
Delta_r
Doxorubicin
YLL008W DRS1
ZShift = 1.07
Z lm Score = −0.22
NG = 0
DB = 0
SM = 0
Delta_AUC
Doxorubicin

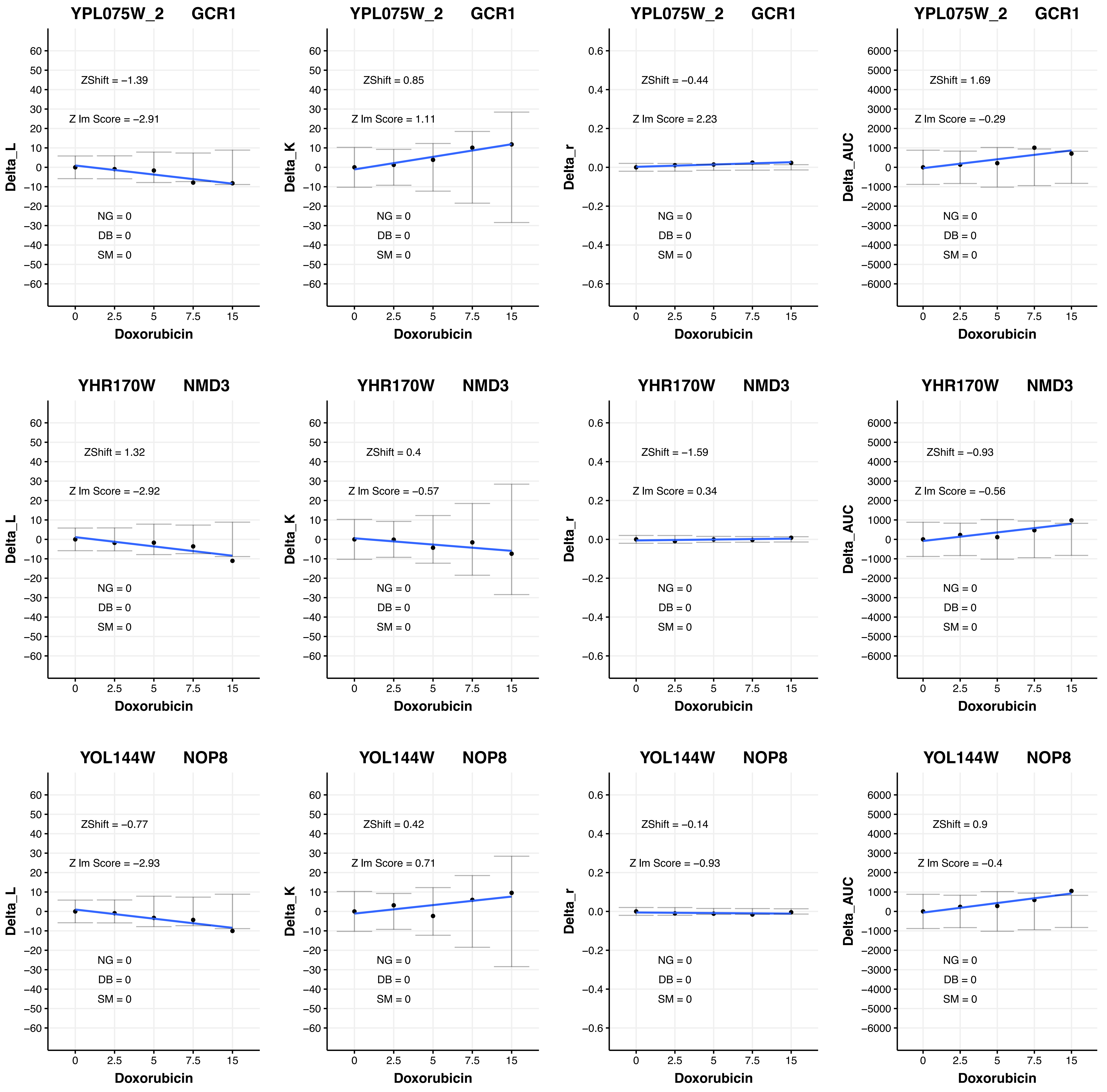

YPL075W_2 GCR1
ZShift = −1.39
Z lm Score = −2.91
NG = 0
DB = 0
SM = 0
Delta_L
Doxorubicin
YPL075W_2 GCR1
ZShift = 0.85
Z lm Score = 1.11
NG = 0
DB = 0
SM = 0
Delta_K
Doxorubicin
YPL075W_2 GCR1
ZShift = −0.44
Z lm Score = 2.23
NG = 0
DB = 0
SM = 0
Delta_r
Doxorubicin
YPL075W_2 GCR1
ZShift = 1.69
Z lm Score = −0.29
NG = 0
DB = 0
SM = 0
Delta_AUC
Doxorubicin
YHR170W NMD3
ZShift = 1.32
Z lm Score = −2.92
NG = 0
DB = 0
SM = 0
Delta_L
Doxorubicin
YHR170W NMD3
ZShift = 0.4
Z lm Score = −0.57
NG = 0
DB = 0
SM = 0
Delta_K
Doxorubicin
YHR170W NMD3
ZShift = −1.59
Z lm Score = 0.34
NG = 0
DB = 0
SM = 0
Delta_r
Doxorubicin
YHR170W NMD3
ZShift = −0.93
Z lm Score = −0.56
NG = 0
DB = 0
SM = 0
Delta_AUC
Doxorubicin
YOL144W NOP8
ZShift = −0.77
Z lm Score = −2.93
NG = 0
DB = 0
SM = 0
Delta_L
Doxorubicin
YOL144W NOP8
ZShift = 0.42
Z lm Score = 0.71
NG = 0
DB = 0
SM = 0
Delta_K
Doxorubicin
YOL144W NOP8
ZShift = −0.14
Z lm Score = −0.93
NG = 0
DB = 0
SM = 0
Delta_r
Doxorubicin
YOL144W NOP8
ZShift = 0.9
Z lm Score = −0.4
NG = 0
DB = 0
SM = 0
Delta_AUC
Doxorubicin

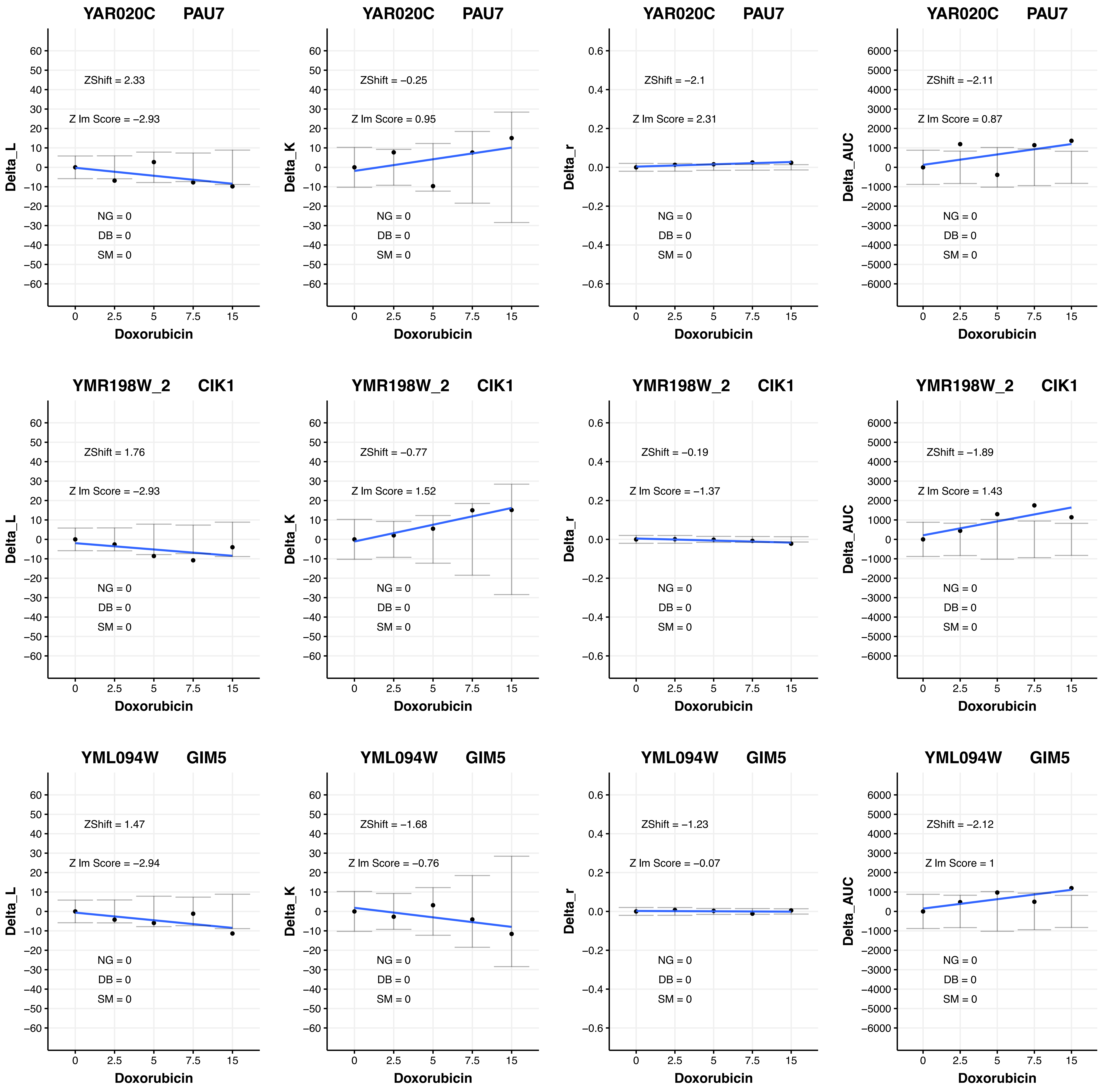
YAR020C PAU7
ZShift = 2.33
Z lm Score = −2.93
NG = 0
DB = 0
SM = 0
Delta_L
Doxorubicin
YAR020C PAU7
ZShift = −0.25
Z lm Score = 0.95
NG = 0
DB = 0
SM = 0
Delta_K
Doxorubicin
YAR020C PAU7
ZShift = −2.1
Z lm Score = 2.31
NG = 0
DB = 0
SM = 0
Delta_r
Doxorubicin
YAR020C PAU7
ZShift = −2.11
Z lm Score = 0.87
NG = 0
DB = 0
SM = 0
Delta_AUC
Doxorubicin
YMR198W_2 CIK1
ZShift = 1.76
Z lm Score = −2.93
NG = 0
DB = 0
SM = 0
Delta_L
Doxorubicin
YMR198W_2 CIK1
ZShift = −0.77
Z lm Score = 1.52
NG = 0
DB = 0
SM = 0
Delta_K
Doxorubicin
YMR198W_2 CIK1
ZShift = −0.19
Z lm Score = −1.37
NG = 0
DB = 0
SM = 0
Delta_r
Doxorubicin
YMR198W_2 CIK1
ZShift = −1.89
Z lm Score = 1.43
NG = 0
DB = 0
SM = 0
Delta_AUC
Doxorubicin
YML094W GIM5
ZShift = 1.47
Z lm Score = −2.94
NG = 0
DB = 0
SM = 0
Delta_L
Doxorubicin
YML094W GIM5
ZShift = −1.68
Z lm Score = −0.76
NG = 0
DB = 0
SM = 0
Delta_K
Doxorubicin
YML094W GIM5
ZShift = −1.23
Z lm Score = −0.07
NG = 0
DB = 0
SM = 0
Delta_r
Doxorubicin
YML094W GIM5
ZShift = −2.12
Z lm Score = 1
NG = 0
DB = 0
SM = 0
Delta_AUC
Doxorubicin

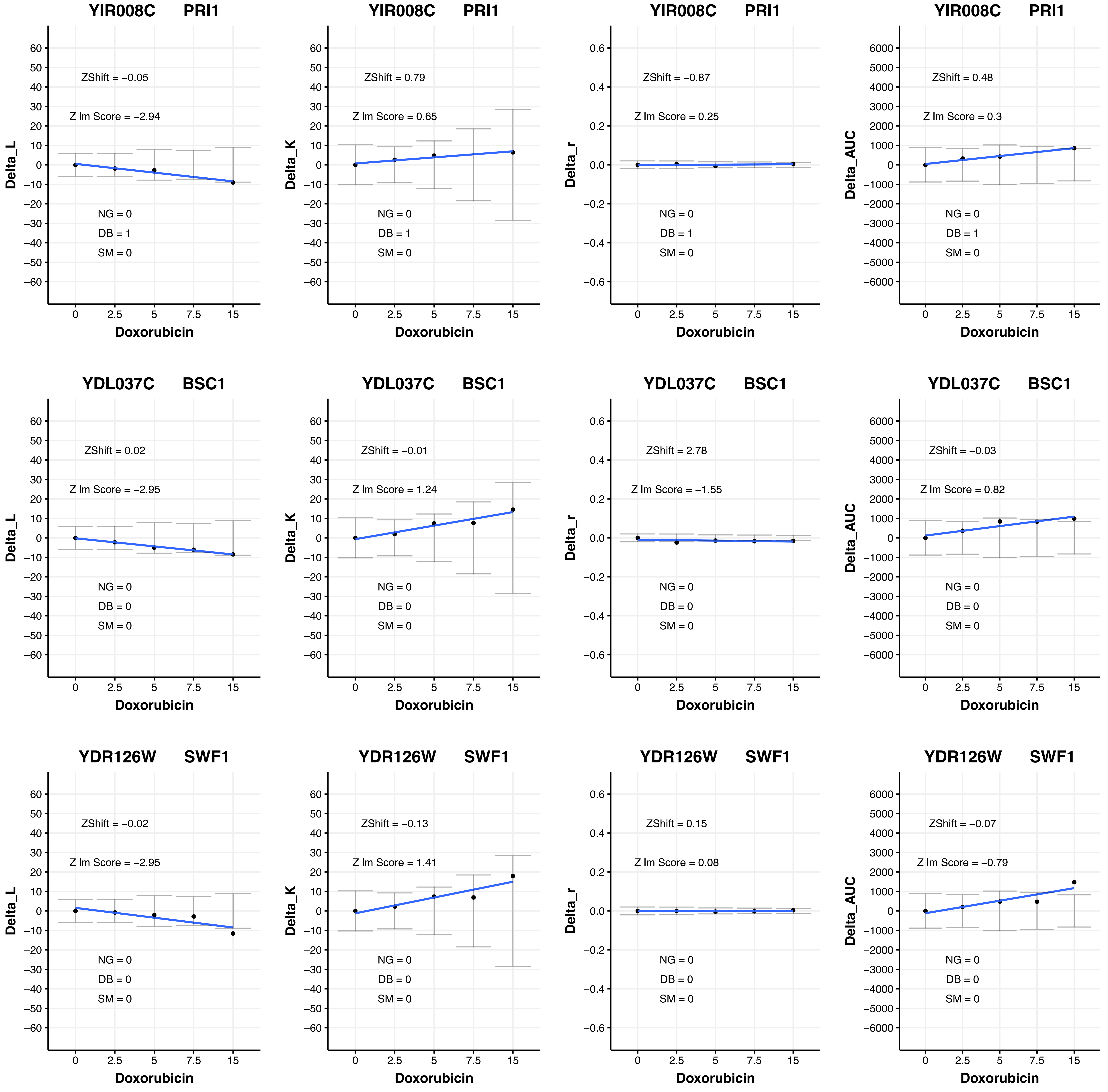
YIR008C PRI1
ZShift = −0.05
Z lm Score = −2.94
NG = 0
DB = 1
SM = 0
Delta_L
Doxorubicin
YIR008C PRI1
ZShift = 0.79
Z lm Score = 0.65
NG = 0
DB = 1
SM = 0
Delta_K
Doxorubicin
YIR008C PRI1
ZShift = −0.87
Z lm Score = 0.25
NG = 0
DB = 1
SM = 0
Delta_r
Doxorubicin
YIR008C PRI1
ZShift = 0.48
Z lm Score = 0.3
NG = 0
DB = 1
SM = 0
Delta_AUC
Doxorubicin
YDL037C BSC1
ZShift = 0.02
Z lm Score = −2.95
NG = 0
DB = 0
SM = 0
Delta_L
Doxorubicin
YDL037C BSC1
ZShift = −0.01
Z lm Score = 1.24
NG = 0
DB = 0
SM = 0
Delta_K
Doxorubicin
YDL037C BSC1
ZShift = 2.78
Z lm Score = −1.55
NG = 0
DB = 0
SM = 0
Delta_r
Doxorubicin
YDL037C BSC1
ZShift = −0.03
Z lm Score = 0.82
NG = 0
DB = 0
SM = 0
Delta_AUC
Doxorubicin
YDR126W SWF1
ZShift = −0.02
Z lm Score = −2.95
NG = 0
DB = 0
SM = 0
Delta_L
Doxorubicin
YDR126W SWF1
ZShift = −0.13
Z lm Score = 1.41
NG = 0
DB = 0
SM = 0
Delta_K
Doxorubicin
YDR126W SWF1
ZShift = 0.15
Z lm Score = 0.08
NG = 0
DB = 0
SM = 0
Delta_r
Doxorubicin
YDR126W SWF1
ZShift = −0.07
Z lm Score = −0.79
NG = 0
DB = 0
SM = 0
Delta_AUC
Doxorubicin

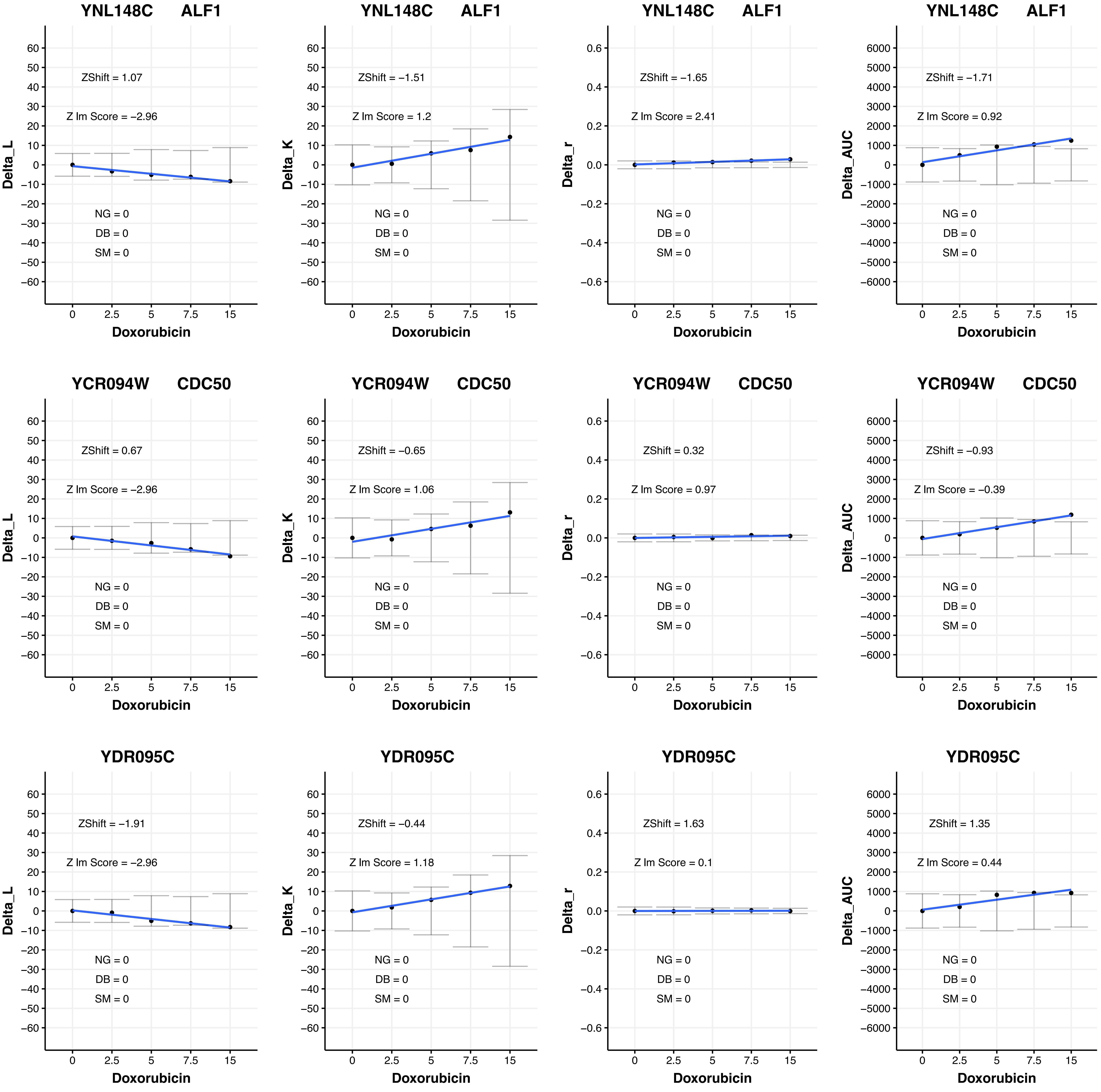

YNL148C ALF1
ZShift = 1.07
Z lm Score = −2.96
NG = 0
DB = 0
SM = 0
Delta_L
Doxorubicin
YNL148C ALF1
ZShift = −1.51
Z lm Score = 1.2
NG = 0
DB = 0
SM = 0
Delta_K
Doxorubicin
YNL148C ALF1
ZShift = −1.65
Z lm Score = 2.41
NG = 0
DB = 0
SM = 0
Delta_r
Doxorubicin
YNL148C ALF1
ZShift = −1.71
Z lm Score = 0.92
NG = 0
DB = 0
SM = 0
Delta_AUC
Doxorubicin
YCR094W CDC50
ZShift = 0.67
Z lm Score = −2.96
NG = 0
DB = 0
SM = 0
Delta_L
Doxorubicin
YCR094W CDC50
ZShift = −0.65
Z lm Score = 1.06
NG = 0
DB = 0
SM = 0
Delta_K
Doxorubicin
YCR094W CDC50
ZShift = 0.32
Z lm Score = 0.97
NG = 0
DB = 0
SM = 0
Delta_r
Doxorubicin
YCR094W CDC50
ZShift = −0.93
Z lm Score = −0.39
NG = 0
DB = 0
SM = 0
Delta_AUC
Doxorubicin
YDR095C
ZShift = −1.91
Z lm Score = −2.96
NG = 0
DB = 0
SM = 0
Delta_L
Doxorubicin
YDR095C
ZShift = −0.44
Z lm Score = 1.18
NG = 0
DB = 0
SM = 0
Delta_K
Doxorubicin
YDR095C
ZShift = 1.63
Z lm Score = 0.1
NG = 0
DB = 0
SM = 0
Delta_r
Doxorubicin
YDR095C
ZShift = 1.35
Z lm Score = 0.44
NG = 0
DB = 0
SM = 0
Delta_AUC
Doxorubicin

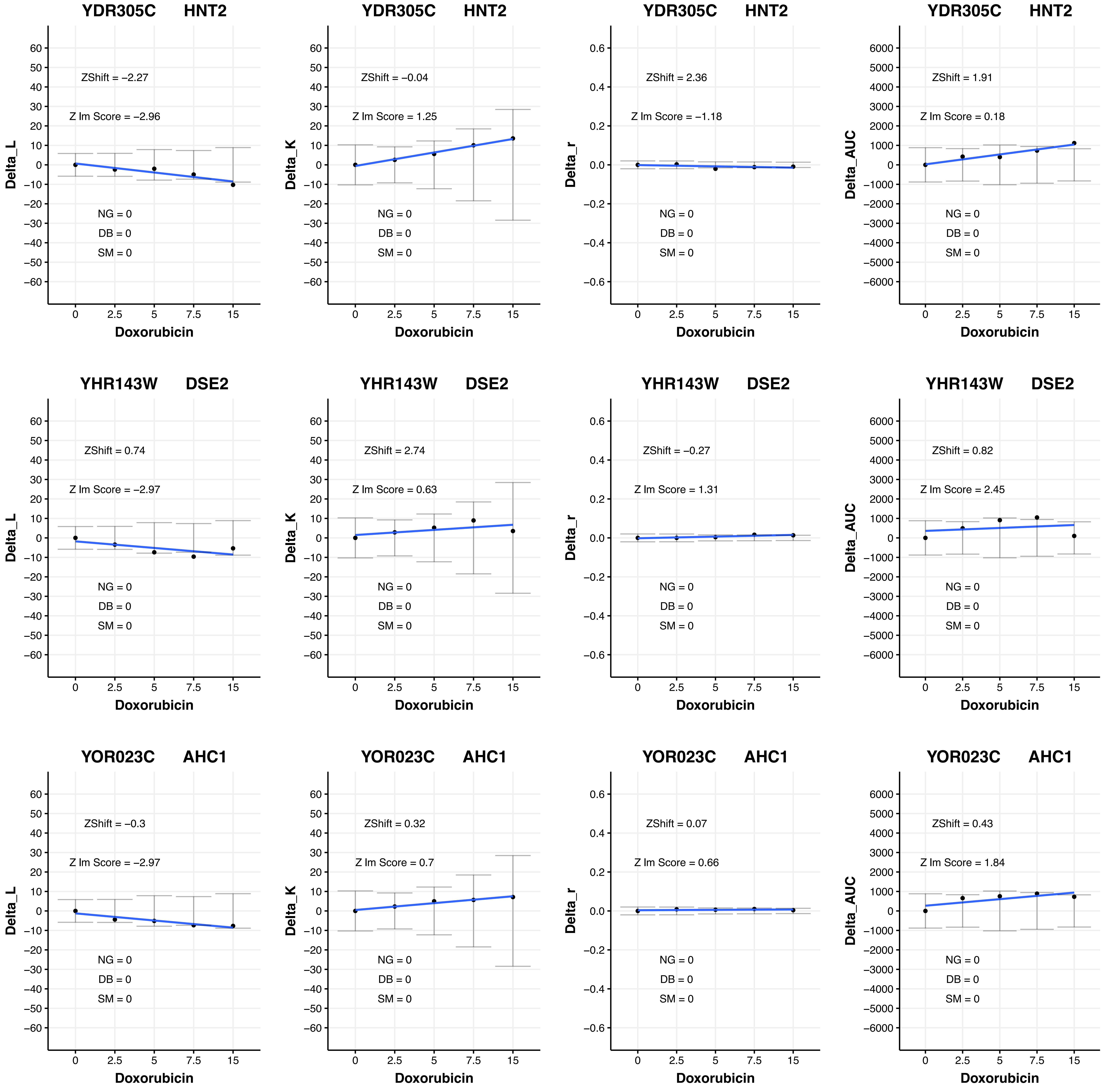

YDR305C HNT2
ZShift = −2.27
Z lm Score = −2.96
NG = 0
DB = 0
SM = 0
Delta_L
Doxorubicin
YDR305C HNT2
ZShift = −0.04
Z lm Score = 1.25
NG = 0
DB = 0
SM = 0
Delta_K
Doxorubicin
YDR305C HNT2
ZShift = 2.36
Z lm Score = −1.18
NG = 0
DB = 0
SM = 0
Delta_r
Doxorubicin
YDR305C HNT2
ZShift = 1.91
Z lm Score = 0.18
NG = 0
DB = 0
SM = 0
Delta_AUC
Doxorubicin
YHR143W DSE2
ZShift = 0.74
Z lm Score = −2.97
NG = 0
DB = 0
SM = 0
Delta_L
Doxorubicin
YHR143W DSE2
ZShift = 2.74
Z lm Score = 0.63
NG = 0
DB = 0
SM = 0
Delta_K
Doxorubicin
YHR143W DSE2
ZShift = −0.27
Z lm Score = 1.31
NG = 0
DB = 0
SM = 0
Delta_r
Doxorubicin
YHR143W DSE2
ZShift = 0.82
Z lm Score = 2.45
NG = 0
DB = 0
SM = 0
Delta_AUC
Doxorubicin
YOR023C AHC1
ZShift = −0.3
Z lm Score = −2.97
NG = 0
DB = 0
SM = 0
Delta_L
Doxorubicin
YOR023C AHC1
ZShift = 0.32
Z lm Score = 0.7
NG = 0
DB = 0
SM = 0
Delta_K
Doxorubicin
YOR023C AHC1
ZShift = 0.07
Z lm Score = 0.66
NG = 0
DB = 0
SM = 0
Delta_r
Doxorubicin
YOR023C AHC1
ZShift = 0.43
Z lm Score = 1.84
NG = 0
DB = 0
SM = 0
Delta_AUC
Doxorubicin

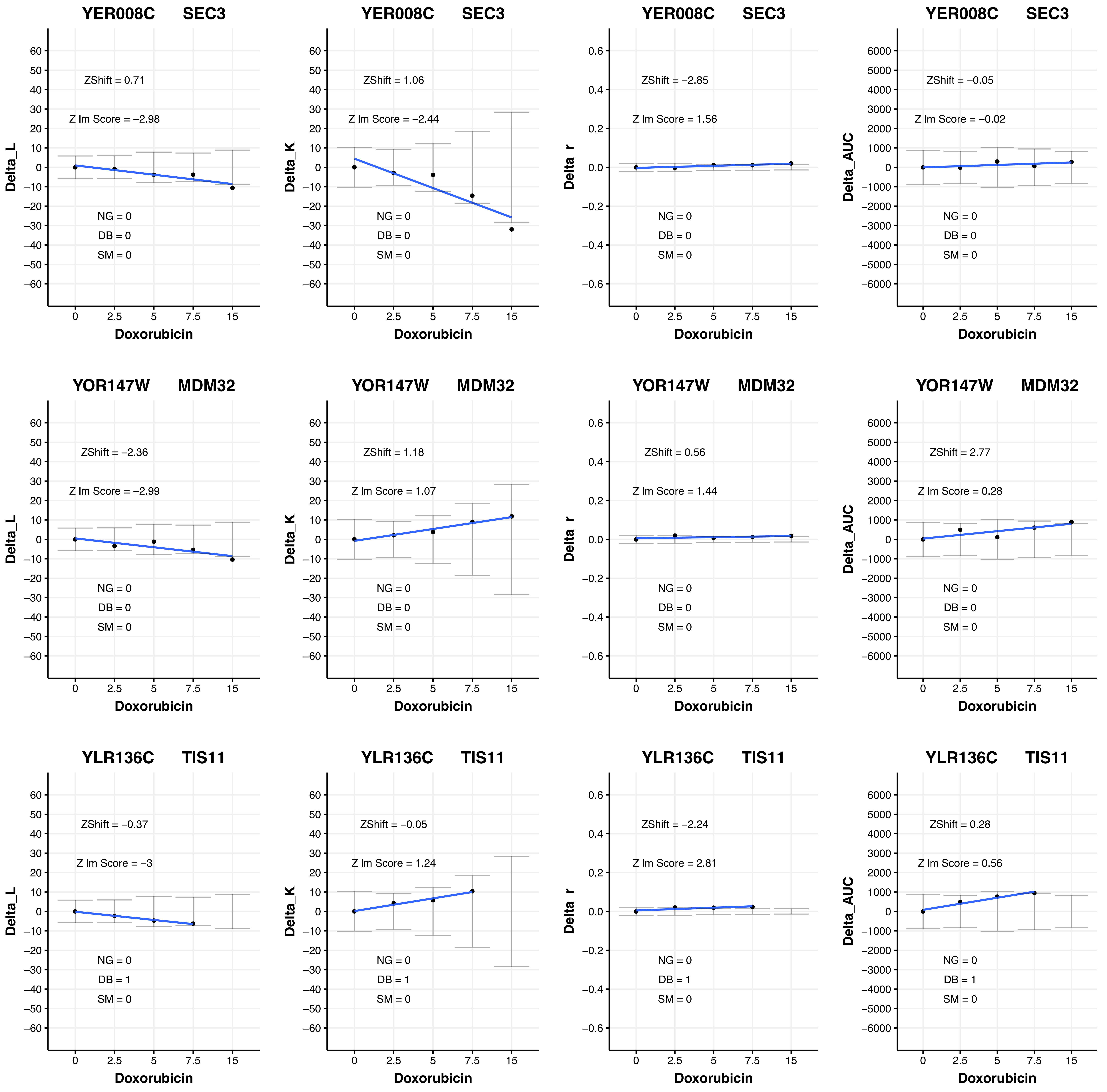

YER008C SEC3
ZShift = 0.71
Z lm Score = −2.98
NG = 0
DB = 0
SM = 0
Delta_L
Doxorubicin
YER008C SEC3
ZShift = 1.06
Z lm Score = −2.44
NG = 0
DB = 0
SM = 0
Delta_K
Doxorubicin
YER008C SEC3
ZShift = −2.85
Z lm Score = 1.56
NG = 0
DB = 0
SM = 0
Delta_r
Doxorubicin
YER008C SEC3
ZShift = −0.05
Z lm Score = −0.02
NG = 0
DB = 0
SM = 0
Delta_AUC
Doxorubicin
YOR147W MDM32
ZShift = −2.36
Z lm Score = −2.99
NG = 0
DB = 0
SM = 0
Delta_L
Doxorubicin
YOR147W MDM32
ZShift = 1.18
Z lm Score = 1.07
NG = 0
DB = 0
SM = 0
Delta_K
Doxorubicin
YOR147W MDM32
ZShift = 0.56
Z lm Score = 1.44
NG = 0
DB = 0
SM = 0
Delta_r
Doxorubicin
YOR147W MDM32
ZShift = 2.77
Z lm Score = 0.28
NG = 0
DB = 0
SM = 0
Delta_AUC
Doxorubicin
YLR136C TIS11
ZShift = −0.37
Z lm Score = −3
NG = 0
DB = 1
SM = 0
Delta_L
Doxorubicin
YLR136C TIS11
ZShift = −0.05
Z lm Score = 1.24
NG = 0
DB = 1
SM = 0
Delta_K
Doxorubicin
YLR136C TIS11
ZShift = −2.24
Z lm Score = 2.81
NG = 0
DB = 1
SM = 0
Delta_r
Doxorubicin
YLR136C TIS11
ZShift = 0.28
Z lm Score = 0.56
NG = 0
DB = 1
SM = 0
Delta_AUC
Doxorubicin

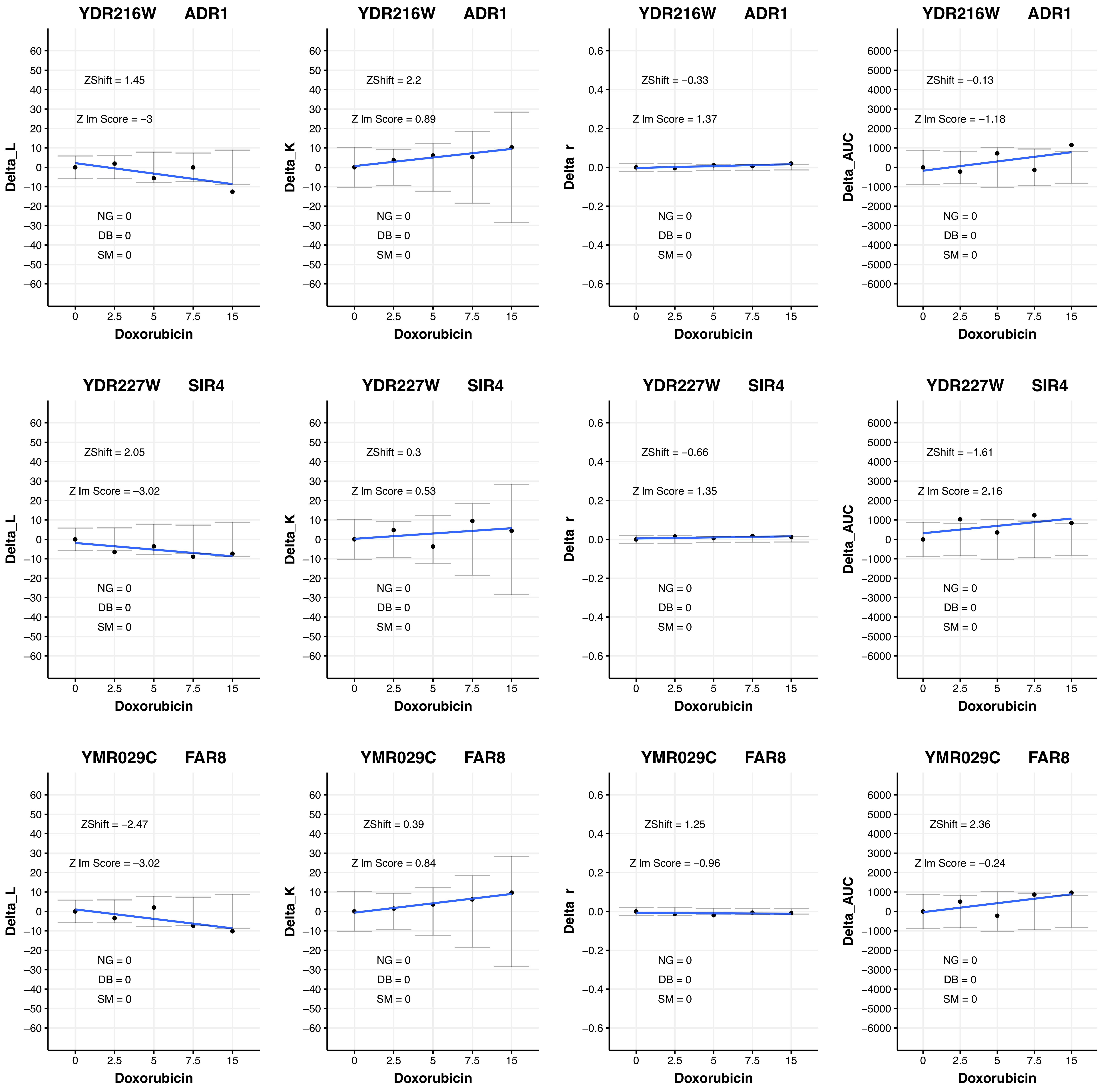
YDR216W ADR1
ZShift = 1.45
Z lm Score = −3
NG = 0
DB = 0
SM = 0
Delta_L
Doxorubicin
YDR216W ADR1
ZShift = 2.2
Z lm Score = 0.89
NG = 0
DB = 0
SM = 0
Delta_K
Doxorubicin
YDR216W ADR1
ZShift = −0.33
Z lm Score = 1.37
NG = 0
DB = 0
SM = 0
Delta_r
Doxorubicin
YDR216W ADR1
ZShift = −0.13
Z lm Score = −1.18
NG = 0
DB = 0
SM = 0
Delta_AUC
Doxorubicin
YDR227W SIR4
ZShift = 2.05
Z lm Score = −3.02
NG = 0
DB = 0
SM = 0
Delta_L
Doxorubicin
YDR227W SIR4
ZShift = 0.3
Z lm Score = 0.53
NG = 0
DB = 0
SM = 0
Delta_K
Doxorubicin
YDR227W SIR4
ZShift = −0.66
Z lm Score = 1.35
NG = 0
DB = 0
SM = 0
Delta_r
Doxorubicin
YDR227W SIR4
ZShift = −1.61
Z lm Score = 2.16
NG = 0
DB = 0
SM = 0
Delta_AUC
Doxorubicin
YMR029C FAR8
ZShift = −2.47
Z lm Score = −3.02
NG = 0
DB = 0
SM = 0
Delta_L
Doxorubicin
YMR029C FAR8
ZShift = 0.39
Z lm Score = 0.84
NG = 0
DB = 0
SM = 0
Delta_K
Doxorubicin
YMR029C FAR8
ZShift = 1.25
Z lm Score = −0.96
NG = 0
DB = 0
SM = 0
Delta_r
Doxorubicin
YMR029C FAR8
ZShift = 2.36
Z lm Score = −0.24
NG = 0
DB = 0
SM = 0
Delta_AUC
Doxorubicin

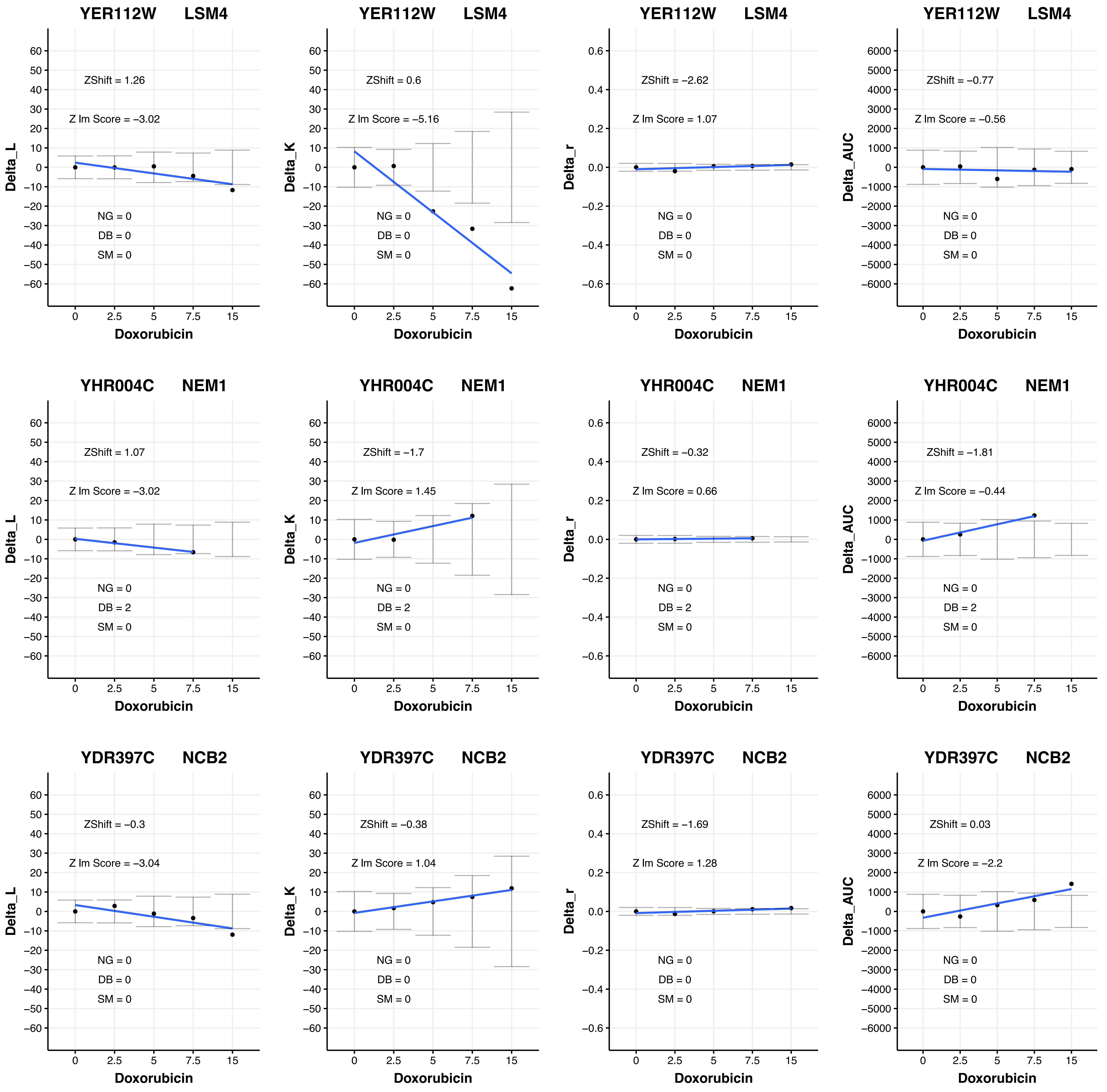

YER112W LSM4
Delta_L
ZShift = 1.26
Z lm Score = −3.02
NG = 0
DB = 0
SM = 0
Doxorubicin
YER112W LSM4
Delta_K
ZShift = 0.6
Z lm Score = −5.16
NG = 0
DB = 0
SM = 0
Doxorubicin
YER112W LSM4
Delta_r
ZShift = −2.62
Z lm Score = 1.07
NG = 0
DB = 0
SM = 0
Doxorubicin
YER112W LSM4
Delta_AUC
ZShift = −0.77
Z lm Score = −0.56
NG = 0
DB = 0
SM = 0
Doxorubicin
YHR004C NEM1
Delta_L
ZShift = 1.07
Z lm Score = −3.02
NG = 0
DB = 2
SM = 0
Doxorubicin
YHR004C NEM1
Delta_K
ZShift = −1.7
Z lm Score = 1.45
NG = 0
DB = 2
SM = 0
Doxorubicin
YHR004C NEM1
Delta_r
ZShift = −0.32
Z lm Score = 0.66
NG = 0
DB = 2
SM = 0
Doxorubicin
YHR004C NEM1
Delta_AUC
ZShift = −1.81
Z lm Score = −0.44
NG = 0
DB = 2
SM = 0
Doxorubicin
YDR397C NCB2
Delta_L
ZShift = −0.3
Z lm Score = −3.04
NG = 0
DB = 0
SM = 0
Doxorubicin
YDR397C NCB2
Delta_K
ZShift = −0.38
Z lm Score = 1.04
NG = 0
DB = 0
SM = 0
Doxorubicin
YDR397C NCB2
Delta_r
ZShift = −1.69
Z lm Score = 1.28
NG = 0
DB = 0
SM = 0
Doxorubicin
YDR397C NCB2
Delta_AUC
ZShift = 0.03
Z lm Score = −2.2
NG = 0
DB = 0
SM = 0
Doxorubicin

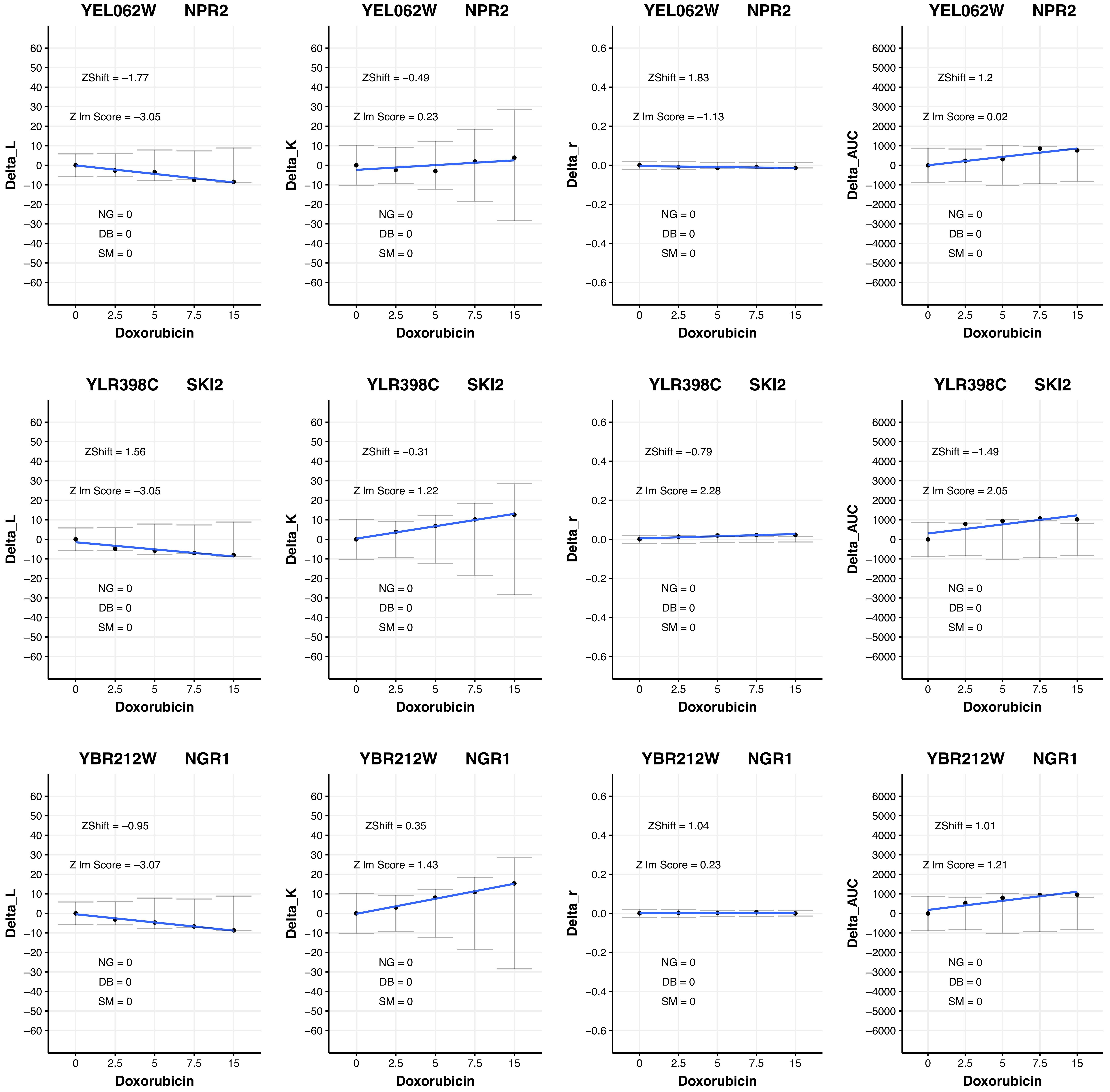
YEL062W NPR2
ZShift = −1.77
Z lm Score = −3.05
NG = 0
DB = 0
SM = 0
Delta_L
Doxorubicin
YEL062W NPR2
ZShift = −0.49
Z lm Score = 0.23
NG = 0
DB = 0
SM = 0
Delta_K
Doxorubicin
YEL062W NPR2
ZShift = 1.83
Z lm Score = −1.13
NG = 0
DB = 0
SM = 0
Delta_r
Doxorubicin
YEL062W NPR2
ZShift = 1.2
Z lm Score = 0.02
NG = 0
DB = 0
SM = 0
Delta_AUC
Doxorubicin
YLR398C SKI2
ZShift = 1.56
Z lm Score = −3.05
NG = 0
DB = 0
SM = 0
Delta_L
Doxorubicin
YLR398C SKI2
ZShift = −0.31
Z lm Score = 1.22
NG = 0
DB = 0
SM = 0
Delta_K
Doxorubicin
YLR398C SKI2
ZShift = −0.79
Z lm Score = 2.28
NG = 0
DB = 0
SM = 0
Delta_r
Doxorubicin
YLR398C SKI2
ZShift = −1.49
Z lm Score = 2.05
NG = 0
DB = 0
SM = 0
Delta_AUC
Doxorubicin
YBR212W NGR1
ZShift = −0.95
Z lm Score = −3.07
NG = 0
DB = 0
SM = 0
Delta_L
Doxorubicin
YBR212W NGR1
ZShift = 0.35
Z lm Score = 1.43
NG = 0
DB = 0
SM = 0
Delta_K
Doxorubicin
YBR212W NGR1
ZShift = 1.04
Z lm Score = 0.23
NG = 0
DB = 0
SM = 0
Delta_r
Doxorubicin
YBR212W NGR1
ZShift = 1.01
Z lm Score = 1.21
NG = 0
DB = 0
SM = 0
Delta_AUC
Doxorubicin

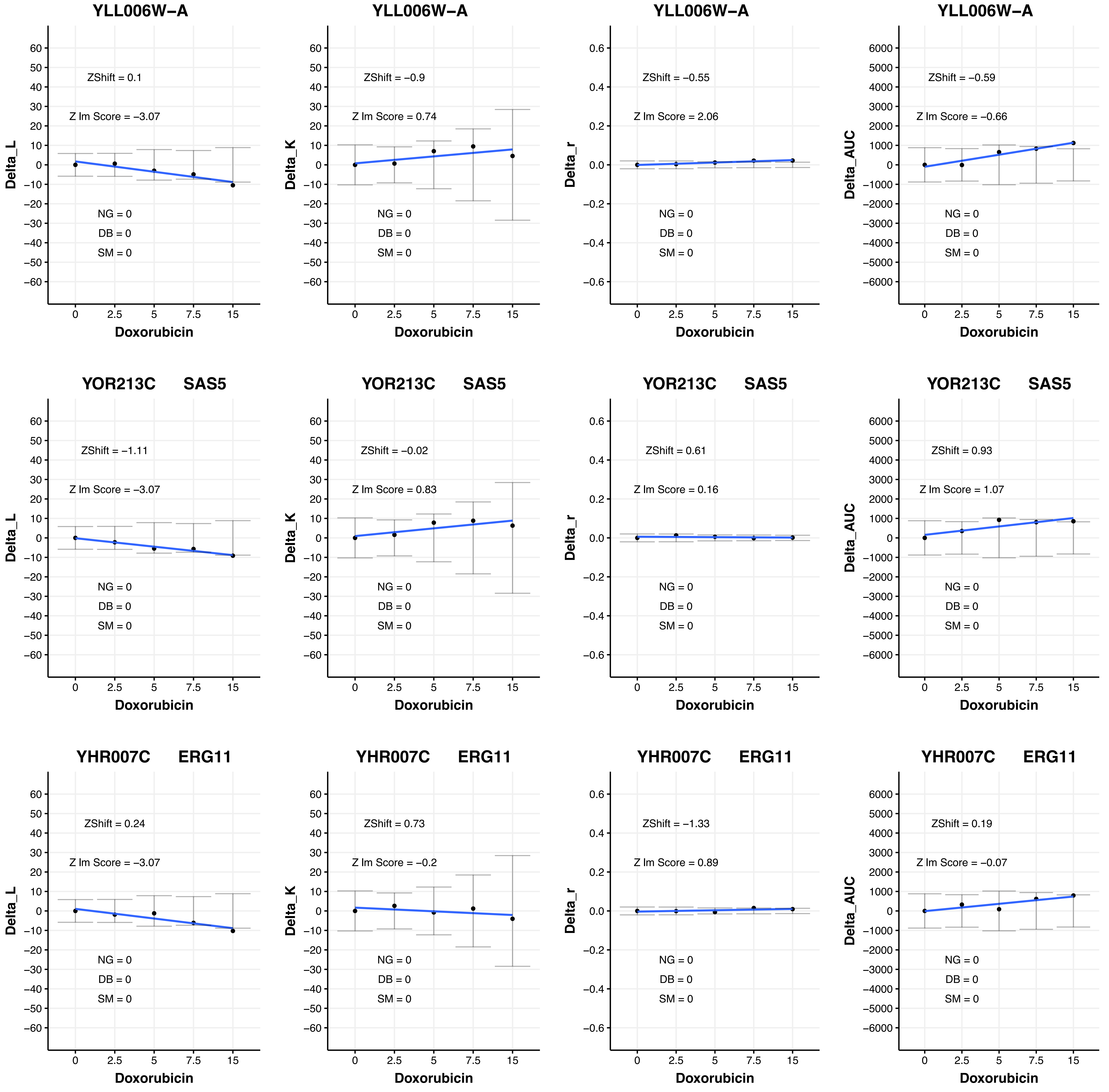
YLL006W−A
ZShift = 0.1
Z lm Score = −3.07
NG = 0
DB = 0
SM = 0
Delta_L
Doxorubicin
YLL006W−A
ZShift = −0.9
Z lm Score = 0.74
NG = 0
DB = 0
SM = 0
Delta_K
Doxorubicin
YLL006W−A
ZShift = −0.55
Z lm Score = 2.06
NG = 0
DB = 0
SM = 0
Delta_r
Doxorubicin
YLL006W−A
ZShift = −0.59
Z lm Score = −0.66
NG = 0
DB = 0
SM = 0
Delta_AUC
Doxorubicin
YOR213C SAS5
ZShift = −1.11
Z lm Score = −3.07
NG = 0
DB = 0
SM = 0
Delta_L
Doxorubicin
YOR213C SAS5
ZShift = −0.02
Z lm Score = 0.83
NG = 0
DB = 0
SM = 0
Delta_K
Doxorubicin
YOR213C SAS5
ZShift = 0.61
Z lm Score = 0.16
NG = 0
DB = 0
SM = 0
Delta_r
Doxorubicin
YOR213C SAS5
ZShift = 0.93
Z lm Score = 1.07
NG = 0
DB = 0
SM = 0
Delta_AUC
Doxorubicin
YHR007C ERG11
ZShift = 0.24
Z lm Score = −3.07
NG = 0
DB = 0
SM = 0
Delta_L
Doxorubicin
YHR007C ERG11
ZShift = 0.73
Z lm Score = −0.2
NG = 0
DB = 0
SM = 0
Delta_K
Doxorubicin
YHR007C ERG11
ZShift = −1.33
Z lm Score = 0.89
NG = 0
DB = 0
SM = 0
Delta_r
Doxorubicin
YHR007C ERG11
ZShift = 0.19
Z lm Score = −0.07
NG = 0
DB = 0
SM = 0
Delta_AUC
Doxorubicin

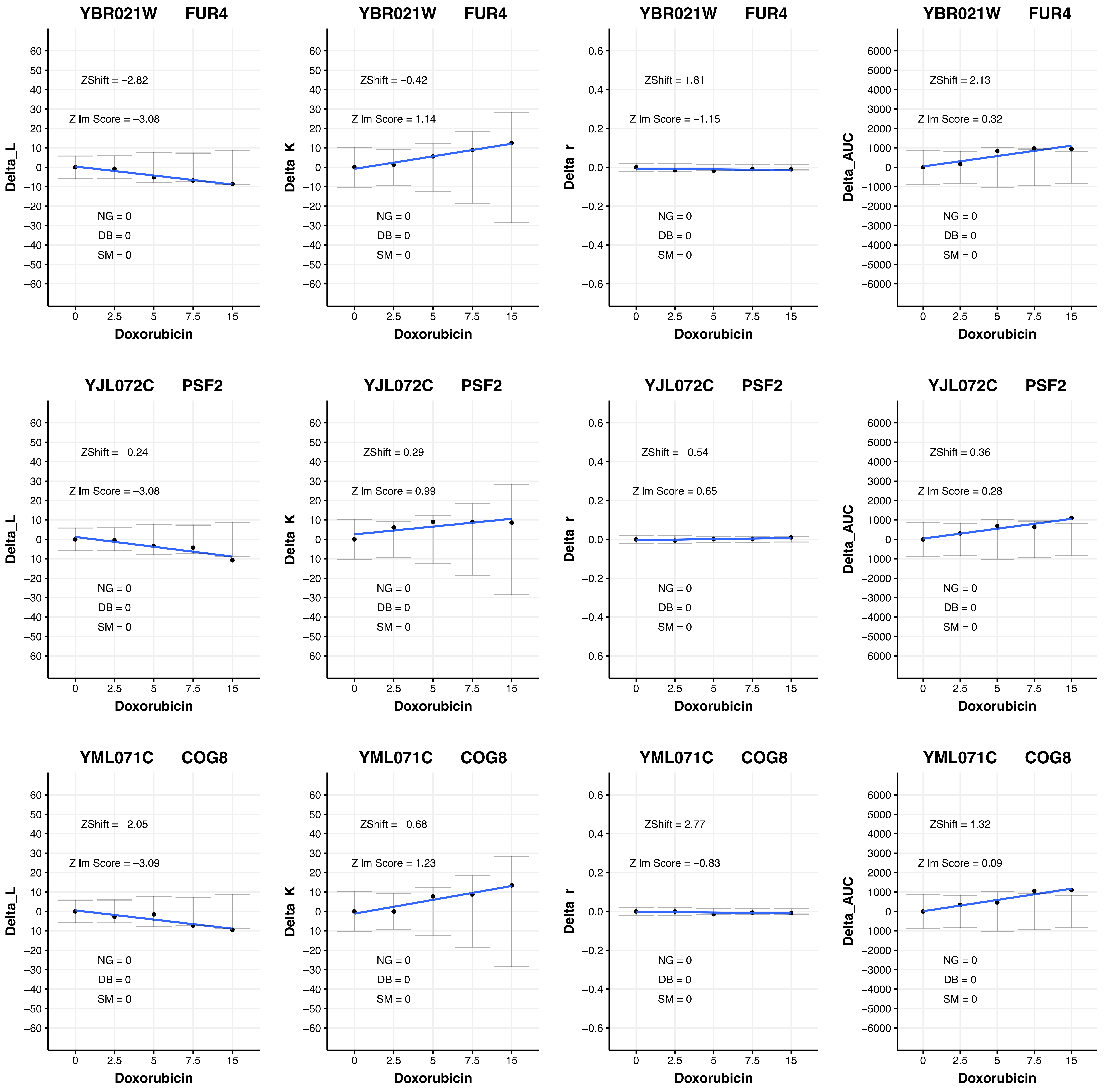
YBR021W FUR4
Delta_L
Doxorubicin
ZShift = −2.82
Z lm Score = −3.08
NG = 0
DB = 0
SM = 0
YBR021W FUR4
Delta_K
Doxorubicin
ZShift = −0.42
Z lm Score = 1.14
NG = 0
DB = 0
SM = 0
YBR021W FUR4
Delta_r
Doxorubicin
ZShift = 1.81
Z lm Score = −1.15
NG = 0
DB = 0
SM = 0
YBR021W FUR4
Delta_AUC
Doxorubicin
ZShift = 2.13
Z lm Score = 0.32
NG = 0
DB = 0
SM = 0
YJL072C PSF2
Delta_L
Doxorubicin
ZShift = −0.24
Z lm Score = −3.08
NG = 0
DB = 0
SM = 0
YJL072C PSF2
Delta_K
Doxorubicin
ZShift = 0.29
Z lm Score = 0.99
NG = 0
DB = 0
SM = 0
YJL072C PSF2
Delta_r
Doxorubicin
ZShift = −0.54
Z lm Score = 0.65
NG = 0
DB = 0
SM = 0
YJL072C PSF2
Delta_AUC
Doxorubicin
ZShift = 0.36
Z lm Score = 0.28
NG = 0
DB = 0
SM = 0
YML071C COG8
Delta_L
Doxorubicin
ZShift = −2.05
Z lm Score = −3.09
NG = 0
DB = 0
SM = 0
YML071C COG8
Delta_K
Doxorubicin
ZShift = −0.68
Z lm Score = 1.23
NG = 0
DB = 0
SM = 0
YML071C COG8
Delta_r
Doxorubicin
ZShift = 2.77
Z lm Score = −0.83
NG = 0
DB = 0
SM = 0
YML071C COG8
Delta_AUC
Doxorubicin
ZShift = 1.32
Z lm Score = 0.09
NG = 0
DB = 0
SM = 0

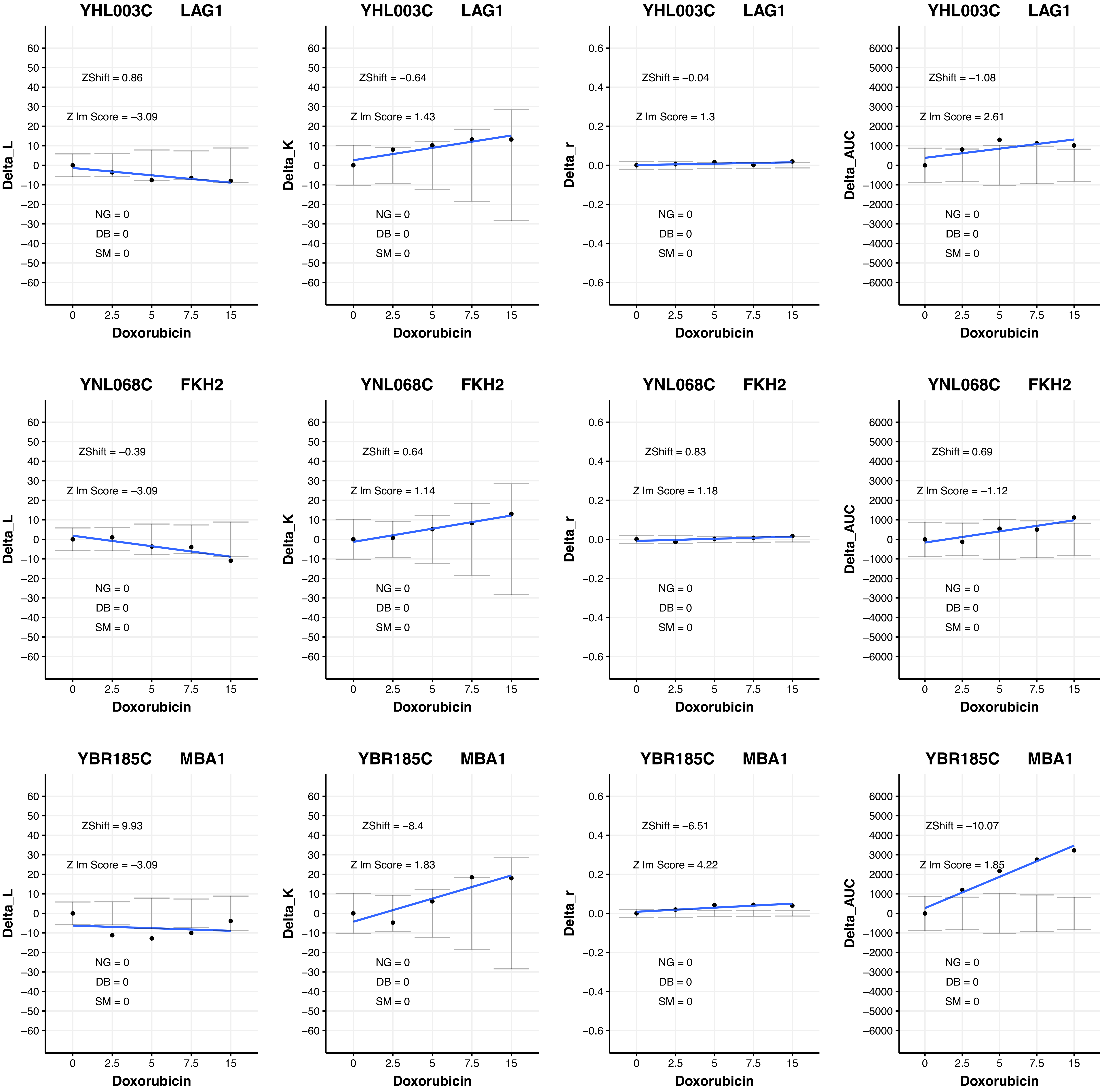
YHL003C LAG1
ZShift = 0.86
Z lm Score = −3.09
NG = 0
DB = 0
SM = 0
Delta_L
Doxorubicin
YHL003C LAG1
ZShift = −0.64
Z lm Score = 1.43
NG = 0
DB = 0
SM = 0
Delta_K
Doxorubicin
YHL003C LAG1
ZShift = −0.04
Z lm Score = 1.3
NG = 0
DB = 0
SM = 0
Delta_r
Doxorubicin
YHL003C LAG1
ZShift = −1.08
Z lm Score = 2.61
NG = 0
DB = 0
SM = 0
Delta_AUC
Doxorubicin
YNL068C FKH2
ZShift = −0.39
Z lm Score = −3.09
NG = 0
DB = 0
SM = 0
Delta_L
Doxorubicin
YNL068C FKH2
ZShift = 0.64
Z lm Score = 1.14
NG = 0
DB = 0
SM = 0
Delta_K
Doxorubicin
YNL068C FKH2
ZShift = 0.83
Z lm Score = 1.18
NG = 0
DB = 0
SM = 0
Delta_r
Doxorubicin
YNL068C FKH2
ZShift = 0.69
Z lm Score = −1.12
NG = 0
DB = 0
SM = 0
Delta_AUC
Doxorubicin
YBR185C MBA1
ZShift = 9.93
Z lm Score = −3.09
NG = 0
DB = 0
SM = 0
Delta_L
Doxorubicin
YBR185C MBA1
ZShift = −8.4
Z lm Score = 1.83
NG = 0
DB = 0
SM = 0
Delta_K
Doxorubicin
YBR185C MBA1
ZShift = −6.51
Z lm Score = 4.22
NG = 0
DB = 0
SM = 0
Delta_r
Doxorubicin
YBR185C MBA1
ZShift = −10.07
Z lm Score = 1.85
NG = 0
DB = 0
SM = 0
Delta_AUC
Doxorubicin

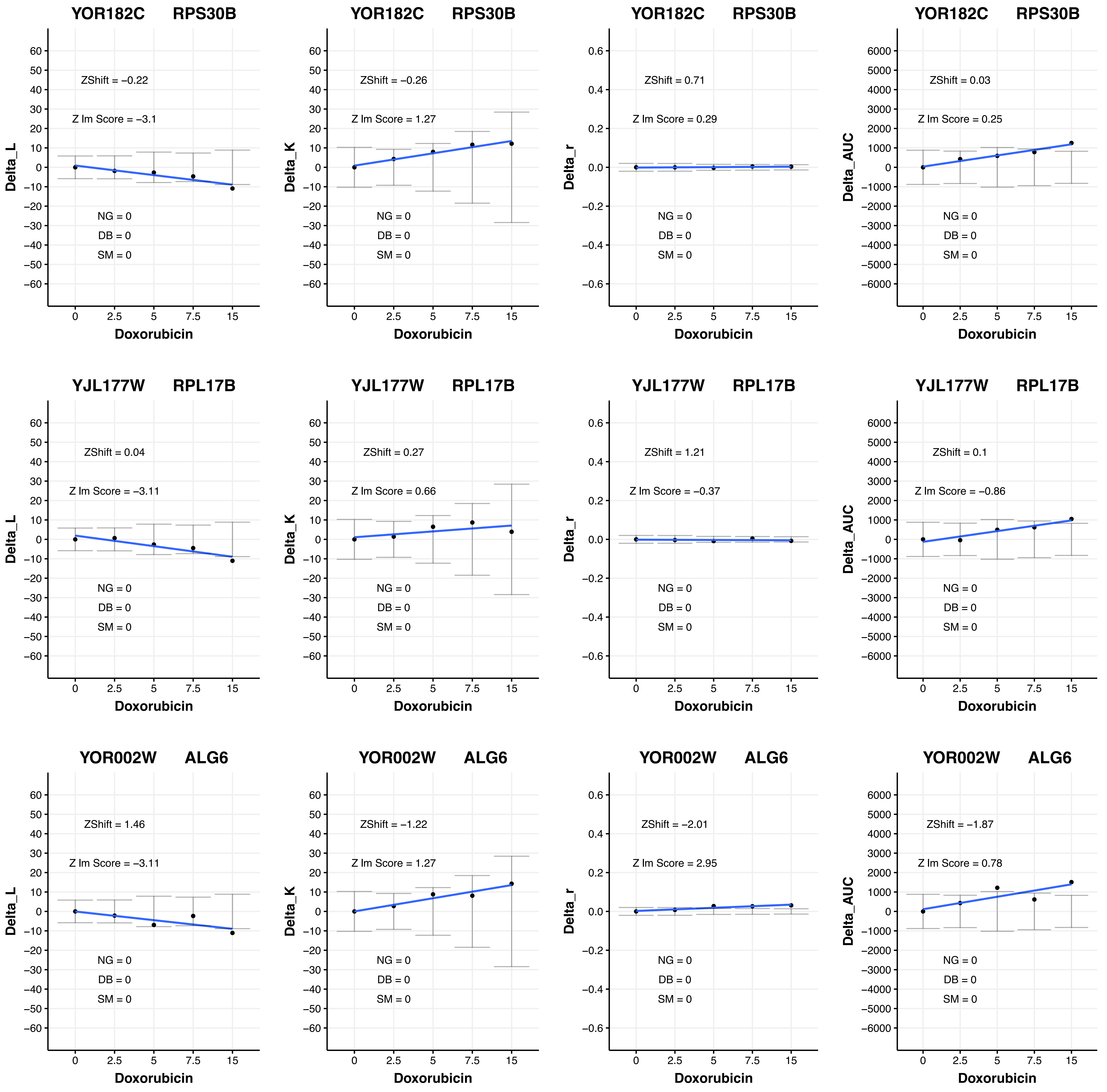
YOR182C RPS30B
ZShift = −0.22
Z lm Score = −3.1
NG = 0
DB = 0
SM = 0
Delta_L
Doxorubicin
YOR182C RPS30B
ZShift = −0.26
Z lm Score = 1.27
NG = 0
DB = 0
SM = 0
Delta_K
Doxorubicin
YOR182C RPS30B
ZShift = 0.71
Z lm Score = 0.29
NG = 0
DB = 0
SM = 0
Delta_r
Doxorubicin
YOR182C RPS30B
ZShift = 0.03
Z lm Score = 0.25
NG = 0
DB = 0
SM = 0
Delta_AUC
Doxorubicin
YJL177W RPL17B
ZShift = 0.04
Z lm Score = −3.11
NG = 0
DB = 0
SM = 0
Delta_L
Doxorubicin
YJL177W RPL17B
ZShift = 0.27
Z lm Score = 0.66
NG = 0
DB = 0
SM = 0
Delta_K
Doxorubicin
YJL177W RPL17B
ZShift = 1.21
Z lm Score = −0.37
NG = 0
DB = 0
SM = 0
Delta_r
Doxorubicin
YJL177W RPL17B
ZShift = 0.1
Z lm Score = −0.86
NG = 0
DB = 0
SM = 0
Delta_AUC
Doxorubicin
YOR002W ALG6
ZShift = 1.46
Z lm Score = −3.11
NG = 0
DB = 0
SM = 0
Delta_L
Doxorubicin
YOR002W ALG6
ZShift = −1.22
Z lm Score = 1.27
NG = 0
DB = 0
SM = 0
Delta_K
Doxorubicin
YOR002W ALG6
ZShift = −2.01
Z lm Score = 2.95
NG = 0
DB = 0
SM = 0
Delta_r
Doxorubicin
YOR002W ALG6
ZShift = −1.87
Z lm Score = 0.78
NG = 0
DB = 0
SM = 0
Delta_AUC
Doxorubicin

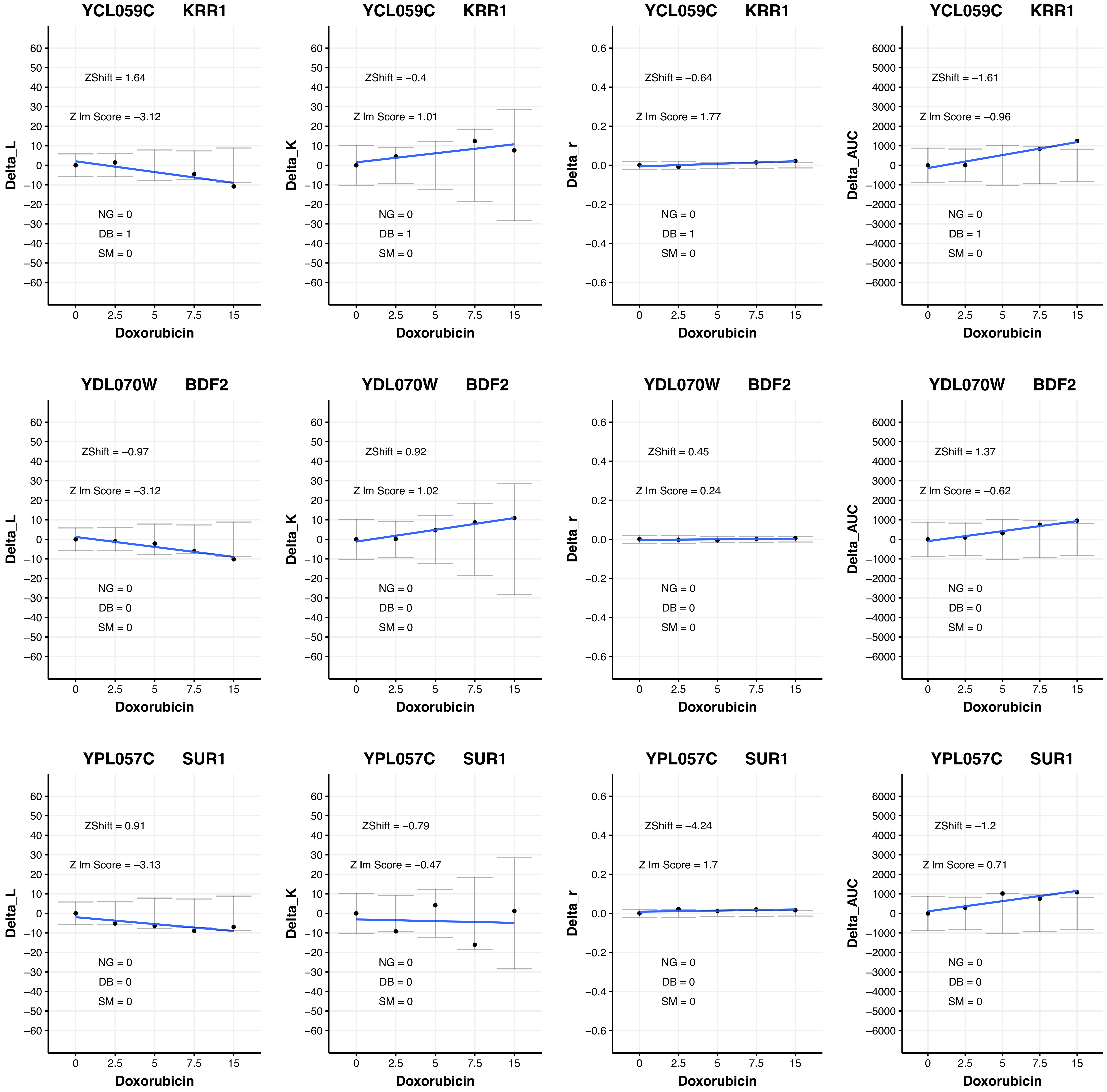
YCL059C KRR1
Delta_L
ZShift = 1.64
Z lm Score = −3.12
NG = 0
DB = 1
SM = 0
Doxorubicin
YCL059C KRR1
Delta_K
ZShift = −0.4
Z lm Score = 1.01
NG = 0
DB = 1
SM = 0
Doxorubicin
YCL059C KRR1
Delta_r
ZShift = −0.64
Z lm Score = 1.77
NG = 0
DB = 1
SM = 0
Doxorubicin
YCL059C KRR1
Delta_AUC
ZShift = −1.61
Z lm Score = −0.96
NG = 0
DB = 1
SM = 0
Doxorubicin
YDL070W BDF2
Delta_L
ZShift = −0.97
Z lm Score = −3.12
NG = 0
DB = 0
SM = 0
Doxorubicin
YDL070W BDF2
Delta_K
ZShift = 0.92
Z lm Score = 1.02
NG = 0
DB = 0
SM = 0
Doxorubicin
YDL070W BDF2
Delta_r
ZShift = 0.45
Z lm Score = 0.24
NG = 0
DB = 0
SM = 0
Doxorubicin
YDL070W BDF2
Delta_AUC
ZShift = 1.37
Z lm Score = −0.62
NG = 0
DB = 0
SM = 0
Doxorubicin
YPL057C SUR1
Delta_L
ZShift = 0.91
Z lm Score = −3.13
NG = 0
DB = 0
SM = 0
Doxorubicin
YPL057C SUR1
Delta_K
ZShift = −0.79
Z lm Score = −0.47
NG = 0
DB = 0
SM = 0
Doxorubicin
YPL057C SUR1
Delta_r
ZShift = −4.24
Z lm Score = 1.7
NG = 0
DB = 0
SM = 0
Doxorubicin
YPL057C SUR1
Delta_AUC
ZShift = −1.2
Z lm Score = 0.71
NG = 0
DB = 0
SM = 0
Doxorubicin

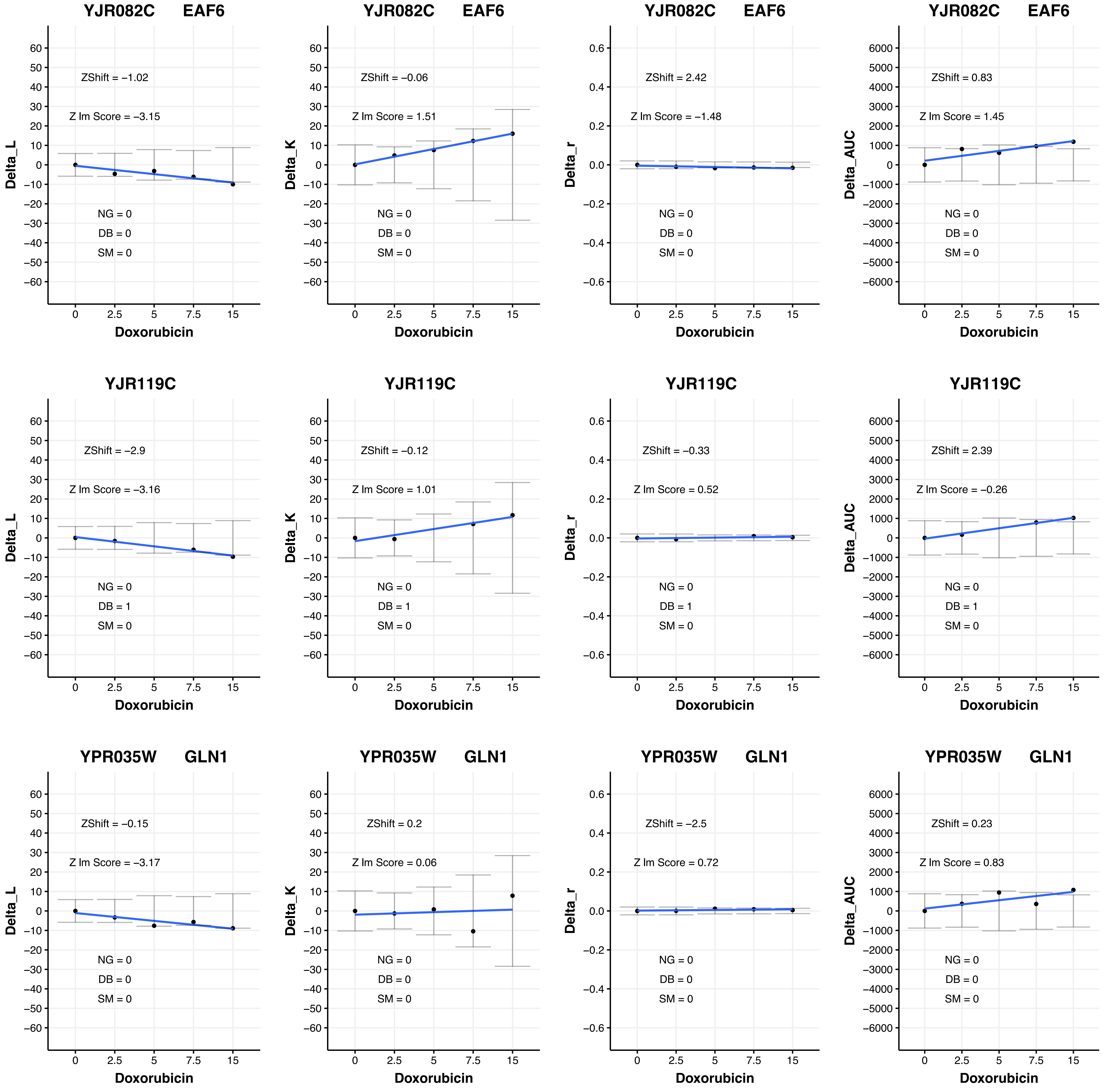
YJR082C EAF6
ZShift = −1.02
Z lm Score = −3.15
NG = 0
DB = 0
SM = 0
Delta_L
Doxorubicin
YJR082C EAF6
ZShift = −0.06
Z lm Score = 1.51
NG = 0
DB = 0
SM = 0
Delta_K
Doxorubicin
YJR082C EAF6
ZShift = 2.42
Z lm Score = −1.48
NG = 0
DB = 0
SM = 0
Delta_r
Doxorubicin
YJR082C EAF6
ZShift = 0.83
Z lm Score = 1.45
NG = 0
DB = 0
SM = 0
Delta_AUC
Doxorubicin
YJR119C
ZShift = −2.9
Z lm Score = −3.16
NG = 0
DB = 1
SM = 0
Delta_L
Doxorubicin
YJR119C
ZShift = −0.12
Z lm Score = 1.01
NG = 0
DB = 1
SM = 0
Delta_K
Doxorubicin
YJR119C
ZShift = −0.33
Z lm Score = 0.52
NG = 0
DB = 1
SM = 0
Delta_r
Doxorubicin
YJR119C
ZShift = 2.39
Z lm Score = −0.26
NG = 0
DB = 1
SM = 0
Delta_AUC
Doxorubicin
YPR035W GLN1
ZShift = −0.15
Z lm Score = −3.17
NG = 0
DB = 0
SM = 0
Delta_L
Doxorubicin
YPR035W GLN1
ZShift = 0.2
Z lm Score = 0.06
NG = 0
DB = 0
SM = 0
Delta_K
Doxorubicin
YPR035W GLN1
ZShift = −2.5
Z lm Score = 0.72
NG = 0
DB = 0
SM = 0
Delta_r
Doxorubicin
YPR035W GLN1
ZShift = 0.23
Z lm Score = 0.83
NG = 0
DB = 0
SM = 0
Delta_AUC
Doxorubicin

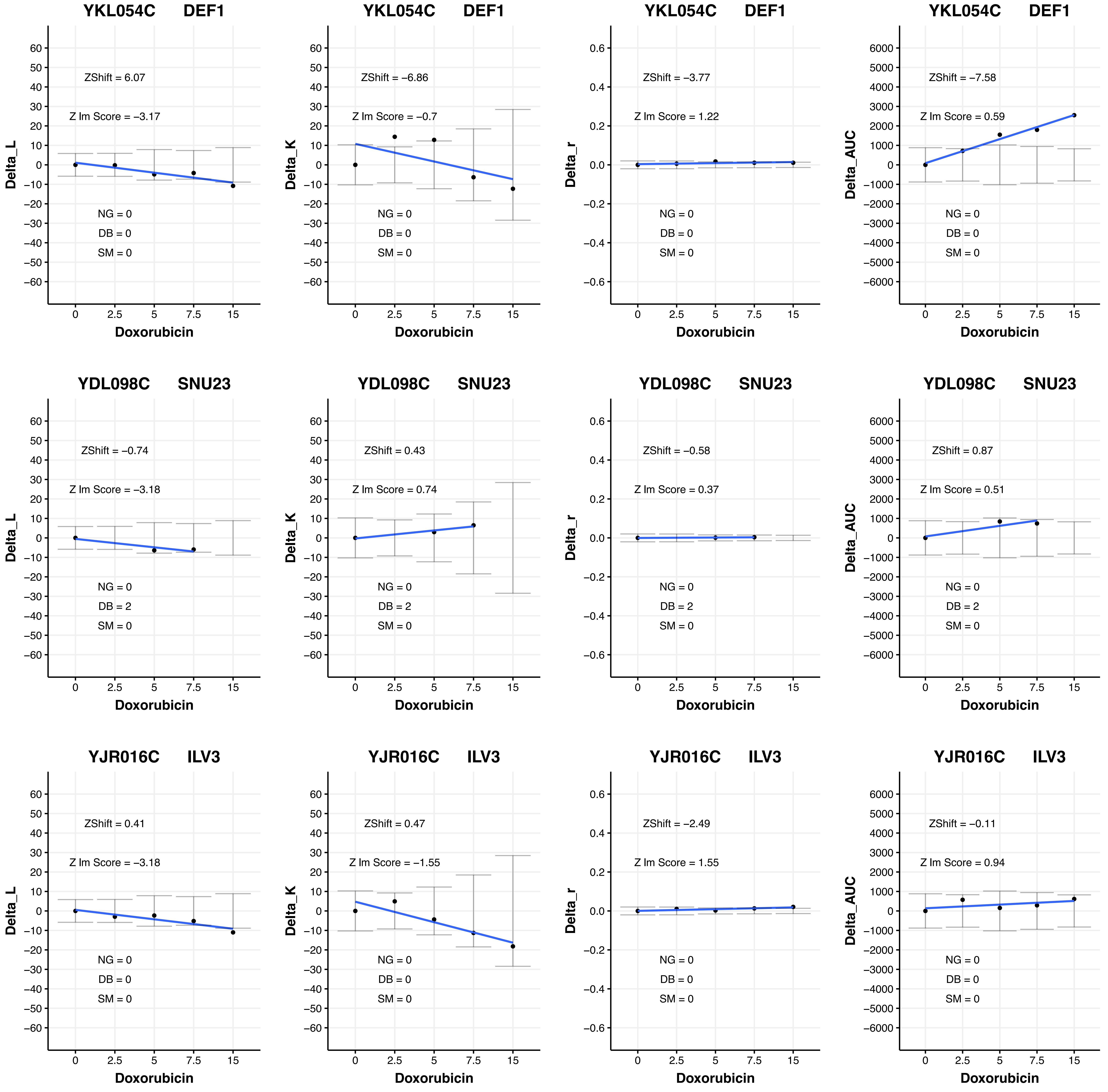
YKL054C DEF1
ZShift = 6.07
Z lm Score = −3.17
NG = 0
DB = 0
SM = 0
Delta_L
Doxorubicin
YKL054C DEF1
ZShift = −6.86
Z lm Score = −0.7
NG = 0
DB = 0
SM = 0
Delta_K
Doxorubicin
YKL054C DEF1
ZShift = −3.77
Z lm Score = 1.22
NG = 0
DB = 0
SM = 0
Delta_r
Doxorubicin
YKL054C DEF1
ZShift = −7.58
Z lm Score = 0.59
NG = 0
DB = 0
SM = 0
Delta_AUC
Doxorubicin
YDL098C SNU23
ZShift = −0.74
Z lm Score = −3.18
NG = 0
DB = 2
SM = 0
Delta_L
Doxorubicin
YDL098C SNU23
ZShift = 0.43
Z lm Score = 0.74
NG = 0
DB = 2
SM = 0
Delta_K
Doxorubicin
YDL098C SNU23
ZShift = −0.58
Z lm Score = 0.37
NG = 0
DB = 2
SM = 0
Delta_r
Doxorubicin
YDL098C SNU23
ZShift = 0.87
Z lm Score = 0.51
NG = 0
DB = 2
SM = 0
Delta_AUC
Doxorubicin
YJR016C ILV3
ZShift = 0.41
Z lm Score = −3.18
NG = 0
DB = 0
SM = 0
Delta_L
Doxorubicin
YJR016C ILV3
ZShift = 0.47
Z lm Score = −1.55
NG = 0
DB = 0
SM = 0
Delta_K
Doxorubicin
YJR016C ILV3
ZShift = −2.49
Z lm Score = 1.55
NG = 0
DB = 0
SM = 0
Delta_r
Doxorubicin
YJR016C ILV3
ZShift = −0.11
Z lm Score = 0.94
NG = 0
DB = 0
SM = 0
Delta_AUC
Doxorubicin

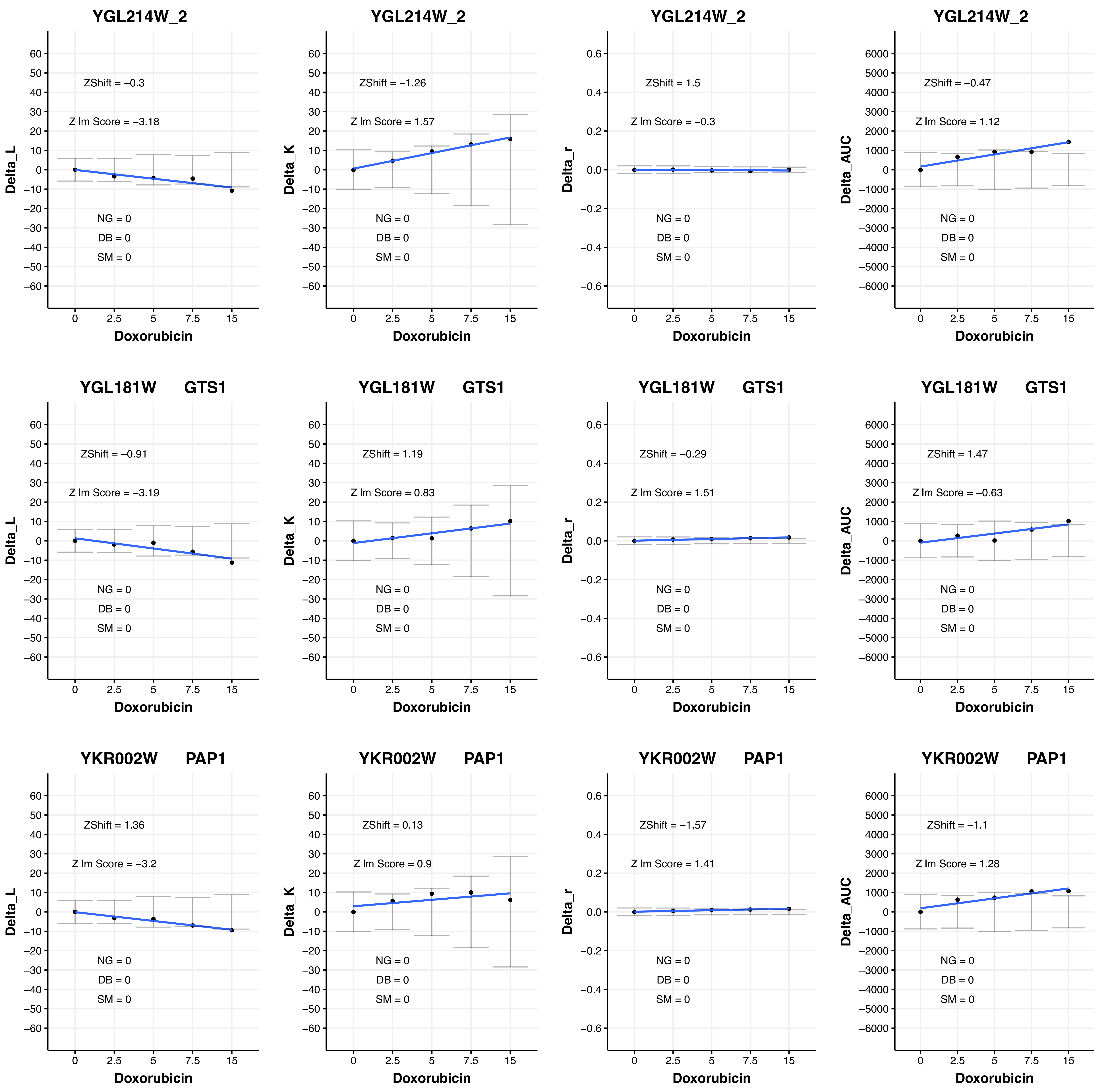

YGL214W_2
ZShift = −0.3
Z Im Score = −3.18
NG = 0
DB = 0
SM = 0
Delta_L
Doxorubicin
YGL214W_2
ZShift = −1.26
Z Im Score = 1.57
NG = 0
DB = 0
SM = 0
Delta_K
Doxorubicin
YGL214W_2
ZShift = 1.5
Z Im Score = −0.3
NG = 0
DB = 0
SM = 0
Delta_r
Doxorubicin
YGL214W_2
ZShift = −0.47
Z Im Score = 1.12
NG = 0
DB = 0
SM = 0
Delta_AUC
Doxorubicin
YGL181W GTS1
ZShift = −0.91
Z Im Score = −3.19
NG = 0
DB = 0
SM = 0
Delta_L
Doxorubicin
YGL181W GTS1
ZShift = 1.19
Z Im Score = 0.83
NG = 0
DB = 0
SM = 0
Delta_K
Doxorubicin
YGL181W GTS1
ZShift = −0.29
Z Im Score = 1.51
NG = 0
DB = 0
SM = 0
Delta_r
Doxorubicin
YGL181W GTS1
ZShift = 1.47
Z Im Score = −0.63
NG = 0
DB = 0
SM = 0
Delta_AUC
Doxorubicin
YKR002W PAP1
ZShift = 1.36
Z Im Score = −3.2
NG = 0
DB = 0
SM = 0
Delta_L
Doxorubicin
YKR002W PAP1
ZShift = 0.13
Z Im Score = 0.9
NG = 0
DB = 0
SM = 0
Delta_K
Doxorubicin
YKR002W PAP1
ZShift = −1.57
Z Im Score = 1.41
NG = 0
DB = 0
SM = 0
Delta_r
Doxorubicin
YKR002W PAP1
ZShift = −1.1
Z Im Score = 1.28
NG = 0
DB = 0
SM = 0
Delta_AUC
Doxorubicin

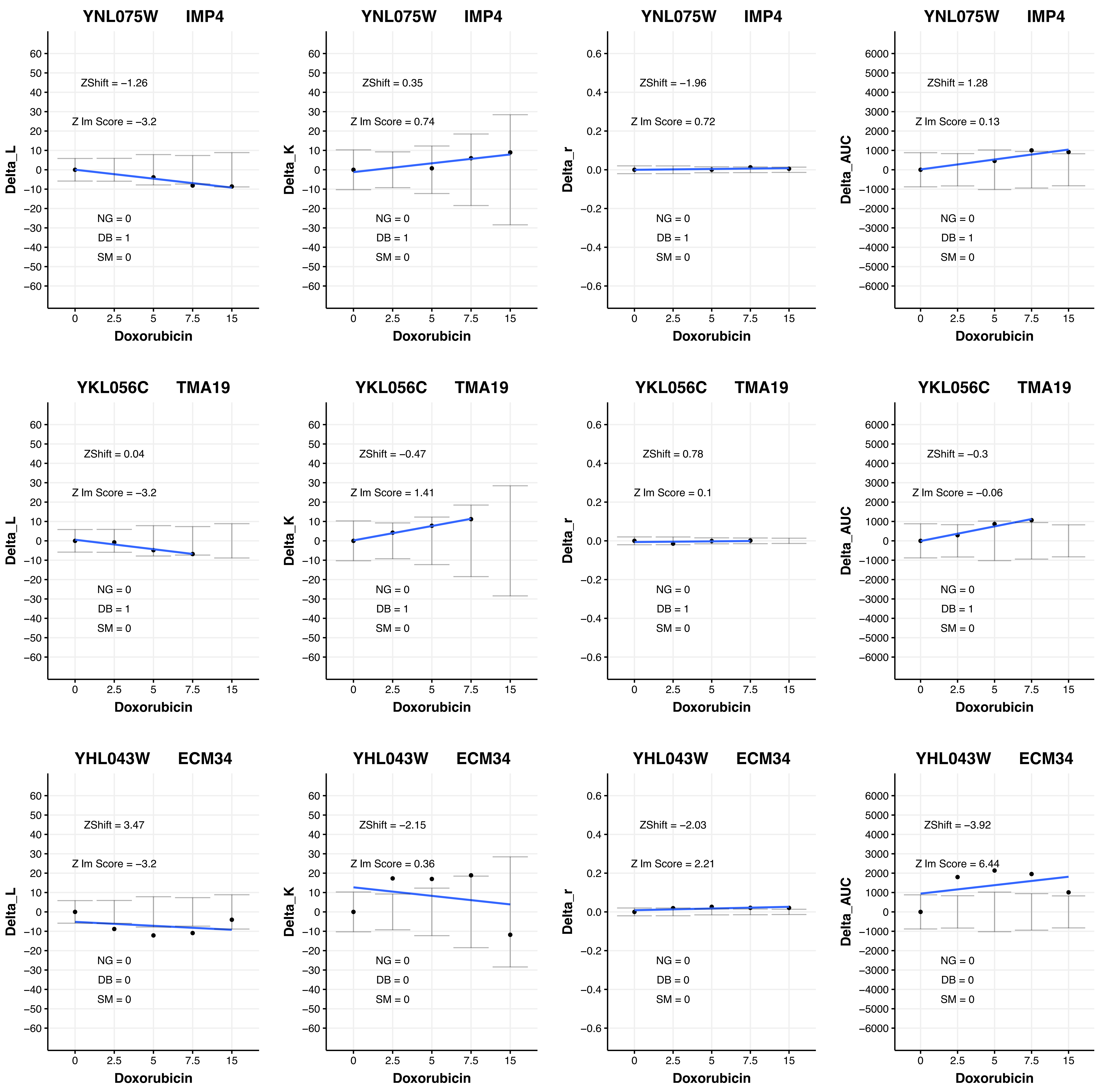
YNL075W IMP4
ZShift = −1.26
Z lm Score = −3.2
NG = 0
DB = 1
SM = 0
Delta_L
Doxorubicin
YNL075W IMP4
ZShift = 0.35
Z lm Score = 0.74
NG = 0
DB = 1
SM = 0
Delta_K
Doxorubicin
YNL075W IMP4
ZShift = −1.96
Z lm Score = 0.72
NG = 0
DB = 1
SM = 0
Delta_r
Doxorubicin
YNL075W IMP4
ZShift = 1.28
Z lm Score = 0.13
NG = 0
DB = 1
SM = 0
Delta_AUC
Doxorubicin
YKL056C TMA19
ZShift = 0.04
Z lm Score = −3.2
NG = 0
DB = 1
SM = 0
Delta_L
Doxorubicin
YKL056C TMA19
ZShift = −0.47
Z lm Score = 1.41
NG = 0
DB = 1
SM = 0
Delta_K
Doxorubicin
YKL056C TMA19
ZShift = 0.78
Z lm Score = 0.1
NG = 0
DB = 1
SM = 0
Delta_r
Doxorubicin
YKL056C TMA19
ZShift = −0.3
Z lm Score = −0.06
NG = 0
DB = 1
SM = 0
Delta_AUC
Doxorubicin
YHL043W ECM34
ZShift = 3.47
Z lm Score = −3.2
NG = 0
DB = 0
SM = 0
Delta_L
Doxorubicin
YHL043W ECM34
ZShift = −2.15
Z lm Score = 0.36
NG = 0
DB = 0
SM = 0
Delta_K
Doxorubicin
YHL043W ECM34
ZShift = −2.03
Z lm Score = 2.21
NG = 0
DB = 0
SM = 0
Delta_r
Doxorubicin
YHL043W ECM34
ZShift = −3.92
Z lm Score = 6.44
NG = 0
DB = 0
SM = 0
Delta_AUC
Doxorubicin

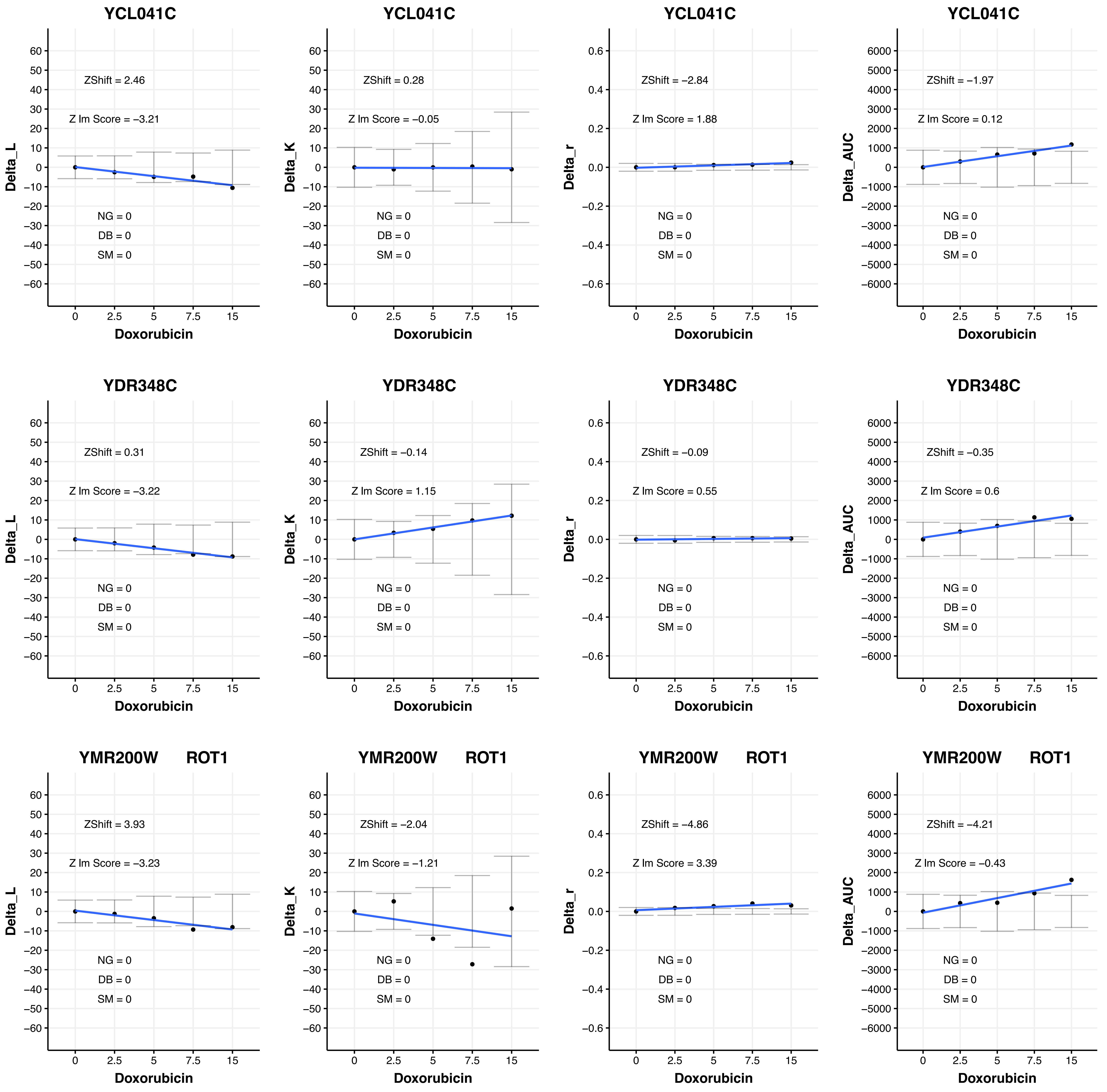

YCL041C
ZShift = 2.46
Z lm Score = −3.21
NG = 0
DB = 0
SM = 0
Delta_L
Doxorubicin
YCL041C
ZShift = 0.28
Z lm Score = −0.05
NG = 0
DB = 0
SM = 0
Delta_K
Doxorubicin
YCL041C
ZShift = −2.84
Z lm Score = 1.88
NG = 0
DB = 0
SM = 0
Delta_r
Doxorubicin
YCL041C
ZShift = −1.97
Z lm Score = 0.12
NG = 0
DB = 0
SM = 0
Delta_AUC
Doxorubicin
YDR348C
ZShift = 0.31
Z lm Score = −3.22
NG = 0
DB = 0
SM = 0
Delta_L
Doxorubicin
YDR348C
ZShift = −0.14
Z lm Score = 1.15
NG = 0
DB = 0
SM = 0
Delta_K
Doxorubicin
YDR348C
ZShift = −0.09
Z lm Score = 0.55
NG = 0
DB = 0
SM = 0
Delta_r
Doxorubicin
YDR348C
ZShift = −0.35
Z lm Score = 0.6
NG = 0
DB = 0
SM = 0
Delta_AUC
Doxorubicin
YMR200W ROT1
ZShift = 3.93
Z lm Score = −3.23
NG = 0
DB = 0
SM = 0
Delta_L
Doxorubicin
YMR200W ROT1
ZShift = −2.04
Z lm Score = −1.21
NG = 0
DB = 0
SM = 0
Delta_K
Doxorubicin
YMR200W ROT1
ZShift = −4.86
Z lm Score = 3.39
NG = 0
DB = 0
SM = 0
Delta_r
Doxorubicin
YMR200W ROT1
ZShift = −4.21
Z lm Score = −0.43
NG = 0
DB = 0
SM = 0
Delta_AUC
Doxorubicin

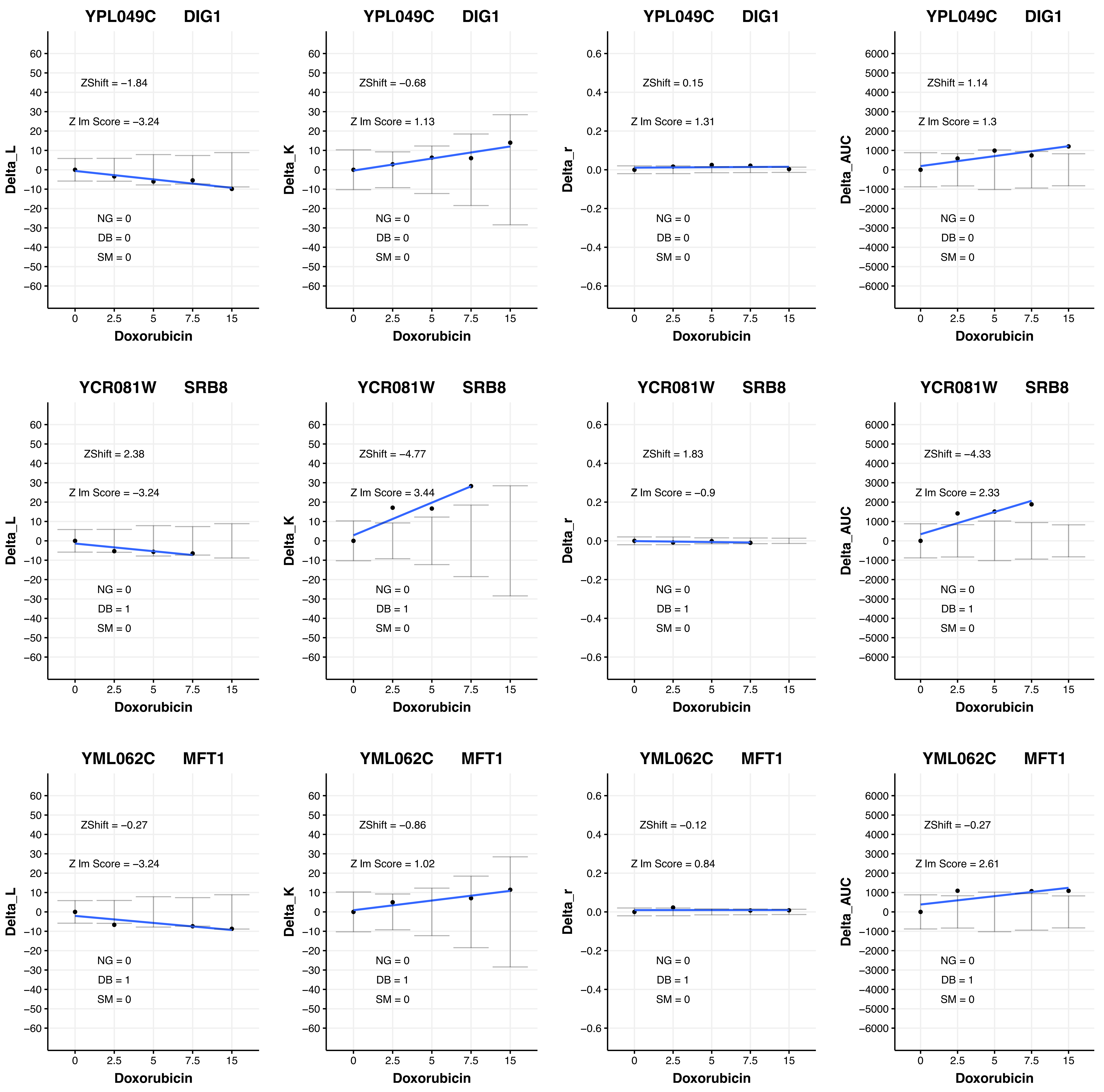
YPL049C DIG1
ZShift = −1.84
Z lm Score = −3.24
NG = 0
DB = 0
SM = 0
Delta_L
Doxorubicin
YPL049C DIG1
ZShift = −0.68
Z lm Score = 1.13
NG = 0
DB = 0
SM = 0
Delta_K
Doxorubicin
YPL049C DIG1
ZShift = 0.15
Z lm Score = 1.31
NG = 0
DB = 0
SM = 0
Delta_r
Doxorubicin
YPL049C DIG1
ZShift = 1.14
Z lm Score = 1.3
NG = 0
DB = 0
SM = 0
Delta_AUC
Doxorubicin
YCR081W SRB8
ZShift = 2.38
Z lm Score = −3.24
NG = 0
DB = 1
SM = 0
Delta_L
Doxorubicin
YCR081W SRB8
ZShift = −4.77
Z lm Score = 3.44
NG = 0
DB = 1
SM = 0
Delta_K
Doxorubicin
YCR081W SRB8
ZShift = 1.83
Z lm Score = −0.9
NG = 0
DB = 1
SM = 0
Delta_r
Doxorubicin
YCR081W SRB8
ZShift = −4.33
Z lm Score = 2.33
NG = 0
DB = 1
SM = 0
Delta_AUC
Doxorubicin
YML062C MFT1
ZShift = −0.27
Z lm Score = −3.24
NG = 0
DB = 1
SM = 0
Delta_L
Doxorubicin
YML062C MFT1
ZShift = −0.86
Z lm Score = 1.02
NG = 0
DB = 1
SM = 0
Delta_K
Doxorubicin
YML062C MFT1
ZShift = −0.12
Z lm Score = 0.84
NG = 0
DB = 1
SM = 0
Delta_r
Doxorubicin
YML062C MFT1
ZShift = −0.27
Z lm Score = 2.61
NG = 0
DB = 1
SM = 0
Delta_AUC
Doxorubicin

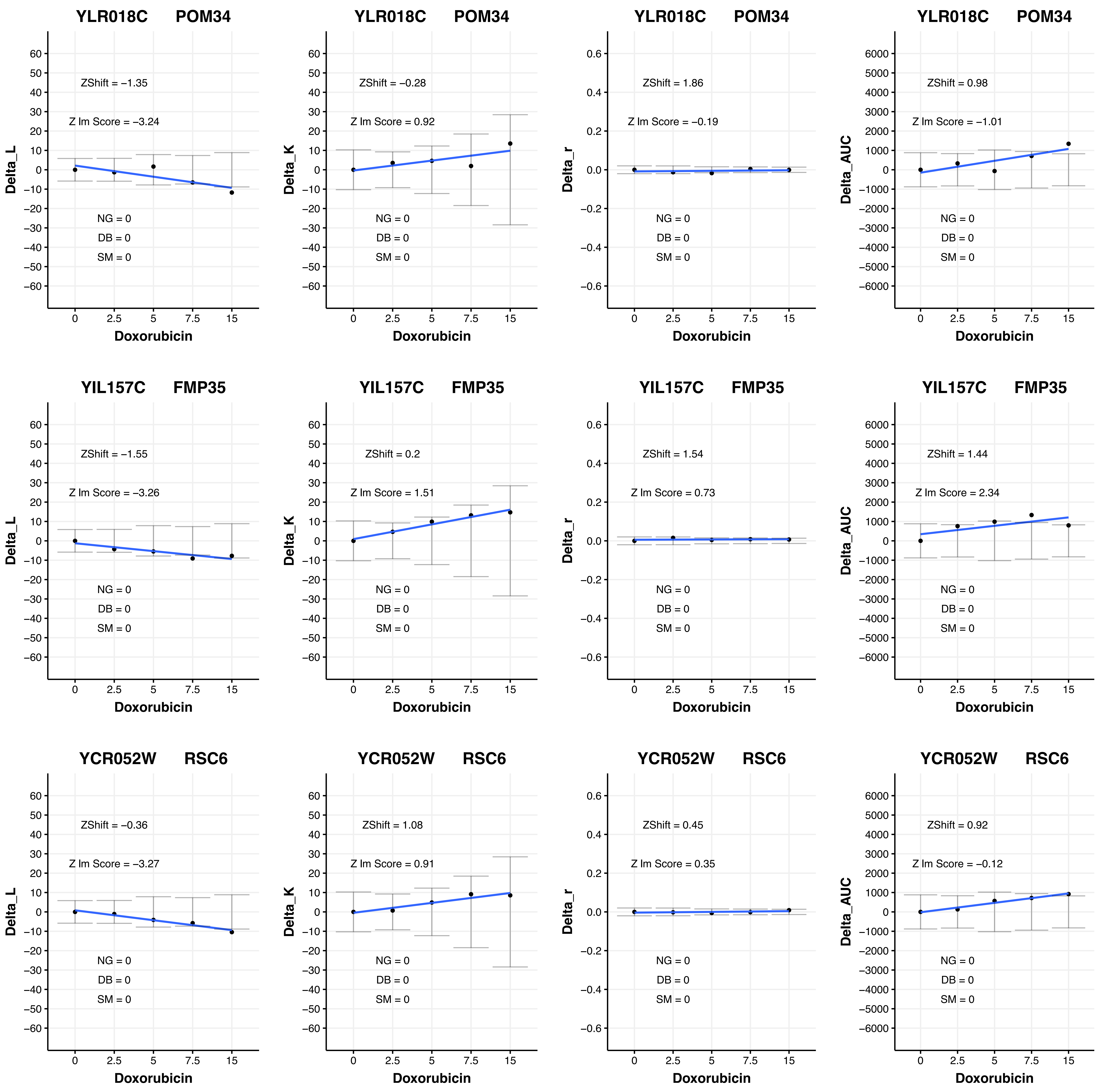
YLR018C POM34
ZShift = −1.35
Z lm Score = −3.24
NG = 0
DB = 0
SM = 0
Delta_L
Doxorubicin
YLR018C POM34
ZShift = −0.28
Z lm Score = 0.92
NG = 0
DB = 0
SM = 0
Delta_K
Doxorubicin
YLR018C POM34
ZShift = 1.86
Z lm Score = −0.19
NG = 0
DB = 0
SM = 0
Delta_r
Doxorubicin
YLR018C POM34
ZShift = 0.98
Z lm Score = −1.01
NG = 0
DB = 0
SM = 0
Delta_AUC
Doxorubicin
YIL157C FMP35
ZShift = −1.55
Z lm Score = −3.26
NG = 0
DB = 0
SM = 0
Delta_L
Doxorubicin
YIL157C FMP35
ZShift = 0.2
Z lm Score = 1.51
NG = 0
DB = 0
SM = 0
Delta_K
Doxorubicin
YIL157C FMP35
ZShift = 1.54
Z lm Score = 0.73
NG = 0
DB = 0
SM = 0
Delta_r
Doxorubicin
YIL157C FMP35
ZShift = 1.44
Z lm Score = 2.34
NG = 0
DB = 0
SM = 0
Delta_AUC
Doxorubicin
YCR052W RSC6
ZShift = −0.36
Z lm Score = −3.27
NG = 0
DB = 0
SM = 0
Delta_L
Doxorubicin
YCR052W RSC6
ZShift = 1.08
Z lm Score = 0.91
NG = 0
DB = 0
SM = 0
Delta_K
Doxorubicin
YCR052W RSC6
ZShift = 0.45
Z lm Score = 0.35
NG = 0
DB = 0
SM = 0
Delta_r
Doxorubicin
YCR052W RSC6
ZShift = 0.92
Z lm Score = −0.12
NG = 0
DB = 0
SM = 0
Delta_AUC
Doxorubicin

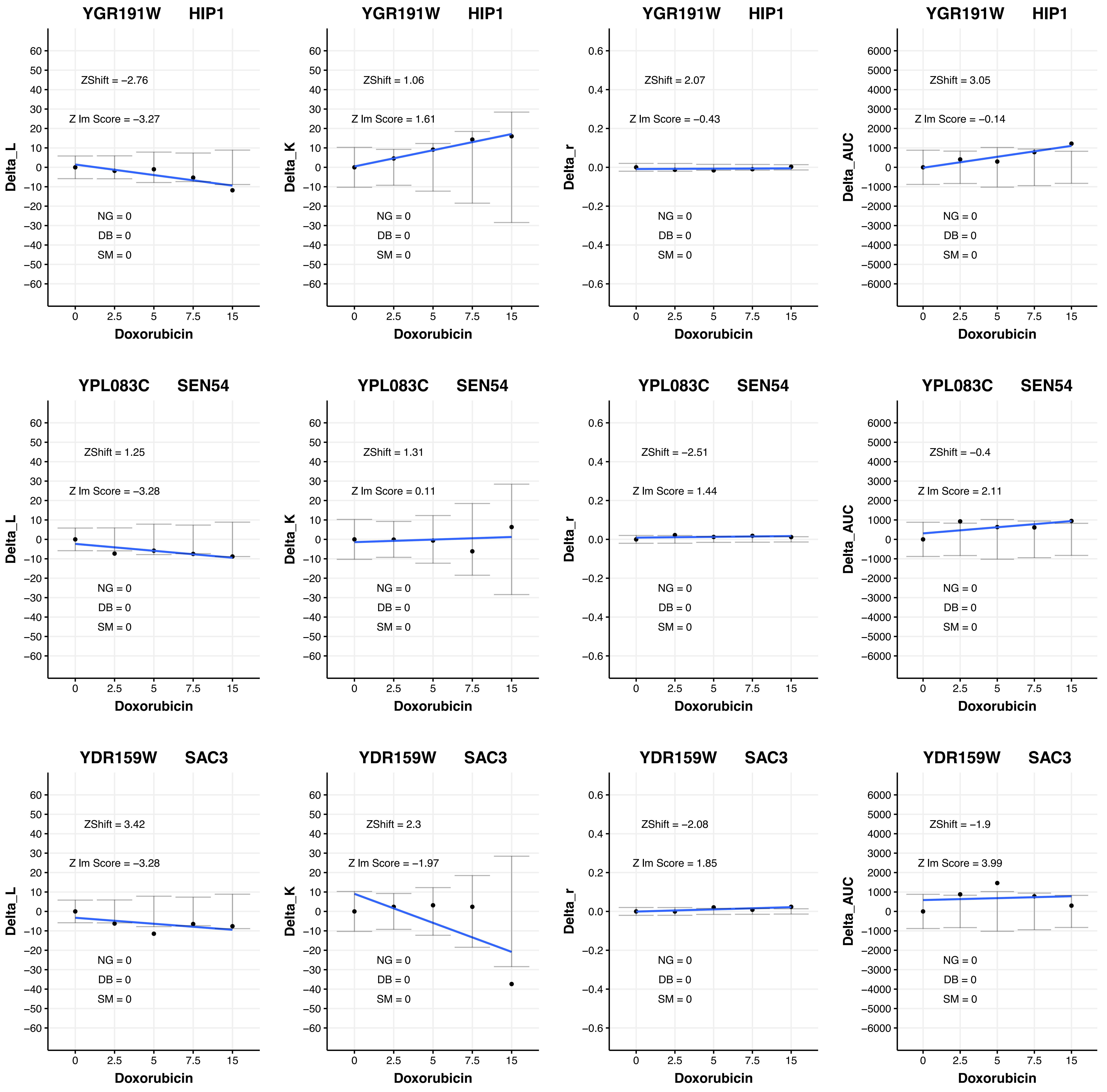
YGR191W HIP1
ZShift = −2.76
Z lm Score = −3.27
NG = 0
DB = 0
SM = 0
Delta_L
Doxorubicin
YGR191W HIP1
ZShift = 1.06
Z lm Score = 1.61
NG = 0
DB = 0
SM = 0
Delta_K
Doxorubicin
YGR191W HIP1
ZShift = 2.07
Z lm Score = −0.43
NG = 0
DB = 0
SM = 0
Delta_r
Doxorubicin
YGR191W HIP1
ZShift = 3.05
Z lm Score = −0.14
NG = 0
DB = 0
SM = 0
Delta_AUC
Doxorubicin
YPL083C SEN54
ZShift = 1.25
Z lm Score = −3.28
NG = 0
DB = 0
SM = 0
Delta_L
Doxorubicin
YPL083C SEN54
ZShift = 1.31
Z lm Score = 0.11
NG = 0
DB = 0
SM = 0
Delta_K
Doxorubicin
YPL083C SEN54
ZShift = −2.51
Z lm Score = 1.44
NG = 0
DB = 0
SM = 0
Delta_r
Doxorubicin
YPL083C SEN54
ZShift = −0.4
Z lm Score = 2.11
NG = 0
DB = 0
SM = 0
Delta_AUC
Doxorubicin
YDR159W SAC3
ZShift = 3.42
Z lm Score = −3.28
NG = 0
DB = 0
SM = 0
Delta_L
Doxorubicin
YDR159W SAC3
ZShift = 2.3
Z lm Score = −1.97
NG = 0
DB = 0
SM = 0
Delta_K
Doxorubicin
YDR159W SAC3
ZShift = −2.08
Z lm Score = 1.85
NG = 0
DB = 0
SM = 0
Delta_r
Doxorubicin
YDR159W SAC3
ZShift = −1.9
Z lm Score = 3.99
NG = 0
DB = 0
SM = 0
Delta_AUC
Doxorubicin

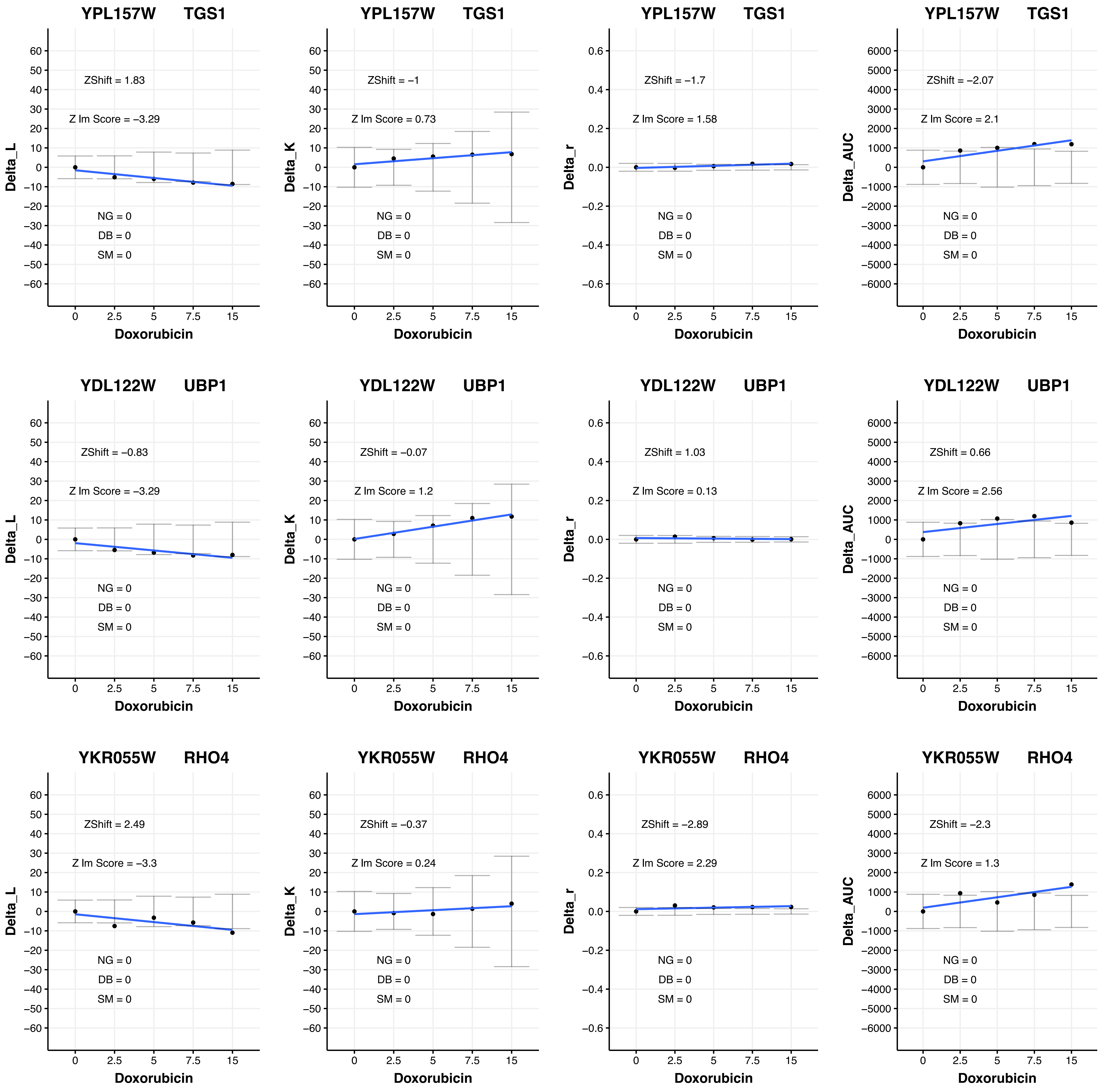
YPL157W TGS1
ZShift = 1.83
Z lm Score = −3.29
NG = 0
DB = 0
SM = 0
Delta_L
Doxorubicin
YPL157W TGS1
ZShift = −1
Z lm Score = 0.73
NG = 0
DB = 0
SM = 0
Delta_K
Doxorubicin
YPL157W TGS1
ZShift = −1.7
Z lm Score = 1.58
NG = 0
DB = 0
SM = 0
Delta_r
Doxorubicin
YPL157W TGS1
ZShift = −2.07
Z lm Score = 2.1
NG = 0
DB = 0
SM = 0
Delta_AUC
Doxorubicin
YDL122W UBP1
ZShift = −0.83
Z lm Score = −3.29
NG = 0
DB = 0
SM = 0
Delta_L
Doxorubicin
YDL122W UBP1
ZShift = −0.07
Z lm Score = 1.2
NG = 0
DB = 0
SM = 0
Delta_K
Doxorubicin
YDL122W UBP1
ZShift = 1.03
Z lm Score = 0.13
NG = 0
DB = 0
SM = 0
Delta_r
Doxorubicin
YDL122W UBP1
ZShift = 0.66
Z lm Score = 2.56
NG = 0
DB = 0
SM = 0
Delta_AUC
Doxorubicin
YKR055W RHO4
ZShift = 2.49
Z lm Score = −3.3
NG = 0
DB = 0
SM = 0
Delta_L
Doxorubicin
YKR055W RHO4
ZShift = −0.37
Z lm Score = 0.24
NG = 0
DB = 0
SM = 0
Delta_K
Doxorubicin
YKR055W RHO4
ZShift = −2.89
Z lm Score = 2.29
NG = 0
DB = 0
SM = 0
Delta_r
Doxorubicin
YKR055W RHO4
ZShift = −2.3
Z lm Score = 1.3
NG = 0
DB = 0
SM = 0
Delta_AUC
Doxorubicin

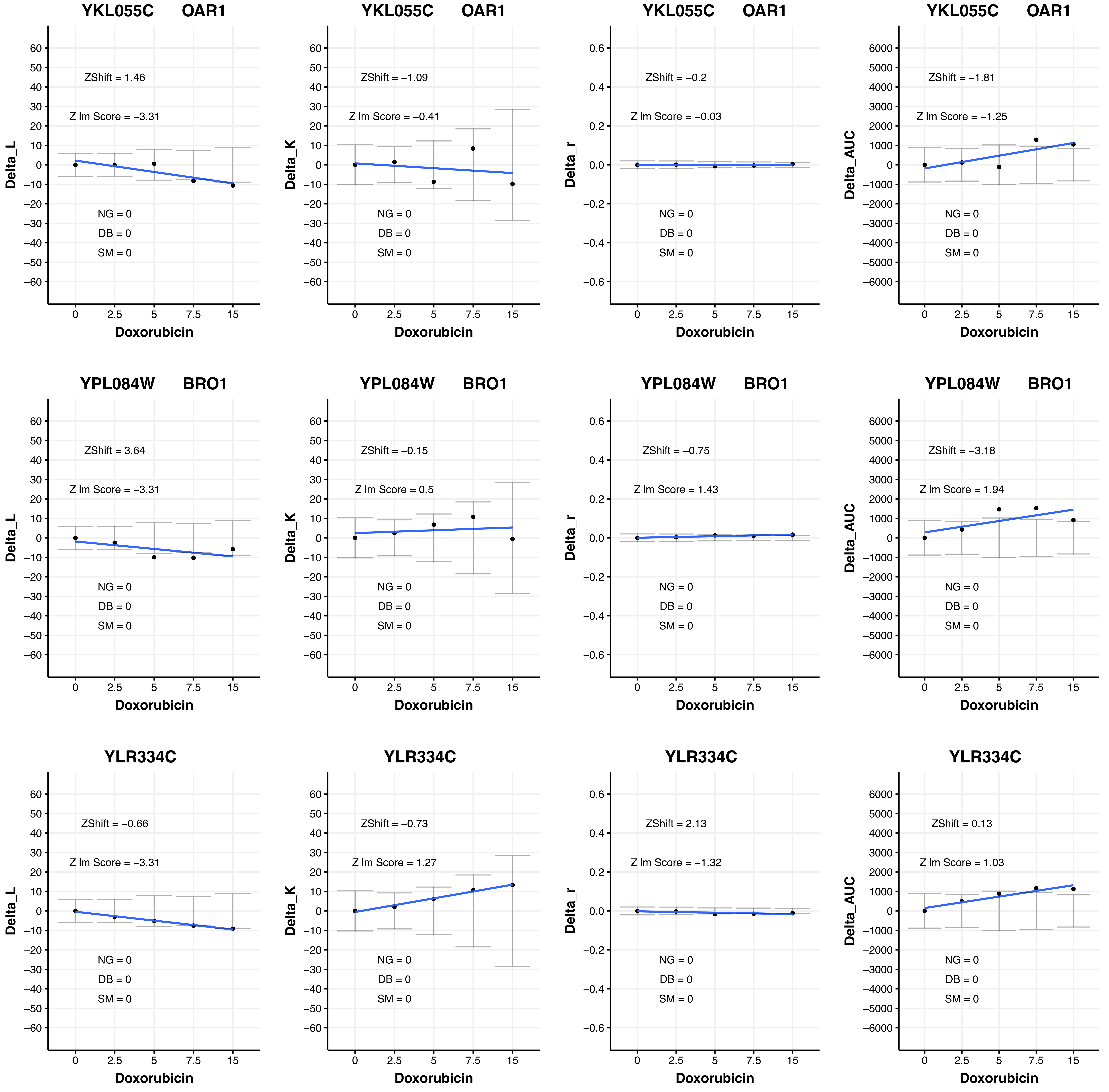
YKL055C OAR1
ZShift = 1.46
Z lm Score = −3.31
NG = 0
DB = 0
SM = 0
Delta_L
Doxorubicin
YKL055C OAR1
ZShift = −1.09
Z lm Score = −0.41
NG = 0
DB = 0
SM = 0
Delta_K
Doxorubicin
YKL055C OAR1
ZShift = −0.2
Z lm Score = −0.03
NG = 0
DB = 0
SM = 0
Delta_r
Doxorubicin
YKL055C OAR1
ZShift = −1.81
Z lm Score = −1.25
NG = 0
DB = 0
SM = 0
Delta_AUC
Doxorubicin
YPL084W BRO1
ZShift = 3.64
Z lm Score = −3.31
NG = 0
DB = 0
SM = 0
Delta_L
Doxorubicin
YPL084W BRO1
ZShift = −0.15
Z lm Score = 0.5
NG = 0
DB = 0
SM = 0
Delta_K
Doxorubicin
YPL084W BRO1
ZShift = −0.75
Z lm Score = 1.43
NG = 0
DB = 0
SM = 0
Delta_r
Doxorubicin
YPL084W BRO1
ZShift = −3.18
Z lm Score = 1.94
NG = 0
DB = 0
SM = 0
Delta_AUC
Doxorubicin
YLR334C
ZShift = −0.66
Z lm Score = −3.31
NG = 0
DB = 0
SM = 0
Delta_L
Doxorubicin
YLR334C
ZShift = −0.73
Z lm Score = 1.27
NG = 0
DB = 0
SM = 0
Delta_K
Doxorubicin
YLR334C
ZShift = 2.13
Z lm Score = −1.32
NG = 0
DB = 0
SM = 0
Delta_r
Doxorubicin
YLR334C
ZShift = 0.13
Z lm Score = 1.03
NG = 0
DB = 0
SM = 0
Delta_AUC
Doxorubicin

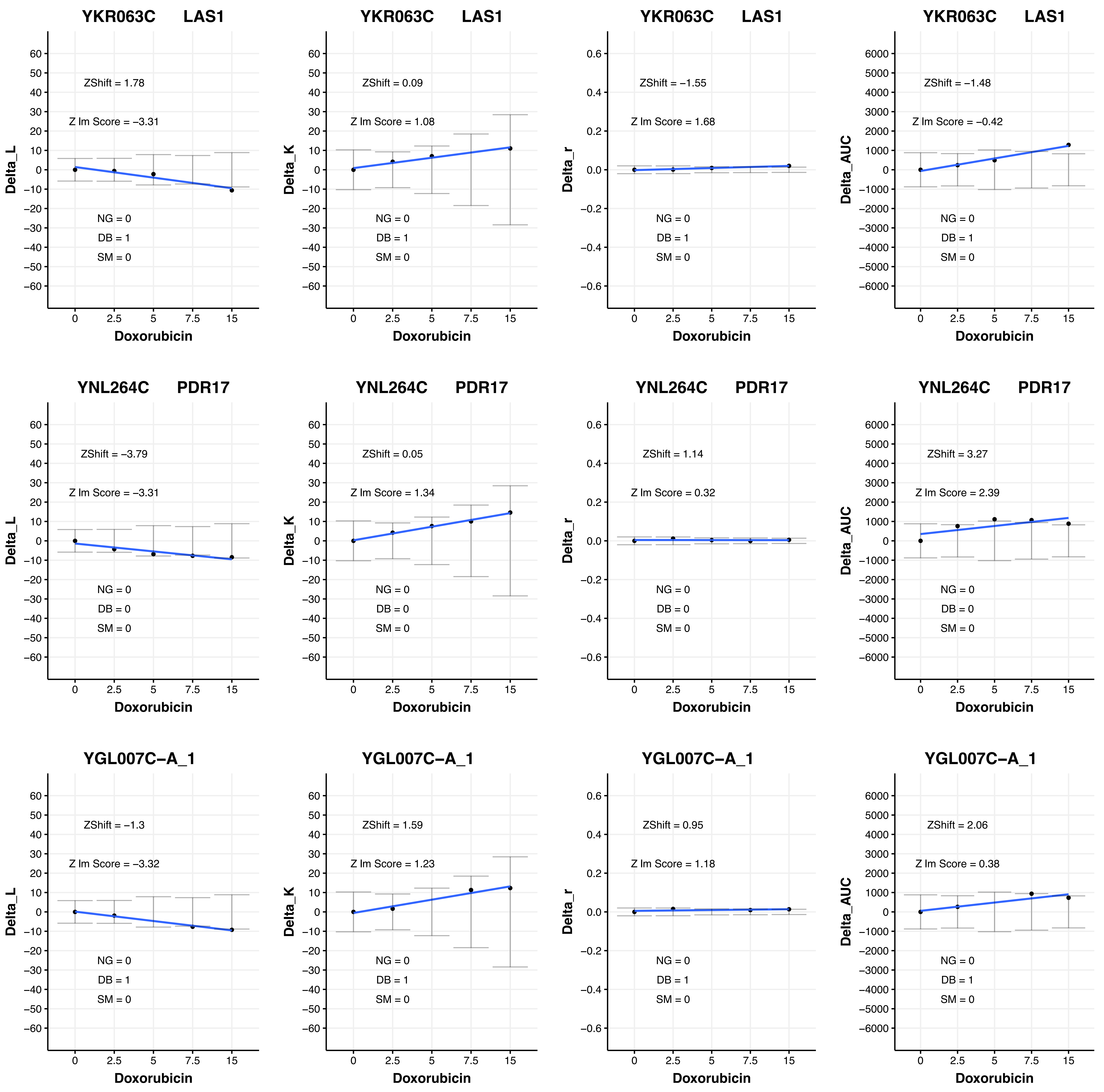
YKR063C LAS1
ZShift = 1.78
Z lm Score = −3.31
NG = 0
DB = 1
SM = 0
Delta_L
Doxorubicin
YKR063C LAS1
ZShift = 0.09
Z lm Score = 1.08
NG = 0
DB = 1
SM = 0
Delta_K
Doxorubicin
YKR063C LAS1
ZShift = −1.55
Z lm Score = 1.68
NG = 0
DB = 1
SM = 0
Delta_r
Doxorubicin
YKR063C LAS1
ZShift = −1.48
Z lm Score = −0.42
NG = 0
DB = 1
SM = 0
Delta_AUC
Doxorubicin
YNL264C PDR17
ZShift = −3.79
Z lm Score = −3.31
NG = 0
DB = 0
SM = 0
Delta_L
Doxorubicin
YNL264C PDR17
ZShift = 0.05
Z lm Score = 1.34
NG = 0
DB = 0
SM = 0
Delta_K
Doxorubicin
YNL264C PDR17
ZShift = 1.14
Z lm Score = 0.32
NG = 0
DB = 0
SM = 0
Delta_r
Doxorubicin
YNL264C PDR17
ZShift = 3.27
Z lm Score = 2.39
NG = 0
DB = 0
SM = 0
Delta_AUC
Doxorubicin
YGL007C−A_1
ZShift = −1.3
Z lm Score = −3.32
NG = 0
DB = 1
SM = 0
Delta_L
Doxorubicin
YGL007C−A_1
ZShift = 1.59
Z lm Score = 1.23
NG = 0
DB = 1
SM = 0
Delta_K
Doxorubicin
YGL007C−A_1
ZShift = 0.95
Z lm Score = 1.18
NG = 0
DB = 1
SM = 0
Delta_r
Doxorubicin
YGL007C−A_1
ZShift = 2.06
Z lm Score = 0.38
NG = 0
DB = 1
SM = 0
Delta_AUC
Doxorubicin

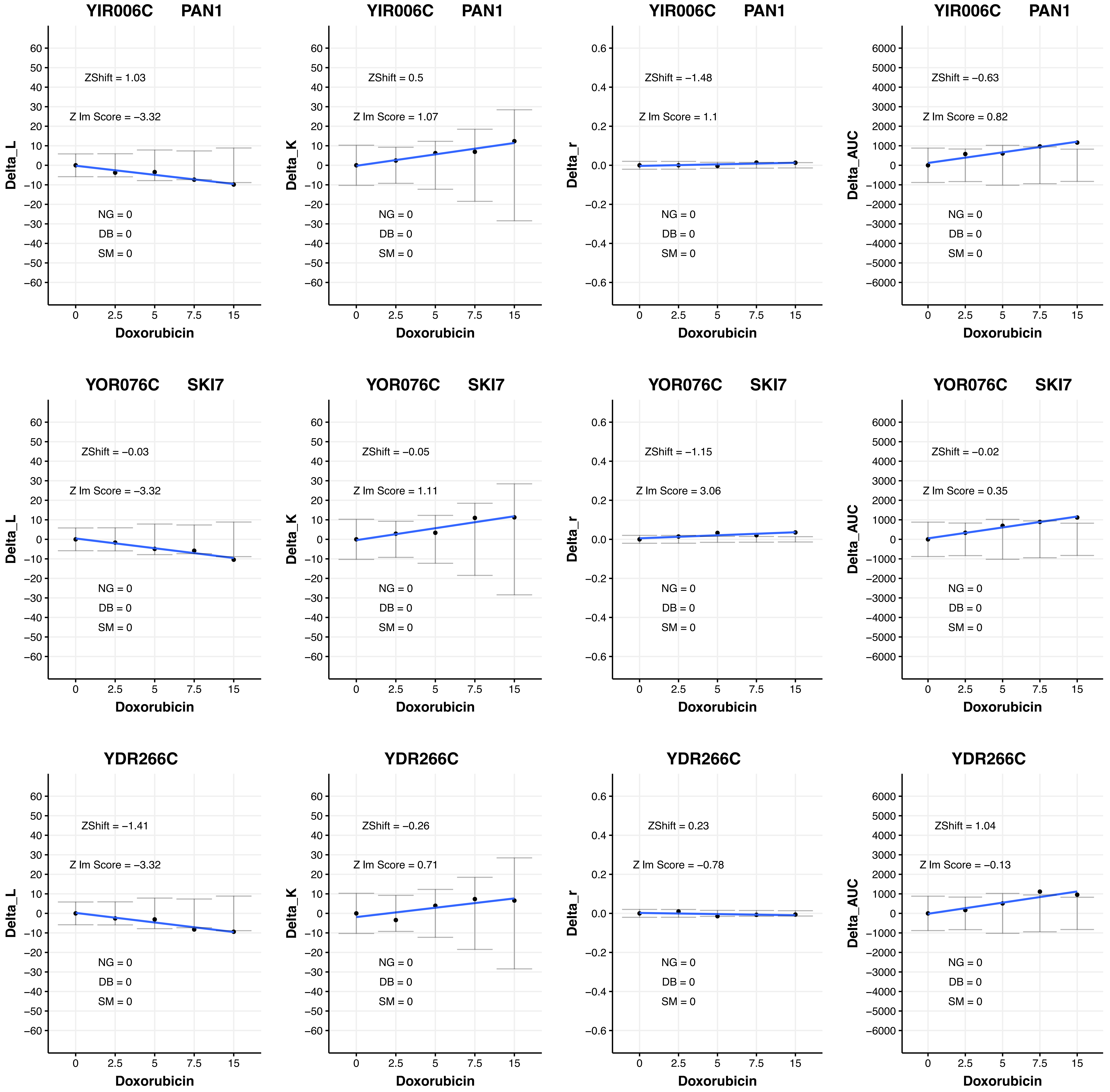
YIR006C PAN1
ZShift = 1.03
Z lm Score = −3.32
NG = 0
DB = 0
SM = 0
Delta_L
Doxorubicin
YIR006C PAN1
ZShift = 0.5
Z lm Score = 1.07
NG = 0
DB = 0
SM = 0
Delta_K
Doxorubicin
YIR006C PAN1
ZShift = −1.48
Z lm Score = 1.1
NG = 0
DB = 0
SM = 0
Delta_r
Doxorubicin
YIR006C PAN1
ZShift = −0.63
Z lm Score = 0.82
NG = 0
DB = 0
SM = 0
Delta_AUC
Doxorubicin
YOR076C SKI7
ZShift = −0.03
Z lm Score = −3.32
NG = 0
DB = 0
SM = 0
Delta_L
Doxorubicin
YOR076C SKI7
ZShift = −0.05
Z lm Score = 1.11
NG = 0
DB = 0
SM = 0
Delta_K
Doxorubicin
YOR076C SKI7
ZShift = −1.15
Z lm Score = 3.06
NG = 0
DB = 0
SM = 0
Delta_r
Doxorubicin
YOR076C SKI7
ZShift = −0.02
Z lm Score = 0.35
NG = 0
DB = 0
SM = 0
Delta_AUC
Doxorubicin
YDR266C
ZShift = −1.41
Z lm Score = −3.32
NG = 0
DB = 0
SM = 0
Delta_L
Doxorubicin
YDR266C
ZShift = −0.26
Z lm Score = 0.71
NG = 0
DB = 0
SM = 0
Delta_K
Doxorubicin
YDR266C
ZShift = 0.23
Z lm Score = −0.78
NG = 0
DB = 0
SM = 0
Delta_r
Doxorubicin
YDR266C
ZShift = 1.04
Z lm Score = −0.13
NG = 0
DB = 0
SM = 0
Delta_AUC
Doxorubicin

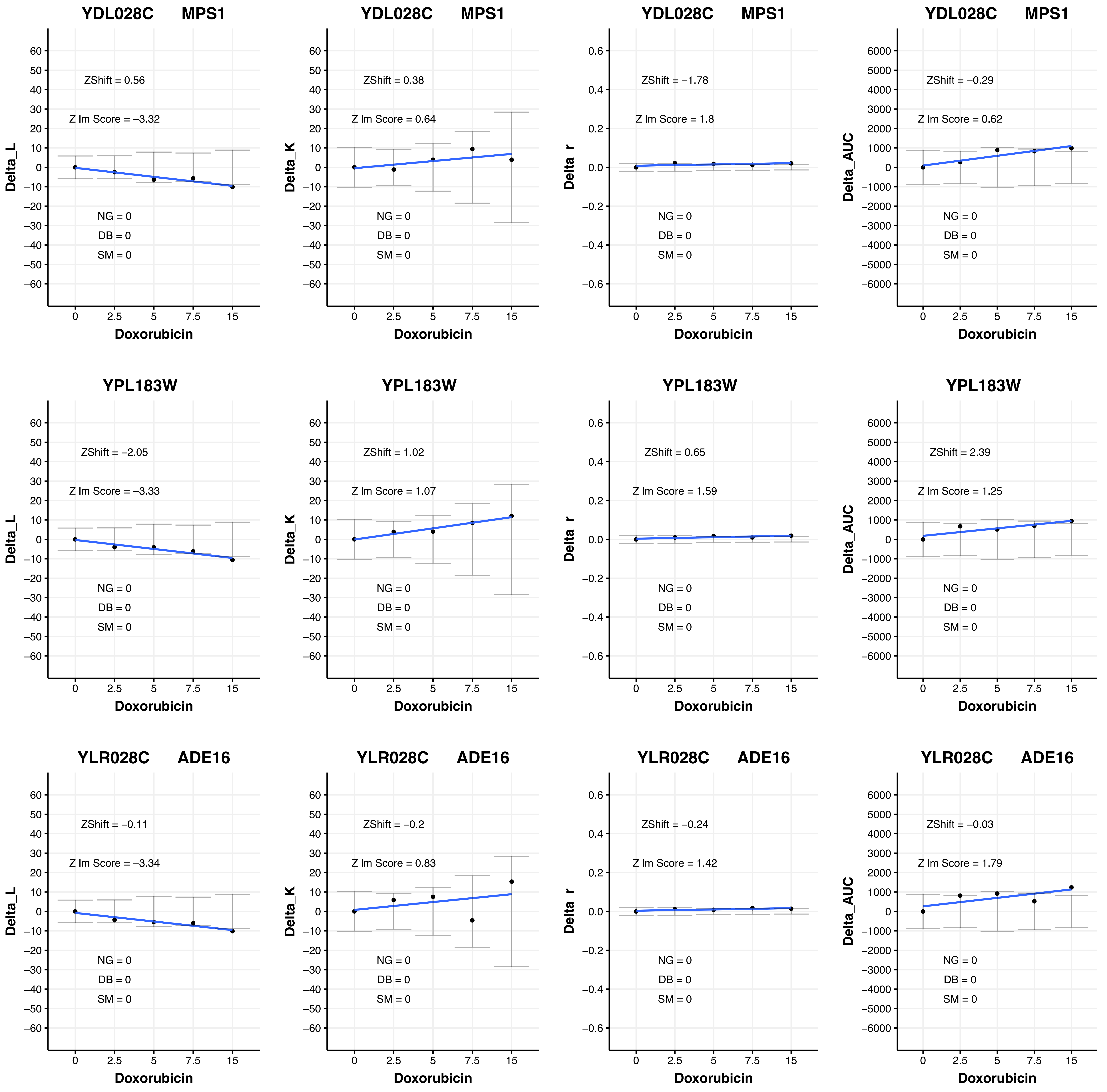

YDL028C MPS1
ZShift = 0.56
Z lm Score = −3.32
NG = 0
DB = 0
SM = 0
Delta_L
Doxorubicin
YDL028C MPS1
ZShift = 0.38
Z lm Score = 0.64
NG = 0
DB = 0
SM = 0
Delta_K
Doxorubicin
YDL028C MPS1
ZShift = −1.78
Z lm Score = 1.8
NG = 0
DB = 0
SM = 0
Delta_r
Doxorubicin
YDL028C MPS1
ZShift = −0.29
Z lm Score = 0.62
NG = 0
DB = 0
SM = 0
Delta_AUC
Doxorubicin
YPL183W
ZShift = −2.05
Z lm Score = −3.33
NG = 0
DB = 0
SM = 0
Delta_L
Doxorubicin
YPL183W
ZShift = 1.02
Z lm Score = 1.07
NG = 0
DB = 0
SM = 0
Delta_K
Doxorubicin
YPL183W
ZShift = 0.65
Z lm Score = 1.59
NG = 0
DB = 0
SM = 0
Delta_r
Doxorubicin
YPL183W
ZShift = 2.39
Z lm Score = 1.25
NG = 0
DB = 0
SM = 0
Delta_AUC
Doxorubicin
YLR028C ADE16
ZShift = −0.11
Z lm Score = −3.34
NG = 0
DB = 0
SM = 0
Delta_L
Doxorubicin
YLR028C ADE16
ZShift = −0.2
Z lm Score = 0.83
NG = 0
DB = 0
SM = 0
Delta_K
Doxorubicin
YLR028C ADE16
ZShift = −0.24
Z lm Score = 1.42
NG = 0
DB = 0
SM = 0
Delta_r
Doxorubicin
YLR028C ADE16
ZShift = −0.03
Z lm Score = 1.79
NG = 0
DB = 0
SM = 0
Delta_AUC
Doxorubicin

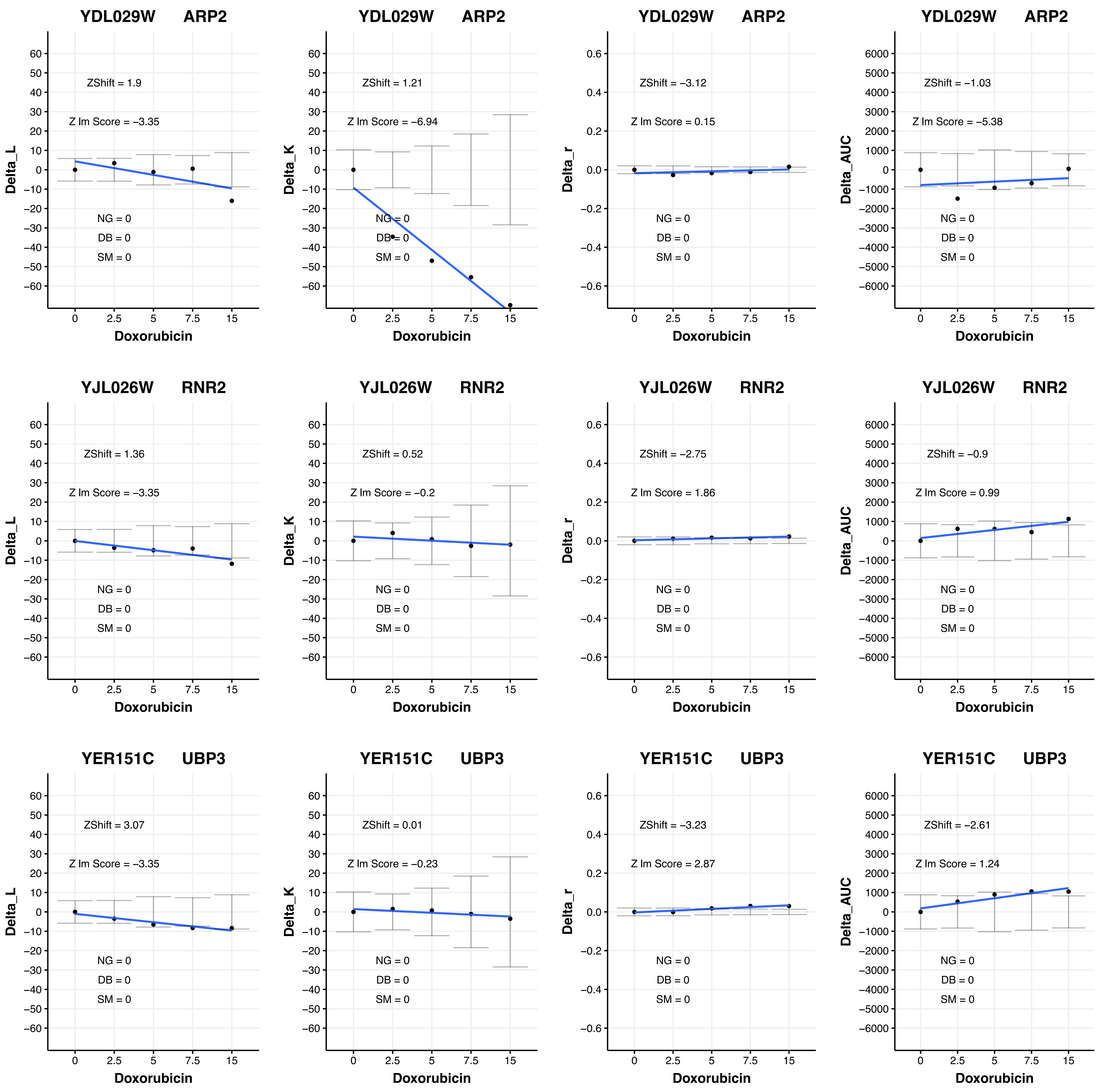
YDL029W ARP2
ZShift = 1.9
Z lm Score = −3.35
NG = 0
DB = 0
SM = 0
Delta_L
Doxorubicin
YDL029W ARP2
ZShift = 1.21
Z lm Score = −6.94
NG = 0
DB = 0
SM = 0
Delta_K
Doxorubicin
YDL029W ARP2
ZShift = −3.12
Z lm Score = 0.15
NG = 0
DB = 0
SM = 0
Delta_r
Doxorubicin
YDL029W ARP2
ZShift = −1.03
Z lm Score = −5.38
NG = 0
DB = 0
SM = 0
Delta_AUC
Doxorubicin
YJL026W RNR2
ZShift = 1.36
Z lm Score = −3.35
NG = 0
DB = 0
SM = 0
Delta_L
Doxorubicin
YJL026W RNR2
ZShift = 0.52
Z lm Score = −0.2
NG = 0
DB = 0
SM = 0
Delta_K
Doxorubicin
YJL026W RNR2
ZShift = −2.75
Z lm Score = 1.86
NG = 0
DB = 0
SM = 0
Delta_r
Doxorubicin
YJL026W RNR2
ZShift = −0.9
Z lm Score = 0.99
NG = 0
DB = 0
SM = 0
Delta_AUC
Doxorubicin
YER151C UBP3
ZShift = 3.07
Z lm Score = −3.35
NG = 0
DB = 0
SM = 0
Delta_L
Doxorubicin
YER151C UBP3
ZShift = 0.01
Z lm Score = −0.23
NG = 0
DB = 0
SM = 0
Delta_K
Doxorubicin
YER151C UBP3
ZShift = −3.23
Z lm Score = 2.87
NG = 0
DB = 0
SM = 0
Delta_r
Doxorubicin
YER151C UBP3
ZShift = −2.61
Z lm Score = 1.24
NG = 0
DB = 0
SM = 0
Delta_AUC
Doxorubicin

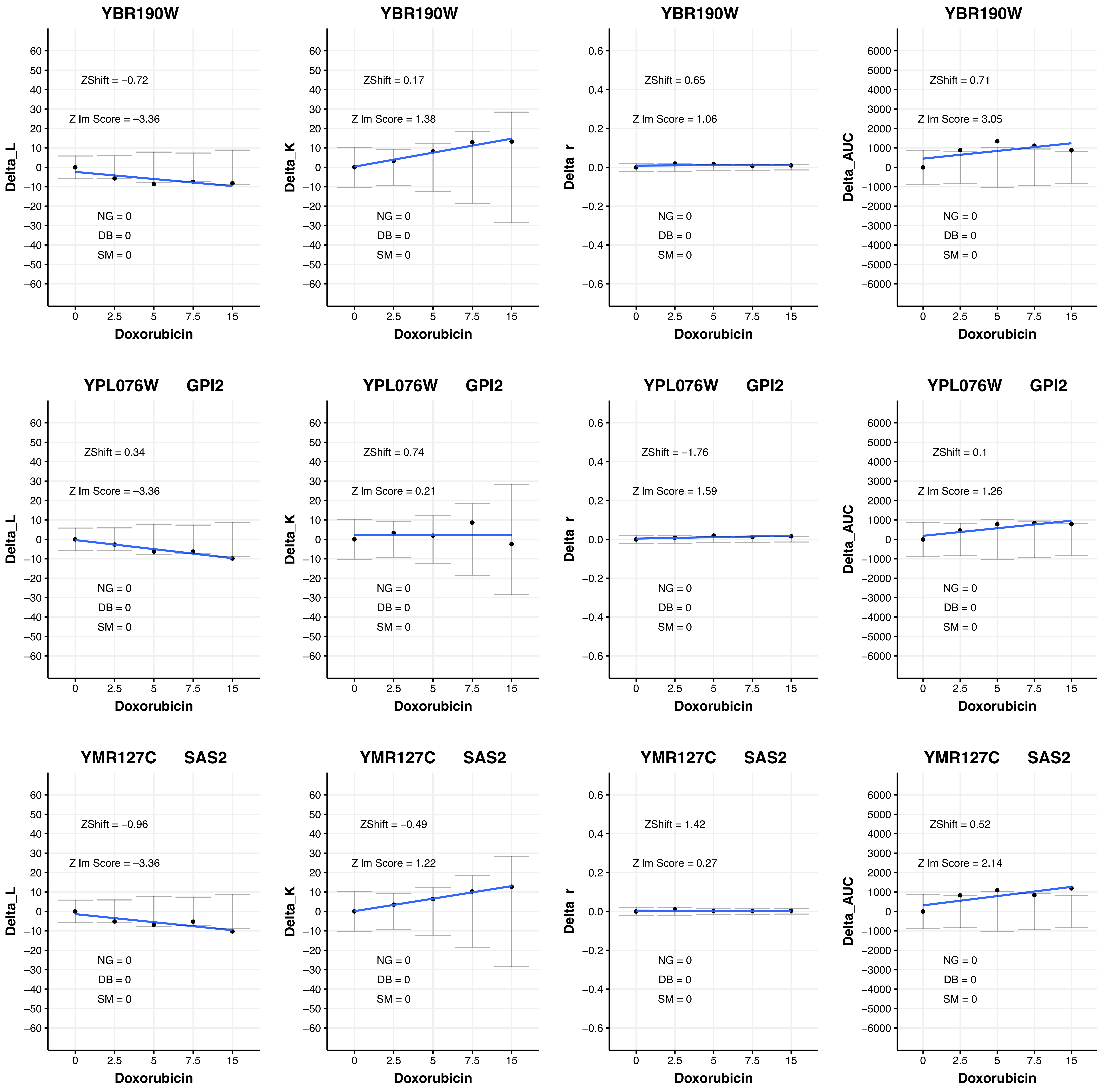
YBR190W
ZShift = −0.72
Z lm Score = −3.36
NG = 0
DB = 0
SM = 0
Delta_L
Doxorubicin
YBR190W
ZShift = 0.17
Z lm Score = 1.38
NG = 0
DB = 0
SM = 0
Delta_K
Doxorubicin
YBR190W
ZShift = 0.65
Z lm Score = 1.06
NG = 0
DB = 0
SM = 0
Delta_r
Doxorubicin
YBR190W
ZShift = 0.71
Z lm Score = 3.05
NG = 0
DB = 0
SM = 0
Delta_AUC
Doxorubicin
YPL076W GPI2
ZShift = 0.34
Z lm Score = −3.36
NG = 0
DB = 0
SM = 0
Delta_L
Doxorubicin
YPL076W GPI2
ZShift = 0.74
Z lm Score = 0.21
NG = 0
DB = 0
SM = 0
Delta_K
Doxorubicin
YPL076W GPI2
ZShift = −1.76
Z lm Score = 1.59
NG = 0
DB = 0
SM = 0
Delta_r
Doxorubicin
YPL076W GPI2
ZShift = 0.1
Z lm Score = 1.26
NG = 0
DB = 0
SM = 0
Delta_AUC
Doxorubicin
YMR127C SAS2
ZShift = −0.96
Z lm Score = −3.36
NG = 0
DB = 0
SM = 0
Delta_L
Doxorubicin
YMR127C SAS2
ZShift = −0.49
Z lm Score = 1.22
NG = 0
DB = 0
SM = 0
Delta_K
Doxorubicin
YMR127C SAS2
ZShift = 1.42
Z lm Score = 0.27
NG = 0
DB = 0
SM = 0
Delta_r
Doxorubicin
YMR127C SAS2
ZShift = 0.52
Z lm Score = 2.14
NG = 0
DB = 0
SM = 0
Delta_AUC
Doxorubicin

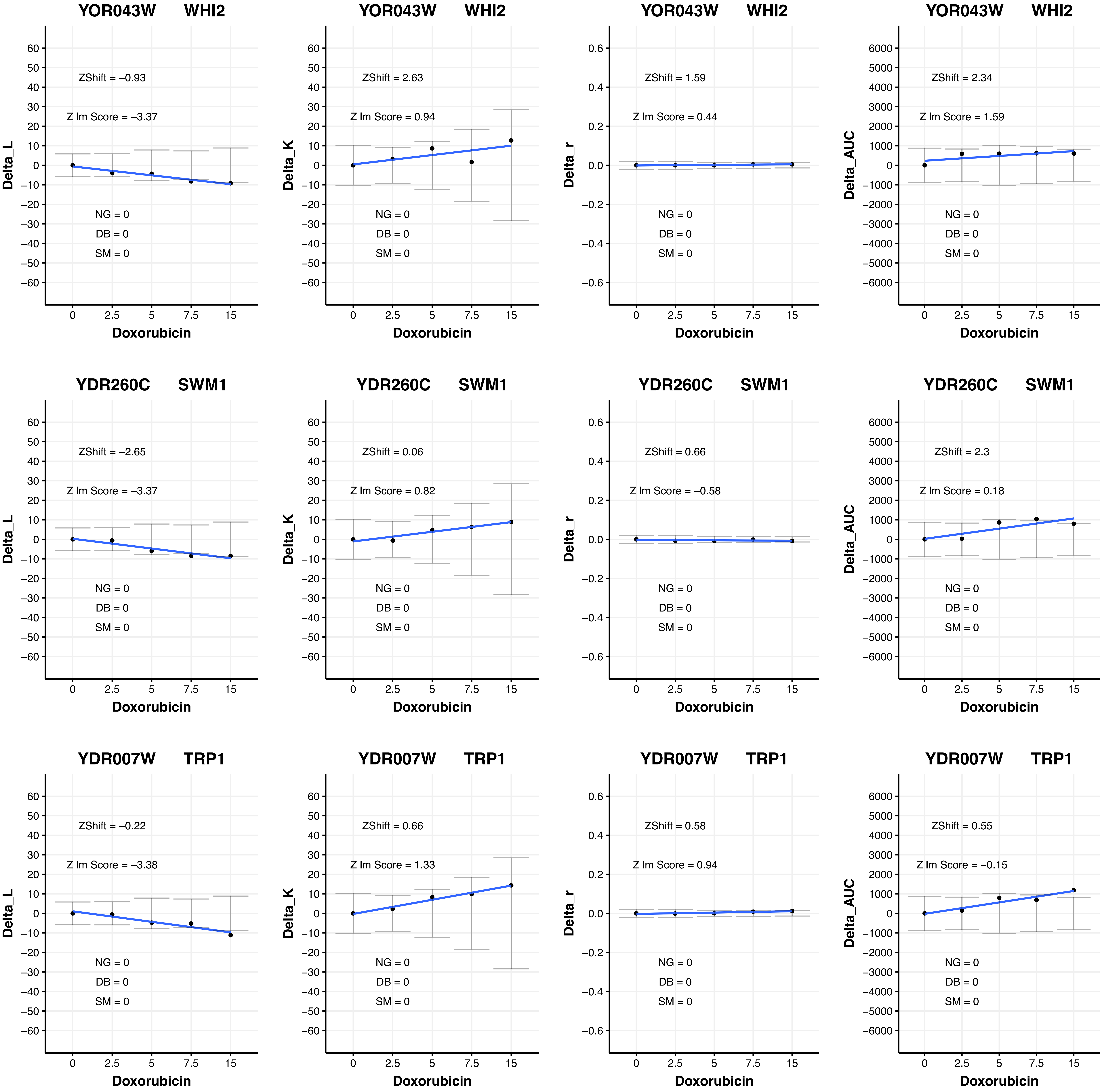
YOR043W WHI2
ZShift = −0.93
Z lm Score = −3.37
NG = 0
DB = 0
SM = 0
Delta_L
Doxorubicin
YOR043W WHI2
ZShift = 2.63
Z lm Score = 0.94
NG = 0
DB = 0
SM = 0
Delta_K
Doxorubicin
YOR043W WHI2
ZShift = 1.59
Z lm Score = 0.44
NG = 0
DB = 0
SM = 0
Delta_r
Doxorubicin
YOR043W WHI2
ZShift = 2.34
Z lm Score = 1.59
NG = 0
DB = 0
SM = 0
Delta_AUC
Doxorubicin
YDR260C SWM1
ZShift = −2.65
Z lm Score = −3.37
NG = 0
DB = 0
SM = 0
Delta_L
Doxorubicin
YDR260C SWM1
ZShift = 0.06
Z lm Score = 0.82
NG = 0
DB = 0
SM = 0
Delta_K
Doxorubicin
YDR260C SWM1
ZShift = 0.66
Z lm Score = −0.58
NG = 0
DB = 0
SM = 0
Delta_r
Doxorubicin
YDR260C SWM1
ZShift = 2.3
Z lm Score = 0.18
NG = 0
DB = 0
SM = 0
Delta_AUC
Doxorubicin
YDR007W TRP1
ZShift = −0.22
Z lm Score = −3.38
NG = 0
DB = 0
SM = 0
Delta_L
Doxorubicin
YDR007W TRP1
ZShift = 0.66
Z lm Score = 1.33
NG = 0
DB = 0
SM = 0
Delta_K
Doxorubicin
YDR007W TRP1
ZShift = 0.58
Z lm Score = 0.94
NG = 0
DB = 0
SM = 0
Delta_r
Doxorubicin
YDR007W TRP1
ZShift = 0.55
Z lm Score = −0.15
NG = 0
DB = 0
SM = 0
Delta_AUC
Doxorubicin

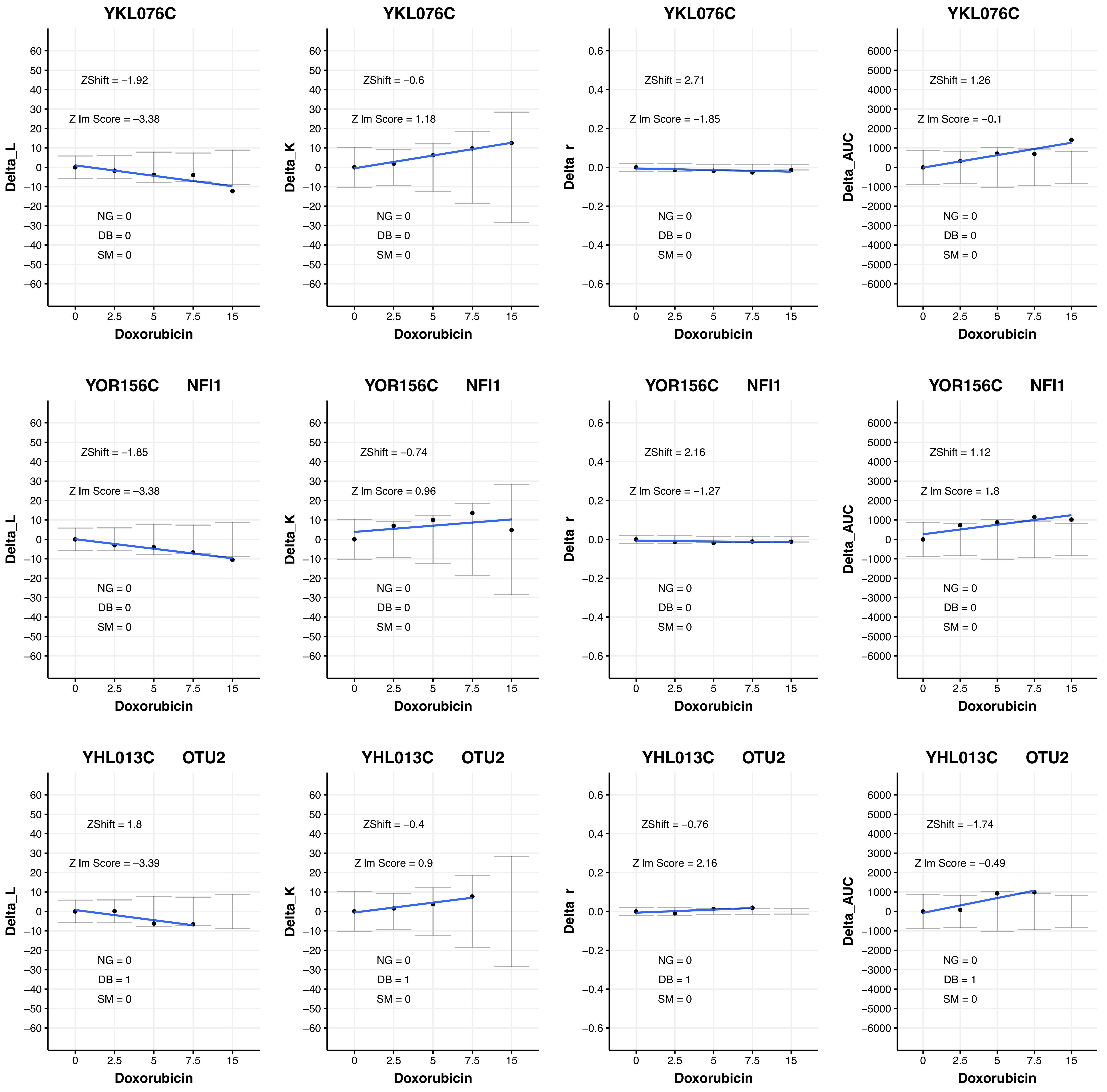
YKL076C
ZShift = −1.92
Z lm Score = −3.38
NG = 0
DB = 0
SM = 0
Delta_L
60 50 40 30 20 10 0 −10 −20 −30 −40 −50 −60
0 2.5 5 7.5 15
Doxorubicin
YKL076C
ZShift = −0.6
Z lm Score = 1.18
NG = 0
DB = 0
SM = 0
Delta_K
60 50 40 30 20 10 0 −10 −20 −30 −40 −50 −60
0 2.5 5 7.5 15
Doxorubicin
YKL076C
ZShift = 2.71
Z lm Score = −1.85
NG = 0
DB = 0
SM = 0
Delta_r
0.6 0.4 0.2 0.0 −0.2 −0.4 −0.6
0 2.5 5 7.5 15
Doxorubicin
YKL076C
ZShift = 1.26
Z lm Score = −0.1
NG = 0
DB = 0
SM = 0
Delta_AUC
6000 5000 4000 3000 2000 1000 0 −1000 −2000 −3000 −4000 −5000 −6000
0 2.5 5 7.5 15
Doxorubicin
YOR156C NFI1
ZShift = −1.85
Z lm Score = −3.38
NG = 0
DB = 0
SM = 0
Delta_L
60 50 40 30 20 10 0 −10 −20 −30 −40 −50 −60
0 2.5 5 7.5 15
Doxorubicin
YOR156C NFI1
ZShift = −0.74
Z lm Score = 0.96
NG = 0
DB = 0
SM = 0
Delta_K
60 50 40 30 20 10 0 −10 −20 −30 −40 −50 −60
0 2.5 5 7.5 15
Doxorubicin
YOR156C NFI1
ZShift = 2.16
Z lm Score = −1.27
NG = 0
DB = 0
SM = 0
Delta_r
0.6 0.4 0.2 0.0 −0.2 −0.4 −0.6
0 2.5 5 7.5 15
Doxorubicin
YOR156C NFI1
ZShift = 1.12
Z lm Score = 1.8
NG = 0
DB = 0
SM = 0
Delta_AUC
6000 5000 4000 3000 2000 1000 0 −1000 −2000 −3000 −4000 −5000 −6000
0 2.5 5 7.5 15
Doxorubicin
YHL013C OTU2
ZShift = 1.8
Z lm Score = −3.39
NG = 0
DB = 1
SM = 0
Delta_L
60 50 40 30 20 10 0 −10 −20 −30 −40 −50 −60
0 2.5 5 7.5 15
Doxorubicin
YHL013C OTU2
ZShift = −0.4
Z lm Score = 0.9
NG = 0
DB = 1
SM = 0
Delta_K
60 50 40 30 20 10 0 −10 −20 −30 −40 −50 −60
0 2.5 5 7.5 15
Doxorubicin
YHL013C OTU2
ZShift = −0.76
Z lm Score = 2.16
NG = 0
DB = 1
SM = 0
Delta_r
0.6 0.4 0.2 0.0 −0.2 −0.4 −0.6
0 2.5 5 7.5 15
Doxorubicin
YHL013C OTU2
ZShift = −1.74
Z lm Score = −0.49
NG = 0
DB = 1
SM = 0
Delta_AUC
6000 5000 4000 3000 2000 1000 0 −1000 −2000 −3000 −4000 −5000 −6000
0 2.5 5 7.5 15
Doxorubicin

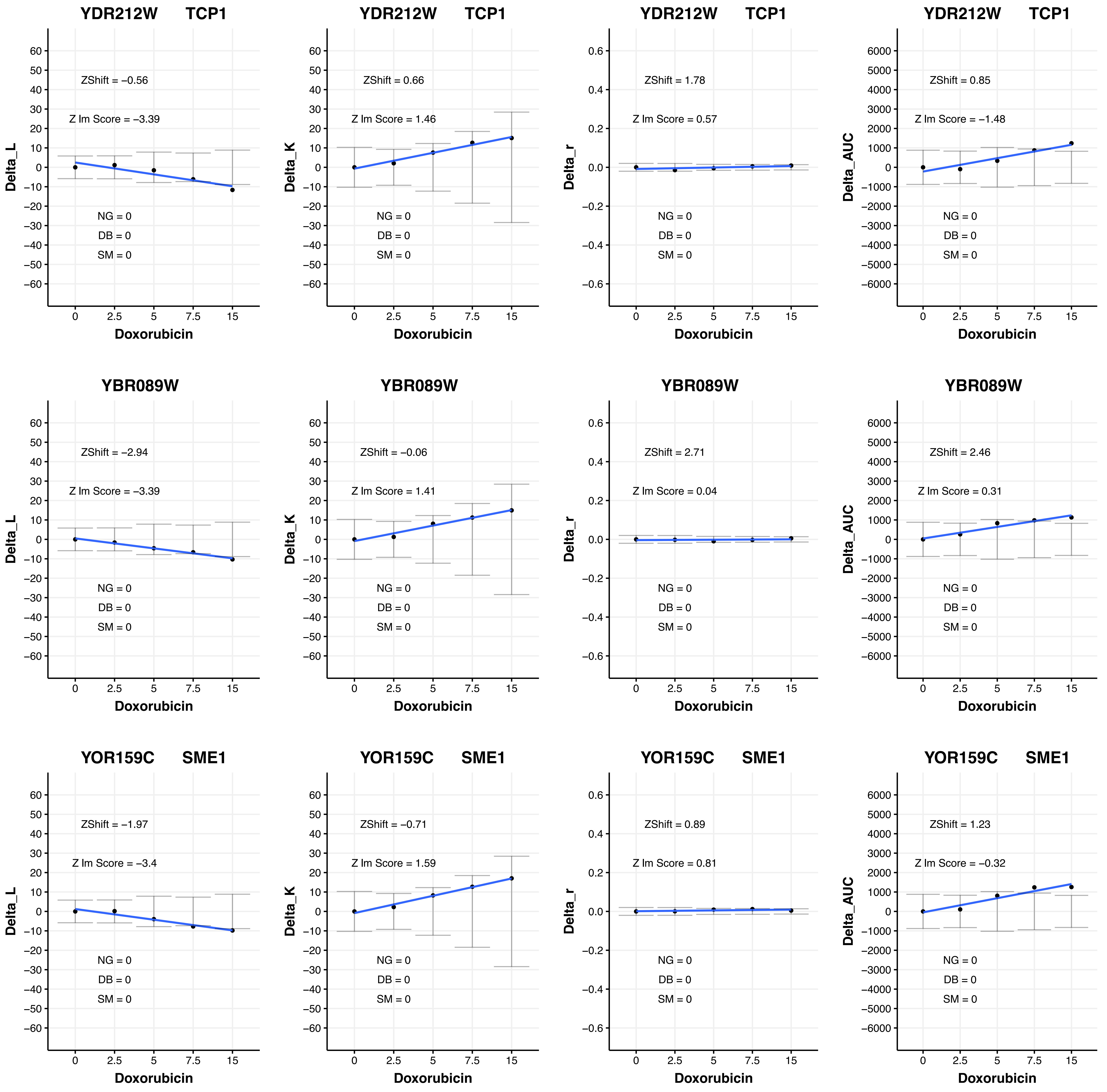
YDR212W TCP1
ZShift = −0.56
Z lm Score = −3.39
NG = 0
DB = 0
SM = 0
Delta_L
Doxorubicin
YDR212W TCP1
ZShift = 0.66
Z lm Score = 1.46
NG = 0
DB = 0
SM = 0
Delta_K
Doxorubicin
YDR212W TCP1
ZShift = 1.78
Z lm Score = 0.57
NG = 0
DB = 0
SM = 0
Delta_r
Doxorubicin
YDR212W TCP1
ZShift = 0.85
Z lm Score = −1.48
NG = 0
DB = 0
SM = 0
Delta_AUC
Doxorubicin
YBR089W
ZShift = −2.94
Z lm Score = −3.39
NG = 0
DB = 0
SM = 0
Delta_L
Doxorubicin
YBR089W
ZShift = −0.06
Z lm Score = 1.41
NG = 0
DB = 0
SM = 0
Delta_K
Doxorubicin
YBR089W
ZShift = 2.71
Z lm Score = 0.04
NG = 0
DB = 0
SM = 0
Delta_r
Doxorubicin
YBR089W
ZShift = 2.46
Z lm Score = 0.31
NG = 0
DB = 0
SM = 0
Delta_AUC
Doxorubicin
YOR159C SME1
ZShift = −1.97
Z lm Score = −3.4
NG = 0
DB = 0
SM = 0
Delta_L
Doxorubicin
YOR159C SME1
ZShift = −0.71
Z lm Score = 1.59
NG = 0
DB = 0
SM = 0
Delta_K
Doxorubicin
YOR159C SME1
ZShift = 0.89
Z lm Score = 0.81
NG = 0
DB = 0
SM = 0
Delta_r
Doxorubicin
YOR159C SME1
ZShift = 1.23
Z lm Score = −0.32
NG = 0
DB = 0
SM = 0
Delta_AUC
Doxorubicin

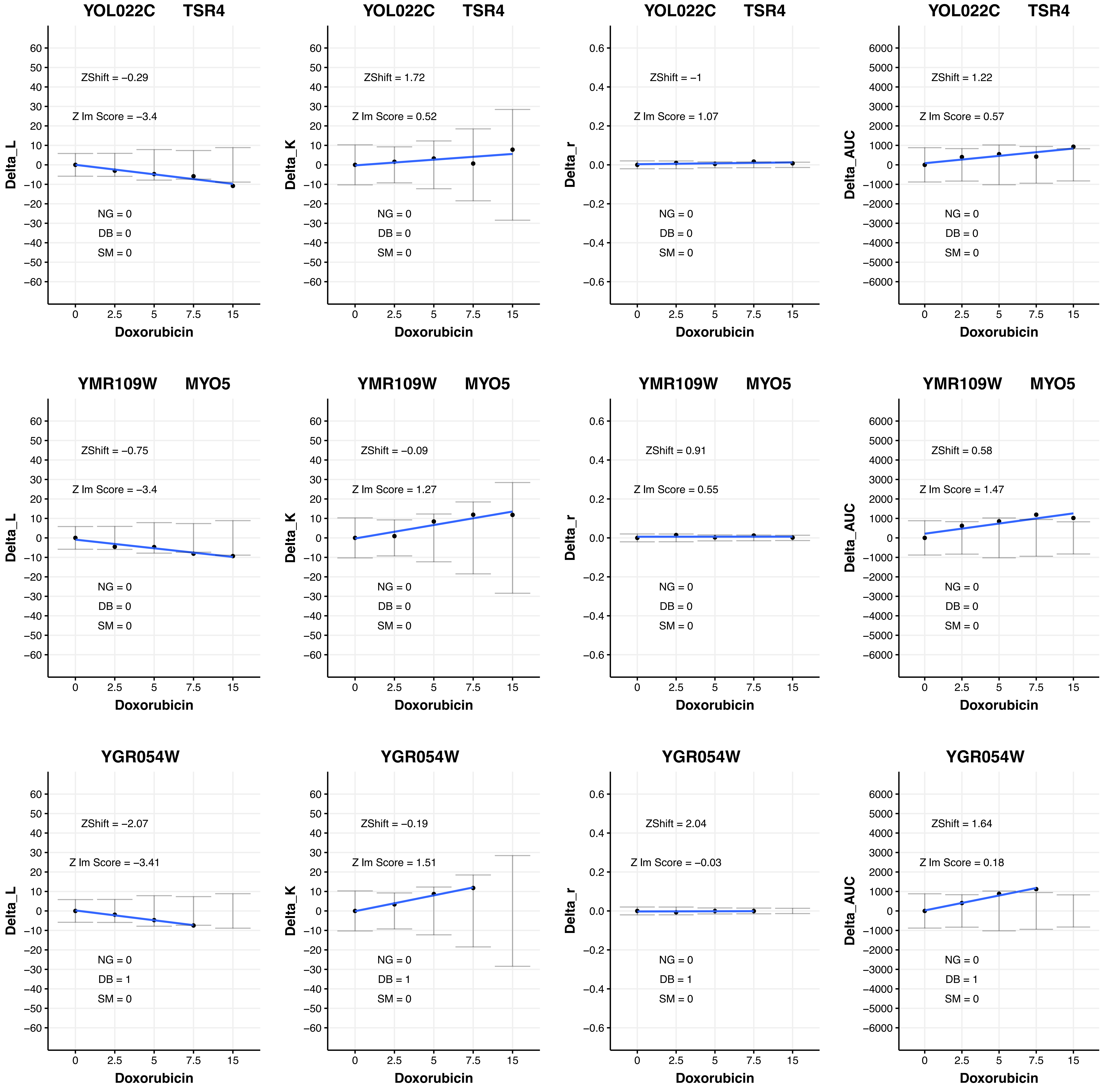
YOL022C TSR4
ZShift = −0.29
Z lm Score = −3.4
NG = 0
DB = 0
SM = 0
Delta_L
Doxorubicin
YOL022C TSR4
ZShift = 1.72
Z lm Score = 0.52
NG = 0
DB = 0
SM = 0
Delta_K
Doxorubicin
YOL022C TSR4
ZShift = −1
Z lm Score = 1.07
NG = 0
DB = 0
SM = 0
Delta_r
Doxorubicin
YOL022C TSR4
ZShift = 1.22
Z lm Score = 0.57
NG = 0
DB = 0
SM = 0
Delta_AUC
Doxorubicin
YMR109W MYO5
ZShift = −0.75
Z lm Score = −3.4
NG = 0
DB = 0
SM = 0
Delta_L
Doxorubicin
YMR109W MYO5
ZShift = −0.09
Z lm Score = 1.27
NG = 0
DB = 0
SM = 0
Delta_K
Doxorubicin
YMR109W MYO5
ZShift = 0.91
Z lm Score = 0.55
NG = 0
DB = 0
SM = 0
Delta_r
Doxorubicin
YMR109W MYO5
ZShift = 0.58
Z lm Score = 1.47
NG = 0
DB = 0
SM = 0
Delta_AUC
Doxorubicin
YGR054W
ZShift = −2.07
Z lm Score = −3.41
NG = 0
DB = 1
SM = 0
Delta_L
Doxorubicin
YGR054W
ZShift = −0.19
Z lm Score = 1.51
NG = 0
DB = 1
SM = 0
Delta_K
Doxorubicin
YGR054W
ZShift = 2.04
Z lm Score = −0.03
NG = 0
DB = 1
SM = 0
Delta_r
Doxorubicin
YGR054W
ZShift = 1.64
Z lm Score = 0.18
NG = 0
DB = 1
SM = 0
Delta_AUC
Doxorubicin

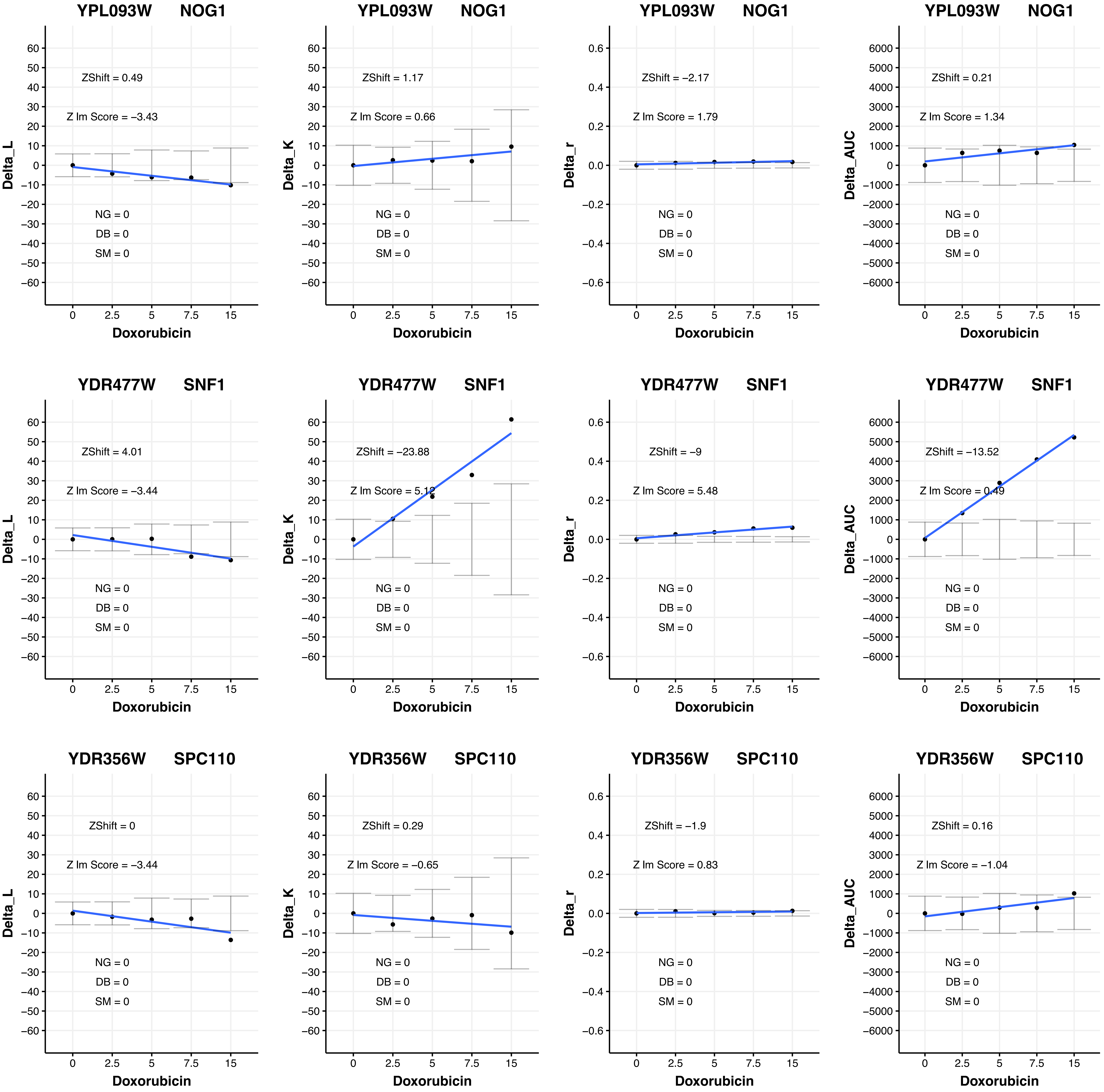
YPL093W NOG1
ZShift = 0.49
Z lm Score = −3.43
NG = 0
DB = 0
SM = 0
Delta_L
Doxorubicin
YPL093W NOG1
ZShift = 1.17
Z lm Score = 0.66
NG = 0
DB = 0
SM = 0
Delta_K
Doxorubicin
YPL093W NOG1
ZShift = −2.17
Z lm Score = 1.79
NG = 0
DB = 0
SM = 0
Delta_r
Doxorubicin
YPL093W NOG1
ZShift = 0.21
Z lm Score = 1.34
NG = 0
DB = 0
SM = 0
Delta_AUC
Doxorubicin
YDR477W SNF1
ZShift = 4.01
Z lm Score = −3.44
NG = 0
DB = 0
SM = 0
Delta_L
Doxorubicin
YDR477W SNF1
ZShift = −23.88
Z lm Score = 5.12
NG = 0
DB = 0
SM = 0
Delta_K
Doxorubicin
YDR477W SNF1
ZShift = −9
Z lm Score = 5.48
NG = 0
DB = 0
SM = 0
Delta_r
Doxorubicin
YDR477W SNF1
ZShift = −13.52
Z lm Score = 0.49
NG = 0
DB = 0
SM = 0
Delta_AUC
Doxorubicin
YDR356W SPC110
ZShift = 0
Z lm Score = −3.44
NG = 0
DB = 0
SM = 0
Delta_L
Doxorubicin
YDR356W SPC110
ZShift = 0.29
Z lm Score = −0.65
NG = 0
DB = 0
SM = 0
Delta_K
Doxorubicin
YDR356W SPC110
ZShift = −1.9
Z lm Score = 0.83
NG = 0
DB = 0
SM = 0
Delta_r
Doxorubicin
YDR356W SPC110
ZShift = 0.16
Z lm Score = −1.04
NG = 0
DB = 0
SM = 0
Delta_AUC
Doxorubicin

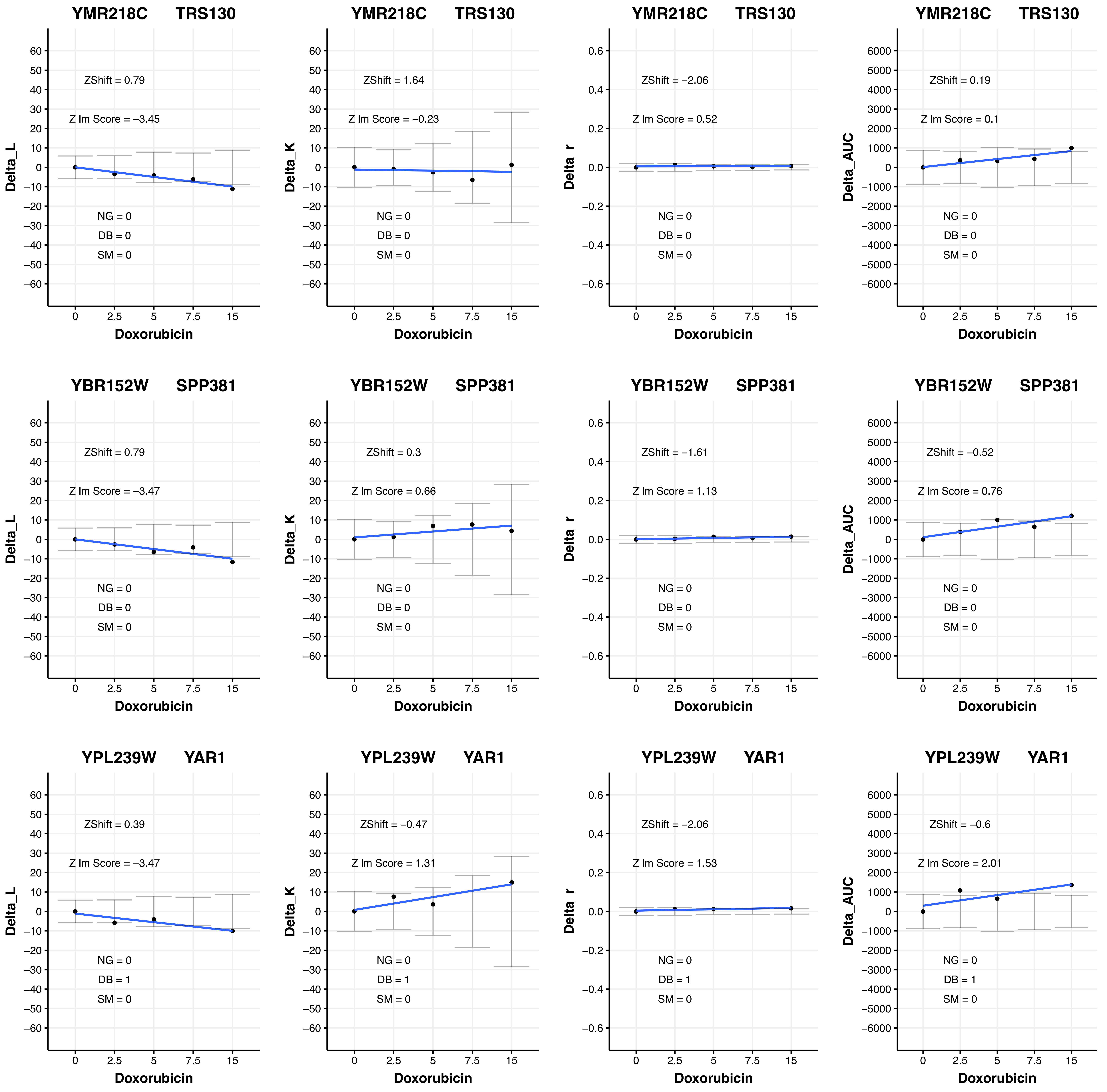

YMR218C TRS130
ZShift = 0.79
Z lm Score = −3.45
NG = 0
DB = 0
SM = 0
Delta_L
Doxorubicin
YMR218C TRS130
ZShift = 1.64
Z lm Score = −0.23
NG = 0
DB = 0
SM = 0
Delta_K
Doxorubicin
YMR218C TRS130
ZShift = −2.06
Z lm Score = 0.52
NG = 0
DB = 0
SM = 0
Delta_r
Doxorubicin
YMR218C TRS130
ZShift = 0.19
Z lm Score = 0.1
NG = 0
DB = 0
SM = 0
Delta_AUC
Doxorubicin
YBR152W SPP381
ZShift = 0.79
Z lm Score = −3.47
NG = 0
DB = 0
SM = 0
Delta_L
Doxorubicin
YBR152W SPP381
ZShift = 0.3
Z lm Score = 0.66
NG = 0
DB = 0
SM = 0
Delta_K
Doxorubicin
YBR152W SPP381
ZShift = −1.61
Z lm Score = 1.13
NG = 0
DB = 0
SM = 0
Delta_r
Doxorubicin
YBR152W SPP381
ZShift = −0.52
Z lm Score = 0.76
NG = 0
DB = 0
SM = 0
Delta_AUC
Doxorubicin
YPL239W YAR1
ZShift = 0.39
Z lm Score = −3.47
NG = 0
DB = 1
SM = 0
Delta_L
Doxorubicin
YPL239W YAR1
ZShift = −0.47
Z lm Score = 1.31
NG = 0
DB = 1
SM = 0
Delta_K
Doxorubicin
YPL239W YAR1
ZShift = −2.06
Z lm Score = 1.53
NG = 0
DB = 1
SM = 0
Delta_r
Doxorubicin
YPL239W YAR1
ZShift = −0.6
Z lm Score = 2.01
NG = 0
DB = 1
SM = 0
Delta_AUC
Doxorubicin

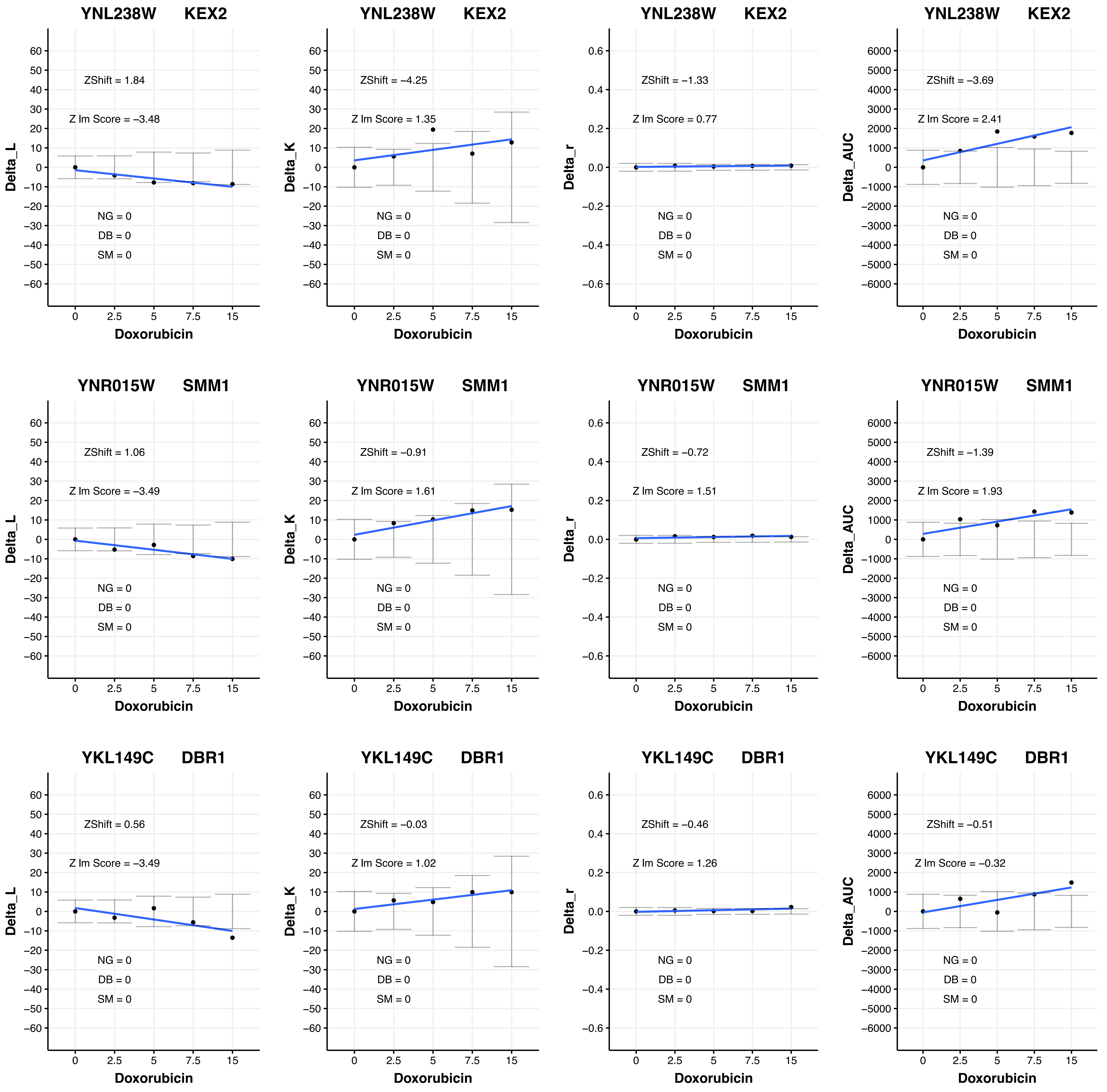
YNL238W KEX2
ZShift = 1.84
Z lm Score = −3.48
NG = 0
DB = 0
SM = 0
Delta_L
Doxorubicin
YNL238W KEX2
ZShift = −4.25
Z lm Score = 1.35
NG = 0
DB = 0
SM = 0
Delta_K
Doxorubicin
YNL238W KEX2
ZShift = −1.33
Z lm Score = 0.77
NG = 0
DB = 0
SM = 0
Delta_r
Doxorubicin
YNL238W KEX2
ZShift = −3.69
Z lm Score = 2.41
NG = 0
DB = 0
SM = 0
Delta_AUC
Doxorubicin
YNR015W SMM1
ZShift = 1.06
Z lm Score = −3.49
NG = 0
DB = 0
SM = 0
Delta_L
Doxorubicin
YNR015W SMM1
ZShift = −0.91
Z lm Score = 1.61
NG = 0
DB = 0
SM = 0
Delta_K
Doxorubicin
YNR015W SMM1
ZShift = −0.72
Z lm Score = 1.51
NG = 0
DB = 0
SM = 0
Delta_r
Doxorubicin
YNR015W SMM1
ZShift = −1.39
Z lm Score = 1.93
NG = 0
DB = 0
SM = 0
Delta_AUC
Doxorubicin
YKL149C DBR1
ZShift = 0.56
Z lm Score = −3.49
NG = 0
DB = 0
SM = 0
Delta_L
Doxorubicin
YKL149C DBR1
ZShift = −0.03
Z lm Score = 1.02
NG = 0
DB = 0
SM = 0
Delta_K
Doxorubicin
YKL149C DBR1
ZShift = −0.46
Z lm Score = 1.26
NG = 0
DB = 0
SM = 0
Delta_r
Doxorubicin
YKL149C DBR1
ZShift = −0.51
Z lm Score = −0.32
NG = 0
DB = 0
SM = 0
Delta_AUC
Doxorubicin

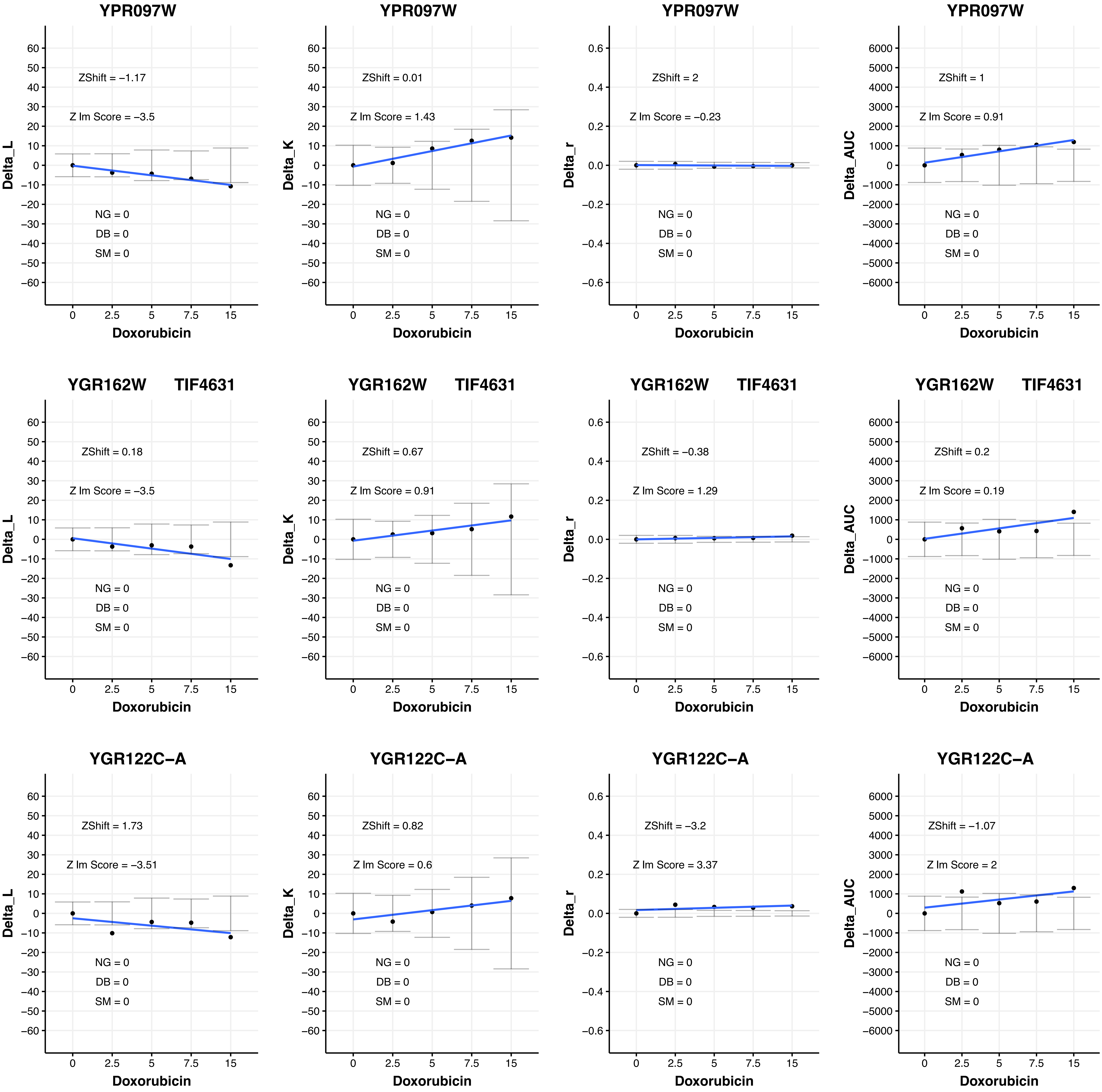
YPR097W
ZShift = −1.17
Z lm Score = −3.5
NG = 0
DB = 0
SM = 0
Delta_L
Doxorubicin
YPR097W
ZShift = 0.01
Z lm Score = 1.43
NG = 0
DB = 0
SM = 0
Delta_K
Doxorubicin
YPR097W
ZShift = 2
Z lm Score = −0.23
NG = 0
DB = 0
SM = 0
Delta_r
Doxorubicin
YPR097W
ZShift = 1
Z lm Score = 0.91
NG = 0
DB = 0
SM = 0
Delta_AUC
Doxorubicin
YGR162W TIF4631
ZShift = 0.18
Z lm Score = −3.5
NG = 0
DB = 0
SM = 0
Delta_L
Doxorubicin
YGR162W TIF4631
ZShift = 0.67
Z lm Score = 0.91
NG = 0
DB = 0
SM = 0
Delta_K
Doxorubicin
YGR162W TIF4631
ZShift = −0.38
Z lm Score = 1.29
NG = 0
DB = 0
SM = 0
Delta_r
Doxorubicin
YGR162W TIF4631
ZShift = 0.2
Z lm Score = 0.19
NG = 0
DB = 0
SM = 0
Delta_AUC
Doxorubicin
YGR122C−A
ZShift = 1.73
Z lm Score = −3.51
NG = 0
DB = 0
SM = 0
Delta_L
Doxorubicin
YGR122C−A
ZShift = 0.82
Z lm Score = 0.6
NG = 0
DB = 0
SM = 0
Delta_K
Doxorubicin
YGR122C−A
ZShift = −3.2
Z lm Score = 3.37
NG = 0
DB = 0
SM = 0
Delta_r
Doxorubicin
YGR122C−A
ZShift = −1.07
Z lm Score = 2
NG = 0
DB = 0
SM = 0
Delta_AUC
Doxorubicin

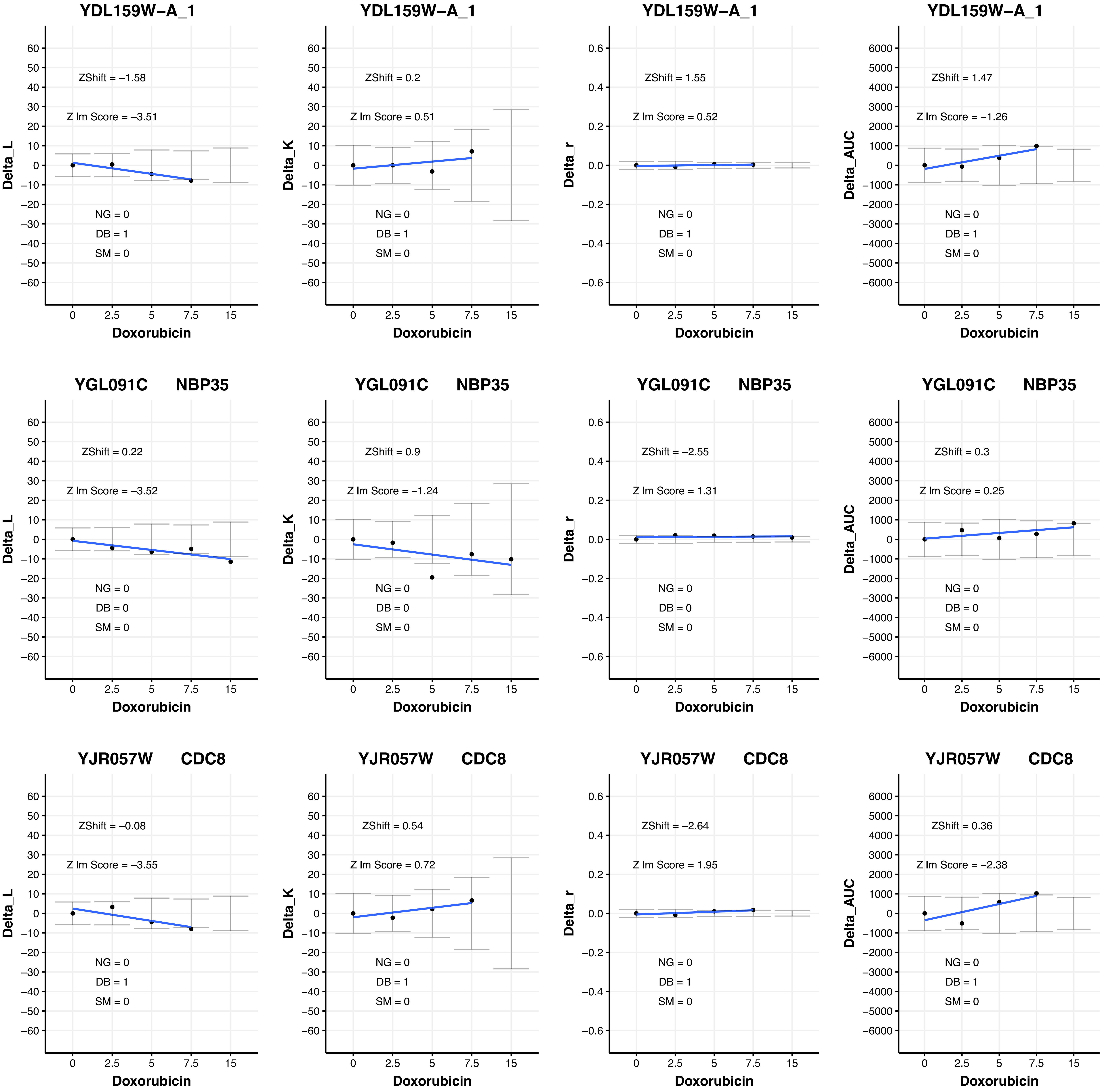

YDL159W−A_1
ZShift = −1.58
Z lm Score = −3.51
NG = 0
DB = 1
SM = 0
Delta_L
Doxorubicin
YDL159W−A_1
ZShift = 0.2
Z lm Score = 0.51
NG = 0
DB = 1
SM = 0
Delta_K
Doxorubicin
YDL159W−A_1
ZShift = 1.55
Z lm Score = 0.52
NG = 0
DB = 1
SM = 0
Delta_r
Doxorubicin
YDL159W−A_1
ZShift = 1.47
Z lm Score = −1.26
NG = 0
DB = 1
SM = 0
Delta_AUC
Doxorubicin
YGL091C NBP35
ZShift = 0.22
Z lm Score = −3.52
NG = 0
DB = 0
SM = 0
Delta_L
Doxorubicin
YGL091C NBP35
ZShift = 0.9
Z lm Score = −1.24
NG = 0
DB = 0
SM = 0
Delta_K
Doxorubicin
YGL091C NBP35
ZShift = −2.55
Z lm Score = 1.31
NG = 0
DB = 0
SM = 0
Delta_r
Doxorubicin
YGL091C NBP35
ZShift = 0.3
Z lm Score = 0.25
NG = 0
DB = 0
SM = 0
Delta_AUC
Doxorubicin
YJR057W CDC8
ZShift = −0.08
Z lm Score = −3.55
NG = 0
DB = 1
SM = 0
Delta_L
Doxorubicin
YJR057W CDC8
ZShift = 0.54
Z lm Score = 0.72
NG = 0
DB = 1
SM = 0
Delta_K
Doxorubicin
YJR057W CDC8
ZShift = −2.64
Z lm Score = 1.95
NG = 0
DB = 1
SM = 0
Delta_r
Doxorubicin
YJR057W CDC8
ZShift = 0.36
Z lm Score = −2.38
NG = 0
DB = 1
SM = 0
Delta_AUC
Doxorubicin

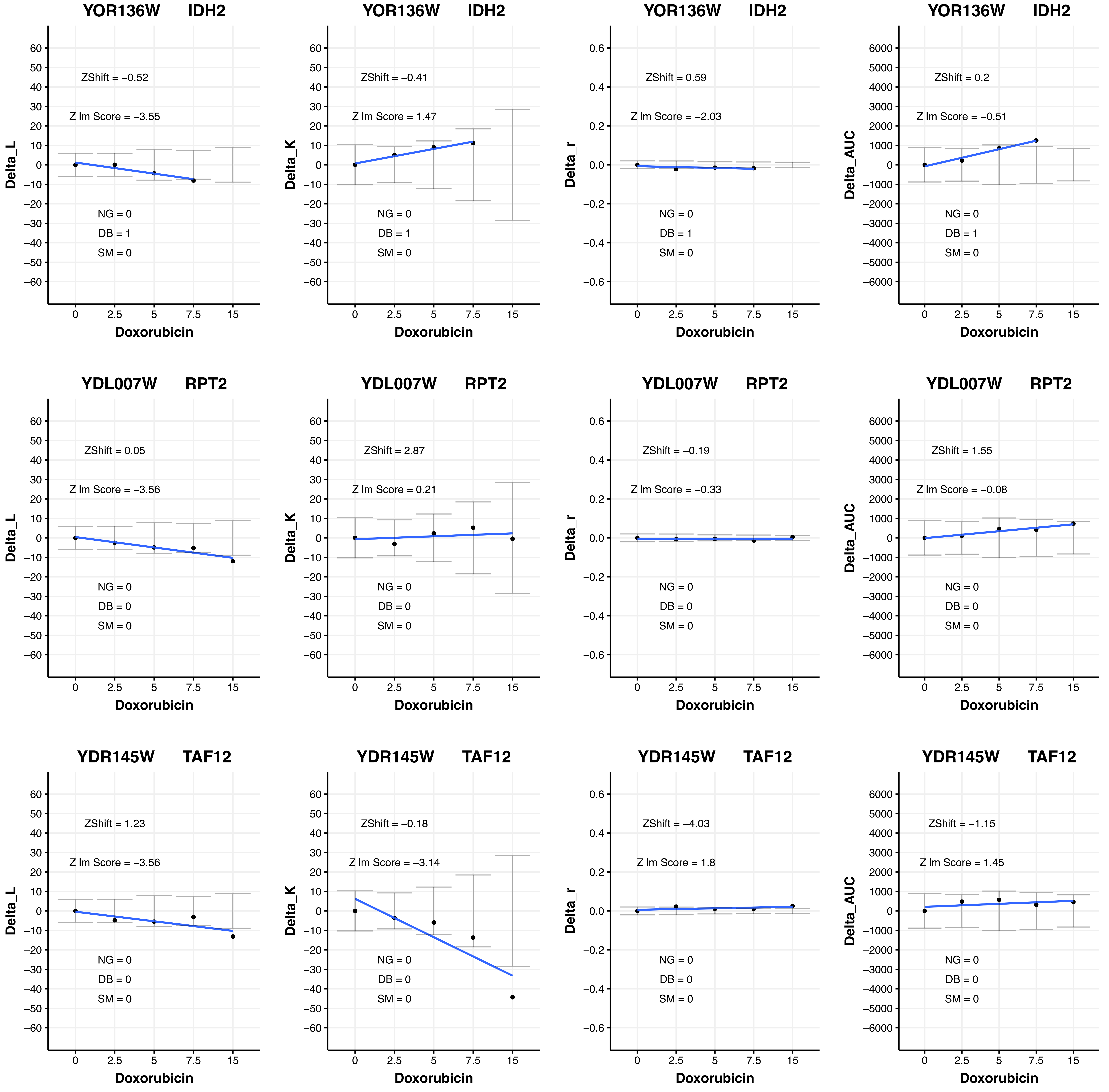
YOR136W IDH2
Delta_L
ZShift = −0.52
Z lm Score = −3.55
NG = 0
DB = 1
SM = 0
Doxorubicin
YOR136W IDH2
Delta_K
ZShift = −0.41
Z lm Score = 1.47
NG = 0
DB = 1
SM = 0
Doxorubicin
YOR136W IDH2
Delta_r
ZShift = 0.59
Z lm Score = −2.03
NG = 0
DB = 1
SM = 0
Doxorubicin
YOR136W IDH2
Delta_AUC
ZShift = 0.2
Z lm Score = −0.51
NG = 0
DB = 1
SM = 0
Doxorubicin
YDL007W RPT2
Delta_L
ZShift = 0.05
Z lm Score = −3.56
NG = 0
DB = 0
SM = 0
Doxorubicin
YDL007W RPT2
Delta_K
ZShift = 2.87
Z lm Score = 0.21
NG = 0
DB = 0
SM = 0
Doxorubicin
YDL007W RPT2
Delta_r
ZShift = −0.19
Z lm Score = −0.33
NG = 0
DB = 0
SM = 0
Doxorubicin
YDL007W RPT2
Delta_AUC
ZShift = 1.55
Z lm Score = −0.08
NG = 0
DB = 0
SM = 0
Doxorubicin
YDR145W TAF12
Delta_L
ZShift = 1.23
Z lm Score = −3.56
NG = 0
DB = 0
SM = 0
Doxorubicin
YDR145W TAF12
Delta_K
ZShift = −0.18
Z lm Score = −3.14
NG = 0
DB = 0
SM = 0
Doxorubicin
YDR145W TAF12
Delta_r
ZShift = −4.03
Z lm Score = 1.8
NG = 0
DB = 0
SM = 0
Doxorubicin
YDR145W TAF12
Delta_AUC
ZShift = −1.15
Z lm Score = 1.45
NG = 0
DB = 0
SM = 0
Doxorubicin

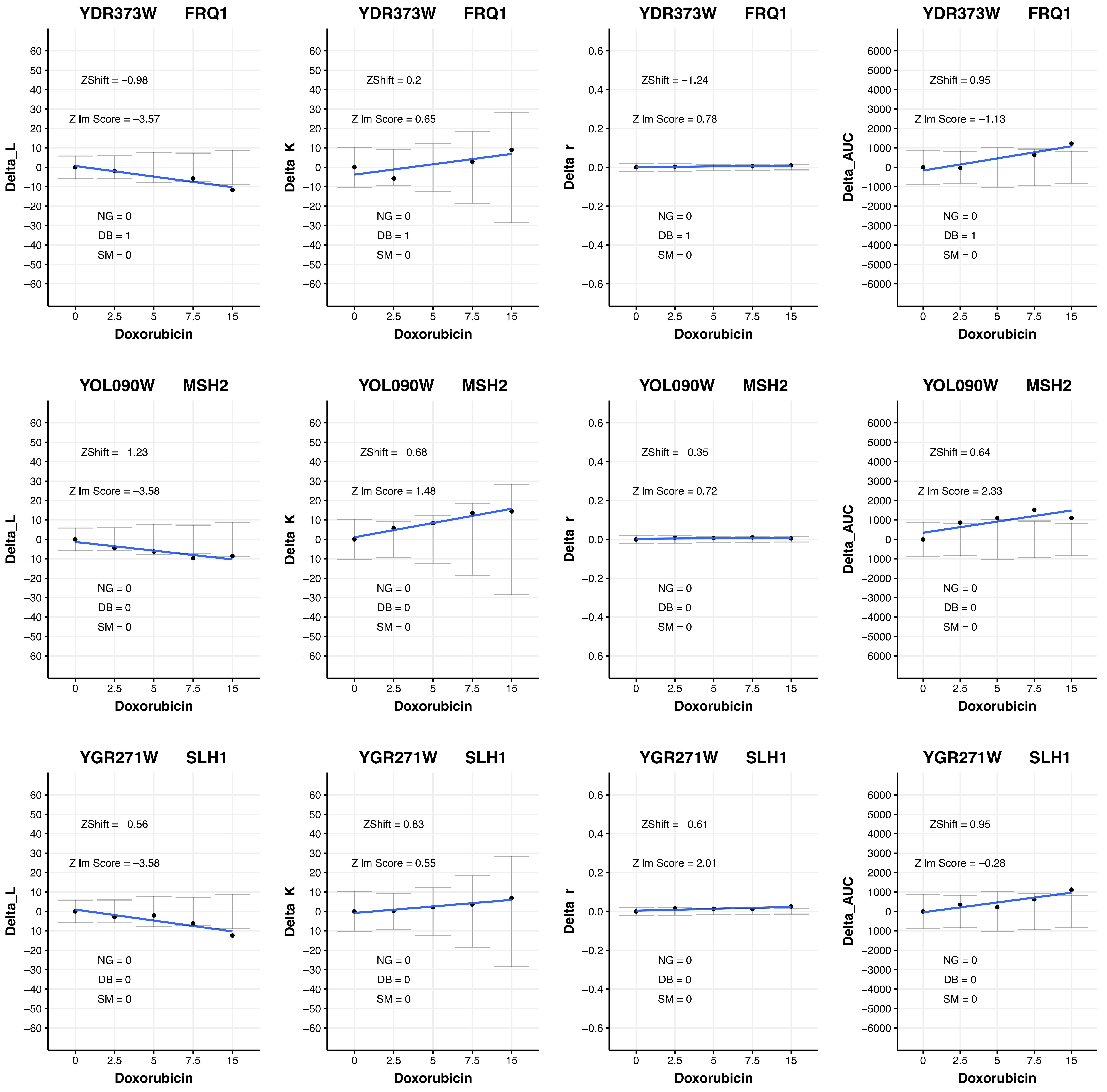
YDR373W FRQ1
ZShift = −0.98
Z lm Score = −3.57
NG = 0
DB = 1
SM = 0
Delta_L
Doxorubicin
YDR373W FRQ1
ZShift = 0.2
Z lm Score = 0.65
NG = 0
DB = 1
SM = 0
Delta_K
Doxorubicin
YDR373W FRQ1
ZShift = −1.24
Z lm Score = 0.78
NG = 0
DB = 1
SM = 0
Delta_r
Doxorubicin
YDR373W FRQ1
ZShift = 0.95
Z lm Score = −1.13
NG = 0
DB = 1
SM = 0
Delta_AUC
Doxorubicin
YOL090W MSH2
ZShift = −1.23
Z lm Score = −3.58
NG = 0
DB = 0
SM = 0
Delta_L
Doxorubicin
YOL090W MSH2
ZShift = −0.68
Z lm Score = 1.48
NG = 0
DB = 0
SM = 0
Delta_K
Doxorubicin
YOL090W MSH2
ZShift = −0.35
Z lm Score = 0.72
NG = 0
DB = 0
SM = 0
Delta_r
Doxorubicin
YOL090W MSH2
ZShift = 0.64
Z lm Score = 2.33
NG = 0
DB = 0
SM = 0
Delta_AUC
Doxorubicin
YGR271W SLH1
ZShift = −0.56
Z lm Score = −3.58
NG = 0
DB = 0
SM = 0
Delta_L
Doxorubicin
YGR271W SLH1
ZShift = 0.83
Z lm Score = 0.55
NG = 0
DB = 0
SM = 0
Delta_K
Doxorubicin
YGR271W SLH1
ZShift = −0.61
Z lm Score = 2.01
NG = 0
DB = 0
SM = 0
Delta_r
Doxorubicin
YGR271W SLH1
ZShift = 0.95
Z lm Score = −0.28
NG = 0
DB = 0
SM = 0
Delta_AUC
Doxorubicin

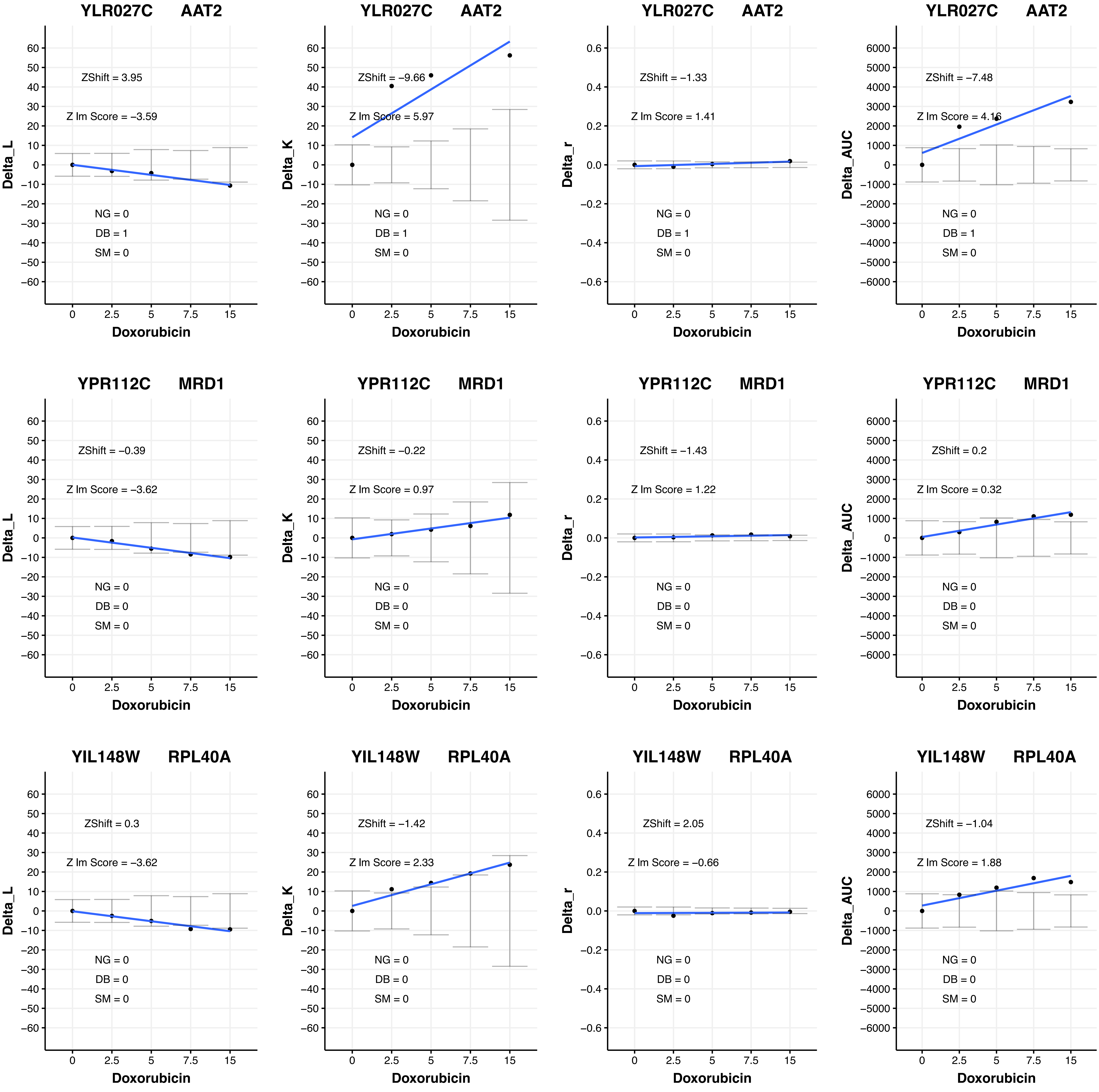

YLR027C AAT2
ZShift = 3.95
Z lm Score = −3.59
NG = 0
DB = 1
SM = 0
Delta_L
Doxorubicin
YLR027C AAT2
ZShift = −9.66
Z lm Score = 5.97
NG = 0
DB = 1
SM = 0
Delta_K
Doxorubicin
YLR027C AAT2
ZShift = −1.33
Z lm Score = 1.41
NG = 0
DB = 1
SM = 0
Delta_r
Doxorubicin
YLR027C AAT2
ZShift = −7.48
Z lm Score = 4.16
NG = 0
DB = 1
SM = 0
Delta_AUC
Doxorubicin
YPR112C MRD1
ZShift = −0.39
Z lm Score = −3.62
NG = 0
DB = 0
SM = 0
Delta_L
Doxorubicin
YPR112C MRD1
ZShift = −0.22
Z lm Score = 0.97
NG = 0
DB = 0
SM = 0
Delta_K
Doxorubicin
YPR112C MRD1
ZShift = −1.43
Z lm Score = 1.22
NG = 0
DB = 0
SM = 0
Delta_r
Doxorubicin
YPR112C MRD1
ZShift = 0.2
Z lm Score = 0.32
NG = 0
DB = 0
SM = 0
Delta_AUC
Doxorubicin
YIL148W RPL40A
ZShift = 0.3
Z lm Score = −3.62
NG = 0
DB = 0
SM = 0
Delta_L
Doxorubicin
YIL148W RPL40A
ZShift = −1.42
Z lm Score = 2.33
NG = 0
DB = 0
SM = 0
Delta_K
Doxorubicin
YIL148W RPL40A
ZShift = 2.05
Z lm Score = −0.66
NG = 0
DB = 0
SM = 0
Delta_r
Doxorubicin
YIL148W RPL40A
ZShift = −1.04
Z lm Score = 1.88
NG = 0
DB = 0
SM = 0
Delta_AUC
Doxorubicin

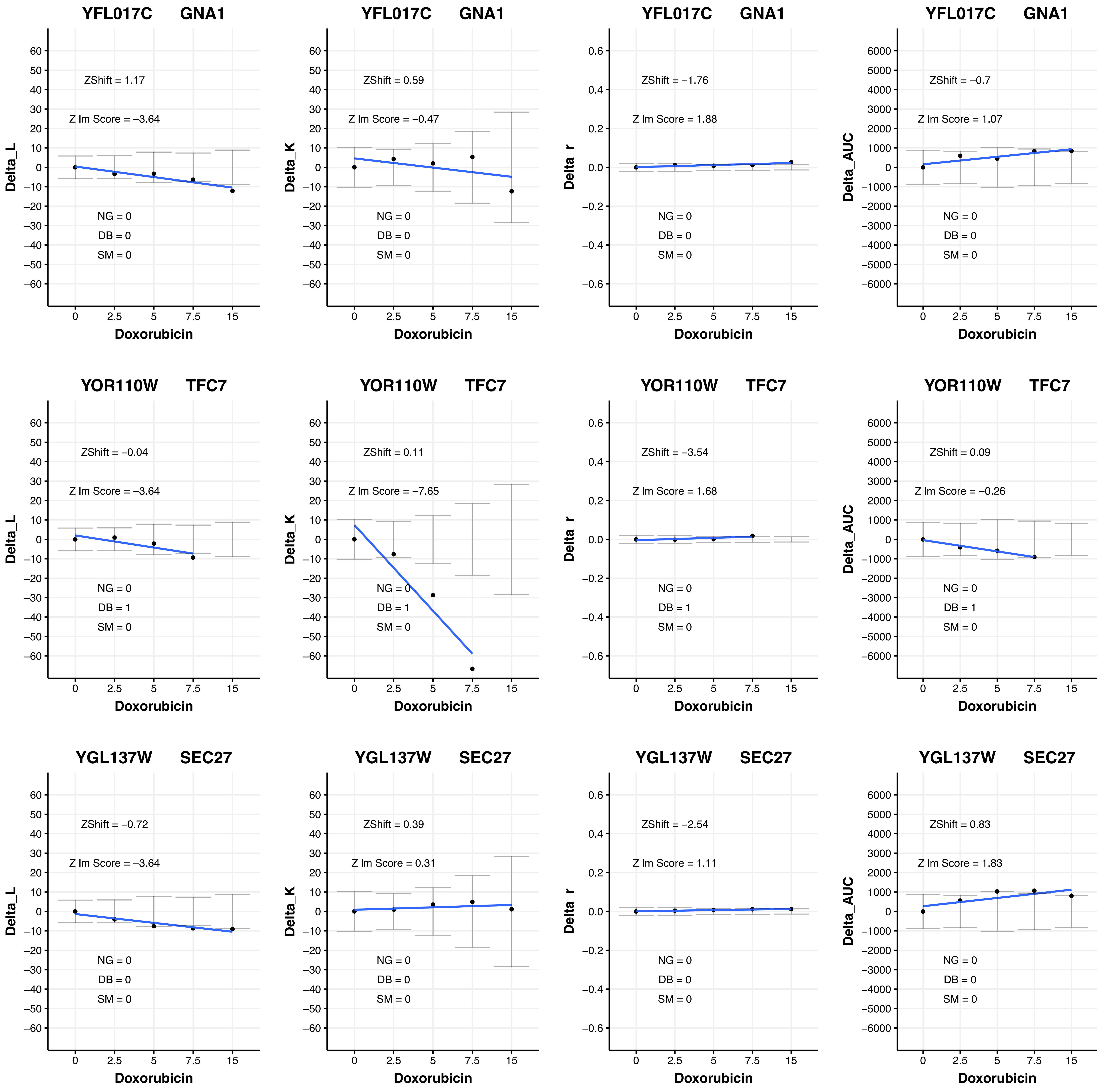
YFL017C GNA1
ZShift = 1.17
Z lm Score = −3.64
NG = 0
DB = 0
SM = 0
Delta_L
Doxorubicin
YFL017C GNA1
ZShift = 0.59
Z lm Score = −0.47
NG = 0
DB = 0
SM = 0
Delta_K
Doxorubicin
YFL017C GNA1
ZShift = −1.76
Z lm Score = 1.88
NG = 0
DB = 0
SM = 0
Delta_r
Doxorubicin
YFL017C GNA1
ZShift = −0.7
Z lm Score = 1.07
NG = 0
DB = 0
SM = 0
Delta_AUC
Doxorubicin
YOR110W TFC7
ZShift = −0.04
Z lm Score = −3.64
NG = 0
DB = 1
SM = 0
Delta_L
Doxorubicin
YOR110W TFC7
ZShift = 0.11
Z lm Score = −7.65
NG = 0
DB = 1
SM = 0
Delta_K
Doxorubicin
YOR110W TFC7
ZShift = −3.54
Z lm Score = 1.68
NG = 0
DB = 1
SM = 0
Delta_r
Doxorubicin
YOR110W TFC7
ZShift = 0.09
Z lm Score = −0.26
NG = 0
DB = 1
SM = 0
Delta_AUC
Doxorubicin
YGL137W SEC27
ZShift = −0.72
Z lm Score = −3.64
NG = 0
DB = 0
SM = 0
Delta_L
Doxorubicin
YGL137W SEC27
ZShift = 0.39
Z lm Score = 0.31
NG = 0
DB = 0
SM = 0
Delta_K
Doxorubicin
YGL137W SEC27
ZShift = −2.54
Z lm Score = 1.11
NG = 0
DB = 0
SM = 0
Delta_r
Doxorubicin
YGL137W SEC27
ZShift = 0.83
Z lm Score = 1.83
NG = 0
DB = 0
SM = 0
Delta_AUC
Doxorubicin

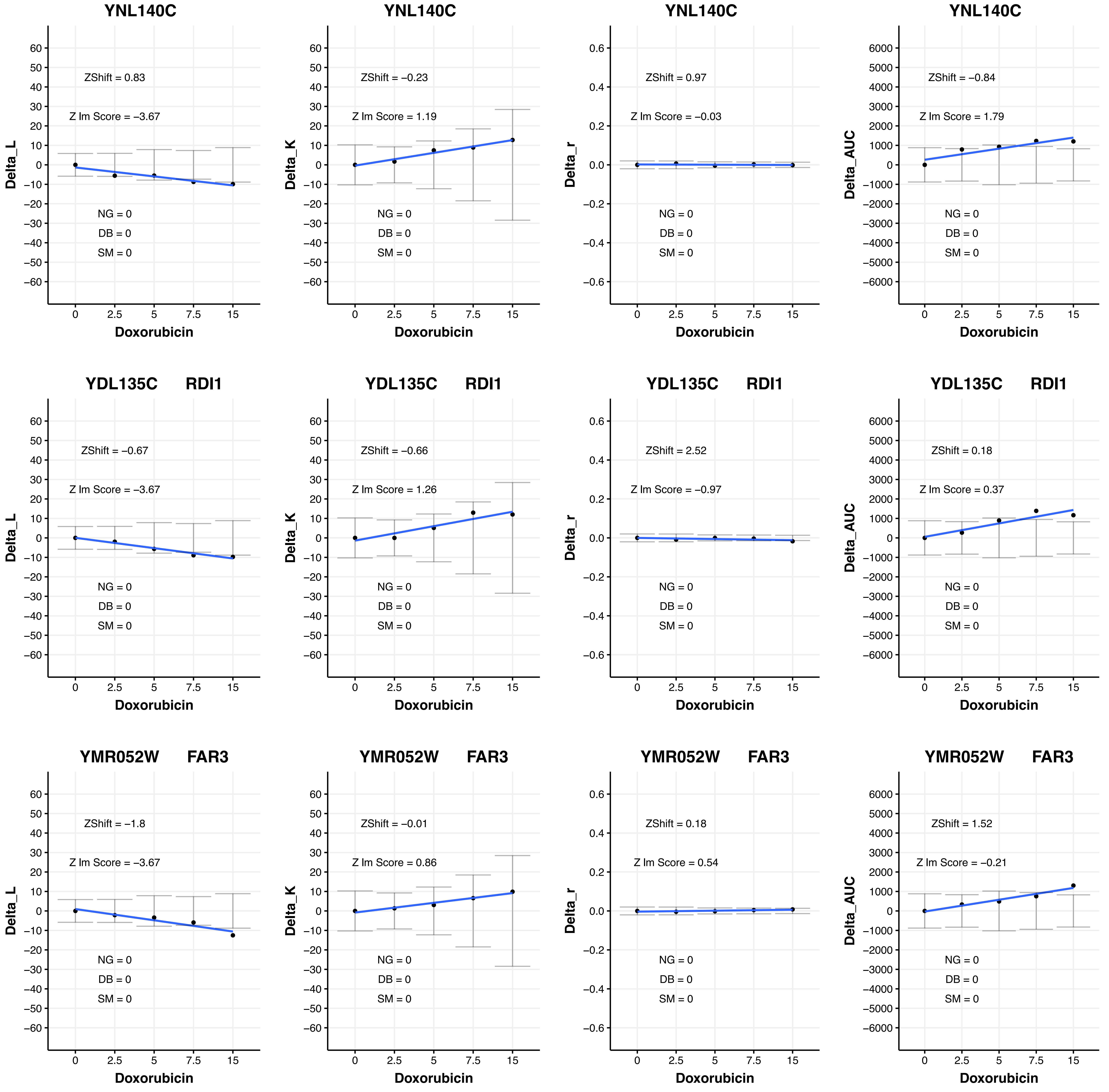

YNL140C
ZShift = 0.83
Z lm Score = −3.67
NG = 0
DB = 0
SM = 0
Delta_L
Doxorubicin
YNL140C
ZShift = −0.23
Z lm Score = 1.19
NG = 0
DB = 0
SM = 0
Delta_K
Doxorubicin
YNL140C
ZShift = 0.97
Z lm Score = −0.03
NG = 0
DB = 0
SM = 0
Delta_r
Doxorubicin
YNL140C
ZShift = −0.84
Z lm Score = 1.79
NG = 0
DB = 0
SM = 0
Delta_AUC
Doxorubicin
YDL135C RDI1
ZShift = −0.67
Z lm Score = −3.67
NG = 0
DB = 0
SM = 0
Delta_L
Doxorubicin
YDL135C RDI1
ZShift = −0.66
Z lm Score = 1.26
NG = 0
DB = 0
SM = 0
Delta_K
Doxorubicin
YDL135C RDI1
ZShift = 2.52
Z lm Score = −0.97
NG = 0
DB = 0
SM = 0
Delta_r
Doxorubicin
YDL135C RDI1
ZShift = 0.18
Z lm Score = 0.37
NG = 0
DB = 0
SM = 0
Delta_AUC
Doxorubicin
YMR052W FAR3
ZShift = −1.8
Z lm Score = −3.67
NG = 0
DB = 0
SM = 0
Delta_L
Doxorubicin
YMR052W FAR3
ZShift = −0.01
Z lm Score = 0.86
NG = 0
DB = 0
SM = 0
Delta_K
Doxorubicin
YMR052W FAR3
ZShift = 0.18
Z lm Score = 0.54
NG = 0
DB = 0
SM = 0
Delta_r
Doxorubicin
YMR052W FAR3
ZShift = 1.52
Z lm Score = −0.21
NG = 0
DB = 0
SM = 0
Delta_AUC
Doxorubicin

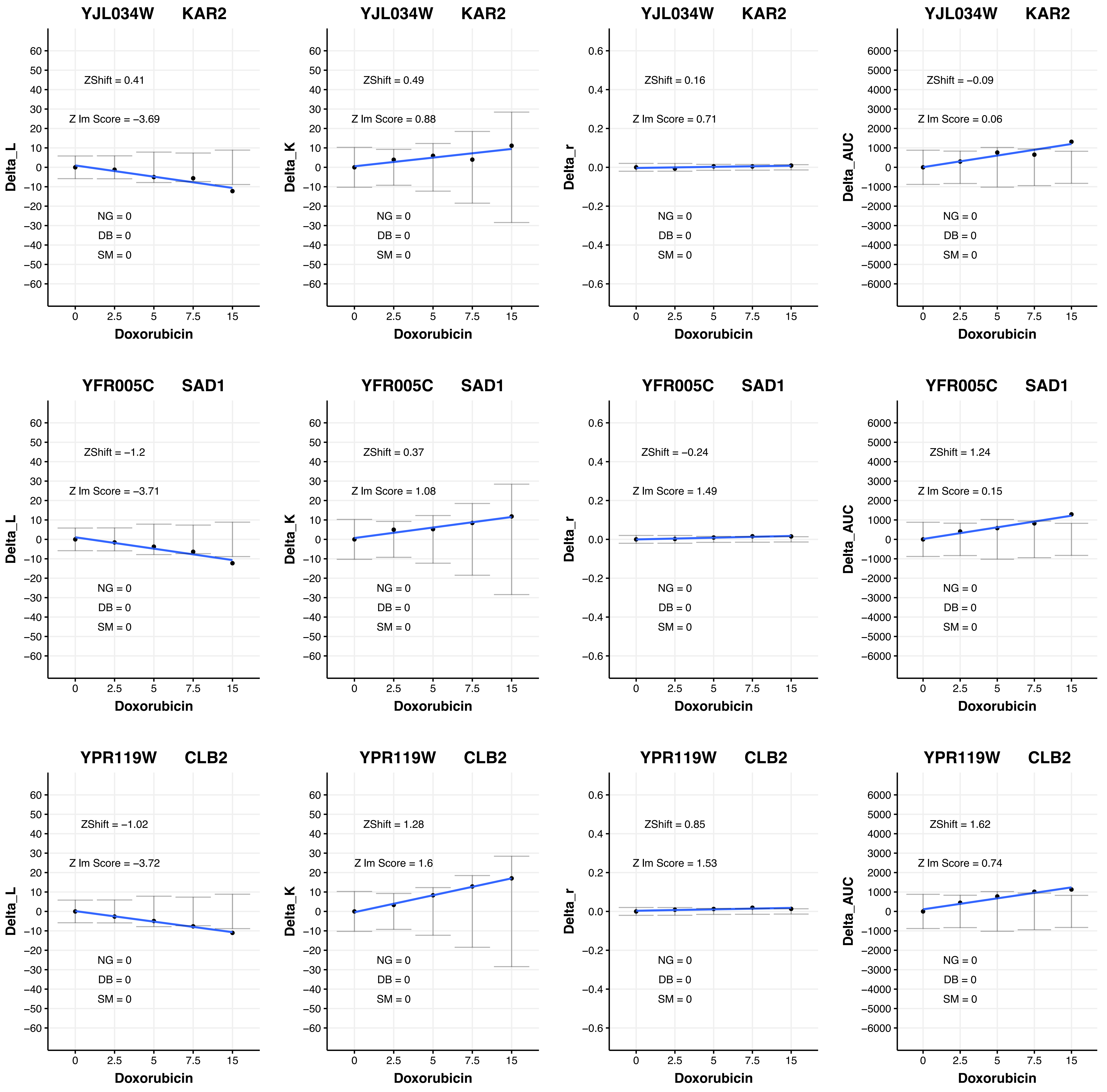

YJL034W KAR2
ZShift = 0.41
Z lm Score = −3.69
NG = 0
DB = 0
SM = 0
Delta_L
Doxorubicin
YJL034W KAR2
ZShift = 0.49
Z lm Score = 0.88
NG = 0
DB = 0
SM = 0
Delta_K
Doxorubicin
YJL034W KAR2
ZShift = 0.16
Z lm Score = 0.71
NG = 0
DB = 0
SM = 0
Delta_r
Doxorubicin
YJL034W KAR2
ZShift = −0.09
Z lm Score = 0.06
NG = 0
DB = 0
SM = 0
Delta_AUC
Doxorubicin
YFR005C SAD1
ZShift = −1.2
Z lm Score = −3.71
NG = 0
DB = 0
SM = 0
Delta_L
Doxorubicin
YFR005C SAD1
ZShift = 0.37
Z lm Score = 1.08
NG = 0
DB = 0
SM = 0
Delta_K
Doxorubicin
YFR005C SAD1
ZShift = −0.24
Z lm Score = 1.49
NG = 0
DB = 0
SM = 0
Delta_r
Doxorubicin
YFR005C SAD1
ZShift = 1.24
Z lm Score = 0.15
NG = 0
DB = 0
SM = 0
Delta_AUC
Doxorubicin
YPR119W CLB2
ZShift = −1.02
Z lm Score = −3.72
NG = 0
DB = 0
SM = 0
Delta_L
Doxorubicin
YPR119W CLB2
ZShift = 1.28
Z lm Score = 1.6
NG = 0
DB = 0
SM = 0
Delta_K
Doxorubicin
YPR119W CLB2
ZShift = 0.85
Z lm Score = 1.53
NG = 0
DB = 0
SM = 0
Delta_r
Doxorubicin
YPR119W CLB2
ZShift = 1.62
Z lm Score = 0.74
NG = 0
DB = 0
SM = 0
Delta_AUC
Doxorubicin

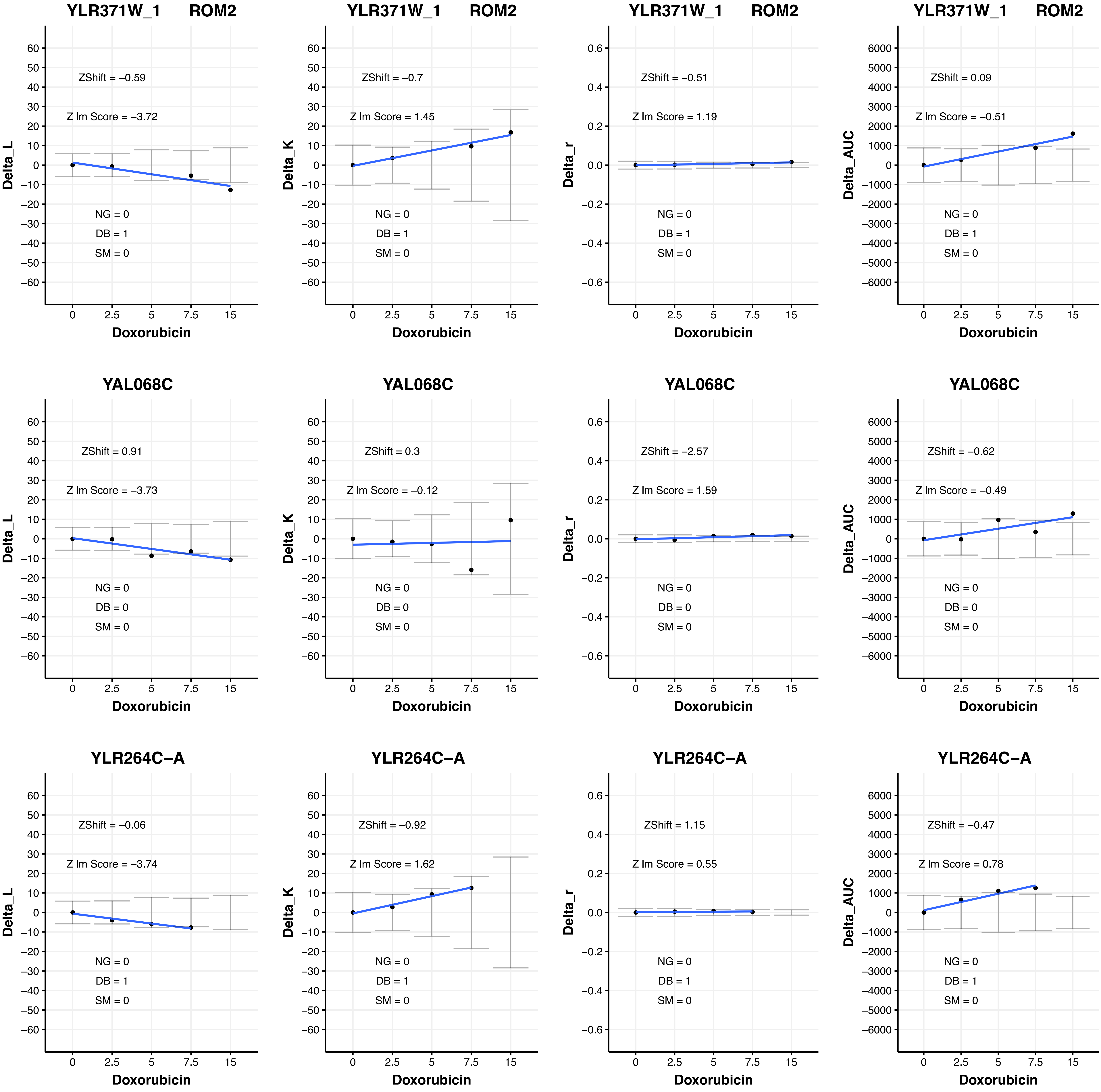

YLR371W_1 ROM2
ZShift = −0.59
Z lm Score = −3.72
NG = 0
DB = 1
SM = 0
Delta_L
Doxorubicin
YLR371W_1 ROM2
ZShift = −0.7
Z lm Score = 1.45
NG = 0
DB = 1
SM = 0
Delta_K
Doxorubicin
YLR371W_1 ROM2
ZShift = −0.51
Z lm Score = 1.19
NG = 0
DB = 1
SM = 0
Delta_r
Doxorubicin
YLR371W_1 ROM2
ZShift = 0.09
Z lm Score = −0.51
NG = 0
DB = 1
SM = 0
Delta_AUC
Doxorubicin
YAL068C
ZShift = 0.91
Z lm Score = −3.73
NG = 0
DB = 0
SM = 0
Delta_L
Doxorubicin
YAL068C
ZShift = 0.3
Z lm Score = −0.12
NG = 0
DB = 0
SM = 0
Delta_K
Doxorubicin
YAL068C
ZShift = −2.57
Z lm Score = 1.59
NG = 0
DB = 0
SM = 0
Delta_r
Doxorubicin
YAL068C
ZShift = −0.62
Z lm Score = −0.49
NG = 0
DB = 0
SM = 0
Delta_AUC
Doxorubicin
YLR264C−A
ZShift = −0.06
Z lm Score = −3.74
NG = 0
DB = 1
SM = 0
Delta_L
Doxorubicin
YLR264C−A
ZShift = −0.92
Z lm Score = 1.62
NG = 0
DB = 1
SM = 0
Delta_K
Doxorubicin
YLR264C−A
ZShift = 1.15
Z lm Score = 0.55
NG = 0
DB = 1
SM = 0
Delta_r
Doxorubicin
YLR264C−A
ZShift = −0.47
Z lm Score = 0.78
NG = 0
DB = 1
SM = 0
Delta_AUC
Doxorubicin

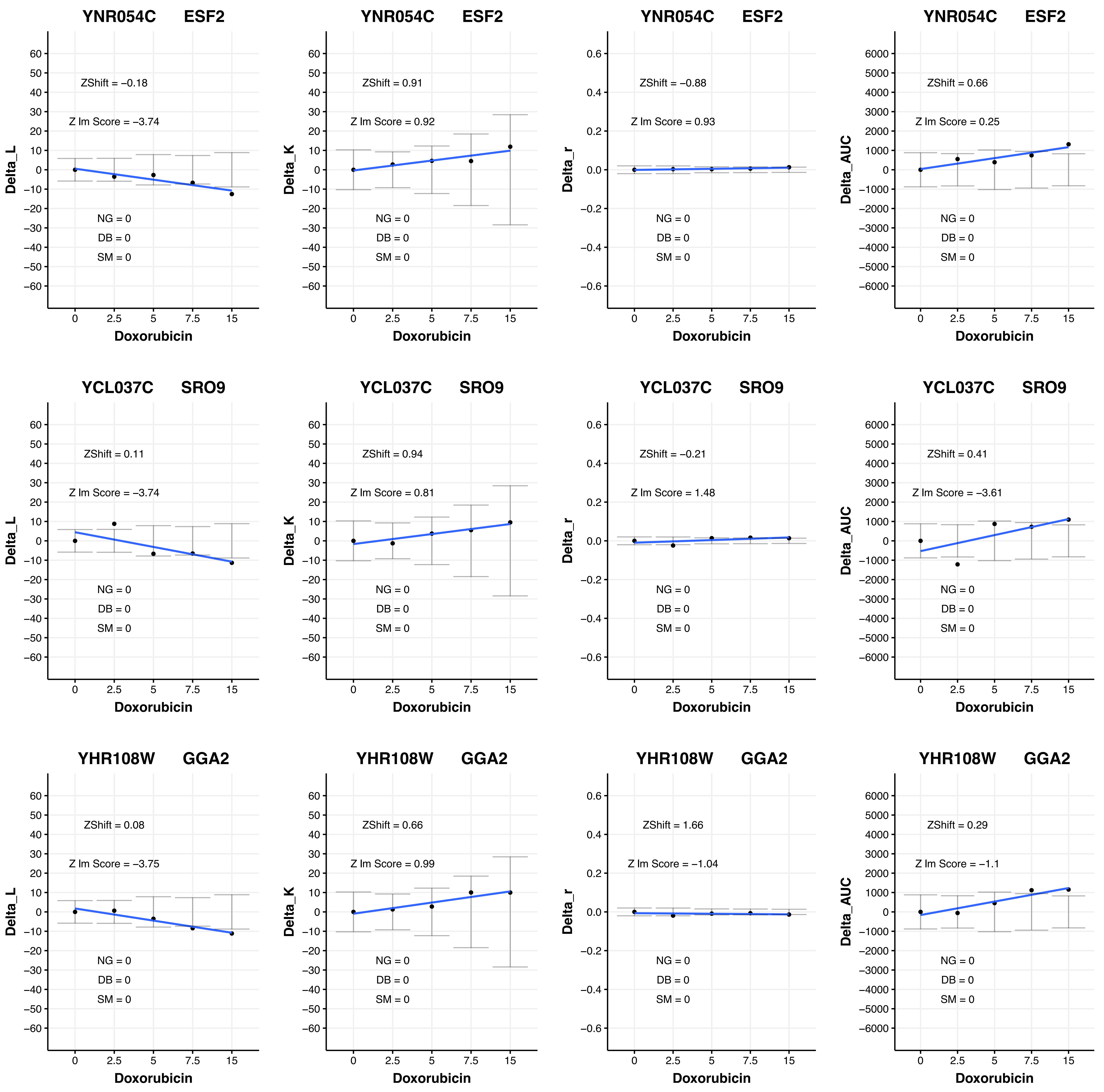
YNR054C ESF2
ZShift = −0.18
Z lm Score = −3.74
NG = 0
DB = 0
SM = 0
Delta_L
Doxorubicin
YNR054C ESF2
ZShift = 0.91
Z lm Score = 0.92
NG = 0
DB = 0
SM = 0
Delta_K
Doxorubicin
YNR054C ESF2
ZShift = −0.88
Z lm Score = 0.93
NG = 0
DB = 0
SM = 0
Delta_r
Doxorubicin
YNR054C ESF2
ZShift = 0.66
Z lm Score = 0.25
NG = 0
DB = 0
SM = 0
Delta_AUC
Doxorubicin
YCL037C SRO9
ZShift = 0.11
Z lm Score = −3.74
NG = 0
DB = 0
SM = 0
Delta_L
Doxorubicin
YCL037C SRO9
ZShift = 0.94
Z lm Score = 0.81
NG = 0
DB = 0
SM = 0
Delta_K
Doxorubicin
YCL037C SRO9
ZShift = −0.21
Z lm Score = 1.48
NG = 0
DB = 0
SM = 0
Delta_r
Doxorubicin
YCL037C SRO9
ZShift = 0.41
Z lm Score = −3.61
NG = 0
DB = 0
SM = 0
Delta_AUC
Doxorubicin
YHR108W GGA2
ZShift = 0.08
Z lm Score = −3.75
NG = 0
DB = 0
SM = 0
Delta_L
Doxorubicin
YHR108W GGA2
ZShift = 0.66
Z lm Score = 0.99
NG = 0
DB = 0
SM = 0
Delta_K
Doxorubicin
YHR108W GGA2
ZShift = 1.66
Z lm Score = −1.04
NG = 0
DB = 0
SM = 0
Delta_r
Doxorubicin
YHR108W GGA2
ZShift = 0.29
Z lm Score = −1.1
NG = 0
DB = 0
SM = 0
Delta_AUC
Doxorubicin

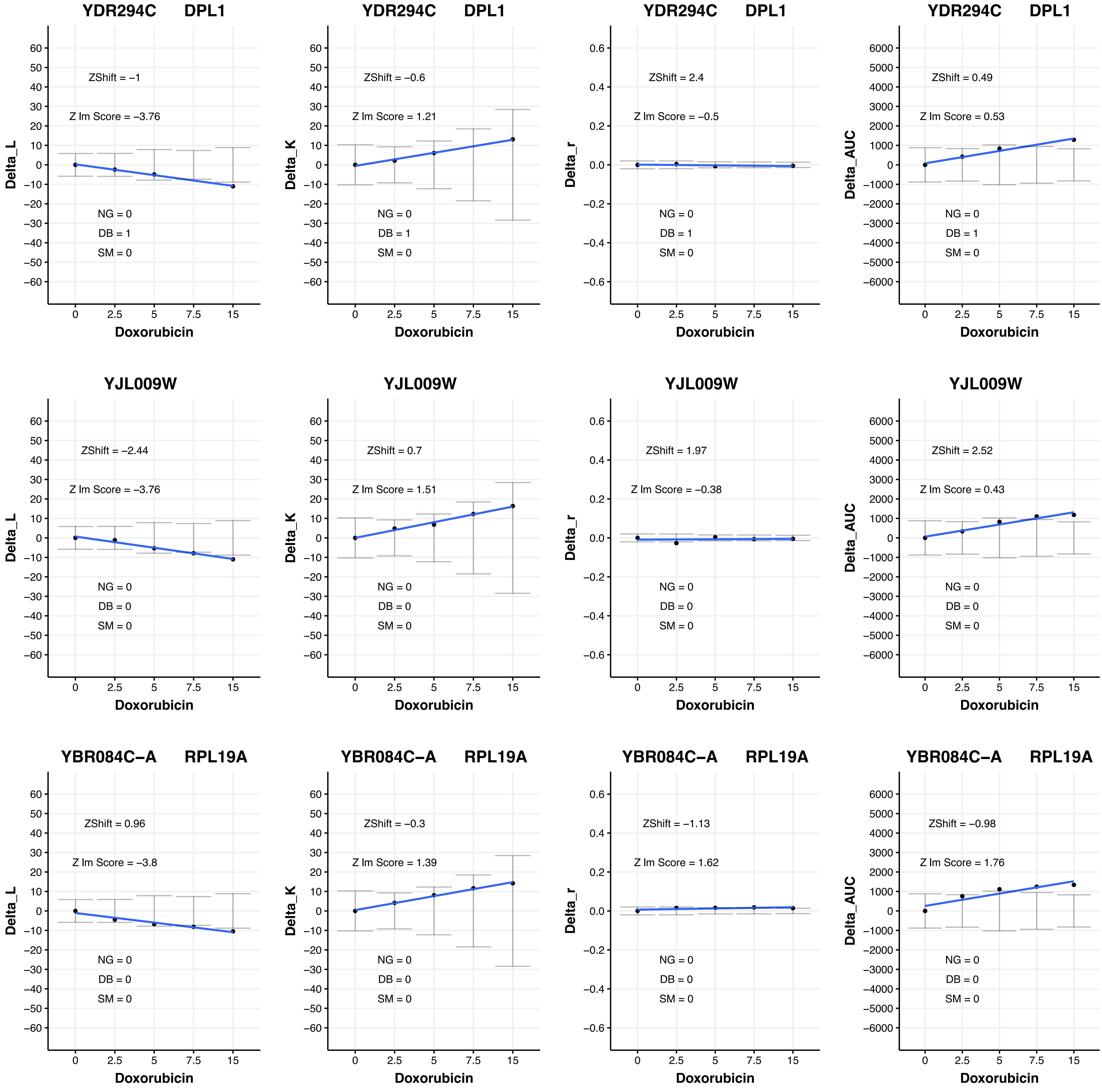
YDR294C DPL1
ZShift = −1
Z lm Score = −3.76
NG = 0
DB = 1
SM = 0
Delta_L
Doxorubicin
YDR294C DPL1
ZShift = −0.6
Z lm Score = 1.21
NG = 0
DB = 1
SM = 0
Delta_K
Doxorubicin
YDR294C DPL1
ZShift = 2.4
Z lm Score = −0.5
NG = 0
DB = 1
SM = 0
Delta_r
Doxorubicin
YDR294C DPL1
ZShift = 0.49
Z lm Score = 0.53
NG = 0
DB = 1
SM = 0
Delta_AUC
Doxorubicin
YJL009W
ZShift = −2.44
Z lm Score = −3.76
NG = 0
DB = 0
SM = 0
Delta_L
Doxorubicin
YJL009W
ZShift = 0.7
Z lm Score = 1.51
NG = 0
DB = 0
SM = 0
Delta_K
Doxorubicin
YJL009W
ZShift = 1.97
Z lm Score = −0.38
NG = 0
DB = 0
SM = 0
Delta_r
Doxorubicin
YJL009W
ZShift = 2.52
Z lm Score = 0.43
NG = 0
DB = 0
SM = 0
Delta_AUC
Doxorubicin
YBR084C−A RPL19A
ZShift = 0.96
Z lm Score = −3.8
NG = 0
DB = 0
SM = 0
Delta_L
Doxorubicin
YBR084C−A RPL19A
ZShift = −0.3
Z lm Score = 1.39
NG = 0
DB = 0
SM = 0
Delta_K
Doxorubicin
YBR084C−A RPL19A
ZShift = −1.13
Z lm Score = 1.62
NG = 0
DB = 0
SM = 0
Delta_r
Doxorubicin
YBR084C−A RPL19A
ZShift = −0.98
Z lm Score = 1.76
NG = 0
DB = 0
SM = 0
Delta_AUC
Doxorubicin

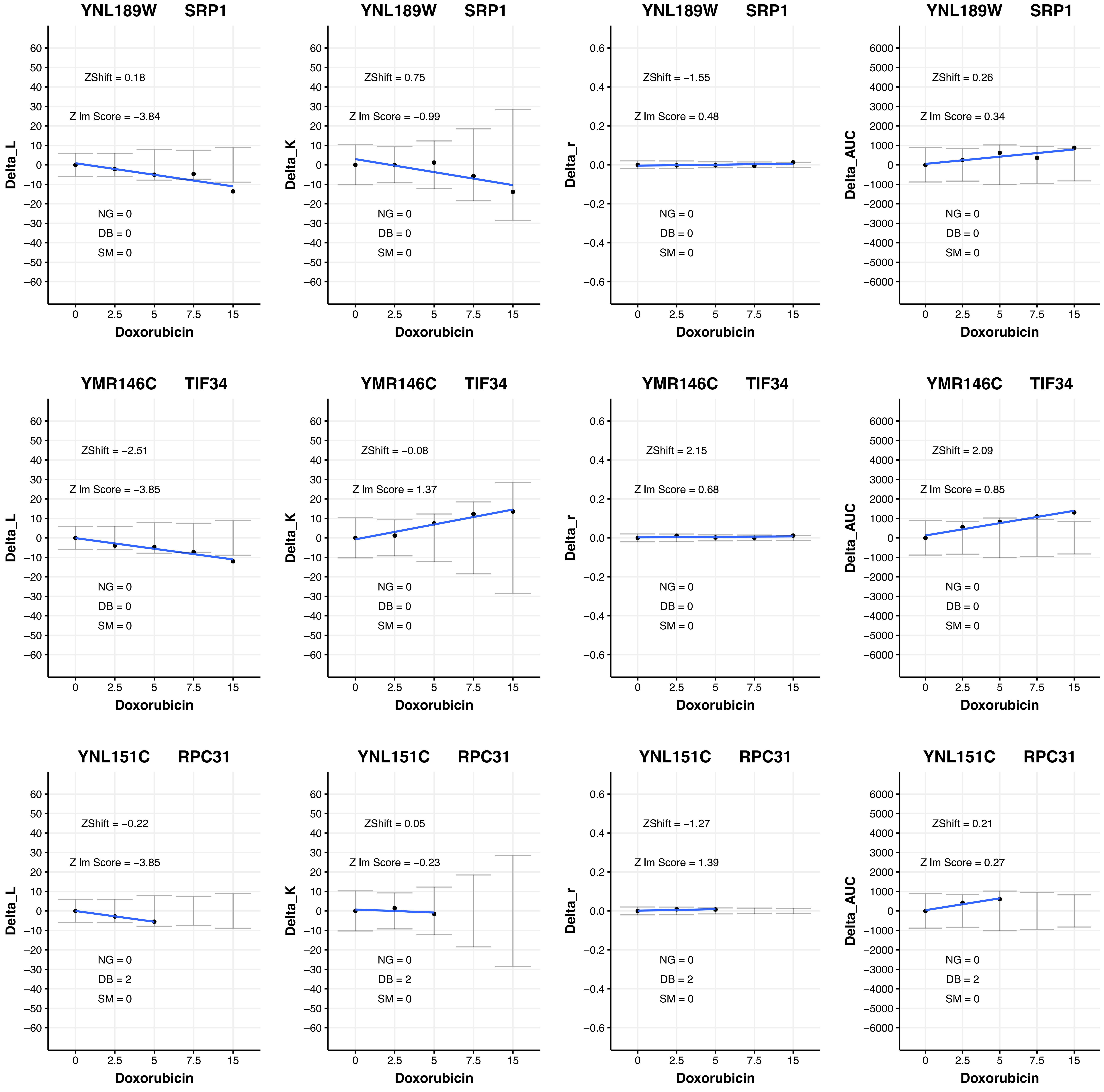

YNL189W SRP1
ZShift = 0.18
Z lm Score = −3.84
NG = 0
DB = 0
SM = 0
Delta_L
Doxorubicin
YNL189W SRP1
ZShift = 0.75
Z lm Score = −0.99
NG = 0
DB = 0
SM = 0
Delta_K
Doxorubicin
YNL189W SRP1
ZShift = −1.55
Z lm Score = 0.48
NG = 0
DB = 0
SM = 0
Delta_r
Doxorubicin
YNL189W SRP1
ZShift = 0.26
Z lm Score = 0.34
NG = 0
DB = 0
SM = 0
Delta_AUC
Doxorubicin
YMR146C TIF34
ZShift = −2.51
Z lm Score = −3.85
NG = 0
DB = 0
SM = 0
Delta_L
Doxorubicin
YMR146C TIF34
ZShift = −0.08
Z lm Score = 1.37
NG = 0
DB = 0
SM = 0
Delta_K
Doxorubicin
YMR146C TIF34
ZShift = 2.15
Z lm Score = 0.68
NG = 0
DB = 0
SM = 0
Delta_r
Doxorubicin
YMR146C TIF34
ZShift = 2.09
Z lm Score = 0.85
NG = 0
DB = 0
SM = 0
Delta_AUC
Doxorubicin
YNL151C RPC31
ZShift = −0.22
Z lm Score = −3.85
NG = 0
DB = 2
SM = 0
Delta_L
Doxorubicin
YNL151C RPC31
ZShift = 0.05
Z lm Score = −0.23
NG = 0
DB = 2
SM = 0
Delta_K
Doxorubicin
YNL151C RPC31
ZShift = −1.27
Z lm Score = 1.39
NG = 0
DB = 2
SM = 0
Delta_r
Doxorubicin
YNL151C RPC31
ZShift = 0.21
Z lm Score = 0.27
NG = 0
DB = 2
SM = 0
Delta_AUC
Doxorubicin

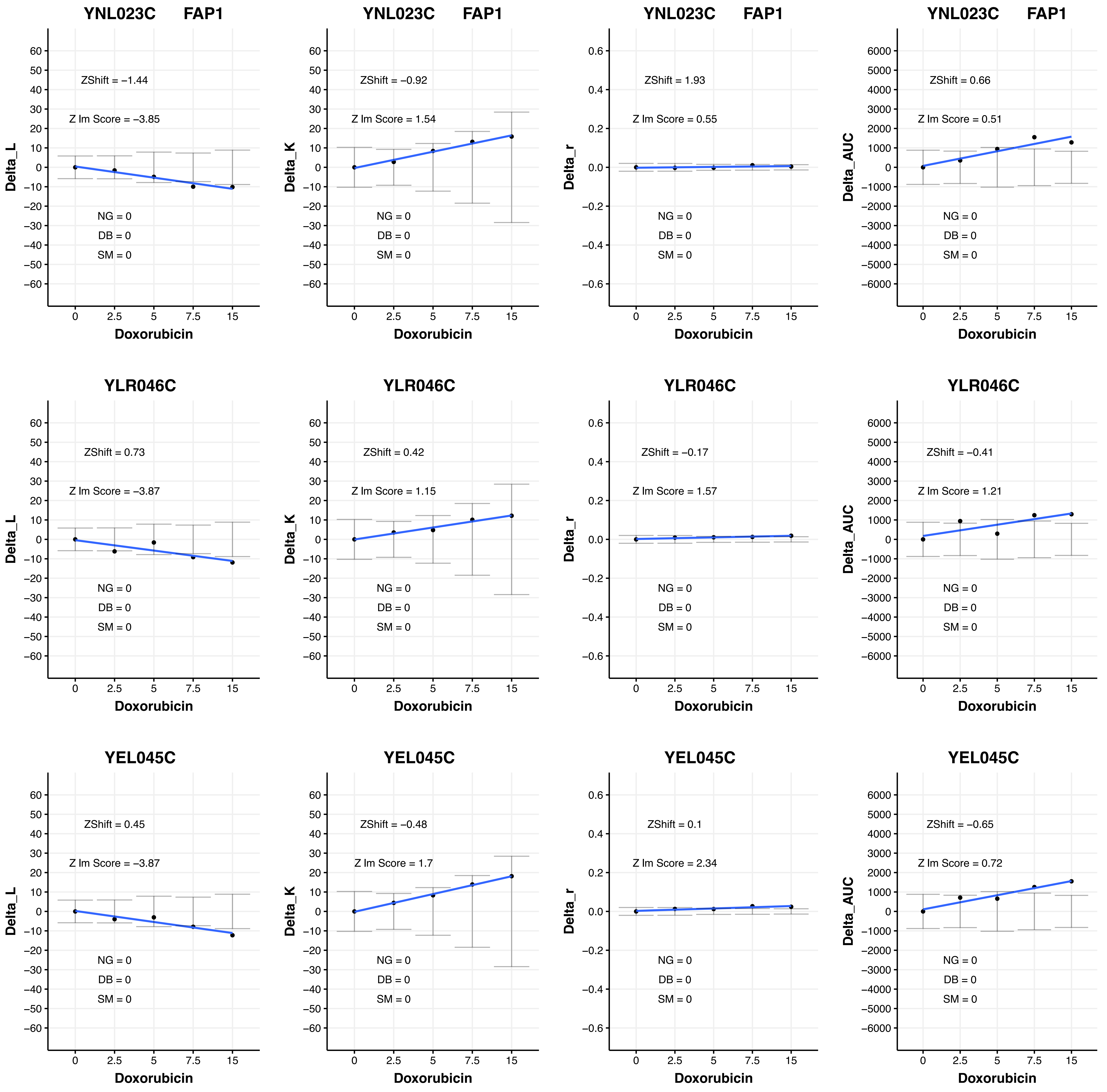

YNL023C FAP1
ZShift = −1.44
Z lm Score = −3.85
NG = 0
DB = 0
SM = 0
Delta_L
Doxorubicin
YNL023C FAP1
ZShift = −0.92
Z lm Score = 1.54
NG = 0
DB = 0
SM = 0
Delta_K
Doxorubicin
YNL023C FAP1
ZShift = 1.93
Z lm Score = 0.55
NG = 0
DB = 0
SM = 0
Delta_r
Doxorubicin
YNL023C FAP1
ZShift = 0.66
Z lm Score = 0.51
NG = 0
DB = 0
SM = 0
Delta_AUC
Doxorubicin
YLR046C
ZShift = 0.73
Z lm Score = −3.87
NG = 0
DB = 0
SM = 0
Delta_L
Doxorubicin
YLR046C
ZShift = 0.42
Z lm Score = 1.15
NG = 0
DB = 0
SM = 0
Delta_K
Doxorubicin
YLR046C
ZShift = −0.17
Z lm Score = 1.57
NG = 0
DB = 0
SM = 0
Delta_r
Doxorubicin
YLR046C
ZShift = −0.41
Z lm Score = 1.21
NG = 0
DB = 0
SM = 0
Delta_AUC
Doxorubicin
YEL045C
ZShift = 0.45
Z lm Score = −3.87
NG = 0
DB = 0
SM = 0
Delta_L
Doxorubicin
YEL045C
ZShift = −0.48
Z lm Score = 1.7
NG = 0
DB = 0
SM = 0
Delta_K
Doxorubicin
YEL045C
ZShift = 0.1
Z lm Score = 2.34
NG = 0
DB = 0
SM = 0
Delta_r
Doxorubicin
YEL045C
ZShift = −0.65
Z lm Score = 0.72
NG = 0
DB = 0
SM = 0
Delta_AUC
Doxorubicin

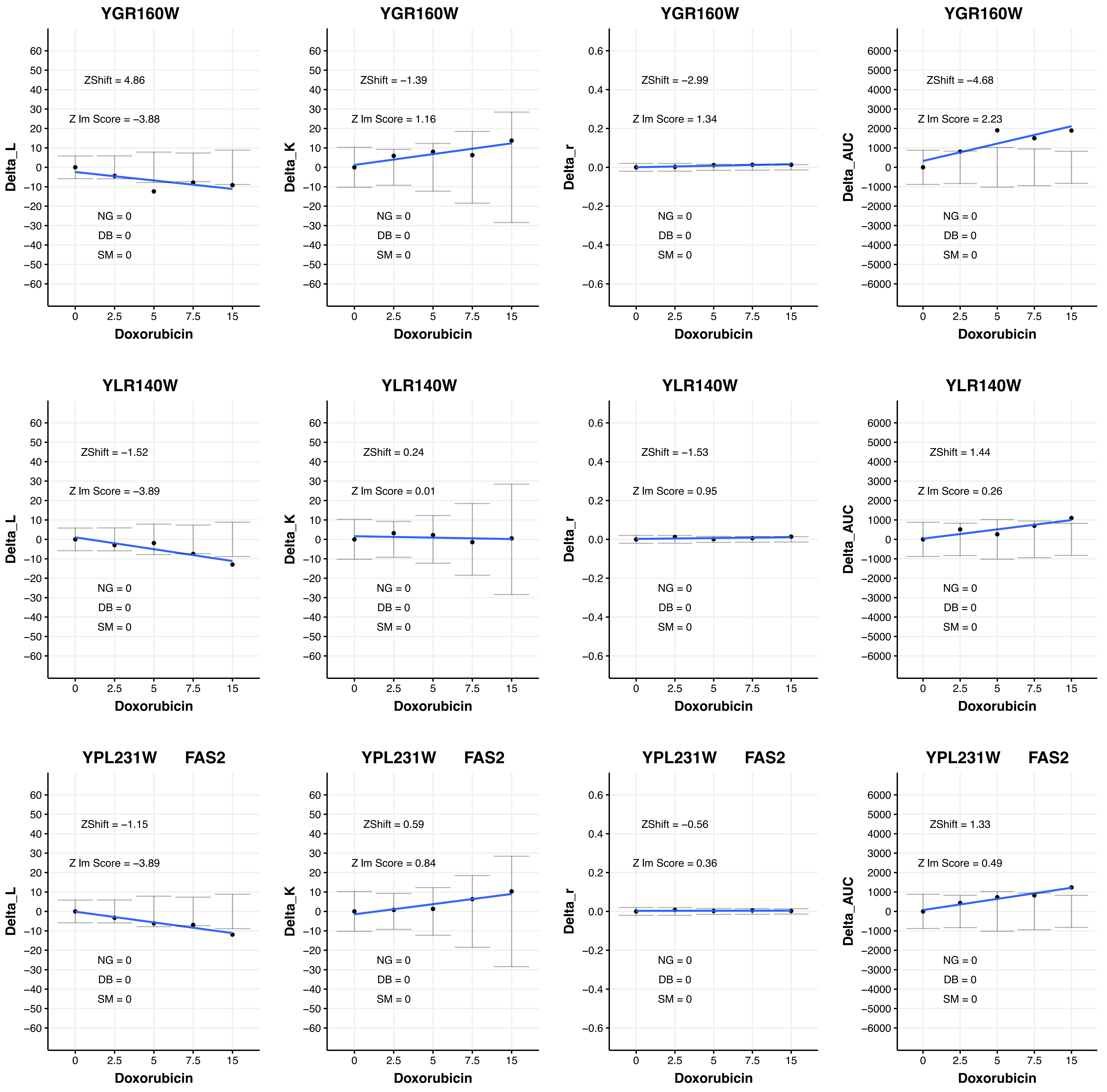
YGR160W
ZShift = 4.86
Z lm Score = −3.88
NG = 0
DB = 0
SM = 0
Delta_L
Doxorubicin
YGR160W
ZShift = −1.39
Z lm Score = 1.16
NG = 0
DB = 0
SM = 0
Delta_K
Doxorubicin
YGR160W
ZShift = −2.99
Z lm Score = 1.34
NG = 0
DB = 0
SM = 0
Delta_r
Doxorubicin
YGR160W
ZShift = −4.68
Z lm Score = 2.23
NG = 0
DB = 0
SM = 0
Delta_AUC
Doxorubicin
YLR140W
ZShift = −1.52
Z lm Score = −3.89
NG = 0
DB = 0
SM = 0
Delta_L
Doxorubicin
YLR140W
ZShift = 0.24
Z lm Score = 0.01
NG = 0
DB = 0
SM = 0
Delta_K
Doxorubicin
YLR140W
ZShift = −1.53
Z lm Score = 0.95
NG = 0
DB = 0
SM = 0
Delta_r
Doxorubicin
YLR140W
ZShift = 1.44
Z lm Score = 0.26
NG = 0
DB = 0
SM = 0
Delta_AUC
Doxorubicin
YPL231W FAS2
ZShift = −1.15
Z lm Score = −3.89
NG = 0
DB = 0
SM = 0
Delta_L
Doxorubicin
YPL231W FAS2
ZShift = 0.59
Z lm Score = 0.84
NG = 0
DB = 0
SM = 0
Delta_K
Doxorubicin
YPL231W FAS2
ZShift = −0.56
Z lm Score = 0.36
NG = 0
DB = 0
SM = 0
Delta_r
Doxorubicin
YPL231W FAS2
ZShift = 1.33
Z lm Score = 0.49
NG = 0
DB = 0
SM = 0
Delta_AUC
Doxorubicin

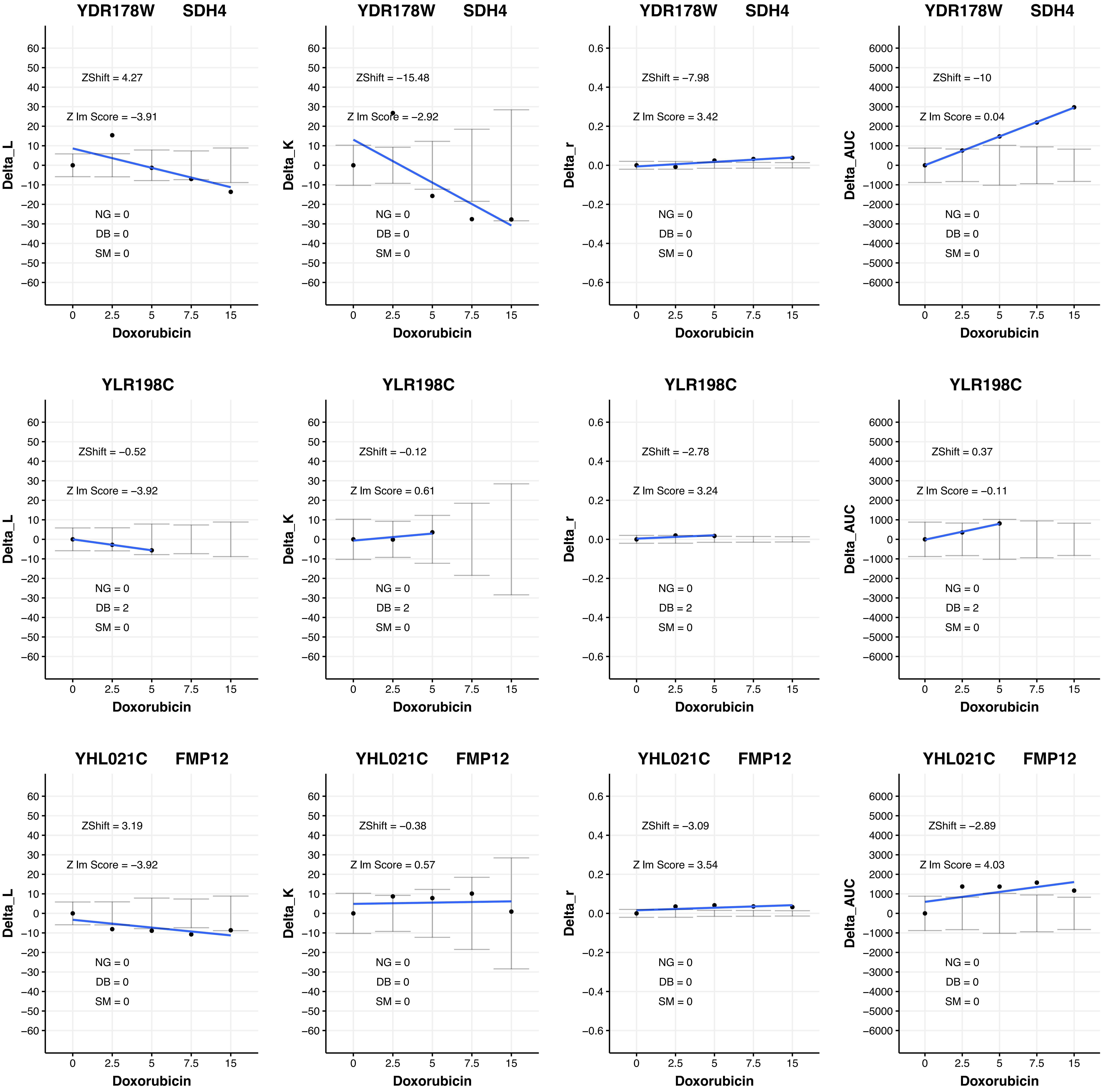
YDR178W SDH4
ZShift = 4.27
Z lm Score = −3.91
NG = 0
DB = 0
SM = 0
Delta_L
Doxorubicin
YDR178W SDH4
ZShift = −15.48
Z lm Score = −2.92
NG = 0
DB = 0
SM = 0
Delta_K
Doxorubicin
YDR178W SDH4
ZShift = −7.98
Z lm Score = 3.42
NG = 0
DB = 0
SM = 0
Delta_r
Doxorubicin
YDR178W SDH4
ZShift = −10
Z lm Score = 0.04
NG = 0
DB = 0
SM = 0
Delta_AUC
Doxorubicin
YLR198C
ZShift = −0.52
Z lm Score = −3.92
NG = 0
DB = 2
SM = 0
Delta_L
Doxorubicin
YLR198C
ZShift = −0.12
Z lm Score = 0.61
NG = 0
DB = 2
SM = 0
Delta_K
Doxorubicin
YLR198C
ZShift = −2.78
Z lm Score = 3.24
NG = 0
DB = 2
SM = 0
Delta_r
Doxorubicin
YLR198C
ZShift = 0.37
Z lm Score = −0.11
NG = 0
DB = 2
SM = 0
Delta_AUC
Doxorubicin
YHL021C FMP12
ZShift = 3.19
Z lm Score = −3.92
NG = 0
DB = 0
SM = 0
Delta_L
Doxorubicin
YHL021C FMP12
ZShift = −0.38
Z lm Score = 0.57
NG = 0
DB = 0
SM = 0
Delta_K
Doxorubicin
YHL021C FMP12
ZShift = −3.09
Z lm Score = 3.54
NG = 0
DB = 0
SM = 0
Delta_r
Doxorubicin
YHL021C FMP12
ZShift = −2.89
Z lm Score = 4.03
NG = 0
DB = 0
SM = 0
Delta_AUC
Doxorubicin

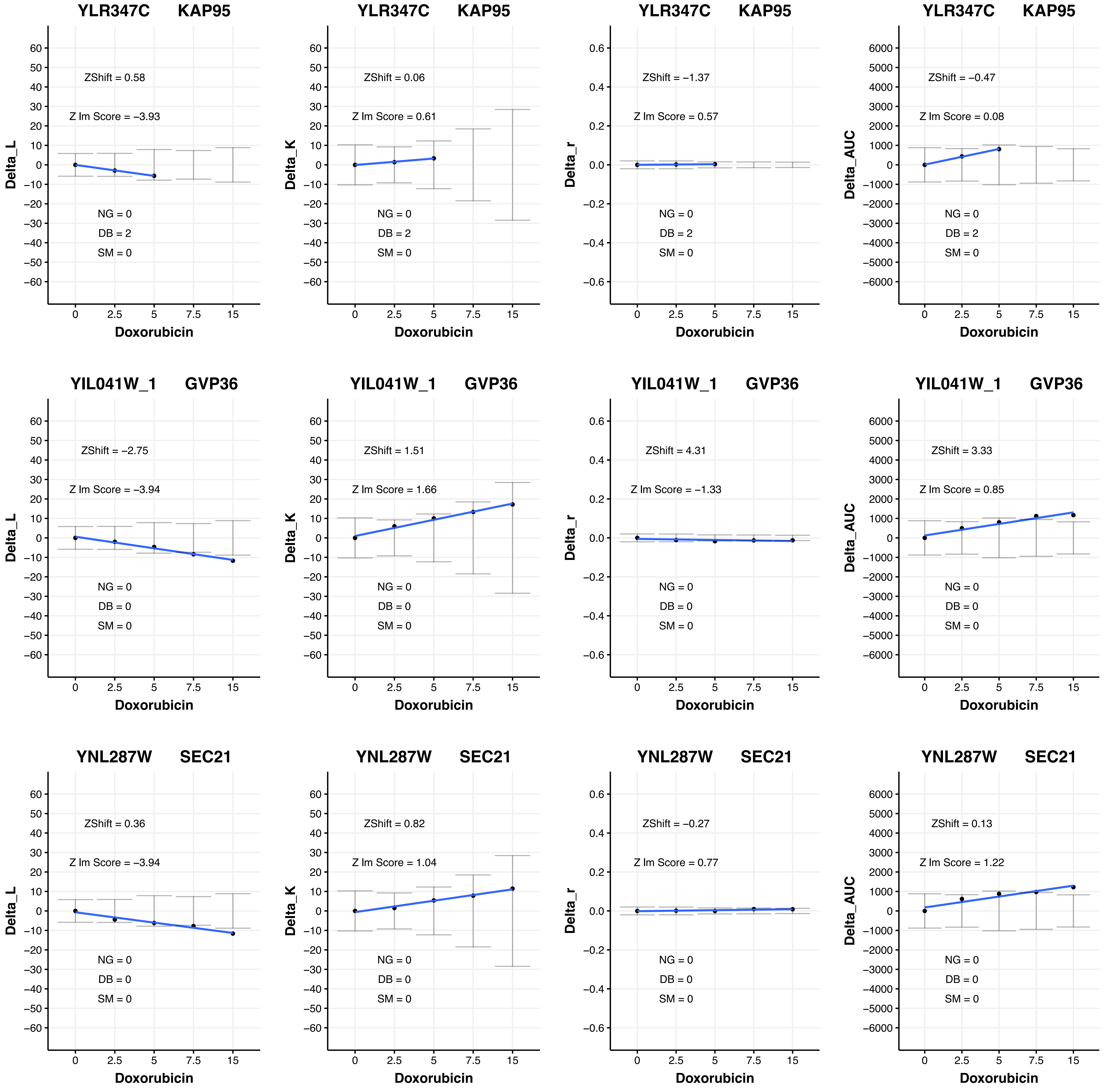
YLR347C KAP95
Delta_L
ZShift = 0.58
Z lm Score = −3.93
NG = 0
DB = 2
SM = 0
Doxorubicin
YLR347C KAP95
Delta_K
ZShift = 0.06
Z lm Score = 0.61
NG = 0
DB = 2
SM = 0
Doxorubicin
YLR347C KAP95
Delta_r
ZShift = −1.37
Z lm Score = 0.57
NG = 0
DB = 2
SM = 0
Doxorubicin
YLR347C KAP95
Delta_AUC
ZShift = −0.47
Z lm Score = 0.08
NG = 0
DB = 2
SM = 0
Doxorubicin
YIL041W_1 GVP36
Delta_L
ZShift = −2.75
Z lm Score = −3.94
NG = 0
DB = 0
SM = 0
Doxorubicin
YIL041W_1 GVP36
Delta_K
ZShift = 1.51
Z lm Score = 1.66
NG = 0
DB = 0
SM = 0
Doxorubicin
YIL041W_1 GVP36
Delta_r
ZShift = 4.31
Z lm Score = −1.33
NG = 0
DB = 0
SM = 0
Doxorubicin
YIL041W_1 GVP36
Delta_AUC
ZShift = 3.33
Z lm Score = 0.85
NG = 0
DB = 0
SM = 0
Doxorubicin
YNL287W SEC21
Delta_L
ZShift = 0.36
Z lm Score = −3.94
NG = 0
DB = 0
SM = 0
Doxorubicin
YNL287W SEC21
Delta_K
ZShift = 0.82
Z lm Score = 1.04
NG = 0
DB = 0
SM = 0
Doxorubicin
YNL287W SEC21
Delta_r
ZShift = −0.27
Z lm Score = 0.77
NG = 0
DB = 0
SM = 0
Doxorubicin
YNL287W SEC21
Delta_AUC
ZShift = 0.13
Z lm Score = 1.22
NG = 0
DB = 0
SM = 0
Doxorubicin

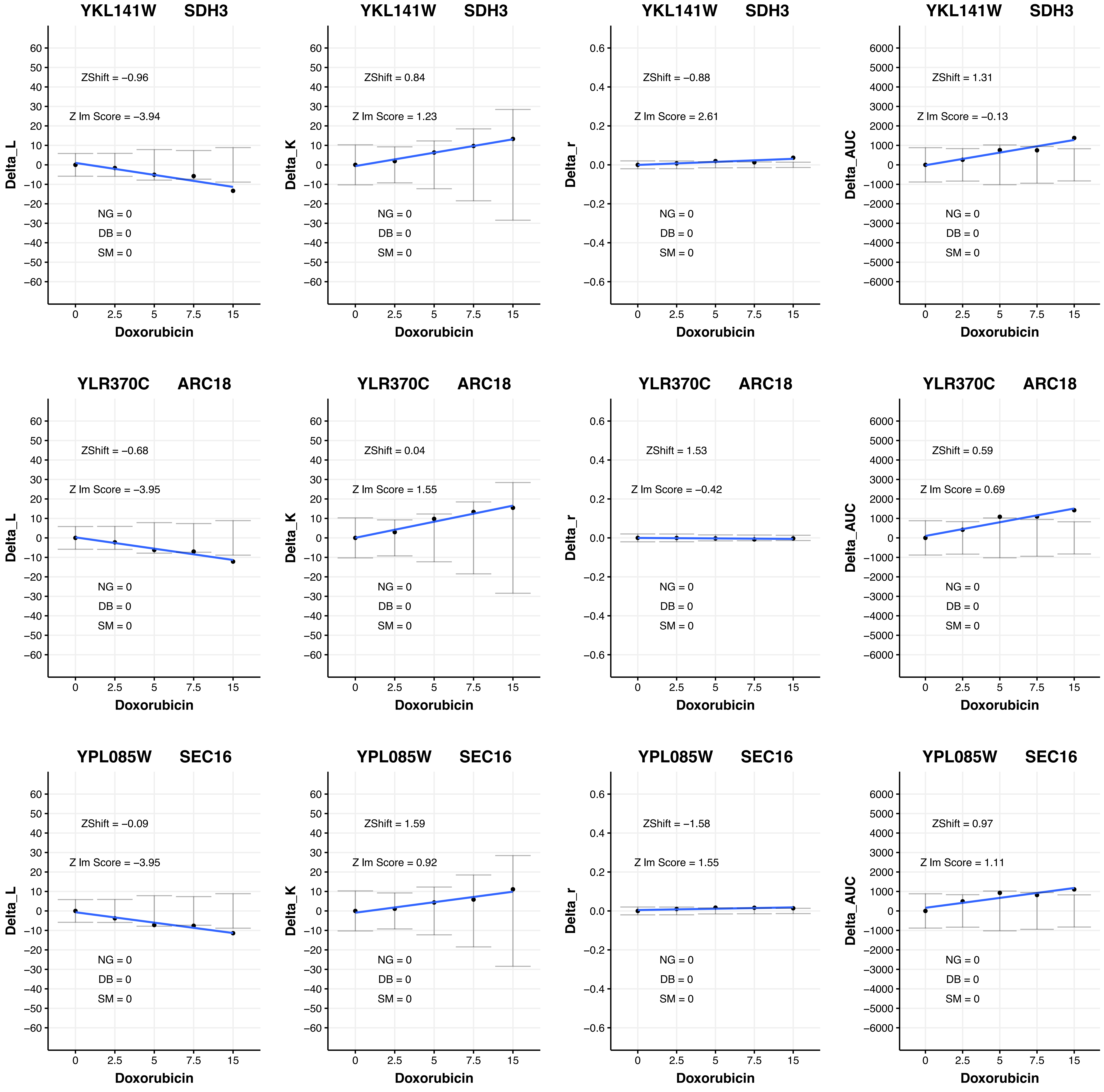
YKL141W SDH3
ZShift = −0.96
Z lm Score = −3.94
NG = 0
DB = 0
SM = 0
Delta_L
Doxorubicin
YKL141W SDH3
ZShift = 0.84
Z lm Score = 1.23
NG = 0
DB = 0
SM = 0
Delta_K
Doxorubicin
YKL141W SDH3
ZShift = −0.88
Z lm Score = 2.61
NG = 0
DB = 0
SM = 0
Delta_r
Doxorubicin
YKL141W SDH3
ZShift = 1.31
Z lm Score = −0.13
NG = 0
DB = 0
SM = 0
Delta_AUC
Doxorubicin
YLR370C ARC18
ZShift = −0.68
Z lm Score = −3.95
NG = 0
DB = 0
SM = 0
Delta_L
Doxorubicin
YLR370C ARC18
ZShift = 0.04
Z lm Score = 1.55
NG = 0
DB = 0
SM = 0
Delta_K
Doxorubicin
YLR370C ARC18
ZShift = 1.53
Z lm Score = −0.42
NG = 0
DB = 0
SM = 0
Delta_r
Doxorubicin
YLR370C ARC18
ZShift = 0.59
Z lm Score = 0.69
NG = 0
DB = 0
SM = 0
Delta_AUC
Doxorubicin
YPL085W SEC16
ZShift = −0.09
Z lm Score = −3.95
NG = 0
DB = 0
SM = 0
Delta_L
Doxorubicin
YPL085W SEC16
ZShift = 1.59
Z lm Score = 0.92
NG = 0
DB = 0
SM = 0
Delta_K
Doxorubicin
YPL085W SEC16
ZShift = −1.58
Z lm Score = 1.55
NG = 0
DB = 0
SM = 0
Delta_r
Doxorubicin
YPL085W SEC16
ZShift = 0.97
Z lm Score = 1.11
NG = 0
DB = 0
SM = 0
Delta_AUC
Doxorubicin

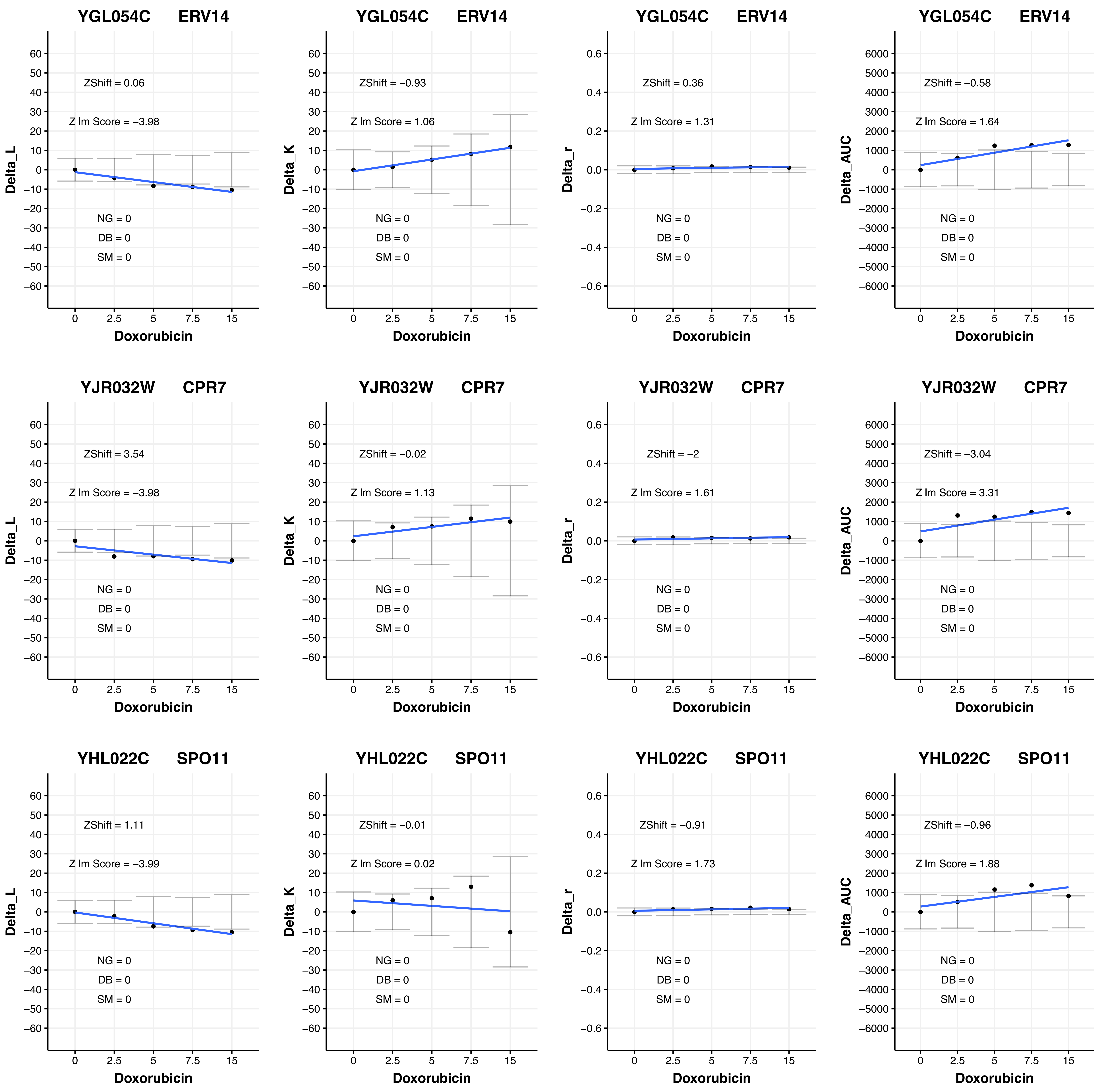

YGL054C ERV14
ZShift = 0.06
Z lm Score = −3.98
NG = 0
DB = 0
SM = 0
Delta_L
Doxorubicin
YGL054C ERV14
ZShift = −0.93
Z lm Score = 1.06
NG = 0
DB = 0
SM = 0
Delta_K
Doxorubicin
YGL054C ERV14
ZShift = 0.36
Z lm Score = 1.31
NG = 0
DB = 0
SM = 0
Delta_r
Doxorubicin
YGL054C ERV14
ZShift = −0.58
Z lm Score = 1.64
NG = 0
DB = 0
SM = 0
Delta_AUC
Doxorubicin
YJR032W CPR7
ZShift = 3.54
Z lm Score = −3.98
NG = 0
DB = 0
SM = 0
Delta_L
Doxorubicin
YJR032W CPR7
ZShift = −0.02
Z lm Score = 1.13
NG = 0
DB = 0
SM = 0
Delta_K
Doxorubicin
YJR032W CPR7
ZShift = −2
Z lm Score = 1.61
NG = 0
DB = 0
SM = 0
Delta_r
Doxorubicin
YJR032W CPR7
ZShift = −3.04
Z lm Score = 3.31
NG = 0
DB = 0
SM = 0
Delta_AUC
Doxorubicin
YHL022C SPO11
ZShift = 1.11
Z lm Score = −3.99
NG = 0
DB = 0
SM = 0
Delta_L
Doxorubicin
YHL022C SPO11
ZShift = −0.01
Z lm Score = 0.02
NG = 0
DB = 0
SM = 0
Delta_K
Doxorubicin
YHL022C SPO11
ZShift = −0.91
Z lm Score = 1.73
NG = 0
DB = 0
SM = 0
Delta_r
Doxorubicin
YHL022C SPO11
ZShift = −0.96
Z lm Score = 1.88
NG = 0
DB = 0
SM = 0
Delta_AUC
Doxorubicin

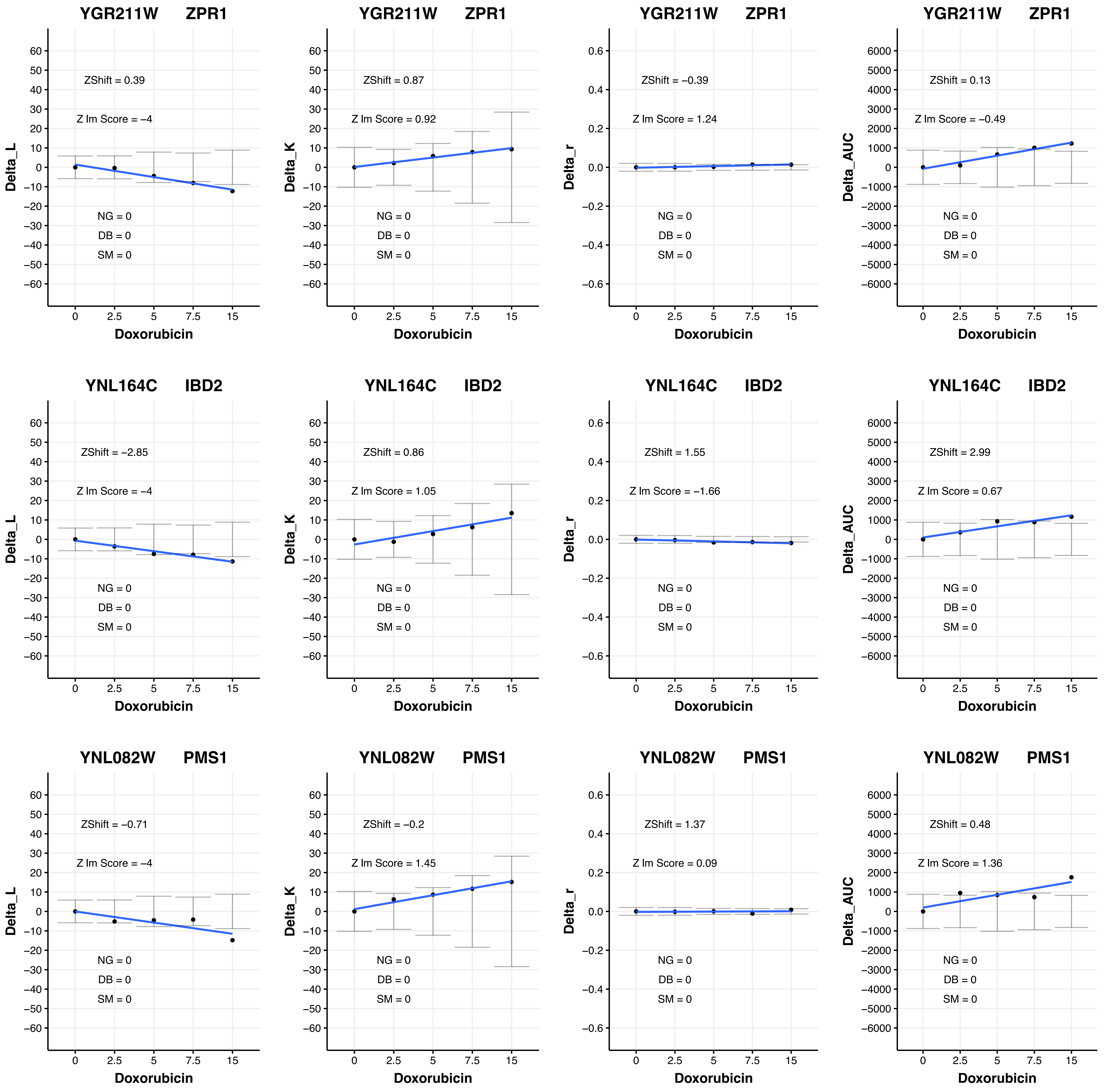
YGR211W ZPR1
ZShift = 0.39
Z lm Score = −4
NG = 0
DB = 0
SM = 0
Delta_L
Doxorubicin
YGR211W ZPR1
ZShift = 0.87
Z lm Score = 0.92
NG = 0
DB = 0
SM = 0
Delta_K
Doxorubicin
YGR211W ZPR1
ZShift = −0.39
Z lm Score = 1.24
NG = 0
DB = 0
SM = 0
Delta_r
Doxorubicin
YGR211W ZPR1
ZShift = 0.13
Z lm Score = −0.49
NG = 0
DB = 0
SM = 0
Delta_AUC
Doxorubicin
YNL164C IBD2
ZShift = −2.85
Z lm Score = −4
NG = 0
DB = 0
SM = 0
Delta_L
Doxorubicin
YNL164C IBD2
ZShift = 0.86
Z lm Score = 1.05
NG = 0
DB = 0
SM = 0
Delta_K
Doxorubicin
YNL164C IBD2
ZShift = 1.55
Z lm Score = −1.66
NG = 0
DB = 0
SM = 0
Delta_r
Doxorubicin
YNL164C IBD2
ZShift = 2.99
Z lm Score = 0.67
NG = 0
DB = 0
SM = 0
Delta_AUC
Doxorubicin
YNL082W PMS1
ZShift = −0.71
Z lm Score = −4
NG = 0
DB = 0
SM = 0
Delta_L
Doxorubicin
YNL082W PMS1
ZShift = −0.2
Z lm Score = 1.45
NG = 0
DB = 0
SM = 0
Delta_K
Doxorubicin
YNL082W PMS1
ZShift = 1.37
Z lm Score = 0.09
NG = 0
DB = 0
SM = 0
Delta_r
Doxorubicin
YNL082W PMS1
ZShift = 0.48
Z lm Score = 1.36
NG = 0
DB = 0
SM = 0
Delta_AUC
Doxorubicin

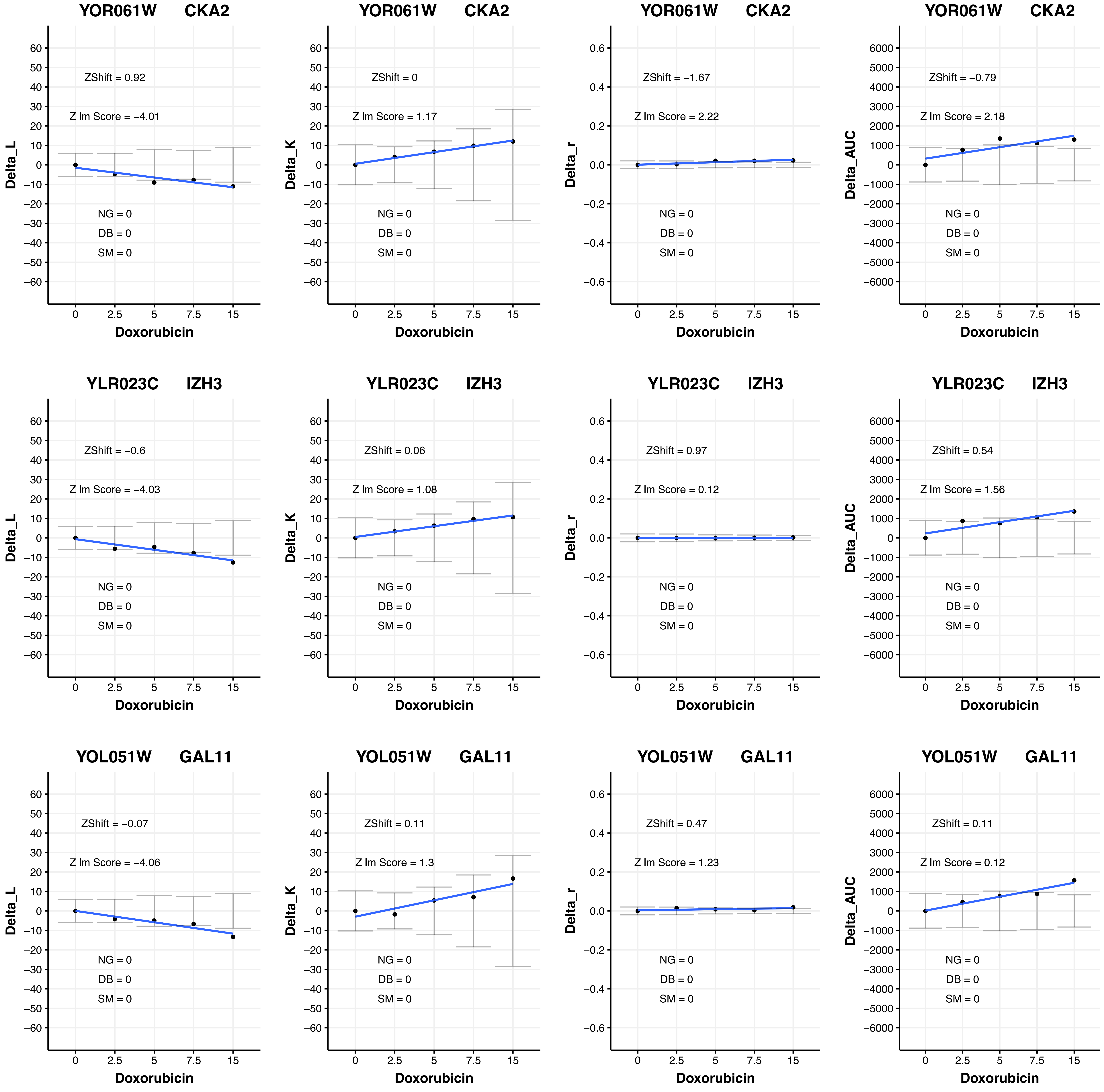
YOR061W CKA2
ZShift = 0.92
Z lm Score = −4.01
NG = 0
DB = 0
SM = 0
Delta_L
Doxorubicin
YOR061W CKA2
ZShift = 0
Z lm Score = 1.17
NG = 0
DB = 0
SM = 0
Delta_K
Doxorubicin
YOR061W CKA2
ZShift = −1.67
Z lm Score = 2.22
NG = 0
DB = 0
SM = 0
Delta_r
Doxorubicin
YOR061W CKA2
ZShift = −0.79
Z lm Score = 2.18
NG = 0
DB = 0
SM = 0
Delta_AUC
Doxorubicin
YLR023C IZH3
ZShift = −0.6
Z lm Score = −4.03
NG = 0
DB = 0
SM = 0
Delta_L
Doxorubicin
YLR023C IZH3
ZShift = 0.06
Z lm Score = 1.08
NG = 0
DB = 0
SM = 0
Delta_K
Doxorubicin
YLR023C IZH3
ZShift = 0.97
Z lm Score = 0.12
NG = 0
DB = 0
SM = 0
Delta_r
Doxorubicin
YLR023C IZH3
ZShift = 0.54
Z lm Score = 1.56
NG = 0
DB = 0
SM = 0
Delta_AUC
Doxorubicin
YOL051W GAL11
ZShift = −0.07
Z lm Score = −4.06
NG = 0
DB = 0
SM = 0
Delta_L
Doxorubicin
YOL051W GAL11
ZShift = 0.11
Z lm Score = 1.3
NG = 0
DB = 0
SM = 0
Delta_K
Doxorubicin
YOL051W GAL11
ZShift = 0.47
Z lm Score = 1.23
NG = 0
DB = 0
SM = 0
Delta_r
Doxorubicin
YOL051W GAL11
ZShift = 0.11
Z lm Score = 0.12
NG = 0
DB = 0
SM = 0
Delta_AUC
Doxorubicin

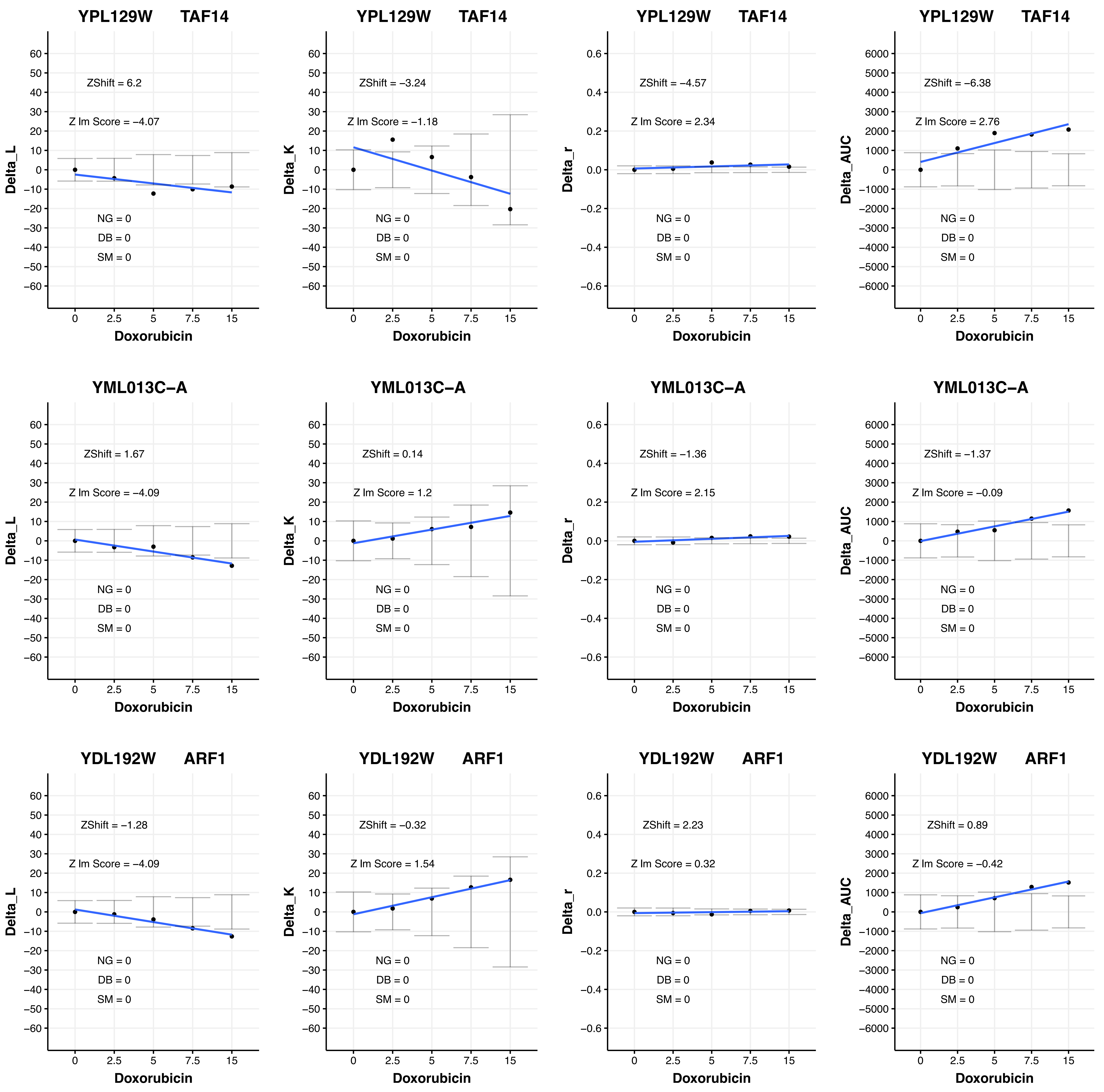

YPL129W TAF14
ZShift = 6.2
Z lm Score = −4.07
NG = 0
DB = 0
SM = 0
Delta_L
Doxorubicin
YPL129W TAF14
ZShift = −3.24
Z lm Score = −1.18
NG = 0
DB = 0
SM = 0
Delta_K
Doxorubicin
YPL129W TAF14
ZShift = −4.57
Z lm Score = 2.34
NG = 0
DB = 0
SM = 0
Delta_r
Doxorubicin
YPL129W TAF14
ZShift = −6.38
Z lm Score = 2.76
NG = 0
DB = 0
SM = 0
Delta_AUC
Doxorubicin
YML013C−A
ZShift = 1.67
Z lm Score = −4.09
NG = 0
DB = 0
SM = 0
Delta_L
Doxorubicin
YML013C−A
ZShift = 0.14
Z lm Score = 1.2
NG = 0
DB = 0
SM = 0
Delta_K
Doxorubicin
YML013C−A
ZShift = −1.36
Z lm Score = 2.15
NG = 0
DB = 0
SM = 0
Delta_r
Doxorubicin
YML013C−A
ZShift = −1.37
Z lm Score = −0.09
NG = 0
DB = 0
SM = 0
Delta_AUC
Doxorubicin
YDL192W ARF1
ZShift = −1.28
Z lm Score = −4.09
NG = 0
DB = 0
SM = 0
Delta_L
Doxorubicin
YDL192W ARF1
ZShift = −0.32
Z lm Score = 1.54
NG = 0
DB = 0
SM = 0
Delta_K
Doxorubicin
YDL192W ARF1
ZShift = 2.23
Z lm Score = 0.32
NG = 0
DB = 0
SM = 0
Delta_r
Doxorubicin
YDL192W ARF1
ZShift = 0.89
Z lm Score = −0.42
NG = 0
DB = 0
SM = 0
Delta_AUC
Doxorubicin

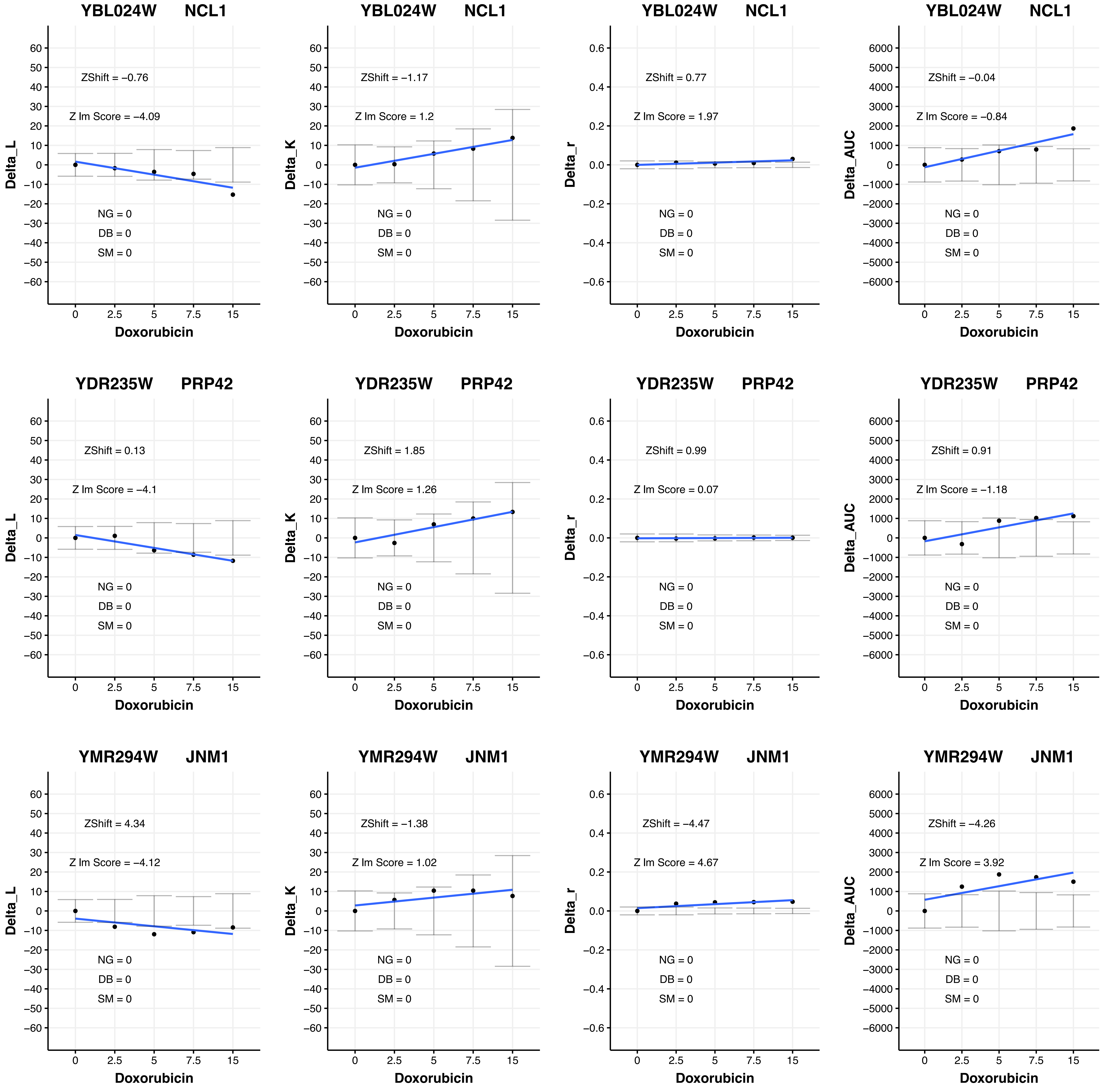

YBL024W NCL1
ZShift = −0.76
Z lm Score = −4.09
NG = 0
DB = 0
SM = 0
Delta_L
Doxorubicin
YBL024W NCL1
ZShift = −1.17
Z lm Score = 1.2
NG = 0
DB = 0
SM = 0
Delta_K
Doxorubicin
YBL024W NCL1
ZShift = 0.77
Z lm Score = 1.97
NG = 0
DB = 0
SM = 0
Delta_r
Doxorubicin
YBL024W NCL1
ZShift = −0.04
Z lm Score = −0.84
NG = 0
DB = 0
SM = 0
Delta_AUC
Doxorubicin
YDR235W PRP42
ZShift = 0.13
Z lm Score = −4.1
NG = 0
DB = 0
SM = 0
Delta_L
Doxorubicin
YDR235W PRP42
ZShift = 1.85
Z lm Score = 1.26
NG = 0
DB = 0
SM = 0
Delta_K
Doxorubicin
YDR235W PRP42
ZShift = 0.99
Z lm Score = 0.07
NG = 0
DB = 0
SM = 0
Delta_r
Doxorubicin
YDR235W PRP42
ZShift = 0.91
Z lm Score = −1.18
NG = 0
DB = 0
SM = 0
Delta_AUC
Doxorubicin
YMR294W JNM1
ZShift = 4.34
Z lm Score = −4.12
NG = 0
DB = 0
SM = 0
Delta_L
Doxorubicin
YMR294W JNM1
ZShift = −1.38
Z lm Score = 1.02
NG = 0
DB = 0
SM = 0
Delta_K
Doxorubicin
YMR294W JNM1
ZShift = −4.47
Z lm Score = 4.67
NG = 0
DB = 0
SM = 0
Delta_r
Doxorubicin
YMR294W JNM1
ZShift = −4.26
Z lm Score = 3.92
NG = 0
DB = 0
SM = 0
Delta_AUC
Doxorubicin

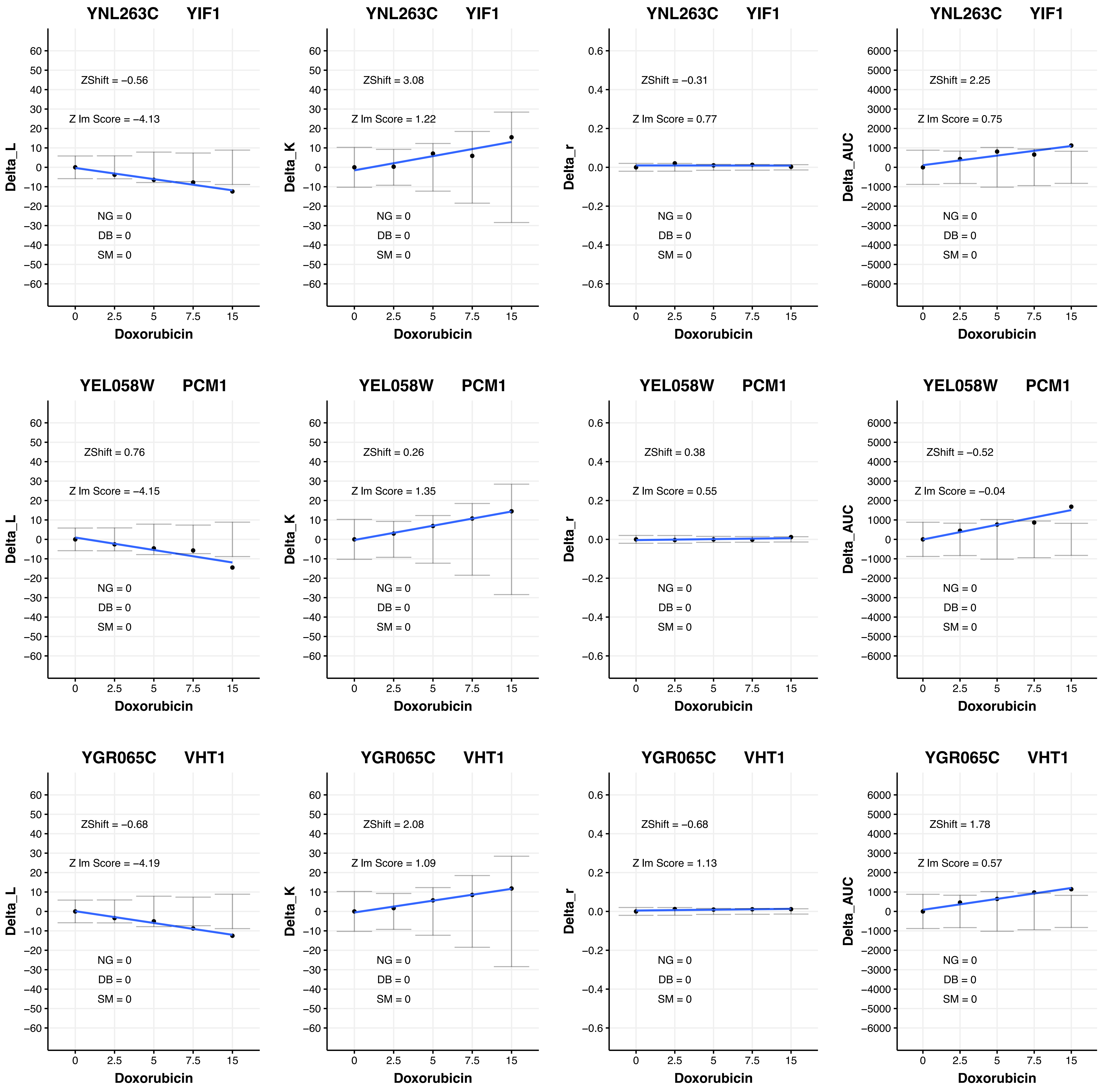
YNL263C YIF1
ZShift = −0.56
Z lm Score = −4.13
NG = 0
DB = 0
SM = 0
Delta_L
Doxorubicin
YNL263C YIF1
ZShift = 3.08
Z lm Score = 1.22
NG = 0
DB = 0
SM = 0
Delta_K
Doxorubicin
YNL263C YIF1
ZShift = −0.31
Z lm Score = 0.77
NG = 0
DB = 0
SM = 0
Delta_r
Doxorubicin
YNL263C YIF1
ZShift = 2.25
Z lm Score = 0.75
NG = 0
DB = 0
SM = 0
Delta_AUC
Doxorubicin
YEL058W PCM1
ZShift = 0.76
Z lm Score = −4.15
NG = 0
DB = 0
SM = 0
Delta_L
Doxorubicin
YEL058W PCM1
ZShift = 0.26
Z lm Score = 1.35
NG = 0
DB = 0
SM = 0
Delta_K
Doxorubicin
YEL058W PCM1
ZShift = 0.38
Z lm Score = 0.55
NG = 0
DB = 0
SM = 0
Delta_r
Doxorubicin
YEL058W PCM1
ZShift = −0.52
Z lm Score = −0.04
NG = 0
DB = 0
SM = 0
Delta_AUC
Doxorubicin
YGR065C VHT1
ZShift = −0.68
Z lm Score = −4.19
NG = 0
DB = 0
SM = 0
Delta_L
Doxorubicin
YGR065C VHT1
ZShift = 2.08
Z lm Score = 1.09
NG = 0
DB = 0
SM = 0
Delta_K
Doxorubicin
YGR065C VHT1
ZShift = −0.68
Z lm Score = 1.13
NG = 0
DB = 0
SM = 0
Delta_r
Doxorubicin
YGR065C VHT1
ZShift = 1.78
Z lm Score = 0.57
NG = 0
DB = 0
SM = 0
Delta_AUC
Doxorubicin

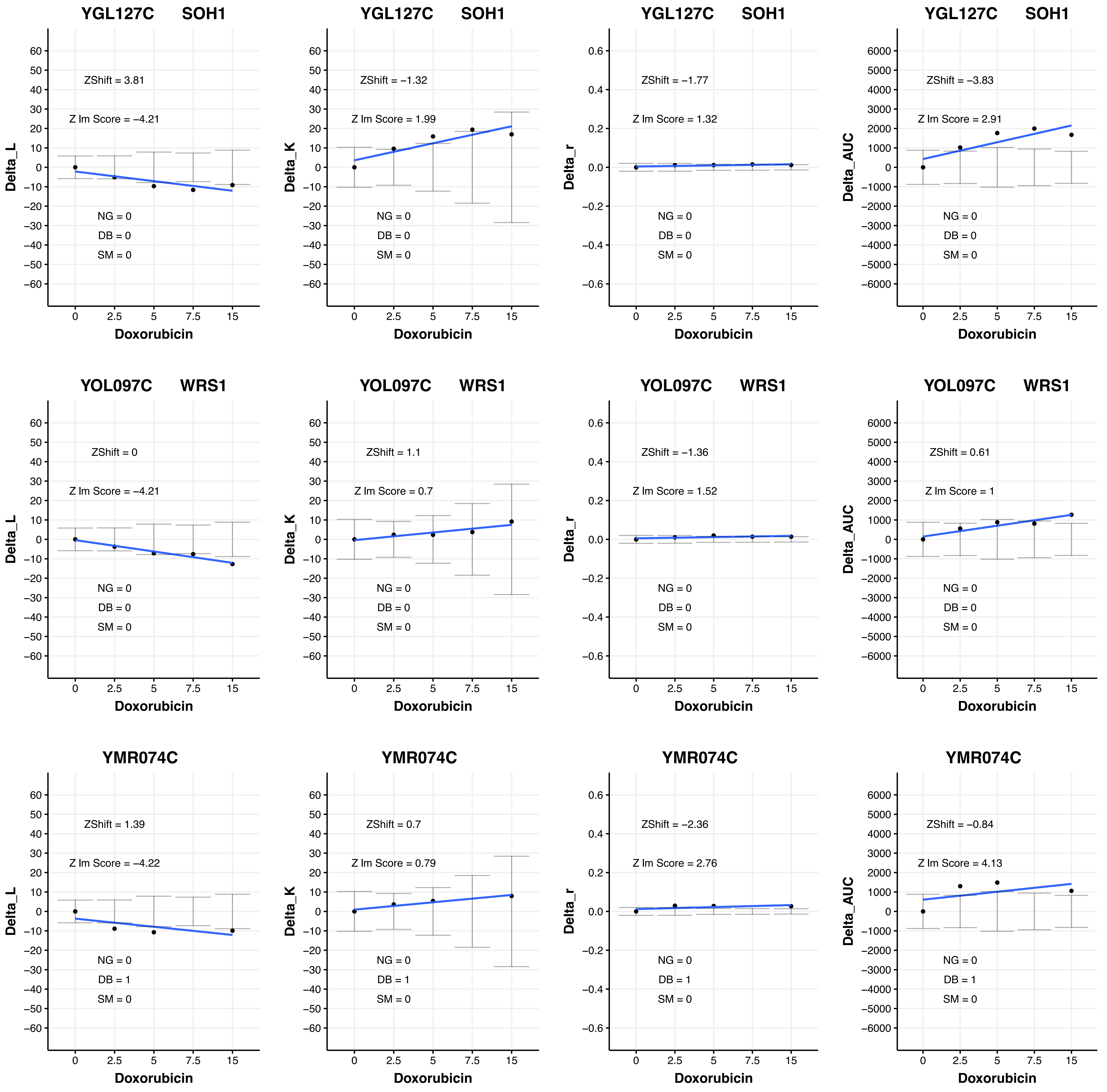
YGL127C SOH1
ZShift = 3.81
Z lm Score = −4.21
NG = 0
DB = 0
SM = 0
Delta_L
Doxorubicin
YGL127C SOH1
ZShift = −1.32
Z lm Score = 1.99
NG = 0
DB = 0
SM = 0
Delta_K
Doxorubicin
YGL127C SOH1
ZShift = −1.77
Z lm Score = 1.32
NG = 0
DB = 0
SM = 0
Delta_r
Doxorubicin
YGL127C SOH1
ZShift = −3.83
Z lm Score = 2.91
NG = 0
DB = 0
SM = 0
Delta_AUC
Doxorubicin
YOL097C WRS1
ZShift = 0
Z lm Score = −4.21
NG = 0
DB = 0
SM = 0
Delta_L
Doxorubicin
YOL097C WRS1
ZShift = 1.1
Z lm Score = 0.7
NG = 0
DB = 0
SM = 0
Delta_K
Doxorubicin
YOL097C WRS1
ZShift = −1.36
Z lm Score = 1.52
NG = 0
DB = 0
SM = 0
Delta_r
Doxorubicin
YOL097C WRS1
ZShift = 0.61
Z lm Score = 1
NG = 0
DB = 0
SM = 0
Delta_AUC
Doxorubicin
YMR074C
ZShift = 1.39
Z lm Score = −4.22
NG = 0
DB = 1
SM = 0
Delta_L
Doxorubicin
YMR074C
ZShift = 0.7
Z lm Score = 0.79
NG = 0
DB = 1
SM = 0
Delta_K
Doxorubicin
YMR074C
ZShift = −2.36
Z lm Score = 2.76
NG = 0
DB = 1
SM = 0
Delta_r
Doxorubicin
YMR074C
ZShift = −0.84
Z lm Score = 4.13
NG = 0
DB = 1
SM = 0
Delta_AUC
Doxorubicin

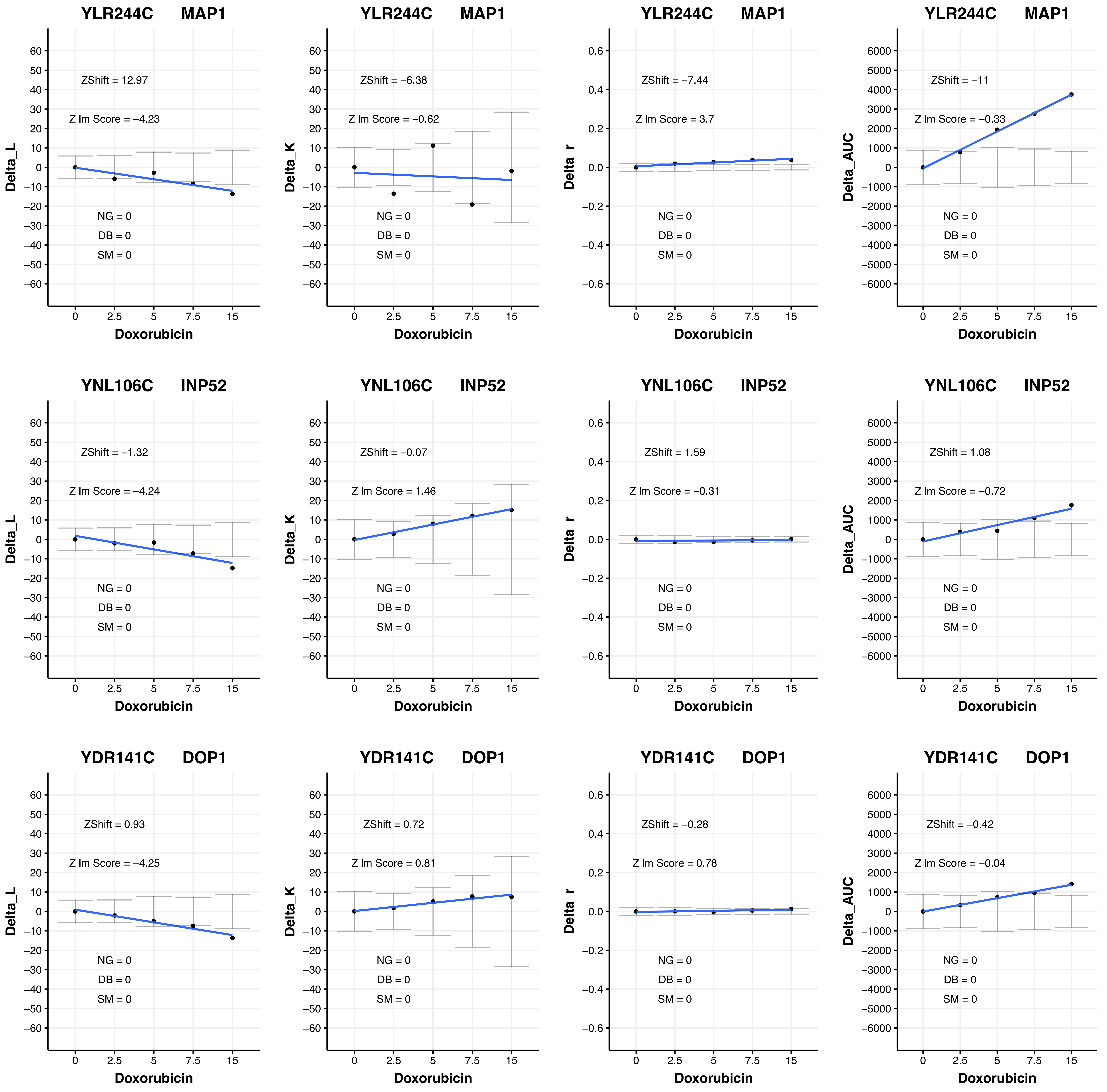

YLR244C MAP1
ZShift = 12.97
Z lm Score = −4.23
NG = 0
DB = 0
SM = 0
Delta_L
Doxorubicin
YLR244C MAP1
ZShift = −6.38
Z lm Score = −0.62
NG = 0
DB = 0
SM = 0
Delta_K
Doxorubicin
YLR244C MAP1
ZShift = −7.44
Z lm Score = 3.7
NG = 0
DB = 0
SM = 0
Delta_r
Doxorubicin
YLR244C MAP1
ZShift = −11
Z lm Score = −0.33
NG = 0
DB = 0
SM = 0
Delta_AUC
Doxorubicin
YNL106C INP52
ZShift = −1.32
Z lm Score = −4.24
NG = 0
DB = 0
SM = 0
Delta_L
Doxorubicin
YNL106C INP52
ZShift = −0.07
Z lm Score = 1.46
NG = 0
DB = 0
SM = 0
Delta_K
Doxorubicin
YNL106C INP52
ZShift = 1.59
Z lm Score = −0.31
NG = 0
DB = 0
SM = 0
Delta_r
Doxorubicin
YNL106C INP52
ZShift = 1.08
Z lm Score = −0.72
NG = 0
DB = 0
SM = 0
Delta_AUC
Doxorubicin
YDR141C DOP1
ZShift = 0.93
Z lm Score = −4.25
NG = 0
DB = 0
SM = 0
Delta_L
Doxorubicin
YDR141C DOP1
ZShift = 0.72
Z lm Score = 0.81
NG = 0
DB = 0
SM = 0
Delta_K
Doxorubicin
YDR141C DOP1
ZShift = −0.28
Z lm Score = 0.78
NG = 0
DB = 0
SM = 0
Delta_r
Doxorubicin
YDR141C DOP1
ZShift = −0.42
Z lm Score = −0.04
NG = 0
DB = 0
SM = 0
Delta_AUC
Doxorubicin

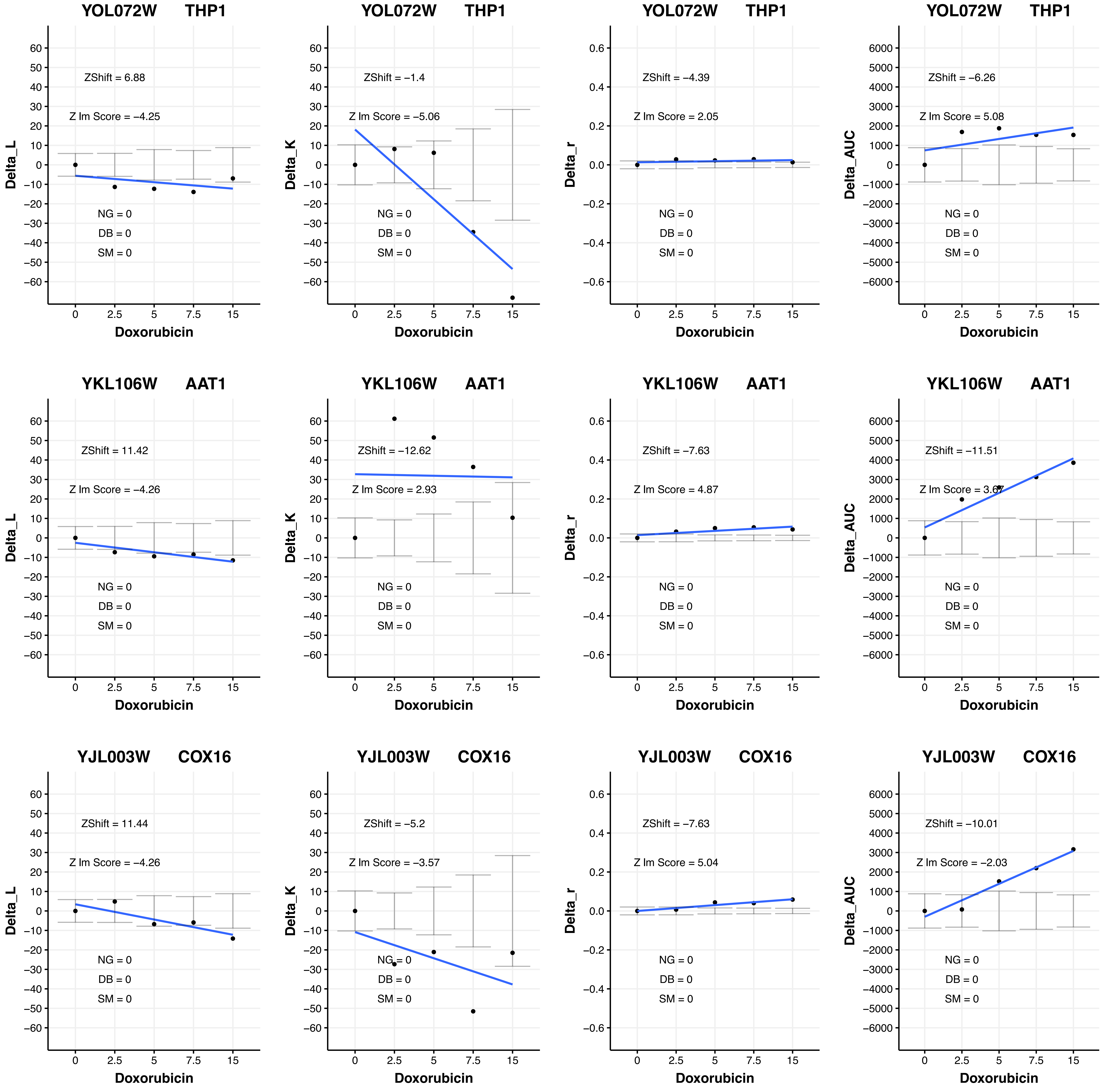
YOL072W THP1
ZShift = 6.88
Z lm Score = −4.25
NG = 0
DB = 0
SM = 0
Delta_L
Doxorubicin
YOL072W THP1
ZShift = −1.4
Z lm Score = −5.06
NG = 0
DB = 0
SM = 0
Delta_K
Doxorubicin
YOL072W THP1
ZShift = −4.39
Z lm Score = 2.05
NG = 0
DB = 0
SM = 0
Delta_r
Doxorubicin
YOL072W THP1
ZShift = −6.26
Z lm Score = 5.08
NG = 0
DB = 0
SM = 0
Delta_AUC
Doxorubicin
YKL106W AAT1
ZShift = 11.42
Z lm Score = −4.26
NG = 0
DB = 0
SM = 0
Delta_L
Doxorubicin
YKL106W AAT1
ZShift = −12.62
Z lm Score = 2.93
NG = 0
DB = 0
SM = 0
Delta_K
Doxorubicin
YKL106W AAT1
ZShift = −7.63
Z lm Score = 4.87
NG = 0
DB = 0
SM = 0
Delta_r
Doxorubicin
YKL106W AAT1
ZShift = −11.51
Z lm Score = 3.67
NG = 0
DB = 0
SM = 0
Delta_AUC
Doxorubicin
YJL003W COX16
ZShift = 11.44
Z lm Score = −4.26
NG = 0
DB = 0
SM = 0
Delta_L
Doxorubicin
YJL003W COX16
ZShift = −5.2
Z lm Score = −3.57
NG = 0
DB = 0
SM = 0
Delta_K
Doxorubicin
YJL003W COX16
ZShift = −7.63
Z lm Score = 5.04
NG = 0
DB = 0
SM = 0
Delta_r
Doxorubicin
YJL003W COX16
ZShift = −10.01
Z lm Score = −2.03
NG = 0
DB = 0
SM = 0
Delta_AUC
Doxorubicin

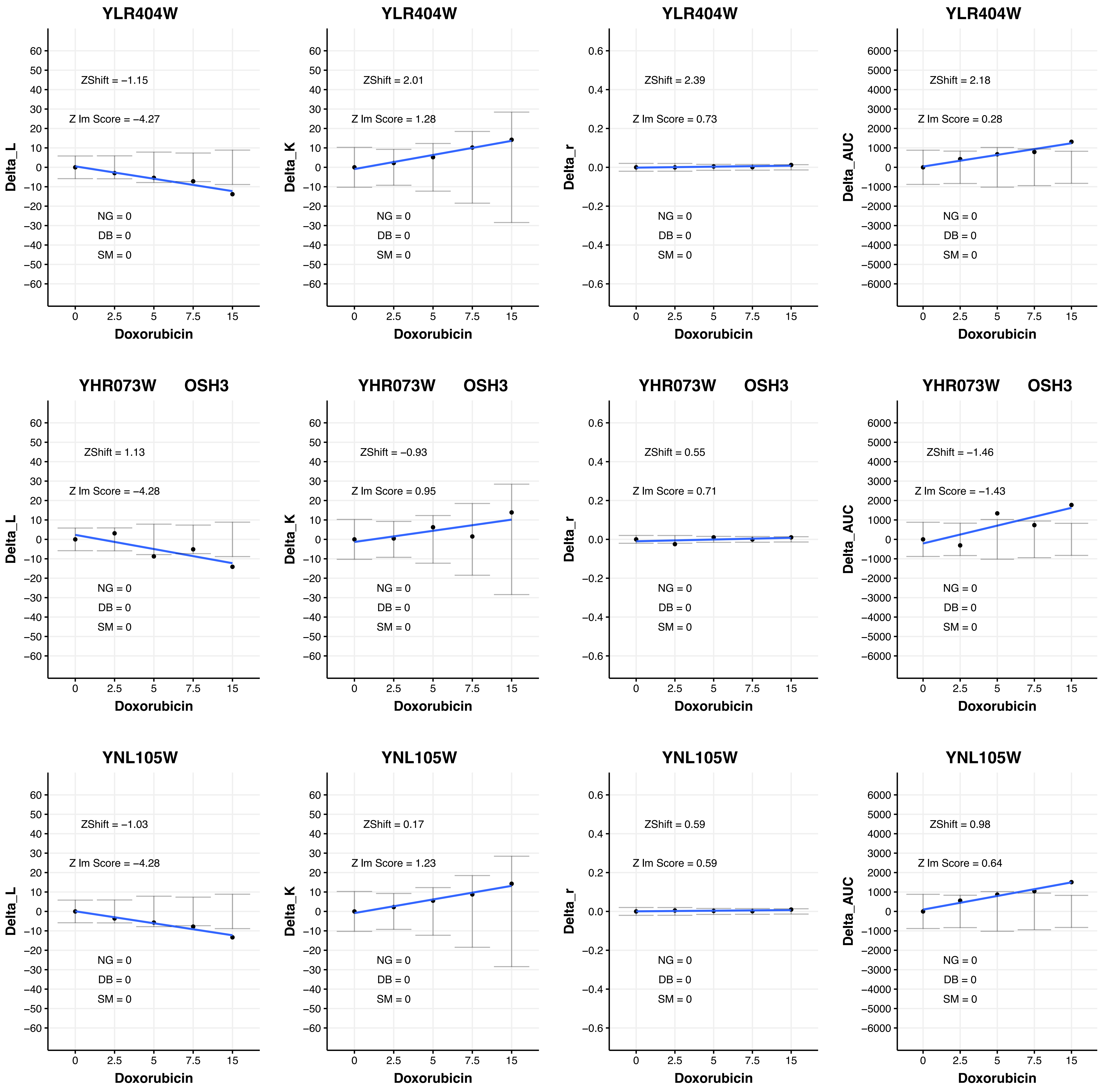

YLR404W
Delta_L
ZShift = −1.15
Z lm Score = −4.27
NG = 0
DB = 0
SM = 0
Doxorubicin
YLR404W
Delta_K
ZShift = 2.01
Z lm Score = 1.28
NG = 0
DB = 0
SM = 0
Doxorubicin
YLR404W
Delta_r
ZShift = 2.39
Z lm Score = 0.73
NG = 0
DB = 0
SM = 0
Doxorubicin
YLR404W
Delta_AUC
ZShift = 2.18
Z lm Score = 0.28
NG = 0
DB = 0
SM = 0
Doxorubicin
YHR073W OSH3
Delta_L
ZShift = 1.13
Z lm Score = −4.28
NG = 0
DB = 0
SM = 0
Doxorubicin
YHR073W OSH3
Delta_K
ZShift = −0.93
Z lm Score = 0.95
NG = 0
DB = 0
SM = 0
Doxorubicin
YHR073W OSH3
Delta_r
ZShift = 0.55
Z lm Score = 0.71
NG = 0
DB = 0
SM = 0
Doxorubicin
YHR073W OSH3
Delta_AUC
ZShift = −1.46
Z lm Score = −1.43
NG = 0
DB = 0
SM = 0
Doxorubicin
YNL105W
Delta_L
ZShift = −1.03
Z lm Score = −4.28
NG = 0
DB = 0
SM = 0
Doxorubicin
YNL105W
Delta_K
ZShift = 0.17
Z lm Score = 1.23
NG = 0
DB = 0
SM = 0
Doxorubicin
YNL105W
Delta_r
ZShift = 0.59
Z lm Score = 0.59
NG = 0
DB = 0
SM = 0
Doxorubicin
YNL105W
Delta_AUC
ZShift = 0.98
Z lm Score = 0.64
NG = 0
DB = 0
SM = 0
Doxorubicin

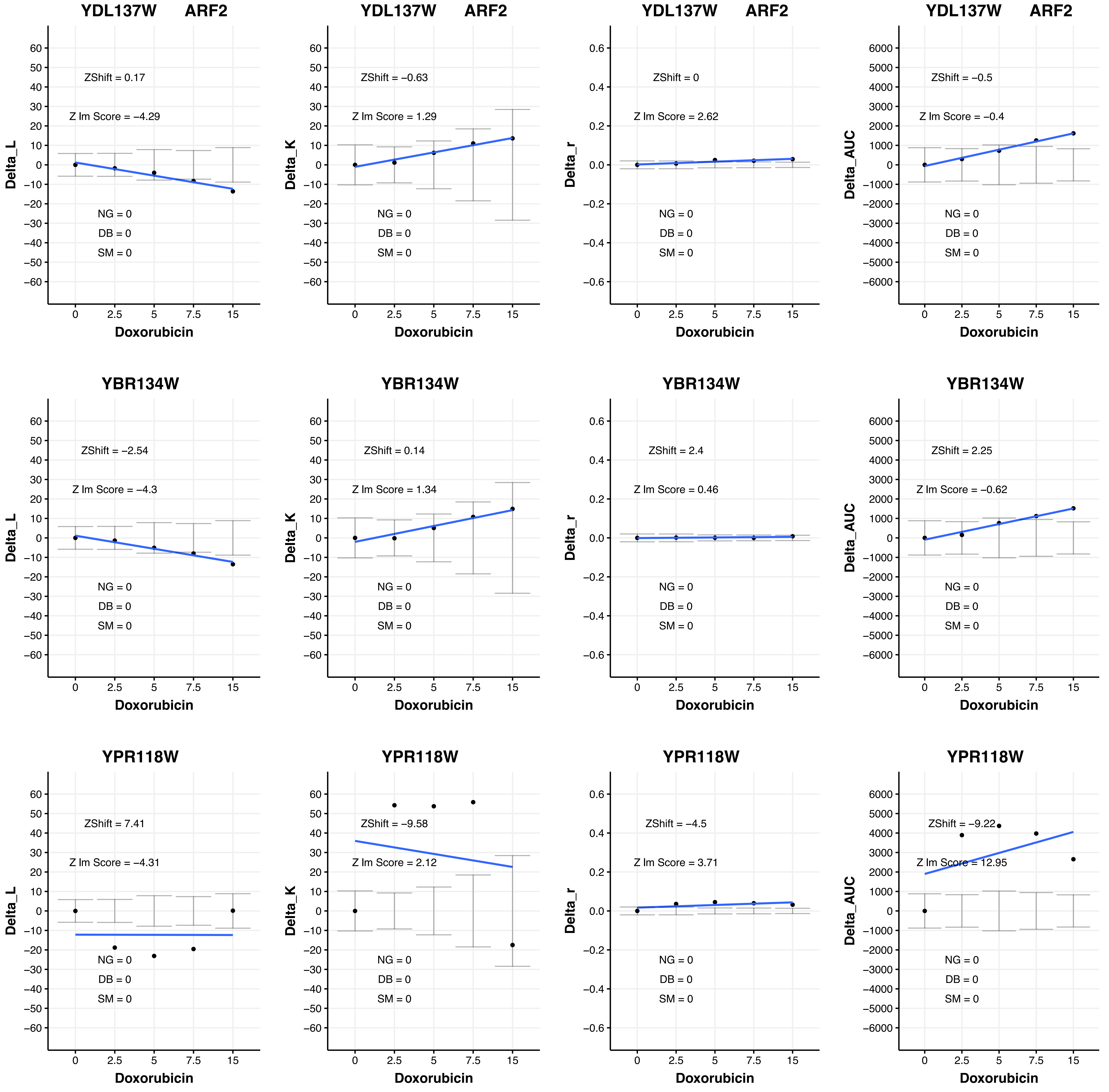

YDL137W ARF2
ZShift = 0.17
Z lm Score = −4.29
NG = 0
DB = 0
SM = 0
Delta_L
Doxorubicin
YDL137W ARF2
ZShift = −0.63
Z lm Score = 1.29
NG = 0
DB = 0
SM = 0
Delta_K
Doxorubicin
YDL137W ARF2
ZShift = 0
Z lm Score = 2.62
NG = 0
DB = 0
SM = 0
Delta_r
Doxorubicin
YDL137W ARF2
ZShift = −0.5
Z lm Score = −0.4
NG = 0
DB = 0
SM = 0
Delta_AUC
Doxorubicin
YBR134W
ZShift = −2.54
Z lm Score = −4.3
NG = 0
DB = 0
SM = 0
Delta_L
Doxorubicin
YBR134W
ZShift = 0.14
Z lm Score = 1.34
NG = 0
DB = 0
SM = 0
Delta_K
Doxorubicin
YBR134W
ZShift = 2.4
Z lm Score = 0.46
NG = 0
DB = 0
SM = 0
Delta_r
Doxorubicin
YBR134W
ZShift = 2.25
Z lm Score = −0.62
NG = 0
DB = 0
SM = 0
Delta_AUC
Doxorubicin
YPR118W
ZShift = 7.41
Z lm Score = −4.31
NG = 0
DB = 0
SM = 0
Delta_L
Doxorubicin
YPR118W
ZShift = −9.58
Z lm Score = 2.12
NG = 0
DB = 0
SM = 0
Delta_K
Doxorubicin
YPR118W
ZShift = −4.5
Z lm Score = 3.71
NG = 0
DB = 0
SM = 0
Delta_r
Doxorubicin
YPR118W
ZShift = −9.22
Z lm Score = 12.95
NG = 0
DB = 0
SM = 0
Delta_AUC
Doxorubicin

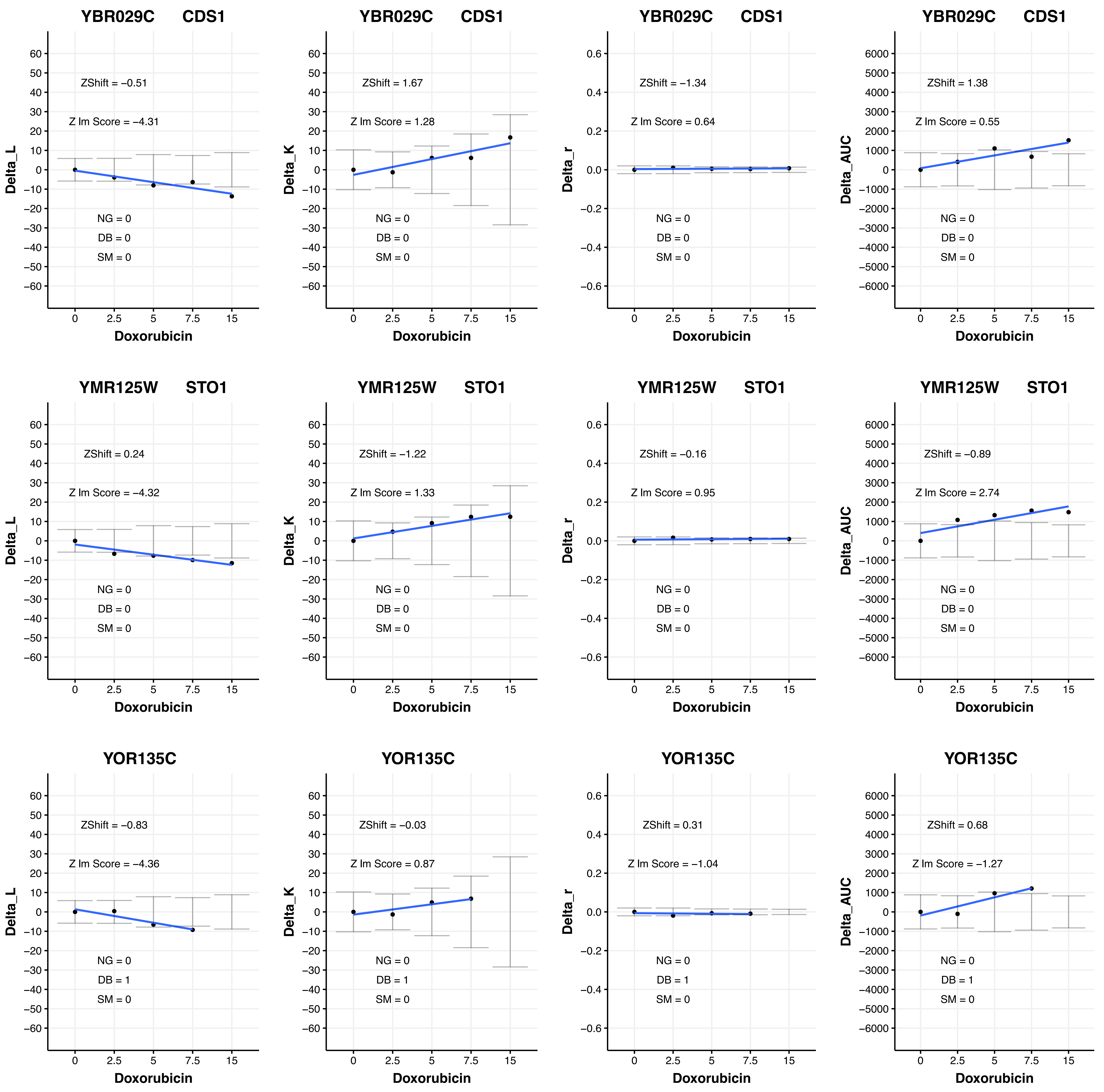
YBR029C CDS1
ZShift = −0.51
Z lm Score = −4.31
NG = 0
DB = 0
SM = 0
Delta_L
Doxorubicin
YBR029C CDS1
ZShift = 1.67
Z lm Score = 1.28
NG = 0
DB = 0
SM = 0
Delta_K
Doxorubicin
YBR029C CDS1
ZShift = −1.34
Z lm Score = 0.64
NG = 0
DB = 0
SM = 0
Delta_r
Doxorubicin
YBR029C CDS1
ZShift = 1.38
Z lm Score = 0.55
NG = 0
DB = 0
SM = 0
Delta_AUC
Doxorubicin
YMR125W STO1
ZShift = 0.24
Z lm Score = −4.32
NG = 0
DB = 0
SM = 0
Delta_L
Doxorubicin
YMR125W STO1
ZShift = −1.22
Z lm Score = 1.33
NG = 0
DB = 0
SM = 0
Delta_K
Doxorubicin
YMR125W STO1
ZShift = −0.16
Z lm Score = 0.95
NG = 0
DB = 0
SM = 0
Delta_r
Doxorubicin
YMR125W STO1
ZShift = −0.89
Z lm Score = 2.74
NG = 0
DB = 0
SM = 0
Delta_AUC
Doxorubicin
YOR135C
ZShift = −0.83
Z lm Score = −4.36
NG = 0
DB = 1
SM = 0
Delta_L
Doxorubicin
YOR135C
ZShift = −0.03
Z lm Score = 0.87
NG = 0
DB = 1
SM = 0
Delta_K
Doxorubicin
YOR135C
ZShift = 0.31
Z lm Score = −1.04
NG = 0
DB = 1
SM = 0
Delta_r
Doxorubicin
YOR135C
ZShift = 0.68
Z lm Score = −1.27
NG = 0
DB = 1
SM = 0
Delta_AUC
Doxorubicin

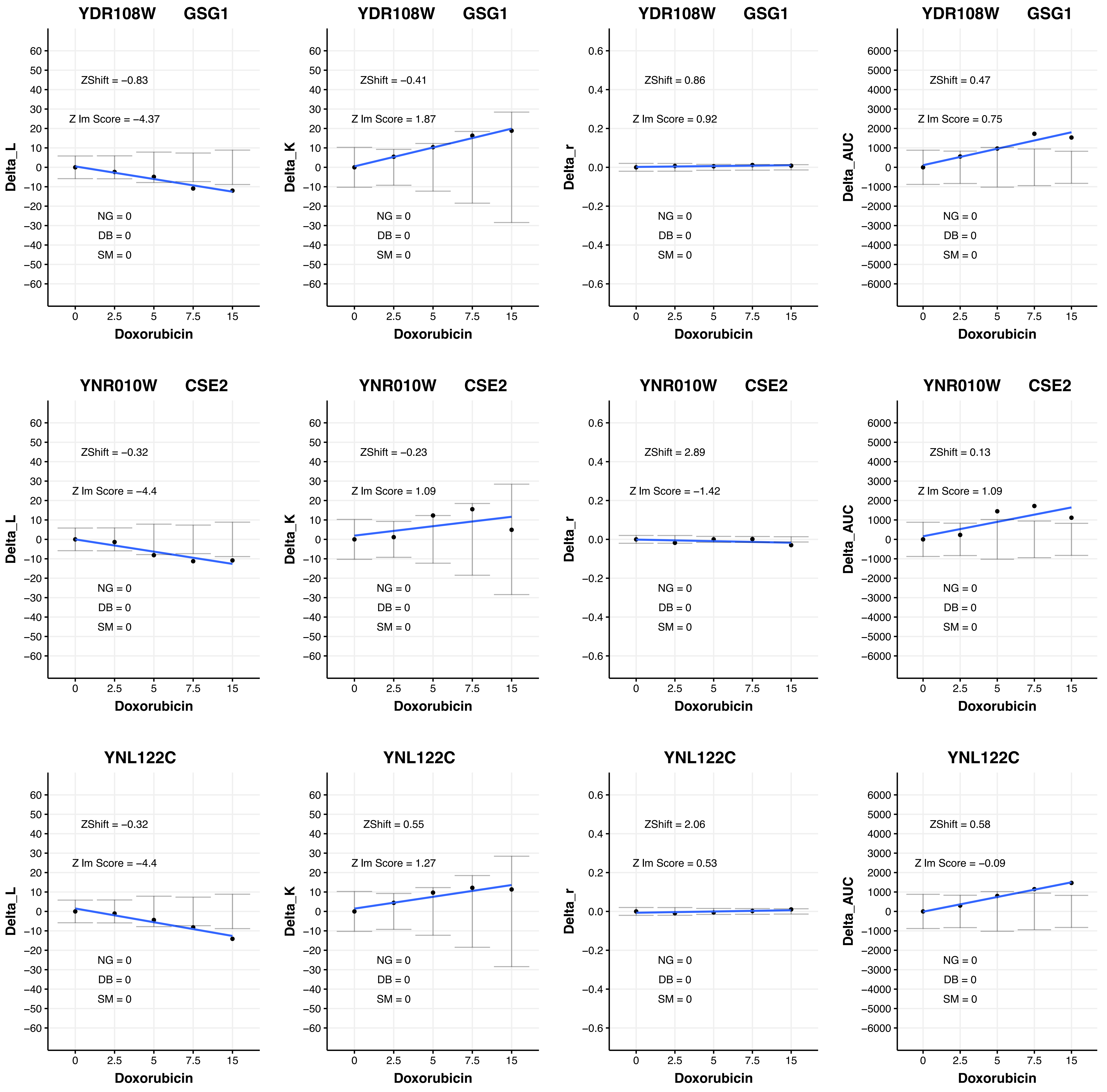

YDR108W GSG1
ZShift = −0.83
Z lm Score = −4.37
NG = 0
DB = 0
SM = 0
Delta_L
Doxorubicin
YDR108W GSG1
ZShift = −0.41
Z lm Score = 1.87
NG = 0
DB = 0
SM = 0
Delta_K
Doxorubicin
YDR108W GSG1
ZShift = 0.86
Z lm Score = 0.92
NG = 0
DB = 0
SM = 0
Delta_r
Doxorubicin
YDR108W GSG1
ZShift = 0.47
Z lm Score = 0.75
NG = 0
DB = 0
SM = 0
Delta_AUC
Doxorubicin
YNR010W CSE2
ZShift = −0.32
Z lm Score = −4.4
NG = 0
DB = 0
SM = 0
Delta_L
Doxorubicin
YNR010W CSE2
ZShift = −0.23
Z lm Score = 1.09
NG = 0
DB = 0
SM = 0
Delta_K
Doxorubicin
YNR010W CSE2
ZShift = 2.89
Z lm Score = −1.42
NG = 0
DB = 0
SM = 0
Delta_r
Doxorubicin
YNR010W CSE2
ZShift = 0.13
Z lm Score = 1.09
NG = 0
DB = 0
SM = 0
Delta_AUC
Doxorubicin
YNL122C
ZShift = −0.32
Z lm Score = −4.4
NG = 0
DB = 0
SM = 0
Delta_L
Doxorubicin
YNL122C
ZShift = 0.55
Z lm Score = 1.27
NG = 0
DB = 0
SM = 0
Delta_K
Doxorubicin
YNL122C
ZShift = 2.06
Z lm Score = 0.53
NG = 0
DB = 0
SM = 0
Delta_r
Doxorubicin
YNL122C
ZShift = 0.58
Z lm Score = −0.09
NG = 0
DB = 0
SM = 0
Delta_AUC
Doxorubicin

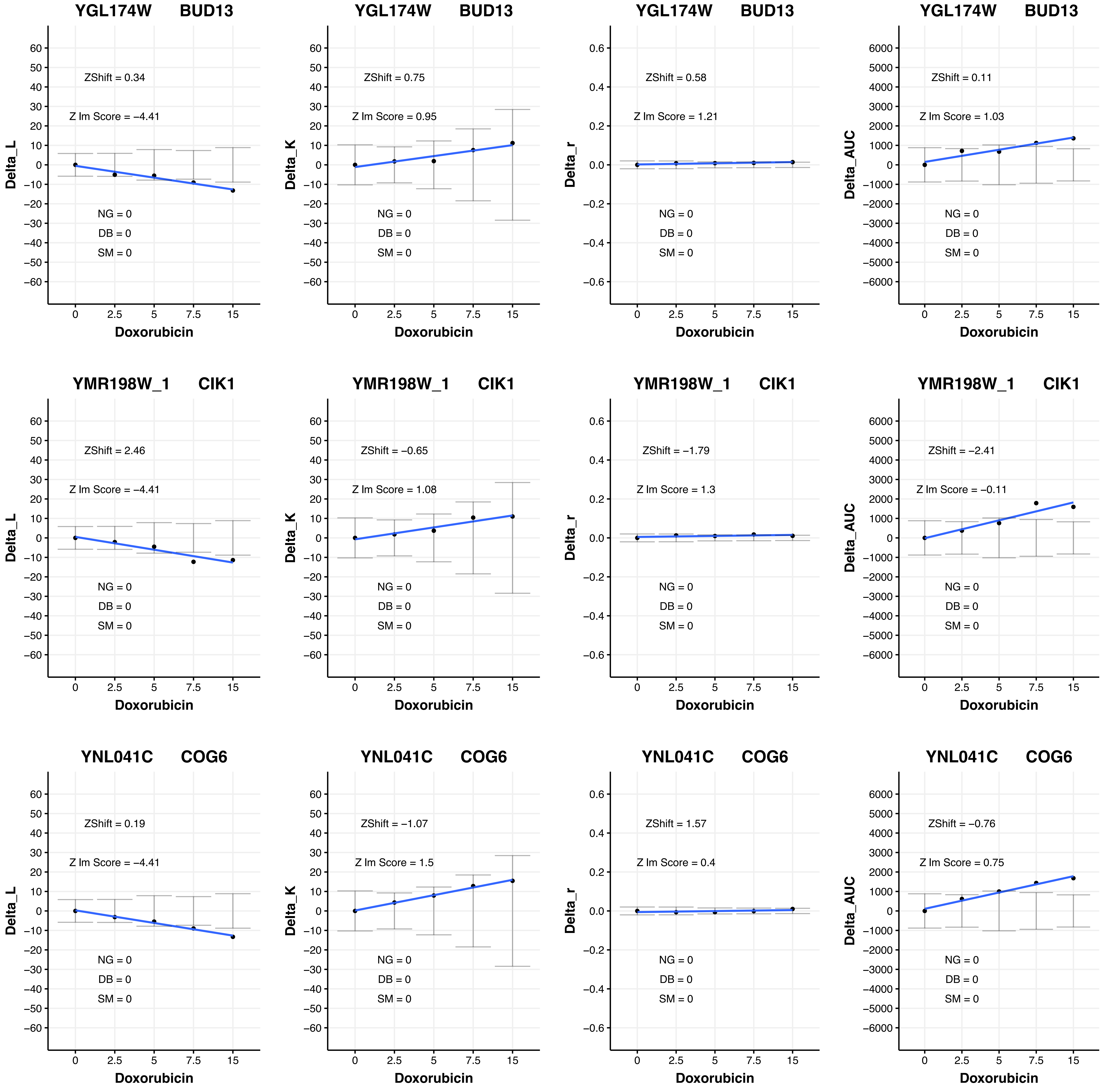

YGL174W BUD13
ZShift = 0.34
Z lm Score = −4.41
NG = 0
DB = 0
SM = 0
Delta_L
Doxorubicin
YGL174W BUD13
ZShift = 0.75
Z lm Score = 0.95
NG = 0
DB = 0
SM = 0
Delta_K
Doxorubicin
YGL174W BUD13
ZShift = 0.58
Z lm Score = 1.21
NG = 0
DB = 0
SM = 0
Delta_r
Doxorubicin
YGL174W BUD13
ZShift = 0.11
Z lm Score = 1.03
NG = 0
DB = 0
SM = 0
Delta_AUC
Doxorubicin
YMR198W_1 CIK1
ZShift = 2.46
Z lm Score = −4.41
NG = 0
DB = 0
SM = 0
Delta_L
Doxorubicin
YMR198W_1 CIK1
ZShift = −0.65
Z lm Score = 1.08
NG = 0
DB = 0
SM = 0
Delta_K
Doxorubicin
YMR198W_1 CIK1
ZShift = −1.79
Z lm Score = 1.3
NG = 0
DB = 0
SM = 0
Delta_r
Doxorubicin
YMR198W_1 CIK1
ZShift = −2.41
Z lm Score = −0.11
NG = 0
DB = 0
SM = 0
Delta_AUC
Doxorubicin
YNL041C COG6
ZShift = 0.19
Z lm Score = −4.41
NG = 0
DB = 0
SM = 0
Delta_L
Doxorubicin
YNL041C COG6
ZShift = −1.07
Z lm Score = 1.5
NG = 0
DB = 0
SM = 0
Delta_K
Doxorubicin
YNL041C COG6
ZShift = 1.57
Z lm Score = 0.4
NG = 0
DB = 0
SM = 0
Delta_r
Doxorubicin
YNL041C COG6
ZShift = −0.76
Z lm Score = 0.75
NG = 0
DB = 0
SM = 0
Delta_AUC
Doxorubicin

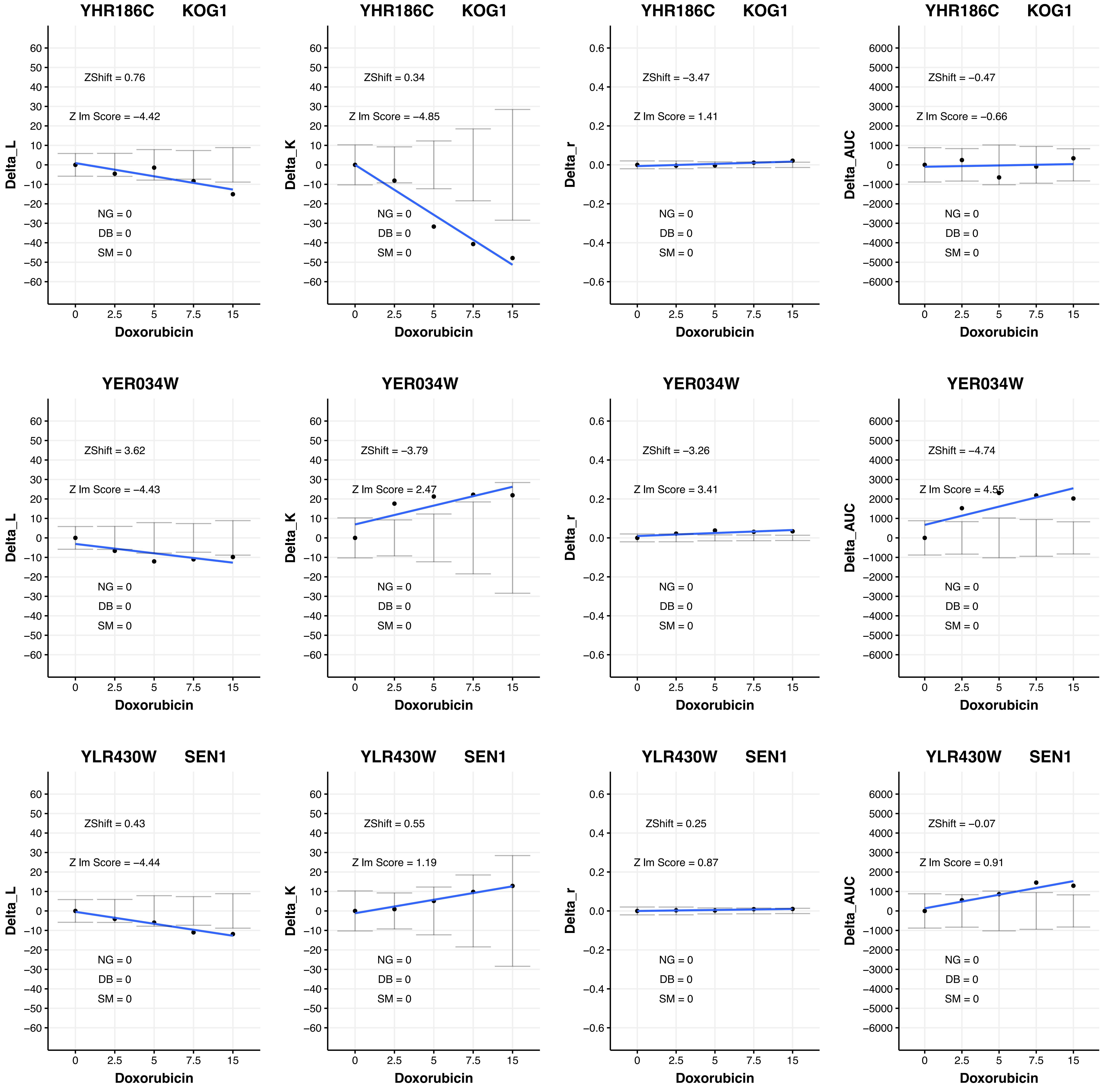
YHR186C KOG1
ZShift = 0.76
Z lm Score = −4.42
NG = 0
DB = 0
SM = 0
Delta_L
Doxorubicin
YHR186C KOG1
ZShift = 0.34
Z lm Score = −4.85
NG = 0
DB = 0
SM = 0
Delta_K
Doxorubicin
YHR186C KOG1
ZShift = −3.47
Z lm Score = 1.41
NG = 0
DB = 0
SM = 0
Delta_r
Doxorubicin
YHR186C KOG1
ZShift = −0.47
Z lm Score = −0.66
NG = 0
DB = 0
SM = 0
Delta_AUC
Doxorubicin
YER034W
ZShift = 3.62
Z lm Score = −4.43
NG = 0
DB = 0
SM = 0
Delta_L
Doxorubicin
YER034W
ZShift = −3.79
Z lm Score = 2.47
NG = 0
DB = 0
SM = 0
Delta_K
Doxorubicin
YER034W
ZShift = −3.26
Z lm Score = 3.41
NG = 0
DB = 0
SM = 0
Delta_r
Doxorubicin
YER034W
ZShift = −4.74
Z lm Score = 4.55
NG = 0
DB = 0
SM = 0
Delta_AUC
Doxorubicin
YLR430W SEN1
ZShift = 0.43
Z lm Score = −4.44
NG = 0
DB = 0
SM = 0
Delta_L
Doxorubicin
YLR430W SEN1
ZShift = 0.55
Z lm Score = 1.19
NG = 0
DB = 0
SM = 0
Delta_K
Doxorubicin
YLR430W SEN1
ZShift = 0.25
Z lm Score = 0.87
NG = 0
DB = 0
SM = 0
Delta_r
Doxorubicin
YLR430W SEN1
ZShift = −0.07
Z lm Score = 0.91
NG = 0
DB = 0
SM = 0
Delta_AUC
Doxorubicin

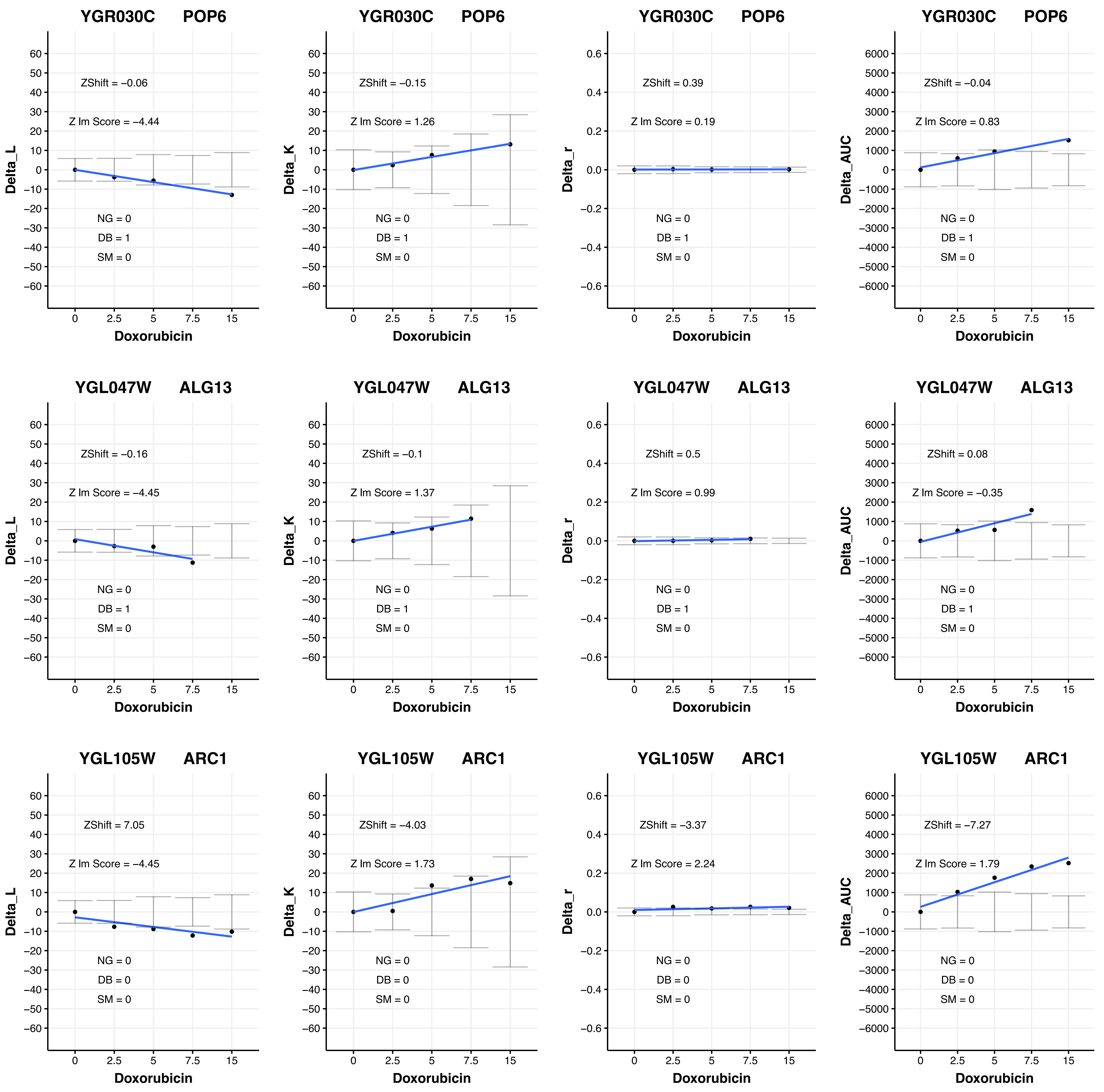

YGR030C POP6
ZShift = −0.06
Z lm Score = −4.44
NG = 0
DB = 1
SM = 0
Delta_L
Doxorubicin
YGR030C POP6
ZShift = −0.15
Z lm Score = 1.26
NG = 0
DB = 1
SM = 0
Delta_K
Doxorubicin
YGR030C POP6
ZShift = 0.39
Z lm Score = 0.19
NG = 0
DB = 1
SM = 0
Delta_r
Doxorubicin
YGR030C POP6
ZShift = −0.04
Z lm Score = 0.83
NG = 0
DB = 1
SM = 0
Delta_AUC
Doxorubicin
YGL047W ALG13
ZShift = −0.16
Z lm Score = −4.45
NG = 0
DB = 1
SM = 0
Delta_L
Doxorubicin
YGL047W ALG13
ZShift = −0.1
Z lm Score = 1.37
NG = 0
DB = 1
SM = 0
Delta_K
Doxorubicin
YGL047W ALG13
ZShift = 0.5
Z lm Score = 0.99
NG = 0
DB = 1
SM = 0
Delta_r
Doxorubicin
YGL047W ALG13
ZShift = 0.08
Z lm Score = −0.35
NG = 0
DB = 1
SM = 0
Delta_AUC
Doxorubicin
YGL105W ARC1
ZShift = 7.05
Z lm Score = −4.45
NG = 0
DB = 0
SM = 0
Delta_L
Doxorubicin
YGL105W ARC1
ZShift = −4.03
Z lm Score = 1.73
NG = 0
DB = 0
SM = 0
Delta_K
Doxorubicin
YGL105W ARC1
ZShift = −3.37
Z lm Score = 2.24
NG = 0
DB = 0
SM = 0
Delta_r
Doxorubicin
YGL105W ARC1
ZShift = −7.27
Z lm Score = 1.79
NG = 0
DB = 0
SM = 0
Delta_AUC
Doxorubicin

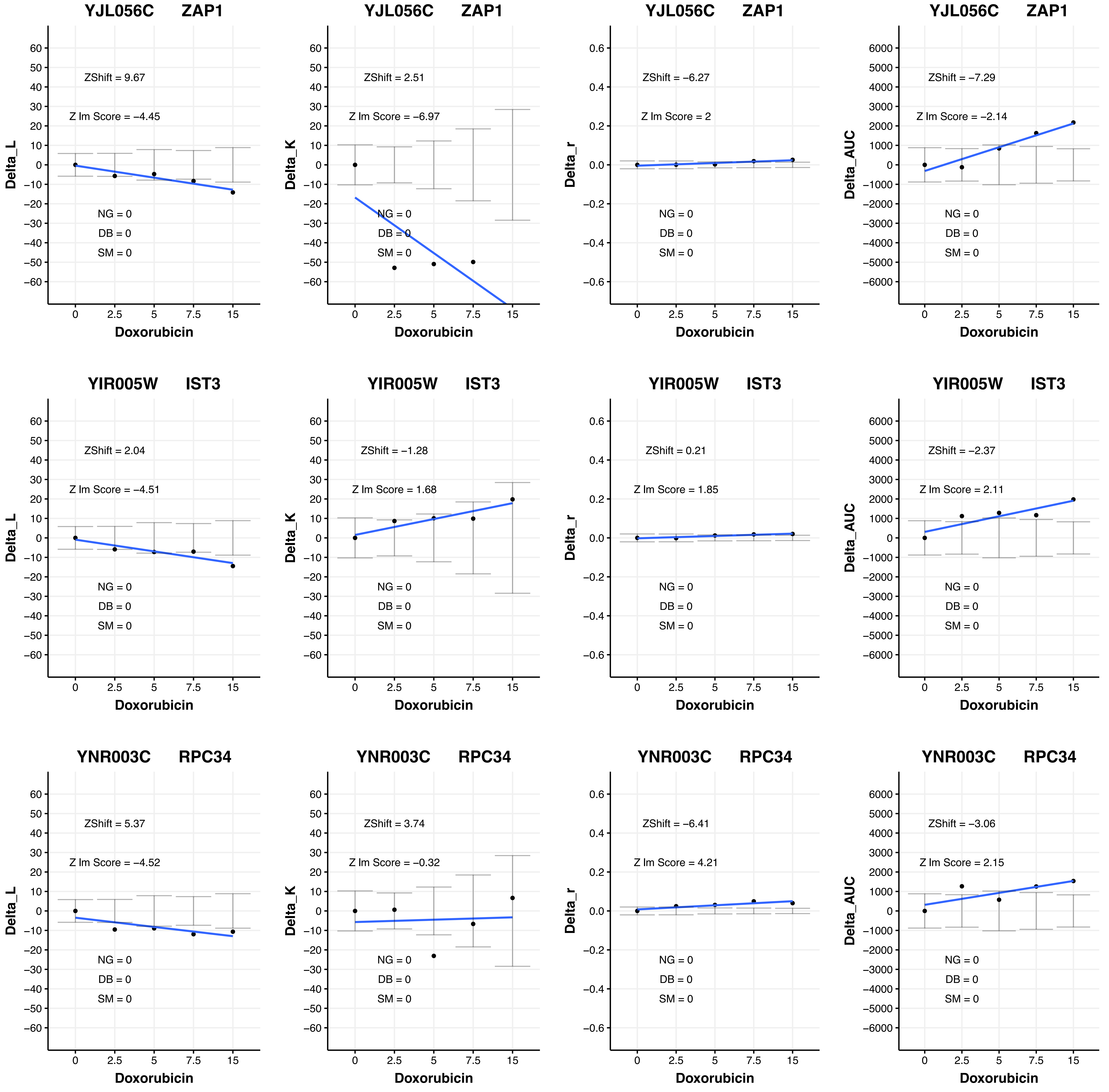
YJL056C ZAP1
ZShift = 9.67
Z lm Score = −4.45
NG = 0
DB = 0
SM = 0
Delta_L
Doxorubicin
YJL056C ZAP1
ZShift = 2.51
Z lm Score = −6.97
NG = 0
DB = 0
SM = 0
Delta_K
Doxorubicin
YJL056C ZAP1
ZShift = −6.27
Z lm Score = 2
NG = 0
DB = 0
SM = 0
Delta_r
Doxorubicin
YJL056C ZAP1
ZShift = −7.29
Z lm Score = −2.14
NG = 0
DB = 0
SM = 0
Delta_AUC
Doxorubicin
YIR005W IST3
ZShift = 2.04
Z lm Score = −4.51
NG = 0
DB = 0
SM = 0
Delta_L
Doxorubicin
YIR005W IST3
ZShift = −1.28
Z lm Score = 1.68
NG = 0
DB = 0
SM = 0
Delta_K
Doxorubicin
YIR005W IST3
ZShift = 0.21
Z lm Score = 1.85
NG = 0
DB = 0
SM = 0
Delta_r
Doxorubicin
YIR005W IST3
ZShift = −2.37
Z lm Score = 2.11
NG = 0
DB = 0
SM = 0
Delta_AUC
Doxorubicin
YNR003C RPC34
ZShift = 5.37
Z lm Score = −4.52
NG = 0
DB = 0
SM = 0
Delta_L
Doxorubicin
YNR003C RPC34
ZShift = 3.74
Z lm Score = −0.32
NG = 0
DB = 0
SM = 0
Delta_K
Doxorubicin
YNR003C RPC34
ZShift = −6.41
Z lm Score = 4.21
NG = 0
DB = 0
SM = 0
Delta_r
Doxorubicin
YNR003C RPC34
ZShift = −3.06
Z lm Score = 2.15
NG = 0
DB = 0
SM = 0
Delta_AUC
Doxorubicin

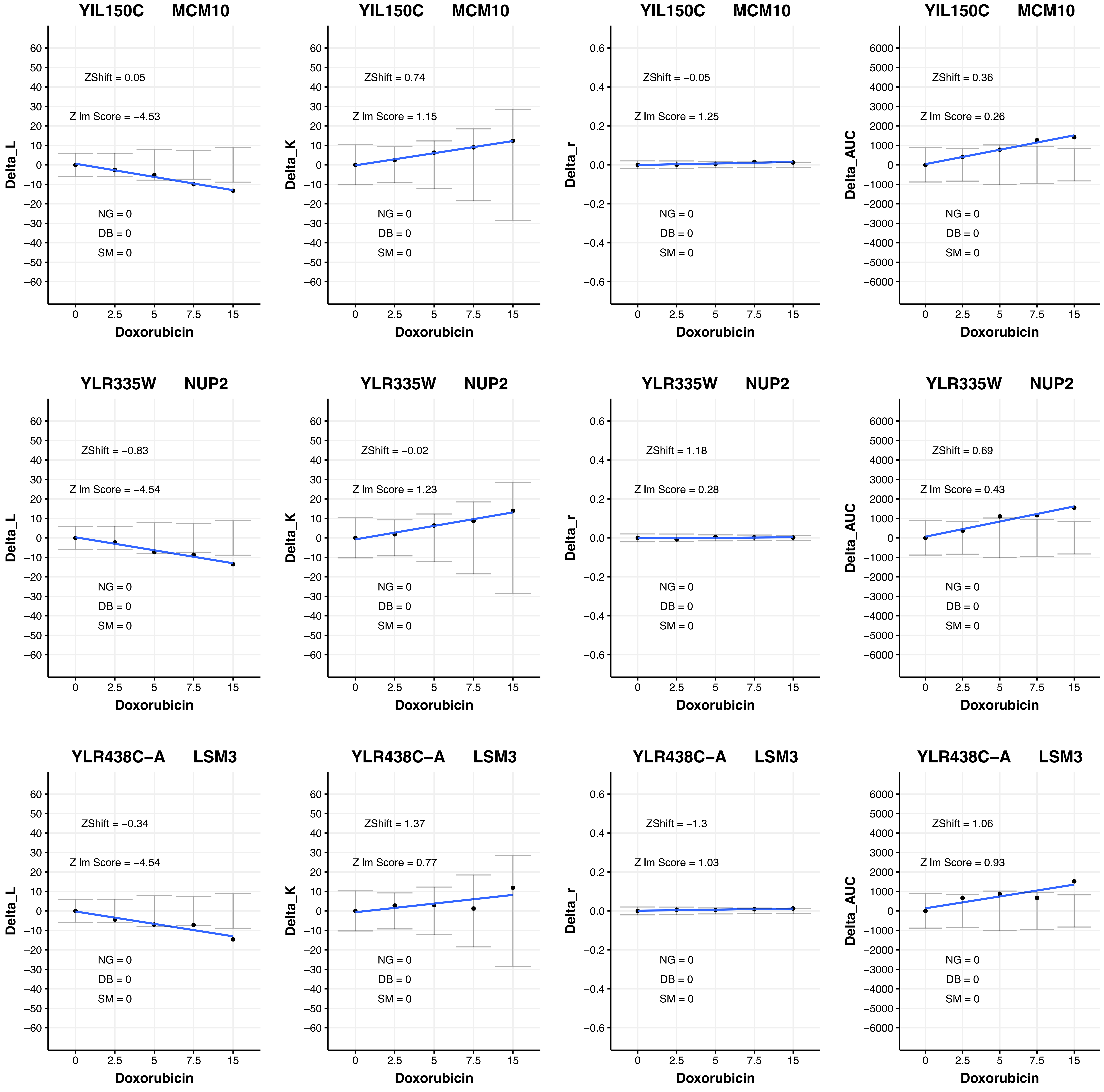
YIL150C MCM10
ZShift = 0.05
Z lm Score = −4.53
NG = 0
DB = 0
SM = 0
Delta_L
Doxorubicin
YIL150C MCM10
ZShift = 0.74
Z lm Score = 1.15
NG = 0
DB = 0
SM = 0
Delta_K
Doxorubicin
YIL150C MCM10
ZShift = −0.05
Z lm Score = 1.25
NG = 0
DB = 0
SM = 0
Delta_r
Doxorubicin
YIL150C MCM10
ZShift = 0.36
Z lm Score = 0.26
NG = 0
DB = 0
SM = 0
Delta_AUC
Doxorubicin
YLR335W NUP2
ZShift = −0.83
Z lm Score = −4.54
NG = 0
DB = 0
SM = 0
Delta_L
Doxorubicin
YLR335W NUP2
ZShift = −0.02
Z lm Score = 1.23
NG = 0
DB = 0
SM = 0
Delta_K
Doxorubicin
YLR335W NUP2
ZShift = 1.18
Z lm Score = 0.28
NG = 0
DB = 0
SM = 0
Delta_r
Doxorubicin
YLR335W NUP2
ZShift = 0.69
Z lm Score = 0.43
NG = 0
DB = 0
SM = 0
Delta_AUC
Doxorubicin
YLR438C−A LSM3
ZShift = −0.34
Z lm Score = −4.54
NG = 0
DB = 0
SM = 0
Delta_L
Doxorubicin
YLR438C−A LSM3
ZShift = 1.37
Z lm Score = 0.77
NG = 0
DB = 0
SM = 0
Delta_K
Doxorubicin
YLR438C−A LSM3
ZShift = −1.3
Z lm Score = 1.03
NG = 0
DB = 0
SM = 0
Delta_r
Doxorubicin
YLR438C−A LSM3
ZShift = 1.06
Z lm Score = 0.93
NG = 0
DB = 0
SM = 0
Delta_AUC
Doxorubicin

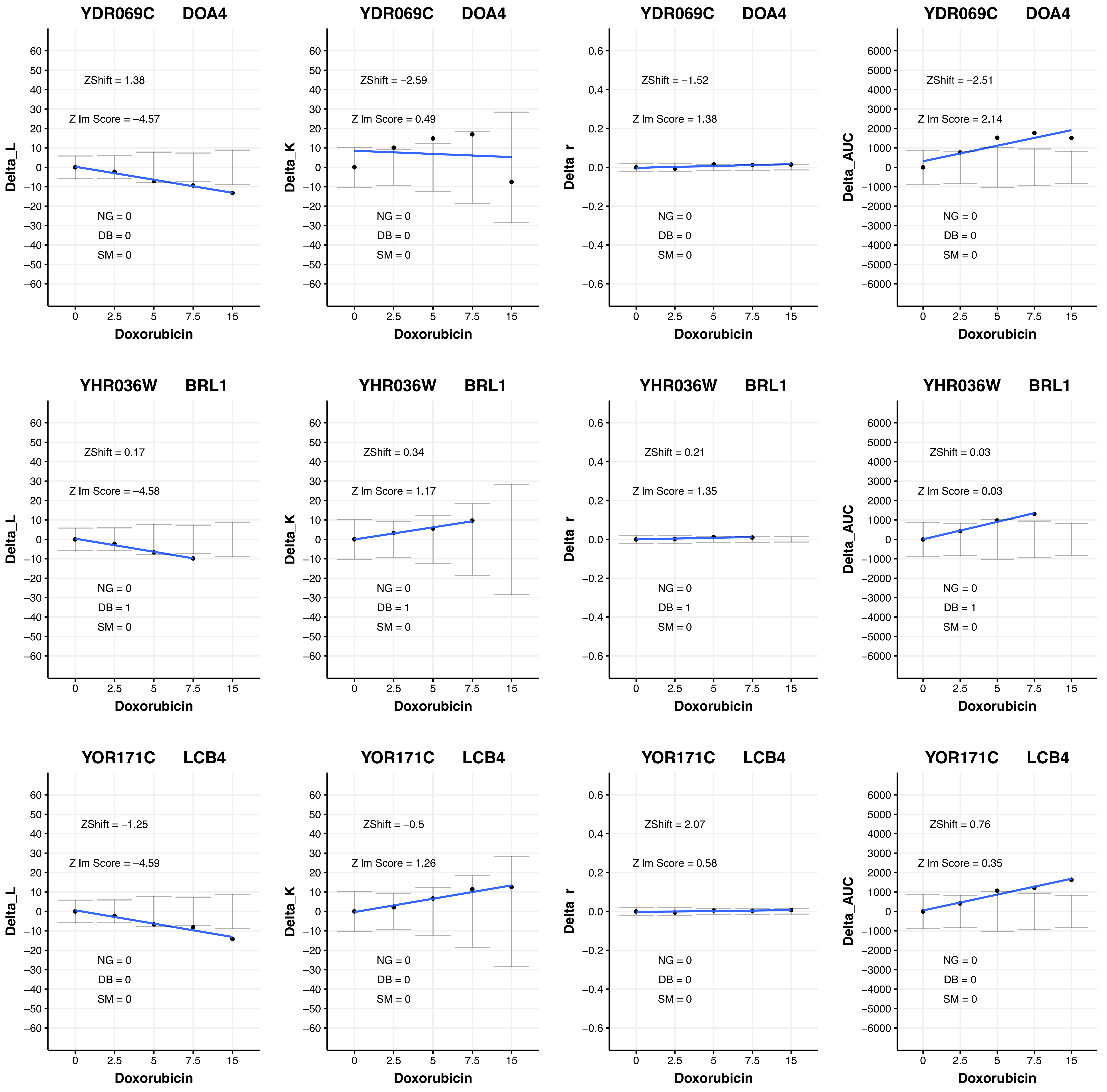
YDR069C DOA4
ZShift = 1.38
Z Im Score = −4.57
NG = 0
DB = 0
SM = 0
Delta_L
Doxorubicin
YDR069C DOA4
ZShift = −2.59
Z Im Score = 0.49
NG = 0
DB = 0
SM = 0
Delta_K
Doxorubicin
YDR069C DOA4
ZShift = −1.52
Z Im Score = 1.38
NG = 0
DB = 0
SM = 0
Delta_r
Doxorubicin
YDR069C DOA4
ZShift = −2.51
Z Im Score = 2.14
NG = 0
DB = 0
SM = 0
Delta_AUC
Doxorubicin
YHR036W BRL1
ZShift = 0.17
Z Im Score = −4.58
NG = 0
DB = 1
SM = 0
Delta_L
Doxorubicin
YHR036W BRL1
ZShift = 0.34
Z Im Score = 1.17
NG = 0
DB = 1
SM = 0
Delta_K
Doxorubicin
YHR036W BRL1
ZShift = 0.21
Z Im Score = 1.35
NG = 0
DB = 1
SM = 0
Delta_r
Doxorubicin
YHR036W BRL1
ZShift = 0.03
Z Im Score = 0.03
NG = 0
DB = 1
SM = 0
Delta_AUC
Doxorubicin
YOR171C LCB4
ZShift = −1.25
Z Im Score = −4.59
NG = 0
DB = 0
SM = 0
Delta_L
Doxorubicin
YOR171C LCB4
ZShift = −0.5
Z Im Score = 1.26
NG = 0
DB = 0
SM = 0
Delta_K
Doxorubicin
YOR171C LCB4
ZShift = 2.07
Z Im Score = 0.58
NG = 0
DB = 0
SM = 0
Delta_r
Doxorubicin
YOR171C LCB4
ZShift = 0.76
Z Im Score = 0.35
NG = 0
DB = 0
SM = 0
Delta_AUC
Doxorubicin

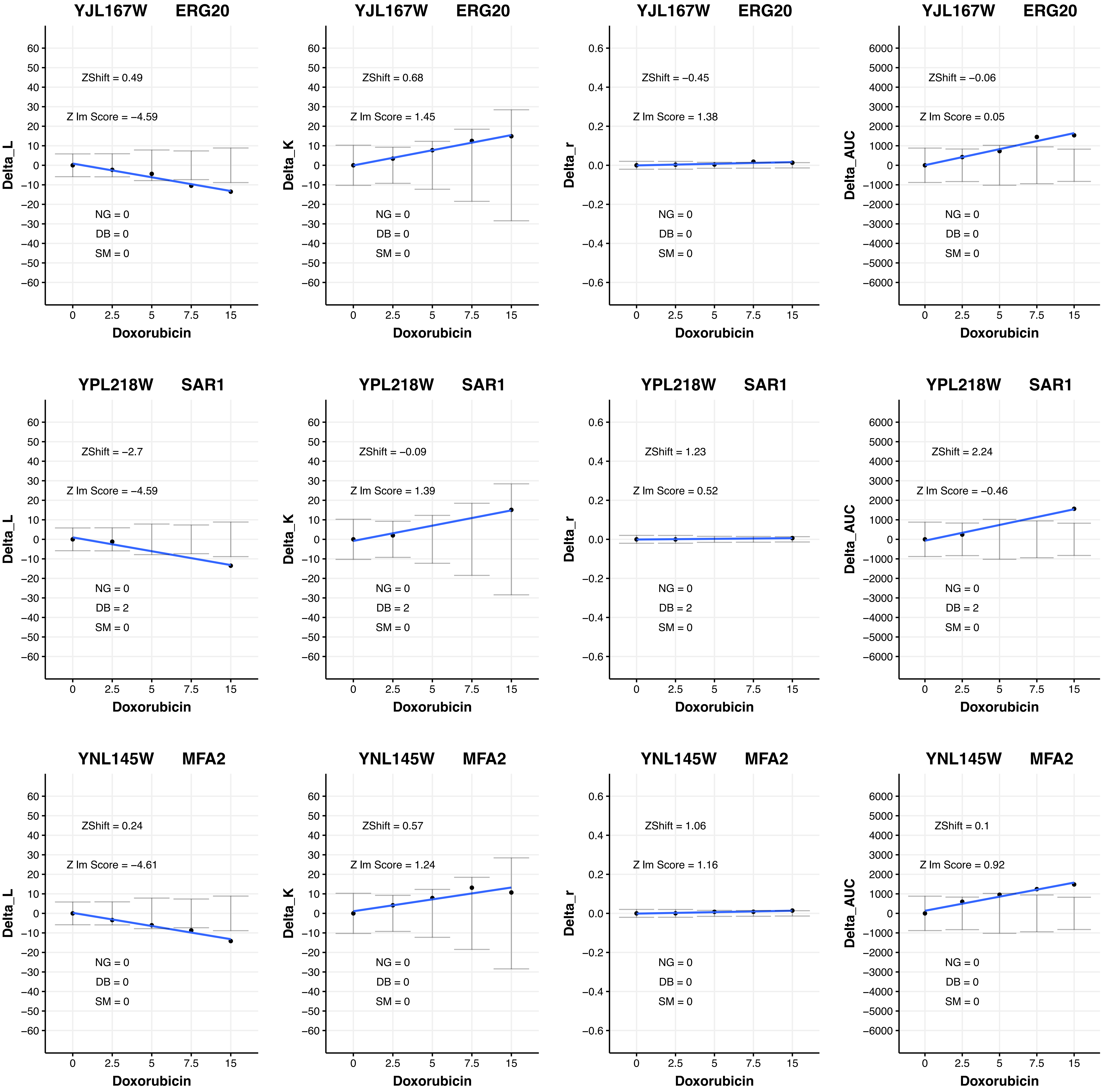
YJL167W ERG20
ZShift = 0.49
Z lm Score = −4.59
NG = 0
DB = 0
SM = 0
Delta_L
Doxorubicin
YJL167W ERG20
ZShift = 0.68
Z lm Score = 1.45
NG = 0
DB = 0
SM = 0
Delta_K
Doxorubicin
YJL167W ERG20
ZShift = −0.45
Z lm Score = 1.38
NG = 0
DB = 0
SM = 0
Delta_r
Doxorubicin
YJL167W ERG20
ZShift = −0.06
Z lm Score = 0.05
NG = 0
DB = 0
SM = 0
Delta_AUC
Doxorubicin
YPL218W SAR1
ZShift = −2.7
Z lm Score = −4.59
NG = 0
DB = 2
SM = 0
Delta_L
Doxorubicin
YPL218W SAR1
ZShift = −0.09
Z lm Score = 1.39
NG = 0
DB = 2
SM = 0
Delta_K
Doxorubicin
YPL218W SAR1
ZShift = 1.23
Z lm Score = 0.52
NG = 0
DB = 2
SM = 0
Delta_r
Doxorubicin
YPL218W SAR1
ZShift = 2.24
Z lm Score = −0.46
NG = 0
DB = 2
SM = 0
Delta_AUC
Doxorubicin
YNL145W MFA2
ZShift = 0.24
Z lm Score = −4.61
NG = 0
DB = 0
SM = 0
Delta_L
Doxorubicin
YNL145W MFA2
ZShift = 0.57
Z lm Score = 1.24
NG = 0
DB = 0
SM = 0
Delta_K
Doxorubicin
YNL145W MFA2
ZShift = 1.06
Z lm Score = 1.16
NG = 0
DB = 0
SM = 0
Delta_r
Doxorubicin
YNL145W MFA2
ZShift = 0.1
Z lm Score = 0.92
NG = 0
DB = 0
SM = 0
Delta_AUC
Doxorubicin

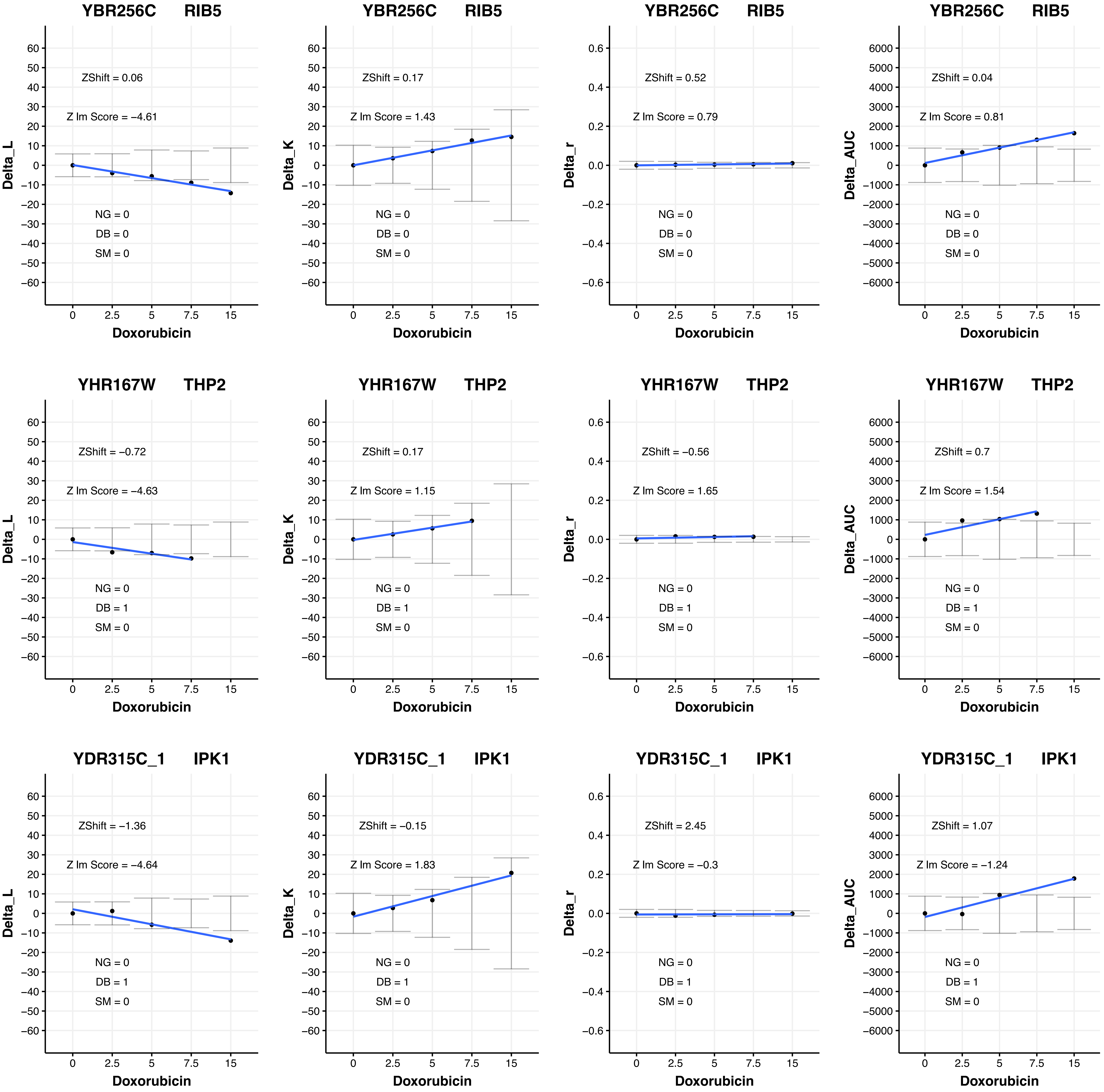
YBR256C RIB5
ZShift = 0.06
Z lm Score = −4.61
NG = 0
DB = 0
SM = 0
Delta_L
Doxorubicin
YBR256C RIB5
ZShift = 0.17
Z lm Score = 1.43
NG = 0
DB = 0
SM = 0
Delta_K
Doxorubicin
YBR256C RIB5
ZShift = 0.52
Z lm Score = 0.79
NG = 0
DB = 0
SM = 0
Delta_r
Doxorubicin
YBR256C RIB5
ZShift = 0.04
Z lm Score = 0.81
NG = 0
DB = 0
SM = 0
Delta_AUC
Doxorubicin
YHR167W THP2
ZShift = −0.72
Z lm Score = −4.63
NG = 0
DB = 1
SM = 0
Delta_L
Doxorubicin
YHR167W THP2
ZShift = 0.17
Z lm Score = 1.15
NG = 0
DB = 1
SM = 0
Delta_K
Doxorubicin
YHR167W THP2
ZShift = −0.56
Z lm Score = 1.65
NG = 0
DB = 1
SM = 0
Delta_r
Doxorubicin
YHR167W THP2
ZShift = 0.7
Z lm Score = 1.54
NG = 0
DB = 1
SM = 0
Delta_AUC
Doxorubicin
YDR315C_1 IPK1
ZShift = −1.36
Z lm Score = −4.64
NG = 0
DB = 1
SM = 0
Delta_L
Doxorubicin
YDR315C_1 IPK1
ZShift = −0.15
Z lm Score = 1.83
NG = 0
DB = 1
SM = 0
Delta_K
Doxorubicin
YDR315C_1 IPK1
ZShift = 2.45
Z lm Score = −0.3
NG = 0
DB = 1
SM = 0
Delta_r
Doxorubicin
YDR315C_1 IPK1
ZShift = 1.07
Z lm Score = −1.24
NG = 0
DB = 1
SM = 0
Delta_AUC
Doxorubicin

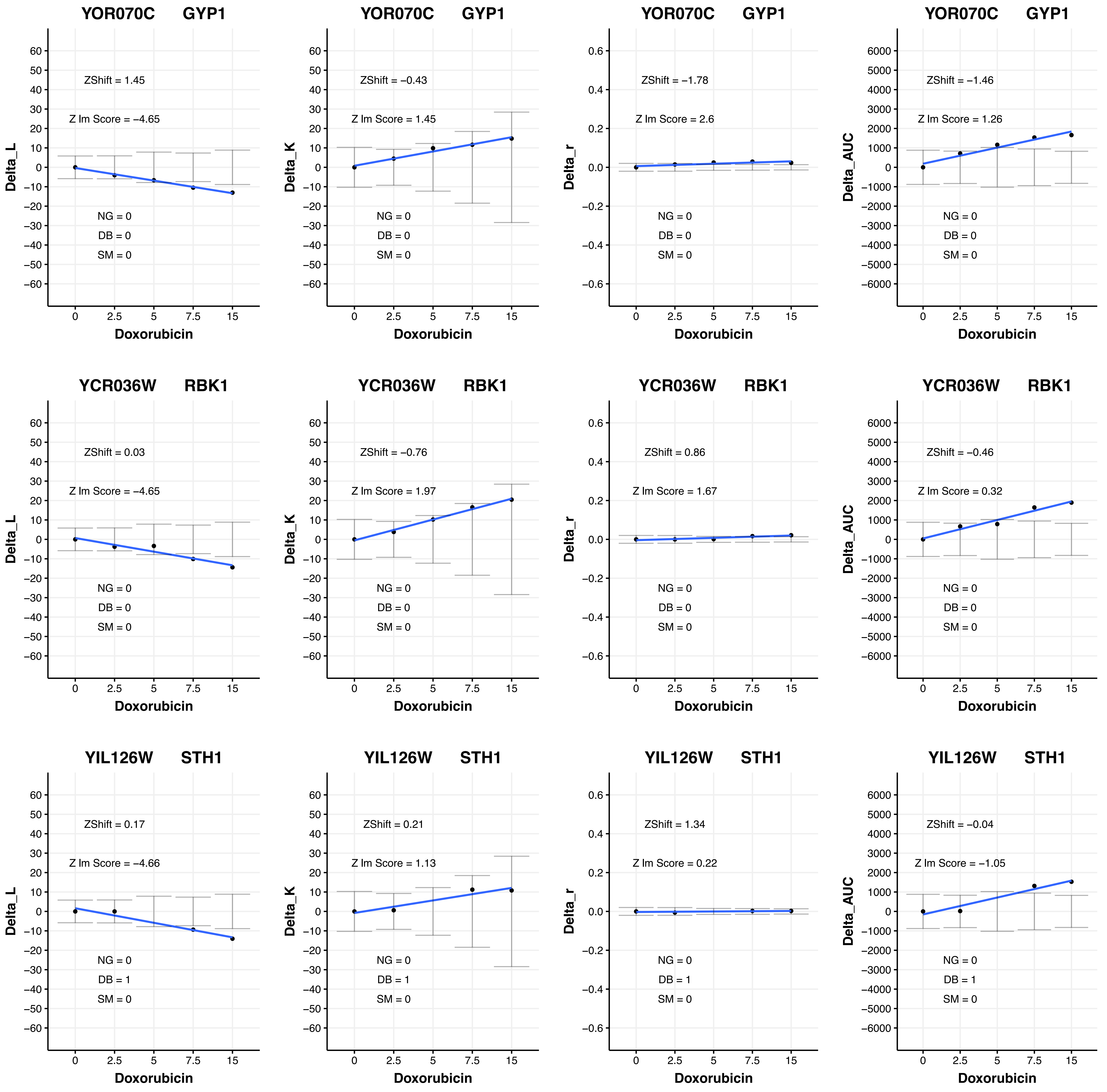
YOR070C GYP1
ZShift = 1.45
Z lm Score = −4.65
NG = 0
DB = 0
SM = 0
Delta_L
Doxorubicin
YOR070C GYP1
ZShift = −0.43
Z lm Score = 1.45
NG = 0
DB = 0
SM = 0
Delta_K
Doxorubicin
YOR070C GYP1
ZShift = −1.78
Z lm Score = 2.6
NG = 0
DB = 0
SM = 0
Delta_r
Doxorubicin
YOR070C GYP1
ZShift = −1.46
Z lm Score = 1.26
NG = 0
DB = 0
SM = 0
Delta_AUC
Doxorubicin
YCR036W RBK1
ZShift = 0.03
Z lm Score = −4.65
NG = 0
DB = 0
SM = 0
Delta_L
Doxorubicin
YCR036W RBK1
ZShift = −0.76
Z lm Score = 1.97
NG = 0
DB = 0
SM = 0
Delta_K
Doxorubicin
YCR036W RBK1
ZShift = 0.86
Z lm Score = 1.67
NG = 0
DB = 0
SM = 0
Delta_r
Doxorubicin
YCR036W RBK1
ZShift = −0.46
Z lm Score = 0.32
NG = 0
DB = 0
SM = 0
Delta_AUC
Doxorubicin
YIL126W STH1
ZShift = 0.17
Z lm Score = −4.66
NG = 0
DB = 1
SM = 0
Delta_L
Doxorubicin
YIL126W STH1
ZShift = 0.21
Z lm Score = 1.13
NG = 0
DB = 1
SM = 0
Delta_K
Doxorubicin
YIL126W STH1
ZShift = 1.34
Z lm Score = 0.22
NG = 0
DB = 1
SM = 0
Delta_r
Doxorubicin
YIL126W STH1
ZShift = −0.04
Z lm Score = −1.05
NG = 0
DB = 1
SM = 0
Delta_AUC
Doxorubicin

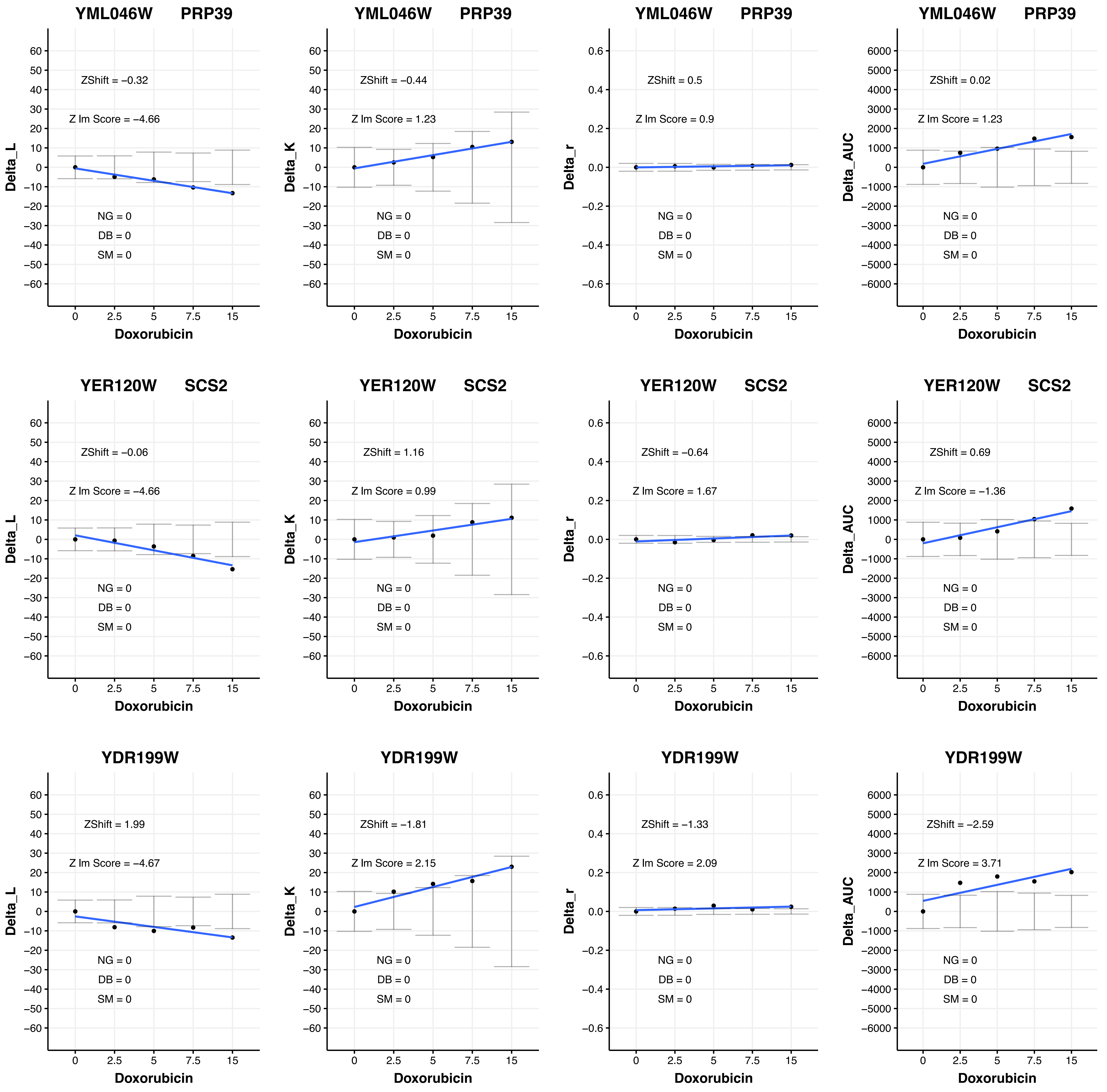
YML046W PRP39
Delta_L
ZShift = −0.32
Z lm Score = −4.66
NG = 0
DB = 0
SM = 0
Doxorubicin
YML046W PRP39
Delta_K
ZShift = −0.44
Z lm Score = 1.23
NG = 0
DB = 0
SM = 0
Doxorubicin
YML046W PRP39
Delta_r
ZShift = 0.5
Z lm Score = 0.9
NG = 0
DB = 0
SM = 0
Doxorubicin
YML046W PRP39
Delta_AUC
ZShift = 0.02
Z lm Score = 1.23
NG = 0
DB = 0
SM = 0
Doxorubicin
YER120W SCS2
Delta_L
ZShift = −0.06
Z lm Score = −4.66
NG = 0
DB = 0
SM = 0
Doxorubicin
YER120W SCS2
Delta_K
ZShift = 1.16
Z lm Score = 0.99
NG = 0
DB = 0
SM = 0
Doxorubicin
YER120W SCS2
Delta_r
ZShift = −0.64
Z lm Score = 1.67
NG = 0
DB = 0
SM = 0
Doxorubicin
YER120W SCS2
Delta_AUC
ZShift = 0.69
Z lm Score = −1.36
NG = 0
DB = 0
SM = 0
Doxorubicin
YDR199W
Delta_L
ZShift = 1.99
Z lm Score = −4.67
NG = 0
DB = 0
SM = 0
Doxorubicin
YDR199W
Delta_K
ZShift = −1.81
Z lm Score = 2.15
NG = 0
DB = 0
SM = 0
Doxorubicin
YDR199W
Delta_r
ZShift = −1.33
Z lm Score = 2.09
NG = 0
DB = 0
SM = 0
Doxorubicin
YDR199W
Delta_AUC
ZShift = −2.59
Z lm Score = 3.71
NG = 0
DB = 0
SM = 0
Doxorubicin

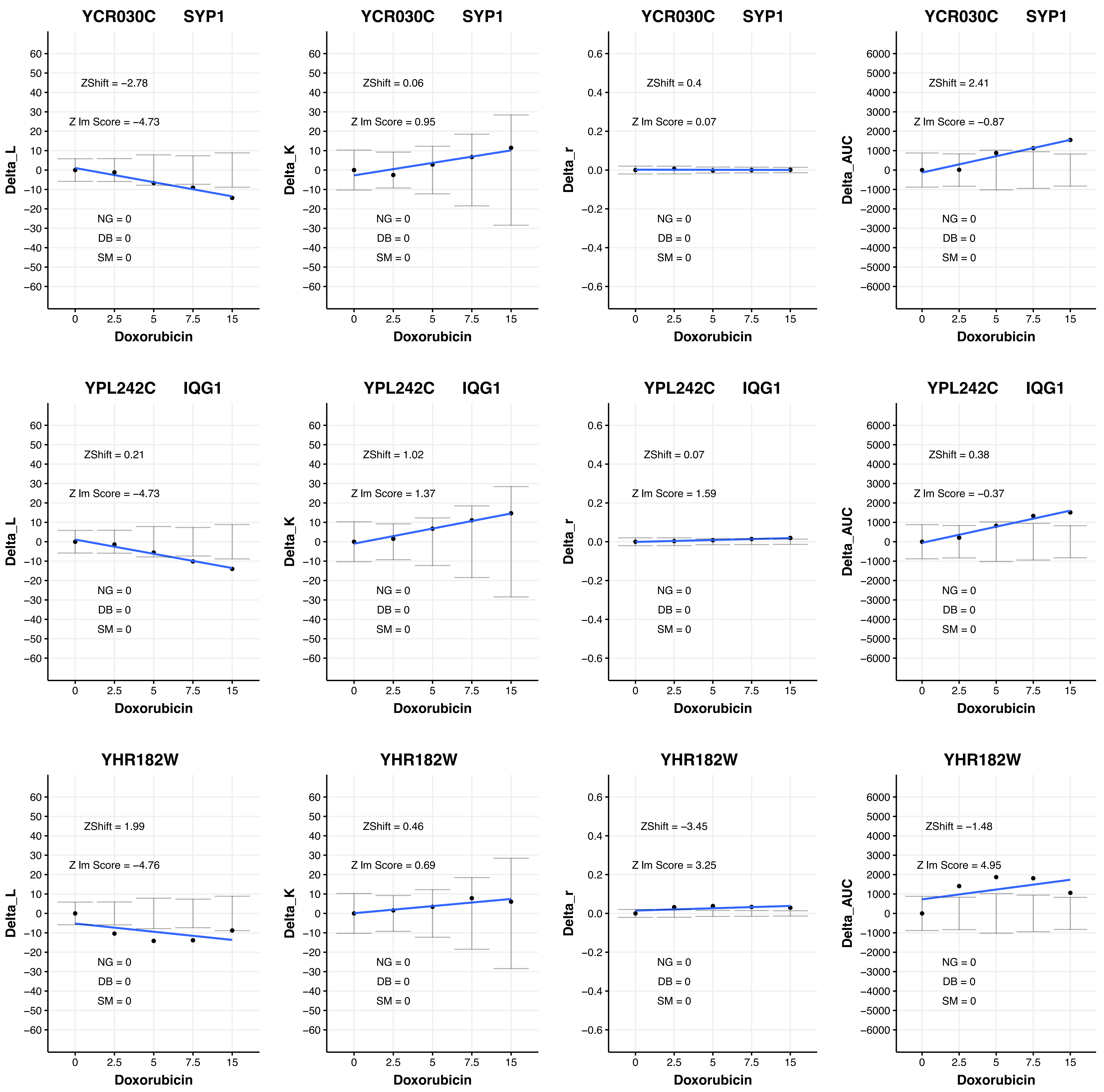
YCR030C SYP1
ZShift = −2.78
Z lm Score = −4.73
NG = 0
DB = 0
SM = 0
Delta_L
Doxorubicin
YCR030C SYP1
ZShift = 0.06
Z lm Score = 0.95
NG = 0
DB = 0
SM = 0
Delta_K
Doxorubicin
YCR030C SYP1
ZShift = 0.4
Z lm Score = 0.07
NG = 0
DB = 0
SM = 0
Delta_r
Doxorubicin
YCR030C SYP1
ZShift = 2.41
Z lm Score = −0.87
NG = 0
DB = 0
SM = 0
Delta_AUC
Doxorubicin
YPL242C IQG1
ZShift = 0.21
Z lm Score = −4.73
NG = 0
DB = 0
SM = 0
Delta_L
Doxorubicin
YPL242C IQG1
ZShift = 1.02
Z lm Score = 1.37
NG = 0
DB = 0
SM = 0
Delta_K
Doxorubicin
YPL242C IQG1
ZShift = 0.07
Z lm Score = 1.59
NG = 0
DB = 0
SM = 0
Delta_r
Doxorubicin
YPL242C IQG1
ZShift = 0.38
Z lm Score = −0.37
NG = 0
DB = 0
SM = 0
Delta_AUC
Doxorubicin
YHR182W
ZShift = 1.99
Z lm Score = −4.76
NG = 0
DB = 0
SM = 0
Delta_L
Doxorubicin
YHR182W
ZShift = 0.46
Z lm Score = 0.69
NG = 0
DB = 0
SM = 0
Delta_K
Doxorubicin
YHR182W
ZShift = −3.45
Z lm Score = 3.25
NG = 0
DB = 0
SM = 0
Delta_r
Doxorubicin
YHR182W
ZShift = −1.48
Z lm Score = 4.95
NG = 0
DB = 0
SM = 0
Delta_AUC
Doxorubicin

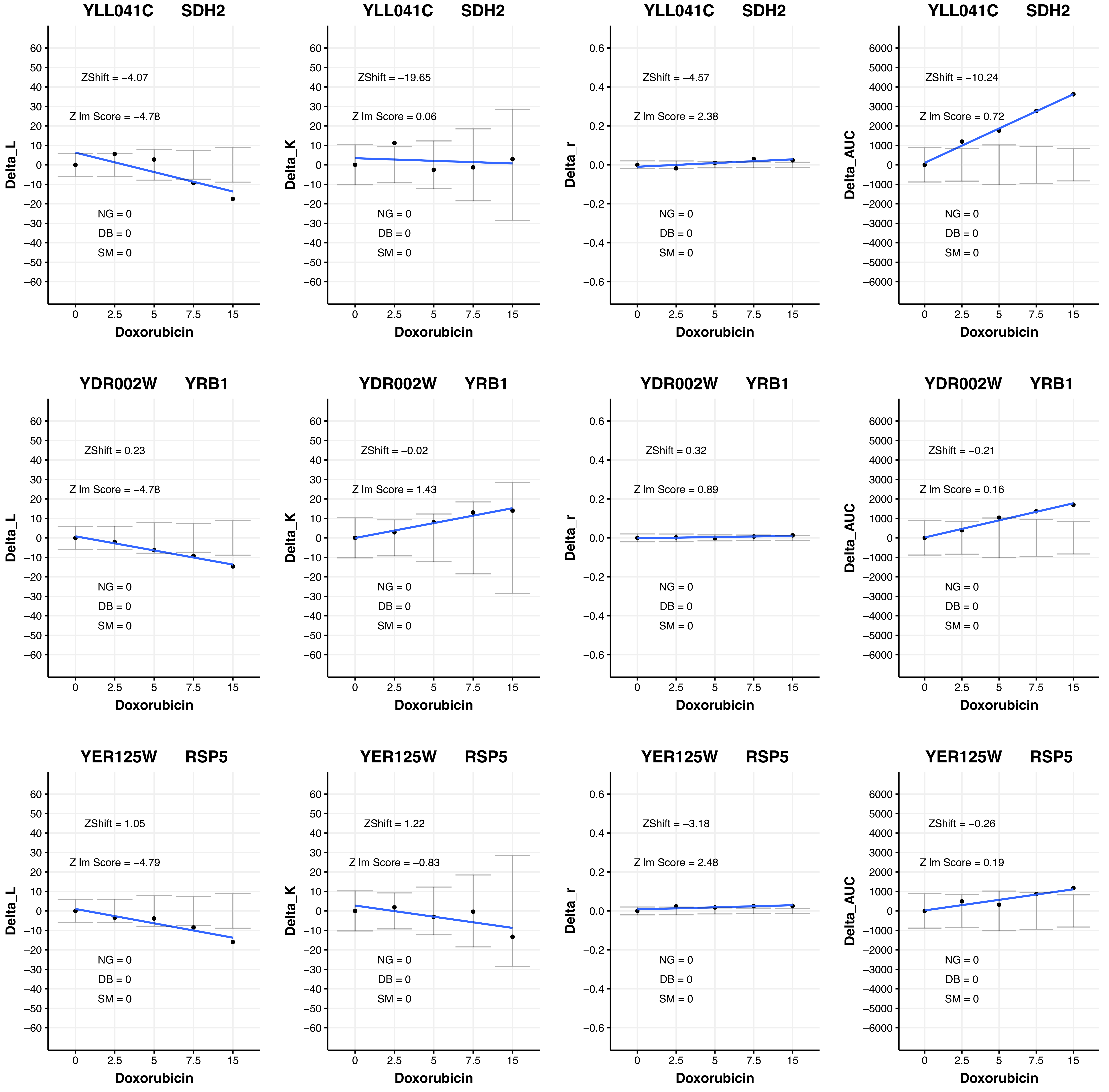
YLL041C SDH2
ZShift = −4.07
Z lm Score = −4.78
NG = 0
DB = 0
SM = 0
Delta_L
Doxorubicin
YLL041C SDH2
ZShift = −19.65
Z lm Score = 0.06
NG = 0
DB = 0
SM = 0
Delta_K
Doxorubicin
YLL041C SDH2
ZShift = −4.57
Z lm Score = 2.38
NG = 0
DB = 0
SM = 0
Delta_r
Doxorubicin
YLL041C SDH2
ZShift = −10.24
Z lm Score = 0.72
NG = 0
DB = 0
SM = 0
Delta_AUC
Doxorubicin
YDR002W YRB1
ZShift = 0.23
Z lm Score = −4.78
NG = 0
DB = 0
SM = 0
Delta_L
Doxorubicin
YDR002W YRB1
ZShift = −0.02
Z lm Score = 1.43
NG = 0
DB = 0
SM = 0
Delta_K
Doxorubicin
YDR002W YRB1
ZShift = 0.32
Z lm Score = 0.89
NG = 0
DB = 0
SM = 0
Delta_r
Doxorubicin
YDR002W YRB1
ZShift = −0.21
Z lm Score = 0.16
NG = 0
DB = 0
SM = 0
Delta_AUC
Doxorubicin
YER125W RSP5
ZShift = 1.05
Z lm Score = −4.79
NG = 0
DB = 0
SM = 0
Delta_L
Doxorubicin
YER125W RSP5
ZShift = 1.22
Z lm Score = −0.83
NG = 0
DB = 0
SM = 0
Delta_K
Doxorubicin
YER125W RSP5
ZShift = −3.18
Z lm Score = 2.48
NG = 0
DB = 0
SM = 0
Delta_r
Doxorubicin
YER125W RSP5
ZShift = −0.26
Z lm Score = 0.19
NG = 0
DB = 0
SM = 0
Delta_AUC
Doxorubicin

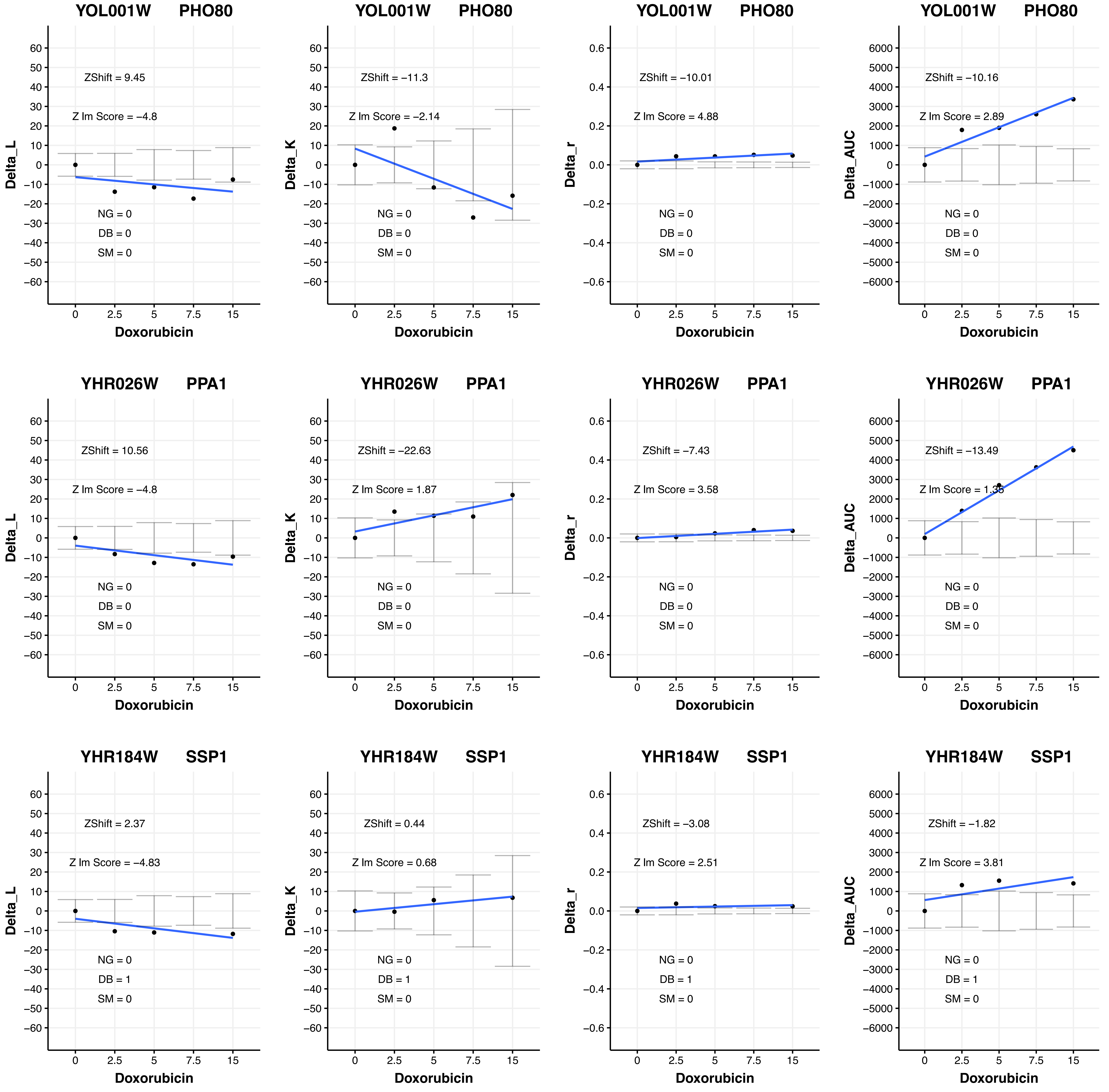
YOL001W PHO80
ZShift = 9.45
Z lm Score = −4.8
NG = 0
DB = 0
SM = 0
Delta_L
Doxorubicin
YOL001W PHO80
ZShift = −11.3
Z lm Score = −2.14
NG = 0
DB = 0
SM = 0
Delta_K
Doxorubicin
YOL001W PHO80
ZShift = −10.01
Z lm Score = 4.88
NG = 0
DB = 0
SM = 0
Delta_r
Doxorubicin
YOL001W PHO80
ZShift = −10.16
Z lm Score = 2.89
NG = 0
DB = 0
SM = 0
Delta_AUC
Doxorubicin
YHR026W PPA1
ZShift = 10.56
Z lm Score = −4.8
NG = 0
DB = 0
SM = 0
Delta_L
Doxorubicin
YHR026W PPA1
ZShift = −22.63
Z lm Score = 1.87
NG = 0
DB = 0
SM = 0
Delta_K
Doxorubicin
YHR026W PPA1
ZShift = −7.43
Z lm Score = 3.58
NG = 0
DB = 0
SM = 0
Delta_r
Doxorubicin
YHR026W PPA1
ZShift = −13.49
Z lm Score = 1.35
NG = 0
DB = 0
SM = 0
Delta_AUC
Doxorubicin
YHR184W SSP1
ZShift = 2.37
Z lm Score = −4.83
NG = 0
DB = 1
SM = 0
Delta_L
Doxorubicin
YHR184W SSP1
ZShift = 0.44
Z lm Score = 0.68
NG = 0
DB = 1
SM = 0
Delta_K
Doxorubicin
YHR184W SSP1
ZShift = −3.08
Z lm Score = 2.51
NG = 0
DB = 1
SM = 0
Delta_r
Doxorubicin
YHR184W SSP1
ZShift = −1.82
Z lm Score = 3.81
NG = 0
DB = 1
SM = 0
Delta_AUC
Doxorubicin

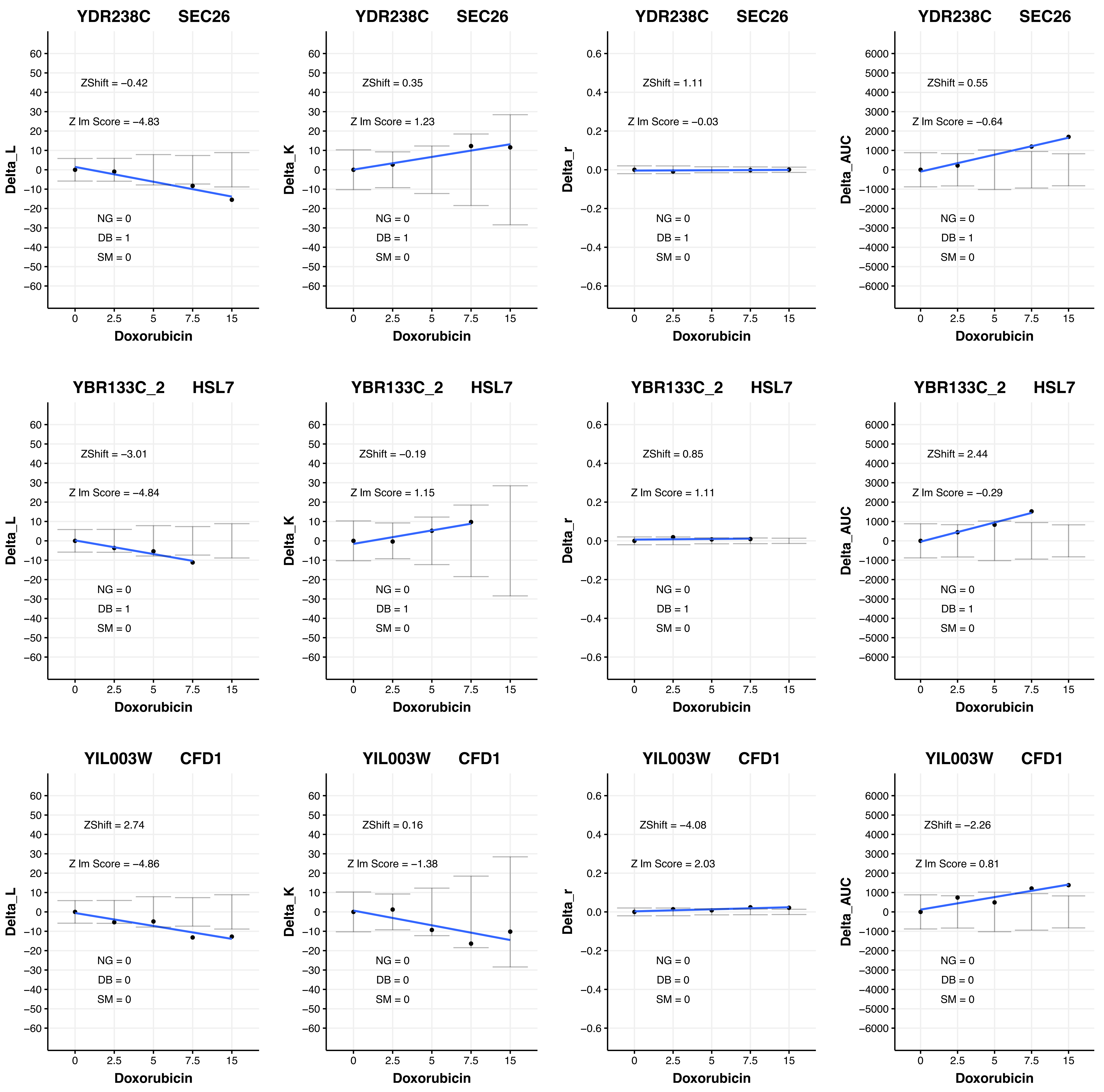
YDR238C SEC26
ZShift = −0.42
Z lm Score = −4.83
NG = 0
DB = 1
SM = 0
Delta_L
Doxorubicin
YDR238C SEC26
ZShift = 0.35
Z lm Score = 1.23
NG = 0
DB = 1
SM = 0
Delta_K
Doxorubicin
YDR238C SEC26
ZShift = 1.11
Z lm Score = −0.03
NG = 0
DB = 1
SM = 0
Delta_r
Doxorubicin
YDR238C SEC26
ZShift = 0.55
Z lm Score = −0.64
NG = 0
DB = 1
SM = 0
Delta_AUC
Doxorubicin
YBR133C_2 HSL7
ZShift = −3.01
Z lm Score = −4.84
NG = 0
DB = 1
SM = 0
Delta_L
Doxorubicin
YBR133C_2 HSL7
ZShift = −0.19
Z lm Score = 1.15
NG = 0
DB = 1
SM = 0
Delta_K
Doxorubicin
YBR133C_2 HSL7
ZShift = 0.85
Z lm Score = 1.11
NG = 0
DB = 1
SM = 0
Delta_r
Doxorubicin
YBR133C_2 HSL7
ZShift = 2.44
Z lm Score = −0.29
NG = 0
DB = 1
SM = 0
Delta_AUC
Doxorubicin
YIL003W CFD1
ZShift = 2.74
Z lm Score = −4.86
NG = 0
DB = 0
SM = 0
Delta_L
Doxorubicin
YIL003W CFD1
ZShift = 0.16
Z lm Score = −1.38
NG = 0
DB = 0
SM = 0
Delta_K
Doxorubicin
YIL003W CFD1
ZShift = −4.08
Z lm Score = 2.03
NG = 0
DB = 0
SM = 0
Delta_r
Doxorubicin
YIL003W CFD1
ZShift = −2.26
Z lm Score = 0.81
NG = 0
DB = 0
SM = 0
Delta_AUC
Doxorubicin

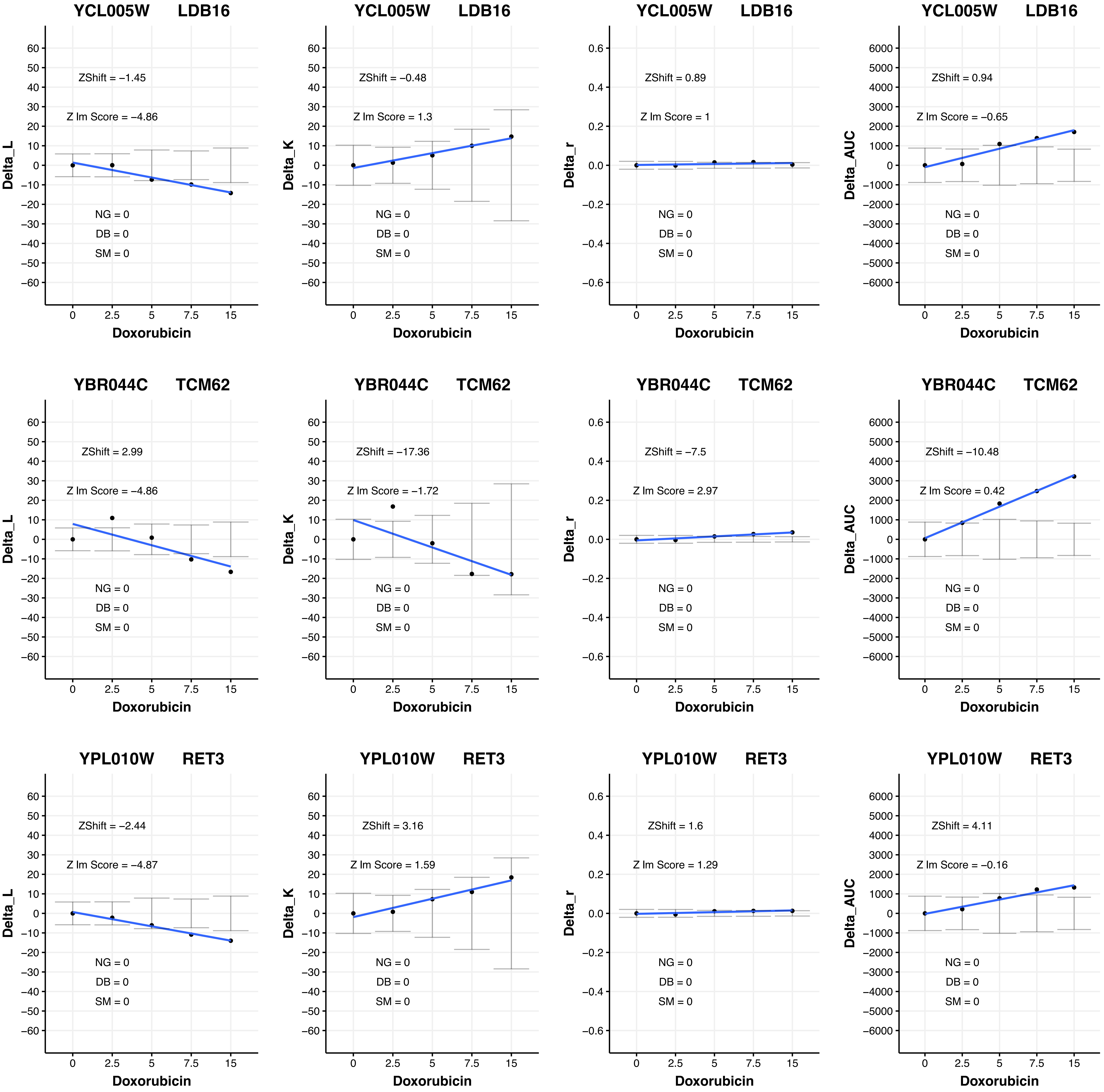

YCL005W LDB16
ZShift = −1.45
Z lm Score = −4.86
NG = 0
DB = 0
SM = 0
Delta_L
Doxorubicin
YCL005W LDB16
ZShift = −0.48
Z lm Score = 1.3
NG = 0
DB = 0
SM = 0
Delta_K
Doxorubicin
YCL005W LDB16
ZShift = 0.89
Z lm Score = 1
NG = 0
DB = 0
SM = 0
Delta_r
Doxorubicin
YCL005W LDB16
ZShift = 0.94
Z lm Score = −0.65
NG = 0
DB = 0
SM = 0
Delta_AUC
Doxorubicin
YBR044C TCM62
ZShift = 2.99
Z lm Score = −4.86
NG = 0
DB = 0
SM = 0
Delta_L
Doxorubicin
YBR044C TCM62
ZShift = −17.36
Z lm Score = −1.72
NG = 0
DB = 0
SM = 0
Delta_K
Doxorubicin
YBR044C TCM62
ZShift = −7.5
Z lm Score = 2.97
NG = 0
DB = 0
SM = 0
Delta_r
Doxorubicin
YBR044C TCM62
ZShift = −10.48
Z lm Score = 0.42
NG = 0
DB = 0
SM = 0
Delta_AUC
Doxorubicin
YPL010W RET3
ZShift = −2.44
Z lm Score = −4.87
NG = 0
DB = 0
SM = 0
Delta_L
Doxorubicin
YPL010W RET3
ZShift = 3.16
Z lm Score = 1.59
NG = 0
DB = 0
SM = 0
Delta_K
Doxorubicin
YPL010W RET3
ZShift = 1.6
Z lm Score = 1.29
NG = 0
DB = 0
SM = 0
Delta_r
Doxorubicin
YPL010W RET3
ZShift = 4.11
Z lm Score = −0.16
NG = 0
DB = 0
SM = 0
Delta_AUC
Doxorubicin

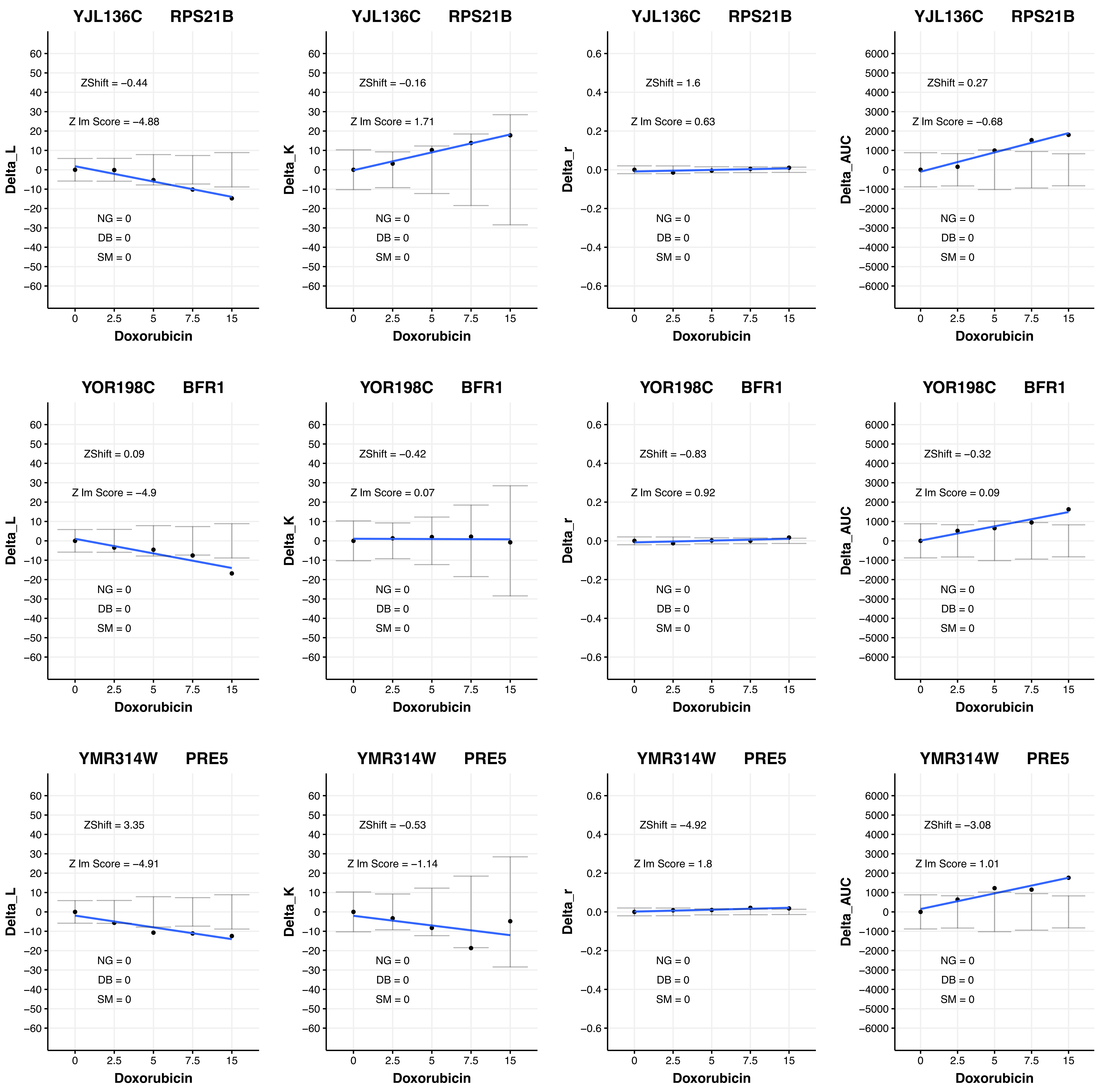

YJL136C RPS21B
ZShift = −0.44
Z lm Score = −4.88
ZShift = −0.16
Z lm Score = 1.71
ZShift = 1.6
Z lm Score = 0.63
ZShift = 0.27
Z lm Score = −0.68
YOR198C BFR1
ZShift = 0.09
Z lm Score = −4.9
ZShift = −0.42
Z lm Score = 0.07
ZShift = −0.83
Z lm Score = 0.92
ZShift = −0.32
Z lm Score = 0.09
YMR314W PRE5
ZShift = 3.35
Z lm Score = −4.91
ZShift = −0.53
Z lm Score = −1.14
ZShift = −4.92
Z lm Score = 1.8
ZShift = −3.08
Z lm Score = 1.01
NG = 0
DB = 0
SM = 0
Delta_L
Delta_K
Delta_r
Delta_AUC
Doxorubicin

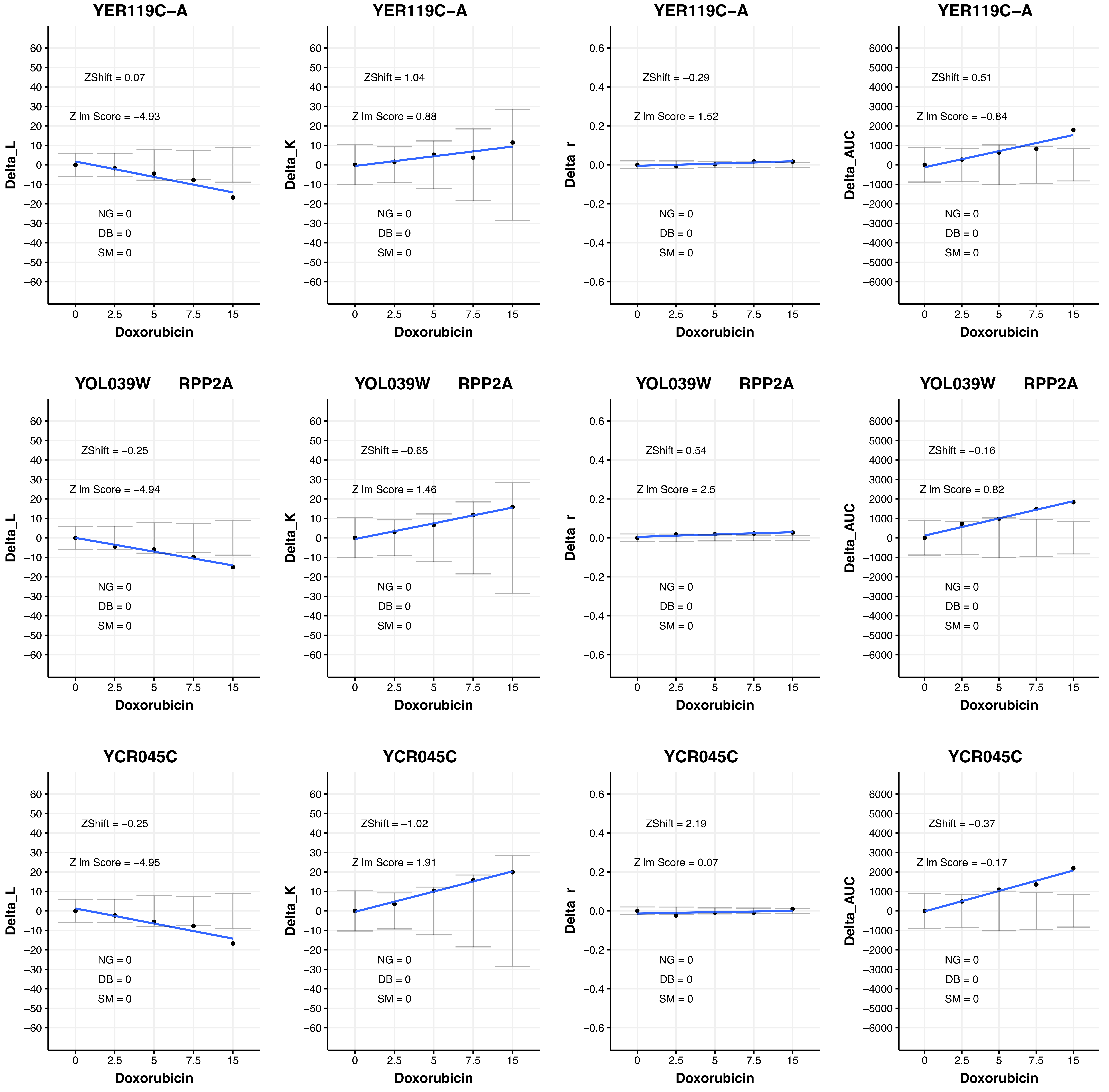

YER119C–A
ZShift = 0.07
Z lm Score = –4.93
NG = 0
DB = 0
SM = 0
Delta_L
Doxorubicin
YER119C–A
ZShift = 1.04
Z lm Score = 0.88
NG = 0
DB = 0
SM = 0
Delta_K
Doxorubicin
YER119C–A
ZShift = –0.29
Z lm Score = 1.52
NG = 0
DB = 0
SM = 0
Delta_r
Doxorubicin
YER119C–A
ZShift = 0.51
Z lm Score = –0.84
NG = 0
DB = 0
SM = 0
Delta_AUC
Doxorubicin
YOL039W RPP2A
ZShift = –0.25
Z lm Score = –4.94
NG = 0
DB = 0
SM = 0
Delta_L
Doxorubicin
YOL039W RPP2A
ZShift = –0.65
Z lm Score = 1.46
NG = 0
DB = 0
SM = 0
Delta_K
Doxorubicin
YOL039W RPP2A
ZShift = 0.54
Z lm Score = 2.5
NG = 0
DB = 0
SM = 0
Delta_r
Doxorubicin
YOL039W RPP2A
ZShift = –0.16
Z lm Score = 0.82
NG = 0
DB = 0
SM = 0
Delta_AUC
Doxorubicin
YCR045C
ZShift = –0.25
Z lm Score = –4.95
NG = 0
DB = 0
SM = 0
Delta_L
Doxorubicin
YCR045C
ZShift = –1.02
Z lm Score = 1.91
NG = 0
DB = 0
SM = 0
Delta_K
Doxorubicin
YCR045C
ZShift = 2.19
Z lm Score = 0.07
NG = 0
DB = 0
SM = 0
Delta_r
Doxorubicin
YCR045C
ZShift = –0.37
Z lm Score = –0.17
NG = 0
DB = 0
SM = 0
Delta_AUC
Doxorubicin

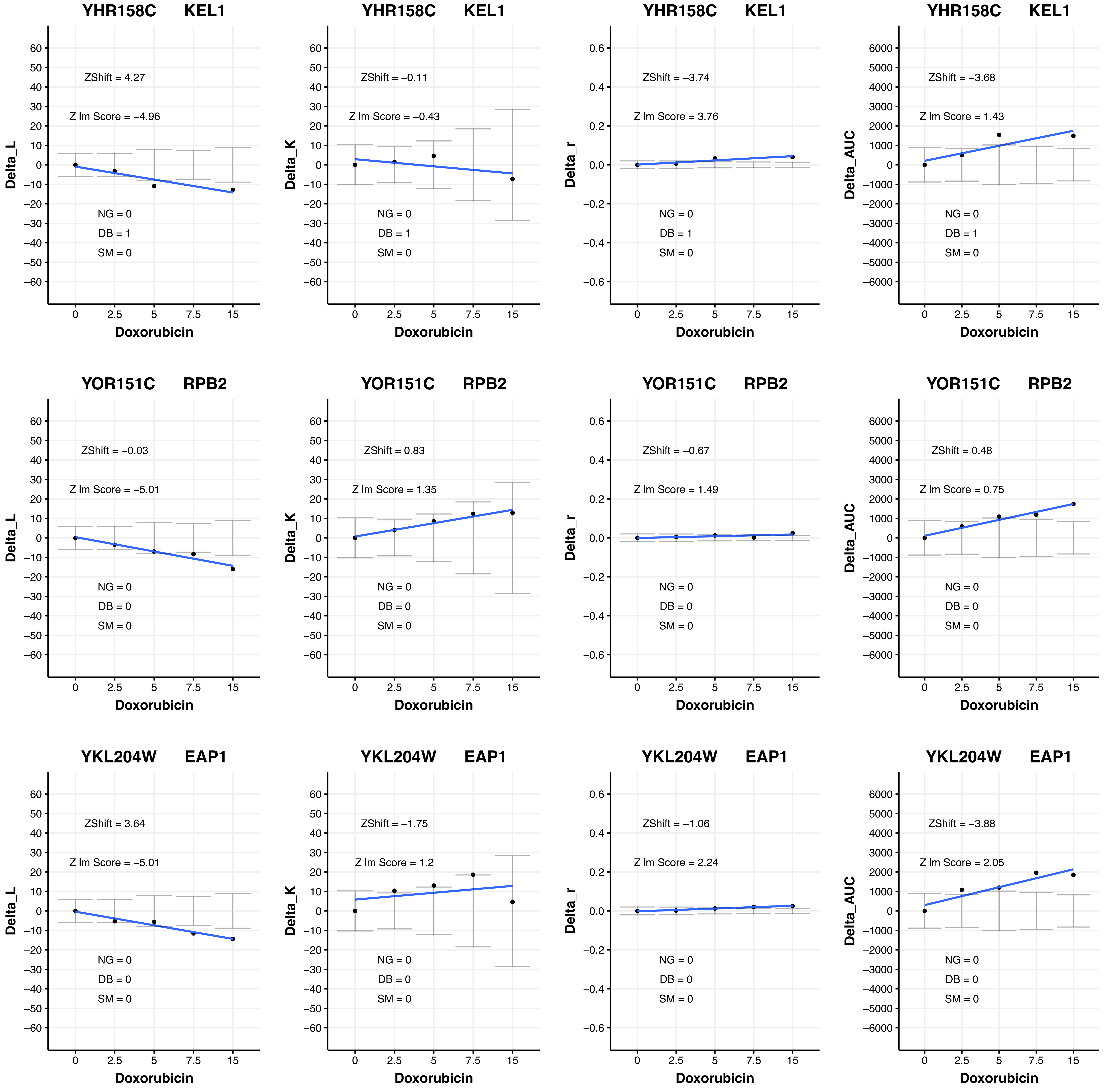
YHR158C KEL1
ZShift = 4.27
Z lm Score = −4.96
NG = 0
DB = 1
SM = 0
Delta_L
Doxorubicin
YHR158C KEL1
ZShift = −0.11
Z lm Score = −0.43
NG = 0
DB = 1
SM = 0
Delta_K
Doxorubicin
YHR158C KEL1
ZShift = −3.74
Z lm Score = 3.76
NG = 0
DB = 1
SM = 0
Delta_r
Doxorubicin
YHR158C KEL1
ZShift = −3.68
Z lm Score = 1.43
NG = 0
DB = 1
SM = 0
Delta_AUC
Doxorubicin
YOR151C RPB2
ZShift = −0.03
Z lm Score = −5.01
NG = 0
DB = 0
SM = 0
Delta_L
Doxorubicin
YOR151C RPB2
ZShift = 0.83
Z lm Score = 1.35
NG = 0
DB = 0
SM = 0
Delta_K
Doxorubicin
YOR151C RPB2
ZShift = −0.67
Z lm Score = 1.49
NG = 0
DB = 0
SM = 0
Delta_r
Doxorubicin
YOR151C RPB2
ZShift = 0.48
Z lm Score = 0.75
NG = 0
DB = 0
SM = 0
Delta_AUC
Doxorubicin
YKL204W EAP1
ZShift = 3.64
Z lm Score = −5.01
NG = 0
DB = 0
SM = 0
Delta_L
Doxorubicin
YKL204W EAP1
ZShift = −1.75
Z lm Score = 1.2
NG = 0
DB = 0
SM = 0
Delta_K
Doxorubicin
YKL204W EAP1
ZShift = −1.06
Z lm Score = 2.24
NG = 0
DB = 0
SM = 0
Delta_r
Doxorubicin
YKL204W EAP1
ZShift = −3.88
Z lm Score = 2.05
NG = 0
DB = 0
SM = 0
Delta_AUC
Doxorubicin

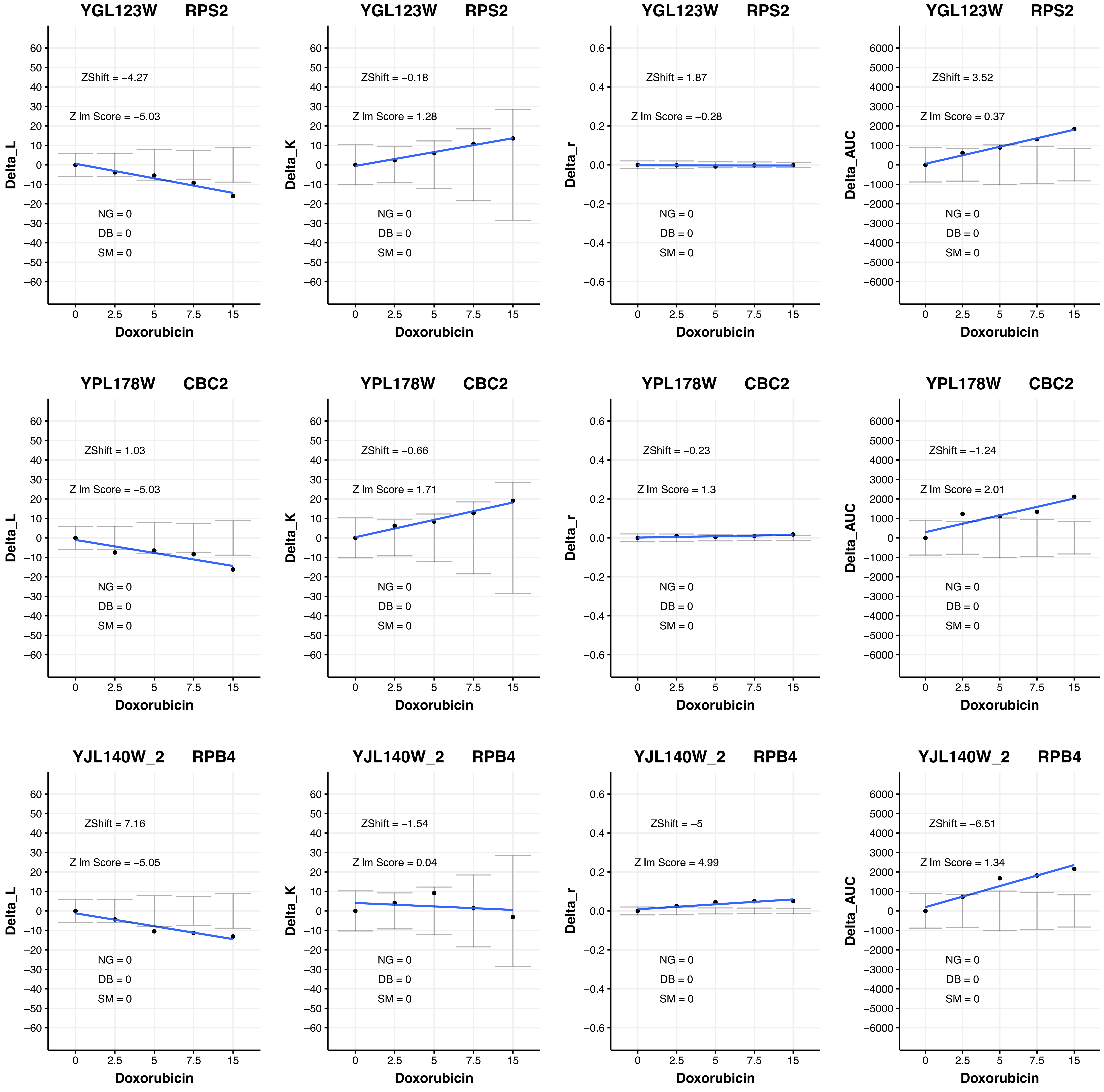
YGL123W RPS2
ZShift = −4.27
Z lm Score = −5.03
NG = 0
DB = 0
SM = 0
Delta_L
YGL123W RPS2
ZShift = −0.18
Z lm Score = 1.28
NG = 0
DB = 0
SM = 0
Delta_K
YGL123W RPS2
ZShift = 1.87
Z lm Score = −0.28
NG = 0
DB = 0
SM = 0
Delta_r
YGL123W RPS2
ZShift = 3.52
Z lm Score = 0.37
NG = 0
DB = 0
SM = 0
Delta_AUC
YPL178W CBC2
ZShift = 1.03
Z lm Score = −5.03
NG = 0
DB = 0
SM = 0
Delta_L
YPL178W CBC2
ZShift = −0.66
Z lm Score = 1.71
NG = 0
DB = 0
SM = 0
Delta_K
YPL178W CBC2
ZShift = −0.23
Z lm Score = 1.3
NG = 0
DB = 0
SM = 0
Delta_r
YPL178W CBC2
ZShift = −1.24
Z lm Score = 2.01
NG = 0
DB = 0
SM = 0
Delta_AUC
YJL140W_2 RPB4
ZShift = 7.16
Z lm Score = −5.05
NG = 0
DB = 0
SM = 0
Delta_L
YJL140W_2 RPB4
ZShift = −1.54
Z lm Score = 0.04
NG = 0
DB = 0
SM = 0
Delta_K
YJL140W_2 RPB4
ZShift = −5
Z lm Score = 4.99
NG = 0
DB = 0
SM = 0
Delta_r
YJL140W_2 RPB4
ZShift = −6.51
Z lm Score = 1.34
NG = 0
DB = 0
SM = 0
Delta_AUC
0 2.5 5 7.5 15
Doxorubicin

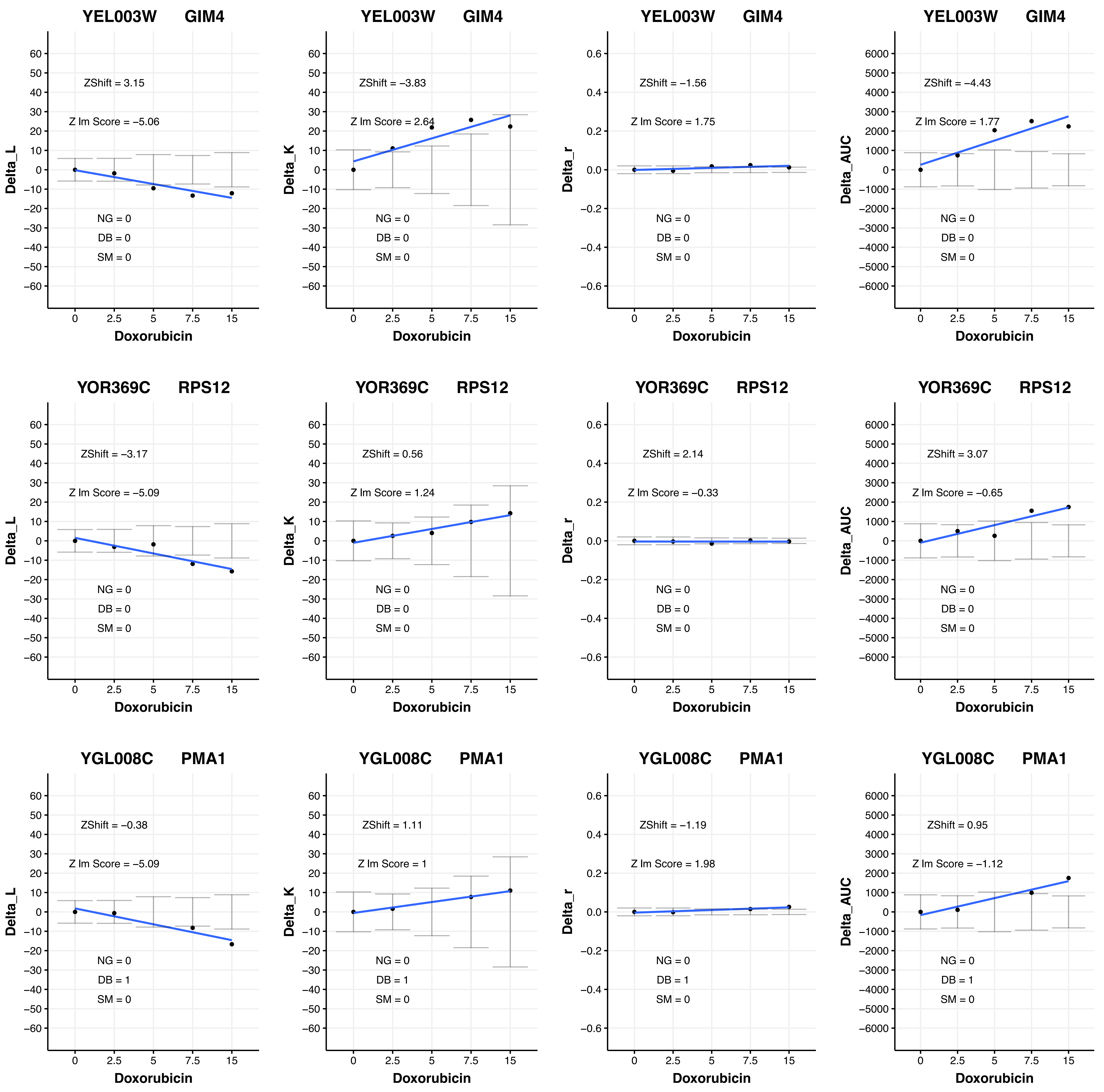
YEL003W GIM4
ZShift = 3.15
Z lm Score = −5.06
NG = 0
DB = 0
SM = 0
Delta_L
Doxorubicin
YEL003W GIM4
ZShift = −3.83
Z lm Score = 2.64
NG = 0
DB = 0
SM = 0
Delta_K
Doxorubicin
YEL003W GIM4
ZShift = −1.56
Z lm Score = 1.75
NG = 0
DB = 0
SM = 0
Delta_r
Doxorubicin
YEL003W GIM4
ZShift = −4.43
Z lm Score = 1.77
NG = 0
DB = 0
SM = 0
Delta_AUC
Doxorubicin
YOR369C RPS12
ZShift = −3.17
Z lm Score = −5.09
NG = 0
DB = 0
SM = 0
Delta_L
Doxorubicin
YOR369C RPS12
ZShift = 0.56
Z lm Score = 1.24
NG = 0
DB = 0
SM = 0
Delta_K
Doxorubicin
YOR369C RPS12
ZShift = 2.14
Z lm Score = −0.33
NG = 0
DB = 0
SM = 0
Delta_r
Doxorubicin
YOR369C RPS12
ZShift = 3.07
Z lm Score = −0.65
NG = 0
DB = 0
SM = 0
Delta_AUC
Doxorubicin
YGL008C PMA1
ZShift = −0.38
Z lm Score = −5.09
NG = 0
DB = 1
SM = 0
Delta_L
Doxorubicin
YGL008C PMA1
ZShift = 1.11
Z lm Score = 1
NG = 0
DB = 1
SM = 0
Delta_K
Doxorubicin
YGL008C PMA1
ZShift = −1.19
Z lm Score = 1.98
NG = 0
DB = 1
SM = 0
Delta_r
Doxorubicin
YGL008C PMA1
ZShift = 0.95
Z lm Score = −1.12
NG = 0
DB = 1
SM = 0
Delta_AUC
Doxorubicin

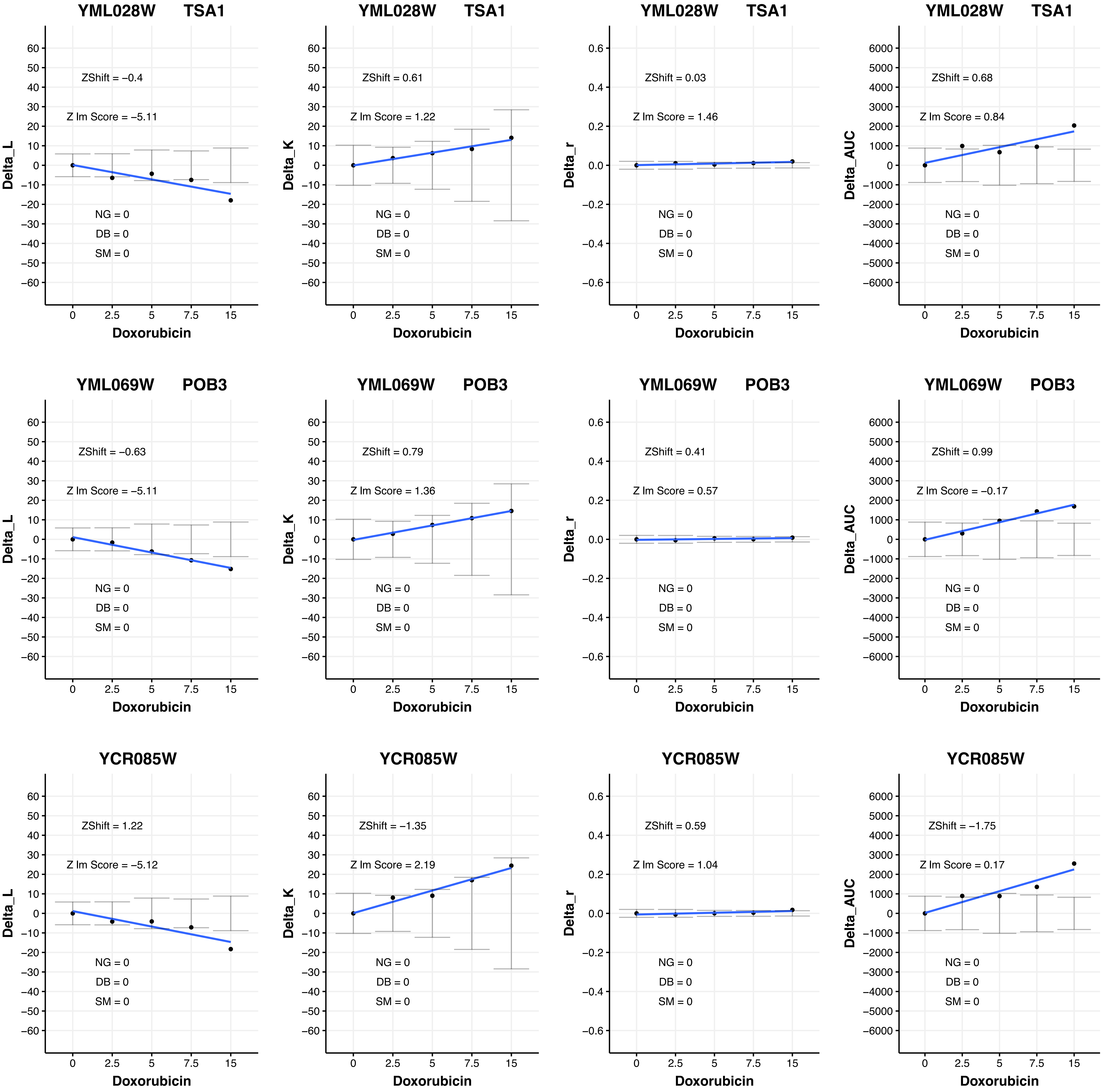
YML028W TSA1
ZShift = −0.4
Z lm Score = −5.11
NG = 0
DB = 0
SM = 0
Delta_L
Doxorubicin
YML028W TSA1
ZShift = 0.61
Z lm Score = 1.22
NG = 0
DB = 0
SM = 0
Delta_K
Doxorubicin
YML028W TSA1
ZShift = 0.03
Z lm Score = 1.46
NG = 0
DB = 0
SM = 0
Delta_r
Doxorubicin
YML028W TSA1
ZShift = 0.68
Z lm Score = 0.84
NG = 0
DB = 0
SM = 0
Delta_AUC
Doxorubicin
YML069W POB3
ZShift = −0.63
Z lm Score = −5.11
NG = 0
DB = 0
SM = 0
Delta_L
Doxorubicin
YML069W POB3
ZShift = 0.79
Z lm Score = 1.36
NG = 0
DB = 0
SM = 0
Delta_K
Doxorubicin
YML069W POB3
ZShift = 0.41
Z lm Score = 0.57
NG = 0
DB = 0
SM = 0
Delta_r
Doxorubicin
YML069W POB3
ZShift = 0.99
Z lm Score = −0.17
NG = 0
DB = 0
SM = 0
Delta_AUC
Doxorubicin
YCR085W
ZShift = 1.22
Z lm Score = −5.12
NG = 0
DB = 0
SM = 0
Delta_L
Doxorubicin
YCR085W
ZShift = −1.35
Z lm Score = 2.19
NG = 0
DB = 0
SM = 0
Delta_K
Doxorubicin
YCR085W
ZShift = 0.59
Z lm Score = 1.04
NG = 0
DB = 0
SM = 0
Delta_r
Doxorubicin
YCR085W
ZShift = −1.75
Z lm Score = 0.17
NG = 0
DB = 0
SM = 0
Delta_AUC
Doxorubicin

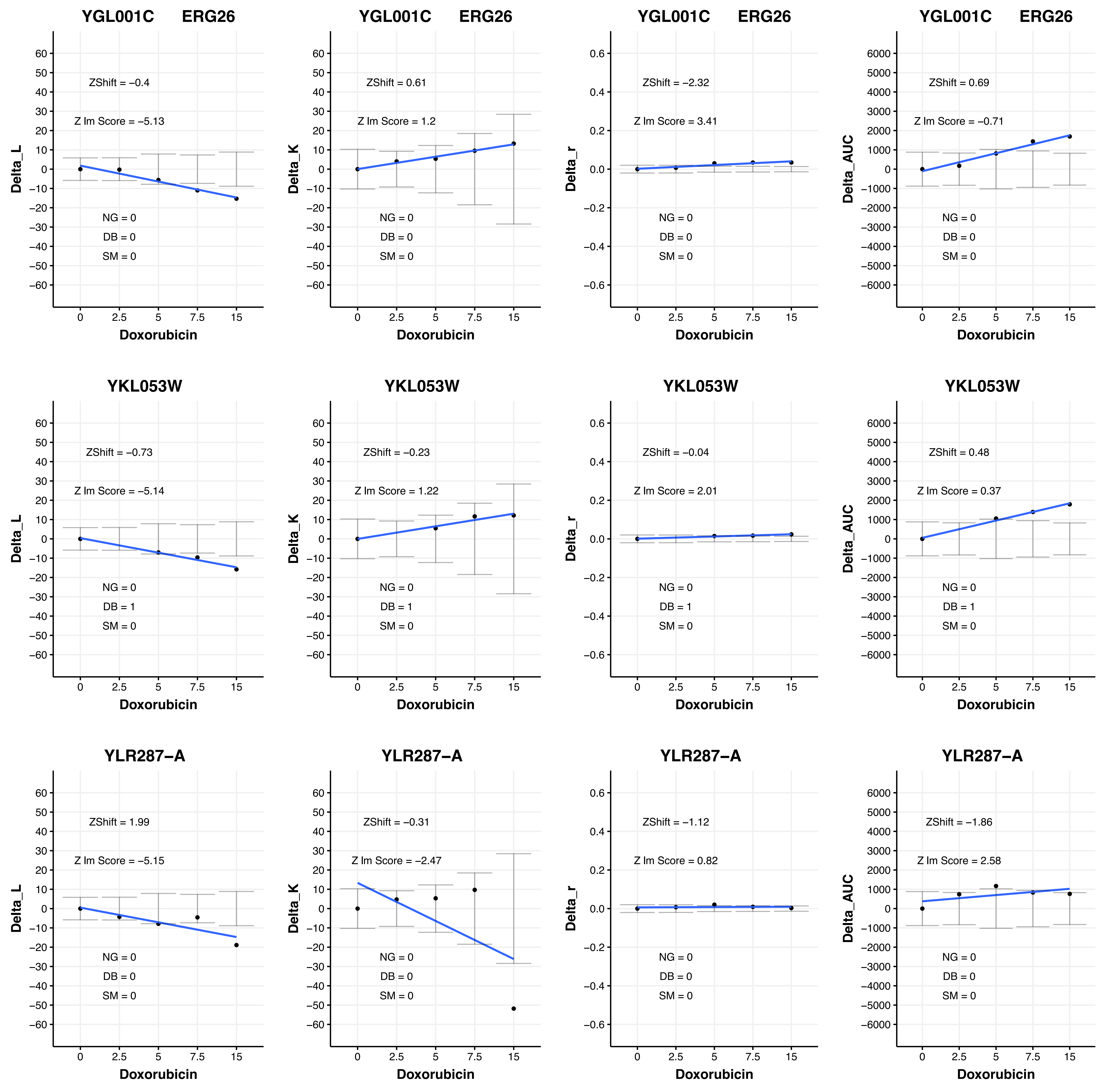
YGL001C ERG26
ZShift = −0.4
Z lm Score = −5.13
NG = 0
DB = 0
SM = 0
Delta_L
Doxorubicin
YGL001C ERG26
ZShift = 0.61
Z lm Score = 1.2
NG = 0
DB = 0
SM = 0
Delta_K
Doxorubicin
YGL001C ERG26
ZShift = −2.32
Z lm Score = 3.41
NG = 0
DB = 0
SM = 0
Delta_r
Doxorubicin
YGL001C ERG26
ZShift = 0.69
Z lm Score = −0.71
NG = 0
DB = 0
SM = 0
Delta_AUC
Doxorubicin
YKL053W
ZShift = −0.73
Z lm Score = −5.14
NG = 0
DB = 1
SM = 0
Delta_L
Doxorubicin
YKL053W
ZShift = −0.23
Z lm Score = 1.22
NG = 0
DB = 1
SM = 0
Delta_K
Doxorubicin
YKL053W
ZShift = −0.04
Z lm Score = 2.01
NG = 0
DB = 1
SM = 0
Delta_r
Doxorubicin
YKL053W
ZShift = 0.48
Z lm Score = 0.37
NG = 0
DB = 1
SM = 0
Delta_AUC
Doxorubicin
YLR287−A
ZShift = 1.99
Z lm Score = −5.15
NG = 0
DB = 0
SM = 0
Delta_L
Doxorubicin
YLR287−A
ZShift = −0.31
Z lm Score = −2.47
NG = 0
DB = 0
SM = 0
Delta_K
Doxorubicin
YLR287−A
ZShift = −1.12
Z lm Score = 0.82
NG = 0
DB = 0
SM = 0
Delta_r
Doxorubicin
YLR287−A
ZShift = −1.86
Z lm Score = 2.58
NG = 0
DB = 0
SM = 0
Delta_AUC
Doxorubicin

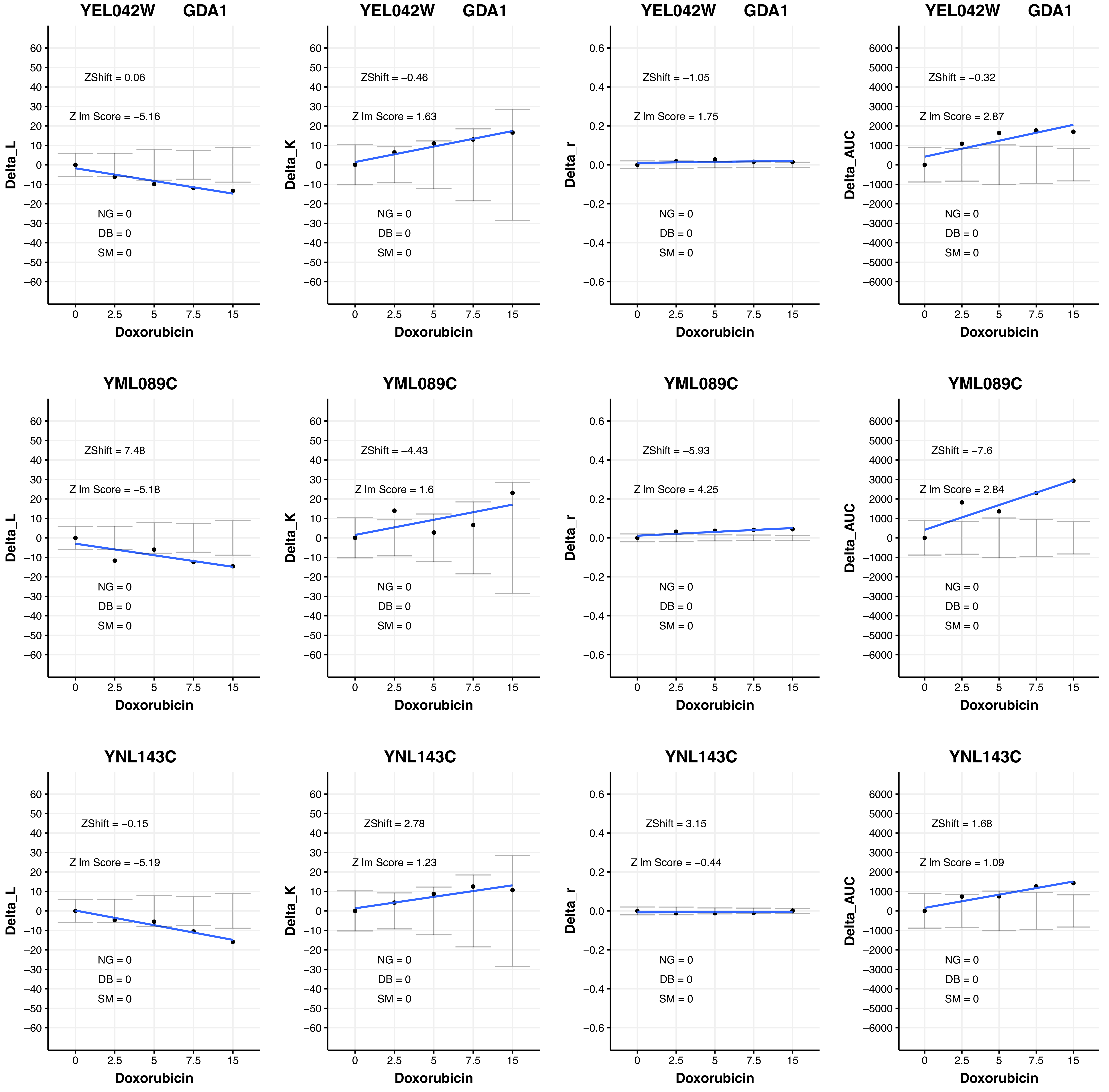
YEL042W GDA1
ZShift = 0.06
Z lm Score = −5.16
NG = 0
DB = 0
SM = 0
Delta_L
Doxorubicin
YEL042W GDA1
ZShift = −0.46
Z lm Score = 1.63
NG = 0
DB = 0
SM = 0
Delta_K
Doxorubicin
YEL042W GDA1
ZShift = −1.05
Z lm Score = 1.75
NG = 0
DB = 0
SM = 0
Delta_r
Doxorubicin
YEL042W GDA1
ZShift = −0.32
Z lm Score = 2.87
NG = 0
DB = 0
SM = 0
Delta_AUC
Doxorubicin
YML089C
ZShift = 7.48
Z lm Score = −5.18
NG = 0
DB = 0
SM = 0
Delta_L
Doxorubicin
YML089C
ZShift = −4.43
Z lm Score = 1.6
NG = 0
DB = 0
SM = 0
Delta_K
Doxorubicin
YML089C
ZShift = −5.93
Z lm Score = 4.25
NG = 0
DB = 0
SM = 0
Delta_r
Doxorubicin
YML089C
ZShift = −7.6
Z lm Score = 2.84
NG = 0
DB = 0
SM = 0
Delta_AUC
Doxorubicin
YNL143C
ZShift = −0.15
Z lm Score = −5.19
NG = 0
DB = 0
SM = 0
Delta_L
Doxorubicin
YNL143C
ZShift = 2.78
Z lm Score = 1.23
NG = 0
DB = 0
SM = 0
Delta_K
Doxorubicin
YNL143C
ZShift = 3.15
Z lm Score = −0.44
NG = 0
DB = 0
SM = 0
Delta_r
Doxorubicin
YNL143C
ZShift = 1.68
Z lm Score = 1.09
NG = 0
DB = 0
SM = 0
Delta_AUC
Doxorubicin

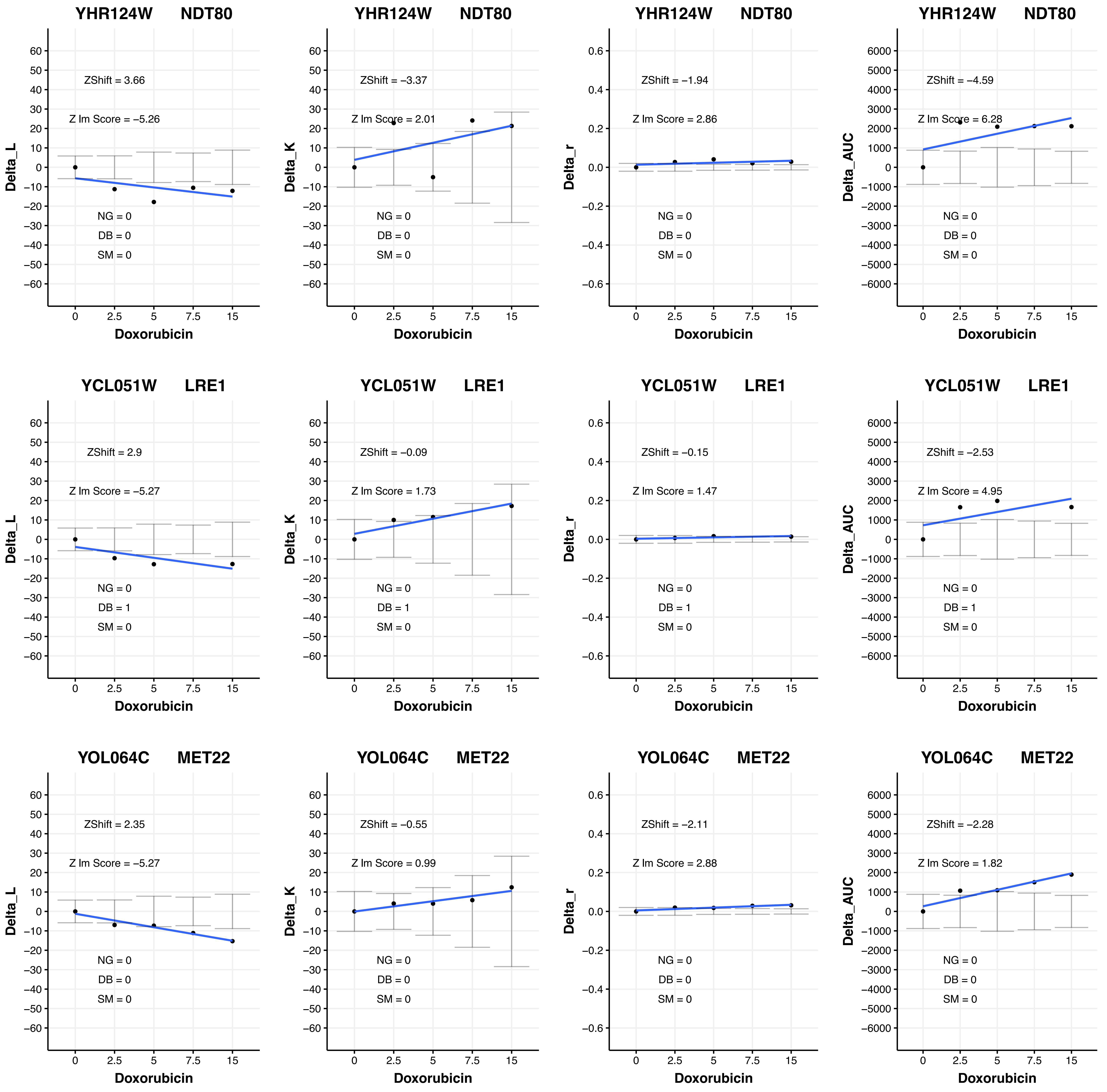
YHR124W NDT80
ZShift = 3.66
Z lm Score = −5.26
NG = 0
DB = 0
SM = 0
Delta_L
Doxorubicin
YHR124W NDT80
ZShift = −3.37
Z lm Score = 2.01
NG = 0
DB = 0
SM = 0
Delta_K
Doxorubicin
YHR124W NDT80
ZShift = −1.94
Z lm Score = 2.86
NG = 0
DB = 0
SM = 0
Delta_r
Doxorubicin
YHR124W NDT80
ZShift = −4.59
Z lm Score = 6.28
NG = 0
DB = 0
SM = 0
Delta_AUC
Doxorubicin
YCL051W LRE1
ZShift = 2.9
Z lm Score = −5.27
NG = 0
DB = 1
SM = 0
Delta_L
Doxorubicin
YCL051W LRE1
ZShift = −0.09
Z lm Score = 1.73
NG = 0
DB = 1
SM = 0
Delta_K
Doxorubicin
YCL051W LRE1
ZShift = −0.15
Z lm Score = 1.47
NG = 0
DB = 1
SM = 0
Delta_r
Doxorubicin
YCL051W LRE1
ZShift = −2.53
Z lm Score = 4.95
NG = 0
DB = 1
SM = 0
Delta_AUC
Doxorubicin
YOL064C MET22
ZShift = 2.35
Z lm Score = −5.27
NG = 0
DB = 0
SM = 0
Delta_L
Doxorubicin
YOL064C MET22
ZShift = −0.55
Z lm Score = 0.99
NG = 0
DB = 0
SM = 0
Delta_K
Doxorubicin
YOL064C MET22
ZShift = −2.11
Z lm Score = 2.88
NG = 0
DB = 0
SM = 0
Delta_r
Doxorubicin
YOL064C MET22
ZShift = −2.28
Z lm Score = 1.82
NG = 0
DB = 0
SM = 0
Delta_AUC
Doxorubicin

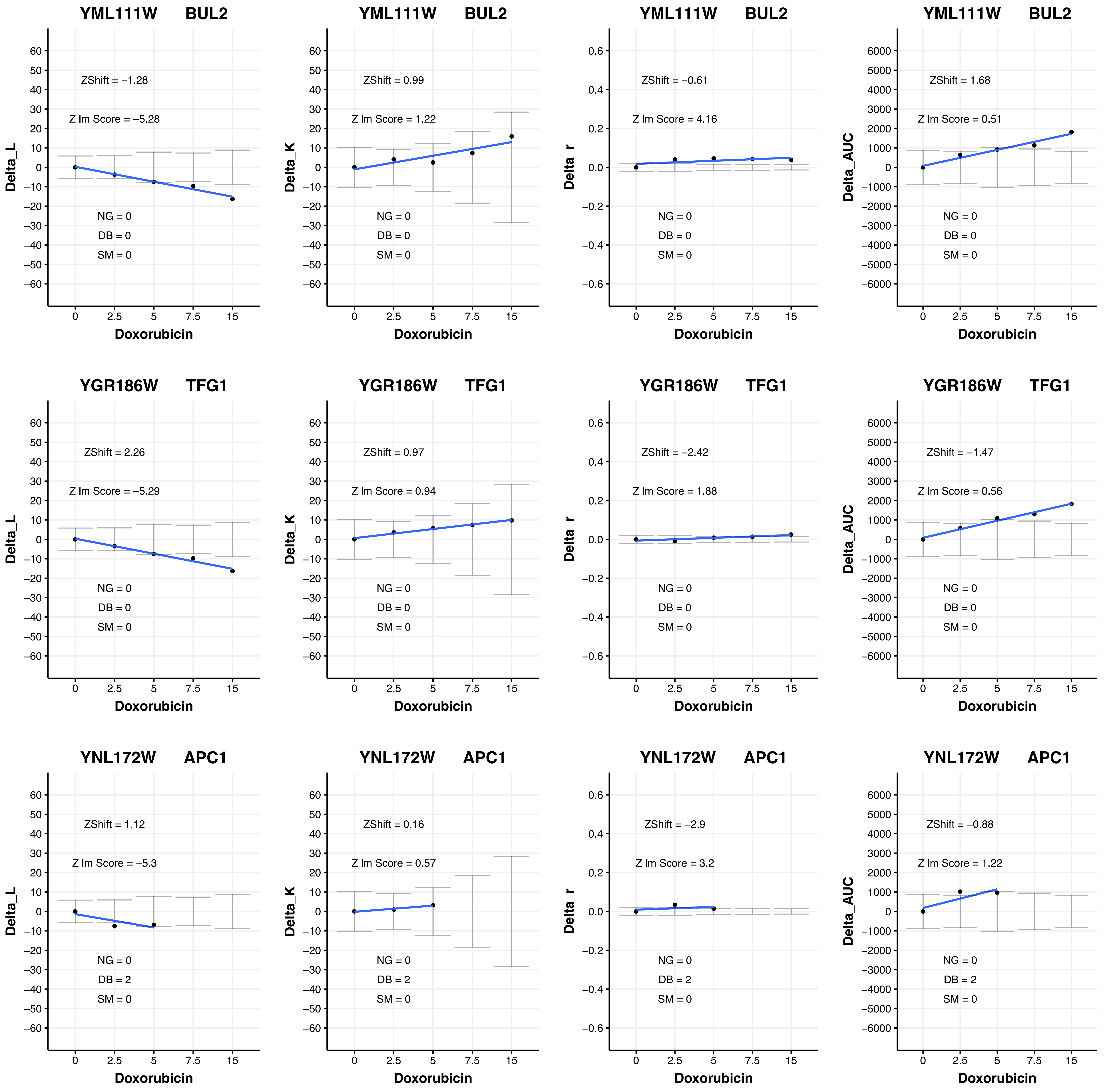
YML111W BUL2
ZShift = −1.28
Z lm Score = −5.28
NG = 0
DB = 0
SM = 0
Delta_L
Doxorubicin
YML111W BUL2
ZShift = 0.99
Z lm Score = 1.22
NG = 0
DB = 0
SM = 0
Delta_K
Doxorubicin
YML111W BUL2
ZShift = −0.61
Z lm Score = 4.16
NG = 0
DB = 0
SM = 0
Delta_r
Doxorubicin
YML111W BUL2
ZShift = 1.68
Z lm Score = 0.51
NG = 0
DB = 0
SM = 0
Delta_AUC
Doxorubicin
YGR186W TFG1
ZShift = 2.26
Z lm Score = −5.29
NG = 0
DB = 0
SM = 0
Delta_L
Doxorubicin
YGR186W TFG1
ZShift = 0.97
Z lm Score = 0.94
NG = 0
DB = 0
SM = 0
Delta_K
Doxorubicin
YGR186W TFG1
ZShift = −2.42
Z lm Score = 1.88
NG = 0
DB = 0
SM = 0
Delta_r
Doxorubicin
YGR186W TFG1
ZShift = −1.47
Z lm Score = 0.56
NG = 0
DB = 0
SM = 0
Delta_AUC
Doxorubicin
YNL172W APC1
ZShift = 1.12
Z lm Score = −5.3
NG = 0
DB = 2
SM = 0
Delta_L
Doxorubicin
YNL172W APC1
ZShift = 0.16
Z lm Score = 0.57
NG = 0
DB = 2
SM = 0
Delta_K
Doxorubicin
YNL172W APC1
ZShift = −2.9
Z lm Score = 3.2
NG = 0
DB = 2
SM = 0
Delta_r
Doxorubicin
YNL172W APC1
ZShift = −0.88
Z lm Score = 1.22
NG = 0
DB = 2
SM = 0
Delta_AUC
Doxorubicin

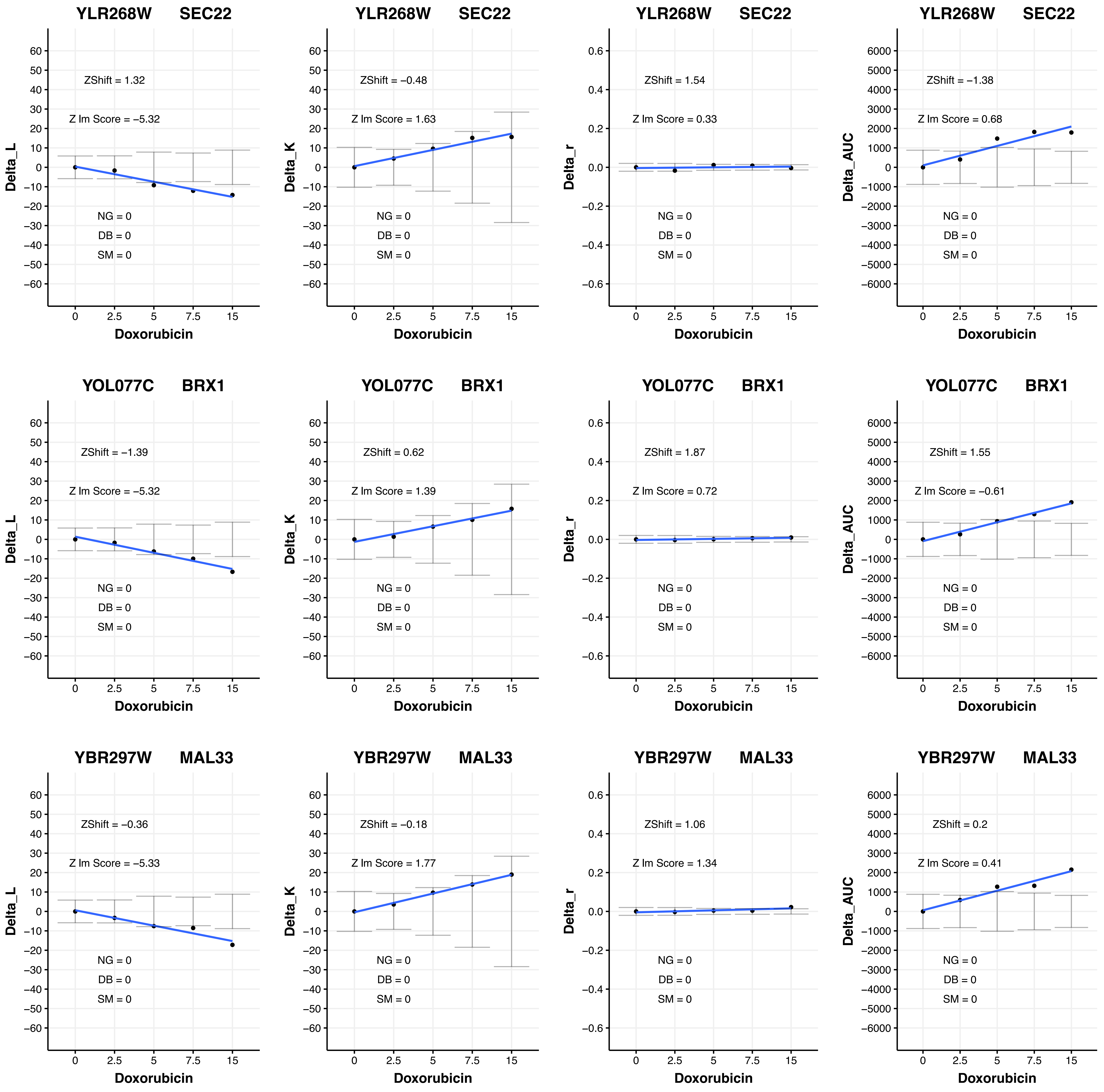

YLR268W SEC22
ZShift = 1.32
Z lm Score = −5.32
NG = 0
DB = 0
SM = 0
Delta_L
Doxorubicin
YLR268W SEC22
ZShift = −0.48
Z lm Score = 1.63
NG = 0
DB = 0
SM = 0
Delta_K
Doxorubicin
YLR268W SEC22
ZShift = 1.54
Z lm Score = 0.33
NG = 0
DB = 0
SM = 0
Delta_r
Doxorubicin
YLR268W SEC22
ZShift = −1.38
Z lm Score = 0.68
NG = 0
DB = 0
SM = 0
Delta_AUC
Doxorubicin
YOL077C BRX1
ZShift = −1.39
Z lm Score = −5.32
NG = 0
DB = 0
SM = 0
Delta_L
Doxorubicin
YOL077C BRX1
ZShift = 0.62
Z lm Score = 1.39
NG = 0
DB = 0
SM = 0
Delta_K
Doxorubicin
YOL077C BRX1
ZShift = 1.87
Z lm Score = 0.72
NG = 0
DB = 0
SM = 0
Delta_r
Doxorubicin
YOL077C BRX1
ZShift = 1.55
Z lm Score = −0.61
NG = 0
DB = 0
SM = 0
Delta_AUC
Doxorubicin
YBR297W MAL33
ZShift = −0.36
Z lm Score = −5.33
NG = 0
DB = 0
SM = 0
Delta_L
Doxorubicin
YBR297W MAL33
ZShift = −0.18
Z lm Score = 1.77
NG = 0
DB = 0
SM = 0
Delta_K
Doxorubicin
YBR297W MAL33
ZShift = 1.06
Z lm Score = 1.34
NG = 0
DB = 0
SM = 0
Delta_r
Doxorubicin
YBR297W MAL33
ZShift = 0.2
Z lm Score = 0.41
NG = 0
DB = 0
SM = 0
Delta_AUC
Doxorubicin

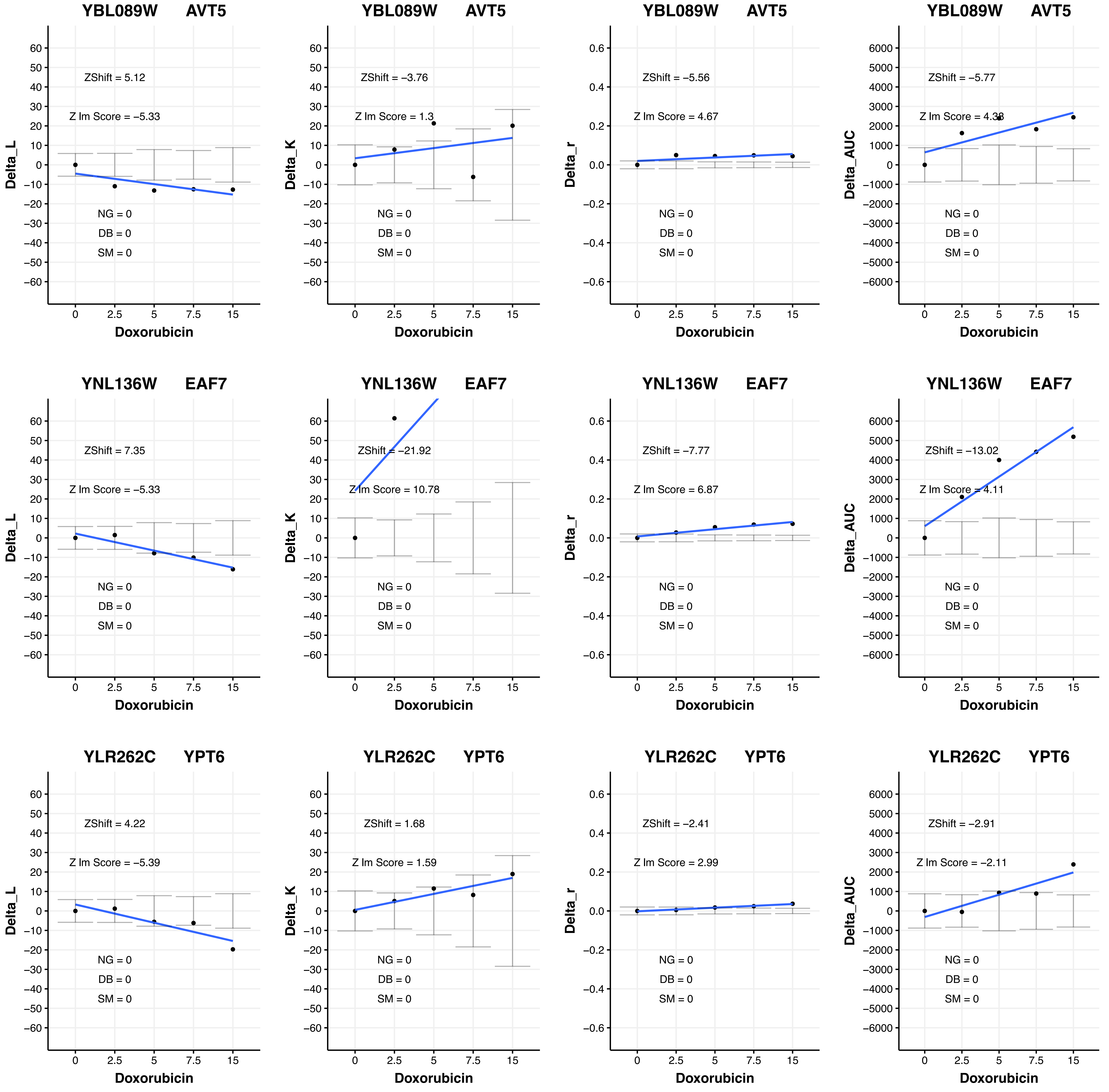
YBL089W AVT5
ZShift = 5.12
Z lm Score = −5.33
NG = 0
DB = 0
SM = 0
Delta_L
Doxorubicin
YBL089W AVT5
ZShift = −3.76
Z lm Score = 1.3
NG = 0
DB = 0
SM = 0
Delta_K
Doxorubicin
YBL089W AVT5
ZShift = −5.56
Z lm Score = 4.67
NG = 0
DB = 0
SM = 0
Delta_r
Doxorubicin
YBL089W AVT5
ZShift = −5.77
Z lm Score = 4.38
NG = 0
DB = 0
SM = 0
Delta_AUC
Doxorubicin
YNL136W EAF7
ZShift = 7.35
Z lm Score = −5.33
NG = 0
DB = 0
SM = 0
Delta_L
Doxorubicin
YNL136W EAF7
ZShift = −21.92
Z lm Score = 10.78
NG = 0
DB = 0
SM = 0
Delta_K
Doxorubicin
YNL136W EAF7
ZShift = −7.77
Z lm Score = 6.87
NG = 0
DB = 0
SM = 0
Delta_r
Doxorubicin
YNL136W EAF7
ZShift = −13.02
Z lm Score = 4.11
NG = 0
DB = 0
SM = 0
Delta_AUC
Doxorubicin
YLR262C YPT6
ZShift = 4.22
Z lm Score = −5.39
NG = 0
DB = 0
SM = 0
Delta_L
Doxorubicin
YLR262C YPT6
ZShift = 1.68
Z lm Score = 1.59
NG = 0
DB = 0
SM = 0
Delta_K
Doxorubicin
YLR262C YPT6
ZShift = −2.41
Z lm Score = 2.99
NG = 0
DB = 0
SM = 0
Delta_r
Doxorubicin
YLR262C YPT6
ZShift = −2.91
Z lm Score = −2.11
NG = 0
DB = 0
SM = 0
Delta_AUC
Doxorubicin

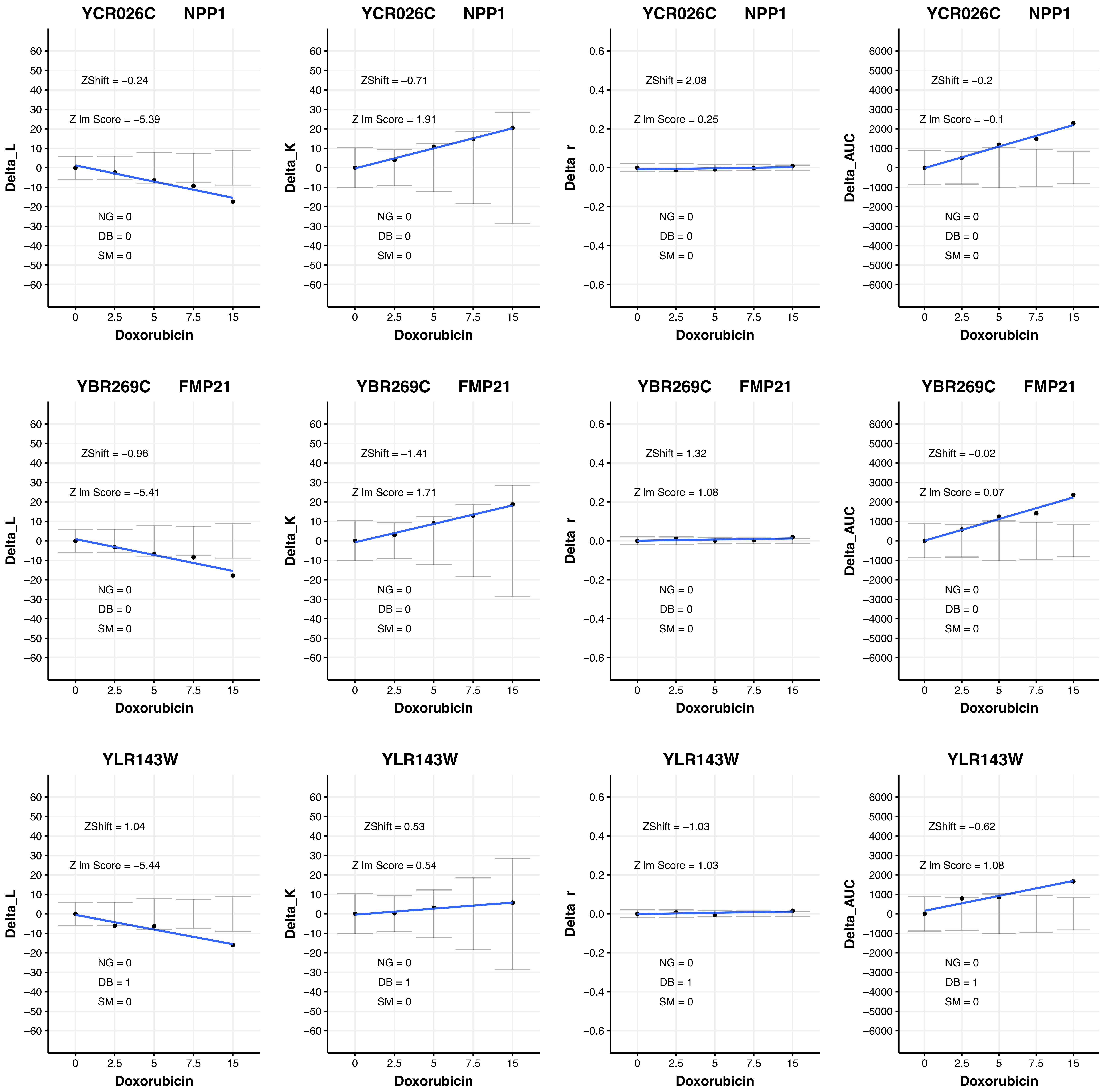
YCR026C NPP1
ZShift = −0.24
Z lm Score = −5.39
NG = 0
DB = 0
SM = 0
Delta_L
Doxorubicin
YCR026C NPP1
ZShift = −0.71
Z lm Score = 1.91
NG = 0
DB = 0
SM = 0
Delta_K
Doxorubicin
YCR026C NPP1
ZShift = 2.08
Z lm Score = 0.25
NG = 0
DB = 0
SM = 0
Delta_r
Doxorubicin
YCR026C NPP1
ZShift = −0.2
Z lm Score = −0.1
NG = 0
DB = 0
SM = 0
Delta_AUC
Doxorubicin
YBR269C FMP21
ZShift = −0.96
Z lm Score = −5.41
NG = 0
DB = 0
SM = 0
Delta_L
Doxorubicin
YBR269C FMP21
ZShift = −1.41
Z lm Score = 1.71
NG = 0
DB = 0
SM = 0
Delta_K
Doxorubicin
YBR269C FMP21
ZShift = 1.32
Z lm Score = 1.08
NG = 0
DB = 0
SM = 0
Delta_r
Doxorubicin
YBR269C FMP21
ZShift = −0.02
Z lm Score = 0.07
NG = 0
DB = 0
SM = 0
Delta_AUC
Doxorubicin
YLR143W
ZShift = 1.04
Z lm Score = −5.44
NG = 0
DB = 1
SM = 0
Delta_L
Doxorubicin
YLR143W
ZShift = 0.53
Z lm Score = 0.54
NG = 0
DB = 1
SM = 0
Delta_K
Doxorubicin
YLR143W
ZShift = −1.03
Z lm Score = 1.03
NG = 0
DB = 1
SM = 0
Delta_r
Doxorubicin
YLR143W
ZShift = −0.62
Z lm Score = 1.08
NG = 0
DB = 1
SM = 0
Delta_AUC
Doxorubicin

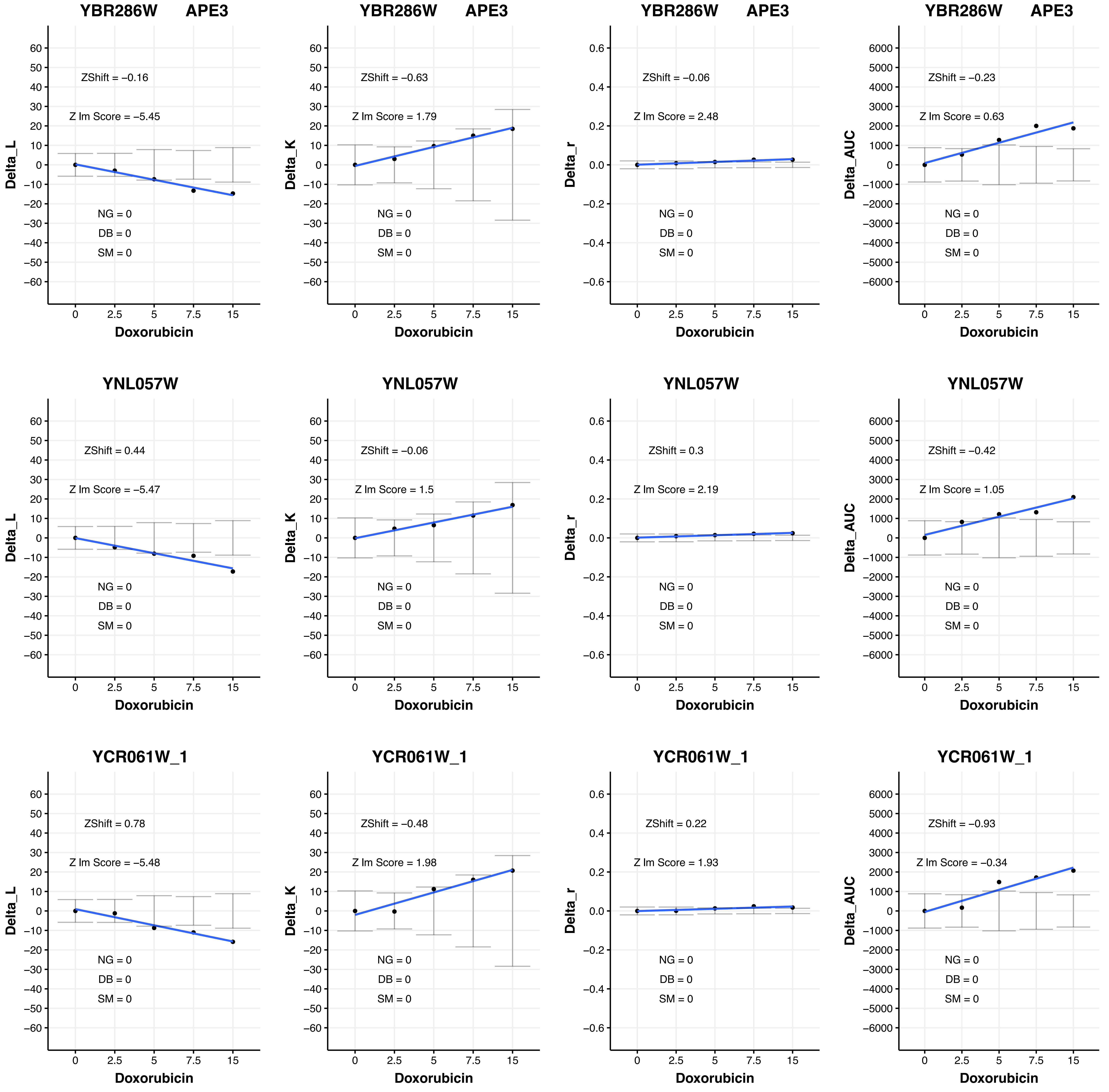
YBR286W APE3
ZShift = −0.16
Z lm Score = −5.45
NG = 0
DB = 0
SM = 0
Delta_L
Doxorubicin
YBR286W APE3
ZShift = −0.63
Z lm Score = 1.79
NG = 0
DB = 0
SM = 0
Delta_K
Doxorubicin
YBR286W APE3
ZShift = −0.06
Z lm Score = 2.48
NG = 0
DB = 0
SM = 0
Delta_r
Doxorubicin
YBR286W APE3
ZShift = −0.23
Z lm Score = 0.63
NG = 0
DB = 0
SM = 0
Delta_AUC
Doxorubicin
YNL057W
ZShift = 0.44
Z lm Score = −5.47
NG = 0
DB = 0
SM = 0
Delta_L
Doxorubicin
YNL057W
ZShift = −0.06
Z lm Score = 1.5
NG = 0
DB = 0
SM = 0
Delta_K
Doxorubicin
YNL057W
ZShift = 0.3
Z lm Score = 2.19
NG = 0
DB = 0
SM = 0
Delta_r
Doxorubicin
YNL057W
ZShift = −0.42
Z lm Score = 1.05
NG = 0
DB = 0
SM = 0
Delta_AUC
Doxorubicin
YCR061W_1
ZShift = 0.78
Z lm Score = −5.48
NG = 0
DB = 0
SM = 0
Delta_L
Doxorubicin
YCR061W_1
ZShift = −0.48
Z lm Score = 1.98
NG = 0
DB = 0
SM = 0
Delta_K
Doxorubicin
YCR061W_1
ZShift = 0.22
Z lm Score = 1.93
NG = 0
DB = 0
SM = 0
Delta_r
Doxorubicin
YCR061W_1
ZShift = −0.93
Z lm Score = −0.34
NG = 0
DB = 0
SM = 0
Delta_AUC
Doxorubicin

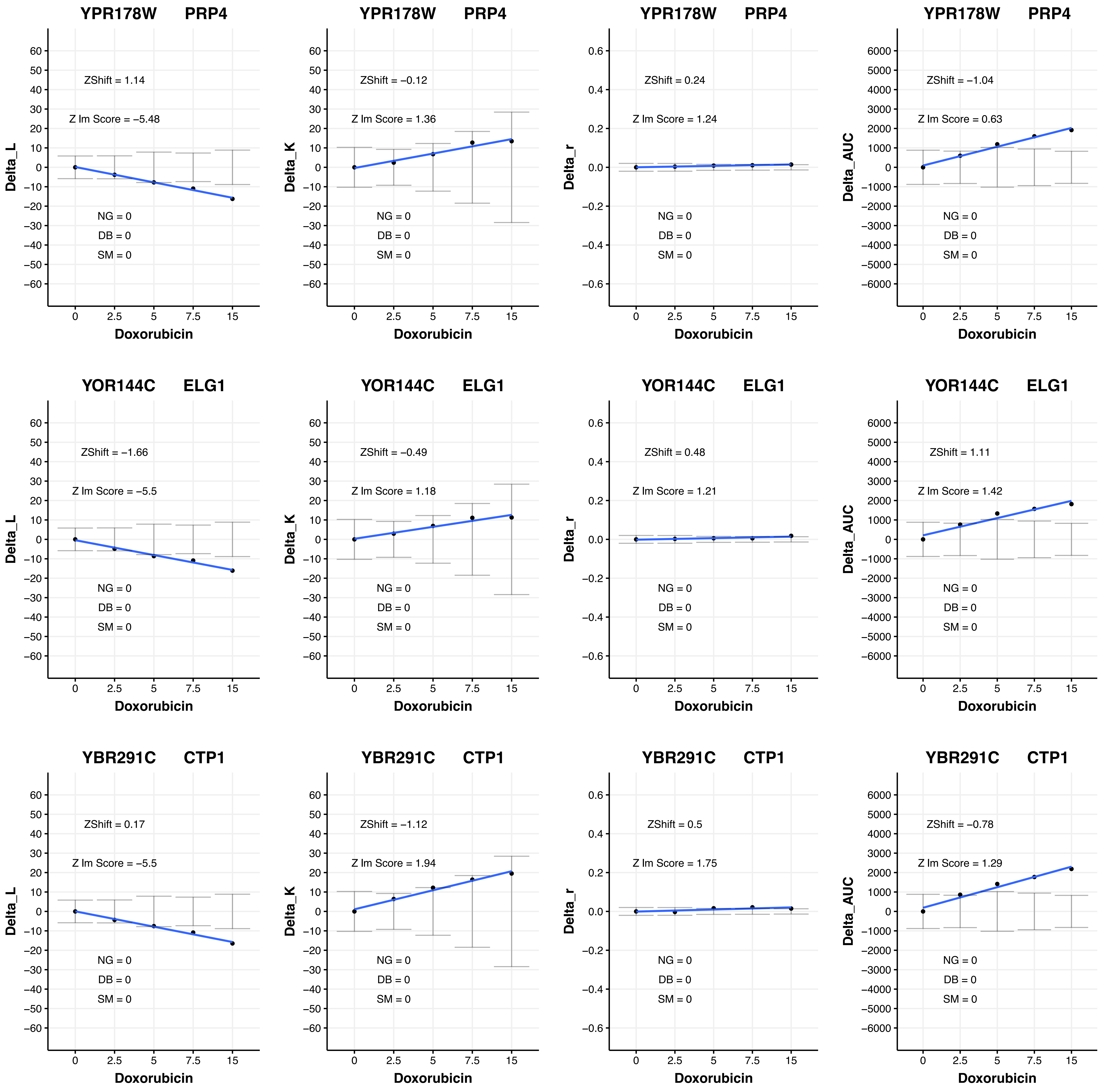
YPR178W PRP4
ZShift = 1.14
Z lm Score = −5.48
NG = 0
DB = 0
SM = 0
Delta_L
Doxorubicin
YPR178W PRP4
ZShift = −0.12
Z lm Score = 1.36
NG = 0
DB = 0
SM = 0
Delta_K
Doxorubicin
YPR178W PRP4
ZShift = 0.24
Z lm Score = 1.24
NG = 0
DB = 0
SM = 0
Delta_r
Doxorubicin
YPR178W PRP4
ZShift = −1.04
Z lm Score = 0.63
NG = 0
DB = 0
SM = 0
Delta_AUC
Doxorubicin
YOR144C ELG1
ZShift = −1.66
Z lm Score = −5.5
NG = 0
DB = 0
SM = 0
Delta_L
Doxorubicin
YOR144C ELG1
ZShift = −0.49
Z lm Score = 1.18
NG = 0
DB = 0
SM = 0
Delta_K
Doxorubicin
YOR144C ELG1
ZShift = 0.48
Z lm Score = 1.21
NG = 0
DB = 0
SM = 0
Delta_r
Doxorubicin
YOR144C ELG1
ZShift = 1.11
Z lm Score = 1.42
NG = 0
DB = 0
SM = 0
Delta_AUC
Doxorubicin
YBR291C CTP1
ZShift = 0.17
Z lm Score = −5.5
NG = 0
DB = 0
SM = 0
Delta_L
Doxorubicin
YBR291C CTP1
ZShift = −1.12
Z lm Score = 1.94
NG = 0
DB = 0
SM = 0
Delta_K
Doxorubicin
YBR291C CTP1
ZShift = 0.5
Z lm Score = 1.75
NG = 0
DB = 0
SM = 0
Delta_r
Doxorubicin
YBR291C CTP1
ZShift = −0.78
Z lm Score = 1.29
NG = 0
DB = 0
SM = 0
Delta_AUC
Doxorubicin

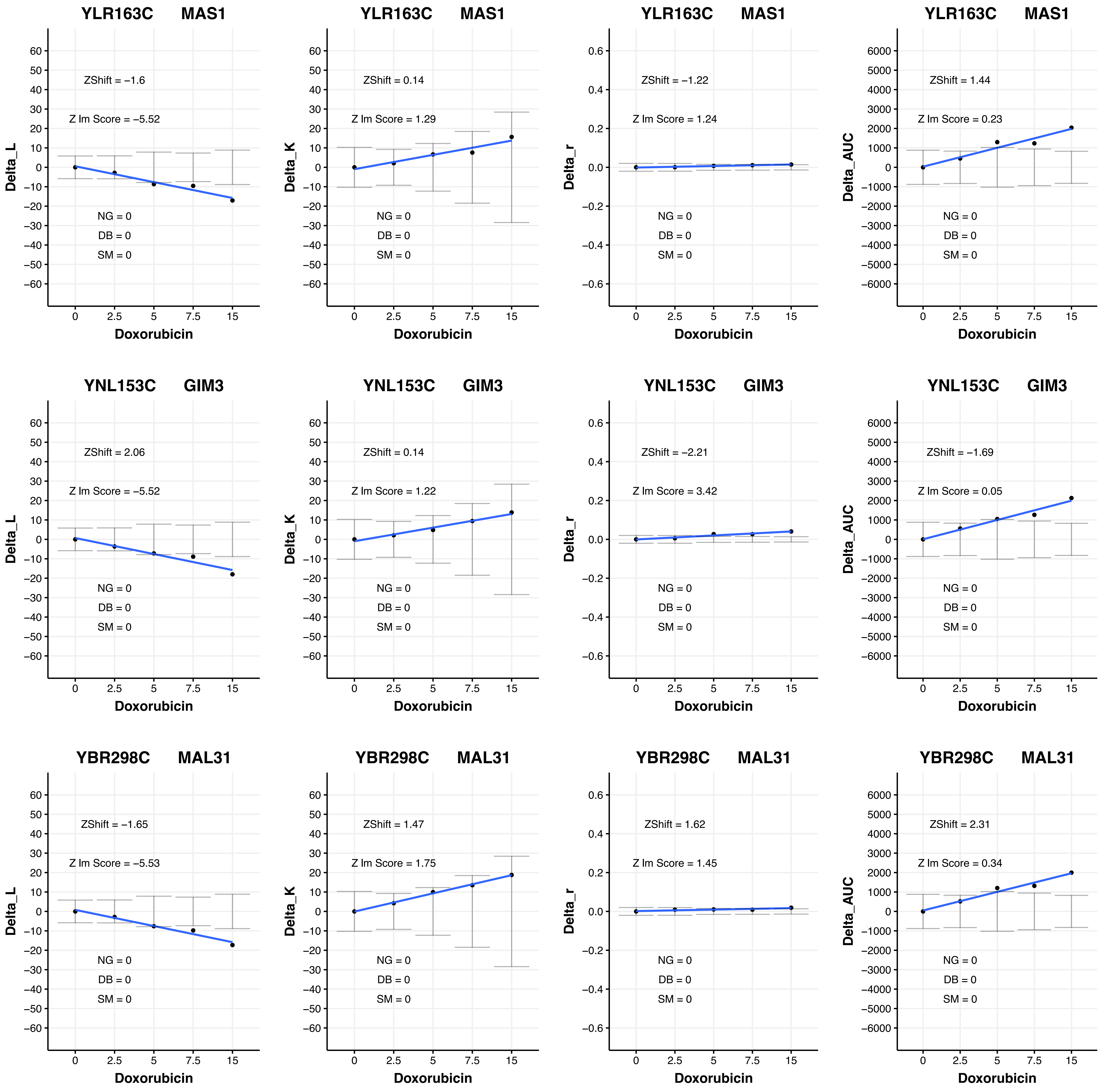
YLR163C MAS1
ZShift = −1.6
Z lm Score = −5.52
NG = 0
DB = 0
SM = 0
Delta_L
Doxorubicin
YLR163C MAS1
ZShift = 0.14
Z lm Score = 1.29
NG = 0
DB = 0
SM = 0
Delta_K
Doxorubicin
YLR163C MAS1
ZShift = −1.22
Z lm Score = 1.24
NG = 0
DB = 0
SM = 0
Delta_r
Doxorubicin
YLR163C MAS1
ZShift = 1.44
Z lm Score = 0.23
NG = 0
DB = 0
SM = 0
Delta_AUC
Doxorubicin
YNL153C GIM3
ZShift = 2.06
Z lm Score = −5.52
NG = 0
DB = 0
SM = 0
Delta_L
Doxorubicin
YNL153C GIM3
ZShift = 0.14
Z lm Score = 1.22
NG = 0
DB = 0
SM = 0
Delta_K
Doxorubicin
YNL153C GIM3
ZShift = −2.21
Z lm Score = 3.42
NG = 0
DB = 0
SM = 0
Delta_r
Doxorubicin
YNL153C GIM3
ZShift = −1.69
Z lm Score = 0.05
NG = 0
DB = 0
SM = 0
Delta_AUC
Doxorubicin
YBR298C MAL31
ZShift = −1.65
Z lm Score = −5.53
NG = 0
DB = 0
SM = 0
Delta_L
Doxorubicin
YBR298C MAL31
ZShift = 1.47
Z lm Score = 1.75
NG = 0
DB = 0
SM = 0
Delta_K
Doxorubicin
YBR298C MAL31
ZShift = 1.62
Z lm Score = 1.45
NG = 0
DB = 0
SM = 0
Delta_r
Doxorubicin
YBR298C MAL31
ZShift = 2.31
Z lm Score = 0.34
NG = 0
DB = 0
SM = 0
Delta_AUC
Doxorubicin

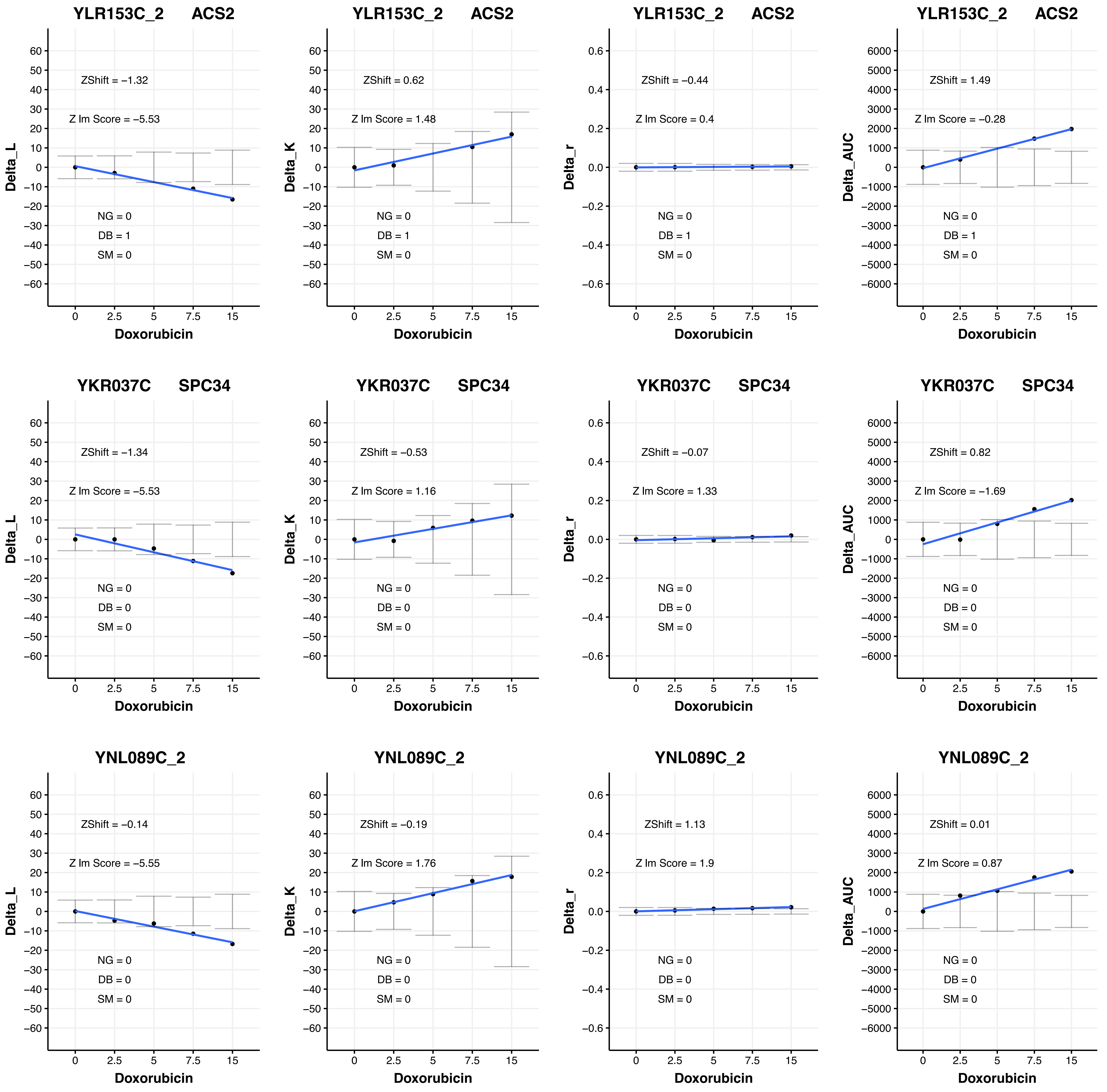
YLR153C_2 ACS2
ZShift = −1.32
Z lm Score = −5.53
NG = 0
DB = 1
SM = 0
Delta_L
Doxorubicin
YLR153C_2 ACS2
ZShift = 0.62
Z lm Score = 1.48
NG = 0
DB = 1
SM = 0
Delta_K
Doxorubicin
YLR153C_2 ACS2
ZShift = −0.44
Z lm Score = 0.4
NG = 0
DB = 1
SM = 0
Delta_r
Doxorubicin
YLR153C_2 ACS2
ZShift = 1.49
Z lm Score = −0.28
NG = 0
DB = 1
SM = 0
Delta_AUC
Doxorubicin
YKR037C SPC34
ZShift = −1.34
Z lm Score = −5.53
NG = 0
DB = 0
SM = 0
Delta_L
Doxorubicin
YKR037C SPC34
ZShift = −0.53
Z lm Score = 1.16
NG = 0
DB = 0
SM = 0
Delta_K
Doxorubicin
YKR037C SPC34
ZShift = −0.07
Z lm Score = 1.33
NG = 0
DB = 0
SM = 0
Delta_r
Doxorubicin
YKR037C SPC34
ZShift = 0.82
Z lm Score = −1.69
NG = 0
DB = 0
SM = 0
Delta_AUC
Doxorubicin
YNL089C_2
ZShift = −0.14
Z lm Score = −5.55
NG = 0
DB = 0
SM = 0
Delta_L
Doxorubicin
YNL089C_2
ZShift = −0.19
Z lm Score = 1.76
NG = 0
DB = 0
SM = 0
Delta_K
Doxorubicin
YNL089C_2
ZShift = 1.13
Z lm Score = 1.9
NG = 0
DB = 0
SM = 0
Delta_r
Doxorubicin
YNL089C_2
ZShift = 0.01
Z lm Score = 0.87
NG = 0
DB = 0
SM = 0
Delta_AUC
Doxorubicin

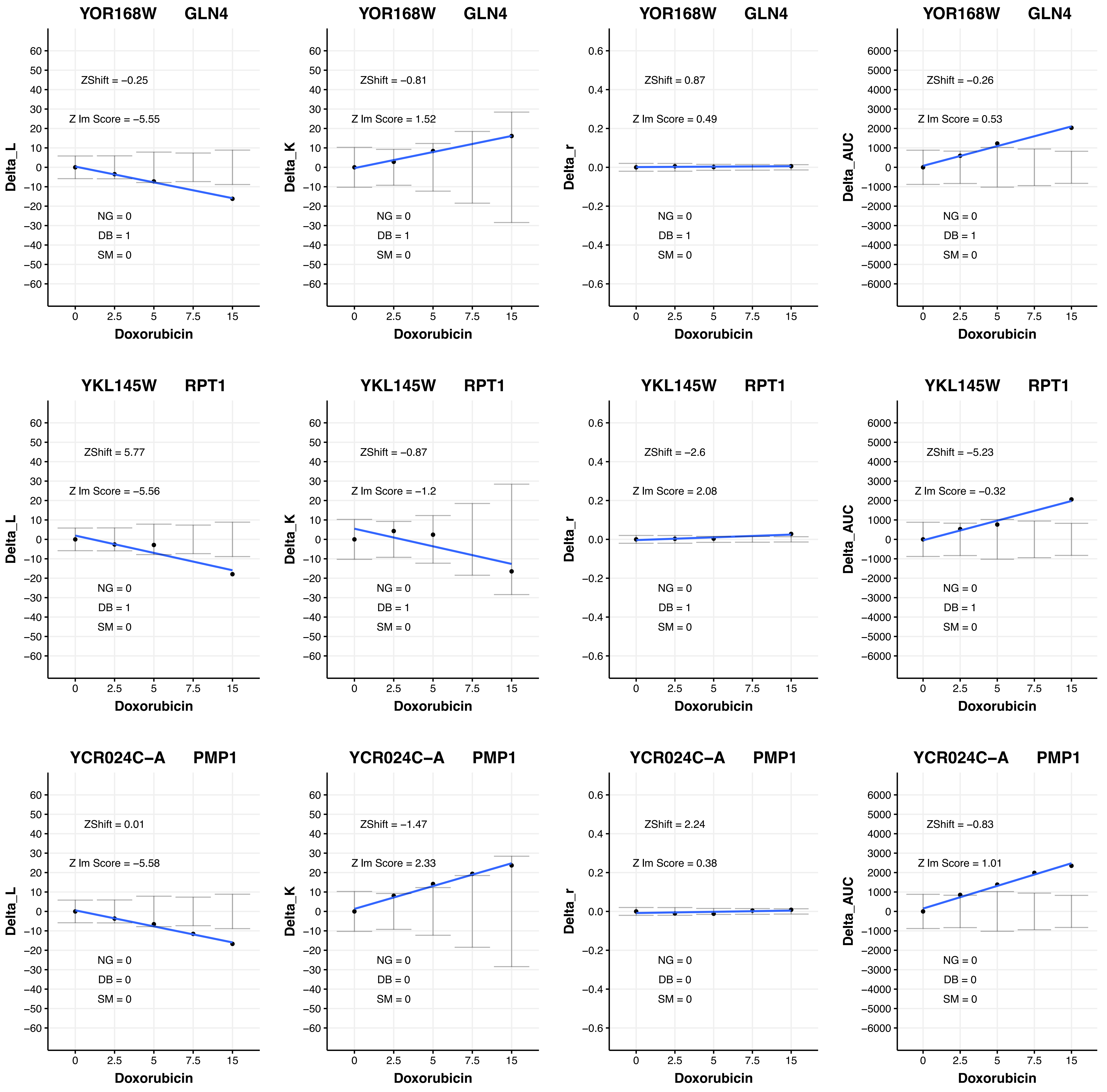
YOR168W GLN4
ZShift = −0.25
Z lm Score = −5.55
NG = 0
DB = 1
SM = 0
Delta_L
Doxorubicin
YOR168W GLN4
ZShift = −0.81
Z lm Score = 1.52
NG = 0
DB = 1
SM = 0
Delta_K
Doxorubicin
YOR168W GLN4
ZShift = 0.87
Z lm Score = 0.49
NG = 0
DB = 1
SM = 0
Delta_r
Doxorubicin
YOR168W GLN4
ZShift = −0.26
Z lm Score = 0.53
NG = 0
DB = 1
SM = 0
Delta_AUC
Doxorubicin
YKL145W RPT1
ZShift = 5.77
Z lm Score = −5.56
NG = 0
DB = 1
SM = 0
Delta_L
Doxorubicin
YKL145W RPT1
ZShift = −0.87
Z lm Score = −1.2
NG = 0
DB = 1
SM = 0
Delta_K
Doxorubicin
YKL145W RPT1
ZShift = −2.6
Z lm Score = 2.08
NG = 0
DB = 1
SM = 0
Delta_r
Doxorubicin
YKL145W RPT1
ZShift = −5.23
Z lm Score = −0.32
NG = 0
DB = 1
SM = 0
Delta_AUC
Doxorubicin
YCR024C−A PMP1
ZShift = 0.01
Z lm Score = −5.58
NG = 0
DB = 0
SM = 0
Delta_L
Doxorubicin
YCR024C−A PMP1
ZShift = −1.47
Z lm Score = 2.33
NG = 0
DB = 0
SM = 0
Delta_K
Doxorubicin
YCR024C−A PMP1
ZShift = 2.24
Z lm Score = 0.38
NG = 0
DB = 0
SM = 0
Delta_r
Doxorubicin
YCR024C−A PMP1
ZShift = −0.83
Z lm Score = 1.01
NG = 0
DB = 0
SM = 0
Delta_AUC
Doxorubicin

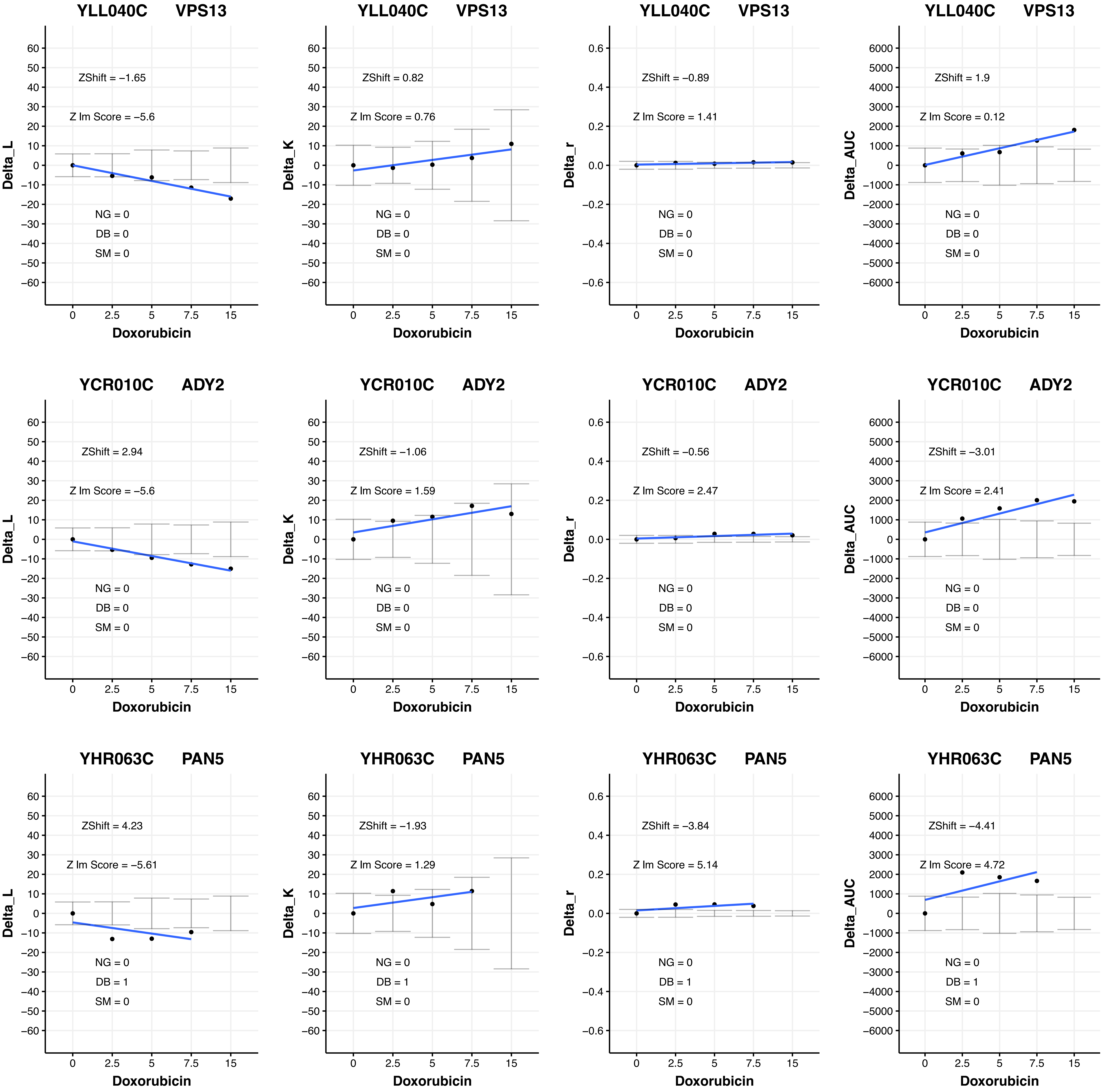
YLL040C VPS13
Delta_L
ZShift = −1.65
Z lm Score = −5.6
NG = 0
DB = 0
SM = 0
Doxorubicin
YLL040C VPS13
Delta_K
ZShift = 0.82
Z lm Score = 0.76
NG = 0
DB = 0
SM = 0
Doxorubicin
YLL040C VPS13
Delta_r
ZShift = −0.89
Z lm Score = 1.41
NG = 0
DB = 0
SM = 0
Doxorubicin
YLL040C VPS13
Delta_AUC
ZShift = 1.9
Z lm Score = 0.12
NG = 0
DB = 0
SM = 0
Doxorubicin
YCR010C ADY2
Delta_L
ZShift = 2.94
Z lm Score = −5.6
NG = 0
DB = 0
SM = 0
Doxorubicin
YCR010C ADY2
Delta_K
ZShift = −1.06
Z lm Score = 1.59
NG = 0
DB = 0
SM = 0
Doxorubicin
YCR010C ADY2
Delta_r
ZShift = −0.56
Z lm Score = 2.47
NG = 0
DB = 0
SM = 0
Doxorubicin
YCR010C ADY2
Delta_AUC
ZShift = −3.01
Z lm Score = 2.41
NG = 0
DB = 0
SM = 0
Doxorubicin
YHR063C PAN5
Delta_L
ZShift = 4.23
Z lm Score = −5.61
NG = 0
DB = 1
SM = 0
Doxorubicin
YHR063C PAN5
Delta_K
ZShift = −1.93
Z lm Score = 1.29
NG = 0
DB = 1
SM = 0
Doxorubicin
YHR063C PAN5
Delta_r
ZShift = −3.84
Z lm Score = 5.14
NG = 0
DB = 1
SM = 0
Doxorubicin
YHR063C PAN5
Delta_AUC
ZShift = −4.41
Z lm Score = 4.72
NG = 0
DB = 1
SM = 0
Doxorubicin

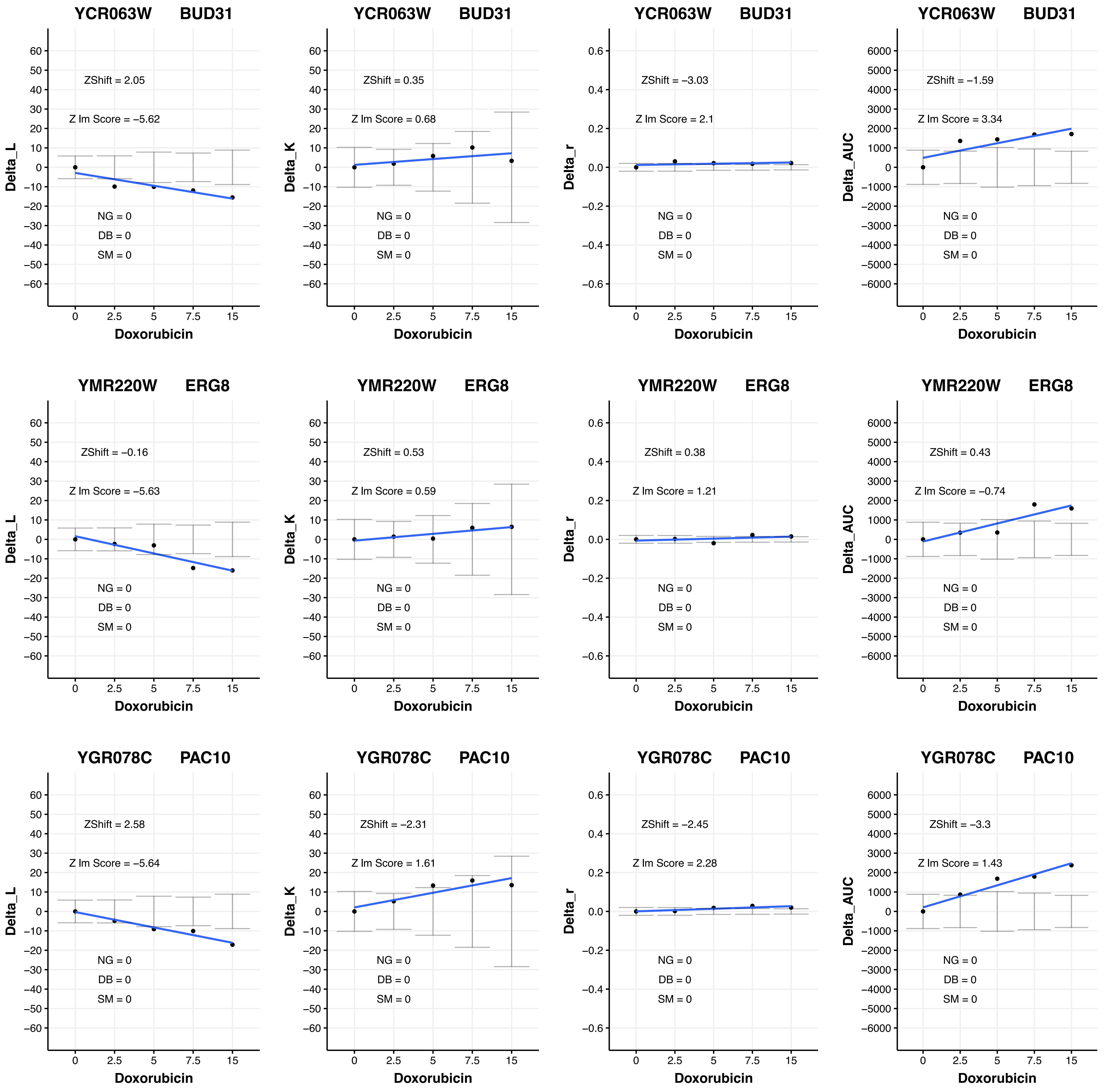
YCR063W BUD31
ZShift = 2.05
Z lm Score = −5.62
NG = 0
DB = 0
SM = 0
Delta_L
Doxorubicin
YCR063W BUD31
ZShift = 0.35
Z lm Score = 0.68
NG = 0
DB = 0
SM = 0
Delta_K
Doxorubicin
YCR063W BUD31
ZShift = −3.03
Z lm Score = 2.1
NG = 0
DB = 0
SM = 0
Delta_r
Doxorubicin
YCR063W BUD31
ZShift = −1.59
Z lm Score = 3.34
NG = 0
DB = 0
SM = 0
Delta_AUC
Doxorubicin
YMR220W ERG8
ZShift = −0.16
Z lm Score = −5.63
NG = 0
DB = 0
SM = 0
Delta_L
Doxorubicin
YMR220W ERG8
ZShift = 0.53
Z lm Score = 0.59
NG = 0
DB = 0
SM = 0
Delta_K
Doxorubicin
YMR220W ERG8
ZShift = 0.38
Z lm Score = 1.21
NG = 0
DB = 0
SM = 0
Delta_r
Doxorubicin
YMR220W ERG8
ZShift = 0.43
Z lm Score = −0.74
NG = 0
DB = 0
SM = 0
Delta_AUC
Doxorubicin
YGR078C PAC10
ZShift = 2.58
Z lm Score = −5.64
NG = 0
DB = 0
SM = 0
Delta_L
Doxorubicin
YGR078C PAC10
ZShift = −2.31
Z lm Score = 1.61
NG = 0
DB = 0
SM = 0
Delta_K
Doxorubicin
YGR078C PAC10
ZShift = −2.45
Z lm Score = 2.28
NG = 0
DB = 0
SM = 0
Delta_r
Doxorubicin
YGR078C PAC10
ZShift = −3.3
Z lm Score = 1.43
NG = 0
DB = 0
SM = 0
Delta_AUC
Doxorubicin

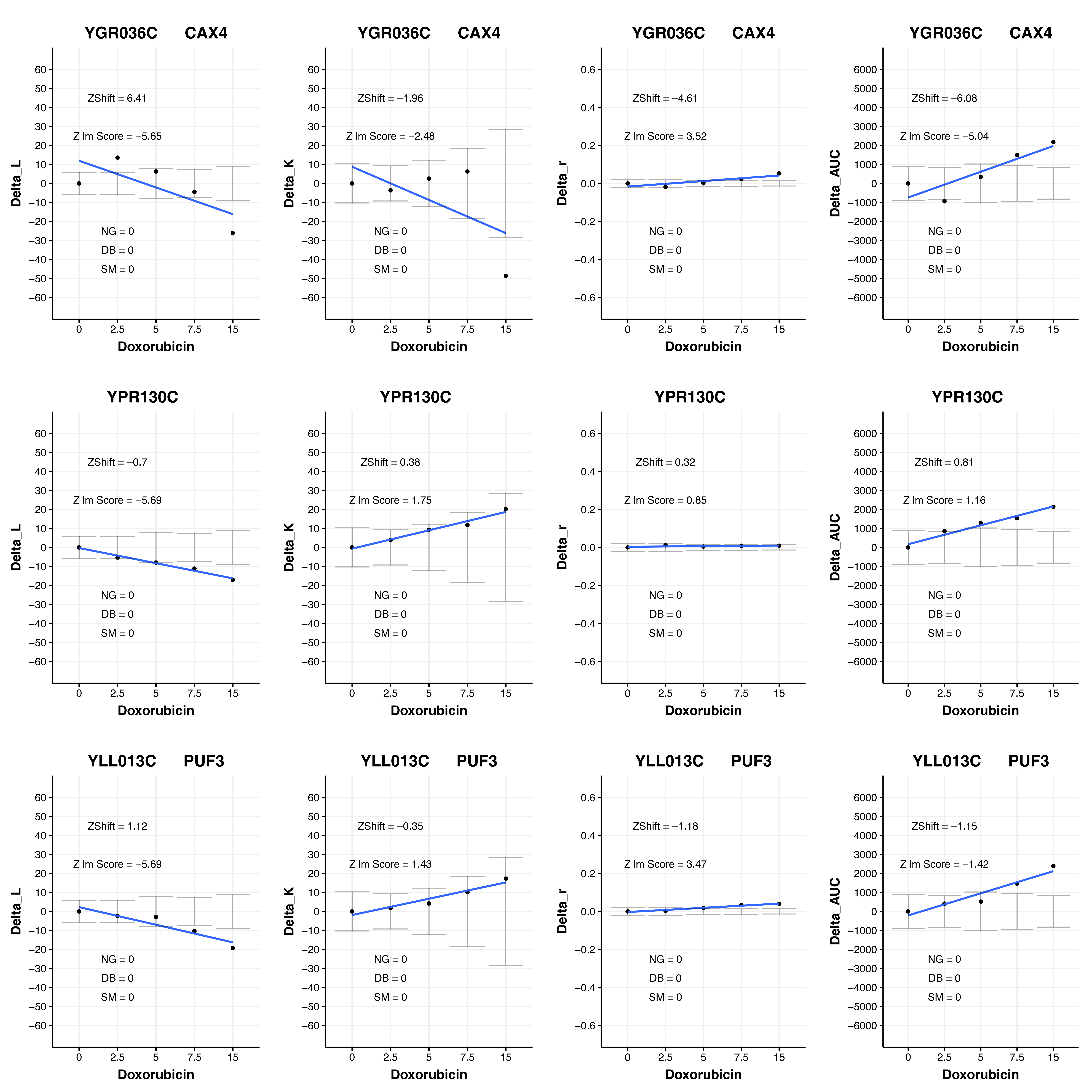
YGR036C CAX4
ZShift = 6.41
Z lm Score = −5.65
NG = 0
DB = 0
SM = 0
Delta_L
Doxorubicin
YGR036C CAX4
ZShift = −1.96
Z lm Score = −2.48
NG = 0
DB = 0
SM = 0
Delta_K
Doxorubicin
YGR036C CAX4
ZShift = −4.61
Z lm Score = 3.52
NG = 0
DB = 0
SM = 0
Delta_r
Doxorubicin
YGR036C CAX4
ZShift = −6.08
Z lm Score = −5.04
NG = 0
DB = 0
SM = 0
Delta_AUC
Doxorubicin
YPR130C
ZShift = −0.7
Z lm Score = −5.69
NG = 0
DB = 0
SM = 0
Delta_L
Doxorubicin
YPR130C
ZShift = 0.38
Z lm Score = 1.75
NG = 0
DB = 0
SM = 0
Delta_K
Doxorubicin
YPR130C
ZShift = 0.32
Z lm Score = 0.85
NG = 0
DB = 0
SM = 0
Delta_r
Doxorubicin
YPR130C
ZShift = 0.81
Z lm Score = 1.16
NG = 0
DB = 0
SM = 0
Delta_AUC
Doxorubicin
YLL013C PUF3
ZShift = 1.12
Z lm Score = −5.69
NG = 0
DB = 0
SM = 0
Delta_L
Doxorubicin
YLL013C PUF3
ZShift = −0.35
Z lm Score = 1.43
NG = 0
DB = 0
SM = 0
Delta_K
Doxorubicin
YLL013C PUF3
ZShift = −1.18
Z lm Score = 3.47
NG = 0
DB = 0
SM = 0
Delta_r
Doxorubicin
YLL013C PUF3
ZShift = −1.15
Z lm Score = −1.42
NG = 0
DB = 0
SM = 0
Delta_AUC
Doxorubicin

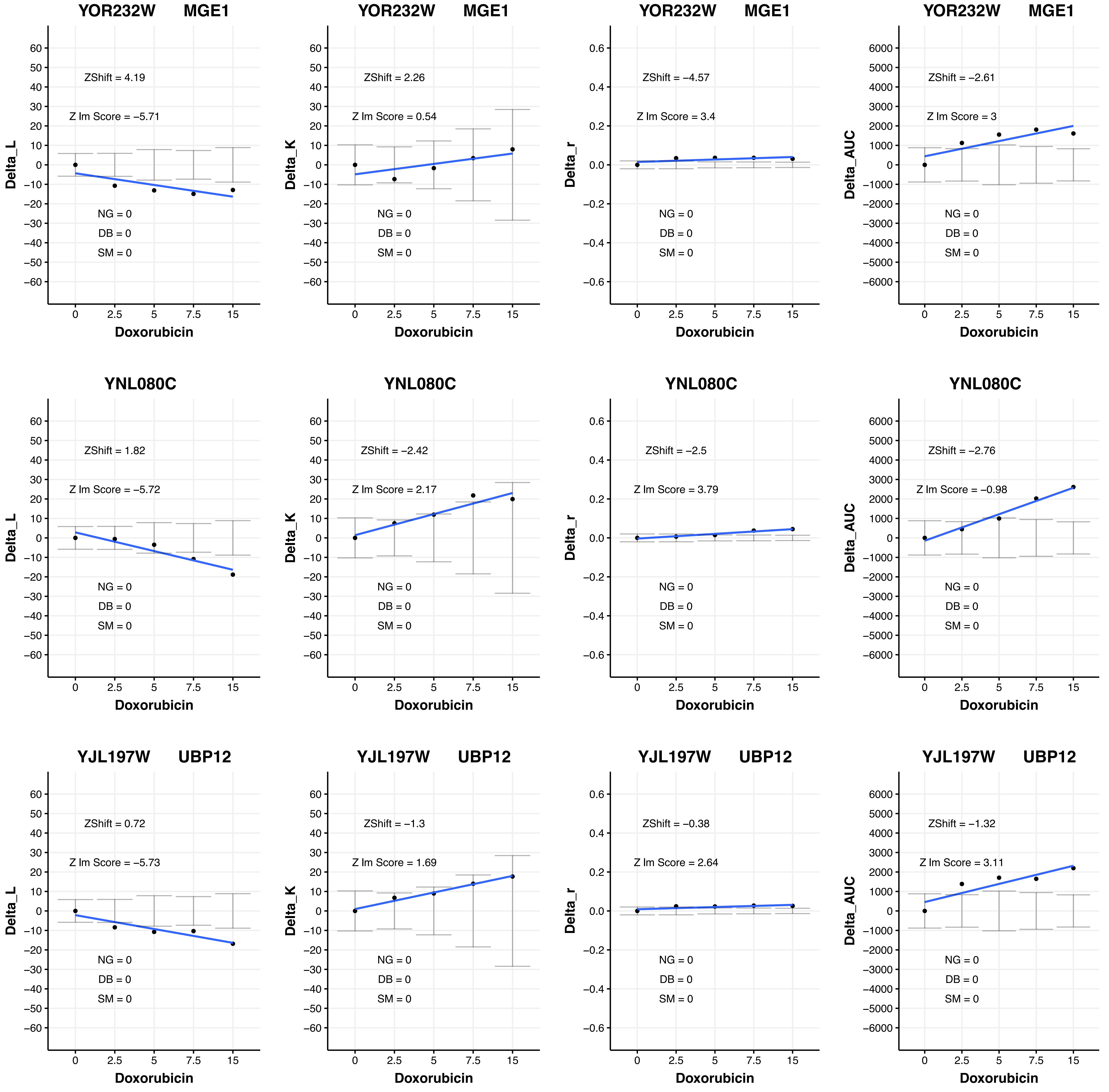
YOR232W MGE1
ZShift = 4.19
Z lm Score = −5.71
NG = 0
DB = 0
SM = 0
Delta_L
Doxorubicin
YOR232W MGE1
ZShift = 2.26
Z lm Score = 0.54
NG = 0
DB = 0
SM = 0
Delta_K
Doxorubicin
YOR232W MGE1
ZShift = −4.57
Z lm Score = 3.4
NG = 0
DB = 0
SM = 0
Delta_r
Doxorubicin
YOR232W MGE1
ZShift = −2.61
Z lm Score = 3
NG = 0
DB = 0
SM = 0
Delta_AUC
Doxorubicin
YNL080C
ZShift = 1.82
Z lm Score = −5.72
NG = 0
DB = 0
SM = 0
Delta_L
Doxorubicin
YNL080C
ZShift = −2.42
Z lm Score = 2.17
NG = 0
DB = 0
SM = 0
Delta_K
Doxorubicin
YNL080C
ZShift = −2.5
Z lm Score = 3.79
NG = 0
DB = 0
SM = 0
Delta_r
Doxorubicin
YNL080C
ZShift = −2.76
Z lm Score = −0.98
NG = 0
DB = 0
SM = 0
Delta_AUC
Doxorubicin
YJL197W UBP12
ZShift = 0.72
Z lm Score = −5.73
NG = 0
DB = 0
SM = 0
Delta_L
Doxorubicin
YJL197W UBP12
ZShift = −1.3
Z lm Score = 1.69
NG = 0
DB = 0
SM = 0
Delta_K
Doxorubicin
YJL197W UBP12
ZShift = −0.38
Z lm Score = 2.64
NG = 0
DB = 0
SM = 0
Delta_r
Doxorubicin
YJL197W UBP12
ZShift = −1.32
Z lm Score = 3.11
NG = 0
DB = 0
SM = 0
Delta_AUC
Doxorubicin

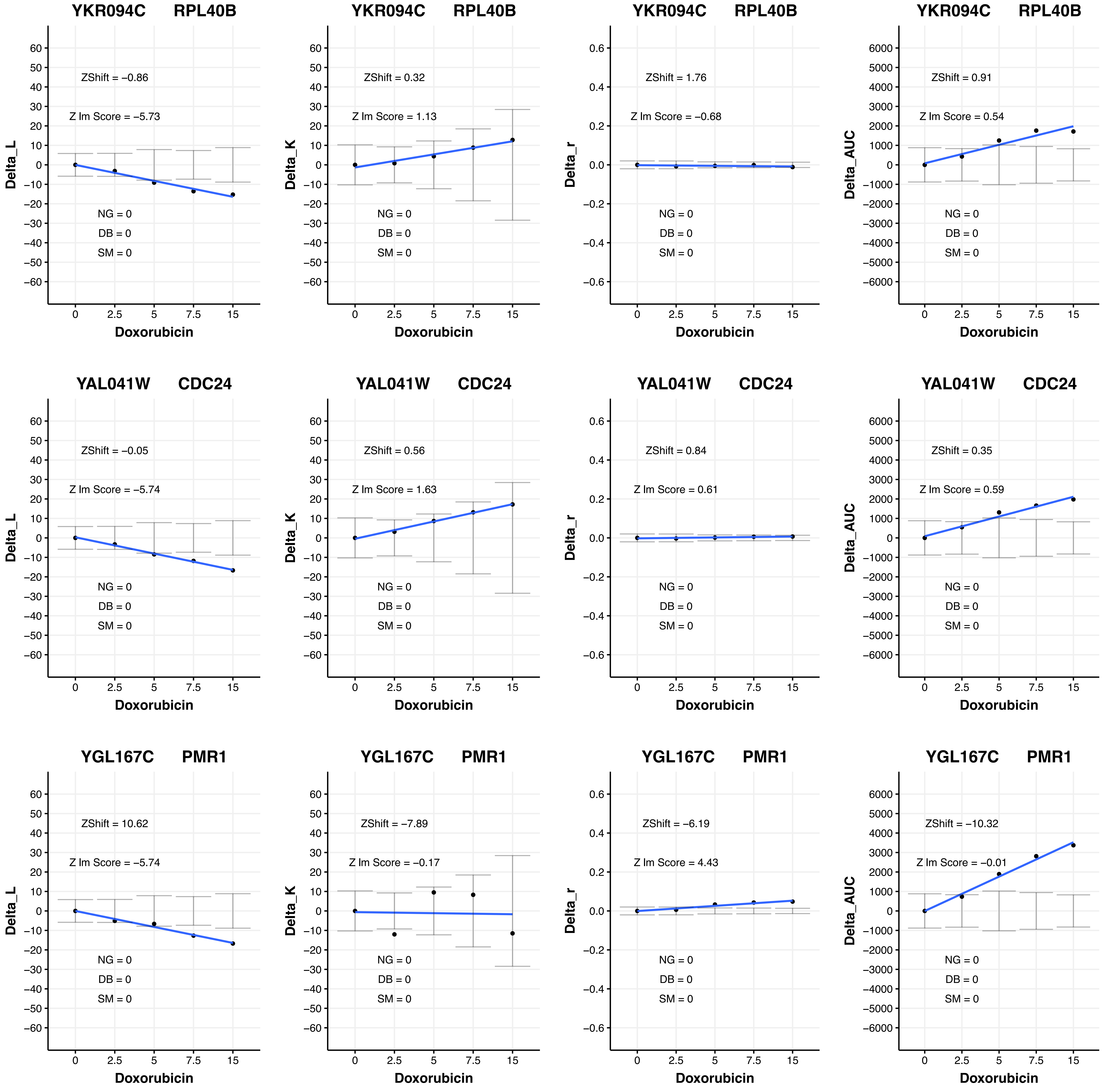
YKR094C RPL40B
ZShift = −0.86
Z lm Score = −5.73
NG = 0
DB = 0
SM = 0
Delta_L
Doxorubicin
YKR094C RPL40B
ZShift = 0.32
Z lm Score = 1.13
NG = 0
DB = 0
SM = 0
Delta_K
Doxorubicin
YKR094C RPL40B
ZShift = 1.76
Z lm Score = −0.68
NG = 0
DB = 0
SM = 0
Delta_r
Doxorubicin
YKR094C RPL40B
ZShift = 0.91
Z lm Score = 0.54
NG = 0
DB = 0
SM = 0
Delta_AUC
Doxorubicin
YAL041W CDC24
ZShift = −0.05
Z lm Score = −5.74
NG = 0
DB = 0
SM = 0
Delta_L
Doxorubicin
YAL041W CDC24
ZShift = 0.56
Z lm Score = 1.63
NG = 0
DB = 0
SM = 0
Delta_K
Doxorubicin
YAL041W CDC24
ZShift = 0.84
Z lm Score = 0.61
NG = 0
DB = 0
SM = 0
Delta_r
Doxorubicin
YAL041W CDC24
ZShift = 0.35
Z lm Score = 0.59
NG = 0
DB = 0
SM = 0
Delta_AUC
Doxorubicin
YGL167C PMR1
ZShift = 10.62
Z lm Score = −5.74
NG = 0
DB = 0
SM = 0
Delta_L
Doxorubicin
YGL167C PMR1
ZShift = −7.89
Z lm Score = −0.17
NG = 0
DB = 0
SM = 0
Delta_K
Doxorubicin
YGL167C PMR1
ZShift = −6.19
Z lm Score = 4.43
NG = 0
DB = 0
SM = 0
Delta_r
Doxorubicin
YGL167C PMR1
ZShift = −10.32
Z lm Score = −0.01
NG = 0
DB = 0
SM = 0
Delta_AUC
Doxorubicin

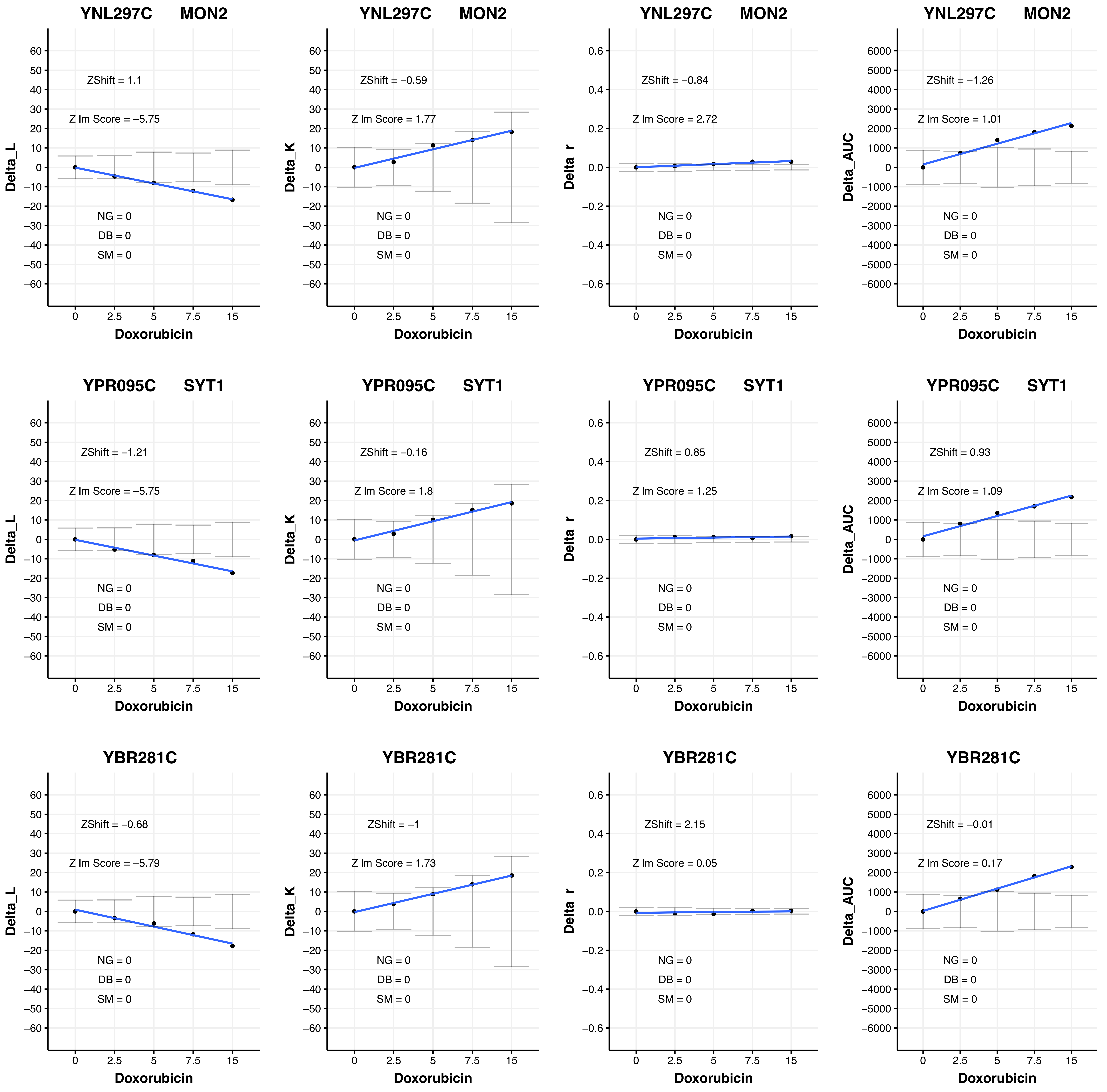
YNL297C MON2
ZShift = 1.1
Z lm Score = −5.75
NG = 0
DB = 0
SM = 0
Delta_L
Doxorubicin
YNL297C MON2
ZShift = −0.59
Z lm Score = 1.77
NG = 0
DB = 0
SM = 0
Delta_K
Doxorubicin
YNL297C MON2
ZShift = −0.84
Z lm Score = 2.72
NG = 0
DB = 0
SM = 0
Delta_r
Doxorubicin
YNL297C MON2
ZShift = −1.26
Z lm Score = 1.01
NG = 0
DB = 0
SM = 0
Delta_AUC
Doxorubicin
YPR095C SYT1
ZShift = −1.21
Z lm Score = −5.75
NG = 0
DB = 0
SM = 0
Delta_L
Doxorubicin
YPR095C SYT1
ZShift = −0.16
Z lm Score = 1.8
NG = 0
DB = 0
SM = 0
Delta_K
Doxorubicin
YPR095C SYT1
ZShift = 0.85
Z lm Score = 1.25
NG = 0
DB = 0
SM = 0
Delta_r
Doxorubicin
YPR095C SYT1
ZShift = 0.93
Z lm Score = 1.09
NG = 0
DB = 0
SM = 0
Delta_AUC
Doxorubicin
YBR281C
ZShift = −0.68
Z lm Score = −5.79
NG = 0
DB = 0
SM = 0
Delta_L
Doxorubicin
YBR281C
ZShift = −1
Z lm Score = 1.73
NG = 0
DB = 0
SM = 0
Delta_K
Doxorubicin
YBR281C
ZShift = 2.15
Z lm Score = 0.05
NG = 0
DB = 0
SM = 0
Delta_r
Doxorubicin
YBR281C
ZShift = −0.01
Z lm Score = 0.17
NG = 0
DB = 0
SM = 0
Delta_AUC
Doxorubicin

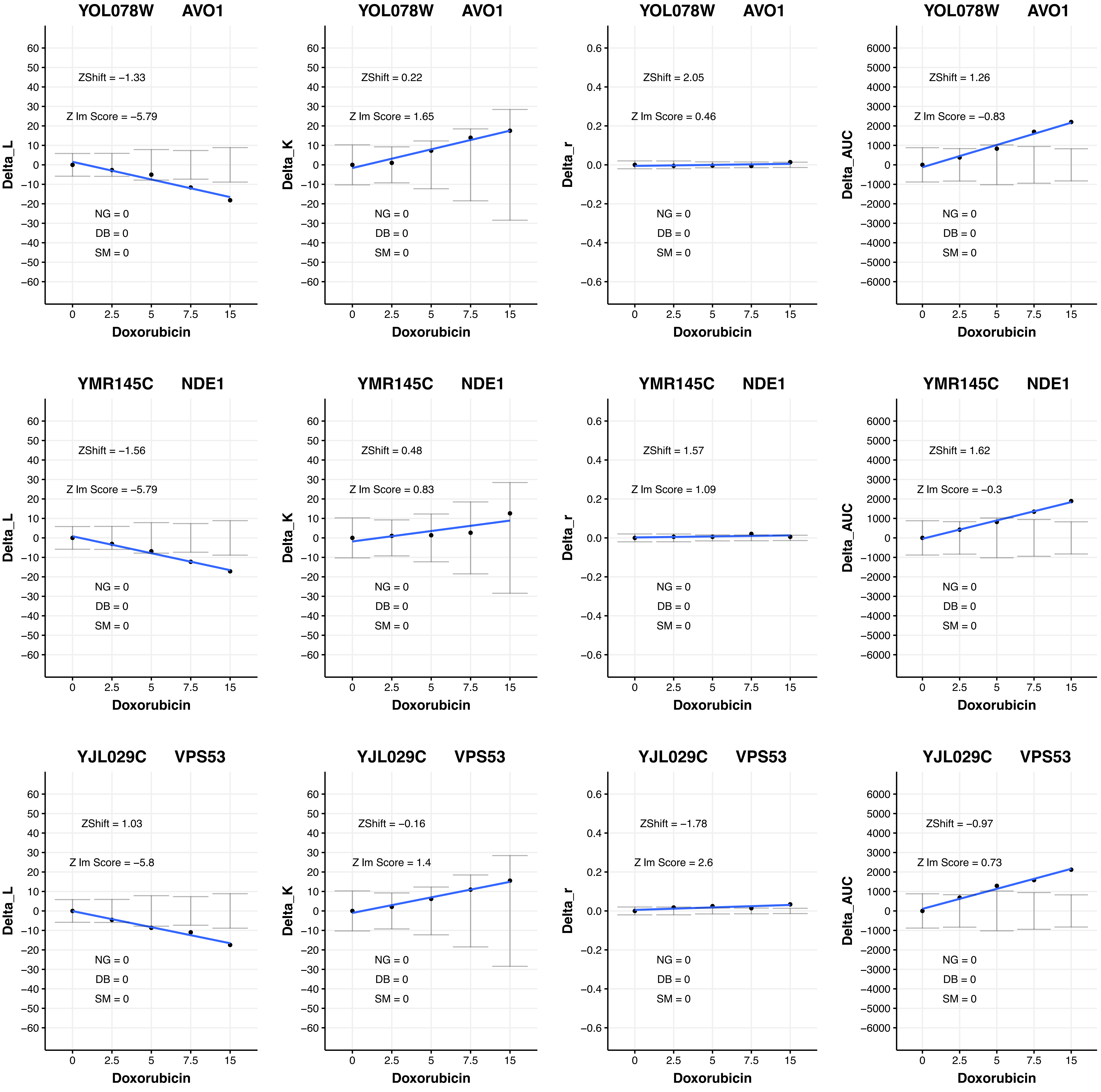

YOL078W AVO1
ZShift = −1.33
Z lm Score = −5.79
NG = 0
DB = 0
SM = 0
Delta_L
Doxorubicin
YOL078W AVO1
ZShift = 0.22
Z lm Score = 1.65
NG = 0
DB = 0
SM = 0
Delta_K
Doxorubicin
YOL078W AVO1
ZShift = 2.05
Z lm Score = 0.46
NG = 0
DB = 0
SM = 0
Delta_r
Doxorubicin
YOL078W AVO1
ZShift = 1.26
Z lm Score = −0.83
NG = 0
DB = 0
SM = 0
Delta_AUC
Doxorubicin
YMR145C NDE1
ZShift = −1.56
Z lm Score = −5.79
NG = 0
DB = 0
SM = 0
Delta_L
Doxorubicin
YMR145C NDE1
ZShift = 0.48
Z lm Score = 0.83
NG = 0
DB = 0
SM = 0
Delta_K
Doxorubicin
YMR145C NDE1
ZShift = 1.57
Z lm Score = 1.09
NG = 0
DB = 0
SM = 0
Delta_r
Doxorubicin
YMR145C NDE1
ZShift = 1.62
Z lm Score = −0.3
NG = 0
DB = 0
SM = 0
Delta_AUC
Doxorubicin
YJL029C VPS53
ZShift = 1.03
Z lm Score = −5.8
NG = 0
DB = 0
SM = 0
Delta_L
Doxorubicin
YJL029C VPS53
ZShift = −0.16
Z lm Score = 1.4
NG = 0
DB = 0
SM = 0
Delta_K
Doxorubicin
YJL029C VPS53
ZShift = −1.78
Z lm Score = 2.6
NG = 0
DB = 0
SM = 0
Delta_r
Doxorubicin
YJL029C VPS53
ZShift = −0.97
Z lm Score = 0.73
NG = 0
DB = 0
SM = 0
Delta_AUC
Doxorubicin

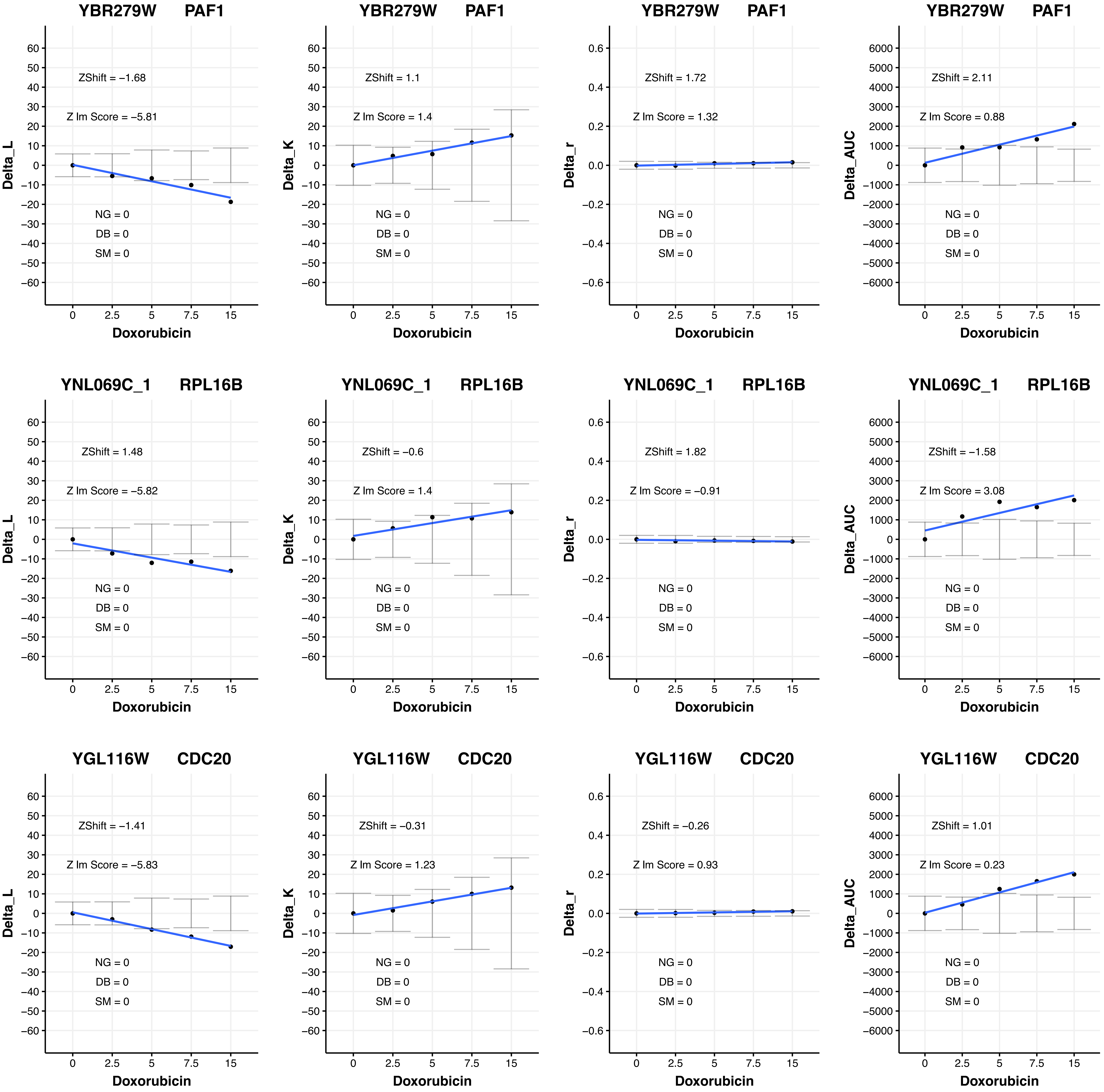
YBR279W PAF1
ZShift = −1.68
Z lm Score = −5.81
NG = 0
DB = 0
SM = 0
Delta_L
Doxorubicin
YBR279W PAF1
ZShift = 1.1
Z lm Score = 1.4
NG = 0
DB = 0
SM = 0
Delta_K
Doxorubicin
YBR279W PAF1
ZShift = 1.72
Z lm Score = 1.32
NG = 0
DB = 0
SM = 0
Delta_r
Doxorubicin
YBR279W PAF1
ZShift = 2.11
Z lm Score = 0.88
NG = 0
DB = 0
SM = 0
Delta_AUC
Doxorubicin
YNL069C_1 RPL16B
ZShift = 1.48
Z lm Score = −5.82
NG = 0
DB = 0
SM = 0
Delta_L
Doxorubicin
YNL069C_1 RPL16B
ZShift = −0.6
Z lm Score = 1.4
NG = 0
DB = 0
SM = 0
Delta_K
Doxorubicin
YNL069C_1 RPL16B
ZShift = 1.82
Z lm Score = −0.91
NG = 0
DB = 0
SM = 0
Delta_r
Doxorubicin
YNL069C_1 RPL16B
ZShift = −1.58
Z lm Score = 3.08
NG = 0
DB = 0
SM = 0
Delta_AUC
Doxorubicin
YGL116W CDC20
ZShift = −1.41
Z lm Score = −5.83
NG = 0
DB = 0
SM = 0
Delta_L
Doxorubicin
YGL116W CDC20
ZShift = −0.31
Z lm Score = 1.23
NG = 0
DB = 0
SM = 0
Delta_K
Doxorubicin
YGL116W CDC20
ZShift = −0.26
Z lm Score = 0.93
NG = 0
DB = 0
SM = 0
Delta_r
Doxorubicin
YGL116W CDC20
ZShift = 1.01
Z lm Score = 0.23
NG = 0
DB = 0
SM = 0
Delta_AUC
Doxorubicin

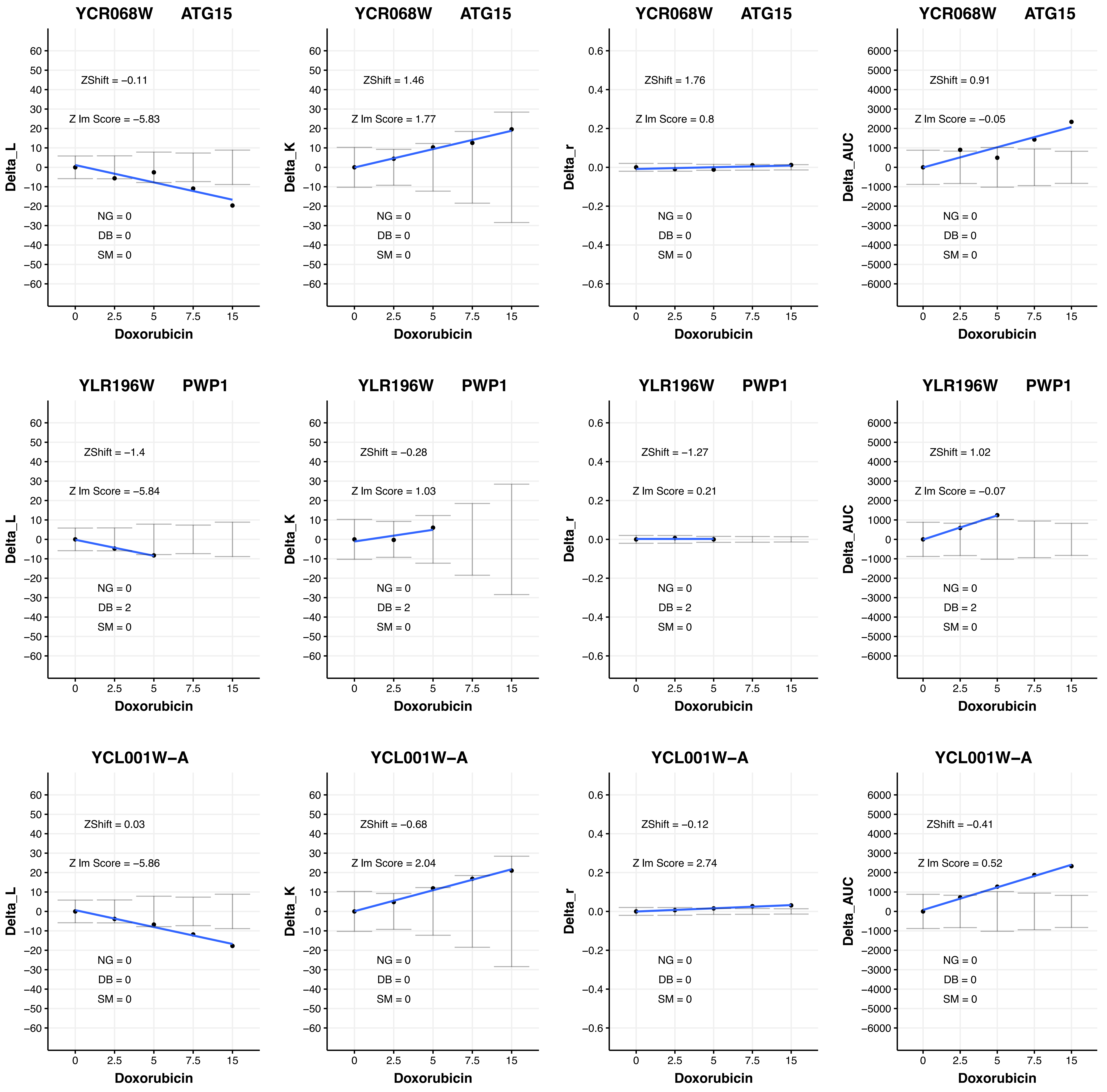
YCR068W ATG15
ZShift = −0.11
Z lm Score = −5.83
NG = 0
DB = 0
SM = 0
Delta_L
Doxorubicin
YCR068W ATG15
ZShift = 1.46
Z lm Score = 1.77
NG = 0
DB = 0
SM = 0
Delta_K
Doxorubicin
YCR068W ATG15
ZShift = 1.76
Z lm Score = 0.8
NG = 0
DB = 0
SM = 0
Delta_r
Doxorubicin
YCR068W ATG15
ZShift = 0.91
Z lm Score = −0.05
NG = 0
DB = 0
SM = 0
Delta_AUC
Doxorubicin
YLR196W PWP1
ZShift = −1.4
Z lm Score = −5.84
NG = 0
DB = 2
SM = 0
Delta_L
Doxorubicin
YLR196W PWP1
ZShift = −0.28
Z lm Score = 1.03
NG = 0
DB = 2
SM = 0
Delta_K
Doxorubicin
YLR196W PWP1
ZShift = −1.27
Z lm Score = 0.21
NG = 0
DB = 2
SM = 0
Delta_r
Doxorubicin
YLR196W PWP1
ZShift = 1.02
Z lm Score = −0.07
NG = 0
DB = 2
SM = 0
Delta_AUC
Doxorubicin
YCL001W−A
ZShift = 0.03
Z lm Score = −5.86
NG = 0
DB = 0
SM = 0
Delta_L
Doxorubicin
YCL001W−A
ZShift = −0.68
Z lm Score = 2.04
NG = 0
DB = 0
SM = 0
Delta_K
Doxorubicin
YCL001W−A
ZShift = −0.12
Z lm Score = 2.74
NG = 0
DB = 0
SM = 0
Delta_r
Doxorubicin
YCL001W−A
ZShift = −0.41
Z lm Score = 0.52
NG = 0
DB = 0
SM = 0
Delta_AUC
Doxorubicin

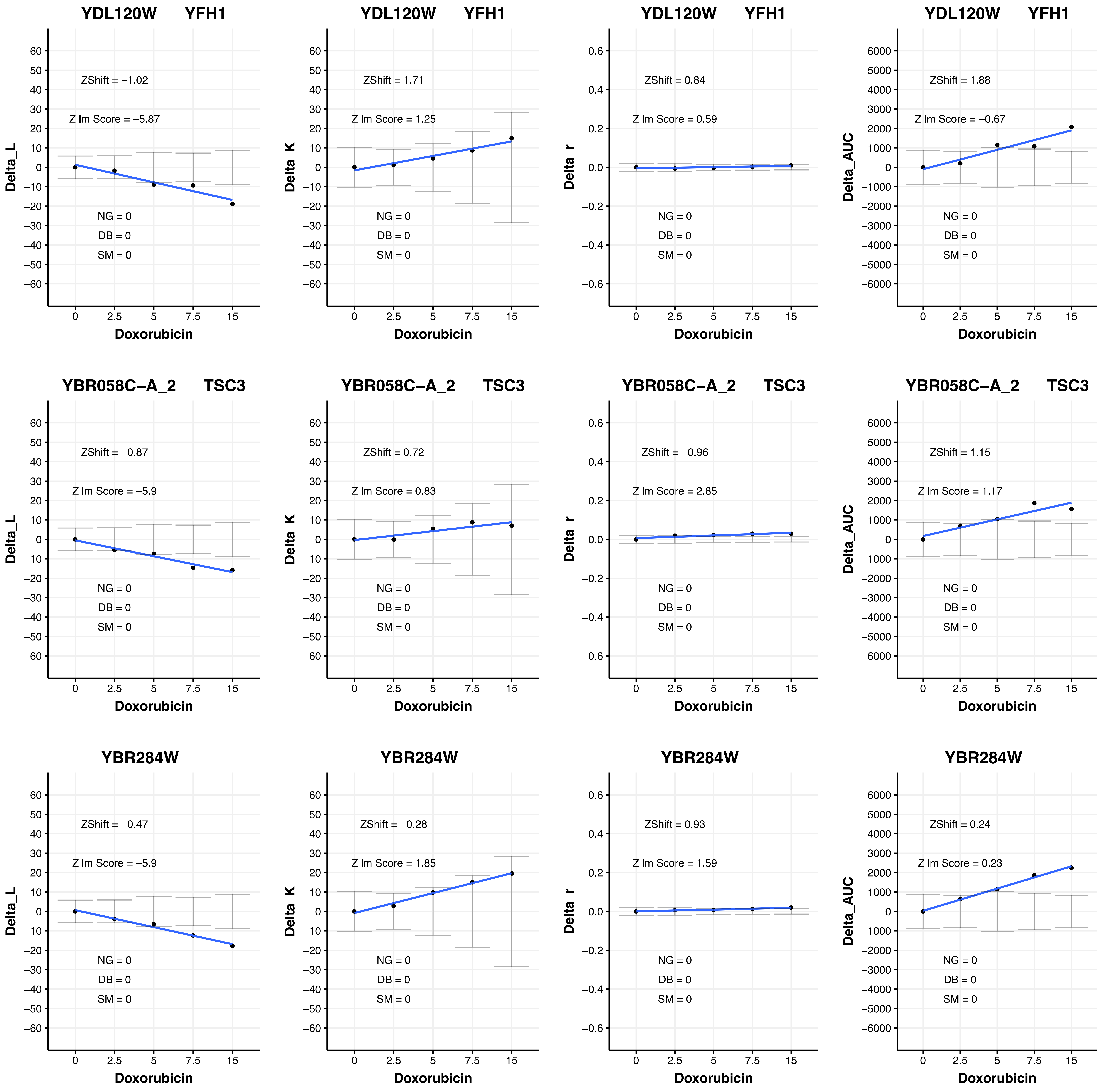
YDL120W YFH1
ZShift = −1.02
Z lm Score = −5.87
NG = 0
DB = 0
SM = 0
Delta_L
Doxorubicin
YDL120W YFH1
ZShift = 1.71
Z lm Score = 1.25
NG = 0
DB = 0
SM = 0
Delta_K
Doxorubicin
YDL120W YFH1
ZShift = 0.84
Z lm Score = 0.59
NG = 0
DB = 0
SM = 0
Delta_r
Doxorubicin
YDL120W YFH1
ZShift = 1.88
Z lm Score = −0.67
NG = 0
DB = 0
SM = 0
Delta_AUC
Doxorubicin
YBR058C−A_2 TSC3
ZShift = −0.87
Z lm Score = −5.9
NG = 0
DB = 0
SM = 0
Delta_L
Doxorubicin
YBR058C−A_2 TSC3
ZShift = 0.72
Z lm Score = 0.83
NG = 0
DB = 0
SM = 0
Delta_K
Doxorubicin
YBR058C−A_2 TSC3
ZShift = −0.96
Z lm Score = 2.85
NG = 0
DB = 0
SM = 0
Delta_r
Doxorubicin
YBR058C−A_2 TSC3
ZShift = 1.15
Z lm Score = 1.17
NG = 0
DB = 0
SM = 0
Delta_AUC
Doxorubicin
YBR284W
ZShift = −0.47
Z lm Score = −5.9
NG = 0
DB = 0
SM = 0
Delta_L
Doxorubicin
YBR284W
ZShift = −0.28
Z lm Score = 1.85
NG = 0
DB = 0
SM = 0
Delta_K
Doxorubicin
YBR284W
ZShift = 0.93
Z lm Score = 1.59
NG = 0
DB = 0
SM = 0
Delta_r
Doxorubicin
YBR284W
ZShift = 0.24
Z lm Score = 0.23
NG = 0
DB = 0
SM = 0
Delta_AUC
Doxorubicin

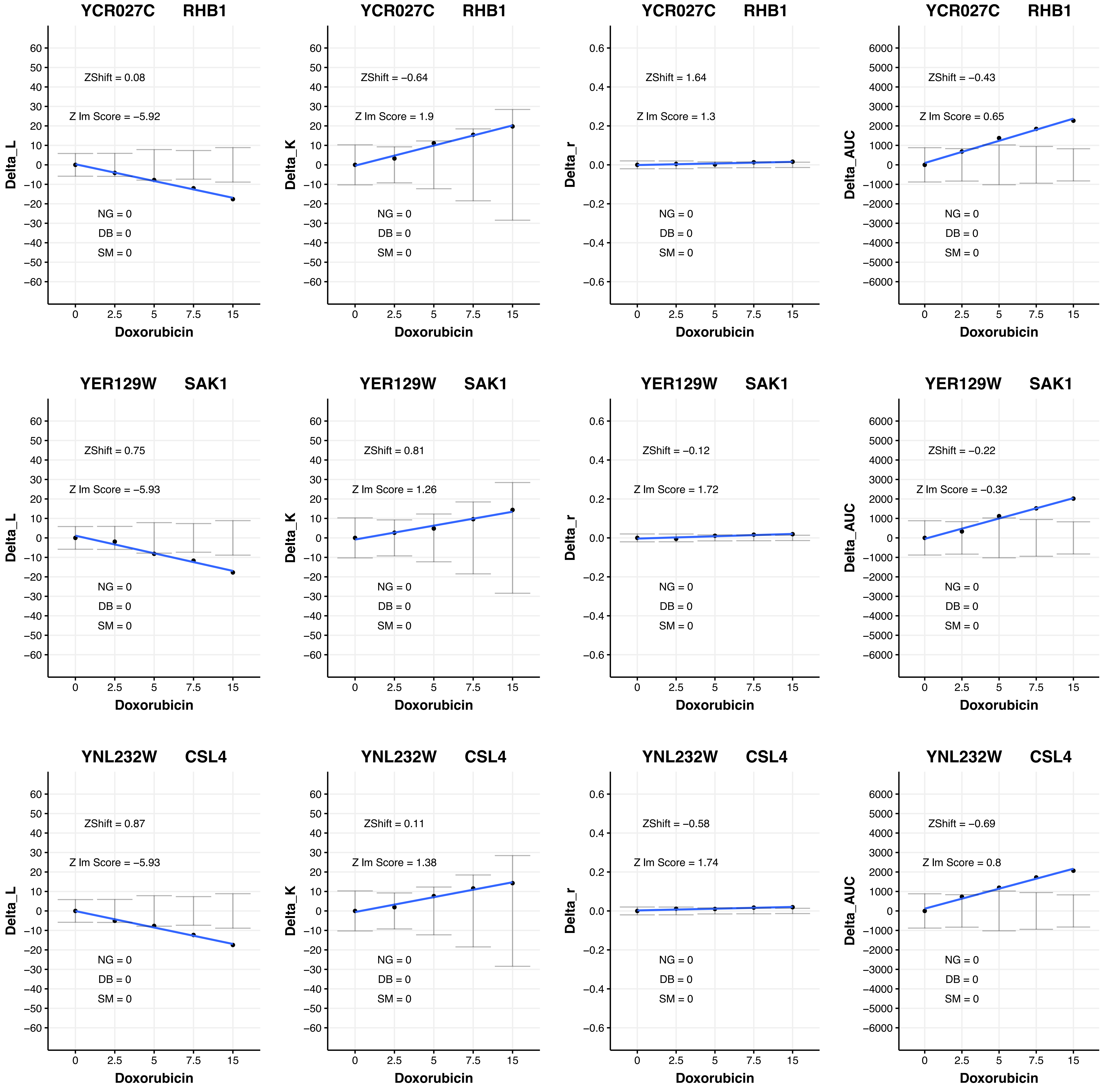

YCR027C RHB1
ZShift = 0.08
Z lm Score = −5.92
NG = 0
DB = 0
SM = 0
Delta_L
Doxorubicin
YCR027C RHB1
ZShift = −0.64
Z lm Score = 1.9
NG = 0
DB = 0
SM = 0
Delta_K
Doxorubicin
YCR027C RHB1
ZShift = 1.64
Z lm Score = 1.3
NG = 0
DB = 0
SM = 0
Delta_r
Doxorubicin
YCR027C RHB1
ZShift = −0.43
Z lm Score = 0.65
NG = 0
DB = 0
SM = 0
Delta_AUC
Doxorubicin
YER129W SAK1
ZShift = 0.75
Z lm Score = −5.93
NG = 0
DB = 0
SM = 0
Delta_L
Doxorubicin
YER129W SAK1
ZShift = 0.81
Z lm Score = 1.26
NG = 0
DB = 0
SM = 0
Delta_K
Doxorubicin
YER129W SAK1
ZShift = −0.12
Z lm Score = 1.72
NG = 0
DB = 0
SM = 0
Delta_r
Doxorubicin
YER129W SAK1
ZShift = −0.22
Z lm Score = −0.32
NG = 0
DB = 0
SM = 0
Delta_AUC
Doxorubicin
YNL232W CSL4
ZShift = 0.87
Z lm Score = −5.93
NG = 0
DB = 0
SM = 0
Delta_L
Doxorubicin
YNL232W CSL4
ZShift = 0.11
Z lm Score = 1.38
NG = 0
DB = 0
SM = 0
Delta_K
Doxorubicin
YNL232W CSL4
ZShift = −0.58
Z lm Score = 1.74
NG = 0
DB = 0
SM = 0
Delta_r
Doxorubicin
YNL232W CSL4
ZShift = −0.69
Z lm Score = 0.8
NG = 0
DB = 0
SM = 0
Delta_AUC
Doxorubicin

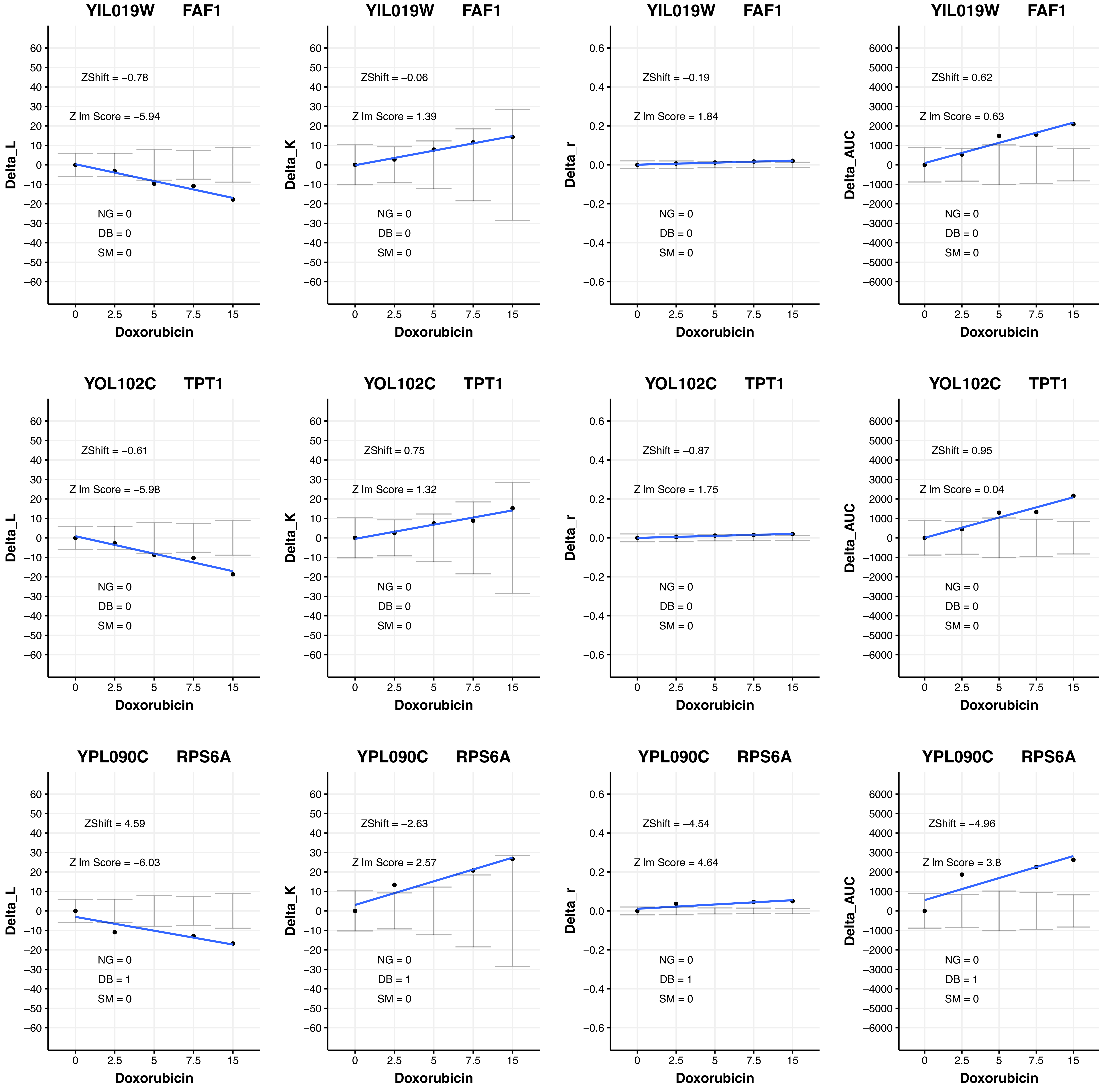
YIL019W FAF1
ZShift = −0.78
Z lm Score = −5.94
NG = 0
DB = 0
SM = 0
Delta_L
Doxorubicin
YIL019W FAF1
ZShift = −0.06
Z lm Score = 1.39
NG = 0
DB = 0
SM = 0
Delta_K
Doxorubicin
YIL019W FAF1
ZShift = −0.19
Z lm Score = 1.84
NG = 0
DB = 0
SM = 0
Delta_r
Doxorubicin
YIL019W FAF1
ZShift = 0.62
Z lm Score = 0.63
NG = 0
DB = 0
SM = 0
Delta_AUC
Doxorubicin
YOL102C TPT1
ZShift = −0.61
Z lm Score = −5.98
NG = 0
DB = 0
SM = 0
Delta_L
Doxorubicin
YOL102C TPT1
ZShift = 0.75
Z lm Score = 1.32
NG = 0
DB = 0
SM = 0
Delta_K
Doxorubicin
YOL102C TPT1
ZShift = −0.87
Z lm Score = 1.75
NG = 0
DB = 0
SM = 0
Delta_r
Doxorubicin
YOL102C TPT1
ZShift = 0.95
Z lm Score = 0.04
NG = 0
DB = 0
SM = 0
Delta_AUC
Doxorubicin
YPL090C RPS6A
ZShift = 4.59
Z lm Score = −6.03
NG = 0
DB = 1
SM = 0
Delta_L
Doxorubicin
YPL090C RPS6A
ZShift = −2.63
Z lm Score = 2.57
NG = 0
DB = 1
SM = 0
Delta_K
Doxorubicin
YPL090C RPS6A
ZShift = −4.54
Z lm Score = 4.64
NG = 0
DB = 1
SM = 0
Delta_r
Doxorubicin
YPL090C RPS6A
ZShift = −4.96
Z lm Score = 3.8
NG = 0
DB = 1
SM = 0
Delta_AUC
Doxorubicin

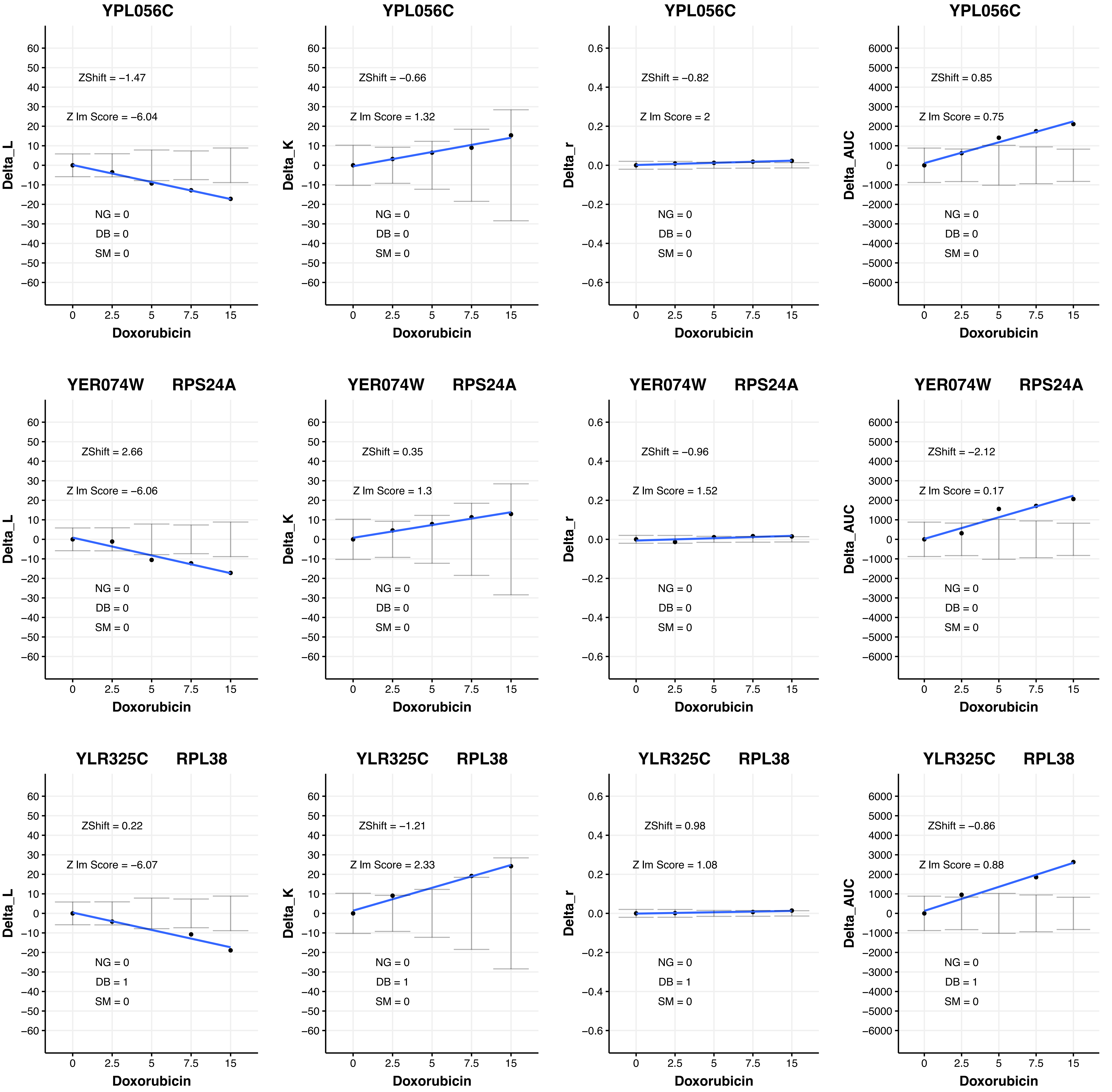
YPL056C
ZShift = −1.47
Z lm Score = −6.04
NG = 0
DB = 0
SM = 0
Delta_L
Doxorubicin
YPL056C
ZShift = −0.66
Z lm Score = 1.32
NG = 0
DB = 0
SM = 0
Delta_K
Doxorubicin
YPL056C
ZShift = −0.82
Z lm Score = 2
NG = 0
DB = 0
SM = 0
Delta_r
Doxorubicin
YPL056C
ZShift = 0.85
Z lm Score = 0.75
NG = 0
DB = 0
SM = 0
Delta_AUC
Doxorubicin
YER074W RPS24A
ZShift = 2.66
Z lm Score = −6.06
NG = 0
DB = 0
SM = 0
Delta_L
Doxorubicin
YER074W RPS24A
ZShift = 0.35
Z lm Score = 1.3
NG = 0
DB = 0
SM = 0
Delta_K
Doxorubicin
YER074W RPS24A
ZShift = −0.96
Z lm Score = 1.52
NG = 0
DB = 0
SM = 0
Delta_r
Doxorubicin
YER074W RPS24A
ZShift = −2.12
Z lm Score = 0.17
NG = 0
DB = 0
SM = 0
Delta_AUC
Doxorubicin
YLR325C RPL38
ZShift = 0.22
Z lm Score = −6.07
NG = 0
DB = 1
SM = 0
Delta_L
Doxorubicin
YLR325C RPL38
ZShift = −1.21
Z lm Score = 2.33
NG = 0
DB = 1
SM = 0
Delta_K
Doxorubicin
YLR325C RPL38
ZShift = 0.98
Z lm Score = 1.08
NG = 0
DB = 1
SM = 0
Delta_r
Doxorubicin
YLR325C RPL38
ZShift = −0.86
Z lm Score = 0.88
NG = 0
DB = 1
SM = 0
Delta_AUC
Doxorubicin

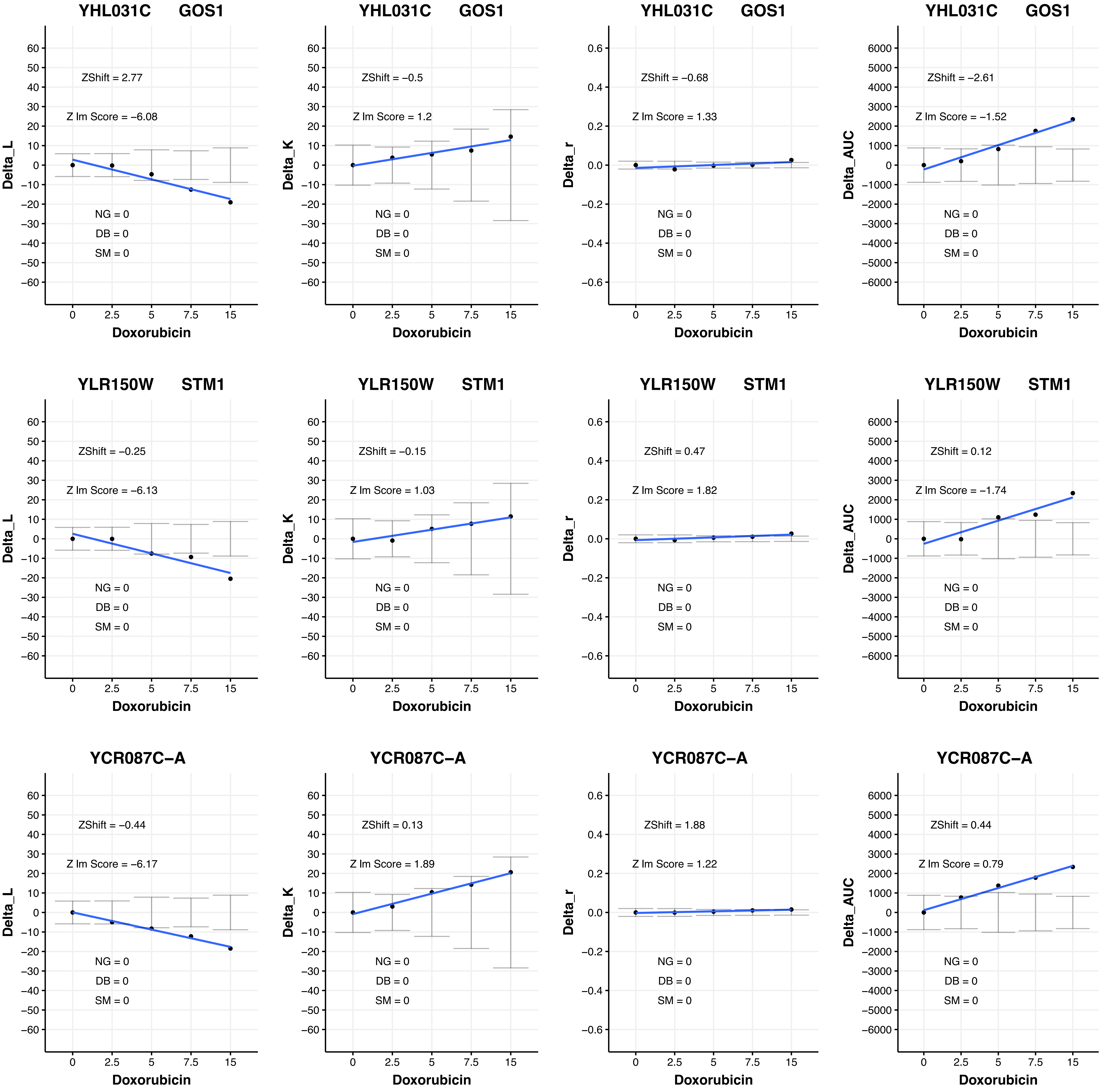
YHL031C GOS1
ZShift = 2.77
Z lm Score = −6.08
NG = 0
DB = 0
SM = 0
Delta_L
Doxorubicin
YHL031C GOS1
ZShift = −0.5
Z lm Score = 1.2
NG = 0
DB = 0
SM = 0
Delta_K
Doxorubicin
YHL031C GOS1
ZShift = −0.68
Z lm Score = 1.33
NG = 0
DB = 0
SM = 0
Delta_r
Doxorubicin
YHL031C GOS1
ZShift = −2.61
Z lm Score = −1.52
NG = 0
DB = 0
SM = 0
Delta_AUC
Doxorubicin
YLR150W STM1
ZShift = −0.25
Z lm Score = −6.13
NG = 0
DB = 0
SM = 0
Delta_L
Doxorubicin
YLR150W STM1
ZShift = −0.15
Z lm Score = 1.03
NG = 0
DB = 0
SM = 0
Delta_K
Doxorubicin
YLR150W STM1
ZShift = 0.47
Z lm Score = 1.82
NG = 0
DB = 0
SM = 0
Delta_r
Doxorubicin
YLR150W STM1
ZShift = 0.12
Z lm Score = −1.74
NG = 0
DB = 0
SM = 0
Delta_AUC
Doxorubicin
YCR087C−A
ZShift = −0.44
Z lm Score = −6.17
NG = 0
DB = 0
SM = 0
Delta_L
Doxorubicin
YCR087C−A
ZShift = 0.13
Z lm Score = 1.89
NG = 0
DB = 0
SM = 0
Delta_K
Doxorubicin
YCR087C−A
ZShift = 1.88
Z lm Score = 1.22
NG = 0
DB = 0
SM = 0
Delta_r
Doxorubicin
YCR087C−A
ZShift = 0.44
Z lm Score = 0.79
NG = 0
DB = 0
SM = 0
Delta_AUC
Doxorubicin

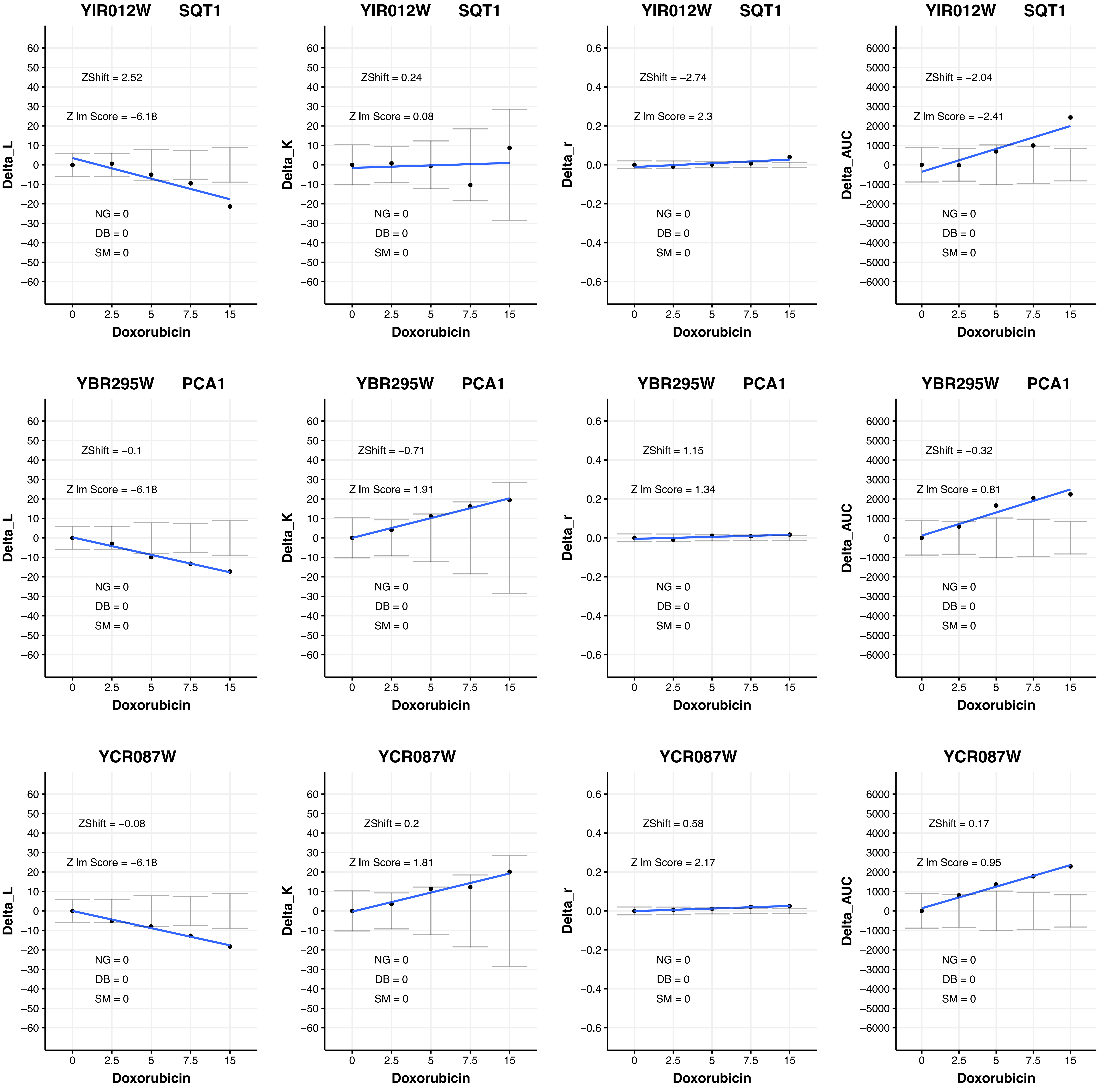
YIR012W SQT1
ZShift = 2.52
Z lm Score = −6.18
NG = 0
DB = 0
SM = 0
Delta_L
Doxorubicin
YIR012W SQT1
ZShift = 0.24
Z lm Score = 0.08
NG = 0
DB = 0
SM = 0
Delta_K
Doxorubicin
YIR012W SQT1
ZShift = −2.74
Z lm Score = 2.3
NG = 0
DB = 0
SM = 0
Delta_r
Doxorubicin
YIR012W SQT1
ZShift = −2.04
Z lm Score = −2.41
NG = 0
DB = 0
SM = 0
Delta_AUC
Doxorubicin
YBR295W PCA1
ZShift = −0.1
Z lm Score = −6.18
NG = 0
DB = 0
SM = 0
Delta_L
Doxorubicin
YBR295W PCA1
ZShift = −0.71
Z lm Score = 1.91
NG = 0
DB = 0
SM = 0
Delta_K
Doxorubicin
YBR295W PCA1
ZShift = 1.15
Z lm Score = 1.34
NG = 0
DB = 0
SM = 0
Delta_r
Doxorubicin
YBR295W PCA1
ZShift = −0.32
Z lm Score = 0.81
NG = 0
DB = 0
SM = 0
Delta_AUC
Doxorubicin
YCR087W
ZShift = −0.08
Z lm Score = −6.18
NG = 0
DB = 0
SM = 0
Delta_L
Doxorubicin
YCR087W
ZShift = 0.2
Z lm Score = 1.81
NG = 0
DB = 0
SM = 0
Delta_K
Doxorubicin
YCR087W
ZShift = 0.58
Z lm Score = 2.17
NG = 0
DB = 0
SM = 0
Delta_r
Doxorubicin
YCR087W
ZShift = 0.17
Z lm Score = 0.95
NG = 0
DB = 0
SM = 0
Delta_AUC
Doxorubicin

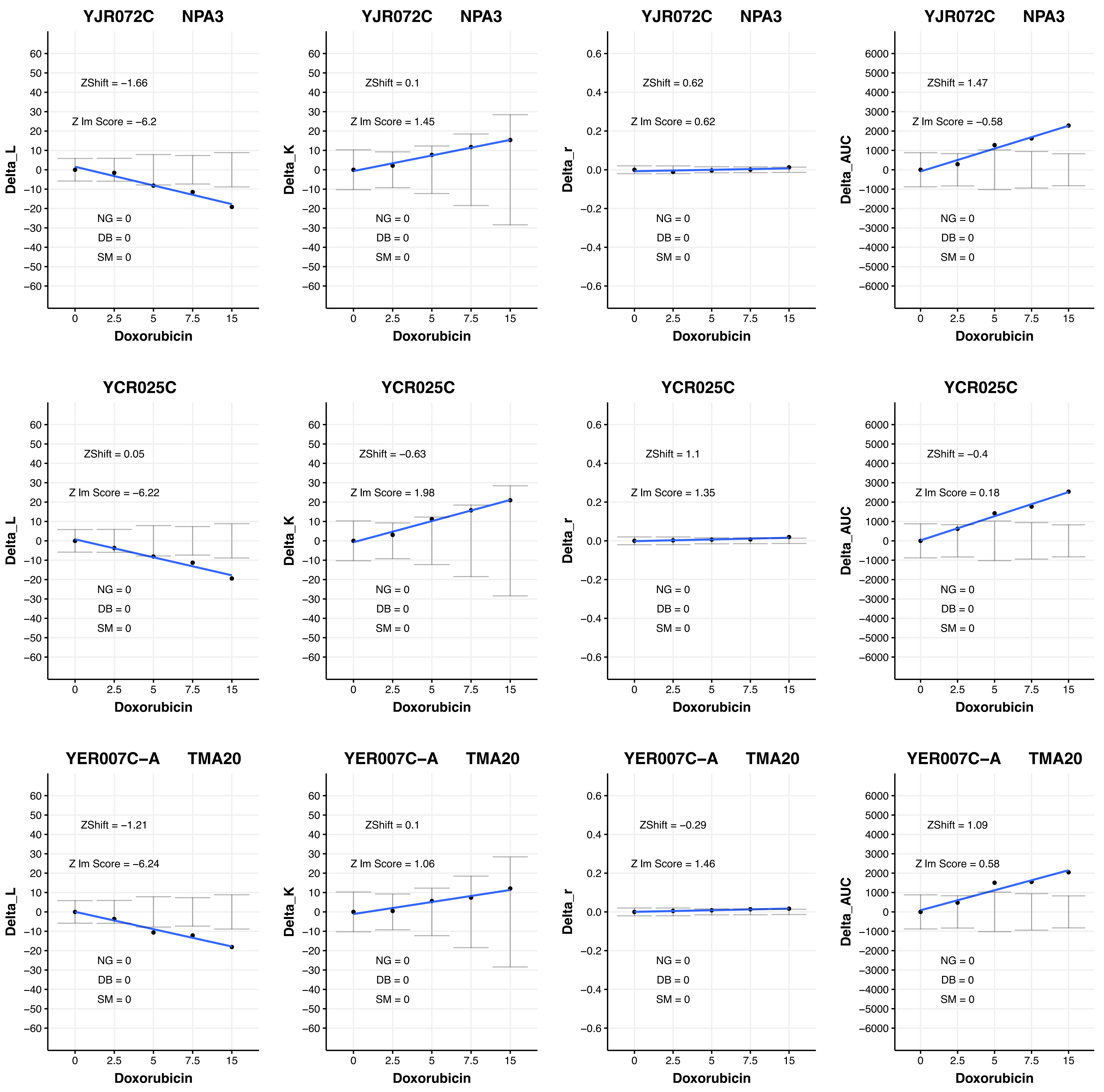

YJR072C NPA3
ZShift = −1.66
Z lm Score = −6.2
NG = 0
DB = 0
SM = 0
Delta_L
Doxorubicin
YJR072C NPA3
ZShift = 0.1
Z lm Score = 1.45
NG = 0
DB = 0
SM = 0
Delta_K
Doxorubicin
YJR072C NPA3
ZShift = 0.62
Z lm Score = 0.62
NG = 0
DB = 0
SM = 0
Delta_r
Doxorubicin
YJR072C NPA3
ZShift = 1.47
Z lm Score = −0.58
NG = 0
DB = 0
SM = 0
Delta_AUC
Doxorubicin
YCR025C
ZShift = 0.05
Z lm Score = −6.22
NG = 0
DB = 0
SM = 0
Delta_L
Doxorubicin
YCR025C
ZShift = −0.63
Z lm Score = 1.98
NG = 0
DB = 0
SM = 0
Delta_K
Doxorubicin
YCR025C
ZShift = 1.1
Z lm Score = 1.35
NG = 0
DB = 0
SM = 0
Delta_r
Doxorubicin
YCR025C
ZShift = −0.4
Z lm Score = 0.18
NG = 0
DB = 0
SM = 0
Delta_AUC
Doxorubicin
YER007C−A TMA20
ZShift = −1.21
Z lm Score = −6.24
NG = 0
DB = 0
SM = 0
Delta_L
Doxorubicin
YER007C−A TMA20
ZShift = 0.1
Z lm Score = 1.06
NG = 0
DB = 0
SM = 0
Delta_K
Doxorubicin
YER007C−A TMA20
ZShift = −0.29
Z lm Score = 1.46
NG = 0
DB = 0
SM = 0
Delta_r
Doxorubicin
YER007C−A TMA20
ZShift = 1.09
Z lm Score = 0.58
NG = 0
DB = 0
SM = 0
Delta_AUC
Doxorubicin

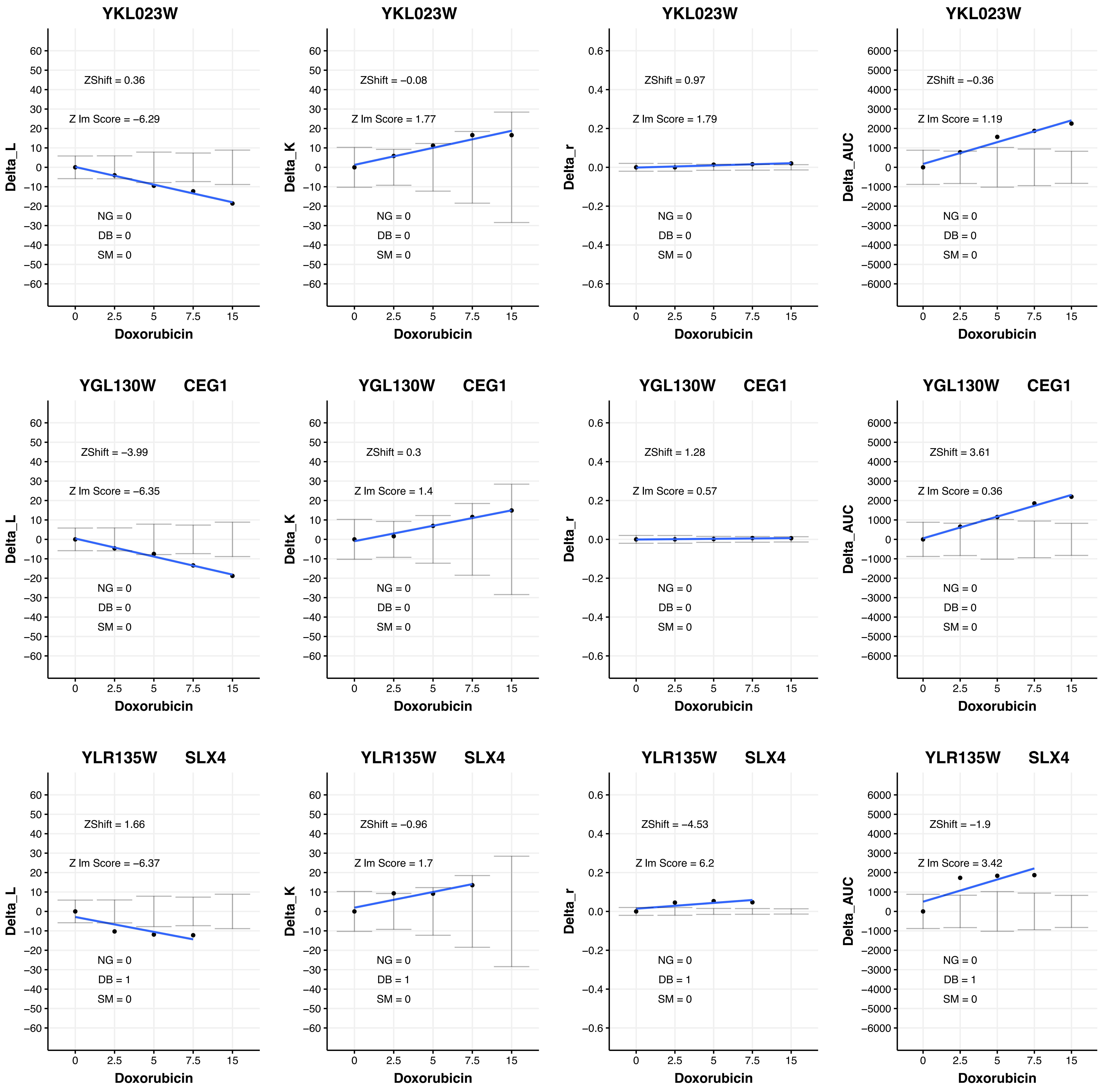
YKL023W
ZShift = 0.36
Z lm Score = −6.29
NG = 0
DB = 0
SM = 0
Delta_L
Doxorubicin
YKL023W
ZShift = −0.08
Z lm Score = 1.77
NG = 0
DB = 0
SM = 0
Delta_K
Doxorubicin
YKL023W
ZShift = 0.97
Z lm Score = 1.79
NG = 0
DB = 0
SM = 0
Delta_r
Doxorubicin
YKL023W
ZShift = −0.36
Z lm Score = 1.19
NG = 0
DB = 0
SM = 0
Delta_AUC
Doxorubicin
YGL130W CEG1
ZShift = −3.99
Z lm Score = −6.35
NG = 0
DB = 0
SM = 0
Delta_L
Doxorubicin
YGL130W CEG1
ZShift = 0.3
Z lm Score = 1.4
NG = 0
DB = 0
SM = 0
Delta_K
Doxorubicin
YGL130W CEG1
ZShift = 1.28
Z lm Score = 0.57
NG = 0
DB = 0
SM = 0
Delta_r
Doxorubicin
YGL130W CEG1
ZShift = 3.61
Z lm Score = 0.36
NG = 0
DB = 0
SM = 0
Delta_AUC
Doxorubicin
YLR135W SLX4
ZShift = 1.66
Z lm Score = −6.37
NG = 0
DB = 1
SM = 0
Delta_L
Doxorubicin
YLR135W SLX4
ZShift = −0.96
Z lm Score = 1.7
NG = 0
DB = 1
SM = 0
Delta_K
Doxorubicin
YLR135W SLX4
ZShift = −4.53
Z lm Score = 6.2
NG = 0
DB = 1
SM = 0
Delta_r
Doxorubicin
YLR135W SLX4
ZShift = −1.9
Z lm Score = 3.42
NG = 0
DB = 1
SM = 0
Delta_AUC
Doxorubicin

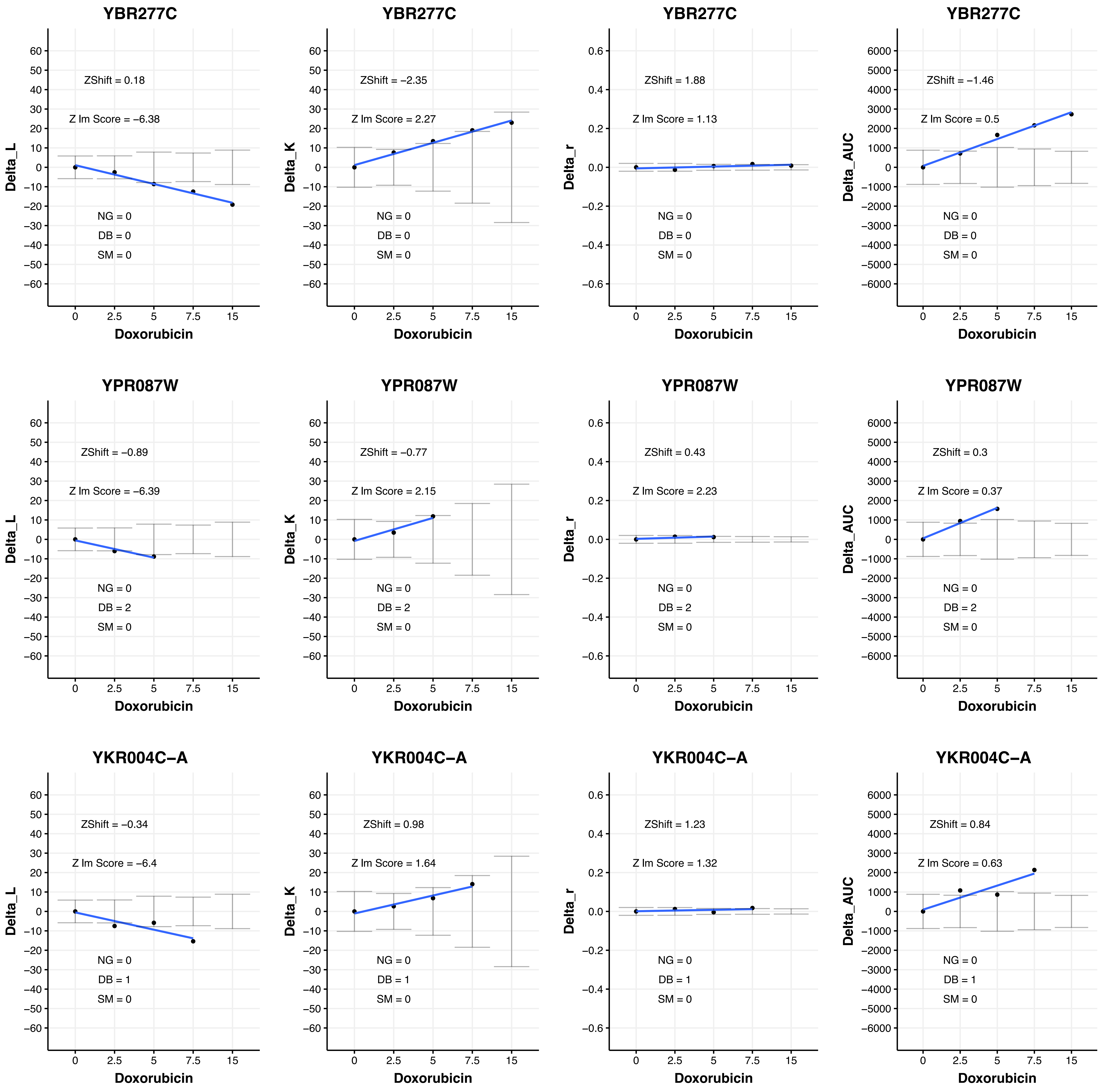
YBR277C
ZShift = 0.18
Z lm Score = −6.38
NG = 0
DB = 0
SM = 0
Delta_L
Doxorubicin
YBR277C
ZShift = −2.35
Z lm Score = 2.27
NG = 0
DB = 0
SM = 0
Delta_K
Doxorubicin
YBR277C
ZShift = 1.88
Z lm Score = 1.13
NG = 0
DB = 0
SM = 0
Delta_r
Doxorubicin
YBR277C
ZShift = −1.46
Z lm Score = 0.5
NG = 0
DB = 0
SM = 0
Delta_AUC
Doxorubicin
YPR087W
ZShift = −0.89
Z lm Score = −6.39
NG = 0
DB = 2
SM = 0
Delta_L
Doxorubicin
YPR087W
ZShift = −0.77
Z lm Score = 2.15
NG = 0
DB = 2
SM = 0
Delta_K
Doxorubicin
YPR087W
ZShift = 0.43
Z lm Score = 2.23
NG = 0
DB = 2
SM = 0
Delta_r
Doxorubicin
YPR087W
ZShift = 0.3
Z lm Score = 0.37
NG = 0
DB = 2
SM = 0
Delta_AUC
Doxorubicin
YKR004C−A
ZShift = −0.34
Z lm Score = −6.4
NG = 0
DB = 1
SM = 0
Delta_L
Doxorubicin
YKR004C−A
ZShift = 0.98
Z lm Score = 1.64
NG = 0
DB = 1
SM = 0
Delta_K
Doxorubicin
YKR004C−A
ZShift = 1.23
Z lm Score = 1.32
NG = 0
DB = 1
SM = 0
Delta_r
Doxorubicin
YKR004C−A
ZShift = 0.84
Z lm Score = 0.63
NG = 0
DB = 1
SM = 0
Delta_AUC
Doxorubicin

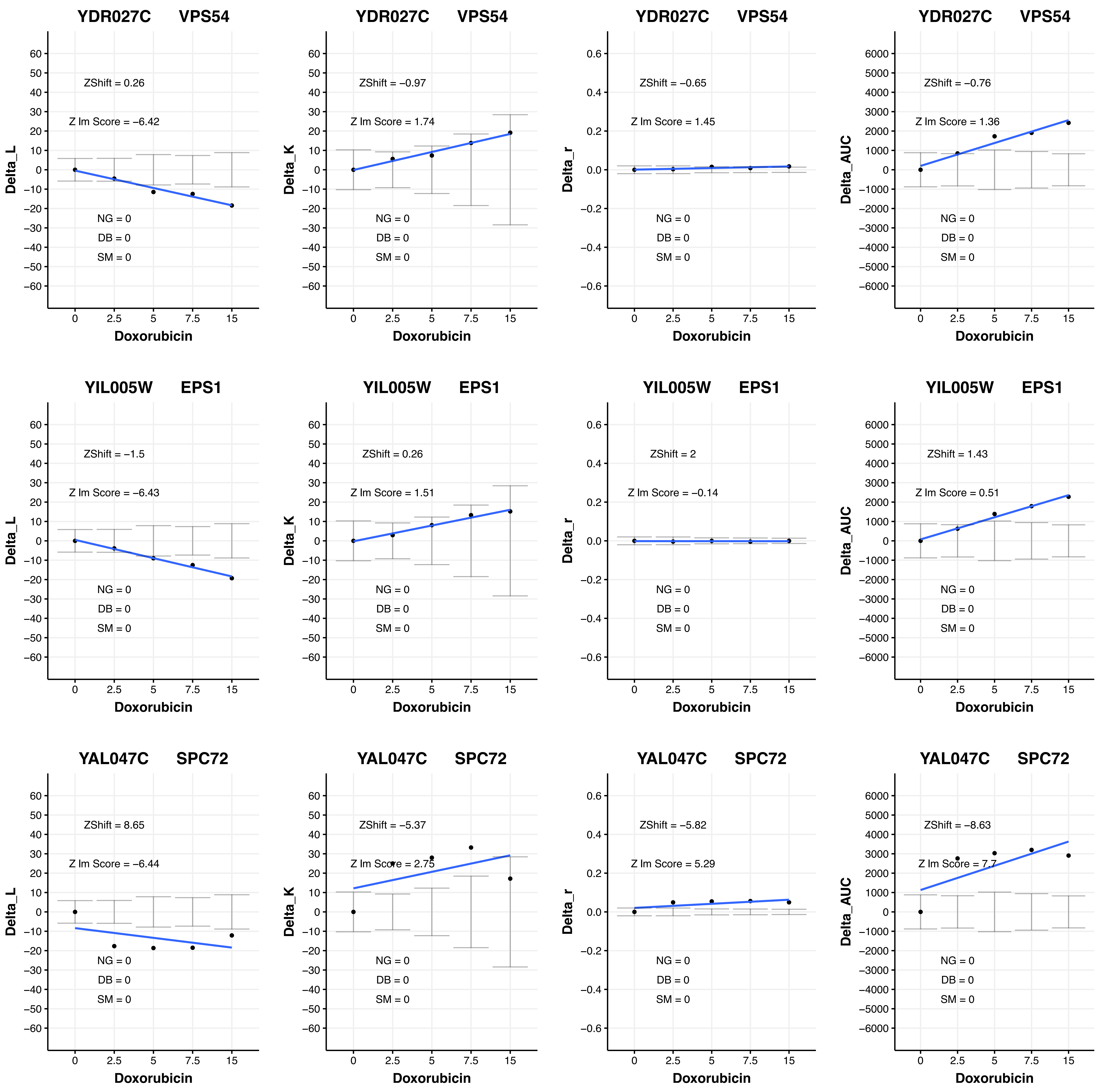
YDR027C VPS54
ZShift = 0.26
Z lm Score = −6.42
NG = 0
DB = 0
SM = 0
Delta_L
Doxorubicin
YDR027C VPS54
ZShift = −0.97
Z lm Score = 1.74
NG = 0
DB = 0
SM = 0
Delta_K
Doxorubicin
YDR027C VPS54
ZShift = −0.65
Z lm Score = 1.45
NG = 0
DB = 0
SM = 0
Delta_r
Doxorubicin
YDR027C VPS54
ZShift = −0.76
Z lm Score = 1.36
NG = 0
DB = 0
SM = 0
Delta_AUC
Doxorubicin
YIL005W EPS1
ZShift = −1.5
Z lm Score = −6.43
NG = 0
DB = 0
SM = 0
Delta_L
Doxorubicin
YIL005W EPS1
ZShift = 0.26
Z lm Score = 1.51
NG = 0
DB = 0
SM = 0
Delta_K
Doxorubicin
YIL005W EPS1
ZShift = 2
Z lm Score = −0.14
NG = 0
DB = 0
SM = 0
Delta_r
Doxorubicin
YIL005W EPS1
ZShift = 1.43
Z lm Score = 0.51
NG = 0
DB = 0
SM = 0
Delta_AUC
Doxorubicin
YAL047C SPC72
ZShift = 8.65
Z lm Score = −6.44
NG = 0
DB = 0
SM = 0
Delta_L
Doxorubicin
YAL047C SPC72
ZShift = −5.37
Z lm Score = 2.75
NG = 0
DB = 0
SM = 0
Delta_K
Doxorubicin
YAL047C SPC72
ZShift = −5.82
Z lm Score = 5.29
NG = 0
DB = 0
SM = 0
Delta_r
Doxorubicin
YAL047C SPC72
ZShift = −8.63
Z lm Score = 7.7
NG = 0
DB = 0
SM = 0
Delta_AUC
Doxorubicin

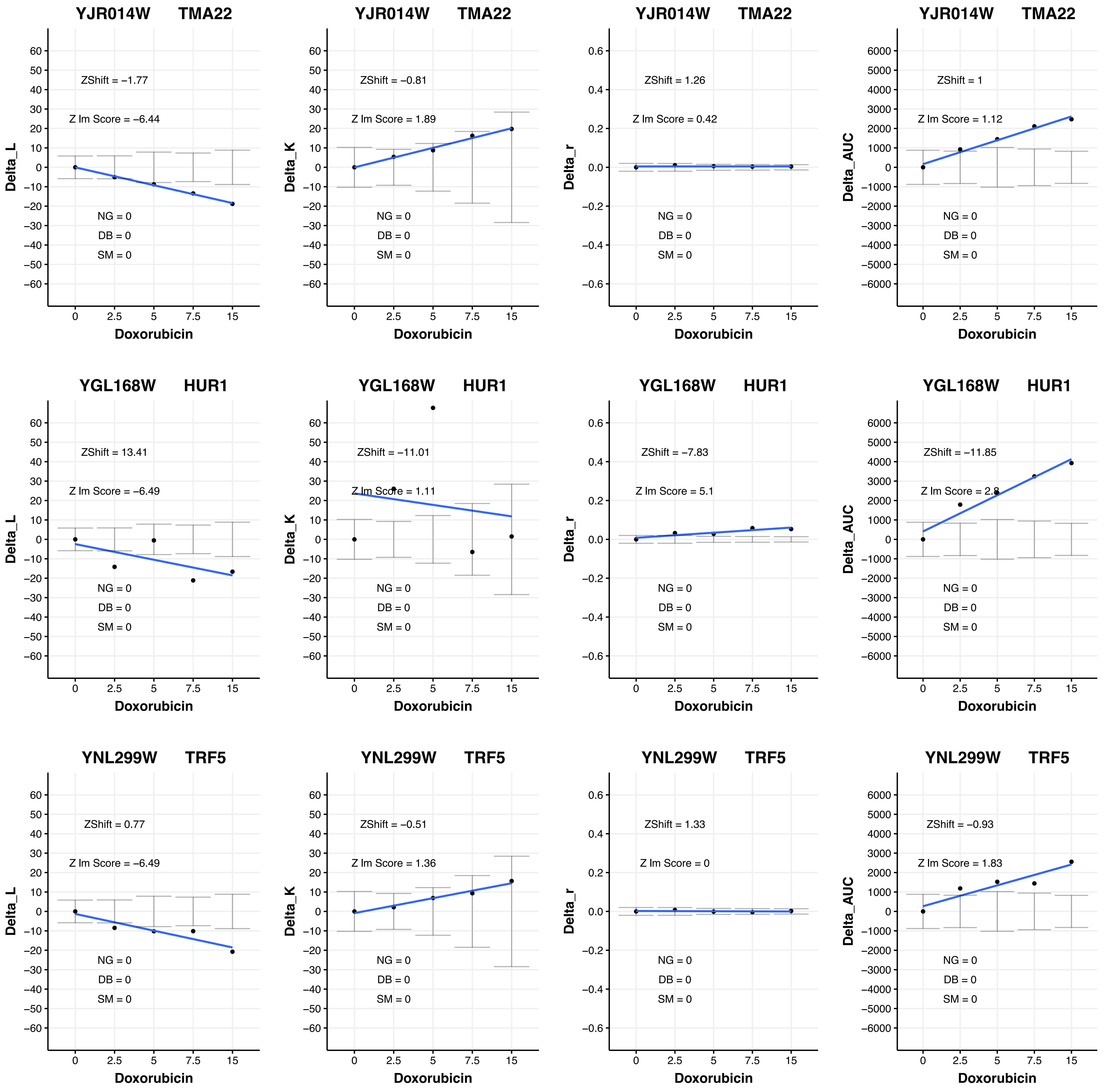

YJR014W TMA22
ZShift = −1.77
Z lm Score = −6.44
NG = 0
DB = 0
SM = 0
Delta_L
Doxorubicin
YJR014W TMA22
ZShift = −0.81
Z lm Score = 1.89
NG = 0
DB = 0
SM = 0
Delta_K
Doxorubicin
YJR014W TMA22
ZShift = 1.26
Z lm Score = 0.42
NG = 0
DB = 0
SM = 0
Delta_r
Doxorubicin
YJR014W TMA22
ZShift = 1
Z lm Score = 1.12
NG = 0
DB = 0
SM = 0
Delta_AUC
Doxorubicin
YGL168W HUR1
ZShift = 13.41
Z lm Score = −6.49
NG = 0
DB = 0
SM = 0
Delta_L
Doxorubicin
YGL168W HUR1
ZShift = −11.01
Z lm Score = 1.11
NG = 0
DB = 0
SM = 0
Delta_K
Doxorubicin
YGL168W HUR1
ZShift = −7.83
Z lm Score = 5.1
NG = 0
DB = 0
SM = 0
Delta_r
Doxorubicin
YGL168W HUR1
ZShift = −11.85
Z lm Score = 2.8
NG = 0
DB = 0
SM = 0
Delta_AUC
Doxorubicin
YNL299W TRF5
ZShift = 0.77
Z lm Score = −6.49
NG = 0
DB = 0
SM = 0
Delta_L
Doxorubicin
YNL299W TRF5
ZShift = −0.51
Z lm Score = 1.36
NG = 0
DB = 0
SM = 0
Delta_K
Doxorubicin
YNL299W TRF5
ZShift = 1.33
Z lm Score = 0
NG = 0
DB = 0
SM = 0
Delta_r
Doxorubicin
YNL299W TRF5
ZShift = −0.93
Z lm Score = 1.83
NG = 0
DB = 0
SM = 0
Delta_AUC
Doxorubicin

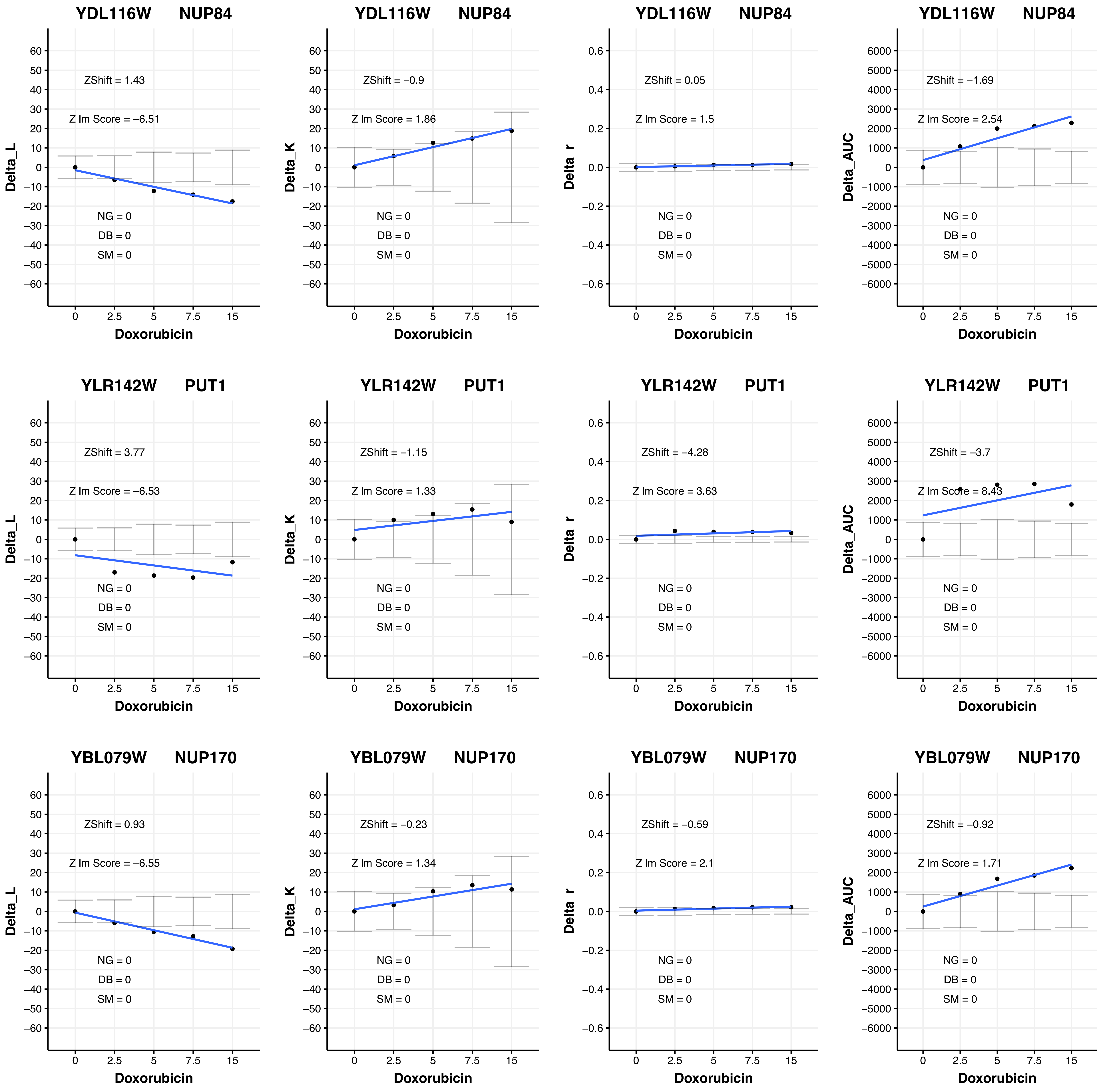
YDL116W NUP84
Delta_L
ZShift = 1.43
Z lm Score = −6.51
NG = 0
DB = 0
SM = 0
Doxorubicin
YDL116W NUP84
Delta_K
ZShift = −0.9
Z lm Score = 1.86
NG = 0
DB = 0
SM = 0
Doxorubicin
YDL116W NUP84
Delta_r
ZShift = 0.05
Z lm Score = 1.5
NG = 0
DB = 0
SM = 0
Doxorubicin
YDL116W NUP84
Delta_AUC
ZShift = −1.69
Z lm Score = 2.54
NG = 0
DB = 0
SM = 0
Doxorubicin
YLR142W PUT1
Delta_L
ZShift = 3.77
Z lm Score = −6.53
NG = 0
DB = 0
SM = 0
Doxorubicin
YLR142W PUT1
Delta_K
ZShift = −1.15
Z lm Score = 1.33
NG = 0
DB = 0
SM = 0
Doxorubicin
YLR142W PUT1
Delta_r
ZShift = −4.28
Z lm Score = 3.63
NG = 0
DB = 0
SM = 0
Doxorubicin
YLR142W PUT1
Delta_AUC
ZShift = −3.7
Z lm Score = 8.43
NG = 0
DB = 0
SM = 0
Doxorubicin
YBL079W NUP170
Delta_L
ZShift = 0.93
Z lm Score = −6.55
NG = 0
DB = 0
SM = 0
Doxorubicin
YBL079W NUP170
Delta_K
ZShift = −0.23
Z lm Score = 1.34
NG = 0
DB = 0
SM = 0
Doxorubicin
YBL079W NUP170
Delta_r
ZShift = −0.59
Z lm Score = 2.1
NG = 0
DB = 0
SM = 0
Doxorubicin
YBL079W NUP170
Delta_AUC
ZShift = −0.92
Z lm Score = 1.71
NG = 0
DB = 0
SM = 0
Doxorubicin

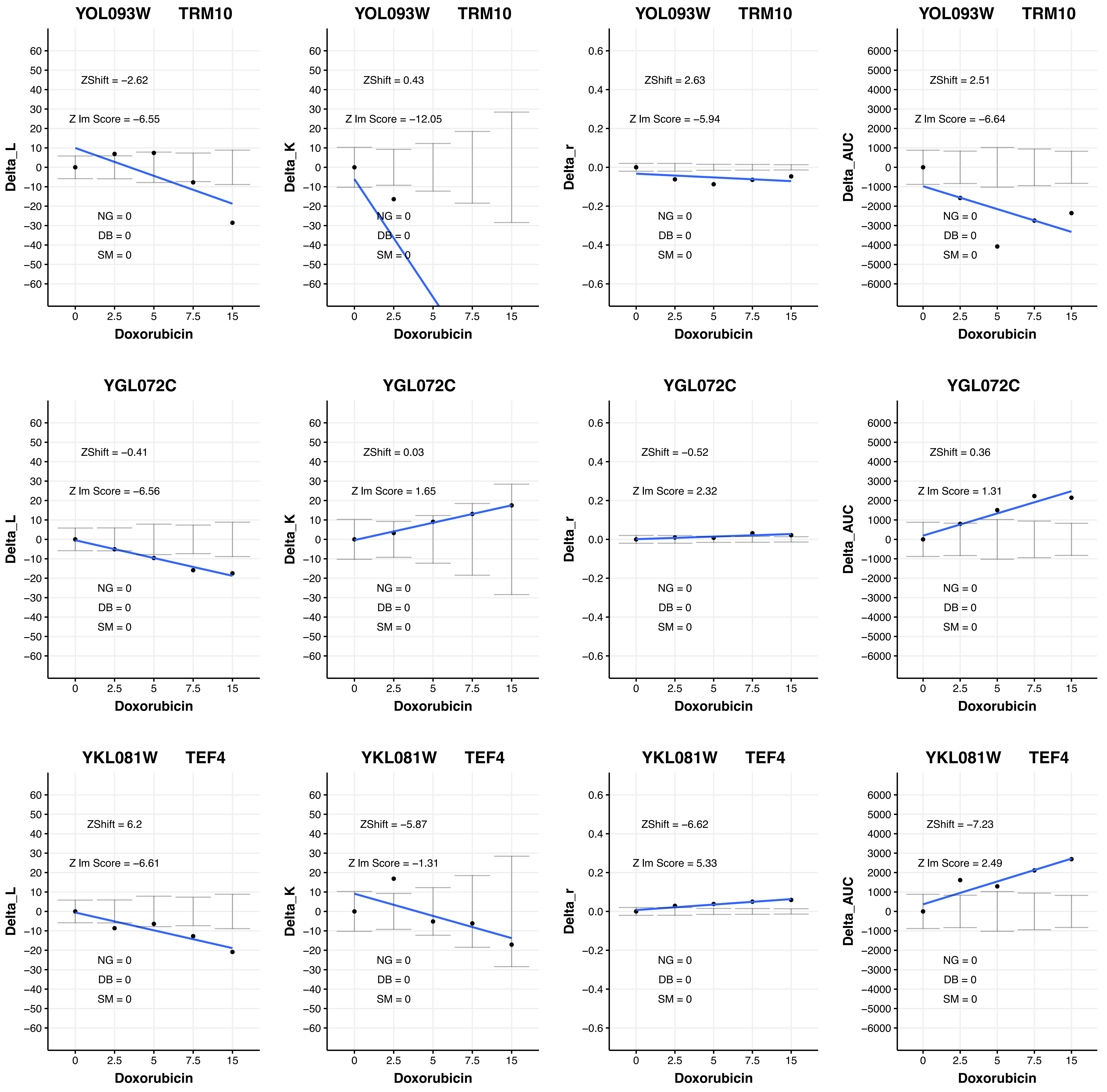

YOL093W TRM10
ZShift = −2.62
Z lm Score = −6.55
ZShift = 0.43
Z lm Score = −12.05
ZShift = 2.63
Z lm Score = −5.94
ZShift = 2.51
Z lm Score = −6.64
YGL072C
ZShift = −0.41
Z lm Score = −6.56
ZShift = 0.03
Z lm Score = 1.65
ZShift = −0.52
Z lm Score = 2.32
ZShift = 0.36
Z lm Score = 1.31
YKL081W TEF4
ZShift = 6.2
Z lm Score = −6.61
ZShift = −5.87
Z lm Score = −1.31
ZShift = −6.62
Z lm Score = 5.33
ZShift = −7.23
Z lm Score = 2.49
NG = 0
DB = 0
SM = 0
Delta_L
Delta_K
Delta_r
Delta_AUC
Doxorubicin

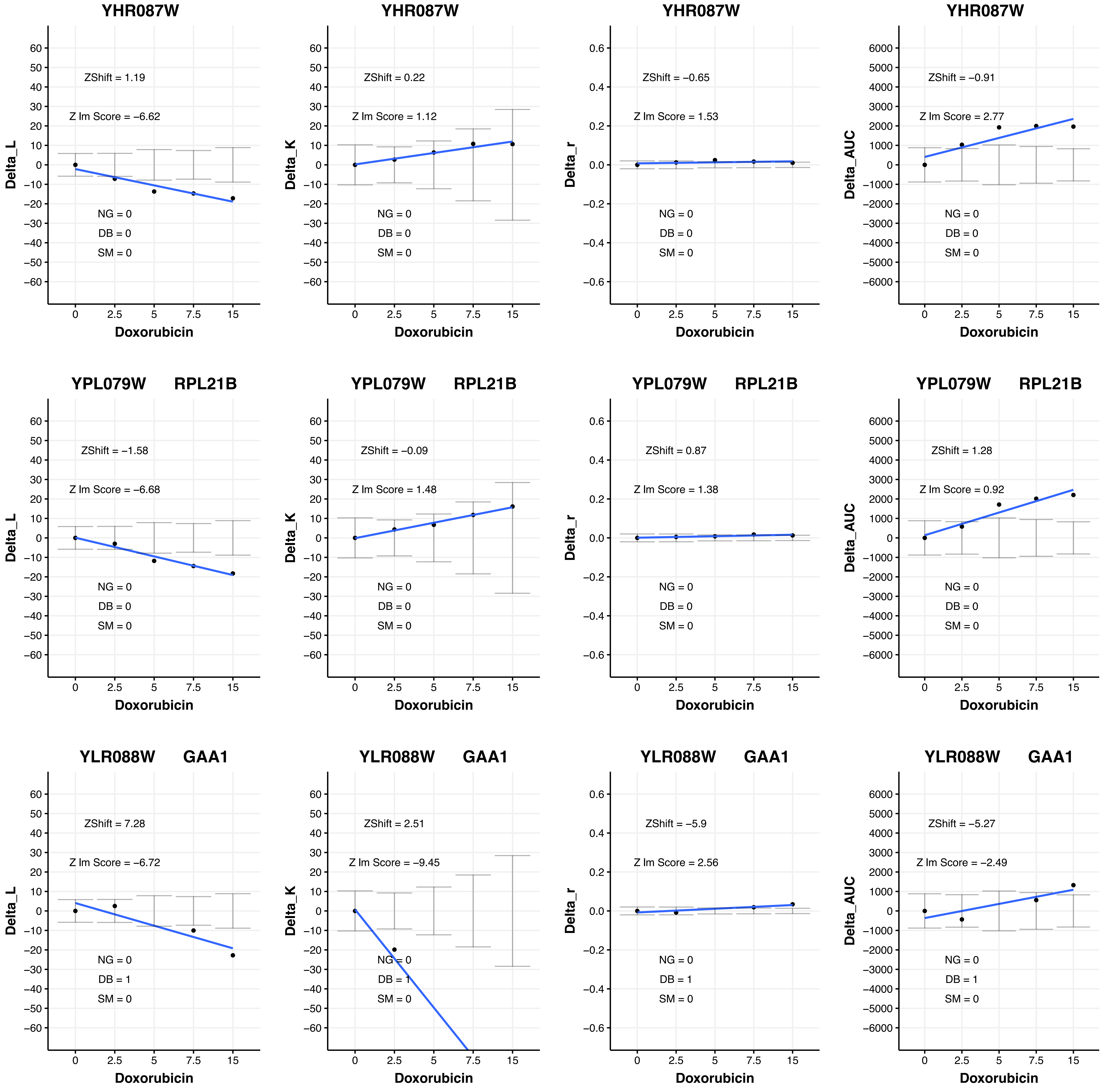
YHR087W
ZShift = 1.19
Z lm Score = −6.62
NG = 0
DB = 0
SM = 0
Delta_L
Doxorubicin
YHR087W
ZShift = 0.22
Z lm Score = 1.12
NG = 0
DB = 0
SM = 0
Delta_K
Doxorubicin
YHR087W
ZShift = −0.65
Z lm Score = 1.53
NG = 0
DB = 0
SM = 0
Delta_r
Doxorubicin
YHR087W
ZShift = −0.91
Z lm Score = 2.77
NG = 0
DB = 0
SM = 0
Delta_AUC
Doxorubicin
YPL079W RPL21B
ZShift = −1.58
Z lm Score = −6.68
NG = 0
DB = 0
SM = 0
Delta_L
Doxorubicin
YPL079W RPL21B
ZShift = −0.09
Z lm Score = 1.48
NG = 0
DB = 0
SM = 0
Delta_K
Doxorubicin
YPL079W RPL21B
ZShift = 0.87
Z lm Score = 1.38
NG = 0
DB = 0
SM = 0
Delta_r
Doxorubicin
YPL079W RPL21B
ZShift = 1.28
Z lm Score = 0.92
NG = 0
DB = 0
SM = 0
Delta_AUC
Doxorubicin
YLR088W GAA1
ZShift = 7.28
Z lm Score = −6.72
NG = 0
DB = 1
SM = 0
Delta_L
Doxorubicin
YLR088W GAA1
ZShift = 2.51
Z lm Score = −9.45
NG = 0
DB = 1
SM = 0
Delta_K
Doxorubicin
YLR088W GAA1
ZShift = −5.9
Z lm Score = 2.56
NG = 0
DB = 1
SM = 0
Delta_r
Doxorubicin
YLR088W GAA1
ZShift = −5.27
Z lm Score = −2.49
NG = 0
DB = 1
SM = 0
Delta_AUC
Doxorubicin

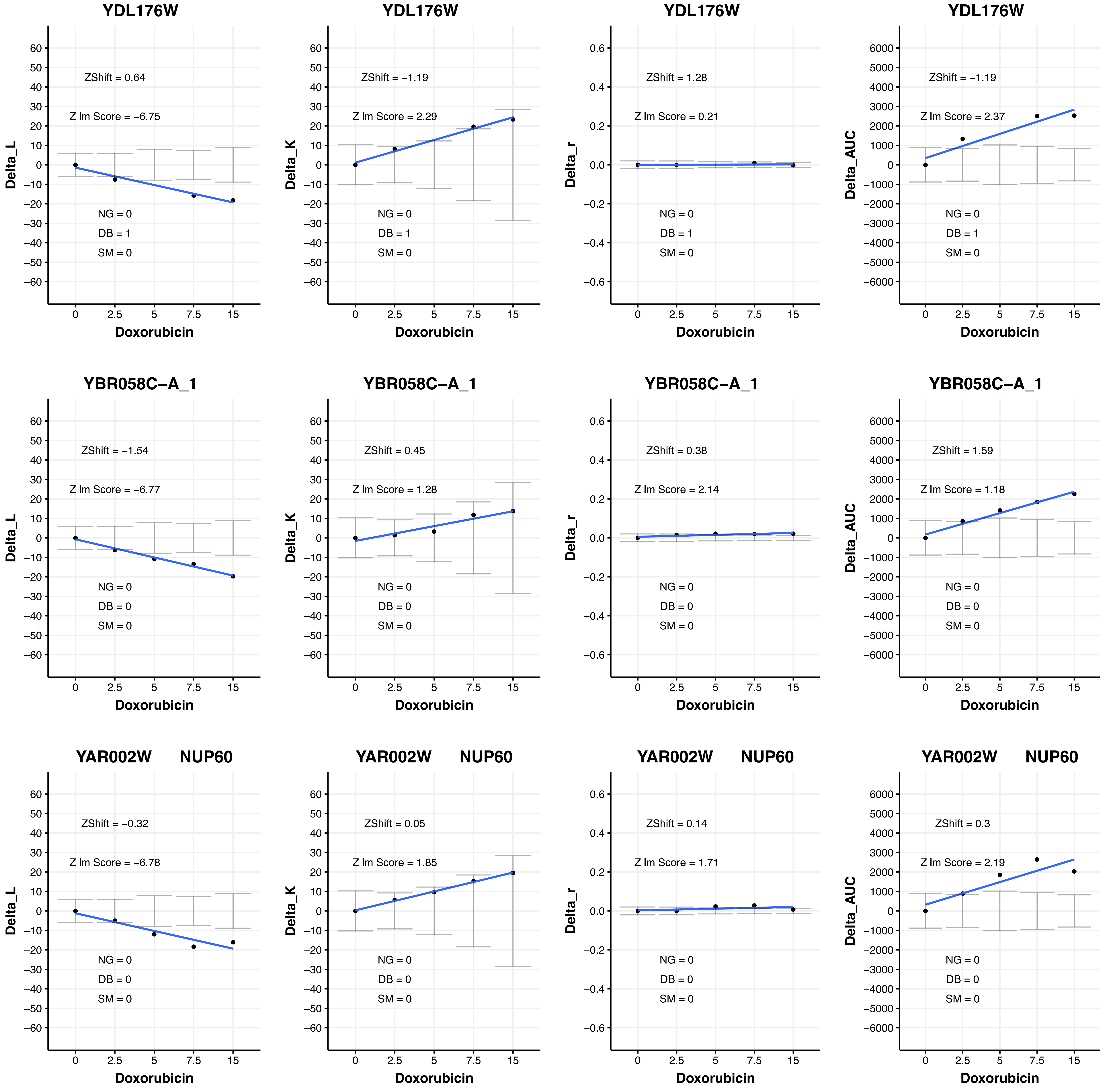
YDL176W
ZShift = 0.64
Z lm Score = −6.75
NG = 0
DB = 1
SM = 0
Delta_L
Doxorubicin
YDL176W
ZShift = −1.19
Z lm Score = 2.29
NG = 0
DB = 1
SM = 0
Delta_K
Doxorubicin
YDL176W
ZShift = 1.28
Z lm Score = 0.21
NG = 0
DB = 1
SM = 0
Delta_r
Doxorubicin
YDL176W
ZShift = −1.19
Z lm Score = 2.37
NG = 0
DB = 1
SM = 0
Delta_AUC
Doxorubicin
YBR058C−A_1
ZShift = −1.54
Z lm Score = −6.77
NG = 0
DB = 0
SM = 0
Delta_L
Doxorubicin
YBR058C−A_1
ZShift = 0.45
Z lm Score = 1.28
NG = 0
DB = 0
SM = 0
Delta_K
Doxorubicin
YBR058C−A_1
ZShift = 0.38
Z lm Score = 2.14
NG = 0
DB = 0
SM = 0
Delta_r
Doxorubicin
YBR058C−A_1
ZShift = 1.59
Z lm Score = 1.18
NG = 0
DB = 0
SM = 0
Delta_AUC
Doxorubicin
YAR002W NUP60
ZShift = −0.32
Z lm Score = −6.78
NG = 0
DB = 0
SM = 0
Delta_L
Doxorubicin
YAR002W NUP60
ZShift = 0.05
Z lm Score = 1.85
NG = 0
DB = 0
SM = 0
Delta_K
Doxorubicin
YAR002W NUP60
ZShift = 0.14
Z lm Score = 1.71
NG = 0
DB = 0
SM = 0
Delta_r
Doxorubicin
YAR002W NUP60
ZShift = 0.3
Z lm Score = 2.19
NG = 0
DB = 0
SM = 0
Delta_AUC
Doxorubicin

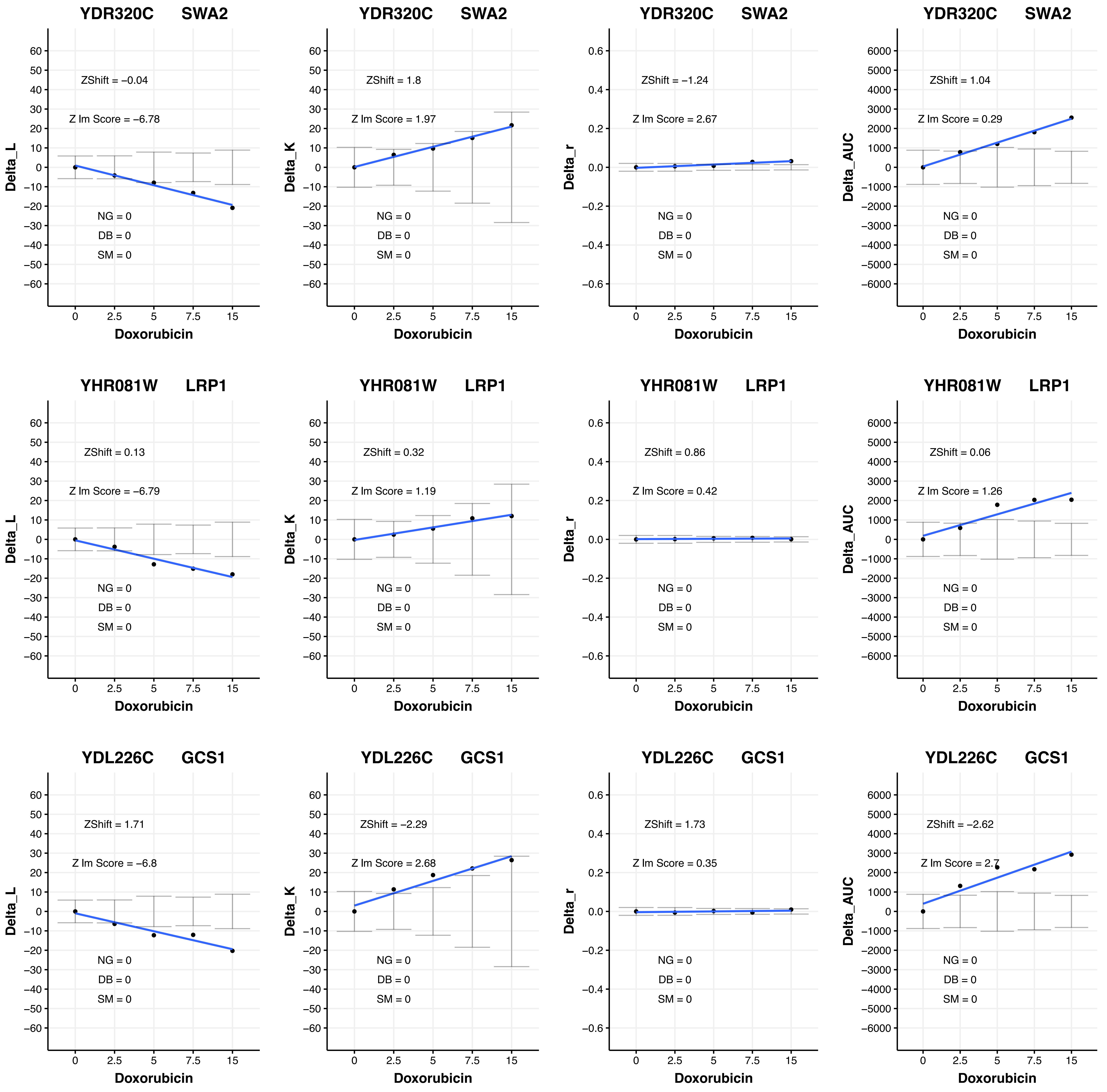
YDR320C SWA2
ZShift = −0.04
Z lm Score = −6.78
NG = 0
DB = 0
SM = 0
Delta_L
Doxorubicin
YDR320C SWA2
ZShift = 1.8
Z lm Score = 1.97
NG = 0
DB = 0
SM = 0
Delta_K
Doxorubicin
YDR320C SWA2
ZShift = −1.24
Z lm Score = 2.67
NG = 0
DB = 0
SM = 0
Delta_r
Doxorubicin
YDR320C SWA2
ZShift = 1.04
Z lm Score = 0.29
NG = 0
DB = 0
SM = 0
Delta_AUC
Doxorubicin
YHR081W LRP1
ZShift = 0.13
Z lm Score = −6.79
NG = 0
DB = 0
SM = 0
Delta_L
Doxorubicin
YHR081W LRP1
ZShift = 0.32
Z lm Score = 1.19
NG = 0
DB = 0
SM = 0
Delta_K
Doxorubicin
YHR081W LRP1
ZShift = 0.86
Z lm Score = 0.42
NG = 0
DB = 0
SM = 0
Delta_r
Doxorubicin
YHR081W LRP1
ZShift = 0.06
Z lm Score = 1.26
NG = 0
DB = 0
SM = 0
Delta_AUC
Doxorubicin
YDL226C GCS1
ZShift = 1.71
Z lm Score = −6.8
NG = 0
DB = 0
SM = 0
Delta_L
Doxorubicin
YDL226C GCS1
ZShift = −2.29
Z lm Score = 2.68
NG = 0
DB = 0
SM = 0
Delta_K
Doxorubicin
YDL226C GCS1
ZShift = 1.73
Z lm Score = 0.35
NG = 0
DB = 0
SM = 0
Delta_r
Doxorubicin
YDL226C GCS1
ZShift = −2.62
Z lm Score = 2.7
NG = 0
DB = 0
SM = 0
Delta_AUC
Doxorubicin

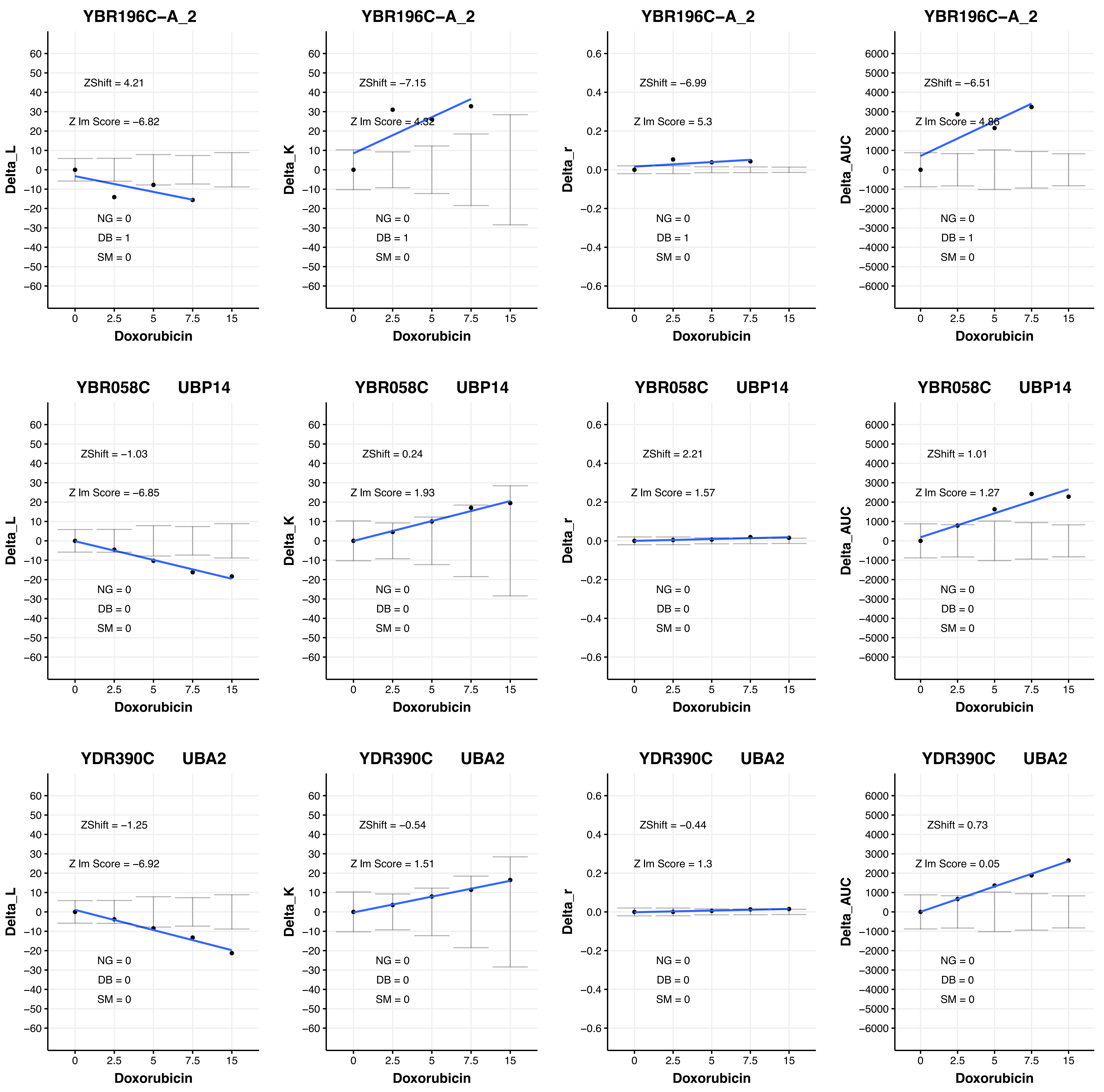
YBR196C−A_2
ZShift = 4.21
Z lm Score = −6.82
NG = 0
DB = 1
SM = 0
Delta_L
Doxorubicin
YBR196C−A_2
ZShift = −7.15
Z lm Score = 4.32
NG = 0
DB = 1
SM = 0
Delta_K
Doxorubicin
YBR196C−A_2
ZShift = −6.99
Z lm Score = 5.3
NG = 0
DB = 1
SM = 0
Delta_r
Doxorubicin
YBR196C−A_2
ZShift = −6.51
Z lm Score = 4.86
NG = 0
DB = 1
SM = 0
Delta_AUC
Doxorubicin
YBR058C UBP14
ZShift = −1.03
Z lm Score = −6.85
NG = 0
DB = 0
SM = 0
Delta_L
Doxorubicin
YBR058C UBP14
ZShift = 0.24
Z lm Score = 1.93
NG = 0
DB = 0
SM = 0
Delta_K
Doxorubicin
YBR058C UBP14
ZShift = 2.21
Z lm Score = 1.57
NG = 0
DB = 0
SM = 0
Delta_r
Doxorubicin
YBR058C UBP14
ZShift = 1.01
Z lm Score = 1.27
NG = 0
DB = 0
SM = 0
Delta_AUC
Doxorubicin
YDR390C UBA2
ZShift = −1.25
Z lm Score = −6.92
NG = 0
DB = 0
SM = 0
Delta_L
Doxorubicin
YDR390C UBA2
ZShift = −0.54
Z lm Score = 1.51
NG = 0
DB = 0
SM = 0
Delta_K
Doxorubicin
YDR390C UBA2
ZShift = −0.44
Z lm Score = 1.3
NG = 0
DB = 0
SM = 0
Delta_r
Doxorubicin
YDR390C UBA2
ZShift = 0.73
Z lm Score = 0.05
NG = 0
DB = 0
SM = 0
Delta_AUC
Doxorubicin

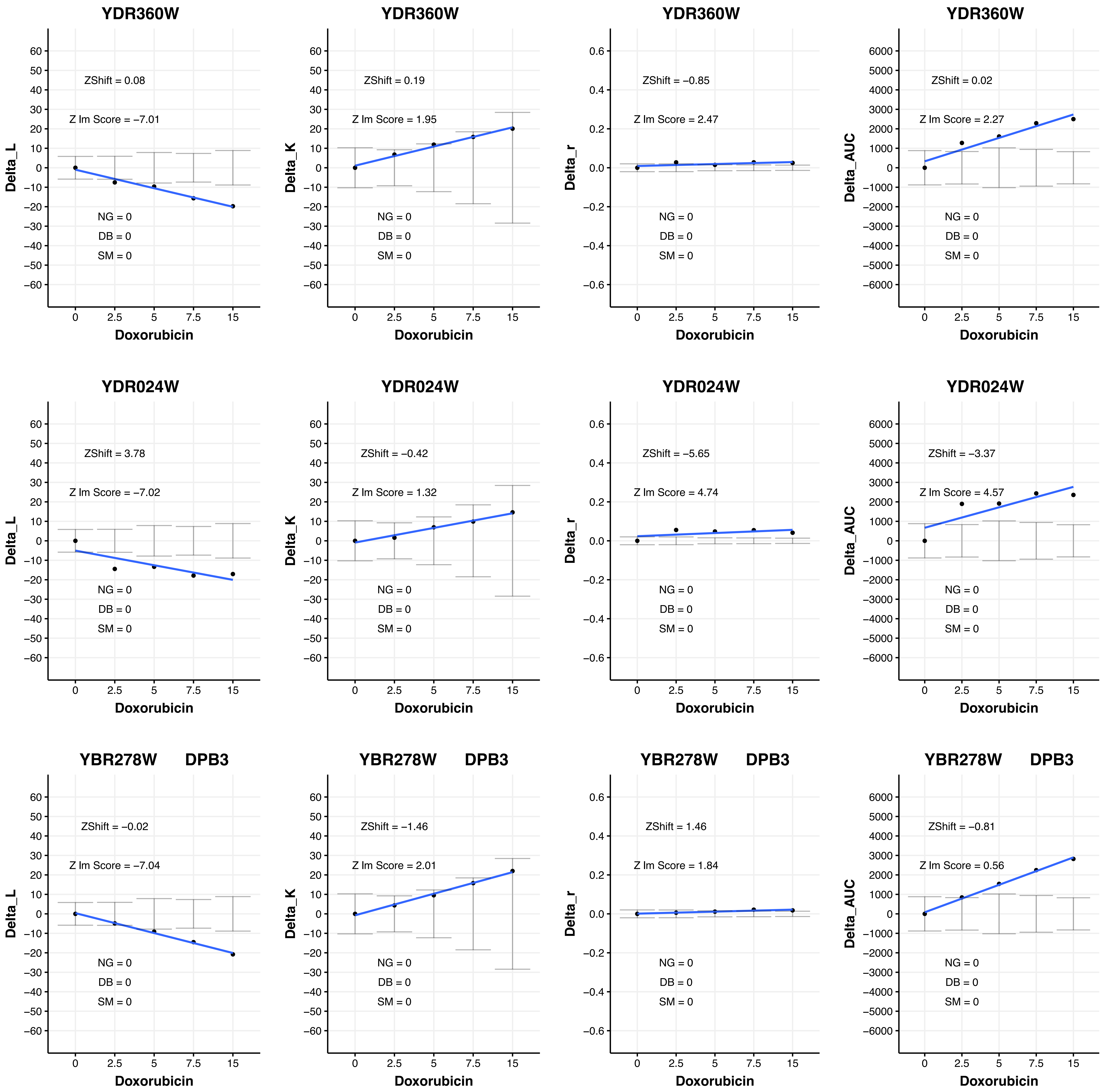
YDR360W
ZShift = 0.08
Z lm Score = −7.01
NG = 0
DB = 0
SM = 0
Delta_L
Doxorubicin
YDR360W
ZShift = 0.19
Z lm Score = 1.95
NG = 0
DB = 0
SM = 0
Delta_K
Doxorubicin
YDR360W
ZShift = −0.85
Z lm Score = 2.47
NG = 0
DB = 0
SM = 0
Delta_r
Doxorubicin
YDR360W
ZShift = 0.02
Z lm Score = 2.27
NG = 0
DB = 0
SM = 0
Delta_AUC
Doxorubicin
YDR024W
ZShift = 3.78
Z lm Score = −7.02
NG = 0
DB = 0
SM = 0
Delta_L
Doxorubicin
YDR024W
ZShift = −0.42
Z lm Score = 1.32
NG = 0
DB = 0
SM = 0
Delta_K
Doxorubicin
YDR024W
ZShift = −5.65
Z lm Score = 4.74
NG = 0
DB = 0
SM = 0
Delta_r
Doxorubicin
YDR024W
ZShift = −3.37
Z lm Score = 4.57
NG = 0
DB = 0
SM = 0
Delta_AUC
Doxorubicin
YBR278W DPB3
ZShift = −0.02
Z lm Score = −7.04
NG = 0
DB = 0
SM = 0
Delta_L
Doxorubicin
YBR278W DPB3
ZShift = −1.46
Z lm Score = 2.01
NG = 0
DB = 0
SM = 0
Delta_K
Doxorubicin
YBR278W DPB3
ZShift = 1.46
Z lm Score = 1.84
NG = 0
DB = 0
SM = 0
Delta_r
Doxorubicin
YBR278W DPB3
ZShift = −0.81
Z lm Score = 0.56
NG = 0
DB = 0
SM = 0
Delta_AUC
Doxorubicin

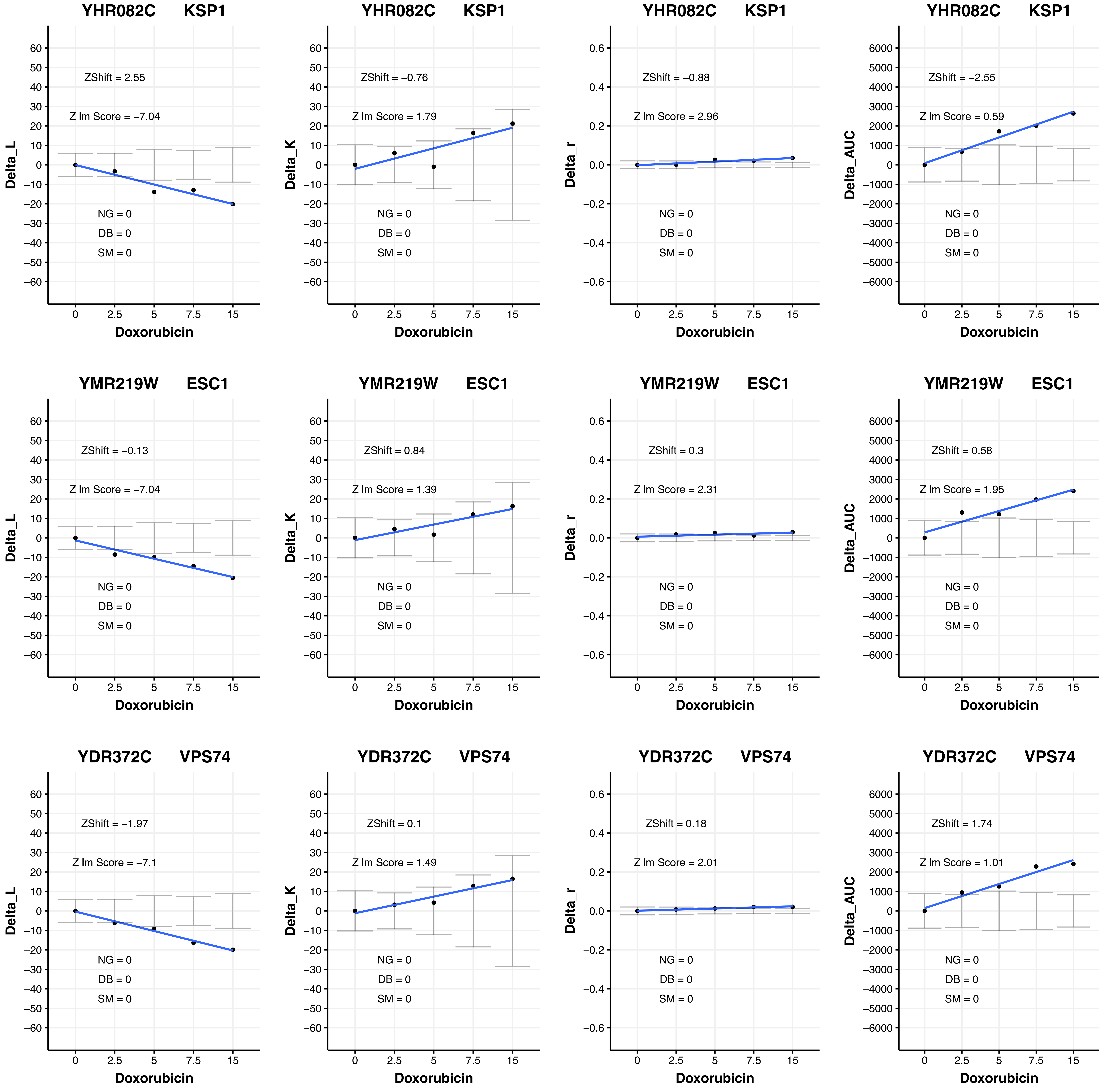

YHR082C KSP1
ZShift = 2.55
Z lm Score = −7.04
NG = 0
DB = 0
SM = 0
Delta_L
Doxorubicin
YHR082C KSP1
ZShift = −0.76
Z lm Score = 1.79
NG = 0
DB = 0
SM = 0
Delta_K
Doxorubicin
YHR082C KSP1
ZShift = −0.88
Z lm Score = 2.96
NG = 0
DB = 0
SM = 0
Delta_r
Doxorubicin
YHR082C KSP1
ZShift = −2.55
Z lm Score = 0.59
NG = 0
DB = 0
SM = 0
Delta_AUC
Doxorubicin
YMR219W ESC1
ZShift = −0.13
Z lm Score = −7.04
NG = 0
DB = 0
SM = 0
Delta_L
Doxorubicin
YMR219W ESC1
ZShift = 0.84
Z lm Score = 1.39
NG = 0
DB = 0
SM = 0
Delta_K
Doxorubicin
YMR219W ESC1
ZShift = 0.3
Z lm Score = 2.31
NG = 0
DB = 0
SM = 0
Delta_r
Doxorubicin
YMR219W ESC1
ZShift = 0.58
Z lm Score = 1.95
NG = 0
DB = 0
SM = 0
Delta_AUC
Doxorubicin
YDR372C VPS74
ZShift = −1.97
Z lm Score = −7.1
NG = 0
DB = 0
SM = 0
Delta_L
Doxorubicin
YDR372C VPS74
ZShift = 0.1
Z lm Score = 1.49
NG = 0
DB = 0
SM = 0
Delta_K
Doxorubicin
YDR372C VPS74
ZShift = 0.18
Z lm Score = 2.01
NG = 0
DB = 0
SM = 0
Delta_r
Doxorubicin
YDR372C VPS74
ZShift = 1.74
Z lm Score = 1.01
NG = 0
DB = 0
SM = 0
Delta_AUC
Doxorubicin

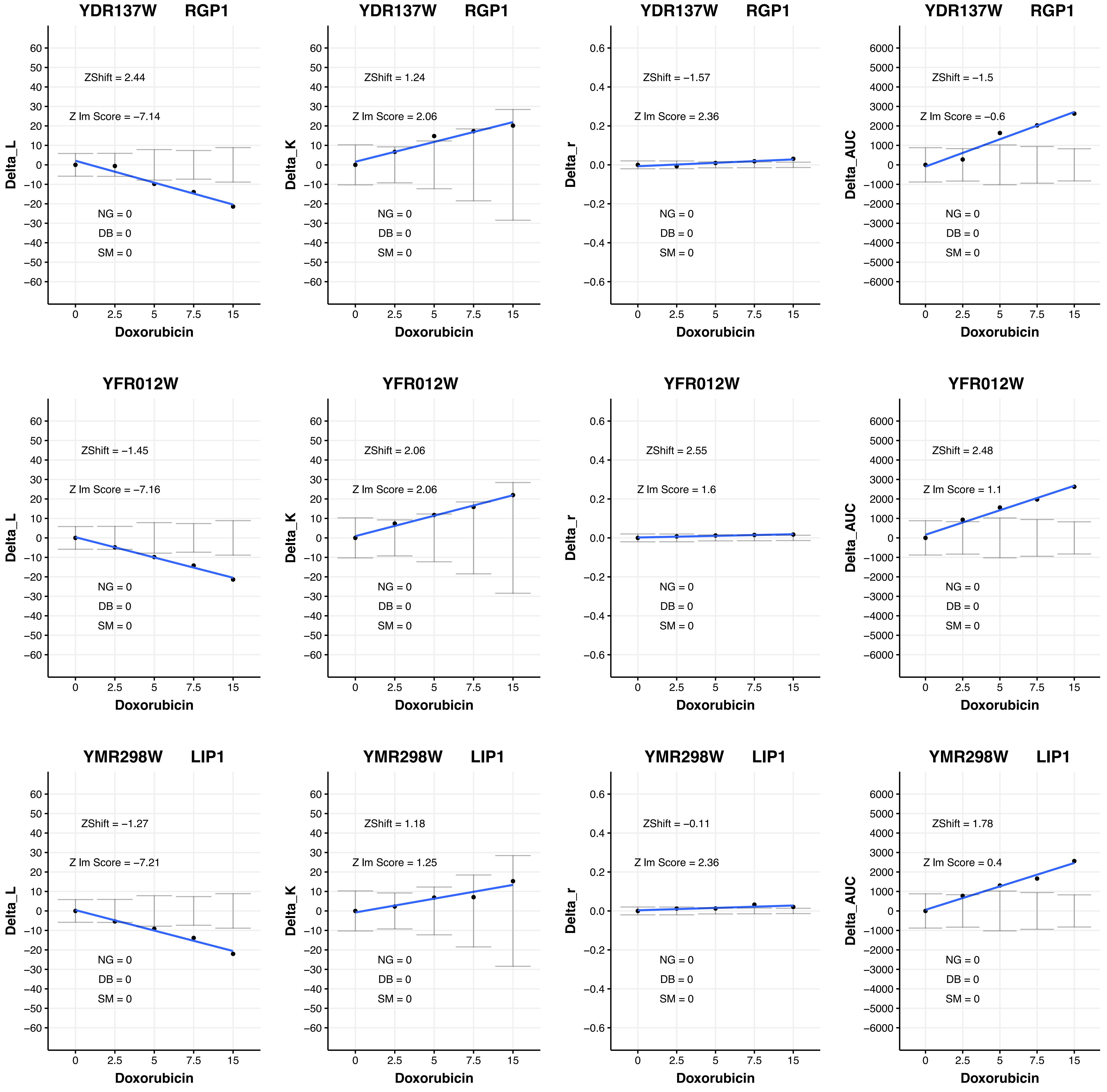
YDR137W RGP1
ZShift = 2.44
Z lm Score = −7.14
NG = 0
DB = 0
SM = 0
Delta_L
Doxorubicin
YDR137W RGP1
ZShift = 1.24
Z lm Score = 2.06
NG = 0
DB = 0
SM = 0
Delta_K
Doxorubicin
YDR137W RGP1
ZShift = −1.57
Z lm Score = 2.36
NG = 0
DB = 0
SM = 0
Delta_r
Doxorubicin
YDR137W RGP1
ZShift = −1.5
Z lm Score = −0.6
NG = 0
DB = 0
SM = 0
Delta_AUC
Doxorubicin
YFR012W
ZShift = −1.45
Z lm Score = −7.16
NG = 0
DB = 0
SM = 0
Delta_L
Doxorubicin
YFR012W
ZShift = 2.06
Z lm Score = 2.06
NG = 0
DB = 0
SM = 0
Delta_K
Doxorubicin
YFR012W
ZShift = 2.55
Z lm Score = 1.6
NG = 0
DB = 0
SM = 0
Delta_r
Doxorubicin
YFR012W
ZShift = 2.48
Z lm Score = 1.1
NG = 0
DB = 0
SM = 0
Delta_AUC
Doxorubicin
YMR298W LIP1
ZShift = −1.27
Z lm Score = −7.21
NG = 0
DB = 0
SM = 0
Delta_L
Doxorubicin
YMR298W LIP1
ZShift = 1.18
Z lm Score = 1.25
NG = 0
DB = 0
SM = 0
Delta_K
Doxorubicin
YMR298W LIP1
ZShift = −0.11
Z lm Score = 2.36
NG = 0
DB = 0
SM = 0
Delta_r
Doxorubicin
YMR298W LIP1
ZShift = 1.78
Z lm Score = 0.4
NG = 0
DB = 0
SM = 0
Delta_AUC
Doxorubicin

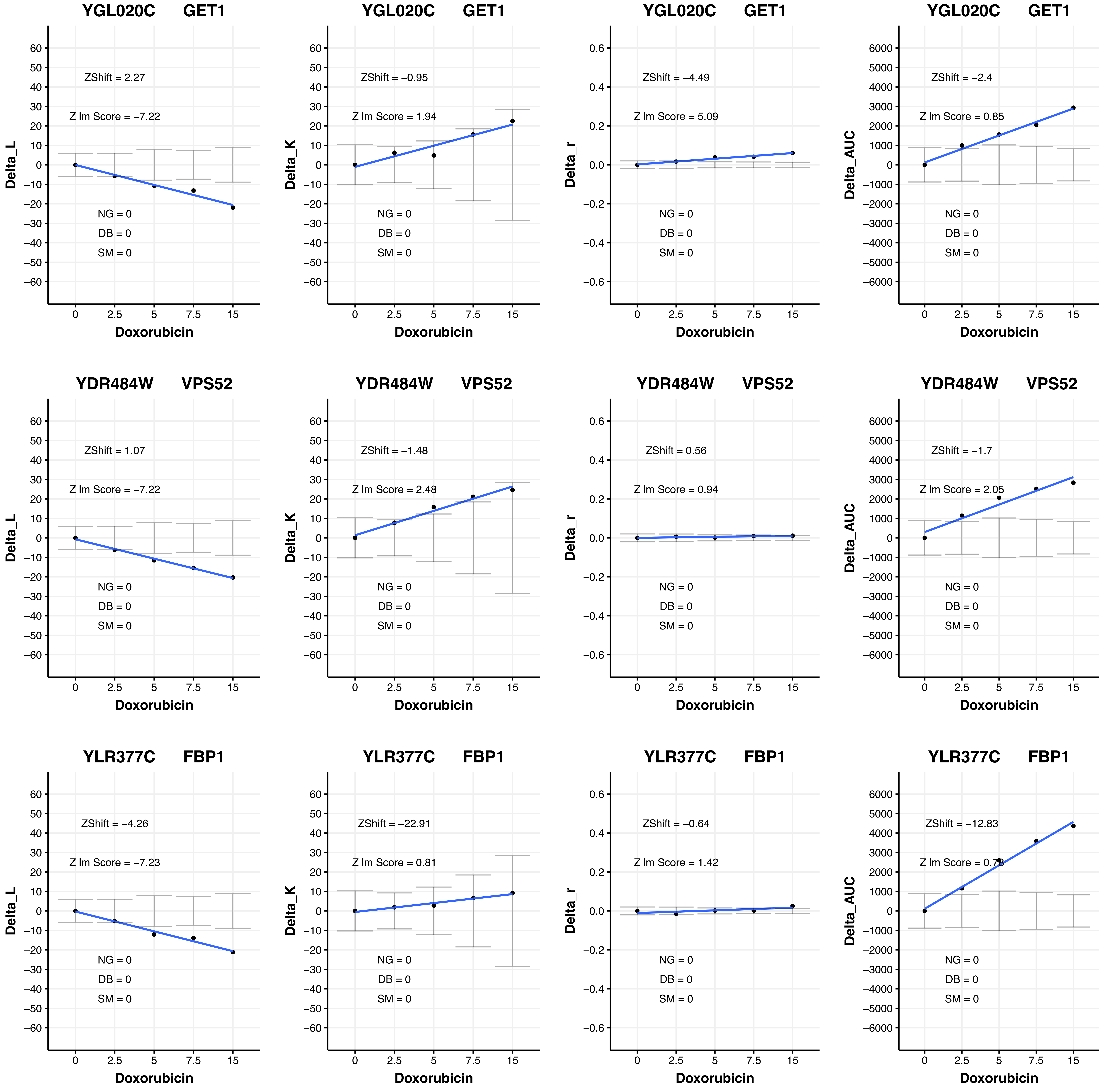

YGL020C GET1
ZShift = 2.27
Z Im Score = −7.22
NG = 0
DB = 0
SM = 0
Delta_L
Doxorubicin
YGL020C GET1
ZShift = −0.95
Z Im Score = 1.94
NG = 0
DB = 0
SM = 0
Delta_K
Doxorubicin
YGL020C GET1
ZShift = −4.49
Z Im Score = 5.09
NG = 0
DB = 0
SM = 0
Delta_r
Doxorubicin
YGL020C GET1
ZShift = −2.4
Z Im Score = 0.85
NG = 0
DB = 0
SM = 0
Delta_AUC
Doxorubicin
YDR484W VPS52
ZShift = 1.07
Z Im Score = −7.22
NG = 0
DB = 0
SM = 0
Delta_L
Doxorubicin
YDR484W VPS52
ZShift = −1.48
Z Im Score = 2.48
NG = 0
DB = 0
SM = 0
Delta_K
Doxorubicin
YDR484W VPS52
ZShift = 0.56
Z Im Score = 0.94
NG = 0
DB = 0
SM = 0
Delta_r
Doxorubicin
YDR484W VPS52
ZShift = −1.7
Z Im Score = 2.05
NG = 0
DB = 0
SM = 0
Delta_AUC
Doxorubicin
YLR377C FBP1
ZShift = −4.26
Z Im Score = −7.23
NG = 0
DB = 0
SM = 0
Delta_L
Doxorubicin
YLR377C FBP1
ZShift = −22.91
Z Im Score = 0.81
NG = 0
DB = 0
SM = 0
Delta_K
Doxorubicin
YLR377C FBP1
ZShift = −0.64
Z Im Score = 1.42
NG = 0
DB = 0
SM = 0
Delta_r
Doxorubicin
YLR377C FBP1
ZShift = −12.83
NG = 0
DB = 0
SM = 0
Delta_AUC
Doxorubicin

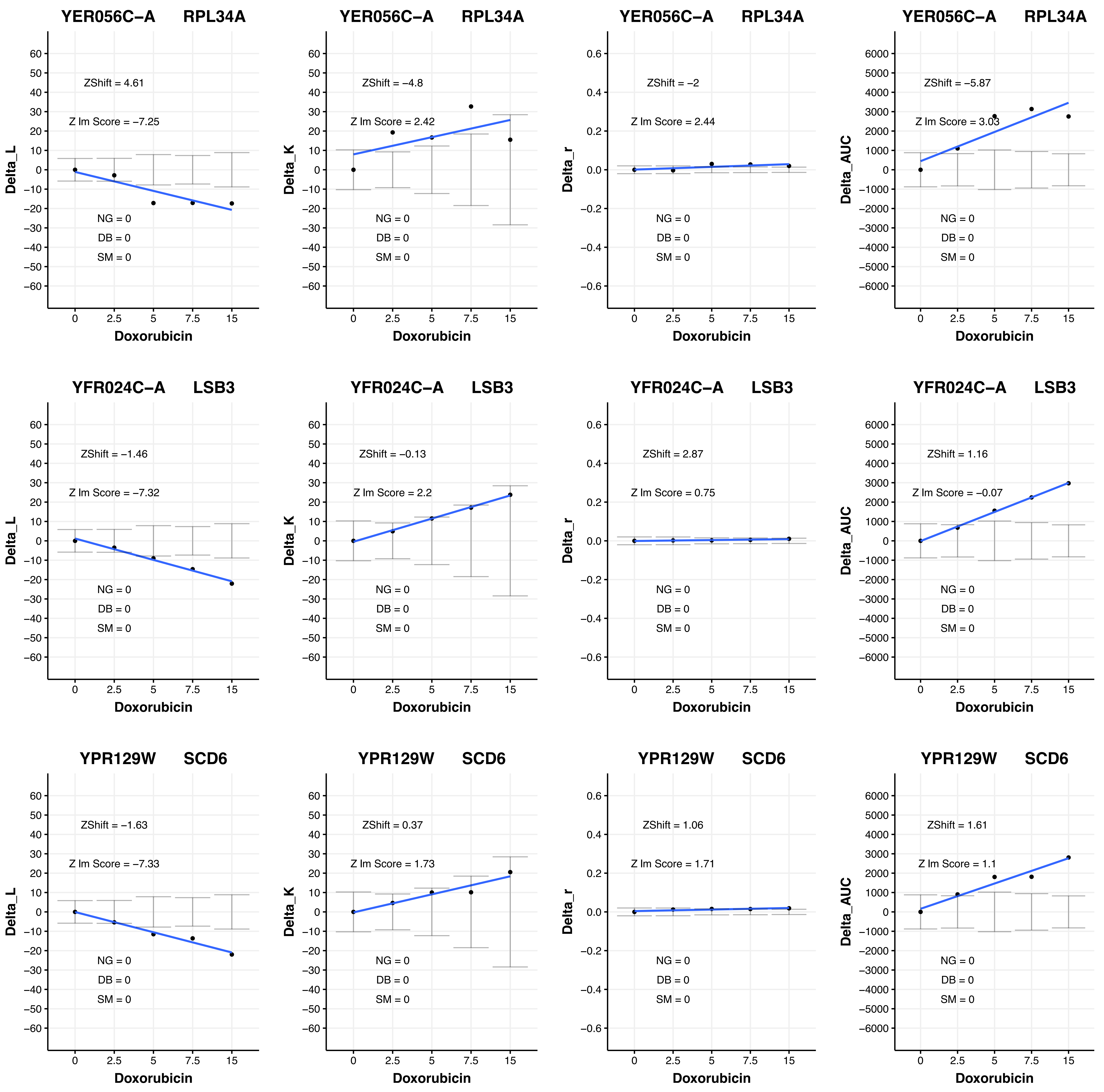

YER056C−A RPL34A
ZShift = 4.61
Z lm Score = −7.25
NG = 0
DB = 0
SM = 0
Delta_L
Doxorubicin
YER056C−A RPL34A
ZShift = −4.8
Z lm Score = 2.42
NG = 0
DB = 0
SM = 0
Delta_K
Doxorubicin
YER056C−A RPL34A
ZShift = −2
Z lm Score = 2.44
NG = 0
DB = 0
SM = 0
Delta_r
Doxorubicin
YER056C−A RPL34A
ZShift = −5.87
Z lm Score = 3.03
NG = 0
DB = 0
SM = 0
Delta_AUC
Doxorubicin
YFR024C−A LSB3
ZShift = −1.46
Z lm Score = −7.32
NG = 0
DB = 0
SM = 0
Delta_L
Doxorubicin
YFR024C−A LSB3
ZShift = −0.13
Z lm Score = 2.2
NG = 0
DB = 0
SM = 0
Delta_K
Doxorubicin
YFR024C−A LSB3
ZShift = 2.87
Z lm Score = 0.75
NG = 0
DB = 0
SM = 0
Delta_r
Doxorubicin
YFR024C−A LSB3
ZShift = 1.16
Z lm Score = −0.07
NG = 0
DB = 0
SM = 0
Delta_AUC
Doxorubicin
YPR129W SCD6
ZShift = −1.63
Z lm Score = −7.33
NG = 0
DB = 0
SM = 0
Delta_L
Doxorubicin
YPR129W SCD6
ZShift = 0.37
Z lm Score = 1.73
NG = 0
DB = 0
SM = 0
Delta_K
Doxorubicin
YPR129W SCD6
ZShift = 1.06
Z lm Score = 1.71
NG = 0
DB = 0
SM = 0
Delta_r
Doxorubicin
YPR129W SCD6
ZShift = 1.61
Z lm Score = 1.1
NG = 0
DB = 0
SM = 0
Delta_AUC
Doxorubicin

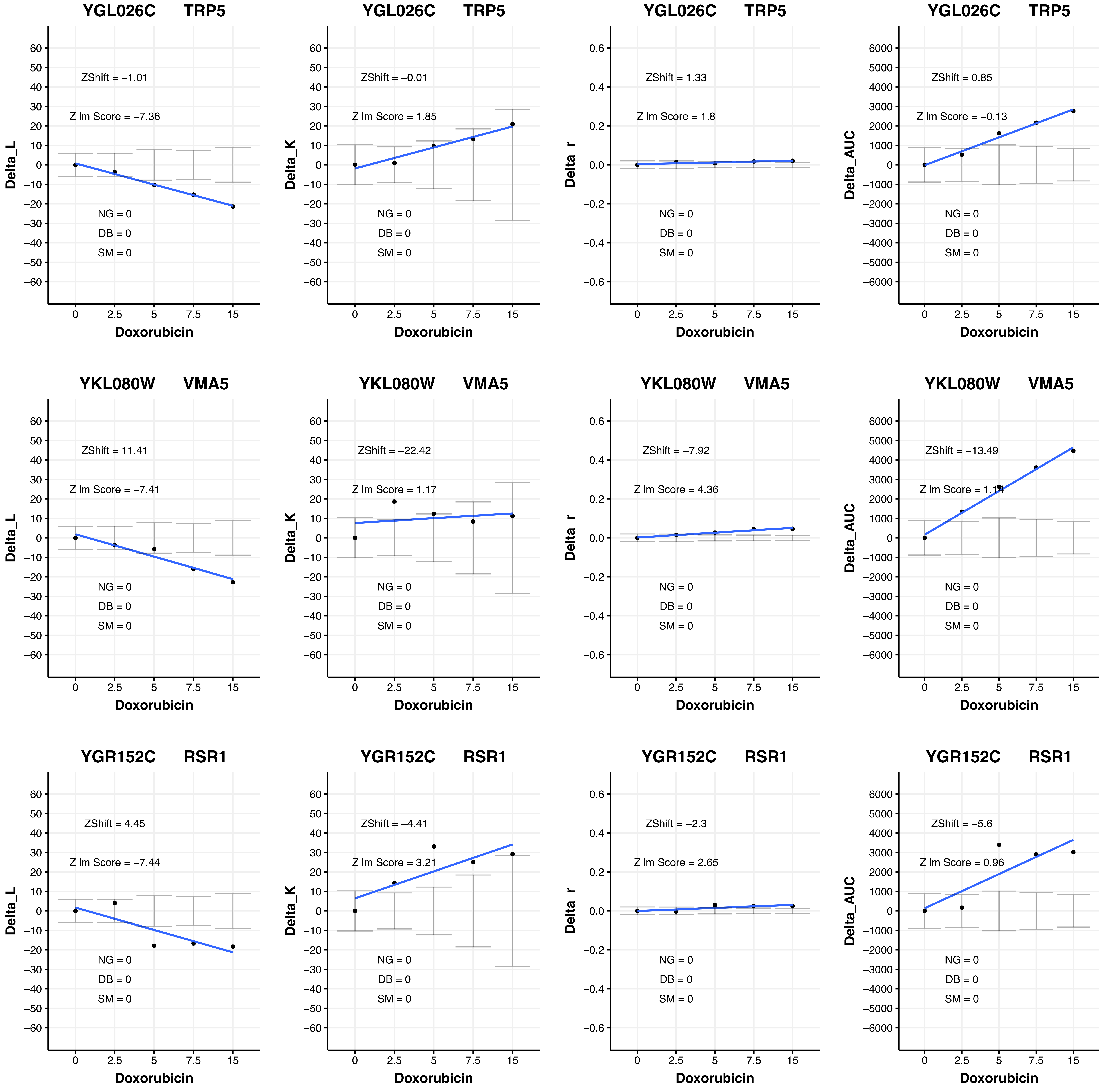
YGL026C TRP5
ZShift = −1.01
Z lm Score = −7.36
NG = 0
DB = 0
SM = 0
Delta_L
Doxorubicin
YGL026C TRP5
ZShift = −0.01
Z lm Score = 1.85
NG = 0
DB = 0
SM = 0
Delta_K
Doxorubicin
YGL026C TRP5
ZShift = 1.33
Z lm Score = 1.8
NG = 0
DB = 0
SM = 0
Delta_r
Doxorubicin
YGL026C TRP5
ZShift = 0.85
Z lm Score = −0.13
NG = 0
DB = 0
SM = 0
Delta_AUC
Doxorubicin
YKL080W VMA5
ZShift = 11.41
Z lm Score = −7.41
NG = 0
DB = 0
SM = 0
Delta_L
Doxorubicin
YKL080W VMA5
ZShift = −22.42
Z lm Score = 1.17
NG = 0
DB = 0
SM = 0
Delta_K
Doxorubicin
YKL080W VMA5
ZShift = −7.92
Z lm Score = 4.36
NG = 0
DB = 0
SM = 0
Delta_r
Doxorubicin
YKL080W VMA5
ZShift = −13.49
Z lm Score = 1.14
NG = 0
DB = 0
SM = 0
Delta_AUC
Doxorubicin
YGR152C RSR1
ZShift = 4.45
Z lm Score = −7.44
NG = 0
DB = 0
SM = 0
Delta_L
Doxorubicin
YGR152C RSR1
ZShift = −4.41
Z lm Score = 3.21
NG = 0
DB = 0
SM = 0
Delta_K
Doxorubicin
YGR152C RSR1
ZShift = −2.3
Z lm Score = 2.65
NG = 0
DB = 0
SM = 0
Delta_r
Doxorubicin
YGR152C RSR1
ZShift = −5.6
Z lm Score = 0.96
NG = 0
DB = 0
SM = 0
Delta_AUC
Doxorubicin

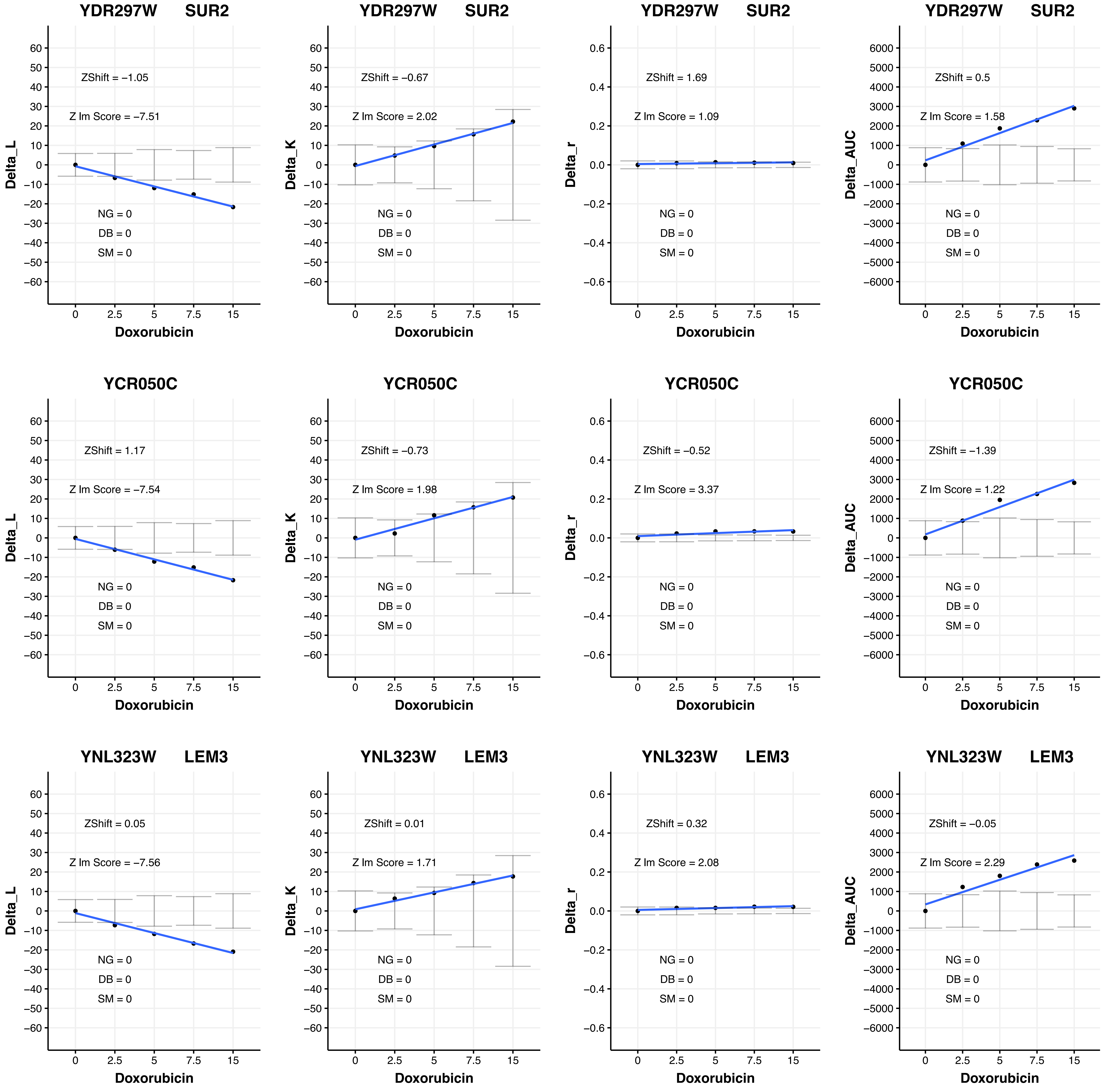
YDR297W SUR2
ZShift = −1.05
Z lm Score = −7.51
NG = 0
DB = 0
SM = 0
Delta_L
Doxorubicin
YDR297W SUR2
ZShift = −0.67
Z lm Score = 2.02
NG = 0
DB = 0
SM = 0
Delta_K
Doxorubicin
YDR297W SUR2
ZShift = 1.69
Z lm Score = 1.09
NG = 0
DB = 0
SM = 0
Delta_r
Doxorubicin
YDR297W SUR2
ZShift = 0.5
Z lm Score = 1.58
NG = 0
DB = 0
SM = 0
Delta_AUC
Doxorubicin
YCR050C
ZShift = 1.17
Z lm Score = −7.54
NG = 0
DB = 0
SM = 0
Delta_L
Doxorubicin
YCR050C
ZShift = −0.73
Z lm Score = 1.98
NG = 0
DB = 0
SM = 0
Delta_K
Doxorubicin
YCR050C
ZShift = −0.52
Z lm Score = 3.37
NG = 0
DB = 0
SM = 0
Delta_r
Doxorubicin
YCR050C
ZShift = −1.39
Z lm Score = 1.22
NG = 0
DB = 0
SM = 0
Delta_AUC
Doxorubicin
YNL323W LEM3
ZShift = 0.05
Z lm Score = −7.56
NG = 0
DB = 0
SM = 0
Delta_L
Doxorubicin
YNL323W LEM3
ZShift = 0.01
Z lm Score = 1.71
NG = 0
DB = 0
SM = 0
Delta_K
Doxorubicin
YNL323W LEM3
ZShift = 0.32
Z lm Score = 2.08
NG = 0
DB = 0
SM = 0
Delta_r
Doxorubicin
YNL323W LEM3
ZShift = −0.05
Z lm Score = 2.29
NG = 0
DB = 0
SM = 0
Delta_AUC
Doxorubicin

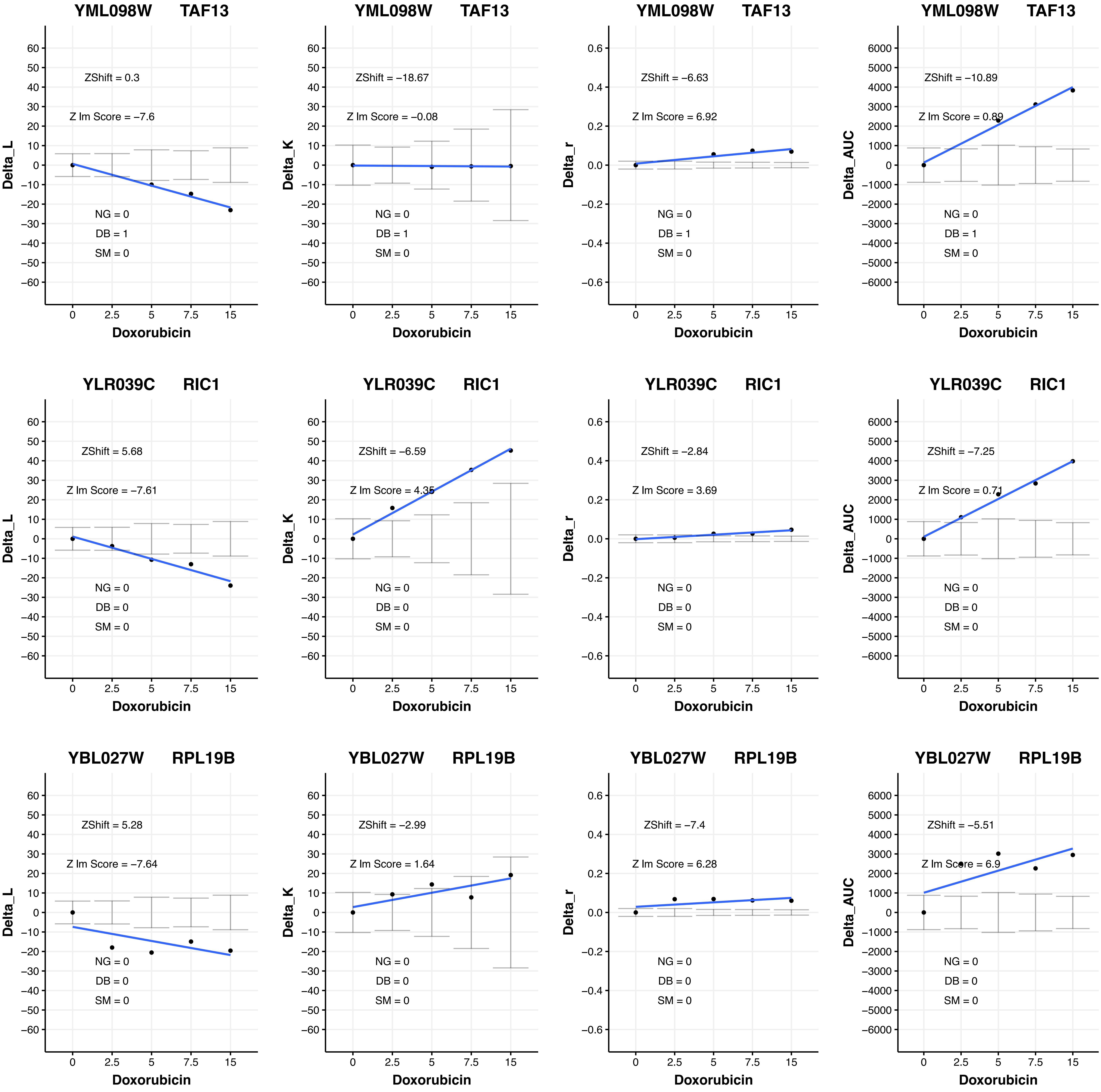
YML098W TAF13
ZShift = 0.3
Z lm Score = −7.6
NG = 0
DB = 1
SM = 0
Delta_L
Doxorubicin
YML098W TAF13
ZShift = −18.67
Z lm Score = −0.08
NG = 0
DB = 1
SM = 0
Delta_K
Doxorubicin
YML098W TAF13
ZShift = −6.63
Z lm Score = 6.92
NG = 0
DB = 1
SM = 0
Delta_r
Doxorubicin
YML098W TAF13
ZShift = −10.89
Z lm Score = 0.89
NG = 0
DB = 1
SM = 0
Delta_AUC
Doxorubicin
YLR039C RIC1
ZShift = 5.68
Z lm Score = −7.61
NG = 0
DB = 0
SM = 0
Delta_L
Doxorubicin
YLR039C RIC1
ZShift = −6.59
Z lm Score = 4.35
NG = 0
DB = 0
SM = 0
Delta_K
Doxorubicin
YLR039C RIC1
ZShift = −2.84
Z lm Score = 3.69
NG = 0
DB = 0
SM = 0
Delta_r
Doxorubicin
YLR039C RIC1
ZShift = −7.25
Z lm Score = 0.71
NG = 0
DB = 0
SM = 0
Delta_AUC
Doxorubicin
YBL027W RPL19B
ZShift = 5.28
Z lm Score = −7.64
NG = 0
DB = 0
SM = 0
Delta_L
Doxorubicin
YBL027W RPL19B
ZShift = −2.99
Z lm Score = 1.64
NG = 0
DB = 0
SM = 0
Delta_K
Doxorubicin
YBL027W RPL19B
ZShift = −7.4
Z lm Score = 6.28
NG = 0
DB = 0
SM = 0
Delta_r
Doxorubicin
YBL027W RPL19B
ZShift = −5.51
Z lm Score = 6.9
NG = 0
DB = 0
SM = 0
Delta_AUC
Doxorubicin

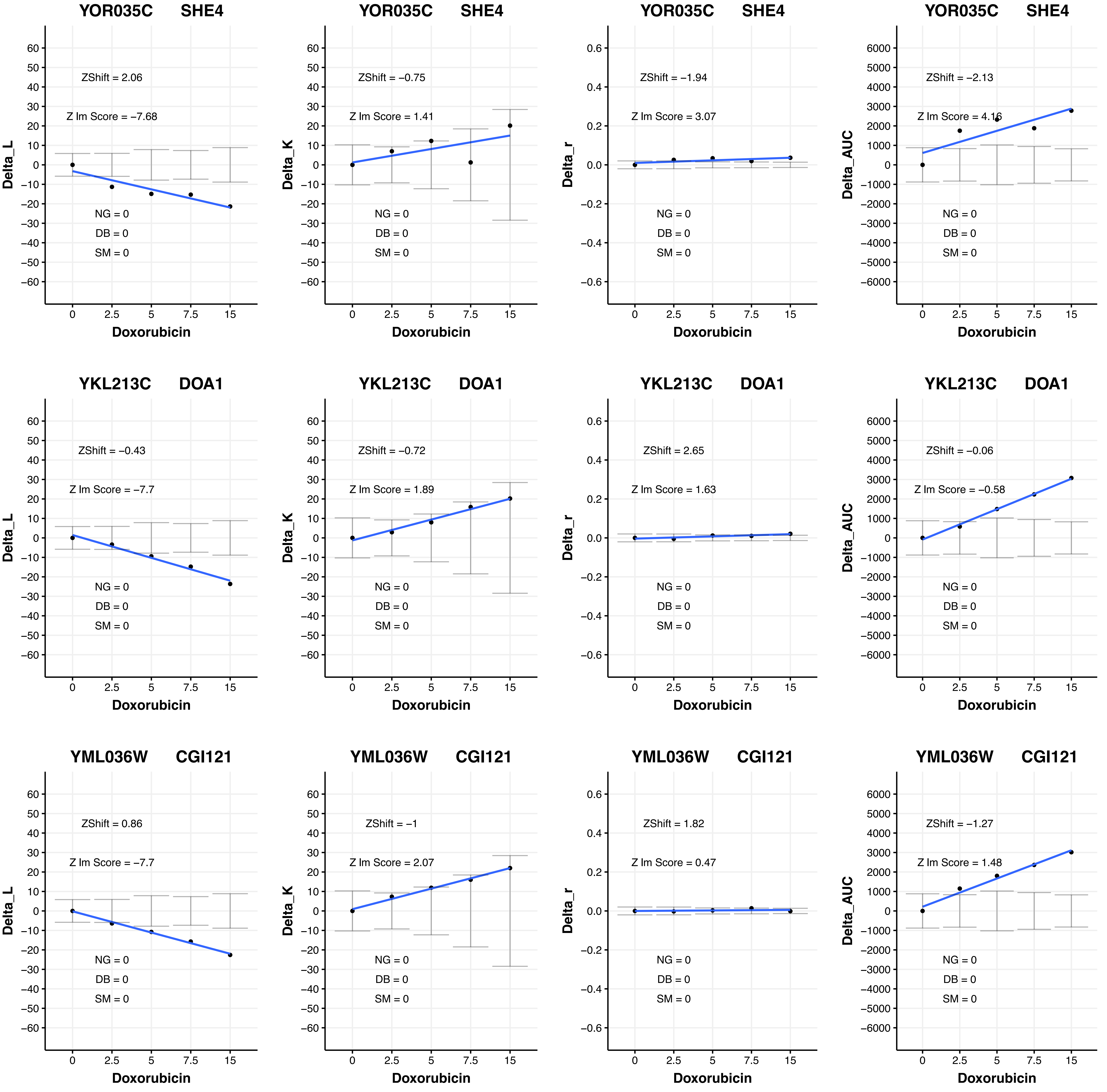

YOR035C SHE4
ZShift = 2.06
Z lm Score = −7.68
NG = 0
DB = 0
SM = 0
Delta_L
Doxorubicin
YOR035C SHE4
ZShift = −0.75
Z lm Score = 1.41
NG = 0
DB = 0
SM = 0
Delta_K
Doxorubicin
YOR035C SHE4
ZShift = −1.94
Z lm Score = 3.07
NG = 0
DB = 0
SM = 0
Delta_r
Doxorubicin
YOR035C SHE4
ZShift = −2.13
Z lm Score = 4.16
NG = 0
DB = 0
SM = 0
Delta_AUC
Doxorubicin
YKL213C DOA1
ZShift = −0.43
Z lm Score = −7.7
NG = 0
DB = 0
SM = 0
Delta_L
Doxorubicin
YKL213C DOA1
ZShift = −0.72
Z lm Score = 1.89
NG = 0
DB = 0
SM = 0
Delta_K
Doxorubicin
YKL213C DOA1
ZShift = 2.65
Z lm Score = 1.63
NG = 0
DB = 0
SM = 0
Delta_r
Doxorubicin
YKL213C DOA1
ZShift = −0.06
Z lm Score = −0.58
NG = 0
DB = 0
SM = 0
Delta_AUC
Doxorubicin
YML036W CGI121
ZShift = 0.86
Z lm Score = −7.7
NG = 0
DB = 0
SM = 0
Delta_L
Doxorubicin
YML036W CGI121
ZShift = −1
Z lm Score = 2.07
NG = 0
DB = 0
SM = 0
Delta_K
Doxorubicin
YML036W CGI121
ZShift = 1.82
Z lm Score = 0.47
NG = 0
DB = 0
SM = 0
Delta_r
Doxorubicin
YML036W CGI121
ZShift = −1.27
Z lm Score = 1.48
NG = 0
DB = 0
SM = 0
Delta_AUC
Doxorubicin

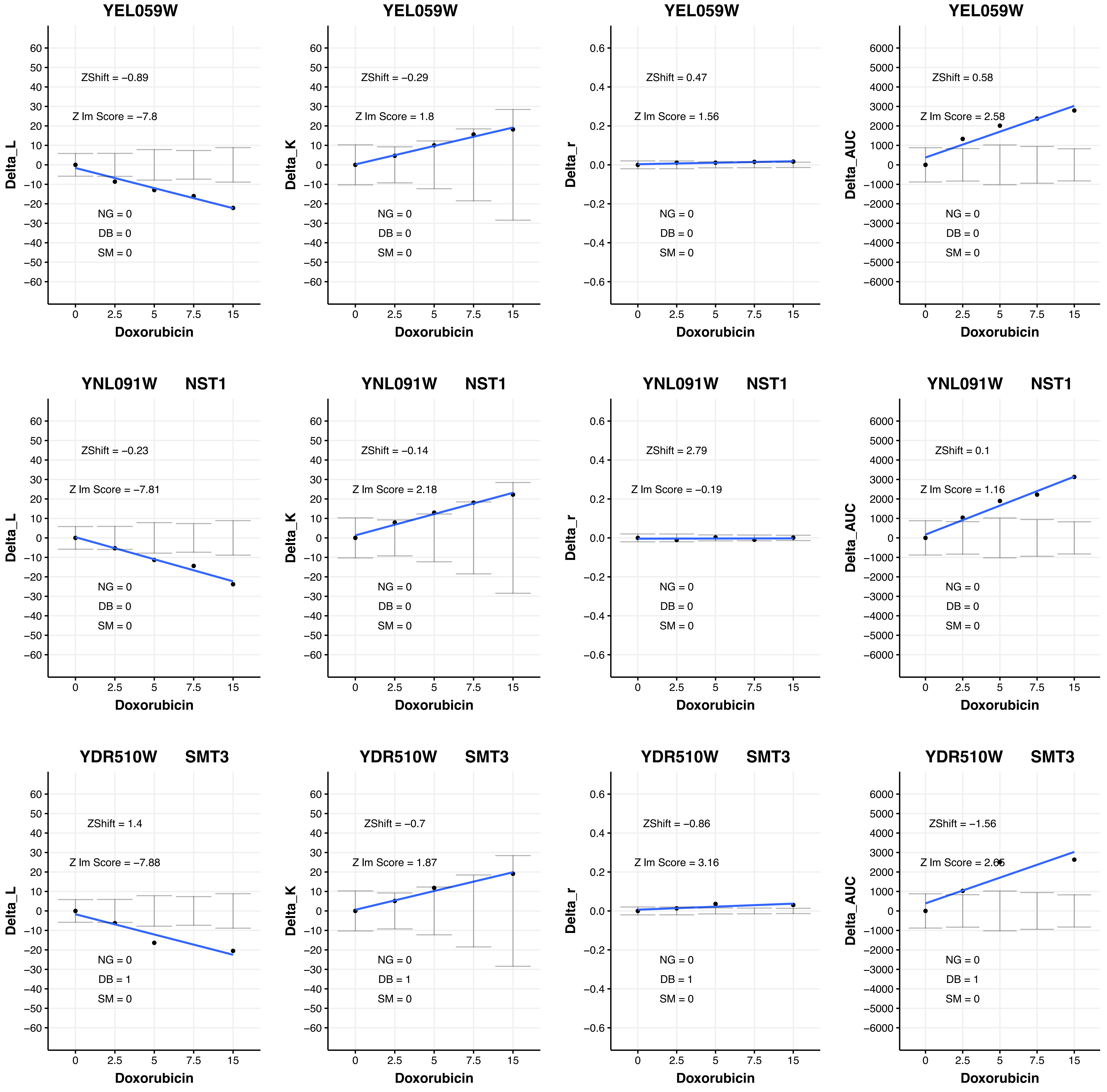
YEL059W
ZShift = −0.89
Z lm Score = −7.8
NG = 0
DB = 0
SM = 0
Delta_L
Doxorubicin
YEL059W
ZShift = −0.29
Z lm Score = 1.8
NG = 0
DB = 0
SM = 0
Delta_K
Doxorubicin
YEL059W
ZShift = 0.47
Z lm Score = 1.56
NG = 0
DB = 0
SM = 0
Delta_r
Doxorubicin
YEL059W
ZShift = 0.58
Z lm Score = 2.58
NG = 0
DB = 0
SM = 0
Delta_AUC
Doxorubicin
YNL091W NST1
ZShift = −0.23
Z lm Score = −7.81
NG = 0
DB = 0
SM = 0
Delta_L
Doxorubicin
YNL091W NST1
ZShift = −0.14
Z lm Score = 2.18
NG = 0
DB = 0
SM = 0
Delta_K
Doxorubicin
YNL091W NST1
ZShift = 2.79
Z lm Score = −0.19
NG = 0
DB = 0
SM = 0
Delta_r
Doxorubicin
YNL091W NST1
ZShift = 0.1
Z lm Score = 1.16
NG = 0
DB = 0
SM = 0
Delta_AUC
Doxorubicin
YDR510W SMT3
ZShift = 1.4
Z lm Score = −7.88
NG = 0
DB = 1
SM = 0
Delta_L
Doxorubicin
YDR510W SMT3
ZShift = −0.7
Z lm Score = 1.87
NG = 0
DB = 1
SM = 0
Delta_K
Doxorubicin
YDR510W SMT3
ZShift = −0.86
Z lm Score = 3.16
NG = 0
DB = 1
SM = 0
Delta_r
Doxorubicin
YDR510W SMT3
ZShift = −1.56
Z lm Score = 2.65
NG = 0
DB = 1
SM = 0
Delta_AUC
Doxorubicin

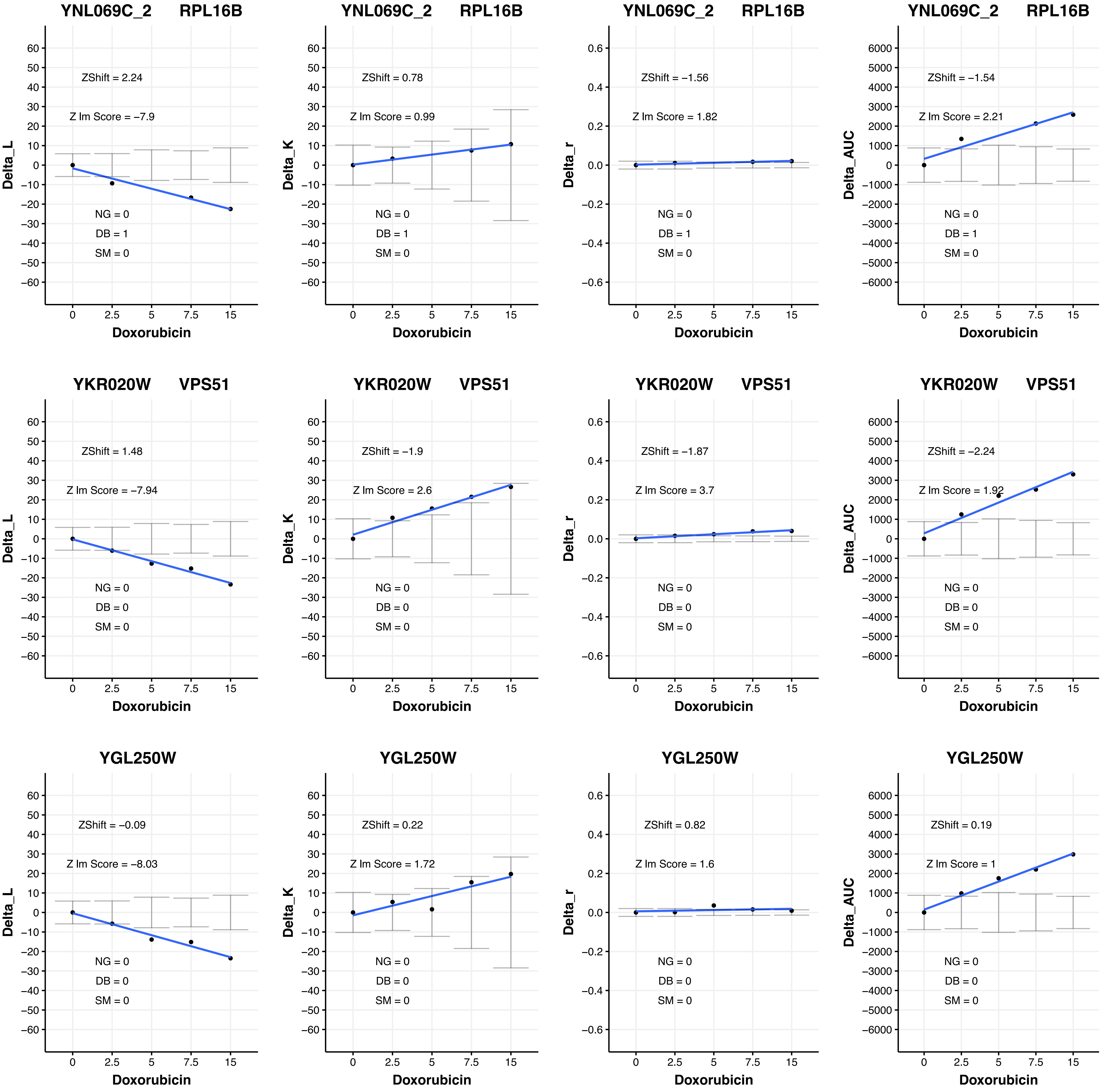
YNL069C_2 RPL16B
ZShift = 2.24
Z lm Score = −7.9
NG = 0
DB = 1
SM = 0
Delta_L
Doxorubicin
YNL069C_2 RPL16B
ZShift = 0.78
Z lm Score = 0.99
NG = 0
DB = 1
SM = 0
Delta_K
Doxorubicin
YNL069C_2 RPL16B
ZShift = −1.56
Z lm Score = 1.82
NG = 0
DB = 1
SM = 0
Delta_r
Doxorubicin
YNL069C_2 RPL16B
ZShift = −1.54
Z lm Score = 2.21
NG = 0
DB = 1
SM = 0
Delta_AUC
Doxorubicin
YKR020W VPS51
ZShift = 1.48
Z lm Score = −7.94
NG = 0
DB = 0
SM = 0
Delta_L
Doxorubicin
YKR020W VPS51
ZShift = −1.9
Z lm Score = 2.6
NG = 0
DB = 0
SM = 0
Delta_K
Doxorubicin
YKR020W VPS51
ZShift = −1.87
Z lm Score = 3.7
NG = 0
DB = 0
SM = 0
Delta_r
Doxorubicin
YKR020W VPS51
ZShift = −2.24
Z lm Score = 1.92
NG = 0
DB = 0
SM = 0
Delta_AUC
Doxorubicin
YGL250W
ZShift = −0.09
Z lm Score = −8.03
NG = 0
DB = 0
SM = 0
Delta_L
Doxorubicin
YGL250W
ZShift = 0.22
Z lm Score = 1.72
NG = 0
DB = 0
SM = 0
Delta_K
Doxorubicin
YGL250W
ZShift = 0.82
Z lm Score = 1.6
NG = 0
DB = 0
SM = 0
Delta_r
Doxorubicin
YGL250W
ZShift = 0.19
Z lm Score = 1
NG = 0
DB = 0
SM = 0
Delta_AUC
Doxorubicin

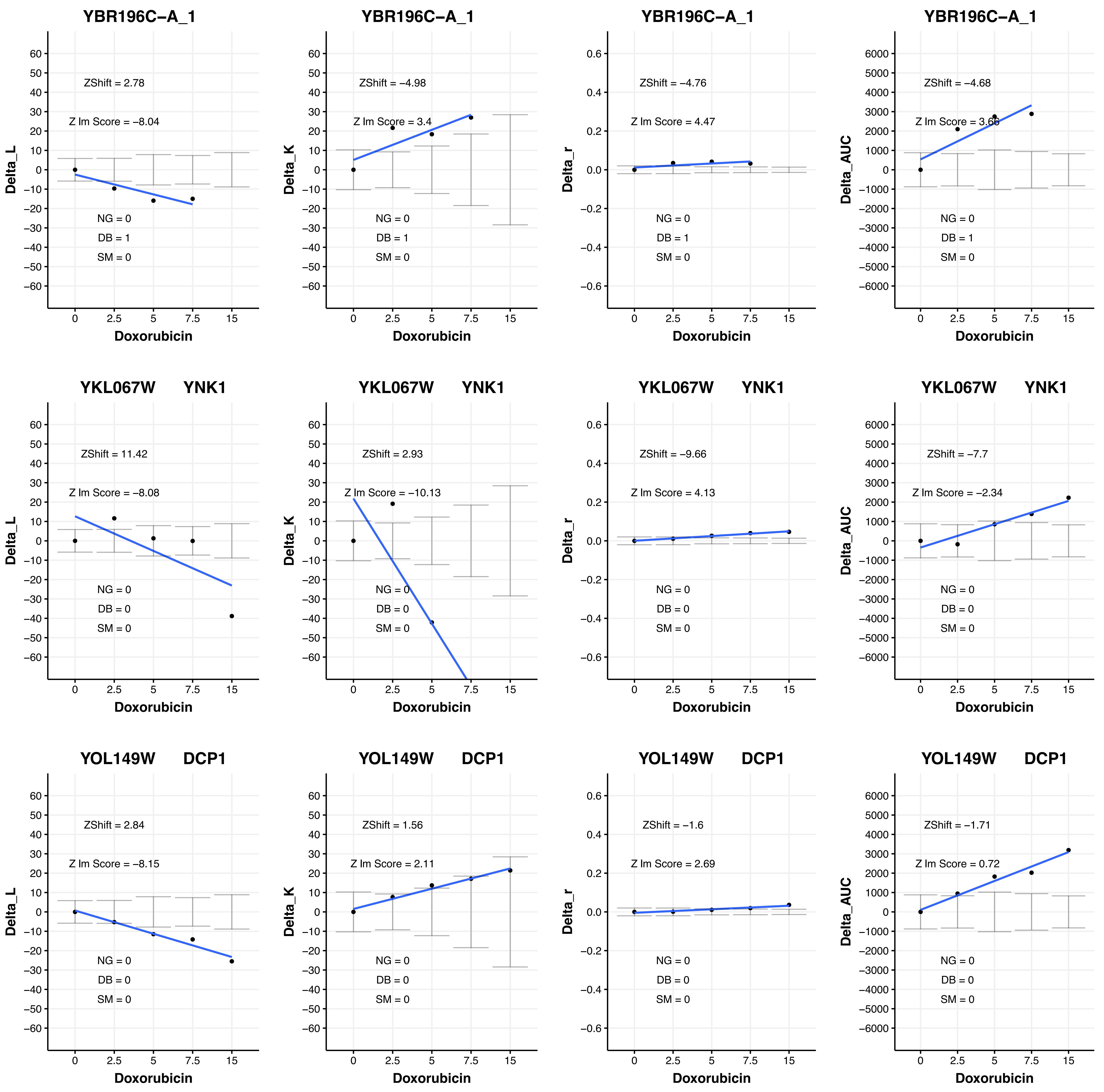
YBR196C−A_1
ZShift = 2.78
Z lm Score = −8.04
NG = 0
DB = 1
SM = 0
Delta_L
Doxorubicin
YBR196C−A_1
ZShift = −4.98
Z lm Score = 3.4
NG = 0
DB = 1
SM = 0
Delta_K
Doxorubicin
YBR196C−A_1
ZShift = −4.76
Z lm Score = 4.47
NG = 0
DB = 1
SM = 0
Delta_r
Doxorubicin
YBR196C−A_1
ZShift = −4.68
Z lm Score = 3.66
NG = 0
DB = 1
SM = 0
Delta_AUC
Doxorubicin
YKL067W YNK1
ZShift = 11.42
Z lm Score = −8.08
NG = 0
DB = 0
SM = 0
Delta_L
Doxorubicin
YKL067W YNK1
ZShift = 2.93
Z lm Score = −10.13
NG = 0
DB = 0
SM = 0
Delta_K
Doxorubicin
YKL067W YNK1
ZShift = −9.66
Z lm Score = 4.13
NG = 0
DB = 0
SM = 0
Delta_r
Doxorubicin
YKL067W YNK1
ZShift = −7.7
Z lm Score = −2.34
NG = 0
DB = 0
SM = 0
Delta_AUC
Doxorubicin
YOL149W DCP1
ZShift = 2.84
Z lm Score = −8.15
NG = 0
DB = 0
SM = 0
Delta_L
Doxorubicin
YOL149W DCP1
ZShift = 1.56
Z lm Score = 2.11
NG = 0
DB = 0
SM = 0
Delta_K
Doxorubicin
YOL149W DCP1
ZShift = −1.6
Z lm Score = 2.69
NG = 0
DB = 0
SM = 0
Delta_r
Doxorubicin
YOL149W DCP1
ZShift = −1.71
Z lm Score = 0.72
NG = 0
DB = 0
SM = 0
Delta_AUC
Doxorubicin

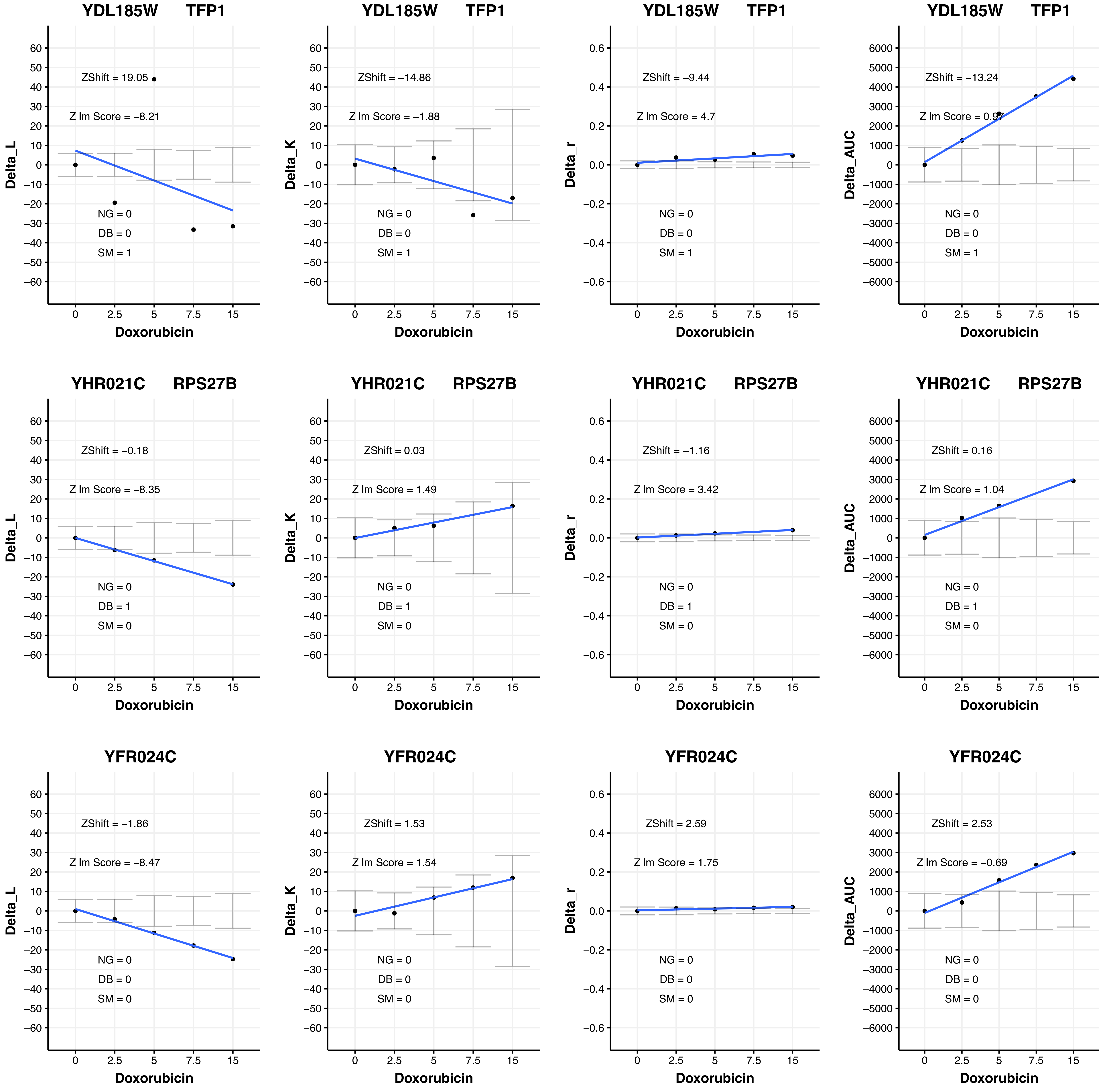
YDL185W TFP1
ZShift = 19.05
Z lm Score = −8.21
NG = 0
DB = 0
SM = 1
Delta_L
Doxorubicin
YDL185W TFP1
ZShift = −14.86
Z lm Score = −1.88
NG = 0
DB = 0
SM = 1
Delta_K
Doxorubicin
YDL185W TFP1
ZShift = −9.44
Z lm Score = 4.7
NG = 0
DB = 0
SM = 1
Delta_r
Doxorubicin
YDL185W TFP1
ZShift = −13.24
Z lm Score = 0.97
NG = 0
DB = 0
SM = 1
Delta_AUC
Doxorubicin
YHR021C RPS27B
ZShift = −0.18
Z lm Score = −8.35
NG = 0
DB = 1
SM = 0
Delta_L
Doxorubicin
YHR021C RPS27B
ZShift = 0.03
Z lm Score = 1.49
NG = 0
DB = 1
SM = 0
Delta_K
Doxorubicin
YHR021C RPS27B
ZShift = −1.16
Z lm Score = 3.42
NG = 0
DB = 1
SM = 0
Delta_r
Doxorubicin
YHR021C RPS27B
ZShift = 0.16
Z lm Score = 1.04
NG = 0
DB = 1
SM = 0
Delta_AUC
Doxorubicin
YFR024C
ZShift = −1.86
Z lm Score = −8.47
NG = 0
DB = 0
SM = 0
Delta_L
Doxorubicin
YFR024C
ZShift = 1.53
Z lm Score = 1.54
NG = 0
DB = 0
SM = 0
Delta_K
Doxorubicin
YFR024C
ZShift = 2.59
Z lm Score = 1.75
NG = 0
DB = 0
SM = 0
Delta_r
Doxorubicin
YFR024C
ZShift = 2.53
Z lm Score = −0.69
NG = 0
DB = 0
SM = 0
Delta_AUC
Doxorubicin

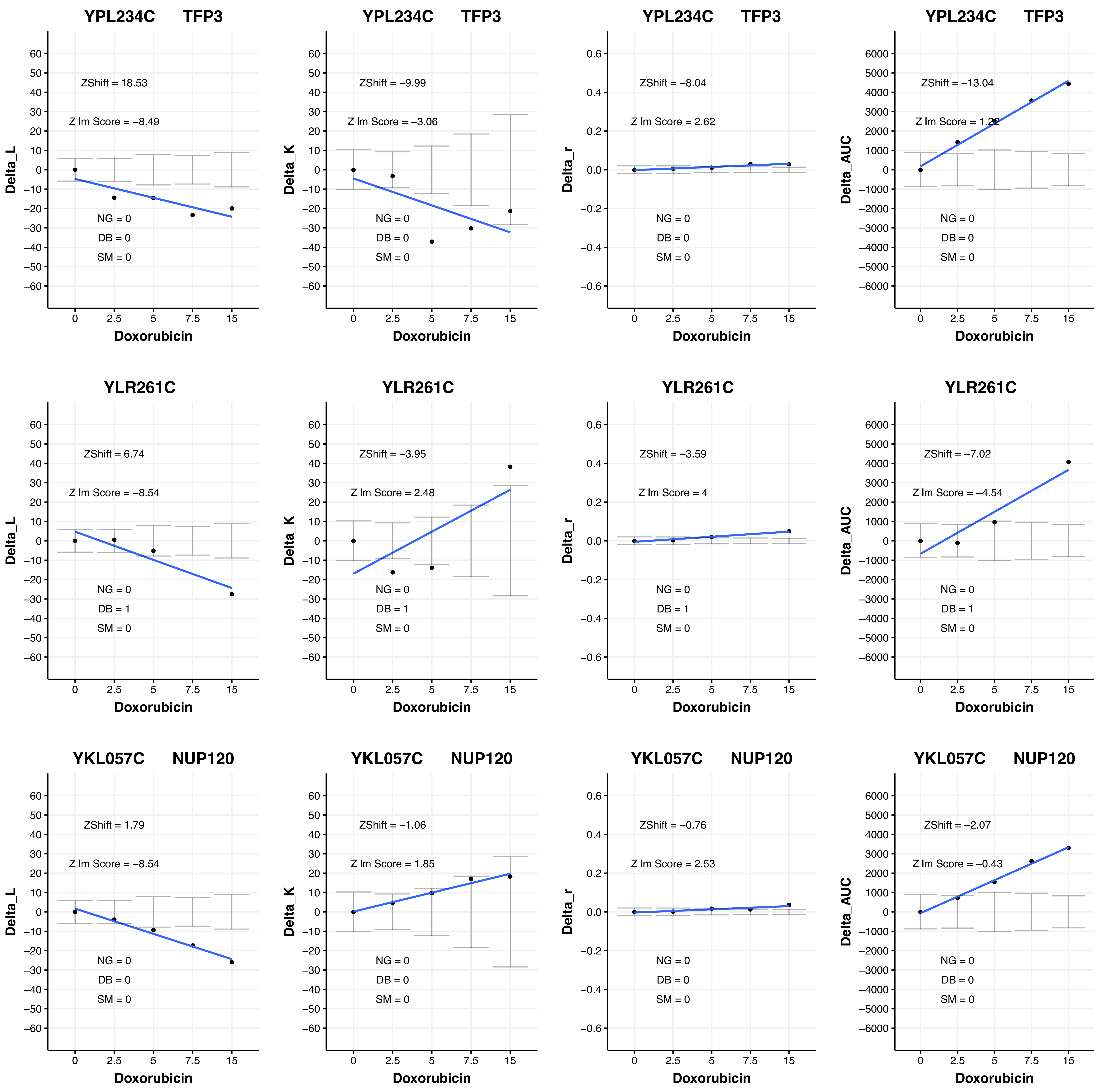
YPL234C TFP3
ZShift = 18.53
Z Im Score = −8.49
NG = 0
DB = 0
SM = 0
Delta_L
Doxorubicin
YPL234C TFP3
ZShift = −9.99
Z Im Score = −3.06
NG = 0
DB = 0
SM = 0
Delta_K
Doxorubicin
YPL234C TFP3
ZShift = −8.04
Z Im Score = 2.62
NG = 0
DB = 0
SM = 0
Delta_r
Doxorubicin
YPL234C TFP3
ZShift = −13.04
Z Im Score = 1.22
NG = 0
DB = 0
SM = 0
Delta_AUC
Doxorubicin
YLR261C
ZShift = 6.74
Z Im Score = −8.54
NG = 0
DB = 1
SM = 0
Delta_L
Doxorubicin
YLR261C
ZShift = −3.95
Z Im Score = 2.48
NG = 0
DB = 1
SM = 0
Delta_K
Doxorubicin
YLR261C
ZShift = −3.59
Z Im Score = 4
NG = 0
DB = 1
SM = 0
Delta_r
Doxorubicin
YLR261C
ZShift = −7.02
Z Im Score = −4.54
NG = 0
DB = 1
SM = 0
Delta_AUC
Doxorubicin
YKL057C NUP120
ZShift = 1.79
Z Im Score = −8.54
NG = 0
DB = 0
SM = 0
Delta_L
Doxorubicin
YKL057C NUP120
ZShift = −1.06
Z Im Score = 1.85
NG = 0
DB = 0
SM = 0
Delta_K
Doxorubicin
YKL057C NUP120
ZShift = −0.76
Z Im Score = 2.53
NG = 0
DB = 0
SM = 0
Delta_r
Doxorubicin
YKL057C NUP120
ZShift = −2.07
Z Im Score = −0.43
NG = 0
DB = 0
SM = 0
Delta_AUC
Doxorubicin

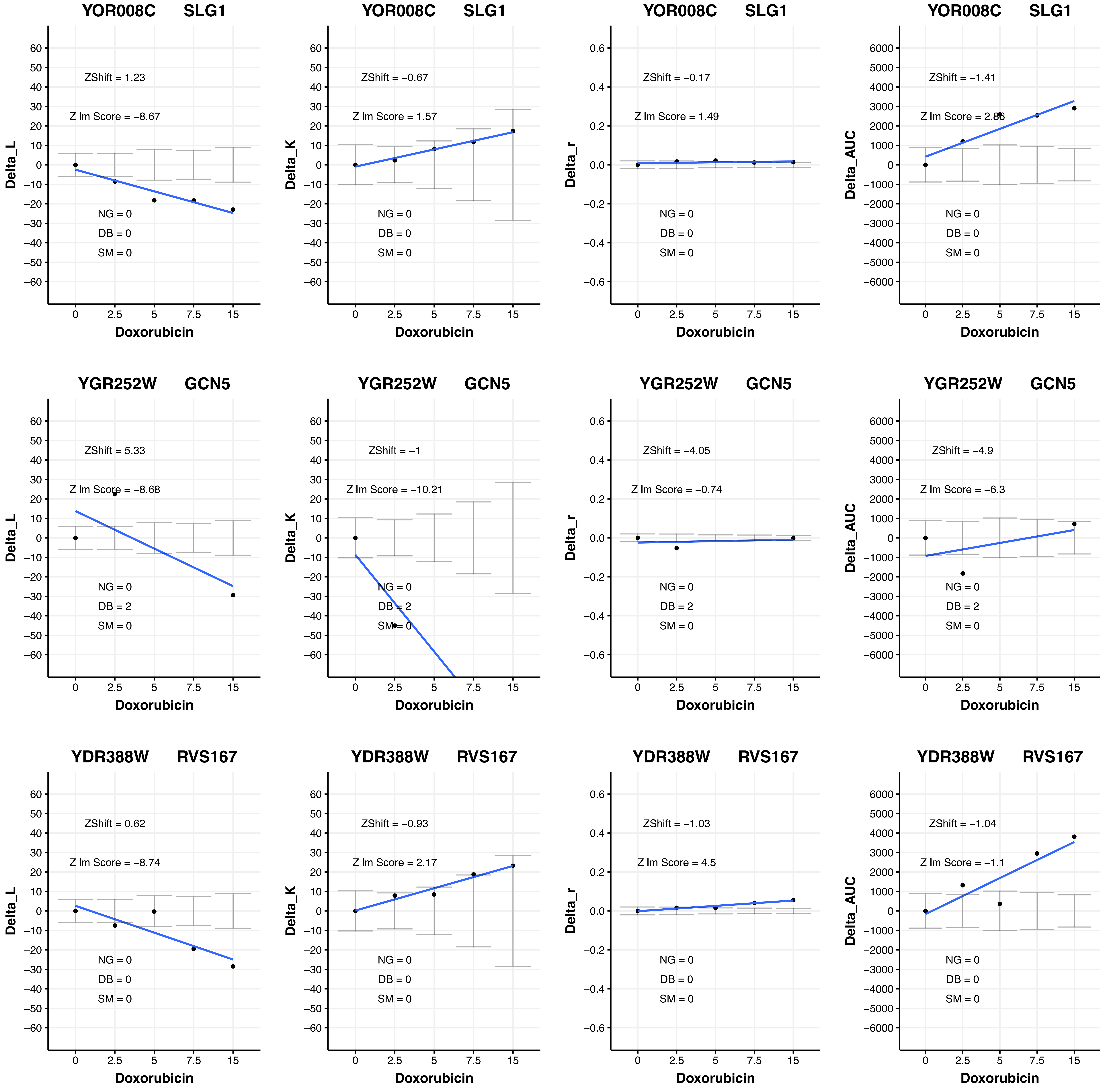
YOR008C SLG1
ZShift = 1.23
Z lm Score = −8.67
NG = 0
DB = 0
SM = 0
Delta_L
Doxorubicin
YOR008C SLG1
ZShift = −0.67
Z lm Score = 1.57
NG = 0
DB = 0
SM = 0
Delta_K
Doxorubicin
YOR008C SLG1
ZShift = −0.17
Z lm Score = 1.49
NG = 0
DB = 0
SM = 0
Delta_r
Doxorubicin
YOR008C SLG1
ZShift = −1.41
Z lm Score = 2.86
NG = 0
DB = 0
SM = 0
Delta_AUC
Doxorubicin
YGR252W GCN5
ZShift = 5.33
Z lm Score = −8.68
NG = 0
DB = 2
SM = 0
Delta_L
Doxorubicin
YGR252W GCN5
ZShift = −1
Z lm Score = −10.21
NG = 0
DB = 2
SM = 0
Delta_K
Doxorubicin
YGR252W GCN5
ZShift = −4.05
Z lm Score = −0.74
NG = 0
DB = 2
SM = 0
Delta_r
Doxorubicin
YGR252W GCN5
ZShift = −4.9
Z lm Score = −6.3
NG = 0
DB = 2
SM = 0
Delta_AUC
Doxorubicin
YDR388W RVS167
ZShift = 0.62
Z lm Score = −8.74
NG = 0
DB = 0
SM = 0
Delta_L
Doxorubicin
YDR388W RVS167
ZShift = −0.93
Z lm Score = 2.17
NG = 0
DB = 0
SM = 0
Delta_K
Doxorubicin
YDR388W RVS167
ZShift = −1.03
Z lm Score = 4.5
NG = 0
DB = 0
SM = 0
Delta_r
Doxorubicin
YDR388W RVS167
ZShift = −1.04
Z lm Score = −1.1
NG = 0
DB = 0
SM = 0
Delta_AUC
Doxorubicin

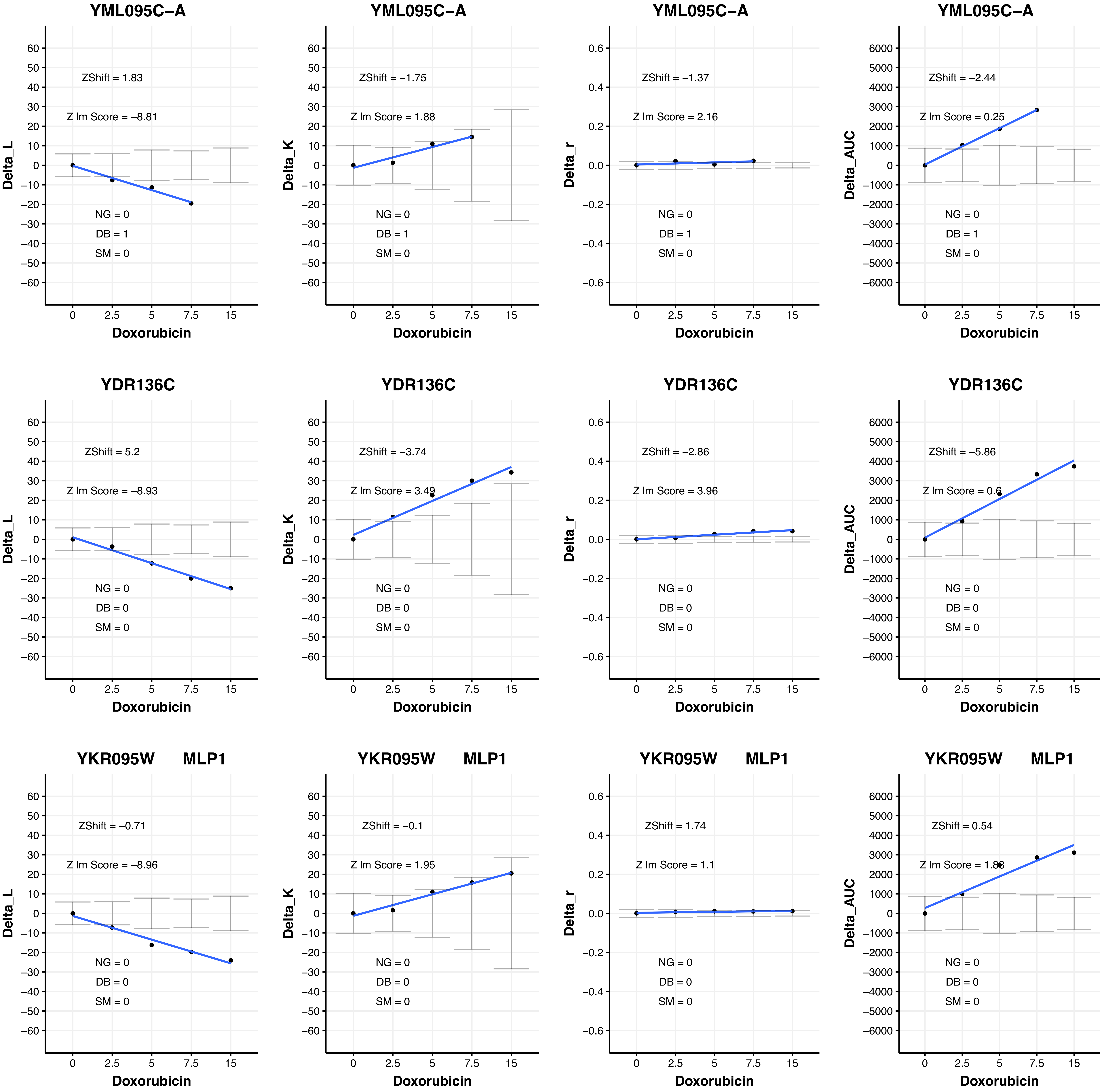

YML095C−A
ZShift = 1.83
Z lm Score = −8.81
NG = 0
DB = 1
SM = 0
Delta_L
Doxorubicin
YML095C−A
ZShift = −1.75
Z lm Score = 1.88
NG = 0
DB = 1
SM = 0
Delta_K
Doxorubicin
YML095C−A
ZShift = −1.37
Z lm Score = 2.16
NG = 0
DB = 1
SM = 0
Delta_r
Doxorubicin
YML095C−A
ZShift = −2.44
Z lm Score = 0.25
NG = 0
DB = 1
SM = 0
Delta_AUC
Doxorubicin
YDR136C
ZShift = 5.2
Z lm Score = −8.93
NG = 0
DB = 0
SM = 0
Delta_L
Doxorubicin
YDR136C
ZShift = −3.74
Z lm Score = 3.49
NG = 0
DB = 0
SM = 0
Delta_K
Doxorubicin
YDR136C
ZShift = −2.86
Z lm Score = 3.96
NG = 0
DB = 0
SM = 0
Delta_r
Doxorubicin
YDR136C
ZShift = −5.86
Z lm Score = 0.6
NG = 0
DB = 0
SM = 0
Delta_AUC
Doxorubicin
YKR095W MLP1
ZShift = −0.71
Z lm Score = −8.96
NG = 0
DB = 0
SM = 0
Delta_L
Doxorubicin
YKR095W MLP1
ZShift = −0.1
Z lm Score = 1.95
NG = 0
DB = 0
SM = 0
Delta_K
Doxorubicin
YKR095W MLP1
ZShift = 1.74
Z lm Score = 1.1
NG = 0
DB = 0
SM = 0
Delta_r
Doxorubicin
YKR095W MLP1
ZShift = 0.54
Z lm Score = 1.88
NG = 0
DB = 0
SM = 0
Delta_AUC
Doxorubicin

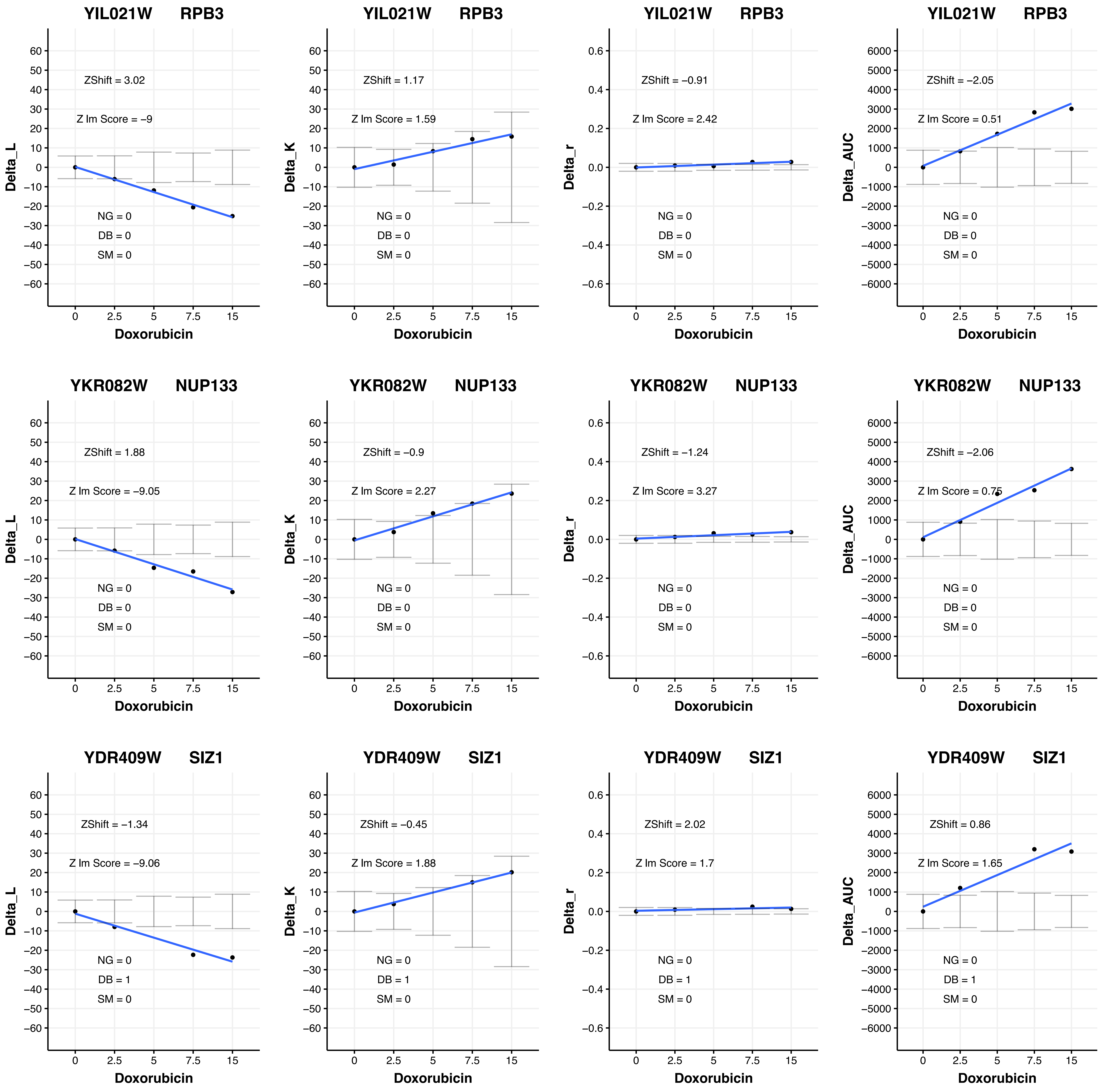

YIL021W RPB3
Delta_L
ZShift = 3.02
Z lm Score = −9
NG = 0
DB = 0
SM = 0
Doxorubicin
YIL021W RPB3
Delta_K
ZShift = 1.17
Z lm Score = 1.59
NG = 0
DB = 0
SM = 0
Doxorubicin
YIL021W RPB3
Delta_r
ZShift = −0.91
Z lm Score = 2.42
NG = 0
DB = 0
SM = 0
Doxorubicin
YIL021W RPB3
Delta_AUC
ZShift = −2.05
Z lm Score = 0.51
NG = 0
DB = 0
SM = 0
Doxorubicin
YKR082W NUP133
Delta_L
ZShift = 1.88
Z lm Score = −9.05
NG = 0
DB = 0
SM = 0
Doxorubicin
YKR082W NUP133
Delta_K
ZShift = −0.9
Z lm Score = 2.27
NG = 0
DB = 0
SM = 0
Doxorubicin
YKR082W NUP133
Delta_r
ZShift = −1.24
Z lm Score = 3.27
NG = 0
DB = 0
SM = 0
Doxorubicin
YKR082W NUP133
Delta_AUC
ZShift = −2.06
Z lm Score = 0.75
NG = 0
DB = 0
SM = 0
Doxorubicin
YDR409W SIZ1
Delta_L
ZShift = −1.34
Z lm Score = −9.06
NG = 0
DB = 1
SM = 0
Doxorubicin
YDR409W SIZ1
Delta_K
ZShift = −0.45
Z lm Score = 1.88
NG = 0
DB = 1
SM = 0
Doxorubicin
YDR409W SIZ1
Delta_r
ZShift = 2.02
Z lm Score = 1.7
NG = 0
DB = 1
SM = 0
Doxorubicin
YDR409W SIZ1
Delta_AUC
ZShift = 0.86
Z lm Score = 1.65
NG = 0
DB = 1
SM = 0
Doxorubicin

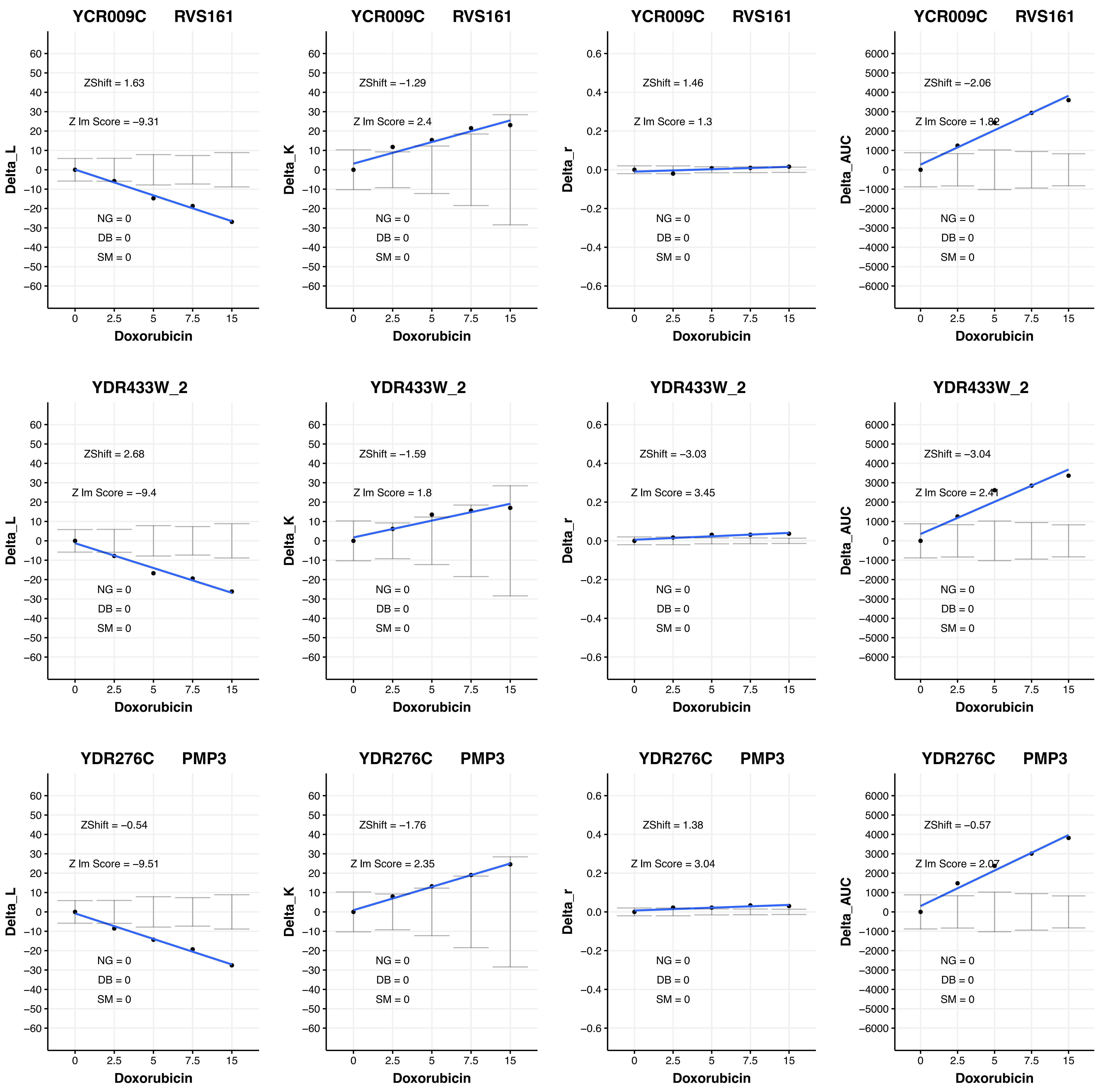
YCR009C RVS161
ZShift = 1.63
Z lm Score = −9.31
NG = 0
DB = 0
SM = 0
Delta_L
Doxorubicin
YCR009C RVS161
ZShift = −1.29
Z lm Score = 2.4
NG = 0
DB = 0
SM = 0
Delta_K
Doxorubicin
YCR009C RVS161
ZShift = 1.46
Z lm Score = 1.3
NG = 0
DB = 0
SM = 0
Delta_r
Doxorubicin
YCR009C RVS161
ZShift = −2.06
Z lm Score = 1.82
NG = 0
DB = 0
SM = 0
Delta_AUC
Doxorubicin
YDR433W_2
ZShift = 2.68
Z lm Score = −9.4
NG = 0
DB = 0
SM = 0
Delta_L
Doxorubicin
YDR433W_2
ZShift = −1.59
Z lm Score = 1.8
NG = 0
DB = 0
SM = 0
Delta_K
Doxorubicin
YDR433W_2
ZShift = −3.03
Z lm Score = 3.45
NG = 0
DB = 0
SM = 0
Delta_r
Doxorubicin
YDR433W_2
ZShift = −3.04
Z lm Score = 2.41
NG = 0
DB = 0
SM = 0
Delta_AUC
Doxorubicin
YDR276C PMP3
ZShift = −0.54
Z lm Score = −9.51
NG = 0
DB = 0
SM = 0
Delta_L
Doxorubicin
YDR276C PMP3
ZShift = −1.76
Z lm Score = 2.35
NG = 0
DB = 0
SM = 0
Delta_K
Doxorubicin
YDR276C PMP3
ZShift = 1.38
Z lm Score = 3.04
NG = 0
DB = 0
SM = 0
Delta_r
Doxorubicin
YDR276C PMP3
ZShift = −0.57
Z lm Score = 2.07
NG = 0
DB = 0
SM = 0
Delta_AUC
Doxorubicin

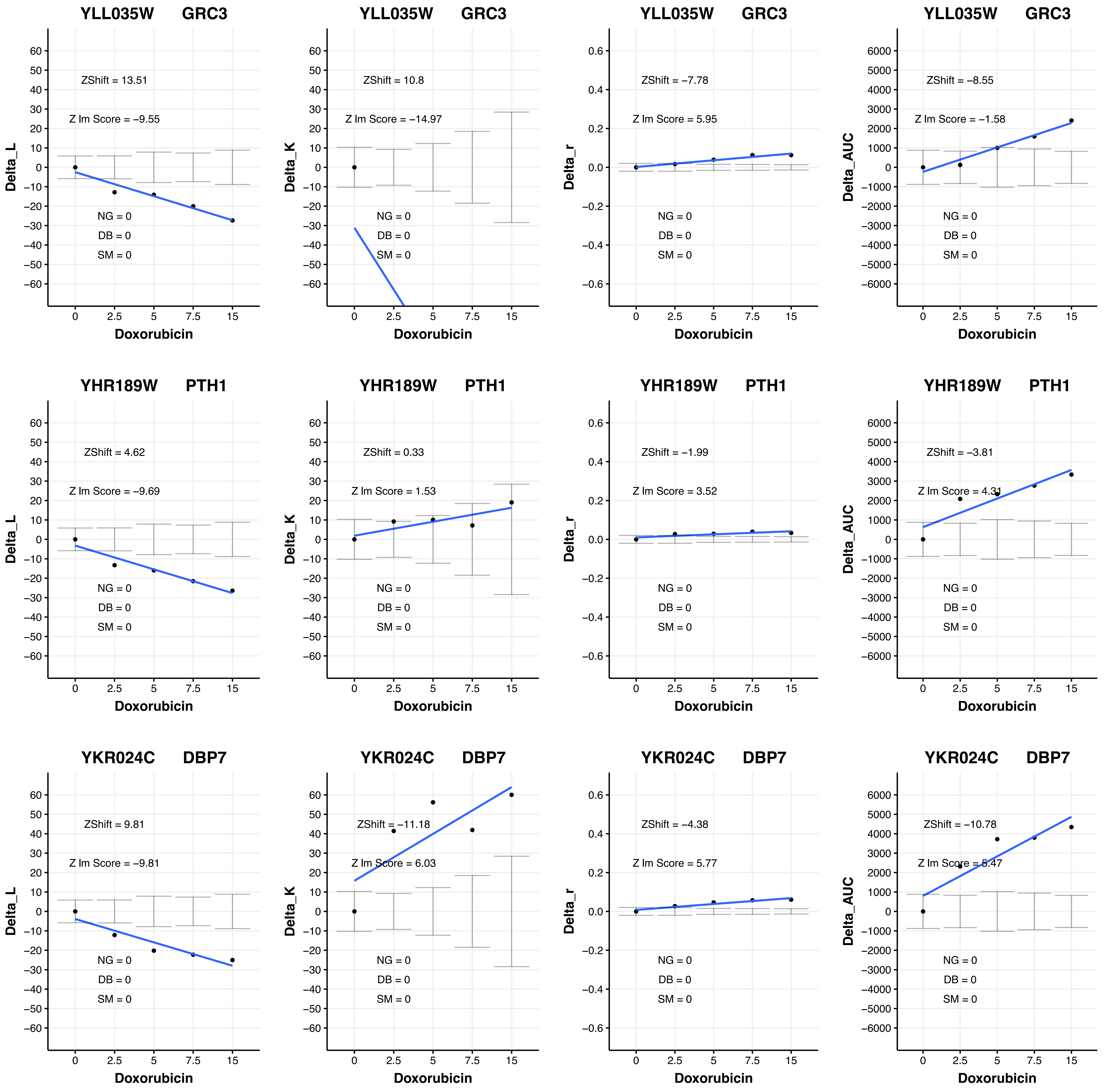

YLL035W GRC3
ZShift = 13.51
Z Im Score = −9.55
NG = 0
DB = 0
SM = 0
Delta_L
Doxorubicin
YLL035W GRC3
ZShift = 10.8
Z Im Score = −14.97
NG = 0
DB = 0
SM = 0
Delta_K
Doxorubicin
YLL035W GRC3
ZShift = −7.78
Z Im Score = 5.95
NG = 0
DB = 0
SM = 0
Delta_r
Doxorubicin
YLL035W GRC3
ZShift = −8.55
Z Im Score = −1.58
NG = 0
DB = 0
SM = 0
Delta_AUC
Doxorubicin
YHR189W PTH1
ZShift = 4.62
Z Im Score = −9.69
NG = 0
DB = 0
SM = 0
Delta_L
Doxorubicin
YHR189W PTH1
ZShift = 0.33
Z Im Score = 1.53
NG = 0
DB = 0
SM = 0
Delta_K
Doxorubicin
YHR189W PTH1
ZShift = −1.99
Z Im Score = 3.52
NG = 0
DB = 0
SM = 0
Delta_r
Doxorubicin
YHR189W PTH1
ZShift = −3.81
Z Im Score = 4.31
NG = 0
DB = 0
SM = 0
Delta_AUC
Doxorubicin
YKR024C DBP7
ZShift = 9.81
Z Im Score = −9.81
NG = 0
DB = 0
SM = 0
Delta_L
Doxorubicin
YKR024C DBP7
ZShift = −11.18
Z Im Score = 6.03
NG = 0
DB = 0
SM = 0
Delta_K
Doxorubicin
YKR024C DBP7
ZShift = −4.38
Z Im Score = 5.77
NG = 0
DB = 0
SM = 0
Delta_r
Doxorubicin
YKR024C DBP7
ZShift = −10.78
Z Im Score = 5.47
NG = 0
DB = 0
SM = 0
Delta_AUC
Doxorubicin

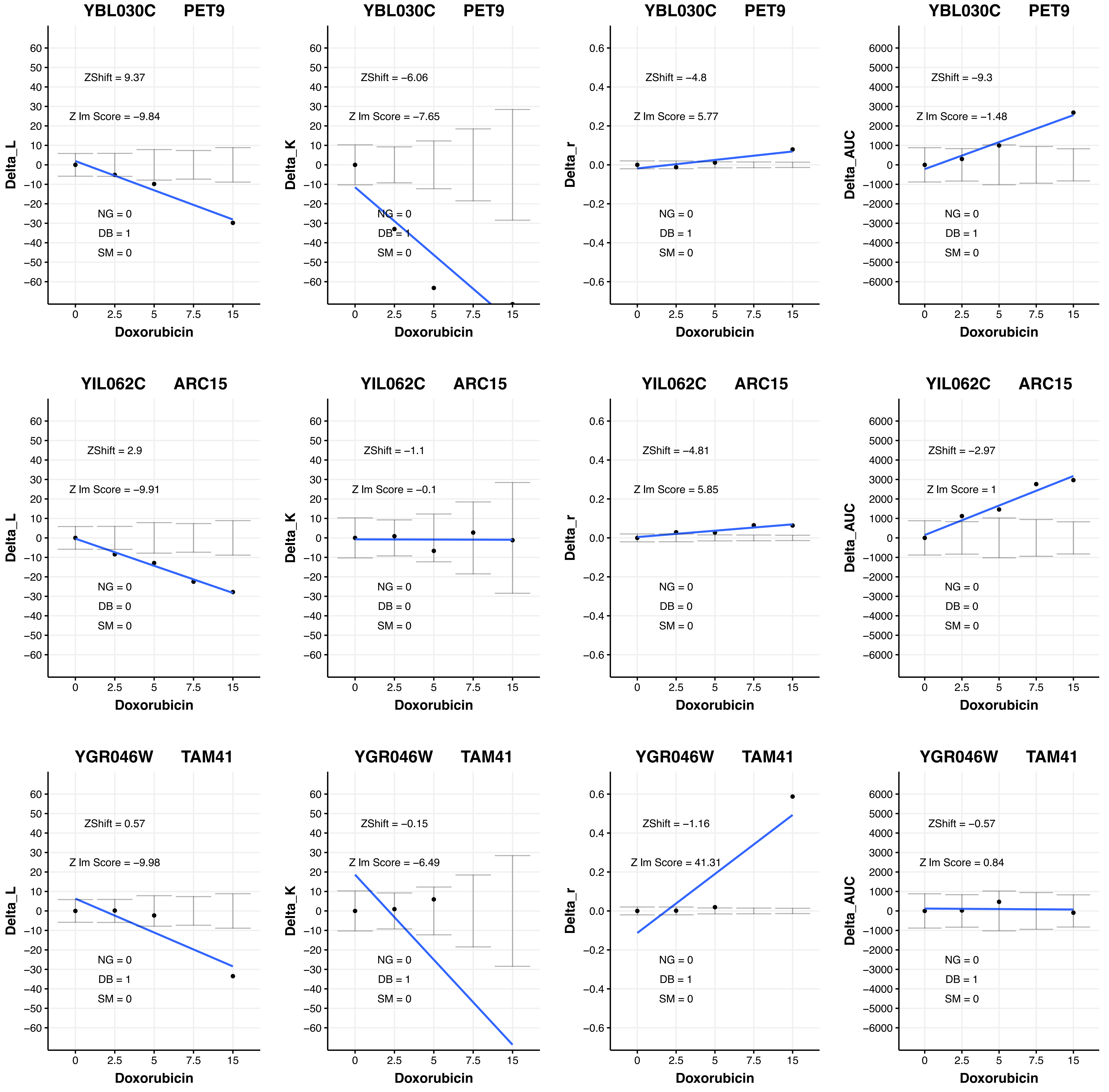
YBL030C PET9
ZShift = 9.37
Z lm Score = −9.84
NG = 0
DB = 1
SM = 0
Delta_L
Doxorubicin
YBL030C PET9
ZShift = −6.06
Z lm Score = −7.65
NG = 0
DB = 1
SM = 0
Delta_K
Doxorubicin
YBL030C PET9
ZShift = −4.8
Z lm Score = 5.77
NG = 0
DB = 1
SM = 0
Delta_r
Doxorubicin
YBL030C PET9
ZShift = −9.3
Z lm Score = −1.48
NG = 0
DB = 1
SM = 0
Delta_AUC
Doxorubicin
YIL062C ARC15
ZShift = 2.9
Z lm Score = −9.91
NG = 0
DB = 0
SM = 0
Delta_L
Doxorubicin
YIL062C ARC15
ZShift = −1.1
Z lm Score = −0.1
NG = 0
DB = 0
SM = 0
Delta_K
Doxorubicin
YIL062C ARC15
ZShift = −4.81
Z lm Score = 5.85
NG = 0
DB = 0
SM = 0
Delta_r
Doxorubicin
YIL062C ARC15
ZShift = −2.97
Z lm Score = 1
NG = 0
DB = 0
SM = 0
Delta_AUC
Doxorubicin
YGR046W TAM41
ZShift = 0.57
Z lm Score = −9.98
NG = 0
DB = 1
SM = 0
Delta_L
Doxorubicin
YGR046W TAM41
ZShift = −0.15
Z lm Score = −6.49
NG = 0
DB = 1
SM = 0
Delta_K
Doxorubicin
YGR046W TAM41
ZShift = −1.16
Z lm Score = 41.31
NG = 0
DB = 1
SM = 0
Delta_r
Doxorubicin
YGR046W TAM41
ZShift = −0.57
Z lm Score = 0.84
NG = 0
DB = 1
SM = 0
Delta_AUC
Doxorubicin

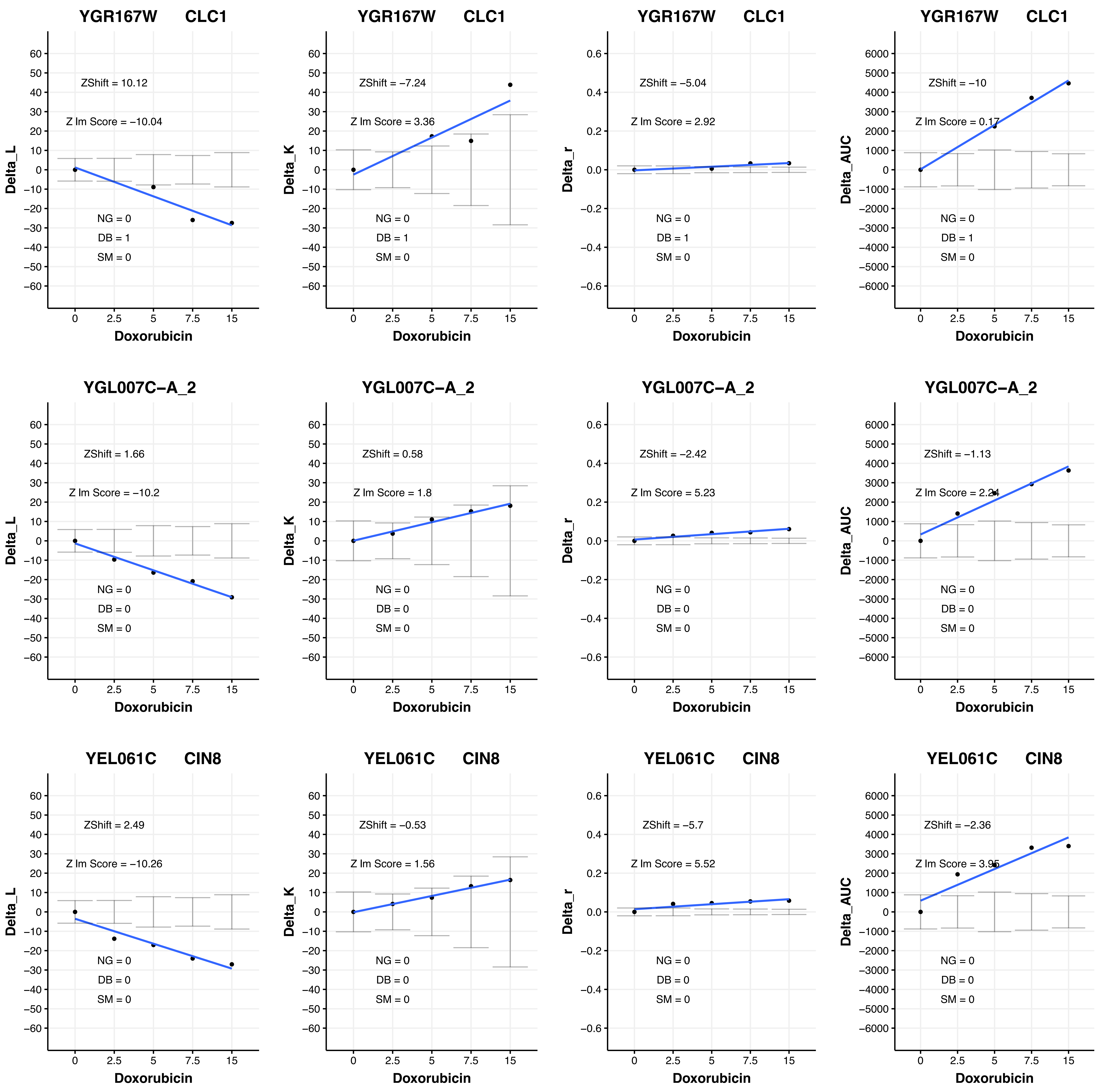
YGR167W CLC1
ZShift = 10.12
Z lm Score = −10.04
NG = 0
DB = 1
SM = 0
Delta_L
Doxorubicin
YGR167W CLC1
ZShift = −7.24
Z lm Score = 3.36
NG = 0
DB = 1
SM = 0
Delta_K
Doxorubicin
YGR167W CLC1
ZShift = −5.04
Z lm Score = 2.92
NG = 0
DB = 1
SM = 0
Delta_r
Doxorubicin
YGR167W CLC1
ZShift = −10
Z lm Score = 0.17
NG = 0
DB = 1
SM = 0
Delta_AUC
Doxorubicin
YGL007C−A_2
ZShift = 1.66
Z lm Score = −10.2
NG = 0
DB = 0
SM = 0
Delta_L
Doxorubicin
YGL007C−A_2
ZShift = 0.58
Z lm Score = 1.8
NG = 0
DB = 0
SM = 0
Delta_K
Doxorubicin
YGL007C−A_2
ZShift = −2.42
Z lm Score = 5.23
NG = 0
DB = 0
SM = 0
Delta_r
Doxorubicin
YGL007C−A_2
ZShift = −1.13
Z lm Score = 2.24
NG = 0
DB = 0
SM = 0
Delta_AUC
Doxorubicin
YEL061C CIN8
ZShift = 2.49
Z lm Score = −10.26
NG = 0
DB = 0
SM = 0
Delta_L
Doxorubicin
YEL061C CIN8
ZShift = −0.53
Z lm Score = 1.56
NG = 0
DB = 0
SM = 0
Delta_K
Doxorubicin
YEL061C CIN8
ZShift = −5.7
Z lm Score = 5.52
NG = 0
DB = 0
SM = 0
Delta_r
Doxorubicin
YEL061C CIN8
ZShift = −2.36
Z lm Score = 3.95
NG = 0
DB = 0
SM = 0
Delta_AUC
Doxorubicin

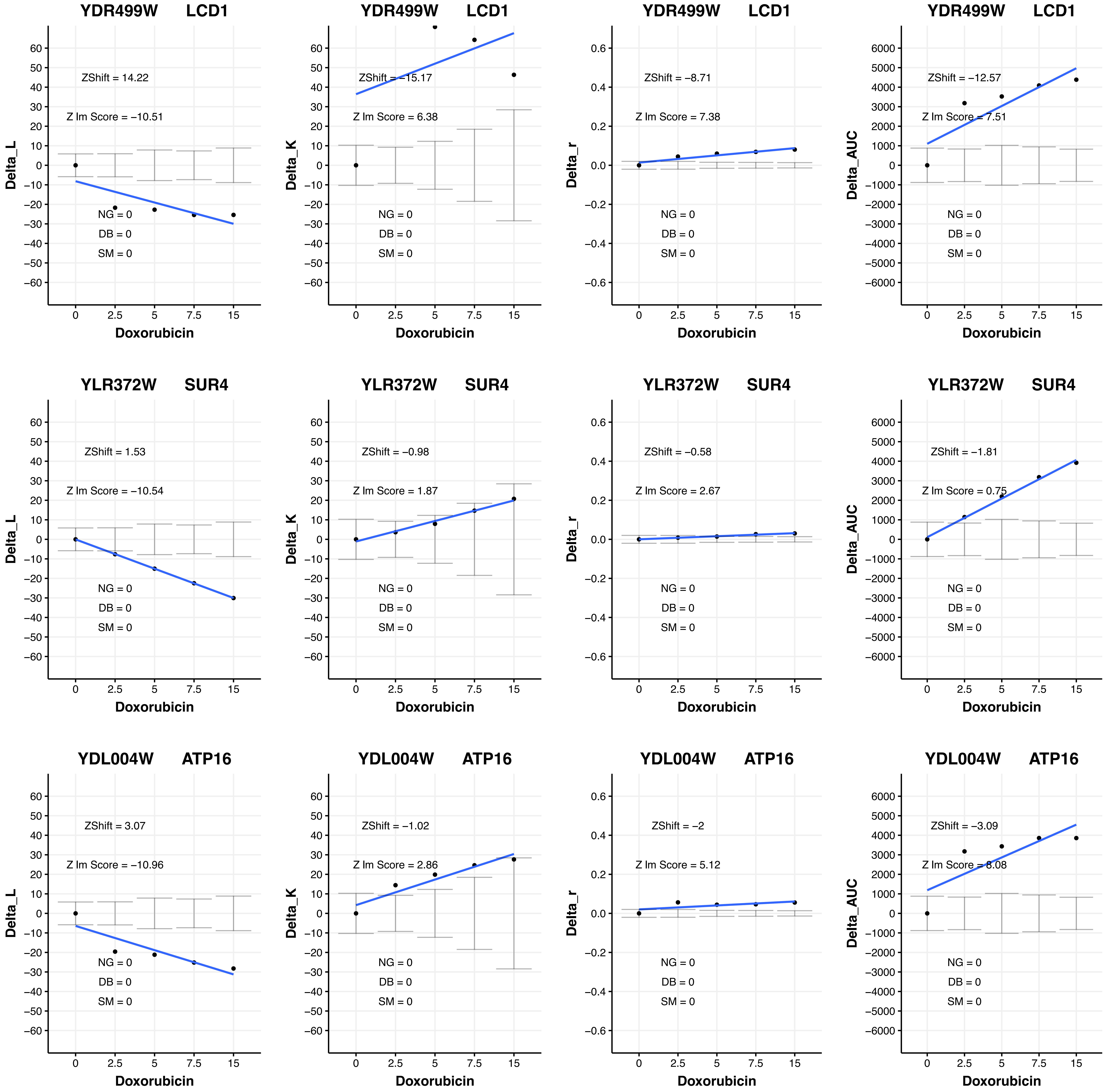
YDR499W LCD1
ZShift = 14.22
Z lm Score = −10.51
NG = 0
DB = 0
SM = 0
Delta_L
Doxorubicin
YDR499W LCD1
ZShift = −15.17
Z lm Score = 6.38
NG = 0
DB = 0
SM = 0
Delta_K
Doxorubicin
YDR499W LCD1
ZShift = −8.71
Z lm Score = 7.38
NG = 0
DB = 0
SM = 0
Delta_r
Doxorubicin
YDR499W LCD1
ZShift = −12.57
Z lm Score = 7.51
NG = 0
DB = 0
SM = 0
Delta_AUC
Doxorubicin
YLR372W SUR4
ZShift = 1.53
Z lm Score = −10.54
NG = 0
DB = 0
SM = 0
Delta_L
Doxorubicin
YLR372W SUR4
ZShift = −0.98
Z lm Score = 1.87
NG = 0
DB = 0
SM = 0
Delta_K
Doxorubicin
YLR372W SUR4
ZShift = −0.58
Z lm Score = 2.67
NG = 0
DB = 0
SM = 0
Delta_r
Doxorubicin
YLR372W SUR4
ZShift = −1.81
Z lm Score = 0.75
NG = 0
DB = 0
SM = 0
Delta_AUC
Doxorubicin
YDL004W ATP16
ZShift = 3.07
Z lm Score = −10.96
NG = 0
DB = 0
SM = 0
Delta_L
Doxorubicin
YDL004W ATP16
ZShift = −1.02
Z lm Score = 2.86
NG = 0
DB = 0
SM = 0
Delta_K
Doxorubicin
YDL004W ATP16
ZShift = −2
Z lm Score = 5.12
NG = 0
DB = 0
SM = 0
Delta_r
Doxorubicin
YDL004W ATP16
ZShift = −3.09
Z lm Score = 8.08
NG = 0
DB = 0
SM = 0
Delta_AUC
Doxorubicin

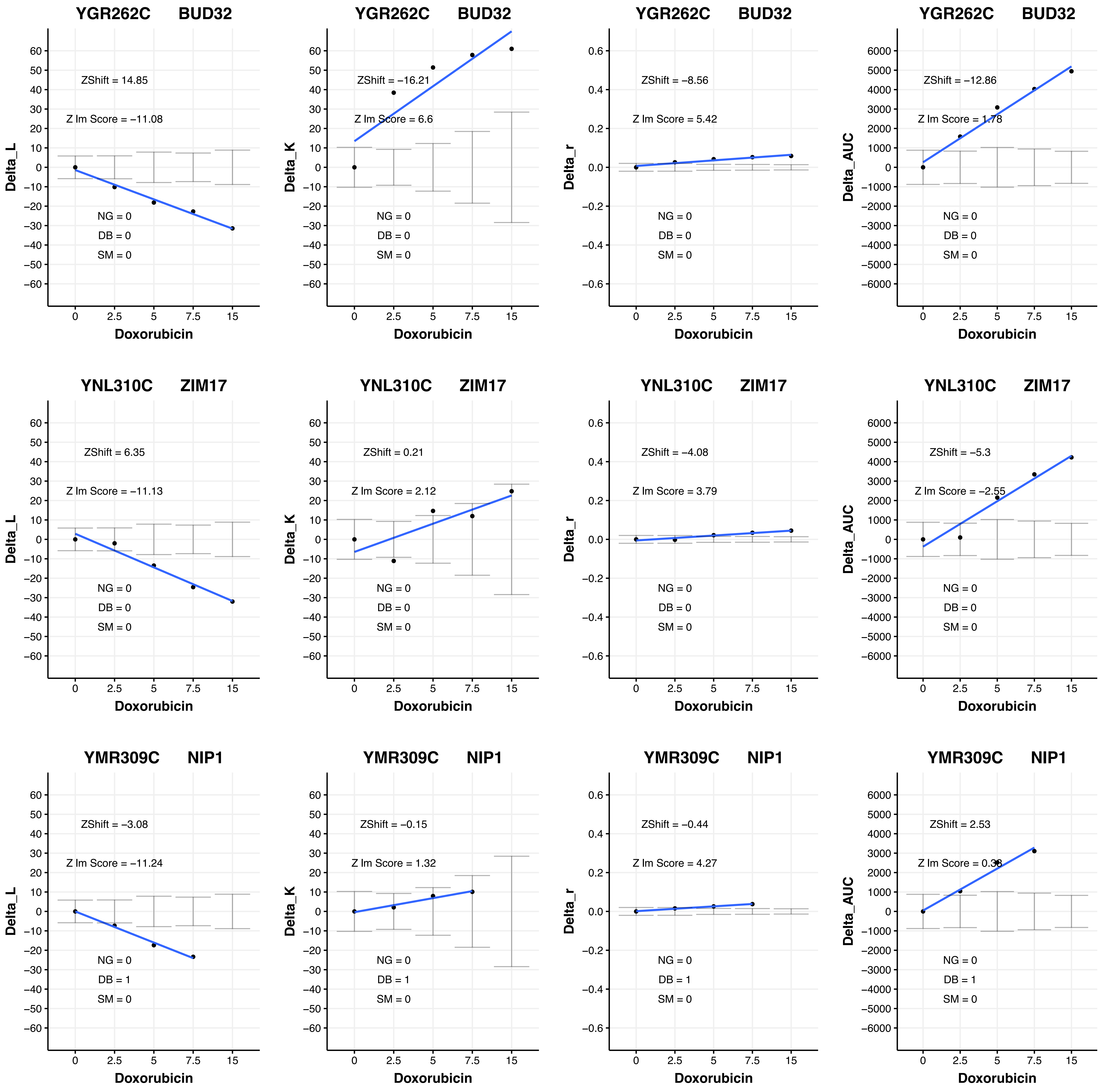
YGR262C BUD32
ZShift = 14.85
Z lm Score = −11.08
NG = 0
DB = 0
SM = 0
Delta_L
Doxorubicin
YGR262C BUD32
ZShift = −16.21
Z lm Score = 6.6
NG = 0
DB = 0
SM = 0
Delta_K
Doxorubicin
YGR262C BUD32
ZShift = −8.56
Z lm Score = 5.42
NG = 0
DB = 0
SM = 0
Delta_r
Doxorubicin
YGR262C BUD32
ZShift = −12.86
Z lm Score = 1.78
NG = 0
DB = 0
SM = 0
Delta_AUC
Doxorubicin
YNL310C ZIM17
ZShift = 6.35
Z lm Score = −11.13
NG = 0
DB = 0
SM = 0
Delta_L
Doxorubicin
YNL310C ZIM17
ZShift = 0.21
Z lm Score = 2.12
NG = 0
DB = 0
SM = 0
Delta_K
Doxorubicin
YNL310C ZIM17
ZShift = −4.08
Z lm Score = 3.79
NG = 0
DB = 0
SM = 0
Delta_r
Doxorubicin
YNL310C ZIM17
ZShift = −5.3
Z lm Score = −2.55
NG = 0
DB = 0
SM = 0
Delta_AUC
Doxorubicin
YMR309C NIP1
ZShift = −3.08
Z lm Score = −11.24
NG = 0
DB = 1
SM = 0
Delta_L
Doxorubicin
YMR309C NIP1
ZShift = −0.15
Z lm Score = 1.32
NG = 0
DB = 1
SM = 0
Delta_K
Doxorubicin
YMR309C NIP1
ZShift = −0.44
Z lm Score = 4.27
NG = 0
DB = 1
SM = 0
Delta_r
Doxorubicin
YMR309C NIP1
ZShift = 2.53
Z lm Score = 0.38
NG = 0
DB = 1
SM = 0
Delta_AUC
Doxorubicin

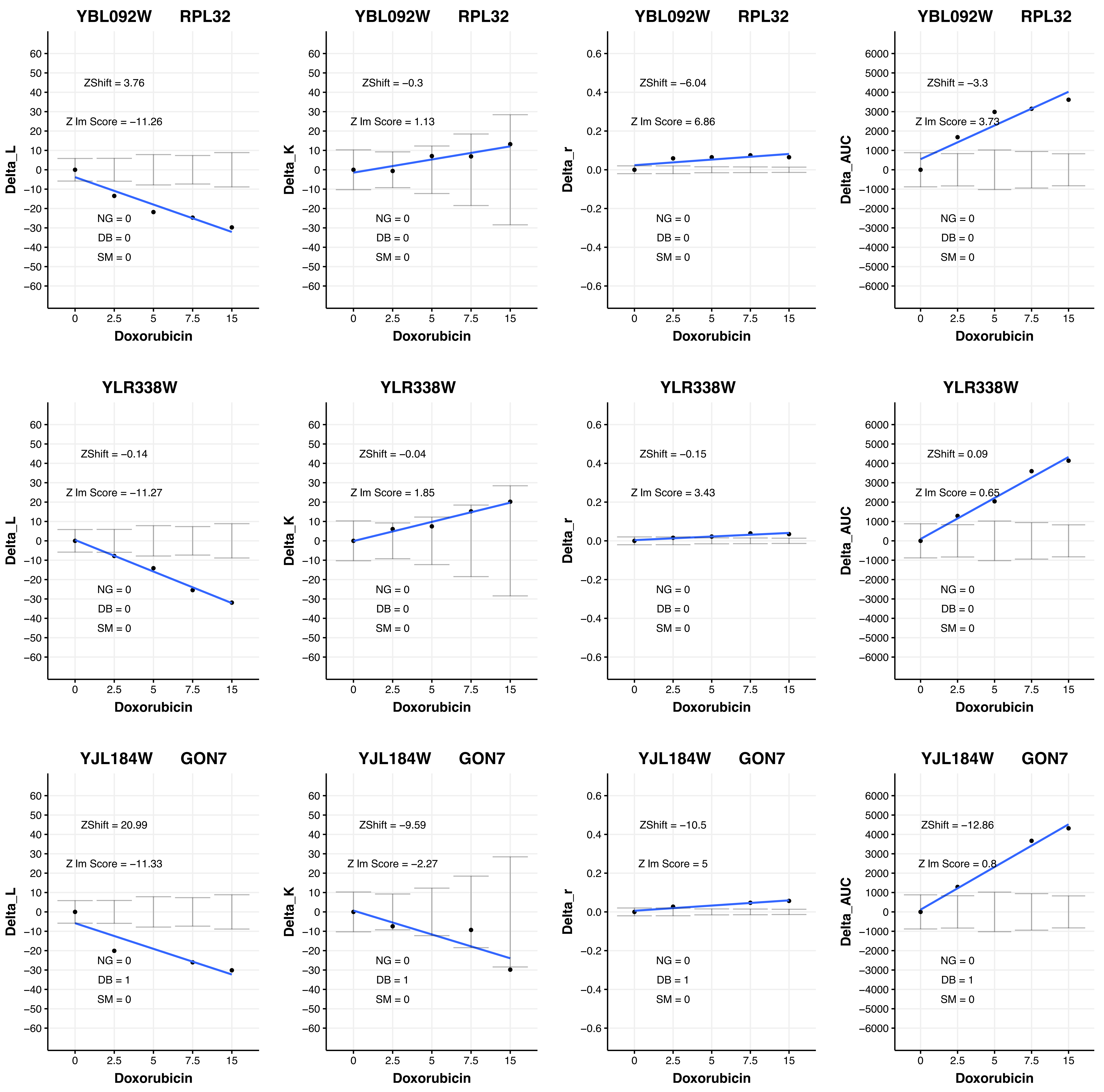
YBL092W RPL32
ZShift = 3.76
Z lm Score = −11.26
NG = 0
DB = 0
SM = 0
Delta_L
Doxorubicin
YBL092W RPL32
ZShift = −0.3
Z lm Score = 1.13
NG = 0
DB = 0
SM = 0
Delta_K
Doxorubicin
YBL092W RPL32
ZShift = −6.04
Z lm Score = 6.86
NG = 0
DB = 0
SM = 0
Delta_r
Doxorubicin
YBL092W RPL32
ZShift = −3.3
Z lm Score = 3.73
NG = 0
DB = 0
SM = 0
Delta_AUC
Doxorubicin
YLR338W
ZShift = −0.14
Z lm Score = −11.27
NG = 0
DB = 0
SM = 0
Delta_L
Doxorubicin
YLR338W
ZShift = −0.04
Z lm Score = 1.85
NG = 0
DB = 0
SM = 0
Delta_K
Doxorubicin
YLR338W
ZShift = −0.15
Z lm Score = 3.43
NG = 0
DB = 0
SM = 0
Delta_r
Doxorubicin
YLR338W
ZShift = 0.09
Z lm Score = 0.65
NG = 0
DB = 0
SM = 0
Delta_AUC
Doxorubicin
YJL184W GON7
ZShift = 20.99
Z lm Score = −11.33
NG = 0
DB = 1
SM = 0
Delta_L
Doxorubicin
YJL184W GON7
ZShift = −9.59
Z lm Score = −2.27
NG = 0
DB = 1
SM = 0
Delta_K
Doxorubicin
YJL184W GON7
ZShift = −10.5
Z lm Score = 5
NG = 0
DB = 1
SM = 0
Delta_r
Doxorubicin
YJL184W GON7
ZShift = −12.86
Z lm Score = 0.8
NG = 0
DB = 1
SM = 0
Delta_AUC
Doxorubicin

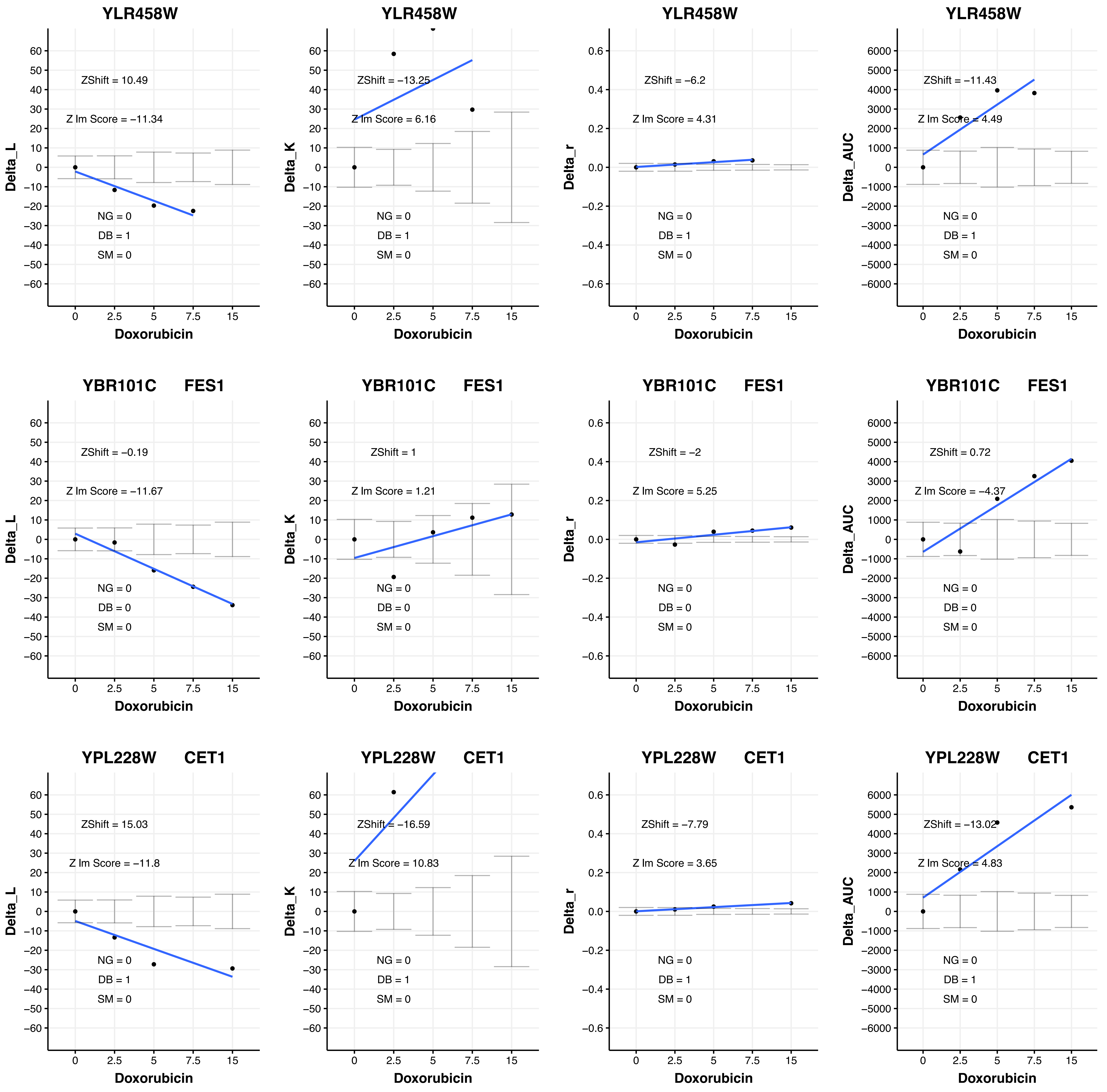
YLR458W
ZShift = 10.49
Z lm Score = −11.34
NG = 0
DB = 1
SM = 0
Delta_L
Doxorubicin
YLR458W
ZShift = −13.25
Z lm Score = 6.16
NG = 0
DB = 1
SM = 0
Delta_K
Doxorubicin
YLR458W
ZShift = −6.2
Z lm Score = 4.31
NG = 0
DB = 1
SM = 0
Delta_r
Doxorubicin
YLR458W
ZShift = −11.43
Z lm Score = 4.49
NG = 0
DB = 1
SM = 0
Delta_AUC
Doxorubicin
YBR101C FES1
ZShift = −0.19
Z lm Score = −11.67
NG = 0
DB = 0
SM = 0
Delta_L
Doxorubicin
YBR101C FES1
ZShift = 1
Z lm Score = 1.21
NG = 0
DB = 0
SM = 0
Delta_K
Doxorubicin
YBR101C FES1
ZShift = −2
Z lm Score = 5.25
NG = 0
DB = 0
SM = 0
Delta_r
Doxorubicin
YBR101C FES1
ZShift = 0.72
Z lm Score = −4.37
NG = 0
DB = 0
SM = 0
Delta_AUC
Doxorubicin
YPL228W CET1
ZShift = 15.03
Z lm Score = −11.8
NG = 0
DB = 1
SM = 0
Delta_L
Doxorubicin
YPL228W CET1
ZShift = −16.59
Z lm Score = 10.83
NG = 0
DB = 1
SM = 0
Delta_K
Doxorubicin
YPL228W CET1
ZShift = −7.79
Z lm Score = 3.65
NG = 0
DB = 1
SM = 0
Delta_r
Doxorubicin
YPL228W CET1
ZShift = −13.02
Z lm Score = 4.83
NG = 0
DB = 1
SM = 0
Delta_AUC
Doxorubicin

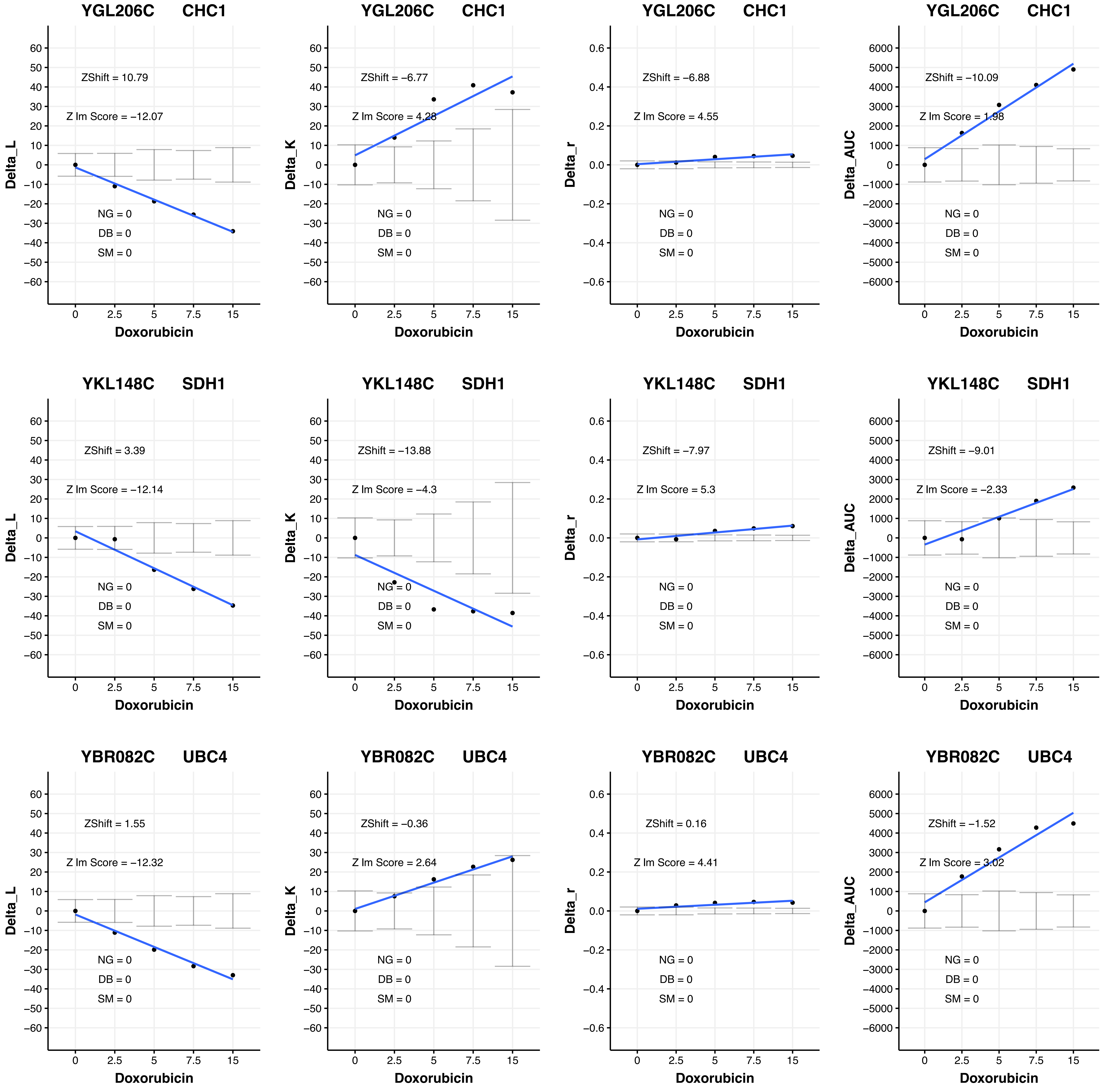
YGL206C CHC1
ZShift = 10.79
Z lm Score = −12.07
NG = 0
DB = 0
SM = 0
Delta_L
Doxorubicin
YGL206C CHC1
ZShift = −6.77
Z lm Score = 4.28
NG = 0
DB = 0
SM = 0
Delta_K
Doxorubicin
YGL206C CHC1
ZShift = −6.88
Z lm Score = 4.55
NG = 0
DB = 0
SM = 0
Delta_r
Doxorubicin
YGL206C CHC1
ZShift = −10.09
Z lm Score = 1.98
NG = 0
DB = 0
SM = 0
Delta_AUC
Doxorubicin
YKL148C SDH1
ZShift = 3.39
Z lm Score = −12.14
NG = 0
DB = 0
SM = 0
Delta_L
Doxorubicin
YKL148C SDH1
ZShift = −13.88
Z lm Score = −4.3
NG = 0
DB = 0
SM = 0
Delta_K
Doxorubicin
YKL148C SDH1
ZShift = −7.97
Z lm Score = 5.3
NG = 0
DB = 0
SM = 0
Delta_r
Doxorubicin
YKL148C SDH1
ZShift = −9.01
Z lm Score = −2.33
NG = 0
DB = 0
SM = 0
Delta_AUC
Doxorubicin
YBR082C UBC4
ZShift = 1.55
Z lm Score = −12.32
NG = 0
DB = 0
SM = 0
Delta_L
Doxorubicin
YBR082C UBC4
ZShift = −0.36
Z lm Score = 2.64
NG = 0
DB = 0
SM = 0
Delta_K
Doxorubicin
YBR082C UBC4
ZShift = 0.16
Z lm Score = 4.41
NG = 0
DB = 0
SM = 0
Delta_r
Doxorubicin
YBR082C UBC4
ZShift = −1.52
Z lm Score = 3.02
NG = 0
DB = 0
SM = 0
Delta_AUC
Doxorubicin

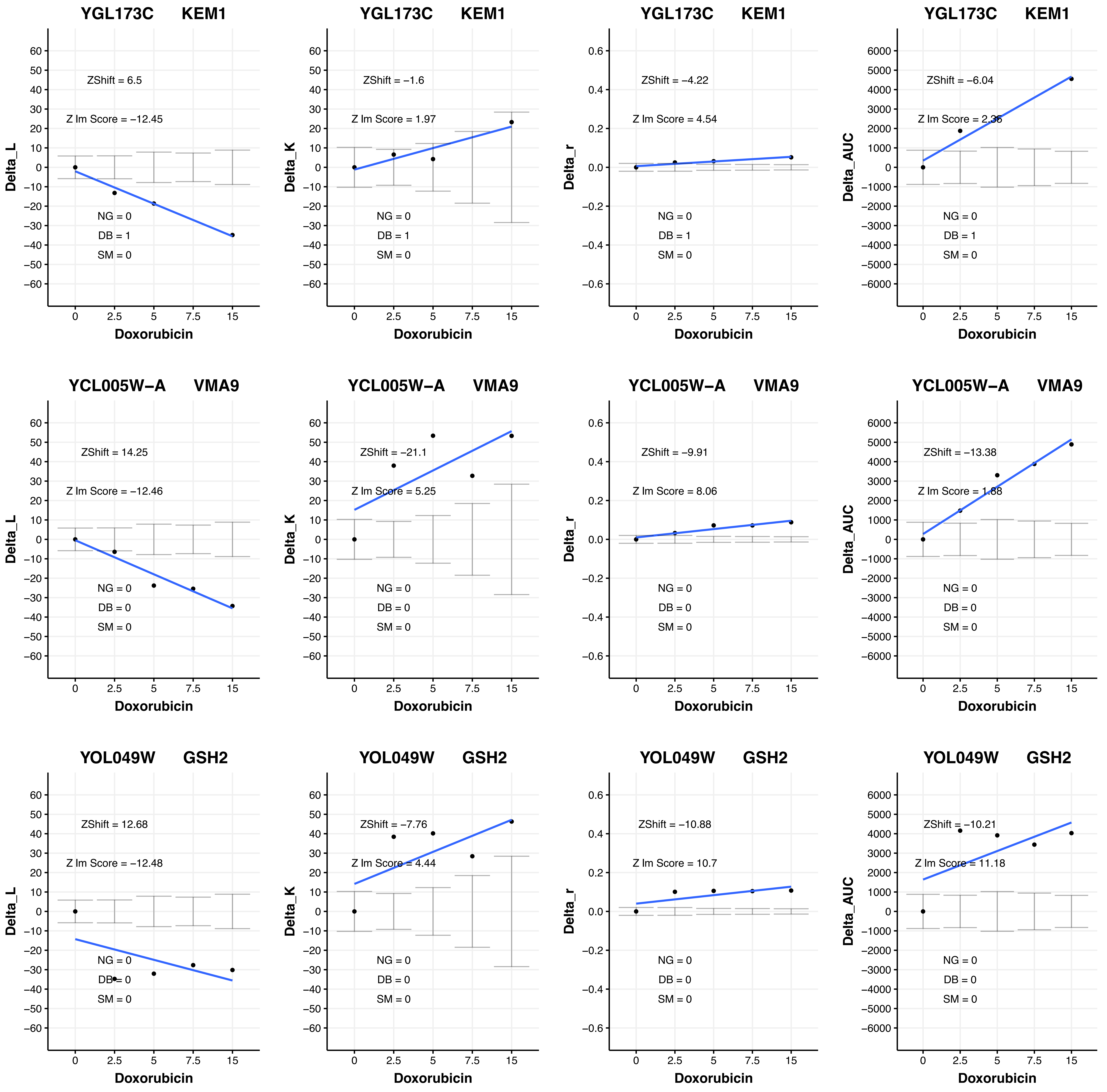
YGL173C KEM1
ZShift = 6.5
Z lm Score = −12.45
NG = 0
DB = 1
SM = 0
Delta_L
Doxorubicin
YGL173C KEM1
ZShift = −1.6
Z lm Score = 1.97
NG = 0
DB = 1
SM = 0
Delta_K
Doxorubicin
YGL173C KEM1
ZShift = −4.22
Z lm Score = 4.54
NG = 0
DB = 1
SM = 0
Delta_r
Doxorubicin
YGL173C KEM1
ZShift = −6.04
Z lm Score = 2.36
NG = 0
DB = 1
SM = 0
Delta_AUC
Doxorubicin
YCL005W−A VMA9
ZShift = 14.25
Z lm Score = −12.46
NG = 0
DB = 0
SM = 0
Delta_L
Doxorubicin
YCL005W−A VMA9
ZShift = −21.1
Z lm Score = 5.25
NG = 0
DB = 0
SM = 0
Delta_K
Doxorubicin
YCL005W−A VMA9
ZShift = −9.91
Z lm Score = 8.06
NG = 0
DB = 0
SM = 0
Delta_r
Doxorubicin
YCL005W−A VMA9
ZShift = −13.38
Z lm Score = 1.88
NG = 0
DB = 0
SM = 0
Delta_AUC
Doxorubicin
YOL049W GSH2
ZShift = 12.68
Z lm Score = −12.48
NG = 0
DB = 0
SM = 0
Delta_L
Doxorubicin
YOL049W GSH2
ZShift = −7.76
Z lm Score = 4.44
NG = 0
DB = 0
SM = 0
Delta_K
Doxorubicin
YOL049W GSH2
ZShift = −10.88
Z lm Score = 10.7
NG = 0
DB = 0
SM = 0
Delta_r
Doxorubicin
YOL049W GSH2
ZShift = −10.21
Z lm Score = 11.18
NG = 0
DB = 0
SM = 0
Delta_AUC
Doxorubicin

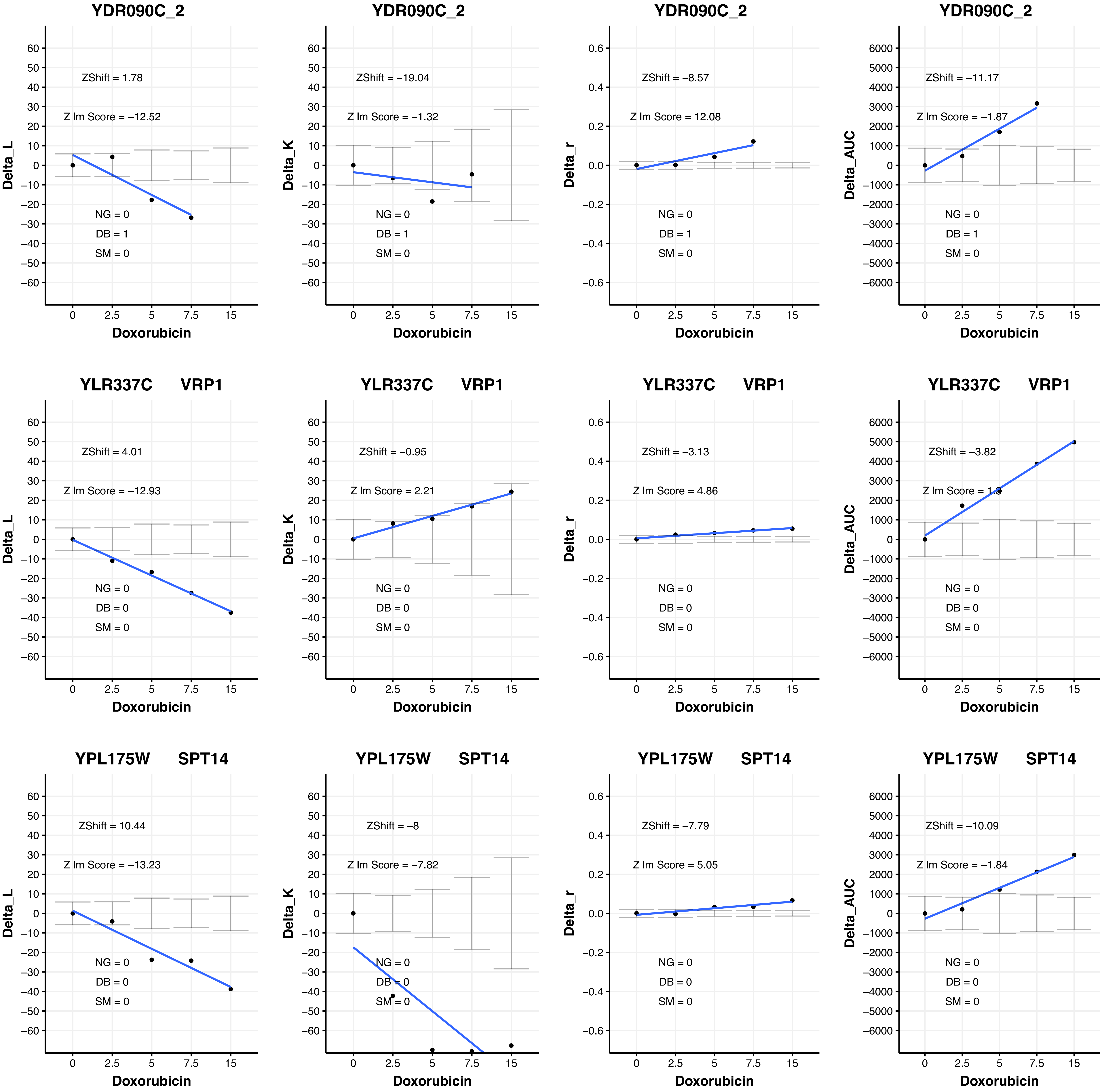
YDR090C_2
ZShift = 1.78
Z lm Score = −12.52
NG = 0
DB = 1
SM = 0
Delta_L
Doxorubicin
YDR090C_2
ZShift = −19.04
Z lm Score = −1.32
NG = 0
DB = 1
SM = 0
Delta_K
Doxorubicin
YDR090C_2
ZShift = −8.57
Z lm Score = 12.08
NG = 0
DB = 1
SM = 0
Delta_r
Doxorubicin
YDR090C_2
ZShift = −11.17
Z lm Score = −1.87
NG = 0
DB = 1
SM = 0
Delta_AUC
Doxorubicin
YLR337C VRP1
ZShift = 4.01
Z lm Score = −12.93
NG = 0
DB = 0
SM = 0
Delta_L
Doxorubicin
YLR337C VRP1
ZShift = −0.95
Z lm Score = 2.21
NG = 0
DB = 0
SM = 0
Delta_K
Doxorubicin
YLR337C VRP1
ZShift = −3.13
Z lm Score = 4.86
NG = 0
DB = 0
SM = 0
Delta_r
Doxorubicin
YLR337C VRP1
ZShift = −3.82
NG = 0
DB = 0
SM = 0
Delta_AUC
Doxorubicin
YPL175W SPT14
ZShift = 10.44
Z lm Score = −13.23
NG = 0
DB = 0
SM = 0
Delta_L
Doxorubicin
YPL175W SPT14
ZShift = −8
Z lm Score = −7.82
NG = 0
DB = 0
SM = 0
Delta_K
Doxorubicin
YPL175W SPT14
ZShift = −7.79
Z lm Score = 5.05
NG = 0
DB = 0
SM = 0
Delta_r
Doxorubicin
YPL175W SPT14
ZShift = −10.09
Z lm Score = −1.84
NG = 0
DB = 0
SM = 0
Delta_AUC
Doxorubicin

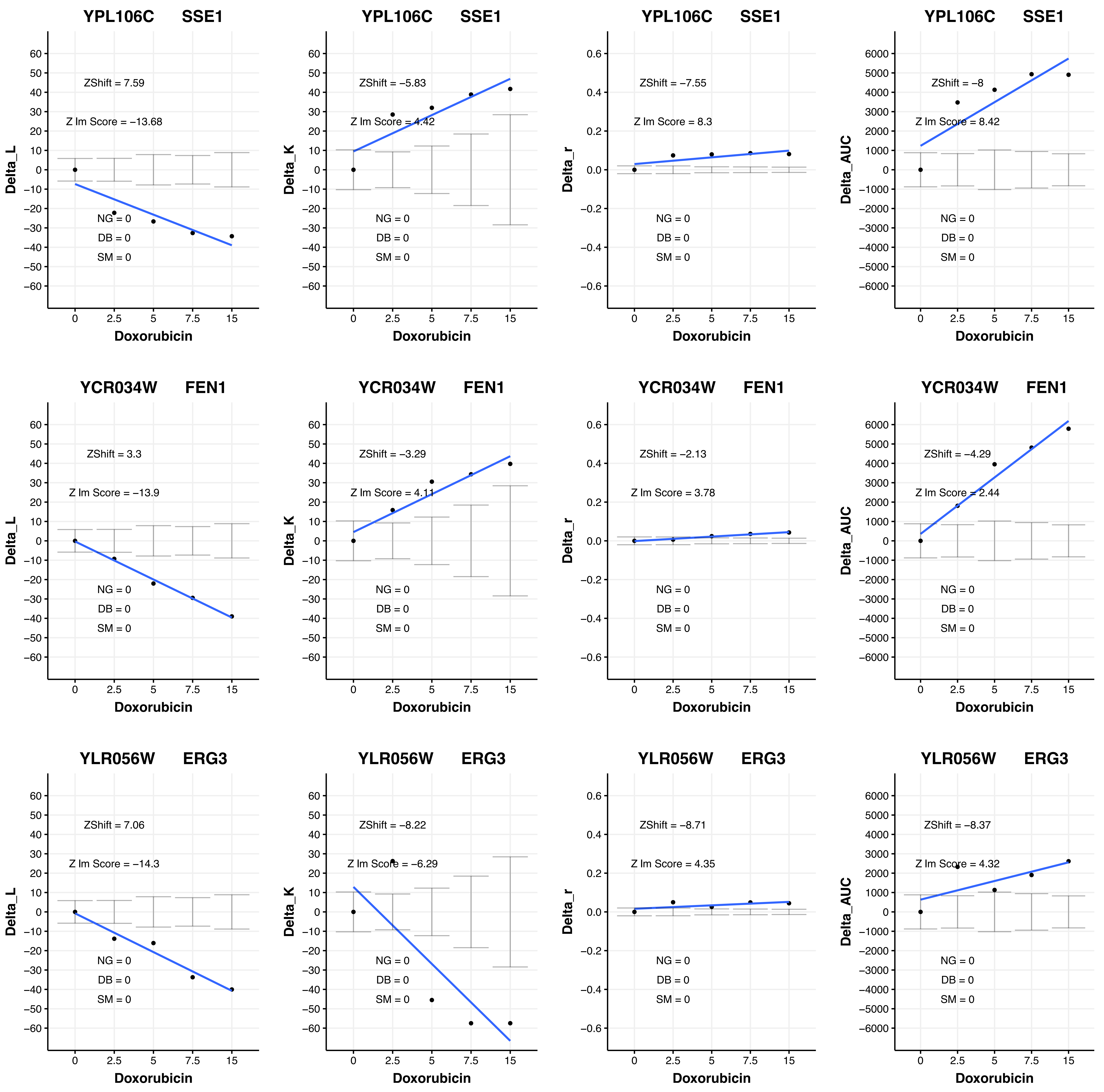

YPL106C SSE1
ZShift = 7.59
Z lm Score = −13.68
NG = 0
DB = 0
SM = 0
Delta_L
Doxorubicin
YPL106C SSE1
ZShift = −5.83
Z lm Score = 4.42
NG = 0
DB = 0
SM = 0
Delta_K
Doxorubicin
YPL106C SSE1
ZShift = −7.55
Z lm Score = 8.3
NG = 0
DB = 0
SM = 0
Delta_r
Doxorubicin
YPL106C SSE1
ZShift = −8
Z lm Score = 8.42
NG = 0
DB = 0
SM = 0
Delta_AUC
Doxorubicin
YCR034W FEN1
ZShift = 3.3
Z lm Score = −13.9
NG = 0
DB = 0
SM = 0
Delta_L
Doxorubicin
YCR034W FEN1
ZShift = −3.29
Z lm Score = 4.11
NG = 0
DB = 0
SM = 0
Delta_K
Doxorubicin
YCR034W FEN1
ZShift = −2.13
Z lm Score = 3.78
NG = 0
DB = 0
SM = 0
Delta_r
Doxorubicin
YCR034W FEN1
ZShift = −4.29
Z lm Score = 2.44
NG = 0
DB = 0
SM = 0
Delta_AUC
Doxorubicin
YLR056W ERG3
ZShift = 7.06
Z lm Score = −14.3
NG = 0
DB = 0
SM = 0
Delta_L
Doxorubicin
YLR056W ERG3
ZShift = −8.22
Z lm Score = −6.29
NG = 0
DB = 0
SM = 0
Delta_K
Doxorubicin
YLR056W ERG3
ZShift = −8.71
Z lm Score = 4.35
NG = 0
DB = 0
SM = 0
Delta_r
Doxorubicin
YLR056W ERG3
ZShift = −8.37
Z lm Score = 4.32
NG = 0
DB = 0
SM = 0
Delta_AUC
Doxorubicin

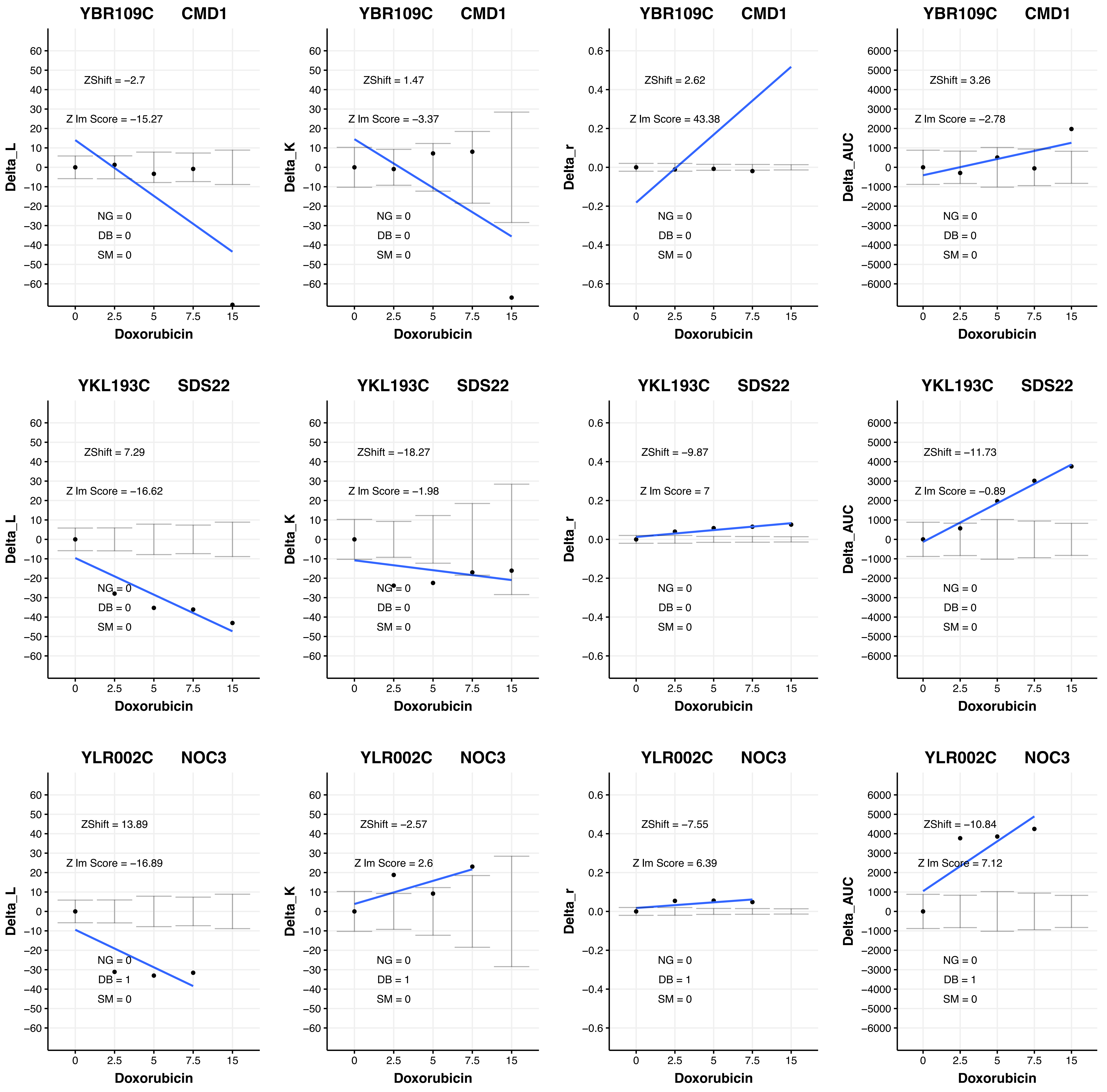
YBR109C CMD1
ZShift = −2.7
Z lm Score = −15.27
NG = 0
DB = 0
SM = 0
Delta_L
Doxorubicin
YBR109C CMD1
ZShift = 1.47
Z lm Score = −3.37
NG = 0
DB = 0
SM = 0
Delta_K
Doxorubicin
YBR109C CMD1
ZShift = 2.62
Z lm Score = 43.38
NG = 0
DB = 0
SM = 0
Delta_r
Doxorubicin
YBR109C CMD1
ZShift = 3.26
Z lm Score = −2.78
NG = 0
DB = 0
SM = 0
Delta_AUC
Doxorubicin
YKL193C SDS22
ZShift = 7.29
Z lm Score = −16.62
NG = 0
DB = 0
SM = 0
Delta_L
Doxorubicin
YKL193C SDS22
ZShift = −18.27
Z lm Score = −1.98
NG = 0
DB = 0
SM = 0
Delta_K
Doxorubicin
YKL193C SDS22
ZShift = −9.87
Z lm Score = 7
NG = 0
DB = 0
SM = 0
Delta_r
Doxorubicin
YKL193C SDS22
ZShift = −11.73
Z lm Score = −0.89
NG = 0
DB = 0
SM = 0
Delta_AUC
Doxorubicin
YLR002C NOC3
ZShift = 13.89
Z lm Score = −16.89
NG = 0
DB = 1
SM = 0
Delta_L
Doxorubicin
YLR002C NOC3
ZShift = −2.57
Z lm Score = 2.6
NG = 0
DB = 1
SM = 0
Delta_K
Doxorubicin
YLR002C NOC3
ZShift = −7.55
Z lm Score = 6.39
NG = 0
DB = 1
SM = 0
Delta_r
Doxorubicin
YLR002C NOC3
ZShift = −10.84
Z lm Score = 7.12
NG = 0
DB = 1
SM = 0
Delta_AUC
Doxorubicin

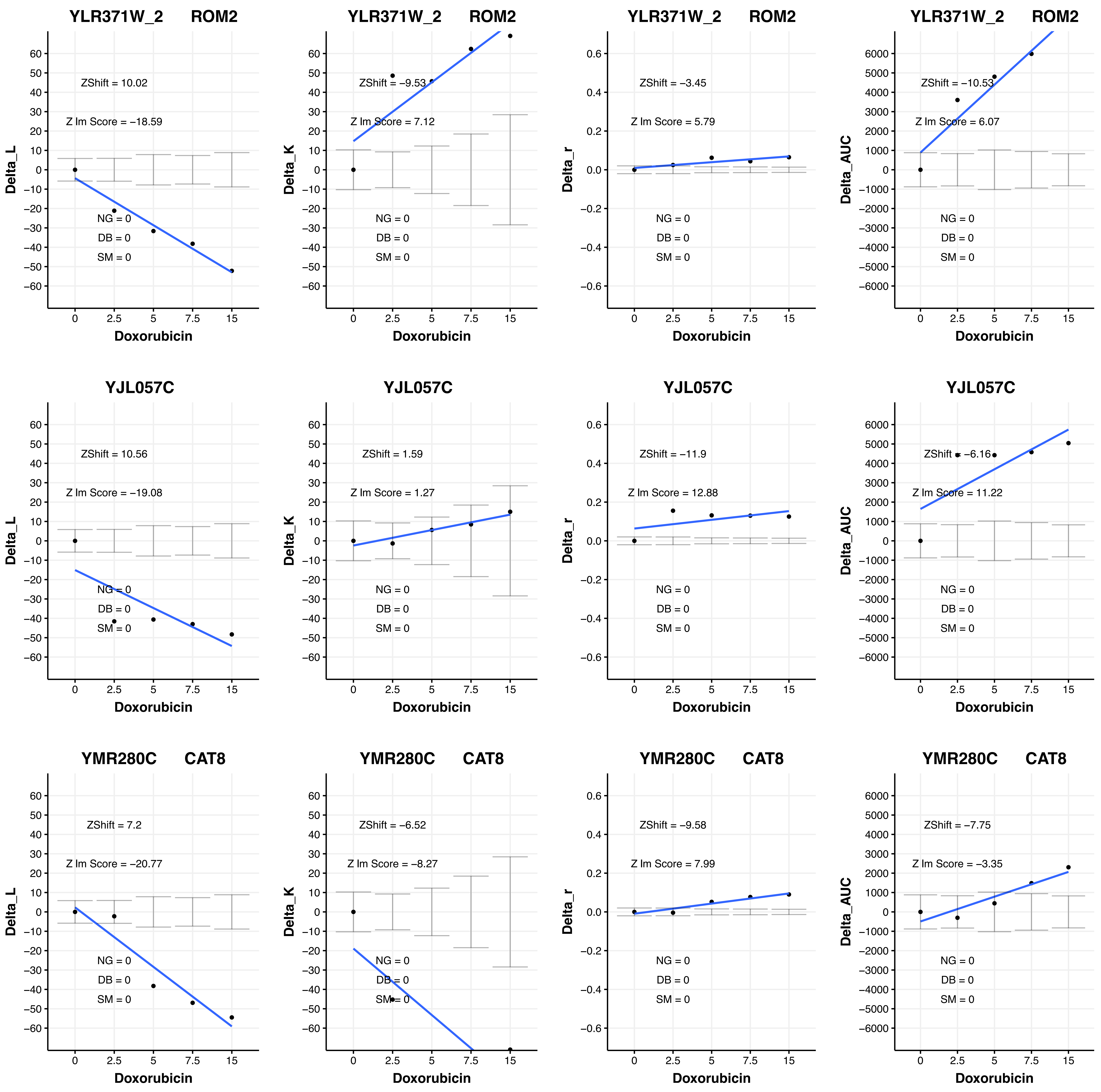
YLR371W_2 ROM2
ZShift = 10.02
Z lm Score = −18.59
NG = 0
DB = 0
SM = 0
Delta_L
Doxorubicin
YLR371W_2 ROM2
ZShift = −9.53
Z lm Score = 7.12
NG = 0
DB = 0
SM = 0
Delta_K
Doxorubicin
YLR371W_2 ROM2
ZShift = −3.45
Z lm Score = 5.79
NG = 0
DB = 0
SM = 0
Delta_r
Doxorubicin
YLR371W_2 ROM2
ZShift = −10.53
Z lm Score = 6.07
NG = 0
DB = 0
SM = 0
Delta_AUC
Doxorubicin
YJL057C
ZShift = 10.56
Z lm Score = −19.08
NG = 0
DB = 0
SM = 0
Delta_L
Doxorubicin
YJL057C
ZShift = 1.59
Z lm Score = 1.27
NG = 0
DB = 0
SM = 0
Delta_K
Doxorubicin
YJL057C
ZShift = −11.9
Z lm Score = 12.88
NG = 0
DB = 0
SM = 0
Delta_r
Doxorubicin
YJL057C
ZShift = −6.16
Z lm Score = 11.22
NG = 0
DB = 0
SM = 0
Delta_AUC
Doxorubicin
YMR280C CAT8
ZShift = 7.2
Z lm Score = −20.77
NG = 0
DB = 0
SM = 0
Delta_L
Doxorubicin
YMR280C CAT8
ZShift = −6.52
Z lm Score = −8.27
NG = 0
DB = 0
SM = 0
Delta_K
Doxorubicin
YMR280C CAT8
ZShift = −9.58
Z lm Score = 7.99
NG = 0
DB = 0
SM = 0
Delta_r
Doxorubicin
YMR280C CAT8
ZShift = −7.75
Z lm Score = −3.35
NG = 0
DB = 0
SM = 0
Delta_AUC
Doxorubicin
