## Additional File 3 for "A yeast phenomic model for the influence of Warburg metabolism on genetic buffering of doxorubicin": C - InteractionPlots_Doxo_HLEG_V.pdf

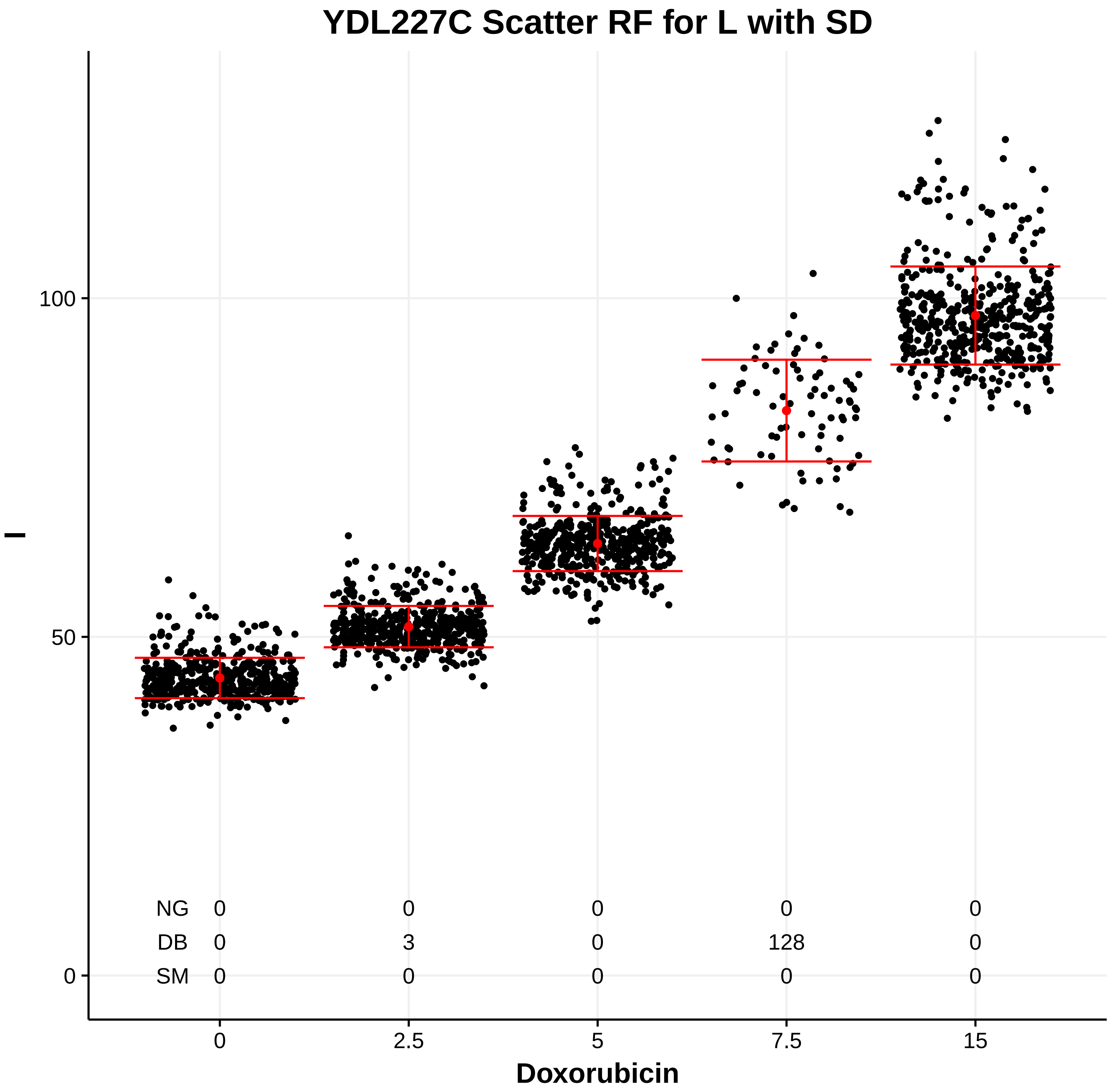
YDL227C Scatter RF for L with SD
l
Doxorubicin
0 2.5 5 7.5 15
NG 0 0 0 0 0
DB 0 3 0 128 0
SM 0 0 0 0 0

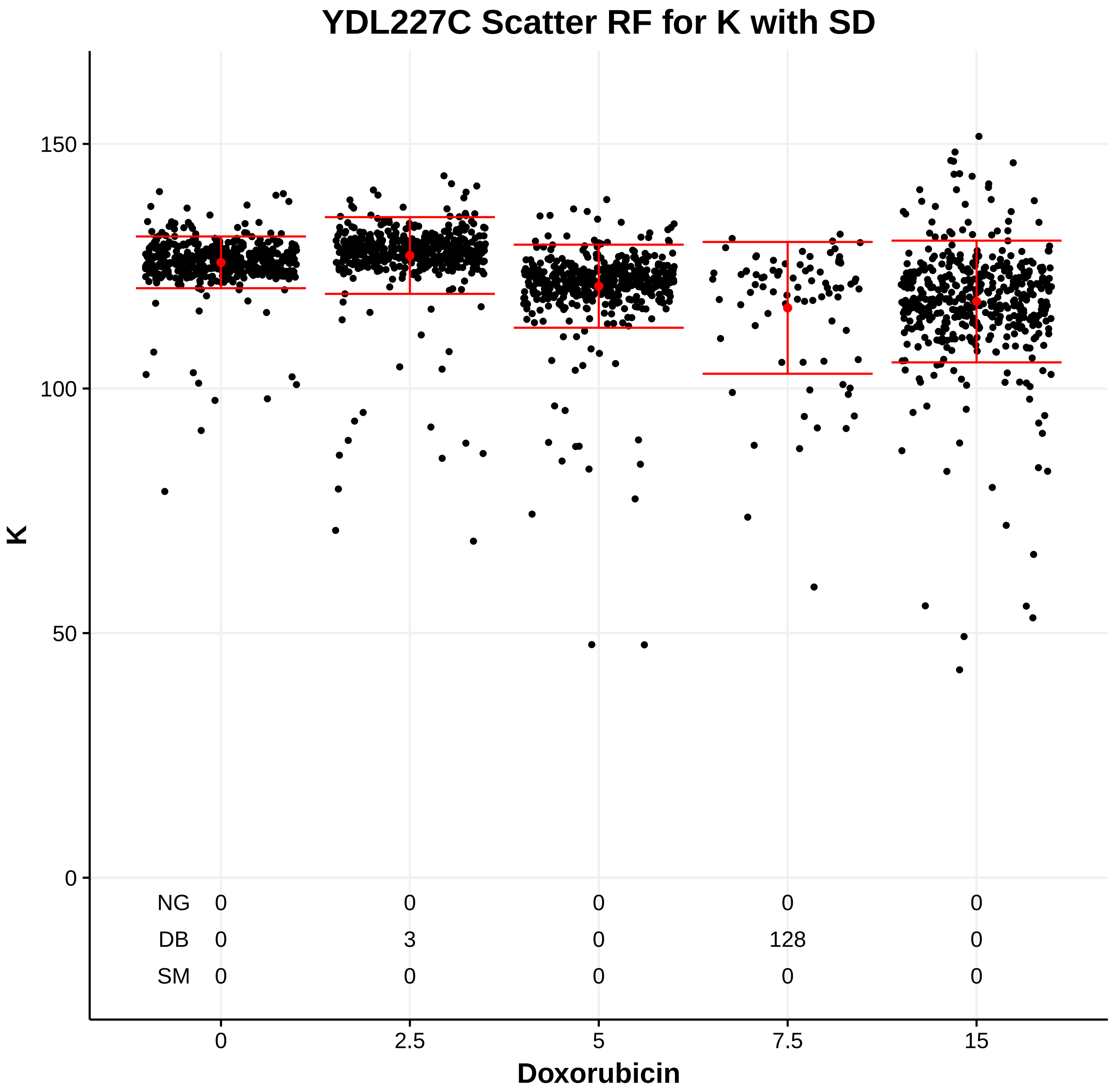
YDL227C Scatter RF for K with SD
K
Doxorubicin
0 2.5 5 7.5 15
NG 0 0 0 0 0
DB 0 3 0 128 0
SM 0 0 0 0 0

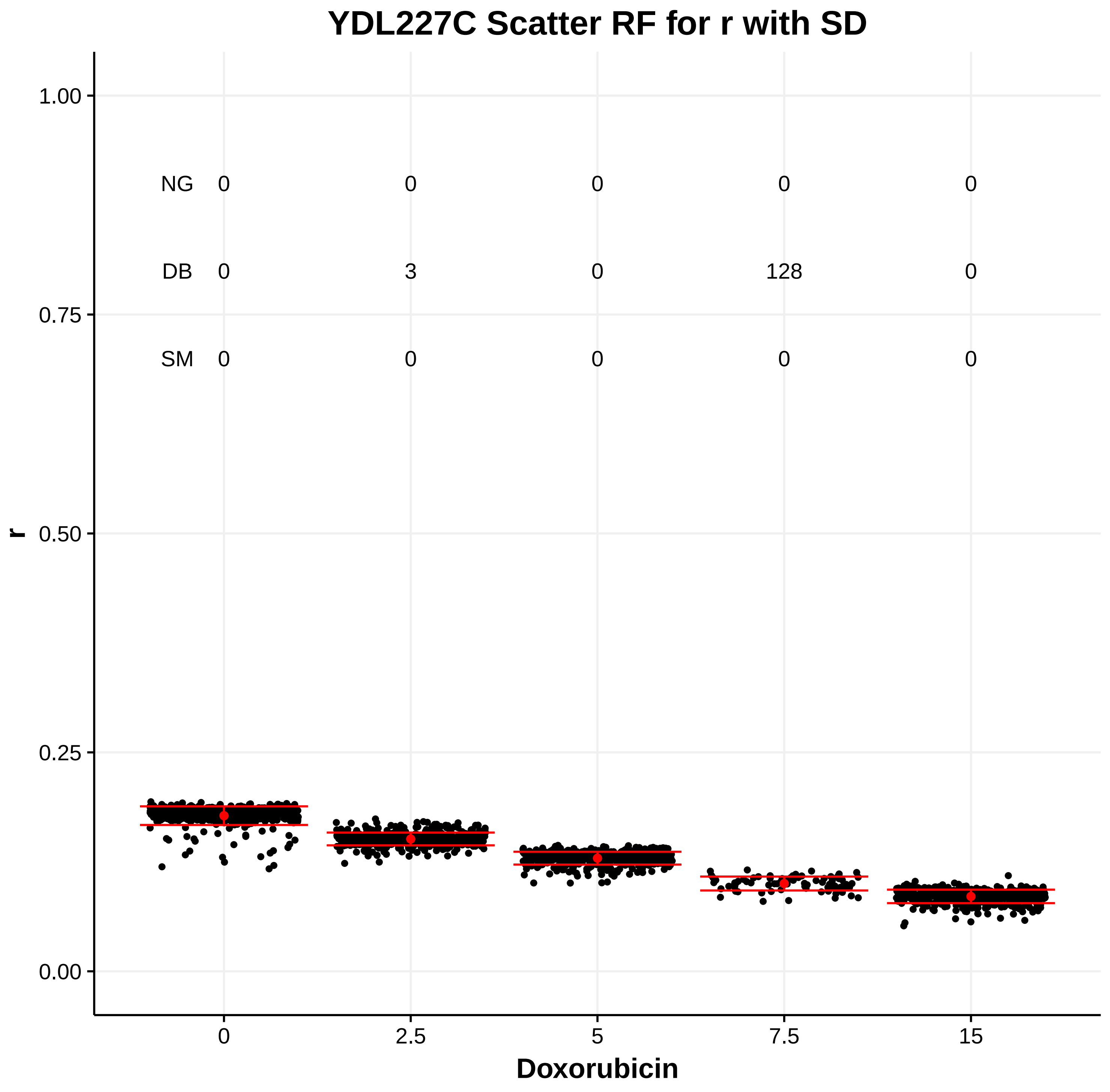
YDL227C Scatter RF for r with SD
r
Doxorubicin
0 2.5 5 7.5 15
NG 0 0 0 0 0
DB 0 3 0 128 0
SM 0 0 0 0 0

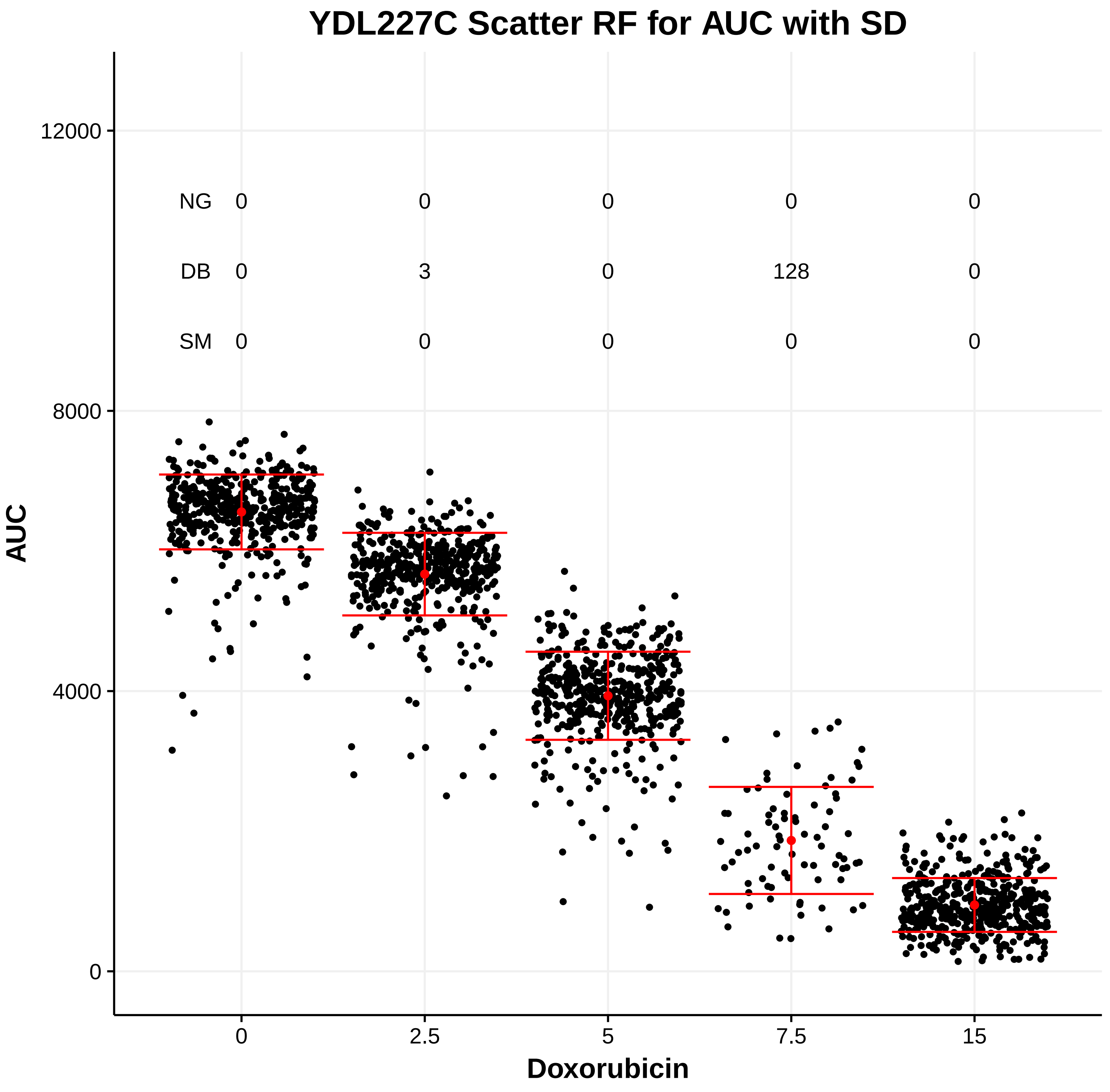
YDL227C Scatter RF for AUC with SD
AUC
Doxorubicin
0 2.5 5 7.5 15
NG 0 0 0 0 0
DB 0 3 0 128 0
SM 0 0 0 0 0

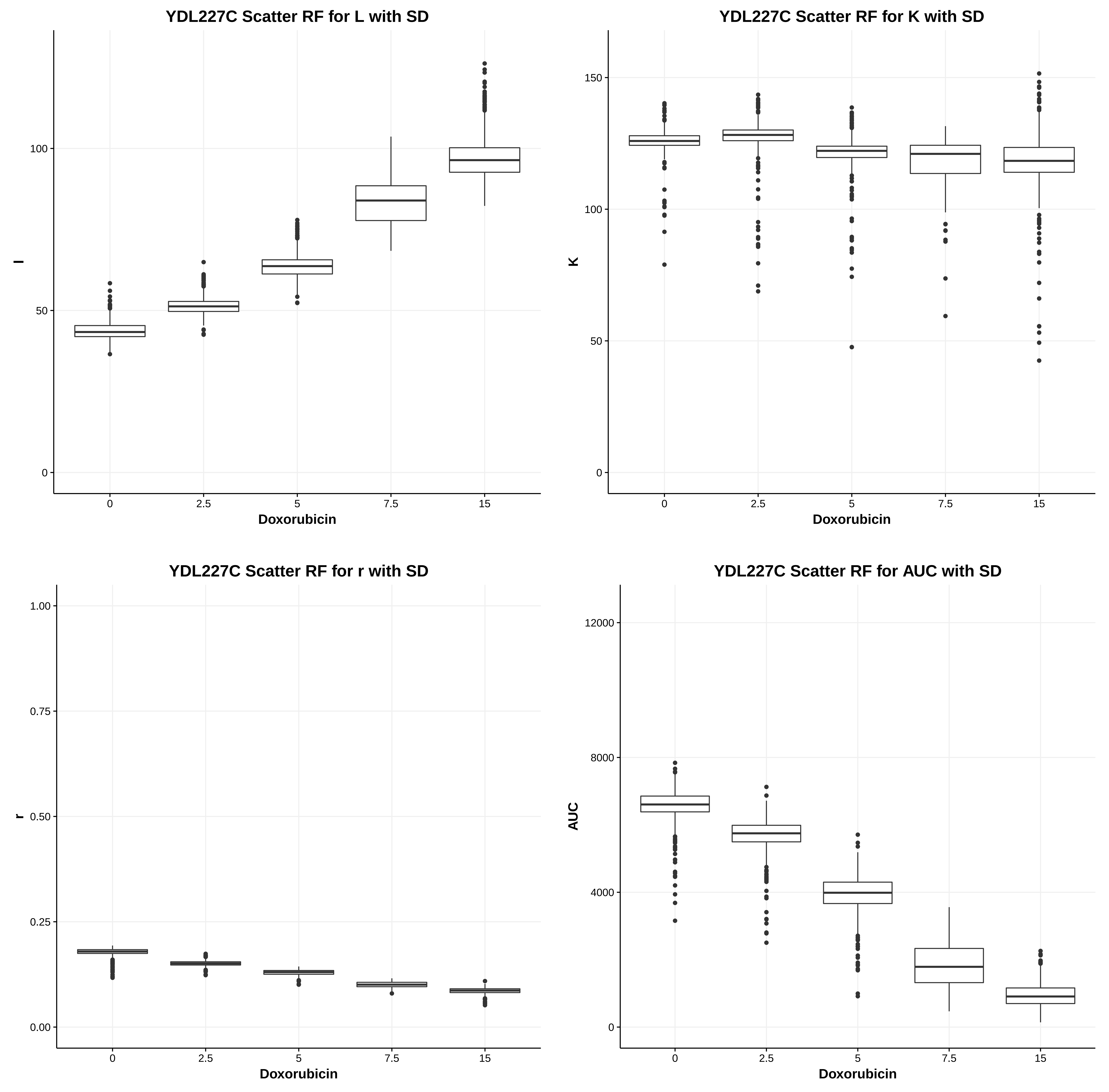

YDL227C Scatter RF for L with SD
l
Doxorubicin
0
50
100
0
2.5
5
7.5
15
YDL227C Scatter RF for K with SD
K
Doxorubicin
0
50
100
150
YDL227C Scatter RF for r with SD
r
Doxorubicin
0.00
0.25
0.50
0.75
1.00
YDL227C Scatter RF for AUC with SD
AUC
Doxorubicin
0
4000
8000
12000

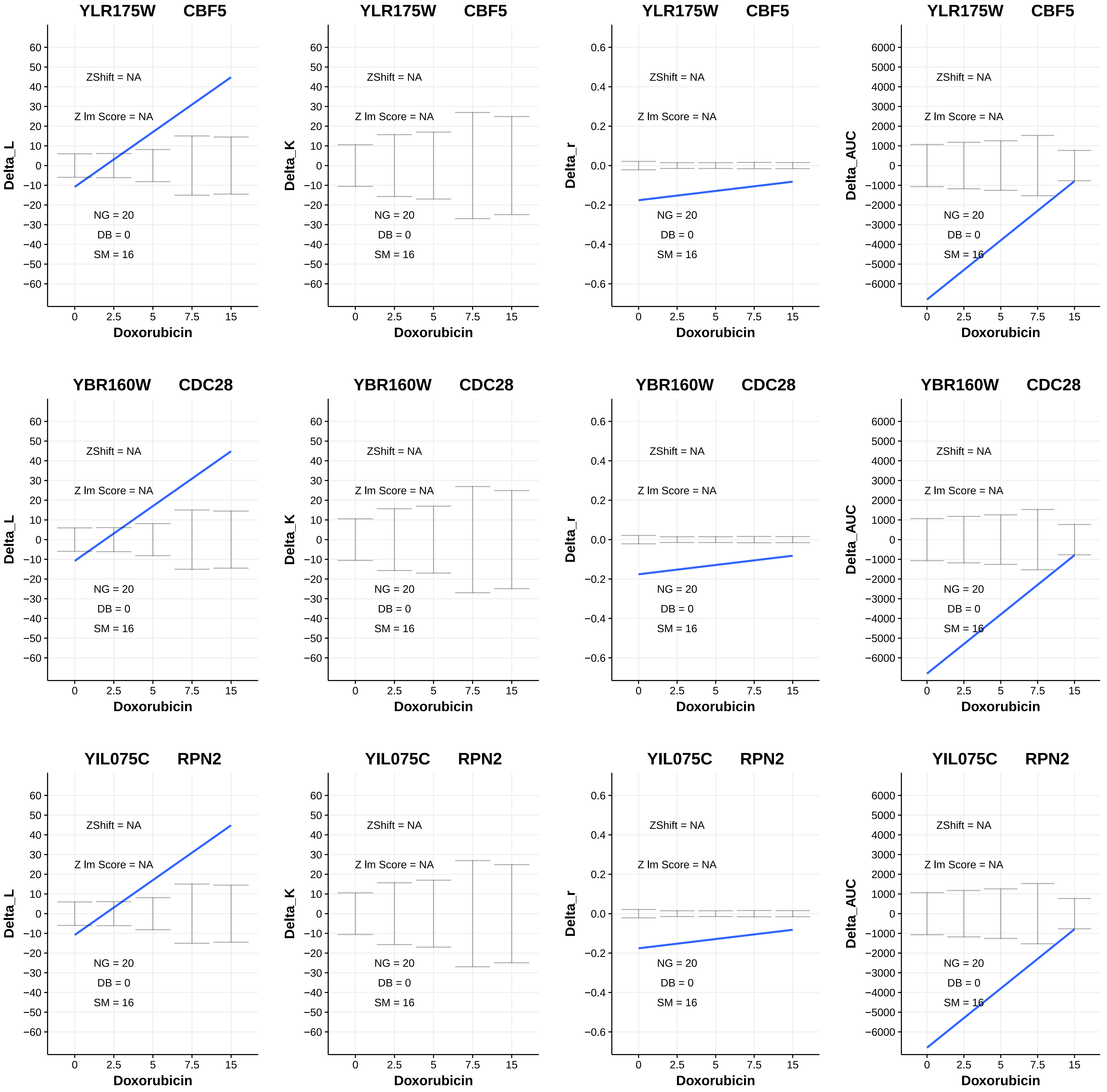
YLR175W CBF5
ZShift = NA
Z lm Score = NA
NG = 20
DB = 0
SM = 16
Delta_L
Delta_K
Delta_r
Delta_AUC
Doxorubicin
0
2.5
5
7.5
15
YBR160W CDC28
ZShift = NA
Z lm Score = NA
NG = 20
DB = 0
SM = 16
Delta_L
Delta_K
Delta_r
Delta_AUC
Doxorubicin
YIL075C RPN2
ZShift = NA
Z lm Score = NA
NG = 20
DB = 0
SM = 16
Delta_L
Delta_K
Delta_r
Delta_AUC
Doxorubicin

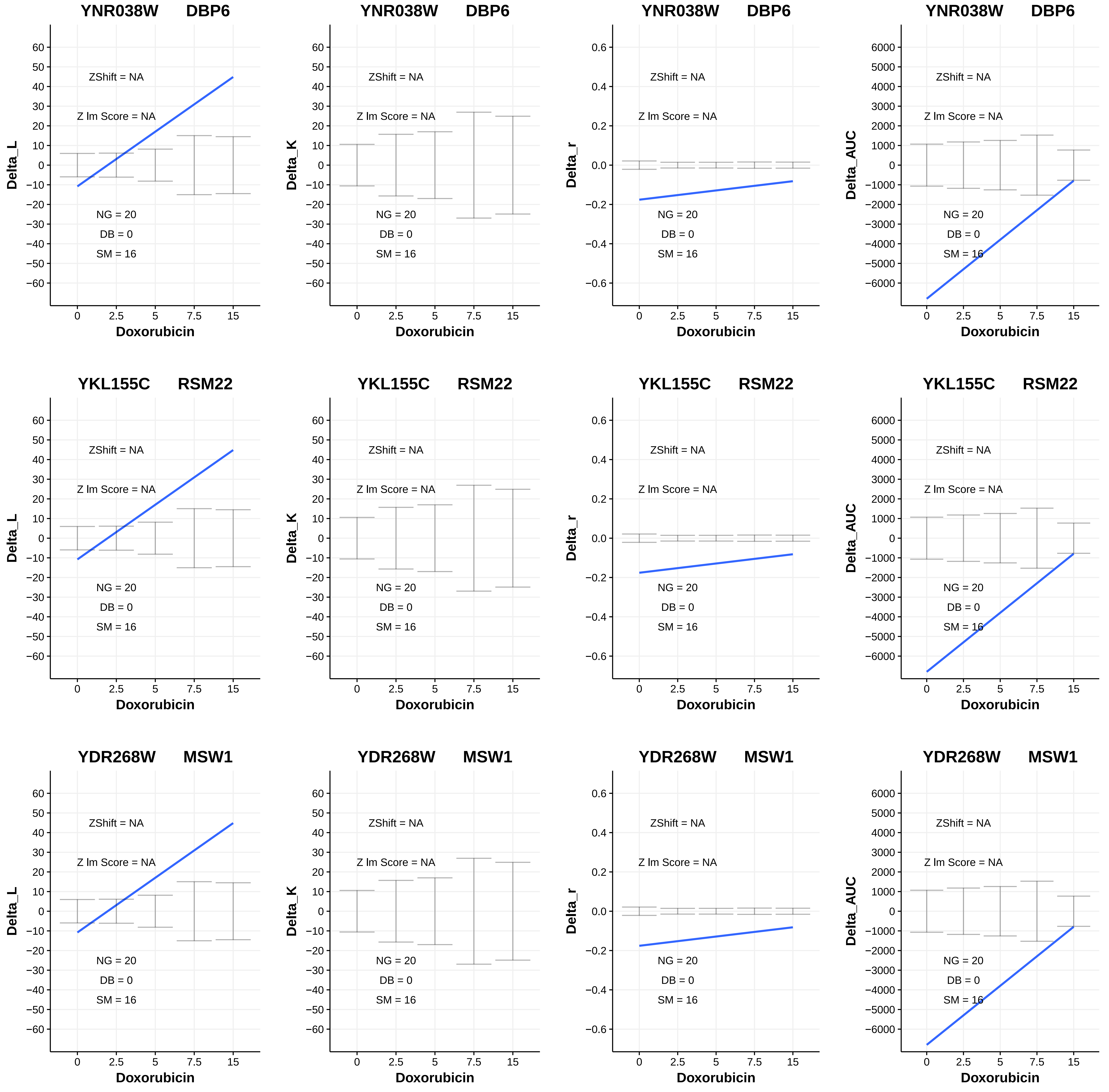
YNR038W DBP6
ZShift = NA
Z lm Score = NA
NG = 20
DB = 0
SM = 16
Delta_L
Doxorubicin
0 2.5 5 7.5 15
60 50 40 30 20 10 0 −10 −20 −30 −40 −50 −60
YNR038W DBP6
ZShift = NA
Z lm Score = NA
NG = 20
DB = 0
SM = 16
Delta_K
Doxorubicin
YNR038W DBP6
ZShift = NA
Z lm Score = NA
NG = 20
DB = 0
SM = 16
Delta_r
Doxorubicin
0.6 0.4 0.2 0.0 −0.2 −0.4 −0.6
YNR038W DBP6
ZShift = NA
Z lm Score = NA
NG = 20
DB = 0
SM = 16
Delta_AUC
Doxorubicin
6000 5000 4000 3000 2000 1000 0 −1000 −2000 −3000 −4000 −5000 −6000
YKL155C RSM22
ZShift = NA
Z lm Score = NA
NG = 20
DB = 0
SM = 16
Delta_L
Doxorubicin
YKL155C RSM22
ZShift = NA
Z lm Score = NA
NG = 20
DB = 0
SM = 16
Delta_K
Doxorubicin
YKL155C RSM22
ZShift = NA
Z lm Score = NA
NG = 20
DB = 0
SM = 16
Delta_r
Doxorubicin
YKL155C RSM22
ZShift = NA
Z lm Score = NA
NG = 20
DB = 0
SM = 16
Delta_AUC
Doxorubicin
YDR268W MSW1
ZShift = NA
Z lm Score = NA
NG = 20
DB = 0
SM = 16
Delta_L
Doxorubicin
YDR268W MSW1
ZShift = NA
Z lm Score = NA
NG = 20
DB = 0
SM = 16
Delta_K
Doxorubicin
YDR268W MSW1
ZShift = NA
Z lm Score = NA
NG = 20
DB = 0
SM = 16
Delta_r
Doxorubicin
YDR268W MSW1
ZShift = NA
Z lm Score = NA
NG = 20
DB = 0
SM = 16
Delta_AUC
Doxorubicin

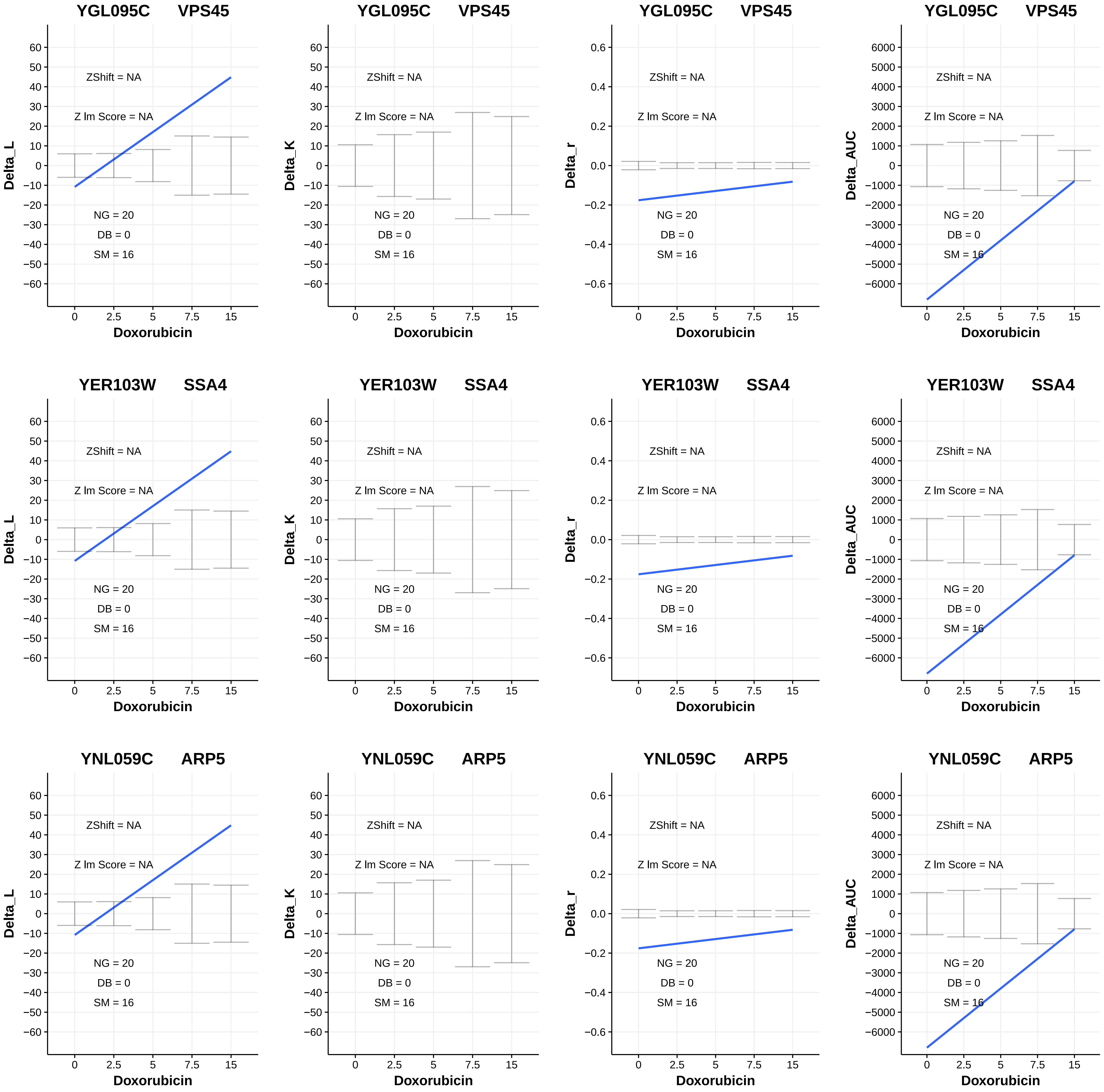
YGL095C VPS45
ZShift = NA
Z lm Score = NA
NG = 20
DB = 0
SM = 16
Delta_L
Delta_K
Delta_r
Delta_AUC
Doxorubicin
0
2.5
5
7.5
15
YER103W SSA4
ZShift = NA
Z lm Score = NA
NG = 20
DB = 0
SM = 16
YNL059C ARP5
ZShift = NA
Z lm Score = NA
NG = 20
DB = 0
SM = 16

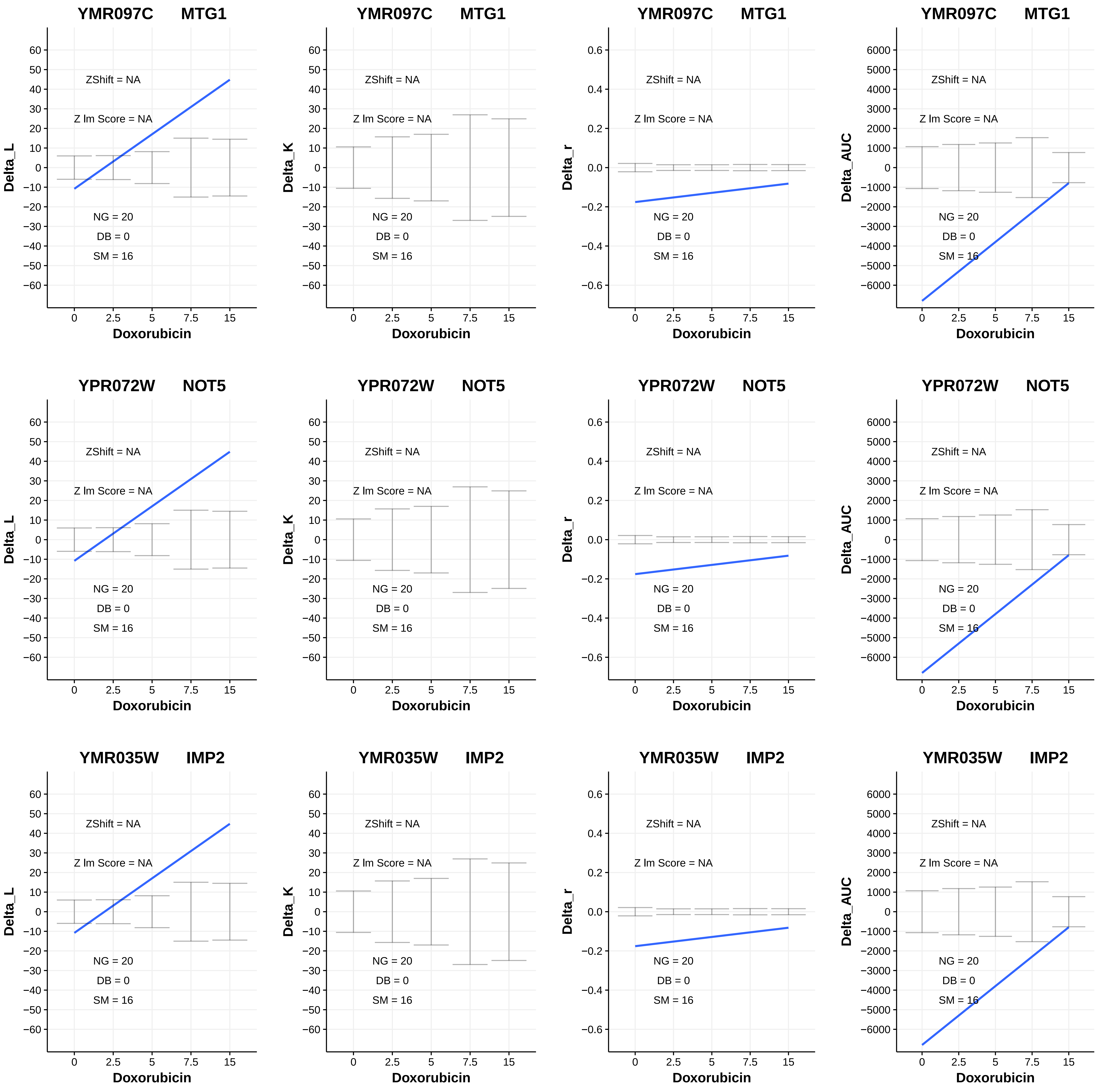
YMR097C MTG1
ZShift = NA
Z lm Score = NA
NG = 20
DB = 0
SM = 16
Delta_L
Doxorubicin
0
2.5
5
7.5
15
YMR097C MTG1
ZShift = NA
Z lm Score = NA
NG = 20
DB = 0
SM = 16
Delta_K
Doxorubicin
YMR097C MTG1
ZShift = NA
Z lm Score = NA
NG = 20
DB = 0
SM = 16
Delta_r
Doxorubicin
YMR097C MTG1
ZShift = NA
Z lm Score = NA
NG = 20
DB = 0
SM = 16
Delta_AUC
Doxorubicin
YPR072W NOT5
ZShift = NA
Z lm Score = NA
NG = 20
DB = 0
SM = 16
Delta_L
Doxorubicin
YPR072W NOT5
ZShift = NA
Z lm Score = NA
NG = 20
DB = 0
SM = 16
Delta_K
Doxorubicin
YPR072W NOT5
ZShift = NA
Z lm Score = NA
NG = 20
DB = 0
SM = 16
Delta_r
Doxorubicin
YPR072W NOT5
ZShift = NA
Z lm Score = NA
NG = 20
DB = 0
SM = 16
Delta_AUC
Doxorubicin
YMR035W IMP2
ZShift = NA
Z lm Score = NA
NG = 20
DB = 0
SM = 16
Delta_L
Doxorubicin
YMR035W IMP2
ZShift = NA
Z lm Score = NA
NG = 20
DB = 0
SM = 16
Delta_K
Doxorubicin
YMR035W IMP2
ZShift = NA
Z lm Score = NA
NG = 20
DB = 0
SM = 16
Delta_r
Doxorubicin
YMR035W IMP2
ZShift = NA
Z lm Score = NA
NG = 20
DB = 0
SM = 16
Delta_AUC
Doxorubicin

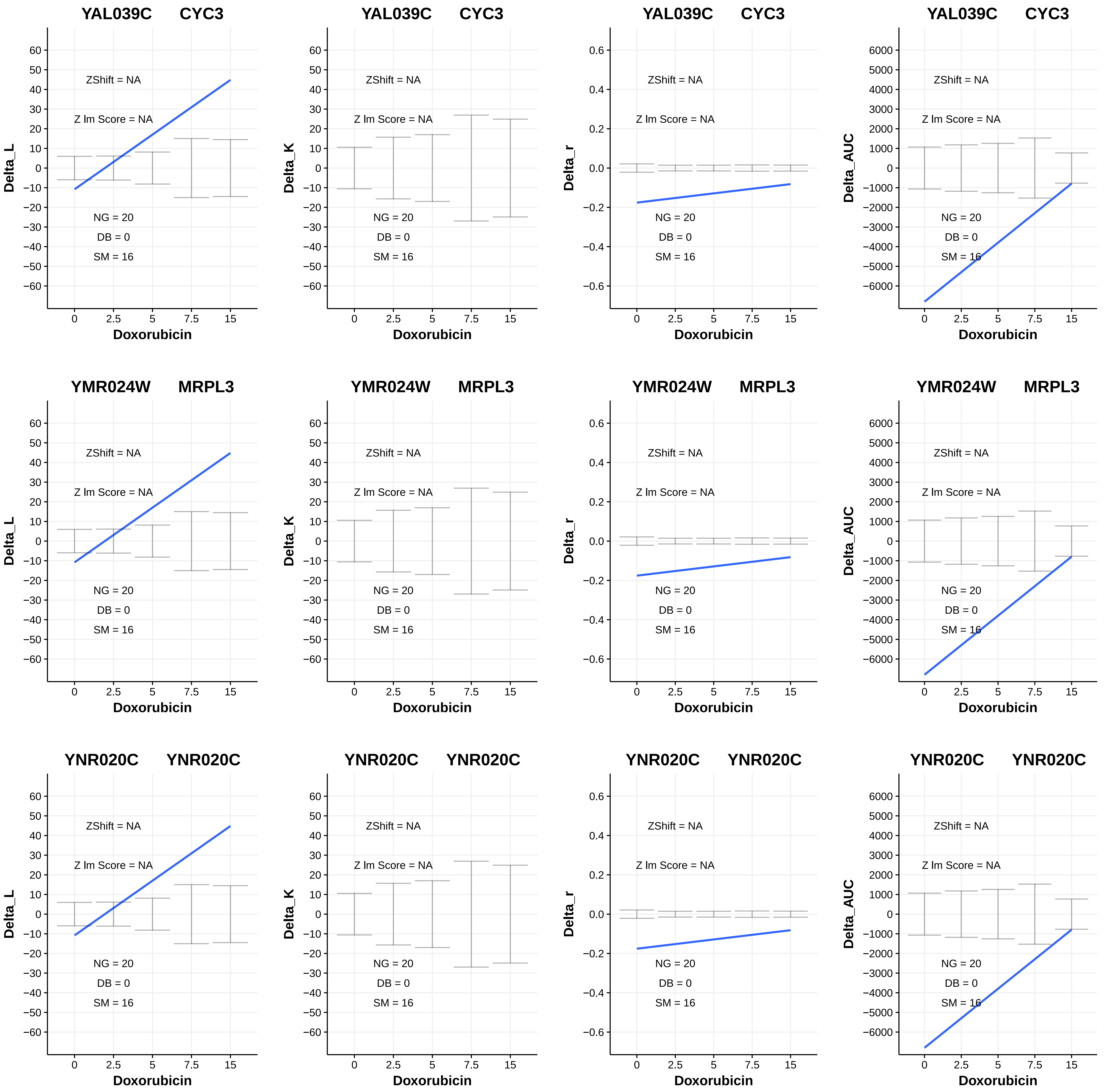
YAL039C CYC3
ZShift = NA
Z lm Score = NA
NG = 20
DB = 0
SM = 16
Delta_L
Delta_K
Delta_r
Delta_AUC
Doxorubicin
YMR024W MRPL3
ZShift = NA
Z lm Score = NA
NG = 20
DB = 0
SM = 16
Delta_L
Delta_K
Delta_r
Delta_AUC
Doxorubicin
YNR020C YNR020C
ZShift = NA
Z lm Score = NA
NG = 20
DB = 0
SM = 16
Delta_L
Delta_K
Delta_r
Delta_AUC
Doxorubicin

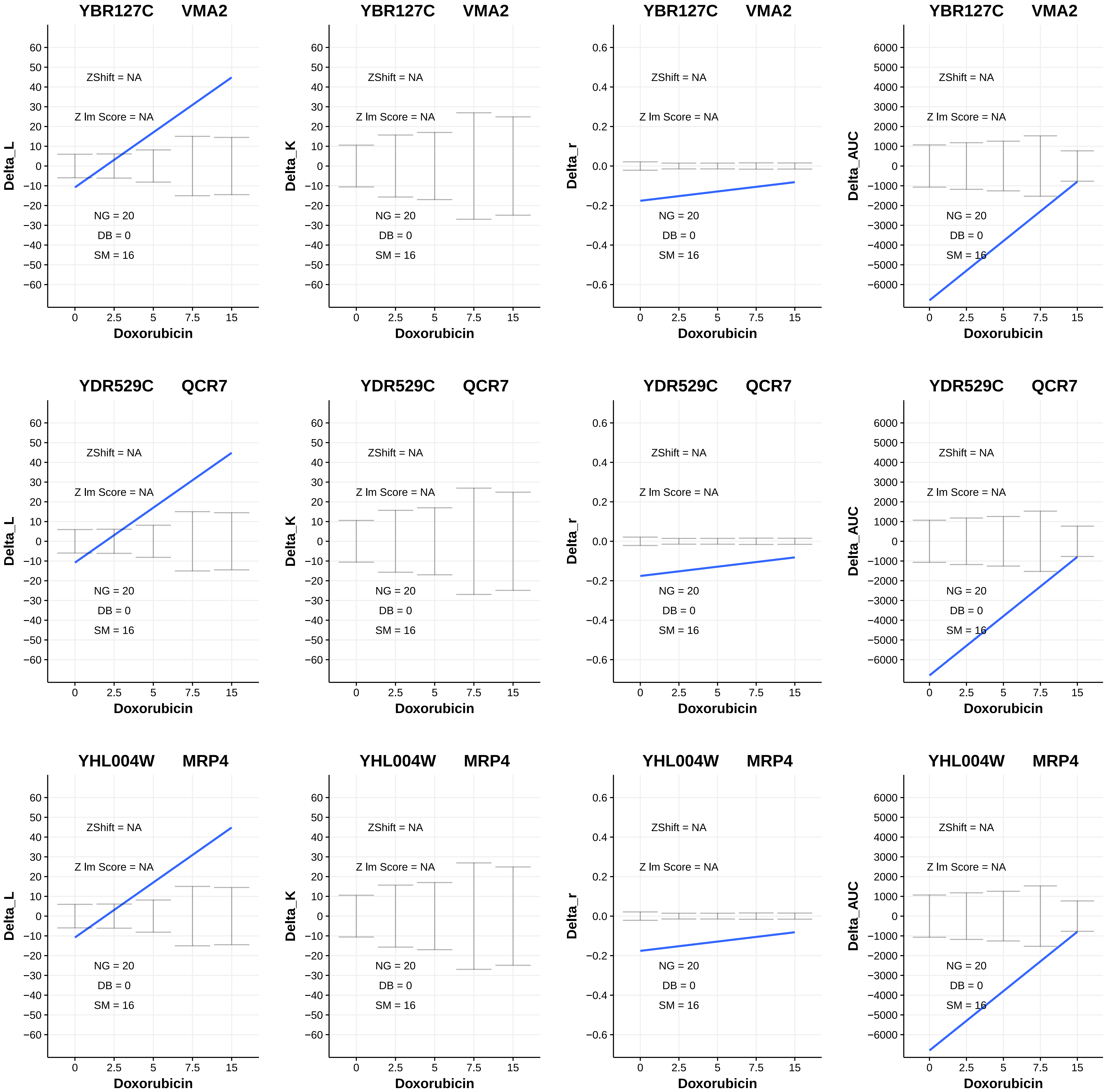
YBR127C VMA2
ZShift = NA
Z lm Score = NA
NG = 20
DB = 0
SM = 16
Delta_L
Delta_K
Delta_r
Delta_AUC
Doxorubicin
0 2.5 5 7.5 15
YDR529C QCR7
ZShift = NA
Z lm Score = NA
NG = 20
DB = 0
SM = 16
Delta_L
Delta_K
Delta_r
Delta_AUC
Doxorubicin
0 2.5 5 7.5 15
YHL004W MRP4
ZShift = NA
Z lm Score = NA
NG = 20
DB = 0
SM = 16
Delta_L
Delta_K
Delta_r
Delta_AUC
Doxorubicin
0 2.5 5 7.5 15
60 50 40 30 20 10 0 −10 −20 −30 −40 −50 −60
0.6 0.4 0.2 0.0 −0.2 −0.4 −0.6
6000 5000 4000 3000 2000 1000 0 −1000 −2000 −3000 −4000 −5000 −6000

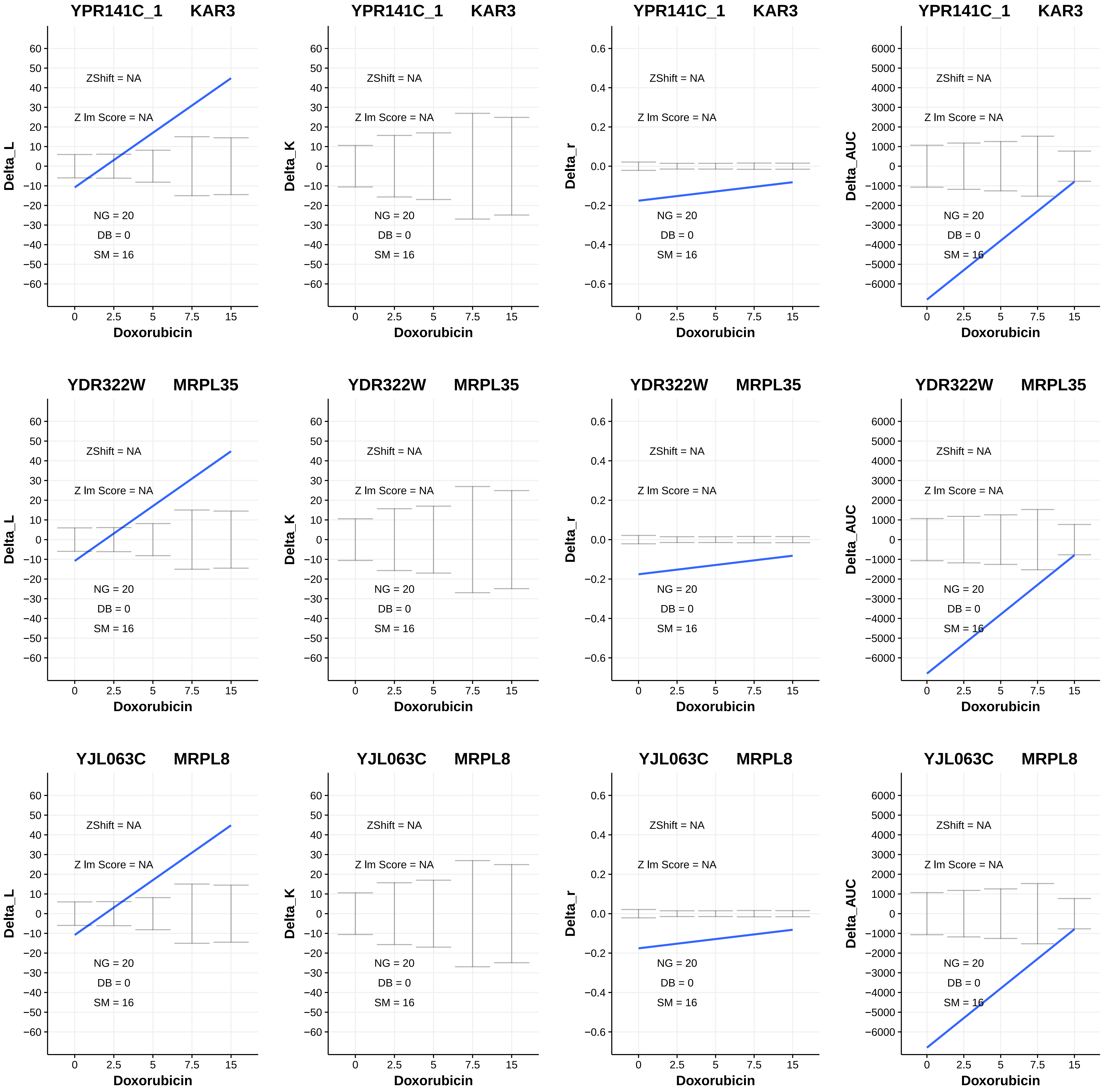
YPR141C_1 KAR3
ZShift = NA
Z lm Score = NA
NG = 20
DB = 0
SM = 16
Delta_L
Delta_K
Delta_r
Delta_AUC
Doxorubicin
0 2.5 5 7.5 15
YDR322W MRPL35
ZShift = NA
Z lm Score = NA
NG = 20
DB = 0
SM = 16
Delta_L
Delta_K
Delta_r
Delta_AUC
Doxorubicin
0 2.5 5 7.5 15
YJL063C MRPL8
ZShift = NA
Z lm Score = NA
NG = 20
DB = 0
SM = 16
Delta_L
Delta_K
Delta_r
Delta_AUC
Doxorubicin
0 2.5 5 7.5 15

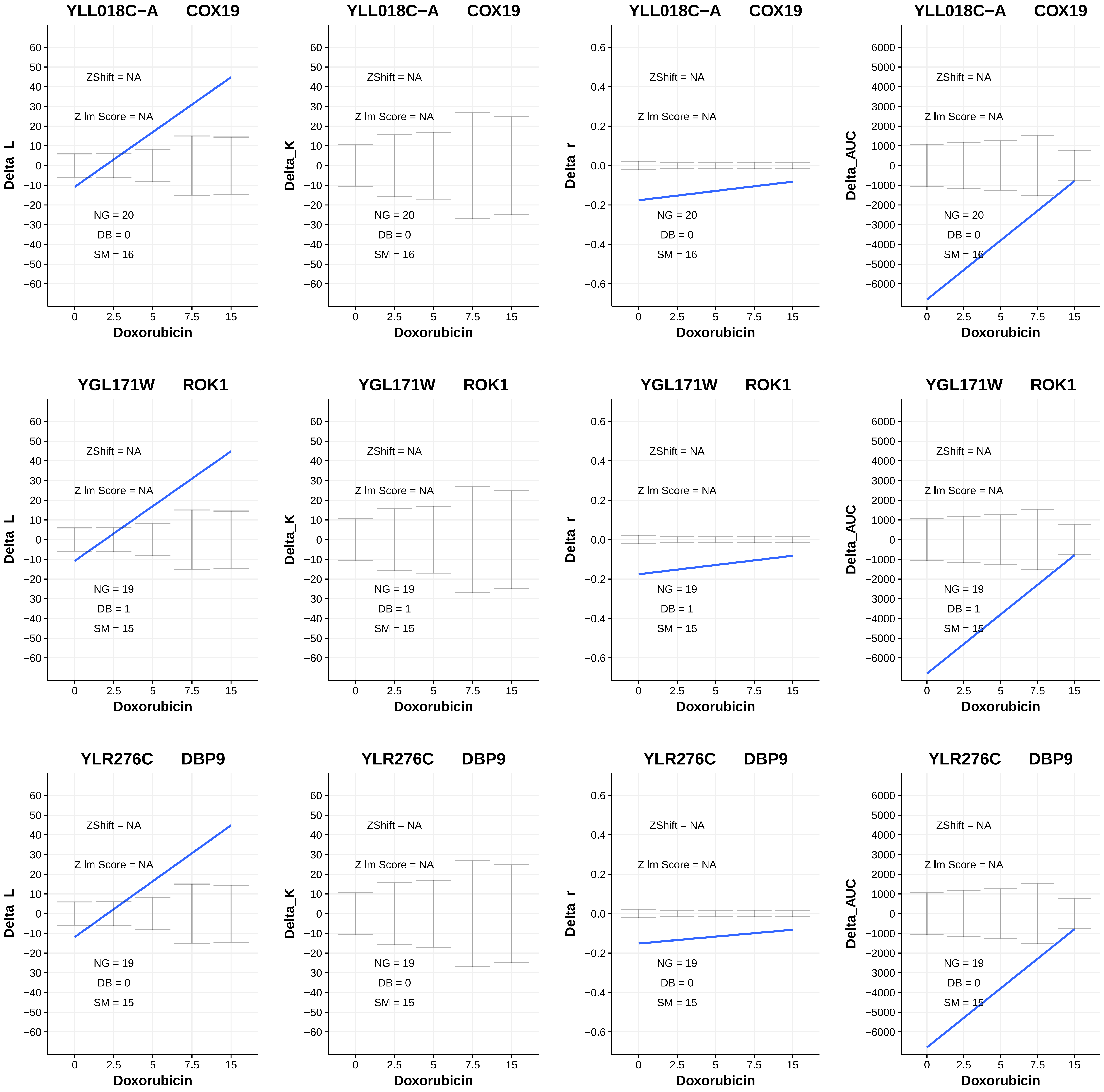

YLL018C−A COX19
ZShift = NA
Z lm Score = NA
NG = 20
DB = 0
SM = 16
Delta_L
Delta_K
Delta_r
Delta_AUC
Doxorubicin
0
2.5
5
7.5
15
YGL171W ROK1
ZShift = NA
Z lm Score = NA
NG = 19
DB = 1
SM = 15
YLR276C DBP9
ZShift = NA
Z lm Score = NA
NG = 19
DB = 0
SM = 15

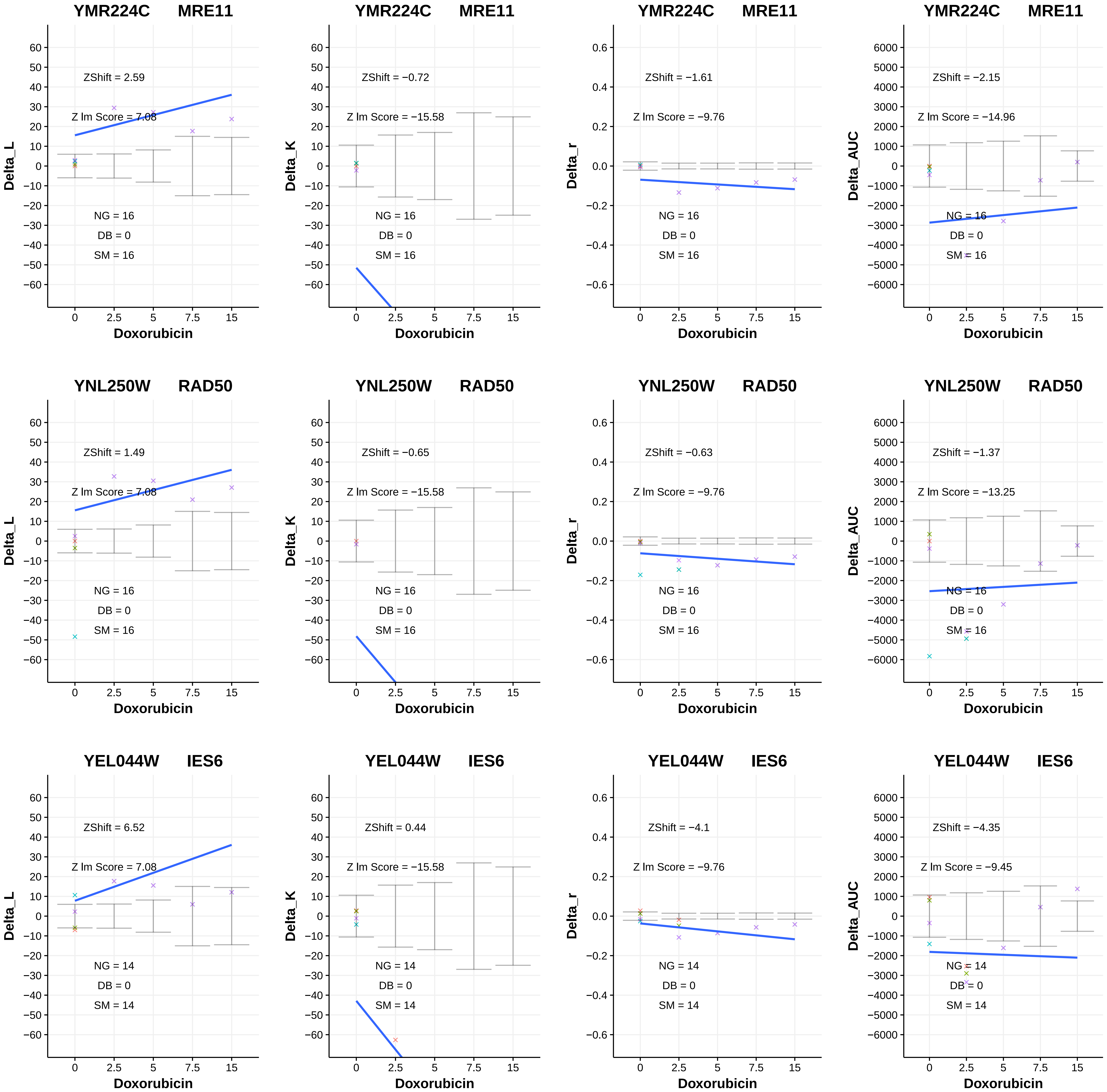

YMR224C MRE11
ZShift = 2.59
Z lm Score = 7.08
NG = 16
DB = 0
SM = 16
Delta_L
Doxorubicin
YMR224C MRE11
ZShift = −0.72
Z lm Score = −15.58
NG = 16
DB = 0
SM = 16
Delta_K
Doxorubicin
YMR224C MRE11
ZShift = −1.61
Z lm Score = −9.76
NG = 16
DB = 0
SM = 16
Delta_r
Doxorubicin
YMR224C MRE11
ZShift = −2.15
Z lm Score = −14.96
NG = 16
DB = 0
SM = 16
Delta_AUC
Doxorubicin
YNL250W RAD50
ZShift = 1.49
Z lm Score = 7.08
NG = 16
DB = 0
SM = 16
Delta_L
Doxorubicin
YNL250W RAD50
ZShift = −0.65
Z lm Score = −15.58
NG = 16
DB = 0
SM = 16
Delta_K
Doxorubicin
YNL250W RAD50
ZShift = −0.63
Z lm Score = −9.76
NG = 16
DB = 0
SM = 16
Delta_r
Doxorubicin
YNL250W RAD50
ZShift = −1.37
Z lm Score = −13.25
NG = 16
DB = 0
SM = 16
Delta_AUC
Doxorubicin
YEL044W IES6
ZShift = 6.52
Z lm Score = 7.08
NG = 14
DB = 0
SM = 14
Delta_L
Doxorubicin
YEL044W IES6
ZShift = 0.44
Z lm Score = −15.58
NG = 14
DB = 0
SM = 14
Delta_K
Doxorubicin
YEL044W IES6
ZShift = −4.1
Z lm Score = −9.76
NG = 14
DB = 0
SM = 14
Delta_r
Doxorubicin
YEL044W IES6
ZShift = −4.35
Z lm Score = −9.45
NG = 14
DB = 0
SM = 14
Delta_AUC
Doxorubicin

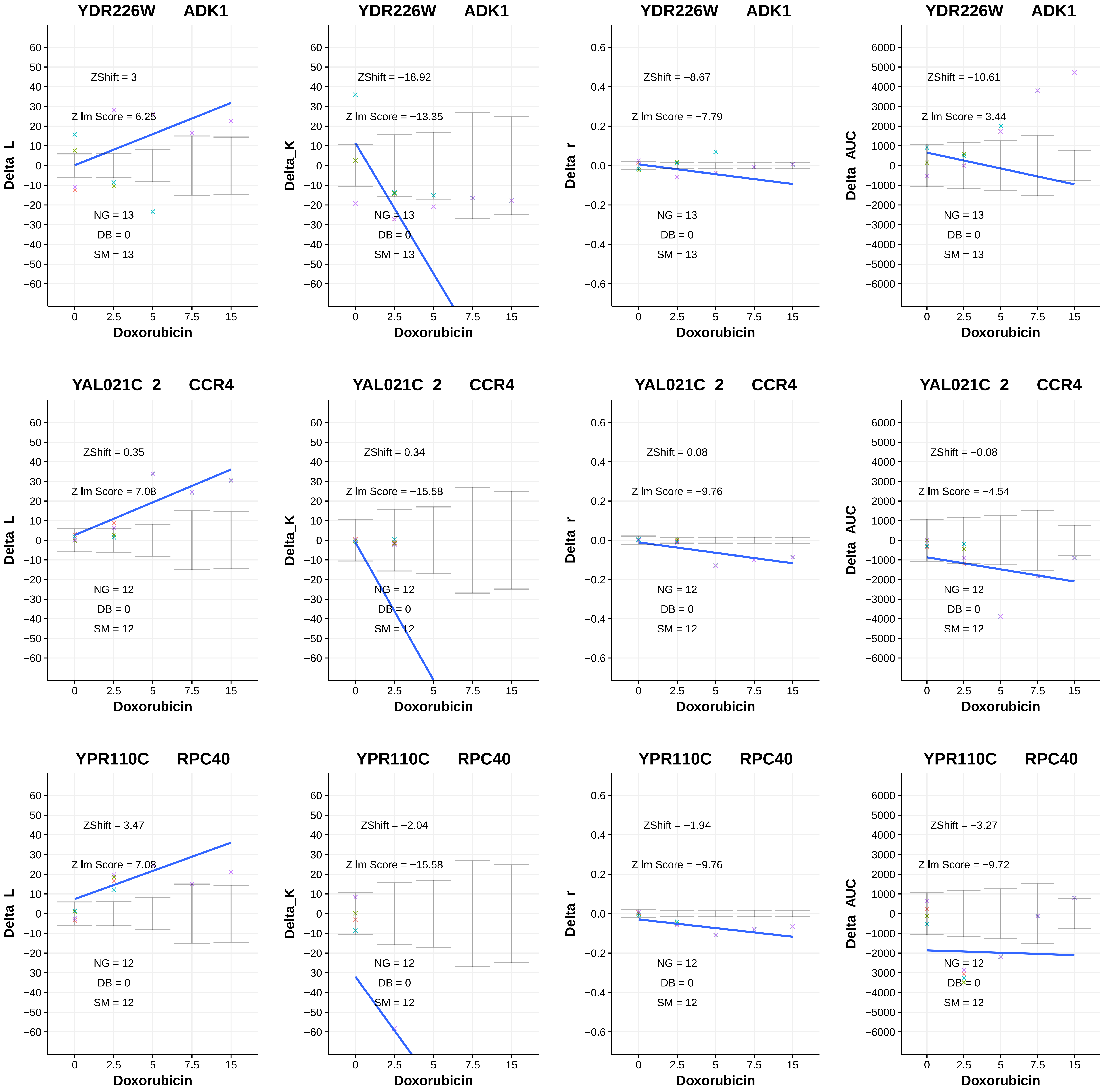
YDR226W ADK1
ZShift = 3
Z lm Score = 6.25
NG = 13
DB = 0
SM = 13
Delta_L
Doxorubicin
YDR226W ADK1
ZShift = −18.92
Z lm Score = −13.35
NG = 13
DB = 0
SM = 13
Delta_K
Doxorubicin
YDR226W ADK1
ZShift = −8.67
Z lm Score = −7.79
NG = 13
DB = 0
SM = 13
Delta_r
Doxorubicin
YDR226W ADK1
ZShift = −10.61
Z lm Score = 3.44
NG = 13
DB = 0
SM = 13
Delta_AUC
Doxorubicin
YAL021C_2 CCR4
ZShift = 0.35
Z lm Score = 7.08
NG = 12
DB = 0
SM = 12
Delta_L
Doxorubicin
YAL021C_2 CCR4
ZShift = 0.34
Z lm Score = −15.58
NG = 12
DB = 0
SM = 12
Delta_K
Doxorubicin
YAL021C_2 CCR4
ZShift = 0.08
Z lm Score = −9.76
NG = 12
DB = 0
SM = 12
Delta_r
Doxorubicin
YAL021C_2 CCR4
ZShift = −0.08
Z lm Score = −4.54
NG = 12
DB = 0
SM = 12
Delta_AUC
Doxorubicin
YPR110C RPC40
ZShift = 3.47
Z lm Score = 7.08
NG = 12
DB = 0
SM = 12
Delta_L
Doxorubicin
YPR110C RPC40
ZShift = −2.04
Z lm Score = −15.58
NG = 12
DB = 0
SM = 12
Delta_K
Doxorubicin
YPR110C RPC40
ZShift = −1.94
Z lm Score = −9.76
NG = 12
DB = 0
SM = 12
Delta_r
Doxorubicin
YPR110C RPC40
ZShift = −3.27
Z lm Score = −9.72
NG = 12
DB = 0
SM = 12
Delta_AUC
Doxorubicin

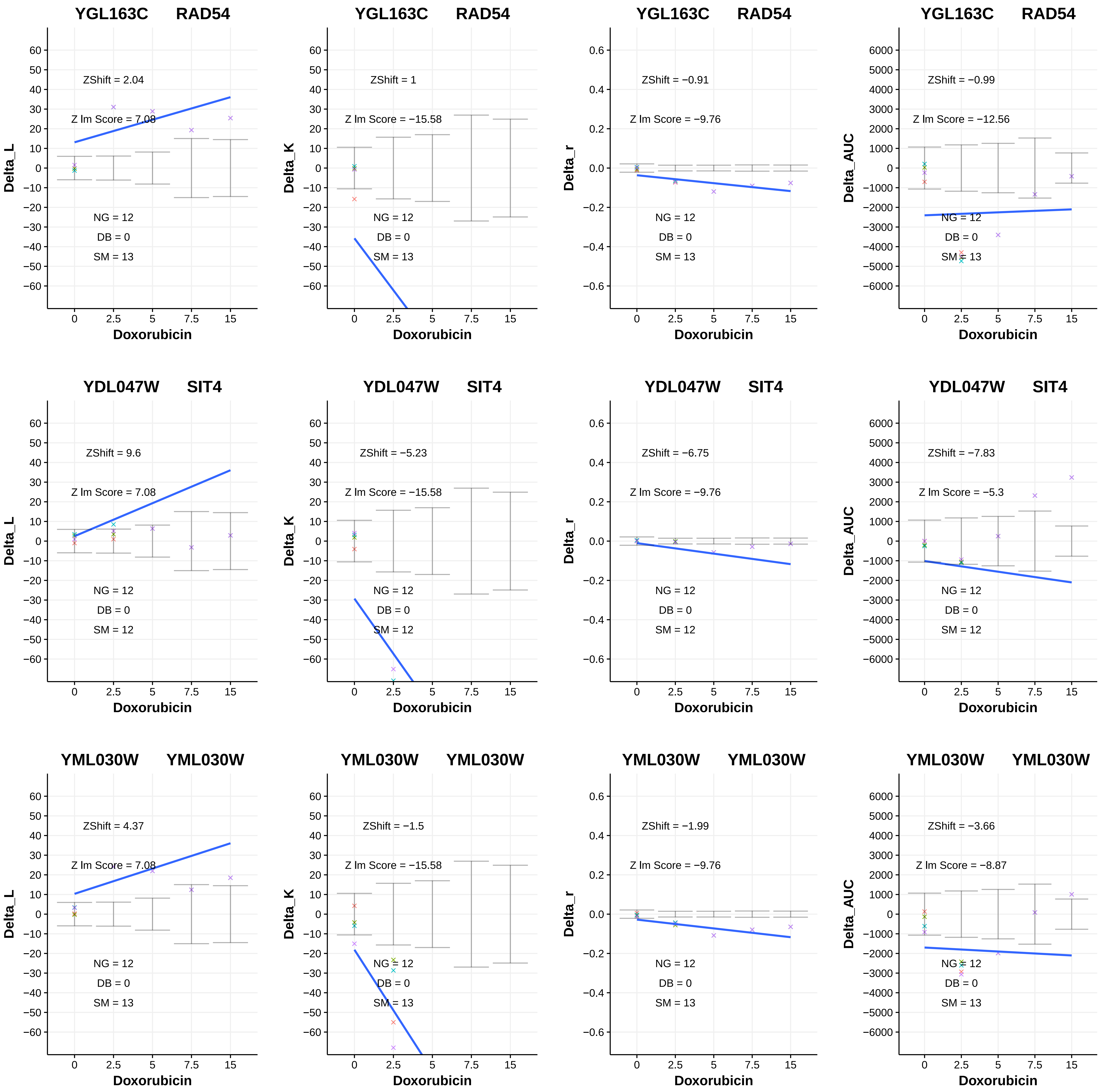
YGL163C RAD54
Delta_L
ZShift = 2.04
Z lm Score = 7.08
NG = 12
DB = 0
SM = 13
Doxorubicin
YGL163C RAD54
Delta_K
ZShift = 1
Z lm Score = −15.58
NG = 12
DB = 0
SM = 13
Doxorubicin
YGL163C RAD54
Delta_r
ZShift = −0.91
Z lm Score = −9.76
NG = 12
DB = 0
SM = 13
Doxorubicin
YGL163C RAD54
Delta_AUC
ZShift = −0.99
Z lm Score = −12.56
NG = 12
DB = 0
SM = 13
Doxorubicin
YDL047W SIT4
Delta_L
ZShift = 9.6
Z lm Score = 7.08
NG = 12
DB = 0
SM = 12
Doxorubicin
YDL047W SIT4
Delta_K
ZShift = −5.23
Z lm Score = −15.58
NG = 12
DB = 0
SM = 12
Doxorubicin
YDL047W SIT4
Delta_r
ZShift = −6.75
Z lm Score = −9.76
NG = 12
DB = 0
SM = 12
Doxorubicin
YDL047W SIT4
Delta_AUC
ZShift = −7.83
Z lm Score = −5.3
NG = 12
DB = 0
SM = 12
Doxorubicin
YML030W YML030W
Delta_L
ZShift = 4.37
Z lm Score = 7.08
NG = 12
DB = 0
SM = 13
Doxorubicin
YML030W YML030W
Delta_K
ZShift = −1.5
Z lm Score = −15.58
NG = 12
DB = 0
SM = 13
Doxorubicin
YML030W YML030W
Delta_r
ZShift = −1.99
Z lm Score = −9.76
NG = 12
DB = 0
SM = 13
Doxorubicin
YML030W YML030W
Delta_AUC
ZShift = −3.66
Z lm Score = −8.87
NG = 12
DB = 0
SM = 13
Doxorubicin

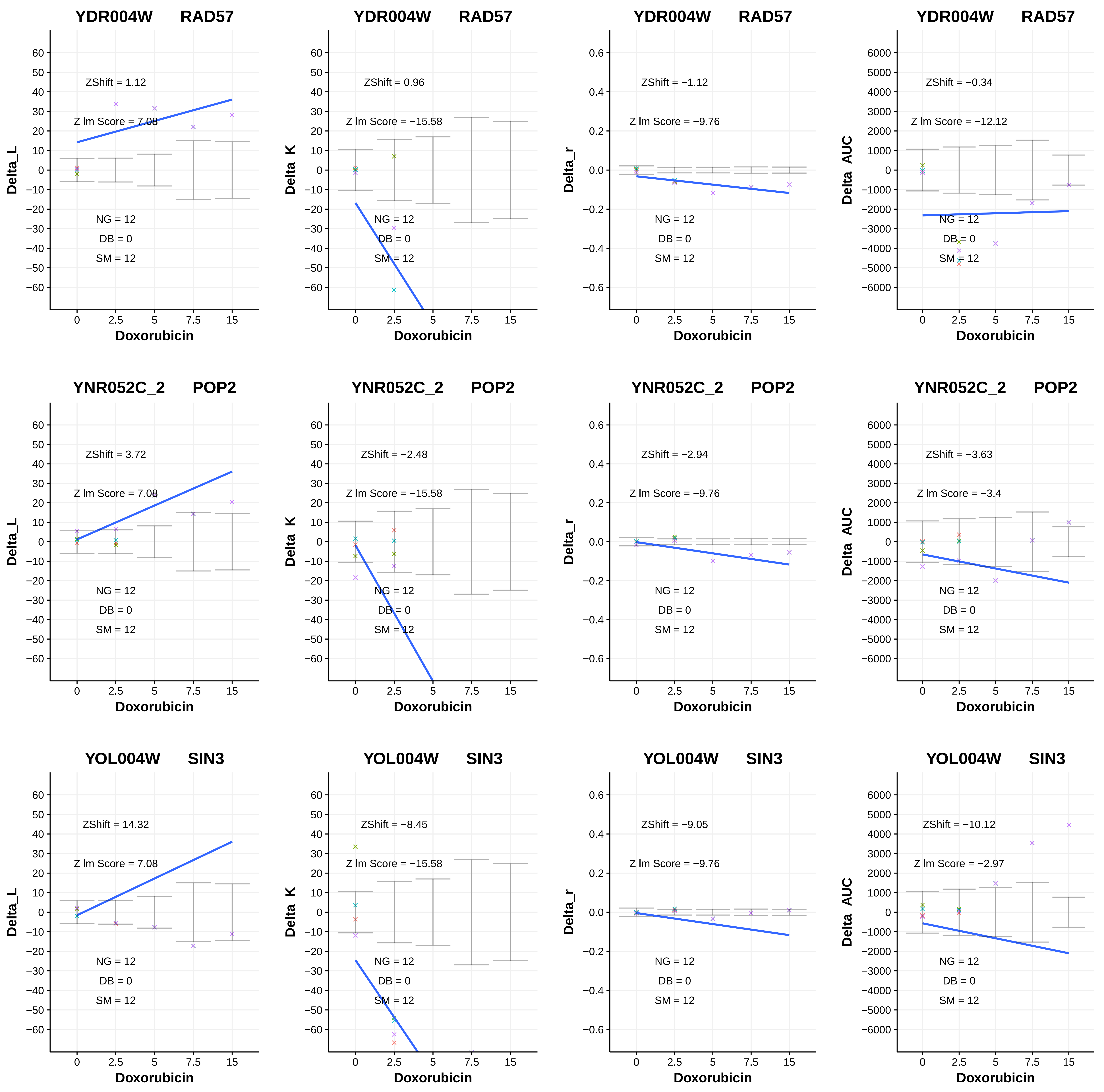
YDR004W RAD57
ZShift = 1.12
Z lm Score = 7.08
NG = 12
DB = 0
SM = 12
Delta_L
Doxorubicin
YDR004W RAD57
ZShift = 0.96
Z lm Score = −15.58
NG = 12
DB = 0
SM = 12
Delta_K
Doxorubicin
YDR004W RAD57
ZShift = −1.12
Z lm Score = −9.76
NG = 12
DB = 0
SM = 12
Delta_r
Doxorubicin
YDR004W RAD57
ZShift = −0.34
Z lm Score = −12.12
NG = 12
DB = 0
SM = 12
Delta_AUC
Doxorubicin
YNR052C_2 POP2
ZShift = 3.72
Z lm Score = 7.08
NG = 12
DB = 0
SM = 12
Delta_L
Doxorubicin
YNR052C_2 POP2
ZShift = −2.48
Z lm Score = −15.58
NG = 12
DB = 0
SM = 12
Delta_K
Doxorubicin
YNR052C_2 POP2
ZShift = −2.94
Z lm Score = −9.76
NG = 12
DB = 0
SM = 12
Delta_r
Doxorubicin
YNR052C_2 POP2
ZShift = −3.63
Z lm Score = −3.4
NG = 12
DB = 0
SM = 12
Delta_AUC
Doxorubicin
YOL004W SIN3
ZShift = 14.32
Z lm Score = 7.08
NG = 12
DB = 0
SM = 12
Delta_L
Doxorubicin
YOL004W SIN3
ZShift = −8.45
Z lm Score = −15.58
NG = 12
DB = 0
SM = 12
Delta_K
Doxorubicin
YOL004W SIN3
ZShift = −9.05
Z lm Score = −9.76
NG = 12
DB = 0
SM = 12
Delta_r
Doxorubicin
YOL004W SIN3
ZShift = −10.12
Z lm Score = −2.97
NG = 12
DB = 0
SM = 12
Delta_AUC
Doxorubicin

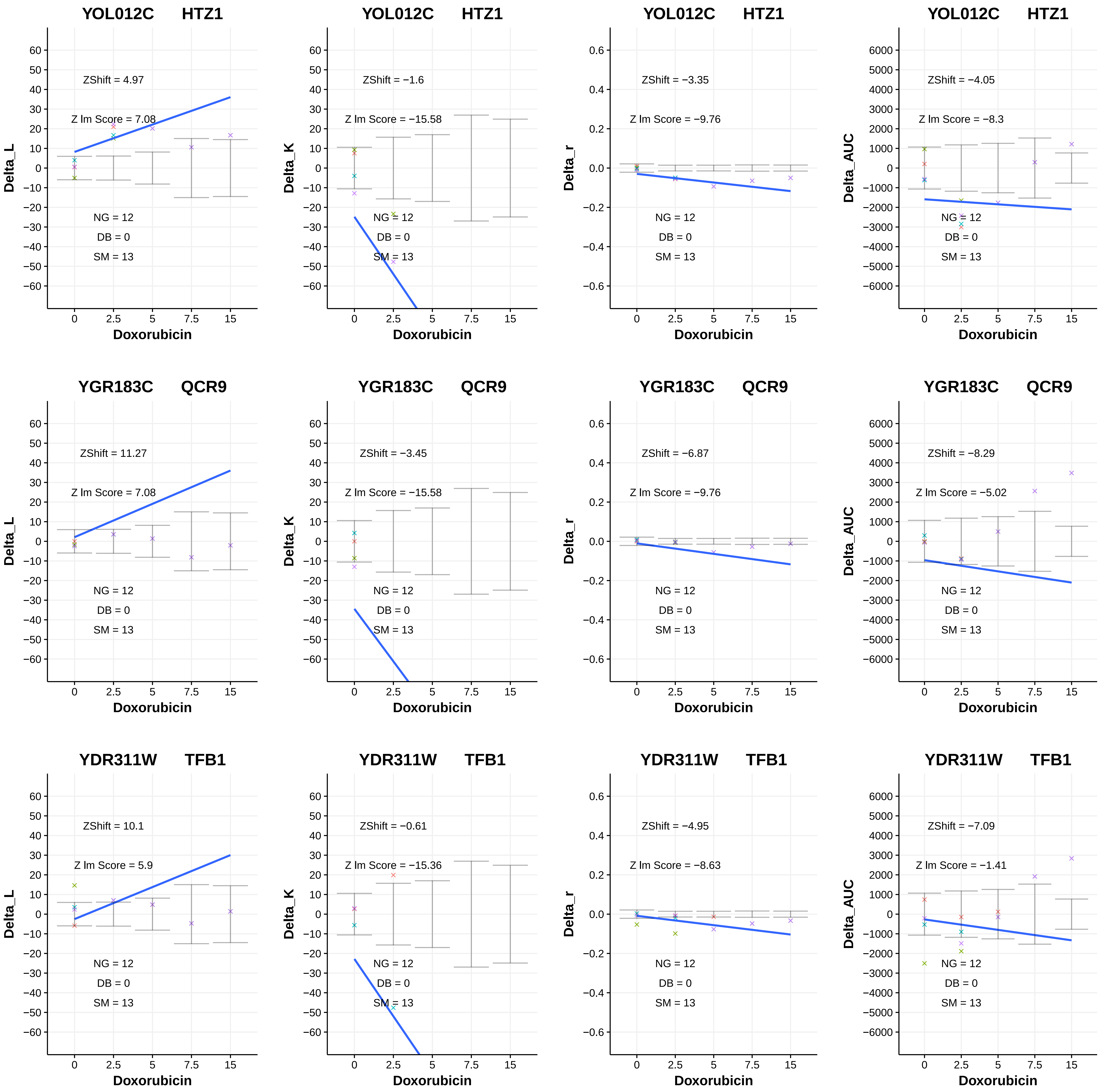
YOL012C HTZ1
ZShift = 4.97
Z lm Score = 7.08
NG = 12
DB = 0
SM = 13
Delta_L
Doxorubicin
YOL012C HTZ1
ZShift = −1.6
Z lm Score = −15.58
NG = 12
DB = 0
SM = 13
Delta_K
Doxorubicin
YOL012C HTZ1
ZShift = −3.35
Z lm Score = −9.76
NG = 12
DB = 0
SM = 13
Delta_r
Doxorubicin
YOL012C HTZ1
ZShift = −4.05
Z lm Score = −8.3
NG = 12
DB = 0
SM = 13
Delta_AUC
Doxorubicin
YGR183C QCR9
ZShift = 11.27
Z lm Score = 7.08
NG = 12
DB = 0
SM = 13
Delta_L
Doxorubicin
YGR183C QCR9
ZShift = −3.45
Z lm Score = −15.58
NG = 12
DB = 0
SM = 13
Delta_K
Doxorubicin
YGR183C QCR9
ZShift = −6.87
Z lm Score = −9.76
NG = 12
DB = 0
SM = 13
Delta_r
Doxorubicin
YGR183C QCR9
ZShift = −8.29
Z lm Score = −5.02
NG = 12
DB = 0
SM = 13
Delta_AUC
Doxorubicin
YDR311W TFB1
ZShift = 10.1
Z lm Score = 5.9
NG = 12
DB = 0
SM = 13
Delta_L
Doxorubicin
YDR311W TFB1
ZShift = −0.61
Z lm Score = −15.36
NG = 12
DB = 0
SM = 13
Delta_K
Doxorubicin
YDR311W TFB1
ZShift = −4.95
Z lm Score = −8.63
NG = 12
DB = 0
SM = 13
Delta_r
Doxorubicin
YDR311W TFB1
ZShift = −7.09
Z lm Score = −1.41
NG = 12
DB = 0
SM = 13
Delta_AUC
Doxorubicin

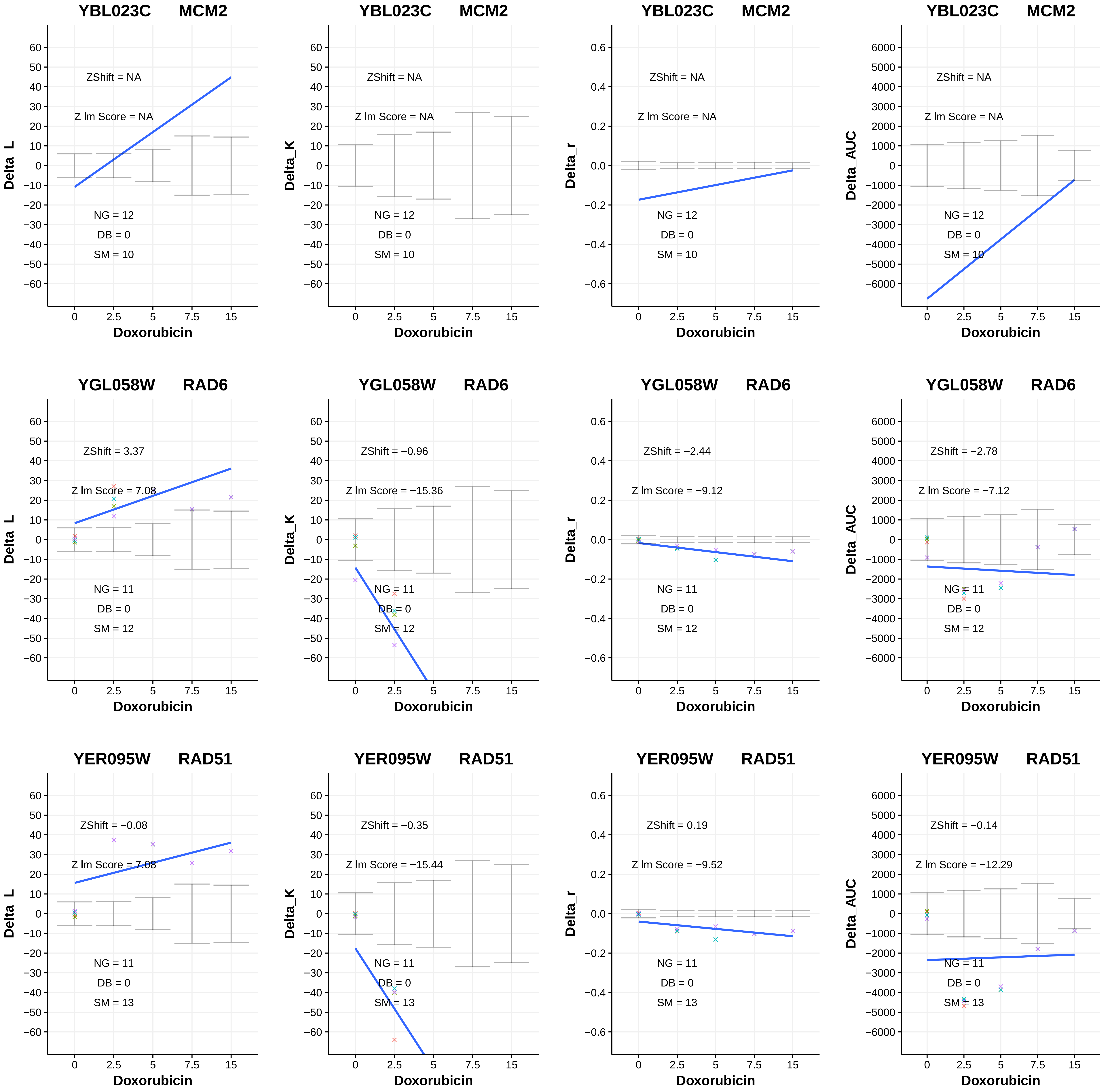

YBL023C MCM2
ZShift = NA
Z lm Score = NA
NG = 12
DB = 0
SM = 10
Delta_L
Doxorubicin
YBL023C MCM2
ZShift = NA
Z lm Score = NA
NG = 12
DB = 0
SM = 10
Delta_K
Doxorubicin
YBL023C MCM2
ZShift = NA
Z lm Score = NA
NG = 12
DB = 0
SM = 10
Delta_r
Doxorubicin
YBL023C MCM2
ZShift = NA
Z lm Score = NA
NG = 12
DB = 0
SM = 10
Delta_AUC
Doxorubicin
YGL058W RAD6
ZShift = 3.37
Z lm Score = 7.08
NG = 11
DB = 0
SM = 12
Delta_L
Doxorubicin
YGL058W RAD6
ZShift = −0.96
Z lm Score = −15.36
NG = 11
DB = 0
SM = 12
Delta_K
Doxorubicin
YGL058W RAD6
ZShift = −2.44
Z lm Score = −9.12
NG = 11
DB = 0
SM = 12
Delta_r
Doxorubicin
YGL058W RAD6
ZShift = −2.78
Z lm Score = −7.12
NG = 11
DB = 0
SM = 12
Delta_AUC
Doxorubicin
YER095W RAD51
ZShift = −0.08
Z lm Score = 7.08
NG = 11
DB = 0
SM = 13
Delta_L
Doxorubicin
YER095W RAD51
ZShift = −0.35
Z lm Score = −15.44
NG = 11
DB = 0
SM = 13
Delta_K
Doxorubicin
YER095W RAD51
ZShift = 0.19
Z lm Score = −9.52
NG = 11
DB = 0
SM = 13
Delta_r
Doxorubicin
YER095W RAD51
ZShift = −0.14
Z lm Score = −12.29
NG = 11
DB = 0
SM = 13
Delta_AUC
Doxorubicin

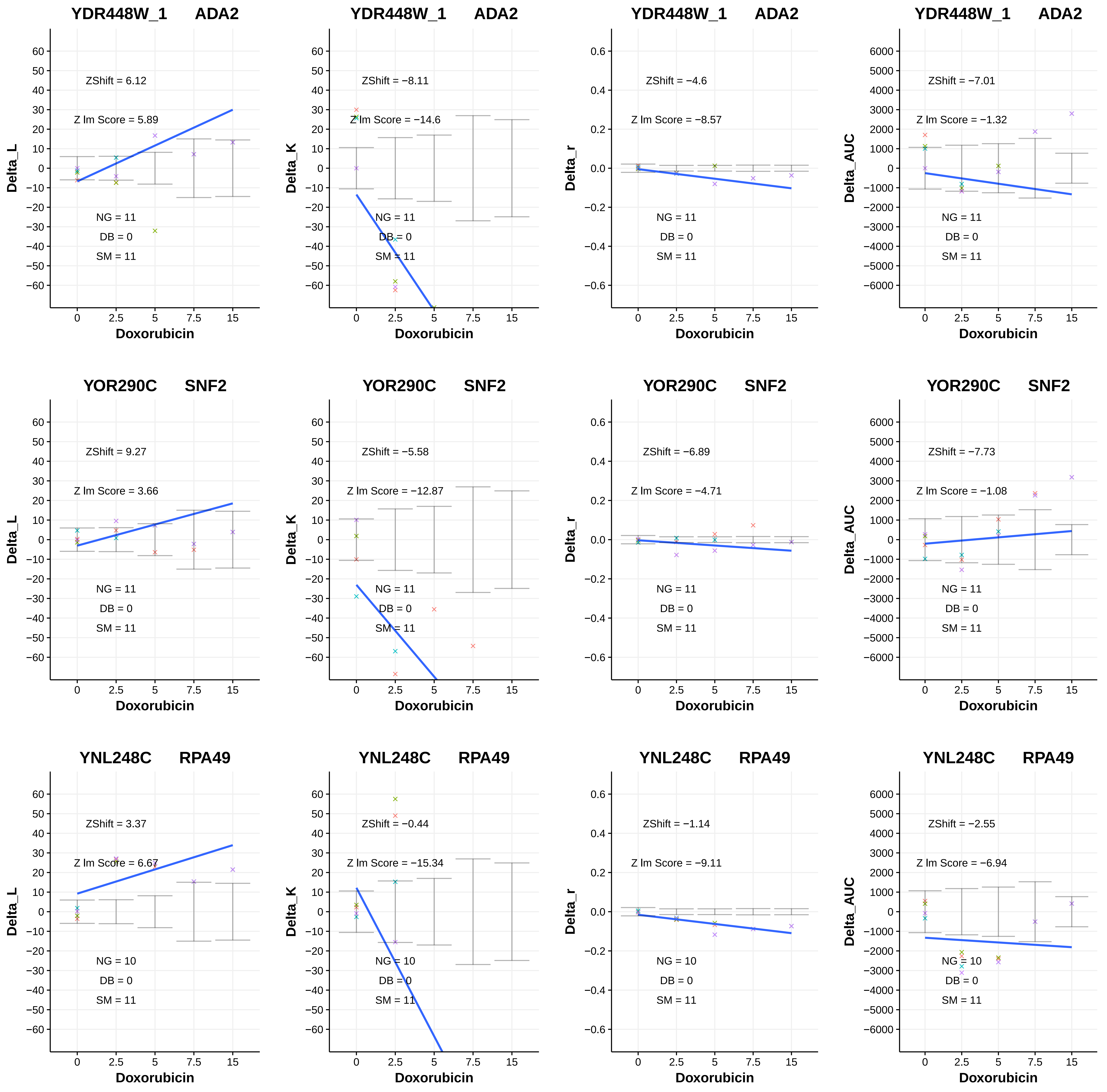

YDR448W_1 ADA2
ZShift = 6.12
Z lm Score = 5.89
NG = 11
DB = 0
SM = 11
Delta_L
Doxorubicin
YDR448W_1 ADA2
ZShift = −8.11
Z lm Score = −14.6
NG = 11
DB = 0
SM = 11
Delta_K
Doxorubicin
YDR448W_1 ADA2
ZShift = −4.6
Z lm Score = −8.57
NG = 11
DB = 0
SM = 11
Delta_r
Doxorubicin
YDR448W_1 ADA2
ZShift = −7.01
Z lm Score = −1.32
NG = 11
DB = 0
SM = 11
Delta_AUC
Doxorubicin
YOR290C SNF2
ZShift = 9.27
Z lm Score = 3.66
NG = 11
DB = 0
SM = 11
Delta_L
Doxorubicin
YOR290C SNF2
ZShift = −5.58
Z lm Score = −12.87
NG = 11
DB = 0
SM = 11
Delta_K
Doxorubicin
YOR290C SNF2
ZShift = −6.89
Z lm Score = −4.71
NG = 11
DB = 0
SM = 11
Delta_r
Doxorubicin
YOR290C SNF2
ZShift = −7.73
Z lm Score = −1.08
NG = 11
DB = 0
SM = 11
Delta_AUC
Doxorubicin
YNL248C RPA49
ZShift = 3.37
Z lm Score = 6.67
NG = 10
DB = 0
SM = 11
Delta_L
Doxorubicin
YNL248C RPA49
ZShift = −0.44
Z lm Score = −15.34
NG = 10
DB = 0
SM = 11
Delta_K
Doxorubicin
YNL248C RPA49
ZShift = −1.14
Z lm Score = −9.11
NG = 10
DB = 0
SM = 11
Delta_r
Doxorubicin
YNL248C RPA49
ZShift = −2.55
Z lm Score = −6.94
NG = 10
DB = 0
SM = 11
Delta_AUC
Doxorubicin

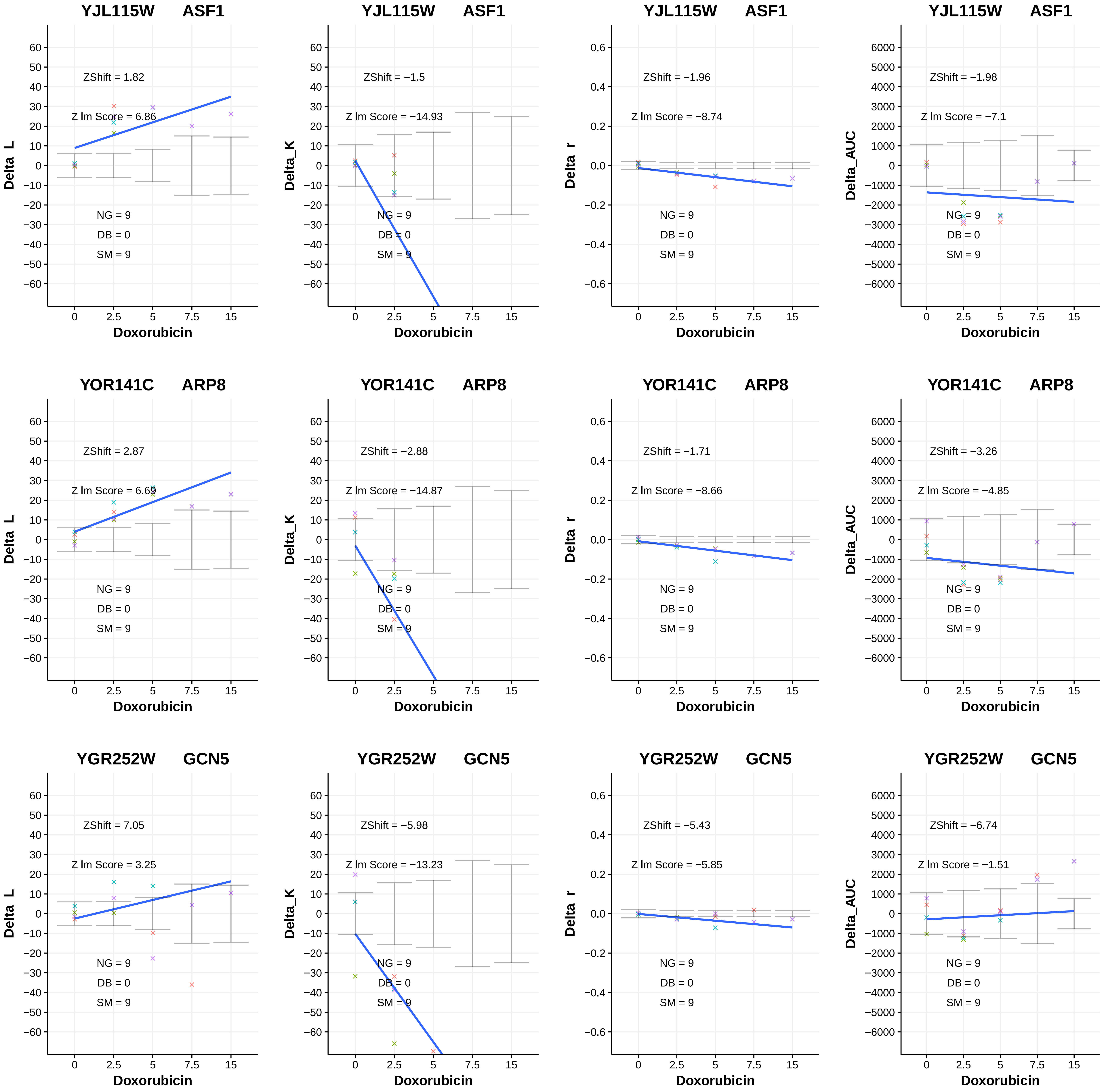
YJL115W ASF1
ZShift = 1.82
Z lm Score = 6.86
NG = 9
DB = 0
SM = 9
Delta_L
Doxorubicin
YJL115W ASF1
ZShift = −1.5
Z lm Score = −14.93
NG = 9
DB = 0
SM = 9
Delta_K
Doxorubicin
YJL115W ASF1
ZShift = −1.96
Z lm Score = −8.74
NG = 9
DB = 0
SM = 9
Delta_r
Doxorubicin
YJL115W ASF1
ZShift = −1.98
Z lm Score = −7.1
NG = 9
DB = 0
SM = 9
Delta_AUC
Doxorubicin
YOR141C ARP8
ZShift = 2.87
Z lm Score = 6.69
NG = 9
DB = 0
SM = 9
Delta_L
Doxorubicin
YOR141C ARP8
ZShift = −2.88
Z lm Score = −14.87
NG = 9
DB = 0
SM = 9
Delta_K
Doxorubicin
YOR141C ARP8
ZShift = −1.71
Z lm Score = −8.66
NG = 9
DB = 0
SM = 9
Delta_r
Doxorubicin
YOR141C ARP8
ZShift = −3.26
Z lm Score = −4.85
NG = 9
DB = 0
SM = 9
Delta_AUC
Doxorubicin
YGR252W GCN5
ZShift = 7.05
Z lm Score = 3.25
NG = 9
DB = 0
SM = 9
Delta_L
Doxorubicin
YGR252W GCN5
ZShift = −5.98
Z lm Score = −13.23
NG = 9
DB = 0
SM = 9
Delta_K
Doxorubicin
YGR252W GCN5
ZShift = −5.43
Z lm Score = −5.85
NG = 9
DB = 0
SM = 9
Delta_r
Doxorubicin
YGR252W GCN5
ZShift = −6.74
Z lm Score = −1.51
NG = 9
DB = 0
SM = 9
Delta_AUC
Doxorubicin

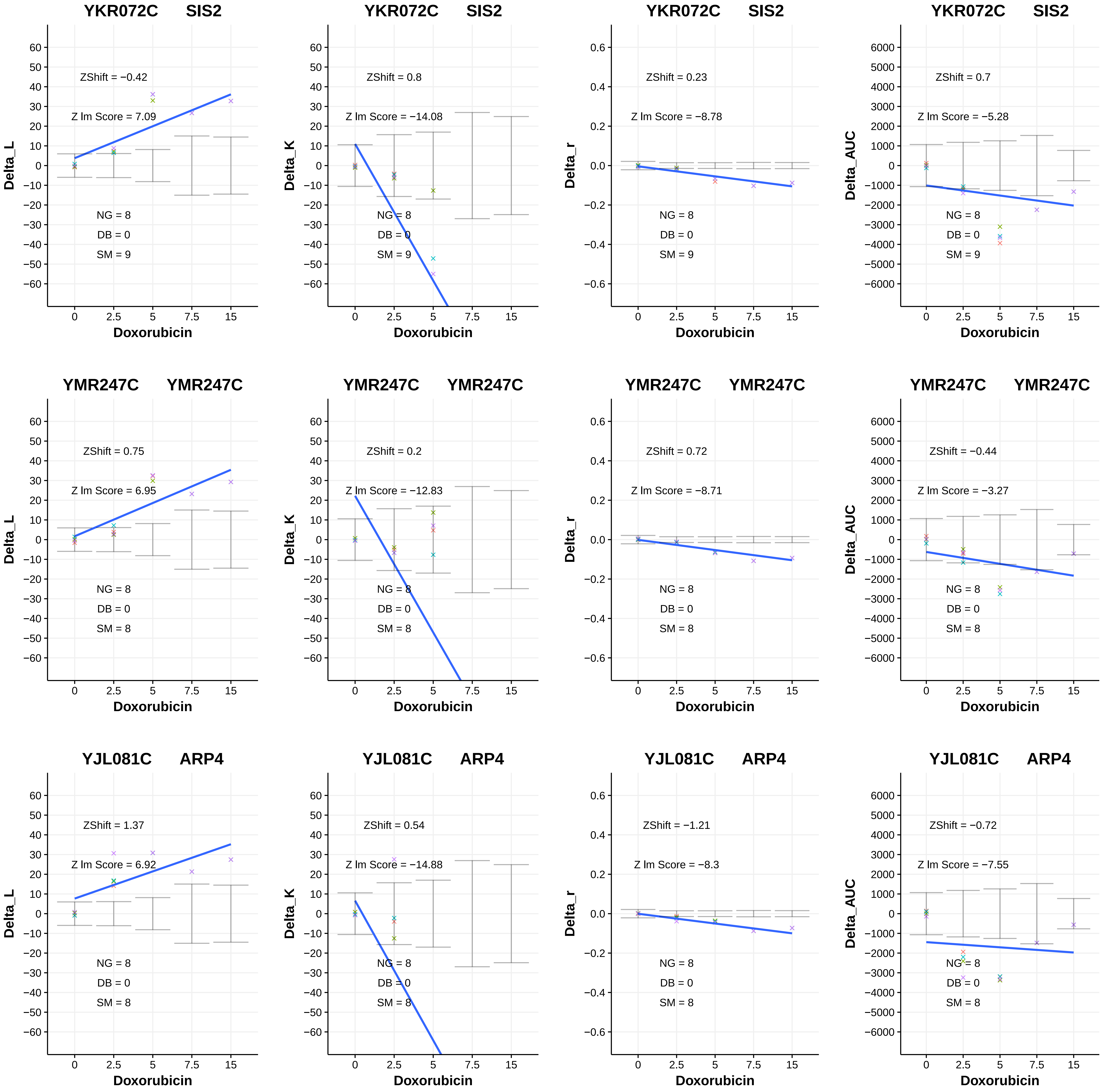

YKR072C SIS2
ZShift = −0.42
Z lm Score = 7.09
NG = 8
DB = 0
SM = 9
Delta_L
Doxorubicin
YKR072C SIS2
ZShift = 0.8
Z lm Score = −14.08
NG = 8
DB = 0
SM = 9
Delta_K
Doxorubicin
YKR072C SIS2
ZShift = 0.23
Z lm Score = −8.78
NG = 8
DB = 0
SM = 9
Delta_r
Doxorubicin
YKR072C SIS2
ZShift = 0.7
Z lm Score = −5.28
NG = 8
DB = 0
SM = 9
Delta_AUC
Doxorubicin
YMR247C YMR247C
ZShift = 0.75
Z lm Score = 6.95
NG = 8
DB = 0
SM = 8
Delta_L
Doxorubicin
YMR247C YMR247C
ZShift = 0.2
Z lm Score = −12.83
NG = 8
DB = 0
SM = 8
Delta_K
Doxorubicin
YMR247C YMR247C
ZShift = 0.72
Z lm Score = −8.71
NG = 8
DB = 0
SM = 8
Delta_r
Doxorubicin
YMR247C YMR247C
ZShift = −0.44
Z lm Score = −3.27
NG = 8
DB = 0
SM = 8
Delta_AUC
Doxorubicin
YJL081C ARP4
ZShift = 1.37
Z lm Score = 6.92
NG = 8
DB = 0
SM = 8
Delta_L
Doxorubicin
YJL081C ARP4
ZShift = 0.54
Z lm Score = −14.88
NG = 8
DB = 0
SM = 8
Delta_K
Doxorubicin
YJL081C ARP4
ZShift = −1.21
Z lm Score = −8.3
NG = 8
DB = 0
SM = 8
Delta_r
Doxorubicin
YJL081C ARP4
ZShift = −0.72
Z lm Score = −7.55
NG = 8
DB = 0
SM = 8
Delta_AUC
Doxorubicin

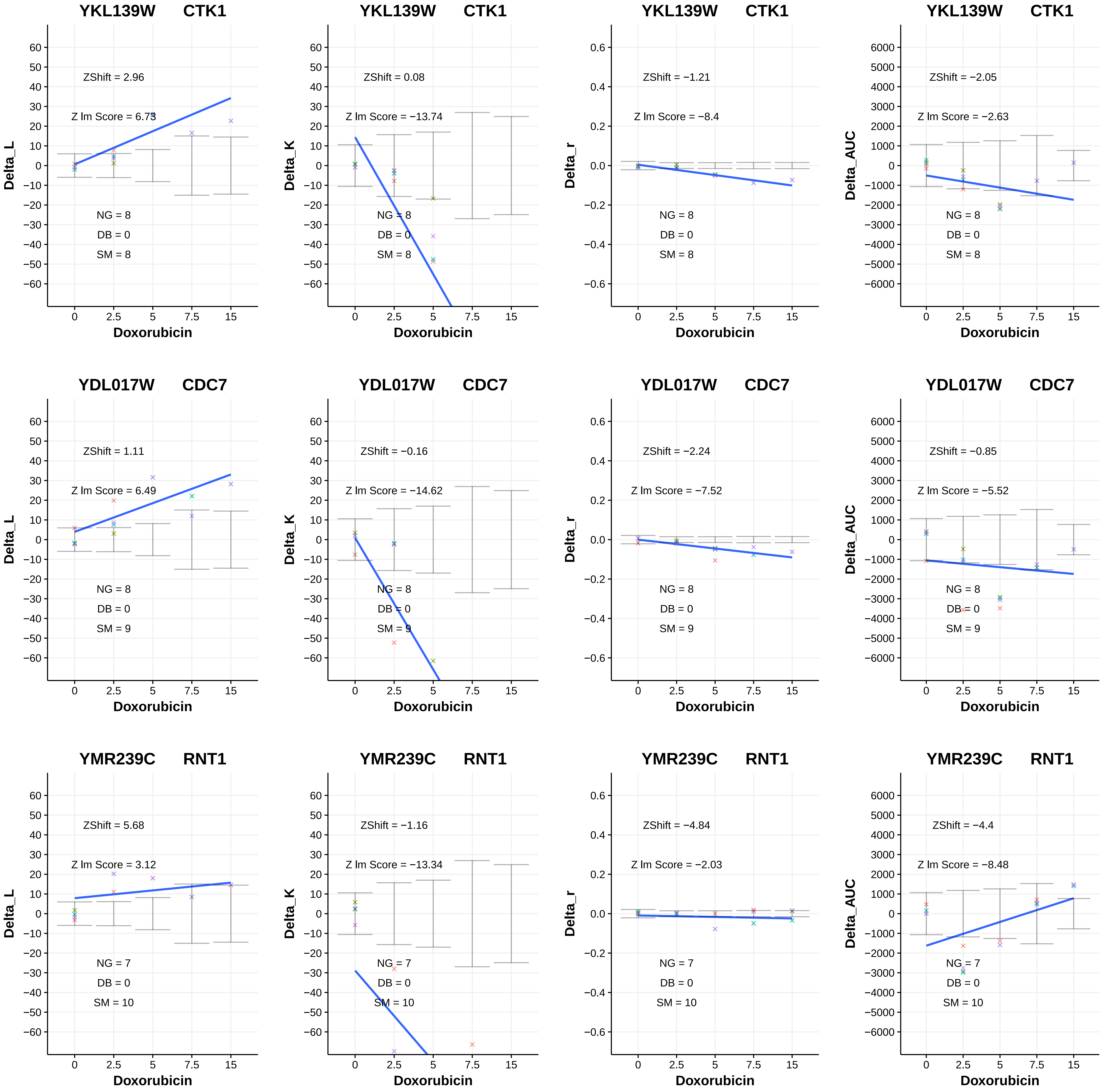

YKL139W CTK1
ZShift = 2.96
Z lm Score = 6.73
NG = 8
DB = 0
SM = 8
Delta_L
Doxorubicin
YKL139W CTK1
ZShift = 0.08
Z lm Score = −13.74
NG = 8
DB = 0
SM = 8
Delta_K
Doxorubicin
YKL139W CTK1
ZShift = −1.21
Z lm Score = −8.4
NG = 8
DB = 0
SM = 8
Delta_r
Doxorubicin
YKL139W CTK1
ZShift = −2.05
Z lm Score = −2.63
NG = 8
DB = 0
SM = 8
Delta_AUC
Doxorubicin
YDL017W CDC7
ZShift = 1.11
Z lm Score = 6.49
NG = 8
DB = 0
SM = 9
Delta_L
Doxorubicin
YDL017W CDC7
ZShift = −0.16
Z lm Score = −14.62
NG = 8
DB = 0
SM = 9
Delta_K
Doxorubicin
YDL017W CDC7
ZShift = −2.24
Z lm Score = −7.52
NG = 8
DB = 0
SM = 9
Delta_r
Doxorubicin
YDL017W CDC7
ZShift = −0.85
Z lm Score = −5.52
NG = 8
DB = 0
SM = 9
Delta_AUC
Doxorubicin
YMR239C RNT1
ZShift = 5.68
Z lm Score = 3.12
NG = 7
DB = 0
SM = 10
Delta_L
Doxorubicin
YMR239C RNT1
ZShift = −1.16
Z lm Score = −13.34
NG = 7
DB = 0
SM = 10
Delta_K
Doxorubicin
YMR239C RNT1
ZShift = −4.84
Z lm Score = −2.03
NG = 7
DB = 0
SM = 10
Delta_r
Doxorubicin
YMR239C RNT1
ZShift = −4.4
Z lm Score = −8.48
NG = 7
DB = 0
SM = 10
Delta_AUC
Doxorubicin

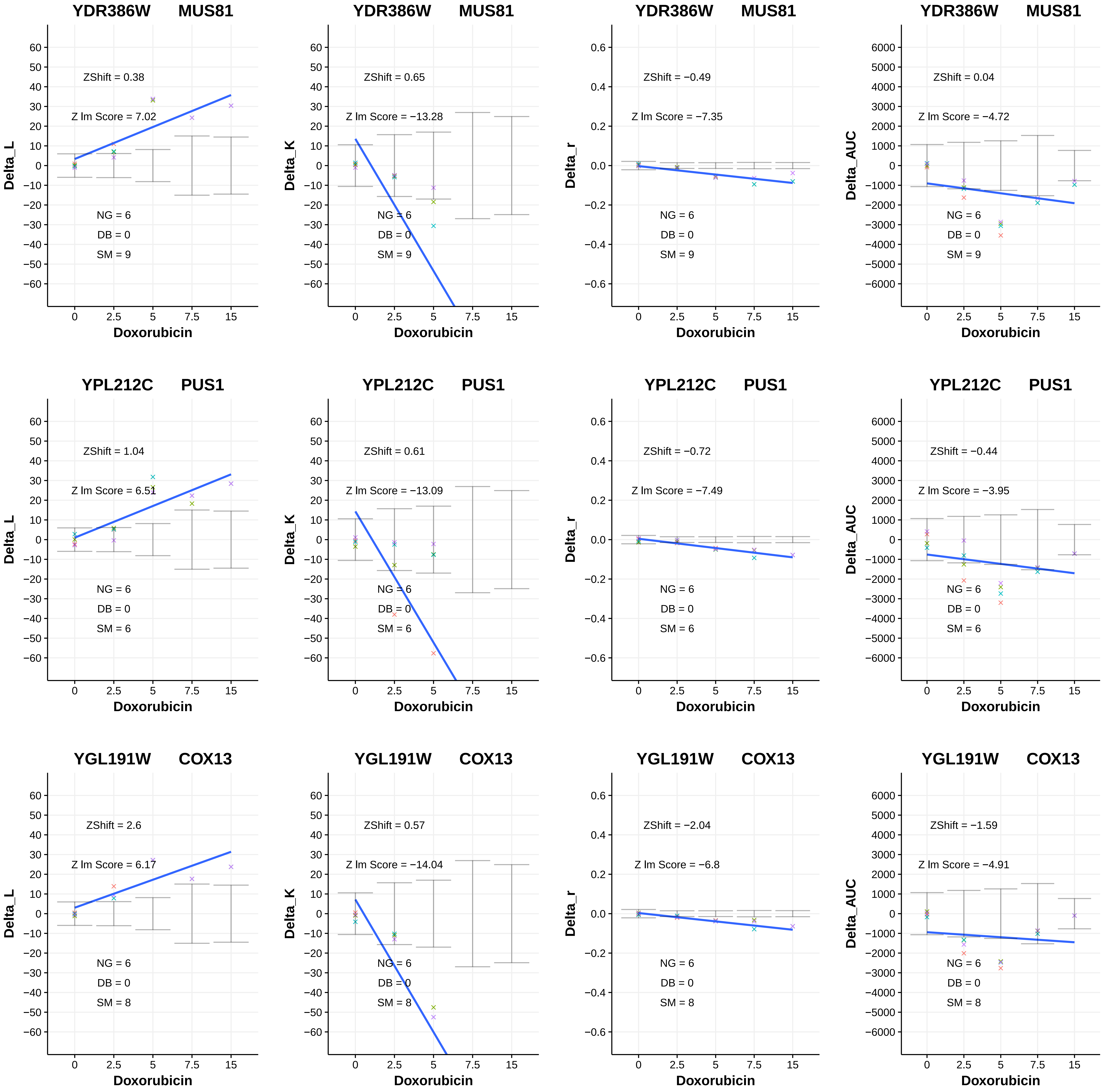
YDR386W MUS81
Delta_L
ZShift = 0.38
Z lm Score = 7.02
NG = 6
DB = 0
SM = 9
Doxorubicin
YDR386W MUS81
Delta_K
ZShift = 0.65
Z lm Score = −13.28
NG = 6
DB = 0
SM = 9
Doxorubicin
YDR386W MUS81
Delta_r
ZShift = −0.49
Z lm Score = −7.35
NG = 6
DB = 0
SM = 9
Doxorubicin
YDR386W MUS81
Delta_AUC
ZShift = 0.04
Z lm Score = −4.72
NG = 6
DB = 0
SM = 9
Doxorubicin
YPL212C PUS1
Delta_L
ZShift = 1.04
Z lm Score = 6.51
NG = 6
DB = 0
SM = 6
Doxorubicin
YPL212C PUS1
Delta_K
ZShift = 0.61
Z lm Score = −13.09
NG = 6
DB = 0
SM = 6
Doxorubicin
YPL212C PUS1
Delta_r
ZShift = −0.72
Z lm Score = −7.49
NG = 6
DB = 0
SM = 6
Doxorubicin
YPL212C PUS1
Delta_AUC
ZShift = −0.44
Z lm Score = −3.95
NG = 6
DB = 0
SM = 6
Doxorubicin
YGL191W COX13
Delta_L
ZShift = 2.6
Z lm Score = 6.17
NG = 6
DB = 0
SM = 8
Doxorubicin
YGL191W COX13
Delta_K
ZShift = 0.57
Z lm Score = −14.04
NG = 6
DB = 0
SM = 8
Doxorubicin
YGL191W COX13
Delta_r
ZShift = −2.04
Z lm Score = −6.8
NG = 6
DB = 0
SM = 8
Doxorubicin
YGL191W COX13
Delta_AUC
ZShift = −1.59
Z lm Score = −4.91
NG = 6
DB = 0
SM = 8
Doxorubicin

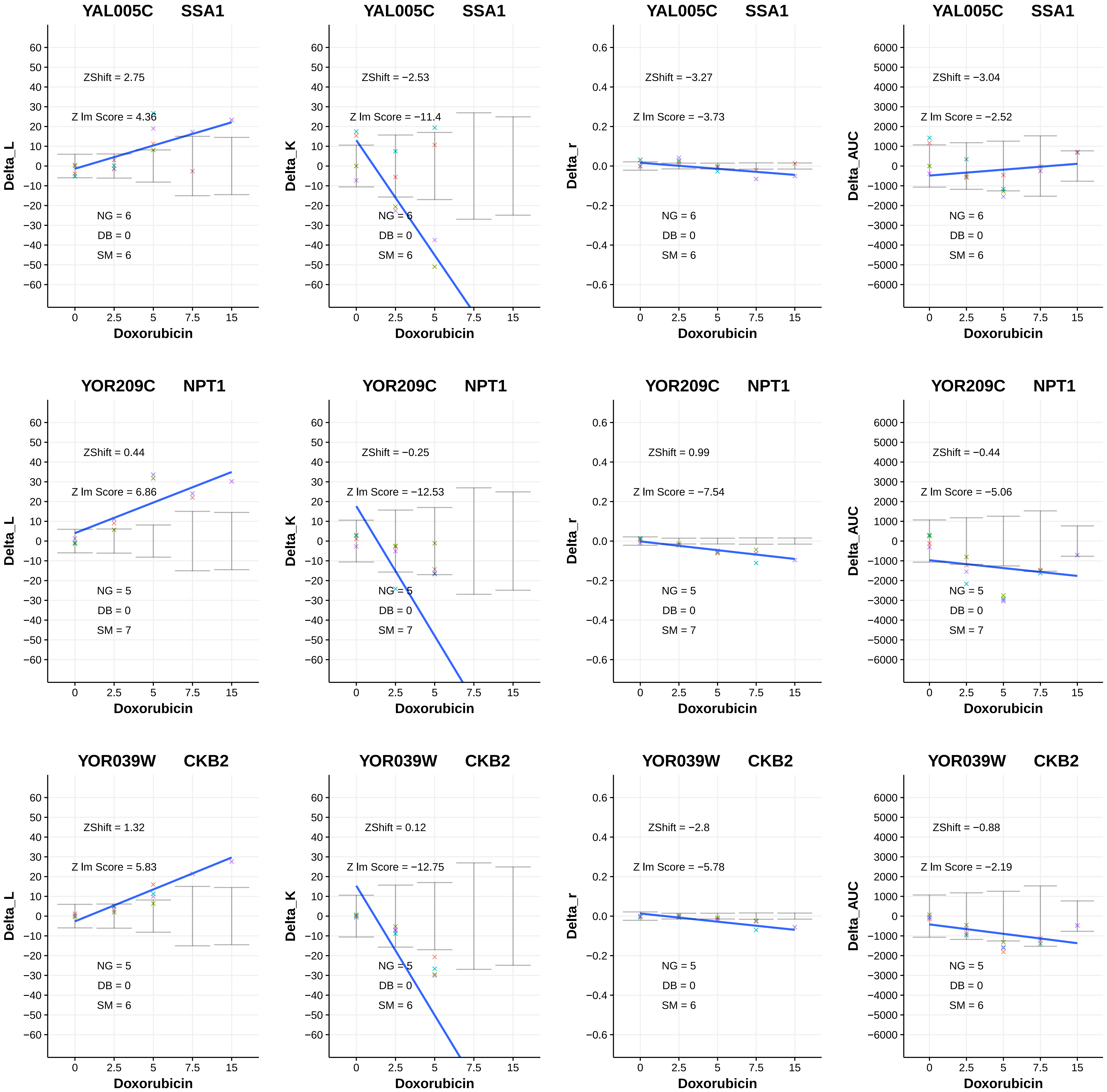
YAL005C SSA1
Delta_L
ZShift = 2.75
Z lm Score = 4.36
NG = 6
DB = 0
SM = 6
Doxorubicin
YAL005C SSA1
Delta_K
ZShift = −2.53
Z lm Score = −11.4
NG = 6
DB = 0
SM = 6
Doxorubicin
YAL005C SSA1
Delta_r
ZShift = −3.27
Z lm Score = −3.73
NG = 6
DB = 0
SM = 6
Doxorubicin
YAL005C SSA1
Delta_AUC
ZShift = −3.04
Z lm Score = −2.52
NG = 6
DB = 0
SM = 6
Doxorubicin
YOR209C NPT1
Delta_L
ZShift = 0.44
Z lm Score = 6.86
NG = 5
DB = 0
SM = 7
Doxorubicin
YOR209C NPT1
Delta_K
ZShift = −0.25
Z lm Score = −12.53
NG = 5
DB = 0
SM = 7
Doxorubicin
YOR209C NPT1
Delta_r
ZShift = 0.99
Z lm Score = −7.54
NG = 5
DB = 0
SM = 7
Doxorubicin
YOR209C NPT1
Delta_AUC
ZShift = −0.44
Z lm Score = −5.06
NG = 5
DB = 0
SM = 7
Doxorubicin
YOR039W CKB2
Delta_L
ZShift = 1.32
Z lm Score = 5.83
NG = 5
DB = 0
SM = 6
Doxorubicin
YOR039W CKB2
Delta_K
ZShift = 0.12
Z lm Score = −12.75
NG = 5
DB = 0
SM = 6
Doxorubicin
YOR039W CKB2
Delta_r
ZShift = −2.8
Z lm Score = −5.78
NG = 5
DB = 0
SM = 6
Doxorubicin
YOR039W CKB2
Delta_AUC
ZShift = −0.88
Z lm Score = −2.19
NG = 5
DB = 0
SM = 6
Doxorubicin

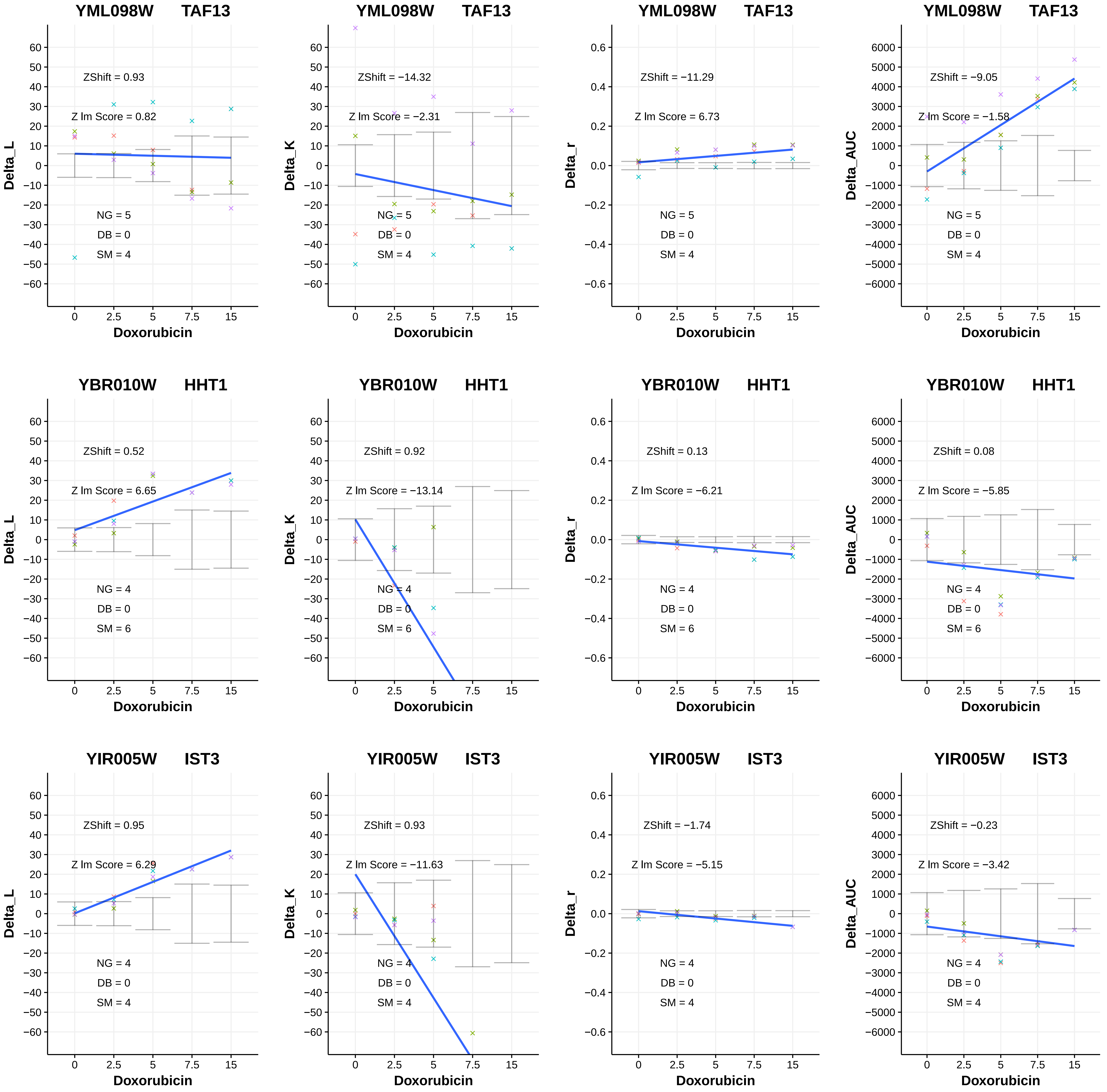
YML098W TAF13
ZShift = 0.93
Z lm Score = 0.82
NG = 5
DB = 0
SM = 4
Delta_L
Doxorubicin
YML098W TAF13
ZShift = −14.32
Z lm Score = −2.31
NG = 5
DB = 0
SM = 4
Delta_K
Doxorubicin
YML098W TAF13
ZShift = −11.29
Z lm Score = 6.73
NG = 5
DB = 0
SM = 4
Delta_r
Doxorubicin
YML098W TAF13
ZShift = −9.05
Z lm Score = −1.58
NG = 5
DB = 0
SM = 4
Delta_AUC
Doxorubicin
YBR010W HHT1
ZShift = 0.52
Z lm Score = 6.65
NG = 4
DB = 0
SM = 6
Delta_L
Doxorubicin
YBR010W HHT1
ZShift = 0.92
Z lm Score = −13.14
NG = 4
DB = 0
SM = 6
Delta_K
Doxorubicin
YBR010W HHT1
ZShift = 0.13
Z lm Score = −6.21
NG = 4
DB = 0
SM = 6
Delta_r
Doxorubicin
YBR010W HHT1
ZShift = 0.08
Z lm Score = −5.85
NG = 4
DB = 0
SM = 6
Delta_AUC
Doxorubicin
YIR005W IST3
ZShift = 0.95
Z lm Score = 6.29
NG = 4
DB = 0
SM = 4
Delta_L
Doxorubicin
YIR005W IST3
ZShift = 0.93
Z lm Score = −11.63
NG = 4
DB = 0
SM = 4
Delta_K
Doxorubicin
YIR005W IST3
ZShift = −1.74
Z lm Score = −5.15
NG = 4
DB = 0
SM = 4
Delta_r
Doxorubicin
YIR005W IST3
ZShift = −0.23
Z lm Score = −3.42
NG = 4
DB = 0
SM = 4
Delta_AUC
Doxorubicin
0 2.5 5 7.5 15

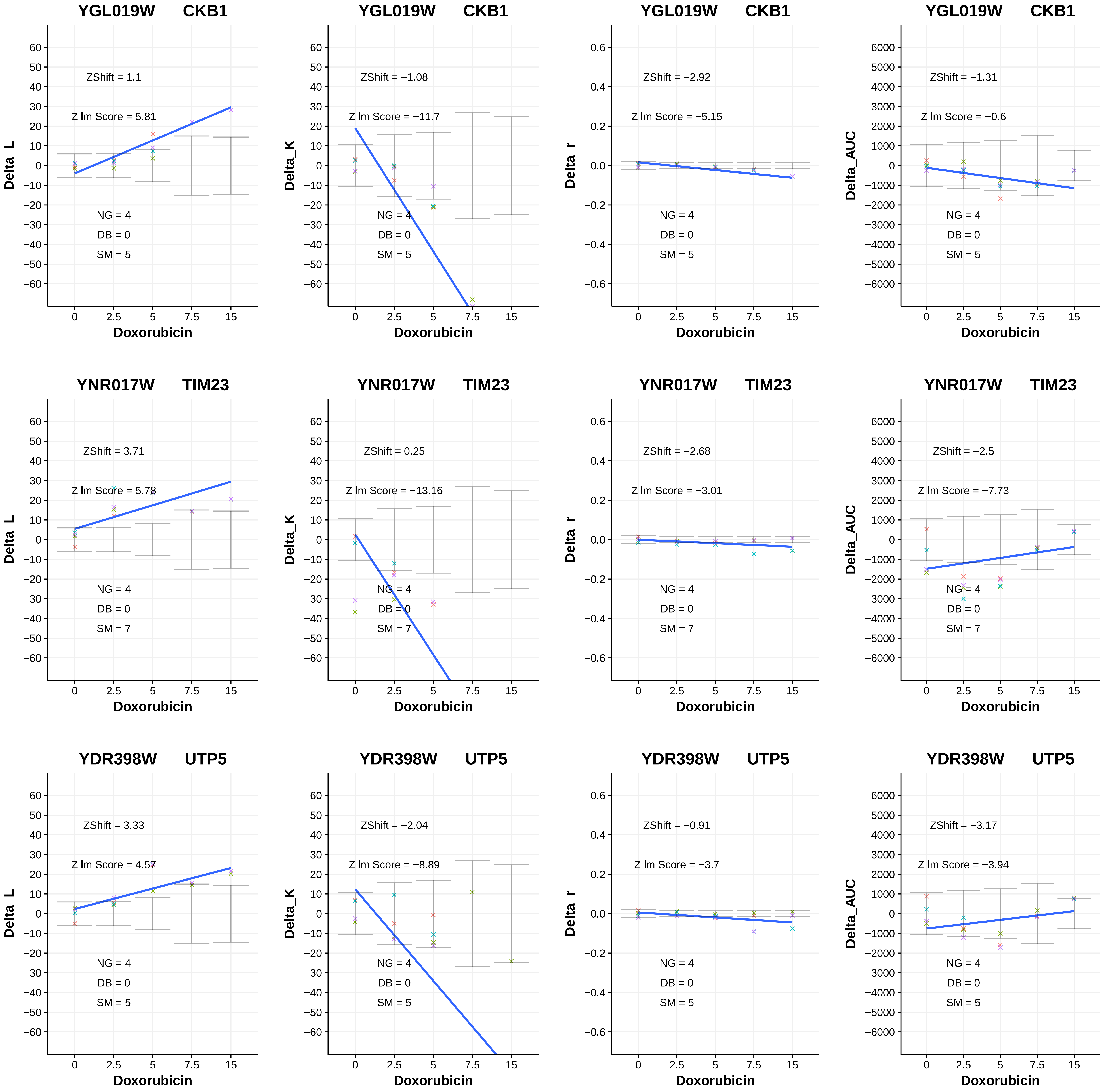
YGL019W CKB1
ZShift = 1.1
Z lm Score = 5.81
NG = 4
DB = 0
SM = 5
Delta_L
Doxorubicin
YGL019W CKB1
ZShift = −1.08
Z lm Score = −11.7
NG = 4
DB = 0
SM = 5
Delta_K
Doxorubicin
YGL019W CKB1
ZShift = −2.92
Z lm Score = −5.15
NG = 4
DB = 0
SM = 5
Delta_r
Doxorubicin
YGL019W CKB1
ZShift = −1.31
Z lm Score = −0.6
NG = 4
DB = 0
SM = 5
Delta_AUC
Doxorubicin
YNR017W TIM23
ZShift = 3.71
Z lm Score = 5.78
NG = 4
DB = 0
SM = 7
Delta_L
Doxorubicin
YNR017W TIM23
ZShift = 0.25
Z lm Score = −13.16
NG = 4
DB = 0
SM = 7
Delta_K
Doxorubicin
YNR017W TIM23
ZShift = −2.68
Z lm Score = −3.01
NG = 4
DB = 0
SM = 7
Delta_r
Doxorubicin
YNR017W TIM23
ZShift = −2.5
Z lm Score = −7.73
NG = 4
DB = 0
SM = 7
Delta_AUC
Doxorubicin
YDR398W UTP5
ZShift = 3.33
Z lm Score = 4.57
NG = 4
DB = 0
SM = 5
Delta_L
Doxorubicin
YDR398W UTP5
ZShift = −2.04
Z lm Score = −8.89
NG = 4
DB = 0
SM = 5
Delta_K
Doxorubicin
YDR398W UTP5
ZShift = −0.91
Z lm Score = −3.7
NG = 4
DB = 0
SM = 5
Delta_r
Doxorubicin
YDR398W UTP5
ZShift = −3.17
Z lm Score = −3.94
NG = 4
DB = 0
SM = 5
Delta_AUC
Doxorubicin

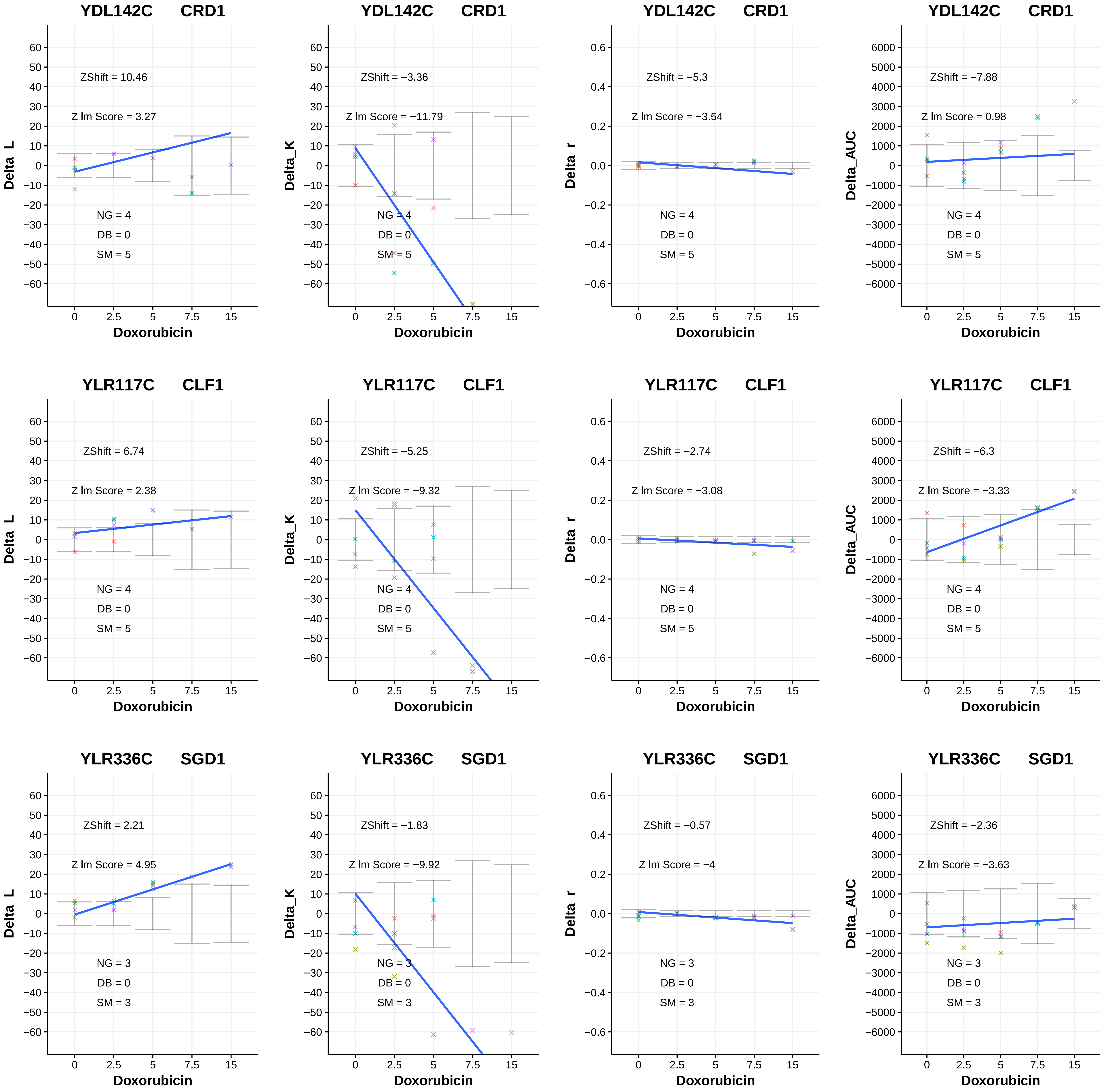
YDL142C CRD1
ZShift = 10.46
Z lm Score = 3.27
NG = 4
DB = 0
SM = 5
Delta_L
Doxorubicin
YDL142C CRD1
ZShift = −3.36
Z lm Score = −11.79
NG = 4
DB = 0
SM = 5
Delta_K
Doxorubicin
YDL142C CRD1
ZShift = −5.3
Z lm Score = −3.54
NG = 4
DB = 0
SM = 5
Delta_r
Doxorubicin
YDL142C CRD1
ZShift = −7.88
Z lm Score = 0.98
NG = 4
DB = 0
SM = 5
Delta_AUC
Doxorubicin
YLR117C CLF1
ZShift = 6.74
Z lm Score = 2.38
NG = 4
DB = 0
SM = 5
Delta_L
Doxorubicin
YLR117C CLF1
ZShift = −5.25
Z lm Score = −9.32
NG = 4
DB = 0
SM = 5
Delta_K
Doxorubicin
YLR117C CLF1
ZShift = −2.74
Z lm Score = −3.08
NG = 4
DB = 0
SM = 5
Delta_r
Doxorubicin
YLR117C CLF1
ZShift = −6.3
Z lm Score = −3.33
NG = 4
DB = 0
SM = 5
Delta_AUC
Doxorubicin
YLR336C SGD1
ZShift = 2.21
Z lm Score = 4.95
NG = 3
DB = 0
SM = 3
Delta_L
Doxorubicin
YLR336C SGD1
ZShift = −1.83
Z lm Score = −9.92
NG = 3
DB = 0
SM = 3
Delta_K
Doxorubicin
YLR336C SGD1
ZShift = −0.57
Z lm Score = −4
NG = 3
DB = 0
SM = 3
Delta_r
Doxorubicin
YLR336C SGD1
ZShift = −2.36
Z lm Score = −3.63
NG = 3
DB = 0
SM = 3
Delta_AUC
Doxorubicin

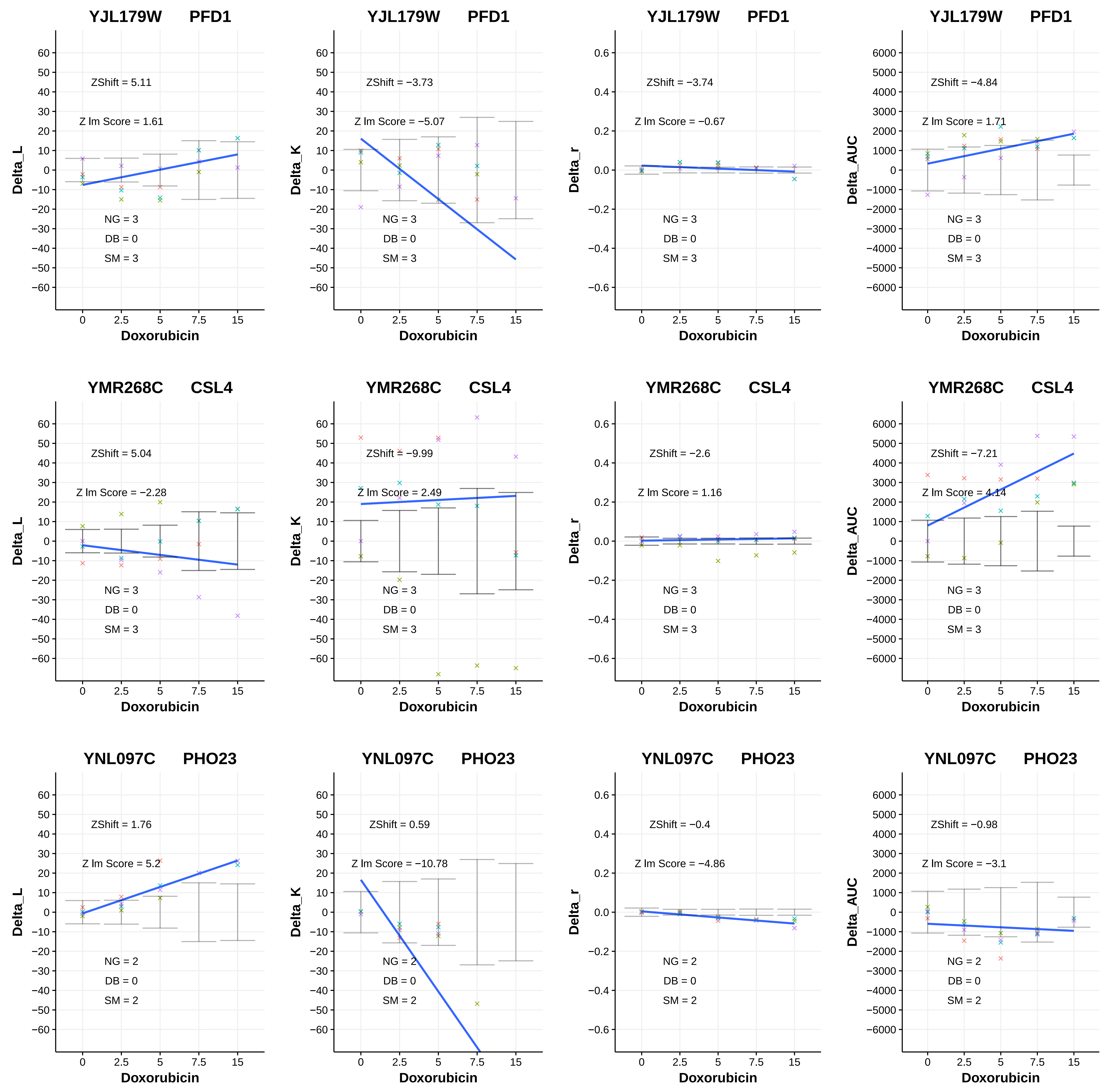
YJL179W PFD1
Delta_L
ZShift = 5.11
Z lm Score = 1.61
NG = 3
DB = 0
SM = 3
Doxorubicin
YJL179W PFD1
Delta_K
ZShift = −3.73
Z lm Score = −5.07
NG = 3
DB = 0
SM = 3
Doxorubicin
YJL179W PFD1
Delta_r
ZShift = −3.74
Z lm Score = −0.67
NG = 3
DB = 0
SM = 3
Doxorubicin
YJL179W PFD1
Delta_AUC
ZShift = −4.84
Z lm Score = 1.71
NG = 3
DB = 0
SM = 3
Doxorubicin
YMR268C CSL4
Delta_L
ZShift = 5.04
Z lm Score = −2.28
NG = 3
DB = 0
SM = 3
Doxorubicin
YMR268C CSL4
Delta_K
ZShift = −9.99
Z lm Score = 2.49
NG = 3
DB = 0
SM = 3
Doxorubicin
YMR268C CSL4
Delta_r
ZShift = −2.6
Z lm Score = 1.16
NG = 3
DB = 0
SM = 3
Doxorubicin
YMR268C CSL4
Delta_AUC
ZShift = −7.21
Z lm Score = 4.14
NG = 3
DB = 0
SM = 3
Doxorubicin
YNL097C PHO23
Delta_L
ZShift = 1.76
Z lm Score = 5.2
NG = 2
DB = 0
SM = 2
Doxorubicin
YNL097C PHO23
Delta_K
ZShift = 0.59
Z lm Score = −10.78
NG = 2
DB = 0
SM = 2
Doxorubicin
YNL097C PHO23
Delta_r
ZShift = −0.4
Z lm Score = −4.86
NG = 2
DB = 0
SM = 2
Doxorubicin
YNL097C PHO23
Delta_AUC
ZShift = −0.98
Z lm Score = −3.1
NG = 2
DB = 0
SM = 2
Doxorubicin

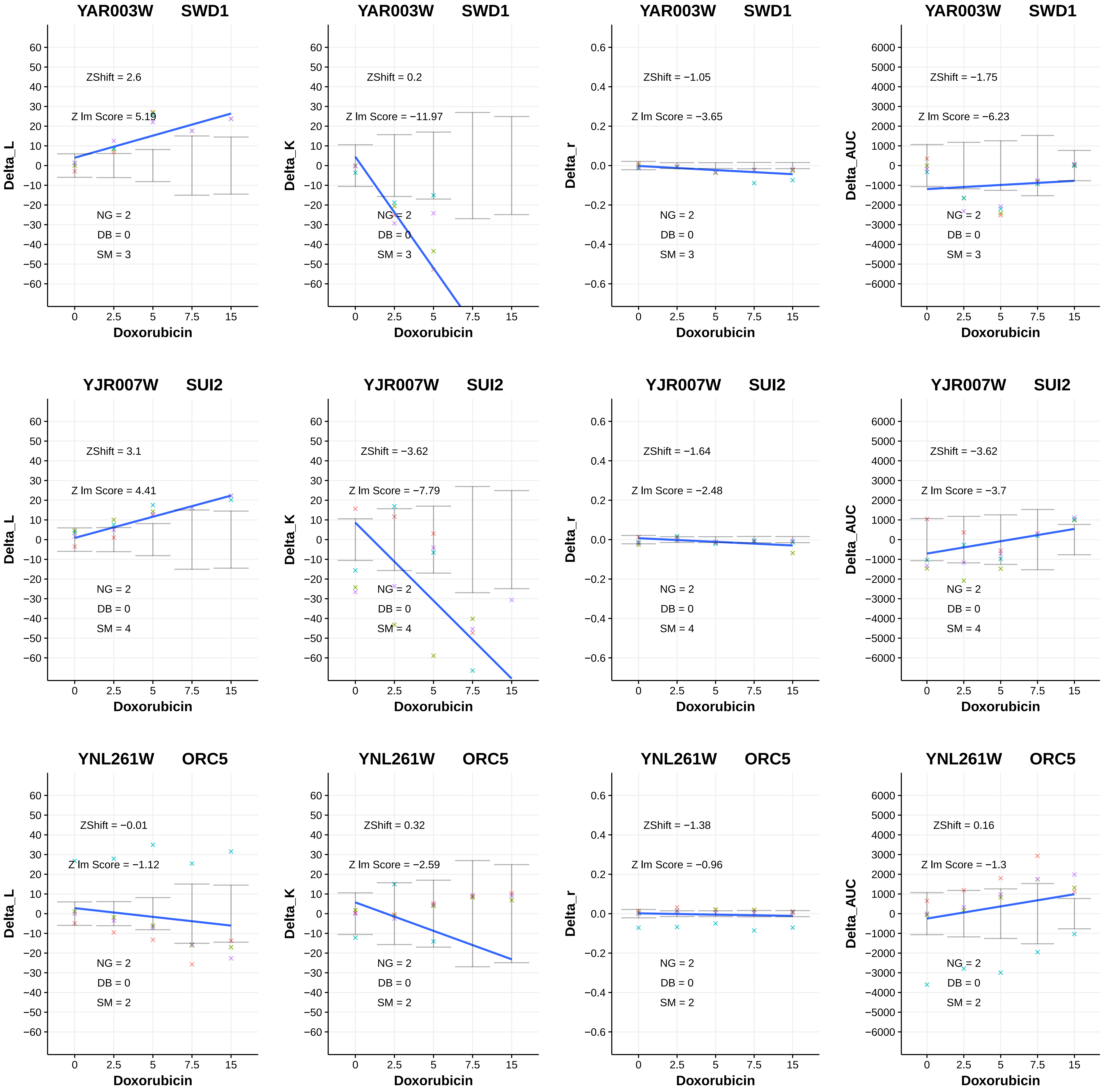

YAR003W SWD1
ZShift = 2.6
Z lm Score = 5.19
NG = 2
DB = 0
SM = 3
Delta_L
Doxorubicin
YAR003W SWD1
ZShift = 0.2
Z lm Score = −11.97
NG = 2
DB = 0
SM = 3
Delta_K
Doxorubicin
YAR003W SWD1
ZShift = −1.05
Z lm Score = −3.65
NG = 2
DB = 0
SM = 3
Delta_r
Doxorubicin
YAR003W SWD1
ZShift = −1.75
Z lm Score = −6.23
NG = 2
DB = 0
SM = 3
Delta_AUC
Doxorubicin
YJR007W SUI2
ZShift = 3.1
Z lm Score = 4.41
NG = 2
DB = 0
SM = 4
Delta_L
Doxorubicin
YJR007W SUI2
ZShift = −3.62
Z lm Score = −7.79
NG = 2
DB = 0
SM = 4
Delta_K
Doxorubicin
YJR007W SUI2
ZShift = −1.64
Z lm Score = −2.48
NG = 2
DB = 0
SM = 4
Delta_r
Doxorubicin
YJR007W SUI2
ZShift = −3.62
Z lm Score = −3.7
NG = 2
DB = 0
SM = 4
Delta_AUC
Doxorubicin
YNL261W ORC5
ZShift = −0.01
Z lm Score = −1.12
NG = 2
DB = 0
SM = 2
Delta_L
Doxorubicin
YNL261W ORC5
ZShift = 0.32
Z lm Score = −2.59
NG = 2
DB = 0
SM = 2
Delta_K
Doxorubicin
YNL261W ORC5
ZShift = −1.38
Z lm Score = −0.96
NG = 2
DB = 0
SM = 2
Delta_r
Doxorubicin
YNL261W ORC5
ZShift = 0.16
Z lm Score = −1.3
NG = 2
DB = 0
SM = 2
Delta_AUC
Doxorubicin

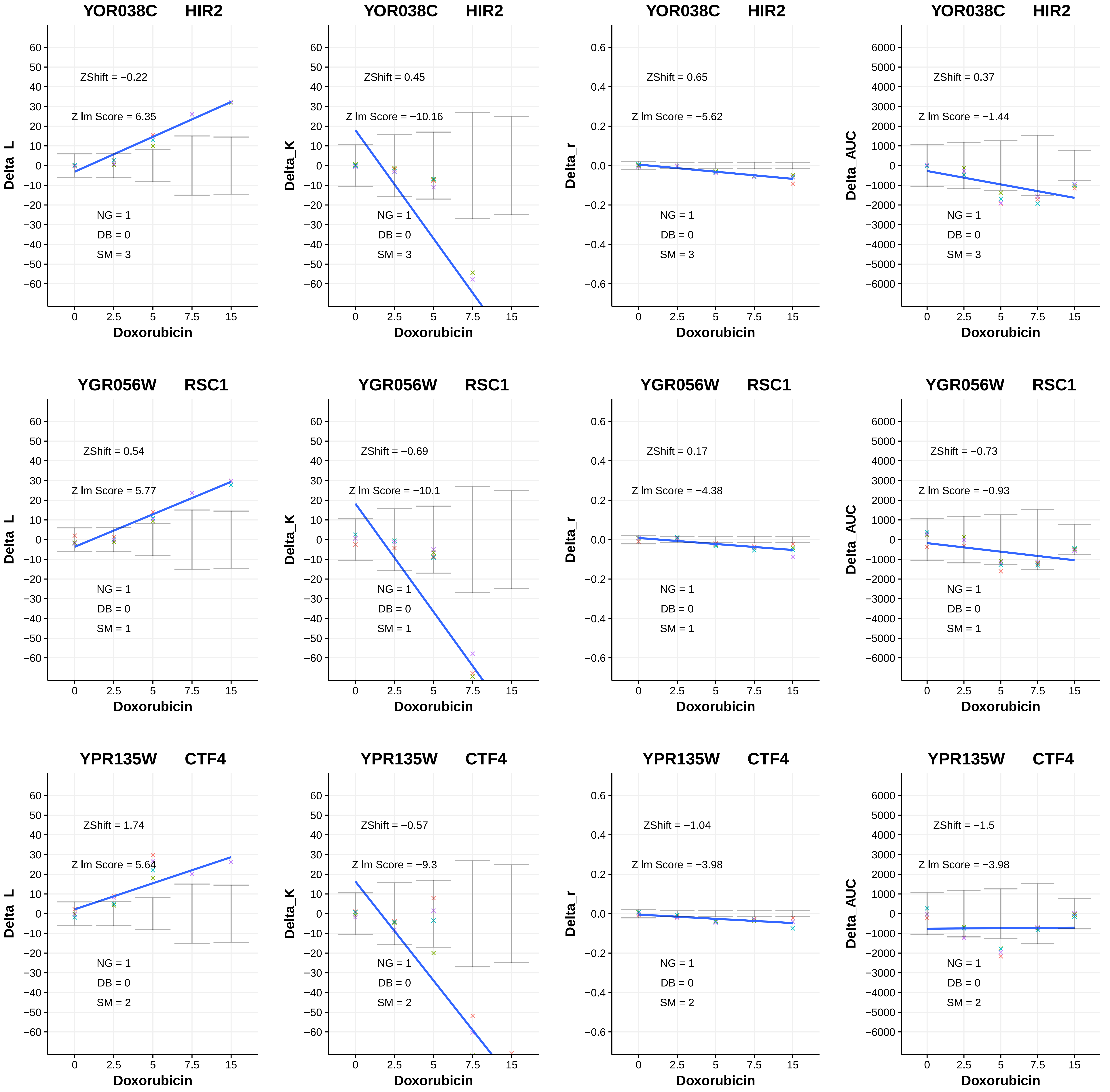
YOR038C HIR2
ZShift = −0.22
Z lm Score = 6.35
NG = 1
DB = 0
SM = 3
Delta_L
Doxorubicin
YOR038C HIR2
ZShift = 0.45
Z lm Score = −10.16
NG = 1
DB = 0
SM = 3
Delta_K
Doxorubicin
YOR038C HIR2
ZShift = 0.65
Z lm Score = −5.62
NG = 1
DB = 0
SM = 3
Delta_r
Doxorubicin
YOR038C HIR2
ZShift = 0.37
Z lm Score = −1.44
NG = 1
DB = 0
SM = 3
Delta_AUC
Doxorubicin
YGR056W RSC1
ZShift = 0.54
Z lm Score = 5.77
NG = 1
DB = 0
SM = 1
Delta_L
Doxorubicin
YGR056W RSC1
ZShift = −0.69
Z lm Score = −10.1
NG = 1
DB = 0
SM = 1
Delta_K
Doxorubicin
YGR056W RSC1
ZShift = 0.17
Z lm Score = −4.38
NG = 1
DB = 0
SM = 1
Delta_r
Doxorubicin
YGR056W RSC1
ZShift = −0.73
Z lm Score = −0.93
NG = 1
DB = 0
SM = 1
Delta_AUC
Doxorubicin
YPR135W CTF4
ZShift = 1.74
Z lm Score = 5.64
NG = 1
DB = 0
SM = 2
Delta_L
Doxorubicin
YPR135W CTF4
ZShift = −0.57
Z lm Score = −9.3
NG = 1
DB = 0
SM = 2
Delta_K
Doxorubicin
YPR135W CTF4
ZShift = −1.04
Z lm Score = −3.98
NG = 1
DB = 0
SM = 2
Delta_r
Doxorubicin
YPR135W CTF4
ZShift = −1.5
Z lm Score = −3.98
NG = 1
DB = 0
SM = 2
Delta_AUC
Doxorubicin

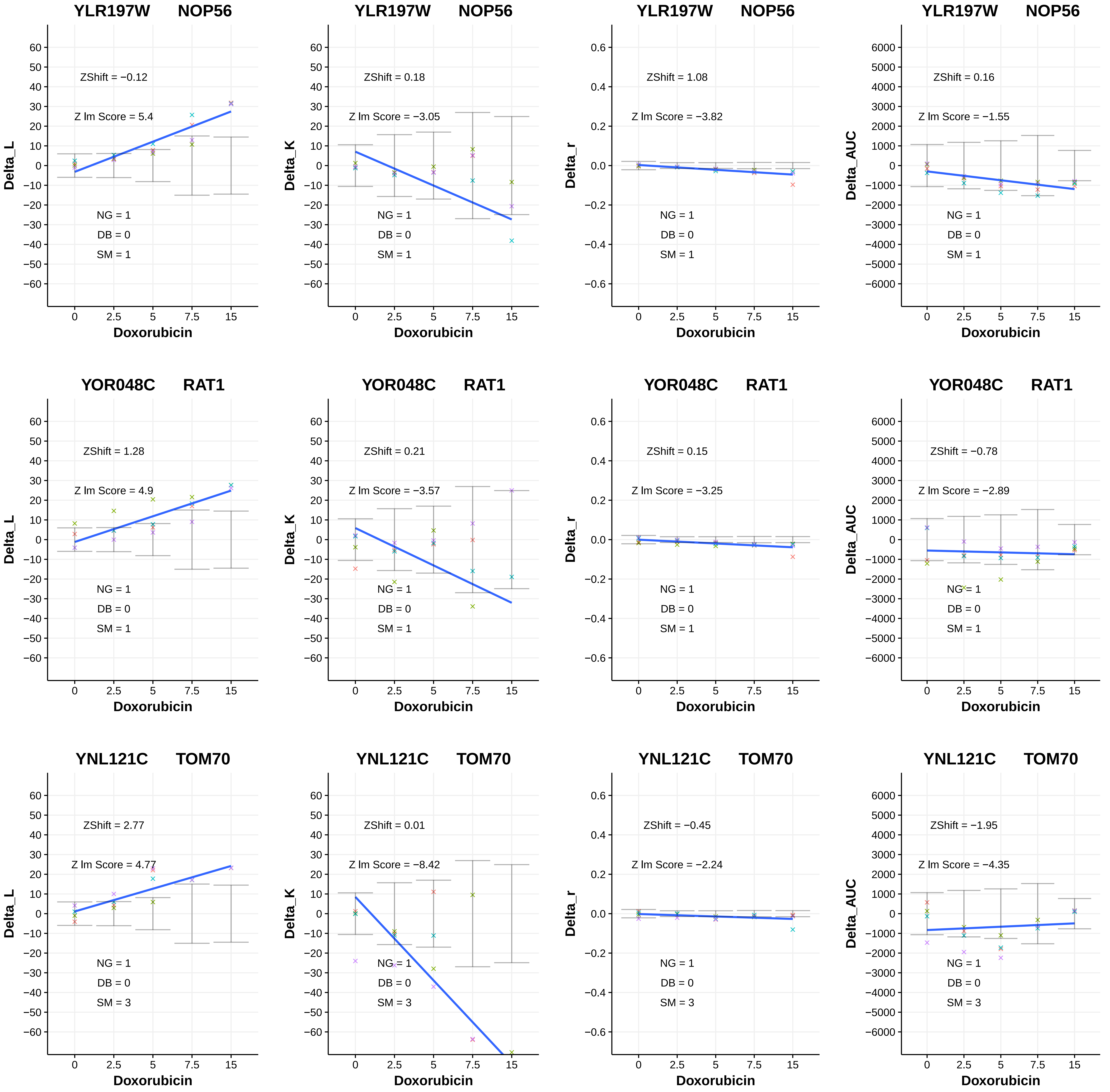
YLR197W NOP56
ZShift = −0.12
Z lm Score = 5.4
NG = 1
DB = 0
SM = 1
Delta_L
Doxorubicin
YLR197W NOP56
ZShift = 0.18
Z lm Score = −3.05
NG = 1
DB = 0
SM = 1
Delta_K
Doxorubicin
YLR197W NOP56
ZShift = 1.08
Z lm Score = −3.82
NG = 1
DB = 0
SM = 1
Delta_r
Doxorubicin
YLR197W NOP56
ZShift = 0.16
Z lm Score = −1.55
NG = 1
DB = 0
SM = 1
Delta_AUC
Doxorubicin
YOR048C RAT1
ZShift = 1.28
Z lm Score = 4.9
NG = 1
DB = 0
SM = 1
Delta_L
Doxorubicin
YOR048C RAT1
ZShift = 0.21
Z lm Score = −3.57
NG = 1
DB = 0
SM = 1
Delta_K
Doxorubicin
YOR048C RAT1
ZShift = 0.15
Z lm Score = −3.25
NG = 1
DB = 0
SM = 1
Delta_r
Doxorubicin
YOR048C RAT1
ZShift = −0.78
Z lm Score = −2.89
NG = 1
DB = 0
SM = 1
Delta_AUC
Doxorubicin
YNL121C TOM70
ZShift = 2.77
Z lm Score = 4.77
NG = 1
DB = 0
SM = 3
Delta_L
Doxorubicin
YNL121C TOM70
ZShift = 0.01
Z lm Score = −8.42
NG = 1
DB = 0
SM = 3
Delta_K
Doxorubicin
YNL121C TOM70
ZShift = −0.45
Z lm Score = −2.24
NG = 1
DB = 0
SM = 3
Delta_r
Doxorubicin
YNL121C TOM70
ZShift = −1.95
Z lm Score = −4.35
NG = 1
DB = 0
SM = 3
Delta_AUC
Doxorubicin

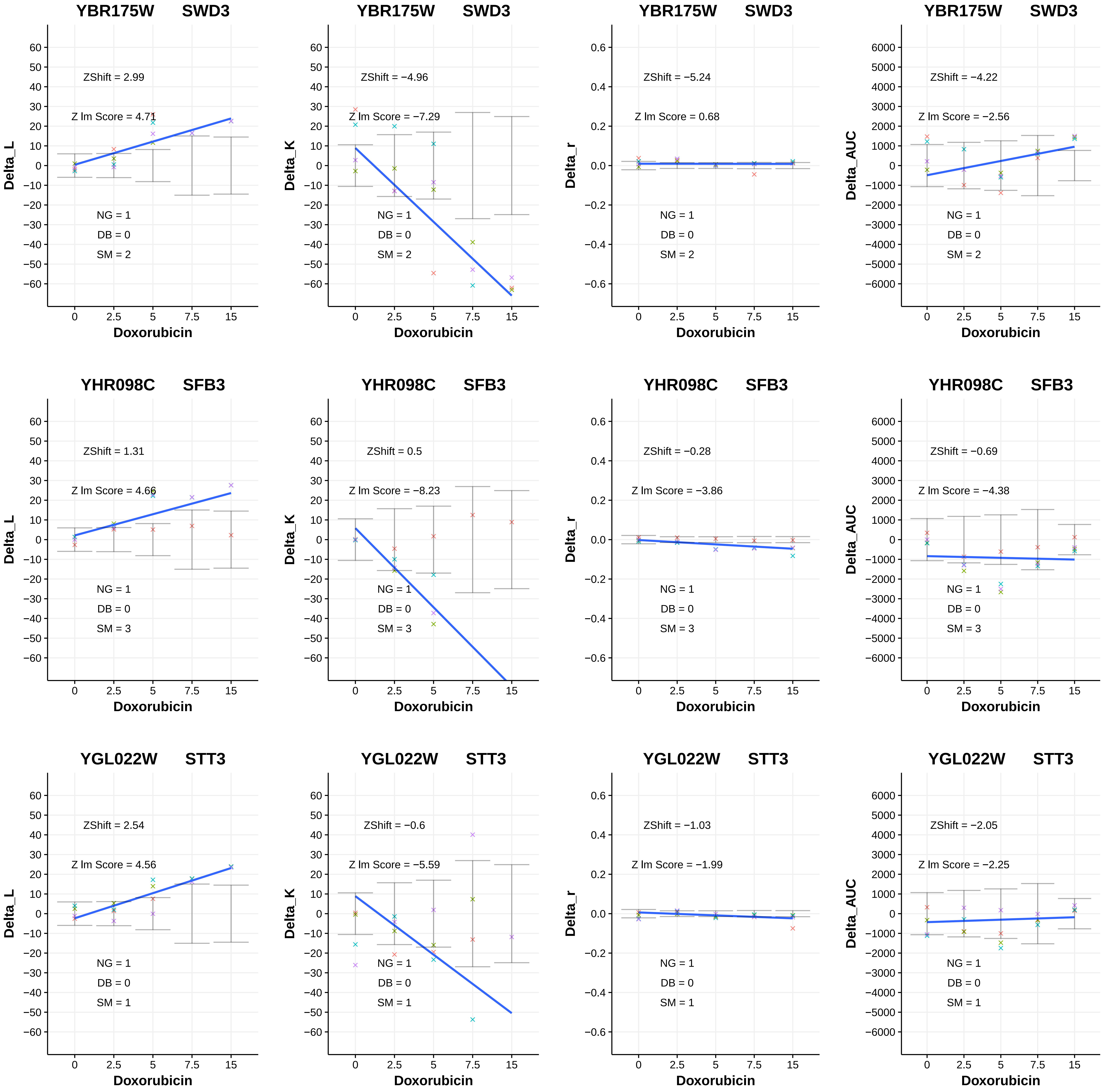

YBR175W SWD3
ZShift = 2.99
Z lm Score = 4.71
NG = 1
DB = 0
SM = 2
Delta_L
Doxorubicin
YBR175W SWD3
ZShift = −4.96
Z lm Score = −7.29
NG = 1
DB = 0
SM = 2
Delta_K
Doxorubicin
YBR175W SWD3
ZShift = −5.24
Z lm Score = 0.68
NG = 1
DB = 0
SM = 2
Delta_r
Doxorubicin
YBR175W SWD3
ZShift = −4.22
Z lm Score = −2.56
NG = 1
DB = 0
SM = 2
Delta_AUC
Doxorubicin
YHR098C SFB3
ZShift = 1.31
Z lm Score = 4.66
NG = 1
DB = 0
SM = 3
Delta_L
Doxorubicin
YHR098C SFB3
ZShift = 0.5
Z lm Score = −8.23
NG = 1
DB = 0
SM = 3
Delta_K
Doxorubicin
YHR098C SFB3
ZShift = −0.28
Z lm Score = −3.86
NG = 1
DB = 0
SM = 3
Delta_r
Doxorubicin
YHR098C SFB3
ZShift = −0.69
Z lm Score = −4.38
NG = 1
DB = 0
SM = 3
Delta_AUC
Doxorubicin
YGL022W STT3
ZShift = 2.54
Z lm Score = 4.56
NG = 1
DB = 0
SM = 1
Delta_L
Doxorubicin
YGL022W STT3
ZShift = −0.6
Z lm Score = −5.59
NG = 1
DB = 0
SM = 1
Delta_K
Doxorubicin
YGL022W STT3
ZShift = −1.03
Z lm Score = −1.99
NG = 1
DB = 0
SM = 1
Delta_r
Doxorubicin
YGL022W STT3
ZShift = −2.05
Z lm Score = −2.25
NG = 1
DB = 0
SM = 1
Delta_AUC
Doxorubicin

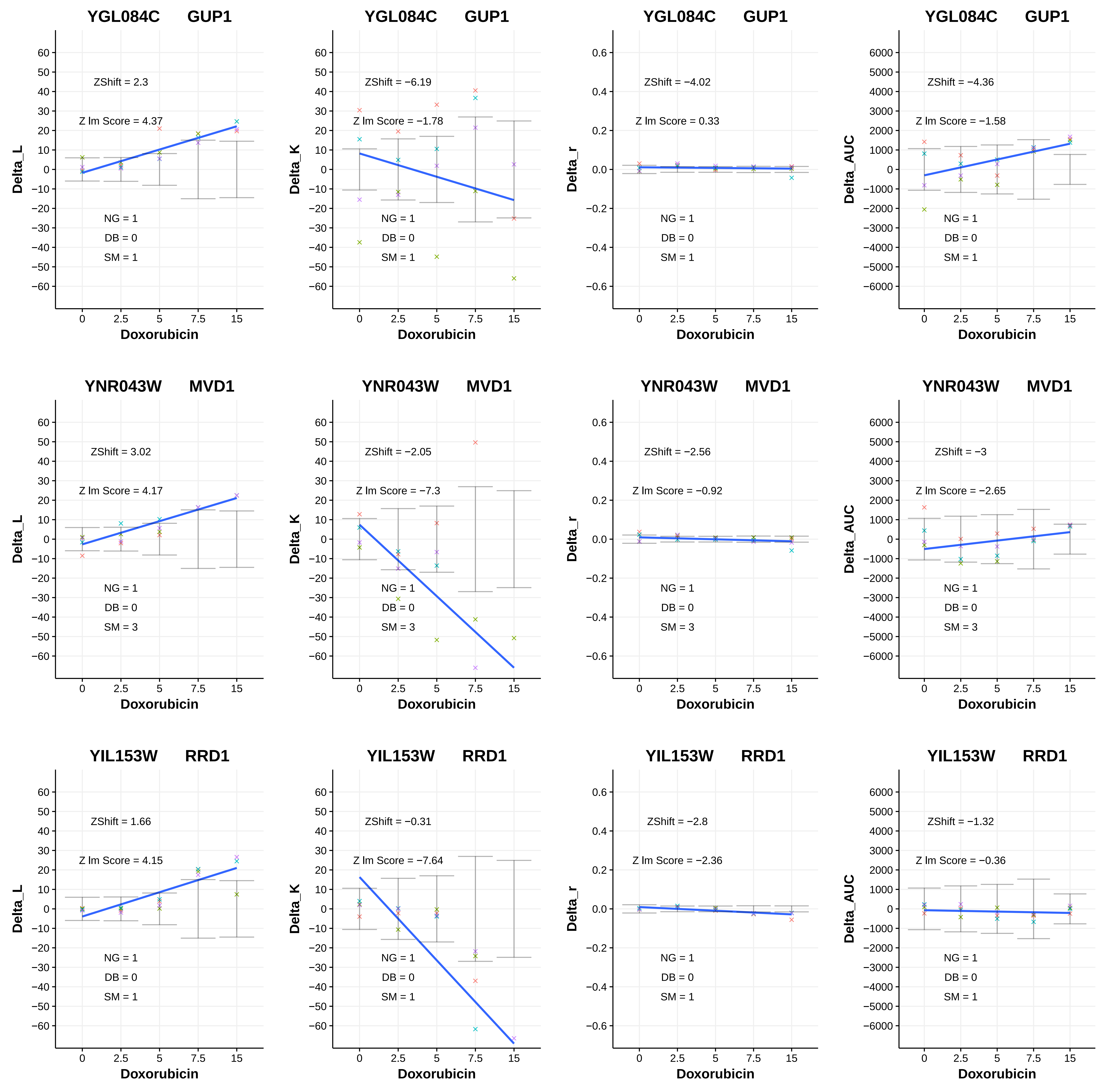

YGL084C GUP1
ZShift = 2.3
Z lm Score = 4.37
NG = 1
DB = 0
SM = 1
Delta_L
Doxorubicin
YGL084C GUP1
ZShift = −6.19
Z lm Score = −1.78
NG = 1
DB = 0
SM = 1
Delta_K
Doxorubicin
YGL084C GUP1
ZShift = −4.02
Z lm Score = 0.33
NG = 1
DB = 0
SM = 1
Delta_r
Doxorubicin
YGL084C GUP1
ZShift = −4.36
Z lm Score = −1.58
NG = 1
DB = 0
SM = 1
Delta_AUC
Doxorubicin
YNR043W MVD1
ZShift = 3.02
Z lm Score = 4.17
NG = 1
DB = 0
SM = 3
Delta_L
Doxorubicin
YNR043W MVD1
ZShift = −2.05
Z lm Score = −7.3
NG = 1
DB = 0
SM = 3
Delta_K
Doxorubicin
YNR043W MVD1
ZShift = −2.56
Z lm Score = −0.92
NG = 1
DB = 0
SM = 3
Delta_r
Doxorubicin
YNR043W MVD1
ZShift = −3
Z lm Score = −2.65
NG = 1
DB = 0
SM = 3
Delta_AUC
Doxorubicin
YIL153W RRD1
ZShift = 1.66
Z lm Score = 4.15
NG = 1
DB = 0
SM = 1
Delta_L
Doxorubicin
YIL153W RRD1
ZShift = −0.31
Z lm Score = −7.64
NG = 1
DB = 0
SM = 1
Delta_K
Doxorubicin
YIL153W RRD1
ZShift = −2.8
Z lm Score = −2.36
NG = 1
DB = 0
SM = 1
Delta_r
Doxorubicin
YIL153W RRD1
ZShift = −1.32
Z lm Score = −0.36
NG = 1
DB = 0
SM = 1
Delta_AUC
Doxorubicin

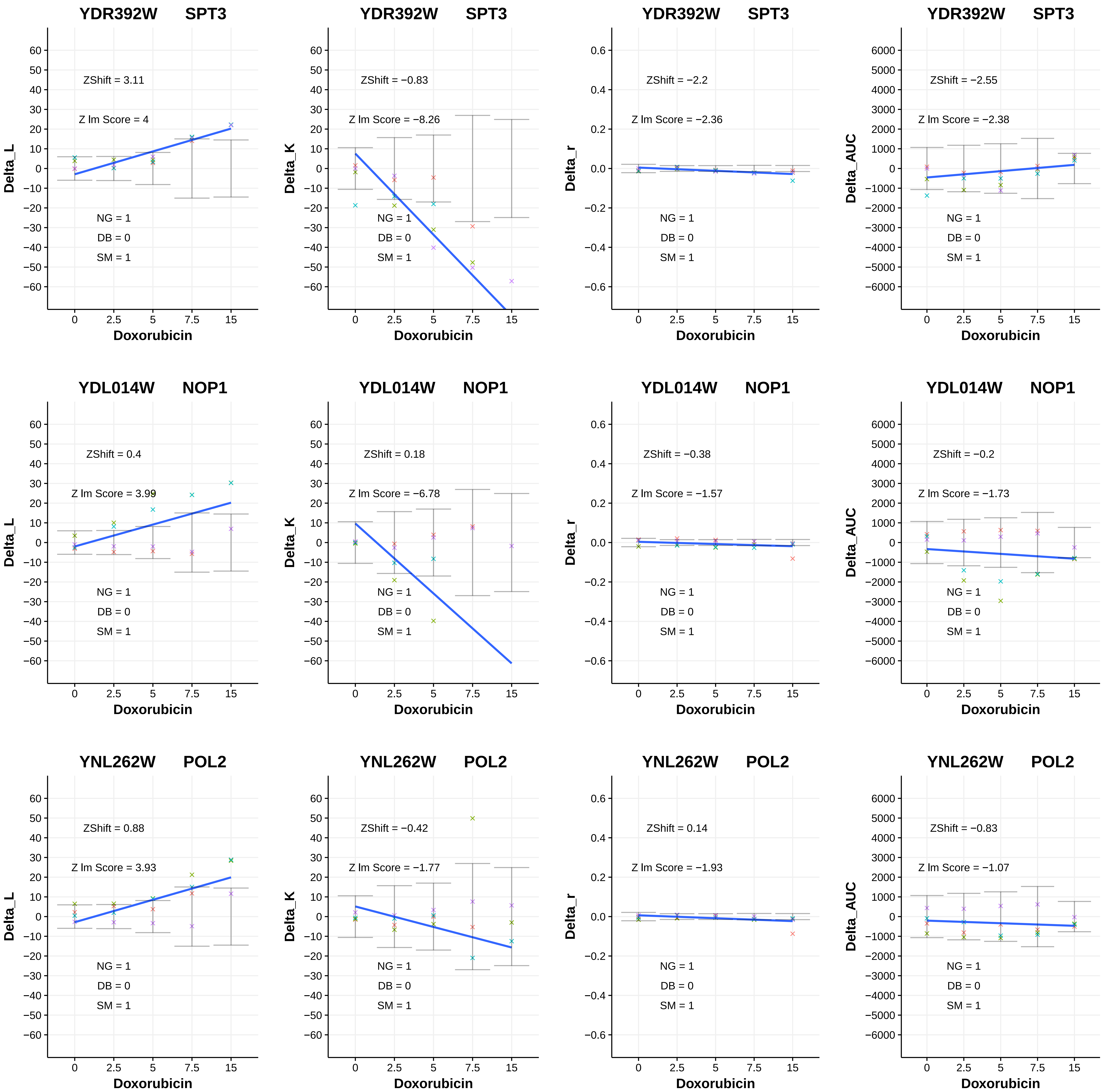
YDR392W SPT3
Delta_L
ZShift = 3.11
Z lm Score = 4
NG = 1
DB = 0
SM = 1
Doxorubicin
YDR392W SPT3
Delta_K
ZShift = −0.83
Z lm Score = −8.26
NG = 1
DB = 0
SM = 1
Doxorubicin
YDR392W SPT3
Delta_r
ZShift = −2.2
Z lm Score = −2.36
NG = 1
DB = 0
SM = 1
Doxorubicin
YDR392W SPT3
Delta_AUC
ZShift = −2.55
Z lm Score = −2.38
NG = 1
DB = 0
SM = 1
Doxorubicin
YDL014W NOP1
Delta_L
ZShift = 0.4
Z lm Score = 3.99
NG = 1
DB = 0
SM = 1
Doxorubicin
YDL014W NOP1
Delta_K
ZShift = 0.18
Z lm Score = −6.78
NG = 1
DB = 0
SM = 1
Doxorubicin
YDL014W NOP1
Delta_r
ZShift = −0.38
Z lm Score = −1.57
NG = 1
DB = 0
SM = 1
Doxorubicin
YDL014W NOP1
Delta_AUC
ZShift = −0.2
Z lm Score = −1.73
NG = 1
DB = 0
SM = 1
Doxorubicin
YNL262W POL2
Delta_L
ZShift = 0.88
Z lm Score = 3.93
NG = 1
DB = 0
SM = 1
Doxorubicin
YNL262W POL2
Delta_K
ZShift = −0.42
Z lm Score = −1.77
NG = 1
DB = 0
SM = 1
Doxorubicin
YNL262W POL2
Delta_r
ZShift = 0.14
Z lm Score = −1.93
NG = 1
DB = 0
SM = 1
Doxorubicin
YNL262W POL2
Delta_AUC
ZShift = −0.83
Z lm Score = −1.07
NG = 1
DB = 0
SM = 1
Doxorubicin

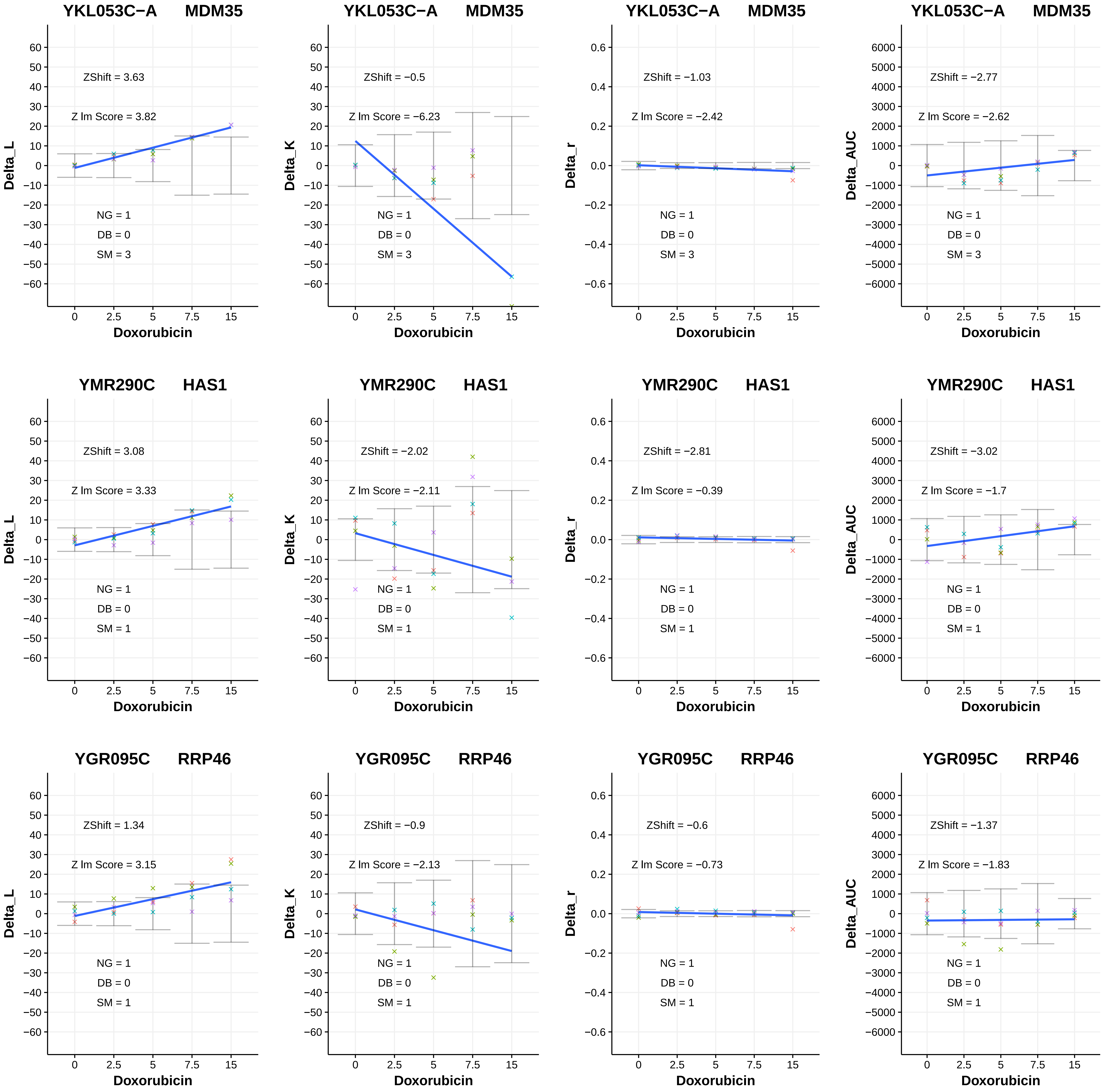
YKL053C−A MDM35
ZShift = 3.63
Z lm Score = 3.82
NG = 1
DB = 0
SM = 3
Delta_L
Doxorubicin
YKL053C−A MDM35
ZShift = −0.5
Z lm Score = −6.23
NG = 1
DB = 0
SM = 3
Delta_K
Doxorubicin
YKL053C−A MDM35
ZShift = −1.03
Z lm Score = −2.42
NG = 1
DB = 0
SM = 3
Delta_r
Doxorubicin
YKL053C−A MDM35
ZShift = −2.77
Z lm Score = −2.62
NG = 1
DB = 0
SM = 3
Delta_AUC
Doxorubicin
YMR290C HAS1
ZShift = 3.08
Z lm Score = 3.33
NG = 1
DB = 0
SM = 1
Delta_L
Doxorubicin
YMR290C HAS1
ZShift = −2.02
Z lm Score = −2.11
NG = 1
DB = 0
SM = 1
Delta_K
Doxorubicin
YMR290C HAS1
ZShift = −2.81
Z lm Score = −0.39
NG = 1
DB = 0
SM = 1
Delta_r
Doxorubicin
YMR290C HAS1
ZShift = −3.02
Z lm Score = −1.7
NG = 1
DB = 0
SM = 1
Delta_AUC
Doxorubicin
YGR095C RRP46
ZShift = 1.34
Z lm Score = 3.15
NG = 1
DB = 0
SM = 1
Delta_L
Doxorubicin
YGR095C RRP46
ZShift = −0.9
Z lm Score = −2.13
NG = 1
DB = 0
SM = 1
Delta_K
Doxorubicin
YGR095C RRP46
ZShift = −0.6
Z lm Score = −0.73
NG = 1
DB = 0
SM = 1
Delta_r
Doxorubicin
YGR095C RRP46
ZShift = −1.37
Z lm Score = −1.83
NG = 1
DB = 0
SM = 1
Delta_AUC
Doxorubicin

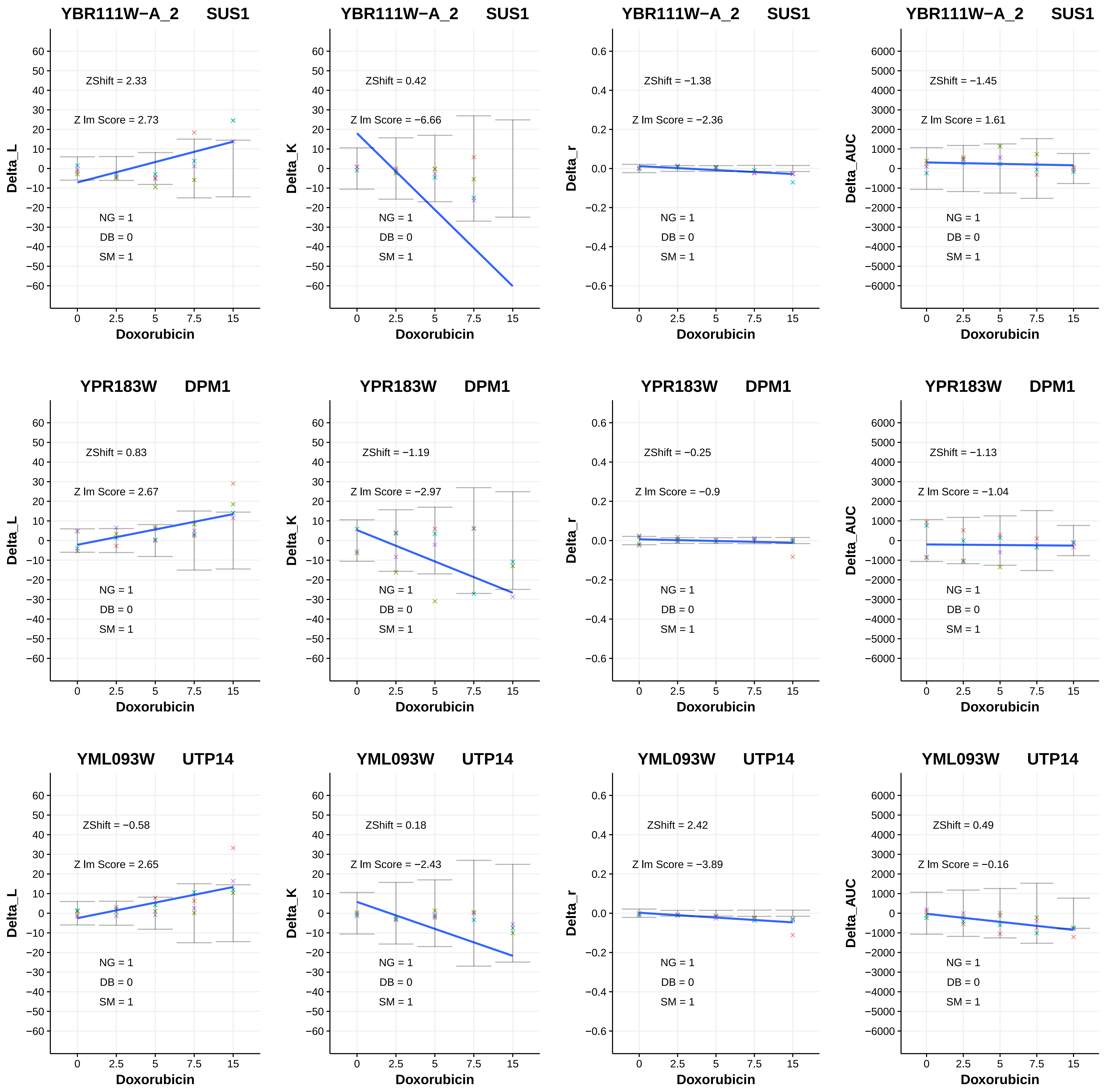
YBR111W−A_2 SUS1
ZShift = 2.33
Z lm Score = 2.73
NG = 1
DB = 0
SM = 1
Delta_L
Doxorubicin
YBR111W−A_2 SUS1
ZShift = 0.42
Z lm Score = −6.66
NG = 1
DB = 0
SM = 1
Delta_K
Doxorubicin
YBR111W−A_2 SUS1
ZShift = −1.38
Z lm Score = −2.36
NG = 1
DB = 0
SM = 1
Delta_r
Doxorubicin
YBR111W−A_2 SUS1
ZShift = −1.45
Z lm Score = 1.61
NG = 1
DB = 0
SM = 1
Delta_AUC
Doxorubicin
YPR183W DPM1
ZShift = 0.83
Z lm Score = 2.67
NG = 1
DB = 0
SM = 1
Delta_L
Doxorubicin
YPR183W DPM1
ZShift = −1.19
Z lm Score = −2.97
NG = 1
DB = 0
SM = 1
Delta_K
Doxorubicin
YPR183W DPM1
ZShift = −0.25
Z lm Score = −0.9
NG = 1
DB = 0
SM = 1
Delta_r
Doxorubicin
YPR183W DPM1
ZShift = −1.13
Z lm Score = −1.04
NG = 1
DB = 0
SM = 1
Delta_AUC
Doxorubicin
YML093W UTP14
ZShift = −0.58
Z lm Score = 2.65
NG = 1
DB = 0
SM = 1
Delta_L
Doxorubicin
YML093W UTP14
ZShift = 0.18
Z lm Score = −2.43
NG = 1
DB = 0
SM = 1
Delta_K
Doxorubicin
YML093W UTP14
ZShift = 2.42
Z lm Score = −3.89
NG = 1
DB = 0
SM = 1
Delta_r
Doxorubicin
YML093W UTP14
ZShift = 0.49
Z lm Score = −0.16
NG = 1
DB = 0
SM = 1
Delta_AUC
Doxorubicin

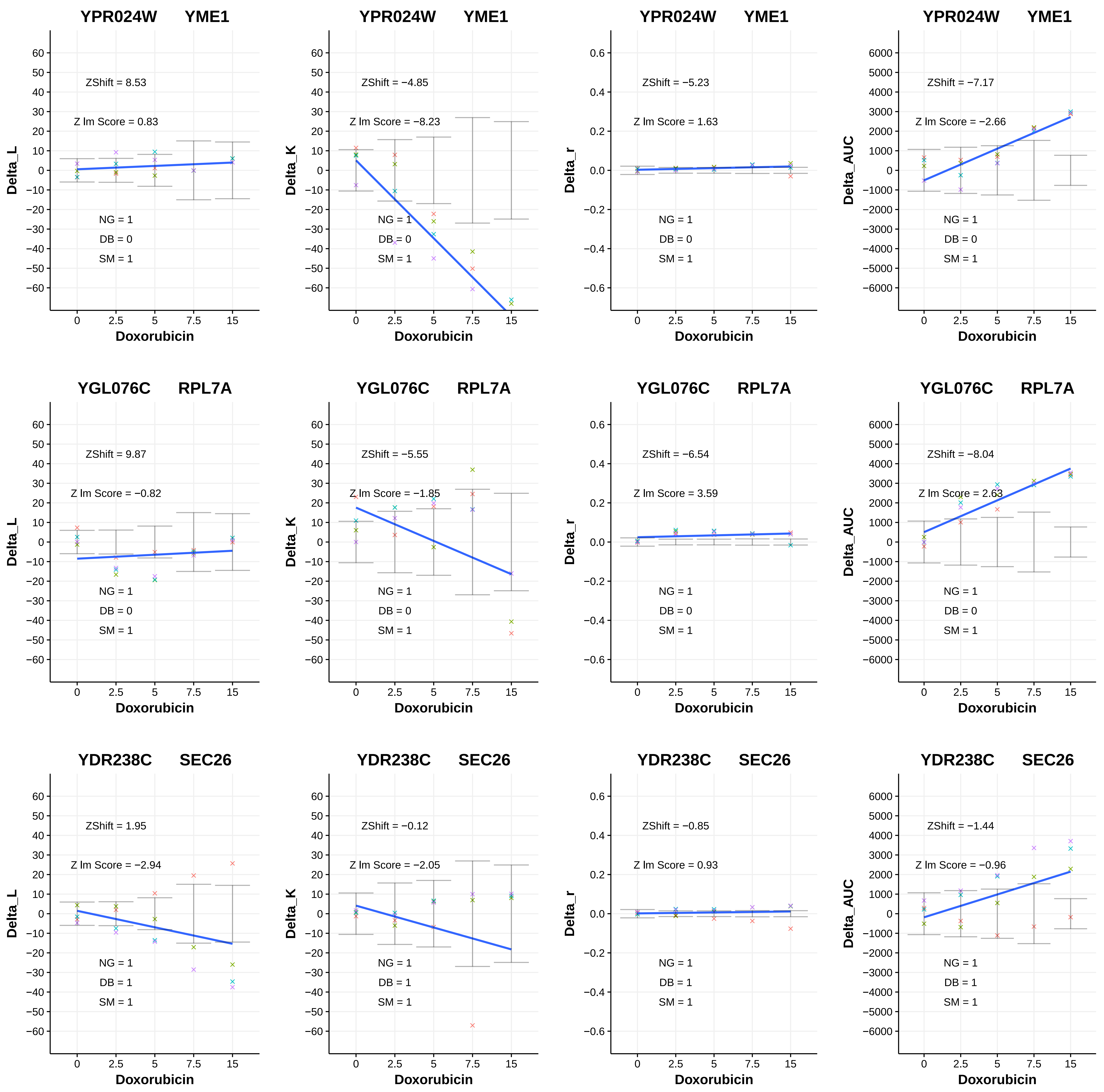

YPR024W YME1
ZShift = 8.53
Z lm Score = 0.83
NG = 1
DB = 0
SM = 1
Delta_L
Doxorubicin
YPR024W YME1
ZShift = −4.85
Z lm Score = −8.23
NG = 1
DB = 0
SM = 1
Delta_K
Doxorubicin
YPR024W YME1
ZShift = −5.23
Z lm Score = 1.63
NG = 1
DB = 0
SM = 1
Delta_r
Doxorubicin
YPR024W YME1
ZShift = −7.17
Z lm Score = −2.66
NG = 1
DB = 0
SM = 1
Delta_AUC
Doxorubicin
YGL076C RPL7A
ZShift = 9.87
Z lm Score = −0.82
NG = 1
DB = 0
SM = 1
Delta_L
Doxorubicin
YGL076C RPL7A
ZShift = −5.55
Z lm Score = −1.85
NG = 1
DB = 0
SM = 1
Delta_K
Doxorubicin
YGL076C RPL7A
ZShift = −6.54
Z lm Score = 3.59
NG = 1
DB = 0
SM = 1
Delta_r
Doxorubicin
YGL076C RPL7A
ZShift = −8.04
Z lm Score = 2.63
NG = 1
DB = 0
SM = 1
Delta_AUC
Doxorubicin
YDR238C SEC26
ZShift = 1.95
Z lm Score = −2.94
NG = 1
DB = 1
SM = 1
Delta_L
Doxorubicin
YDR238C SEC26
ZShift = −0.12
Z lm Score = −2.05
NG = 1
DB = 1
SM = 1
Delta_K
Doxorubicin
YDR238C SEC26
ZShift = −0.85
Z lm Score = 0.93
NG = 1
DB = 1
SM = 1
Delta_r
Doxorubicin
YDR238C SEC26
ZShift = −1.44
Z lm Score = −0.96
NG = 1
DB = 1
SM = 1
Delta_AUC
Doxorubicin

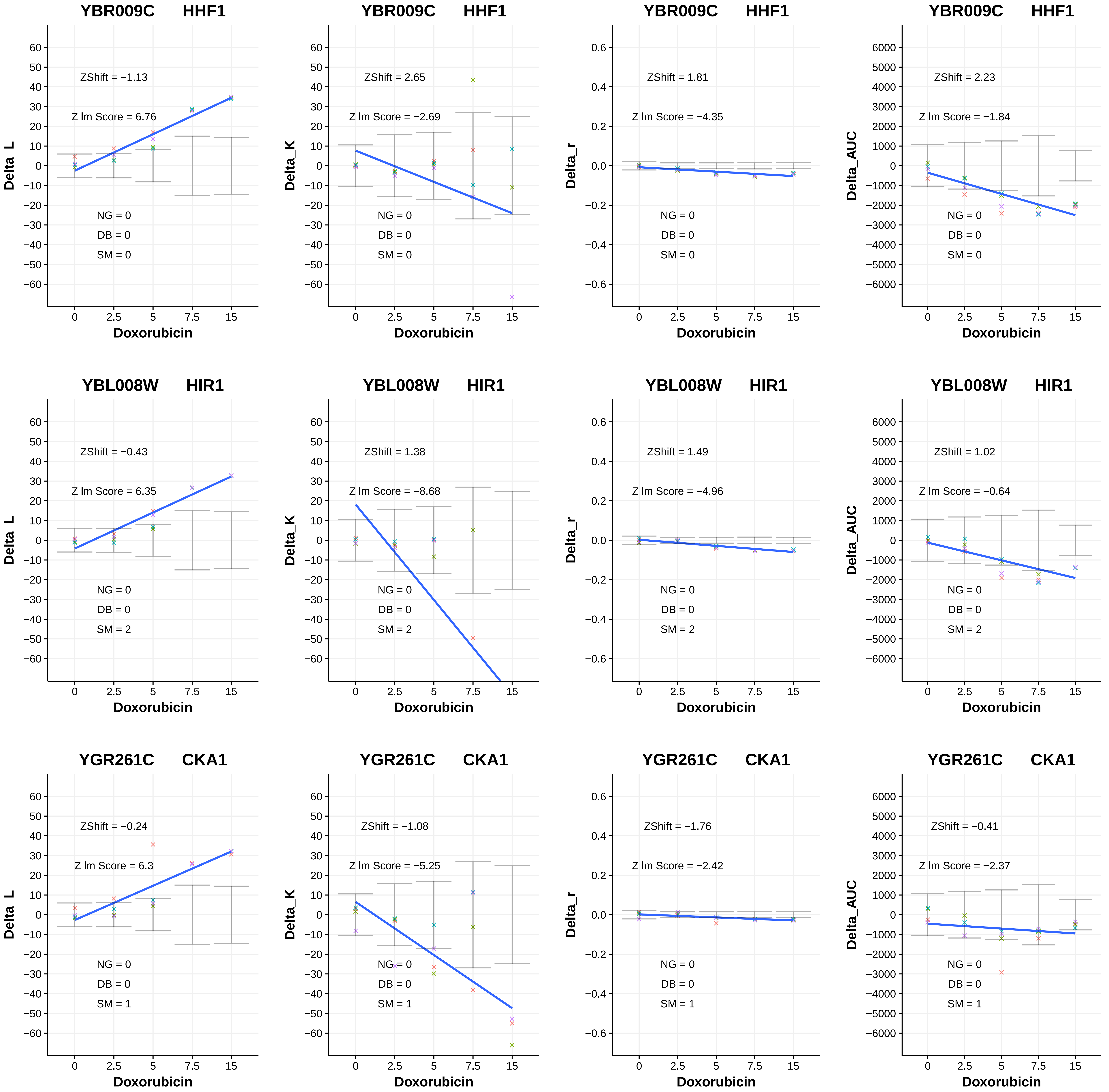
YBR009C HHF1
ZShift = −1.13
Z lm Score = 6.76
NG = 0
DB = 0
SM = 0
Delta_L
Doxorubicin
YBR009C HHF1
ZShift = 2.65
Z lm Score = −2.69
NG = 0
DB = 0
SM = 0
Delta_K
Doxorubicin
YBR009C HHF1
ZShift = 1.81
Z lm Score = −4.35
NG = 0
DB = 0
SM = 0
Delta_r
Doxorubicin
YBR009C HHF1
ZShift = 2.23
Z lm Score = −1.84
NG = 0
DB = 0
SM = 0
Delta_AUC
Doxorubicin
YBL008W HIR1
ZShift = −0.43
Z lm Score = 6.35
NG = 0
DB = 0
SM = 2
Delta_L
Doxorubicin
YBL008W HIR1
ZShift = 1.38
Z lm Score = −8.68
NG = 0
DB = 0
SM = 2
Delta_K
Doxorubicin
YBL008W HIR1
ZShift = 1.49
Z lm Score = −4.96
NG = 0
DB = 0
SM = 2
Delta_r
Doxorubicin
YBL008W HIR1
ZShift = 1.02
Z lm Score = −0.64
NG = 0
DB = 0
SM = 2
Delta_AUC
Doxorubicin
YGR261C CKA1
ZShift = −0.24
Z lm Score = 6.3
NG = 0
DB = 0
SM = 1
Delta_L
Doxorubicin
YGR261C CKA1
ZShift = −1.08
Z lm Score = −5.25
NG = 0
DB = 0
SM = 1
Delta_K
Doxorubicin
YGR261C CKA1
ZShift = −1.76
Z lm Score = −2.42
NG = 0
DB = 0
SM = 1
Delta_r
Doxorubicin
YGR261C CKA1
ZShift = −0.41
Z lm Score = −2.37
NG = 0
DB = 0
SM = 1
Delta_AUC
Doxorubicin

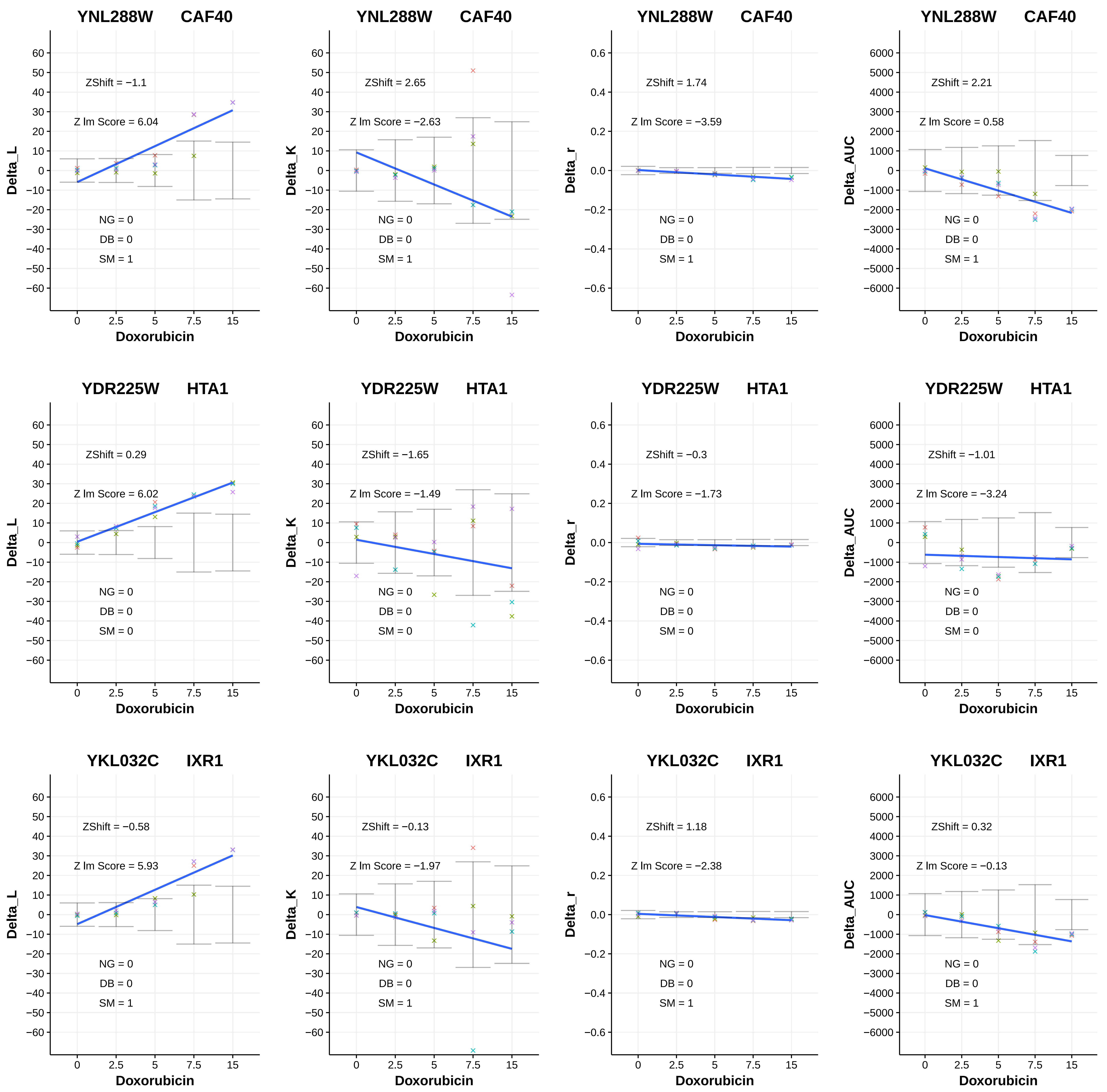
YNL288W CAF40
ZShift = −1.1
Z lm Score = 6.04
NG = 0
DB = 0
SM = 1
Delta_L
Doxorubicin
YNL288W CAF40
ZShift = 2.65
Z lm Score = −2.63
NG = 0
DB = 0
SM = 1
Delta_K
Doxorubicin
YNL288W CAF40
ZShift = 1.74
Z lm Score = −3.59
NG = 0
DB = 0
SM = 1
Delta_r
Doxorubicin
YNL288W CAF40
ZShift = 2.21
Z lm Score = 0.58
NG = 0
DB = 0
SM = 1
Delta_AUC
Doxorubicin
YDR225W HTA1
ZShift = 0.29
Z lm Score = 6.02
NG = 0
DB = 0
SM = 0
Delta_L
Doxorubicin
YDR225W HTA1
ZShift = −1.65
Z lm Score = −1.49
NG = 0
DB = 0
SM = 0
Delta_K
Doxorubicin
YDR225W HTA1
ZShift = −0.3
Z lm Score = −1.73
NG = 0
DB = 0
SM = 0
Delta_r
Doxorubicin
YDR225W HTA1
ZShift = −1.01
Z lm Score = −3.24
NG = 0
DB = 0
SM = 0
Delta_AUC
Doxorubicin
YKL032C IXR1
ZShift = −0.58
Z lm Score = 5.93
NG = 0
DB = 0
SM = 1
Delta_L
Doxorubicin
YKL032C IXR1
ZShift = −0.13
Z lm Score = −1.97
NG = 0
DB = 0
SM = 1
Delta_K
Doxorubicin
YKL032C IXR1
ZShift = 1.18
Z lm Score = −2.38
NG = 0
DB = 0
SM = 1
Delta_r
Doxorubicin
YKL032C IXR1
ZShift = 0.32
Z lm Score = −0.13
NG = 0
DB = 0
SM = 1
Delta_AUC
Doxorubicin

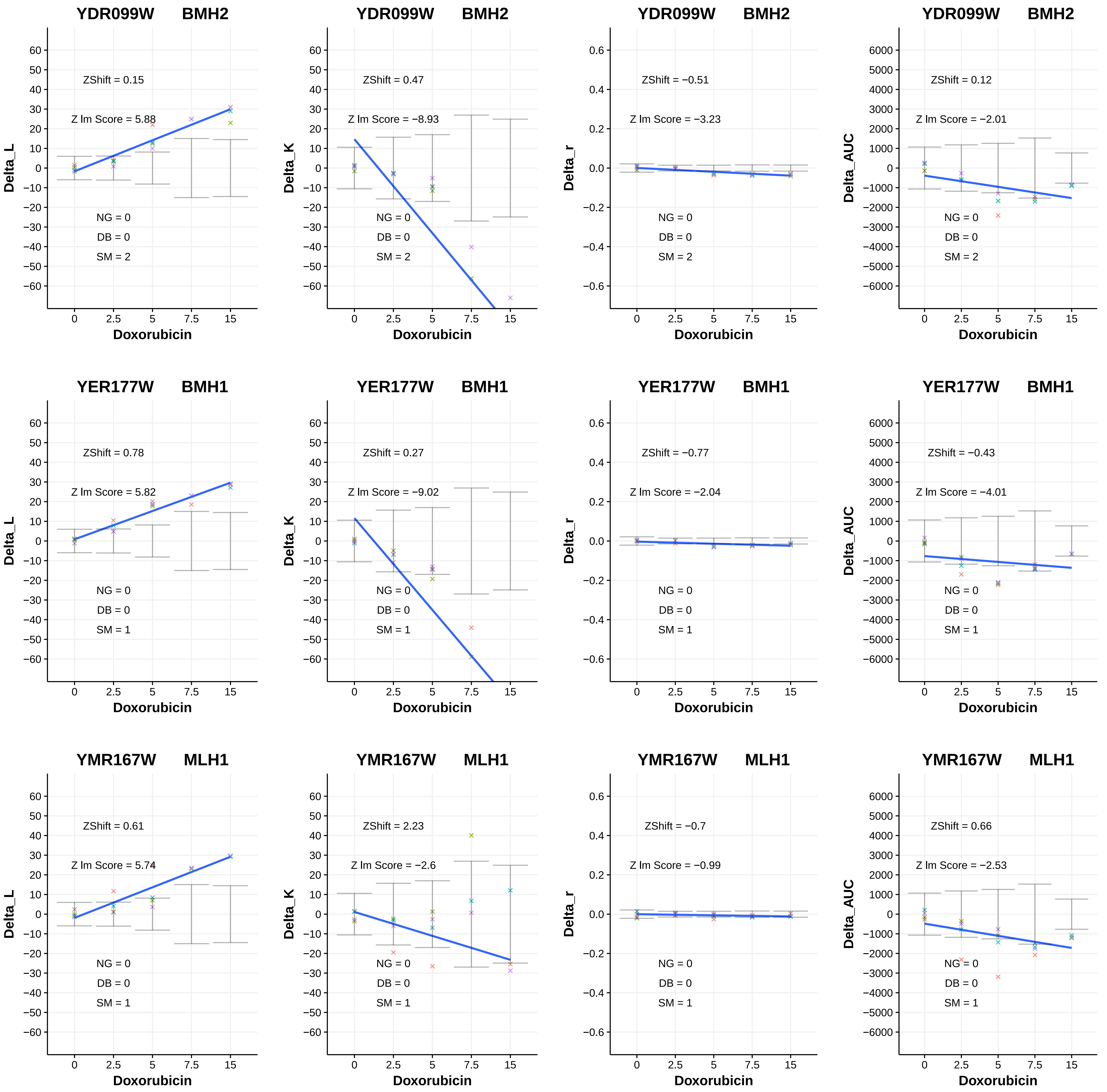

YDR099W BMH2
ZShift = 0.15
Z lm Score = 5.88
NG = 0
DB = 0
SM = 2
Delta_L
Doxorubicin
YDR099W BMH2
ZShift = 0.47
Z lm Score = −8.93
NG = 0
DB = 0
SM = 2
Delta_K
Doxorubicin
YDR099W BMH2
ZShift = −0.51
Z lm Score = −3.23
NG = 0
DB = 0
SM = 2
Delta_r
Doxorubicin
YDR099W BMH2
ZShift = 0.12
Z lm Score = −2.01
NG = 0
DB = 0
SM = 2
Delta_AUC
Doxorubicin
YER177W BMH1
ZShift = 0.78
Z lm Score = 5.82
NG = 0
DB = 0
SM = 1
Delta_L
Doxorubicin
YER177W BMH1
ZShift = 0.27
Z lm Score = −9.02
NG = 0
DB = 0
SM = 1
Delta_K
Doxorubicin
YER177W BMH1
ZShift = −0.77
Z lm Score = −2.04
NG = 0
DB = 0
SM = 1
Delta_r
Doxorubicin
YER177W BMH1
ZShift = −0.43
Z lm Score = −4.01
NG = 0
DB = 0
SM = 1
Delta_AUC
Doxorubicin
YMR167W MLH1
ZShift = 0.61
Z lm Score = 5.74
NG = 0
DB = 0
SM = 1
Delta_L
Doxorubicin
YMR167W MLH1
ZShift = 2.23
Z lm Score = −2.6
NG = 0
DB = 0
SM = 1
Delta_K
Doxorubicin
YMR167W MLH1
ZShift = −0.7
Z lm Score = −0.99
NG = 0
DB = 0
SM = 1
Delta_r
Doxorubicin
YMR167W MLH1
ZShift = 0.66
Z lm Score = −2.53
NG = 0
DB = 0
SM = 1
Delta_AUC
Doxorubicin

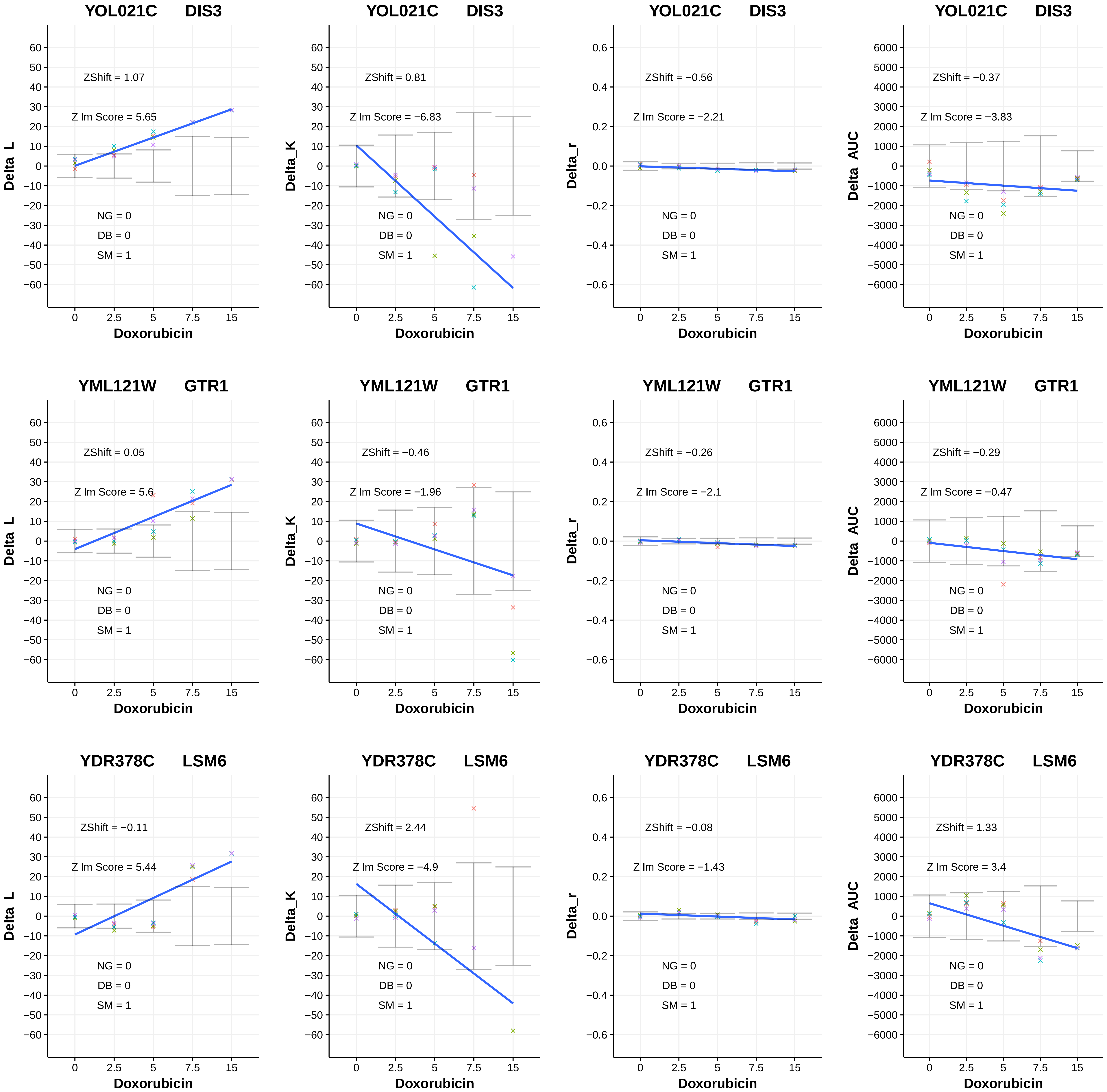

YOL021C DIS3
Delta_L
ZShift = 1.07
Z lm Score = 5.65
NG = 0
DB = 0
SM = 1
Doxorubicin
YOL021C DIS3
Delta_K
ZShift = 0.81
Z lm Score = −6.83
NG = 0
DB = 0
SM = 1
Doxorubicin
YOL021C DIS3
Delta_r
ZShift = −0.56
Z lm Score = −2.21
NG = 0
DB = 0
SM = 1
Doxorubicin
YOL021C DIS3
Delta_AUC
ZShift = −0.37
Z lm Score = −3.83
NG = 0
DB = 0
SM = 1
Doxorubicin
YML121W GTR1
Delta_L
ZShift = 0.05
Z lm Score = 5.6
NG = 0
DB = 0
SM = 1
Doxorubicin
YML121W GTR1
Delta_K
ZShift = −0.46
Z lm Score = −1.96
NG = 0
DB = 0
SM = 1
Doxorubicin
YML121W GTR1
Delta_r
ZShift = −0.26
Z lm Score = −2.1
NG = 0
DB = 0
SM = 1
Doxorubicin
YML121W GTR1
Delta_AUC
ZShift = −0.29
Z lm Score = −0.47
NG = 0
DB = 0
SM = 1
Doxorubicin
YDR378C LSM6
Delta_L
ZShift = −0.11
Z lm Score = 5.44
NG = 0
DB = 0
SM = 1
Doxorubicin
YDR378C LSM6
Delta_K
ZShift = 2.44
Z lm Score = −4.9
NG = 0
DB = 0
SM = 1
Doxorubicin
YDR378C LSM6
Delta_r
ZShift = −0.08
Z lm Score = −1.43
NG = 0
DB = 0
SM = 1
Doxorubicin
YDR378C LSM6
Delta_AUC
ZShift = 1.33
Z lm Score = 3.4
NG = 0
DB = 0
SM = 1
Doxorubicin

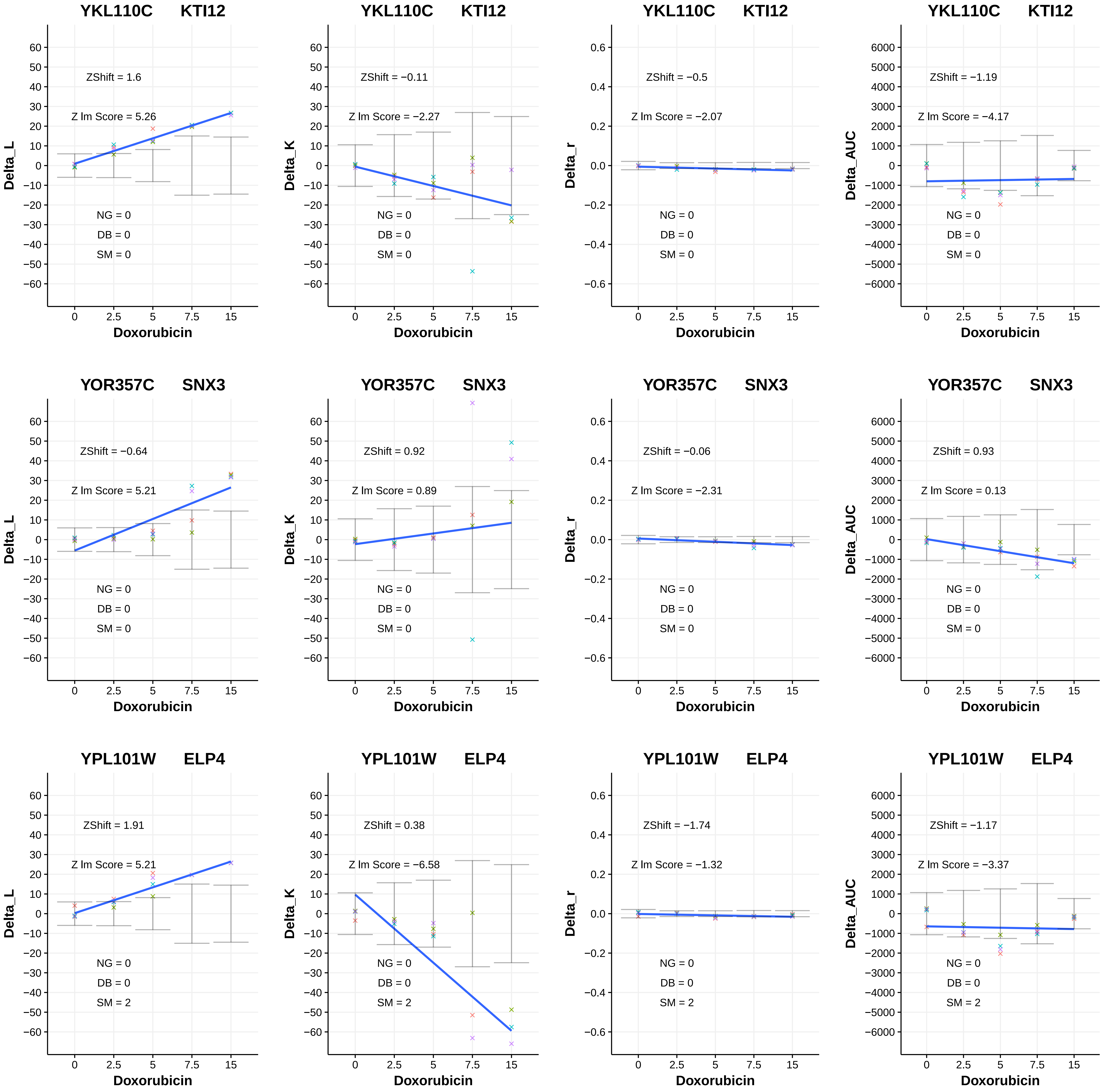

YKL110C KTI12
Delta_L
ZShift = 1.6
Z lm Score = 5.26
NG = 0
DB = 0
SM = 0
Doxorubicin
YKL110C KTI12
Delta_K
ZShift = −0.11
Z lm Score = −2.27
NG = 0
DB = 0
SM = 0
Doxorubicin
YKL110C KTI12
Delta_r
ZShift = −0.5
Z lm Score = −2.07
NG = 0
DB = 0
SM = 0
Doxorubicin
YKL110C KTI12
Delta_AUC
ZShift = −1.19
Z lm Score = −4.17
NG = 0
DB = 0
SM = 0
Doxorubicin
YOR357C SNX3
Delta_L
ZShift = −0.64
Z lm Score = 5.21
NG = 0
DB = 0
SM = 0
Doxorubicin
YOR357C SNX3
Delta_K
ZShift = 0.92
Z lm Score = 0.89
NG = 0
DB = 0
SM = 0
Doxorubicin
YOR357C SNX3
Delta_r
ZShift = −0.06
Z lm Score = −2.31
NG = 0
DB = 0
SM = 0
Doxorubicin
YOR357C SNX3
Delta_AUC
ZShift = 0.93
Z lm Score = 0.13
NG = 0
DB = 0
SM = 0
Doxorubicin
YPL101W ELP4
Delta_L
ZShift = 1.91
Z lm Score = 5.21
NG = 0
DB = 0
SM = 2
Doxorubicin
YPL101W ELP4
Delta_K
ZShift = 0.38
Z lm Score = −6.58
NG = 0
DB = 0
SM = 2
Doxorubicin
YPL101W ELP4
Delta_r
ZShift = −1.74
Z lm Score = −1.32
NG = 0
DB = 0
SM = 2
Doxorubicin
YPL101W ELP4
Delta_AUC
ZShift = −1.17
Z lm Score = −3.37
NG = 0
DB = 0
SM = 2
Doxorubicin

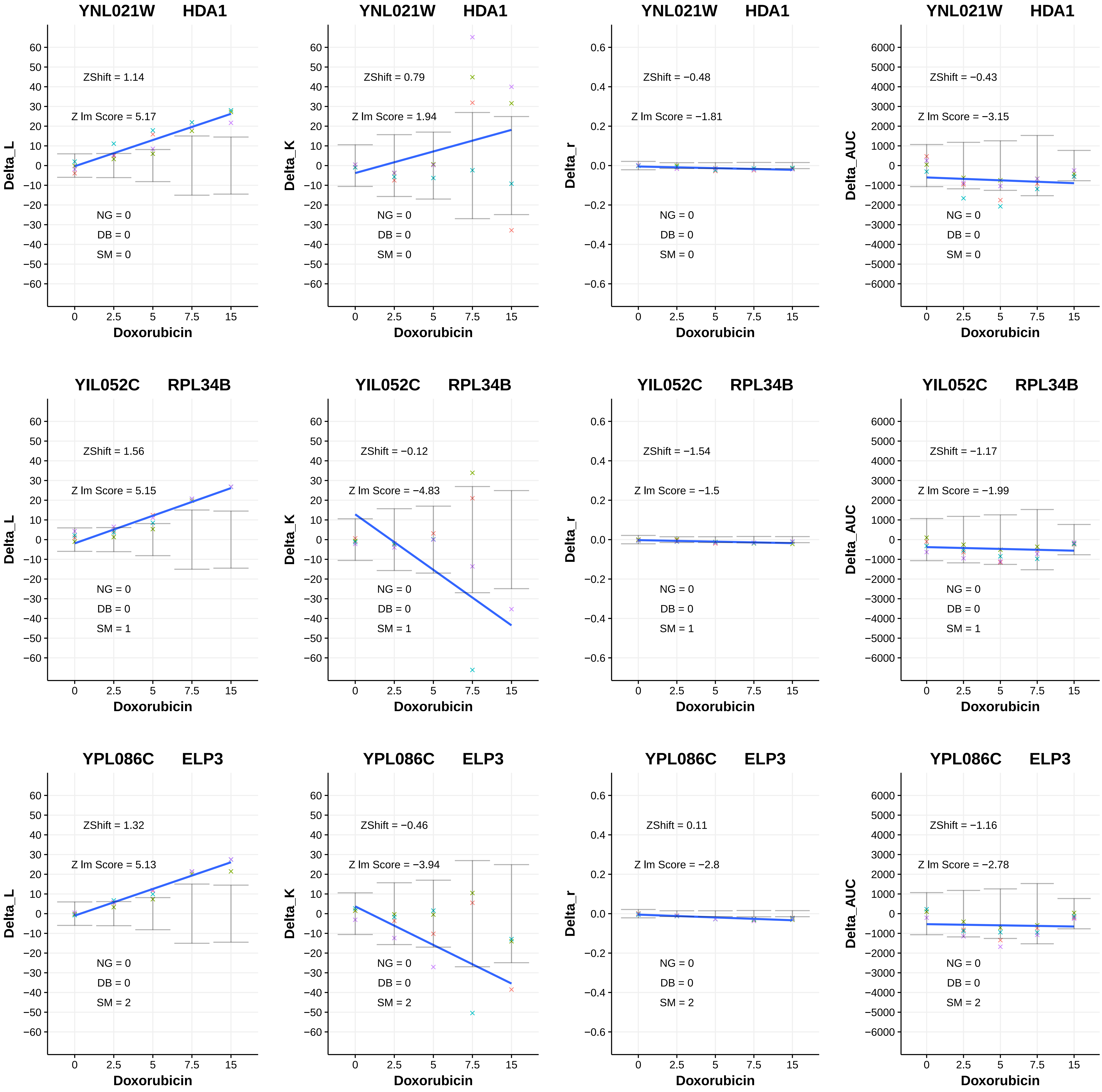

YNL021W HDA1
ZShift = 1.14
Z lm Score = 5.17
NG = 0
DB = 0
SM = 0
Delta_L
Doxorubicin
YNL021W HDA1
ZShift = 0.79
Z lm Score = 1.94
NG = 0
DB = 0
SM = 0
Delta_K
Doxorubicin
YNL021W HDA1
ZShift = −0.48
Z lm Score = −1.81
NG = 0
DB = 0
SM = 0
Delta_r
Doxorubicin
YNL021W HDA1
ZShift = −0.43
Z lm Score = −3.15
NG = 0
DB = 0
SM = 0
Delta_AUC
Doxorubicin
YIL052C RPL34B
ZShift = 1.56
Z lm Score = 5.15
NG = 0
DB = 0
SM = 1
Delta_L
Doxorubicin
YIL052C RPL34B
ZShift = −0.12
Z lm Score = −4.83
NG = 0
DB = 0
SM = 1
Delta_K
Doxorubicin
YIL052C RPL34B
ZShift = −1.54
Z lm Score = −1.5
NG = 0
DB = 0
SM = 1
Delta_r
Doxorubicin
YIL052C RPL34B
ZShift = −1.17
Z lm Score = −1.99
NG = 0
DB = 0
SM = 1
Delta_AUC
Doxorubicin
YPL086C ELP3
ZShift = 1.32
Z lm Score = 5.13
NG = 0
DB = 0
SM = 2
Delta_L
Doxorubicin
YPL086C ELP3
ZShift = −0.46
Z lm Score = −3.94
NG = 0
DB = 0
SM = 2
Delta_K
Doxorubicin
YPL086C ELP3
ZShift = 0.11
Z lm Score = −2.8
NG = 0
DB = 0
SM = 2
Delta_r
Doxorubicin
YPL086C ELP3
ZShift = −1.16
Z lm Score = −2.78
NG = 0
DB = 0
SM = 2
Delta_AUC
Doxorubicin

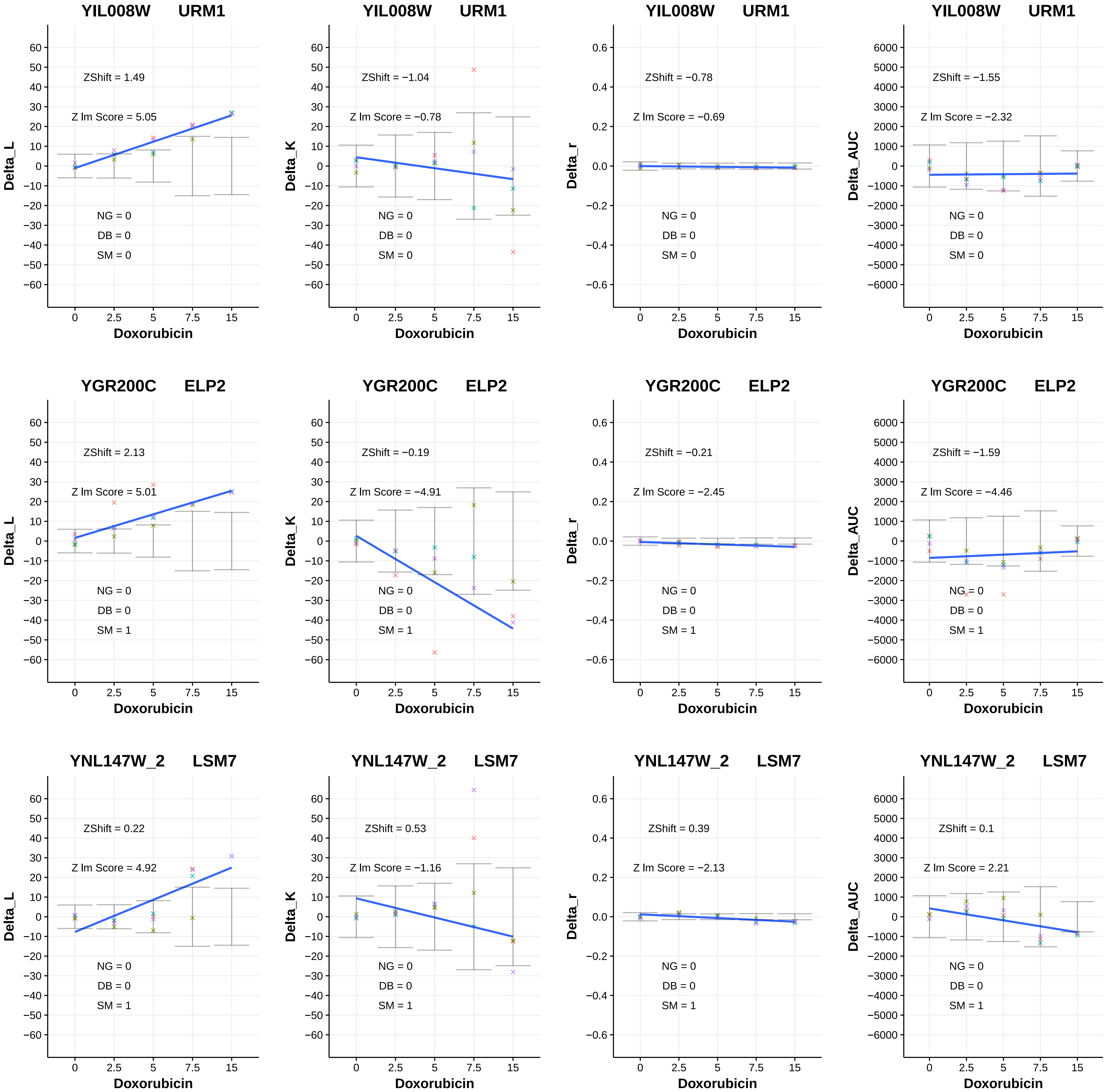
YIL008W URM1
ZShift = 1.49
Z lm Score = 5.05
NG = 0
DB = 0
SM = 0
Delta_L
Doxorubicin
YIL008W URM1
ZShift = −1.04
Z lm Score = −0.78
NG = 0
DB = 0
SM = 0
Delta_K
Doxorubicin
YIL008W URM1
ZShift = −0.78
Z lm Score = −0.69
NG = 0
DB = 0
SM = 0
Delta_r
Doxorubicin
YIL008W URM1
ZShift = −1.55
Z lm Score = −2.32
NG = 0
DB = 0
SM = 0
Delta_AUC
Doxorubicin
YGR200C ELP2
ZShift = 2.13
Z lm Score = 5.01
NG = 0
DB = 0
SM = 1
Delta_L
Doxorubicin
YGR200C ELP2
ZShift = −0.19
Z lm Score = −4.91
NG = 0
DB = 0
SM = 1
Delta_K
Doxorubicin
YGR200C ELP2
ZShift = −0.21
Z lm Score = −2.45
NG = 0
DB = 0
SM = 1
Delta_r
Doxorubicin
YGR200C ELP2
ZShift = −1.59
Z lm Score = −4.46
NG = 0
DB = 0
SM = 1
Delta_AUC
Doxorubicin
YNL147W_2 LSM7
ZShift = 0.22
Z lm Score = 4.92
NG = 0
DB = 0
SM = 1
Delta_L
Doxorubicin
YNL147W_2 LSM7
ZShift = 0.53
Z lm Score = −1.16
NG = 0
DB = 0
SM = 1
Delta_K
Doxorubicin
YNL147W_2 LSM7
ZShift = 0.39
Z lm Score = −2.13
NG = 0
DB = 0
SM = 1
Delta_r
Doxorubicin
YNL147W_2 LSM7
ZShift = 0.1
Z lm Score = 2.21
NG = 0
DB = 0
SM = 1
Delta_AUC
Doxorubicin

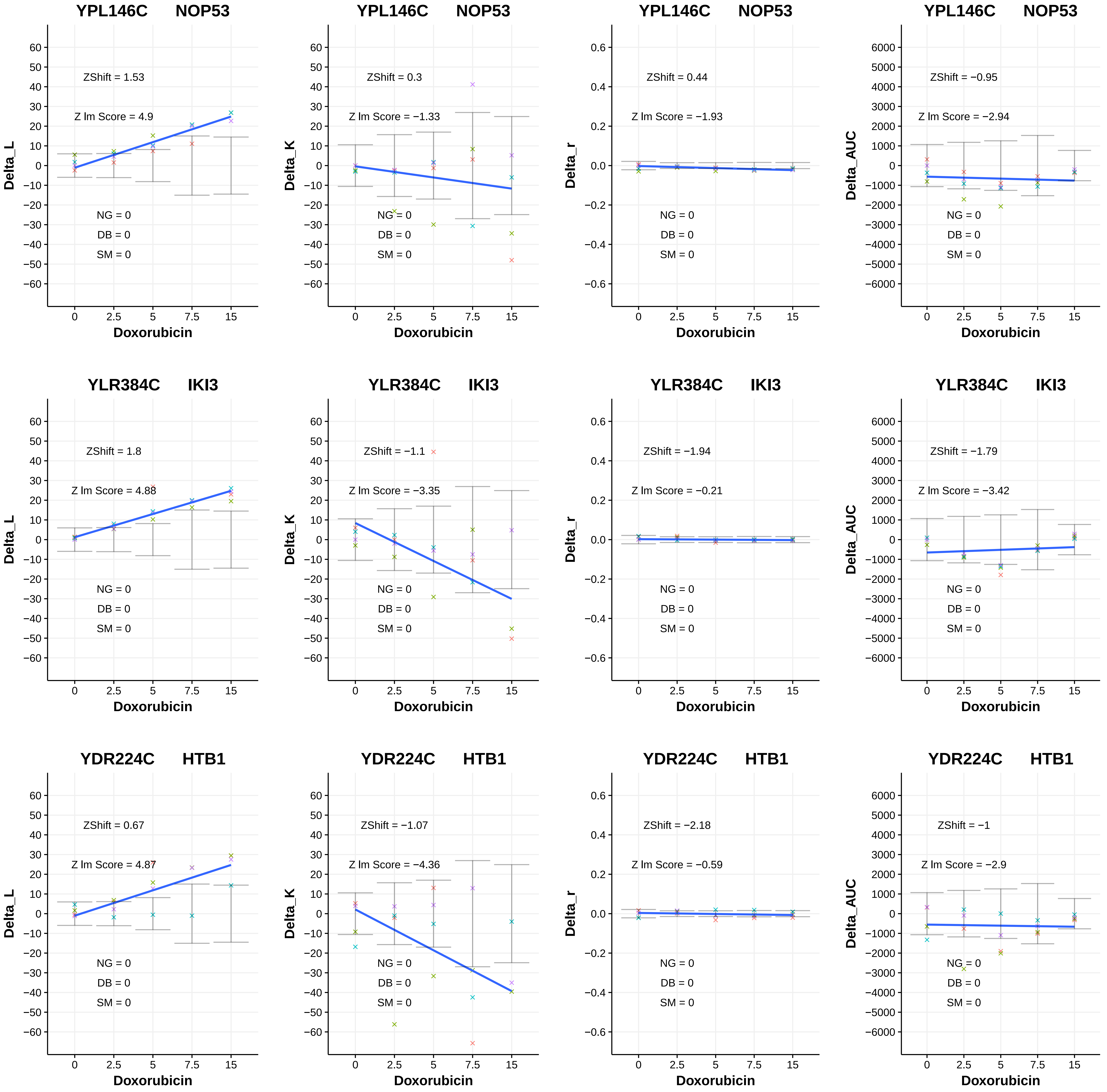
YPL146C NOP53
ZShift = 1.53
Z lm Score = 4.9
NG = 0
DB = 0
SM = 0
Delta_L
Doxorubicin
YPL146C NOP53
ZShift = 0.3
Z lm Score = −1.33
NG = 0
DB = 0
SM = 0
Delta_K
Doxorubicin
YPL146C NOP53
ZShift = 0.44
Z lm Score = −1.93
NG = 0
DB = 0
SM = 0
Delta_r
Doxorubicin
YPL146C NOP53
ZShift = −0.95
Z lm Score = −2.94
NG = 0
DB = 0
SM = 0
Delta_AUC
Doxorubicin
YLR384C IKI3
ZShift = 1.8
Z lm Score = 4.88
NG = 0
DB = 0
SM = 0
Delta_L
Doxorubicin
YLR384C IKI3
ZShift = −1.1
Z lm Score = −3.35
NG = 0
DB = 0
SM = 0
Delta_K
Doxorubicin
YLR384C IKI3
ZShift = −1.94
Z lm Score = −0.21
NG = 0
DB = 0
SM = 0
Delta_r
Doxorubicin
YLR384C IKI3
ZShift = −1.79
Z lm Score = −3.42
NG = 0
DB = 0
SM = 0
Delta_AUC
Doxorubicin
YDR224C HTB1
ZShift = 0.67
Z lm Score = 4.87
NG = 0
DB = 0
SM = 0
Delta_L
Doxorubicin
YDR224C HTB1
ZShift = −1.07
Z lm Score = −4.36
NG = 0
DB = 0
SM = 0
Delta_K
Doxorubicin
YDR224C HTB1
ZShift = −2.18
Z lm Score = −0.59
NG = 0
DB = 0
SM = 0
Delta_r
Doxorubicin
YDR224C HTB1
ZShift = −1
Z lm Score = −2.9
NG = 0
DB = 0
SM = 0
Delta_AUC
Doxorubicin

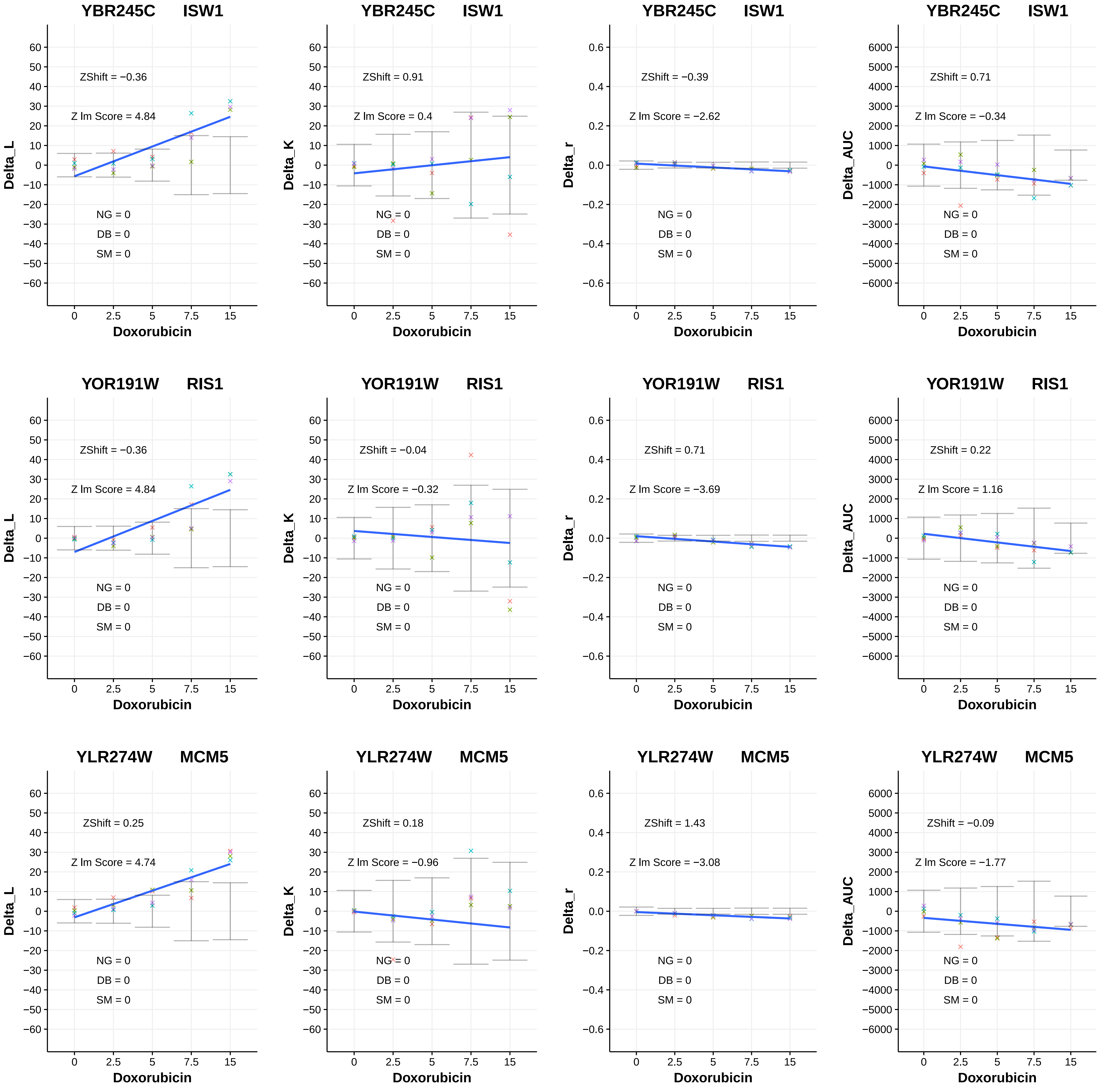
YBR245C ISW1
ZShift = −0.36
Z lm Score = 4.84
NG = 0
DB = 0
SM = 0
Delta_L
Doxorubicin
YBR245C ISW1
ZShift = 0.91
Z lm Score = 0.4
NG = 0
DB = 0
SM = 0
Delta_K
Doxorubicin
YBR245C ISW1
ZShift = −0.39
Z lm Score = −2.62
NG = 0
DB = 0
SM = 0
Delta_r
Doxorubicin
YBR245C ISW1
ZShift = 0.71
Z lm Score = −0.34
NG = 0
DB = 0
SM = 0
Delta_AUC
Doxorubicin
YOR191W RIS1
ZShift = −0.36
Z lm Score = 4.84
NG = 0
DB = 0
SM = 0
Delta_L
Doxorubicin
YOR191W RIS1
ZShift = −0.04
Z lm Score = −0.32
NG = 0
DB = 0
SM = 0
Delta_K
Doxorubicin
YOR191W RIS1
ZShift = 0.71
Z lm Score = −3.69
NG = 0
DB = 0
SM = 0
Delta_r
Doxorubicin
YOR191W RIS1
ZShift = 0.22
Z lm Score = 1.16
NG = 0
DB = 0
SM = 0
Delta_AUC
Doxorubicin
YLR274W MCM5
ZShift = 0.25
Z lm Score = 4.74
NG = 0
DB = 0
SM = 0
Delta_L
Doxorubicin
YLR274W MCM5
ZShift = 0.18
Z lm Score = −0.96
NG = 0
DB = 0
SM = 0
Delta_K
Doxorubicin
YLR274W MCM5
ZShift = 1.43
Z lm Score = −3.08
NG = 0
DB = 0
SM = 0
Delta_r
Doxorubicin
YLR274W MCM5
ZShift = −0.09
Z lm Score = −1.77
NG = 0
DB = 0
SM = 0
Delta_AUC
Doxorubicin

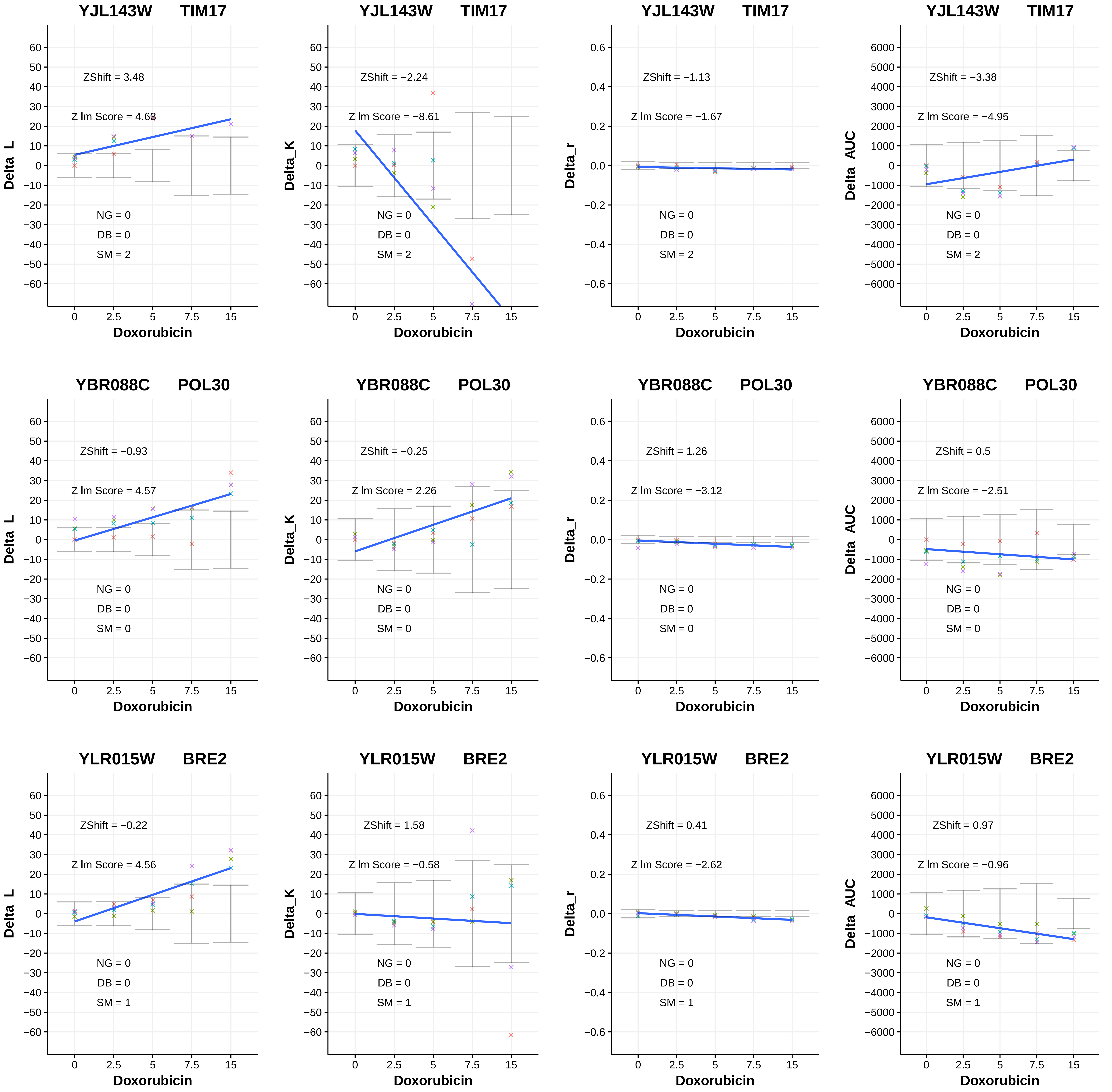
YJL143W TIM17
ZShift = 3.48
Z lm Score = 4.63
NG = 0
DB = 0
SM = 2
Delta_L
Doxorubicin
YJL143W TIM17
ZShift = −2.24
Z lm Score = −8.61
NG = 0
DB = 0
SM = 2
Delta_K
Doxorubicin
YJL143W TIM17
ZShift = −1.13
Z lm Score = −1.67
NG = 0
DB = 0
SM = 2
Delta_r
Doxorubicin
YJL143W TIM17
ZShift = −3.38
Z lm Score = −4.95
NG = 0
DB = 0
SM = 2
Delta_AUC
Doxorubicin
YBR088C POL30
ZShift = −0.93
Z lm Score = 4.57
NG = 0
DB = 0
SM = 0
Delta_L
Doxorubicin
YBR088C POL30
ZShift = −0.25
Z lm Score = 2.26
NG = 0
DB = 0
SM = 0
Delta_K
Doxorubicin
YBR088C POL30
ZShift = 1.26
Z lm Score = −3.12
NG = 0
DB = 0
SM = 0
Delta_r
Doxorubicin
YBR088C POL30
ZShift = 0.5
Z lm Score = −2.51
NG = 0
DB = 0
SM = 0
Delta_AUC
Doxorubicin
YLR015W BRE2
ZShift = −0.22
Z lm Score = 4.56
NG = 0
DB = 0
SM = 1
Delta_L
Doxorubicin
YLR015W BRE2
ZShift = 1.58
Z lm Score = −0.58
NG = 0
DB = 0
SM = 1
Delta_K
Doxorubicin
YLR015W BRE2
ZShift = 0.41
Z lm Score = −2.62
NG = 0
DB = 0
SM = 1
Delta_r
Doxorubicin
YLR015W BRE2
ZShift = 0.97
Z lm Score = −0.96
NG = 0
DB = 0
SM = 1
Delta_AUC
Doxorubicin

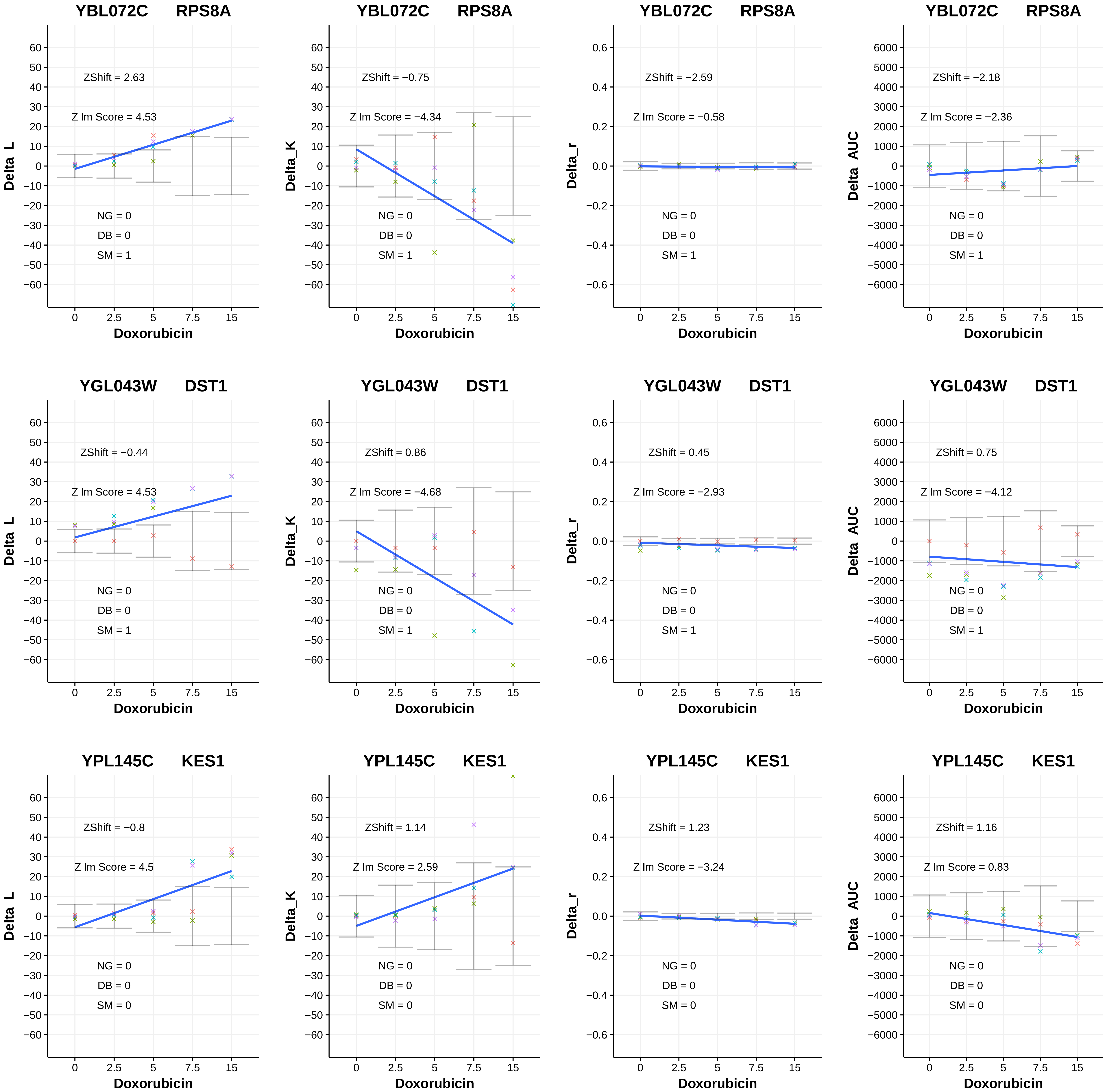

YBL072C RPS8A
ZShift = 2.63
Z lm Score = 4.53
NG = 0
DB = 0
SM = 1
Delta_L
Doxorubicin
YBL072C RPS8A
ZShift = −0.75
Z lm Score = −4.34
NG = 0
DB = 0
SM = 1
Delta_K
Doxorubicin
YBL072C RPS8A
ZShift = −2.59
Z lm Score = −0.58
NG = 0
DB = 0
SM = 1
Delta_r
Doxorubicin
YBL072C RPS8A
ZShift = −2.18
Z lm Score = −2.36
NG = 0
DB = 0
SM = 1
Delta_AUC
Doxorubicin
YGL043W DST1
ZShift = −0.44
Z lm Score = 4.53
NG = 0
DB = 0
SM = 1
Delta_L
Doxorubicin
YGL043W DST1
ZShift = 0.86
Z lm Score = −4.68
NG = 0
DB = 0
SM = 1
Delta_K
Doxorubicin
YGL043W DST1
ZShift = 0.45
Z lm Score = −2.93
NG = 0
DB = 0
SM = 1
Delta_r
Doxorubicin
YGL043W DST1
ZShift = 0.75
Z lm Score = −4.12
NG = 0
DB = 0
SM = 1
Delta_AUC
Doxorubicin
YPL145C KES1
ZShift = −0.8
Z lm Score = 4.5
NG = 0
DB = 0
SM = 0
Delta_L
Doxorubicin
YPL145C KES1
ZShift = 1.14
Z lm Score = 2.59
NG = 0
DB = 0
SM = 0
Delta_K
Doxorubicin
YPL145C KES1
ZShift = 1.23
Z lm Score = −3.24
NG = 0
DB = 0
SM = 0
Delta_r
Doxorubicin
YPL145C KES1
ZShift = 1.16
Z lm Score = 0.83
NG = 0
DB = 0
SM = 0
Delta_AUC
Doxorubicin

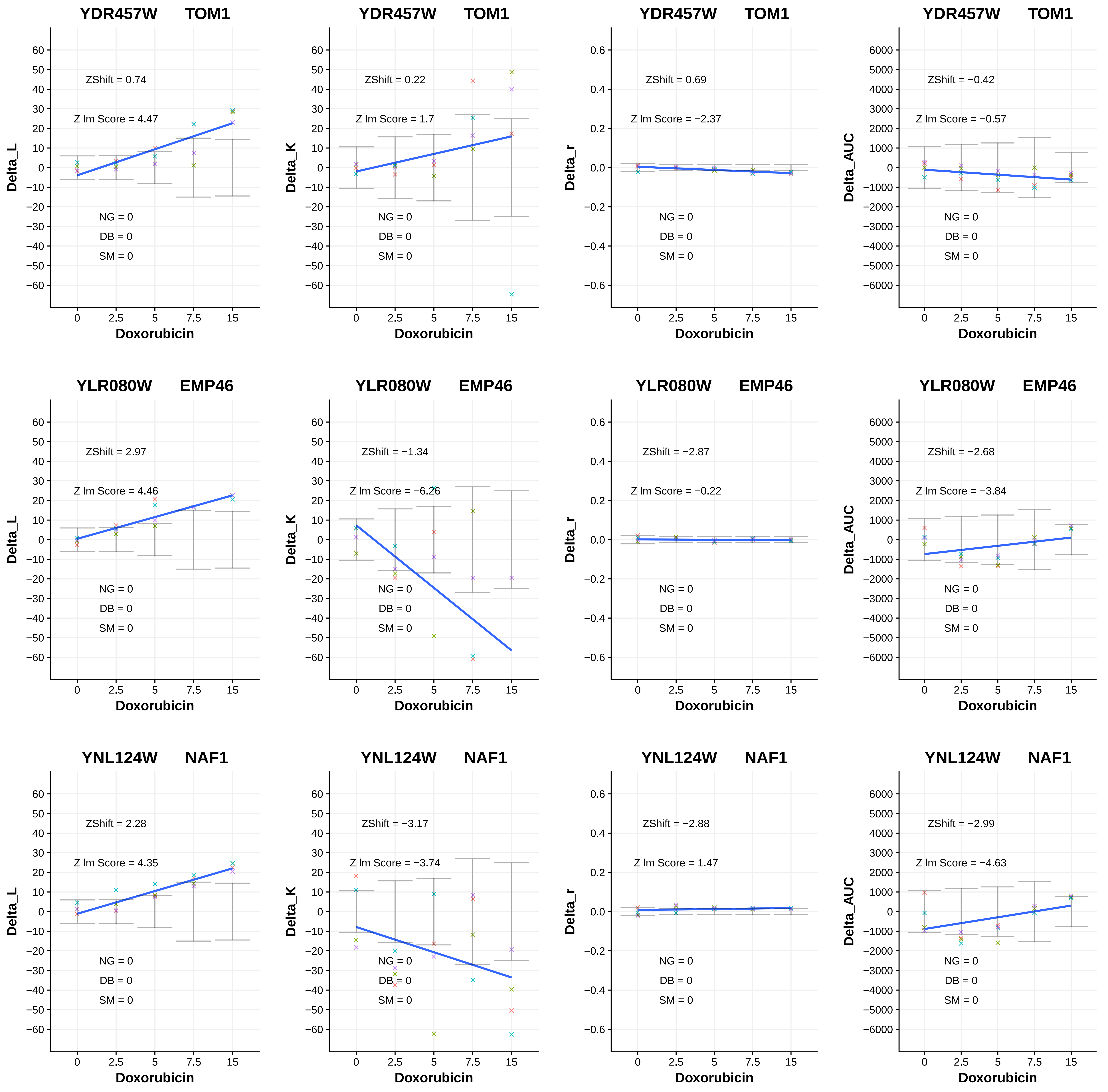

YDR457W TOM1
ZShift = 0.74
Z lm Score = 4.47
NG = 0
DB = 0
SM = 0
Delta_L
Doxorubicin
YDR457W TOM1
ZShift = 0.22
Z lm Score = 1.7
NG = 0
DB = 0
SM = 0
Delta_K
Doxorubicin
YDR457W TOM1
ZShift = 0.69
Z lm Score = −2.37
NG = 0
DB = 0
SM = 0
Delta_r
Doxorubicin
YDR457W TOM1
ZShift = −0.42
Z lm Score = −0.57
NG = 0
DB = 0
SM = 0
Delta_AUC
Doxorubicin
YLR080W EMP46
ZShift = 2.97
Z lm Score = 4.46
NG = 0
DB = 0
SM = 0
Delta_L
Doxorubicin
YLR080W EMP46
ZShift = −1.34
Z lm Score = −6.26
NG = 0
DB = 0
SM = 0
Delta_K
Doxorubicin
YLR080W EMP46
ZShift = −2.87
Z lm Score = −0.22
NG = 0
DB = 0
SM = 0
Delta_r
Doxorubicin
YLR080W EMP46
ZShift = −2.68
Z lm Score = −3.84
NG = 0
DB = 0
SM = 0
Delta_AUC
Doxorubicin
YNL124W NAF1
ZShift = 2.28
Z lm Score = 4.35
NG = 0
DB = 0
SM = 0
Delta_L
Doxorubicin
YNL124W NAF1
ZShift = −3.17
Z lm Score = −3.74
NG = 0
DB = 0
SM = 0
Delta_K
Doxorubicin
YNL124W NAF1
ZShift = −2.88
Z lm Score = 1.47
NG = 0
DB = 0
SM = 0
Delta_r
Doxorubicin
YNL124W NAF1
ZShift = −2.99
Z lm Score = −4.63
NG = 0
DB = 0
SM = 0
Delta_AUC
Doxorubicin

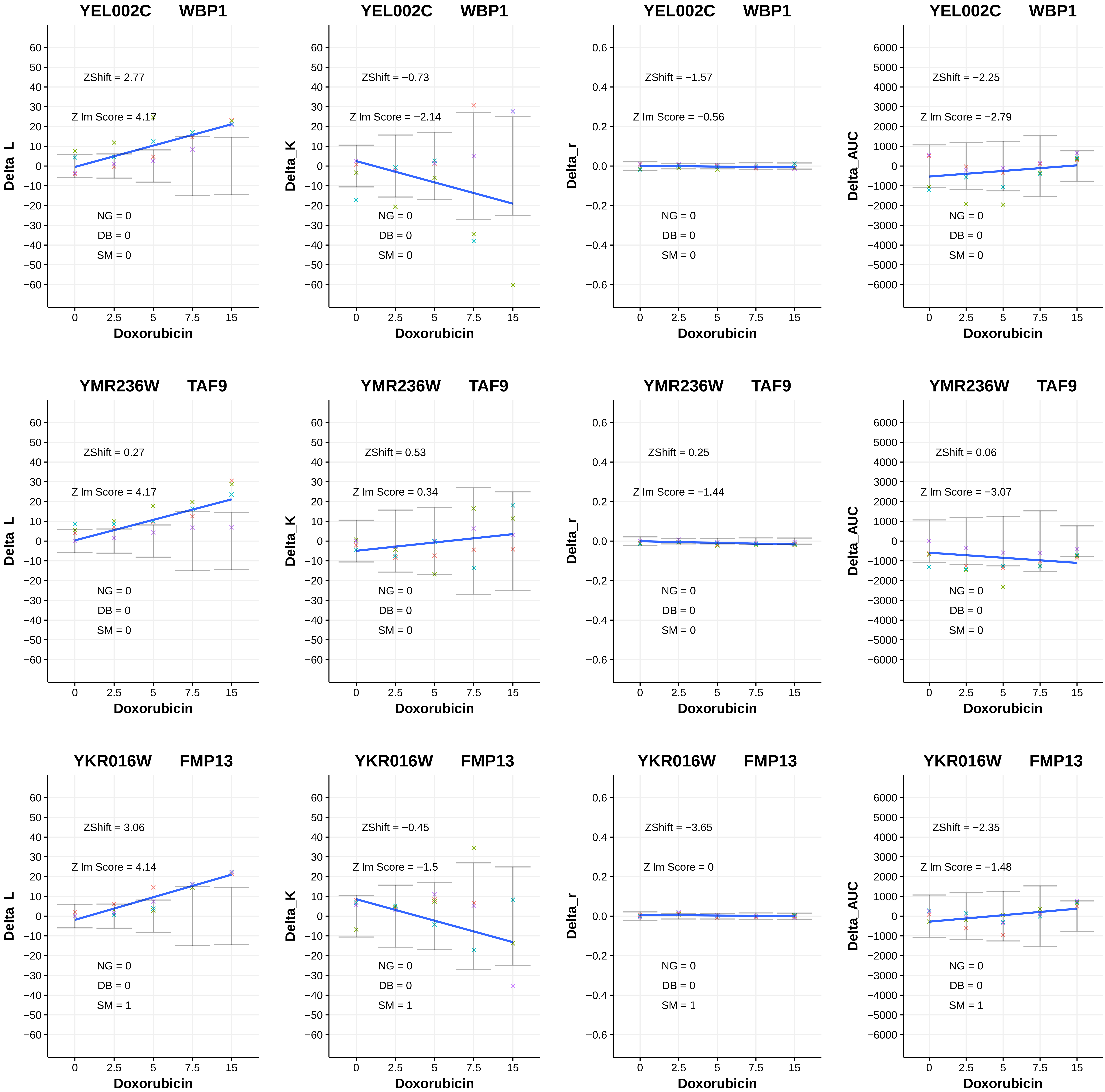
YEL002C WBP1
ZShift = 2.77
Z lm Score = 4.17
NG = 0
DB = 0
SM = 0
Delta_L
Doxorubicin
YEL002C WBP1
ZShift = −0.73
Z lm Score = −2.14
NG = 0
DB = 0
SM = 0
Delta_K
Doxorubicin
YEL002C WBP1
ZShift = −1.57
Z lm Score = −0.56
NG = 0
DB = 0
SM = 0
Delta_r
Doxorubicin
YEL002C WBP1
ZShift = −2.25
Z lm Score = −2.79
NG = 0
DB = 0
SM = 0
Delta_AUC
Doxorubicin
YMR236W TAF9
ZShift = 0.27
Z lm Score = 4.17
NG = 0
DB = 0
SM = 0
Delta_L
Doxorubicin
YMR236W TAF9
ZShift = 0.53
Z lm Score = 0.34
NG = 0
DB = 0
SM = 0
Delta_K
Doxorubicin
YMR236W TAF9
ZShift = 0.25
Z lm Score = −1.44
NG = 0
DB = 0
SM = 0
Delta_r
Doxorubicin
YMR236W TAF9
ZShift = 0.06
Z lm Score = −3.07
NG = 0
DB = 0
SM = 0
Delta_AUC
Doxorubicin
YKR016W FMP13
ZShift = 3.06
Z lm Score = 4.14
NG = 0
DB = 0
SM = 1
Delta_L
Doxorubicin
YKR016W FMP13
ZShift = −0.45
Z lm Score = −1.5
NG = 0
DB = 0
SM = 1
Delta_K
Doxorubicin
YKR016W FMP13
ZShift = −3.65
Z lm Score = 0
NG = 0
DB = 0
SM = 1
Delta_r
Doxorubicin
YKR016W FMP13
ZShift = −2.35
Z lm Score = −1.48
NG = 0
DB = 0
SM = 1
Delta_AUC
Doxorubicin

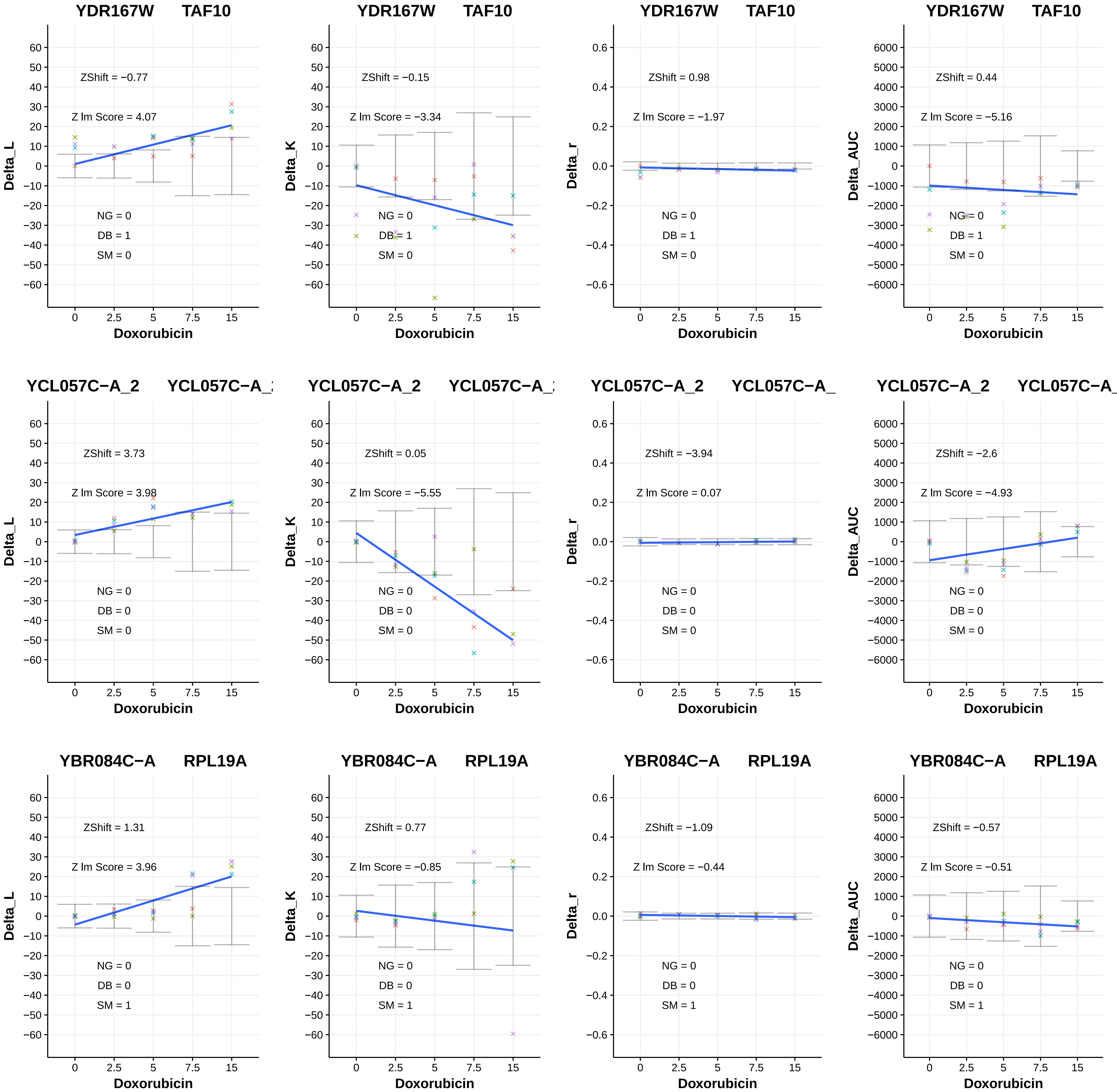
YDR167W TAF10
Delta_L
ZShift = −0.77
Z lm Score = 4.07
NG = 0
DB = 1
SM = 0
Doxorubicin
YDR167W TAF10
Delta_K
ZShift = −0.15
Z lm Score = −3.34
NG = 0
DB = 1
SM = 0
Doxorubicin
YDR167W TAF10
Delta_r
ZShift = 0.98
Z lm Score = −1.97
NG = 0
DB = 1
SM = 0
Doxorubicin
YDR167W TAF10
Delta_AUC
ZShift = 0.44
Z lm Score = −5.16
NG = 0
DB = 1
SM = 0
Doxorubicin
YCL057C−A_2 YCL057C−A_
Delta_L
ZShift = 3.73
Z lm Score = 3.98
NG = 0
DB = 0
SM = 0
Doxorubicin
YCL057C−A_2 YCL057C−A_
Delta_K
ZShift = 0.05
Z lm Score = −5.55
NG = 0
DB = 0
SM = 0
Doxorubicin
YCL057C−A_2 YCL057C−A_
Delta_r
ZShift = −3.94
Z lm Score = 0.07
NG = 0
DB = 0
SM = 0
Doxorubicin
YCL057C−A_2 YCL057C−A_
Delta_AUC
ZShift = −2.6
Z lm Score = −4.93
NG = 0
DB = 0
SM = 0
Doxorubicin
YBR084C−A RPL19A
Delta_L
ZShift = 1.31
Z lm Score = 3.96
NG = 0
DB = 0
SM = 1
Doxorubicin
YBR084C−A RPL19A
Delta_K
ZShift = 0.77
Z lm Score = −0.85
NG = 0
DB = 0
SM = 1
Doxorubicin
YBR084C−A RPL19A
Delta_r
ZShift = −1.09
Z lm Score = −0.44
NG = 0
DB = 0
SM = 1
Doxorubicin
YBR084C−A RPL19A
Delta_AUC
ZShift = −0.57
Z lm Score = −0.51
NG = 0
DB = 0
SM = 1
Doxorubicin

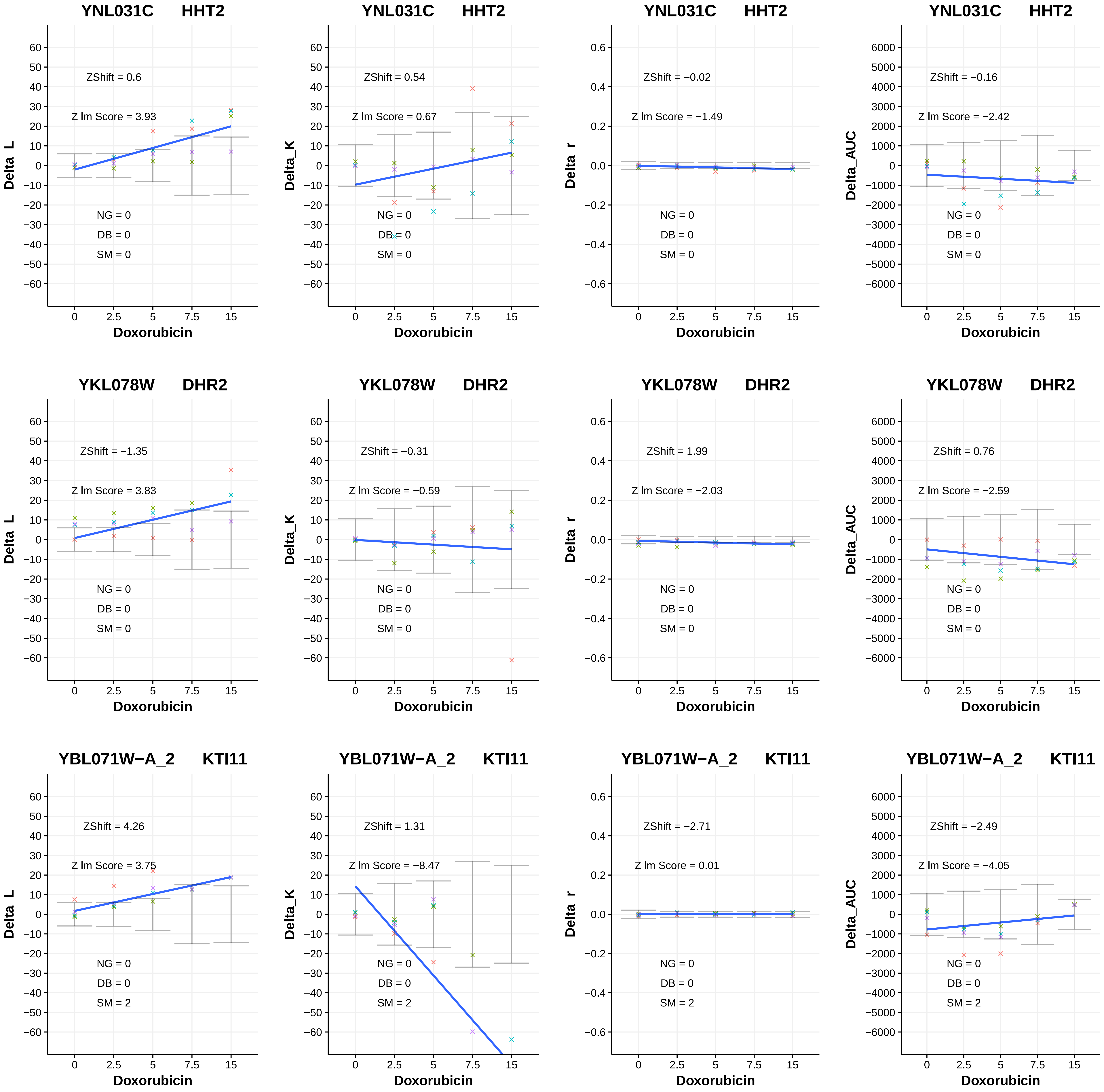
YNL031C HHT2
ZShift = 0.6
Z lm Score = 3.93
NG = 0
DB = 0
SM = 0
Delta_L
Doxorubicin
YNL031C HHT2
ZShift = 0.54
Z lm Score = 0.67
NG = 0
DB = 0
SM = 0
Delta_K
Doxorubicin
YNL031C HHT2
ZShift = −0.02
Z lm Score = −1.49
NG = 0
DB = 0
SM = 0
Delta_r
Doxorubicin
YNL031C HHT2
ZShift = −0.16
Z lm Score = −2.42
NG = 0
DB = 0
SM = 0
Delta_AUC
Doxorubicin
YKL078W DHR2
ZShift = −1.35
Z lm Score = 3.83
NG = 0
DB = 0
SM = 0
Delta_L
Doxorubicin
YKL078W DHR2
ZShift = −0.31
Z lm Score = −0.59
NG = 0
DB = 0
SM = 0
Delta_K
Doxorubicin
YKL078W DHR2
ZShift = 1.99
Z lm Score = −2.03
NG = 0
DB = 0
SM = 0
Delta_r
Doxorubicin
YKL078W DHR2
ZShift = 0.76
Z lm Score = −2.59
NG = 0
DB = 0
SM = 0
Delta_AUC
Doxorubicin
YBL071W−A_2 KTI11
ZShift = 4.26
Z lm Score = 3.75
NG = 0
DB = 0
SM = 2
Delta_L
Doxorubicin
YBL071W−A_2 KTI11
ZShift = 1.31
Z lm Score = −8.47
NG = 0
DB = 0
SM = 2
Delta_K
Doxorubicin
YBL071W−A_2 KTI11
ZShift = −2.71
Z lm Score = 0.01
NG = 0
DB = 0
SM = 2
Delta_r
Doxorubicin
YBL071W−A_2 KTI11
ZShift = −2.49
Z lm Score = −4.05
NG = 0
DB = 0
SM = 2
Delta_AUC
Doxorubicin

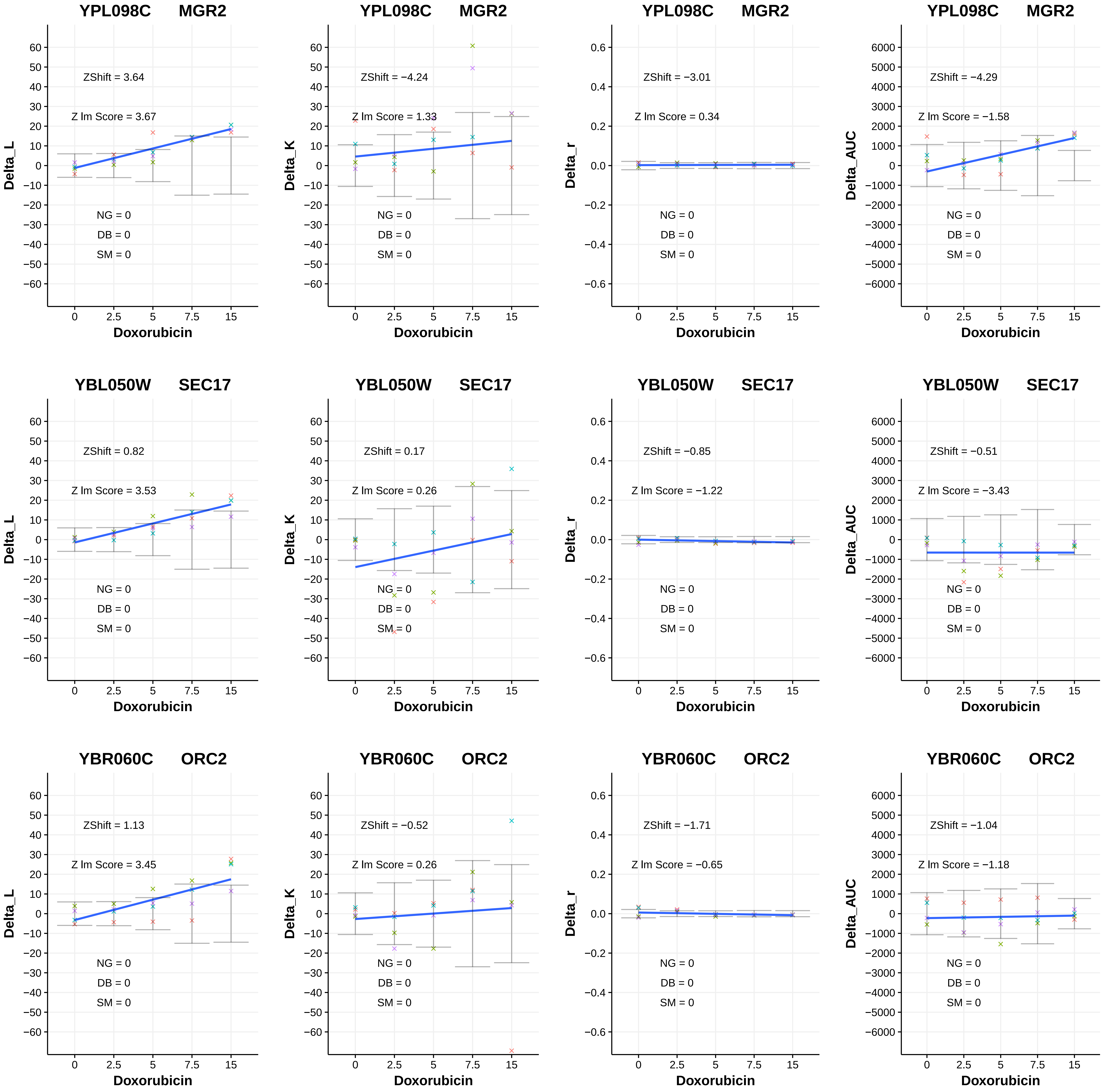

YPL098C MGR2
ZShift = 3.64
Z lm Score = 3.67
NG = 0
DB = 0
SM = 0
Delta_L
Doxorubicin
YPL098C MGR2
ZShift = −4.24
Z lm Score = 1.33
NG = 0
DB = 0
SM = 0
Delta_K
Doxorubicin
YPL098C MGR2
ZShift = −3.01
Z lm Score = 0.34
NG = 0
DB = 0
SM = 0
Delta_r
Doxorubicin
YPL098C MGR2
ZShift = −4.29
Z lm Score = −1.58
NG = 0
DB = 0
SM = 0
Delta_AUC
Doxorubicin
YBL050W SEC17
ZShift = 0.82
Z lm Score = 3.53
NG = 0
DB = 0
SM = 0
Delta_L
Doxorubicin
YBL050W SEC17
ZShift = 0.17
Z lm Score = 0.26
NG = 0
DB = 0
SM = 0
Delta_K
Doxorubicin
YBL050W SEC17
ZShift = −0.85
Z lm Score = −1.22
NG = 0
DB = 0
SM = 0
Delta_r
Doxorubicin
YBL050W SEC17
ZShift = −0.51
Z lm Score = −3.43
NG = 0
DB = 0
SM = 0
Delta_AUC
Doxorubicin
YBR060C ORC2
ZShift = 1.13
Z lm Score = 3.45
NG = 0
DB = 0
SM = 0
Delta_L
Doxorubicin
YBR060C ORC2
ZShift = −0.52
Z lm Score = 0.26
NG = 0
DB = 0
SM = 0
Delta_K
Doxorubicin
YBR060C ORC2
ZShift = −1.71
Z lm Score = −0.65
NG = 0
DB = 0
SM = 0
Delta_r
Doxorubicin
YBR060C ORC2
ZShift = −1.04
Z lm Score = −1.18
NG = 0
DB = 0
SM = 0
Delta_AUC
Doxorubicin

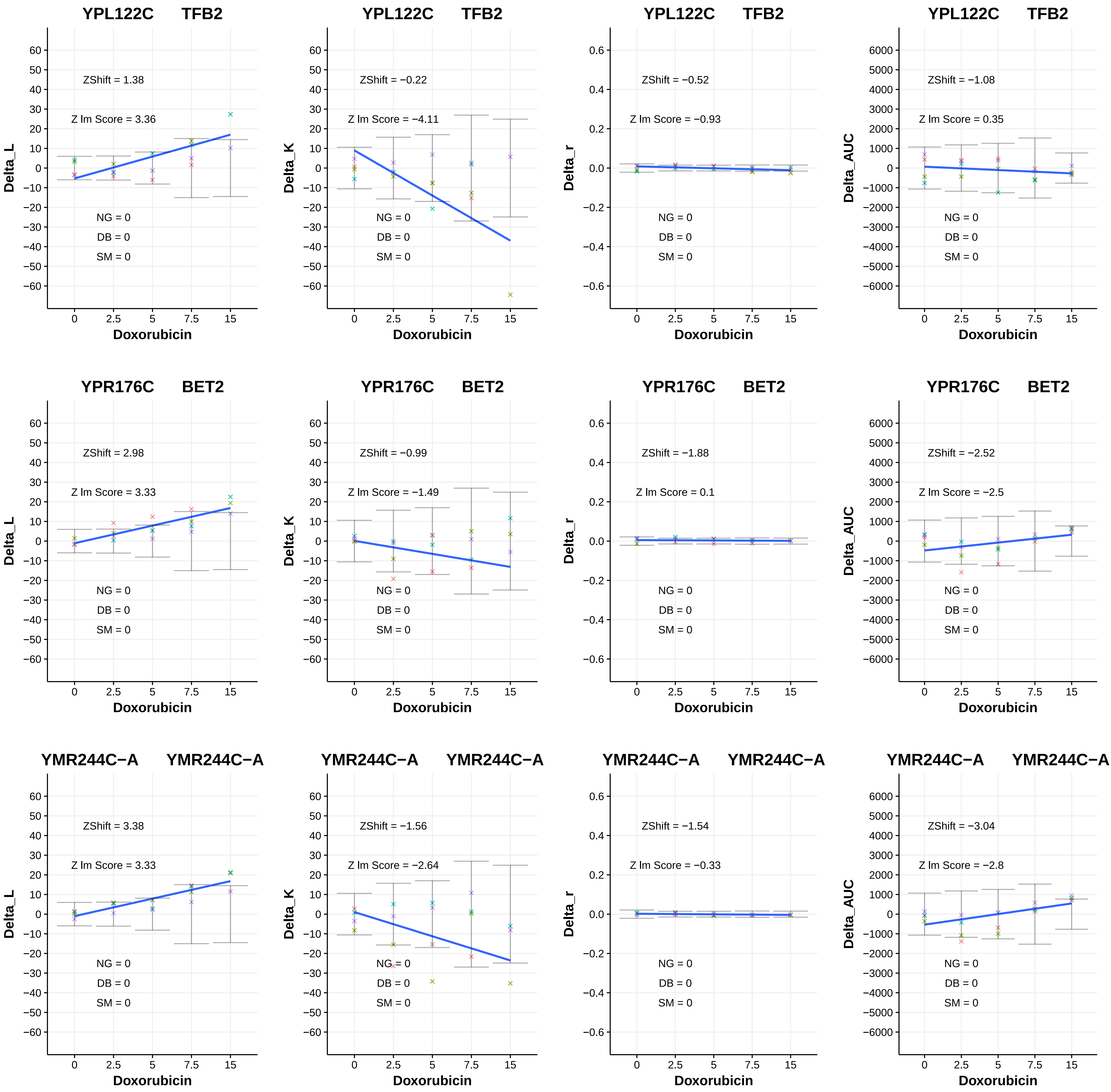
YPL122C TFB2
ZShift = 1.38
Z lm Score = 3.36
NG = 0
DB = 0
SM = 0
Delta_L
Doxorubicin
YPL122C TFB2
ZShift = −0.22
Z lm Score = −4.11
NG = 0
DB = 0
SM = 0
Delta_K
Doxorubicin
YPL122C TFB2
ZShift = −0.52
Z lm Score = −0.93
NG = 0
DB = 0
SM = 0
Delta_r
Doxorubicin
YPL122C TFB2
ZShift = −1.08
Z lm Score = 0.35
NG = 0
DB = 0
SM = 0
Delta_AUC
Doxorubicin
YPR176C BET2
ZShift = 2.98
Z lm Score = 3.33
NG = 0
DB = 0
SM = 0
Delta_L
Doxorubicin
YPR176C BET2
ZShift = −0.99
Z lm Score = −1.49
NG = 0
DB = 0
SM = 0
Delta_K
Doxorubicin
YPR176C BET2
ZShift = −1.88
Z lm Score = 0.1
NG = 0
DB = 0
SM = 0
Delta_r
Doxorubicin
YPR176C BET2
ZShift = −2.52
Z lm Score = −2.5
NG = 0
DB = 0
SM = 0
Delta_AUC
Doxorubicin
YMR244C−A YMR244C−A
ZShift = 3.38
Z lm Score = 3.33
NG = 0
DB = 0
SM = 0
Delta_L
Doxorubicin
YMR244C−A YMR244C−A
ZShift = −1.56
Z lm Score = −2.64
NG = 0
DB = 0
SM = 0
Delta_K
Doxorubicin
YMR244C−A YMR244C−A
ZShift = −1.54
Z lm Score = −0.33
NG = 0
DB = 0
SM = 0
Delta_r
Doxorubicin
YMR244C−A YMR244C−A
ZShift = −3.04
Z lm Score = −2.8
NG = 0
DB = 0
SM = 0
Delta_AUC
Doxorubicin

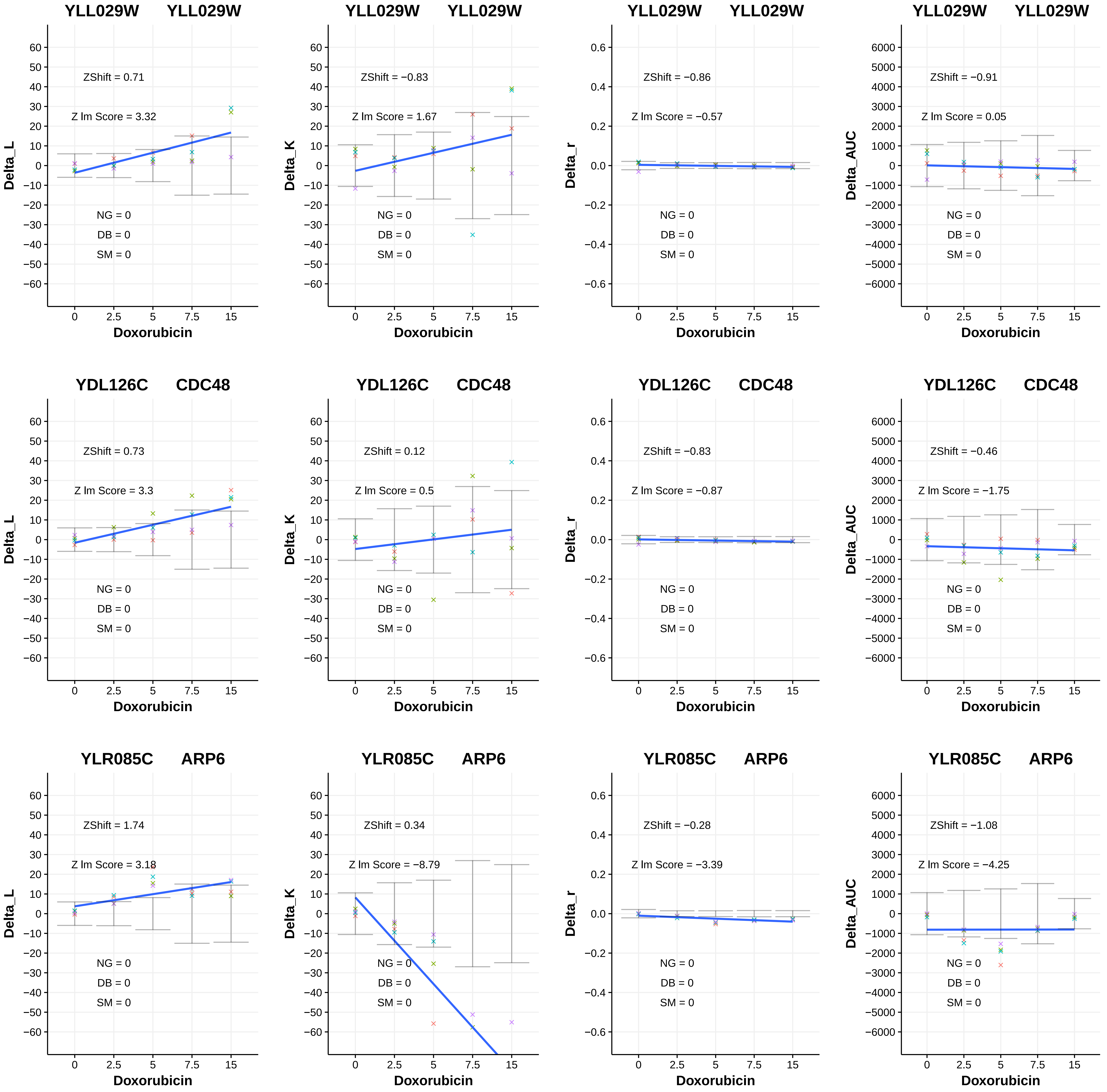

YLL029W YLL029W
ZShift = 0.71
Z lm Score = 3.32
NG = 0
DB = 0
SM = 0
Delta_L
Doxorubicin
YLL029W YLL029W
ZShift = −0.83
Z lm Score = 1.67
NG = 0
DB = 0
SM = 0
Delta_K
Doxorubicin
YLL029W YLL029W
ZShift = −0.86
Z lm Score = −0.57
NG = 0
DB = 0
SM = 0
Delta_r
Doxorubicin
YLL029W YLL029W
ZShift = −0.91
Z lm Score = 0.05
NG = 0
DB = 0
SM = 0
Delta_AUC
Doxorubicin
YDL126C CDC48
ZShift = 0.73
Z lm Score = 3.3
NG = 0
DB = 0
SM = 0
Delta_L
Doxorubicin
YDL126C CDC48
ZShift = 0.12
Z lm Score = 0.5
NG = 0
DB = 0
SM = 0
Delta_K
Doxorubicin
YDL126C CDC48
ZShift = −0.83
Z lm Score = −0.87
NG = 0
DB = 0
SM = 0
Delta_r
Doxorubicin
YDL126C CDC48
ZShift = −0.46
Z lm Score = −1.75
NG = 0
DB = 0
SM = 0
Delta_AUC
Doxorubicin
YLR085C ARP6
ZShift = 1.74
Z lm Score = 3.18
NG = 0
DB = 0
SM = 0
Delta_L
Doxorubicin
YLR085C ARP6
ZShift = 0.34
Z lm Score = −8.79
NG = 0
DB = 0
SM = 0
Delta_K
Doxorubicin
YLR085C ARP6
ZShift = −0.28
Z lm Score = −3.39
NG = 0
DB = 0
SM = 0
Delta_r
Doxorubicin
YLR085C ARP6
ZShift = −1.08
Z lm Score = −4.25
NG = 0
DB = 0
SM = 0
Delta_AUC
Doxorubicin

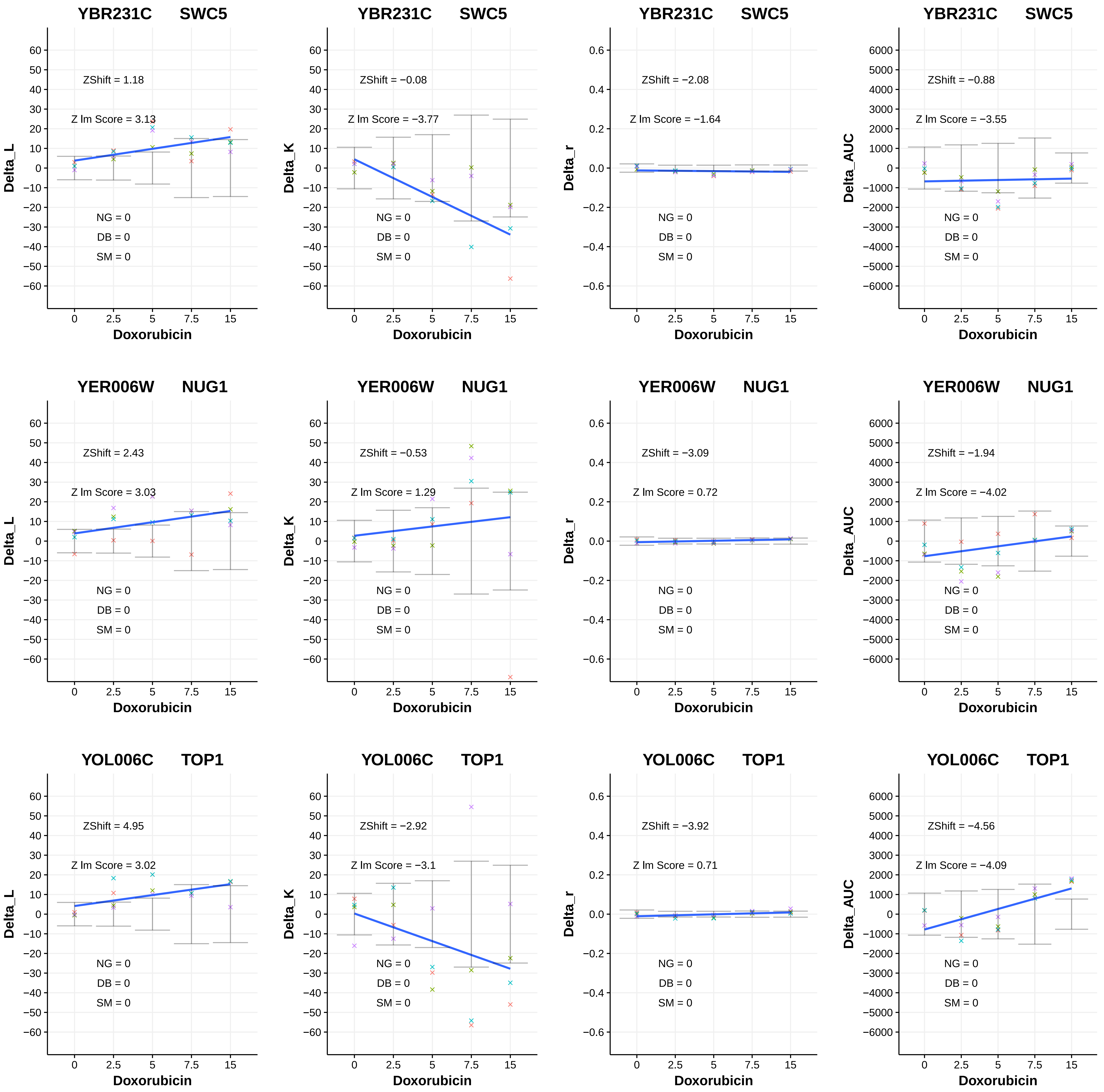
YBR231C SWC5
ZShift = 1.18
Z lm Score = 3.13
NG = 0
DB = 0
SM = 0
Delta_L
Doxorubicin
YBR231C SWC5
ZShift = −0.08
Z lm Score = −3.77
NG = 0
DB = 0
SM = 0
Delta_K
Doxorubicin
YBR231C SWC5
ZShift = −2.08
Z lm Score = −1.64
NG = 0
DB = 0
SM = 0
Delta_r
Doxorubicin
YBR231C SWC5
ZShift = −0.88
Z lm Score = −3.55
NG = 0
DB = 0
SM = 0
Delta_AUC
Doxorubicin
YER006W NUG1
ZShift = 2.43
Z lm Score = 3.03
NG = 0
DB = 0
SM = 0
Delta_L
Doxorubicin
YER006W NUG1
ZShift = −0.53
Z lm Score = 1.29
NG = 0
DB = 0
SM = 0
Delta_K
Doxorubicin
YER006W NUG1
ZShift = −3.09
Z lm Score = 0.72
NG = 0
DB = 0
SM = 0
Delta_r
Doxorubicin
YER006W NUG1
ZShift = −1.94
Z lm Score = −4.02
NG = 0
DB = 0
SM = 0
Delta_AUC
Doxorubicin
YOL006C TOP1
ZShift = 4.95
Z lm Score = 3.02
NG = 0
DB = 0
SM = 0
Delta_L
Doxorubicin
YOL006C TOP1
ZShift = −2.92
Z lm Score = −3.1
NG = 0
DB = 0
SM = 0
Delta_K
Doxorubicin
YOL006C TOP1
ZShift = −3.92
Z lm Score = 0.71
NG = 0
DB = 0
SM = 0
Delta_r
Doxorubicin
YOL006C TOP1
ZShift = −4.56
Z lm Score = −4.09
NG = 0
DB = 0
SM = 0
Delta_AUC
Doxorubicin

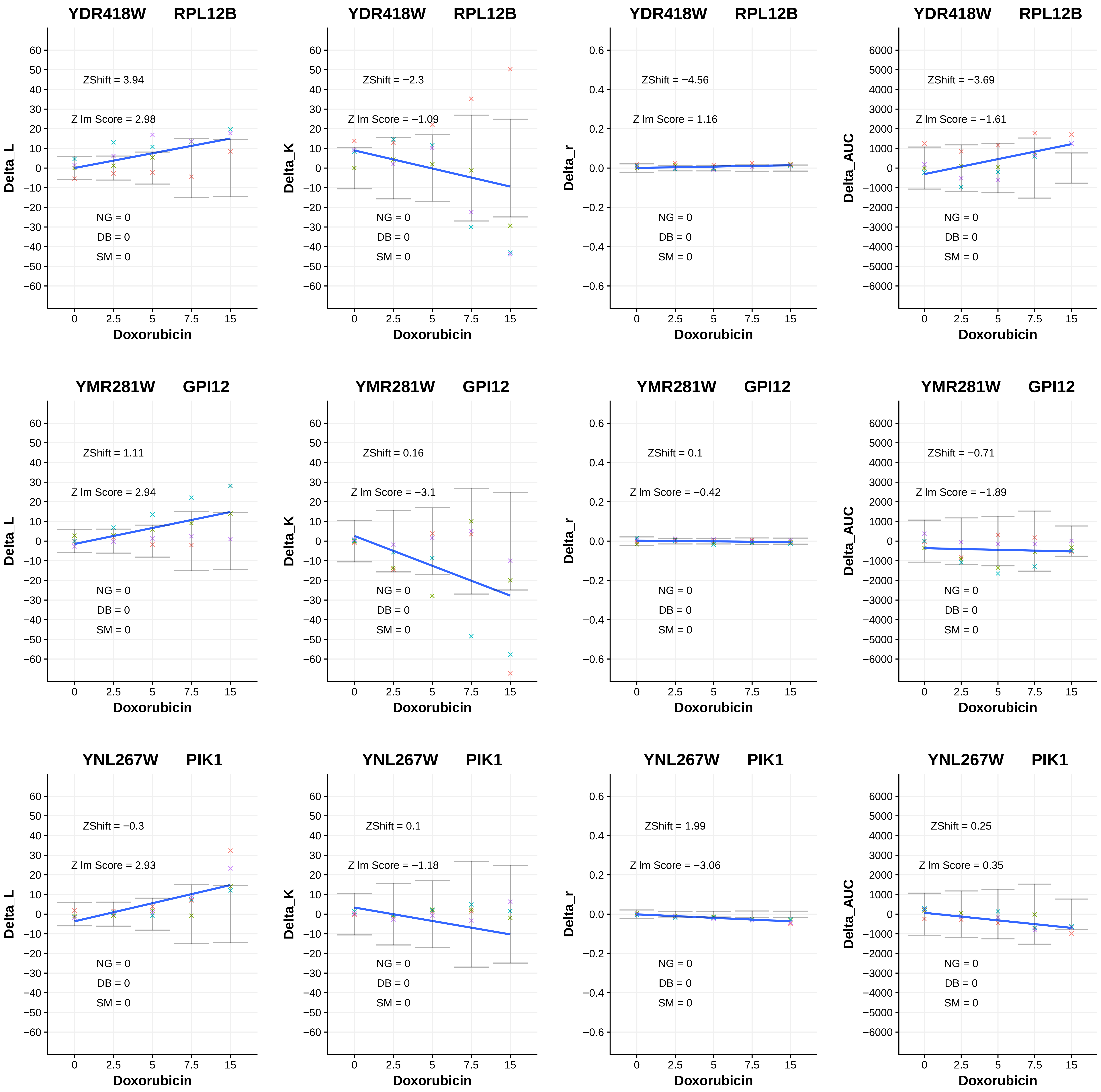
YDR418W RPL12B
ZShift = 3.94
Z lm Score = 2.98
NG = 0
DB = 0
SM = 0
Delta_L
Doxorubicin
YDR418W RPL12B
ZShift = −2.3
Z lm Score = −1.09
NG = 0
DB = 0
SM = 0
Delta_K
Doxorubicin
YDR418W RPL12B
ZShift = −4.56
Z lm Score = 1.16
NG = 0
DB = 0
SM = 0
Delta_r
Doxorubicin
YDR418W RPL12B
ZShift = −3.69
Z lm Score = −1.61
NG = 0
DB = 0
SM = 0
Delta_AUC
Doxorubicin
YMR281W GPI12
ZShift = 1.11
Z lm Score = 2.94
NG = 0
DB = 0
SM = 0
Delta_L
Doxorubicin
YMR281W GPI12
ZShift = 0.16
Z lm Score = −3.1
NG = 0
DB = 0
SM = 0
Delta_K
Doxorubicin
YMR281W GPI12
ZShift = 0.1
Z lm Score = −0.42
NG = 0
DB = 0
SM = 0
Delta_r
Doxorubicin
YMR281W GPI12
ZShift = −0.71
Z lm Score = −1.89
NG = 0
DB = 0
SM = 0
Delta_AUC
Doxorubicin
YNL267W PIK1
ZShift = −0.3
Z lm Score = 2.93
NG = 0
DB = 0
SM = 0
Delta_L
Doxorubicin
YNL267W PIK1
ZShift = 0.1
Z lm Score = −1.18
NG = 0
DB = 0
SM = 0
Delta_K
Doxorubicin
YNL267W PIK1
ZShift = 1.99
Z lm Score = −3.06
NG = 0
DB = 0
SM = 0
Delta_r
Doxorubicin
YNL267W PIK1
ZShift = 0.25
Z lm Score = 0.35
NG = 0
DB = 0
SM = 0
Delta_AUC
Doxorubicin

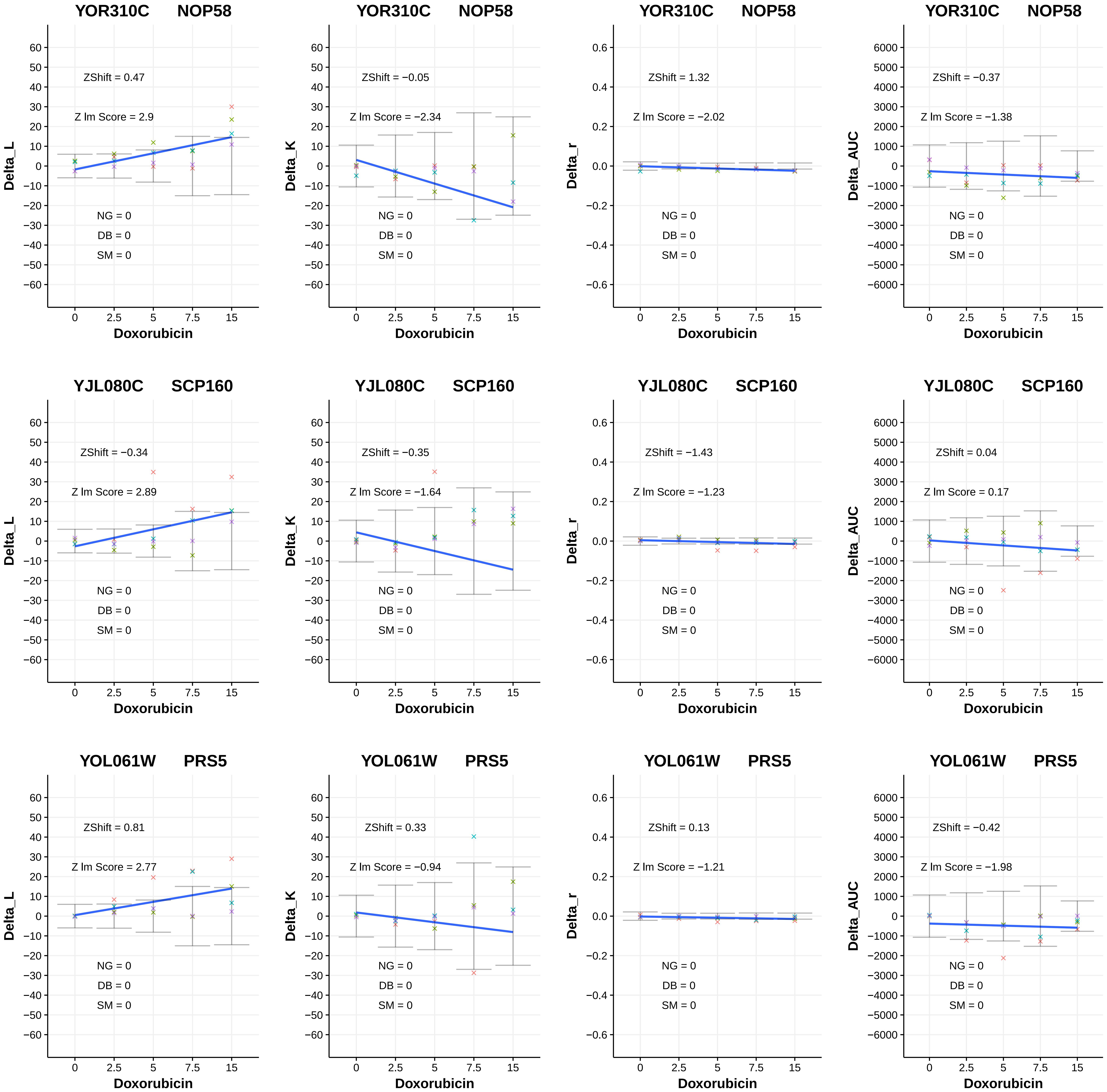

YOR310C NOP58
ZShift = 0.47
Z lm Score = 2.9
NG = 0
DB = 0
SM = 0
Delta_L
Doxorubicin
YOR310C NOP58
ZShift = −0.05
Z lm Score = −2.34
NG = 0
DB = 0
SM = 0
Delta_K
Doxorubicin
YOR310C NOP58
ZShift = 1.32
Z lm Score = −2.02
NG = 0
DB = 0
SM = 0
Delta_r
Doxorubicin
YOR310C NOP58
ZShift = −0.37
Z lm Score = −1.38
NG = 0
DB = 0
SM = 0
Delta_AUC
Doxorubicin
YJL080C SCP160
ZShift = −0.34
Z lm Score = 2.89
NG = 0
DB = 0
SM = 0
Delta_L
Doxorubicin
YJL080C SCP160
ZShift = −0.35
Z lm Score = −1.64
NG = 0
DB = 0
SM = 0
Delta_K
Doxorubicin
YJL080C SCP160
ZShift = −1.43
Z lm Score = −1.23
NG = 0
DB = 0
SM = 0
Delta_r
Doxorubicin
YJL080C SCP160
ZShift = 0.04
Z lm Score = 0.17
NG = 0
DB = 0
SM = 0
Delta_AUC
Doxorubicin
YOL061W PRS5
ZShift = 0.81
Z lm Score = 2.77
NG = 0
DB = 0
SM = 0
Delta_L
Doxorubicin
YOL061W PRS5
ZShift = 0.33
Z lm Score = −0.94
NG = 0
DB = 0
SM = 0
Delta_K
Doxorubicin
YOL061W PRS5
ZShift = 0.13
Z lm Score = −1.21
NG = 0
DB = 0
SM = 0
Delta_r
Doxorubicin
YOL061W PRS5
ZShift = −0.42
Z lm Score = −1.98
NG = 0
DB = 0
SM = 0
Delta_AUC
Doxorubicin

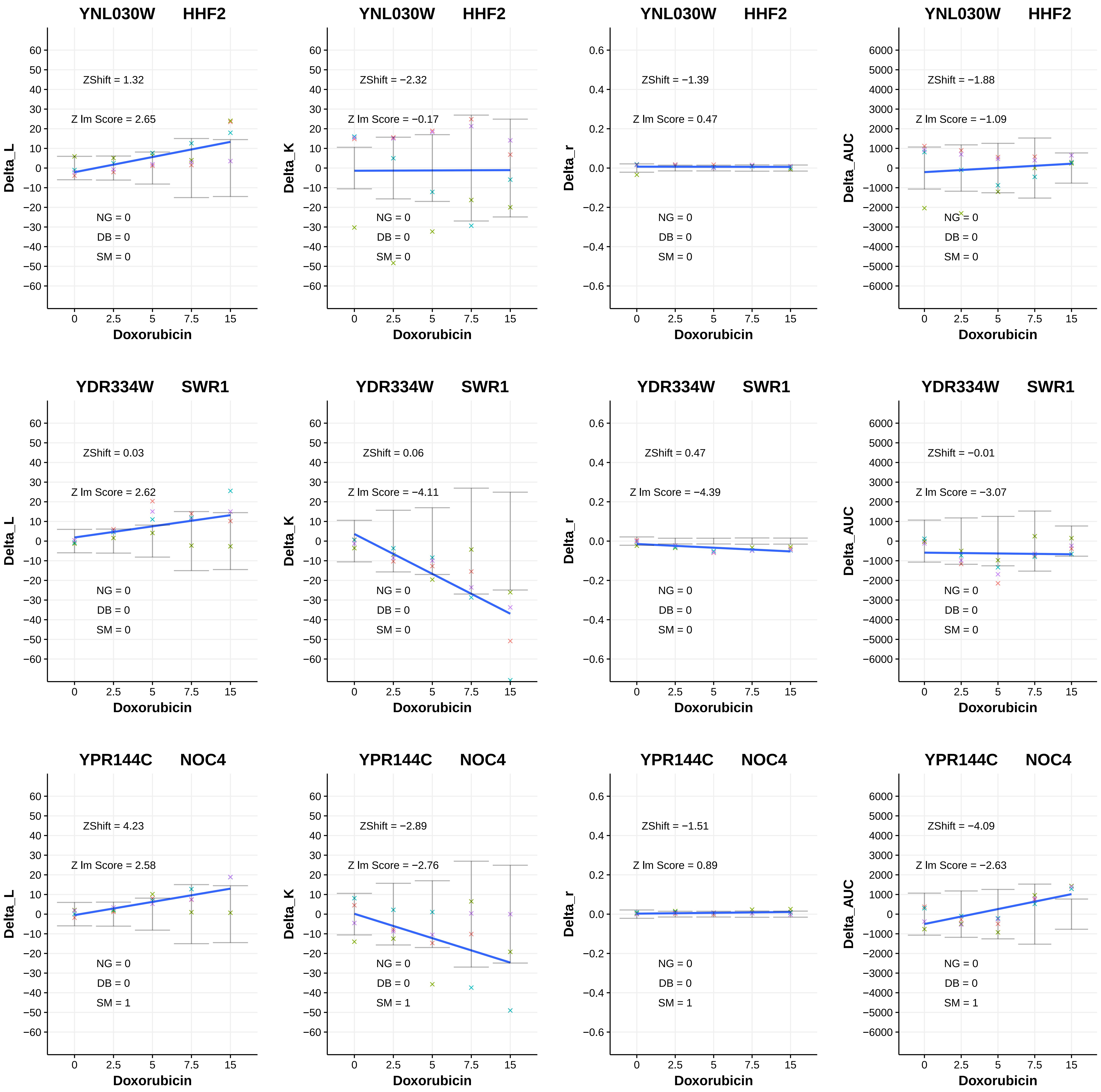
YNL030W HHF2
ZShift = 1.32
Z lm Score = 2.65
NG = 0
DB = 0
SM = 0
Delta_L
Doxorubicin
YNL030W HHF2
ZShift = −2.32
Z lm Score = −0.17
NG = 0
DB = 0
SM = 0
Delta_K
Doxorubicin
YNL030W HHF2
ZShift = −1.39
Z lm Score = 0.47
NG = 0
DB = 0
SM = 0
Delta_r
Doxorubicin
YNL030W HHF2
ZShift = −1.88
Z lm Score = −1.09
NG = 0
DB = 0
SM = 0
Delta_AUC
Doxorubicin
YDR334W SWR1
ZShift = 0.03
Z lm Score = 2.62
NG = 0
DB = 0
SM = 0
Delta_L
Doxorubicin
YDR334W SWR1
ZShift = 0.06
Z lm Score = −4.11
NG = 0
DB = 0
SM = 0
Delta_K
Doxorubicin
YDR334W SWR1
ZShift = 0.47
Z lm Score = −4.39
NG = 0
DB = 0
SM = 0
Delta_r
Doxorubicin
YDR334W SWR1
ZShift = −0.01
Z lm Score = −3.07
NG = 0
DB = 0
SM = 0
Delta_AUC
Doxorubicin
YPR144C NOC4
ZShift = 4.23
Z lm Score = 2.58
NG = 0
DB = 0
SM = 1
Delta_L
Doxorubicin
YPR144C NOC4
ZShift = −2.89
Z lm Score = −2.76
NG = 0
DB = 0
SM = 1
Delta_K
Doxorubicin
YPR144C NOC4
ZShift = −1.51
Z lm Score = 0.89
NG = 0
DB = 0
SM = 1
Delta_r
Doxorubicin
YPR144C NOC4
ZShift = −4.09
Z lm Score = −2.63
NG = 0
DB = 0
SM = 1
Delta_AUC
Doxorubicin

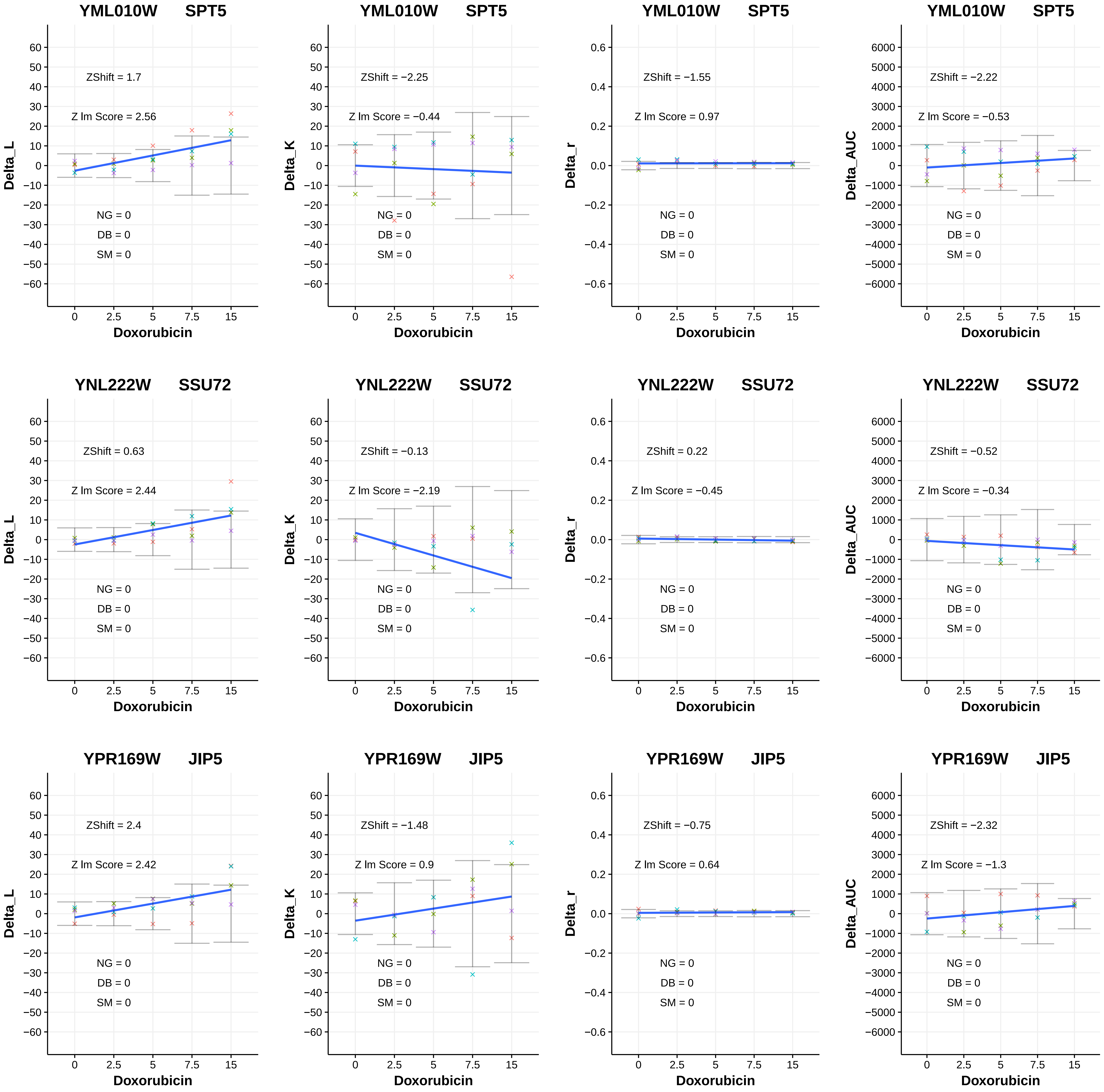

YML010W SPT5
ZShift = 1.7
Z lm Score = 2.56
NG = 0
DB = 0
SM = 0
Delta_L
Doxorubicin
YML010W SPT5
ZShift = −2.25
Z lm Score = −0.44
NG = 0
DB = 0
SM = 0
Delta_K
Doxorubicin
YML010W SPT5
ZShift = −1.55
Z lm Score = 0.97
NG = 0
DB = 0
SM = 0
Delta_r
Doxorubicin
YML010W SPT5
ZShift = −2.22
Z lm Score = −0.53
NG = 0
DB = 0
SM = 0
Delta_AUC
Doxorubicin
YNL222W SSU72
ZShift = 0.63
Z lm Score = 2.44
NG = 0
DB = 0
SM = 0
Delta_L
Doxorubicin
YNL222W SSU72
ZShift = −0.13
Z lm Score = −2.19
NG = 0
DB = 0
SM = 0
Delta_K
Doxorubicin
YNL222W SSU72
ZShift = 0.22
Z lm Score = −0.45
NG = 0
DB = 0
SM = 0
Delta_r
Doxorubicin
YNL222W SSU72
ZShift = −0.52
Z lm Score = −0.34
NG = 0
DB = 0
SM = 0
Delta_AUC
Doxorubicin
YPR169W JIP5
ZShift = 2.4
Z lm Score = 2.42
NG = 0
DB = 0
SM = 0
Delta_L
Doxorubicin
YPR169W JIP5
ZShift = −1.48
Z lm Score = 0.9
NG = 0
DB = 0
SM = 0
Delta_K
Doxorubicin
YPR169W JIP5
ZShift = −0.75
Z lm Score = 0.64
NG = 0
DB = 0
SM = 0
Delta_r
Doxorubicin
YPR169W JIP5
ZShift = −2.32
Z lm Score = −1.3
NG = 0
DB = 0
SM = 0
Delta_AUC
Doxorubicin

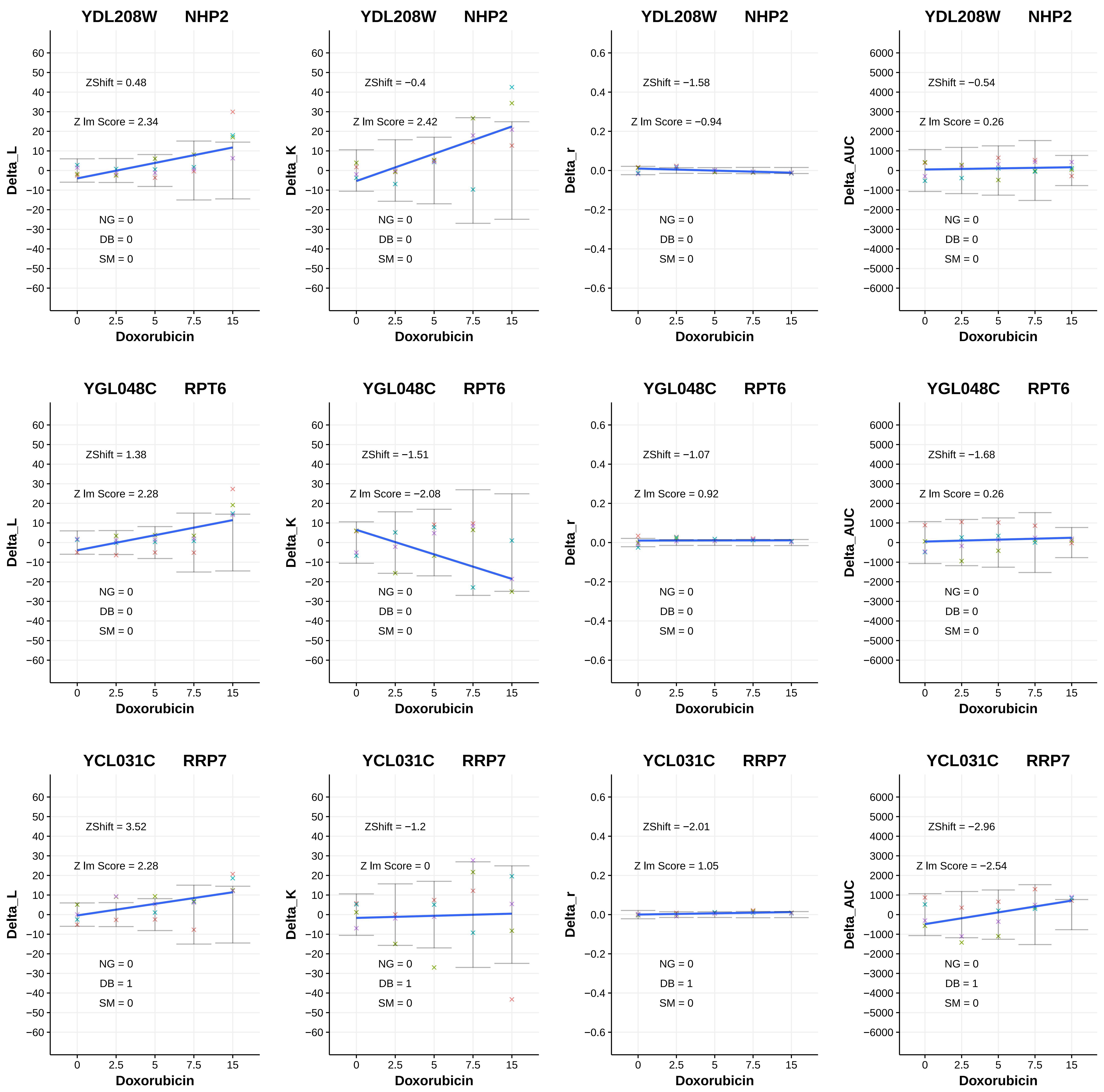

YDL208W NHP2
ZShift = 0.48
Z lm Score = 2.34
NG = 0
DB = 0
SM = 0
Delta_L
Doxorubicin
YDL208W NHP2
ZShift = −0.4
Z lm Score = 2.42
NG = 0
DB = 0
SM = 0
Delta_K
Doxorubicin
YDL208W NHP2
ZShift = −1.58
Z lm Score = −0.94
NG = 0
DB = 0
SM = 0
Delta_r
Doxorubicin
YDL208W NHP2
ZShift = −0.54
Z lm Score = 0.26
NG = 0
DB = 0
SM = 0
Delta_AUC
Doxorubicin
YGL048C RPT6
ZShift = 1.38
Z lm Score = 2.28
NG = 0
DB = 0
SM = 0
Delta_L
Doxorubicin
YGL048C RPT6
ZShift = −1.51
Z lm Score = −2.08
NG = 0
DB = 0
SM = 0
Delta_K
Doxorubicin
YGL048C RPT6
ZShift = −1.07
Z lm Score = 0.92
NG = 0
DB = 0
SM = 0
Delta_r
Doxorubicin
YGL048C RPT6
ZShift = −1.68
Z lm Score = 0.26
NG = 0
DB = 0
SM = 0
Delta_AUC
Doxorubicin
YCL031C RRP7
ZShift = 3.52
Z lm Score = 2.28
NG = 0
DB = 1
SM = 0
Delta_L
Doxorubicin
YCL031C RRP7
ZShift = −1.2
Z lm Score = 0
NG = 0
DB = 1
SM = 0
Delta_K
Doxorubicin
YCL031C RRP7
ZShift = −2.01
Z lm Score = 1.05
NG = 0
DB = 1
SM = 0
Delta_r
Doxorubicin
YCL031C RRP7
ZShift = −2.96
Z lm Score = −2.54
NG = 0
DB = 1
SM = 0
Delta_AUC
Doxorubicin

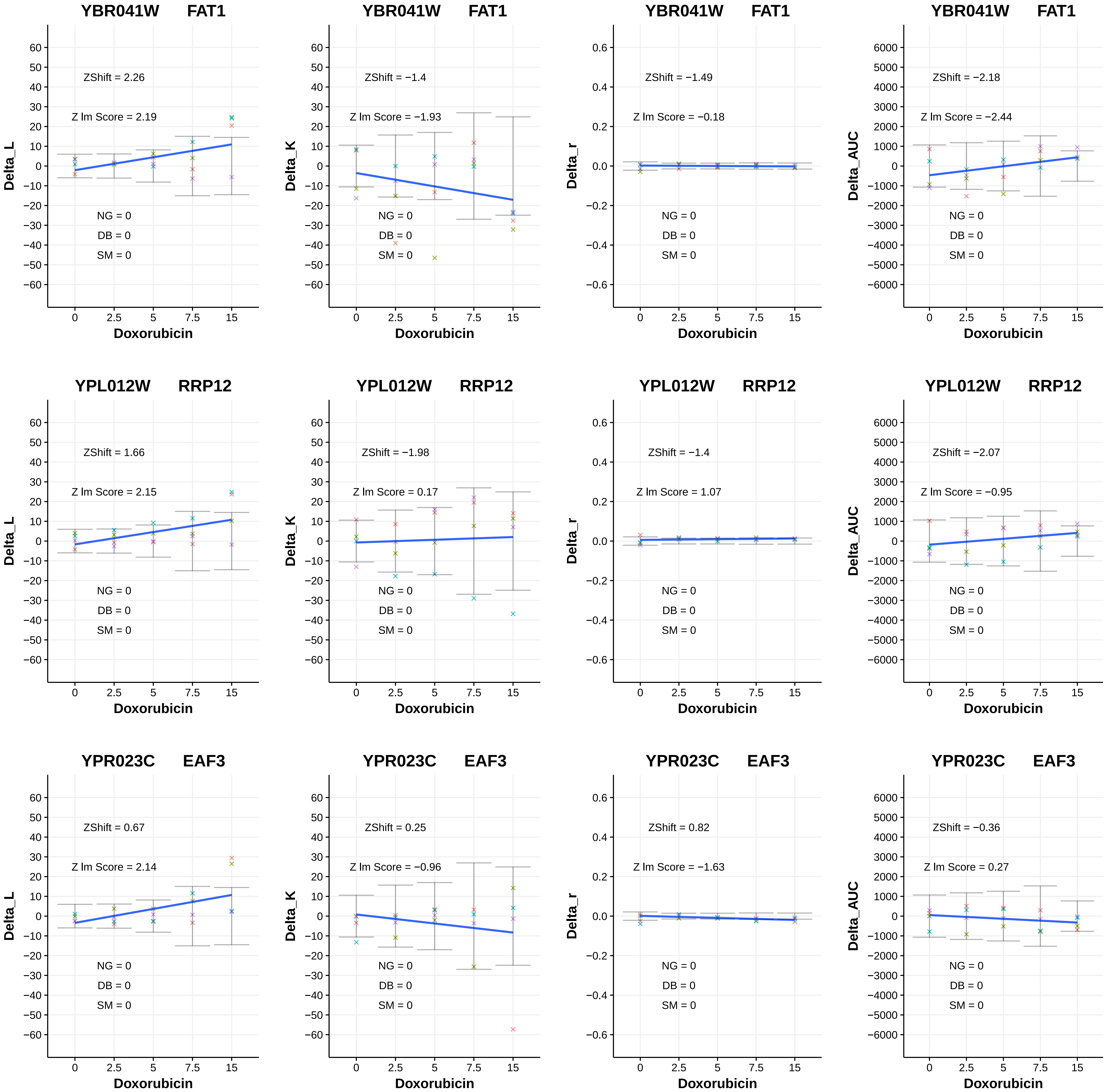
YBR041W FAT1
ZShift = 2.26
Z lm Score = 2.19
NG = 0
DB = 0
SM = 0
Delta_L
Doxorubicin
YBR041W FAT1
ZShift = −1.4
Z lm Score = −1.93
NG = 0
DB = 0
SM = 0
Delta_K
Doxorubicin
YBR041W FAT1
ZShift = −1.49
Z lm Score = −0.18
NG = 0
DB = 0
SM = 0
Delta_r
Doxorubicin
YBR041W FAT1
ZShift = −2.18
Z lm Score = −2.44
NG = 0
DB = 0
SM = 0
Delta_AUC
Doxorubicin
YPL012W RRP12
ZShift = 1.66
Z lm Score = 2.15
NG = 0
DB = 0
SM = 0
Delta_L
Doxorubicin
YPL012W RRP12
ZShift = −1.98
Z lm Score = 0.17
NG = 0
DB = 0
SM = 0
Delta_K
Doxorubicin
YPL012W RRP12
ZShift = −1.4
Z lm Score = 1.07
NG = 0
DB = 0
SM = 0
Delta_r
Doxorubicin
YPL012W RRP12
ZShift = −2.07
Z lm Score = −0.95
NG = 0
DB = 0
SM = 0
Delta_AUC
Doxorubicin
YPR023C EAF3
ZShift = 0.67
Z lm Score = 2.14
NG = 0
DB = 0
SM = 0
Delta_L
Doxorubicin
YPR023C EAF3
ZShift = 0.25
Z lm Score = −0.96
NG = 0
DB = 0
SM = 0
Delta_K
Doxorubicin
YPR023C EAF3
ZShift = 0.82
Z lm Score = −1.63
NG = 0
DB = 0
SM = 0
Delta_r
Doxorubicin
YPR023C EAF3
ZShift = −0.36
Z lm Score = 0.27
NG = 0
DB = 0
SM = 0
Delta_AUC
Doxorubicin

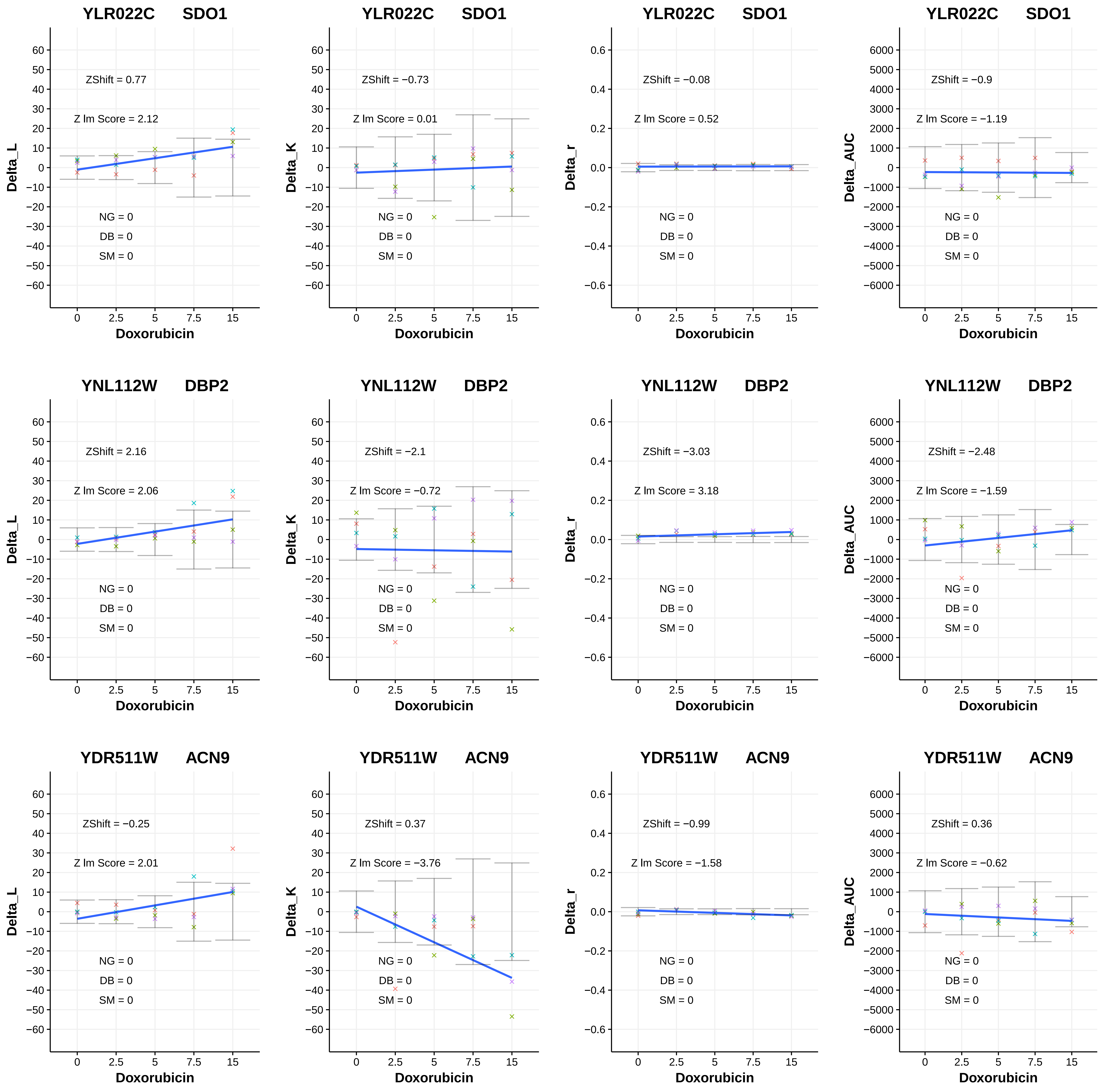

YLR022C SDO1
ZShift = 0.77
Z lm Score = 2.12
NG = 0
DB = 0
SM = 0
Delta_L
Doxorubicin
YLR022C SDO1
ZShift = −0.73
Z lm Score = 0.01
NG = 0
DB = 0
SM = 0
Delta_K
Doxorubicin
YLR022C SDO1
ZShift = −0.08
Z lm Score = 0.52
NG = 0
DB = 0
SM = 0
Delta_r
Doxorubicin
YLR022C SDO1
ZShift = −0.9
Z lm Score = −1.19
NG = 0
DB = 0
SM = 0
Delta_AUC
Doxorubicin
YNL112W DBP2
ZShift = 2.16
Z lm Score = 2.06
NG = 0
DB = 0
SM = 0
Delta_L
Doxorubicin
YNL112W DBP2
ZShift = −2.1
Z lm Score = −0.72
NG = 0
DB = 0
SM = 0
Delta_K
Doxorubicin
YNL112W DBP2
ZShift = −3.03
Z lm Score = 3.18
NG = 0
DB = 0
SM = 0
Delta_r
Doxorubicin
YNL112W DBP2
ZShift = −2.48
Z lm Score = −1.59
NG = 0
DB = 0
SM = 0
Delta_AUC
Doxorubicin
YDR511W ACN9
ZShift = −0.25
Z lm Score = 2.01
NG = 0
DB = 0
SM = 0
Delta_L
Doxorubicin
YDR511W ACN9
ZShift = 0.37
Z lm Score = −3.76
NG = 0
DB = 0
SM = 0
Delta_K
Doxorubicin
YDR511W ACN9
ZShift = −0.99
Z lm Score = −1.58
NG = 0
DB = 0
SM = 0
Delta_r
Doxorubicin
YDR511W ACN9
ZShift = 0.36
Z lm Score = −0.62
NG = 0
DB = 0
SM = 0
Delta_AUC
Doxorubicin

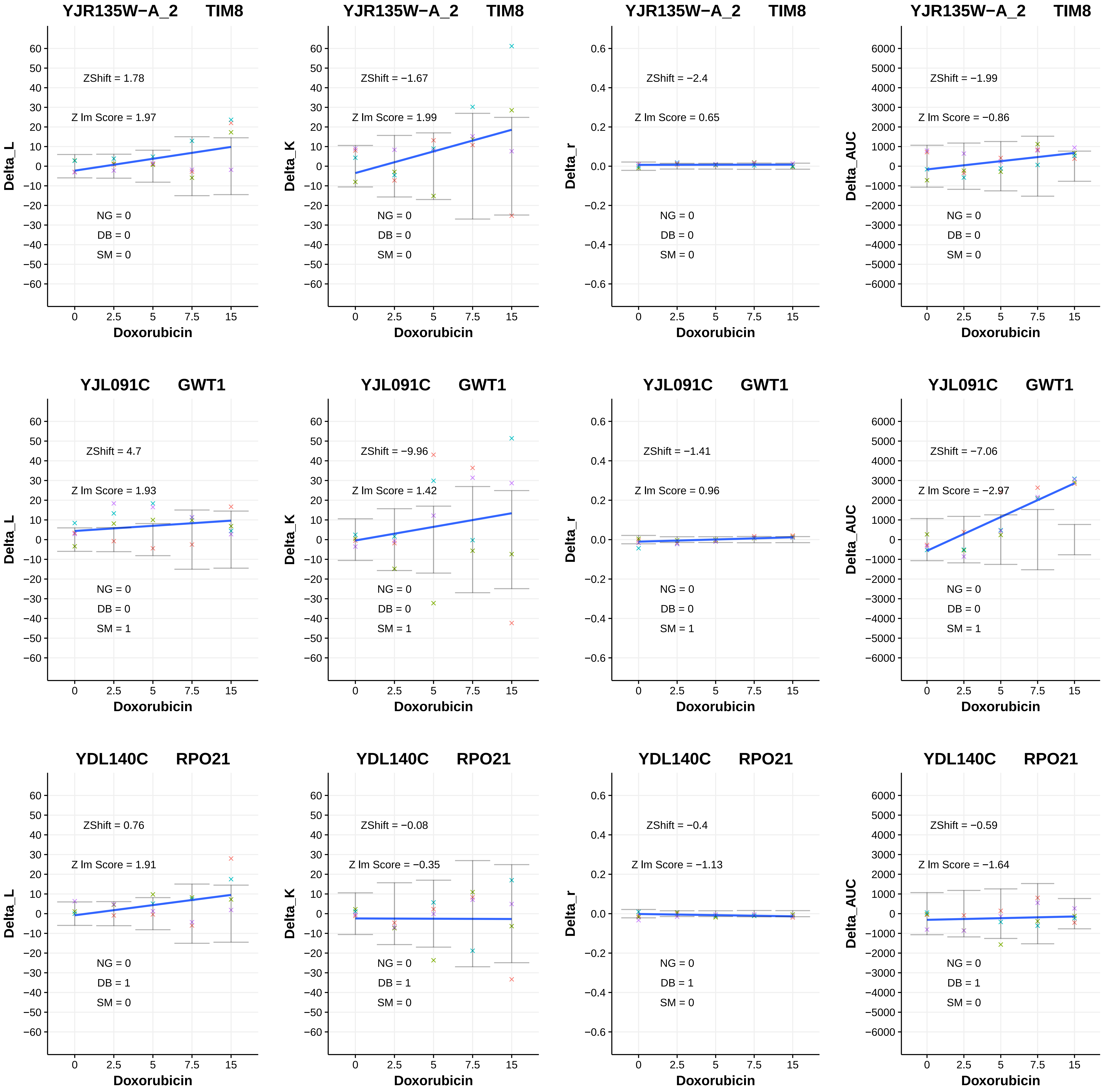

YJR135W−A_2 TIM8
ZShift = 1.78
Z lm Score = 1.97
NG = 0
DB = 0
SM = 0
Delta_L
Doxorubicin
YJR135W−A_2 TIM8
ZShift = −1.67
Z lm Score = 1.99
NG = 0
DB = 0
SM = 0
Delta_K
Doxorubicin
YJR135W−A_2 TIM8
ZShift = −2.4
Z lm Score = 0.65
NG = 0
DB = 0
SM = 0
Delta_r
Doxorubicin
YJR135W−A_2 TIM8
ZShift = −1.99
Z lm Score = −0.86
NG = 0
DB = 0
SM = 0
Delta_AUC
Doxorubicin
YJL091C GWT1
ZShift = 4.7
Z lm Score = 1.93
NG = 0
DB = 0
SM = 1
Delta_L
Doxorubicin
YJL091C GWT1
ZShift = −9.96
Z lm Score = 1.42
NG = 0
DB = 0
SM = 1
Delta_K
Doxorubicin
YJL091C GWT1
ZShift = −1.41
Z lm Score = 0.96
NG = 0
DB = 0
SM = 1
Delta_r
Doxorubicin
YJL091C GWT1
ZShift = −7.06
Z lm Score = −2.97
NG = 0
DB = 0
SM = 1
Delta_AUC
Doxorubicin
YDL140C RPO21
ZShift = 0.76
Z lm Score = 1.91
NG = 0
DB = 1
SM = 0
Delta_L
Doxorubicin
YDL140C RPO21
ZShift = −0.08
Z lm Score = −0.35
NG = 0
DB = 1
SM = 0
Delta_K
Doxorubicin
YDL140C RPO21
ZShift = −0.4
Z lm Score = −1.13
NG = 0
DB = 1
SM = 0
Delta_r
Doxorubicin
YDL140C RPO21
ZShift = −0.59
Z lm Score = −1.64
NG = 0
DB = 1
SM = 0
Delta_AUC
Doxorubicin

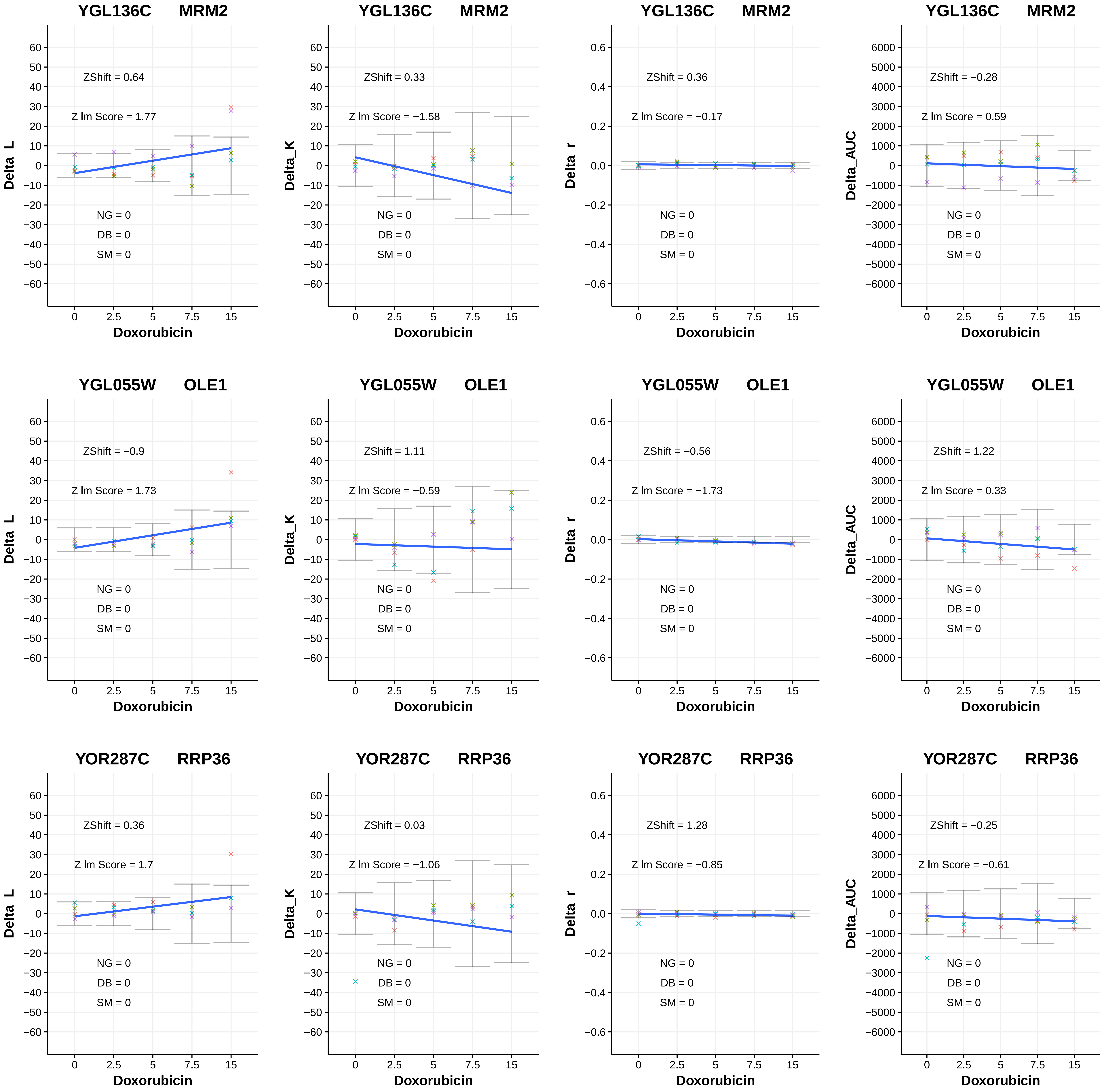

YGL136C MRM2
ZShift = 0.64
Z lm Score = 1.77
NG = 0
DB = 0
SM = 0
Delta_L
Doxorubicin
YGL136C MRM2
ZShift = 0.33
Z lm Score = −1.58
NG = 0
DB = 0
SM = 0
Delta_K
Doxorubicin
YGL136C MRM2
ZShift = 0.36
Z lm Score = −0.17
NG = 0
DB = 0
SM = 0
Delta_r
Doxorubicin
YGL136C MRM2
ZShift = −0.28
Z lm Score = 0.59
NG = 0
DB = 0
SM = 0
Delta_AUC
Doxorubicin
YGL055W OLE1
ZShift = −0.9
Z lm Score = 1.73
NG = 0
DB = 0
SM = 0
Delta_L
Doxorubicin
YGL055W OLE1
ZShift = 1.11
Z lm Score = −0.59
NG = 0
DB = 0
SM = 0
Delta_K
Doxorubicin
YGL055W OLE1
ZShift = −0.56
Z lm Score = −1.73
NG = 0
DB = 0
SM = 0
Delta_r
Doxorubicin
YGL055W OLE1
ZShift = 1.22
Z lm Score = 0.33
NG = 0
DB = 0
SM = 0
Delta_AUC
Doxorubicin
YOR287C RRP36
ZShift = 0.36
Z lm Score = 1.7
NG = 0
DB = 0
SM = 0
Delta_L
Doxorubicin
YOR287C RRP36
ZShift = 0.03
Z lm Score = −1.06
NG = 0
DB = 0
SM = 0
Delta_K
Doxorubicin
YOR287C RRP36
ZShift = 1.28
Z lm Score = −0.85
NG = 0
DB = 0
SM = 0
Delta_r
Doxorubicin
YOR287C RRP36
ZShift = −0.25
Z lm Score = −0.61
NG = 0
DB = 0
SM = 0
Delta_AUC
Doxorubicin

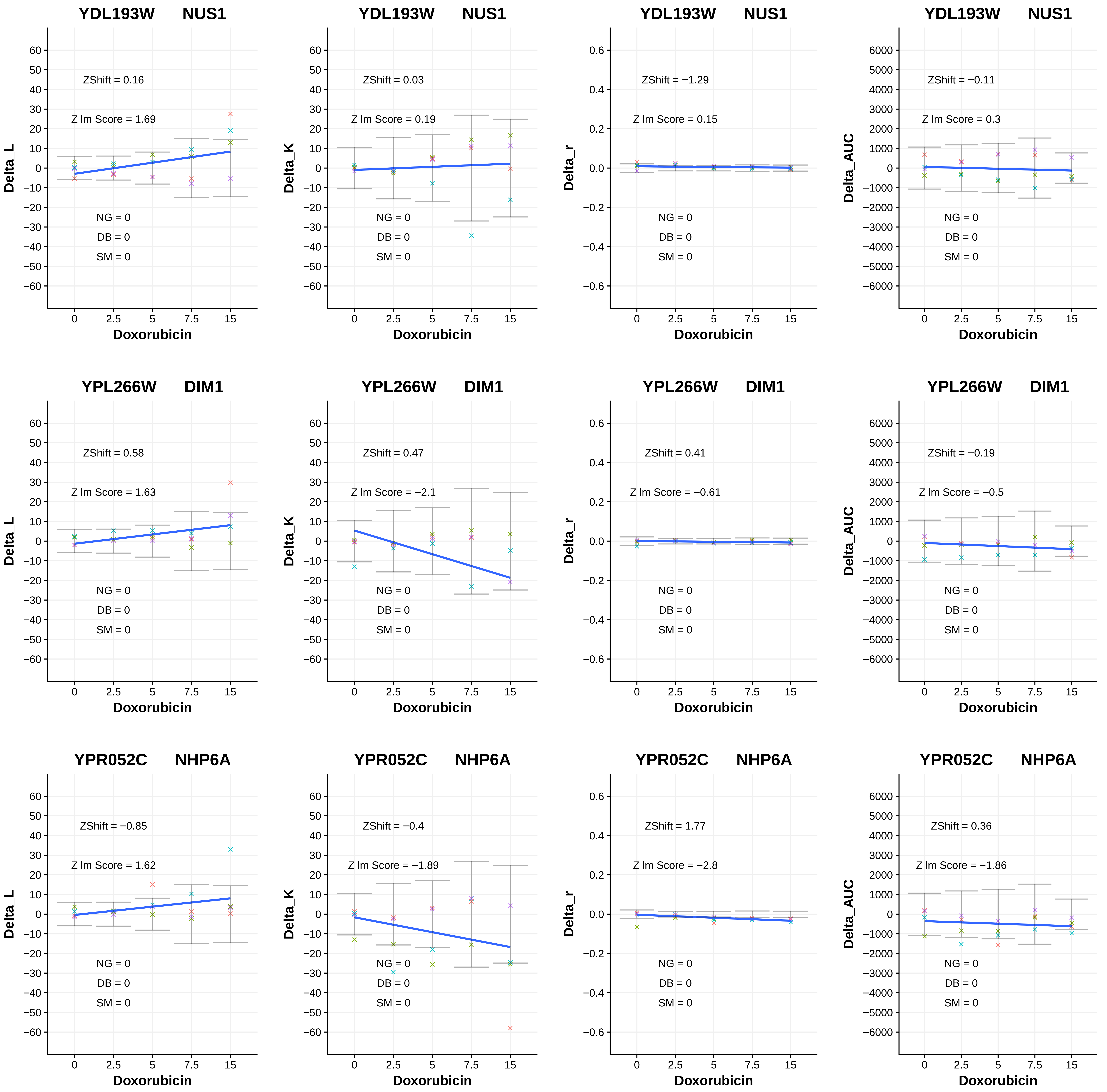
YDL193W NUS1
ZShift = 0.16
Z lm Score = 1.69
NG = 0
DB = 0
SM = 0
Delta_L
Doxorubicin
YDL193W NUS1
ZShift = 0.03
Z lm Score = 0.19
NG = 0
DB = 0
SM = 0
Delta_K
Doxorubicin
YDL193W NUS1
ZShift = −1.29
Z lm Score = 0.15
NG = 0
DB = 0
SM = 0
Delta_r
Doxorubicin
YDL193W NUS1
ZShift = −0.11
Z lm Score = 0.3
NG = 0
DB = 0
SM = 0
Delta_AUC
Doxorubicin
YPL266W DIM1
ZShift = 0.58
Z lm Score = 1.63
NG = 0
DB = 0
SM = 0
Delta_L
Doxorubicin
YPL266W DIM1
ZShift = 0.47
Z lm Score = −2.1
NG = 0
DB = 0
SM = 0
Delta_K
Doxorubicin
YPL266W DIM1
ZShift = 0.41
Z lm Score = −0.61
NG = 0
DB = 0
SM = 0
Delta_r
Doxorubicin
YPL266W DIM1
ZShift = −0.19
Z lm Score = −0.5
NG = 0
DB = 0
SM = 0
Delta_AUC
Doxorubicin
YPR052C NHP6A
ZShift = −0.85
Z lm Score = 1.62
NG = 0
DB = 0
SM = 0
Delta_L
Doxorubicin
YPR052C NHP6A
ZShift = −0.4
Z lm Score = −1.89
NG = 0
DB = 0
SM = 0
Delta_K
Doxorubicin
YPR052C NHP6A
ZShift = 1.77
Z lm Score = −2.8
NG = 0
DB = 0
SM = 0
Delta_r
Doxorubicin
YPR052C NHP6A
ZShift = 0.36
Z lm Score = −1.86
NG = 0
DB = 0
SM = 0
Delta_AUC
Doxorubicin

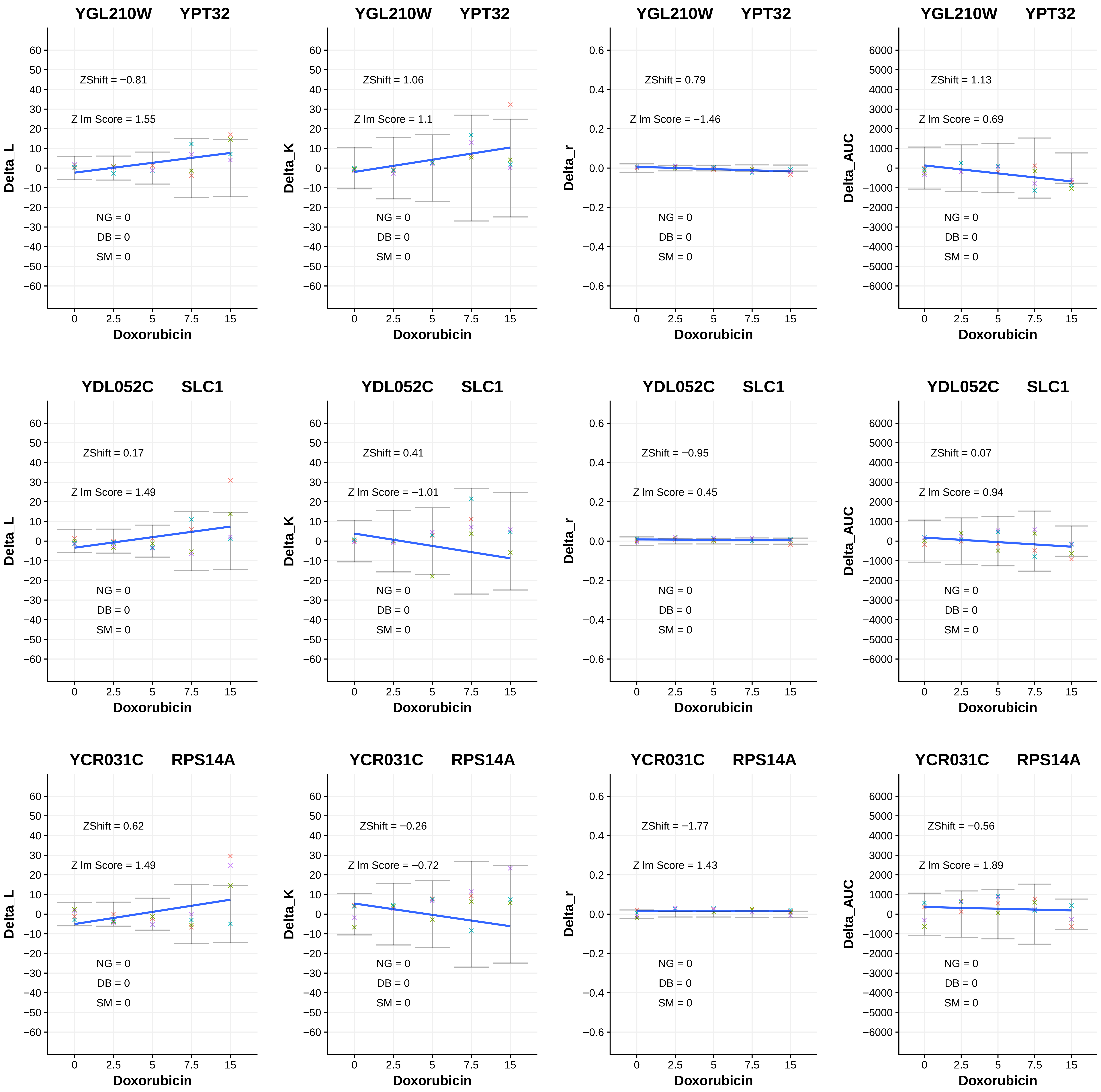

YGL210W YPT32
ZShift = −0.81
Z lm Score = 1.55
NG = 0
DB = 0
SM = 0
Delta_L
Doxorubicin
YGL210W YPT32
ZShift = 1.06
Z lm Score = 1.1
NG = 0
DB = 0
SM = 0
Delta_K
Doxorubicin
YGL210W YPT32
ZShift = 0.79
Z lm Score = −1.46
NG = 0
DB = 0
SM = 0
Delta_r
Doxorubicin
YGL210W YPT32
ZShift = 1.13
Z lm Score = 0.69
NG = 0
DB = 0
SM = 0
Delta_AUC
Doxorubicin
YDL052C SLC1
ZShift = 0.17
Z lm Score = 1.49
NG = 0
DB = 0
SM = 0
Delta_L
Doxorubicin
YDL052C SLC1
ZShift = 0.41
Z lm Score = −1.01
NG = 0
DB = 0
SM = 0
Delta_K
Doxorubicin
YDL052C SLC1
ZShift = −0.95
Z lm Score = 0.45
NG = 0
DB = 0
SM = 0
Delta_r
Doxorubicin
YDL052C SLC1
ZShift = 0.07
Z lm Score = 0.94
NG = 0
DB = 0
SM = 0
Delta_AUC
Doxorubicin
YCR031C RPS14A
ZShift = 0.62
Z lm Score = 1.49
NG = 0
DB = 0
SM = 0
Delta_L
Doxorubicin
YCR031C RPS14A
ZShift = −0.26
Z lm Score = −0.72
NG = 0
DB = 0
SM = 0
Delta_K
Doxorubicin
YCR031C RPS14A
ZShift = −1.77
Z lm Score = 1.43
NG = 0
DB = 0
SM = 0
Delta_r
Doxorubicin
YCR031C RPS14A
ZShift = −0.56
Z lm Score = 1.89
NG = 0
DB = 0
SM = 0
Delta_AUC
Doxorubicin

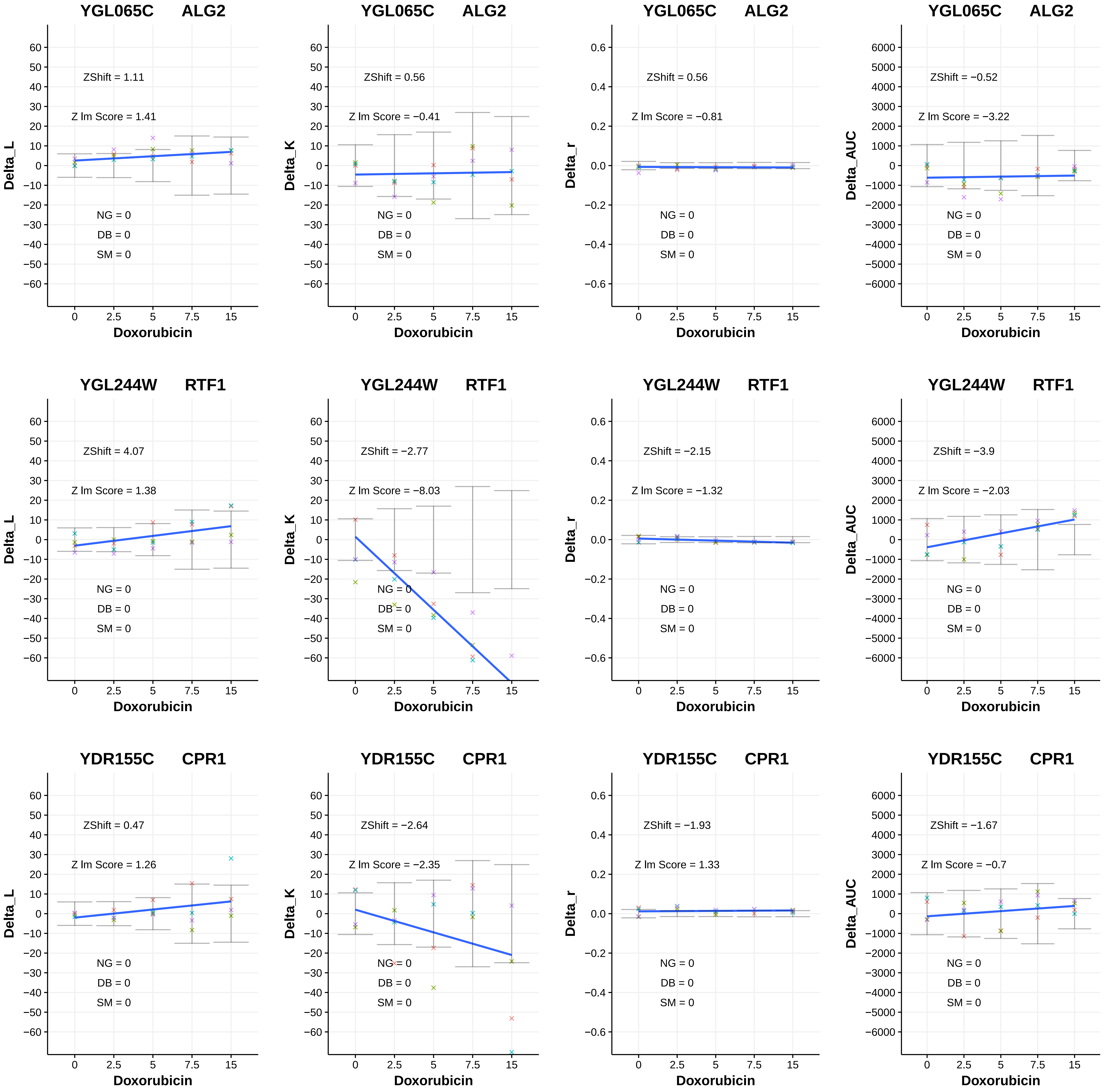

YGL065C ALG2
ZShift = 1.11
Z lm Score = 1.41
NG = 0
DB = 0
SM = 0
Delta_L
Doxorubicin
YGL065C ALG2
ZShift = 0.56
Z lm Score = −0.41
NG = 0
DB = 0
SM = 0
Delta_K
Doxorubicin
YGL065C ALG2
ZShift = 0.56
Z lm Score = −0.81
NG = 0
DB = 0
SM = 0
Delta_r
Doxorubicin
YGL065C ALG2
ZShift = −0.52
Z lm Score = −3.22
NG = 0
DB = 0
SM = 0
Delta_AUC
Doxorubicin
YGL244W RTF1
ZShift = 4.07
Z lm Score = 1.38
NG = 0
DB = 0
SM = 0
Delta_L
Doxorubicin
YGL244W RTF1
ZShift = −2.77
Z lm Score = −8.03
NG = 0
DB = 0
SM = 0
Delta_K
Doxorubicin
YGL244W RTF1
ZShift = −2.15
Z lm Score = −1.32
NG = 0
DB = 0
SM = 0
Delta_r
Doxorubicin
YGL244W RTF1
ZShift = −3.9
Z lm Score = −2.03
NG = 0
DB = 0
SM = 0
Delta_AUC
Doxorubicin
YDR155C CPR1
ZShift = 0.47
Z lm Score = 1.26
NG = 0
DB = 0
SM = 0
Delta_L
Doxorubicin
YDR155C CPR1
ZShift = −2.64
Z lm Score = −2.35
NG = 0
DB = 0
SM = 0
Delta_K
Doxorubicin
YDR155C CPR1
ZShift = −1.93
Z lm Score = 1.33
NG = 0
DB = 0
SM = 0
Delta_r
Doxorubicin
YDR155C CPR1
ZShift = −1.67
Z lm Score = −0.7
NG = 0
DB = 0
SM = 0
Delta_AUC
Doxorubicin

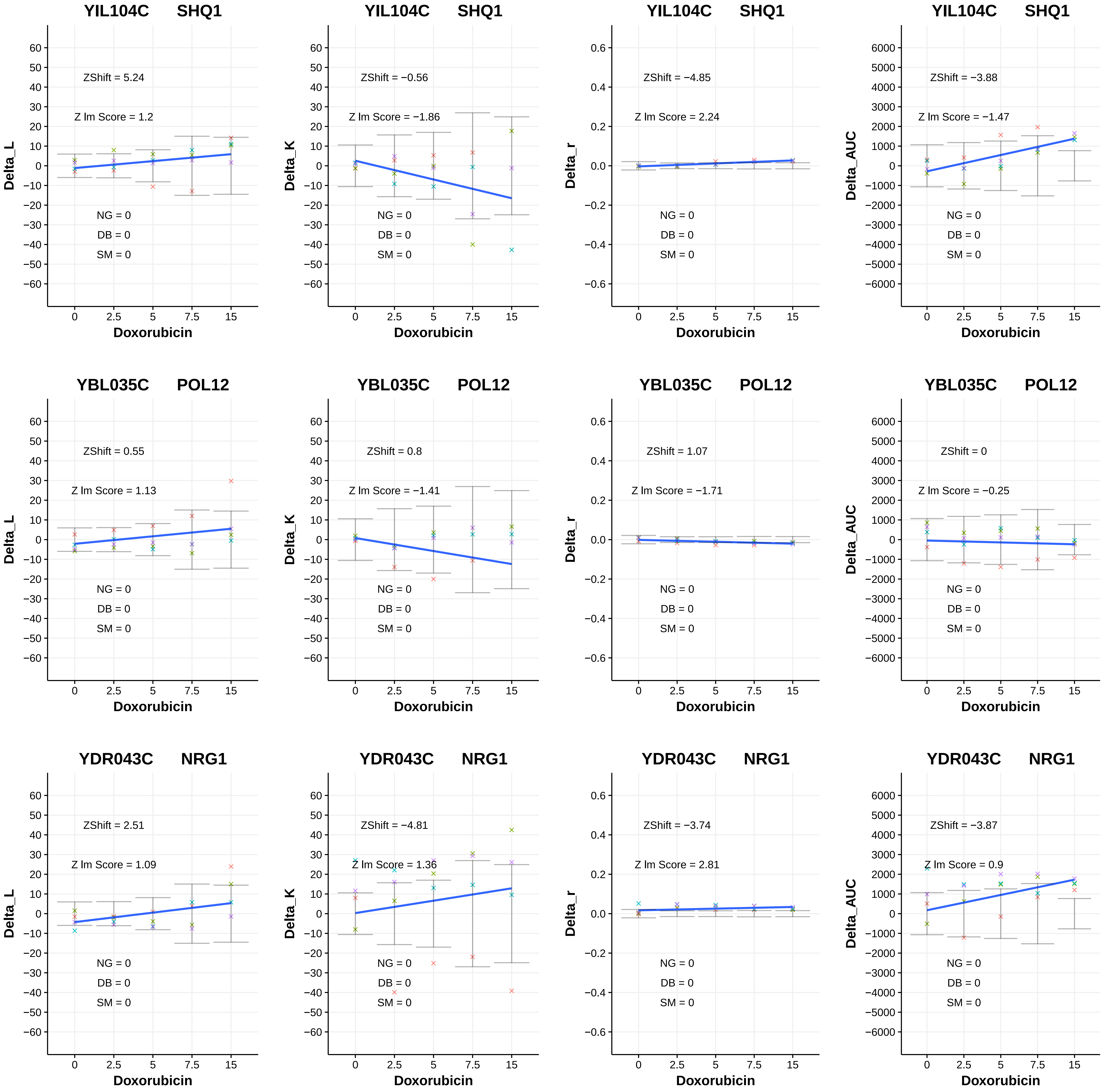

YIL104C SHQ1
ZShift = 5.24
Z lm Score = 1.2
NG = 0
DB = 0
SM = 0
Delta_L
Doxorubicin
YIL104C SHQ1
ZShift = −0.56
Z lm Score = −1.86
NG = 0
DB = 0
SM = 0
Delta_K
Doxorubicin
YIL104C SHQ1
ZShift = −4.85
Z lm Score = 2.24
NG = 0
DB = 0
SM = 0
Delta_r
Doxorubicin
YIL104C SHQ1
ZShift = −3.88
Z lm Score = −1.47
NG = 0
DB = 0
SM = 0
Delta_AUC
Doxorubicin
YBL035C POL12
ZShift = 0.55
Z lm Score = 1.13
NG = 0
DB = 0
SM = 0
Delta_L
Doxorubicin
YBL035C POL12
ZShift = 0.8
Z lm Score = −1.41
NG = 0
DB = 0
SM = 0
Delta_K
Doxorubicin
YBL035C POL12
ZShift = 1.07
Z lm Score = −1.71
NG = 0
DB = 0
SM = 0
Delta_r
Doxorubicin
YBL035C POL12
ZShift = 0
Z lm Score = −0.25
NG = 0
DB = 0
SM = 0
Delta_AUC
Doxorubicin
YDR043C NRG1
ZShift = 2.51
Z lm Score = 1.09
NG = 0
DB = 0
SM = 0
Delta_L
Doxorubicin
YDR043C NRG1
ZShift = −4.81
Z lm Score = 1.36
NG = 0
DB = 0
SM = 0
Delta_K
Doxorubicin
YDR043C NRG1
ZShift = −3.74
Z lm Score = 2.81
NG = 0
DB = 0
SM = 0
Delta_r
Doxorubicin
YDR043C NRG1
ZShift = −3.87
Z lm Score = 0.9
NG = 0
DB = 0
SM = 0
Delta_AUC
Doxorubicin

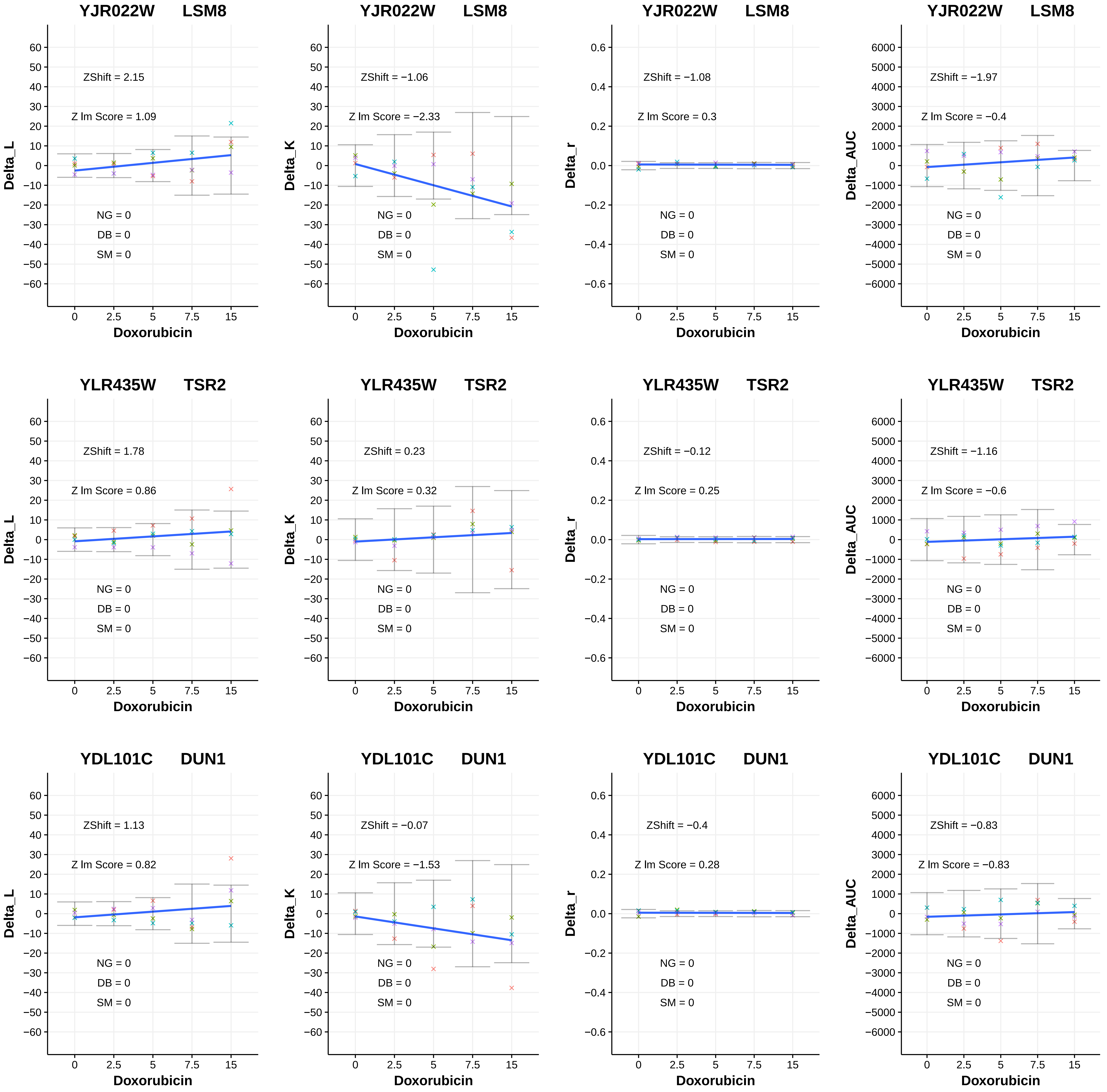
YJR022W LSM8
ZShift = 2.15
Z lm Score = 1.09
NG = 0
DB = 0
SM = 0
Delta_L
Doxorubicin
YJR022W LSM8
ZShift = −1.06
Z lm Score = −2.33
NG = 0
DB = 0
SM = 0
Delta_K
Doxorubicin
YJR022W LSM8
ZShift = −1.08
Z lm Score = 0.3
NG = 0
DB = 0
SM = 0
Delta_r
Doxorubicin
YJR022W LSM8
ZShift = −1.97
Z lm Score = −0.4
NG = 0
DB = 0
SM = 0
Delta_AUC
Doxorubicin
YLR435W TSR2
ZShift = 1.78
Z lm Score = 0.86
NG = 0
DB = 0
SM = 0
Delta_L
Doxorubicin
YLR435W TSR2
ZShift = 0.23
Z lm Score = 0.32
NG = 0
DB = 0
SM = 0
Delta_K
Doxorubicin
YLR435W TSR2
ZShift = −0.12
Z lm Score = 0.25
NG = 0
DB = 0
SM = 0
Delta_r
Doxorubicin
YLR435W TSR2
ZShift = −1.16
Z lm Score = −0.6
NG = 0
DB = 0
SM = 0
Delta_AUC
Doxorubicin
YDL101C DUN1
ZShift = 1.13
Z lm Score = 0.82
NG = 0
DB = 0
SM = 0
Delta_L
Doxorubicin
YDL101C DUN1
ZShift = −0.07
Z lm Score = −1.53
NG = 0
DB = 0
SM = 0
Delta_K
Doxorubicin
YDL101C DUN1
ZShift = −0.4
Z lm Score = 0.28
NG = 0
DB = 0
SM = 0
Delta_r
Doxorubicin
YDL101C DUN1
ZShift = −0.83
Z lm Score = −0.83
NG = 0
DB = 0
SM = 0
Delta_AUC
Doxorubicin

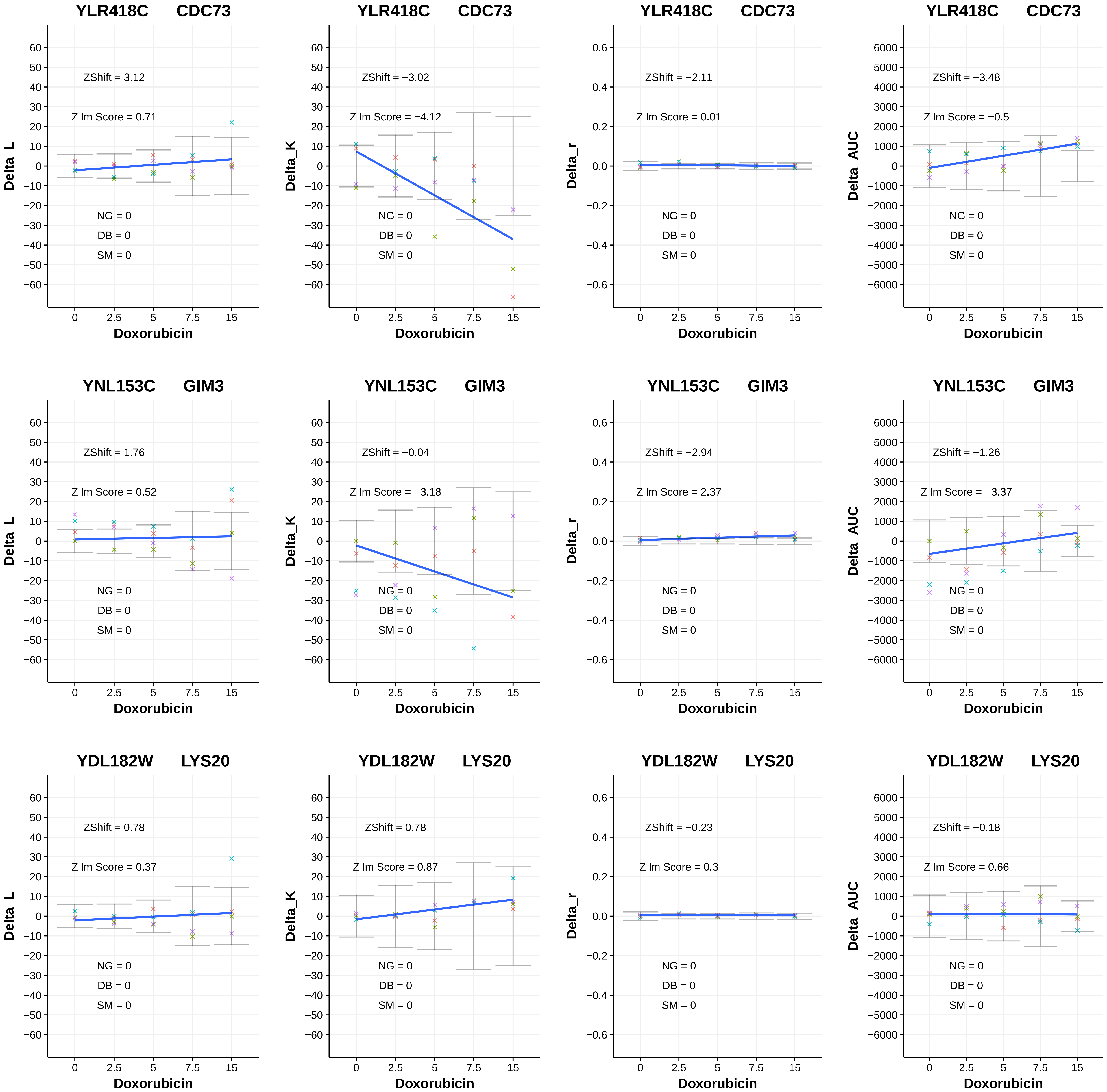

YLR418C CDC73
ZShift = 3.12
Z lm Score = 0.71
NG = 0
DB = 0
SM = 0
Delta_L
Doxorubicin
YLR418C CDC73
ZShift = −3.02
Z lm Score = −4.12
NG = 0
DB = 0
SM = 0
Delta_K
Doxorubicin
YLR418C CDC73
ZShift = −2.11
Z lm Score = 0.01
NG = 0
DB = 0
SM = 0
Delta_r
Doxorubicin
YLR418C CDC73
ZShift = −3.48
Z lm Score = −0.5
NG = 0
DB = 0
SM = 0
Delta_AUC
Doxorubicin
YNL153C GIM3
ZShift = 1.76
Z lm Score = 0.52
NG = 0
DB = 0
SM = 0
Delta_L
Doxorubicin
YNL153C GIM3
ZShift = −0.04
Z lm Score = −3.18
NG = 0
DB = 0
SM = 0
Delta_K
Doxorubicin
YNL153C GIM3
ZShift = −2.94
Z lm Score = 2.37
NG = 0
DB = 0
SM = 0
Delta_r
Doxorubicin
YNL153C GIM3
ZShift = −1.26
Z lm Score = −3.37
NG = 0
DB = 0
SM = 0
Delta_AUC
Doxorubicin
YDL182W LYS20
ZShift = 0.78
Z lm Score = 0.37
NG = 0
DB = 0
SM = 0
Delta_L
Doxorubicin
YDL182W LYS20
ZShift = 0.78
Z lm Score = 0.87
NG = 0
DB = 0
SM = 0
Delta_K
Doxorubicin
YDL182W LYS20
ZShift = −0.23
Z lm Score = 0.3
NG = 0
DB = 0
SM = 0
Delta_r
Doxorubicin
YDL182W LYS20
ZShift = −0.18
Z lm Score = 0.66
NG = 0
DB = 0
SM = 0
Delta_AUC
Doxorubicin

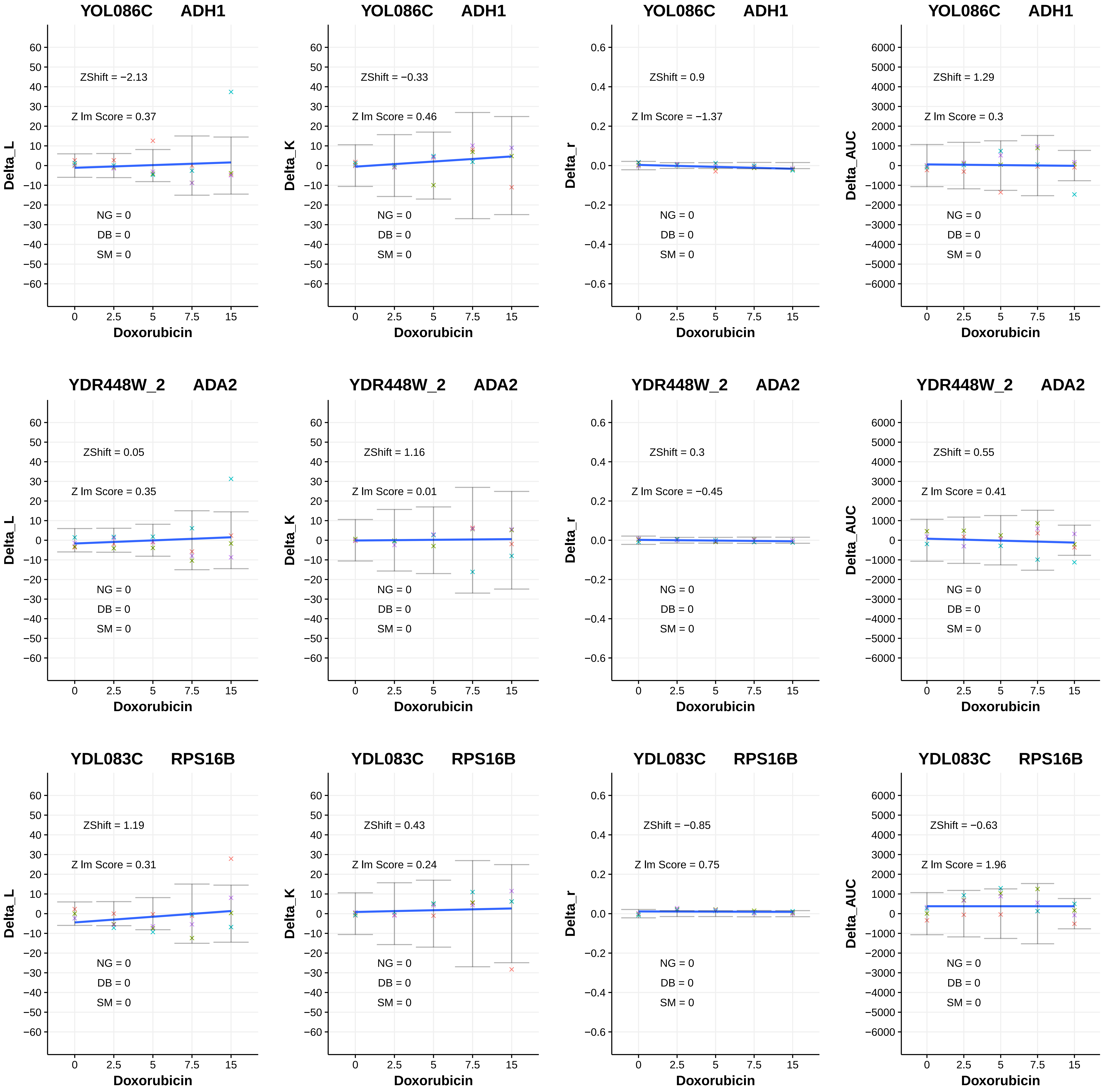
YOL086C ADH1
ZShift = −2.13
Z lm Score = 0.37
NG = 0
DB = 0
SM = 0
Delta_L
Doxorubicin
YOL086C ADH1
ZShift = −0.33
Z lm Score = 0.46
NG = 0
DB = 0
SM = 0
Delta_K
Doxorubicin
YOL086C ADH1
ZShift = 0.9
Z lm Score = −1.37
NG = 0
DB = 0
SM = 0
Delta_r
Doxorubicin
YOL086C ADH1
ZShift = 1.29
Z lm Score = 0.3
NG = 0
DB = 0
SM = 0
Delta_AUC
Doxorubicin
YDR448W_2 ADA2
ZShift = 0.05
Z lm Score = 0.35
NG = 0
DB = 0
SM = 0
Delta_L
Doxorubicin
YDR448W_2 ADA2
ZShift = 1.16
Z lm Score = 0.01
NG = 0
DB = 0
SM = 0
Delta_K
Doxorubicin
YDR448W_2 ADA2
ZShift = 0.3
Z lm Score = −0.45
NG = 0
DB = 0
SM = 0
Delta_r
Doxorubicin
YDR448W_2 ADA2
ZShift = 0.55
Z lm Score = 0.41
NG = 0
DB = 0
SM = 0
Delta_AUC
Doxorubicin
YDL083C RPS16B
ZShift = 1.19
Z lm Score = 0.31
NG = 0
DB = 0
SM = 0
Delta_L
Doxorubicin
YDL083C RPS16B
ZShift = 0.43
Z lm Score = 0.24
NG = 0
DB = 0
SM = 0
Delta_K
Doxorubicin
YDL083C RPS16B
ZShift = −0.85
Z lm Score = 0.75
NG = 0
DB = 0
SM = 0
Delta_r
Doxorubicin
YDL083C RPS16B
ZShift = −0.63
Z lm Score = 1.96
NG = 0
DB = 0
SM = 0
Delta_AUC
Doxorubicin

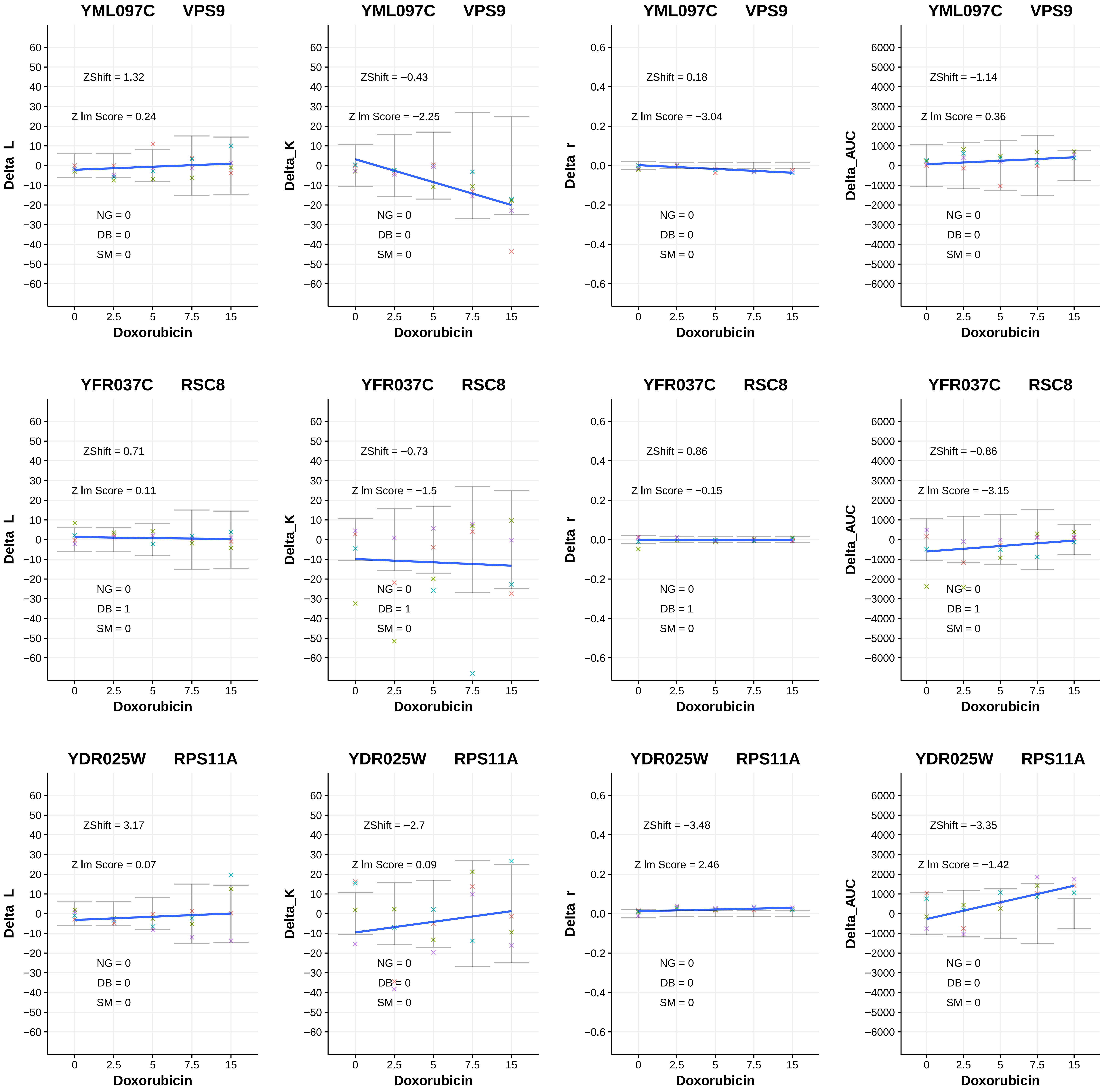

YML097C VPS9
ZShift = 1.32
Z lm Score = 0.24
NG = 0
DB = 0
SM = 0
Delta_L
Doxorubicin
YML097C VPS9
ZShift = −0.43
Z lm Score = −2.25
NG = 0
DB = 0
SM = 0
Delta_K
Doxorubicin
YML097C VPS9
ZShift = 0.18
Z lm Score = −3.04
NG = 0
DB = 0
SM = 0
Delta_r
Doxorubicin
YML097C VPS9
ZShift = −1.14
Z lm Score = 0.36
NG = 0
DB = 0
SM = 0
Delta_AUC
Doxorubicin
YFR037C RSC8
ZShift = 0.71
Z lm Score = 0.11
NG = 0
DB = 1
SM = 0
Delta_L
Doxorubicin
YFR037C RSC8
ZShift = −0.73
Z lm Score = −1.5
NG = 0
DB = 1
SM = 0
Delta_K
Doxorubicin
YFR037C RSC8
ZShift = 0.86
Z lm Score = −0.15
NG = 0
DB = 1
SM = 0
Delta_r
Doxorubicin
YFR037C RSC8
ZShift = −0.86
Z lm Score = −3.15
NG = 0
DB = 1
SM = 0
Delta_AUC
Doxorubicin
YDR025W RPS11A
ZShift = 3.17
Z lm Score = 0.07
NG = 0
DB = 0
SM = 0
Delta_L
Doxorubicin
YDR025W RPS11A
ZShift = −2.7
Z lm Score = 0.09
NG = 0
DB = 0
SM = 0
Delta_K
Doxorubicin
YDR025W RPS11A
ZShift = −3.48
Z lm Score = 2.46
NG = 0
DB = 0
SM = 0
Delta_r
Doxorubicin
YDR025W RPS11A
ZShift = −3.35
Z lm Score = −1.42
NG = 0
DB = 0
SM = 0
Delta_AUC
Doxorubicin

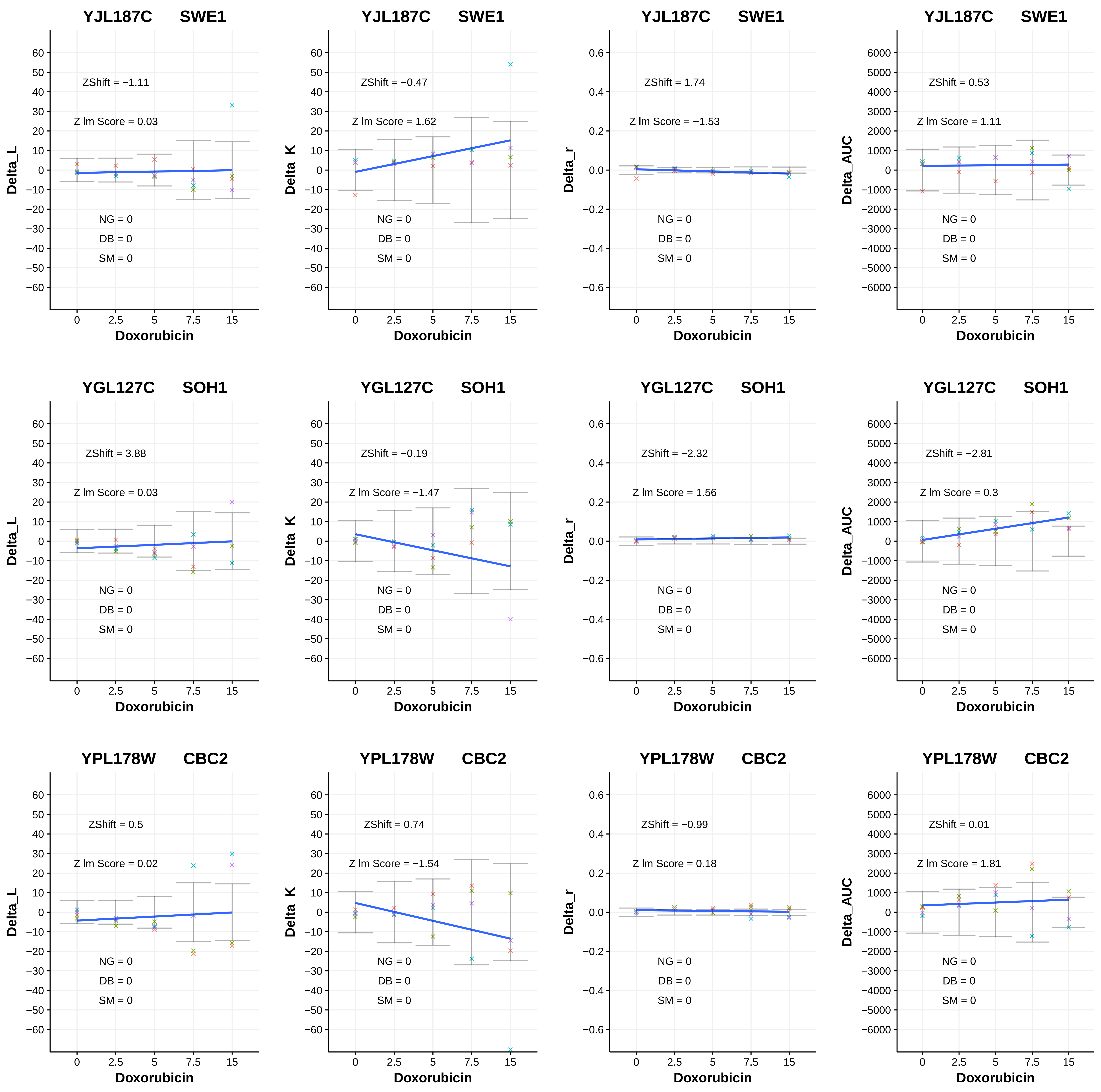
YJL187C SWE1
ZShift = −1.11
Z lm Score = 0.03
NG = 0
DB = 0
SM = 0
Delta_L
Doxorubicin
YJL187C SWE1
ZShift = −0.47
Z lm Score = 1.62
NG = 0
DB = 0
SM = 0
Delta_K
Doxorubicin
YJL187C SWE1
ZShift = 1.74
Z lm Score = −1.53
NG = 0
DB = 0
SM = 0
Delta_r
Doxorubicin
YJL187C SWE1
ZShift = 0.53
Z lm Score = 1.11
NG = 0
DB = 0
SM = 0
Delta_AUC
Doxorubicin
YGL127C SOH1
ZShift = 3.88
Z lm Score = 0.03
NG = 0
DB = 0
SM = 0
Delta_L
Doxorubicin
YGL127C SOH1
ZShift = −0.19
Z lm Score = −1.47
NG = 0
DB = 0
SM = 0
Delta_K
Doxorubicin
YGL127C SOH1
ZShift = −2.32
Z lm Score = 1.56
NG = 0
DB = 0
SM = 0
Delta_r
Doxorubicin
YGL127C SOH1
ZShift = −2.81
Z lm Score = 0.3
NG = 0
DB = 0
SM = 0
Delta_AUC
Doxorubicin
YPL178W CBC2
ZShift = 0.5
Z lm Score = 0.02
NG = 0
DB = 0
SM = 0
Delta_L
Doxorubicin
YPL178W CBC2
ZShift = 0.74
Z lm Score = −1.54
NG = 0
DB = 0
SM = 0
Delta_K
Doxorubicin
YPL178W CBC2
ZShift = −0.99
Z lm Score = 0.18
NG = 0
DB = 0
SM = 0
Delta_r
Doxorubicin
YPL178W CBC2
ZShift = 0.01
Z lm Score = 1.81
NG = 0
DB = 0
SM = 0
Delta_AUC
Doxorubicin

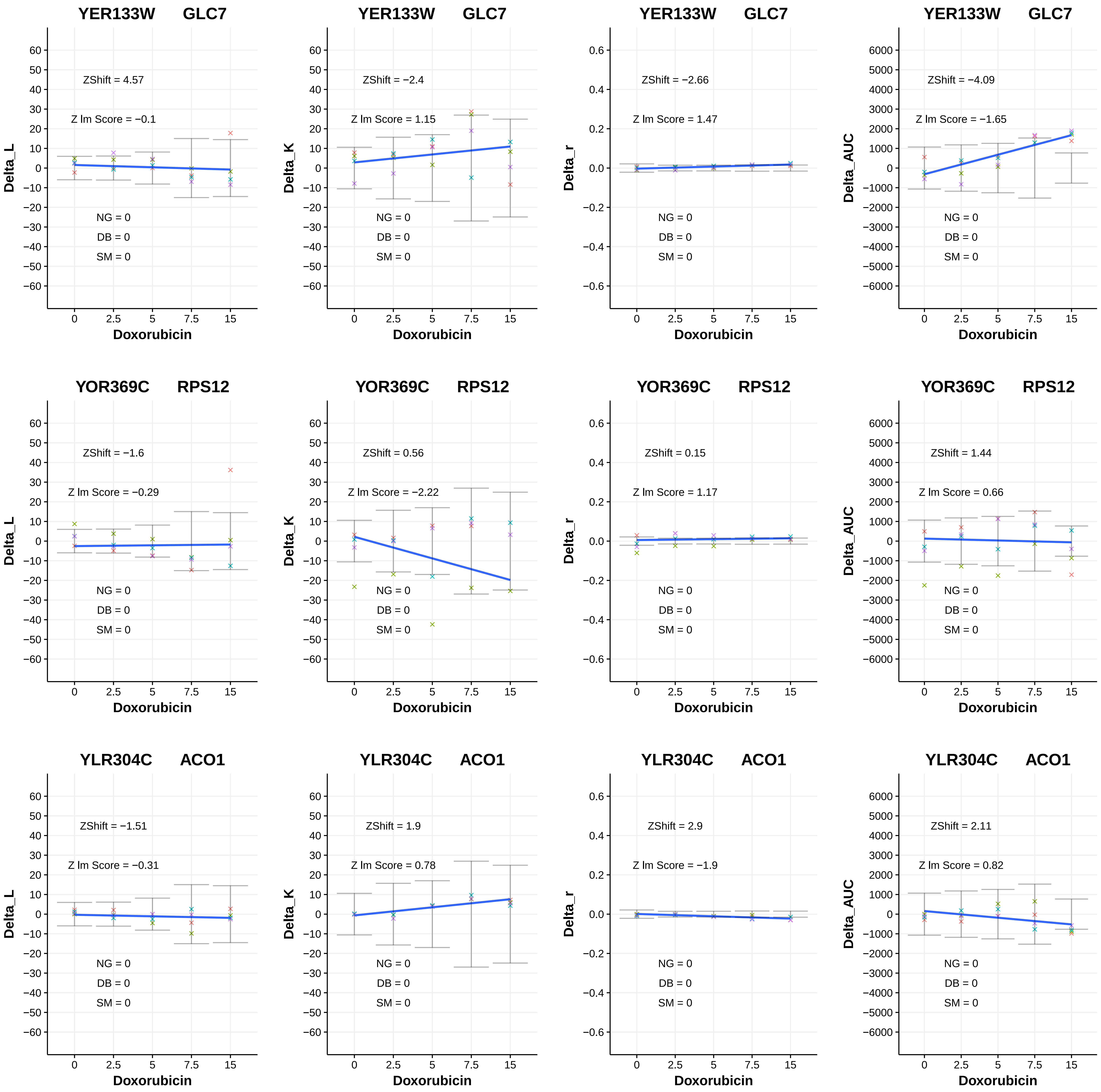

YER133W GLC7
ZShift = 4.57
Z lm Score = −0.1
NG = 0
DB = 0
SM = 0
Delta_L
Doxorubicin
YER133W GLC7
ZShift = −2.4
Z lm Score = 1.15
NG = 0
DB = 0
SM = 0
Delta_K
Doxorubicin
YER133W GLC7
ZShift = −2.66
Z lm Score = 1.47
NG = 0
DB = 0
SM = 0
Delta_r
Doxorubicin
YER133W GLC7
ZShift = −4.09
Z lm Score = −1.65
NG = 0
DB = 0
SM = 0
Delta_AUC
Doxorubicin
YOR369C RPS12
ZShift = −1.6
Z lm Score = −0.29
NG = 0
DB = 0
SM = 0
Delta_L
Doxorubicin
YOR369C RPS12
ZShift = 0.56
Z lm Score = −2.22
NG = 0
DB = 0
SM = 0
Delta_K
Doxorubicin
YOR369C RPS12
ZShift = 0.15
Z lm Score = 1.17
NG = 0
DB = 0
SM = 0
Delta_r
Doxorubicin
YOR369C RPS12
ZShift = 1.44
Z lm Score = 0.66
NG = 0
DB = 0
SM = 0
Delta_AUC
Doxorubicin
YLR304C ACO1
ZShift = −1.51
Z lm Score = −0.31
NG = 0
DB = 0
SM = 0
Delta_L
Doxorubicin
YLR304C ACO1
ZShift = 1.9
Z lm Score = 0.78
NG = 0
DB = 0
SM = 0
Delta_K
Doxorubicin
YLR304C ACO1
ZShift = 2.9
Z lm Score = −1.9
NG = 0
DB = 0
SM = 0
Delta_r
Doxorubicin
YLR304C ACO1
ZShift = 2.11
Z lm Score = 0.82
NG = 0
DB = 0
SM = 0
Delta_AUC
Doxorubicin

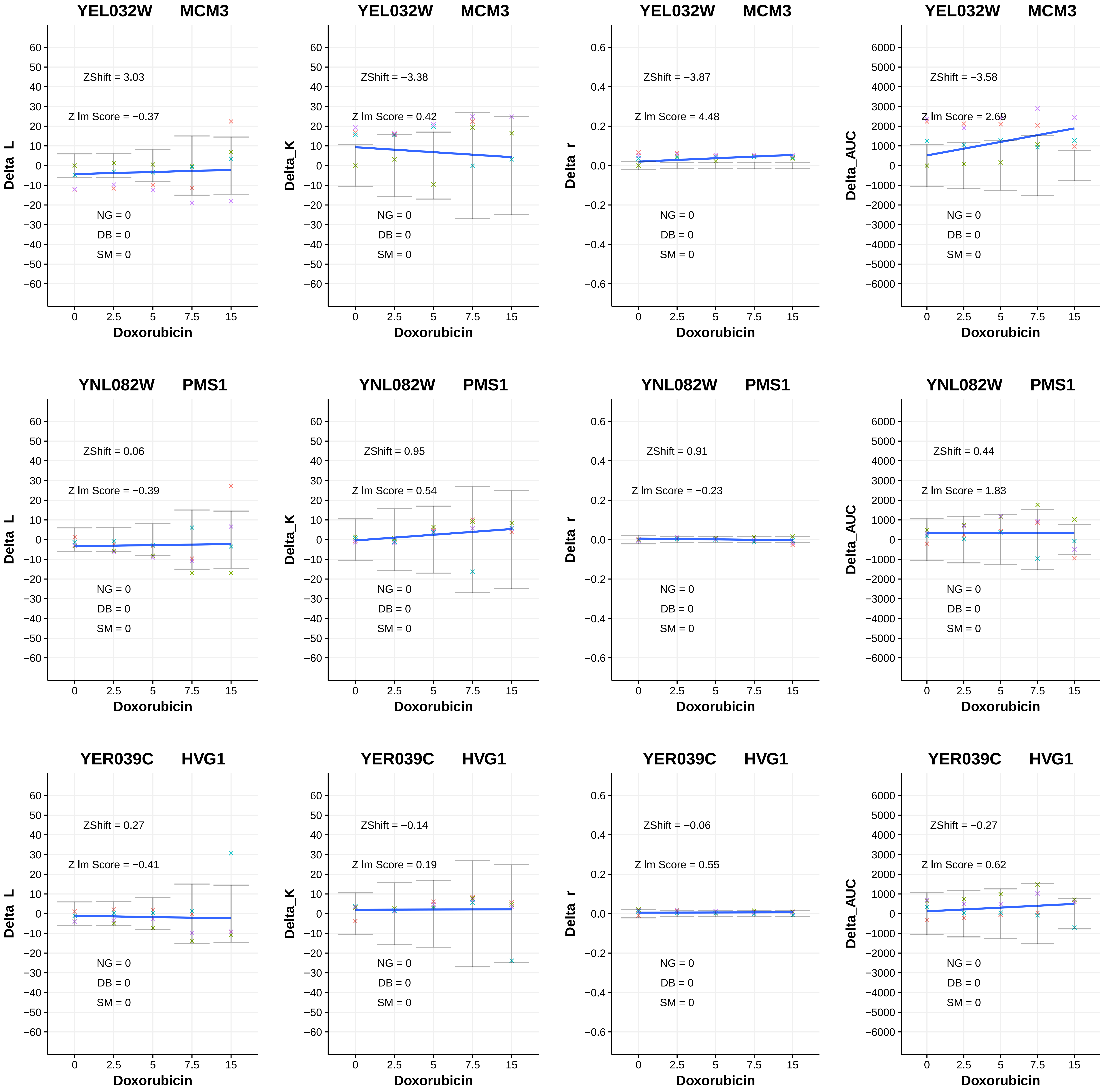
YEL032W MCM3
ZShift = 3.03
Z lm Score = −0.37
NG = 0
DB = 0
SM = 0
Delta_L
60 50 40 30 20 10 0 −10 −20 −30 −40 −50 −60
0 2.5 5 7.5 15
Doxorubicin
YEL032W MCM3
ZShift = −3.38
Z lm Score = 0.42
NG = 0
DB = 0
SM = 0
Delta_K
60 50 40 30 20 10 0 −10 −20 −30 −40 −50 −60
0 2.5 5 7.5 15
Doxorubicin
YEL032W MCM3
ZShift = −3.87
Z lm Score = 4.48
NG = 0
DB = 0
SM = 0
Delta_r
0.6 0.4 0.2 0.0 −0.2 −0.4 −0.6
0 2.5 5 7.5 15
Doxorubicin
YEL032W MCM3
ZShift = −3.58
Z lm Score = 2.69
NG = 0
DB = 0
SM = 0
Delta_AUC
6000 5000 4000 3000 2000 1000 0 −1000 −2000 −3000 −4000 −5000 −6000
0 2.5 5 7.5 15
Doxorubicin
YNL082W PMS1
ZShift = 0.06
Z lm Score = −0.39
NG = 0
DB = 0
SM = 0
Delta_L
60 50 40 30 20 10 0 −10 −20 −30 −40 −50 −60
0 2.5 5 7.5 15
Doxorubicin
YNL082W PMS1
ZShift = 0.95
Z lm Score = 0.54
NG = 0
DB = 0
SM = 0
Delta_K
60 50 40 30 20 10 0 −10 −20 −30 −40 −50 −60
0 2.5 5 7.5 15
Doxorubicin
YNL082W PMS1
ZShift = 0.91
Z lm Score = −0.23
NG = 0
DB = 0
SM = 0
Delta_r
0.6 0.4 0.2 0.0 −0.2 −0.4 −0.6
0 2.5 5 7.5 15
Doxorubicin
YNL082W PMS1
ZShift = 0.44
Z lm Score = 1.83
NG = 0
DB = 0
SM = 0
Delta_AUC
6000 5000 4000 3000 2000 1000 0 −1000 −2000 −3000 −4000 −5000 −6000
0 2.5 5 7.5 15
Doxorubicin
YER039C HVG1
ZShift = 0.27
Z lm Score = −0.41
NG = 0
DB = 0
SM = 0
Delta_L
60 50 40 30 20 10 0 −10 −20 −30 −40 −50 −60
0 2.5 5 7.5 15
Doxorubicin
YER039C HVG1
ZShift = −0.14
Z lm Score = 0.19
NG = 0
DB = 0
SM = 0
Delta_K
60 50 40 30 20 10 0 −10 −20 −30 −40 −50 −60
0 2.5 5 7.5 15
Doxorubicin
YER039C HVG1
ZShift = −0.06
Z lm Score = 0.55
NG = 0
DB = 0
SM = 0
Delta_r
0.6 0.4 0.2 0.0 −0.2 −0.4 −0.6
0 2.5 5 7.5 15
Doxorubicin
YER039C HVG1
ZShift = −0.27
Z lm Score = 0.62
NG = 0
DB = 0
SM = 0
Delta_AUC
6000 5000 4000 3000 2000 1000 0 −1000 −2000 −3000 −4000 −5000 −6000
0 2.5 5 7.5 15
Doxorubicin

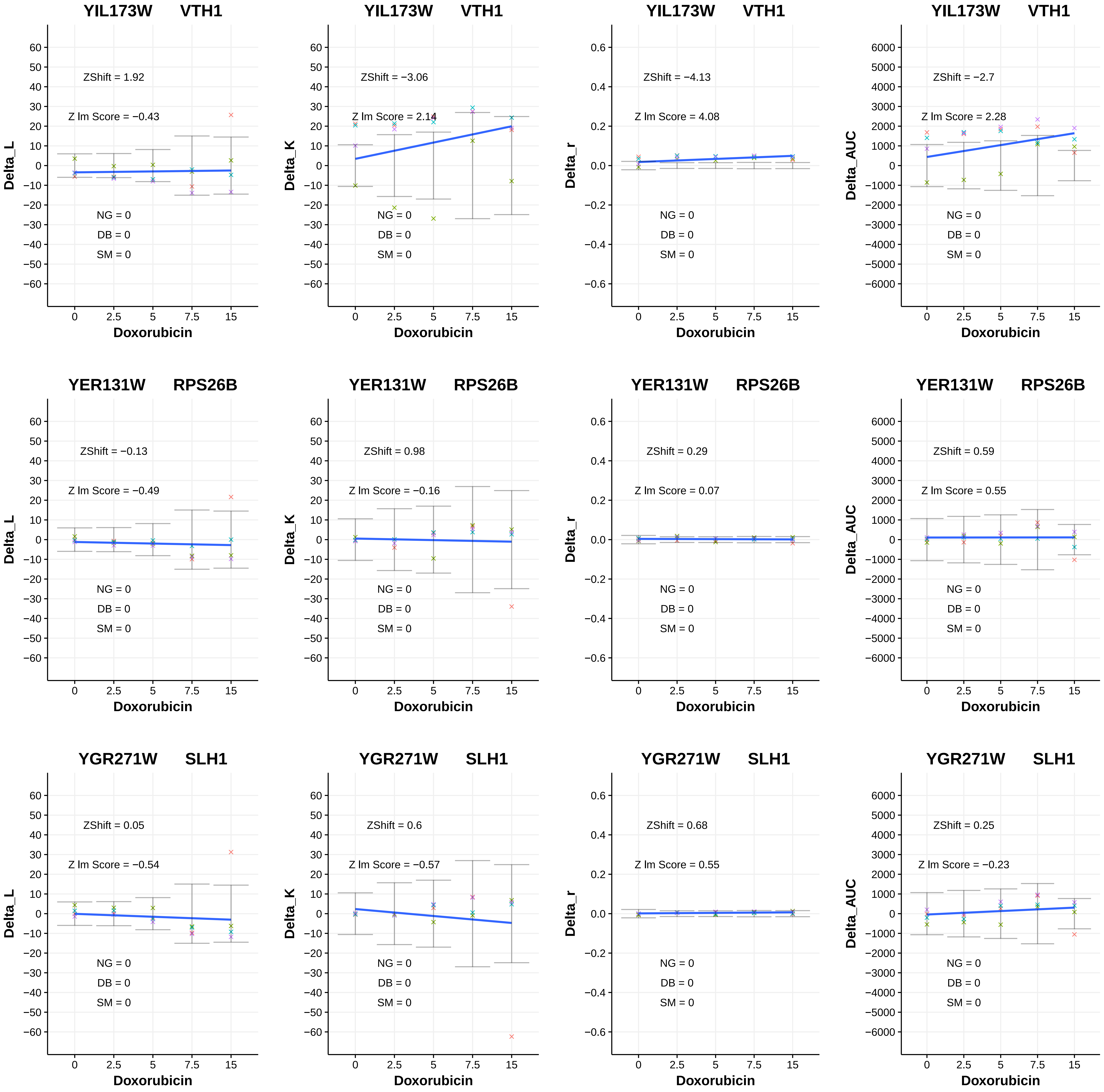
YIL173W VTH1
ZShift = 1.92
Z lm Score = −0.43
NG = 0
DB = 0
SM = 0
Delta_L
Doxorubicin
YIL173W VTH1
ZShift = −3.06
Z lm Score = 2.14
NG = 0
DB = 0
SM = 0
Delta_K
Doxorubicin
YIL173W VTH1
ZShift = −4.13
Z lm Score = 4.08
NG = 0
DB = 0
SM = 0
Delta_r
Doxorubicin
YIL173W VTH1
ZShift = −2.7
Z lm Score = 2.28
NG = 0
DB = 0
SM = 0
Delta_AUC
Doxorubicin
YER131W RPS26B
ZShift = −0.13
Z lm Score = −0.49
NG = 0
DB = 0
SM = 0
Delta_L
Doxorubicin
YER131W RPS26B
ZShift = 0.98
Z lm Score = −0.16
NG = 0
DB = 0
SM = 0
Delta_K
Doxorubicin
YER131W RPS26B
ZShift = 0.29
Z lm Score = 0.07
NG = 0
DB = 0
SM = 0
Delta_r
Doxorubicin
YER131W RPS26B
ZShift = 0.59
Z lm Score = 0.55
NG = 0
DB = 0
SM = 0
Delta_AUC
Doxorubicin
YGR271W SLH1
ZShift = 0.05
Z lm Score = −0.54
NG = 0
DB = 0
SM = 0
Delta_L
Doxorubicin
YGR271W SLH1
ZShift = 0.6
Z lm Score = −0.57
NG = 0
DB = 0
SM = 0
Delta_K
Doxorubicin
YGR271W SLH1
ZShift = 0.68
Z lm Score = 0.55
NG = 0
DB = 0
SM = 0
Delta_r
Doxorubicin
YGR271W SLH1
ZShift = 0.25
Z lm Score = −0.23
NG = 0
DB = 0
SM = 0
Delta_AUC
Doxorubicin

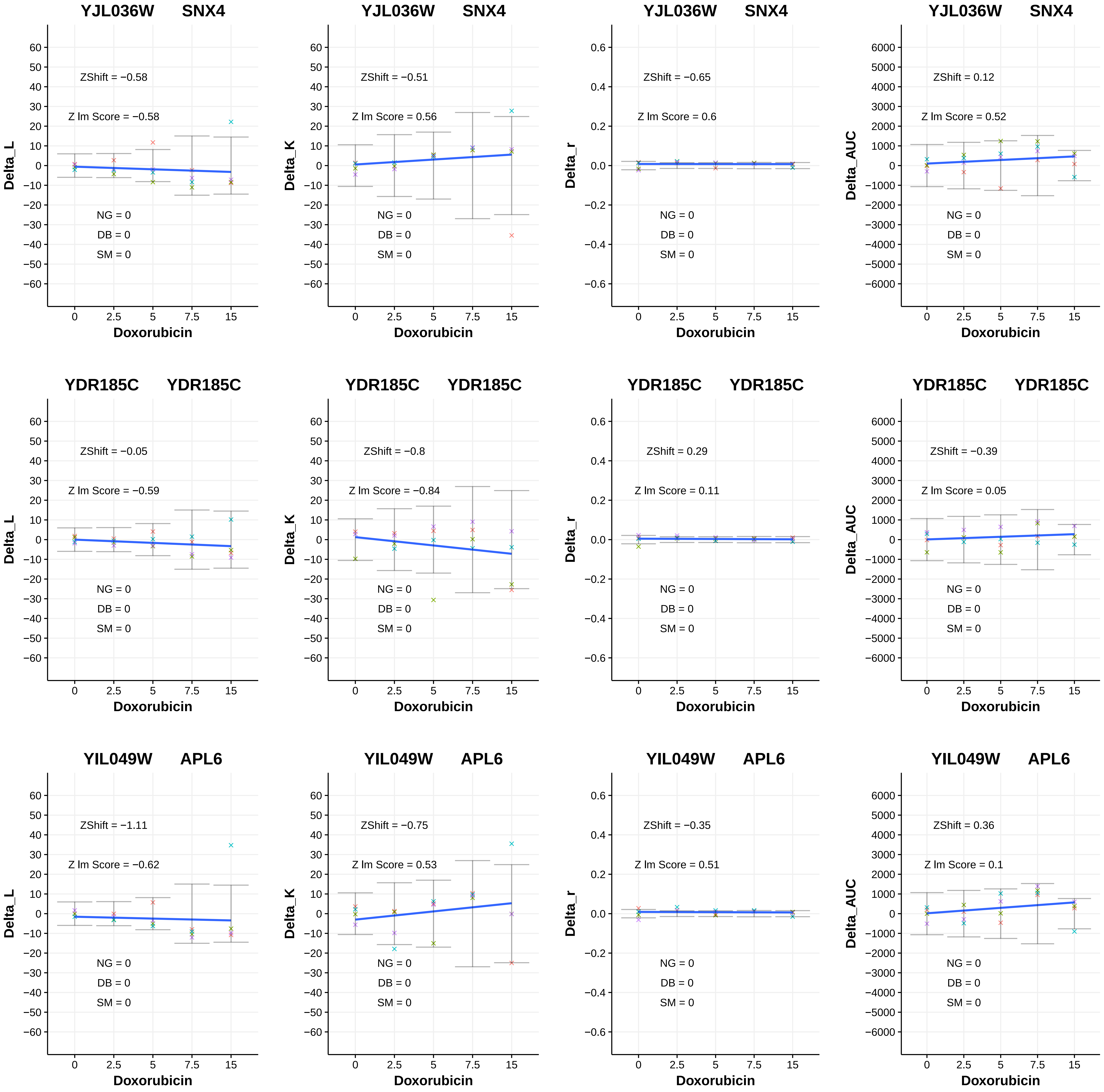

YJL036W SNX4
Delta_L
ZShift = −0.58
Z lm Score = −0.58
NG = 0
DB = 0
SM = 0
Doxorubicin
YJL036W SNX4
Delta_K
ZShift = −0.51
Z lm Score = 0.56
NG = 0
DB = 0
SM = 0
Doxorubicin
YJL036W SNX4
Delta_r
ZShift = −0.65
Z lm Score = 0.6
NG = 0
DB = 0
SM = 0
Doxorubicin
YJL036W SNX4
Delta_AUC
ZShift = 0.12
Z lm Score = 0.52
NG = 0
DB = 0
SM = 0
Doxorubicin
YDR185C YDR185C
Delta_L
ZShift = −0.05
Z lm Score = −0.59
NG = 0
DB = 0
SM = 0
Doxorubicin
YDR185C YDR185C
Delta_K
ZShift = −0.8
Z lm Score = −0.84
NG = 0
DB = 0
SM = 0
Doxorubicin
YDR185C YDR185C
Delta_r
ZShift = 0.29
Z lm Score = 0.11
NG = 0
DB = 0
SM = 0
Doxorubicin
YDR185C YDR185C
Delta_AUC
ZShift = −0.39
Z lm Score = 0.05
NG = 0
DB = 0
SM = 0
Doxorubicin
YIL049W APL6
Delta_L
ZShift = −1.11
Z lm Score = −0.62
NG = 0
DB = 0
SM = 0
Doxorubicin
YIL049W APL6
Delta_K
ZShift = −0.75
Z lm Score = 0.53
NG = 0
DB = 0
SM = 0
Doxorubicin
YIL049W APL6
Delta_r
ZShift = −0.35
Z lm Score = 0.51
NG = 0
DB = 0
SM = 0
Doxorubicin
YIL049W APL6
Delta_AUC
ZShift = 0.36
Z lm Score = 0.1
NG = 0
DB = 0
SM = 0
Doxorubicin

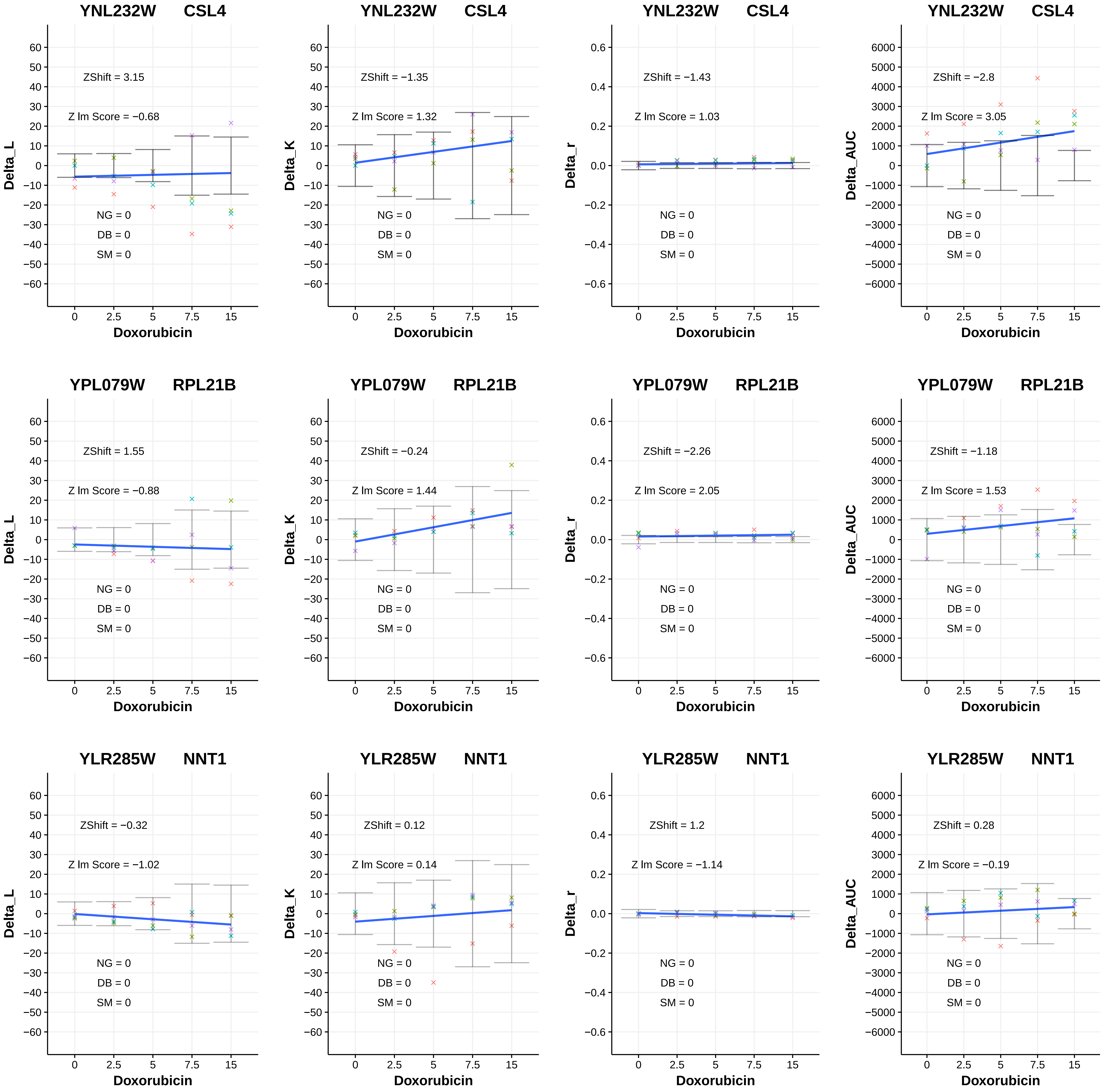
YNL232W CSL4
ZShift = 3.15
Z lm Score = −0.68
NG = 0
DB = 0
SM = 0
Delta_L
Doxorubicin
YNL232W CSL4
ZShift = −1.35
Z lm Score = 1.32
NG = 0
DB = 0
SM = 0
Delta_K
Doxorubicin
YNL232W CSL4
ZShift = −1.43
Z lm Score = 1.03
NG = 0
DB = 0
SM = 0
Delta_r
Doxorubicin
YNL232W CSL4
ZShift = −2.8
Z lm Score = 3.05
NG = 0
DB = 0
SM = 0
Delta_AUC
Doxorubicin
YPL079W RPL21B
ZShift = 1.55
Z lm Score = −0.88
NG = 0
DB = 0
SM = 0
Delta_L
Doxorubicin
YPL079W RPL21B
ZShift = −0.24
Z lm Score = 1.44
NG = 0
DB = 0
SM = 0
Delta_K
Doxorubicin
YPL079W RPL21B
ZShift = −2.26
Z lm Score = 2.05
NG = 0
DB = 0
SM = 0
Delta_r
Doxorubicin
YPL079W RPL21B
ZShift = −1.18
Z lm Score = 1.53
NG = 0
DB = 0
SM = 0
Delta_AUC
Doxorubicin
YLR285W NNT1
ZShift = −0.32
Z lm Score = −1.02
NG = 0
DB = 0
SM = 0
Delta_L
Doxorubicin
YLR285W NNT1
ZShift = 0.12
Z lm Score = 0.14
NG = 0
DB = 0
SM = 0
Delta_K
Doxorubicin
YLR285W NNT1
ZShift = 1.2
Z lm Score = −1.14
NG = 0
DB = 0
SM = 0
Delta_r
Doxorubicin
YLR285W NNT1
ZShift = 0.28
Z lm Score = −0.19
NG = 0
DB = 0
SM = 0
Delta_AUC
Doxorubicin

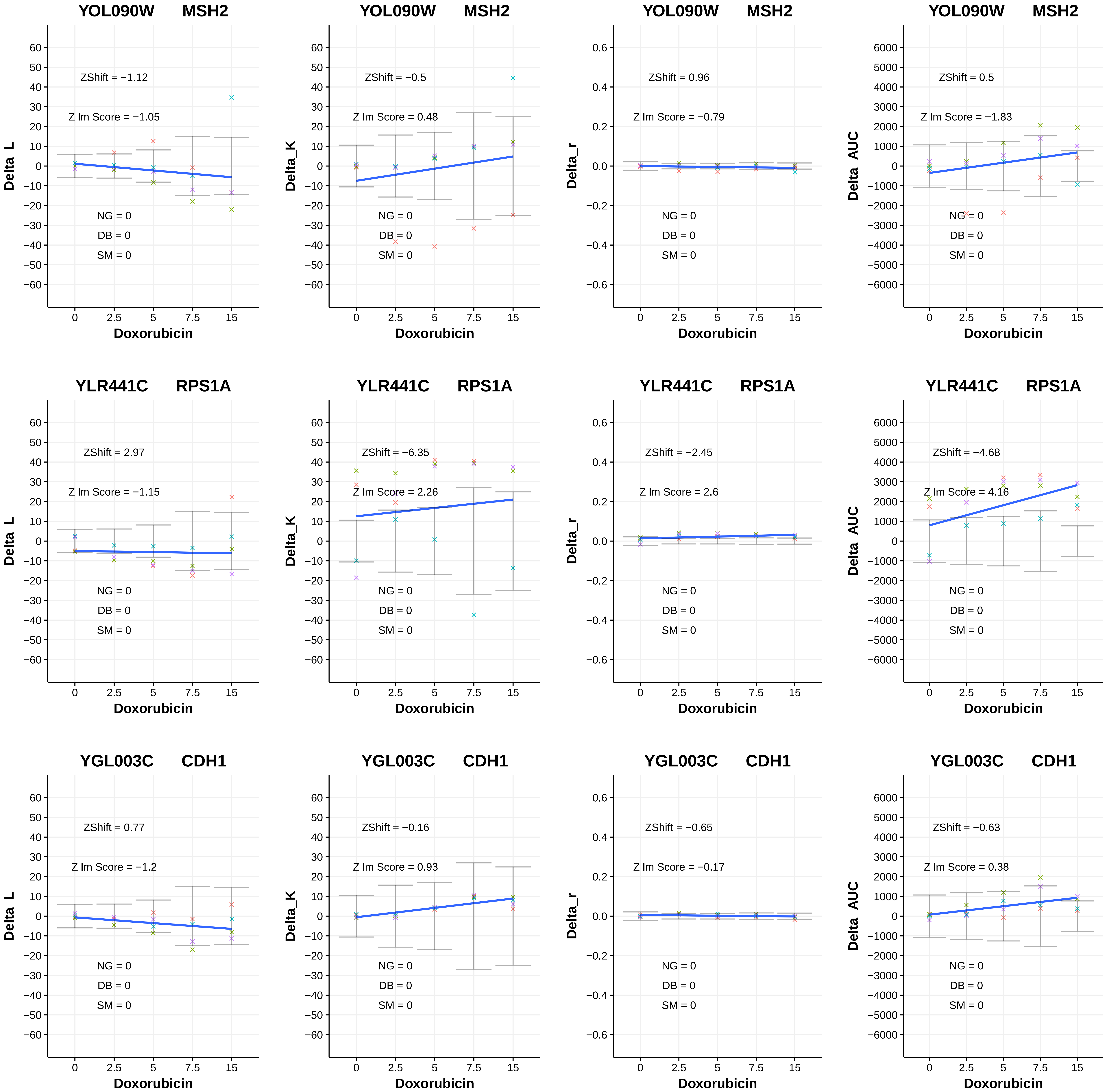

YOL090W MSH2
ZShift = −1.12
Z lm Score = −1.05
NG = 0
DB = 0
SM = 0
Delta_L
Doxorubicin
YOL090W MSH2
ZShift = −0.5
Z lm Score = 0.48
NG = 0
DB = 0
SM = 0
Delta_K
Doxorubicin
YOL090W MSH2
ZShift = 0.96
Z lm Score = −0.79
NG = 0
DB = 0
SM = 0
Delta_r
Doxorubicin
YOL090W MSH2
ZShift = 0.5
Z lm Score = −1.83
NG = 0
DB = 0
SM = 0
Delta_AUC
Doxorubicin
YLR441C RPS1A
ZShift = 2.97
Z lm Score = −1.15
NG = 0
DB = 0
SM = 0
Delta_L
Doxorubicin
YLR441C RPS1A
ZShift = −6.35
Z lm Score = 2.26
NG = 0
DB = 0
SM = 0
Delta_K
Doxorubicin
YLR441C RPS1A
ZShift = −2.45
Z lm Score = 2.6
NG = 0
DB = 0
SM = 0
Delta_r
Doxorubicin
YLR441C RPS1A
ZShift = −4.68
Z lm Score = 4.16
NG = 0
DB = 0
SM = 0
Delta_AUC
Doxorubicin
YGL003C CDH1
ZShift = 0.77
Z lm Score = −1.2
NG = 0
DB = 0
SM = 0
Delta_L
Doxorubicin
YGL003C CDH1
ZShift = −0.16
Z lm Score = 0.93
NG = 0
DB = 0
SM = 0
Delta_K
Doxorubicin
YGL003C CDH1
ZShift = −0.65
Z lm Score = −0.17
NG = 0
DB = 0
SM = 0
Delta_r
Doxorubicin
YGL003C CDH1
ZShift = −0.63
Z lm Score = 0.38
NG = 0
DB = 0
SM = 0
Delta_AUC
Doxorubicin

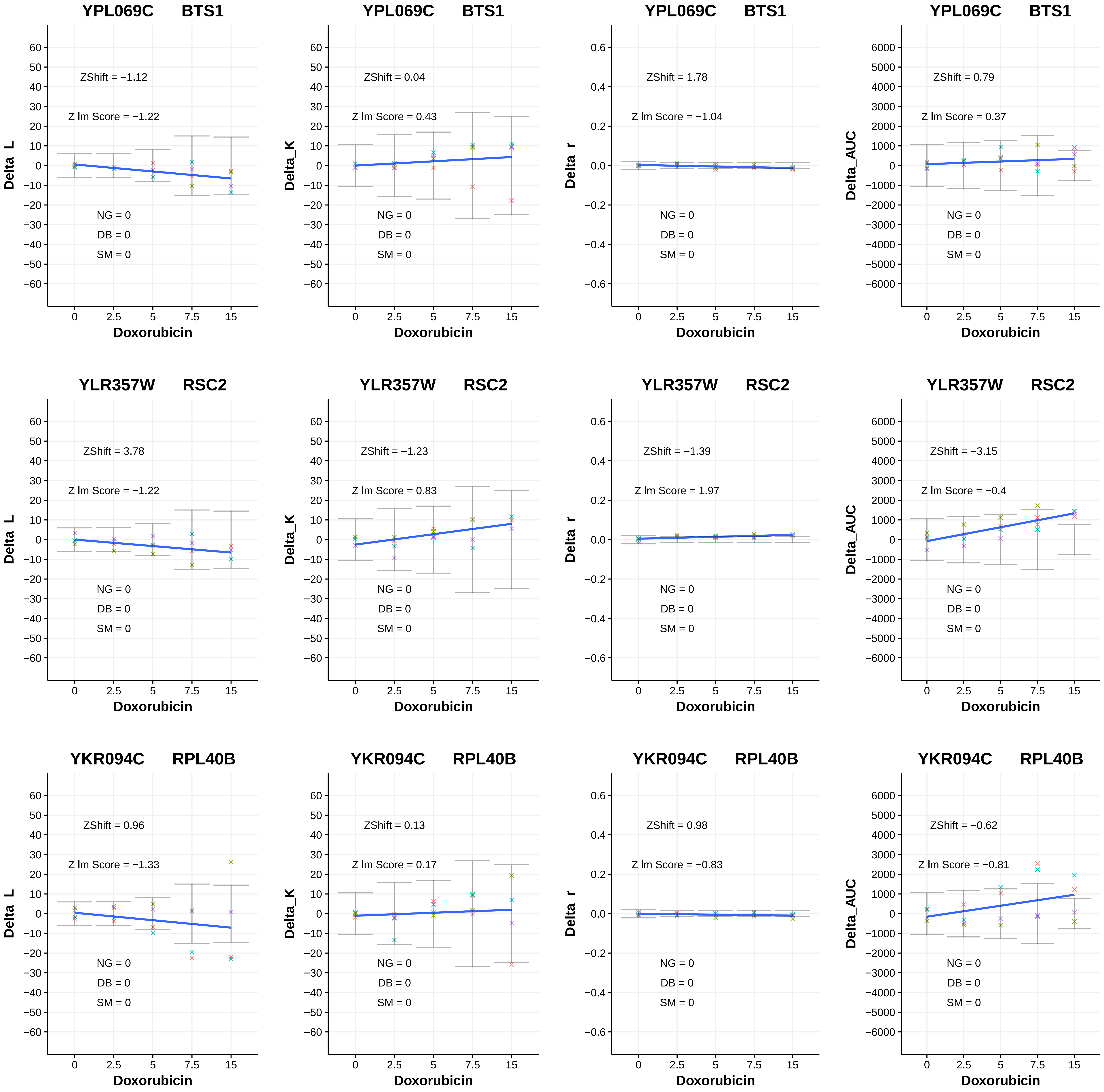
YPL069C BTS1
ZShift = −1.12
Z lm Score = −1.22
NG = 0
DB = 0
SM = 0
Delta_L
Doxorubicin
YPL069C BTS1
ZShift = 0.04
Z lm Score = 0.43
NG = 0
DB = 0
SM = 0
Delta_K
Doxorubicin
YPL069C BTS1
ZShift = 1.78
Z lm Score = −1.04
NG = 0
DB = 0
SM = 0
Delta_r
Doxorubicin
YPL069C BTS1
ZShift = 0.79
Z lm Score = 0.37
NG = 0
DB = 0
SM = 0
Delta_AUC
Doxorubicin
YLR357W RSC2
ZShift = 3.78
Z lm Score = −1.22
NG = 0
DB = 0
SM = 0
Delta_L
Doxorubicin
YLR357W RSC2
ZShift = −1.23
Z lm Score = 0.83
NG = 0
DB = 0
SM = 0
Delta_K
Doxorubicin
YLR357W RSC2
ZShift = −1.39
Z lm Score = 1.97
NG = 0
DB = 0
SM = 0
Delta_r
Doxorubicin
YLR357W RSC2
ZShift = −3.15
Z lm Score = −0.4
NG = 0
DB = 0
SM = 0
Delta_AUC
Doxorubicin
YKR094C RPL40B
ZShift = 0.96
Z lm Score = −1.33
NG = 0
DB = 0
SM = 0
Delta_L
Doxorubicin
YKR094C RPL40B
ZShift = 0.13
Z lm Score = 0.17
NG = 0
DB = 0
SM = 0
Delta_K
Doxorubicin
YKR094C RPL40B
ZShift = 0.98
Z lm Score = −0.83
NG = 0
DB = 0
SM = 0
Delta_r
Doxorubicin
YKR094C RPL40B
ZShift = −0.62
Z lm Score = −0.81
NG = 0
DB = 0
SM = 0
Delta_AUC
Doxorubicin
0
2.5
5
7.5
15

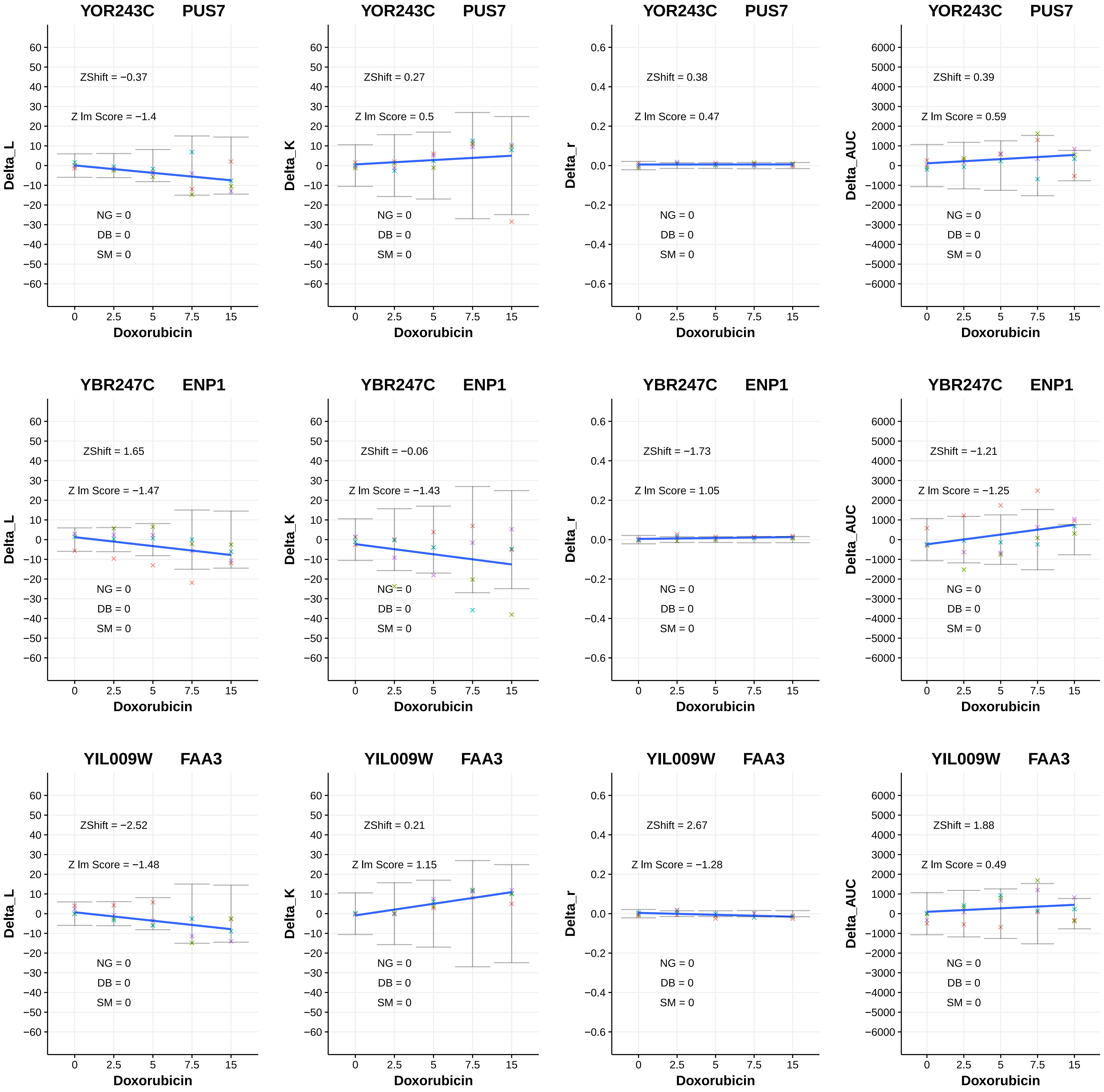

YOR243C PUS7
ZShift = −0.37
Z lm Score = −1.4
NG = 0
DB = 0
SM = 0
Delta_L
Doxorubicin
YOR243C PUS7
ZShift = 0.27
Z lm Score = 0.5
NG = 0
DB = 0
SM = 0
Delta_K
Doxorubicin
YOR243C PUS7
ZShift = 0.38
Z lm Score = 0.47
NG = 0
DB = 0
SM = 0
Delta_r
Doxorubicin
YOR243C PUS7
ZShift = 0.39
Z lm Score = 0.59
NG = 0
DB = 0
SM = 0
Delta_AUC
Doxorubicin
YBR247C ENP1
ZShift = 1.65
Z lm Score = −1.47
NG = 0
DB = 0
SM = 0
Delta_L
Doxorubicin
YBR247C ENP1
ZShift = −0.06
Z lm Score = −1.43
NG = 0
DB = 0
SM = 0
Delta_K
Doxorubicin
YBR247C ENP1
ZShift = −1.73
Z lm Score = 1.05
NG = 0
DB = 0
SM = 0
Delta_r
Doxorubicin
YBR247C ENP1
ZShift = −1.21
Z lm Score = −1.25
NG = 0
DB = 0
SM = 0
Delta_AUC
Doxorubicin
YIL009W FAA3
ZShift = −2.52
Z lm Score = −1.48
NG = 0
DB = 0
SM = 0
Delta_L
Doxorubicin
YIL009W FAA3
ZShift = 0.21
Z lm Score = 1.15
NG = 0
DB = 0
SM = 0
Delta_K
Doxorubicin
YIL009W FAA3
ZShift = 2.67
Z lm Score = −1.28
NG = 0
DB = 0
SM = 0
Delta_r
Doxorubicin
YIL009W FAA3
ZShift = 1.88
Z lm Score = 0.49
NG = 0
DB = 0
SM = 0
Delta_AUC
Doxorubicin

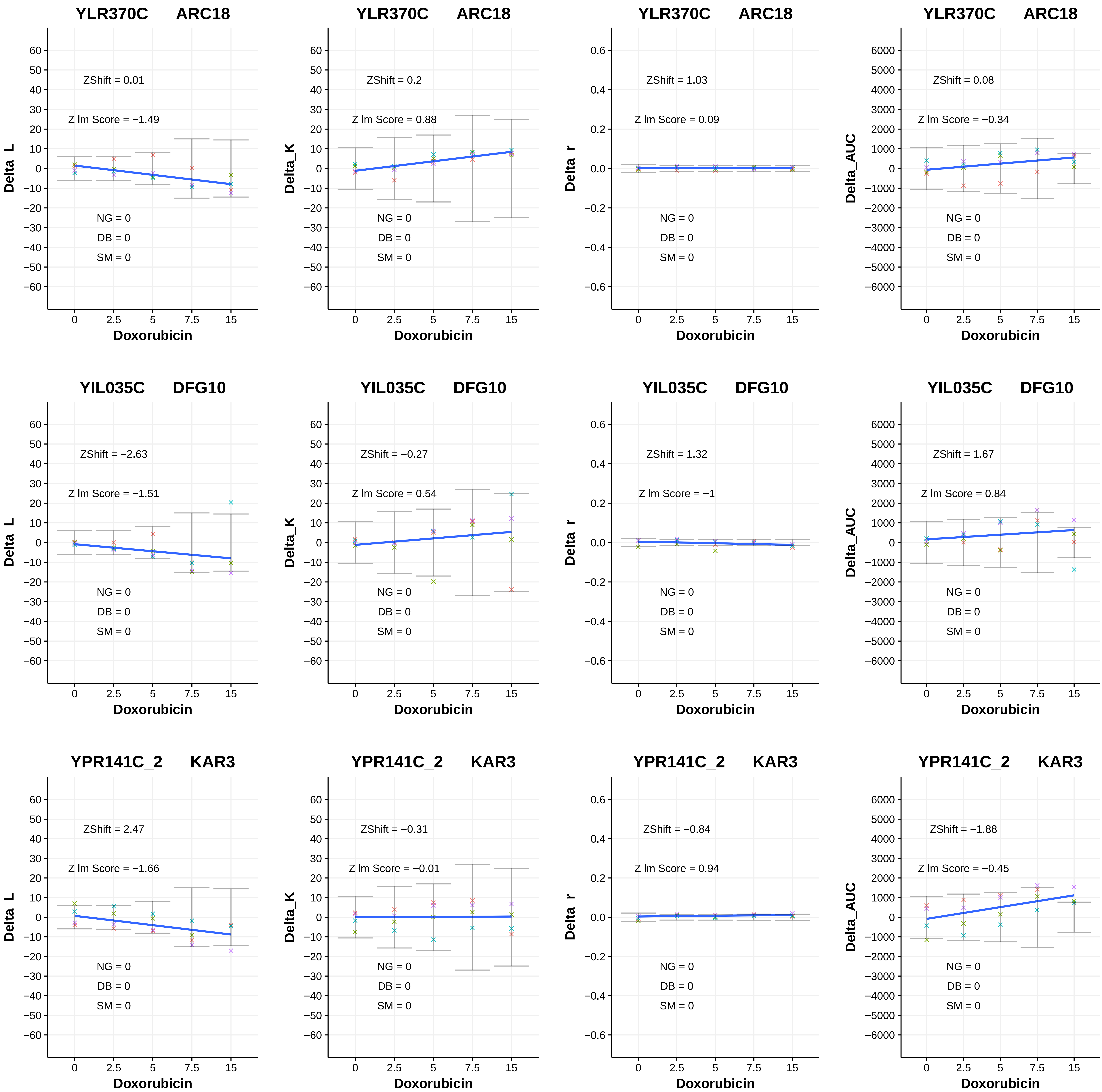
YLR370C ARC18
ZShift = 0.01
Z lm Score = −1.49
NG = 0
DB = 0
SM = 0
Delta_L
Doxorubicin
YLR370C ARC18
ZShift = 0.2
Z lm Score = 0.88
NG = 0
DB = 0
SM = 0
Delta_K
Doxorubicin
YLR370C ARC18
ZShift = 1.03
Z lm Score = 0.09
NG = 0
DB = 0
SM = 0
Delta_r
Doxorubicin
YLR370C ARC18
ZShift = 0.08
Z lm Score = −0.34
NG = 0
DB = 0
SM = 0
Delta_AUC
Doxorubicin
YIL035C DFG10
ZShift = −2.63
Z lm Score = −1.51
NG = 0
DB = 0
SM = 0
Delta_L
Doxorubicin
YIL035C DFG10
ZShift = −0.27
Z lm Score = 0.54
NG = 0
DB = 0
SM = 0
Delta_K
Doxorubicin
YIL035C DFG10
ZShift = 1.32
Z lm Score = −1
NG = 0
DB = 0
SM = 0
Delta_r
Doxorubicin
YIL035C DFG10
ZShift = 1.67
Z lm Score = 0.84
NG = 0
DB = 0
SM = 0
Delta_AUC
Doxorubicin
YPR141C_2 KAR3
ZShift = 2.47
Z lm Score = −1.66
NG = 0
DB = 0
SM = 0
Delta_L
Doxorubicin
YPR141C_2 KAR3
ZShift = −0.31
Z lm Score = −0.01
NG = 0
DB = 0
SM = 0
Delta_K
Doxorubicin
YPR141C_2 KAR3
ZShift = −0.84
Z lm Score = 0.94
NG = 0
DB = 0
SM = 0
Delta_r
Doxorubicin
YPR141C_2 KAR3
ZShift = −1.88
Z lm Score = −0.45
NG = 0
DB = 0
SM = 0
Delta_AUC
Doxorubicin

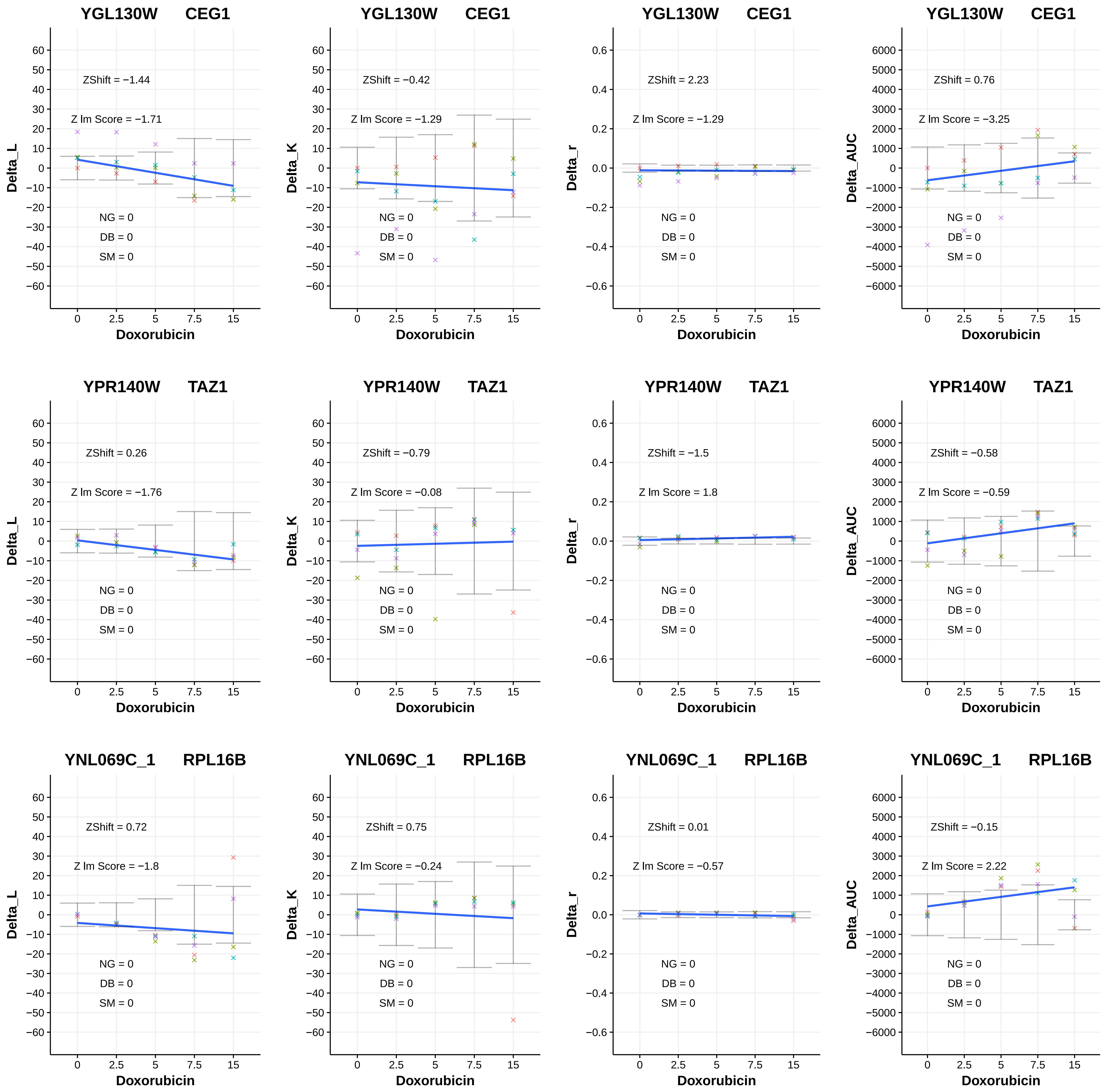
YGL130W CEG1
ZShift = −1.44
Z lm Score = −1.71
NG = 0
DB = 0
SM = 0
Delta_L
Doxorubicin
YGL130W CEG1
ZShift = −0.42
Z lm Score = −1.29
NG = 0
DB = 0
SM = 0
Delta_K
Doxorubicin
YGL130W CEG1
ZShift = 2.23
Z lm Score = −1.29
NG = 0
DB = 0
SM = 0
Delta_r
Doxorubicin
YGL130W CEG1
ZShift = 0.76
Z lm Score = −3.25
NG = 0
DB = 0
SM = 0
Delta_AUC
Doxorubicin
YPR140W TAZ1
ZShift = 0.26
Z lm Score = −1.76
NG = 0
DB = 0
SM = 0
Delta_L
Doxorubicin
YPR140W TAZ1
ZShift = −0.79
Z lm Score = −0.08
NG = 0
DB = 0
SM = 0
Delta_K
Doxorubicin
YPR140W TAZ1
ZShift = −1.5
Z lm Score = 1.8
NG = 0
DB = 0
SM = 0
Delta_r
Doxorubicin
YPR140W TAZ1
ZShift = −0.58
Z lm Score = −0.59
NG = 0
DB = 0
SM = 0
Delta_AUC
Doxorubicin
YNL069C_1 RPL16B
ZShift = 0.72
Z lm Score = −1.8
NG = 0
DB = 0
SM = 0
Delta_L
Doxorubicin
YNL069C_1 RPL16B
ZShift = 0.75
Z lm Score = −0.24
NG = 0
DB = 0
SM = 0
Delta_K
Doxorubicin
YNL069C_1 RPL16B
ZShift = 0.01
Z lm Score = −0.57
NG = 0
DB = 0
SM = 0
Delta_r
Doxorubicin
YNL069C_1 RPL16B
ZShift = −0.15
Z lm Score = 2.22
NG = 0
DB = 0
SM = 0
Delta_AUC
Doxorubicin

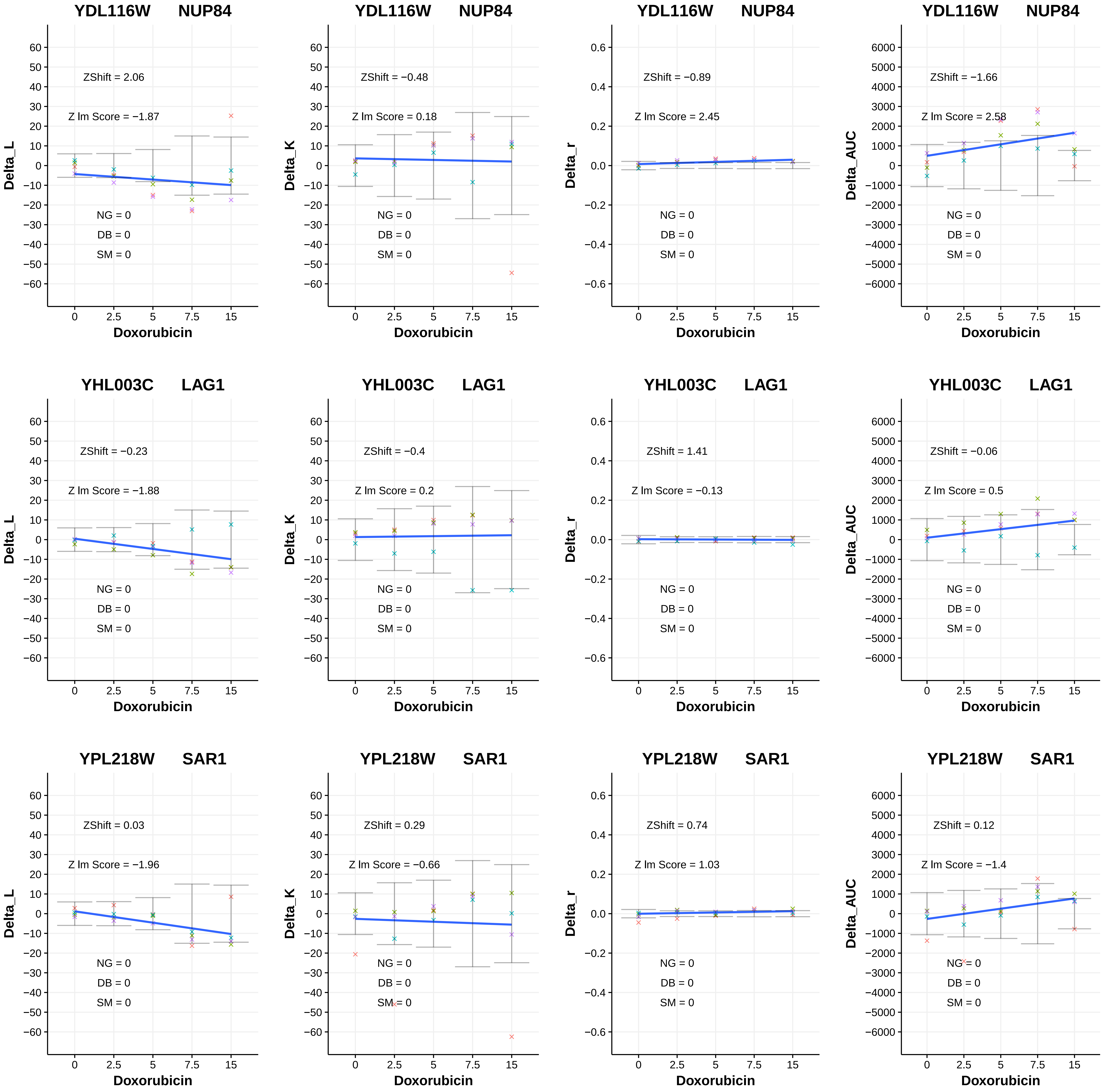
YDL116W NUP84
ZShift = 2.06
Z lm Score = −1.87
NG = 0
DB = 0
SM = 0
Delta_L
Doxorubicin
YDL116W NUP84
ZShift = −0.48
Z lm Score = 0.18
NG = 0
DB = 0
SM = 0
Delta_K
Doxorubicin
YDL116W NUP84
ZShift = −0.89
Z lm Score = 2.45
NG = 0
DB = 0
SM = 0
Delta_r
Doxorubicin
YDL116W NUP84
ZShift = −1.66
Z lm Score = 2.58
NG = 0
DB = 0
SM = 0
Delta_AUC
Doxorubicin
YHL003C LAG1
ZShift = −0.23
Z lm Score = −1.88
NG = 0
DB = 0
SM = 0
Delta_L
Doxorubicin
YHL003C LAG1
ZShift = −0.4
Z lm Score = 0.2
NG = 0
DB = 0
SM = 0
Delta_K
Doxorubicin
YHL003C LAG1
ZShift = 1.41
Z lm Score = −0.13
NG = 0
DB = 0
SM = 0
Delta_r
Doxorubicin
YHL003C LAG1
ZShift = −0.06
Z lm Score = 0.5
NG = 0
DB = 0
SM = 0
Delta_AUC
Doxorubicin
YPL218W SAR1
ZShift = 0.03
Z lm Score = −1.96
NG = 0
DB = 0
SM = 0
Delta_L
Doxorubicin
YPL218W SAR1
ZShift = 0.29
Z lm Score = −0.66
NG = 0
DB = 0
SM = 0
Delta_K
Doxorubicin
YPL218W SAR1
ZShift = 0.74
Z lm Score = 1.03
NG = 0
DB = 0
SM = 0
Delta_r
Doxorubicin
YPL218W SAR1
ZShift = 0.12
Z lm Score = −1.4
NG = 0
DB = 0
SM = 0
Delta_AUC
Doxorubicin

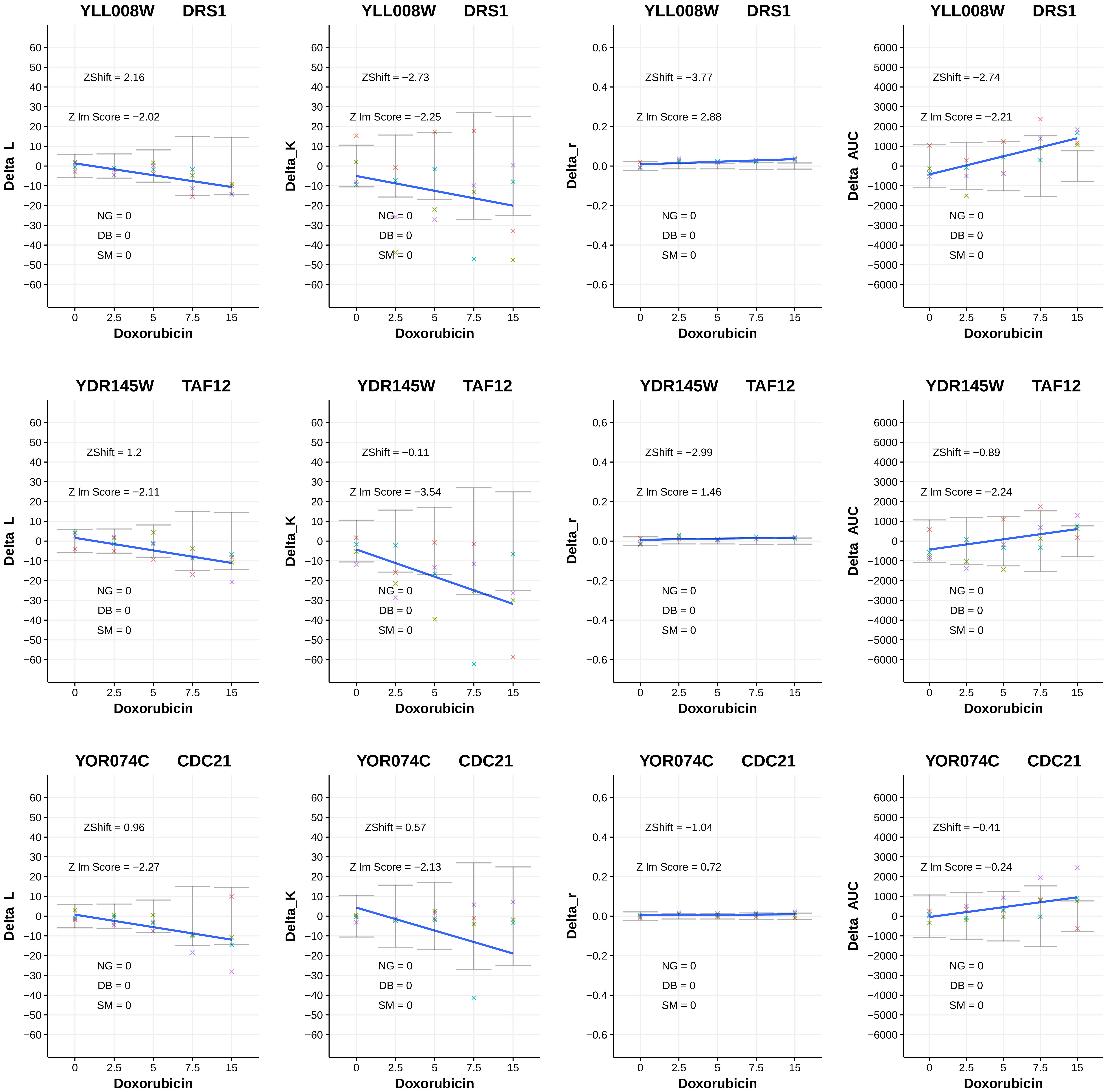
YLL008W DRS1
ZShift = 2.16
Z lm Score = −2.02
NG = 0
DB = 0
SM = 0
Delta_L
Doxorubicin
YLL008W DRS1
ZShift = −2.73
Z lm Score = −2.25
NG = 0
DB = 0
SM = 0
Delta_K
Doxorubicin
YLL008W DRS1
ZShift = −3.77
Z lm Score = 2.88
NG = 0
DB = 0
SM = 0
Delta_r
Doxorubicin
YLL008W DRS1
ZShift = −2.74
Z lm Score = −2.21
NG = 0
DB = 0
SM = 0
Delta_AUC
Doxorubicin
YDR145W TAF12
ZShift = 1.2
Z lm Score = −2.11
NG = 0
DB = 0
SM = 0
Delta_L
Doxorubicin
YDR145W TAF12
ZShift = −0.11
Z lm Score = −3.54
NG = 0
DB = 0
SM = 0
Delta_K
Doxorubicin
YDR145W TAF12
ZShift = −2.99
Z lm Score = 1.46
NG = 0
DB = 0
SM = 0
Delta_r
Doxorubicin
YDR145W TAF12
ZShift = −0.89
Z lm Score = −2.24
NG = 0
DB = 0
SM = 0
Delta_AUC
Doxorubicin
YOR074C CDC21
ZShift = 0.96
Z lm Score = −2.27
NG = 0
DB = 0
SM = 0
Delta_L
Doxorubicin
YOR074C CDC21
ZShift = 0.57
Z lm Score = −2.13
NG = 0
DB = 0
SM = 0
Delta_K
Doxorubicin
YOR074C CDC21
ZShift = −1.04
Z lm Score = 0.72
NG = 0
DB = 0
SM = 0
Delta_r
Doxorubicin
YOR074C CDC21
ZShift = −0.41
Z lm Score = −0.24
NG = 0
DB = 0
SM = 0
Delta_AUC
Doxorubicin

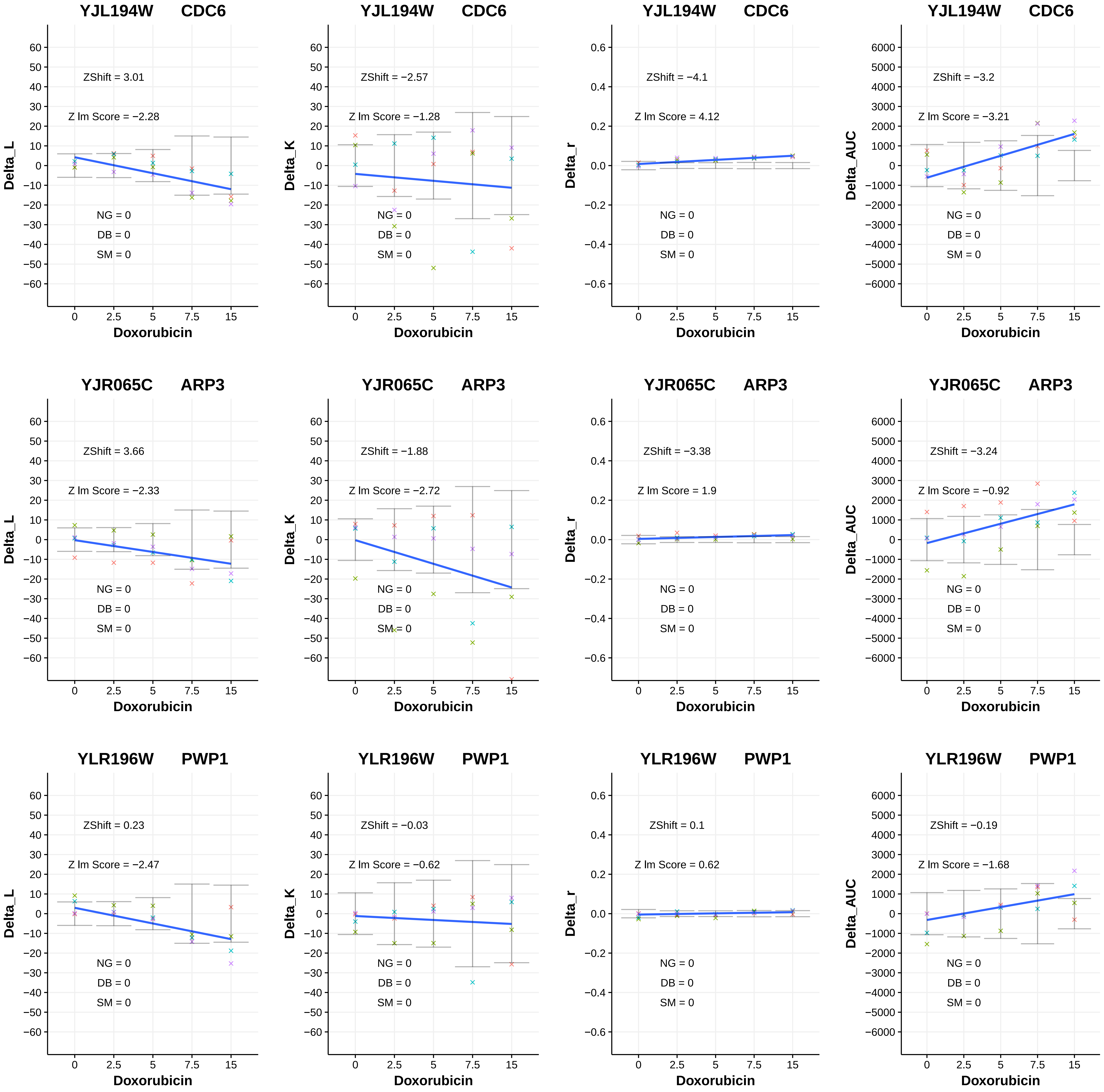

YJL194W CDC6
ZShift = 3.01
Z lm Score = −2.28
NG = 0
DB = 0
SM = 0
Delta_L
Doxorubicin
YJL194W CDC6
ZShift = −2.57
Z lm Score = −1.28
NG = 0
DB = 0
SM = 0
Delta_K
Doxorubicin
YJL194W CDC6
ZShift = −4.1
Z lm Score = 4.12
NG = 0
DB = 0
SM = 0
Delta_r
Doxorubicin
YJL194W CDC6
ZShift = −3.2
Z lm Score = −3.21
NG = 0
DB = 0
SM = 0
Delta_AUC
Doxorubicin
YJR065C ARP3
ZShift = 3.66
Z lm Score = −2.33
NG = 0
DB = 0
SM = 0
Delta_L
Doxorubicin
YJR065C ARP3
ZShift = −1.88
Z lm Score = −2.72
NG = 0
DB = 0
SM = 0
Delta_K
Doxorubicin
YJR065C ARP3
ZShift = −3.38
Z lm Score = 1.9
NG = 0
DB = 0
SM = 0
Delta_r
Doxorubicin
YJR065C ARP3
ZShift = −3.24
Z lm Score = −0.92
NG = 0
DB = 0
SM = 0
Delta_AUC
Doxorubicin
YLR196W PWP1
ZShift = 0.23
Z lm Score = −2.47
NG = 0
DB = 0
SM = 0
Delta_L
Doxorubicin
YLR196W PWP1
ZShift = −0.03
Z lm Score = −0.62
NG = 0
DB = 0
SM = 0
Delta_K
Doxorubicin
YLR196W PWP1
ZShift = 0.1
Z lm Score = 0.62
NG = 0
DB = 0
SM = 0
Delta_r
Doxorubicin
YLR196W PWP1
ZShift = −0.19
Z lm Score = −1.68
NG = 0
DB = 0
SM = 0
Delta_AUC
Doxorubicin

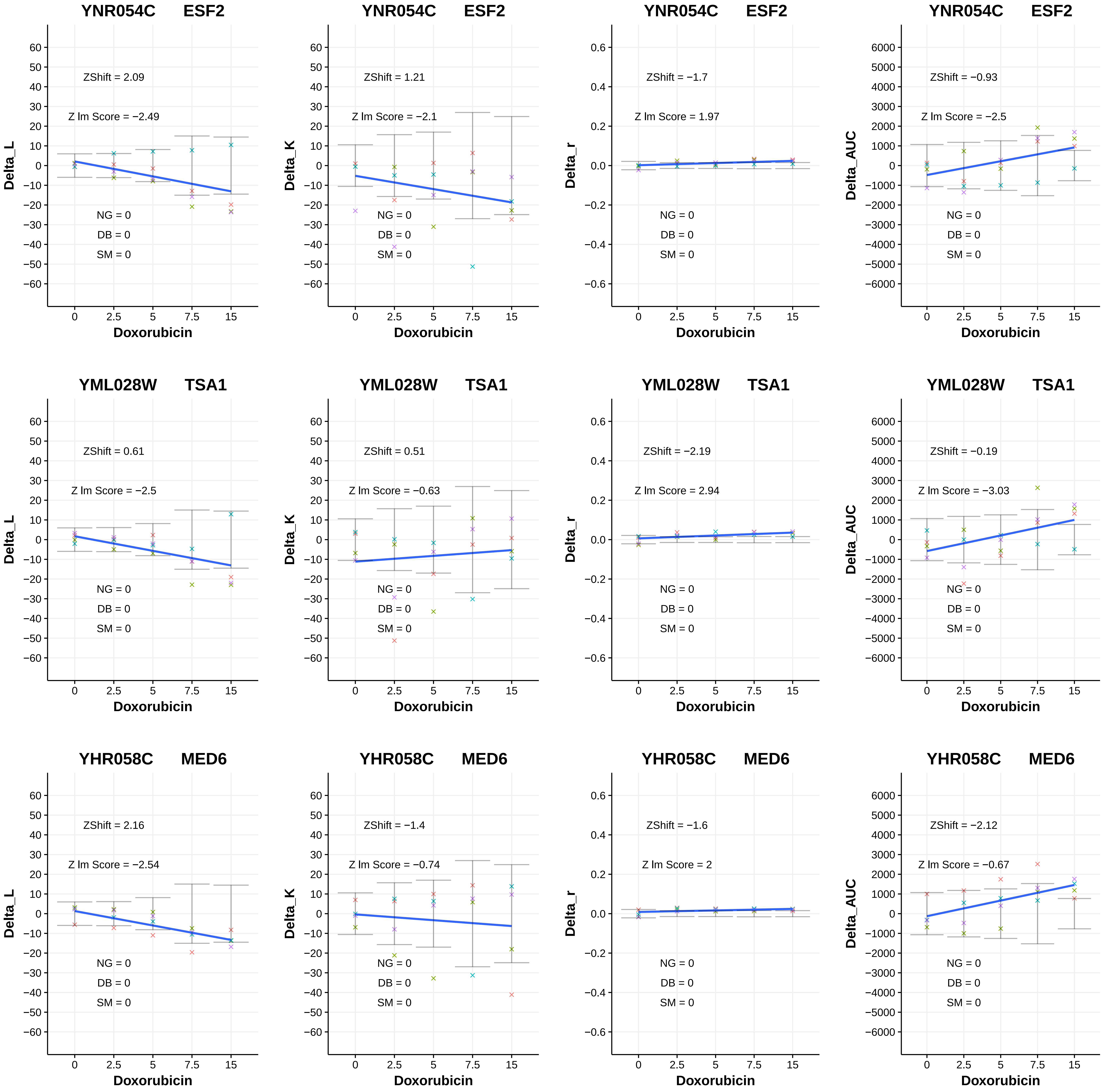
YNR054C ESF2
Delta_L
ZShift = 2.09
Z lm Score = −2.49
NG = 0
DB = 0
SM = 0
Doxorubicin
YNR054C ESF2
Delta_K
ZShift = 1.21
Z lm Score = −2.1
NG = 0
DB = 0
SM = 0
Doxorubicin
YNR054C ESF2
Delta_r
ZShift = −1.7
Z lm Score = 1.97
NG = 0
DB = 0
SM = 0
Doxorubicin
YNR054C ESF2
Delta_AUC
ZShift = −0.93
Z lm Score = −2.5
NG = 0
DB = 0
SM = 0
Doxorubicin
YML028W TSA1
Delta_L
ZShift = 0.61
Z lm Score = −2.5
NG = 0
DB = 0
SM = 0
Doxorubicin
YML028W TSA1
Delta_K
ZShift = 0.51
Z lm Score = −0.63
NG = 0
DB = 0
SM = 0
Doxorubicin
YML028W TSA1
Delta_r
ZShift = −2.19
Z lm Score = 2.94
NG = 0
DB = 0
SM = 0
Doxorubicin
YML028W TSA1
Delta_AUC
ZShift = −0.19
Z lm Score = −3.03
NG = 0
DB = 0
SM = 0
Doxorubicin
YHR058C MED6
Delta_L
ZShift = 2.16
Z lm Score = −2.54
NG = 0
DB = 0
SM = 0
Doxorubicin
YHR058C MED6
Delta_K
ZShift = −1.4
Z lm Score = −0.74
NG = 0
DB = 0
SM = 0
Doxorubicin
YHR058C MED6
Delta_r
ZShift = −1.6
Z lm Score = 2
NG = 0
DB = 0
SM = 0
Doxorubicin
YHR058C MED6
Delta_AUC
ZShift = −2.12
Z lm Score = −0.67
NG = 0
DB = 0
SM = 0
Doxorubicin

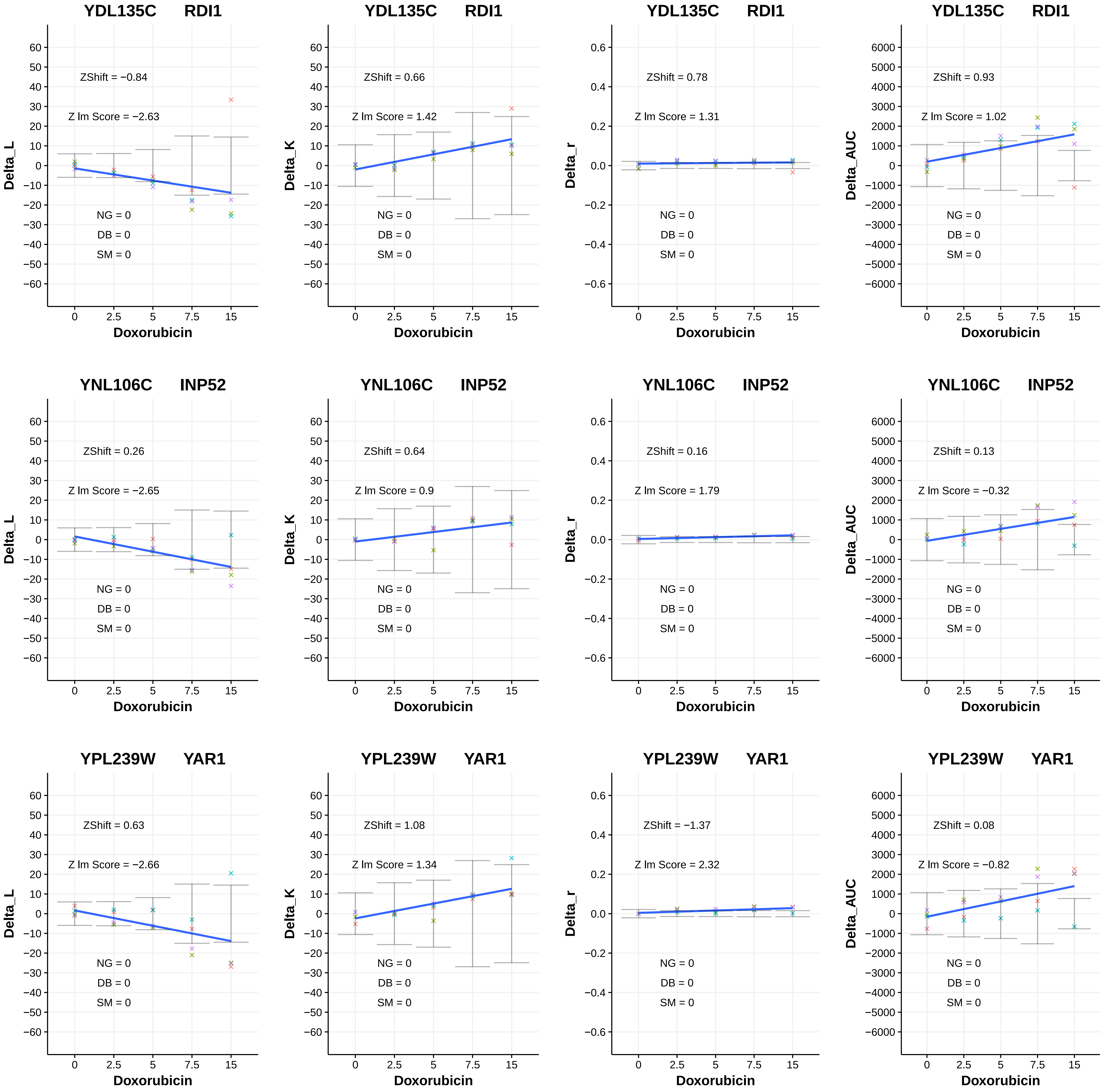
YDL135C RDI1
ZShift = −0.84
Z lm Score = −2.63
NG = 0
DB = 0
SM = 0
Delta_L
Doxorubicin
YDL135C RDI1
ZShift = 0.66
Z lm Score = 1.42
NG = 0
DB = 0
SM = 0
Delta_K
Doxorubicin
YDL135C RDI1
ZShift = 0.78
Z lm Score = 1.31
NG = 0
DB = 0
SM = 0
Delta_r
Doxorubicin
YDL135C RDI1
ZShift = 0.93
Z lm Score = 1.02
NG = 0
DB = 0
SM = 0
Delta_AUC
Doxorubicin
YNL106C INP52
ZShift = 0.26
Z lm Score = −2.65
NG = 0
DB = 0
SM = 0
Delta_L
Doxorubicin
YNL106C INP52
ZShift = 0.64
Z lm Score = 0.9
NG = 0
DB = 0
SM = 0
Delta_K
Doxorubicin
YNL106C INP52
ZShift = 0.16
Z lm Score = 1.79
NG = 0
DB = 0
SM = 0
Delta_r
Doxorubicin
YNL106C INP52
ZShift = 0.13
Z lm Score = −0.32
NG = 0
DB = 0
SM = 0
Delta_AUC
Doxorubicin
YPL239W YAR1
ZShift = 0.63
Z lm Score = −2.66
NG = 0
DB = 0
SM = 0
Delta_L
Doxorubicin
YPL239W YAR1
ZShift = 1.08
Z lm Score = 1.34
NG = 0
DB = 0
SM = 0
Delta_K
Doxorubicin
YPL239W YAR1
ZShift = −1.37
Z lm Score = 2.32
NG = 0
DB = 0
SM = 0
Delta_r
Doxorubicin
YPL239W YAR1
ZShift = 0.08
Z lm Score = −0.82
NG = 0
DB = 0
SM = 0
Delta_AUC
Doxorubicin

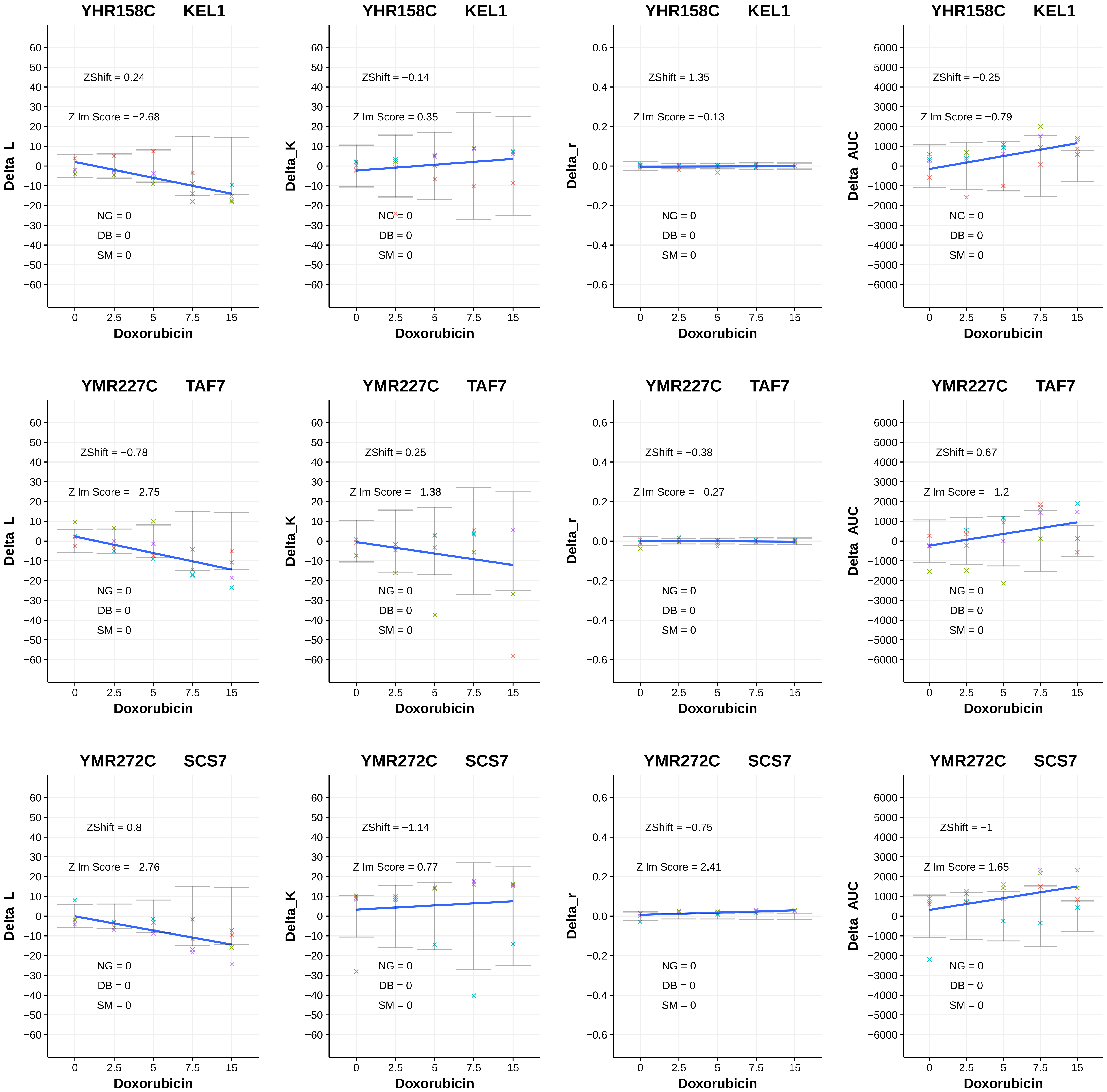
YHR158C KEL1
ZShift = 0.24
Z lm Score = −2.68
NG = 0
DB = 0
SM = 0
Delta_L
Doxorubicin
YHR158C KEL1
ZShift = −0.14
Z lm Score = 0.35
NG = 0
DB = 0
SM = 0
Delta_K
Doxorubicin
YHR158C KEL1
ZShift = 1.35
Z lm Score = −0.13
NG = 0
DB = 0
SM = 0
Delta_r
Doxorubicin
YHR158C KEL1
ZShift = −0.25
Z lm Score = −0.79
NG = 0
DB = 0
SM = 0
Delta_AUC
Doxorubicin
YMR227C TAF7
ZShift = −0.78
Z lm Score = −2.75
NG = 0
DB = 0
SM = 0
Delta_L
Doxorubicin
YMR227C TAF7
ZShift = 0.25
Z lm Score = −1.38
NG = 0
DB = 0
SM = 0
Delta_K
Doxorubicin
YMR227C TAF7
ZShift = −0.38
Z lm Score = −0.27
NG = 0
DB = 0
SM = 0
Delta_r
Doxorubicin
YMR227C TAF7
ZShift = 0.67
Z lm Score = −1.2
NG = 0
DB = 0
SM = 0
Delta_AUC
Doxorubicin
YMR272C SCS7
ZShift = 0.8
Z lm Score = −2.76
NG = 0
DB = 0
SM = 0
Delta_L
Doxorubicin
YMR272C SCS7
ZShift = −1.14
Z lm Score = 0.77
NG = 0
DB = 0
SM = 0
Delta_K
Doxorubicin
YMR272C SCS7
ZShift = −0.75
Z lm Score = 2.41
NG = 0
DB = 0
SM = 0
Delta_r
Doxorubicin
YMR272C SCS7
ZShift = −1
Z lm Score = 1.65
NG = 0
DB = 0
SM = 0
Delta_AUC
Doxorubicin

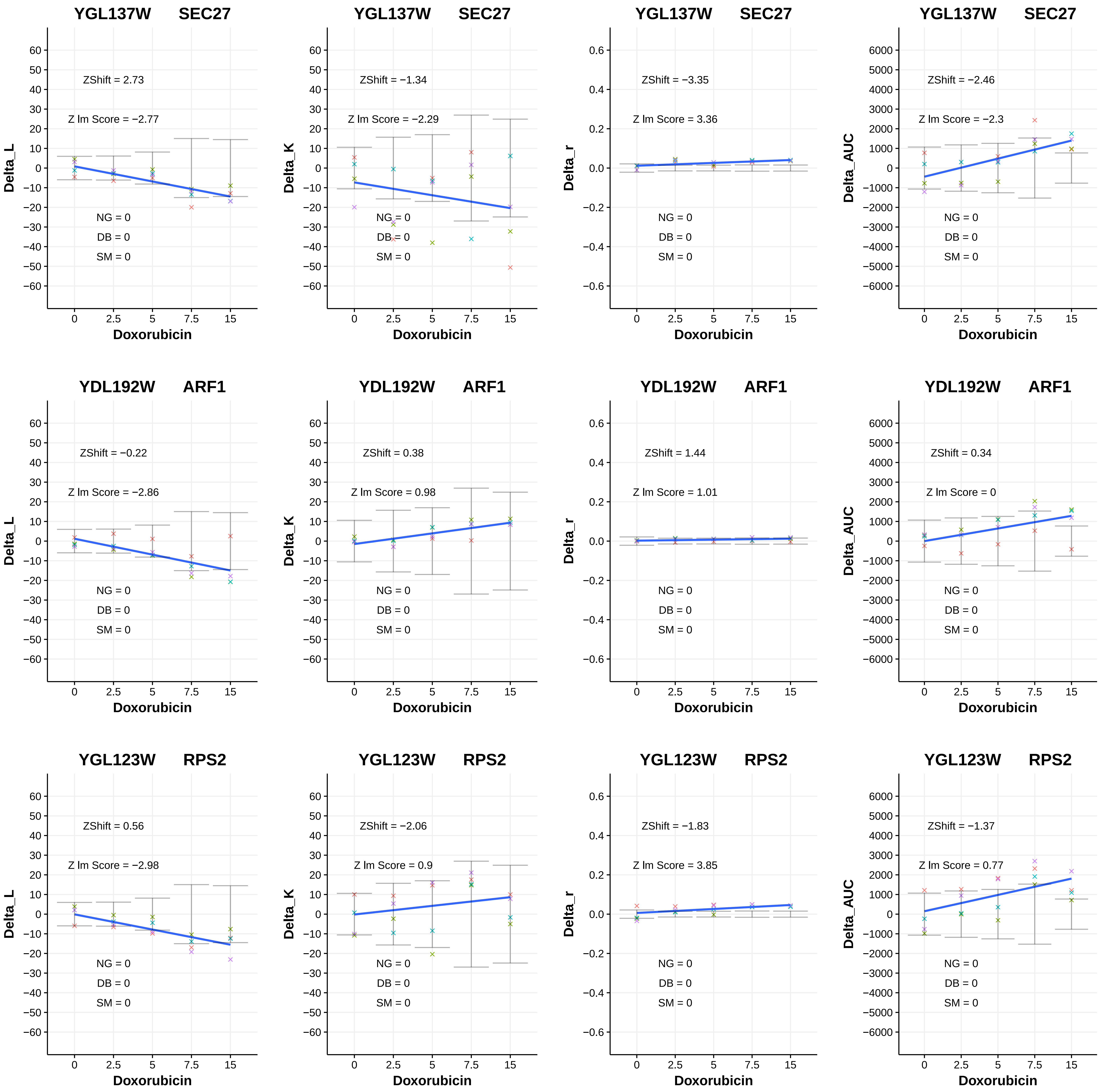
YGL137W SEC27
ZShift = 2.73
Z lm Score = −2.77
NG = 0
DB = 0
SM = 0
Delta_L
Doxorubicin
YGL137W SEC27
ZShift = −1.34
Z lm Score = −2.29
NG = 0
DB = 0
SM = 0
Delta_K
Doxorubicin
YGL137W SEC27
ZShift = −3.35
Z lm Score = 3.36
NG = 0
DB = 0
SM = 0
Delta_r
Doxorubicin
YGL137W SEC27
ZShift = −2.46
Z lm Score = −2.3
NG = 0
DB = 0
SM = 0
Delta_AUC
Doxorubicin
YDL192W ARF1
ZShift = −0.22
Z lm Score = −2.86
NG = 0
DB = 0
SM = 0
Delta_L
Doxorubicin
YDL192W ARF1
ZShift = 0.38
Z lm Score = 0.98
NG = 0
DB = 0
SM = 0
Delta_K
Doxorubicin
YDL192W ARF1
ZShift = 1.44
Z lm Score = 1.01
NG = 0
DB = 0
SM = 0
Delta_r
Doxorubicin
YDL192W ARF1
ZShift = 0.34
Z lm Score = 0
NG = 0
DB = 0
SM = 0
Delta_AUC
Doxorubicin
YGL123W RPS2
ZShift = 0.56
Z lm Score = −2.98
NG = 0
DB = 0
SM = 0
Delta_L
Doxorubicin
YGL123W RPS2
ZShift = −2.06
Z lm Score = 0.9
NG = 0
DB = 0
SM = 0
Delta_K
Doxorubicin
YGL123W RPS2
ZShift = −1.83
Z lm Score = 3.85
NG = 0
DB = 0
SM = 0
Delta_r
Doxorubicin
YGL123W RPS2
ZShift = −1.37
Z lm Score = 0.77
NG = 0
DB = 0
SM = 0
Delta_AUC
Doxorubicin

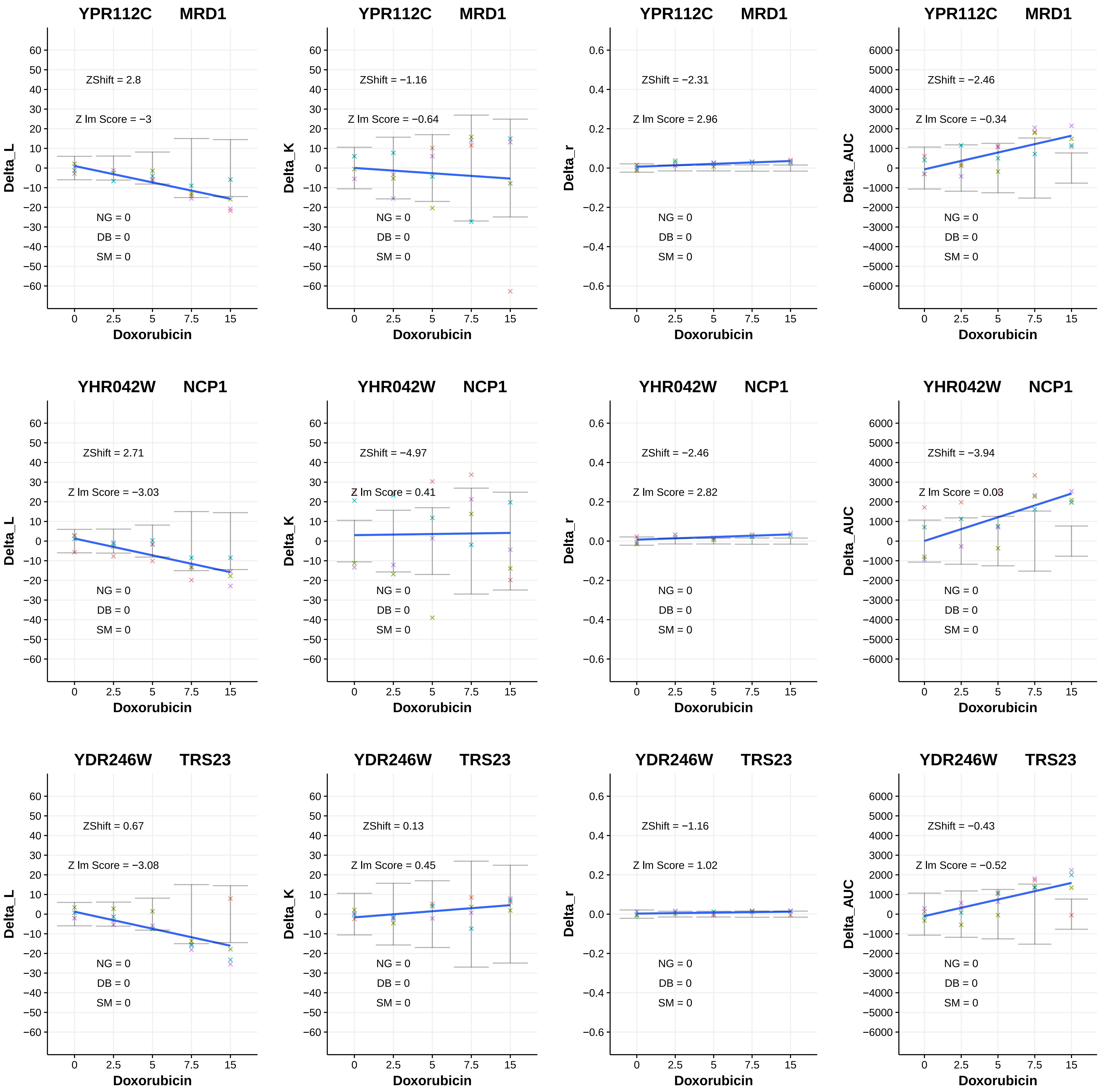
YPR112C MRD1
ZShift = 2.8
Z lm Score = −3
NG = 0
DB = 0
SM = 0
Delta_L
Doxorubicin
YPR112C MRD1
ZShift = −1.16
Z lm Score = −0.64
NG = 0
DB = 0
SM = 0
Delta_K
Doxorubicin
YPR112C MRD1
ZShift = −2.31
Z lm Score = 2.96
NG = 0
DB = 0
SM = 0
Delta_r
Doxorubicin
YPR112C MRD1
ZShift = −2.46
Z lm Score = −0.34
NG = 0
DB = 0
SM = 0
Delta_AUC
Doxorubicin
YHR042W NCP1
ZShift = 2.71
Z lm Score = −3.03
NG = 0
DB = 0
SM = 0
Delta_L
Doxorubicin
YHR042W NCP1
ZShift = −4.97
Z lm Score = 0.41
NG = 0
DB = 0
SM = 0
Delta_K
Doxorubicin
YHR042W NCP1
ZShift = −2.46
Z lm Score = 2.82
NG = 0
DB = 0
SM = 0
Delta_r
Doxorubicin
YHR042W NCP1
ZShift = −3.94
Z lm Score = 0.03
NG = 0
DB = 0
SM = 0
Delta_AUC
Doxorubicin
YDR246W TRS23
ZShift = 0.67
Z lm Score = −3.08
NG = 0
DB = 0
SM = 0
Delta_L
Doxorubicin
YDR246W TRS23
ZShift = 0.13
Z lm Score = 0.45
NG = 0
DB = 0
SM = 0
Delta_K
Doxorubicin
YDR246W TRS23
ZShift = −1.16
Z lm Score = 1.02
NG = 0
DB = 0
SM = 0
Delta_r
Doxorubicin
YDR246W TRS23
ZShift = −0.43
Z lm Score = −0.52
NG = 0
DB = 0
SM = 0
Delta_AUC
Doxorubicin

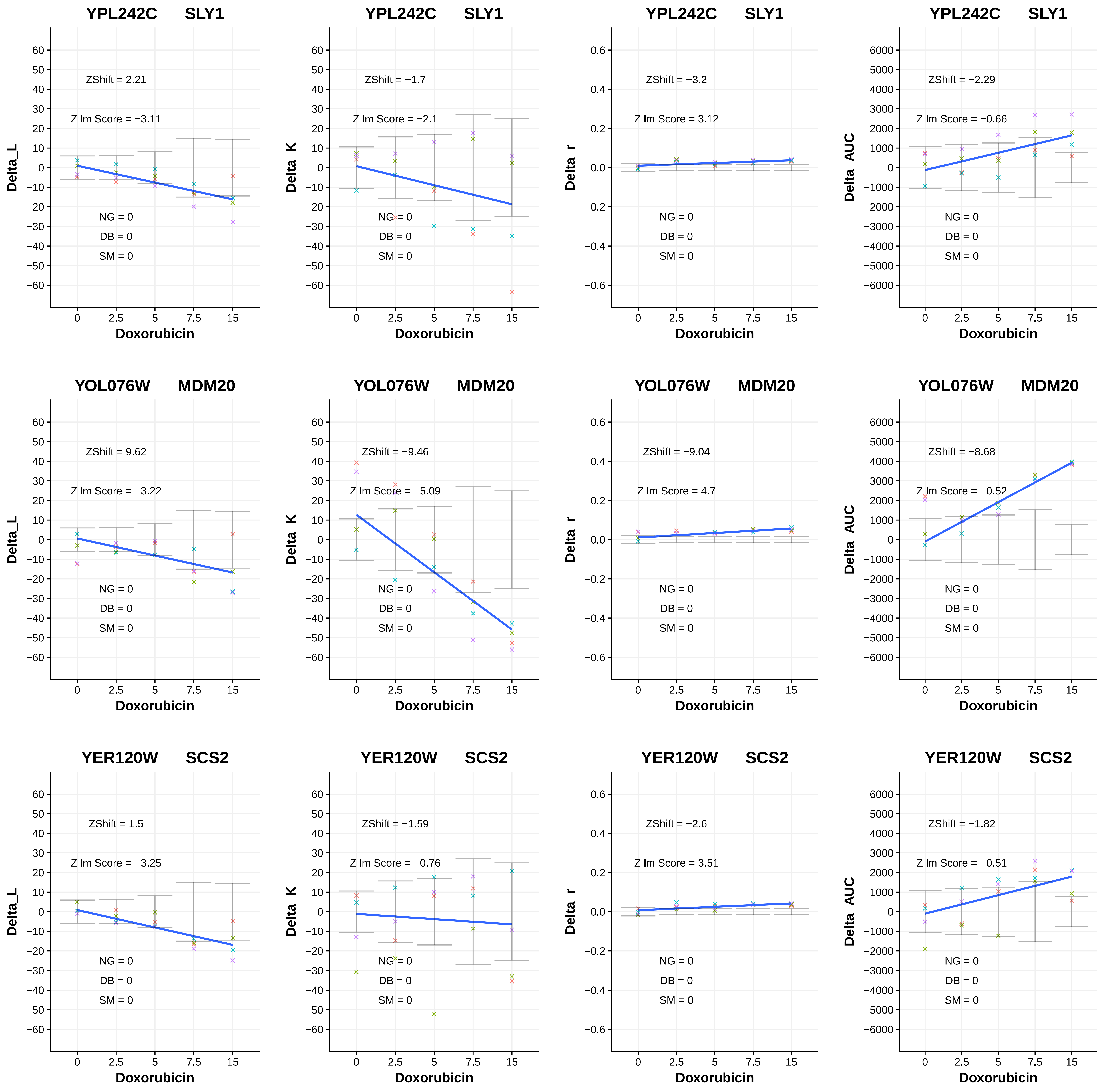

YPL242C SLY1
ZShift = 2.21
Z lm Score = −3.11
NG = 0
DB = 0
SM = 0
Delta_L
Doxorubicin
YPL242C SLY1
ZShift = −1.7
Z lm Score = −2.1
NG = 0
DB = 0
SM = 0
Delta_K
Doxorubicin
YPL242C SLY1
ZShift = −3.2
Z lm Score = 3.12
NG = 0
DB = 0
SM = 0
Delta_r
Doxorubicin
YPL242C SLY1
ZShift = −2.29
Z lm Score = −0.66
NG = 0
DB = 0
SM = 0
Delta_AUC
Doxorubicin
YOL076W MDM20
ZShift = 9.62
Z lm Score = −3.22
NG = 0
DB = 0
SM = 0
Delta_L
Doxorubicin
YOL076W MDM20
ZShift = −9.46
Z lm Score = −5.09
NG = 0
DB = 0
SM = 0
Delta_K
Doxorubicin
YOL076W MDM20
ZShift = −9.04
Z lm Score = 4.7
NG = 0
DB = 0
SM = 0
Delta_r
Doxorubicin
YOL076W MDM20
ZShift = −8.68
Z lm Score = −0.52
NG = 0
DB = 0
SM = 0
Delta_AUC
Doxorubicin
YER120W SCS2
ZShift = 1.5
Z lm Score = −3.25
NG = 0
DB = 0
SM = 0
Delta_L
Doxorubicin
YER120W SCS2
ZShift = −1.59
Z lm Score = −0.76
NG = 0
DB = 0
SM = 0
Delta_K
Doxorubicin
YER120W SCS2
ZShift = −2.6
Z lm Score = 3.51
NG = 0
DB = 0
SM = 0
Delta_r
Doxorubicin
YER120W SCS2
ZShift = −1.82
Z lm Score = −0.51
NG = 0
DB = 0
SM = 0
Delta_AUC
Doxorubicin

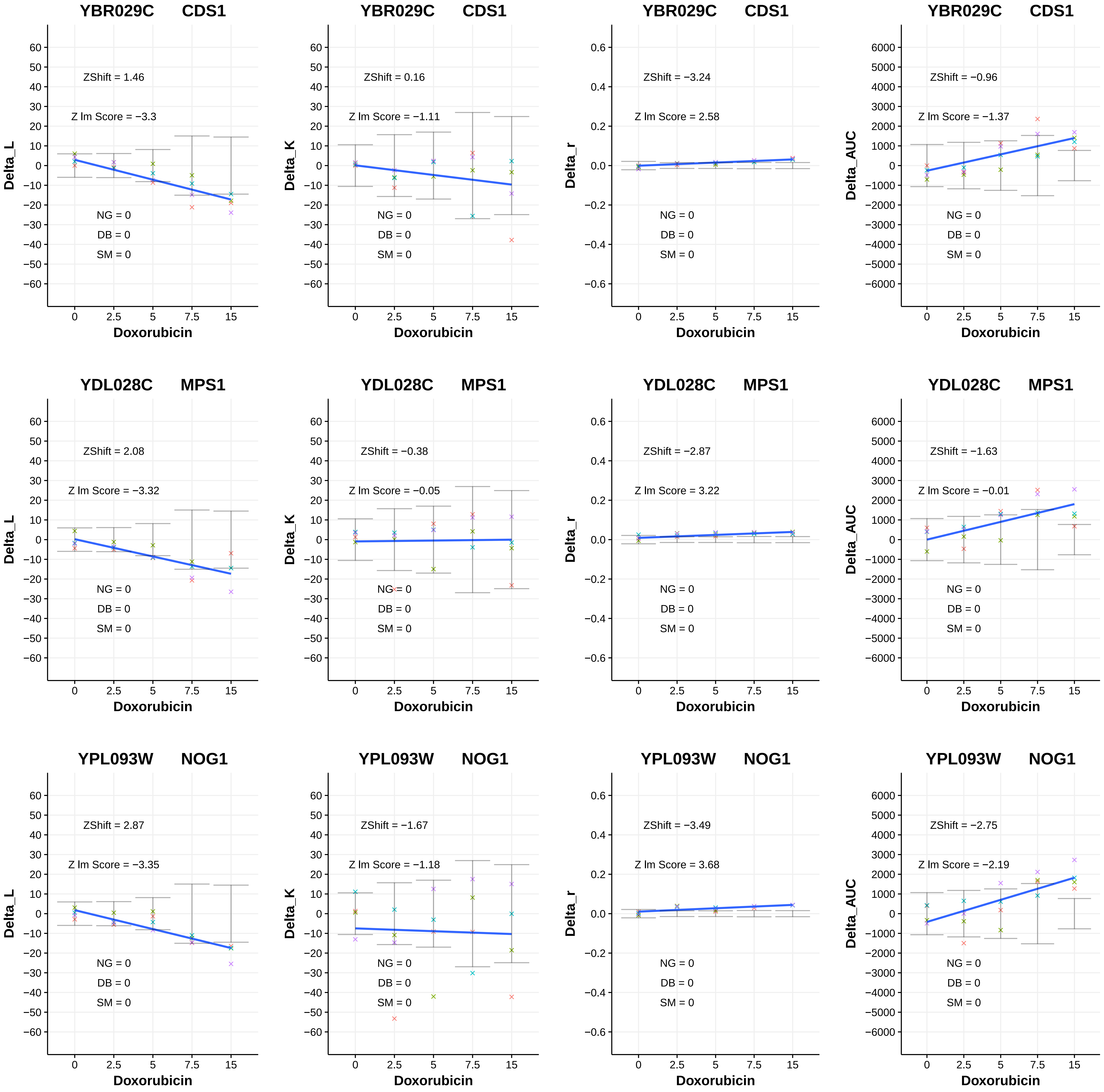

YBR029C CDS1
ZShift = 1.46
Z lm Score = −3.3
NG = 0
DB = 0
SM = 0
Delta_L
Doxorubicin
YBR029C CDS1
ZShift = 0.16
Z lm Score = −1.11
NG = 0
DB = 0
SM = 0
Delta_K
Doxorubicin
YBR029C CDS1
ZShift = −3.24
Z lm Score = 2.58
NG = 0
DB = 0
SM = 0
Delta_r
Doxorubicin
YBR029C CDS1
ZShift = −0.96
Z lm Score = −1.37
NG = 0
DB = 0
SM = 0
Delta_AUC
Doxorubicin
YDL028C MPS1
ZShift = 2.08
Z lm Score = −3.32
NG = 0
DB = 0
SM = 0
Delta_L
Doxorubicin
YDL028C MPS1
ZShift = −0.38
Z lm Score = −0.05
NG = 0
DB = 0
SM = 0
Delta_K
Doxorubicin
YDL028C MPS1
ZShift = −2.87
Z lm Score = 3.22
NG = 0
DB = 0
SM = 0
Delta_r
Doxorubicin
YDL028C MPS1
ZShift = −1.63
Z lm Score = −0.01
NG = 0
DB = 0
SM = 0
Delta_AUC
Doxorubicin
YPL093W NOG1
ZShift = 2.87
Z lm Score = −3.35
NG = 0
DB = 0
SM = 0
Delta_L
Doxorubicin
YPL093W NOG1
ZShift = −1.67
Z lm Score = −1.18
NG = 0
DB = 0
SM = 0
Delta_K
Doxorubicin
YPL093W NOG1
ZShift = −3.49
Z lm Score = 3.68
NG = 0
DB = 0
SM = 0
Delta_r
Doxorubicin
YPL093W NOG1
ZShift = −2.75
Z lm Score = −2.19
NG = 0
DB = 0
SM = 0
Delta_AUC
Doxorubicin

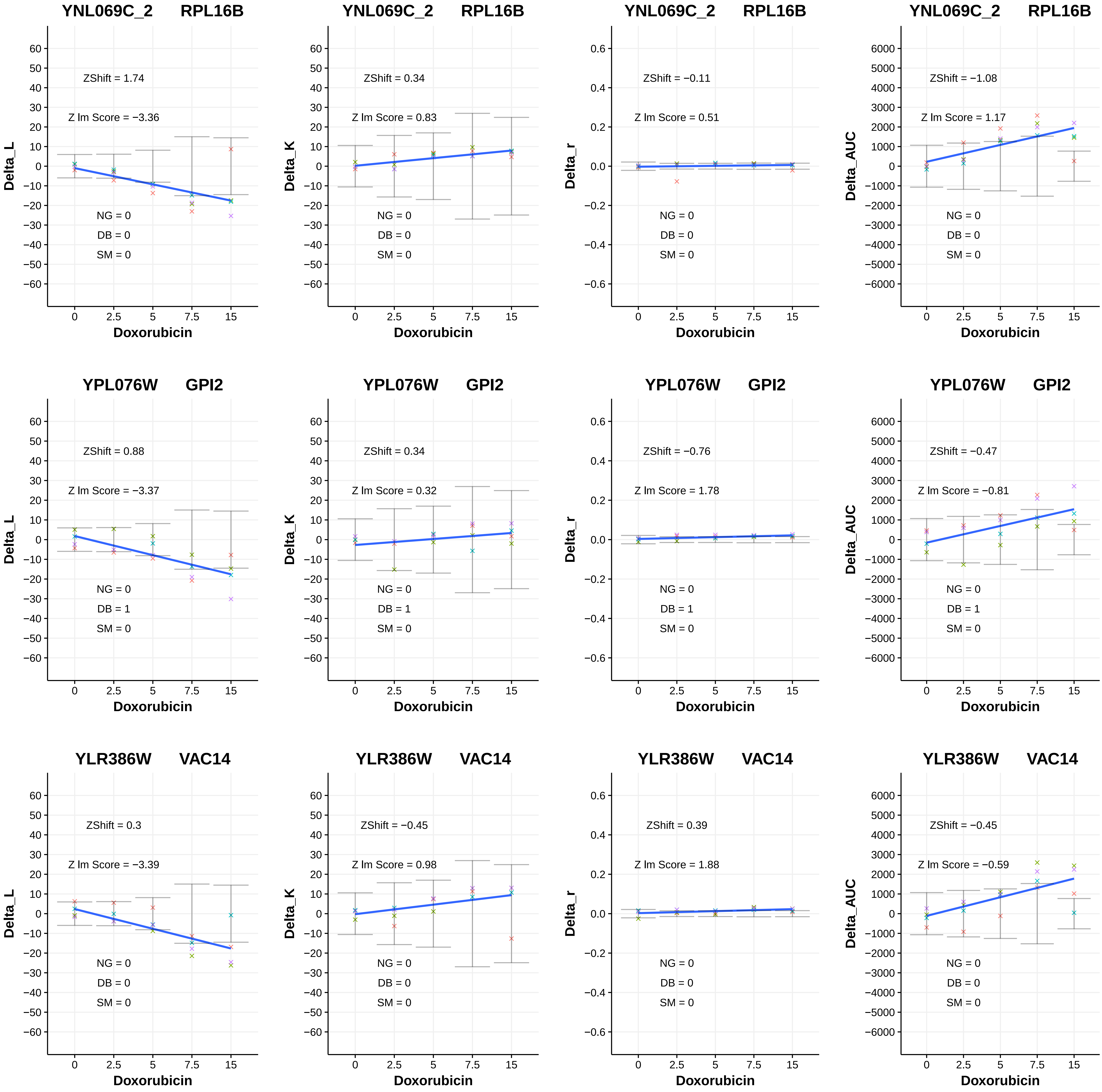

YNL069C_2 RPL16B
ZShift = 1.74
Z lm Score = −3.36
NG = 0
DB = 0
SM = 0
Delta_L
Doxorubicin
YNL069C_2 RPL16B
ZShift = 0.34
Z lm Score = 0.83
NG = 0
DB = 0
SM = 0
Delta_K
Doxorubicin
YNL069C_2 RPL16B
ZShift = −0.11
Z lm Score = 0.51
NG = 0
DB = 0
SM = 0
Delta_r
Doxorubicin
YNL069C_2 RPL16B
ZShift = −1.08
Z lm Score = 1.17
NG = 0
DB = 0
SM = 0
Delta_AUC
Doxorubicin
YPL076W GPI2
ZShift = 0.88
Z lm Score = −3.37
NG = 0
DB = 1
SM = 0
Delta_L
Doxorubicin
YPL076W GPI2
ZShift = 0.34
Z lm Score = 0.32
NG = 0
DB = 1
SM = 0
Delta_K
Doxorubicin
YPL076W GPI2
ZShift = −0.76
Z lm Score = 1.78
NG = 0
DB = 1
SM = 0
Delta_r
Doxorubicin
YPL076W GPI2
ZShift = −0.47
Z lm Score = −0.81
NG = 0
DB = 1
SM = 0
Delta_AUC
Doxorubicin
YLR386W VAC14
ZShift = 0.3
Z lm Score = −3.39
NG = 0
DB = 0
SM = 0
Delta_L
Doxorubicin
YLR386W VAC14
ZShift = −0.45
Z lm Score = 0.98
NG = 0
DB = 0
SM = 0
Delta_K
Doxorubicin
YLR386W VAC14
ZShift = 0.39
Z lm Score = 1.88
NG = 0
DB = 0
SM = 0
Delta_r
Doxorubicin
YLR386W VAC14
ZShift = −0.45
Z lm Score = −0.59
NG = 0
DB = 0
SM = 0
Delta_AUC
Doxorubicin

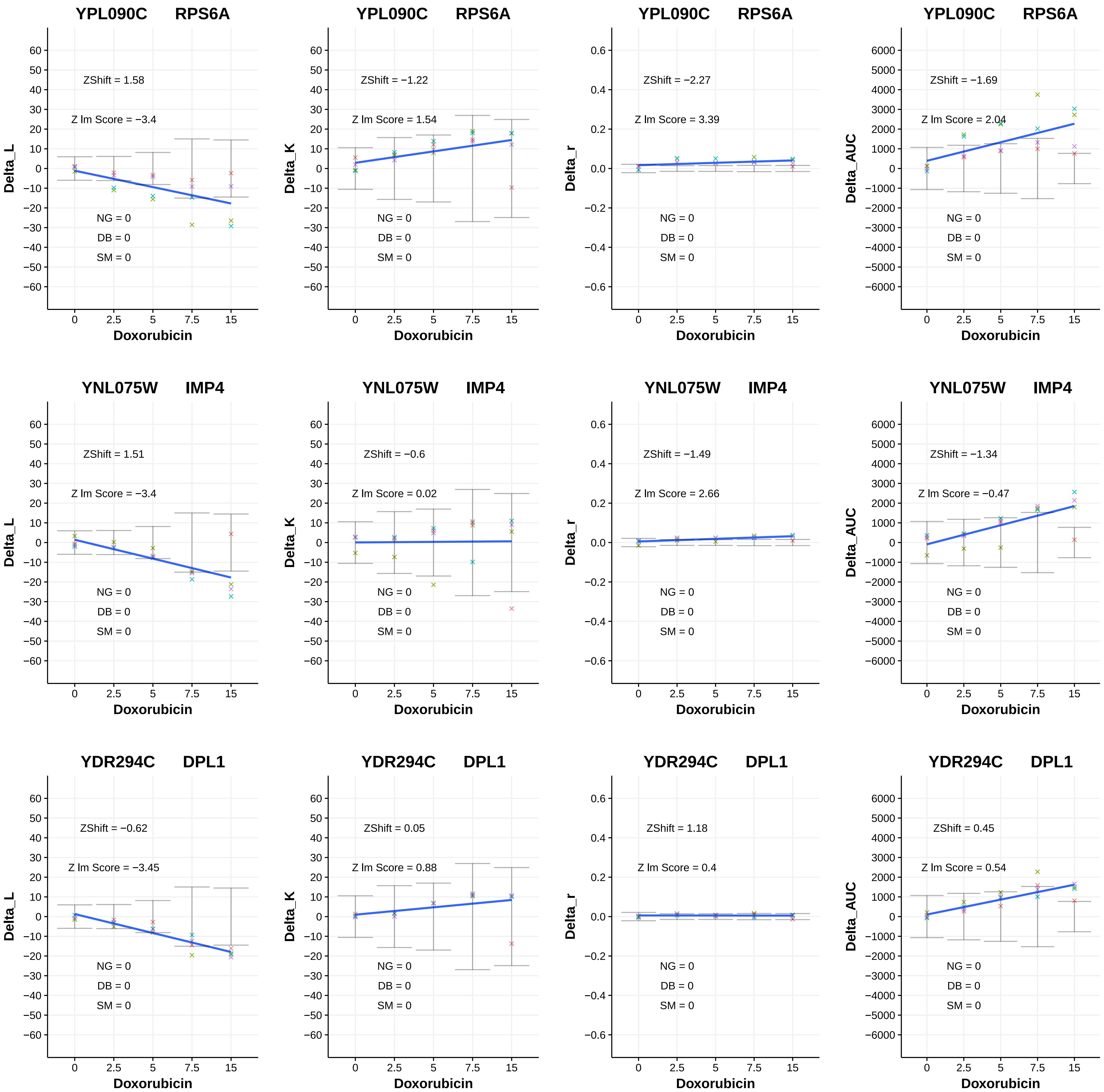

YPL090C RPS6A
ZShift = 1.58
Z lm Score = −3.4
NG = 0
DB = 0
SM = 0
Delta_L
Doxorubicin
YPL090C RPS6A
ZShift = −1.22
Z lm Score = 1.54
NG = 0
DB = 0
SM = 0
Delta_K
Doxorubicin
YPL090C RPS6A
ZShift = −2.27
Z lm Score = 3.39
NG = 0
DB = 0
SM = 0
Delta_r
Doxorubicin
YPL090C RPS6A
ZShift = −1.69
Z lm Score = 2.04
NG = 0
DB = 0
SM = 0
Delta_AUC
Doxorubicin
YNL075W IMP4
ZShift = 1.51
Z lm Score = −3.4
NG = 0
DB = 0
SM = 0
Delta_L
Doxorubicin
YNL075W IMP4
ZShift = −0.6
Z lm Score = 0.02
NG = 0
DB = 0
SM = 0
Delta_K
Doxorubicin
YNL075W IMP4
ZShift = −1.49
Z lm Score = 2.66
NG = 0
DB = 0
SM = 0
Delta_r
Doxorubicin
YNL075W IMP4
ZShift = −1.34
Z lm Score = −0.47
NG = 0
DB = 0
SM = 0
Delta_AUC
Doxorubicin
YDR294C DPL1
ZShift = −0.62
Z lm Score = −3.45
NG = 0
DB = 0
SM = 0
Delta_L
Doxorubicin
YDR294C DPL1
ZShift = 0.05
Z lm Score = 0.88
NG = 0
DB = 0
SM = 0
Delta_K
Doxorubicin
YDR294C DPL1
ZShift = 1.18
Z lm Score = 0.4
NG = 0
DB = 0
SM = 0
Delta_r
Doxorubicin
YDR294C DPL1
ZShift = 0.45
Z lm Score = 0.54
NG = 0
DB = 0
SM = 0
Delta_AUC
Doxorubicin

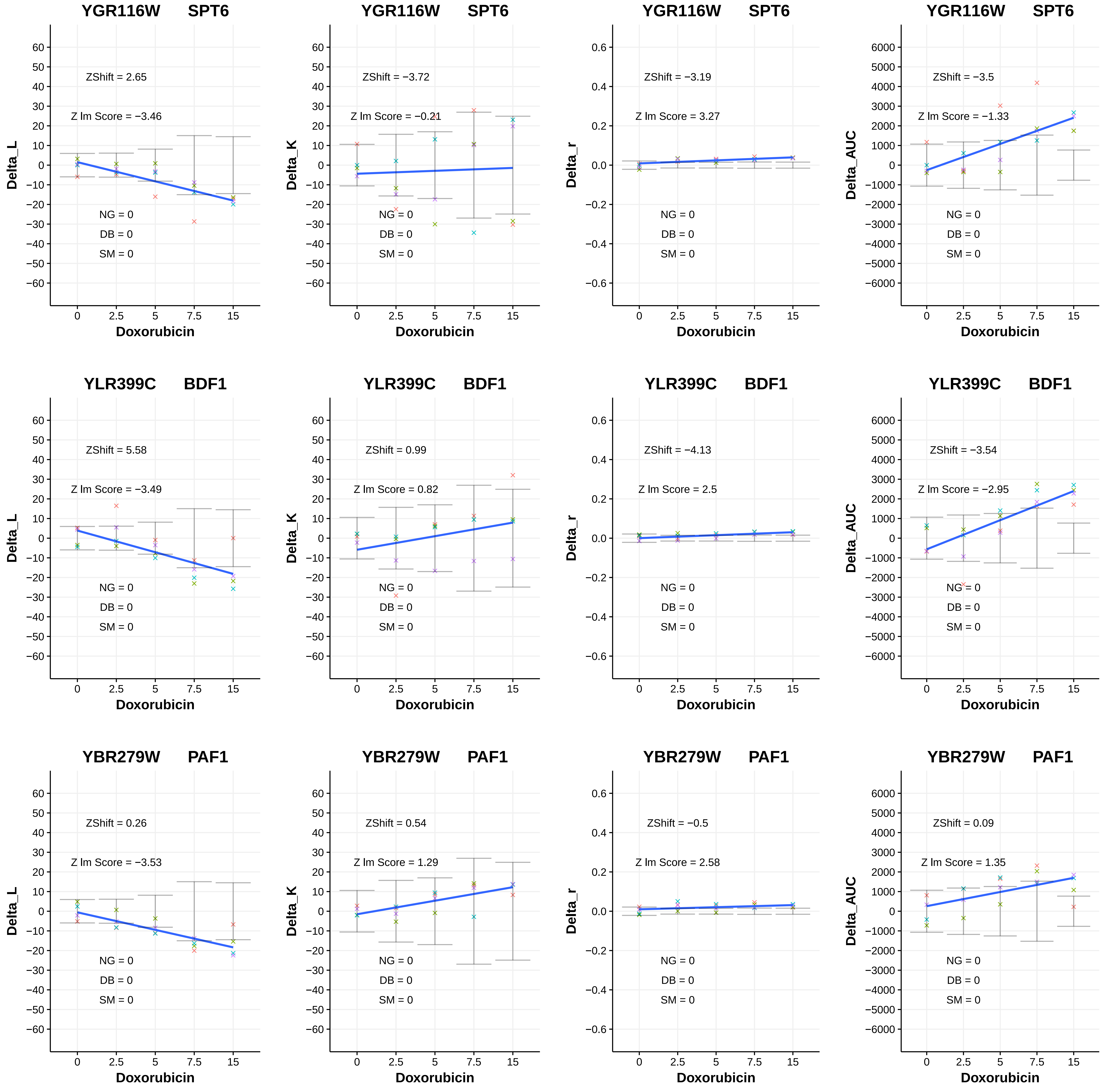

YGR116W SPT6
ZShift = 2.65
Z lm Score = −3.46
NG = 0
DB = 0
SM = 0
Delta_L
Doxorubicin
YGR116W SPT6
ZShift = −3.72
Z lm Score = −0.21
NG = 0
DB = 0
SM = 0
Delta_K
Doxorubicin
YGR116W SPT6
ZShift = −3.19
Z lm Score = 3.27
NG = 0
DB = 0
SM = 0
Delta_r
Doxorubicin
YGR116W SPT6
ZShift = −3.5
Z lm Score = −1.33
NG = 0
DB = 0
SM = 0
Delta_AUC
Doxorubicin
YLR399C BDF1
ZShift = 5.58
Z lm Score = −3.49
NG = 0
DB = 0
SM = 0
Delta_L
Doxorubicin
YLR399C BDF1
ZShift = 0.99
Z lm Score = 0.82
NG = 0
DB = 0
SM = 0
Delta_K
Doxorubicin
YLR399C BDF1
ZShift = −4.13
Z lm Score = 2.5
NG = 0
DB = 0
SM = 0
Delta_r
Doxorubicin
YLR399C BDF1
ZShift = −3.54
Z lm Score = −2.95
NG = 0
DB = 0
SM = 0
Delta_AUC
Doxorubicin
YBR279W PAF1
ZShift = 0.26
Z lm Score = −3.53
NG = 0
DB = 0
SM = 0
Delta_L
Doxorubicin
YBR279W PAF1
ZShift = 0.54
Z lm Score = 1.29
NG = 0
DB = 0
SM = 0
Delta_K
Doxorubicin
YBR279W PAF1
ZShift = −0.5
Z lm Score = 2.58
NG = 0
DB = 0
SM = 0
Delta_r
Doxorubicin
YBR279W PAF1
ZShift = 0.09
Z lm Score = 1.35
NG = 0
DB = 0
SM = 0
Delta_AUC
Doxorubicin

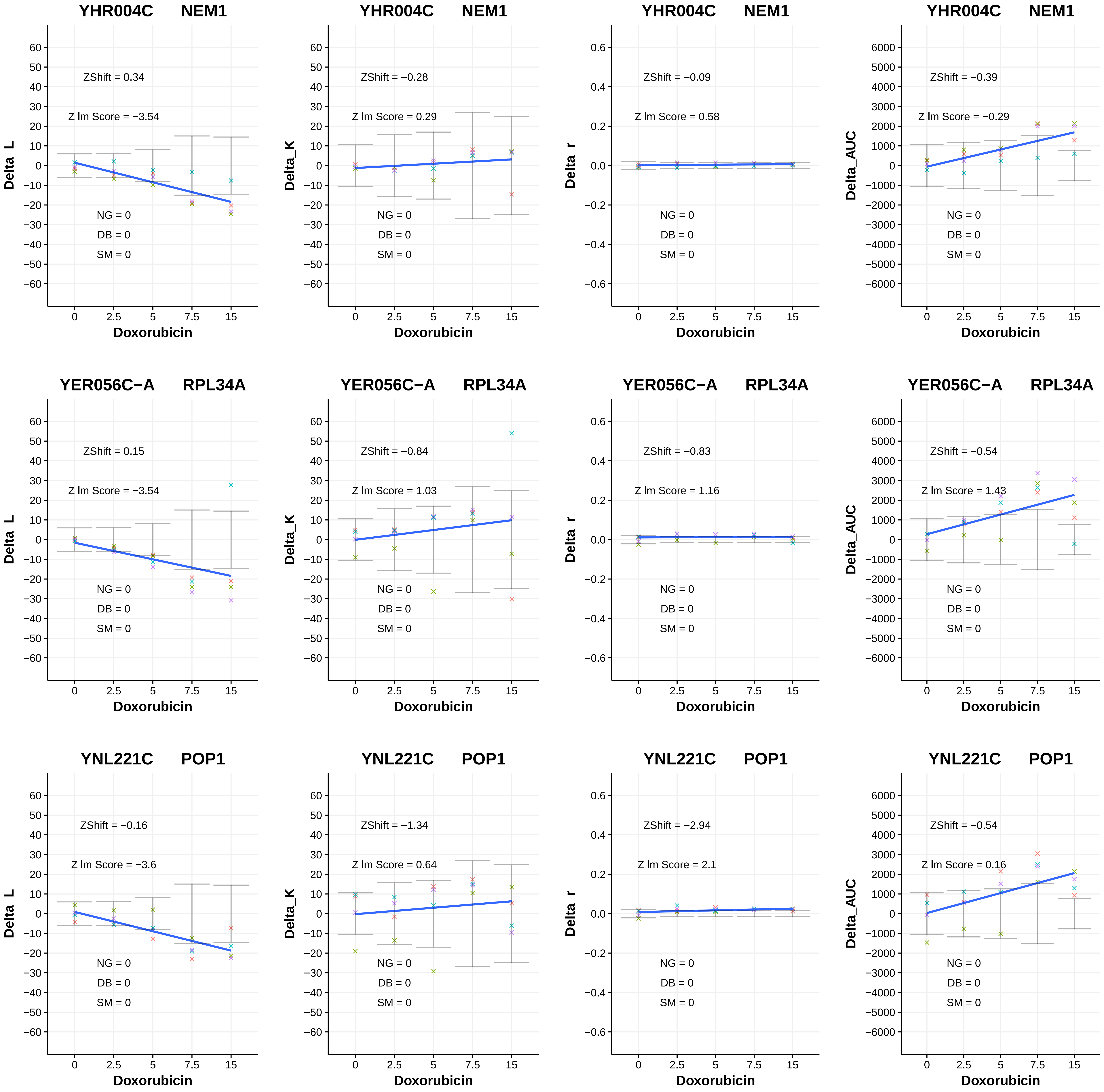

YHR004C NEM1
ZShift = 0.34
Z lm Score = −3.54
NG = 0
DB = 0
SM = 0
Delta_L
Doxorubicin
YHR004C NEM1
ZShift = −0.28
Z lm Score = 0.29
NG = 0
DB = 0
SM = 0
Delta_K
Doxorubicin
YHR004C NEM1
ZShift = −0.09
Z lm Score = 0.58
NG = 0
DB = 0
SM = 0
Delta_r
Doxorubicin
YHR004C NEM1
ZShift = −0.39
Z lm Score = −0.29
NG = 0
DB = 0
SM = 0
Delta_AUC
Doxorubicin
YER056C−A RPL34A
ZShift = 0.15
Z lm Score = −3.54
NG = 0
DB = 0
SM = 0
Delta_L
Doxorubicin
YER056C−A RPL34A
ZShift = −0.84
Z lm Score = 1.03
NG = 0
DB = 0
SM = 0
Delta_K
Doxorubicin
YER056C−A RPL34A
ZShift = −0.83
Z lm Score = 1.16
NG = 0
DB = 0
SM = 0
Delta_r
Doxorubicin
YER056C−A RPL34A
ZShift = −0.54
Z lm Score = 1.43
NG = 0
DB = 0
SM = 0
Delta_AUC
Doxorubicin
YNL221C POP1
ZShift = −0.16
Z lm Score = −3.6
NG = 0
DB = 0
SM = 0
Delta_L
Doxorubicin
YNL221C POP1
ZShift = −1.34
Z lm Score = 0.64
NG = 0
DB = 0
SM = 0
Delta_K
Doxorubicin
YNL221C POP1
ZShift = −2.94
Z lm Score = 2.1
NG = 0
DB = 0
SM = 0
Delta_r
Doxorubicin
YNL221C POP1
ZShift = −0.54
Z lm Score = 0.16
NG = 0
DB = 0
SM = 0
Delta_AUC
Doxorubicin

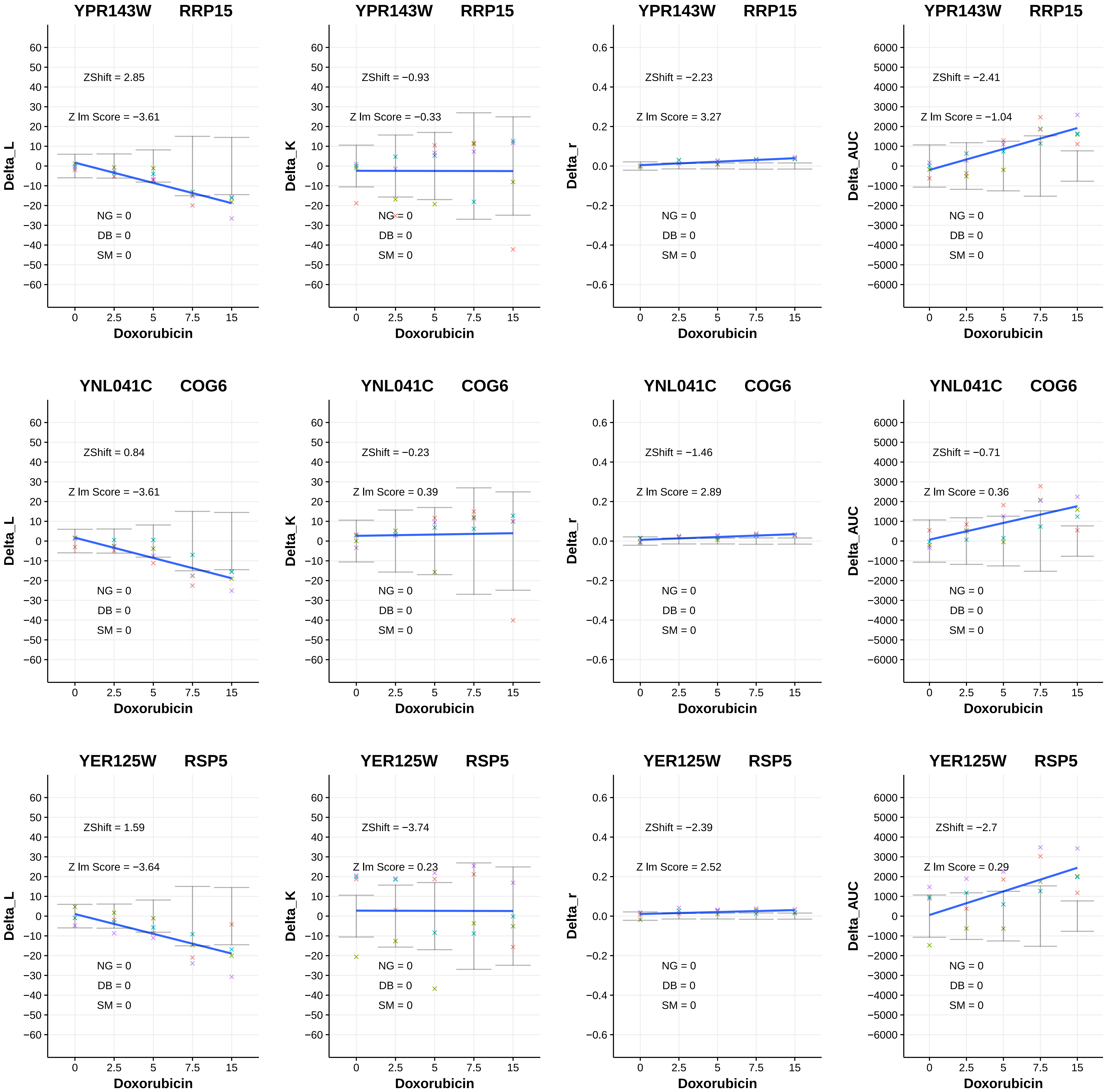

YPR143W RRP15
ZShift = 2.85
Z lm Score = −3.61
NG = 0
DB = 0
SM = 0
Delta_L
Doxorubicin
YPR143W RRP15
ZShift = −0.93
Z lm Score = −0.33
NG = 0
DB = 0
SM = 0
Delta_K
Doxorubicin
YPR143W RRP15
ZShift = −2.23
Z lm Score = 3.27
NG = 0
DB = 0
SM = 0
Delta_r
Doxorubicin
YPR143W RRP15
ZShift = −2.41
Z lm Score = −1.04
NG = 0
DB = 0
SM = 0
Delta_AUC
Doxorubicin
YNL041C COG6
ZShift = 0.84
Z lm Score = −3.61
NG = 0
DB = 0
SM = 0
Delta_L
Doxorubicin
YNL041C COG6
ZShift = −0.23
Z lm Score = 0.39
NG = 0
DB = 0
SM = 0
Delta_K
Doxorubicin
YNL041C COG6
ZShift = −1.46
Z lm Score = 2.89
NG = 0
DB = 0
SM = 0
Delta_r
Doxorubicin
YNL041C COG6
ZShift = −0.71
Z lm Score = 0.36
NG = 0
DB = 0
SM = 0
Delta_AUC
Doxorubicin
YER125W RSP5
ZShift = 1.59
Z lm Score = −3.64
NG = 0
DB = 0
SM = 0
Delta_L
Doxorubicin
YER125W RSP5
ZShift = −3.74
Z lm Score = 0.23
NG = 0
DB = 0
SM = 0
Delta_K
Doxorubicin
YER125W RSP5
ZShift = −2.39
Z lm Score = 2.52
NG = 0
DB = 0
SM = 0
Delta_r
Doxorubicin
YER125W RSP5
ZShift = −2.7
Z lm Score = 0.29
NG = 0
DB = 0
SM = 0
Delta_AUC
Doxorubicin

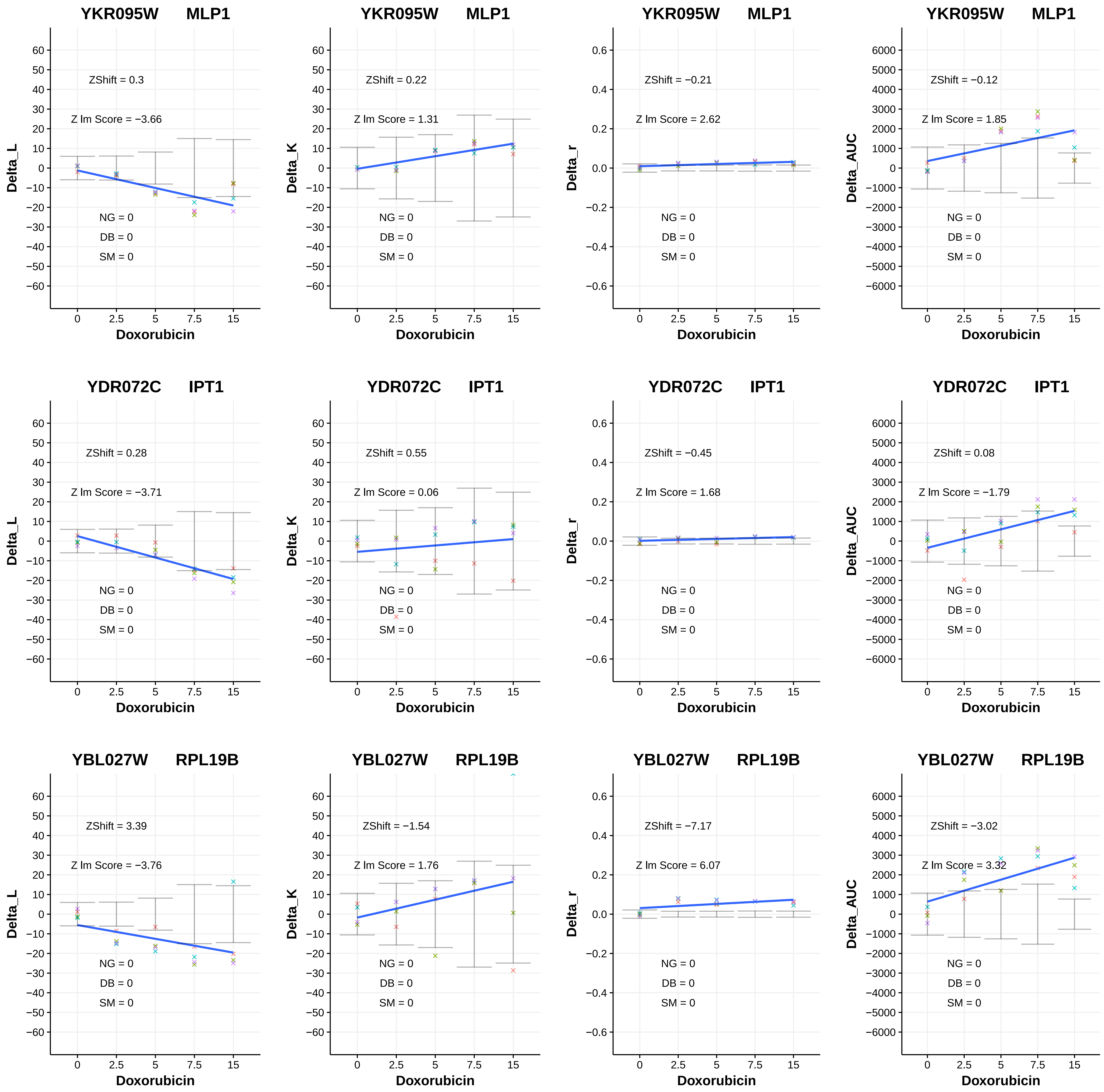
YKR095W MLP1
ZShift = 0.3
Z lm Score = −3.66
NG = 0
DB = 0
SM = 0
Delta_L
Doxorubicin
YKR095W MLP1
ZShift = 0.22
Z lm Score = 1.31
NG = 0
DB = 0
SM = 0
Delta_K
Doxorubicin
YKR095W MLP1
ZShift = −0.21
Z lm Score = 2.62
NG = 0
DB = 0
SM = 0
Delta_r
Doxorubicin
YKR095W MLP1
ZShift = −0.12
Z lm Score = 1.85
NG = 0
DB = 0
SM = 0
Delta_AUC
Doxorubicin
YDR072C IPT1
ZShift = 0.28
Z lm Score = −3.71
NG = 0
DB = 0
SM = 0
Delta_L
Doxorubicin
YDR072C IPT1
ZShift = 0.55
Z lm Score = 0.06
NG = 0
DB = 0
SM = 0
Delta_K
Doxorubicin
YDR072C IPT1
ZShift = −0.45
Z lm Score = 1.68
NG = 0
DB = 0
SM = 0
Delta_r
Doxorubicin
YDR072C IPT1
ZShift = 0.08
Z lm Score = −1.79
NG = 0
DB = 0
SM = 0
Delta_AUC
Doxorubicin
YBL027W RPL19B
ZShift = 3.39
Z lm Score = −3.76
NG = 0
DB = 0
SM = 0
Delta_L
Doxorubicin
YBL027W RPL19B
ZShift = −1.54
Z lm Score = 1.76
NG = 0
DB = 0
SM = 0
Delta_K
Doxorubicin
YBL027W RPL19B
ZShift = −7.17
Z lm Score = 6.07
NG = 0
DB = 0
SM = 0
Delta_r
Doxorubicin
YBL027W RPL19B
ZShift = −3.02
Z lm Score = 3.32
NG = 0
DB = 0
SM = 0
Delta_AUC
Doxorubicin

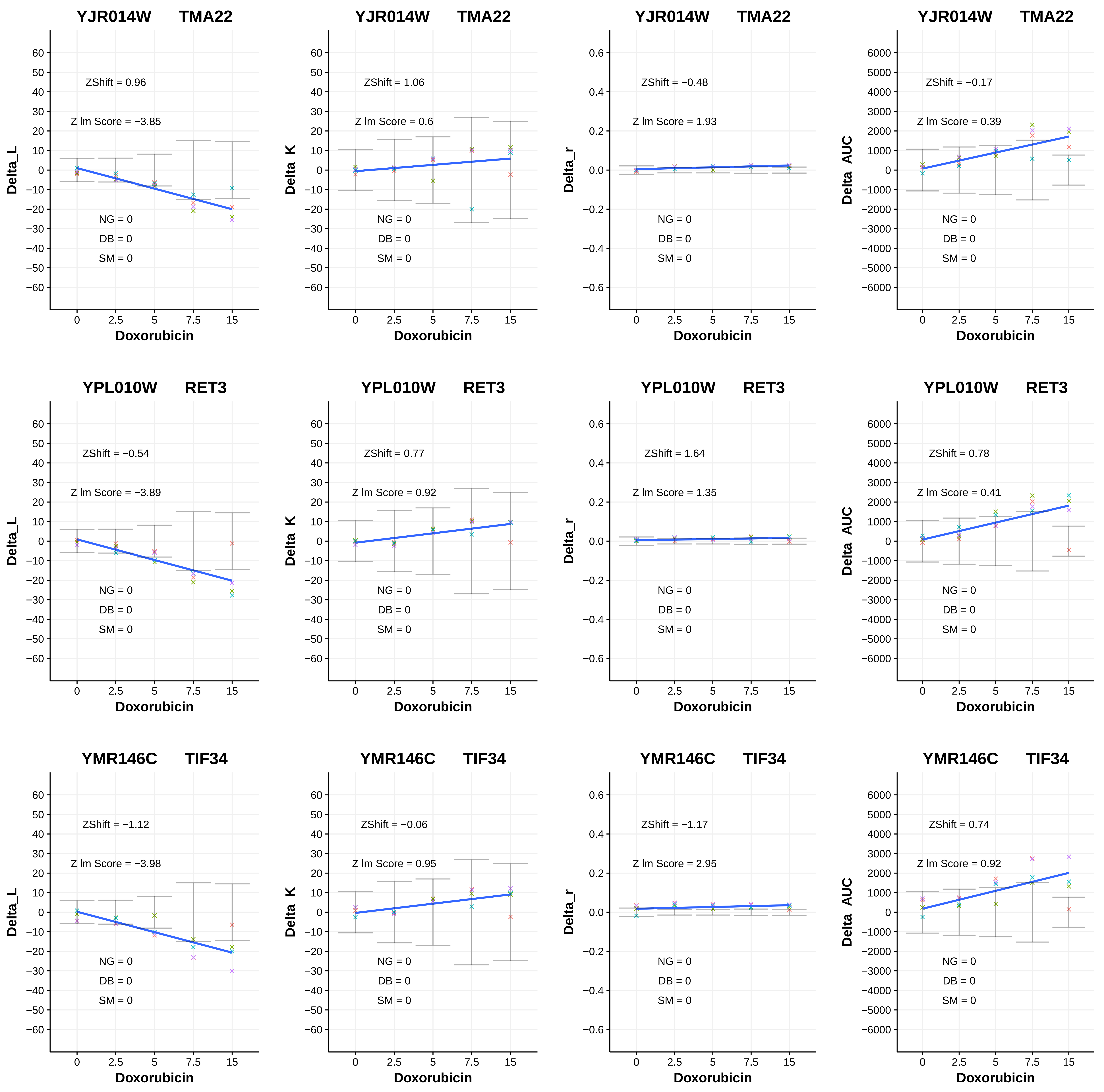

YJR014W TMA22
ZShift = 0.96
Z lm Score = −3.85
NG = 0
DB = 0
SM = 0
Delta_L
Doxorubicin
YJR014W TMA22
ZShift = 1.06
Z lm Score = 0.6
NG = 0
DB = 0
SM = 0
Delta_K
Doxorubicin
YJR014W TMA22
ZShift = −0.48
Z lm Score = 1.93
NG = 0
DB = 0
SM = 0
Delta_r
Doxorubicin
YJR014W TMA22
ZShift = −0.17
Z lm Score = 0.39
NG = 0
DB = 0
SM = 0
Delta_AUC
Doxorubicin
YPL010W RET3
ZShift = −0.54
Z lm Score = −3.89
NG = 0
DB = 0
SM = 0
Delta_L
Doxorubicin
YPL010W RET3
ZShift = 0.77
Z lm Score = 0.92
NG = 0
DB = 0
SM = 0
Delta_K
Doxorubicin
YPL010W RET3
ZShift = 1.64
Z lm Score = 1.35
NG = 0
DB = 0
SM = 0
Delta_r
Doxorubicin
YPL010W RET3
ZShift = 0.78
Z lm Score = 0.41
NG = 0
DB = 0
SM = 0
Delta_AUC
Doxorubicin
YMR146C TIF34
ZShift = −1.12
Z lm Score = −3.98
NG = 0
DB = 0
SM = 0
Delta_L
Doxorubicin
YMR146C TIF34
ZShift = −0.06
Z lm Score = 0.95
NG = 0
DB = 0
SM = 0
Delta_K
Doxorubicin
YMR146C TIF34
ZShift = −1.17
Z lm Score = 2.95
NG = 0
DB = 0
SM = 0
Delta_r
Doxorubicin
YMR146C TIF34
ZShift = 0.74
Z lm Score = 0.92
NG = 0
DB = 0
SM = 0
Delta_AUC
Doxorubicin

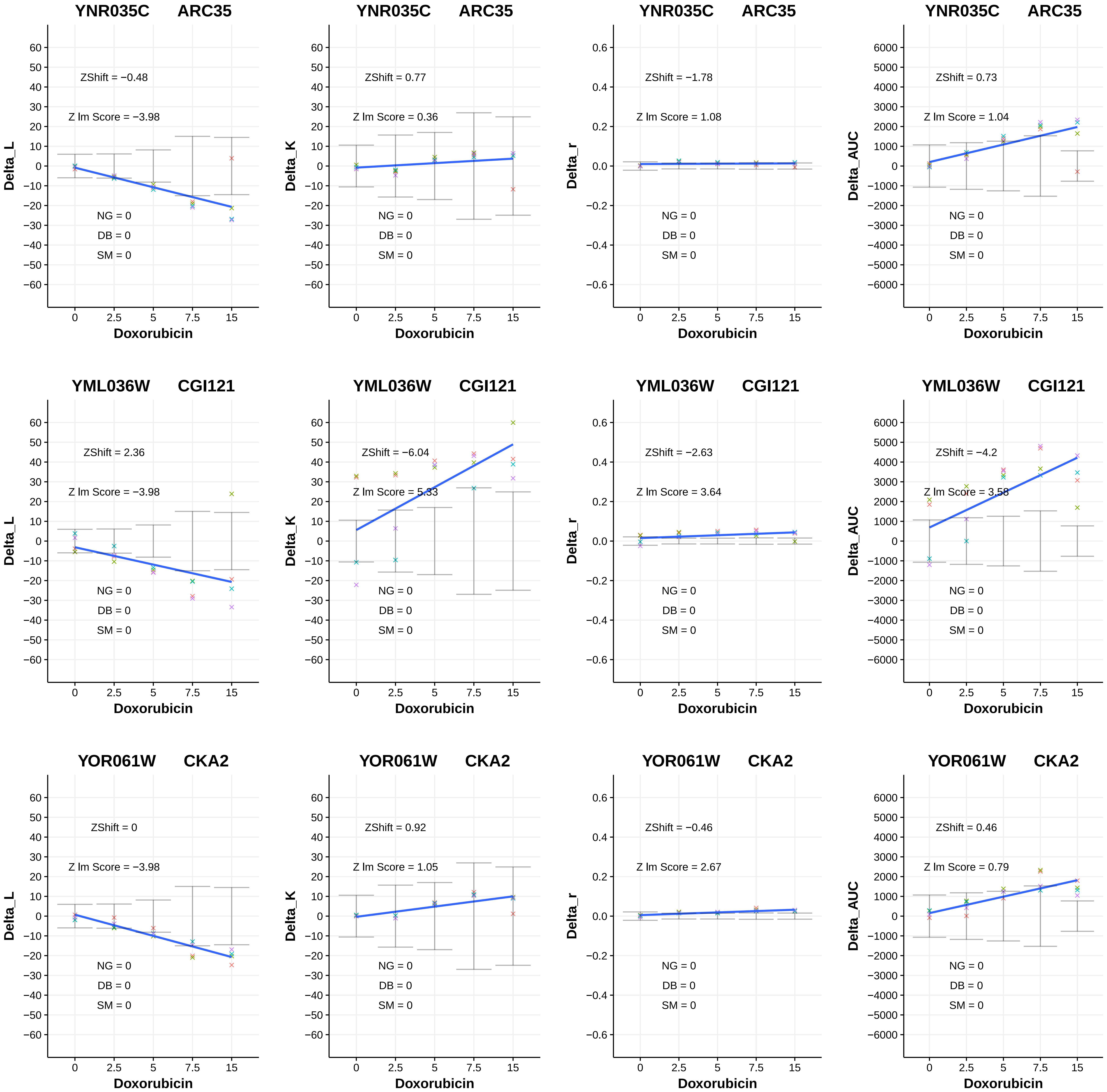

YNR035C ARC35
ZShift = −0.48
Z lm Score = −3.98
NG = 0
DB = 0
SM = 0
Delta_L
Doxorubicin
YNR035C ARC35
ZShift = 0.77
Z lm Score = 0.36
NG = 0
DB = 0
SM = 0
Delta_K
Doxorubicin
YNR035C ARC35
ZShift = −1.78
Z lm Score = 1.08
NG = 0
DB = 0
SM = 0
Delta_r
Doxorubicin
YNR035C ARC35
ZShift = 0.73
Z lm Score = 1.04
NG = 0
DB = 0
SM = 0
Delta_AUC
Doxorubicin
YML036W CGI121
ZShift = 2.36
Z lm Score = −3.98
NG = 0
DB = 0
SM = 0
Delta_L
Doxorubicin
YML036W CGI121
ZShift = −6.04
Z lm Score = 5.33
NG = 0
DB = 0
SM = 0
Delta_K
Doxorubicin
YML036W CGI121
ZShift = −2.63
Z lm Score = 3.64
NG = 0
DB = 0
SM = 0
Delta_r
Doxorubicin
YML036W CGI121
ZShift = −4.2
Z lm Score = 3.58
NG = 0
DB = 0
SM = 0
Delta_AUC
Doxorubicin
YOR061W CKA2
ZShift = 0
Z lm Score = −3.98
NG = 0
DB = 0
SM = 0
Delta_L
Doxorubicin
YOR061W CKA2
ZShift = 0.92
Z lm Score = 1.05
NG = 0
DB = 0
SM = 0
Delta_K
Doxorubicin
YOR061W CKA2
ZShift = −0.46
Z lm Score = 2.67
NG = 0
DB = 0
SM = 0
Delta_r
Doxorubicin
YOR061W CKA2
ZShift = 0.46
Z lm Score = 0.79
NG = 0
DB = 0
SM = 0
Delta_AUC
Doxorubicin

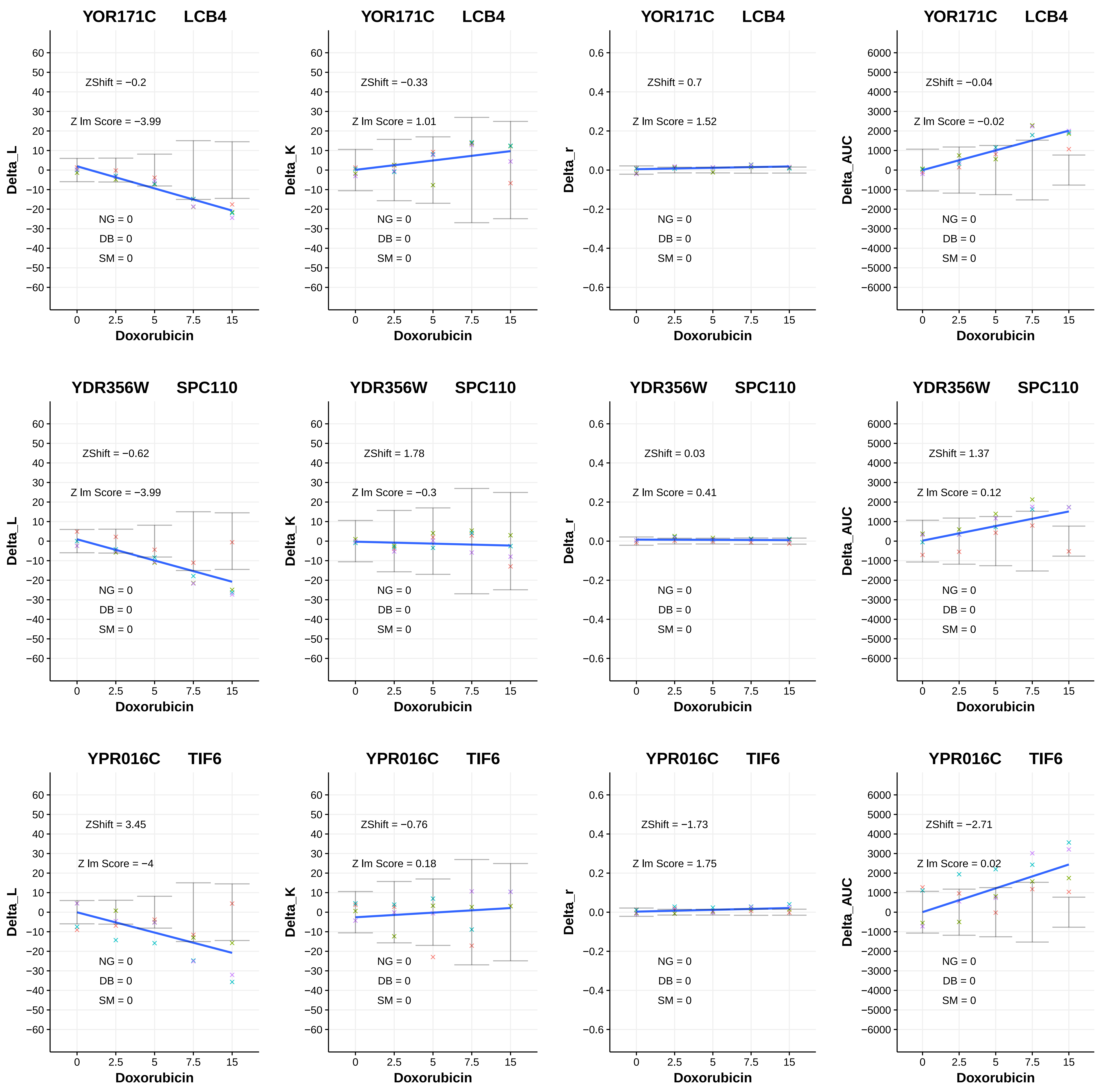

YOR171C LCB4
ZShift = −0.2
Z lm Score = −3.99
NG = 0
DB = 0
SM = 0
Delta_L
Doxorubicin
YOR171C LCB4
ZShift = −0.33
Z lm Score = 1.01
NG = 0
DB = 0
SM = 0
Delta_K
Doxorubicin
YOR171C LCB4
ZShift = 0.7
Z lm Score = 1.52
NG = 0
DB = 0
SM = 0
Delta_r
Doxorubicin
YOR171C LCB4
ZShift = −0.04
Z lm Score = −0.02
NG = 0
DB = 0
SM = 0
Delta_AUC
Doxorubicin
YDR356W SPC110
ZShift = −0.62
Z lm Score = −3.99
NG = 0
DB = 0
SM = 0
Delta_L
Doxorubicin
YDR356W SPC110
ZShift = 1.78
Z lm Score = −0.3
NG = 0
DB = 0
SM = 0
Delta_K
Doxorubicin
YDR356W SPC110
ZShift = 0.03
Z lm Score = 0.41
NG = 0
DB = 0
SM = 0
Delta_r
Doxorubicin
YDR356W SPC110
ZShift = 1.37
Z lm Score = 0.12
NG = 0
DB = 0
SM = 0
Delta_AUC
Doxorubicin
YPR016C TIF6
ZShift = 3.45
Z lm Score = −4
NG = 0
DB = 0
SM = 0
Delta_L
Doxorubicin
YPR016C TIF6
ZShift = −0.76
Z lm Score = 0.18
NG = 0
DB = 0
SM = 0
Delta_K
Doxorubicin
YPR016C TIF6
ZShift = −1.73
Z lm Score = 1.75
NG = 0
DB = 0
SM = 0
Delta_r
Doxorubicin
YPR016C TIF6
ZShift = −2.71
Z lm Score = 0.02
NG = 0
DB = 0
SM = 0
Delta_AUC
Doxorubicin

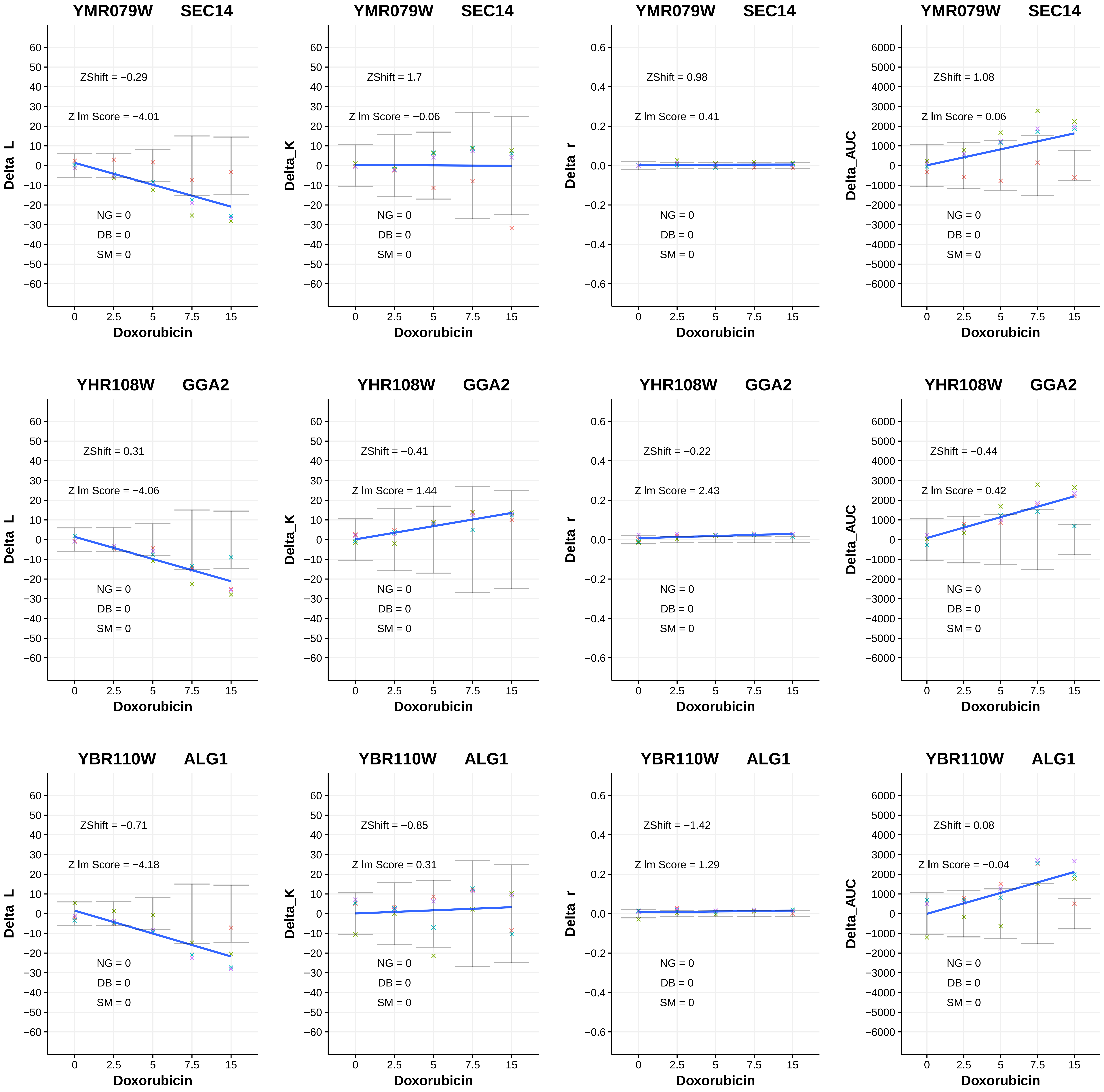

YMR079W SEC14
ZShift = −0.29
Z lm Score = −4.01
NG = 0
DB = 0
SM = 0
Delta_L
Doxorubicin
YMR079W SEC14
ZShift = 1.7
Z lm Score = −0.06
NG = 0
DB = 0
SM = 0
Delta_K
Doxorubicin
YMR079W SEC14
ZShift = 0.98
Z lm Score = 0.41
NG = 0
DB = 0
SM = 0
Delta_r
Doxorubicin
YMR079W SEC14
ZShift = 1.08
Z lm Score = 0.06
NG = 0
DB = 0
SM = 0
Delta_AUC
Doxorubicin
YHR108W GGA2
ZShift = 0.31
Z lm Score = −4.06
NG = 0
DB = 0
SM = 0
Delta_L
Doxorubicin
YHR108W GGA2
ZShift = −0.41
Z lm Score = 1.44
NG = 0
DB = 0
SM = 0
Delta_K
Doxorubicin
YHR108W GGA2
ZShift = −0.22
Z lm Score = 2.43
NG = 0
DB = 0
SM = 0
Delta_r
Doxorubicin
YHR108W GGA2
ZShift = −0.44
Z lm Score = 0.42
NG = 0
DB = 0
SM = 0
Delta_AUC
Doxorubicin
YBR110W ALG1
ZShift = −0.71
Z lm Score = −4.18
NG = 0
DB = 0
SM = 0
Delta_L
Doxorubicin
YBR110W ALG1
ZShift = −0.85
Z lm Score = 0.31
NG = 0
DB = 0
SM = 0
Delta_K
Doxorubicin
YBR110W ALG1
ZShift = −1.42
Z lm Score = 1.29
NG = 0
DB = 0
SM = 0
Delta_r
Doxorubicin
YBR110W ALG1
ZShift = 0.08
Z lm Score = −0.04
NG = 0
DB = 0
SM = 0
Delta_AUC
Doxorubicin

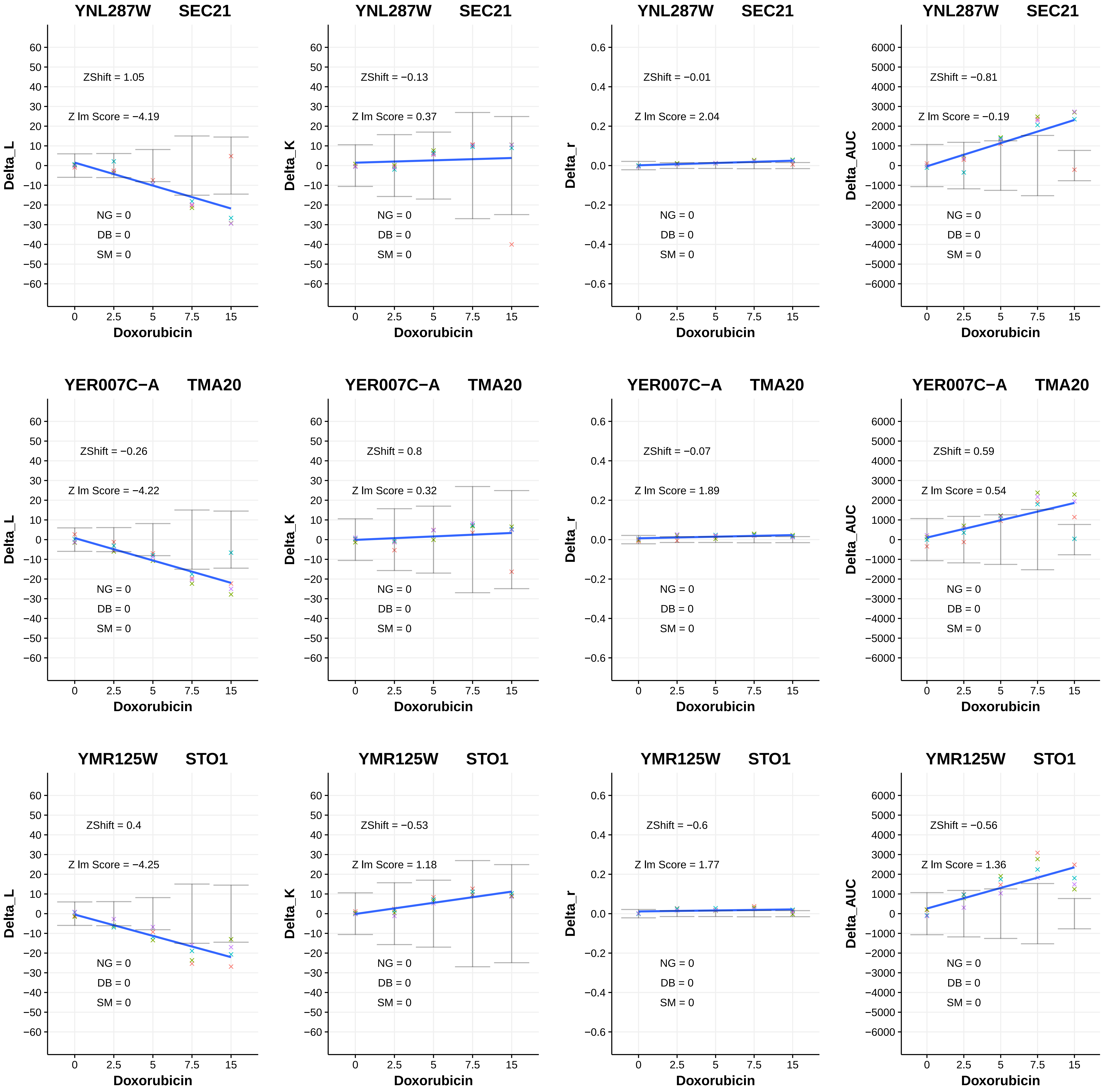

YNL287W SEC21
ZShift = 1.05
Z lm Score = −4.19
NG = 0
DB = 0
SM = 0
Delta_L
Doxorubicin
YNL287W SEC21
ZShift = −0.13
Z lm Score = 0.37
NG = 0
DB = 0
SM = 0
Delta_K
Doxorubicin
YNL287W SEC21
ZShift = −0.01
Z lm Score = 2.04
NG = 0
DB = 0
SM = 0
Delta_r
Doxorubicin
YNL287W SEC21
ZShift = −0.81
Z lm Score = −0.19
NG = 0
DB = 0
SM = 0
Delta_AUC
Doxorubicin
YER007C−A TMA20
ZShift = −0.26
Z lm Score = −4.22
NG = 0
DB = 0
SM = 0
Delta_L
Doxorubicin
YER007C−A TMA20
ZShift = 0.8
Z lm Score = 0.32
NG = 0
DB = 0
SM = 0
Delta_K
Doxorubicin
YER007C−A TMA20
ZShift = −0.07
Z lm Score = 1.89
NG = 0
DB = 0
SM = 0
Delta_r
Doxorubicin
YER007C−A TMA20
ZShift = 0.59
Z lm Score = 0.54
NG = 0
DB = 0
SM = 0
Delta_AUC
Doxorubicin
YMR125W STO1
ZShift = 0.4
Z lm Score = −4.25
NG = 0
DB = 0
SM = 0
Delta_L
Doxorubicin
YMR125W STO1
ZShift = −0.53
Z lm Score = 1.18
NG = 0
DB = 0
SM = 0
Delta_K
Doxorubicin
YMR125W STO1
ZShift = −0.6
Z lm Score = 1.77
NG = 0
DB = 0
SM = 0
Delta_r
Doxorubicin
YMR125W STO1
ZShift = −0.56
Z lm Score = 1.36
NG = 0
DB = 0
SM = 0
Delta_AUC
Doxorubicin

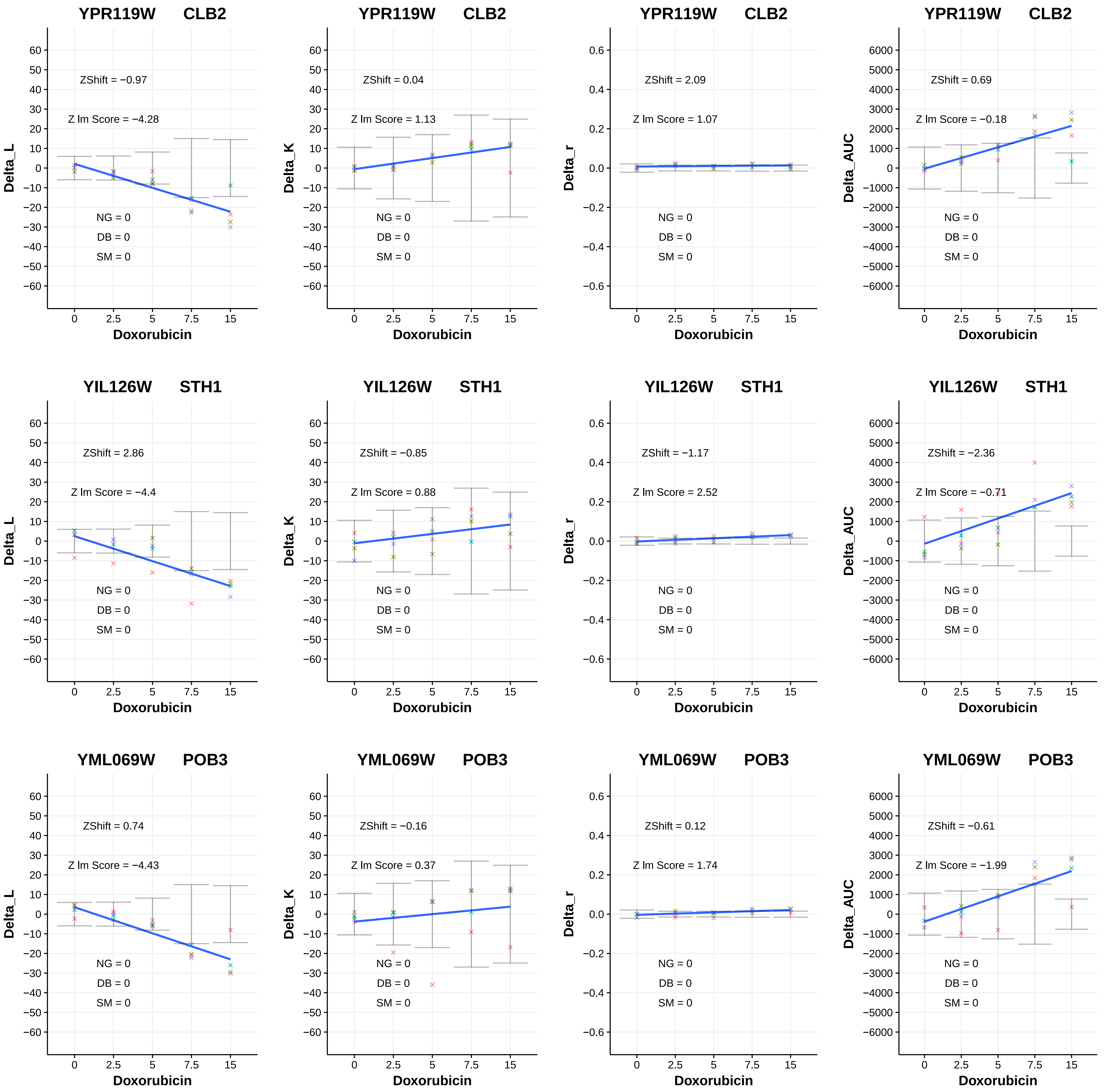

YPR119W CLB2
ZShift = −0.97
Z lm Score = −4.28
NG = 0
DB = 0
SM = 0
Delta_L
Doxorubicin
YPR119W CLB2
ZShift = 0.04
Z lm Score = 1.13
NG = 0
DB = 0
SM = 0
Delta_K
Doxorubicin
YPR119W CLB2
ZShift = 2.09
Z lm Score = 1.07
NG = 0
DB = 0
SM = 0
Delta_r
Doxorubicin
YPR119W CLB2
ZShift = 0.69
Z lm Score = −0.18
NG = 0
DB = 0
SM = 0
Delta_AUC
Doxorubicin
YIL126W STH1
ZShift = 2.86
Z lm Score = −4.4
NG = 0
DB = 0
SM = 0
Delta_L
Doxorubicin
YIL126W STH1
ZShift = −0.85
Z lm Score = 0.88
NG = 0
DB = 0
SM = 0
Delta_K
Doxorubicin
YIL126W STH1
ZShift = −1.17
Z lm Score = 2.52
NG = 0
DB = 0
SM = 0
Delta_r
Doxorubicin
YIL126W STH1
ZShift = −2.36
Z lm Score = −0.71
NG = 0
DB = 0
SM = 0
Delta_AUC
Doxorubicin
YML069W POB3
ZShift = 0.74
Z lm Score = −4.43
NG = 0
DB = 0
SM = 0
Delta_L
Doxorubicin
YML069W POB3
ZShift = −0.16
Z lm Score = 0.37
NG = 0
DB = 0
SM = 0
Delta_K
Doxorubicin
YML069W POB3
ZShift = 0.12
Z lm Score = 1.74
NG = 0
DB = 0
SM = 0
Delta_r
Doxorubicin
YML069W POB3
ZShift = −0.61
Z lm Score = −1.99
NG = 0
DB = 0
SM = 0
Delta_AUC
Doxorubicin

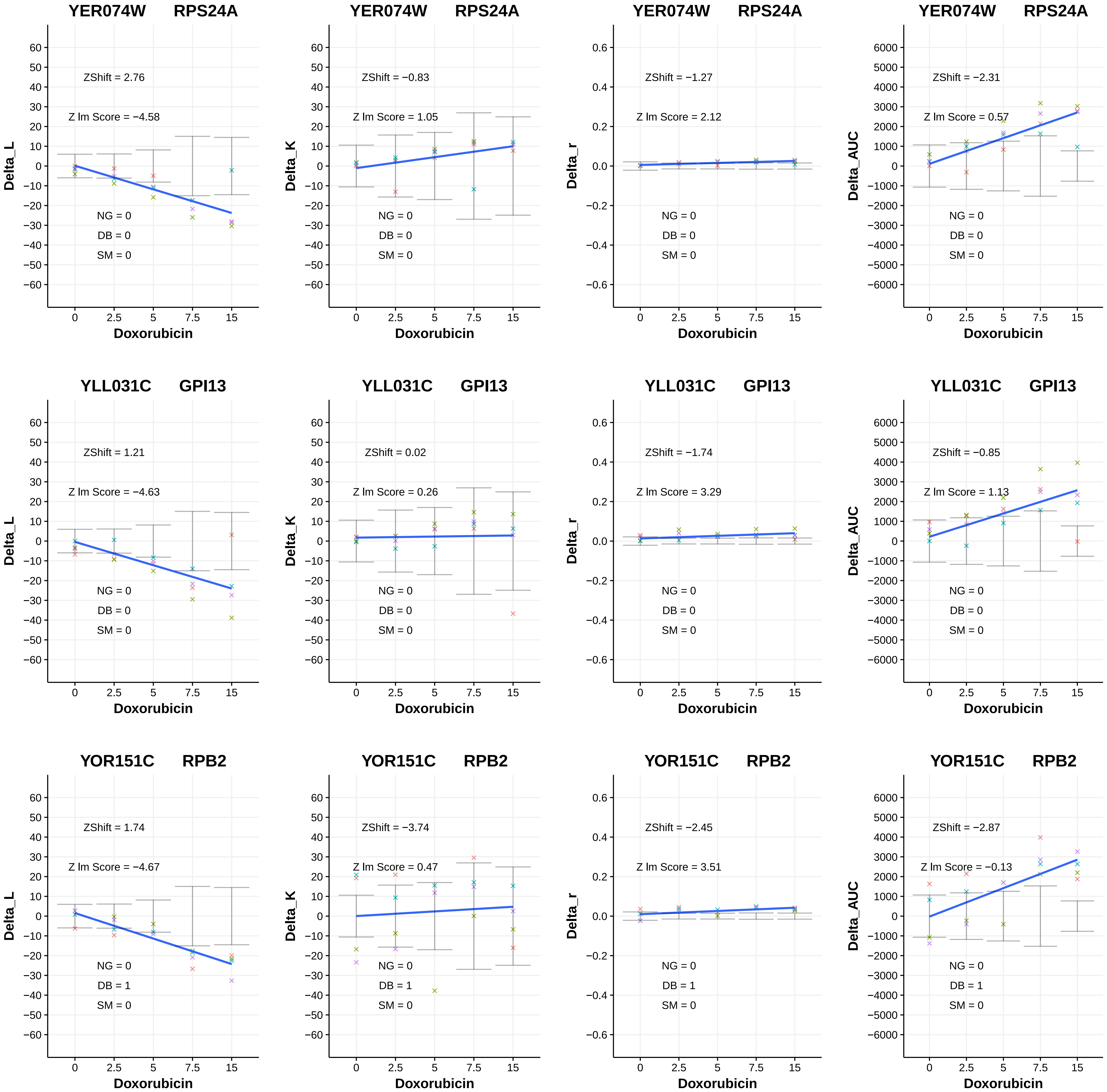

YER074W RPS24A
ZShift = 2.76
Z lm Score = −4.58
NG = 0
DB = 0
SM = 0
Delta_L
Doxorubicin
YER074W RPS24A
ZShift = −0.83
Z lm Score = 1.05
NG = 0
DB = 0
SM = 0
Delta_K
Doxorubicin
YER074W RPS24A
ZShift = −1.27
Z lm Score = 2.12
NG = 0
DB = 0
SM = 0
Delta_r
Doxorubicin
YER074W RPS24A
ZShift = −2.31
Z lm Score = 0.57
NG = 0
DB = 0
SM = 0
Delta_AUC
Doxorubicin
YLL031C GPI13
ZShift = 1.21
Z lm Score = −4.63
NG = 0
DB = 0
SM = 0
Delta_L
Doxorubicin
YLL031C GPI13
ZShift = 0.02
Z lm Score = 0.26
NG = 0
DB = 0
SM = 0
Delta_K
Doxorubicin
YLL031C GPI13
ZShift = −1.74
Z lm Score = 3.29
NG = 0
DB = 0
SM = 0
Delta_r
Doxorubicin
YLL031C GPI13
ZShift = −0.85
Z lm Score = 1.13
NG = 0
DB = 0
SM = 0
Delta_AUC
Doxorubicin
YOR151C RPB2
ZShift = 1.74
Z lm Score = −4.67
NG = 0
DB = 1
SM = 0
Delta_L
Doxorubicin
YOR151C RPB2
ZShift = −3.74
Z lm Score = 0.47
NG = 0
DB = 1
SM = 0
Delta_K
Doxorubicin
YOR151C RPB2
ZShift = −2.45
Z lm Score = 3.51
NG = 0
DB = 1
SM = 0
Delta_r
Doxorubicin
YOR151C RPB2
ZShift = −2.87
Z lm Score = −0.13
NG = 0
DB = 1
SM = 0
Delta_AUC
Doxorubicin

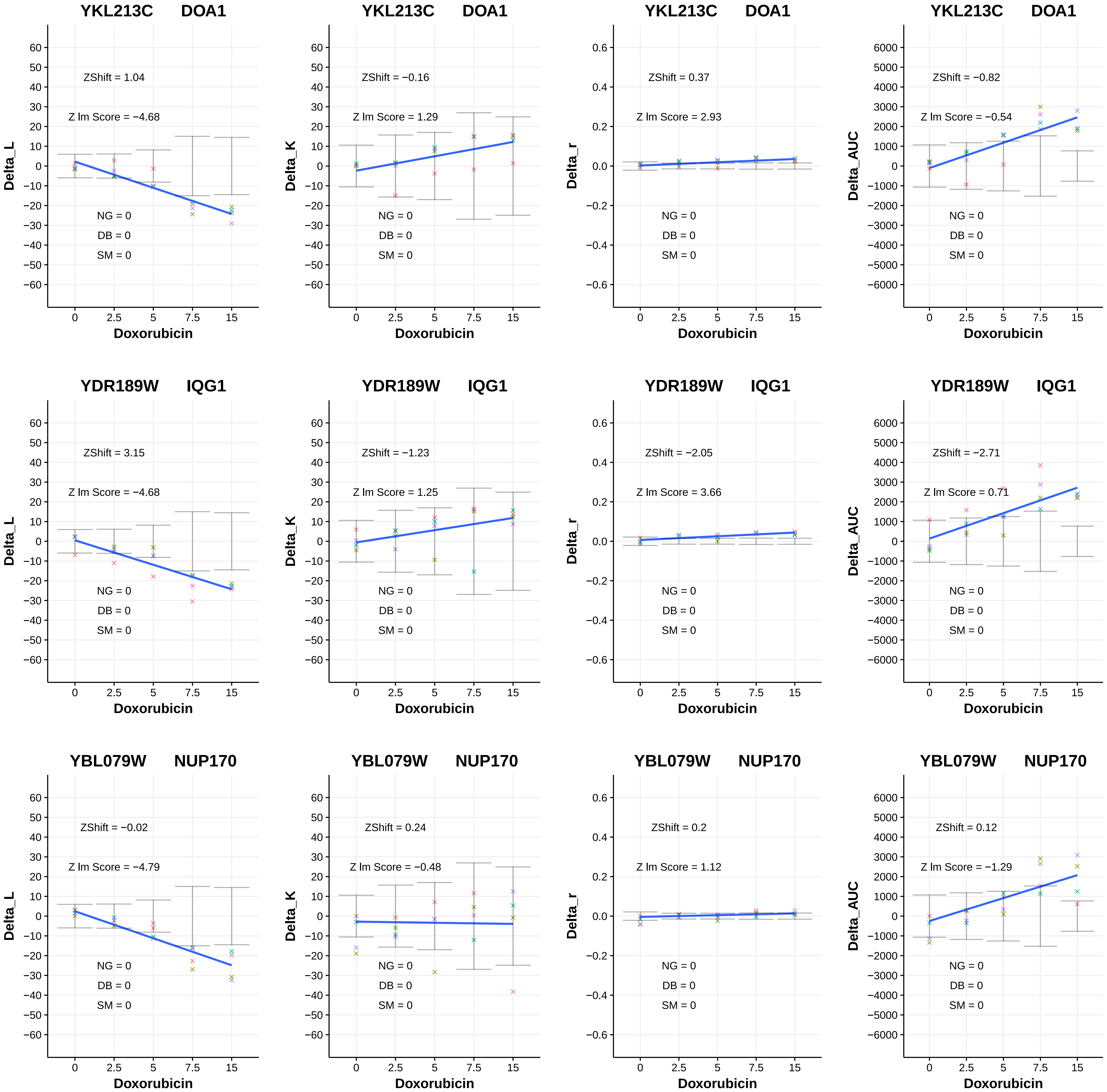
YKL213C DOA1
ZShift = 1.04
Z lm Score = −4.68
NG = 0
DB = 0
SM = 0
Delta_L
Doxorubicin
YKL213C DOA1
ZShift = −0.16
Z lm Score = 1.29
NG = 0
DB = 0
SM = 0
Delta_K
Doxorubicin
YKL213C DOA1
ZShift = 0.37
Z lm Score = 2.93
NG = 0
DB = 0
SM = 0
Delta_r
Doxorubicin
YKL213C DOA1
ZShift = −0.82
Z lm Score = −0.54
NG = 0
DB = 0
SM = 0
Delta_AUC
Doxorubicin
YDR189W IQG1
ZShift = 3.15
Z lm Score = −4.68
NG = 0
DB = 0
SM = 0
Delta_L
Doxorubicin
YDR189W IQG1
ZShift = −1.23
Z lm Score = 1.25
NG = 0
DB = 0
SM = 0
Delta_K
Doxorubicin
YDR189W IQG1
ZShift = −2.05
Z lm Score = 3.66
NG = 0
DB = 0
SM = 0
Delta_r
Doxorubicin
YDR189W IQG1
ZShift = −2.71
Z lm Score = 0.71
NG = 0
DB = 0
SM = 0
Delta_AUC
Doxorubicin
YBL079W NUP170
ZShift = −0.02
Z lm Score = −4.79
NG = 0
DB = 0
SM = 0
Delta_L
Doxorubicin
YBL079W NUP170
ZShift = 0.24
Z lm Score = −0.48
NG = 0
DB = 0
SM = 0
Delta_K
Doxorubicin
YBL079W NUP170
ZShift = 0.2
Z lm Score = 1.12
NG = 0
DB = 0
SM = 0
Delta_r
Doxorubicin
YBL079W NUP170
ZShift = 0.12
Z lm Score = −1.29
NG = 0
DB = 0
SM = 0
Delta_AUC
Doxorubicin

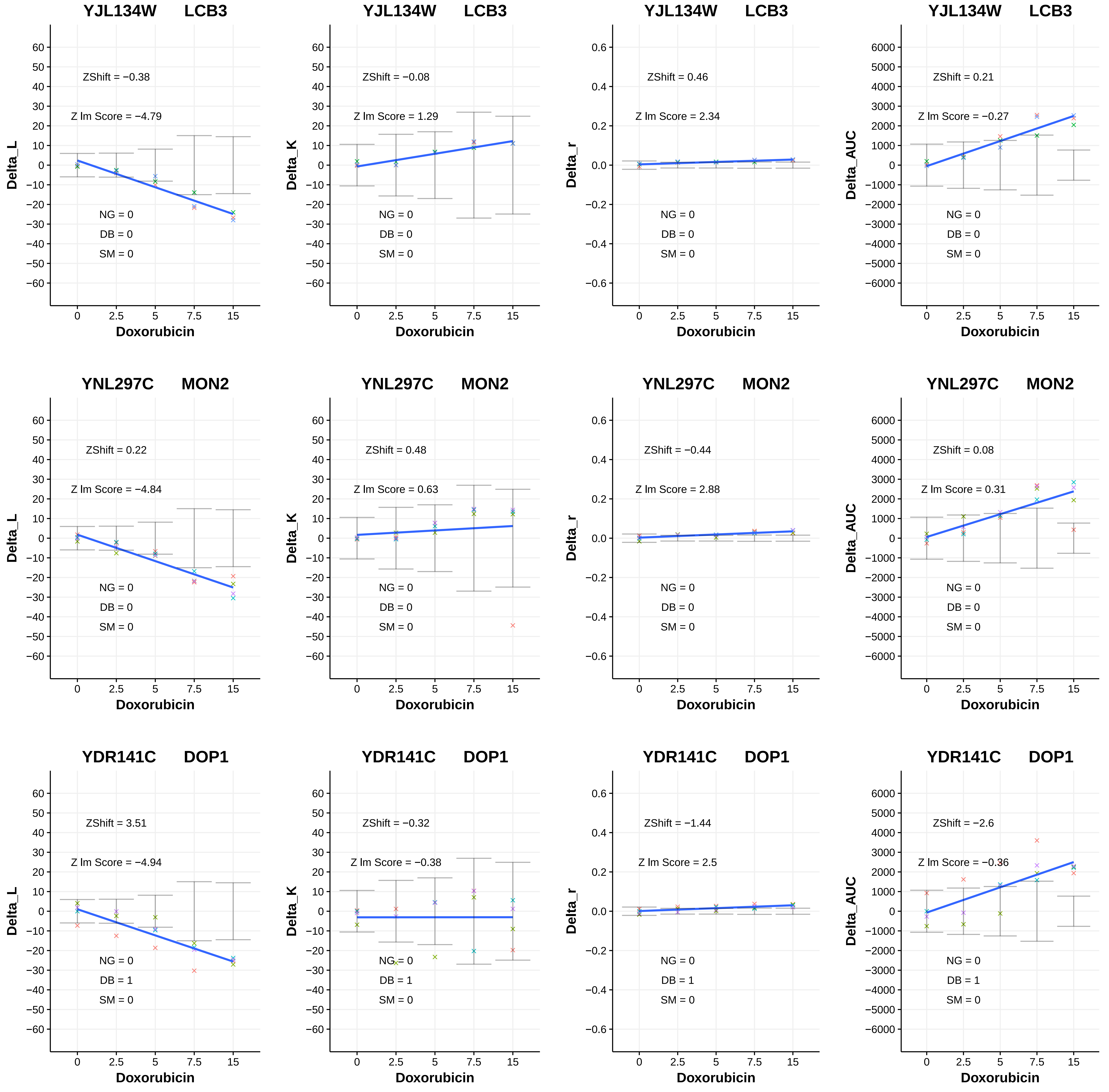
YJL134W LCB3
ZShift = −0.38
Z lm Score = −4.79
NG = 0
DB = 0
SM = 0
Delta_L
Doxorubicin
YJL134W LCB3
ZShift = −0.08
Z lm Score = 1.29
NG = 0
DB = 0
SM = 0
Delta_K
Doxorubicin
YJL134W LCB3
ZShift = 0.46
Z lm Score = 2.34
NG = 0
DB = 0
SM = 0
Delta_r
Doxorubicin
YJL134W LCB3
ZShift = 0.21
Z lm Score = −0.27
NG = 0
DB = 0
SM = 0
Delta_AUC
Doxorubicin
YNL297C MON2
ZShift = 0.22
Z lm Score = −4.84
NG = 0
DB = 0
SM = 0
Delta_L
Doxorubicin
YNL297C MON2
ZShift = 0.48
Z lm Score = 0.63
NG = 0
DB = 0
SM = 0
Delta_K
Doxorubicin
YNL297C MON2
ZShift = −0.44
Z lm Score = 2.88
NG = 0
DB = 0
SM = 0
Delta_r
Doxorubicin
YNL297C MON2
ZShift = 0.08
Z lm Score = 0.31
NG = 0
DB = 0
SM = 0
Delta_AUC
Doxorubicin
YDR141C DOP1
ZShift = 3.51
Z lm Score = −4.94
NG = 0
DB = 1
SM = 0
Delta_L
Doxorubicin
YDR141C DOP1
ZShift = −0.32
Z lm Score = −0.38
NG = 0
DB = 1
SM = 0
Delta_K
Doxorubicin
YDR141C DOP1
ZShift = −1.44
Z lm Score = 2.5
NG = 0
DB = 1
SM = 0
Delta_r
Doxorubicin
YDR141C DOP1
ZShift = −2.6
Z lm Score = −0.36
NG = 0
DB = 1
SM = 0
Delta_AUC
Doxorubicin

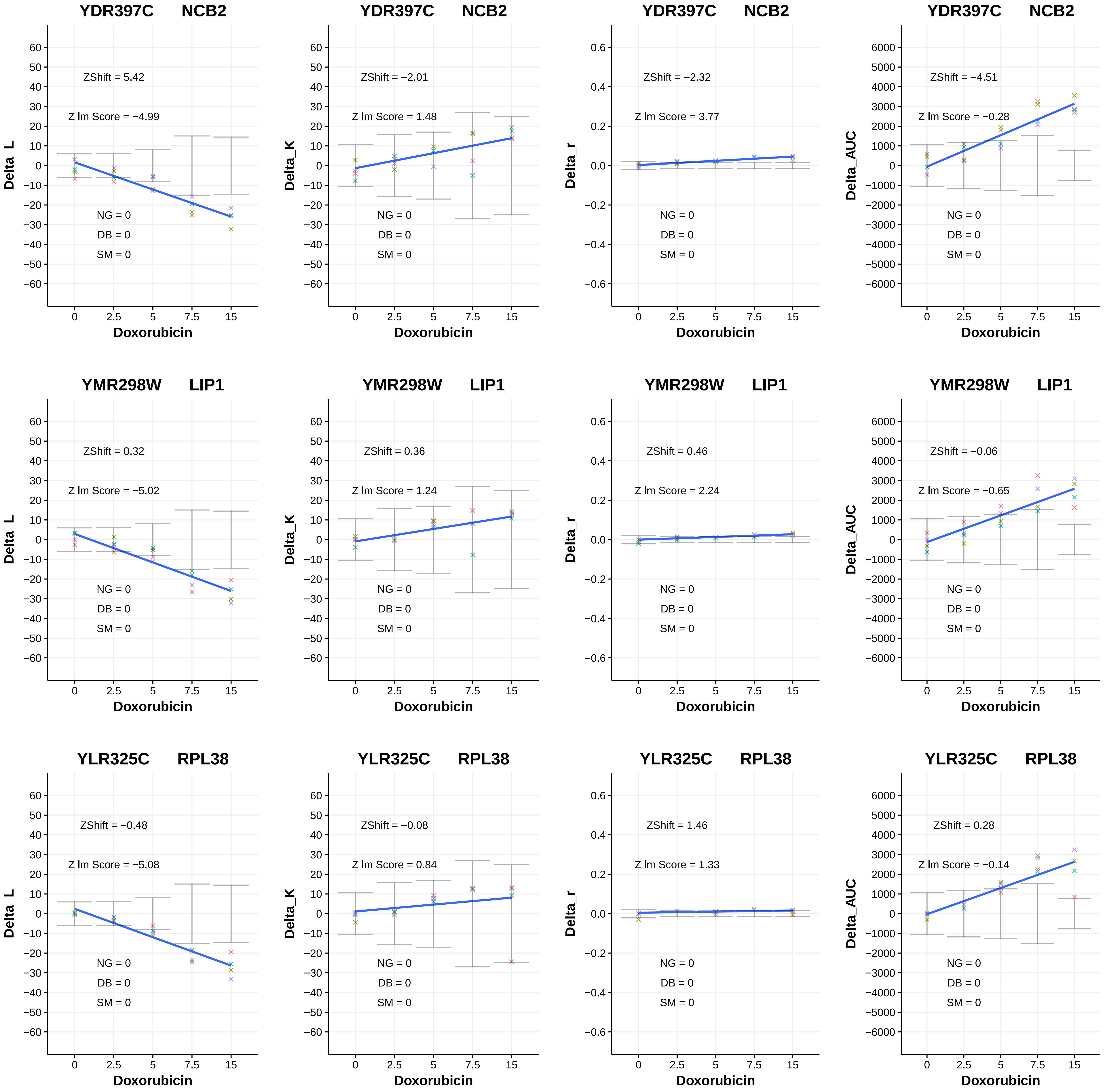
YDR397C NCB2
ZShift = 5.42
Z lm Score = −4.99
NG = 0
DB = 0
SM = 0
Delta_L
Doxorubicin
YDR397C NCB2
ZShift = −2.01
Z lm Score = 1.48
NG = 0
DB = 0
SM = 0
Delta_K
Doxorubicin
YDR397C NCB2
ZShift = −2.32
Z lm Score = 3.77
NG = 0
DB = 0
SM = 0
Delta_r
Doxorubicin
YDR397C NCB2
ZShift = −4.51
Z lm Score = −0.28
NG = 0
DB = 0
SM = 0
Delta_AUC
Doxorubicin
YMR298W LIP1
ZShift = 0.32
Z lm Score = −5.02
NG = 0
DB = 0
SM = 0
Delta_L
Doxorubicin
YMR298W LIP1
ZShift = 0.36
Z lm Score = 1.24
NG = 0
DB = 0
SM = 0
Delta_K
Doxorubicin
YMR298W LIP1
ZShift = 0.46
Z lm Score = 2.24
NG = 0
DB = 0
SM = 0
Delta_r
Doxorubicin
YMR298W LIP1
ZShift = −0.06
Z lm Score = −0.65
NG = 0
DB = 0
SM = 0
Delta_AUC
Doxorubicin
YLR325C RPL38
ZShift = −0.48
Z lm Score = −5.08
NG = 0
DB = 0
SM = 0
Delta_L
Doxorubicin
YLR325C RPL38
ZShift = −0.08
Z lm Score = 0.84
NG = 0
DB = 0
SM = 0
Delta_K
Doxorubicin
YLR325C RPL38
ZShift = 1.46
Z lm Score = 1.33
NG = 0
DB = 0
SM = 0
Delta_r
Doxorubicin
YLR325C RPL38
ZShift = 0.28
Z lm Score = −0.14
NG = 0
DB = 0
SM = 0
Delta_AUC
Doxorubicin

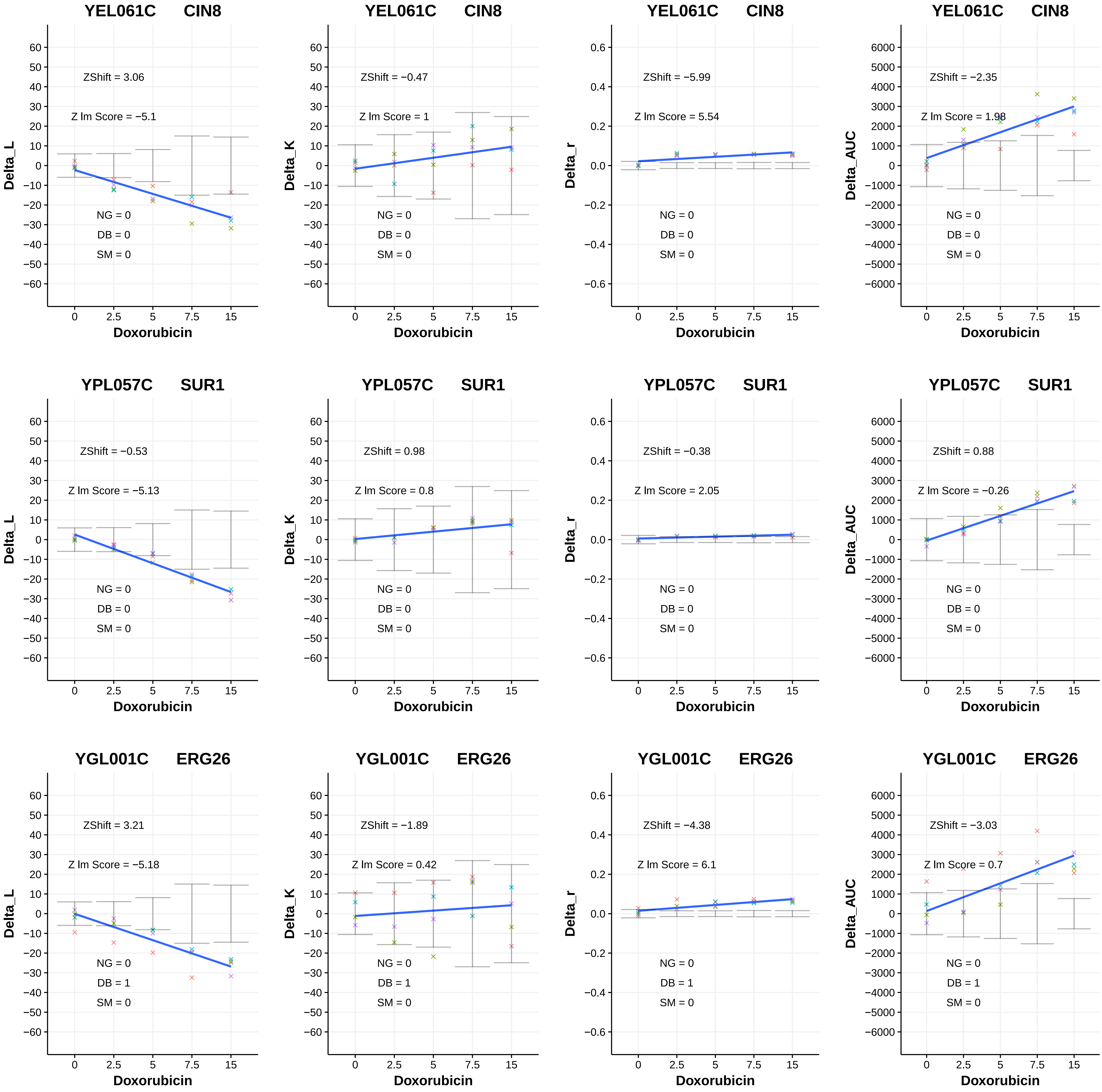

YEL061C CIN8
ZShift = 3.06
Z lm Score = −5.1
NG = 0
DB = 0
SM = 0
Delta_L
Doxorubicin
YEL061C CIN8
ZShift = −0.47
Z lm Score = 1
NG = 0
DB = 0
SM = 0
Delta_K
Doxorubicin
YEL061C CIN8
ZShift = −5.99
Z lm Score = 5.54
NG = 0
DB = 0
SM = 0
Delta_r
Doxorubicin
YEL061C CIN8
ZShift = −2.35
Z lm Score = 1.98
NG = 0
DB = 0
SM = 0
Delta_AUC
Doxorubicin
YPL057C SUR1
ZShift = −0.53
Z lm Score = −5.13
NG = 0
DB = 0
SM = 0
Delta_L
Doxorubicin
YPL057C SUR1
ZShift = 0.98
Z lm Score = 0.8
NG = 0
DB = 0
SM = 0
Delta_K
Doxorubicin
YPL057C SUR1
ZShift = −0.38
Z lm Score = 2.05
NG = 0
DB = 0
SM = 0
Delta_r
Doxorubicin
YPL057C SUR1
ZShift = 0.88
Z lm Score = −0.26
NG = 0
DB = 0
SM = 0
Delta_AUC
Doxorubicin
YGL001C ERG26
ZShift = 3.21
Z lm Score = −5.18
NG = 0
DB = 1
SM = 0
Delta_L
Doxorubicin
YGL001C ERG26
ZShift = −1.89
Z lm Score = 0.42
NG = 0
DB = 1
SM = 0
Delta_K
Doxorubicin
YGL001C ERG26
ZShift = −4.38
Z lm Score = 6.1
NG = 0
DB = 1
SM = 0
Delta_r
Doxorubicin
YGL001C ERG26
ZShift = −3.03
Z lm Score = 0.7
NG = 0
DB = 1
SM = 0
Delta_AUC
Doxorubicin

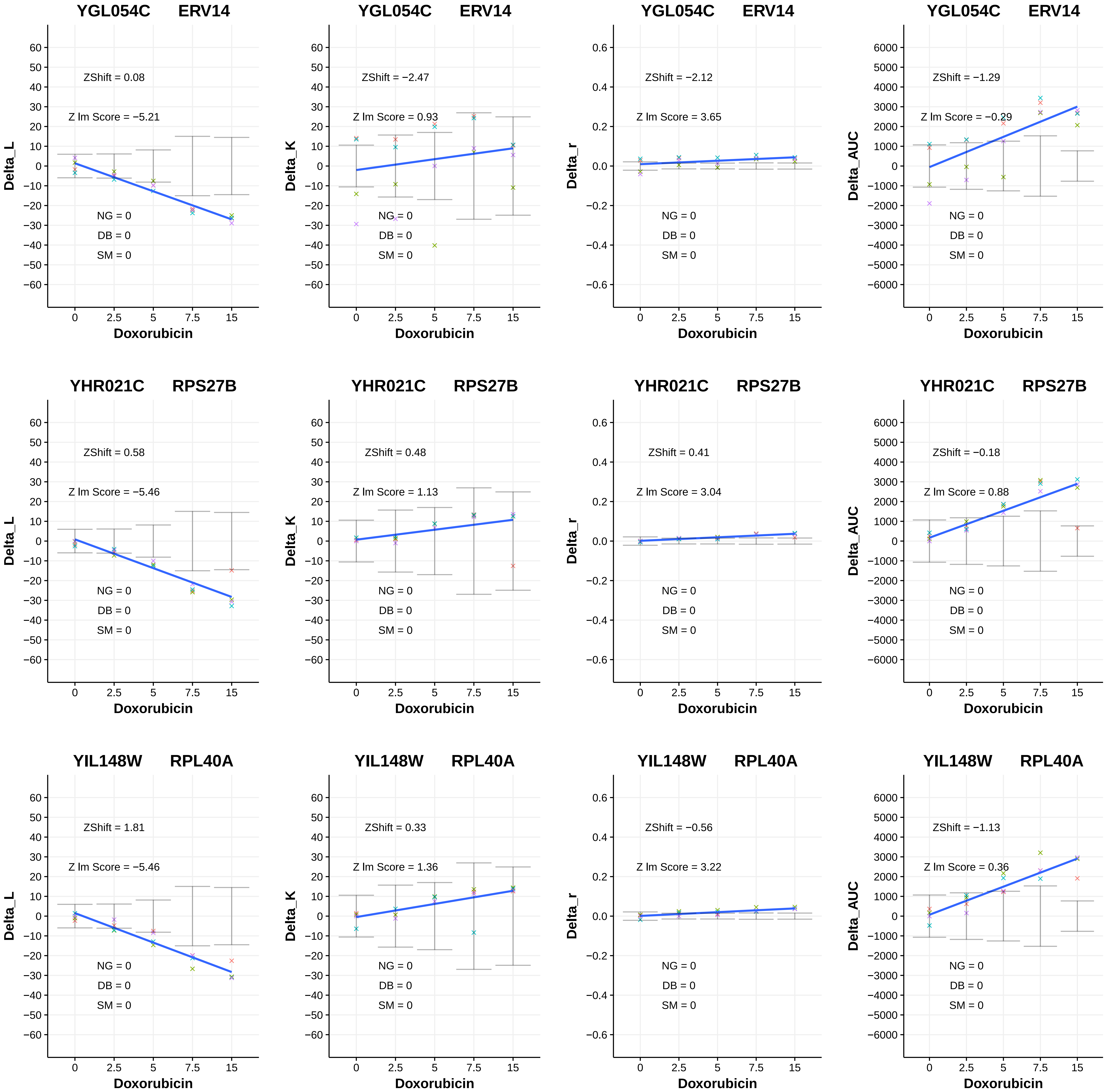

YGL054C ERV14
ZShift = 0.08
Z lm Score = −5.21
NG = 0
DB = 0
SM = 0
Delta_L
Doxorubicin
YGL054C ERV14
ZShift = −2.47
Z lm Score = 0.93
NG = 0
DB = 0
SM = 0
Delta_K
Doxorubicin
YGL054C ERV14
ZShift = −2.12
Z lm Score = 3.65
NG = 0
DB = 0
SM = 0
Delta_r
Doxorubicin
YGL054C ERV14
ZShift = −1.29
Z lm Score = −0.29
NG = 0
DB = 0
SM = 0
Delta_AUC
Doxorubicin
YHR021C RPS27B
ZShift = 0.58
Z lm Score = −5.46
NG = 0
DB = 0
SM = 0
Delta_L
Doxorubicin
YHR021C RPS27B
ZShift = 0.48
Z lm Score = 1.13
NG = 0
DB = 0
SM = 0
Delta_K
Doxorubicin
YHR021C RPS27B
ZShift = 0.41
Z lm Score = 3.04
NG = 0
DB = 0
SM = 0
Delta_r
Doxorubicin
YHR021C RPS27B
ZShift = −0.18
Z lm Score = 0.88
NG = 0
DB = 0
SM = 0
Delta_AUC
Doxorubicin
YIL148W RPL40A
ZShift = 1.81
Z lm Score = −5.46
NG = 0
DB = 0
SM = 0
Delta_L
Doxorubicin
YIL148W RPL40A
ZShift = 0.33
Z lm Score = 1.36
NG = 0
DB = 0
SM = 0
Delta_K
Doxorubicin
YIL148W RPL40A
ZShift = −0.56
Z lm Score = 3.22
NG = 0
DB = 0
SM = 0
Delta_r
Doxorubicin
YIL148W RPL40A
ZShift = −1.13
Z lm Score = 0.36
NG = 0
DB = 0
SM = 0
Delta_AUC
Doxorubicin

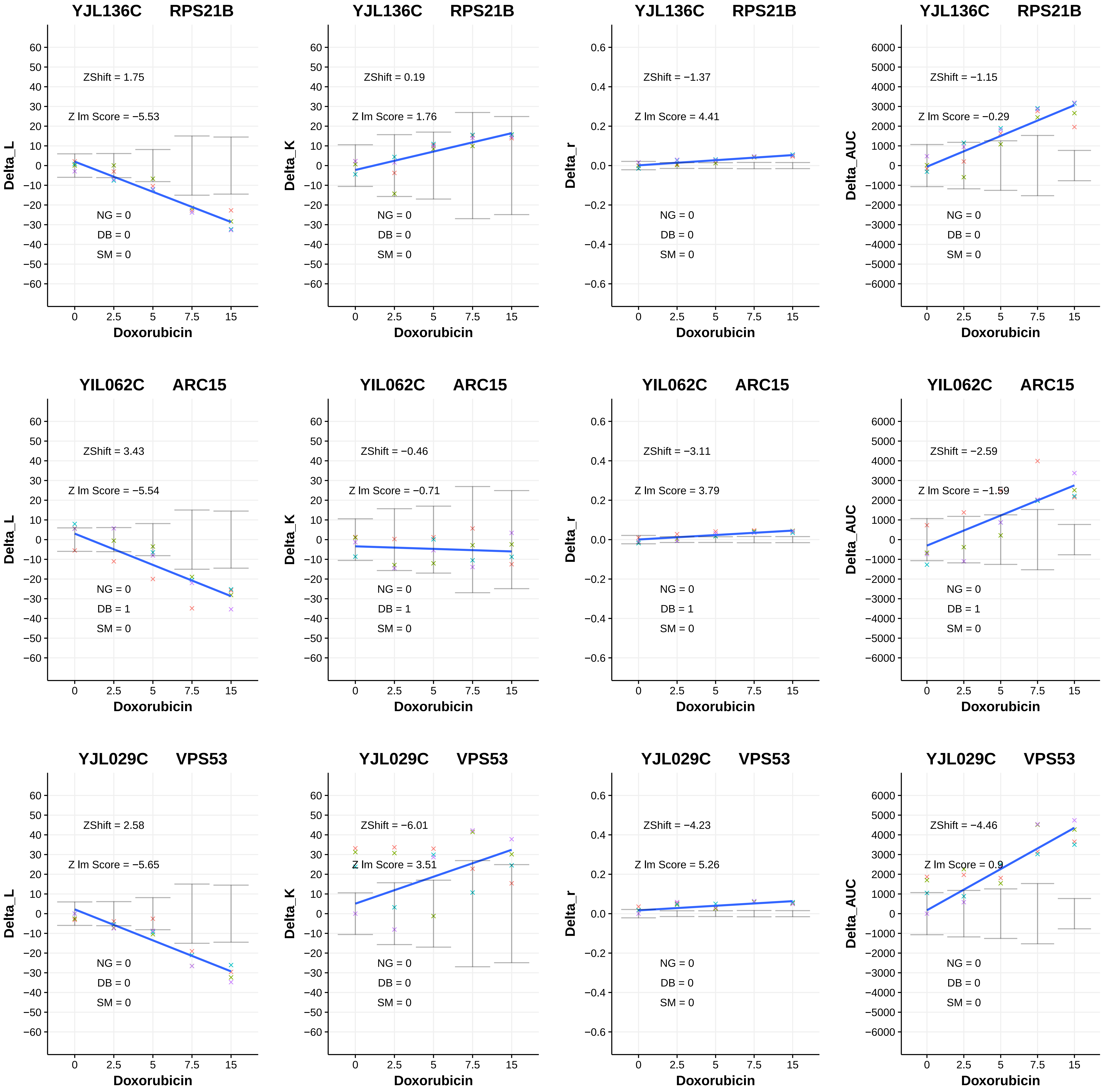

YJL136C RPS21B
ZShift = 1.75
Z lm Score = −5.53
NG = 0
DB = 0
SM = 0
Delta_L
Doxorubicin
YJL136C RPS21B
ZShift = 0.19
Z lm Score = 1.76
NG = 0
DB = 0
SM = 0
Delta_K
Doxorubicin
YJL136C RPS21B
ZShift = −1.37
Z lm Score = 4.41
NG = 0
DB = 0
SM = 0
Delta_r
Doxorubicin
YJL136C RPS21B
ZShift = −1.15
Z lm Score = −0.29
NG = 0
DB = 0
SM = 0
Delta_AUC
Doxorubicin
YIL062C ARC15
ZShift = 3.43
Z lm Score = −5.54
NG = 0
DB = 1
SM = 0
Delta_L
Doxorubicin
YIL062C ARC15
ZShift = −0.46
Z lm Score = −0.71
NG = 0
DB = 1
SM = 0
Delta_K
Doxorubicin
YIL062C ARC15
ZShift = −3.11
Z lm Score = 3.79
NG = 0
DB = 1
SM = 0
Delta_r
Doxorubicin
YIL062C ARC15
ZShift = −2.59
Z lm Score = −1.59
NG = 0
DB = 1
SM = 0
Delta_AUC
Doxorubicin
YJL029C VPS53
ZShift = 2.58
Z lm Score = −5.65
NG = 0
DB = 0
SM = 0
Delta_L
Doxorubicin
YJL029C VPS53
ZShift = −6.01
Z lm Score = 3.51
NG = 0
DB = 0
SM = 0
Delta_K
Doxorubicin
YJL029C VPS53
ZShift = −4.23
Z lm Score = 5.26
NG = 0
DB = 0
SM = 0
Delta_r
Doxorubicin
YJL029C VPS53
ZShift = −4.46
Z lm Score = 0.9
NG = 0
DB = 0
SM = 0
Delta_AUC
Doxorubicin

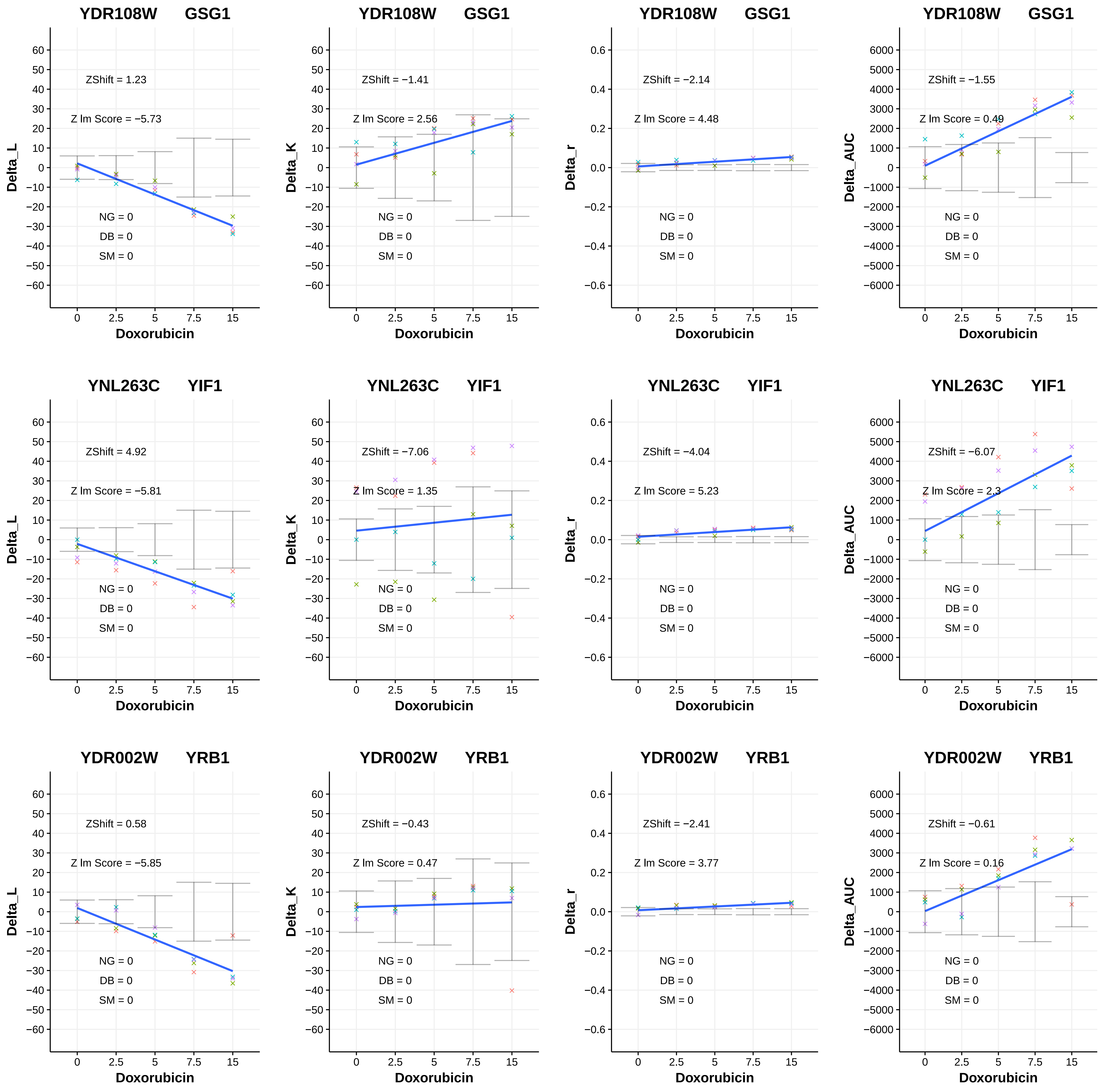
YDR108W GSG1
ZShift = 1.23
Z lm Score = −5.73
NG = 0
DB = 0
SM = 0
Delta_L
Doxorubicin
YDR108W GSG1
ZShift = −1.41
Z lm Score = 2.56
NG = 0
DB = 0
SM = 0
Delta_K
Doxorubicin
YDR108W GSG1
ZShift = −2.14
Z lm Score = 4.48
NG = 0
DB = 0
SM = 0
Delta_r
Doxorubicin
YDR108W GSG1
ZShift = −1.55
Z lm Score = 0.49
NG = 0
DB = 0
SM = 0
Delta_AUC
Doxorubicin
YNL263C YIF1
ZShift = 4.92
Z lm Score = −5.81
NG = 0
DB = 0
SM = 0
Delta_L
Doxorubicin
YNL263C YIF1
ZShift = −7.06
Z lm Score = 1.35
NG = 0
DB = 0
SM = 0
Delta_K
Doxorubicin
YNL263C YIF1
ZShift = −4.04
Z lm Score = 5.23
NG = 0
DB = 0
SM = 0
Delta_r
Doxorubicin
YNL263C YIF1
ZShift = −6.07
Z lm Score = 2.3
NG = 0
DB = 0
SM = 0
Delta_AUC
Doxorubicin
YDR002W YRB1
ZShift = 0.58
Z lm Score = −5.85
NG = 0
DB = 0
SM = 0
Delta_L
Doxorubicin
YDR002W YRB1
ZShift = −0.43
Z lm Score = 0.47
NG = 0
DB = 0
SM = 0
Delta_K
Doxorubicin
YDR002W YRB1
ZShift = −2.41
Z lm Score = 3.77
NG = 0
DB = 0
SM = 0
Delta_r
Doxorubicin
YDR002W YRB1
ZShift = −0.61
Z lm Score = 0.16
NG = 0
DB = 0
SM = 0
Delta_AUC
Doxorubicin

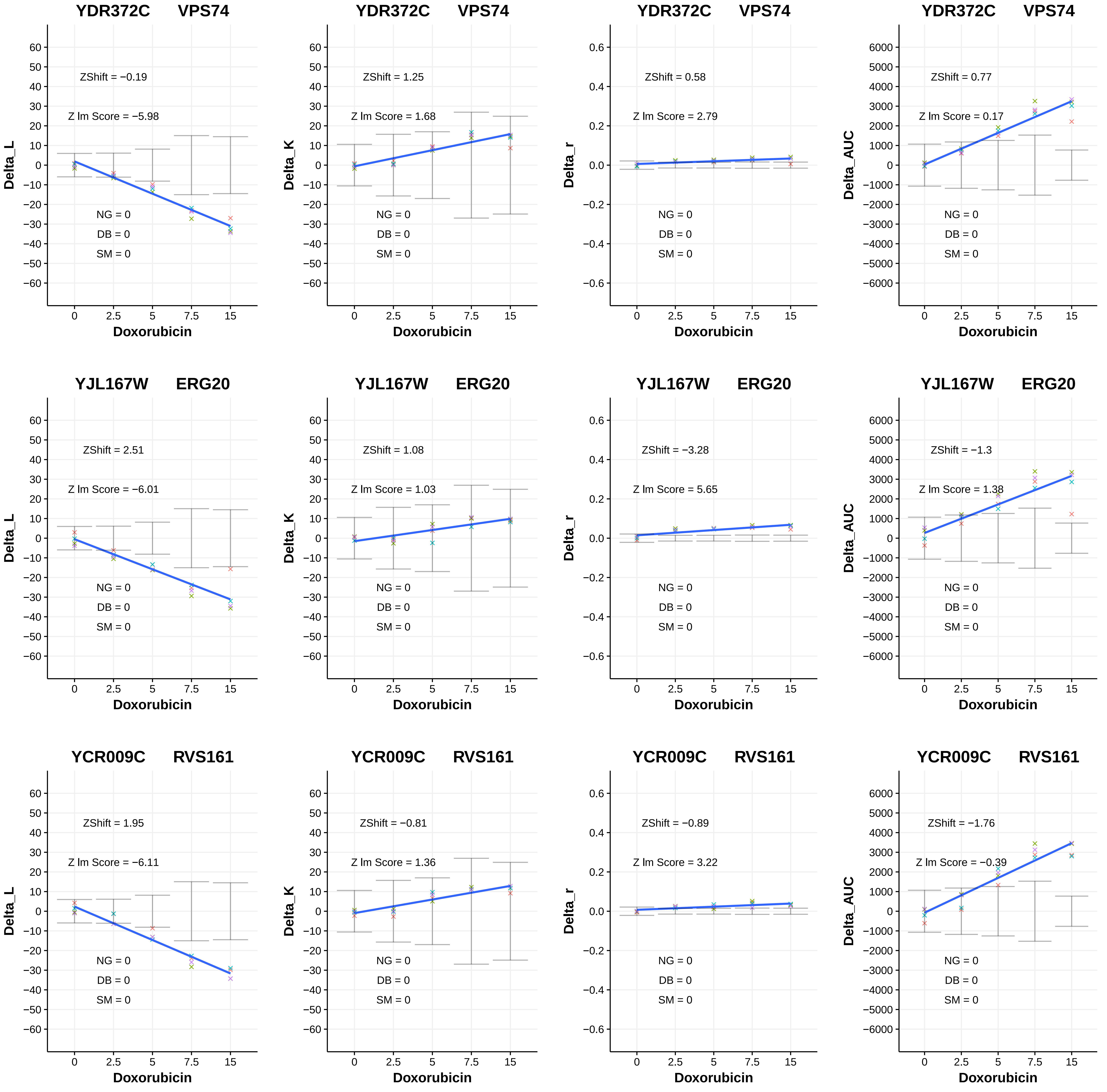
YDR372C VPS74
ZShift = −0.19
Z lm Score = −5.98
NG = 0
DB = 0
SM = 0
Delta_L
Doxorubicin
YDR372C VPS74
ZShift = 1.25
Z lm Score = 1.68
NG = 0
DB = 0
SM = 0
Delta_K
Doxorubicin
YDR372C VPS74
ZShift = 0.58
Z lm Score = 2.79
NG = 0
DB = 0
SM = 0
Delta_r
Doxorubicin
YDR372C VPS74
ZShift = 0.77
Z lm Score = 0.17
NG = 0
DB = 0
SM = 0
Delta_AUC
Doxorubicin
YJL167W ERG20
ZShift = 2.51
Z lm Score = −6.01
NG = 0
DB = 0
SM = 0
Delta_L
Doxorubicin
YJL167W ERG20
ZShift = 1.08
Z lm Score = 1.03
NG = 0
DB = 0
SM = 0
Delta_K
Doxorubicin
YJL167W ERG20
ZShift = −3.28
Z lm Score = 5.65
NG = 0
DB = 0
SM = 0
Delta_r
Doxorubicin
YJL167W ERG20
ZShift = −1.3
Z lm Score = 1.38
NG = 0
DB = 0
SM = 0
Delta_AUC
Doxorubicin
YCR009C RVS161
ZShift = 1.95
Z lm Score = −6.11
NG = 0
DB = 0
SM = 0
Delta_L
Doxorubicin
YCR009C RVS161
ZShift = −0.81
Z lm Score = 1.36
NG = 0
DB = 0
SM = 0
Delta_K
Doxorubicin
YCR009C RVS161
ZShift = −0.89
Z lm Score = 3.22
NG = 0
DB = 0
SM = 0
Delta_r
Doxorubicin
YCR009C RVS161
ZShift = −1.76
Z lm Score = −0.39
NG = 0
DB = 0
SM = 0
Delta_AUC
Doxorubicin

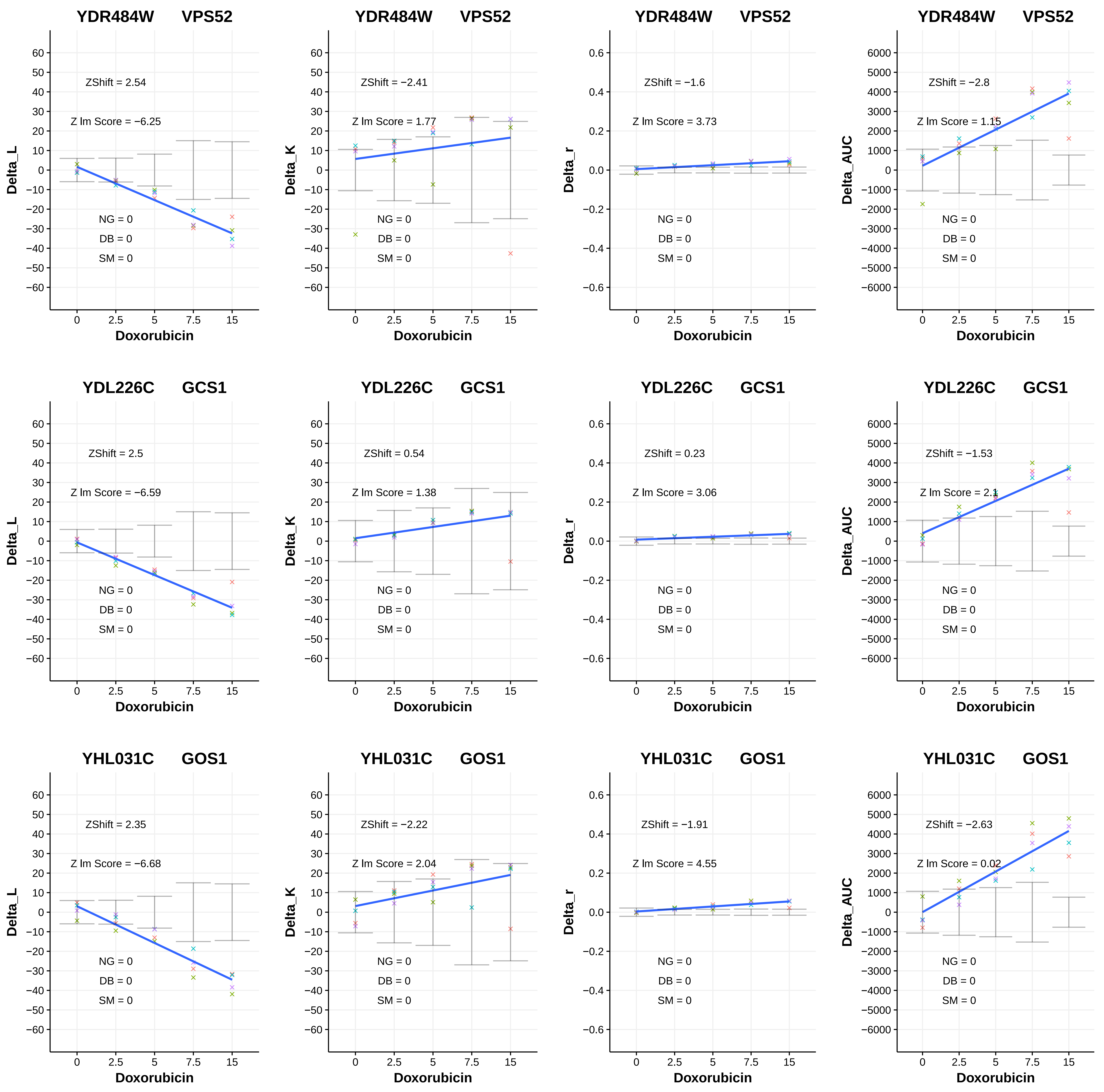
YDR484W VPS52
ZShift = 2.54
Z lm Score = −6.25
NG = 0
DB = 0
SM = 0
Delta_L
YDR484W VPS52
ZShift = −2.41
Z lm Score = 1.77
NG = 0
DB = 0
SM = 0
Delta_K
YDR484W VPS52
ZShift = −1.6
Z lm Score = 3.73
NG = 0
DB = 0
SM = 0
Delta_r
YDR484W VPS52
ZShift = −2.8
Z lm Score = 1.15
NG = 0
DB = 0
SM = 0
Delta_AUC
YDL226C GCS1
ZShift = 2.5
Z lm Score = −6.59
NG = 0
DB = 0
SM = 0
Delta_L
YDL226C GCS1
ZShift = 0.54
Z lm Score = 1.38
NG = 0
DB = 0
SM = 0
Delta_K
YDL226C GCS1
ZShift = 0.23
Z lm Score = 3.06
NG = 0
DB = 0
SM = 0
Delta_r
YDL226C GCS1
ZShift = −1.53
Z lm Score = 2.1
NG = 0
DB = 0
SM = 0
Delta_AUC
YHL031C GOS1
ZShift = 2.35
Z lm Score = −6.68
NG = 0
DB = 0
SM = 0
Delta_L
YHL031C GOS1
ZShift = −2.22
Z lm Score = 2.04
NG = 0
DB = 0
SM = 0
Delta_K
YHL031C GOS1
ZShift = −1.91
Z lm Score = 4.55
NG = 0
DB = 0
SM = 0
Delta_r
YHL031C GOS1
ZShift = −2.63
Z lm Score = 0.02
NG = 0
DB = 0
SM = 0
Delta_AUC
Doxorubicin
0
2.5
5
7.5
15

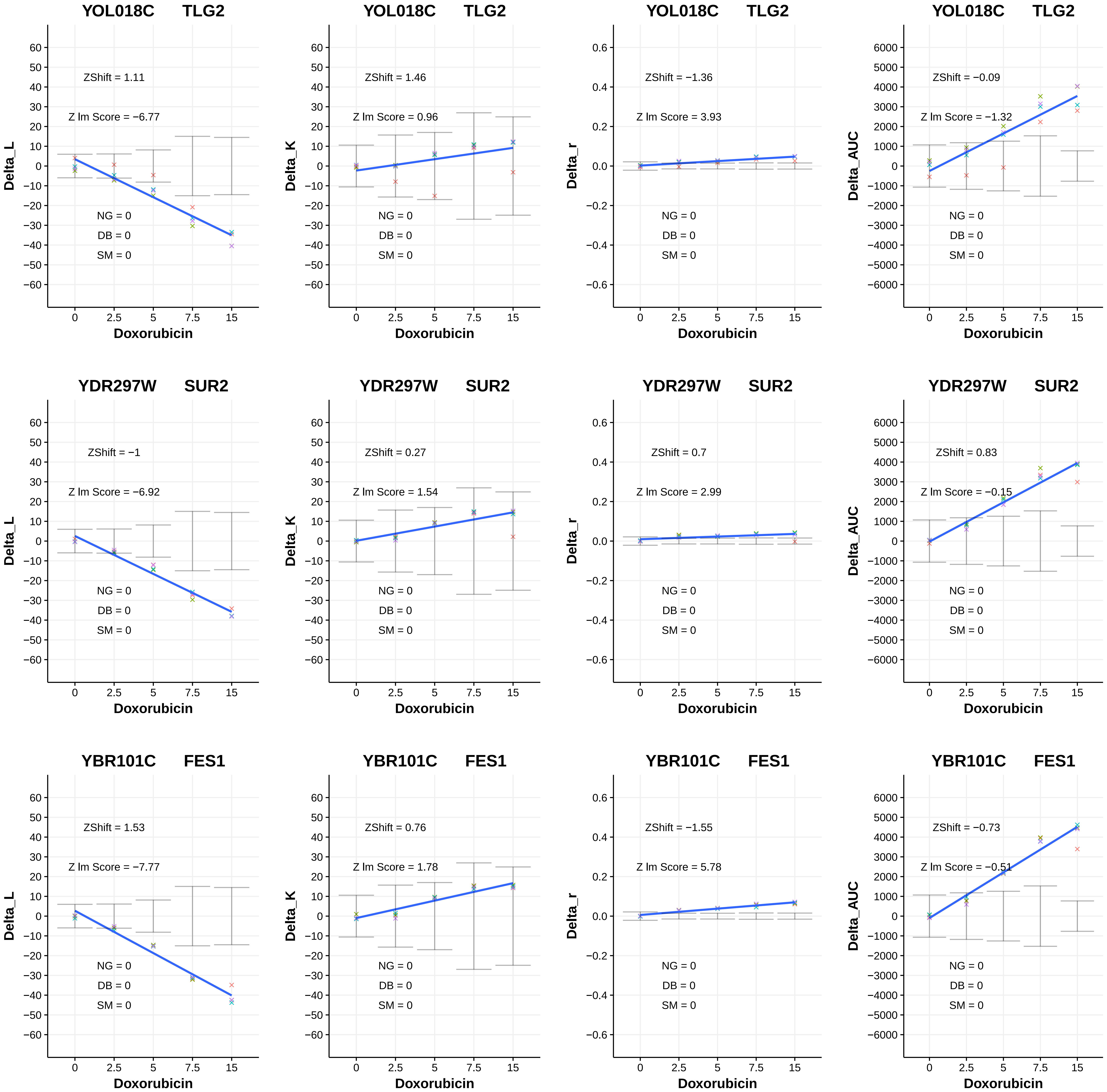
YOL018C TLG2
ZShift = 1.11
Z lm Score = −6.77
NG = 0
DB = 0
SM = 0
Delta_L
Doxorubicin
YOL018C TLG2
ZShift = 1.46
Z lm Score = 0.96
NG = 0
DB = 0
SM = 0
Delta_K
Doxorubicin
YOL018C TLG2
ZShift = −1.36
Z lm Score = 3.93
NG = 0
DB = 0
SM = 0
Delta_r
Doxorubicin
YOL018C TLG2
ZShift = −0.09
Z lm Score = −1.32
NG = 0
DB = 0
SM = 0
Delta_AUC
Doxorubicin
YDR297W SUR2
ZShift = −1
Z lm Score = −6.92
NG = 0
DB = 0
SM = 0
Delta_L
Doxorubicin
YDR297W SUR2
ZShift = 0.27
Z lm Score = 1.54
NG = 0
DB = 0
SM = 0
Delta_K
Doxorubicin
YDR297W SUR2
ZShift = 0.7
Z lm Score = 2.99
NG = 0
DB = 0
SM = 0
Delta_r
Doxorubicin
YDR297W SUR2
ZShift = 0.83
Z lm Score = −0.15
NG = 0
DB = 0
SM = 0
Delta_AUC
Doxorubicin
YBR101C FES1
ZShift = 1.53
Z lm Score = −7.77
NG = 0
DB = 0
SM = 0
Delta_L
Doxorubicin
YBR101C FES1
ZShift = 0.76
Z lm Score = 1.78
NG = 0
DB = 0
SM = 0
Delta_K
Doxorubicin
YBR101C FES1
ZShift = −1.55
Z lm Score = 5.78
NG = 0
DB = 0
SM = 0
Delta_r
Doxorubicin
YBR101C FES1
ZShift = −0.73
Z lm Score = −0.51
NG = 0
DB = 0
SM = 0
Delta_AUC
Doxorubicin

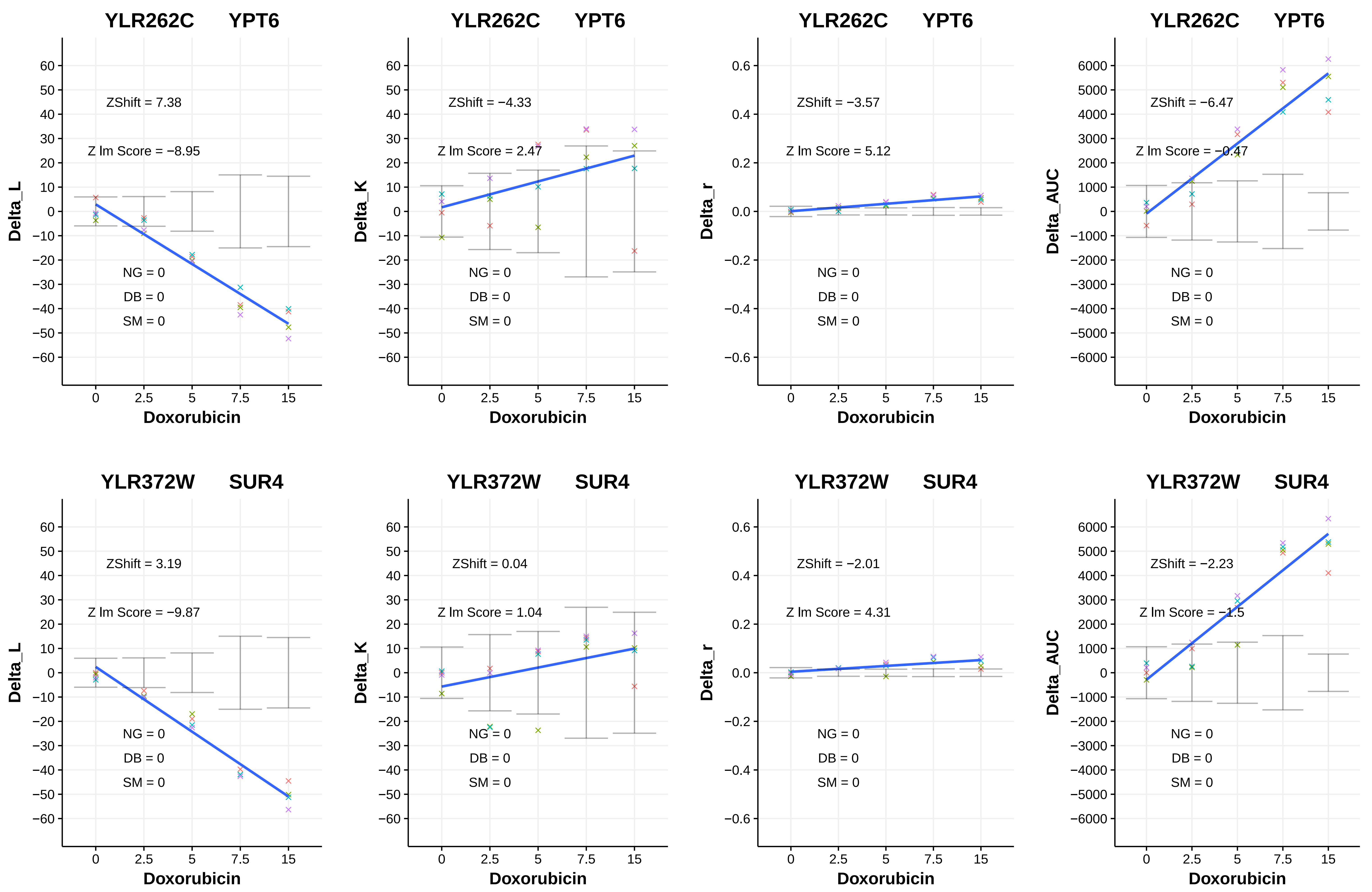
YLR262C YPT6
Delta_L
ZShift = 7.38
Z lm Score = −8.95
NG = 0
DB = 0
SM = 0
Doxorubicin
YLR262C YPT6
Delta_K
ZShift = −4.33
Z lm Score = 2.47
NG = 0
DB = 0
SM = 0
Doxorubicin
YLR262C YPT6
Delta_r
ZShift = −3.57
Z lm Score = 5.12
NG = 0
DB = 0
SM = 0
Doxorubicin
YLR262C YPT6
Delta_AUC
ZShift = −6.47
Z lm Score = −0.47
NG = 0
DB = 0
SM = 0
Doxorubicin
YLR372W SUR4
Delta_L
ZShift = 3.19
Z lm Score = −9.87
NG = 0
DB = 0
SM = 0
Doxorubicin
YLR372W SUR4
Delta_K
ZShift = 0.04
Z lm Score = 1.04
NG = 0
DB = 0
SM = 0
Doxorubicin
YLR372W SUR4
Delta_r
ZShift = −2.01
Z lm Score = 4.31
NG = 0
DB = 0
SM = 0
Doxorubicin
YLR372W SUR4
Delta_AUC
ZShift = −2.23
Z lm Score = −1.5
NG = 0
DB = 0
SM = 0
Doxorubicin
