## Additional File 3 for "A yeast phenomic model for the influence of Warburg metabolism on genetic buffering of doxorubicin": D - RF_InteractionPlots_Doxo_HLEG_V.pdf

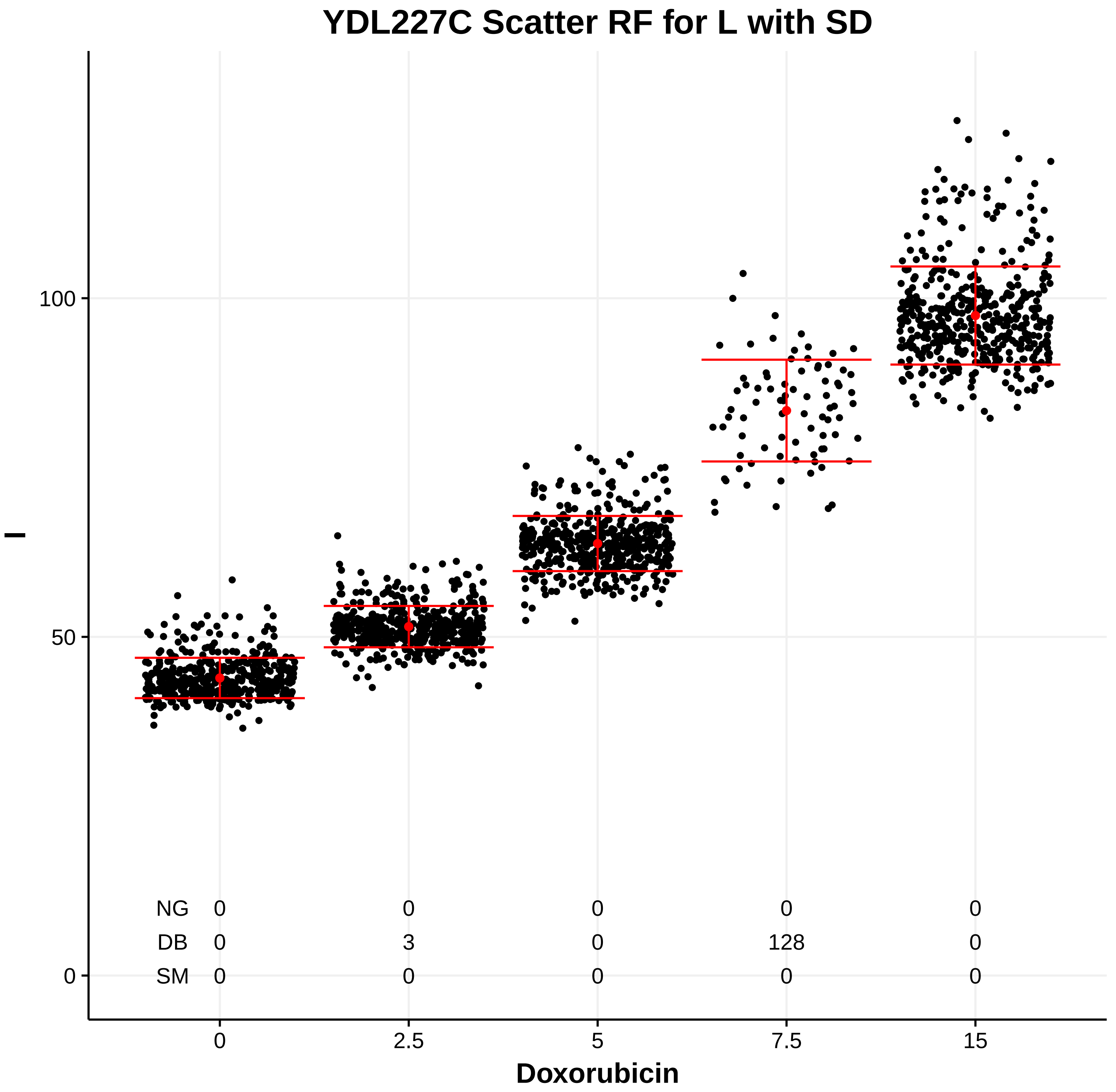
YDL227C Scatter RF for L with SD
l
0
50
100
NG 0 0 0 0 0
DB 0 3 0 128 0
SM 0 0 0 0 0
0 2.5 5 7.5 15
Doxorubicin

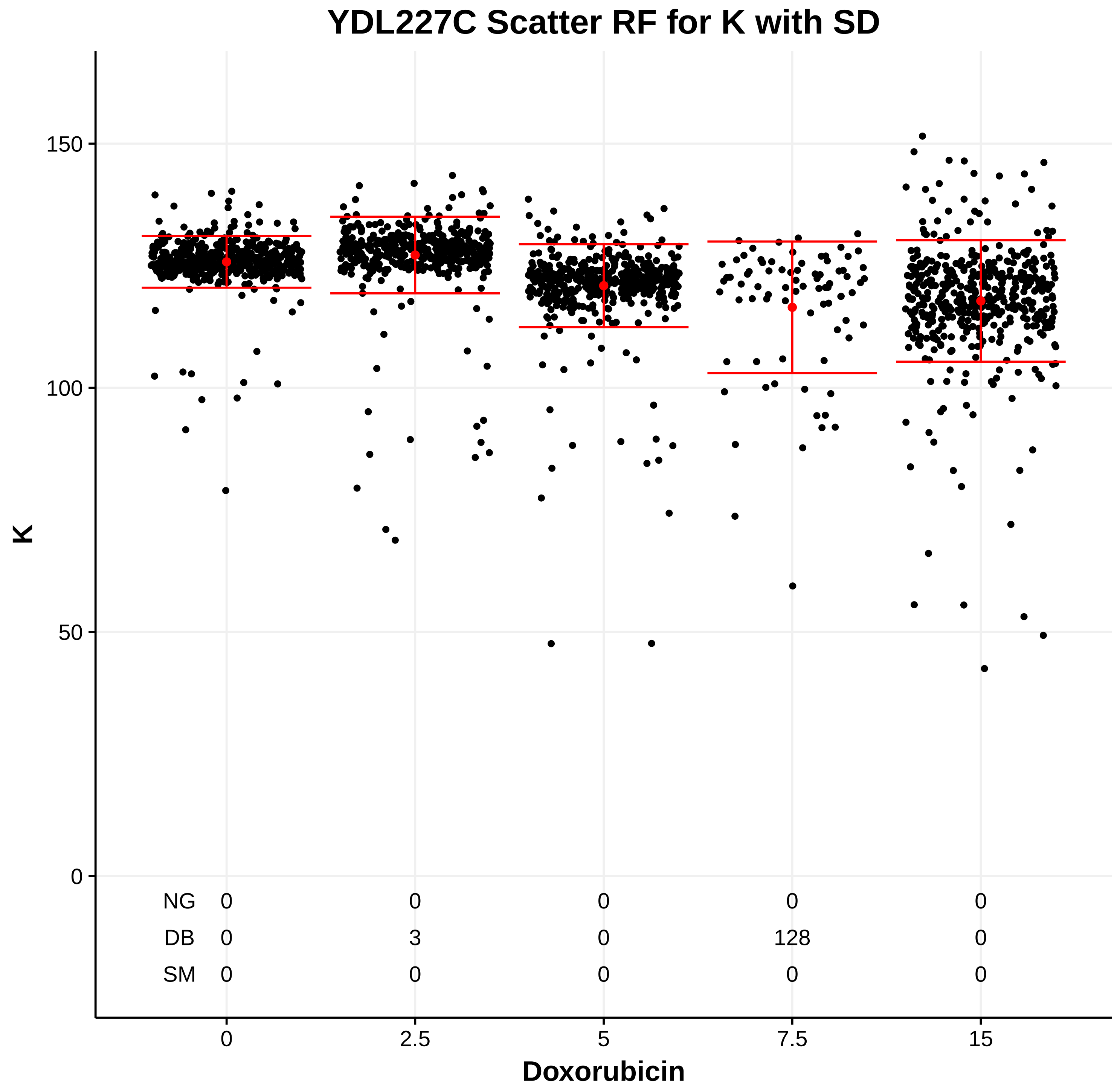
YDL227C Scatter RF for K with SD
K
0
50
100
150
NG 0 0 0 0 0
DB 0 3 0 128 0
SM 0 0 0 0 0
0 2.5 5 7.5 15
Doxorubicin

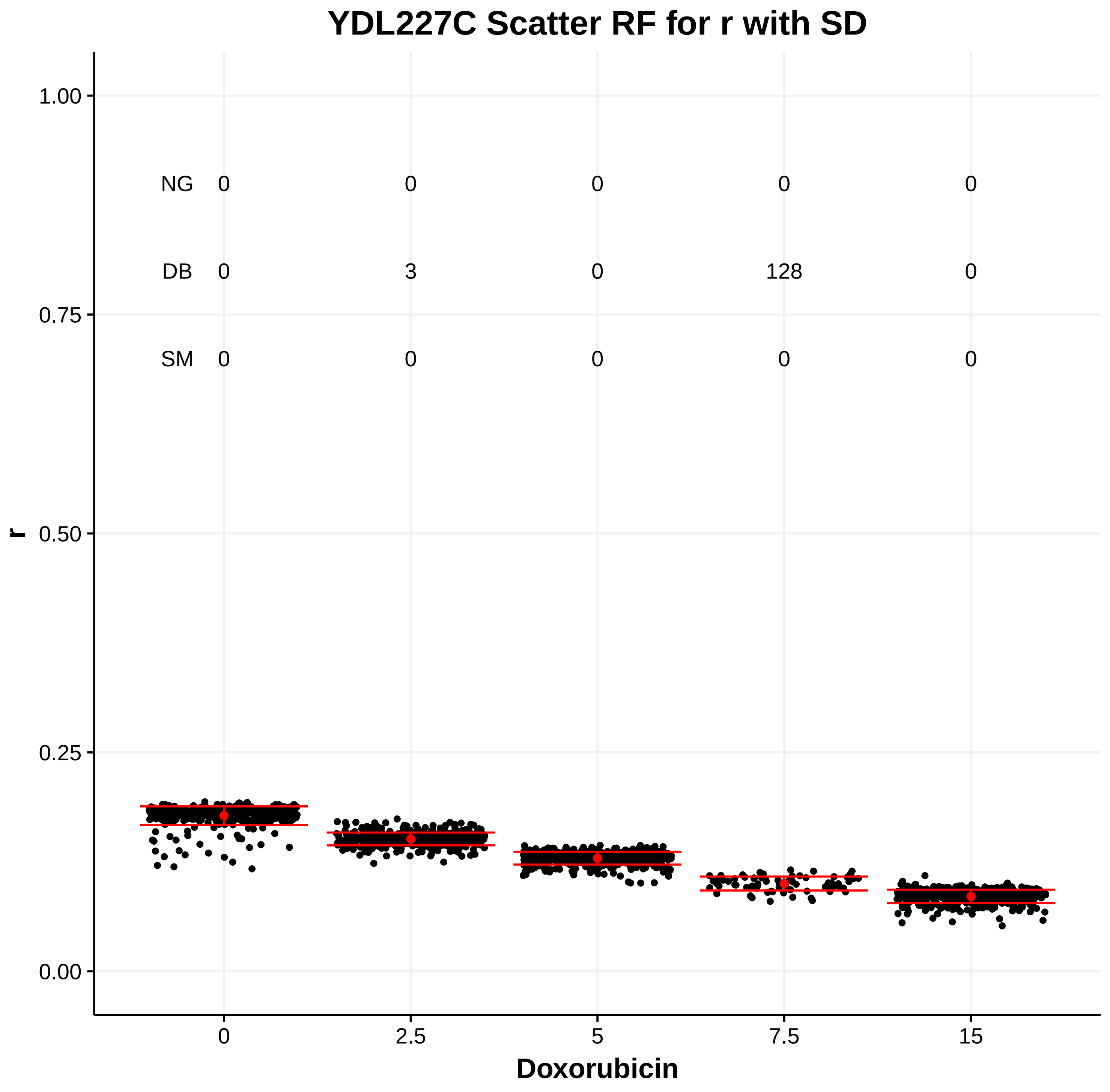
YDL227C Scatter RF for r with SD
r
0.00
0.25
0.50
0.75
1.00
NG 0 0 0 0 0
DB 0 3 0 128 0
SM 0 0 0 0 0
0 2.5 5 7.5 15
Doxorubicin

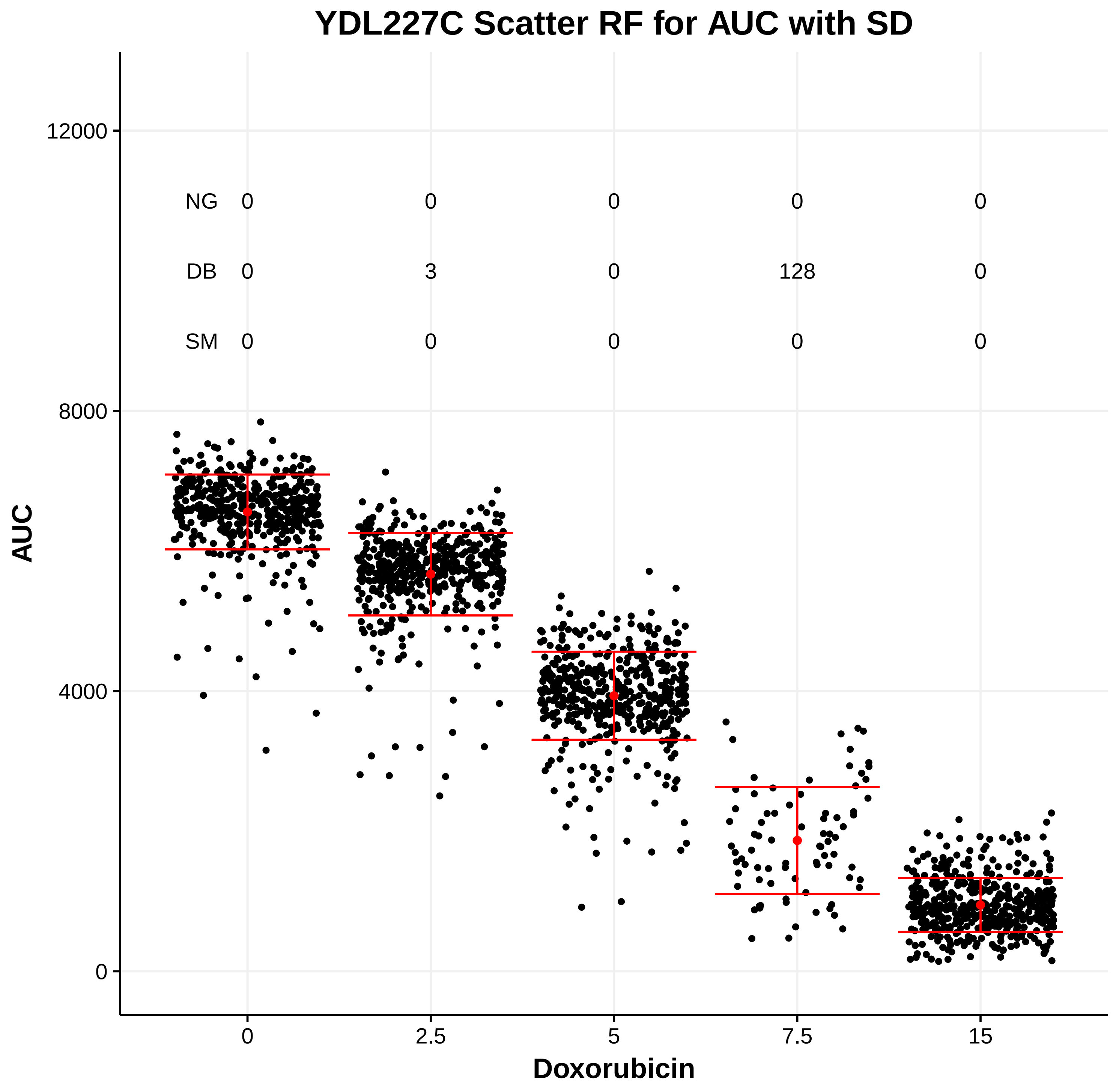
YDL227C Scatter RF for AUC with SD
AUC
0
4000
8000
12000
NG 0 0 0 0 0
DB 0 3 0 128 0
SM 0 0 0 0 0
0 2.5 5 7.5 15
Doxorubicin

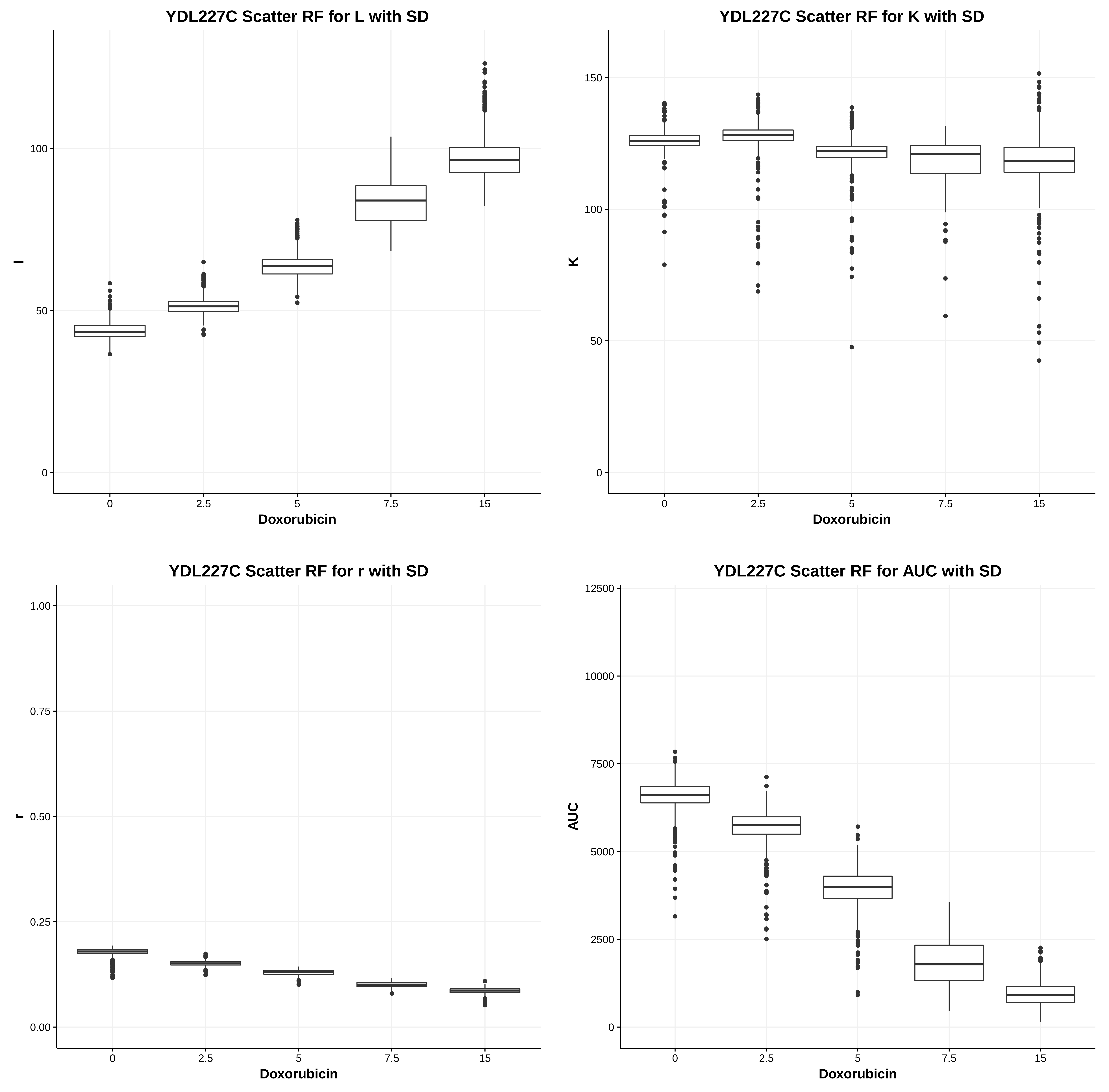
YDL227C Scatter RF for L with SD
l
Doxorubicin
0
50
100
2.5
5
7.5
15
YDL227C Scatter RF for K with SD
K
Doxorubicin
150
YDL227C Scatter RF for r with SD
r
0.00
0.25
0.50
0.75
1.00
YDL227C Scatter RF for AUC with SD
AUC
2500
5000
7500
10000
12500

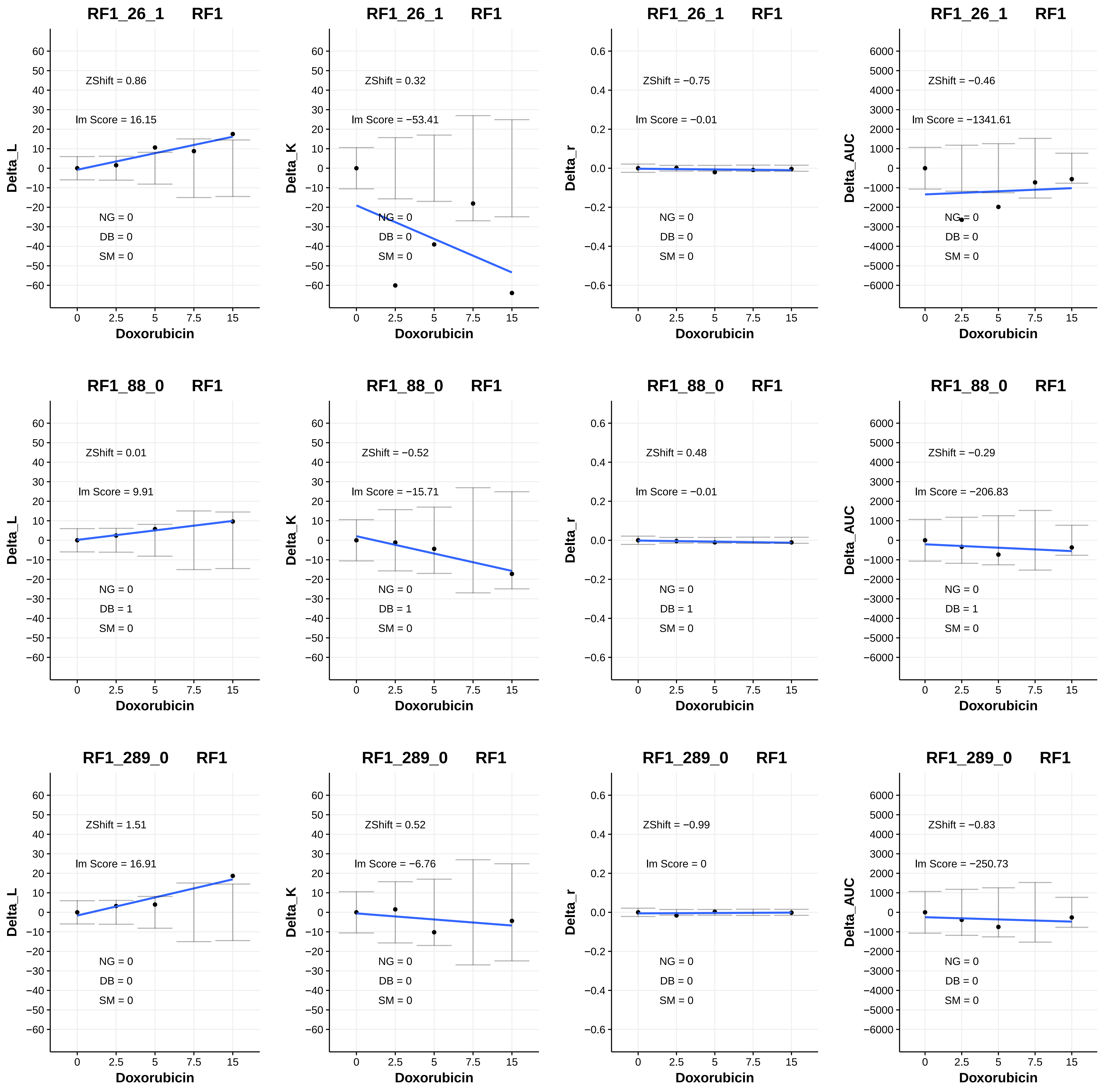
RF1_26_1 RF1
ZShift = 0.86
Im Score = 16.15
NG = 0
DB = 0
SM = 0
Delta_L
Doxorubicin
RF1_26_1 RF1
ZShift = 0.32
Im Score = −53.41
NG = 0
DB = 0
SM = 0
Delta_K
Doxorubicin
RF1_26_1 RF1
ZShift = −0.75
Im Score = −0.01
NG = 0
DB = 0
SM = 0
Delta_r
Doxorubicin
RF1_26_1 RF1
ZShift = −0.46
Im Score = −1341.61
NG = 0
DB = 0
SM = 0
Delta_AUC
Doxorubicin
RF1_88_0 RF1
ZShift = 0.01
Im Score = 9.91
NG = 0
DB = 1
SM = 0
Delta_L
Doxorubicin
RF1_88_0 RF1
ZShift = −0.52
Im Score = −15.71
NG = 0
DB = 1
SM = 0
Delta_K
Doxorubicin
RF1_88_0 RF1
ZShift = 0.48
Im Score = −0.01
NG = 0
DB = 1
SM = 0
Delta_r
Doxorubicin
RF1_88_0 RF1
ZShift = −0.29
Im Score = −206.83
NG = 0
DB = 1
SM = 0
Delta_AUC
Doxorubicin
RF1_289_0 RF1
ZShift = 1.51
Im Score = 16.91
NG = 0
DB = 0
SM = 0
Delta_L
Doxorubicin
RF1_289_0 RF1
ZShift = 0.52
Im Score = −6.76
NG = 0
DB = 0
SM = 0
Delta_K
Doxorubicin
RF1_289_0 RF1
ZShift = −0.99
Im Score = 0
NG = 0
DB = 0
SM = 0
Delta_r
Doxorubicin
RF1_289_0 RF1
ZShift = −0.83
Im Score = −250.73
NG = 0
DB = 0
SM = 0
Delta_AUC
Doxorubicin

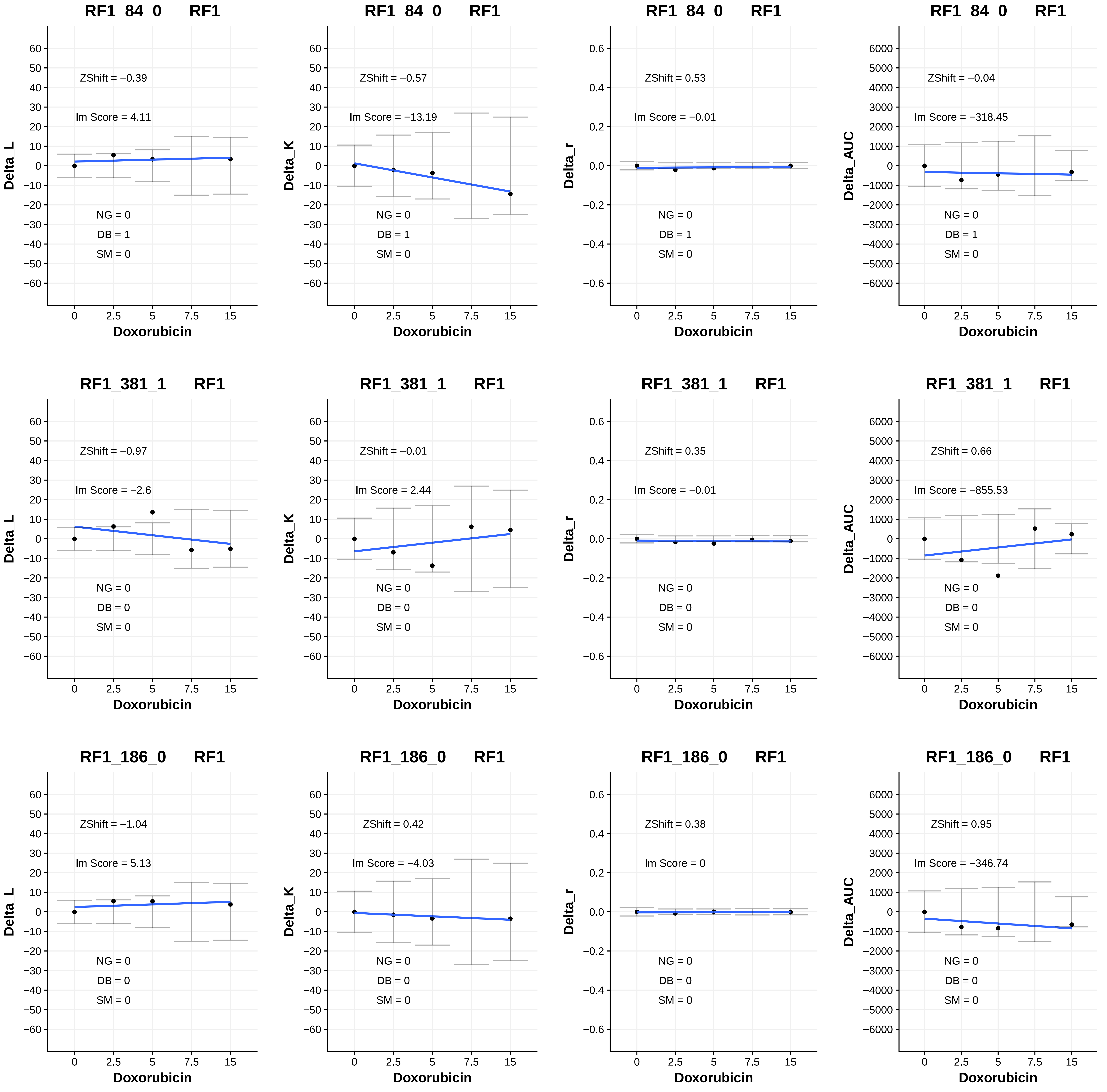

RF1_84_0 RF1
ZShift = −0.39
Im Score = 4.11
NG = 0
DB = 1
SM = 0
Delta_L
Doxorubicin
RF1_84_0 RF1
ZShift = −0.57
Im Score = −13.19
NG = 0
DB = 1
SM = 0
Delta_K
Doxorubicin
RF1_84_0 RF1
ZShift = 0.53
Im Score = −0.01
NG = 0
DB = 1
SM = 0
Delta_r
Doxorubicin
RF1_84_0 RF1
ZShift = −0.04
Im Score = −318.45
NG = 0
DB = 1
SM = 0
Delta_AUC
Doxorubicin
RF1_381_1 RF1
ZShift = −0.97
Im Score = −2.6
NG = 0
DB = 0
SM = 0
Delta_L
Doxorubicin
RF1_381_1 RF1
ZShift = −0.01
Im Score = 2.44
NG = 0
DB = 0
SM = 0
Delta_K
Doxorubicin
RF1_381_1 RF1
ZShift = 0.35
Im Score = −0.01
NG = 0
DB = 0
SM = 0
Delta_r
Doxorubicin
RF1_381_1 RF1
ZShift = 0.66
Im Score = −855.53
NG = 0
DB = 0
SM = 0
Delta_AUC
Doxorubicin
RF1_186_0 RF1
ZShift = −1.04
Im Score = 5.13
NG = 0
DB = 0
SM = 0
Delta_L
Doxorubicin
RF1_186_0 RF1
ZShift = 0.42
Im Score = −4.03
NG = 0
DB = 0
SM = 0
Delta_K
Doxorubicin
RF1_186_0 RF1
ZShift = 0.38
Im Score = 0
NG = 0
DB = 0
SM = 0
Delta_r
Doxorubicin
RF1_186_0 RF1
ZShift = 0.95
Im Score = −346.74
NG = 0
DB = 0
SM = 0
Delta_AUC
Doxorubicin

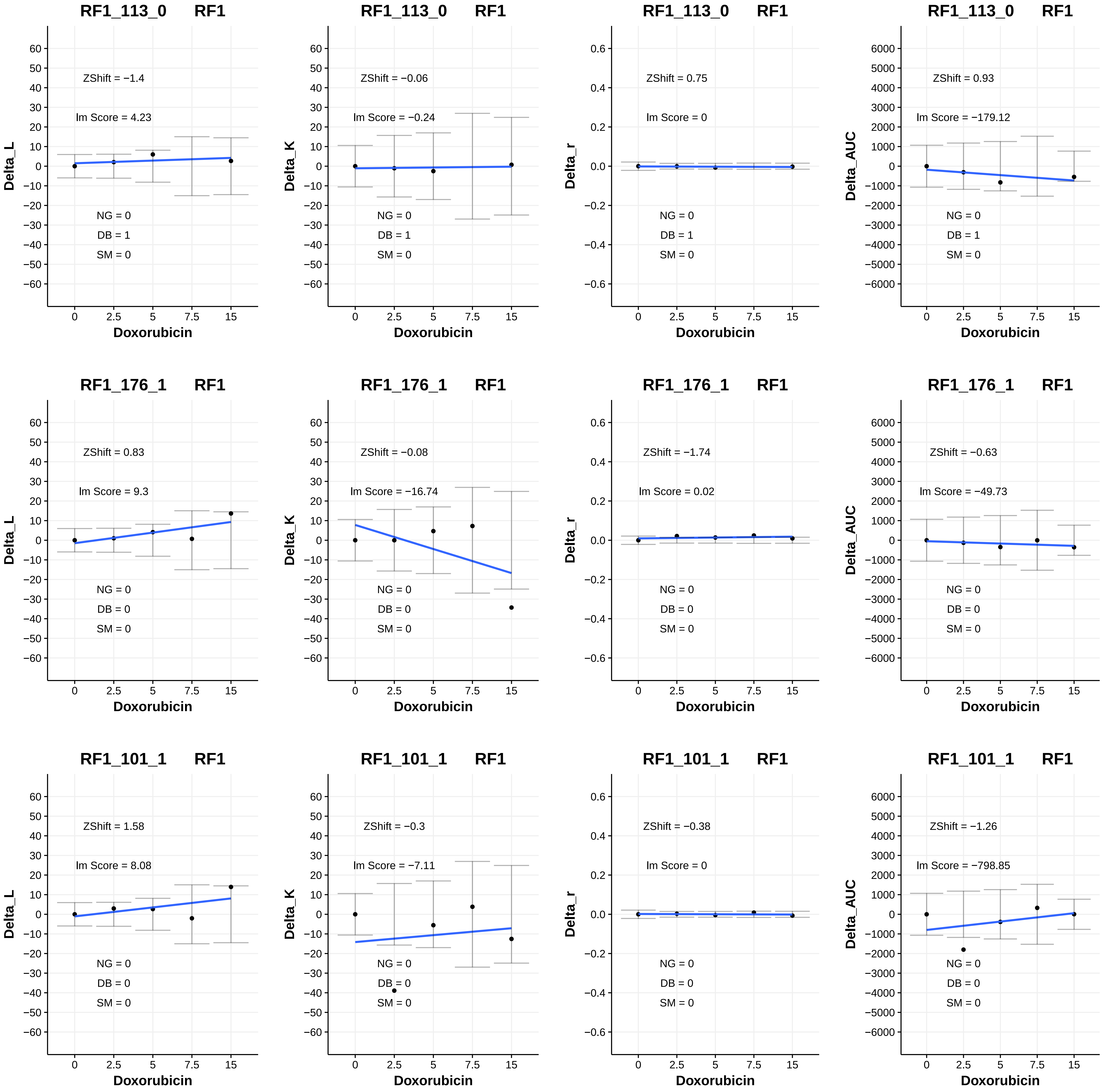

RF1_113_0 RF1
ZShift = −1.4
lm Score = 4.23
NG = 0
DB = 1
SM = 0
Delta_L
Doxorubicin
RF1_113_0 RF1
ZShift = −0.06
lm Score = −0.24
NG = 0
DB = 1
SM = 0
Delta_K
Doxorubicin
RF1_113_0 RF1
ZShift = 0.75
lm Score = 0
NG = 0
DB = 1
SM = 0
Delta_r
Doxorubicin
RF1_113_0 RF1
ZShift = 0.93
lm Score = −179.12
NG = 0
DB = 1
SM = 0
Delta_AUC
Doxorubicin
RF1_176_1 RF1
ZShift = 0.83
lm Score = 9.3
NG = 0
DB = 0
SM = 0
Delta_L
Doxorubicin
RF1_176_1 RF1
ZShift = −0.08
lm Score = −16.74
NG = 0
DB = 0
SM = 0
Delta_K
Doxorubicin
RF1_176_1 RF1
ZShift = −1.74
lm Score = 0.02
NG = 0
DB = 0
SM = 0
Delta_r
Doxorubicin
RF1_176_1 RF1
ZShift = −0.63
lm Score = −49.73
NG = 0
DB = 0
SM = 0
Delta_AUC
Doxorubicin
RF1_101_1 RF1
ZShift = 1.58
lm Score = 8.08
NG = 0
DB = 0
SM = 0
Delta_L
Doxorubicin
RF1_101_1 RF1
ZShift = −0.3
lm Score = −7.11
NG = 0
DB = 0
SM = 0
Delta_K
Doxorubicin
RF1_101_1 RF1
ZShift = −0.38
lm Score = 0
NG = 0
DB = 0
SM = 0
Delta_r
Doxorubicin
RF1_101_1 RF1
ZShift = −1.26
lm Score = −798.85
NG = 0
DB = 0
SM = 0
Delta_AUC
Doxorubicin

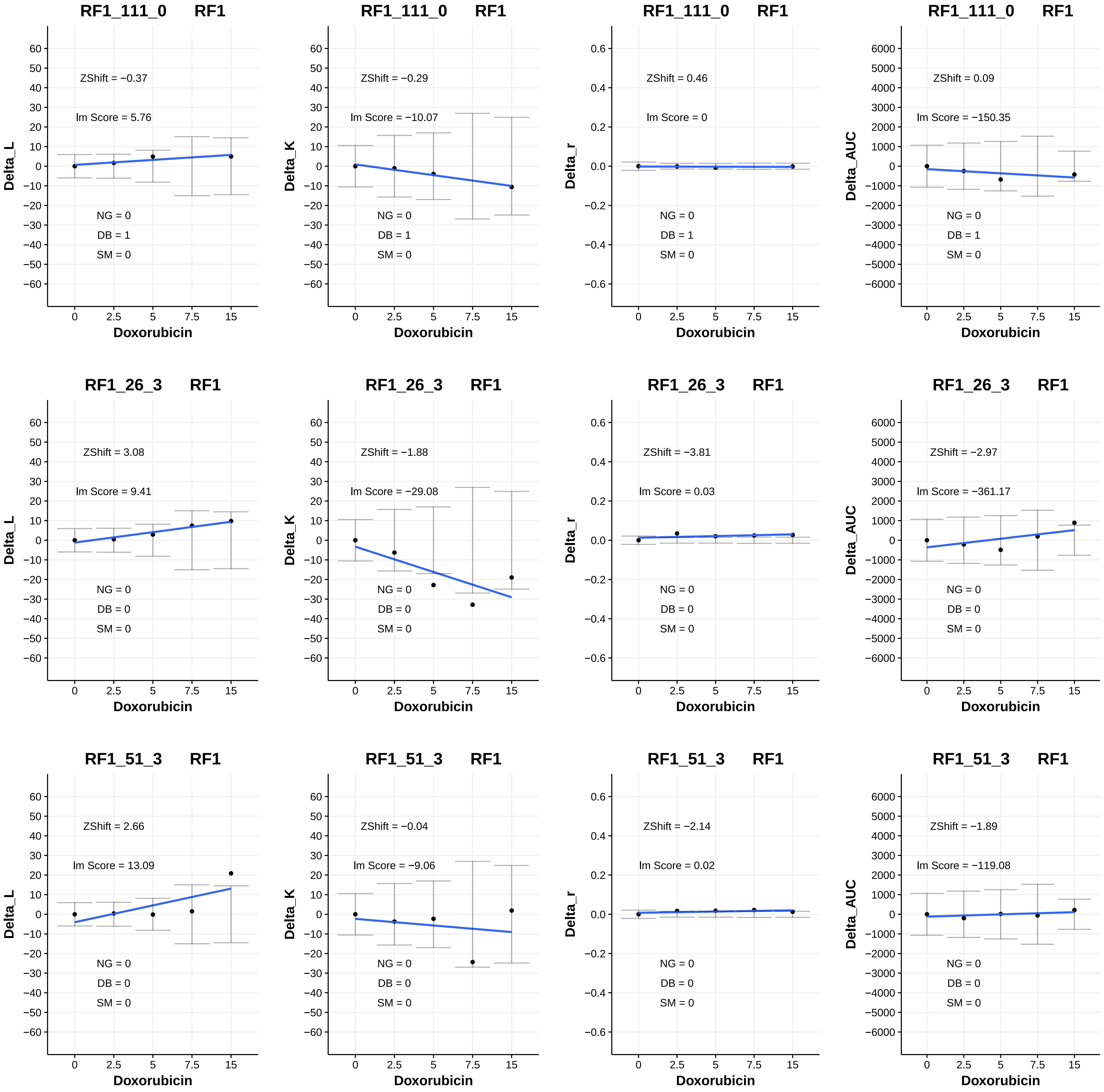
RF1_111_0 RF1
Delta_L
ZShift = −0.37
lm Score = 5.76
NG = 0
DB = 1
SM = 0
Doxorubicin
RF1_111_0 RF1
Delta_K
ZShift = −0.29
lm Score = −10.07
NG = 0
DB = 1
SM = 0
Doxorubicin
RF1_111_0 RF1
Delta_r
ZShift = 0.46
lm Score = 0
NG = 0
DB = 1
SM = 0
Doxorubicin
RF1_111_0 RF1
Delta_AUC
ZShift = 0.09
lm Score = −150.35
NG = 0
DB = 1
SM = 0
Doxorubicin
RF1_26_3 RF1
Delta_L
ZShift = 3.08
lm Score = 9.41
NG = 0
DB = 0
SM = 0
Doxorubicin
RF1_26_3 RF1
Delta_K
ZShift = −1.88
lm Score = −29.08
NG = 0
DB = 0
SM = 0
Doxorubicin
RF1_26_3 RF1
Delta_r
ZShift = −3.81
lm Score = 0.03
NG = 0
DB = 0
SM = 0
Doxorubicin
RF1_26_3 RF1
Delta_AUC
ZShift = −2.97
lm Score = −361.17
NG = 0
DB = 0
SM = 0
Doxorubicin
RF1_51_3 RF1
Delta_L
ZShift = 2.66
lm Score = 13.09
NG = 0
DB = 0
SM = 0
Doxorubicin
RF1_51_3 RF1
Delta_K
ZShift = −0.04
lm Score = −9.06
NG = 0
DB = 0
SM = 0
Doxorubicin
RF1_51_3 RF1
Delta_r
ZShift = −2.14
lm Score = 0.02
NG = 0
DB = 0
SM = 0
Doxorubicin
RF1_51_3 RF1
Delta_AUC
ZShift = −1.89
lm Score = −119.08
NG = 0
DB = 0
SM = 0
Doxorubicin

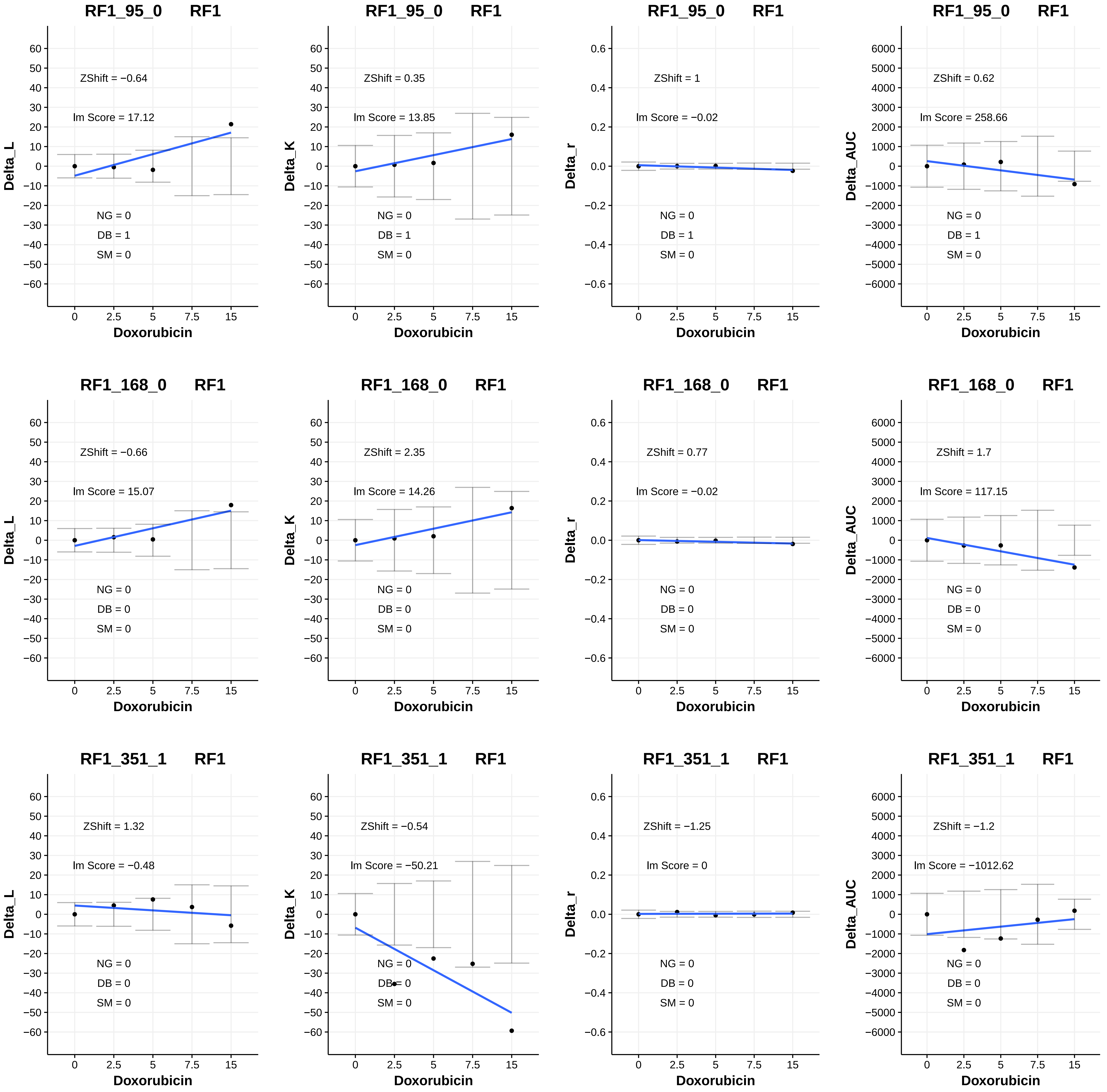
RF1_95_0 RF1
ZShift = −0.64
lm Score = 17.12
NG = 0
DB = 1
SM = 0
Delta_L
Doxorubicin
RF1_95_0 RF1
ZShift = 0.35
lm Score = 13.85
NG = 0
DB = 1
SM = 0
Delta_K
Doxorubicin
RF1_95_0 RF1
ZShift = 1
lm Score = −0.02
NG = 0
DB = 1
SM = 0
Delta_r
Doxorubicin
RF1_95_0 RF1
ZShift = 0.62
lm Score = 258.66
NG = 0
DB = 1
SM = 0
Delta_AUC
Doxorubicin
RF1_168_0 RF1
ZShift = −0.66
lm Score = 15.07
NG = 0
DB = 0
SM = 0
Delta_L
Doxorubicin
RF1_168_0 RF1
ZShift = 2.35
lm Score = 14.26
NG = 0
DB = 0
SM = 0
Delta_K
Doxorubicin
RF1_168_0 RF1
ZShift = 0.77
lm Score = −0.02
NG = 0
DB = 0
SM = 0
Delta_r
Doxorubicin
RF1_168_0 RF1
ZShift = 1.7
lm Score = 117.15
NG = 0
DB = 0
SM = 0
Delta_AUC
Doxorubicin
RF1_351_1 RF1
ZShift = 1.32
lm Score = −0.48
NG = 0
DB = 0
SM = 0
Delta_L
Doxorubicin
RF1_351_1 RF1
ZShift = −0.54
lm Score = −50.21
NG = 0
DB = 0
SM = 0
Delta_K
Doxorubicin
RF1_351_1 RF1
ZShift = −1.25
lm Score = 0
NG = 0
DB = 0
SM = 0
Delta_r
Doxorubicin
RF1_351_1 RF1
ZShift = −1.2
lm Score = −1012.62
NG = 0
DB = 0
SM = 0
Delta_AUC
Doxorubicin

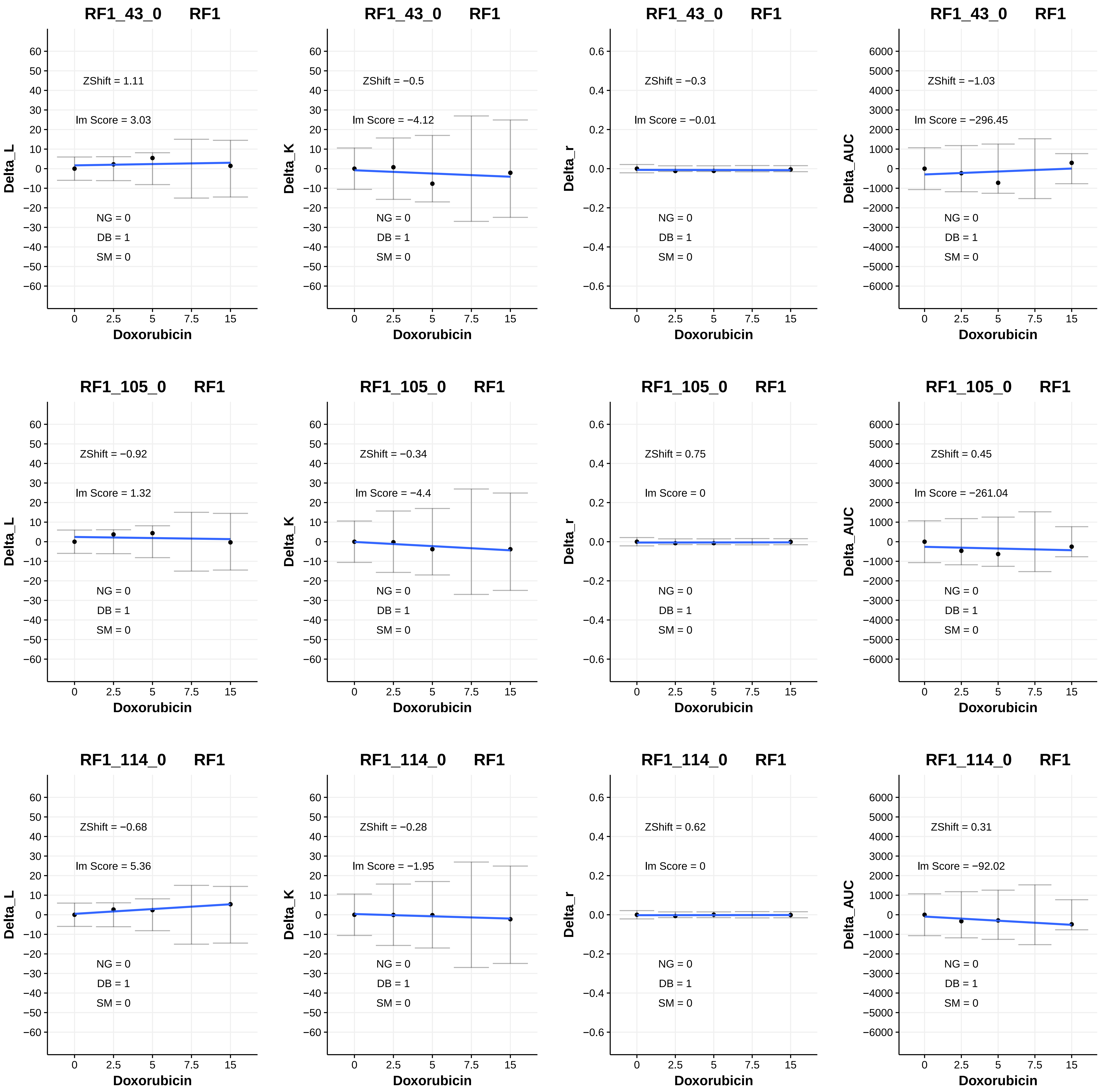
RF1_43_0 RF1
ZShift = 1.11
lm Score = 3.03
NG = 0
DB = 1
SM = 0
Delta_L
Doxorubicin
RF1_43_0 RF1
ZShift = −0.5
lm Score = −4.12
NG = 0
DB = 1
SM = 0
Delta_K
Doxorubicin
RF1_43_0 RF1
ZShift = −0.3
lm Score = −0.01
NG = 0
DB = 1
SM = 0
Delta_r
Doxorubicin
RF1_43_0 RF1
ZShift = −1.03
lm Score = −296.45
NG = 0
DB = 1
SM = 0
Delta_AUC
Doxorubicin
RF1_105_0 RF1
ZShift = −0.92
lm Score = 1.32
NG = 0
DB = 1
SM = 0
Delta_L
Doxorubicin
RF1_105_0 RF1
ZShift = −0.34
lm Score = −4.4
NG = 0
DB = 1
SM = 0
Delta_K
Doxorubicin
RF1_105_0 RF1
ZShift = 0.75
lm Score = 0
NG = 0
DB = 1
SM = 0
Delta_r
Doxorubicin
RF1_105_0 RF1
ZShift = 0.45
lm Score = −261.04
NG = 0
DB = 1
SM = 0
Delta_AUC
Doxorubicin
RF1_114_0 RF1
ZShift = −0.68
lm Score = 5.36
NG = 0
DB = 1
SM = 0
Delta_L
Doxorubicin
RF1_114_0 RF1
ZShift = −0.28
lm Score = −1.95
NG = 0
DB = 1
SM = 0
Delta_K
Doxorubicin
RF1_114_0 RF1
ZShift = 0.62
lm Score = 0
NG = 0
DB = 1
SM = 0
Delta_r
Doxorubicin
RF1_114_0 RF1
ZShift = 0.31
lm Score = −92.02
NG = 0
DB = 1
SM = 0
Delta_AUC
Doxorubicin

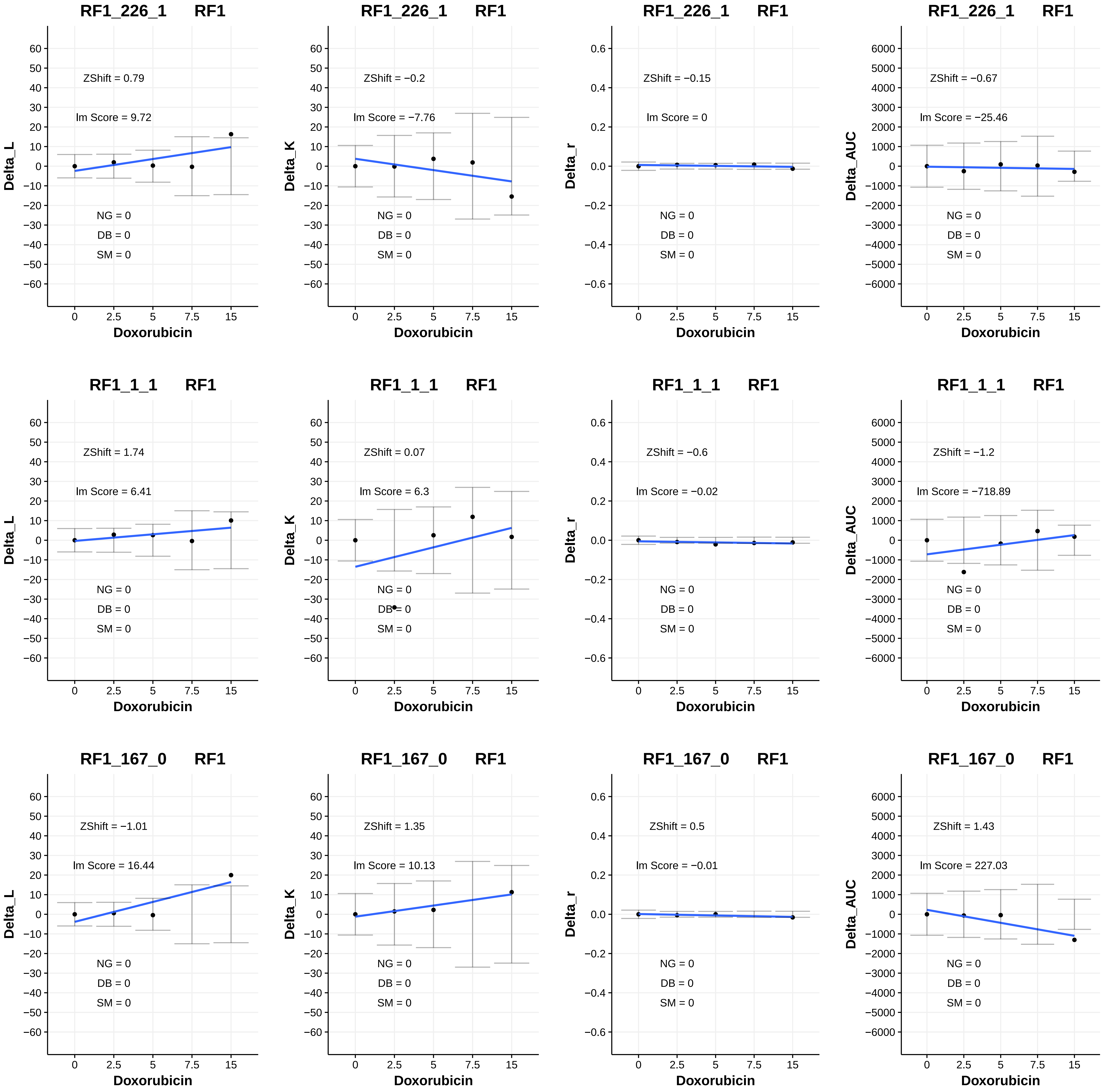
RF1_226_1 RF1
ZShift = 0.79
lm Score = 9.72
NG = 0
DB = 0
SM = 0
Delta_L
Doxorubicin
RF1_226_1 RF1
ZShift = −0.2
lm Score = −7.76
NG = 0
DB = 0
SM = 0
Delta_K
Doxorubicin
RF1_226_1 RF1
ZShift = −0.15
lm Score = 0
NG = 0
DB = 0
SM = 0
Delta_r
Doxorubicin
RF1_226_1 RF1
ZShift = −0.67
lm Score = −25.46
NG = 0
DB = 0
SM = 0
Delta_AUC
Doxorubicin
RF1_1_1 RF1
ZShift = 1.74
lm Score = 6.41
NG = 0
DB = 0
SM = 0
Delta_L
Doxorubicin
RF1_1_1 RF1
ZShift = 0.07
lm Score = 6.3
NG = 0
DB = 0
SM = 0
Delta_K
Doxorubicin
RF1_1_1 RF1
ZShift = −0.6
lm Score = −0.02
NG = 0
DB = 0
SM = 0
Delta_r
Doxorubicin
RF1_1_1 RF1
ZShift = −1.2
lm Score = −718.89
NG = 0
DB = 0
SM = 0
Delta_AUC
Doxorubicin
RF1_167_0 RF1
ZShift = −1.01
lm Score = 16.44
NG = 0
DB = 0
SM = 0
Delta_L
Doxorubicin
RF1_167_0 RF1
ZShift = 1.35
lm Score = 10.13
NG = 0
DB = 0
SM = 0
Delta_K
Doxorubicin
RF1_167_0 RF1
ZShift = 0.5
lm Score = −0.01
NG = 0
DB = 0
SM = 0
Delta_r
Doxorubicin
RF1_167_0 RF1
ZShift = 1.43
lm Score = 227.03
NG = 0
DB = 0
SM = 0
Delta_AUC
Doxorubicin

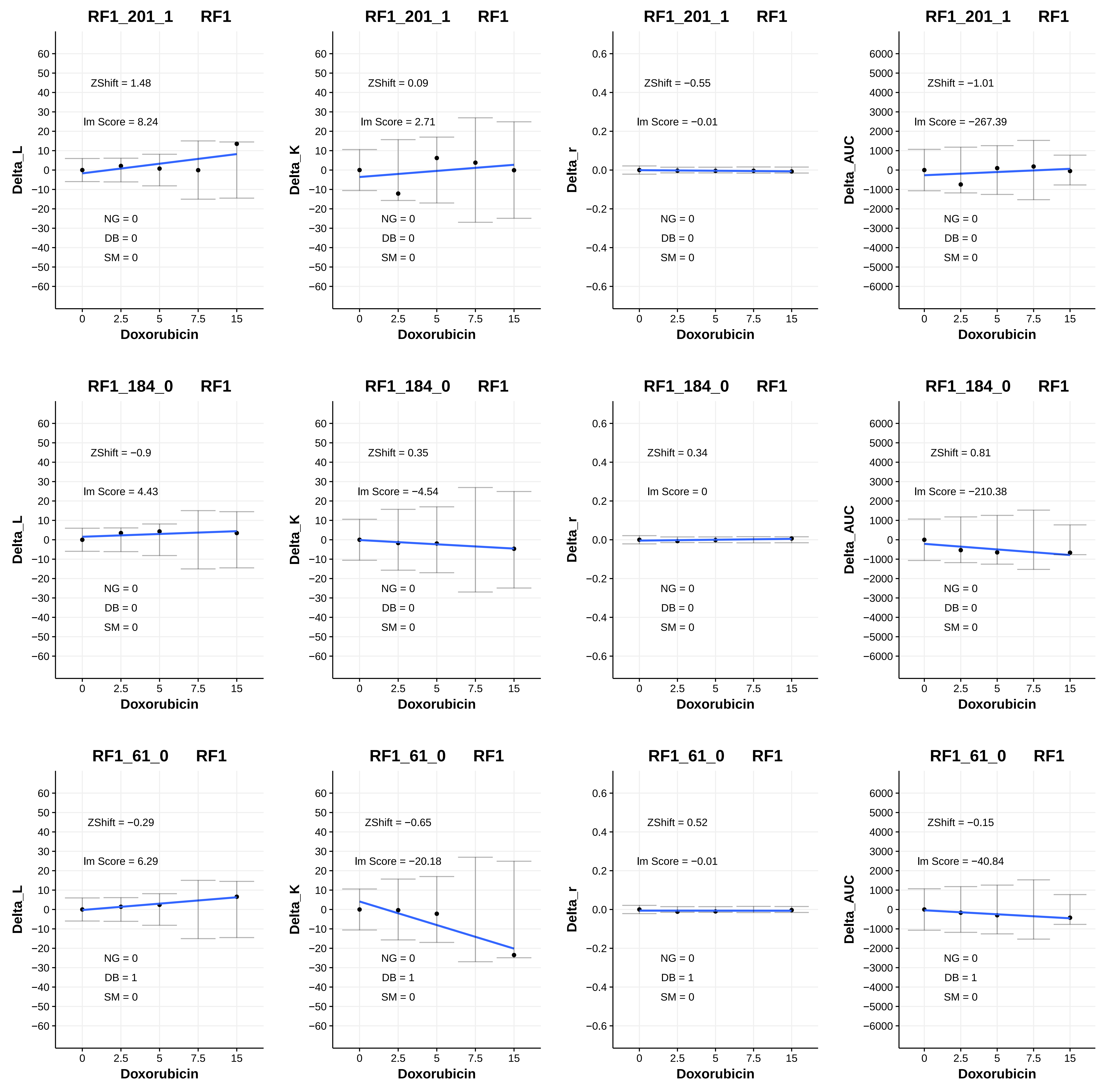

RF1_201_1 RF1
ZShift = 1.48
lm Score = 8.24
NG = 0
DB = 0
SM = 0
Delta_L
Doxorubicin
RF1_201_1 RF1
ZShift = 0.09
lm Score = 2.71
NG = 0
DB = 0
SM = 0
Delta_K
Doxorubicin
RF1_201_1 RF1
ZShift = −0.55
lm Score = −0.01
NG = 0
DB = 0
SM = 0
Delta_r
Doxorubicin
RF1_201_1 RF1
ZShift = −1.01
lm Score = −267.39
NG = 0
DB = 0
SM = 0
Delta_AUC
Doxorubicin
RF1_184_0 RF1
ZShift = −0.9
lm Score = 4.43
NG = 0
DB = 0
SM = 0
Delta_L
Doxorubicin
RF1_184_0 RF1
ZShift = 0.35
lm Score = −4.54
NG = 0
DB = 0
SM = 0
Delta_K
Doxorubicin
RF1_184_0 RF1
ZShift = 0.34
lm Score = 0
NG = 0
DB = 0
SM = 0
Delta_r
Doxorubicin
RF1_184_0 RF1
ZShift = 0.81
lm Score = −210.38
NG = 0
DB = 0
SM = 0
Delta_AUC
Doxorubicin
RF1_61_0 RF1
ZShift = −0.29
lm Score = 6.29
NG = 0
DB = 1
SM = 0
Delta_L
Doxorubicin
RF1_61_0 RF1
ZShift = −0.65
lm Score = −20.18
NG = 0
DB = 1
SM = 0
Delta_K
Doxorubicin
RF1_61_0 RF1
ZShift = 0.52
lm Score = −0.01
NG = 0
DB = 1
SM = 0
Delta_r
Doxorubicin
RF1_61_0 RF1
ZShift = −0.15
lm Score = −40.84
NG = 0
DB = 1
SM = 0
Delta_AUC
Doxorubicin

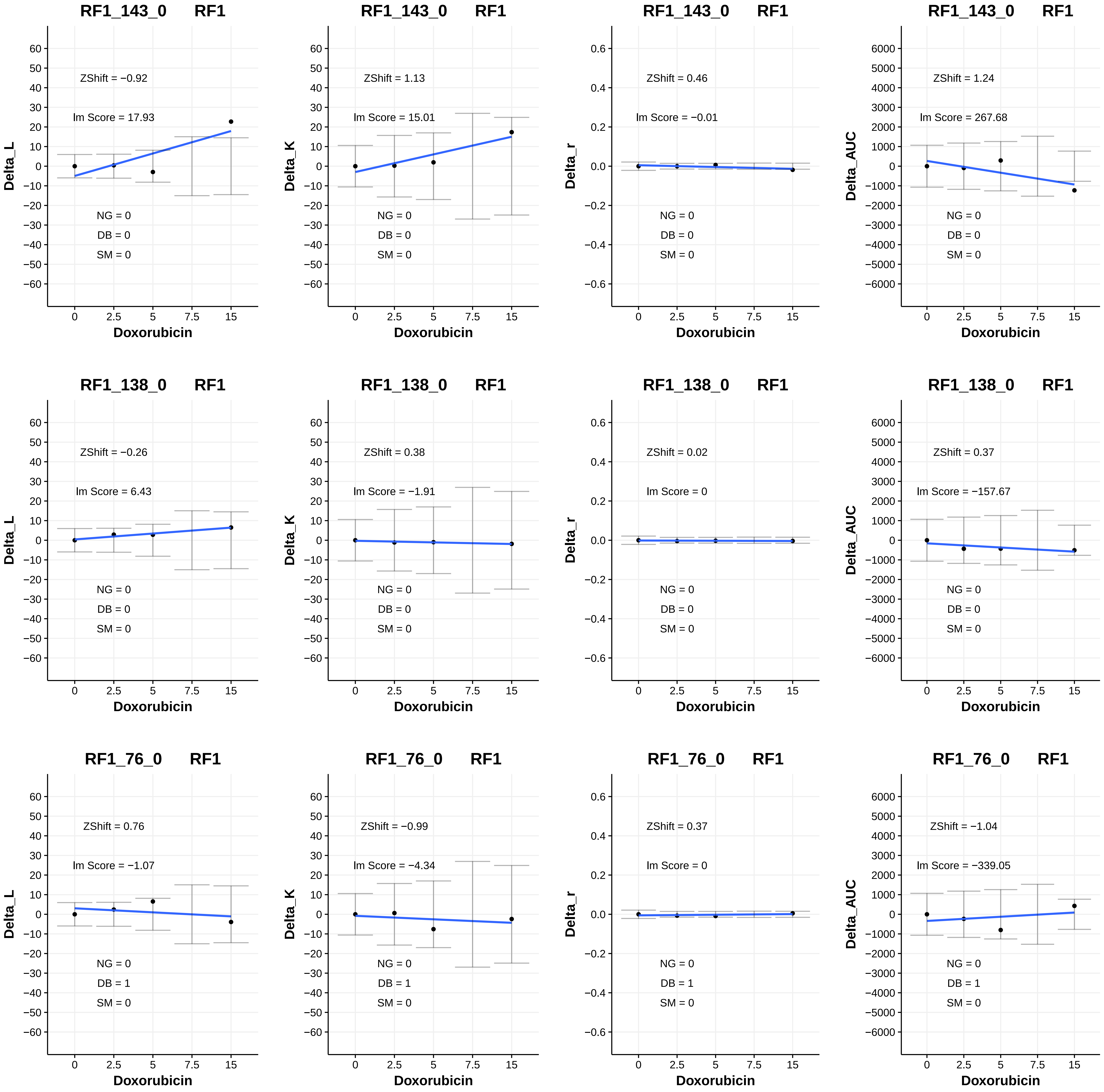

RF1_143_0 RF1
ZShift = −0.92
lm Score = 17.93
NG = 0
DB = 0
SM = 0
Delta_L
Doxorubicin
RF1_143_0 RF1
ZShift = 1.13
lm Score = 15.01
NG = 0
DB = 0
SM = 0
Delta_K
Doxorubicin
RF1_143_0 RF1
ZShift = 0.46
lm Score = −0.01
NG = 0
DB = 0
SM = 0
Delta_r
Doxorubicin
RF1_143_0 RF1
ZShift = 1.24
lm Score = 267.68
NG = 0
DB = 0
SM = 0
Delta_AUC
Doxorubicin
RF1_138_0 RF1
ZShift = −0.26
lm Score = 6.43
NG = 0
DB = 0
SM = 0
Delta_L
Doxorubicin
RF1_138_0 RF1
ZShift = 0.38
lm Score = −1.91
NG = 0
DB = 0
SM = 0
Delta_K
Doxorubicin
RF1_138_0 RF1
ZShift = 0.02
lm Score = 0
NG = 0
DB = 0
SM = 0
Delta_r
Doxorubicin
RF1_138_0 RF1
ZShift = 0.37
lm Score = −157.67
NG = 0
DB = 0
SM = 0
Delta_AUC
Doxorubicin
RF1_76_0 RF1
ZShift = 0.76
lm Score = −1.07
NG = 0
DB = 1
SM = 0
Delta_L
Doxorubicin
RF1_76_0 RF1
ZShift = −0.99
lm Score = −4.34
NG = 0
DB = 1
SM = 0
Delta_K
Doxorubicin
RF1_76_0 RF1
ZShift = 0.37
lm Score = 0
NG = 0
DB = 1
SM = 0
Delta_r
Doxorubicin
RF1_76_0 RF1
ZShift = −1.04
lm Score = −339.05
NG = 0
DB = 1
SM = 0
Delta_AUC
Doxorubicin

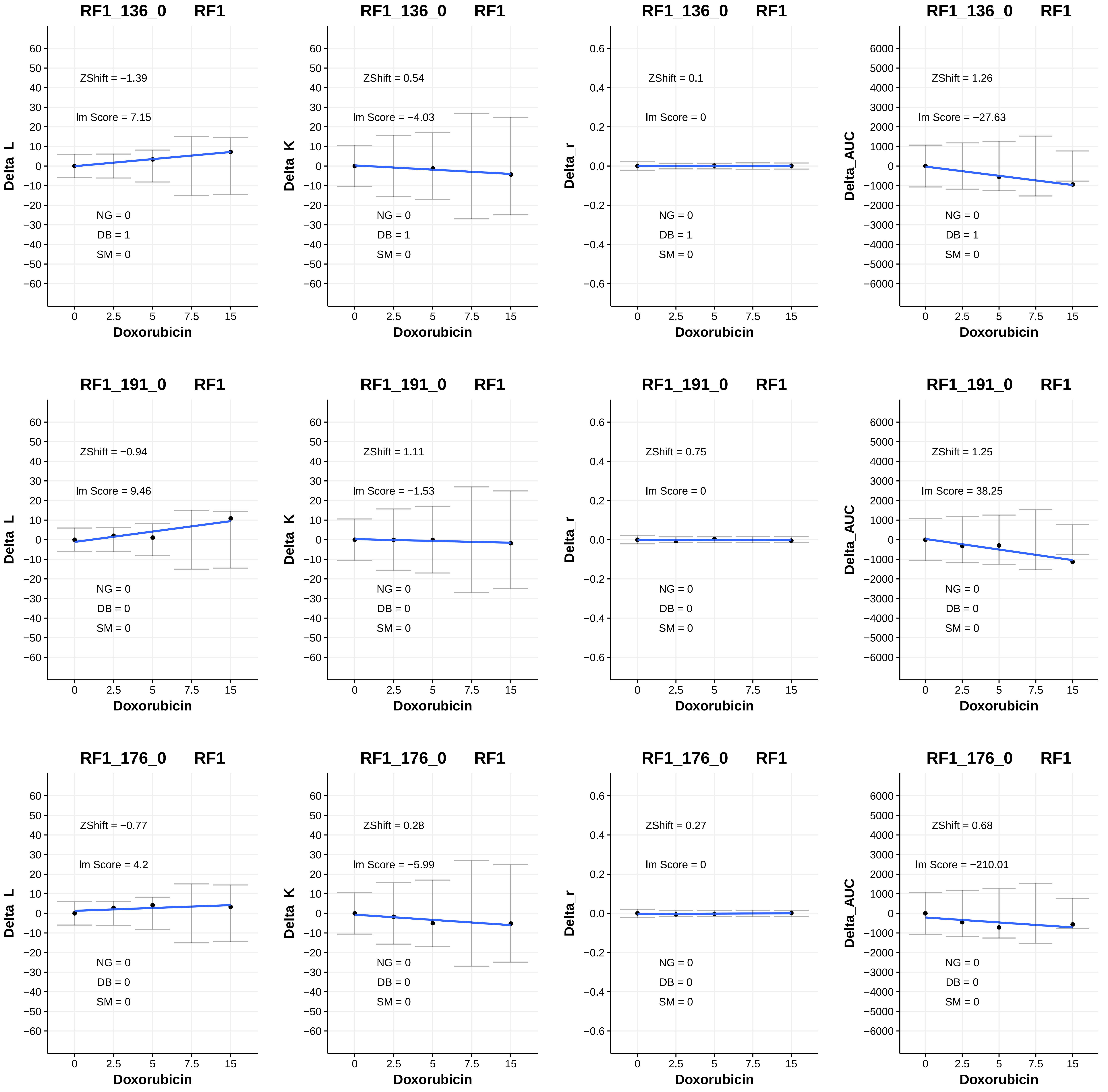
RF1_136_0 RF1
ZShift = −1.39
lm Score = 7.15
NG = 0
DB = 1
SM = 0
Delta_L
Doxorubicin
RF1_136_0 RF1
ZShift = 0.54
lm Score = −4.03
NG = 0
DB = 1
SM = 0
Delta_K
Doxorubicin
RF1_136_0 RF1
ZShift = 0.1
lm Score = 0
NG = 0
DB = 1
SM = 0
Delta_r
Doxorubicin
RF1_136_0 RF1
ZShift = 1.26
lm Score = −27.63
NG = 0
DB = 1
SM = 0
Delta_AUC
Doxorubicin
RF1_191_0 RF1
ZShift = −0.94
lm Score = 9.46
NG = 0
DB = 0
SM = 0
Delta_L
Doxorubicin
RF1_191_0 RF1
ZShift = 1.11
lm Score = −1.53
NG = 0
DB = 0
SM = 0
Delta_K
Doxorubicin
RF1_191_0 RF1
ZShift = 0.75
lm Score = 0
NG = 0
DB = 0
SM = 0
Delta_r
Doxorubicin
RF1_191_0 RF1
ZShift = 1.25
lm Score = 38.25
NG = 0
DB = 0
SM = 0
Delta_AUC
Doxorubicin
RF1_176_0 RF1
ZShift = −0.77
lm Score = 4.2
NG = 0
DB = 0
SM = 0
Delta_L
Doxorubicin
RF1_176_0 RF1
ZShift = 0.28
lm Score = −5.99
NG = 0
DB = 0
SM = 0
Delta_K
Doxorubicin
RF1_176_0 RF1
ZShift = 0.27
lm Score = 0
NG = 0
DB = 0
SM = 0
Delta_r
Doxorubicin
RF1_176_0 RF1
ZShift = 0.68
lm Score = −210.01
NG = 0
DB = 0
SM = 0
Delta_AUC
Doxorubicin

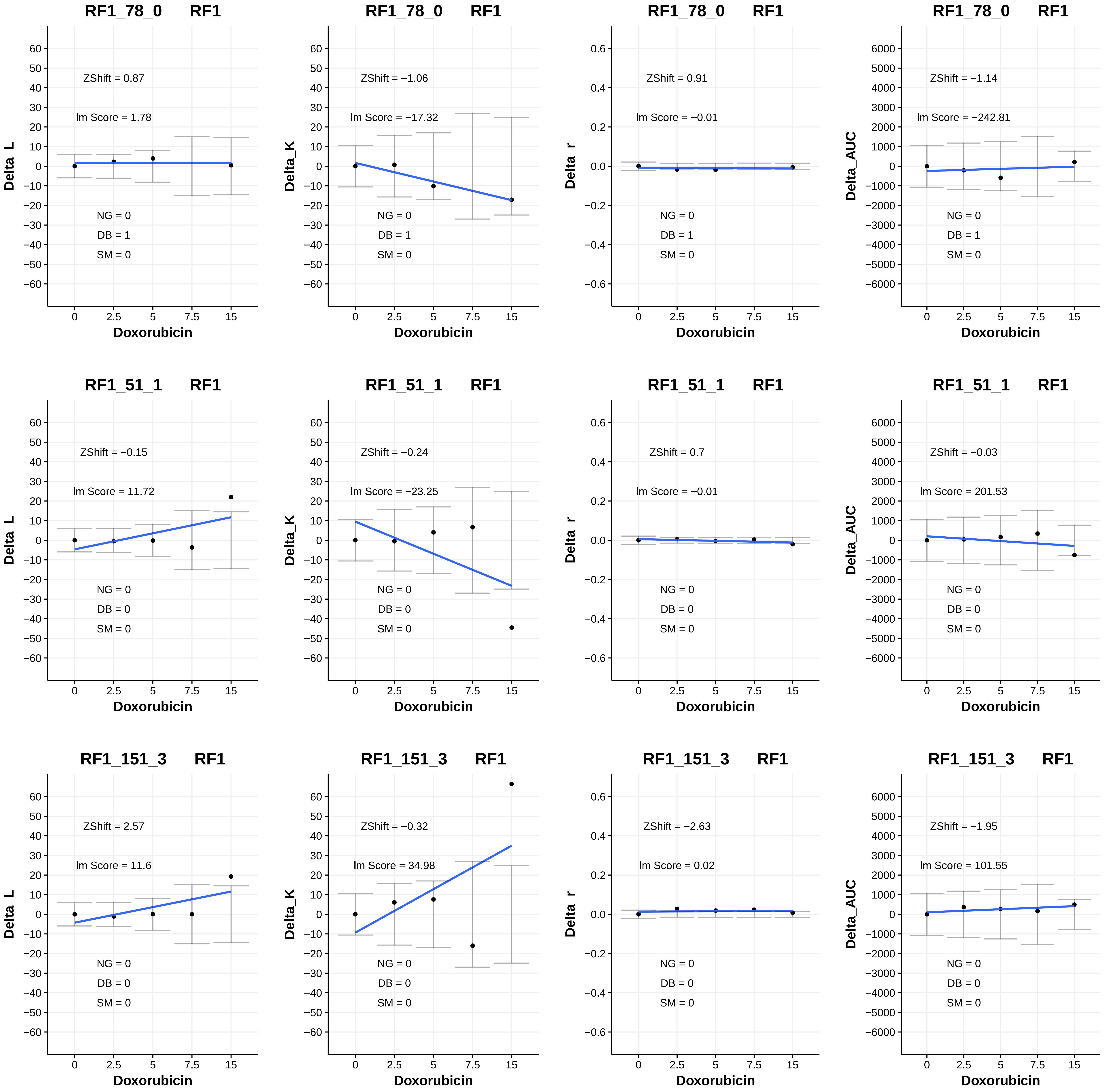
RF1_78_0 RF1
ZShift = 0.87
Im Score = 1.78
NG = 0
DB = 1
SM = 0
Delta_L
Doxorubicin
RF1_78_0 RF1
ZShift = −1.06
Im Score = −17.32
NG = 0
DB = 1
SM = 0
Delta_K
Doxorubicin
RF1_78_0 RF1
ZShift = 0.91
Im Score = −0.01
NG = 0
DB = 1
SM = 0
Delta_r
Doxorubicin
RF1_78_0 RF1
ZShift = −1.14
Im Score = −242.81
NG = 0
DB = 1
SM = 0
Delta_AUC
Doxorubicin
RF1_51_1 RF1
ZShift = −0.15
Im Score = 11.72
NG = 0
DB = 0
SM = 0
Delta_L
Doxorubicin
RF1_51_1 RF1
ZShift = −0.24
Im Score = −23.25
NG = 0
DB = 0
SM = 0
Delta_K
Doxorubicin
RF1_51_1 RF1
ZShift = 0.7
Im Score = −0.01
NG = 0
DB = 0
SM = 0
Delta_r
Doxorubicin
RF1_51_1 RF1
ZShift = −0.03
Im Score = 201.53
NG = 0
DB = 0
SM = 0
Delta_AUC
Doxorubicin
RF1_151_3 RF1
ZShift = 2.57
Im Score = 11.6
NG = 0
DB = 0
SM = 0
Delta_L
Doxorubicin
RF1_151_3 RF1
ZShift = −0.32
Im Score = 34.98
NG = 0
DB = 0
SM = 0
Delta_K
Doxorubicin
RF1_151_3 RF1
ZShift = −2.63
Im Score = 0.02
NG = 0
DB = 0
SM = 0
Delta_r
Doxorubicin
RF1_151_3 RF1
ZShift = −1.95
Im Score = 101.55
NG = 0
DB = 0
SM = 0
Delta_AUC
Doxorubicin

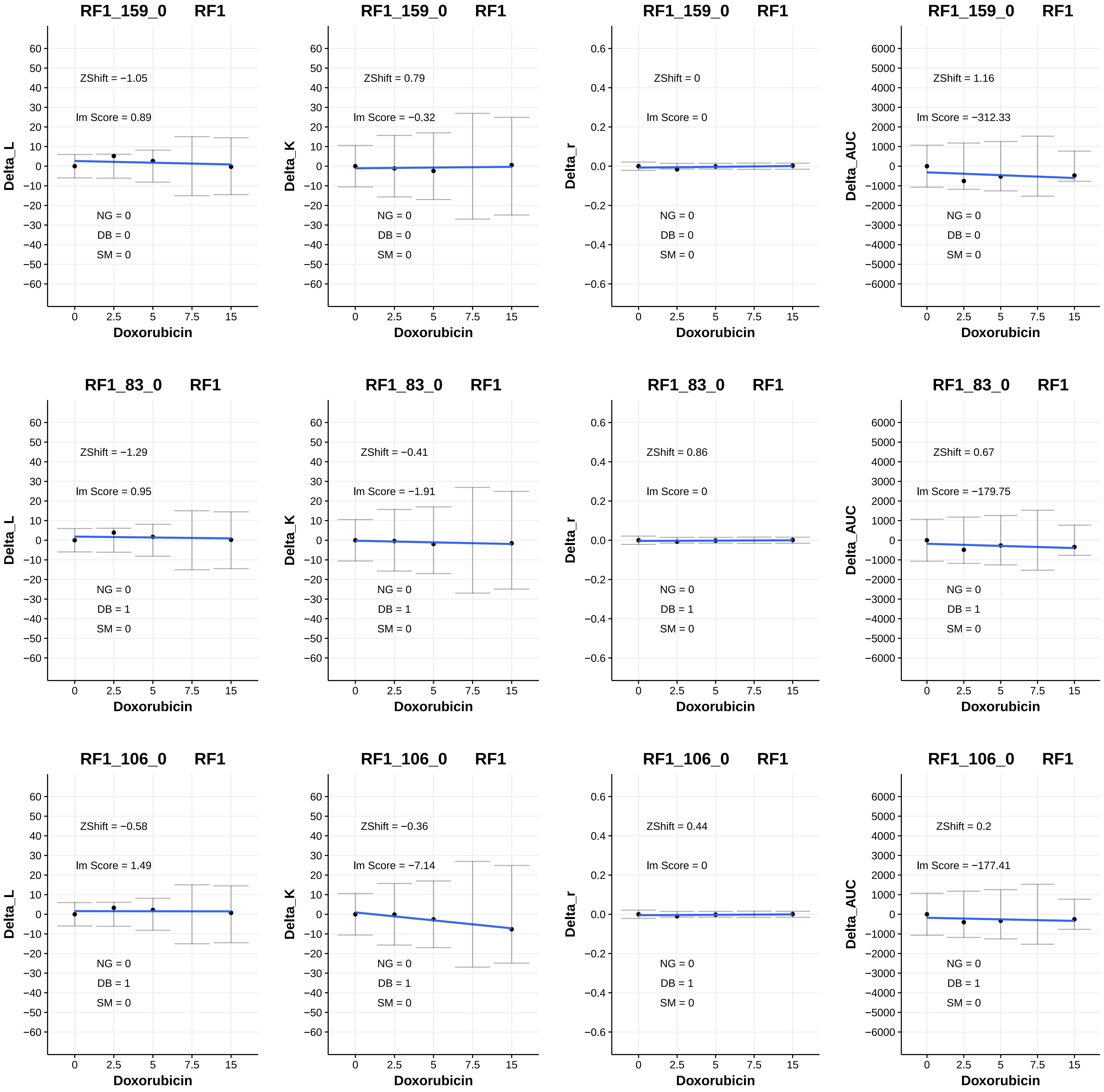

RF1_159_0 RF1
ZShift = −1.05
lm Score = 0.89
NG = 0
DB = 0
SM = 0
Delta_L
Doxorubicin
RF1_159_0 RF1
ZShift = 0.79
lm Score = −0.32
NG = 0
DB = 0
SM = 0
Delta_K
Doxorubicin
RF1_159_0 RF1
ZShift = 0
lm Score = 0
NG = 0
DB = 0
SM = 0
Delta_r
Doxorubicin
RF1_159_0 RF1
ZShift = 1.16
lm Score = −312.33
NG = 0
DB = 0
SM = 0
Delta_AUC
Doxorubicin
RF1_83_0 RF1
ZShift = −1.29
lm Score = 0.95
NG = 0
DB = 1
SM = 0
Delta_L
Doxorubicin
RF1_83_0 RF1
ZShift = −0.41
lm Score = −1.91
NG = 0
DB = 1
SM = 0
Delta_K
Doxorubicin
RF1_83_0 RF1
ZShift = 0.86
lm Score = 0
NG = 0
DB = 1
SM = 0
Delta_r
Doxorubicin
RF1_83_0 RF1
ZShift = 0.67
lm Score = −179.75
NG = 0
DB = 1
SM = 0
Delta_AUC
Doxorubicin
RF1_106_0 RF1
ZShift = −0.58
lm Score = 1.49
NG = 0
DB = 1
SM = 0
Delta_L
Doxorubicin
RF1_106_0 RF1
ZShift = −0.36
lm Score = −7.14
NG = 0
DB = 1
SM = 0
Delta_K
Doxorubicin
RF1_106_0 RF1
ZShift = 0.44
lm Score = 0
NG = 0
DB = 1
SM = 0
Delta_r
Doxorubicin
RF1_106_0 RF1
ZShift = 0.2
lm Score = −177.41
NG = 0
DB = 1
SM = 0
Delta_AUC
Doxorubicin

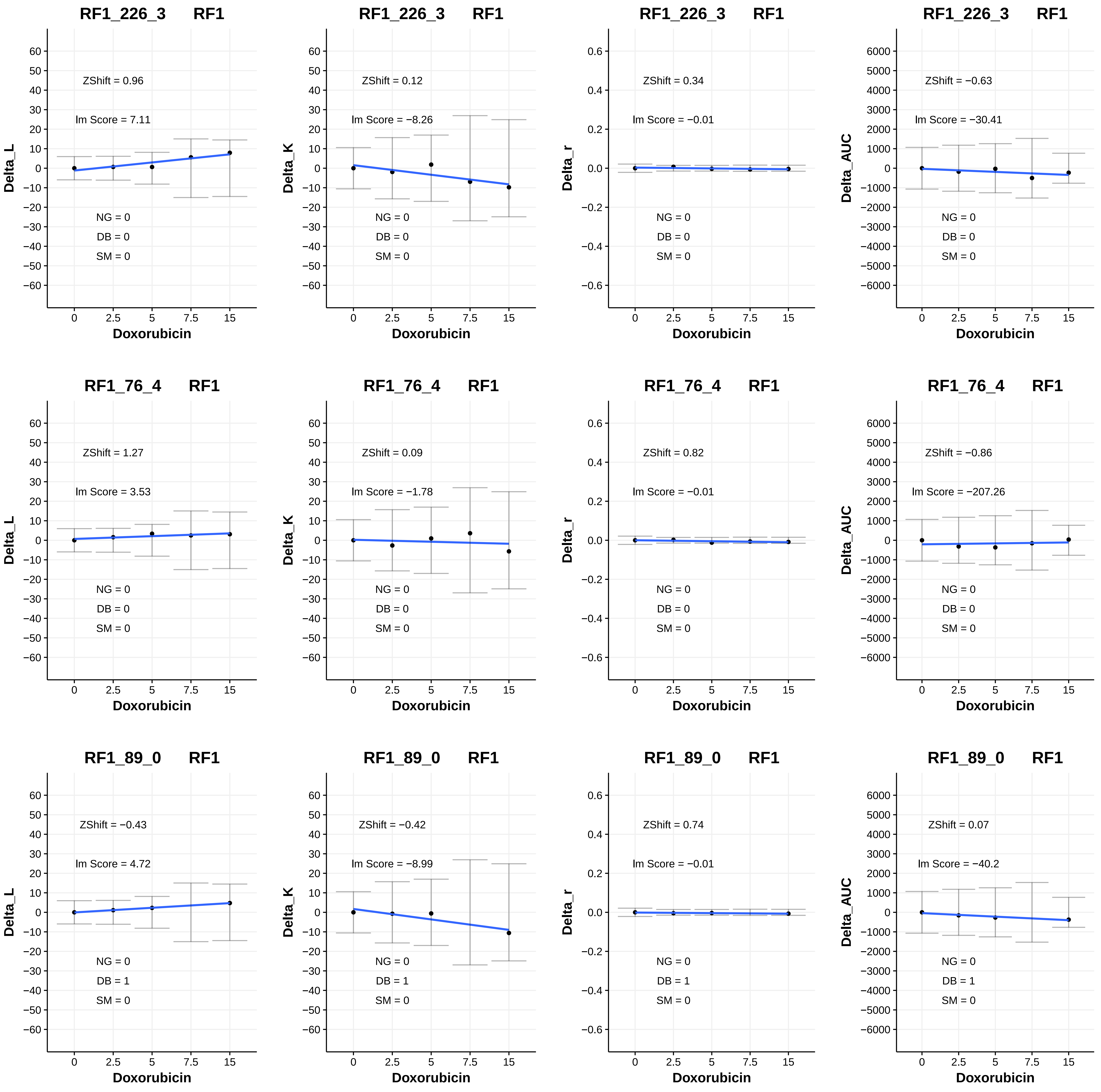
RF1_226_3 RF1
ZShift = 0.96
Im Score = 7.11
NG = 0
DB = 0
SM = 0
Delta_L
Doxorubicin
RF1_226_3 RF1
ZShift = 0.12
Im Score = −8.26
NG = 0
DB = 0
SM = 0
Delta_K
Doxorubicin
RF1_226_3 RF1
ZShift = 0.34
Im Score = −0.01
NG = 0
DB = 0
SM = 0
Delta_r
Doxorubicin
RF1_226_3 RF1
ZShift = −0.63
Im Score = −30.41
NG = 0
DB = 0
SM = 0
Delta_AUC
Doxorubicin
RF1_76_4 RF1
ZShift = 1.27
Im Score = 3.53
NG = 0
DB = 0
SM = 0
Delta_L
Doxorubicin
RF1_76_4 RF1
ZShift = 0.09
Im Score = −1.78
NG = 0
DB = 0
SM = 0
Delta_K
Doxorubicin
RF1_76_4 RF1
ZShift = 0.82
Im Score = −0.01
NG = 0
DB = 0
SM = 0
Delta_r
Doxorubicin
RF1_76_4 RF1
ZShift = −0.86
Im Score = −207.26
NG = 0
DB = 0
SM = 0
Delta_AUC
Doxorubicin
RF1_89_0 RF1
ZShift = −0.43
Im Score = 4.72
NG = 0
DB = 1
SM = 0
Delta_L
Doxorubicin
RF1_89_0 RF1
ZShift = −0.42
Im Score = −8.99
NG = 0
DB = 1
SM = 0
Delta_K
Doxorubicin
RF1_89_0 RF1
ZShift = 0.74
Im Score = −0.01
NG = 0
DB = 1
SM = 0
Delta_r
Doxorubicin
RF1_89_0 RF1
ZShift = 0.07
Im Score = −40.2
NG = 0
DB = 1
SM = 0
Delta_AUC
Doxorubicin

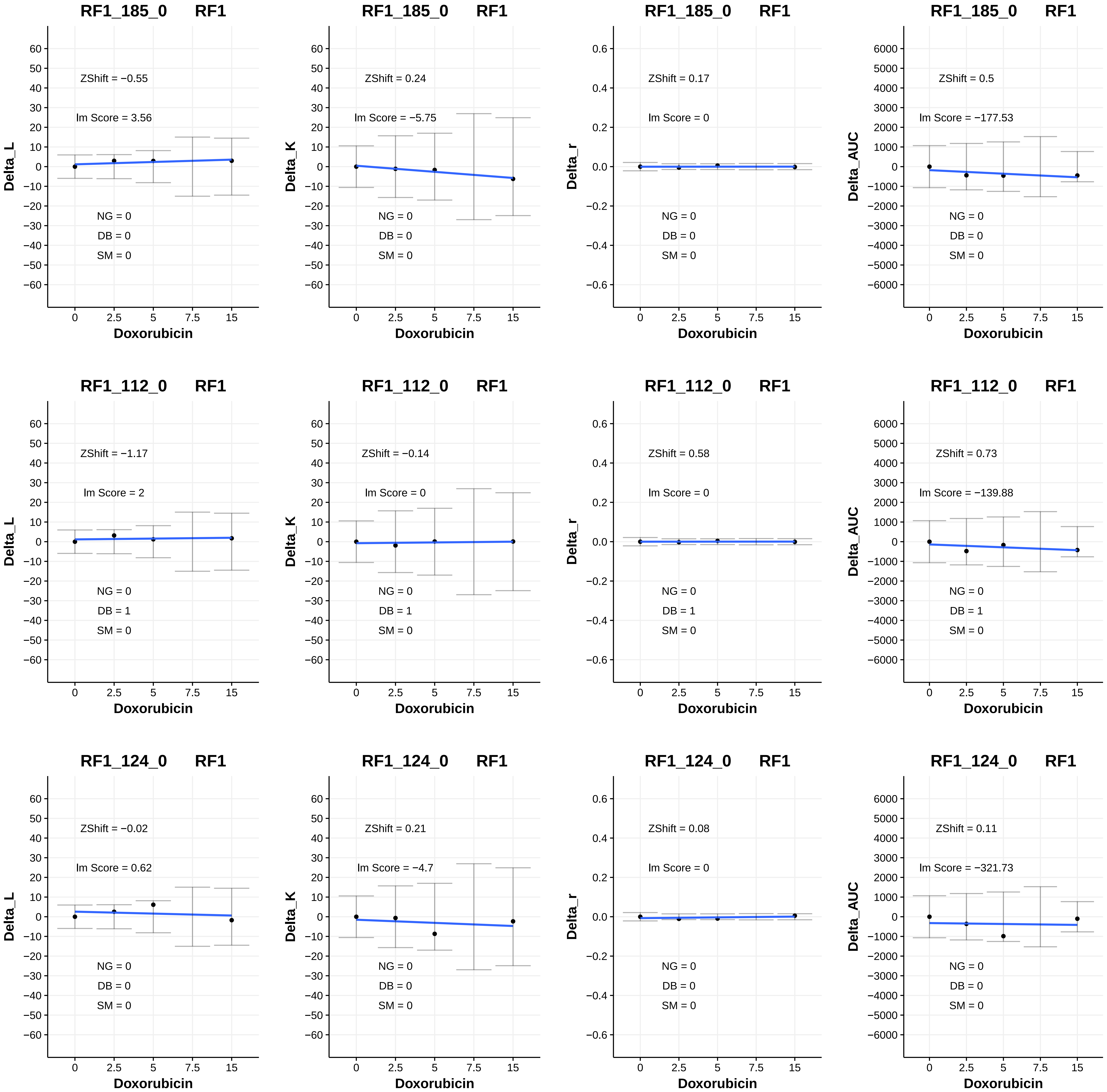
RF1_185_0 RF1
ZShift = −0.55
lm Score = 3.56
NG = 0
DB = 0
SM = 0
Delta_L
Doxorubicin
RF1_185_0 RF1
ZShift = 0.24
lm Score = −5.75
NG = 0
DB = 0
SM = 0
Delta_K
Doxorubicin
RF1_185_0 RF1
ZShift = 0.17
lm Score = 0
NG = 0
DB = 0
SM = 0
Delta_r
Doxorubicin
RF1_185_0 RF1
ZShift = 0.5
lm Score = −177.53
NG = 0
DB = 0
SM = 0
Delta_AUC
Doxorubicin
RF1_112_0 RF1
ZShift = −1.17
lm Score = 2
NG = 0
DB = 1
SM = 0
Delta_L
Doxorubicin
RF1_112_0 RF1
ZShift = −0.14
lm Score = 0
NG = 0
DB = 1
SM = 0
Delta_K
Doxorubicin
RF1_112_0 RF1
ZShift = 0.58
lm Score = 0
NG = 0
DB = 1
SM = 0
Delta_r
Doxorubicin
RF1_112_0 RF1
ZShift = 0.73
lm Score = −139.88
NG = 0
DB = 1
SM = 0
Delta_AUC
Doxorubicin
RF1_124_0 RF1
ZShift = −0.02
lm Score = 0.62
NG = 0
DB = 0
SM = 0
Delta_L
Doxorubicin
RF1_124_0 RF1
ZShift = 0.21
lm Score = −4.7
NG = 0
DB = 0
SM = 0
Delta_K
Doxorubicin
RF1_124_0 RF1
ZShift = 0.08
lm Score = 0
NG = 0
DB = 0
SM = 0
Delta_r
Doxorubicin
RF1_124_0 RF1
ZShift = 0.11
lm Score = −321.73
NG = 0
DB = 0
SM = 0
Delta_AUC
Doxorubicin

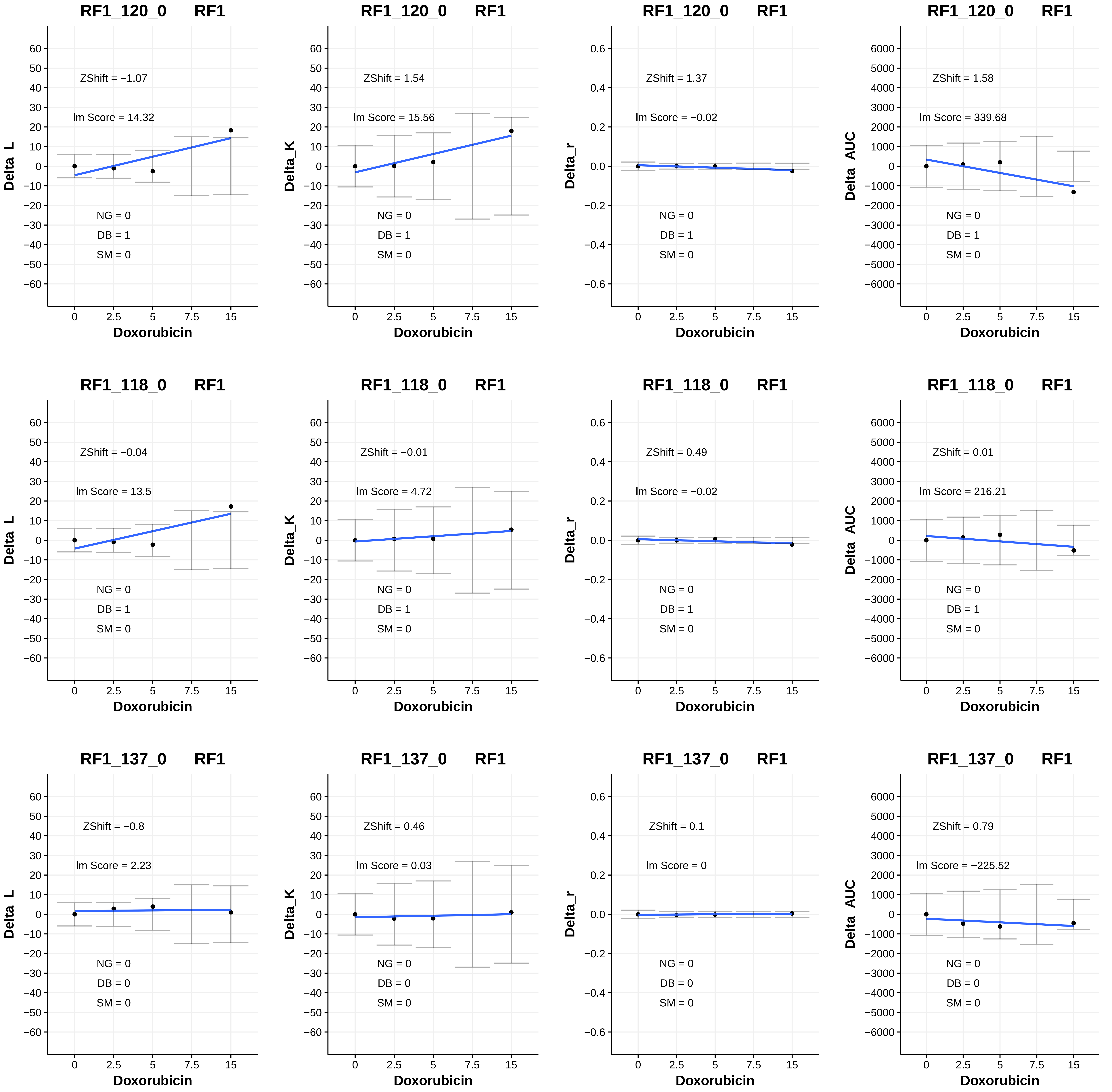
RF1_120_0 RF1
ZShift = −1.07
lm Score = 14.32
NG = 0
DB = 1
SM = 0
Delta_L
Doxorubicin
RF1_120_0 RF1
ZShift = 1.54
lm Score = 15.56
NG = 0
DB = 1
SM = 0
Delta_K
Doxorubicin
RF1_120_0 RF1
ZShift = 1.37
lm Score = −0.02
NG = 0
DB = 1
SM = 0
Delta_r
Doxorubicin
RF1_120_0 RF1
ZShift = 1.58
lm Score = 339.68
NG = 0
DB = 1
SM = 0
Delta_AUC
Doxorubicin
RF1_118_0 RF1
ZShift = −0.04
lm Score = 13.5
NG = 0
DB = 1
SM = 0
Delta_L
Doxorubicin
RF1_118_0 RF1
ZShift = −0.01
lm Score = 4.72
NG = 0
DB = 1
SM = 0
Delta_K
Doxorubicin
RF1_118_0 RF1
ZShift = 0.49
lm Score = −0.02
NG = 0
DB = 1
SM = 0
Delta_r
Doxorubicin
RF1_118_0 RF1
ZShift = 0.01
lm Score = 216.21
NG = 0
DB = 1
SM = 0
Delta_AUC
Doxorubicin
RF1_137_0 RF1
ZShift = −0.8
lm Score = 2.23
NG = 0
DB = 0
SM = 0
Delta_L
Doxorubicin
RF1_137_0 RF1
ZShift = 0.46
lm Score = 0.03
NG = 0
DB = 0
SM = 0
Delta_K
Doxorubicin
RF1_137_0 RF1
ZShift = 0.1
lm Score = 0
NG = 0
DB = 0
SM = 0
Delta_r
Doxorubicin
RF1_137_0 RF1
ZShift = 0.79
lm Score = −225.52
NG = 0
DB = 0
SM = 0
Delta_AUC
Doxorubicin

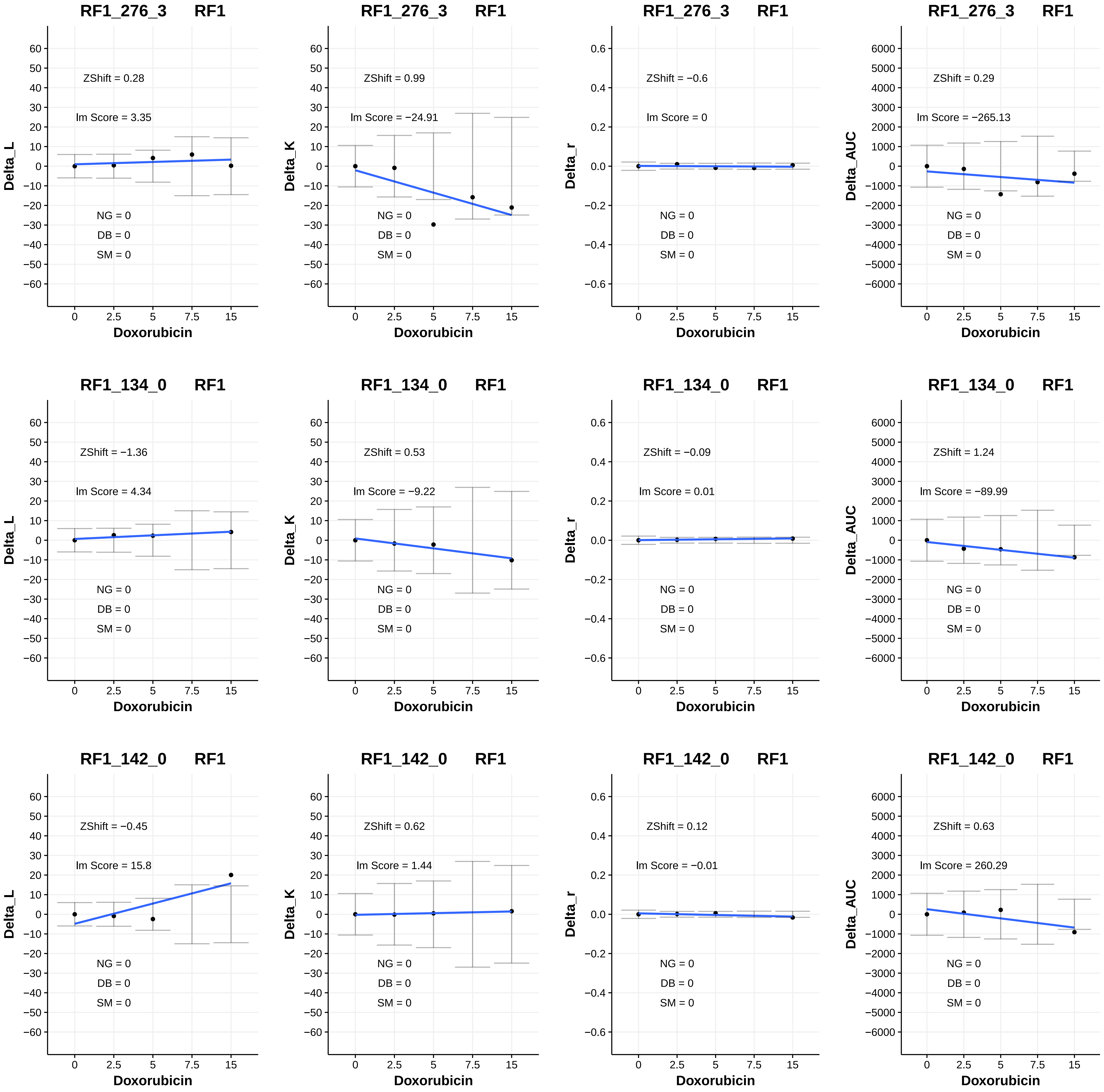

RF1_276_3 RF1
ZShift = 0.28
lm Score = 3.35
NG = 0
DB = 0
SM = 0
Delta_L
Doxorubicin
RF1_276_3 RF1
ZShift = 0.99
lm Score = −24.91
NG = 0
DB = 0
SM = 0
Delta_K
Doxorubicin
RF1_276_3 RF1
ZShift = −0.6
lm Score = 0
NG = 0
DB = 0
SM = 0
Delta_r
Doxorubicin
RF1_276_3 RF1
ZShift = 0.29
lm Score = −265.13
NG = 0
DB = 0
SM = 0
Delta_AUC
Doxorubicin
RF1_134_0 RF1
ZShift = −1.36
lm Score = 4.34
NG = 0
DB = 0
SM = 0
Delta_L
Doxorubicin
RF1_134_0 RF1
ZShift = 0.53
lm Score = −9.22
NG = 0
DB = 0
SM = 0
Delta_K
Doxorubicin
RF1_134_0 RF1
ZShift = −0.09
lm Score = 0.01
NG = 0
DB = 0
SM = 0
Delta_r
Doxorubicin
RF1_134_0 RF1
ZShift = 1.24
lm Score = −89.99
NG = 0
DB = 0
SM = 0
Delta_AUC
Doxorubicin
RF1_142_0 RF1
ZShift = −0.45
lm Score = 15.8
NG = 0
DB = 0
SM = 0
Delta_L
Doxorubicin
RF1_142_0 RF1
ZShift = 0.62
lm Score = 1.44
NG = 0
DB = 0
SM = 0
Delta_K
Doxorubicin
RF1_142_0 RF1
ZShift = 0.12
lm Score = −0.01
NG = 0
DB = 0
SM = 0
Delta_r
Doxorubicin
RF1_142_0 RF1
ZShift = 0.63
lm Score = 260.29
NG = 0
DB = 0
SM = 0
Delta_AUC
Doxorubicin

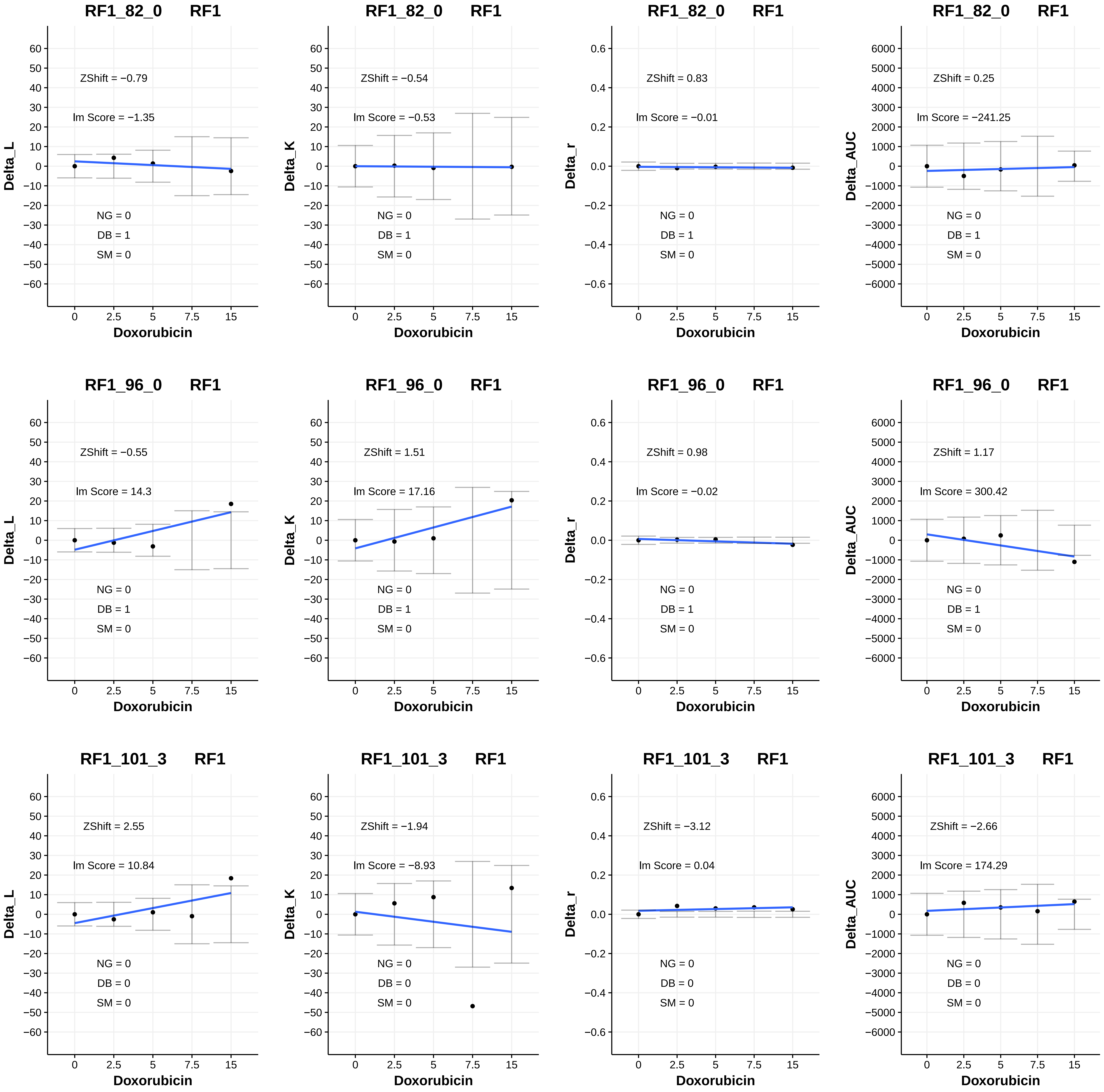
RF1_82_0 RF1
ZShift = −0.79
lm Score = −1.35
NG = 0
DB = 1
SM = 0
Delta_L
Doxorubicin
RF1_82_0 RF1
ZShift = −0.54
lm Score = −0.53
NG = 0
DB = 1
SM = 0
Delta_K
Doxorubicin
RF1_82_0 RF1
ZShift = 0.83
lm Score = −0.01
NG = 0
DB = 1
SM = 0
Delta_r
Doxorubicin
RF1_82_0 RF1
ZShift = 0.25
lm Score = −241.25
NG = 0
DB = 1
SM = 0
Delta_AUC
Doxorubicin
RF1_96_0 RF1
ZShift = −0.55
lm Score = 14.3
NG = 0
DB = 1
SM = 0
Delta_L
Doxorubicin
RF1_96_0 RF1
ZShift = 1.51
lm Score = 17.16
NG = 0
DB = 1
SM = 0
Delta_K
Doxorubicin
RF1_96_0 RF1
ZShift = 0.98
lm Score = −0.02
NG = 0
DB = 1
SM = 0
Delta_r
Doxorubicin
RF1_96_0 RF1
ZShift = 1.17
lm Score = 300.42
NG = 0
DB = 1
SM = 0
Delta_AUC
Doxorubicin
RF1_101_3 RF1
ZShift = 2.55
lm Score = 10.84
NG = 0
DB = 0
SM = 0
Delta_L
Doxorubicin
RF1_101_3 RF1
ZShift = −1.94
lm Score = −8.93
NG = 0
DB = 0
SM = 0
Delta_K
Doxorubicin
RF1_101_3 RF1
ZShift = −3.12
lm Score = 0.04
NG = 0
DB = 0
SM = 0
Delta_r
Doxorubicin
RF1_101_3 RF1
ZShift = −2.66
lm Score = 174.29
NG = 0
DB = 0
SM = 0
Delta_AUC
Doxorubicin

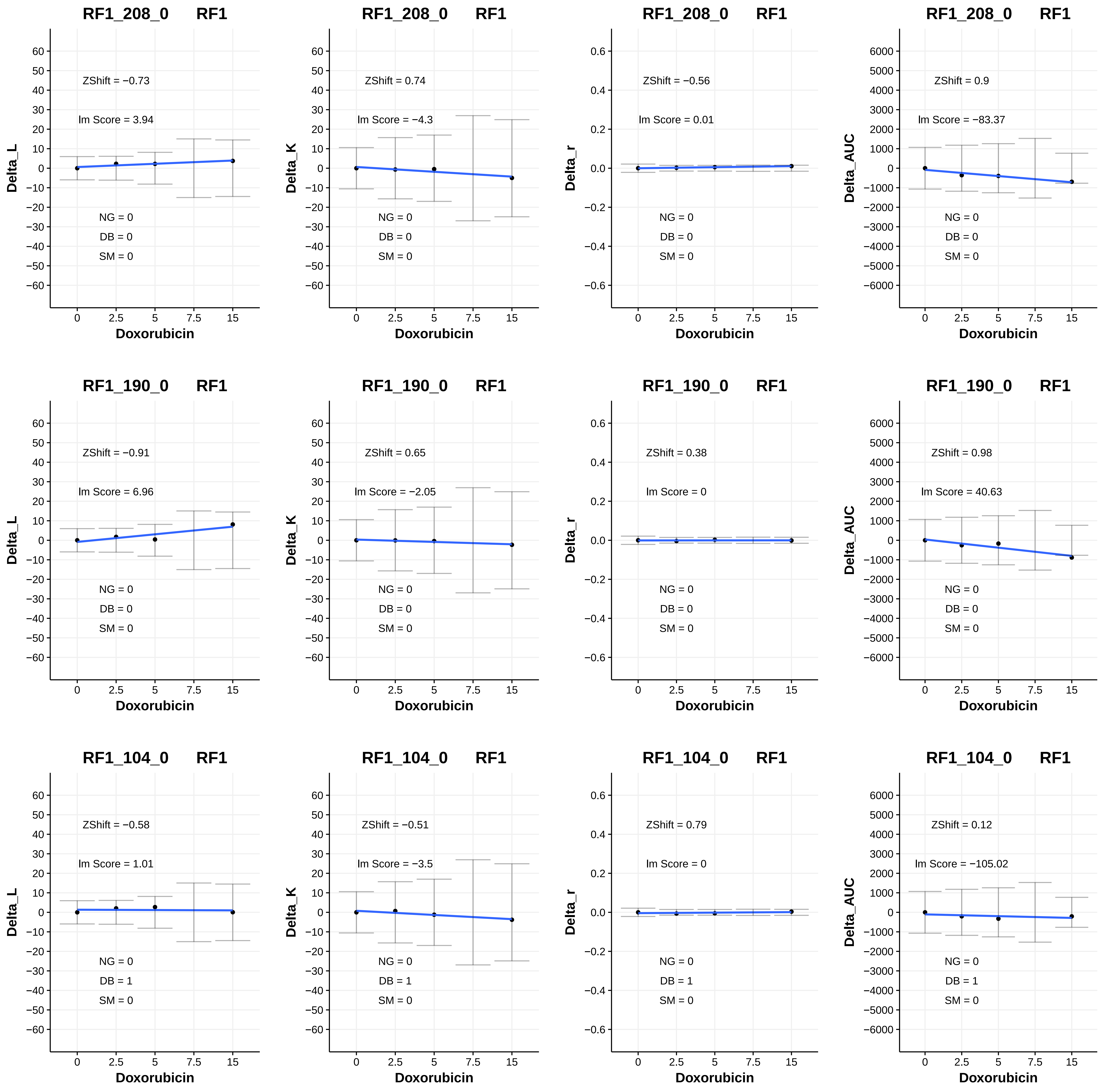
RF1_208_0 RF1
ZShift = −0.73
lm Score = 3.94
NG = 0
DB = 0
SM = 0
Delta_L
Doxorubicin
RF1_208_0 RF1
ZShift = 0.74
lm Score = −4.3
NG = 0
DB = 0
SM = 0
Delta_K
Doxorubicin
RF1_208_0 RF1
ZShift = −0.56
lm Score = 0.01
NG = 0
DB = 0
SM = 0
Delta_r
Doxorubicin
RF1_208_0 RF1
ZShift = 0.9
lm Score = −83.37
NG = 0
DB = 0
SM = 0
Delta_AUC
Doxorubicin
RF1_190_0 RF1
ZShift = −0.91
lm Score = 6.96
NG = 0
DB = 0
SM = 0
Delta_L
Doxorubicin
RF1_190_0 RF1
ZShift = 0.65
lm Score = −2.05
NG = 0
DB = 0
SM = 0
Delta_K
Doxorubicin
RF1_190_0 RF1
ZShift = 0.38
lm Score = 0
NG = 0
DB = 0
SM = 0
Delta_r
Doxorubicin
RF1_190_0 RF1
ZShift = 0.98
lm Score = 40.63
NG = 0
DB = 0
SM = 0
Delta_AUC
Doxorubicin
RF1_104_0 RF1
ZShift = −0.58
lm Score = 1.01
NG = 0
DB = 1
SM = 0
Delta_L
Doxorubicin
RF1_104_0 RF1
ZShift = −0.51
lm Score = −3.5
NG = 0
DB = 1
SM = 0
Delta_K
Doxorubicin
RF1_104_0 RF1
ZShift = 0.79
lm Score = 0
NG = 0
DB = 1
SM = 0
Delta_r
Doxorubicin
RF1_104_0 RF1
ZShift = 0.12
lm Score = −105.02
NG = 0
DB = 1
SM = 0
Delta_AUC
Doxorubicin

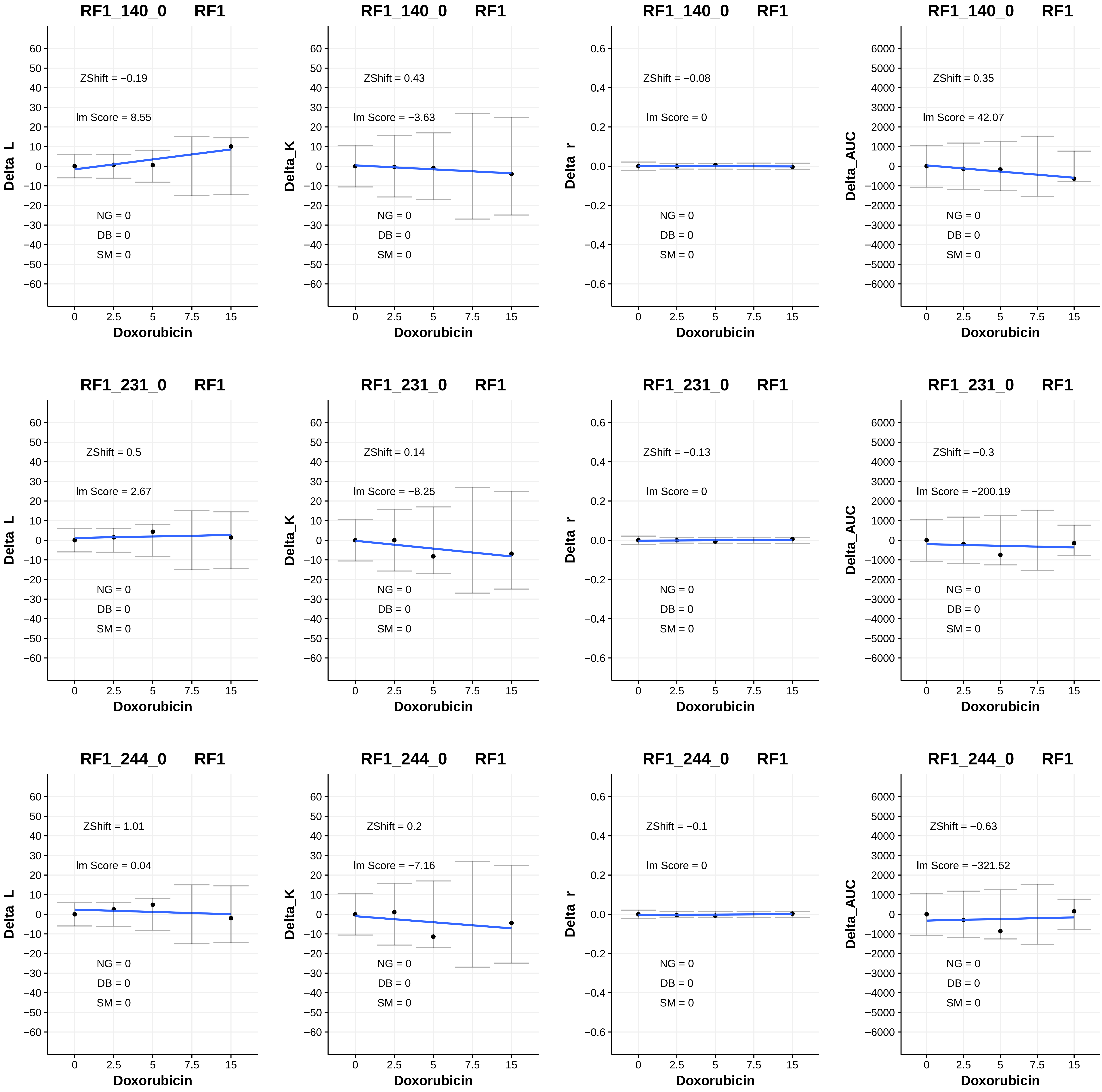
RF1_140_0 RF1
ZShift = −0.19
Im Score = 8.55
NG = 0
DB = 0
SM = 0
Delta_L
Doxorubicin
RF1_140_0 RF1
ZShift = 0.43
Im Score = −3.63
NG = 0
DB = 0
SM = 0
Delta_K
Doxorubicin
RF1_140_0 RF1
ZShift = −0.08
Im Score = 0
NG = 0
DB = 0
SM = 0
Delta_r
Doxorubicin
RF1_140_0 RF1
ZShift = 0.35
Im Score = 42.07
NG = 0
DB = 0
SM = 0
Delta_AUC
Doxorubicin
RF1_231_0 RF1
ZShift = 0.5
Im Score = 2.67
NG = 0
DB = 0
SM = 0
Delta_L
Doxorubicin
RF1_231_0 RF1
ZShift = 0.14
Im Score = −8.25
NG = 0
DB = 0
SM = 0
Delta_K
Doxorubicin
RF1_231_0 RF1
ZShift = −0.13
Im Score = 0
NG = 0
DB = 0
SM = 0
Delta_r
Doxorubicin
RF1_231_0 RF1
ZShift = −0.3
Im Score = −200.19
NG = 0
DB = 0
SM = 0
Delta_AUC
Doxorubicin
RF1_244_0 RF1
ZShift = 1.01
Im Score = 0.04
NG = 0
DB = 0
SM = 0
Delta_L
Doxorubicin
RF1_244_0 RF1
ZShift = 0.2
Im Score = −7.16
NG = 0
DB = 0
SM = 0
Delta_K
Doxorubicin
RF1_244_0 RF1
ZShift = −0.1
Im Score = 0
NG = 0
DB = 0
SM = 0
Delta_r
Doxorubicin
RF1_244_0 RF1
ZShift = −0.63
Im Score = −321.52
NG = 0
DB = 0
SM = 0
Delta_AUC
Doxorubicin

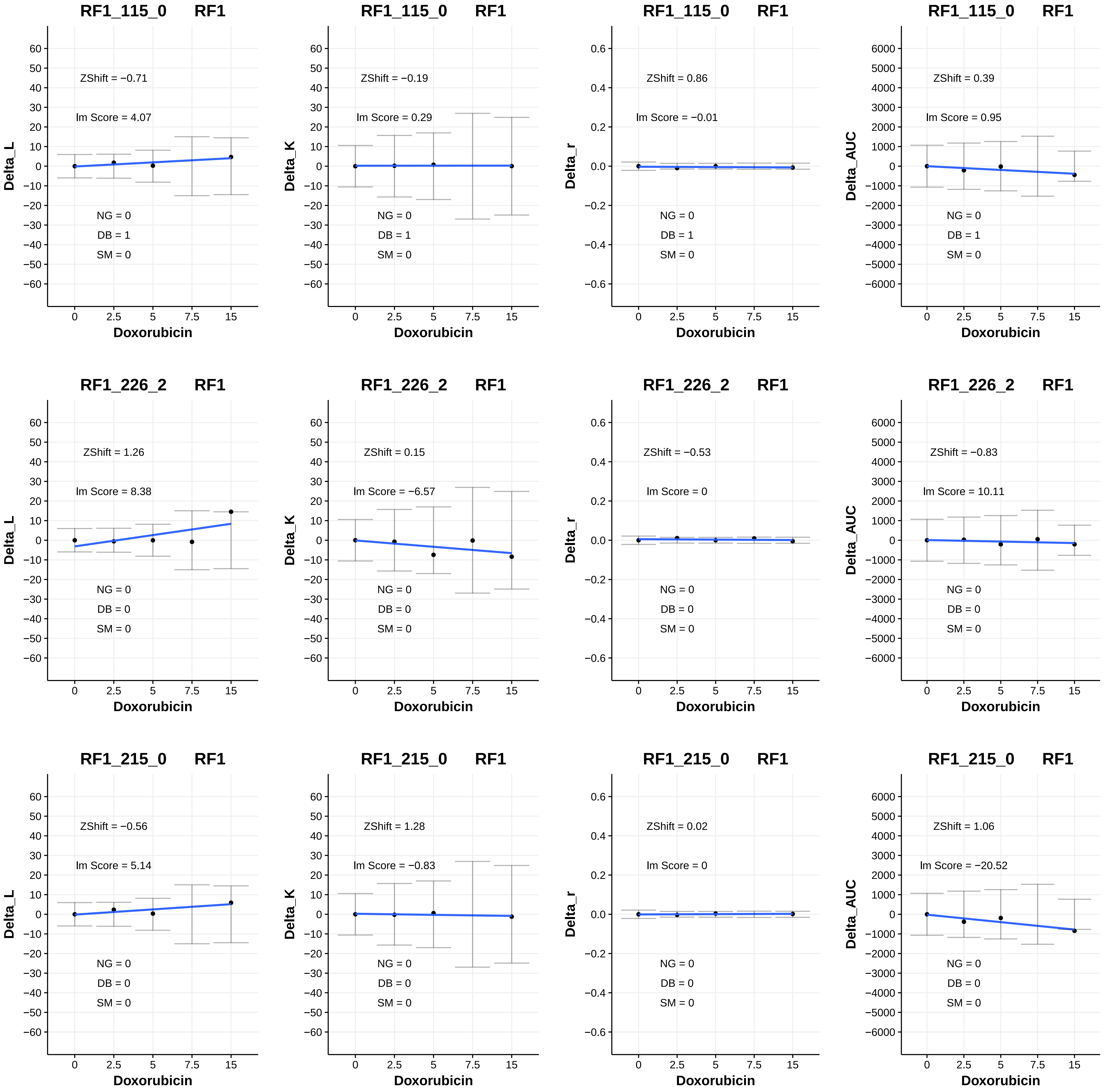
RF1_115_0 RF1
ZShift = −0.71
lm Score = 4.07
NG = 0
DB = 1
SM = 0
Delta_L
Doxorubicin
RF1_115_0 RF1
ZShift = −0.19
lm Score = 0.29
NG = 0
DB = 1
SM = 0
Delta_K
Doxorubicin
RF1_115_0 RF1
ZShift = 0.86
lm Score = −0.01
NG = 0
DB = 1
SM = 0
Delta_r
Doxorubicin
RF1_115_0 RF1
ZShift = 0.39
lm Score = 0.95
NG = 0
DB = 1
SM = 0
Delta_AUC
Doxorubicin
RF1_226_2 RF1
ZShift = 1.26
lm Score = 8.38
NG = 0
DB = 0
SM = 0
Delta_L
Doxorubicin
RF1_226_2 RF1
ZShift = 0.15
lm Score = −6.57
NG = 0
DB = 0
SM = 0
Delta_K
Doxorubicin
RF1_226_2 RF1
ZShift = −0.53
lm Score = 0
NG = 0
DB = 0
SM = 0
Delta_r
Doxorubicin
RF1_226_2 RF1
ZShift = −0.83
lm Score = 10.11
NG = 0
DB = 0
SM = 0
Delta_AUC
Doxorubicin
RF1_215_0 RF1
ZShift = −0.56
lm Score = 5.14
NG = 0
DB = 0
SM = 0
Delta_L
Doxorubicin
RF1_215_0 RF1
ZShift = 1.28
lm Score = −0.83
NG = 0
DB = 0
SM = 0
Delta_K
Doxorubicin
RF1_215_0 RF1
ZShift = 0.02
lm Score = 0
NG = 0
DB = 0
SM = 0
Delta_r
Doxorubicin
RF1_215_0 RF1
ZShift = 1.06
lm Score = −20.52
NG = 0
DB = 0
SM = 0
Delta_AUC
Doxorubicin

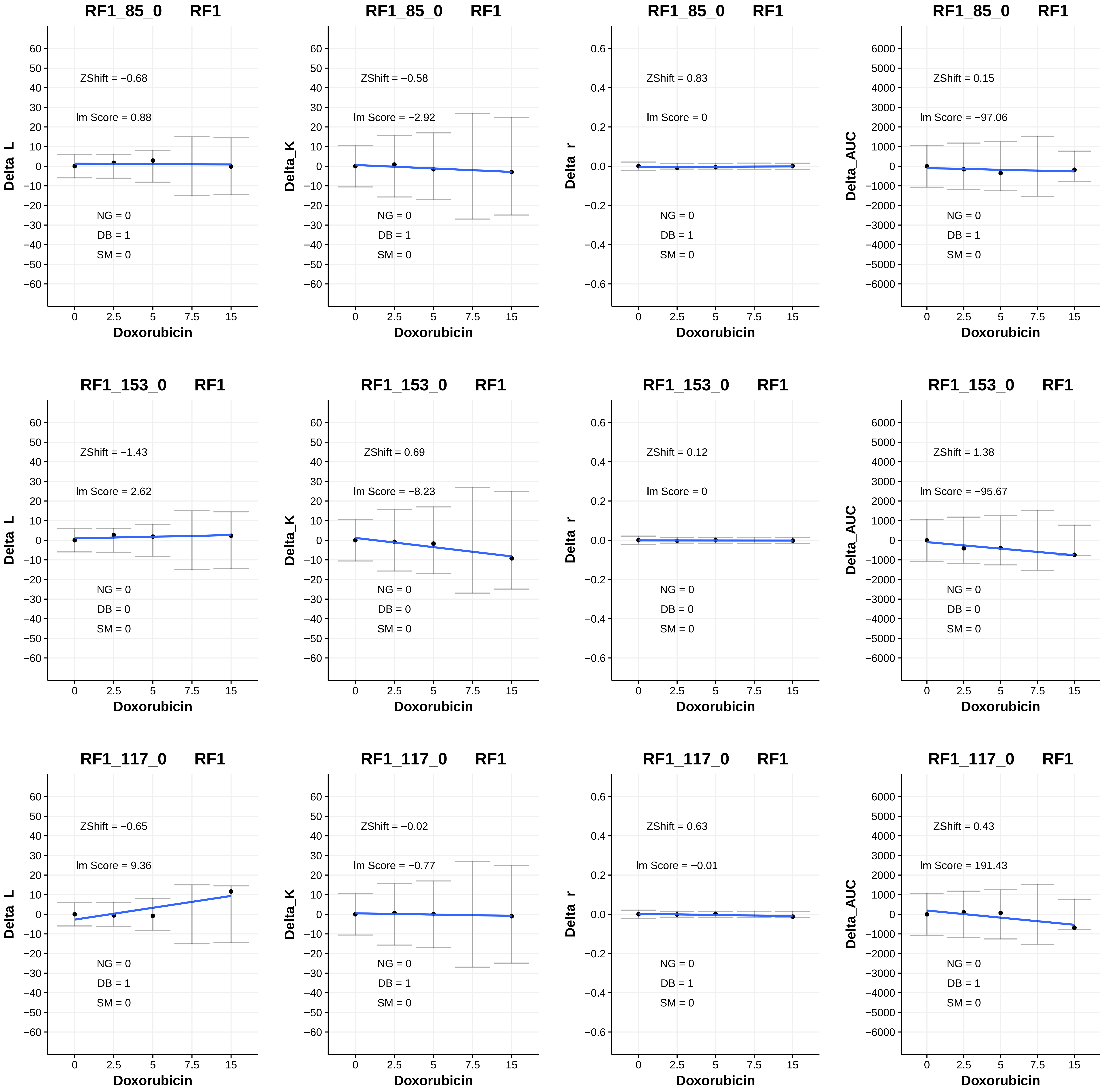

RF1_85_0 RF1
ZShift = −0.68
Im Score = 0.88
NG = 0
DB = 1
SM = 0
Delta_L
Doxorubicin
RF1_85_0 RF1
ZShift = −0.58
Im Score = −2.92
NG = 0
DB = 1
SM = 0
Delta_K
Doxorubicin
RF1_85_0 RF1
ZShift = 0.83
Im Score = 0
NG = 0
DB = 1
SM = 0
Delta_r
Doxorubicin
RF1_85_0 RF1
ZShift = 0.15
Im Score = −97.06
NG = 0
DB = 1
SM = 0
Delta_AUC
Doxorubicin
RF1_153_0 RF1
ZShift = −1.43
Im Score = 2.62
NG = 0
DB = 0
SM = 0
Delta_L
Doxorubicin
RF1_153_0 RF1
ZShift = 0.69
Im Score = −8.23
NG = 0
DB = 0
SM = 0
Delta_K
Doxorubicin
RF1_153_0 RF1
ZShift = 0.12
Im Score = 0
NG = 0
DB = 0
SM = 0
Delta_r
Doxorubicin
RF1_153_0 RF1
ZShift = 1.38
Im Score = −95.67
NG = 0
DB = 0
SM = 0
Delta_AUC
Doxorubicin
RF1_117_0 RF1
ZShift = −0.65
Im Score = 9.36
NG = 0
DB = 1
SM = 0
Delta_L
Doxorubicin
RF1_117_0 RF1
ZShift = −0.02
Im Score = −0.77
NG = 0
DB = 1
SM = 0
Delta_K
Doxorubicin
RF1_117_0 RF1
ZShift = 0.63
Im Score = −0.01
NG = 0
DB = 1
SM = 0
Delta_r
Doxorubicin
RF1_117_0 RF1
ZShift = 0.43
Im Score = 191.43
NG = 0
DB = 1
SM = 0
Delta_AUC
Doxorubicin

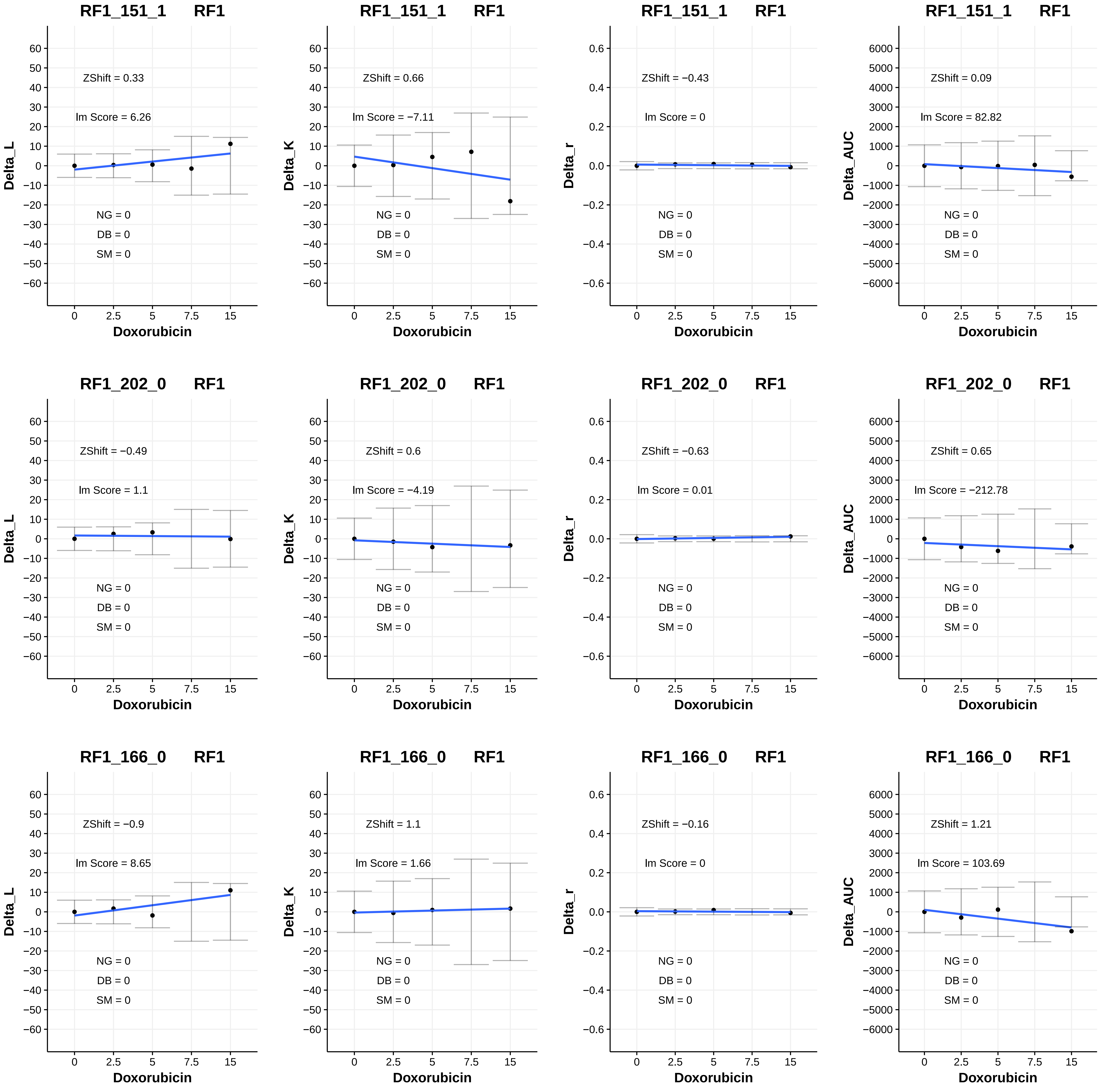

RF1_151_1 RF1
ZShift = 0.33
lm Score = 6.26
NG = 0
DB = 0
SM = 0
Delta_L
Doxorubicin
RF1_151_1 RF1
ZShift = 0.66
lm Score = −7.11
NG = 0
DB = 0
SM = 0
Delta_K
Doxorubicin
RF1_151_1 RF1
ZShift = −0.43
lm Score = 0
NG = 0
DB = 0
SM = 0
Delta_r
Doxorubicin
RF1_151_1 RF1
ZShift = 0.09
lm Score = 82.82
NG = 0
DB = 0
SM = 0
Delta_AUC
Doxorubicin
RF1_202_0 RF1
ZShift = −0.49
lm Score = 1.1
NG = 0
DB = 0
SM = 0
Delta_L
Doxorubicin
RF1_202_0 RF1
ZShift = 0.6
lm Score = −4.19
NG = 0
DB = 0
SM = 0
Delta_K
Doxorubicin
RF1_202_0 RF1
ZShift = −0.63
lm Score = 0.01
NG = 0
DB = 0
SM = 0
Delta_r
Doxorubicin
RF1_202_0 RF1
ZShift = 0.65
lm Score = −212.78
NG = 0
DB = 0
SM = 0
Delta_AUC
Doxorubicin
RF1_166_0 RF1
ZShift = −0.9
lm Score = 8.65
NG = 0
DB = 0
SM = 0
Delta_L
Doxorubicin
RF1_166_0 RF1
ZShift = 1.1
lm Score = 1.66
NG = 0
DB = 0
SM = 0
Delta_K
Doxorubicin
RF1_166_0 RF1
ZShift = −0.16
lm Score = 0
NG = 0
DB = 0
SM = 0
Delta_r
Doxorubicin
RF1_166_0 RF1
ZShift = 1.21
lm Score = 103.69
NG = 0
DB = 0
SM = 0
Delta_AUC
Doxorubicin

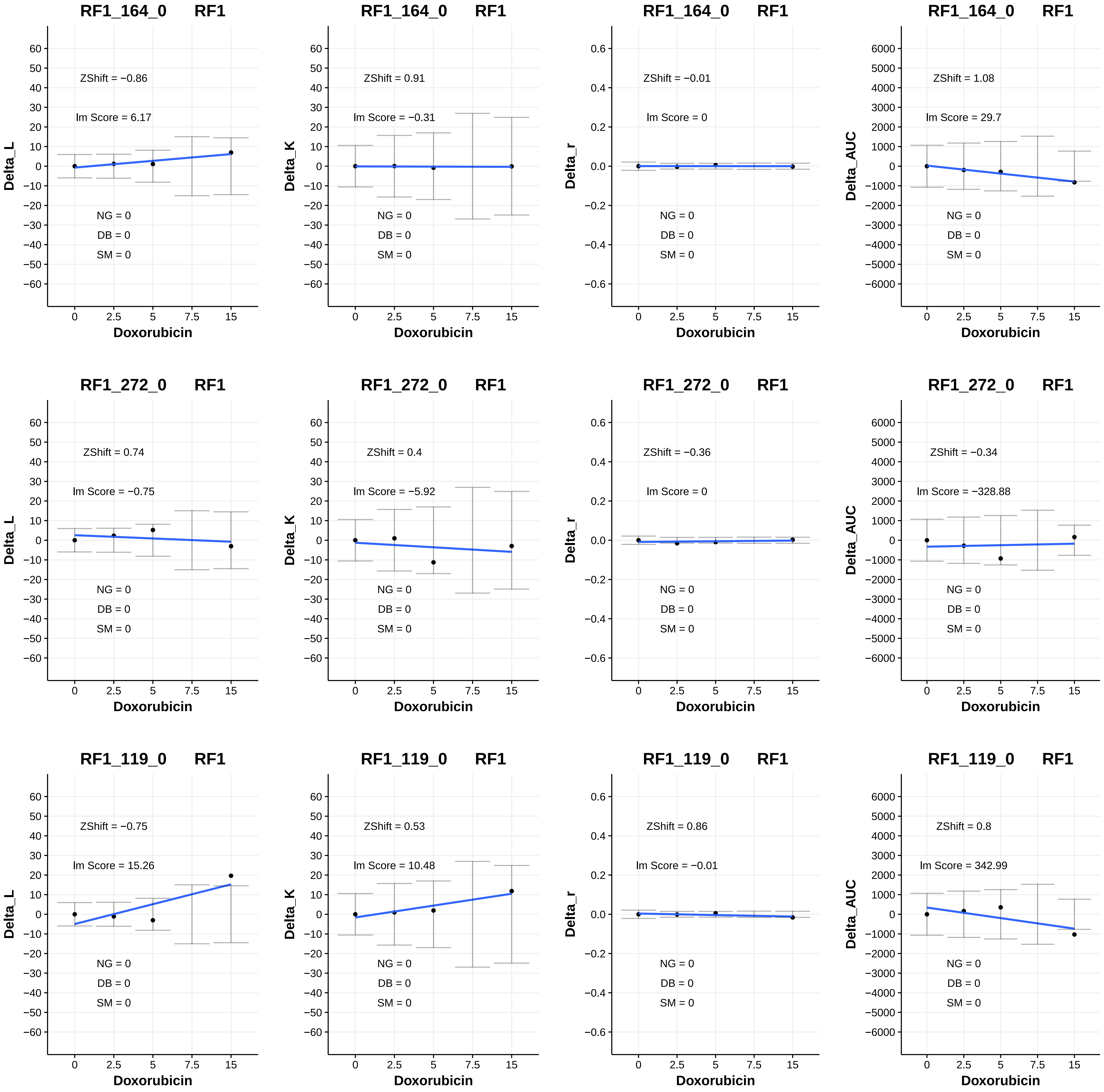

RF1_164_0 RF1
ZShift = −0.86
Im Score = 6.17
NG = 0
DB = 0
SM = 0
Delta_L
Doxorubicin
RF1_164_0 RF1
ZShift = 0.91
Im Score = −0.31
NG = 0
DB = 0
SM = 0
Delta_K
Doxorubicin
RF1_164_0 RF1
ZShift = −0.01
Im Score = 0
NG = 0
DB = 0
SM = 0
Delta_r
Doxorubicin
RF1_164_0 RF1
ZShift = 1.08
Im Score = 29.7
NG = 0
DB = 0
SM = 0
Delta_AUC
Doxorubicin
RF1_272_0 RF1
ZShift = 0.74
Im Score = −0.75
NG = 0
DB = 0
SM = 0
Delta_L
Doxorubicin
RF1_272_0 RF1
ZShift = 0.4
Im Score = −5.92
NG = 0
DB = 0
SM = 0
Delta_K
Doxorubicin
RF1_272_0 RF1
ZShift = −0.36
Im Score = 0
NG = 0
DB = 0
SM = 0
Delta_r
Doxorubicin
RF1_272_0 RF1
ZShift = −0.34
Im Score = −328.88
NG = 0
DB = 0
SM = 0
Delta_AUC
Doxorubicin
RF1_119_0 RF1
ZShift = −0.75
Im Score = 15.26
NG = 0
DB = 0
SM = 0
Delta_L
Doxorubicin
RF1_119_0 RF1
ZShift = 0.53
Im Score = 10.48
NG = 0
DB = 0
SM = 0
Delta_K
Doxorubicin
RF1_119_0 RF1
ZShift = 0.86
Im Score = −0.01
NG = 0
DB = 0
SM = 0
Delta_r
Doxorubicin
RF1_119_0 RF1
ZShift = 0.8
Im Score = 342.99
NG = 0
DB = 0
SM = 0
Delta_AUC
Doxorubicin

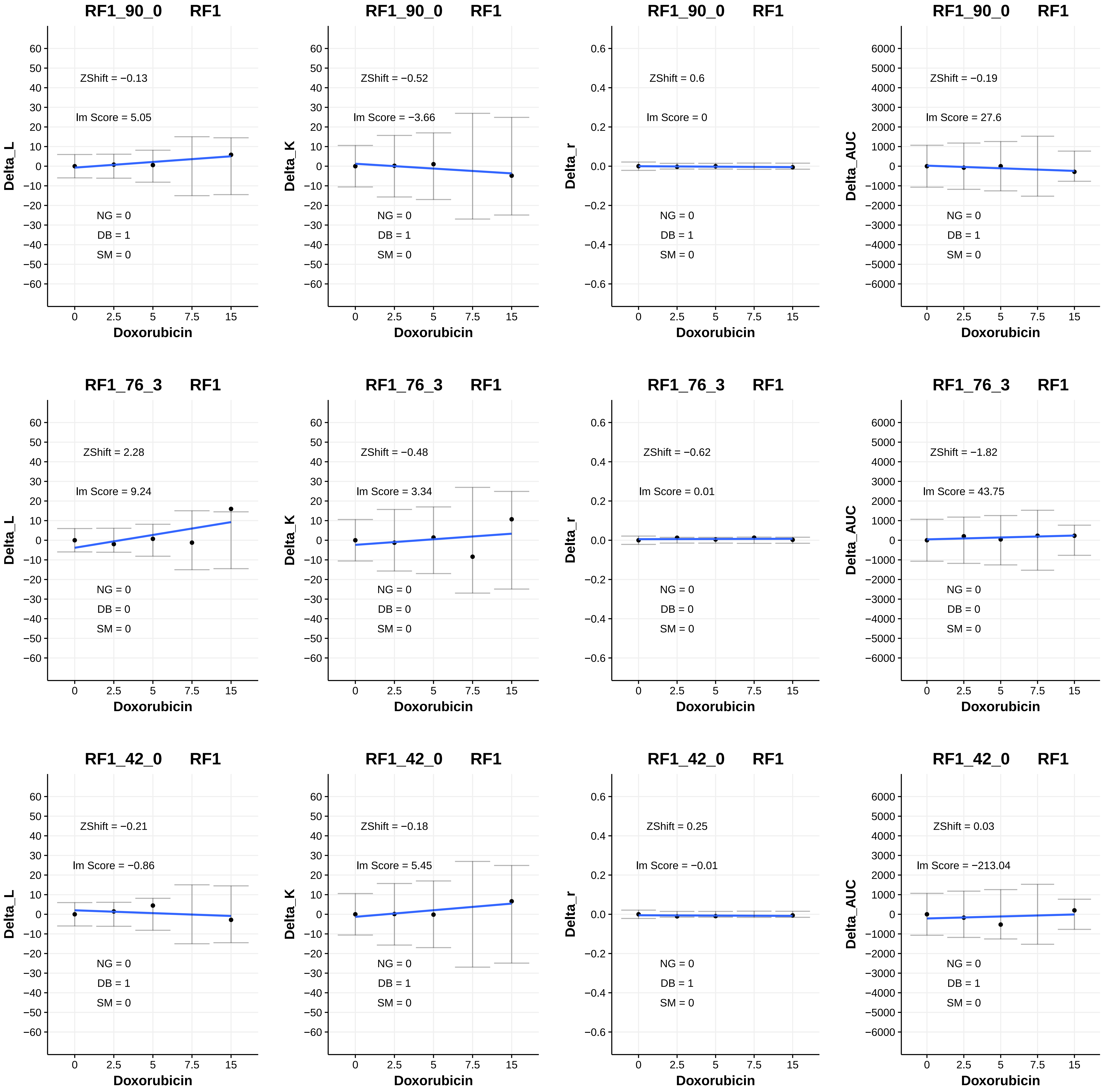
RF1_90_0 RF1
ZShift = −0.13
lm Score = 5.05
NG = 0
DB = 1
SM = 0
Delta_L
Doxorubicin
RF1_90_0 RF1
ZShift = −0.52
lm Score = −3.66
NG = 0
DB = 1
SM = 0
Delta_K
Doxorubicin
RF1_90_0 RF1
ZShift = 0.6
lm Score = 0
NG = 0
DB = 1
SM = 0
Delta_r
Doxorubicin
RF1_90_0 RF1
ZShift = −0.19
lm Score = 27.6
NG = 0
DB = 1
SM = 0
Delta_AUC
Doxorubicin
RF1_76_3 RF1
ZShift = 2.28
lm Score = 9.24
NG = 0
DB = 0
SM = 0
Delta_L
Doxorubicin
RF1_76_3 RF1
ZShift = −0.48
lm Score = 3.34
NG = 0
DB = 0
SM = 0
Delta_K
Doxorubicin
RF1_76_3 RF1
ZShift = −0.62
lm Score = 0.01
NG = 0
DB = 0
SM = 0
Delta_r
Doxorubicin
RF1_76_3 RF1
ZShift = −1.82
lm Score = 43.75
NG = 0
DB = 0
SM = 0
Delta_AUC
Doxorubicin
RF1_42_0 RF1
ZShift = −0.21
lm Score = −0.86
NG = 0
DB = 1
SM = 0
Delta_L
Doxorubicin
RF1_42_0 RF1
ZShift = −0.18
lm Score = 5.45
NG = 0
DB = 1
SM = 0
Delta_K
Doxorubicin
RF1_42_0 RF1
ZShift = 0.25
lm Score = −0.01
NG = 0
DB = 1
SM = 0
Delta_r
Doxorubicin
RF1_42_0 RF1
ZShift = 0.03
lm Score = −213.04
NG = 0
DB = 1
SM = 0
Delta_AUC
Doxorubicin

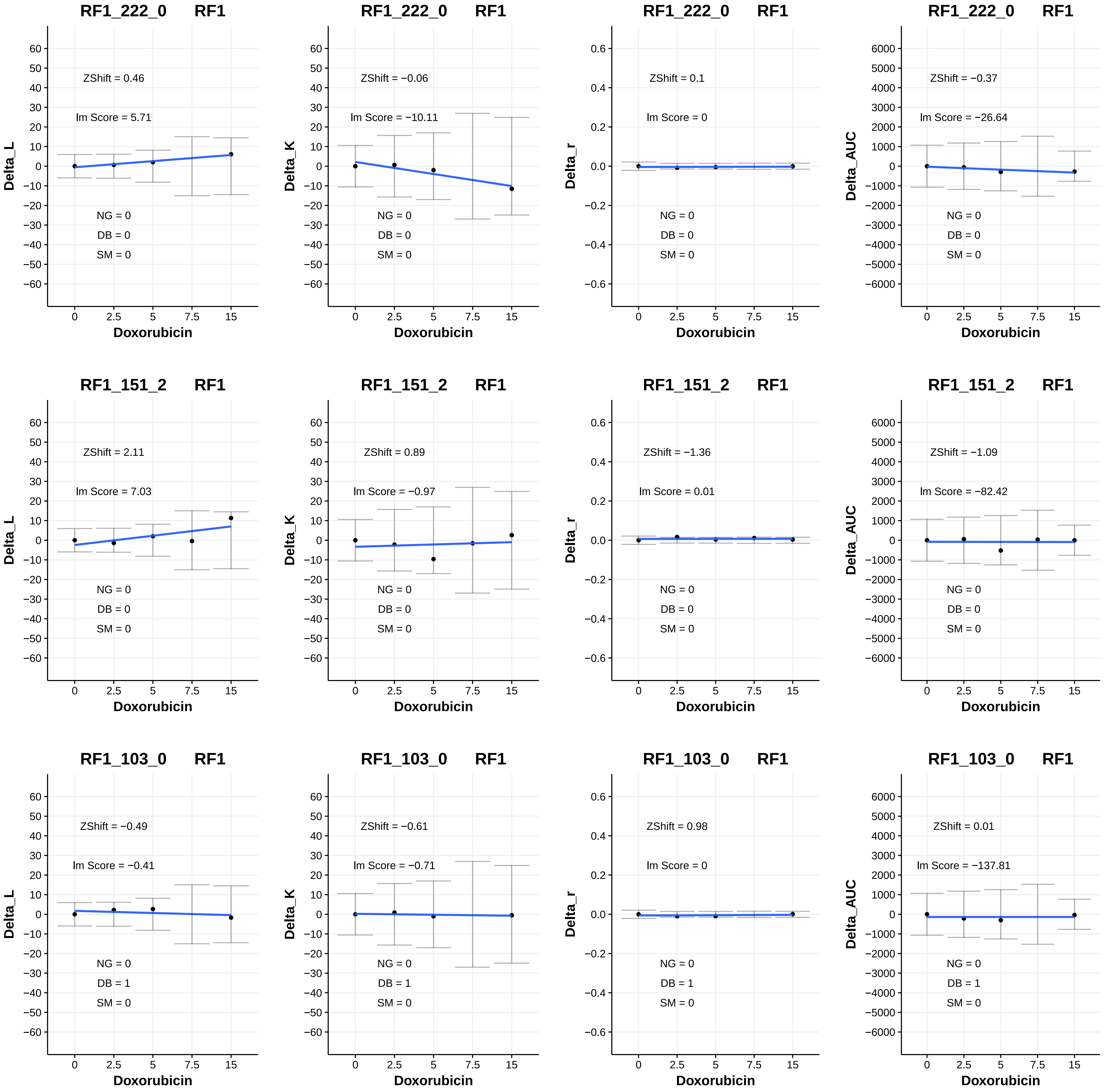

RF1_222_0 RF1
ZShift = 0.46
lm Score = 5.71
NG = 0
DB = 0
SM = 0
Delta_L
Doxorubicin
RF1_222_0 RF1
ZShift = −0.06
lm Score = −10.11
NG = 0
DB = 0
SM = 0
Delta_K
Doxorubicin
RF1_222_0 RF1
ZShift = 0.1
lm Score = 0
NG = 0
DB = 0
SM = 0
Delta_r
Doxorubicin
RF1_222_0 RF1
ZShift = −0.37
lm Score = −26.64
NG = 0
DB = 0
SM = 0
Delta_AUC
Doxorubicin
RF1_151_2 RF1
ZShift = 2.11
lm Score = 7.03
NG = 0
DB = 0
SM = 0
Delta_L
Doxorubicin
RF1_151_2 RF1
ZShift = 0.89
lm Score = −0.97
NG = 0
DB = 0
SM = 0
Delta_K
Doxorubicin
RF1_151_2 RF1
ZShift = −1.36
lm Score = 0.01
NG = 0
DB = 0
SM = 0
Delta_r
Doxorubicin
RF1_151_2 RF1
ZShift = −1.09
lm Score = −82.42
NG = 0
DB = 0
SM = 0
Delta_AUC
Doxorubicin
RF1_103_0 RF1
ZShift = −0.49
lm Score = −0.41
NG = 0
DB = 1
SM = 0
Delta_L
Doxorubicin
RF1_103_0 RF1
ZShift = −0.61
lm Score = −0.71
NG = 0
DB = 1
SM = 0
Delta_K
Doxorubicin
RF1_103_0 RF1
ZShift = 0.98
lm Score = 0
NG = 0
DB = 1
SM = 0
Delta_r
Doxorubicin
RF1_103_0 RF1
ZShift = 0.01
lm Score = −137.81
NG = 0
DB = 1
SM = 0
Delta_AUC
Doxorubicin

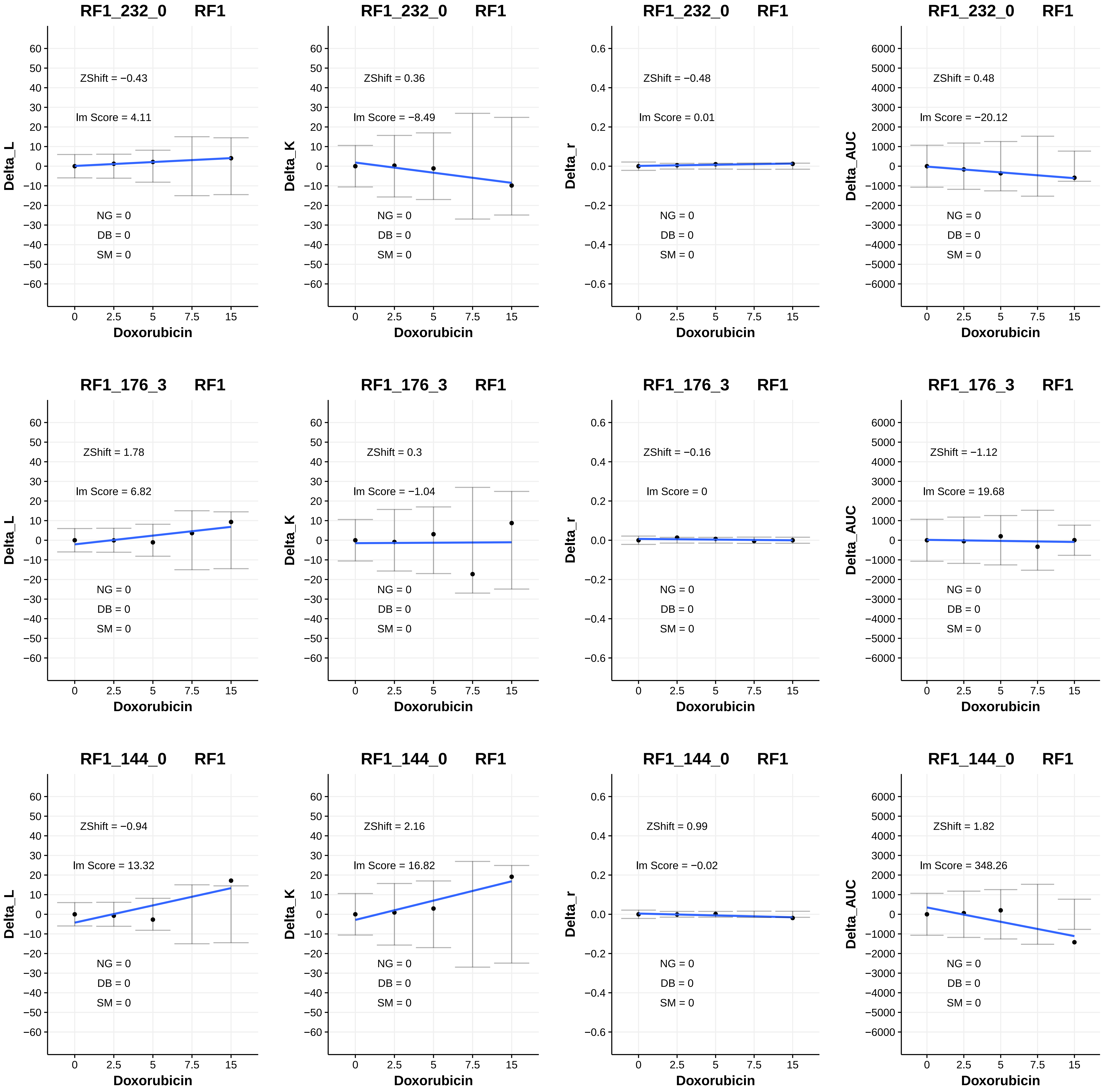

RF1_232_0 RF1
ZShift = −0.43
lm Score = 4.11
NG = 0
DB = 0
SM = 0
Delta_L
Doxorubicin
RF1_232_0 RF1
ZShift = 0.36
lm Score = −8.49
NG = 0
DB = 0
SM = 0
Delta_K
Doxorubicin
RF1_232_0 RF1
ZShift = −0.48
lm Score = 0.01
NG = 0
DB = 0
SM = 0
Delta_r
Doxorubicin
RF1_232_0 RF1
ZShift = 0.48
lm Score = −20.12
NG = 0
DB = 0
SM = 0
Delta_AUC
Doxorubicin
RF1_176_3 RF1
ZShift = 1.78
lm Score = 6.82
NG = 0
DB = 0
SM = 0
Delta_L
Doxorubicin
RF1_176_3 RF1
ZShift = 0.3
lm Score = −1.04
NG = 0
DB = 0
SM = 0
Delta_K
Doxorubicin
RF1_176_3 RF1
ZShift = −0.16
lm Score = 0
NG = 0
DB = 0
SM = 0
Delta_r
Doxorubicin
RF1_176_3 RF1
ZShift = −1.12
lm Score = 19.68
NG = 0
DB = 0
SM = 0
Delta_AUC
Doxorubicin
RF1_144_0 RF1
ZShift = −0.94
lm Score = 13.32
NG = 0
DB = 0
SM = 0
Delta_L
Doxorubicin
RF1_144_0 RF1
ZShift = 2.16
lm Score = 16.82
NG = 0
DB = 0
SM = 0
Delta_K
Doxorubicin
RF1_144_0 RF1
ZShift = 0.99
lm Score = −0.02
NG = 0
DB = 0
SM = 0
Delta_r
Doxorubicin
RF1_144_0 RF1
ZShift = 1.82
lm Score = 348.26
NG = 0
DB = 0
SM = 0
Delta_AUC
Doxorubicin

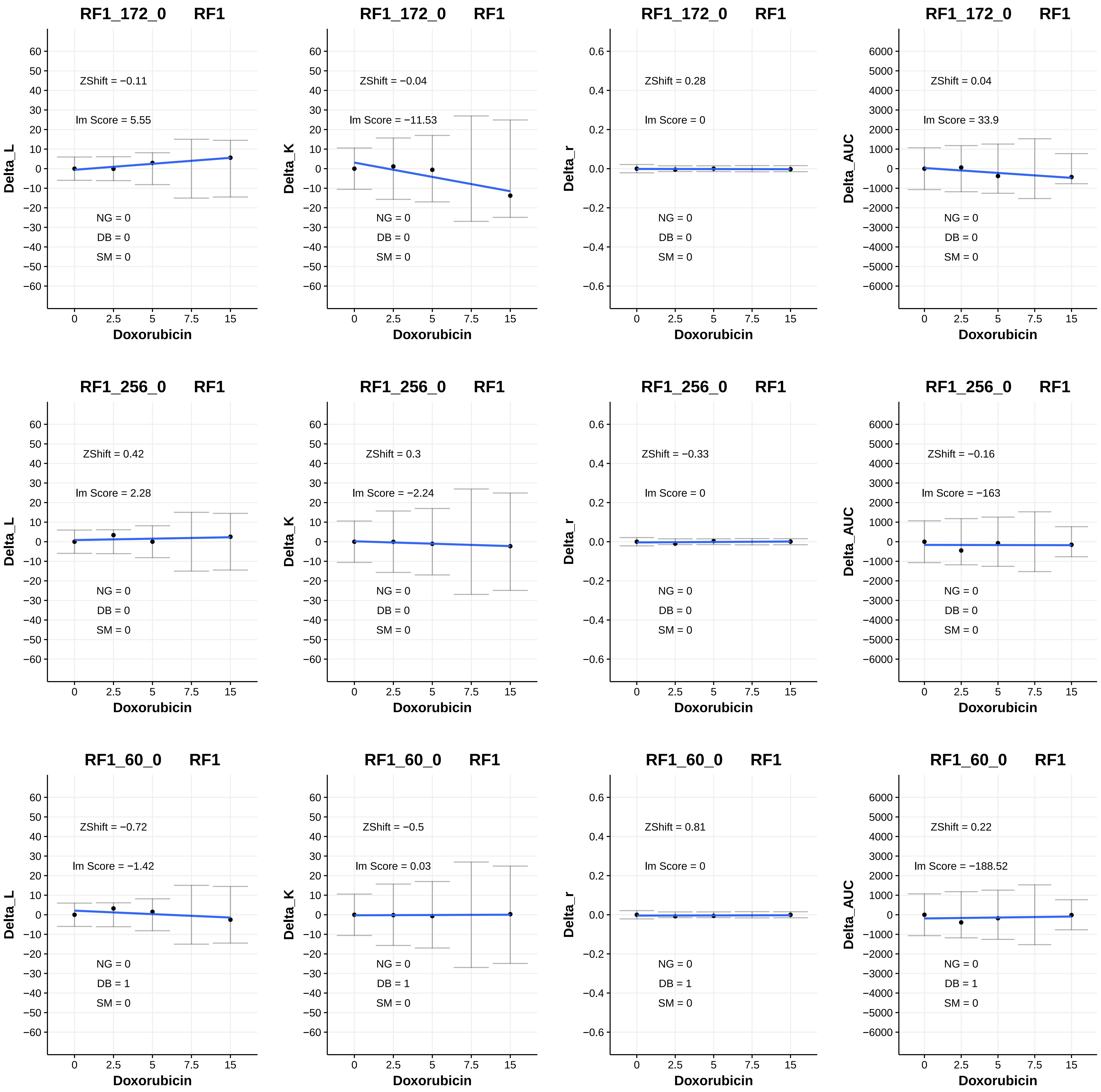

RF1_172_0 RF1
ZShift = −0.11
Im Score = 5.55
NG = 0
DB = 0
SM = 0
Delta_L
Doxorubicin
RF1_172_0 RF1
ZShift = −0.04
Im Score = −11.53
NG = 0
DB = 0
SM = 0
Delta_K
Doxorubicin
RF1_172_0 RF1
ZShift = 0.28
Im Score = 0
NG = 0
DB = 0
SM = 0
Delta_r
Doxorubicin
RF1_172_0 RF1
ZShift = 0.04
Im Score = 33.9
NG = 0
DB = 0
SM = 0
Delta_AUC
Doxorubicin
RF1_256_0 RF1
ZShift = 0.42
Im Score = 2.28
NG = 0
DB = 0
SM = 0
Delta_L
Doxorubicin
RF1_256_0 RF1
ZShift = 0.3
Im Score = −2.24
NG = 0
DB = 0
SM = 0
Delta_K
Doxorubicin
RF1_256_0 RF1
ZShift = −0.33
Im Score = 0
NG = 0
DB = 0
SM = 0
Delta_r
Doxorubicin
RF1_256_0 RF1
ZShift = −0.16
Im Score = −163
NG = 0
DB = 0
SM = 0
Delta_AUC
Doxorubicin
RF1_60_0 RF1
ZShift = −0.72
Im Score = −1.42
NG = 0
DB = 1
SM = 0
Delta_L
Doxorubicin
RF1_60_0 RF1
ZShift = −0.5
Im Score = 0.03
NG = 0
DB = 1
SM = 0
Delta_K
Doxorubicin
RF1_60_0 RF1
ZShift = 0.81
Im Score = 0
NG = 0
DB = 1
SM = 0
Delta_r
Doxorubicin
RF1_60_0 RF1
ZShift = 0.22
Im Score = −188.52
NG = 0
DB = 1
SM = 0
Delta_AUC
Doxorubicin

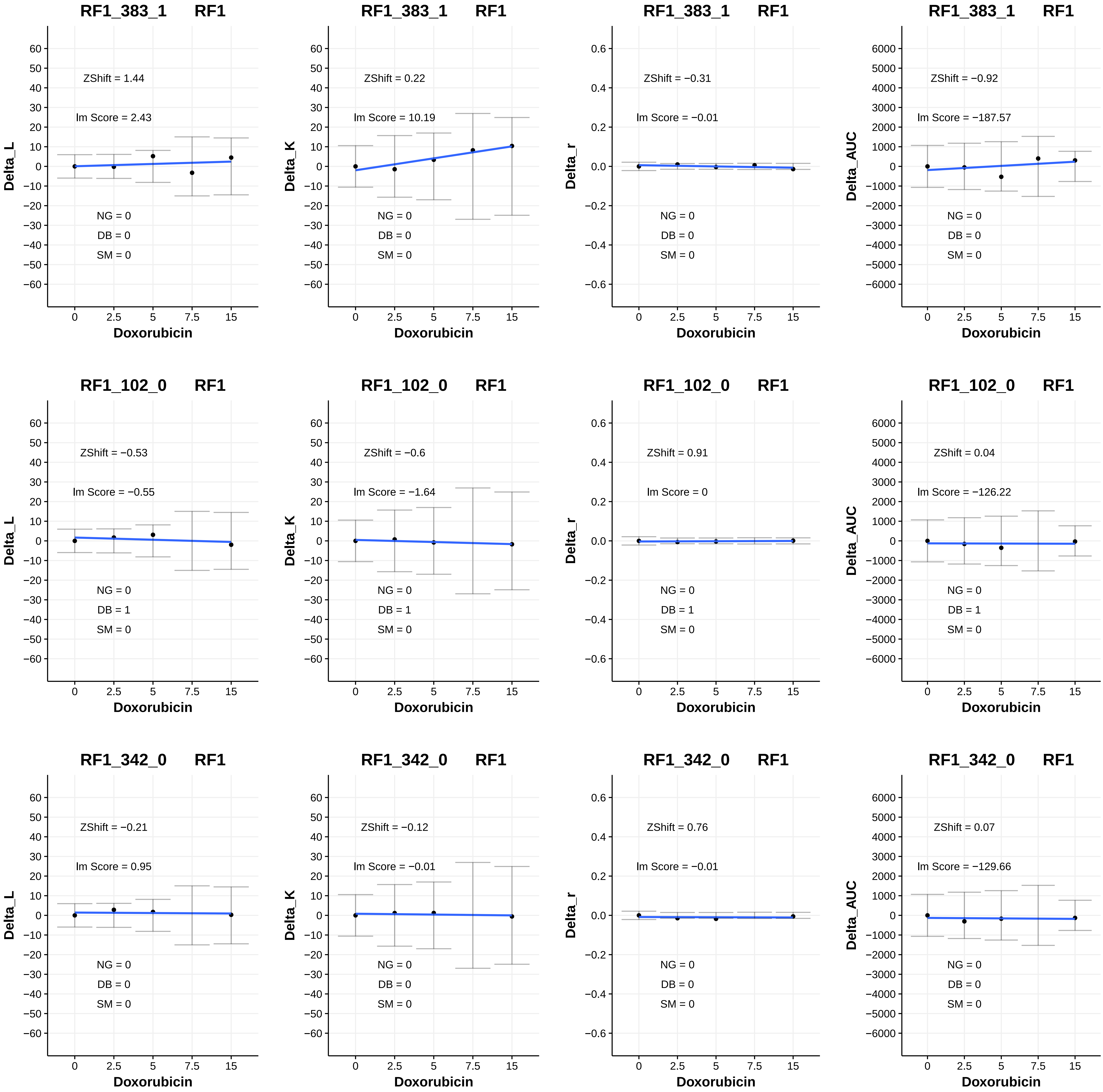

RF1_383_1 RF1
ZShift = 1.44
lm Score = 2.43
NG = 0
DB = 0
SM = 0
Delta_L
Doxorubicin
RF1_383_1 RF1
ZShift = 0.22
lm Score = 10.19
NG = 0
DB = 0
SM = 0
Delta_K
Doxorubicin
RF1_383_1 RF1
ZShift = −0.31
lm Score = −0.01
NG = 0
DB = 0
SM = 0
Delta_r
Doxorubicin
RF1_383_1 RF1
ZShift = −0.92
lm Score = −187.57
NG = 0
DB = 0
SM = 0
Delta_AUC
Doxorubicin
RF1_102_0 RF1
ZShift = −0.53
lm Score = −0.55
NG = 0
DB = 1
SM = 0
Delta_L
Doxorubicin
RF1_102_0 RF1
ZShift = −0.6
lm Score = −1.64
NG = 0
DB = 1
SM = 0
Delta_K
Doxorubicin
RF1_102_0 RF1
ZShift = 0.91
lm Score = 0
NG = 0
DB = 1
SM = 0
Delta_r
Doxorubicin
RF1_102_0 RF1
ZShift = 0.04
lm Score = −126.22
NG = 0
DB = 1
SM = 0
Delta_AUC
Doxorubicin
RF1_342_0 RF1
ZShift = −0.21
lm Score = 0.95
NG = 0
DB = 0
SM = 0
Delta_L
Doxorubicin
RF1_342_0 RF1
ZShift = −0.12
lm Score = −0.01
NG = 0
DB = 0
SM = 0
Delta_K
Doxorubicin
RF1_342_0 RF1
ZShift = 0.76
lm Score = −0.01
NG = 0
DB = 0
SM = 0
Delta_r
Doxorubicin
RF1_342_0 RF1
ZShift = 0.07
lm Score = −129.66
NG = 0
DB = 0
SM = 0
Delta_AUC
Doxorubicin

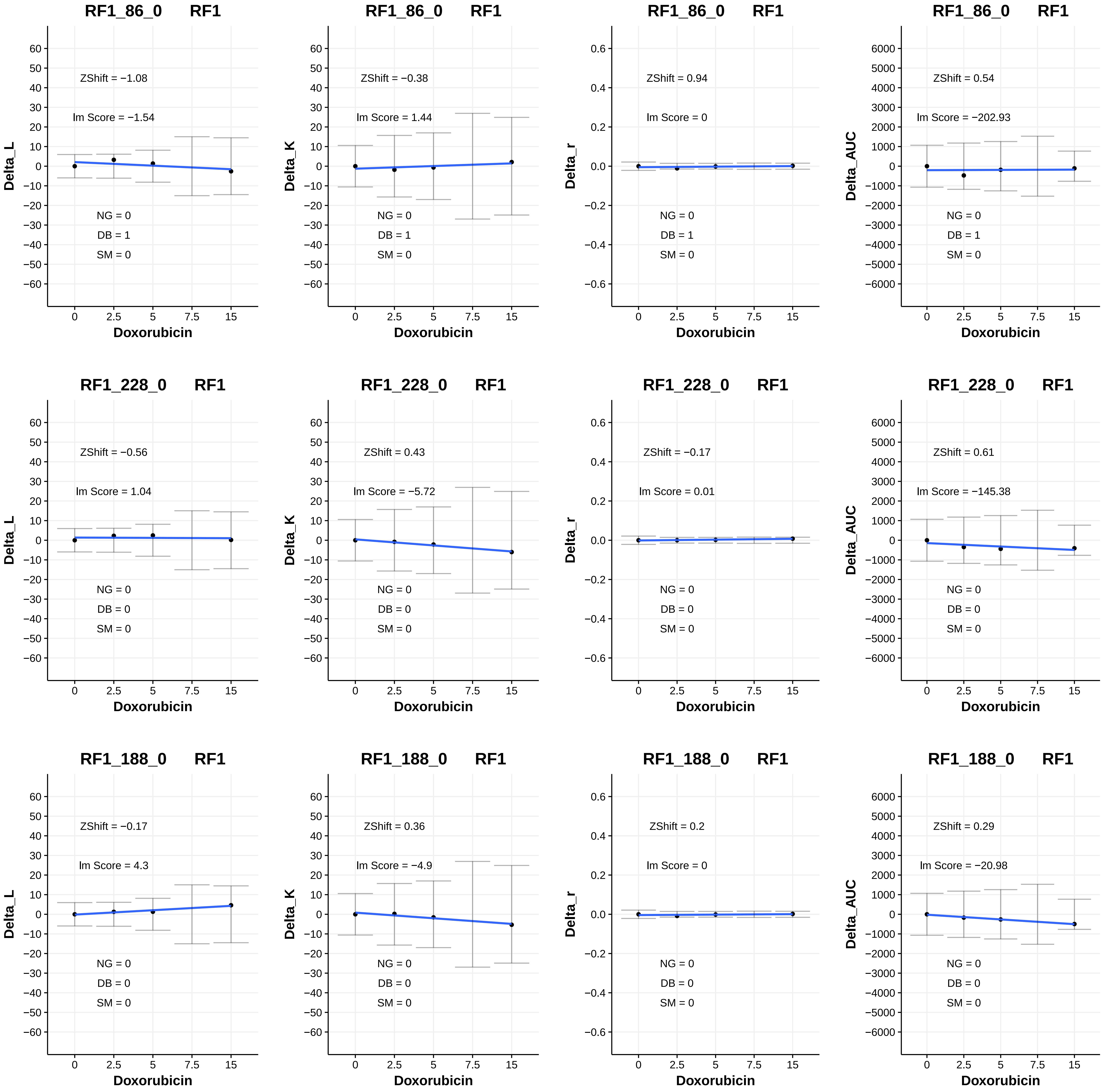
RF1_86_0 RF1
ZShift = −1.08
lm Score = −1.54
NG = 0
DB = 1
SM = 0
Delta_L
Doxorubicin
RF1_86_0 RF1
ZShift = −0.38
lm Score = 1.44
NG = 0
DB = 1
SM = 0
Delta_K
Doxorubicin
RF1_86_0 RF1
ZShift = 0.94
lm Score = 0
NG = 0
DB = 1
SM = 0
Delta_r
Doxorubicin
RF1_86_0 RF1
ZShift = 0.54
lm Score = −202.93
NG = 0
DB = 1
SM = 0
Delta_AUC
Doxorubicin
RF1_228_0 RF1
ZShift = −0.56
lm Score = 1.04
NG = 0
DB = 0
SM = 0
Delta_L
Doxorubicin
RF1_228_0 RF1
ZShift = 0.43
lm Score = −5.72
NG = 0
DB = 0
SM = 0
Delta_K
Doxorubicin
RF1_228_0 RF1
ZShift = −0.17
lm Score = 0.01
NG = 0
DB = 0
SM = 0
Delta_r
Doxorubicin
RF1_228_0 RF1
ZShift = 0.61
lm Score = −145.38
NG = 0
DB = 0
SM = 0
Delta_AUC
Doxorubicin
RF1_188_0 RF1
ZShift = −0.17
lm Score = 4.3
NG = 0
DB = 0
SM = 0
Delta_L
Doxorubicin
RF1_188_0 RF1
ZShift = 0.36
lm Score = −4.9
NG = 0
DB = 0
SM = 0
Delta_K
Doxorubicin
RF1_188_0 RF1
ZShift = 0.2
lm Score = 0
NG = 0
DB = 0
SM = 0
Delta_r
Doxorubicin
RF1_188_0 RF1
ZShift = 0.29
lm Score = −20.98
NG = 0
DB = 0
SM = 0
Delta_AUC
Doxorubicin

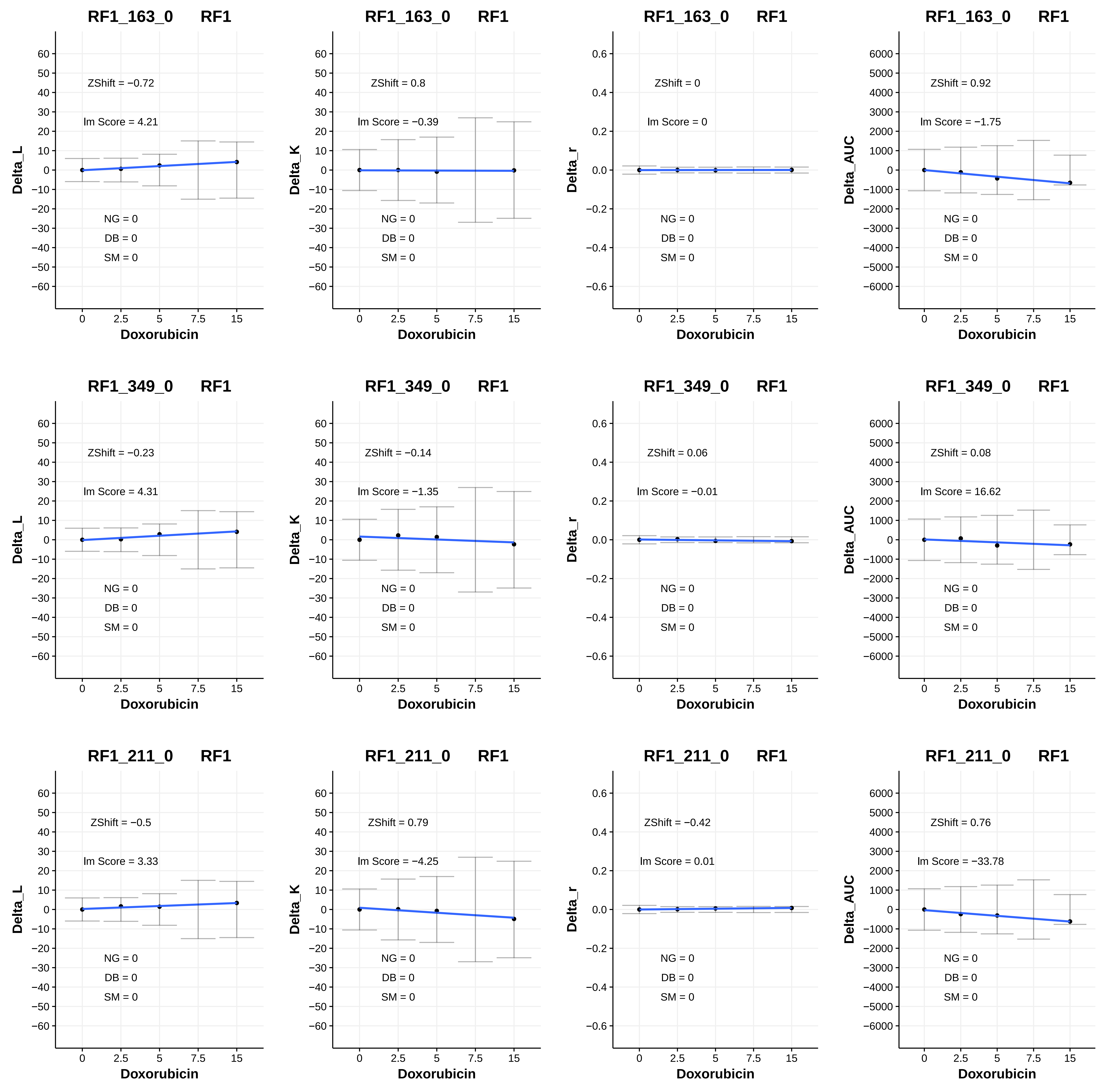

RF1_163_0 RF1
ZShift = −0.72
lm Score = 4.21
NG = 0
DB = 0
SM = 0
Delta_L
Doxorubicin
RF1_163_0 RF1
ZShift = 0.8
lm Score = −0.39
NG = 0
DB = 0
SM = 0
Delta_K
Doxorubicin
RF1_163_0 RF1
ZShift = 0
lm Score = 0
NG = 0
DB = 0
SM = 0
Delta_r
Doxorubicin
RF1_163_0 RF1
ZShift = 0.92
lm Score = −1.75
NG = 0
DB = 0
SM = 0
Delta_AUC
Doxorubicin
RF1_349_0 RF1
ZShift = −0.23
lm Score = 4.31
NG = 0
DB = 0
SM = 0
Delta_L
Doxorubicin
RF1_349_0 RF1
ZShift = −0.14
lm Score = −1.35
NG = 0
DB = 0
SM = 0
Delta_K
Doxorubicin
RF1_349_0 RF1
ZShift = 0.06
lm Score = −0.01
NG = 0
DB = 0
SM = 0
Delta_r
Doxorubicin
RF1_349_0 RF1
ZShift = 0.08
lm Score = 16.62
NG = 0
DB = 0
SM = 0
Delta_AUC
Doxorubicin
RF1_211_0 RF1
ZShift = −0.5
lm Score = 3.33
NG = 0
DB = 0
SM = 0
Delta_L
Doxorubicin
RF1_211_0 RF1
ZShift = 0.79
lm Score = −4.25
NG = 0
DB = 0
SM = 0
Delta_K
Doxorubicin
RF1_211_0 RF1
ZShift = −0.42
lm Score = 0.01
NG = 0
DB = 0
SM = 0
Delta_r
Doxorubicin
RF1_211_0 RF1
ZShift = 0.76
lm Score = −33.78
NG = 0
DB = 0
SM = 0
Delta_AUC
Doxorubicin

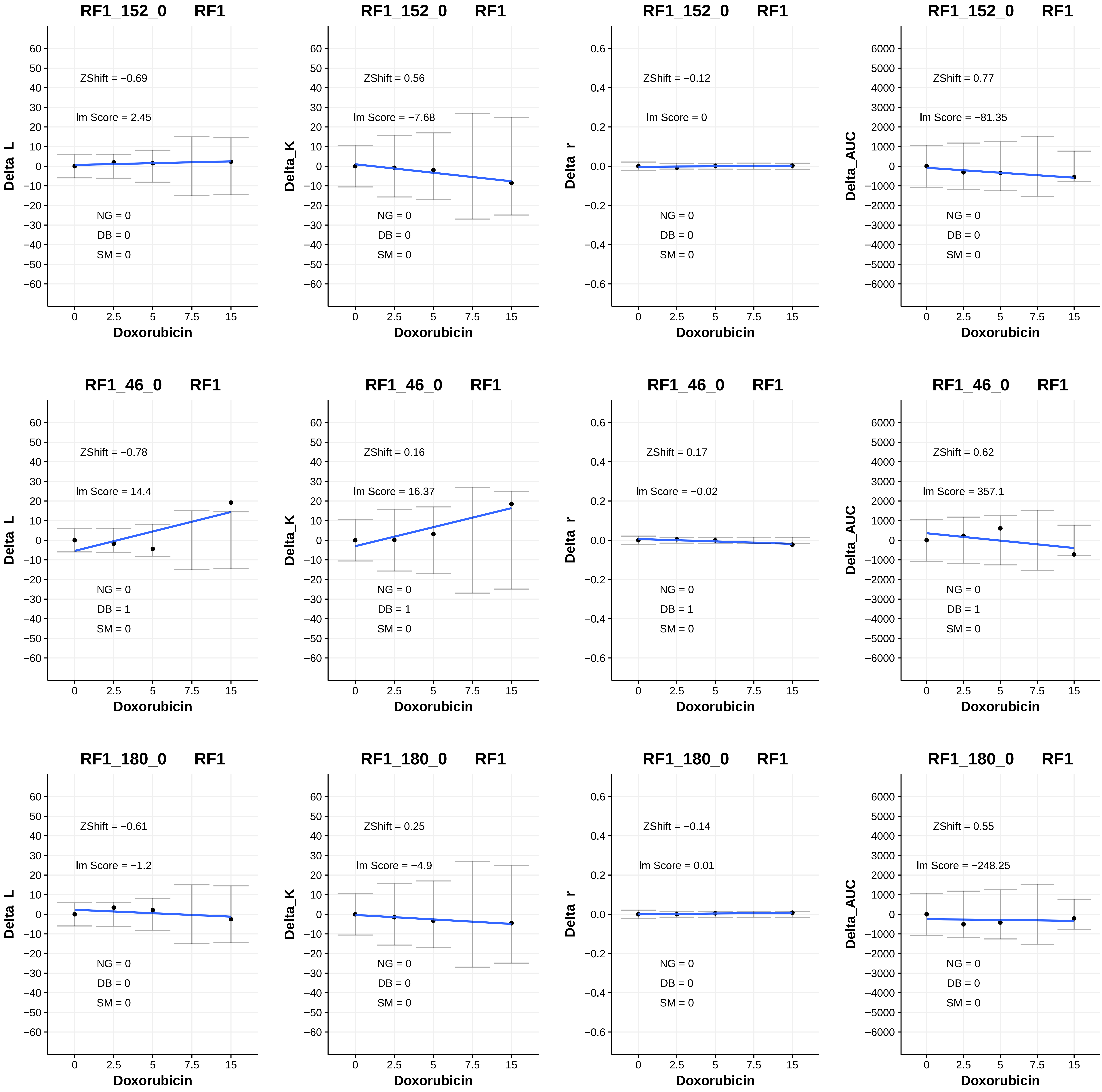
RF1_152_0 RF1
Delta_L
ZShift = −0.69
lm Score = 2.45
NG = 0
DB = 0
SM = 0
Doxorubicin
RF1_152_0 RF1
Delta_K
ZShift = 0.56
lm Score = −7.68
NG = 0
DB = 0
SM = 0
Doxorubicin
RF1_152_0 RF1
Delta_r
ZShift = −0.12
lm Score = 0
NG = 0
DB = 0
SM = 0
Doxorubicin
RF1_152_0 RF1
Delta_AUC
ZShift = 0.77
lm Score = −81.35
NG = 0
DB = 0
SM = 0
Doxorubicin
RF1_46_0 RF1
Delta_L
ZShift = −0.78
lm Score = 14.4
NG = 0
DB = 1
SM = 0
Doxorubicin
RF1_46_0 RF1
Delta_K
ZShift = 0.16
lm Score = 16.37
NG = 0
DB = 1
SM = 0
Doxorubicin
RF1_46_0 RF1
Delta_r
ZShift = 0.17
lm Score = −0.02
NG = 0
DB = 1
SM = 0
Doxorubicin
RF1_46_0 RF1
Delta_AUC
ZShift = 0.62
lm Score = 357.1
NG = 0
DB = 1
SM = 0
Doxorubicin
RF1_180_0 RF1
Delta_L
ZShift = −0.61
lm Score = −1.2
NG = 0
DB = 0
SM = 0
Doxorubicin
RF1_180_0 RF1
Delta_K
ZShift = 0.25
lm Score = −4.9
NG = 0
DB = 0
SM = 0
Doxorubicin
RF1_180_0 RF1
Delta_r
ZShift = −0.14
lm Score = 0.01
NG = 0
DB = 0
SM = 0
Doxorubicin
RF1_180_0 RF1
Delta_AUC
ZShift = 0.55
lm Score = −248.25
NG = 0
DB = 0
SM = 0
Doxorubicin

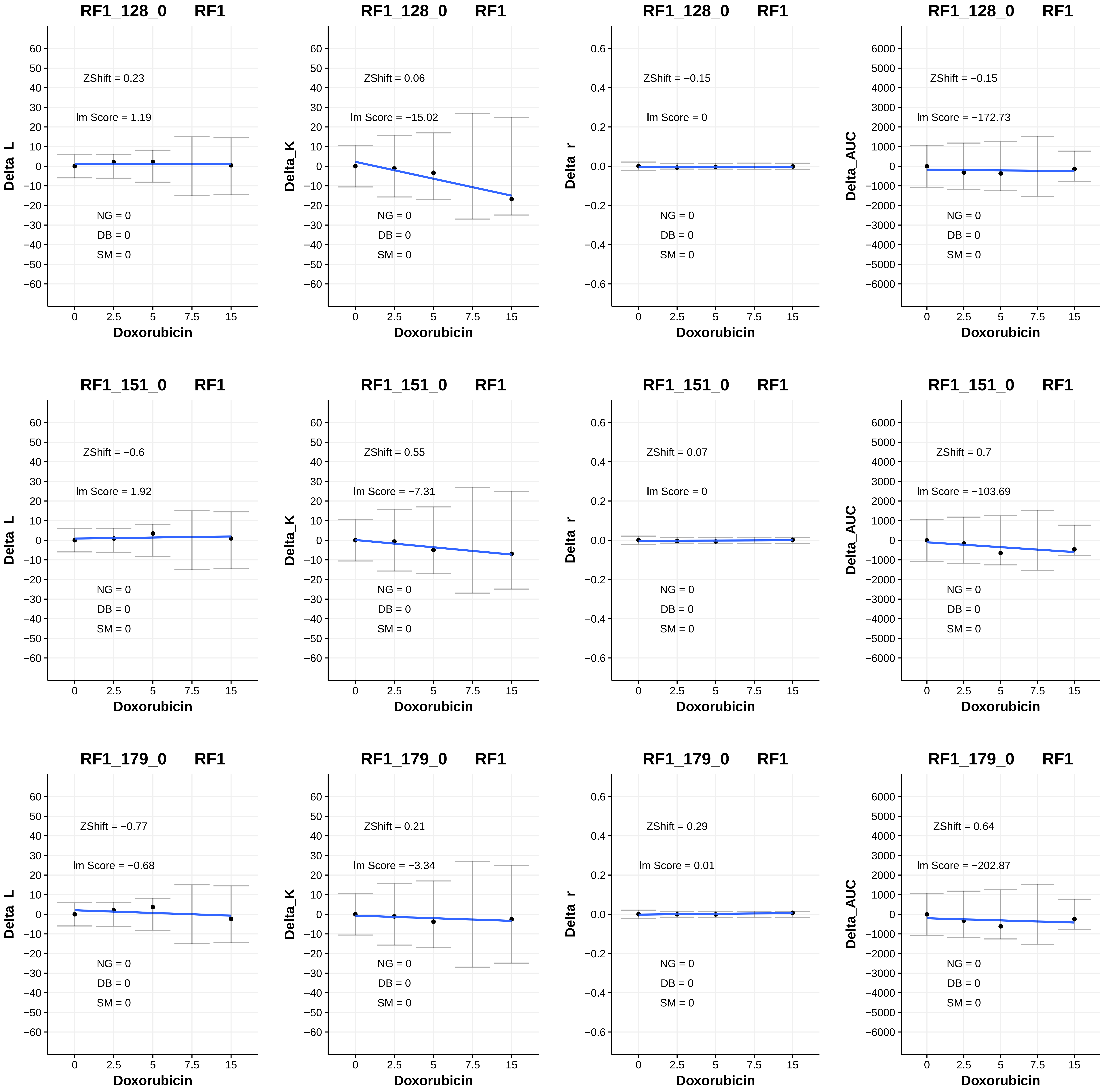

RF1_128_0 RF1
ZShift = 0.23
Im Score = 1.19
NG = 0
DB = 0
SM = 0
Delta_L
Doxorubicin
RF1_128_0 RF1
ZShift = 0.06
Im Score = −15.02
NG = 0
DB = 0
SM = 0
Delta_K
Doxorubicin
RF1_128_0 RF1
ZShift = −0.15
Im Score = 0
NG = 0
DB = 0
SM = 0
Delta_r
Doxorubicin
RF1_128_0 RF1
ZShift = −0.15
Im Score = −172.73
NG = 0
DB = 0
SM = 0
Delta_AUC
Doxorubicin
RF1_151_0 RF1
ZShift = −0.6
Im Score = 1.92
NG = 0
DB = 0
SM = 0
Delta_L
Doxorubicin
RF1_151_0 RF1
ZShift = 0.55
Im Score = −7.31
NG = 0
DB = 0
SM = 0
Delta_K
Doxorubicin
RF1_151_0 RF1
ZShift = 0.07
Im Score = 0
NG = 0
DB = 0
SM = 0
Delta_r
Doxorubicin
RF1_151_0 RF1
ZShift = 0.7
Im Score = −103.69
NG = 0
DB = 0
SM = 0
Delta_AUC
Doxorubicin
RF1_179_0 RF1
ZShift = −0.77
Im Score = −0.68
NG = 0
DB = 0
SM = 0
Delta_L
Doxorubicin
RF1_179_0 RF1
ZShift = 0.21
Im Score = −3.34
NG = 0
DB = 0
SM = 0
Delta_K
Doxorubicin
RF1_179_0 RF1
ZShift = 0.29
Im Score = 0.01
NG = 0
DB = 0
SM = 0
Delta_r
Doxorubicin
RF1_179_0 RF1
ZShift = 0.64
Im Score = −202.87
NG = 0
DB = 0
SM = 0
Delta_AUC
Doxorubicin

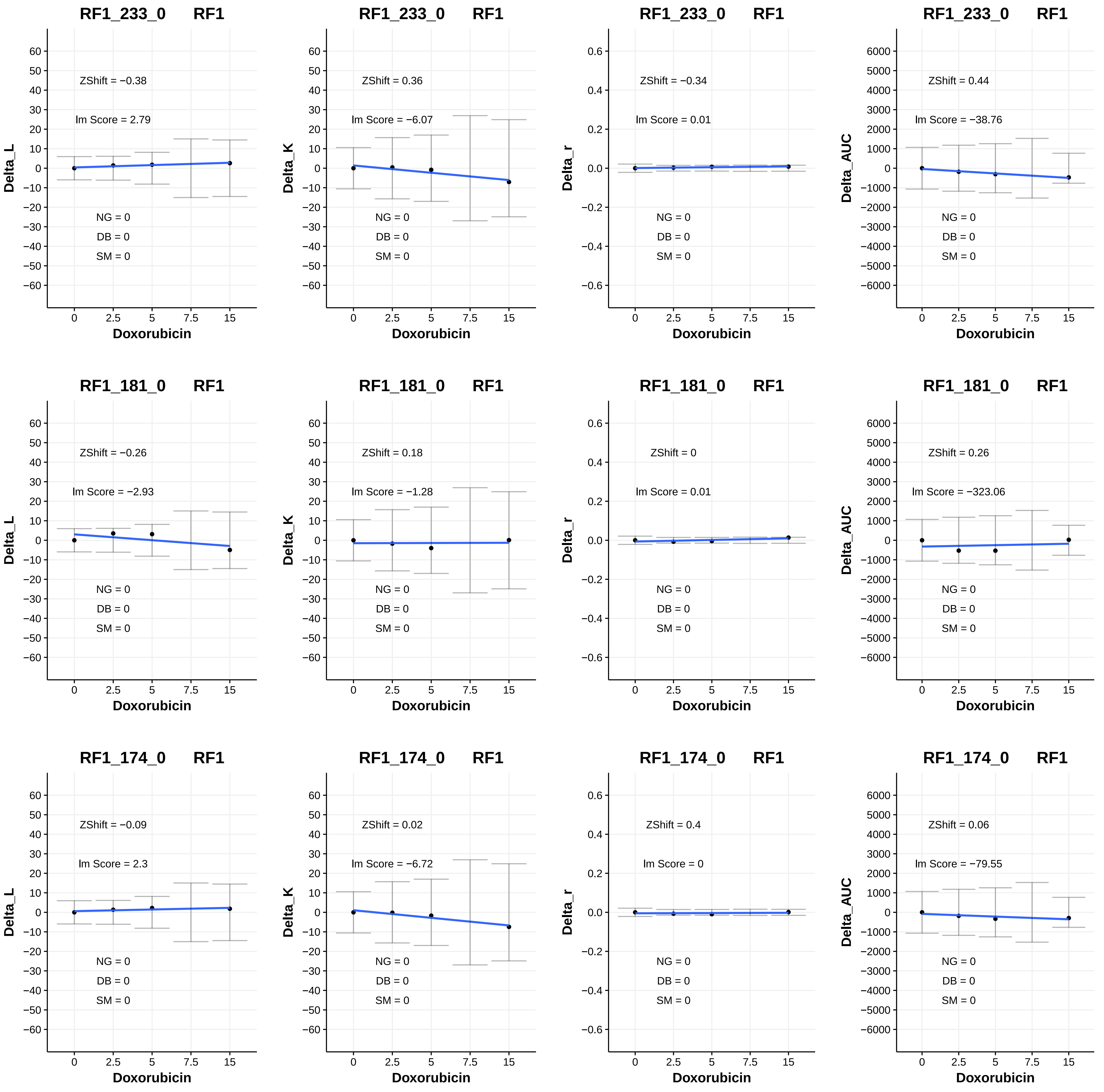
RF1_233_0 RF1
ZShift = −0.38
lm Score = 2.79
NG = 0
DB = 0
SM = 0
Delta_L
Doxorubicin
RF1_233_0 RF1
ZShift = 0.36
lm Score = −6.07
NG = 0
DB = 0
SM = 0
Delta_K
Doxorubicin
RF1_233_0 RF1
ZShift = −0.34
lm Score = 0.01
NG = 0
DB = 0
SM = 0
Delta_r
Doxorubicin
RF1_233_0 RF1
ZShift = 0.44
lm Score = −38.76
NG = 0
DB = 0
SM = 0
Delta_AUC
Doxorubicin
RF1_181_0 RF1
ZShift = −0.26
lm Score = −2.93
NG = 0
DB = 0
SM = 0
Delta_L
Doxorubicin
RF1_181_0 RF1
ZShift = 0.18
lm Score = −1.28
NG = 0
DB = 0
SM = 0
Delta_K
Doxorubicin
RF1_181_0 RF1
ZShift = 0
lm Score = 0.01
NG = 0
DB = 0
SM = 0
Delta_r
Doxorubicin
RF1_181_0 RF1
ZShift = 0.26
lm Score = −323.06
NG = 0
DB = 0
SM = 0
Delta_AUC
Doxorubicin
RF1_174_0 RF1
ZShift = −0.09
lm Score = 2.3
NG = 0
DB = 0
SM = 0
Delta_L
Doxorubicin
RF1_174_0 RF1
ZShift = 0.02
lm Score = −6.72
NG = 0
DB = 0
SM = 0
Delta_K
Doxorubicin
RF1_174_0 RF1
ZShift = 0.4
lm Score = 0
NG = 0
DB = 0
SM = 0
Delta_r
Doxorubicin
RF1_174_0 RF1
ZShift = 0.06
lm Score = −79.55
NG = 0
DB = 0
SM = 0
Delta_AUC
Doxorubicin

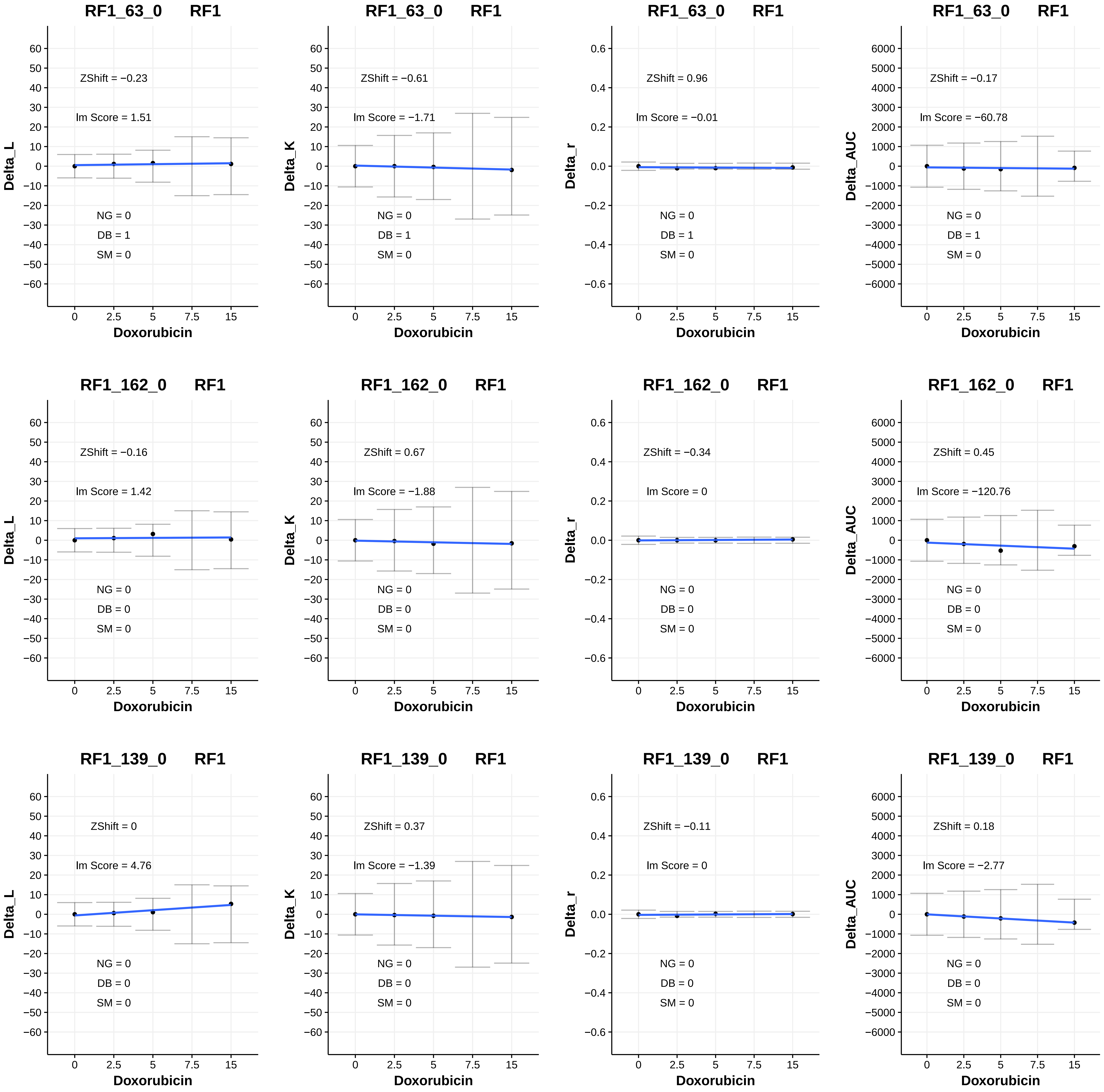

RF1_63_0 RF1
ZShift = −0.23
Im Score = 1.51
NG = 0
DB = 1
SM = 0
Delta_L
Doxorubicin
RF1_63_0 RF1
ZShift = −0.61
Im Score = −1.71
NG = 0
DB = 1
SM = 0
Delta_K
Doxorubicin
RF1_63_0 RF1
ZShift = 0.96
Im Score = −0.01
NG = 0
DB = 1
SM = 0
Delta_r
Doxorubicin
RF1_63_0 RF1
ZShift = −0.17
Im Score = −60.78
NG = 0
DB = 1
SM = 0
Delta_AUC
Doxorubicin
RF1_162_0 RF1
ZShift = −0.16
Im Score = 1.42
NG = 0
DB = 0
SM = 0
Delta_L
Doxorubicin
RF1_162_0 RF1
ZShift = 0.67
Im Score = −1.88
NG = 0
DB = 0
SM = 0
Delta_K
Doxorubicin
RF1_162_0 RF1
ZShift = −0.34
Im Score = 0
NG = 0
DB = 0
SM = 0
Delta_r
Doxorubicin
RF1_162_0 RF1
ZShift = 0.45
Im Score = −120.76
NG = 0
DB = 0
SM = 0
Delta_AUC
Doxorubicin
RF1_139_0 RF1
ZShift = 0
Im Score = 4.76
NG = 0
DB = 0
SM = 0
Delta_L
Doxorubicin
RF1_139_0 RF1
ZShift = 0.37
Im Score = −1.39
NG = 0
DB = 0
SM = 0
Delta_K
Doxorubicin
RF1_139_0 RF1
ZShift = −0.11
Im Score = 0
NG = 0
DB = 0
SM = 0
Delta_r
Doxorubicin
RF1_139_0 RF1
ZShift = 0.18
Im Score = −2.77
NG = 0
DB = 0
SM = 0
Delta_AUC
Doxorubicin

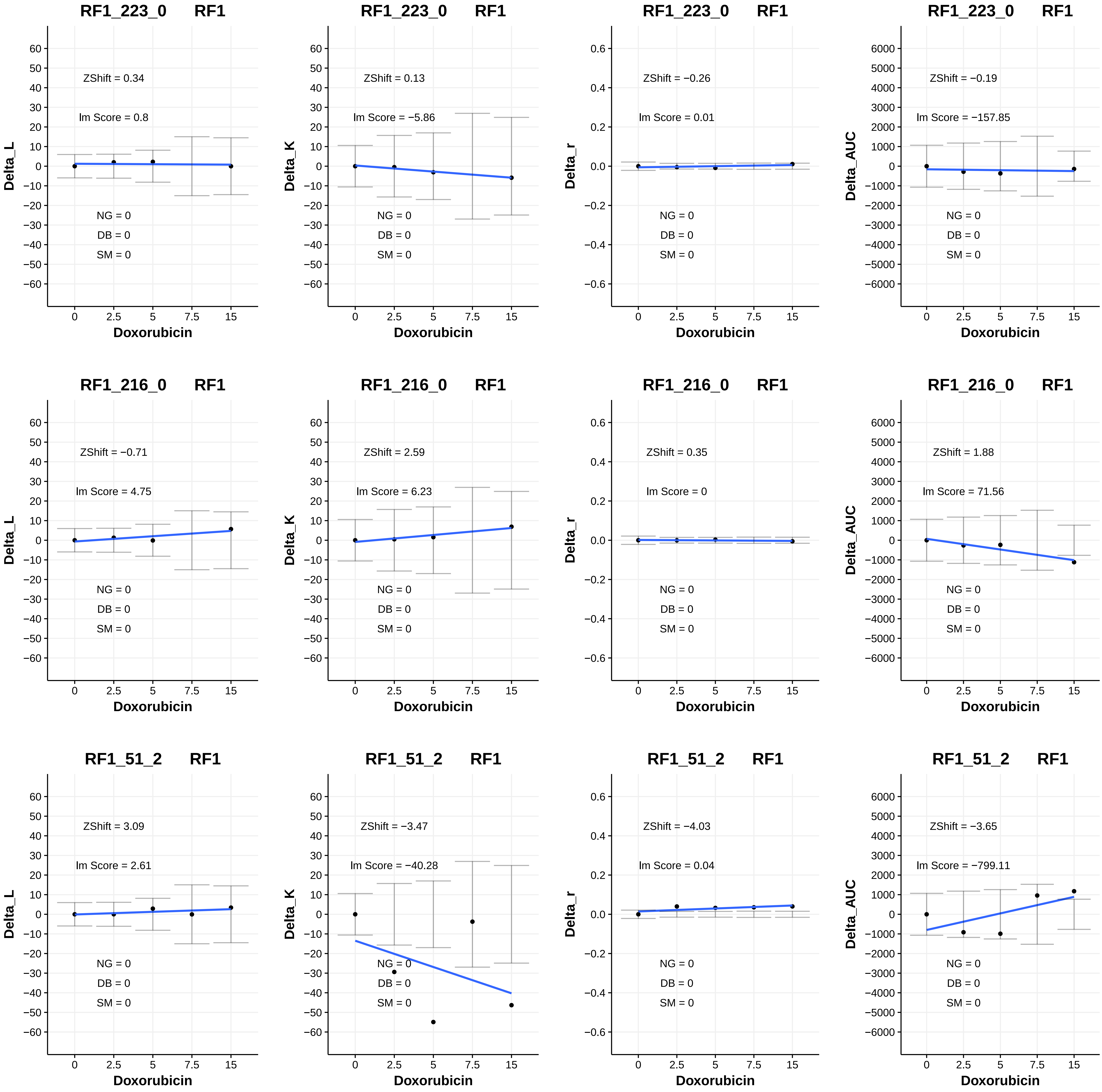
RF1_223_0 RF1
ZShift = 0.34
lm Score = 0.8
NG = 0
DB = 0
SM = 0
Delta_L
Doxorubicin
RF1_223_0 RF1
ZShift = 0.13
lm Score = −5.86
NG = 0
DB = 0
SM = 0
Delta_K
Doxorubicin
RF1_223_0 RF1
ZShift = −0.26
lm Score = 0.01
NG = 0
DB = 0
SM = 0
Delta_r
Doxorubicin
RF1_223_0 RF1
ZShift = −0.19
lm Score = −157.85
NG = 0
DB = 0
SM = 0
Delta_AUC
Doxorubicin
RF1_216_0 RF1
ZShift = −0.71
lm Score = 4.75
NG = 0
DB = 0
SM = 0
Delta_L
Doxorubicin
RF1_216_0 RF1
ZShift = 2.59
lm Score = 6.23
NG = 0
DB = 0
SM = 0
Delta_K
Doxorubicin
RF1_216_0 RF1
ZShift = 0.35
lm Score = 0
NG = 0
DB = 0
SM = 0
Delta_r
Doxorubicin
RF1_216_0 RF1
ZShift = 1.88
lm Score = 71.56
NG = 0
DB = 0
SM = 0
Delta_AUC
Doxorubicin
RF1_51_2 RF1
ZShift = 3.09
lm Score = 2.61
NG = 0
DB = 0
SM = 0
Delta_L
Doxorubicin
RF1_51_2 RF1
ZShift = −3.47
lm Score = −40.28
NG = 0
DB = 0
SM = 0
Delta_K
Doxorubicin
RF1_51_2 RF1
ZShift = −4.03
lm Score = 0.04
NG = 0
DB = 0
SM = 0
Delta_r
Doxorubicin
RF1_51_2 RF1
ZShift = −3.65
lm Score = −799.11
NG = 0
DB = 0
SM = 0
Delta_AUC
Doxorubicin

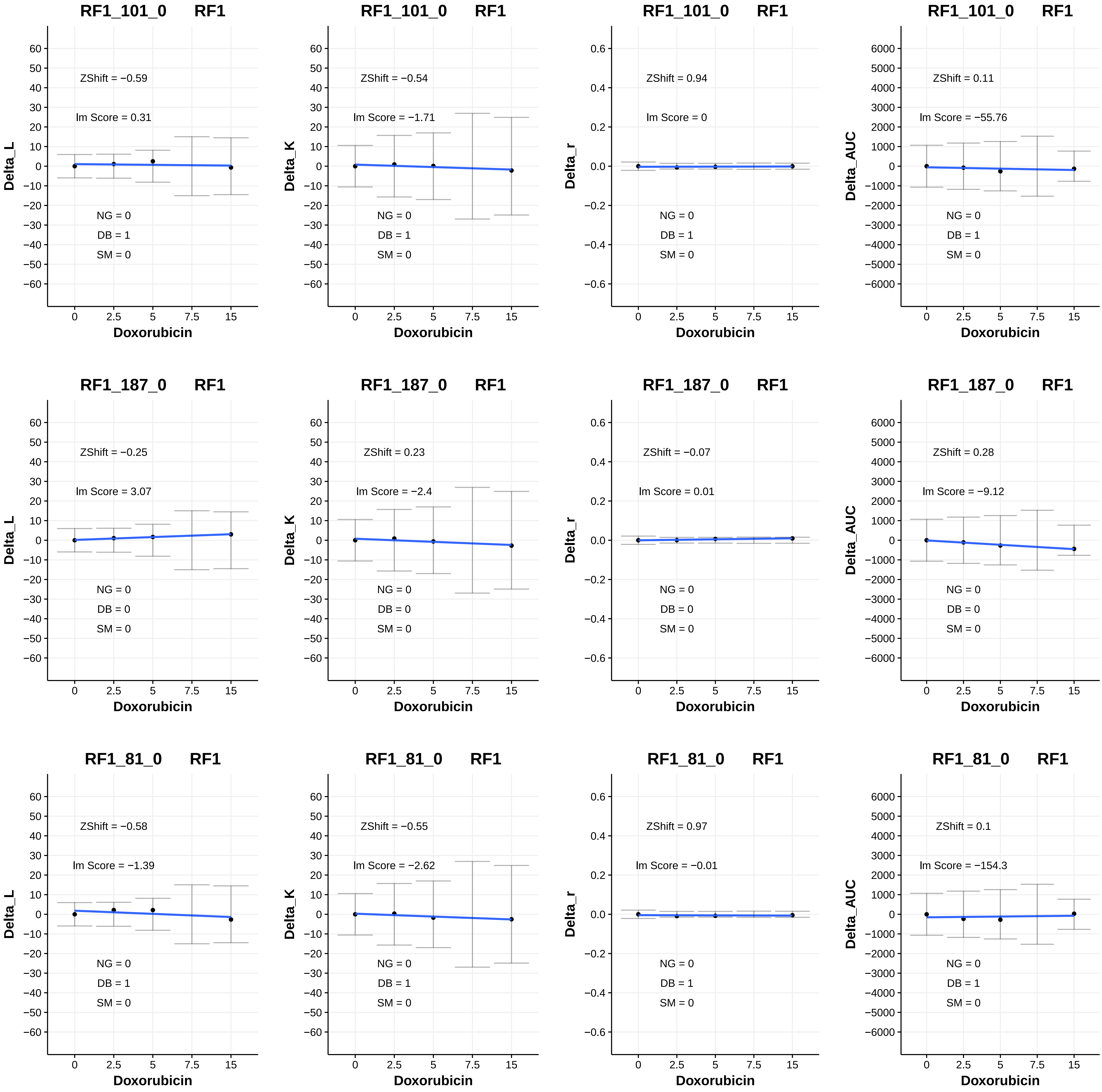

RF1_101_0 RF1
ZShift = −0.59
lm Score = 0.31
NG = 0
DB = 1
SM = 0
Delta_L
Doxorubicin
RF1_101_0 RF1
ZShift = −0.54
lm Score = −1.71
NG = 0
DB = 1
SM = 0
Delta_K
Doxorubicin
RF1_101_0 RF1
ZShift = 0.94
lm Score = 0
NG = 0
DB = 1
SM = 0
Delta_r
Doxorubicin
RF1_101_0 RF1
ZShift = 0.11
lm Score = −55.76
NG = 0
DB = 1
SM = 0
Delta_AUC
Doxorubicin
RF1_187_0 RF1
ZShift = −0.25
lm Score = 3.07
NG = 0
DB = 0
SM = 0
Delta_L
Doxorubicin
RF1_187_0 RF1
ZShift = 0.23
lm Score = −2.4
NG = 0
DB = 0
SM = 0
Delta_K
Doxorubicin
RF1_187_0 RF1
ZShift = −0.07
lm Score = 0.01
NG = 0
DB = 0
SM = 0
Delta_r
Doxorubicin
RF1_187_0 RF1
ZShift = 0.28
lm Score = −9.12
NG = 0
DB = 0
SM = 0
Delta_AUC
Doxorubicin
RF1_81_0 RF1
ZShift = −0.58
lm Score = −1.39
NG = 0
DB = 1
SM = 0
Delta_L
Doxorubicin
RF1_81_0 RF1
ZShift = −0.55
lm Score = −2.62
NG = 0
DB = 1
SM = 0
Delta_K
Doxorubicin
RF1_81_0 RF1
ZShift = 0.97
lm Score = −0.01
NG = 0
DB = 1
SM = 0
Delta_r
Doxorubicin
RF1_81_0 RF1
ZShift = 0.1
lm Score = −154.3
NG = 0
DB = 1
SM = 0
Delta_AUC
Doxorubicin

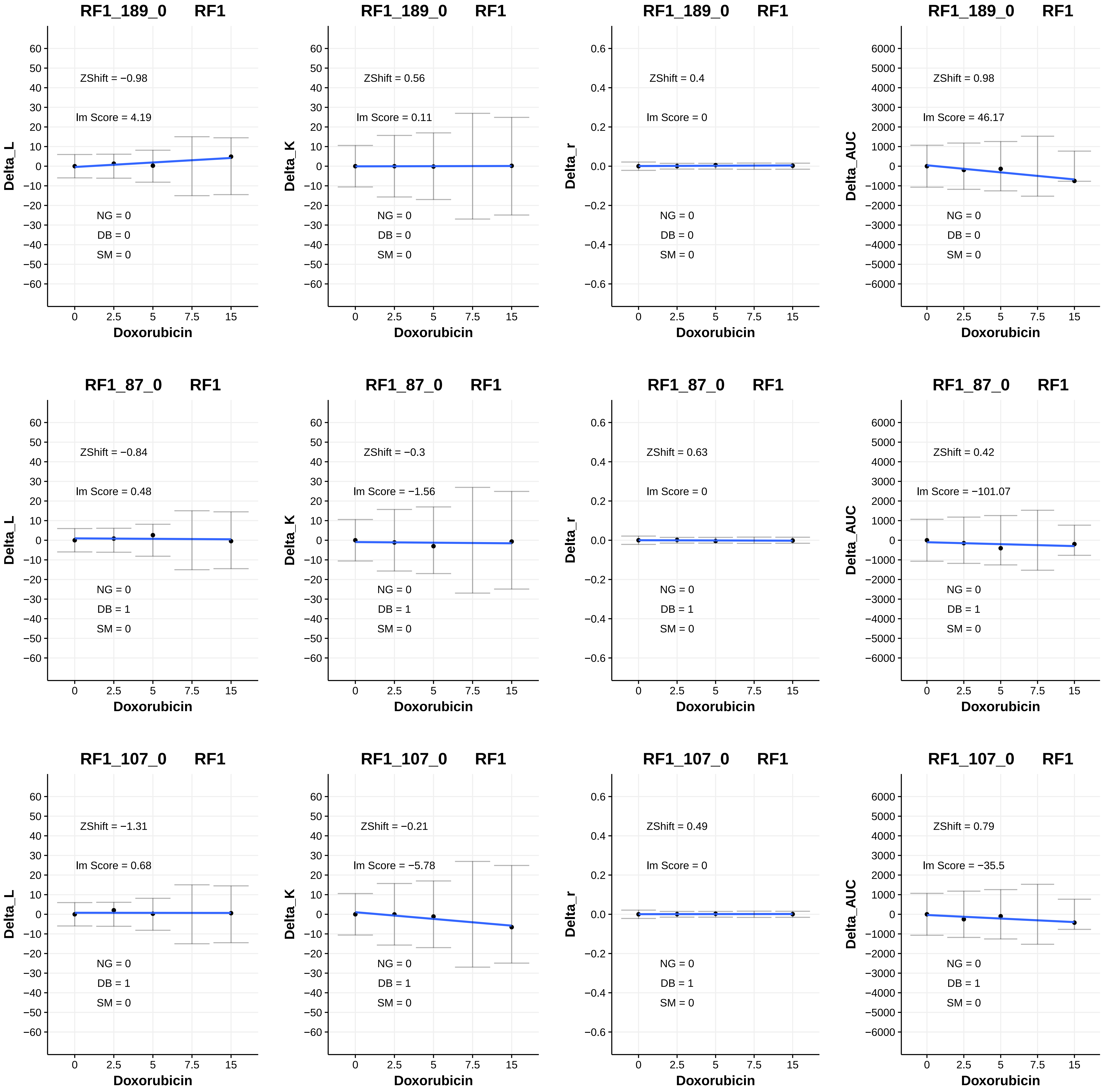
RF1_189_0 RF1
ZShift = −0.98
lm Score = 4.19
NG = 0
DB = 0
SM = 0
Delta_L
Doxorubicin
RF1_189_0 RF1
ZShift = 0.56
lm Score = 0.11
NG = 0
DB = 0
SM = 0
Delta_K
Doxorubicin
RF1_189_0 RF1
ZShift = 0.4
lm Score = 0
NG = 0
DB = 0
SM = 0
Delta_r
Doxorubicin
RF1_189_0 RF1
ZShift = 0.98
lm Score = 46.17
NG = 0
DB = 0
SM = 0
Delta_AUC
Doxorubicin
RF1_87_0 RF1
ZShift = −0.84
lm Score = 0.48
NG = 0
DB = 1
SM = 0
Delta_L
Doxorubicin
RF1_87_0 RF1
ZShift = −0.3
lm Score = −1.56
NG = 0
DB = 1
SM = 0
Delta_K
Doxorubicin
RF1_87_0 RF1
ZShift = 0.63
lm Score = 0
NG = 0
DB = 1
SM = 0
Delta_r
Doxorubicin
RF1_87_0 RF1
ZShift = 0.42
lm Score = −101.07
NG = 0
DB = 1
SM = 0
Delta_AUC
Doxorubicin
RF1_107_0 RF1
ZShift = −1.31
lm Score = 0.68
NG = 0
DB = 1
SM = 0
Delta_L
Doxorubicin
RF1_107_0 RF1
ZShift = −0.21
lm Score = −5.78
NG = 0
DB = 1
SM = 0
Delta_K
Doxorubicin
RF1_107_0 RF1
ZShift = 0.49
lm Score = 0
NG = 0
DB = 1
SM = 0
Delta_r
Doxorubicin
RF1_107_0 RF1
ZShift = 0.79
lm Score = −35.5
NG = 0
DB = 1
SM = 0
Delta_AUC
Doxorubicin

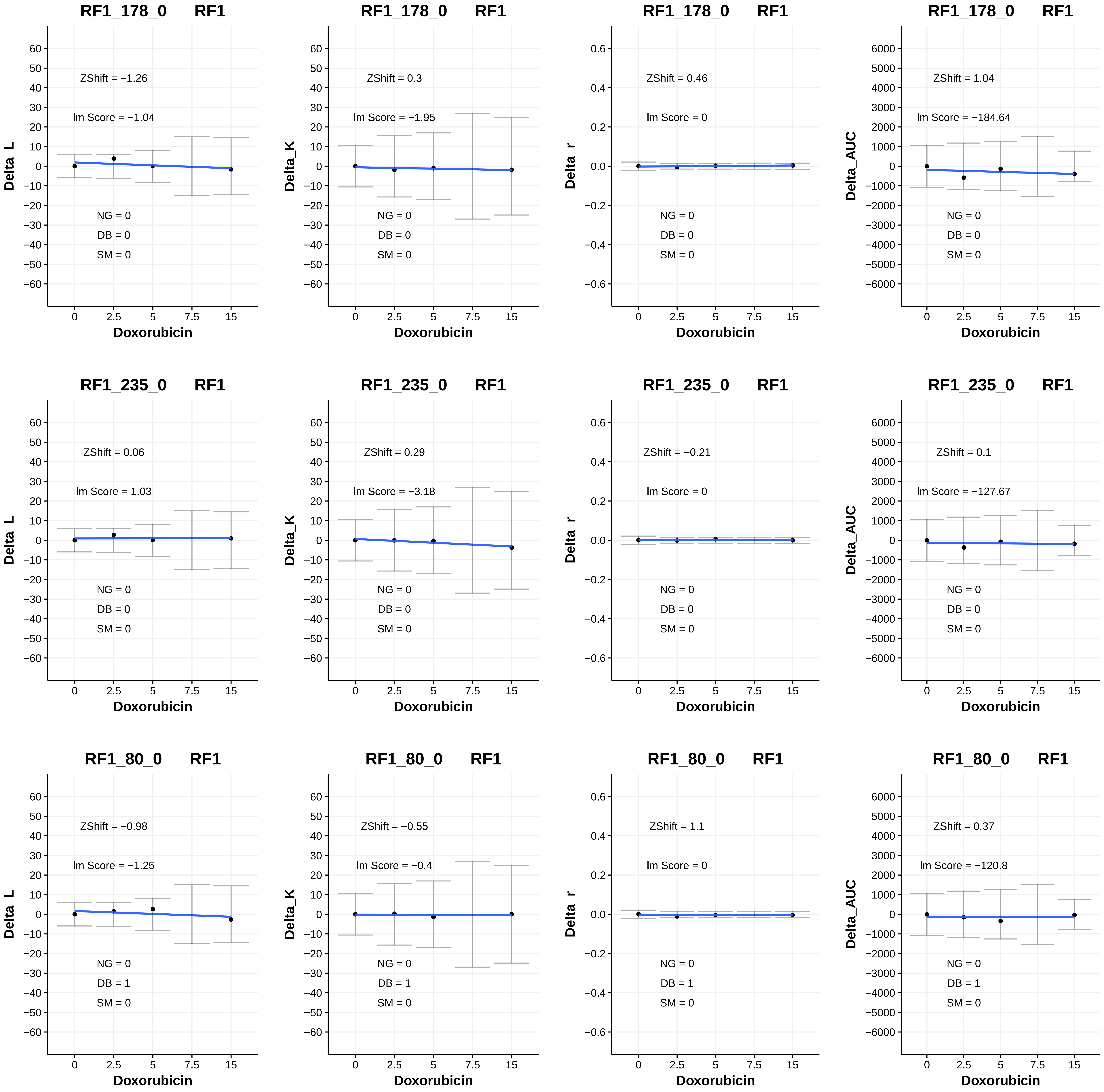
RF1_178_0 RF1
ZShift = −1.26
lm Score = −1.04
NG = 0
DB = 0
SM = 0
Delta_L
Doxorubicin
RF1_178_0 RF1
ZShift = 0.3
lm Score = −1.95
NG = 0
DB = 0
SM = 0
Delta_K
Doxorubicin
RF1_178_0 RF1
ZShift = 0.46
lm Score = 0
NG = 0
DB = 0
SM = 0
Delta_r
Doxorubicin
RF1_178_0 RF1
ZShift = 1.04
lm Score = −184.64
NG = 0
DB = 0
SM = 0
Delta_AUC
Doxorubicin
RF1_235_0 RF1
ZShift = 0.06
lm Score = 1.03
NG = 0
DB = 0
SM = 0
Delta_L
Doxorubicin
RF1_235_0 RF1
ZShift = 0.29
lm Score = −3.18
NG = 0
DB = 0
SM = 0
Delta_K
Doxorubicin
RF1_235_0 RF1
ZShift = −0.21
lm Score = 0
NG = 0
DB = 0
SM = 0
Delta_r
Doxorubicin
RF1_235_0 RF1
ZShift = 0.1
lm Score = −127.67
NG = 0
DB = 0
SM = 0
Delta_AUC
Doxorubicin
RF1_80_0 RF1
ZShift = −0.98
lm Score = −1.25
NG = 0
DB = 1
SM = 0
Delta_L
Doxorubicin
RF1_80_0 RF1
ZShift = −0.55
lm Score = −0.4
NG = 0
DB = 1
SM = 0
Delta_K
Doxorubicin
RF1_80_0 RF1
ZShift = 1.1
lm Score = 0
NG = 0
DB = 1
SM = 0
Delta_r
Doxorubicin
RF1_80_0 RF1
ZShift = 0.37
lm Score = −120.8
NG = 0
DB = 1
SM = 0
Delta_AUC
Doxorubicin

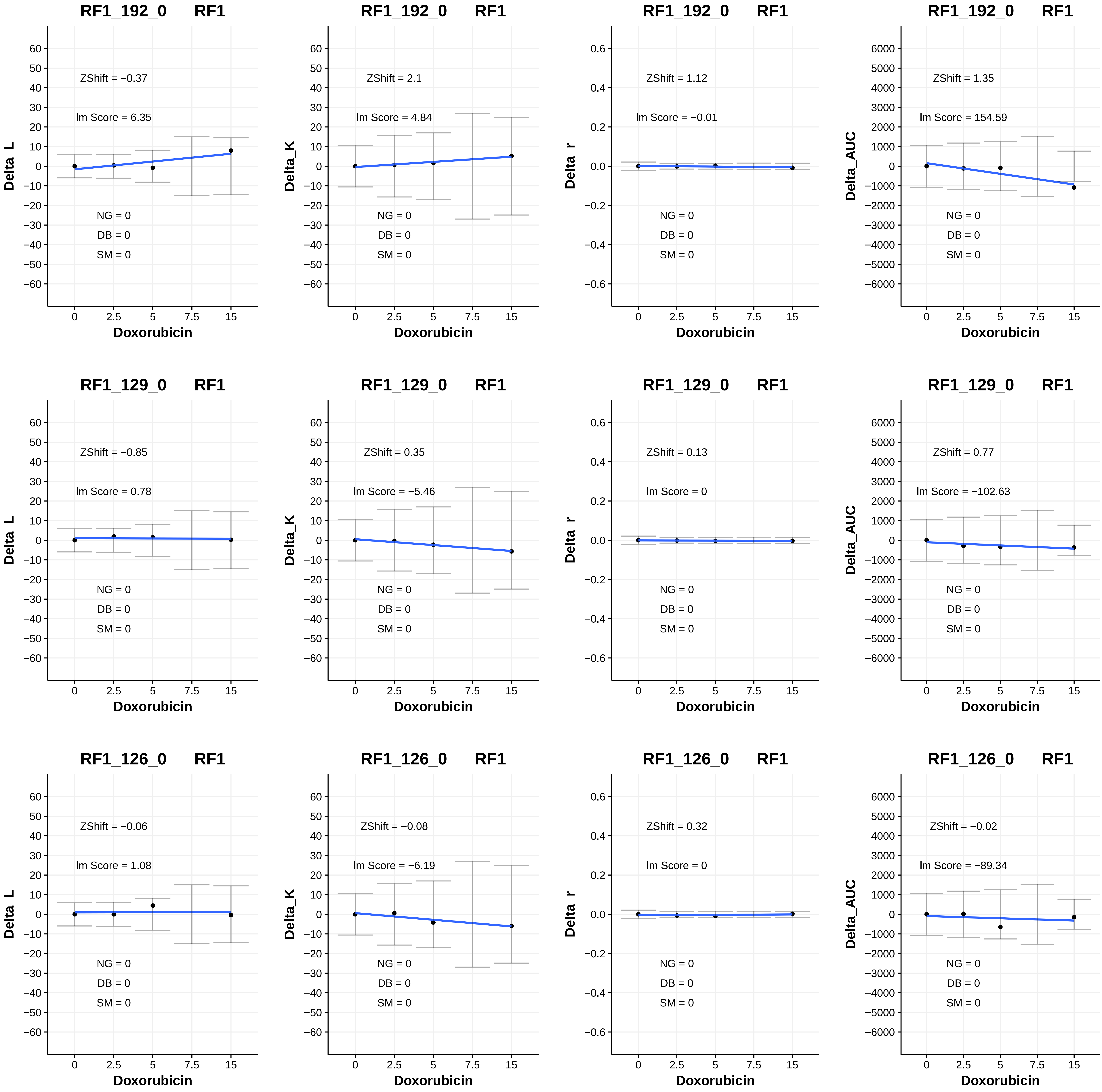
RF1_192_0 RF1
ZShift = −0.37
Im Score = 6.35
NG = 0
DB = 0
SM = 0
Delta_L
Doxorubicin
RF1_192_0 RF1
ZShift = 2.1
Im Score = 4.84
NG = 0
DB = 0
SM = 0
Delta_K
Doxorubicin
RF1_192_0 RF1
ZShift = 1.12
Im Score = −0.01
NG = 0
DB = 0
SM = 0
Delta_r
Doxorubicin
RF1_192_0 RF1
ZShift = 1.35
Im Score = 154.59
NG = 0
DB = 0
SM = 0
Delta_AUC
Doxorubicin
RF1_129_0 RF1
ZShift = −0.85
Im Score = 0.78
NG = 0
DB = 0
SM = 0
Delta_L
Doxorubicin
RF1_129_0 RF1
ZShift = 0.35
Im Score = −5.46
NG = 0
DB = 0
SM = 0
Delta_K
Doxorubicin
RF1_129_0 RF1
ZShift = 0.13
Im Score = 0
NG = 0
DB = 0
SM = 0
Delta_r
Doxorubicin
RF1_129_0 RF1
ZShift = 0.77
Im Score = −102.63
NG = 0
DB = 0
SM = 0
Delta_AUC
Doxorubicin
RF1_126_0 RF1
ZShift = −0.06
Im Score = 1.08
NG = 0
DB = 0
SM = 0
Delta_L
Doxorubicin
RF1_126_0 RF1
ZShift = −0.08
Im Score = −6.19
NG = 0
DB = 0
SM = 0
Delta_K
Doxorubicin
RF1_126_0 RF1
ZShift = 0.32
Im Score = 0
NG = 0
DB = 0
SM = 0
Delta_r
Doxorubicin
RF1_126_0 RF1
ZShift = −0.02
Im Score = −89.34
NG = 0
DB = 0
SM = 0
Delta_AUC
Doxorubicin

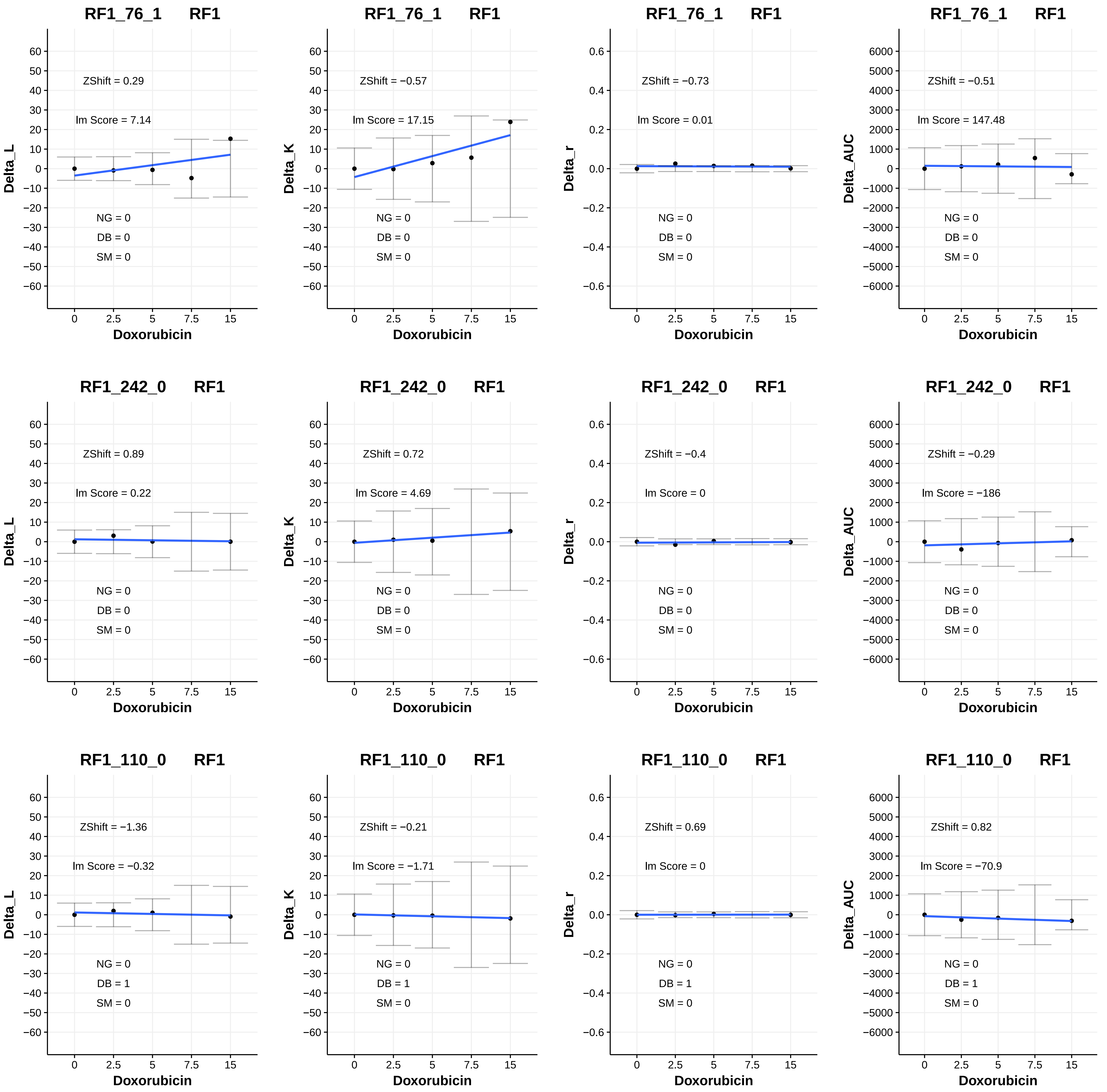

RF1_76_1 RF1
ZShift = 0.29
lm Score = 7.14
NG = 0
DB = 0
SM = 0
Delta_L
Doxorubicin
RF1_76_1 RF1
ZShift = −0.57
lm Score = 17.15
NG = 0
DB = 0
SM = 0
Delta_K
Doxorubicin
RF1_76_1 RF1
ZShift = −0.73
lm Score = 0.01
NG = 0
DB = 0
SM = 0
Delta_r
Doxorubicin
RF1_76_1 RF1
ZShift = −0.51
lm Score = 147.48
NG = 0
DB = 0
SM = 0
Delta_AUC
Doxorubicin
RF1_242_0 RF1
ZShift = 0.89
lm Score = 0.22
NG = 0
DB = 0
SM = 0
Delta_L
Doxorubicin
RF1_242_0 RF1
ZShift = 0.72
lm Score = 4.69
NG = 0
DB = 0
SM = 0
Delta_K
Doxorubicin
RF1_242_0 RF1
ZShift = −0.4
lm Score = 0
NG = 0
DB = 0
SM = 0
Delta_r
Doxorubicin
RF1_242_0 RF1
ZShift = −0.29
lm Score = −186
NG = 0
DB = 0
SM = 0
Delta_AUC
Doxorubicin
RF1_110_0 RF1
ZShift = −1.36
lm Score = −0.32
NG = 0
DB = 1
SM = 0
Delta_L
Doxorubicin
RF1_110_0 RF1
ZShift = −0.21
lm Score = −1.71
NG = 0
DB = 1
SM = 0
Delta_K
Doxorubicin
RF1_110_0 RF1
ZShift = 0.69
lm Score = 0
NG = 0
DB = 1
SM = 0
Delta_r
Doxorubicin
RF1_110_0 RF1
ZShift = 0.82
lm Score = −70.9
NG = 0
DB = 1
SM = 0
Delta_AUC
Doxorubicin

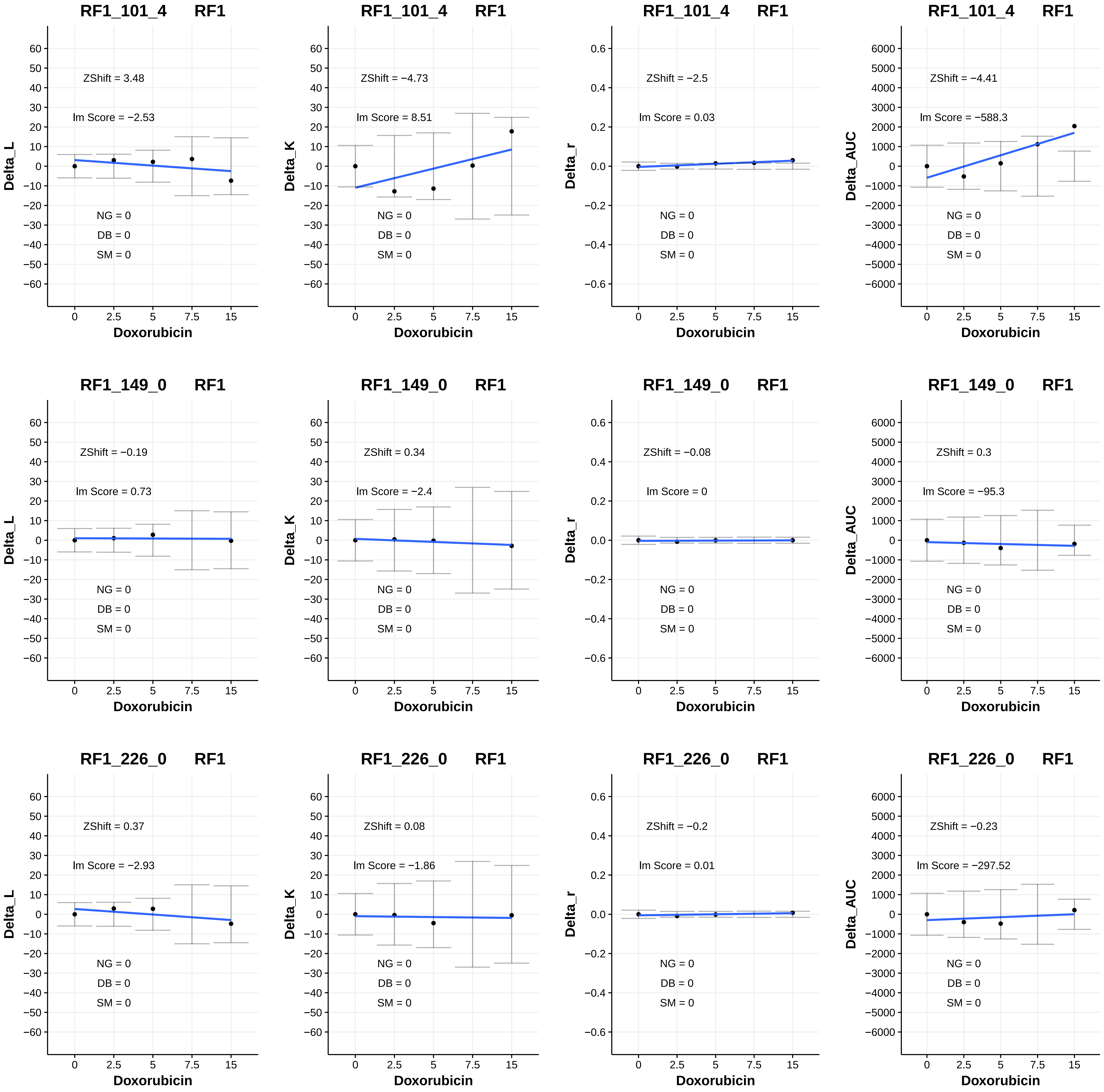
RF1_101_4 RF1
ZShift = 3.48
lm Score = −2.53
NG = 0
DB = 0
SM = 0
Delta_L
Doxorubicin
RF1_101_4 RF1
ZShift = −4.73
lm Score = 8.51
NG = 0
DB = 0
SM = 0
Delta_K
Doxorubicin
RF1_101_4 RF1
ZShift = −2.5
lm Score = 0.03
NG = 0
DB = 0
SM = 0
Delta_r
Doxorubicin
RF1_101_4 RF1
ZShift = −4.41
lm Score = −588.3
NG = 0
DB = 0
SM = 0
Delta_AUC
Doxorubicin
RF1_149_0 RF1
ZShift = −0.19
lm Score = 0.73
NG = 0
DB = 0
SM = 0
Delta_L
Doxorubicin
RF1_149_0 RF1
ZShift = 0.34
lm Score = −2.4
NG = 0
DB = 0
SM = 0
Delta_K
Doxorubicin
RF1_149_0 RF1
ZShift = −0.08
lm Score = 0
NG = 0
DB = 0
SM = 0
Delta_r
Doxorubicin
RF1_149_0 RF1
ZShift = 0.3
lm Score = −95.3
NG = 0
DB = 0
SM = 0
Delta_AUC
Doxorubicin
RF1_226_0 RF1
ZShift = 0.37
lm Score = −2.93
NG = 0
DB = 0
SM = 0
Delta_L
Doxorubicin
RF1_226_0 RF1
ZShift = 0.08
lm Score = −1.86
NG = 0
DB = 0
SM = 0
Delta_K
Doxorubicin
RF1_226_0 RF1
ZShift = −0.2
lm Score = 0.01
NG = 0
DB = 0
SM = 0
Delta_r
Doxorubicin
RF1_226_0 RF1
ZShift = −0.23
lm Score = −297.52
NG = 0
DB = 0
SM = 0
Delta_AUC
Doxorubicin

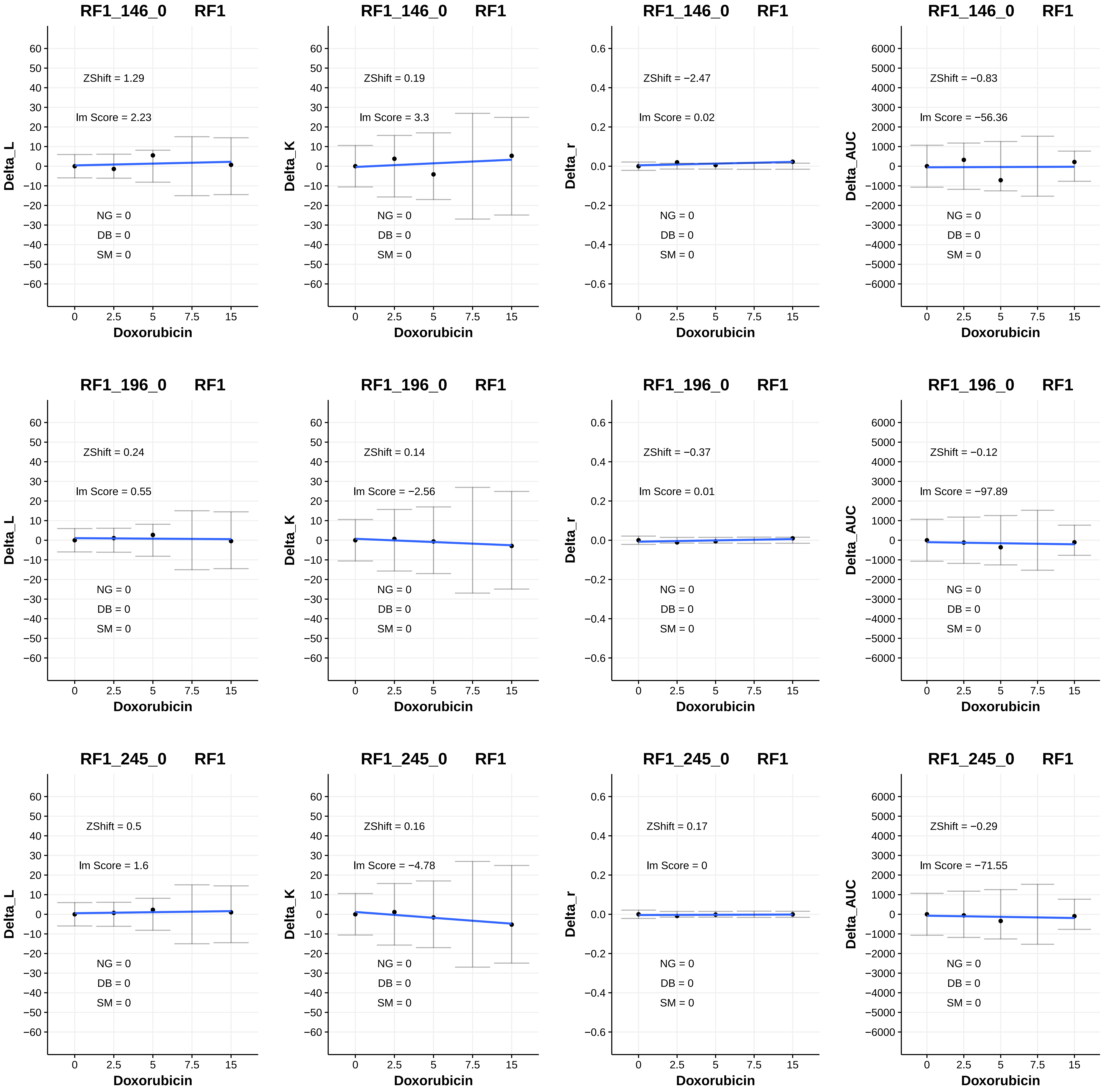
RF1_146_0 RF1
ZShift = 1.29
lm Score = 2.23
NG = 0
DB = 0
SM = 0
Delta_L
Doxorubicin
RF1_146_0 RF1
ZShift = 0.19
lm Score = 3.3
NG = 0
DB = 0
SM = 0
Delta_K
Doxorubicin
RF1_146_0 RF1
ZShift = −2.47
lm Score = 0.02
NG = 0
DB = 0
SM = 0
Delta_r
Doxorubicin
RF1_146_0 RF1
ZShift = −0.83
lm Score = −56.36
NG = 0
DB = 0
SM = 0
Delta_AUC
Doxorubicin
RF1_196_0 RF1
ZShift = 0.24
lm Score = 0.55
NG = 0
DB = 0
SM = 0
Delta_L
Doxorubicin
RF1_196_0 RF1
ZShift = 0.14
lm Score = −2.56
NG = 0
DB = 0
SM = 0
Delta_K
Doxorubicin
RF1_196_0 RF1
ZShift = −0.37
lm Score = 0.01
NG = 0
DB = 0
SM = 0
Delta_r
Doxorubicin
RF1_196_0 RF1
ZShift = −0.12
lm Score = −97.89
NG = 0
DB = 0
SM = 0
Delta_AUC
Doxorubicin
RF1_245_0 RF1
ZShift = 0.5
lm Score = 1.6
NG = 0
DB = 0
SM = 0
Delta_L
Doxorubicin
RF1_245_0 RF1
ZShift = 0.16
lm Score = −4.78
NG = 0
DB = 0
SM = 0
Delta_K
Doxorubicin
RF1_245_0 RF1
ZShift = 0.17
lm Score = 0
NG = 0
DB = 0
SM = 0
Delta_r
Doxorubicin
RF1_245_0 RF1
ZShift = −0.29
lm Score = −71.55
NG = 0
DB = 0
SM = 0
Delta_AUC
Doxorubicin

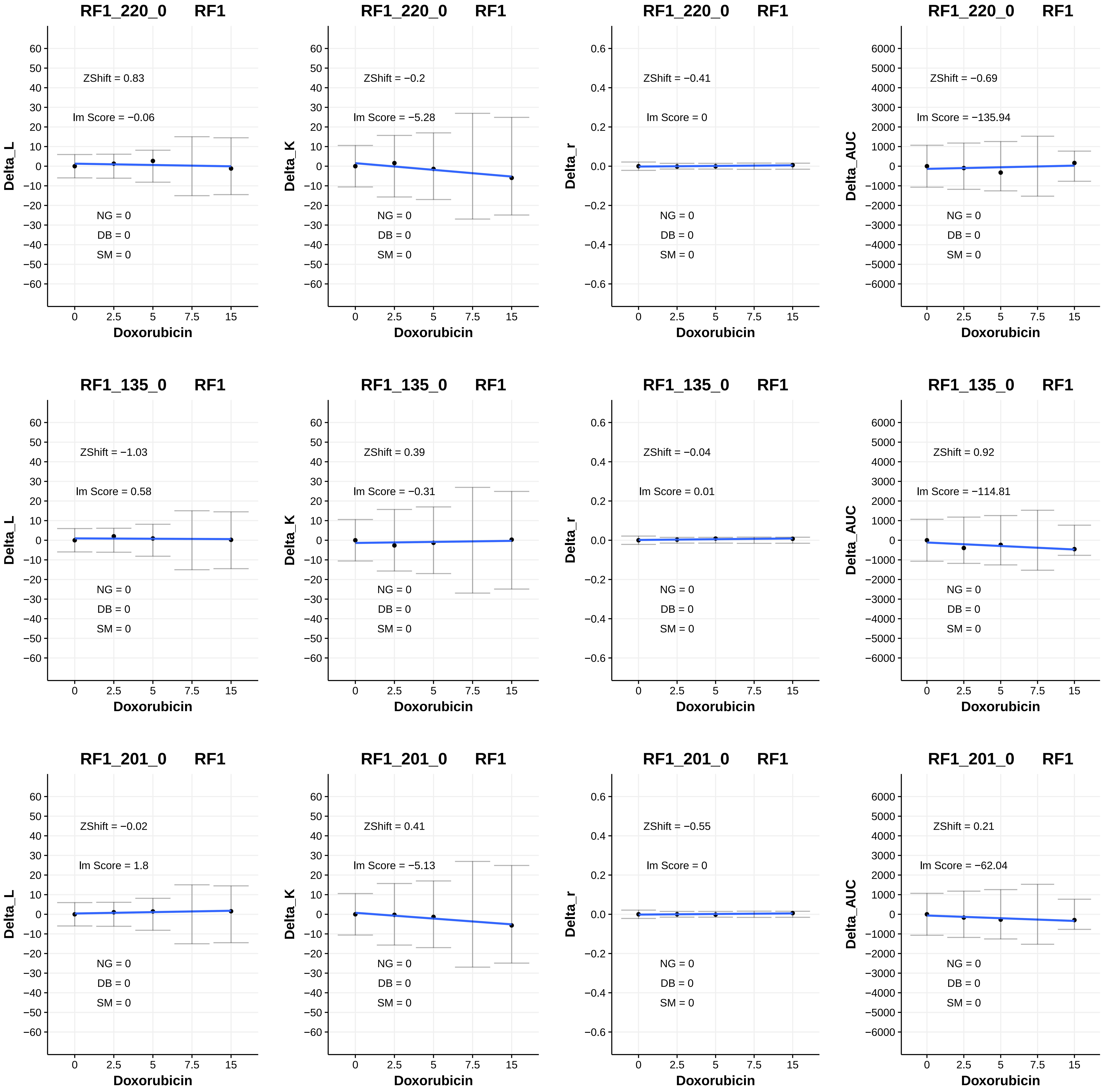
RF1_220_0 RF1
ZShift = 0.83
lm Score = −0.06
NG = 0
DB = 0
SM = 0
Delta_L
Doxorubicin
RF1_220_0 RF1
ZShift = −0.2
lm Score = −5.28
NG = 0
DB = 0
SM = 0
Delta_K
Doxorubicin
RF1_220_0 RF1
ZShift = −0.41
lm Score = 0
NG = 0
DB = 0
SM = 0
Delta_r
Doxorubicin
RF1_220_0 RF1
ZShift = −0.69
lm Score = −135.94
NG = 0
DB = 0
SM = 0
Delta_AUC
Doxorubicin
RF1_135_0 RF1
ZShift = −1.03
lm Score = 0.58
NG = 0
DB = 0
SM = 0
Delta_L
Doxorubicin
RF1_135_0 RF1
ZShift = 0.39
lm Score = −0.31
NG = 0
DB = 0
SM = 0
Delta_K
Doxorubicin
RF1_135_0 RF1
ZShift = −0.04
lm Score = 0.01
NG = 0
DB = 0
SM = 0
Delta_r
Doxorubicin
RF1_135_0 RF1
ZShift = 0.92
lm Score = −114.81
NG = 0
DB = 0
SM = 0
Delta_AUC
Doxorubicin
RF1_201_0 RF1
ZShift = −0.02
lm Score = 1.8
NG = 0
DB = 0
SM = 0
Delta_L
Doxorubicin
RF1_201_0 RF1
ZShift = 0.41
lm Score = −5.13
NG = 0
DB = 0
SM = 0
Delta_K
Doxorubicin
RF1_201_0 RF1
ZShift = −0.55
lm Score = 0
NG = 0
DB = 0
SM = 0
Delta_r
Doxorubicin
RF1_201_0 RF1
ZShift = 0.21
lm Score = −62.04
NG = 0
DB = 0
SM = 0
Delta_AUC
Doxorubicin

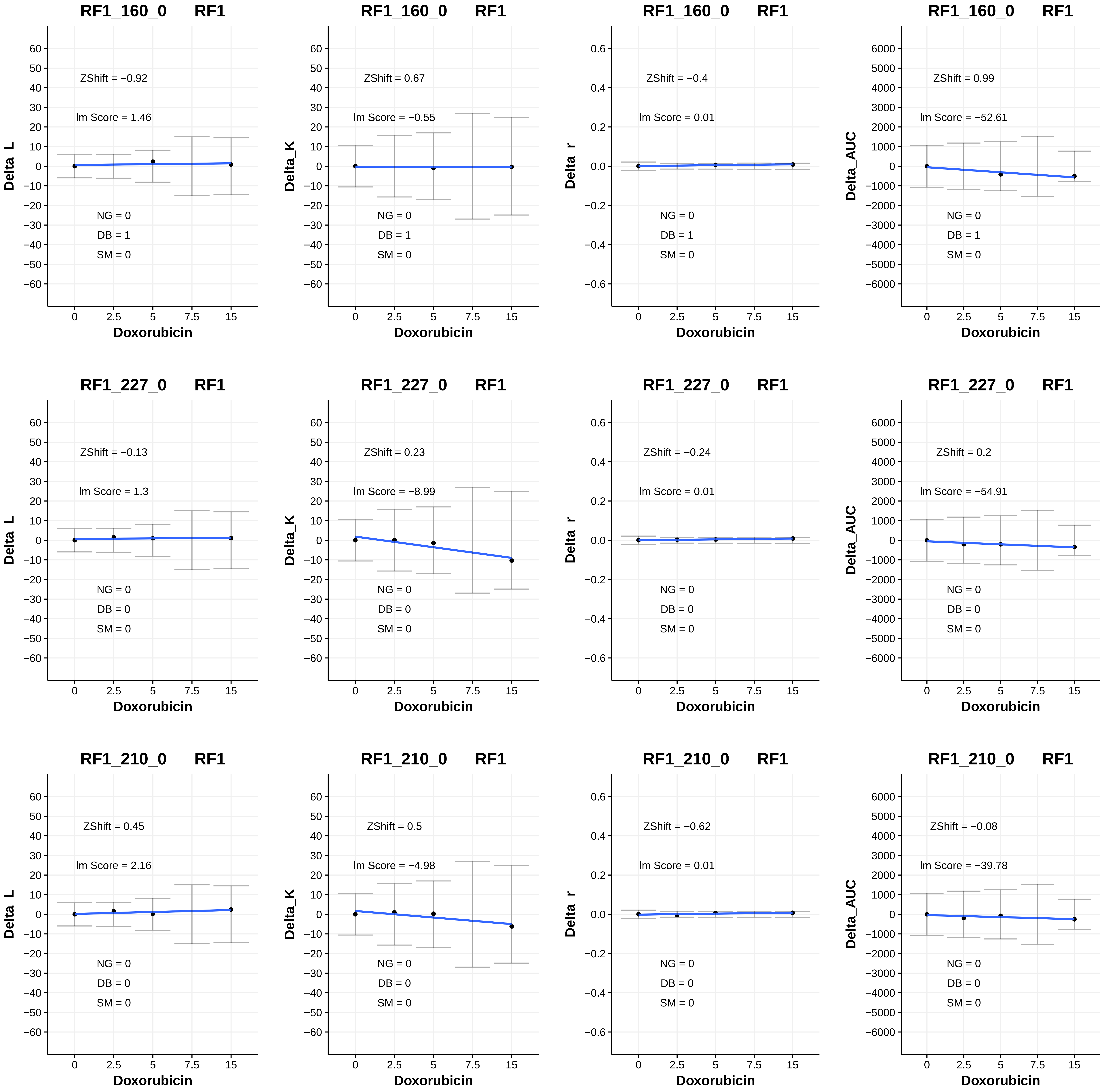
RF1_160_0 RF1
ZShift = −0.92
lm Score = 1.46
NG = 0
DB = 1
SM = 0
Delta_L
Doxorubicin
RF1_160_0 RF1
ZShift = 0.67
lm Score = −0.55
NG = 0
DB = 1
SM = 0
Delta_K
Doxorubicin
RF1_160_0 RF1
ZShift = −0.4
lm Score = 0.01
NG = 0
DB = 1
SM = 0
Delta_r
Doxorubicin
RF1_160_0 RF1
ZShift = 0.99
lm Score = −52.61
NG = 0
DB = 1
SM = 0
Delta_AUC
Doxorubicin
RF1_227_0 RF1
ZShift = −0.13
lm Score = 1.3
NG = 0
DB = 0
SM = 0
Delta_L
Doxorubicin
RF1_227_0 RF1
ZShift = 0.23
lm Score = −8.99
NG = 0
DB = 0
SM = 0
Delta_K
Doxorubicin
RF1_227_0 RF1
ZShift = −0.24
lm Score = 0.01
NG = 0
DB = 0
SM = 0
Delta_r
Doxorubicin
RF1_227_0 RF1
ZShift = 0.2
lm Score = −54.91
NG = 0
DB = 0
SM = 0
Delta_AUC
Doxorubicin
RF1_210_0 RF1
ZShift = 0.45
lm Score = 2.16
NG = 0
DB = 0
SM = 0
Delta_L
Doxorubicin
RF1_210_0 RF1
ZShift = 0.5
lm Score = −4.98
NG = 0
DB = 0
SM = 0
Delta_K
Doxorubicin
RF1_210_0 RF1
ZShift = −0.62
lm Score = 0.01
NG = 0
DB = 0
SM = 0
Delta_r
Doxorubicin
RF1_210_0 RF1
ZShift = −0.08
lm Score = −39.78
NG = 0
DB = 0
SM = 0
Delta_AUC
Doxorubicin

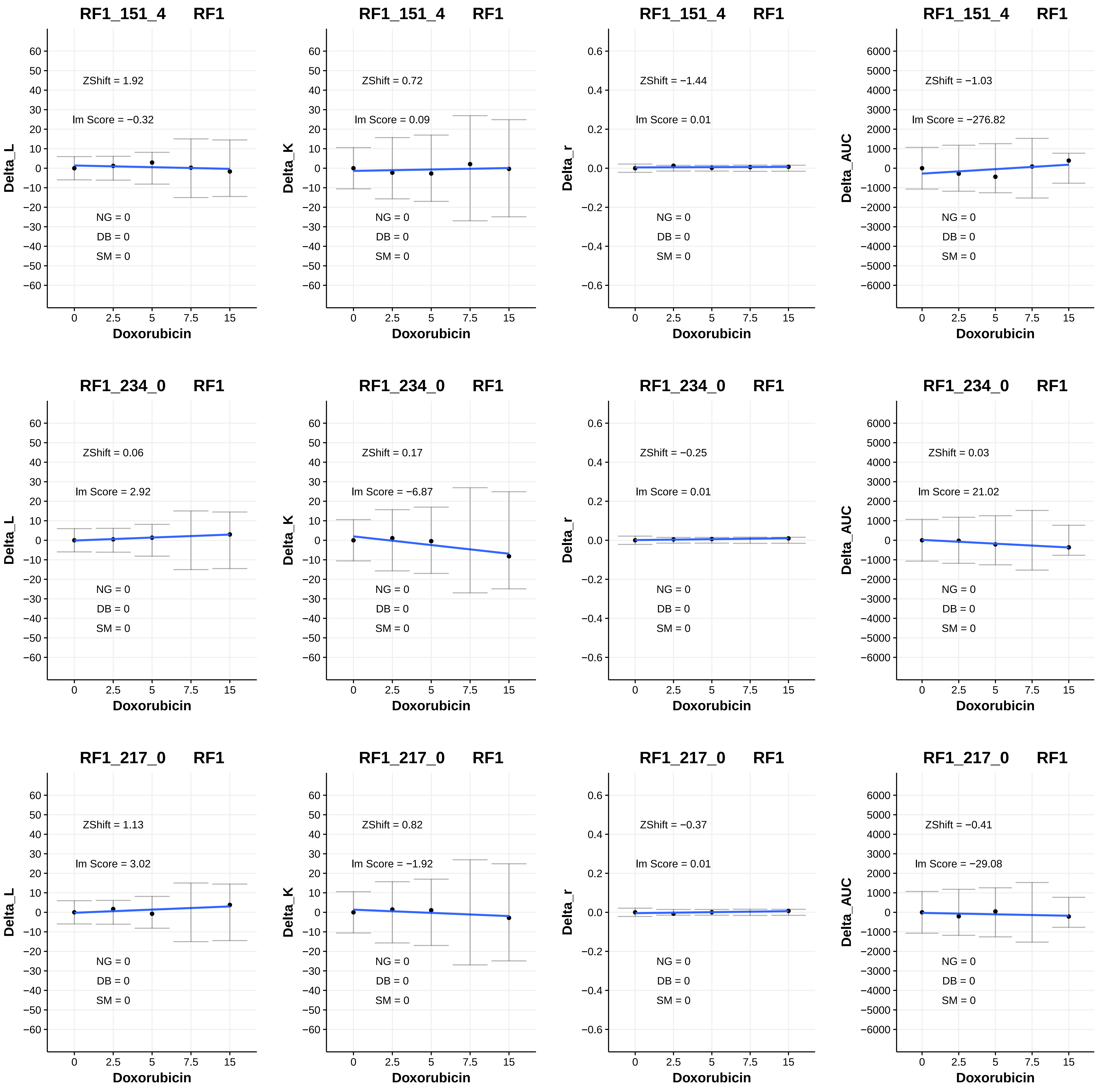

RF1_151_4 RF1
ZShift = 1.92
lm Score = −0.32
NG = 0
DB = 0
SM = 0
Delta_L
Doxorubicin
RF1_151_4 RF1
ZShift = 0.72
lm Score = 0.09
NG = 0
DB = 0
SM = 0
Delta_K
Doxorubicin
RF1_151_4 RF1
ZShift = −1.44
lm Score = 0.01
NG = 0
DB = 0
SM = 0
Delta_r
Doxorubicin
RF1_151_4 RF1
ZShift = −1.03
lm Score = −276.82
NG = 0
DB = 0
SM = 0
Delta_AUC
Doxorubicin
RF1_234_0 RF1
ZShift = 0.06
lm Score = 2.92
NG = 0
DB = 0
SM = 0
Delta_L
Doxorubicin
RF1_234_0 RF1
ZShift = 0.17
lm Score = −6.87
NG = 0
DB = 0
SM = 0
Delta_K
Doxorubicin
RF1_234_0 RF1
ZShift = −0.25
lm Score = 0.01
NG = 0
DB = 0
SM = 0
Delta_r
Doxorubicin
RF1_234_0 RF1
ZShift = 0.03
lm Score = 21.02
NG = 0
DB = 0
SM = 0
Delta_AUC
Doxorubicin
RF1_217_0 RF1
ZShift = 1.13
lm Score = 3.02
NG = 0
DB = 0
SM = 0
Delta_L
Doxorubicin
RF1_217_0 RF1
ZShift = 0.82
lm Score = −1.92
NG = 0
DB = 0
SM = 0
Delta_K
Doxorubicin
RF1_217_0 RF1
ZShift = −0.37
lm Score = 0.01
NG = 0
DB = 0
SM = 0
Delta_r
Doxorubicin
RF1_217_0 RF1
ZShift = −0.41
lm Score = −29.08
NG = 0
DB = 0
SM = 0
Delta_AUC
Doxorubicin

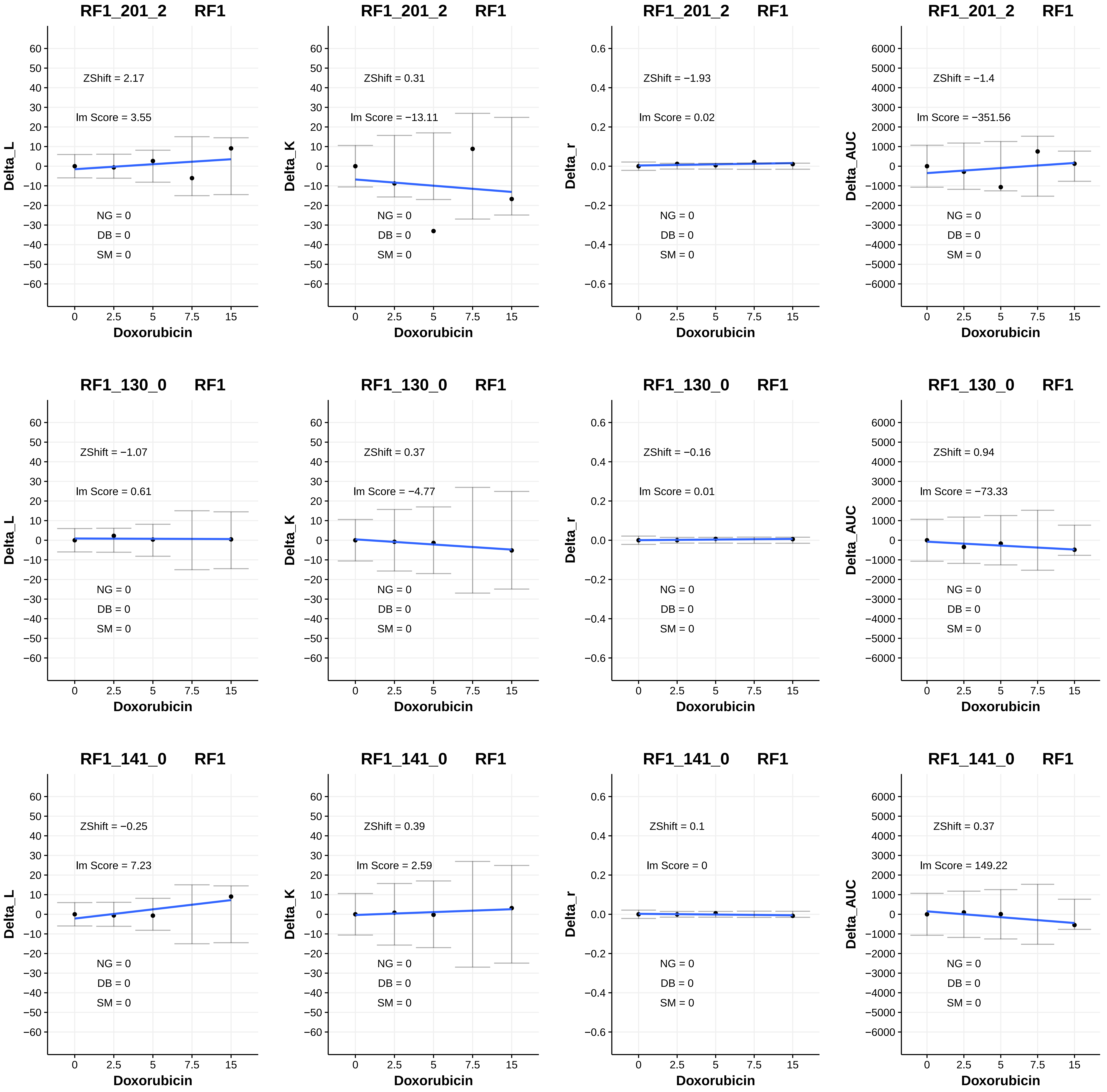

RF1_201_2 RF1
ZShift = 2.17
lm Score = 3.55
NG = 0
DB = 0
SM = 0
Delta_L
Doxorubicin
RF1_201_2 RF1
ZShift = 0.31
lm Score = −13.11
NG = 0
DB = 0
SM = 0
Delta_K
Doxorubicin
RF1_201_2 RF1
ZShift = −1.93
lm Score = 0.02
NG = 0
DB = 0
SM = 0
Delta_r
Doxorubicin
RF1_201_2 RF1
ZShift = −1.4
lm Score = −351.56
NG = 0
DB = 0
SM = 0
Delta_AUC
Doxorubicin
RF1_130_0 RF1
ZShift = −1.07
lm Score = 0.61
NG = 0
DB = 0
SM = 0
Delta_L
Doxorubicin
RF1_130_0 RF1
ZShift = 0.37
lm Score = −4.77
NG = 0
DB = 0
SM = 0
Delta_K
Doxorubicin
RF1_130_0 RF1
ZShift = −0.16
lm Score = 0.01
NG = 0
DB = 0
SM = 0
Delta_r
Doxorubicin
RF1_130_0 RF1
ZShift = 0.94
lm Score = −73.33
NG = 0
DB = 0
SM = 0
Delta_AUC
Doxorubicin
RF1_141_0 RF1
ZShift = −0.25
lm Score = 7.23
NG = 0
DB = 0
SM = 0
Delta_L
Doxorubicin
RF1_141_0 RF1
ZShift = 0.39
lm Score = 2.59
NG = 0
DB = 0
SM = 0
Delta_K
Doxorubicin
RF1_141_0 RF1
ZShift = 0.1
lm Score = 0
NG = 0
DB = 0
SM = 0
Delta_r
Doxorubicin
RF1_141_0 RF1
ZShift = 0.37
lm Score = 149.22
NG = 0
DB = 0
SM = 0
Delta_AUC
Doxorubicin

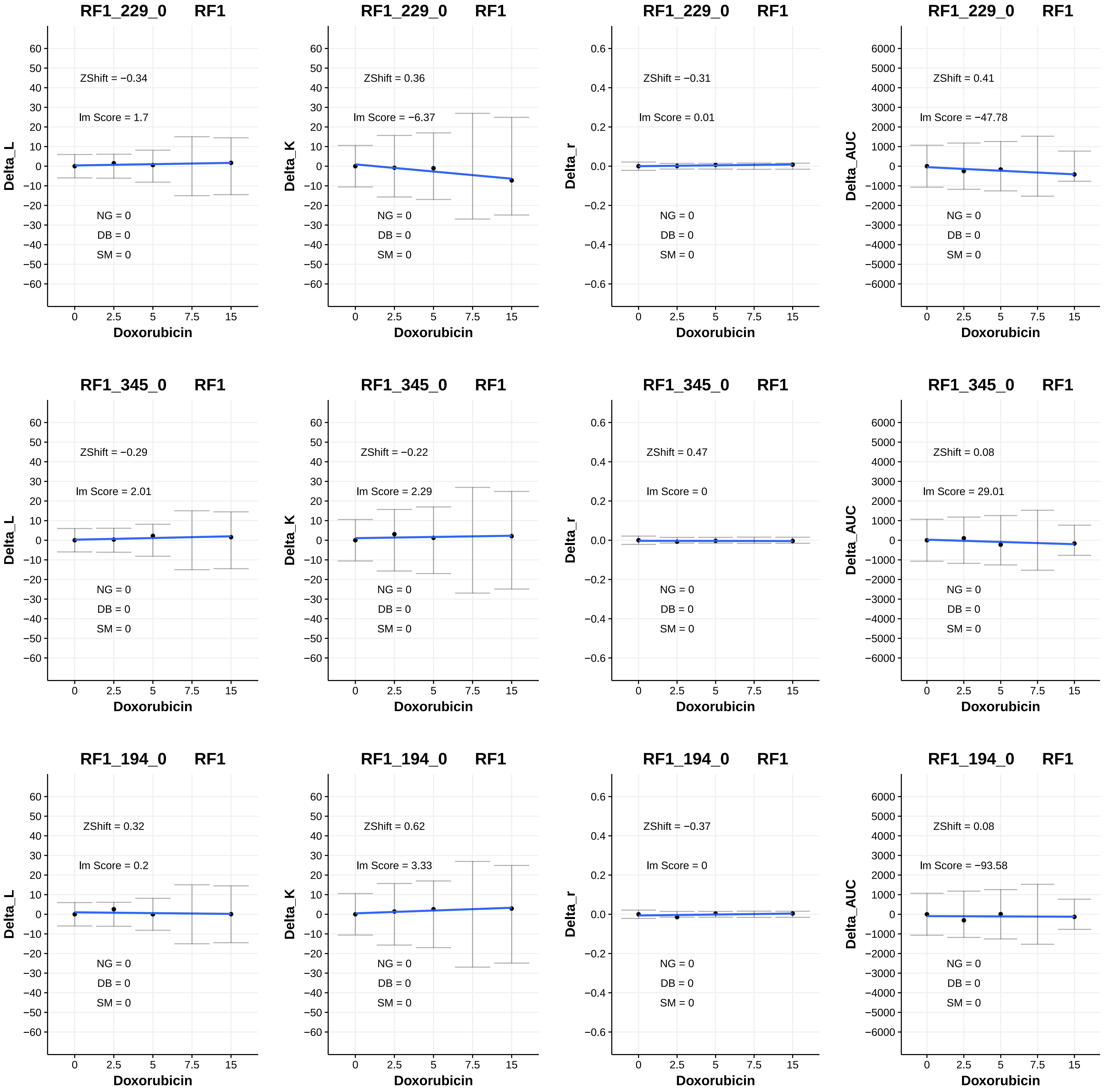
RF1_229_0 RF1
ZShift = −0.34
lm Score = 1.7
NG = 0
DB = 0
SM = 0
Delta_L
Doxorubicin
RF1_229_0 RF1
ZShift = 0.36
lm Score = −6.37
NG = 0
DB = 0
SM = 0
Delta_K
Doxorubicin
RF1_229_0 RF1
ZShift = −0.31
lm Score = 0.01
NG = 0
DB = 0
SM = 0
Delta_r
Doxorubicin
RF1_229_0 RF1
ZShift = 0.41
lm Score = −47.78
NG = 0
DB = 0
SM = 0
Delta_AUC
Doxorubicin
RF1_345_0 RF1
ZShift = −0.29
lm Score = 2.01
NG = 0
DB = 0
SM = 0
Delta_L
Doxorubicin
RF1_345_0 RF1
ZShift = −0.22
lm Score = 2.29
NG = 0
DB = 0
SM = 0
Delta_K
Doxorubicin
RF1_345_0 RF1
ZShift = 0.47
lm Score = 0
NG = 0
DB = 0
SM = 0
Delta_r
Doxorubicin
RF1_345_0 RF1
ZShift = 0.08
lm Score = 29.01
NG = 0
DB = 0
SM = 0
Delta_AUC
Doxorubicin
RF1_194_0 RF1
ZShift = 0.32
lm Score = 0.2
NG = 0
DB = 0
SM = 0
Delta_L
Doxorubicin
RF1_194_0 RF1
ZShift = 0.62
lm Score = 3.33
NG = 0
DB = 0
SM = 0
Delta_K
Doxorubicin
RF1_194_0 RF1
ZShift = −0.37
lm Score = 0
NG = 0
DB = 0
SM = 0
Delta_r
Doxorubicin
RF1_194_0 RF1
ZShift = 0.08
lm Score = −93.58
NG = 0
DB = 0
SM = 0
Delta_AUC
Doxorubicin

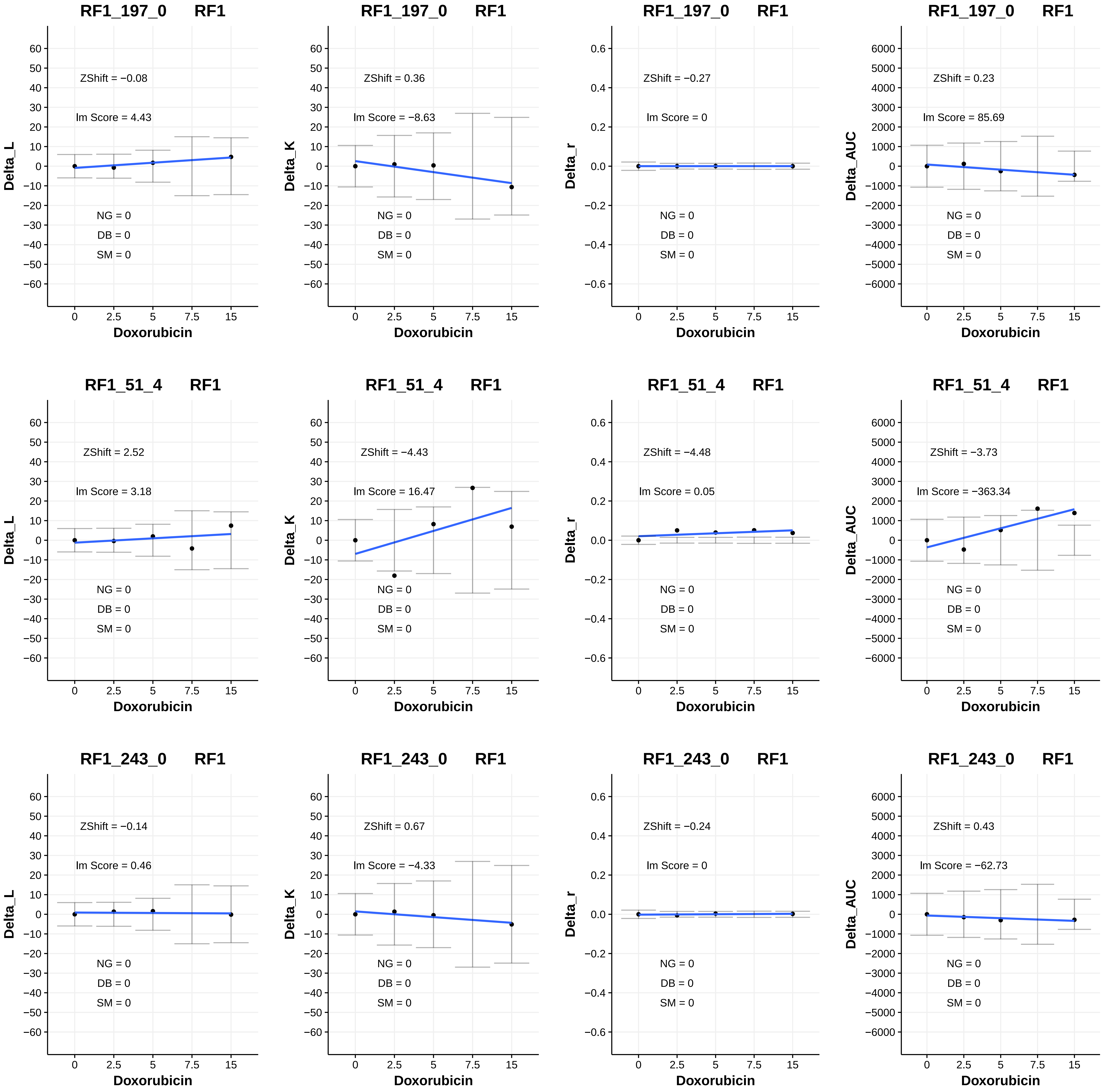
RF1_197_0 RF1
ZShift = −0.08
lm Score = 4.43
NG = 0
DB = 0
SM = 0
Delta_L
Doxorubicin
RF1_197_0 RF1
ZShift = 0.36
lm Score = −8.63
NG = 0
DB = 0
SM = 0
Delta_K
Doxorubicin
RF1_197_0 RF1
ZShift = −0.27
lm Score = 0
NG = 0
DB = 0
SM = 0
Delta_r
Doxorubicin
RF1_197_0 RF1
ZShift = 0.23
lm Score = 85.69
NG = 0
DB = 0
SM = 0
Delta_AUC
Doxorubicin
RF1_51_4 RF1
ZShift = 2.52
lm Score = 3.18
NG = 0
DB = 0
SM = 0
Delta_L
Doxorubicin
RF1_51_4 RF1
ZShift = −4.43
lm Score = 16.47
NG = 0
DB = 0
SM = 0
Delta_K
Doxorubicin
RF1_51_4 RF1
ZShift = −4.48
lm Score = 0.05
NG = 0
DB = 0
SM = 0
Delta_r
Doxorubicin
RF1_51_4 RF1
ZShift = −3.73
lm Score = −363.34
NG = 0
DB = 0
SM = 0
Delta_AUC
Doxorubicin
RF1_243_0 RF1
ZShift = −0.14
lm Score = 0.46
NG = 0
DB = 0
SM = 0
Delta_L
Doxorubicin
RF1_243_0 RF1
ZShift = 0.67
lm Score = −4.33
NG = 0
DB = 0
SM = 0
Delta_K
Doxorubicin
RF1_243_0 RF1
ZShift = −0.24
lm Score = 0
NG = 0
DB = 0
SM = 0
Delta_r
Doxorubicin
RF1_243_0 RF1
ZShift = 0.43
lm Score = −62.73
NG = 0
DB = 0
SM = 0
Delta_AUC
Doxorubicin

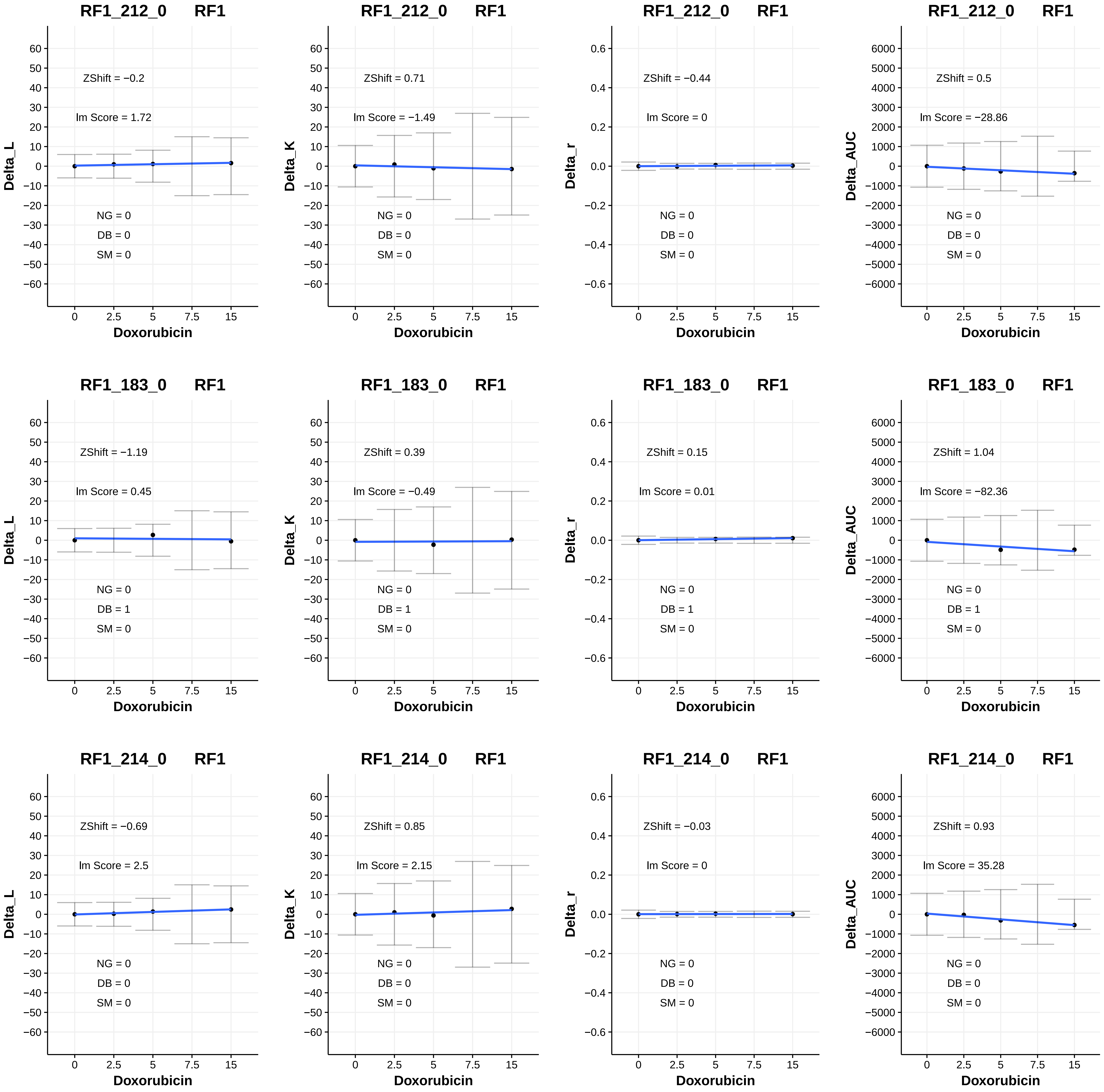
RF1_212_0 RF1
ZShift = −0.2
lm Score = 1.72
NG = 0
DB = 0
SM = 0
Delta_L
Doxorubicin
RF1_212_0 RF1
ZShift = 0.71
lm Score = −1.49
NG = 0
DB = 0
SM = 0
Delta_K
Doxorubicin
RF1_212_0 RF1
ZShift = −0.44
lm Score = 0
NG = 0
DB = 0
SM = 0
Delta_r
Doxorubicin
RF1_212_0 RF1
ZShift = 0.5
lm Score = −28.86
NG = 0
DB = 0
SM = 0
Delta_AUC
Doxorubicin
RF1_183_0 RF1
ZShift = −1.19
lm Score = 0.45
NG = 0
DB = 1
SM = 0
Delta_L
Doxorubicin
RF1_183_0 RF1
ZShift = 0.39
lm Score = −0.49
NG = 0
DB = 1
SM = 0
Delta_K
Doxorubicin
RF1_183_0 RF1
ZShift = 0.15
lm Score = 0.01
NG = 0
DB = 1
SM = 0
Delta_r
Doxorubicin
RF1_183_0 RF1
ZShift = 1.04
lm Score = −82.36
NG = 0
DB = 1
SM = 0
Delta_AUC
Doxorubicin
RF1_214_0 RF1
ZShift = −0.69
lm Score = 2.5
NG = 0
DB = 0
SM = 0
Delta_L
Doxorubicin
RF1_214_0 RF1
ZShift = 0.85
lm Score = 2.15
NG = 0
DB = 0
SM = 0
Delta_K
Doxorubicin
RF1_214_0 RF1
ZShift = −0.03
lm Score = 0
NG = 0
DB = 0
SM = 0
Delta_r
Doxorubicin
RF1_214_0 RF1
ZShift = 0.93
lm Score = 35.28
NG = 0
DB = 0
SM = 0
Delta_AUC
Doxorubicin

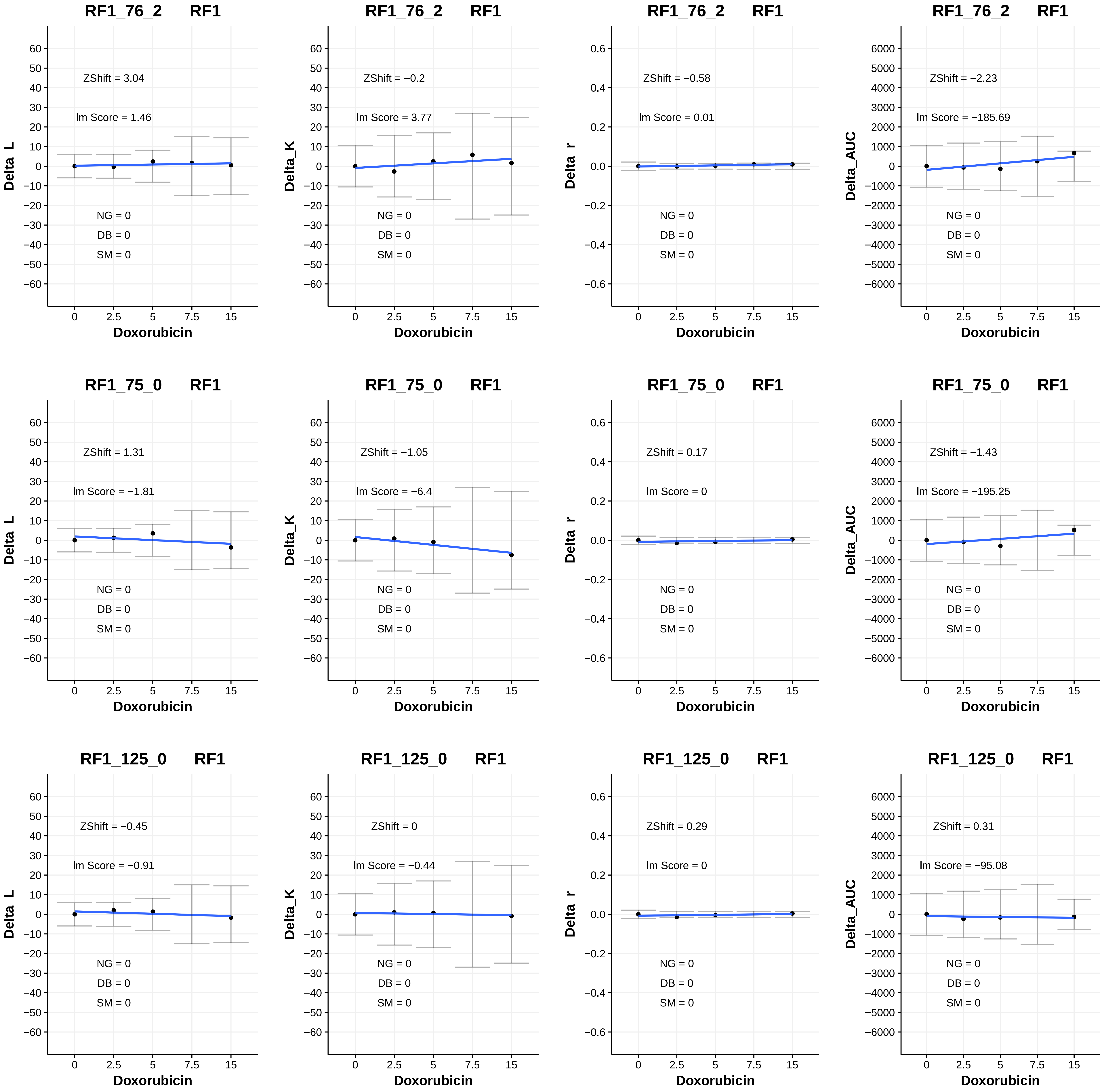
RF1_76_2 RF1
ZShift = 3.04
lm Score = 1.46
NG = 0
DB = 0
SM = 0
Delta_L
Doxorubicin
RF1_76_2 RF1
ZShift = −0.2
lm Score = 3.77
NG = 0
DB = 0
SM = 0
Delta_K
Doxorubicin
RF1_76_2 RF1
ZShift = −0.58
lm Score = 0.01
NG = 0
DB = 0
SM = 0
Delta_r
Doxorubicin
RF1_76_2 RF1
ZShift = −2.23
lm Score = −185.69
NG = 0
DB = 0
SM = 0
Delta_AUC
Doxorubicin
RF1_75_0 RF1
ZShift = 1.31
lm Score = −1.81
NG = 0
DB = 0
SM = 0
Delta_L
Doxorubicin
RF1_75_0 RF1
ZShift = −1.05
lm Score = −6.4
NG = 0
DB = 0
SM = 0
Delta_K
Doxorubicin
RF1_75_0 RF1
ZShift = 0.17
lm Score = 0
NG = 0
DB = 0
SM = 0
Delta_r
Doxorubicin
RF1_75_0 RF1
ZShift = −1.43
lm Score = −195.25
NG = 0
DB = 0
SM = 0
Delta_AUC
Doxorubicin
RF1_125_0 RF1
ZShift = −0.45
lm Score = −0.91
NG = 0
DB = 0
SM = 0
Delta_L
Doxorubicin
RF1_125_0 RF1
ZShift = 0
lm Score = −0.44
NG = 0
DB = 0
SM = 0
Delta_K
Doxorubicin
RF1_125_0 RF1
ZShift = 0.29
lm Score = 0
NG = 0
DB = 0
SM = 0
Delta_r
Doxorubicin
RF1_125_0 RF1
ZShift = 0.31
lm Score = −95.08
NG = 0
DB = 0
SM = 0
Delta_AUC
Doxorubicin

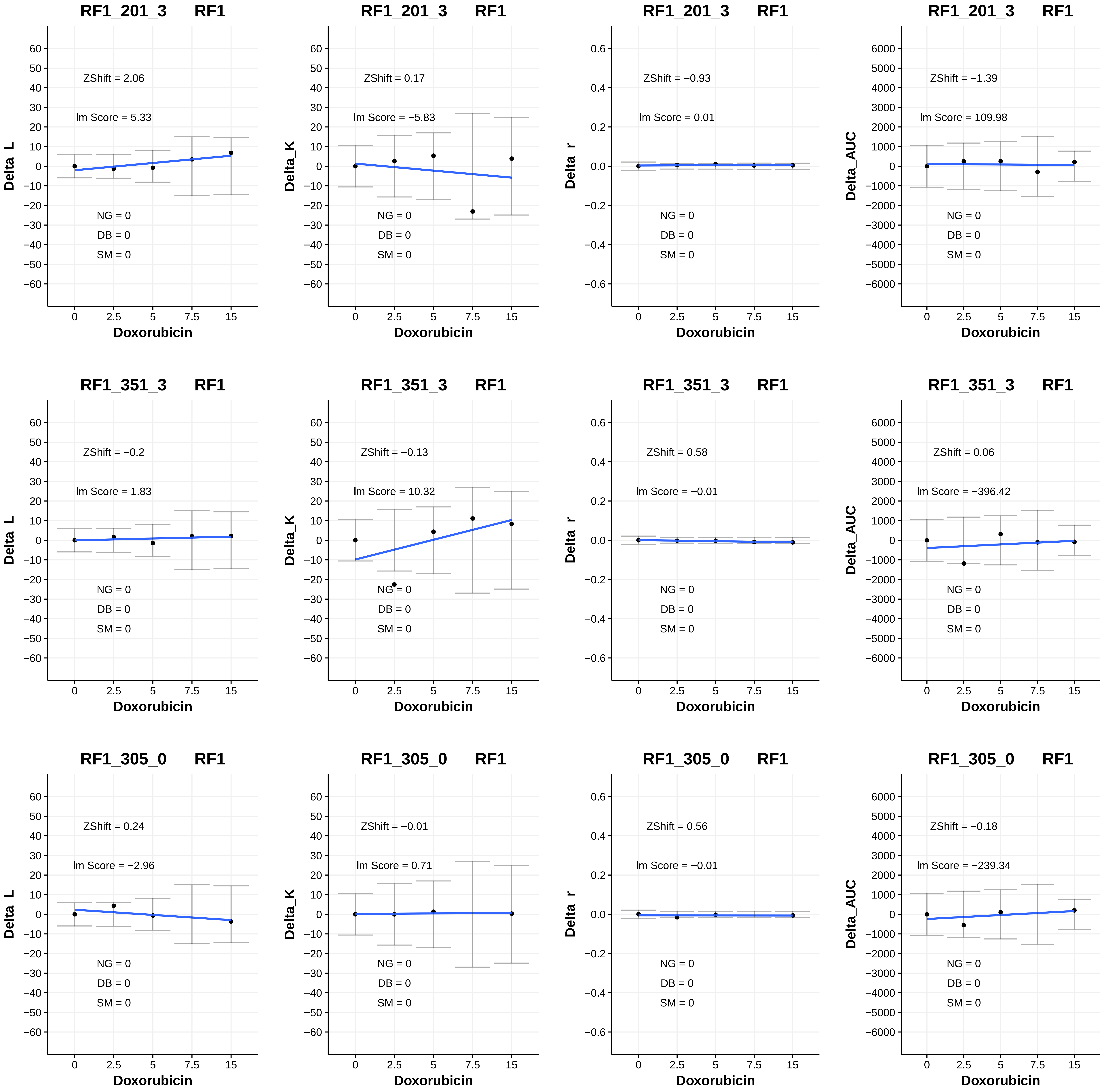
RF1_201_3 RF1
ZShift = 2.06
Im Score = 5.33
NG = 0
DB = 0
SM = 0
Delta_L
Doxorubicin
RF1_201_3 RF1
ZShift = 0.17
Im Score = −5.83
NG = 0
DB = 0
SM = 0
Delta_K
Doxorubicin
RF1_201_3 RF1
ZShift = −0.93
Im Score = 0.01
NG = 0
DB = 0
SM = 0
Delta_r
Doxorubicin
RF1_201_3 RF1
ZShift = −1.39
Im Score = 109.98
NG = 0
DB = 0
SM = 0
Delta_AUC
Doxorubicin
RF1_351_3 RF1
ZShift = −0.2
Im Score = 1.83
NG = 0
DB = 0
SM = 0
Delta_L
Doxorubicin
RF1_351_3 RF1
ZShift = −0.13
Im Score = 10.32
NG = 0
DB = 0
SM = 0
Delta_K
Doxorubicin
RF1_351_3 RF1
ZShift = 0.58
Im Score = −0.01
NG = 0
DB = 0
SM = 0
Delta_r
Doxorubicin
RF1_351_3 RF1
ZShift = 0.06
Im Score = −396.42
NG = 0
DB = 0
SM = 0
Delta_AUC
Doxorubicin
RF1_305_0 RF1
ZShift = 0.24
Im Score = −2.96
NG = 0
DB = 0
SM = 0
Delta_L
Doxorubicin
RF1_305_0 RF1
ZShift = −0.01
Im Score = 0.71
NG = 0
DB = 0
SM = 0
Delta_K
Doxorubicin
RF1_305_0 RF1
ZShift = 0.56
Im Score = −0.01
NG = 0
DB = 0
SM = 0
Delta_r
Doxorubicin
RF1_305_0 RF1
ZShift = −0.18
Im Score = −239.34
NG = 0
DB = 0
SM = 0
Delta_AUC
Doxorubicin

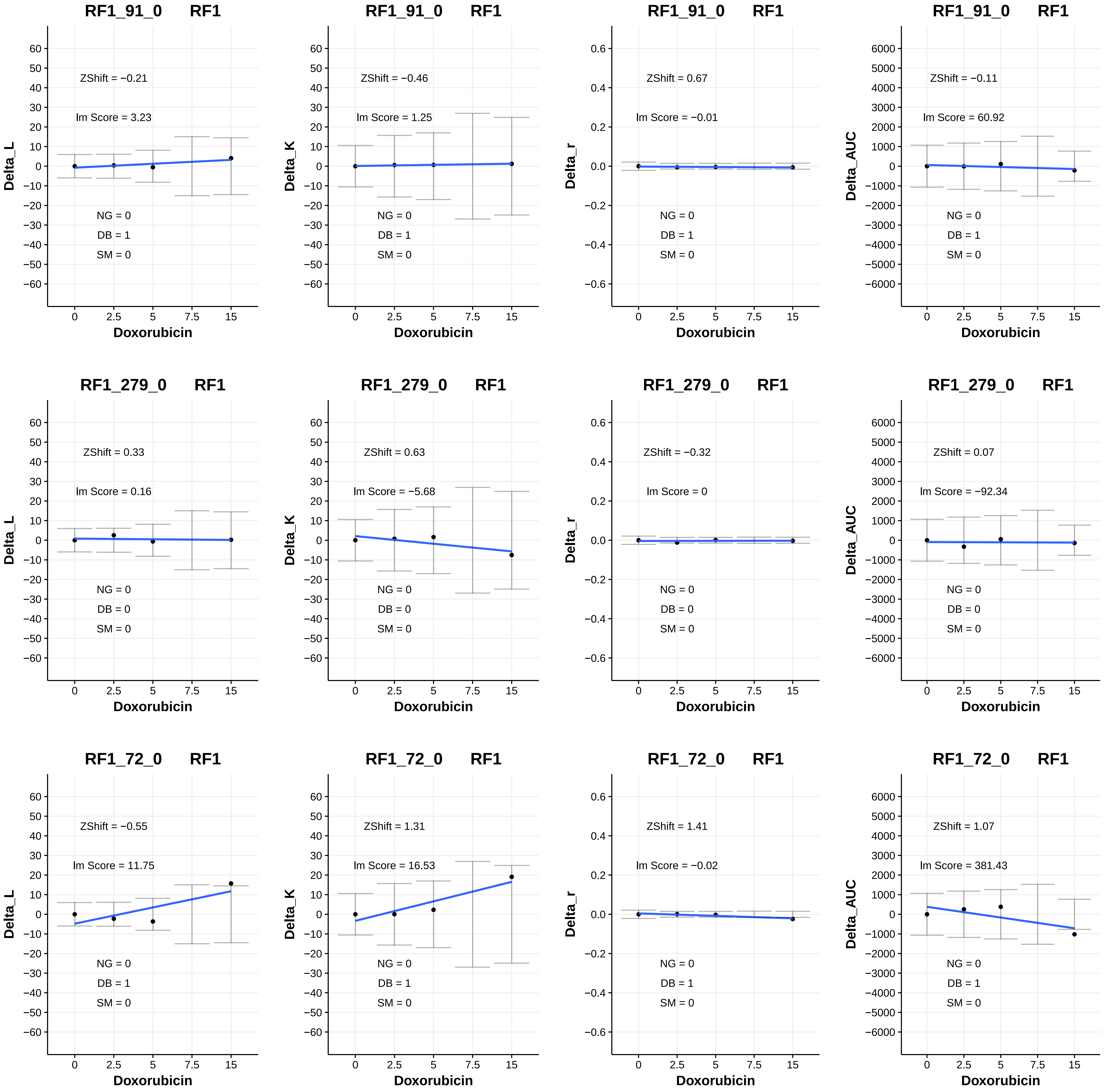
RF1_91_0 RF1
ZShift = −0.21
lm Score = 3.23
NG = 0
DB = 1
SM = 0
Delta_L
Doxorubicin
RF1_91_0 RF1
ZShift = −0.46
lm Score = 1.25
NG = 0
DB = 1
SM = 0
Delta_K
Doxorubicin
RF1_91_0 RF1
ZShift = 0.67
lm Score = −0.01
NG = 0
DB = 1
SM = 0
Delta_r
Doxorubicin
RF1_91_0 RF1
ZShift = −0.11
lm Score = 60.92
NG = 0
DB = 1
SM = 0
Delta_AUC
Doxorubicin
RF1_279_0 RF1
ZShift = 0.33
lm Score = 0.16
NG = 0
DB = 0
SM = 0
Delta_L
Doxorubicin
RF1_279_0 RF1
ZShift = 0.63
lm Score = −5.68
NG = 0
DB = 0
SM = 0
Delta_K
Doxorubicin
RF1_279_0 RF1
ZShift = −0.32
lm Score = 0
NG = 0
DB = 0
SM = 0
Delta_r
Doxorubicin
RF1_279_0 RF1
ZShift = 0.07
lm Score = −92.34
NG = 0
DB = 0
SM = 0
Delta_AUC
Doxorubicin
RF1_72_0 RF1
ZShift = −0.55
lm Score = 11.75
NG = 0
DB = 1
SM = 0
Delta_L
Doxorubicin
RF1_72_0 RF1
ZShift = 1.31
lm Score = 16.53
NG = 0
DB = 1
SM = 0
Delta_K
Doxorubicin
RF1_72_0 RF1
ZShift = 1.41
lm Score = −0.02
NG = 0
DB = 1
SM = 0
Delta_r
Doxorubicin
RF1_72_0 RF1
ZShift = 1.07
lm Score = 381.43
NG = 0
DB = 1
SM = 0
Delta_AUC
Doxorubicin

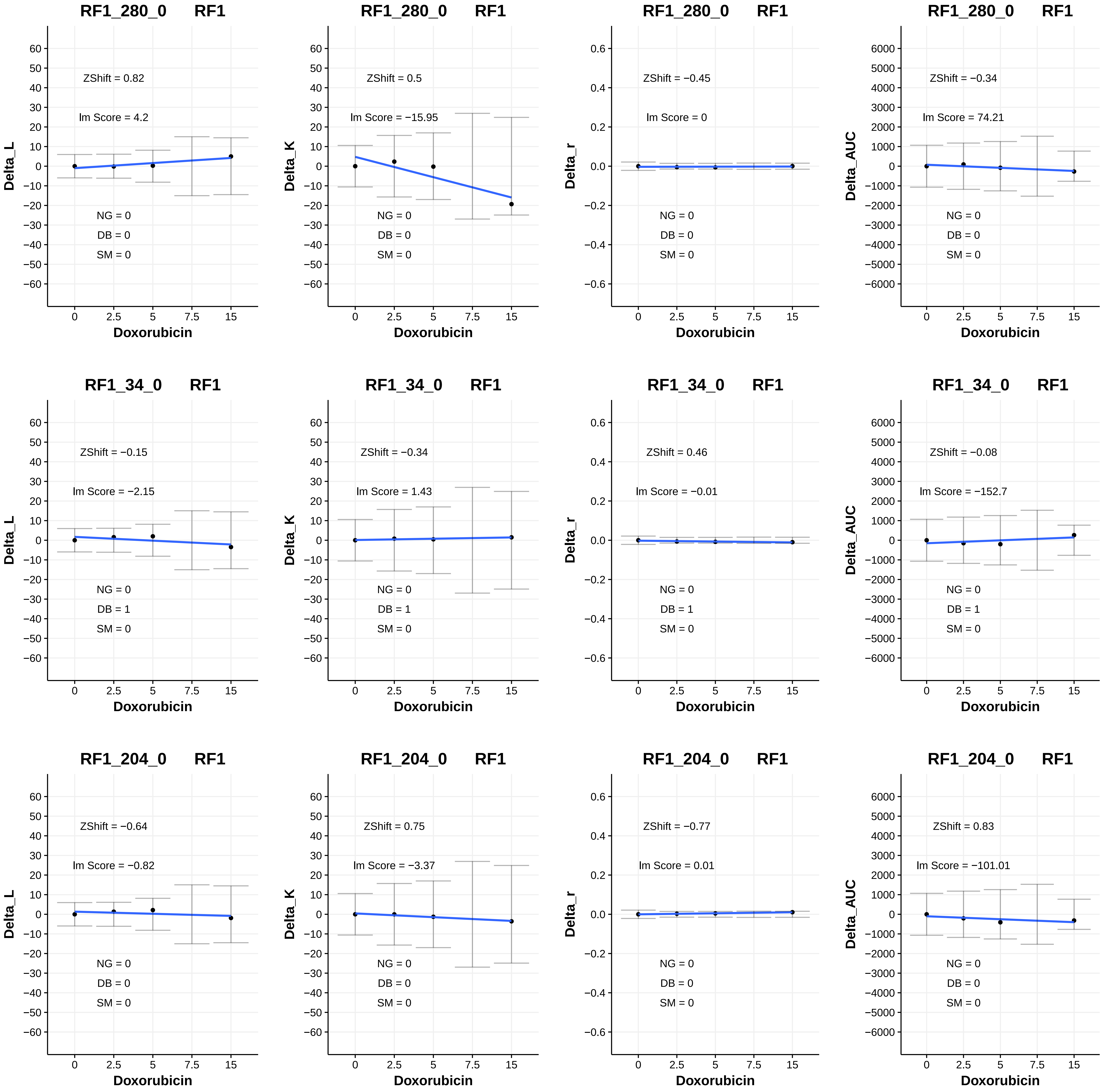

RF1_280_0 RF1
ZShift = 0.82
lm Score = 4.2
NG = 0
DB = 0
SM = 0
Delta_L
Doxorubicin
RF1_280_0 RF1
ZShift = 0.5
lm Score = −15.95
NG = 0
DB = 0
SM = 0
Delta_K
Doxorubicin
RF1_280_0 RF1
ZShift = −0.45
lm Score = 0
NG = 0
DB = 0
SM = 0
Delta_r
Doxorubicin
RF1_280_0 RF1
ZShift = −0.34
lm Score = 74.21
NG = 0
DB = 0
SM = 0
Delta_AUC
Doxorubicin
RF1_34_0 RF1
ZShift = −0.15
lm Score = −2.15
NG = 0
DB = 1
SM = 0
Delta_L
Doxorubicin
RF1_34_0 RF1
ZShift = −0.34
lm Score = 1.43
NG = 0
DB = 1
SM = 0
Delta_K
Doxorubicin
RF1_34_0 RF1
ZShift = 0.46
lm Score = −0.01
NG = 0
DB = 1
SM = 0
Delta_r
Doxorubicin
RF1_34_0 RF1
ZShift = −0.08
lm Score = −152.7
NG = 0
DB = 1
SM = 0
Delta_AUC
Doxorubicin
RF1_204_0 RF1
ZShift = −0.64
lm Score = −0.82
NG = 0
DB = 0
SM = 0
Delta_L
Doxorubicin
RF1_204_0 RF1
ZShift = 0.75
lm Score = −3.37
NG = 0
DB = 0
SM = 0
Delta_K
Doxorubicin
RF1_204_0 RF1
ZShift = −0.77
lm Score = 0.01
NG = 0
DB = 0
SM = 0
Delta_r
Doxorubicin
RF1_204_0 RF1
ZShift = 0.83
lm Score = −101.01
NG = 0
DB = 0
SM = 0
Delta_AUC
Doxorubicin

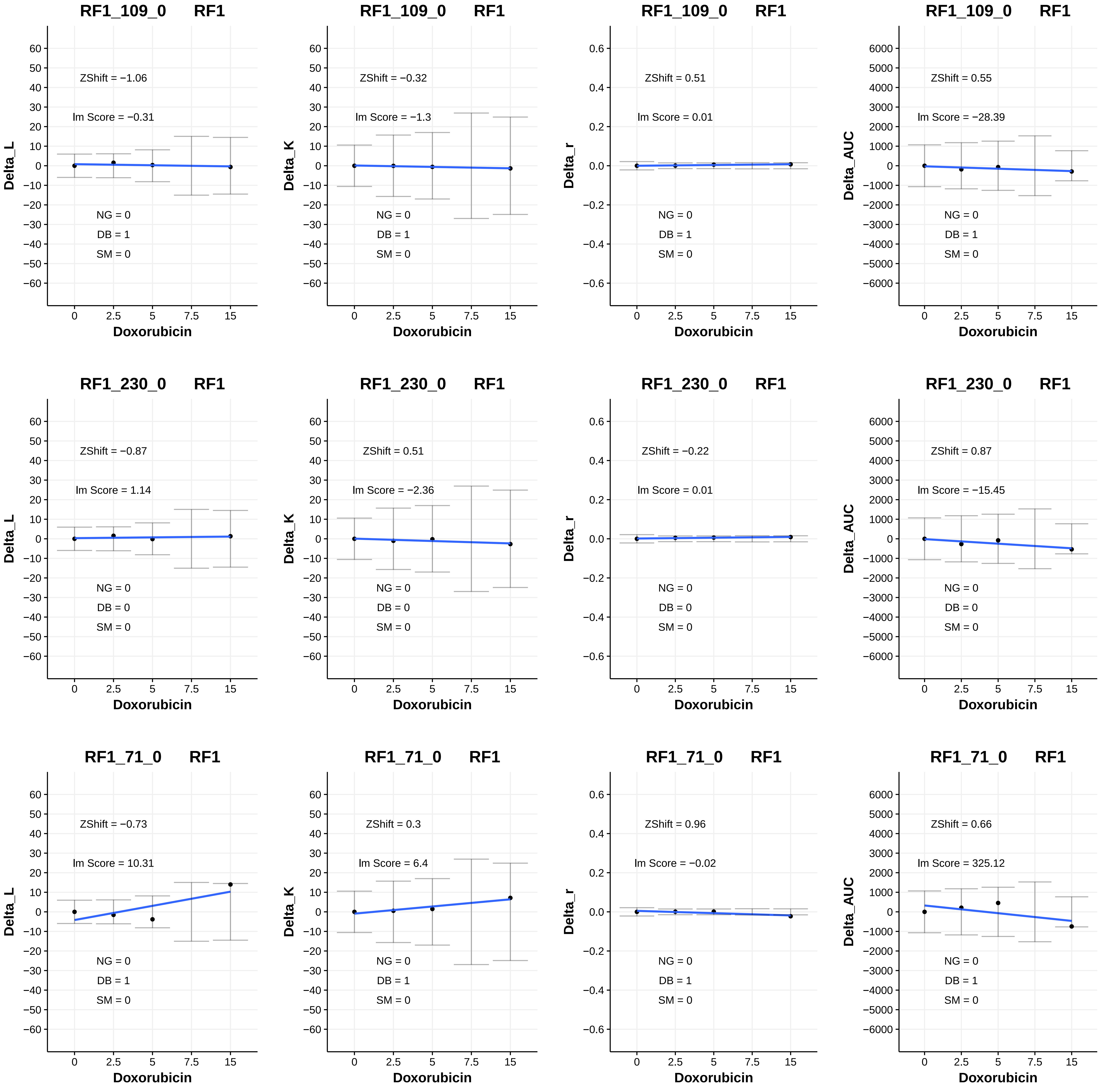
RF1_109_0 RF1
ZShift = −1.06
lm Score = −0.31
NG = 0
DB = 1
SM = 0
Delta_L
Doxorubicin
RF1_109_0 RF1
ZShift = −0.32
lm Score = −1.3
NG = 0
DB = 1
SM = 0
Delta_K
Doxorubicin
RF1_109_0 RF1
ZShift = 0.51
lm Score = 0.01
NG = 0
DB = 1
SM = 0
Delta_r
Doxorubicin
RF1_109_0 RF1
ZShift = 0.55
lm Score = −28.39
NG = 0
DB = 1
SM = 0
Delta_AUC
Doxorubicin
RF1_230_0 RF1
ZShift = −0.87
lm Score = 1.14
NG = 0
DB = 0
SM = 0
Delta_L
Doxorubicin
RF1_230_0 RF1
ZShift = 0.51
lm Score = −2.36
NG = 0
DB = 0
SM = 0
Delta_K
Doxorubicin
RF1_230_0 RF1
ZShift = −0.22
lm Score = 0.01
NG = 0
DB = 0
SM = 0
Delta_r
Doxorubicin
RF1_230_0 RF1
ZShift = 0.87
lm Score = −15.45
NG = 0
DB = 0
SM = 0
Delta_AUC
Doxorubicin
RF1_71_0 RF1
ZShift = −0.73
lm Score = 10.31
NG = 0
DB = 1
SM = 0
Delta_L
Doxorubicin
RF1_71_0 RF1
ZShift = 0.3
lm Score = 6.4
NG = 0
DB = 1
SM = 0
Delta_K
Doxorubicin
RF1_71_0 RF1
ZShift = 0.96
lm Score = −0.02
NG = 0
DB = 1
SM = 0
Delta_r
Doxorubicin
RF1_71_0 RF1
ZShift = 0.66
lm Score = 325.12
NG = 0
DB = 1
SM = 0
Delta_AUC
Doxorubicin

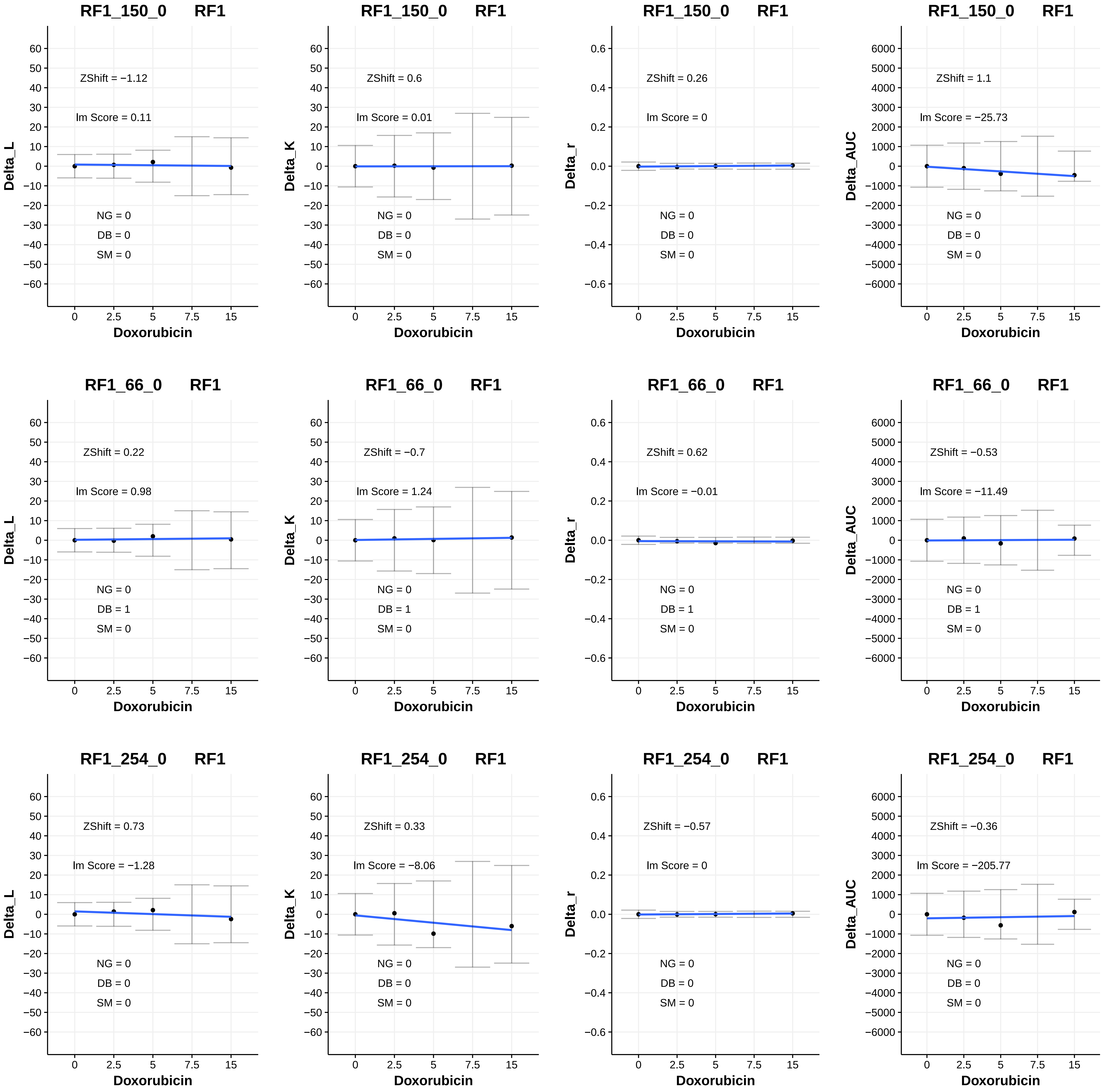
RF1_150_0 RF1
ZShift = −1.12
lm Score = 0.11
NG = 0
DB = 0
SM = 0
Delta_L
Doxorubicin
RF1_150_0 RF1
ZShift = 0.6
lm Score = 0.01
NG = 0
DB = 0
SM = 0
Delta_K
Doxorubicin
RF1_150_0 RF1
ZShift = 0.26
lm Score = 0
NG = 0
DB = 0
SM = 0
Delta_r
Doxorubicin
RF1_150_0 RF1
ZShift = 1.1
lm Score = −25.73
NG = 0
DB = 0
SM = 0
Delta_AUC
Doxorubicin
RF1_66_0 RF1
ZShift = 0.22
lm Score = 0.98
NG = 0
DB = 1
SM = 0
Delta_L
Doxorubicin
RF1_66_0 RF1
ZShift = −0.7
lm Score = 1.24
NG = 0
DB = 1
SM = 0
Delta_K
Doxorubicin
RF1_66_0 RF1
ZShift = 0.62
lm Score = −0.01
NG = 0
DB = 1
SM = 0
Delta_r
Doxorubicin
RF1_66_0 RF1
ZShift = −0.53
lm Score = −11.49
NG = 0
DB = 1
SM = 0
Delta_AUC
Doxorubicin
RF1_254_0 RF1
ZShift = 0.73
lm Score = −1.28
NG = 0
DB = 0
SM = 0
Delta_L
Doxorubicin
RF1_254_0 RF1
ZShift = 0.33
lm Score = −8.06
NG = 0
DB = 0
SM = 0
Delta_K
Doxorubicin
RF1_254_0 RF1
ZShift = −0.57
lm Score = 0
NG = 0
DB = 0
SM = 0
Delta_r
Doxorubicin
RF1_254_0 RF1
ZShift = −0.36
lm Score = −205.77
NG = 0
DB = 0
SM = 0
Delta_AUC
Doxorubicin

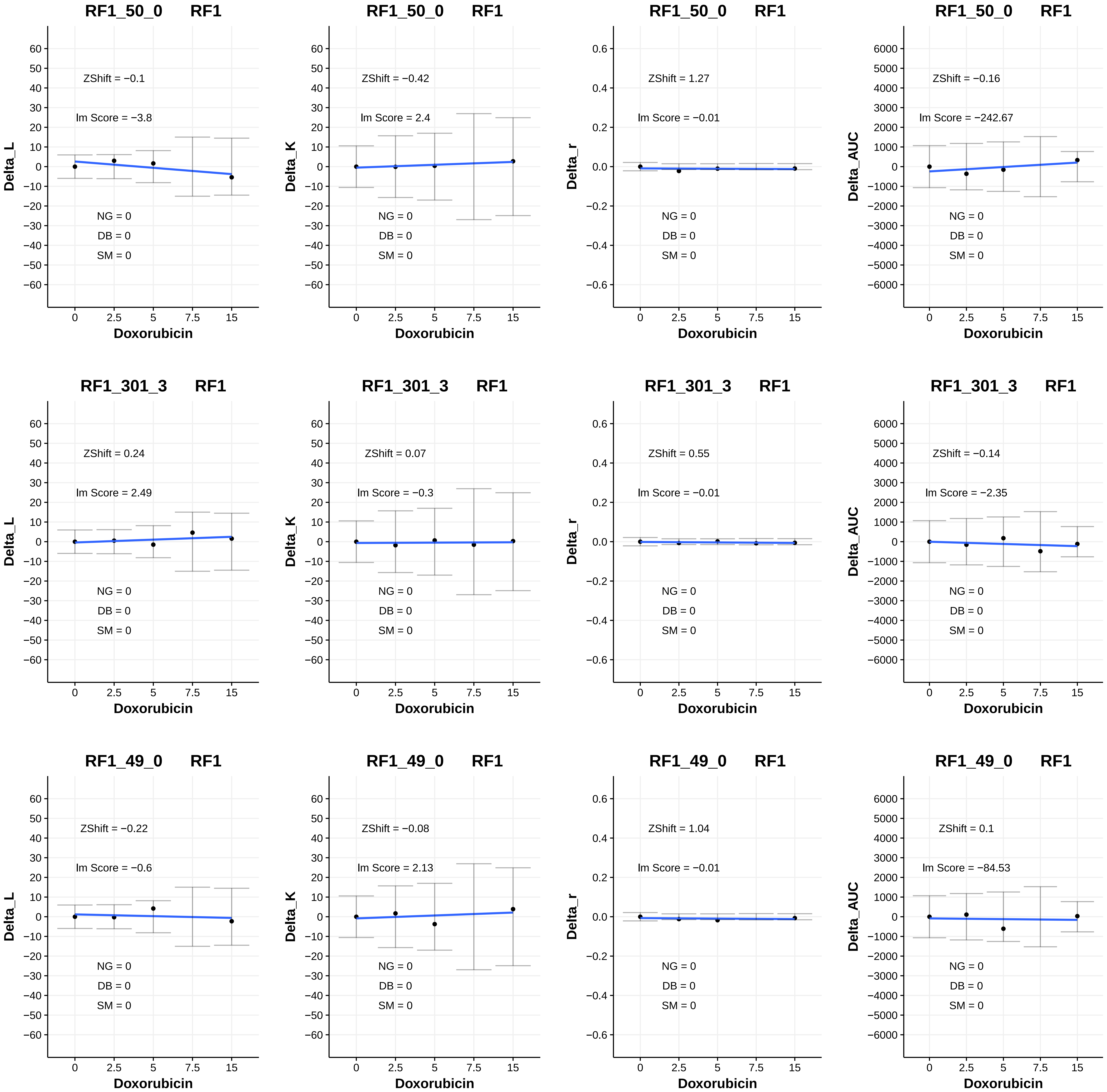
RF1_50_0 RF1
ZShift = −0.1
Im Score = −3.8
NG = 0
DB = 0
SM = 0
Delta_L
Doxorubicin
RF1_50_0 RF1
ZShift = −0.42
Im Score = 2.4
NG = 0
DB = 0
SM = 0
Delta_K
Doxorubicin
RF1_50_0 RF1
ZShift = 1.27
Im Score = −0.01
NG = 0
DB = 0
SM = 0
Delta_r
Doxorubicin
RF1_50_0 RF1
ZShift = −0.16
Im Score = −242.67
NG = 0
DB = 0
SM = 0
Delta_AUC
Doxorubicin
RF1_301_3 RF1
ZShift = 0.24
Im Score = 2.49
NG = 0
DB = 0
SM = 0
Delta_L
Doxorubicin
RF1_301_3 RF1
ZShift = 0.07
Im Score = −0.3
NG = 0
DB = 0
SM = 0
Delta_K
Doxorubicin
RF1_301_3 RF1
ZShift = 0.55
Im Score = −0.01
NG = 0
DB = 0
SM = 0
Delta_r
Doxorubicin
RF1_301_3 RF1
ZShift = −0.14
Im Score = −2.35
NG = 0
DB = 0
SM = 0
Delta_AUC
Doxorubicin
RF1_49_0 RF1
ZShift = −0.22
Im Score = −0.6
NG = 0
DB = 0
SM = 0
Delta_L
Doxorubicin
RF1_49_0 RF1
ZShift = −0.08
Im Score = 2.13
NG = 0
DB = 0
SM = 0
Delta_K
Doxorubicin
RF1_49_0 RF1
ZShift = 1.04
Im Score = −0.01
NG = 0
DB = 0
SM = 0
Delta_r
Doxorubicin
RF1_49_0 RF1
ZShift = 0.1
Im Score = −84.53
NG = 0
DB = 0
SM = 0
Delta_AUC
Doxorubicin

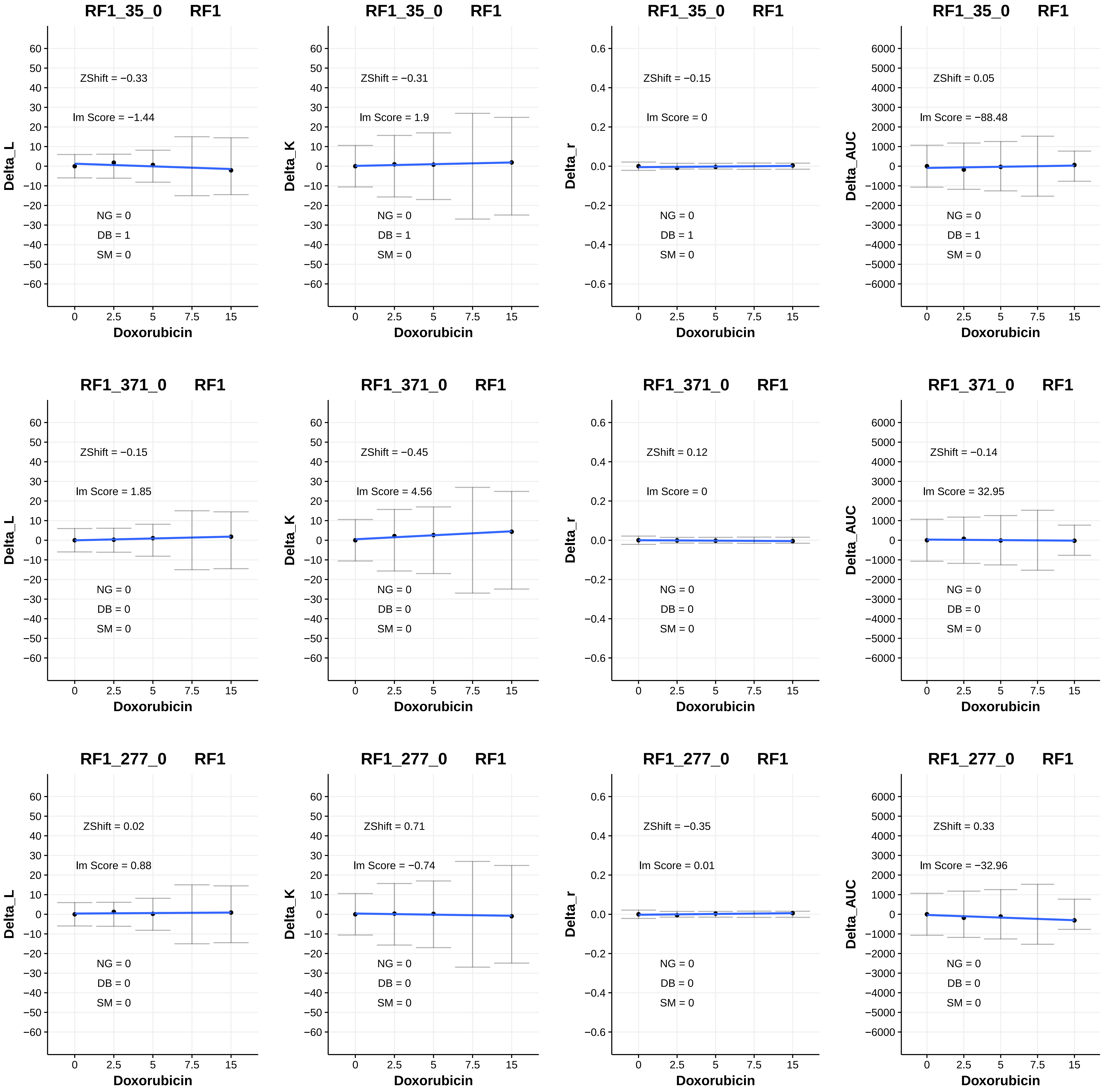

RF1_35_0 RF1
ZShift = −0.33
Im Score = −1.44
NG = 0
DB = 1
SM = 0
Delta_L
Doxorubicin
RF1_35_0 RF1
ZShift = −0.31
Im Score = 1.9
NG = 0
DB = 1
SM = 0
Delta_K
Doxorubicin
RF1_35_0 RF1
ZShift = −0.15
Im Score = 0
NG = 0
DB = 1
SM = 0
Delta_r
Doxorubicin
RF1_35_0 RF1
ZShift = 0.05
Im Score = −88.48
NG = 0
DB = 1
SM = 0
Delta_AUC
Doxorubicin
RF1_371_0 RF1
ZShift = −0.15
Im Score = 1.85
NG = 0
DB = 0
SM = 0
Delta_L
Doxorubicin
RF1_371_0 RF1
ZShift = −0.45
Im Score = 4.56
NG = 0
DB = 0
SM = 0
Delta_K
Doxorubicin
RF1_371_0 RF1
ZShift = 0.12
Im Score = 0
NG = 0
DB = 0
SM = 0
Delta_r
Doxorubicin
RF1_371_0 RF1
ZShift = −0.14
Im Score = 32.95
NG = 0
DB = 0
SM = 0
Delta_AUC
Doxorubicin
RF1_277_0 RF1
ZShift = 0.02
Im Score = 0.88
NG = 0
DB = 0
SM = 0
Delta_L
Doxorubicin
RF1_277_0 RF1
ZShift = 0.71
Im Score = −0.74
NG = 0
DB = 0
SM = 0
Delta_K
Doxorubicin
RF1_277_0 RF1
ZShift = −0.35
Im Score = 0.01
NG = 0
DB = 0
SM = 0
Delta_r
Doxorubicin
RF1_277_0 RF1
ZShift = 0.33
Im Score = −32.96
NG = 0
DB = 0
SM = 0
Delta_AUC
Doxorubicin

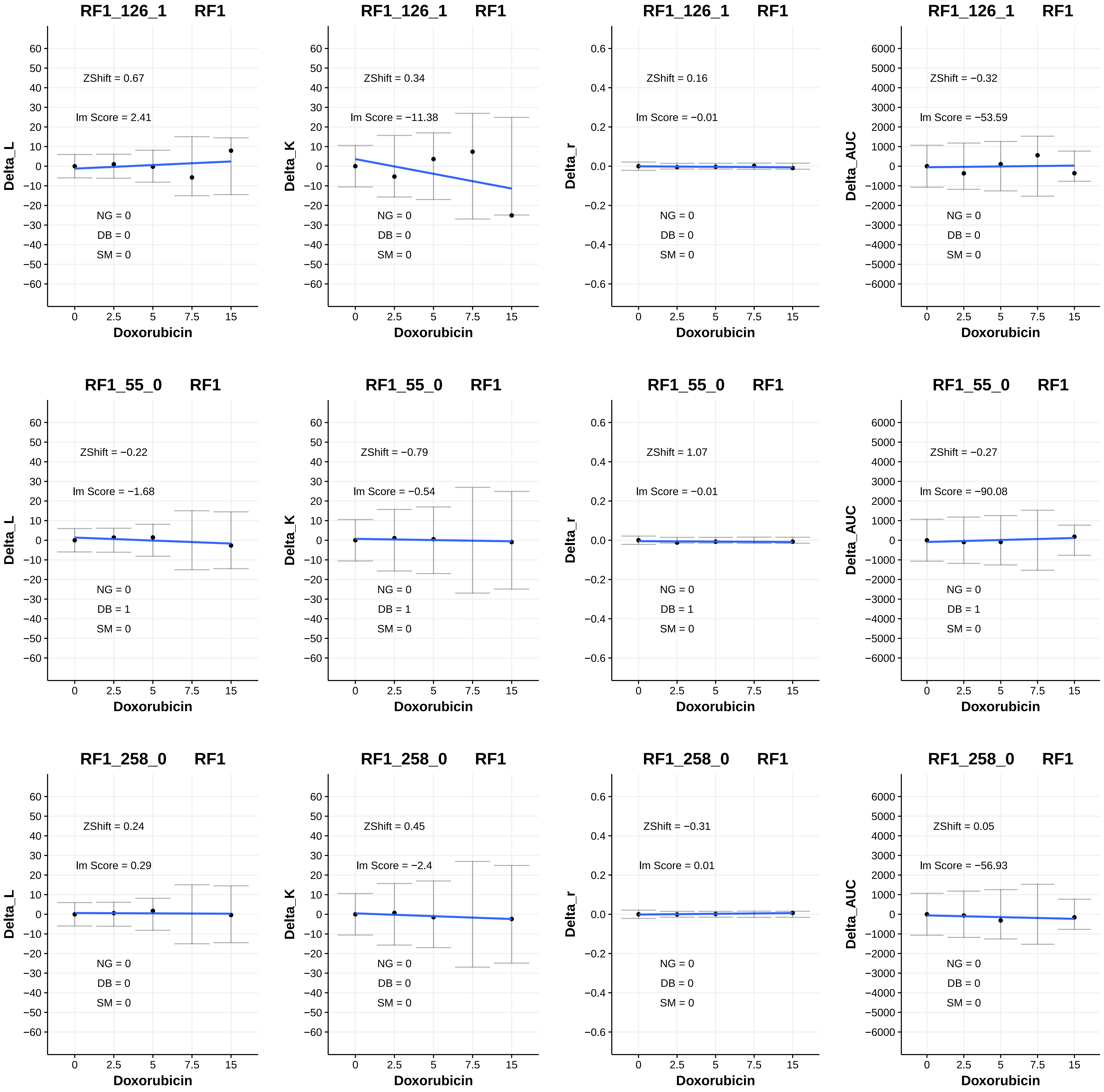
RF1_126_1 RF1
ZShift = 0.67
lm Score = 2.41
NG = 0
DB = 0
SM = 0
Delta_L
Doxorubicin
RF1_126_1 RF1
ZShift = 0.34
lm Score = −11.38
NG = 0
DB = 0
SM = 0
Delta_K
Doxorubicin
RF1_126_1 RF1
ZShift = 0.16
lm Score = −0.01
NG = 0
DB = 0
SM = 0
Delta_r
Doxorubicin
RF1_126_1 RF1
ZShift = −0.32
lm Score = −53.59
NG = 0
DB = 0
SM = 0
Delta_AUC
Doxorubicin
RF1_55_0 RF1
ZShift = −0.22
lm Score = −1.68
NG = 0
DB = 1
SM = 0
Delta_L
Doxorubicin
RF1_55_0 RF1
ZShift = −0.79
lm Score = −0.54
NG = 0
DB = 1
SM = 0
Delta_K
Doxorubicin
RF1_55_0 RF1
ZShift = 1.07
lm Score = −0.01
NG = 0
DB = 1
SM = 0
Delta_r
Doxorubicin
RF1_55_0 RF1
ZShift = −0.27
lm Score = −90.08
NG = 0
DB = 1
SM = 0
Delta_AUC
Doxorubicin
RF1_258_0 RF1
ZShift = 0.24
lm Score = 0.29
NG = 0
DB = 0
SM = 0
Delta_L
Doxorubicin
RF1_258_0 RF1
ZShift = 0.45
lm Score = −2.4
NG = 0
DB = 0
SM = 0
Delta_K
Doxorubicin
RF1_258_0 RF1
ZShift = −0.31
lm Score = 0.01
NG = 0
DB = 0
SM = 0
Delta_r
Doxorubicin
RF1_258_0 RF1
ZShift = 0.05
lm Score = −56.93
NG = 0
DB = 0
SM = 0
Delta_AUC
Doxorubicin

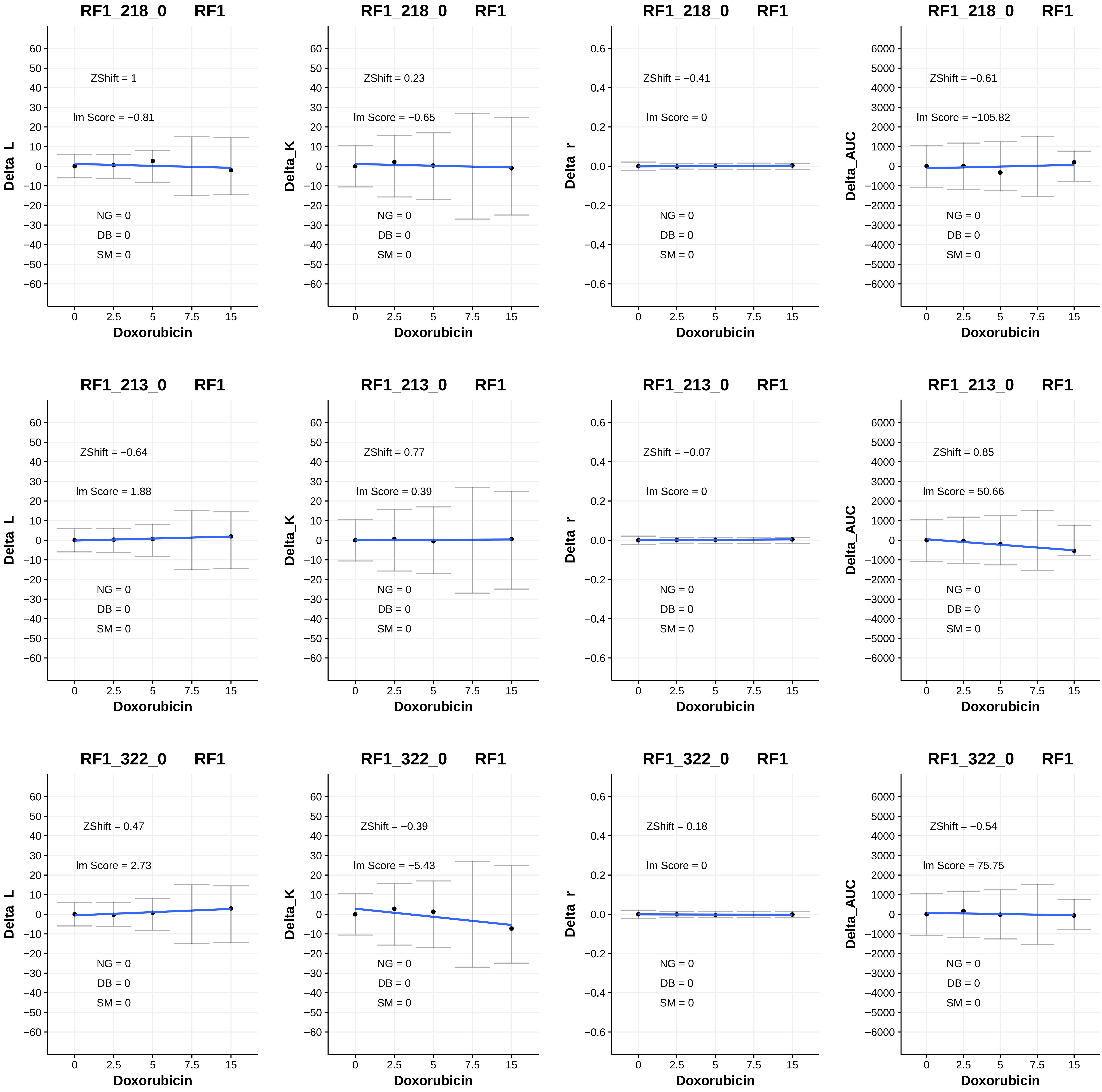
RF1_218_0 RF1
ZShift = 1
lm Score = −0.81
NG = 0
DB = 0
SM = 0
Delta_L
Doxorubicin
RF1_218_0 RF1
ZShift = 0.23
lm Score = −0.65
NG = 0
DB = 0
SM = 0
Delta_K
Doxorubicin
RF1_218_0 RF1
ZShift = −0.41
lm Score = 0
NG = 0
DB = 0
SM = 0
Delta_r
Doxorubicin
RF1_218_0 RF1
ZShift = −0.61
lm Score = −105.82
NG = 0
DB = 0
SM = 0
Delta_AUC
Doxorubicin
RF1_213_0 RF1
ZShift = −0.64
lm Score = 1.88
NG = 0
DB = 0
SM = 0
Delta_L
Doxorubicin
RF1_213_0 RF1
ZShift = 0.77
lm Score = 0.39
NG = 0
DB = 0
SM = 0
Delta_K
Doxorubicin
RF1_213_0 RF1
ZShift = −0.07
lm Score = 0
NG = 0
DB = 0
SM = 0
Delta_r
Doxorubicin
RF1_213_0 RF1
ZShift = 0.85
lm Score = 50.66
NG = 0
DB = 0
SM = 0
Delta_AUC
Doxorubicin
RF1_322_0 RF1
ZShift = 0.47
lm Score = 2.73
NG = 0
DB = 0
SM = 0
Delta_L
Doxorubicin
RF1_322_0 RF1
ZShift = −0.39
lm Score = −5.43
NG = 0
DB = 0
SM = 0
Delta_K
Doxorubicin
RF1_322_0 RF1
ZShift = 0.18
lm Score = 0
NG = 0
DB = 0
SM = 0
Delta_r
Doxorubicin
RF1_322_0 RF1
ZShift = −0.54
lm Score = 75.75
NG = 0
DB = 0
SM = 0
Delta_AUC
Doxorubicin

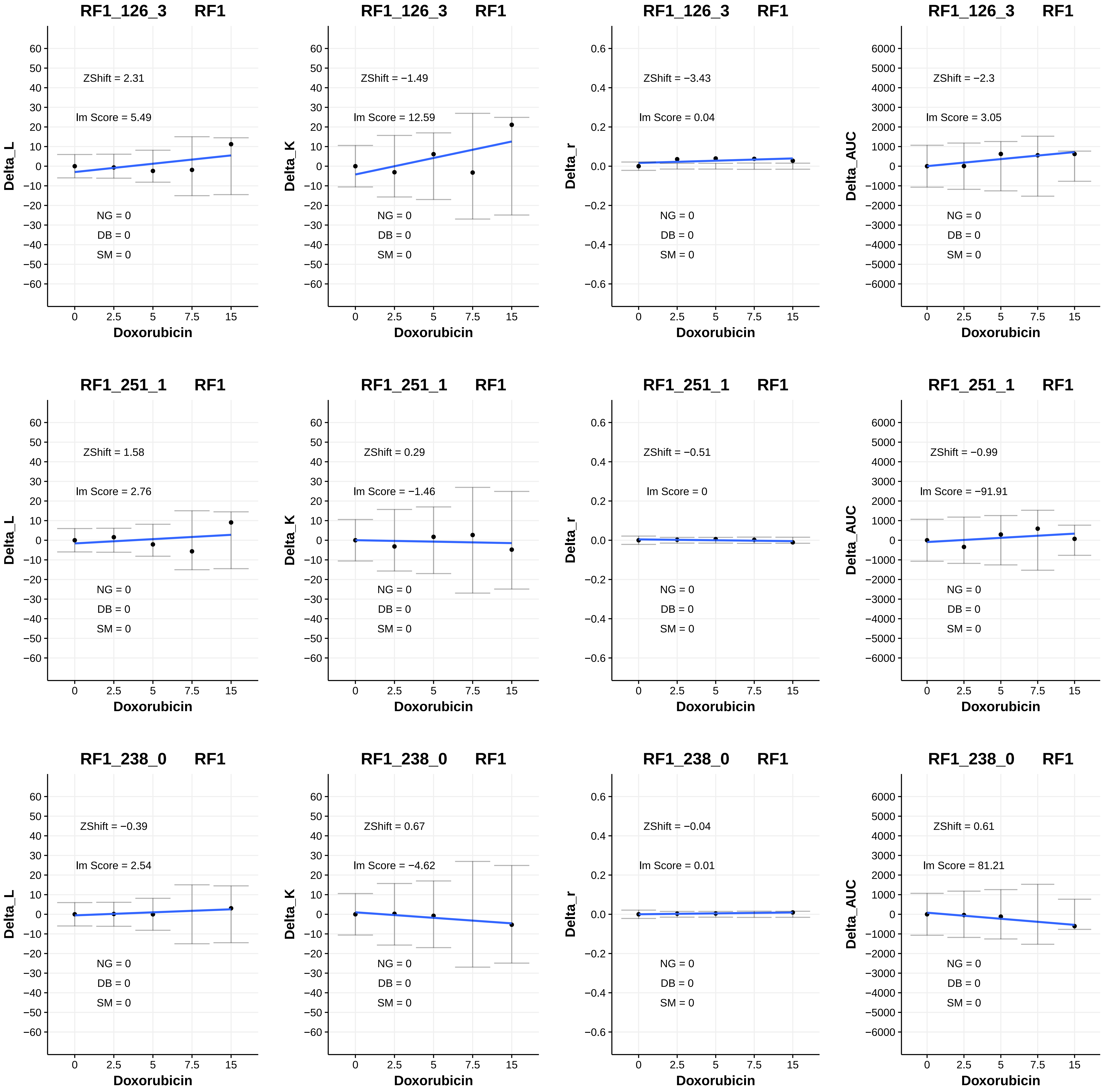
RF1_126_3 RF1
Delta_L
ZShift = 2.31
lm Score = 5.49
NG = 0
DB = 0
SM = 0
Doxorubicin
RF1_126_3 RF1
Delta_K
ZShift = −1.49
lm Score = 12.59
NG = 0
DB = 0
SM = 0
Doxorubicin
RF1_126_3 RF1
Delta_r
ZShift = −3.43
lm Score = 0.04
NG = 0
DB = 0
SM = 0
Doxorubicin
RF1_126_3 RF1
Delta_AUC
ZShift = −2.3
lm Score = 3.05
NG = 0
DB = 0
SM = 0
Doxorubicin
RF1_251_1 RF1
Delta_L
ZShift = 1.58
lm Score = 2.76
NG = 0
DB = 0
SM = 0
Doxorubicin
RF1_251_1 RF1
Delta_K
ZShift = 0.29
lm Score = −1.46
NG = 0
DB = 0
SM = 0
Doxorubicin
RF1_251_1 RF1
Delta_r
ZShift = −0.51
lm Score = 0
NG = 0
DB = 0
SM = 0
Doxorubicin
RF1_251_1 RF1
Delta_AUC
ZShift = −0.99
lm Score = −91.91
NG = 0
DB = 0
SM = 0
Doxorubicin
RF1_238_0 RF1
Delta_L
ZShift = −0.39
lm Score = 2.54
NG = 0
DB = 0
SM = 0
Doxorubicin
RF1_238_0 RF1
Delta_K
ZShift = 0.67
lm Score = −4.62
NG = 0
DB = 0
SM = 0
Doxorubicin
RF1_238_0 RF1
Delta_r
ZShift = −0.04
lm Score = 0.01
NG = 0
DB = 0
SM = 0
Doxorubicin
RF1_238_0 RF1
Delta_AUC
ZShift = 0.61
lm Score = 81.21
NG = 0
DB = 0
SM = 0
Doxorubicin

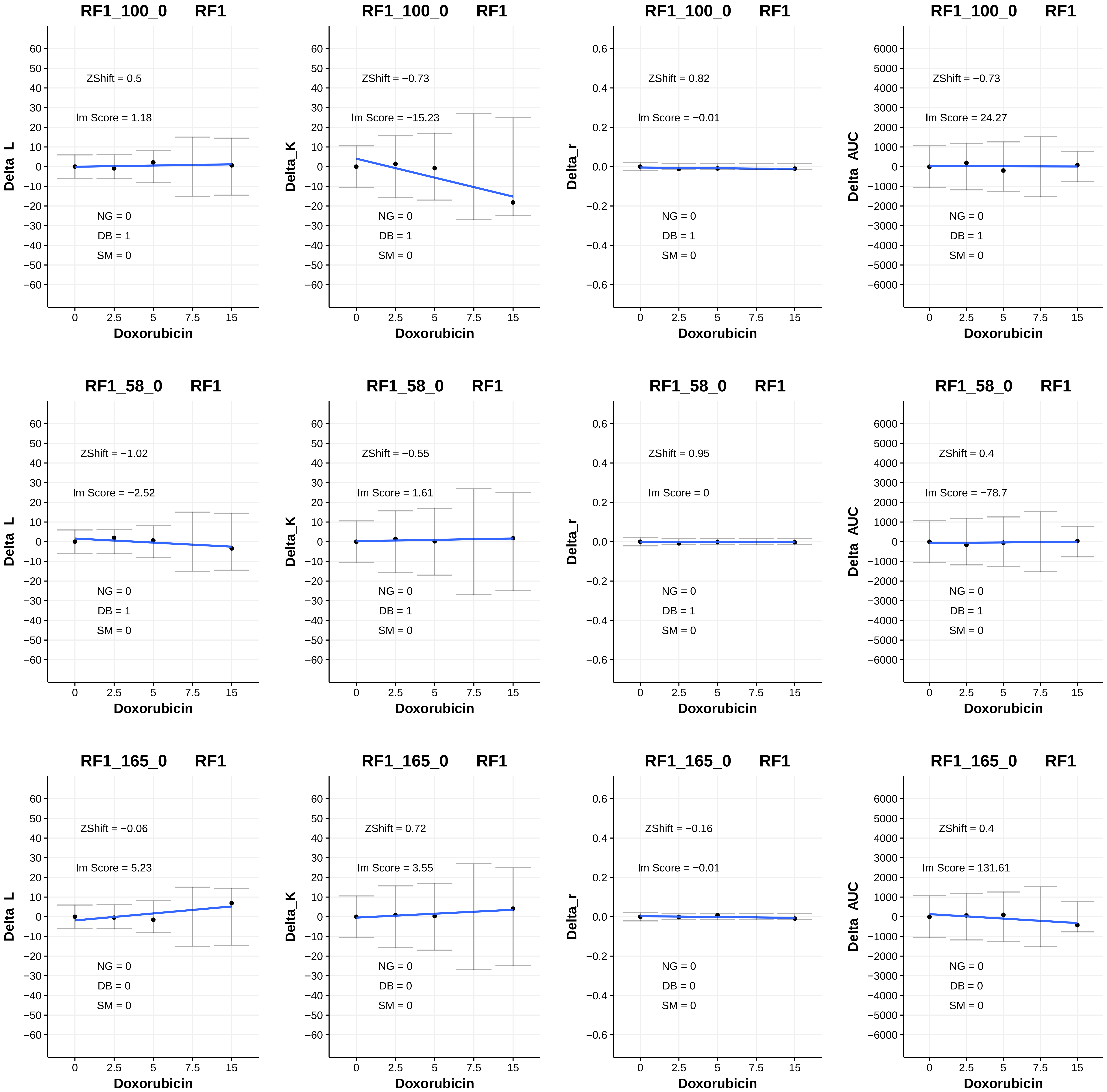
RF1_100_0 RF1
ZShift = 0.5
lm Score = 1.18
NG = 0
DB = 1
SM = 0
Delta_L
Doxorubicin
RF1_100_0 RF1
ZShift = −0.73
lm Score = −15.23
NG = 0
DB = 1
SM = 0
Delta_K
Doxorubicin
RF1_100_0 RF1
ZShift = 0.82
lm Score = −0.01
NG = 0
DB = 1
SM = 0
Delta_r
Doxorubicin
RF1_100_0 RF1
ZShift = −0.73
lm Score = 24.27
NG = 0
DB = 1
SM = 0
Delta_AUC
Doxorubicin
RF1_58_0 RF1
ZShift = −1.02
lm Score = −2.52
NG = 0
DB = 1
SM = 0
Delta_L
Doxorubicin
RF1_58_0 RF1
ZShift = −0.55
lm Score = 1.61
NG = 0
DB = 1
SM = 0
Delta_K
Doxorubicin
RF1_58_0 RF1
ZShift = 0.95
lm Score = 0
NG = 0
DB = 1
SM = 0
Delta_r
Doxorubicin
RF1_58_0 RF1
ZShift = 0.4
lm Score = −78.7
NG = 0
DB = 1
SM = 0
Delta_AUC
Doxorubicin
RF1_165_0 RF1
ZShift = −0.06
lm Score = 5.23
NG = 0
DB = 0
SM = 0
Delta_L
Doxorubicin
RF1_165_0 RF1
ZShift = 0.72
lm Score = 3.55
NG = 0
DB = 0
SM = 0
Delta_K
Doxorubicin
RF1_165_0 RF1
ZShift = −0.16
lm Score = −0.01
NG = 0
DB = 0
SM = 0
Delta_r
Doxorubicin
RF1_165_0 RF1
ZShift = 0.4
lm Score = 131.61
NG = 0
DB = 0
SM = 0
Delta_AUC
Doxorubicin

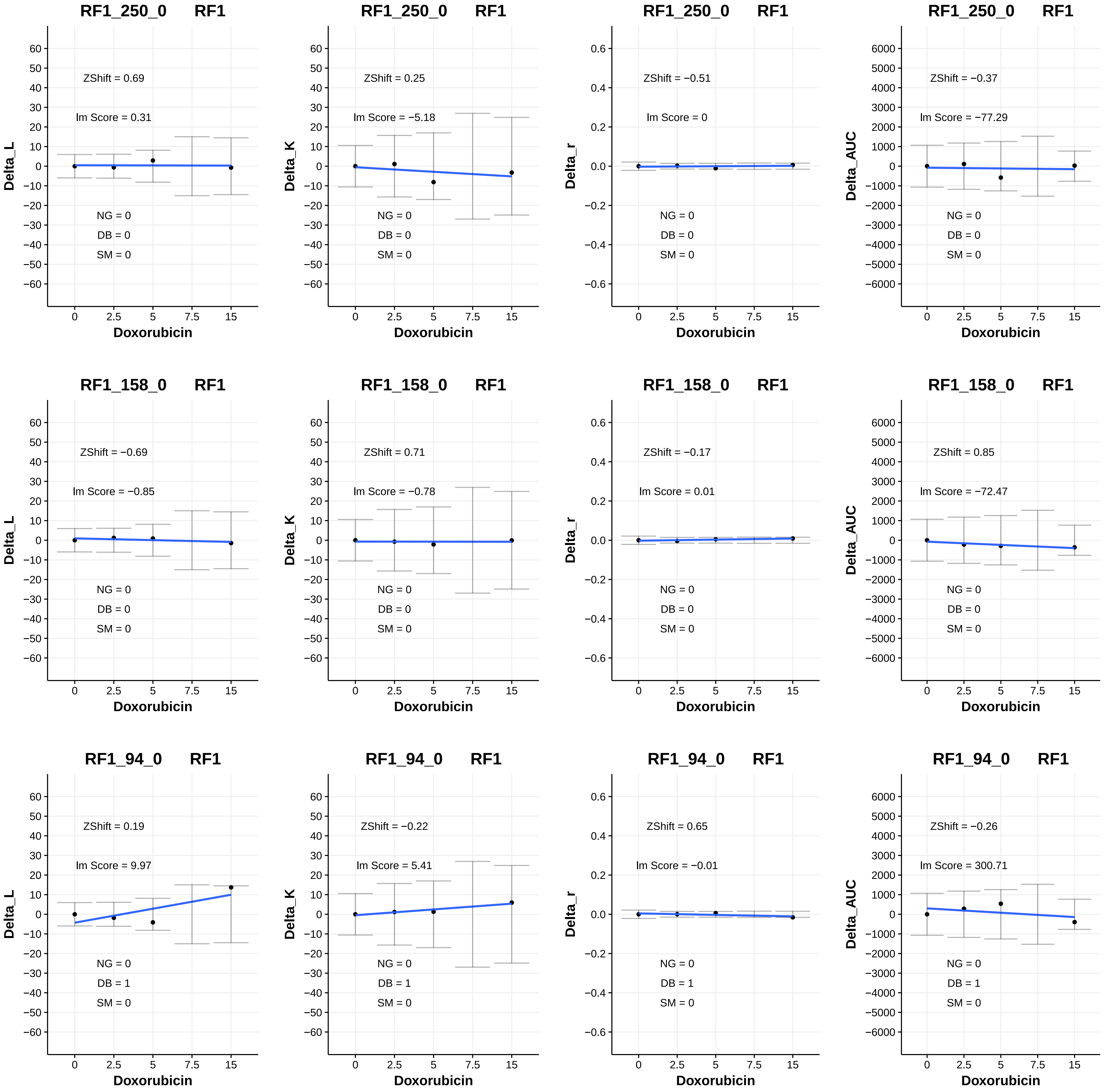

RF1_250_0 RF1
ZShift = 0.69
lm Score = 0.31
NG = 0
DB = 0
SM = 0
Delta_L
Doxorubicin
RF1_250_0 RF1
ZShift = 0.25
lm Score = −5.18
NG = 0
DB = 0
SM = 0
Delta_K
Doxorubicin
RF1_250_0 RF1
ZShift = −0.51
lm Score = 0
NG = 0
DB = 0
SM = 0
Delta_r
Doxorubicin
RF1_250_0 RF1
ZShift = −0.37
lm Score = −77.29
NG = 0
DB = 0
SM = 0
Delta_AUC
Doxorubicin
RF1_158_0 RF1
ZShift = −0.69
lm Score = −0.85
NG = 0
DB = 0
SM = 0
Delta_L
Doxorubicin
RF1_158_0 RF1
ZShift = 0.71
lm Score = −0.78
NG = 0
DB = 0
SM = 0
Delta_K
Doxorubicin
RF1_158_0 RF1
ZShift = −0.17
lm Score = 0.01
NG = 0
DB = 0
SM = 0
Delta_r
Doxorubicin
RF1_158_0 RF1
ZShift = 0.85
lm Score = −72.47
NG = 0
DB = 0
SM = 0
Delta_AUC
Doxorubicin
RF1_94_0 RF1
ZShift = 0.19
lm Score = 9.97
NG = 0
DB = 1
SM = 0
Delta_L
Doxorubicin
RF1_94_0 RF1
ZShift = −0.22
lm Score = 5.41
NG = 0
DB = 1
SM = 0
Delta_K
Doxorubicin
RF1_94_0 RF1
ZShift = 0.65
lm Score = −0.01
NG = 0
DB = 1
SM = 0
Delta_r
Doxorubicin
RF1_94_0 RF1
ZShift = −0.26
lm Score = 300.71
NG = 0
DB = 1
SM = 0
Delta_AUC
Doxorubicin

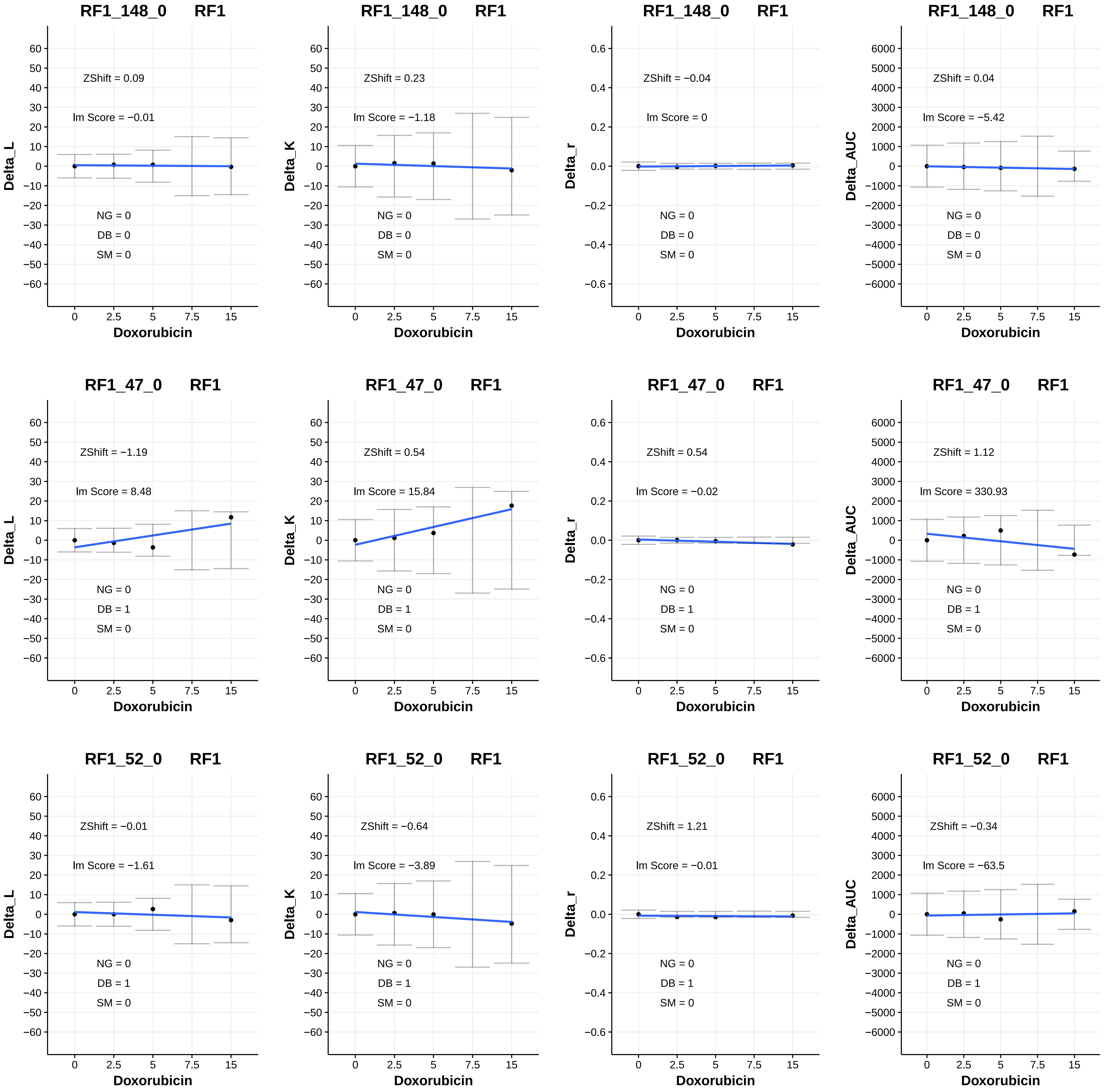
RF1_148_0 RF1
ZShift = 0.09
lm Score = −0.01
NG = 0
DB = 0
SM = 0
Delta_L
Doxorubicin
RF1_148_0 RF1
ZShift = 0.23
lm Score = −1.18
NG = 0
DB = 0
SM = 0
Delta_K
Doxorubicin
RF1_148_0 RF1
ZShift = −0.04
lm Score = 0
NG = 0
DB = 0
SM = 0
Delta_r
Doxorubicin
RF1_148_0 RF1
ZShift = 0.04
lm Score = −5.42
NG = 0
DB = 0
SM = 0
Delta_AUC
Doxorubicin
RF1_47_0 RF1
ZShift = −1.19
lm Score = 8.48
NG = 0
DB = 1
SM = 0
Delta_L
Doxorubicin
RF1_47_0 RF1
ZShift = 0.54
lm Score = 15.84
NG = 0
DB = 1
SM = 0
Delta_K
Doxorubicin
RF1_47_0 RF1
ZShift = 0.54
lm Score = −0.02
NG = 0
DB = 1
SM = 0
Delta_r
Doxorubicin
RF1_47_0 RF1
ZShift = 1.12
lm Score = 330.93
NG = 0
DB = 1
SM = 0
Delta_AUC
Doxorubicin
RF1_52_0 RF1
ZShift = −0.01
lm Score = −1.61
NG = 0
DB = 1
SM = 0
Delta_L
Doxorubicin
RF1_52_0 RF1
ZShift = −0.64
lm Score = −3.89
NG = 0
DB = 1
SM = 0
Delta_K
Doxorubicin
RF1_52_0 RF1
ZShift = 1.21
lm Score = −0.01
NG = 0
DB = 1
SM = 0
Delta_r
Doxorubicin
RF1_52_0 RF1
ZShift = −0.34
lm Score = −63.5
NG = 0
DB = 1
SM = 0
Delta_AUC
Doxorubicin

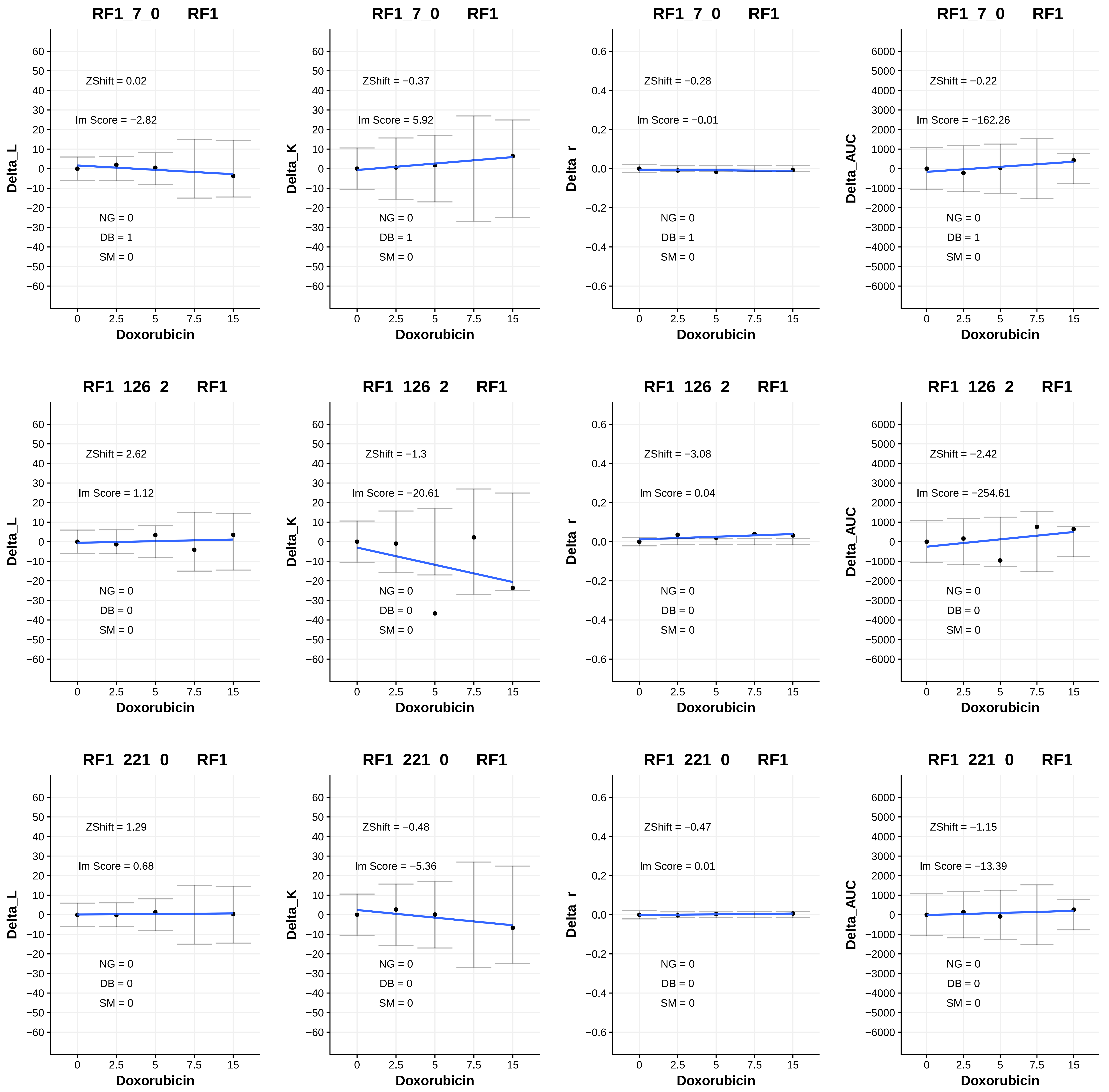
RF1_7_0 RF1
ZShift = 0.02
Im Score = −2.82
NG = 0
DB = 1
SM = 0
Delta_L
Doxorubicin
RF1_7_0 RF1
ZShift = −0.37
Im Score = 5.92
NG = 0
DB = 1
SM = 0
Delta_K
Doxorubicin
RF1_7_0 RF1
ZShift = −0.28
Im Score = −0.01
NG = 0
DB = 1
SM = 0
Delta_r
Doxorubicin
RF1_7_0 RF1
ZShift = −0.22
Im Score = −162.26
NG = 0
DB = 1
SM = 0
Delta_AUC
Doxorubicin
RF1_126_2 RF1
ZShift = 2.62
Im Score = 1.12
NG = 0
DB = 0
SM = 0
Delta_L
Doxorubicin
RF1_126_2 RF1
ZShift = −1.3
Im Score = −20.61
NG = 0
DB = 0
SM = 0
Delta_K
Doxorubicin
RF1_126_2 RF1
ZShift = −3.08
Im Score = 0.04
NG = 0
DB = 0
SM = 0
Delta_r
Doxorubicin
RF1_126_2 RF1
ZShift = −2.42
Im Score = −254.61
NG = 0
DB = 0
SM = 0
Delta_AUC
Doxorubicin
RF1_221_0 RF1
ZShift = 1.29
Im Score = 0.68
NG = 0
DB = 0
SM = 0
Delta_L
Doxorubicin
RF1_221_0 RF1
ZShift = −0.48
Im Score = −5.36
NG = 0
DB = 0
SM = 0
Delta_K
Doxorubicin
RF1_221_0 RF1
ZShift = −0.47
Im Score = 0.01
NG = 0
DB = 0
SM = 0
Delta_r
Doxorubicin
RF1_221_0 RF1
ZShift = −1.15
Im Score = −13.39
NG = 0
DB = 0
SM = 0
Delta_AUC
Doxorubicin

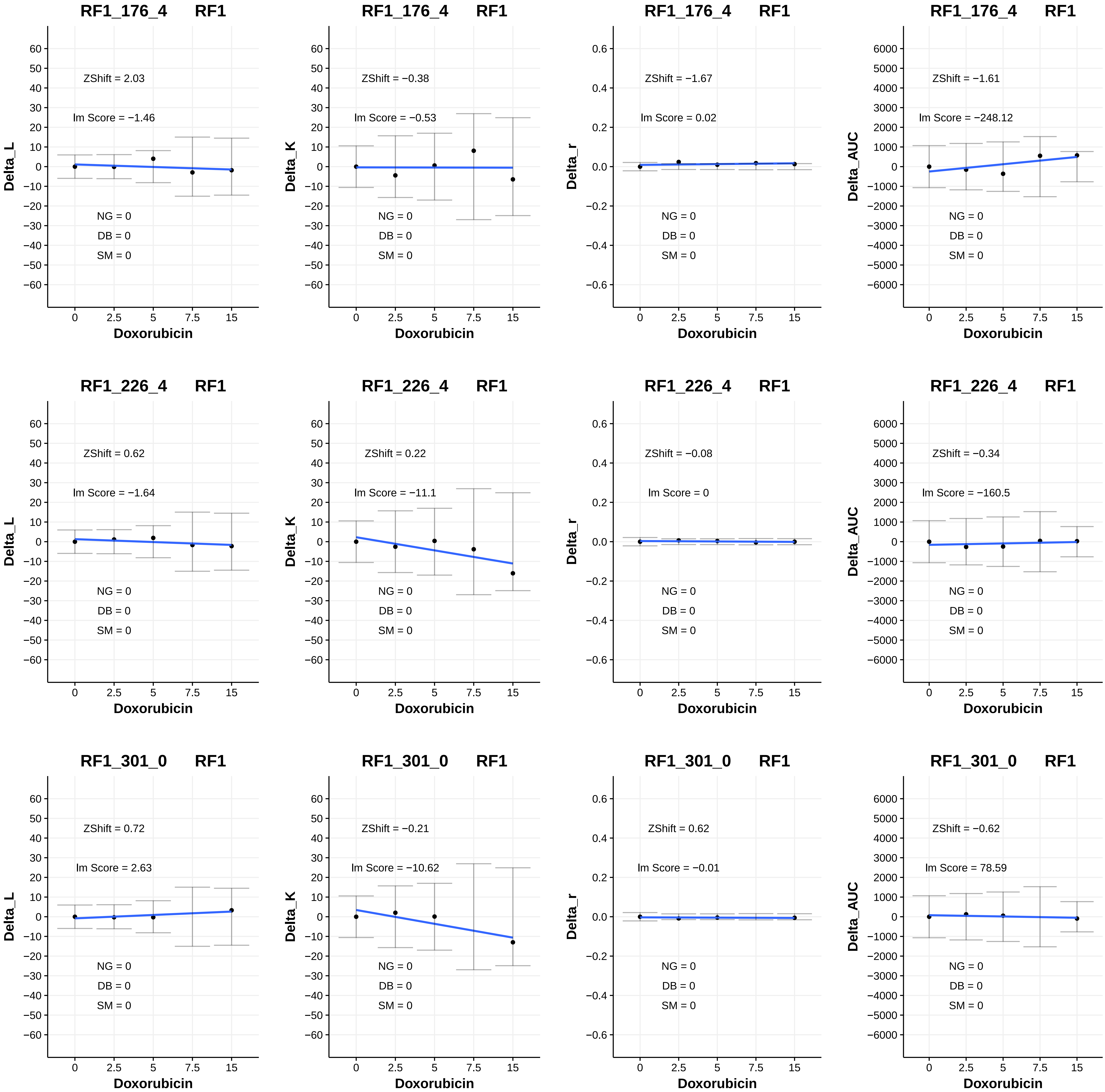
RF1_176_4 RF1
ZShift = 2.03
Im Score = −1.46
NG = 0
DB = 0
SM = 0
Delta_L
Doxorubicin
RF1_176_4 RF1
ZShift = −0.38
Im Score = −0.53
NG = 0
DB = 0
SM = 0
Delta_K
Doxorubicin
RF1_176_4 RF1
ZShift = −1.67
Im Score = 0.02
NG = 0
DB = 0
SM = 0
Delta_r
Doxorubicin
RF1_176_4 RF1
ZShift = −1.61
Im Score = −248.12
NG = 0
DB = 0
SM = 0
Delta_AUC
Doxorubicin
RF1_226_4 RF1
ZShift = 0.62
Im Score = −1.64
NG = 0
DB = 0
SM = 0
Delta_L
Doxorubicin
RF1_226_4 RF1
ZShift = 0.22
Im Score = −11.1
NG = 0
DB = 0
SM = 0
Delta_K
Doxorubicin
RF1_226_4 RF1
ZShift = −0.08
Im Score = 0
NG = 0
DB = 0
SM = 0
Delta_r
Doxorubicin
RF1_226_4 RF1
ZShift = −0.34
Im Score = −160.5
NG = 0
DB = 0
SM = 0
Delta_AUC
Doxorubicin
RF1_301_0 RF1
ZShift = 0.72
Im Score = 2.63
NG = 0
DB = 0
SM = 0
Delta_L
Doxorubicin
RF1_301_0 RF1
ZShift = −0.21
Im Score = −10.62
NG = 0
DB = 0
SM = 0
Delta_K
Doxorubicin
RF1_301_0 RF1
ZShift = 0.62
Im Score = −0.01
NG = 0
DB = 0
SM = 0
Delta_r
Doxorubicin
RF1_301_0 RF1
ZShift = −0.62
Im Score = 78.59
NG = 0
DB = 0
SM = 0
Delta_AUC
Doxorubicin

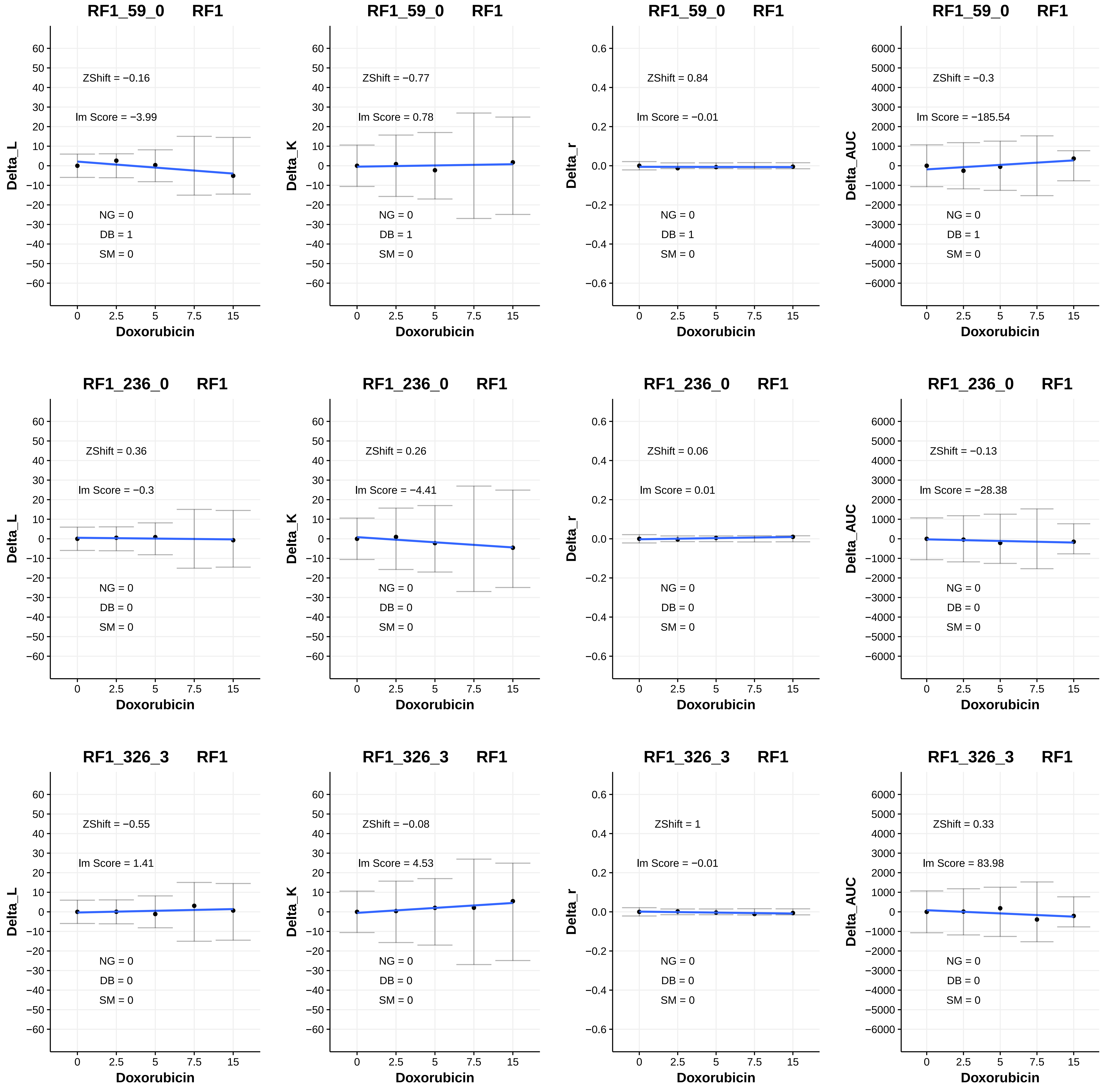

RF1_59_0 RF1
ZShift = −0.16
Im Score = −3.99
NG = 0
DB = 1
SM = 0
Delta_L
Doxorubicin
RF1_59_0 RF1
ZShift = −0.77
Im Score = 0.78
NG = 0
DB = 1
SM = 0
Delta_K
Doxorubicin
RF1_59_0 RF1
ZShift = 0.84
Im Score = −0.01
NG = 0
DB = 1
SM = 0
Delta_r
Doxorubicin
RF1_59_0 RF1
ZShift = −0.3
Im Score = −185.54
NG = 0
DB = 1
SM = 0
Delta_AUC
Doxorubicin
RF1_236_0 RF1
ZShift = 0.36
Im Score = −0.3
NG = 0
DB = 0
SM = 0
Delta_L
Doxorubicin
RF1_236_0 RF1
ZShift = 0.26
Im Score = −4.41
NG = 0
DB = 0
SM = 0
Delta_K
Doxorubicin
RF1_236_0 RF1
ZShift = 0.06
Im Score = 0.01
NG = 0
DB = 0
SM = 0
Delta_r
Doxorubicin
RF1_236_0 RF1
ZShift = −0.13
Im Score = −28.38
NG = 0
DB = 0
SM = 0
Delta_AUC
Doxorubicin
RF1_326_3 RF1
ZShift = −0.55
Im Score = 1.41
NG = 0
DB = 0
SM = 0
Delta_L
Doxorubicin
RF1_326_3 RF1
ZShift = −0.08
Im Score = 4.53
NG = 0
DB = 0
SM = 0
Delta_K
Doxorubicin
RF1_326_3 RF1
ZShift = 1
Im Score = −0.01
NG = 0
DB = 0
SM = 0
Delta_r
Doxorubicin
RF1_326_3 RF1
ZShift = 0.33
Im Score = 83.98
NG = 0
DB = 0
SM = 0
Delta_AUC
Doxorubicin

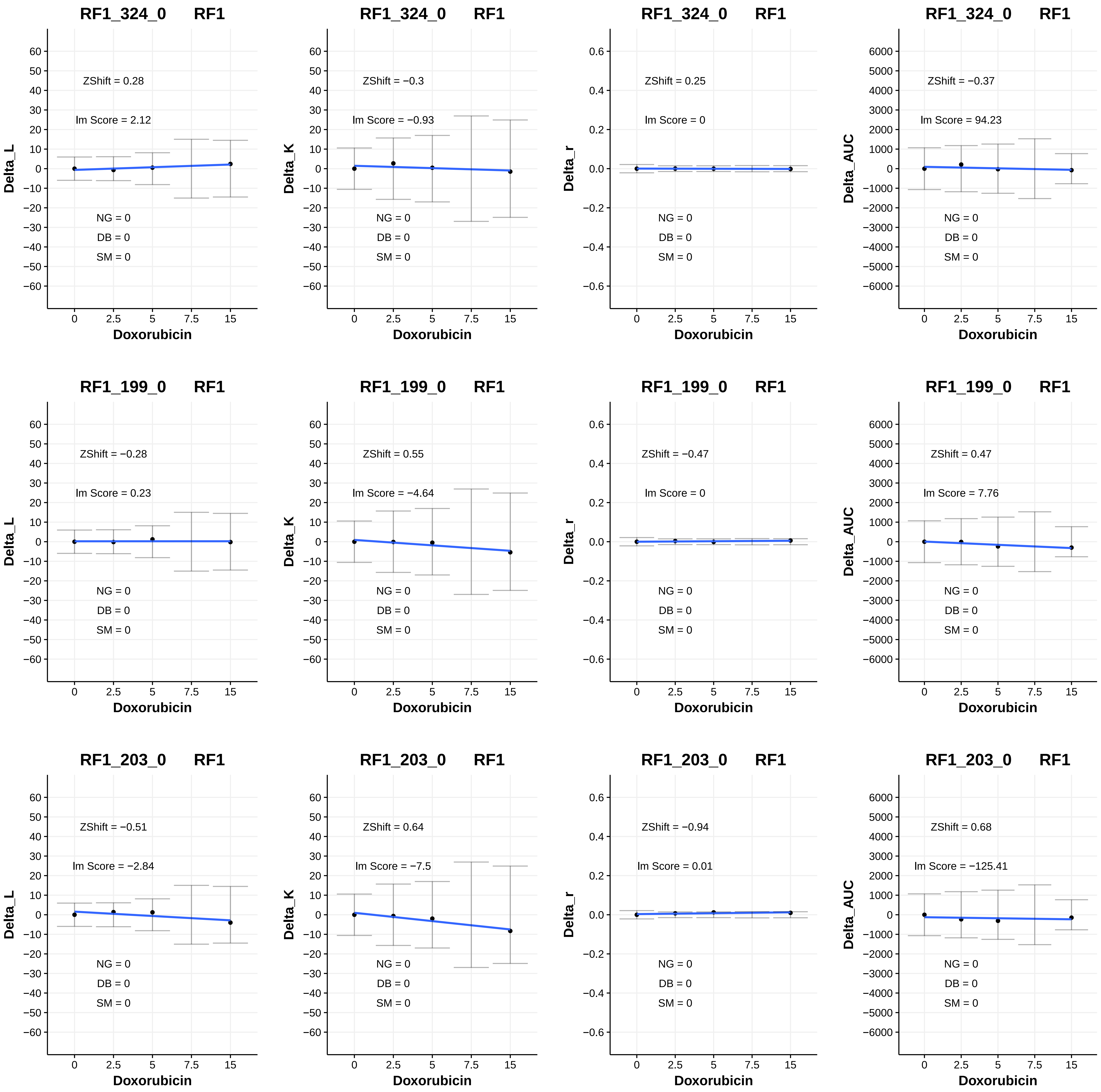
RF1_324_0 RF1
ZShift = 0.28
lm Score = 2.12
NG = 0
DB = 0
SM = 0
Delta_L
Doxorubicin
RF1_324_0 RF1
ZShift = −0.3
lm Score = −0.93
NG = 0
DB = 0
SM = 0
Delta_K
Doxorubicin
RF1_324_0 RF1
ZShift = 0.25
lm Score = 0
NG = 0
DB = 0
SM = 0
Delta_r
Doxorubicin
RF1_324_0 RF1
ZShift = −0.37
lm Score = 94.23
NG = 0
DB = 0
SM = 0
Delta_AUC
Doxorubicin
RF1_199_0 RF1
ZShift = −0.28
lm Score = 0.23
NG = 0
DB = 0
SM = 0
Delta_L
Doxorubicin
RF1_199_0 RF1
ZShift = 0.55
lm Score = −4.64
NG = 0
DB = 0
SM = 0
Delta_K
Doxorubicin
RF1_199_0 RF1
ZShift = −0.47
lm Score = 0
NG = 0
DB = 0
SM = 0
Delta_r
Doxorubicin
RF1_199_0 RF1
ZShift = 0.47
lm Score = 7.76
NG = 0
DB = 0
SM = 0
Delta_AUC
Doxorubicin
RF1_203_0 RF1
ZShift = −0.51
lm Score = −2.84
NG = 0
DB = 0
SM = 0
Delta_L
Doxorubicin
RF1_203_0 RF1
ZShift = 0.64
lm Score = −7.5
NG = 0
DB = 0
SM = 0
Delta_K
Doxorubicin
RF1_203_0 RF1
ZShift = −0.94
lm Score = 0.01
NG = 0
DB = 0
SM = 0
Delta_r
Doxorubicin
RF1_203_0 RF1
ZShift = 0.68
lm Score = −125.41
NG = 0
DB = 0
SM = 0
Delta_AUC
Doxorubicin

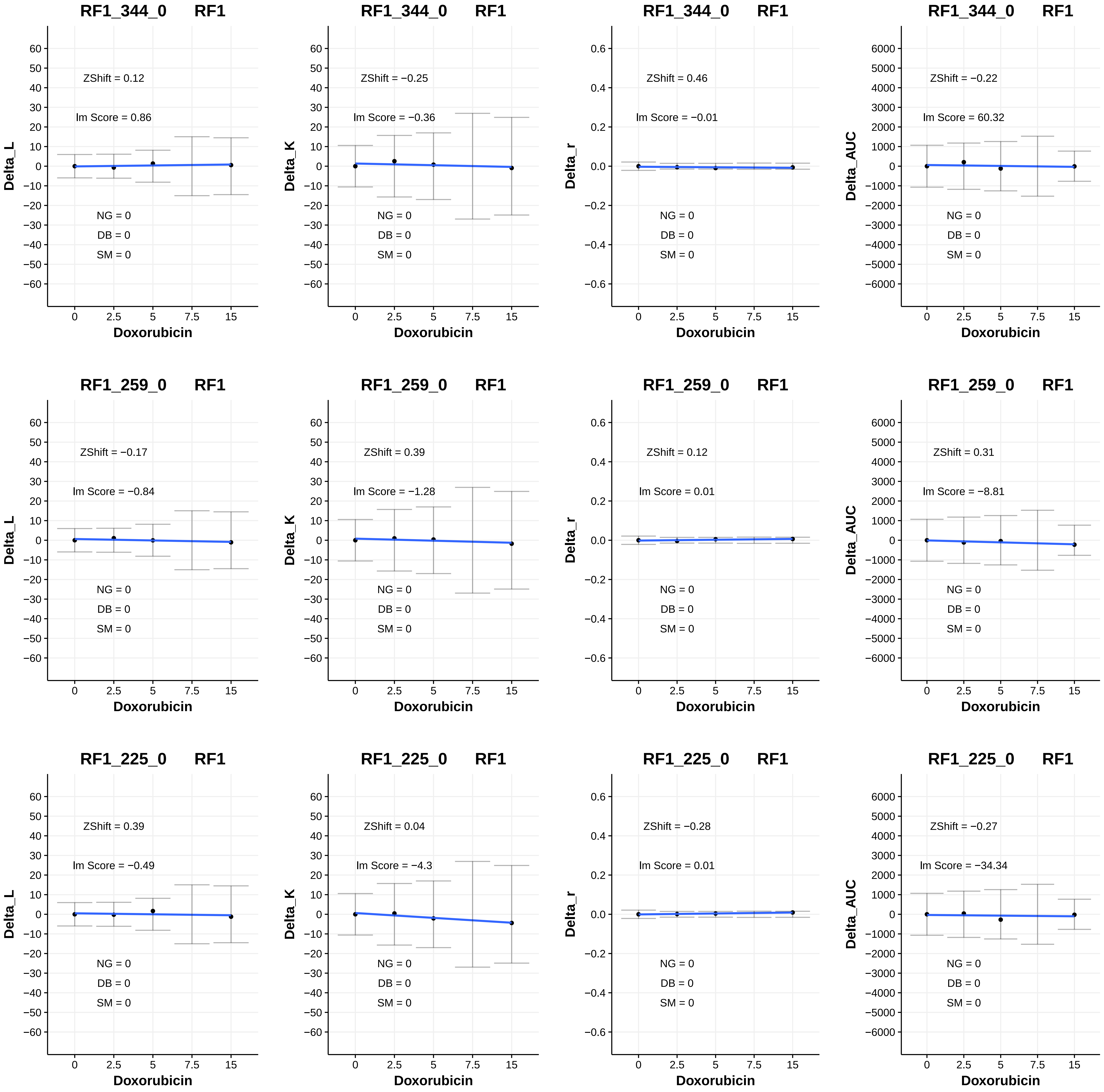

RF1_344_0 RF1
ZShift = 0.12
Im Score = 0.86
NG = 0
DB = 0
SM = 0
Delta_L
Doxorubicin
RF1_344_0 RF1
ZShift = −0.25
Im Score = −0.36
NG = 0
DB = 0
SM = 0
Delta_K
Doxorubicin
RF1_344_0 RF1
ZShift = 0.46
Im Score = −0.01
NG = 0
DB = 0
SM = 0
Delta_r
Doxorubicin
RF1_344_0 RF1
ZShift = −0.22
Im Score = 60.32
NG = 0
DB = 0
SM = 0
Delta_AUC
Doxorubicin
RF1_259_0 RF1
ZShift = −0.17
Im Score = −0.84
NG = 0
DB = 0
SM = 0
Delta_L
Doxorubicin
RF1_259_0 RF1
ZShift = 0.39
Im Score = −1.28
NG = 0
DB = 0
SM = 0
Delta_K
Doxorubicin
RF1_259_0 RF1
ZShift = 0.12
Im Score = 0.01
NG = 0
DB = 0
SM = 0
Delta_r
Doxorubicin
RF1_259_0 RF1
ZShift = 0.31
Im Score = −8.81
NG = 0
DB = 0
SM = 0
Delta_AUC
Doxorubicin
RF1_225_0 RF1
ZShift = 0.39
Im Score = −0.49
NG = 0
DB = 0
SM = 0
Delta_L
Doxorubicin
RF1_225_0 RF1
ZShift = 0.04
Im Score = −4.3
NG = 0
DB = 0
SM = 0
Delta_K
Doxorubicin
RF1_225_0 RF1
ZShift = −0.28
Im Score = 0.01
NG = 0
DB = 0
SM = 0
Delta_r
Doxorubicin
RF1_225_0 RF1
ZShift = −0.27
Im Score = −34.34
NG = 0
DB = 0
SM = 0
Delta_AUC
Doxorubicin

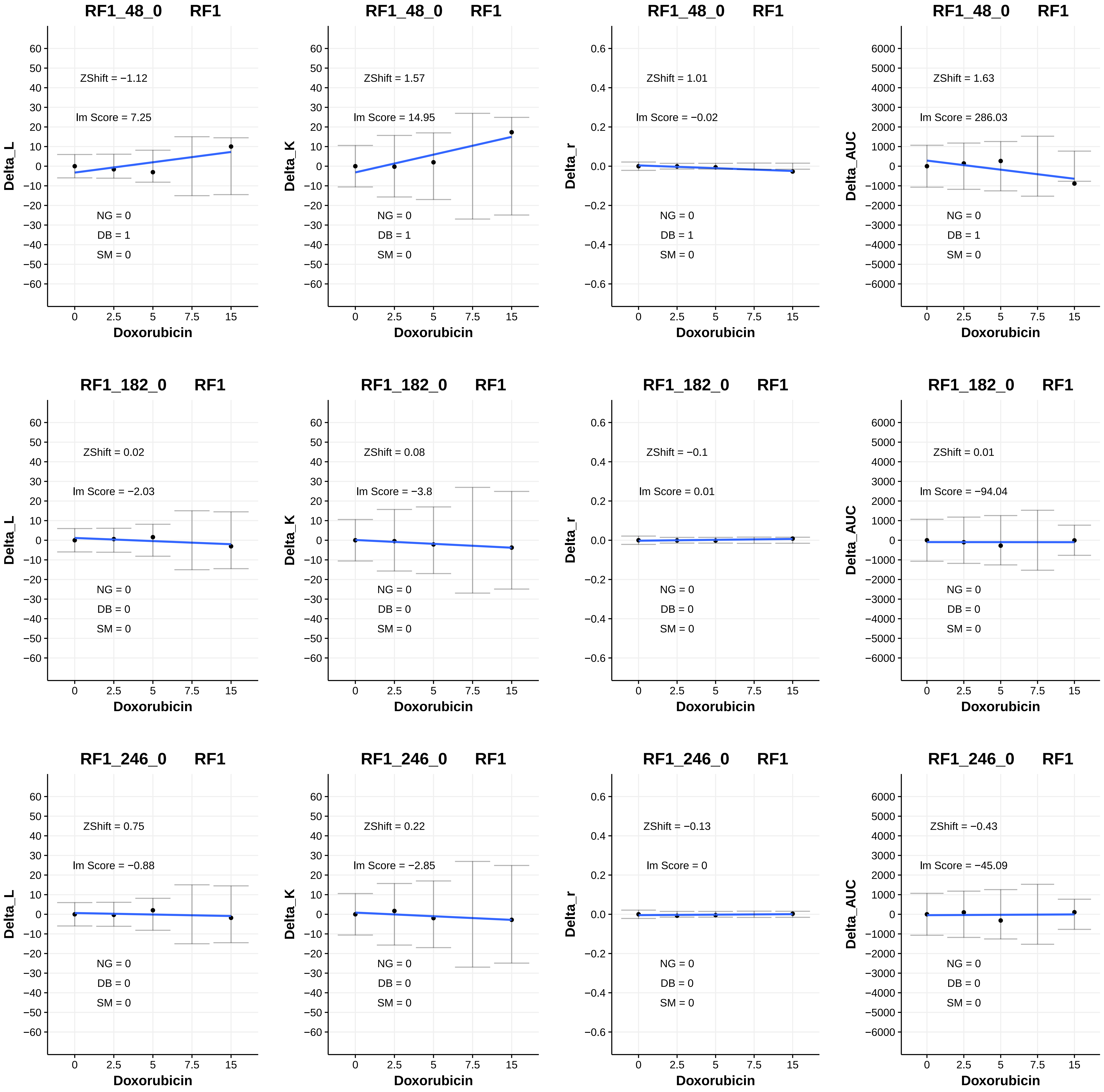

RF1_48_0 RF1
ZShift = −1.12
lm Score = 7.25
NG = 0
DB = 1
SM = 0
Delta_L
Doxorubicin
RF1_48_0 RF1
ZShift = 1.57
lm Score = 14.95
NG = 0
DB = 1
SM = 0
Delta_K
Doxorubicin
RF1_48_0 RF1
ZShift = 1.01
lm Score = −0.02
NG = 0
DB = 1
SM = 0
Delta_r
Doxorubicin
RF1_48_0 RF1
ZShift = 1.63
lm Score = 286.03
NG = 0
DB = 1
SM = 0
Delta_AUC
Doxorubicin
RF1_182_0 RF1
ZShift = 0.02
lm Score = −2.03
NG = 0
DB = 0
SM = 0
Delta_L
Doxorubicin
RF1_182_0 RF1
ZShift = 0.08
lm Score = −3.8
NG = 0
DB = 0
SM = 0
Delta_K
Doxorubicin
RF1_182_0 RF1
ZShift = −0.1
lm Score = 0.01
NG = 0
DB = 0
SM = 0
Delta_r
Doxorubicin
RF1_182_0 RF1
ZShift = 0.01
lm Score = −94.04
NG = 0
DB = 0
SM = 0
Delta_AUC
Doxorubicin
RF1_246_0 RF1
ZShift = 0.75
lm Score = −0.88
NG = 0
DB = 0
SM = 0
Delta_L
Doxorubicin
RF1_246_0 RF1
ZShift = 0.22
lm Score = −2.85
NG = 0
DB = 0
SM = 0
Delta_K
Doxorubicin
RF1_246_0 RF1
ZShift = −0.13
lm Score = 0
NG = 0
DB = 0
SM = 0
Delta_r
Doxorubicin
RF1_246_0 RF1
ZShift = −0.43
lm Score = −45.09
NG = 0
DB = 0
SM = 0
Delta_AUC
Doxorubicin

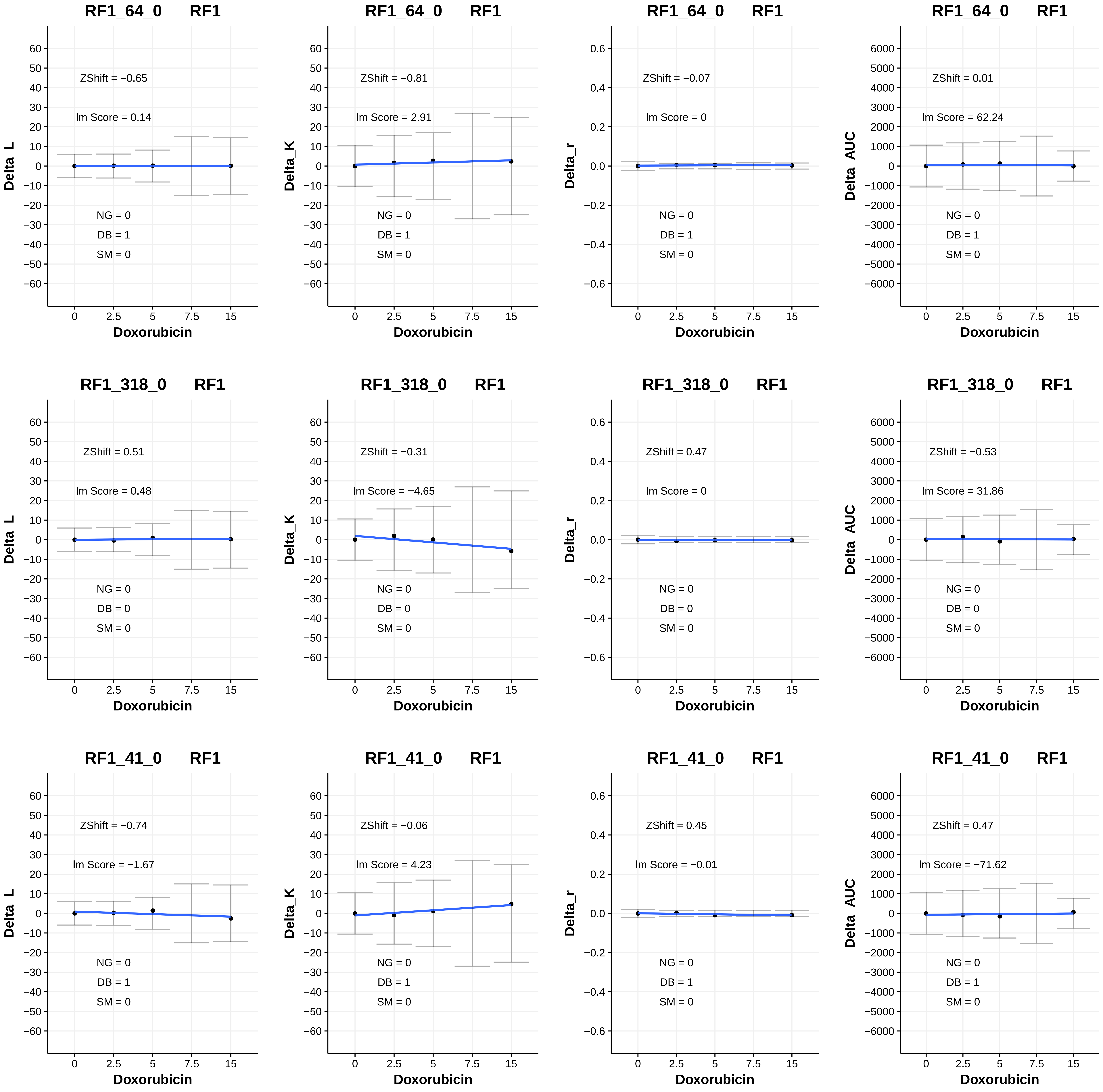
RF1_64_0 RF1
ZShift = −0.65
lm Score = 0.14
NG = 0
DB = 1
SM = 0
Delta_L
Doxorubicin
RF1_64_0 RF1
ZShift = −0.81
lm Score = 2.91
NG = 0
DB = 1
SM = 0
Delta_K
Doxorubicin
RF1_64_0 RF1
ZShift = −0.07
lm Score = 0
NG = 0
DB = 1
SM = 0
Delta_r
Doxorubicin
RF1_64_0 RF1
ZShift = 0.01
lm Score = 62.24
NG = 0
DB = 1
SM = 0
Delta_AUC
Doxorubicin
RF1_318_0 RF1
ZShift = 0.51
lm Score = 0.48
NG = 0
DB = 0
SM = 0
Delta_L
Doxorubicin
RF1_318_0 RF1
ZShift = −0.31
lm Score = −4.65
NG = 0
DB = 0
SM = 0
Delta_K
Doxorubicin
RF1_318_0 RF1
ZShift = 0.47
lm Score = 0
NG = 0
DB = 0
SM = 0
Delta_r
Doxorubicin
RF1_318_0 RF1
ZShift = −0.53
lm Score = 31.86
NG = 0
DB = 0
SM = 0
Delta_AUC
Doxorubicin
RF1_41_0 RF1
ZShift = −0.74
lm Score = −1.67
NG = 0
DB = 1
SM = 0
Delta_L
Doxorubicin
RF1_41_0 RF1
ZShift = −0.06
lm Score = 4.23
NG = 0
DB = 1
SM = 0
Delta_K
Doxorubicin
RF1_41_0 RF1
ZShift = 0.45
lm Score = −0.01
NG = 0
DB = 1
SM = 0
Delta_r
Doxorubicin
RF1_41_0 RF1
ZShift = 0.47
lm Score = −71.62
NG = 0
DB = 1
SM = 0
Delta_AUC
Doxorubicin

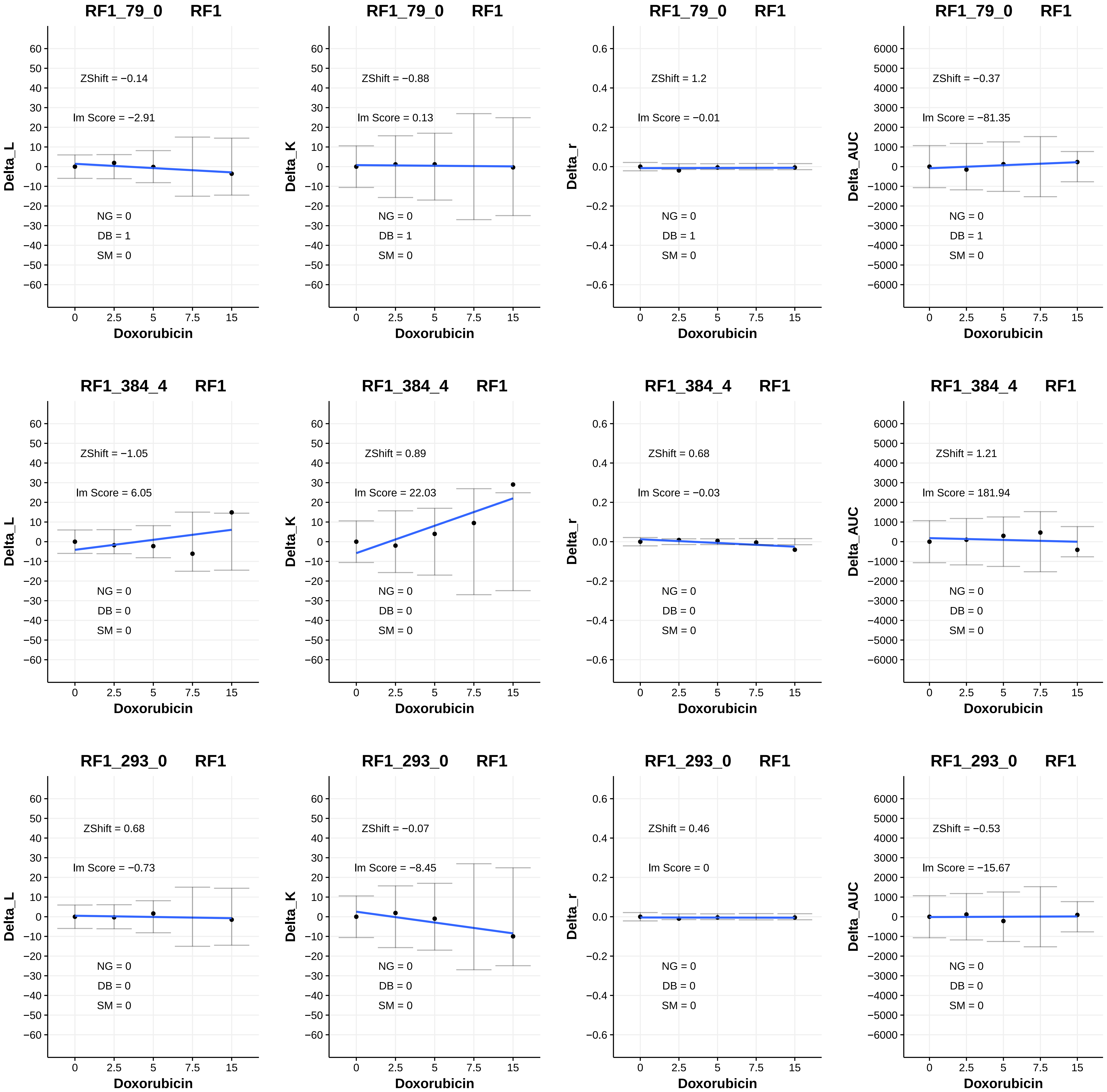

RF1_79_0 RF1
ZShift = −0.14
lm Score = −2.91
NG = 0
DB = 1
SM = 0
Delta_L
Doxorubicin
RF1_79_0 RF1
ZShift = −0.88
lm Score = 0.13
NG = 0
DB = 1
SM = 0
Delta_K
Doxorubicin
RF1_79_0 RF1
ZShift = 1.2
lm Score = −0.01
NG = 0
DB = 1
SM = 0
Delta_r
Doxorubicin
RF1_79_0 RF1
ZShift = −0.37
lm Score = −81.35
NG = 0
DB = 1
SM = 0
Delta_AUC
Doxorubicin
RF1_384_4 RF1
ZShift = −1.05
lm Score = 6.05
NG = 0
DB = 0
SM = 0
Delta_L
Doxorubicin
RF1_384_4 RF1
ZShift = 0.89
lm Score = 22.03
NG = 0
DB = 0
SM = 0
Delta_K
Doxorubicin
RF1_384_4 RF1
ZShift = 0.68
lm Score = −0.03
NG = 0
DB = 0
SM = 0
Delta_r
Doxorubicin
RF1_384_4 RF1
ZShift = 1.21
lm Score = 181.94
NG = 0
DB = 0
SM = 0
Delta_AUC
Doxorubicin
RF1_293_0 RF1
ZShift = 0.68
lm Score = −0.73
NG = 0
DB = 0
SM = 0
Delta_L
Doxorubicin
RF1_293_0 RF1
ZShift = −0.07
lm Score = −8.45
NG = 0
DB = 0
SM = 0
Delta_K
Doxorubicin
RF1_293_0 RF1
ZShift = 0.46
lm Score = 0
NG = 0
DB = 0
SM = 0
Delta_r
Doxorubicin
RF1_293_0 RF1
ZShift = −0.53
lm Score = −15.67
NG = 0
DB = 0
SM = 0
Delta_AUC
Doxorubicin

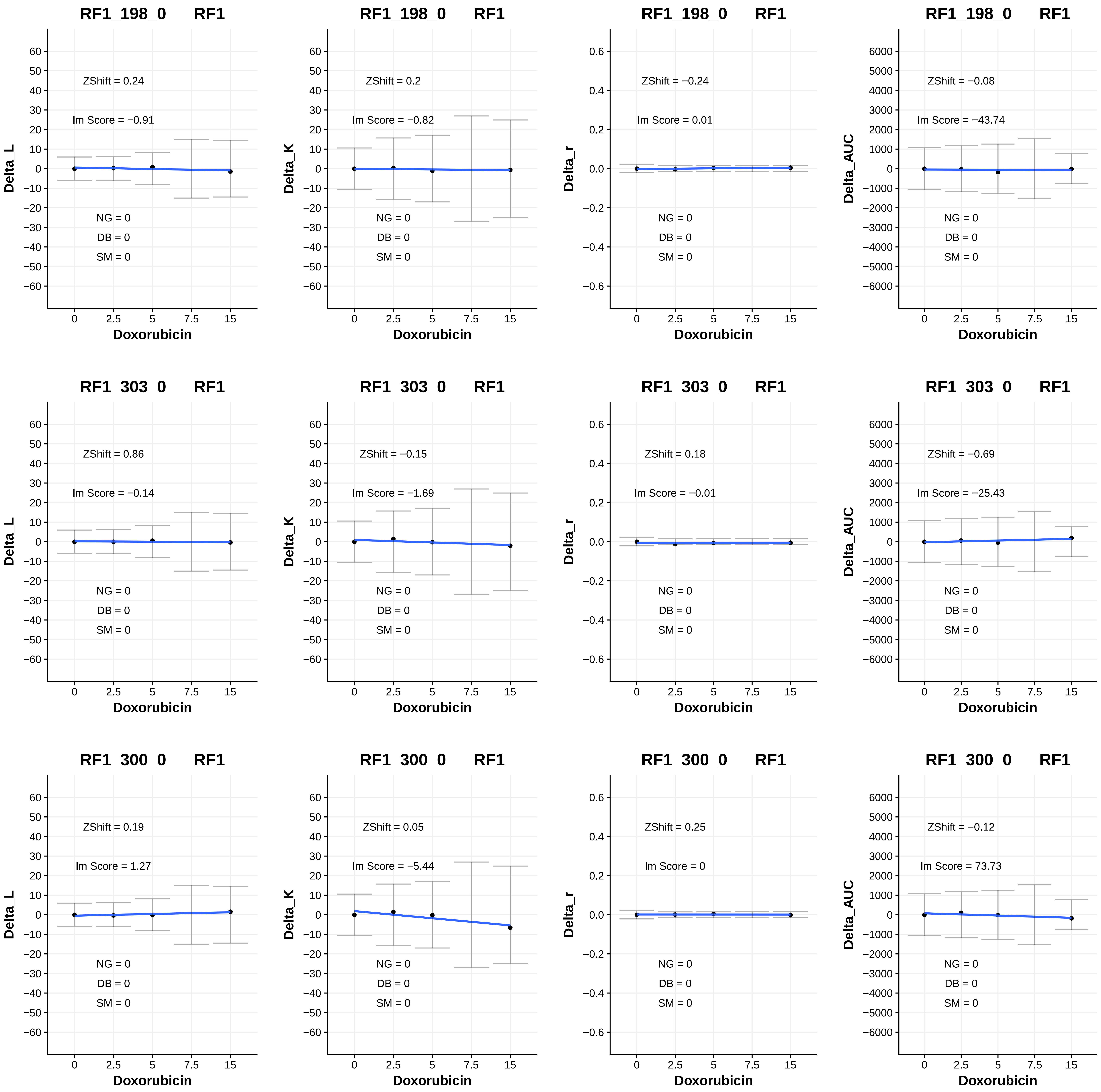
RF1_198_0 RF1
ZShift = 0.24
Im Score = −0.91
NG = 0
DB = 0
SM = 0
Delta_L
Doxorubicin
RF1_198_0 RF1
ZShift = 0.2
Im Score = −0.82
NG = 0
DB = 0
SM = 0
Delta_K
Doxorubicin
RF1_198_0 RF1
ZShift = −0.24
Im Score = 0.01
NG = 0
DB = 0
SM = 0
Delta_r
Doxorubicin
RF1_198_0 RF1
ZShift = −0.08
Im Score = −43.74
NG = 0
DB = 0
SM = 0
Delta_AUC
Doxorubicin
RF1_303_0 RF1
ZShift = 0.86
Im Score = −0.14
NG = 0
DB = 0
SM = 0
Delta_L
Doxorubicin
RF1_303_0 RF1
ZShift = −0.15
Im Score = −1.69
NG = 0
DB = 0
SM = 0
Delta_K
Doxorubicin
RF1_303_0 RF1
ZShift = 0.18
Im Score = −0.01
NG = 0
DB = 0
SM = 0
Delta_r
Doxorubicin
RF1_303_0 RF1
ZShift = −0.69
Im Score = −25.43
NG = 0
DB = 0
SM = 0
Delta_AUC
Doxorubicin
RF1_300_0 RF1
ZShift = 0.19
Im Score = 1.27
NG = 0
DB = 0
SM = 0
Delta_L
Doxorubicin
RF1_300_0 RF1
ZShift = 0.05
Im Score = −5.44
NG = 0
DB = 0
SM = 0
Delta_K
Doxorubicin
RF1_300_0 RF1
ZShift = 0.25
Im Score = 0
NG = 0
DB = 0
SM = 0
Delta_r
Doxorubicin
RF1_300_0 RF1
ZShift = −0.12
Im Score = 73.73
NG = 0
DB = 0
SM = 0
Delta_AUC
Doxorubicin

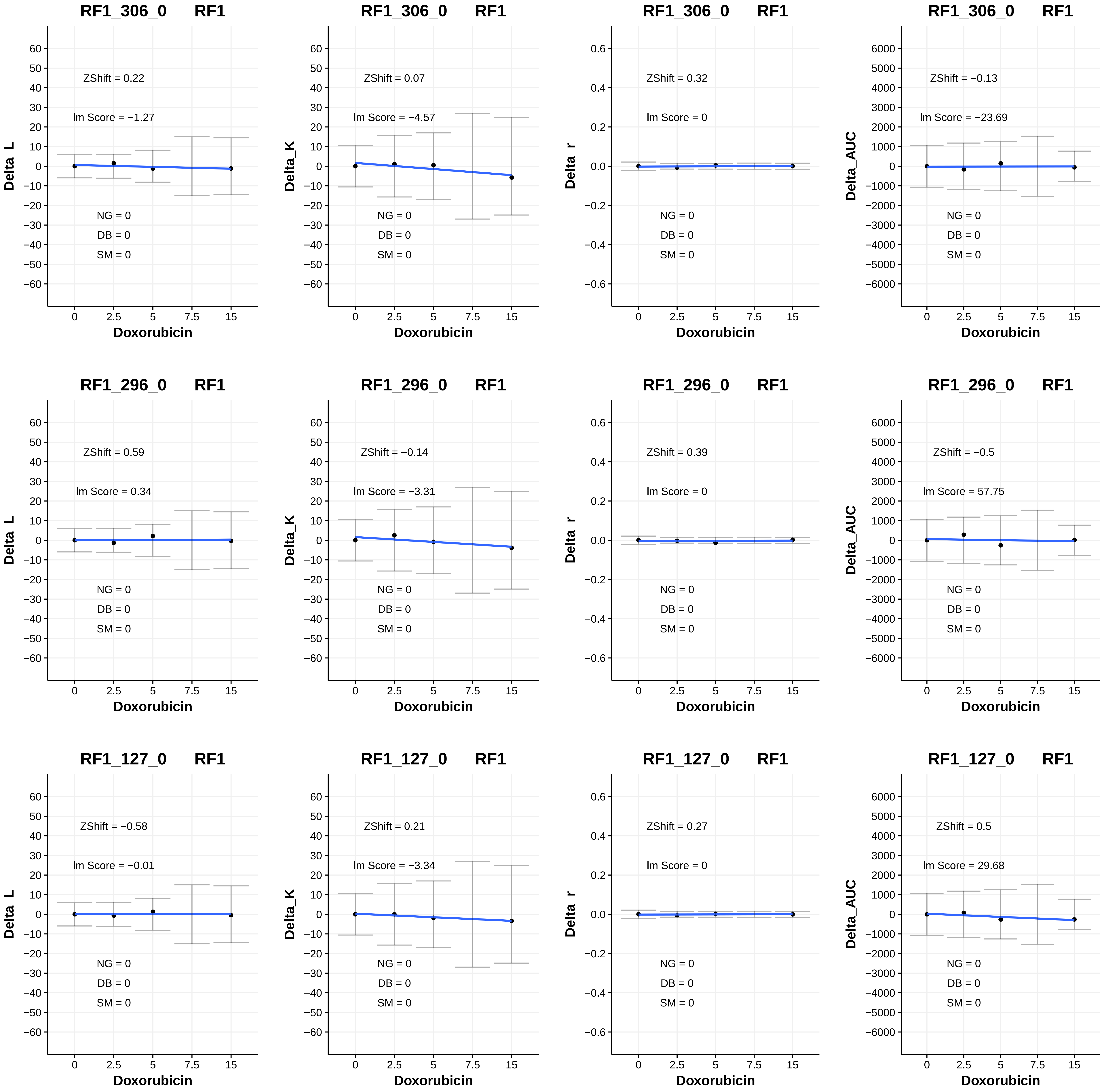

RF1_306_0 RF1
ZShift = 0.22
lm Score = −1.27
NG = 0
DB = 0
SM = 0
Delta_L
Doxorubicin
RF1_306_0 RF1
ZShift = 0.07
lm Score = −4.57
NG = 0
DB = 0
SM = 0
Delta_K
Doxorubicin
RF1_306_0 RF1
ZShift = 0.32
lm Score = 0
NG = 0
DB = 0
SM = 0
Delta_r
Doxorubicin
RF1_306_0 RF1
ZShift = −0.13
lm Score = −23.69
NG = 0
DB = 0
SM = 0
Delta_AUC
Doxorubicin
RF1_296_0 RF1
ZShift = 0.59
lm Score = 0.34
NG = 0
DB = 0
SM = 0
Delta_L
Doxorubicin
RF1_296_0 RF1
ZShift = −0.14
lm Score = −3.31
NG = 0
DB = 0
SM = 0
Delta_K
Doxorubicin
RF1_296_0 RF1
ZShift = 0.39
lm Score = 0
NG = 0
DB = 0
SM = 0
Delta_r
Doxorubicin
RF1_296_0 RF1
ZShift = −0.5
lm Score = 57.75
NG = 0
DB = 0
SM = 0
Delta_AUC
Doxorubicin
RF1_127_0 RF1
ZShift = −0.58
lm Score = −0.01
NG = 0
DB = 0
SM = 0
Delta_L
Doxorubicin
RF1_127_0 RF1
ZShift = 0.21
lm Score = −3.34
NG = 0
DB = 0
SM = 0
Delta_K
Doxorubicin
RF1_127_0 RF1
ZShift = 0.27
lm Score = 0
NG = 0
DB = 0
SM = 0
Delta_r
Doxorubicin
RF1_127_0 RF1
ZShift = 0.5
lm Score = 29.68
NG = 0
DB = 0
SM = 0
Delta_AUC
Doxorubicin

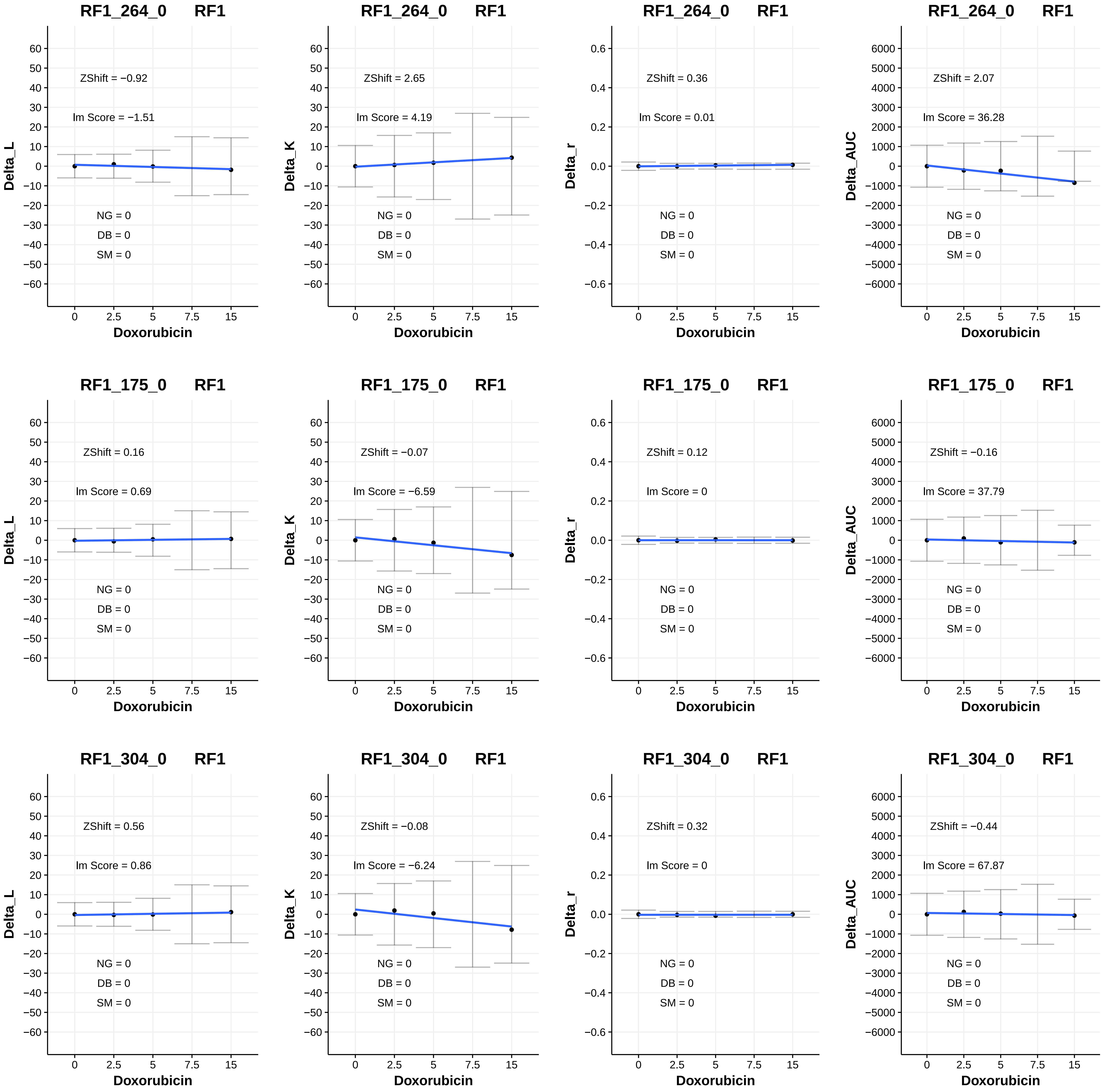

RF1_264_0 RF1
ZShift = −0.92
lm Score = −1.51
NG = 0
DB = 0
SM = 0
Delta_L
Doxorubicin
RF1_264_0 RF1
ZShift = 2.65
lm Score = 4.19
NG = 0
DB = 0
SM = 0
Delta_K
Doxorubicin
RF1_264_0 RF1
ZShift = 0.36
lm Score = 0.01
NG = 0
DB = 0
SM = 0
Delta_r
Doxorubicin
RF1_264_0 RF1
ZShift = 2.07
lm Score = 36.28
NG = 0
DB = 0
SM = 0
Delta_AUC
Doxorubicin
RF1_175_0 RF1
ZShift = 0.16
lm Score = 0.69
NG = 0
DB = 0
SM = 0
Delta_L
Doxorubicin
RF1_175_0 RF1
ZShift = −0.07
lm Score = −6.59
NG = 0
DB = 0
SM = 0
Delta_K
Doxorubicin
RF1_175_0 RF1
ZShift = 0.12
lm Score = 0
NG = 0
DB = 0
SM = 0
Delta_r
Doxorubicin
RF1_175_0 RF1
ZShift = −0.16
lm Score = 37.79
NG = 0
DB = 0
SM = 0
Delta_AUC
Doxorubicin
RF1_304_0 RF1
ZShift = 0.56
lm Score = 0.86
NG = 0
DB = 0
SM = 0
Delta_L
Doxorubicin
RF1_304_0 RF1
ZShift = −0.08
lm Score = −6.24
NG = 0
DB = 0
SM = 0
Delta_K
Doxorubicin
RF1_304_0 RF1
ZShift = 0.32
lm Score = 0
NG = 0
DB = 0
SM = 0
Delta_r
Doxorubicin
RF1_304_0 RF1
ZShift = −0.44
lm Score = 67.87
NG = 0
DB = 0
SM = 0
Delta_AUC
Doxorubicin

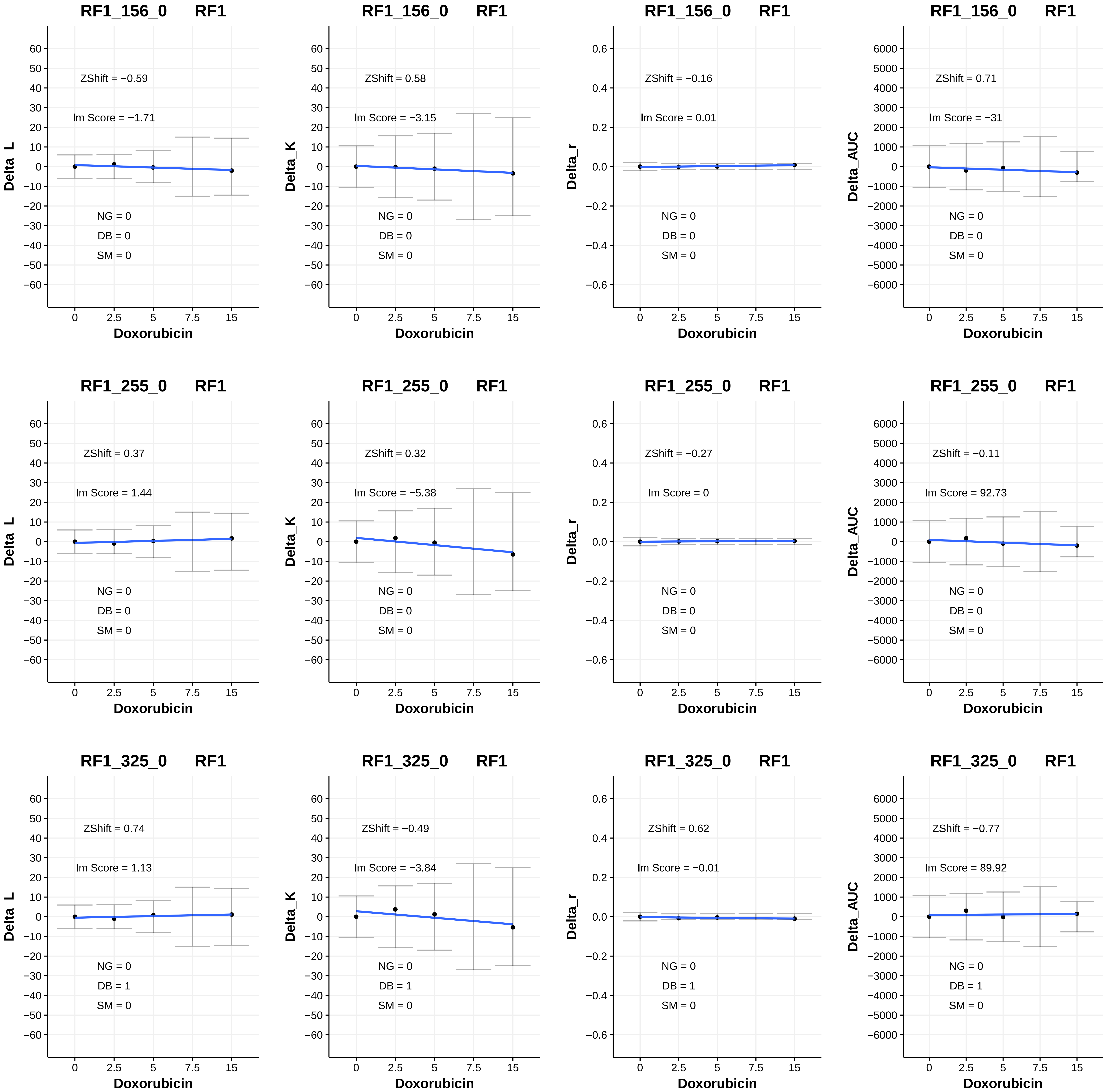
RF1_156_0 RF1
ZShift = −0.59
lm Score = −1.71
NG = 0
DB = 0
SM = 0
Delta_L
Doxorubicin
RF1_156_0 RF1
ZShift = 0.58
lm Score = −3.15
NG = 0
DB = 0
SM = 0
Delta_K
Doxorubicin
RF1_156_0 RF1
ZShift = −0.16
lm Score = 0.01
NG = 0
DB = 0
SM = 0
Delta_r
Doxorubicin
RF1_156_0 RF1
ZShift = 0.71
lm Score = −31
NG = 0
DB = 0
SM = 0
Delta_AUC
Doxorubicin
RF1_255_0 RF1
ZShift = 0.37
lm Score = 1.44
NG = 0
DB = 0
SM = 0
Delta_L
Doxorubicin
RF1_255_0 RF1
ZShift = 0.32
lm Score = −5.38
NG = 0
DB = 0
SM = 0
Delta_K
Doxorubicin
RF1_255_0 RF1
ZShift = −0.27
lm Score = 0
NG = 0
DB = 0
SM = 0
Delta_r
Doxorubicin
RF1_255_0 RF1
ZShift = −0.11
lm Score = 92.73
NG = 0
DB = 0
SM = 0
Delta_AUC
Doxorubicin
RF1_325_0 RF1
ZShift = 0.74
lm Score = 1.13
NG = 0
DB = 1
SM = 0
Delta_L
Doxorubicin
RF1_325_0 RF1
ZShift = −0.49
lm Score = −3.84
NG = 0
DB = 1
SM = 0
Delta_K
Doxorubicin
RF1_325_0 RF1
ZShift = 0.62
lm Score = −0.01
NG = 0
DB = 1
SM = 0
Delta_r
Doxorubicin
RF1_325_0 RF1
ZShift = −0.77
lm Score = 89.92
NG = 0
DB = 1
SM = 0
Delta_AUC
Doxorubicin

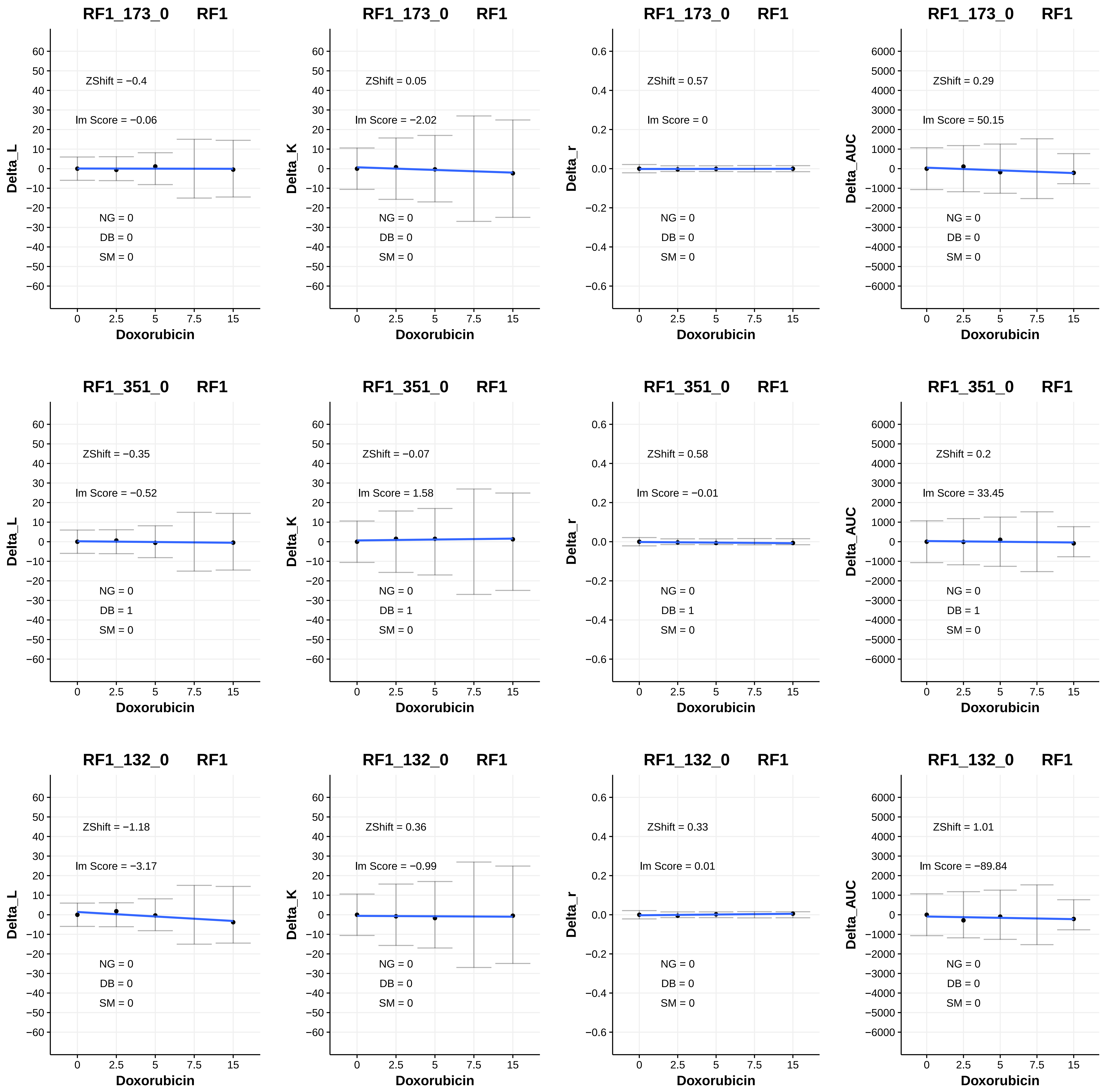
RF1_173_0 RF1
ZShift = −0.4
lm Score = −0.06
NG = 0
DB = 0
SM = 0
Delta_L
Doxorubicin
RF1_173_0 RF1
ZShift = 0.05
lm Score = −2.02
NG = 0
DB = 0
SM = 0
Delta_K
Doxorubicin
RF1_173_0 RF1
ZShift = 0.57
lm Score = 0
NG = 0
DB = 0
SM = 0
Delta_r
Doxorubicin
RF1_173_0 RF1
ZShift = 0.29
lm Score = 50.15
NG = 0
DB = 0
SM = 0
Delta_AUC
Doxorubicin
RF1_351_0 RF1
ZShift = −0.35
lm Score = −0.52
NG = 0
DB = 1
SM = 0
Delta_L
Doxorubicin
RF1_351_0 RF1
ZShift = −0.07
lm Score = 1.58
NG = 0
DB = 1
SM = 0
Delta_K
Doxorubicin
RF1_351_0 RF1
ZShift = 0.58
lm Score = −0.01
NG = 0
DB = 1
SM = 0
Delta_r
Doxorubicin
RF1_351_0 RF1
ZShift = 0.2
lm Score = 33.45
NG = 0
DB = 1
SM = 0
Delta_AUC
Doxorubicin
RF1_132_0 RF1
ZShift = −1.18
lm Score = −3.17
NG = 0
DB = 0
SM = 0
Delta_L
Doxorubicin
RF1_132_0 RF1
ZShift = 0.36
lm Score = −0.99
NG = 0
DB = 0
SM = 0
Delta_K
Doxorubicin
RF1_132_0 RF1
ZShift = 0.33
lm Score = 0.01
NG = 0
DB = 0
SM = 0
Delta_r
Doxorubicin
RF1_132_0 RF1
ZShift = 1.01
lm Score = −89.84
NG = 0
DB = 0
SM = 0
Delta_AUC
Doxorubicin

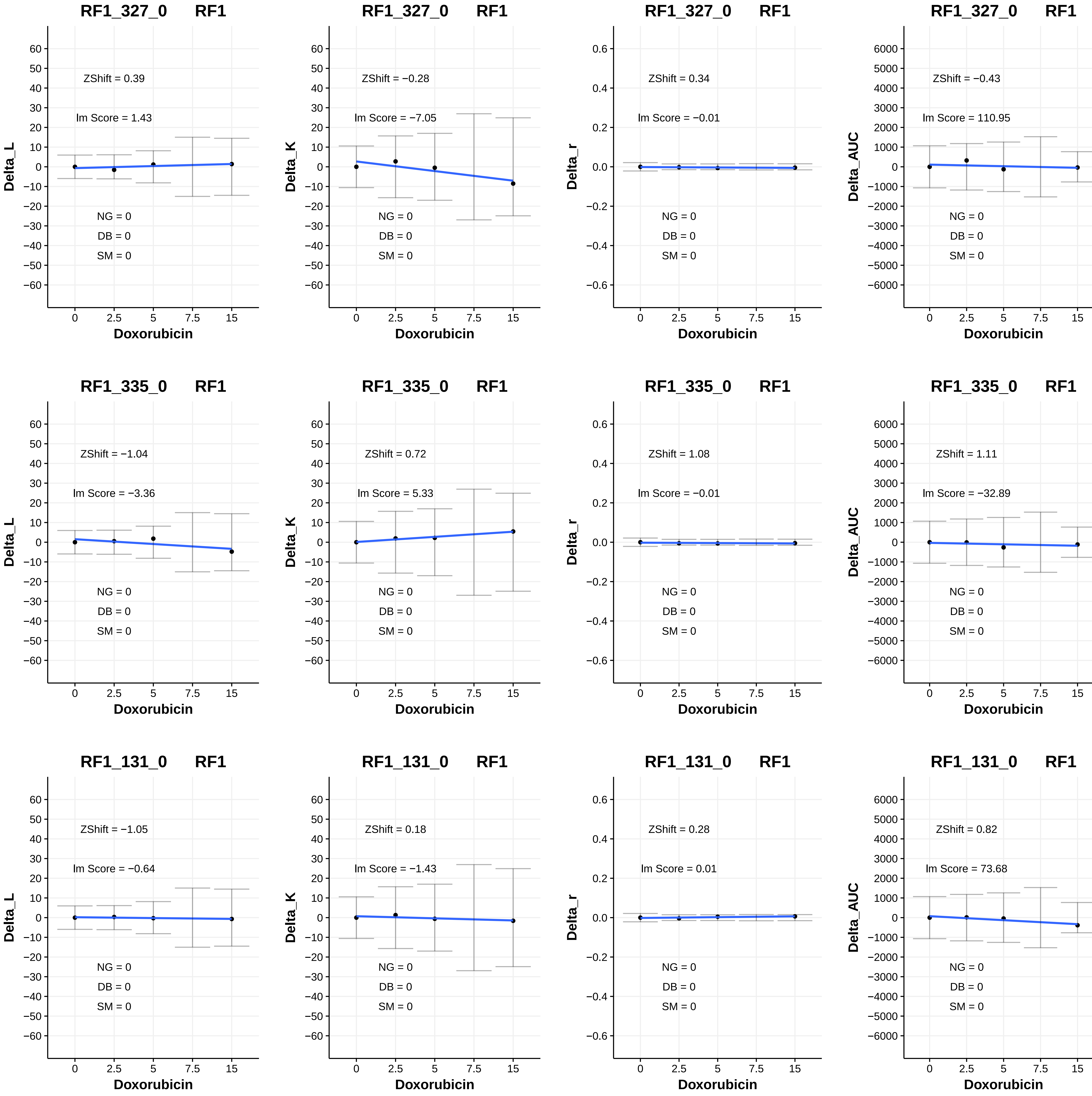
RF1_327_0 RF1
ZShift = 0.39
lm Score = 1.43
NG = 0
DB = 0
SM = 0
Delta_L
RF1_327_0 RF1
ZShift = −0.28
lm Score = −7.05
NG = 0
DB = 0
SM = 0
Delta_K
RF1_327_0 RF1
ZShift = 0.34
lm Score = −0.01
NG = 0
DB = 0
SM = 0
Delta_r
RF1_327_0 RF1
ZShift = −0.43
lm Score = 110.95
NG = 0
DB = 0
SM = 0
Delta_AUC
RF1_335_0 RF1
ZShift = −1.04
lm Score = −3.36
NG = 0
DB = 0
SM = 0
Delta_L
RF1_335_0 RF1
ZShift = 0.72
lm Score = 5.33
NG = 0
DB = 0
SM = 0
Delta_K
RF1_335_0 RF1
ZShift = 1.08
lm Score = −0.01
NG = 0
DB = 0
SM = 0
Delta_r
RF1_335_0 RF1
ZShift = 1.11
lm Score = −32.89
NG = 0
DB = 0
SM = 0
Delta_AUC
RF1_131_0 RF1
ZShift = −1.05
lm Score = −0.64
NG = 0
DB = 0
SM = 0
Delta_L
RF1_131_0 RF1
ZShift = 0.18
lm Score = −1.43
NG = 0
DB = 0
SM = 0
Delta_K
RF1_131_0 RF1
ZShift = 0.28
lm Score = 0.01
NG = 0
DB = 0
SM = 0
Delta_r
RF1_131_0 RF1
ZShift = 0.82
lm Score = 73.68
NG = 0
DB = 0
SM = 0
Delta_AUC
Doxorubicin
0
2.5
5
7.5
15

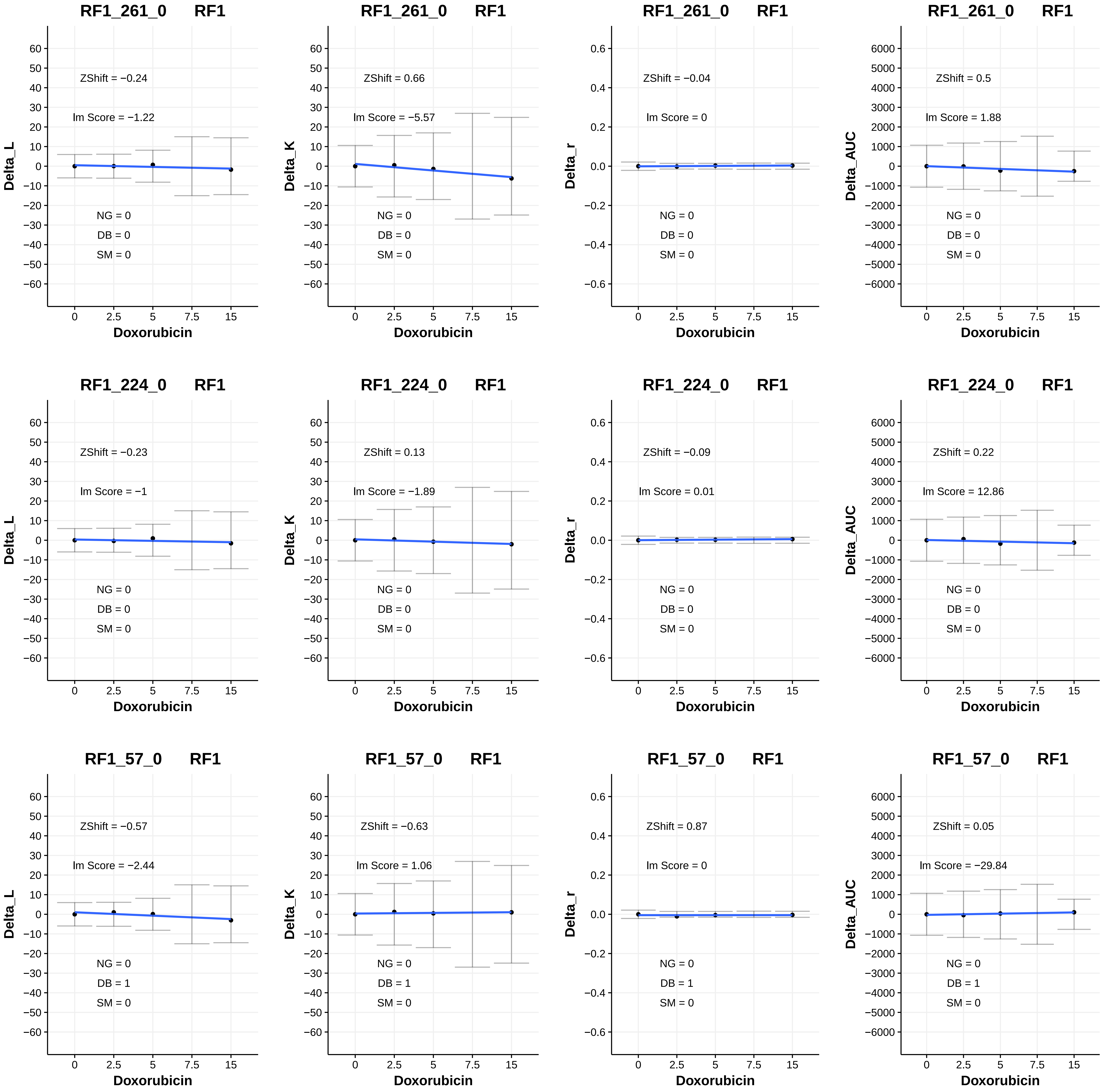

RF1_261_0 RF1
ZShift = −0.24
lm Score = −1.22
NG = 0
DB = 0
SM = 0
Delta_L
Doxorubicin
RF1_261_0 RF1
ZShift = 0.66
lm Score = −5.57
NG = 0
DB = 0
SM = 0
Delta_K
Doxorubicin
RF1_261_0 RF1
ZShift = −0.04
lm Score = 0
NG = 0
DB = 0
SM = 0
Delta_r
Doxorubicin
RF1_261_0 RF1
ZShift = 0.5
lm Score = 1.88
NG = 0
DB = 0
SM = 0
Delta_AUC
Doxorubicin
RF1_224_0 RF1
ZShift = −0.23
lm Score = −1
NG = 0
DB = 0
SM = 0
Delta_L
Doxorubicin
RF1_224_0 RF1
ZShift = 0.13
lm Score = −1.89
NG = 0
DB = 0
SM = 0
Delta_K
Doxorubicin
RF1_224_0 RF1
ZShift = −0.09
lm Score = 0.01
NG = 0
DB = 0
SM = 0
Delta_r
Doxorubicin
RF1_224_0 RF1
ZShift = 0.22
lm Score = 12.86
NG = 0
DB = 0
SM = 0
Delta_AUC
Doxorubicin
RF1_57_0 RF1
ZShift = −0.57
lm Score = −2.44
NG = 0
DB = 1
SM = 0
Delta_L
Doxorubicin
RF1_57_0 RF1
ZShift = −0.63
lm Score = 1.06
NG = 0
DB = 1
SM = 0
Delta_K
Doxorubicin
RF1_57_0 RF1
ZShift = 0.87
lm Score = 0
NG = 0
DB = 1
SM = 0
Delta_r
Doxorubicin
RF1_57_0 RF1
ZShift = 0.05
lm Score = −29.84
NG = 0
DB = 1
SM = 0
Delta_AUC
Doxorubicin

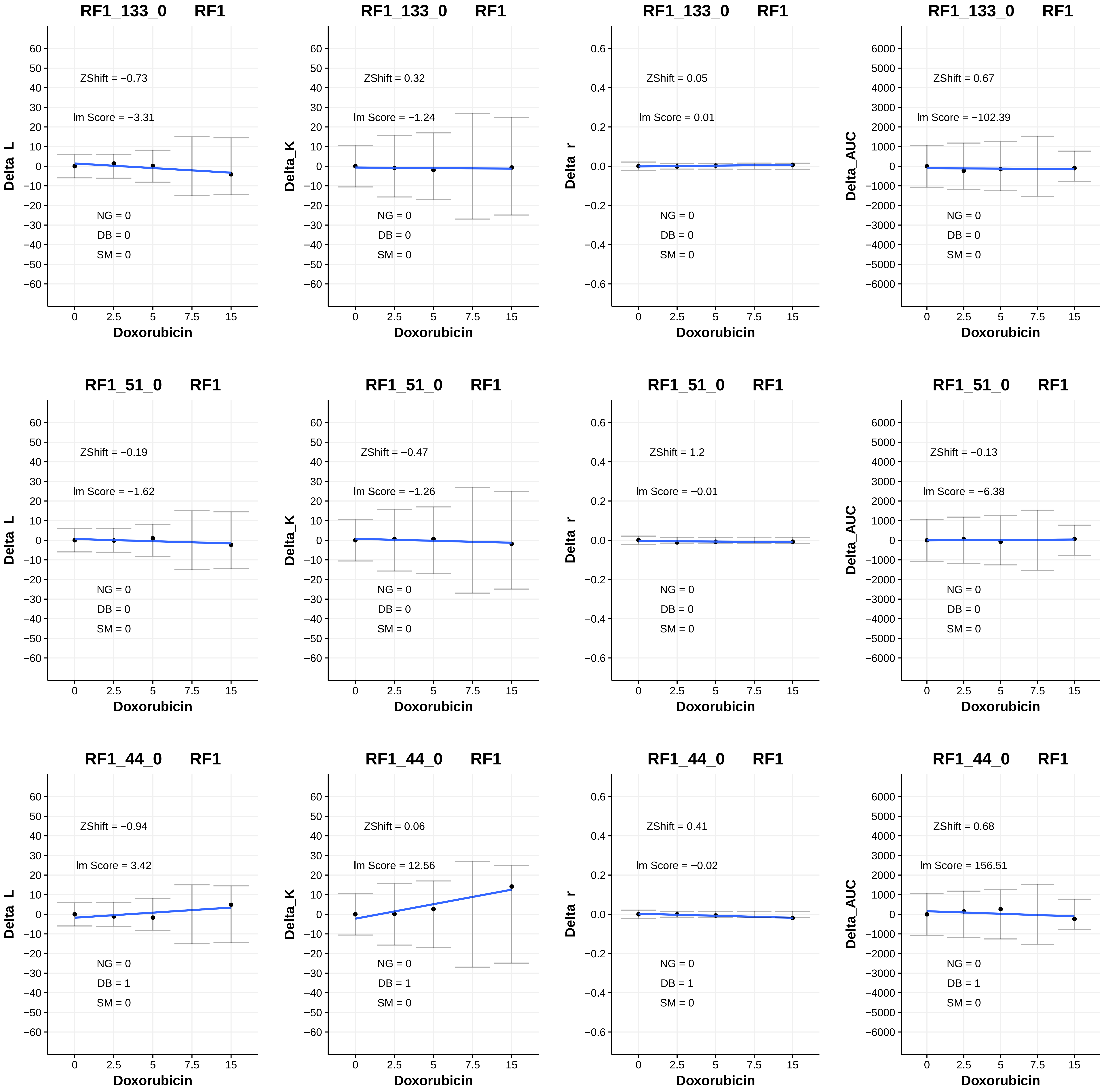

RF1_133_0 RF1
ZShift = −0.73
lm Score = −3.31
NG = 0
DB = 0
SM = 0
Delta_L
Doxorubicin
RF1_133_0 RF1
ZShift = 0.32
lm Score = −1.24
NG = 0
DB = 0
SM = 0
Delta_K
Doxorubicin
RF1_133_0 RF1
ZShift = 0.05
lm Score = 0.01
NG = 0
DB = 0
SM = 0
Delta_r
Doxorubicin
RF1_133_0 RF1
ZShift = 0.67
lm Score = −102.39
NG = 0
DB = 0
SM = 0
Delta_AUC
Doxorubicin
RF1_51_0 RF1
ZShift = −0.19
lm Score = −1.62
NG = 0
DB = 0
SM = 0
Delta_L
Doxorubicin
RF1_51_0 RF1
ZShift = −0.47
lm Score = −1.26
NG = 0
DB = 0
SM = 0
Delta_K
Doxorubicin
RF1_51_0 RF1
ZShift = 1.2
lm Score = −0.01
NG = 0
DB = 0
SM = 0
Delta_r
Doxorubicin
RF1_51_0 RF1
ZShift = −0.13
lm Score = −6.38
NG = 0
DB = 0
SM = 0
Delta_AUC
Doxorubicin
RF1_44_0 RF1
ZShift = −0.94
lm Score = 3.42
NG = 0
DB = 1
SM = 0
Delta_L
Doxorubicin
RF1_44_0 RF1
ZShift = 0.06
lm Score = 12.56
NG = 0
DB = 1
SM = 0
Delta_K
Doxorubicin
RF1_44_0 RF1
ZShift = 0.41
lm Score = −0.02
NG = 0
DB = 1
SM = 0
Delta_r
Doxorubicin
RF1_44_0 RF1
ZShift = 0.68
lm Score = 156.51
NG = 0
DB = 1
SM = 0
Delta_AUC
Doxorubicin

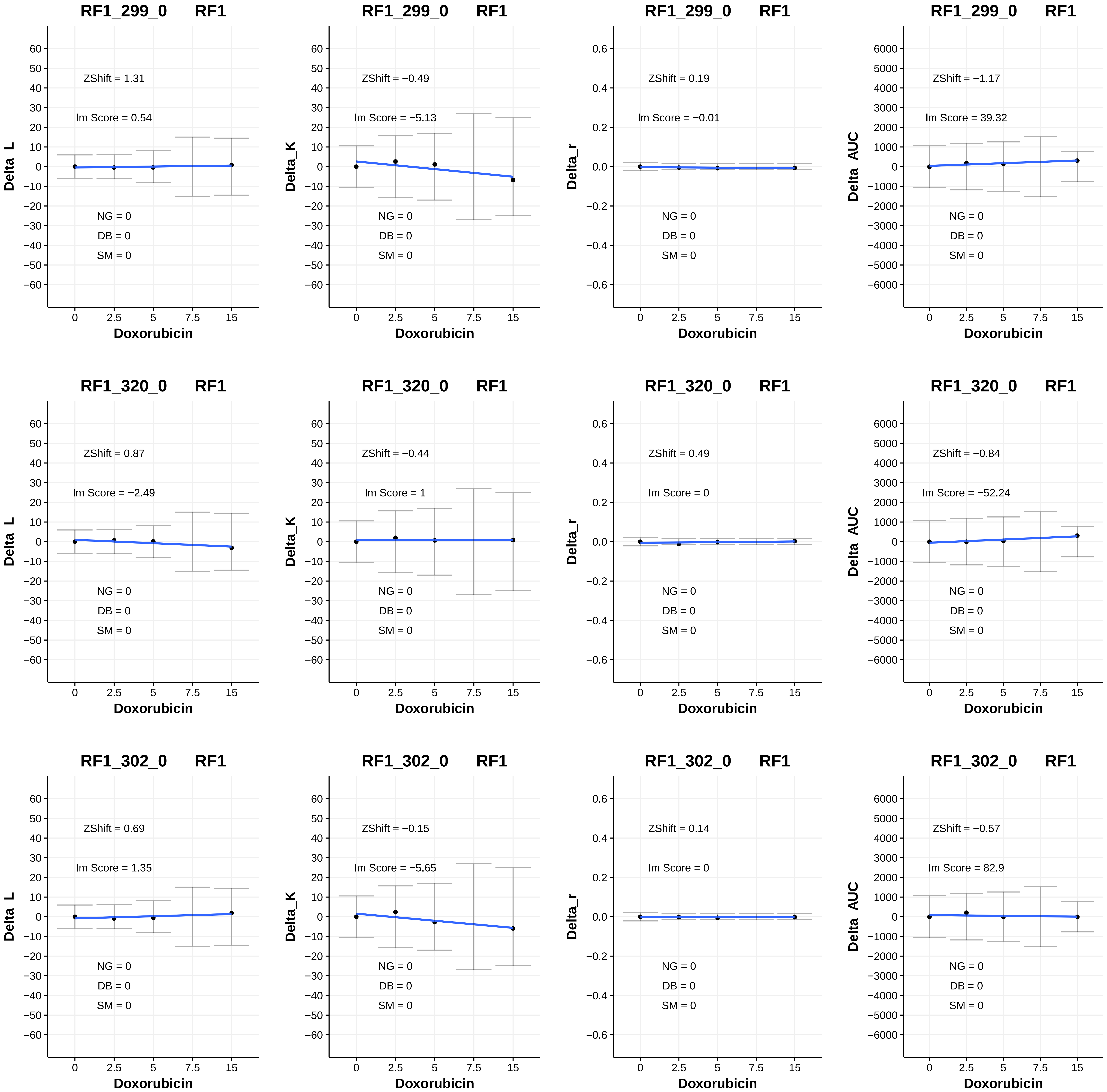
RF1_299_0 RF1
ZShift = 1.31
Im Score = 0.54
NG = 0
DB = 0
SM = 0
Delta_L
Doxorubicin
RF1_299_0 RF1
ZShift = −0.49
Im Score = −5.13
NG = 0
DB = 0
SM = 0
Delta_K
Doxorubicin
RF1_299_0 RF1
ZShift = 0.19
Im Score = −0.01
NG = 0
DB = 0
SM = 0
Delta_r
Doxorubicin
RF1_299_0 RF1
ZShift = −1.17
Im Score = 39.32
NG = 0
DB = 0
SM = 0
Delta_AUC
Doxorubicin
RF1_320_0 RF1
ZShift = 0.87
Im Score = −2.49
NG = 0
DB = 0
SM = 0
Delta_L
Doxorubicin
RF1_320_0 RF1
ZShift = −0.44
Im Score = 1
NG = 0
DB = 0
SM = 0
Delta_K
Doxorubicin
RF1_320_0 RF1
ZShift = 0.49
Im Score = 0
NG = 0
DB = 0
SM = 0
Delta_r
Doxorubicin
RF1_320_0 RF1
ZShift = −0.84
Im Score = −52.24
NG = 0
DB = 0
SM = 0
Delta_AUC
Doxorubicin
RF1_302_0 RF1
ZShift = 0.69
Im Score = 1.35
NG = 0
DB = 0
SM = 0
Delta_L
Doxorubicin
RF1_302_0 RF1
ZShift = −0.15
Im Score = −5.65
NG = 0
DB = 0
SM = 0
Delta_K
Doxorubicin
RF1_302_0 RF1
ZShift = 0.14
Im Score = 0
NG = 0
DB = 0
SM = 0
Delta_r
Doxorubicin
RF1_302_0 RF1
ZShift = −0.57
Im Score = 82.9
NG = 0
DB = 0
SM = 0
Delta_AUC
Doxorubicin

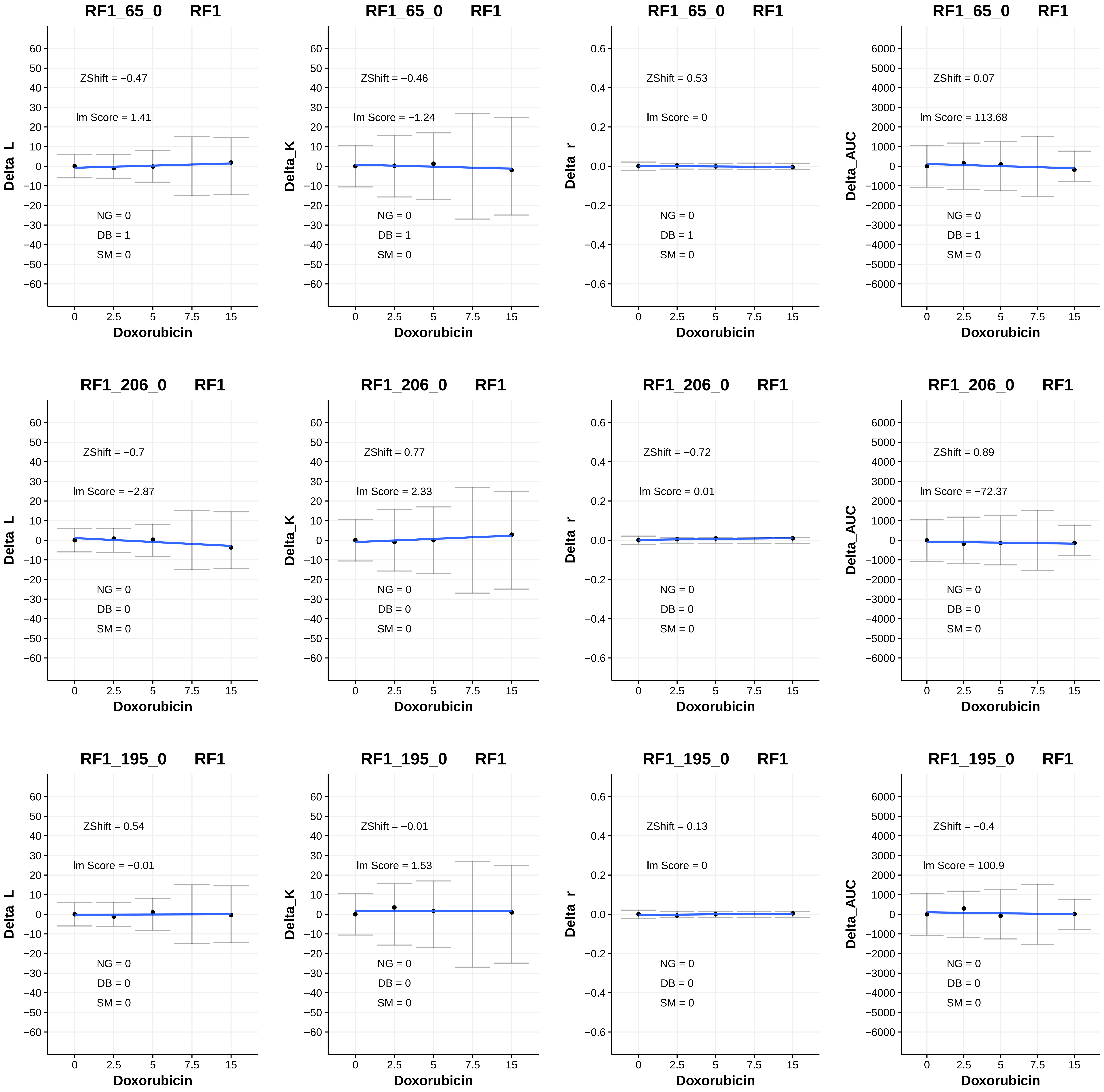

RF1_65_0 RF1
ZShift = −0.47
lm Score = 1.41
NG = 0
DB = 1
SM = 0
Delta_L
Doxorubicin
RF1_65_0 RF1
ZShift = −0.46
lm Score = −1.24
NG = 0
DB = 1
SM = 0
Delta_K
Doxorubicin
RF1_65_0 RF1
ZShift = 0.53
lm Score = 0
NG = 0
DB = 1
SM = 0
Delta_r
Doxorubicin
RF1_65_0 RF1
ZShift = 0.07
lm Score = 113.68
NG = 0
DB = 1
SM = 0
Delta_AUC
Doxorubicin
RF1_206_0 RF1
ZShift = −0.7
lm Score = −2.87
NG = 0
DB = 0
SM = 0
Delta_L
Doxorubicin
RF1_206_0 RF1
ZShift = 0.77
lm Score = 2.33
NG = 0
DB = 0
SM = 0
Delta_K
Doxorubicin
RF1_206_0 RF1
ZShift = −0.72
lm Score = 0.01
NG = 0
DB = 0
SM = 0
Delta_r
Doxorubicin
RF1_206_0 RF1
ZShift = 0.89
lm Score = −72.37
NG = 0
DB = 0
SM = 0
Delta_AUC
Doxorubicin
RF1_195_0 RF1
ZShift = 0.54
lm Score = −0.01
NG = 0
DB = 0
SM = 0
Delta_L
Doxorubicin
RF1_195_0 RF1
ZShift = −0.01
lm Score = 1.53
NG = 0
DB = 0
SM = 0
Delta_K
Doxorubicin
RF1_195_0 RF1
ZShift = 0.13
lm Score = 0
NG = 0
DB = 0
SM = 0
Delta_r
Doxorubicin
RF1_195_0 RF1
ZShift = −0.4
lm Score = 100.9
NG = 0
DB = 0
SM = 0
Delta_AUC
Doxorubicin

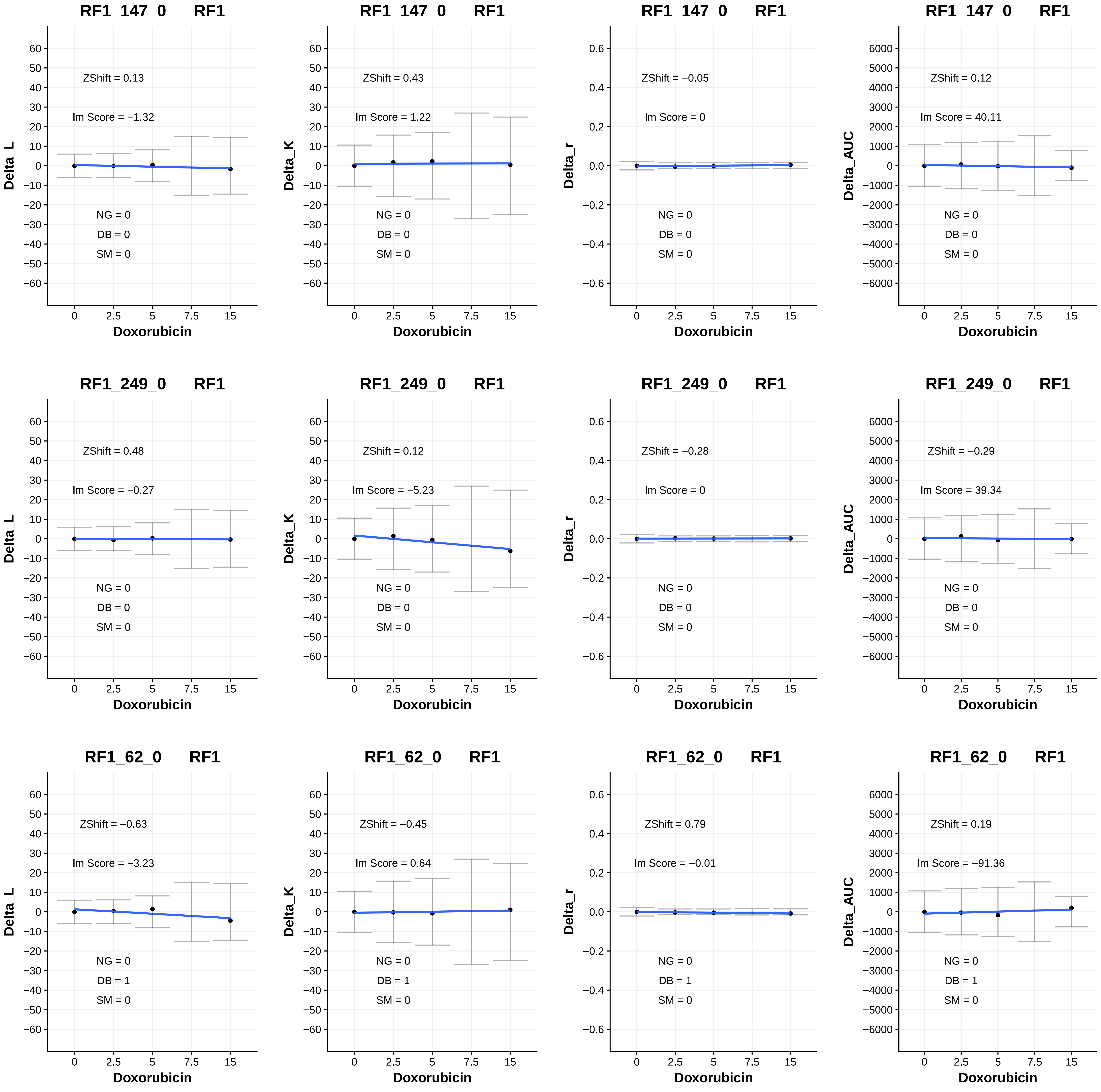

RF1_147_0 RF1
ZShift = 0.13
lm Score = −1.32
NG = 0
DB = 0
SM = 0
Delta_L
Doxorubicin
RF1_147_0 RF1
ZShift = 0.43
lm Score = 1.22
NG = 0
DB = 0
SM = 0
Delta_K
Doxorubicin
RF1_147_0 RF1
ZShift = −0.05
lm Score = 0
NG = 0
DB = 0
SM = 0
Delta_r
Doxorubicin
RF1_147_0 RF1
ZShift = 0.12
lm Score = 40.11
NG = 0
DB = 0
SM = 0
Delta_AUC
Doxorubicin
RF1_249_0 RF1
ZShift = 0.48
lm Score = −0.27
NG = 0
DB = 0
SM = 0
Delta_L
Doxorubicin
RF1_249_0 RF1
ZShift = 0.12
lm Score = −5.23
NG = 0
DB = 0
SM = 0
Delta_K
Doxorubicin
RF1_249_0 RF1
ZShift = −0.28
lm Score = 0
NG = 0
DB = 0
SM = 0
Delta_r
Doxorubicin
RF1_249_0 RF1
ZShift = −0.29
lm Score = 39.34
NG = 0
DB = 0
SM = 0
Delta_AUC
Doxorubicin
RF1_62_0 RF1
ZShift = −0.63
lm Score = −3.23
NG = 0
DB = 1
SM = 0
Delta_L
Doxorubicin
RF1_62_0 RF1
ZShift = −0.45
lm Score = 0.64
NG = 0
DB = 1
SM = 0
Delta_K
Doxorubicin
RF1_62_0 RF1
ZShift = 0.79
lm Score = −0.01
NG = 0
DB = 1
SM = 0
Delta_r
Doxorubicin
RF1_62_0 RF1
ZShift = 0.19
lm Score = −91.36
NG = 0
DB = 1
SM = 0
Delta_AUC
Doxorubicin

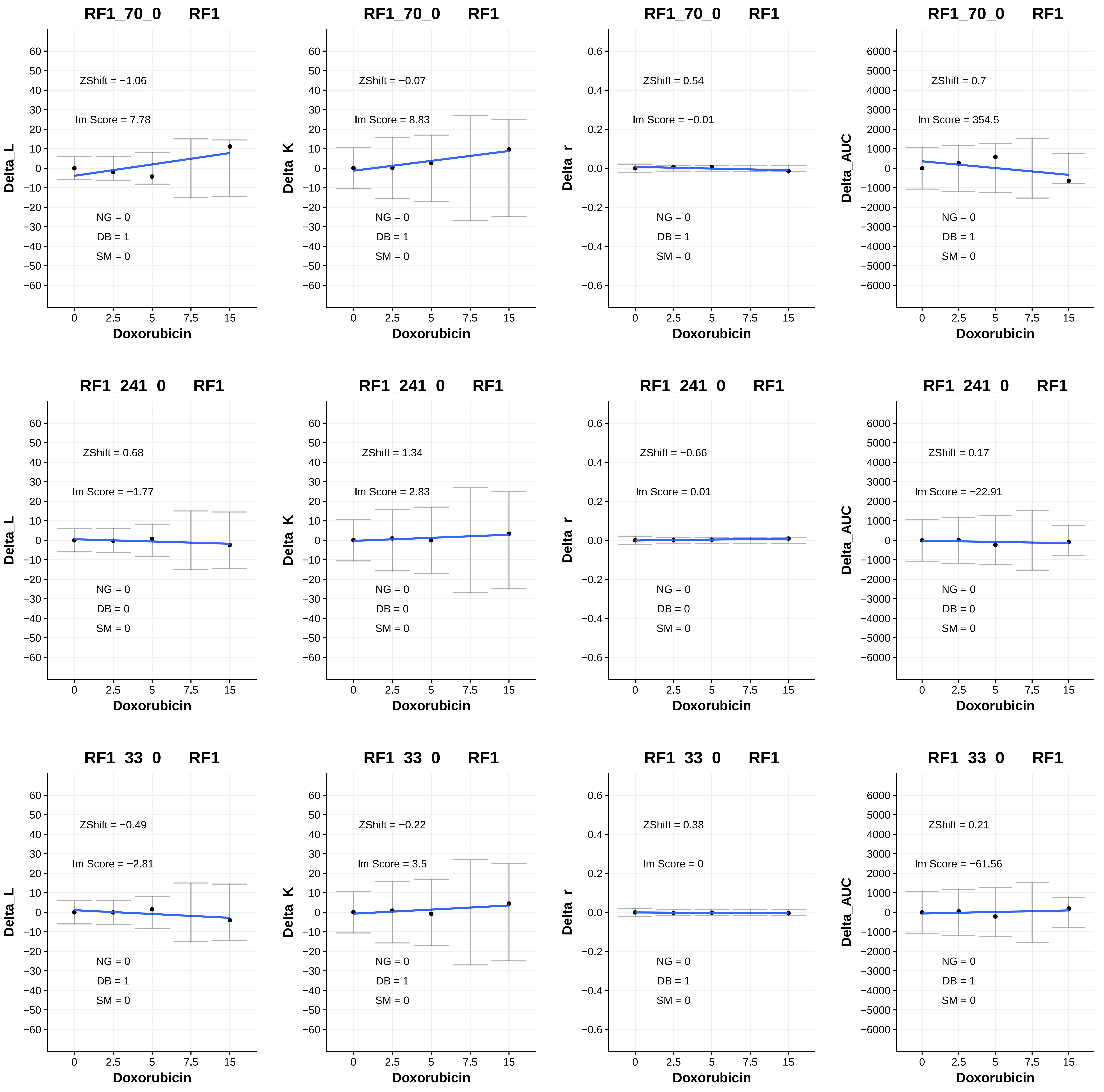
RF1_70_0 RF1
ZShift = −1.06
lm Score = 7.78
NG = 0
DB = 1
SM = 0
Delta_L
Doxorubicin
RF1_70_0 RF1
ZShift = −0.07
lm Score = 8.83
NG = 0
DB = 1
SM = 0
Delta_K
Doxorubicin
RF1_70_0 RF1
ZShift = 0.54
lm Score = −0.01
NG = 0
DB = 1
SM = 0
Delta_r
Doxorubicin
RF1_70_0 RF1
ZShift = 0.7
lm Score = 354.5
NG = 0
DB = 1
SM = 0
Delta_AUC
Doxorubicin
RF1_241_0 RF1
ZShift = 0.68
lm Score = −1.77
NG = 0
DB = 0
SM = 0
Delta_L
Doxorubicin
RF1_241_0 RF1
ZShift = 1.34
lm Score = 2.83
NG = 0
DB = 0
SM = 0
Delta_K
Doxorubicin
RF1_241_0 RF1
ZShift = −0.66
lm Score = 0.01
NG = 0
DB = 0
SM = 0
Delta_r
Doxorubicin
RF1_241_0 RF1
ZShift = 0.17
lm Score = −22.91
NG = 0
DB = 0
SM = 0
Delta_AUC
Doxorubicin
RF1_33_0 RF1
ZShift = −0.49
lm Score = −2.81
NG = 0
DB = 1
SM = 0
Delta_L
Doxorubicin
RF1_33_0 RF1
ZShift = −0.22
lm Score = 3.5
NG = 0
DB = 1
SM = 0
Delta_K
Doxorubicin
RF1_33_0 RF1
ZShift = 0.38
lm Score = 0
NG = 0
DB = 1
SM = 0
Delta_r
Doxorubicin
RF1_33_0 RF1
ZShift = 0.21
lm Score = −61.56
NG = 0
DB = 1
SM = 0
Delta_AUC
Doxorubicin
0 2.5 5 7.5 15

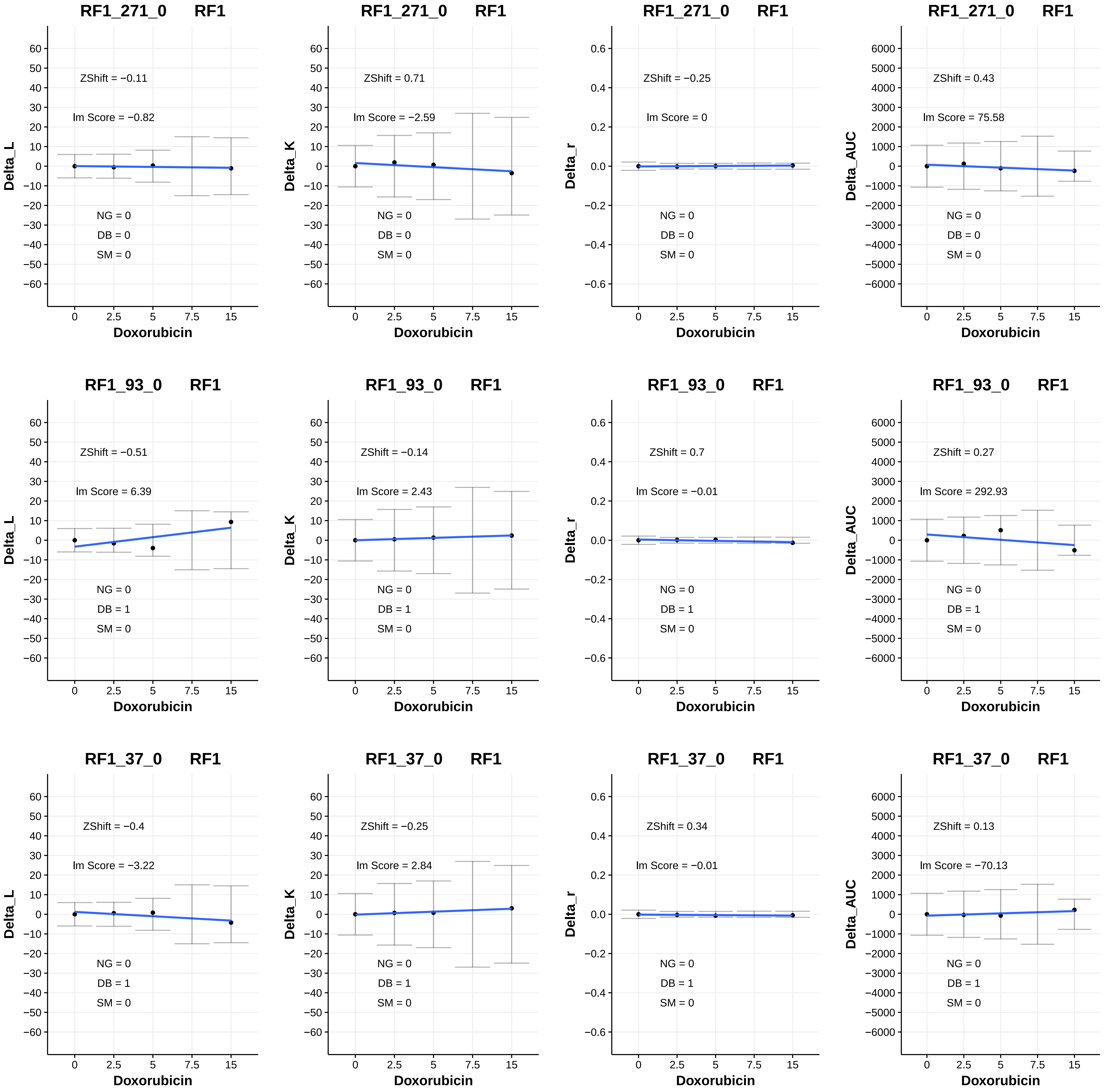
RF1_271_0 RF1
ZShift = −0.11
lm Score = −0.82
NG = 0
DB = 0
SM = 0
Delta_L
Doxorubicin
RF1_271_0 RF1
ZShift = 0.71
lm Score = −2.59
NG = 0
DB = 0
SM = 0
Delta_K
Doxorubicin
RF1_271_0 RF1
ZShift = −0.25
lm Score = 0
NG = 0
DB = 0
SM = 0
Delta_r
Doxorubicin
RF1_271_0 RF1
ZShift = 0.43
lm Score = 75.58
NG = 0
DB = 0
SM = 0
Delta_AUC
Doxorubicin
RF1_93_0 RF1
ZShift = −0.51
lm Score = 6.39
NG = 0
DB = 1
SM = 0
Delta_L
Doxorubicin
RF1_93_0 RF1
ZShift = −0.14
lm Score = 2.43
NG = 0
DB = 1
SM = 0
Delta_K
Doxorubicin
RF1_93_0 RF1
ZShift = 0.7
lm Score = −0.01
NG = 0
DB = 1
SM = 0
Delta_r
Doxorubicin
RF1_93_0 RF1
ZShift = 0.27
lm Score = 292.93
NG = 0
DB = 1
SM = 0
Delta_AUC
Doxorubicin
RF1_37_0 RF1
ZShift = −0.4
lm Score = −3.22
NG = 0
DB = 1
SM = 0
Delta_L
Doxorubicin
RF1_37_0 RF1
ZShift = −0.25
lm Score = 2.84
NG = 0
DB = 1
SM = 0
Delta_K
Doxorubicin
RF1_37_0 RF1
ZShift = 0.34
lm Score = −0.01
NG = 0
DB = 1
SM = 0
Delta_r
Doxorubicin
RF1_37_0 RF1
ZShift = 0.13
lm Score = −70.13
NG = 0
DB = 1
SM = 0
Delta_AUC
Doxorubicin

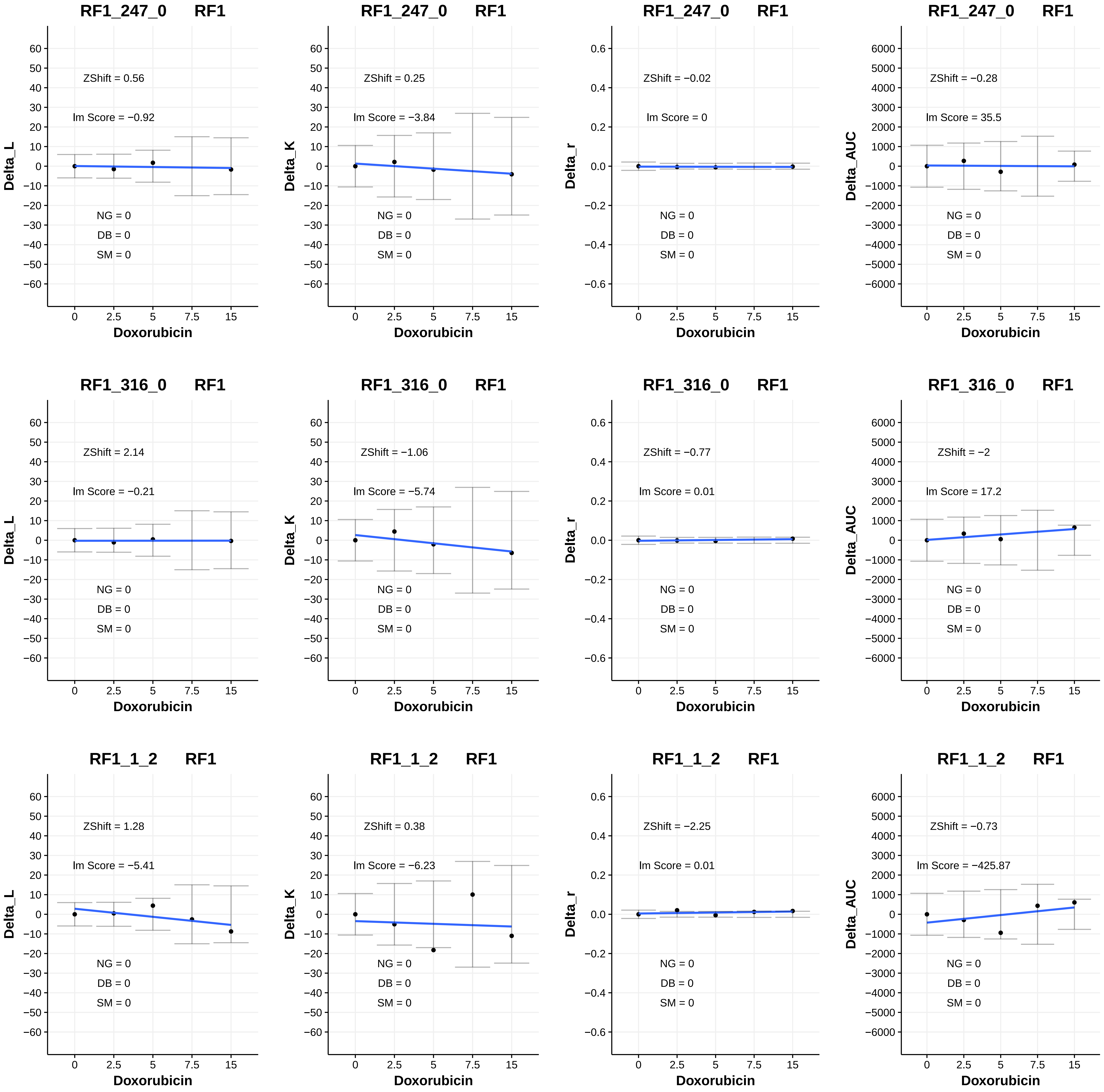
RF1_247_0 RF1
ZShift = 0.56
Im Score = −0.92
NG = 0
DB = 0
SM = 0
Delta_L
Doxorubicin
RF1_247_0 RF1
ZShift = 0.25
Im Score = −3.84
NG = 0
DB = 0
SM = 0
Delta_K
Doxorubicin
RF1_247_0 RF1
ZShift = −0.02
Im Score = 0
NG = 0
DB = 0
SM = 0
Delta_r
Doxorubicin
RF1_247_0 RF1
ZShift = −0.28
Im Score = 35.5
NG = 0
DB = 0
SM = 0
Delta_AUC
Doxorubicin
RF1_316_0 RF1
ZShift = 2.14
Im Score = −0.21
NG = 0
DB = 0
SM = 0
Delta_L
Doxorubicin
RF1_316_0 RF1
ZShift = −1.06
Im Score = −5.74
NG = 0
DB = 0
SM = 0
Delta_K
Doxorubicin
RF1_316_0 RF1
ZShift = −0.77
Im Score = 0.01
NG = 0
DB = 0
SM = 0
Delta_r
Doxorubicin
RF1_316_0 RF1
ZShift = −2
Im Score = 17.2
NG = 0
DB = 0
SM = 0
Delta_AUC
Doxorubicin
RF1_1_2 RF1
ZShift = 1.28
Im Score = −5.41
NG = 0
DB = 0
SM = 0
Delta_L
Doxorubicin
RF1_1_2 RF1
ZShift = 0.38
Im Score = −6.23
NG = 0
DB = 0
SM = 0
Delta_K
Doxorubicin
RF1_1_2 RF1
ZShift = −2.25
Im Score = 0.01
NG = 0
DB = 0
SM = 0
Delta_r
Doxorubicin
RF1_1_2 RF1
ZShift = −0.73
Im Score = −425.87
NG = 0
DB = 0
SM = 0
Delta_AUC
Doxorubicin

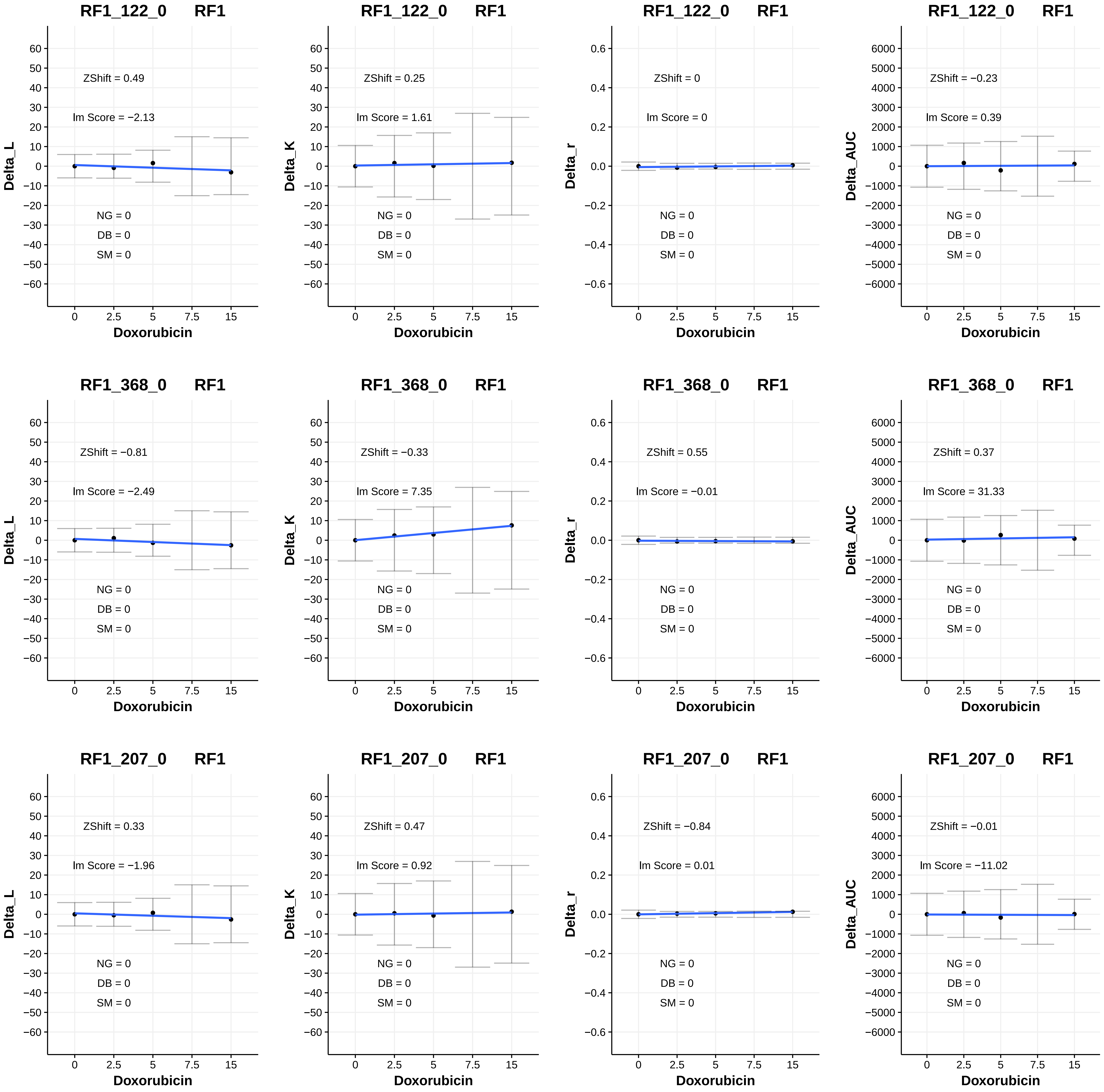
RF1_122_0 RF1
ZShift = 0.49
Im Score = −2.13
NG = 0
DB = 0
SM = 0
Delta_L
Doxorubicin
RF1_122_0 RF1
ZShift = 0.25
Im Score = 1.61
NG = 0
DB = 0
SM = 0
Delta_K
Doxorubicin
RF1_122_0 RF1
ZShift = 0
Im Score = 0
NG = 0
DB = 0
SM = 0
Delta_r
Doxorubicin
RF1_122_0 RF1
ZShift = −0.23
Im Score = 0.39
NG = 0
DB = 0
SM = 0
Delta_AUC
Doxorubicin
RF1_368_0 RF1
ZShift = −0.81
Im Score = −2.49
NG = 0
DB = 0
SM = 0
Delta_L
Doxorubicin
RF1_368_0 RF1
ZShift = −0.33
Im Score = 7.35
NG = 0
DB = 0
SM = 0
Delta_K
Doxorubicin
RF1_368_0 RF1
ZShift = 0.55
Im Score = −0.01
NG = 0
DB = 0
SM = 0
Delta_r
Doxorubicin
RF1_368_0 RF1
ZShift = 0.37
Im Score = 31.33
NG = 0
DB = 0
SM = 0
Delta_AUC
Doxorubicin
RF1_207_0 RF1
ZShift = 0.33
Im Score = −1.96
NG = 0
DB = 0
SM = 0
Delta_L
Doxorubicin
RF1_207_0 RF1
ZShift = 0.47
Im Score = 0.92
NG = 0
DB = 0
SM = 0
Delta_K
Doxorubicin
RF1_207_0 RF1
ZShift = −0.84
Im Score = 0.01
NG = 0
DB = 0
SM = 0
Delta_r
Doxorubicin
RF1_207_0 RF1
ZShift = −0.01
Im Score = −11.02
NG = 0
DB = 0
SM = 0
Delta_AUC
Doxorubicin

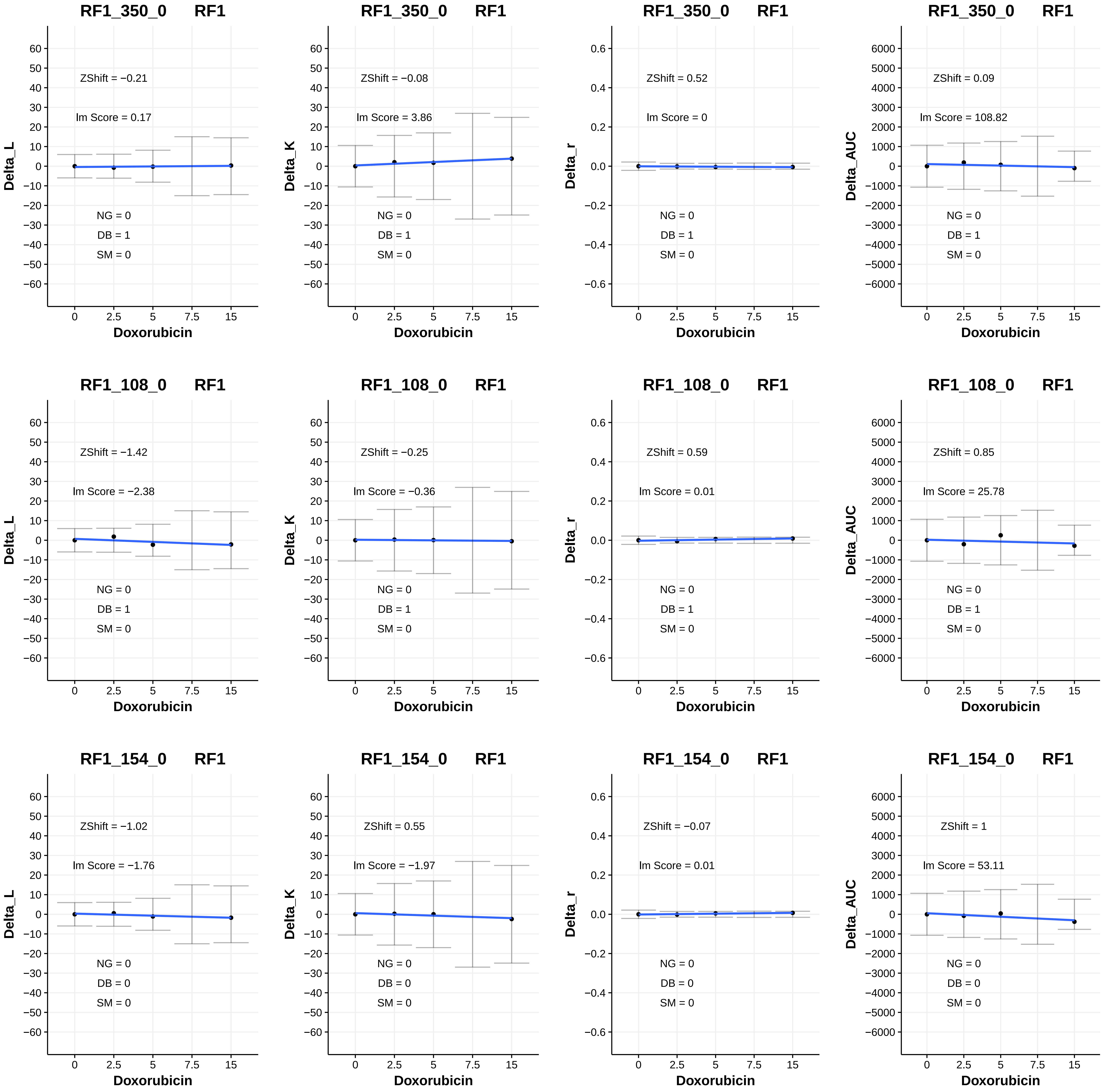
RF1_350_0 RF1
ZShift = −0.21
lm Score = 0.17
NG = 0
DB = 1
SM = 0
Delta_L
Doxorubicin
RF1_350_0 RF1
ZShift = −0.08
lm Score = 3.86
NG = 0
DB = 1
SM = 0
Delta_K
Doxorubicin
RF1_350_0 RF1
ZShift = 0.52
lm Score = 0
NG = 0
DB = 1
SM = 0
Delta_r
Doxorubicin
RF1_350_0 RF1
ZShift = 0.09
lm Score = 108.82
NG = 0
DB = 1
SM = 0
Delta_AUC
Doxorubicin
RF1_108_0 RF1
ZShift = −1.42
lm Score = −2.38
NG = 0
DB = 1
SM = 0
Delta_L
Doxorubicin
RF1_108_0 RF1
ZShift = −0.25
lm Score = −0.36
NG = 0
DB = 1
SM = 0
Delta_K
Doxorubicin
RF1_108_0 RF1
ZShift = 0.59
lm Score = 0.01
NG = 0
DB = 1
SM = 0
Delta_r
Doxorubicin
RF1_108_0 RF1
ZShift = 0.85
lm Score = 25.78
NG = 0
DB = 1
SM = 0
Delta_AUC
Doxorubicin
RF1_154_0 RF1
ZShift = −1.02
lm Score = −1.76
NG = 0
DB = 0
SM = 0
Delta_L
Doxorubicin
RF1_154_0 RF1
ZShift = 0.55
lm Score = −1.97
NG = 0
DB = 0
SM = 0
Delta_K
Doxorubicin
RF1_154_0 RF1
ZShift = −0.07
lm Score = 0.01
NG = 0
DB = 0
SM = 0
Delta_r
Doxorubicin
RF1_154_0 RF1
ZShift = 1
lm Score = 53.11
NG = 0
DB = 0
SM = 0
Delta_AUC
Doxorubicin

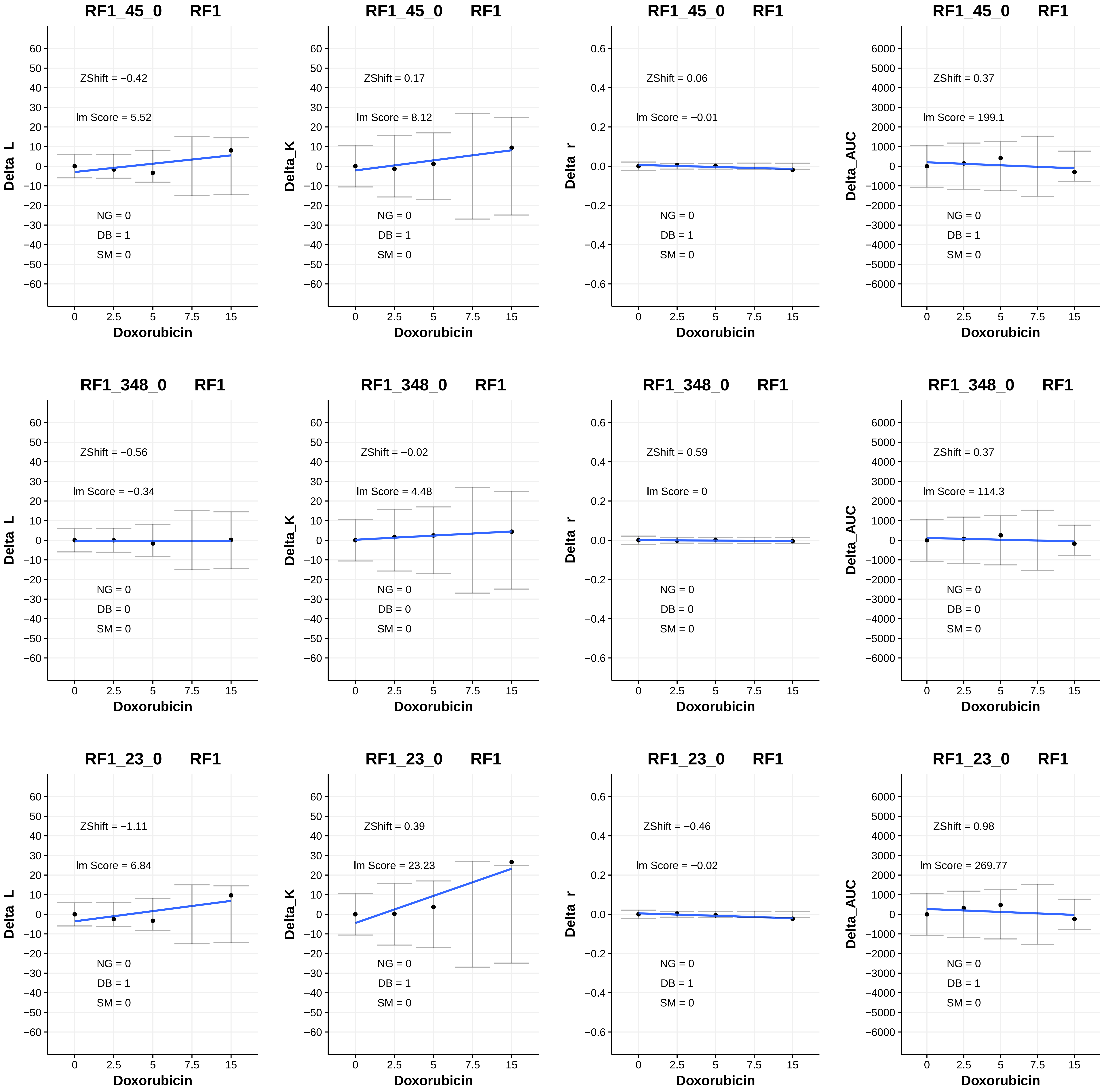

RF1_45_0 RF1
ZShift = −0.42
lm Score = 5.52
NG = 0
DB = 1
SM = 0
Delta_L
Doxorubicin
RF1_45_0 RF1
ZShift = 0.17
lm Score = 8.12
NG = 0
DB = 1
SM = 0
Delta_K
Doxorubicin
RF1_45_0 RF1
ZShift = 0.06
lm Score = −0.01
NG = 0
DB = 1
SM = 0
Delta_r
Doxorubicin
RF1_45_0 RF1
ZShift = 0.37
lm Score = 199.1
NG = 0
DB = 1
SM = 0
Delta_AUC
Doxorubicin
RF1_348_0 RF1
ZShift = −0.56
lm Score = −0.34
NG = 0
DB = 0
SM = 0
Delta_L
Doxorubicin
RF1_348_0 RF1
ZShift = −0.02
lm Score = 4.48
NG = 0
DB = 0
SM = 0
Delta_K
Doxorubicin
RF1_348_0 RF1
ZShift = 0.59
lm Score = 0
NG = 0
DB = 0
SM = 0
Delta_r
Doxorubicin
RF1_348_0 RF1
ZShift = 0.37
lm Score = 114.3
NG = 0
DB = 0
SM = 0
Delta_AUC
Doxorubicin
RF1_23_0 RF1
ZShift = −1.11
lm Score = 6.84
NG = 0
DB = 1
SM = 0
Delta_L
Doxorubicin
RF1_23_0 RF1
ZShift = 0.39
lm Score = 23.23
NG = 0
DB = 1
SM = 0
Delta_K
Doxorubicin
RF1_23_0 RF1
ZShift = −0.46
lm Score = −0.02
NG = 0
DB = 1
SM = 0
Delta_r
Doxorubicin
RF1_23_0 RF1
ZShift = 0.98
lm Score = 269.77
NG = 0
DB = 1
SM = 0
Delta_AUC
Doxorubicin

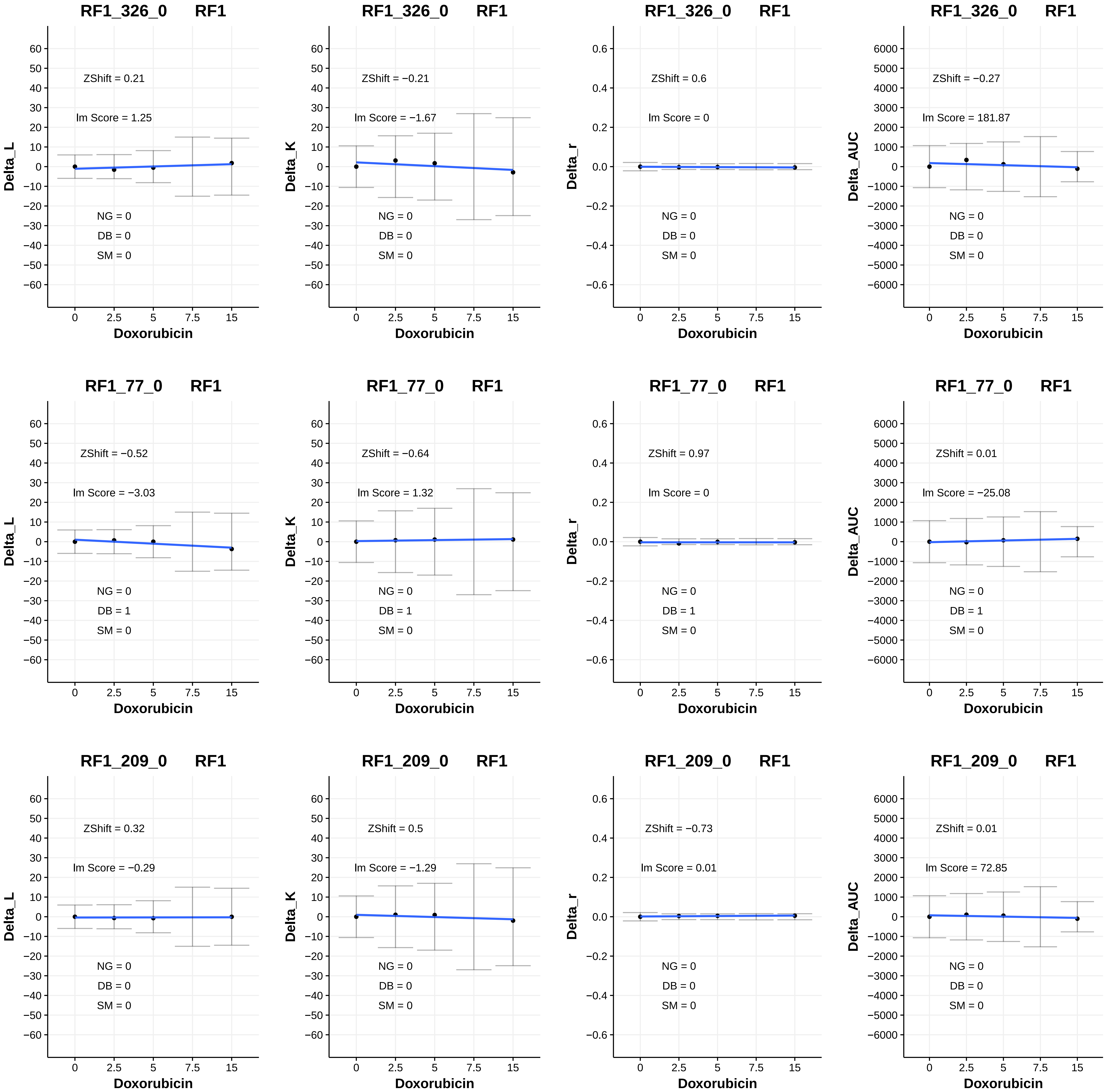
RF1_326_0 RF1
ZShift = 0.21
lm Score = 1.25
NG = 0
DB = 0
SM = 0
Delta_L
Doxorubicin
0 2.5 5 7.5 15
RF1_326_0 RF1
ZShift = −0.21
lm Score = −1.67
NG = 0
DB = 0
SM = 0
Delta_K
Doxorubicin
RF1_326_0 RF1
ZShift = 0.6
lm Score = 0
NG = 0
DB = 0
SM = 0
Delta_r
Doxorubicin
RF1_326_0 RF1
ZShift = −0.27
lm Score = 181.87
NG = 0
DB = 0
SM = 0
Delta_AUC
Doxorubicin
RF1_77_0 RF1
ZShift = −0.52
lm Score = −3.03
NG = 0
DB = 1
SM = 0
Delta_L
Doxorubicin
RF1_77_0 RF1
ZShift = −0.64
lm Score = 1.32
NG = 0
DB = 1
SM = 0
Delta_K
Doxorubicin
RF1_77_0 RF1
ZShift = 0.97
lm Score = 0
NG = 0
DB = 1
SM = 0
Delta_r
Doxorubicin
RF1_77_0 RF1
ZShift = 0.01
lm Score = −25.08
NG = 0
DB = 1
SM = 0
Delta_AUC
Doxorubicin
RF1_209_0 RF1
ZShift = 0.32
lm Score = −0.29
NG = 0
DB = 0
SM = 0
Delta_L
Doxorubicin
RF1_209_0 RF1
ZShift = 0.5
lm Score = −1.29
NG = 0
DB = 0
SM = 0
Delta_K
Doxorubicin
RF1_209_0 RF1
ZShift = −0.73
lm Score = 0.01
NG = 0
DB = 0
SM = 0
Delta_r
Doxorubicin
RF1_209_0 RF1
ZShift = 0.01
lm Score = 72.85
NG = 0
DB = 0
SM = 0
Delta_AUC
Doxorubicin

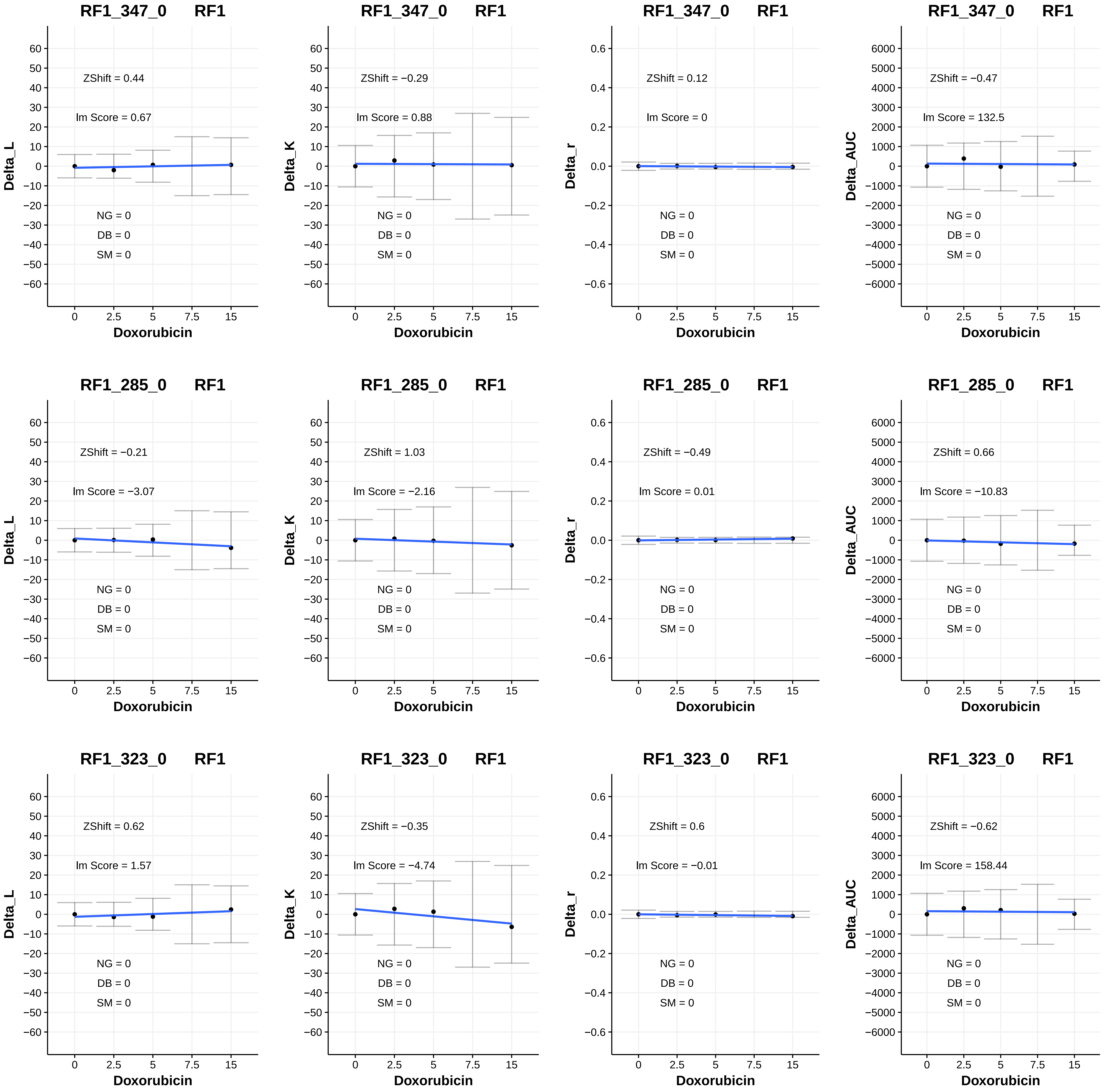
RF1_347_0 RF1
ZShift = 0.44
lm Score = 0.67
NG = 0
DB = 0
SM = 0
Delta_L
Doxorubicin
RF1_347_0 RF1
ZShift = −0.29
lm Score = 0.88
NG = 0
DB = 0
SM = 0
Delta_K
Doxorubicin
RF1_347_0 RF1
ZShift = 0.12
lm Score = 0
NG = 0
DB = 0
SM = 0
Delta_r
Doxorubicin
RF1_347_0 RF1
ZShift = −0.47
lm Score = 132.5
NG = 0
DB = 0
SM = 0
Delta_AUC
Doxorubicin
RF1_285_0 RF1
ZShift = −0.21
lm Score = −3.07
NG = 0
DB = 0
SM = 0
Delta_L
Doxorubicin
RF1_285_0 RF1
ZShift = 1.03
lm Score = −2.16
NG = 0
DB = 0
SM = 0
Delta_K
Doxorubicin
RF1_285_0 RF1
ZShift = −0.49
lm Score = 0.01
NG = 0
DB = 0
SM = 0
Delta_r
Doxorubicin
RF1_285_0 RF1
ZShift = 0.66
lm Score = −10.83
NG = 0
DB = 0
SM = 0
Delta_AUC
Doxorubicin
RF1_323_0 RF1
ZShift = 0.62
lm Score = 1.57
NG = 0
DB = 0
SM = 0
Delta_L
Doxorubicin
RF1_323_0 RF1
ZShift = −0.35
lm Score = −4.74
NG = 0
DB = 0
SM = 0
Delta_K
Doxorubicin
RF1_323_0 RF1
ZShift = 0.6
lm Score = −0.01
NG = 0
DB = 0
SM = 0
Delta_r
Doxorubicin
RF1_323_0 RF1
ZShift = −0.62
lm Score = 158.44
NG = 0
DB = 0
SM = 0
Delta_AUC
Doxorubicin

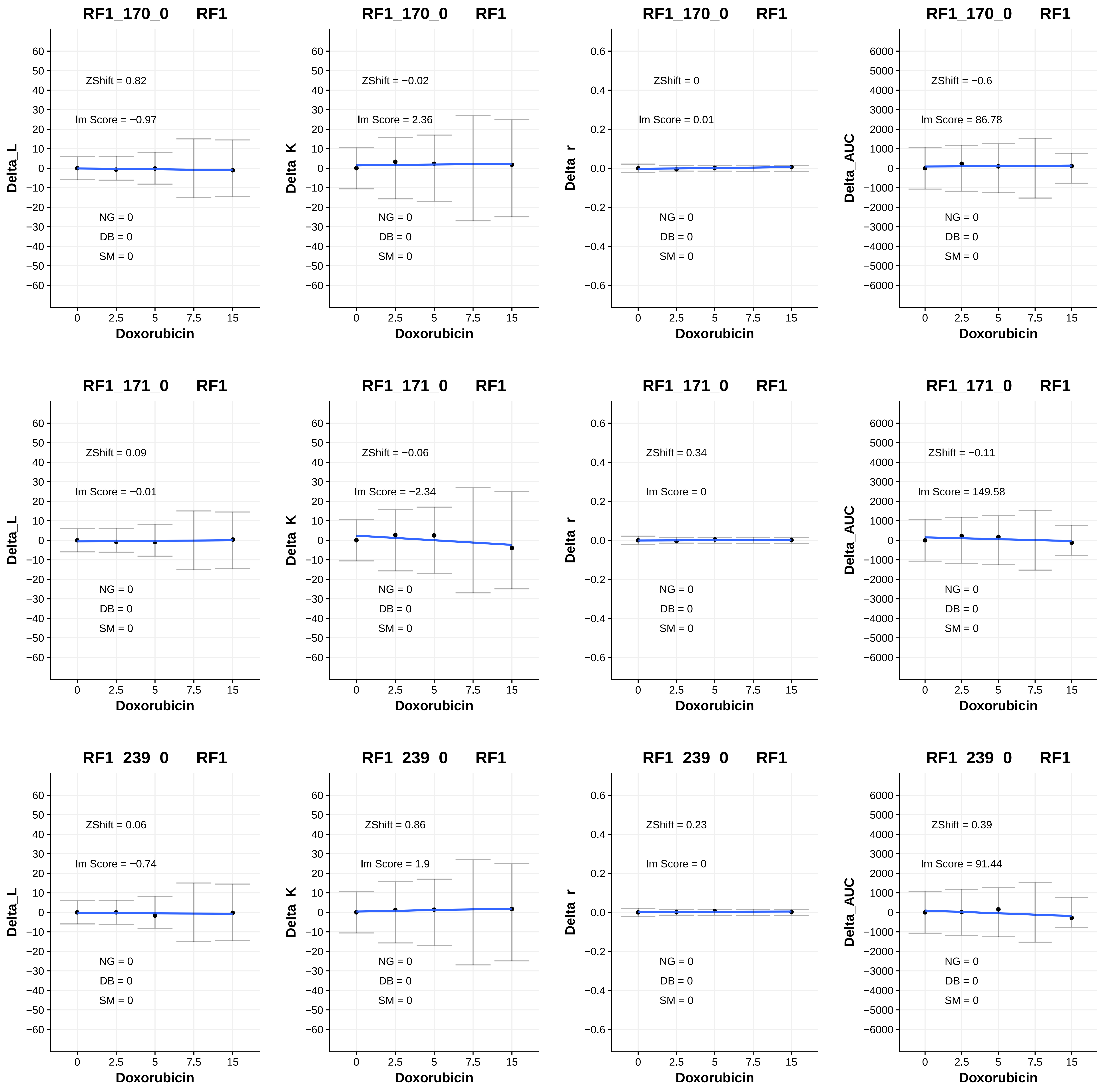
RF1_170_0 RF1
ZShift = 0.82
lm Score = −0.97
NG = 0
DB = 0
SM = 0
Delta_L
Doxorubicin
RF1_170_0 RF1
ZShift = −0.02
lm Score = 2.36
NG = 0
DB = 0
SM = 0
Delta_K
Doxorubicin
RF1_170_0 RF1
ZShift = 0
lm Score = 0.01
NG = 0
DB = 0
SM = 0
Delta_r
Doxorubicin
RF1_170_0 RF1
ZShift = −0.6
lm Score = 86.78
NG = 0
DB = 0
SM = 0
Delta_AUC
Doxorubicin
RF1_171_0 RF1
ZShift = 0.09
lm Score = −0.01
NG = 0
DB = 0
SM = 0
Delta_L
Doxorubicin
RF1_171_0 RF1
ZShift = −0.06
lm Score = −2.34
NG = 0
DB = 0
SM = 0
Delta_K
Doxorubicin
RF1_171_0 RF1
ZShift = 0.34
lm Score = 0
NG = 0
DB = 0
SM = 0
Delta_r
Doxorubicin
RF1_171_0 RF1
ZShift = −0.11
lm Score = 149.58
NG = 0
DB = 0
SM = 0
Delta_AUC
Doxorubicin
RF1_239_0 RF1
ZShift = 0.06
lm Score = −0.74
NG = 0
DB = 0
SM = 0
Delta_L
Doxorubicin
RF1_239_0 RF1
ZShift = 0.86
lm Score = 1.9
NG = 0
DB = 0
SM = 0
Delta_K
Doxorubicin
RF1_239_0 RF1
ZShift = 0.23
lm Score = 0
NG = 0
DB = 0
SM = 0
Delta_r
Doxorubicin
RF1_239_0 RF1
ZShift = 0.39
lm Score = 91.44
NG = 0
DB = 0
SM = 0
Delta_AUC
Doxorubicin

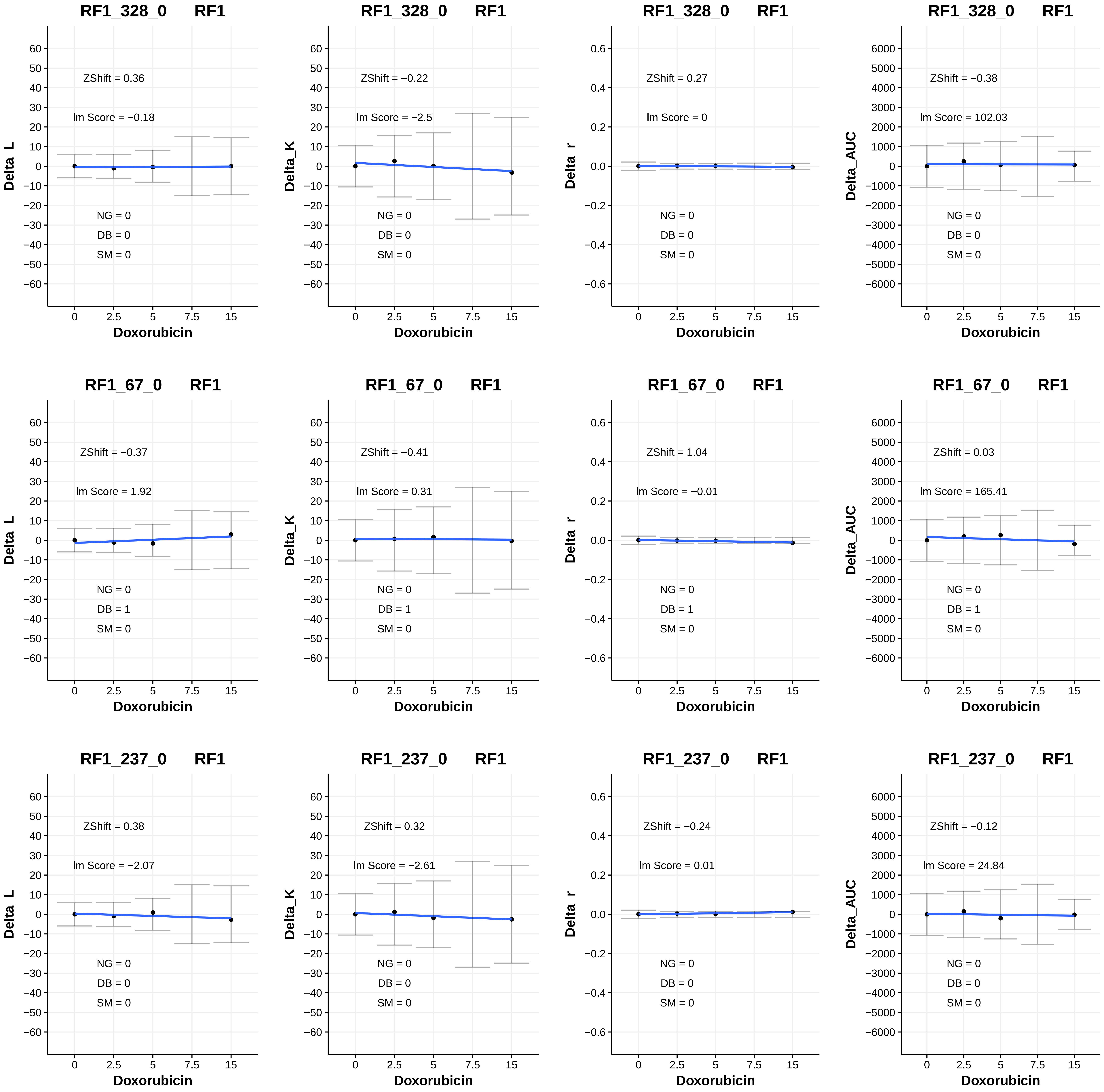
RF1_328_0 RF1
ZShift = 0.36
lm Score = −0.18
NG = 0
DB = 0
SM = 0
Delta_L
Doxorubicin
RF1_328_0 RF1
ZShift = −0.22
lm Score = −2.5
NG = 0
DB = 0
SM = 0
Delta_K
Doxorubicin
RF1_328_0 RF1
ZShift = 0.27
lm Score = 0
NG = 0
DB = 0
SM = 0
Delta_r
Doxorubicin
RF1_328_0 RF1
ZShift = −0.38
lm Score = 102.03
NG = 0
DB = 0
SM = 0
Delta_AUC
Doxorubicin
RF1_67_0 RF1
ZShift = −0.37
lm Score = 1.92
NG = 0
DB = 1
SM = 0
Delta_L
Doxorubicin
RF1_67_0 RF1
ZShift = −0.41
lm Score = 0.31
NG = 0
DB = 1
SM = 0
Delta_K
Doxorubicin
RF1_67_0 RF1
ZShift = 1.04
lm Score = −0.01
NG = 0
DB = 1
SM = 0
Delta_r
Doxorubicin
RF1_67_0 RF1
ZShift = 0.03
lm Score = 165.41
NG = 0
DB = 1
SM = 0
Delta_AUC
Doxorubicin
RF1_237_0 RF1
ZShift = 0.38
lm Score = −2.07
NG = 0
DB = 0
SM = 0
Delta_L
Doxorubicin
RF1_237_0 RF1
ZShift = 0.32
lm Score = −2.61
NG = 0
DB = 0
SM = 0
Delta_K
Doxorubicin
RF1_237_0 RF1
ZShift = −0.24
lm Score = 0.01
NG = 0
DB = 0
SM = 0
Delta_r
Doxorubicin
RF1_237_0 RF1
ZShift = −0.12
lm Score = 24.84
NG = 0
DB = 0
SM = 0
Delta_AUC
Doxorubicin

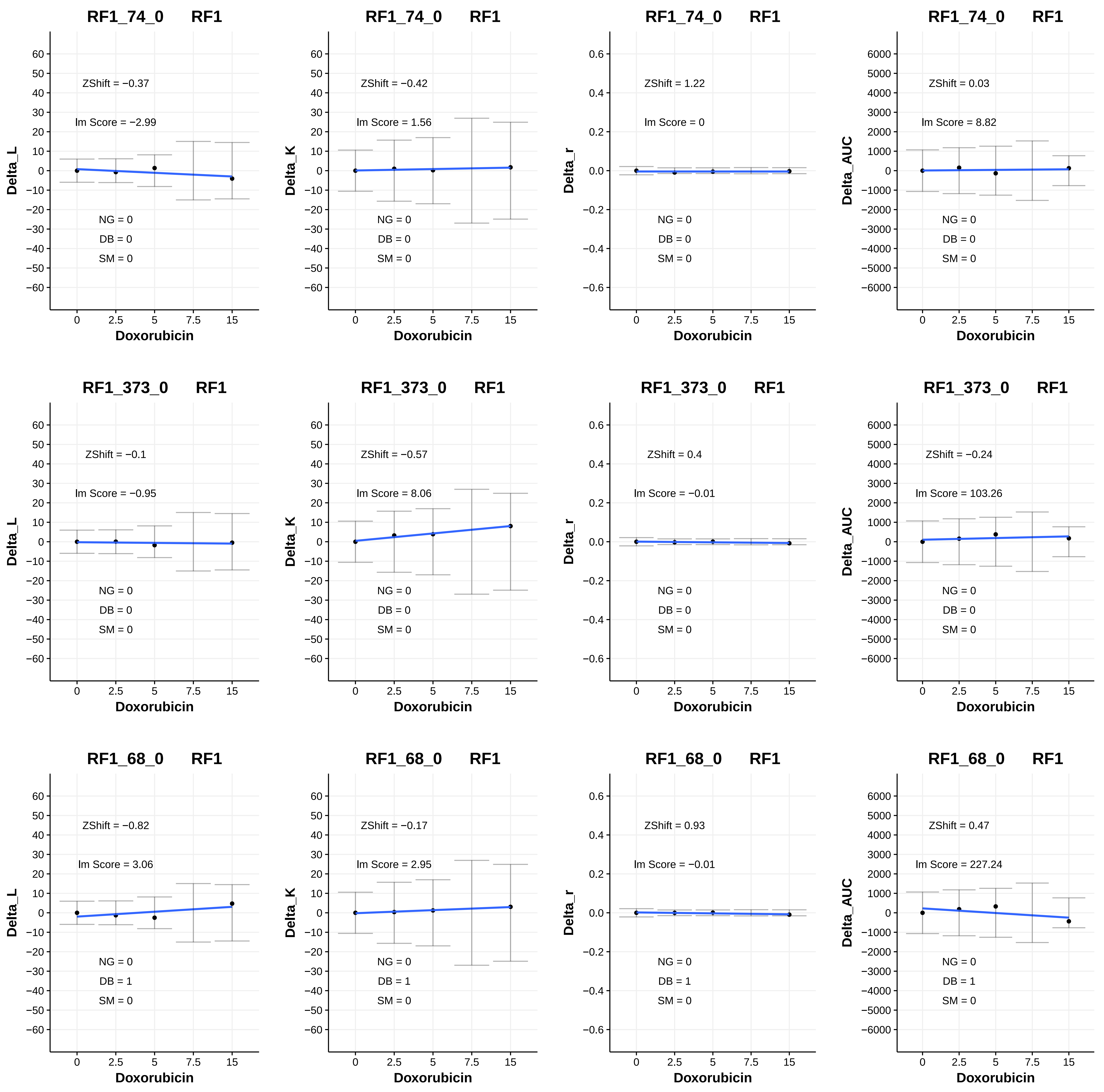

RF1_74_0 RF1
ZShift = −0.37
lm Score = −2.99
NG = 0
DB = 0
SM = 0
Delta_L
Doxorubicin
RF1_74_0 RF1
ZShift = −0.42
lm Score = 1.56
NG = 0
DB = 0
SM = 0
Delta_K
Doxorubicin
RF1_74_0 RF1
ZShift = 1.22
lm Score = 0
NG = 0
DB = 0
SM = 0
Delta_r
Doxorubicin
RF1_74_0 RF1
ZShift = 0.03
lm Score = 8.82
NG = 0
DB = 0
SM = 0
Delta_AUC
Doxorubicin
RF1_373_0 RF1
ZShift = −0.1
lm Score = −0.95
NG = 0
DB = 0
SM = 0
Delta_L
Doxorubicin
RF1_373_0 RF1
ZShift = −0.57
lm Score = 8.06
NG = 0
DB = 0
SM = 0
Delta_K
Doxorubicin
RF1_373_0 RF1
ZShift = 0.4
lm Score = −0.01
NG = 0
DB = 0
SM = 0
Delta_r
Doxorubicin
RF1_373_0 RF1
ZShift = −0.24
lm Score = 103.26
NG = 0
DB = 0
SM = 0
Delta_AUC
Doxorubicin
RF1_68_0 RF1
ZShift = −0.82
lm Score = 3.06
NG = 0
DB = 1
SM = 0
Delta_L
Doxorubicin
RF1_68_0 RF1
ZShift = −0.17
lm Score = 2.95
NG = 0
DB = 1
SM = 0
Delta_K
Doxorubicin
RF1_68_0 RF1
ZShift = 0.93
lm Score = −0.01
NG = 0
DB = 1
SM = 0
Delta_r
Doxorubicin
RF1_68_0 RF1
ZShift = 0.47
lm Score = 227.24
NG = 0
DB = 1
SM = 0
Delta_AUC
Doxorubicin
0
2.5
5
7.5
15

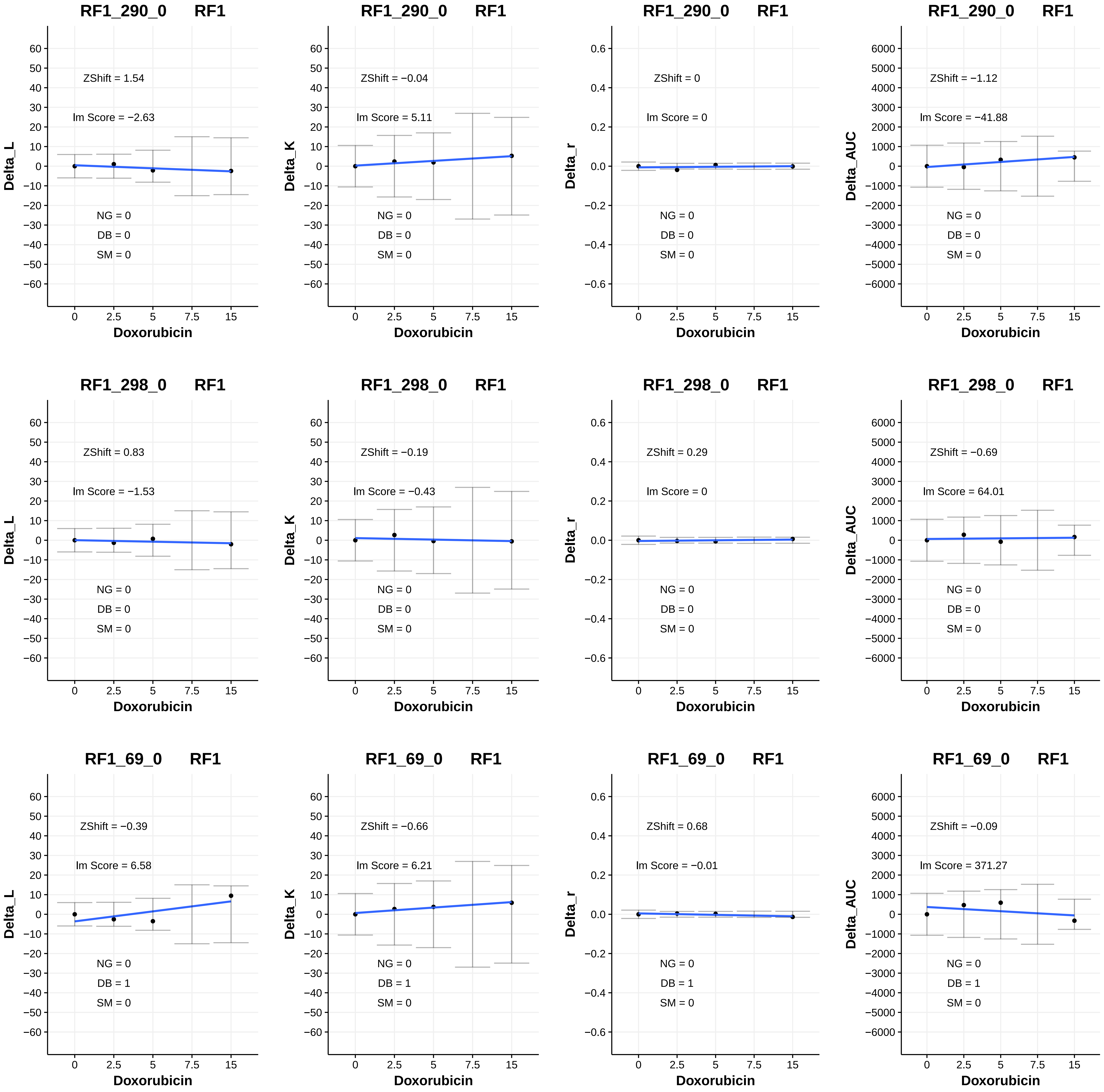
RF1_290_0 RF1
ZShift = 1.54
Im Score = −2.63
NG = 0
DB = 0
SM = 0
Delta_L
Doxorubicin
RF1_290_0 RF1
ZShift = −0.04
Im Score = 5.11
NG = 0
DB = 0
SM = 0
Delta_K
Doxorubicin
RF1_290_0 RF1
ZShift = 0
Im Score = 0
NG = 0
DB = 0
SM = 0
Delta_r
Doxorubicin
RF1_290_0 RF1
ZShift = −1.12
Im Score = −41.88
NG = 0
DB = 0
SM = 0
Delta_AUC
Doxorubicin
RF1_298_0 RF1
ZShift = 0.83
Im Score = −1.53
NG = 0
DB = 0
SM = 0
Delta_L
Doxorubicin
RF1_298_0 RF1
ZShift = −0.19
Im Score = −0.43
NG = 0
DB = 0
SM = 0
Delta_K
Doxorubicin
RF1_298_0 RF1
ZShift = 0.29
Im Score = 0
NG = 0
DB = 0
SM = 0
Delta_r
Doxorubicin
RF1_298_0 RF1
ZShift = −0.69
Im Score = 64.01
NG = 0
DB = 0
SM = 0
Delta_AUC
Doxorubicin
RF1_69_0 RF1
ZShift = −0.39
Im Score = 6.58
NG = 0
DB = 1
SM = 0
Delta_L
Doxorubicin
RF1_69_0 RF1
ZShift = −0.66
Im Score = 6.21
NG = 0
DB = 1
SM = 0
Delta_K
Doxorubicin
RF1_69_0 RF1
ZShift = 0.68
Im Score = −0.01
NG = 0
DB = 1
SM = 0
Delta_r
Doxorubicin
RF1_69_0 RF1
ZShift = −0.09
Im Score = 371.27
NG = 0
DB = 1
SM = 0
Delta_AUC
Doxorubicin

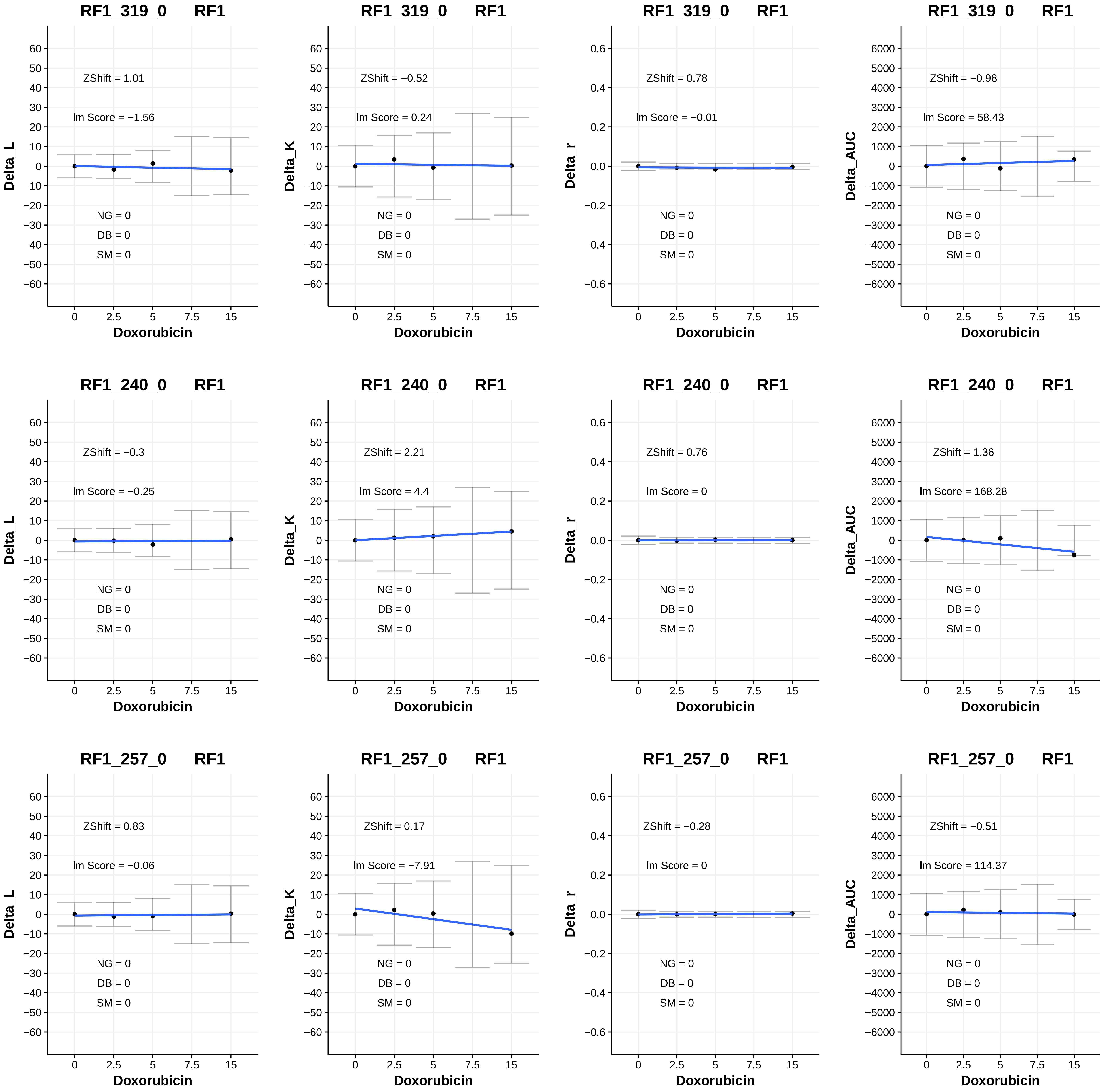

RF1_319_0 RF1
ZShift = 1.01
lm Score = −1.56
NG = 0
DB = 0
SM = 0
Delta_L
Doxorubicin
RF1_319_0 RF1
ZShift = −0.52
lm Score = 0.24
NG = 0
DB = 0
SM = 0
Delta_K
Doxorubicin
RF1_319_0 RF1
ZShift = 0.78
lm Score = −0.01
NG = 0
DB = 0
SM = 0
Delta_r
Doxorubicin
RF1_319_0 RF1
ZShift = −0.98
lm Score = 58.43
NG = 0
DB = 0
SM = 0
Delta_AUC
Doxorubicin
RF1_240_0 RF1
ZShift = −0.3
lm Score = −0.25
NG = 0
DB = 0
SM = 0
Delta_L
Doxorubicin
RF1_240_0 RF1
ZShift = 2.21
lm Score = 4.4
NG = 0
DB = 0
SM = 0
Delta_K
Doxorubicin
RF1_240_0 RF1
ZShift = 0.76
lm Score = 0
NG = 0
DB = 0
SM = 0
Delta_r
Doxorubicin
RF1_240_0 RF1
ZShift = 1.36
lm Score = 168.28
NG = 0
DB = 0
SM = 0
Delta_AUC
Doxorubicin
RF1_257_0 RF1
ZShift = 0.83
lm Score = −0.06
NG = 0
DB = 0
SM = 0
Delta_L
Doxorubicin
RF1_257_0 RF1
ZShift = 0.17
lm Score = −7.91
NG = 0
DB = 0
SM = 0
Delta_K
Doxorubicin
RF1_257_0 RF1
ZShift = −0.28
lm Score = 0
NG = 0
DB = 0
SM = 0
Delta_r
Doxorubicin
RF1_257_0 RF1
ZShift = −0.51
lm Score = 114.37
NG = 0
DB = 0
SM = 0
Delta_AUC
Doxorubicin

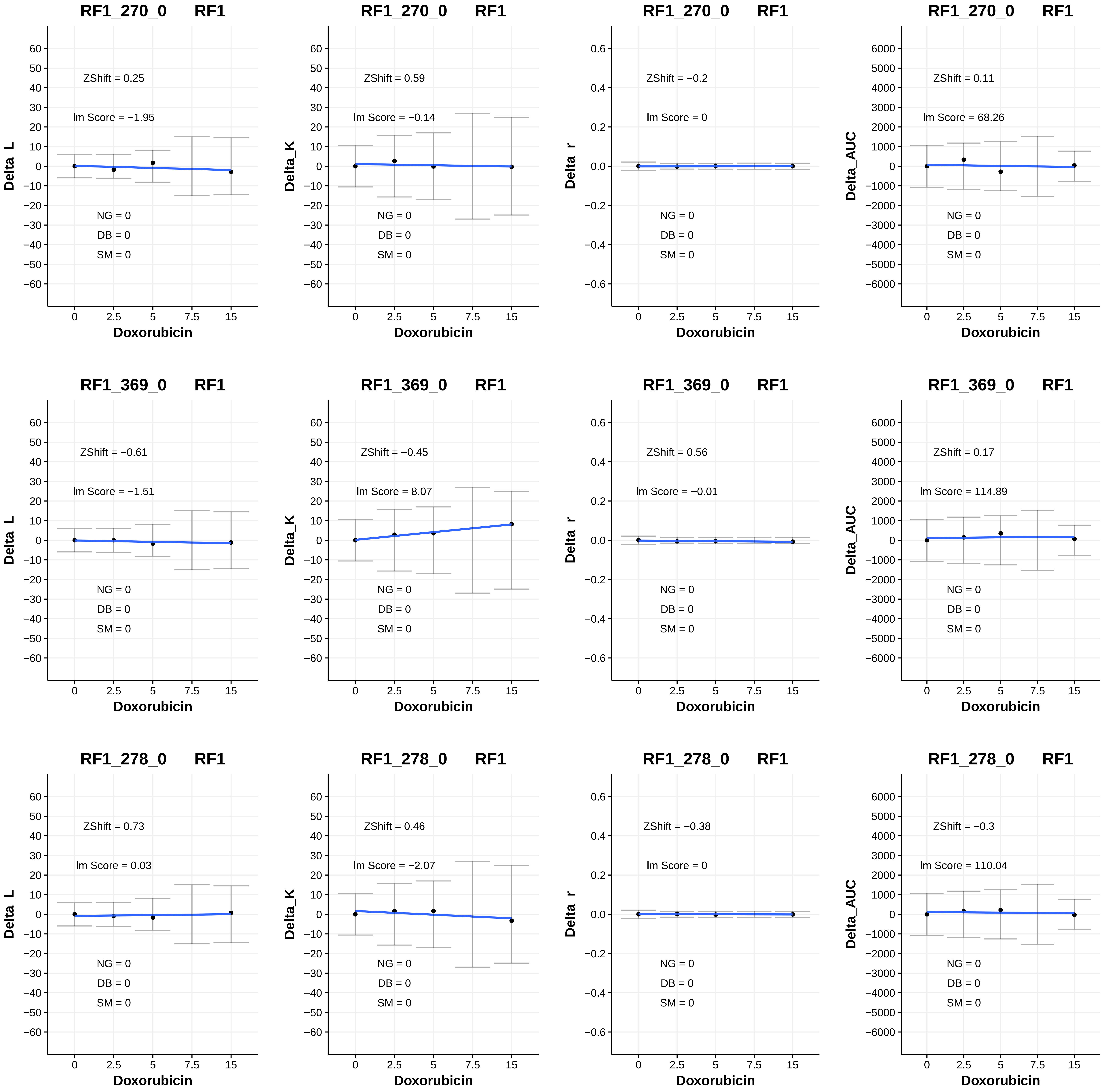
RF1_270_0 RF1
ZShift = 0.25
lm Score = −1.95
NG = 0
DB = 0
SM = 0
Delta_L
Doxorubicin
RF1_270_0 RF1
ZShift = 0.59
lm Score = −0.14
NG = 0
DB = 0
SM = 0
Delta_K
Doxorubicin
RF1_270_0 RF1
ZShift = −0.2
lm Score = 0
NG = 0
DB = 0
SM = 0
Delta_r
Doxorubicin
RF1_270_0 RF1
ZShift = 0.11
lm Score = 68.26
NG = 0
DB = 0
SM = 0
Delta_AUC
Doxorubicin
RF1_369_0 RF1
ZShift = −0.61
lm Score = −1.51
NG = 0
DB = 0
SM = 0
Delta_L
Doxorubicin
RF1_369_0 RF1
ZShift = −0.45
lm Score = 8.07
NG = 0
DB = 0
SM = 0
Delta_K
Doxorubicin
RF1_369_0 RF1
ZShift = 0.56
lm Score = −0.01
NG = 0
DB = 0
SM = 0
Delta_r
Doxorubicin
RF1_369_0 RF1
ZShift = 0.17
lm Score = 114.89
NG = 0
DB = 0
SM = 0
Delta_AUC
Doxorubicin
RF1_278_0 RF1
ZShift = 0.73
lm Score = 0.03
NG = 0
DB = 0
SM = 0
Delta_L
Doxorubicin
RF1_278_0 RF1
ZShift = 0.46
lm Score = −2.07
NG = 0
DB = 0
SM = 0
Delta_K
Doxorubicin
RF1_278_0 RF1
ZShift = −0.38
lm Score = 0
NG = 0
DB = 0
SM = 0
Delta_r
Doxorubicin
RF1_278_0 RF1
ZShift = −0.3
lm Score = 110.04
NG = 0
DB = 0
SM = 0
Delta_AUC
Doxorubicin

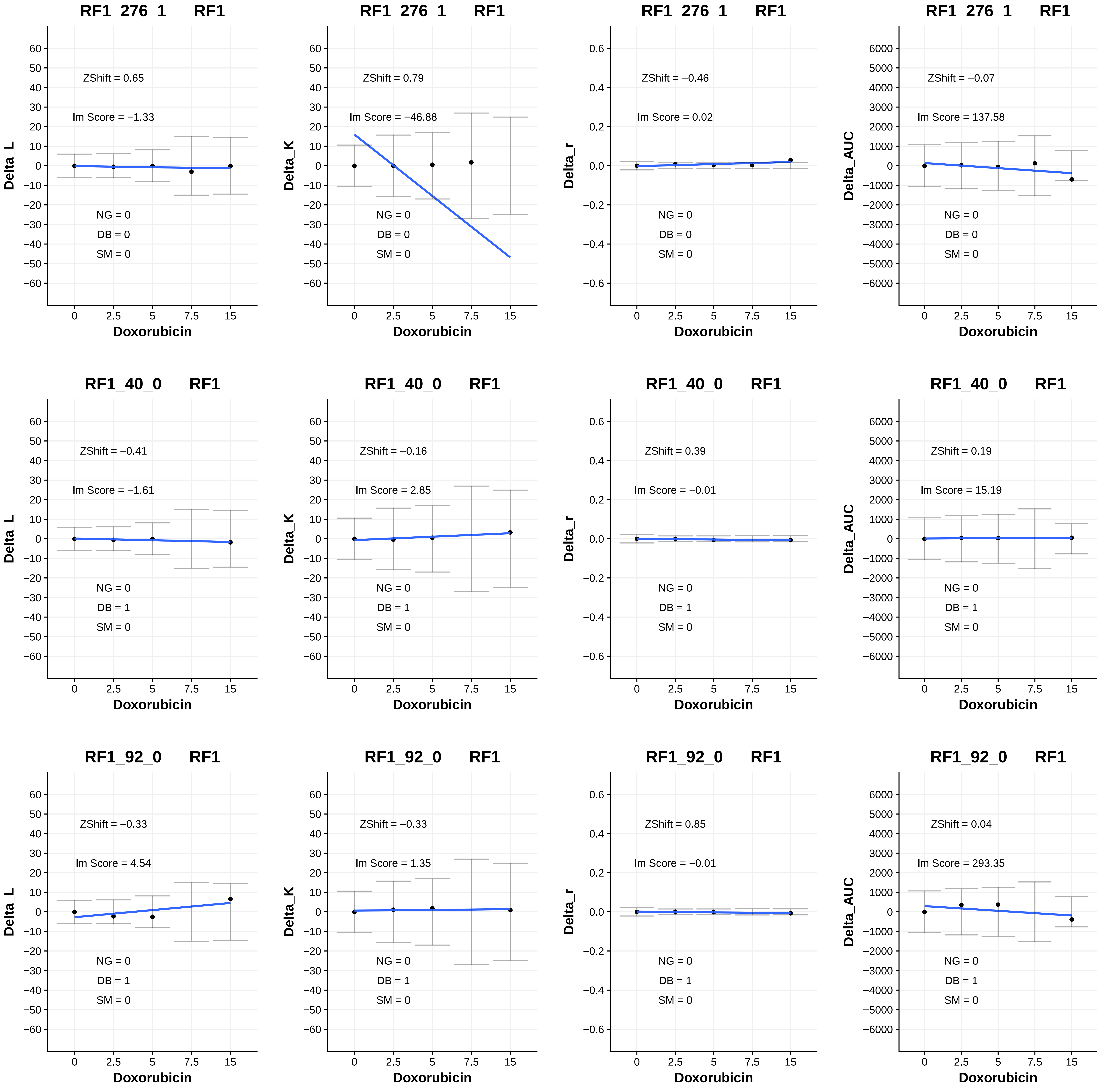

RF1_276_1 RF1
ZShift = 0.65
lm Score = −1.33
NG = 0
DB = 0
SM = 0
Delta_L
Doxorubicin
RF1_276_1 RF1
ZShift = 0.79
lm Score = −46.88
NG = 0
DB = 0
SM = 0
Delta_K
Doxorubicin
RF1_276_1 RF1
ZShift = −0.46
lm Score = 0.02
NG = 0
DB = 0
SM = 0
Delta_r
Doxorubicin
RF1_276_1 RF1
ZShift = −0.07
lm Score = 137.58
NG = 0
DB = 0
SM = 0
Delta_AUC
Doxorubicin
RF1_40_0 RF1
ZShift = −0.41
lm Score = −1.61
NG = 0
DB = 1
SM = 0
Delta_L
Doxorubicin
RF1_40_0 RF1
ZShift = −0.16
lm Score = 2.85
NG = 0
DB = 1
SM = 0
Delta_K
Doxorubicin
RF1_40_0 RF1
ZShift = 0.39
lm Score = −0.01
NG = 0
DB = 1
SM = 0
Delta_r
Doxorubicin
RF1_40_0 RF1
ZShift = 0.19
lm Score = 15.19
NG = 0
DB = 1
SM = 0
Delta_AUC
Doxorubicin
RF1_92_0 RF1
ZShift = −0.33
lm Score = 4.54
NG = 0
DB = 1
SM = 0
Delta_L
Doxorubicin
RF1_92_0 RF1
ZShift = −0.33
lm Score = 1.35
NG = 0
DB = 1
SM = 0
Delta_K
Doxorubicin
RF1_92_0 RF1
ZShift = 0.85
lm Score = −0.01
NG = 0
DB = 1
SM = 0
Delta_r
Doxorubicin
RF1_92_0 RF1
ZShift = 0.04
lm Score = 293.35
NG = 0
DB = 1
SM = 0
Delta_AUC
Doxorubicin

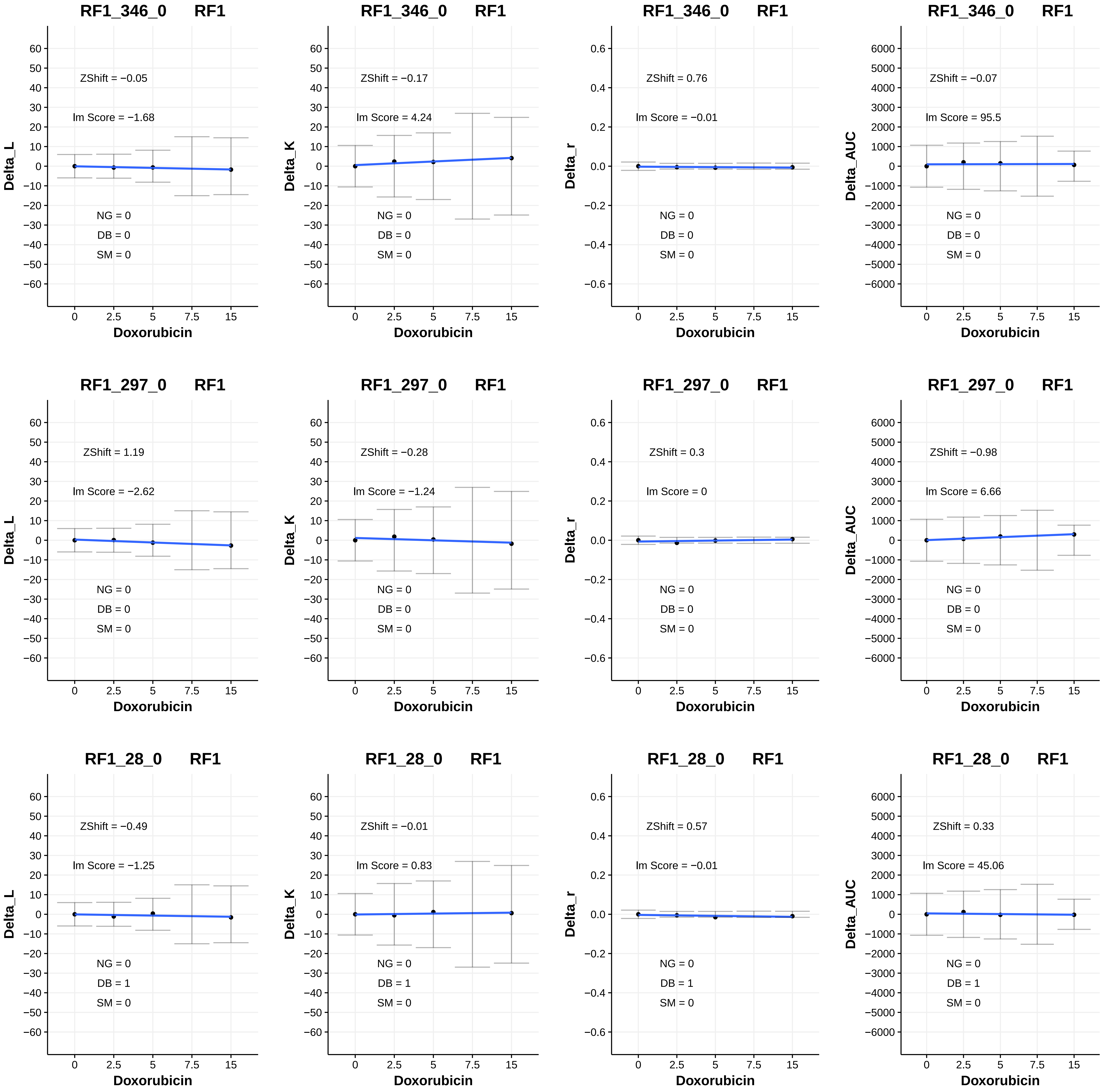

RF1_346_0 RF1
ZShift = −0.05
Im Score = −1.68
NG = 0
DB = 0
SM = 0
Delta_L
Doxorubicin
RF1_346_0 RF1
ZShift = −0.17
Im Score = 4.24
NG = 0
DB = 0
SM = 0
Delta_K
Doxorubicin
RF1_346_0 RF1
ZShift = 0.76
Im Score = −0.01
NG = 0
DB = 0
SM = 0
Delta_r
Doxorubicin
RF1_346_0 RF1
ZShift = −0.07
Im Score = 95.5
NG = 0
DB = 0
SM = 0
Delta_AUC
Doxorubicin
RF1_297_0 RF1
ZShift = 1.19
Im Score = −2.62
NG = 0
DB = 0
SM = 0
Delta_L
Doxorubicin
RF1_297_0 RF1
ZShift = −0.28
Im Score = −1.24
NG = 0
DB = 0
SM = 0
Delta_K
Doxorubicin
RF1_297_0 RF1
ZShift = 0.3
Im Score = 0
NG = 0
DB = 0
SM = 0
Delta_r
Doxorubicin
RF1_297_0 RF1
ZShift = −0.98
Im Score = 6.66
NG = 0
DB = 0
SM = 0
Delta_AUC
Doxorubicin
RF1_28_0 RF1
ZShift = −0.49
Im Score = −1.25
NG = 0
DB = 1
SM = 0
Delta_L
Doxorubicin
RF1_28_0 RF1
ZShift = −0.01
Im Score = 0.83
NG = 0
DB = 1
SM = 0
Delta_K
Doxorubicin
RF1_28_0 RF1
ZShift = 0.57
Im Score = −0.01
NG = 0
DB = 1
SM = 0
Delta_r
Doxorubicin
RF1_28_0 RF1
ZShift = 0.33
Im Score = 45.06
NG = 0
DB = 1
SM = 0
Delta_AUC
Doxorubicin
0
2.5
5
7.5
15

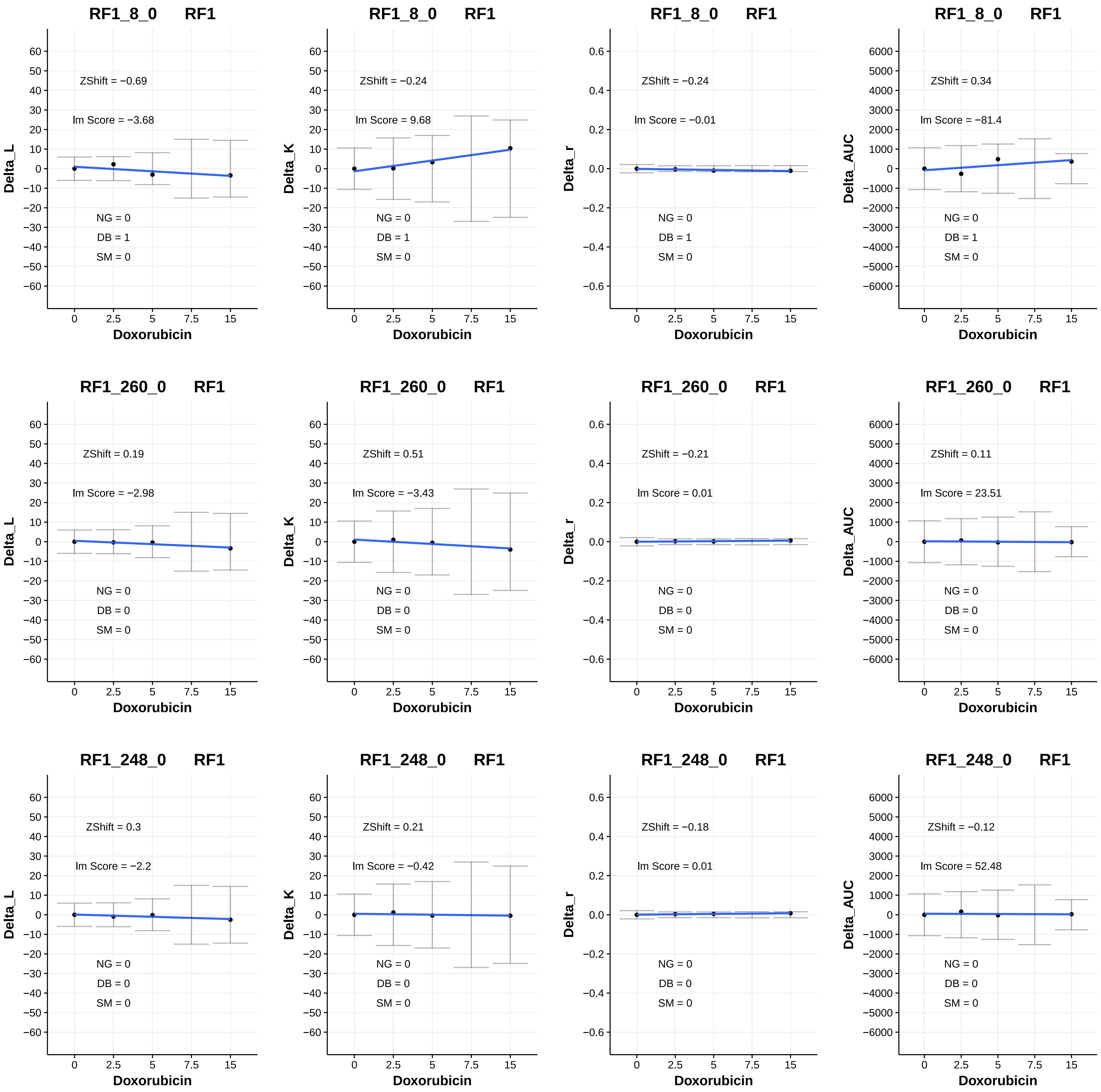

RF1_8_0 RF1
Delta_L
ZShift = −0.69
lm Score = −3.68
NG = 0
DB = 1
SM = 0
Doxorubicin
RF1_8_0 RF1
Delta_K
ZShift = −0.24
lm Score = 9.68
NG = 0
DB = 1
SM = 0
Doxorubicin
RF1_8_0 RF1
Delta_r
ZShift = −0.24
lm Score = −0.01
NG = 0
DB = 1
SM = 0
Doxorubicin
RF1_8_0 RF1
Delta_AUC
ZShift = 0.34
lm Score = −81.4
NG = 0
DB = 1
SM = 0
Doxorubicin
RF1_260_0 RF1
Delta_L
ZShift = 0.19
lm Score = −2.98
NG = 0
DB = 0
SM = 0
Doxorubicin
RF1_260_0 RF1
Delta_K
ZShift = 0.51
lm Score = −3.43
NG = 0
DB = 0
SM = 0
Doxorubicin
RF1_260_0 RF1
Delta_r
ZShift = −0.21
lm Score = 0.01
NG = 0
DB = 0
SM = 0
Doxorubicin
RF1_260_0 RF1
Delta_AUC
ZShift = 0.11
lm Score = 23.51
NG = 0
DB = 0
SM = 0
Doxorubicin
RF1_248_0 RF1
Delta_L
ZShift = 0.3
lm Score = −2.2
NG = 0
DB = 0
SM = 0
Doxorubicin
RF1_248_0 RF1
Delta_K
ZShift = 0.21
lm Score = −0.42
NG = 0
DB = 0
SM = 0
Doxorubicin
RF1_248_0 RF1
Delta_r
ZShift = −0.18
lm Score = 0.01
NG = 0
DB = 0
SM = 0
Doxorubicin
RF1_248_0 RF1
Delta_AUC
ZShift = −0.12
lm Score = 52.48
NG = 0
DB = 0
SM = 0
Doxorubicin

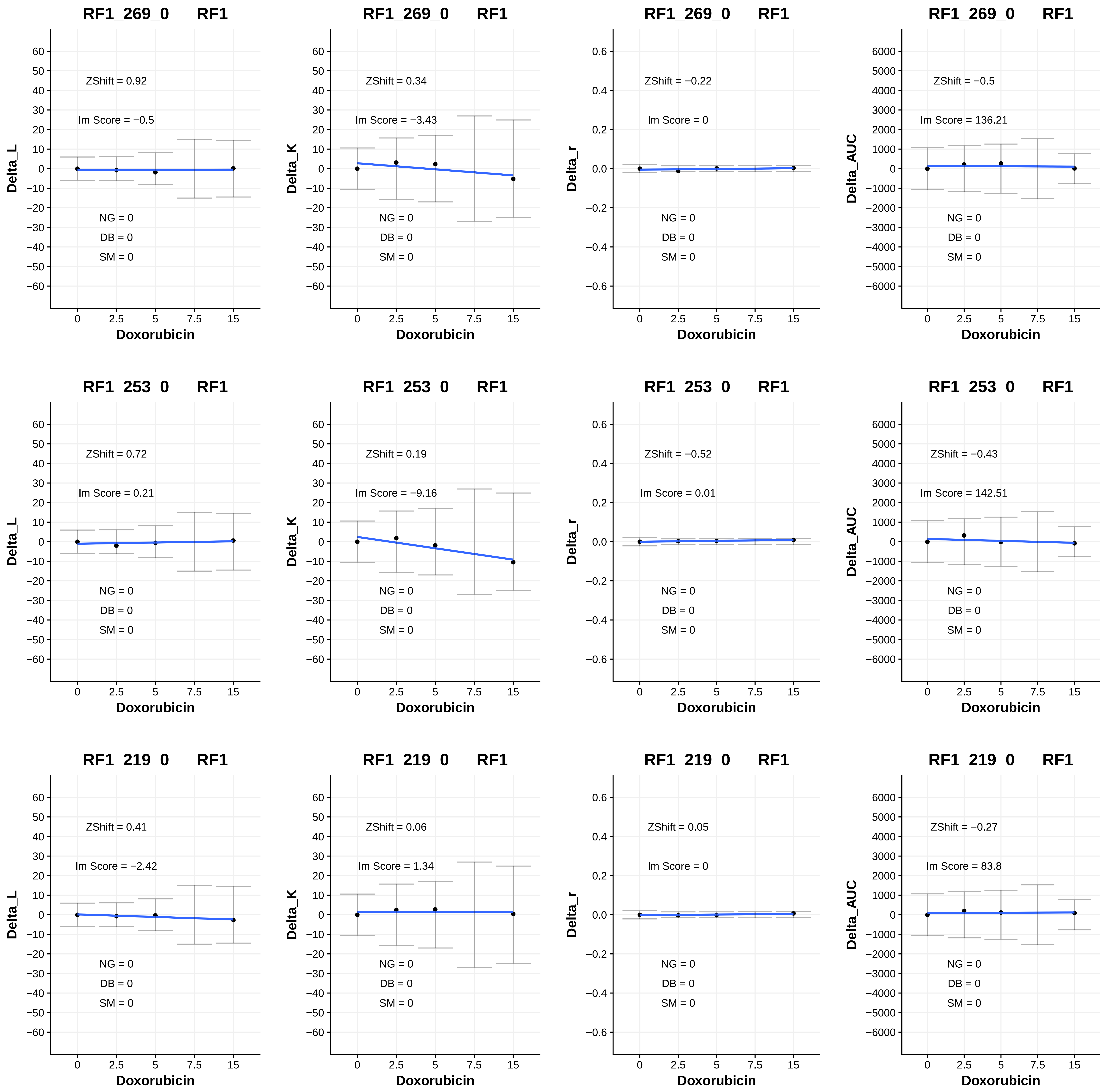
RF1_269_0 RF1
ZShift = 0.92
Im Score = −0.5
NG = 0
DB = 0
SM = 0
Delta_L
Doxorubicin
RF1_269_0 RF1
ZShift = 0.34
Im Score = −3.43
NG = 0
DB = 0
SM = 0
Delta_K
Doxorubicin
RF1_269_0 RF1
ZShift = −0.22
Im Score = 0
NG = 0
DB = 0
SM = 0
Delta_r
Doxorubicin
RF1_269_0 RF1
ZShift = −0.5
Im Score = 136.21
NG = 0
DB = 0
SM = 0
Delta_AUC
Doxorubicin
RF1_253_0 RF1
ZShift = 0.72
Im Score = 0.21
NG = 0
DB = 0
SM = 0
Delta_L
Doxorubicin
RF1_253_0 RF1
ZShift = 0.19
Im Score = −9.16
NG = 0
DB = 0
SM = 0
Delta_K
Doxorubicin
RF1_253_0 RF1
ZShift = −0.52
Im Score = 0.01
NG = 0
DB = 0
SM = 0
Delta_r
Doxorubicin
RF1_253_0 RF1
ZShift = −0.43
Im Score = 142.51
NG = 0
DB = 0
SM = 0
Delta_AUC
Doxorubicin
RF1_219_0 RF1
ZShift = 0.41
Im Score = −2.42
NG = 0
DB = 0
SM = 0
Delta_L
Doxorubicin
RF1_219_0 RF1
ZShift = 0.06
Im Score = 1.34
NG = 0
DB = 0
SM = 0
Delta_K
Doxorubicin
RF1_219_0 RF1
ZShift = 0.05
Im Score = 0
NG = 0
DB = 0
SM = 0
Delta_r
Doxorubicin
RF1_219_0 RF1
ZShift = −0.27
Im Score = 83.8
NG = 0
DB = 0
SM = 0
Delta_AUC
Doxorubicin

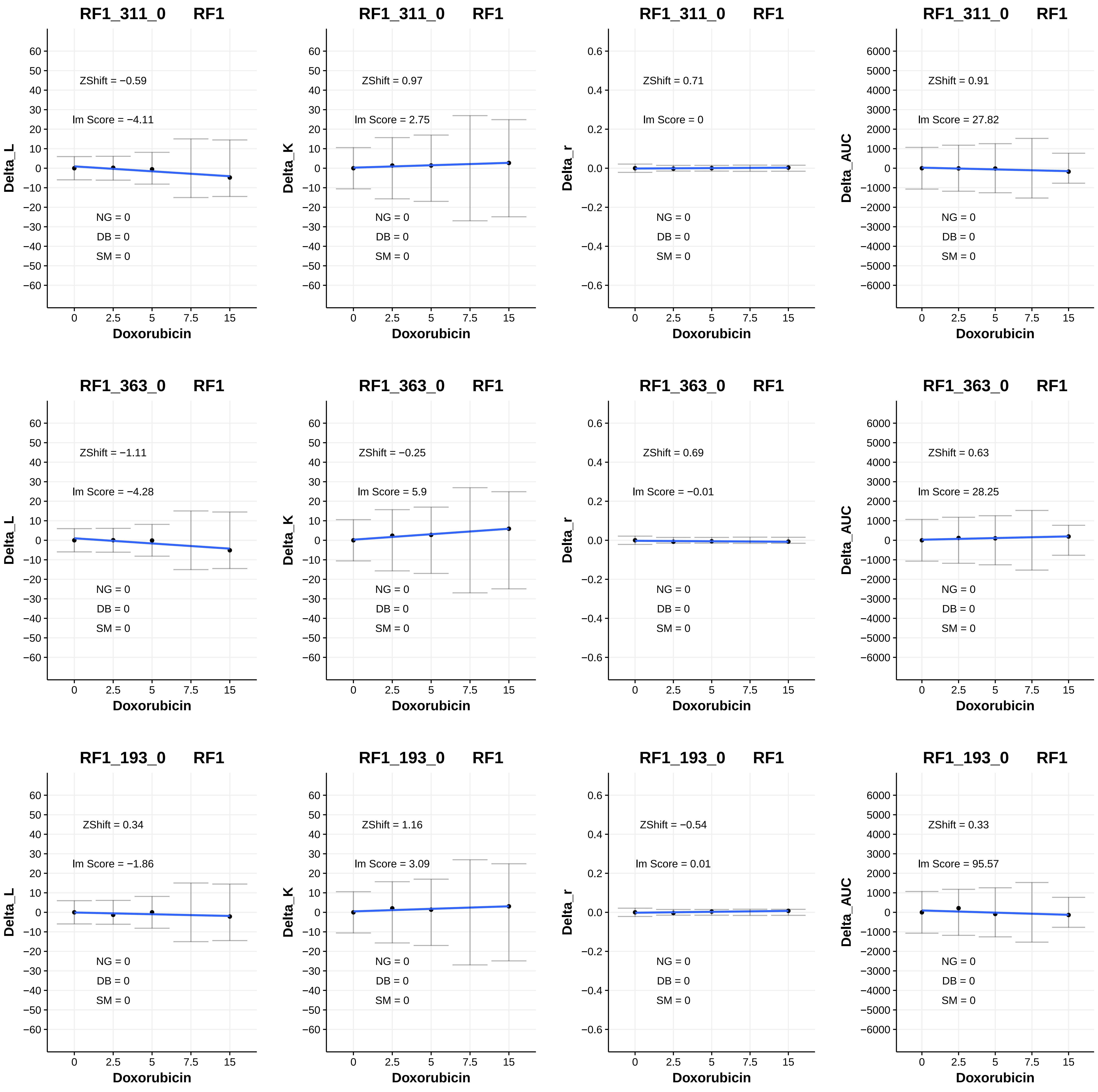
RF1_311_0 RF1
ZShift = −0.59
Im Score = −4.11
NG = 0
DB = 0
SM = 0
Delta_L
Doxorubicin
RF1_311_0 RF1
ZShift = 0.97
Im Score = 2.75
NG = 0
DB = 0
SM = 0
Delta_K
Doxorubicin
RF1_311_0 RF1
ZShift = 0.71
Im Score = 0
NG = 0
DB = 0
SM = 0
Delta_r
Doxorubicin
RF1_311_0 RF1
ZShift = 0.91
Im Score = 27.82
NG = 0
DB = 0
SM = 0
Delta_AUC
Doxorubicin
RF1_363_0 RF1
ZShift = −1.11
Im Score = −4.28
NG = 0
DB = 0
SM = 0
Delta_L
Doxorubicin
RF1_363_0 RF1
ZShift = −0.25
Im Score = 5.9
NG = 0
DB = 0
SM = 0
Delta_K
Doxorubicin
RF1_363_0 RF1
ZShift = 0.69
Im Score = −0.01
NG = 0
DB = 0
SM = 0
Delta_r
Doxorubicin
RF1_363_0 RF1
ZShift = 0.63
Im Score = 28.25
NG = 0
DB = 0
SM = 0
Delta_AUC
Doxorubicin
RF1_193_0 RF1
ZShift = 0.34
Im Score = −1.86
NG = 0
DB = 0
SM = 0
Delta_L
Doxorubicin
RF1_193_0 RF1
ZShift = 1.16
Im Score = 3.09
NG = 0
DB = 0
SM = 0
Delta_K
Doxorubicin
RF1_193_0 RF1
ZShift = −0.54
Im Score = 0.01
NG = 0
DB = 0
SM = 0
Delta_r
Doxorubicin
RF1_193_0 RF1
ZShift = 0.33
Im Score = 95.57
NG = 0
DB = 0
SM = 0
Delta_AUC
Doxorubicin

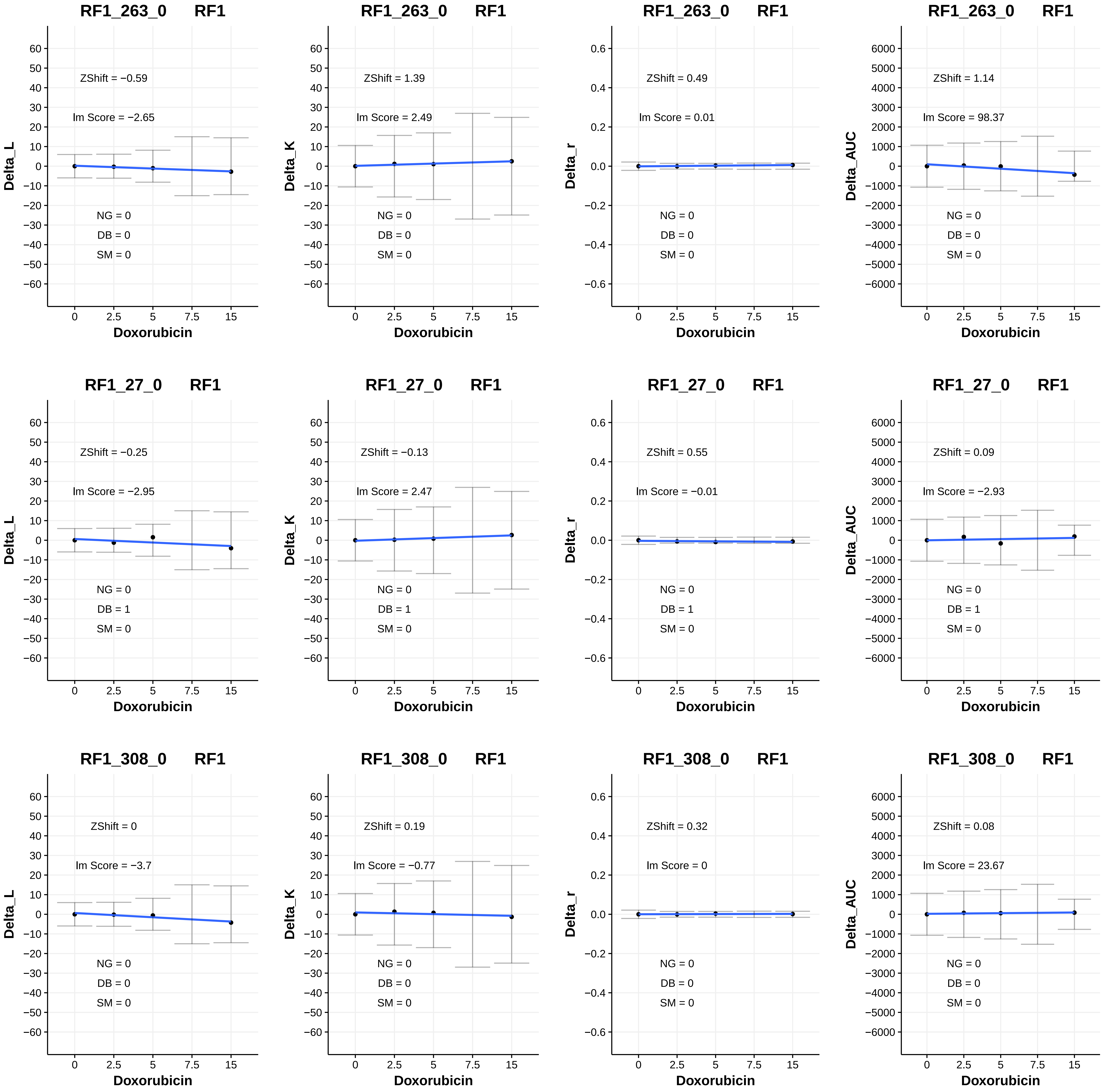
RF1_263_0 RF1
ZShift = −0.59
Im Score = −2.65
NG = 0
DB = 0
SM = 0
Delta_L
Doxorubicin
RF1_263_0 RF1
ZShift = 1.39
Im Score = 2.49
NG = 0
DB = 0
SM = 0
Delta_K
Doxorubicin
RF1_263_0 RF1
ZShift = 0.49
Im Score = 0.01
NG = 0
DB = 0
SM = 0
Delta_r
Doxorubicin
RF1_263_0 RF1
ZShift = 1.14
Im Score = 98.37
NG = 0
DB = 0
SM = 0
Delta_AUC
Doxorubicin
RF1_27_0 RF1
ZShift = −0.25
Im Score = −2.95
NG = 0
DB = 1
SM = 0
Delta_L
Doxorubicin
RF1_27_0 RF1
ZShift = −0.13
Im Score = 2.47
NG = 0
DB = 1
SM = 0
Delta_K
Doxorubicin
RF1_27_0 RF1
ZShift = 0.55
Im Score = −0.01
NG = 0
DB = 1
SM = 0
Delta_r
Doxorubicin
RF1_27_0 RF1
ZShift = 0.09
Im Score = −2.93
NG = 0
DB = 1
SM = 0
Delta_AUC
Doxorubicin
RF1_308_0 RF1
ZShift = 0
Im Score = −3.7
NG = 0
DB = 0
SM = 0
Delta_L
Doxorubicin
RF1_308_0 RF1
ZShift = 0.19
Im Score = −0.77
NG = 0
DB = 0
SM = 0
Delta_K
Doxorubicin
RF1_308_0 RF1
ZShift = 0.32
Im Score = 0
NG = 0
DB = 0
SM = 0
Delta_r
Doxorubicin
RF1_308_0 RF1
ZShift = 0.08
Im Score = 23.67
NG = 0
DB = 0
SM = 0
Delta_AUC
Doxorubicin

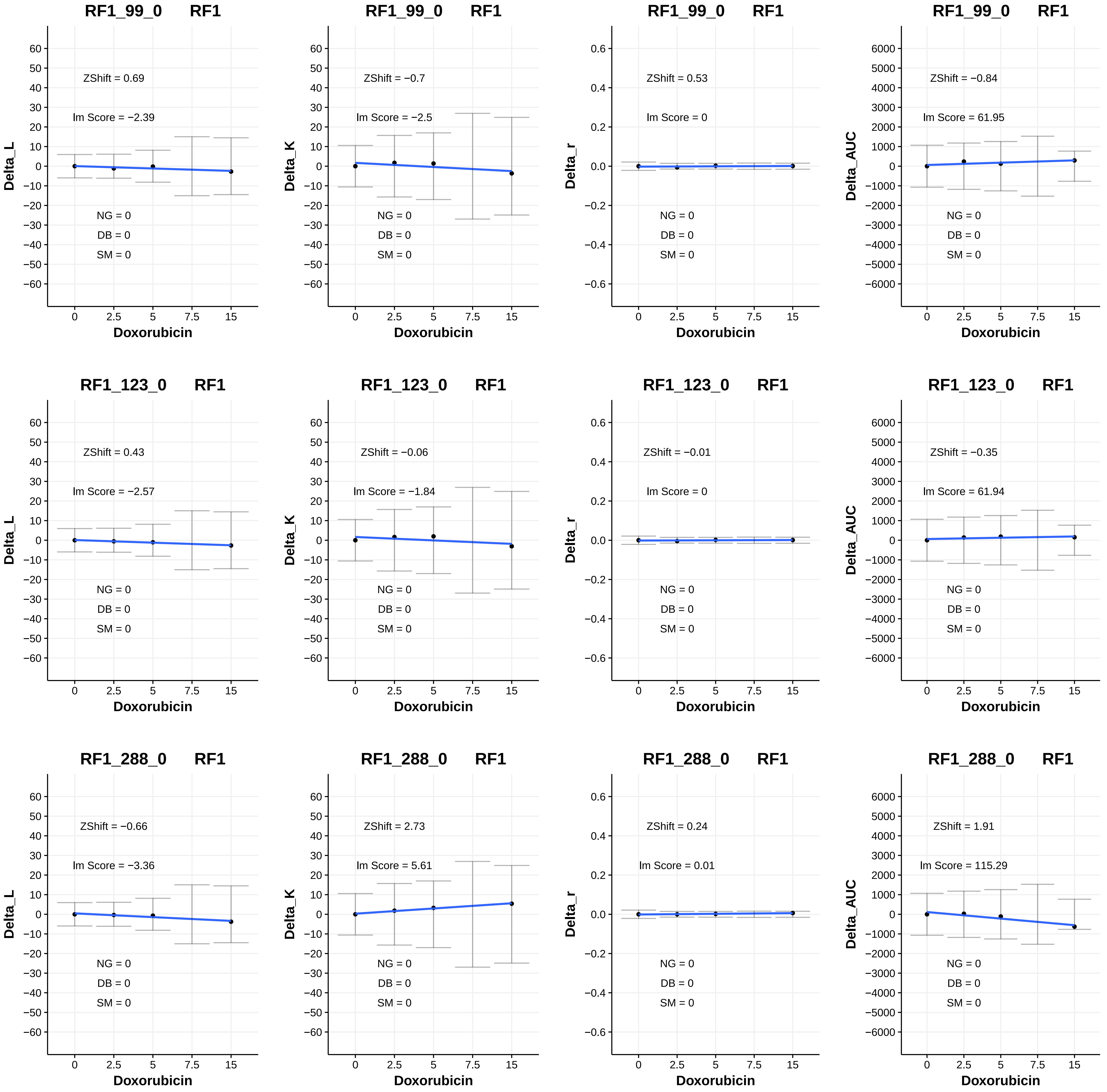

RF1_99_0 RF1
ZShift = 0.69
Im Score = −2.39
NG = 0
DB = 0
SM = 0
Delta_L
Doxorubicin
RF1_99_0 RF1
ZShift = −0.7
Im Score = −2.5
NG = 0
DB = 0
SM = 0
Delta_K
Doxorubicin
RF1_99_0 RF1
ZShift = 0.53
Im Score = 0
NG = 0
DB = 0
SM = 0
Delta_r
Doxorubicin
RF1_99_0 RF1
ZShift = −0.84
Im Score = 61.95
NG = 0
DB = 0
SM = 0
Delta_AUC
Doxorubicin
RF1_123_0 RF1
ZShift = 0.43
Im Score = −2.57
NG = 0
DB = 0
SM = 0
Delta_L
Doxorubicin
RF1_123_0 RF1
ZShift = −0.06
Im Score = −1.84
NG = 0
DB = 0
SM = 0
Delta_K
Doxorubicin
RF1_123_0 RF1
ZShift = −0.01
Im Score = 0
NG = 0
DB = 0
SM = 0
Delta_r
Doxorubicin
RF1_123_0 RF1
ZShift = −0.35
Im Score = 61.94
NG = 0
DB = 0
SM = 0
Delta_AUC
Doxorubicin
RF1_288_0 RF1
ZShift = −0.66
Im Score = −3.36
NG = 0
DB = 0
SM = 0
Delta_L
Doxorubicin
RF1_288_0 RF1
ZShift = 2.73
Im Score = 5.61
NG = 0
DB = 0
SM = 0
Delta_K
Doxorubicin
RF1_288_0 RF1
ZShift = 0.24
Im Score = 0.01
NG = 0
DB = 0
SM = 0
Delta_r
Doxorubicin
RF1_288_0 RF1
ZShift = 1.91
Im Score = 115.29
NG = 0
DB = 0
SM = 0
Delta_AUC
Doxorubicin

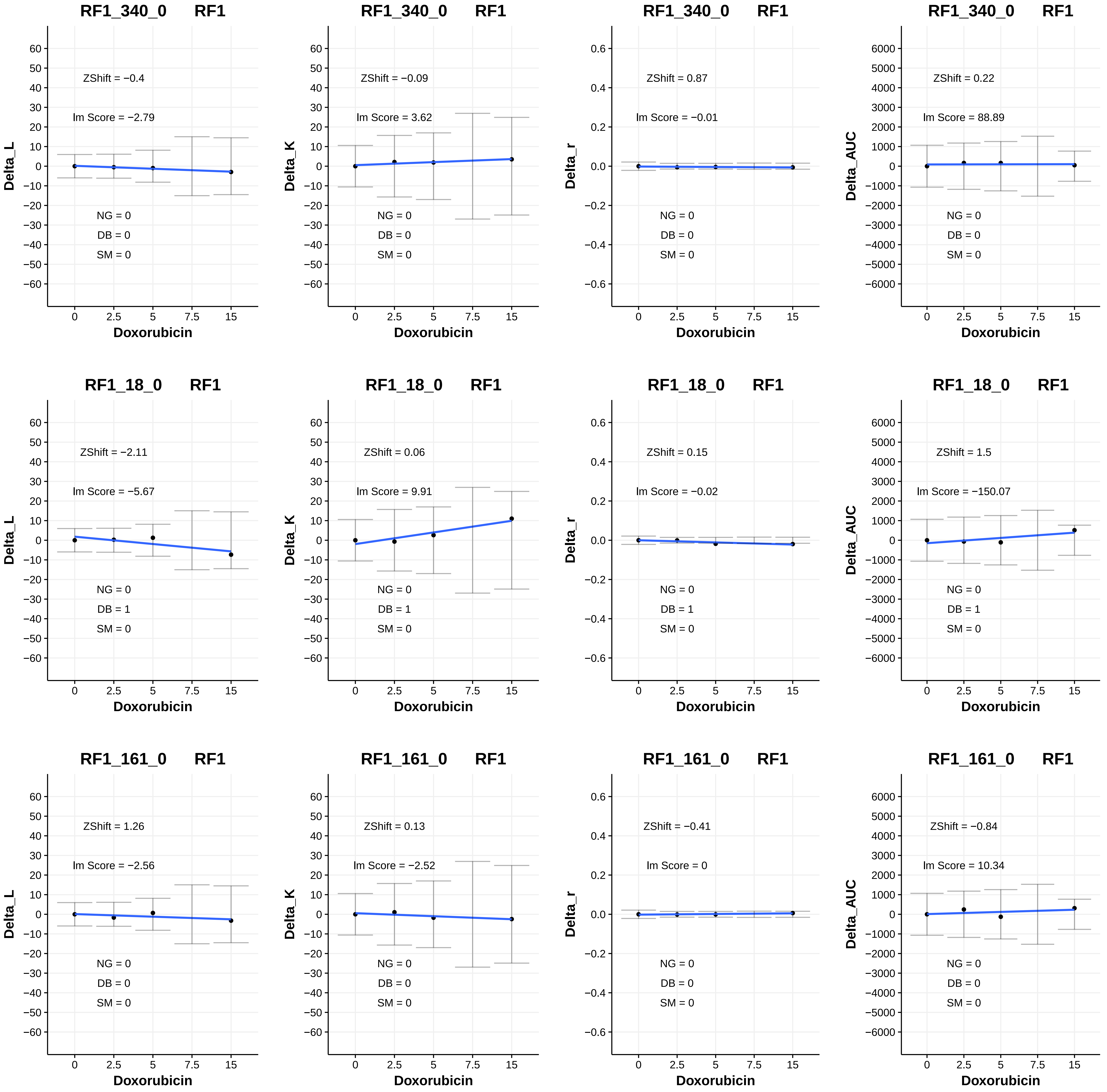
RF1_340_0 RF1
ZShift = −0.4
Im Score = −2.79
NG = 0
DB = 0
SM = 0
Delta_L
Doxorubicin
RF1_340_0 RF1
ZShift = −0.09
Im Score = 3.62
NG = 0
DB = 0
SM = 0
Delta_K
Doxorubicin
RF1_340_0 RF1
ZShift = 0.87
Im Score = −0.01
NG = 0
DB = 0
SM = 0
Delta_r
Doxorubicin
RF1_340_0 RF1
ZShift = 0.22
Im Score = 88.89
NG = 0
DB = 0
SM = 0
Delta_AUC
Doxorubicin
RF1_18_0 RF1
ZShift = −2.11
Im Score = −5.67
NG = 0
DB = 1
SM = 0
Delta_L
Doxorubicin
RF1_18_0 RF1
ZShift = 0.06
Im Score = 9.91
NG = 0
DB = 1
SM = 0
Delta_K
Doxorubicin
RF1_18_0 RF1
ZShift = 0.15
Im Score = −0.02
NG = 0
DB = 1
SM = 0
Delta_r
Doxorubicin
RF1_18_0 RF1
ZShift = 1.5
Im Score = −150.07
NG = 0
DB = 1
SM = 0
Delta_AUC
Doxorubicin
RF1_161_0 RF1
ZShift = 1.26
Im Score = −2.56
NG = 0
DB = 0
SM = 0
Delta_L
Doxorubicin
RF1_161_0 RF1
ZShift = 0.13
Im Score = −2.52
NG = 0
DB = 0
SM = 0
Delta_K
Doxorubicin
RF1_161_0 RF1
ZShift = −0.41
Im Score = 0
NG = 0
DB = 0
SM = 0
Delta_r
Doxorubicin
RF1_161_0 RF1
ZShift = −0.84
Im Score = 10.34
NG = 0
DB = 0
SM = 0
Delta_AUC
Doxorubicin

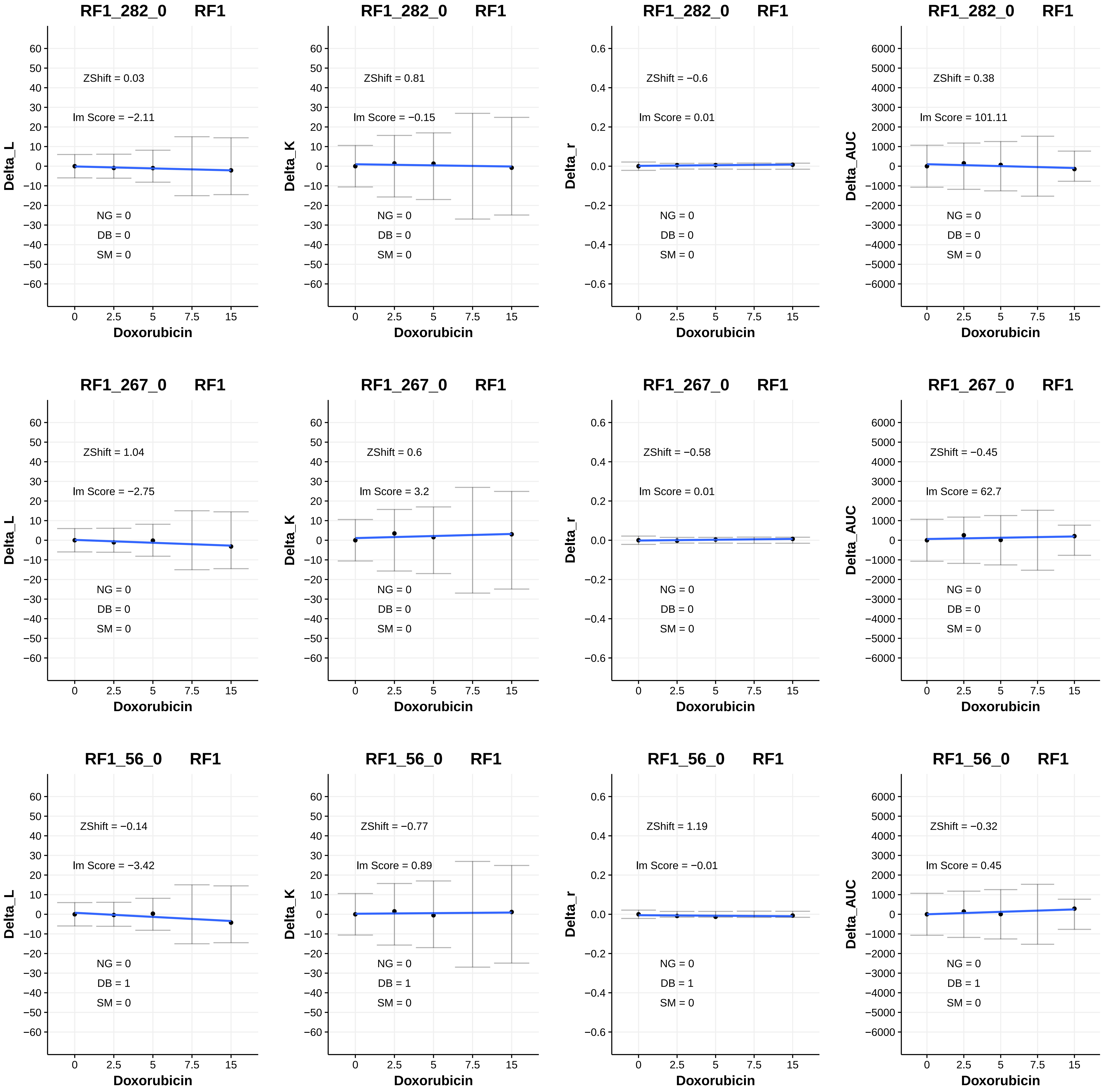
RF1_282_0 RF1
ZShift = 0.03
lm Score = −2.11
NG = 0
DB = 0
SM = 0
Delta_L
Doxorubicin
RF1_282_0 RF1
ZShift = 0.81
lm Score = −0.15
NG = 0
DB = 0
SM = 0
Delta_K
Doxorubicin
RF1_282_0 RF1
ZShift = −0.6
lm Score = 0.01
NG = 0
DB = 0
SM = 0
Delta_r
Doxorubicin
RF1_282_0 RF1
ZShift = 0.38
lm Score = 101.11
NG = 0
DB = 0
SM = 0
Delta_AUC
Doxorubicin
RF1_267_0 RF1
ZShift = 1.04
lm Score = −2.75
NG = 0
DB = 0
SM = 0
Delta_L
Doxorubicin
RF1_267_0 RF1
ZShift = 0.6
lm Score = 3.2
NG = 0
DB = 0
SM = 0
Delta_K
Doxorubicin
RF1_267_0 RF1
ZShift = −0.58
lm Score = 0.01
NG = 0
DB = 0
SM = 0
Delta_r
Doxorubicin
RF1_267_0 RF1
ZShift = −0.45
lm Score = 62.7
NG = 0
DB = 0
SM = 0
Delta_AUC
Doxorubicin
RF1_56_0 RF1
ZShift = −0.14
lm Score = −3.42
NG = 0
DB = 1
SM = 0
Delta_L
Doxorubicin
RF1_56_0 RF1
ZShift = −0.77
lm Score = 0.89
NG = 0
DB = 1
SM = 0
Delta_K
Doxorubicin
RF1_56_0 RF1
ZShift = 1.19
lm Score = −0.01
NG = 0
DB = 1
SM = 0
Delta_r
Doxorubicin
RF1_56_0 RF1
ZShift = −0.32
lm Score = 0.45
NG = 0
DB = 1
SM = 0
Delta_AUC
Doxorubicin

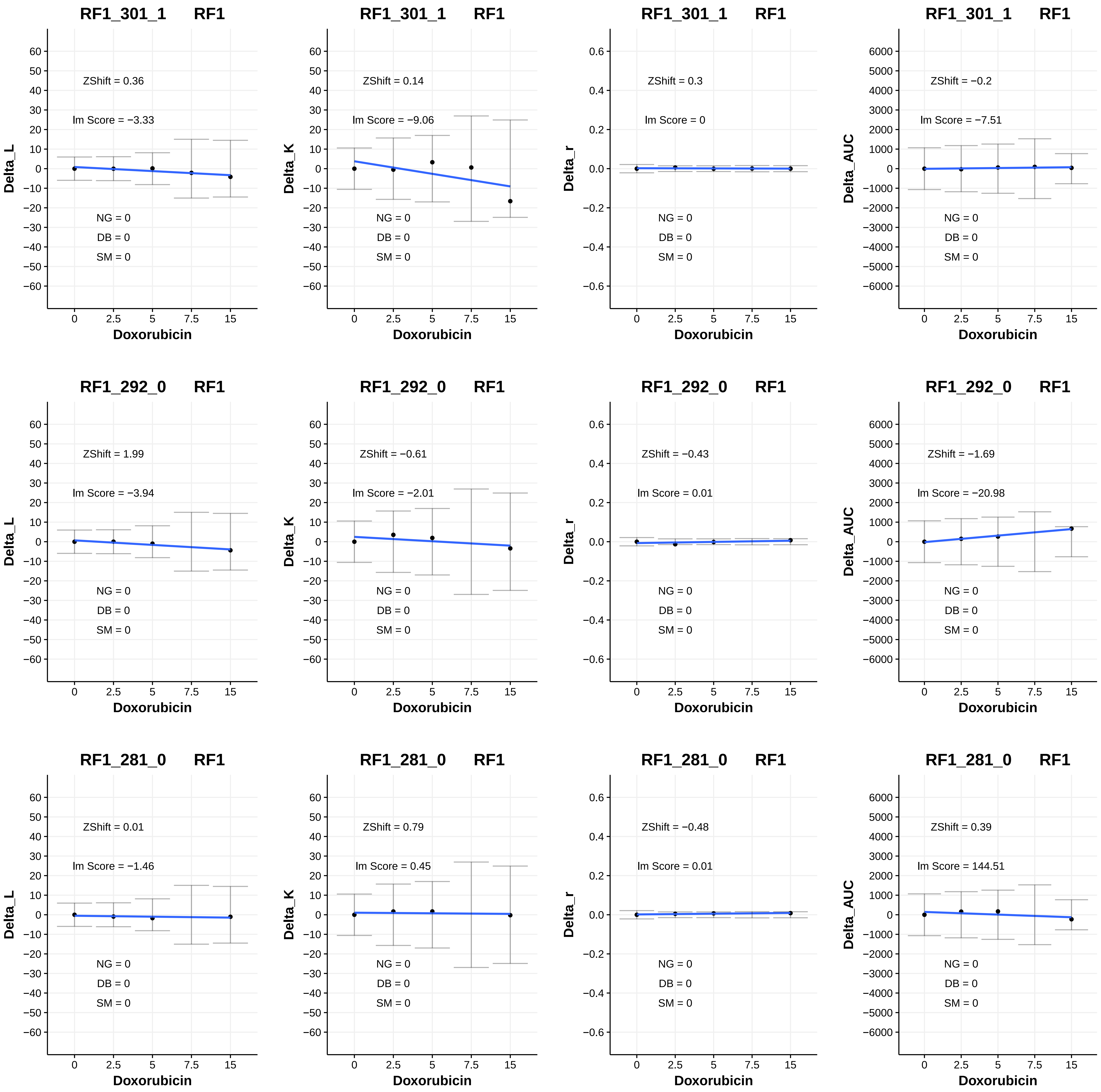

RF1_301_1 RF1
ZShift = 0.36
lm Score = −3.33
NG = 0
DB = 0
SM = 0
Delta_L
Doxorubicin
RF1_301_1 RF1
ZShift = 0.14
lm Score = −9.06
NG = 0
DB = 0
SM = 0
Delta_K
Doxorubicin
RF1_301_1 RF1
ZShift = 0.3
lm Score = 0
NG = 0
DB = 0
SM = 0
Delta_r
Doxorubicin
RF1_301_1 RF1
ZShift = −0.2
lm Score = −7.51
NG = 0
DB = 0
SM = 0
Delta_AUC
Doxorubicin
RF1_292_0 RF1
ZShift = 1.99
lm Score = −3.94
NG = 0
DB = 0
SM = 0
Delta_L
Doxorubicin
RF1_292_0 RF1
ZShift = −0.61
lm Score = −2.01
NG = 0
DB = 0
SM = 0
Delta_K
Doxorubicin
RF1_292_0 RF1
ZShift = −0.43
lm Score = 0.01
NG = 0
DB = 0
SM = 0
Delta_r
Doxorubicin
RF1_292_0 RF1
ZShift = −1.69
lm Score = −20.98
NG = 0
DB = 0
SM = 0
Delta_AUC
Doxorubicin
RF1_281_0 RF1
ZShift = 0.01
lm Score = −1.46
NG = 0
DB = 0
SM = 0
Delta_L
Doxorubicin
RF1_281_0 RF1
ZShift = 0.79
lm Score = 0.45
NG = 0
DB = 0
SM = 0
Delta_K
Doxorubicin
RF1_281_0 RF1
ZShift = −0.48
lm Score = 0.01
NG = 0
DB = 0
SM = 0
Delta_r
Doxorubicin
RF1_281_0 RF1
ZShift = 0.39
lm Score = 144.51
NG = 0
DB = 0
SM = 0
Delta_AUC
Doxorubicin

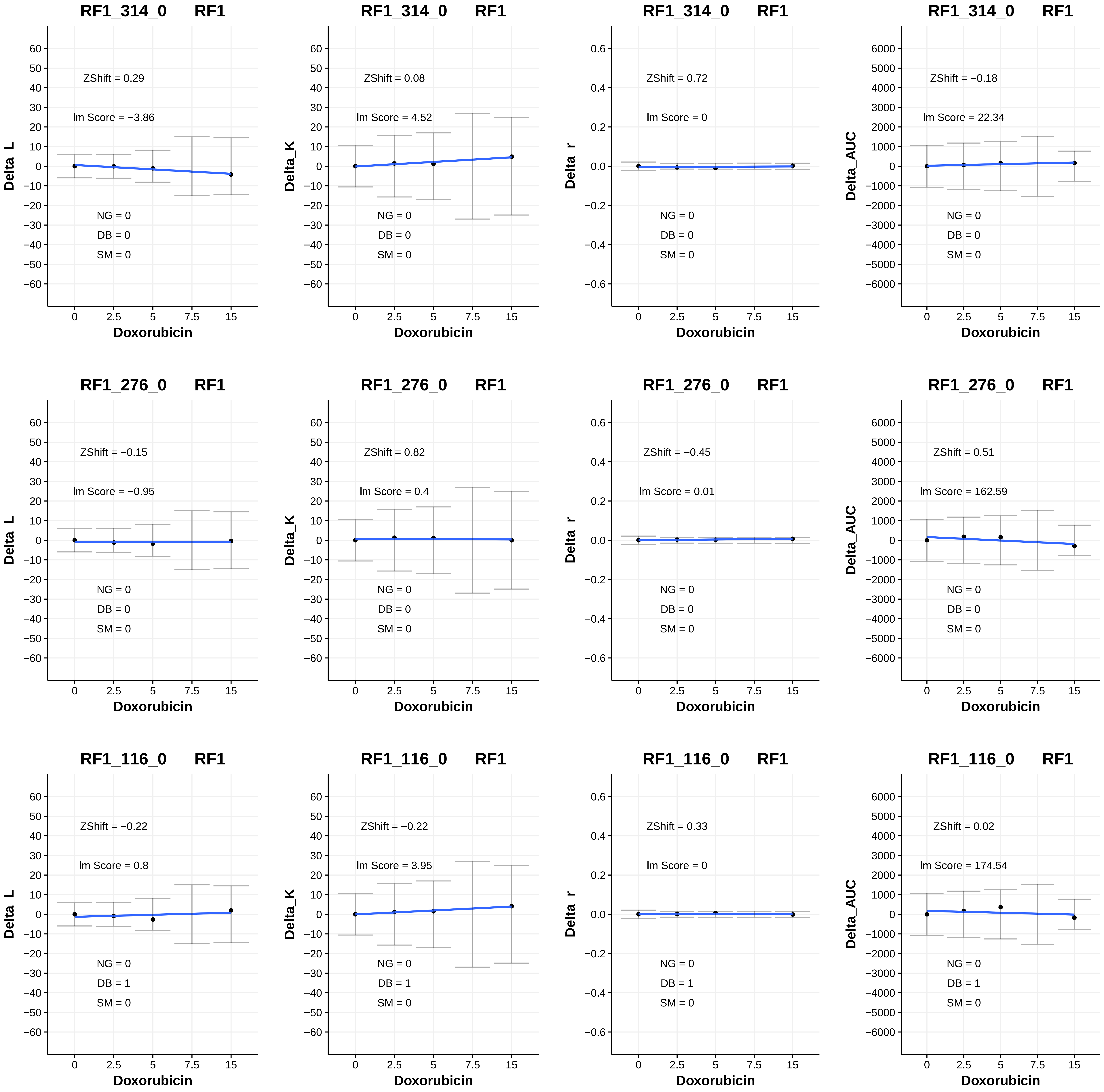

RF1_314_0 RF1
ZShift = 0.29
lm Score = −3.86
NG = 0
DB = 0
SM = 0
Delta_L
Doxorubicin
RF1_314_0 RF1
ZShift = 0.08
lm Score = 4.52
NG = 0
DB = 0
SM = 0
Delta_K
Doxorubicin
RF1_314_0 RF1
ZShift = 0.72
lm Score = 0
NG = 0
DB = 0
SM = 0
Delta_r
Doxorubicin
RF1_314_0 RF1
ZShift = −0.18
lm Score = 22.34
NG = 0
DB = 0
SM = 0
Delta_AUC
Doxorubicin
RF1_276_0 RF1
ZShift = −0.15
lm Score = −0.95
NG = 0
DB = 0
SM = 0
Delta_L
Doxorubicin
RF1_276_0 RF1
ZShift = 0.82
lm Score = 0.4
NG = 0
DB = 0
SM = 0
Delta_K
Doxorubicin
RF1_276_0 RF1
ZShift = −0.45
lm Score = 0.01
NG = 0
DB = 0
SM = 0
Delta_r
Doxorubicin
RF1_276_0 RF1
ZShift = 0.51
lm Score = 162.59
NG = 0
DB = 0
SM = 0
Delta_AUC
Doxorubicin
RF1_116_0 RF1
ZShift = −0.22
lm Score = 0.8
NG = 0
DB = 1
SM = 0
Delta_L
Doxorubicin
RF1_116_0 RF1
ZShift = −0.22
lm Score = 3.95
NG = 0
DB = 1
SM = 0
Delta_K
Doxorubicin
RF1_116_0 RF1
ZShift = 0.33
lm Score = 0
NG = 0
DB = 1
SM = 0
Delta_r
Doxorubicin
RF1_116_0 RF1
ZShift = 0.02
lm Score = 174.54
NG = 0
DB = 1
SM = 0
Delta_AUC
Doxorubicin

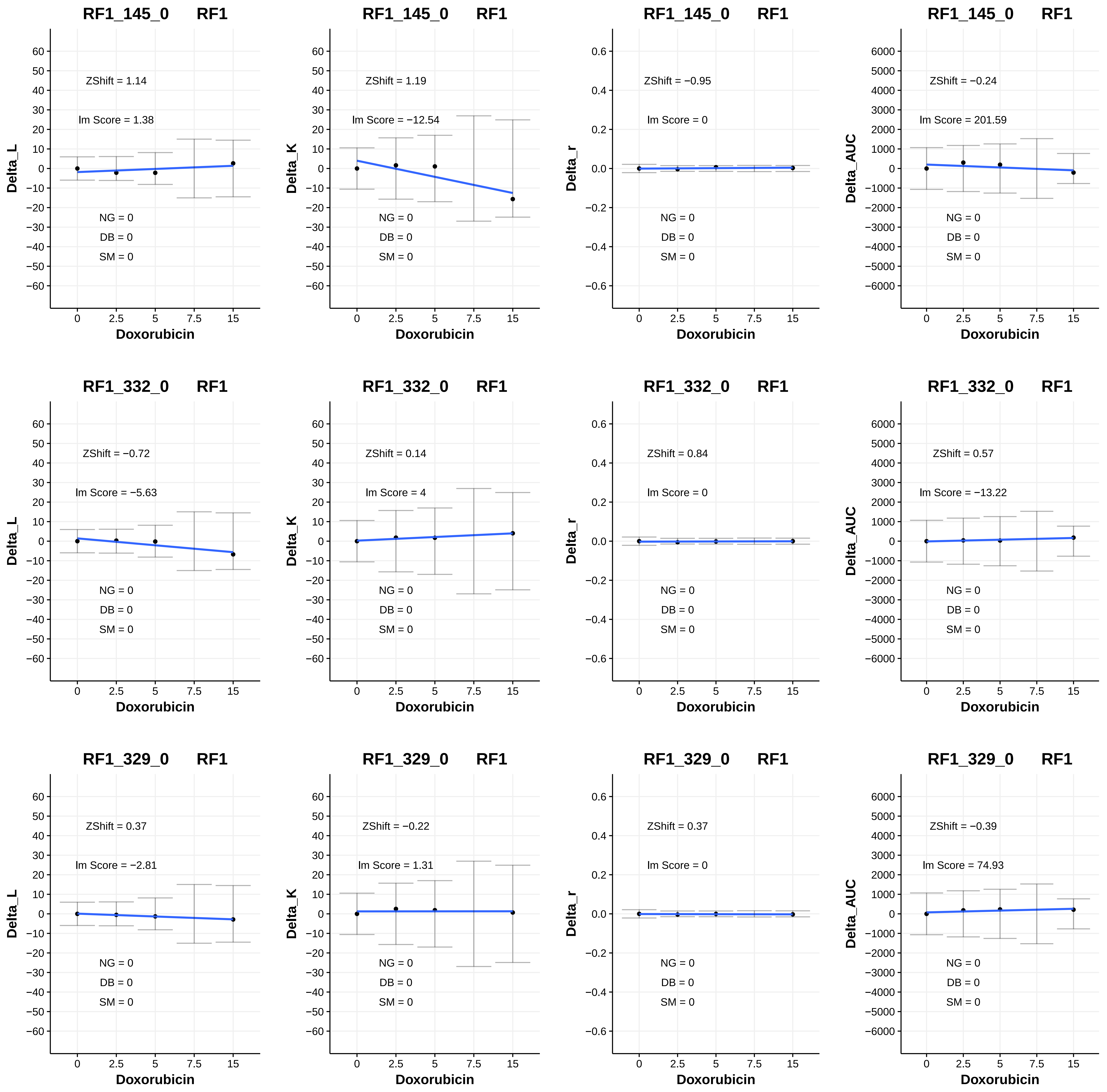
RF1_145_0 RF1
Delta_L
ZShift = 1.14
lm Score = 1.38
NG = 0
DB = 0
SM = 0
Doxorubicin
RF1_145_0 RF1
Delta_K
ZShift = 1.19
lm Score = −12.54
NG = 0
DB = 0
SM = 0
Doxorubicin
RF1_145_0 RF1
Delta_r
ZShift = −0.95
lm Score = 0
NG = 0
DB = 0
SM = 0
Doxorubicin
RF1_145_0 RF1
Delta_AUC
ZShift = −0.24
lm Score = 201.59
NG = 0
DB = 0
SM = 0
Doxorubicin
RF1_332_0 RF1
Delta_L
ZShift = −0.72
lm Score = −5.63
NG = 0
DB = 0
SM = 0
Doxorubicin
RF1_332_0 RF1
Delta_K
ZShift = 0.14
lm Score = 4
NG = 0
DB = 0
SM = 0
Doxorubicin
RF1_332_0 RF1
Delta_r
ZShift = 0.84
lm Score = 0
NG = 0
DB = 0
SM = 0
Doxorubicin
RF1_332_0 RF1
Delta_AUC
ZShift = 0.57
lm Score = −13.22
NG = 0
DB = 0
SM = 0
Doxorubicin
RF1_329_0 RF1
Delta_L
ZShift = 0.37
lm Score = −2.81
NG = 0
DB = 0
SM = 0
Doxorubicin
RF1_329_0 RF1
Delta_K
ZShift = −0.22
lm Score = 1.31
NG = 0
DB = 0
SM = 0
Doxorubicin
RF1_329_0 RF1
Delta_r
ZShift = 0.37
lm Score = 0
NG = 0
DB = 0
SM = 0
Doxorubicin
RF1_329_0 RF1
Delta_AUC
ZShift = −0.39
lm Score = 74.93
NG = 0
DB = 0
SM = 0
Doxorubicin

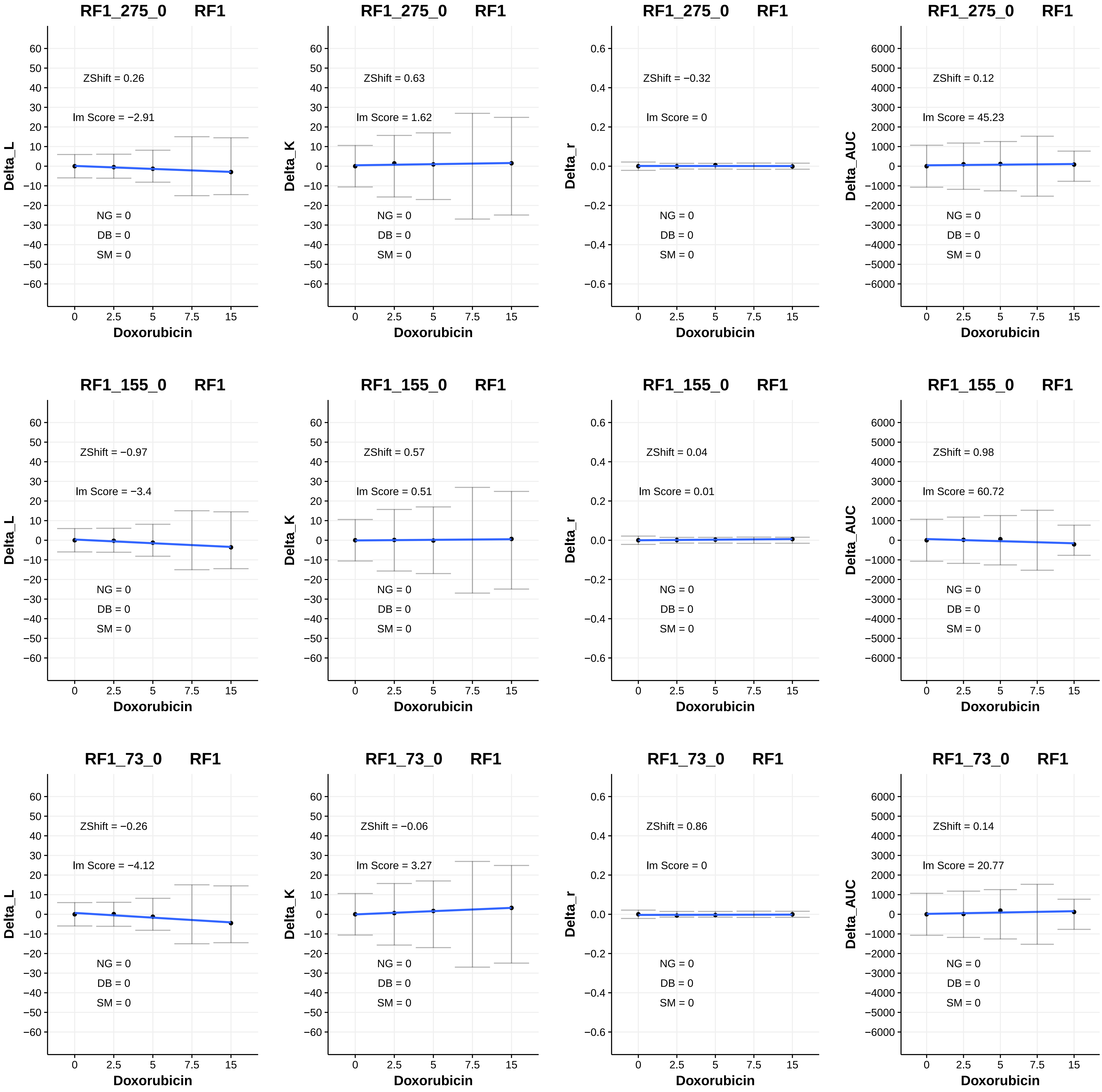
RF1_275_0 RF1
Delta_L
ZShift = 0.26
lm Score = −2.91
NG = 0
DB = 0
SM = 0
Doxorubicin
RF1_275_0 RF1
Delta_K
ZShift = 0.63
lm Score = 1.62
NG = 0
DB = 0
SM = 0
Doxorubicin
RF1_275_0 RF1
Delta_r
ZShift = −0.32
lm Score = 0
NG = 0
DB = 0
SM = 0
Doxorubicin
RF1_275_0 RF1
Delta_AUC
ZShift = 0.12
lm Score = 45.23
NG = 0
DB = 0
SM = 0
Doxorubicin
RF1_155_0 RF1
Delta_L
ZShift = −0.97
lm Score = −3.4
NG = 0
DB = 0
SM = 0
Doxorubicin
RF1_155_0 RF1
Delta_K
ZShift = 0.57
lm Score = 0.51
NG = 0
DB = 0
SM = 0
Doxorubicin
RF1_155_0 RF1
Delta_r
ZShift = 0.04
lm Score = 0.01
NG = 0
DB = 0
SM = 0
Doxorubicin
RF1_155_0 RF1
Delta_AUC
ZShift = 0.98
lm Score = 60.72
NG = 0
DB = 0
SM = 0
Doxorubicin
RF1_73_0 RF1
Delta_L
ZShift = −0.26
lm Score = −4.12
NG = 0
DB = 0
SM = 0
Doxorubicin
RF1_73_0 RF1
Delta_K
ZShift = −0.06
lm Score = 3.27
NG = 0
DB = 0
SM = 0
Doxorubicin
RF1_73_0 RF1
Delta_r
ZShift = 0.86
lm Score = 0
NG = 0
DB = 0
SM = 0
Doxorubicin
RF1_73_0 RF1
Delta_AUC
ZShift = 0.14
lm Score = 20.77
NG = 0
DB = 0
SM = 0
Doxorubicin

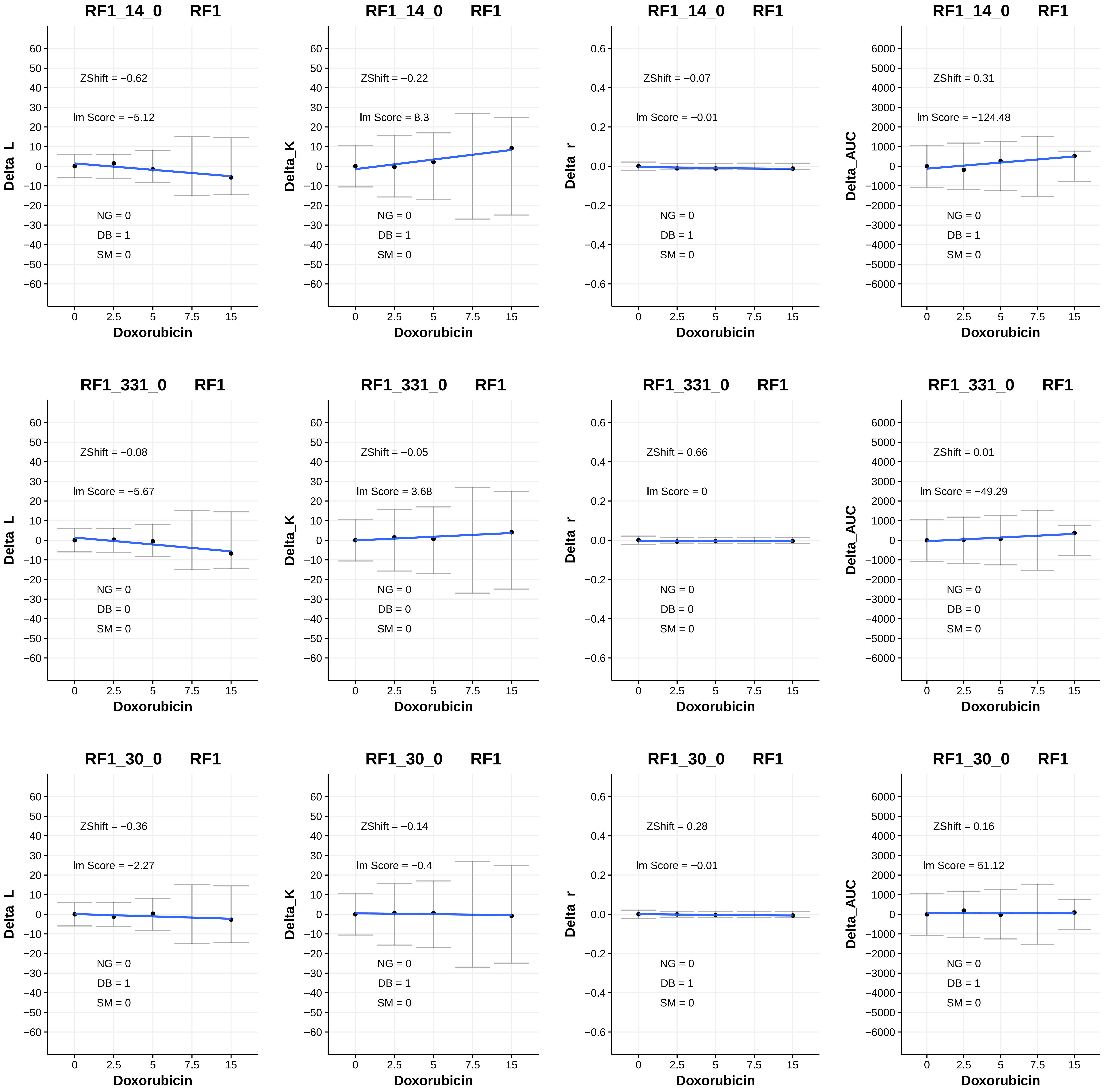
RF1_14_0 RF1
ZShift = −0.62
lm Score = −5.12
NG = 0
DB = 1
SM = 0
Delta_L
Doxorubicin
RF1_14_0 RF1
ZShift = −0.22
lm Score = 8.3
NG = 0
DB = 1
SM = 0
Delta_K
Doxorubicin
RF1_14_0 RF1
ZShift = −0.07
lm Score = −0.01
NG = 0
DB = 1
SM = 0
Delta_r
Doxorubicin
RF1_14_0 RF1
ZShift = 0.31
lm Score = −124.48
NG = 0
DB = 1
SM = 0
Delta_AUC
Doxorubicin
RF1_331_0 RF1
ZShift = −0.08
lm Score = −5.67
NG = 0
DB = 0
SM = 0
Delta_L
Doxorubicin
RF1_331_0 RF1
ZShift = −0.05
lm Score = 3.68
NG = 0
DB = 0
SM = 0
Delta_K
Doxorubicin
RF1_331_0 RF1
ZShift = 0.66
lm Score = 0
NG = 0
DB = 0
SM = 0
Delta_r
Doxorubicin
RF1_331_0 RF1
ZShift = 0.01
lm Score = −49.29
NG = 0
DB = 0
SM = 0
Delta_AUC
Doxorubicin
RF1_30_0 RF1
ZShift = −0.36
lm Score = −2.27
NG = 0
DB = 1
SM = 0
Delta_L
Doxorubicin
RF1_30_0 RF1
ZShift = −0.14
lm Score = −0.4
NG = 0
DB = 1
SM = 0
Delta_K
Doxorubicin
RF1_30_0 RF1
ZShift = 0.28
lm Score = −0.01
NG = 0
DB = 1
SM = 0
Delta_r
Doxorubicin
RF1_30_0 RF1
ZShift = 0.16
lm Score = 51.12
NG = 0
DB = 1
SM = 0
Delta_AUC
Doxorubicin

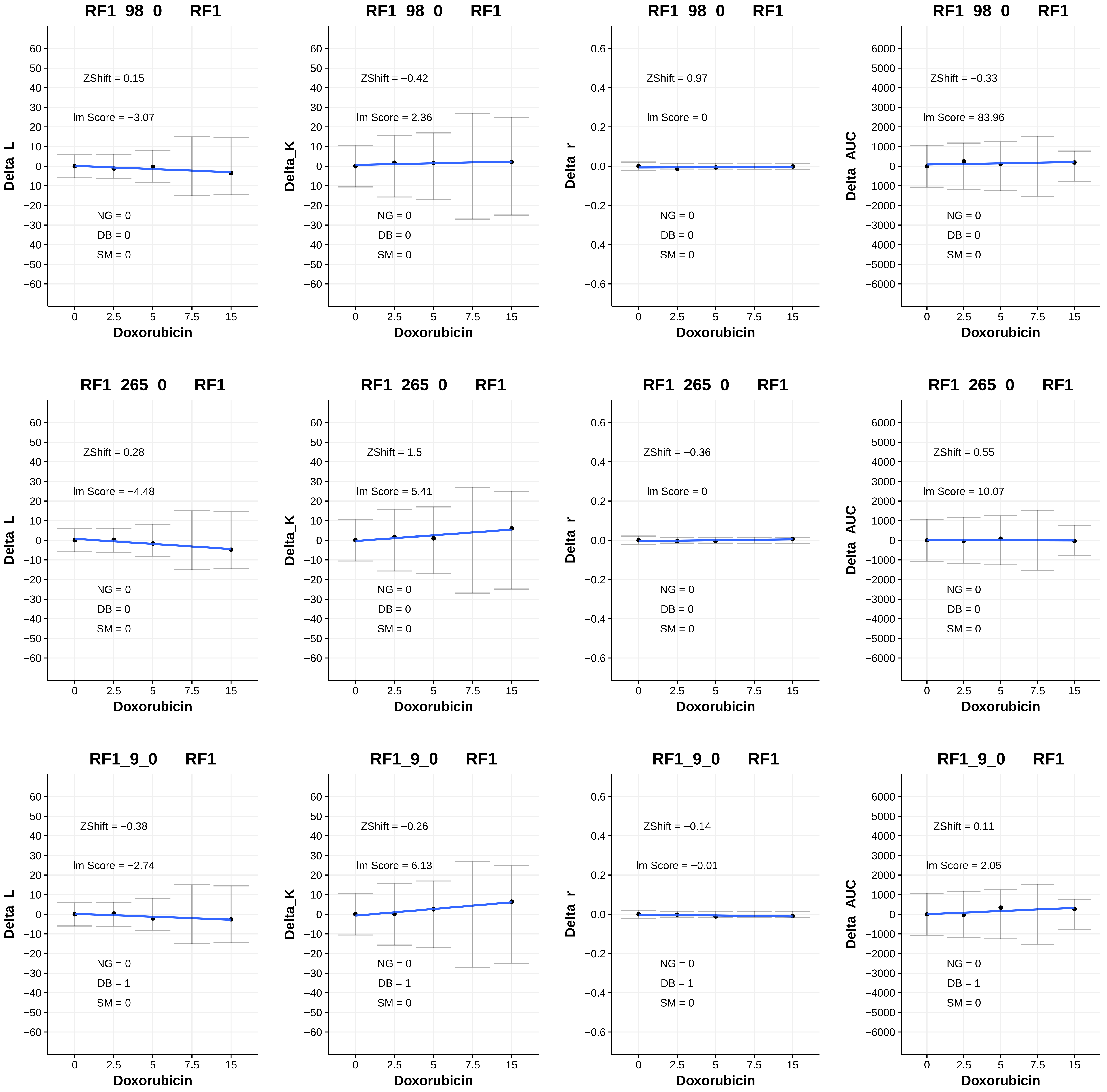
RF1_98_0 RF1
ZShift = 0.15
lm Score = −3.07
NG = 0
DB = 0
SM = 0
Delta_L
Doxorubicin
RF1_98_0 RF1
ZShift = −0.42
lm Score = 2.36
NG = 0
DB = 0
SM = 0
Delta_K
Doxorubicin
RF1_98_0 RF1
ZShift = 0.97
lm Score = 0
NG = 0
DB = 0
SM = 0
Delta_r
Doxorubicin
RF1_98_0 RF1
ZShift = −0.33
lm Score = 83.96
NG = 0
DB = 0
SM = 0
Delta_AUC
Doxorubicin
RF1_265_0 RF1
ZShift = 0.28
lm Score = −4.48
NG = 0
DB = 0
SM = 0
Delta_L
Doxorubicin
RF1_265_0 RF1
ZShift = 1.5
lm Score = 5.41
NG = 0
DB = 0
SM = 0
Delta_K
Doxorubicin
RF1_265_0 RF1
ZShift = −0.36
lm Score = 0
NG = 0
DB = 0
SM = 0
Delta_r
Doxorubicin
RF1_265_0 RF1
ZShift = 0.55
lm Score = 10.07
NG = 0
DB = 0
SM = 0
Delta_AUC
Doxorubicin
RF1_9_0 RF1
ZShift = −0.38
lm Score = −2.74
NG = 0
DB = 1
SM = 0
Delta_L
Doxorubicin
RF1_9_0 RF1
ZShift = −0.26
lm Score = 6.13
NG = 0
DB = 1
SM = 0
Delta_K
Doxorubicin
RF1_9_0 RF1
ZShift = −0.14
lm Score = −0.01
NG = 0
DB = 1
SM = 0
Delta_r
Doxorubicin
RF1_9_0 RF1
ZShift = 0.11
lm Score = 2.05
NG = 0
DB = 1
SM = 0
Delta_AUC
Doxorubicin

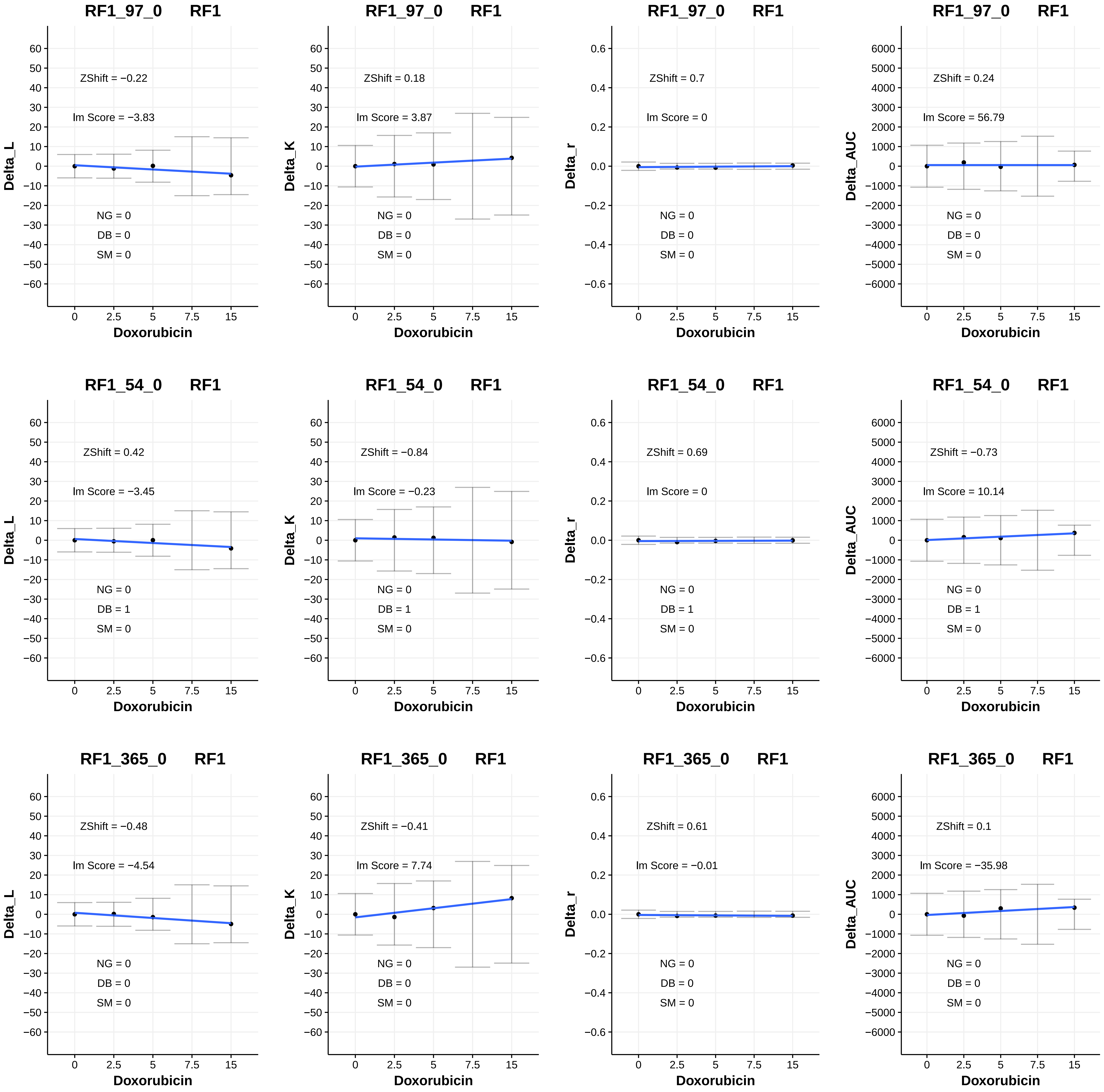

RF1_97_0 RF1
ZShift = −0.22
Im Score = −3.83
NG = 0
DB = 0
SM = 0
Delta_L
Doxorubicin
RF1_97_0 RF1
ZShift = 0.18
Im Score = 3.87
NG = 0
DB = 0
SM = 0
Delta_K
Doxorubicin
RF1_97_0 RF1
ZShift = 0.7
Im Score = 0
NG = 0
DB = 0
SM = 0
Delta_r
Doxorubicin
RF1_97_0 RF1
ZShift = 0.24
Im Score = 56.79
NG = 0
DB = 0
SM = 0
Delta_AUC
Doxorubicin
RF1_54_0 RF1
ZShift = 0.42
Im Score = −3.45
NG = 0
DB = 1
SM = 0
Delta_L
Doxorubicin
RF1_54_0 RF1
ZShift = −0.84
Im Score = −0.23
NG = 0
DB = 1
SM = 0
Delta_K
Doxorubicin
RF1_54_0 RF1
ZShift = 0.69
Im Score = 0
NG = 0
DB = 1
SM = 0
Delta_r
Doxorubicin
RF1_54_0 RF1
ZShift = −0.73
Im Score = 10.14
NG = 0
DB = 1
SM = 0
Delta_AUC
Doxorubicin
RF1_365_0 RF1
ZShift = −0.48
Im Score = −4.54
NG = 0
DB = 0
SM = 0
Delta_L
Doxorubicin
RF1_365_0 RF1
ZShift = −0.41
Im Score = 7.74
NG = 0
DB = 0
SM = 0
Delta_K
Doxorubicin
RF1_365_0 RF1
ZShift = 0.61
Im Score = −0.01
NG = 0
DB = 0
SM = 0
Delta_r
Doxorubicin
RF1_365_0 RF1
ZShift = 0.1
Im Score = −35.98
NG = 0
DB = 0
SM = 0
Delta_AUC
Doxorubicin

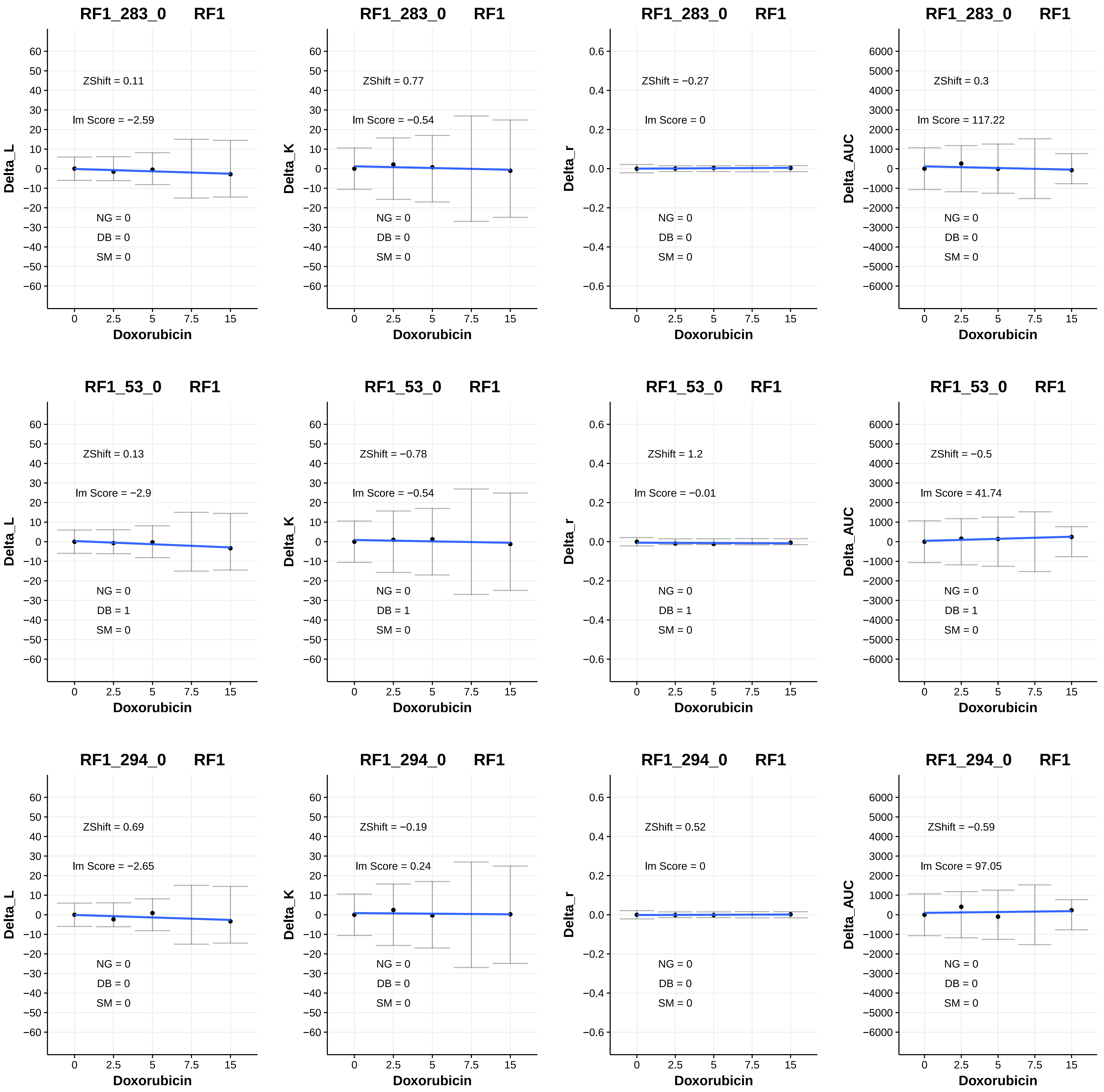

RF1_283_0 RF1
ZShift = 0.11
Im Score = −2.59
NG = 0
DB = 0
SM = 0
Delta_L
Doxorubicin
RF1_283_0 RF1
ZShift = 0.77
Im Score = −0.54
NG = 0
DB = 0
SM = 0
Delta_K
Doxorubicin
RF1_283_0 RF1
ZShift = −0.27
Im Score = 0
NG = 0
DB = 0
SM = 0
Delta_r
Doxorubicin
RF1_283_0 RF1
ZShift = 0.3
Im Score = 117.22
NG = 0
DB = 0
SM = 0
Delta_AUC
Doxorubicin
RF1_53_0 RF1
ZShift = 0.13
Im Score = −2.9
NG = 0
DB = 1
SM = 0
Delta_L
Doxorubicin
RF1_53_0 RF1
ZShift = −0.78
Im Score = −0.54
NG = 0
DB = 1
SM = 0
Delta_K
Doxorubicin
RF1_53_0 RF1
ZShift = 1.2
Im Score = −0.01
NG = 0
DB = 1
SM = 0
Delta_r
Doxorubicin
RF1_53_0 RF1
ZShift = −0.5
Im Score = 41.74
NG = 0
DB = 1
SM = 0
Delta_AUC
Doxorubicin
RF1_294_0 RF1
ZShift = 0.69
Im Score = −2.65
NG = 0
DB = 0
SM = 0
Delta_L
Doxorubicin
RF1_294_0 RF1
ZShift = −0.19
Im Score = 0.24
NG = 0
DB = 0
SM = 0
Delta_K
Doxorubicin
RF1_294_0 RF1
ZShift = 0.52
Im Score = 0
NG = 0
DB = 0
SM = 0
Delta_r
Doxorubicin
RF1_294_0 RF1
ZShift = −0.59
Im Score = 97.05
NG = 0
DB = 0
SM = 0
Delta_AUC
Doxorubicin

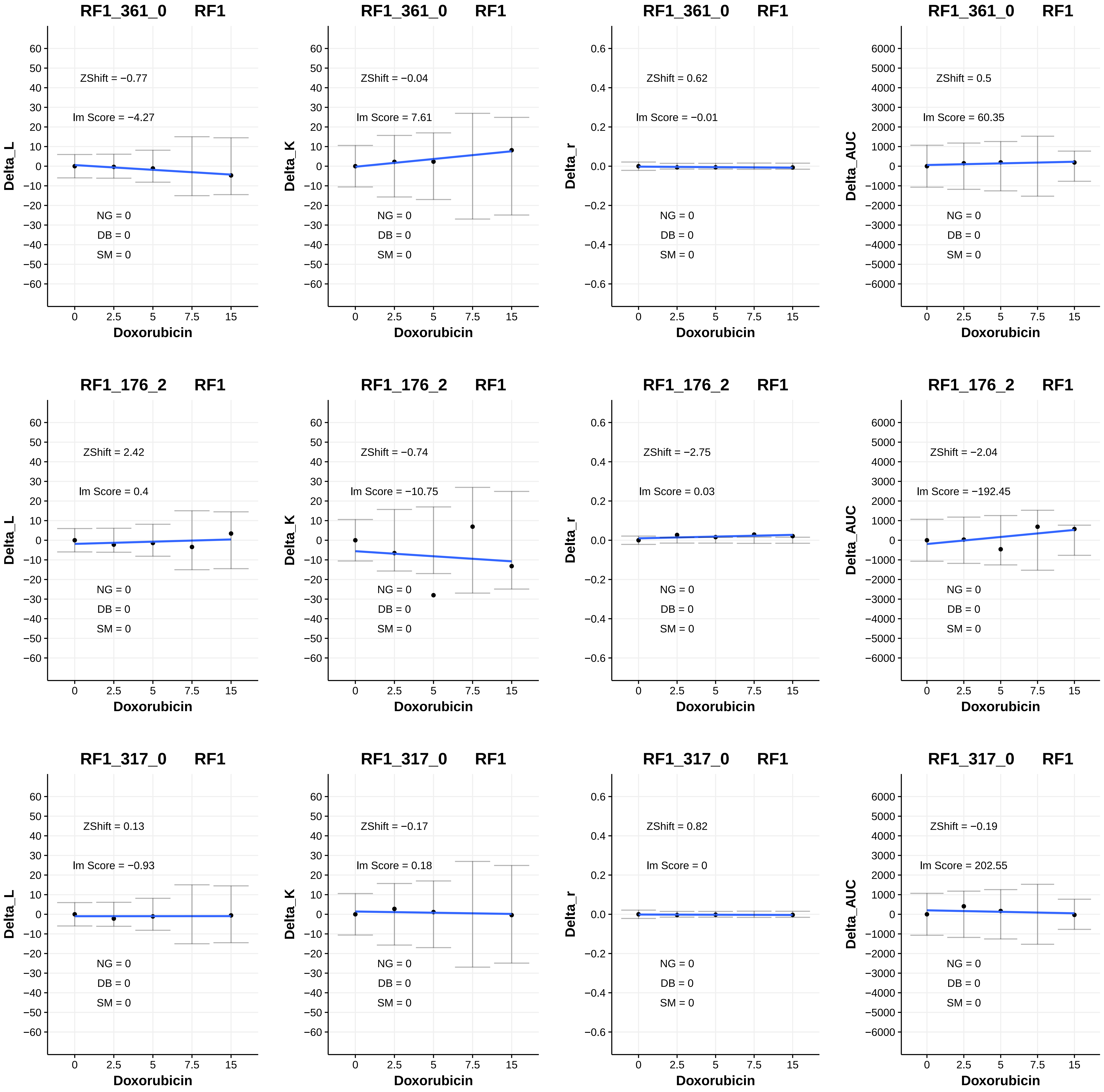

RF1_361_0 RF1
ZShift = −0.77
lm Score = −4.27
NG = 0
DB = 0
SM = 0
Delta_L
Doxorubicin
RF1_361_0 RF1
ZShift = −0.04
lm Score = 7.61
NG = 0
DB = 0
SM = 0
Delta_K
Doxorubicin
RF1_361_0 RF1
ZShift = 0.62
lm Score = −0.01
NG = 0
DB = 0
SM = 0
Delta_r
Doxorubicin
RF1_361_0 RF1
ZShift = 0.5
lm Score = 60.35
NG = 0
DB = 0
SM = 0
Delta_AUC
Doxorubicin
RF1_176_2 RF1
ZShift = 2.42
lm Score = 0.4
NG = 0
DB = 0
SM = 0
Delta_L
Doxorubicin
RF1_176_2 RF1
ZShift = −0.74
lm Score = −10.75
NG = 0
DB = 0
SM = 0
Delta_K
Doxorubicin
RF1_176_2 RF1
ZShift = −2.75
lm Score = 0.03
NG = 0
DB = 0
SM = 0
Delta_r
Doxorubicin
RF1_176_2 RF1
ZShift = −2.04
lm Score = −192.45
NG = 0
DB = 0
SM = 0
Delta_AUC
Doxorubicin
RF1_317_0 RF1
ZShift = 0.13
lm Score = −0.93
NG = 0
DB = 0
SM = 0
Delta_L
Doxorubicin
RF1_317_0 RF1
ZShift = −0.17
lm Score = 0.18
NG = 0
DB = 0
SM = 0
Delta_K
Doxorubicin
RF1_317_0 RF1
ZShift = 0.82
lm Score = 0
NG = 0
DB = 0
SM = 0
Delta_r
Doxorubicin
RF1_317_0 RF1
ZShift = −0.19
lm Score = 202.55
NG = 0
DB = 0
SM = 0
Delta_AUC
Doxorubicin

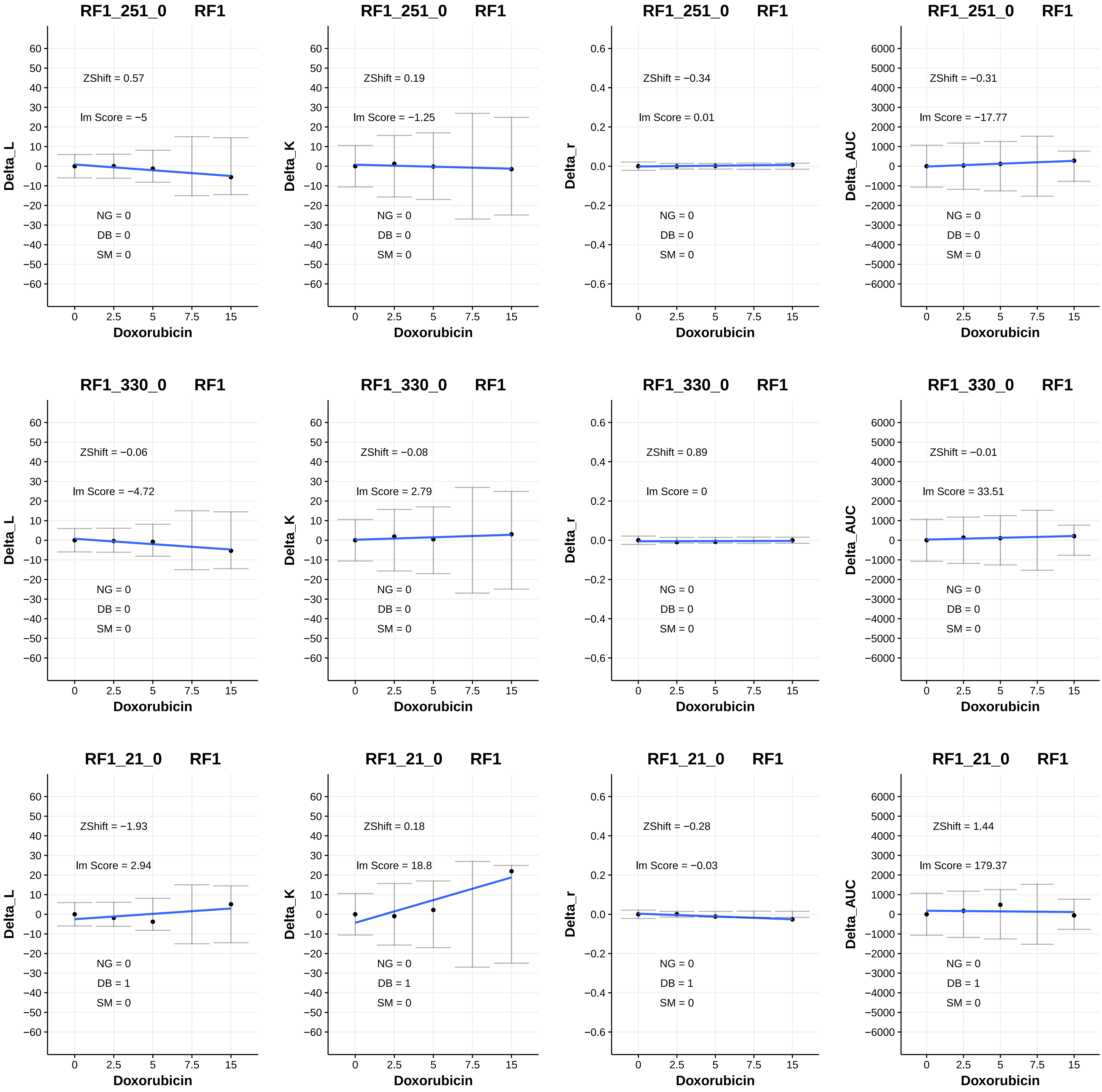
RF1_251_0 RF1
ZShift = 0.57
lm Score = −5
NG = 0
DB = 0
SM = 0
Delta_L
Doxorubicin
RF1_251_0 RF1
ZShift = 0.19
lm Score = −1.25
NG = 0
DB = 0
SM = 0
Delta_K
Doxorubicin
RF1_251_0 RF1
ZShift = −0.34
lm Score = 0.01
NG = 0
DB = 0
SM = 0
Delta_r
Doxorubicin
RF1_251_0 RF1
ZShift = −0.31
lm Score = −17.77
NG = 0
DB = 0
SM = 0
Delta_AUC
Doxorubicin
RF1_330_0 RF1
ZShift = −0.06
lm Score = −4.72
NG = 0
DB = 0
SM = 0
Delta_L
Doxorubicin
RF1_330_0 RF1
ZShift = −0.08
lm Score = 2.79
NG = 0
DB = 0
SM = 0
Delta_K
Doxorubicin
RF1_330_0 RF1
ZShift = 0.89
lm Score = 0
NG = 0
DB = 0
SM = 0
Delta_r
Doxorubicin
RF1_330_0 RF1
ZShift = −0.01
lm Score = 33.51
NG = 0
DB = 0
SM = 0
Delta_AUC
Doxorubicin
RF1_21_0 RF1
ZShift = −1.93
lm Score = 2.94
NG = 0
DB = 1
SM = 0
Delta_L
Doxorubicin
RF1_21_0 RF1
ZShift = 0.18
lm Score = 18.8
NG = 0
DB = 1
SM = 0
Delta_K
Doxorubicin
RF1_21_0 RF1
ZShift = −0.28
lm Score = −0.03
NG = 0
DB = 1
SM = 0
Delta_r
Doxorubicin
RF1_21_0 RF1
ZShift = 1.44
lm Score = 179.37
NG = 0
DB = 1
SM = 0
Delta_AUC
Doxorubicin

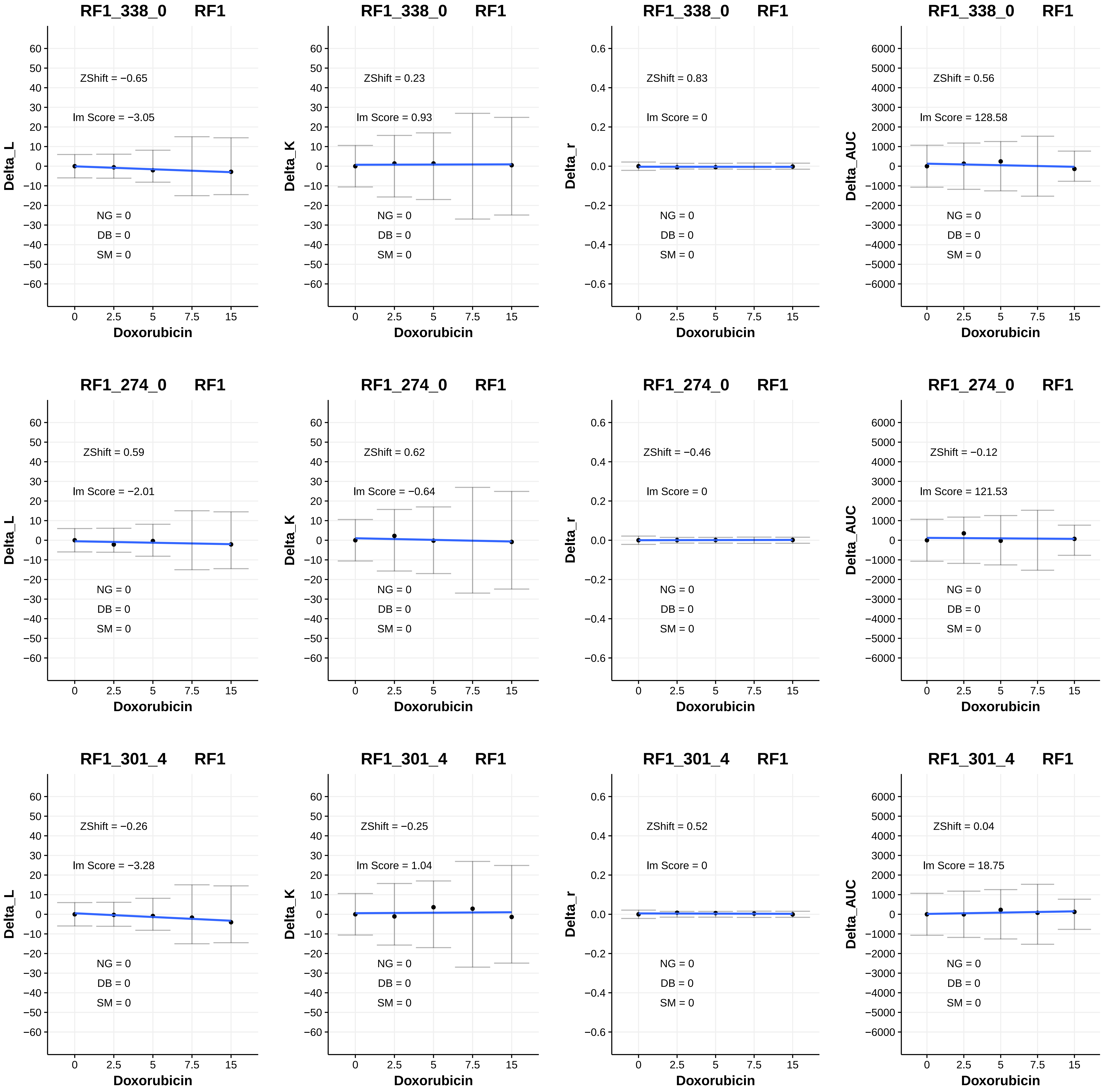
RF1_338_0 RF1
ZShift = −0.65
lm Score = −3.05
NG = 0
DB = 0
SM = 0
Delta_L
Doxorubicin
RF1_338_0 RF1
ZShift = 0.23
lm Score = 0.93
NG = 0
DB = 0
SM = 0
Delta_K
Doxorubicin
RF1_338_0 RF1
ZShift = 0.83
lm Score = 0
NG = 0
DB = 0
SM = 0
Delta_r
Doxorubicin
RF1_338_0 RF1
ZShift = 0.56
lm Score = 128.58
NG = 0
DB = 0
SM = 0
Delta_AUC
Doxorubicin
RF1_274_0 RF1
ZShift = 0.59
lm Score = −2.01
NG = 0
DB = 0
SM = 0
Delta_L
Doxorubicin
RF1_274_0 RF1
ZShift = 0.62
lm Score = −0.64
NG = 0
DB = 0
SM = 0
Delta_K
Doxorubicin
RF1_274_0 RF1
ZShift = −0.46
lm Score = 0
NG = 0
DB = 0
SM = 0
Delta_r
Doxorubicin
RF1_274_0 RF1
ZShift = −0.12
lm Score = 121.53
NG = 0
DB = 0
SM = 0
Delta_AUC
Doxorubicin
RF1_301_4 RF1
ZShift = −0.26
lm Score = −3.28
NG = 0
DB = 0
SM = 0
Delta_L
Doxorubicin
RF1_301_4 RF1
ZShift = −0.25
lm Score = 1.04
NG = 0
DB = 0
SM = 0
Delta_K
Doxorubicin
RF1_301_4 RF1
ZShift = 0.52
lm Score = 0
NG = 0
DB = 0
SM = 0
Delta_r
Doxorubicin
RF1_301_4 RF1
ZShift = 0.04
lm Score = 18.75
NG = 0
DB = 0
SM = 0
Delta_AUC
Doxorubicin

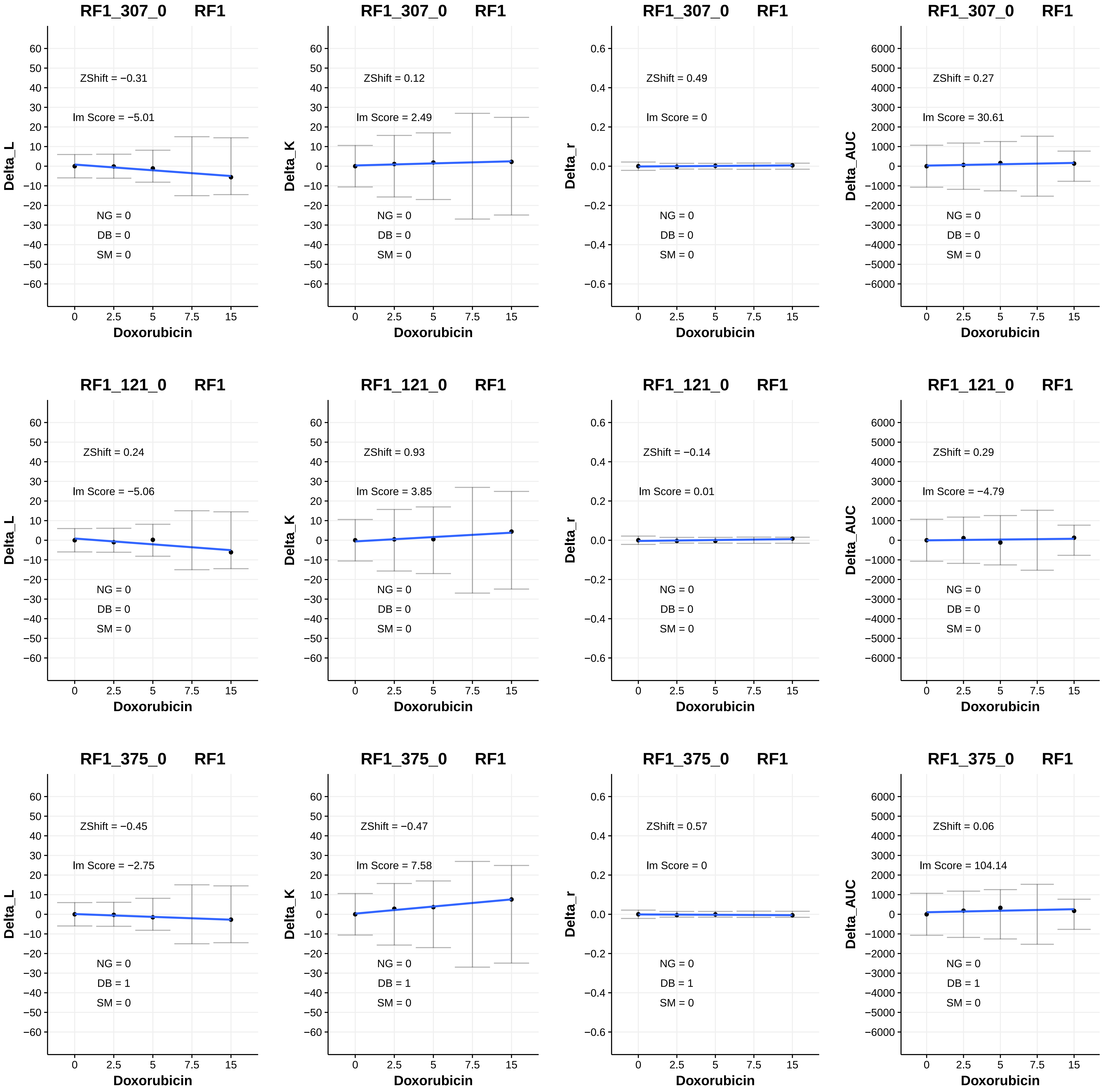

RF1_307_0 RF1
ZShift = −0.31
Im Score = −5.01
NG = 0
DB = 0
SM = 0
Delta_L
Doxorubicin
RF1_307_0 RF1
ZShift = 0.12
Im Score = 2.49
NG = 0
DB = 0
SM = 0
Delta_K
Doxorubicin
RF1_307_0 RF1
ZShift = 0.49
Im Score = 0
NG = 0
DB = 0
SM = 0
Delta_r
Doxorubicin
RF1_307_0 RF1
ZShift = 0.27
Im Score = 30.61
NG = 0
DB = 0
SM = 0
Delta_AUC
Doxorubicin
RF1_121_0 RF1
ZShift = 0.24
Im Score = −5.06
NG = 0
DB = 0
SM = 0
Delta_L
Doxorubicin
RF1_121_0 RF1
ZShift = 0.93
Im Score = 3.85
NG = 0
DB = 0
SM = 0
Delta_K
Doxorubicin
RF1_121_0 RF1
ZShift = −0.14
Im Score = 0.01
NG = 0
DB = 0
SM = 0
Delta_r
Doxorubicin
RF1_121_0 RF1
ZShift = 0.29
Im Score = −4.79
NG = 0
DB = 0
SM = 0
Delta_AUC
Doxorubicin
RF1_375_0 RF1
ZShift = −0.45
Im Score = −2.75
NG = 0
DB = 1
SM = 0
Delta_L
Doxorubicin
RF1_375_0 RF1
ZShift = −0.47
Im Score = 7.58
NG = 0
DB = 1
SM = 0
Delta_K
Doxorubicin
RF1_375_0 RF1
ZShift = 0.57
Im Score = 0
NG = 0
DB = 1
SM = 0
Delta_r
Doxorubicin
RF1_375_0 RF1
ZShift = 0.06
Im Score = 104.14
NG = 0
DB = 1
SM = 0
Delta_AUC
Doxorubicin

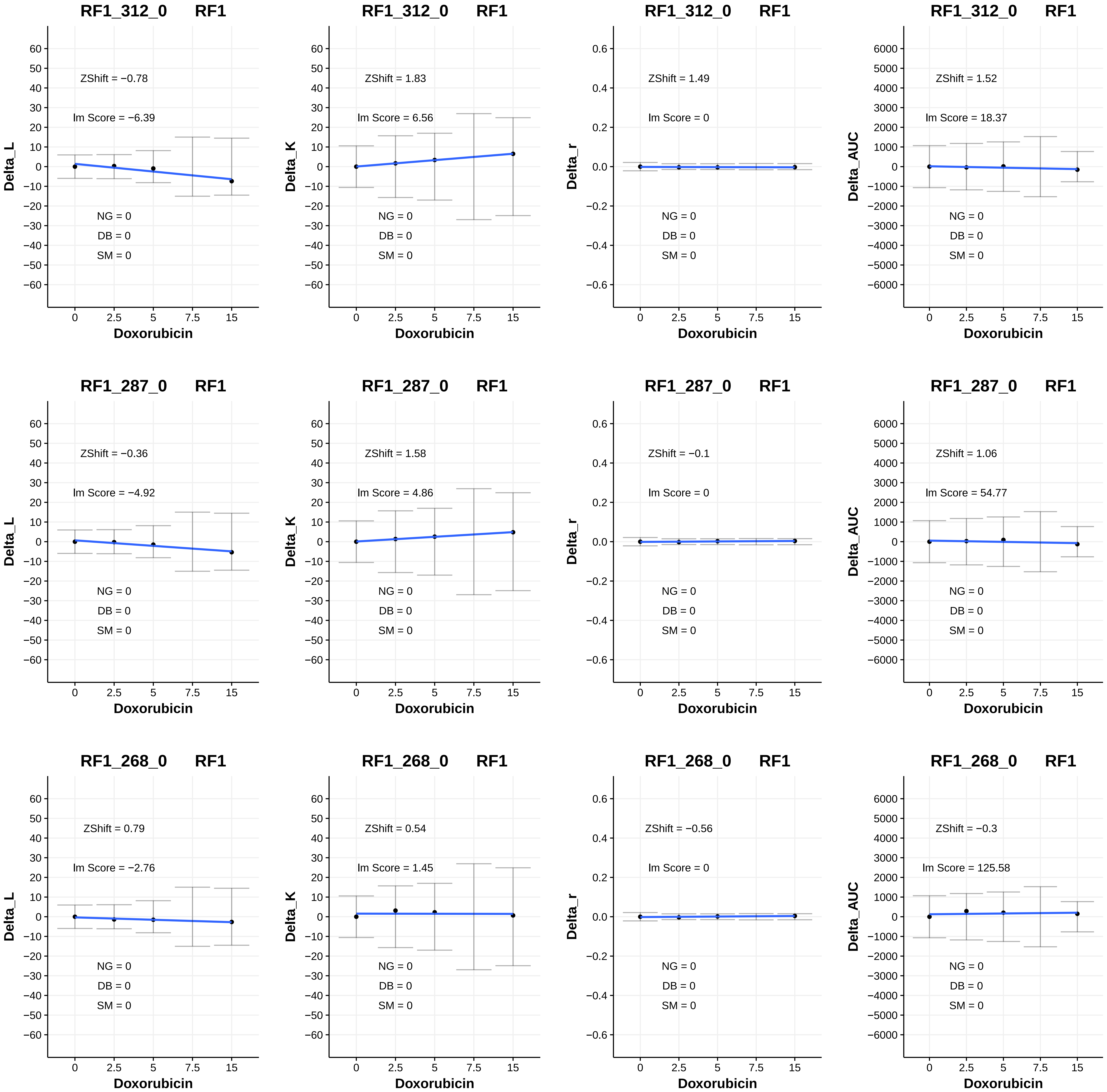
RF1_312_0 RF1
ZShift = −0.78
Im Score = −6.39
NG = 0
DB = 0
SM = 0
Delta_L
Doxorubicin
RF1_312_0 RF1
ZShift = 1.83
Im Score = 6.56
NG = 0
DB = 0
SM = 0
Delta_K
Doxorubicin
RF1_312_0 RF1
ZShift = 1.49
Im Score = 0
NG = 0
DB = 0
SM = 0
Delta_r
Doxorubicin
RF1_312_0 RF1
ZShift = 1.52
Im Score = 18.37
NG = 0
DB = 0
SM = 0
Delta_AUC
Doxorubicin
RF1_287_0 RF1
ZShift = −0.36
Im Score = −4.92
NG = 0
DB = 0
SM = 0
Delta_L
Doxorubicin
RF1_287_0 RF1
ZShift = 1.58
Im Score = 4.86
NG = 0
DB = 0
SM = 0
Delta_K
Doxorubicin
RF1_287_0 RF1
ZShift = −0.1
Im Score = 0
NG = 0
DB = 0
SM = 0
Delta_r
Doxorubicin
RF1_287_0 RF1
ZShift = 1.06
Im Score = 54.77
NG = 0
DB = 0
SM = 0
Delta_AUC
Doxorubicin
RF1_268_0 RF1
ZShift = 0.79
Im Score = −2.76
NG = 0
DB = 0
SM = 0
Delta_L
Doxorubicin
RF1_268_0 RF1
ZShift = 0.54
Im Score = 1.45
NG = 0
DB = 0
SM = 0
Delta_K
Doxorubicin
RF1_268_0 RF1
ZShift = −0.56
Im Score = 0
NG = 0
DB = 0
SM = 0
Delta_r
Doxorubicin
RF1_268_0 RF1
ZShift = −0.3
Im Score = 125.58
NG = 0
DB = 0
SM = 0
Delta_AUC
Doxorubicin

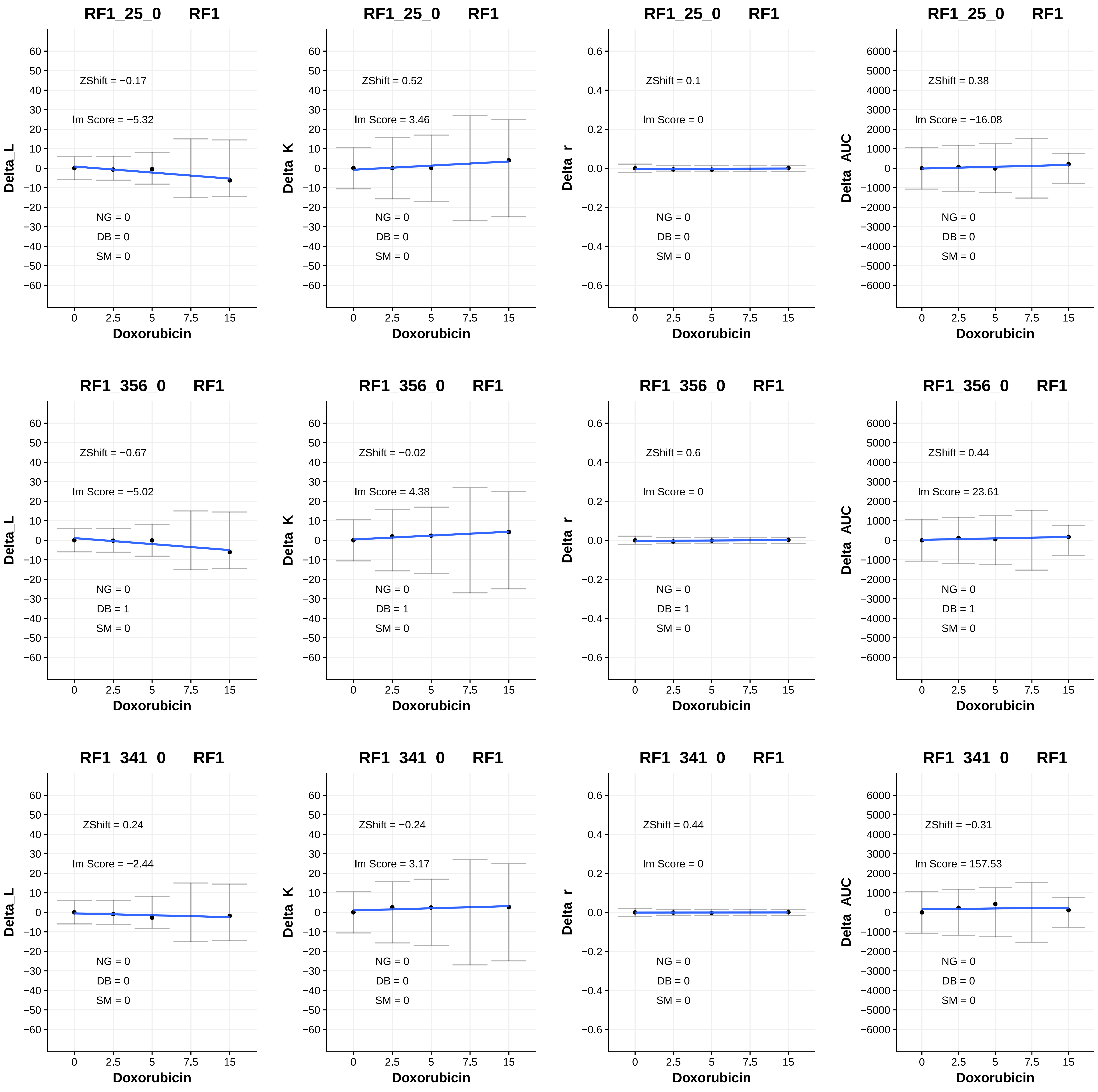
RF1_25_0 RF1
ZShift = −0.17
lm Score = −5.32
NG = 0
DB = 0
SM = 0
Delta_L
Doxorubicin
RF1_25_0 RF1
ZShift = 0.52
lm Score = 3.46
NG = 0
DB = 0
SM = 0
Delta_K
Doxorubicin
RF1_25_0 RF1
ZShift = 0.1
lm Score = 0
NG = 0
DB = 0
SM = 0
Delta_r
Doxorubicin
RF1_25_0 RF1
ZShift = 0.38
lm Score = −16.08
NG = 0
DB = 0
SM = 0
Delta_AUC
Doxorubicin
RF1_356_0 RF1
ZShift = −0.67
lm Score = −5.02
NG = 0
DB = 1
SM = 0
Delta_L
Doxorubicin
RF1_356_0 RF1
ZShift = −0.02
lm Score = 4.38
NG = 0
DB = 1
SM = 0
Delta_K
Doxorubicin
RF1_356_0 RF1
ZShift = 0.6
lm Score = 0
NG = 0
DB = 1
SM = 0
Delta_r
Doxorubicin
RF1_356_0 RF1
ZShift = 0.44
lm Score = 23.61
NG = 0
DB = 1
SM = 0
Delta_AUC
Doxorubicin
RF1_341_0 RF1
ZShift = 0.24
lm Score = −2.44
NG = 0
DB = 0
SM = 0
Delta_L
Doxorubicin
RF1_341_0 RF1
ZShift = −0.24
lm Score = 3.17
NG = 0
DB = 0
SM = 0
Delta_K
Doxorubicin
RF1_341_0 RF1
ZShift = 0.44
lm Score = 0
NG = 0
DB = 0
SM = 0
Delta_r
Doxorubicin
RF1_341_0 RF1
ZShift = −0.31
lm Score = 157.53
NG = 0
DB = 0
SM = 0
Delta_AUC
Doxorubicin

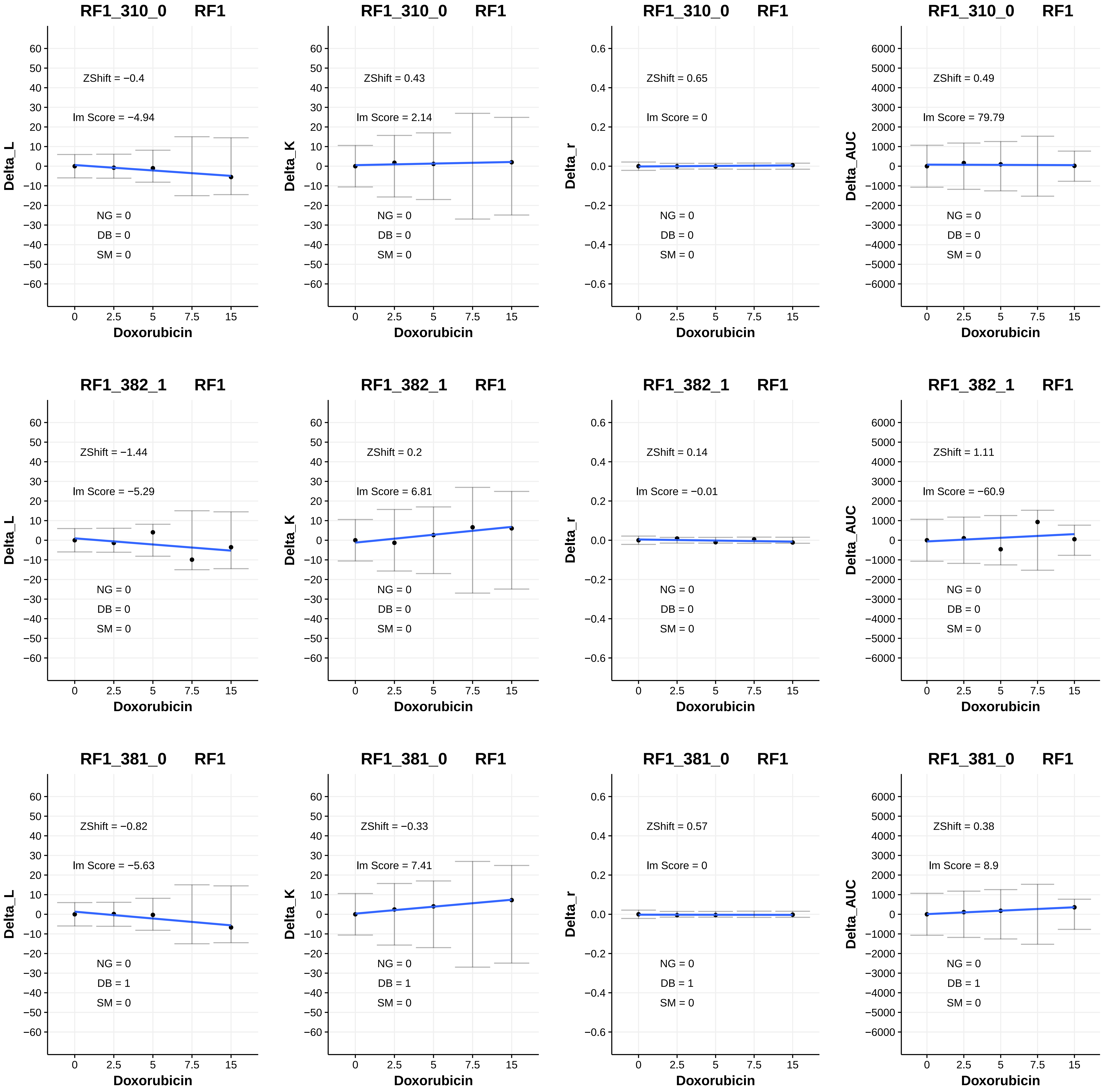

RF1_310_0 RF1
ZShift = −0.4
Im Score = −4.94
NG = 0
DB = 0
SM = 0
Delta_L
Doxorubicin
RF1_310_0 RF1
ZShift = 0.43
Im Score = 2.14
NG = 0
DB = 0
SM = 0
Delta_K
Doxorubicin
RF1_310_0 RF1
ZShift = 0.65
Im Score = 0
NG = 0
DB = 0
SM = 0
Delta_r
Doxorubicin
RF1_310_0 RF1
ZShift = 0.49
Im Score = 79.79
NG = 0
DB = 0
SM = 0
Delta_AUC
Doxorubicin
RF1_382_1 RF1
ZShift = −1.44
Im Score = −5.29
NG = 0
DB = 0
SM = 0
Delta_L
Doxorubicin
RF1_382_1 RF1
ZShift = 0.2
Im Score = 6.81
NG = 0
DB = 0
SM = 0
Delta_K
Doxorubicin
RF1_382_1 RF1
ZShift = 0.14
Im Score = −0.01
NG = 0
DB = 0
SM = 0
Delta_r
Doxorubicin
RF1_382_1 RF1
ZShift = 1.11
Im Score = −60.9
NG = 0
DB = 0
SM = 0
Delta_AUC
Doxorubicin
RF1_381_0 RF1
ZShift = −0.82
Im Score = −5.63
NG = 0
DB = 1
SM = 0
Delta_L
Doxorubicin
RF1_381_0 RF1
ZShift = −0.33
Im Score = 7.41
NG = 0
DB = 1
SM = 0
Delta_K
Doxorubicin
RF1_381_0 RF1
ZShift = 0.57
Im Score = 0
NG = 0
DB = 1
SM = 0
Delta_r
Doxorubicin
RF1_381_0 RF1
ZShift = 0.38
Im Score = 8.9
NG = 0
DB = 1
SM = 0
Delta_AUC
Doxorubicin

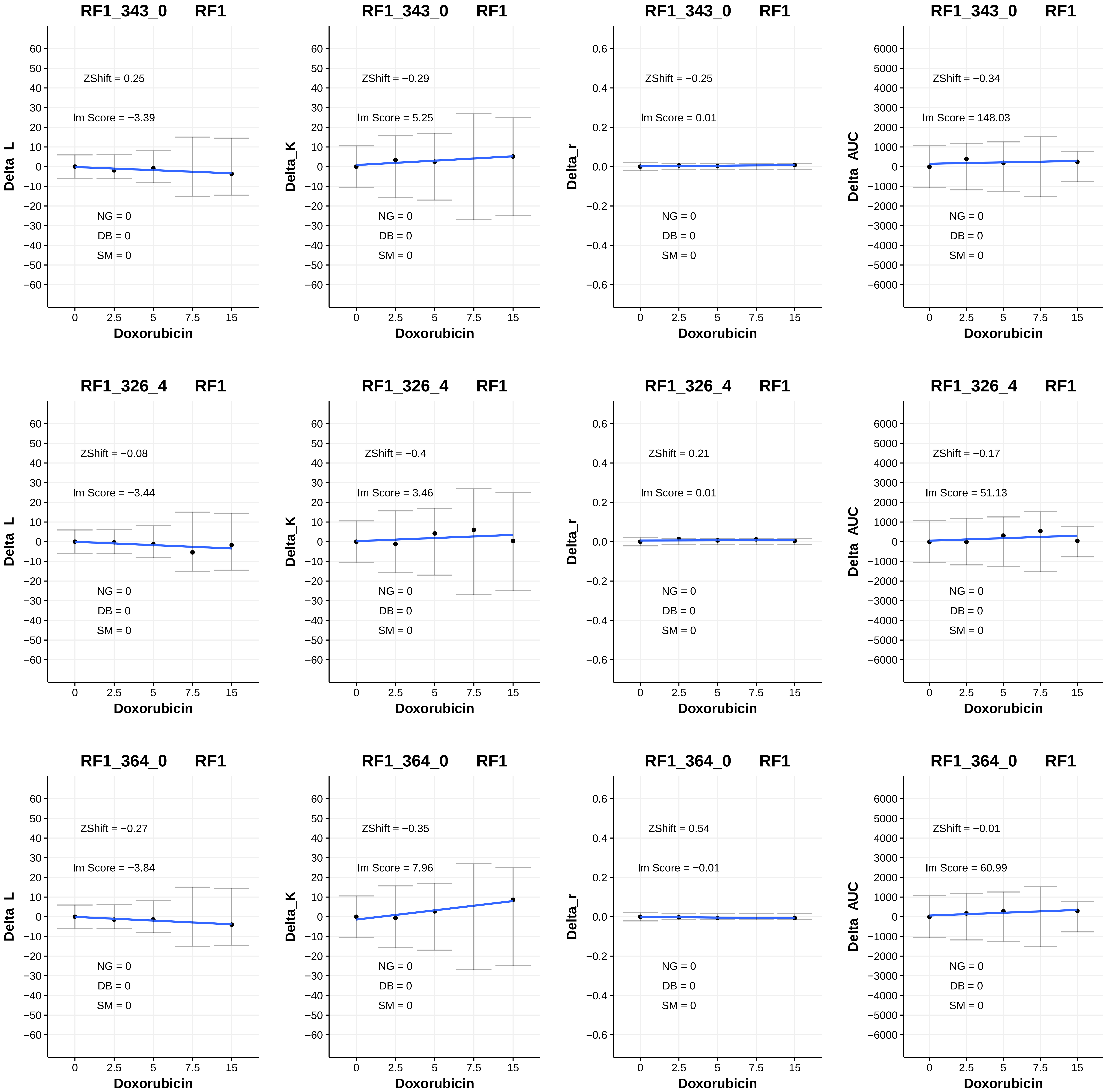
RF1_343_0 RF1
Delta_L
ZShift = 0.25
lm Score = −3.39
NG = 0
DB = 0
SM = 0
Doxorubicin
RF1_343_0 RF1
Delta_K
ZShift = −0.29
lm Score = 5.25
NG = 0
DB = 0
SM = 0
Doxorubicin
RF1_343_0 RF1
Delta_r
ZShift = −0.25
lm Score = 0.01
NG = 0
DB = 0
SM = 0
Doxorubicin
RF1_343_0 RF1
Delta_AUC
ZShift = −0.34
lm Score = 148.03
NG = 0
DB = 0
SM = 0
Doxorubicin
RF1_326_4 RF1
Delta_L
ZShift = −0.08
lm Score = −3.44
NG = 0
DB = 0
SM = 0
Doxorubicin
RF1_326_4 RF1
Delta_K
ZShift = −0.4
lm Score = 3.46
NG = 0
DB = 0
SM = 0
Doxorubicin
RF1_326_4 RF1
Delta_r
ZShift = 0.21
lm Score = 0.01
NG = 0
DB = 0
SM = 0
Doxorubicin
RF1_326_4 RF1
Delta_AUC
ZShift = −0.17
lm Score = 51.13
NG = 0
DB = 0
SM = 0
Doxorubicin
RF1_364_0 RF1
Delta_L
ZShift = −0.27
lm Score = −3.84
NG = 0
DB = 0
SM = 0
Doxorubicin
RF1_364_0 RF1
Delta_K
ZShift = −0.35
lm Score = 7.96
NG = 0
DB = 0
SM = 0
Doxorubicin
RF1_364_0 RF1
Delta_r
ZShift = 0.54
lm Score = −0.01
NG = 0
DB = 0
SM = 0
Doxorubicin
RF1_364_0 RF1
Delta_AUC
ZShift = −0.01
lm Score = 60.99
NG = 0
DB = 0
SM = 0
Doxorubicin

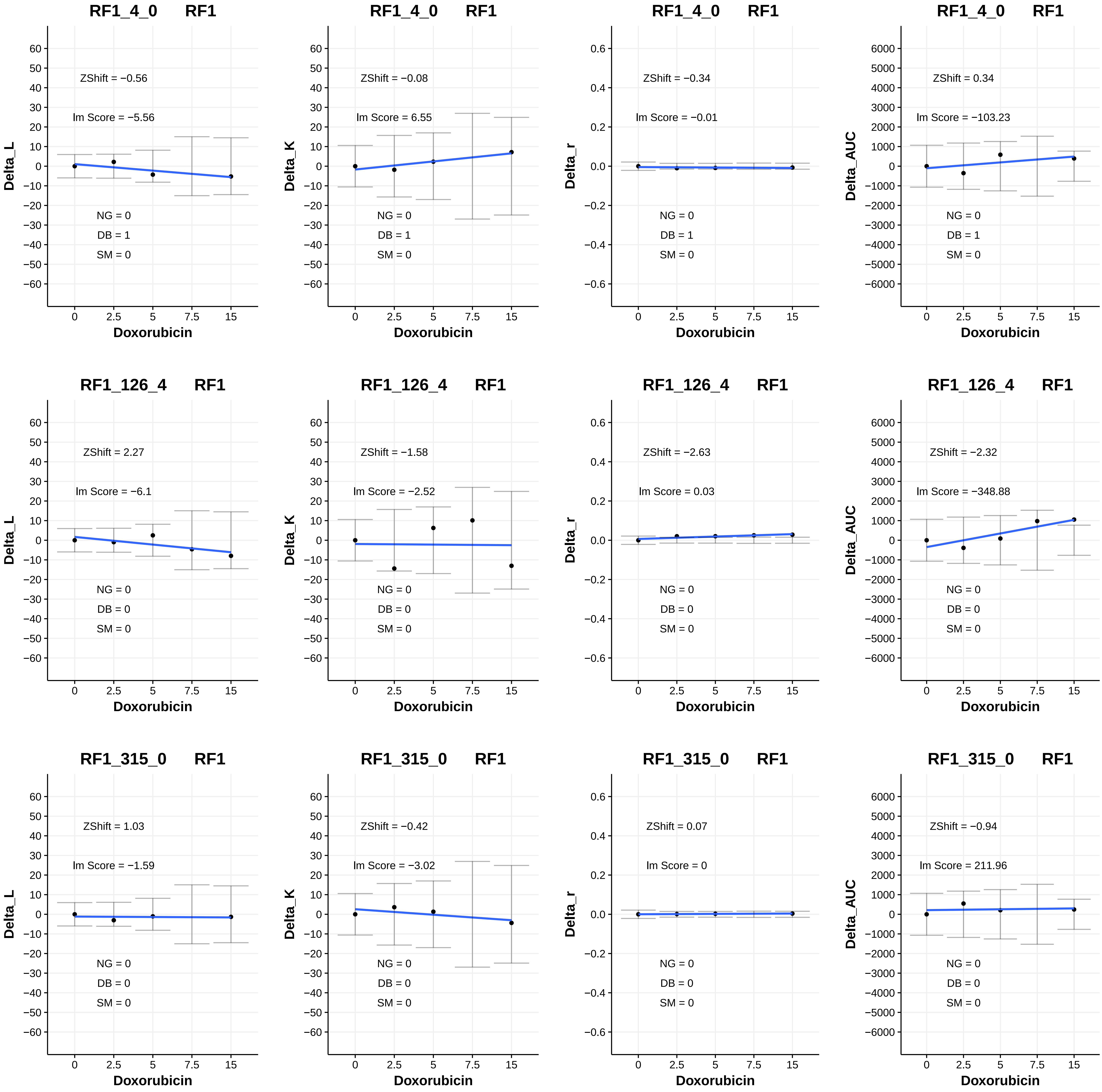

RF1_4_0 RF1
ZShift = −0.56
Im Score = −5.56
NG = 0
DB = 1
SM = 0
Delta_L
Doxorubicin
RF1_4_0 RF1
ZShift = −0.08
Im Score = 6.55
NG = 0
DB = 1
SM = 0
Delta_K
Doxorubicin
RF1_4_0 RF1
ZShift = −0.34
Im Score = −0.01
NG = 0
DB = 1
SM = 0
Delta_r
Doxorubicin
RF1_4_0 RF1
ZShift = 0.34
Im Score = −103.23
NG = 0
DB = 1
SM = 0
Delta_AUC
Doxorubicin
RF1_126_4 RF1
ZShift = 2.27
Im Score = −6.1
NG = 0
DB = 0
SM = 0
Delta_L
Doxorubicin
RF1_126_4 RF1
ZShift = −1.58
Im Score = −2.52
NG = 0
DB = 0
SM = 0
Delta_K
Doxorubicin
RF1_126_4 RF1
ZShift = −2.63
Im Score = 0.03
NG = 0
DB = 0
SM = 0
Delta_r
Doxorubicin
RF1_126_4 RF1
ZShift = −2.32
Im Score = −348.88
NG = 0
DB = 0
SM = 0
Delta_AUC
Doxorubicin
RF1_315_0 RF1
ZShift = 1.03
Im Score = −1.59
NG = 0
DB = 0
SM = 0
Delta_L
Doxorubicin
RF1_315_0 RF1
ZShift = −0.42
Im Score = −3.02
NG = 0
DB = 0
SM = 0
Delta_K
Doxorubicin
RF1_315_0 RF1
ZShift = 0.07
Im Score = 0
NG = 0
DB = 0
SM = 0
Delta_r
Doxorubicin
RF1_315_0 RF1
ZShift = −0.94
Im Score = 211.96
NG = 0
DB = 0
SM = 0
Delta_AUC
Doxorubicin

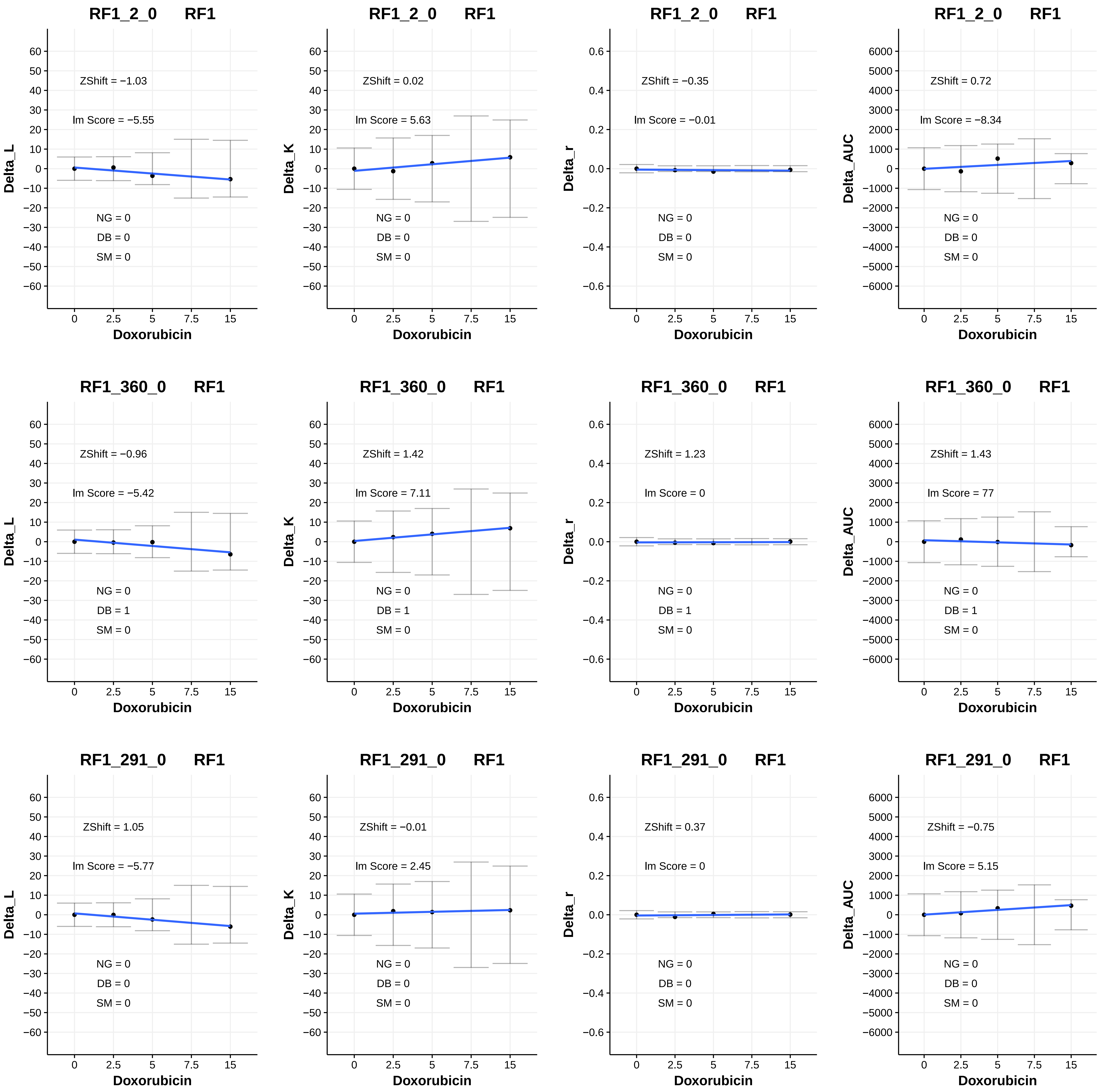

RF1_2_0 RF1
ZShift = −1.03
Im Score = −5.55
NG = 0
DB = 0
SM = 0
Delta_L
Doxorubicin
RF1_2_0 RF1
ZShift = 0.02
Im Score = 5.63
NG = 0
DB = 0
SM = 0
Delta_K
Doxorubicin
RF1_2_0 RF1
ZShift = −0.35
Im Score = −0.01
NG = 0
DB = 0
SM = 0
Delta_r
Doxorubicin
RF1_2_0 RF1
ZShift = 0.72
Im Score = −8.34
NG = 0
DB = 0
SM = 0
Delta_AUC
Doxorubicin
RF1_360_0 RF1
ZShift = −0.96
Im Score = −5.42
NG = 0
DB = 1
SM = 0
Delta_L
Doxorubicin
RF1_360_0 RF1
ZShift = 1.42
Im Score = 7.11
NG = 0
DB = 1
SM = 0
Delta_K
Doxorubicin
RF1_360_0 RF1
ZShift = 1.23
Im Score = 0
NG = 0
DB = 1
SM = 0
Delta_r
Doxorubicin
RF1_360_0 RF1
ZShift = 1.43
Im Score = 77
NG = 0
DB = 1
SM = 0
Delta_AUC
Doxorubicin
RF1_291_0 RF1
ZShift = 1.05
Im Score = −5.77
NG = 0
DB = 0
SM = 0
Delta_L
Doxorubicin
RF1_291_0 RF1
ZShift = −0.01
Im Score = 2.45
NG = 0
DB = 0
SM = 0
Delta_K
Doxorubicin
RF1_291_0 RF1
ZShift = 0.37
Im Score = 0
NG = 0
DB = 0
SM = 0
Delta_r
Doxorubicin
RF1_291_0 RF1
ZShift = −0.75
Im Score = 5.15
NG = 0
DB = 0
SM = 0
Delta_AUC
Doxorubicin

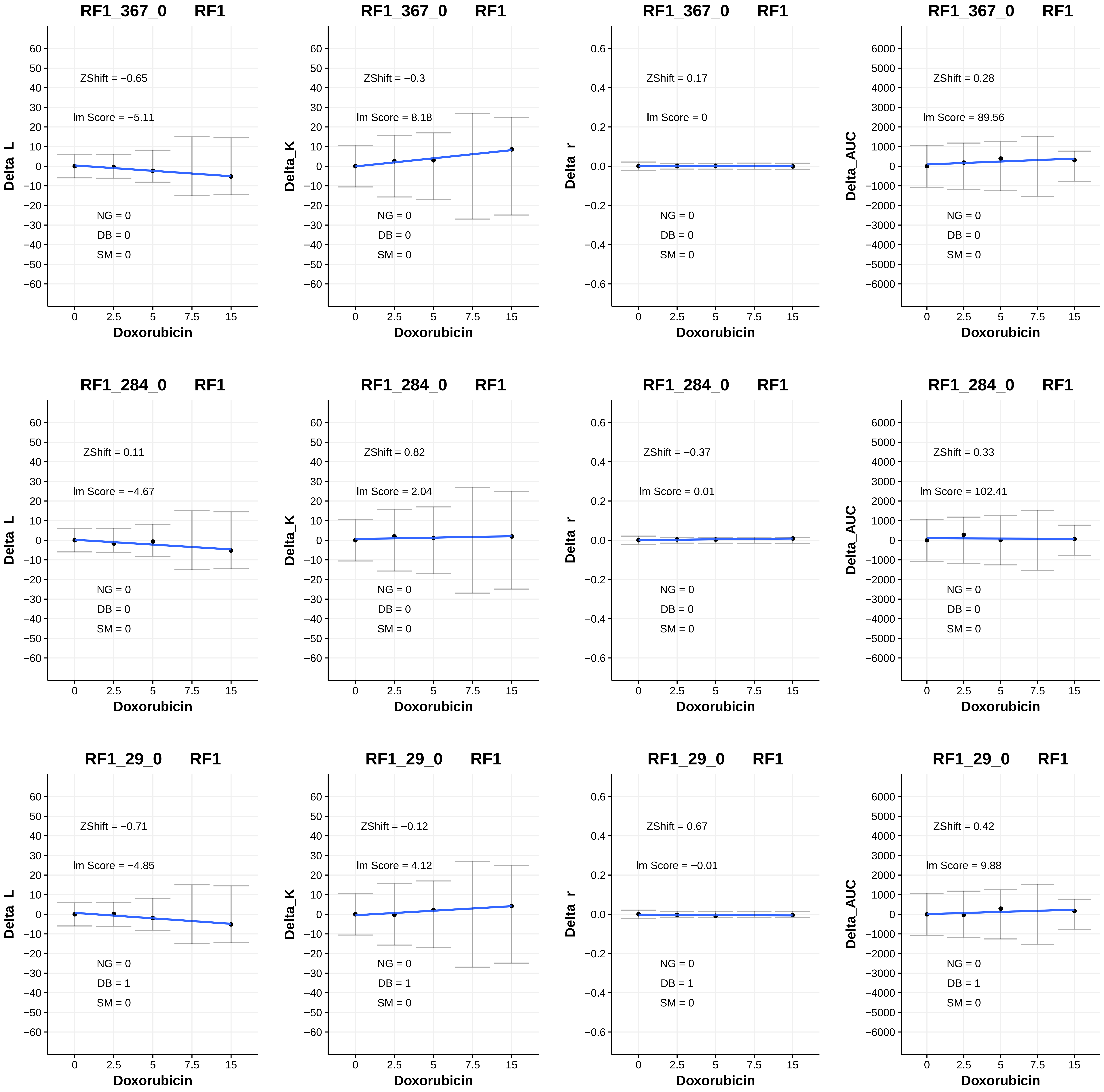
RF1_367_0 RF1
ZShift = −0.65
lm Score = −5.11
NG = 0
DB = 0
SM = 0
Delta_L
Doxorubicin
RF1_367_0 RF1
ZShift = −0.3
lm Score = 8.18
NG = 0
DB = 0
SM = 0
Delta_K
Doxorubicin
RF1_367_0 RF1
ZShift = 0.17
lm Score = 0
NG = 0
DB = 0
SM = 0
Delta_r
Doxorubicin
RF1_367_0 RF1
ZShift = 0.28
lm Score = 89.56
NG = 0
DB = 0
SM = 0
Delta_AUC
Doxorubicin
RF1_284_0 RF1
ZShift = 0.11
lm Score = −4.67
NG = 0
DB = 0
SM = 0
Delta_L
Doxorubicin
RF1_284_0 RF1
ZShift = 0.82
lm Score = 2.04
NG = 0
DB = 0
SM = 0
Delta_K
Doxorubicin
RF1_284_0 RF1
ZShift = −0.37
lm Score = 0.01
NG = 0
DB = 0
SM = 0
Delta_r
Doxorubicin
RF1_284_0 RF1
ZShift = 0.33
lm Score = 102.41
NG = 0
DB = 0
SM = 0
Delta_AUC
Doxorubicin
RF1_29_0 RF1
ZShift = −0.71
lm Score = −4.85
NG = 0
DB = 1
SM = 0
Delta_L
Doxorubicin
RF1_29_0 RF1
ZShift = −0.12
lm Score = 4.12
NG = 0
DB = 1
SM = 0
Delta_K
Doxorubicin
RF1_29_0 RF1
ZShift = 0.67
lm Score = −0.01
NG = 0
DB = 1
SM = 0
Delta_r
Doxorubicin
RF1_29_0 RF1
ZShift = 0.42
lm Score = 9.88
NG = 0
DB = 1
SM = 0
Delta_AUC
Doxorubicin

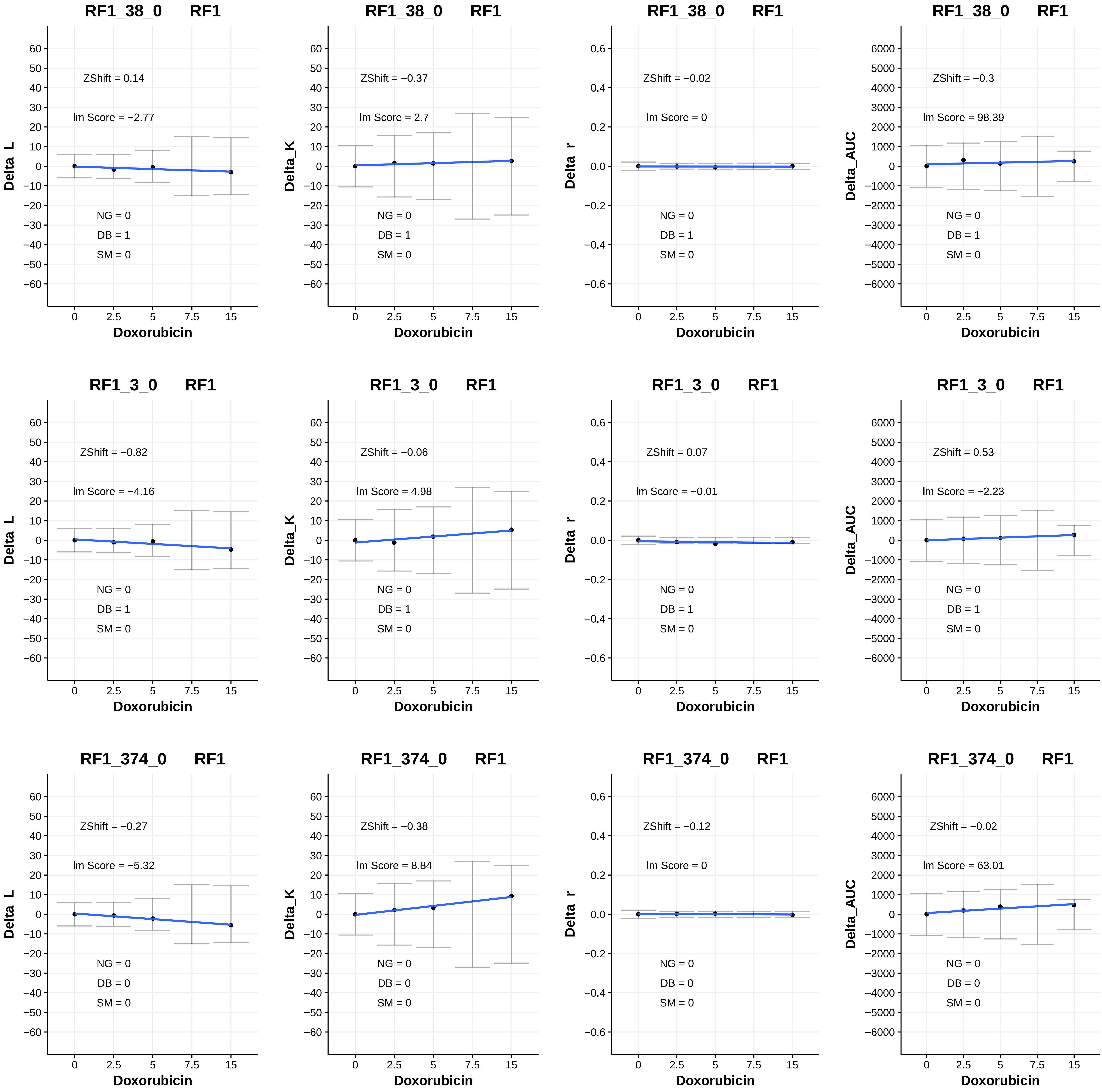

RF1_38_0 RF1
ZShift = 0.14
lm Score = −2.77
NG = 0
DB = 1
SM = 0
Delta_L
Doxorubicin
RF1_38_0 RF1
ZShift = −0.37
lm Score = 2.7
NG = 0
DB = 1
SM = 0
Delta_K
Doxorubicin
RF1_38_0 RF1
ZShift = −0.02
lm Score = 0
NG = 0
DB = 1
SM = 0
Delta_r
Doxorubicin
RF1_38_0 RF1
ZShift = −0.3
lm Score = 98.39
NG = 0
DB = 1
SM = 0
Delta_AUC
Doxorubicin
RF1_3_0 RF1
ZShift = −0.82
lm Score = −4.16
NG = 0
DB = 1
SM = 0
Delta_L
Doxorubicin
RF1_3_0 RF1
ZShift = −0.06
lm Score = 4.98
NG = 0
DB = 1
SM = 0
Delta_K
Doxorubicin
RF1_3_0 RF1
ZShift = 0.07
lm Score = −0.01
NG = 0
DB = 1
SM = 0
Delta_r
Doxorubicin
RF1_3_0 RF1
ZShift = 0.53
lm Score = −2.23
NG = 0
DB = 1
SM = 0
Delta_AUC
Doxorubicin
RF1_374_0 RF1
ZShift = −0.27
lm Score = −5.32
NG = 0
DB = 0
SM = 0
Delta_L
Doxorubicin
RF1_374_0 RF1
ZShift = −0.38
lm Score = 8.84
NG = 0
DB = 0
SM = 0
Delta_K
Doxorubicin
RF1_374_0 RF1
ZShift = −0.12
lm Score = 0
NG = 0
DB = 0
SM = 0
Delta_r
Doxorubicin
RF1_374_0 RF1
ZShift = −0.02
lm Score = 63.01
NG = 0
DB = 0
SM = 0
Delta_AUC
Doxorubicin

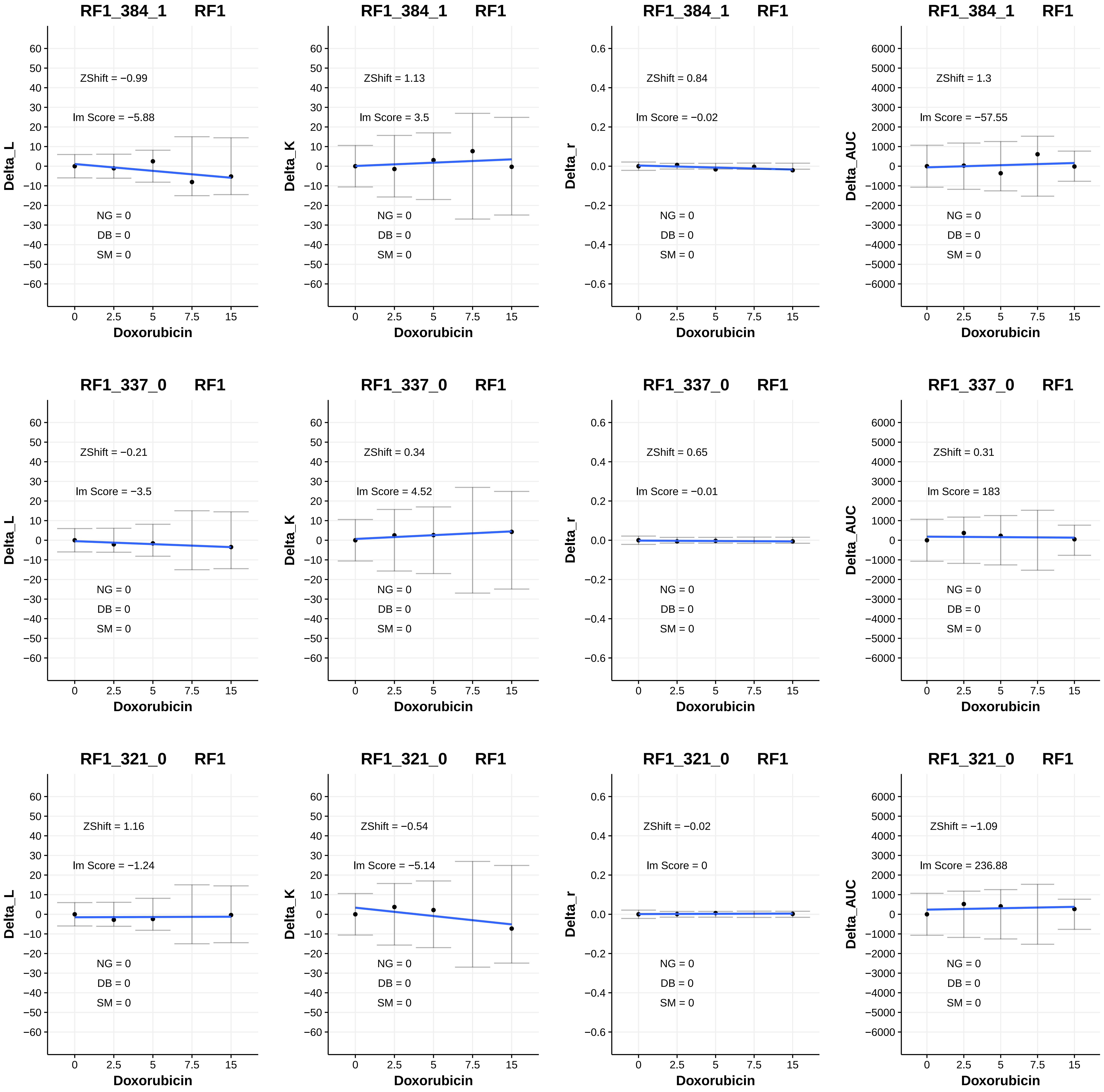

RF1_384_1 RF1
ZShift = −0.99
lm Score = −5.88
NG = 0
DB = 0
SM = 0
Delta_L
Doxorubicin
RF1_384_1 RF1
ZShift = 1.13
lm Score = 3.5
NG = 0
DB = 0
SM = 0
Delta_K
Doxorubicin
RF1_384_1 RF1
ZShift = 0.84
lm Score = −0.02
NG = 0
DB = 0
SM = 0
Delta_r
Doxorubicin
RF1_384_1 RF1
ZShift = 1.3
lm Score = −57.55
NG = 0
DB = 0
SM = 0
Delta_AUC
Doxorubicin
RF1_337_0 RF1
ZShift = −0.21
lm Score = −3.5
NG = 0
DB = 0
SM = 0
Delta_L
Doxorubicin
RF1_337_0 RF1
ZShift = 0.34
lm Score = 4.52
NG = 0
DB = 0
SM = 0
Delta_K
Doxorubicin
RF1_337_0 RF1
ZShift = 0.65
lm Score = −0.01
NG = 0
DB = 0
SM = 0
Delta_r
Doxorubicin
RF1_337_0 RF1
ZShift = 0.31
lm Score = 183
NG = 0
DB = 0
SM = 0
Delta_AUC
Doxorubicin
RF1_321_0 RF1
ZShift = 1.16
lm Score = −1.24
NG = 0
DB = 0
SM = 0
Delta_L
Doxorubicin
RF1_321_0 RF1
ZShift = −0.54
lm Score = −5.14
NG = 0
DB = 0
SM = 0
Delta_K
Doxorubicin
RF1_321_0 RF1
ZShift = −0.02
lm Score = 0
NG = 0
DB = 0
SM = 0
Delta_r
Doxorubicin
RF1_321_0 RF1
ZShift = −1.09
lm Score = 236.88
NG = 0
DB = 0
SM = 0
Delta_AUC
Doxorubicin
0
2.5
5
7.5
15

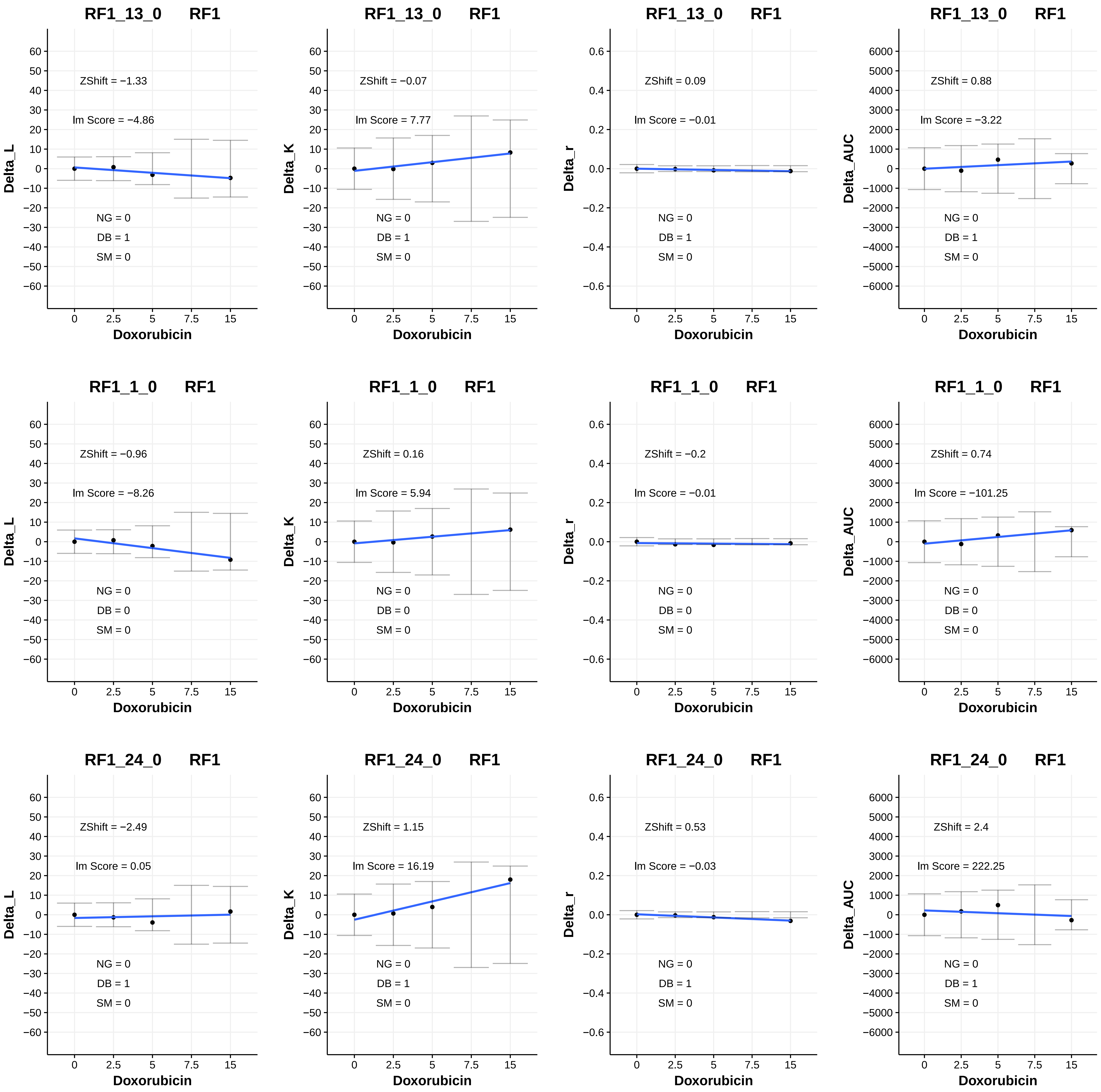
RF1_13_0 RF1
ZShift = −1.33
Im Score = −4.86
NG = 0
DB = 1
SM = 0
Delta_L
Doxorubicin
RF1_13_0 RF1
ZShift = −0.07
Im Score = 7.77
NG = 0
DB = 1
SM = 0
Delta_K
Doxorubicin
RF1_13_0 RF1
ZShift = 0.09
Im Score = −0.01
NG = 0
DB = 1
SM = 0
Delta_r
Doxorubicin
RF1_13_0 RF1
ZShift = 0.88
Im Score = −3.22
NG = 0
DB = 1
SM = 0
Delta_AUC
Doxorubicin
RF1_1_0 RF1
ZShift = −0.96
Im Score = −8.26
NG = 0
DB = 0
SM = 0
Delta_L
Doxorubicin
RF1_1_0 RF1
ZShift = 0.16
Im Score = 5.94
NG = 0
DB = 0
SM = 0
Delta_K
Doxorubicin
RF1_1_0 RF1
ZShift = −0.2
Im Score = −0.01
NG = 0
DB = 0
SM = 0
Delta_r
Doxorubicin
RF1_1_0 RF1
ZShift = 0.74
Im Score = −101.25
NG = 0
DB = 0
SM = 0
Delta_AUC
Doxorubicin
RF1_24_0 RF1
ZShift = −2.49
Im Score = 0.05
NG = 0
DB = 1
SM = 0
Delta_L
Doxorubicin
RF1_24_0 RF1
ZShift = 1.15
Im Score = 16.19
NG = 0
DB = 1
SM = 0
Delta_K
Doxorubicin
RF1_24_0 RF1
ZShift = 0.53
Im Score = −0.03
NG = 0
DB = 1
SM = 0
Delta_r
Doxorubicin
RF1_24_0 RF1
ZShift = 2.4
Im Score = 222.25
NG = 0
DB = 1
SM = 0
Delta_AUC
Doxorubicin

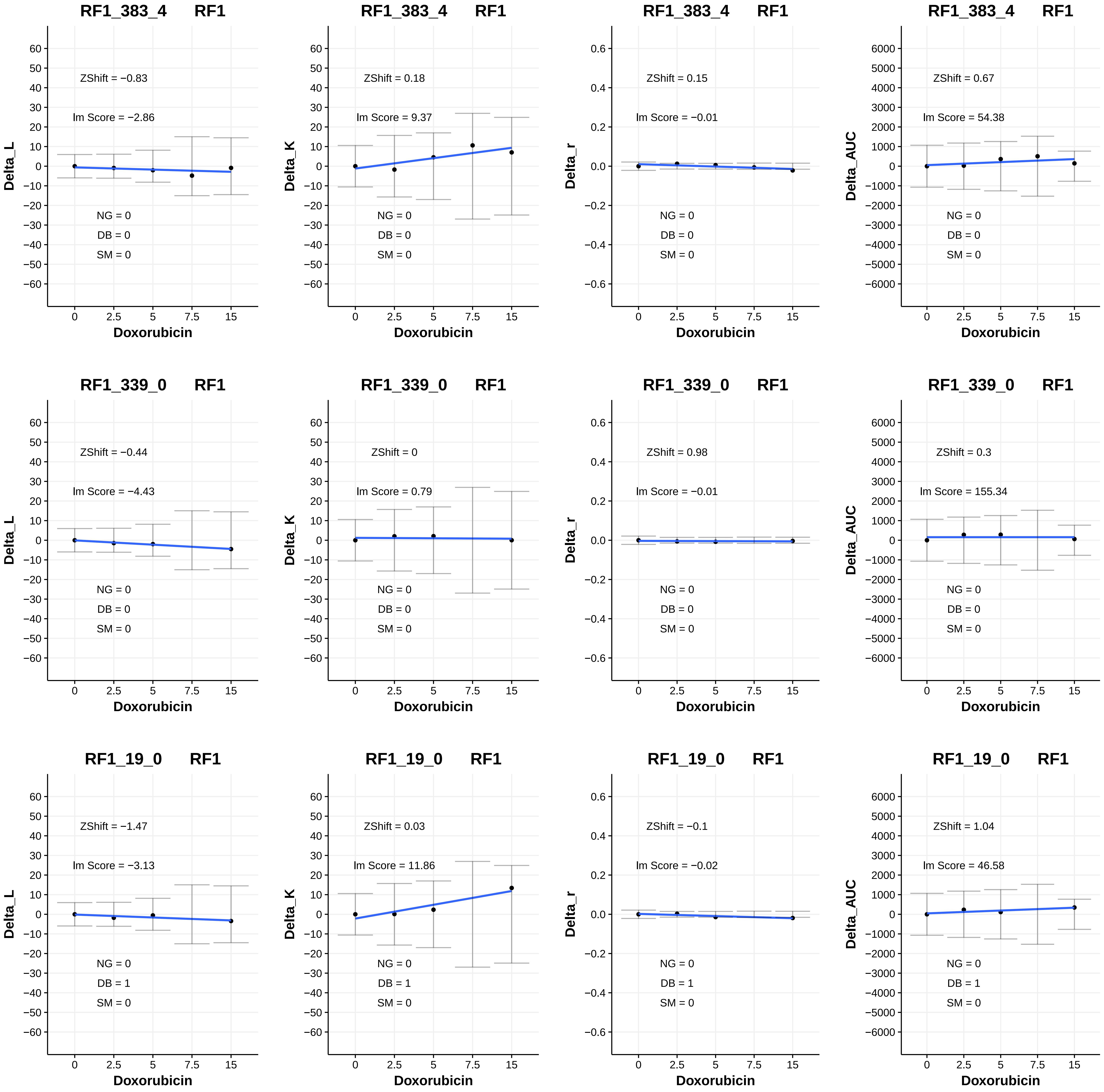

RF1_383_4 RF1
ZShift = −0.83
lm Score = −2.86
NG = 0
DB = 0
SM = 0
Delta_L
Doxorubicin
RF1_383_4 RF1
ZShift = 0.18
lm Score = 9.37
NG = 0
DB = 0
SM = 0
Delta_K
Doxorubicin
RF1_383_4 RF1
ZShift = 0.15
lm Score = −0.01
NG = 0
DB = 0
SM = 0
Delta_r
Doxorubicin
RF1_383_4 RF1
ZShift = 0.67
lm Score = 54.38
NG = 0
DB = 0
SM = 0
Delta_AUC
Doxorubicin
RF1_339_0 RF1
ZShift = −0.44
lm Score = −4.43
NG = 0
DB = 0
SM = 0
Delta_L
Doxorubicin
RF1_339_0 RF1
ZShift = 0
lm Score = 0.79
NG = 0
DB = 0
SM = 0
Delta_K
Doxorubicin
RF1_339_0 RF1
ZShift = 0.98
lm Score = −0.01
NG = 0
DB = 0
SM = 0
Delta_r
Doxorubicin
RF1_339_0 RF1
ZShift = 0.3
lm Score = 155.34
NG = 0
DB = 0
SM = 0
Delta_AUC
Doxorubicin
RF1_19_0 RF1
ZShift = −1.47
lm Score = −3.13
NG = 0
DB = 1
SM = 0
Delta_L
Doxorubicin
RF1_19_0 RF1
ZShift = 0.03
lm Score = 11.86
NG = 0
DB = 1
SM = 0
Delta_K
Doxorubicin
RF1_19_0 RF1
ZShift = −0.1
lm Score = −0.02
NG = 0
DB = 1
SM = 0
Delta_r
Doxorubicin
RF1_19_0 RF1
ZShift = 1.04
lm Score = 46.58
NG = 0
DB = 1
SM = 0
Delta_AUC
Doxorubicin

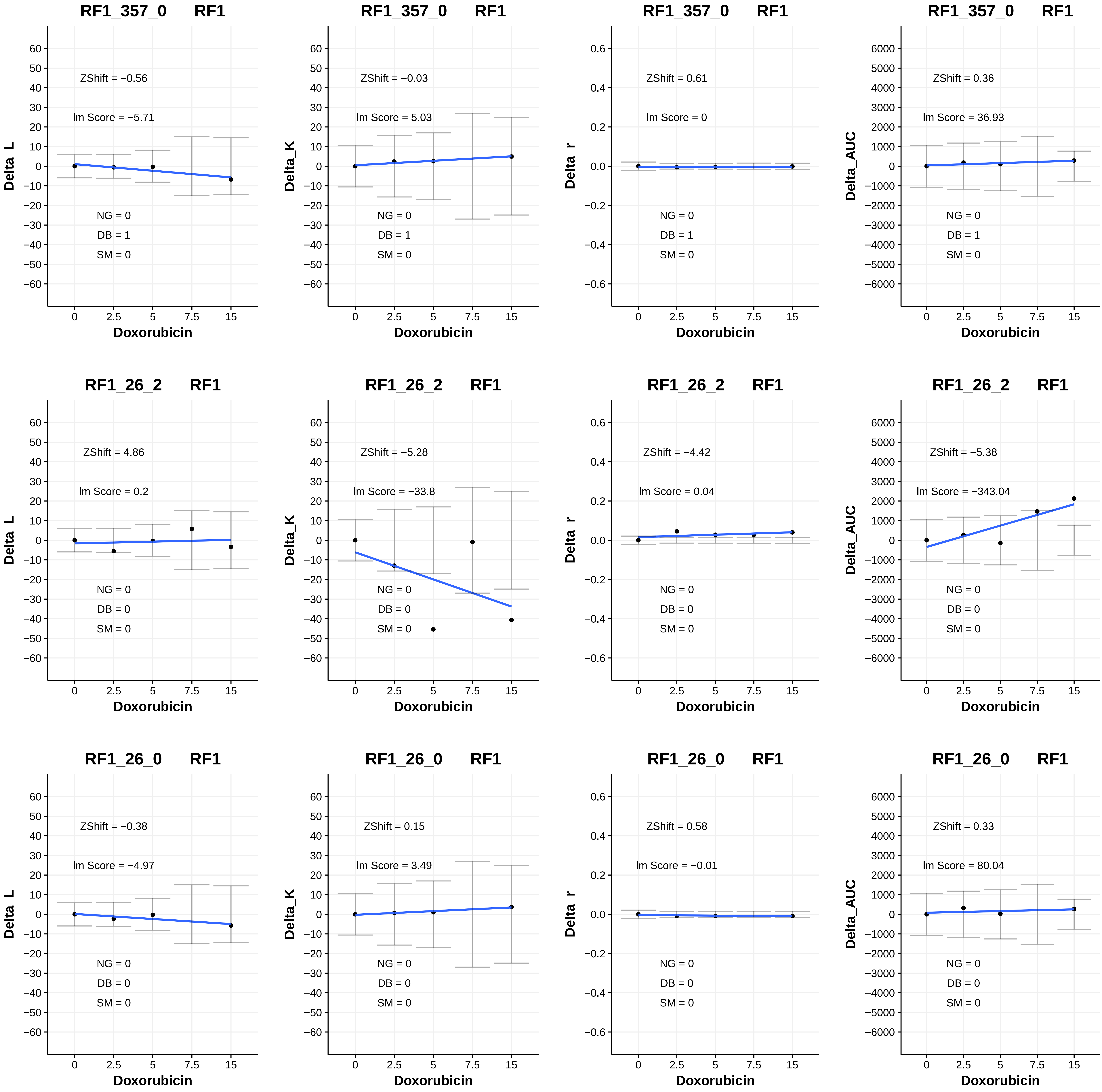
RF1_357_0 RF1
ZShift = −0.56
Im Score = −5.71
NG = 0
DB = 1
SM = 0
Delta_L
Doxorubicin
RF1_357_0 RF1
ZShift = −0.03
Im Score = 5.03
NG = 0
DB = 1
SM = 0
Delta_K
Doxorubicin
RF1_357_0 RF1
ZShift = 0.61
Im Score = 0
NG = 0
DB = 1
SM = 0
Delta_r
Doxorubicin
RF1_357_0 RF1
ZShift = 0.36
Im Score = 36.93
NG = 0
DB = 1
SM = 0
Delta_AUC
Doxorubicin
RF1_26_2 RF1
ZShift = 4.86
Im Score = 0.2
NG = 0
DB = 0
SM = 0
Delta_L
Doxorubicin
RF1_26_2 RF1
ZShift = −5.28
Im Score = −33.8
NG = 0
DB = 0
SM = 0
Delta_K
Doxorubicin
RF1_26_2 RF1
ZShift = −4.42
Im Score = 0.04
NG = 0
DB = 0
SM = 0
Delta_r
Doxorubicin
RF1_26_2 RF1
ZShift = −5.38
Im Score = −343.04
NG = 0
DB = 0
SM = 0
Delta_AUC
Doxorubicin
RF1_26_0 RF1
ZShift = −0.38
Im Score = −4.97
NG = 0
DB = 0
SM = 0
Delta_L
Doxorubicin
RF1_26_0 RF1
ZShift = 0.15
Im Score = 3.49
NG = 0
DB = 0
SM = 0
Delta_K
Doxorubicin
RF1_26_0 RF1
ZShift = 0.58
Im Score = −0.01
NG = 0
DB = 0
SM = 0
Delta_r
Doxorubicin
RF1_26_0 RF1
ZShift = 0.33
Im Score = 80.04
NG = 0
DB = 0
SM = 0
Delta_AUC
Doxorubicin

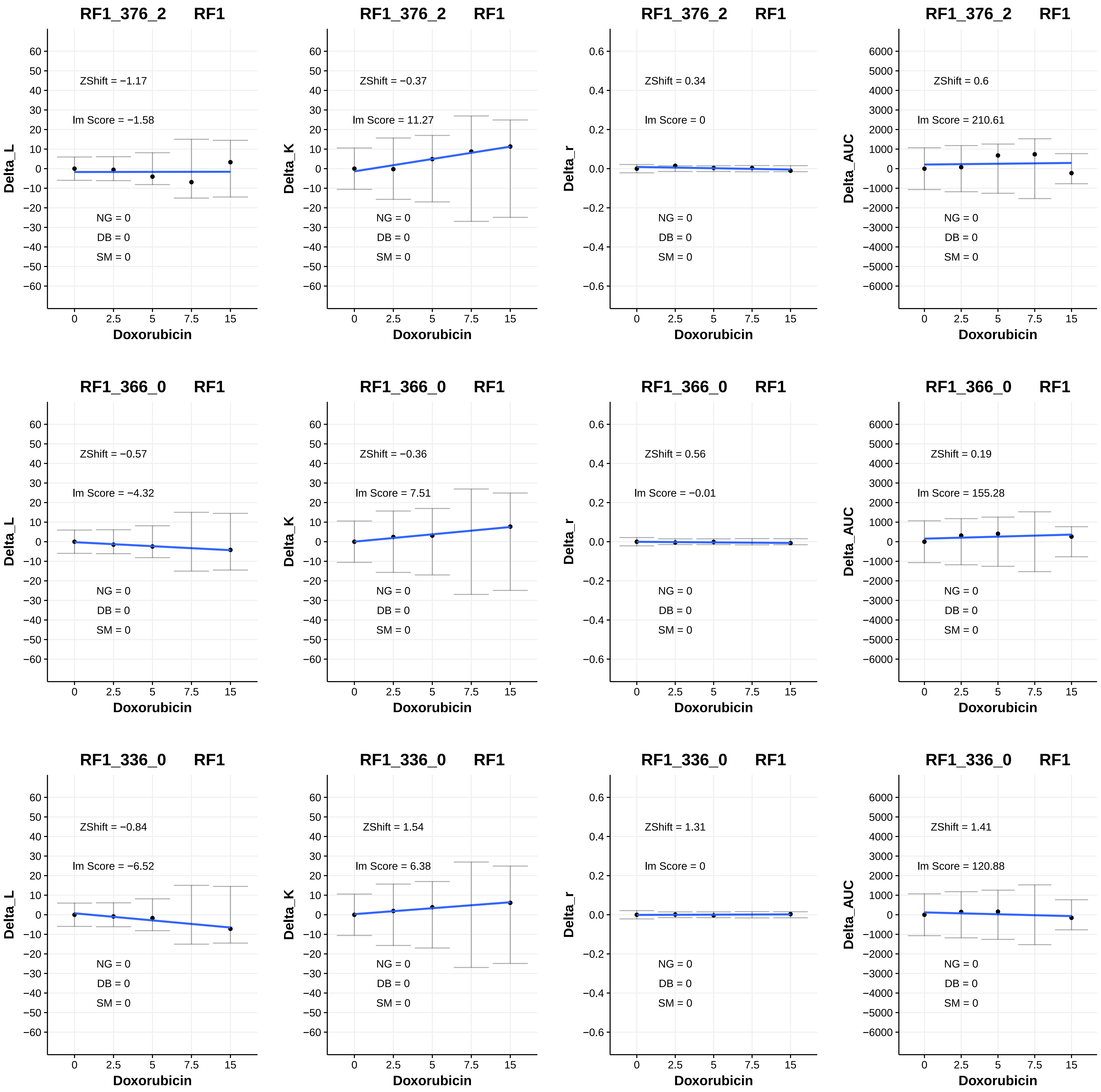
RF1_376_2 RF1
ZShift = −1.17
lm Score = −1.58
NG = 0
DB = 0
SM = 0
Delta_L
Doxorubicin
RF1_376_2 RF1
ZShift = −0.37
lm Score = 11.27
NG = 0
DB = 0
SM = 0
Delta_K
Doxorubicin
RF1_376_2 RF1
ZShift = 0.34
lm Score = 0
NG = 0
DB = 0
SM = 0
Delta_r
Doxorubicin
RF1_376_2 RF1
ZShift = 0.6
lm Score = 210.61
NG = 0
DB = 0
SM = 0
Delta_AUC
Doxorubicin
RF1_366_0 RF1
ZShift = −0.57
lm Score = −4.32
NG = 0
DB = 0
SM = 0
Delta_L
Doxorubicin
RF1_366_0 RF1
ZShift = −0.36
lm Score = 7.51
NG = 0
DB = 0
SM = 0
Delta_K
Doxorubicin
RF1_366_0 RF1
ZShift = 0.56
lm Score = −0.01
NG = 0
DB = 0
SM = 0
Delta_r
Doxorubicin
RF1_366_0 RF1
ZShift = 0.19
lm Score = 155.28
NG = 0
DB = 0
SM = 0
Delta_AUC
Doxorubicin
RF1_336_0 RF1
ZShift = −0.84
lm Score = −6.52
NG = 0
DB = 0
SM = 0
Delta_L
Doxorubicin
RF1_336_0 RF1
ZShift = 1.54
lm Score = 6.38
NG = 0
DB = 0
SM = 0
Delta_K
Doxorubicin
RF1_336_0 RF1
ZShift = 1.31
lm Score = 0
NG = 0
DB = 0
SM = 0
Delta_r
Doxorubicin
RF1_336_0 RF1
ZShift = 1.41
lm Score = 120.88
NG = 0
DB = 0
SM = 0
Delta_AUC
Doxorubicin

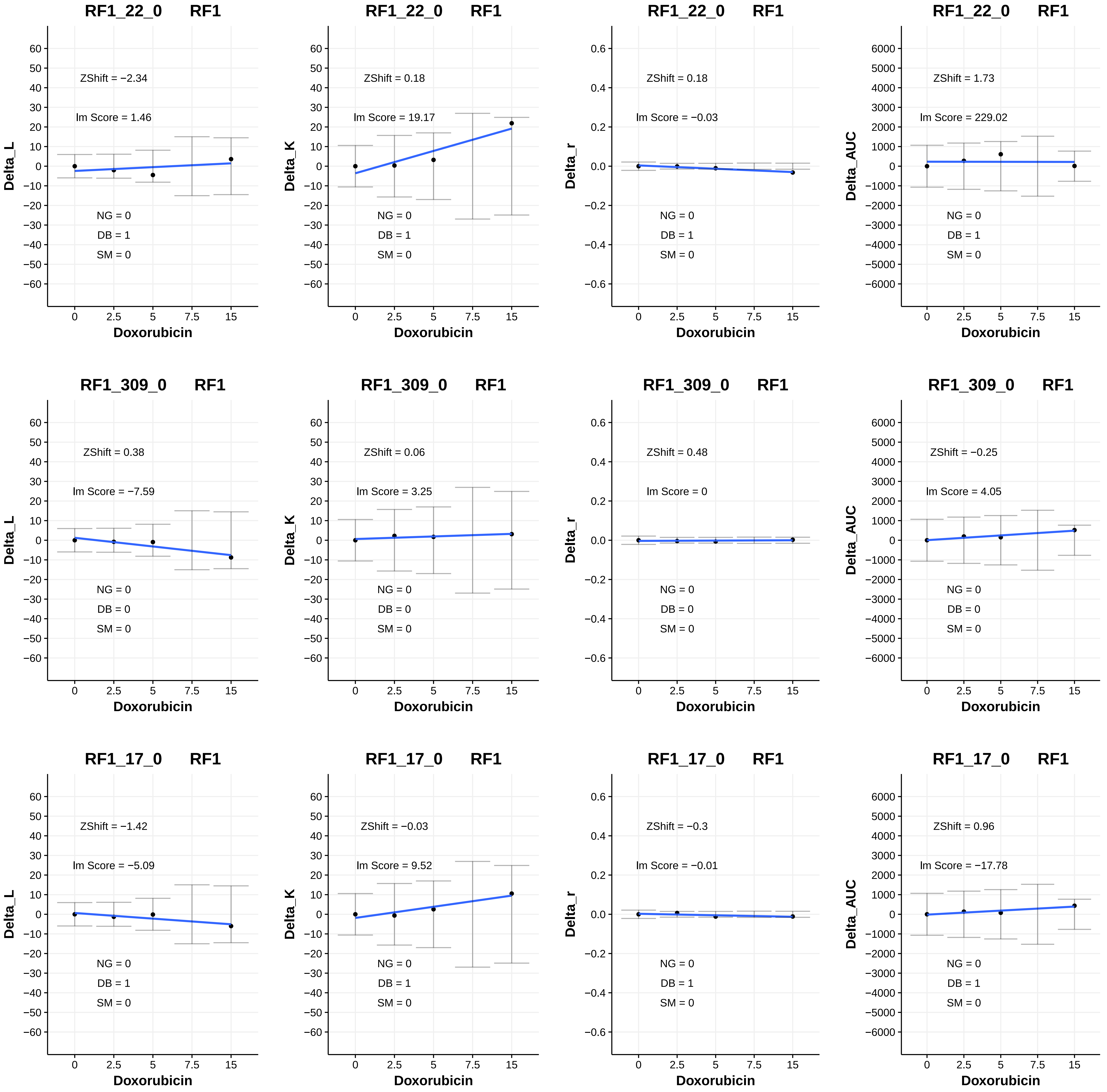
RF1_22_0 RF1
ZShift = −2.34
lm Score = 1.46
NG = 0
DB = 1
SM = 0
Delta_L
Doxorubicin
RF1_22_0 RF1
ZShift = 0.18
lm Score = 19.17
NG = 0
DB = 1
SM = 0
Delta_K
Doxorubicin
RF1_22_0 RF1
ZShift = 0.18
lm Score = −0.03
NG = 0
DB = 1
SM = 0
Delta_r
Doxorubicin
RF1_22_0 RF1
ZShift = 1.73
lm Score = 229.02
NG = 0
DB = 1
SM = 0
Delta_AUC
Doxorubicin
RF1_309_0 RF1
ZShift = 0.38
lm Score = −7.59
NG = 0
DB = 0
SM = 0
Delta_L
Doxorubicin
RF1_309_0 RF1
ZShift = 0.06
lm Score = 3.25
NG = 0
DB = 0
SM = 0
Delta_K
Doxorubicin
RF1_309_0 RF1
ZShift = 0.48
lm Score = 0
NG = 0
DB = 0
SM = 0
Delta_r
Doxorubicin
RF1_309_0 RF1
ZShift = −0.25
lm Score = 4.05
NG = 0
DB = 0
SM = 0
Delta_AUC
Doxorubicin
RF1_17_0 RF1
ZShift = −1.42
lm Score = −5.09
NG = 0
DB = 1
SM = 0
Delta_L
Doxorubicin
RF1_17_0 RF1
ZShift = −0.03
lm Score = 9.52
NG = 0
DB = 1
SM = 0
Delta_K
Doxorubicin
RF1_17_0 RF1
ZShift = −0.3
lm Score = −0.01
NG = 0
DB = 1
SM = 0
Delta_r
Doxorubicin
RF1_17_0 RF1
ZShift = 0.96
lm Score = −17.78
NG = 0
DB = 1
SM = 0
Delta_AUC
Doxorubicin

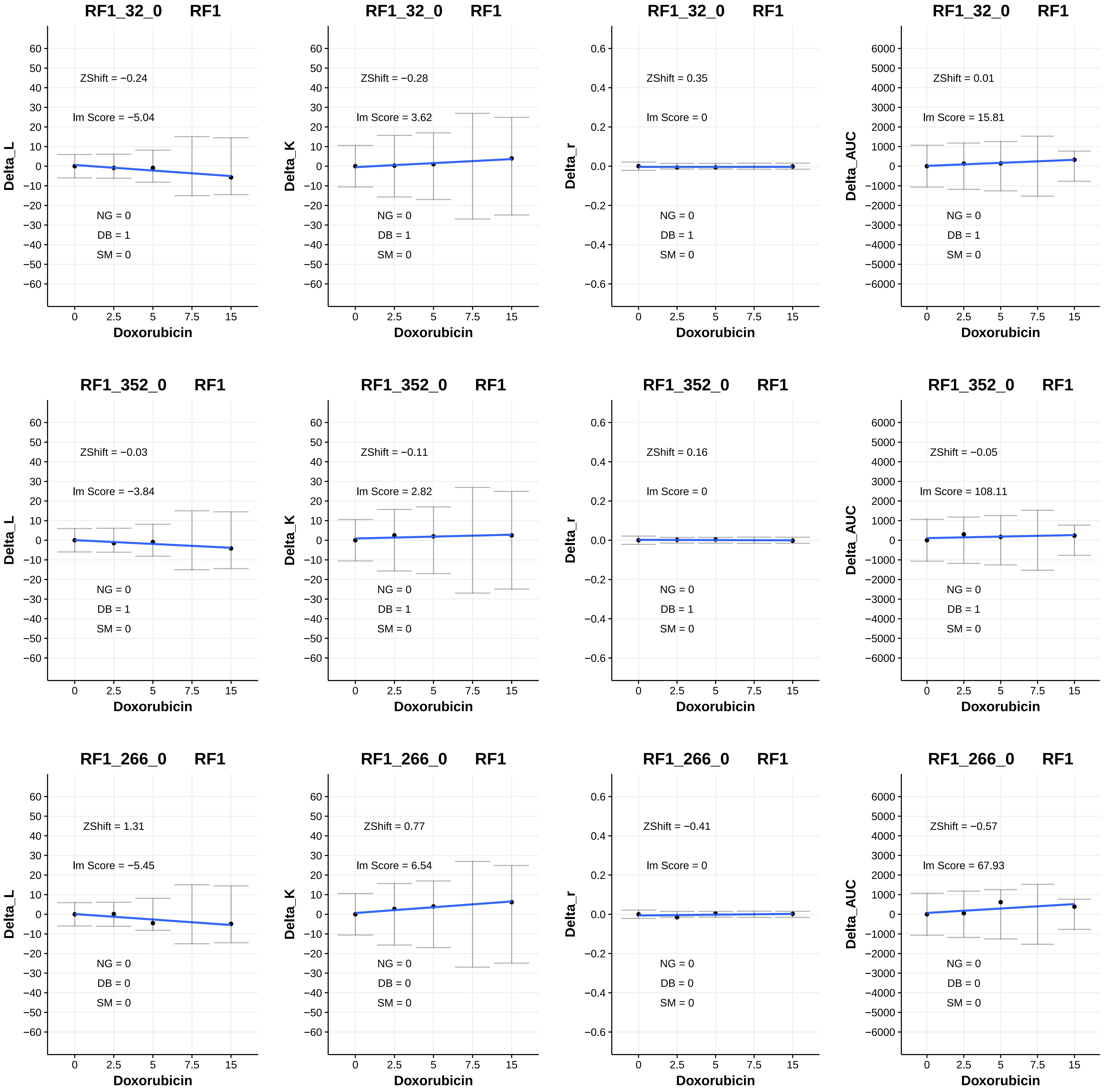

RF1_32_0 RF1
ZShift = −0.24
Im Score = −5.04
NG = 0
DB = 1
SM = 0
Delta_L
Doxorubicin
RF1_32_0 RF1
ZShift = −0.28
Im Score = 3.62
NG = 0
DB = 1
SM = 0
Delta_K
Doxorubicin
RF1_32_0 RF1
ZShift = 0.35
Im Score = 0
NG = 0
DB = 1
SM = 0
Delta_r
Doxorubicin
RF1_32_0 RF1
ZShift = 0.01
Im Score = 15.81
NG = 0
DB = 1
SM = 0
Delta_AUC
Doxorubicin
RF1_352_0 RF1
ZShift = −0.03
Im Score = −3.84
NG = 0
DB = 1
SM = 0
Delta_L
Doxorubicin
RF1_352_0 RF1
ZShift = −0.11
Im Score = 2.82
NG = 0
DB = 1
SM = 0
Delta_K
Doxorubicin
RF1_352_0 RF1
ZShift = 0.16
Im Score = 0
NG = 0
DB = 1
SM = 0
Delta_r
Doxorubicin
RF1_352_0 RF1
ZShift = −0.05
Im Score = 108.11
NG = 0
DB = 1
SM = 0
Delta_AUC
Doxorubicin
RF1_266_0 RF1
ZShift = 1.31
Im Score = −5.45
NG = 0
DB = 0
SM = 0
Delta_L
Doxorubicin
RF1_266_0 RF1
ZShift = 0.77
Im Score = 6.54
NG = 0
DB = 0
SM = 0
Delta_K
Doxorubicin
RF1_266_0 RF1
ZShift = −0.41
Im Score = 0
NG = 0
DB = 0
SM = 0
Delta_r
Doxorubicin
RF1_266_0 RF1
ZShift = −0.57
Im Score = 67.93
NG = 0
DB = 0
SM = 0
Delta_AUC
Doxorubicin

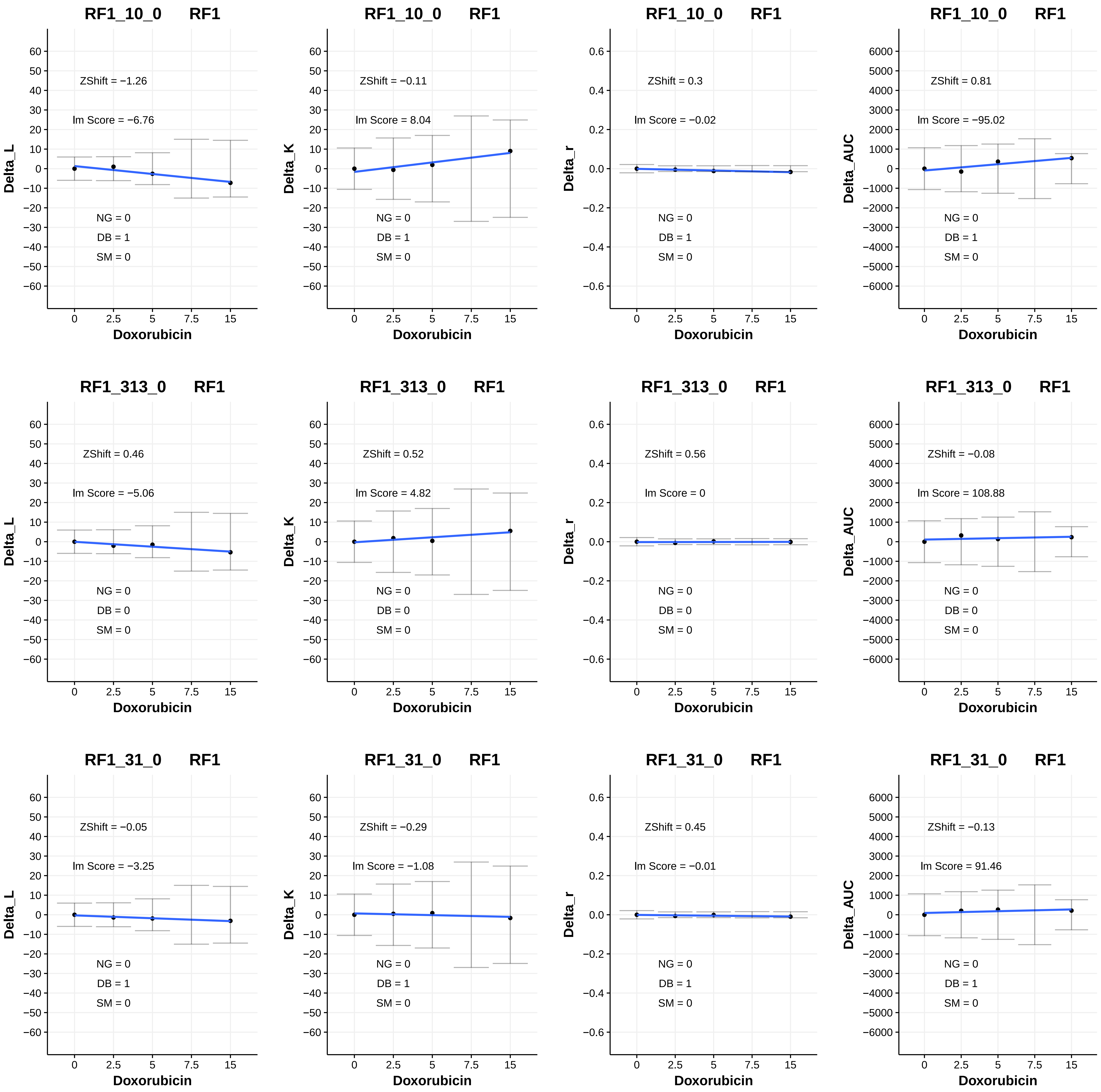

RF1_10_0 RF1
ZShift = −1.26
Im Score = −6.76
NG = 0
DB = 1
SM = 0
Delta_L
Doxorubicin
RF1_10_0 RF1
ZShift = −0.11
Im Score = 8.04
NG = 0
DB = 1
SM = 0
Delta_K
Doxorubicin
RF1_10_0 RF1
ZShift = 0.3
Im Score = −0.02
NG = 0
DB = 1
SM = 0
Delta_r
Doxorubicin
RF1_10_0 RF1
ZShift = 0.81
Im Score = −95.02
NG = 0
DB = 1
SM = 0
Delta_AUC
Doxorubicin
RF1_313_0 RF1
ZShift = 0.46
Im Score = −5.06
NG = 0
DB = 0
SM = 0
Delta_L
Doxorubicin
RF1_313_0 RF1
ZShift = 0.52
Im Score = 4.82
NG = 0
DB = 0
SM = 0
Delta_K
Doxorubicin
RF1_313_0 RF1
ZShift = 0.56
Im Score = 0
NG = 0
DB = 0
SM = 0
Delta_r
Doxorubicin
RF1_313_0 RF1
ZShift = −0.08
Im Score = 108.88
NG = 0
DB = 0
SM = 0
Delta_AUC
Doxorubicin
RF1_31_0 RF1
ZShift = −0.05
Im Score = −3.25
NG = 0
DB = 1
SM = 0
Delta_L
Doxorubicin
RF1_31_0 RF1
ZShift = −0.29
Im Score = −1.08
NG = 0
DB = 1
SM = 0
Delta_K
Doxorubicin
RF1_31_0 RF1
ZShift = 0.45
Im Score = −0.01
NG = 0
DB = 1
SM = 0
Delta_r
Doxorubicin
RF1_31_0 RF1
ZShift = −0.13
Im Score = 91.46
NG = 0
DB = 1
SM = 0
Delta_AUC
Doxorubicin

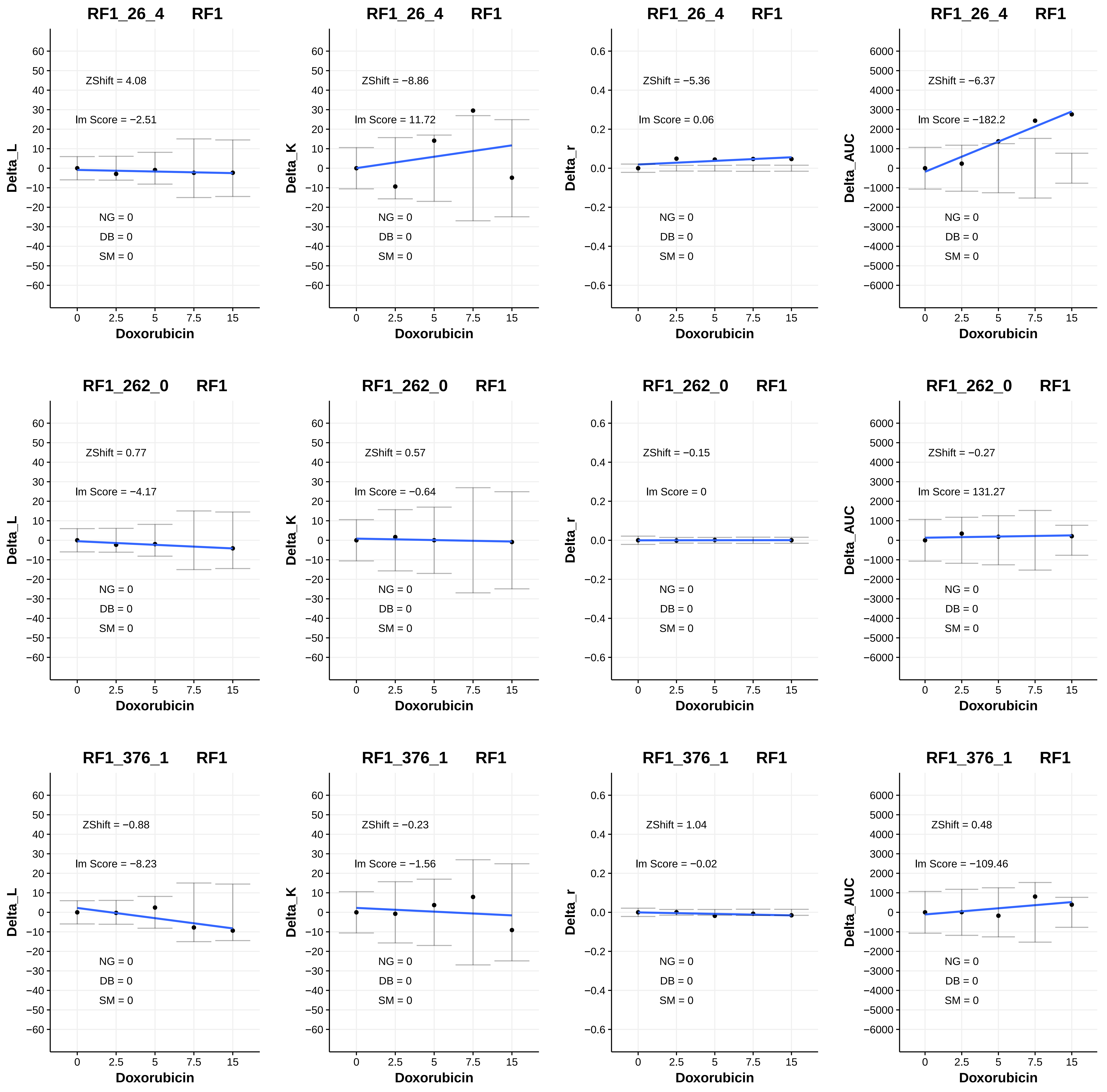
RF1_26_4 RF1
ZShift = 4.08
lm Score = −2.51
NG = 0
DB = 0
SM = 0
Delta_L
Doxorubicin
RF1_26_4 RF1
ZShift = −8.86
lm Score = 11.72
NG = 0
DB = 0
SM = 0
Delta_K
Doxorubicin
RF1_26_4 RF1
ZShift = −5.36
lm Score = 0.06
NG = 0
DB = 0
SM = 0
Delta_r
Doxorubicin
RF1_26_4 RF1
ZShift = −6.37
lm Score = −182.2
NG = 0
DB = 0
SM = 0
Delta_AUC
Doxorubicin
RF1_262_0 RF1
ZShift = 0.77
lm Score = −4.17
NG = 0
DB = 0
SM = 0
Delta_L
Doxorubicin
RF1_262_0 RF1
ZShift = 0.57
lm Score = −0.64
NG = 0
DB = 0
SM = 0
Delta_K
Doxorubicin
RF1_262_0 RF1
ZShift = −0.15
lm Score = 0
NG = 0
DB = 0
SM = 0
Delta_r
Doxorubicin
RF1_262_0 RF1
ZShift = −0.27
lm Score = 131.27
NG = 0
DB = 0
SM = 0
Delta_AUC
Doxorubicin
RF1_376_1 RF1
ZShift = −0.88
lm Score = −8.23
NG = 0
DB = 0
SM = 0
Delta_L
Doxorubicin
RF1_376_1 RF1
ZShift = −0.23
lm Score = −1.56
NG = 0
DB = 0
SM = 0
Delta_K
Doxorubicin
RF1_376_1 RF1
ZShift = 1.04
lm Score = −0.02
NG = 0
DB = 0
SM = 0
Delta_r
Doxorubicin
RF1_376_1 RF1
ZShift = 0.48
lm Score = −109.46
NG = 0
DB = 0
SM = 0
Delta_AUC
Doxorubicin

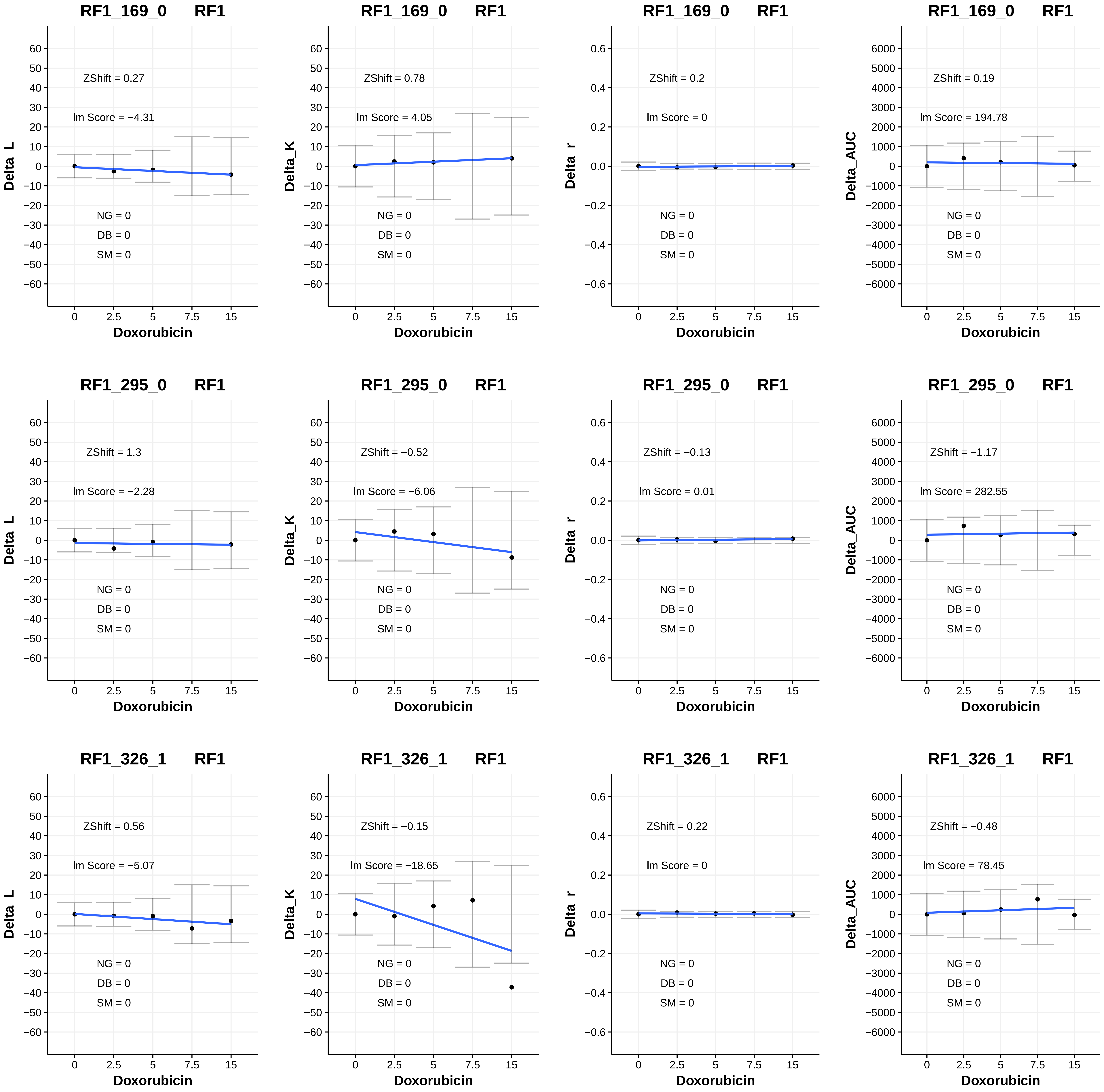
RF1_169_0 RF1
ZShift = 0.27
lm Score = −4.31
NG = 0
DB = 0
SM = 0
Delta_L
Doxorubicin
RF1_169_0 RF1
ZShift = 0.78
lm Score = 4.05
NG = 0
DB = 0
SM = 0
Delta_K
Doxorubicin
RF1_169_0 RF1
ZShift = 0.2
lm Score = 0
NG = 0
DB = 0
SM = 0
Delta_r
Doxorubicin
RF1_169_0 RF1
ZShift = 0.19
lm Score = 194.78
NG = 0
DB = 0
SM = 0
Delta_AUC
Doxorubicin
RF1_295_0 RF1
ZShift = 1.3
lm Score = −2.28
NG = 0
DB = 0
SM = 0
Delta_L
Doxorubicin
RF1_295_0 RF1
ZShift = −0.52
lm Score = −6.06
NG = 0
DB = 0
SM = 0
Delta_K
Doxorubicin
RF1_295_0 RF1
ZShift = −0.13
lm Score = 0.01
NG = 0
DB = 0
SM = 0
Delta_r
Doxorubicin
RF1_295_0 RF1
ZShift = −1.17
lm Score = 282.55
NG = 0
DB = 0
SM = 0
Delta_AUC
Doxorubicin
RF1_326_1 RF1
ZShift = 0.56
lm Score = −5.07
NG = 0
DB = 0
SM = 0
Delta_L
Doxorubicin
RF1_326_1 RF1
ZShift = −0.15
lm Score = −18.65
NG = 0
DB = 0
SM = 0
Delta_K
Doxorubicin
RF1_326_1 RF1
ZShift = 0.22
lm Score = 0
NG = 0
DB = 0
SM = 0
Delta_r
Doxorubicin
RF1_326_1 RF1
ZShift = −0.48
lm Score = 78.45
NG = 0
DB = 0
SM = 0
Delta_AUC
Doxorubicin

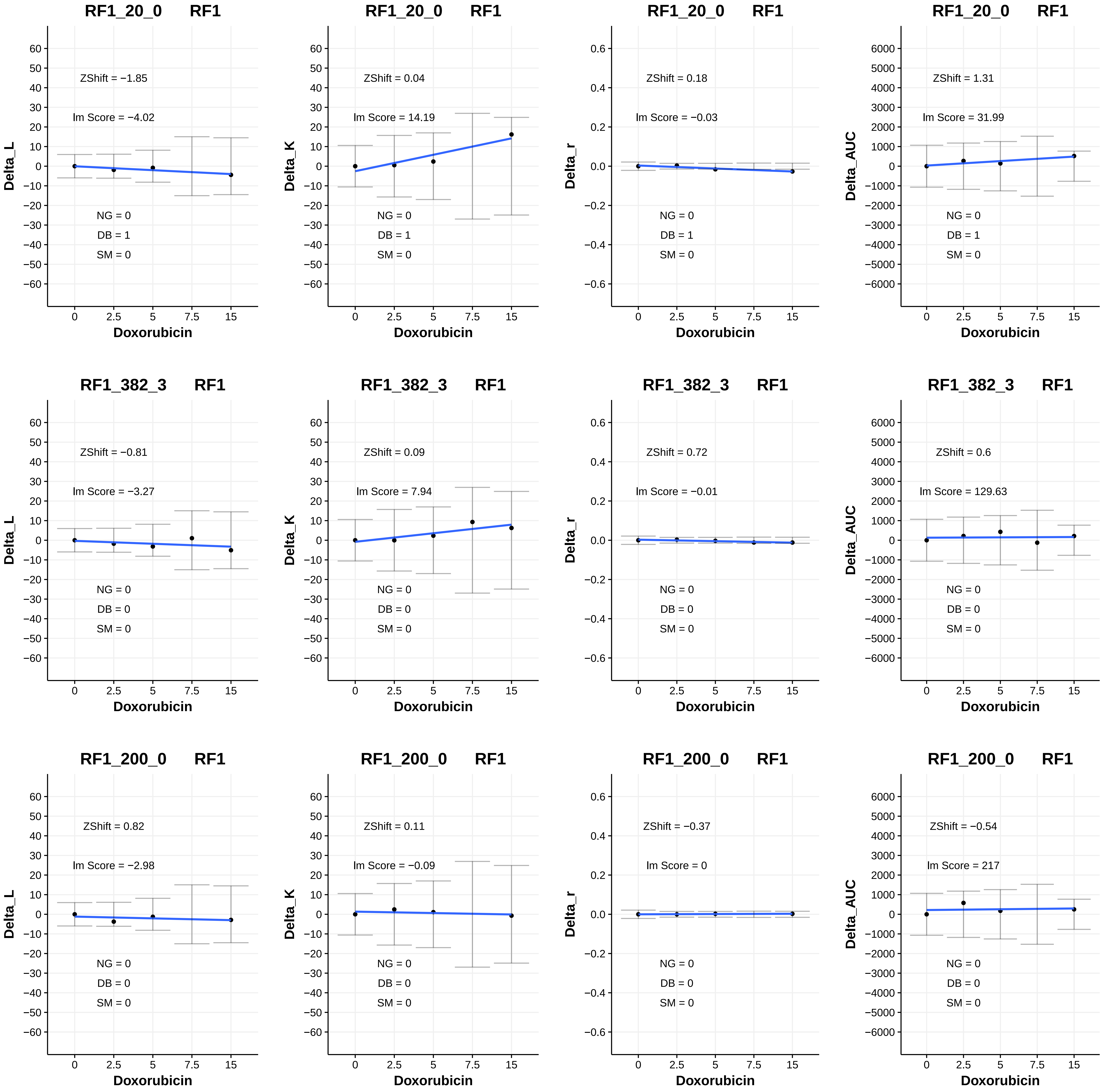
RF1_20_0 RF1
ZShift = −1.85
lm Score = −4.02
NG = 0
DB = 1
SM = 0
Delta_L
Doxorubicin
RF1_20_0 RF1
ZShift = 0.04
lm Score = 14.19
NG = 0
DB = 1
SM = 0
Delta_K
Doxorubicin
RF1_20_0 RF1
ZShift = 0.18
lm Score = −0.03
NG = 0
DB = 1
SM = 0
Delta_r
Doxorubicin
RF1_20_0 RF1
ZShift = 1.31
lm Score = 31.99
NG = 0
DB = 1
SM = 0
Delta_AUC
Doxorubicin
RF1_382_3 RF1
ZShift = −0.81
lm Score = −3.27
NG = 0
DB = 0
SM = 0
Delta_L
Doxorubicin
RF1_382_3 RF1
ZShift = 0.09
lm Score = 7.94
NG = 0
DB = 0
SM = 0
Delta_K
Doxorubicin
RF1_382_3 RF1
ZShift = 0.72
lm Score = −0.01
NG = 0
DB = 0
SM = 0
Delta_r
Doxorubicin
RF1_382_3 RF1
ZShift = 0.6
lm Score = 129.63
NG = 0
DB = 0
SM = 0
Delta_AUC
Doxorubicin
RF1_200_0 RF1
ZShift = 0.82
lm Score = −2.98
NG = 0
DB = 0
SM = 0
Delta_L
Doxorubicin
RF1_200_0 RF1
ZShift = 0.11
lm Score = −0.09
NG = 0
DB = 0
SM = 0
Delta_K
Doxorubicin
RF1_200_0 RF1
ZShift = −0.37
lm Score = 0
NG = 0
DB = 0
SM = 0
Delta_r
Doxorubicin
RF1_200_0 RF1
ZShift = −0.54
lm Score = 217
NG = 0
DB = 0
SM = 0
Delta_AUC
Doxorubicin

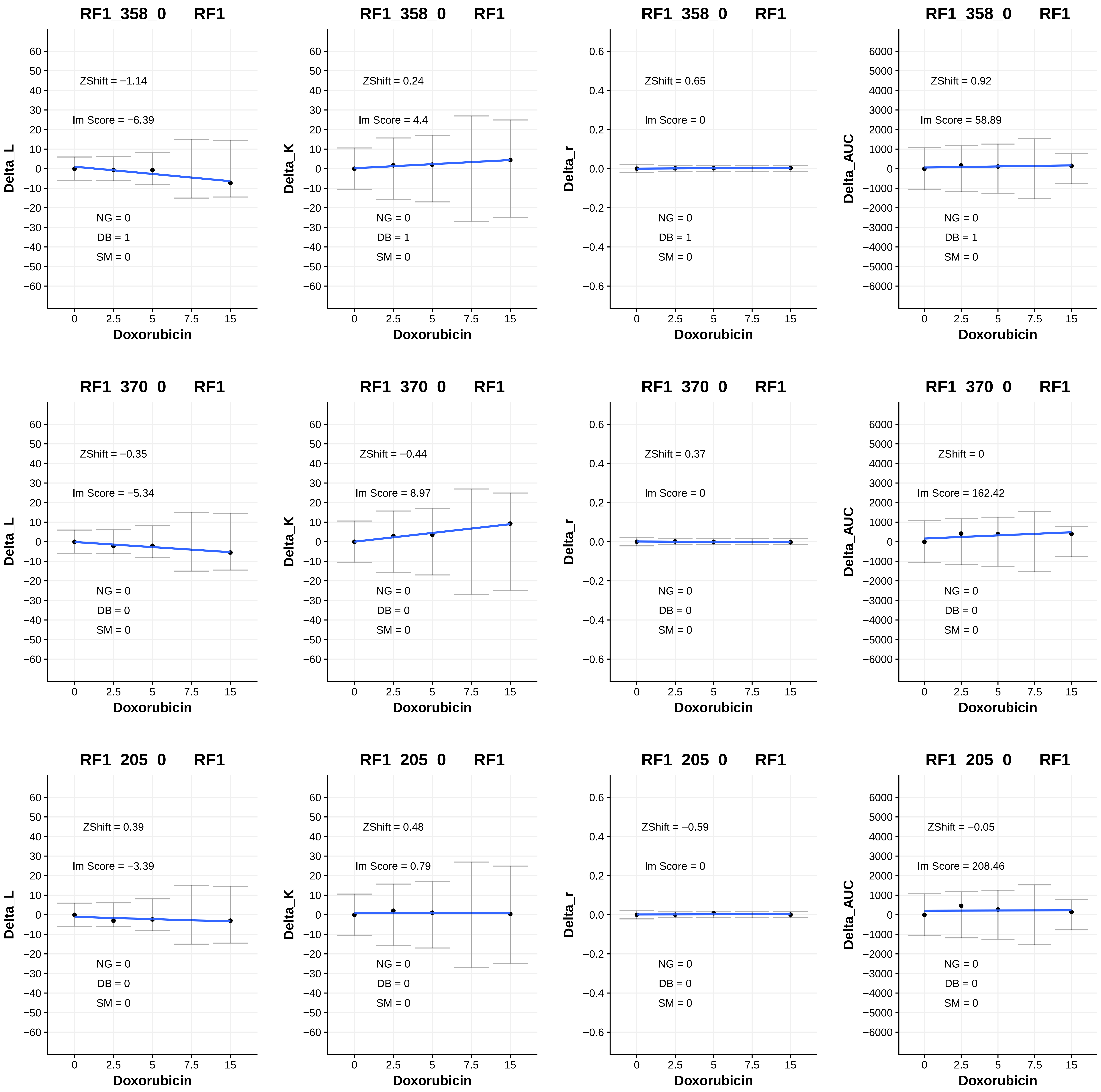
RF1_358_0 RF1
ZShift = −1.14
lm Score = −6.39
NG = 0
DB = 1
SM = 0
Delta_L
Doxorubicin
RF1_358_0 RF1
ZShift = 0.24
lm Score = 4.4
NG = 0
DB = 1
SM = 0
Delta_K
Doxorubicin
RF1_358_0 RF1
ZShift = 0.65
lm Score = 0
NG = 0
DB = 1
SM = 0
Delta_r
Doxorubicin
RF1_358_0 RF1
ZShift = 0.92
lm Score = 58.89
NG = 0
DB = 1
SM = 0
Delta_AUC
Doxorubicin
RF1_370_0 RF1
ZShift = −0.35
lm Score = −5.34
NG = 0
DB = 0
SM = 0
Delta_L
Doxorubicin
RF1_370_0 RF1
ZShift = −0.44
lm Score = 8.97
NG = 0
DB = 0
SM = 0
Delta_K
Doxorubicin
RF1_370_0 RF1
ZShift = 0.37
lm Score = 0
NG = 0
DB = 0
SM = 0
Delta_r
Doxorubicin
RF1_370_0 RF1
ZShift = 0
lm Score = 162.42
NG = 0
DB = 0
SM = 0
Delta_AUC
Doxorubicin
RF1_205_0 RF1
ZShift = 0.39
lm Score = −3.39
NG = 0
DB = 0
SM = 0
Delta_L
Doxorubicin
RF1_205_0 RF1
ZShift = 0.48
lm Score = 0.79
NG = 0
DB = 0
SM = 0
Delta_K
Doxorubicin
RF1_205_0 RF1
ZShift = −0.59
lm Score = 0
NG = 0
DB = 0
SM = 0
Delta_r
Doxorubicin
RF1_205_0 RF1
ZShift = −0.05
lm Score = 208.46
NG = 0
DB = 0
SM = 0
Delta_AUC
Doxorubicin

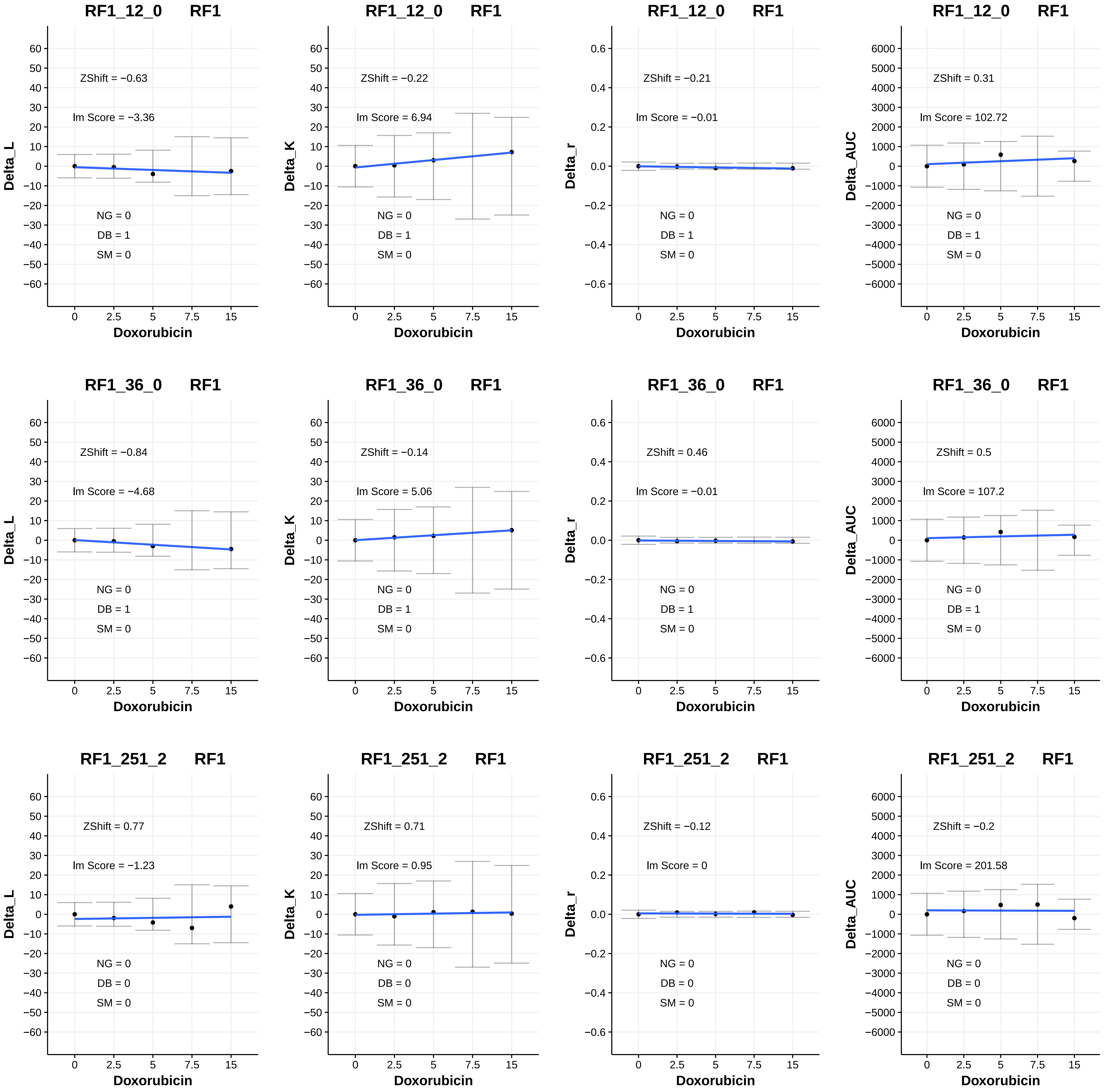
RF1_12_0 RF1
ZShift = −0.63
lm Score = −3.36
NG = 0
DB = 1
SM = 0
Delta_L
Doxorubicin
RF1_12_0 RF1
ZShift = −0.22
lm Score = 6.94
NG = 0
DB = 1
SM = 0
Delta_K
Doxorubicin
RF1_12_0 RF1
ZShift = −0.21
lm Score = −0.01
NG = 0
DB = 1
SM = 0
Delta_r
Doxorubicin
RF1_12_0 RF1
ZShift = 0.31
lm Score = 102.72
NG = 0
DB = 1
SM = 0
Delta_AUC
Doxorubicin
RF1_36_0 RF1
ZShift = −0.84
lm Score = −4.68
NG = 0
DB = 1
SM = 0
Delta_L
Doxorubicin
RF1_36_0 RF1
ZShift = −0.14
lm Score = 5.06
NG = 0
DB = 1
SM = 0
Delta_K
Doxorubicin
RF1_36_0 RF1
ZShift = 0.46
lm Score = −0.01
NG = 0
DB = 1
SM = 0
Delta_r
Doxorubicin
RF1_36_0 RF1
ZShift = 0.5
lm Score = 107.2
NG = 0
DB = 1
SM = 0
Delta_AUC
Doxorubicin
RF1_251_2 RF1
ZShift = 0.77
lm Score = −1.23
NG = 0
DB = 0
SM = 0
Delta_L
Doxorubicin
RF1_251_2 RF1
ZShift = 0.71
lm Score = 0.95
NG = 0
DB = 0
SM = 0
Delta_K
Doxorubicin
RF1_251_2 RF1
ZShift = −0.12
lm Score = 0
NG = 0
DB = 0
SM = 0
Delta_r
Doxorubicin
RF1_251_2 RF1
ZShift = −0.2
lm Score = 201.58
NG = 0
DB = 0
SM = 0
Delta_AUC
Doxorubicin

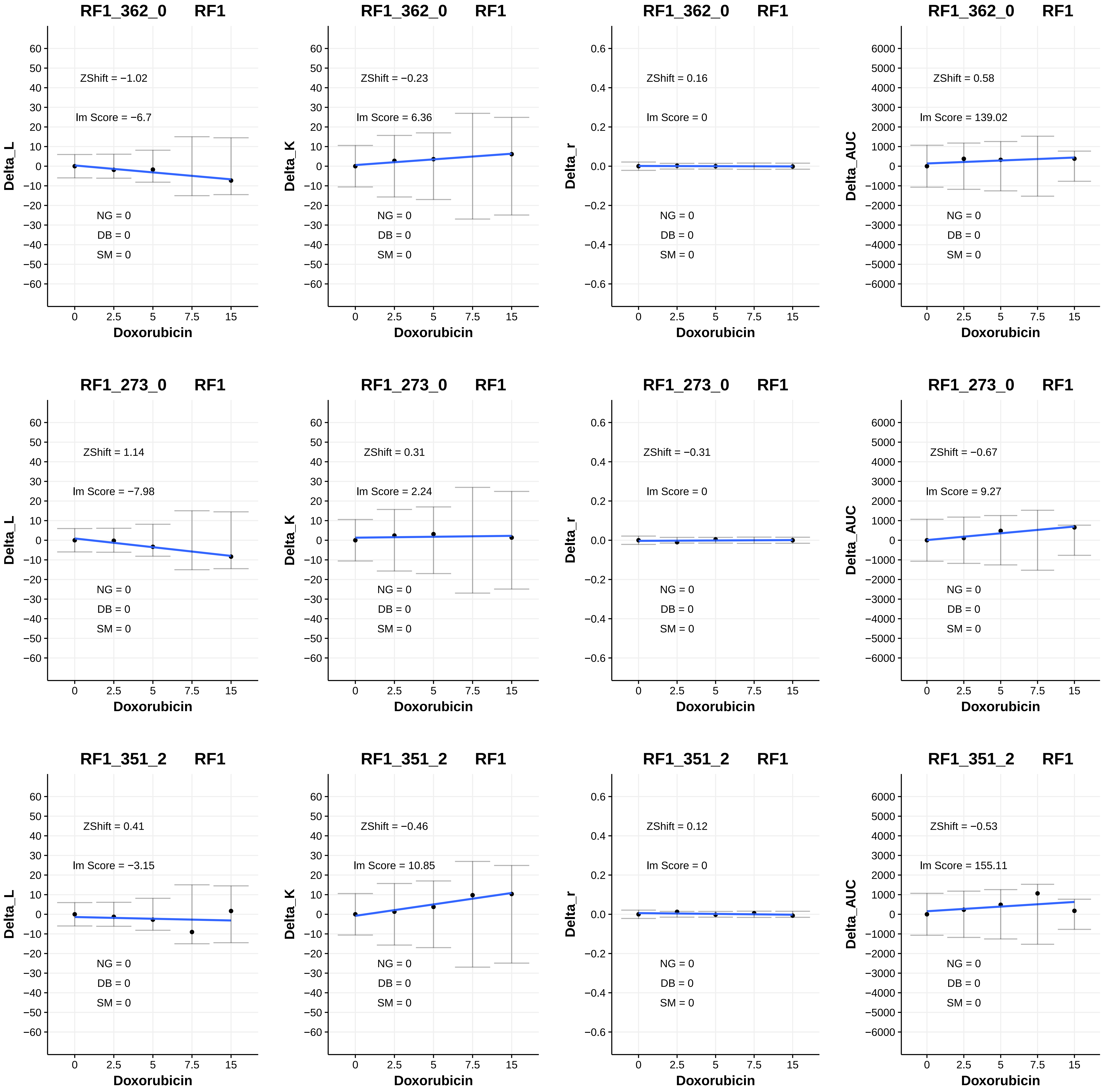
RF1_362_0 RF1
ZShift = −1.02
Im Score = −6.7
NG = 0
DB = 0
SM = 0
Delta_L
Doxorubicin
RF1_362_0 RF1
ZShift = −0.23
Im Score = 6.36
NG = 0
DB = 0
SM = 0
Delta_K
Doxorubicin
RF1_362_0 RF1
ZShift = 0.16
Im Score = 0
NG = 0
DB = 0
SM = 0
Delta_r
Doxorubicin
RF1_362_0 RF1
ZShift = 0.58
Im Score = 139.02
NG = 0
DB = 0
SM = 0
Delta_AUC
Doxorubicin
RF1_273_0 RF1
ZShift = 1.14
Im Score = −7.98
NG = 0
DB = 0
SM = 0
Delta_L
Doxorubicin
RF1_273_0 RF1
ZShift = 0.31
Im Score = 2.24
NG = 0
DB = 0
SM = 0
Delta_K
Doxorubicin
RF1_273_0 RF1
ZShift = −0.31
Im Score = 0
NG = 0
DB = 0
SM = 0
Delta_r
Doxorubicin
RF1_273_0 RF1
ZShift = −0.67
Im Score = 9.27
NG = 0
DB = 0
SM = 0
Delta_AUC
Doxorubicin
RF1_351_2 RF1
ZShift = 0.41
Im Score = −3.15
NG = 0
DB = 0
SM = 0
Delta_L
Doxorubicin
RF1_351_2 RF1
ZShift = −0.46
Im Score = 10.85
NG = 0
DB = 0
SM = 0
Delta_K
Doxorubicin
RF1_351_2 RF1
ZShift = 0.12
Im Score = 0
NG = 0
DB = 0
SM = 0
Delta_r
Doxorubicin
RF1_351_2 RF1
ZShift = −0.53
Im Score = 155.11
NG = 0
DB = 0
SM = 0
Delta_AUC
Doxorubicin

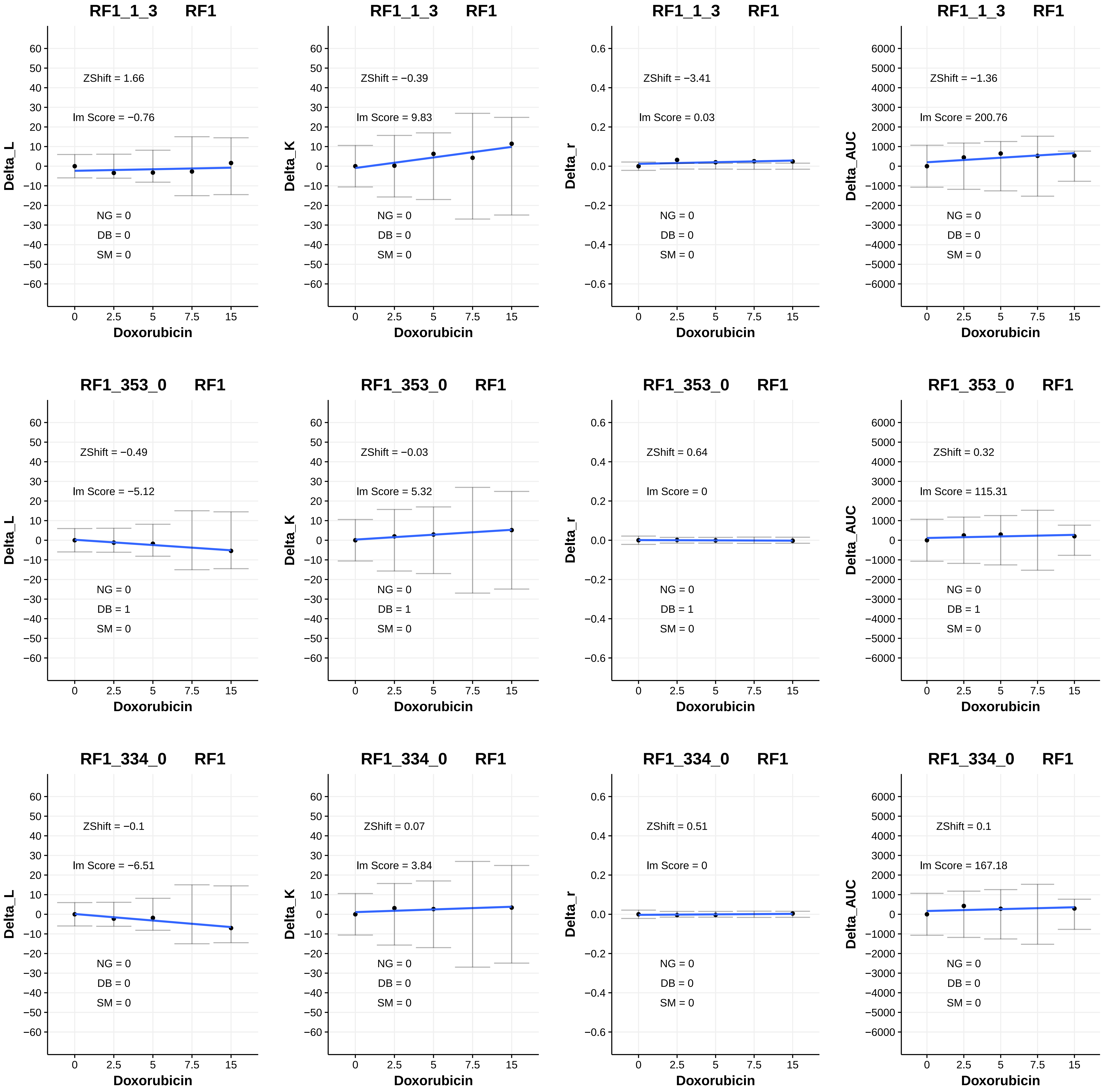

RF1_1_3 RF1
ZShift = 1.66
lm Score = −0.76
NG = 0
DB = 0
SM = 0
Delta_L
Doxorubicin
RF1_1_3 RF1
ZShift = −0.39
lm Score = 9.83
NG = 0
DB = 0
SM = 0
Delta_K
Doxorubicin
RF1_1_3 RF1
ZShift = −3.41
lm Score = 0.03
NG = 0
DB = 0
SM = 0
Delta_r
Doxorubicin
RF1_1_3 RF1
ZShift = −1.36
lm Score = 200.76
NG = 0
DB = 0
SM = 0
Delta_AUC
Doxorubicin
RF1_353_0 RF1
ZShift = −0.49
lm Score = −5.12
NG = 0
DB = 1
SM = 0
Delta_L
Doxorubicin
RF1_353_0 RF1
ZShift = −0.03
lm Score = 5.32
NG = 0
DB = 1
SM = 0
Delta_K
Doxorubicin
RF1_353_0 RF1
ZShift = 0.64
lm Score = 0
NG = 0
DB = 1
SM = 0
Delta_r
Doxorubicin
RF1_353_0 RF1
ZShift = 0.32
lm Score = 115.31
NG = 0
DB = 1
SM = 0
Delta_AUC
Doxorubicin
RF1_334_0 RF1
ZShift = −0.1
lm Score = −6.51
NG = 0
DB = 0
SM = 0
Delta_L
Doxorubicin
RF1_334_0 RF1
ZShift = 0.07
lm Score = 3.84
NG = 0
DB = 0
SM = 0
Delta_K
Doxorubicin
RF1_334_0 RF1
ZShift = 0.51
lm Score = 0
NG = 0
DB = 0
SM = 0
Delta_r
Doxorubicin
RF1_334_0 RF1
ZShift = 0.1
lm Score = 167.18
NG = 0
DB = 0
SM = 0
Delta_AUC
Doxorubicin

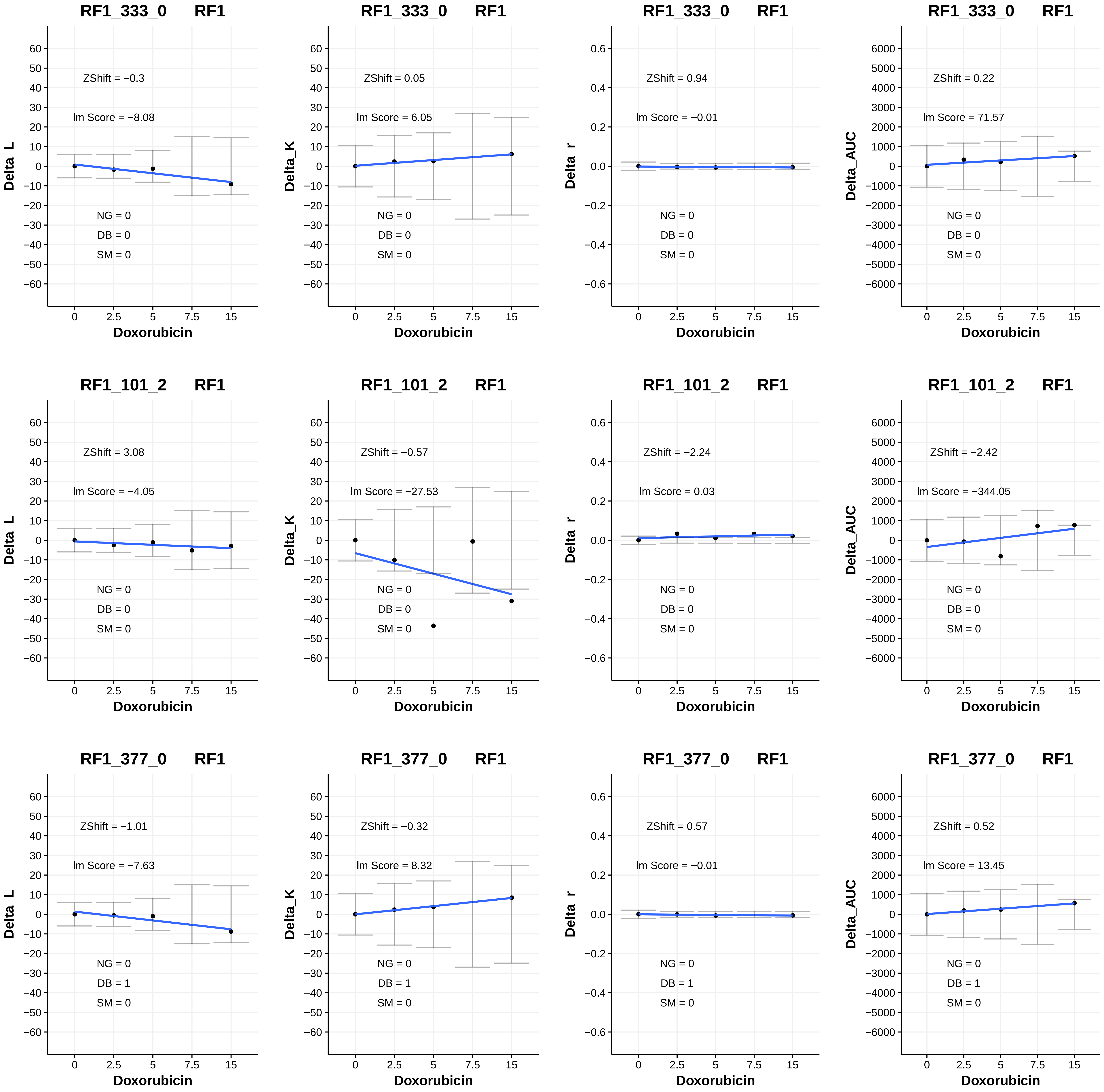
RF1_333_0 RF1
Delta_L
ZShift = −0.3
lm Score = −8.08
NG = 0
DB = 0
SM = 0
Doxorubicin
RF1_333_0 RF1
Delta_K
ZShift = 0.05
lm Score = 6.05
NG = 0
DB = 0
SM = 0
Doxorubicin
RF1_333_0 RF1
Delta_r
ZShift = 0.94
lm Score = −0.01
NG = 0
DB = 0
SM = 0
Doxorubicin
RF1_333_0 RF1
Delta_AUC
ZShift = 0.22
lm Score = 71.57
NG = 0
DB = 0
SM = 0
Doxorubicin
RF1_101_2 RF1
Delta_L
ZShift = 3.08
lm Score = −4.05
NG = 0
DB = 0
SM = 0
Doxorubicin
RF1_101_2 RF1
Delta_K
ZShift = −0.57
lm Score = −27.53
NG = 0
DB = 0
SM = 0
Doxorubicin
RF1_101_2 RF1
Delta_r
ZShift = −2.24
lm Score = 0.03
NG = 0
DB = 0
SM = 0
Doxorubicin
RF1_101_2 RF1
Delta_AUC
ZShift = −2.42
lm Score = −344.05
NG = 0
DB = 0
SM = 0
Doxorubicin
RF1_377_0 RF1
Delta_L
ZShift = −1.01
lm Score = −7.63
NG = 0
DB = 1
SM = 0
Doxorubicin
RF1_377_0 RF1
Delta_K
ZShift = −0.32
lm Score = 8.32
NG = 0
DB = 1
SM = 0
Doxorubicin
RF1_377_0 RF1
Delta_r
ZShift = 0.57
lm Score = −0.01
NG = 0
DB = 1
SM = 0
Doxorubicin
RF1_377_0 RF1
Delta_AUC
ZShift = 0.52
lm Score = 13.45
NG = 0
DB = 1
SM = 0
Doxorubicin

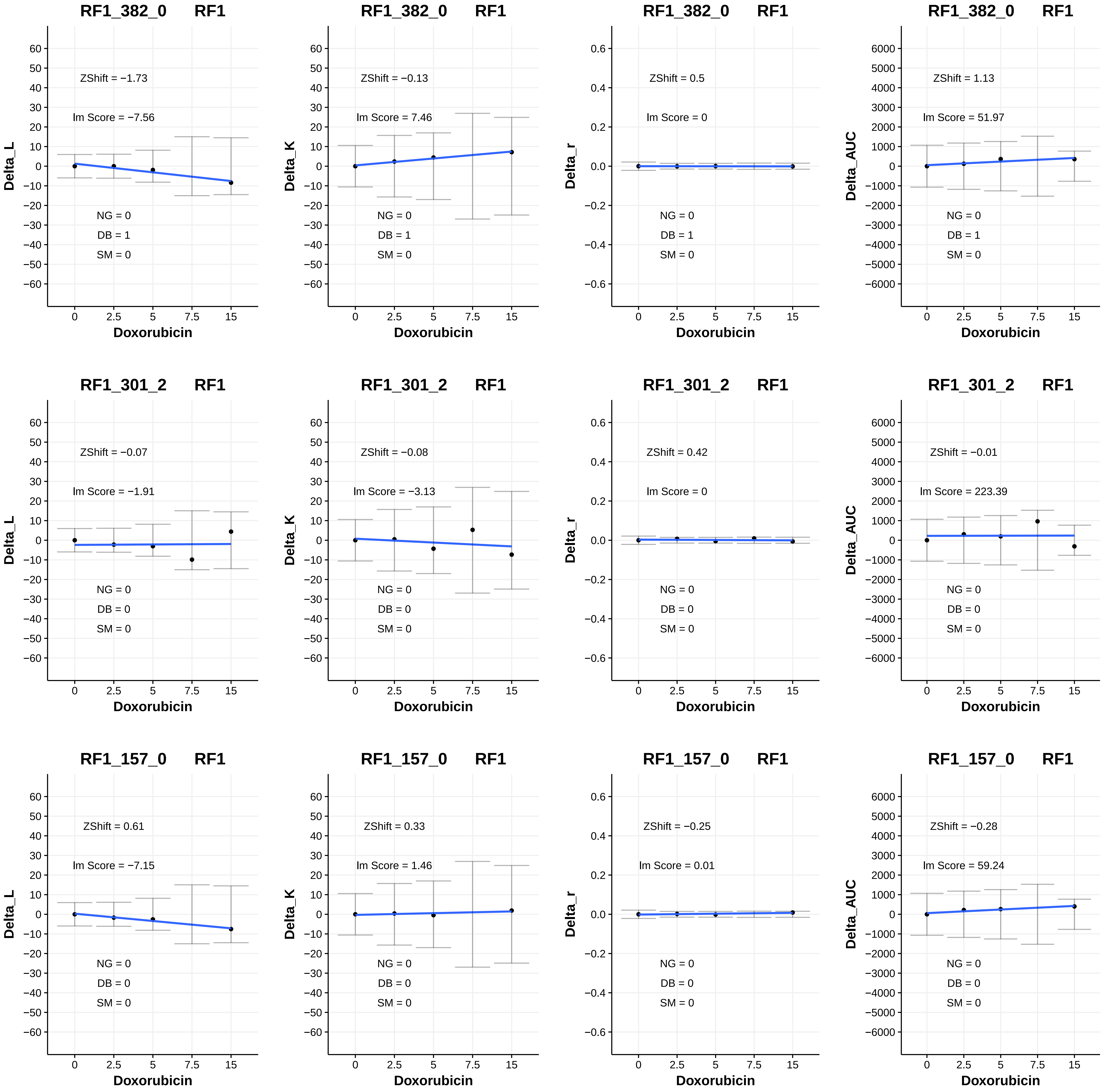
RF1_382_0 RF1
ZShift = −1.73
lm Score = −7.56
NG = 0
DB = 1
SM = 0
Delta_L
Doxorubicin
RF1_382_0 RF1
ZShift = −0.13
lm Score = 7.46
NG = 0
DB = 1
SM = 0
Delta_K
Doxorubicin
RF1_382_0 RF1
ZShift = 0.5
lm Score = 0
NG = 0
DB = 1
SM = 0
Delta_r
Doxorubicin
RF1_382_0 RF1
ZShift = 1.13
lm Score = 51.97
NG = 0
DB = 1
SM = 0
Delta_AUC
Doxorubicin
RF1_301_2 RF1
ZShift = −0.07
lm Score = −1.91
NG = 0
DB = 0
SM = 0
Delta_L
Doxorubicin
RF1_301_2 RF1
ZShift = −0.08
lm Score = −3.13
NG = 0
DB = 0
SM = 0
Delta_K
Doxorubicin
RF1_301_2 RF1
ZShift = 0.42
lm Score = 0
NG = 0
DB = 0
SM = 0
Delta_r
Doxorubicin
RF1_301_2 RF1
ZShift = −0.01
lm Score = 223.39
NG = 0
DB = 0
SM = 0
Delta_AUC
Doxorubicin
RF1_157_0 RF1
ZShift = 0.61
lm Score = −7.15
NG = 0
DB = 0
SM = 0
Delta_L
Doxorubicin
RF1_157_0 RF1
ZShift = 0.33
lm Score = 1.46
NG = 0
DB = 0
SM = 0
Delta_K
Doxorubicin
RF1_157_0 RF1
ZShift = −0.25
lm Score = 0.01
NG = 0
DB = 0
SM = 0
Delta_r
Doxorubicin
RF1_157_0 RF1
ZShift = −0.28
lm Score = 59.24
NG = 0
DB = 0
SM = 0
Delta_AUC
Doxorubicin

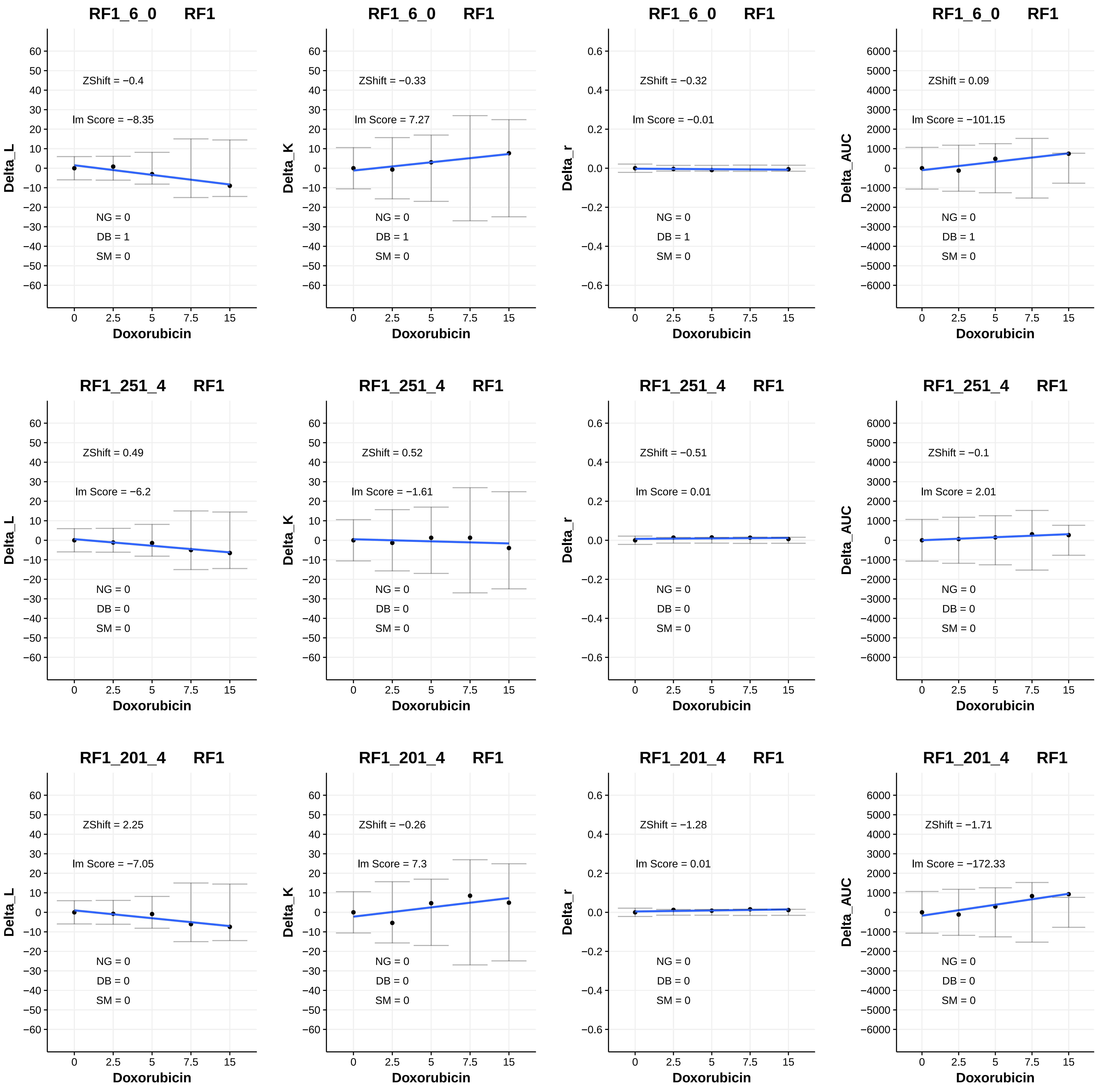

RF1_6_0 RF1
ZShift = −0.4
Im Score = −8.35
NG = 0
DB = 1
SM = 0
Delta_L
Doxorubicin
RF1_6_0 RF1
ZShift = −0.33
Im Score = 7.27
NG = 0
DB = 1
SM = 0
Delta_K
Doxorubicin
RF1_6_0 RF1
ZShift = −0.32
Im Score = −0.01
NG = 0
DB = 1
SM = 0
Delta_r
Doxorubicin
RF1_6_0 RF1
ZShift = 0.09
Im Score = −101.15
NG = 0
DB = 1
SM = 0
Delta_AUC
Doxorubicin
RF1_251_4 RF1
ZShift = 0.49
Im Score = −6.2
NG = 0
DB = 0
SM = 0
Delta_L
Doxorubicin
RF1_251_4 RF1
ZShift = 0.52
Im Score = −1.61
NG = 0
DB = 0
SM = 0
Delta_K
Doxorubicin
RF1_251_4 RF1
ZShift = −0.51
Im Score = 0.01
NG = 0
DB = 0
SM = 0
Delta_r
Doxorubicin
RF1_251_4 RF1
ZShift = −0.1
Im Score = 2.01
NG = 0
DB = 0
SM = 0
Delta_AUC
Doxorubicin
RF1_201_4 RF1
ZShift = 2.25
Im Score = −7.05
NG = 0
DB = 0
SM = 0
Delta_L
Doxorubicin
RF1_201_4 RF1
ZShift = −0.26
Im Score = 7.3
NG = 0
DB = 0
SM = 0
Delta_K
Doxorubicin
RF1_201_4 RF1
ZShift = −1.28
Im Score = 0.01
NG = 0
DB = 0
SM = 0
Delta_r
Doxorubicin
RF1_201_4 RF1
ZShift = −1.71
Im Score = −172.33
NG = 0
DB = 0
SM = 0
Delta_AUC
Doxorubicin

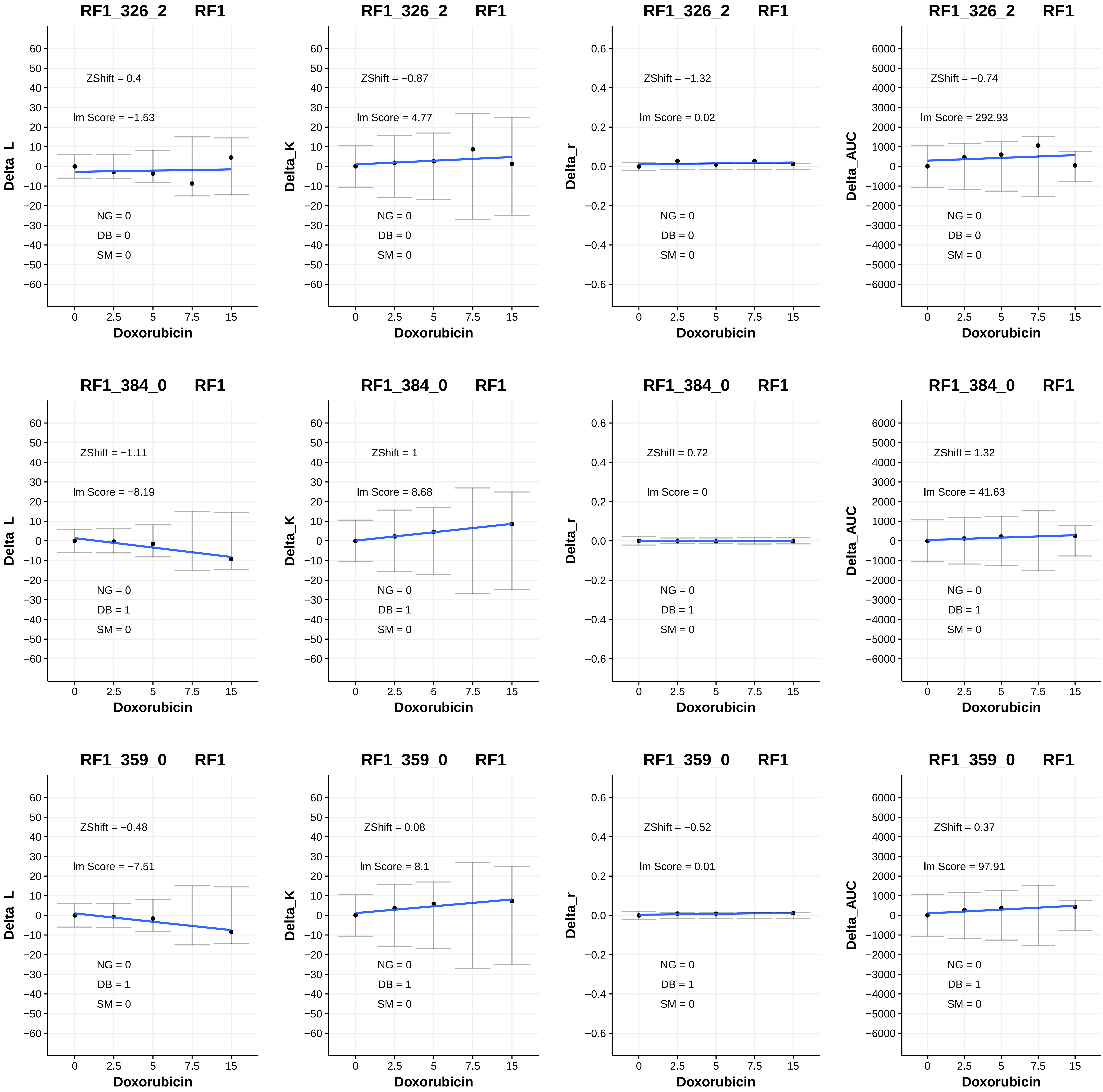

RF1_326_2 RF1
ZShift = 0.4
lm Score = −1.53
NG = 0
DB = 0
SM = 0
Delta_L
Doxorubicin
RF1_326_2 RF1
ZShift = −0.87
lm Score = 4.77
NG = 0
DB = 0
SM = 0
Delta_K
Doxorubicin
RF1_326_2 RF1
ZShift = −1.32
lm Score = 0.02
NG = 0
DB = 0
SM = 0
Delta_r
Doxorubicin
RF1_326_2 RF1
ZShift = −0.74
lm Score = 292.93
NG = 0
DB = 0
SM = 0
Delta_AUC
Doxorubicin
RF1_384_0 RF1
ZShift = −1.11
lm Score = −8.19
NG = 0
DB = 1
SM = 0
Delta_L
Doxorubicin
RF1_384_0 RF1
ZShift = 1
lm Score = 8.68
NG = 0
DB = 1
SM = 0
Delta_K
Doxorubicin
RF1_384_0 RF1
ZShift = 0.72
lm Score = 0
NG = 0
DB = 1
SM = 0
Delta_r
Doxorubicin
RF1_384_0 RF1
ZShift = 1.32
lm Score = 41.63
NG = 0
DB = 1
SM = 0
Delta_AUC
Doxorubicin
RF1_359_0 RF1
ZShift = −0.48
lm Score = −7.51
NG = 0
DB = 1
SM = 0
Delta_L
Doxorubicin
RF1_359_0 RF1
ZShift = 0.08
lm Score = 8.1
NG = 0
DB = 1
SM = 0
Delta_K
Doxorubicin
RF1_359_0 RF1
ZShift = −0.52
lm Score = 0.01
NG = 0
DB = 1
SM = 0
Delta_r
Doxorubicin
RF1_359_0 RF1
ZShift = 0.37
lm Score = 97.91
NG = 0
DB = 1
SM = 0
Delta_AUC
Doxorubicin

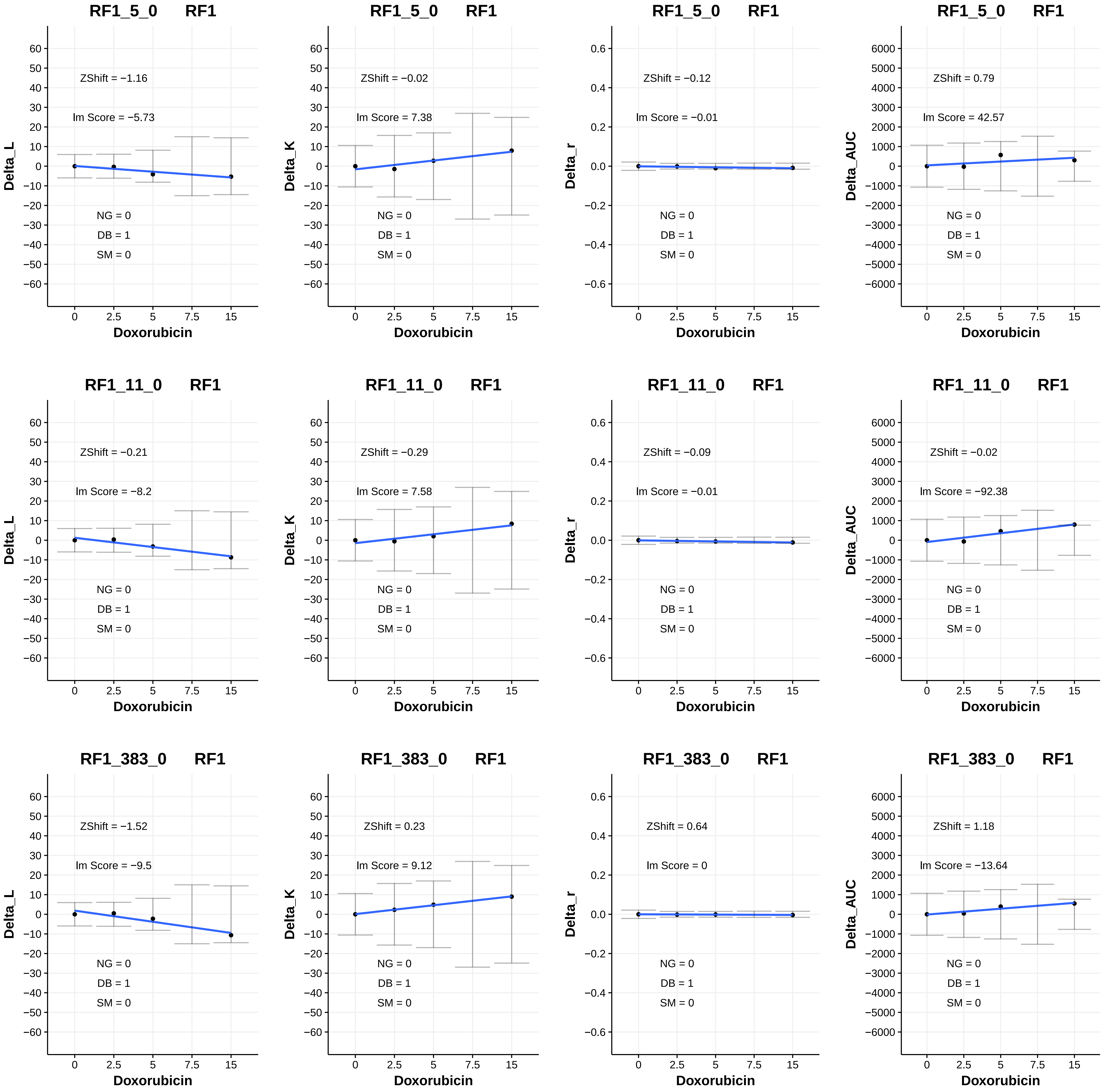

RF1_5_0 RF1
ZShift = −1.16
Im Score = −5.73
NG = 0
DB = 1
SM = 0
Delta_L
Doxorubicin
RF1_5_0 RF1
ZShift = −0.02
Im Score = 7.38
NG = 0
DB = 1
SM = 0
Delta_K
Doxorubicin
RF1_5_0 RF1
ZShift = −0.12
Im Score = −0.01
NG = 0
DB = 1
SM = 0
Delta_r
Doxorubicin
RF1_5_0 RF1
ZShift = 0.79
Im Score = 42.57
NG = 0
DB = 1
SM = 0
Delta_AUC
Doxorubicin
RF1_11_0 RF1
ZShift = −0.21
Im Score = −8.2
NG = 0
DB = 1
SM = 0
Delta_L
Doxorubicin
RF1_11_0 RF1
ZShift = −0.29
Im Score = 7.58
NG = 0
DB = 1
SM = 0
Delta_K
Doxorubicin
RF1_11_0 RF1
ZShift = −0.09
Im Score = −0.01
NG = 0
DB = 1
SM = 0
Delta_r
Doxorubicin
RF1_11_0 RF1
ZShift = −0.02
Im Score = −92.38
NG = 0
DB = 1
SM = 0
Delta_AUC
Doxorubicin
RF1_383_0 RF1
ZShift = −1.52
Im Score = −9.5
NG = 0
DB = 1
SM = 0
Delta_L
Doxorubicin
RF1_383_0 RF1
ZShift = 0.23
Im Score = 9.12
NG = 0
DB = 1
SM = 0
Delta_K
Doxorubicin
RF1_383_0 RF1
ZShift = 0.64
Im Score = 0
NG = 0
DB = 1
SM = 0
Delta_r
Doxorubicin
RF1_383_0 RF1
ZShift = 1.18
Im Score = −13.64
NG = 0
DB = 1
SM = 0
Delta_AUC
Doxorubicin

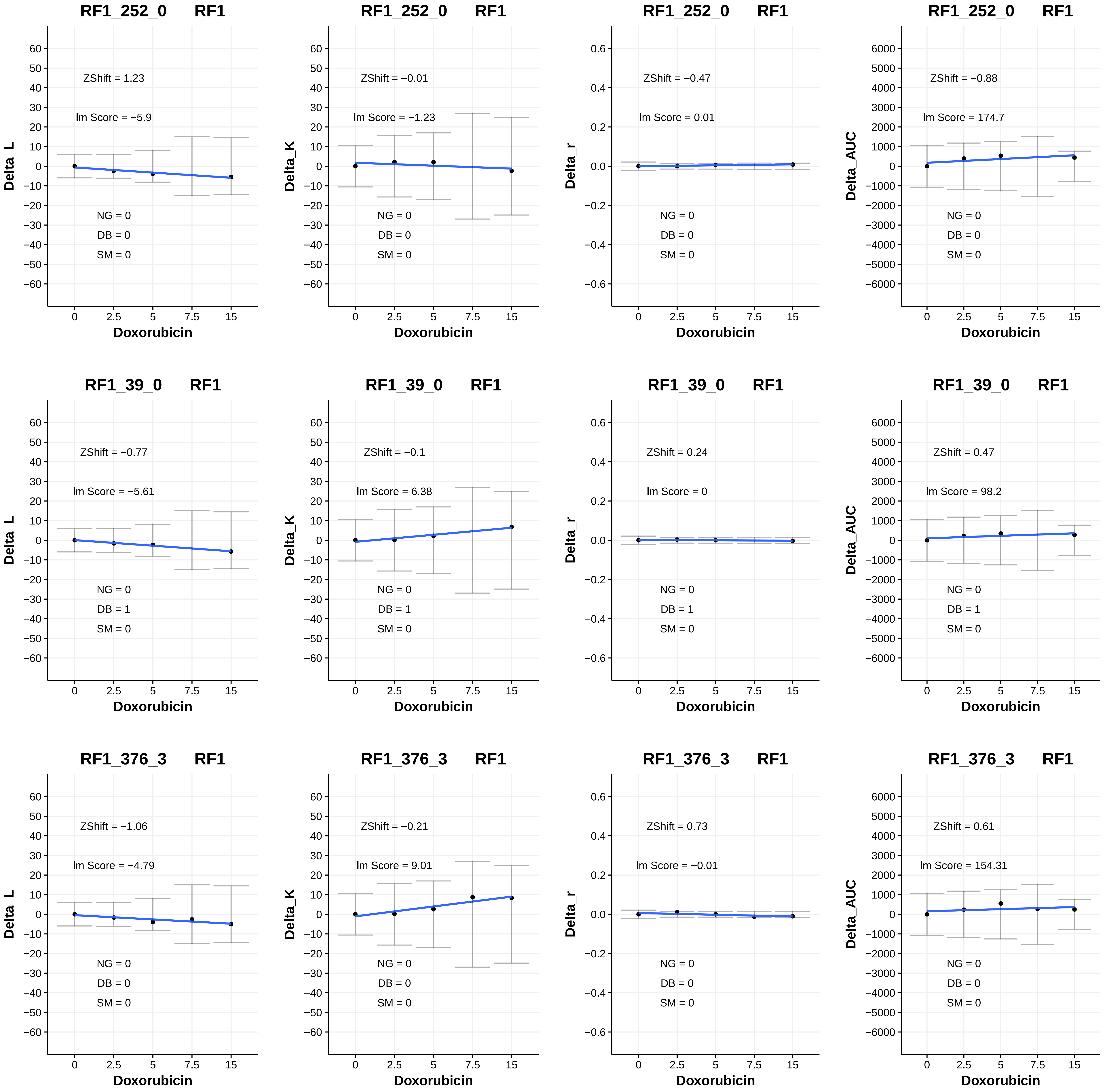

RF1_252_0 RF1
Delta_L
ZShift = 1.23
lm Score = −5.9
NG = 0
DB = 0
SM = 0
Doxorubicin
RF1_252_0 RF1
Delta_K
ZShift = −0.01
lm Score = −1.23
NG = 0
DB = 0
SM = 0
Doxorubicin
RF1_252_0 RF1
Delta_r
ZShift = −0.47
lm Score = 0.01
NG = 0
DB = 0
SM = 0
Doxorubicin
RF1_252_0 RF1
Delta_AUC
ZShift = −0.88
lm Score = 174.7
NG = 0
DB = 0
SM = 0
Doxorubicin
RF1_39_0 RF1
Delta_L
ZShift = −0.77
lm Score = −5.61
NG = 0
DB = 1
SM = 0
Doxorubicin
RF1_39_0 RF1
Delta_K
ZShift = −0.1
lm Score = 6.38
NG = 0
DB = 1
SM = 0
Doxorubicin
RF1_39_0 RF1
Delta_r
ZShift = 0.24
lm Score = 0
NG = 0
DB = 1
SM = 0
Doxorubicin
RF1_39_0 RF1
Delta_AUC
ZShift = 0.47
lm Score = 98.2
NG = 0
DB = 1
SM = 0
Doxorubicin
RF1_376_3 RF1
Delta_L
ZShift = −1.06
lm Score = −4.79
NG = 0
DB = 0
SM = 0
Doxorubicin
RF1_376_3 RF1
Delta_K
ZShift = −0.21
lm Score = 9.01
NG = 0
DB = 0
SM = 0
Doxorubicin
RF1_376_3 RF1
Delta_r
ZShift = 0.73
lm Score = −0.01
NG = 0
DB = 0
SM = 0
Doxorubicin
RF1_376_3 RF1
Delta_AUC
ZShift = 0.61
lm Score = 154.31
NG = 0
DB = 0
SM = 0
Doxorubicin

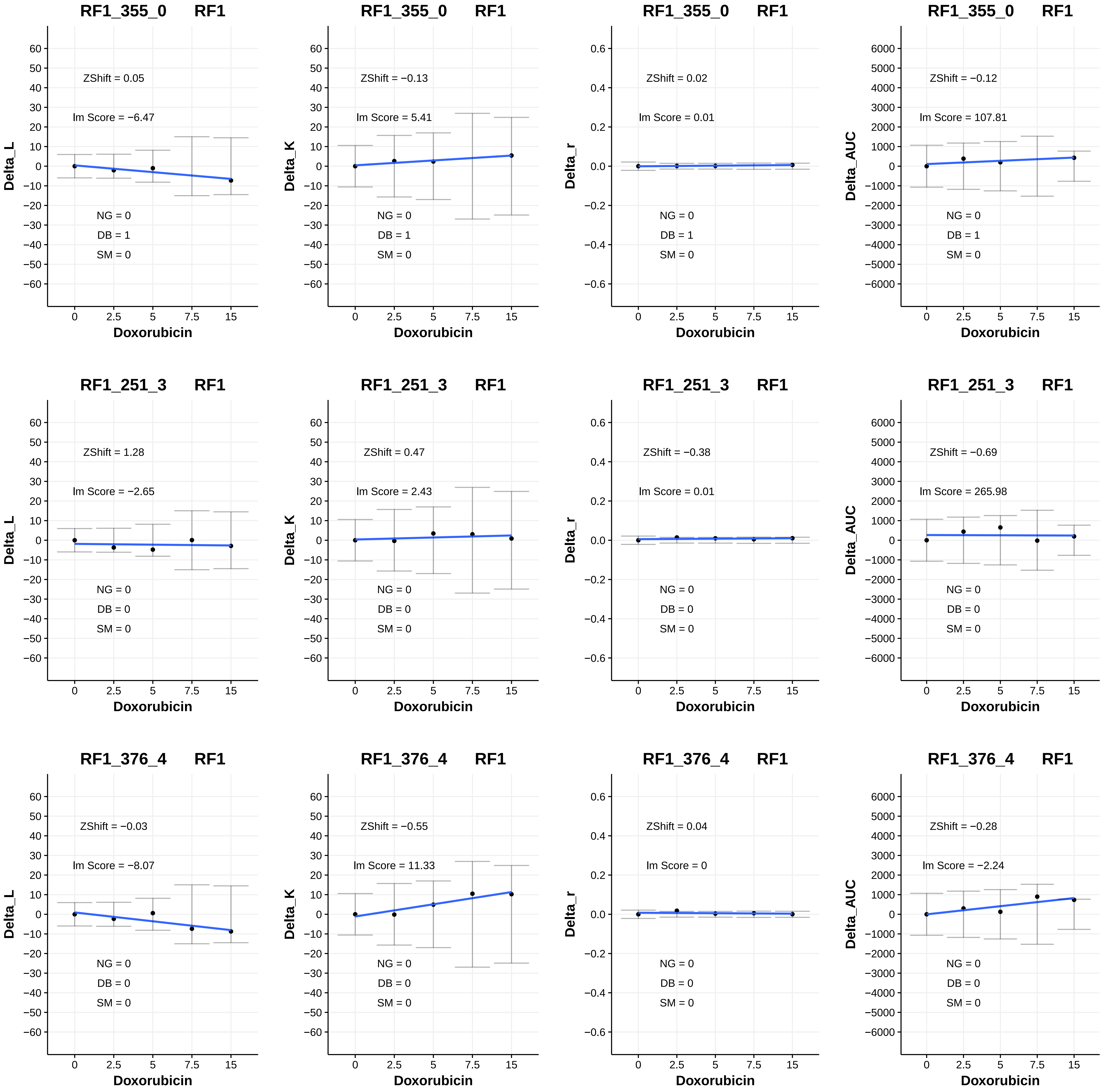
RF1_355_0 RF1
ZShift = 0.05
Im Score = −6.47
NG = 0
DB = 1
SM = 0
Delta_L
Doxorubicin
RF1_355_0 RF1
ZShift = −0.13
Im Score = 5.41
NG = 0
DB = 1
SM = 0
Delta_K
Doxorubicin
RF1_355_0 RF1
ZShift = 0.02
Im Score = 0.01
NG = 0
DB = 1
SM = 0
Delta_r
Doxorubicin
RF1_355_0 RF1
ZShift = −0.12
Im Score = 107.81
NG = 0
DB = 1
SM = 0
Delta_AUC
Doxorubicin
RF1_251_3 RF1
ZShift = 1.28
Im Score = −2.65
NG = 0
DB = 0
SM = 0
Delta_L
Doxorubicin
RF1_251_3 RF1
ZShift = 0.47
Im Score = 2.43
NG = 0
DB = 0
SM = 0
Delta_K
Doxorubicin
RF1_251_3 RF1
ZShift = −0.38
Im Score = 0.01
NG = 0
DB = 0
SM = 0
Delta_r
Doxorubicin
RF1_251_3 RF1
ZShift = −0.69
Im Score = 265.98
NG = 0
DB = 0
SM = 0
Delta_AUC
Doxorubicin
RF1_376_4 RF1
ZShift = −0.03
Im Score = −8.07
NG = 0
DB = 0
SM = 0
Delta_L
Doxorubicin
RF1_376_4 RF1
ZShift = −0.55
Im Score = 11.33
NG = 0
DB = 0
SM = 0
Delta_K
Doxorubicin
RF1_376_4 RF1
ZShift = 0.04
Im Score = 0
NG = 0
DB = 0
SM = 0
Delta_r
Doxorubicin
RF1_376_4 RF1
ZShift = −0.28
Im Score = −2.24
NG = 0
DB = 0
SM = 0
Delta_AUC
Doxorubicin

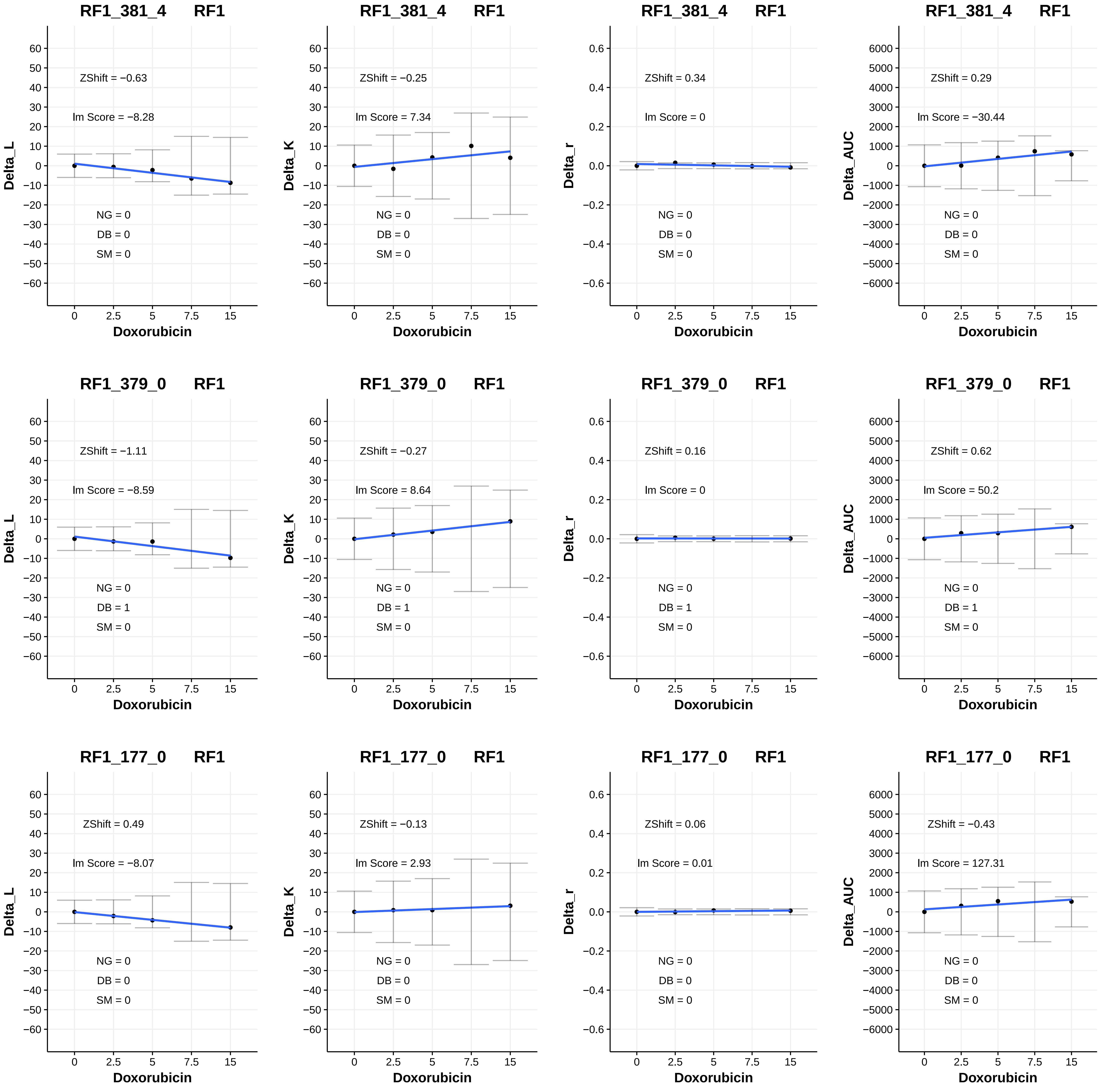
RF1_381_4 RF1
Delta_L
ZShift = −0.63
lm Score = −8.28
NG = 0
DB = 0
SM = 0
Doxorubicin
RF1_381_4 RF1
Delta_K
ZShift = −0.25
lm Score = 7.34
NG = 0
DB = 0
SM = 0
Doxorubicin
RF1_381_4 RF1
Delta_r
ZShift = 0.34
lm Score = 0
NG = 0
DB = 0
SM = 0
Doxorubicin
RF1_381_4 RF1
Delta_AUC
ZShift = 0.29
lm Score = −30.44
NG = 0
DB = 0
SM = 0
Doxorubicin
RF1_379_0 RF1
Delta_L
ZShift = −1.11
lm Score = −8.59
NG = 0
DB = 1
SM = 0
Doxorubicin
RF1_379_0 RF1
Delta_K
ZShift = −0.27
lm Score = 8.64
NG = 0
DB = 1
SM = 0
Doxorubicin
RF1_379_0 RF1
Delta_r
ZShift = 0.16
lm Score = 0
NG = 0
DB = 1
SM = 0
Doxorubicin
RF1_379_0 RF1
Delta_AUC
ZShift = 0.62
lm Score = 50.2
NG = 0
DB = 1
SM = 0
Doxorubicin
RF1_177_0 RF1
Delta_L
ZShift = 0.49
lm Score = −8.07
NG = 0
DB = 0
SM = 0
Doxorubicin
RF1_177_0 RF1
Delta_K
ZShift = −0.13
lm Score = 2.93
NG = 0
DB = 0
SM = 0
Doxorubicin
RF1_177_0 RF1
Delta_r
ZShift = 0.06
lm Score = 0.01
NG = 0
DB = 0
SM = 0
Doxorubicin
RF1_177_0 RF1
Delta_AUC
ZShift = −0.43
lm Score = 127.31
NG = 0
DB = 0
SM = 0
Doxorubicin

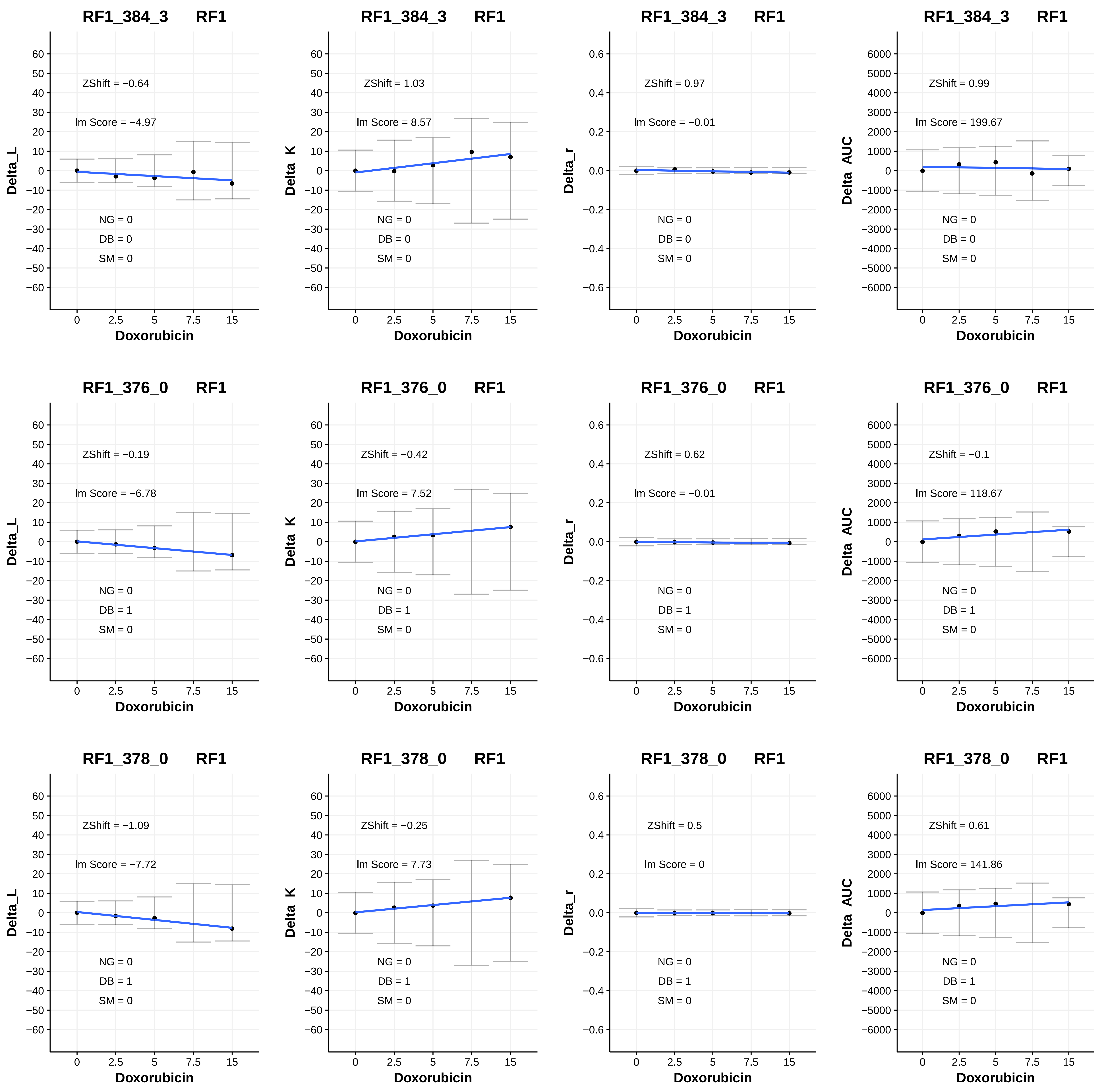
RF1_384_3 RF1
ZShift = −0.64
lm Score = −4.97
NG = 0
DB = 0
SM = 0
Delta_L
Doxorubicin
RF1_384_3 RF1
ZShift = 1.03
lm Score = 8.57
NG = 0
DB = 0
SM = 0
Delta_K
Doxorubicin
RF1_384_3 RF1
ZShift = 0.97
lm Score = −0.01
NG = 0
DB = 0
SM = 0
Delta_r
Doxorubicin
RF1_384_3 RF1
ZShift = 0.99
lm Score = 199.67
NG = 0
DB = 0
SM = 0
Delta_AUC
Doxorubicin
RF1_376_0 RF1
ZShift = −0.19
lm Score = −6.78
NG = 0
DB = 1
SM = 0
Delta_L
Doxorubicin
RF1_376_0 RF1
ZShift = −0.42
lm Score = 7.52
NG = 0
DB = 1
SM = 0
Delta_K
Doxorubicin
RF1_376_0 RF1
ZShift = 0.62
lm Score = −0.01
NG = 0
DB = 1
SM = 0
Delta_r
Doxorubicin
RF1_376_0 RF1
ZShift = −0.1
lm Score = 118.67
NG = 0
DB = 1
SM = 0
Delta_AUC
Doxorubicin
RF1_378_0 RF1
ZShift = −1.09
lm Score = −7.72
NG = 0
DB = 1
SM = 0
Delta_L
Doxorubicin
RF1_378_0 RF1
ZShift = −0.25
lm Score = 7.73
NG = 0
DB = 1
SM = 0
Delta_K
Doxorubicin
RF1_378_0 RF1
ZShift = 0.5
lm Score = 0
NG = 0
DB = 1
SM = 0
Delta_r
Doxorubicin
RF1_378_0 RF1
ZShift = 0.61
lm Score = 141.86
NG = 0
DB = 1
SM = 0
Delta_AUC
Doxorubicin

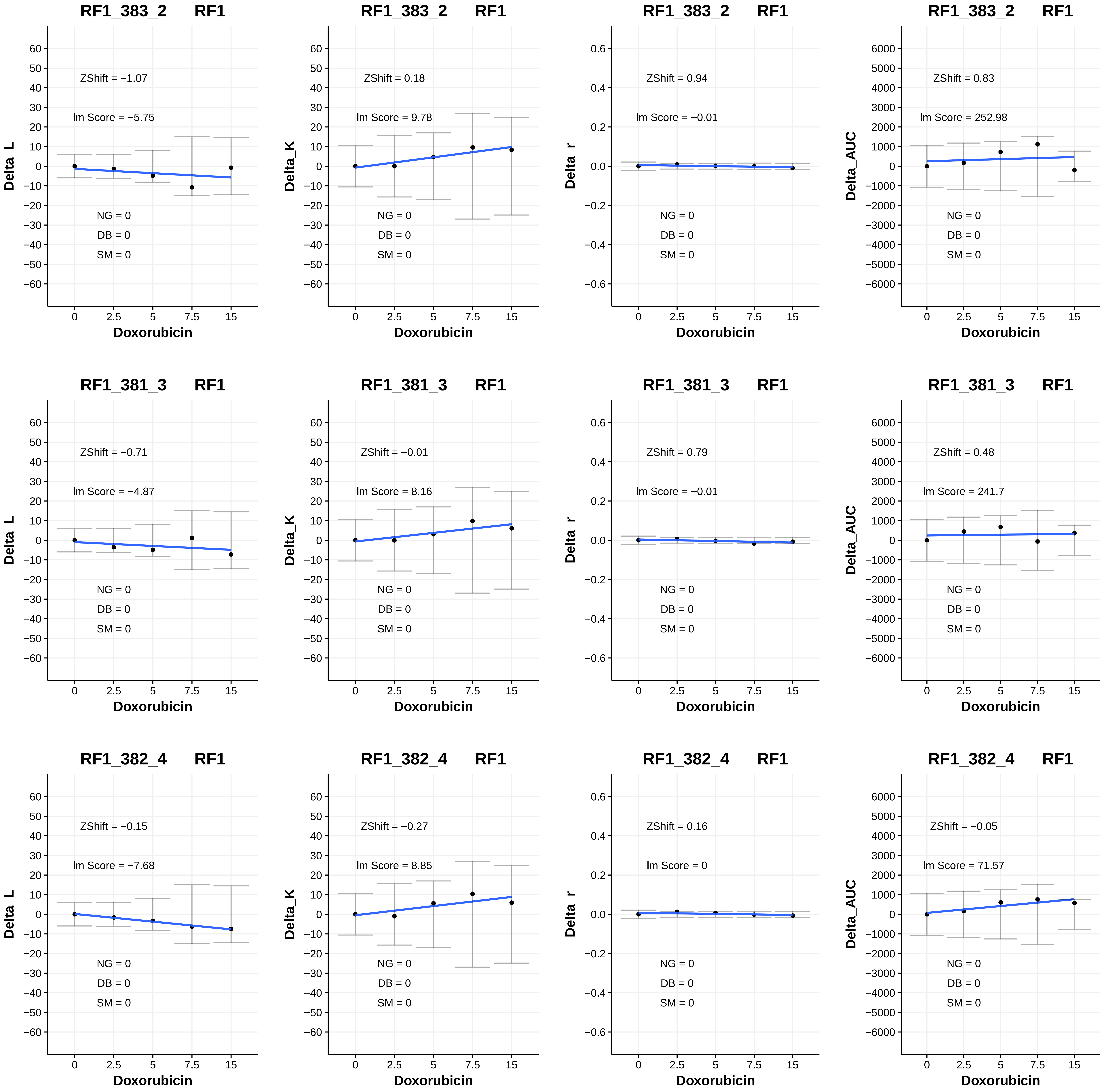
RF1_383_2 RF1
ZShift = −1.07
lm Score = −5.75
NG = 0
DB = 0
SM = 0
Delta_L
Doxorubicin
RF1_383_2 RF1
ZShift = 0.18
lm Score = 9.78
NG = 0
DB = 0
SM = 0
Delta_K
Doxorubicin
RF1_383_2 RF1
ZShift = 0.94
lm Score = −0.01
NG = 0
DB = 0
SM = 0
Delta_r
Doxorubicin
RF1_383_2 RF1
ZShift = 0.83
lm Score = 252.98
NG = 0
DB = 0
SM = 0
Delta_AUC
Doxorubicin
RF1_381_3 RF1
ZShift = −0.71
lm Score = −4.87
NG = 0
DB = 0
SM = 0
Delta_L
Doxorubicin
RF1_381_3 RF1
ZShift = −0.01
lm Score = 8.16
NG = 0
DB = 0
SM = 0
Delta_K
Doxorubicin
RF1_381_3 RF1
ZShift = 0.79
lm Score = −0.01
NG = 0
DB = 0
SM = 0
Delta_r
Doxorubicin
RF1_381_3 RF1
ZShift = 0.48
lm Score = 241.7
NG = 0
DB = 0
SM = 0
Delta_AUC
Doxorubicin
RF1_382_4 RF1
ZShift = −0.15
lm Score = −7.68
NG = 0
DB = 0
SM = 0
Delta_L
Doxorubicin
RF1_382_4 RF1
ZShift = −0.27
lm Score = 8.85
NG = 0
DB = 0
SM = 0
Delta_K
Doxorubicin
RF1_382_4 RF1
ZShift = 0.16
lm Score = 0
NG = 0
DB = 0
SM = 0
Delta_r
Doxorubicin
RF1_382_4 RF1
ZShift = −0.05
lm Score = 71.57
NG = 0
DB = 0
SM = 0
Delta_AUC
Doxorubicin

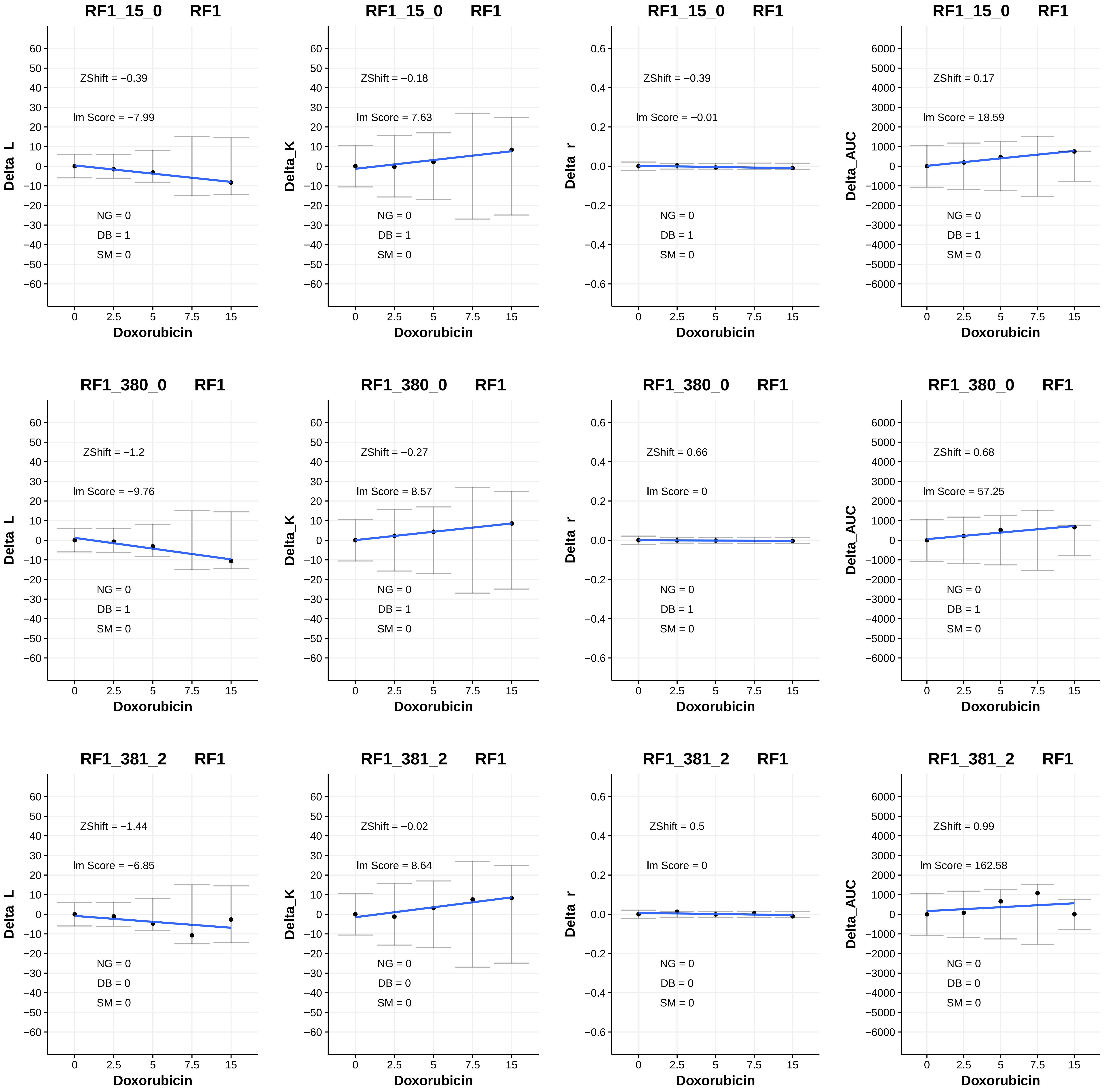
RF1_15_0 RF1
ZShift = −0.39
lm Score = −7.99
NG = 0
DB = 1
SM = 0
Delta_L
Doxorubicin
RF1_15_0 RF1
ZShift = −0.18
lm Score = 7.63
NG = 0
DB = 1
SM = 0
Delta_K
Doxorubicin
RF1_15_0 RF1
ZShift = −0.39
lm Score = −0.01
NG = 0
DB = 1
SM = 0
Delta_r
Doxorubicin
RF1_15_0 RF1
ZShift = 0.17
lm Score = 18.59
NG = 0
DB = 1
SM = 0
Delta_AUC
Doxorubicin
RF1_380_0 RF1
ZShift = −1.2
lm Score = −9.76
NG = 0
DB = 1
SM = 0
Delta_L
Doxorubicin
RF1_380_0 RF1
ZShift = −0.27
lm Score = 8.57
NG = 0
DB = 1
SM = 0
Delta_K
Doxorubicin
RF1_380_0 RF1
ZShift = 0.66
lm Score = 0
NG = 0
DB = 1
SM = 0
Delta_r
Doxorubicin
RF1_380_0 RF1
ZShift = 0.68
lm Score = 57.25
NG = 0
DB = 1
SM = 0
Delta_AUC
Doxorubicin
RF1_381_2 RF1
ZShift = −1.44
lm Score = −6.85
NG = 0
DB = 0
SM = 0
Delta_L
Doxorubicin
RF1_381_2 RF1
ZShift = −0.02
lm Score = 8.64
NG = 0
DB = 0
SM = 0
Delta_K
Doxorubicin
RF1_381_2 RF1
ZShift = 0.5
lm Score = 0
NG = 0
DB = 0
SM = 0
Delta_r
Doxorubicin
RF1_381_2 RF1
ZShift = 0.99
lm Score = 162.58
NG = 0
DB = 0
SM = 0
Delta_AUC
Doxorubicin

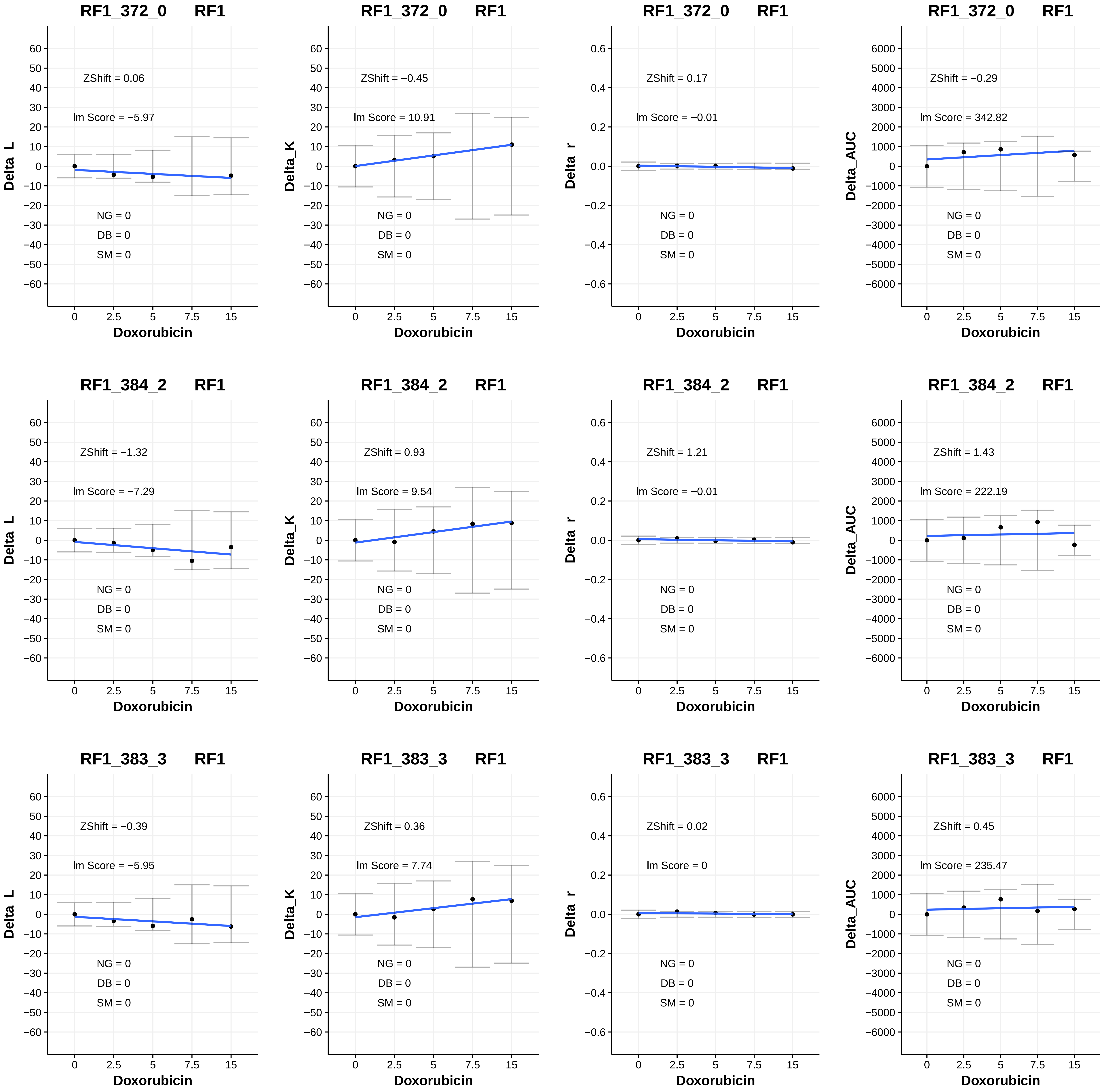
RF1_372_0 RF1
ZShift = 0.06
lm Score = −5.97
NG = 0
DB = 0
SM = 0
Delta_L
Doxorubicin
RF1_372_0 RF1
ZShift = −0.45
lm Score = 10.91
NG = 0
DB = 0
SM = 0
Delta_K
Doxorubicin
RF1_372_0 RF1
ZShift = 0.17
lm Score = −0.01
NG = 0
DB = 0
SM = 0
Delta_r
Doxorubicin
RF1_372_0 RF1
ZShift = −0.29
lm Score = 342.82
NG = 0
DB = 0
SM = 0
Delta_AUC
Doxorubicin
RF1_384_2 RF1
ZShift = −1.32
lm Score = −7.29
NG = 0
DB = 0
SM = 0
Delta_L
Doxorubicin
RF1_384_2 RF1
ZShift = 0.93
lm Score = 9.54
NG = 0
DB = 0
SM = 0
Delta_K
Doxorubicin
RF1_384_2 RF1
ZShift = 1.21
lm Score = −0.01
NG = 0
DB = 0
SM = 0
Delta_r
Doxorubicin
RF1_384_2 RF1
ZShift = 1.43
lm Score = 222.19
NG = 0
DB = 0
SM = 0
Delta_AUC
Doxorubicin
RF1_383_3 RF1
ZShift = −0.39
lm Score = −5.95
NG = 0
DB = 0
SM = 0
Delta_L
Doxorubicin
RF1_383_3 RF1
ZShift = 0.36
lm Score = 7.74
NG = 0
DB = 0
SM = 0
Delta_K
Doxorubicin
RF1_383_3 RF1
ZShift = 0.02
lm Score = 0
NG = 0
DB = 0
SM = 0
Delta_r
Doxorubicin
RF1_383_3 RF1
ZShift = 0.45
lm Score = 235.47
NG = 0
DB = 0
SM = 0
Delta_AUC
Doxorubicin

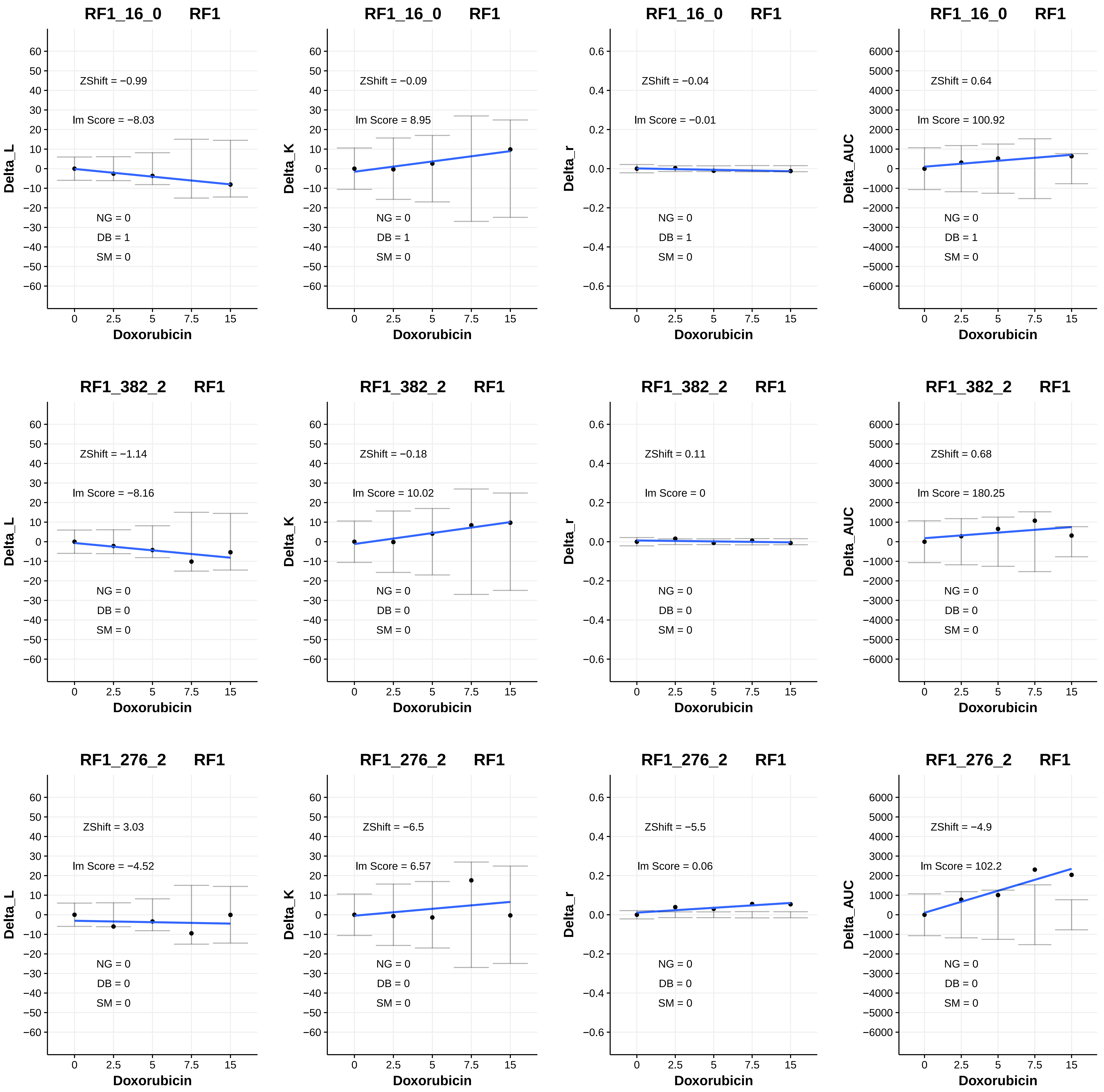

RF1_16_0 RF1
ZShift = −0.99
Im Score = −8.03
NG = 0
DB = 1
SM = 0
Delta_L
Doxorubicin
RF1_16_0 RF1
ZShift = −0.09
Im Score = 8.95
NG = 0
DB = 1
SM = 0
Delta_K
Doxorubicin
RF1_16_0 RF1
ZShift = −0.04
Im Score = −0.01
NG = 0
DB = 1
SM = 0
Delta_r
Doxorubicin
RF1_16_0 RF1
ZShift = 0.64
Im Score = 100.92
NG = 0
DB = 1
SM = 0
Delta_AUC
Doxorubicin
RF1_382_2 RF1
ZShift = −1.14
Im Score = −8.16
NG = 0
DB = 0
SM = 0
Delta_L
Doxorubicin
RF1_382_2 RF1
ZShift = −0.18
Im Score = 10.02
NG = 0
DB = 0
SM = 0
Delta_K
Doxorubicin
RF1_382_2 RF1
ZShift = 0.11
Im Score = 0
NG = 0
DB = 0
SM = 0
Delta_r
Doxorubicin
RF1_382_2 RF1
ZShift = 0.68
Im Score = 180.25
NG = 0
DB = 0
SM = 0
Delta_AUC
Doxorubicin
RF1_276_2 RF1
ZShift = 3.03
Im Score = −4.52
NG = 0
DB = 0
SM = 0
Delta_L
Doxorubicin
RF1_276_2 RF1
ZShift = −6.5
Im Score = 6.57
NG = 0
DB = 0
SM = 0
Delta_K
Doxorubicin
RF1_276_2 RF1
ZShift = −5.5
Im Score = 0.06
NG = 0
DB = 0
SM = 0
Delta_r
Doxorubicin
RF1_276_2 RF1
ZShift = −4.9
Im Score = 102.2
NG = 0
DB = 0
SM = 0
Delta_AUC
Doxorubicin

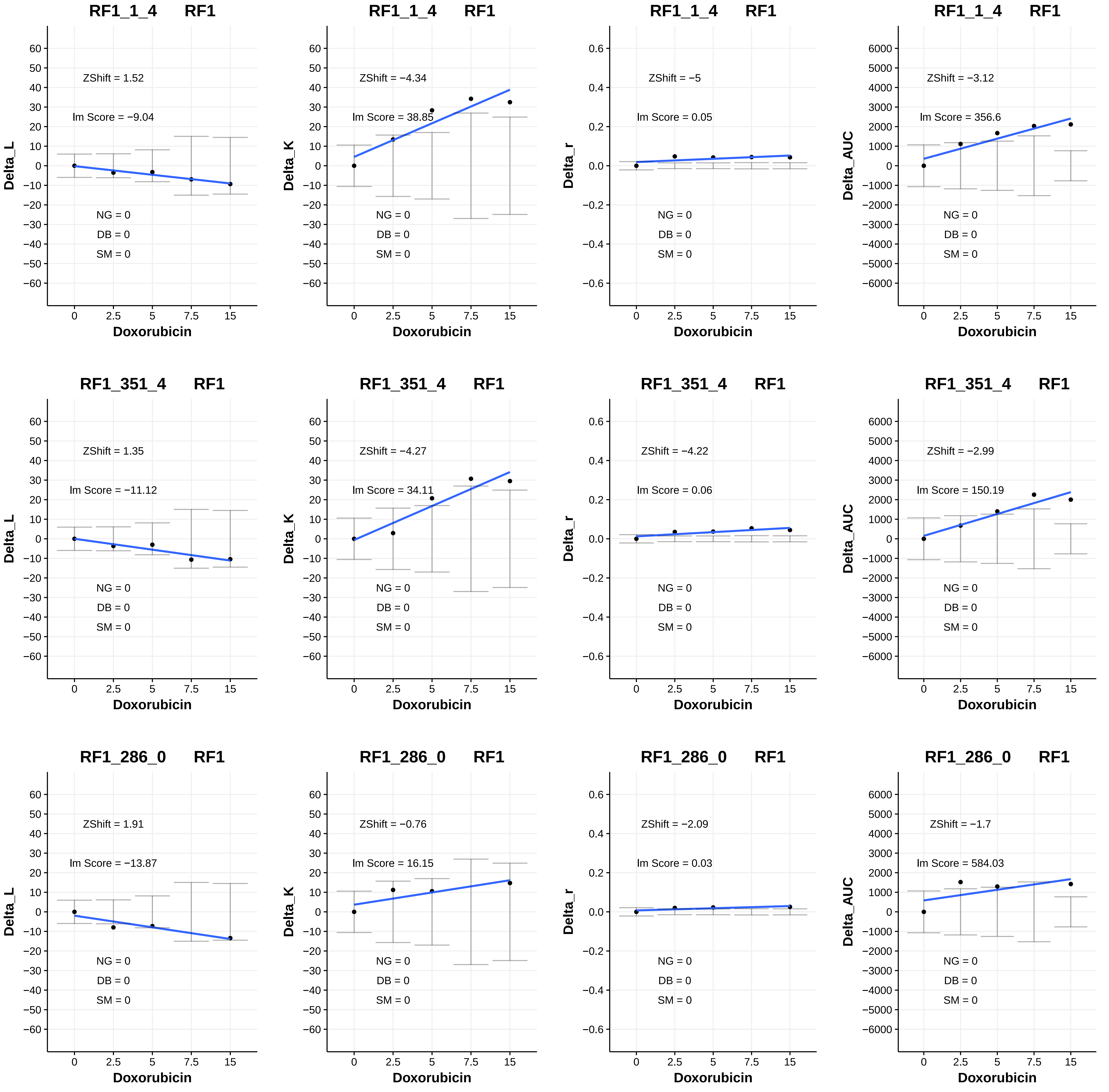

RF1_1_4 RF1
ZShift = 1.52
lm Score = −9.04
NG = 0
DB = 0
SM = 0
Delta_L
Doxorubicin
RF1_1_4 RF1
ZShift = −4.34
lm Score = 38.85
NG = 0
DB = 0
SM = 0
Delta_K
Doxorubicin
RF1_1_4 RF1
ZShift = −5
lm Score = 0.05
NG = 0
DB = 0
SM = 0
Delta_r
Doxorubicin
RF1_1_4 RF1
ZShift = −3.12
lm Score = 356.6
NG = 0
DB = 0
SM = 0
Delta_AUC
Doxorubicin
RF1_351_4 RF1
ZShift = 1.35
lm Score = −11.12
NG = 0
DB = 0
SM = 0
Delta_L
Doxorubicin
RF1_351_4 RF1
ZShift = −4.27
lm Score = 34.11
NG = 0
DB = 0
SM = 0
Delta_K
Doxorubicin
RF1_351_4 RF1
ZShift = −4.22
lm Score = 0.06
NG = 0
DB = 0
SM = 0
Delta_r
Doxorubicin
RF1_351_4 RF1
ZShift = −2.99
lm Score = 150.19
NG = 0
DB = 0
SM = 0
Delta_AUC
Doxorubicin
RF1_286_0 RF1
ZShift = 1.91
lm Score = −13.87
NG = 0
DB = 0
SM = 0
Delta_L
Doxorubicin
RF1_286_0 RF1
ZShift = −0.76
lm Score = 16.15
NG = 0
DB = 0
SM = 0
Delta_K
Doxorubicin
RF1_286_0 RF1
ZShift = −2.09
lm Score = 0.03
NG = 0
DB = 0
SM = 0
Delta_r
Doxorubicin
RF1_286_0 RF1
ZShift = −1.7
lm Score = 584.03
NG = 0
DB = 0
SM = 0
Delta_AUC
Doxorubicin

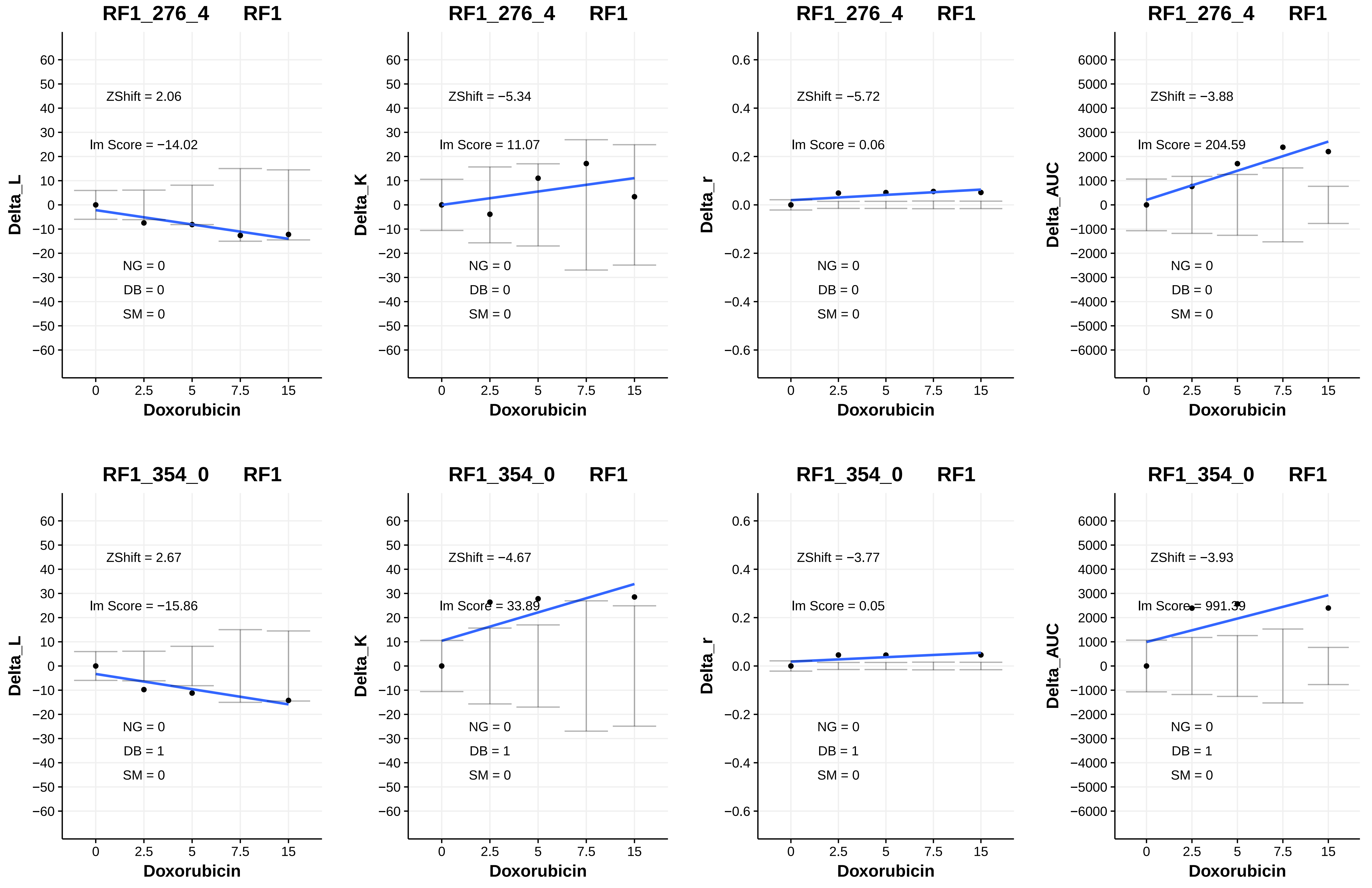
RF1_276_4 RF1
ZShift = 2.06
lm Score = −14.02
NG = 0
DB = 0
SM = 0
Delta_L
Doxorubicin
RF1_276_4 RF1
ZShift = −5.34
lm Score = 11.07
NG = 0
DB = 0
SM = 0
Delta_K
Doxorubicin
RF1_276_4 RF1
ZShift = −5.72
lm Score = 0.06
NG = 0
DB = 0
SM = 0
Delta_r
Doxorubicin
RF1_276_4 RF1
ZShift = −3.88
lm Score = 204.59
NG = 0
DB = 0
SM = 0
Delta_AUC
Doxorubicin
RF1_354_0 RF1
ZShift = 2.67
lm Score = −15.86
NG = 0
DB = 1
SM = 0
Delta_L
Doxorubicin
RF1_354_0 RF1
ZShift = −4.67
lm Score = 33.89
NG = 0
DB = 1
SM = 0
Delta_K
Doxorubicin
RF1_354_0 RF1
ZShift = −3.77
lm Score = 0.05
NG = 0
DB = 1
SM = 0
Delta_r
Doxorubicin
RF1_354_0 RF1
ZShift = −3.93
lm Score = 991.39
NG = 0
DB = 1
SM = 0
Delta_AUC
Doxorubicin
