## Additional File 4 for "A yeast phenomic model for the influence of Warburg metabolism on genetic buffering of doxorubicin": A - InteractionPlots_Doxo_HLD.compressed.pdf

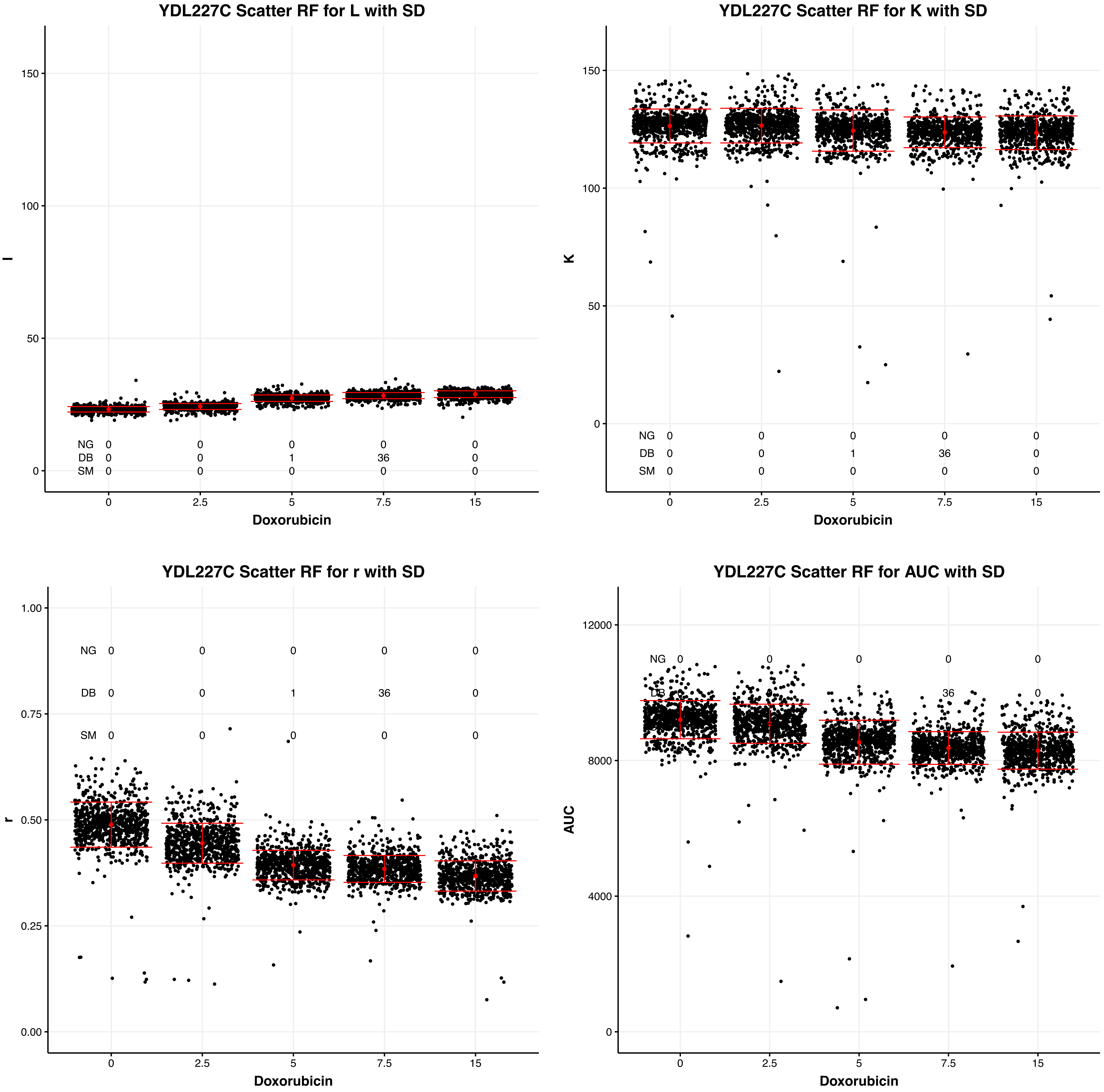

YDL227C Scatter RF for L with SD
l
Doxorubicin
0
2.5
5
7.5
15
NG 0 0 0 0 0
DB 0 0 1 36 0
SM 0 0 0 0 0
YDL227C Scatter RF for K with SD
K
Doxorubicin
NG 0 0 0 0 0
DB 0 0 1 36 0
SM 0 0 0 0 0
YDL227C Scatter RF for r with SD
r
Doxorubicin
NG 0 0 0 0 0
DB 0 0 1 36 0
SM 0 0 0 0 0
YDL227C Scatter RF for AUC with SD
AUC
Doxorubicin
NG 0 0 0 0 0
DB 0 0 1 36 0

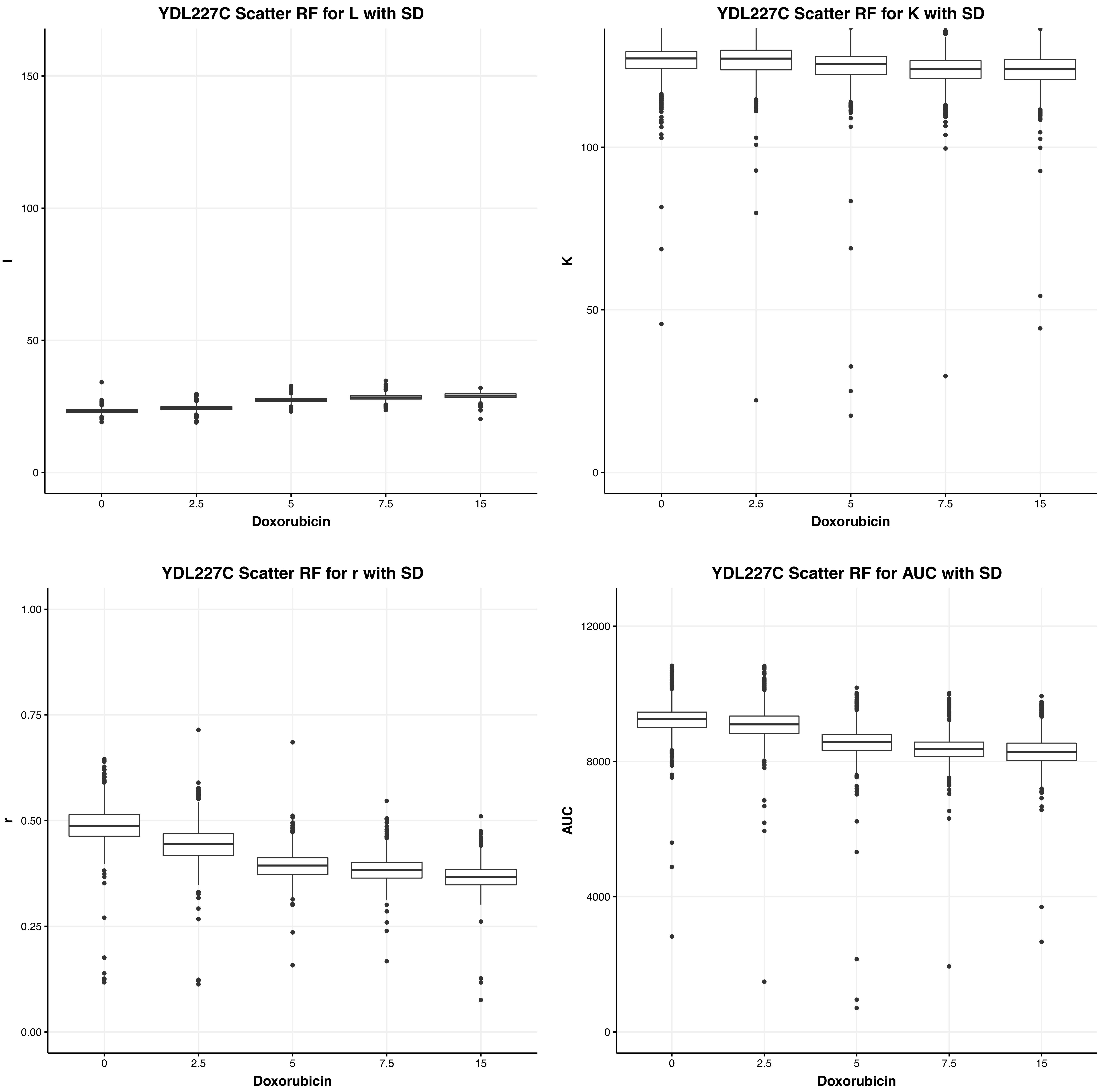

YDL227C Scatter RF for L with SD
l
Doxorubicin
YDL227C Scatter RF for K with SD
K
Doxorubicin
YDL227C Scatter RF for r with SD
r
Doxorubicin
YDL227C Scatter RF for AUC with SD
AUC
Doxorubicin

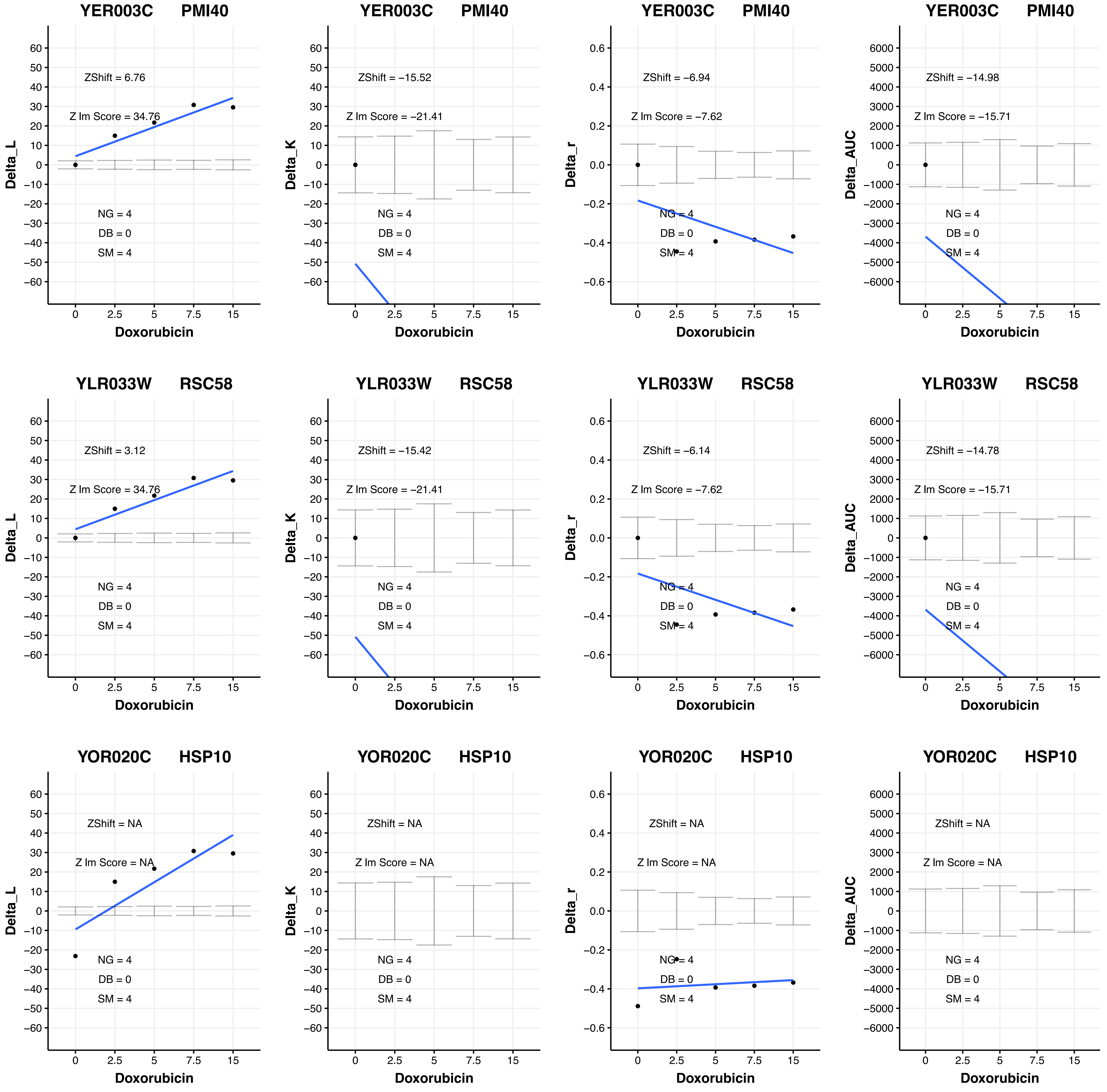
YER003C PMI40
ZShift = 6.76
Z lm Score = 34.76
NG = 4
DB = 0
SM = 4
Delta_L
Doxorubicin
YER003C PMI40
ZShift = −15.52
Z lm Score = −21.41
NG = 4
DB = 0
SM = 4
Delta_K
Doxorubicin
YER003C PMI40
ZShift = −6.94
Z lm Score = −7.62
NG = 4
DB = 0
SM = 4
Delta_r
Doxorubicin
YER003C PMI40
ZShift = −14.98
Z lm Score = −15.71
NG = 4
DB = 0
SM = 4
Delta_AUC
Doxorubicin
YLR033W RSC58
ZShift = 3.12
Z lm Score = 34.76
NG = 4
DB = 0
SM = 4
Delta_L
Doxorubicin
YLR033W RSC58
ZShift = −15.42
Z lm Score = −21.41
NG = 4
DB = 0
SM = 4
Delta_K
Doxorubicin
YLR033W RSC58
ZShift = −6.14
Z lm Score = −7.62
NG = 4
DB = 0
SM = 4
Delta_r
Doxorubicin
YLR033W RSC58
ZShift = −14.78
Z lm Score = −15.71
NG = 4
DB = 0
SM = 4
Delta_AUC
Doxorubicin
YOR020C HSP10
ZShift = NA
Z lm Score = NA
NG = 4
DB = 0
SM = 4
Delta_L
Doxorubicin
YOR020C HSP10
ZShift = NA
Z lm Score = NA
NG = 4
DB = 0
SM = 4
Delta_K
Doxorubicin
YOR020C HSP10
ZShift = NA
Z lm Score = NA
NG = 4
DB = 0
SM = 4
Delta_r
Doxorubicin
YOR020C HSP10
ZShift = NA
Z lm Score = NA
NG = 4
DB = 0
SM = 4
Delta_AUC
Doxorubicin

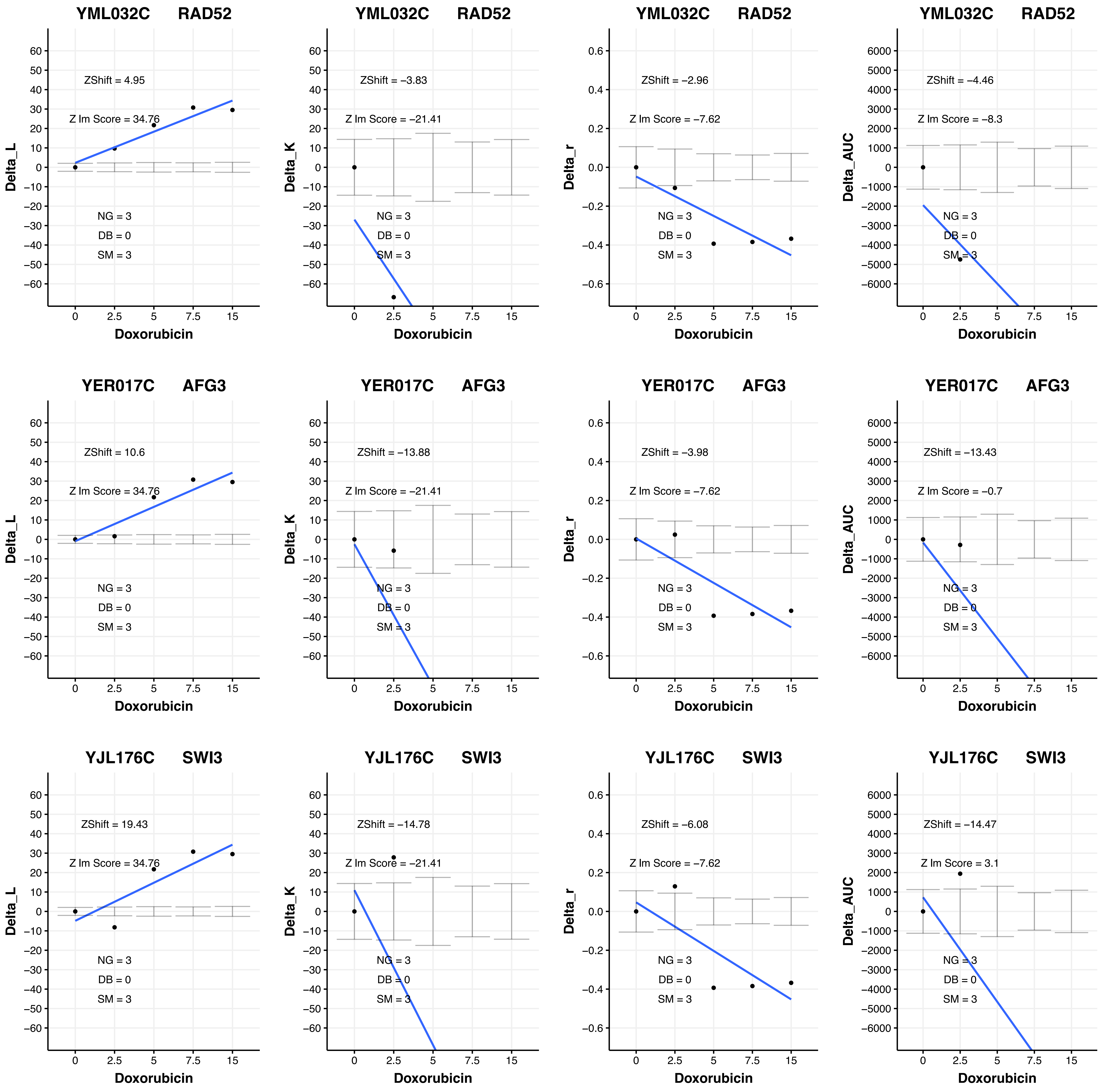
YML032C RAD52
ZShift = 4.95
Z lm Score = 34.76
NG = 3
DB = 0
SM = 3
Delta_L
Doxorubicin
YML032C RAD52
ZShift = −3.83
Z lm Score = −21.41
Delta_K
YML032C RAD52
ZShift = −2.96
Z lm Score = −7.62
Delta_r
YML032C RAD52
ZShift = −4.46
Z lm Score = −8.3
Delta_AUC
YER017C AFG3
ZShift = 10.6
Z lm Score = 34.76
YER017C AFG3
ZShift = −13.88
Z lm Score = −21.41
YER017C AFG3
ZShift = −3.98
Z lm Score = −7.62
YER017C AFG3
ZShift = −13.43
Z lm Score = −0.7
YJL176C SWI3
ZShift = 19.43
Z lm Score = 34.76
YJL176C SWI3
ZShift = −14.78
Z lm Score = −21.41
YJL176C SWI3
ZShift = −6.08
Z lm Score = −7.62
YJL176C SWI3
ZShift = −14.47
Z lm Score = 3.1

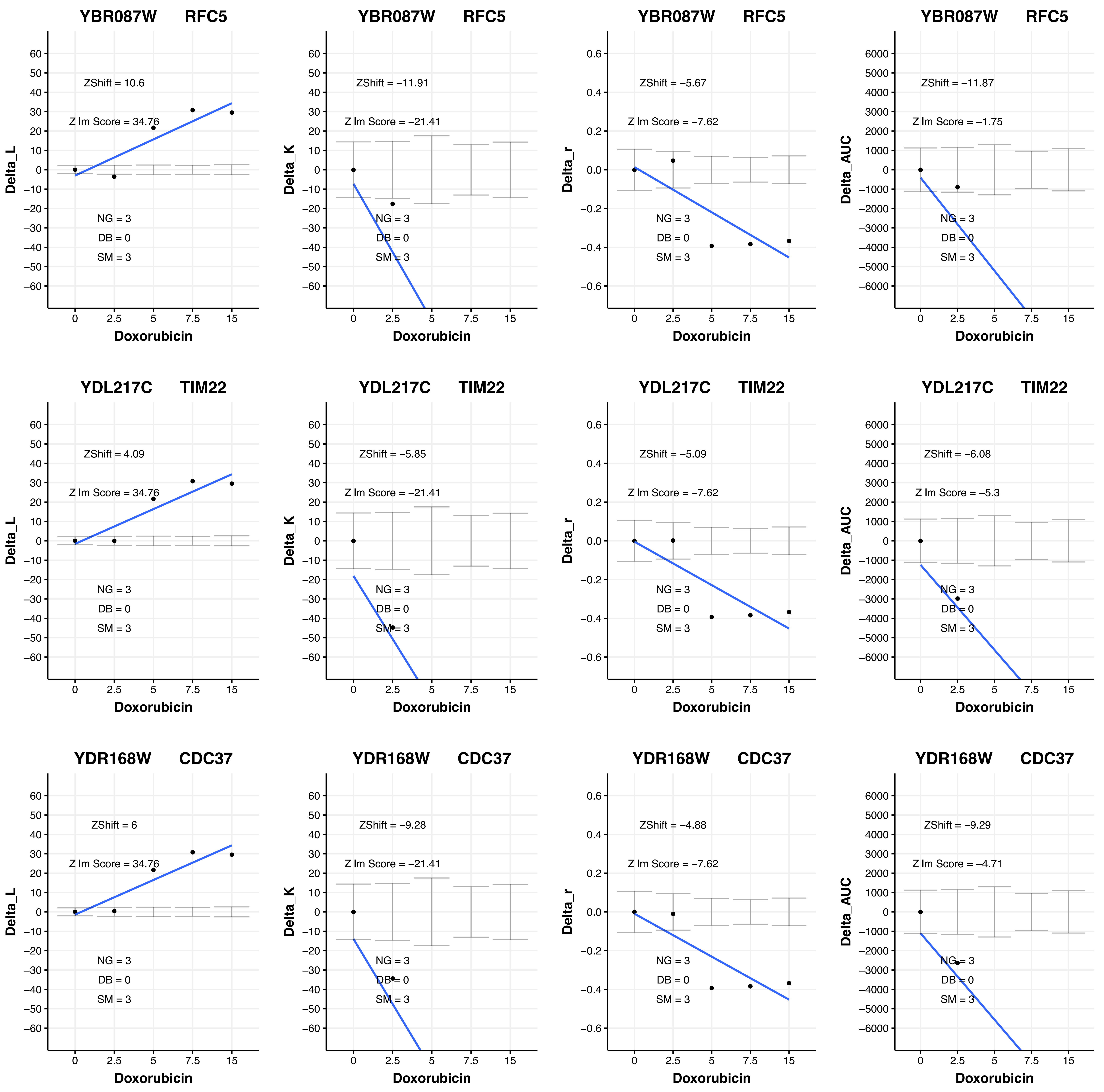

YBR087W RFC5
ZShift = 10.6
Z lm Score = 34.76
NG = 3
DB = 0
SM = 3
Delta_L
Doxorubicin
YBR087W RFC5
ZShift = −11.91
Z lm Score = −21.41
NG = 3
DB = 0
SM = 3
Delta_K
Doxorubicin
YBR087W RFC5
ZShift = −5.67
Z lm Score = −7.62
NG = 3
DB = 0
SM = 3
Delta_r
Doxorubicin
YBR087W RFC5
ZShift = −11.87
Z lm Score = −1.75
NG = 3
DB = 0
SM = 3
Delta_AUC
Doxorubicin
YDL217C TIM22
ZShift = 4.09
Z lm Score = 34.76
NG = 3
DB = 0
SM = 3
Delta_L
Doxorubicin
YDL217C TIM22
ZShift = −5.85
Z lm Score = −21.41
NG = 3
DB = 0
SM = 3
Delta_K
Doxorubicin
YDL217C TIM22
ZShift = −5.09
Z lm Score = −7.62
NG = 3
DB = 0
SM = 3
Delta_r
Doxorubicin
YDL217C TIM22
ZShift = −6.08
Z lm Score = −5.3
NG = 3
DB = 0
SM = 3
Delta_AUC
Doxorubicin
YDR168W CDC37
ZShift = 6
Z lm Score = 34.76
NG = 3
DB = 0
SM = 3
Delta_L
Doxorubicin
YDR168W CDC37
ZShift = −9.28
Z lm Score = −21.41
NG = 3
DB = 0
SM = 3
Delta_K
Doxorubicin
YDR168W CDC37
ZShift = −4.88
Z lm Score = −7.62
NG = 3
DB = 0
SM = 3
Delta_r
Doxorubicin
YDR168W CDC37
ZShift = −9.29
Z lm Score = −4.71
NG = 3
DB = 0
SM = 3
Delta_AUC
Doxorubicin

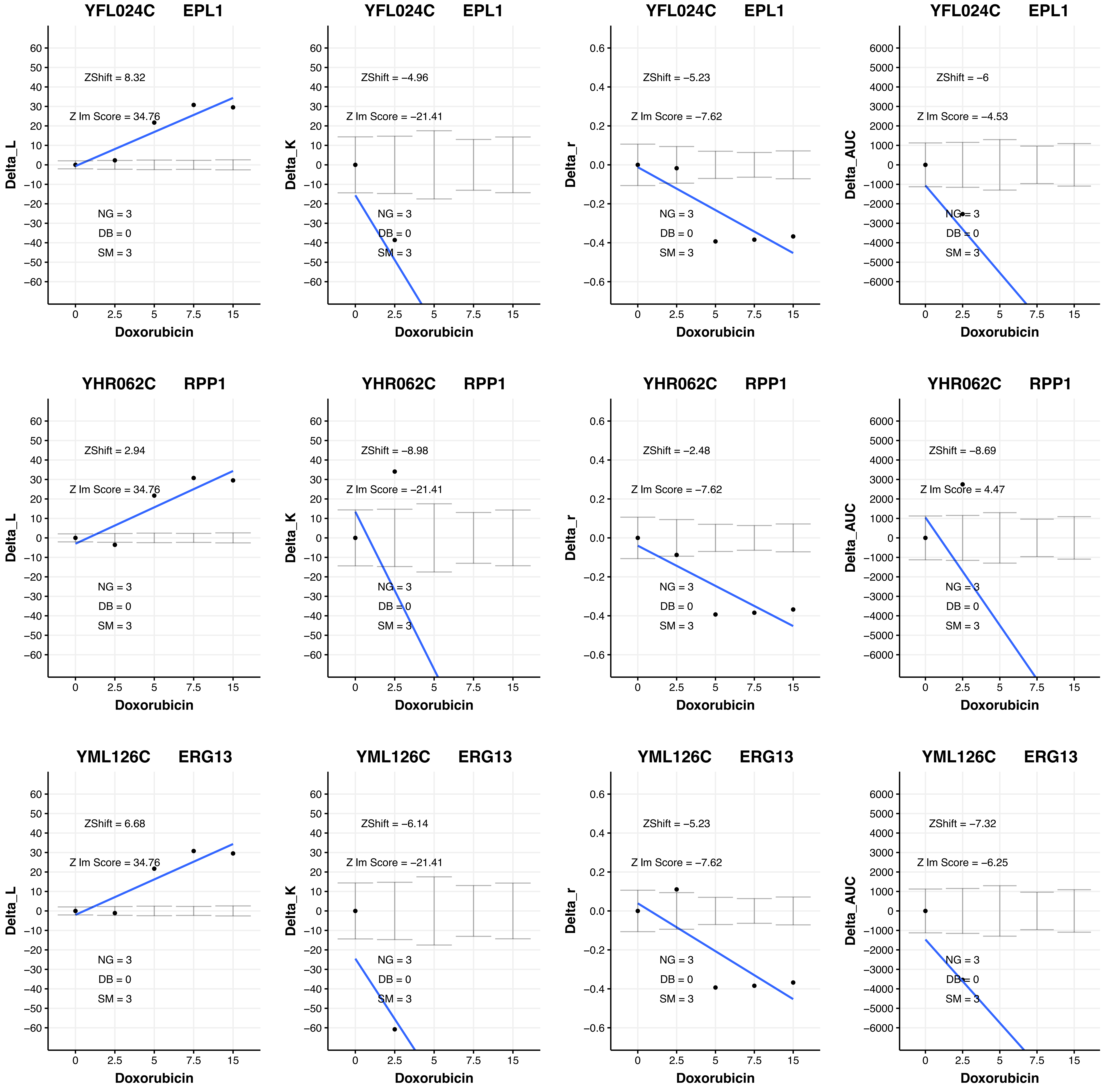
YFL024C EPL1
ZShift = 8.32
Z lm Score = 34.76
NG = 3
DB = 0
SM = 3
Delta_L
Doxorubicin
YFL024C EPL1
ZShift = −4.96
Z lm Score = −21.41
NG = 3
DB = 0
SM = 3
Delta_K
Doxorubicin
YFL024C EPL1
ZShift = −5.23
Z lm Score = −7.62
NG = 3
DB = 0
SM = 3
Delta_r
Doxorubicin
YFL024C EPL1
ZShift = −6
Z lm Score = −4.53
NG = 3
DB = 0
SM = 3
Delta_AUC
Doxorubicin
YHR062C RPP1
ZShift = 2.94
Z lm Score = 34.76
NG = 3
DB = 0
SM = 3
Delta_L
Doxorubicin
YHR062C RPP1
ZShift = −8.98
Z lm Score = −21.41
NG = 3
DB = 0
SM = 3
Delta_K
Doxorubicin
YHR062C RPP1
ZShift = −2.48
Z lm Score = −7.62
NG = 3
DB = 0
SM = 3
Delta_r
Doxorubicin
YHR062C RPP1
ZShift = −8.69
Z lm Score = 4.47
NG = 3
DB = 0
SM = 3
Delta_AUC
Doxorubicin
YML126C ERG13
ZShift = 6.68
Z lm Score = 34.76
NG = 3
DB = 0
SM = 3
Delta_L
Doxorubicin
YML126C ERG13
ZShift = −6.14
Z lm Score = −21.41
NG = 3
DB = 0
SM = 3
Delta_K
Doxorubicin
YML126C ERG13
ZShift = −5.23
Z lm Score = −7.62
NG = 3
DB = 0
SM = 3
Delta_r
Doxorubicin
YML126C ERG13
ZShift = −7.32
Z lm Score = −6.25
NG = 3
DB = 0
SM = 3
Delta_AUC
Doxorubicin

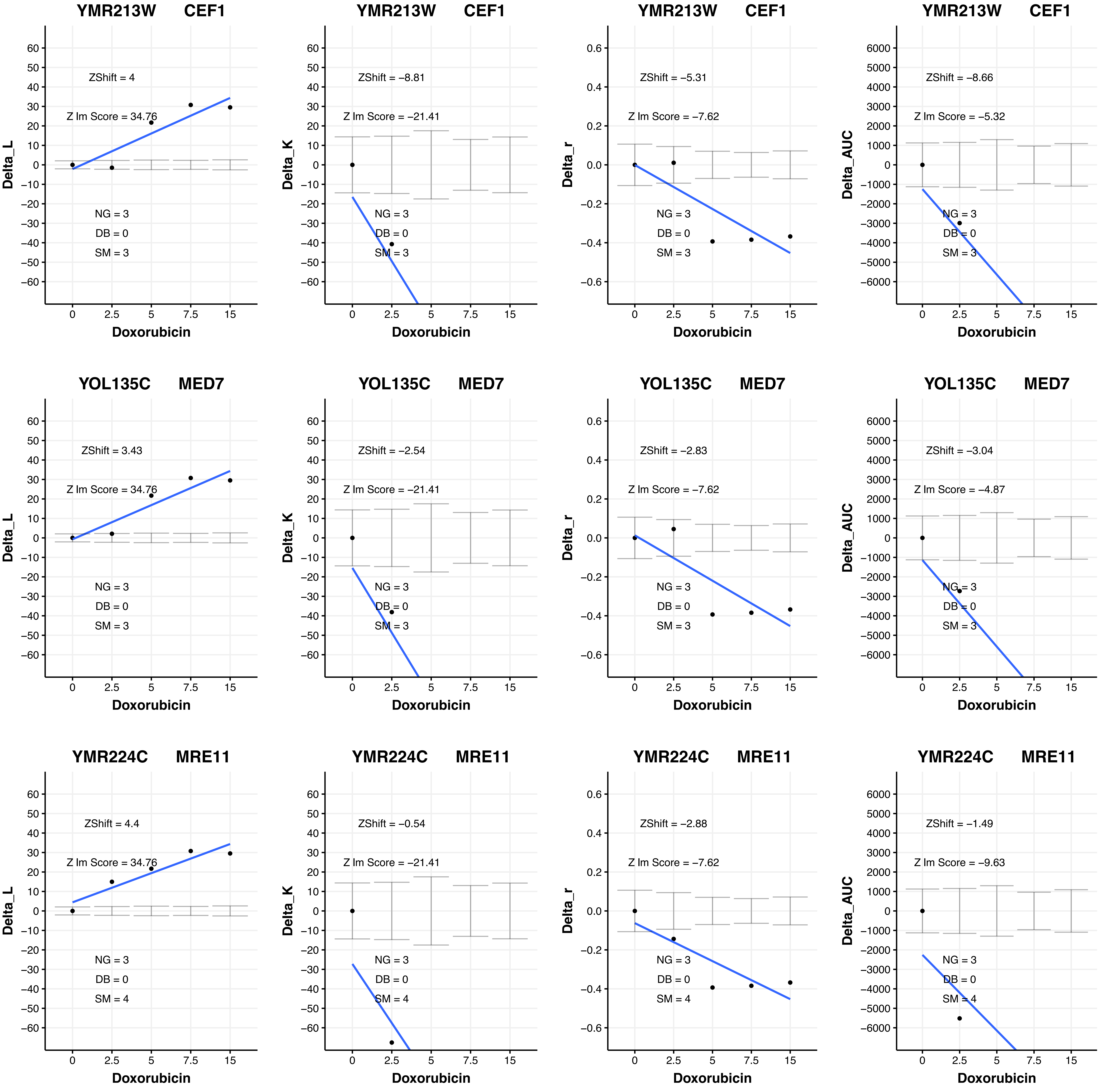
YMR213W CEF1
Delta_L
ZShift = 4
Z lm Score = 34.76
NG = 3
DB = 0
SM = 3
Doxorubicin
YMR213W CEF1
Delta_K
ZShift = −8.81
Z lm Score = −21.41
NG = 3
DB = 0
SM = 3
Doxorubicin
YMR213W CEF1
Delta_r
ZShift = −5.31
Z lm Score = −7.62
NG = 3
DB = 0
SM = 3
Doxorubicin
YMR213W CEF1
Delta_AUC
ZShift = −8.66
Z lm Score = −5.32
NG = 3
DB = 0
SM = 3
Doxorubicin
YOL135C MED7
Delta_L
ZShift = 3.43
Z lm Score = 34.76
NG = 3
DB = 0
SM = 3
Doxorubicin
YOL135C MED7
Delta_K
ZShift = −2.54
Z lm Score = −21.41
NG = 3
DB = 0
SM = 3
Doxorubicin
YOL135C MED7
Delta_r
ZShift = −2.83
Z lm Score = −7.62
NG = 3
DB = 0
SM = 3
Doxorubicin
YOL135C MED7
Delta_AUC
ZShift = −3.04
Z lm Score = −4.87
NG = 3
DB = 0
SM = 3
Doxorubicin
YMR224C MRE11
Delta_L
ZShift = 4.4
Z lm Score = 34.76
NG = 3
DB = 0
SM = 4
Doxorubicin
YMR224C MRE11
Delta_K
ZShift = −0.54
Z lm Score = −21.41
NG = 3
DB = 0
SM = 4
Doxorubicin
YMR224C MRE11
Delta_r
ZShift = −2.88
Z lm Score = −7.62
NG = 3
DB = 0
SM = 4
Doxorubicin
YMR224C MRE11
Delta_AUC
ZShift = −1.49
Z lm Score = −9.63
NG = 3
DB = 0
SM = 4
Doxorubicin

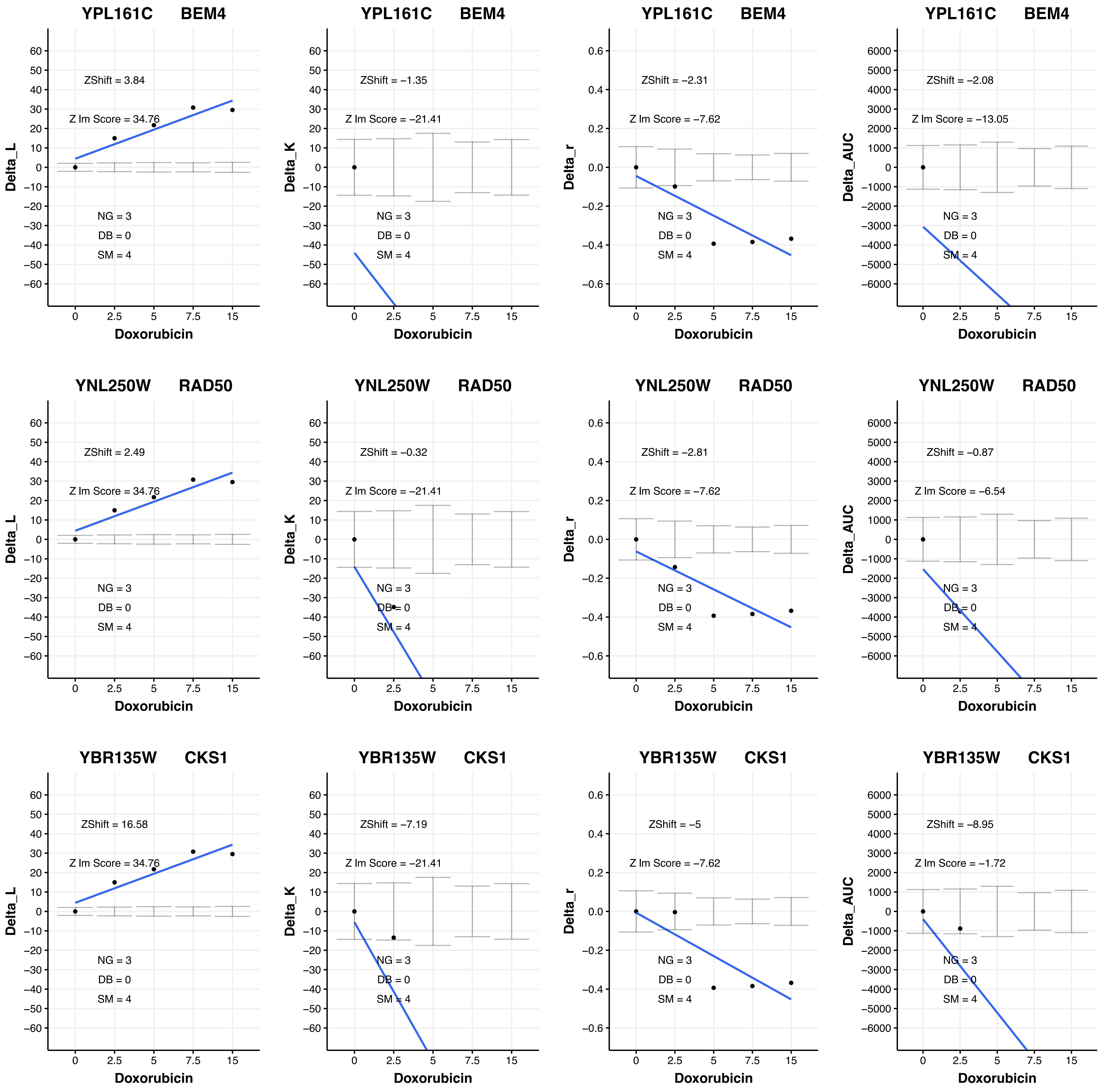
YPL161C BEM4
ZShift = 3.84
Z lm Score = 34.76
NG = 3
DB = 0
SM = 4
Delta_L
Doxorubicin
YPL161C BEM4
ZShift = −1.35
Z lm Score = −21.41
NG = 3
DB = 0
SM = 4
Delta_K
Doxorubicin
YPL161C BEM4
ZShift = −2.31
Z lm Score = −7.62
NG = 3
DB = 0
SM = 4
Delta_r
Doxorubicin
YPL161C BEM4
ZShift = −2.08
Z lm Score = −13.05
NG = 3
DB = 0
SM = 4
Delta_AUC
Doxorubicin
YNL250W RAD50
ZShift = 2.49
Z lm Score = 34.76
NG = 3
DB = 0
SM = 4
Delta_L
Doxorubicin
YNL250W RAD50
ZShift = −0.32
Z lm Score = −21.41
NG = 3
DB = 0
SM = 4
Delta_K
Doxorubicin
YNL250W RAD50
ZShift = −2.81
Z lm Score = −7.62
NG = 3
DB = 0
SM = 4
Delta_r
Doxorubicin
YNL250W RAD50
ZShift = −0.87
Z lm Score = −6.54
NG = 3
DB = 0
SM = 4
Delta_AUC
Doxorubicin
YBR135W CKS1
ZShift = 16.58
Z lm Score = 34.76
NG = 3
DB = 0
SM = 4
Delta_L
Doxorubicin
YBR135W CKS1
ZShift = −7.19
Z lm Score = −21.41
NG = 3
DB = 0
SM = 4
Delta_K
Doxorubicin
YBR135W CKS1
ZShift = −5
Z lm Score = −7.62
NG = 3
DB = 0
SM = 4
Delta_r
Doxorubicin
YBR135W CKS1
ZShift = −8.95
Z lm Score = −1.72
NG = 3
DB = 0
SM = 4
Delta_AUC
Doxorubicin

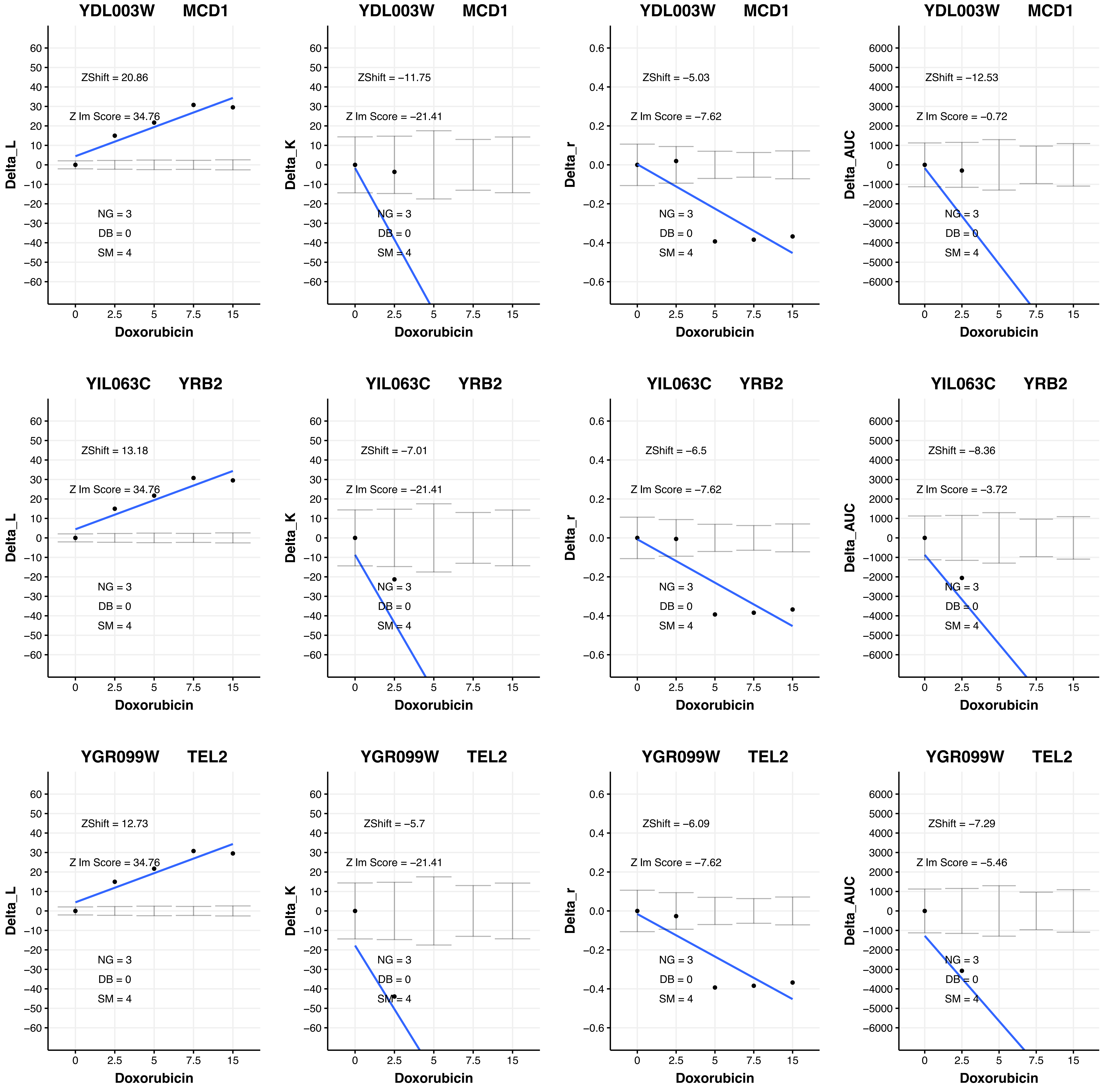
YDL003W MCD1
ZShift = 20.86
Z lm Score = 34.76
NG = 3
DB = 0
SM = 4
Delta_L
Doxorubicin
YDL003W MCD1
ZShift = −11.75
Z lm Score = −21.41
NG = 3
DB = 0
SM = 4
Delta_K
Doxorubicin
YDL003W MCD1
ZShift = −5.03
Z lm Score = −7.62
NG = 3
DB = 0
SM = 4
Delta_r
Doxorubicin
YDL003W MCD1
ZShift = −12.53
Z lm Score = −0.72
NG = 3
DB = 0
SM = 4
Delta_AUC
Doxorubicin
YIL063C YRB2
ZShift = 13.18
Z lm Score = 34.76
NG = 3
DB = 0
SM = 4
Delta_L
Doxorubicin
YIL063C YRB2
ZShift = −7.01
Z lm Score = −21.41
NG = 3
DB = 0
SM = 4
Delta_K
Doxorubicin
YIL063C YRB2
ZShift = −6.5
Z lm Score = −7.62
NG = 3
DB = 0
SM = 4
Delta_r
Doxorubicin
YIL063C YRB2
ZShift = −8.36
Z lm Score = −3.72
NG = 3
DB = 0
SM = 4
Delta_AUC
Doxorubicin
YGR099W TEL2
ZShift = 12.73
Z lm Score = 34.76
NG = 3
DB = 0
SM = 4
Delta_L
Doxorubicin
YGR099W TEL2
ZShift = −5.7
Z lm Score = −21.41
NG = 3
DB = 0
SM = 4
Delta_K
Doxorubicin
YGR099W TEL2
ZShift = −6.09
Z lm Score = −7.62
NG = 3
DB = 0
SM = 4
Delta_r
Doxorubicin
YGR099W TEL2
ZShift = −7.29
Z lm Score = −5.46
NG = 3
DB = 0
SM = 4
Delta_AUC
Doxorubicin

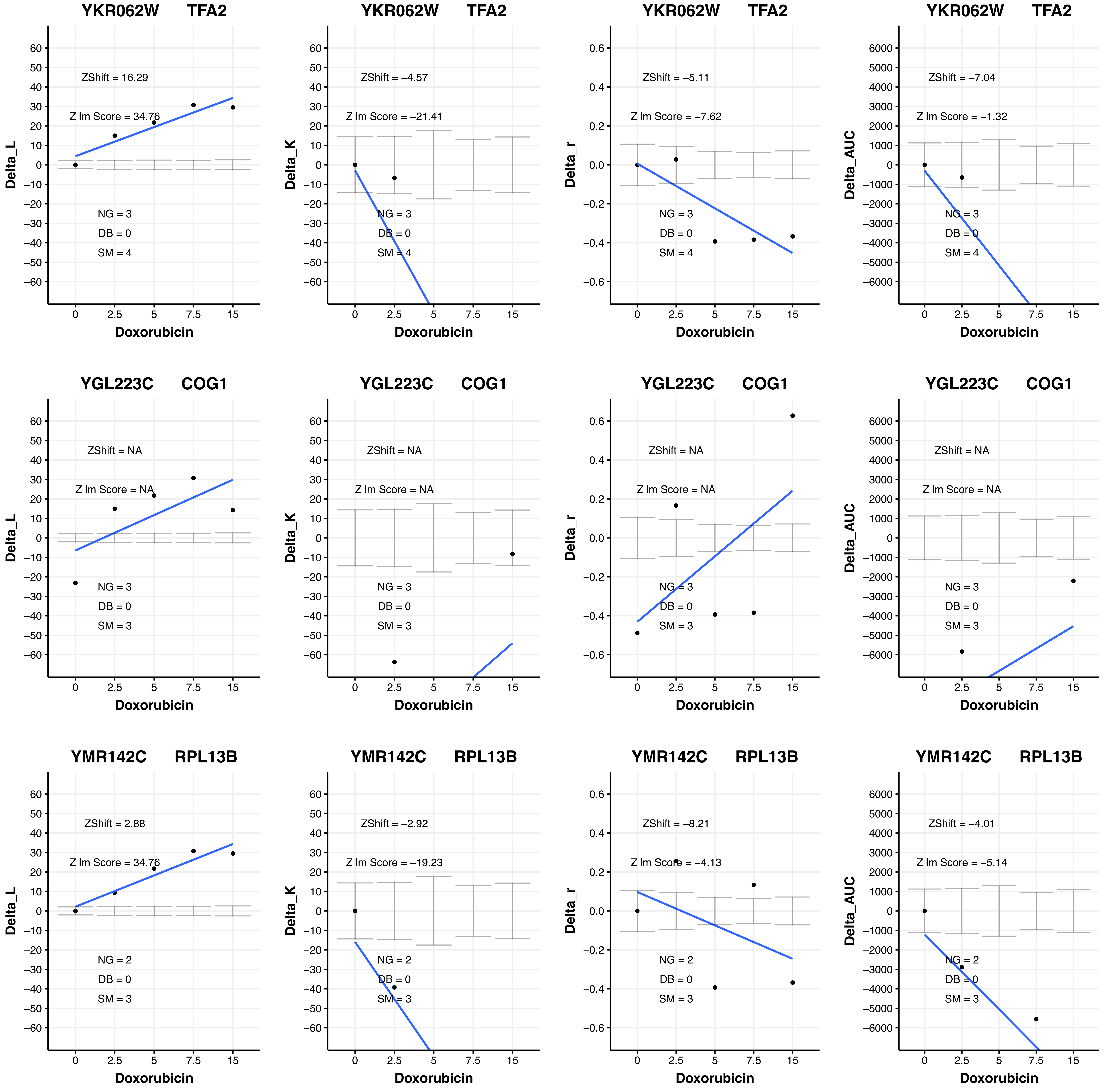
YKR062W TFA2
ZShift = 16.29
Z lm Score = 34.76
NG = 3
DB = 0
SM = 4
Delta_L
Doxorubicin
YKR062W TFA2
ZShift = −4.57
Z lm Score = −21.41
NG = 3
DB = 0
SM = 4
Delta_K
Doxorubicin
YKR062W TFA2
ZShift = −5.11
Z lm Score = −7.62
NG = 3
DB = 0
SM = 4
Delta_r
Doxorubicin
YKR062W TFA2
ZShift = −7.04
Z lm Score = −1.32
NG = 3
DB = 0
SM = 4
Delta_AUC
Doxorubicin
YGL223C COG1
ZShift = NA
Z lm Score = NA
NG = 3
DB = 0
SM = 3
Delta_L
Doxorubicin
YGL223C COG1
ZShift = NA
Z lm Score = NA
NG = 3
DB = 0
SM = 3
Delta_K
Doxorubicin
YGL223C COG1
ZShift = NA
Z lm Score = NA
NG = 3
DB = 0
SM = 3
Delta_r
Doxorubicin
YGL223C COG1
ZShift = NA
Z lm Score = NA
NG = 3
DB = 0
SM = 3
Delta_AUC
Doxorubicin
YMR142C RPL13B
ZShift = 2.88
Z lm Score = 34.76
NG = 2
DB = 0
SM = 3
Delta_L
Doxorubicin
YMR142C RPL13B
ZShift = −2.92
Z lm Score = −19.23
NG = 2
DB = 0
SM = 3
Delta_K
Doxorubicin
YMR142C RPL13B
ZShift = −8.21
Z lm Score = −4.13
NG = 2
DB = 0
SM = 3
Delta_r
Doxorubicin
YMR142C RPL13B
ZShift = −4.01
Z lm Score = −5.14
NG = 2
DB = 0
SM = 3
Delta_AUC
Doxorubicin

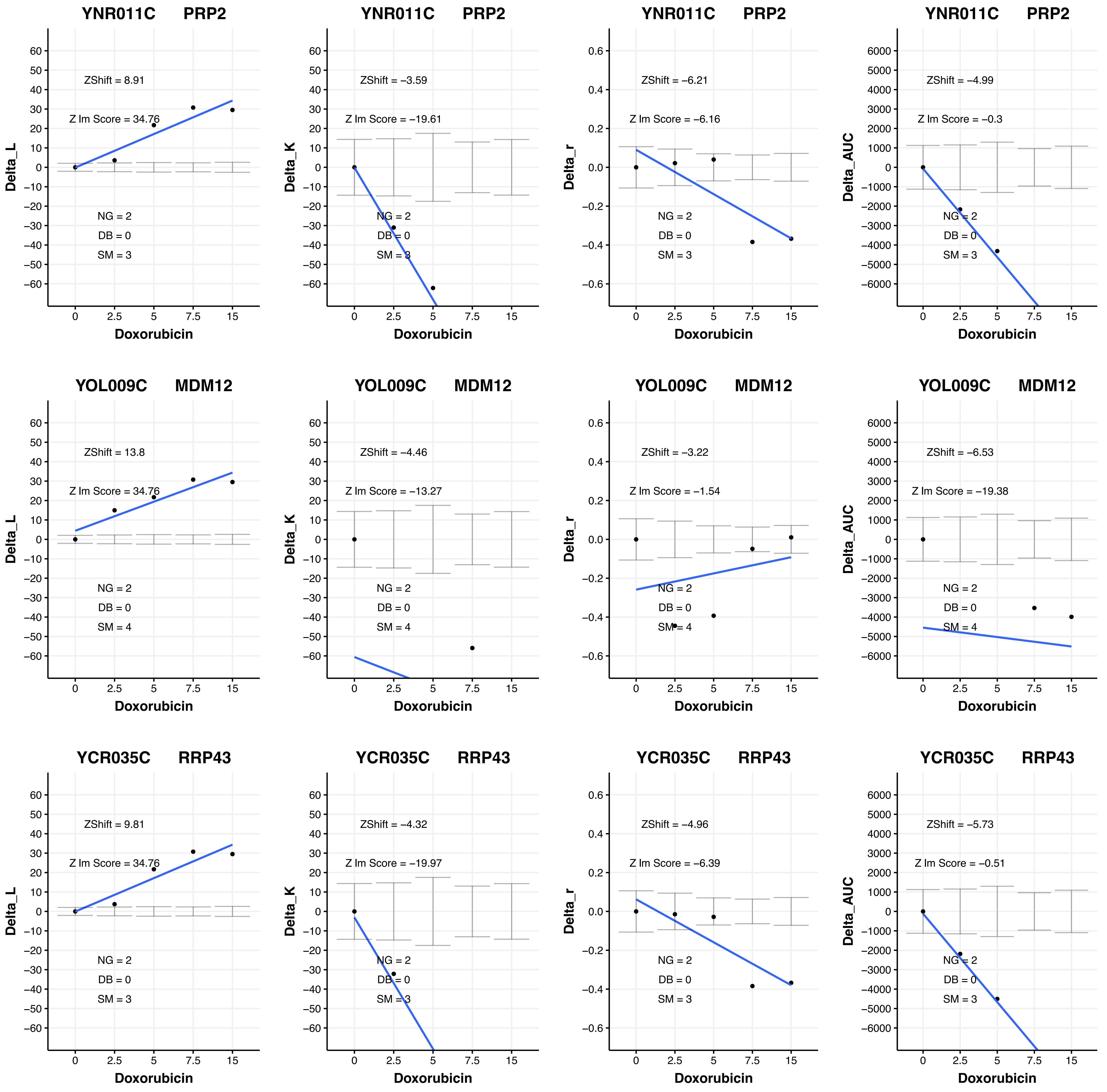
YNR011C PRP2
ZShift = 8.91
Z lm Score = 34.76
NG = 2
DB = 0
SM = 3
Delta_L
Doxorubicin
YNR011C PRP2
ZShift = −3.59
Z lm Score = −19.61
NG = 2
DB = 0
SM = 3
Delta_K
Doxorubicin
YNR011C PRP2
ZShift = −6.21
Z lm Score = −6.16
NG = 2
DB = 0
SM = 3
Delta_r
Doxorubicin
YNR011C PRP2
ZShift = −4.99
Z lm Score = −0.3
NG = 2
DB = 0
SM = 3
Delta_AUC
Doxorubicin
YOL009C MDM12
ZShift = 13.8
Z lm Score = 34.76
NG = 2
DB = 0
SM = 4
Delta_L
Doxorubicin
YOL009C MDM12
ZShift = −4.46
Z lm Score = −13.27
NG = 2
DB = 0
SM = 4
Delta_K
Doxorubicin
YOL009C MDM12
ZShift = −3.22
Z lm Score = −1.54
NG = 2
DB = 0
SM = 4
Delta_r
Doxorubicin
YOL009C MDM12
ZShift = −6.53
Z lm Score = −19.38
NG = 2
DB = 0
SM = 4
Delta_AUC
Doxorubicin
YCR035C RRP43
ZShift = 9.81
Z lm Score = 34.76
NG = 2
DB = 0
SM = 3
Delta_L
Doxorubicin
YCR035C RRP43
ZShift = −4.32
Z lm Score = −19.97
NG = 2
DB = 0
SM = 3
Delta_K
Doxorubicin
YCR035C RRP43
ZShift = −4.96
Z lm Score = −6.39
NG = 2
DB = 0
SM = 3
Delta_r
Doxorubicin
YCR035C RRP43
ZShift = −5.73
Z lm Score = −0.51
NG = 2
DB = 0
SM = 3
Delta_AUC
Doxorubicin

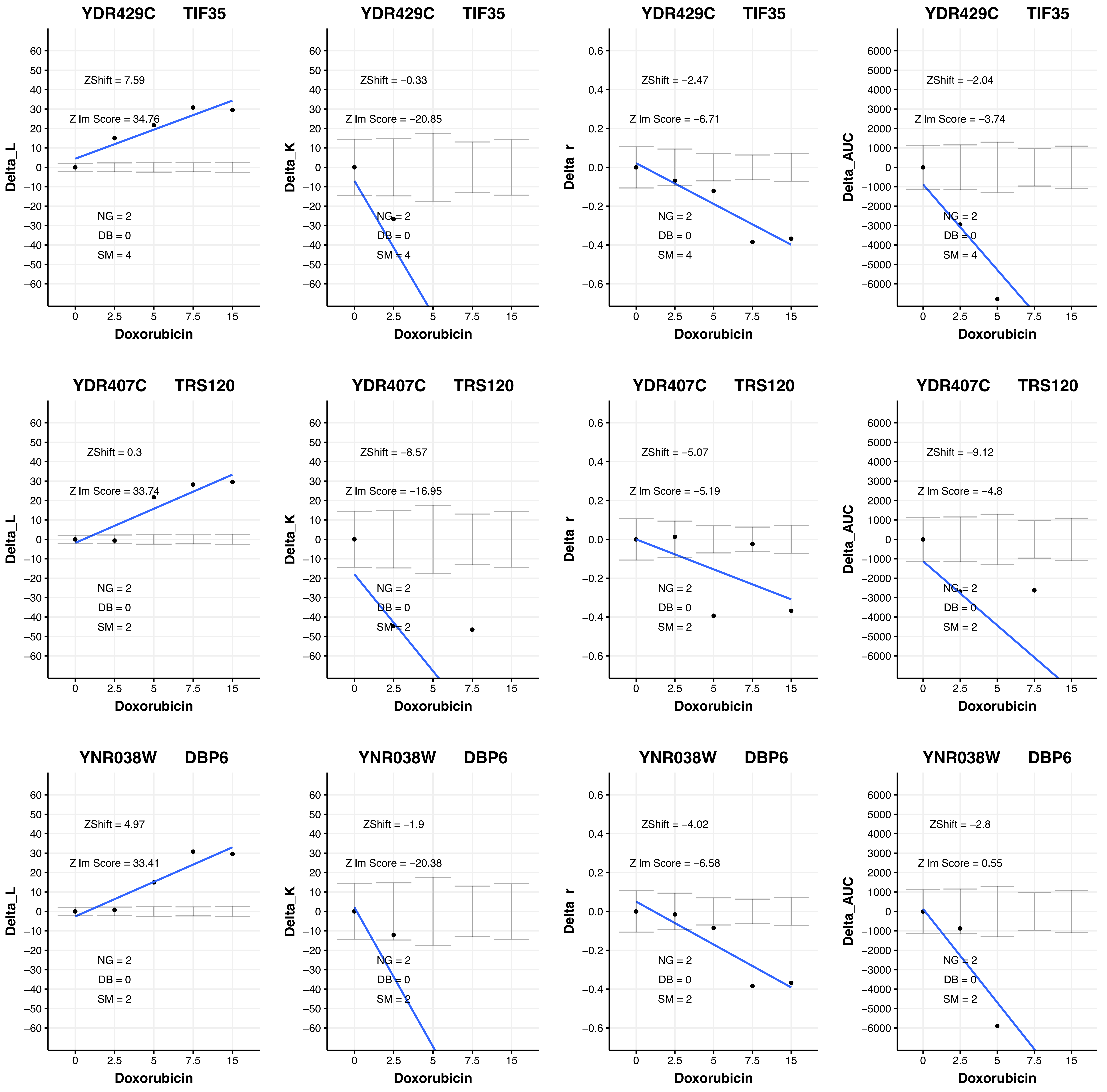
YDR429C TIF35
ZShift = 7.59
Z lm Score = 34.76
NG = 2
DB = 0
SM = 4
Delta_L
Doxorubicin
YDR429C TIF35
ZShift = −0.33
Z lm Score = −20.85
NG = 2
DB = 0
SM = 4
Delta_K
Doxorubicin
YDR429C TIF35
ZShift = −2.47
Z lm Score = −6.71
NG = 2
DB = 0
SM = 4
Delta_r
Doxorubicin
YDR429C TIF35
ZShift = −2.04
Z lm Score = −3.74
NG = 2
DB = 0
SM = 4
Delta_AUC
Doxorubicin
YDR407C TRS120
ZShift = 0.3
Z lm Score = 33.74
NG = 2
DB = 0
SM = 2
Delta_L
Doxorubicin
YDR407C TRS120
ZShift = −8.57
Z lm Score = −16.95
NG = 2
DB = 0
SM = 2
Delta_K
Doxorubicin
YDR407C TRS120
ZShift = −5.07
Z lm Score = −5.19
NG = 2
DB = 0
SM = 2
Delta_r
Doxorubicin
YDR407C TRS120
ZShift = −9.12
Z lm Score = −4.8
NG = 2
DB = 0
SM = 2
Delta_AUC
Doxorubicin
YNR038W DBP6
ZShift = 4.97
Z lm Score = 33.41
NG = 2
DB = 0
SM = 2
Delta_L
Doxorubicin
YNR038W DBP6
ZShift = −1.9
Z lm Score = −20.38
NG = 2
DB = 0
SM = 2
Delta_K
Doxorubicin
YNR038W DBP6
ZShift = −4.02
Z lm Score = −6.58
NG = 2
DB = 0
SM = 2
Delta_r
Doxorubicin
YNR038W DBP6
ZShift = −2.8
Z lm Score = 0.55
NG = 2
DB = 0
SM = 2
Delta_AUC
Doxorubicin

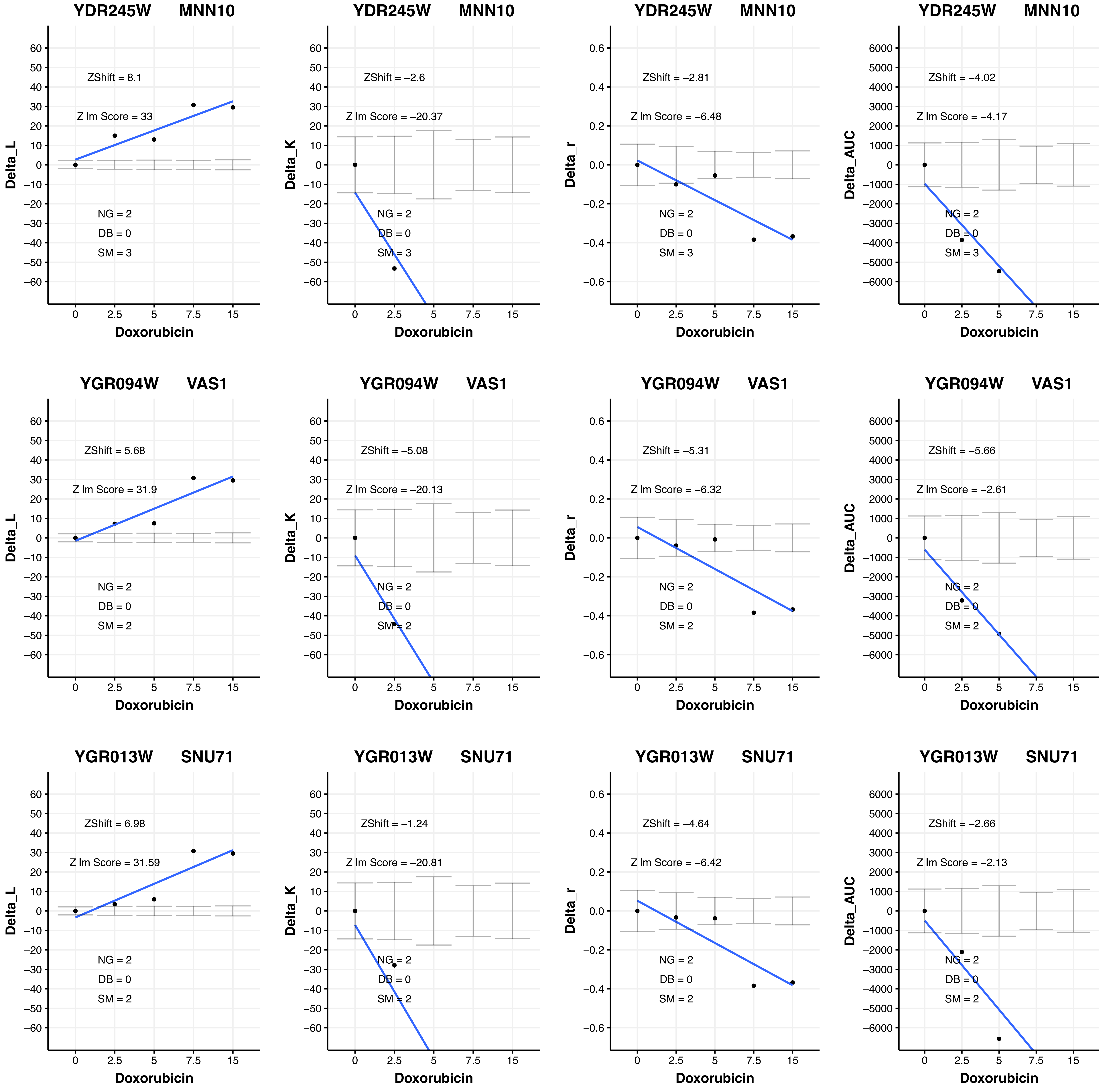

YDR245W MNN10
ZShift = 8.1
Z lm Score = 33
NG = 2
DB = 0
SM = 3
Delta_L
Doxorubicin
YDR245W MNN10
ZShift = −2.6
Z lm Score = −20.37
NG = 2
DB = 0
SM = 3
Delta_K
Doxorubicin
YDR245W MNN10
ZShift = −2.81
Z lm Score = −6.48
NG = 2
DB = 0
SM = 3
Delta_r
Doxorubicin
YDR245W MNN10
ZShift = −4.02
Z lm Score = −4.17
NG = 2
DB = 0
SM = 3
Delta_AUC
Doxorubicin
YGR094W VAS1
ZShift = 5.68
Z lm Score = 31.9
NG = 2
DB = 0
SM = 2
Delta_L
Doxorubicin
YGR094W VAS1
ZShift = −5.08
Z lm Score = −20.13
NG = 2
DB = 0
SM = 2
Delta_K
Doxorubicin
YGR094W VAS1
ZShift = −5.31
Z lm Score = −6.32
NG = 2
DB = 0
SM = 2
Delta_r
Doxorubicin
YGR094W VAS1
ZShift = −5.66
Z lm Score = −2.61
NG = 2
DB = 0
SM = 2
Delta_AUC
Doxorubicin
YGR013W SNU71
ZShift = 6.98
Z lm Score = 31.59
NG = 2
DB = 0
SM = 2
Delta_L
Doxorubicin
YGR013W SNU71
ZShift = −1.24
Z lm Score = −20.81
NG = 2
DB = 0
SM = 2
Delta_K
Doxorubicin
YGR013W SNU71
ZShift = −4.64
Z lm Score = −6.42
NG = 2
DB = 0
SM = 2
Delta_r
Doxorubicin
YGR013W SNU71
ZShift = −2.66
Z lm Score = −2.13
NG = 2
DB = 0
SM = 2
Delta_AUC
Doxorubicin

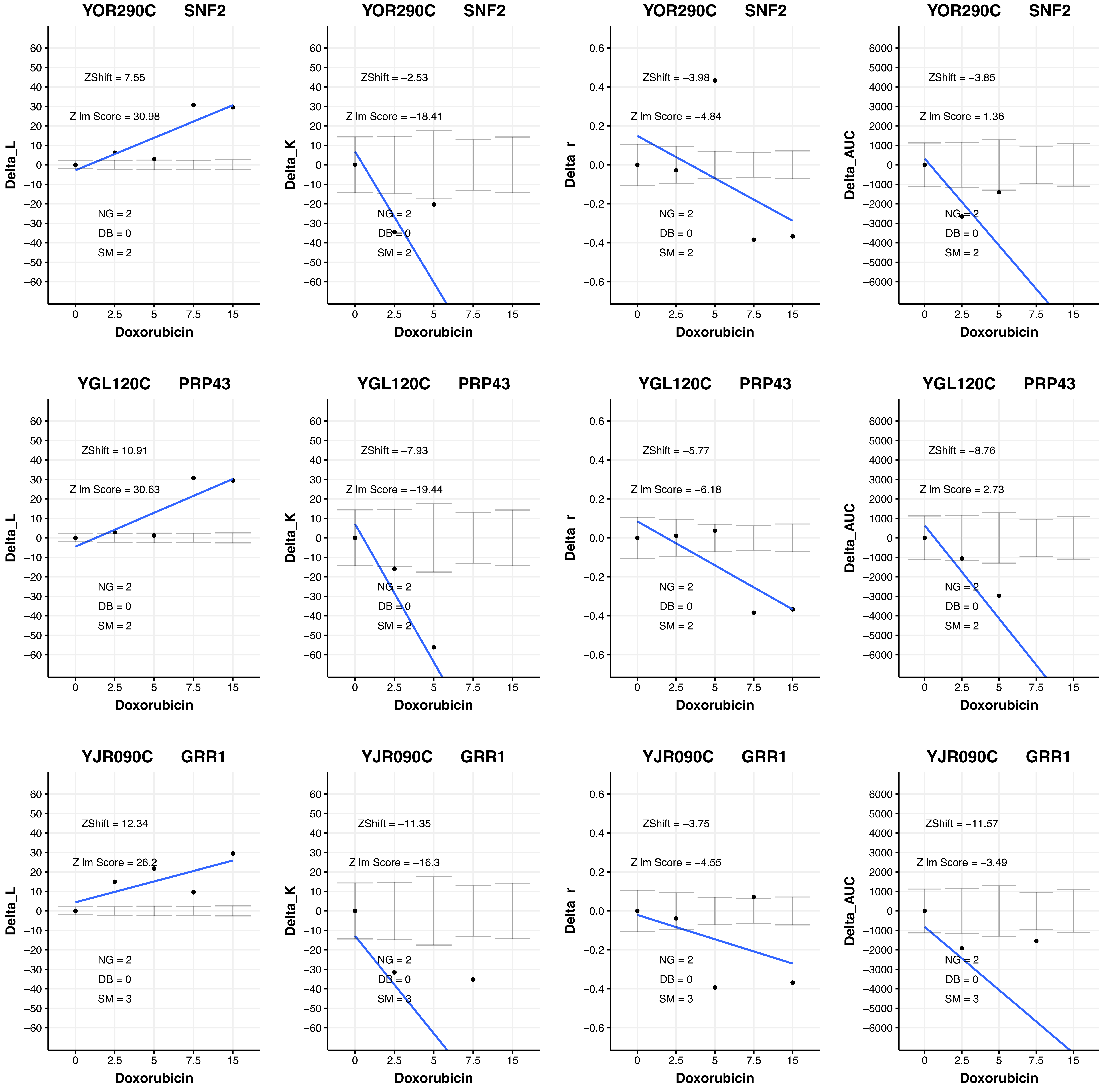
YOR290C SNF2
ZShift = 7.55
Z lm Score = 30.98
NG = 2
DB = 0
SM = 2
Delta_L
Doxorubicin
YOR290C SNF2
ZShift = −2.53
Z lm Score = −18.41
NG = 2
DB = 0
SM = 2
Delta_K
Doxorubicin
YOR290C SNF2
ZShift = −3.98
Z lm Score = −4.84
NG = 2
DB = 0
SM = 2
Delta_r
Doxorubicin
YOR290C SNF2
ZShift = −3.85
Z lm Score = 1.36
NG = 2
DB = 0
SM = 2
Delta_AUC
Doxorubicin
YGL120C PRP43
ZShift = 10.91
Z lm Score = 30.63
NG = 2
DB = 0
SM = 2
Delta_L
Doxorubicin
YGL120C PRP43
ZShift = −7.93
Z lm Score = −19.44
NG = 2
DB = 0
SM = 2
Delta_K
Doxorubicin
YGL120C PRP43
ZShift = −5.77
Z lm Score = −6.18
NG = 2
DB = 0
SM = 2
Delta_r
Doxorubicin
YGL120C PRP43
ZShift = −8.76
Z lm Score = 2.73
NG = 2
DB = 0
SM = 2
Delta_AUC
Doxorubicin
YJR090C GRR1
ZShift = 12.34
Z lm Score = 26.2
NG = 2
DB = 0
SM = 3
Delta_L
Doxorubicin
YJR090C GRR1
ZShift = −11.35
Z lm Score = −16.3
NG = 2
DB = 0
SM = 3
Delta_K
Doxorubicin
YJR090C GRR1
ZShift = −3.75
Z lm Score = −4.55
NG = 2
DB = 0
SM = 3
Delta_r
Doxorubicin
YJR090C GRR1
ZShift = −11.57
Z lm Score = −3.49
NG = 2
DB = 0
SM = 3
Delta_AUC
Doxorubicin

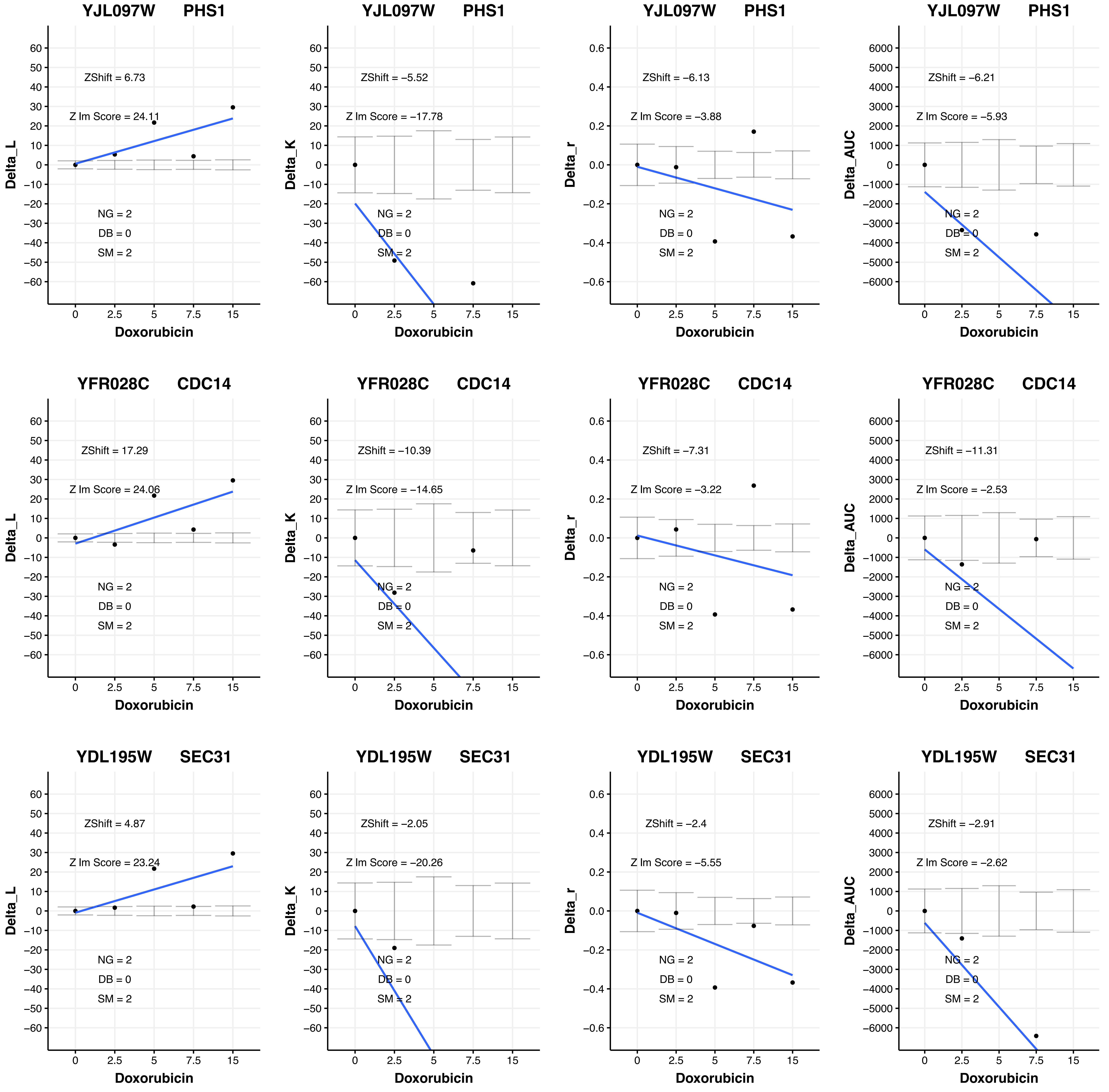
YJL097W PHS1
ZShift = 6.73
Z lm Score = 24.11
NG = 2
DB = 0
SM = 2
Delta_L
Doxorubicin
YJL097W PHS1
ZShift = −5.52
Z lm Score = −17.78
NG = 2
DB = 0
SM = 2
Delta_K
Doxorubicin
YJL097W PHS1
ZShift = −6.13
Z lm Score = −3.88
NG = 2
DB = 0
SM = 2
Delta_r
Doxorubicin
YJL097W PHS1
ZShift = −6.21
Z lm Score = −5.93
NG = 2
DB = 0
SM = 2
Delta_AUC
Doxorubicin
YFR028C CDC14
ZShift = 17.29
Z lm Score = 24.06
NG = 2
DB = 0
SM = 2
Delta_L
Doxorubicin
YFR028C CDC14
ZShift = −10.39
Z lm Score = −14.65
NG = 2
DB = 0
SM = 2
Delta_K
Doxorubicin
YFR028C CDC14
ZShift = −7.31
Z lm Score = −3.22
NG = 2
DB = 0
SM = 2
Delta_r
Doxorubicin
YFR028C CDC14
ZShift = −11.31
Z lm Score = −2.53
NG = 2
DB = 0
SM = 2
Delta_AUC
Doxorubicin
YDL195W SEC31
ZShift = 4.87
Z lm Score = 23.24
NG = 2
DB = 0
SM = 2
Delta_L
Doxorubicin
YDL195W SEC31
ZShift = −2.05
Z lm Score = −20.26
NG = 2
DB = 0
SM = 2
Delta_K
Doxorubicin
YDL195W SEC31
ZShift = −2.4
Z lm Score = −5.55
NG = 2
DB = 0
SM = 2
Delta_r
Doxorubicin
YDL195W SEC31
ZShift = −2.91
Z lm Score = −2.62
NG = 2
DB = 0
SM = 2
Delta_AUC
Doxorubicin

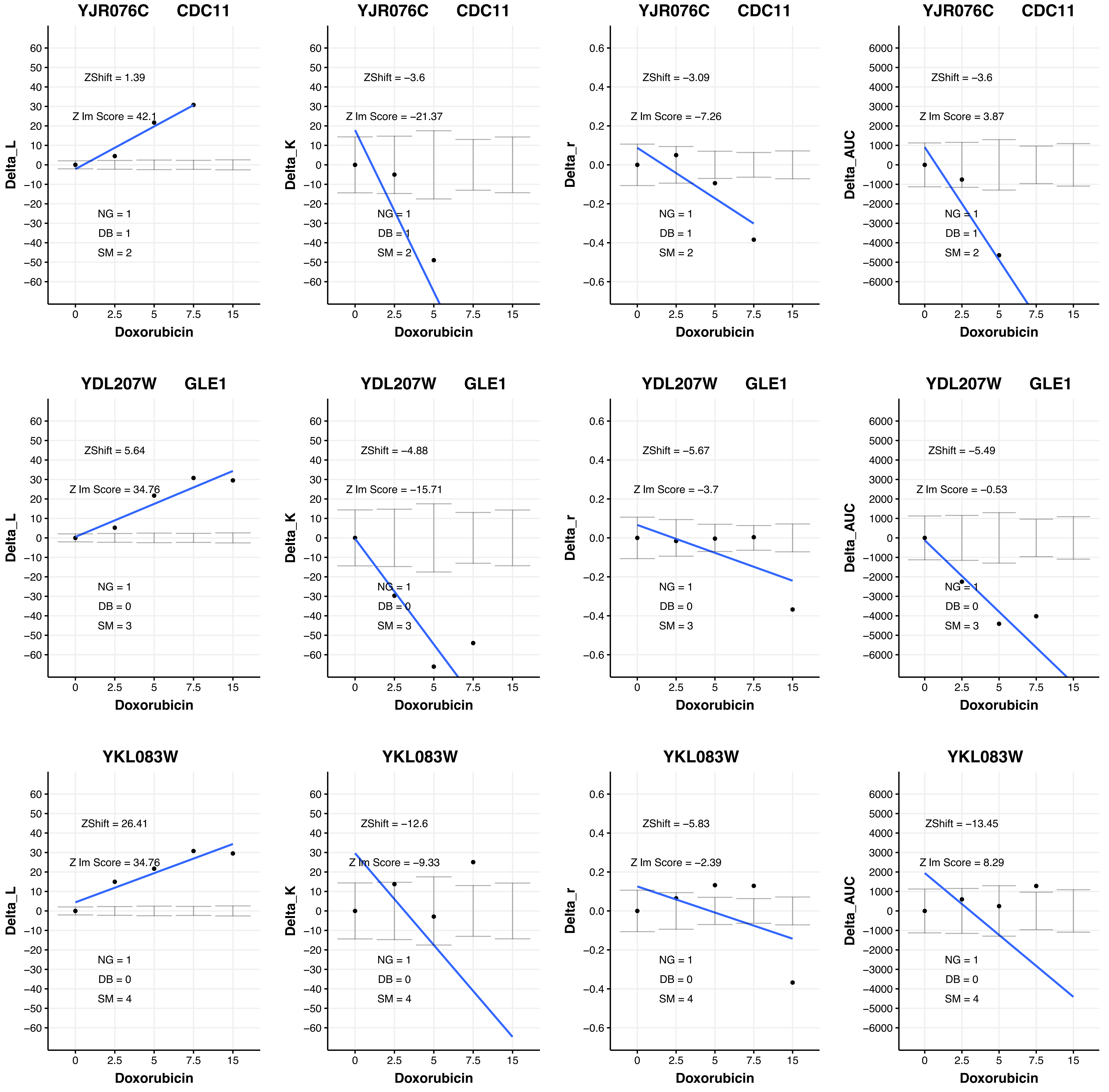

YJR076C CDC11
ZShift = 1.39
Z lm Score = 42.1
NG = 1
DB = 1
SM = 2
Delta_L
Doxorubicin
YJR076C CDC11
ZShift = −3.6
Z lm Score = −21.37
NG = 1
DB = 1
SM = 2
Delta_K
Doxorubicin
YJR076C CDC11
ZShift = −3.09
Z lm Score = −7.26
NG = 1
DB = 1
SM = 2
Delta_r
Doxorubicin
YJR076C CDC11
ZShift = −3.6
Z lm Score = 3.87
NG = 1
DB = 1
SM = 2
Delta_AUC
Doxorubicin
YDL207W GLE1
ZShift = 5.64
Z lm Score = 34.76
NG = 1
DB = 0
SM = 3
Delta_L
Doxorubicin
YDL207W GLE1
ZShift = −4.88
Z lm Score = −15.71
NG = 1
DB = 0
SM = 3
Delta_K
Doxorubicin
YDL207W GLE1
ZShift = −5.67
Z lm Score = −3.7
NG = 1
DB = 0
SM = 3
Delta_r
Doxorubicin
YDL207W GLE1
ZShift = −5.49
Z lm Score = −0.53
NG = 1
DB = 0
SM = 3
Delta_AUC
Doxorubicin
YKL083W
ZShift = 26.41
Z lm Score = 34.76
NG = 1
DB = 0
SM = 4
Delta_L
Doxorubicin
YKL083W
ZShift = −12.6
Z lm Score = −9.33
NG = 1
DB = 0
SM = 4
Delta_K
Doxorubicin
YKL083W
ZShift = −5.83
Z lm Score = −2.39
NG = 1
DB = 0
SM = 4
Delta_r
Doxorubicin
YKL083W
ZShift = −13.45
Z lm Score = 8.29
NG = 1
DB = 0
SM = 4
Delta_AUC
Doxorubicin

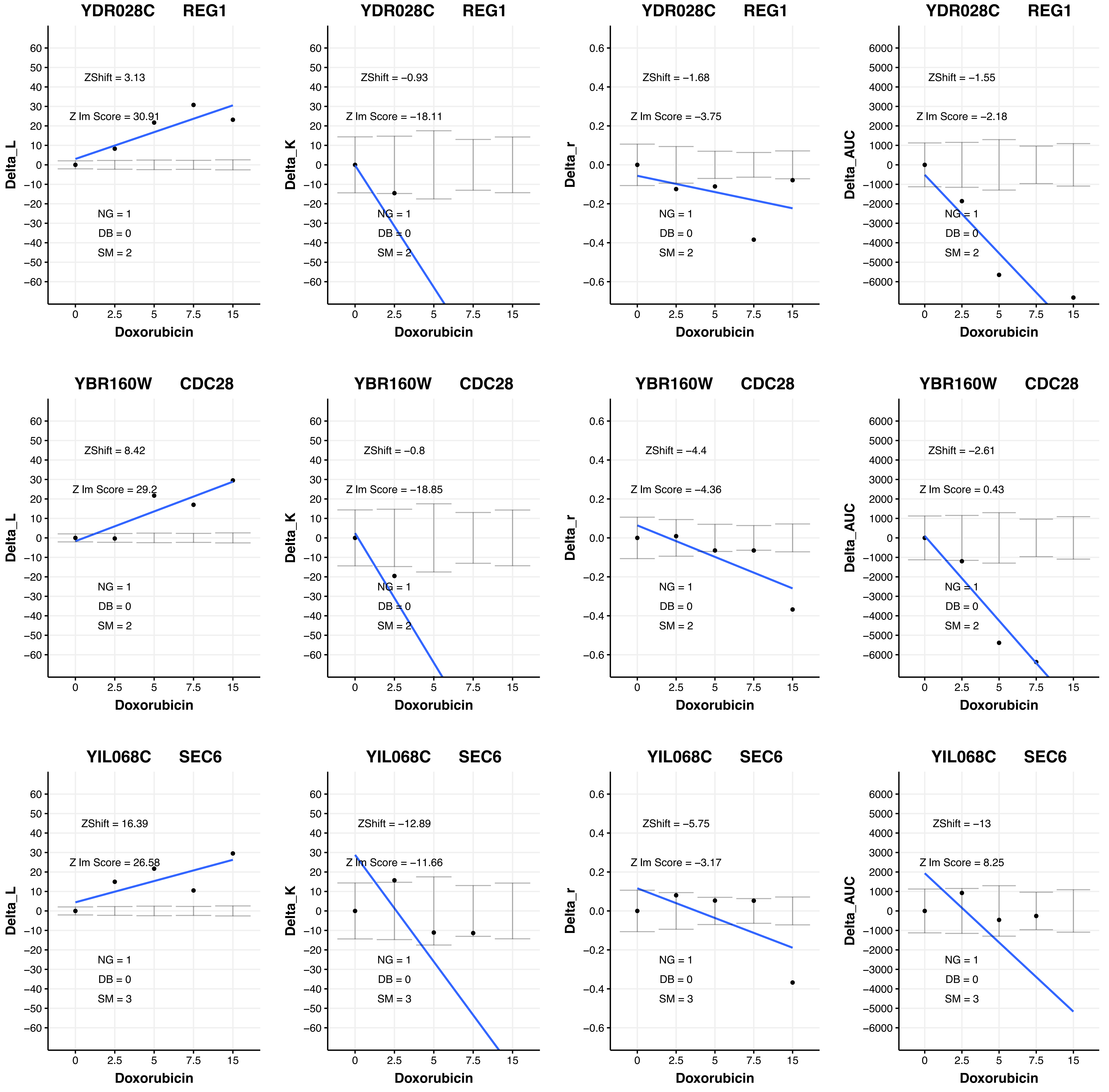
YDR028C REG1
ZShift = 3.13
Z lm Score = 30.91
NG = 1
DB = 0
SM = 2
Delta_L
Doxorubicin
YDR028C REG1
ZShift = −0.93
Z lm Score = −18.11
NG = 1
DB = 0
SM = 2
Delta_K
Doxorubicin
YDR028C REG1
ZShift = −1.68
Z lm Score = −3.75
NG = 1
DB = 0
SM = 2
Delta_r
Doxorubicin
YDR028C REG1
ZShift = −1.55
Z lm Score = −2.18
NG = 1
DB = 0
SM = 2
Delta_AUC
Doxorubicin
YBR160W CDC28
ZShift = 8.42
Z lm Score = 29.2
NG = 1
DB = 0
SM = 2
Delta_L
Doxorubicin
YBR160W CDC28
ZShift = −0.8
Z lm Score = −18.85
NG = 1
DB = 0
SM = 2
Delta_K
Doxorubicin
YBR160W CDC28
ZShift = −4.4
Z lm Score = −4.36
NG = 1
DB = 0
SM = 2
Delta_r
Doxorubicin
YBR160W CDC28
ZShift = −2.61
Z lm Score = 0.43
NG = 1
DB = 0
SM = 2
Delta_AUC
Doxorubicin
YIL068C SEC6
ZShift = 16.39
Z lm Score = 26.58
NG = 1
DB = 0
SM = 3
Delta_L
Doxorubicin
YIL068C SEC6
ZShift = −12.89
Z lm Score = −11.66
NG = 1
DB = 0
SM = 3
Delta_K
Doxorubicin
YIL068C SEC6
ZShift = −5.75
Z lm Score = −3.17
NG = 1
DB = 0
SM = 3
Delta_r
Doxorubicin
YIL068C SEC6
ZShift = −13
Z lm Score = 8.25
NG = 1
DB = 0
SM = 3
Delta_AUC
Doxorubicin

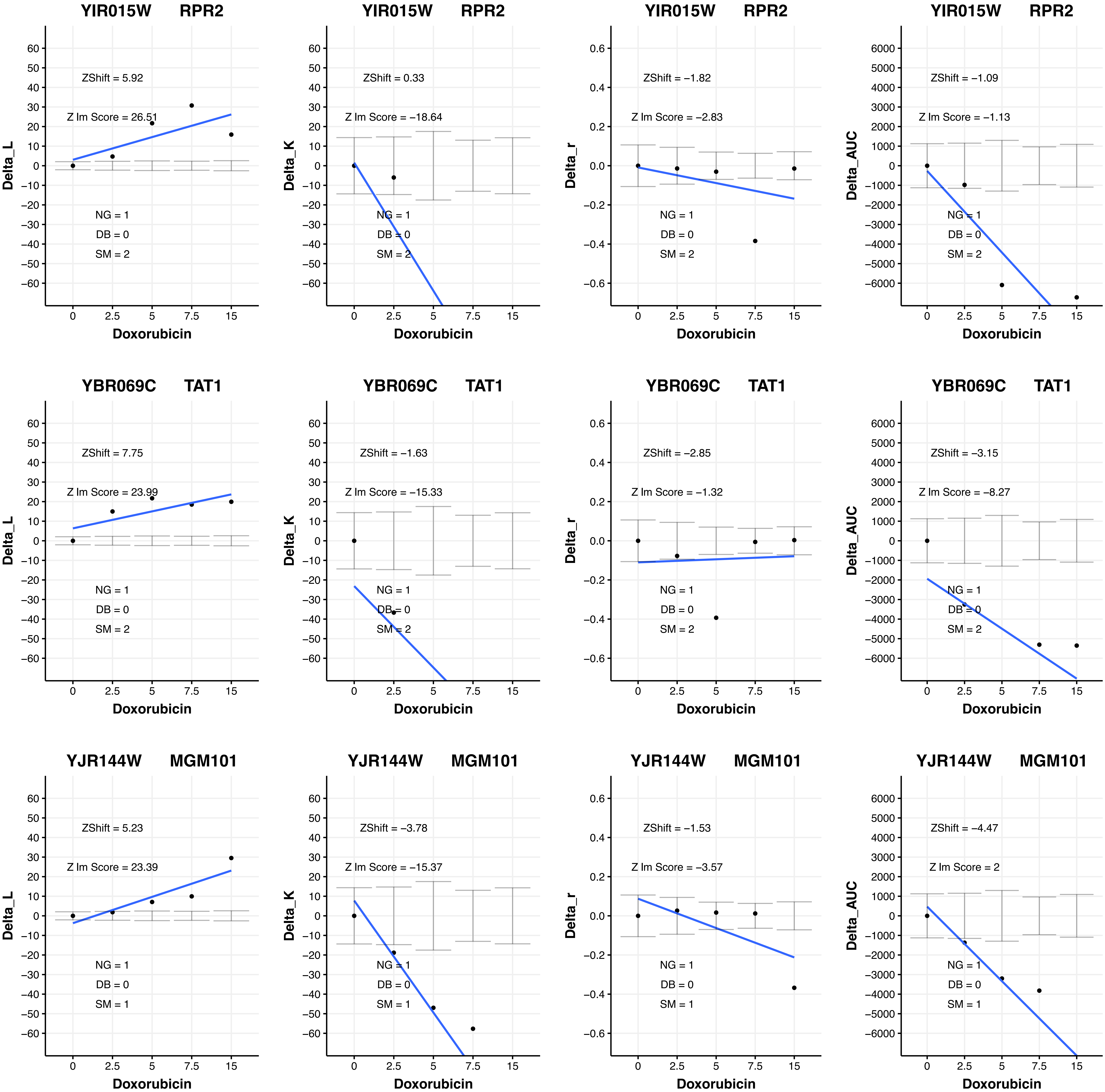
YIR015W RPR2
ZShift = 5.92
Z lm Score = 26.51
NG = 1
DB = 0
SM = 2
Delta_L
Doxorubicin
YIR015W RPR2
ZShift = 0.33
Z lm Score = −18.64
NG = 1
DB = 0
SM = 2
Delta_K
Doxorubicin
YIR015W RPR2
ZShift = −1.82
Z lm Score = −2.83
NG = 1
DB = 0
SM = 2
Delta_r
Doxorubicin
YIR015W RPR2
ZShift = −1.09
Z lm Score = −1.13
NG = 1
DB = 0
SM = 2
Delta_AUC
Doxorubicin
YBR069C TAT1
ZShift = 7.75
Z lm Score = 23.99
NG = 1
DB = 0
SM = 2
Delta_L
Doxorubicin
YBR069C TAT1
ZShift = −1.63
Z lm Score = −15.33
NG = 1
DB = 0
SM = 2
Delta_K
Doxorubicin
YBR069C TAT1
ZShift = −2.85
Z lm Score = −1.32
NG = 1
DB = 0
SM = 2
Delta_r
Doxorubicin
YBR069C TAT1
ZShift = −3.15
Z lm Score = −8.27
NG = 1
DB = 0
SM = 2
Delta_AUC
Doxorubicin
YJR144W MGM101
ZShift = 5.23
Z lm Score = 23.39
NG = 1
DB = 0
SM = 1
Delta_L
Doxorubicin
YJR144W MGM101
ZShift = −3.78
Z lm Score = −15.37
NG = 1
DB = 0
SM = 1
Delta_K
Doxorubicin
YJR144W MGM101
ZShift = −1.53
Z lm Score = −3.57
NG = 1
DB = 0
SM = 1
Delta_r
Doxorubicin
YJR144W MGM101
ZShift = −4.47
Z lm Score = 2
NG = 1
DB = 0
SM = 1
Delta_AUC
Doxorubicin

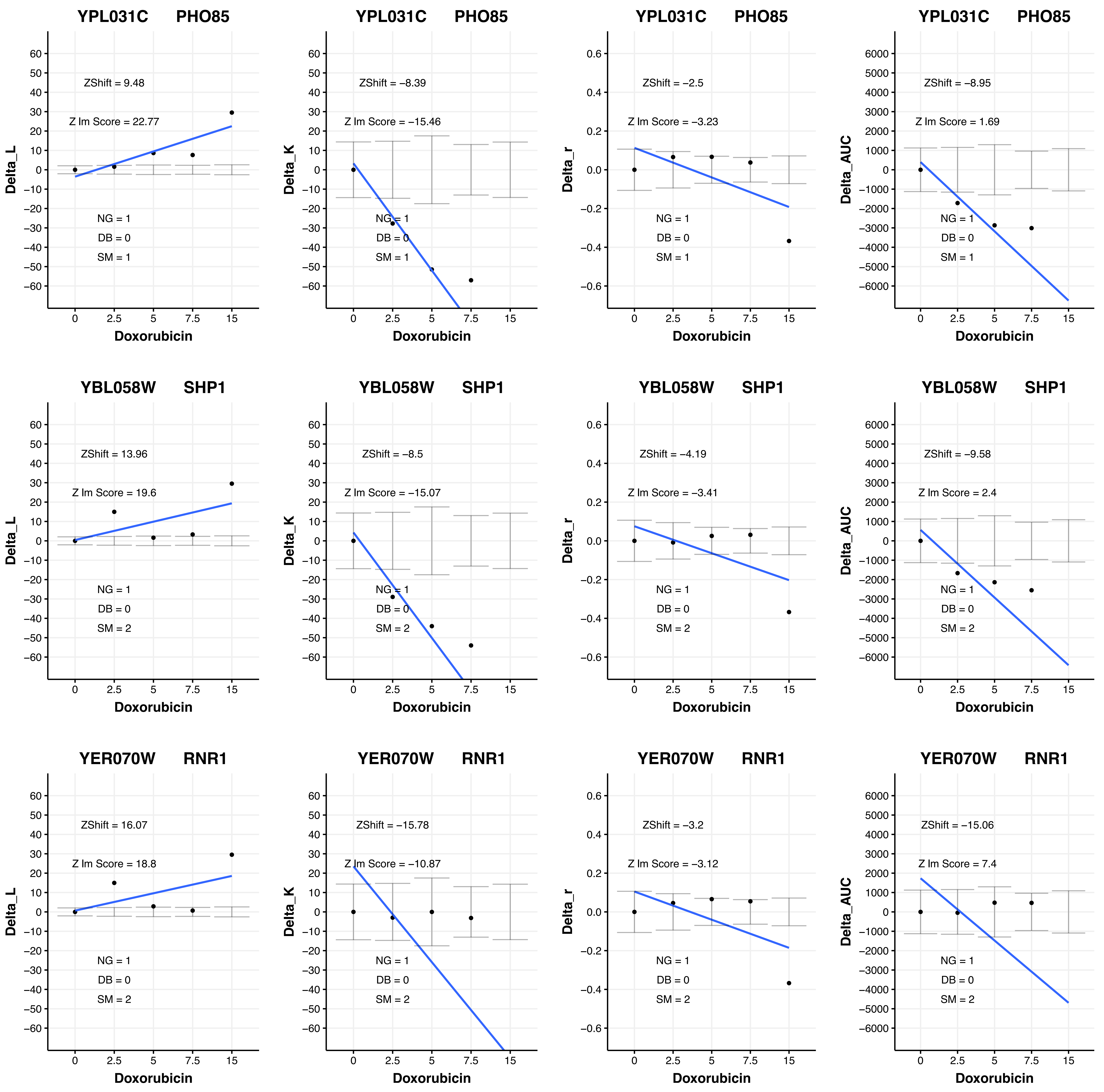

YPL031C PHO85
ZShift = 9.48
Z lm Score = 22.77
NG = 1
DB = 0
SM = 1
Delta_L
Doxorubicin
YPL031C PHO85
ZShift = −8.39
Z lm Score = −15.46
NG = 1
DB = 0
SM = 1
Delta_K
Doxorubicin
YPL031C PHO85
ZShift = −2.5
Z lm Score = −3.23
NG = 1
DB = 0
SM = 1
Delta_r
Doxorubicin
YPL031C PHO85
ZShift = −8.95
Z lm Score = 1.69
NG = 1
DB = 0
SM = 1
Delta_AUC
Doxorubicin
YBL058W SHP1
ZShift = 13.96
Z lm Score = 19.6
NG = 1
DB = 0
SM = 2
Delta_L
Doxorubicin
YBL058W SHP1
ZShift = −8.5
Z lm Score = −15.07
NG = 1
DB = 0
SM = 2
Delta_K
Doxorubicin
YBL058W SHP1
ZShift = −4.19
Z lm Score = −3.41
NG = 1
DB = 0
SM = 2
Delta_r
Doxorubicin
YBL058W SHP1
ZShift = −9.58
Z lm Score = 2.4
NG = 1
DB = 0
SM = 2
Delta_AUC
Doxorubicin
YER070W RNR1
ZShift = 16.07
Z lm Score = 18.8
NG = 1
DB = 0
SM = 2
Delta_L
Doxorubicin
YER070W RNR1
ZShift = −15.78
Z lm Score = −10.87
NG = 1
DB = 0
SM = 2
Delta_K
Doxorubicin
YER070W RNR1
ZShift = −3.2
Z lm Score = −3.12
NG = 1
DB = 0
SM = 2
Delta_r
Doxorubicin
YER070W RNR1
ZShift = −15.06
Z lm Score = 7.4
NG = 1
DB = 0
SM = 2
Delta_AUC
Doxorubicin

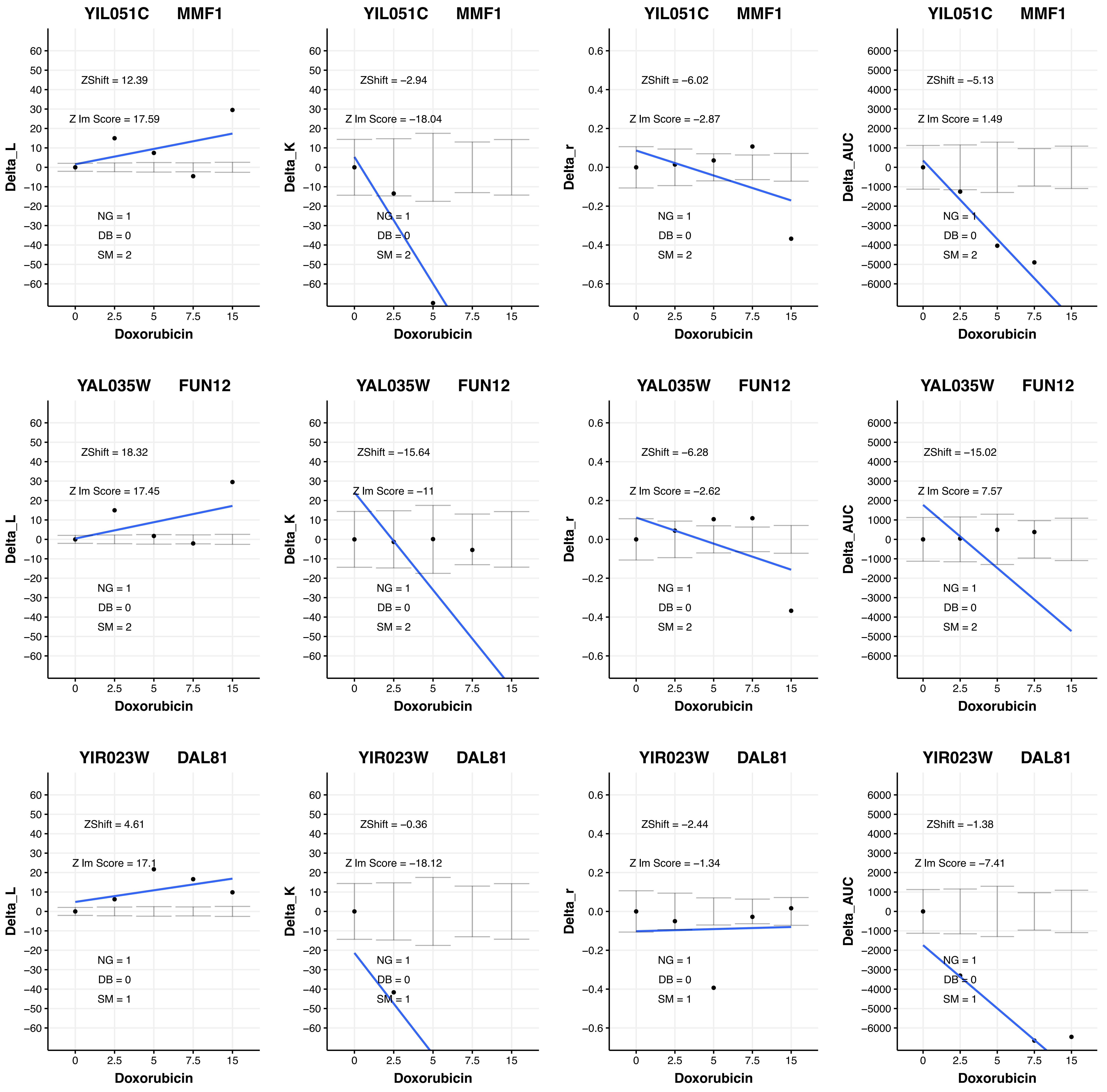
YIL051C MMF1
ZShift = 12.39
Z lm Score = 17.59
NG = 1
DB = 0
SM = 2
Delta_L
Doxorubicin
YIL051C MMF1
ZShift = −2.94
Z lm Score = −18.04
NG = 1
DB = 0
SM = 2
Delta_K
Doxorubicin
YIL051C MMF1
ZShift = −6.02
Z lm Score = −2.87
NG = 1
DB = 0
SM = 2
Delta_r
Doxorubicin
YIL051C MMF1
ZShift = −5.13
Z lm Score = 1.49
NG = 1
DB = 0
SM = 2
Delta_AUC
Doxorubicin
YAL035W FUN12
ZShift = 18.32
Z lm Score = 17.45
NG = 1
DB = 0
SM = 2
Delta_L
Doxorubicin
YAL035W FUN12
ZShift = −15.64
Z lm Score = −11
NG = 1
DB = 0
SM = 2
Delta_K
Doxorubicin
YAL035W FUN12
ZShift = −6.28
Z lm Score = −2.62
NG = 1
DB = 0
SM = 2
Delta_r
Doxorubicin
YAL035W FUN12
ZShift = −15.02
Z lm Score = 7.57
NG = 1
DB = 0
SM = 2
Delta_AUC
Doxorubicin
YIR023W DAL81
ZShift = 4.61
Z lm Score = 17.1
NG = 1
DB = 0
SM = 1
Delta_L
Doxorubicin
YIR023W DAL81
ZShift = −0.36
Z lm Score = −18.12
NG = 1
DB = 0
SM = 1
Delta_K
Doxorubicin
YIR023W DAL81
ZShift = −2.44
Z lm Score = −1.34
NG = 1
DB = 0
SM = 1
Delta_r
Doxorubicin
YIR023W DAL81
ZShift = −1.38
Z lm Score = −7.41
NG = 1
DB = 0
SM = 1
Delta_AUC
Doxorubicin

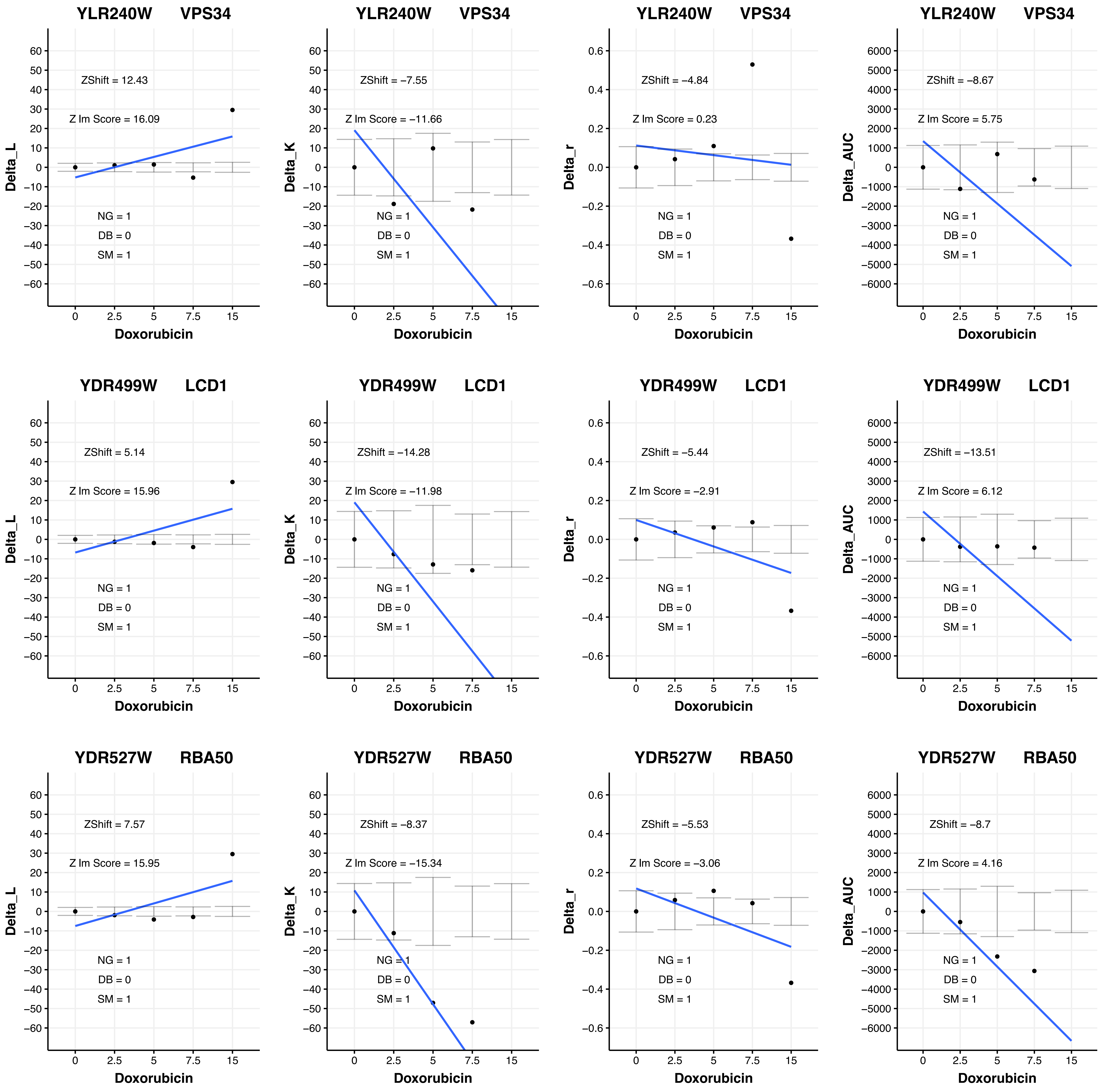
YLR240W VPS34
ZShift = 12.43
Z lm Score = 16.09
NG = 1
DB = 0
SM = 1
Delta_L
Doxorubicin
YLR240W VPS34
ZShift = −7.55
Z lm Score = −11.66
NG = 1
DB = 0
SM = 1
Delta_K
Doxorubicin
YLR240W VPS34
ZShift = −4.84
Z lm Score = 0.23
NG = 1
DB = 0
SM = 1
Delta_r
Doxorubicin
YLR240W VPS34
ZShift = −8.67
Z lm Score = 5.75
NG = 1
DB = 0
SM = 1
Delta_AUC
Doxorubicin
YDR499W LCD1
ZShift = 5.14
Z lm Score = 15.96
NG = 1
DB = 0
SM = 1
Delta_L
Doxorubicin
YDR499W LCD1
ZShift = −14.28
Z lm Score = −11.98
NG = 1
DB = 0
SM = 1
Delta_K
Doxorubicin
YDR499W LCD1
ZShift = −5.44
Z lm Score = −2.91
NG = 1
DB = 0
SM = 1
Delta_r
Doxorubicin
YDR499W LCD1
ZShift = −13.51
Z lm Score = 6.12
NG = 1
DB = 0
SM = 1
Delta_AUC
Doxorubicin
YDR527W RBA50
ZShift = 7.57
Z lm Score = 15.95
NG = 1
DB = 0
SM = 1
Delta_L
Doxorubicin
YDR527W RBA50
ZShift = −8.37
Z lm Score = −15.34
NG = 1
DB = 0
SM = 1
Delta_K
Doxorubicin
YDR527W RBA50
ZShift = −5.53
Z lm Score = −3.06
NG = 1
DB = 0
SM = 1
Delta_r
Doxorubicin
YDR527W RBA50
ZShift = −8.7
Z lm Score = 4.16
NG = 1
DB = 0
SM = 1
Delta_AUC
Doxorubicin

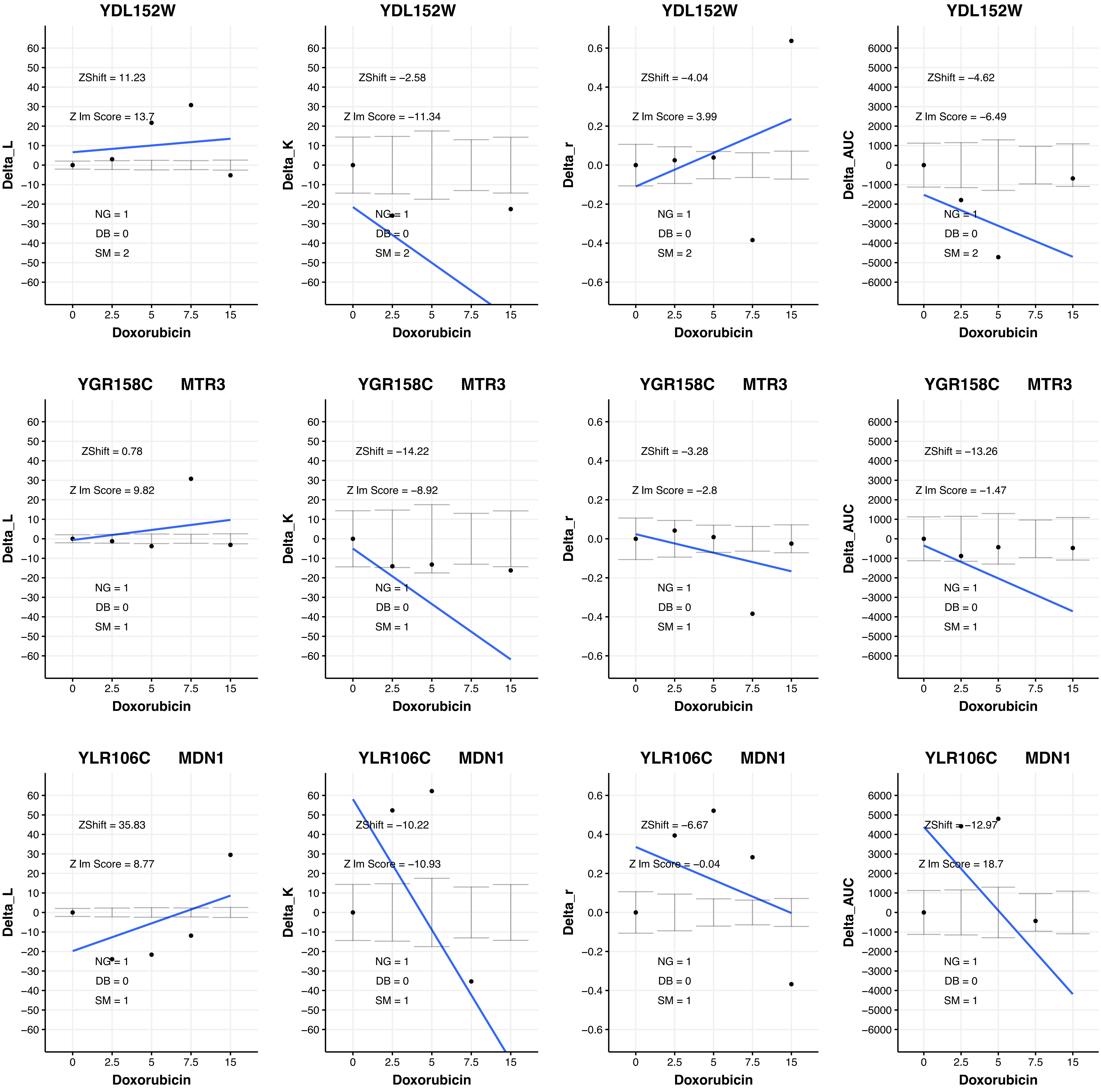
YDL152W
ZShift = 11.23
Z lm Score = 13.7
NG = 1
DB = 0
SM = 2
Delta_L
Doxorubicin
YDL152W
ZShift = −2.58
Z lm Score = −11.34
NG = 1
DB = 0
SM = 2
Delta_K
Doxorubicin
YDL152W
ZShift = −4.04
Z lm Score = 3.99
NG = 1
DB = 0
SM = 2
Delta_r
Doxorubicin
YDL152W
ZShift = −4.62
Z lm Score = −6.49
NG = 1
DB = 0
SM = 2
Delta_AUC
Doxorubicin
YGR158C MTR3
ZShift = 0.78
Z lm Score = 9.82
NG = 1
DB = 0
SM = 1
Delta_L
Doxorubicin
YGR158C MTR3
ZShift = −14.22
Z lm Score = −8.92
NG = 1
DB = 0
SM = 1
Delta_K
Doxorubicin
YGR158C MTR3
ZShift = −3.28
Z lm Score = −2.8
NG = 1
DB = 0
SM = 1
Delta_r
Doxorubicin
YGR158C MTR3
ZShift = −13.26
Z lm Score = −1.47
NG = 1
DB = 0
SM = 1
Delta_AUC
Doxorubicin
YLR106C MDN1
ZShift = 35.83
Z lm Score = 8.77
NG = 1
DB = 0
SM = 1
Delta_L
Doxorubicin
YLR106C MDN1
ZShift = −10.22
Z lm Score = −10.93
NG = 1
DB = 0
SM = 1
Delta_K
Doxorubicin
YLR106C MDN1
ZShift = −6.67
Z lm Score = −0.04
NG = 1
DB = 0
SM = 1
Delta_r
Doxorubicin
YLR106C MDN1
ZShift = −12.97
Z lm Score = 18.7
NG = 1
DB = 0
SM = 1
Delta_AUC
Doxorubicin

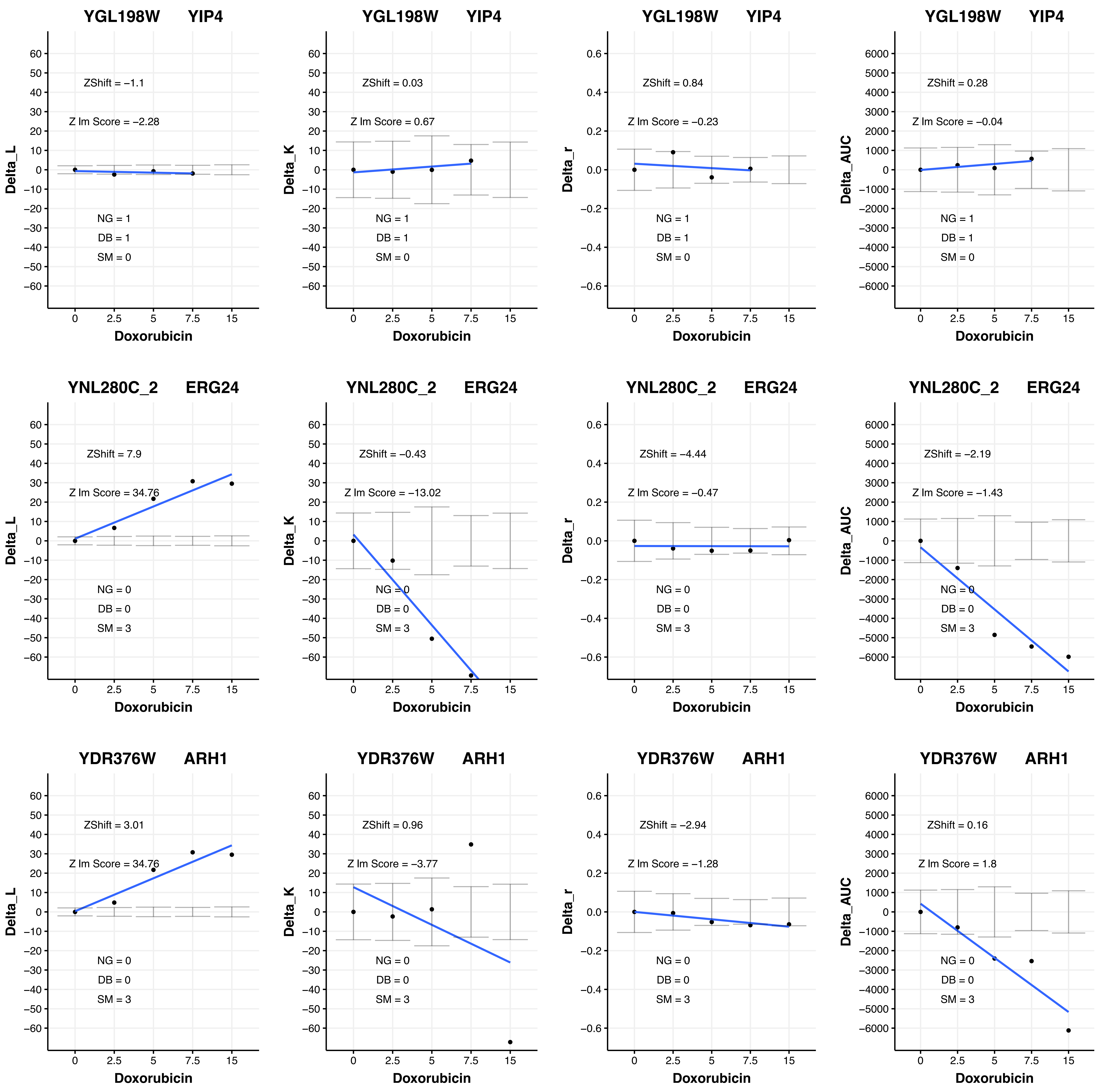
YGL198W YIP4
ZShift = −1.1
Z lm Score = −2.28
NG = 1
DB = 1
SM = 0
Delta_L
Doxorubicin
YGL198W YIP4
ZShift = 0.03
Z lm Score = 0.67
NG = 1
DB = 1
SM = 0
Delta_K
Doxorubicin
YGL198W YIP4
ZShift = 0.84
Z lm Score = −0.23
NG = 1
DB = 1
SM = 0
Delta_r
Doxorubicin
YGL198W YIP4
ZShift = 0.28
Z lm Score = −0.04
NG = 1
DB = 1
SM = 0
Delta_AUC
Doxorubicin
YNL280C_2 ERG24
ZShift = 7.9
Z lm Score = 34.76
NG = 0
DB = 0
SM = 3
Delta_L
Doxorubicin
YNL280C_2 ERG24
ZShift = −0.43
Z lm Score = −13.02
NG = 0
DB = 0
SM = 3
Delta_K
Doxorubicin
YNL280C_2 ERG24
ZShift = −4.44
Z lm Score = −0.47
NG = 0
DB = 0
SM = 3
Delta_r
Doxorubicin
YNL280C_2 ERG24
ZShift = −2.19
Z lm Score = −1.43
NG = 0
DB = 0
SM = 3
Delta_AUC
Doxorubicin
YDR376W ARH1
ZShift = 3.01
Z lm Score = 34.76
NG = 0
DB = 0
SM = 3
Delta_L
Doxorubicin
YDR376W ARH1
ZShift = 0.96
Z lm Score = −3.77
NG = 0
DB = 0
SM = 3
Delta_K
Doxorubicin
YDR376W ARH1
ZShift = −2.94
Z lm Score = −1.28
NG = 0
DB = 0
SM = 3
Delta_r
Doxorubicin
YDR376W ARH1
ZShift = 0.16
Z lm Score = 1.8
NG = 0
DB = 0
SM = 3
Delta_AUC
Doxorubicin

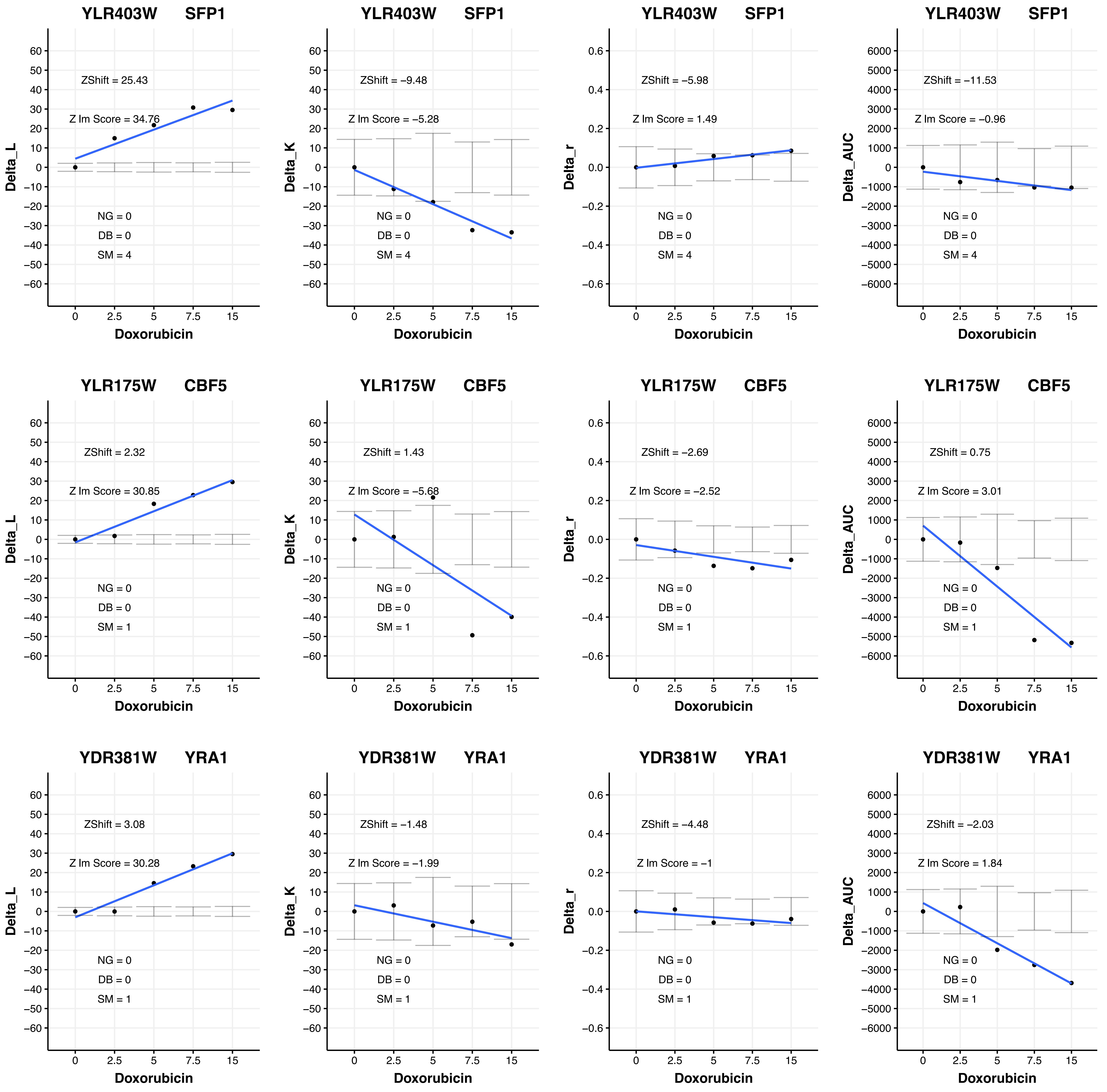

YLR403W SFP1
ZShift = 25.43
Z lm Score = 34.76
NG = 0
DB = 0
SM = 4
Delta_L
Doxorubicin
YLR403W SFP1
ZShift = −9.48
Z lm Score = −5.28
NG = 0
DB = 0
SM = 4
Delta_K
Doxorubicin
YLR403W SFP1
ZShift = −5.98
Z lm Score = 1.49
NG = 0
DB = 0
SM = 4
Delta_r
Doxorubicin
YLR403W SFP1
ZShift = −11.53
Z lm Score = −0.96
NG = 0
DB = 0
SM = 4
Delta_AUC
Doxorubicin
YLR175W CBF5
ZShift = 2.32
Z lm Score = 30.85
NG = 0
DB = 0
SM = 1
Delta_L
Doxorubicin
YLR175W CBF5
ZShift = 1.43
Z lm Score = −5.68
NG = 0
DB = 0
SM = 1
Delta_K
Doxorubicin
YLR175W CBF5
ZShift = −2.69
Z lm Score = −2.52
NG = 0
DB = 0
SM = 1
Delta_r
Doxorubicin
YLR175W CBF5
ZShift = 0.75
Z lm Score = 3.01
NG = 0
DB = 0
SM = 1
Delta_AUC
Doxorubicin
YDR381W YRA1
ZShift = 3.08
Z lm Score = 30.28
NG = 0
DB = 0
SM = 1
Delta_L
Doxorubicin
YDR381W YRA1
ZShift = −1.48
Z lm Score = −1.99
NG = 0
DB = 0
SM = 1
Delta_K
Doxorubicin
YDR381W YRA1
ZShift = −4.48
Z lm Score = −1
NG = 0
DB = 0
SM = 1
Delta_r
Doxorubicin
YDR381W YRA1
ZShift = −2.03
Z lm Score = 1.84
NG = 0
DB = 0
SM = 1
Delta_AUC
Doxorubicin

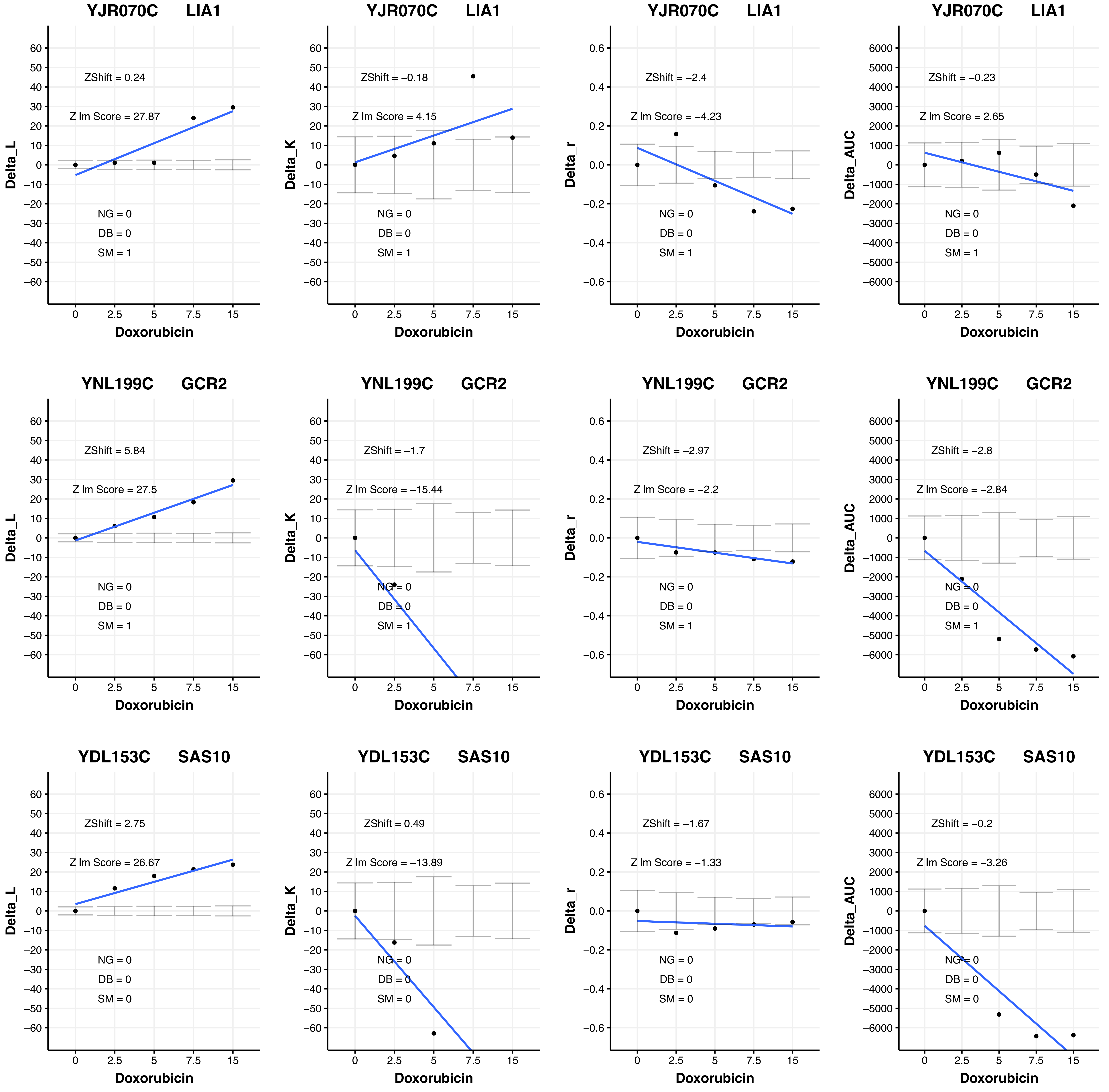
YJR070C LIA1
ZShift = 0.24
Z lm Score = 27.87
NG = 0
DB = 0
SM = 1
Delta_L
Doxorubicin
YJR070C LIA1
ZShift = −0.18
Z lm Score = 4.15
NG = 0
DB = 0
SM = 1
Delta_K
Doxorubicin
YJR070C LIA1
ZShift = −2.4
Z lm Score = −4.23
NG = 0
DB = 0
SM = 1
Delta_r
Doxorubicin
YJR070C LIA1
ZShift = −0.23
Z lm Score = 2.65
NG = 0
DB = 0
SM = 1
Delta_AUC
Doxorubicin
YNL199C GCR2
ZShift = 5.84
Z lm Score = 27.5
NG = 0
DB = 0
SM = 1
Delta_L
Doxorubicin
YNL199C GCR2
ZShift = −1.7
Z lm Score = −15.44
NG = 0
DB = 0
SM = 1
Delta_K
Doxorubicin
YNL199C GCR2
ZShift = −2.97
Z lm Score = −2.2
NG = 0
DB = 0
SM = 1
Delta_r
Doxorubicin
YNL199C GCR2
ZShift = −2.8
Z lm Score = −2.84
NG = 0
DB = 0
SM = 1
Delta_AUC
Doxorubicin
YDL153C SAS10
ZShift = 2.75
Z lm Score = 26.67
NG = 0
DB = 0
SM = 0
Delta_L
Doxorubicin
YDL153C SAS10
ZShift = 0.49
Z lm Score = −13.89
NG = 0
DB = 0
SM = 0
Delta_K
Doxorubicin
YDL153C SAS10
ZShift = −1.67
Z lm Score = −1.33
NG = 0
DB = 0
SM = 0
Delta_r
Doxorubicin
YDL153C SAS10
ZShift = −0.2
Z lm Score = −3.26
NG = 0
DB = 0
SM = 0
Delta_AUC
Doxorubicin

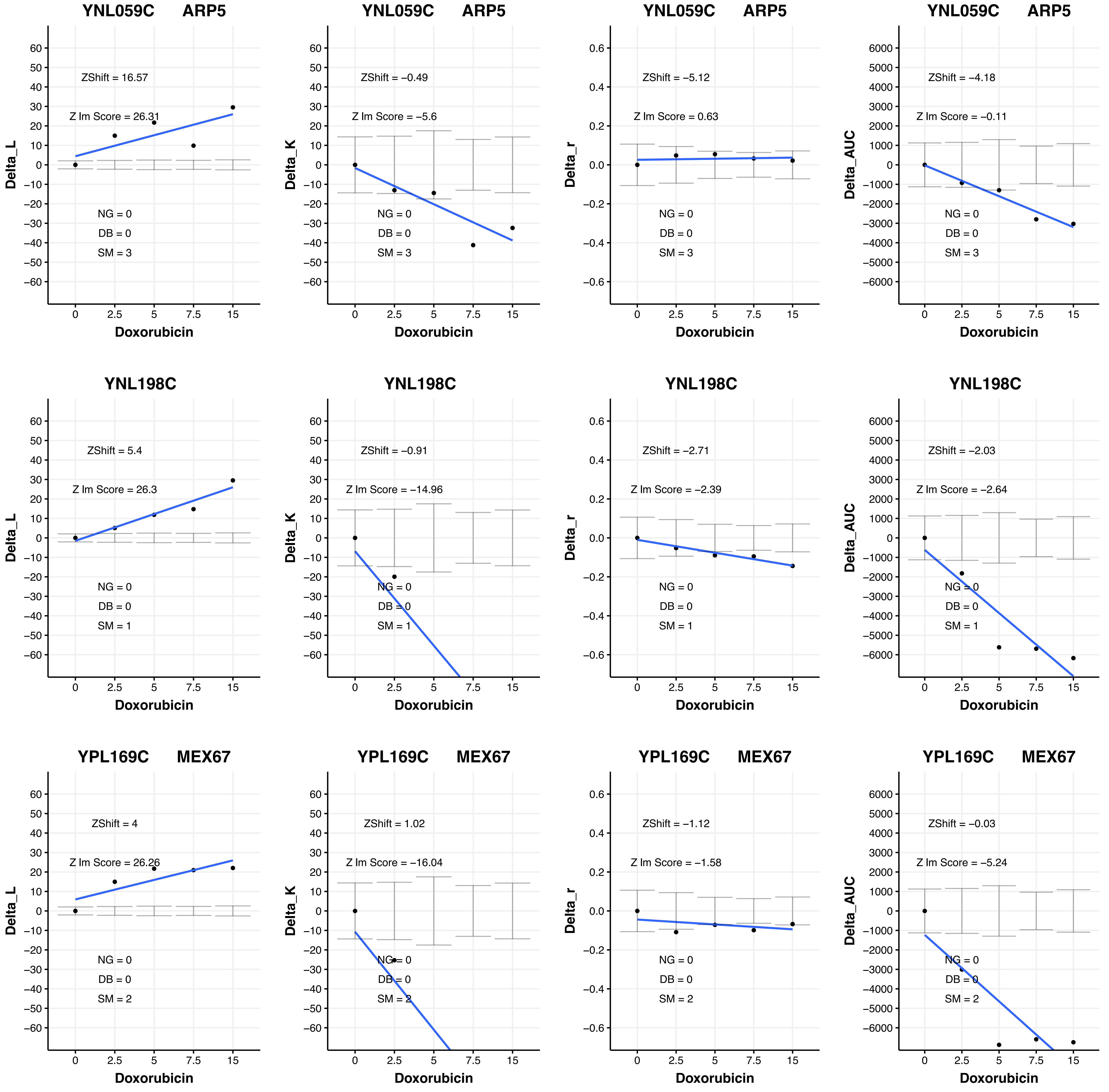
YNL059C ARP5
ZShift = 16.57
Z lm Score = 26.31
NG = 0
DB = 0
SM = 3
Delta_L
Doxorubicin
YNL059C ARP5
ZShift = −0.49
Z lm Score = −5.6
NG = 0
DB = 0
SM = 3
Delta_K
Doxorubicin
YNL059C ARP5
ZShift = −5.12
Z lm Score = 0.63
NG = 0
DB = 0
SM = 3
Delta_r
Doxorubicin
YNL059C ARP5
ZShift = −4.18
Z lm Score = −0.11
NG = 0
DB = 0
SM = 3
Delta_AUC
Doxorubicin
YNL198C
ZShift = 5.4
Z lm Score = 26.3
NG = 0
DB = 0
SM = 1
Delta_L
Doxorubicin
YNL198C
ZShift = −0.91
Z lm Score = −14.96
NG = 0
DB = 0
SM = 1
Delta_K
Doxorubicin
YNL198C
ZShift = −2.71
Z lm Score = −2.39
NG = 0
DB = 0
SM = 1
Delta_r
Doxorubicin
YNL198C
ZShift = −2.03
Z lm Score = −2.64
NG = 0
DB = 0
SM = 1
Delta_AUC
Doxorubicin
YPL169C MEX67
ZShift = 4
Z lm Score = 26.26
NG = 0
DB = 0
SM = 2
Delta_L
Doxorubicin
YPL169C MEX67
ZShift = 1.02
Z lm Score = −16.04
NG = 0
DB = 0
SM = 2
Delta_K
Doxorubicin
YPL169C MEX67
ZShift = −1.12
Z lm Score = −1.58
NG = 0
DB = 0
SM = 2
Delta_r
Doxorubicin
YPL169C MEX67
ZShift = −0.03
Z lm Score = −5.24
NG = 0
DB = 0
SM = 2
Delta_AUC
Doxorubicin

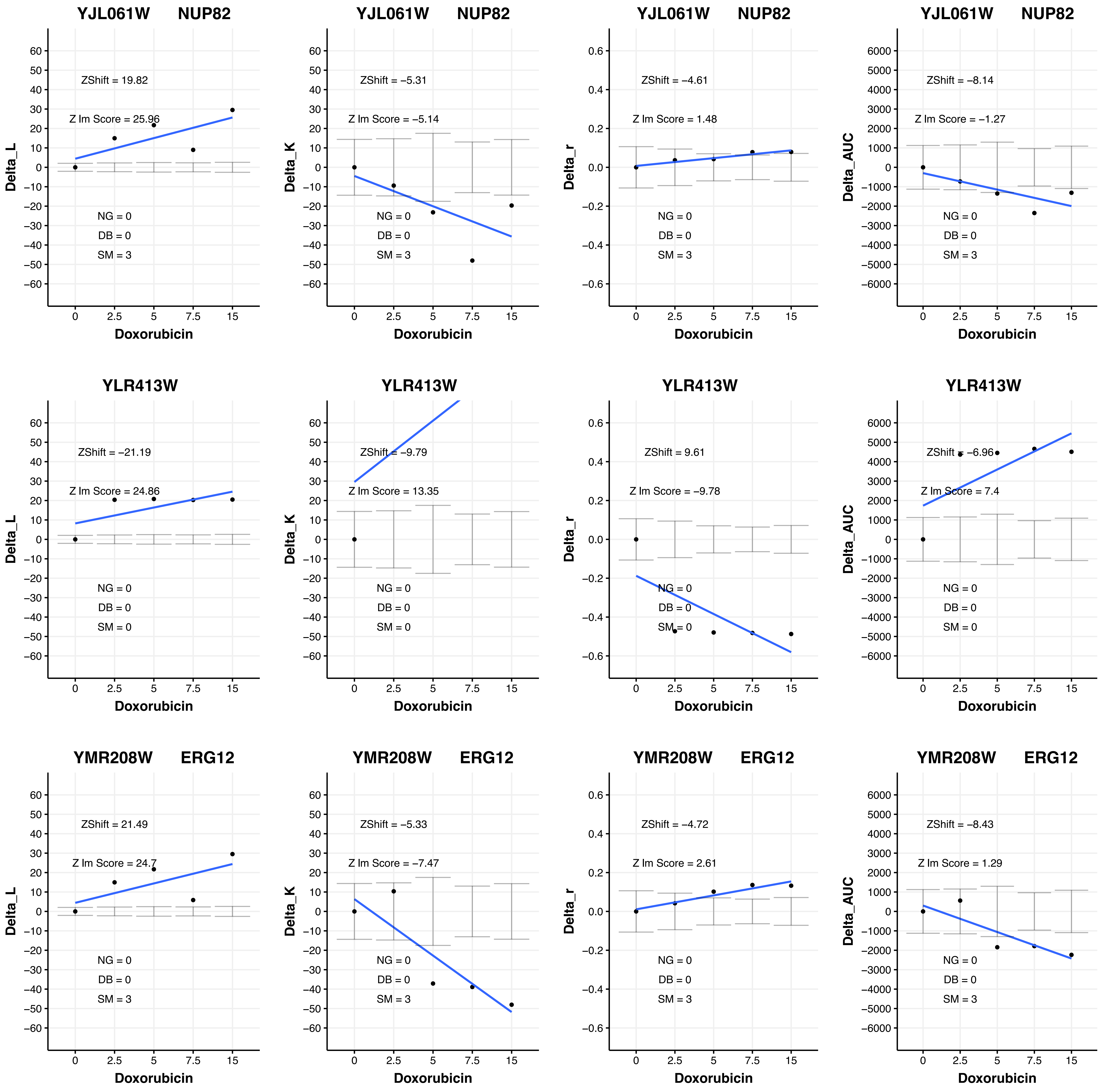

YJL061W NUP82
ZShift = 19.82
Z lm Score = 25.96
NG = 0
DB = 0
SM = 3
Delta_L
Doxorubicin
YJL061W NUP82
ZShift = −5.31
Z lm Score = −5.14
NG = 0
DB = 0
SM = 3
Delta_K
Doxorubicin
YJL061W NUP82
ZShift = −4.61
Z lm Score = 1.48
NG = 0
DB = 0
SM = 3
Delta_r
Doxorubicin
YJL061W NUP82
ZShift = −8.14
Z lm Score = −1.27
NG = 0
DB = 0
SM = 3
Delta_AUC
Doxorubicin
YLR413W
ZShift = −21.19
Z lm Score = 24.86
NG = 0
DB = 0
SM = 0
Delta_L
Doxorubicin
YLR413W
ZShift = −9.79
Z lm Score = 13.35
NG = 0
DB = 0
SM = 0
Delta_K
Doxorubicin
YLR413W
ZShift = 9.61
Z lm Score = −9.78
NG = 0
DB = 0
SM = 0
Delta_r
Doxorubicin
YLR413W
ZShift = −6.96
Z lm Score = 7.4
NG = 0
DB = 0
SM = 0
Delta_AUC
Doxorubicin
YMR208W ERG12
ZShift = 21.49
Z lm Score = 24.7
NG = 0
DB = 0
SM = 3
Delta_L
Doxorubicin
YMR208W ERG12
ZShift = −5.33
Z lm Score = −7.47
NG = 0
DB = 0
SM = 3
Delta_K
Doxorubicin
YMR208W ERG12
ZShift = −4.72
Z lm Score = 2.61
NG = 0
DB = 0
SM = 3
Delta_r
Doxorubicin
YMR208W ERG12
ZShift = −8.43
Z lm Score = 1.29
NG = 0
DB = 0
SM = 3
Delta_AUC
Doxorubicin

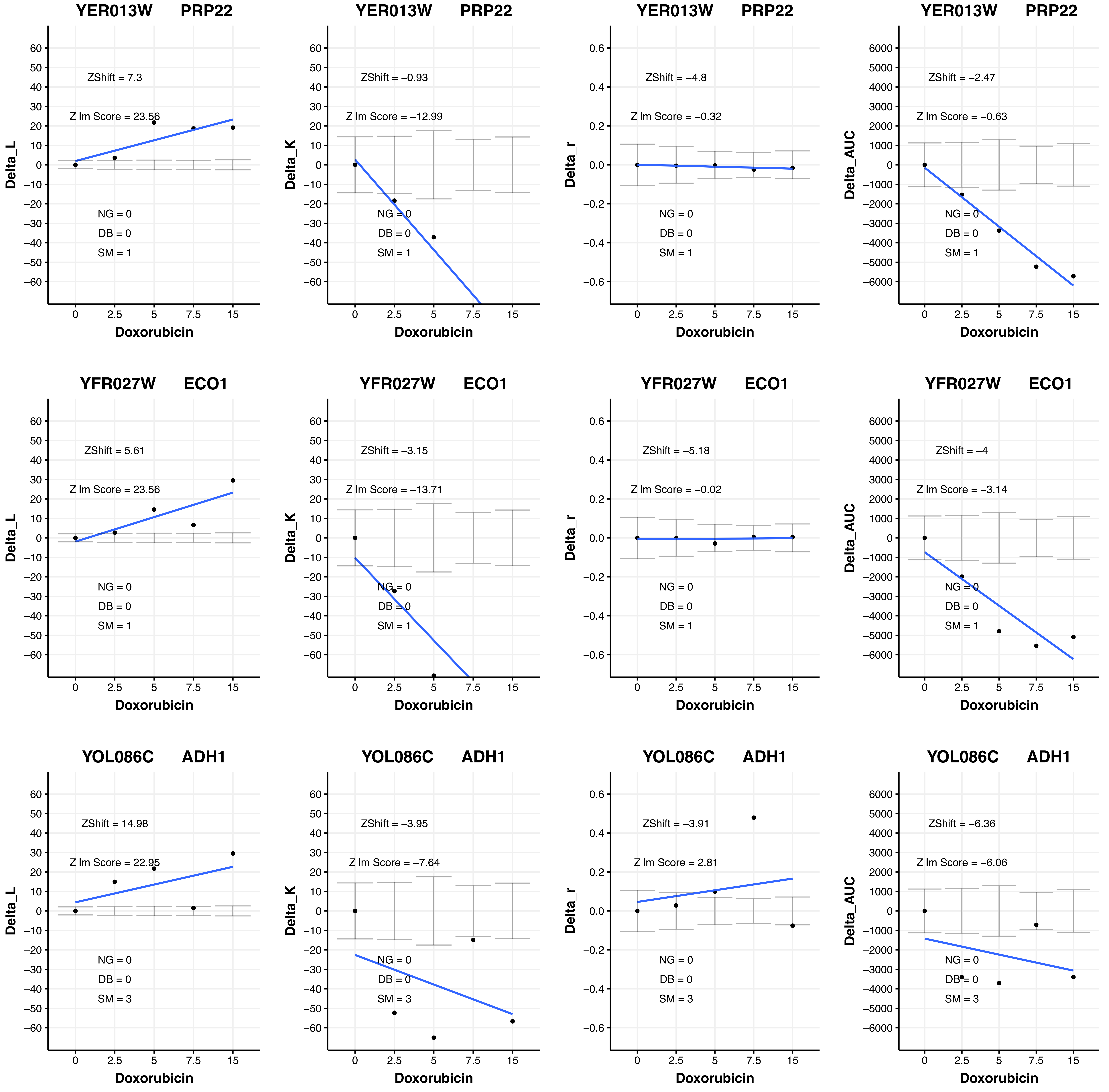

YER013W PRP22
ZShift = 7.3
Z lm Score = 23.56
NG = 0
DB = 0
SM = 1
Delta_L
Doxorubicin
YER013W PRP22
ZShift = −0.93
Z lm Score = −12.99
NG = 0
DB = 0
SM = 1
Delta_K
Doxorubicin
YER013W PRP22
ZShift = −4.8
Z lm Score = −0.32
NG = 0
DB = 0
SM = 1
Delta_r
Doxorubicin
YER013W PRP22
ZShift = −2.47
Z lm Score = −0.63
NG = 0
DB = 0
SM = 1
Delta_AUC
Doxorubicin
YFR027W ECO1
ZShift = 5.61
Z lm Score = 23.56
NG = 0
DB = 0
SM = 1
Delta_L
Doxorubicin
YFR027W ECO1
ZShift = −3.15
Z lm Score = −13.71
NG = 0
DB = 0
SM = 1
Delta_K
Doxorubicin
YFR027W ECO1
ZShift = −5.18
Z lm Score = −0.02
NG = 0
DB = 0
SM = 1
Delta_r
Doxorubicin
YFR027W ECO1
ZShift = −4
Z lm Score = −3.14
NG = 0
DB = 0
SM = 1
Delta_AUC
Doxorubicin
YOL086C ADH1
ZShift = 14.98
Z lm Score = 22.95
NG = 0
DB = 0
SM = 3
Delta_L
Doxorubicin
YOL086C ADH1
ZShift = −3.95
Z lm Score = −7.64
NG = 0
DB = 0
SM = 3
Delta_K
Doxorubicin
YOL086C ADH1
ZShift = −3.91
Z lm Score = 2.81
NG = 0
DB = 0
SM = 3
Delta_r
Doxorubicin
YOL086C ADH1
ZShift = −6.36
Z lm Score = −6.06
NG = 0
DB = 0
SM = 3
Delta_AUC
Doxorubicin

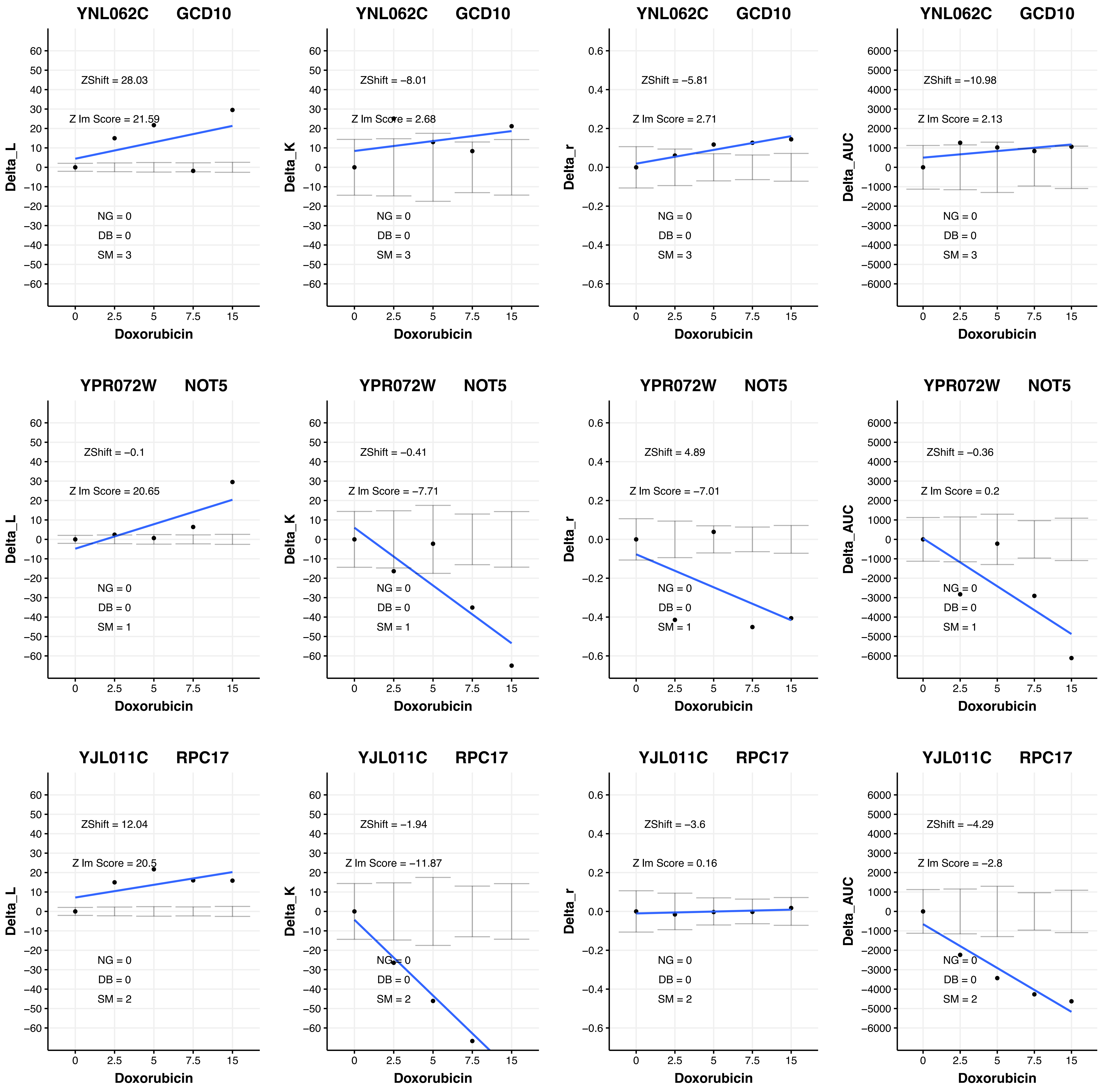
YNL062C GCD10
ZShift = 28.03
Z lm Score = 21.59
NG = 0
DB = 0
SM = 3
Delta_L
Doxorubicin
YNL062C GCD10
ZShift = −8.01
Z lm Score = 2.68
NG = 0
DB = 0
SM = 3
Delta_K
Doxorubicin
YNL062C GCD10
ZShift = −5.81
Z lm Score = 2.71
NG = 0
DB = 0
SM = 3
Delta_r
Doxorubicin
YNL062C GCD10
ZShift = −10.98
Z lm Score = 2.13
NG = 0
DB = 0
SM = 3
Delta_AUC
Doxorubicin
YPR072W NOT5
ZShift = −0.1
Z lm Score = 20.65
NG = 0
DB = 0
SM = 1
Delta_L
Doxorubicin
YPR072W NOT5
ZShift = −0.41
Z lm Score = −7.71
NG = 0
DB = 0
SM = 1
Delta_K
Doxorubicin
YPR072W NOT5
ZShift = 4.89
Z lm Score = −7.01
NG = 0
DB = 0
SM = 1
Delta_r
Doxorubicin
YPR072W NOT5
ZShift = −0.36
Z lm Score = 0.2
NG = 0
DB = 0
SM = 1
Delta_AUC
Doxorubicin
YJL011C RPC17
ZShift = 12.04
Z lm Score = 20.5
NG = 0
DB = 0
SM = 2
Delta_L
Doxorubicin
YJL011C RPC17
ZShift = −1.94
Z lm Score = −11.87
NG = 0
DB = 0
SM = 2
Delta_K
Doxorubicin
YJL011C RPC17
ZShift = −3.6
Z lm Score = 0.16
NG = 0
DB = 0
SM = 2
Delta_r
Doxorubicin
YJL011C RPC17
ZShift = −4.29
Z lm Score = −2.8
NG = 0
DB = 0
SM = 2
Delta_AUC
Doxorubicin

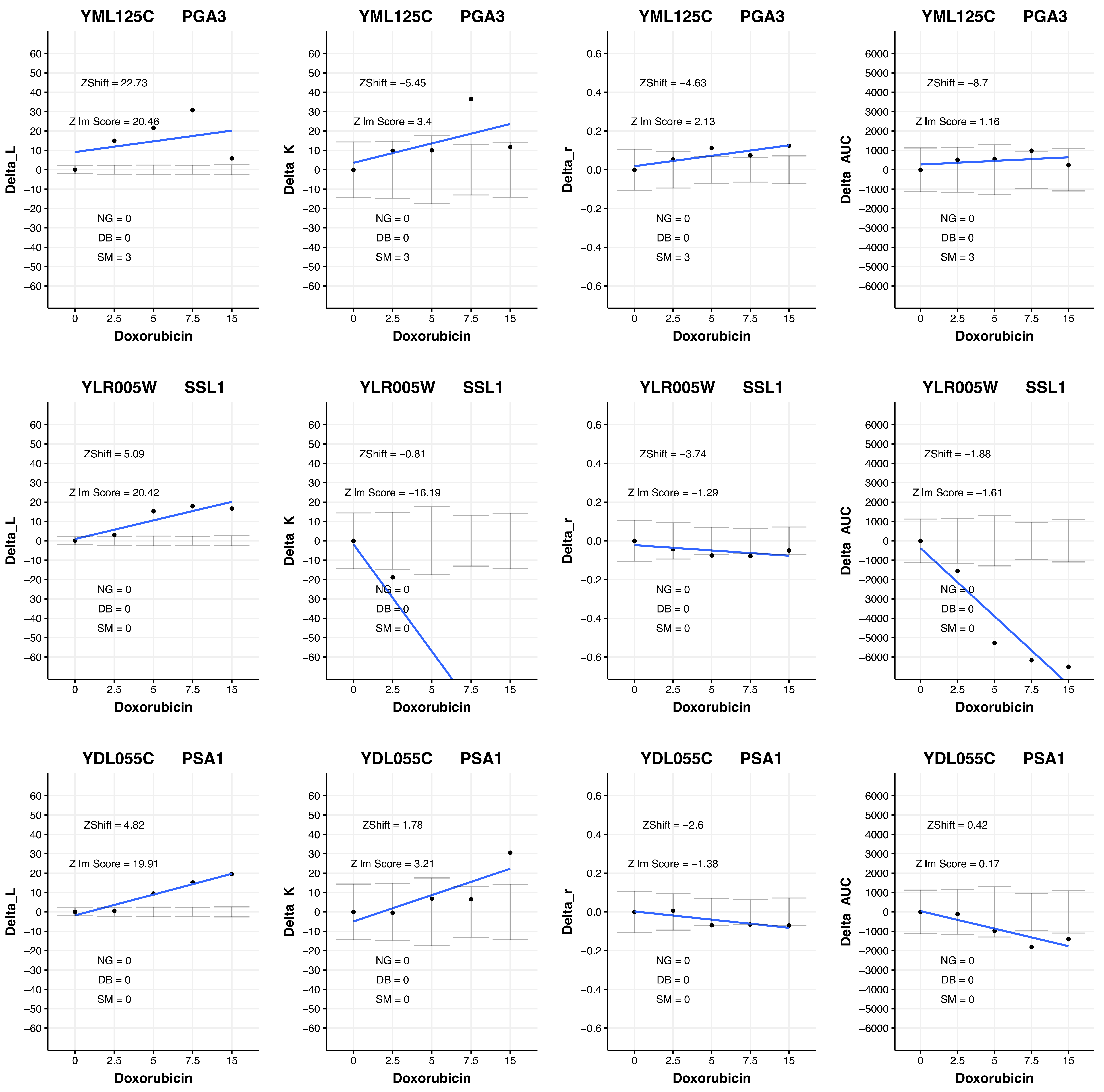
YML125C PGA3
ZShift = 22.73
Z lm Score = 20.46
NG = 0
DB = 0
SM = 3
Delta_L
Doxorubicin
YML125C PGA3
ZShift = −5.45
Z lm Score = 3.4
NG = 0
DB = 0
SM = 3
Delta_K
Doxorubicin
YML125C PGA3
ZShift = −4.63
Z lm Score = 2.13
NG = 0
DB = 0
SM = 3
Delta_r
Doxorubicin
YML125C PGA3
ZShift = −8.7
Z lm Score = 1.16
NG = 0
DB = 0
SM = 3
Delta_AUC
Doxorubicin
YLR005W SSL1
ZShift = 5.09
Z lm Score = 20.42
NG = 0
DB = 0
SM = 0
Delta_L
Doxorubicin
YLR005W SSL1
ZShift = −0.81
Z lm Score = −16.19
NG = 0
DB = 0
SM = 0
Delta_K
Doxorubicin
YLR005W SSL1
ZShift = −3.74
Z lm Score = −1.29
NG = 0
DB = 0
SM = 0
Delta_r
Doxorubicin
YLR005W SSL1
ZShift = −1.88
Z lm Score = −1.61
NG = 0
DB = 0
SM = 0
Delta_AUC
Doxorubicin
YDL055C PSA1
ZShift = 4.82
Z lm Score = 19.91
NG = 0
DB = 0
SM = 0
Delta_L
Doxorubicin
YDL055C PSA1
ZShift = 1.78
Z lm Score = 3.21
NG = 0
DB = 0
SM = 0
Delta_K
Doxorubicin
YDL055C PSA1
ZShift = −2.6
Z lm Score = −1.38
NG = 0
DB = 0
SM = 0
Delta_r
Doxorubicin
YDL055C PSA1
ZShift = 0.42
Z lm Score = 0.17
NG = 0
DB = 0
SM = 0
Delta_AUC
Doxorubicin

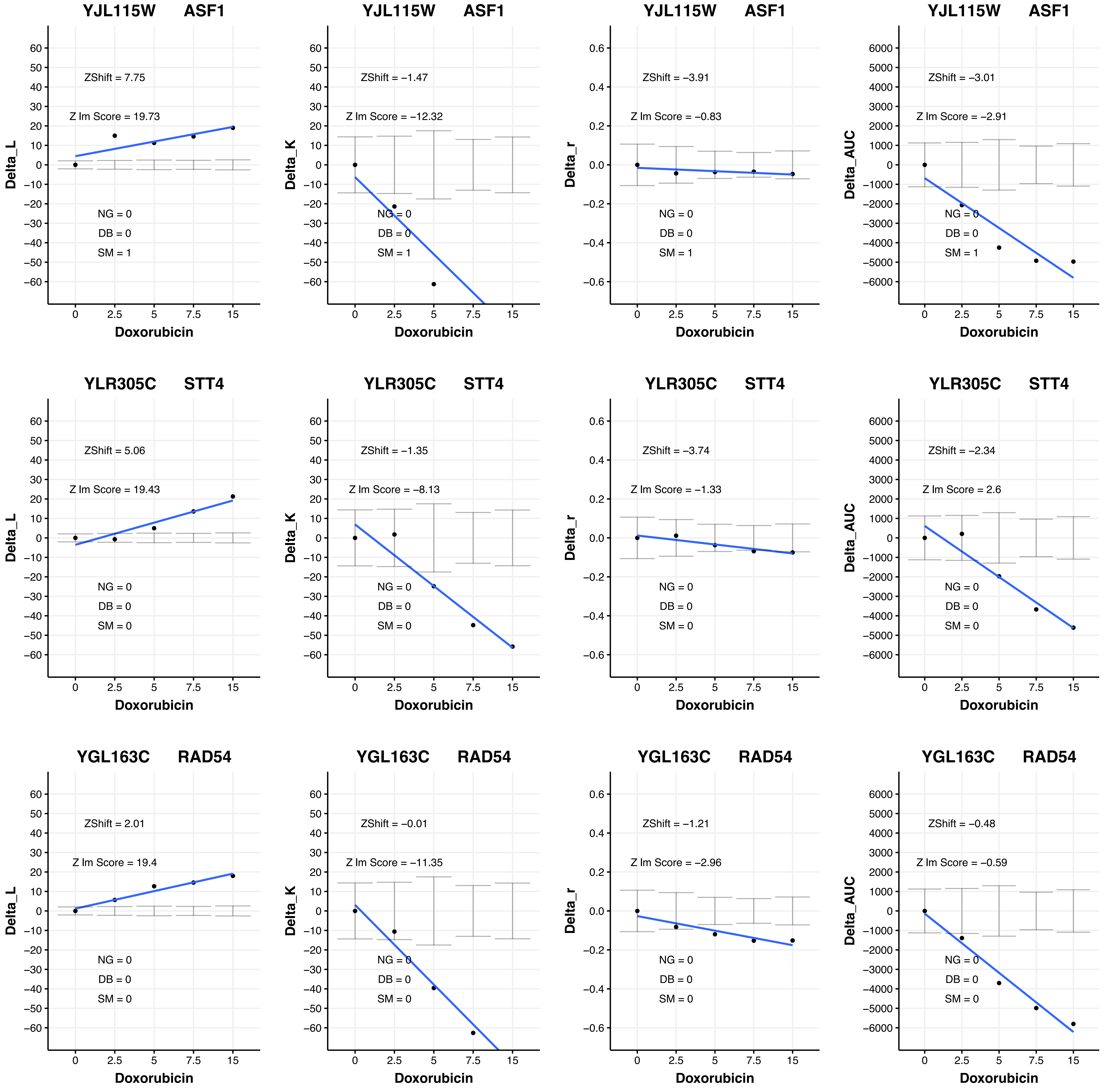
YJL115W ASF1
ZShift = 7.75
Z lm Score = 19.73
NG = 0
DB = 0
SM = 1
Delta_L
Doxorubicin
ZShift = −1.47
Z lm Score = −12.32
Delta_K
ZShift = −3.91
Z lm Score = −0.83
Delta_r
ZShift = −3.01
Z lm Score = −2.91
Delta_AUC
YLR305C STT4
ZShift = 5.06
Z lm Score = 19.43
SM = 0
ZShift = −1.35
Z lm Score = −8.13
ZShift = −3.74
Z lm Score = −1.33
ZShift = −2.34
Z lm Score = 2.6
YGL163C RAD54
ZShift = 2.01
Z lm Score = 19.4
ZShift = −0.01
Z lm Score = −11.35
ZShift = −1.21
Z lm Score = −2.96
ZShift = −0.48
Z lm Score = −0.59

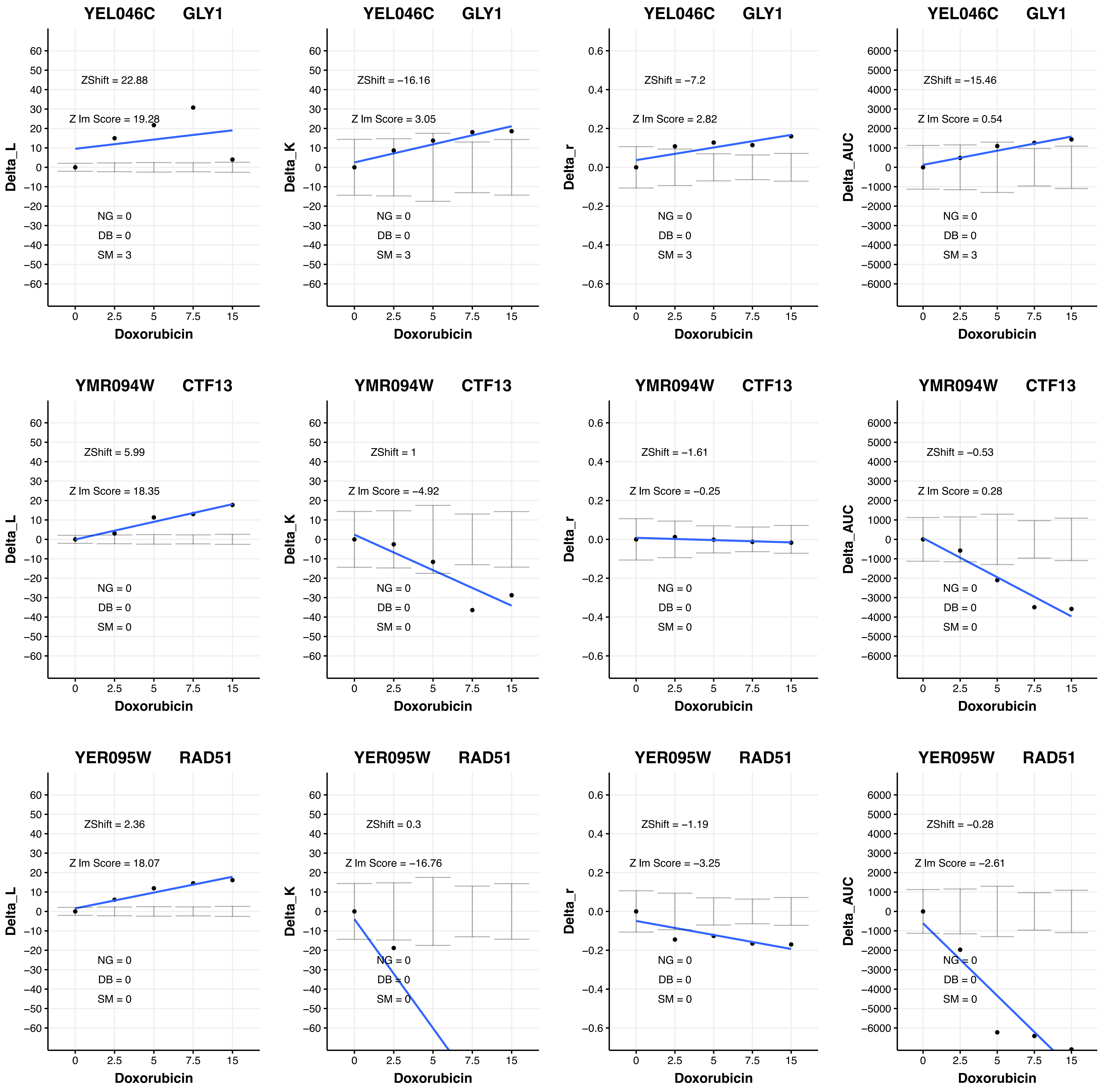
YEL046C GLY1
ZShift = 22.88
Z lm Score = 19.28
NG = 0
DB = 0
SM = 3
Delta_L
Doxorubicin
YEL046C GLY1
ZShift = −16.16
Z lm Score = 3.05
NG = 0
DB = 0
SM = 3
Delta_K
Doxorubicin
YEL046C GLY1
ZShift = −7.2
Z lm Score = 2.82
NG = 0
DB = 0
SM = 3
Delta_r
Doxorubicin
YEL046C GLY1
ZShift = −15.46
Z lm Score = 0.54
NG = 0
DB = 0
SM = 3
Delta_AUC
Doxorubicin
YMR094W CTF13
ZShift = 5.99
Z lm Score = 18.35
NG = 0
DB = 0
SM = 0
Delta_L
Doxorubicin
YMR094W CTF13
ZShift = 1
Z lm Score = −4.92
NG = 0
DB = 0
SM = 0
Delta_K
Doxorubicin
YMR094W CTF13
ZShift = −1.61
Z lm Score = −0.25
NG = 0
DB = 0
SM = 0
Delta_r
Doxorubicin
YMR094W CTF13
ZShift = −0.53
Z lm Score = 0.28
NG = 0
DB = 0
SM = 0
Delta_AUC
Doxorubicin
YER095W RAD51
ZShift = 2.36
Z lm Score = 18.07
NG = 0
DB = 0
SM = 0
Delta_L
Doxorubicin
YER095W RAD51
ZShift = 0.3
Z lm Score = −16.76
NG = 0
DB = 0
SM = 0
Delta_K
Doxorubicin
YER095W RAD51
ZShift = −1.19
Z lm Score = −3.25
NG = 0
DB = 0
SM = 0
Delta_r
Doxorubicin
YER095W RAD51
ZShift = −0.28
Z lm Score = −2.61
NG = 0
DB = 0
SM = 0
Delta_AUC
Doxorubicin

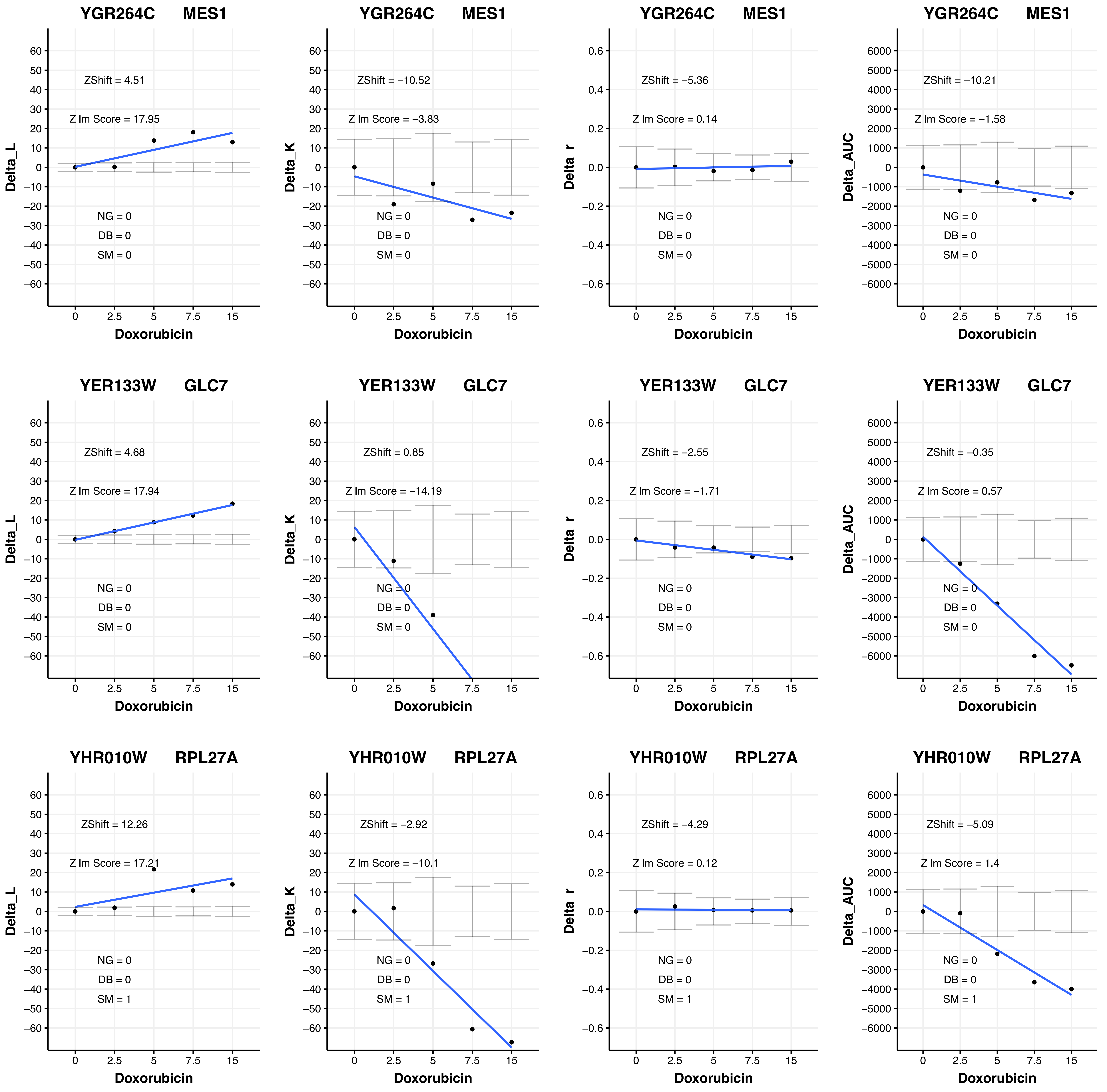

YGR264C MES1
ZShift = 4.51
Z lm Score = 17.95
NG = 0
DB = 0
SM = 0
Delta_L
Doxorubicin
YGR264C MES1
ZShift = −10.52
Z lm Score = −3.83
NG = 0
DB = 0
SM = 0
Delta_K
Doxorubicin
YGR264C MES1
ZShift = −5.36
Z lm Score = 0.14
NG = 0
DB = 0
SM = 0
Delta_r
Doxorubicin
YGR264C MES1
ZShift = −10.21
Z lm Score = −1.58
NG = 0
DB = 0
SM = 0
Delta_AUC
Doxorubicin
YER133W GLC7
ZShift = 4.68
Z lm Score = 17.94
NG = 0
DB = 0
SM = 0
Delta_L
Doxorubicin
YER133W GLC7
ZShift = 0.85
Z lm Score = −14.19
NG = 0
DB = 0
SM = 0
Delta_K
Doxorubicin
YER133W GLC7
ZShift = −2.55
Z lm Score = −1.71
NG = 0
DB = 0
SM = 0
Delta_r
Doxorubicin
YER133W GLC7
ZShift = −0.35
Z lm Score = 0.57
NG = 0
DB = 0
SM = 0
Delta_AUC
Doxorubicin
YHR010W RPL27A
ZShift = 12.26
Z lm Score = 17.21
NG = 0
DB = 0
SM = 1
Delta_L
Doxorubicin
YHR010W RPL27A
ZShift = −2.92
Z lm Score = −10.1
NG = 0
DB = 0
SM = 1
Delta_K
Doxorubicin
YHR010W RPL27A
ZShift = −4.29
Z lm Score = 0.12
NG = 0
DB = 0
SM = 1
Delta_r
Doxorubicin
YHR010W RPL27A
ZShift = −5.09
Z lm Score = 1.4
NG = 0
DB = 0
SM = 1
Delta_AUC
Doxorubicin

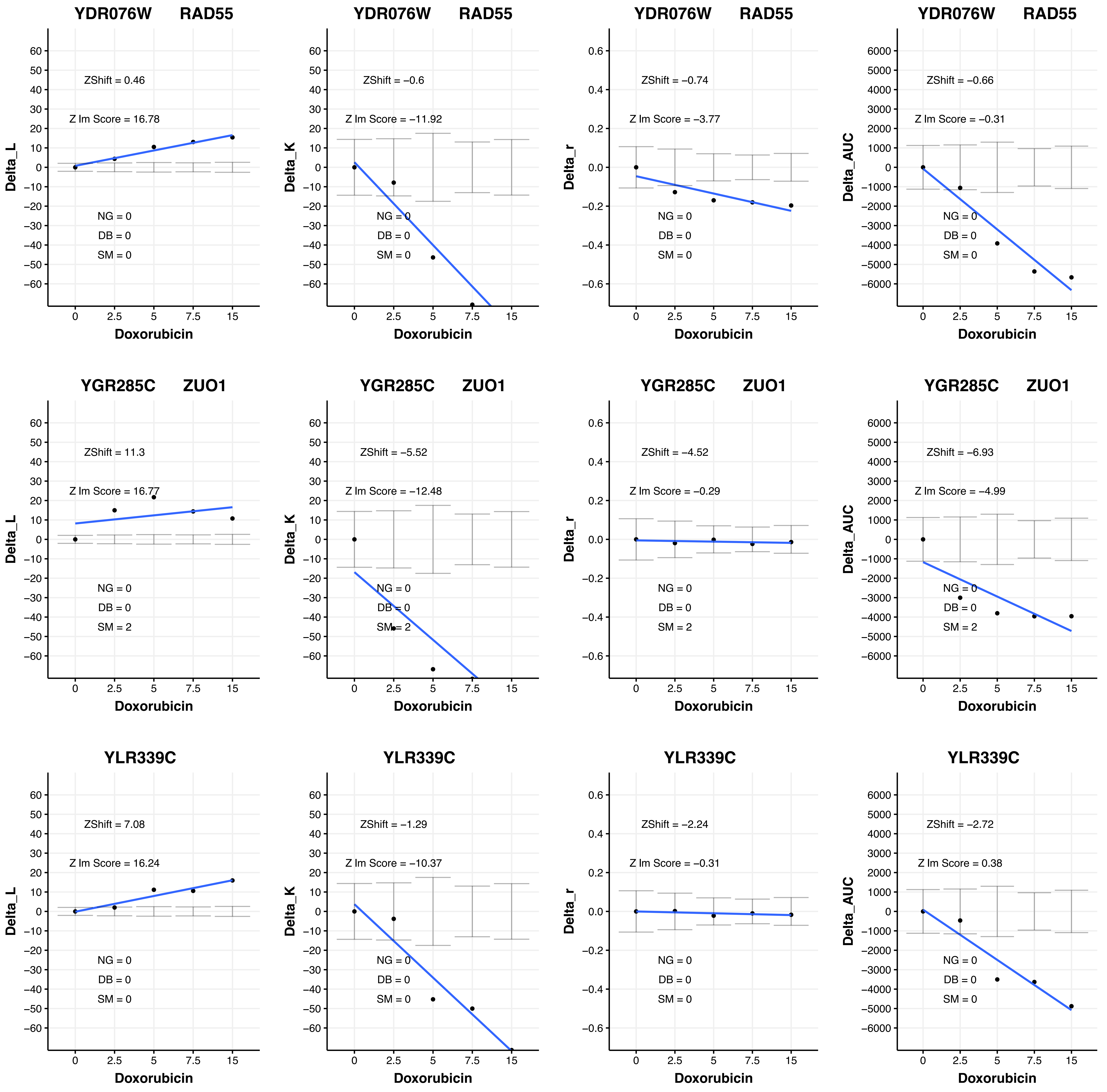
YDR076W RAD55
ZShift = 0.46
Z lm Score = 16.78
Delta_L
NG = 0
DB = 0
SM = 0
Doxorubicin
YDR076W RAD55
ZShift = −0.6
Z lm Score = −11.92
Delta_K
NG = 0
DB = 0
SM = 0
Doxorubicin
YDR076W RAD55
ZShift = −0.74
Z lm Score = −3.77
Delta_r
NG = 0
DB = 0
SM = 0
Doxorubicin
YDR076W RAD55
ZShift = −0.66
Z lm Score = −0.31
Delta_AUC
NG = 0
DB = 0
SM = 0
Doxorubicin
YGR285C ZUO1
ZShift = 11.3
Z lm Score = 16.77
Delta_L
NG = 0
DB = 0
SM = 2
Doxorubicin
YGR285C ZUO1
ZShift = −5.52
Z lm Score = −12.48
Delta_K
NG = 0
DB = 0
SM = 2
Doxorubicin
YGR285C ZUO1
ZShift = −4.52
Z lm Score = −0.29
Delta_r
NG = 0
DB = 0
SM = 2
Doxorubicin
YGR285C ZUO1
ZShift = −6.93
Z lm Score = −4.99
Delta_AUC
NG = 0
DB = 0
SM = 2
Doxorubicin
YLR339C
ZShift = 7.08
Z lm Score = 16.24
Delta_L
NG = 0
DB = 0
SM = 0
Doxorubicin
YLR339C
ZShift = −1.29
Z lm Score = −10.37
Delta_K
NG = 0
DB = 0
SM = 0
Doxorubicin
YLR339C
ZShift = −2.24
Z lm Score = −0.31
Delta_r
NG = 0
DB = 0
SM = 0
Doxorubicin
YLR339C
ZShift = −2.72
Z lm Score = 0.38
Delta_AUC
NG = 0
DB = 0
SM = 0
Doxorubicin

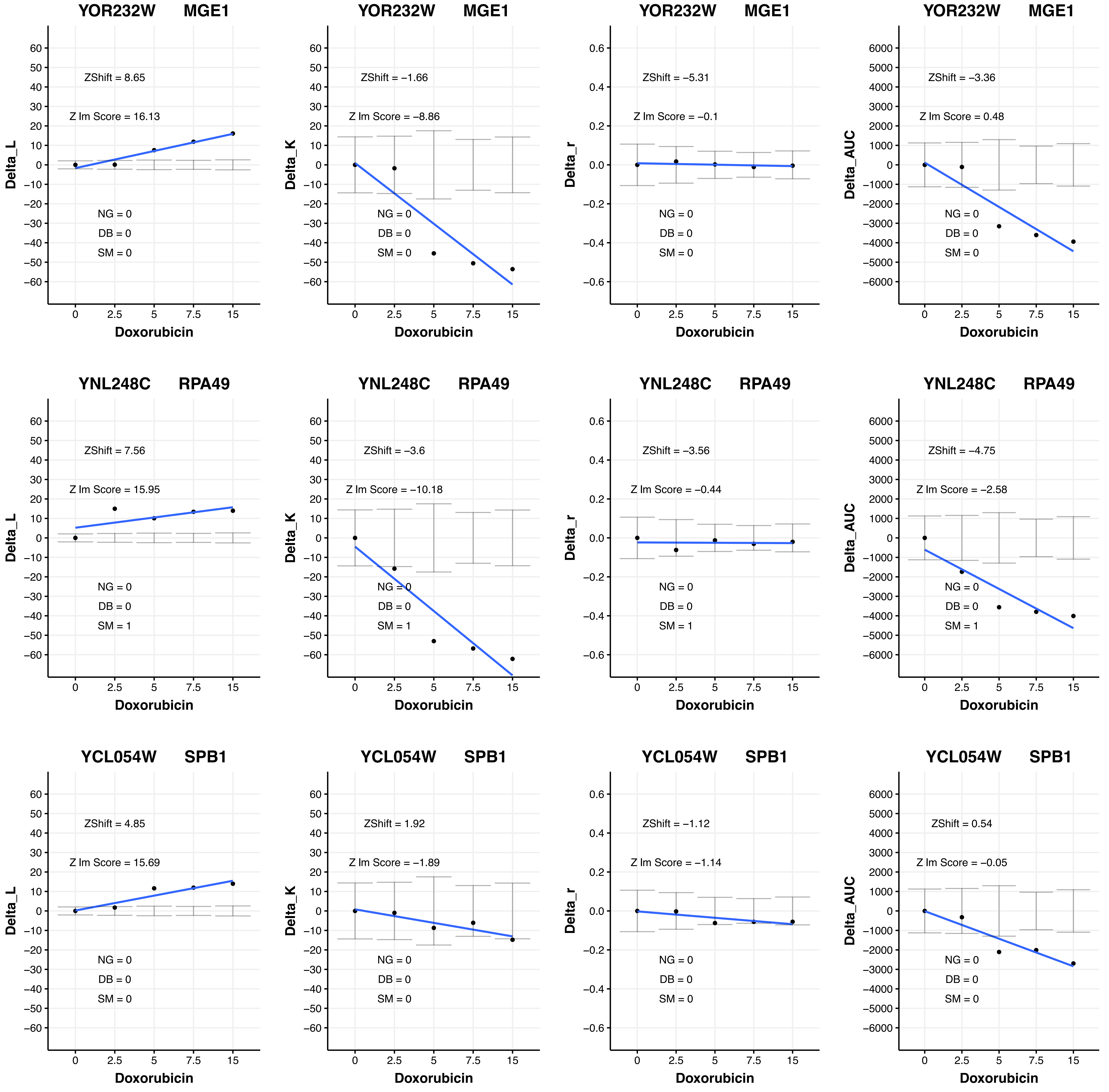

YOR232W MGE1
ZShift = 8.65
Z lm Score = 16.13
NG = 0
DB = 0
SM = 0
Delta_L
Doxorubicin
YOR232W MGE1
ZShift = −1.66
Z lm Score = −8.86
NG = 0
DB = 0
SM = 0
Delta_K
Doxorubicin
YOR232W MGE1
ZShift = −5.31
Z lm Score = −0.1
NG = 0
DB = 0
SM = 0
Delta_r
Doxorubicin
YOR232W MGE1
ZShift = −3.36
Z lm Score = 0.48
NG = 0
DB = 0
SM = 0
Delta_AUC
Doxorubicin
YNL248C RPA49
ZShift = 7.56
Z lm Score = 15.95
NG = 0
DB = 0
SM = 1
Delta_L
Doxorubicin
YNL248C RPA49
ZShift = −3.6
Z lm Score = −10.18
NG = 0
DB = 0
SM = 1
Delta_K
Doxorubicin
YNL248C RPA49
ZShift = −3.56
Z lm Score = −0.44
NG = 0
DB = 0
SM = 1
Delta_r
Doxorubicin
YNL248C RPA49
ZShift = −4.75
Z lm Score = −2.58
NG = 0
DB = 0
SM = 1
Delta_AUC
Doxorubicin
YCL054W SPB1
ZShift = 4.85
Z lm Score = 15.69
NG = 0
DB = 0
SM = 0
Delta_L
Doxorubicin
YCL054W SPB1
ZShift = 1.92
Z lm Score = −1.89
NG = 0
DB = 0
SM = 0
Delta_K
Doxorubicin
YCL054W SPB1
ZShift = −1.12
Z lm Score = −1.14
NG = 0
DB = 0
SM = 0
Delta_r
Doxorubicin
YCL054W SPB1
ZShift = 0.54
Z lm Score = −0.05
NG = 0
DB = 0
SM = 0
Delta_AUC
Doxorubicin

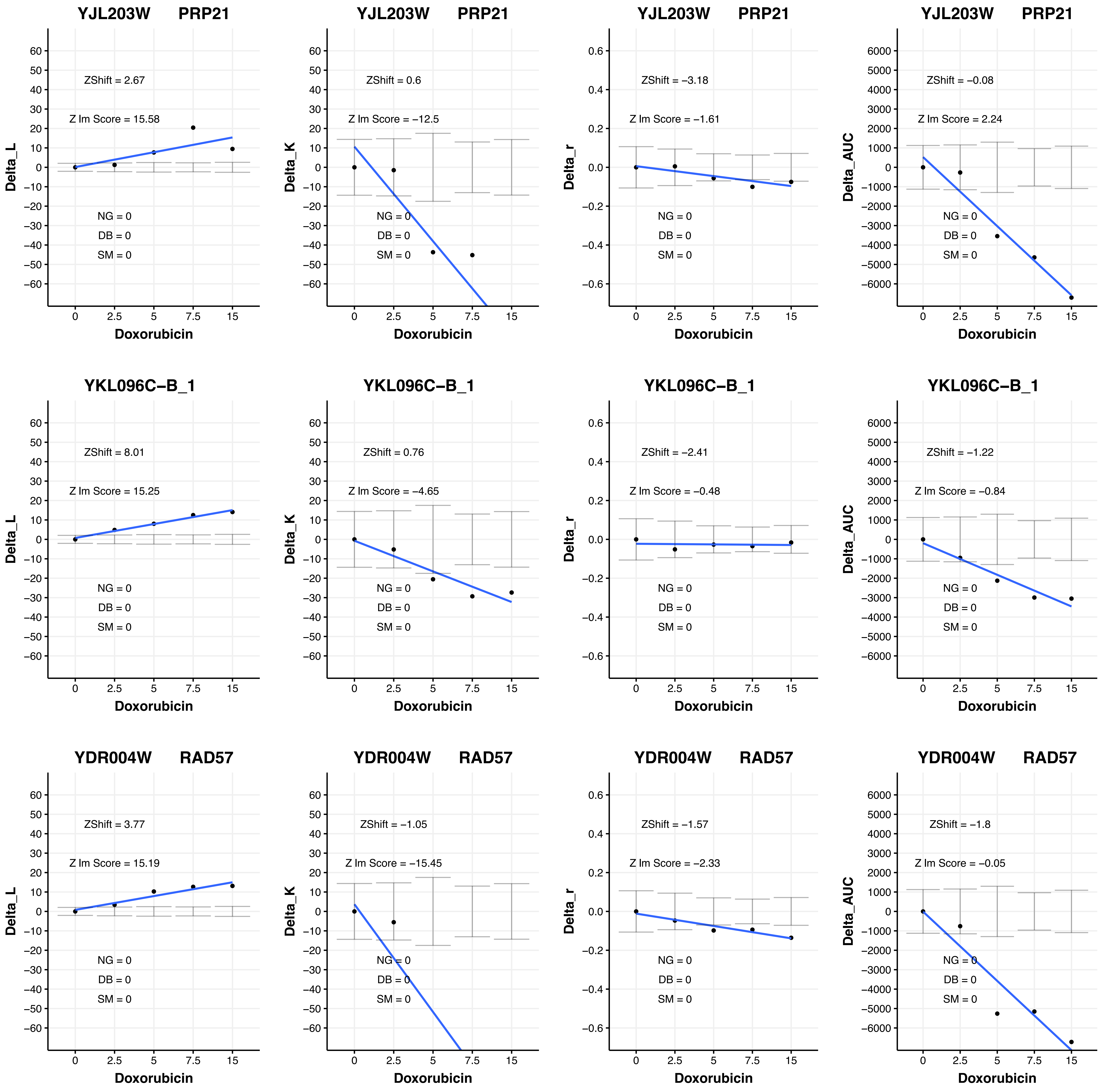
YJL203W PRP21
ZShift = 2.67
Z lm Score = 15.58
NG = 0
DB = 0
SM = 0
Delta_L
Doxorubicin
YJL203W PRP21
ZShift = 0.6
Z lm Score = −12.5
NG = 0
DB = 0
SM = 0
Delta_K
Doxorubicin
YJL203W PRP21
ZShift = −3.18
Z lm Score = −1.61
NG = 0
DB = 0
SM = 0
Delta_r
Doxorubicin
YJL203W PRP21
ZShift = −0.08
Z lm Score = 2.24
NG = 0
DB = 0
SM = 0
Delta_AUC
Doxorubicin
YKL096C−B_1
ZShift = 8.01
Z lm Score = 15.25
NG = 0
DB = 0
SM = 0
Delta_L
Doxorubicin
YKL096C−B_1
ZShift = 0.76
Z lm Score = −4.65
NG = 0
DB = 0
SM = 0
Delta_K
Doxorubicin
YKL096C−B_1
ZShift = −2.41
Z lm Score = −0.48
NG = 0
DB = 0
SM = 0
Delta_r
Doxorubicin
YKL096C−B_1
ZShift = −1.22
Z lm Score = −0.84
NG = 0
DB = 0
SM = 0
Delta_AUC
Doxorubicin
YDR004W RAD57
ZShift = 3.77
Z lm Score = 15.19
NG = 0
DB = 0
SM = 0
Delta_L
Doxorubicin
YDR004W RAD57
ZShift = −1.05
Z lm Score = −15.45
NG = 0
DB = 0
SM = 0
Delta_K
Doxorubicin
YDR004W RAD57
ZShift = −1.57
Z lm Score = −2.33
NG = 0
DB = 0
SM = 0
Delta_r
Doxorubicin
YDR004W RAD57
ZShift = −1.8
Z lm Score = −0.05
NG = 0
DB = 0
SM = 0
Delta_AUC
Doxorubicin

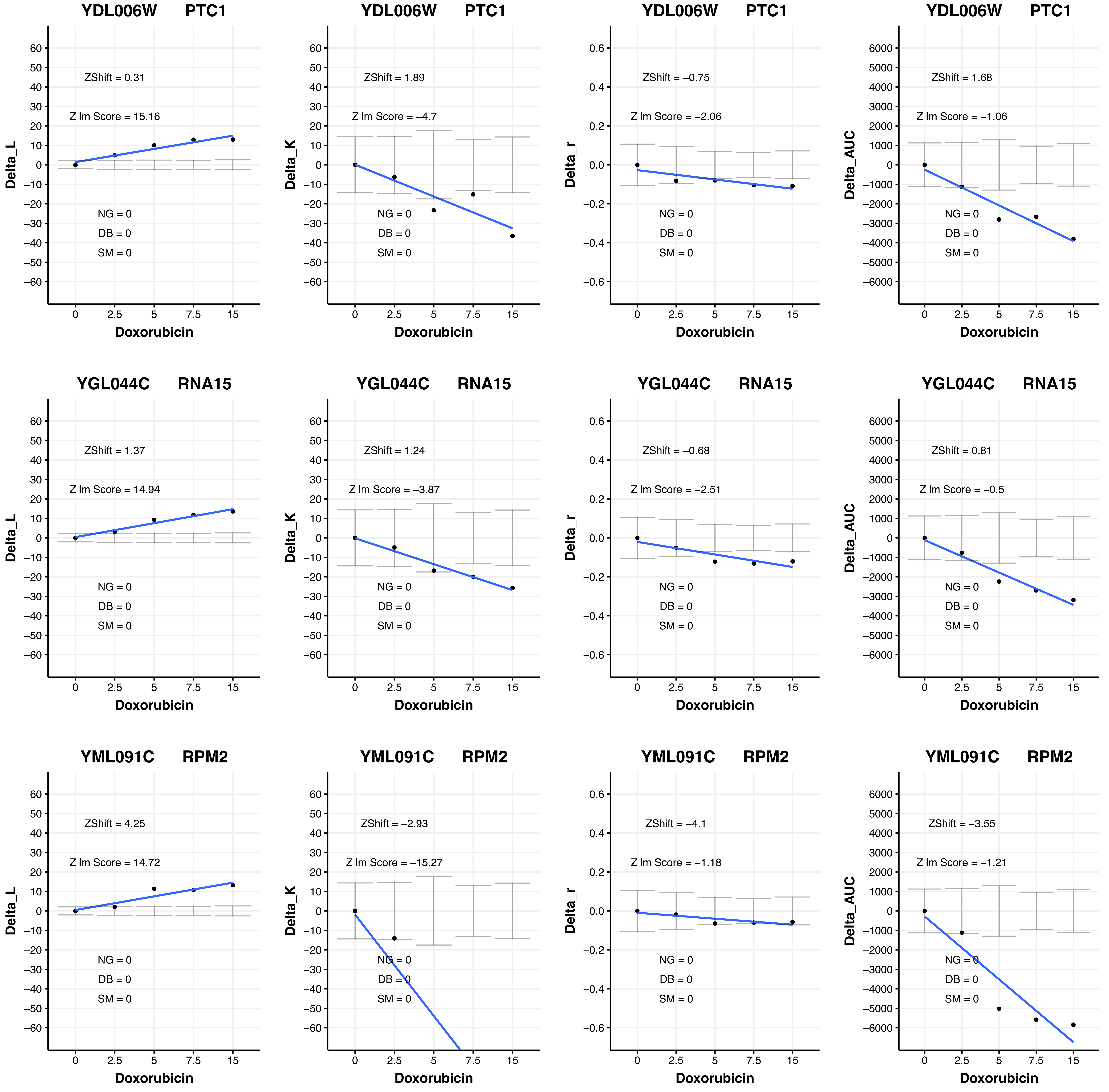

YDL006W PTC1
ZShift = 0.31
Z lm Score = 15.16
NG = 0
DB = 0
SM = 0
Delta_L
Doxorubicin
0 2.5 5 7.5 15
60 50 40 30 20 10 0 −10 −20 −30 −40 −50 −60
YDL006W PTC1
ZShift = 1.89
Z lm Score = −4.7
NG = 0
DB = 0
SM = 0
Delta_K
Doxorubicin
0 2.5 5 7.5 15
60 50 40 30 20 10 0 −10 −20 −30 −40 −50 −60
YDL006W PTC1
ZShift = −0.75
Z lm Score = −2.06
NG = 0
DB = 0
SM = 0
Delta_r
Doxorubicin
0 2.5 5 7.5 15
0.6 0.4 0.2 0.0 −0.2 −0.4 −0.6
YDL006W PTC1
ZShift = 1.68
Z lm Score = −1.06
NG = 0
DB = 0
SM = 0
Delta_AUC
Doxorubicin
0 2.5 5 7.5 15
6000 5000 4000 3000 2000 1000 0 −1000 −2000 −3000 −4000 −5000 −6000
YGL044C RNA15
ZShift = 1.37
Z lm Score = 14.94
NG = 0
DB = 0
SM = 0
Delta_L
Doxorubicin
0 2.5 5 7.5 15
60 50 40 30 20 10 0 −10 −20 −30 −40 −50 −60
YGL044C RNA15
ZShift = 1.24
Z lm Score = −3.87
NG = 0
DB = 0
SM = 0
Delta_K
Doxorubicin
0 2.5 5 7.5 15
60 50 40 30 20 10 0 −10 −20 −30 −40 −50 −60
YGL044C RNA15
ZShift = −0.68
Z lm Score = −2.51
NG = 0
DB = 0
SM = 0
Delta_r
Doxorubicin
0 2.5 5 7.5 15
0.6 0.4 0.2 0.0 −0.2 −0.4 −0.6
YGL044C RNA15
ZShift = 0.81
Z lm Score = −0.5
NG = 0
DB = 0
SM = 0
Delta_AUC
Doxorubicin
0 2.5 5 7.5 15
6000 5000 4000 3000 2000 1000 0 −1000 −2000 −3000 −4000 −5000 −6000
YML091C RPM2
ZShift = 4.25
Z lm Score = 14.72
NG = 0
DB = 0
SM = 0
Delta_L
Doxorubicin
0 2.5 5 7.5 15
60 50 40 30 20 10 0 −10 −20 −30 −40 −50 −60
YML091C RPM2
ZShift = −2.93
Z lm Score = −15.27
NG = 0
DB = 0
SM = 0
Delta_K
Doxorubicin
0 2.5 5 7.5 15
60 50 40 30 20 10 0 −10 −20 −30 −40 −50 −60
YML091C RPM2
ZShift = −4.1
Z lm Score = −1.18
NG = 0
DB = 0
SM = 0
Delta_r
Doxorubicin
0 2.5 5 7.5 15
0.6 0.4 0.2 0.0 −0.2 −0.4 −0.6
YML091C RPM2
ZShift = −3.55
Z lm Score = −1.21
NG = 0
DB = 0
SM = 0
Delta_AUC
Doxorubicin
0 2.5 5 7.5 15
6000 5000 4000 3000 2000 1000 0 −1000 −2000 −3000 −4000 −5000 −6000

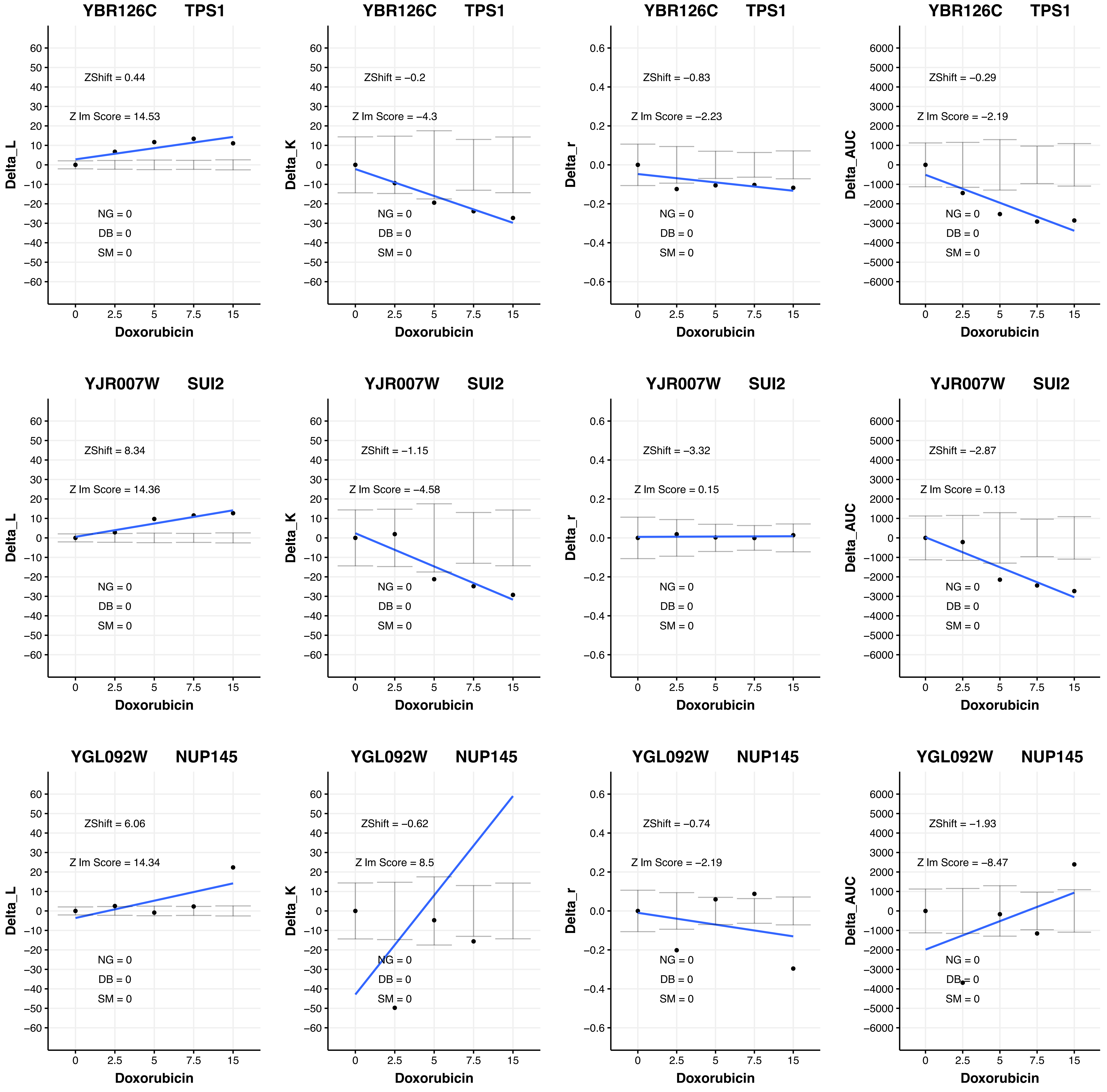
YBR126C TPS1
ZShift = 0.44
Z lm Score = 14.53
NG = 0
DB = 0
SM = 0
Delta_L
Doxorubicin
YBR126C TPS1
ZShift = −0.2
Z lm Score = −4.3
NG = 0
DB = 0
SM = 0
Delta_K
Doxorubicin
YBR126C TPS1
ZShift = −0.83
Z lm Score = −2.23
NG = 0
DB = 0
SM = 0
Delta_r
Doxorubicin
YBR126C TPS1
ZShift = −0.29
Z lm Score = −2.19
NG = 0
DB = 0
SM = 0
Delta_AUC
Doxorubicin
YJR007W SUI2
ZShift = 8.34
Z lm Score = 14.36
NG = 0
DB = 0
SM = 0
Delta_L
Doxorubicin
YJR007W SUI2
ZShift = −1.15
Z lm Score = −4.58
NG = 0
DB = 0
SM = 0
Delta_K
Doxorubicin
YJR007W SUI2
ZShift = −3.32
Z lm Score = 0.15
NG = 0
DB = 0
SM = 0
Delta_r
Doxorubicin
YJR007W SUI2
ZShift = −2.87
Z lm Score = 0.13
NG = 0
DB = 0
SM = 0
Delta_AUC
Doxorubicin
YGL092W NUP145
ZShift = 6.06
Z lm Score = 14.34
NG = 0
DB = 0
SM = 0
Delta_L
Doxorubicin
YGL092W NUP145
ZShift = −0.62
Z lm Score = 8.5
NG = 0
DB = 0
SM = 0
Delta_K
Doxorubicin
YGL092W NUP145
ZShift = −0.74
Z lm Score = −2.19
NG = 0
DB = 0
SM = 0
Delta_r
Doxorubicin
YGL092W NUP145
ZShift = −1.93
Z lm Score = −8.47
NG = 0
DB = 0
SM = 0
Delta_AUC
Doxorubicin

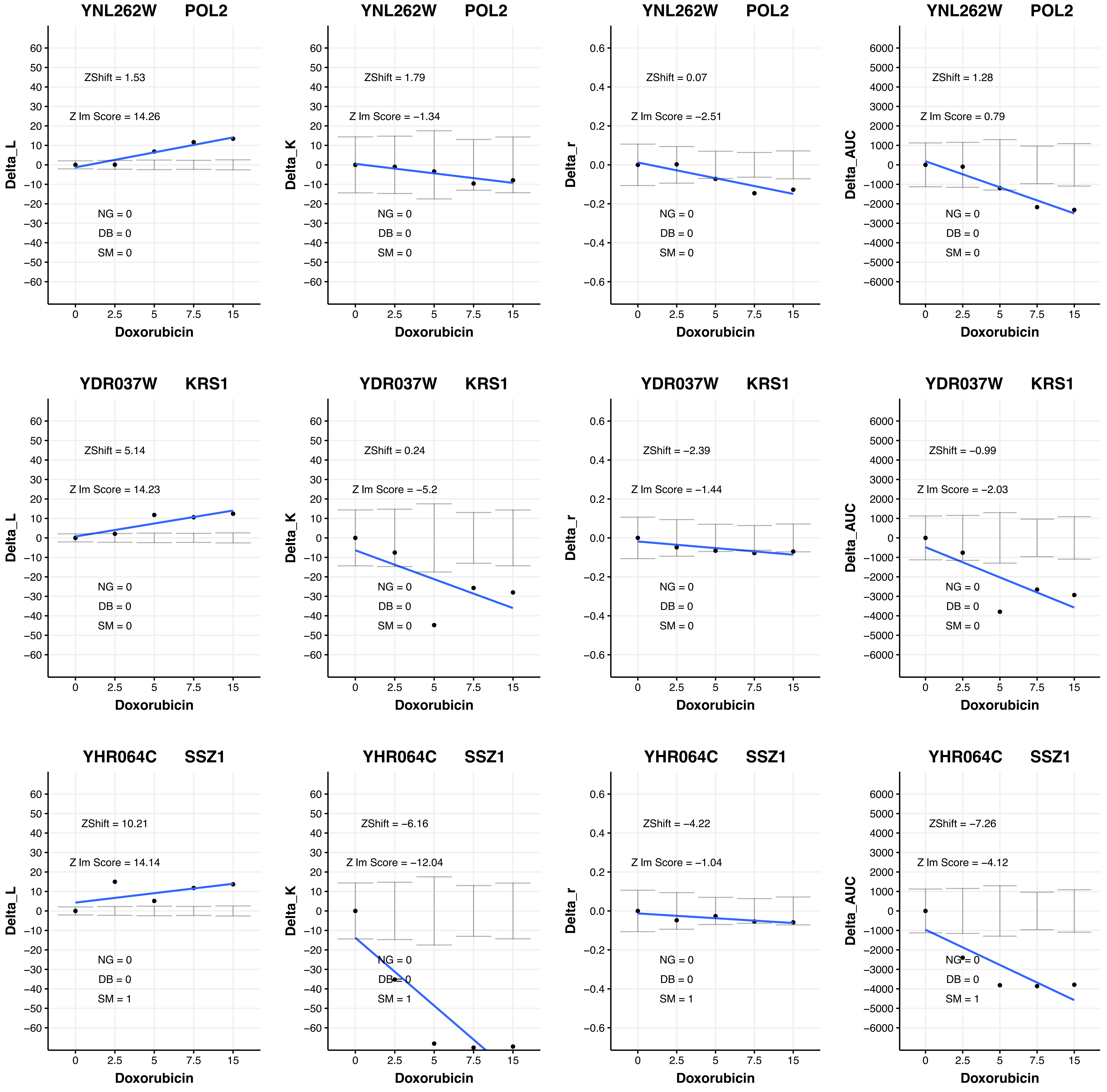
YNL262W POL2
ZShift = 1.53
Z lm Score = 14.26
NG = 0
DB = 0
SM = 0
Delta_L
Doxorubicin
YNL262W POL2
ZShift = 1.79
Z lm Score = −1.34
NG = 0
DB = 0
SM = 0
Delta_K
Doxorubicin
YNL262W POL2
ZShift = 0.07
Z lm Score = −2.51
NG = 0
DB = 0
SM = 0
Delta_r
Doxorubicin
YNL262W POL2
ZShift = 1.28
Z lm Score = 0.79
NG = 0
DB = 0
SM = 0
Delta_AUC
Doxorubicin
YDR037W KRS1
ZShift = 5.14
Z lm Score = 14.23
NG = 0
DB = 0
SM = 0
Delta_L
Doxorubicin
YDR037W KRS1
ZShift = 0.24
Z lm Score = −5.2
NG = 0
DB = 0
SM = 0
Delta_K
Doxorubicin
YDR037W KRS1
ZShift = −2.39
Z lm Score = −1.44
NG = 0
DB = 0
SM = 0
Delta_r
Doxorubicin
YDR037W KRS1
ZShift = −0.99
Z lm Score = −2.03
NG = 0
DB = 0
SM = 0
Delta_AUC
Doxorubicin
YHR064C SSZ1
ZShift = 10.21
Z lm Score = 14.14
NG = 0
DB = 0
SM = 1
Delta_L
Doxorubicin
YHR064C SSZ1
ZShift = −6.16
Z lm Score = −12.04
NG = 0
DB = 0
SM = 1
Delta_K
Doxorubicin
YHR064C SSZ1
ZShift = −4.22
Z lm Score = −1.04
NG = 0
DB = 0
SM = 1
Delta_r
Doxorubicin
YHR064C SSZ1
ZShift = −7.26
Z lm Score = −4.12
NG = 0
DB = 0
SM = 1
Delta_AUC
Doxorubicin

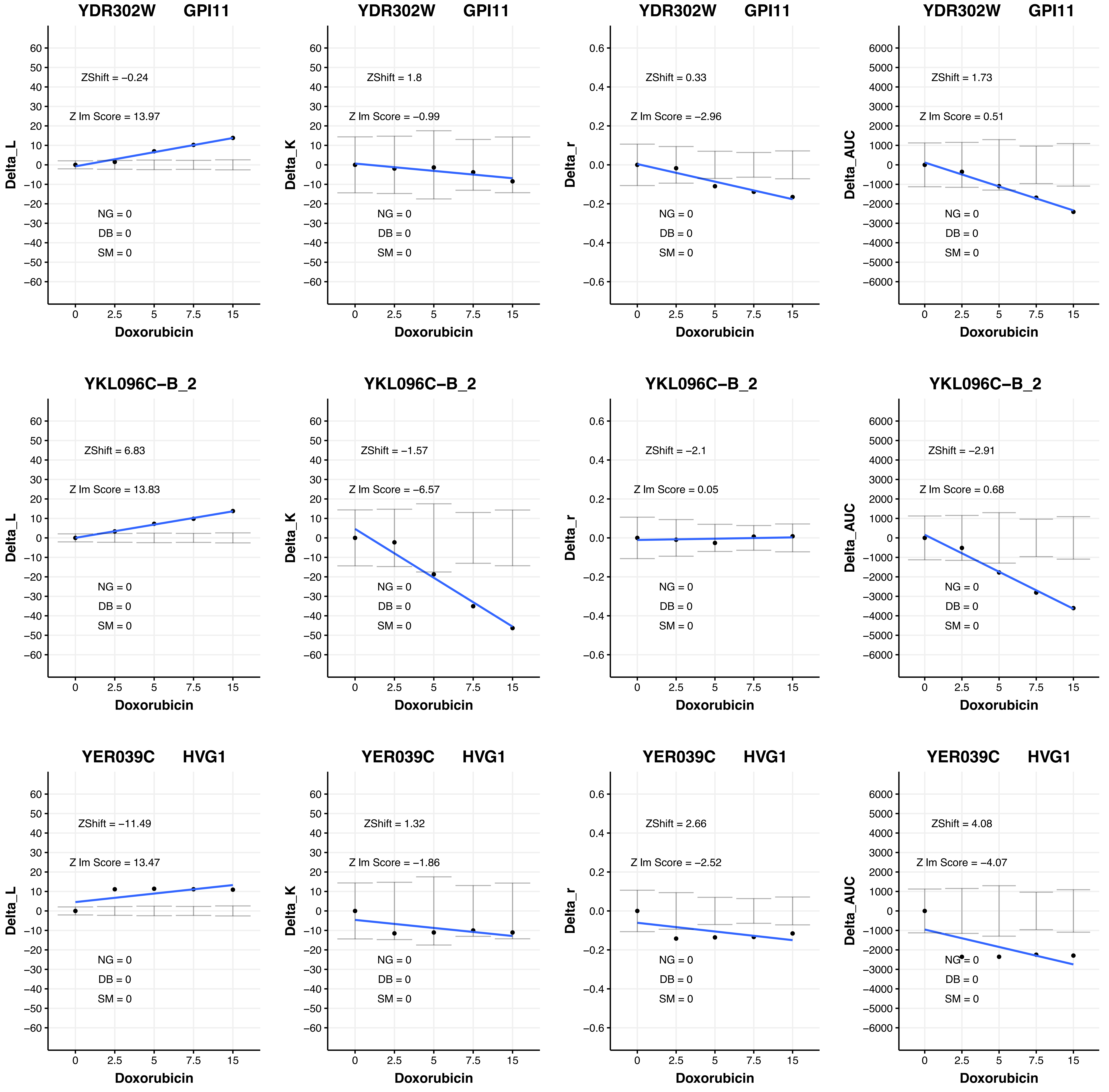

YDR302W GPI11
ZShift = −0.24
Z lm Score = 13.97
NG = 0
DB = 0
SM = 0
Delta_L
Doxorubicin
YDR302W GPI11
ZShift = 1.8
Z lm Score = −0.99
NG = 0
DB = 0
SM = 0
Delta_K
Doxorubicin
YDR302W GPI11
ZShift = 0.33
Z lm Score = −2.96
NG = 0
DB = 0
SM = 0
Delta_r
Doxorubicin
YDR302W GPI11
ZShift = 1.73
Z lm Score = 0.51
NG = 0
DB = 0
SM = 0
Delta_AUC
Doxorubicin
YKL096C−B_2
ZShift = 6.83
Z lm Score = 13.83
NG = 0
DB = 0
SM = 0
Delta_L
Doxorubicin
YKL096C−B_2
ZShift = −1.57
Z lm Score = −6.57
NG = 0
DB = 0
SM = 0
Delta_K
Doxorubicin
YKL096C−B_2
ZShift = −2.1
Z lm Score = 0.05
NG = 0
DB = 0
SM = 0
Delta_r
Doxorubicin
YKL096C−B_2
ZShift = −2.91
Z lm Score = 0.68
NG = 0
DB = 0
SM = 0
Delta_AUC
Doxorubicin
YER039C HVG1
ZShift = −11.49
Z lm Score = 13.47
NG = 0
DB = 0
SM = 0
Delta_L
Doxorubicin
YER039C HVG1
ZShift = 1.32
Z lm Score = −1.86
NG = 0
DB = 0
SM = 0
Delta_K
Doxorubicin
YER039C HVG1
ZShift = 2.66
Z lm Score = −2.52
NG = 0
DB = 0
SM = 0
Delta_r
Doxorubicin
YER039C HVG1
ZShift = 4.08
Z lm Score = −4.07
NG = 0
DB = 0
SM = 0
Delta_AUC
Doxorubicin

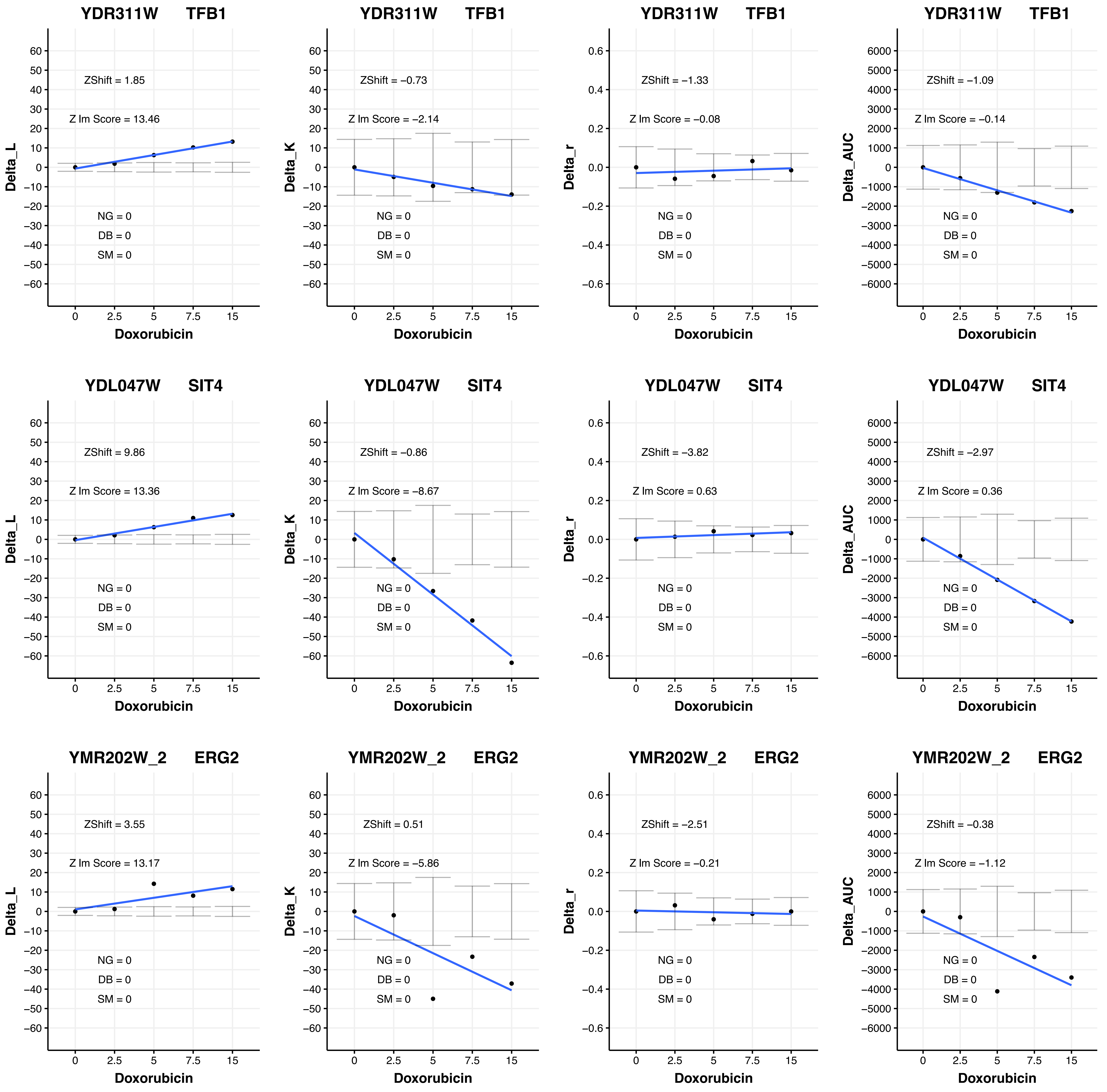

YDR311W TFB1
ZShift = 1.85
Z lm Score = 13.46
NG = 0
DB = 0
SM = 0
Delta_L
Doxorubicin
YDR311W TFB1
ZShift = −0.73
Z lm Score = −2.14
NG = 0
DB = 0
SM = 0
Delta_K
Doxorubicin
YDR311W TFB1
ZShift = −1.33
Z lm Score = −0.08
NG = 0
DB = 0
SM = 0
Delta_r
Doxorubicin
YDR311W TFB1
ZShift = −1.09
Z lm Score = −0.14
NG = 0
DB = 0
SM = 0
Delta_AUC
Doxorubicin
YDL047W SIT4
ZShift = 9.86
Z lm Score = 13.36
NG = 0
DB = 0
SM = 0
Delta_L
Doxorubicin
YDL047W SIT4
ZShift = −0.86
Z lm Score = −8.67
NG = 0
DB = 0
SM = 0
Delta_K
Doxorubicin
YDL047W SIT4
ZShift = −3.82
Z lm Score = 0.63
NG = 0
DB = 0
SM = 0
Delta_r
Doxorubicin
YDL047W SIT4
ZShift = −2.97
Z lm Score = 0.36
NG = 0
DB = 0
SM = 0
Delta_AUC
Doxorubicin
YMR202W_2 ERG2
ZShift = 3.55
Z lm Score = 13.17
NG = 0
DB = 0
SM = 0
Delta_L
Doxorubicin
YMR202W_2 ERG2
ZShift = 0.51
Z lm Score = −5.86
NG = 0
DB = 0
SM = 0
Delta_K
Doxorubicin
YMR202W_2 ERG2
ZShift = −2.51
Z lm Score = −0.21
NG = 0
DB = 0
SM = 0
Delta_r
Doxorubicin
YMR202W_2 ERG2
ZShift = −0.38
Z lm Score = −1.12
NG = 0
DB = 0
SM = 0
Delta_AUC
Doxorubicin

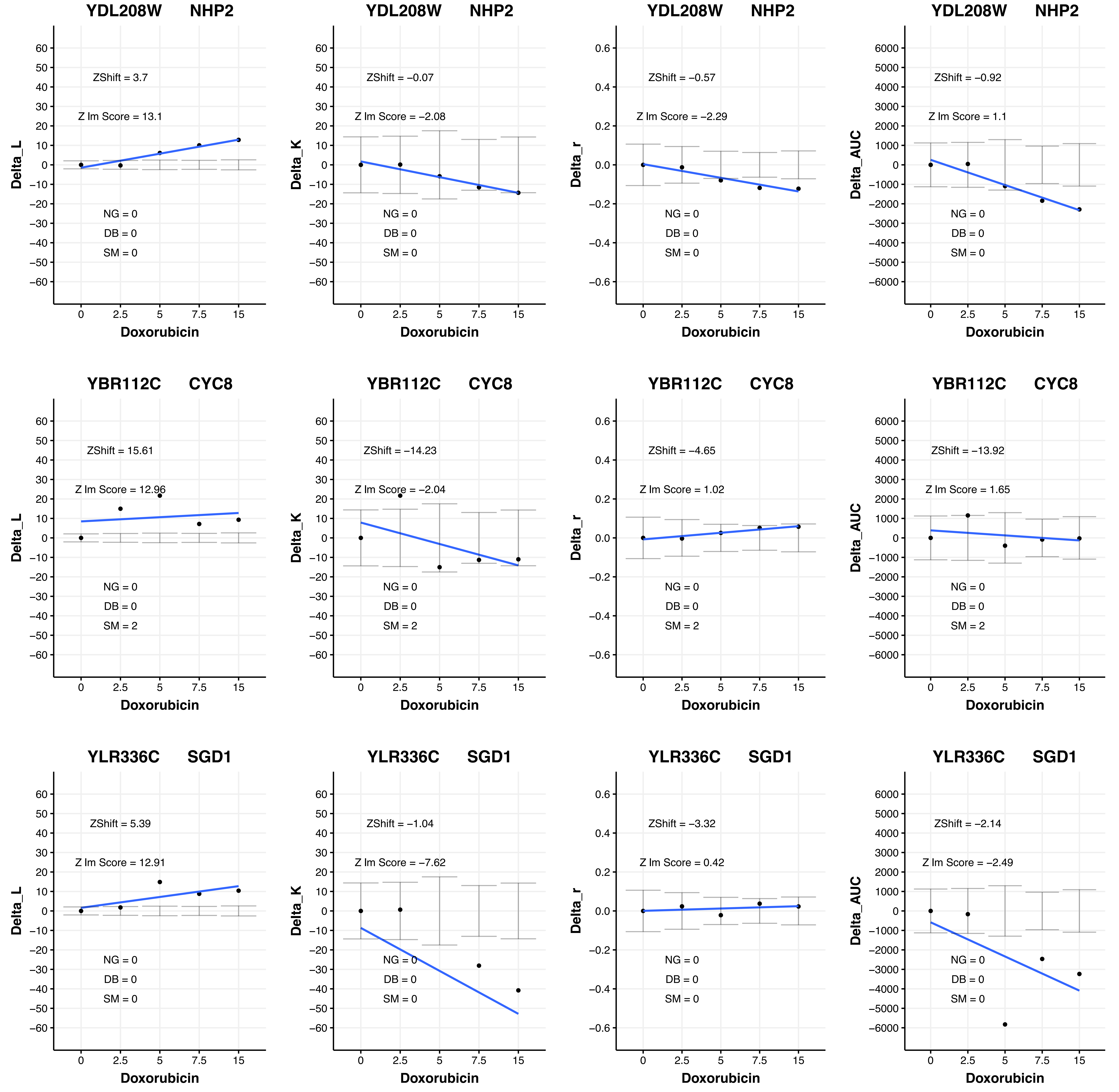
YDL208W NHP2
ZShift = 3.7
Z lm Score = 13.1
NG = 0
DB = 0
SM = 0
Delta_L
Doxorubicin
YDL208W NHP2
ZShift = −0.07
Z lm Score = −2.08
NG = 0
DB = 0
SM = 0
Delta_K
Doxorubicin
YDL208W NHP2
ZShift = −0.57
Z lm Score = −2.29
NG = 0
DB = 0
SM = 0
Delta_r
Doxorubicin
YDL208W NHP2
ZShift = −0.92
Z lm Score = 1.1
NG = 0
DB = 0
SM = 0
Delta_AUC
Doxorubicin
YBR112C CYC8
ZShift = 15.61
Z lm Score = 12.96
NG = 0
DB = 0
SM = 2
Delta_L
Doxorubicin
YBR112C CYC8
ZShift = −14.23
Z lm Score = −2.04
NG = 0
DB = 0
SM = 2
Delta_K
Doxorubicin
YBR112C CYC8
ZShift = −4.65
Z lm Score = 1.02
NG = 0
DB = 0
SM = 2
Delta_r
Doxorubicin
YBR112C CYC8
ZShift = −13.92
Z lm Score = 1.65
NG = 0
DB = 0
SM = 2
Delta_AUC
Doxorubicin
YLR336C SGD1
ZShift = 5.39
Z lm Score = 12.91
NG = 0
DB = 0
SM = 0
Delta_L
Doxorubicin
YLR336C SGD1
ZShift = −1.04
Z lm Score = −7.62
NG = 0
DB = 0
SM = 0
Delta_K
Doxorubicin
YLR336C SGD1
ZShift = −3.32
Z lm Score = 0.42
NG = 0
DB = 0
SM = 0
Delta_r
Doxorubicin
YLR336C SGD1
ZShift = −2.14
Z lm Score = −2.49
NG = 0
DB = 0
SM = 0
Delta_AUC
Doxorubicin

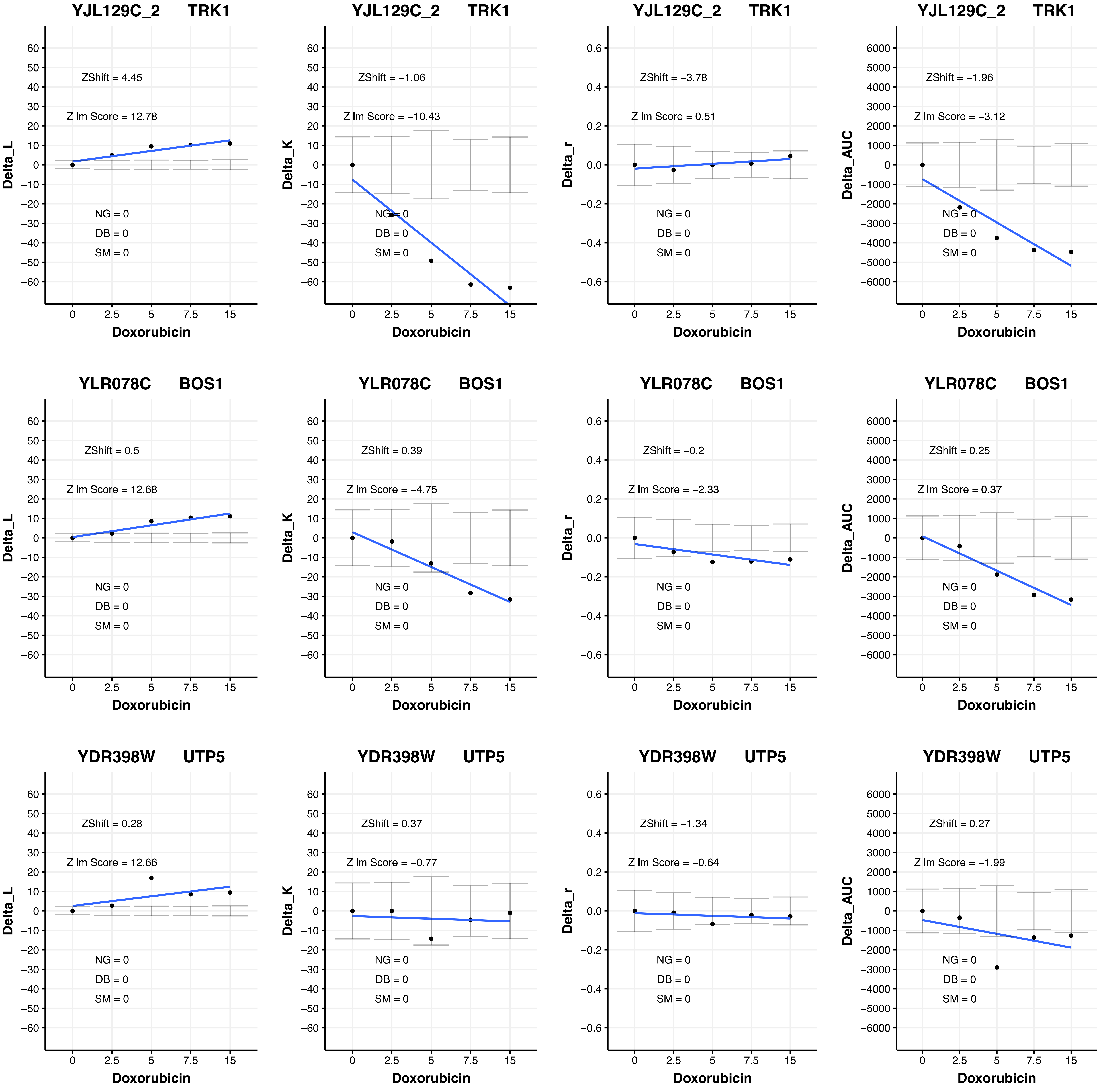
YJL129C_2 TRK1
ZShift = 4.45
Z lm Score = 12.78
NG = 0
DB = 0
SM = 0
Delta_L
Doxorubicin
YJL129C_2 TRK1
ZShift = −1.06
Z lm Score = −10.43
NG = 0
DB = 0
SM = 0
Delta_K
Doxorubicin
YJL129C_2 TRK1
ZShift = −3.78
Z lm Score = 0.51
NG = 0
DB = 0
SM = 0
Delta_r
Doxorubicin
YJL129C_2 TRK1
ZShift = −1.96
Z lm Score = −3.12
NG = 0
DB = 0
SM = 0
Delta_AUC
Doxorubicin
YLR078C BOS1
ZShift = 0.5
Z lm Score = 12.68
NG = 0
DB = 0
SM = 0
Delta_L
Doxorubicin
YLR078C BOS1
ZShift = 0.39
Z lm Score = −4.75
NG = 0
DB = 0
SM = 0
Delta_K
Doxorubicin
YLR078C BOS1
ZShift = −0.2
Z lm Score = −2.33
NG = 0
DB = 0
SM = 0
Delta_r
Doxorubicin
YLR078C BOS1
ZShift = 0.25
Z lm Score = 0.37
NG = 0
DB = 0
SM = 0
Delta_AUC
Doxorubicin
YDR398W UTP5
ZShift = 0.28
Z lm Score = 12.66
NG = 0
DB = 0
SM = 0
Delta_L
Doxorubicin
YDR398W UTP5
ZShift = 0.37
Z lm Score = −0.77
NG = 0
DB = 0
SM = 0
Delta_K
Doxorubicin
YDR398W UTP5
ZShift = −1.34
Z lm Score = −0.64
NG = 0
DB = 0
SM = 0
Delta_r
Doxorubicin
YDR398W UTP5
ZShift = 0.27
Z lm Score = −1.99
NG = 0
DB = 0
SM = 0
Delta_AUC
Doxorubicin

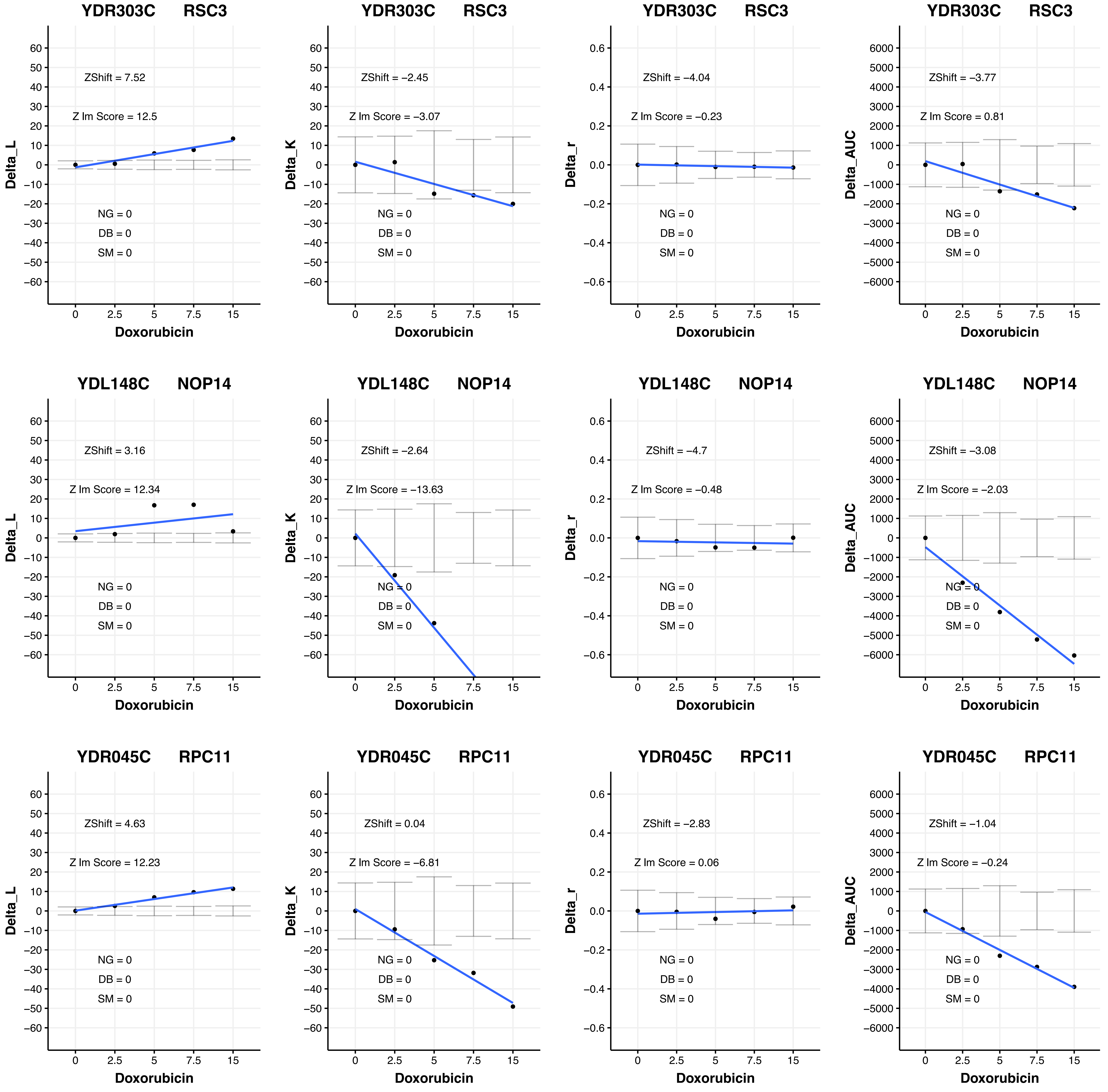
YDR303C RSC3
ZShift = 7.52
Z lm Score = 12.5
NG = 0
DB = 0
SM = 0
Delta_L
Doxorubicin
YDR303C RSC3
ZShift = −2.45
Z lm Score = −3.07
NG = 0
DB = 0
SM = 0
Delta_K
Doxorubicin
YDR303C RSC3
ZShift = −4.04
Z lm Score = −0.23
NG = 0
DB = 0
SM = 0
Delta_r
Doxorubicin
YDR303C RSC3
ZShift = −3.77
Z lm Score = 0.81
NG = 0
DB = 0
SM = 0
Delta_AUC
Doxorubicin
YDL148C NOP14
ZShift = 3.16
Z lm Score = 12.34
NG = 0
DB = 0
SM = 0
Delta_L
Doxorubicin
YDL148C NOP14
ZShift = −2.64
Z lm Score = −13.63
NG = 0
DB = 0
SM = 0
Delta_K
Doxorubicin
YDL148C NOP14
ZShift = −4.7
Z lm Score = −0.48
NG = 0
DB = 0
SM = 0
Delta_r
Doxorubicin
YDL148C NOP14
ZShift = −3.08
Z lm Score = −2.03
NG = 0
DB = 0
SM = 0
Delta_AUC
Doxorubicin
YDR045C RPC11
ZShift = 4.63
Z lm Score = 12.23
NG = 0
DB = 0
SM = 0
Delta_L
Doxorubicin
YDR045C RPC11
ZShift = 0.04
Z lm Score = −6.81
NG = 0
DB = 0
SM = 0
Delta_K
Doxorubicin
YDR045C RPC11
ZShift = −2.83
Z lm Score = 0.06
NG = 0
DB = 0
SM = 0
Delta_r
Doxorubicin
YDR045C RPC11
ZShift = −1.04
Z lm Score = −0.24
NG = 0
DB = 0
SM = 0
Delta_AUC
Doxorubicin

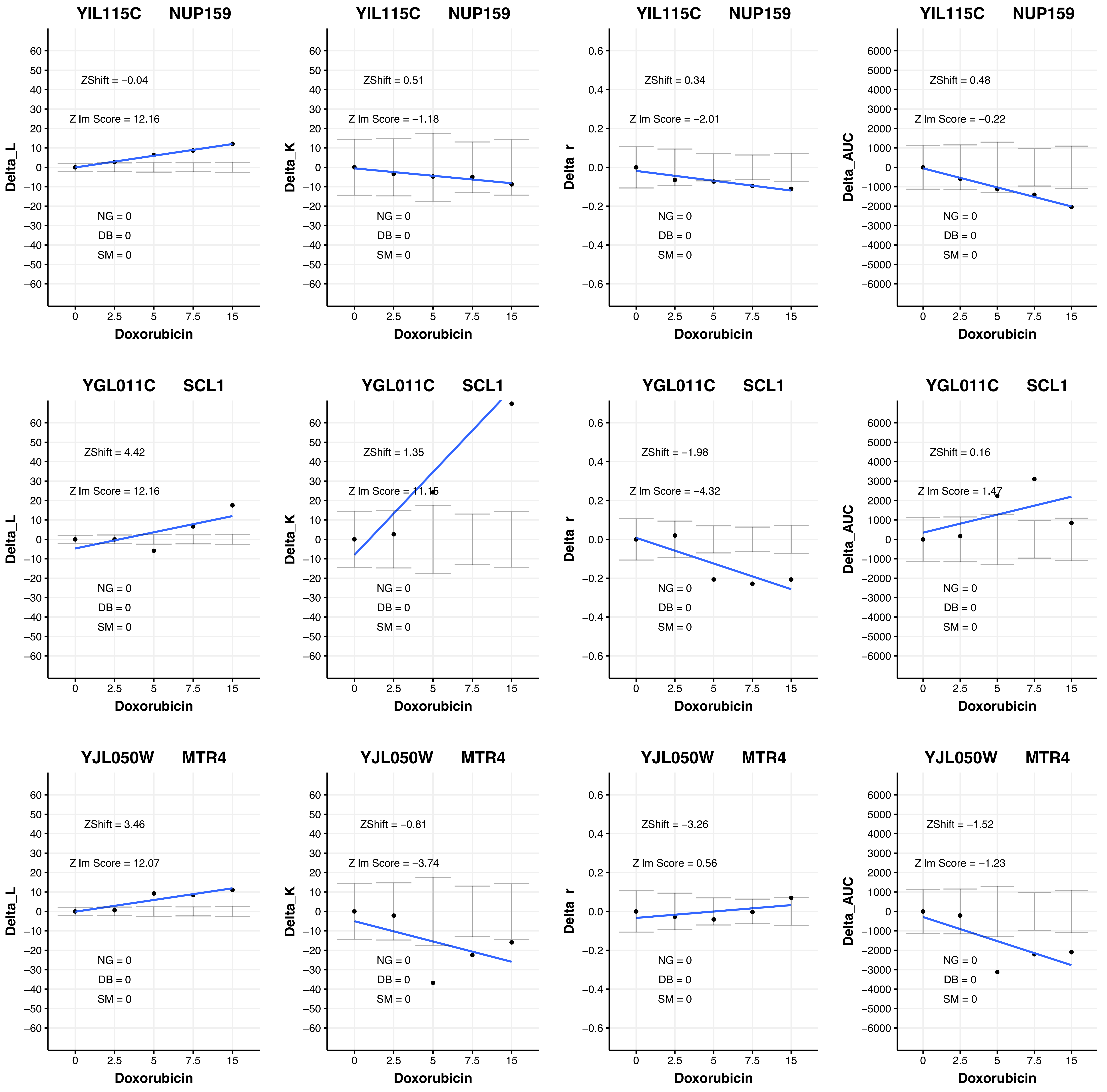

YIL115C NUP159
ZShift = −0.04
Z lm Score = 12.16
NG = 0
DB = 0
SM = 0
Delta_L
Doxorubicin
YIL115C NUP159
ZShift = 0.51
Z lm Score = −1.18
NG = 0
DB = 0
SM = 0
Delta_K
Doxorubicin
YIL115C NUP159
ZShift = 0.34
Z lm Score = −2.01
NG = 0
DB = 0
SM = 0
Delta_r
Doxorubicin
YIL115C NUP159
ZShift = 0.48
Z lm Score = −0.22
NG = 0
DB = 0
SM = 0
Delta_AUC
Doxorubicin
YGL011C SCL1
ZShift = 4.42
Z lm Score = 12.16
NG = 0
DB = 0
SM = 0
Delta_L
Doxorubicin
YGL011C SCL1
ZShift = 1.35
Z lm Score = 11.15
NG = 0
DB = 0
SM = 0
Delta_K
Doxorubicin
YGL011C SCL1
ZShift = −1.98
Z lm Score = −4.32
NG = 0
DB = 0
SM = 0
Delta_r
Doxorubicin
YGL011C SCL1
ZShift = 0.16
Z lm Score = 1.47
NG = 0
DB = 0
SM = 0
Delta_AUC
Doxorubicin
YJL050W MTR4
ZShift = 3.46
Z lm Score = 12.07
NG = 0
DB = 0
SM = 0
Delta_L
Doxorubicin
YJL050W MTR4
ZShift = −0.81
Z lm Score = −3.74
NG = 0
DB = 0
SM = 0
Delta_K
Doxorubicin
YJL050W MTR4
ZShift = −3.26
Z lm Score = 0.56
NG = 0
DB = 0
SM = 0
Delta_r
Doxorubicin
YJL050W MTR4
ZShift = −1.52
Z lm Score = −1.23
NG = 0
DB = 0
SM = 0
Delta_AUC
Doxorubicin

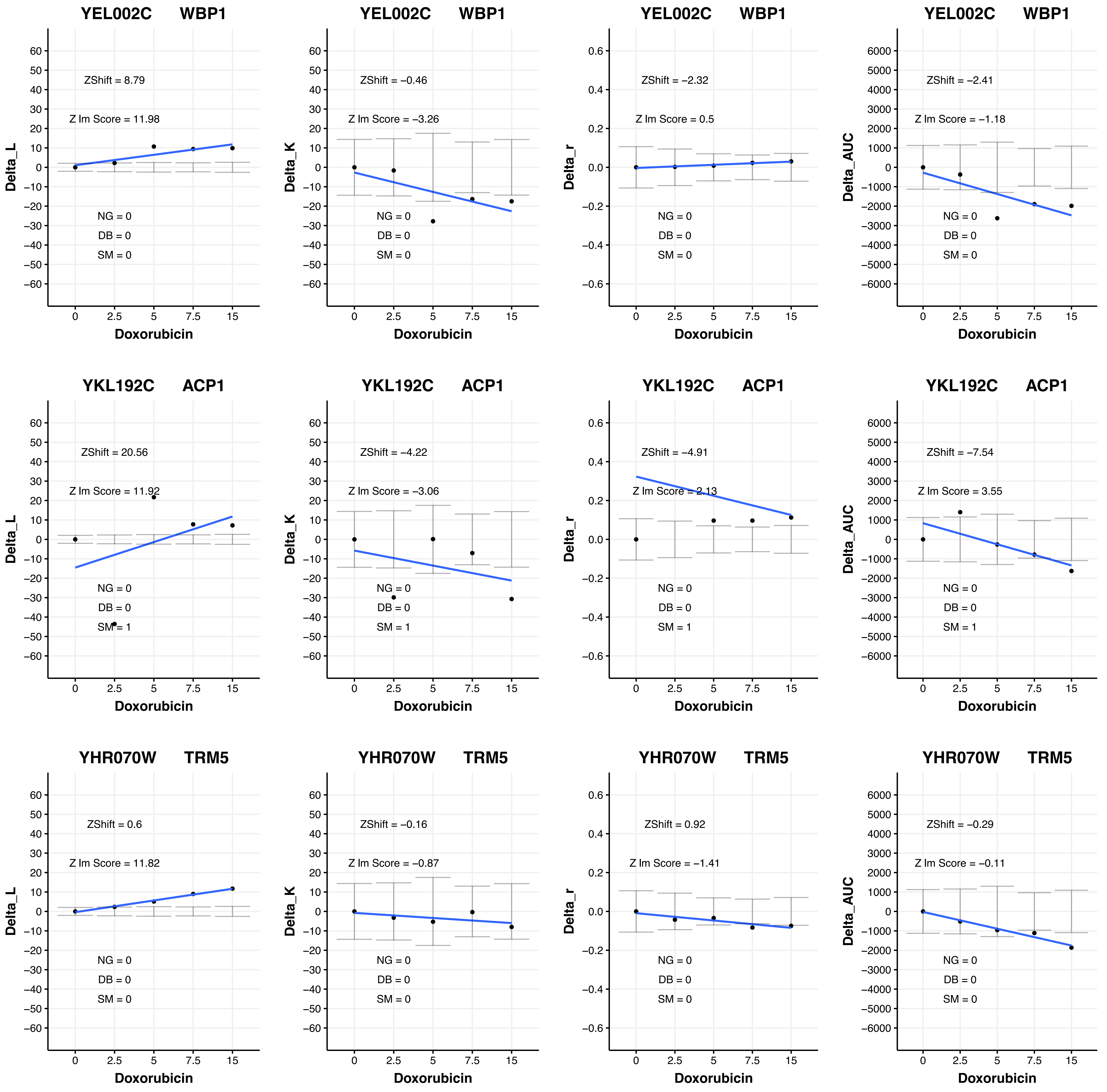

YEL002C WBP1
ZShift = 8.79
Z lm Score = 11.98
NG = 0
DB = 0
SM = 0
Delta_L
Doxorubicin
YEL002C WBP1
ZShift = −0.46
Z lm Score = −3.26
NG = 0
DB = 0
SM = 0
Delta_K
Doxorubicin
YEL002C WBP1
ZShift = −2.32
Z lm Score = 0.5
NG = 0
DB = 0
SM = 0
Delta_r
Doxorubicin
YEL002C WBP1
ZShift = −2.41
Z lm Score = −1.18
NG = 0
DB = 0
SM = 0
Delta_AUC
Doxorubicin
YKL192C ACP1
ZShift = 20.56
Z lm Score = 11.92
NG = 0
DB = 0
SM = 1
Delta_L
Doxorubicin
YKL192C ACP1
ZShift = −4.22
Z lm Score = −3.06
NG = 0
DB = 0
SM = 1
Delta_K
Doxorubicin
YKL192C ACP1
ZShift = −4.91
Z lm Score = 2.13
NG = 0
DB = 0
SM = 1
Delta_r
Doxorubicin
YKL192C ACP1
ZShift = −7.54
Z lm Score = 3.55
NG = 0
DB = 0
SM = 1
Delta_AUC
Doxorubicin
YHR070W TRM5
ZShift = 0.6
Z lm Score = 11.82
NG = 0
DB = 0
SM = 0
Delta_L
Doxorubicin
YHR070W TRM5
ZShift = −0.16
Z lm Score = −0.87
NG = 0
DB = 0
SM = 0
Delta_K
Doxorubicin
YHR070W TRM5
ZShift = 0.92
Z lm Score = −1.41
NG = 0
DB = 0
SM = 0
Delta_r
Doxorubicin
YHR070W TRM5
ZShift = −0.29
Z lm Score = −0.11
NG = 0
DB = 0
SM = 0
Delta_AUC
Doxorubicin

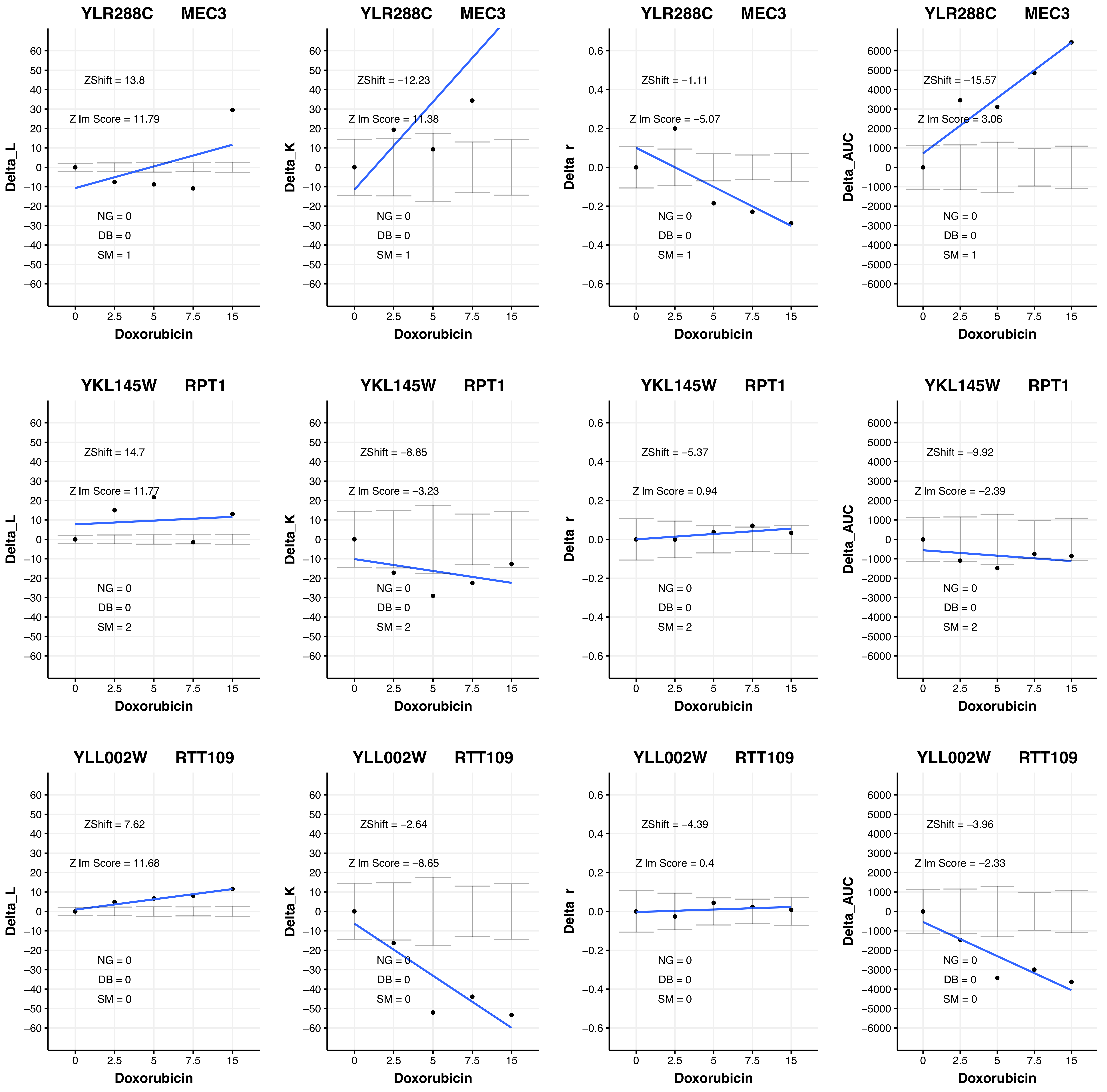

YLR288C MEC3
ZShift = 13.8
Z lm Score = 11.79
NG = 0
DB = 0
SM = 1
Delta_L
Doxorubicin
YLR288C MEC3
ZShift = −12.23
Z lm Score = 11.38
NG = 0
DB = 0
SM = 1
Delta_K
Doxorubicin
YLR288C MEC3
ZShift = −1.11
Z lm Score = −5.07
NG = 0
DB = 0
SM = 1
Delta_r
Doxorubicin
YLR288C MEC3
ZShift = −15.57
Z lm Score = 3.06
NG = 0
DB = 0
SM = 1
Delta_AUC
Doxorubicin
YKL145W RPT1
ZShift = 14.7
Z lm Score = 11.77
NG = 0
DB = 0
SM = 2
Delta_L
Doxorubicin
YKL145W RPT1
ZShift = −8.85
Z lm Score = −3.23
NG = 0
DB = 0
SM = 2
Delta_K
Doxorubicin
YKL145W RPT1
ZShift = −5.37
Z lm Score = 0.94
NG = 0
DB = 0
SM = 2
Delta_r
Doxorubicin
YKL145W RPT1
ZShift = −9.92
Z lm Score = −2.39
NG = 0
DB = 0
SM = 2
Delta_AUC
Doxorubicin
YLL002W RTT109
ZShift = 7.62
Z lm Score = 11.68
NG = 0
DB = 0
SM = 0
Delta_L
Doxorubicin
YLL002W RTT109
ZShift = −2.64
Z lm Score = −8.65
NG = 0
DB = 0
SM = 0
Delta_K
Doxorubicin
YLL002W RTT109
ZShift = −4.39
Z lm Score = 0.4
NG = 0
DB = 0
SM = 0
Delta_r
Doxorubicin
YLL002W RTT109
ZShift = −3.96
Z lm Score = −2.33
NG = 0
DB = 0
SM = 0
Delta_AUC
Doxorubicin

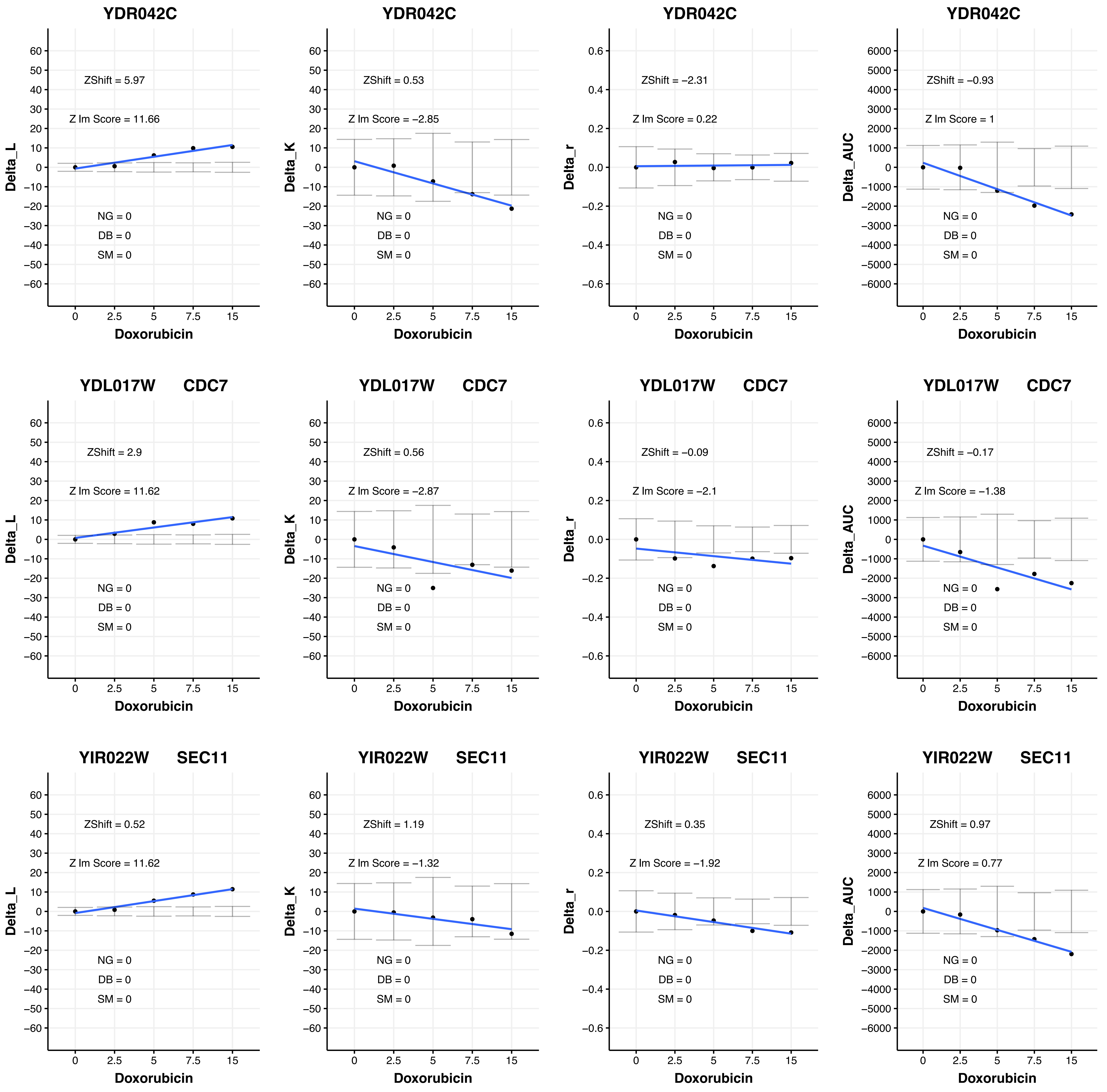
YDR042C
ZShift = 5.97
Z lm Score = 11.66
NG = 0
DB = 0
SM = 0
Delta_L
Doxorubicin
YDR042C
ZShift = 0.53
Z lm Score = −2.85
NG = 0
DB = 0
SM = 0
Delta_K
Doxorubicin
YDR042C
ZShift = −2.31
Z lm Score = 0.22
NG = 0
DB = 0
SM = 0
Delta_r
Doxorubicin
YDR042C
ZShift = −0.93
Z lm Score = 1
NG = 0
DB = 0
SM = 0
Delta_AUC
Doxorubicin
YDL017W CDC7
ZShift = 2.9
Z lm Score = 11.62
NG = 0
DB = 0
SM = 0
Delta_L
Doxorubicin
YDL017W CDC7
ZShift = 0.56
Z lm Score = −2.87
NG = 0
DB = 0
SM = 0
Delta_K
Doxorubicin
YDL017W CDC7
ZShift = −0.09
Z lm Score = −2.1
NG = 0
DB = 0
SM = 0
Delta_r
Doxorubicin
YDL017W CDC7
ZShift = −0.17
Z lm Score = −1.38
NG = 0
DB = 0
SM = 0
Delta_AUC
Doxorubicin
YIR022W SEC11
ZShift = 0.52
Z lm Score = 11.62
NG = 0
DB = 0
SM = 0
Delta_L
Doxorubicin
YIR022W SEC11
ZShift = 1.19
Z lm Score = −1.32
NG = 0
DB = 0
SM = 0
Delta_K
Doxorubicin
YIR022W SEC11
ZShift = 0.35
Z lm Score = −1.92
NG = 0
DB = 0
SM = 0
Delta_r
Doxorubicin
YIR022W SEC11
ZShift = 0.97
Z lm Score = 0.77
NG = 0
DB = 0
SM = 0
Delta_AUC
Doxorubicin

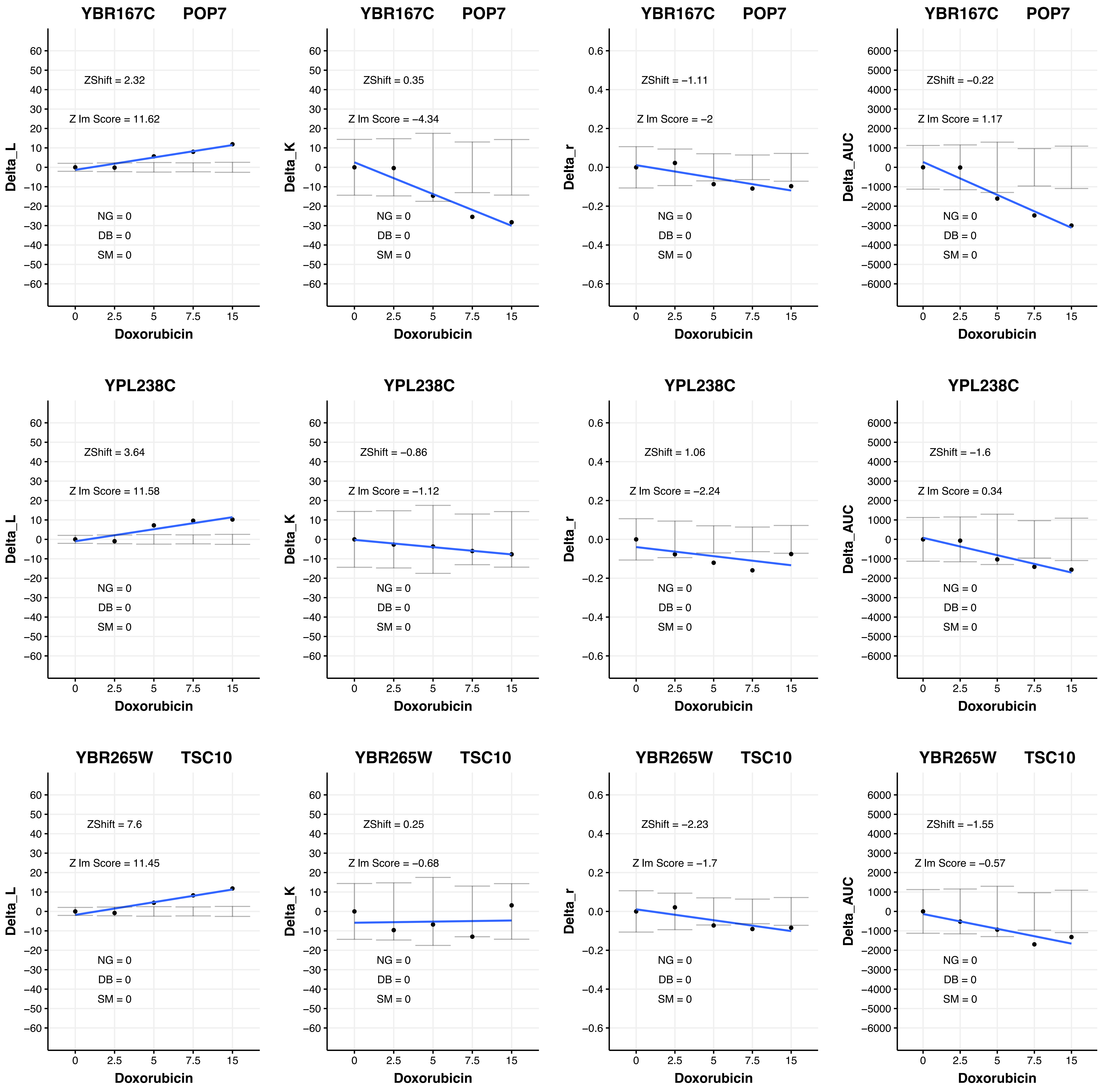

YBR167C POP7
ZShift = 2.32
Z lm Score = 11.62
NG = 0
DB = 0
SM = 0
Delta_L
Doxorubicin
YBR167C POP7
ZShift = 0.35
Z lm Score = −4.34
NG = 0
DB = 0
SM = 0
Delta_K
Doxorubicin
YBR167C POP7
ZShift = −1.11
Z lm Score = −2
NG = 0
DB = 0
SM = 0
Delta_r
Doxorubicin
YBR167C POP7
ZShift = −0.22
Z lm Score = 1.17
NG = 0
DB = 0
SM = 0
Delta_AUC
Doxorubicin
YPL238C
ZShift = 3.64
Z lm Score = 11.58
NG = 0
DB = 0
SM = 0
Delta_L
Doxorubicin
YPL238C
ZShift = −0.86
Z lm Score = −1.12
NG = 0
DB = 0
SM = 0
Delta_K
Doxorubicin
YPL238C
ZShift = 1.06
Z lm Score = −2.24
NG = 0
DB = 0
SM = 0
Delta_r
Doxorubicin
YPL238C
ZShift = −1.6
Z lm Score = 0.34
NG = 0
DB = 0
SM = 0
Delta_AUC
Doxorubicin
YBR265W TSC10
ZShift = 7.6
Z lm Score = 11.45
NG = 0
DB = 0
SM = 0
Delta_L
Doxorubicin
YBR265W TSC10
ZShift = 0.25
Z lm Score = −0.68
NG = 0
DB = 0
SM = 0
Delta_K
Doxorubicin
YBR265W TSC10
ZShift = −2.23
Z lm Score = −1.7
NG = 0
DB = 0
SM = 0
Delta_r
Doxorubicin
YBR265W TSC10
ZShift = −1.55
Z lm Score = −0.57
NG = 0
DB = 0
SM = 0
Delta_AUC
Doxorubicin

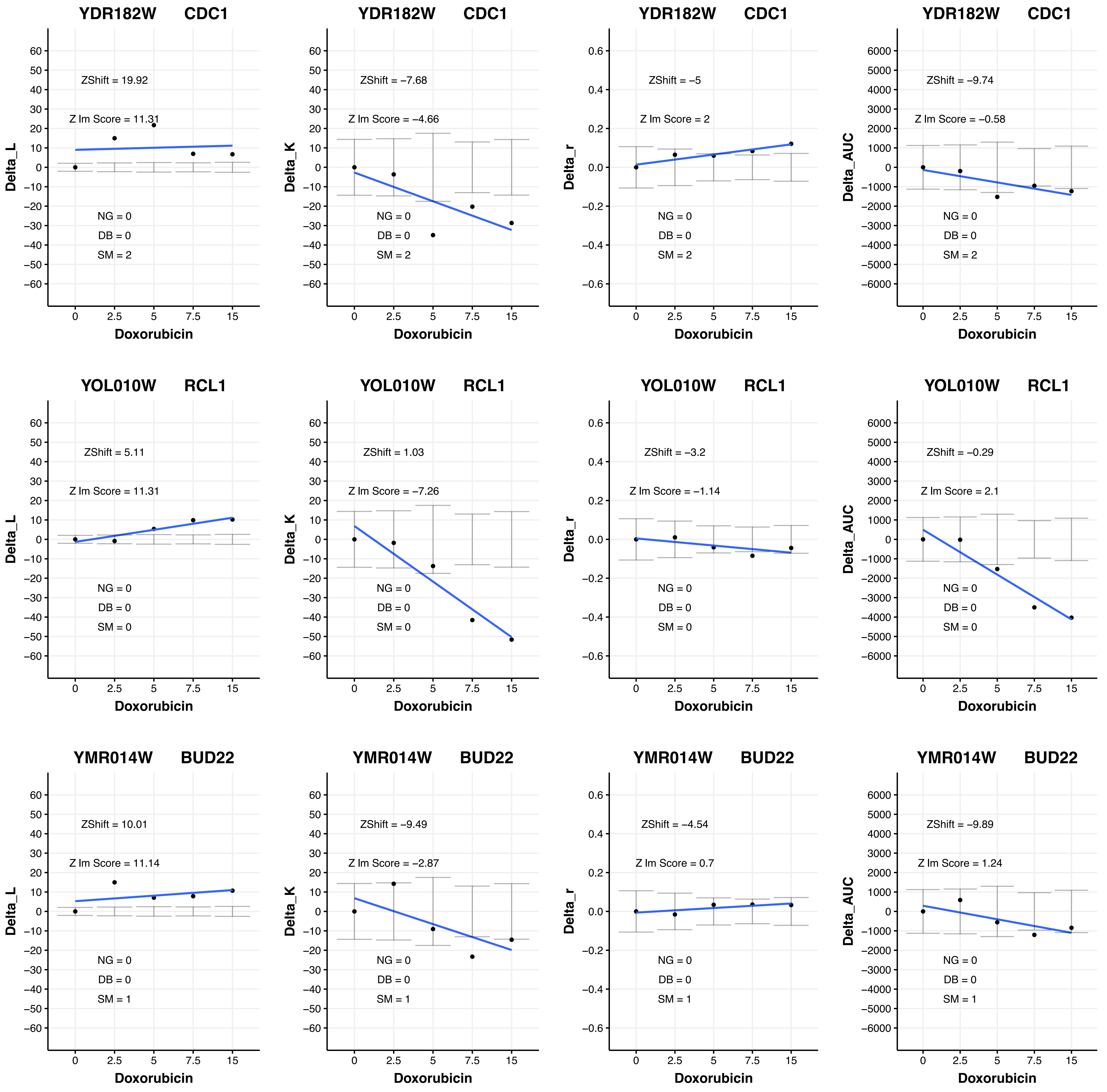
YDR182W CDC1
ZShift = 19.92
Z lm Score = 11.31
NG = 0
DB = 0
SM = 2
Delta_L
Doxorubicin
YDR182W CDC1
ZShift = −7.68
Z lm Score = −4.66
NG = 0
DB = 0
SM = 2
Delta_K
Doxorubicin
YDR182W CDC1
ZShift = −5
Z lm Score = 2
NG = 0
DB = 0
SM = 2
Delta_r
Doxorubicin
YDR182W CDC1
ZShift = −9.74
Z lm Score = −0.58
NG = 0
DB = 0
SM = 2
Delta_AUC
Doxorubicin
YOL010W RCL1
ZShift = 5.11
Z lm Score = 11.31
NG = 0
DB = 0
SM = 0
Delta_L
Doxorubicin
YOL010W RCL1
ZShift = 1.03
Z lm Score = −7.26
NG = 0
DB = 0
SM = 0
Delta_K
Doxorubicin
YOL010W RCL1
ZShift = −3.2
Z lm Score = −1.14
NG = 0
DB = 0
SM = 0
Delta_r
Doxorubicin
YOL010W RCL1
ZShift = −0.29
Z lm Score = 2.1
NG = 0
DB = 0
SM = 0
Delta_AUC
Doxorubicin
YMR014W BUD22
ZShift = 10.01
Z lm Score = 11.14
NG = 0
DB = 0
SM = 1
Delta_L
Doxorubicin
YMR014W BUD22
ZShift = −9.49
Z lm Score = −2.87
NG = 0
DB = 0
SM = 1
Delta_K
Doxorubicin
YMR014W BUD22
ZShift = −4.54
Z lm Score = 0.7
NG = 0
DB = 0
SM = 1
Delta_r
Doxorubicin
YMR014W BUD22
ZShift = −9.89
Z lm Score = 1.24
NG = 0
DB = 0
SM = 1
Delta_AUC
Doxorubicin

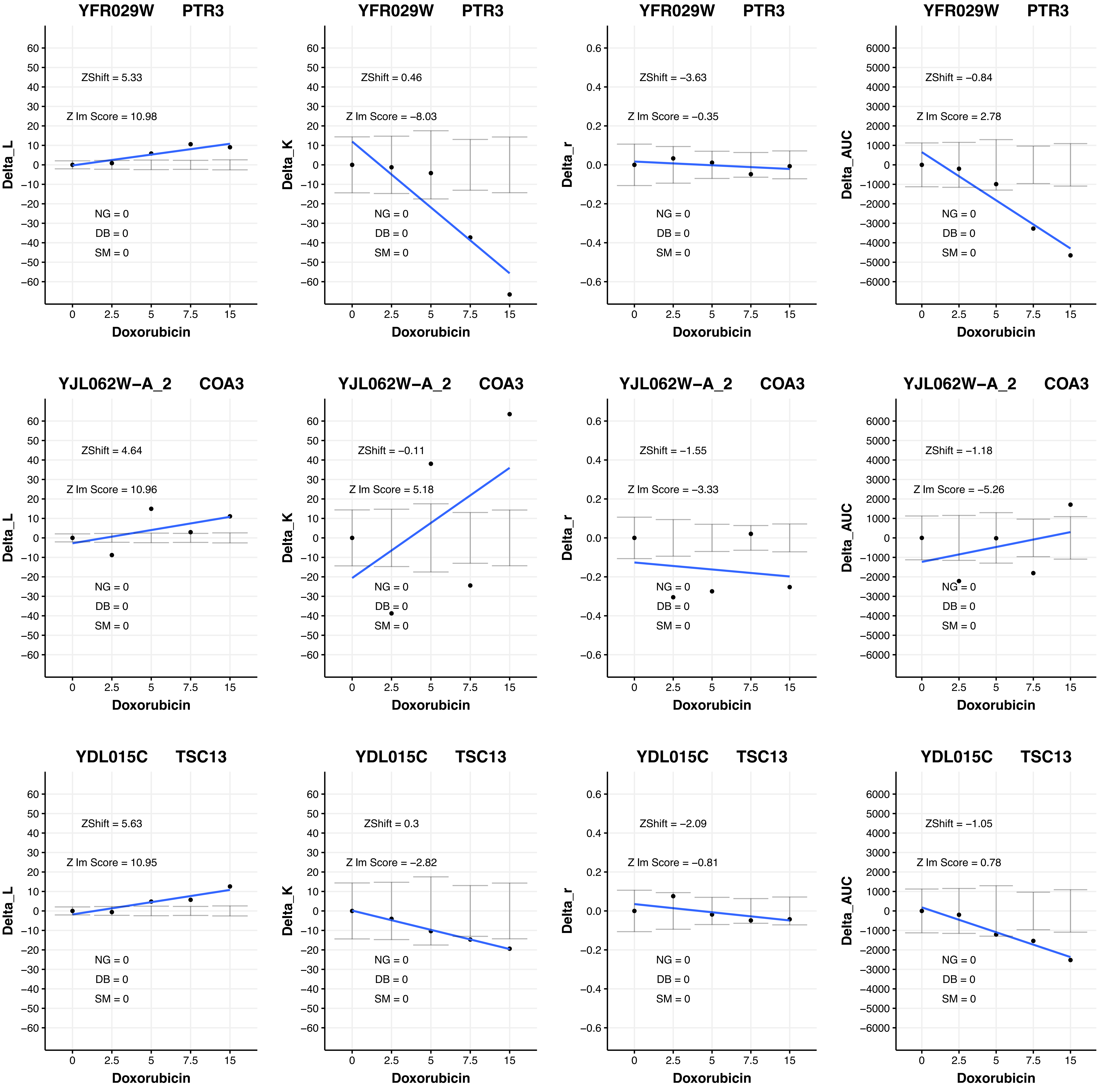
YFR029W PTR3
ZShift = 5.33
Z lm Score = 10.98
NG = 0
DB = 0
SM = 0
Delta_L
Doxorubicin
YFR029W PTR3
ZShift = 0.46
Z lm Score = −8.03
NG = 0
DB = 0
SM = 0
Delta_K
Doxorubicin
YFR029W PTR3
ZShift = −3.63
Z lm Score = −0.35
NG = 0
DB = 0
SM = 0
Delta_r
Doxorubicin
YFR029W PTR3
ZShift = −0.84
Z lm Score = 2.78
NG = 0
DB = 0
SM = 0
Delta_AUC
Doxorubicin
YJL062W−A_2 COA3
ZShift = 4.64
Z lm Score = 10.96
NG = 0
DB = 0
SM = 0
Delta_L
Doxorubicin
YJL062W−A_2 COA3
ZShift = −0.11
Z lm Score = 5.18
NG = 0
DB = 0
SM = 0
Delta_K
Doxorubicin
YJL062W−A_2 COA3
ZShift = −1.55
Z lm Score = −3.33
NG = 0
DB = 0
SM = 0
Delta_r
Doxorubicin
YJL062W−A_2 COA3
ZShift = −1.18
Z lm Score = −5.26
NG = 0
DB = 0
SM = 0
Delta_AUC
Doxorubicin
YDL015C TSC13
ZShift = 5.63
Z lm Score = 10.95
NG = 0
DB = 0
SM = 0
Delta_L
Doxorubicin
YDL015C TSC13
ZShift = 0.3
Z lm Score = −2.82
NG = 0
DB = 0
SM = 0
Delta_K
Doxorubicin
YDL015C TSC13
ZShift = −2.09
Z lm Score = −0.81
NG = 0
DB = 0
SM = 0
Delta_r
Doxorubicin
YDL015C TSC13
ZShift = −1.05
Z lm Score = 0.78
NG = 0
DB = 0
SM = 0
Delta_AUC
Doxorubicin

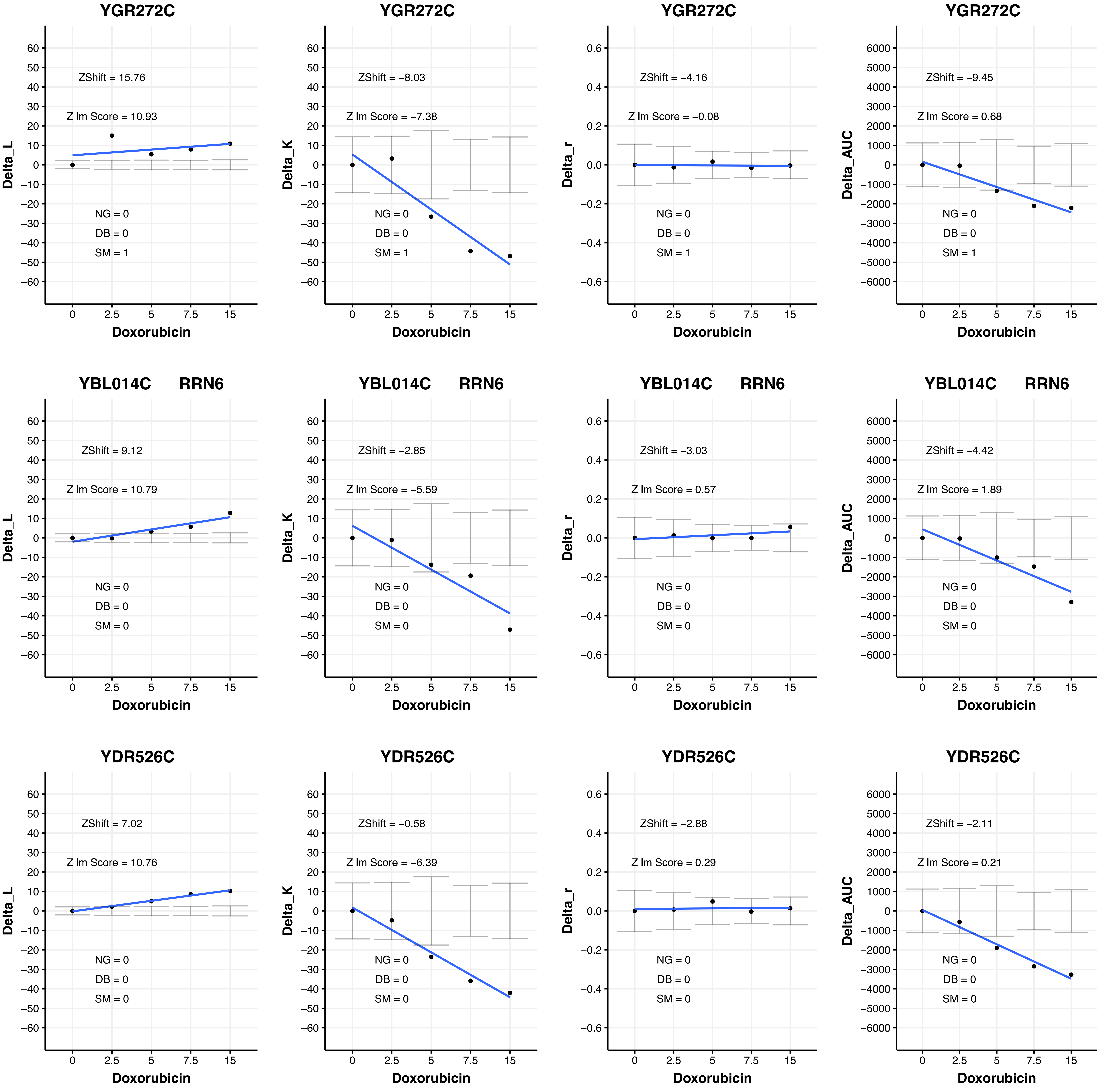
YGR272C
ZShift = 15.76
Z lm Score = 10.93
NG = 0
DB = 0
SM = 1
Delta_L
Doxorubicin
YGR272C
ZShift = −8.03
Z lm Score = −7.38
NG = 0
DB = 0
SM = 1
Delta_K
Doxorubicin
YGR272C
ZShift = −4.16
Z lm Score = −0.08
NG = 0
DB = 0
SM = 1
Delta_r
Doxorubicin
YGR272C
ZShift = −9.45
Z lm Score = 0.68
NG = 0
DB = 0
SM = 1
Delta_AUC
Doxorubicin
YBL014C RRN6
ZShift = 9.12
Z lm Score = 10.79
NG = 0
DB = 0
SM = 0
Delta_L
Doxorubicin
YBL014C RRN6
ZShift = −2.85
Z lm Score = −5.59
NG = 0
DB = 0
SM = 0
Delta_K
Doxorubicin
YBL014C RRN6
ZShift = −3.03
Z lm Score = 0.57
NG = 0
DB = 0
SM = 0
Delta_r
Doxorubicin
YBL014C RRN6
ZShift = −4.42
Z lm Score = 1.89
NG = 0
DB = 0
SM = 0
Delta_AUC
Doxorubicin
YDR526C
ZShift = 7.02
Z lm Score = 10.76
NG = 0
DB = 0
SM = 0
Delta_L
Doxorubicin
YDR526C
ZShift = −0.58
Z lm Score = −6.39
NG = 0
DB = 0
SM = 0
Delta_K
Doxorubicin
YDR526C
ZShift = −2.88
Z lm Score = 0.29
NG = 0
DB = 0
SM = 0
Delta_r
Doxorubicin
YDR526C
ZShift = −2.11
Z lm Score = 0.21
NG = 0
DB = 0
SM = 0
Delta_AUC
Doxorubicin

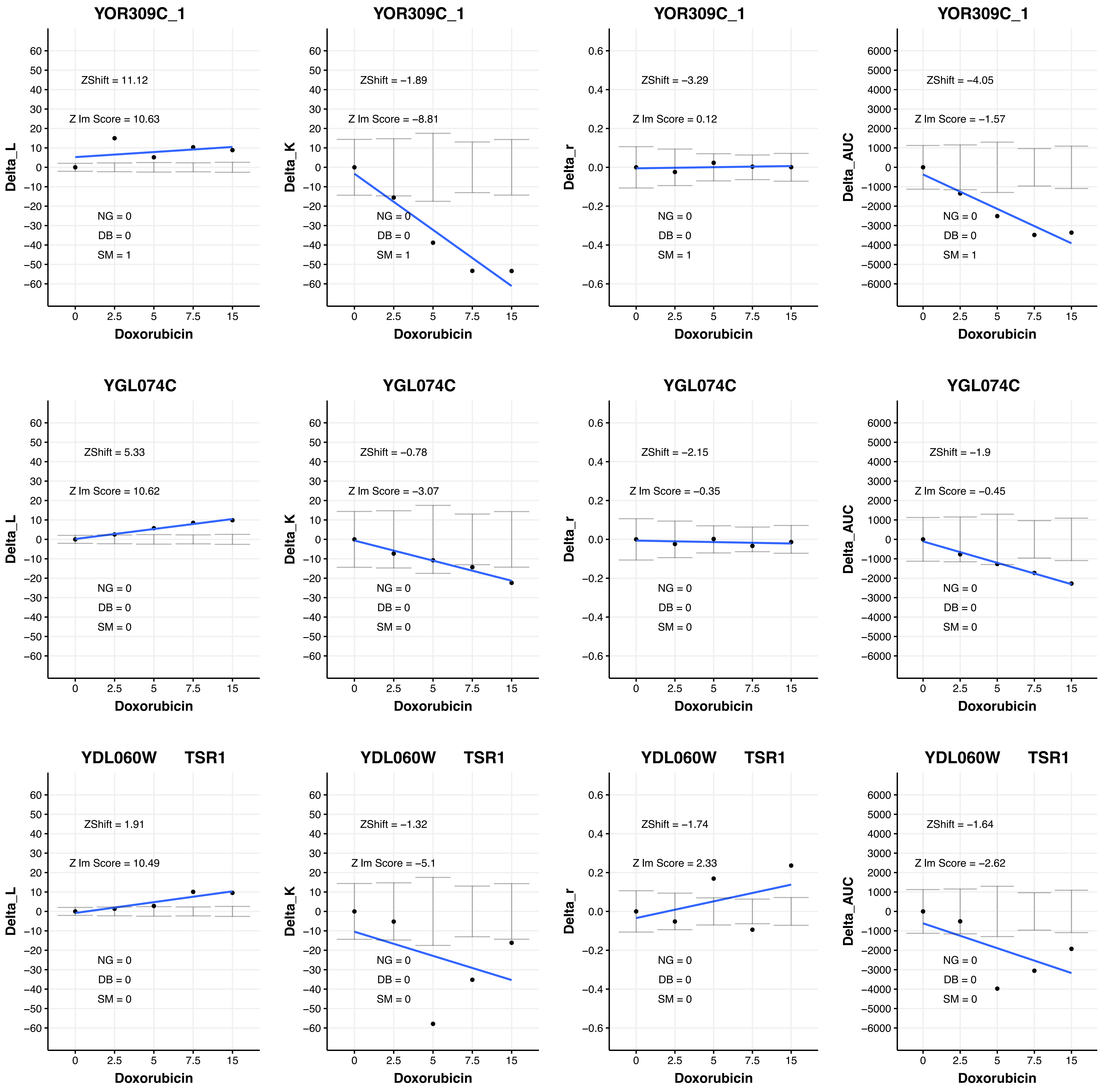
YOR309C_1
ZShift = 11.12
Z lm Score = 10.63
NG = 0
DB = 0
SM = 1
Delta_L
Doxorubicin
YOR309C_1
ZShift = −1.89
Z lm Score = −8.81
NG = 0
DB = 0
SM = 1
Delta_K
Doxorubicin
YOR309C_1
ZShift = −3.29
Z lm Score = 0.12
NG = 0
DB = 0
SM = 1
Delta_r
Doxorubicin
YOR309C_1
ZShift = −4.05
Z lm Score = −1.57
NG = 0
DB = 0
SM = 1
Delta_AUC
Doxorubicin
YGL074C
ZShift = 5.33
Z lm Score = 10.62
NG = 0
DB = 0
SM = 0
Delta_L
Doxorubicin
YGL074C
ZShift = −0.78
Z lm Score = −3.07
NG = 0
DB = 0
SM = 0
Delta_K
Doxorubicin
YGL074C
ZShift = −2.15
Z lm Score = −0.35
NG = 0
DB = 0
SM = 0
Delta_r
Doxorubicin
YGL074C
ZShift = −1.9
Z lm Score = −0.45
NG = 0
DB = 0
SM = 0
Delta_AUC
Doxorubicin
YDL060W TSR1
ZShift = 1.91
Z lm Score = 10.49
NG = 0
DB = 0
SM = 0
Delta_L
Doxorubicin
YDL060W TSR1
ZShift = −1.32
Z lm Score = −5.1
NG = 0
DB = 0
SM = 0
Delta_K
Doxorubicin
YDL060W TSR1
ZShift = −1.74
Z lm Score = 2.33
NG = 0
DB = 0
SM = 0
Delta_r
Doxorubicin
YDL060W TSR1
ZShift = −1.64
Z lm Score = −2.62
NG = 0
DB = 0
SM = 0
Delta_AUC
Doxorubicin

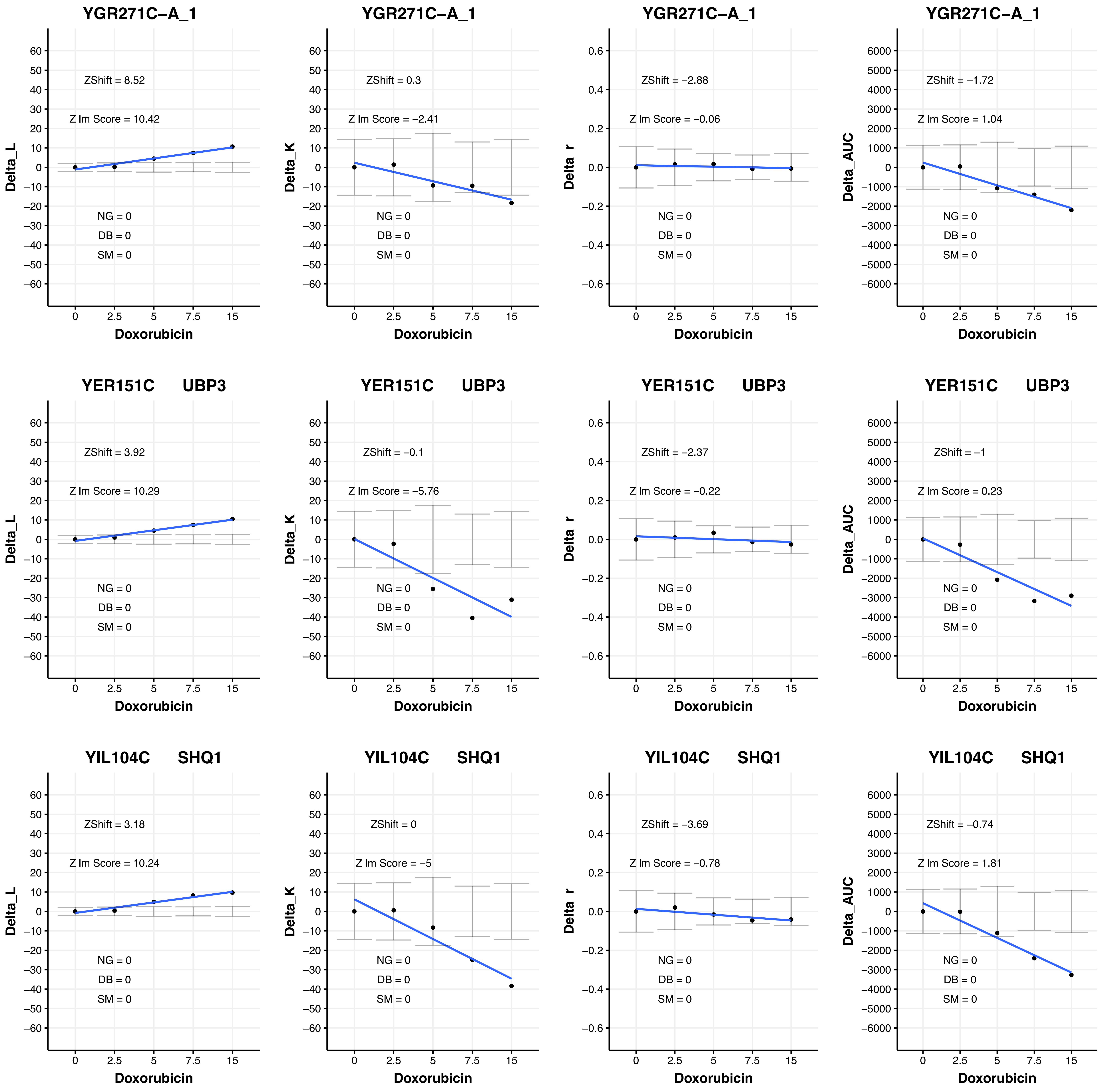
YGR271C−A_1
ZShift = 8.52
Z lm Score = 10.42
NG = 0
DB = 0
SM = 0
Delta_L
Doxorubicin
YGR271C−A_1
ZShift = 0.3
Z lm Score = −2.41
NG = 0
DB = 0
SM = 0
Delta_K
Doxorubicin
YGR271C−A_1
ZShift = −2.88
Z lm Score = −0.06
NG = 0
DB = 0
SM = 0
Delta_r
Doxorubicin
YGR271C−A_1
ZShift = −1.72
Z lm Score = 1.04
NG = 0
DB = 0
SM = 0
Delta_AUC
Doxorubicin
YER151C UBP3
ZShift = 3.92
Z lm Score = 10.29
NG = 0
DB = 0
SM = 0
Delta_L
Doxorubicin
YER151C UBP3
ZShift = −0.1
Z lm Score = −5.76
NG = 0
DB = 0
SM = 0
Delta_K
Doxorubicin
YER151C UBP3
ZShift = −2.37
Z lm Score = −0.22
NG = 0
DB = 0
SM = 0
Delta_r
Doxorubicin
YER151C UBP3
ZShift = −1
Z lm Score = 0.23
NG = 0
DB = 0
SM = 0
Delta_AUC
Doxorubicin
YIL104C SHQ1
ZShift = 3.18
Z lm Score = 10.24
NG = 0
DB = 0
SM = 0
Delta_L
Doxorubicin
YIL104C SHQ1
ZShift = 0
Z lm Score = −5
NG = 0
DB = 0
SM = 0
Delta_K
Doxorubicin
YIL104C SHQ1
ZShift = −3.69
Z lm Score = −0.78
NG = 0
DB = 0
SM = 0
Delta_r
Doxorubicin
YIL104C SHQ1
ZShift = −0.74
Z lm Score = 1.81
NG = 0
DB = 0
SM = 0
Delta_AUC
Doxorubicin

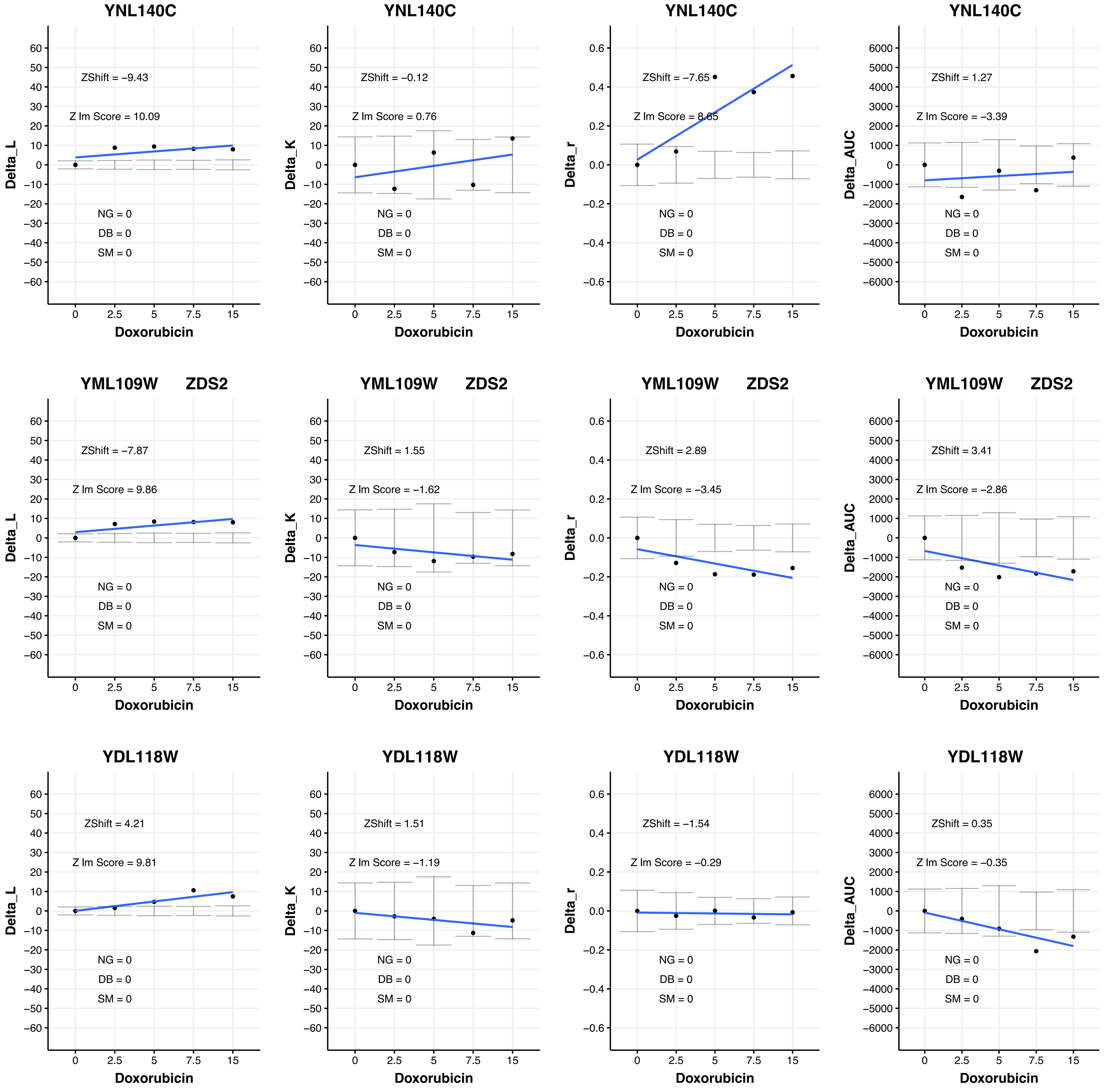
YNL140C
ZShift = −9.43
Z lm Score = 10.09
NG = 0
DB = 0
SM = 0
Delta_L
Doxorubicin
YNL140C
ZShift = −0.12
Z lm Score = 0.76
NG = 0
DB = 0
SM = 0
Delta_K
Doxorubicin
YNL140C
ZShift = −7.65
Z lm Score = 8.65
NG = 0
DB = 0
SM = 0
Delta_r
Doxorubicin
YNL140C
ZShift = 1.27
Z lm Score = −3.39
NG = 0
DB = 0
SM = 0
Delta_AUC
Doxorubicin
YML109W ZDS2
ZShift = −7.87
Z lm Score = 9.86
NG = 0
DB = 0
SM = 0
Delta_L
Doxorubicin
YML109W ZDS2
ZShift = 1.55
Z lm Score = −1.62
NG = 0
DB = 0
SM = 0
Delta_K
Doxorubicin
YML109W ZDS2
ZShift = 2.89
Z lm Score = −3.45
NG = 0
DB = 0
SM = 0
Delta_r
Doxorubicin
YML109W ZDS2
ZShift = 3.41
Z lm Score = −2.86
NG = 0
DB = 0
SM = 0
Delta_AUC
Doxorubicin
YDL118W
ZShift = 4.21
Z lm Score = 9.81
NG = 0
DB = 0
SM = 0
Delta_L
Doxorubicin
YDL118W
ZShift = 1.51
Z lm Score = −1.19
NG = 0
DB = 0
SM = 0
Delta_K
Doxorubicin
YDL118W
ZShift = −1.54
Z lm Score = −0.29
NG = 0
DB = 0
SM = 0
Delta_r
Doxorubicin
YDL118W
ZShift = 0.35
Z lm Score = −0.35
NG = 0
DB = 0
SM = 0
Delta_AUC
Doxorubicin

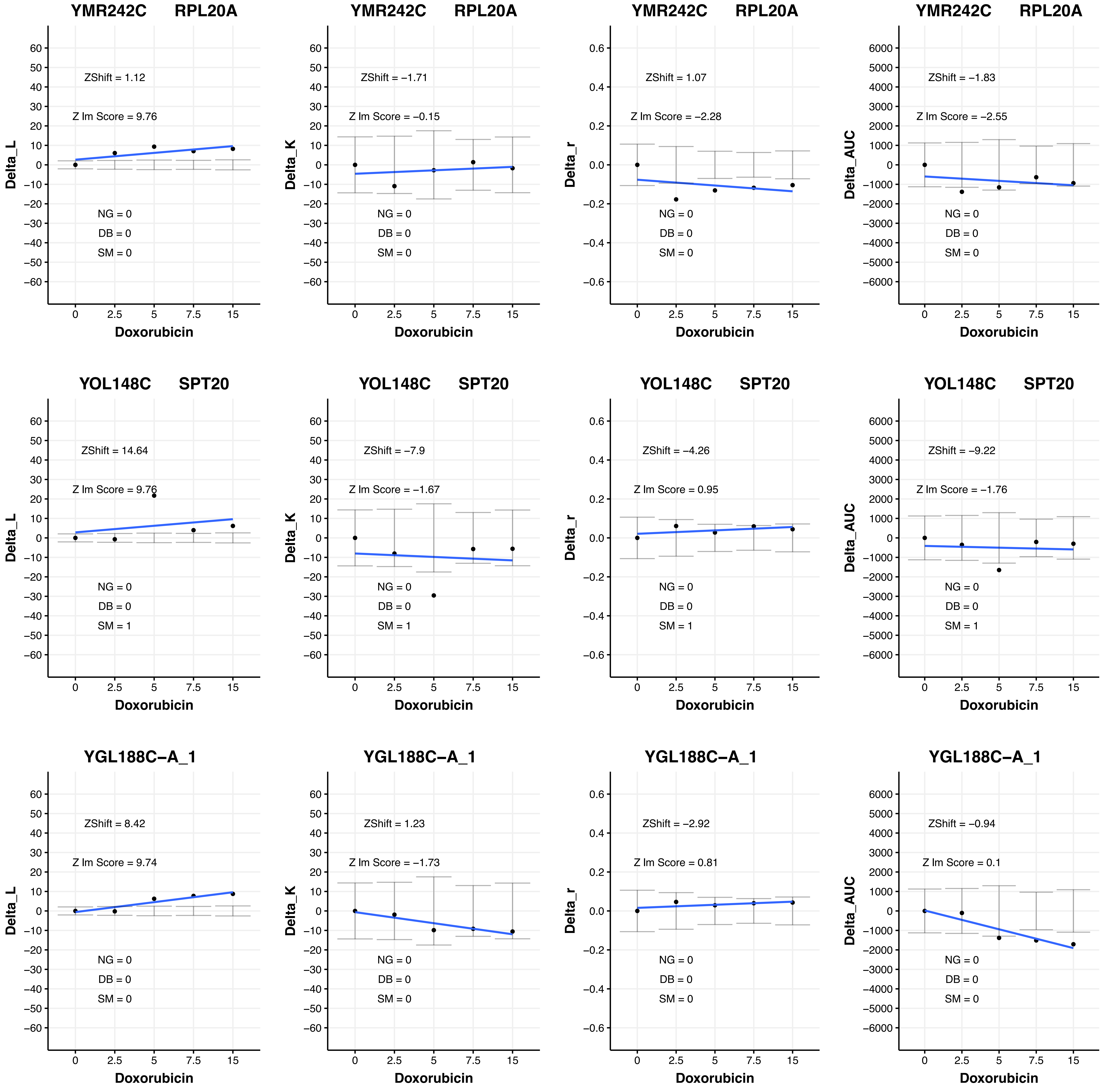

YMR242C RPL20A
ZShift = 1.12
Z lm Score = 9.76
NG = 0
DB = 0
SM = 0
Delta_L
Doxorubicin
YMR242C RPL20A
ZShift = −1.71
Z lm Score = −0.15
NG = 0
DB = 0
SM = 0
Delta_K
Doxorubicin
YMR242C RPL20A
ZShift = 1.07
Z lm Score = −2.28
NG = 0
DB = 0
SM = 0
Delta_r
Doxorubicin
YMR242C RPL20A
ZShift = −1.83
Z lm Score = −2.55
NG = 0
DB = 0
SM = 0
Delta_AUC
Doxorubicin
YOL148C SPT20
ZShift = 14.64
Z lm Score = 9.76
NG = 0
DB = 0
SM = 1
Delta_L
Doxorubicin
YOL148C SPT20
ZShift = −7.9
Z lm Score = −1.67
NG = 0
DB = 0
SM = 1
Delta_K
Doxorubicin
YOL148C SPT20
ZShift = −4.26
Z lm Score = 0.95
NG = 0
DB = 0
SM = 1
Delta_r
Doxorubicin
YOL148C SPT20
ZShift = −9.22
Z lm Score = −1.76
NG = 0
DB = 0
SM = 1
Delta_AUC
Doxorubicin
YGL188C−A_1
ZShift = 8.42
Z lm Score = 9.74
NG = 0
DB = 0
SM = 0
Delta_L
Doxorubicin
YGL188C−A_1
ZShift = 1.23
Z lm Score = −1.73
NG = 0
DB = 0
SM = 0
Delta_K
Doxorubicin
YGL188C−A_1
ZShift = −2.92
Z lm Score = 0.81
NG = 0
DB = 0
SM = 0
Delta_r
Doxorubicin
YGL188C−A_1
ZShift = −0.94
Z lm Score = 0.1
NG = 0
DB = 0
SM = 0
Delta_AUC
Doxorubicin

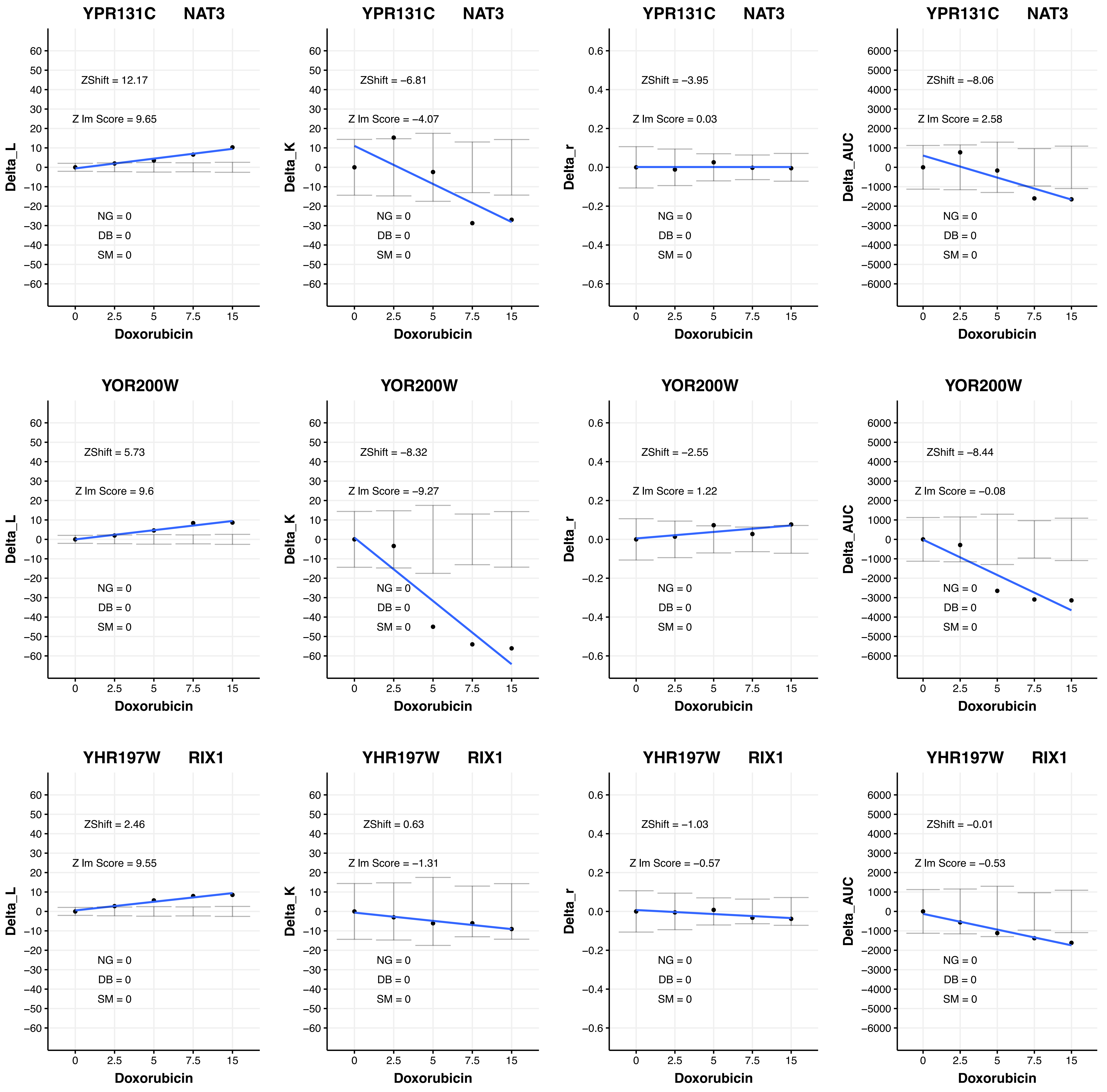
YPR131C NAT3
ZShift = 12.17
Z lm Score = 9.65
NG = 0
DB = 0
SM = 0
Delta_L
Doxorubicin
YPR131C NAT3
ZShift = −6.81
Z lm Score = −4.07
NG = 0
DB = 0
SM = 0
Delta_K
Doxorubicin
YPR131C NAT3
ZShift = −3.95
Z lm Score = 0.03
NG = 0
DB = 0
SM = 0
Delta_r
Doxorubicin
YPR131C NAT3
ZShift = −8.06
Z lm Score = 2.58
NG = 0
DB = 0
SM = 0
Delta_AUC
Doxorubicin
YOR200W
ZShift = 5.73
Z lm Score = 9.6
NG = 0
DB = 0
SM = 0
Delta_L
Doxorubicin
YOR200W
ZShift = −8.32
Z lm Score = −9.27
NG = 0
DB = 0
SM = 0
Delta_K
Doxorubicin
YOR200W
ZShift = −2.55
Z lm Score = 1.22
NG = 0
DB = 0
SM = 0
Delta_r
Doxorubicin
YOR200W
ZShift = −8.44
Z lm Score = −0.08
NG = 0
DB = 0
SM = 0
Delta_AUC
Doxorubicin
YHR197W RIX1
ZShift = 2.46
Z lm Score = 9.55
NG = 0
DB = 0
SM = 0
Delta_L
Doxorubicin
YHR197W RIX1
ZShift = 0.63
Z lm Score = −1.31
NG = 0
DB = 0
SM = 0
Delta_K
Doxorubicin
YHR197W RIX1
ZShift = −1.03
Z lm Score = −0.57
NG = 0
DB = 0
SM = 0
Delta_r
Doxorubicin
YHR197W RIX1
ZShift = −0.01
Z lm Score = −0.53
NG = 0
DB = 0
SM = 0
Delta_AUC
Doxorubicin

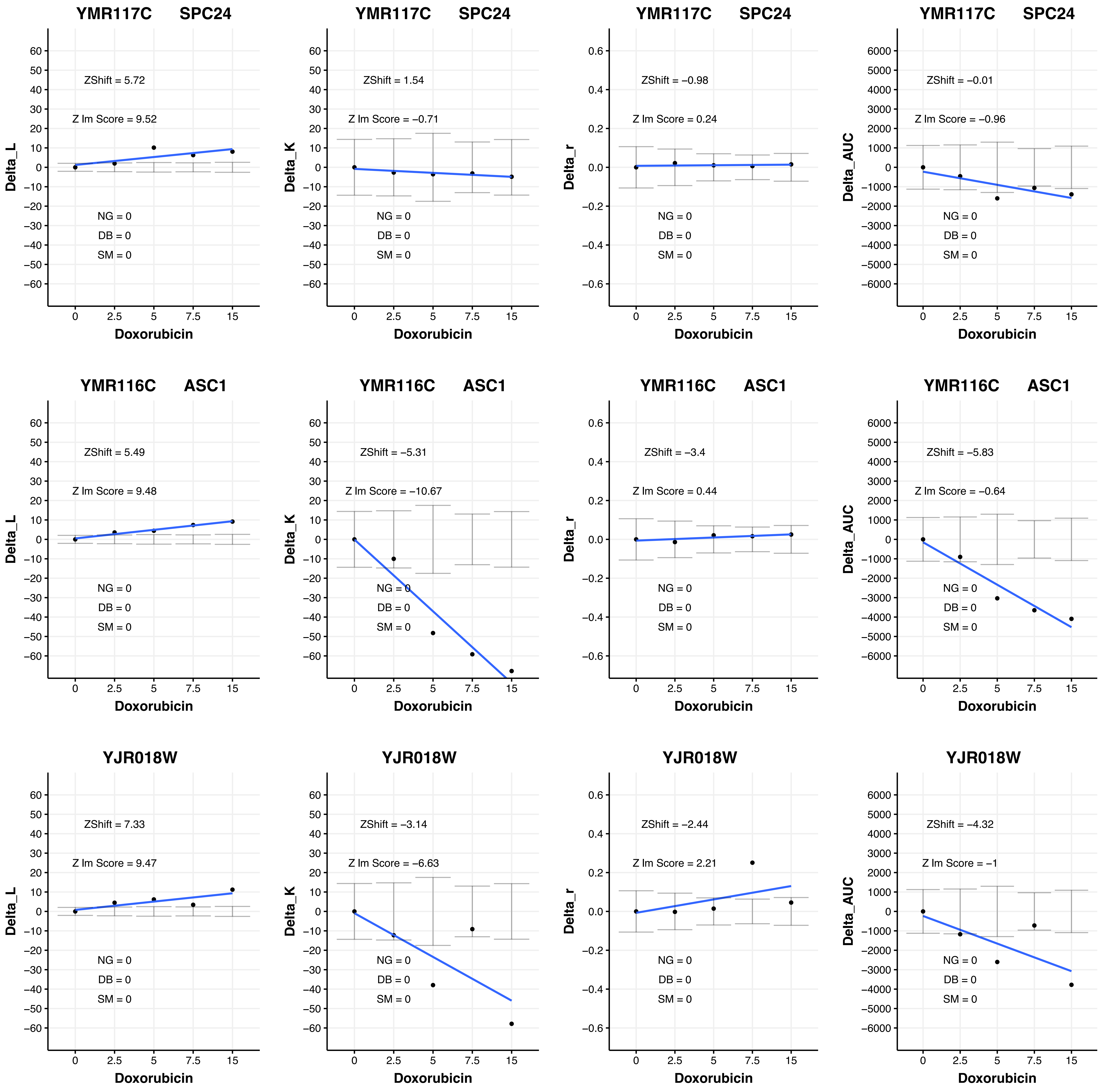
YMR117C SPC24
ZShift = 5.72
Z lm Score = 9.52
NG = 0
DB = 0
SM = 0
Delta_L
Doxorubicin
YMR117C SPC24
ZShift = 1.54
Z lm Score = −0.71
NG = 0
DB = 0
SM = 0
Delta_K
Doxorubicin
YMR117C SPC24
ZShift = −0.98
Z lm Score = 0.24
NG = 0
DB = 0
SM = 0
Delta_r
Doxorubicin
YMR117C SPC24
ZShift = −0.01
Z lm Score = −0.96
NG = 0
DB = 0
SM = 0
Delta_AUC
Doxorubicin
YMR116C ASC1
ZShift = 5.49
Z lm Score = 9.48
NG = 0
DB = 0
SM = 0
Delta_L
Doxorubicin
YMR116C ASC1
ZShift = −5.31
Z lm Score = −10.67
NG = 0
DB = 0
SM = 0
Delta_K
Doxorubicin
YMR116C ASC1
ZShift = −3.4
Z lm Score = 0.44
NG = 0
DB = 0
SM = 0
Delta_r
Doxorubicin
YMR116C ASC1
ZShift = −5.83
Z lm Score = −0.64
NG = 0
DB = 0
SM = 0
Delta_AUC
Doxorubicin
YJR018W
ZShift = 7.33
Z lm Score = 9.47
NG = 0
DB = 0
SM = 0
Delta_L
Doxorubicin
YJR018W
ZShift = −3.14
Z lm Score = −6.63
NG = 0
DB = 0
SM = 0
Delta_K
Doxorubicin
YJR018W
ZShift = −2.44
Z lm Score = 2.21
NG = 0
DB = 0
SM = 0
Delta_r
Doxorubicin
YJR018W
ZShift = −4.32
Z lm Score = −1
NG = 0
DB = 0
SM = 0
Delta_AUC
Doxorubicin

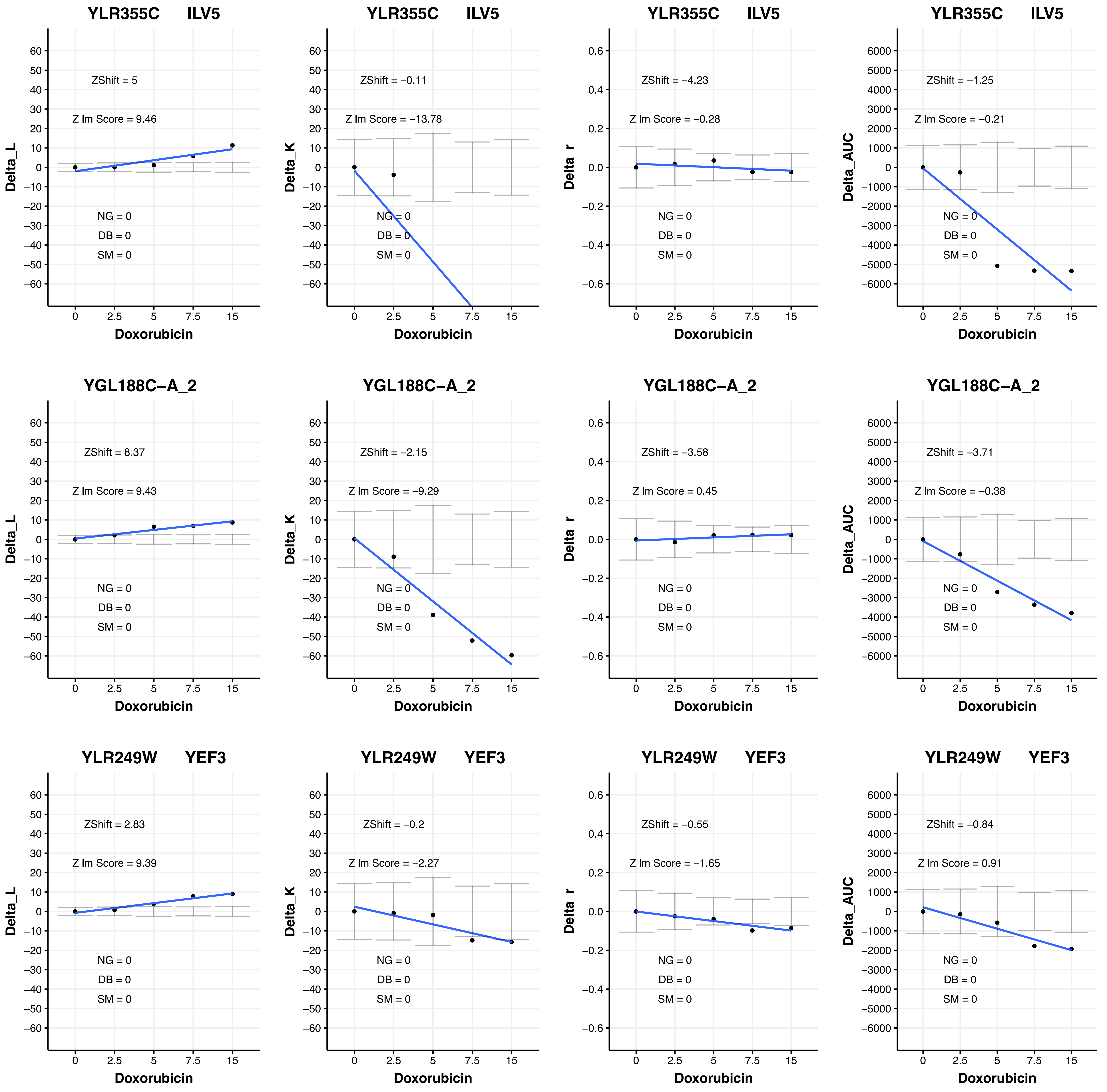
YLR355C ILV5
ZShift = 5
Z lm Score = 9.46
NG = 0
DB = 0
SM = 0
Delta_L
Doxorubicin
YLR355C ILV5
ZShift = −0.11
Z lm Score = −13.78
NG = 0
DB = 0
SM = 0
Delta_K
Doxorubicin
YLR355C ILV5
ZShift = −4.23
Z lm Score = −0.28
NG = 0
DB = 0
SM = 0
Delta_r
Doxorubicin
YLR355C ILV5
ZShift = −1.25
Z lm Score = −0.21
NG = 0
DB = 0
SM = 0
Delta_AUC
Doxorubicin
YGL188C−A_2
ZShift = 8.37
Z lm Score = 9.43
NG = 0
DB = 0
SM = 0
Delta_L
Doxorubicin
YGL188C−A_2
ZShift = −2.15
Z lm Score = −9.29
NG = 0
DB = 0
SM = 0
Delta_K
Doxorubicin
YGL188C−A_2
ZShift = −3.58
Z lm Score = 0.45
NG = 0
DB = 0
SM = 0
Delta_r
Doxorubicin
YGL188C−A_2
ZShift = −3.71
Z lm Score = −0.38
NG = 0
DB = 0
SM = 0
Delta_AUC
Doxorubicin
YLR249W YEF3
ZShift = 2.83
Z lm Score = 9.39
NG = 0
DB = 0
SM = 0
Delta_L
Doxorubicin
YLR249W YEF3
ZShift = −0.2
Z lm Score = −2.27
NG = 0
DB = 0
SM = 0
Delta_K
Doxorubicin
YLR249W YEF3
ZShift = −0.55
Z lm Score = −1.65
NG = 0
DB = 0
SM = 0
Delta_r
Doxorubicin
YLR249W YEF3
ZShift = −0.84
Z lm Score = 0.91
NG = 0
DB = 0
SM = 0
Delta_AUC
Doxorubicin

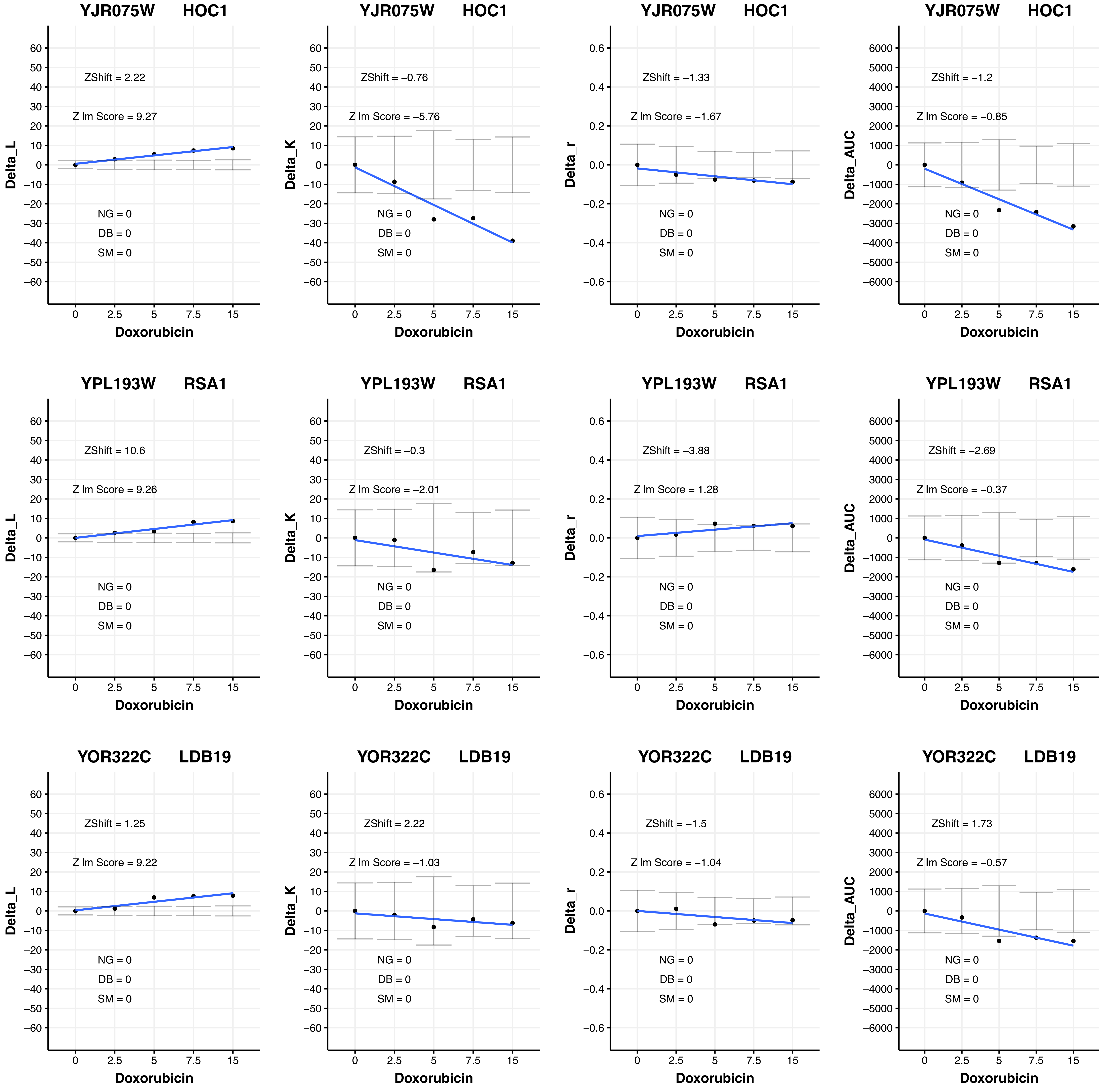
YJR075W HOC1
ZShift = 2.22
Z lm Score = 9.27
NG = 0
DB = 0
SM = 0
Delta_L
Doxorubicin
YJR075W HOC1
ZShift = −0.76
Z lm Score = −5.76
NG = 0
DB = 0
SM = 0
Delta_K
Doxorubicin
YJR075W HOC1
ZShift = −1.33
Z lm Score = −1.67
NG = 0
DB = 0
SM = 0
Delta_r
Doxorubicin
YJR075W HOC1
ZShift = −1.2
Z lm Score = −0.85
NG = 0
DB = 0
SM = 0
Delta_AUC
Doxorubicin
YPL193W RSA1
ZShift = 10.6
Z lm Score = 9.26
NG = 0
DB = 0
SM = 0
Delta_L
Doxorubicin
YPL193W RSA1
ZShift = −0.3
Z lm Score = −2.01
NG = 0
DB = 0
SM = 0
Delta_K
Doxorubicin
YPL193W RSA1
ZShift = −3.88
Z lm Score = 1.28
NG = 0
DB = 0
SM = 0
Delta_r
Doxorubicin
YPL193W RSA1
ZShift = −2.69
Z lm Score = −0.37
NG = 0
DB = 0
SM = 0
Delta_AUC
Doxorubicin
YOR322C LDB19
ZShift = 1.25
Z lm Score = 9.22
NG = 0
DB = 0
SM = 0
Delta_L
Doxorubicin
YOR322C LDB19
ZShift = 2.22
Z lm Score = −1.03
NG = 0
DB = 0
SM = 0
Delta_K
Doxorubicin
YOR322C LDB19
ZShift = −1.5
Z lm Score = −1.04
NG = 0
DB = 0
SM = 0
Delta_r
Doxorubicin
YOR322C LDB19
ZShift = 1.73
Z lm Score = −0.57
NG = 0
DB = 0
SM = 0
Delta_AUC
Doxorubicin

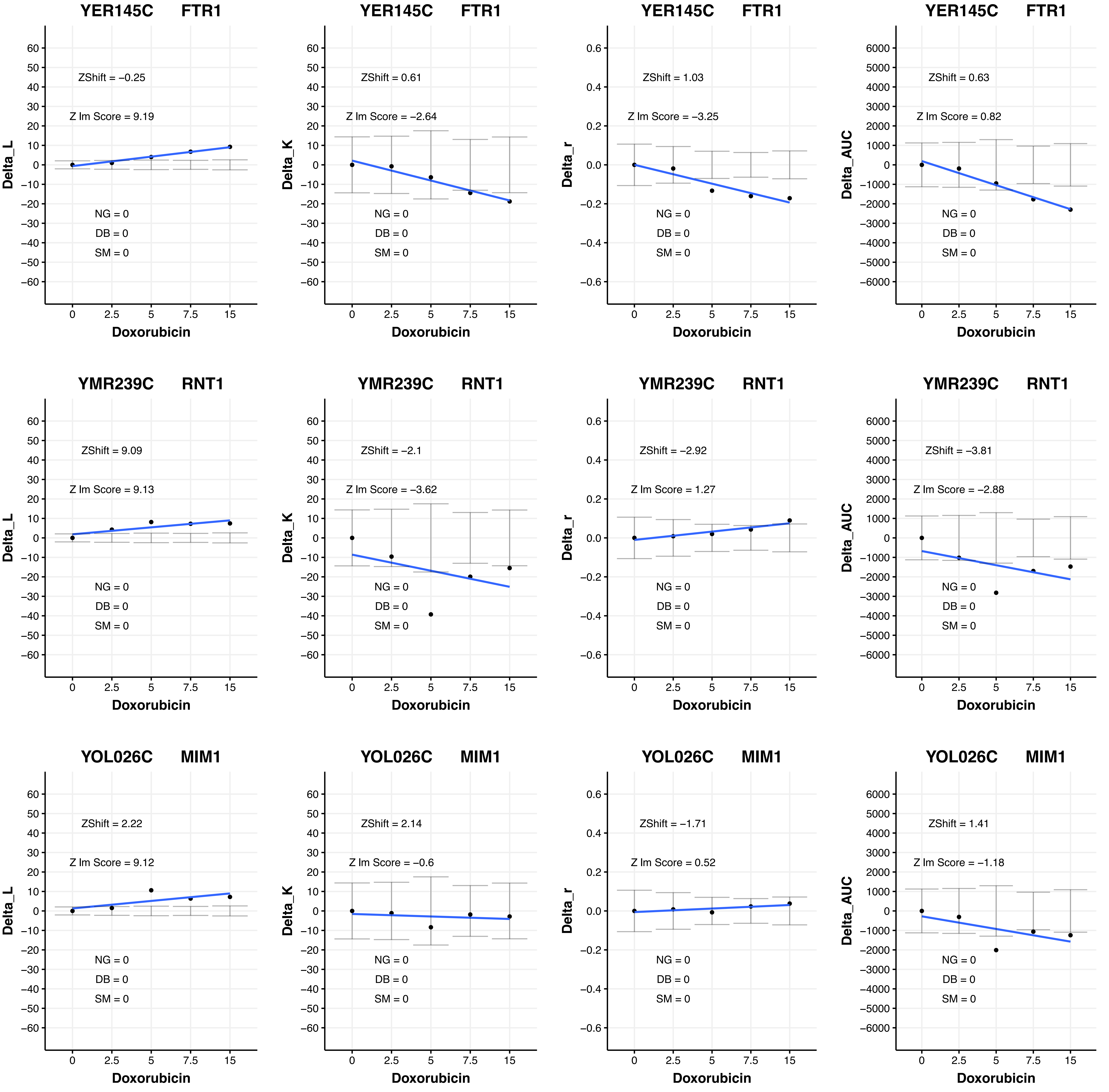
YER145C FTR1
ZShift = −0.25
Z lm Score = 9.19
NG = 0
DB = 0
SM = 0
Delta_L
Doxorubicin
YER145C FTR1
ZShift = 0.61
Z lm Score = −2.64
NG = 0
DB = 0
SM = 0
Delta_K
Doxorubicin
YER145C FTR1
ZShift = 1.03
Z lm Score = −3.25
NG = 0
DB = 0
SM = 0
Delta_r
Doxorubicin
YER145C FTR1
ZShift = 0.63
Z lm Score = 0.82
NG = 0
DB = 0
SM = 0
Delta_AUC
Doxorubicin
YMR239C RNT1
ZShift = 9.09
Z lm Score = 9.13
NG = 0
DB = 0
SM = 0
Delta_L
Doxorubicin
YMR239C RNT1
ZShift = −2.1
Z lm Score = −3.62
NG = 0
DB = 0
SM = 0
Delta_K
Doxorubicin
YMR239C RNT1
ZShift = −2.92
Z lm Score = 1.27
NG = 0
DB = 0
SM = 0
Delta_r
Doxorubicin
YMR239C RNT1
ZShift = −3.81
Z lm Score = −2.88
NG = 0
DB = 0
SM = 0
Delta_AUC
Doxorubicin
YOL026C MIM1
ZShift = 2.22
Z lm Score = 9.12
NG = 0
DB = 0
SM = 0
Delta_L
Doxorubicin
YOL026C MIM1
ZShift = 2.14
Z lm Score = −0.6
NG = 0
DB = 0
SM = 0
Delta_K
Doxorubicin
YOL026C MIM1
ZShift = −1.71
Z lm Score = 0.52
NG = 0
DB = 0
SM = 0
Delta_r
Doxorubicin
YOL026C MIM1
ZShift = 1.41
Z lm Score = −1.18
NG = 0
DB = 0
SM = 0
Delta_AUC
Doxorubicin

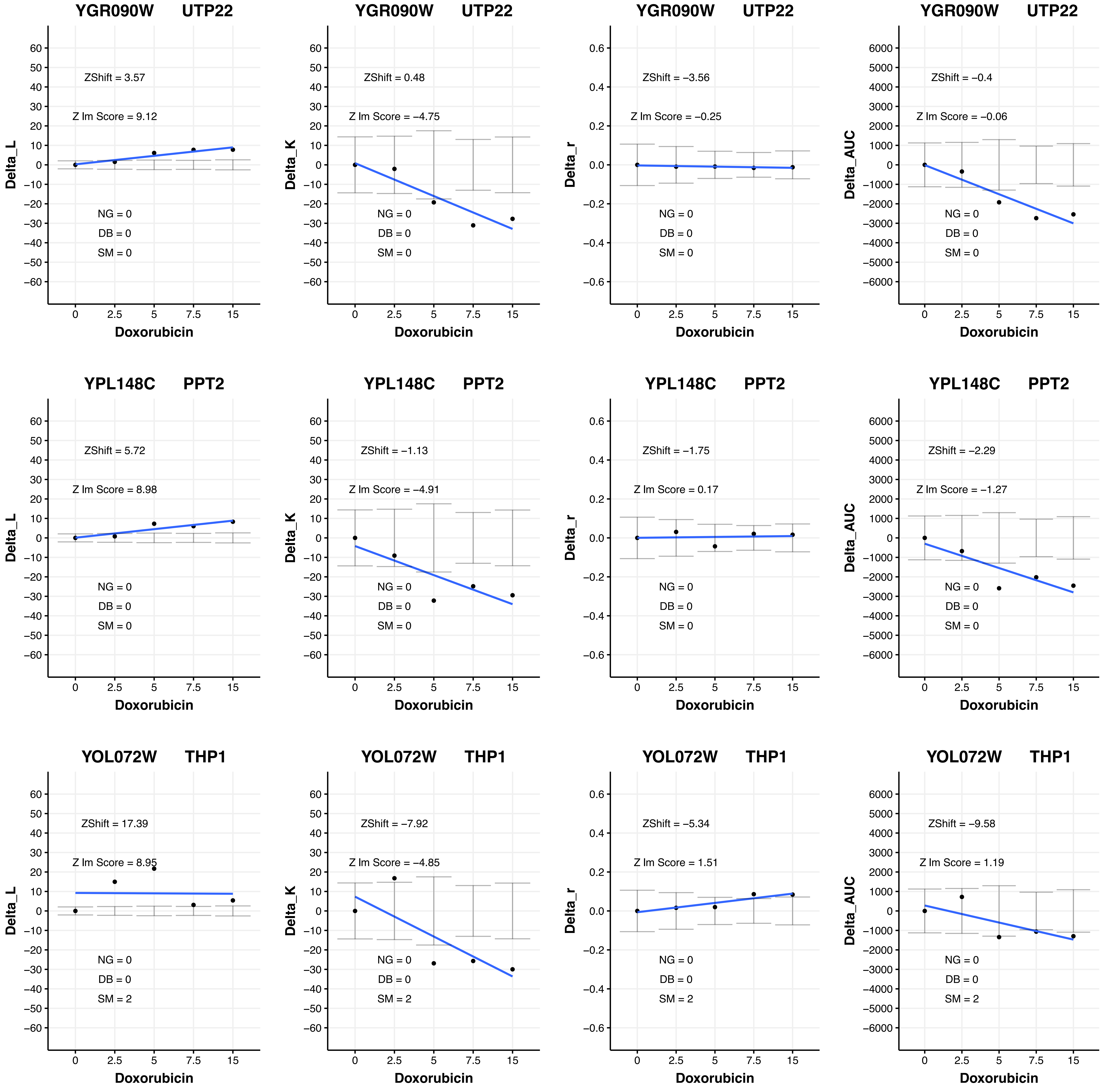
YGR090W UTP22
ZShift = 3.57
Z lm Score = 9.12
NG = 0
DB = 0
SM = 0
Delta_L
Doxorubicin
YGR090W UTP22
ZShift = 0.48
Z lm Score = −4.75
NG = 0
DB = 0
SM = 0
Delta_K
Doxorubicin
YGR090W UTP22
ZShift = −3.56
Z lm Score = −0.25
NG = 0
DB = 0
SM = 0
Delta_r
Doxorubicin
YGR090W UTP22
ZShift = −0.4
Z lm Score = −0.06
NG = 0
DB = 0
SM = 0
Delta_AUC
Doxorubicin
YPL148C PPT2
ZShift = 5.72
Z lm Score = 8.98
NG = 0
DB = 0
SM = 0
Delta_L
Doxorubicin
YPL148C PPT2
ZShift = −1.13
Z lm Score = −4.91
NG = 0
DB = 0
SM = 0
Delta_K
Doxorubicin
YPL148C PPT2
ZShift = −1.75
Z lm Score = 0.17
NG = 0
DB = 0
SM = 0
Delta_r
Doxorubicin
YPL148C PPT2
ZShift = −2.29
Z lm Score = −1.27
NG = 0
DB = 0
SM = 0
Delta_AUC
Doxorubicin
YOL072W THP1
ZShift = 17.39
Z lm Score = 8.95
NG = 0
DB = 0
SM = 2
Delta_L
Doxorubicin
YOL072W THP1
ZShift = −7.92
Z lm Score = −4.85
NG = 0
DB = 0
SM = 2
Delta_K
Doxorubicin
YOL072W THP1
ZShift = −5.34
Z lm Score = 1.51
NG = 0
DB = 0
SM = 2
Delta_r
Doxorubicin
YOL072W THP1
ZShift = −9.58
Z lm Score = 1.19
NG = 0
DB = 0
SM = 2
Delta_AUC
Doxorubicin

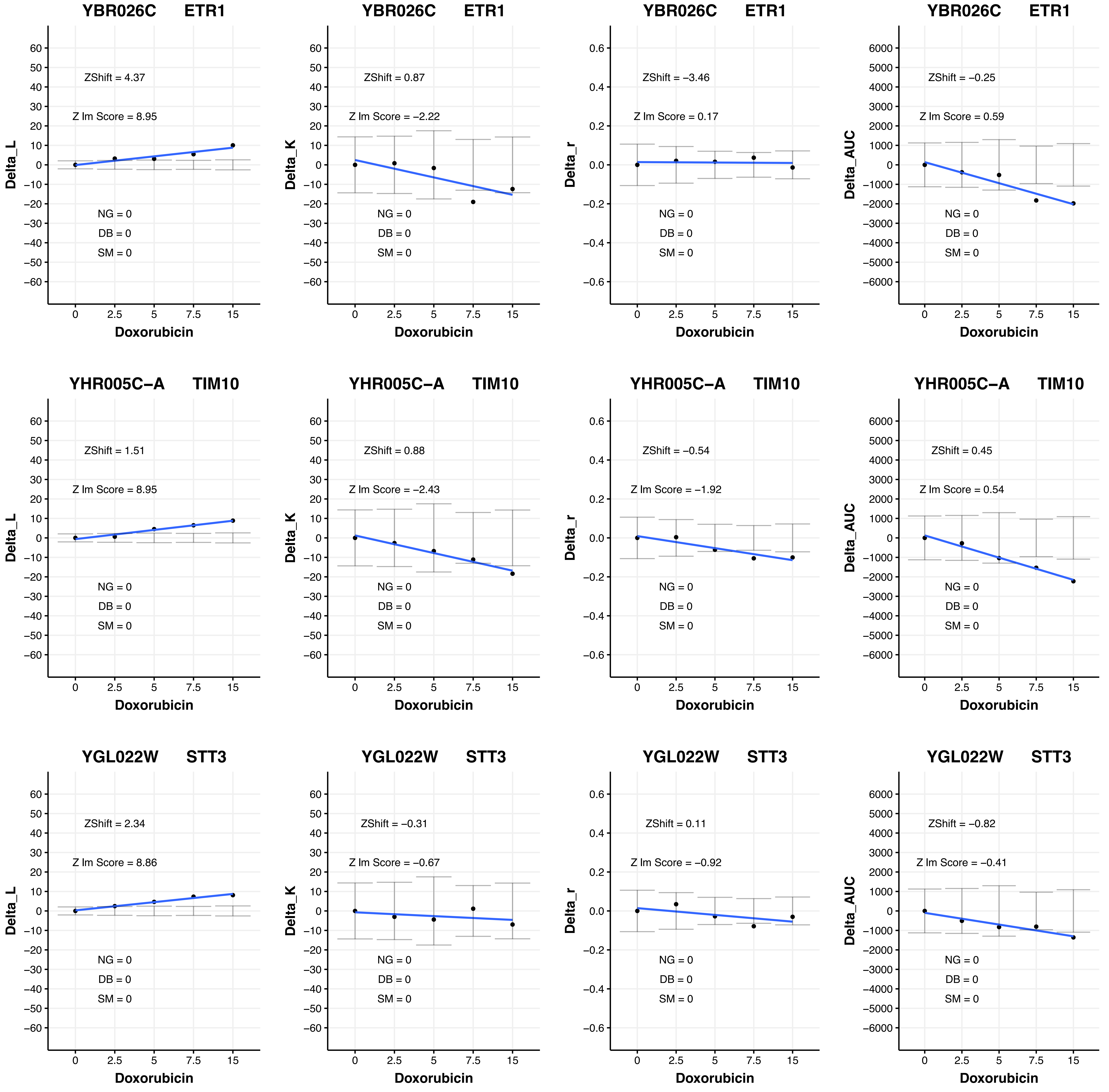
YBR026C ETR1
ZShift = 4.37
Z lm Score = 8.95
NG = 0
DB = 0
SM = 0
Delta_L
Doxorubicin
YBR026C ETR1
ZShift = 0.87
Z lm Score = −2.22
NG = 0
DB = 0
SM = 0
Delta_K
Doxorubicin
YBR026C ETR1
ZShift = −3.46
Z lm Score = 0.17
NG = 0
DB = 0
SM = 0
Delta_r
Doxorubicin
YBR026C ETR1
ZShift = −0.25
Z lm Score = 0.59
NG = 0
DB = 0
SM = 0
Delta_AUC
Doxorubicin
YHR005C−A TIM10
ZShift = 1.51
Z lm Score = 8.95
NG = 0
DB = 0
SM = 0
Delta_L
Doxorubicin
YHR005C−A TIM10
ZShift = 0.88
Z lm Score = −2.43
NG = 0
DB = 0
SM = 0
Delta_K
Doxorubicin
YHR005C−A TIM10
ZShift = −0.54
Z lm Score = −1.92
NG = 0
DB = 0
SM = 0
Delta_r
Doxorubicin
YHR005C−A TIM10
ZShift = 0.45
Z lm Score = 0.54
NG = 0
DB = 0
SM = 0
Delta_AUC
Doxorubicin
YGL022W STT3
ZShift = 2.34
Z lm Score = 8.86
NG = 0
DB = 0
SM = 0
Delta_L
Doxorubicin
YGL022W STT3
ZShift = −0.31
Z lm Score = −0.67
NG = 0
DB = 0
SM = 0
Delta_K
Doxorubicin
YGL022W STT3
ZShift = 0.11
Z lm Score = −0.92
NG = 0
DB = 0
SM = 0
Delta_r
Doxorubicin
YGL022W STT3
ZShift = −0.82
Z lm Score = −0.41
NG = 0
DB = 0
SM = 0
Delta_AUC
Doxorubicin

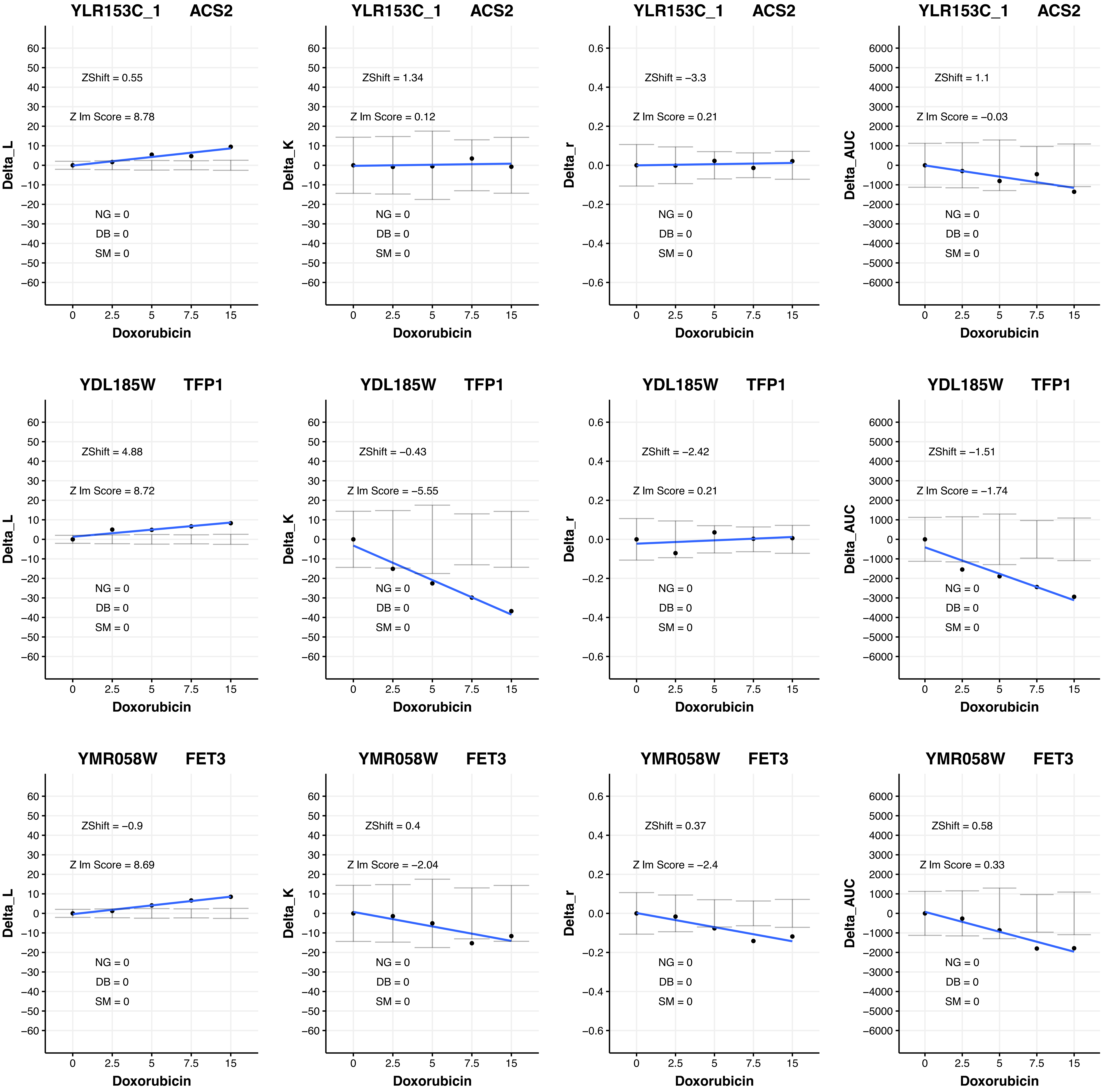
YLR153C_1 ACS2
ZShift = 0.55
Z lm Score = 8.78
NG = 0
DB = 0
SM = 0
Delta_L
Doxorubicin
YLR153C_1 ACS2
ZShift = 1.34
Z lm Score = 0.12
NG = 0
DB = 0
SM = 0
Delta_K
Doxorubicin
YLR153C_1 ACS2
ZShift = −3.3
Z lm Score = 0.21
NG = 0
DB = 0
SM = 0
Delta_r
Doxorubicin
YLR153C_1 ACS2
ZShift = 1.1
Z lm Score = −0.03
NG = 0
DB = 0
SM = 0
Delta_AUC
Doxorubicin
YDL185W TFP1
ZShift = 4.88
Z lm Score = 8.72
NG = 0
DB = 0
SM = 0
Delta_L
Doxorubicin
YDL185W TFP1
ZShift = −0.43
Z lm Score = −5.55
NG = 0
DB = 0
SM = 0
Delta_K
Doxorubicin
YDL185W TFP1
ZShift = −2.42
Z lm Score = 0.21
NG = 0
DB = 0
SM = 0
Delta_r
Doxorubicin
YDL185W TFP1
ZShift = −1.51
Z lm Score = −1.74
NG = 0
DB = 0
SM = 0
Delta_AUC
Doxorubicin
YMR058W FET3
ZShift = −0.9
Z lm Score = 8.69
NG = 0
DB = 0
SM = 0
Delta_L
Doxorubicin
YMR058W FET3
ZShift = 0.4
Z lm Score = −2.04
NG = 0
DB = 0
SM = 0
Delta_K
Doxorubicin
YMR058W FET3
ZShift = 0.37
Z lm Score = −2.4
NG = 0
DB = 0
SM = 0
Delta_r
Doxorubicin
YMR058W FET3
ZShift = 0.58
Z lm Score = 0.33
NG = 0
DB = 0
SM = 0
Delta_AUC
Doxorubicin

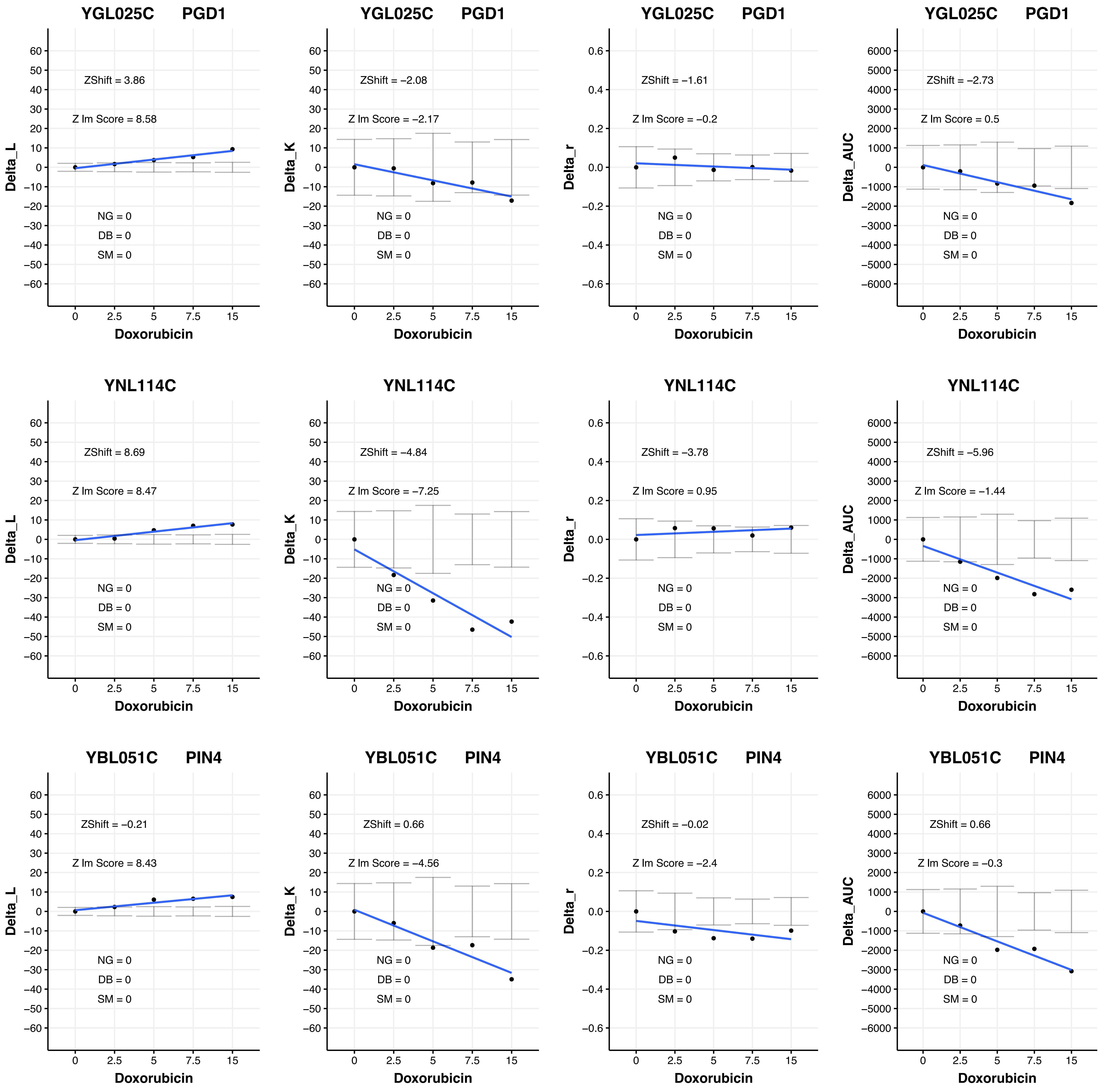

YGL025C PGD1
ZShift = 3.86
Z lm Score = 8.58
NG = 0
DB = 0
SM = 0
Delta_L
Doxorubicin
YGL025C PGD1
ZShift = −2.08
Z lm Score = −2.17
NG = 0
DB = 0
SM = 0
Delta_K
Doxorubicin
YGL025C PGD1
ZShift = −1.61
Z lm Score = −0.2
NG = 0
DB = 0
SM = 0
Delta_r
Doxorubicin
YGL025C PGD1
ZShift = −2.73
Z lm Score = 0.5
NG = 0
DB = 0
SM = 0
Delta_AUC
Doxorubicin
YNL114C
ZShift = 8.69
Z lm Score = 8.47
NG = 0
DB = 0
SM = 0
Delta_L
Doxorubicin
YNL114C
ZShift = −4.84
Z lm Score = −7.25
NG = 0
DB = 0
SM = 0
Delta_K
Doxorubicin
YNL114C
ZShift = −3.78
Z lm Score = 0.95
NG = 0
DB = 0
SM = 0
Delta_r
Doxorubicin
YNL114C
ZShift = −5.96
Z lm Score = −1.44
NG = 0
DB = 0
SM = 0
Delta_AUC
Doxorubicin
YBL051C PIN4
ZShift = −0.21
Z lm Score = 8.43
NG = 0
DB = 0
SM = 0
Delta_L
Doxorubicin
YBL051C PIN4
ZShift = 0.66
Z lm Score = −4.56
NG = 0
DB = 0
SM = 0
Delta_K
Doxorubicin
YBL051C PIN4
ZShift = −0.02
Z lm Score = −2.4
NG = 0
DB = 0
SM = 0
Delta_r
Doxorubicin
YBL051C PIN4
ZShift = 0.66
Z lm Score = −0.3
NG = 0
DB = 0
SM = 0
Delta_AUC
Doxorubicin

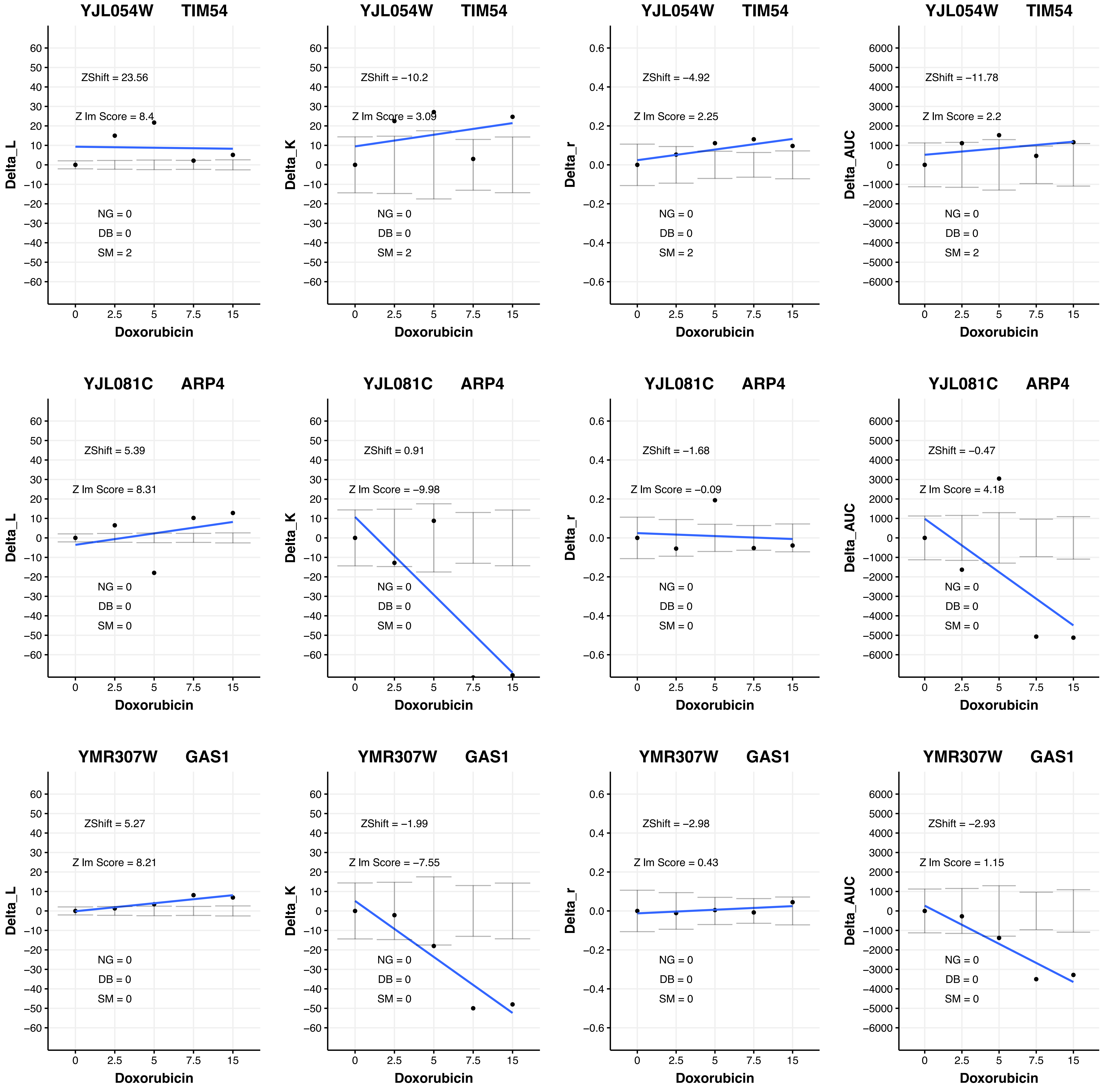

YJL054W TIM54
ZShift = 23.56
Z lm Score = 8.4
NG = 0
DB = 0
SM = 2
Delta_L
Doxorubicin
YJL054W TIM54
ZShift = −10.2
Z lm Score = 3.09
NG = 0
DB = 0
SM = 2
Delta_K
Doxorubicin
YJL054W TIM54
ZShift = −4.92
Z lm Score = 2.25
NG = 0
DB = 0
SM = 2
Delta_r
Doxorubicin
YJL054W TIM54
ZShift = −11.78
Z lm Score = 2.2
NG = 0
DB = 0
SM = 2
Delta_AUC
Doxorubicin
YJL081C ARP4
ZShift = 5.39
Z lm Score = 8.31
NG = 0
DB = 0
SM = 0
Delta_L
Doxorubicin
YJL081C ARP4
ZShift = 0.91
Z lm Score = −9.98
NG = 0
DB = 0
SM = 0
Delta_K
Doxorubicin
YJL081C ARP4
ZShift = −1.68
Z lm Score = −0.09
NG = 0
DB = 0
SM = 0
Delta_r
Doxorubicin
YJL081C ARP4
ZShift = −0.47
Z lm Score = 4.18
NG = 0
DB = 0
SM = 0
Delta_AUC
Doxorubicin
YMR307W GAS1
ZShift = 5.27
Z lm Score = 8.21
NG = 0
DB = 0
SM = 0
Delta_L
Doxorubicin
YMR307W GAS1
ZShift = −1.99
Z lm Score = −7.55
NG = 0
DB = 0
SM = 0
Delta_K
Doxorubicin
YMR307W GAS1
ZShift = −2.98
Z lm Score = 0.43
NG = 0
DB = 0
SM = 0
Delta_r
Doxorubicin
YMR307W GAS1
ZShift = −2.93
Z lm Score = 1.15
NG = 0
DB = 0
SM = 0
Delta_AUC
Doxorubicin

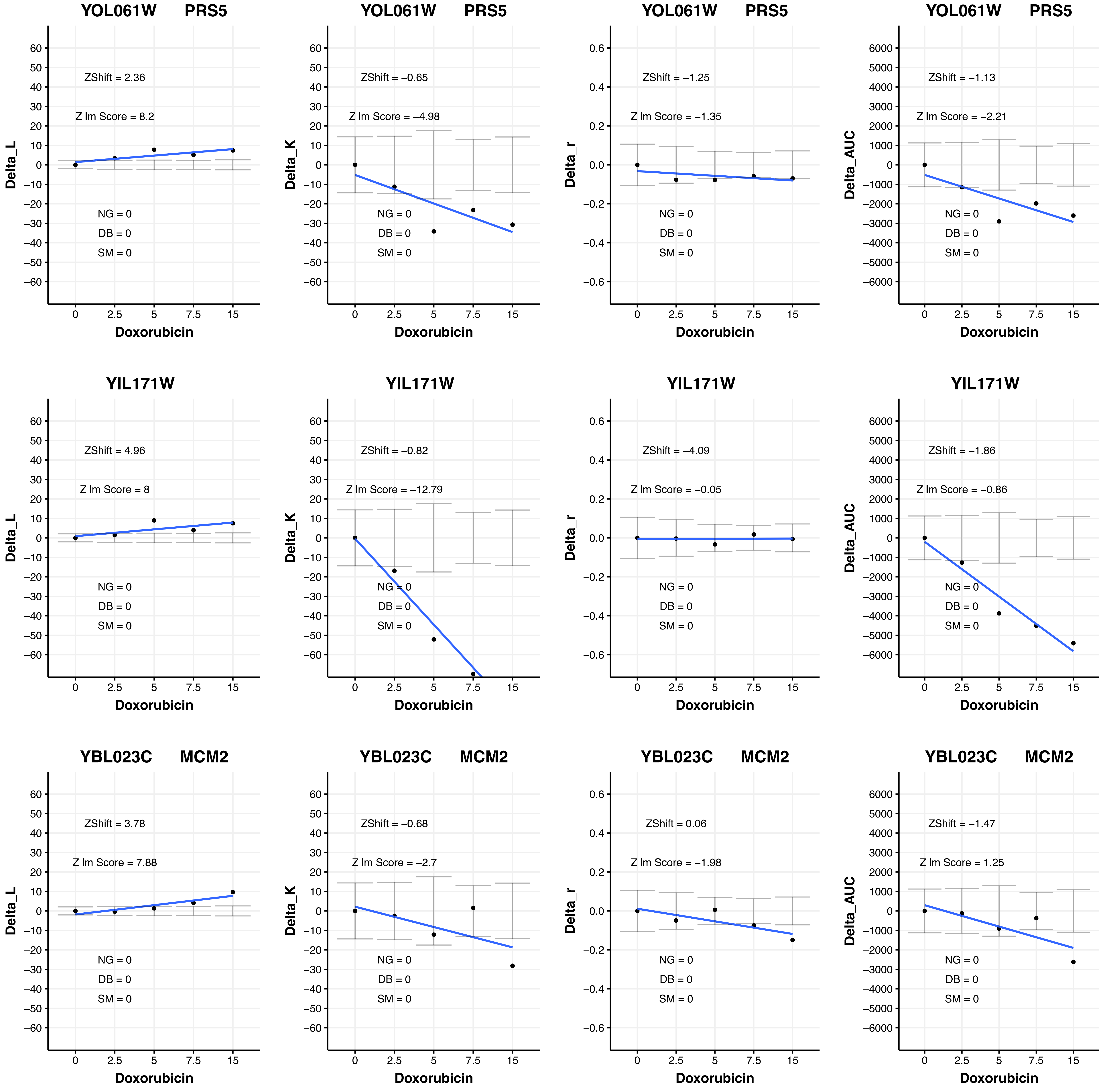

YOL061W PRS5
ZShift = 2.36
Z lm Score = 8.2
NG = 0
DB = 0
SM = 0
Delta_L
Doxorubicin
YOL061W PRS5
ZShift = −0.65
Z lm Score = −4.98
NG = 0
DB = 0
SM = 0
Delta_K
Doxorubicin
YOL061W PRS5
ZShift = −1.25
Z lm Score = −1.35
NG = 0
DB = 0
SM = 0
Delta_r
Doxorubicin
YOL061W PRS5
ZShift = −1.13
Z lm Score = −2.21
NG = 0
DB = 0
SM = 0
Delta_AUC
Doxorubicin
YIL171W
ZShift = 4.96
Z lm Score = 8
NG = 0
DB = 0
SM = 0
Delta_L
Doxorubicin
YIL171W
ZShift = −0.82
Z lm Score = −12.79
NG = 0
DB = 0
SM = 0
Delta_K
Doxorubicin
YIL171W
ZShift = −4.09
Z lm Score = −0.05
NG = 0
DB = 0
SM = 0
Delta_r
Doxorubicin
YIL171W
ZShift = −1.86
Z lm Score = −0.86
NG = 0
DB = 0
SM = 0
Delta_AUC
Doxorubicin
YBL023C MCM2
ZShift = 3.78
Z lm Score = 7.88
NG = 0
DB = 0
SM = 0
Delta_L
Doxorubicin
YBL023C MCM2
ZShift = −0.68
Z lm Score = −2.7
NG = 0
DB = 0
SM = 0
Delta_K
Doxorubicin
YBL023C MCM2
ZShift = 0.06
Z lm Score = −1.98
NG = 0
DB = 0
SM = 0
Delta_r
Doxorubicin
YBL023C MCM2
ZShift = −1.47
Z lm Score = 1.25
NG = 0
DB = 0
SM = 0
Delta_AUC
Doxorubicin

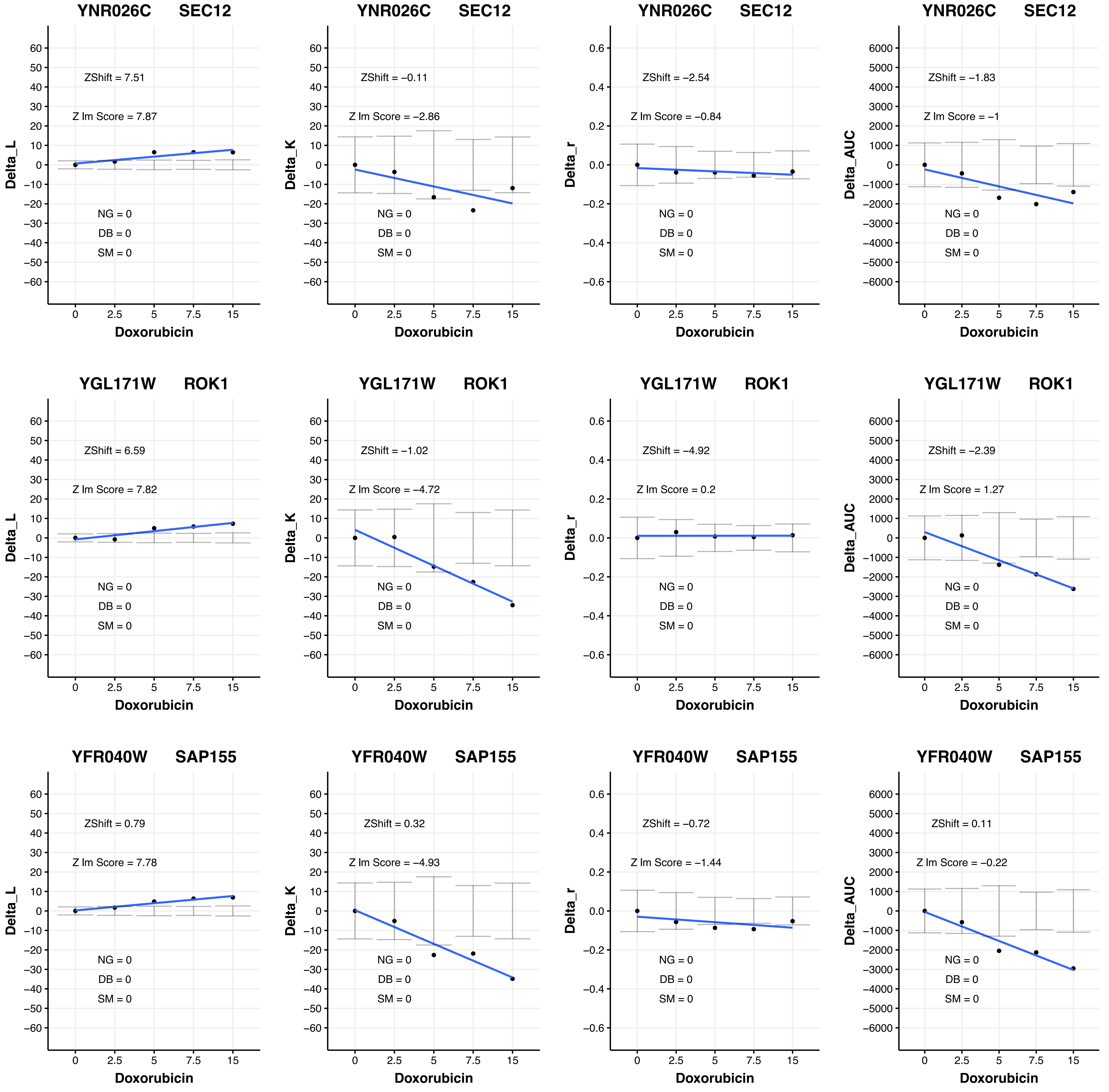
YNR026C SEC12
ZShift = 7.51
Z lm Score = 7.87
NG = 0
DB = 0
SM = 0
Delta_L
Doxorubicin
YNR026C SEC12
ZShift = −0.11
Z lm Score = −2.86
NG = 0
DB = 0
SM = 0
Delta_K
Doxorubicin
YNR026C SEC12
ZShift = −2.54
Z lm Score = −0.84
NG = 0
DB = 0
SM = 0
Delta_r
Doxorubicin
YNR026C SEC12
ZShift = −1.83
Z lm Score = −1
NG = 0
DB = 0
SM = 0
Delta_AUC
Doxorubicin
YGL171W ROK1
ZShift = 6.59
Z lm Score = 7.82
NG = 0
DB = 0
SM = 0
Delta_L
Doxorubicin
YGL171W ROK1
ZShift = −1.02
Z lm Score = −4.72
NG = 0
DB = 0
SM = 0
Delta_K
Doxorubicin
YGL171W ROK1
ZShift = −4.92
Z lm Score = 0.2
NG = 0
DB = 0
SM = 0
Delta_r
Doxorubicin
YGL171W ROK1
ZShift = −2.39
Z lm Score = 1.27
NG = 0
DB = 0
SM = 0
Delta_AUC
Doxorubicin
YFR040W SAP155
ZShift = 0.79
Z lm Score = 7.78
NG = 0
DB = 0
SM = 0
Delta_L
Doxorubicin
YFR040W SAP155
ZShift = 0.32
Z lm Score = −4.93
NG = 0
DB = 0
SM = 0
Delta_K
Doxorubicin
YFR040W SAP155
ZShift = −0.72
Z lm Score = −1.44
NG = 0
DB = 0
SM = 0
Delta_r
Doxorubicin
YFR040W SAP155
ZShift = 0.11
Z lm Score = −0.22
NG = 0
DB = 0
SM = 0
Delta_AUC
Doxorubicin

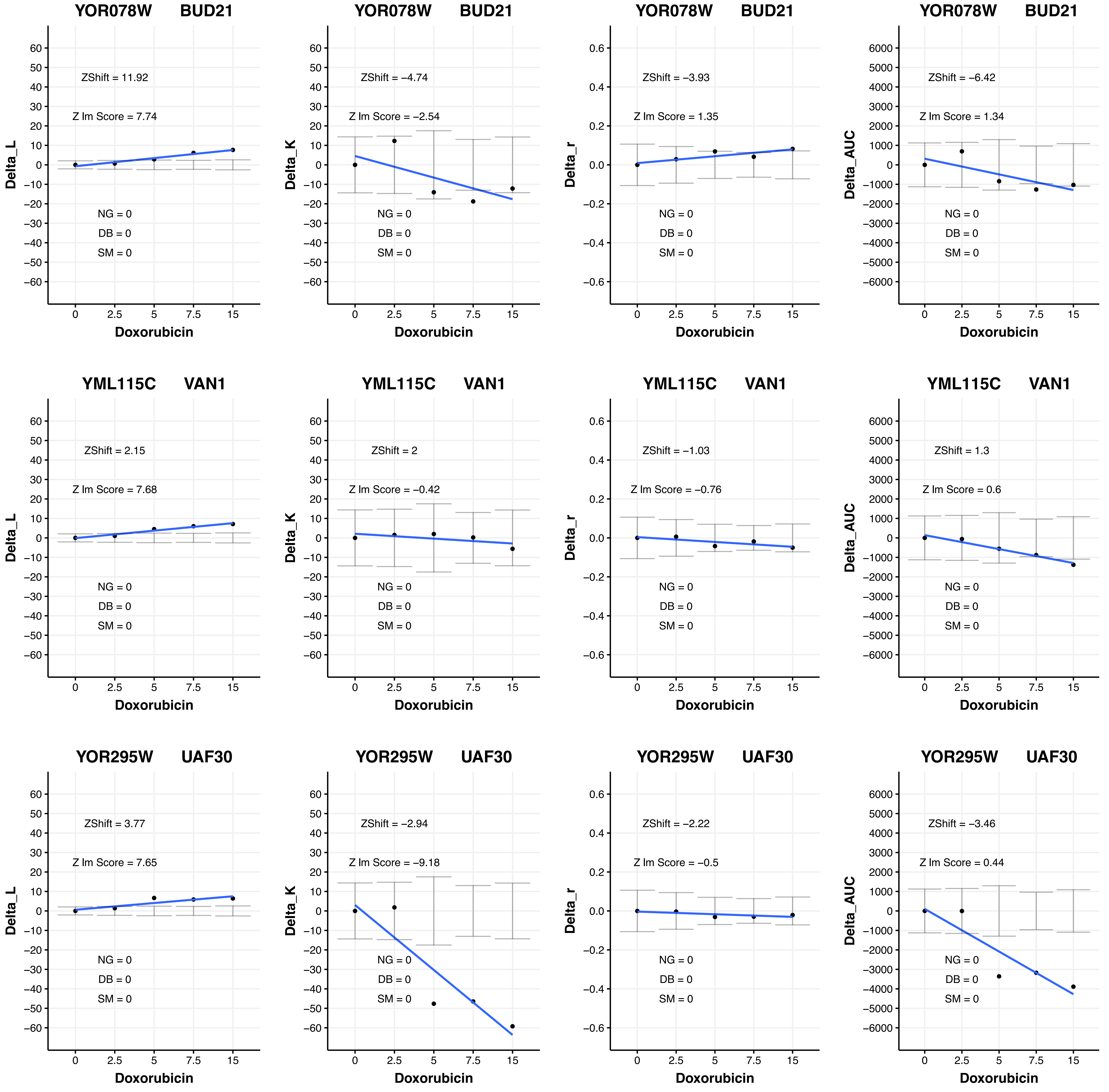
YOR078W BUD21
ZShift = 11.92
Z lm Score = 7.74
NG = 0
DB = 0
SM = 0
Delta_L
Doxorubicin
YOR078W BUD21
ZShift = −4.74
Z lm Score = −2.54
NG = 0
DB = 0
SM = 0
Delta_K
Doxorubicin
YOR078W BUD21
ZShift = −3.93
Z lm Score = 1.35
NG = 0
DB = 0
SM = 0
Delta_r
Doxorubicin
YOR078W BUD21
ZShift = −6.42
Z lm Score = 1.34
NG = 0
DB = 0
SM = 0
Delta_AUC
Doxorubicin
YML115C VAN1
ZShift = 2.15
Z lm Score = 7.68
NG = 0
DB = 0
SM = 0
Delta_L
Doxorubicin
YML115C VAN1
ZShift = 2
Z lm Score = −0.42
NG = 0
DB = 0
SM = 0
Delta_K
Doxorubicin
YML115C VAN1
ZShift = −1.03
Z lm Score = −0.76
NG = 0
DB = 0
SM = 0
Delta_r
Doxorubicin
YML115C VAN1
ZShift = 1.3
Z lm Score = 0.6
NG = 0
DB = 0
SM = 0
Delta_AUC
Doxorubicin
YOR295W UAF30
ZShift = 3.77
Z lm Score = 7.65
NG = 0
DB = 0
SM = 0
Delta_L
Doxorubicin
YOR295W UAF30
ZShift = −2.94
Z lm Score = −9.18
NG = 0
DB = 0
SM = 0
Delta_K
Doxorubicin
YOR295W UAF30
ZShift = −2.22
Z lm Score = −0.5
NG = 0
DB = 0
SM = 0
Delta_r
Doxorubicin
YOR295W UAF30
ZShift = −3.46
Z lm Score = 0.44
NG = 0
DB = 0
SM = 0
Delta_AUC
Doxorubicin

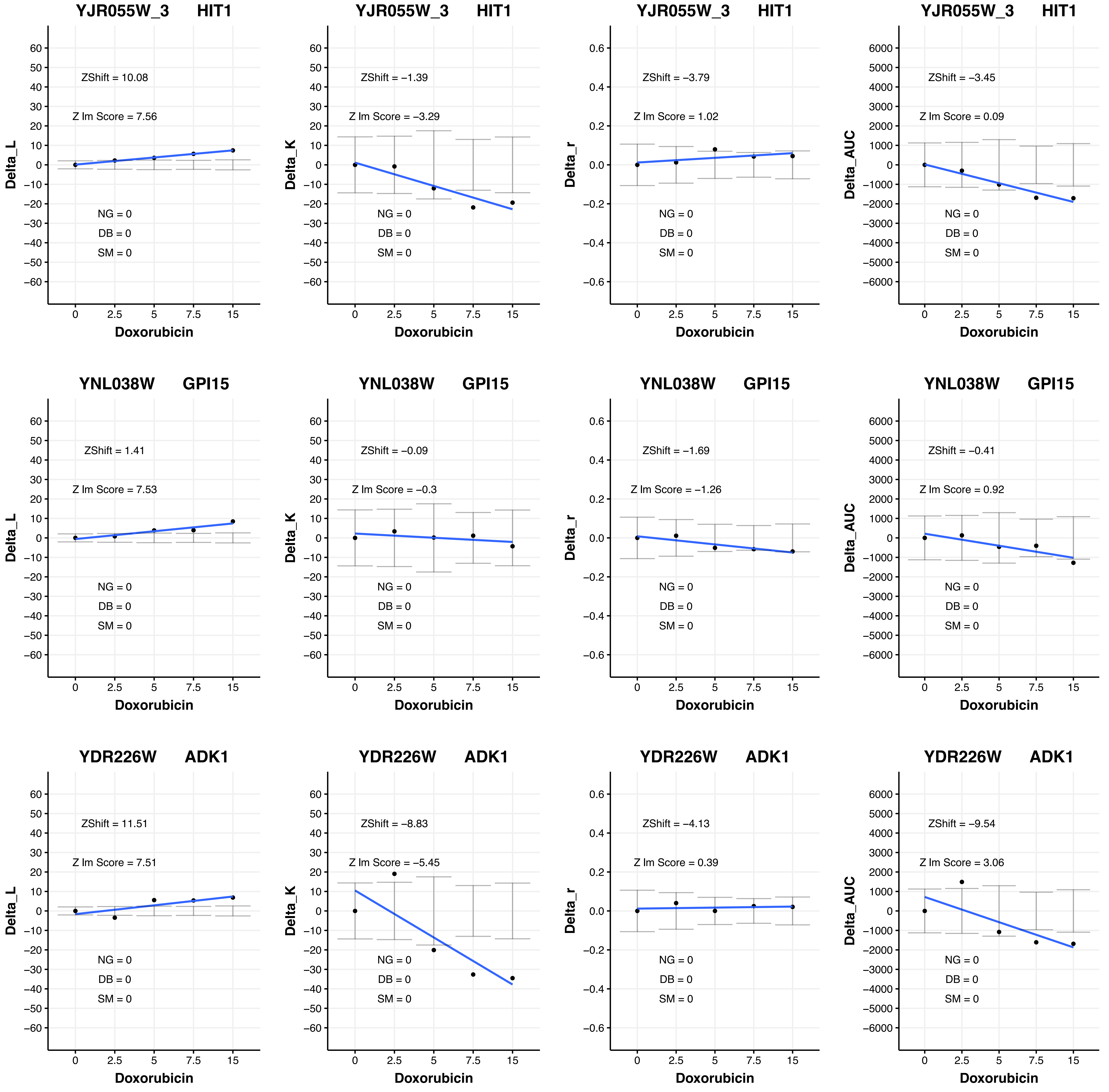
YJR055W_3 HIT1
ZShift = 10.08
Z lm Score = 7.56
NG = 0
DB = 0
SM = 0
Delta_L
Doxorubicin
YJR055W_3 HIT1
ZShift = −1.39
Z lm Score = −3.29
NG = 0
DB = 0
SM = 0
Delta_K
Doxorubicin
YJR055W_3 HIT1
ZShift = −3.79
Z lm Score = 1.02
NG = 0
DB = 0
SM = 0
Delta_r
Doxorubicin
YJR055W_3 HIT1
ZShift = −3.45
Z lm Score = 0.09
NG = 0
DB = 0
SM = 0
Delta_AUC
Doxorubicin
YNL038W GPI15
ZShift = 1.41
Z lm Score = 7.53
NG = 0
DB = 0
SM = 0
Delta_L
Doxorubicin
YNL038W GPI15
ZShift = −0.09
Z lm Score = −0.3
NG = 0
DB = 0
SM = 0
Delta_K
Doxorubicin
YNL038W GPI15
ZShift = −1.69
Z lm Score = −1.26
NG = 0
DB = 0
SM = 0
Delta_r
Doxorubicin
YNL038W GPI15
ZShift = −0.41
Z lm Score = 0.92
NG = 0
DB = 0
SM = 0
Delta_AUC
Doxorubicin
YDR226W ADK1
ZShift = 11.51
Z lm Score = 7.51
NG = 0
DB = 0
SM = 0
Delta_L
Doxorubicin
YDR226W ADK1
ZShift = −8.83
Z lm Score = −5.45
NG = 0
DB = 0
SM = 0
Delta_K
Doxorubicin
YDR226W ADK1
ZShift = −4.13
Z lm Score = 0.39
NG = 0
DB = 0
SM = 0
Delta_r
Doxorubicin
YDR226W ADK1
ZShift = −9.54
Z lm Score = 3.06
NG = 0
DB = 0
SM = 0
Delta_AUC
Doxorubicin

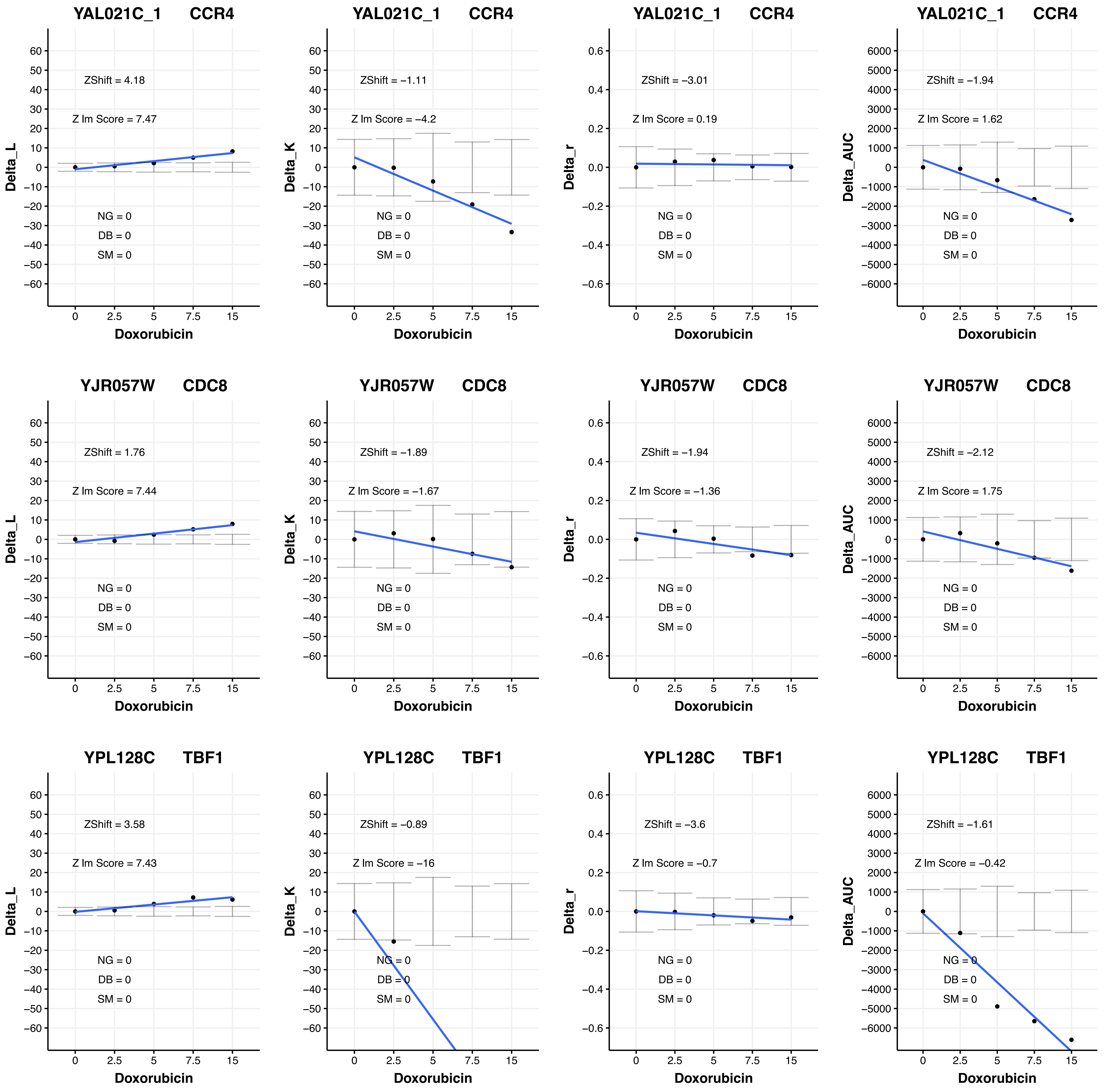

YAL021C_1 CCR4
ZShift = 4.18
Z lm Score = 7.47
NG = 0
DB = 0
SM = 0
Delta_L
Doxorubicin
YAL021C_1 CCR4
ZShift = −1.11
Z lm Score = −4.2
NG = 0
DB = 0
SM = 0
Delta_K
Doxorubicin
YAL021C_1 CCR4
ZShift = −3.01
Z lm Score = 0.19
NG = 0
DB = 0
SM = 0
Delta_r
Doxorubicin
YAL021C_1 CCR4
ZShift = −1.94
Z lm Score = 1.62
NG = 0
DB = 0
SM = 0
Delta_AUC
Doxorubicin
YJR057W CDC8
ZShift = 1.76
Z lm Score = 7.44
NG = 0
DB = 0
SM = 0
Delta_L
Doxorubicin
YJR057W CDC8
ZShift = −1.89
Z lm Score = −1.67
NG = 0
DB = 0
SM = 0
Delta_K
Doxorubicin
YJR057W CDC8
ZShift = −1.94
Z lm Score = −1.36
NG = 0
DB = 0
SM = 0
Delta_r
Doxorubicin
YJR057W CDC8
ZShift = −2.12
Z lm Score = 1.75
NG = 0
DB = 0
SM = 0
Delta_AUC
Doxorubicin
YPL128C TBF1
ZShift = 3.58
Z lm Score = 7.43
NG = 0
DB = 0
SM = 0
Delta_L
Doxorubicin
YPL128C TBF1
ZShift = −0.89
Z lm Score = −16
NG = 0
DB = 0
SM = 0
Delta_K
Doxorubicin
YPL128C TBF1
ZShift = −3.6
Z lm Score = −0.7
NG = 0
DB = 0
SM = 0
Delta_r
Doxorubicin
YPL128C TBF1
ZShift = −1.61
Z lm Score = −0.42
NG = 0
DB = 0
SM = 0
Delta_AUC
Doxorubicin

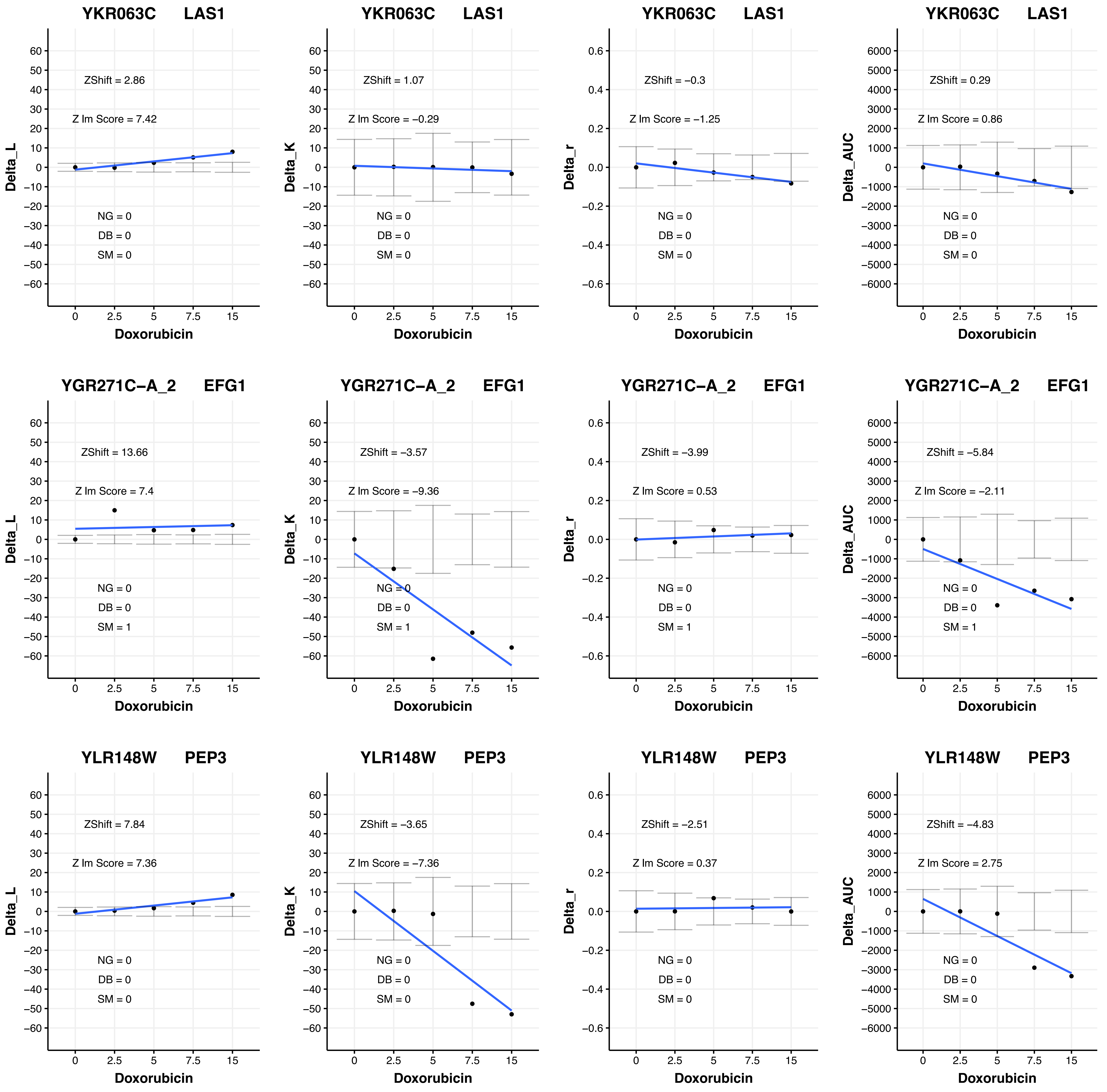
YKR063C LAS1
Delta_L
ZShift = 2.86
Z lm Score = 7.42
NG = 0
DB = 0
SM = 0
Doxorubicin
YKR063C LAS1
Delta_K
ZShift = 1.07
Z lm Score = −0.29
NG = 0
DB = 0
SM = 0
Doxorubicin
YKR063C LAS1
Delta_r
ZShift = −0.3
Z lm Score = −1.25
NG = 0
DB = 0
SM = 0
Doxorubicin
YKR063C LAS1
Delta_AUC
ZShift = 0.29
Z lm Score = 0.86
NG = 0
DB = 0
SM = 0
Doxorubicin
YGR271C−A_2 EFG1
Delta_L
ZShift = 13.66
Z lm Score = 7.4
NG = 0
DB = 0
SM = 1
Doxorubicin
YGR271C−A_2 EFG1
Delta_K
ZShift = −3.57
Z lm Score = −9.36
NG = 0
DB = 0
SM = 1
Doxorubicin
YGR271C−A_2 EFG1
Delta_r
ZShift = −3.99
Z lm Score = 0.53
NG = 0
DB = 0
SM = 1
Doxorubicin
YGR271C−A_2 EFG1
Delta_AUC
ZShift = −5.84
Z lm Score = −2.11
NG = 0
DB = 0
SM = 1
Doxorubicin
YLR148W PEP3
Delta_L
ZShift = 7.84
Z lm Score = 7.36
NG = 0
DB = 0
SM = 0
Doxorubicin
YLR148W PEP3
Delta_K
ZShift = −3.65
Z lm Score = −7.36
NG = 0
DB = 0
SM = 0
Doxorubicin
YLR148W PEP3
Delta_r
ZShift = −2.51
Z lm Score = 0.37
NG = 0
DB = 0
SM = 0
Doxorubicin
YLR148W PEP3
Delta_AUC
ZShift = −4.83
Z lm Score = 2.75
NG = 0
DB = 0
SM = 0
Doxorubicin

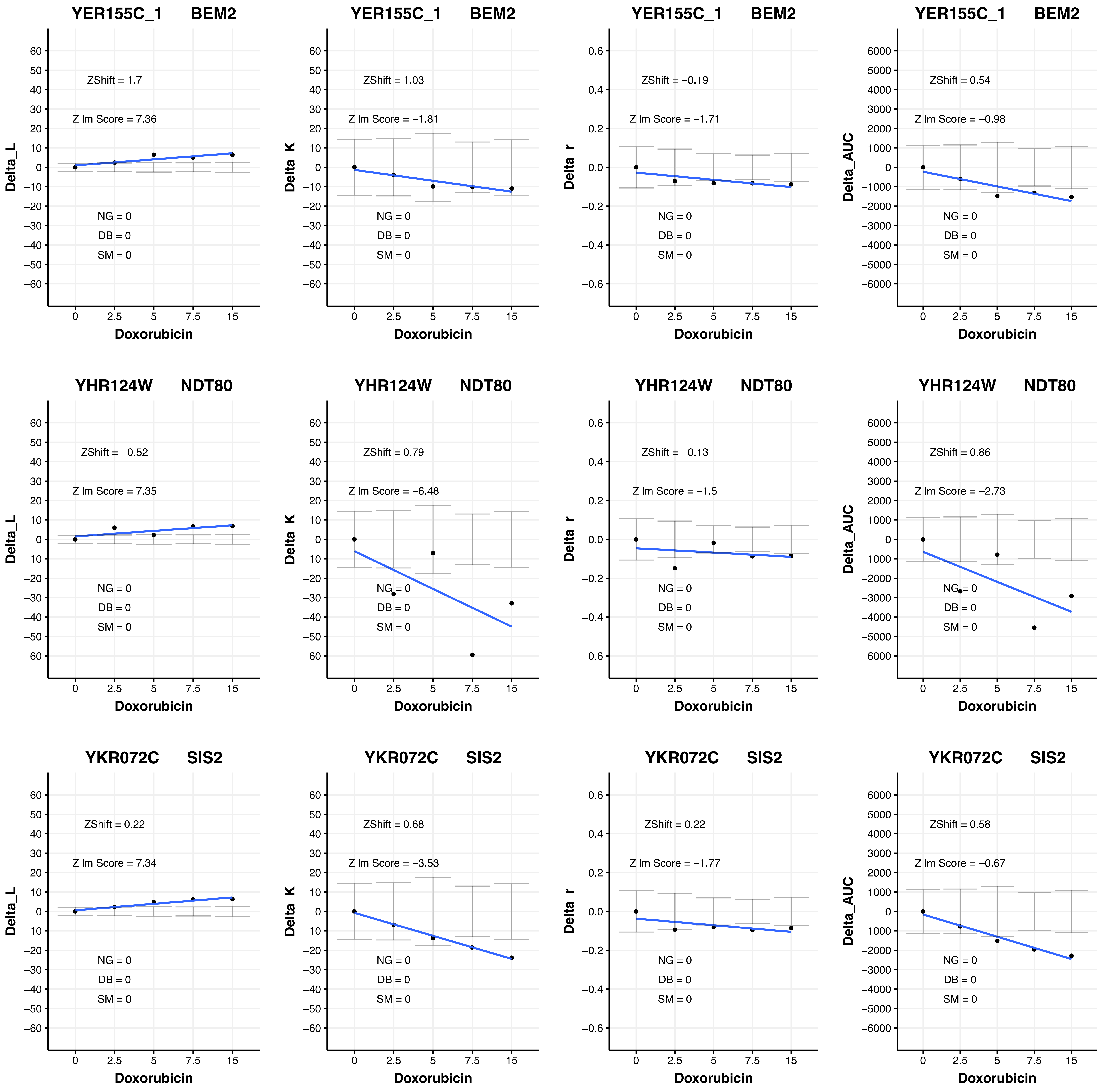
YER155C_1 BEM2
ZShift = 1.7
Z lm Score = 7.36
NG = 0
DB = 0
SM = 0
Delta_L
Doxorubicin
YER155C_1 BEM2
ZShift = 1.03
Z lm Score = −1.81
NG = 0
DB = 0
SM = 0
Delta_K
Doxorubicin
YER155C_1 BEM2
ZShift = −0.19
Z lm Score = −1.71
NG = 0
DB = 0
SM = 0
Delta_r
Doxorubicin
YER155C_1 BEM2
ZShift = 0.54
Z lm Score = −0.98
NG = 0
DB = 0
SM = 0
Delta_AUC
Doxorubicin
YHR124W NDT80
ZShift = −0.52
Z lm Score = 7.35
NG = 0
DB = 0
SM = 0
Delta_L
Doxorubicin
YHR124W NDT80
ZShift = 0.79
Z lm Score = −6.48
NG = 0
DB = 0
SM = 0
Delta_K
Doxorubicin
YHR124W NDT80
ZShift = −0.13
Z lm Score = −1.5
NG = 0
DB = 0
SM = 0
Delta_r
Doxorubicin
YHR124W NDT80
ZShift = 0.86
Z lm Score = −2.73
NG = 0
DB = 0
SM = 0
Delta_AUC
Doxorubicin
YKR072C SIS2
ZShift = 0.22
Z lm Score = 7.34
NG = 0
DB = 0
SM = 0
Delta_L
Doxorubicin
YKR072C SIS2
ZShift = 0.68
Z lm Score = −3.53
NG = 0
DB = 0
SM = 0
Delta_K
Doxorubicin
YKR072C SIS2
ZShift = 0.22
Z lm Score = −1.77
NG = 0
DB = 0
SM = 0
Delta_r
Doxorubicin
YKR072C SIS2
ZShift = 0.58
Z lm Score = −0.67
NG = 0
DB = 0
SM = 0
Delta_AUC
Doxorubicin

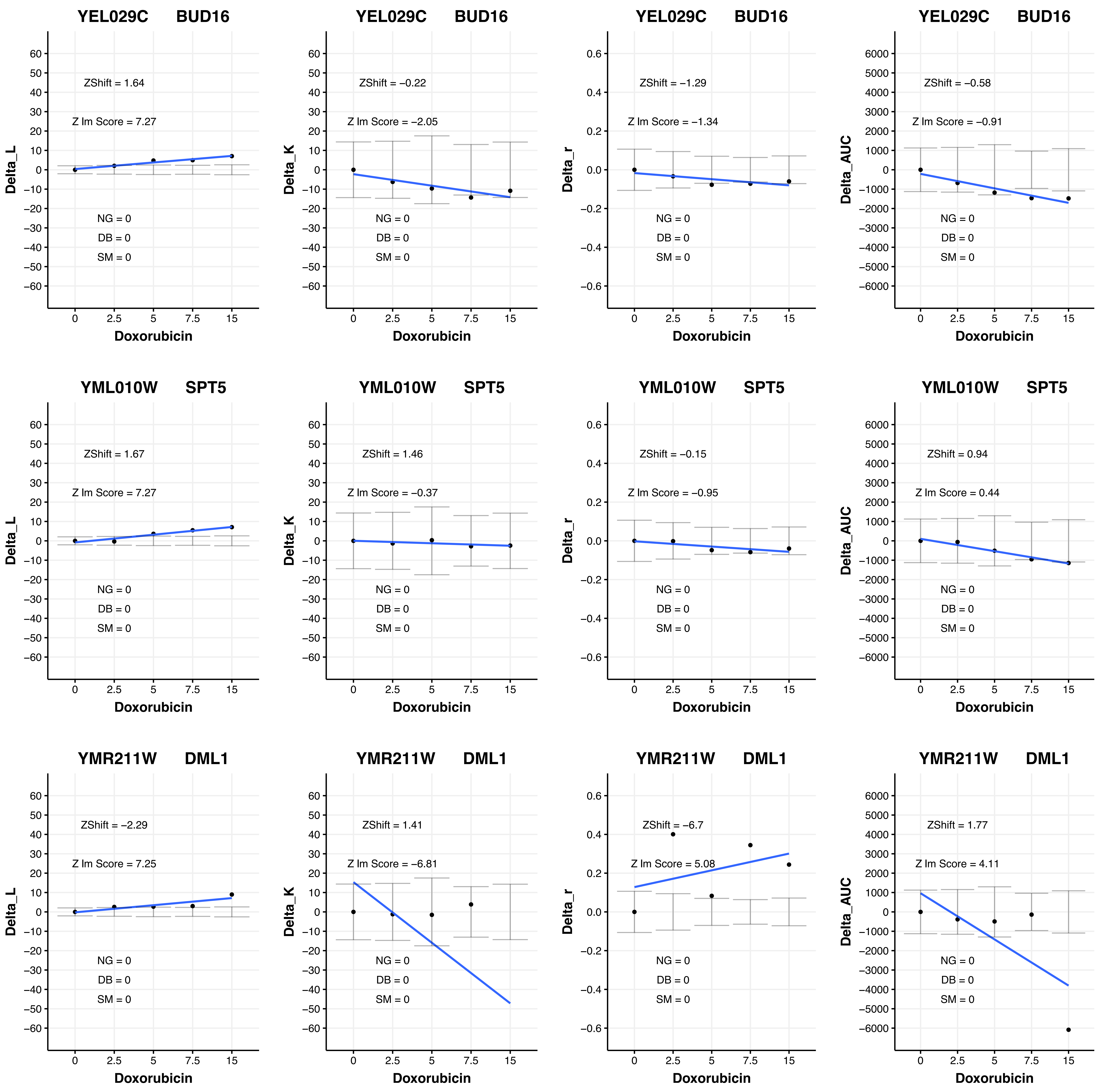
YEL029C BUD16
ZShift = 1.64
Z lm Score = 7.27
NG = 0
DB = 0
SM = 0
Delta_L
Doxorubicin
YEL029C BUD16
ZShift = −0.22
Z lm Score = −2.05
NG = 0
DB = 0
SM = 0
Delta_K
Doxorubicin
YEL029C BUD16
ZShift = −1.29
Z lm Score = −1.34
NG = 0
DB = 0
SM = 0
Delta_r
Doxorubicin
YEL029C BUD16
ZShift = −0.58
Z lm Score = −0.91
NG = 0
DB = 0
SM = 0
Delta_AUC
Doxorubicin
YML010W SPT5
ZShift = 1.67
Z lm Score = 7.27
NG = 0
DB = 0
SM = 0
Delta_L
Doxorubicin
YML010W SPT5
ZShift = 1.46
Z lm Score = −0.37
NG = 0
DB = 0
SM = 0
Delta_K
Doxorubicin
YML010W SPT5
ZShift = −0.15
Z lm Score = −0.95
NG = 0
DB = 0
SM = 0
Delta_r
Doxorubicin
YML010W SPT5
ZShift = 0.94
Z lm Score = 0.44
NG = 0
DB = 0
SM = 0
Delta_AUC
Doxorubicin
YMR211W DML1
ZShift = −2.29
Z lm Score = 7.25
NG = 0
DB = 0
SM = 0
Delta_L
Doxorubicin
YMR211W DML1
ZShift = 1.41
Z lm Score = −6.81
NG = 0
DB = 0
SM = 0
Delta_K
Doxorubicin
YMR211W DML1
ZShift = −6.7
Z lm Score = 5.08
NG = 0
DB = 0
SM = 0
Delta_r
Doxorubicin
YMR211W DML1
ZShift = 1.77
Z lm Score = 4.11
NG = 0
DB = 0
SM = 0
Delta_AUC
Doxorubicin

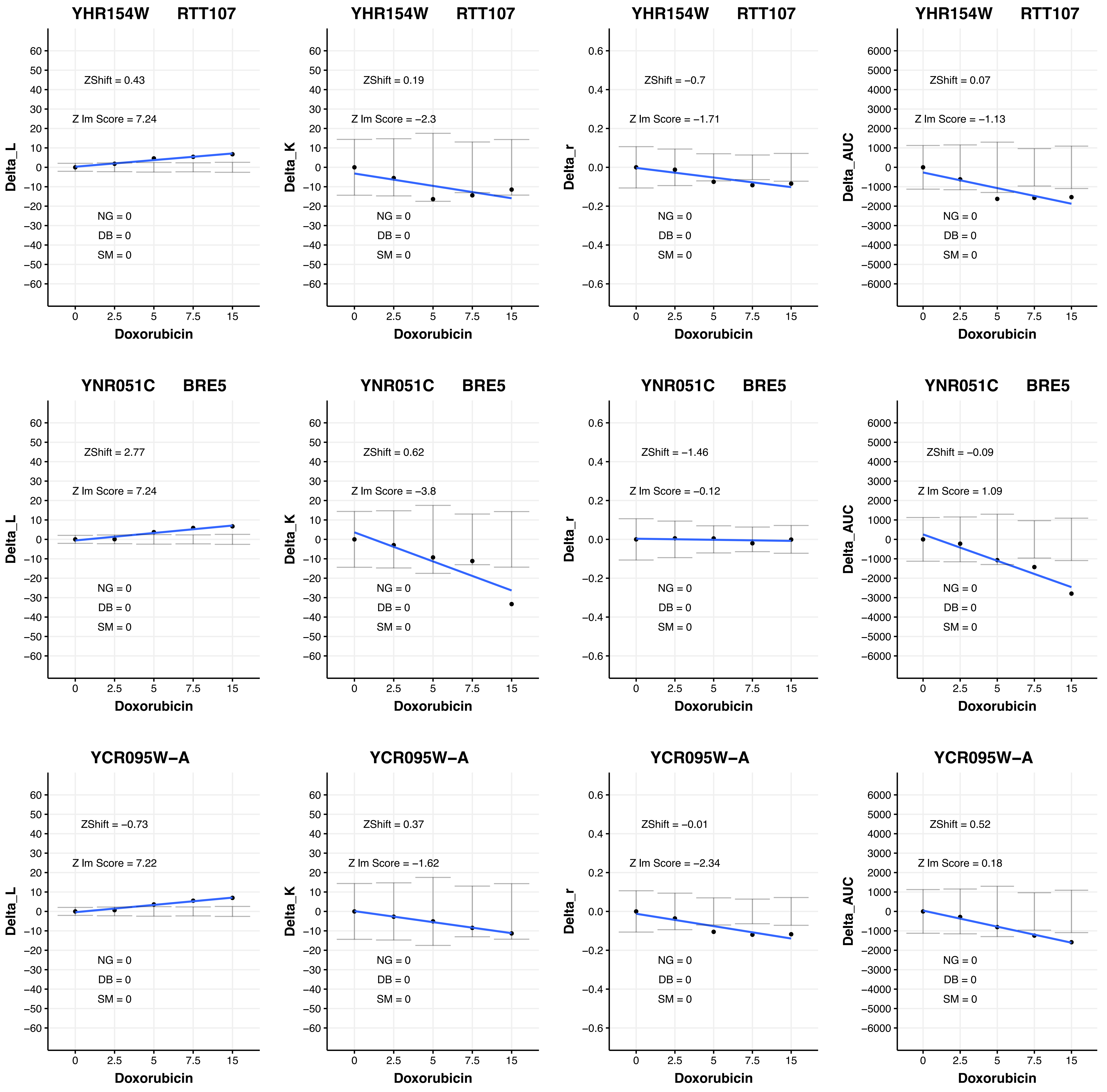
YHR154W RTT107
ZShift = 0.43
Z lm Score = 7.24
NG = 0
DB = 0
SM = 0
Delta_L
Doxorubicin
YHR154W RTT107
ZShift = 0.19
Z lm Score = −2.3
NG = 0
DB = 0
SM = 0
Delta_K
Doxorubicin
YHR154W RTT107
ZShift = −0.7
Z lm Score = −1.71
NG = 0
DB = 0
SM = 0
Delta_r
Doxorubicin
YHR154W RTT107
ZShift = 0.07
Z lm Score = −1.13
NG = 0
DB = 0
SM = 0
Delta_AUC
Doxorubicin
YNR051C BRE5
ZShift = 2.77
Z lm Score = 7.24
NG = 0
DB = 0
SM = 0
Delta_L
Doxorubicin
YNR051C BRE5
ZShift = 0.62
Z lm Score = −3.8
NG = 0
DB = 0
SM = 0
Delta_K
Doxorubicin
YNR051C BRE5
ZShift = −1.46
Z lm Score = −0.12
NG = 0
DB = 0
SM = 0
Delta_r
Doxorubicin
YNR051C BRE5
ZShift = −0.09
Z lm Score = 1.09
NG = 0
DB = 0
SM = 0
Delta_AUC
Doxorubicin
YCR095W−A
ZShift = −0.73
Z lm Score = 7.22
NG = 0
DB = 0
SM = 0
Delta_L
Doxorubicin
YCR095W−A
ZShift = 0.37
Z lm Score = −1.62
NG = 0
DB = 0
SM = 0
Delta_K
Doxorubicin
YCR095W−A
ZShift = −0.01
Z lm Score = −2.34
NG = 0
DB = 0
SM = 0
Delta_r
Doxorubicin
YCR095W−A
ZShift = 0.52
Z lm Score = 0.18
NG = 0
DB = 0
SM = 0
Delta_AUC
Doxorubicin

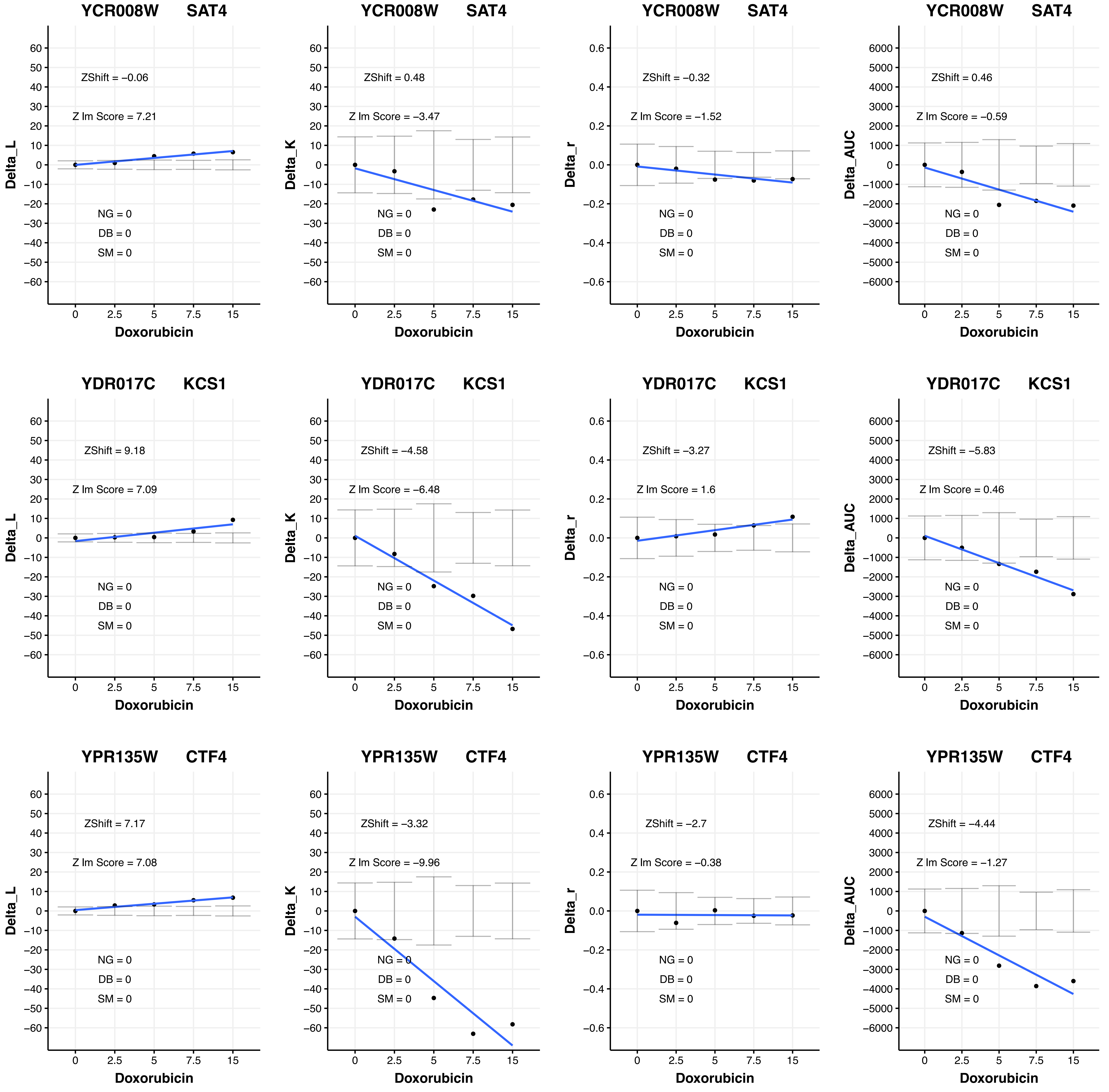
YCR008W SAT4
ZShift = −0.06
Z lm Score = 7.21
NG = 0
DB = 0
SM = 0
Delta_L
Doxorubicin
YCR008W SAT4
ZShift = 0.48
Z lm Score = −3.47
NG = 0
DB = 0
SM = 0
Delta_K
Doxorubicin
YCR008W SAT4
ZShift = −0.32
Z lm Score = −1.52
NG = 0
DB = 0
SM = 0
Delta_r
Doxorubicin
YCR008W SAT4
ZShift = 0.46
Z lm Score = −0.59
NG = 0
DB = 0
SM = 0
Delta_AUC
Doxorubicin
YDR017C KCS1
ZShift = 9.18
Z lm Score = 7.09
NG = 0
DB = 0
SM = 0
Delta_L
Doxorubicin
YDR017C KCS1
ZShift = −4.58
Z lm Score = −6.48
NG = 0
DB = 0
SM = 0
Delta_K
Doxorubicin
YDR017C KCS1
ZShift = −3.27
Z lm Score = 1.6
NG = 0
DB = 0
SM = 0
Delta_r
Doxorubicin
YDR017C KCS1
ZShift = −5.83
Z lm Score = 0.46
NG = 0
DB = 0
SM = 0
Delta_AUC
Doxorubicin
YPR135W CTF4
ZShift = 7.17
Z lm Score = 7.08
NG = 0
DB = 0
SM = 0
Delta_L
Doxorubicin
YPR135W CTF4
ZShift = −3.32
Z lm Score = −9.96
NG = 0
DB = 0
SM = 0
Delta_K
Doxorubicin
YPR135W CTF4
ZShift = −2.7
Z lm Score = −0.38
NG = 0
DB = 0
SM = 0
Delta_r
Doxorubicin
YPR135W CTF4
ZShift = −4.44
Z lm Score = −1.27
NG = 0
DB = 0
SM = 0
Delta_AUC
Doxorubicin

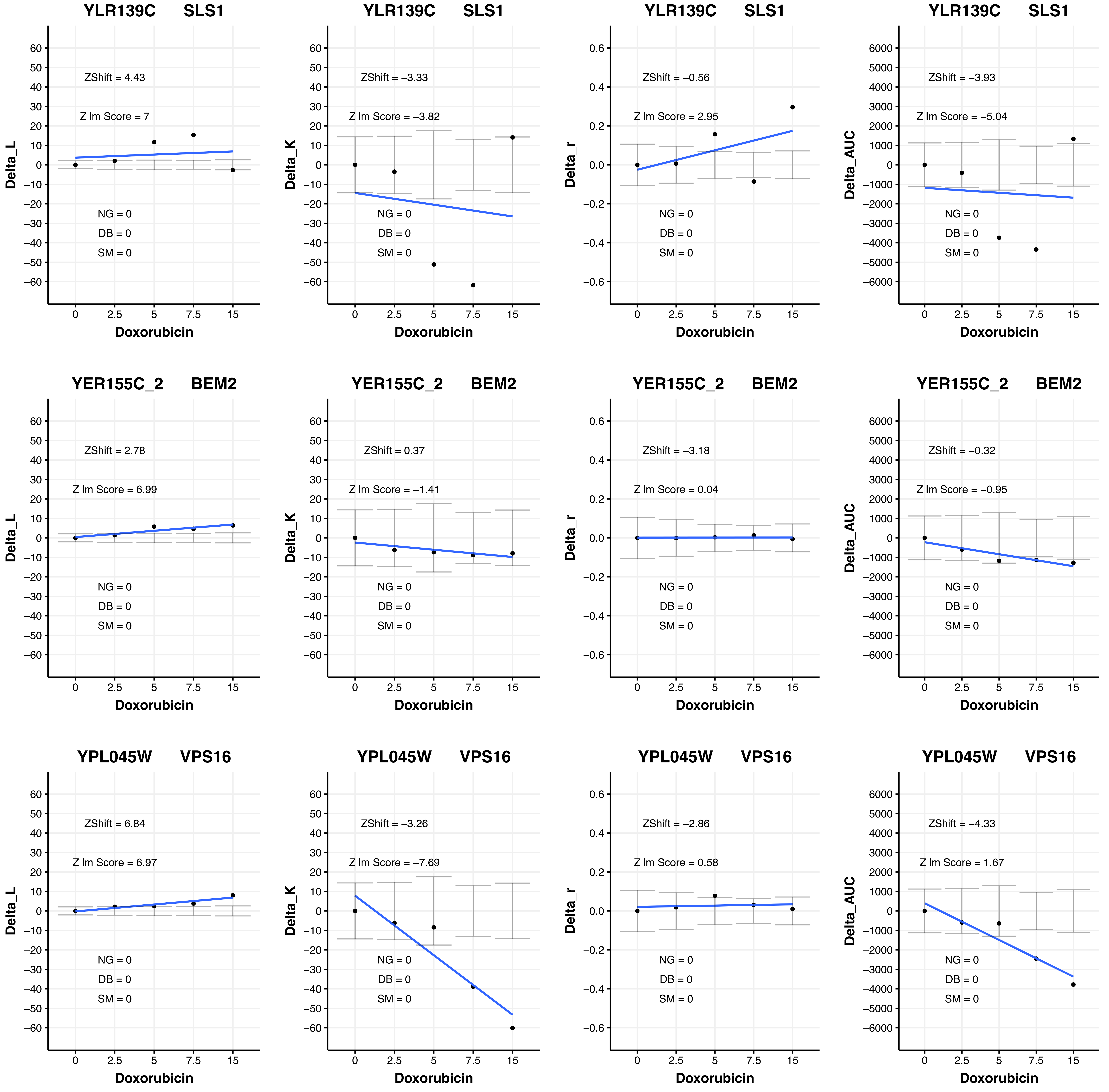
YLR139C SLS1
ZShift = 4.43
Z lm Score = 7
NG = 0
DB = 0
SM = 0
Delta_L
Doxorubicin
YLR139C SLS1
ZShift = −3.33
Z lm Score = −3.82
NG = 0
DB = 0
SM = 0
Delta_K
Doxorubicin
YLR139C SLS1
ZShift = −0.56
Z lm Score = 2.95
NG = 0
DB = 0
SM = 0
Delta_r
Doxorubicin
YLR139C SLS1
ZShift = −3.93
Z lm Score = −5.04
NG = 0
DB = 0
SM = 0
Delta_AUC
Doxorubicin
YER155C_2 BEM2
ZShift = 2.78
Z lm Score = 6.99
NG = 0
DB = 0
SM = 0
Delta_L
Doxorubicin
YER155C_2 BEM2
ZShift = 0.37
Z lm Score = −1.41
NG = 0
DB = 0
SM = 0
Delta_K
Doxorubicin
YER155C_2 BEM2
ZShift = −3.18
Z lm Score = 0.04
NG = 0
DB = 0
SM = 0
Delta_r
Doxorubicin
YER155C_2 BEM2
ZShift = −0.32
Z lm Score = −0.95
NG = 0
DB = 0
SM = 0
Delta_AUC
Doxorubicin
YPL045W VPS16
ZShift = 6.84
Z lm Score = 6.97
NG = 0
DB = 0
SM = 0
Delta_L
Doxorubicin
YPL045W VPS16
ZShift = −3.26
Z lm Score = −7.69
NG = 0
DB = 0
SM = 0
Delta_K
Doxorubicin
YPL045W VPS16
ZShift = −2.86
Z lm Score = 0.58
NG = 0
DB = 0
SM = 0
Delta_r
Doxorubicin
YPL045W VPS16
ZShift = −4.33
Z lm Score = 1.67
NG = 0
DB = 0
SM = 0
Delta_AUC
Doxorubicin

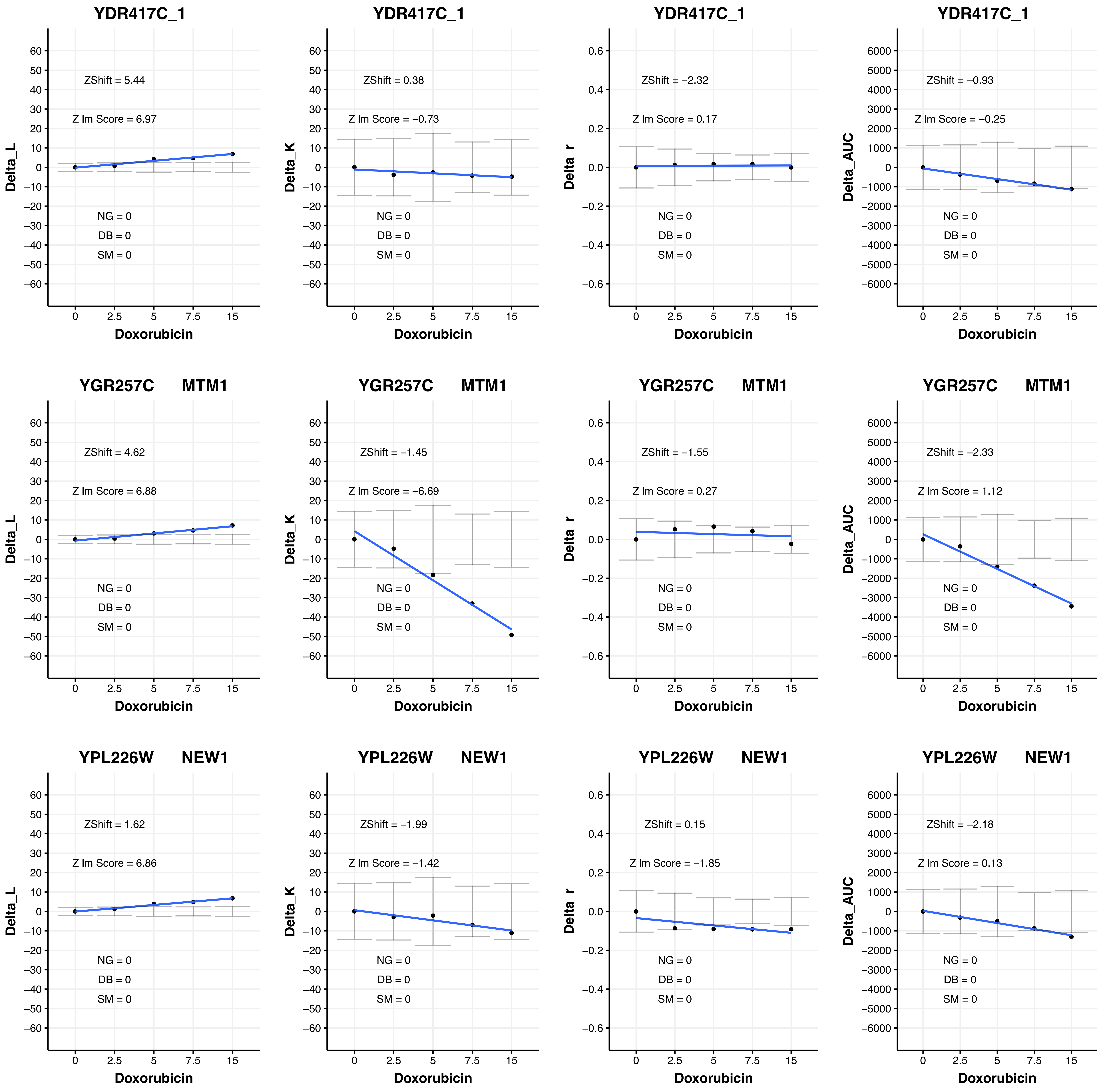

YDR417C_1
ZShift = 5.44
Z lm Score = 6.97
NG = 0
DB = 0
SM = 0
Delta_L
Doxorubicin
YDR417C_1
ZShift = 0.38
Z lm Score = −0.73
NG = 0
DB = 0
SM = 0
Delta_K
Doxorubicin
YDR417C_1
ZShift = −2.32
Z lm Score = 0.17
NG = 0
DB = 0
SM = 0
Delta_r
Doxorubicin
YDR417C_1
ZShift = −0.93
Z lm Score = −0.25
NG = 0
DB = 0
SM = 0
Delta_AUC
Doxorubicin
YGR257C MTM1
ZShift = 4.62
Z lm Score = 6.88
NG = 0
DB = 0
SM = 0
Delta_L
Doxorubicin
YGR257C MTM1
ZShift = −1.45
Z lm Score = −6.69
NG = 0
DB = 0
SM = 0
Delta_K
Doxorubicin
YGR257C MTM1
ZShift = −1.55
Z lm Score = 0.27
NG = 0
DB = 0
SM = 0
Delta_r
Doxorubicin
YGR257C MTM1
ZShift = −2.33
Z lm Score = 1.12
NG = 0
DB = 0
SM = 0
Delta_AUC
Doxorubicin
YPL226W NEW1
ZShift = 1.62
Z lm Score = 6.86
NG = 0
DB = 0
SM = 0
Delta_L
Doxorubicin
YPL226W NEW1
ZShift = −1.99
Z lm Score = −1.42
NG = 0
DB = 0
SM = 0
Delta_K
Doxorubicin
YPL226W NEW1
ZShift = 0.15
Z lm Score = −1.85
NG = 0
DB = 0
SM = 0
Delta_r
Doxorubicin
YPL226W NEW1
ZShift = −2.18
Z lm Score = 0.13
NG = 0
DB = 0
SM = 0
Delta_AUC
Doxorubicin

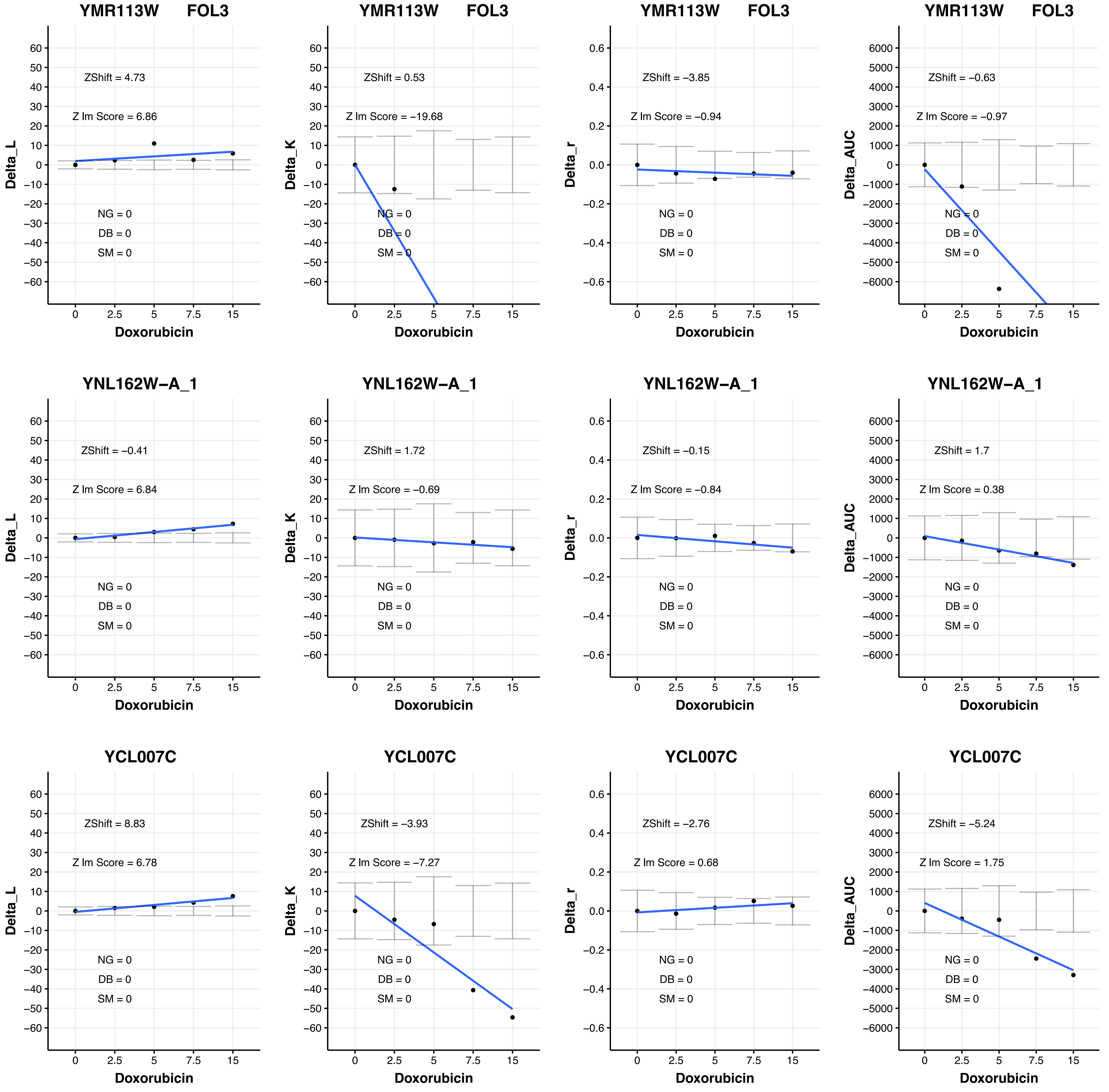

YMR113W FOL3
ZShift = 4.73
Z lm Score = 6.86
NG = 0
DB = 0
SM = 0
Delta_L
Doxorubicin
YMR113W FOL3
ZShift = 0.53
Z lm Score = −19.68
NG = 0
DB = 0
SM = 0
Delta_K
Doxorubicin
YMR113W FOL3
ZShift = −3.85
Z lm Score = −0.94
NG = 0
DB = 0
SM = 0
Delta_r
Doxorubicin
YMR113W FOL3
ZShift = −0.63
Z lm Score = −0.97
NG = 0
DB = 0
SM = 0
Delta_AUC
Doxorubicin
YNL162W−A_1
ZShift = −0.41
Z lm Score = 6.84
NG = 0
DB = 0
SM = 0
Delta_L
Doxorubicin
YNL162W−A_1
ZShift = 1.72
Z lm Score = −0.69
NG = 0
DB = 0
SM = 0
Delta_K
Doxorubicin
YNL162W−A_1
ZShift = −0.15
Z lm Score = −0.84
NG = 0
DB = 0
SM = 0
Delta_r
Doxorubicin
YNL162W−A_1
ZShift = 1.7
Z lm Score = 0.38
NG = 0
DB = 0
SM = 0
Delta_AUC
Doxorubicin
YCL007C
ZShift = 8.83
Z lm Score = 6.78
NG = 0
DB = 0
SM = 0
Delta_L
Doxorubicin
YCL007C
ZShift = −3.93
Z lm Score = −7.27
NG = 0
DB = 0
SM = 0
Delta_K
Doxorubicin
YCL007C
ZShift = −2.76
Z lm Score = 0.68
NG = 0
DB = 0
SM = 0
Delta_r
Doxorubicin
YCL007C
ZShift = −5.24
Z lm Score = 1.75
NG = 0
DB = 0
SM = 0
Delta_AUC
Doxorubicin

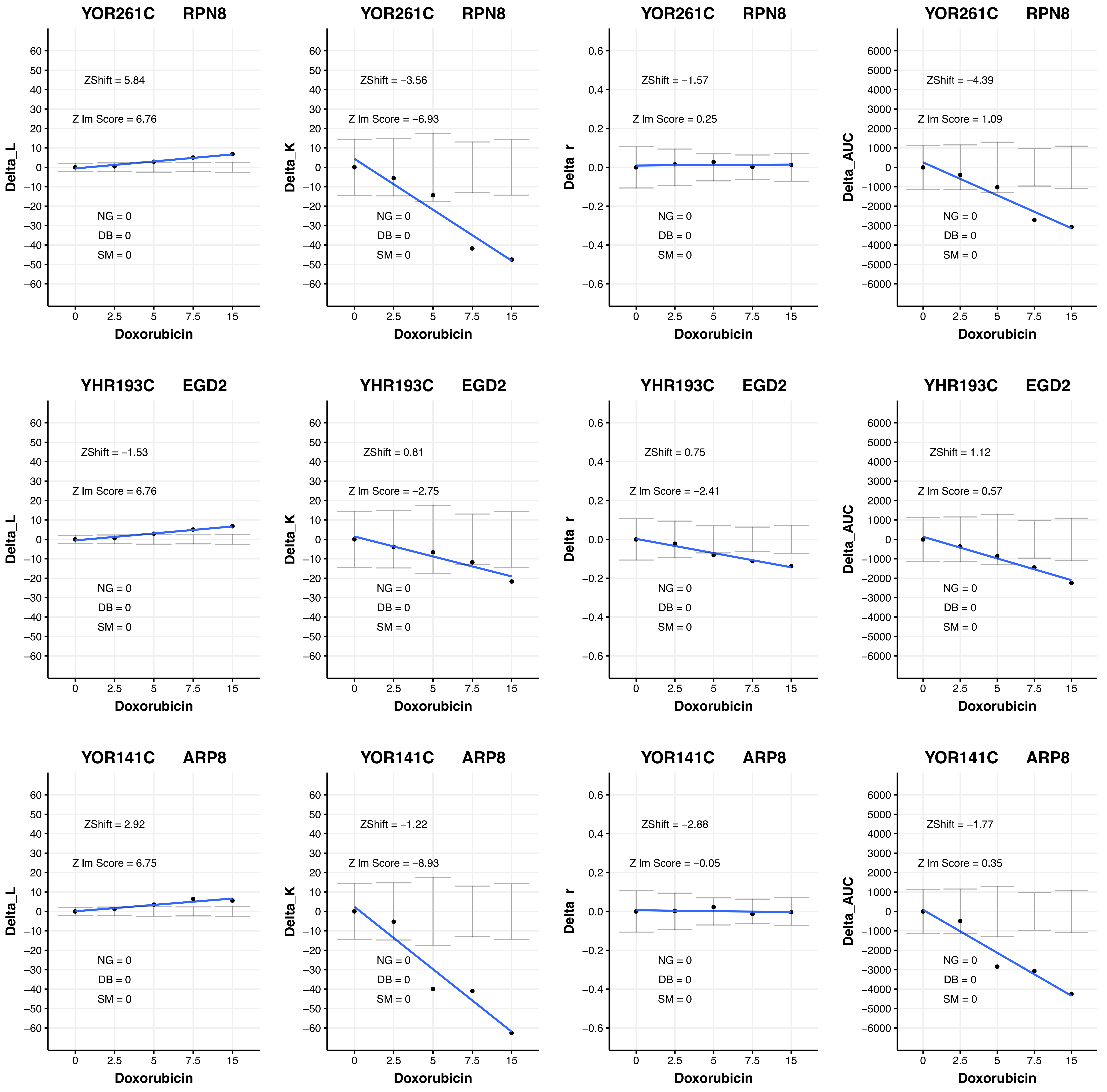
YOR261C RPN8
ZShift = 5.84
Z lm Score = 6.76
Delta_L
YOR261C RPN8
ZShift = −3.56
Z lm Score = −6.93
Delta_K
YOR261C RPN8
ZShift = −1.57
Z lm Score = 0.25
Delta_r
YOR261C RPN8
ZShift = −4.39
Z lm Score = 1.09
Delta_AUC
YHR193C EGD2
ZShift = −1.53
Z lm Score = 6.76
Delta_L
YHR193C EGD2
ZShift = 0.81
Z lm Score = −2.75
Delta_K
YHR193C EGD2
ZShift = 0.75
Z lm Score = −2.41
Delta_r
YHR193C EGD2
ZShift = 1.12
Z lm Score = 0.57
Delta_AUC
YOR141C ARP8
ZShift = 2.92
Z lm Score = 6.75
Delta_L
YOR141C ARP8
ZShift = −1.22
Z lm Score = −8.93
Delta_K
YOR141C ARP8
ZShift = −2.88
Z lm Score = −0.05
Delta_r
YOR141C ARP8
ZShift = −1.77
Z lm Score = 0.35
Delta_AUC
NG = 0
DB = 0
SM = 0
Doxorubicin
0
2.5
5
7.5
15

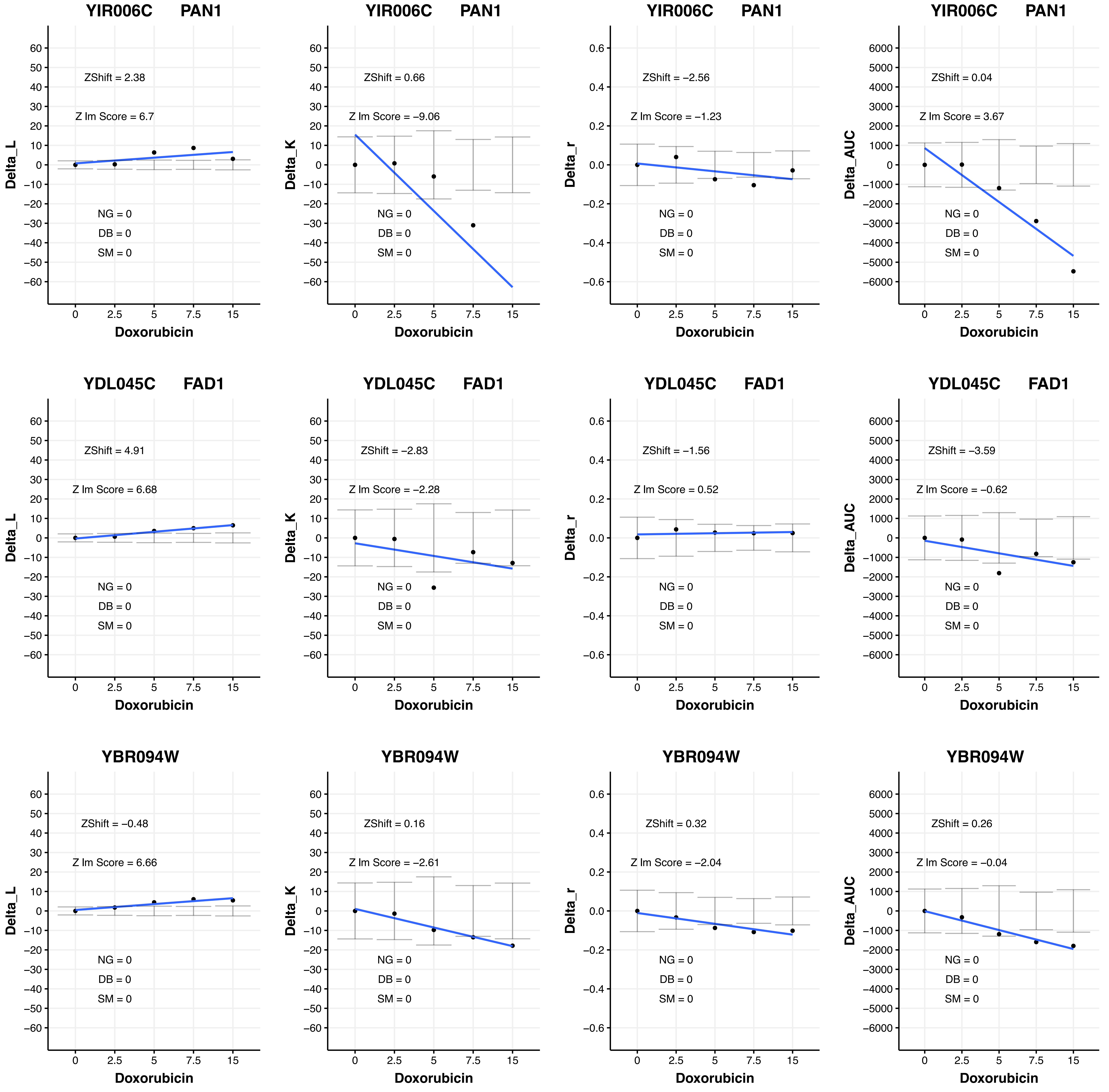
YIR006C PAN1
ZShift = 2.38
Z lm Score = 6.7
NG = 0
DB = 0
SM = 0
Delta_L
Doxorubicin
YIR006C PAN1
ZShift = 0.66
Z lm Score = −9.06
NG = 0
DB = 0
SM = 0
Delta_K
Doxorubicin
YIR006C PAN1
ZShift = −2.56
Z lm Score = −1.23
NG = 0
DB = 0
SM = 0
Delta_r
Doxorubicin
YIR006C PAN1
ZShift = 0.04
Z lm Score = 3.67
NG = 0
DB = 0
SM = 0
Delta_AUC
Doxorubicin
YDL045C FAD1
ZShift = 4.91
Z lm Score = 6.68
NG = 0
DB = 0
SM = 0
Delta_L
Doxorubicin
YDL045C FAD1
ZShift = −2.83
Z lm Score = −2.28
NG = 0
DB = 0
SM = 0
Delta_K
Doxorubicin
YDL045C FAD1
ZShift = −1.56
Z lm Score = 0.52
NG = 0
DB = 0
SM = 0
Delta_r
Doxorubicin
YDL045C FAD1
ZShift = −3.59
Z lm Score = −0.62
NG = 0
DB = 0
SM = 0
Delta_AUC
Doxorubicin
YBR094W
ZShift = −0.48
Z lm Score = 6.66
NG = 0
DB = 0
SM = 0
Delta_L
Doxorubicin
YBR094W
ZShift = 0.16
Z lm Score = −2.61
NG = 0
DB = 0
SM = 0
Delta_K
Doxorubicin
YBR094W
ZShift = 0.32
Z lm Score = −2.04
NG = 0
DB = 0
SM = 0
Delta_r
Doxorubicin
YBR094W
ZShift = 0.26
Z lm Score = −0.04
NG = 0
DB = 0
SM = 0
Delta_AUC
Doxorubicin

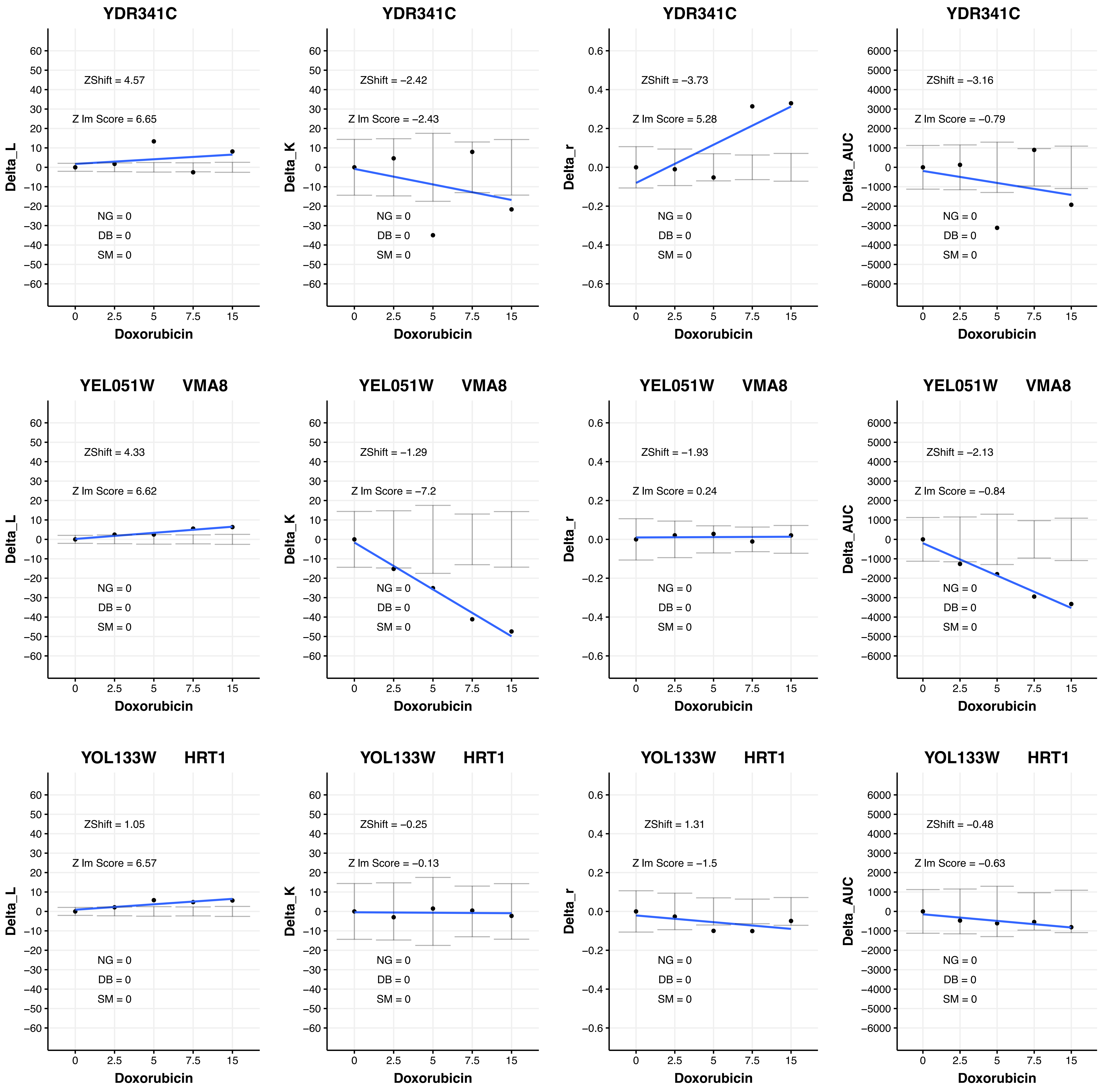
YDR341C
ZShift = 4.57
Z lm Score = 6.65
NG = 0
DB = 0
SM = 0
Delta_L
Doxorubicin
YDR341C
ZShift = −2.42
Z lm Score = −2.43
NG = 0
DB = 0
SM = 0
Delta_K
Doxorubicin
YDR341C
ZShift = −3.73
Z lm Score = 5.28
NG = 0
DB = 0
SM = 0
Delta_r
Doxorubicin
YDR341C
ZShift = −3.16
Z lm Score = −0.79
NG = 0
DB = 0
SM = 0
Delta_AUC
Doxorubicin
YEL051W VMA8
ZShift = 4.33
Z lm Score = 6.62
NG = 0
DB = 0
SM = 0
Delta_L
Doxorubicin
YEL051W VMA8
ZShift = −1.29
Z lm Score = −7.2
NG = 0
DB = 0
SM = 0
Delta_K
Doxorubicin
YEL051W VMA8
ZShift = −1.93
Z lm Score = 0.24
NG = 0
DB = 0
SM = 0
Delta_r
Doxorubicin
YEL051W VMA8
ZShift = −2.13
Z lm Score = −0.84
NG = 0
DB = 0
SM = 0
Delta_AUC
Doxorubicin
YOL133W HRT1
ZShift = 1.05
Z lm Score = 6.57
NG = 0
DB = 0
SM = 0
Delta_L
Doxorubicin
YOL133W HRT1
ZShift = −0.25
Z lm Score = −0.13
NG = 0
DB = 0
SM = 0
Delta_K
Doxorubicin
YOL133W HRT1
ZShift = 1.31
Z lm Score = −1.5
NG = 0
DB = 0
SM = 0
Delta_r
Doxorubicin
YOL133W HRT1
ZShift = −0.48
Z lm Score = −0.63
NG = 0
DB = 0
SM = 0
Delta_AUC
Doxorubicin

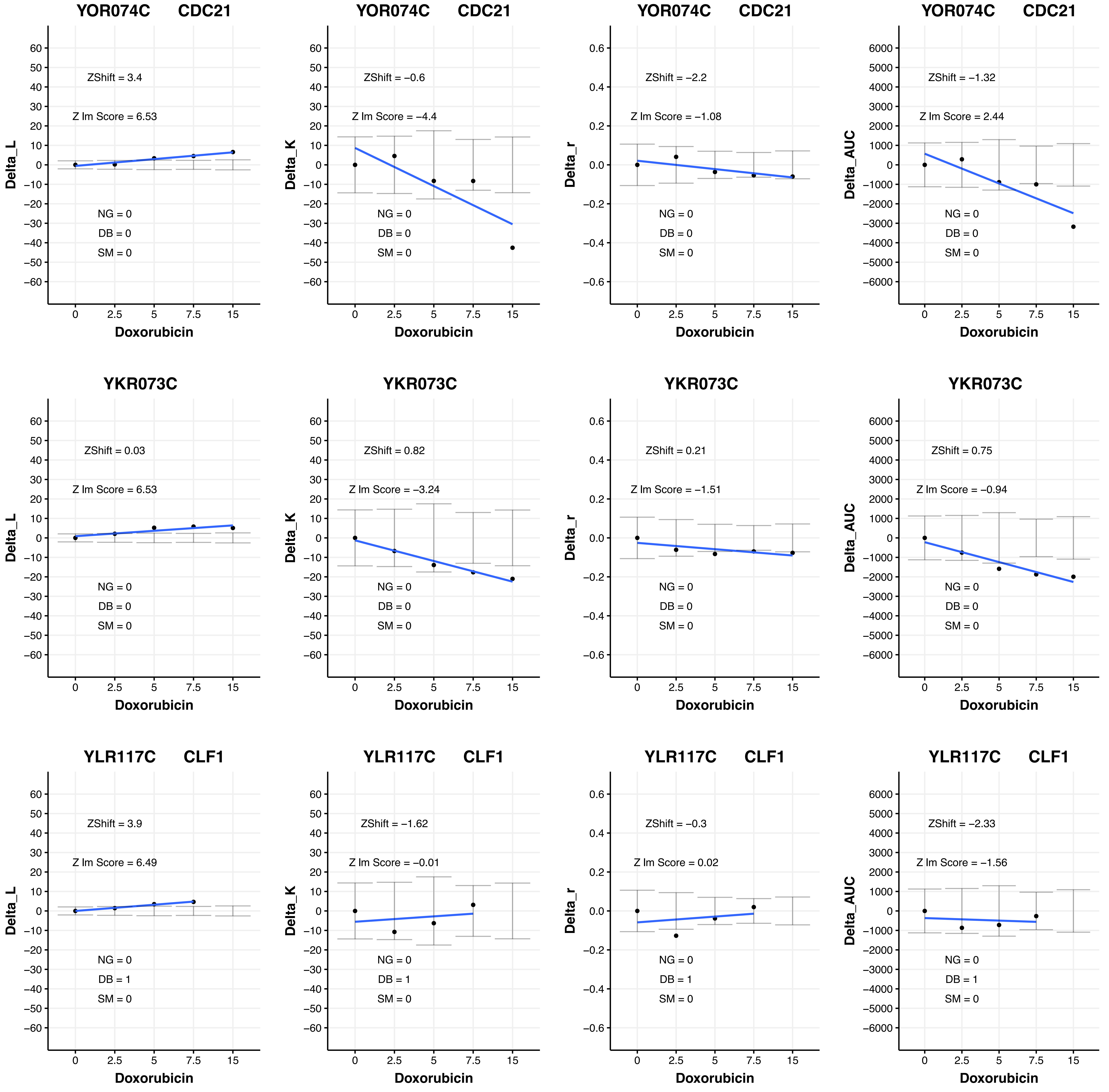
YOR074C CDC21
ZShift = 3.4
Z lm Score = 6.53
NG = 0
DB = 0
SM = 0
Delta_L
Doxorubicin
YOR074C CDC21
ZShift = −0.6
Z lm Score = −4.4
NG = 0
DB = 0
SM = 0
Delta_K
Doxorubicin
YOR074C CDC21
ZShift = −2.2
Z lm Score = −1.08
NG = 0
DB = 0
SM = 0
Delta_r
Doxorubicin
YOR074C CDC21
ZShift = −1.32
Z lm Score = 2.44
NG = 0
DB = 0
SM = 0
Delta_AUC
Doxorubicin
YKR073C
ZShift = 0.03
Z lm Score = 6.53
NG = 0
DB = 0
SM = 0
Delta_L
Doxorubicin
YKR073C
ZShift = 0.82
Z lm Score = −3.24
NG = 0
DB = 0
SM = 0
Delta_K
Doxorubicin
YKR073C
ZShift = 0.21
Z lm Score = −1.51
NG = 0
DB = 0
SM = 0
Delta_r
Doxorubicin
YKR073C
ZShift = 0.75
Z lm Score = −0.94
NG = 0
DB = 0
SM = 0
Delta_AUC
Doxorubicin
YLR117C CLF1
ZShift = 3.9
Z lm Score = 6.49
NG = 0
DB = 1
SM = 0
Delta_L
Doxorubicin
YLR117C CLF1
ZShift = −1.62
Z lm Score = −0.01
NG = 0
DB = 1
SM = 0
Delta_K
Doxorubicin
YLR117C CLF1
ZShift = −0.3
Z lm Score = 0.02
NG = 0
DB = 1
SM = 0
Delta_r
Doxorubicin
YLR117C CLF1
ZShift = −2.33
Z lm Score = −1.56
NG = 0
DB = 1
SM = 0
Delta_AUC
Doxorubicin

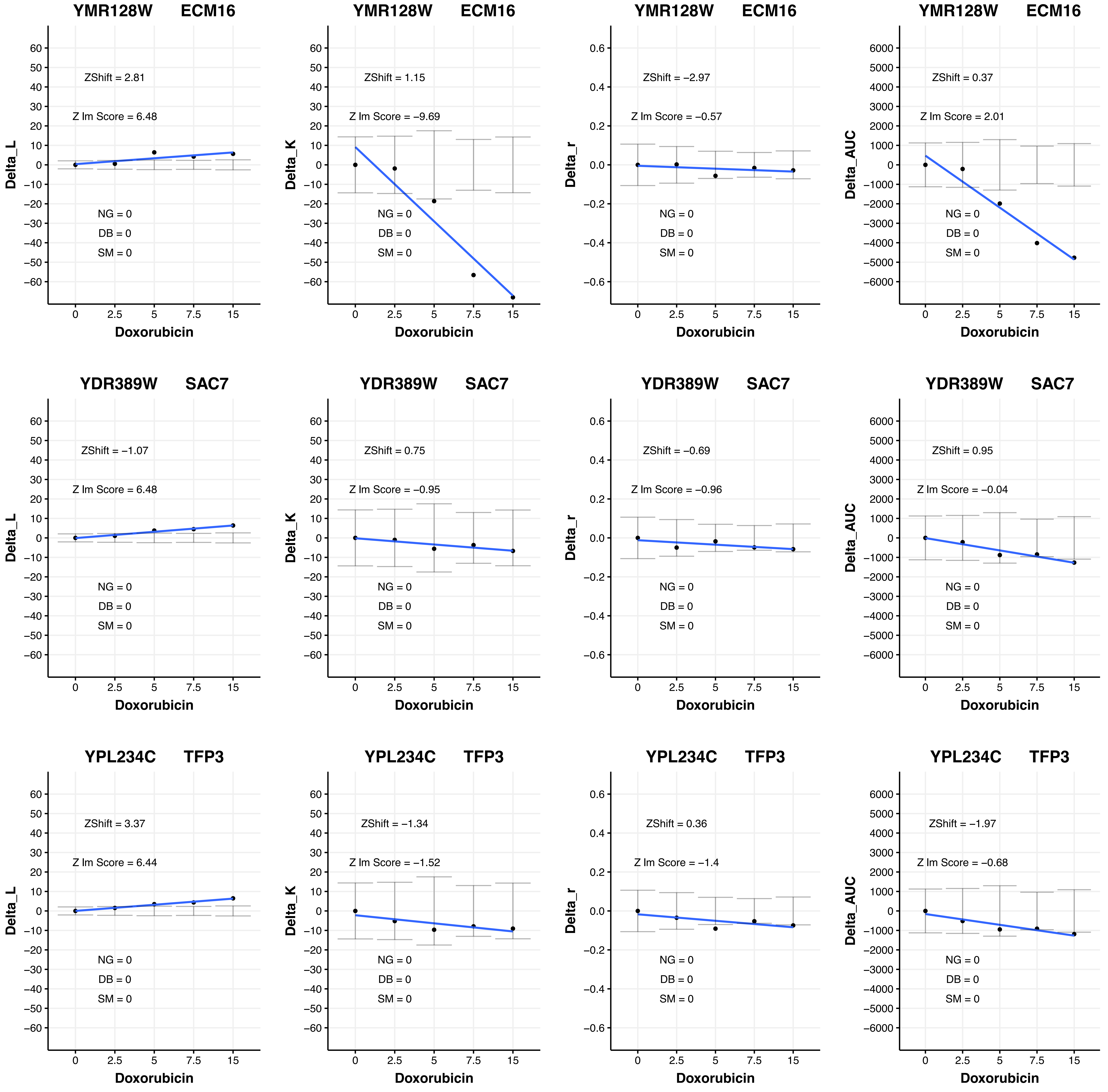
YMR128W ECM16
ZShift = 2.81
Z lm Score = 6.48
NG = 0
DB = 0
SM = 0
Delta_L
Doxorubicin
YMR128W ECM16
ZShift = 1.15
Z lm Score = −9.69
NG = 0
DB = 0
SM = 0
Delta_K
Doxorubicin
YMR128W ECM16
ZShift = −2.97
Z lm Score = −0.57
NG = 0
DB = 0
SM = 0
Delta_r
Doxorubicin
YMR128W ECM16
ZShift = 0.37
Z lm Score = 2.01
NG = 0
DB = 0
SM = 0
Delta_AUC
Doxorubicin
YDR389W SAC7
ZShift = −1.07
Z lm Score = 6.48
NG = 0
DB = 0
SM = 0
Delta_L
Doxorubicin
YDR389W SAC7
ZShift = 0.75
Z lm Score = −0.95
NG = 0
DB = 0
SM = 0
Delta_K
Doxorubicin
YDR389W SAC7
ZShift = −0.69
Z lm Score = −0.96
NG = 0
DB = 0
SM = 0
Delta_r
Doxorubicin
YDR389W SAC7
ZShift = 0.95
Z lm Score = −0.04
NG = 0
DB = 0
SM = 0
Delta_AUC
Doxorubicin
YPL234C TFP3
ZShift = 3.37
Z lm Score = 6.44
NG = 0
DB = 0
SM = 0
Delta_L
Doxorubicin
YPL234C TFP3
ZShift = −1.34
Z lm Score = −1.52
NG = 0
DB = 0
SM = 0
Delta_K
Doxorubicin
YPL234C TFP3
ZShift = 0.36
Z lm Score = −1.4
NG = 0
DB = 0
SM = 0
Delta_r
Doxorubicin
YPL234C TFP3
ZShift = −1.97
Z lm Score = −0.68
NG = 0
DB = 0
SM = 0
Delta_AUC
Doxorubicin

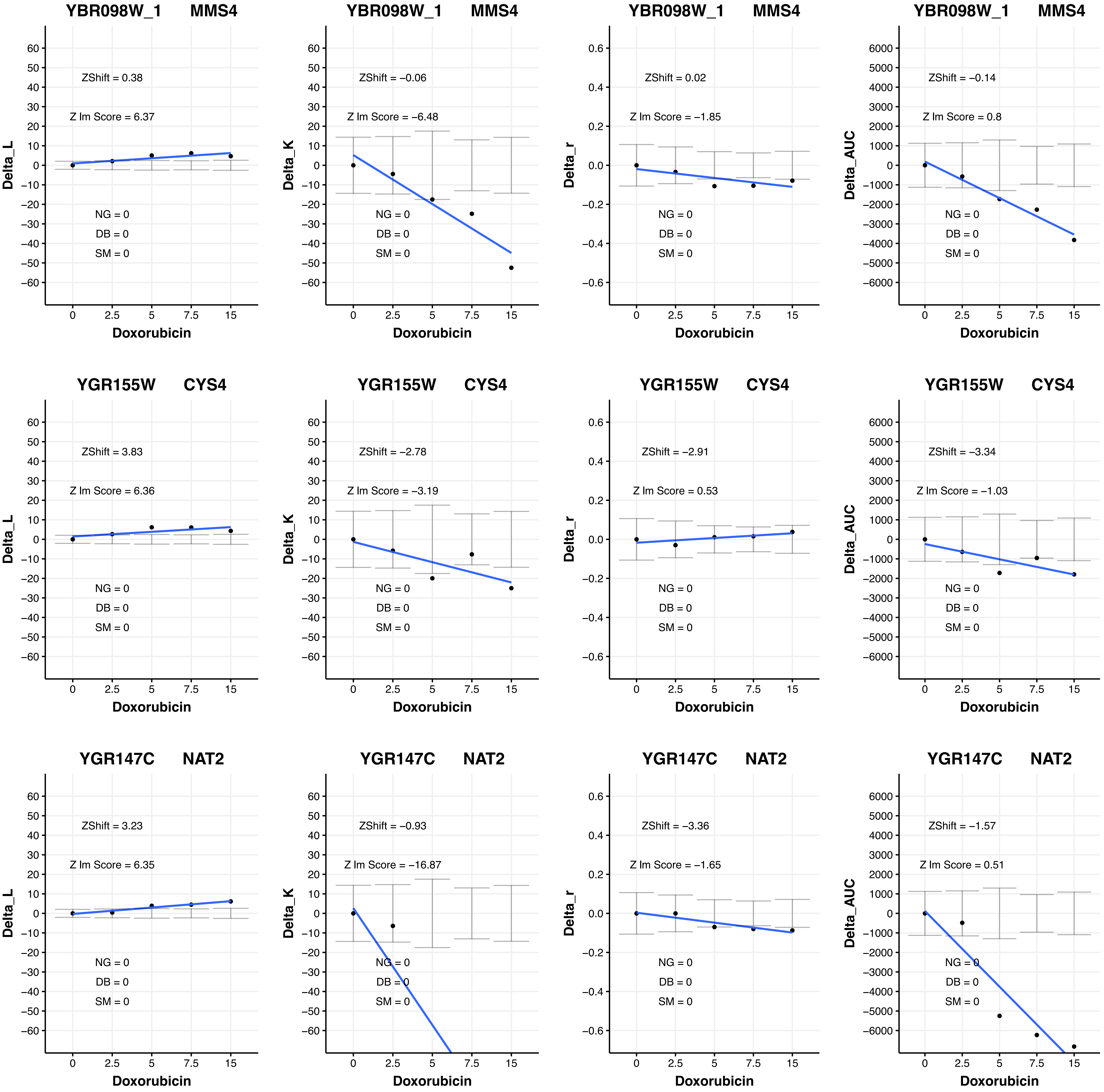
YBR098W_1 MMS4
ZShift = 0.38
Z lm Score = 6.37
NG = 0
DB = 0
SM = 0
Delta_L
Doxorubicin
YBR098W_1 MMS4
ZShift = −0.06
Z lm Score = −6.48
NG = 0
DB = 0
SM = 0
Delta_K
Doxorubicin
YBR098W_1 MMS4
ZShift = 0.02
Z lm Score = −1.85
NG = 0
DB = 0
SM = 0
Delta_r
Doxorubicin
YBR098W_1 MMS4
ZShift = −0.14
Z lm Score = 0.8
NG = 0
DB = 0
SM = 0
Delta_AUC
Doxorubicin
YGR155W CYS4
ZShift = 3.83
Z lm Score = 6.36
NG = 0
DB = 0
SM = 0
Delta_L
Doxorubicin
YGR155W CYS4
ZShift = −2.78
Z lm Score = −3.19
NG = 0
DB = 0
SM = 0
Delta_K
Doxorubicin
YGR155W CYS4
ZShift = −2.91
Z lm Score = 0.53
NG = 0
DB = 0
SM = 0
Delta_r
Doxorubicin
YGR155W CYS4
ZShift = −3.34
Z lm Score = −1.03
NG = 0
DB = 0
SM = 0
Delta_AUC
Doxorubicin
YGR147C NAT2
ZShift = 3.23
Z lm Score = 6.35
NG = 0
DB = 0
SM = 0
Delta_L
Doxorubicin
YGR147C NAT2
ZShift = −0.93
Z lm Score = −16.87
NG = 0
DB = 0
SM = 0
Delta_K
Doxorubicin
YGR147C NAT2
ZShift = −3.36
Z lm Score = −1.65
NG = 0
DB = 0
SM = 0
Delta_r
Doxorubicin
YGR147C NAT2
ZShift = −1.57
Z lm Score = 0.51
NG = 0
DB = 0
SM = 0
Delta_AUC
Doxorubicin

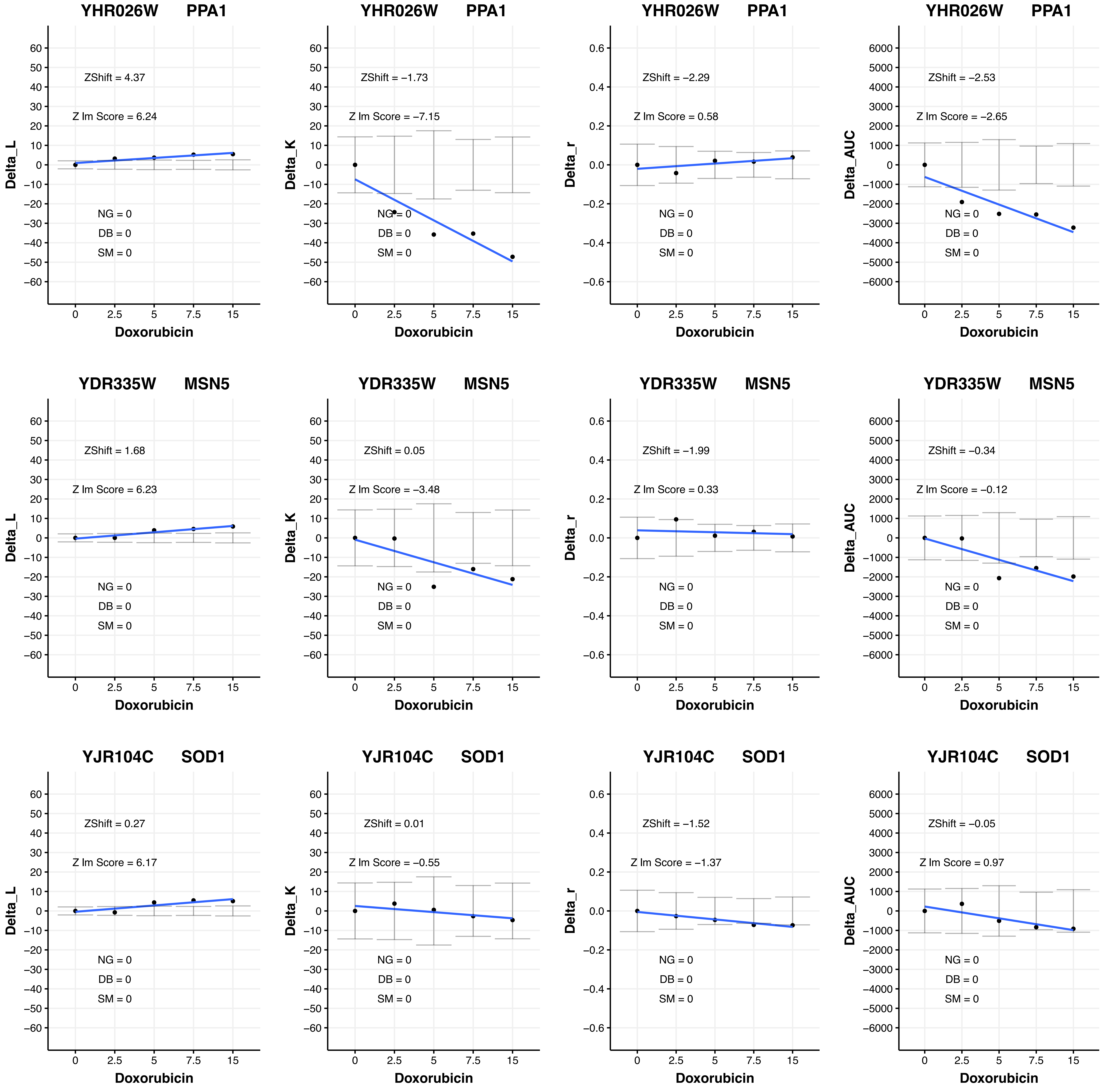
YHR026W PPA1
ZShift = 4.37
Z lm Score = 6.24
NG = 0
DB = 0
SM = 0
Delta_L
Doxorubicin
YHR026W PPA1
ZShift = −1.73
Z lm Score = −7.15
NG = 0
DB = 0
SM = 0
Delta_K
Doxorubicin
YHR026W PPA1
ZShift = −2.29
Z lm Score = 0.58
NG = 0
DB = 0
SM = 0
Delta_r
Doxorubicin
YHR026W PPA1
ZShift = −2.53
Z lm Score = −2.65
NG = 0
DB = 0
SM = 0
Delta_AUC
Doxorubicin
YDR335W MSN5
ZShift = 1.68
Z lm Score = 6.23
NG = 0
DB = 0
SM = 0
Delta_L
Doxorubicin
YDR335W MSN5
ZShift = 0.05
Z lm Score = −3.48
NG = 0
DB = 0
SM = 0
Delta_K
Doxorubicin
YDR335W MSN5
ZShift = −1.99
Z lm Score = 0.33
NG = 0
DB = 0
SM = 0
Delta_r
Doxorubicin
YDR335W MSN5
ZShift = −0.34
Z lm Score = −0.12
NG = 0
DB = 0
SM = 0
Delta_AUC
Doxorubicin
YJR104C SOD1
ZShift = 0.27
Z lm Score = 6.17
NG = 0
DB = 0
SM = 0
Delta_L
Doxorubicin
YJR104C SOD1
ZShift = 0.01
Z lm Score = −0.55
NG = 0
DB = 0
SM = 0
Delta_K
Doxorubicin
YJR104C SOD1
ZShift = −1.52
Z lm Score = −1.37
NG = 0
DB = 0
SM = 0
Delta_r
Doxorubicin
YJR104C SOD1
ZShift = −0.05
Z lm Score = 0.97
NG = 0
DB = 0
SM = 0
Delta_AUC
Doxorubicin

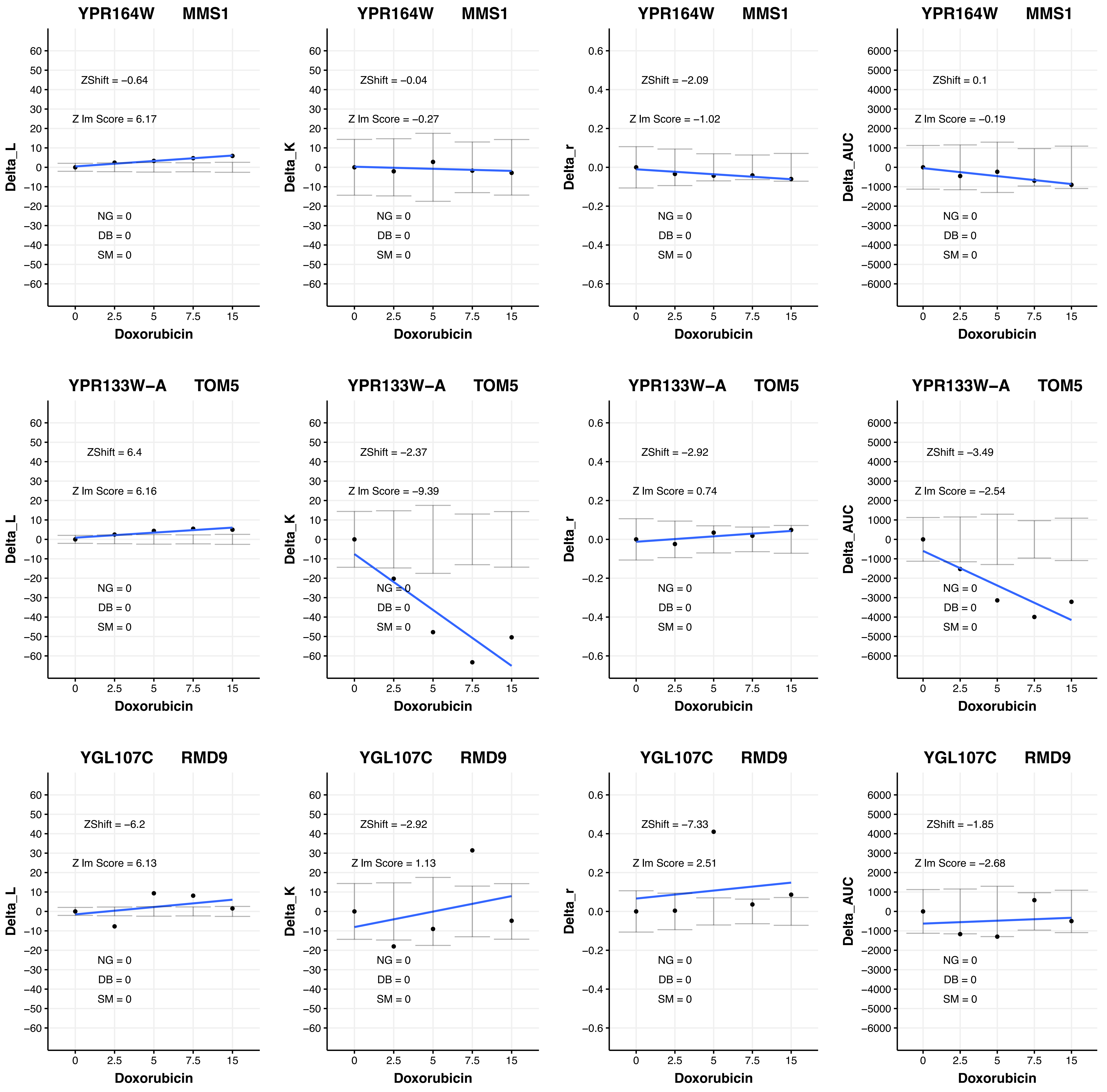

YPR164W MMS1
ZShift = −0.64
Z lm Score = 6.17
Delta_L
NG = 0
DB = 0
SM = 0
Doxorubicin
YPR164W MMS1
ZShift = −0.04
Z lm Score = −0.27
Delta_K
NG = 0
DB = 0
SM = 0
Doxorubicin
YPR164W MMS1
ZShift = −2.09
Z lm Score = −1.02
Delta_r
NG = 0
DB = 0
SM = 0
Doxorubicin
YPR164W MMS1
ZShift = 0.1
Z lm Score = −0.19
Delta_AUC
NG = 0
DB = 0
SM = 0
Doxorubicin
YPR133W−A TOM5
ZShift = 6.4
Z lm Score = 6.16
Delta_L
NG = 0
DB = 0
SM = 0
Doxorubicin
YPR133W−A TOM5
ZShift = −2.37
Z lm Score = −9.39
Delta_K
NG = 0
DB = 0
SM = 0
Doxorubicin
YPR133W−A TOM5
ZShift = −2.92
Z lm Score = 0.74
Delta_r
NG = 0
DB = 0
SM = 0
Doxorubicin
YPR133W−A TOM5
ZShift = −3.49
Z lm Score = −2.54
Delta_AUC
NG = 0
DB = 0
SM = 0
Doxorubicin
YGL107C RMD9
ZShift = −6.2
Z lm Score = 6.13
Delta_L
NG = 0
DB = 0
SM = 0
Doxorubicin
YGL107C RMD9
ZShift = −2.92
Z lm Score = 1.13
Delta_K
NG = 0
DB = 0
SM = 0
Doxorubicin
YGL107C RMD9
ZShift = −7.33
Z lm Score = 2.51
Delta_r
NG = 0
DB = 0
SM = 0
Doxorubicin
YGL107C RMD9
ZShift = −1.85
Z lm Score = −2.68
Delta_AUC
NG = 0
DB = 0
SM = 0
Doxorubicin

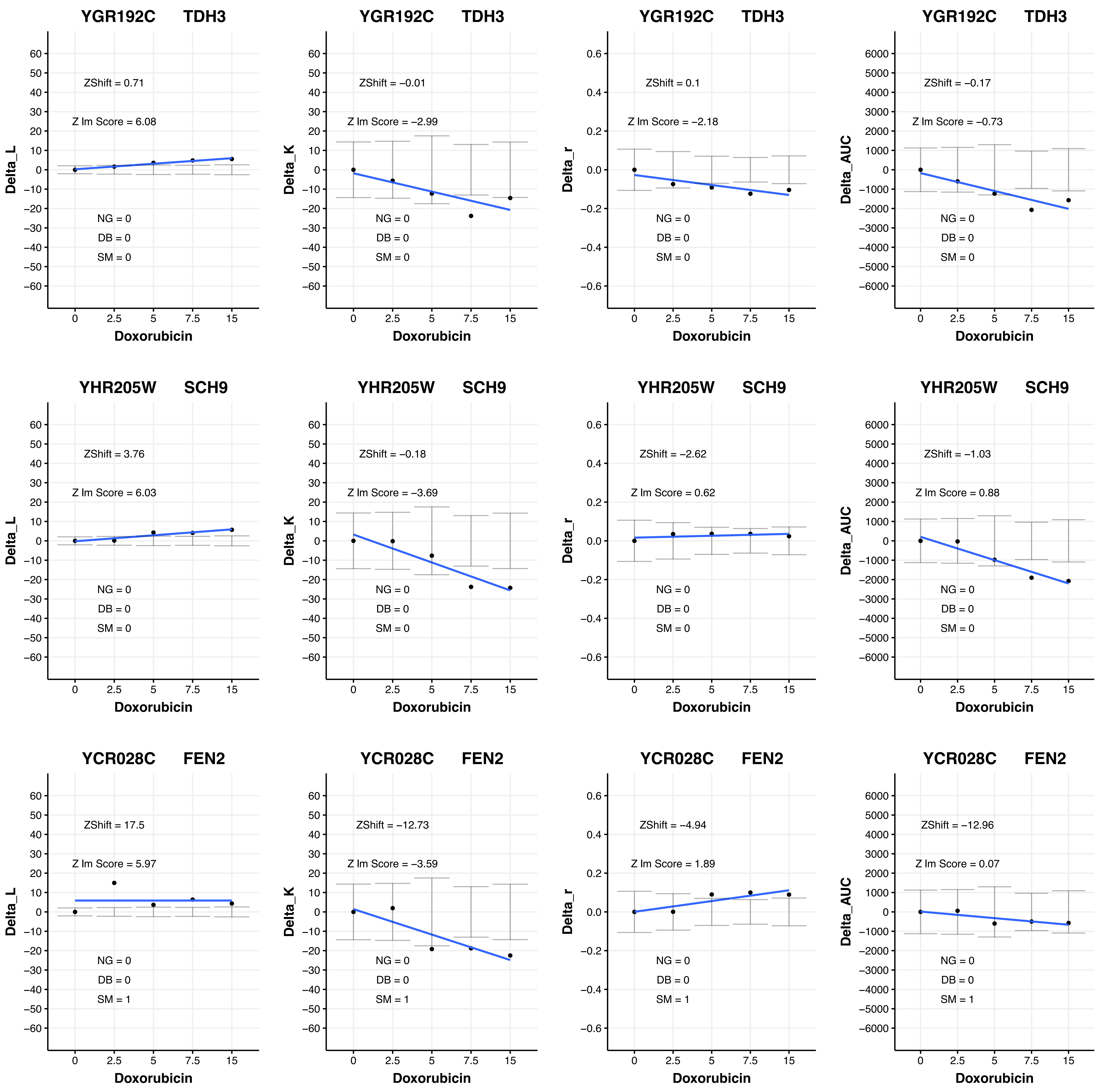

YGR192C TDH3
ZShift = 0.71
Z lm Score = 6.08
NG = 0
DB = 0
SM = 0
Delta_L
Doxorubicin
YGR192C TDH3
ZShift = −0.01
Z lm Score = −2.99
NG = 0
DB = 0
SM = 0
Delta_K
Doxorubicin
YGR192C TDH3
ZShift = 0.1
Z lm Score = −2.18
NG = 0
DB = 0
SM = 0
Delta_r
Doxorubicin
YGR192C TDH3
ZShift = −0.17
Z lm Score = −0.73
NG = 0
DB = 0
SM = 0
Delta_AUC
Doxorubicin
YHR205W SCH9
ZShift = 3.76
Z lm Score = 6.03
NG = 0
DB = 0
SM = 0
Delta_L
Doxorubicin
YHR205W SCH9
ZShift = −0.18
Z lm Score = −3.69
NG = 0
DB = 0
SM = 0
Delta_K
Doxorubicin
YHR205W SCH9
ZShift = −2.62
Z lm Score = 0.62
NG = 0
DB = 0
SM = 0
Delta_r
Doxorubicin
YHR205W SCH9
ZShift = −1.03
Z lm Score = 0.88
NG = 0
DB = 0
SM = 0
Delta_AUC
Doxorubicin
YCR028C FEN2
ZShift = 17.5
Z lm Score = 5.97
NG = 0
DB = 0
SM = 1
Delta_L
Doxorubicin
YCR028C FEN2
ZShift = −12.73
Z lm Score = −3.59
NG = 0
DB = 0
SM = 1
Delta_K
Doxorubicin
YCR028C FEN2
ZShift = −4.94
Z lm Score = 1.89
NG = 0
DB = 0
SM = 1
Delta_r
Doxorubicin
YCR028C FEN2
ZShift = −12.96
Z lm Score = 0.07
NG = 0
DB = 0
SM = 1
Delta_AUC
Doxorubicin

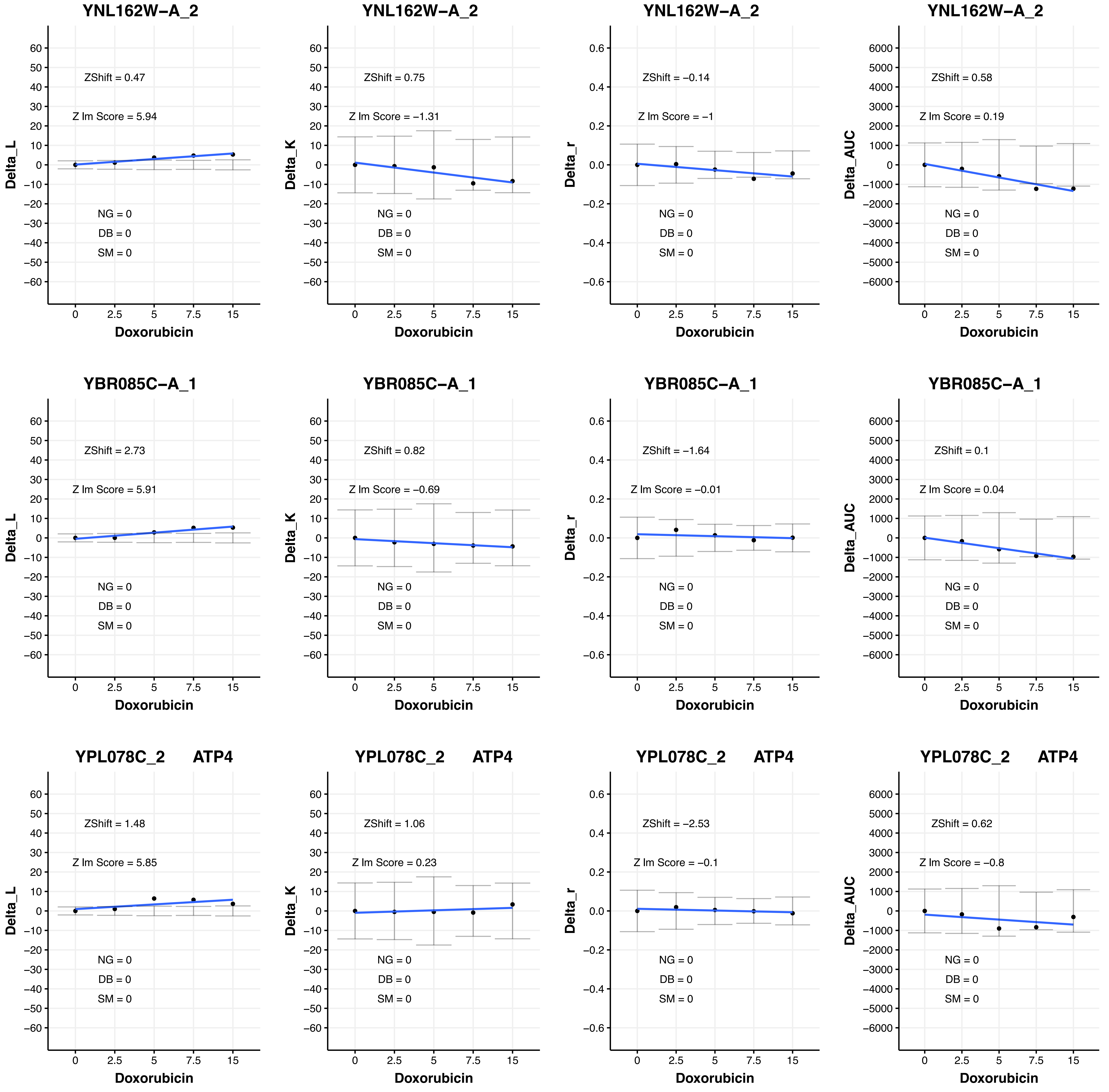

YNL162W−A_2
ZShift = 0.47
Z lm Score = 5.94
NG = 0
DB = 0
SM = 0
Delta_L
Doxorubicin
YNL162W−A_2
ZShift = 0.75
Z lm Score = −1.31
NG = 0
DB = 0
SM = 0
Delta_K
Doxorubicin
YNL162W−A_2
ZShift = −0.14
Z lm Score = −1
NG = 0
DB = 0
SM = 0
Delta_r
Doxorubicin
YNL162W−A_2
ZShift = 0.58
Z lm Score = 0.19
NG = 0
DB = 0
SM = 0
Delta_AUC
Doxorubicin
YBR085C−A_1
ZShift = 2.73
Z lm Score = 5.91
NG = 0
DB = 0
SM = 0
Delta_L
Doxorubicin
YBR085C−A_1
ZShift = 0.82
Z lm Score = −0.69
NG = 0
DB = 0
SM = 0
Delta_K
Doxorubicin
YBR085C−A_1
ZShift = −1.64
Z lm Score = −0.01
NG = 0
DB = 0
SM = 0
Delta_r
Doxorubicin
YBR085C−A_1
ZShift = 0.1
Z lm Score = 0.04
NG = 0
DB = 0
SM = 0
Delta_AUC
Doxorubicin
YPL078C_2 ATP4
ZShift = 1.48
Z lm Score = 5.85
NG = 0
DB = 0
SM = 0
Delta_L
Doxorubicin
YPL078C_2 ATP4
ZShift = 1.06
Z lm Score = 0.23
NG = 0
DB = 0
SM = 0
Delta_K
Doxorubicin
YPL078C_2 ATP4
ZShift = −2.53
Z lm Score = −0.1
NG = 0
DB = 0
SM = 0
Delta_r
Doxorubicin
YPL078C_2 ATP4
ZShift = 0.62
Z lm Score = −0.8
NG = 0
DB = 0
SM = 0
Delta_AUC
Doxorubicin

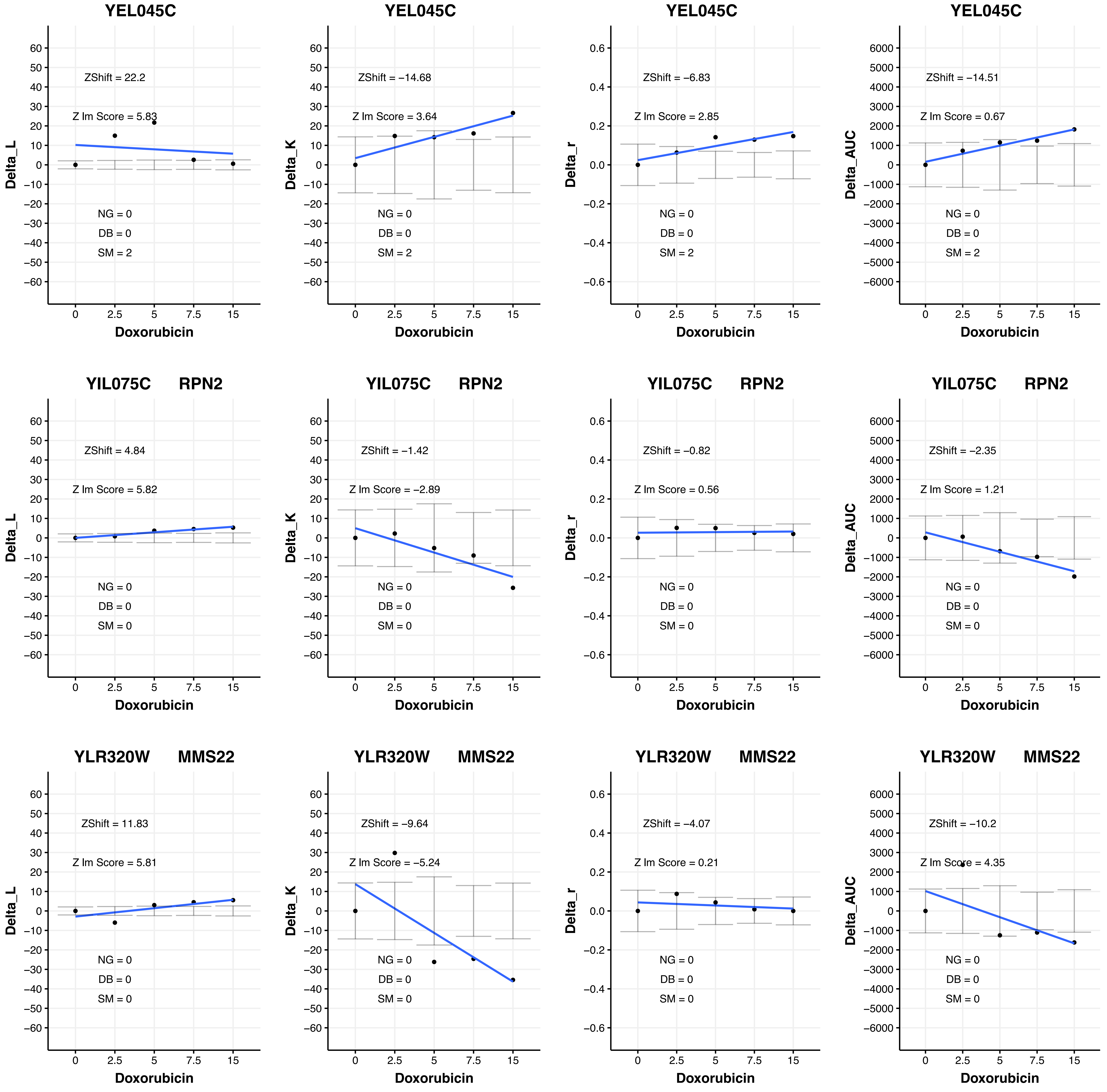
YEL045C
ZShift = 22.2
Z lm Score = 5.83
NG = 0
DB = 0
SM = 2
Delta_L
Doxorubicin
YEL045C
ZShift = −14.68
Z lm Score = 3.64
NG = 0
DB = 0
SM = 2
Delta_K
Doxorubicin
YEL045C
ZShift = −6.83
Z lm Score = 2.85
NG = 0
DB = 0
SM = 2
Delta_r
Doxorubicin
YEL045C
ZShift = −14.51
Z lm Score = 0.67
NG = 0
DB = 0
SM = 2
Delta_AUC
Doxorubicin
YIL075C RPN2
ZShift = 4.84
Z lm Score = 5.82
NG = 0
DB = 0
SM = 0
Delta_L
Doxorubicin
YIL075C RPN2
ZShift = −1.42
Z lm Score = −2.89
NG = 0
DB = 0
SM = 0
Delta_K
Doxorubicin
YIL075C RPN2
ZShift = −0.82
Z lm Score = 0.56
NG = 0
DB = 0
SM = 0
Delta_r
Doxorubicin
YIL075C RPN2
ZShift = −2.35
Z lm Score = 1.21
NG = 0
DB = 0
SM = 0
Delta_AUC
Doxorubicin
YLR320W MMS22
ZShift = 11.83
Z lm Score = 5.81
NG = 0
DB = 0
SM = 0
Delta_L
Doxorubicin
YLR320W MMS22
ZShift = −9.64
Z lm Score = −5.24
NG = 0
DB = 0
SM = 0
Delta_K
Doxorubicin
YLR320W MMS22
ZShift = −4.07
Z lm Score = 0.21
NG = 0
DB = 0
SM = 0
Delta_r
Doxorubicin
YLR320W MMS22
ZShift = −10.2
Z lm Score = 4.35
NG = 0
DB = 0
SM = 0
Delta_AUC
Doxorubicin

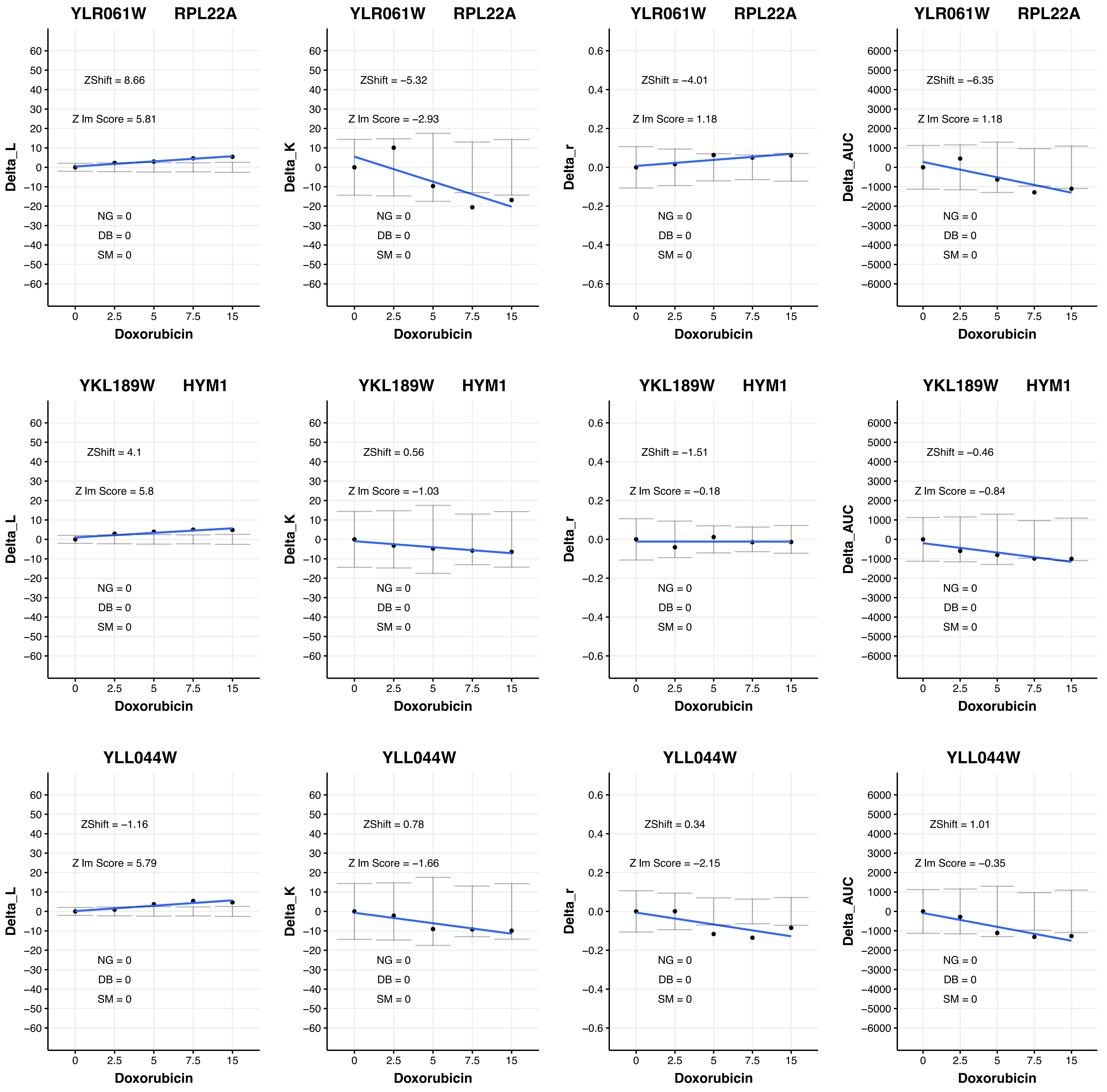

YLR061W RPL22A
ZShift = 8.66
Z lm Score = 5.81
NG = 0
DB = 0
SM = 0
Delta_L
Doxorubicin
YLR061W RPL22A
ZShift = −5.32
Z lm Score = −2.93
NG = 0
DB = 0
SM = 0
Delta_K
Doxorubicin
YLR061W RPL22A
ZShift = −4.01
Z lm Score = 1.18
NG = 0
DB = 0
SM = 0
Delta_r
Doxorubicin
YLR061W RPL22A
ZShift = −6.35
Z lm Score = 1.18
NG = 0
DB = 0
SM = 0
Delta_AUC
Doxorubicin
YKL189W HYM1
ZShift = 4.1
Z lm Score = 5.8
NG = 0
DB = 0
SM = 0
Delta_L
Doxorubicin
YKL189W HYM1
ZShift = 0.56
Z lm Score = −1.03
NG = 0
DB = 0
SM = 0
Delta_K
Doxorubicin
YKL189W HYM1
ZShift = −1.51
Z lm Score = −0.18
NG = 0
DB = 0
SM = 0
Delta_r
Doxorubicin
YKL189W HYM1
ZShift = −0.46
Z lm Score = −0.84
NG = 0
DB = 0
SM = 0
Delta_AUC
Doxorubicin
YLL044W
ZShift = −1.16
Z lm Score = 5.79
NG = 0
DB = 0
SM = 0
Delta_L
Doxorubicin
YLL044W
ZShift = 0.78
Z lm Score = −1.66
NG = 0
DB = 0
SM = 0
Delta_K
Doxorubicin
YLL044W
ZShift = 0.34
Z lm Score = −2.15
NG = 0
DB = 0
SM = 0
Delta_r
Doxorubicin
YLL044W
ZShift = 1.01
Z lm Score = −0.35
NG = 0
DB = 0
SM = 0
Delta_AUC
Doxorubicin

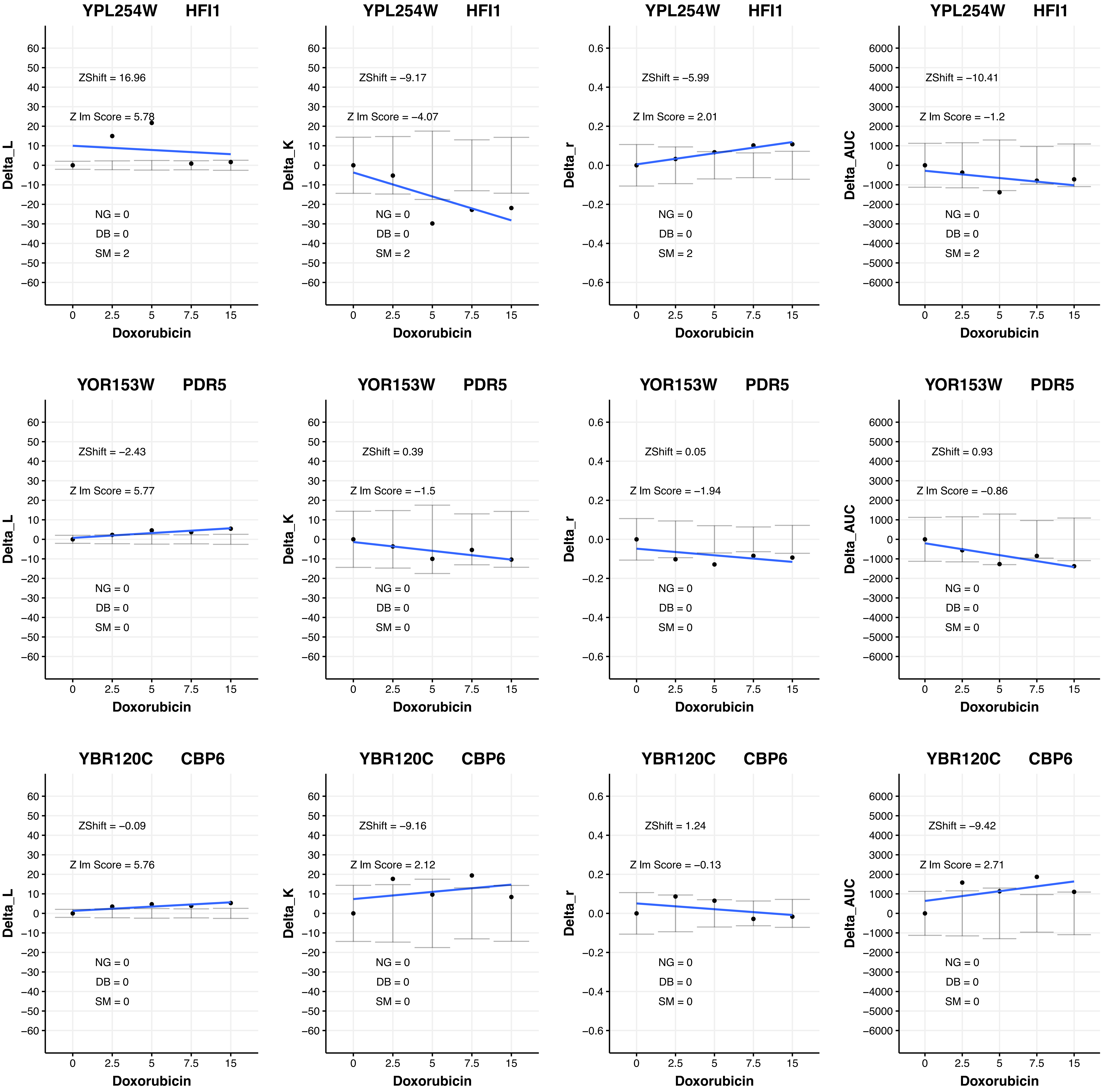
YPL254W HFI1
ZShift = 16.96
Z lm Score = 5.78
NG = 0
DB = 0
SM = 2
Delta_L
Doxorubicin
YPL254W HFI1
ZShift = −9.17
Z lm Score = −4.07
NG = 0
DB = 0
SM = 2
Delta_K
Doxorubicin
YPL254W HFI1
ZShift = −5.99
Z lm Score = 2.01
NG = 0
DB = 0
SM = 2
Delta_r
Doxorubicin
YPL254W HFI1
ZShift = −10.41
Z lm Score = −1.2
NG = 0
DB = 0
SM = 2
Delta_AUC
Doxorubicin
YOR153W PDR5
ZShift = −2.43
Z lm Score = 5.77
NG = 0
DB = 0
SM = 0
Delta_L
Doxorubicin
YOR153W PDR5
ZShift = 0.39
Z lm Score = −1.5
NG = 0
DB = 0
SM = 0
Delta_K
Doxorubicin
YOR153W PDR5
ZShift = 0.05
Z lm Score = −1.94
NG = 0
DB = 0
SM = 0
Delta_r
Doxorubicin
YOR153W PDR5
ZShift = 0.93
Z lm Score = −0.86
NG = 0
DB = 0
SM = 0
Delta_AUC
Doxorubicin
YBR120C CBP6
ZShift = −0.09
Z lm Score = 5.76
NG = 0
DB = 0
SM = 0
Delta_L
Doxorubicin
YBR120C CBP6
ZShift = −9.16
Z lm Score = 2.12
NG = 0
DB = 0
SM = 0
Delta_K
Doxorubicin
YBR120C CBP6
ZShift = 1.24
Z lm Score = −0.13
NG = 0
DB = 0
SM = 0
Delta_r
Doxorubicin
YBR120C CBP6
ZShift = −9.42
Z lm Score = 2.71
NG = 0
DB = 0
SM = 0
Delta_AUC
Doxorubicin

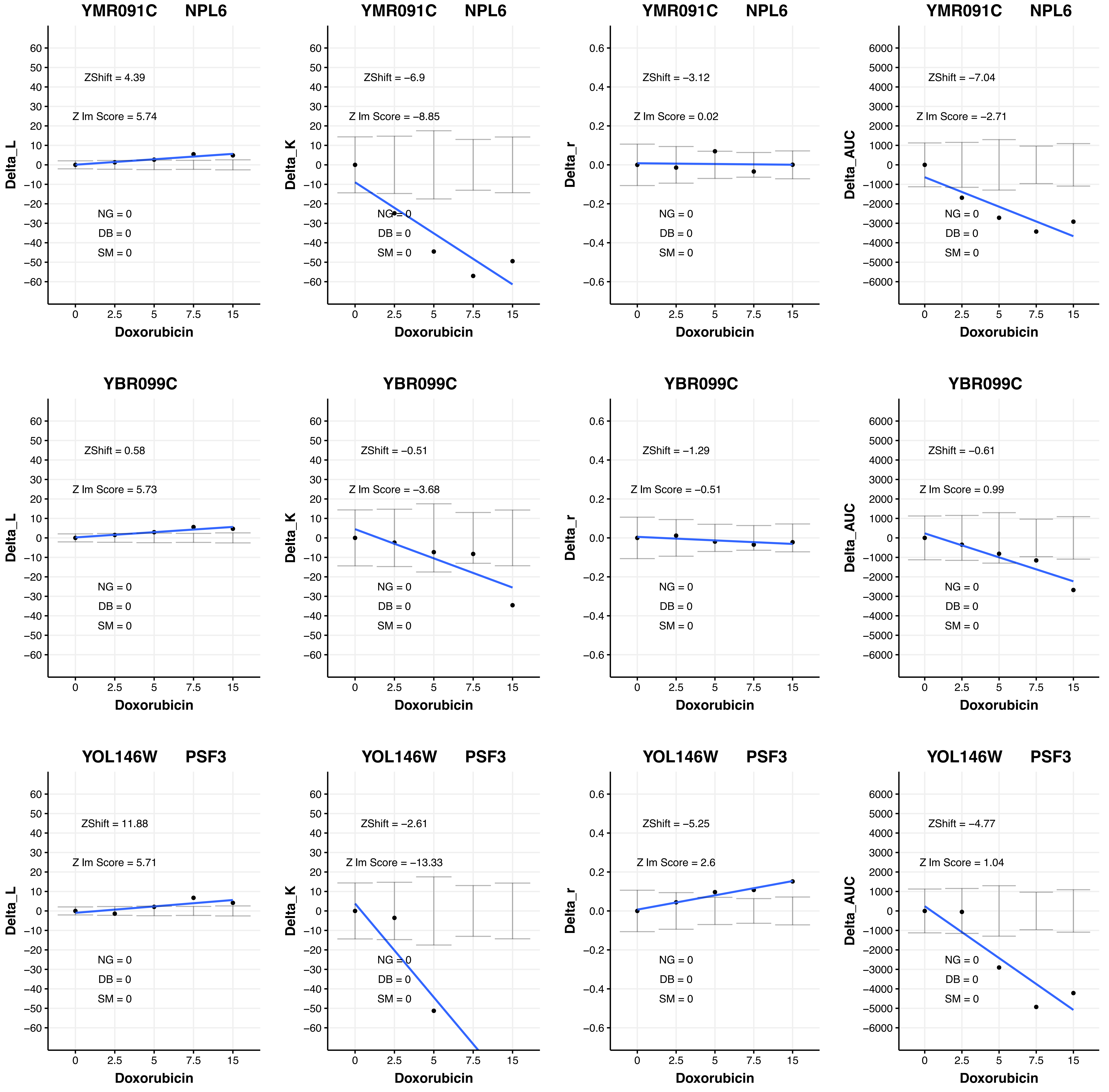
YMR091C NPL6
ZShift = 4.39
Z lm Score = 5.74
NG = 0
DB = 0
SM = 0
Delta_L
Doxorubicin
YMR091C NPL6
ZShift = −6.9
Z lm Score = −8.85
NG = 0
DB = 0
SM = 0
Delta_K
Doxorubicin
YMR091C NPL6
ZShift = −3.12
Z lm Score = 0.02
NG = 0
DB = 0
SM = 0
Delta_r
Doxorubicin
YMR091C NPL6
ZShift = −7.04
Z lm Score = −2.71
NG = 0
DB = 0
SM = 0
Delta_AUC
Doxorubicin
YBR099C
ZShift = 0.58
Z lm Score = 5.73
NG = 0
DB = 0
SM = 0
Delta_L
Doxorubicin
YBR099C
ZShift = −0.51
Z lm Score = −3.68
NG = 0
DB = 0
SM = 0
Delta_K
Doxorubicin
YBR099C
ZShift = −1.29
Z lm Score = −0.51
NG = 0
DB = 0
SM = 0
Delta_r
Doxorubicin
YBR099C
ZShift = −0.61
Z lm Score = 0.99
NG = 0
DB = 0
SM = 0
Delta_AUC
Doxorubicin
YOL146W PSF3
ZShift = 11.88
Z lm Score = 5.71
NG = 0
DB = 0
SM = 0
Delta_L
Doxorubicin
YOL146W PSF3
ZShift = −2.61
Z lm Score = −13.33
NG = 0
DB = 0
SM = 0
Delta_K
Doxorubicin
YOL146W PSF3
ZShift = −5.25
Z lm Score = 2.6
NG = 0
DB = 0
SM = 0
Delta_r
Doxorubicin
YOL146W PSF3
ZShift = −4.77
Z lm Score = 1.04
NG = 0
DB = 0
SM = 0
Delta_AUC
Doxorubicin

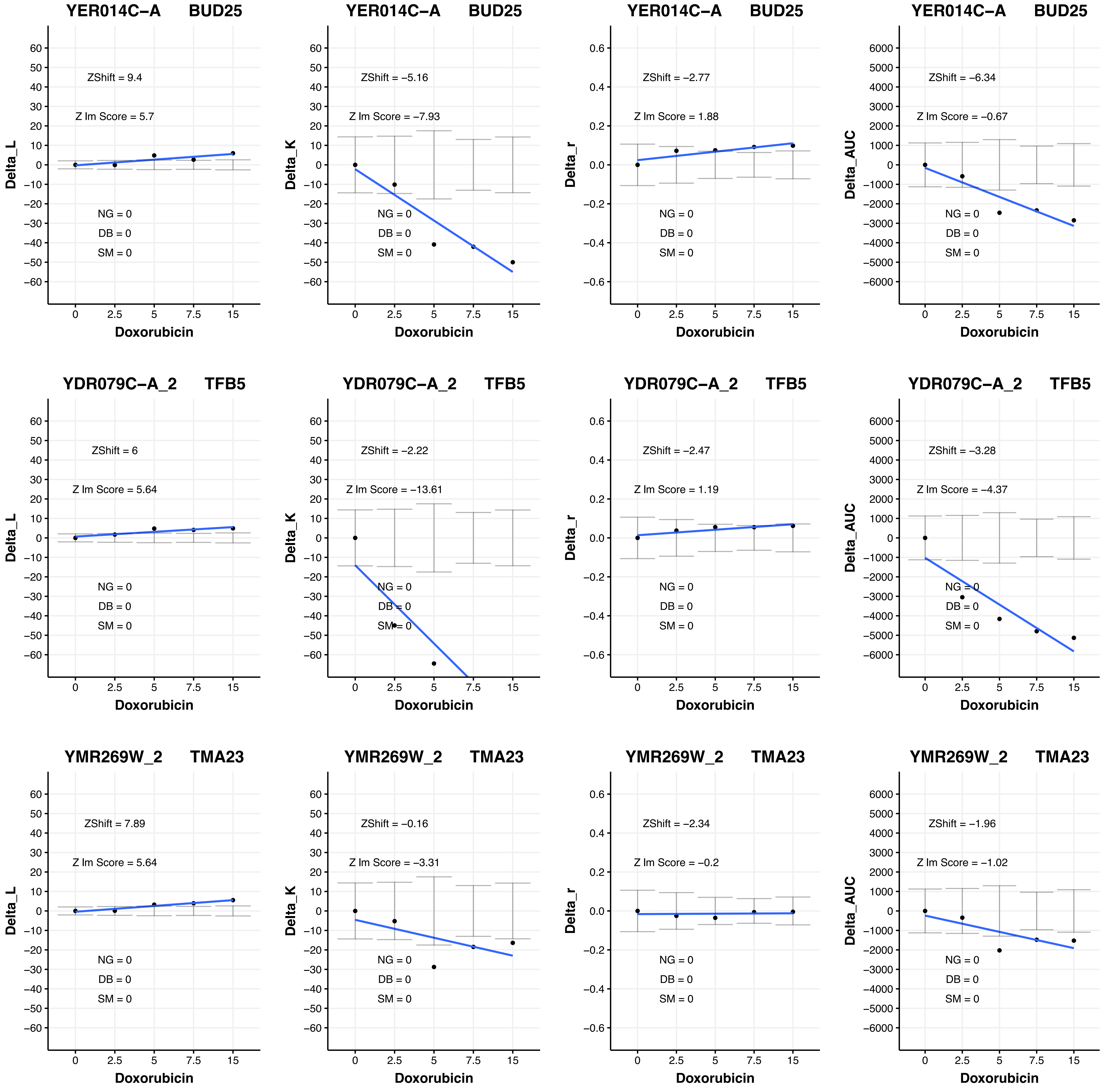

YER014C−A BUD25
ZShift = 9.4
Z lm Score = 5.7
NG = 0
DB = 0
SM = 0
Delta_L
Doxorubicin
YER014C−A BUD25
ZShift = −5.16
Z lm Score = −7.93
NG = 0
DB = 0
SM = 0
Delta_K
Doxorubicin
YER014C−A BUD25
ZShift = −2.77
Z lm Score = 1.88
NG = 0
DB = 0
SM = 0
Delta_r
Doxorubicin
YER014C−A BUD25
ZShift = −6.34
Z lm Score = −0.67
NG = 0
DB = 0
SM = 0
Delta_AUC
Doxorubicin
YDR079C−A_2 TFB5
ZShift = 6
Z lm Score = 5.64
NG = 0
DB = 0
SM = 0
Delta_L
Doxorubicin
YDR079C−A_2 TFB5
ZShift = −2.22
Z lm Score = −13.61
NG = 0
DB = 0
SM = 0
Delta_K
Doxorubicin
YDR079C−A_2 TFB5
ZShift = −2.47
Z lm Score = 1.19
NG = 0
DB = 0
SM = 0
Delta_r
Doxorubicin
YDR079C−A_2 TFB5
ZShift = −3.28
Z lm Score = −4.37
NG = 0
DB = 0
SM = 0
Delta_AUC
Doxorubicin
YMR269W_2 TMA23
ZShift = 7.89
Z lm Score = 5.64
NG = 0
DB = 0
SM = 0
Delta_L
Doxorubicin
YMR269W_2 TMA23
ZShift = −0.16
Z lm Score = −3.31
NG = 0
DB = 0
SM = 0
Delta_K
Doxorubicin
YMR269W_2 TMA23
ZShift = −2.34
Z lm Score = −0.2
NG = 0
DB = 0
SM = 0
Delta_r
Doxorubicin
YMR269W_2 TMA23
ZShift = −1.96
Z lm Score = −1.02
NG = 0
DB = 0
SM = 0
Delta_AUC
Doxorubicin

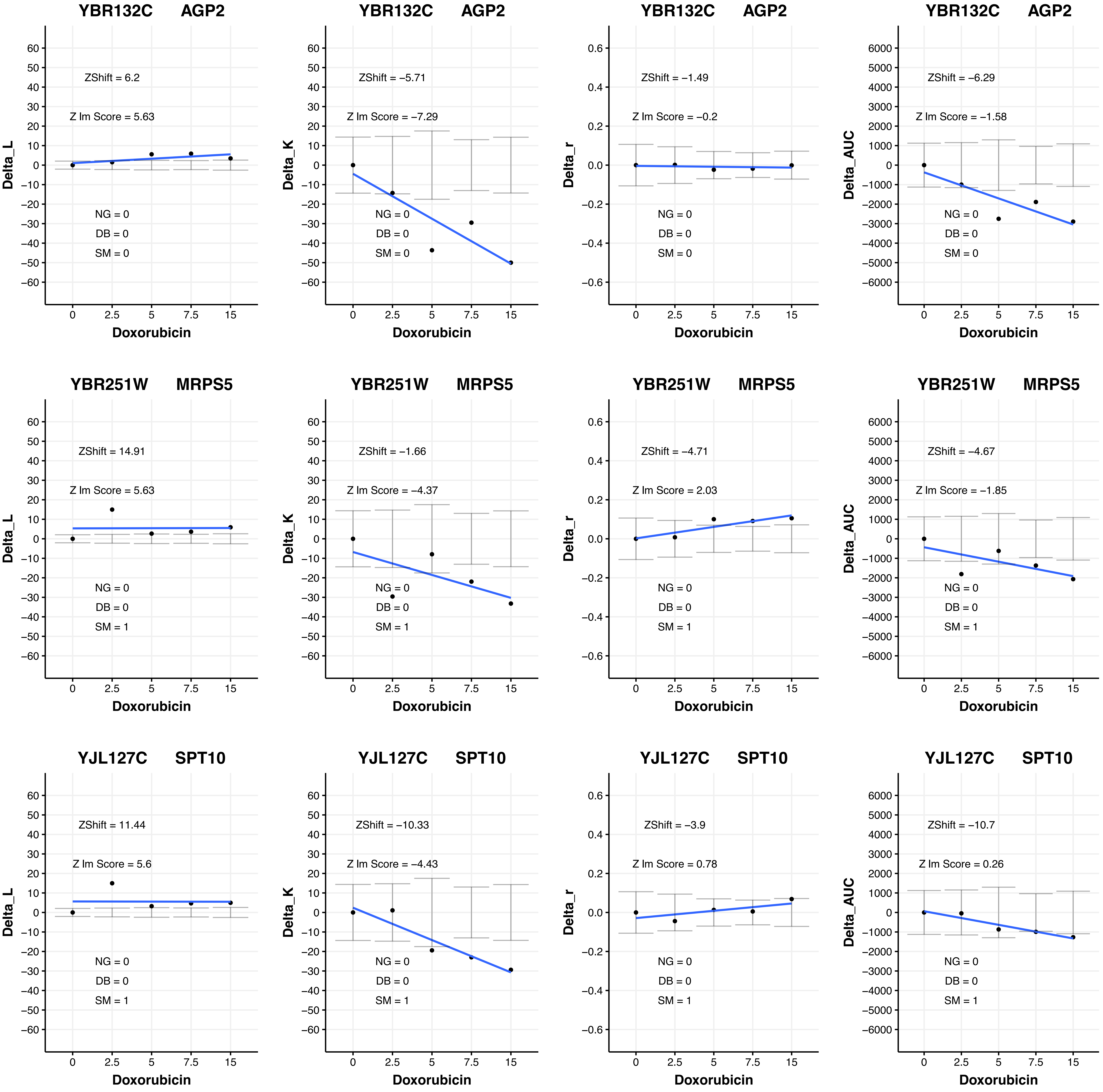
YBR132C AGP2
ZShift = 6.2
Z lm Score = 5.63
NG = 0
DB = 0
SM = 0
Delta_L
Doxorubicin
YBR132C AGP2
ZShift = −5.71
Z lm Score = −7.29
NG = 0
DB = 0
SM = 0
Delta_K
Doxorubicin
YBR132C AGP2
ZShift = −1.49
Z lm Score = −0.2
NG = 0
DB = 0
SM = 0
Delta_r
Doxorubicin
YBR132C AGP2
ZShift = −6.29
Z lm Score = −1.58
NG = 0
DB = 0
SM = 0
Delta_AUC
Doxorubicin
YBR251W MRPS5
ZShift = 14.91
Z lm Score = 5.63
NG = 0
DB = 0
SM = 1
Delta_L
Doxorubicin
YBR251W MRPS5
ZShift = −1.66
Z lm Score = −4.37
NG = 0
DB = 0
SM = 1
Delta_K
Doxorubicin
YBR251W MRPS5
ZShift = −4.71
Z lm Score = 2.03
NG = 0
DB = 0
SM = 1
Delta_r
Doxorubicin
YBR251W MRPS5
ZShift = −4.67
Z lm Score = −1.85
NG = 0
DB = 0
SM = 1
Delta_AUC
Doxorubicin
YJL127C SPT10
ZShift = 11.44
Z lm Score = 5.6
NG = 0
DB = 0
SM = 1
Delta_L
Doxorubicin
YJL127C SPT10
ZShift = −10.33
Z lm Score = −4.43
NG = 0
DB = 0
SM = 1
Delta_K
Doxorubicin
YJL127C SPT10
ZShift = −3.9
Z lm Score = 0.78
NG = 0
DB = 0
SM = 1
Delta_r
Doxorubicin
YJL127C SPT10
ZShift = −10.7
Z lm Score = 0.26
NG = 0
DB = 0
SM = 1
Delta_AUC
Doxorubicin

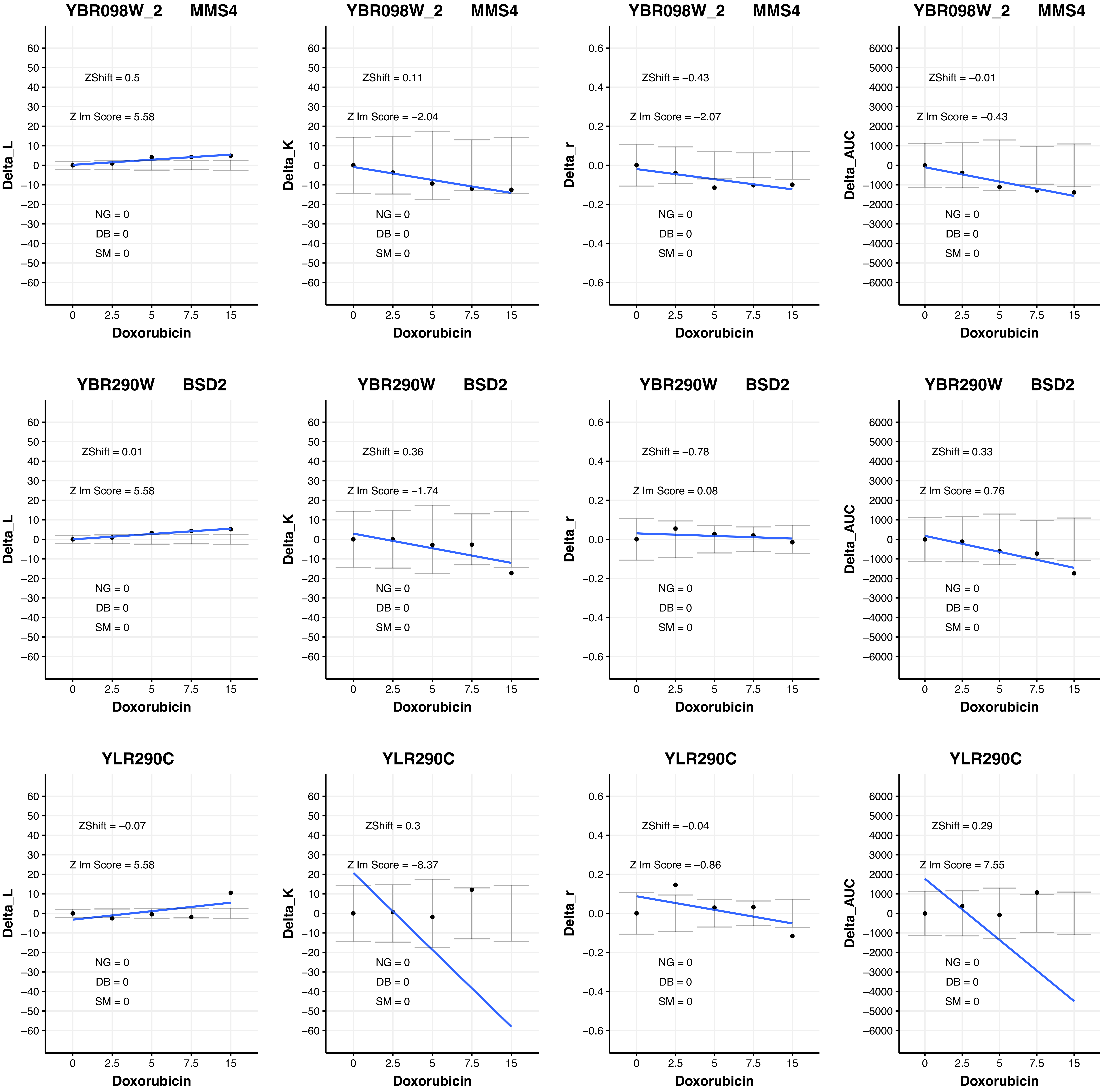

YBR098W_2 MMS4
ZShift = 0.5
Z lm Score = 5.58
NG = 0
DB = 0
SM = 0
Delta_L
Doxorubicin
YBR098W_2 MMS4
ZShift = 0.11
Z lm Score = −2.04
NG = 0
DB = 0
SM = 0
Delta_K
Doxorubicin
YBR098W_2 MMS4
ZShift = −0.43
Z lm Score = −2.07
NG = 0
DB = 0
SM = 0
Delta_r
Doxorubicin
YBR098W_2 MMS4
ZShift = −0.01
Z lm Score = −0.43
NG = 0
DB = 0
SM = 0
Delta_AUC
Doxorubicin
YBR290W BSD2
ZShift = 0.01
Z lm Score = 5.58
NG = 0
DB = 0
SM = 0
Delta_L
Doxorubicin
YBR290W BSD2
ZShift = 0.36
Z lm Score = −1.74
NG = 0
DB = 0
SM = 0
Delta_K
Doxorubicin
YBR290W BSD2
ZShift = −0.78
Z lm Score = 0.08
NG = 0
DB = 0
SM = 0
Delta_r
Doxorubicin
YBR290W BSD2
ZShift = 0.33
Z lm Score = 0.76
NG = 0
DB = 0
SM = 0
Delta_AUC
Doxorubicin
YLR290C
ZShift = −0.07
Z lm Score = 5.58
NG = 0
DB = 0
SM = 0
Delta_L
Doxorubicin
YLR290C
ZShift = 0.3
Z lm Score = −8.37
NG = 0
DB = 0
SM = 0
Delta_K
Doxorubicin
YLR290C
ZShift = −0.04
Z lm Score = −0.86
NG = 0
DB = 0
SM = 0
Delta_r
Doxorubicin
YLR290C
ZShift = 0.29
Z lm Score = 7.55
NG = 0
DB = 0
SM = 0
Delta_AUC
Doxorubicin

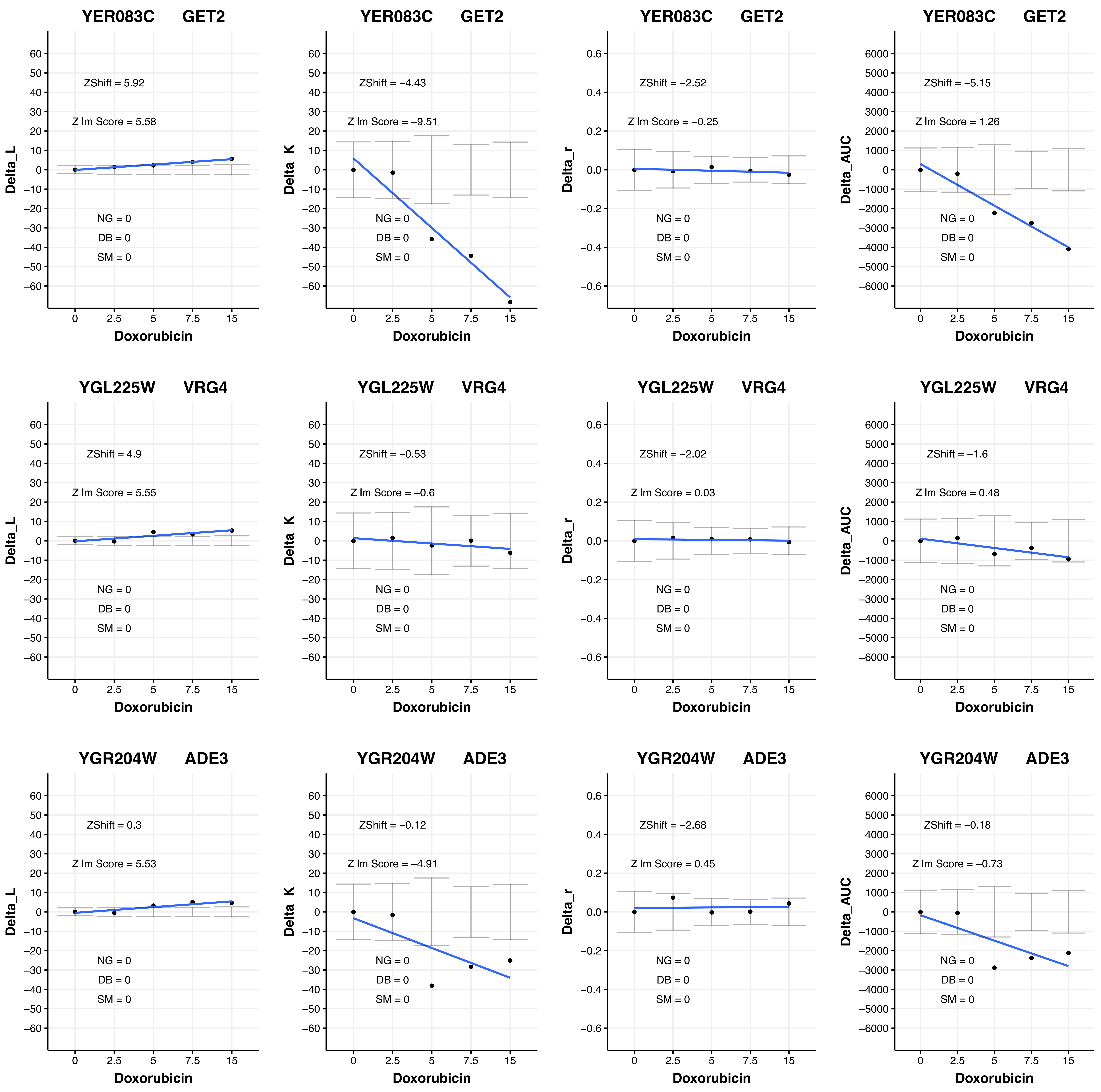
YER083C GET2
ZShift = 5.92
Z lm Score = 5.58
NG = 0
DB = 0
SM = 0
Delta_L
Doxorubicin
YER083C GET2
ZShift = −4.43
Z lm Score = −9.51
NG = 0
DB = 0
SM = 0
Delta_K
Doxorubicin
YER083C GET2
ZShift = −2.52
Z lm Score = −0.25
NG = 0
DB = 0
SM = 0
Delta_r
Doxorubicin
YER083C GET2
ZShift = −5.15
Z lm Score = 1.26
NG = 0
DB = 0
SM = 0
Delta_AUC
Doxorubicin
YGL225W VRG4
ZShift = 4.9
Z lm Score = 5.55
NG = 0
DB = 0
SM = 0
Delta_L
Doxorubicin
YGL225W VRG4
ZShift = −0.53
Z lm Score = −0.6
NG = 0
DB = 0
SM = 0
Delta_K
Doxorubicin
YGL225W VRG4
ZShift = −2.02
Z lm Score = 0.03
NG = 0
DB = 0
SM = 0
Delta_r
Doxorubicin
YGL225W VRG4
ZShift = −1.6
Z lm Score = 0.48
NG = 0
DB = 0
SM = 0
Delta_AUC
Doxorubicin
YGR204W ADE3
ZShift = 0.3
Z lm Score = 5.53
NG = 0
DB = 0
SM = 0
Delta_L
Doxorubicin
YGR204W ADE3
ZShift = −0.12
Z lm Score = −4.91
NG = 0
DB = 0
SM = 0
Delta_K
Doxorubicin
YGR204W ADE3
ZShift = −2.68
Z lm Score = 0.45
NG = 0
DB = 0
SM = 0
Delta_r
Doxorubicin
YGR204W ADE3
ZShift = −0.18
Z lm Score = −0.73
NG = 0
DB = 0
SM = 0
Delta_AUC
Doxorubicin

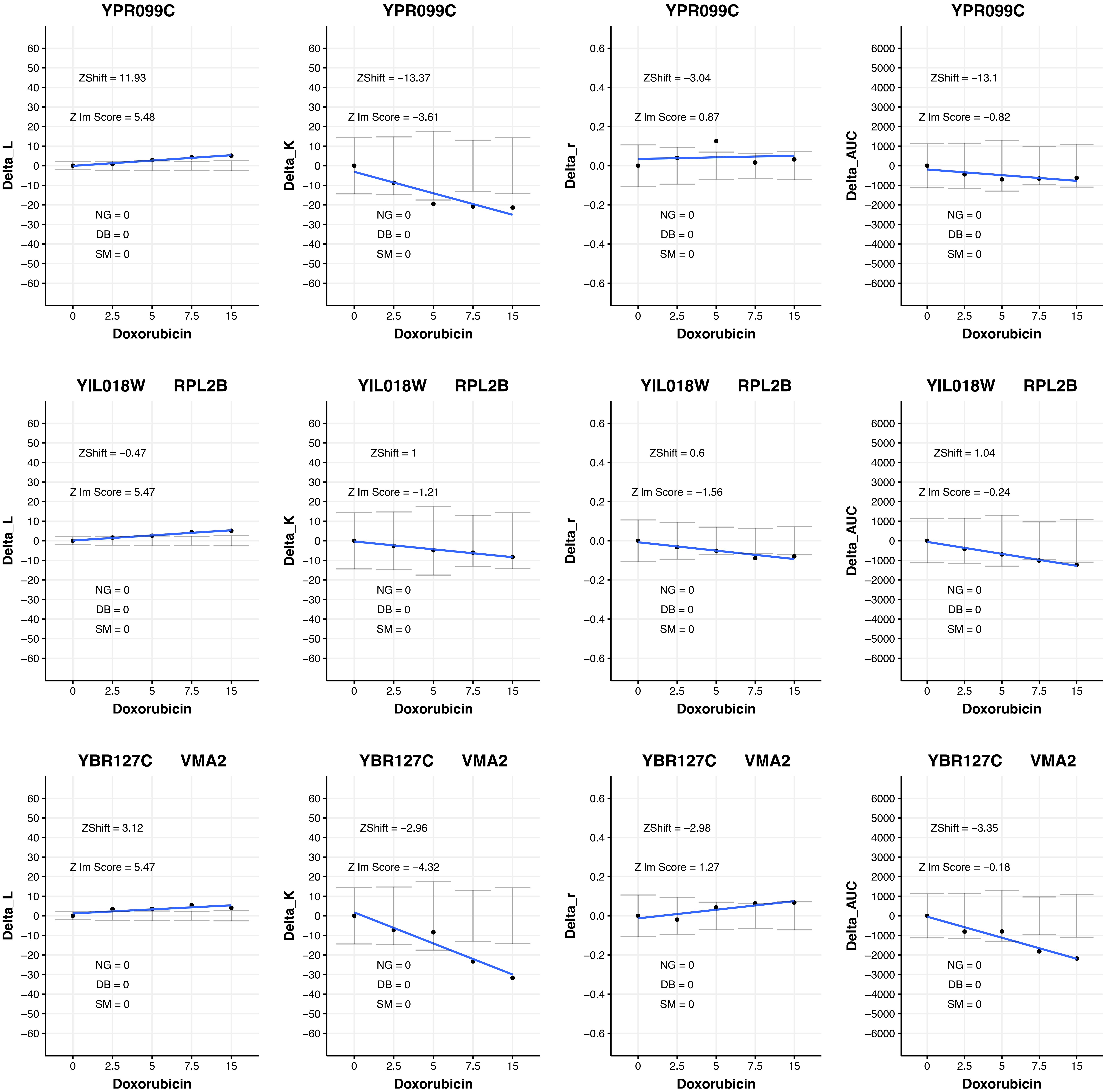
YPR099C
ZShift = 11.93
Z lm Score = 5.48
NG = 0
DB = 0
SM = 0
Delta_L
Doxorubicin
YPR099C
ZShift = −13.37
Z lm Score = −3.61
NG = 0
DB = 0
SM = 0
Delta_K
Doxorubicin
YPR099C
ZShift = −3.04
Z lm Score = 0.87
NG = 0
DB = 0
SM = 0
Delta_r
Doxorubicin
YPR099C
ZShift = −13.1
Z lm Score = −0.82
NG = 0
DB = 0
SM = 0
Delta_AUC
Doxorubicin
YIL018W RPL2B
ZShift = −0.47
Z lm Score = 5.47
NG = 0
DB = 0
SM = 0
Delta_L
Doxorubicin
YIL018W RPL2B
ZShift = 1
Z lm Score = −1.21
NG = 0
DB = 0
SM = 0
Delta_K
Doxorubicin
YIL018W RPL2B
ZShift = 0.6
Z lm Score = −1.56
NG = 0
DB = 0
SM = 0
Delta_r
Doxorubicin
YIL018W RPL2B
ZShift = 1.04
Z lm Score = −0.24
NG = 0
DB = 0
SM = 0
Delta_AUC
Doxorubicin
YBR127C VMA2
ZShift = 3.12
Z lm Score = 5.47
NG = 0
DB = 0
SM = 0
Delta_L
Doxorubicin
YBR127C VMA2
ZShift = −2.96
Z lm Score = −4.32
NG = 0
DB = 0
SM = 0
Delta_K
Doxorubicin
YBR127C VMA2
ZShift = −2.98
Z lm Score = 1.27
NG = 0
DB = 0
SM = 0
Delta_r
Doxorubicin
YBR127C VMA2
ZShift = −3.35
Z lm Score = −0.18
NG = 0
DB = 0
SM = 0
Delta_AUC
Doxorubicin

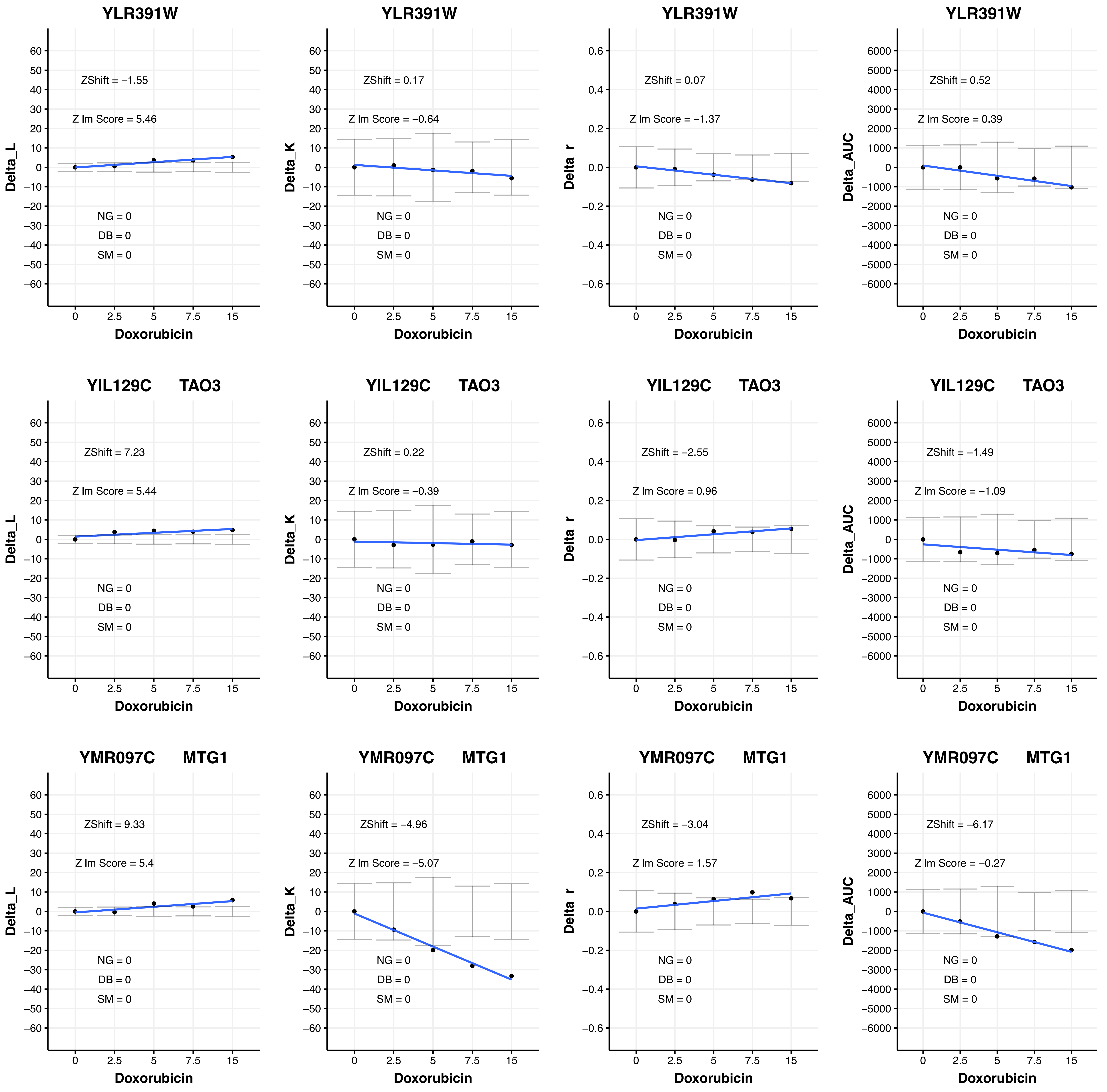

YLR391W
ZShift = −1.55
Z lm Score = 5.46
NG = 0
DB = 0
SM = 0
Delta_L
Doxorubicin
YLR391W
ZShift = 0.17
Z lm Score = −0.64
NG = 0
DB = 0
SM = 0
Delta_K
Doxorubicin
YLR391W
ZShift = 0.07
Z lm Score = −1.37
NG = 0
DB = 0
SM = 0
Delta_r
Doxorubicin
YLR391W
ZShift = 0.52
Z lm Score = 0.39
NG = 0
DB = 0
SM = 0
Delta_AUC
Doxorubicin
YIL129C TAO3
ZShift = 7.23
Z lm Score = 5.44
NG = 0
DB = 0
SM = 0
Delta_L
Doxorubicin
YIL129C TAO3
ZShift = 0.22
Z lm Score = −0.39
NG = 0
DB = 0
SM = 0
Delta_K
Doxorubicin
YIL129C TAO3
ZShift = −2.55
Z lm Score = 0.96
NG = 0
DB = 0
SM = 0
Delta_r
Doxorubicin
YIL129C TAO3
ZShift = −1.49
Z lm Score = −1.09
NG = 0
DB = 0
SM = 0
Delta_AUC
Doxorubicin
YMR097C MTG1
ZShift = 9.33
Z lm Score = 5.4
NG = 0
DB = 0
SM = 0
Delta_L
Doxorubicin
YMR097C MTG1
ZShift = −4.96
Z lm Score = −5.07
NG = 0
DB = 0
SM = 0
Delta_K
Doxorubicin
YMR097C MTG1
ZShift = −3.04
Z lm Score = 1.57
NG = 0
DB = 0
SM = 0
Delta_r
Doxorubicin
YMR097C MTG1
ZShift = −6.17
Z lm Score = −0.27
NG = 0
DB = 0
SM = 0
Delta_AUC
Doxorubicin

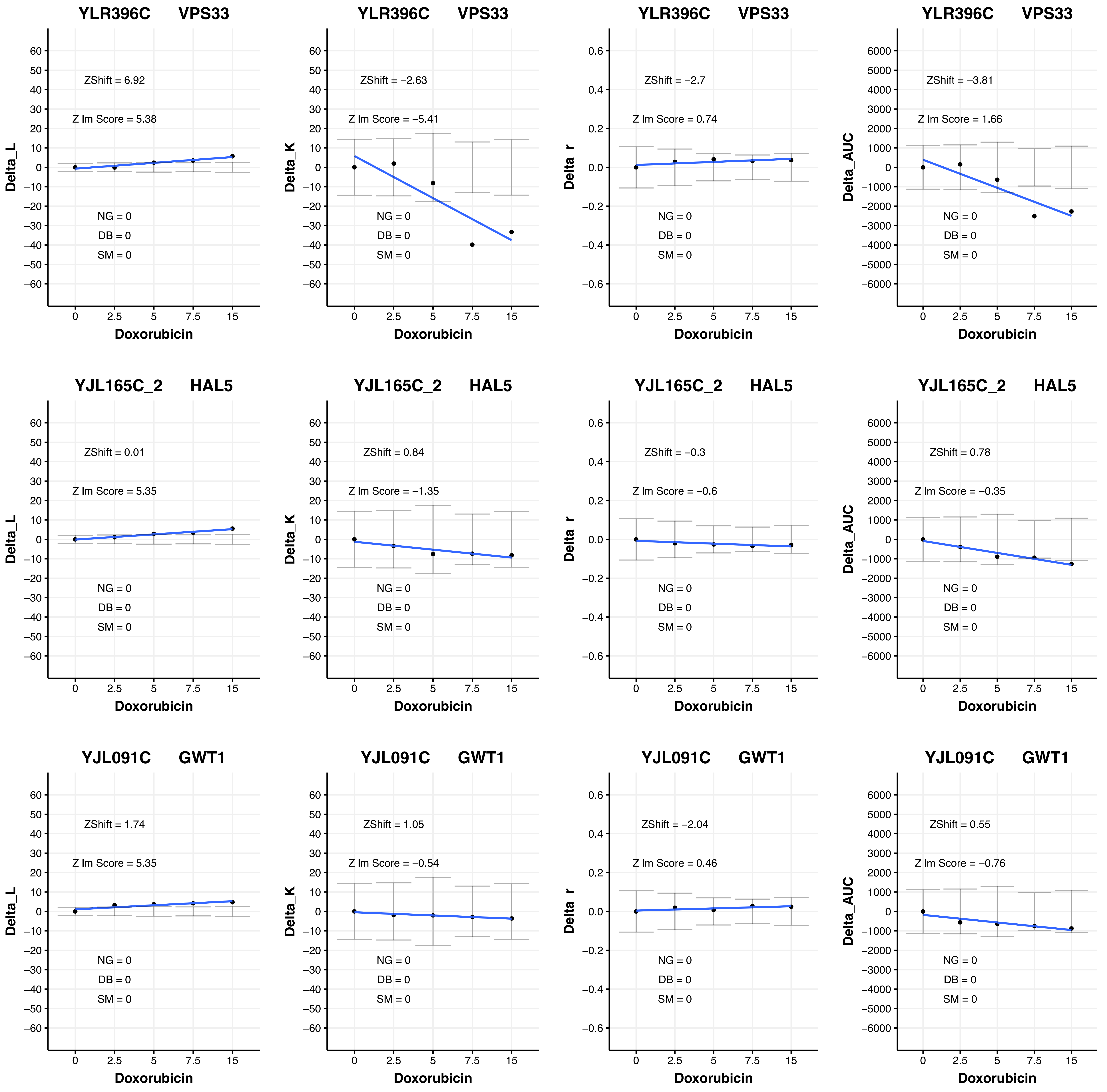

YLR396C VPS33
ZShift = 6.92
Z lm Score = 5.38
NG = 0
DB = 0
SM = 0
Delta_L
Doxorubicin
YLR396C VPS33
ZShift = −2.63
Z lm Score = −5.41
NG = 0
DB = 0
SM = 0
Delta_K
Doxorubicin
YLR396C VPS33
ZShift = −2.7
Z lm Score = 0.74
NG = 0
DB = 0
SM = 0
Delta_r
Doxorubicin
YLR396C VPS33
ZShift = −3.81
Z lm Score = 1.66
NG = 0
DB = 0
SM = 0
Delta_AUC
Doxorubicin
YJL165C_2 HAL5
ZShift = 0.01
Z lm Score = 5.35
NG = 0
DB = 0
SM = 0
Delta_L
Doxorubicin
YJL165C_2 HAL5
ZShift = 0.84
Z lm Score = −1.35
NG = 0
DB = 0
SM = 0
Delta_K
Doxorubicin
YJL165C_2 HAL5
ZShift = −0.3
Z lm Score = −0.6
NG = 0
DB = 0
SM = 0
Delta_r
Doxorubicin
YJL165C_2 HAL5
ZShift = 0.78
Z lm Score = −0.35
NG = 0
DB = 0
SM = 0
Delta_AUC
Doxorubicin
YJL091C GWT1
ZShift = 1.74
Z lm Score = 5.35
NG = 0
DB = 0
SM = 0
Delta_L
Doxorubicin
YJL091C GWT1
ZShift = 1.05
Z lm Score = −0.54
NG = 0
DB = 0
SM = 0
Delta_K
Doxorubicin
YJL091C GWT1
ZShift = −2.04
Z lm Score = 0.46
NG = 0
DB = 0
SM = 0
Delta_r
Doxorubicin
YJL091C GWT1
ZShift = 0.55
Z lm Score = −0.76
NG = 0
DB = 0
SM = 0
Delta_AUC
Doxorubicin

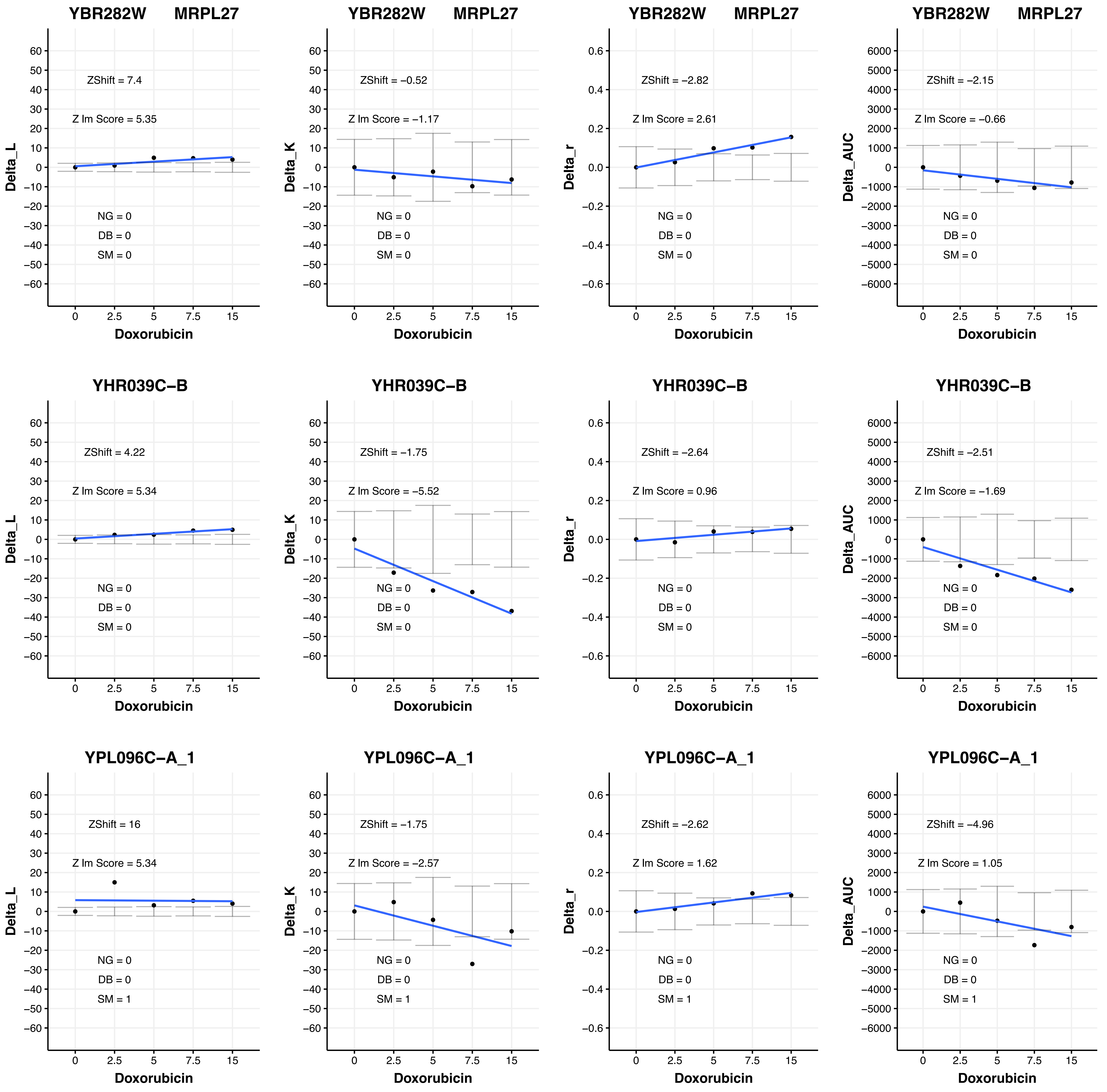

YBR282W MRPL27
ZShift = 7.4
Z lm Score = 5.35
NG = 0
DB = 0
SM = 0
Delta_L
Doxorubicin
YBR282W MRPL27
ZShift = −0.52
Z lm Score = −1.17
NG = 0
DB = 0
SM = 0
Delta_K
Doxorubicin
YBR282W MRPL27
ZShift = −2.82
Z lm Score = 2.61
NG = 0
DB = 0
SM = 0
Delta_r
Doxorubicin
YBR282W MRPL27
ZShift = −2.15
Z lm Score = −0.66
NG = 0
DB = 0
SM = 0
Delta_AUC
Doxorubicin
YHR039C−B
ZShift = 4.22
Z lm Score = 5.34
NG = 0
DB = 0
SM = 0
Delta_L
Doxorubicin
YHR039C−B
ZShift = −1.75
Z lm Score = −5.52
NG = 0
DB = 0
SM = 0
Delta_K
Doxorubicin
YHR039C−B
ZShift = −2.64
Z lm Score = 0.96
NG = 0
DB = 0
SM = 0
Delta_r
Doxorubicin
YHR039C−B
ZShift = −2.51
Z lm Score = −1.69
NG = 0
DB = 0
SM = 0
Delta_AUC
Doxorubicin
YPL096C−A_1
ZShift = 16
Z lm Score = 5.34
NG = 0
DB = 0
SM = 1
Delta_L
Doxorubicin
YPL096C−A_1
ZShift = −1.75
Z lm Score = −2.57
NG = 0
DB = 0
SM = 1
Delta_K
Doxorubicin
YPL096C−A_1
ZShift = −2.62
Z lm Score = 1.62
NG = 0
DB = 0
SM = 1
Delta_r
Doxorubicin
YPL096C−A_1
ZShift = −4.96
Z lm Score = 1.05
NG = 0
DB = 0
SM = 1
Delta_AUC
Doxorubicin
0 2.5 5 7.5 15

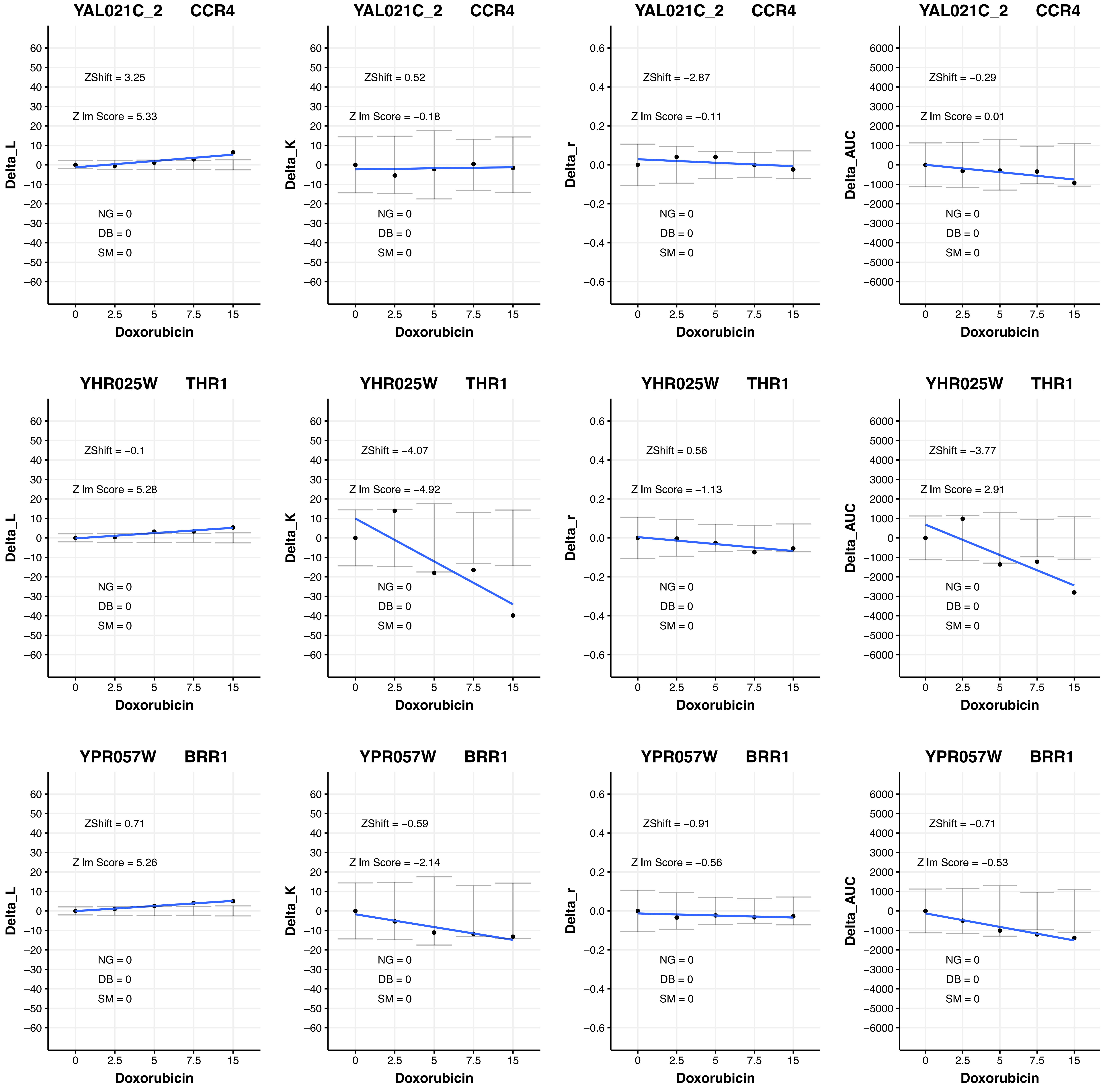
YAL021C_2 CCR4
ZShift = 3.25
Z lm Score = 5.33
NG = 0
DB = 0
SM = 0
Delta_L
Doxorubicin
YAL021C_2 CCR4
ZShift = 0.52
Z lm Score = −0.18
NG = 0
DB = 0
SM = 0
Delta_K
Doxorubicin
YAL021C_2 CCR4
ZShift = −2.87
Z lm Score = −0.11
NG = 0
DB = 0
SM = 0
Delta_r
Doxorubicin
YAL021C_2 CCR4
ZShift = −0.29
Z lm Score = 0.01
NG = 0
DB = 0
SM = 0
Delta_AUC
Doxorubicin
YHR025W THR1
ZShift = −0.1
Z lm Score = 5.28
NG = 0
DB = 0
SM = 0
Delta_L
Doxorubicin
YHR025W THR1
ZShift = −4.07
Z lm Score = −4.92
NG = 0
DB = 0
SM = 0
Delta_K
Doxorubicin
YHR025W THR1
ZShift = 0.56
Z lm Score = −1.13
NG = 0
DB = 0
SM = 0
Delta_r
Doxorubicin
YHR025W THR1
ZShift = −3.77
Z lm Score = 2.91
NG = 0
DB = 0
SM = 0
Delta_AUC
Doxorubicin
YPR057W BRR1
ZShift = 0.71
Z lm Score = 5.26
NG = 0
DB = 0
SM = 0
Delta_L
Doxorubicin
YPR057W BRR1
ZShift = −0.59
Z lm Score = −2.14
NG = 0
DB = 0
SM = 0
Delta_K
Doxorubicin
YPR057W BRR1
ZShift = −0.91
Z lm Score = −0.56
NG = 0
DB = 0
SM = 0
Delta_r
Doxorubicin
YPR057W BRR1
ZShift = −0.71
Z lm Score = −0.53
NG = 0
DB = 0
SM = 0
Delta_AUC
Doxorubicin

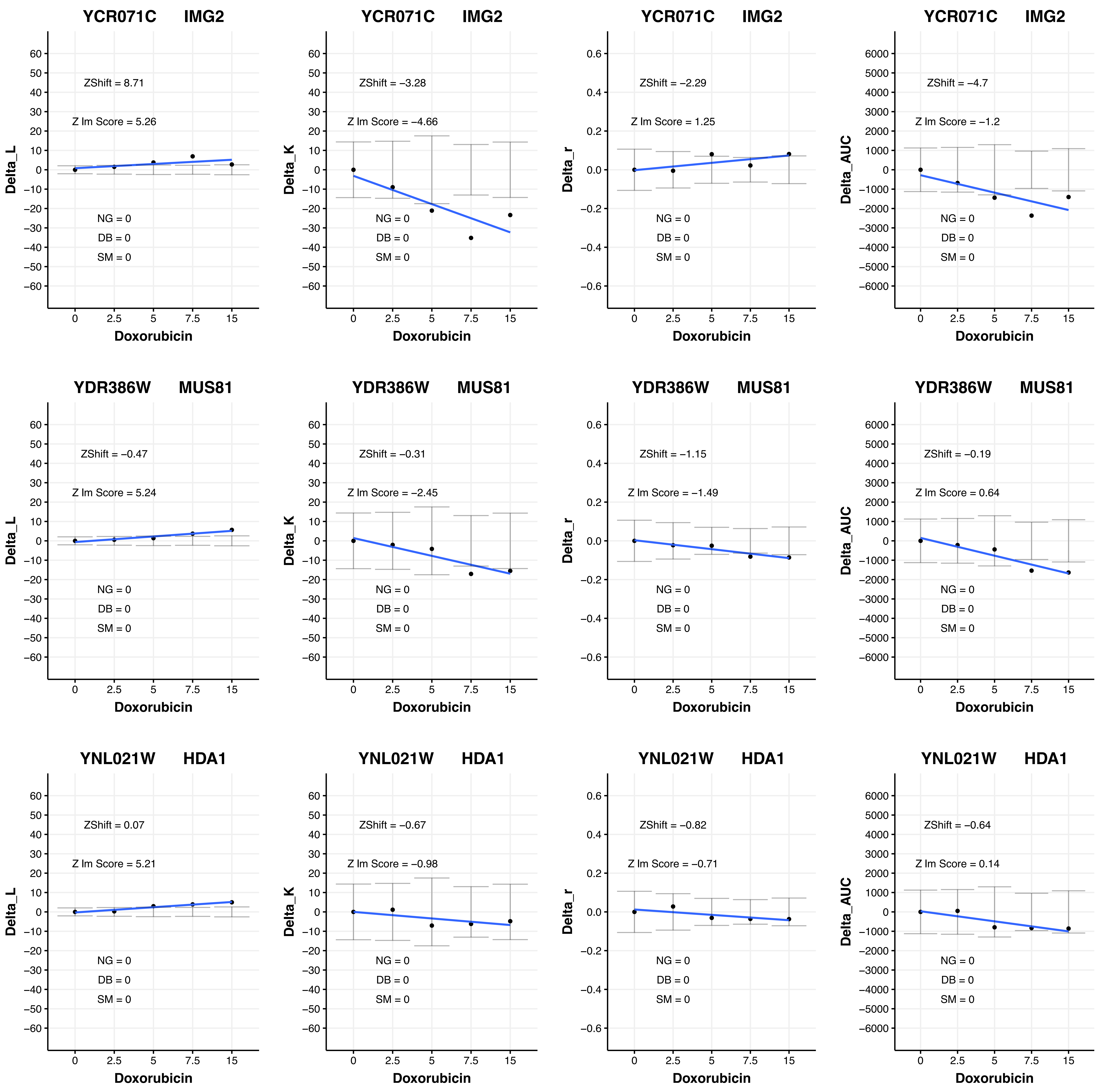
YCR071C IMG2
ZShift = 8.71
Z lm Score = 5.26
NG = 0
DB = 0
SM = 0
Delta_L
Doxorubicin
YCR071C IMG2
ZShift = −3.28
Z lm Score = −4.66
NG = 0
DB = 0
SM = 0
Delta_K
Doxorubicin
YCR071C IMG2
ZShift = −2.29
Z lm Score = 1.25
NG = 0
DB = 0
SM = 0
Delta_r
Doxorubicin
YCR071C IMG2
ZShift = −4.7
Z lm Score = −1.2
NG = 0
DB = 0
SM = 0
Delta_AUC
Doxorubicin
YDR386W MUS81
ZShift = −0.47
Z lm Score = 5.24
NG = 0
DB = 0
SM = 0
Delta_L
Doxorubicin
YDR386W MUS81
ZShift = −0.31
Z lm Score = −2.45
NG = 0
DB = 0
SM = 0
Delta_K
Doxorubicin
YDR386W MUS81
ZShift = −1.15
Z lm Score = −1.49
NG = 0
DB = 0
SM = 0
Delta_r
Doxorubicin
YDR386W MUS81
ZShift = −0.19
Z lm Score = 0.64
NG = 0
DB = 0
SM = 0
Delta_AUC
Doxorubicin
YNL021W HDA1
ZShift = 0.07
Z lm Score = 5.21
NG = 0
DB = 0
SM = 0
Delta_L
Doxorubicin
YNL021W HDA1
ZShift = −0.67
Z lm Score = −0.98
NG = 0
DB = 0
SM = 0
Delta_K
Doxorubicin
YNL021W HDA1
ZShift = −0.82
Z lm Score = −0.71
NG = 0
DB = 0
SM = 0
Delta_r
Doxorubicin
YNL021W HDA1
ZShift = −0.64
Z lm Score = 0.14
NG = 0
DB = 0
SM = 0
Delta_AUC
Doxorubicin

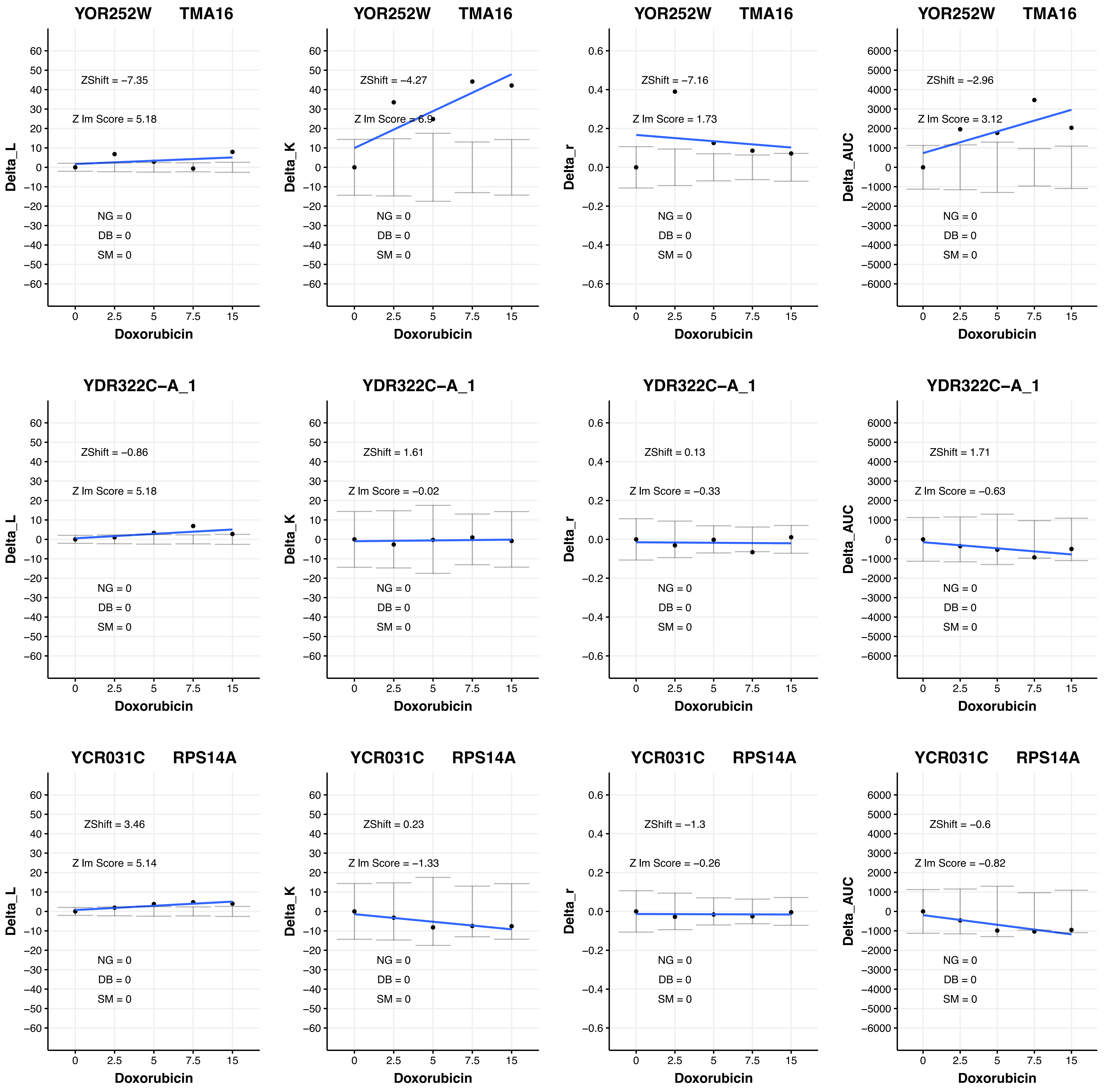

YOR252W TMA16
ZShift = −7.35
Z lm Score = 5.18
NG = 0
DB = 0
SM = 0
Delta_L
Doxorubicin
YOR252W TMA16
ZShift = −4.27
Z lm Score = 6.9
NG = 0
DB = 0
SM = 0
Delta_K
Doxorubicin
YOR252W TMA16
ZShift = −7.16
Z lm Score = 1.73
NG = 0
DB = 0
SM = 0
Delta_r
Doxorubicin
YOR252W TMA16
ZShift = −2.96
Z lm Score = 3.12
NG = 0
DB = 0
SM = 0
Delta_AUC
Doxorubicin
YDR322C−A_1
ZShift = −0.86
Z lm Score = 5.18
NG = 0
DB = 0
SM = 0
Delta_L
Doxorubicin
YDR322C−A_1
ZShift = 1.61
Z lm Score = −0.02
NG = 0
DB = 0
SM = 0
Delta_K
Doxorubicin
YDR322C−A_1
ZShift = 0.13
Z lm Score = −0.33
NG = 0
DB = 0
SM = 0
Delta_r
Doxorubicin
YDR322C−A_1
ZShift = 1.71
Z lm Score = −0.63
NG = 0
DB = 0
SM = 0
Delta_AUC
Doxorubicin
YCR031C RPS14A
ZShift = 3.46
Z lm Score = 5.14
NG = 0
DB = 0
SM = 0
Delta_L
Doxorubicin
YCR031C RPS14A
ZShift = 0.23
Z lm Score = −1.33
NG = 0
DB = 0
SM = 0
Delta_K
Doxorubicin
YCR031C RPS14A
ZShift = −1.3
Z lm Score = −0.26
NG = 0
DB = 0
SM = 0
Delta_r
Doxorubicin
YCR031C RPS14A
ZShift = −0.6
Z lm Score = −0.82
NG = 0
DB = 0
SM = 0
Delta_AUC
Doxorubicin

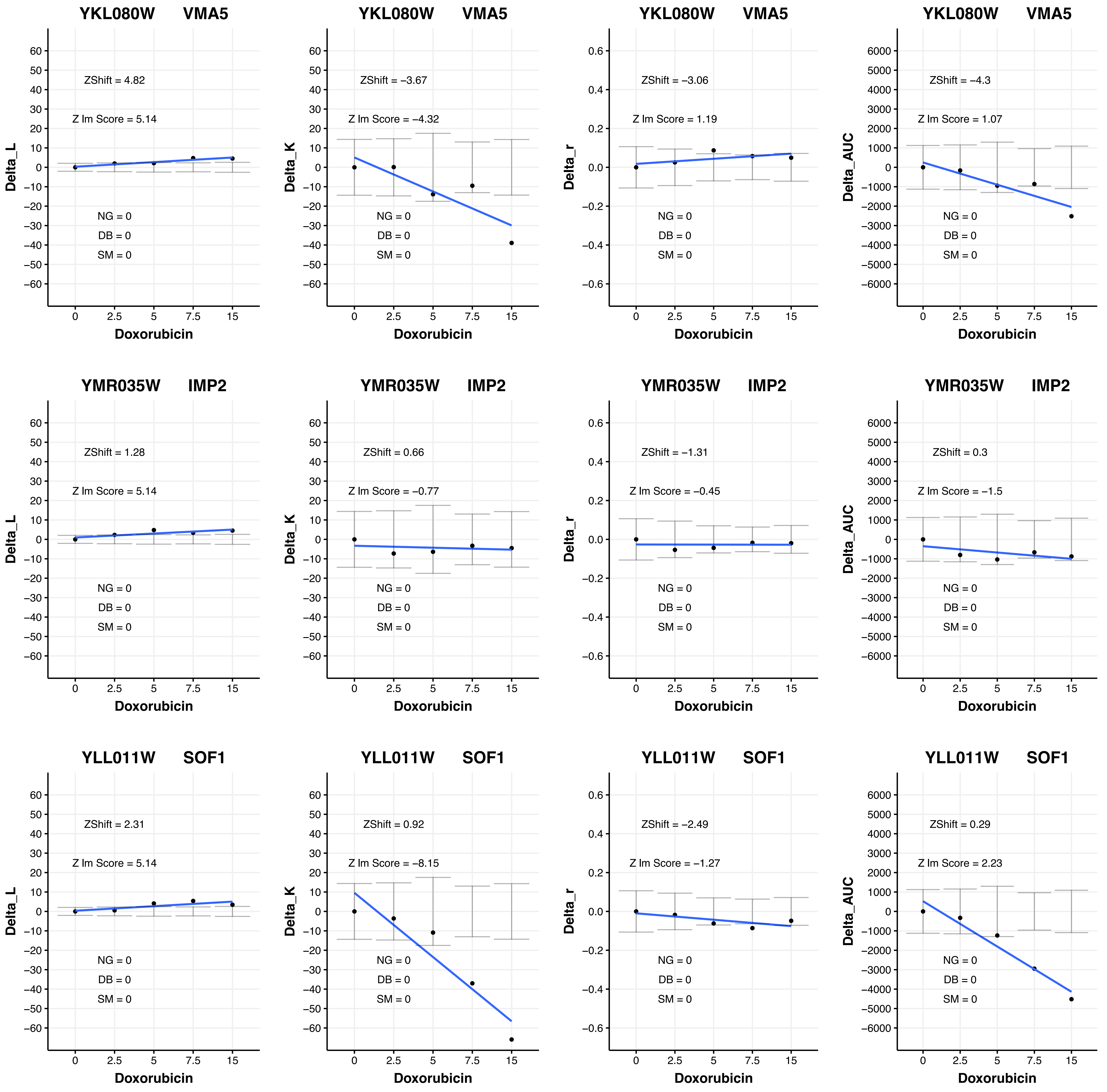

YKL080W VMA5
ZShift = 4.82
Z lm Score = 5.14
NG = 0
DB = 0
SM = 0
Delta_L
Doxorubicin
YKL080W VMA5
ZShift = −3.67
Z lm Score = −4.32
NG = 0
DB = 0
SM = 0
Delta_K
Doxorubicin
YKL080W VMA5
ZShift = −3.06
Z lm Score = 1.19
NG = 0
DB = 0
SM = 0
Delta_r
Doxorubicin
YKL080W VMA5
ZShift = −4.3
Z lm Score = 1.07
NG = 0
DB = 0
SM = 0
Delta_AUC
Doxorubicin
YMR035W IMP2
ZShift = 1.28
Z lm Score = 5.14
NG = 0
DB = 0
SM = 0
Delta_L
Doxorubicin
YMR035W IMP2
ZShift = 0.66
Z lm Score = −0.77
NG = 0
DB = 0
SM = 0
Delta_K
Doxorubicin
YMR035W IMP2
ZShift = −1.31
Z lm Score = −0.45
NG = 0
DB = 0
SM = 0
Delta_r
Doxorubicin
YMR035W IMP2
ZShift = 0.3
Z lm Score = −1.5
NG = 0
DB = 0
SM = 0
Delta_AUC
Doxorubicin
YLL011W SOF1
ZShift = 2.31
Z lm Score = 5.14
NG = 0
DB = 0
SM = 0
Delta_L
Doxorubicin
YLL011W SOF1
ZShift = 0.92
Z lm Score = −8.15
NG = 0
DB = 0
SM = 0
Delta_K
Doxorubicin
YLL011W SOF1
ZShift = −2.49
Z lm Score = −1.27
NG = 0
DB = 0
SM = 0
Delta_r
Doxorubicin
YLL011W SOF1
ZShift = 0.29
Z lm Score = 2.23
NG = 0
DB = 0
SM = 0
Delta_AUC
Doxorubicin

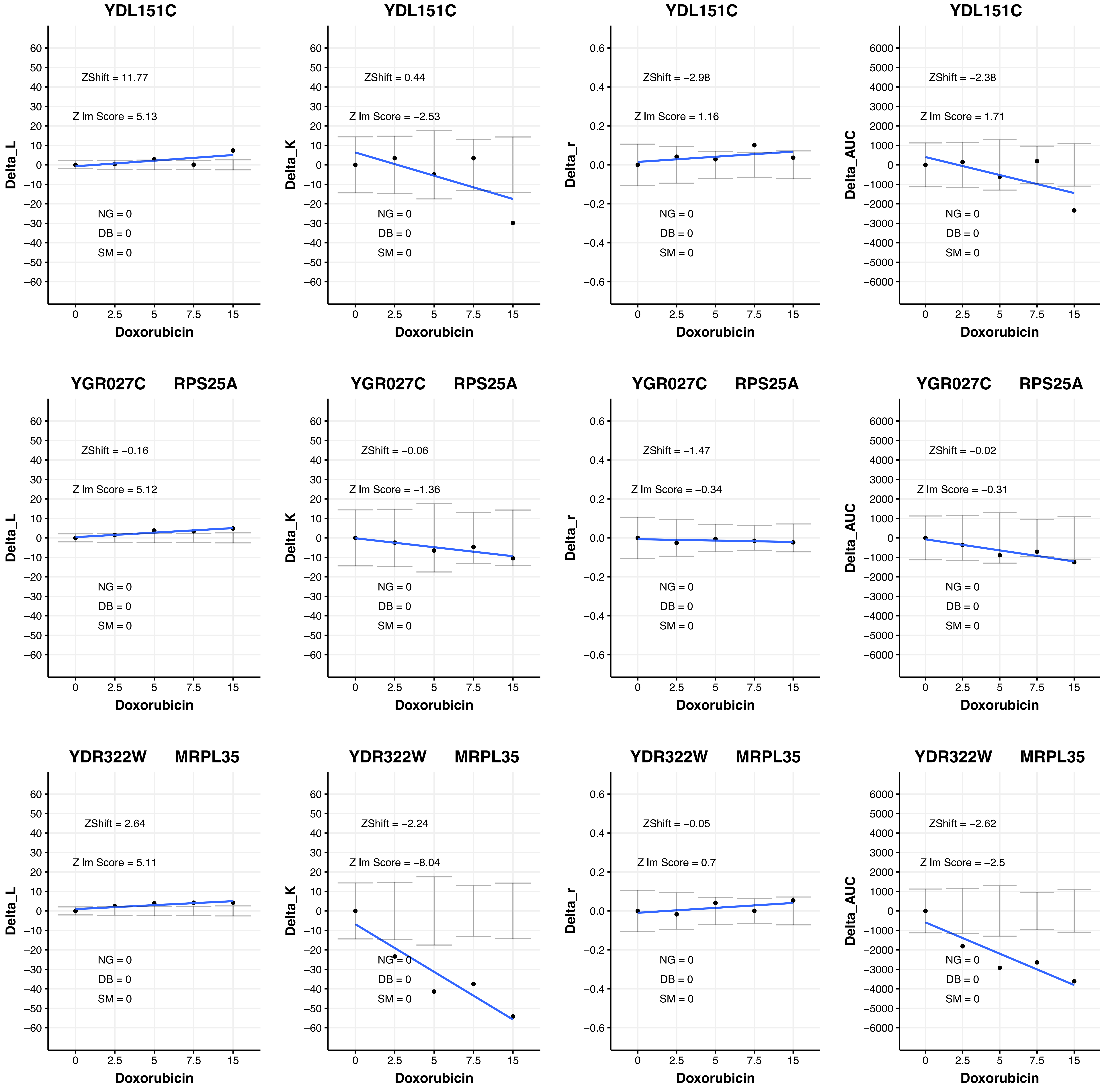

YDL151C
ZShift = 11.77
Z lm Score = 5.13
NG = 0
DB = 0
SM = 0
Delta_L
Doxorubicin
YDL151C
ZShift = 0.44
Z lm Score = −2.53
NG = 0
DB = 0
SM = 0
Delta_K
Doxorubicin
YDL151C
ZShift = −2.98
Z lm Score = 1.16
NG = 0
DB = 0
SM = 0
Delta_r
Doxorubicin
YDL151C
ZShift = −2.38
Z lm Score = 1.71
NG = 0
DB = 0
SM = 0
Delta_AUC
Doxorubicin
YGR027C RPS25A
ZShift = −0.16
Z lm Score = 5.12
NG = 0
DB = 0
SM = 0
Delta_L
Doxorubicin
YGR027C RPS25A
ZShift = −0.06
Z lm Score = −1.36
NG = 0
DB = 0
SM = 0
Delta_K
Doxorubicin
YGR027C RPS25A
ZShift = −1.47
Z lm Score = −0.34
NG = 0
DB = 0
SM = 0
Delta_r
Doxorubicin
YGR027C RPS25A
ZShift = −0.02
Z lm Score = −0.31
NG = 0
DB = 0
SM = 0
Delta_AUC
Doxorubicin
YDR322W MRPL35
ZShift = 2.64
Z lm Score = 5.11
NG = 0
DB = 0
SM = 0
Delta_L
Doxorubicin
YDR322W MRPL35
ZShift = −2.24
Z lm Score = −8.04
NG = 0
DB = 0
SM = 0
Delta_K
Doxorubicin
YDR322W MRPL35
ZShift = −0.05
Z lm Score = 0.7
NG = 0
DB = 0
SM = 0
Delta_r
Doxorubicin
YDR322W MRPL35
ZShift = −2.62
Z lm Score = −2.5
NG = 0
DB = 0
SM = 0
Delta_AUC
Doxorubicin

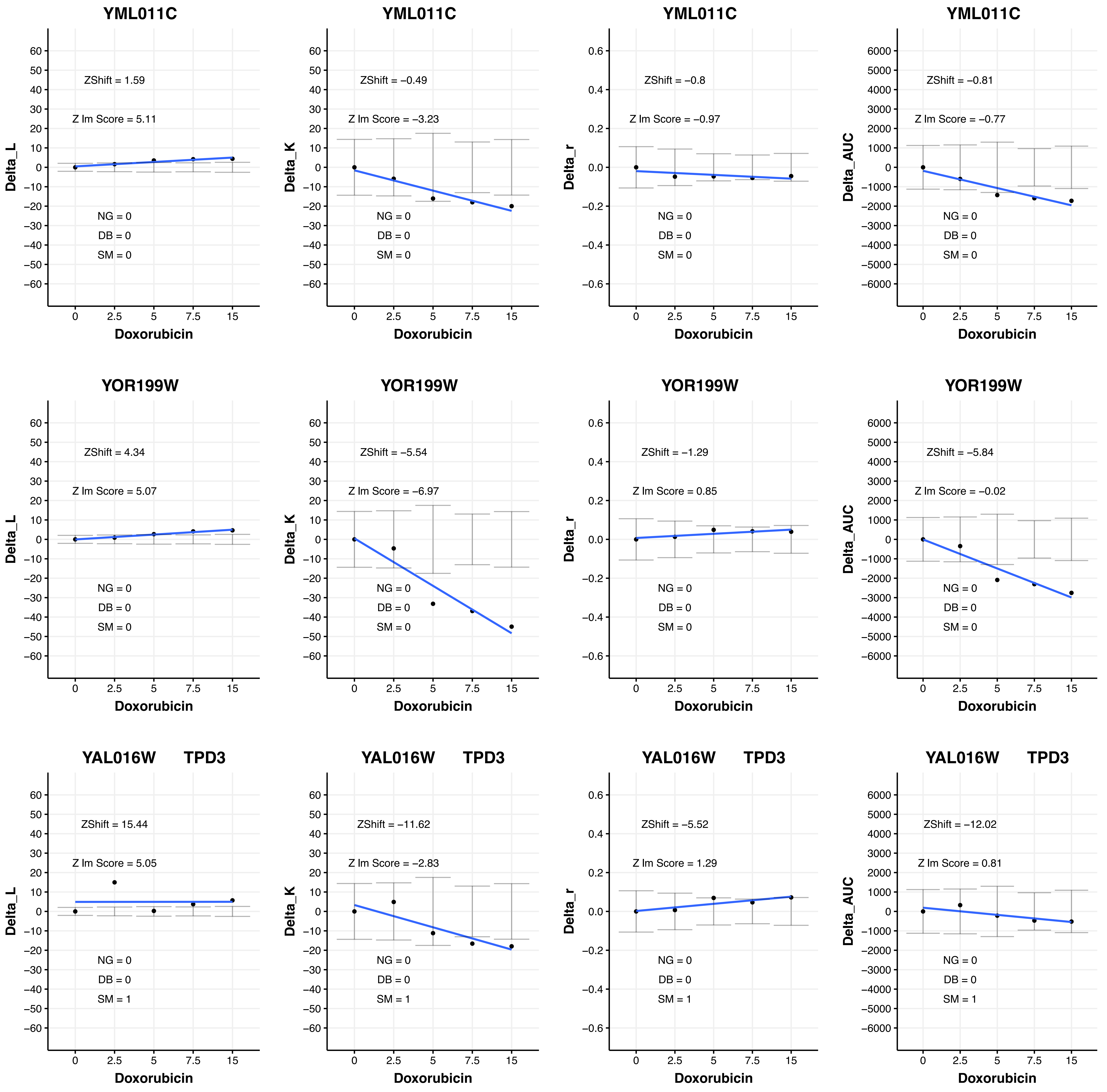

YML011C
ZShift = 1.59
Z lm Score = 5.11
NG = 0
DB = 0
SM = 0
Delta_L
Doxorubicin
YML011C
ZShift = −0.49
Z lm Score = −3.23
NG = 0
DB = 0
SM = 0
Delta_K
Doxorubicin
YML011C
ZShift = −0.8
Z lm Score = −0.97
NG = 0
DB = 0
SM = 0
Delta_r
Doxorubicin
YML011C
ZShift = −0.81
Z lm Score = −0.77
NG = 0
DB = 0
SM = 0
Delta_AUC
Doxorubicin
YOR199W
ZShift = 4.34
Z lm Score = 5.07
NG = 0
DB = 0
SM = 0
Delta_L
Doxorubicin
YOR199W
ZShift = −5.54
Z lm Score = −6.97
NG = 0
DB = 0
SM = 0
Delta_K
Doxorubicin
YOR199W
ZShift = −1.29
Z lm Score = 0.85
NG = 0
DB = 0
SM = 0
Delta_r
Doxorubicin
YOR199W
ZShift = −5.84
Z lm Score = −0.02
NG = 0
DB = 0
SM = 0
Delta_AUC
Doxorubicin
YAL016W TPD3
ZShift = 15.44
Z lm Score = 5.05
NG = 0
DB = 0
SM = 1
Delta_L
Doxorubicin
YAL016W TPD3
ZShift = −11.62
Z lm Score = −2.83
NG = 0
DB = 0
SM = 1
Delta_K
Doxorubicin
YAL016W TPD3
ZShift = −5.52
Z lm Score = 1.29
NG = 0
DB = 0
SM = 1
Delta_r
Doxorubicin
YAL016W TPD3
ZShift = −12.02
Z lm Score = 0.81
NG = 0
DB = 0
SM = 1
Delta_AUC
Doxorubicin

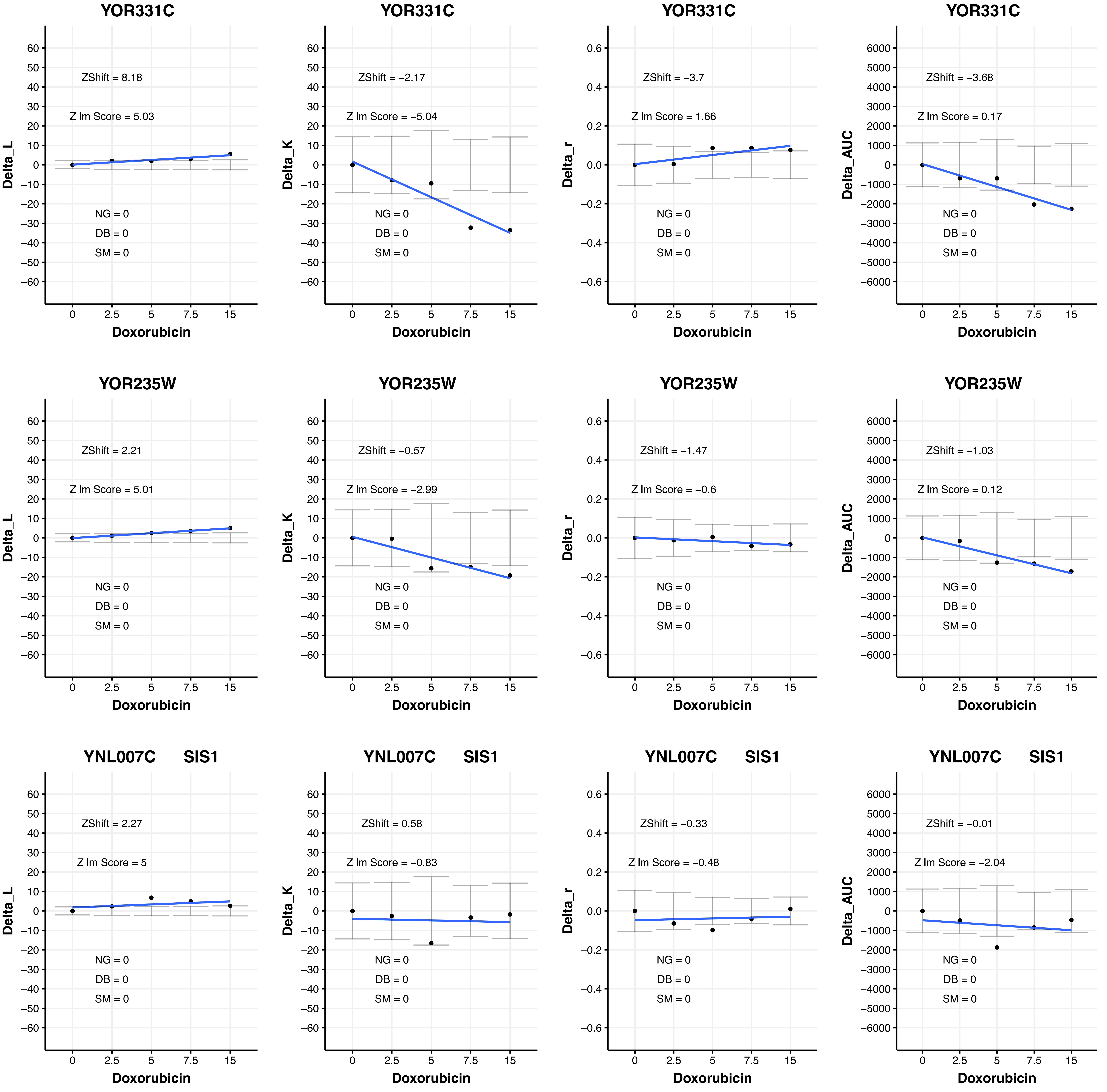
YOR331C
ZShift = 8.18
Z lm Score = 5.03
NG = 0
DB = 0
SM = 0
Delta_L
Doxorubicin
YOR331C
ZShift = −2.17
Z lm Score = −5.04
NG = 0
DB = 0
SM = 0
Delta_K
Doxorubicin
YOR331C
ZShift = −3.7
Z lm Score = 1.66
NG = 0
DB = 0
SM = 0
Delta_r
Doxorubicin
YOR331C
ZShift = −3.68
Z lm Score = 0.17
NG = 0
DB = 0
SM = 0
Delta_AUC
Doxorubicin
YOR235W
ZShift = 2.21
Z lm Score = 5.01
NG = 0
DB = 0
SM = 0
Delta_L
Doxorubicin
YOR235W
ZShift = −0.57
Z lm Score = −2.99
NG = 0
DB = 0
SM = 0
Delta_K
Doxorubicin
YOR235W
ZShift = −1.47
Z lm Score = −0.6
NG = 0
DB = 0
SM = 0
Delta_r
Doxorubicin
YOR235W
ZShift = −1.03
Z lm Score = 0.12
NG = 0
DB = 0
SM = 0
Delta_AUC
Doxorubicin
YNL007C SIS1
ZShift = 2.27
Z lm Score = 5
NG = 0
DB = 0
SM = 0
Delta_L
Doxorubicin
YNL007C SIS1
ZShift = 0.58
Z lm Score = −0.83
NG = 0
DB = 0
SM = 0
Delta_K
Doxorubicin
YNL007C SIS1
ZShift = −0.33
Z lm Score = −0.48
NG = 0
DB = 0
SM = 0
Delta_r
Doxorubicin
YNL007C SIS1
ZShift = −0.01
Z lm Score = −2.04
NG = 0
DB = 0
SM = 0
Delta_AUC
Doxorubicin

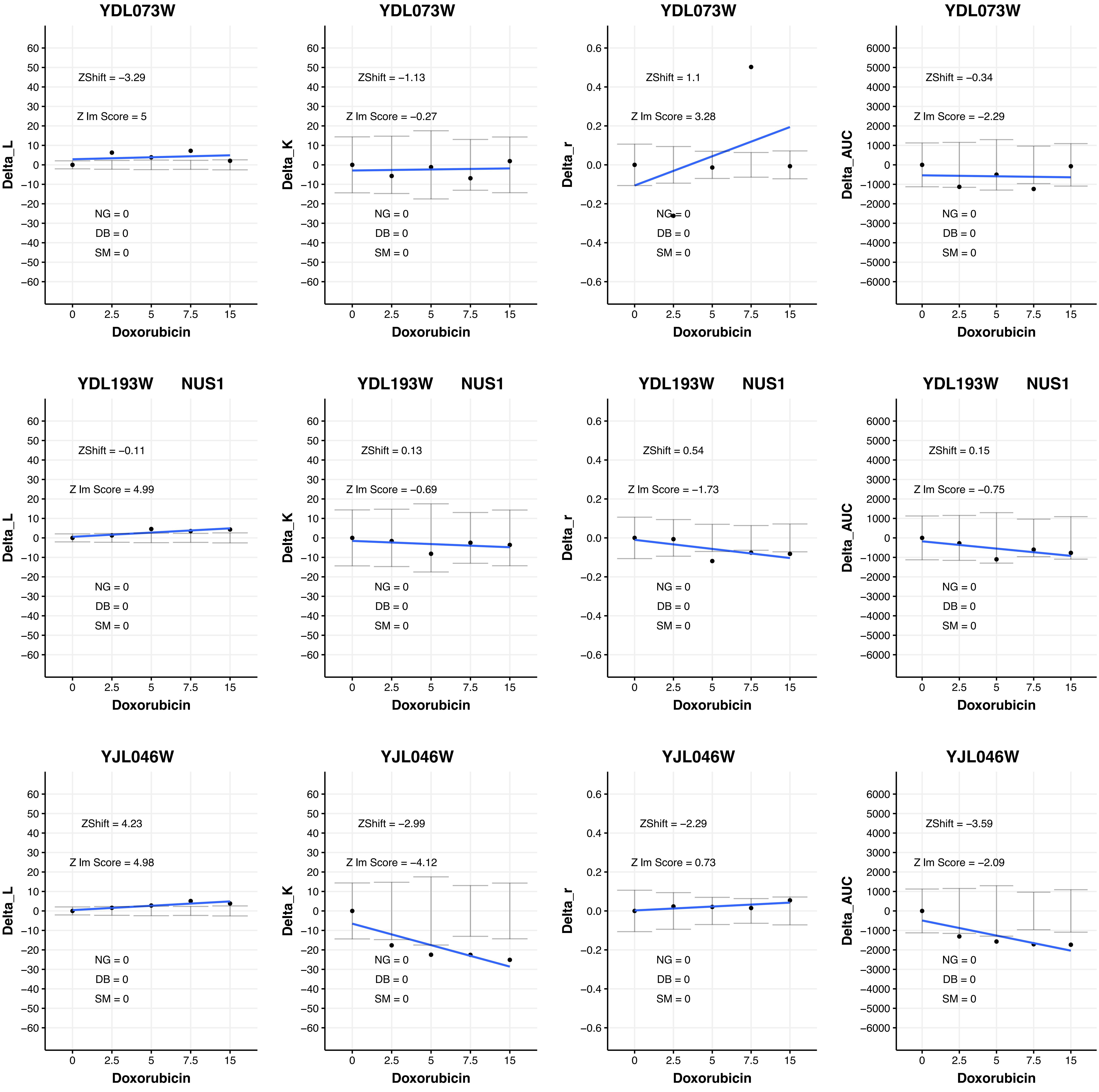
YDL073W
ZShift = −3.29
Z lm Score = 5
NG = 0
DB = 0
SM = 0
Delta_L
Doxorubicin
YDL073W
ZShift = −1.13
Z lm Score = −0.27
NG = 0
DB = 0
SM = 0
Delta_K
Doxorubicin
YDL073W
ZShift = 1.1
Z lm Score = 3.28
NG = 0
DB = 0
SM = 0
Delta_r
Doxorubicin
YDL073W
ZShift = −0.34
Z lm Score = −2.29
NG = 0
DB = 0
SM = 0
Delta_AUC
Doxorubicin
YDL193W NUS1
ZShift = −0.11
Z lm Score = 4.99
NG = 0
DB = 0
SM = 0
Delta_L
Doxorubicin
YDL193W NUS1
ZShift = 0.13
Z lm Score = −0.69
NG = 0
DB = 0
SM = 0
Delta_K
Doxorubicin
YDL193W NUS1
ZShift = 0.54
Z lm Score = −1.73
NG = 0
DB = 0
SM = 0
Delta_r
Doxorubicin
YDL193W NUS1
ZShift = 0.15
Z lm Score = −0.75
NG = 0
DB = 0
SM = 0
Delta_AUC
Doxorubicin
YJL046W
ZShift = 4.23
Z lm Score = 4.98
NG = 0
DB = 0
SM = 0
Delta_L
Doxorubicin
YJL046W
ZShift = −2.99
Z lm Score = −4.12
NG = 0
DB = 0
SM = 0
Delta_K
Doxorubicin
YJL046W
ZShift = −2.29
Z lm Score = 0.73
NG = 0
DB = 0
SM = 0
Delta_r
Doxorubicin
YJL046W
ZShift = −3.59
Z lm Score = −2.09
NG = 0
DB = 0
SM = 0
Delta_AUC
Doxorubicin

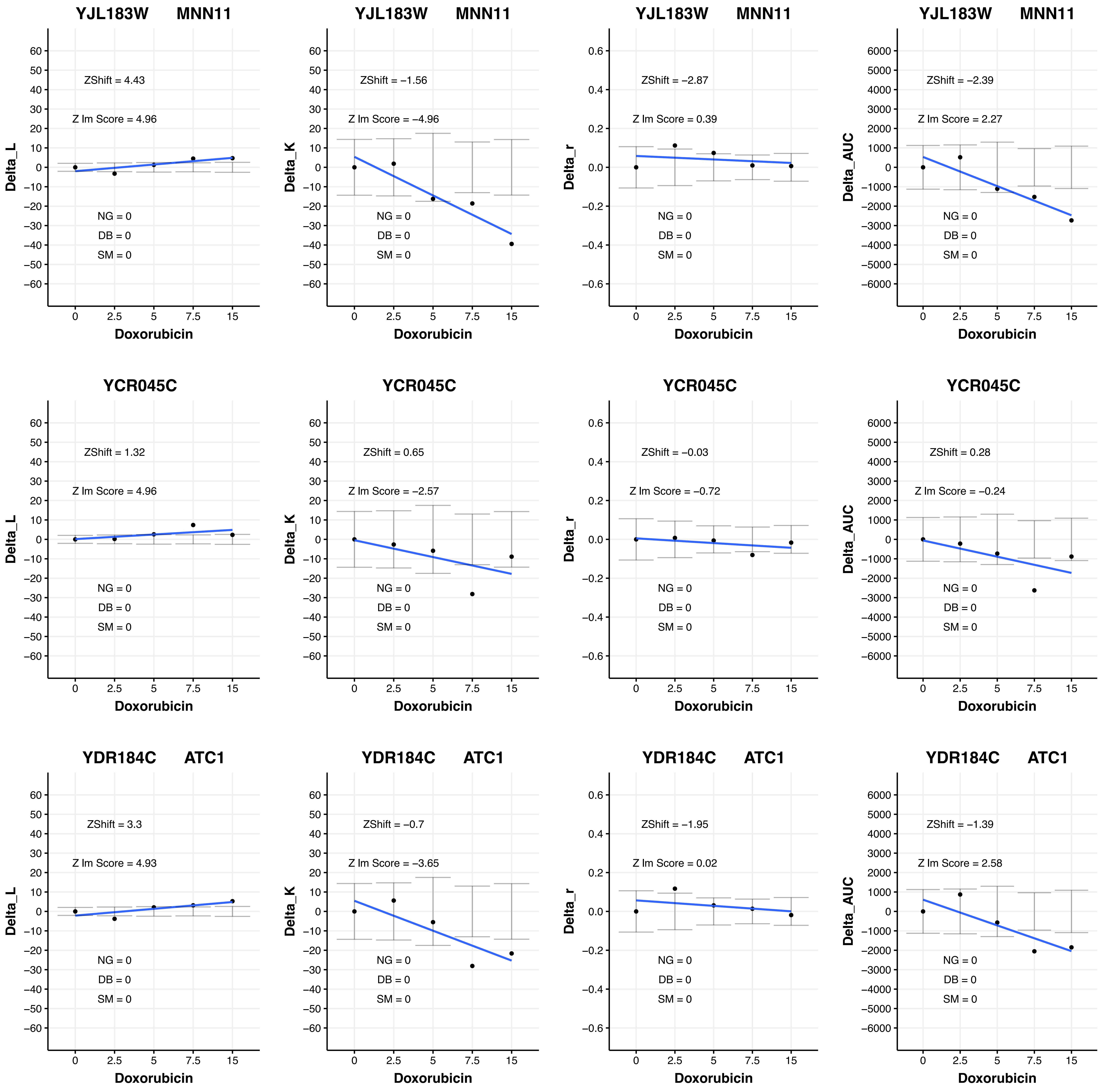
YJL183W MNN11
ZShift = 4.43
Z lm Score = 4.96
NG = 0
DB = 0
SM = 0
Delta_L
Doxorubicin
YJL183W MNN11
ZShift = −1.56
Z lm Score = −4.96
NG = 0
DB = 0
SM = 0
Delta_K
Doxorubicin
YJL183W MNN11
ZShift = −2.87
Z lm Score = 0.39
NG = 0
DB = 0
SM = 0
Delta_r
Doxorubicin
YJL183W MNN11
ZShift = −2.39
Z lm Score = 2.27
NG = 0
DB = 0
SM = 0
Delta_AUC
Doxorubicin
YCR045C
ZShift = 1.32
Z lm Score = 4.96
NG = 0
DB = 0
SM = 0
Delta_L
Doxorubicin
YCR045C
ZShift = 0.65
Z lm Score = −2.57
NG = 0
DB = 0
SM = 0
Delta_K
Doxorubicin
YCR045C
ZShift = −0.03
Z lm Score = −0.72
NG = 0
DB = 0
SM = 0
Delta_r
Doxorubicin
YCR045C
ZShift = 0.28
Z lm Score = −0.24
NG = 0
DB = 0
SM = 0
Delta_AUC
Doxorubicin
YDR184C ATC1
ZShift = 3.3
Z lm Score = 4.93
NG = 0
DB = 0
SM = 0
Delta_L
Doxorubicin
YDR184C ATC1
ZShift = −0.7
Z lm Score = −3.65
NG = 0
DB = 0
SM = 0
Delta_K
Doxorubicin
YDR184C ATC1
ZShift = −1.95
Z lm Score = 0.02
NG = 0
DB = 0
SM = 0
Delta_r
Doxorubicin
YDR184C ATC1
ZShift = −1.39
Z lm Score = 2.58
NG = 0
DB = 0
SM = 0
Delta_AUC
Doxorubicin

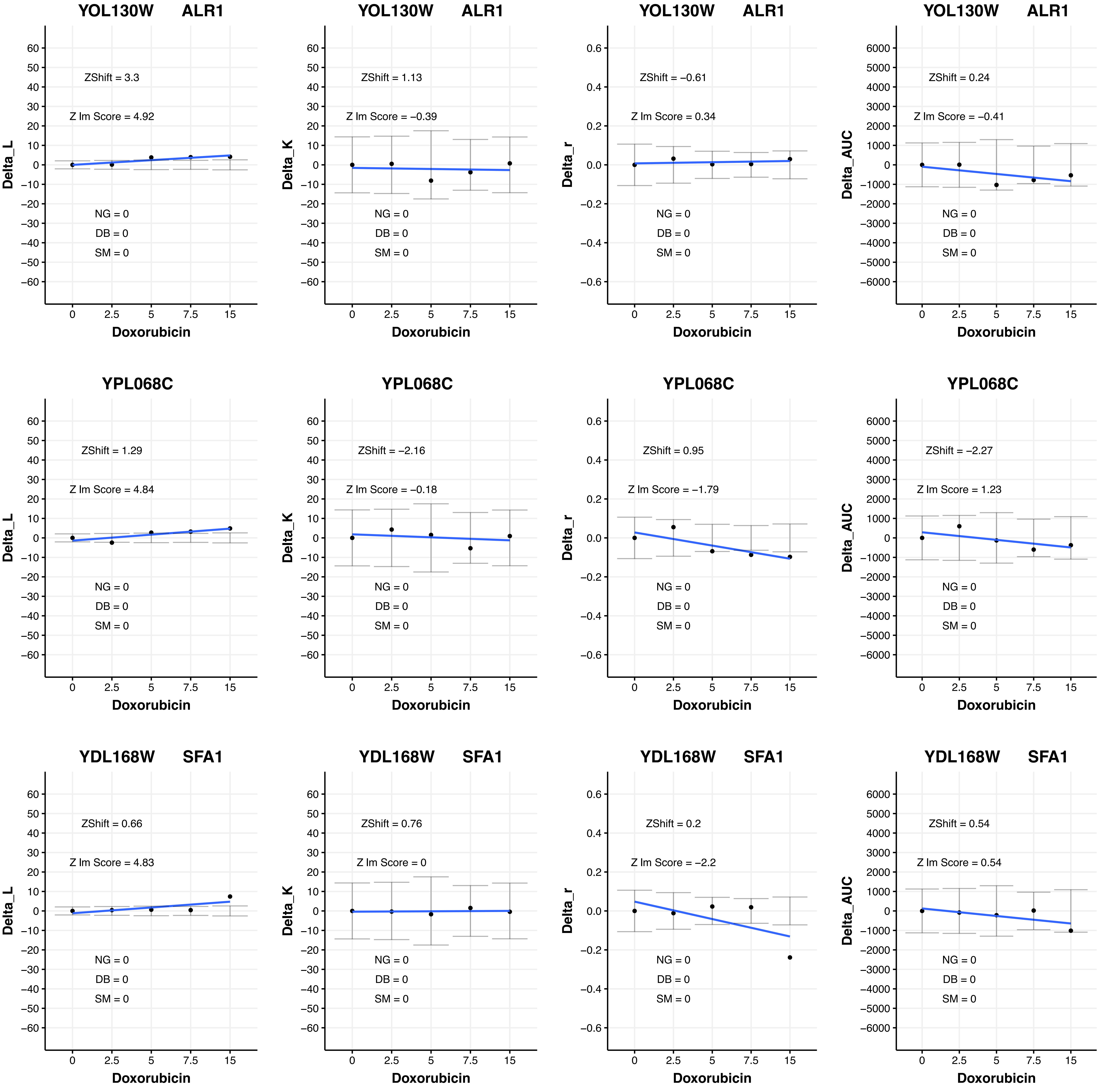
YOL130W ALR1
ZShift = 3.3
Z lm Score = 4.92
NG = 0
DB = 0
SM = 0
Delta_L
Doxorubicin
YOL130W ALR1
ZShift = 1.13
Z lm Score = −0.39
NG = 0
DB = 0
SM = 0
Delta_K
Doxorubicin
YOL130W ALR1
ZShift = −0.61
Z lm Score = 0.34
NG = 0
DB = 0
SM = 0
Delta_r
Doxorubicin
YOL130W ALR1
ZShift = 0.24
Z lm Score = −0.41
NG = 0
DB = 0
SM = 0
Delta_AUC
Doxorubicin
YPL068C
ZShift = 1.29
Z lm Score = 4.84
NG = 0
DB = 0
SM = 0
Delta_L
Doxorubicin
YPL068C
ZShift = −2.16
Z lm Score = −0.18
NG = 0
DB = 0
SM = 0
Delta_K
Doxorubicin
YPL068C
ZShift = 0.95
Z lm Score = −1.79
NG = 0
DB = 0
SM = 0
Delta_r
Doxorubicin
YPL068C
ZShift = −2.27
Z lm Score = 1.23
NG = 0
DB = 0
SM = 0
Delta_AUC
Doxorubicin
YDL168W SFA1
ZShift = 0.66
Z lm Score = 4.83
NG = 0
DB = 0
SM = 0
Delta_L
Doxorubicin
YDL168W SFA1
ZShift = 0.76
Z lm Score = 0
NG = 0
DB = 0
SM = 0
Delta_K
Doxorubicin
YDL168W SFA1
ZShift = 0.2
Z lm Score = −2.2
NG = 0
DB = 0
SM = 0
Delta_r
Doxorubicin
YDL168W SFA1
ZShift = 0.54
Z lm Score = 0.54
NG = 0
DB = 0
SM = 0
Delta_AUC
Doxorubicin

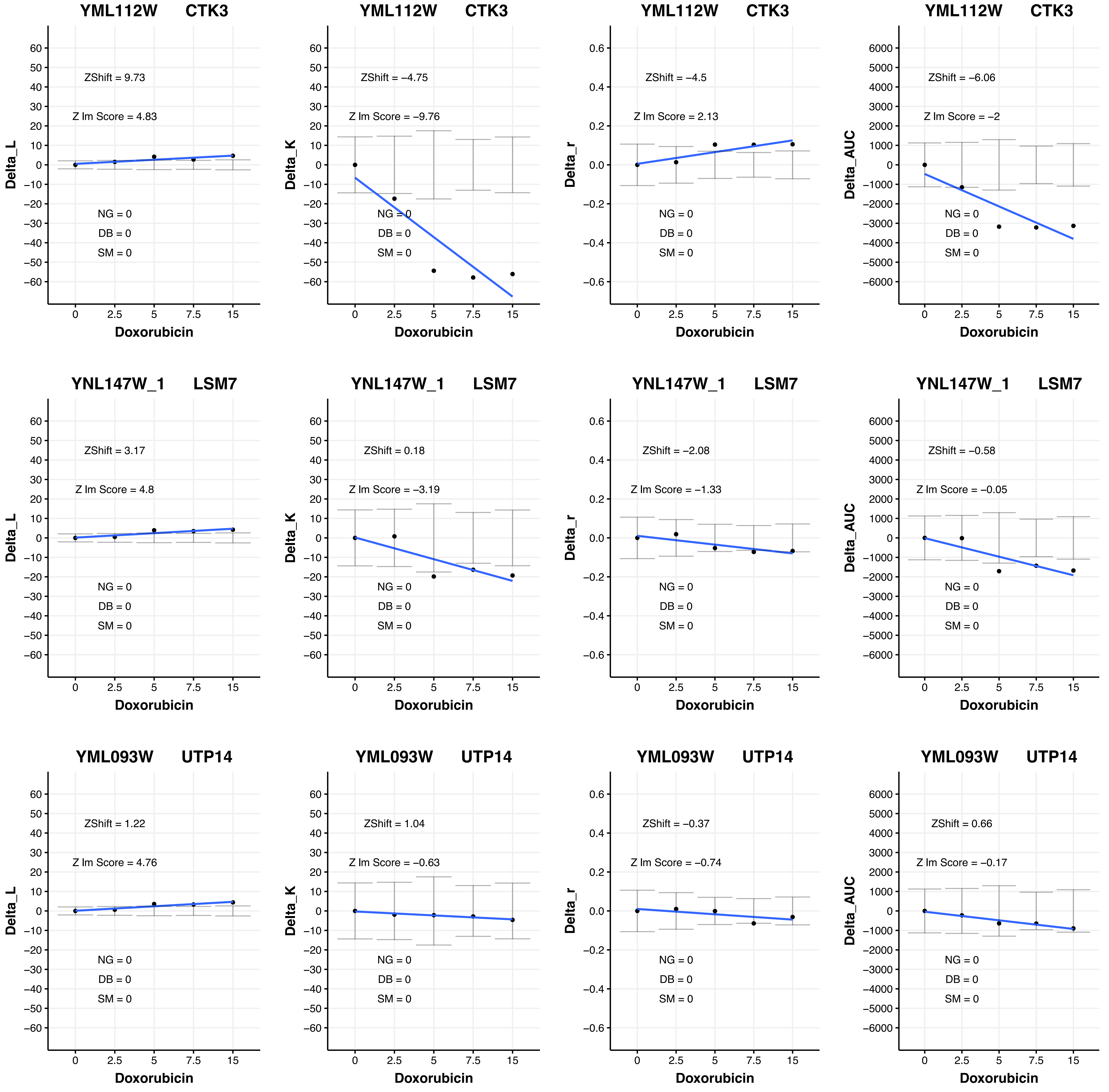
YML112W CTK3
ZShift = 9.73
Z lm Score = 4.83
NG = 0
DB = 0
SM = 0
Delta_L
Doxorubicin
YML112W CTK3
ZShift = −4.75
Z lm Score = −9.76
NG = 0
DB = 0
SM = 0
Delta_K
Doxorubicin
YML112W CTK3
ZShift = −4.5
Z lm Score = 2.13
NG = 0
DB = 0
SM = 0
Delta_r
Doxorubicin
YML112W CTK3
ZShift = −6.06
Z lm Score = −2
NG = 0
DB = 0
SM = 0
Delta_AUC
Doxorubicin
YNL147W_1 LSM7
ZShift = 3.17
Z lm Score = 4.8
NG = 0
DB = 0
SM = 0
Delta_L
Doxorubicin
YNL147W_1 LSM7
ZShift = 0.18
Z lm Score = −3.19
NG = 0
DB = 0
SM = 0
Delta_K
Doxorubicin
YNL147W_1 LSM7
ZShift = −2.08
Z lm Score = −1.33
NG = 0
DB = 0
SM = 0
Delta_r
Doxorubicin
YNL147W_1 LSM7
ZShift = −0.58
Z lm Score = −0.05
NG = 0
DB = 0
SM = 0
Delta_AUC
Doxorubicin
YML093W UTP14
ZShift = 1.22
Z lm Score = 4.76
NG = 0
DB = 0
SM = 0
Delta_L
Doxorubicin
YML093W UTP14
ZShift = 1.04
Z lm Score = −0.63
NG = 0
DB = 0
SM = 0
Delta_K
Doxorubicin
YML093W UTP14
ZShift = −0.37
Z lm Score = −0.74
NG = 0
DB = 0
SM = 0
Delta_r
Doxorubicin
YML093W UTP14
ZShift = 0.66
Z lm Score = −0.17
NG = 0
DB = 0
SM = 0
Delta_AUC
Doxorubicin

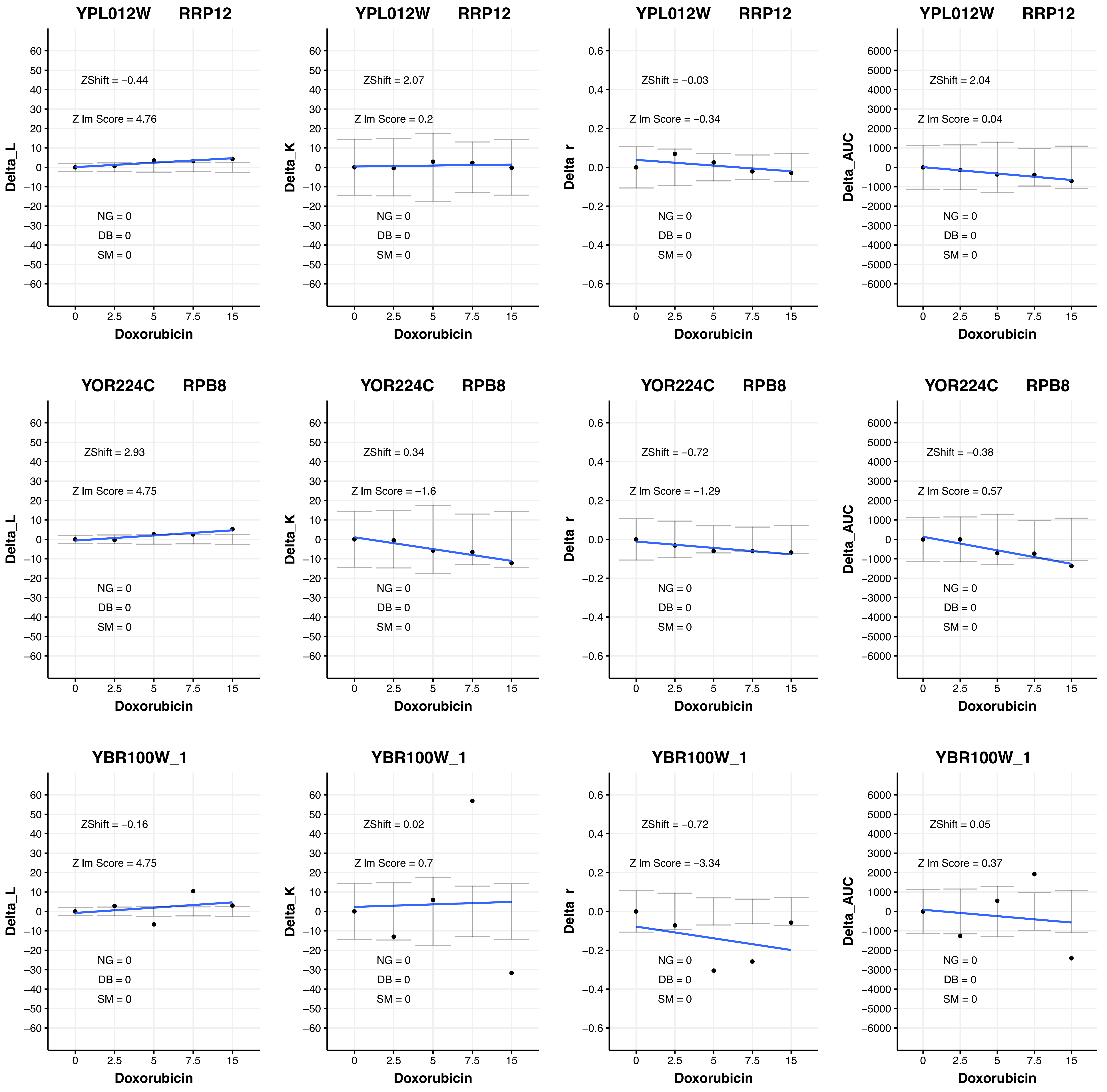

YPL012W RRP12
ZShift = −0.44
Z lm Score = 4.76
NG = 0
DB = 0
SM = 0
Delta_L
Doxorubicin
YPL012W RRP12
ZShift = 2.07
Z lm Score = 0.2
NG = 0
DB = 0
SM = 0
Delta_K
Doxorubicin
YPL012W RRP12
ZShift = −0.03
Z lm Score = −0.34
NG = 0
DB = 0
SM = 0
Delta_r
Doxorubicin
YPL012W RRP12
ZShift = 2.04
Z lm Score = 0.04
NG = 0
DB = 0
SM = 0
Delta_AUC
Doxorubicin
YOR224C RPB8
ZShift = 2.93
Z lm Score = 4.75
NG = 0
DB = 0
SM = 0
Delta_L
Doxorubicin
YOR224C RPB8
ZShift = 0.34
Z lm Score = −1.6
NG = 0
DB = 0
SM = 0
Delta_K
Doxorubicin
YOR224C RPB8
ZShift = −0.72
Z lm Score = −1.29
NG = 0
DB = 0
SM = 0
Delta_r
Doxorubicin
YOR224C RPB8
ZShift = −0.38
Z lm Score = 0.57
NG = 0
DB = 0
SM = 0
Delta_AUC
Doxorubicin
YBR100W_1
ZShift = −0.16
Z lm Score = 4.75
NG = 0
DB = 0
SM = 0
Delta_L
Doxorubicin
YBR100W_1
ZShift = 0.02
Z lm Score = 0.7
NG = 0
DB = 0
SM = 0
Delta_K
Doxorubicin
YBR100W_1
ZShift = −0.72
Z lm Score = −3.34
NG = 0
DB = 0
SM = 0
Delta_r
Doxorubicin
YBR100W_1
ZShift = 0.05
Z lm Score = 0.37
NG = 0
DB = 0
SM = 0
Delta_AUC
Doxorubicin

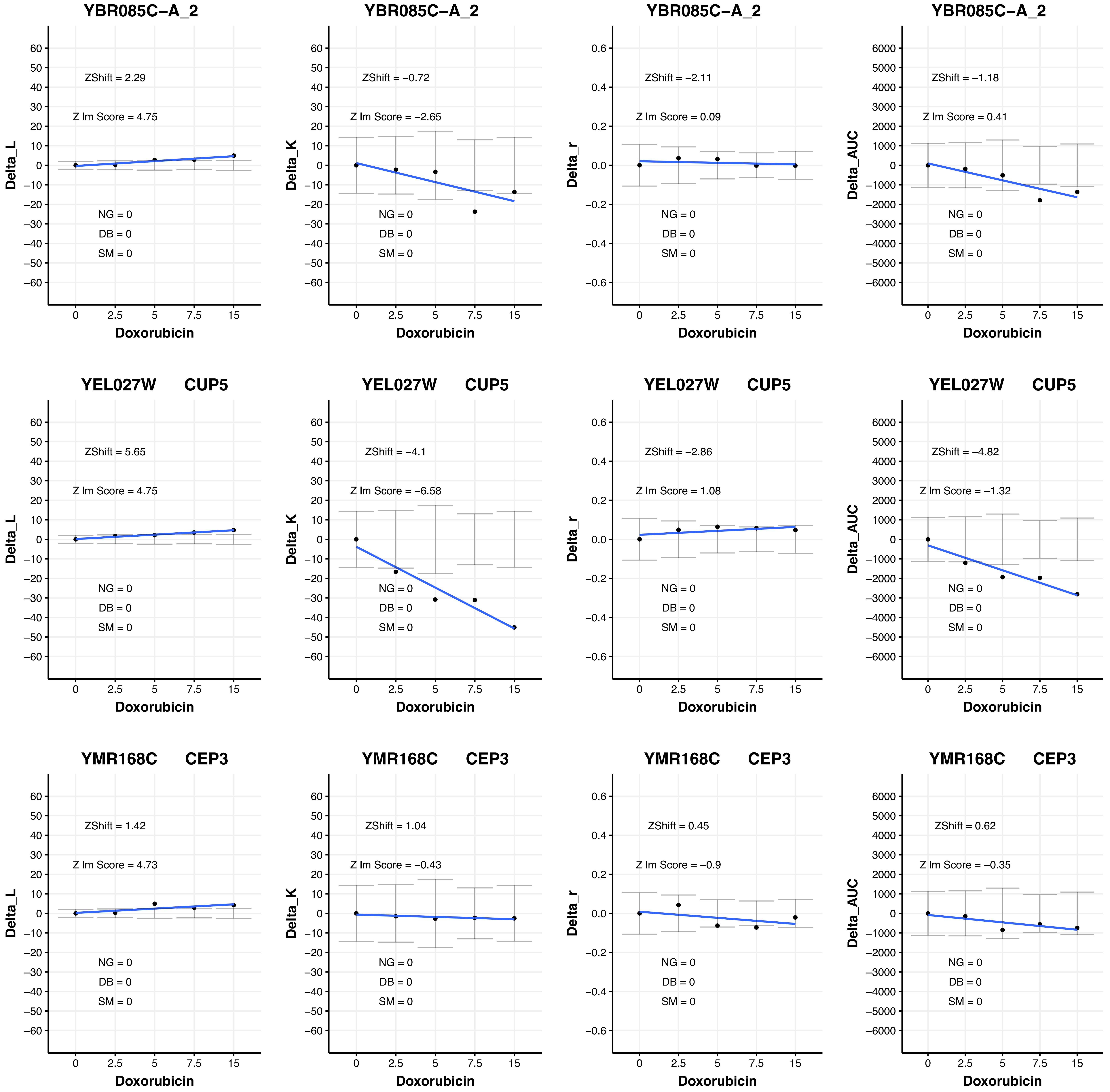
YBR085C−A_2
ZShift = 2.29
Z lm Score = 4.75
NG = 0
DB = 0
SM = 0
Delta_L
Doxorubicin
YBR085C−A_2
ZShift = −0.72
Z lm Score = −2.65
NG = 0
DB = 0
SM = 0
Delta_K
Doxorubicin
YBR085C−A_2
ZShift = −2.11
Z lm Score = 0.09
NG = 0
DB = 0
SM = 0
Delta_r
Doxorubicin
YBR085C−A_2
ZShift = −1.18
Z lm Score = 0.41
NG = 0
DB = 0
SM = 0
Delta_AUC
Doxorubicin
YEL027W CUP5
ZShift = 5.65
Z lm Score = 4.75
NG = 0
DB = 0
SM = 0
Delta_L
Doxorubicin
YEL027W CUP5
ZShift = −4.1
Z lm Score = −6.58
NG = 0
DB = 0
SM = 0
Delta_K
Doxorubicin
YEL027W CUP5
ZShift = −2.86
Z lm Score = 1.08
NG = 0
DB = 0
SM = 0
Delta_r
Doxorubicin
YEL027W CUP5
ZShift = −4.82
Z lm Score = −1.32
NG = 0
DB = 0
SM = 0
Delta_AUC
Doxorubicin
YMR168C CEP3
ZShift = 1.42
Z lm Score = 4.73
NG = 0
DB = 0
SM = 0
Delta_L
Doxorubicin
YMR168C CEP3
ZShift = 1.04
Z lm Score = −0.43
NG = 0
DB = 0
SM = 0
Delta_K
Doxorubicin
YMR168C CEP3
ZShift = 0.45
Z lm Score = −0.9
NG = 0
DB = 0
SM = 0
Delta_r
Doxorubicin
YMR168C CEP3
ZShift = 0.62
Z lm Score = −0.35
NG = 0
DB = 0
SM = 0
Delta_AUC
Doxorubicin

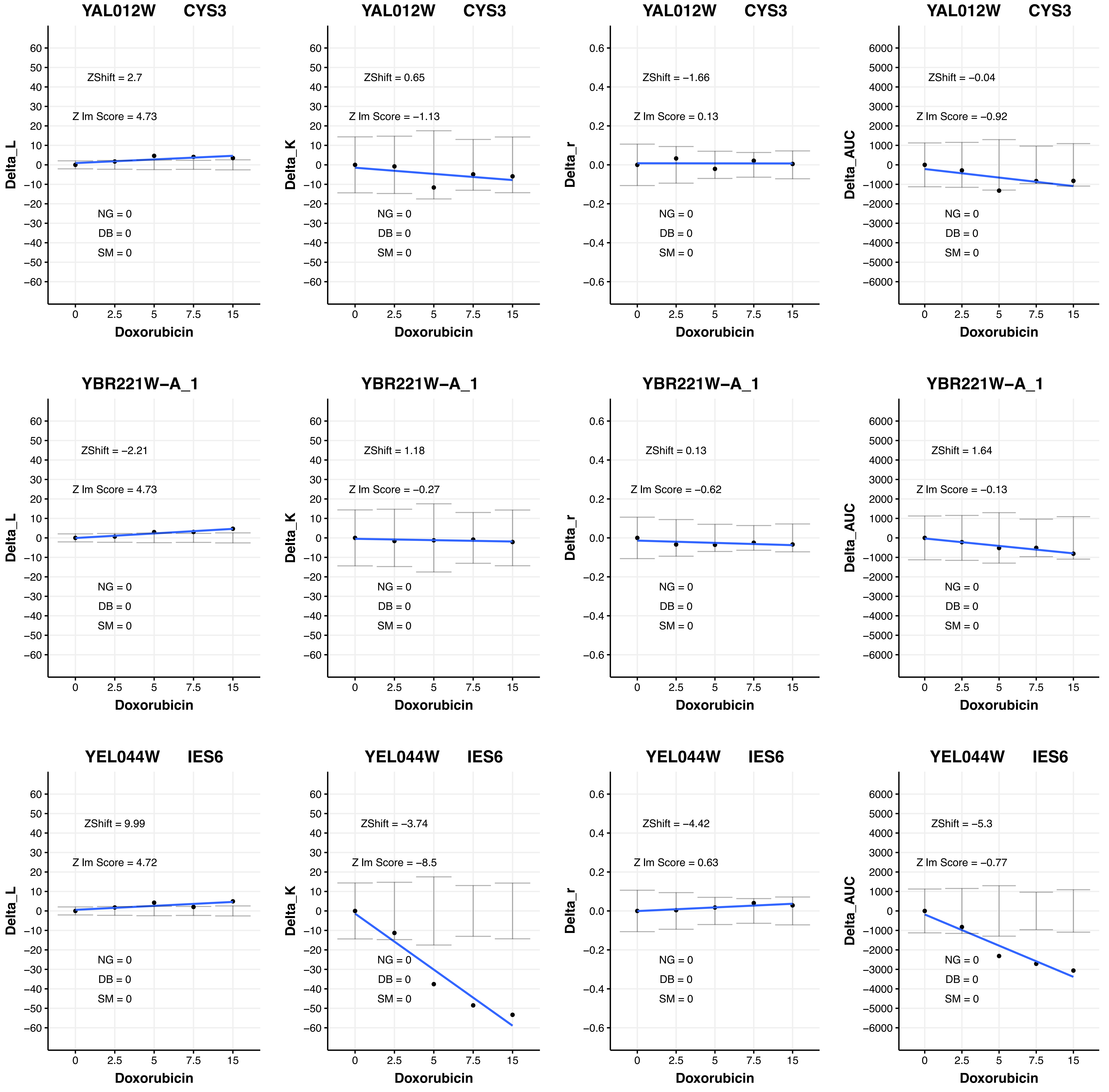
YAL012W CYS3
ZShift = 2.7
Z lm Score = 4.73
NG = 0
DB = 0
SM = 0
Delta_L
Doxorubicin
YAL012W CYS3
ZShift = 0.65
Z lm Score = −1.13
NG = 0
DB = 0
SM = 0
Delta_K
Doxorubicin
YAL012W CYS3
ZShift = −1.66
Z lm Score = 0.13
NG = 0
DB = 0
SM = 0
Delta_r
Doxorubicin
YAL012W CYS3
ZShift = −0.04
Z lm Score = −0.92
NG = 0
DB = 0
SM = 0
Delta_AUC
Doxorubicin
YBR221W−A_1
ZShift = −2.21
Z lm Score = 4.73
NG = 0
DB = 0
SM = 0
Delta_L
Doxorubicin
YBR221W−A_1
ZShift = 1.18
Z lm Score = −0.27
NG = 0
DB = 0
SM = 0
Delta_K
Doxorubicin
YBR221W−A_1
ZShift = 0.13
Z lm Score = −0.62
NG = 0
DB = 0
SM = 0
Delta_r
Doxorubicin
YBR221W−A_1
ZShift = 1.64
Z lm Score = −0.13
NG = 0
DB = 0
SM = 0
Delta_AUC
Doxorubicin
YEL044W IES6
ZShift = 9.99
Z lm Score = 4.72
NG = 0
DB = 0
SM = 0
Delta_L
Doxorubicin
YEL044W IES6
ZShift = −3.74
Z lm Score = −8.5
NG = 0
DB = 0
SM = 0
Delta_K
Doxorubicin
YEL044W IES6
ZShift = −4.42
Z lm Score = 0.63
NG = 0
DB = 0
SM = 0
Delta_r
Doxorubicin
YEL044W IES6
ZShift = −5.3
Z lm Score = −0.77
NG = 0
DB = 0
SM = 0
Delta_AUC
Doxorubicin

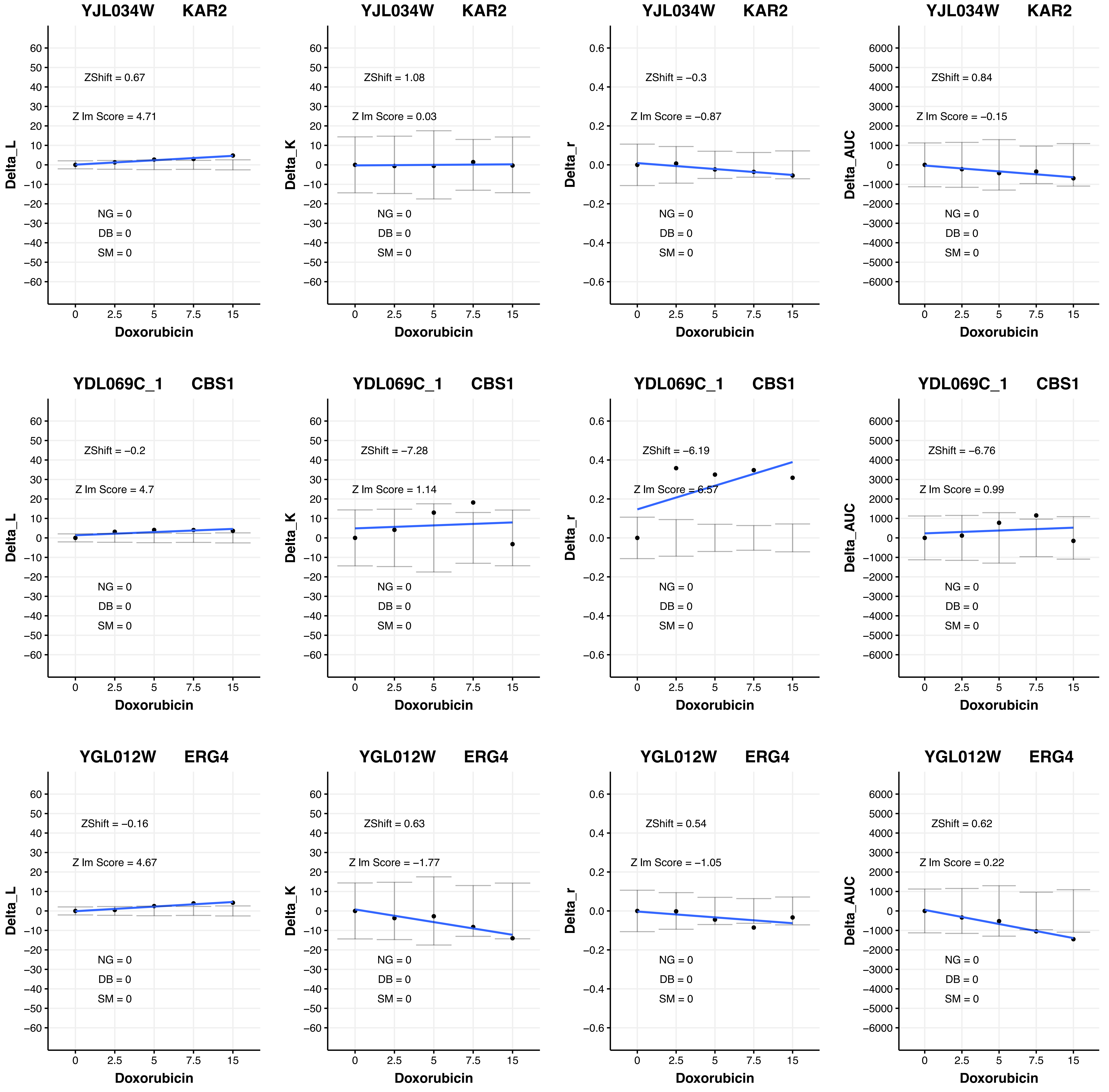
YJL034W KAR2
ZShift = 0.67
Z lm Score = 4.71
NG = 0
DB = 0
SM = 0
Delta_L
Doxorubicin
YJL034W KAR2
ZShift = 1.08
Z lm Score = 0.03
NG = 0
DB = 0
SM = 0
Delta_K
Doxorubicin
YJL034W KAR2
ZShift = −0.3
Z lm Score = −0.87
NG = 0
DB = 0
SM = 0
Delta_r
Doxorubicin
YJL034W KAR2
ZShift = 0.84
Z lm Score = −0.15
NG = 0
DB = 0
SM = 0
Delta_AUC
Doxorubicin
YDL069C_1 CBS1
ZShift = −0.2
Z lm Score = 4.7
NG = 0
DB = 0
SM = 0
Delta_L
Doxorubicin
YDL069C_1 CBS1
ZShift = −7.28
Z lm Score = 1.14
NG = 0
DB = 0
SM = 0
Delta_K
Doxorubicin
YDL069C_1 CBS1
ZShift = −6.19
Z lm Score = 6.57
NG = 0
DB = 0
SM = 0
Delta_r
Doxorubicin
YDL069C_1 CBS1
ZShift = −6.76
Z lm Score = 0.99
NG = 0
DB = 0
SM = 0
Delta_AUC
Doxorubicin
YGL012W ERG4
ZShift = −0.16
Z lm Score = 4.67
NG = 0
DB = 0
SM = 0
Delta_L
Doxorubicin
YGL012W ERG4
ZShift = 0.63
Z lm Score = −1.77
NG = 0
DB = 0
SM = 0
Delta_K
Doxorubicin
YGL012W ERG4
ZShift = 0.54
Z lm Score = −1.05
NG = 0
DB = 0
SM = 0
Delta_r
Doxorubicin
YGL012W ERG4
ZShift = 0.62
Z lm Score = 0.22
NG = 0
DB = 0
SM = 0
Delta_AUC
Doxorubicin

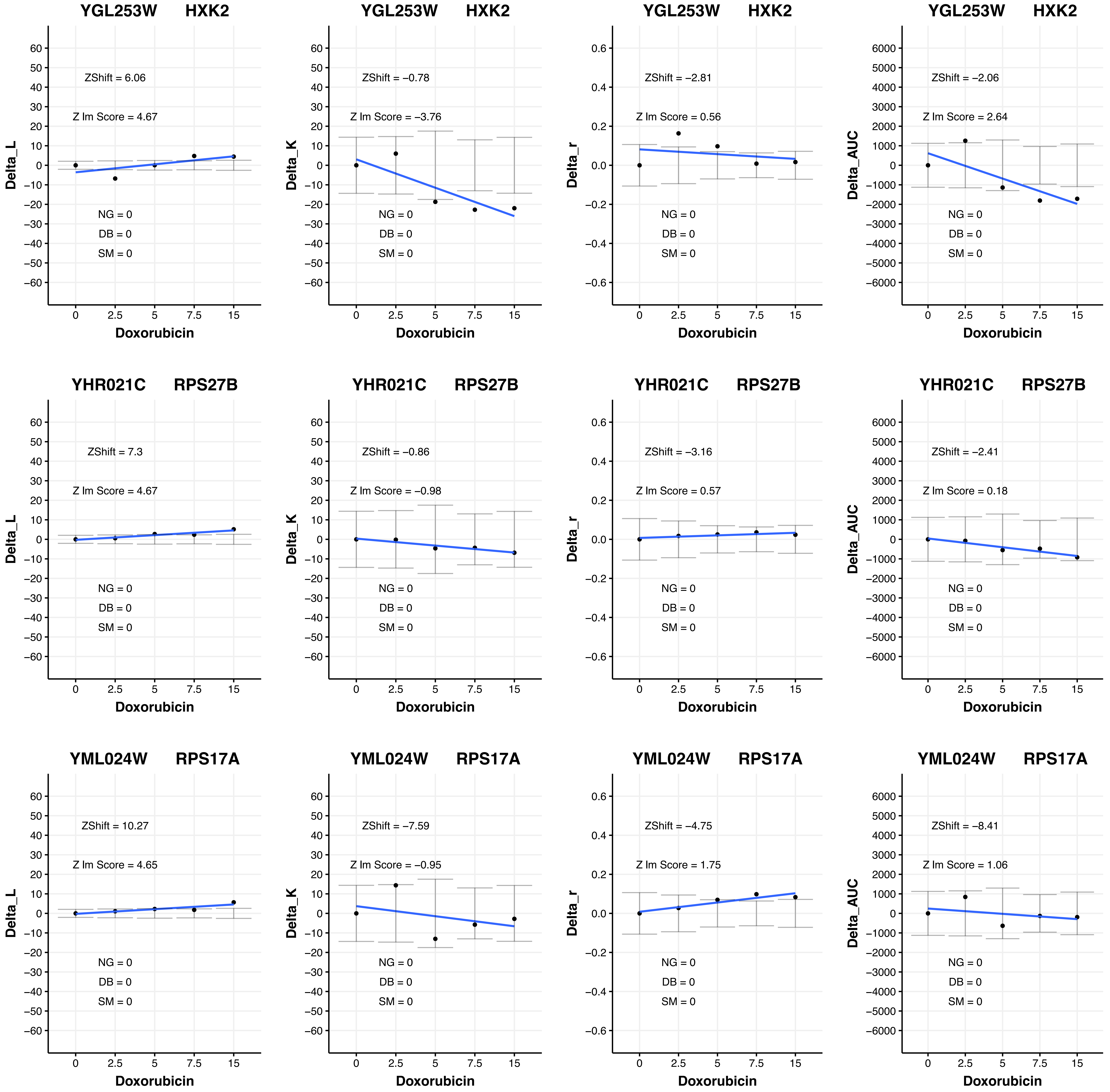

YGL253W HXK2
ZShift = 6.06
Z lm Score = 4.67
NG = 0
DB = 0
SM = 0
Delta_L
Doxorubicin
YGL253W HXK2
ZShift = −0.78
Z lm Score = −3.76
NG = 0
DB = 0
SM = 0
Delta_K
Doxorubicin
YGL253W HXK2
ZShift = −2.81
Z lm Score = 0.56
NG = 0
DB = 0
SM = 0
Delta_r
Doxorubicin
YGL253W HXK2
ZShift = −2.06
Z lm Score = 2.64
NG = 0
DB = 0
SM = 0
Delta_AUC
Doxorubicin
YHR021C RPS27B
ZShift = 7.3
Z lm Score = 4.67
NG = 0
DB = 0
SM = 0
Delta_L
Doxorubicin
YHR021C RPS27B
ZShift = −0.86
Z lm Score = −0.98
NG = 0
DB = 0
SM = 0
Delta_K
Doxorubicin
YHR021C RPS27B
ZShift = −3.16
Z lm Score = 0.57
NG = 0
DB = 0
SM = 0
Delta_r
Doxorubicin
YHR021C RPS27B
ZShift = −2.41
Z lm Score = 0.18
NG = 0
DB = 0
SM = 0
Delta_AUC
Doxorubicin
YML024W RPS17A
ZShift = 10.27
Z lm Score = 4.65
NG = 0
DB = 0
SM = 0
Delta_L
Doxorubicin
YML024W RPS17A
ZShift = −7.59
Z lm Score = −0.95
NG = 0
DB = 0
SM = 0
Delta_K
Doxorubicin
YML024W RPS17A
ZShift = −4.75
Z lm Score = 1.75
NG = 0
DB = 0
SM = 0
Delta_r
Doxorubicin
YML024W RPS17A
ZShift = −8.41
Z lm Score = 1.06
NG = 0
DB = 0
SM = 0
Delta_AUC
Doxorubicin

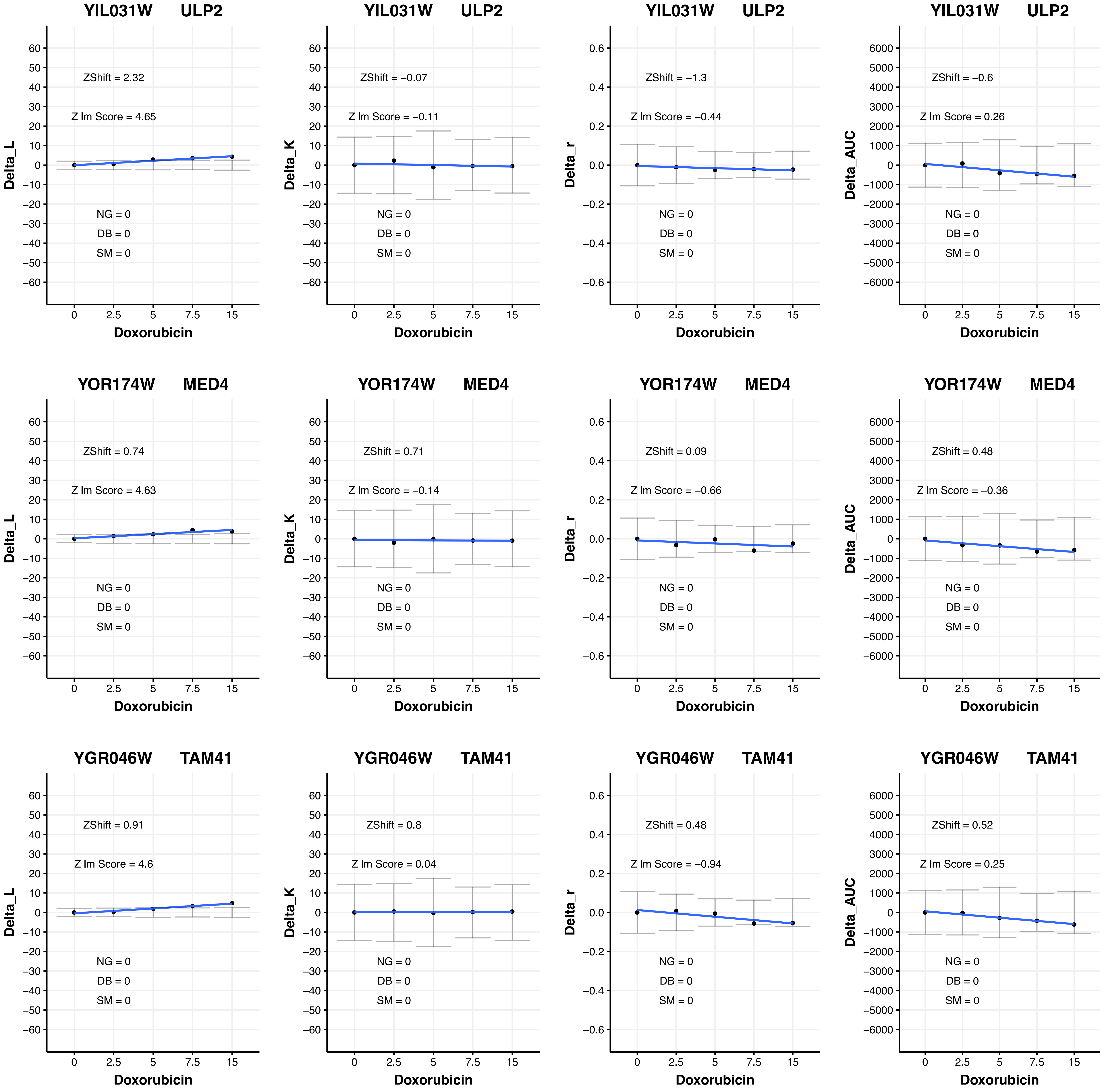
YIL031W ULP2
ZShift = 2.32
Z lm Score = 4.65
NG = 0
DB = 0
SM = 0
Delta_L
Doxorubicin
YIL031W ULP2
ZShift = −0.07
Z lm Score = −0.11
NG = 0
DB = 0
SM = 0
Delta_K
Doxorubicin
YIL031W ULP2
ZShift = −1.3
Z lm Score = −0.44
NG = 0
DB = 0
SM = 0
Delta_r
Doxorubicin
YIL031W ULP2
ZShift = −0.6
Z lm Score = 0.26
NG = 0
DB = 0
SM = 0
Delta_AUC
Doxorubicin
YOR174W MED4
ZShift = 0.74
Z lm Score = 4.63
NG = 0
DB = 0
SM = 0
Delta_L
Doxorubicin
YOR174W MED4
ZShift = 0.71
Z lm Score = −0.14
NG = 0
DB = 0
SM = 0
Delta_K
Doxorubicin
YOR174W MED4
ZShift = 0.09
Z lm Score = −0.66
NG = 0
DB = 0
SM = 0
Delta_r
Doxorubicin
YOR174W MED4
ZShift = 0.48
Z lm Score = −0.36
NG = 0
DB = 0
SM = 0
Delta_AUC
Doxorubicin
YGR046W TAM41
ZShift = 0.91
Z lm Score = 4.6
NG = 0
DB = 0
SM = 0
Delta_L
Doxorubicin
YGR046W TAM41
ZShift = 0.8
Z lm Score = 0.04
NG = 0
DB = 0
SM = 0
Delta_K
Doxorubicin
YGR046W TAM41
ZShift = 0.48
Z lm Score = −0.94
NG = 0
DB = 0
SM = 0
Delta_r
Doxorubicin
YGR046W TAM41
ZShift = 0.52
Z lm Score = 0.25
NG = 0
DB = 0
SM = 0
Delta_AUC
Doxorubicin

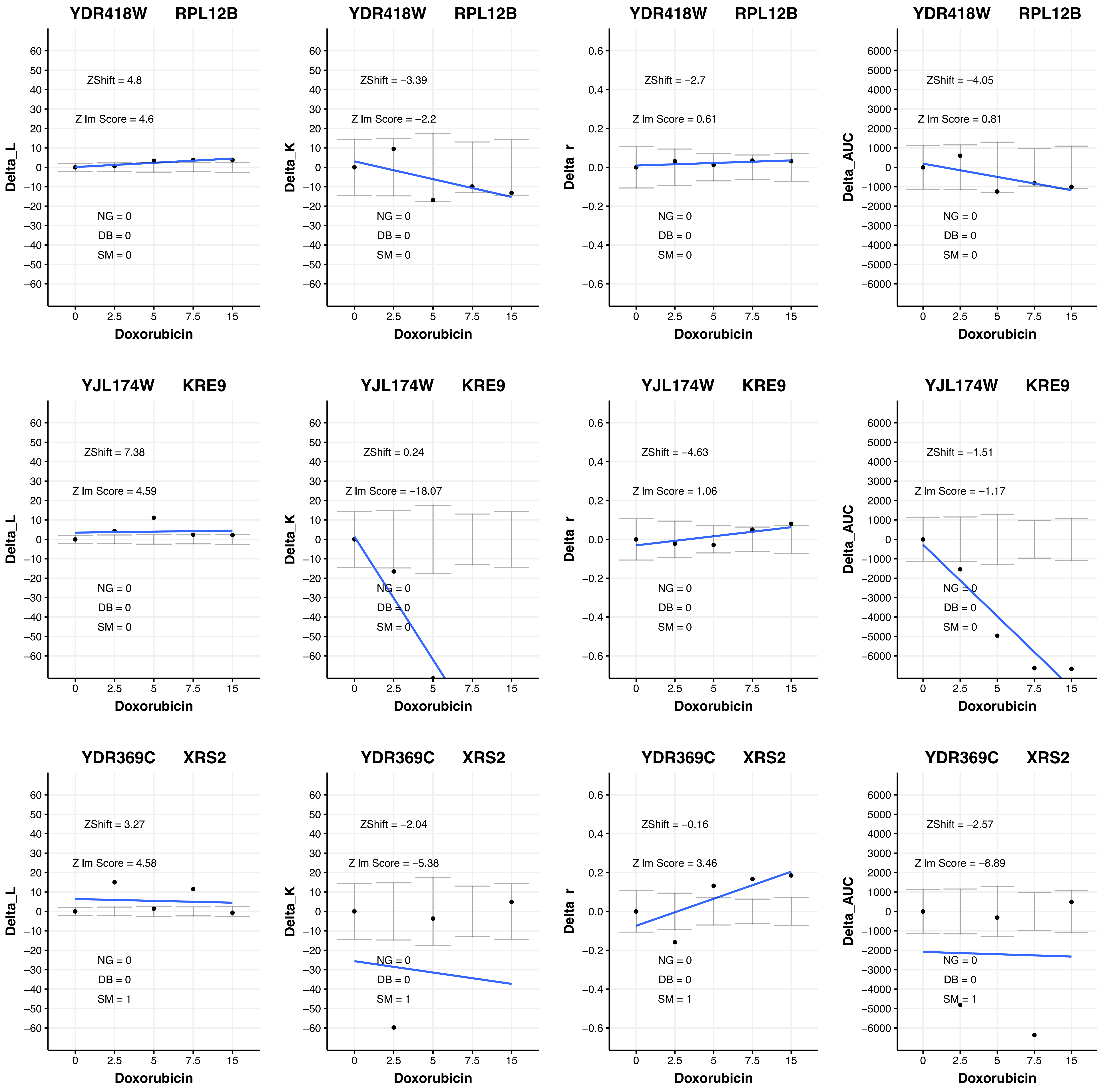

YDR418W RPL12B
ZShift = 4.8
Z lm Score = 4.6
NG = 0
DB = 0
SM = 0
Delta_L
Doxorubicin
YDR418W RPL12B
ZShift = −3.39
Z lm Score = −2.2
NG = 0
DB = 0
SM = 0
Delta_K
Doxorubicin
YDR418W RPL12B
ZShift = −2.7
Z lm Score = 0.61
NG = 0
DB = 0
SM = 0
Delta_r
Doxorubicin
YDR418W RPL12B
ZShift = −4.05
Z lm Score = 0.81
NG = 0
DB = 0
SM = 0
Delta_AUC
Doxorubicin
YJL174W KRE9
ZShift = 7.38
Z lm Score = 4.59
NG = 0
DB = 0
SM = 0
Delta_L
Doxorubicin
YJL174W KRE9
ZShift = 0.24
Z lm Score = −18.07
NG = 0
DB = 0
SM = 0
Delta_K
Doxorubicin
YJL174W KRE9
ZShift = −4.63
Z lm Score = 1.06
NG = 0
DB = 0
SM = 0
Delta_r
Doxorubicin
YJL174W KRE9
ZShift = −1.51
Z lm Score = −1.17
NG = 0
DB = 0
SM = 0
Delta_AUC
Doxorubicin
YDR369C XRS2
ZShift = 3.27
Z lm Score = 4.58
NG = 0
DB = 0
SM = 1
Delta_L
Doxorubicin
YDR369C XRS2
ZShift = −2.04
Z lm Score = −5.38
NG = 0
DB = 0
SM = 1
Delta_K
Doxorubicin
YDR369C XRS2
ZShift = −0.16
Z lm Score = 3.46
NG = 0
DB = 0
SM = 1
Delta_r
Doxorubicin
YDR369C XRS2
ZShift = −2.57
Z lm Score = −8.89
NG = 0
DB = 0
SM = 1
Delta_AUC
Doxorubicin

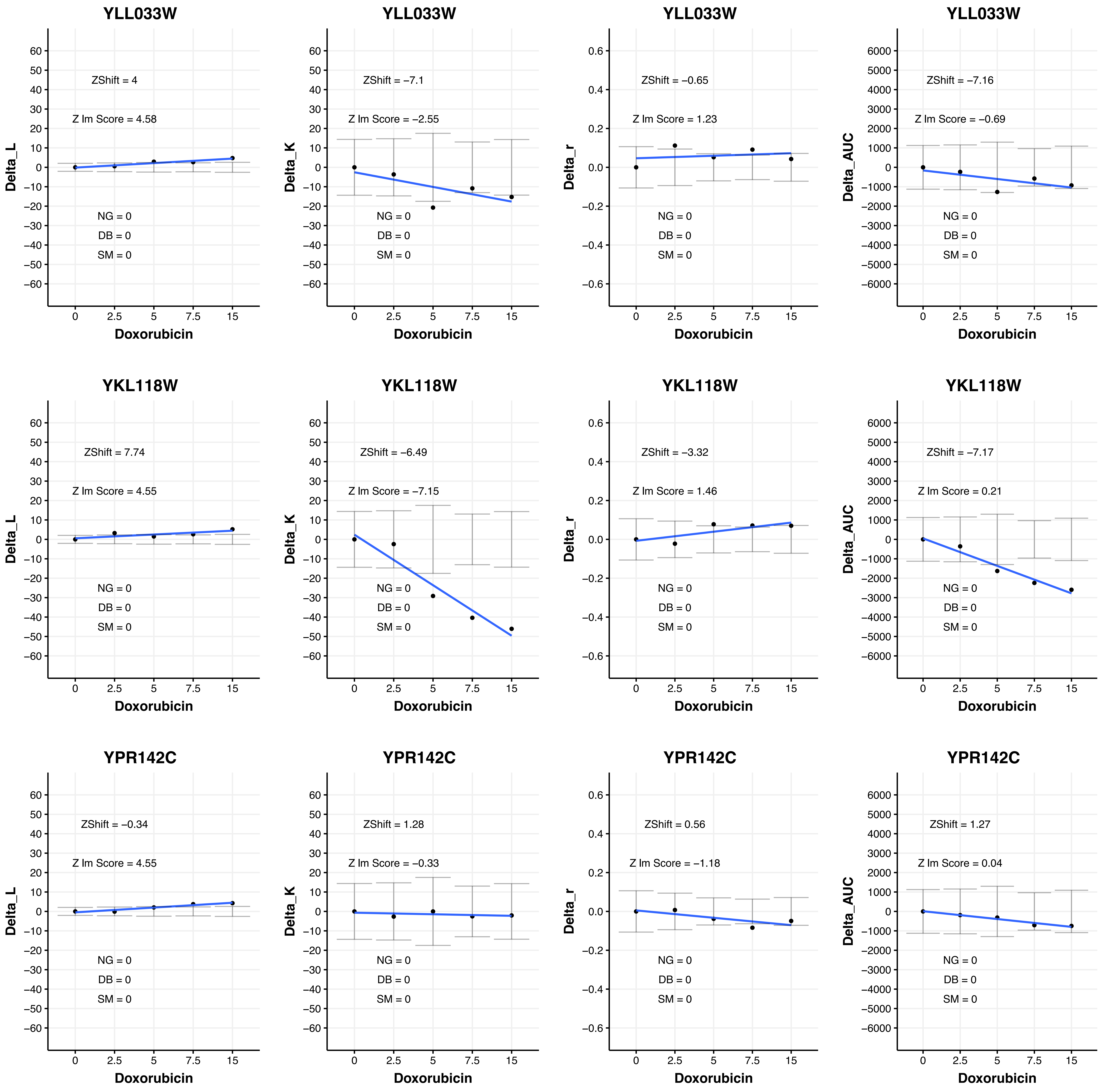
YLL033W
ZShift = 4
Z lm Score = 4.58
NG = 0
DB = 0
SM = 0
Delta_L
Doxorubicin
YLL033W
ZShift = −7.1
Z lm Score = −2.55
NG = 0
DB = 0
SM = 0
Delta_K
Doxorubicin
YLL033W
ZShift = −0.65
Z lm Score = 1.23
NG = 0
DB = 0
SM = 0
Delta_r
Doxorubicin
YLL033W
ZShift = −7.16
Z lm Score = −0.69
NG = 0
DB = 0
SM = 0
Delta_AUC
Doxorubicin
YKL118W
ZShift = 7.74
Z lm Score = 4.55
NG = 0
DB = 0
SM = 0
Delta_L
Doxorubicin
YKL118W
ZShift = −6.49
Z lm Score = −7.15
NG = 0
DB = 0
SM = 0
Delta_K
Doxorubicin
YKL118W
ZShift = −3.32
Z lm Score = 1.46
NG = 0
DB = 0
SM = 0
Delta_r
Doxorubicin
YKL118W
ZShift = −7.17
Z lm Score = 0.21
NG = 0
DB = 0
SM = 0
Delta_AUC
Doxorubicin
YPR142C
ZShift = −0.34
Z lm Score = 4.55
NG = 0
DB = 0
SM = 0
Delta_L
Doxorubicin
YPR142C
ZShift = 1.28
Z lm Score = −0.33
NG = 0
DB = 0
SM = 0
Delta_K
Doxorubicin
YPR142C
ZShift = 0.56
Z lm Score = −1.18
NG = 0
DB = 0
SM = 0
Delta_r
Doxorubicin
YPR142C
ZShift = 1.27
Z lm Score = 0.04
NG = 0
DB = 0
SM = 0
Delta_AUC
Doxorubicin

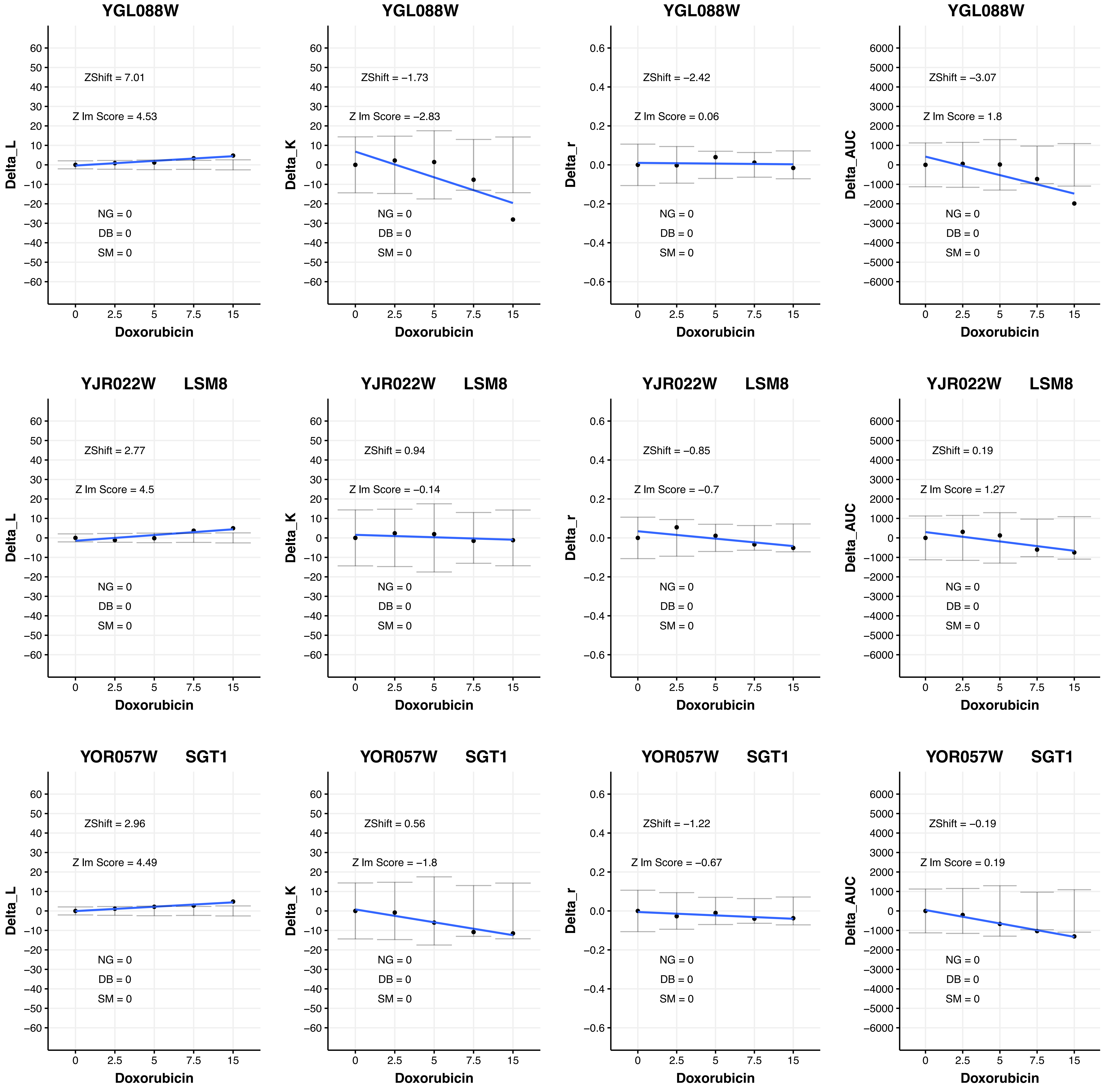
YGL088W
ZShift = 7.01
Z lm Score = 4.53
NG = 0
DB = 0
SM = 0
Delta_L
Doxorubicin
YGL088W
ZShift = −1.73
Z lm Score = −2.83
NG = 0
DB = 0
SM = 0
Delta_K
Doxorubicin
YGL088W
ZShift = −2.42
Z lm Score = 0.06
NG = 0
DB = 0
SM = 0
Delta_r
Doxorubicin
YGL088W
ZShift = −3.07
Z lm Score = 1.8
NG = 0
DB = 0
SM = 0
Delta_AUC
Doxorubicin
YJR022W LSM8
ZShift = 2.77
Z lm Score = 4.5
NG = 0
DB = 0
SM = 0
Delta_L
Doxorubicin
YJR022W LSM8
ZShift = 0.94
Z lm Score = −0.14
NG = 0
DB = 0
SM = 0
Delta_K
Doxorubicin
YJR022W LSM8
ZShift = −0.85
Z lm Score = −0.7
NG = 0
DB = 0
SM = 0
Delta_r
Doxorubicin
YJR022W LSM8
ZShift = 0.19
Z lm Score = 1.27
NG = 0
DB = 0
SM = 0
Delta_AUC
Doxorubicin
YOR057W SGT1
ZShift = 2.96
Z lm Score = 4.49
NG = 0
DB = 0
SM = 0
Delta_L
Doxorubicin
YOR057W SGT1
ZShift = 0.56
Z lm Score = −1.8
NG = 0
DB = 0
SM = 0
Delta_K
Doxorubicin
YOR057W SGT1
ZShift = −1.22
Z lm Score = −0.67
NG = 0
DB = 0
SM = 0
Delta_r
Doxorubicin
YOR057W SGT1
ZShift = −0.19
Z lm Score = 0.19
NG = 0
DB = 0
SM = 0
Delta_AUC
Doxorubicin

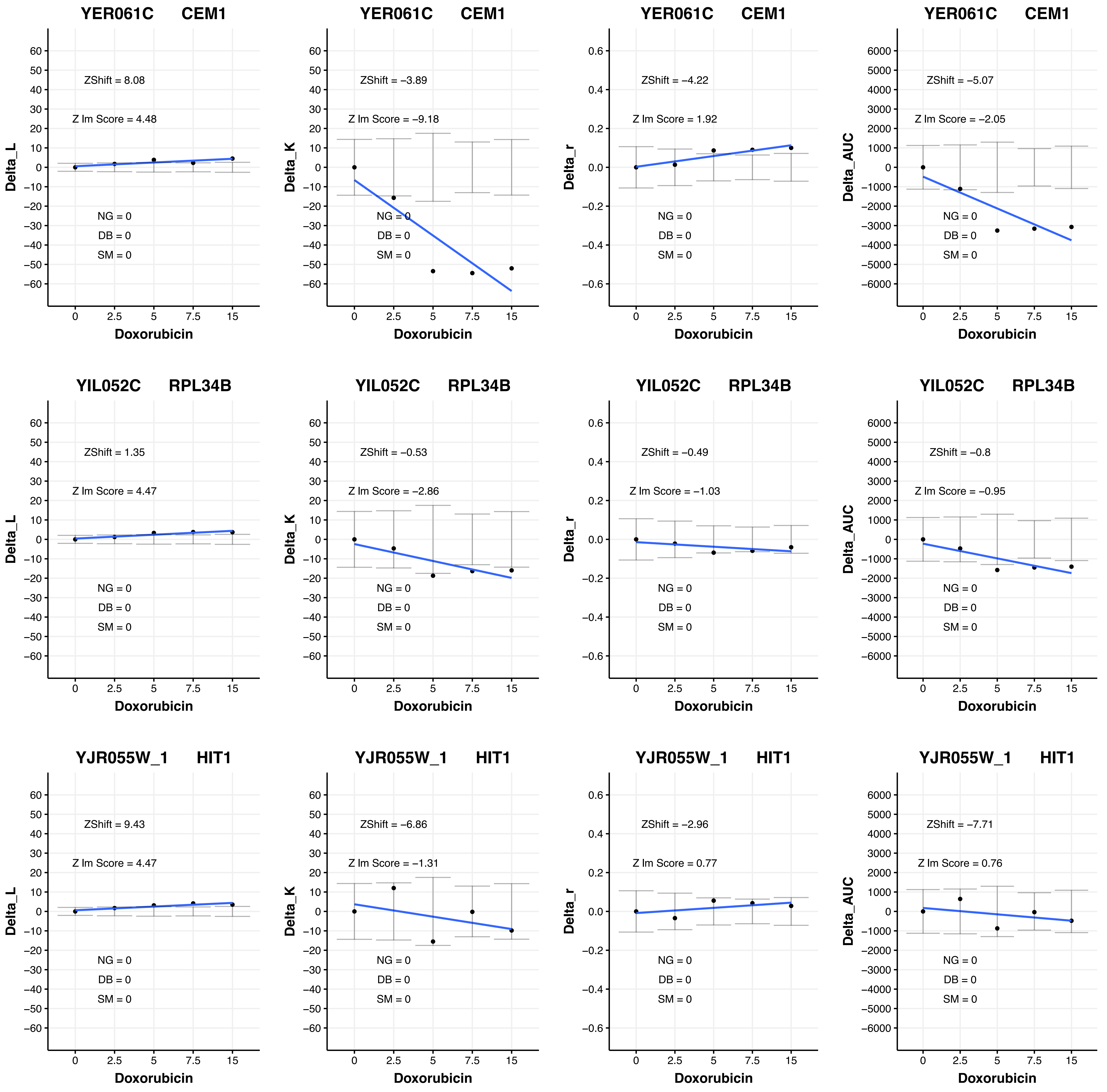
YER061C CEM1
ZShift = 8.08
Z lm Score = 4.48
NG = 0
DB = 0
SM = 0
Delta_L
Doxorubicin
YER061C CEM1
ZShift = −3.89
Z lm Score = −9.18
NG = 0
DB = 0
SM = 0
Delta_K
Doxorubicin
YER061C CEM1
ZShift = −4.22
Z lm Score = 1.92
NG = 0
DB = 0
SM = 0
Delta_r
Doxorubicin
YER061C CEM1
ZShift = −5.07
Z lm Score = −2.05
NG = 0
DB = 0
SM = 0
Delta_AUC
Doxorubicin
YIL052C RPL34B
ZShift = 1.35
Z lm Score = 4.47
NG = 0
DB = 0
SM = 0
Delta_L
Doxorubicin
YIL052C RPL34B
ZShift = −0.53
Z lm Score = −2.86
NG = 0
DB = 0
SM = 0
Delta_K
Doxorubicin
YIL052C RPL34B
ZShift = −0.49
Z lm Score = −1.03
NG = 0
DB = 0
SM = 0
Delta_r
Doxorubicin
YIL052C RPL34B
ZShift = −0.8
Z lm Score = −0.95
NG = 0
DB = 0
SM = 0
Delta_AUC
Doxorubicin
YJR055W_1 HIT1
ZShift = 9.43
Z lm Score = 4.47
NG = 0
DB = 0
SM = 0
Delta_L
Doxorubicin
YJR055W_1 HIT1
ZShift = −6.86
Z lm Score = −1.31
NG = 0
DB = 0
SM = 0
Delta_K
Doxorubicin
YJR055W_1 HIT1
ZShift = −2.96
Z lm Score = 0.77
NG = 0
DB = 0
SM = 0
Delta_r
Doxorubicin
YJR055W_1 HIT1
ZShift = −7.71
Z lm Score = 0.76
NG = 0
DB = 0
SM = 0
Delta_AUC
Doxorubicin

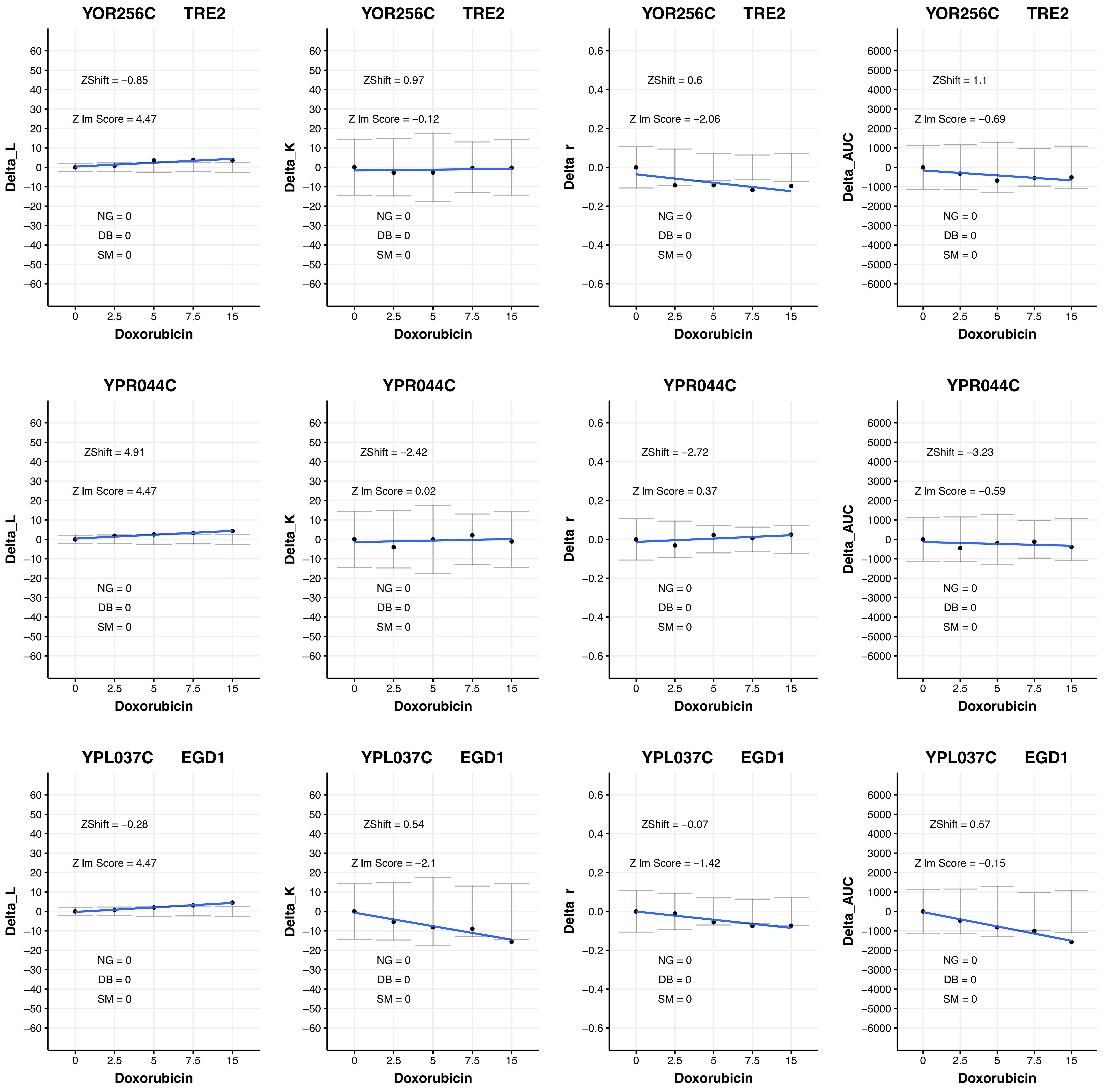
YOR256C TRE2
ZShift = −0.85
Z lm Score = 4.47
NG = 0
DB = 0
SM = 0
Delta_L
Doxorubicin
YOR256C TRE2
ZShift = 0.97
Z lm Score = −0.12
NG = 0
DB = 0
SM = 0
Delta_K
Doxorubicin
YOR256C TRE2
ZShift = 0.6
Z lm Score = −2.06
NG = 0
DB = 0
SM = 0
Delta_r
Doxorubicin
YOR256C TRE2
ZShift = 1.1
Z lm Score = −0.69
NG = 0
DB = 0
SM = 0
Delta_AUC
Doxorubicin
YPR044C
ZShift = 4.91
Z lm Score = 4.47
NG = 0
DB = 0
SM = 0
Delta_L
Doxorubicin
YPR044C
ZShift = −2.42
Z lm Score = 0.02
NG = 0
DB = 0
SM = 0
Delta_K
Doxorubicin
YPR044C
ZShift = −2.72
Z lm Score = 0.37
NG = 0
DB = 0
SM = 0
Delta_r
Doxorubicin
YPR044C
ZShift = −3.23
Z lm Score = −0.59
NG = 0
DB = 0
SM = 0
Delta_AUC
Doxorubicin
YPL037C EGD1
ZShift = −0.28
Z lm Score = 4.47
NG = 0
DB = 0
SM = 0
Delta_L
Doxorubicin
YPL037C EGD1
ZShift = 0.54
Z lm Score = −2.1
NG = 0
DB = 0
SM = 0
Delta_K
Doxorubicin
YPL037C EGD1
ZShift = −0.07
Z lm Score = −1.42
NG = 0
DB = 0
SM = 0
Delta_r
Doxorubicin
YPL037C EGD1
ZShift = 0.57
Z lm Score = −0.15
NG = 0
DB = 0
SM = 0
Delta_AUC
Doxorubicin

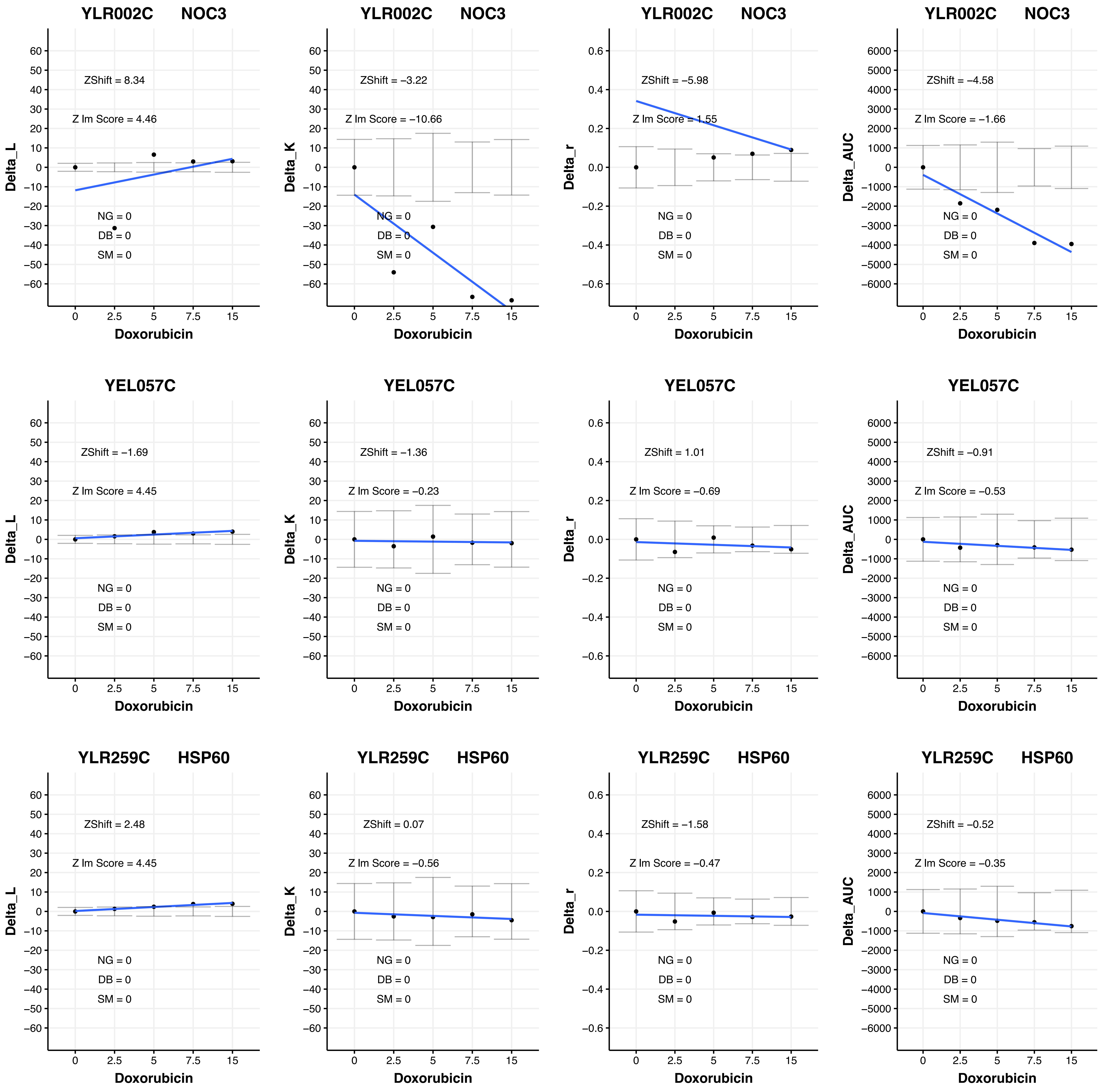

YLR002C NOC3
ZShift = 8.34
Z lm Score = 4.46
NG = 0
DB = 0
SM = 0
Delta_L
Doxorubicin
YLR002C NOC3
ZShift = −3.22
Z lm Score = −10.66
NG = 0
DB = 0
SM = 0
Delta_K
Doxorubicin
YLR002C NOC3
ZShift = −5.98
Z lm Score = 1.55
NG = 0
DB = 0
SM = 0
Delta_r
Doxorubicin
YLR002C NOC3
ZShift = −4.58
Z lm Score = −1.66
NG = 0
DB = 0
SM = 0
Delta_AUC
Doxorubicin
YEL057C
ZShift = −1.69
Z lm Score = 4.45
NG = 0
DB = 0
SM = 0
Delta_L
Doxorubicin
YEL057C
ZShift = −1.36
Z lm Score = −0.23
NG = 0
DB = 0
SM = 0
Delta_K
Doxorubicin
YEL057C
ZShift = 1.01
Z lm Score = −0.69
NG = 0
DB = 0
SM = 0
Delta_r
Doxorubicin
YEL057C
ZShift = −0.91
Z lm Score = −0.53
NG = 0
DB = 0
SM = 0
Delta_AUC
Doxorubicin
YLR259C HSP60
ZShift = 2.48
Z lm Score = 4.45
NG = 0
DB = 0
SM = 0
Delta_L
Doxorubicin
YLR259C HSP60
ZShift = 0.07
Z lm Score = −0.56
NG = 0
DB = 0
SM = 0
Delta_K
Doxorubicin
YLR259C HSP60
ZShift = −1.58
Z lm Score = −0.47
NG = 0
DB = 0
SM = 0
Delta_r
Doxorubicin
YLR259C HSP60
ZShift = −0.52
Z lm Score = −0.35
NG = 0
DB = 0
SM = 0
Delta_AUC
Doxorubicin

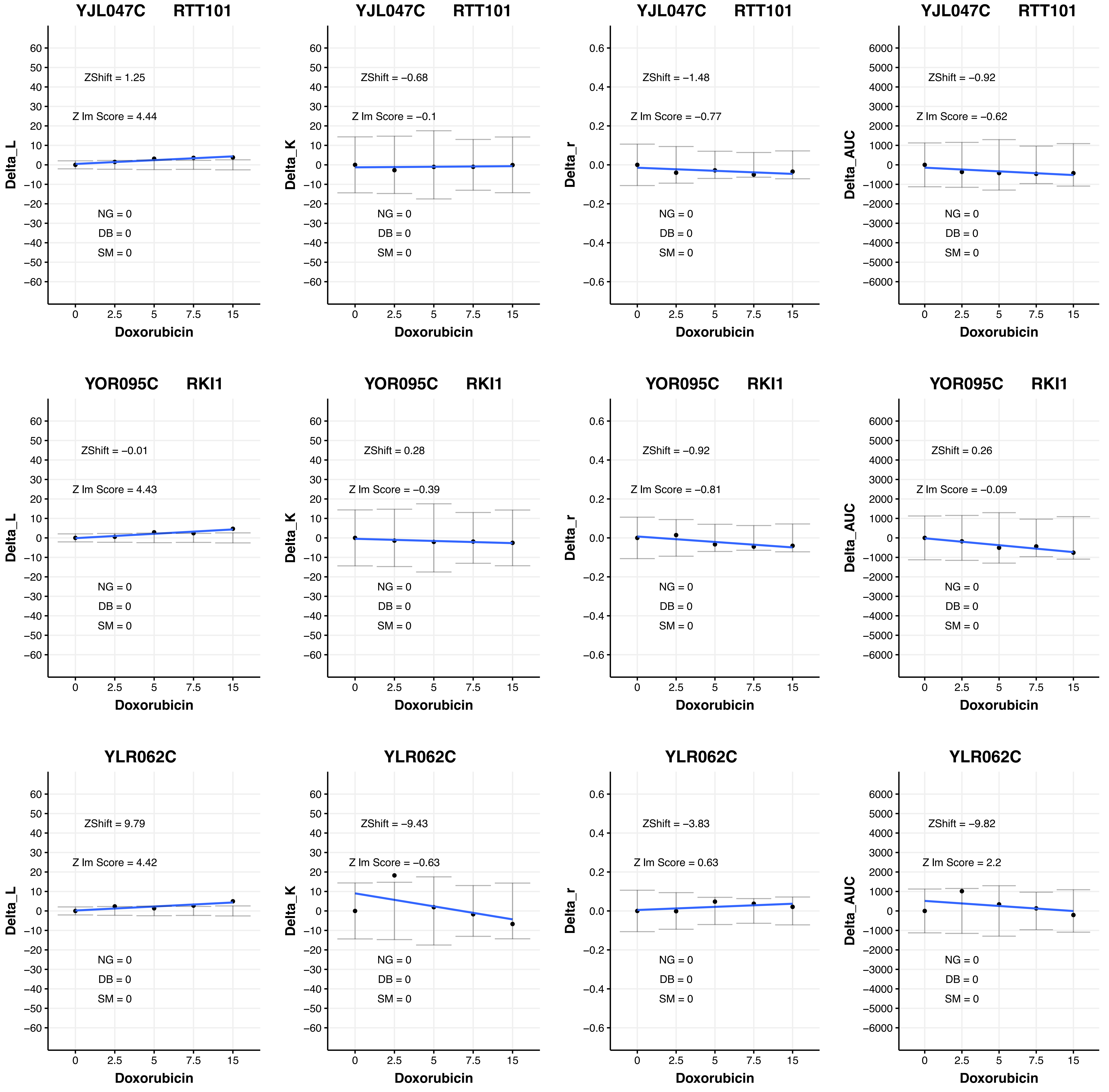
YJL047C RTT101
ZShift = 1.25
Z lm Score = 4.44
NG = 0
DB = 0
SM = 0
Delta_L
Doxorubicin
YJL047C RTT101
ZShift = −0.68
Z lm Score = −0.1
NG = 0
DB = 0
SM = 0
Delta_K
Doxorubicin
YJL047C RTT101
ZShift = −1.48
Z lm Score = −0.77
NG = 0
DB = 0
SM = 0
Delta_r
Doxorubicin
YJL047C RTT101
ZShift = −0.92
Z lm Score = −0.62
NG = 0
DB = 0
SM = 0
Delta_AUC
Doxorubicin
YOR095C RKI1
ZShift = −0.01
Z lm Score = 4.43
NG = 0
DB = 0
SM = 0
Delta_L
Doxorubicin
YOR095C RKI1
ZShift = 0.28
Z lm Score = −0.39
NG = 0
DB = 0
SM = 0
Delta_K
Doxorubicin
YOR095C RKI1
ZShift = −0.92
Z lm Score = −0.81
NG = 0
DB = 0
SM = 0
Delta_r
Doxorubicin
YOR095C RKI1
ZShift = 0.26
Z lm Score = −0.09
NG = 0
DB = 0
SM = 0
Delta_AUC
Doxorubicin
YLR062C
ZShift = 9.79
Z lm Score = 4.42
NG = 0
DB = 0
SM = 0
Delta_L
Doxorubicin
YLR062C
ZShift = −9.43
Z lm Score = −0.63
NG = 0
DB = 0
SM = 0
Delta_K
Doxorubicin
YLR062C
ZShift = −3.83
Z lm Score = 0.63
NG = 0
DB = 0
SM = 0
Delta_r
Doxorubicin
YLR062C
ZShift = −9.82
Z lm Score = 2.2
NG = 0
DB = 0
SM = 0
Delta_AUC
Doxorubicin

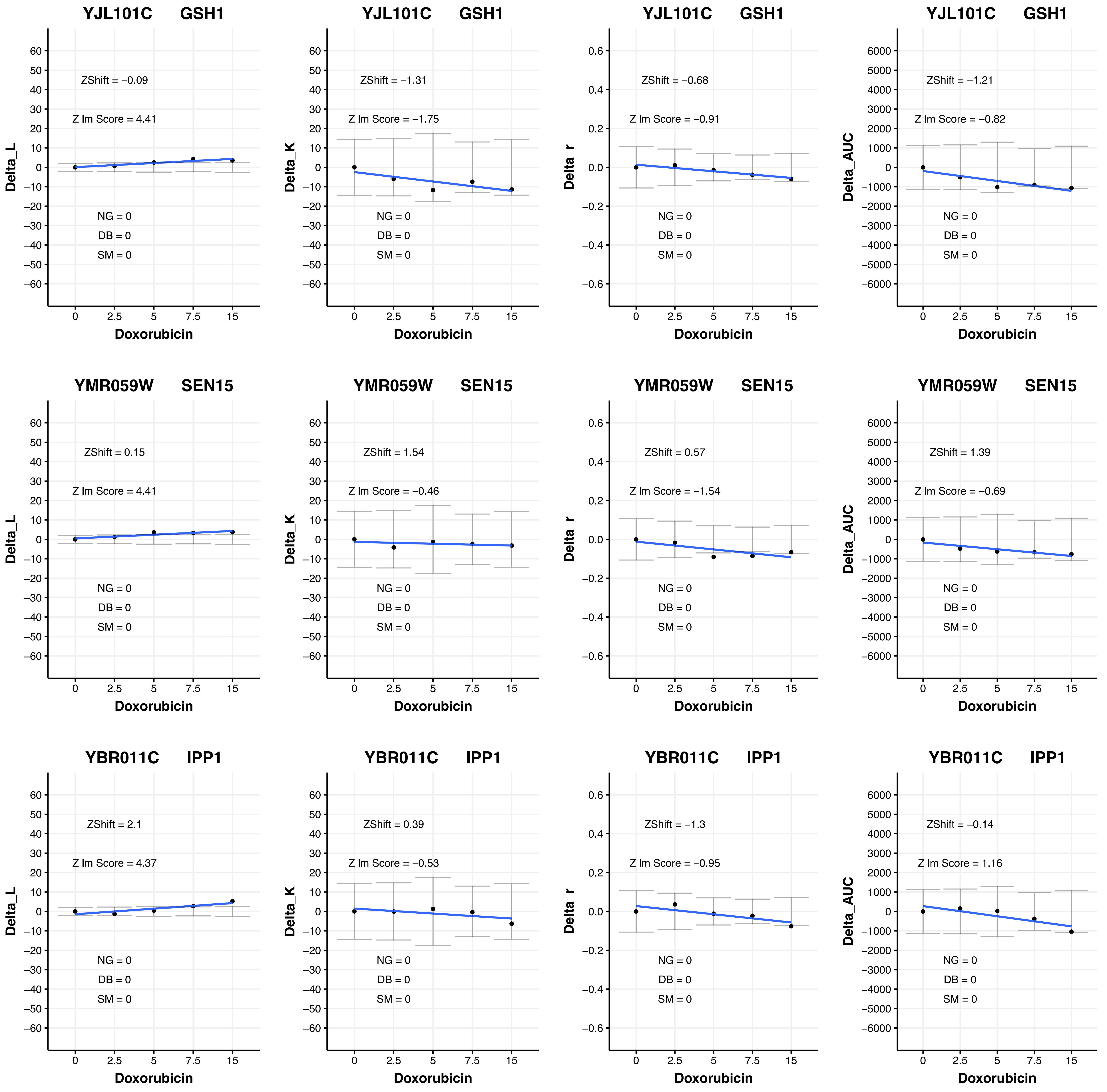

YJL101C GSH1
ZShift = −0.09
Z lm Score = 4.41
NG = 0
DB = 0
SM = 0
Delta_L
Doxorubicin
YJL101C GSH1
ZShift = −1.31
Z lm Score = −1.75
NG = 0
DB = 0
SM = 0
Delta_K
Doxorubicin
YJL101C GSH1
ZShift = −0.68
Z lm Score = −0.91
NG = 0
DB = 0
SM = 0
Delta_r
Doxorubicin
YJL101C GSH1
ZShift = −1.21
Z lm Score = −0.82
NG = 0
DB = 0
SM = 0
Delta_AUC
Doxorubicin
YMR059W SEN15
ZShift = 0.15
Z lm Score = 4.41
NG = 0
DB = 0
SM = 0
Delta_L
Doxorubicin
YMR059W SEN15
ZShift = 1.54
Z lm Score = −0.46
NG = 0
DB = 0
SM = 0
Delta_K
Doxorubicin
YMR059W SEN15
ZShift = 0.57
Z lm Score = −1.54
NG = 0
DB = 0
SM = 0
Delta_r
Doxorubicin
YMR059W SEN15
ZShift = 1.39
Z lm Score = −0.69
NG = 0
DB = 0
SM = 0
Delta_AUC
Doxorubicin
YBR011C IPP1
ZShift = 2.1
Z lm Score = 4.37
NG = 0
DB = 0
SM = 0
Delta_L
Doxorubicin
YBR011C IPP1
ZShift = 0.39
Z lm Score = −0.53
NG = 0
DB = 0
SM = 0
Delta_K
Doxorubicin
YBR011C IPP1
ZShift = −1.3
Z lm Score = −0.95
NG = 0
DB = 0
SM = 0
Delta_r
Doxorubicin
YBR011C IPP1
ZShift = −0.14
Z lm Score = 1.16
NG = 0
DB = 0
SM = 0
Delta_AUC
Doxorubicin

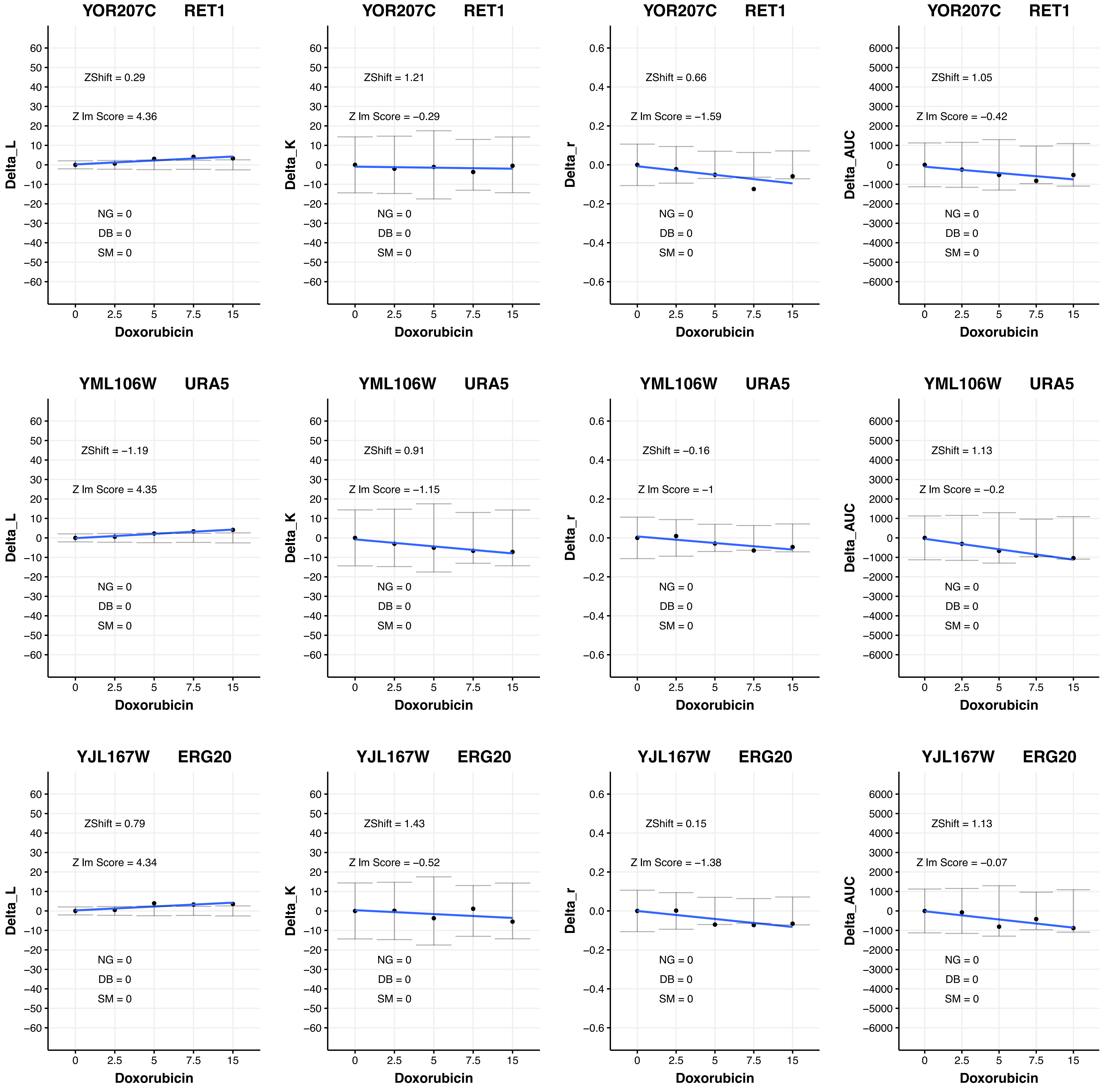
YOR207C RET1
ZShift = 0.29
Z lm Score = 4.36
NG = 0
DB = 0
SM = 0
Delta_L
Doxorubicin
YOR207C RET1
ZShift = 1.21
Z lm Score = −0.29
NG = 0
DB = 0
SM = 0
Delta_K
Doxorubicin
YOR207C RET1
ZShift = 0.66
Z lm Score = −1.59
NG = 0
DB = 0
SM = 0
Delta_r
Doxorubicin
YOR207C RET1
ZShift = 1.05
Z lm Score = −0.42
NG = 0
DB = 0
SM = 0
Delta_AUC
Doxorubicin
YML106W URA5
ZShift = −1.19
Z lm Score = 4.35
NG = 0
DB = 0
SM = 0
Delta_L
Doxorubicin
YML106W URA5
ZShift = 0.91
Z lm Score = −1.15
NG = 0
DB = 0
SM = 0
Delta_K
Doxorubicin
YML106W URA5
ZShift = −0.16
Z lm Score = −1
NG = 0
DB = 0
SM = 0
Delta_r
Doxorubicin
YML106W URA5
ZShift = 1.13
Z lm Score = −0.2
NG = 0
DB = 0
SM = 0
Delta_AUC
Doxorubicin
YJL167W ERG20
ZShift = 0.79
Z lm Score = 4.34
NG = 0
DB = 0
SM = 0
Delta_L
Doxorubicin
YJL167W ERG20
ZShift = 1.43
Z lm Score = −0.52
NG = 0
DB = 0
SM = 0
Delta_K
Doxorubicin
YJL167W ERG20
ZShift = 0.15
Z lm Score = −1.38
NG = 0
DB = 0
SM = 0
Delta_r
Doxorubicin
YJL167W ERG20
ZShift = 1.13
Z lm Score = −0.07
NG = 0
DB = 0
SM = 0
Delta_AUC
Doxorubicin

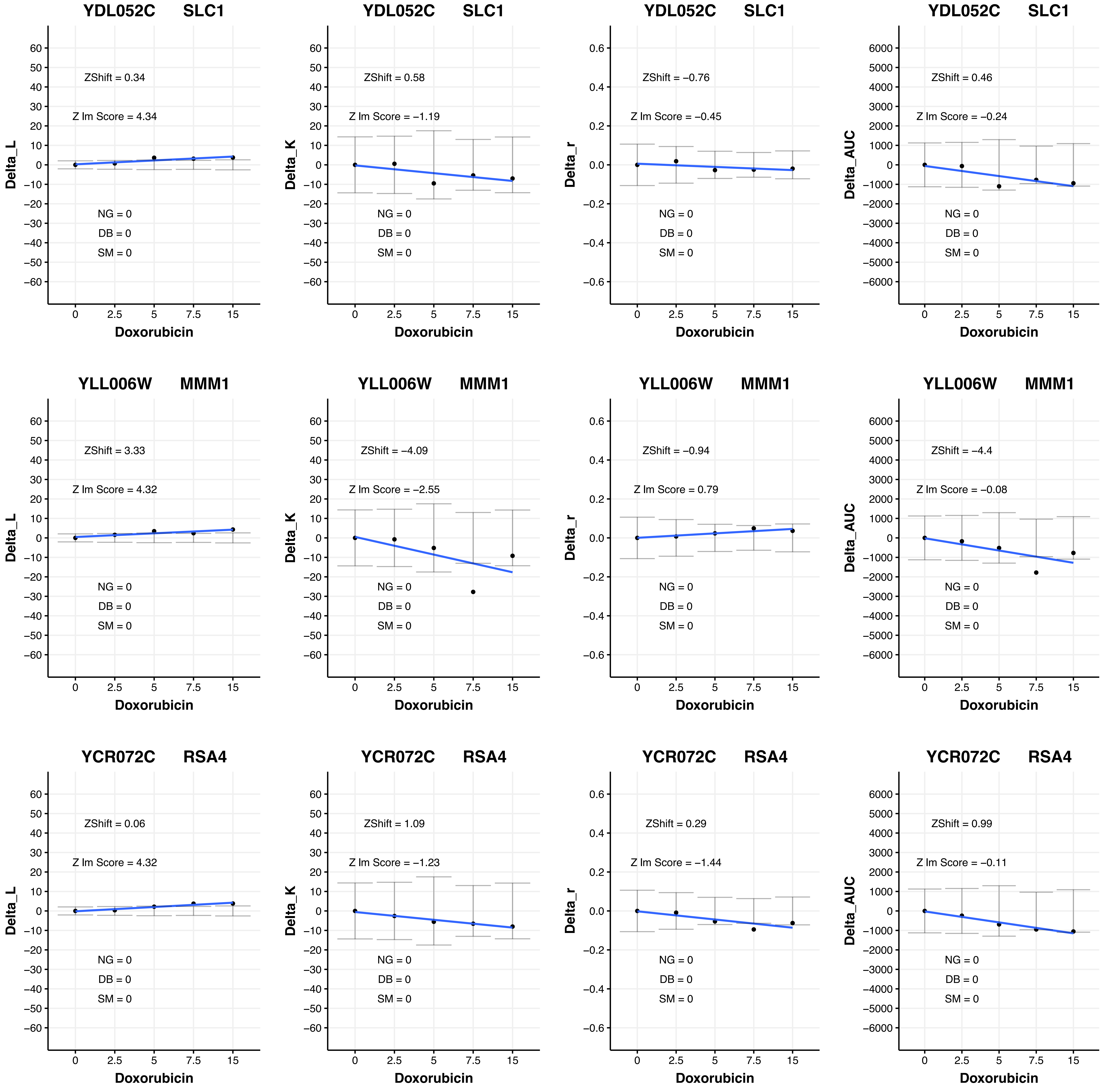
YDL052C SLC1
ZShift = 0.34
Z lm Score = 4.34
NG = 0
DB = 0
SM = 0
Delta_L
Doxorubicin
YDL052C SLC1
ZShift = 0.58
Z lm Score = −1.19
NG = 0
DB = 0
SM = 0
Delta_K
Doxorubicin
YDL052C SLC1
ZShift = −0.76
Z lm Score = −0.45
NG = 0
DB = 0
SM = 0
Delta_r
Doxorubicin
YDL052C SLC1
ZShift = 0.46
Z lm Score = −0.24
NG = 0
DB = 0
SM = 0
Delta_AUC
Doxorubicin
YLL006W MMM1
ZShift = 3.33
Z lm Score = 4.32
NG = 0
DB = 0
SM = 0
Delta_L
Doxorubicin
YLL006W MMM1
ZShift = −4.09
Z lm Score = −2.55
NG = 0
DB = 0
SM = 0
Delta_K
Doxorubicin
YLL006W MMM1
ZShift = −0.94
Z lm Score = 0.79
NG = 0
DB = 0
SM = 0
Delta_r
Doxorubicin
YLL006W MMM1
ZShift = −4.4
Z lm Score = −0.08
NG = 0
DB = 0
SM = 0
Delta_AUC
Doxorubicin
YCR072C RSA4
ZShift = 0.06
Z lm Score = 4.32
NG = 0
DB = 0
SM = 0
Delta_L
Doxorubicin
YCR072C RSA4
ZShift = 1.09
Z lm Score = −1.23
NG = 0
DB = 0
SM = 0
Delta_K
Doxorubicin
YCR072C RSA4
ZShift = 0.29
Z lm Score = −1.44
NG = 0
DB = 0
SM = 0
Delta_r
Doxorubicin
YCR072C RSA4
ZShift = 0.99
Z lm Score = −0.11
NG = 0
DB = 0
SM = 0
Delta_AUC
Doxorubicin

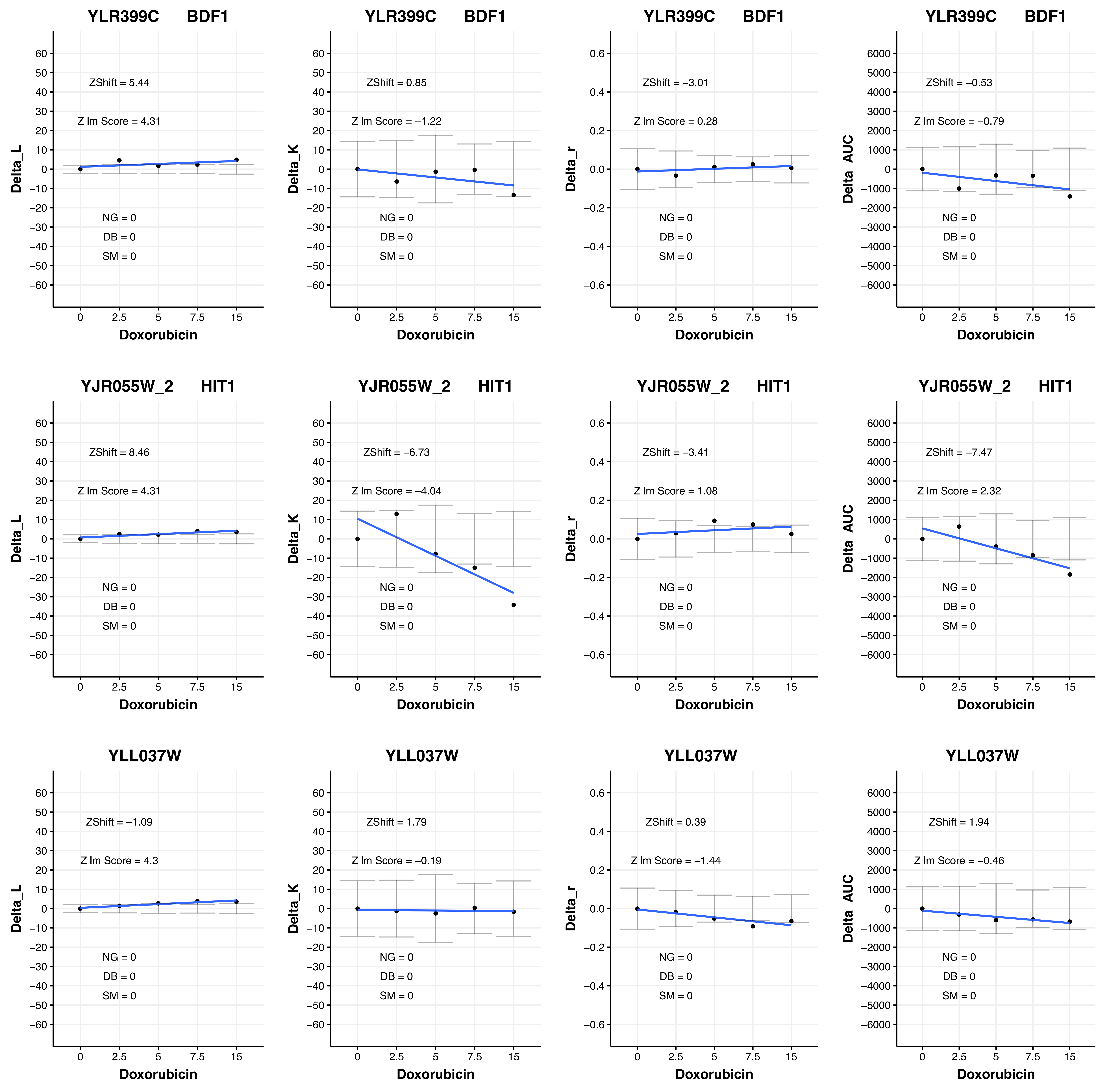

YLR399C BDF1
ZShift = 5.44
Z lm Score = 4.31
NG = 0
DB = 0
SM = 0
Delta_L
Doxorubicin
YLR399C BDF1
ZShift = 0.85
Z lm Score = −1.22
NG = 0
DB = 0
SM = 0
Delta_K
Doxorubicin
YLR399C BDF1
ZShift = −3.01
Z lm Score = 0.28
NG = 0
DB = 0
SM = 0
Delta_r
Doxorubicin
YLR399C BDF1
ZShift = −0.53
Z lm Score = −0.79
NG = 0
DB = 0
SM = 0
Delta_AUC
Doxorubicin
YJR055W_2 HIT1
ZShift = 8.46
Z lm Score = 4.31
NG = 0
DB = 0
SM = 0
Delta_L
Doxorubicin
YJR055W_2 HIT1
ZShift = −6.73
Z lm Score = −4.04
NG = 0
DB = 0
SM = 0
Delta_K
Doxorubicin
YJR055W_2 HIT1
ZShift = −3.41
Z lm Score = 1.08
NG = 0
DB = 0
SM = 0
Delta_r
Doxorubicin
YJR055W_2 HIT1
ZShift = −7.47
Z lm Score = 2.32
NG = 0
DB = 0
SM = 0
Delta_AUC
Doxorubicin
YLL037W
ZShift = −1.09
Z lm Score = 4.3
NG = 0
DB = 0
SM = 0
Delta_L
Doxorubicin
YLL037W
ZShift = 1.79
Z lm Score = −0.19
NG = 0
DB = 0
SM = 0
Delta_K
Doxorubicin
YLL037W
ZShift = 0.39
Z lm Score = −1.44
NG = 0
DB = 0
SM = 0
Delta_r
Doxorubicin
YLL037W
ZShift = 1.94
Z lm Score = −0.46
NG = 0
DB = 0
SM = 0
Delta_AUC
Doxorubicin

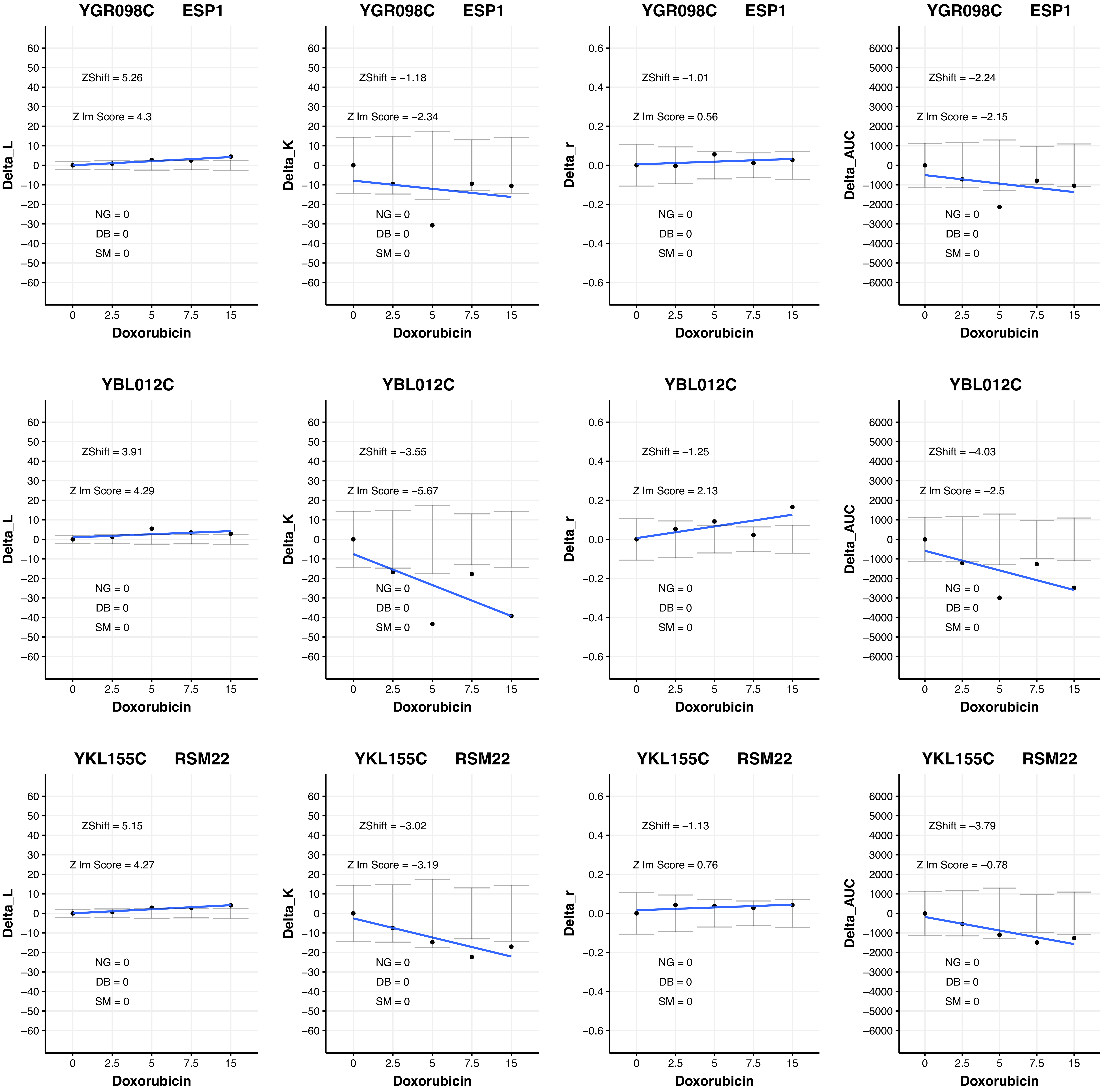
YGR098C ESP1
ZShift = 5.26
Z lm Score = 4.3
NG = 0
DB = 0
SM = 0
Delta_L
Doxorubicin
YGR098C ESP1
ZShift = −1.18
Z lm Score = −2.34
NG = 0
DB = 0
SM = 0
Delta_K
Doxorubicin
YGR098C ESP1
ZShift = −1.01
Z lm Score = 0.56
NG = 0
DB = 0
SM = 0
Delta_r
Doxorubicin
YGR098C ESP1
ZShift = −2.24
Z lm Score = −2.15
NG = 0
DB = 0
SM = 0
Delta_AUC
Doxorubicin
YBL012C
ZShift = 3.91
Z lm Score = 4.29
NG = 0
DB = 0
SM = 0
Delta_L
Doxorubicin
YBL012C
ZShift = −3.55
Z lm Score = −5.67
NG = 0
DB = 0
SM = 0
Delta_K
Doxorubicin
YBL012C
ZShift = −1.25
Z lm Score = 2.13
NG = 0
DB = 0
SM = 0
Delta_r
Doxorubicin
YBL012C
ZShift = −4.03
Z lm Score = −2.5
NG = 0
DB = 0
SM = 0
Delta_AUC
Doxorubicin
YKL155C RSM22
ZShift = 5.15
Z lm Score = 4.27
NG = 0
DB = 0
SM = 0
Delta_L
Doxorubicin
YKL155C RSM22
ZShift = −3.02
Z lm Score = −3.19
NG = 0
DB = 0
SM = 0
Delta_K
Doxorubicin
YKL155C RSM22
ZShift = −1.13
Z lm Score = 0.76
NG = 0
DB = 0
SM = 0
Delta_r
Doxorubicin
YKL155C RSM22
ZShift = −3.79
Z lm Score = −0.78
NG = 0
DB = 0
SM = 0
Delta_AUC
Doxorubicin

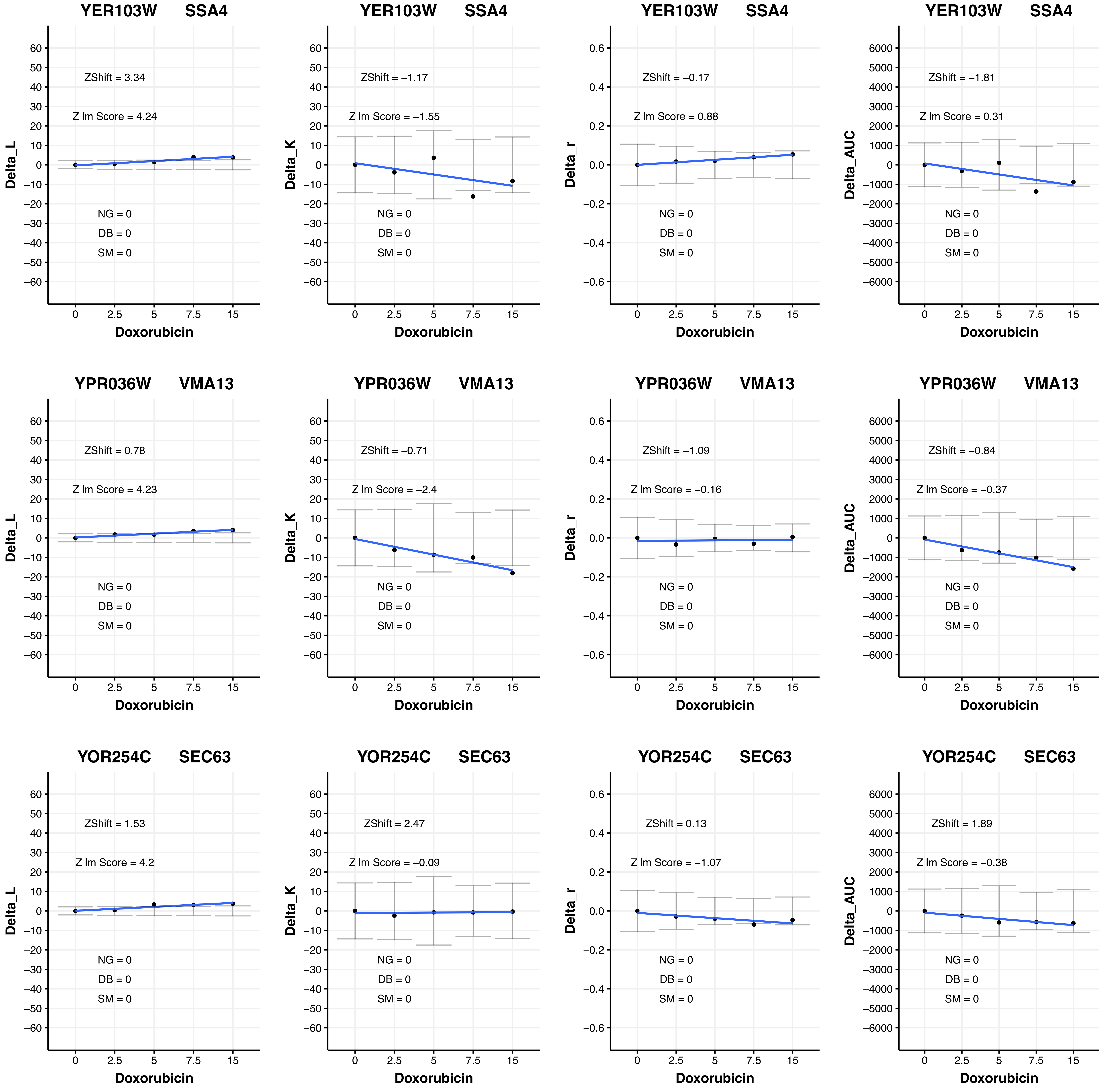
YER103W SSA4
ZShift = 3.34
Z lm Score = 4.24
NG = 0
DB = 0
SM = 0
Delta_L
Doxorubicin
YER103W SSA4
ZShift = −1.17
Z lm Score = −1.55
NG = 0
DB = 0
SM = 0
Delta_K
Doxorubicin
YER103W SSA4
ZShift = −0.17
Z lm Score = 0.88
NG = 0
DB = 0
SM = 0
Delta_r
Doxorubicin
YER103W SSA4
ZShift = −1.81
Z lm Score = 0.31
NG = 0
DB = 0
SM = 0
Delta_AUC
Doxorubicin
YPR036W VMA13
ZShift = 0.78
Z lm Score = 4.23
NG = 0
DB = 0
SM = 0
Delta_L
Doxorubicin
YPR036W VMA13
ZShift = −0.71
Z lm Score = −2.4
NG = 0
DB = 0
SM = 0
Delta_K
Doxorubicin
YPR036W VMA13
ZShift = −1.09
Z lm Score = −0.16
NG = 0
DB = 0
SM = 0
Delta_r
Doxorubicin
YPR036W VMA13
ZShift = −0.84
Z lm Score = −0.37
NG = 0
DB = 0
SM = 0
Delta_AUC
Doxorubicin
YOR254C SEC63
ZShift = 1.53
Z lm Score = 4.2
NG = 0
DB = 0
SM = 0
Delta_L
Doxorubicin
YOR254C SEC63
ZShift = 2.47
Z lm Score = −0.09
NG = 0
DB = 0
SM = 0
Delta_K
Doxorubicin
YOR254C SEC63
ZShift = 0.13
Z lm Score = −1.07
NG = 0
DB = 0
SM = 0
Delta_r
Doxorubicin
YOR254C SEC63
ZShift = 1.89
Z lm Score = −0.38
NG = 0
DB = 0
SM = 0
Delta_AUC
Doxorubicin

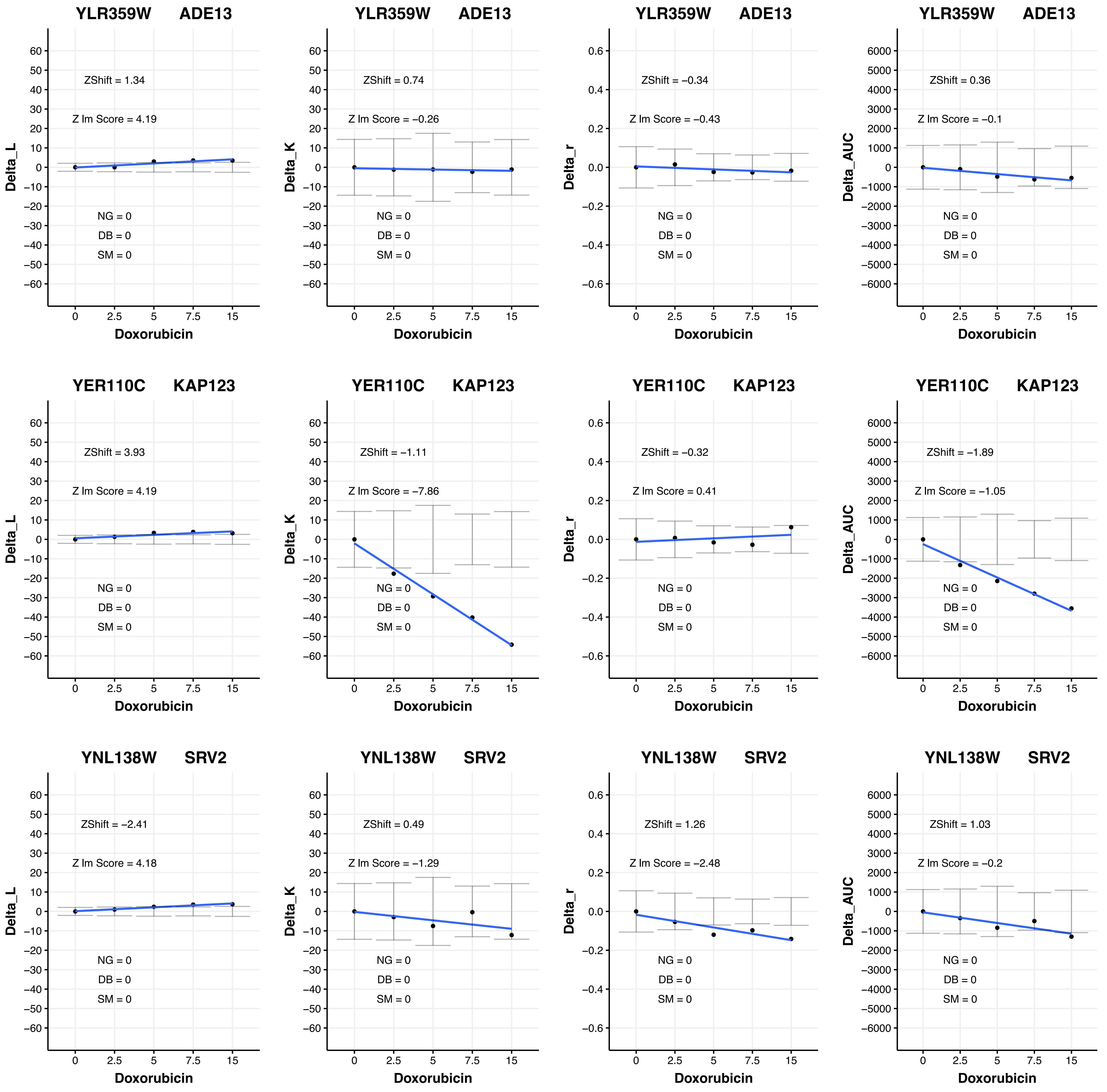
YLR359W ADE13
ZShift = 1.34
Z lm Score = 4.19
NG = 0
DB = 0
SM = 0
Delta_L
Doxorubicin
YLR359W ADE13
ZShift = 0.74
Z lm Score = −0.26
NG = 0
DB = 0
SM = 0
Delta_K
Doxorubicin
YLR359W ADE13
ZShift = −0.34
Z lm Score = −0.43
NG = 0
DB = 0
SM = 0
Delta_r
Doxorubicin
YLR359W ADE13
ZShift = 0.36
Z lm Score = −0.1
NG = 0
DB = 0
SM = 0
Delta_AUC
Doxorubicin
YER110C KAP123
ZShift = 3.93
Z lm Score = 4.19
NG = 0
DB = 0
SM = 0
Delta_L
Doxorubicin
YER110C KAP123
ZShift = −1.11
Z lm Score = −7.86
NG = 0
DB = 0
SM = 0
Delta_K
Doxorubicin
YER110C KAP123
ZShift = −0.32
Z lm Score = 0.41
NG = 0
DB = 0
SM = 0
Delta_r
Doxorubicin
YER110C KAP123
ZShift = −1.89
Z lm Score = −1.05
NG = 0
DB = 0
SM = 0
Delta_AUC
Doxorubicin
YNL138W SRV2
ZShift = −2.41
Z lm Score = 4.18
NG = 0
DB = 0
SM = 0
Delta_L
Doxorubicin
YNL138W SRV2
ZShift = 0.49
Z lm Score = −1.29
NG = 0
DB = 0
SM = 0
Delta_K
Doxorubicin
YNL138W SRV2
ZShift = 1.26
Z lm Score = −2.48
NG = 0
DB = 0
SM = 0
Delta_r
Doxorubicin
YNL138W SRV2
ZShift = 1.03
Z lm Score = −0.2
NG = 0
DB = 0
SM = 0
Delta_AUC
Doxorubicin

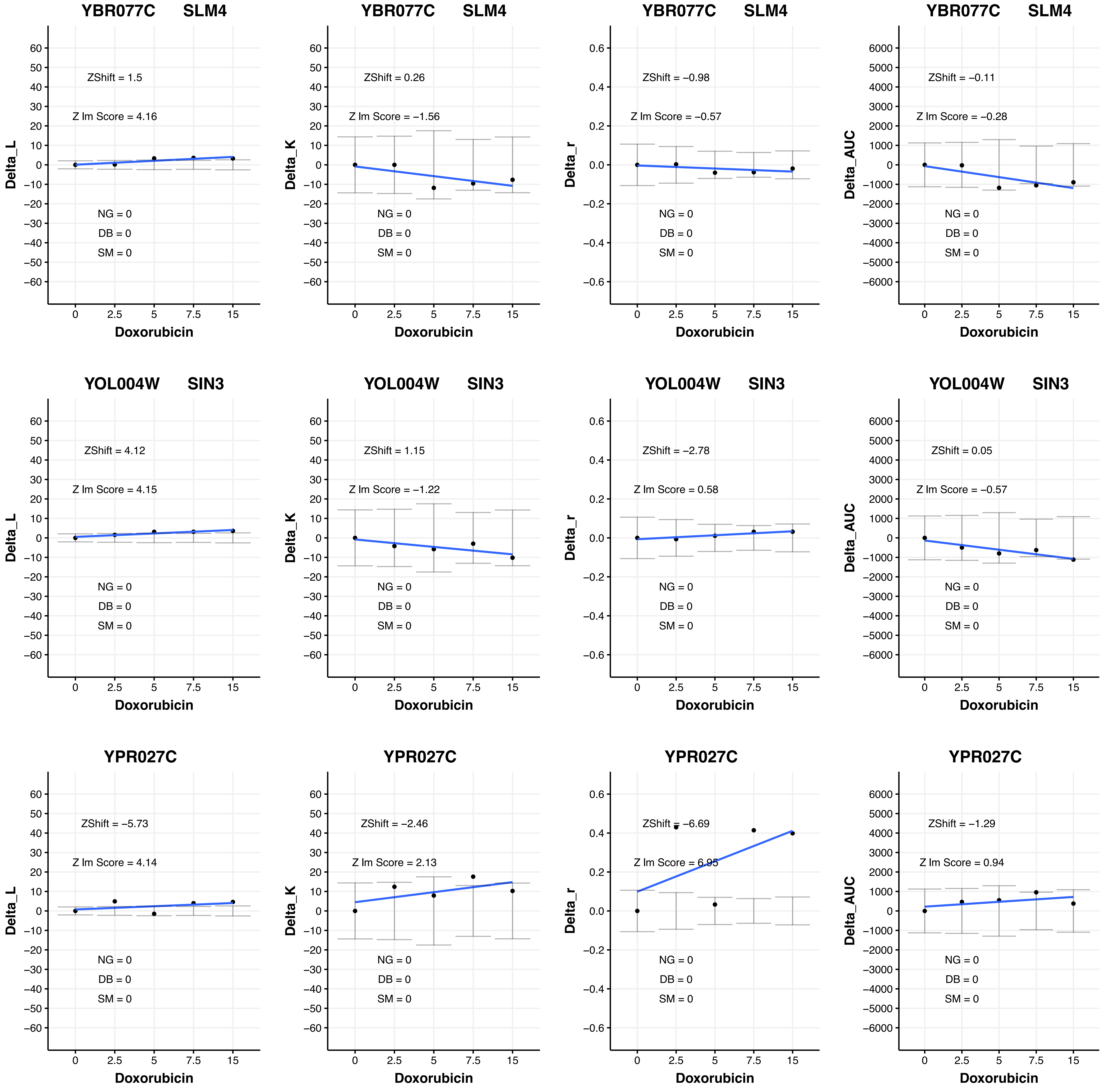

YBR077C SLM4
ZShift = 1.5
Z lm Score = 4.16
NG = 0
DB = 0
SM = 0
Delta_L
Doxorubicin
YBR077C SLM4
ZShift = 0.26
Z lm Score = −1.56
NG = 0
DB = 0
SM = 0
Delta_K
Doxorubicin
YBR077C SLM4
ZShift = −0.98
Z lm Score = −0.57
NG = 0
DB = 0
SM = 0
Delta_r
Doxorubicin
YBR077C SLM4
ZShift = −0.11
Z lm Score = −0.28
NG = 0
DB = 0
SM = 0
Delta_AUC
Doxorubicin
YOL004W SIN3
ZShift = 4.12
Z lm Score = 4.15
NG = 0
DB = 0
SM = 0
Delta_L
Doxorubicin
YOL004W SIN3
ZShift = 1.15
Z lm Score = −1.22
NG = 0
DB = 0
SM = 0
Delta_K
Doxorubicin
YOL004W SIN3
ZShift = −2.78
Z lm Score = 0.58
NG = 0
DB = 0
SM = 0
Delta_r
Doxorubicin
YOL004W SIN3
ZShift = 0.05
Z lm Score = −0.57
NG = 0
DB = 0
SM = 0
Delta_AUC
Doxorubicin
YPR027C
ZShift = −5.73
Z lm Score = 4.14
NG = 0
DB = 0
SM = 0
Delta_L
Doxorubicin
YPR027C
ZShift = −2.46
Z lm Score = 2.13
NG = 0
DB = 0
SM = 0
Delta_K
Doxorubicin
YPR027C
ZShift = −6.69
Z lm Score = 6.95
NG = 0
DB = 0
SM = 0
Delta_r
Doxorubicin
YPR027C
ZShift = −1.29
Z lm Score = 0.94
NG = 0
DB = 0
SM = 0
Delta_AUC
Doxorubicin

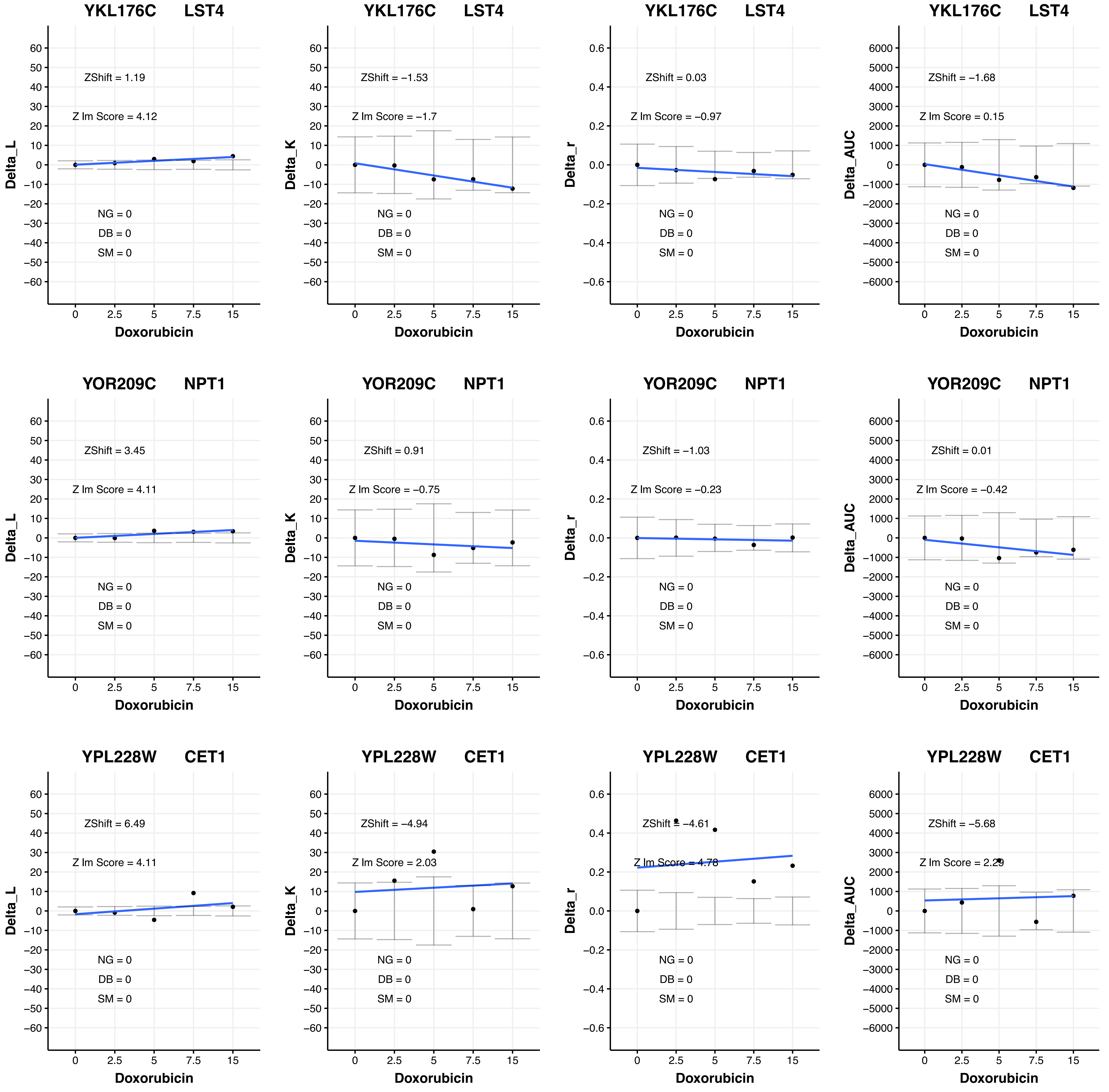

YKL176C LST4
ZShift = 1.19
Z lm Score = 4.12
NG = 0
DB = 0
SM = 0
Delta_L
YKL176C LST4
ZShift = −1.53
Z lm Score = −1.7
NG = 0
DB = 0
SM = 0
Delta_K
YKL176C LST4
ZShift = 0.03
Z lm Score = −0.97
NG = 0
DB = 0
SM = 0
Delta_r
YKL176C LST4
ZShift = −1.68
Z lm Score = 0.15
NG = 0
DB = 0
SM = 0
Delta_AUC
YOR209C NPT1
ZShift = 3.45
Z lm Score = 4.11
NG = 0
DB = 0
SM = 0
Delta_L
YOR209C NPT1
ZShift = 0.91
Z lm Score = −0.75
NG = 0
DB = 0
SM = 0
Delta_K
YOR209C NPT1
ZShift = −1.03
Z lm Score = −0.23
NG = 0
DB = 0
SM = 0
Delta_r
YOR209C NPT1
ZShift = 0.01
Z lm Score = −0.42
NG = 0
DB = 0
SM = 0
Delta_AUC
YPL228W CET1
ZShift = 6.49
Z lm Score = 4.11
NG = 0
DB = 0
SM = 0
Delta_L
YPL228W CET1
ZShift = −4.94
Z lm Score = 2.03
NG = 0
DB = 0
SM = 0
Delta_K
YPL228W CET1
ZShift = −4.61
Z lm Score = 4.78
NG = 0
DB = 0
SM = 0
Delta_r
YPL228W CET1
ZShift = −5.68
Z lm Score = 2.29
NG = 0
DB = 0
SM = 0
Delta_AUC
Doxorubicin
0
2.5
5
7.5
15

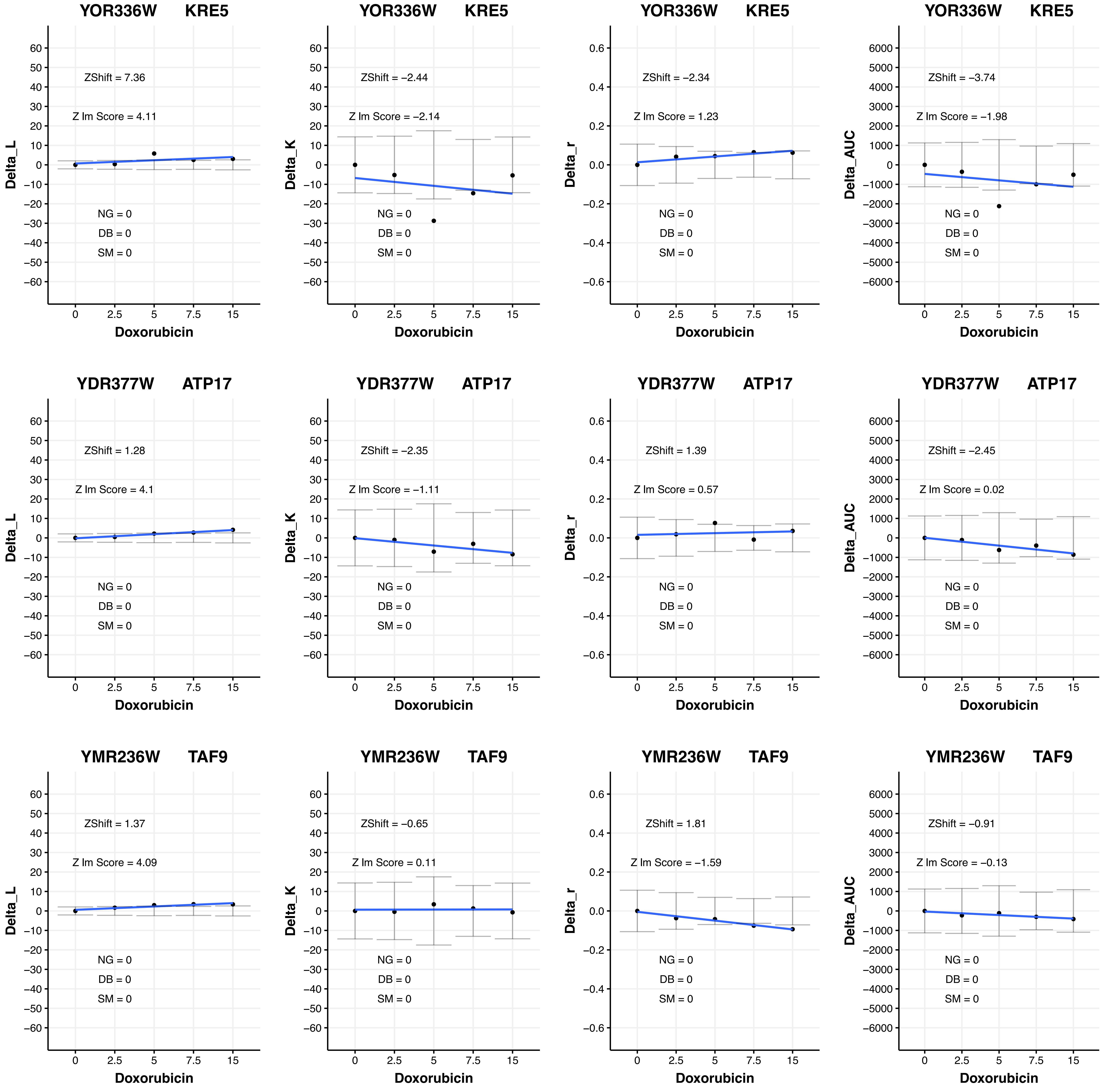
YOR336W KRE5
ZShift = 7.36
Z lm Score = 4.11
NG = 0
DB = 0
SM = 0
Delta_L
Doxorubicin
YOR336W KRE5
ZShift = −2.44
Z lm Score = −2.14
NG = 0
DB = 0
SM = 0
Delta_K
Doxorubicin
YOR336W KRE5
ZShift = −2.34
Z lm Score = 1.23
NG = 0
DB = 0
SM = 0
Delta_r
Doxorubicin
YOR336W KRE5
ZShift = −3.74
Z lm Score = −1.98
NG = 0
DB = 0
SM = 0
Delta_AUC
Doxorubicin
YDR377W ATP17
ZShift = 1.28
Z lm Score = 4.1
NG = 0
DB = 0
SM = 0
Delta_L
Doxorubicin
YDR377W ATP17
ZShift = −2.35
Z lm Score = −1.11
NG = 0
DB = 0
SM = 0
Delta_K
Doxorubicin
YDR377W ATP17
ZShift = 1.39
Z lm Score = 0.57
NG = 0
DB = 0
SM = 0
Delta_r
Doxorubicin
YDR377W ATP17
ZShift = −2.45
Z lm Score = 0.02
NG = 0
DB = 0
SM = 0
Delta_AUC
Doxorubicin
YMR236W TAF9
ZShift = 1.37
Z lm Score = 4.09
NG = 0
DB = 0
SM = 0
Delta_L
Doxorubicin
YMR236W TAF9
ZShift = −0.65
Z lm Score = 0.11
NG = 0
DB = 0
SM = 0
Delta_K
Doxorubicin
YMR236W TAF9
ZShift = 1.81
Z lm Score = −1.59
NG = 0
DB = 0
SM = 0
Delta_r
Doxorubicin
YMR236W TAF9
ZShift = −0.91
Z lm Score = −0.13
NG = 0
DB = 0
SM = 0
Delta_AUC
Doxorubicin

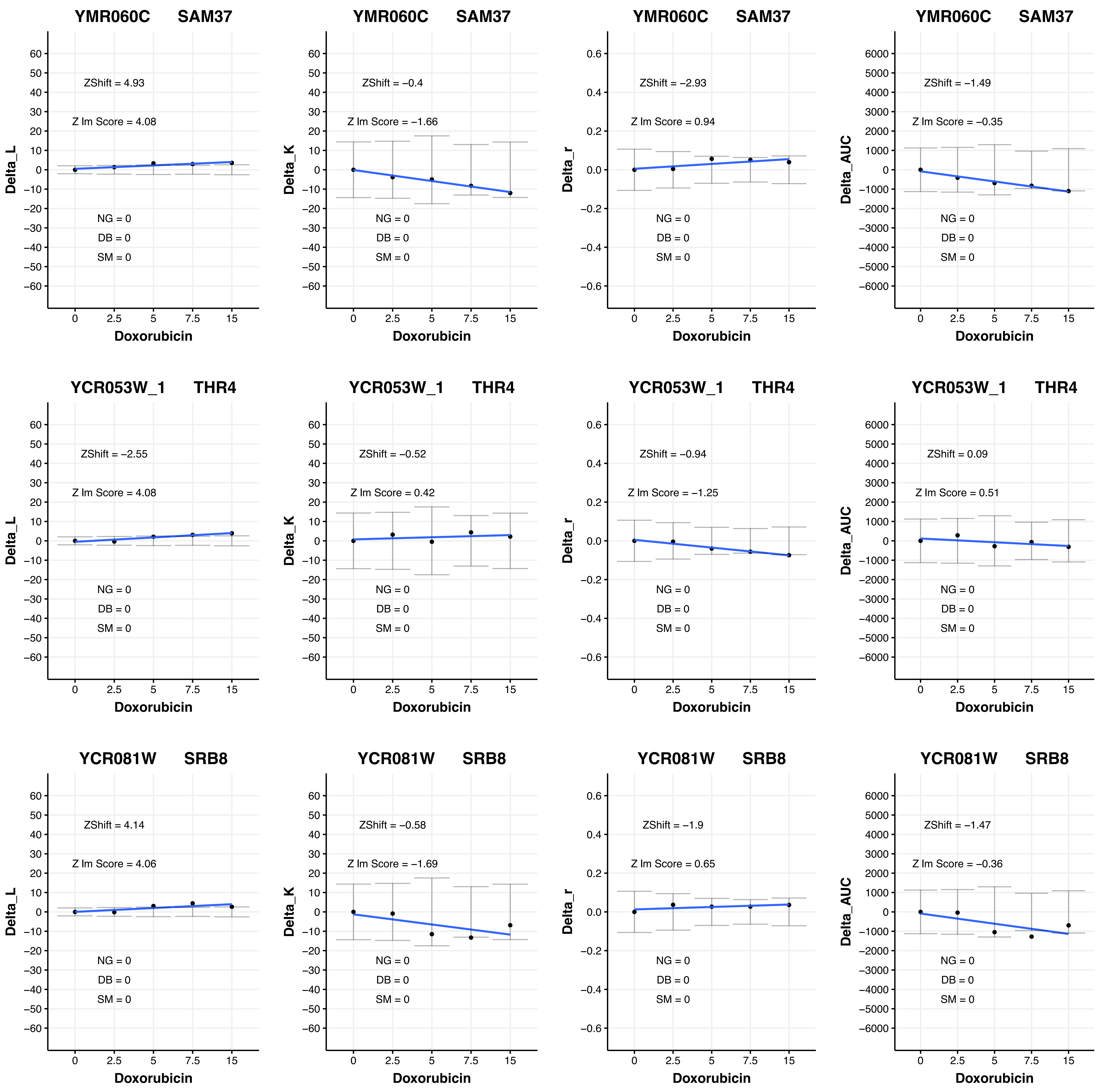

YMR060C SAM37
ZShift = 4.93
Z lm Score = 4.08
NG = 0
DB = 0
SM = 0
Delta_L
Doxorubicin
YMR060C SAM37
ZShift = −0.4
Z lm Score = −1.66
NG = 0
DB = 0
SM = 0
Delta_K
Doxorubicin
YMR060C SAM37
ZShift = −2.93
Z lm Score = 0.94
NG = 0
DB = 0
SM = 0
Delta_r
Doxorubicin
YMR060C SAM37
ZShift = −1.49
Z lm Score = −0.35
NG = 0
DB = 0
SM = 0
Delta_AUC
Doxorubicin
YCR053W_1 THR4
ZShift = −2.55
Z lm Score = 4.08
NG = 0
DB = 0
SM = 0
Delta_L
Doxorubicin
YCR053W_1 THR4
ZShift = −0.52
Z lm Score = 0.42
NG = 0
DB = 0
SM = 0
Delta_K
Doxorubicin
YCR053W_1 THR4
ZShift = −0.94
Z lm Score = −1.25
NG = 0
DB = 0
SM = 0
Delta_r
Doxorubicin
YCR053W_1 THR4
ZShift = 0.09
Z lm Score = 0.51
NG = 0
DB = 0
SM = 0
Delta_AUC
Doxorubicin
YCR081W SRB8
ZShift = 4.14
Z lm Score = 4.06
NG = 0
DB = 0
SM = 0
Delta_L
Doxorubicin
YCR081W SRB8
ZShift = −0.58
Z lm Score = −1.69
NG = 0
DB = 0
SM = 0
Delta_K
Doxorubicin
YCR081W SRB8
ZShift = −1.9
Z lm Score = 0.65
NG = 0
DB = 0
SM = 0
Delta_r
Doxorubicin
YCR081W SRB8
ZShift = −1.47
Z lm Score = −0.36
NG = 0
DB = 0
SM = 0
Delta_AUC
Doxorubicin

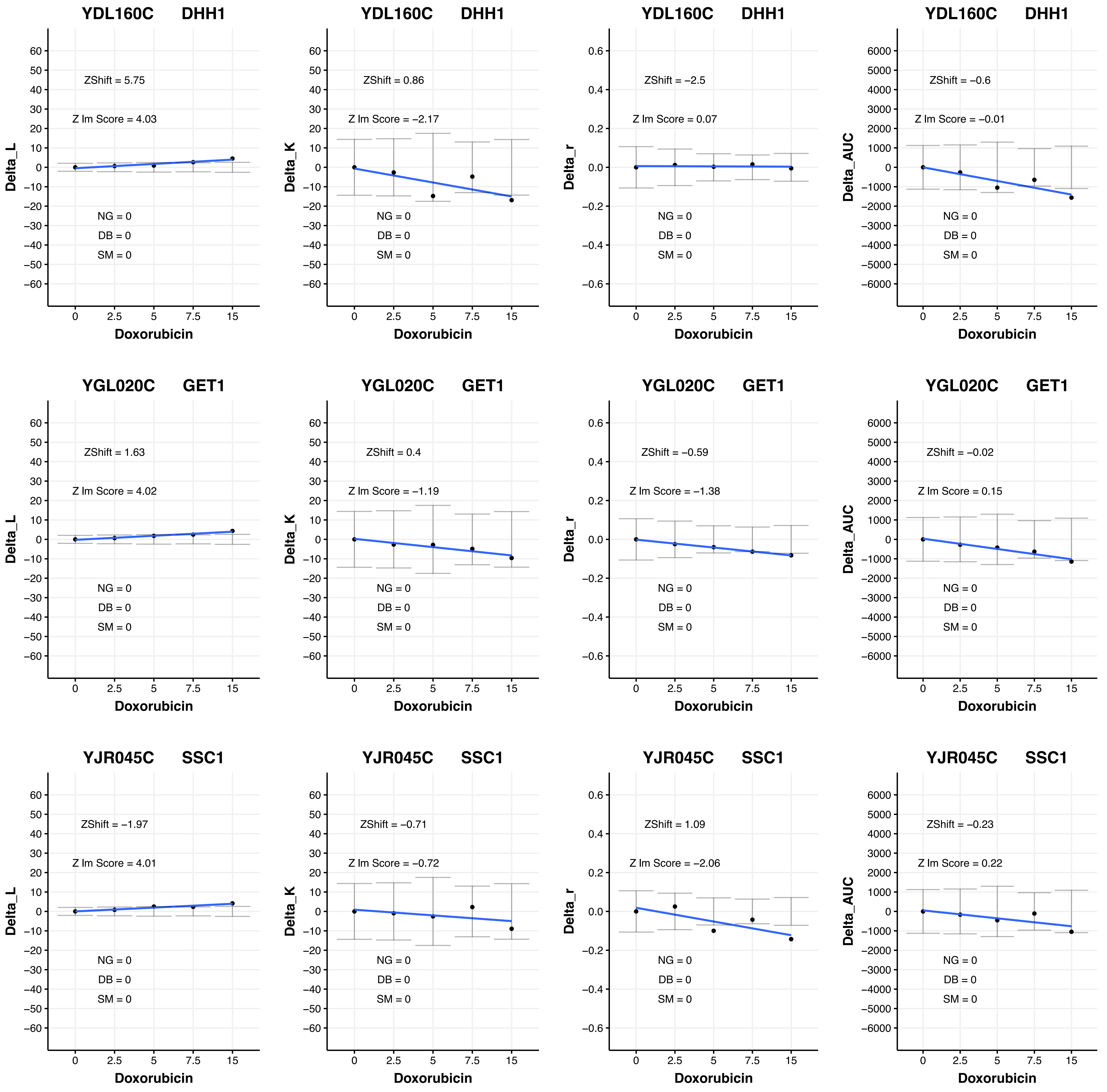
YDL160C DHH1
ZShift = 5.75
Z lm Score = 4.03
NG = 0
DB = 0
SM = 0
Delta_L
Doxorubicin
YDL160C DHH1
ZShift = 0.86
Z lm Score = −2.17
NG = 0
DB = 0
SM = 0
Delta_K
Doxorubicin
YDL160C DHH1
ZShift = −2.5
Z lm Score = 0.07
NG = 0
DB = 0
SM = 0
Delta_r
Doxorubicin
YDL160C DHH1
ZShift = −0.6
Z lm Score = −0.01
NG = 0
DB = 0
SM = 0
Delta_AUC
Doxorubicin
YGL020C GET1
ZShift = 1.63
Z lm Score = 4.02
NG = 0
DB = 0
SM = 0
Delta_L
Doxorubicin
YGL020C GET1
ZShift = 0.4
Z lm Score = −1.19
NG = 0
DB = 0
SM = 0
Delta_K
Doxorubicin
YGL020C GET1
ZShift = −0.59
Z lm Score = −1.38
NG = 0
DB = 0
SM = 0
Delta_r
Doxorubicin
YGL020C GET1
ZShift = −0.02
Z lm Score = 0.15
NG = 0
DB = 0
SM = 0
Delta_AUC
Doxorubicin
YJR045C SSC1
ZShift = −1.97
Z lm Score = 4.01
NG = 0
DB = 0
SM = 0
Delta_L
Doxorubicin
YJR045C SSC1
ZShift = −0.71
Z lm Score = −0.72
NG = 0
DB = 0
SM = 0
Delta_K
Doxorubicin
YJR045C SSC1
ZShift = 1.09
Z lm Score = −2.06
NG = 0
DB = 0
SM = 0
Delta_r
Doxorubicin
YJR045C SSC1
ZShift = −0.23
Z lm Score = 0.22
NG = 0
DB = 0
SM = 0
Delta_AUC
Doxorubicin

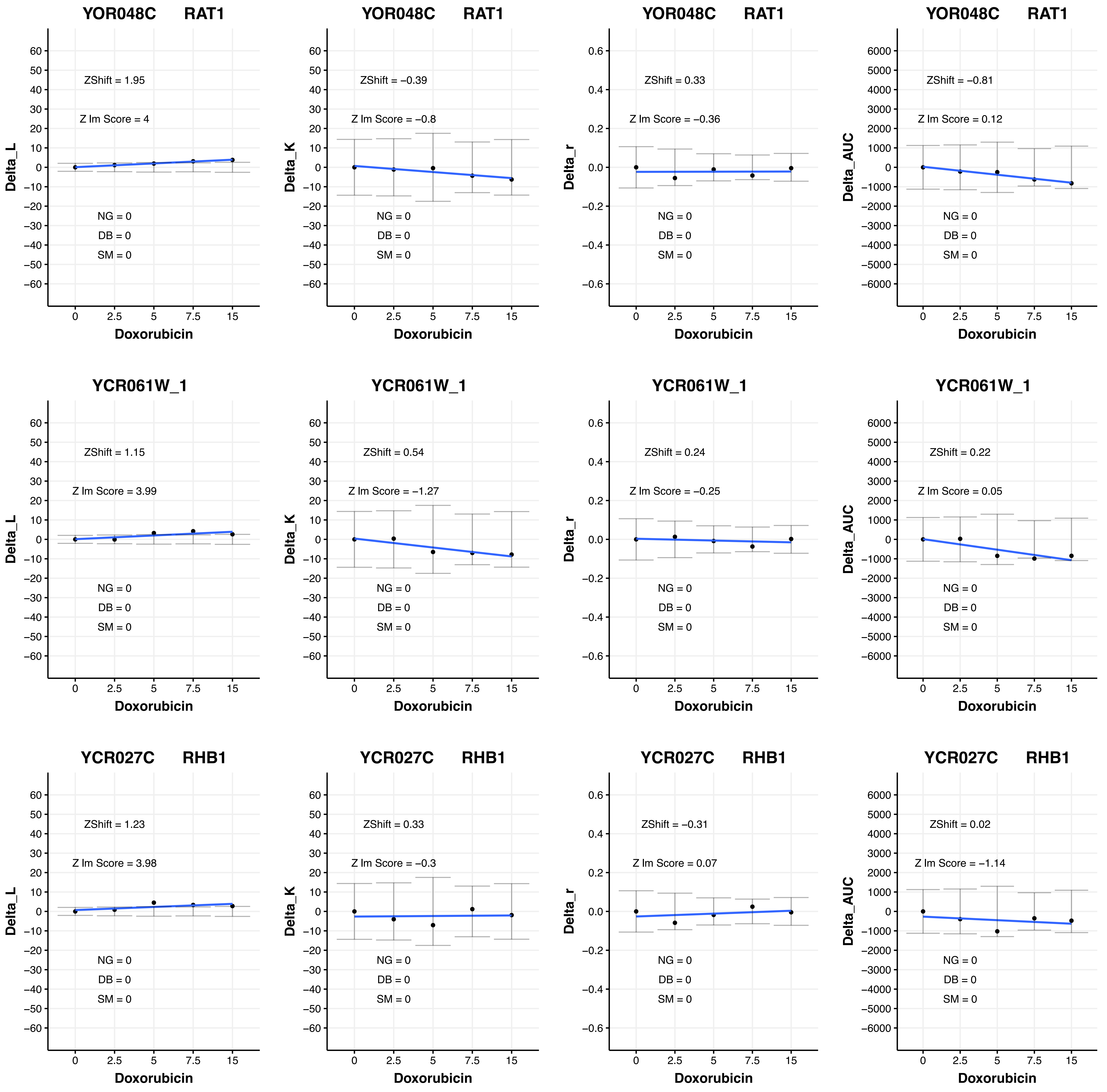

YOR048C RAT1
ZShift = 1.95
Z lm Score = 4
NG = 0
DB = 0
SM = 0
Delta_L
Doxorubicin
YOR048C RAT1
ZShift = −0.39
Z lm Score = −0.8
NG = 0
DB = 0
SM = 0
Delta_K
Doxorubicin
YOR048C RAT1
ZShift = 0.33
Z lm Score = −0.36
NG = 0
DB = 0
SM = 0
Delta_r
Doxorubicin
YOR048C RAT1
ZShift = −0.81
Z lm Score = 0.12
NG = 0
DB = 0
SM = 0
Delta_AUC
Doxorubicin
YCR061W_1
ZShift = 1.15
Z lm Score = 3.99
NG = 0
DB = 0
SM = 0
Delta_L
Doxorubicin
YCR061W_1
ZShift = 0.54
Z lm Score = −1.27
NG = 0
DB = 0
SM = 0
Delta_K
Doxorubicin
YCR061W_1
ZShift = 0.24
Z lm Score = −0.25
NG = 0
DB = 0
SM = 0
Delta_r
Doxorubicin
YCR061W_1
ZShift = 0.22
Z lm Score = 0.05
NG = 0
DB = 0
SM = 0
Delta_AUC
Doxorubicin
YCR027C RHB1
ZShift = 1.23
Z lm Score = 3.98
NG = 0
DB = 0
SM = 0
Delta_L
Doxorubicin
YCR027C RHB1
ZShift = 0.33
Z lm Score = −0.3
NG = 0
DB = 0
SM = 0
Delta_K
Doxorubicin
YCR027C RHB1
ZShift = −0.31
Z lm Score = 0.07
NG = 0
DB = 0
SM = 0
Delta_r
Doxorubicin
YCR027C RHB1
ZShift = 0.02
Z lm Score = −1.14
NG = 0
DB = 0
SM = 0
Delta_AUC
Doxorubicin

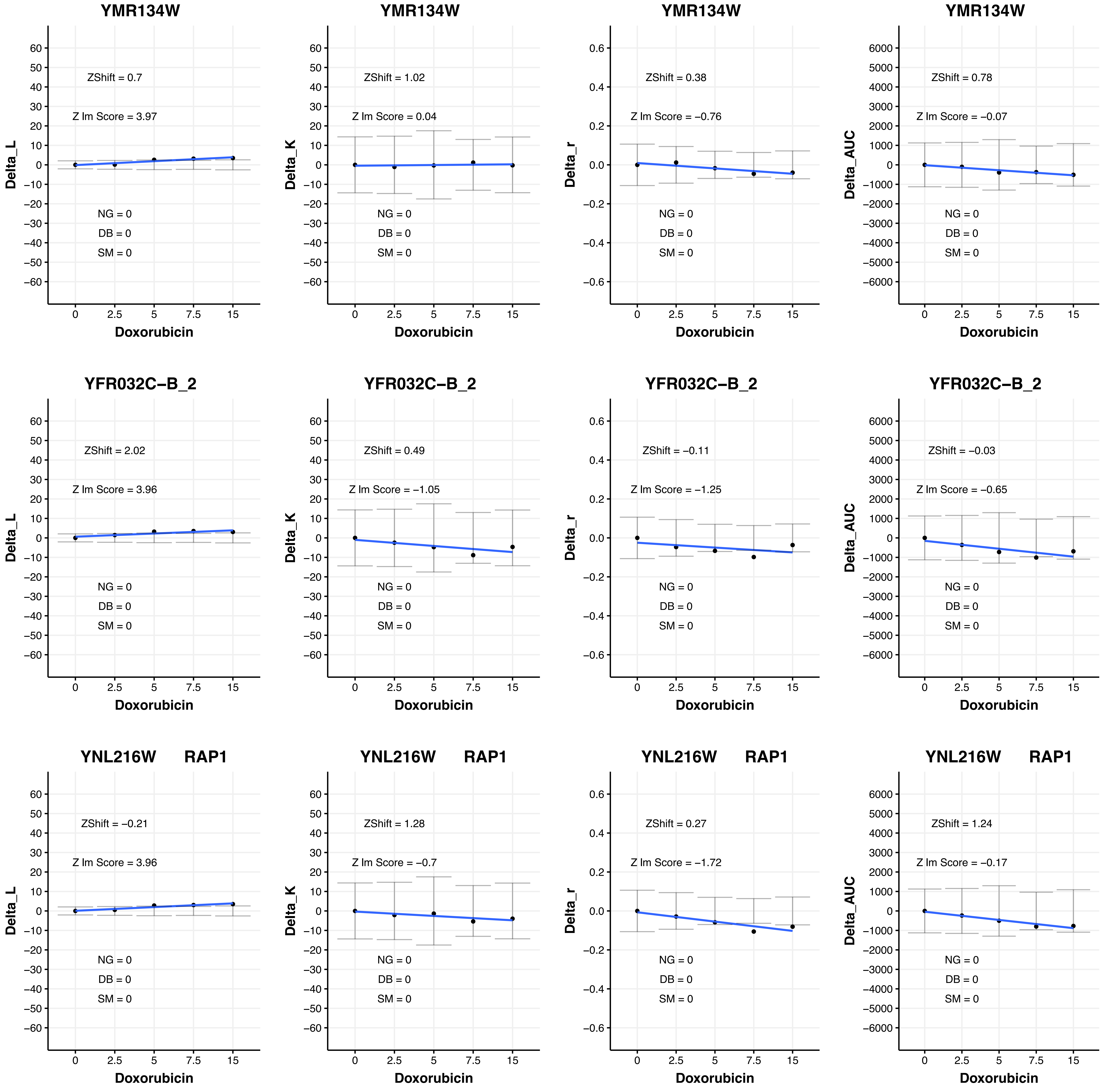
YMR134W
ZShift = 0.7
Z lm Score = 3.97
NG = 0
DB = 0
SM = 0
Delta_L
Doxorubicin
YMR134W
ZShift = 1.02
Z lm Score = 0.04
NG = 0
DB = 0
SM = 0
Delta_K
Doxorubicin
YMR134W
ZShift = 0.38
Z lm Score = −0.76
NG = 0
DB = 0
SM = 0
Delta_r
Doxorubicin
YMR134W
ZShift = 0.78
Z lm Score = −0.07
NG = 0
DB = 0
SM = 0
Delta_AUC
Doxorubicin
YFR032C−B_2
ZShift = 2.02
Z lm Score = 3.96
NG = 0
DB = 0
SM = 0
Delta_L
Doxorubicin
YFR032C−B_2
ZShift = 0.49
Z lm Score = −1.05
NG = 0
DB = 0
SM = 0
Delta_K
Doxorubicin
YFR032C−B_2
ZShift = −0.11
Z lm Score = −1.25
NG = 0
DB = 0
SM = 0
Delta_r
Doxorubicin
YFR032C−B_2
ZShift = −0.03
Z lm Score = −0.65
NG = 0
DB = 0
SM = 0
Delta_AUC
Doxorubicin
YNL216W RAP1
ZShift = −0.21
Z lm Score = 3.96
NG = 0
DB = 0
SM = 0
Delta_L
Doxorubicin
YNL216W RAP1
ZShift = 1.28
Z lm Score = −0.7
NG = 0
DB = 0
SM = 0
Delta_K
Doxorubicin
YNL216W RAP1
ZShift = 0.27
Z lm Score = −1.72
NG = 0
DB = 0
SM = 0
Delta_r
Doxorubicin
YNL216W RAP1
ZShift = 1.24
Z lm Score = −0.17
NG = 0
DB = 0
SM = 0
Delta_AUC
Doxorubicin

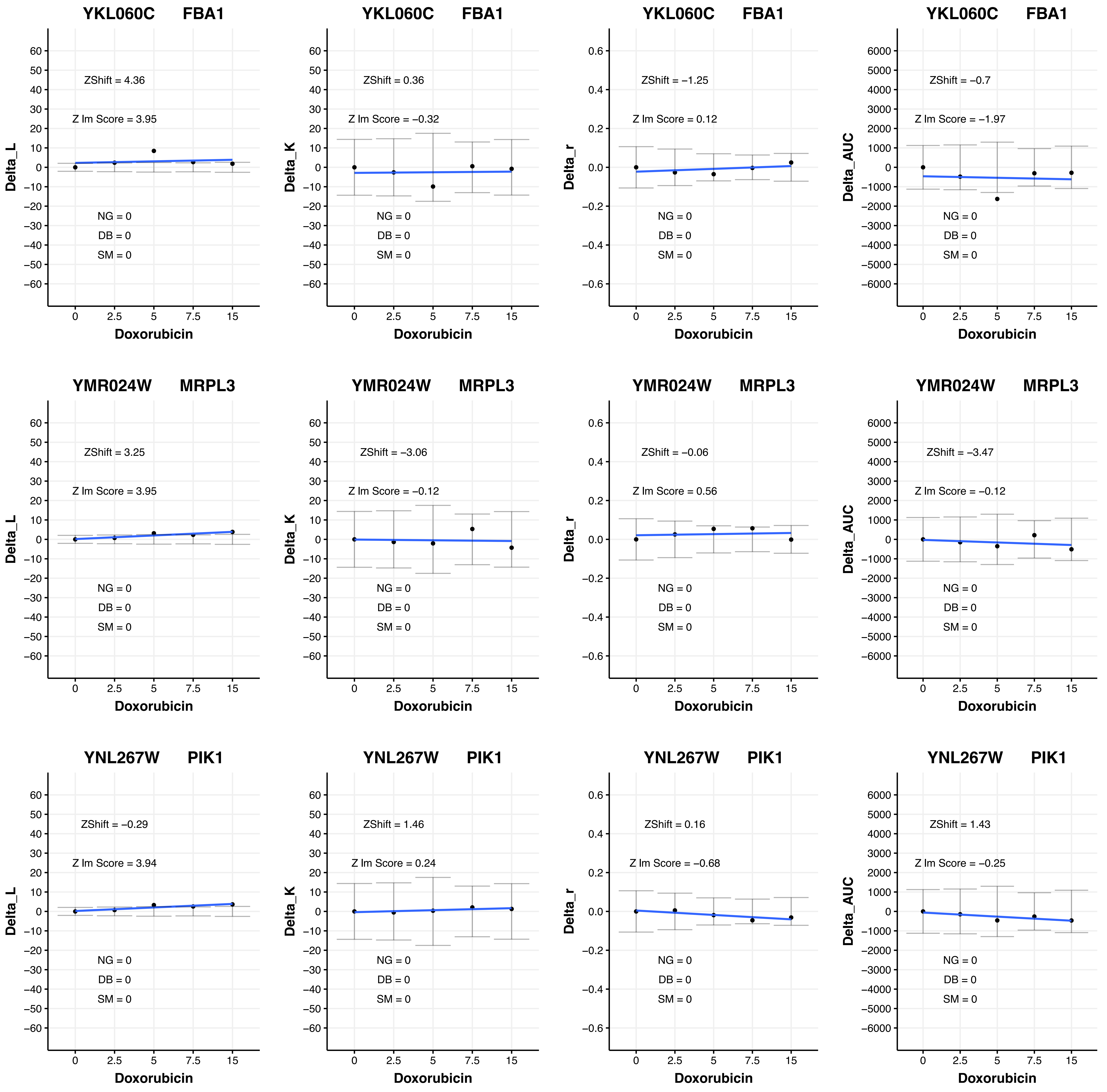
YKL060C FBA1
ZShift = 4.36
Z lm Score = 3.95
NG = 0
DB = 0
SM = 0
Delta_L
Doxorubicin
YKL060C FBA1
ZShift = 0.36
Z lm Score = −0.32
NG = 0
DB = 0
SM = 0
Delta_K
Doxorubicin
YKL060C FBA1
ZShift = −1.25
Z lm Score = 0.12
NG = 0
DB = 0
SM = 0
Delta_r
Doxorubicin
YKL060C FBA1
ZShift = −0.7
Z lm Score = −1.97
NG = 0
DB = 0
SM = 0
Delta_AUC
Doxorubicin
YMR024W MRPL3
ZShift = 3.25
Z lm Score = 3.95
NG = 0
DB = 0
SM = 0
Delta_L
Doxorubicin
YMR024W MRPL3
ZShift = −3.06
Z lm Score = −0.12
NG = 0
DB = 0
SM = 0
Delta_K
Doxorubicin
YMR024W MRPL3
ZShift = −0.06
Z lm Score = 0.56
NG = 0
DB = 0
SM = 0
Delta_r
Doxorubicin
YMR024W MRPL3
ZShift = −3.47
Z lm Score = −0.12
NG = 0
DB = 0
SM = 0
Delta_AUC
Doxorubicin
YNL267W PIK1
ZShift = −0.29
Z lm Score = 3.94
NG = 0
DB = 0
SM = 0
Delta_L
Doxorubicin
YNL267W PIK1
ZShift = 1.46
Z lm Score = 0.24
NG = 0
DB = 0
SM = 0
Delta_K
Doxorubicin
YNL267W PIK1
ZShift = 0.16
Z lm Score = −0.68
NG = 0
DB = 0
SM = 0
Delta_r
Doxorubicin
YNL267W PIK1
ZShift = 1.43
Z lm Score = −0.25
NG = 0
DB = 0
SM = 0
Delta_AUC
Doxorubicin

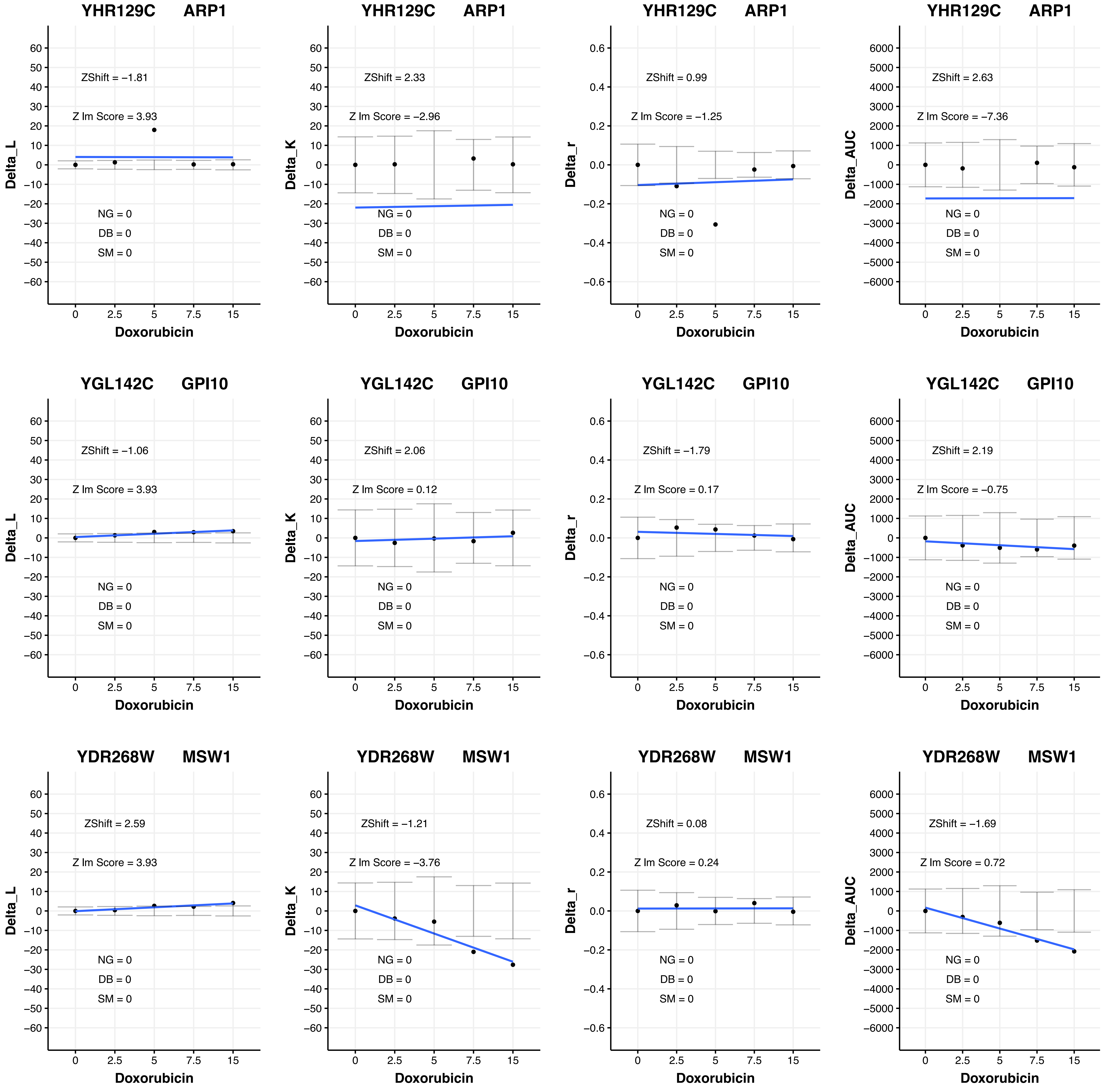

YHR129C ARP1
ZShift = −1.81
Z lm Score = 3.93
NG = 0
DB = 0
SM = 0
Delta_L
Doxorubicin
YHR129C ARP1
ZShift = 2.33
Z lm Score = −2.96
NG = 0
DB = 0
SM = 0
Delta_K
Doxorubicin
YHR129C ARP1
ZShift = 0.99
Z lm Score = −1.25
NG = 0
DB = 0
SM = 0
Delta_r
Doxorubicin
YHR129C ARP1
ZShift = 2.63
Z lm Score = −7.36
NG = 0
DB = 0
SM = 0
Delta_AUC
Doxorubicin
YGL142C GPI10
ZShift = −1.06
Z lm Score = 3.93
NG = 0
DB = 0
SM = 0
Delta_L
Doxorubicin
YGL142C GPI10
ZShift = 2.06
Z lm Score = 0.12
NG = 0
DB = 0
SM = 0
Delta_K
Doxorubicin
YGL142C GPI10
ZShift = −1.79
Z lm Score = 0.17
NG = 0
DB = 0
SM = 0
Delta_r
Doxorubicin
YGL142C GPI10
ZShift = 2.19
Z lm Score = −0.75
NG = 0
DB = 0
SM = 0
Delta_AUC
Doxorubicin
YDR268W MSW1
ZShift = 2.59
Z lm Score = 3.93
NG = 0
DB = 0
SM = 0
Delta_L
Doxorubicin
YDR268W MSW1
ZShift = −1.21
Z lm Score = −3.76
NG = 0
DB = 0
SM = 0
Delta_K
Doxorubicin
YDR268W MSW1
ZShift = 0.08
Z lm Score = 0.24
NG = 0
DB = 0
SM = 0
Delta_r
Doxorubicin
YDR268W MSW1
ZShift = −1.69
Z lm Score = 0.72
NG = 0
DB = 0
SM = 0
Delta_AUC
Doxorubicin

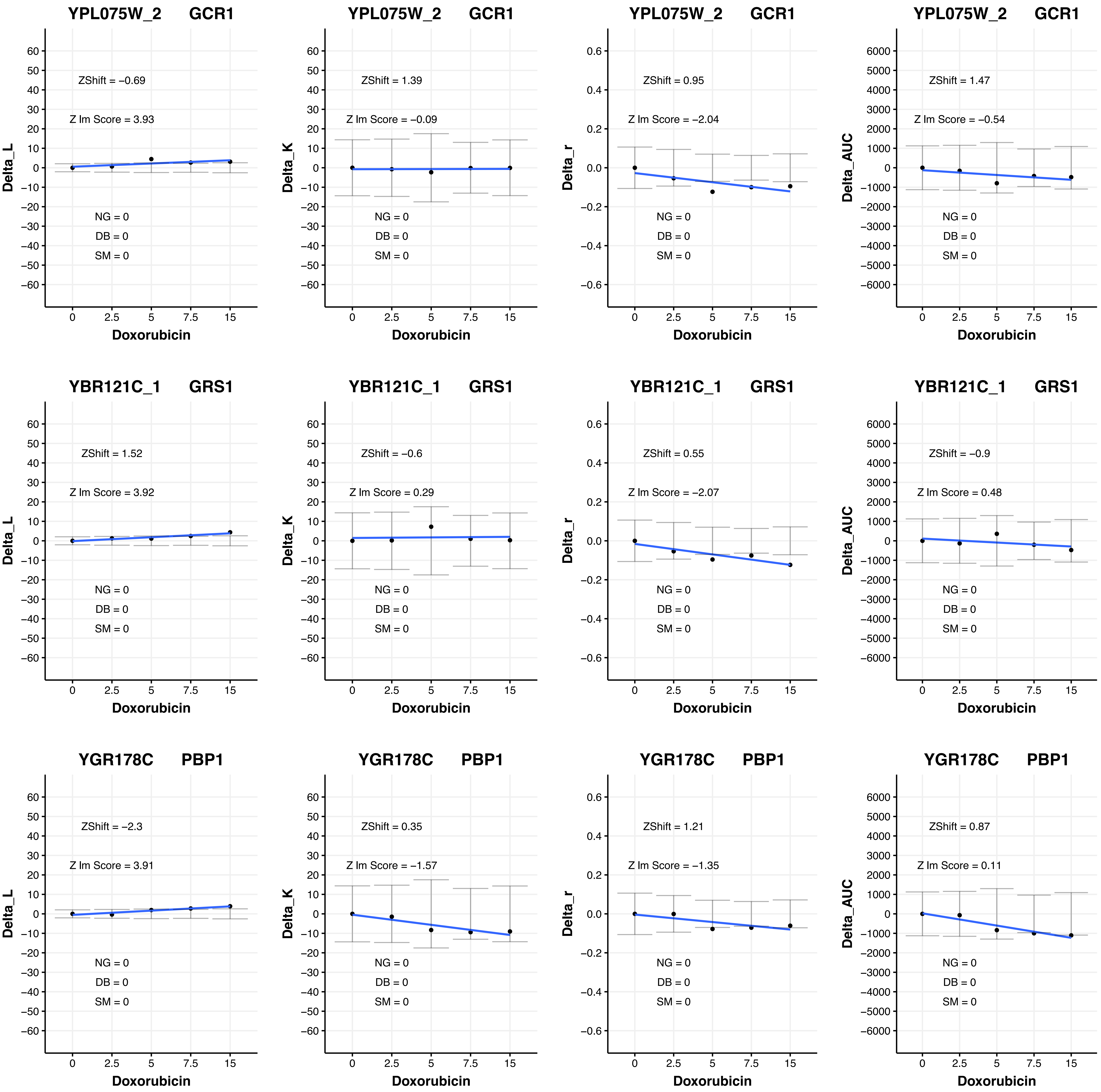

YPL075W_2 GCR1
ZShift = −0.69
Z lm Score = 3.93
NG = 0
DB = 0
SM = 0
Delta_L
Doxorubicin
YPL075W_2 GCR1
ZShift = 1.39
Z lm Score = −0.09
NG = 0
DB = 0
SM = 0
Delta_K
Doxorubicin
YPL075W_2 GCR1
ZShift = 0.95
Z lm Score = −2.04
NG = 0
DB = 0
SM = 0
Delta_r
Doxorubicin
YPL075W_2 GCR1
ZShift = 1.47
Z lm Score = −0.54
NG = 0
DB = 0
SM = 0
Delta_AUC
Doxorubicin
YBR121C_1 GRS1
ZShift = 1.52
Z lm Score = 3.92
NG = 0
DB = 0
SM = 0
Delta_L
Doxorubicin
YBR121C_1 GRS1
ZShift = −0.6
Z lm Score = 0.29
NG = 0
DB = 0
SM = 0
Delta_K
Doxorubicin
YBR121C_1 GRS1
ZShift = 0.55
Z lm Score = −2.07
NG = 0
DB = 0
SM = 0
Delta_r
Doxorubicin
YBR121C_1 GRS1
ZShift = −0.9
Z lm Score = 0.48
NG = 0
DB = 0
SM = 0
Delta_AUC
Doxorubicin
YGR178C PBP1
ZShift = −2.3
Z lm Score = 3.91
NG = 0
DB = 0
SM = 0
Delta_L
Doxorubicin
YGR178C PBP1
ZShift = 0.35
Z lm Score = −1.57
NG = 0
DB = 0
SM = 0
Delta_K
Doxorubicin
YGR178C PBP1
ZShift = 1.21
Z lm Score = −1.35
NG = 0
DB = 0
SM = 0
Delta_r
Doxorubicin
YGR178C PBP1
ZShift = 0.87
Z lm Score = 0.11
NG = 0
DB = 0
SM = 0
Delta_AUC
Doxorubicin

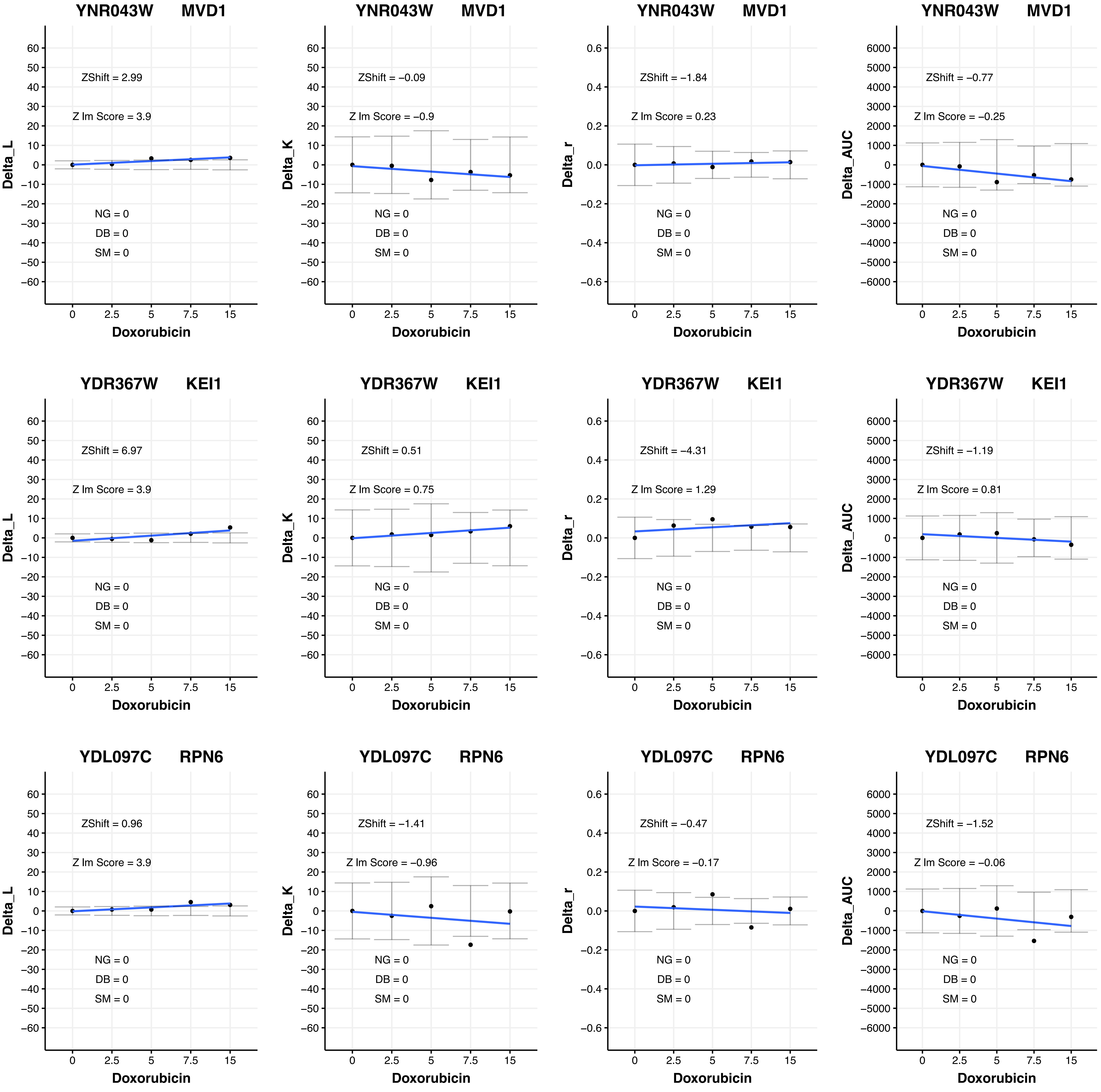
YNR043W MVD1
ZShift = 2.99
Z lm Score = 3.9
Delta_L
NG = 0
DB = 0
SM = 0
Doxorubicin
YNR043W MVD1
ZShift = −0.09
Z lm Score = −0.9
Delta_K
NG = 0
DB = 0
SM = 0
Doxorubicin
YNR043W MVD1
ZShift = −1.84
Z lm Score = 0.23
Delta_r
NG = 0
DB = 0
SM = 0
Doxorubicin
YNR043W MVD1
ZShift = −0.77
Z lm Score = −0.25
Delta_AUC
NG = 0
DB = 0
SM = 0
Doxorubicin
YDR367W KEI1
ZShift = 6.97
Z lm Score = 3.9
Delta_L
NG = 0
DB = 0
SM = 0
Doxorubicin
YDR367W KEI1
ZShift = 0.51
Z lm Score = 0.75
Delta_K
NG = 0
DB = 0
SM = 0
Doxorubicin
YDR367W KEI1
ZShift = −4.31
Z lm Score = 1.29
Delta_r
NG = 0
DB = 0
SM = 0
Doxorubicin
YDR367W KEI1
ZShift = −1.19
Z lm Score = 0.81
Delta_AUC
NG = 0
DB = 0
SM = 0
Doxorubicin
YDL097C RPN6
ZShift = 0.96
Z lm Score = 3.9
Delta_L
NG = 0
DB = 0
SM = 0
Doxorubicin
YDL097C RPN6
ZShift = −1.41
Z lm Score = −0.96
Delta_K
NG = 0
DB = 0
SM = 0
Doxorubicin
YDL097C RPN6
ZShift = −0.47
Z lm Score = −0.17
Delta_r
NG = 0
DB = 0
SM = 0
Doxorubicin
YDL097C RPN6
ZShift = −1.52
Z lm Score = −0.06
Delta_AUC
NG = 0
DB = 0
SM = 0
Doxorubicin

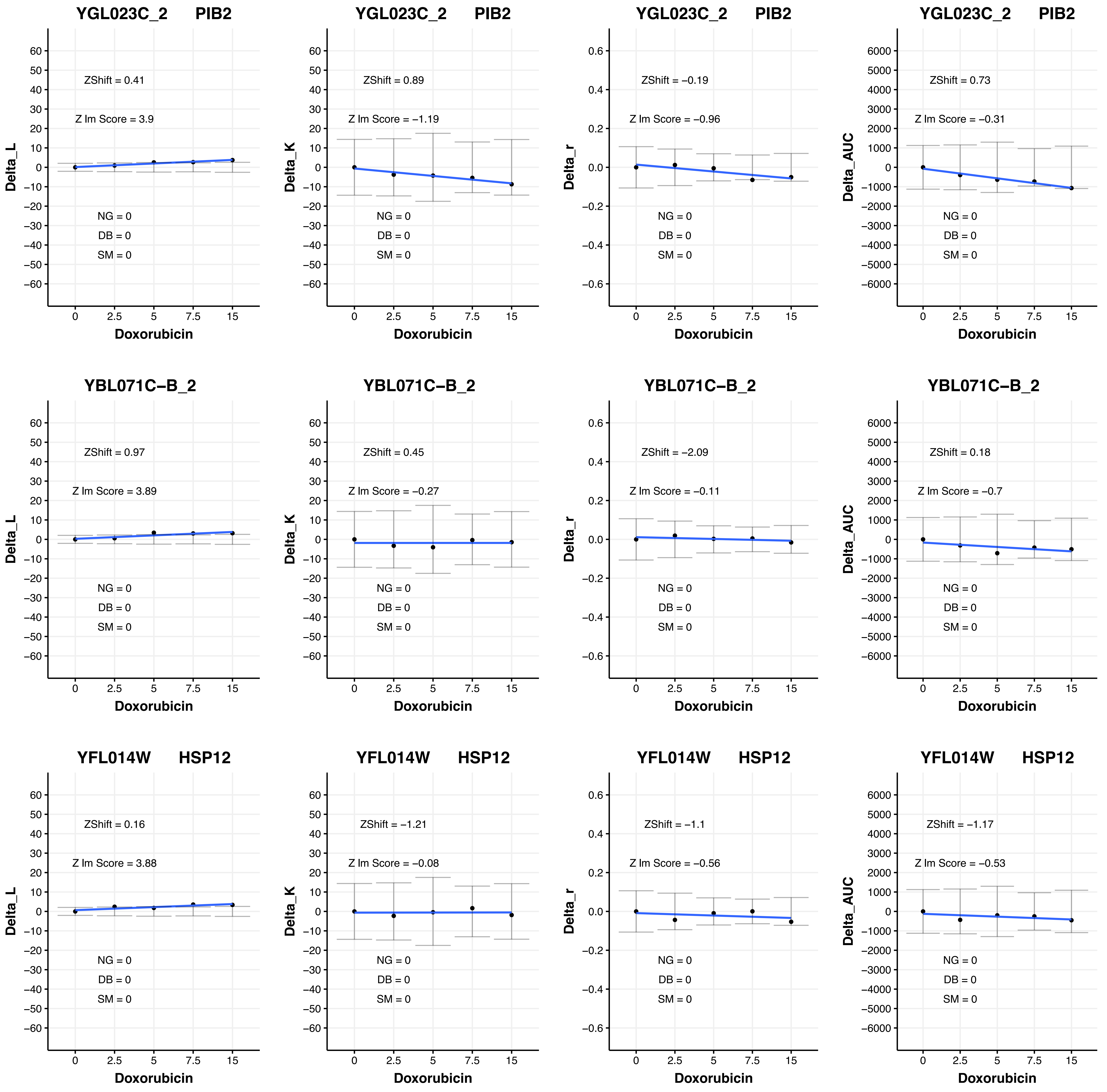

YGL023C_2 PIB2
ZShift = 0.41
Z lm Score = 3.9
NG = 0
DB = 0
SM = 0
Delta_L
Doxorubicin
YGL023C_2 PIB2
ZShift = 0.89
Z lm Score = −1.19
Delta_K
YGL023C_2 PIB2
ZShift = −0.19
Z lm Score = −0.96
Delta_r
YGL023C_2 PIB2
ZShift = 0.73
Z lm Score = −0.31
Delta_AUC
YBL071C−B_2
ZShift = 0.97
Z lm Score = 3.89
YBL071C−B_2
ZShift = 0.45
Z lm Score = −0.27
YBL071C−B_2
ZShift = −2.09
Z lm Score = −0.11
YBL071C−B_2
ZShift = 0.18
Z lm Score = −0.7
YFL014W HSP12
ZShift = 0.16
Z lm Score = 3.88
YFL014W HSP12
ZShift = −1.21
Z lm Score = −0.08
YFL014W HSP12
ZShift = −1.1
Z lm Score = −0.56
YFL014W HSP12
ZShift = −1.17
Z lm Score = −0.53

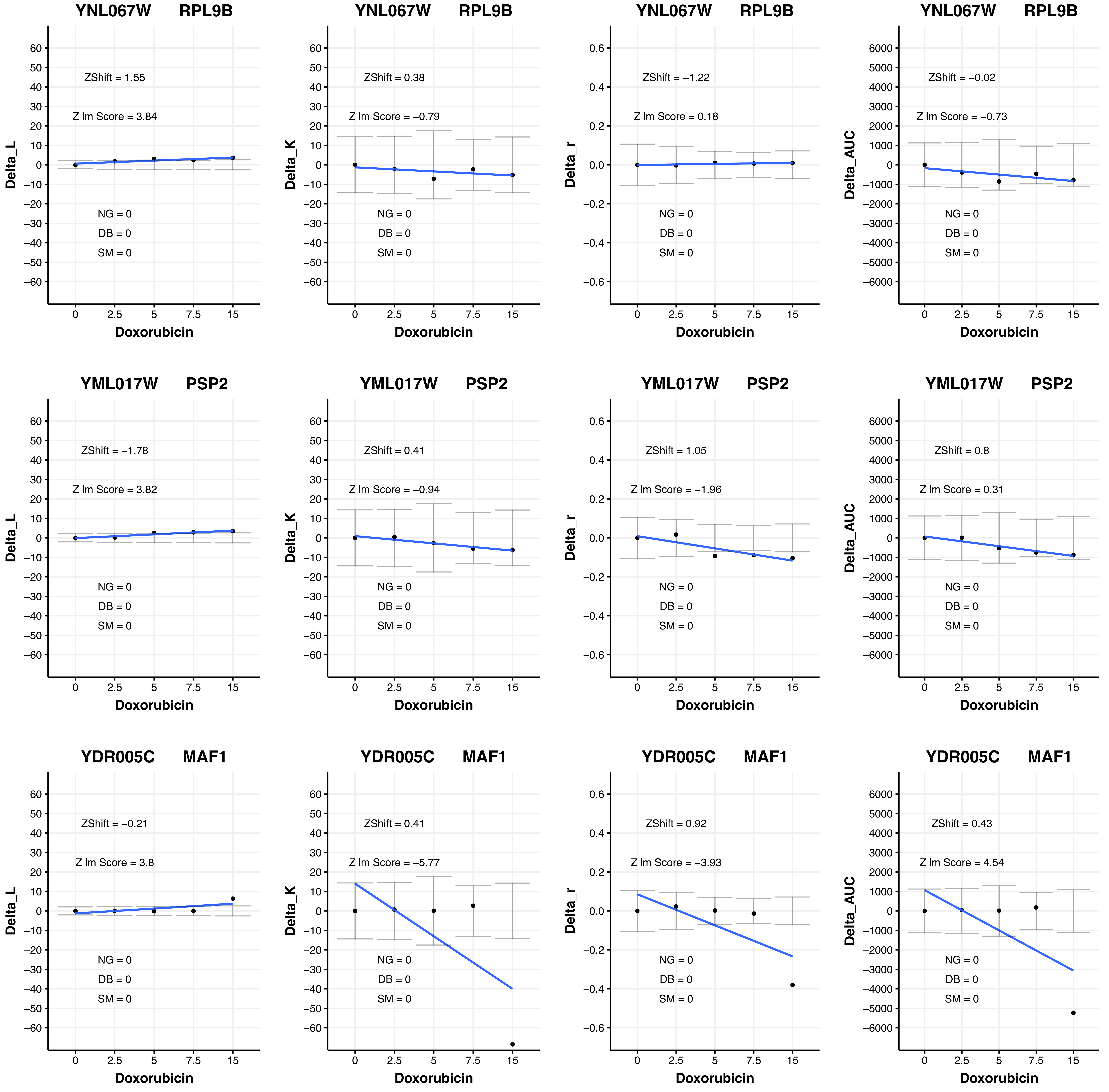

YNL067W RPL9B
ZShift = 1.55
Z lm Score = 3.84
NG = 0
DB = 0
SM = 0
Delta_L
Doxorubicin
YNL067W RPL9B
ZShift = 0.38
Z lm Score = −0.79
NG = 0
DB = 0
SM = 0
Delta_K
Doxorubicin
YNL067W RPL9B
ZShift = −1.22
Z lm Score = 0.18
NG = 0
DB = 0
SM = 0
Delta_r
Doxorubicin
YNL067W RPL9B
ZShift = −0.02
Z lm Score = −0.73
NG = 0
DB = 0
SM = 0
Delta_AUC
Doxorubicin
YML017W PSP2
ZShift = −1.78
Z lm Score = 3.82
NG = 0
DB = 0
SM = 0
Delta_L
Doxorubicin
YML017W PSP2
ZShift = 0.41
Z lm Score = −0.94
NG = 0
DB = 0
SM = 0
Delta_K
Doxorubicin
YML017W PSP2
ZShift = 1.05
Z lm Score = −1.96
NG = 0
DB = 0
SM = 0
Delta_r
Doxorubicin
YML017W PSP2
ZShift = 0.8
Z lm Score = 0.31
NG = 0
DB = 0
SM = 0
Delta_AUC
Doxorubicin
YDR005C MAF1
ZShift = −0.21
Z lm Score = 3.8
NG = 0
DB = 0
SM = 0
Delta_L
Doxorubicin
YDR005C MAF1
ZShift = 0.41
Z lm Score = −5.77
NG = 0
DB = 0
SM = 0
Delta_K
Doxorubicin
YDR005C MAF1
ZShift = 0.92
Z lm Score = −3.93
NG = 0
DB = 0
SM = 0
Delta_r
Doxorubicin
YDR005C MAF1
ZShift = 0.43
Z lm Score = 4.54
NG = 0
DB = 0
SM = 0
Delta_AUC
Doxorubicin

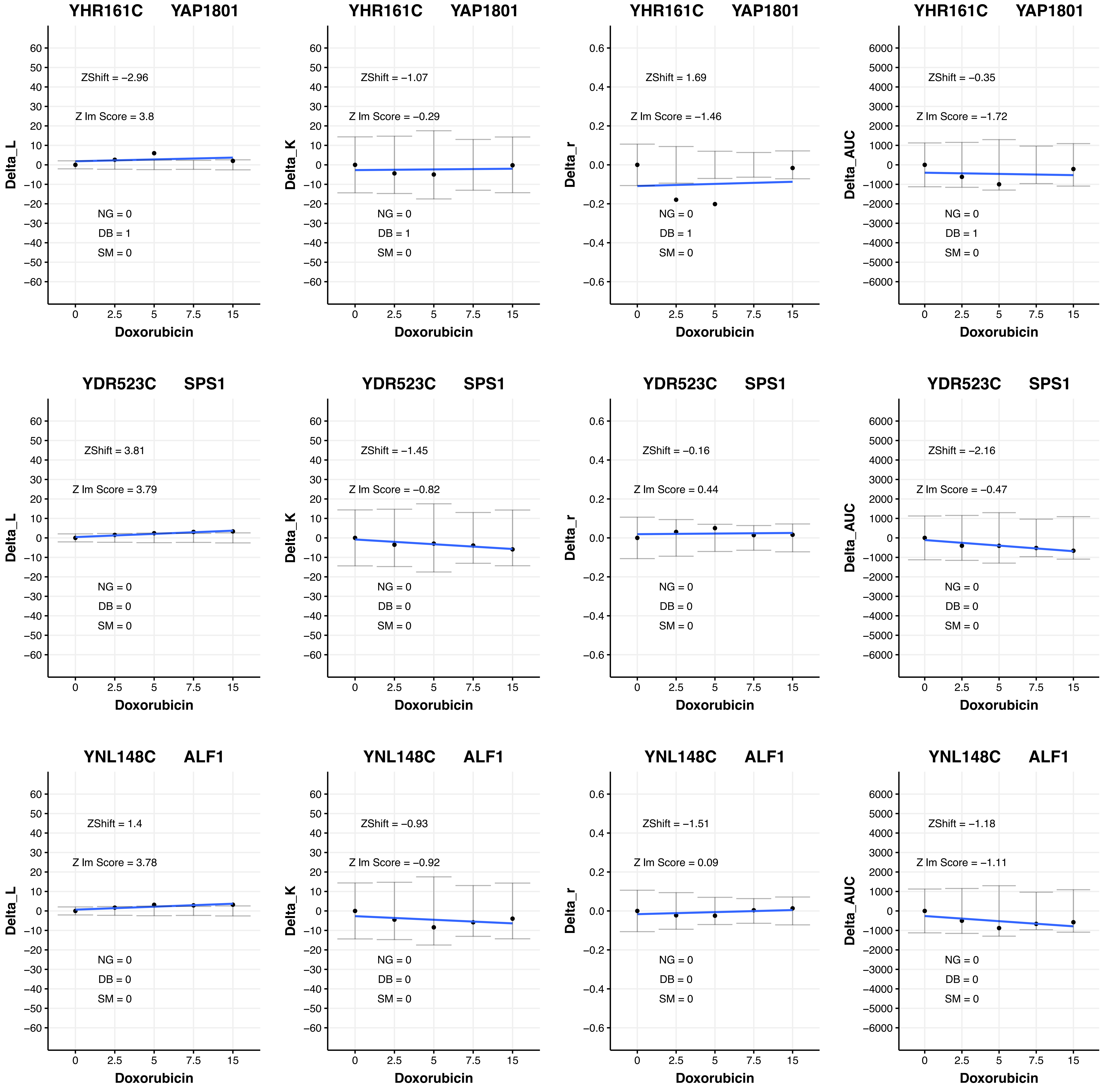

YHR161C YAP1801
ZShift = −2.96
Z lm Score = 3.8
NG = 0
DB = 1
SM = 0
Delta_L
Doxorubicin
YHR161C YAP1801
ZShift = −1.07
Z lm Score = −0.29
NG = 0
DB = 1
SM = 0
Delta_K
Doxorubicin
YHR161C YAP1801
ZShift = 1.69
Z lm Score = −1.46
NG = 0
DB = 1
SM = 0
Delta_r
Doxorubicin
YHR161C YAP1801
ZShift = −0.35
Z lm Score = −1.72
NG = 0
DB = 1
SM = 0
Delta_AUC
Doxorubicin
YDR523C SPS1
ZShift = 3.81
Z lm Score = 3.79
NG = 0
DB = 0
SM = 0
Delta_L
Doxorubicin
YDR523C SPS1
ZShift = −1.45
Z lm Score = −0.82
NG = 0
DB = 0
SM = 0
Delta_K
Doxorubicin
YDR523C SPS1
ZShift = −0.16
Z lm Score = 0.44
NG = 0
DB = 0
SM = 0
Delta_r
Doxorubicin
YDR523C SPS1
ZShift = −2.16
Z lm Score = −0.47
NG = 0
DB = 0
SM = 0
Delta_AUC
Doxorubicin
YNL148C ALF1
ZShift = 1.4
Z lm Score = 3.78
NG = 0
DB = 0
SM = 0
Delta_L
Doxorubicin
YNL148C ALF1
ZShift = −0.93
Z lm Score = −0.92
NG = 0
DB = 0
SM = 0
Delta_K
Doxorubicin
YNL148C ALF1
ZShift = −1.51
Z lm Score = 0.09
NG = 0
DB = 0
SM = 0
Delta_r
Doxorubicin
YNL148C ALF1
ZShift = −1.18
Z lm Score = −1.11
NG = 0
DB = 0
SM = 0
Delta_AUC
Doxorubicin

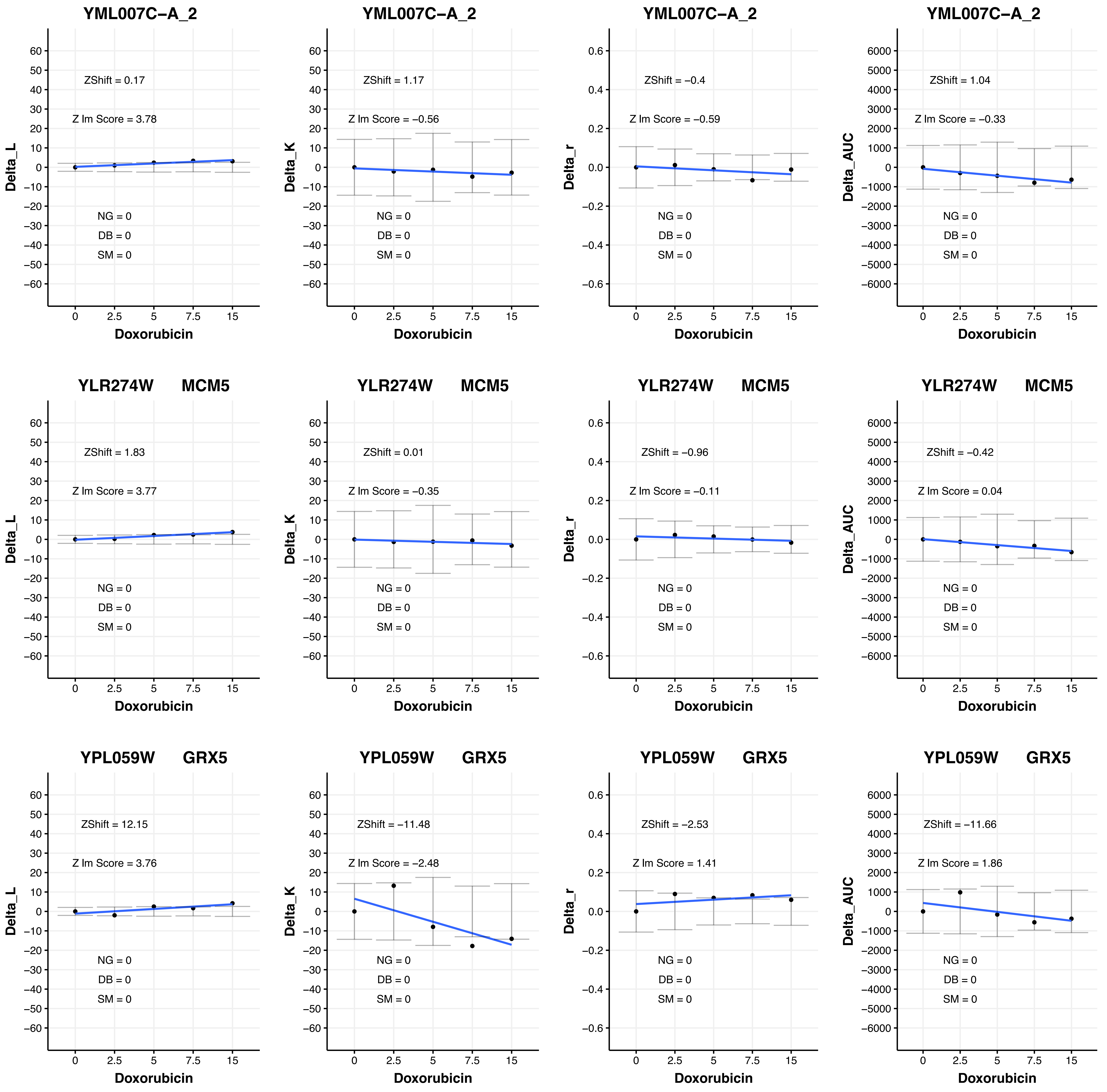

YML007C−A_2
ZShift = 0.17
Z lm Score = 3.78
NG = 0
DB = 0
SM = 0
Delta_L
Doxorubicin
YML007C−A_2
ZShift = 1.17
Z lm Score = −0.56
NG = 0
DB = 0
SM = 0
Delta_K
Doxorubicin
YML007C−A_2
ZShift = −0.4
Z lm Score = −0.59
NG = 0
DB = 0
SM = 0
Delta_r
Doxorubicin
YML007C−A_2
ZShift = 1.04
Z lm Score = −0.33
NG = 0
DB = 0
SM = 0
Delta_AUC
Doxorubicin
YLR274W MCM5
ZShift = 1.83
Z lm Score = 3.77
NG = 0
DB = 0
SM = 0
Delta_L
Doxorubicin
YLR274W MCM5
ZShift = 0.01
Z lm Score = −0.35
NG = 0
DB = 0
SM = 0
Delta_K
Doxorubicin
YLR274W MCM5
ZShift = −0.96
Z lm Score = −0.11
NG = 0
DB = 0
SM = 0
Delta_r
Doxorubicin
YLR274W MCM5
ZShift = −0.42
Z lm Score = 0.04
NG = 0
DB = 0
SM = 0
Delta_AUC
Doxorubicin
YPL059W GRX5
ZShift = 12.15
Z lm Score = 3.76
NG = 0
DB = 0
SM = 0
Delta_L
Doxorubicin
YPL059W GRX5
ZShift = −11.48
Z lm Score = −2.48
NG = 0
DB = 0
SM = 0
Delta_K
Doxorubicin
YPL059W GRX5
ZShift = −2.53
Z lm Score = 1.41
NG = 0
DB = 0
SM = 0
Delta_r
Doxorubicin
YPL059W GRX5
ZShift = −11.66
Z lm Score = 1.86
NG = 0
DB = 0
SM = 0
Delta_AUC
Doxorubicin

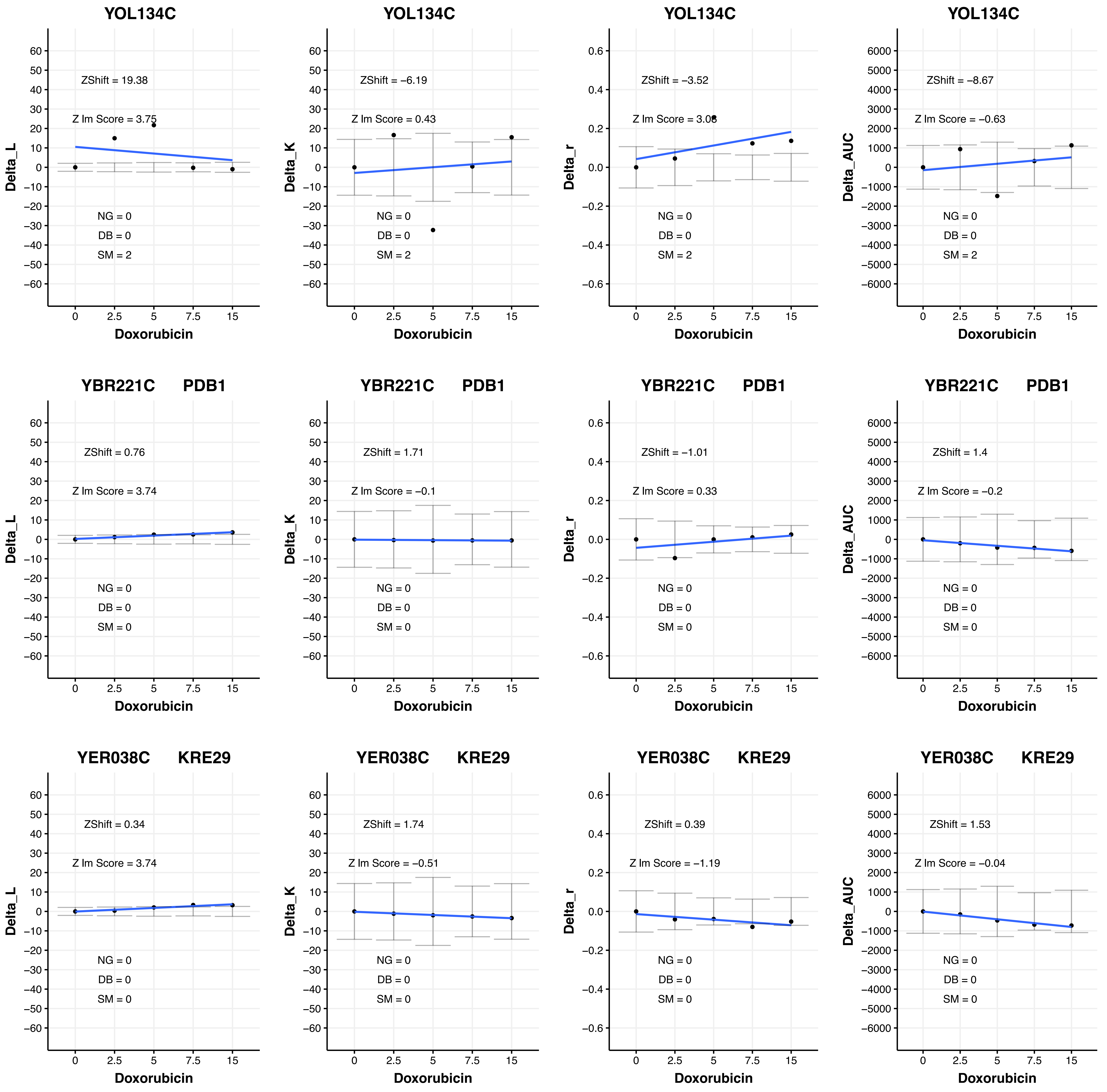
YOL134C
ZShift = 19.38
Z lm Score = 3.75
NG = 0
DB = 0
SM = 2
Delta_L
Doxorubicin
YOL134C
ZShift = −6.19
Z lm Score = 0.43
NG = 0
DB = 0
SM = 2
Delta_K
Doxorubicin
YOL134C
ZShift = −3.52
Z lm Score = 3.08
NG = 0
DB = 0
SM = 2
Delta_r
Doxorubicin
YOL134C
ZShift = −8.67
Z lm Score = −0.63
NG = 0
DB = 0
SM = 2
Delta_AUC
Doxorubicin
YBR221C PDB1
ZShift = 0.76
Z lm Score = 3.74
NG = 0
DB = 0
SM = 0
Delta_L
Doxorubicin
YBR221C PDB1
ZShift = 1.71
Z lm Score = −0.1
NG = 0
DB = 0
SM = 0
Delta_K
Doxorubicin
YBR221C PDB1
ZShift = −1.01
Z lm Score = 0.33
NG = 0
DB = 0
SM = 0
Delta_r
Doxorubicin
YBR221C PDB1
ZShift = 1.4
Z lm Score = −0.2
NG = 0
DB = 0
SM = 0
Delta_AUC
Doxorubicin
YER038C KRE29
ZShift = 0.34
Z lm Score = 3.74
NG = 0
DB = 0
SM = 0
Delta_L
Doxorubicin
YER038C KRE29
ZShift = 1.74
Z lm Score = −0.51
NG = 0
DB = 0
SM = 0
Delta_K
Doxorubicin
YER038C KRE29
ZShift = 0.39
Z lm Score = −1.19
NG = 0
DB = 0
SM = 0
Delta_r
Doxorubicin
YER038C KRE29
ZShift = 1.53
Z lm Score = −0.04
NG = 0
DB = 0
SM = 0
Delta_AUC
Doxorubicin

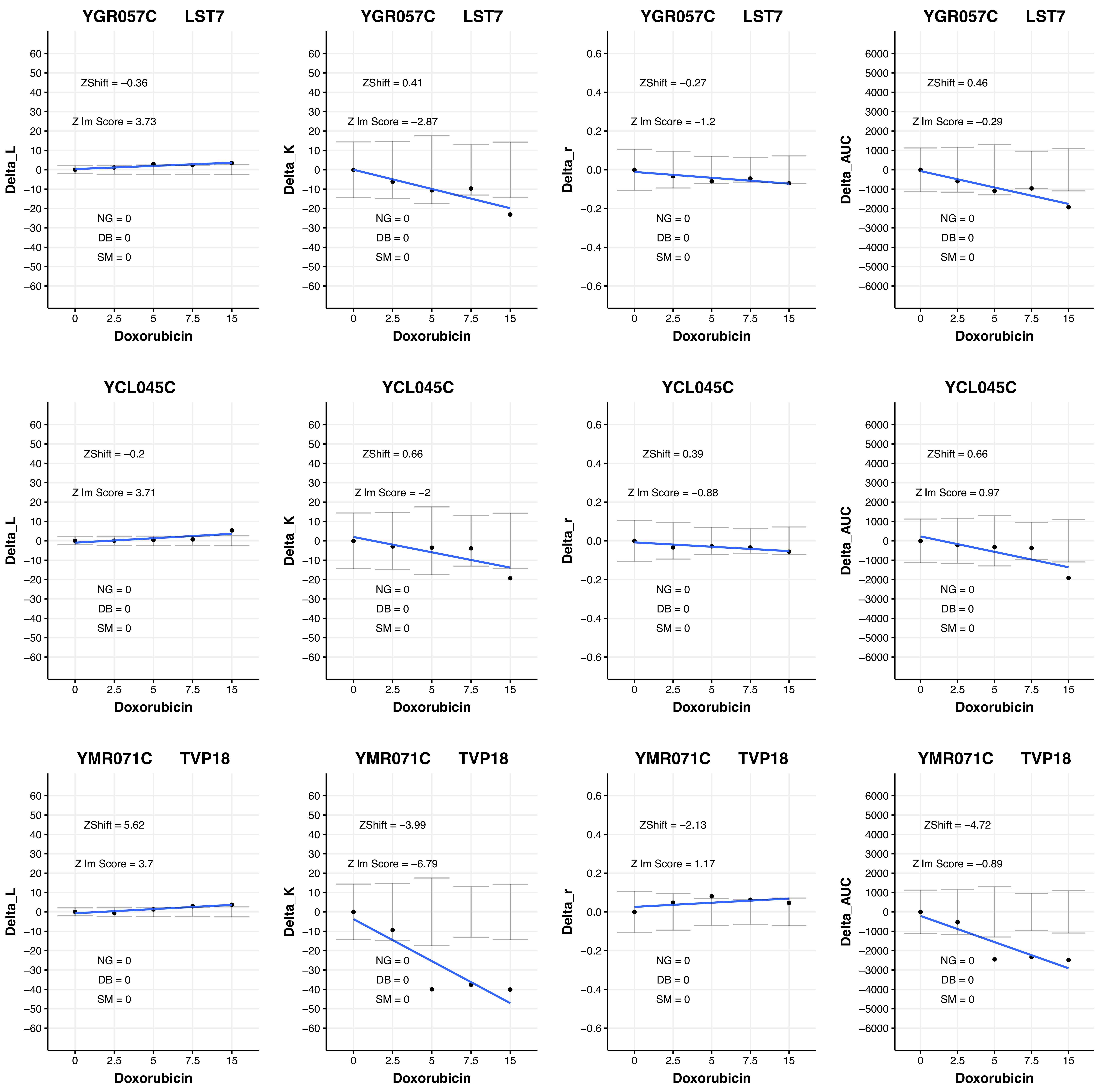
YGR057C LST7
ZShift = −0.36
Z lm Score = 3.73
NG = 0
DB = 0
SM = 0
Delta_L
Doxorubicin
YGR057C LST7
ZShift = 0.41
Z lm Score = −2.87
NG = 0
DB = 0
SM = 0
Delta_K
Doxorubicin
YGR057C LST7
ZShift = −0.27
Z lm Score = −1.2
NG = 0
DB = 0
SM = 0
Delta_r
Doxorubicin
YGR057C LST7
ZShift = 0.46
Z lm Score = −0.29
NG = 0
DB = 0
SM = 0
Delta_AUC
Doxorubicin
YCL045C
ZShift = −0.2
Z lm Score = 3.71
NG = 0
DB = 0
SM = 0
Delta_L
Doxorubicin
YCL045C
ZShift = 0.66
Z lm Score = −2
NG = 0
DB = 0
SM = 0
Delta_K
Doxorubicin
YCL045C
ZShift = 0.39
Z lm Score = −0.88
NG = 0
DB = 0
SM = 0
Delta_r
Doxorubicin
YCL045C
ZShift = 0.66
Z lm Score = 0.97
NG = 0
DB = 0
SM = 0
Delta_AUC
Doxorubicin
YMR071C TVP18
ZShift = 5.62
Z lm Score = 3.7
NG = 0
DB = 0
SM = 0
Delta_L
Doxorubicin
YMR071C TVP18
ZShift = −3.99
Z lm Score = −6.79
NG = 0
DB = 0
SM = 0
Delta_K
Doxorubicin
YMR071C TVP18
ZShift = −2.13
Z lm Score = 1.17
NG = 0
DB = 0
SM = 0
Delta_r
Doxorubicin
YMR071C TVP18
ZShift = −4.72
Z lm Score = −0.89
NG = 0
DB = 0
SM = 0
Delta_AUC
Doxorubicin

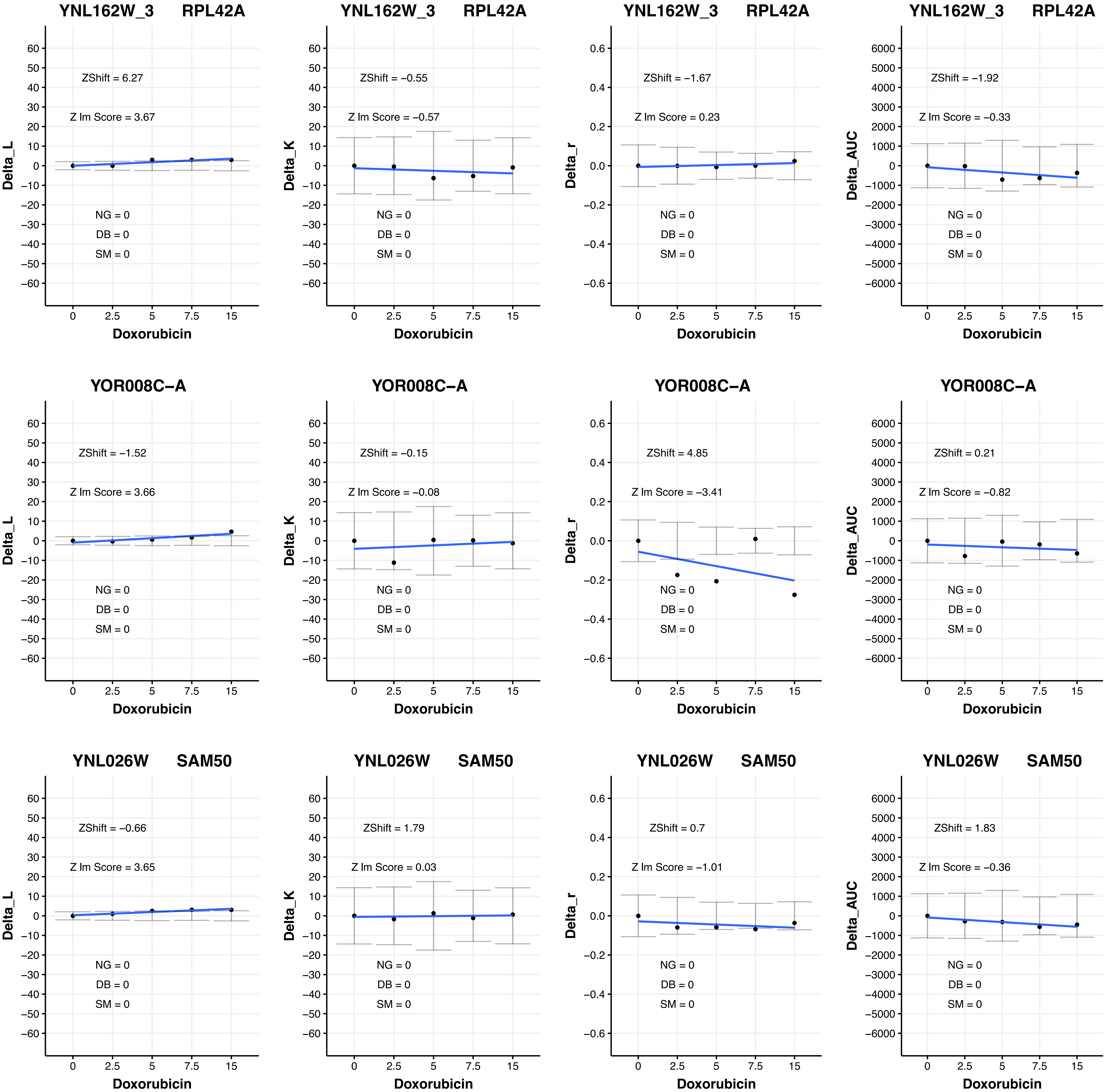
YNL162W_3 RPL42A
ZShift = 6.27
Z lm Score = 3.67
NG = 0
DB = 0
SM = 0
Delta_L
Doxorubicin
YNL162W_3 RPL42A
ZShift = −0.55
Z lm Score = −0.57
NG = 0
DB = 0
SM = 0
Delta_K
Doxorubicin
YNL162W_3 RPL42A
ZShift = −1.67
Z lm Score = 0.23
NG = 0
DB = 0
SM = 0
Delta_r
Doxorubicin
YNL162W_3 RPL42A
ZShift = −1.92
Z lm Score = −0.33
NG = 0
DB = 0
SM = 0
Delta_AUC
Doxorubicin
YOR008C−A
ZShift = −1.52
Z lm Score = 3.66
NG = 0
DB = 0
SM = 0
Delta_L
Doxorubicin
YOR008C−A
ZShift = −0.15
Z lm Score = −0.08
NG = 0
DB = 0
SM = 0
Delta_K
Doxorubicin
YOR008C−A
ZShift = 4.85
Z lm Score = −3.41
NG = 0
DB = 0
SM = 0
Delta_r
Doxorubicin
YOR008C−A
ZShift = 0.21
Z lm Score = −0.82
NG = 0
DB = 0
SM = 0
Delta_AUC
Doxorubicin
YNL026W SAM50
ZShift = −0.66
Z lm Score = 3.65
NG = 0
DB = 0
SM = 0
Delta_L
Doxorubicin
YNL026W SAM50
ZShift = 1.79
Z lm Score = 0.03
NG = 0
DB = 0
SM = 0
Delta_K
Doxorubicin
YNL026W SAM50
ZShift = 0.7
Z lm Score = −1.01
NG = 0
DB = 0
SM = 0
Delta_r
Doxorubicin
YNL026W SAM50
ZShift = 1.83
Z lm Score = −0.36
NG = 0
DB = 0
SM = 0
Delta_AUC
Doxorubicin

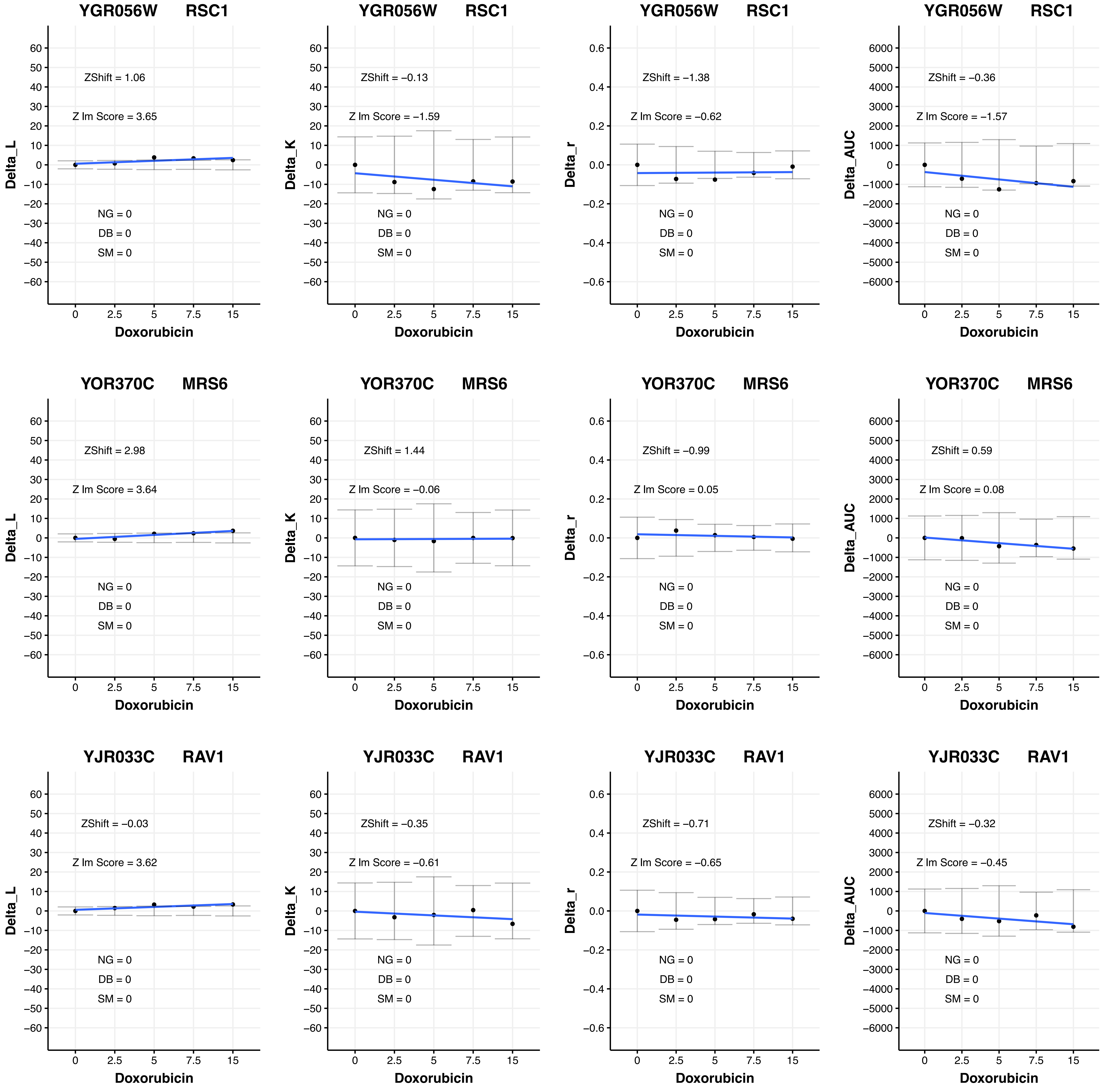

YGR056W RSC1
Delta_L
ZShift = 1.06
Z lm Score = 3.65
NG = 0
DB = 0
SM = 0
Doxorubicin
YGR056W RSC1
Delta_K
ZShift = −0.13
Z lm Score = −1.59
NG = 0
DB = 0
SM = 0
Doxorubicin
YGR056W RSC1
Delta_r
ZShift = −1.38
Z lm Score = −0.62
NG = 0
DB = 0
SM = 0
Doxorubicin
YGR056W RSC1
Delta_AUC
ZShift = −0.36
Z lm Score = −1.57
NG = 0
DB = 0
SM = 0
Doxorubicin
YOR370C MRS6
Delta_L
ZShift = 2.98
Z lm Score = 3.64
NG = 0
DB = 0
SM = 0
Doxorubicin
YOR370C MRS6
Delta_K
ZShift = 1.44
Z lm Score = −0.06
NG = 0
DB = 0
SM = 0
Doxorubicin
YOR370C MRS6
Delta_r
ZShift = −0.99
Z lm Score = 0.05
NG = 0
DB = 0
SM = 0
Doxorubicin
YOR370C MRS6
Delta_AUC
ZShift = 0.59
Z lm Score = 0.08
NG = 0
DB = 0
SM = 0
Doxorubicin
YJR033C RAV1
Delta_L
ZShift = −0.03
Z lm Score = 3.62
NG = 0
DB = 0
SM = 0
Doxorubicin
YJR033C RAV1
Delta_K
ZShift = −0.35
Z lm Score = −0.61
NG = 0
DB = 0
SM = 0
Doxorubicin
YJR033C RAV1
Delta_r
ZShift = −0.71
Z lm Score = −0.65
NG = 0
DB = 0
SM = 0
Doxorubicin
YJR033C RAV1
Delta_AUC
ZShift = −0.32
Z lm Score = −0.45
NG = 0
DB = 0
SM = 0
Doxorubicin

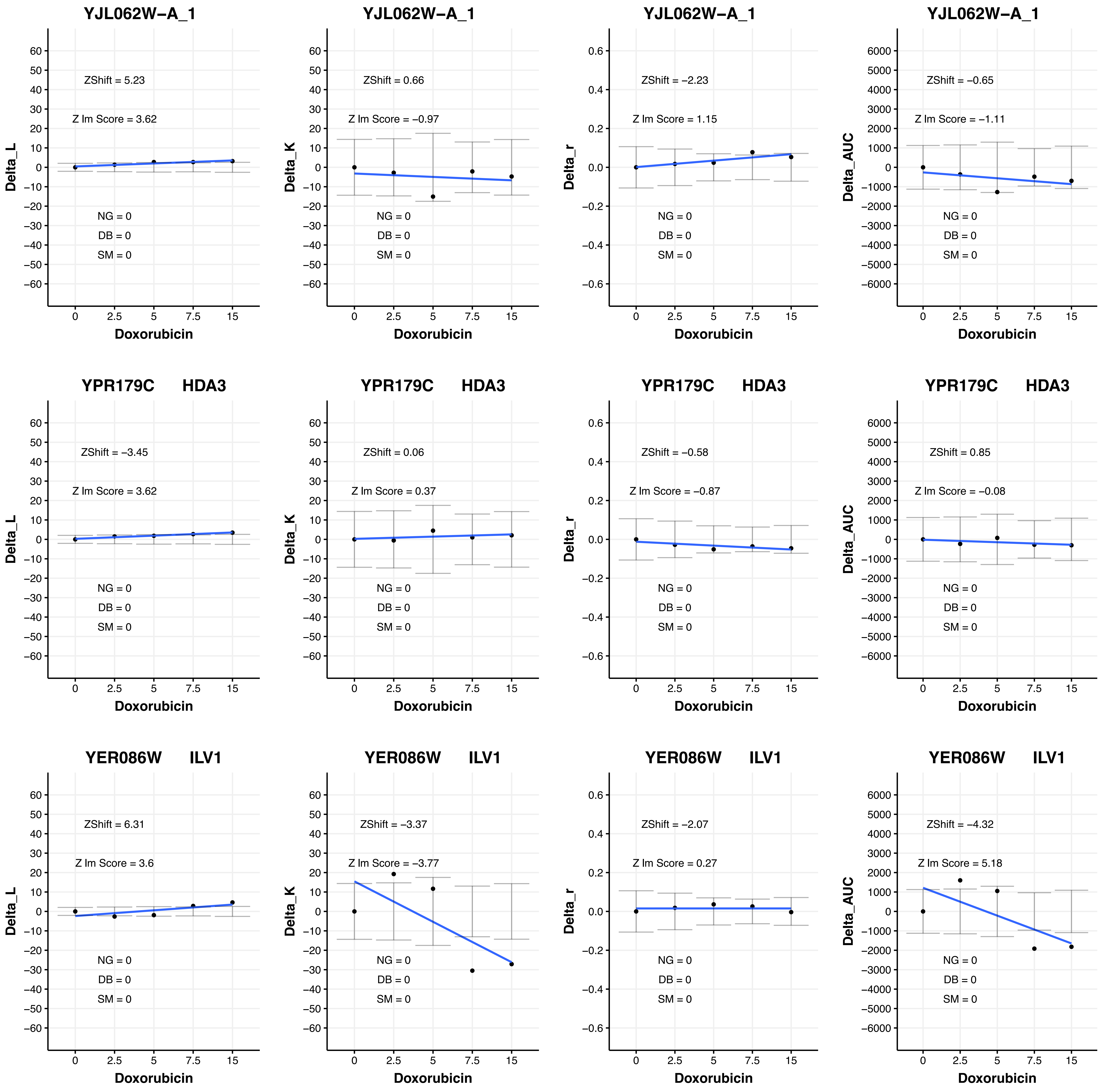
YJL062W-A_1
ZShift = 5.23
Z lm Score = 3.62
NG = 0
DB = 0
SM = 0
Delta_L
Doxorubicin
YJL062W-A_1
ZShift = 0.66
Z lm Score = −0.97
NG = 0
DB = 0
SM = 0
Delta_K
Doxorubicin
YJL062W-A_1
ZShift = −2.23
Z lm Score = 1.15
NG = 0
DB = 0
SM = 0
Delta_r
Doxorubicin
YJL062W-A_1
ZShift = −0.65
Z lm Score = −1.11
NG = 0
DB = 0
SM = 0
Delta_AUC
Doxorubicin
YPR179C HDA3
ZShift = −3.45
Z lm Score = 3.62
NG = 0
DB = 0
SM = 0
Delta_L
Doxorubicin
YPR179C HDA3
ZShift = 0.06
Z lm Score = 0.37
NG = 0
DB = 0
SM = 0
Delta_K
Doxorubicin
YPR179C HDA3
ZShift = −0.58
Z lm Score = −0.87
NG = 0
DB = 0
SM = 0
Delta_r
Doxorubicin
YPR179C HDA3
ZShift = 0.85
Z lm Score = −0.08
NG = 0
DB = 0
SM = 0
Delta_AUC
Doxorubicin
YER086W ILV1
ZShift = 6.31
Z lm Score = 3.6
NG = 0
DB = 0
SM = 0
Delta_L
Doxorubicin
YER086W ILV1
ZShift = −3.37
Z lm Score = −3.77
NG = 0
DB = 0
SM = 0
Delta_K
Doxorubicin
YER086W ILV1
ZShift = −2.07
Z lm Score = 0.27
NG = 0
DB = 0
SM = 0
Delta_r
Doxorubicin
YER086W ILV1
ZShift = −4.32
Z lm Score = 5.18
NG = 0
DB = 0
SM = 0
Delta_AUC
Doxorubicin

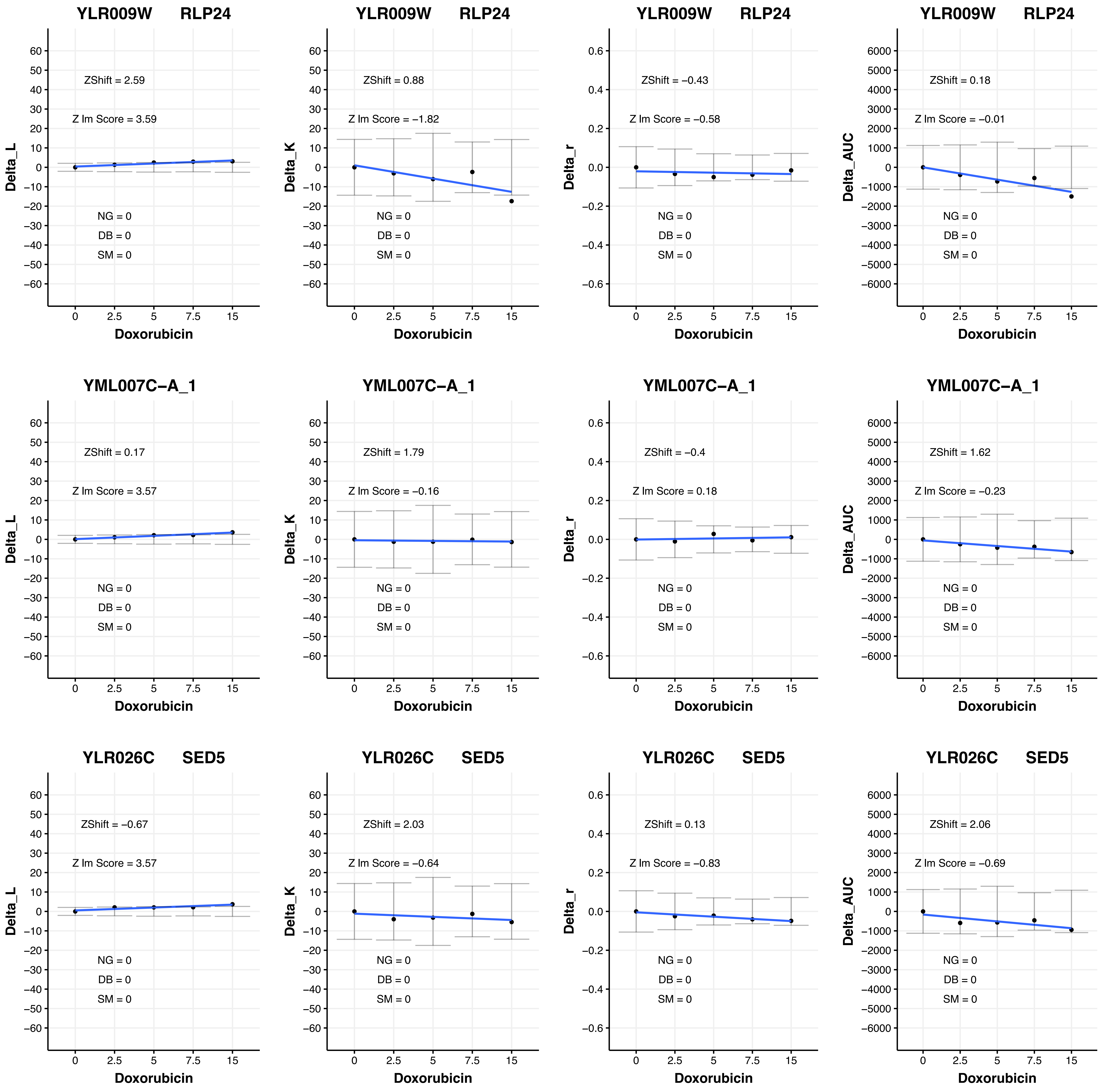
YLR009W RLP24
ZShift = 2.59
Z lm Score = 3.59
NG = 0
DB = 0
SM = 0
Delta_L
Doxorubicin
YLR009W RLP24
ZShift = 0.88
Z lm Score = −1.82
NG = 0
DB = 0
SM = 0
Delta_K
Doxorubicin
YLR009W RLP24
ZShift = −0.43
Z lm Score = −0.58
NG = 0
DB = 0
SM = 0
Delta_r
Doxorubicin
YLR009W RLP24
ZShift = 0.18
Z lm Score = −0.01
NG = 0
DB = 0
SM = 0
Delta_AUC
Doxorubicin
YML007C−A_1
ZShift = 0.17
Z lm Score = 3.57
NG = 0
DB = 0
SM = 0
Delta_L
Doxorubicin
YML007C−A_1
ZShift = 1.79
Z lm Score = −0.16
NG = 0
DB = 0
SM = 0
Delta_K
Doxorubicin
YML007C−A_1
ZShift = −0.4
Z lm Score = 0.18
NG = 0
DB = 0
SM = 0
Delta_r
Doxorubicin
YML007C−A_1
ZShift = 1.62
Z lm Score = −0.23
NG = 0
DB = 0
SM = 0
Delta_AUC
Doxorubicin
YLR026C SED5
ZShift = −0.67
Z lm Score = 3.57
NG = 0
DB = 0
SM = 0
Delta_L
Doxorubicin
YLR026C SED5
ZShift = 2.03
Z lm Score = −0.64
NG = 0
DB = 0
SM = 0
Delta_K
Doxorubicin
YLR026C SED5
ZShift = 0.13
Z lm Score = −0.83
NG = 0
DB = 0
SM = 0
Delta_r
Doxorubicin
YLR026C SED5
ZShift = 2.06
Z lm Score = −0.69
NG = 0
DB = 0
SM = 0
Delta_AUC
Doxorubicin

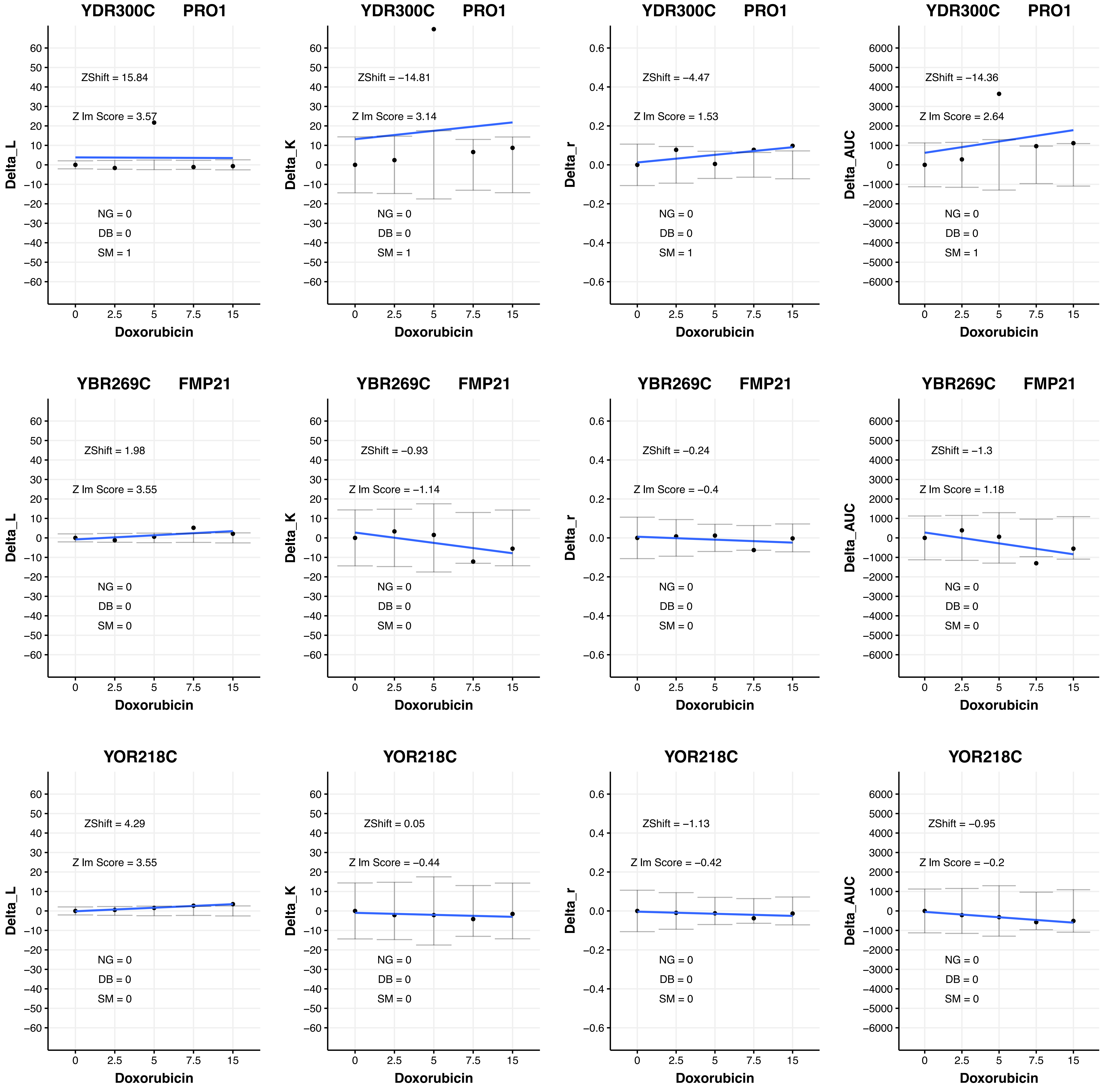

YDR300C PRO1
ZShift = 15.84
Z lm Score = 3.57
NG = 0
DB = 0
SM = 1
Delta_L
Doxorubicin
YDR300C PRO1
ZShift = −14.81
Z lm Score = 3.14
NG = 0
DB = 0
SM = 1
Delta_K
Doxorubicin
YDR300C PRO1
ZShift = −4.47
Z lm Score = 1.53
NG = 0
DB = 0
SM = 1
Delta_r
Doxorubicin
YDR300C PRO1
ZShift = −14.36
Z lm Score = 2.64
NG = 0
DB = 0
SM = 1
Delta_AUC
Doxorubicin
YBR269C FMP21
ZShift = 1.98
Z lm Score = 3.55
NG = 0
DB = 0
SM = 0
Delta_L
Doxorubicin
YBR269C FMP21
ZShift = −0.93
Z lm Score = −1.14
NG = 0
DB = 0
SM = 0
Delta_K
Doxorubicin
YBR269C FMP21
ZShift = −0.24
Z lm Score = −0.4
NG = 0
DB = 0
SM = 0
Delta_r
Doxorubicin
YBR269C FMP21
ZShift = −1.3
Z lm Score = 1.18
NG = 0
DB = 0
SM = 0
Delta_AUC
Doxorubicin
YOR218C
ZShift = 4.29
Z lm Score = 3.55
NG = 0
DB = 0
SM = 0
Delta_L
Doxorubicin
YOR218C
ZShift = 0.05
Z lm Score = −0.44
NG = 0
DB = 0
SM = 0
Delta_K
Doxorubicin
YOR218C
ZShift = −1.13
Z lm Score = −0.42
NG = 0
DB = 0
SM = 0
Delta_r
Doxorubicin
YOR218C
ZShift = −0.95
Z lm Score = −0.2
NG = 0
DB = 0
SM = 0
Delta_AUC
Doxorubicin

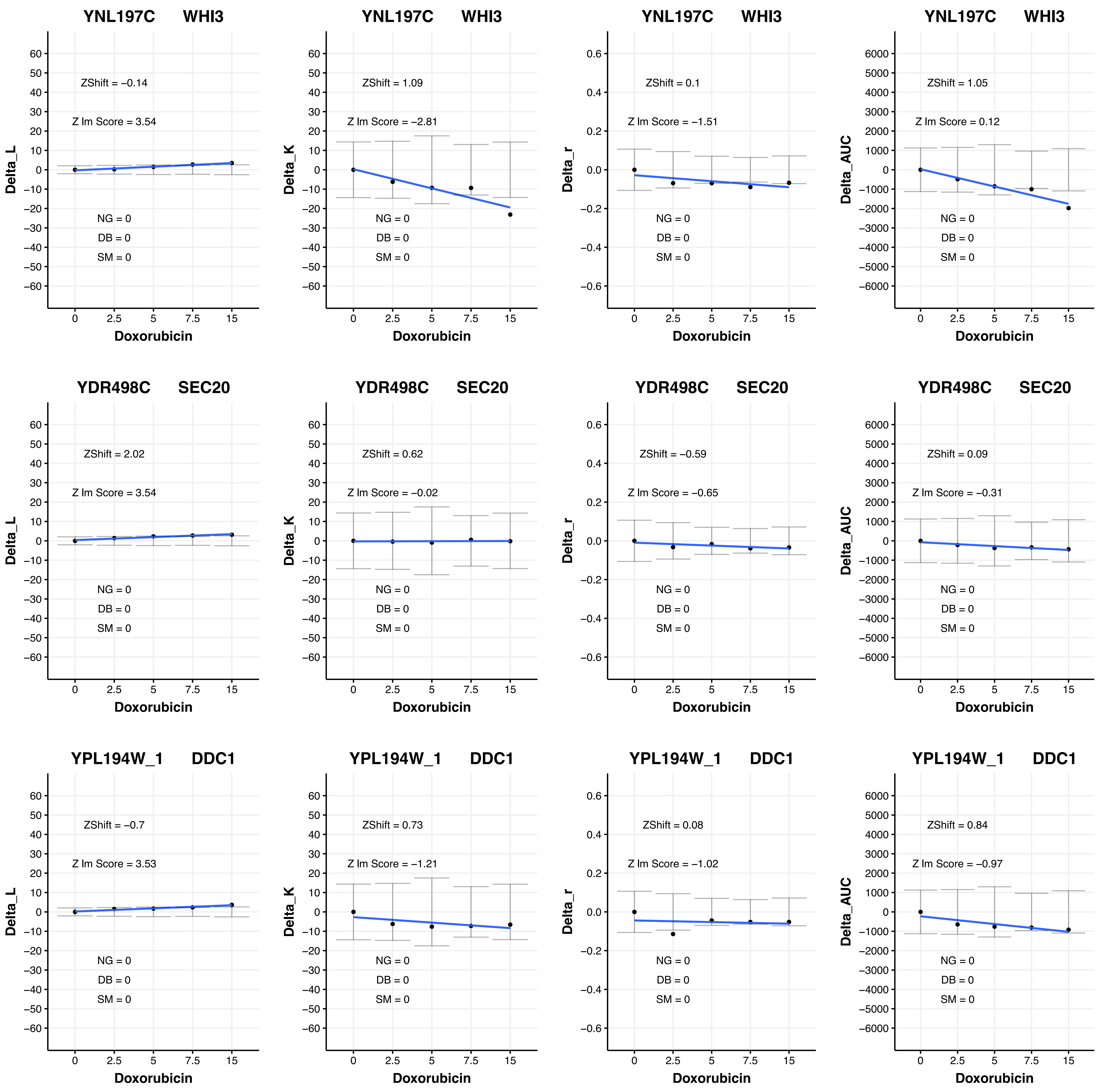
YNL197C WHI3
ZShift = −0.14
Z lm Score = 3.54
NG = 0
DB = 0
SM = 0
Delta_L
Doxorubicin
YNL197C WHI3
ZShift = 1.09
Z lm Score = −2.81
NG = 0
DB = 0
SM = 0
Delta_K
Doxorubicin
YNL197C WHI3
ZShift = 0.1
Z lm Score = −1.51
NG = 0
DB = 0
SM = 0
Delta_r
Doxorubicin
YNL197C WHI3
ZShift = 1.05
Z lm Score = 0.12
NG = 0
DB = 0
SM = 0
Delta_AUC
Doxorubicin
YDR498C SEC20
ZShift = 2.02
Z lm Score = 3.54
NG = 0
DB = 0
SM = 0
Delta_L
Doxorubicin
YDR498C SEC20
ZShift = 0.62
Z lm Score = −0.02
NG = 0
DB = 0
SM = 0
Delta_K
Doxorubicin
YDR498C SEC20
ZShift = −0.59
Z lm Score = −0.65
NG = 0
DB = 0
SM = 0
Delta_r
Doxorubicin
YDR498C SEC20
ZShift = 0.09
Z lm Score = −0.31
NG = 0
DB = 0
SM = 0
Delta_AUC
Doxorubicin
YPL194W_1 DDC1
ZShift = −0.7
Z lm Score = 3.53
NG = 0
DB = 0
SM = 0
Delta_L
Doxorubicin
YPL194W_1 DDC1
ZShift = 0.73
Z lm Score = −1.21
NG = 0
DB = 0
SM = 0
Delta_K
Doxorubicin
YPL194W_1 DDC1
ZShift = 0.08
Z lm Score = −1.02
NG = 0
DB = 0
SM = 0
Delta_r
Doxorubicin
YPL194W_1 DDC1
ZShift = 0.84
Z lm Score = −0.97
NG = 0
DB = 0
SM = 0
Delta_AUC
Doxorubicin

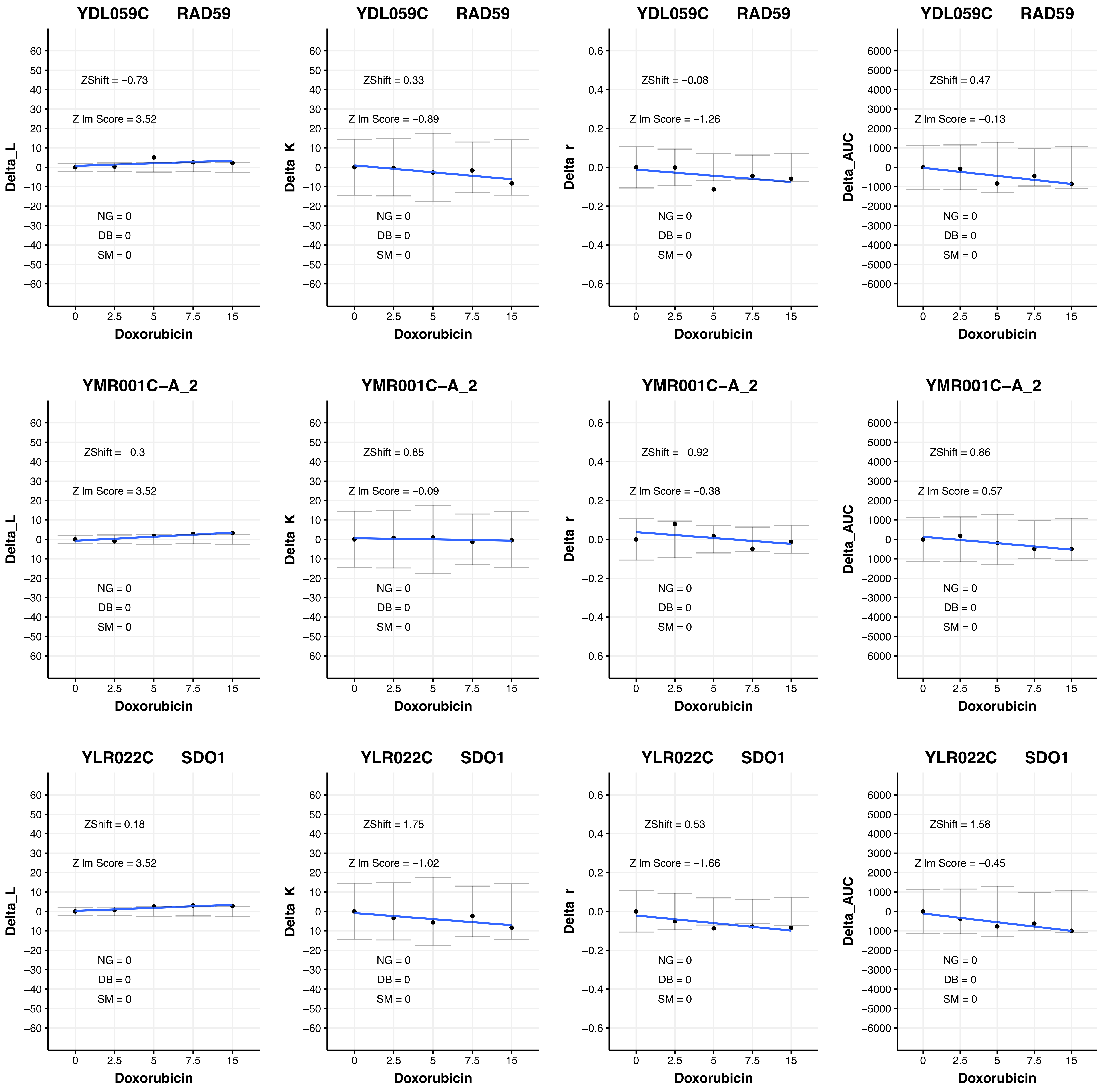
YDL059C RAD59
ZShift = −0.73
Z lm Score = 3.52
NG = 0
DB = 0
SM = 0
Delta_L
Doxorubicin
YDL059C RAD59
ZShift = 0.33
Z lm Score = −0.89
NG = 0
DB = 0
SM = 0
Delta_K
Doxorubicin
YDL059C RAD59
ZShift = −0.08
Z lm Score = −1.26
NG = 0
DB = 0
SM = 0
Delta_r
Doxorubicin
YDL059C RAD59
ZShift = 0.47
Z lm Score = −0.13
NG = 0
DB = 0
SM = 0
Delta_AUC
Doxorubicin
YMR001C−A_2
ZShift = −0.3
Z lm Score = 3.52
NG = 0
DB = 0
SM = 0
Delta_L
Doxorubicin
YMR001C−A_2
ZShift = 0.85
Z lm Score = −0.09
NG = 0
DB = 0
SM = 0
Delta_K
Doxorubicin
YMR001C−A_2
ZShift = −0.92
Z lm Score = −0.38
NG = 0
DB = 0
SM = 0
Delta_r
Doxorubicin
YMR001C−A_2
ZShift = 0.86
Z lm Score = 0.57
NG = 0
DB = 0
SM = 0
Delta_AUC
Doxorubicin
YLR022C SDO1
ZShift = 0.18
Z lm Score = 3.52
NG = 0
DB = 0
SM = 0
Delta_L
Doxorubicin
YLR022C SDO1
ZShift = 1.75
Z lm Score = −1.02
NG = 0
DB = 0
SM = 0
Delta_K
Doxorubicin
YLR022C SDO1
ZShift = 0.53
Z lm Score = −1.66
NG = 0
DB = 0
SM = 0
Delta_r
Doxorubicin
YLR022C SDO1
ZShift = 1.58
Z lm Score = −0.45
NG = 0
DB = 0
SM = 0
Delta_AUC
Doxorubicin

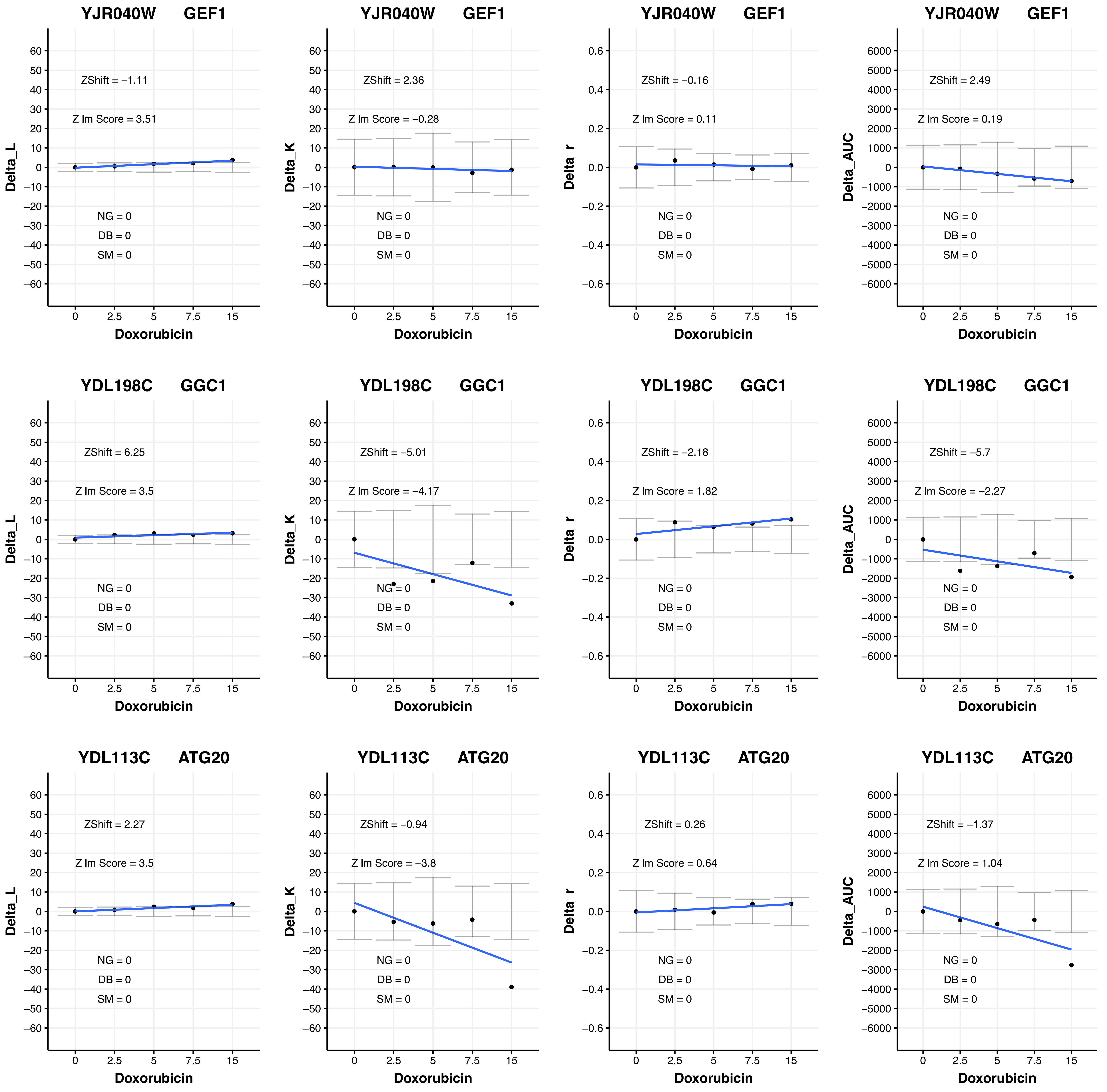
YJR040W GEF1
ZShift = −1.11
Z lm Score = 3.51
NG = 0
DB = 0
SM = 0
Delta_L
Doxorubicin
YJR040W GEF1
ZShift = 2.36
Z lm Score = −0.28
NG = 0
DB = 0
SM = 0
Delta_K
Doxorubicin
YJR040W GEF1
ZShift = −0.16
Z lm Score = 0.11
NG = 0
DB = 0
SM = 0
Delta_r
Doxorubicin
YJR040W GEF1
ZShift = 2.49
Z lm Score = 0.19
NG = 0
DB = 0
SM = 0
Delta_AUC
Doxorubicin
YDL198C GGC1
ZShift = 6.25
Z lm Score = 3.5
NG = 0
DB = 0
SM = 0
Delta_L
Doxorubicin
YDL198C GGC1
ZShift = −5.01
Z lm Score = −4.17
NG = 0
DB = 0
SM = 0
Delta_K
Doxorubicin
YDL198C GGC1
ZShift = −2.18
Z lm Score = 1.82
NG = 0
DB = 0
SM = 0
Delta_r
Doxorubicin
YDL198C GGC1
ZShift = −5.7
Z lm Score = −2.27
NG = 0
DB = 0
SM = 0
Delta_AUC
Doxorubicin
YDL113C ATG20
ZShift = 2.27
Z lm Score = 3.5
NG = 0
DB = 0
SM = 0
Delta_L
Doxorubicin
YDL113C ATG20
ZShift = −0.94
Z lm Score = −3.8
NG = 0
DB = 0
SM = 0
Delta_K
Doxorubicin
YDL113C ATG20
ZShift = 0.26
Z lm Score = 0.64
NG = 0
DB = 0
SM = 0
Delta_r
Doxorubicin
YDL113C ATG20
ZShift = −1.37
Z lm Score = 1.04
NG = 0
DB = 0
SM = 0
Delta_AUC
Doxorubicin

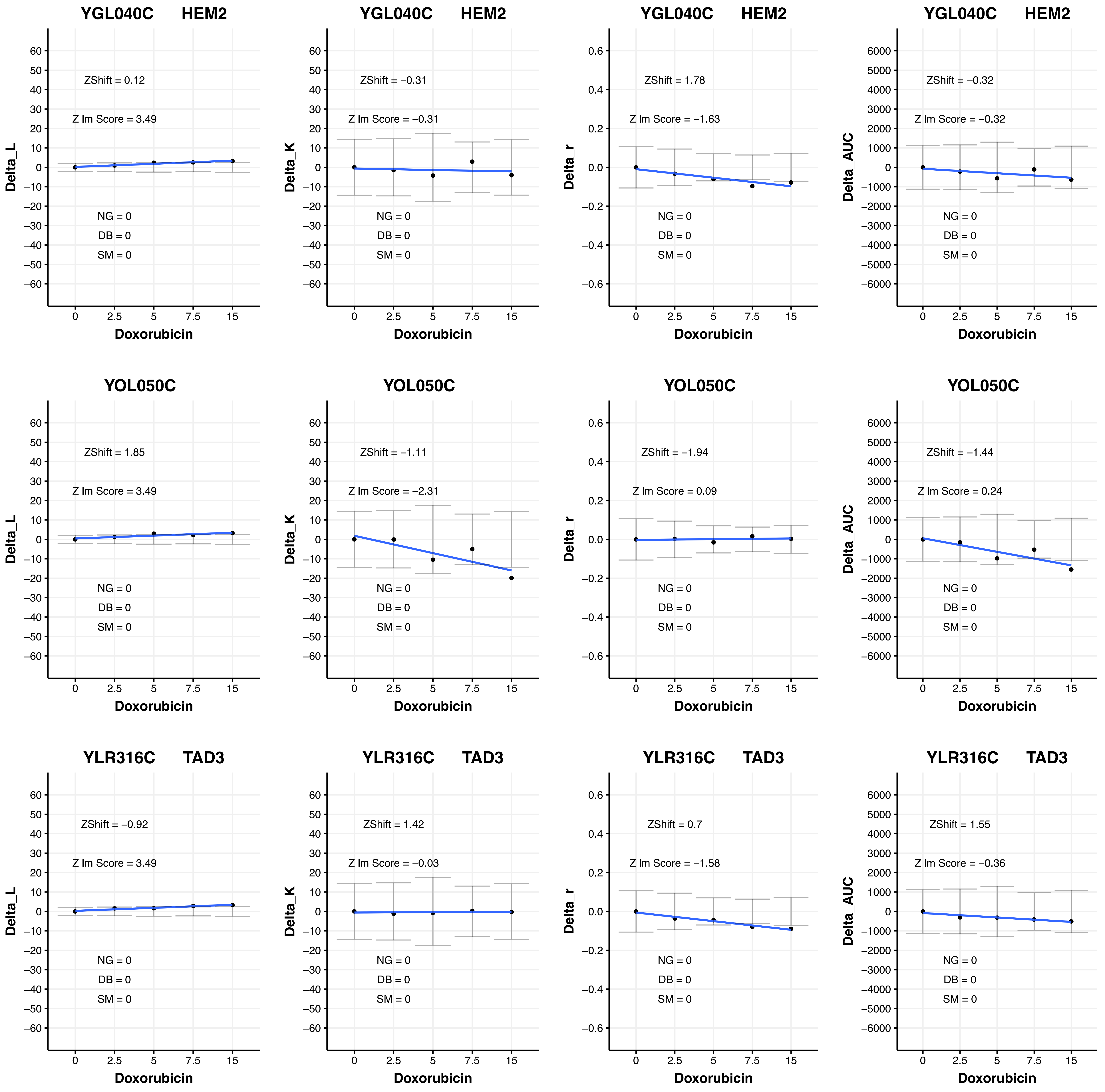
YGL040C HEM2
ZShift = 0.12
Z lm Score = 3.49
NG = 0
DB = 0
SM = 0
Delta_L
Doxorubicin
YGL040C HEM2
ZShift = −0.31
Z lm Score = −0.31
NG = 0
DB = 0
SM = 0
Delta_K
Doxorubicin
YGL040C HEM2
ZShift = 1.78
Z lm Score = −1.63
NG = 0
DB = 0
SM = 0
Delta_r
Doxorubicin
YGL040C HEM2
ZShift = −0.32
Z lm Score = −0.32
NG = 0
DB = 0
SM = 0
Delta_AUC
Doxorubicin
YOL050C
ZShift = 1.85
Z lm Score = 3.49
NG = 0
DB = 0
SM = 0
Delta_L
Doxorubicin
YOL050C
ZShift = −1.11
Z lm Score = −2.31
NG = 0
DB = 0
SM = 0
Delta_K
Doxorubicin
YOL050C
ZShift = −1.94
Z lm Score = 0.09
NG = 0
DB = 0
SM = 0
Delta_r
Doxorubicin
YOL050C
ZShift = −1.44
Z lm Score = 0.24
NG = 0
DB = 0
SM = 0
Delta_AUC
Doxorubicin
YLR316C TAD3
ZShift = −0.92
Z lm Score = 3.49
NG = 0
DB = 0
SM = 0
Delta_L
Doxorubicin
YLR316C TAD3
ZShift = 1.42
Z lm Score = −0.03
NG = 0
DB = 0
SM = 0
Delta_K
Doxorubicin
YLR316C TAD3
ZShift = 0.7
Z lm Score = −1.58
NG = 0
DB = 0
SM = 0
Delta_r
Doxorubicin
YLR316C TAD3
ZShift = 1.55
Z lm Score = −0.36
NG = 0
DB = 0
SM = 0
Delta_AUC
Doxorubicin

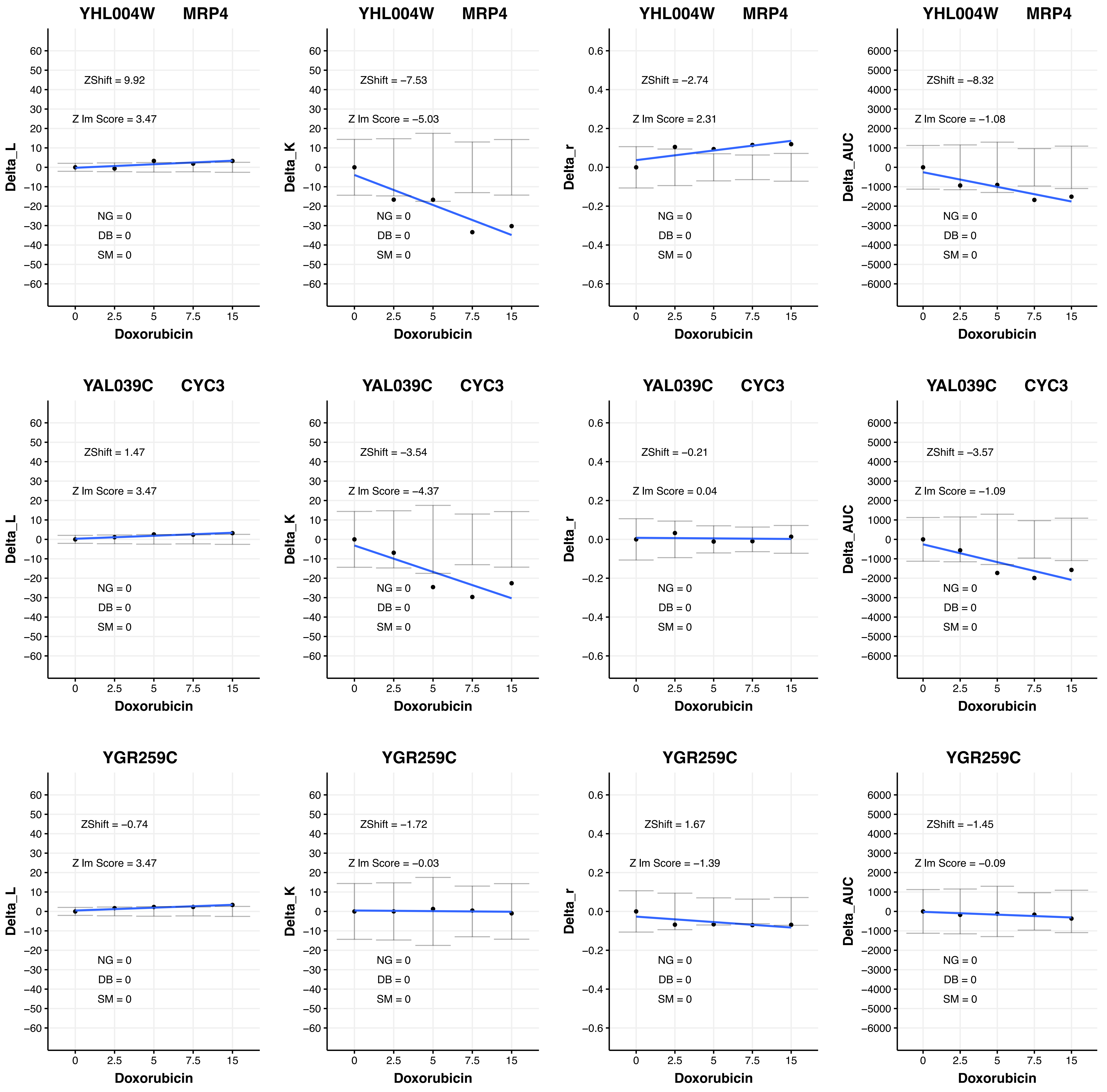
YHL004W MRP4
ZShift = 9.92
Z lm Score = 3.47
NG = 0
DB = 0
SM = 0
Delta_L
Doxorubicin
YHL004W MRP4
ZShift = −7.53
Z lm Score = −5.03
NG = 0
DB = 0
SM = 0
Delta_K
Doxorubicin
YHL004W MRP4
ZShift = −2.74
Z lm Score = 2.31
NG = 0
DB = 0
SM = 0
Delta_r
Doxorubicin
YHL004W MRP4
ZShift = −8.32
Z lm Score = −1.08
NG = 0
DB = 0
SM = 0
Delta_AUC
Doxorubicin
YAL039C CYC3
ZShift = 1.47
Z lm Score = 3.47
NG = 0
DB = 0
SM = 0
Delta_L
Doxorubicin
YAL039C CYC3
ZShift = −3.54
Z lm Score = −4.37
NG = 0
DB = 0
SM = 0
Delta_K
Doxorubicin
YAL039C CYC3
ZShift = −0.21
Z lm Score = 0.04
NG = 0
DB = 0
SM = 0
Delta_r
Doxorubicin
YAL039C CYC3
ZShift = −3.57
Z lm Score = −1.09
NG = 0
DB = 0
SM = 0
Delta_AUC
Doxorubicin
YGR259C
ZShift = −0.74
Z lm Score = 3.47
NG = 0
DB = 0
SM = 0
Delta_L
Doxorubicin
YGR259C
ZShift = −1.72
Z lm Score = −0.03
NG = 0
DB = 0
SM = 0
Delta_K
Doxorubicin
YGR259C
ZShift = 1.67
Z lm Score = −1.39
NG = 0
DB = 0
SM = 0
Delta_r
Doxorubicin
YGR259C
ZShift = −1.45
Z lm Score = −0.09
NG = 0
DB = 0
SM = 0
Delta_AUC
Doxorubicin

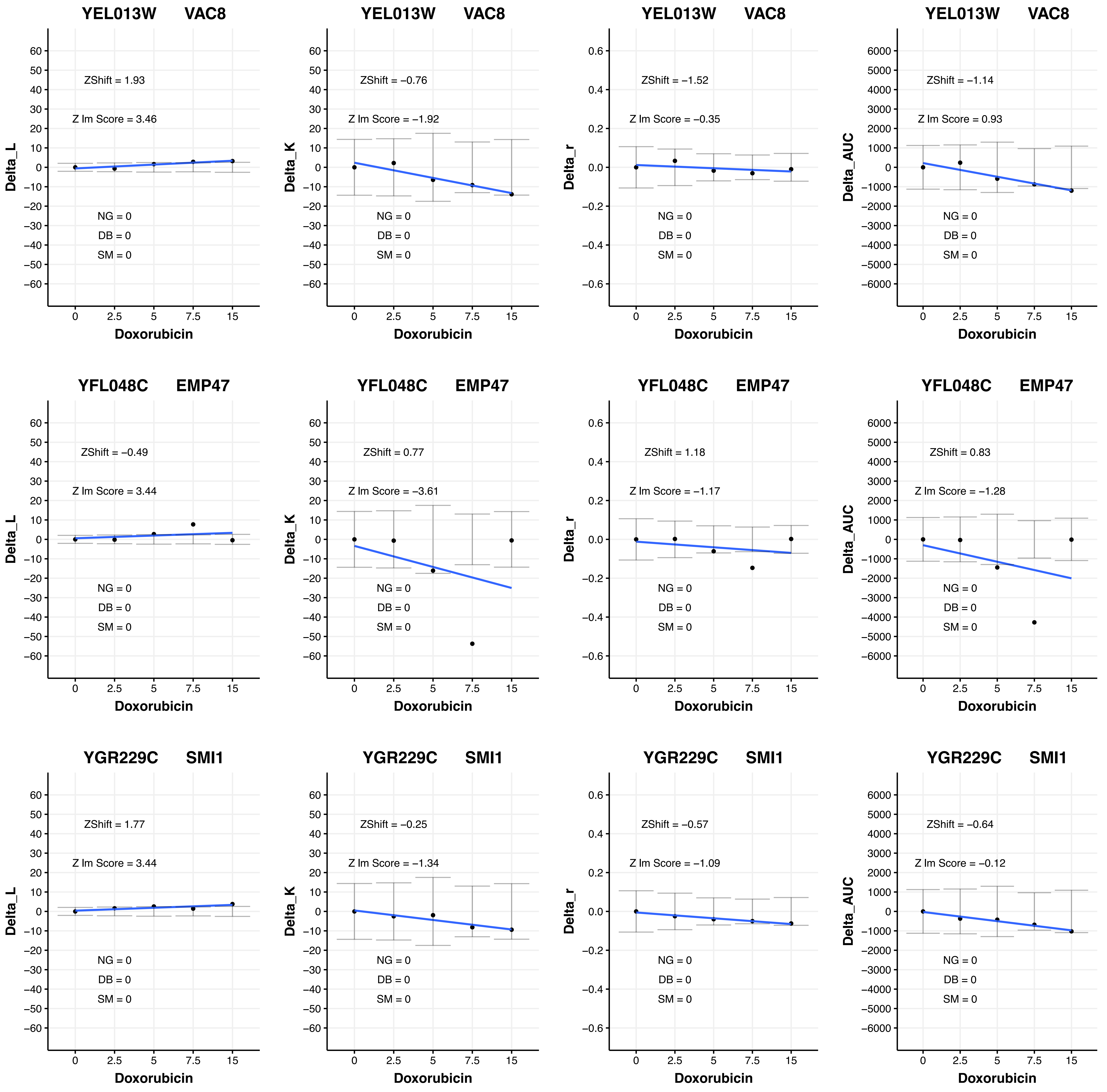

YEL013W VAC8
ZShift = 1.93
Z lm Score = 3.46
NG = 0
DB = 0
SM = 0
Delta_L
Doxorubicin
YEL013W VAC8
ZShift = −0.76
Z lm Score = −1.92
NG = 0
DB = 0
SM = 0
Delta_K
Doxorubicin
YEL013W VAC8
ZShift = −1.52
Z lm Score = −0.35
NG = 0
DB = 0
SM = 0
Delta_r
Doxorubicin
YEL013W VAC8
ZShift = −1.14
Z lm Score = 0.93
NG = 0
DB = 0
SM = 0
Delta_AUC
Doxorubicin
YFL048C EMP47
ZShift = −0.49
Z lm Score = 3.44
NG = 0
DB = 0
SM = 0
Delta_L
Doxorubicin
YFL048C EMP47
ZShift = 0.77
Z lm Score = −3.61
NG = 0
DB = 0
SM = 0
Delta_K
Doxorubicin
YFL048C EMP47
ZShift = 1.18
Z lm Score = −1.17
NG = 0
DB = 0
SM = 0
Delta_r
Doxorubicin
YFL048C EMP47
ZShift = 0.83
Z lm Score = −1.28
NG = 0
DB = 0
SM = 0
Delta_AUC
Doxorubicin
YGR229C SMI1
ZShift = 1.77
Z lm Score = 3.44
NG = 0
DB = 0
SM = 0
Delta_L
Doxorubicin
YGR229C SMI1
ZShift = −0.25
Z lm Score = −1.34
NG = 0
DB = 0
SM = 0
Delta_K
Doxorubicin
YGR229C SMI1
ZShift = −0.57
Z lm Score = −1.09
NG = 0
DB = 0
SM = 0
Delta_r
Doxorubicin
YGR229C SMI1
ZShift = −0.64
Z lm Score = −0.12
NG = 0
DB = 0
SM = 0
Delta_AUC
Doxorubicin

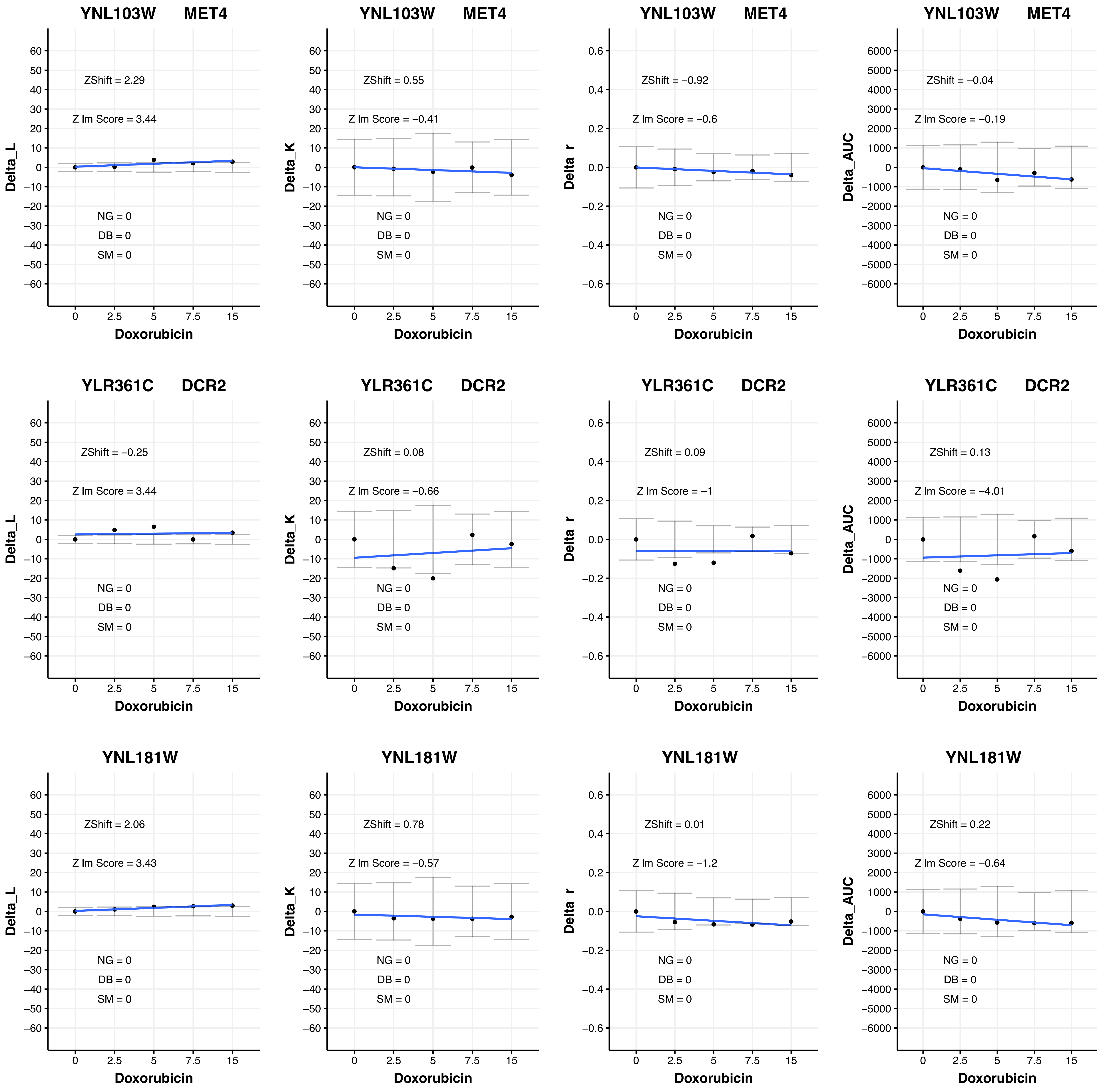

YNL103W MET4
ZShift = 2.29
Z lm Score = 3.44
NG = 0
DB = 0
SM = 0
Delta_L
Doxorubicin
YNL103W MET4
ZShift = 0.55
Z lm Score = −0.41
NG = 0
DB = 0
SM = 0
Delta_K
Doxorubicin
YNL103W MET4
ZShift = −0.92
Z lm Score = −0.6
NG = 0
DB = 0
SM = 0
Delta_r
Doxorubicin
YNL103W MET4
ZShift = −0.04
Z lm Score = −0.19
NG = 0
DB = 0
SM = 0
Delta_AUC
Doxorubicin
YLR361C DCR2
ZShift = −0.25
Z lm Score = 3.44
NG = 0
DB = 0
SM = 0
Delta_L
Doxorubicin
YLR361C DCR2
ZShift = 0.08
Z lm Score = −0.66
NG = 0
DB = 0
SM = 0
Delta_K
Doxorubicin
YLR361C DCR2
ZShift = 0.09
Z lm Score = −1
NG = 0
DB = 0
SM = 0
Delta_r
Doxorubicin
YLR361C DCR2
ZShift = 0.13
Z lm Score = −4.01
NG = 0
DB = 0
SM = 0
Delta_AUC
Doxorubicin
YNL181W
ZShift = 2.06
Z lm Score = 3.43
NG = 0
DB = 0
SM = 0
Delta_L
Doxorubicin
YNL181W
ZShift = 0.78
Z lm Score = −0.57
NG = 0
DB = 0
SM = 0
Delta_K
Doxorubicin
YNL181W
ZShift = 0.01
Z lm Score = −1.2
NG = 0
DB = 0
SM = 0
Delta_r
Doxorubicin
YNL181W
ZShift = 0.22
Z lm Score = −0.64
NG = 0
DB = 0
SM = 0
Delta_AUC
Doxorubicin

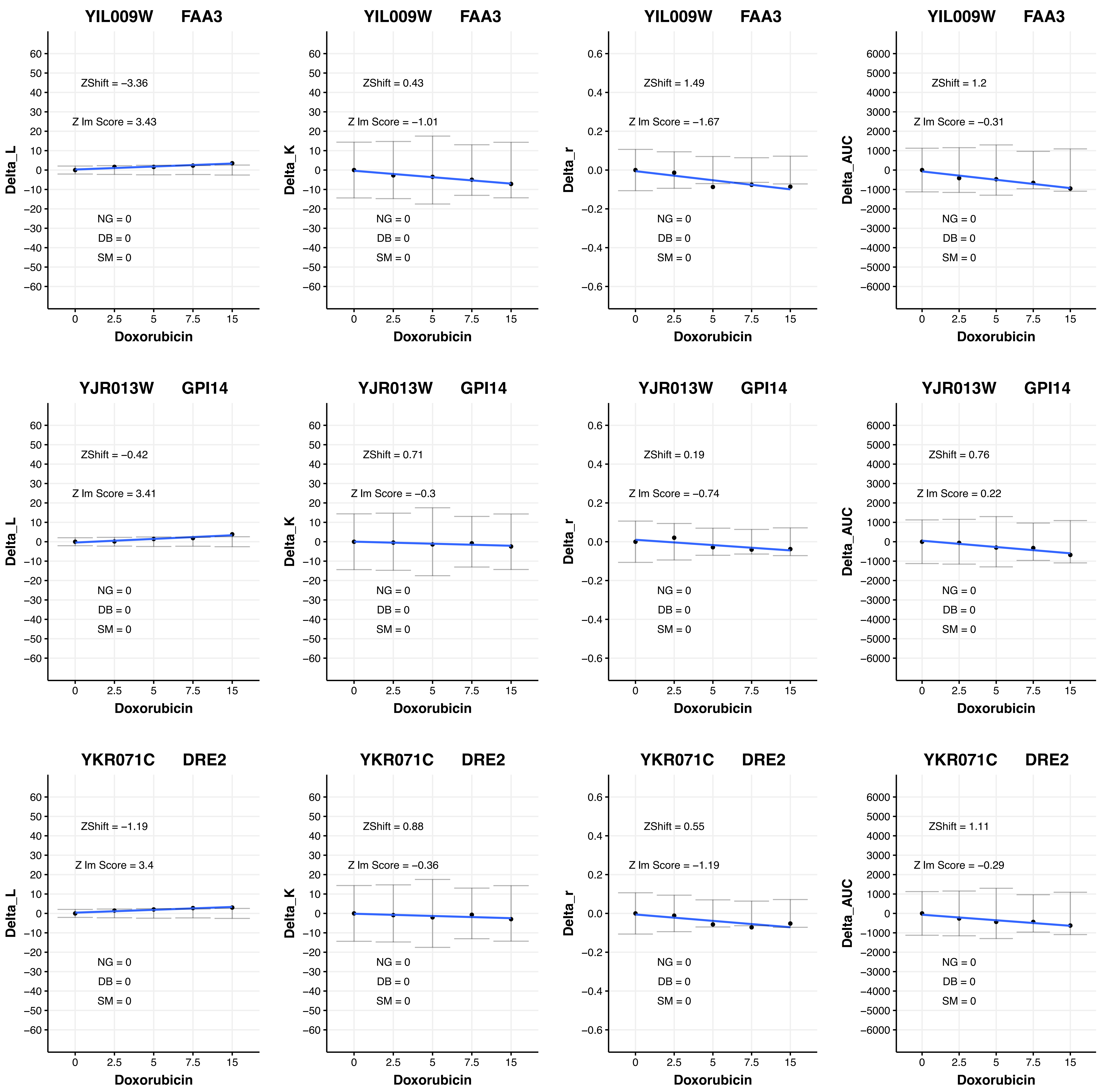

YIL009W FAA3
ZShift = −3.36
Z lm Score = 3.43
NG = 0
DB = 0
SM = 0
Delta_L
Doxorubicin
YIL009W FAA3
ZShift = 0.43
Z lm Score = −1.01
NG = 0
DB = 0
SM = 0
Delta_K
Doxorubicin
YIL009W FAA3
ZShift = 1.49
Z lm Score = −1.67
NG = 0
DB = 0
SM = 0
Delta_r
Doxorubicin
YIL009W FAA3
ZShift = 1.2
Z lm Score = −0.31
NG = 0
DB = 0
SM = 0
Delta_AUC
Doxorubicin
YJR013W GPI14
ZShift = −0.42
Z lm Score = 3.41
NG = 0
DB = 0
SM = 0
Delta_L
Doxorubicin
YJR013W GPI14
ZShift = 0.71
Z lm Score = −0.3
NG = 0
DB = 0
SM = 0
Delta_K
Doxorubicin
YJR013W GPI14
ZShift = 0.19
Z lm Score = −0.74
NG = 0
DB = 0
SM = 0
Delta_r
Doxorubicin
YJR013W GPI14
ZShift = 0.76
Z lm Score = 0.22
NG = 0
DB = 0
SM = 0
Delta_AUC
Doxorubicin
YKR071C DRE2
ZShift = −1.19
Z lm Score = 3.4
NG = 0
DB = 0
SM = 0
Delta_L
Doxorubicin
YKR071C DRE2
ZShift = 0.88
Z lm Score = −0.36
NG = 0
DB = 0
SM = 0
Delta_K
Doxorubicin
YKR071C DRE2
ZShift = 0.55
Z lm Score = −1.19
NG = 0
DB = 0
SM = 0
Delta_r
Doxorubicin
YKR071C DRE2
ZShift = 1.11
Z lm Score = −0.29
NG = 0
DB = 0
SM = 0
Delta_AUC
Doxorubicin

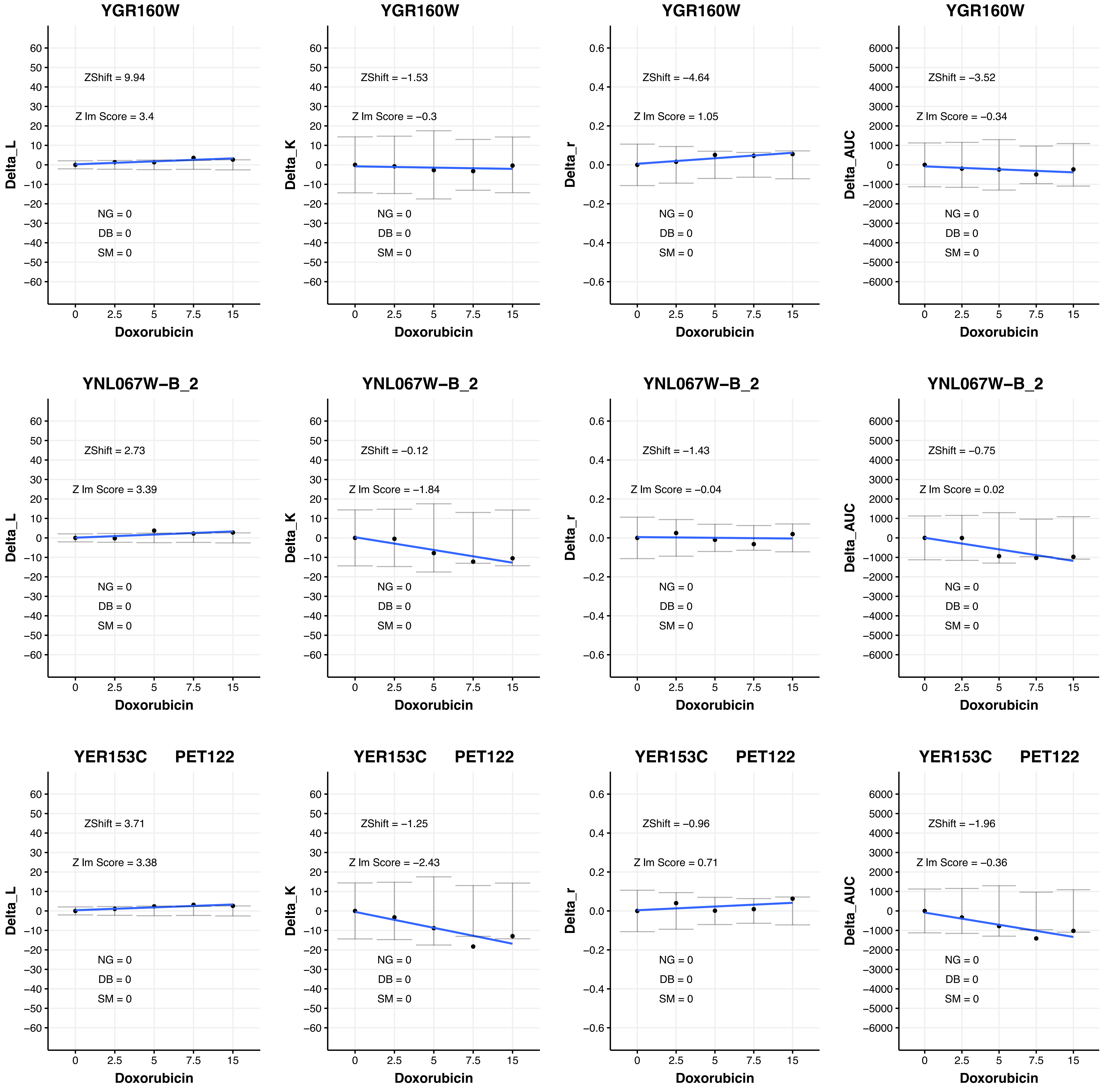

YGR160W
ZShift = 9.94
Z lm Score = 3.4
NG = 0
DB = 0
SM = 0
Delta_L
Doxorubicin
YGR160W
ZShift = −1.53
Z lm Score = −0.3
NG = 0
DB = 0
SM = 0
Delta_K
Doxorubicin
YGR160W
ZShift = −4.64
Z lm Score = 1.05
NG = 0
DB = 0
SM = 0
Delta_r
Doxorubicin
YGR160W
ZShift = −3.52
Z lm Score = −0.34
NG = 0
DB = 0
SM = 0
Delta_AUC
Doxorubicin
YNL067W−B_2
ZShift = 2.73
Z lm Score = 3.39
NG = 0
DB = 0
SM = 0
Delta_L
Doxorubicin
YNL067W−B_2
ZShift = −0.12
Z lm Score = −1.84
NG = 0
DB = 0
SM = 0
Delta_K
Doxorubicin
YNL067W−B_2
ZShift = −1.43
Z lm Score = −0.04
NG = 0
DB = 0
SM = 0
Delta_r
Doxorubicin
YNL067W−B_2
ZShift = −0.75
Z lm Score = 0.02
NG = 0
DB = 0
SM = 0
Delta_AUC
Doxorubicin
YER153C PET122
ZShift = 3.71
Z lm Score = 3.38
NG = 0
DB = 0
SM = 0
Delta_L
Doxorubicin
YER153C PET122
ZShift = −1.25
Z lm Score = −2.43
NG = 0
DB = 0
SM = 0
Delta_K
Doxorubicin
YER153C PET122
ZShift = −0.96
Z lm Score = 0.71
NG = 0
DB = 0
SM = 0
Delta_r
Doxorubicin
YER153C PET122
ZShift = −1.96
Z lm Score = −0.36
NG = 0
DB = 0
SM = 0
Delta_AUC
Doxorubicin

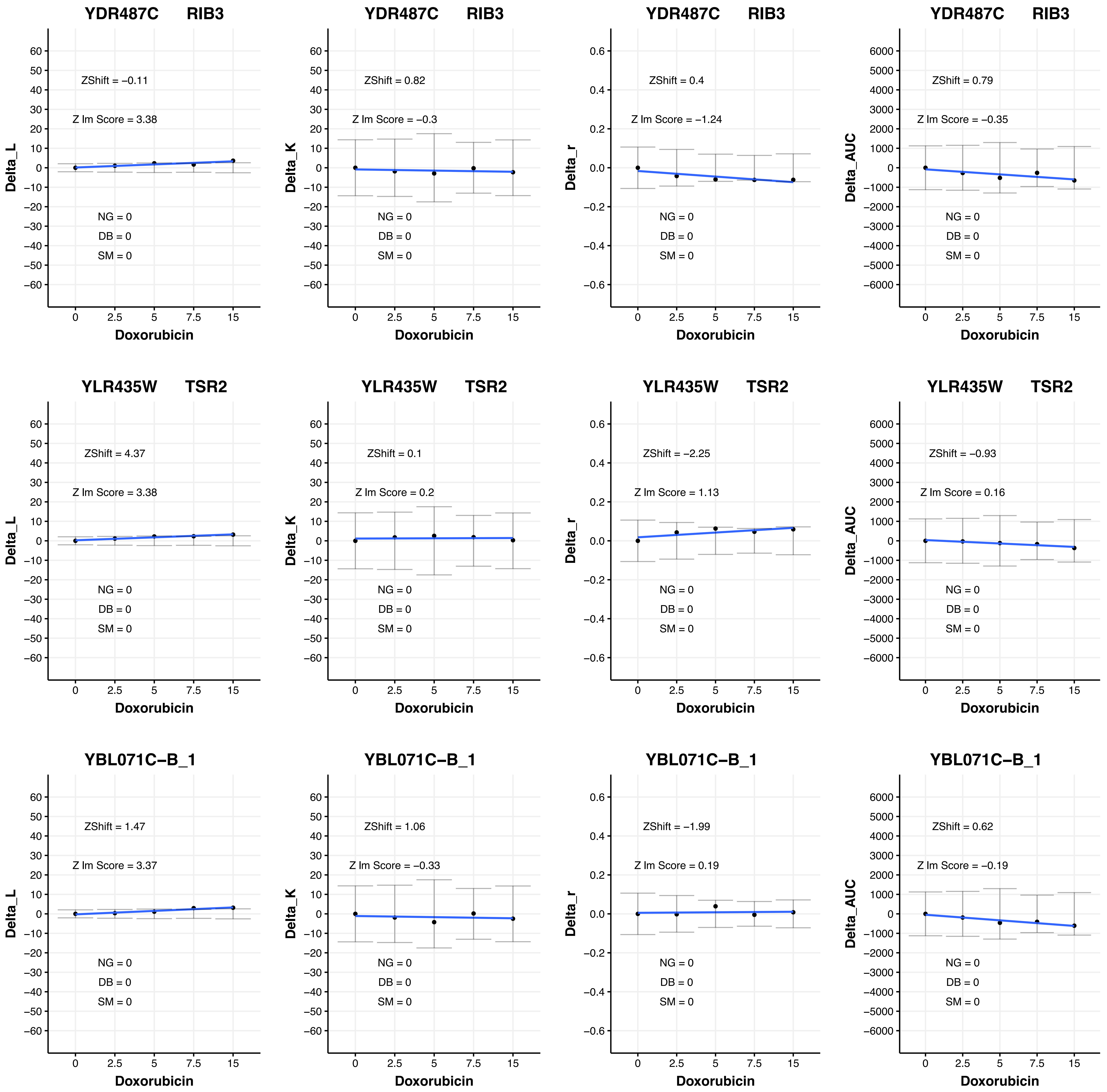
YDR487C RIB3
ZShift = −0.11
Z lm Score = 3.38
NG = 0
DB = 0
SM = 0
Delta_L
Doxorubicin
YDR487C RIB3
ZShift = 0.82
Z lm Score = −0.3
NG = 0
DB = 0
SM = 0
Delta_K
Doxorubicin
YDR487C RIB3
ZShift = 0.4
Z lm Score = −1.24
NG = 0
DB = 0
SM = 0
Delta_r
Doxorubicin
YDR487C RIB3
ZShift = 0.79
Z lm Score = −0.35
NG = 0
DB = 0
SM = 0
Delta_AUC
Doxorubicin
YLR435W TSR2
ZShift = 4.37
Z lm Score = 3.38
NG = 0
DB = 0
SM = 0
Delta_L
Doxorubicin
YLR435W TSR2
ZShift = 0.1
Z lm Score = 0.2
NG = 0
DB = 0
SM = 0
Delta_K
Doxorubicin
YLR435W TSR2
ZShift = −2.25
Z lm Score = 1.13
NG = 0
DB = 0
SM = 0
Delta_r
Doxorubicin
YLR435W TSR2
ZShift = −0.93
Z lm Score = 0.16
NG = 0
DB = 0
SM = 0
Delta_AUC
Doxorubicin
YBL071C−B_1
ZShift = 1.47
Z lm Score = 3.37
NG = 0
DB = 0
SM = 0
Delta_L
Doxorubicin
YBL071C−B_1
ZShift = 1.06
Z lm Score = −0.33
NG = 0
DB = 0
SM = 0
Delta_K
Doxorubicin
YBL071C−B_1
ZShift = −1.99
Z lm Score = 0.19
NG = 0
DB = 0
SM = 0
Delta_r
Doxorubicin
YBL071C−B_1
ZShift = 0.62
Z lm Score = −0.19
NG = 0
DB = 0
SM = 0
Delta_AUC
Doxorubicin

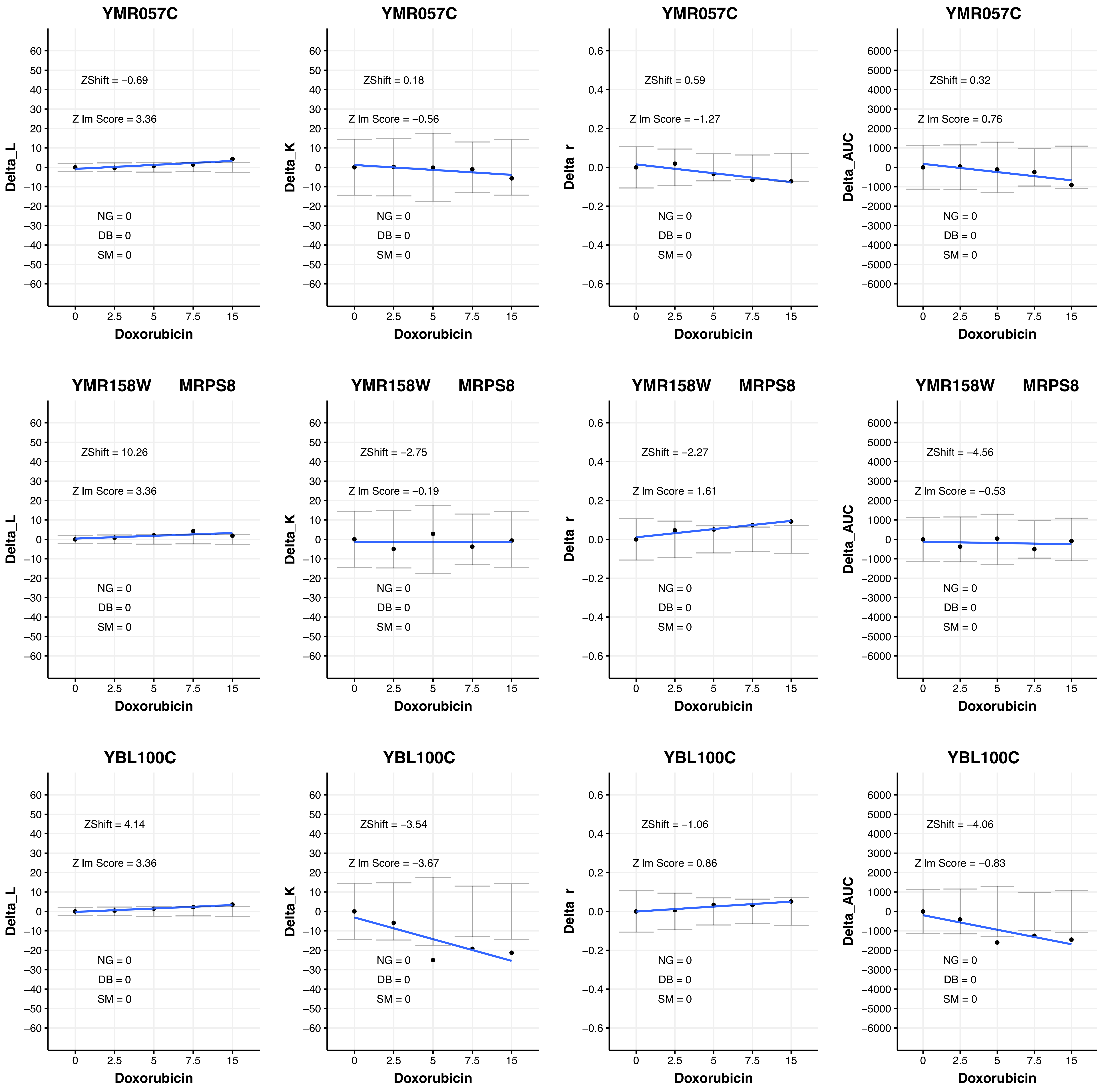

YMR057C
ZShift = −0.69
Z lm Score = 3.36
NG = 0
DB = 0
SM = 0
Delta_L
Doxorubicin
YMR057C
ZShift = 0.18
Z lm Score = −0.56
NG = 0
DB = 0
SM = 0
Delta_K
Doxorubicin
YMR057C
ZShift = 0.59
Z lm Score = −1.27
NG = 0
DB = 0
SM = 0
Delta_r
Doxorubicin
YMR057C
ZShift = 0.32
Z lm Score = 0.76
NG = 0
DB = 0
SM = 0
Delta_AUC
Doxorubicin
YMR158W MRPS8
ZShift = 10.26
Z lm Score = 3.36
NG = 0
DB = 0
SM = 0
Delta_L
Doxorubicin
YMR158W MRPS8
ZShift = −2.75
Z lm Score = −0.19
NG = 0
DB = 0
SM = 0
Delta_K
Doxorubicin
YMR158W MRPS8
ZShift = −2.27
Z lm Score = 1.61
NG = 0
DB = 0
SM = 0
Delta_r
Doxorubicin
YMR158W MRPS8
ZShift = −4.56
Z lm Score = −0.53
NG = 0
DB = 0
SM = 0
Delta_AUC
Doxorubicin
YBL100C
ZShift = 4.14
Z lm Score = 3.36
NG = 0
DB = 0
SM = 0
Delta_L
Doxorubicin
YBL100C
ZShift = −3.54
Z lm Score = −3.67
NG = 0
DB = 0
SM = 0
Delta_K
Doxorubicin
YBL100C
ZShift = −1.06
Z lm Score = 0.86
NG = 0
DB = 0
SM = 0
Delta_r
Doxorubicin
YBL100C
ZShift = −4.06
Z lm Score = −0.83
NG = 0
DB = 0
SM = 0
Delta_AUC
Doxorubicin

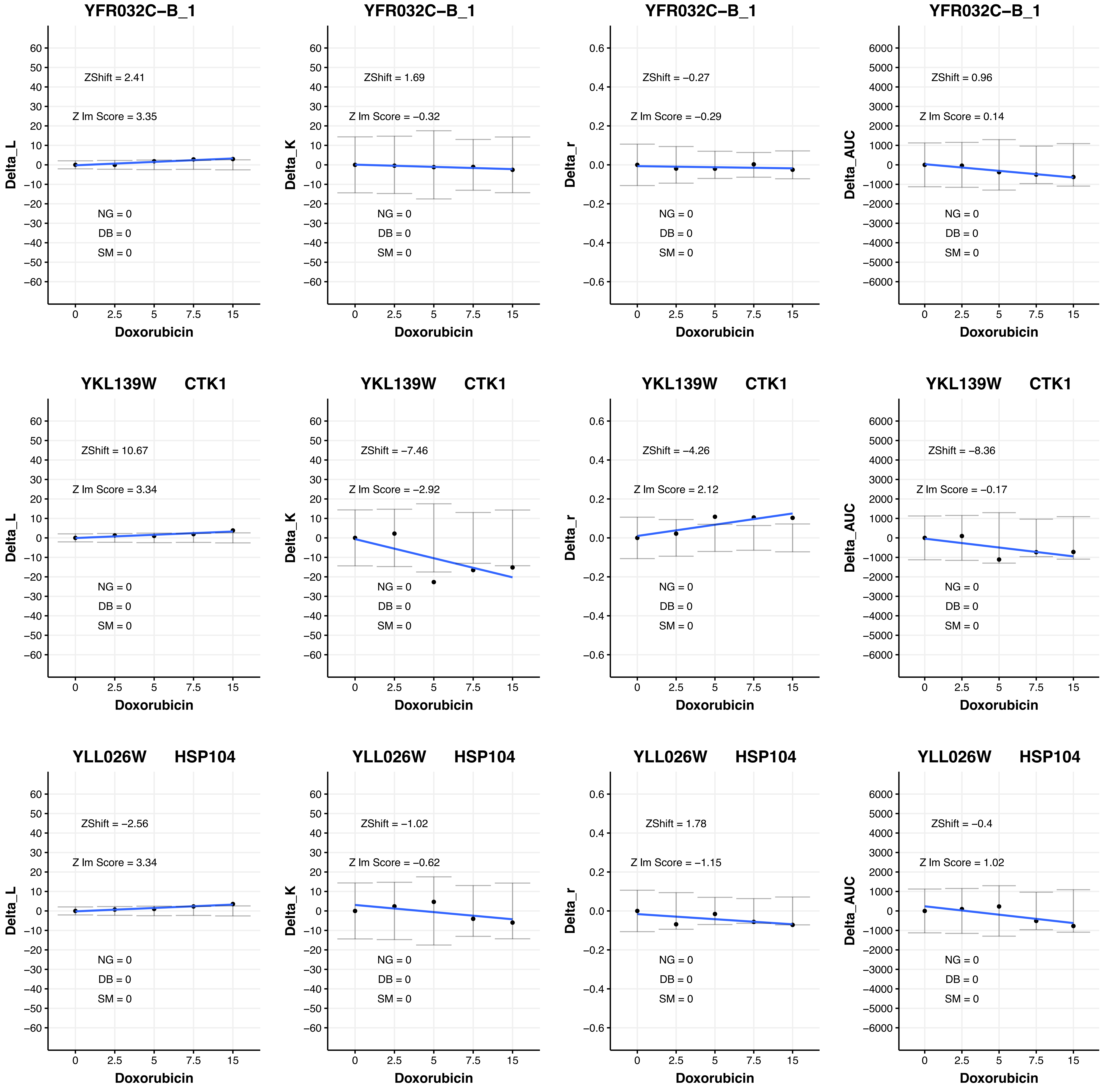
YFR032C−B_1
ZShift = 2.41
Z lm Score = 3.35
NG = 0
DB = 0
SM = 0
Delta_L
Doxorubicin
YFR032C−B_1
ZShift = 1.69
Z lm Score = −0.32
NG = 0
DB = 0
SM = 0
Delta_K
Doxorubicin
YFR032C−B_1
ZShift = −0.27
Z lm Score = −0.29
NG = 0
DB = 0
SM = 0
Delta_r
Doxorubicin
YFR032C−B_1
ZShift = 0.96
Z lm Score = 0.14
NG = 0
DB = 0
SM = 0
Delta_AUC
Doxorubicin
YKL139W CTK1
ZShift = 10.67
Z lm Score = 3.34
NG = 0
DB = 0
SM = 0
Delta_L
Doxorubicin
YKL139W CTK1
ZShift = −7.46
Z lm Score = −2.92
NG = 0
DB = 0
SM = 0
Delta_K
Doxorubicin
YKL139W CTK1
ZShift = −4.26
Z lm Score = 2.12
NG = 0
DB = 0
SM = 0
Delta_r
Doxorubicin
YKL139W CTK1
ZShift = −8.36
Z lm Score = −0.17
NG = 0
DB = 0
SM = 0
Delta_AUC
Doxorubicin
YLL026W HSP104
ZShift = −2.56
Z lm Score = 3.34
NG = 0
DB = 0
SM = 0
Delta_L
Doxorubicin
YLL026W HSP104
ZShift = −1.02
Z lm Score = −0.62
NG = 0
DB = 0
SM = 0
Delta_K
Doxorubicin
YLL026W HSP104
ZShift = 1.78
Z lm Score = −1.15
NG = 0
DB = 0
SM = 0
Delta_r
Doxorubicin
YLL026W HSP104
ZShift = −0.4
Z lm Score = 1.02
NG = 0
DB = 0
SM = 0
Delta_AUC
Doxorubicin

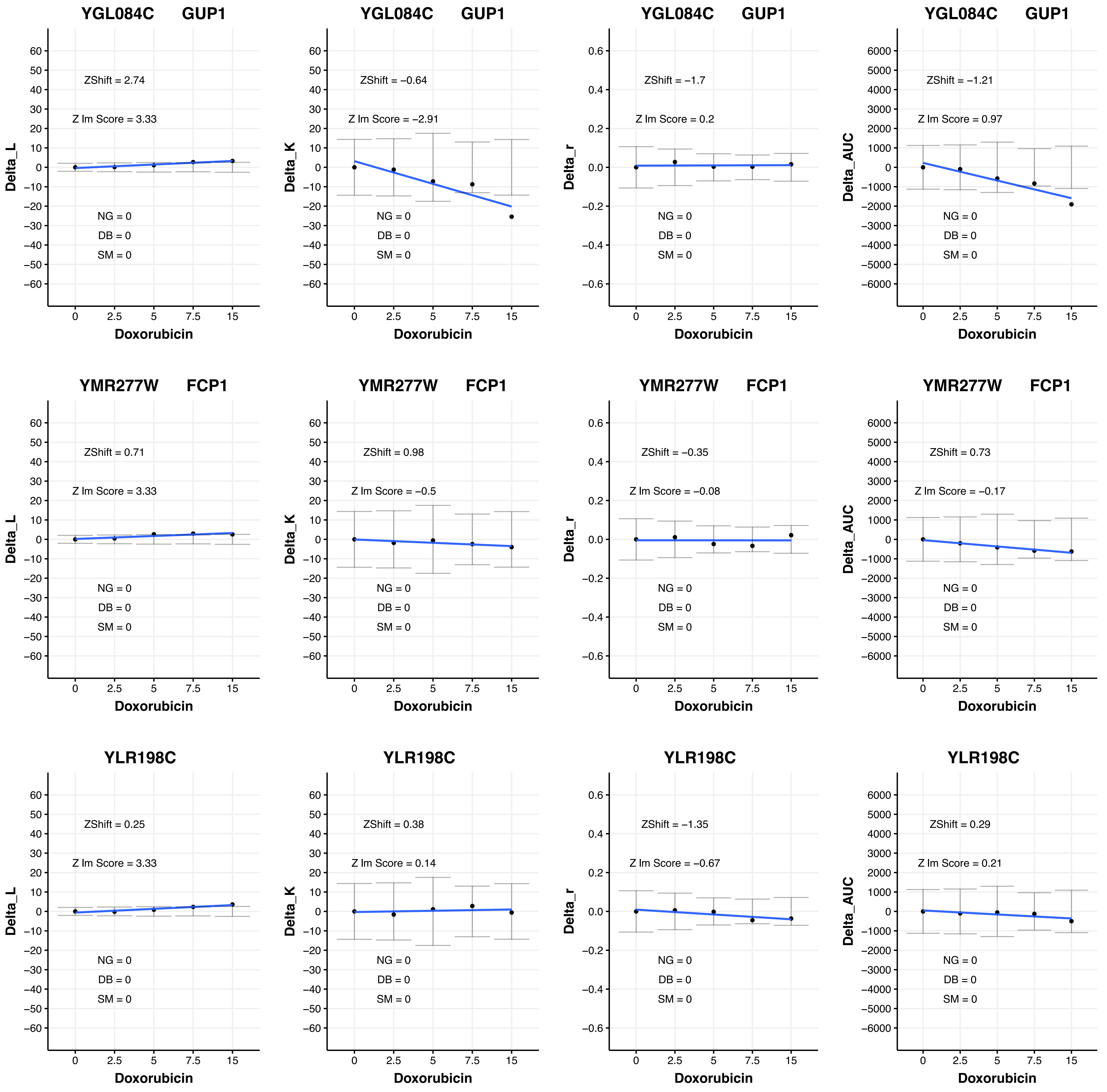
YGL084C GUP1
ZShift = 2.74
Z lm Score = 3.33
NG = 0
DB = 0
SM = 0
Delta_L
Doxorubicin
YGL084C GUP1
ZShift = −0.64
Z lm Score = −2.91
NG = 0
DB = 0
SM = 0
Delta_K
Doxorubicin
YGL084C GUP1
ZShift = −1.7
Z lm Score = 0.2
NG = 0
DB = 0
SM = 0
Delta_r
Doxorubicin
YGL084C GUP1
ZShift = −1.21
Z lm Score = 0.97
NG = 0
DB = 0
SM = 0
Delta_AUC
Doxorubicin
YMR277W FCP1
ZShift = 0.71
Z lm Score = 3.33
NG = 0
DB = 0
SM = 0
Delta_L
Doxorubicin
YMR277W FCP1
ZShift = 0.98
Z lm Score = −0.5
NG = 0
DB = 0
SM = 0
Delta_K
Doxorubicin
YMR277W FCP1
ZShift = −0.35
Z lm Score = −0.08
NG = 0
DB = 0
SM = 0
Delta_r
Doxorubicin
YMR277W FCP1
ZShift = 0.73
Z lm Score = −0.17
NG = 0
DB = 0
SM = 0
Delta_AUC
Doxorubicin
YLR198C
ZShift = 0.25
Z lm Score = 3.33
NG = 0
DB = 0
SM = 0
Delta_L
Doxorubicin
YLR198C
ZShift = 0.38
Z lm Score = 0.14
NG = 0
DB = 0
SM = 0
Delta_K
Doxorubicin
YLR198C
ZShift = −1.35
Z lm Score = −0.67
NG = 0
DB = 0
SM = 0
Delta_r
Doxorubicin
YLR198C
ZShift = 0.29
Z lm Score = 0.21
NG = 0
DB = 0
SM = 0
Delta_AUC
Doxorubicin

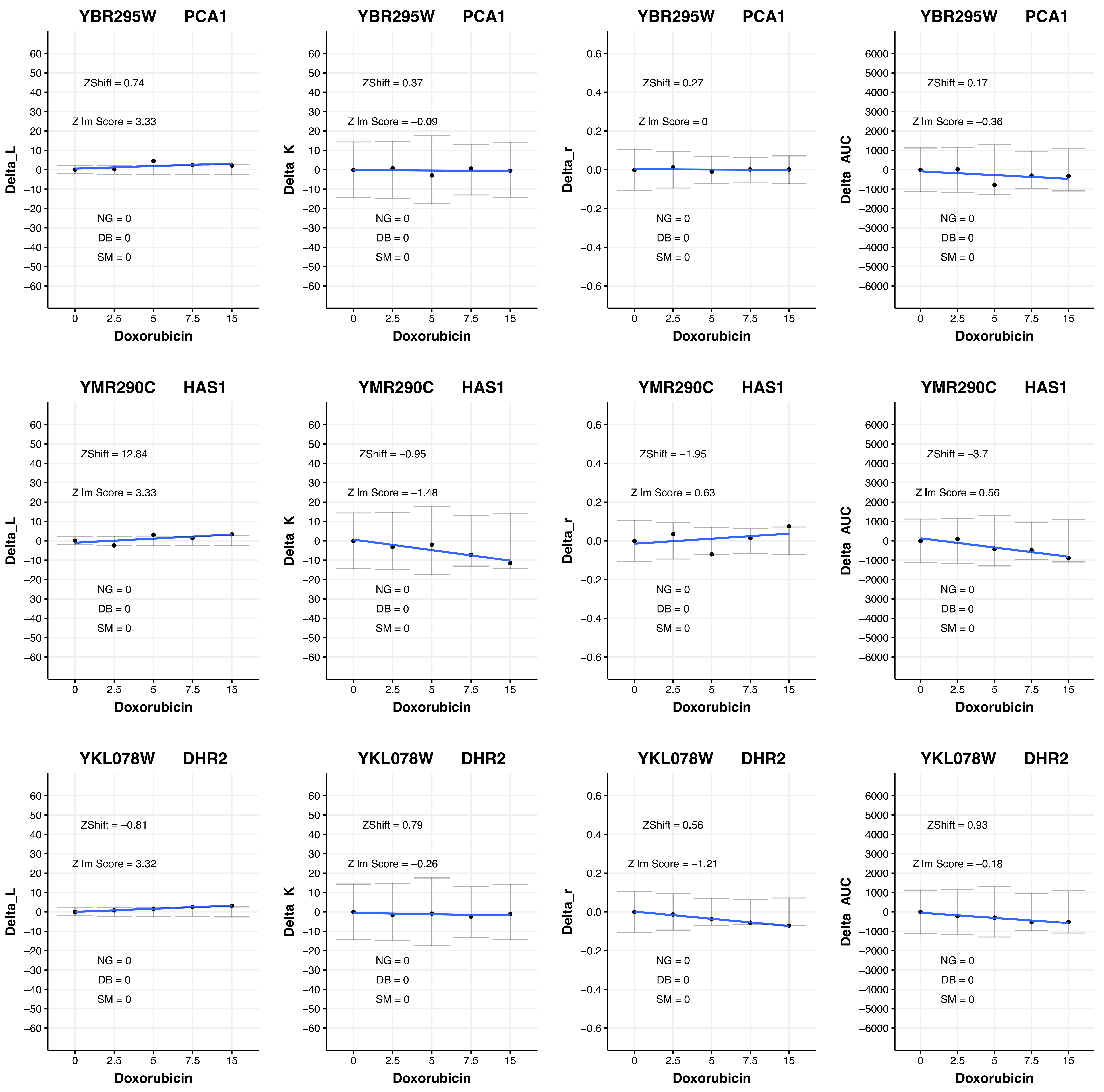
YBR295W PCA1
ZShift = 0.74
Z lm Score = 3.33
NG = 0
DB = 0
SM = 0
Delta_L
Doxorubicin
YBR295W PCA1
ZShift = 0.37
Z lm Score = −0.09
NG = 0
DB = 0
SM = 0
Delta_K
Doxorubicin
YBR295W PCA1
ZShift = 0.27
Z lm Score = 0
NG = 0
DB = 0
SM = 0
Delta_r
Doxorubicin
YBR295W PCA1
ZShift = 0.17
Z lm Score = −0.36
NG = 0
DB = 0
SM = 0
Delta_AUC
Doxorubicin
YMR290C HAS1
ZShift = 12.84
Z lm Score = 3.33
NG = 0
DB = 0
SM = 0
Delta_L
Doxorubicin
YMR290C HAS1
ZShift = −0.95
Z lm Score = −1.48
NG = 0
DB = 0
SM = 0
Delta_K
Doxorubicin
YMR290C HAS1
ZShift = −1.95
Z lm Score = 0.63
NG = 0
DB = 0
SM = 0
Delta_r
Doxorubicin
YMR290C HAS1
ZShift = −3.7
Z lm Score = 0.56
NG = 0
DB = 0
SM = 0
Delta_AUC
Doxorubicin
YKL078W DHR2
ZShift = −0.81
Z lm Score = 3.32
NG = 0
DB = 0
SM = 0
Delta_L
Doxorubicin
YKL078W DHR2
ZShift = 0.79
Z lm Score = −0.26
NG = 0
DB = 0
SM = 0
Delta_K
Doxorubicin
YKL078W DHR2
ZShift = 0.56
Z lm Score = −1.21
NG = 0
DB = 0
SM = 0
Delta_r
Doxorubicin
YKL078W DHR2
ZShift = 0.93
Z lm Score = −0.18
NG = 0
DB = 0
SM = 0
Delta_AUC
Doxorubicin

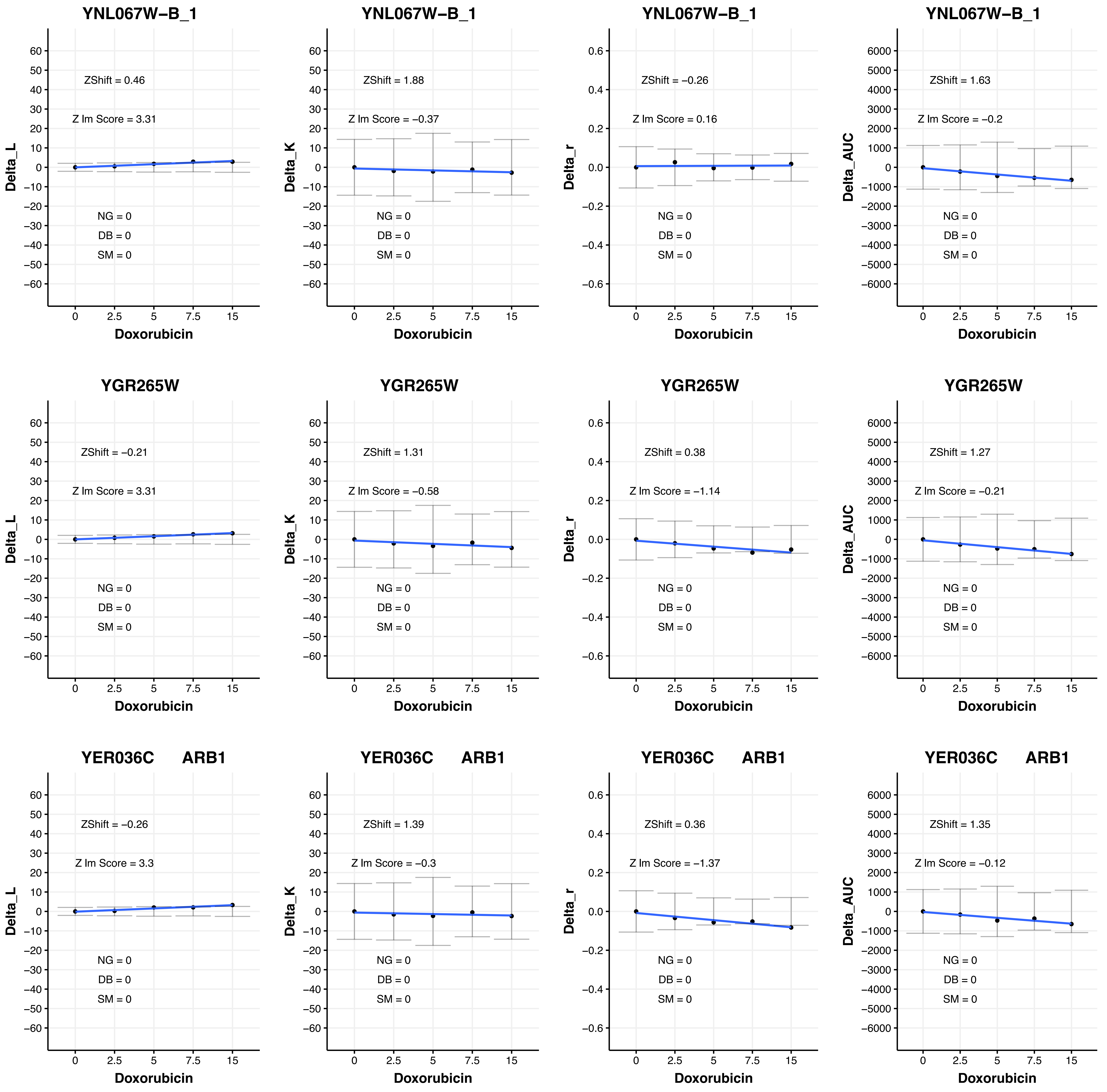
YNL067W−B_1
ZShift = 0.46
Z lm Score = 3.31
NG = 0
DB = 0
SM = 0
Delta_L
Doxorubicin
YNL067W−B_1
ZShift = 1.88
Z lm Score = −0.37
NG = 0
DB = 0
SM = 0
Delta_K
Doxorubicin
YNL067W−B_1
ZShift = −0.26
Z lm Score = 0.16
NG = 0
DB = 0
SM = 0
Delta_r
Doxorubicin
YNL067W−B_1
ZShift = 1.63
Z lm Score = −0.2
NG = 0
DB = 0
SM = 0
Delta_AUC
Doxorubicin
YGR265W
ZShift = −0.21
Z lm Score = 3.31
NG = 0
DB = 0
SM = 0
Delta_L
Doxorubicin
YGR265W
ZShift = 1.31
Z lm Score = −0.58
NG = 0
DB = 0
SM = 0
Delta_K
Doxorubicin
YGR265W
ZShift = 0.38
Z lm Score = −1.14
NG = 0
DB = 0
SM = 0
Delta_r
Doxorubicin
YGR265W
ZShift = 1.27
Z lm Score = −0.21
NG = 0
DB = 0
SM = 0
Delta_AUC
Doxorubicin
YER036C ARB1
ZShift = −0.26
Z lm Score = 3.3
NG = 0
DB = 0
SM = 0
Delta_L
Doxorubicin
YER036C ARB1
ZShift = 1.39
Z lm Score = −0.3
NG = 0
DB = 0
SM = 0
Delta_K
Doxorubicin
YER036C ARB1
ZShift = 0.36
Z lm Score = −1.37
NG = 0
DB = 0
SM = 0
Delta_r
Doxorubicin
YER036C ARB1
ZShift = 1.35
Z lm Score = −0.12
NG = 0
DB = 0
SM = 0
Delta_AUC
Doxorubicin

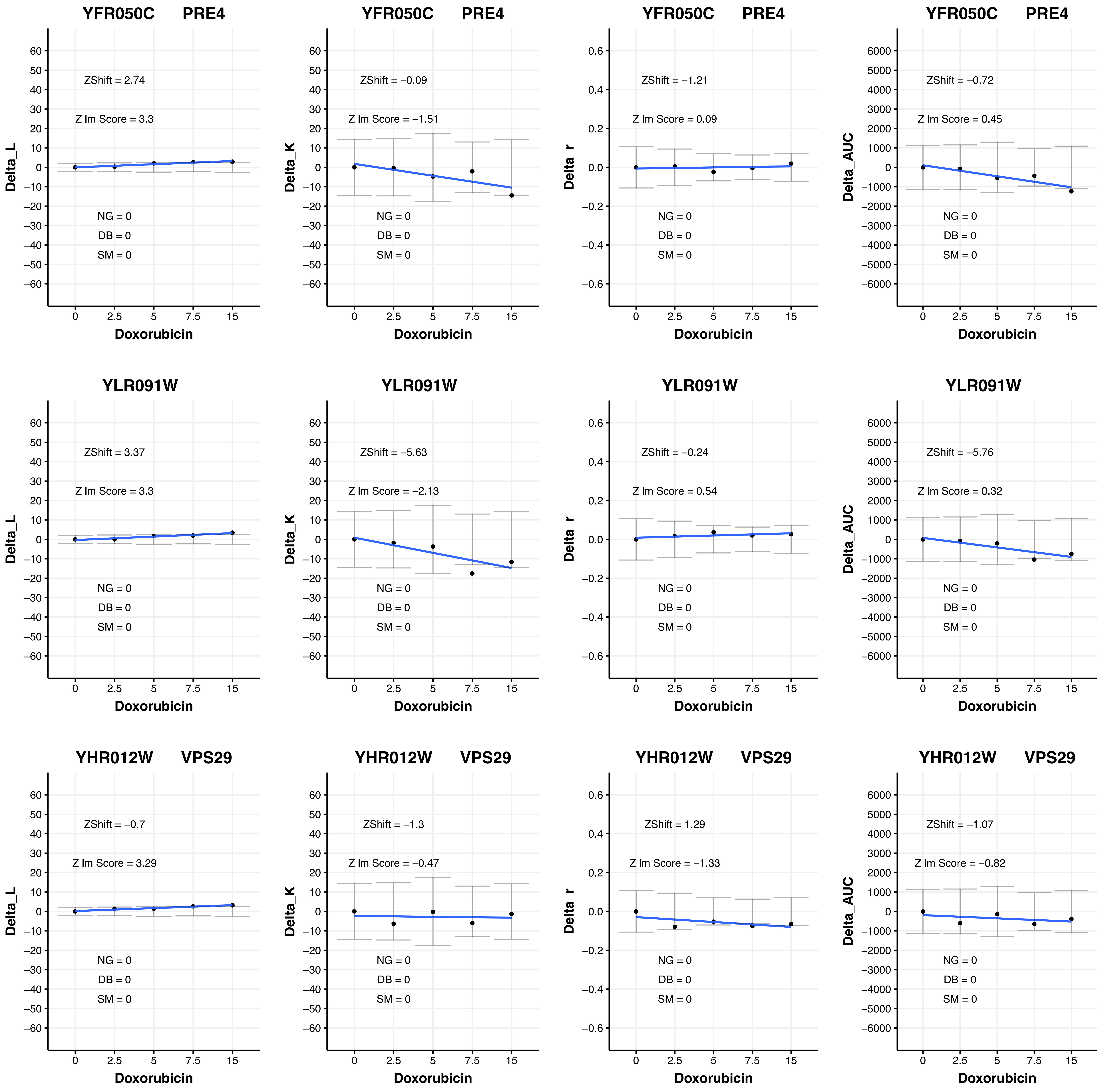
YFR050C PRE4
ZShift = 2.74
Z lm Score = 3.3
NG = 0
DB = 0
SM = 0
Delta_L
Doxorubicin
YFR050C PRE4
ZShift = −0.09
Z lm Score = −1.51
NG = 0
DB = 0
SM = 0
Delta_K
Doxorubicin
YFR050C PRE4
ZShift = −1.21
Z lm Score = 0.09
NG = 0
DB = 0
SM = 0
Delta_r
Doxorubicin
YFR050C PRE4
ZShift = −0.72
Z lm Score = 0.45
NG = 0
DB = 0
SM = 0
Delta_AUC
Doxorubicin
YLR091W
ZShift = 3.37
Z lm Score = 3.3
NG = 0
DB = 0
SM = 0
Delta_L
Doxorubicin
YLR091W
ZShift = −5.63
Z lm Score = −2.13
NG = 0
DB = 0
SM = 0
Delta_K
Doxorubicin
YLR091W
ZShift = −0.24
Z lm Score = 0.54
NG = 0
DB = 0
SM = 0
Delta_r
Doxorubicin
YLR091W
ZShift = −5.76
Z lm Score = 0.32
NG = 0
DB = 0
SM = 0
Delta_AUC
Doxorubicin
YHR012W VPS29
ZShift = −0.7
Z lm Score = 3.29
NG = 0
DB = 0
SM = 0
Delta_L
Doxorubicin
YHR012W VPS29
ZShift = −1.3
Z lm Score = −0.47
NG = 0
DB = 0
SM = 0
Delta_K
Doxorubicin
YHR012W VPS29
ZShift = 1.29
Z lm Score = −1.33
NG = 0
DB = 0
SM = 0
Delta_r
Doxorubicin
YHR012W VPS29
ZShift = −1.07
Z lm Score = −0.82
NG = 0
DB = 0
SM = 0
Delta_AUC
Doxorubicin

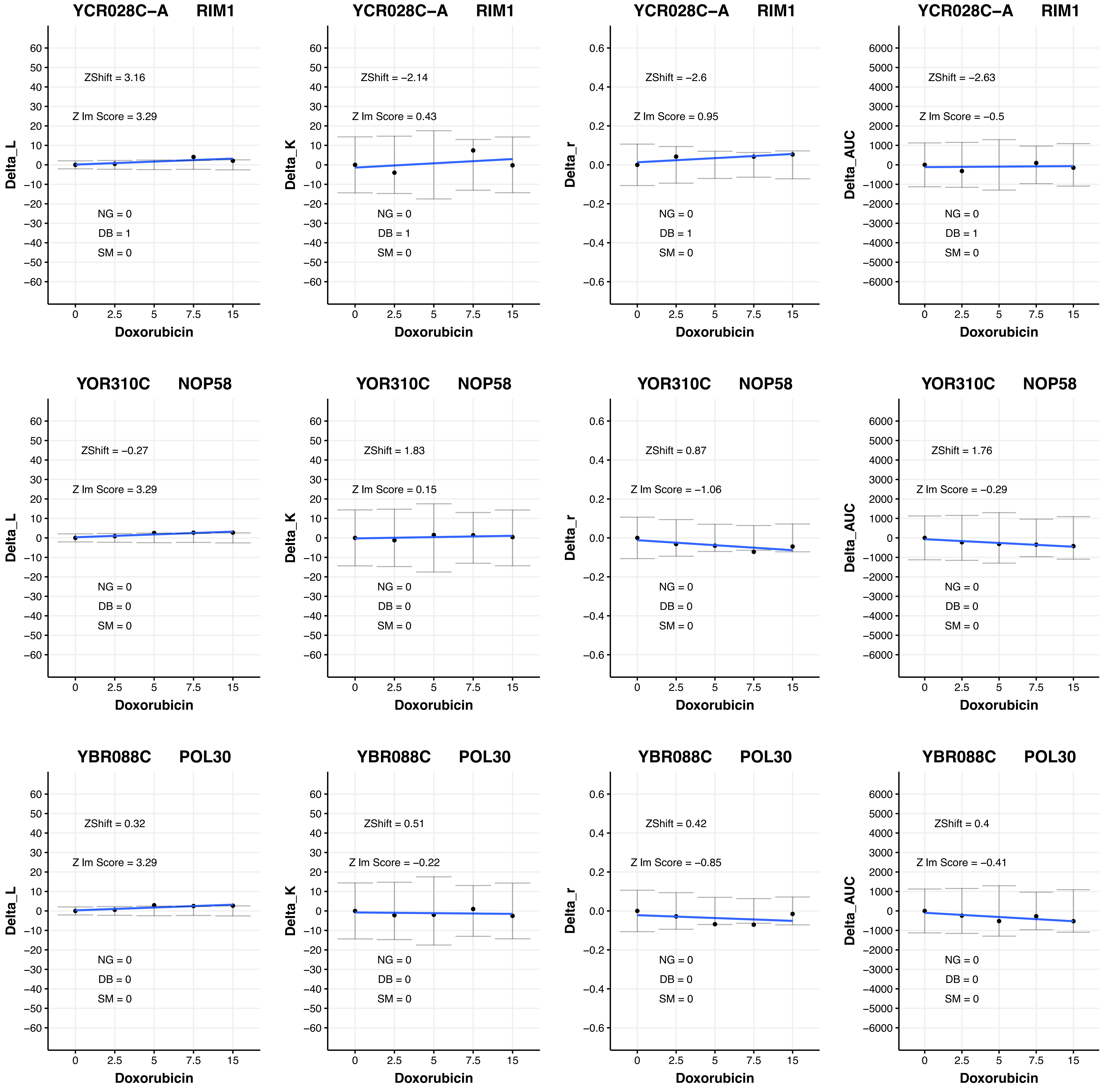

YCR028C−A RIM1
ZShift = 3.16
Z lm Score = 3.29
NG = 0
DB = 1
SM = 0
Delta_L
Doxorubicin
YCR028C−A RIM1
ZShift = −2.14
Z lm Score = 0.43
NG = 0
DB = 1
SM = 0
Delta_K
Doxorubicin
YCR028C−A RIM1
ZShift = −2.6
Z lm Score = 0.95
NG = 0
DB = 1
SM = 0
Delta_r
Doxorubicin
YCR028C−A RIM1
ZShift = −2.63
Z lm Score = −0.5
NG = 0
DB = 1
SM = 0
Delta_AUC
Doxorubicin
YOR310C NOP58
ZShift = −0.27
Z lm Score = 3.29
NG = 0
DB = 0
SM = 0
Delta_L
Doxorubicin
YOR310C NOP58
ZShift = 1.83
Z lm Score = 0.15
NG = 0
DB = 0
SM = 0
Delta_K
Doxorubicin
YOR310C NOP58
ZShift = 0.87
Z lm Score = −1.06
NG = 0
DB = 0
SM = 0
Delta_r
Doxorubicin
YOR310C NOP58
ZShift = 1.76
Z lm Score = −0.29
NG = 0
DB = 0
SM = 0
Delta_AUC
Doxorubicin
YBR088C POL30
ZShift = 0.32
Z lm Score = 3.29
NG = 0
DB = 0
SM = 0
Delta_L
Doxorubicin
YBR088C POL30
ZShift = 0.51
Z lm Score = −0.22
NG = 0
DB = 0
SM = 0
Delta_K
Doxorubicin
YBR088C POL30
ZShift = 0.42
Z lm Score = −0.85
NG = 0
DB = 0
SM = 0
Delta_r
Doxorubicin
YBR088C POL30
ZShift = 0.4
Z lm Score = −0.41
NG = 0
DB = 0
SM = 0
Delta_AUC
Doxorubicin

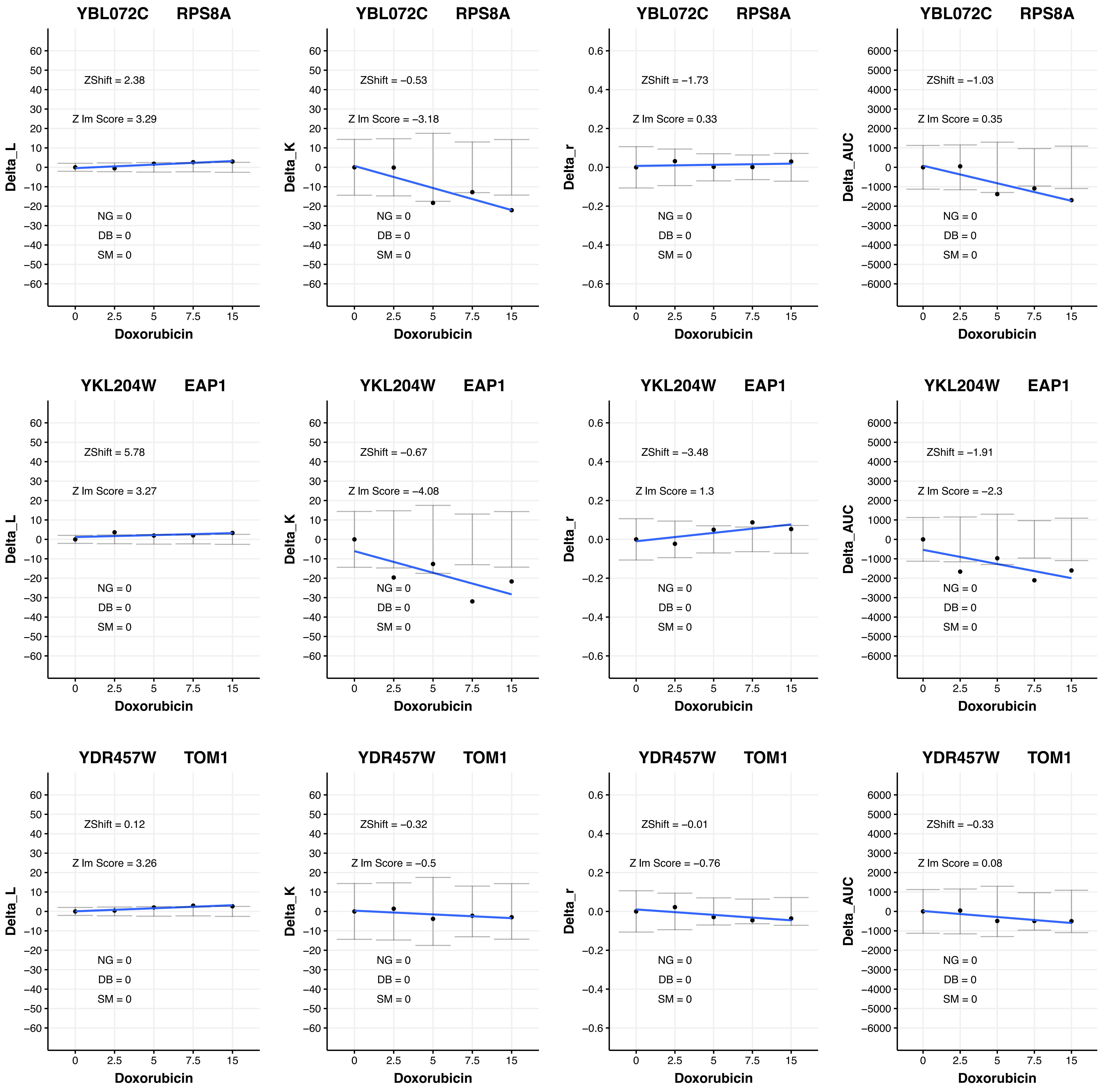
YBL072C RPS8A
ZShift = 2.38
Z lm Score = 3.29
NG = 0
DB = 0
SM = 0
Delta_L
Doxorubicin
YBL072C RPS8A
ZShift = −0.53
Z lm Score = −3.18
NG = 0
DB = 0
SM = 0
Delta_K
Doxorubicin
YBL072C RPS8A
ZShift = −1.73
Z lm Score = 0.33
NG = 0
DB = 0
SM = 0
Delta_r
Doxorubicin
YBL072C RPS8A
ZShift = −1.03
Z lm Score = 0.35
NG = 0
DB = 0
SM = 0
Delta_AUC
Doxorubicin
YKL204W EAP1
ZShift = 5.78
Z lm Score = 3.27
NG = 0
DB = 0
SM = 0
Delta_L
Doxorubicin
YKL204W EAP1
ZShift = −0.67
Z lm Score = −4.08
NG = 0
DB = 0
SM = 0
Delta_K
Doxorubicin
YKL204W EAP1
ZShift = −3.48
Z lm Score = 1.3
NG = 0
DB = 0
SM = 0
Delta_r
Doxorubicin
YKL204W EAP1
ZShift = −1.91
Z lm Score = −2.3
NG = 0
DB = 0
SM = 0
Delta_AUC
Doxorubicin
YDR457W TOM1
ZShift = 0.12
Z lm Score = 3.26
NG = 0
DB = 0
SM = 0
Delta_L
Doxorubicin
YDR457W TOM1
ZShift = −0.32
Z lm Score = −0.5
NG = 0
DB = 0
SM = 0
Delta_K
Doxorubicin
YDR457W TOM1
ZShift = −0.01
Z lm Score = −0.76
NG = 0
DB = 0
SM = 0
Delta_r
Doxorubicin
YDR457W TOM1
ZShift = −0.33
Z lm Score = 0.08
NG = 0
DB = 0
SM = 0
Delta_AUC
Doxorubicin

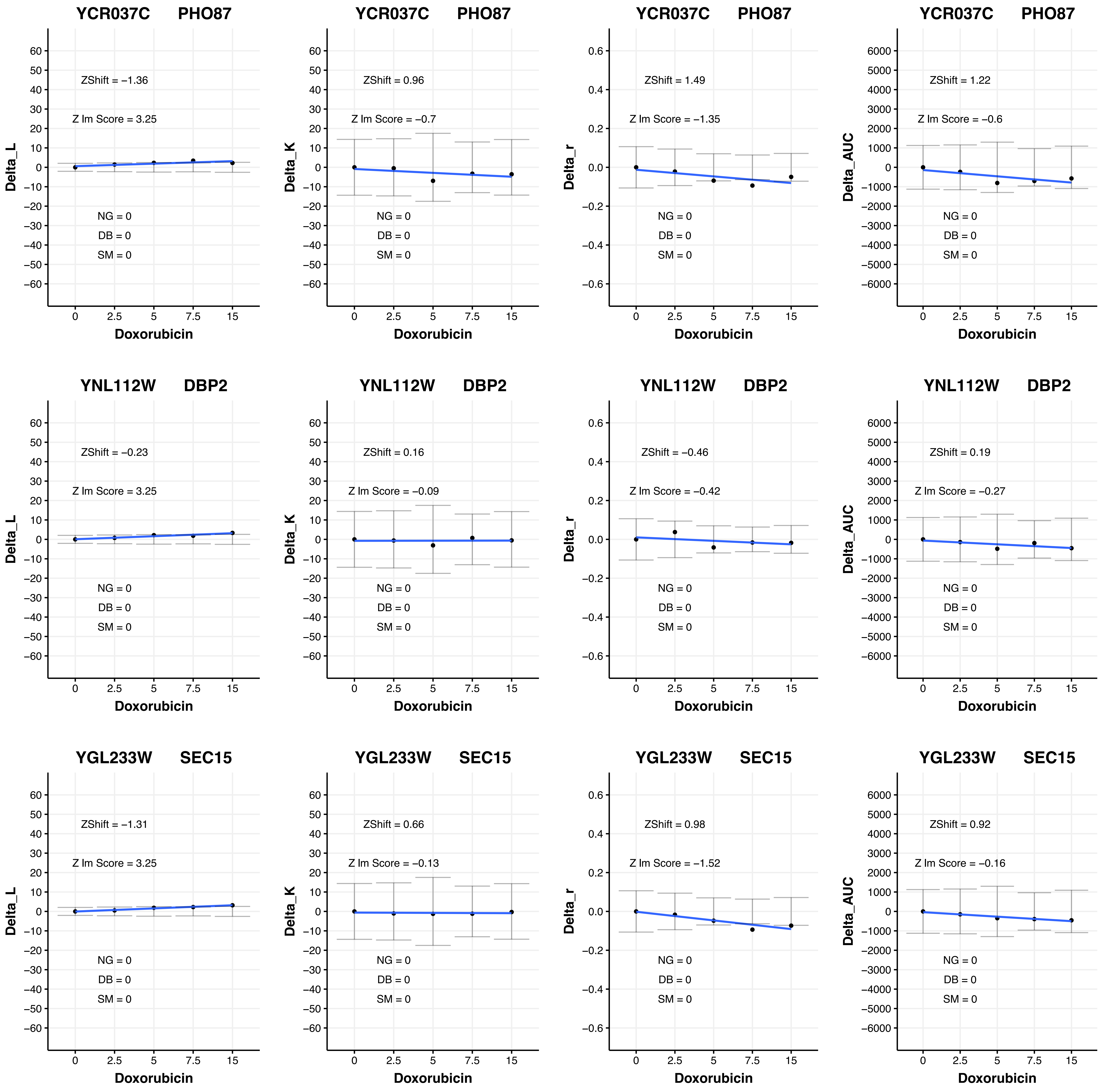

YCR037C PHO87
ZShift = −1.36
Z lm Score = 3.25
NG = 0
DB = 0
SM = 0
Delta_L
Doxorubicin
YCR037C PHO87
ZShift = 0.96
Z lm Score = −0.7
NG = 0
DB = 0
SM = 0
Delta_K
Doxorubicin
YCR037C PHO87
ZShift = 1.49
Z lm Score = −1.35
NG = 0
DB = 0
SM = 0
Delta_r
Doxorubicin
YCR037C PHO87
ZShift = 1.22
Z lm Score = −0.6
NG = 0
DB = 0
SM = 0
Delta_AUC
Doxorubicin
YNL112W DBP2
ZShift = −0.23
Z lm Score = 3.25
NG = 0
DB = 0
SM = 0
Delta_L
Doxorubicin
YNL112W DBP2
ZShift = 0.16
Z lm Score = −0.09
NG = 0
DB = 0
SM = 0
Delta_K
Doxorubicin
YNL112W DBP2
ZShift = −0.46
Z lm Score = −0.42
NG = 0
DB = 0
SM = 0
Delta_r
Doxorubicin
YNL112W DBP2
ZShift = 0.19
Z lm Score = −0.27
NG = 0
DB = 0
SM = 0
Delta_AUC
Doxorubicin
YGL233W SEC15
ZShift = −1.31
Z lm Score = 3.25
NG = 0
DB = 0
SM = 0
Delta_L
Doxorubicin
YGL233W SEC15
ZShift = 0.66
Z lm Score = −0.13
NG = 0
DB = 0
SM = 0
Delta_K
Doxorubicin
YGL233W SEC15
ZShift = 0.98
Z lm Score = −1.52
NG = 0
DB = 0
SM = 0
Delta_r
Doxorubicin
YGL233W SEC15
ZShift = 0.92
Z lm Score = −0.16
NG = 0
DB = 0
SM = 0
Delta_AUC
Doxorubicin

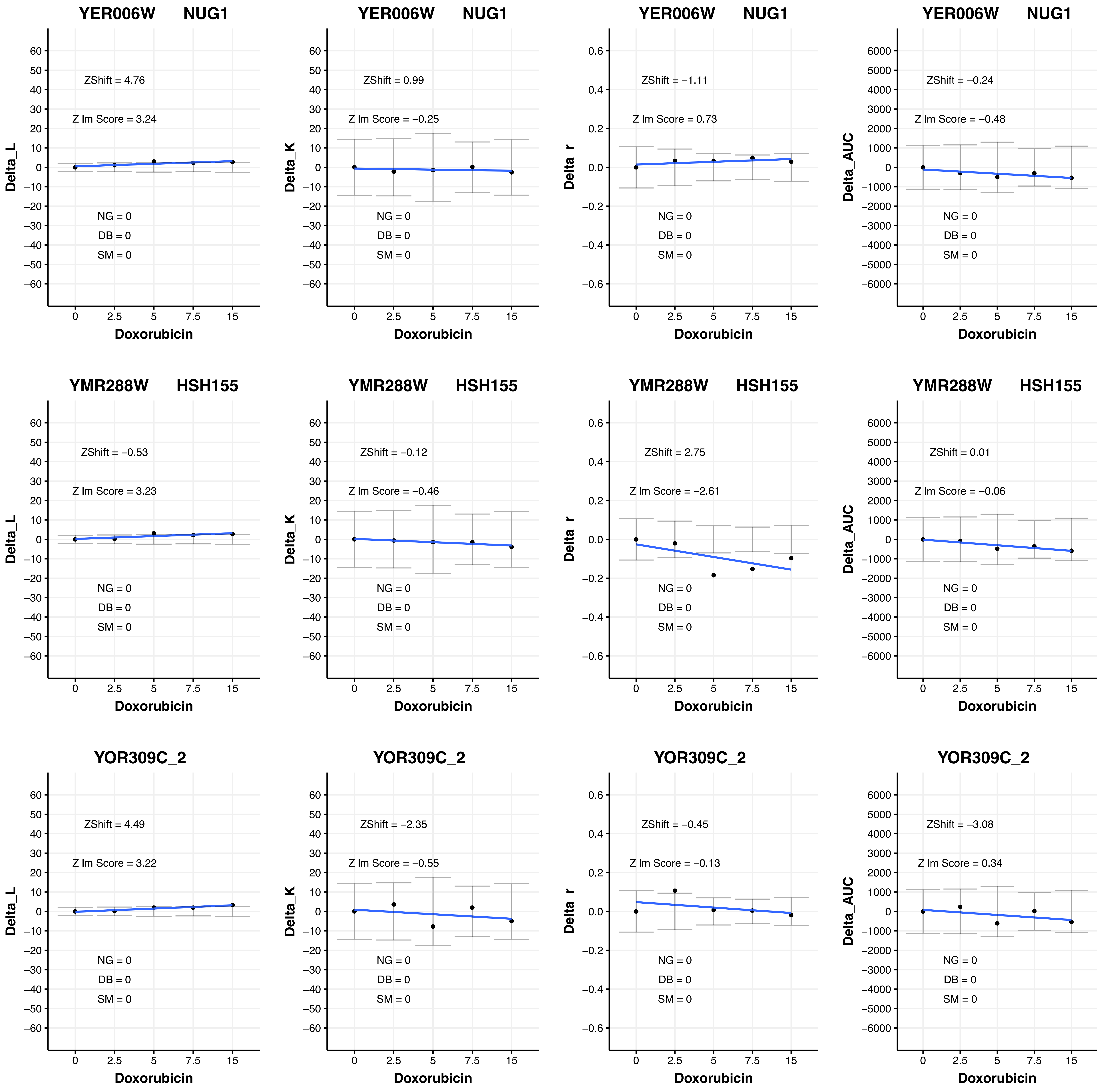
YER006W NUG1
ZShift = 4.76
Z lm Score = 3.24
Delta_L
NG = 0
DB = 0
SM = 0
Doxorubicin
YER006W NUG1
ZShift = 0.99
Z lm Score = −0.25
Delta_K
NG = 0
DB = 0
SM = 0
Doxorubicin
YER006W NUG1
ZShift = −1.11
Z lm Score = 0.73
Delta_r
NG = 0
DB = 0
SM = 0
Doxorubicin
YER006W NUG1
ZShift = −0.24
Z lm Score = −0.48
Delta_AUC
NG = 0
DB = 0
SM = 0
Doxorubicin
YMR288W HSH155
ZShift = −0.53
Z lm Score = 3.23
Delta_L
NG = 0
DB = 0
SM = 0
Doxorubicin
YMR288W HSH155
ZShift = −0.12
Z lm Score = −0.46
Delta_K
NG = 0
DB = 0
SM = 0
Doxorubicin
YMR288W HSH155
ZShift = 2.75
Z lm Score = −2.61
Delta_r
NG = 0
DB = 0
SM = 0
Doxorubicin
YMR288W HSH155
ZShift = 0.01
Z lm Score = −0.06
Delta_AUC
NG = 0
DB = 0
SM = 0
Doxorubicin
YOR309C_2
ZShift = 4.49
Z lm Score = 3.22
Delta_L
NG = 0
DB = 0
SM = 0
Doxorubicin
YOR309C_2
ZShift = −2.35
Z lm Score = −0.55
Delta_K
NG = 0
DB = 0
SM = 0
Doxorubicin
YOR309C_2
ZShift = −0.45
Z lm Score = −0.13
Delta_r
NG = 0
DB = 0
SM = 0
Doxorubicin
YOR309C_2
ZShift = −3.08
Z lm Score = 0.34
Delta_AUC
NG = 0
DB = 0
SM = 0
Doxorubicin

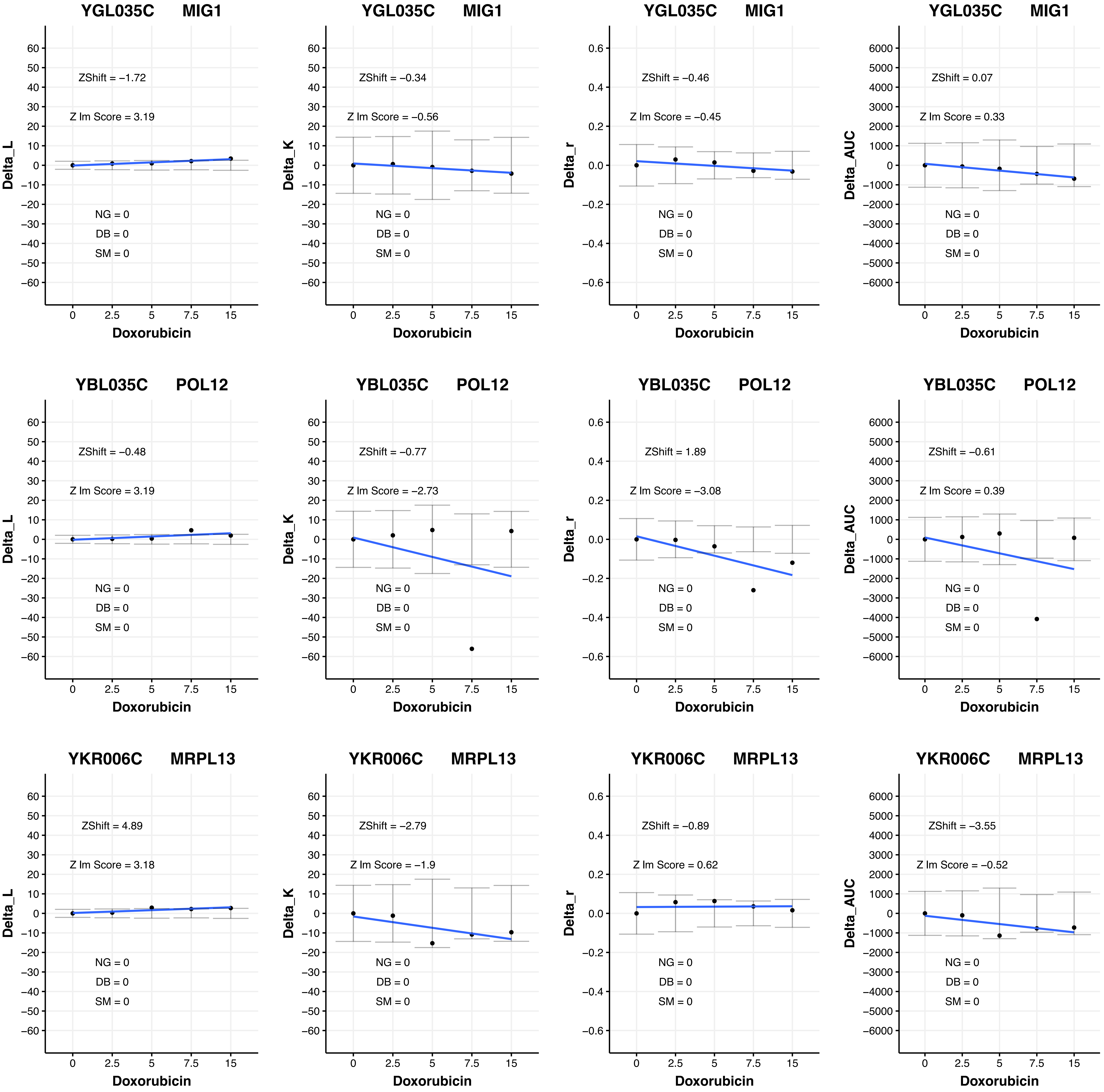
YGL035C MIG1
ZShift = −1.72
Z lm Score = 3.19
NG = 0
DB = 0
SM = 0
Delta_L
Doxorubicin
YGL035C MIG1
ZShift = −0.34
Z lm Score = −0.56
NG = 0
DB = 0
SM = 0
Delta_K
Doxorubicin
YGL035C MIG1
ZShift = −0.46
Z lm Score = −0.45
NG = 0
DB = 0
SM = 0
Delta_r
Doxorubicin
YGL035C MIG1
ZShift = 0.07
Z lm Score = 0.33
NG = 0
DB = 0
SM = 0
Delta_AUC
Doxorubicin
YBL035C POL12
ZShift = −0.48
Z lm Score = 3.19
NG = 0
DB = 0
SM = 0
Delta_L
Doxorubicin
YBL035C POL12
ZShift = −0.77
Z lm Score = −2.73
NG = 0
DB = 0
SM = 0
Delta_K
Doxorubicin
YBL035C POL12
ZShift = 1.89
Z lm Score = −3.08
NG = 0
DB = 0
SM = 0
Delta_r
Doxorubicin
YBL035C POL12
ZShift = −0.61
Z lm Score = 0.39
NG = 0
DB = 0
SM = 0
Delta_AUC
Doxorubicin
YKR006C MRPL13
ZShift = 4.89
Z lm Score = 3.18
NG = 0
DB = 0
SM = 0
Delta_L
Doxorubicin
YKR006C MRPL13
ZShift = −2.79
Z lm Score = −1.9
NG = 0
DB = 0
SM = 0
Delta_K
Doxorubicin
YKR006C MRPL13
ZShift = −0.89
Z lm Score = 0.62
NG = 0
DB = 0
SM = 0
Delta_r
Doxorubicin
YKR006C MRPL13
ZShift = −3.55
Z lm Score = −0.52
NG = 0
DB = 0
SM = 0
Delta_AUC
Doxorubicin

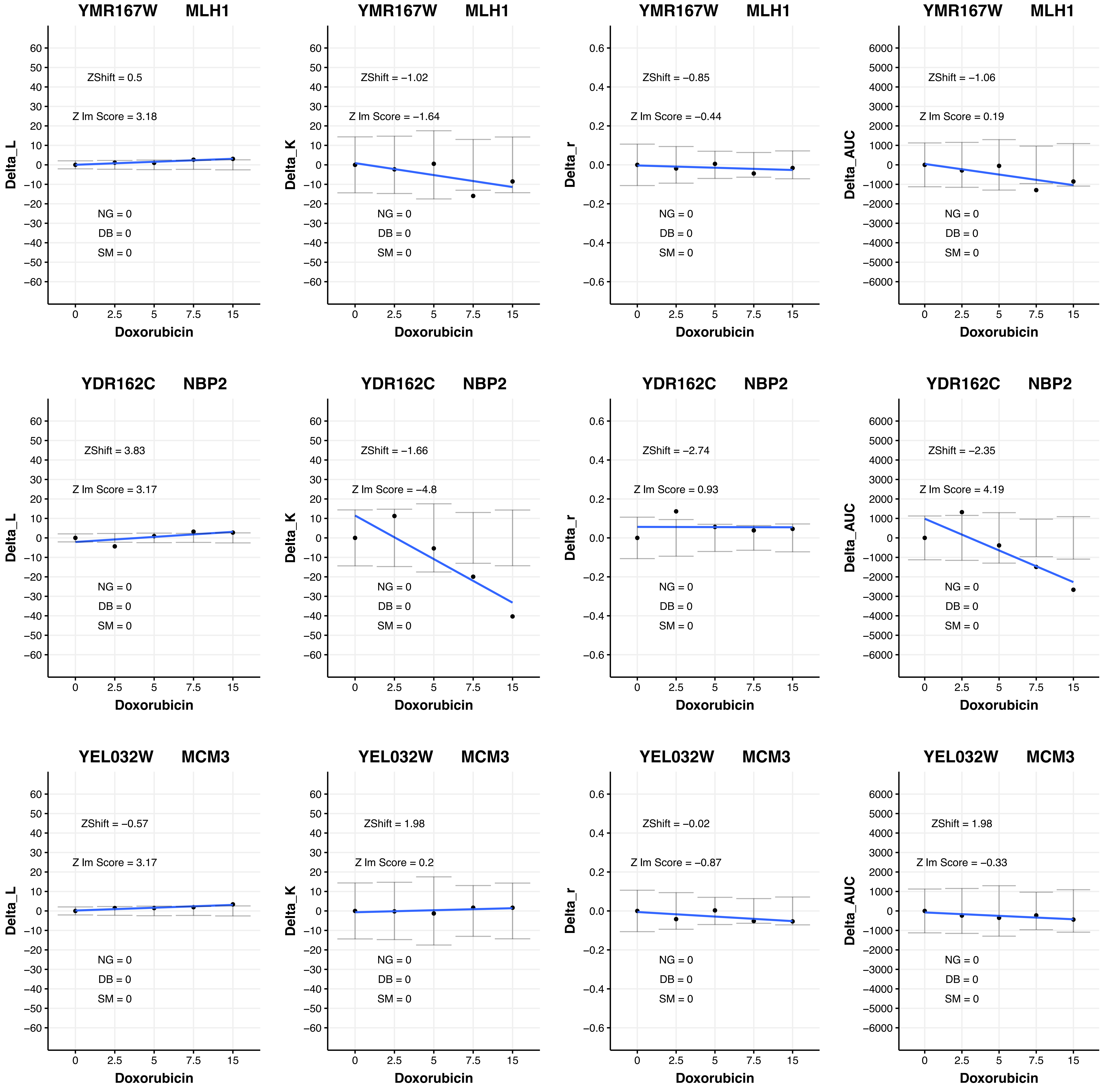
YMR167W MLH1
ZShift = 0.5
Z lm Score = 3.18
NG = 0
DB = 0
SM = 0
Delta_L
Doxorubicin
YMR167W MLH1
ZShift = −1.02
Z lm Score = −1.64
NG = 0
DB = 0
SM = 0
Delta_K
Doxorubicin
YMR167W MLH1
ZShift = −0.85
Z lm Score = −0.44
NG = 0
DB = 0
SM = 0
Delta_r
Doxorubicin
YMR167W MLH1
ZShift = −1.06
Z lm Score = 0.19
NG = 0
DB = 0
SM = 0
Delta_AUC
Doxorubicin
YDR162C NBP2
ZShift = 3.83
Z lm Score = 3.17
NG = 0
DB = 0
SM = 0
Delta_L
Doxorubicin
YDR162C NBP2
ZShift = −1.66
Z lm Score = −4.8
NG = 0
DB = 0
SM = 0
Delta_K
Doxorubicin
YDR162C NBP2
ZShift = −2.74
Z lm Score = 0.93
NG = 0
DB = 0
SM = 0
Delta_r
Doxorubicin
YDR162C NBP2
ZShift = −2.35
Z lm Score = 4.19
NG = 0
DB = 0
SM = 0
Delta_AUC
Doxorubicin
YEL032W MCM3
ZShift = −0.57
Z lm Score = 3.17
NG = 0
DB = 0
SM = 0
Delta_L
Doxorubicin
YEL032W MCM3
ZShift = 1.98
Z lm Score = 0.2
NG = 0
DB = 0
SM = 0
Delta_K
Doxorubicin
YEL032W MCM3
ZShift = −0.02
Z lm Score = −0.87
NG = 0
DB = 0
SM = 0
Delta_r
Doxorubicin
YEL032W MCM3
ZShift = 1.98
Z lm Score = −0.33
NG = 0
DB = 0
SM = 0
Delta_AUC
Doxorubicin

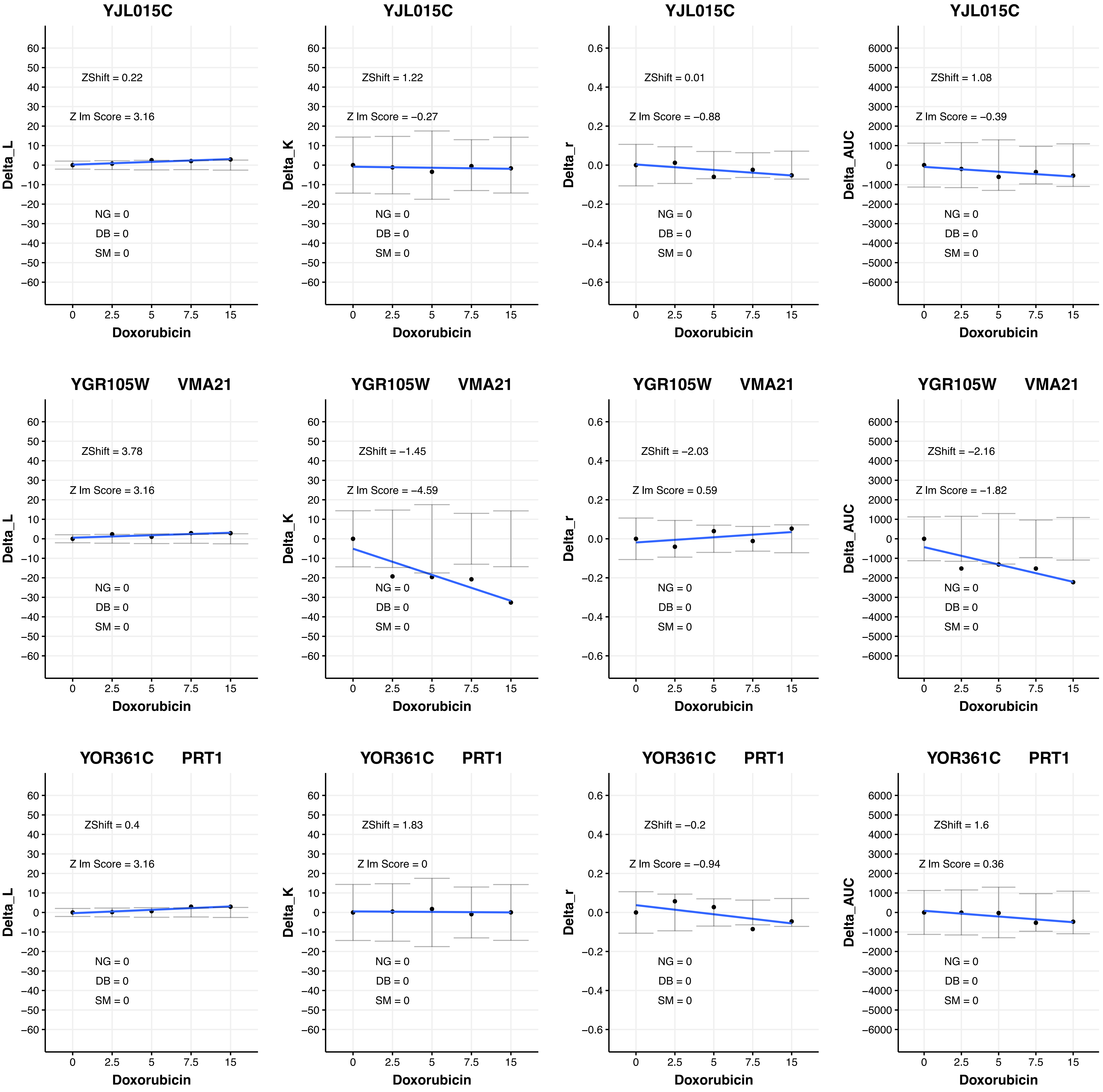
YJL015C
Delta_L
ZShift = 0.22
Z lm Score = 3.16
NG = 0
DB = 0
SM = 0
Doxorubicin
YJL015C
Delta_K
ZShift = 1.22
Z lm Score = −0.27
NG = 0
DB = 0
SM = 0
Doxorubicin
YJL015C
Delta_r
ZShift = 0.01
Z lm Score = −0.88
NG = 0
DB = 0
SM = 0
Doxorubicin
YJL015C
Delta_AUC
ZShift = 1.08
Z lm Score = −0.39
NG = 0
DB = 0
SM = 0
Doxorubicin
YGR105W VMA21
Delta_L
ZShift = 3.78
Z lm Score = 3.16
NG = 0
DB = 0
SM = 0
Doxorubicin
YGR105W VMA21
Delta_K
ZShift = −1.45
Z lm Score = −4.59
NG = 0
DB = 0
SM = 0
Doxorubicin
YGR105W VMA21
Delta_r
ZShift = −2.03
Z lm Score = 0.59
NG = 0
DB = 0
SM = 0
Doxorubicin
YGR105W VMA21
Delta_AUC
ZShift = −2.16
Z lm Score = −1.82
NG = 0
DB = 0
SM = 0
Doxorubicin
YOR361C PRT1
Delta_L
ZShift = 0.4
Z lm Score = 3.16
NG = 0
DB = 0
SM = 0
Doxorubicin
YOR361C PRT1
Delta_K
ZShift = 1.83
Z lm Score = 0
NG = 0
DB = 0
SM = 0
Doxorubicin
YOR361C PRT1
Delta_r
ZShift = −0.2
Z lm Score = −0.94
NG = 0
DB = 0
SM = 0
Doxorubicin
YOR361C PRT1
Delta_AUC
ZShift = 1.6
Z lm Score = 0.36
NG = 0
DB = 0
SM = 0
Doxorubicin

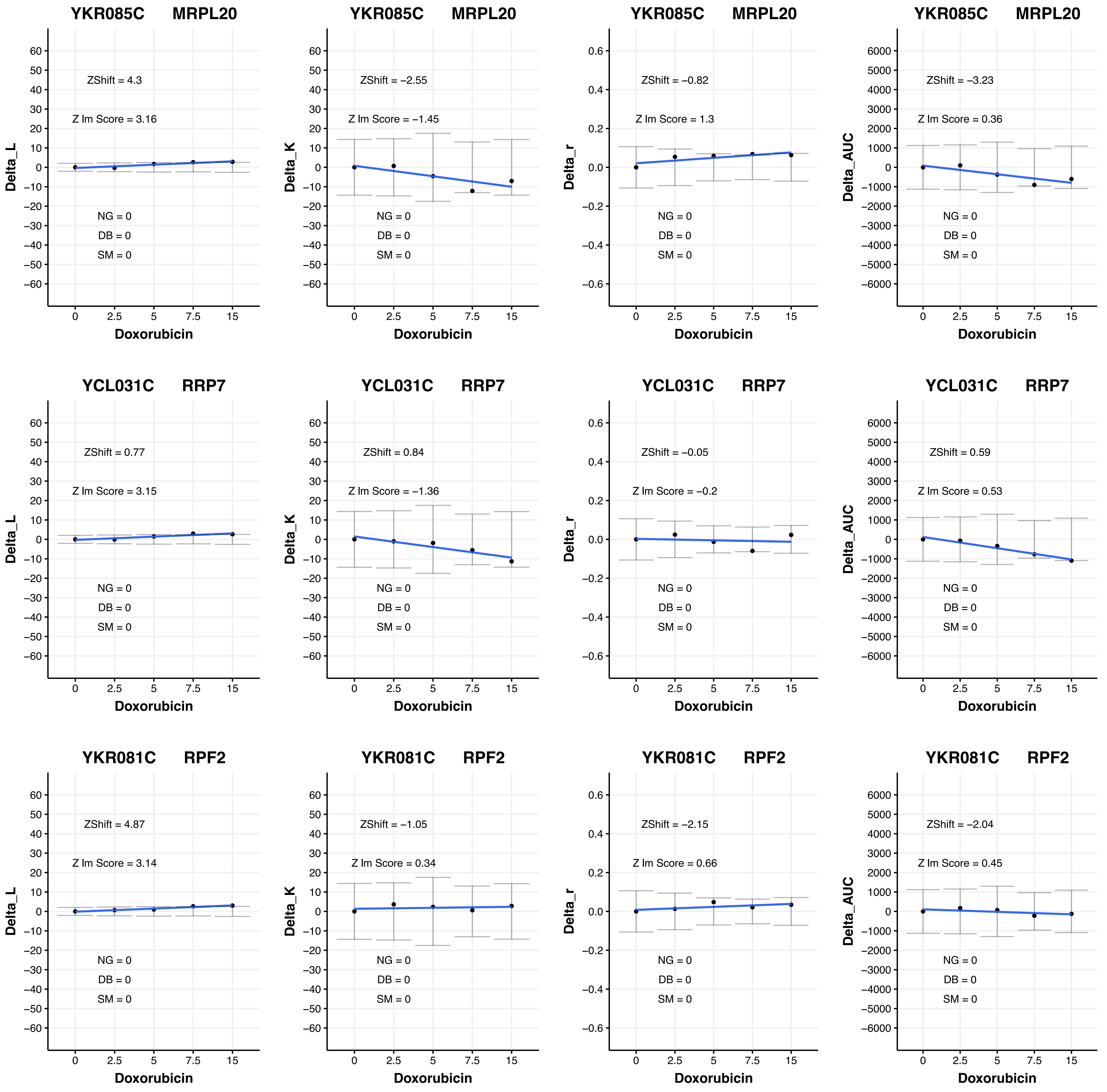

YKR085C MRPL20
Delta_L
ZShift = 4.3
Z lm Score = 3.16
NG = 0
DB = 0
SM = 0
Doxorubicin
YKR085C MRPL20
Delta_K
ZShift = −2.55
Z lm Score = −1.45
NG = 0
DB = 0
SM = 0
Doxorubicin
YKR085C MRPL20
Delta_r
ZShift = −0.82
Z lm Score = 1.3
NG = 0
DB = 0
SM = 0
Doxorubicin
YKR085C MRPL20
Delta_AUC
ZShift = −3.23
Z lm Score = 0.36
NG = 0
DB = 0
SM = 0
Doxorubicin
YCL031C RRP7
Delta_L
ZShift = 0.77
Z lm Score = 3.15
NG = 0
DB = 0
SM = 0
Doxorubicin
YCL031C RRP7
Delta_K
ZShift = 0.84
Z lm Score = −1.36
NG = 0
DB = 0
SM = 0
Doxorubicin
YCL031C RRP7
Delta_r
ZShift = −0.05
Z lm Score = −0.2
NG = 0
DB = 0
SM = 0
Doxorubicin
YCL031C RRP7
Delta_AUC
ZShift = 0.59
Z lm Score = 0.53
NG = 0
DB = 0
SM = 0
Doxorubicin
YKR081C RPF2
Delta_L
ZShift = 4.87
Z lm Score = 3.14
NG = 0
DB = 0
SM = 0
Doxorubicin
YKR081C RPF2
Delta_K
ZShift = −1.05
Z lm Score = 0.34
NG = 0
DB = 0
SM = 0
Doxorubicin
YKR081C RPF2
Delta_r
ZShift = −2.15
Z lm Score = 0.66
NG = 0
DB = 0
SM = 0
Doxorubicin
YKR081C RPF2
Delta_AUC
ZShift = −2.04
Z lm Score = 0.45
NG = 0
DB = 0
SM = 0
Doxorubicin

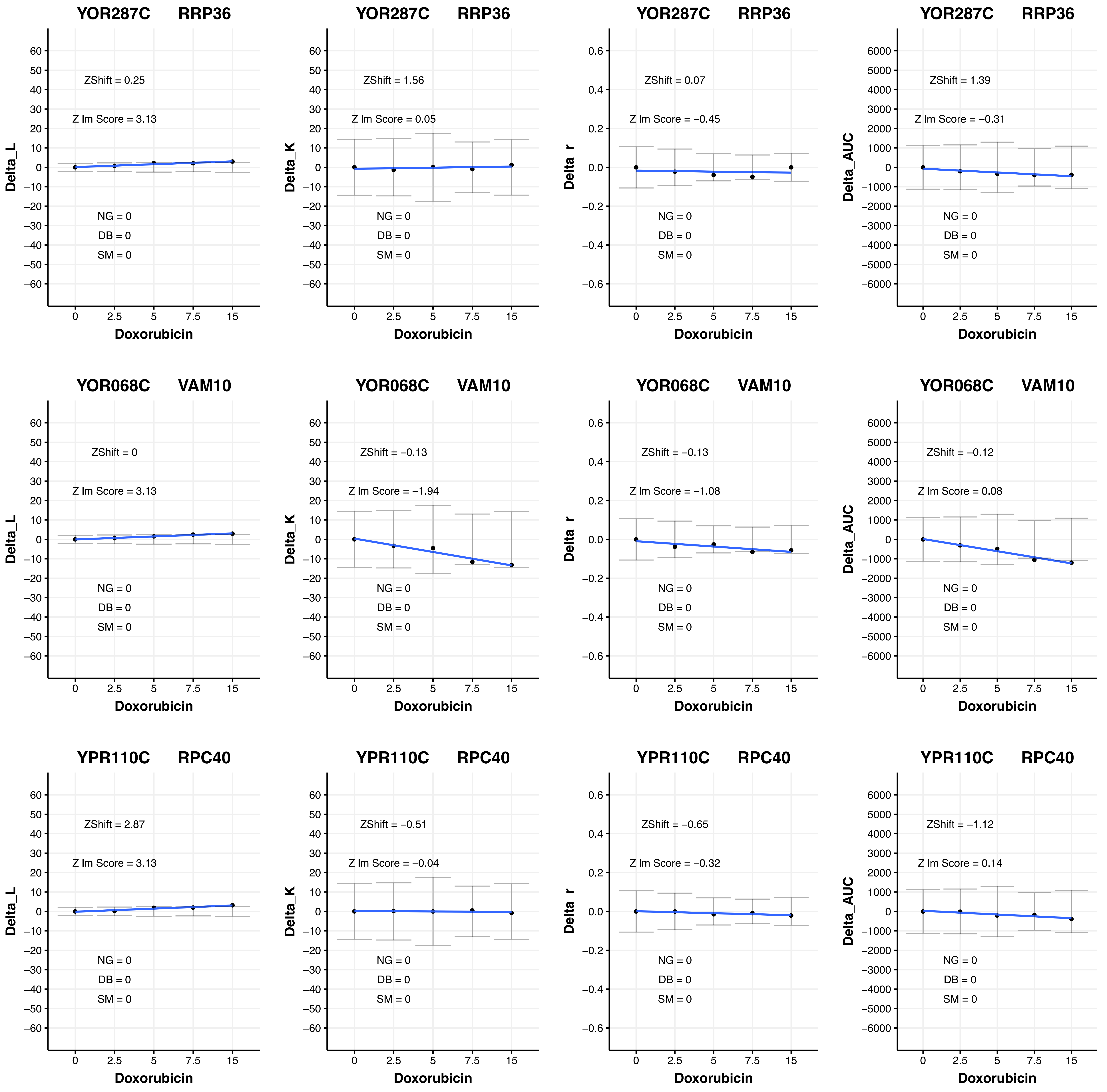
YOR287C RRP36
ZShift = 0.25
Z lm Score = 3.13
NG = 0
DB = 0
SM = 0
Delta_L
Doxorubicin
YOR287C RRP36
ZShift = 1.56
Z lm Score = 0.05
NG = 0
DB = 0
SM = 0
Delta_K
Doxorubicin
YOR287C RRP36
ZShift = 0.07
Z lm Score = −0.45
NG = 0
DB = 0
SM = 0
Delta_r
Doxorubicin
YOR287C RRP36
ZShift = 1.39
Z lm Score = −0.31
NG = 0
DB = 0
SM = 0
Delta_AUC
Doxorubicin
YOR068C VAM10
ZShift = 0
Z lm Score = 3.13
NG = 0
DB = 0
SM = 0
Delta_L
Doxorubicin
YOR068C VAM10
ZShift = −0.13
Z lm Score = −1.94
NG = 0
DB = 0
SM = 0
Delta_K
Doxorubicin
YOR068C VAM10
ZShift = −0.13
Z lm Score = −1.08
NG = 0
DB = 0
SM = 0
Delta_r
Doxorubicin
YOR068C VAM10
ZShift = −0.12
Z lm Score = 0.08
NG = 0
DB = 0
SM = 0
Delta_AUC
Doxorubicin
YPR110C RPC40
ZShift = 2.87
Z lm Score = 3.13
NG = 0
DB = 0
SM = 0
Delta_L
Doxorubicin
YPR110C RPC40
ZShift = −0.51
Z lm Score = −0.04
NG = 0
DB = 0
SM = 0
Delta_K
Doxorubicin
YPR110C RPC40
ZShift = −0.65
Z lm Score = −0.32
NG = 0
DB = 0
SM = 0
Delta_r
Doxorubicin
YPR110C RPC40
ZShift = −1.12
Z lm Score = 0.14
NG = 0
DB = 0
SM = 0
Delta_AUC
Doxorubicin

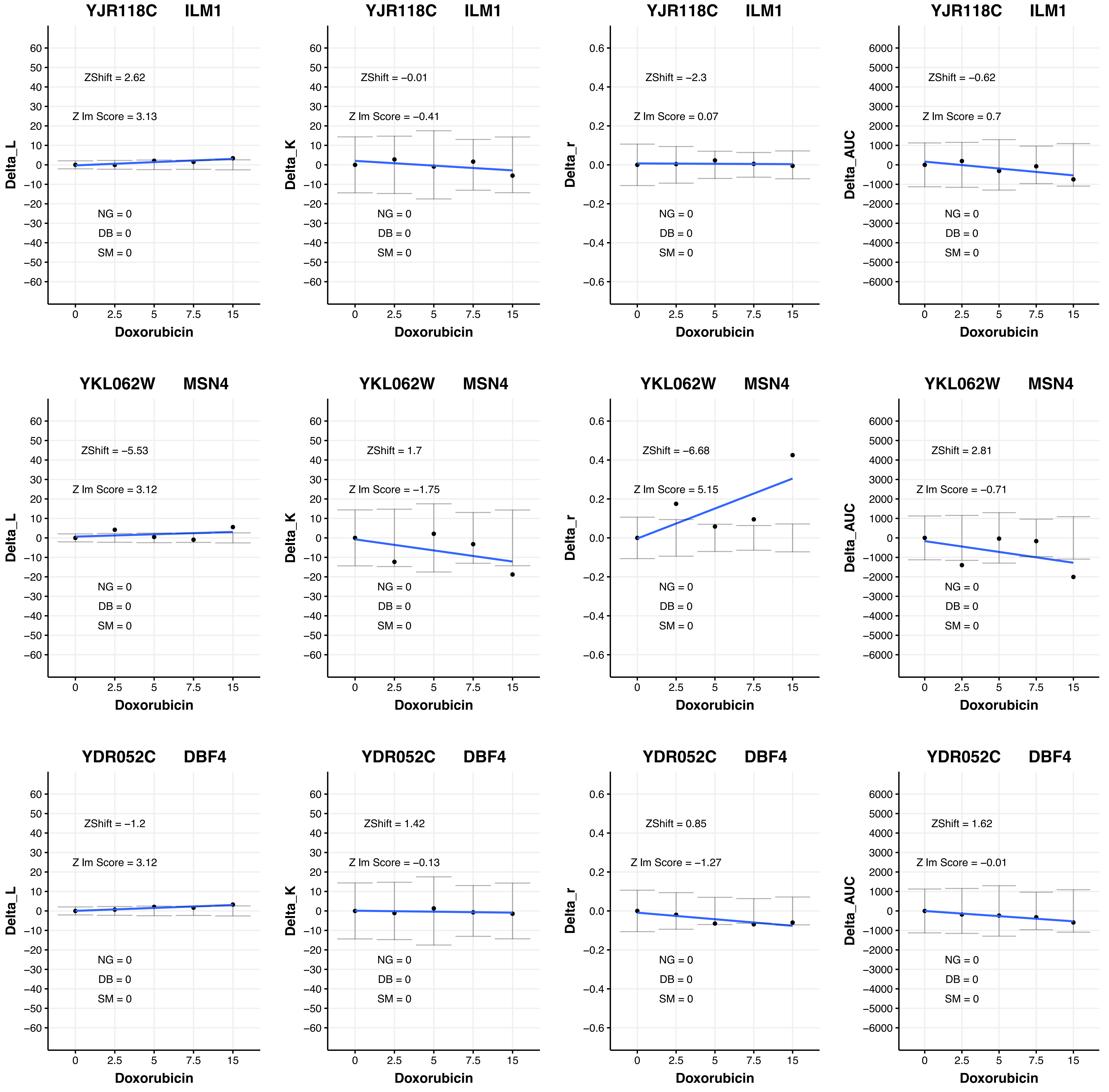

YJR118C ILM1
ZShift = 2.62
Z lm Score = 3.13
NG = 0
DB = 0
SM = 0
Delta_L
Doxorubicin
YJR118C ILM1
ZShift = −0.01
Z lm Score = −0.41
NG = 0
DB = 0
SM = 0
Delta_K
Doxorubicin
YJR118C ILM1
ZShift = −2.3
Z lm Score = 0.07
NG = 0
DB = 0
SM = 0
Delta_r
Doxorubicin
YJR118C ILM1
ZShift = −0.62
Z lm Score = 0.7
NG = 0
DB = 0
SM = 0
Delta_AUC
Doxorubicin
YKL062W MSN4
ZShift = −5.53
Z lm Score = 3.12
NG = 0
DB = 0
SM = 0
Delta_L
Doxorubicin
YKL062W MSN4
ZShift = 1.7
Z lm Score = −1.75
NG = 0
DB = 0
SM = 0
Delta_K
Doxorubicin
YKL062W MSN4
ZShift = −6.68
Z lm Score = 5.15
NG = 0
DB = 0
SM = 0
Delta_r
Doxorubicin
YKL062W MSN4
ZShift = 2.81
Z lm Score = −0.71
NG = 0
DB = 0
SM = 0
Delta_AUC
Doxorubicin
YDR052C DBF4
ZShift = −1.2
Z lm Score = 3.12
NG = 0
DB = 0
SM = 0
Delta_L
Doxorubicin
YDR052C DBF4
ZShift = 1.42
Z lm Score = −0.13
NG = 0
DB = 0
SM = 0
Delta_K
Doxorubicin
YDR052C DBF4
ZShift = 0.85
Z lm Score = −1.27
NG = 0
DB = 0
SM = 0
Delta_r
Doxorubicin
YDR052C DBF4
ZShift = 1.62
Z lm Score = −0.01
NG = 0
DB = 0
SM = 0
Delta_AUC
Doxorubicin

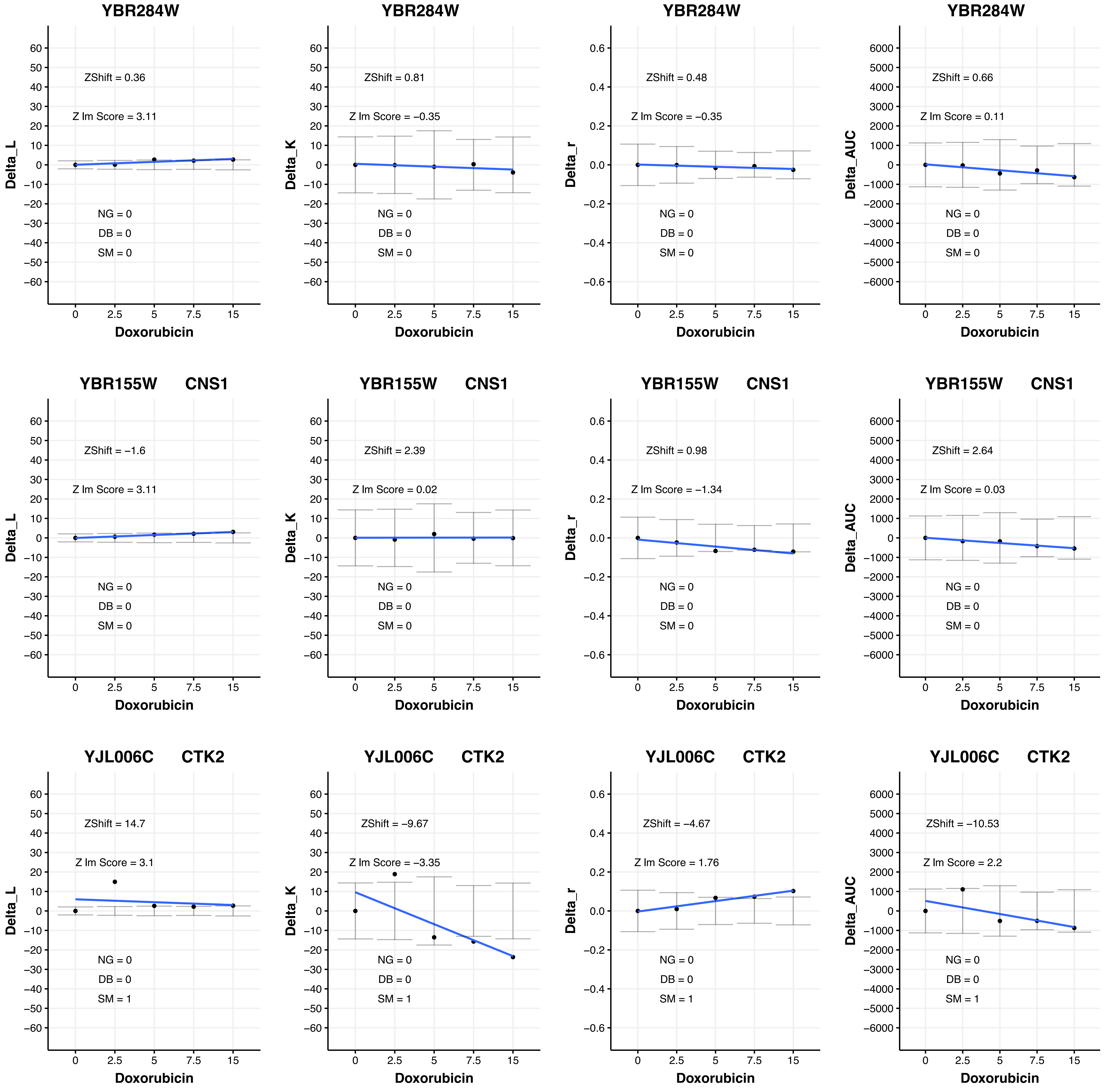
YBR284W
ZShift = 0.36
Z lm Score = 3.11
NG = 0
DB = 0
SM = 0
Delta_L
Doxorubicin
YBR284W
ZShift = 0.81
Z lm Score = −0.35
NG = 0
DB = 0
SM = 0
Delta_K
Doxorubicin
YBR284W
ZShift = 0.48
Z lm Score = −0.35
NG = 0
DB = 0
SM = 0
Delta_r
Doxorubicin
YBR284W
ZShift = 0.66
Z lm Score = 0.11
NG = 0
DB = 0
SM = 0
Delta_AUC
Doxorubicin
YBR155W CNS1
ZShift = −1.6
Z lm Score = 3.11
NG = 0
DB = 0
SM = 0
Delta_L
Doxorubicin
YBR155W CNS1
ZShift = 2.39
Z lm Score = 0.02
NG = 0
DB = 0
SM = 0
Delta_K
Doxorubicin
YBR155W CNS1
ZShift = 0.98
Z lm Score = −1.34
NG = 0
DB = 0
SM = 0
Delta_r
Doxorubicin
YBR155W CNS1
ZShift = 2.64
Z lm Score = 0.03
NG = 0
DB = 0
SM = 0
Delta_AUC
Doxorubicin
YJL006C CTK2
ZShift = 14.7
Z lm Score = 3.1
NG = 0
DB = 0
SM = 1
Delta_L
Doxorubicin
YJL006C CTK2
ZShift = −9.67
Z lm Score = −3.35
NG = 0
DB = 0
SM = 1
Delta_K
Doxorubicin
YJL006C CTK2
ZShift = −4.67
Z lm Score = 1.76
NG = 0
DB = 0
SM = 1
Delta_r
Doxorubicin
YJL006C CTK2
ZShift = −10.53
Z lm Score = 2.2
NG = 0
DB = 0
SM = 1
Delta_AUC
Doxorubicin

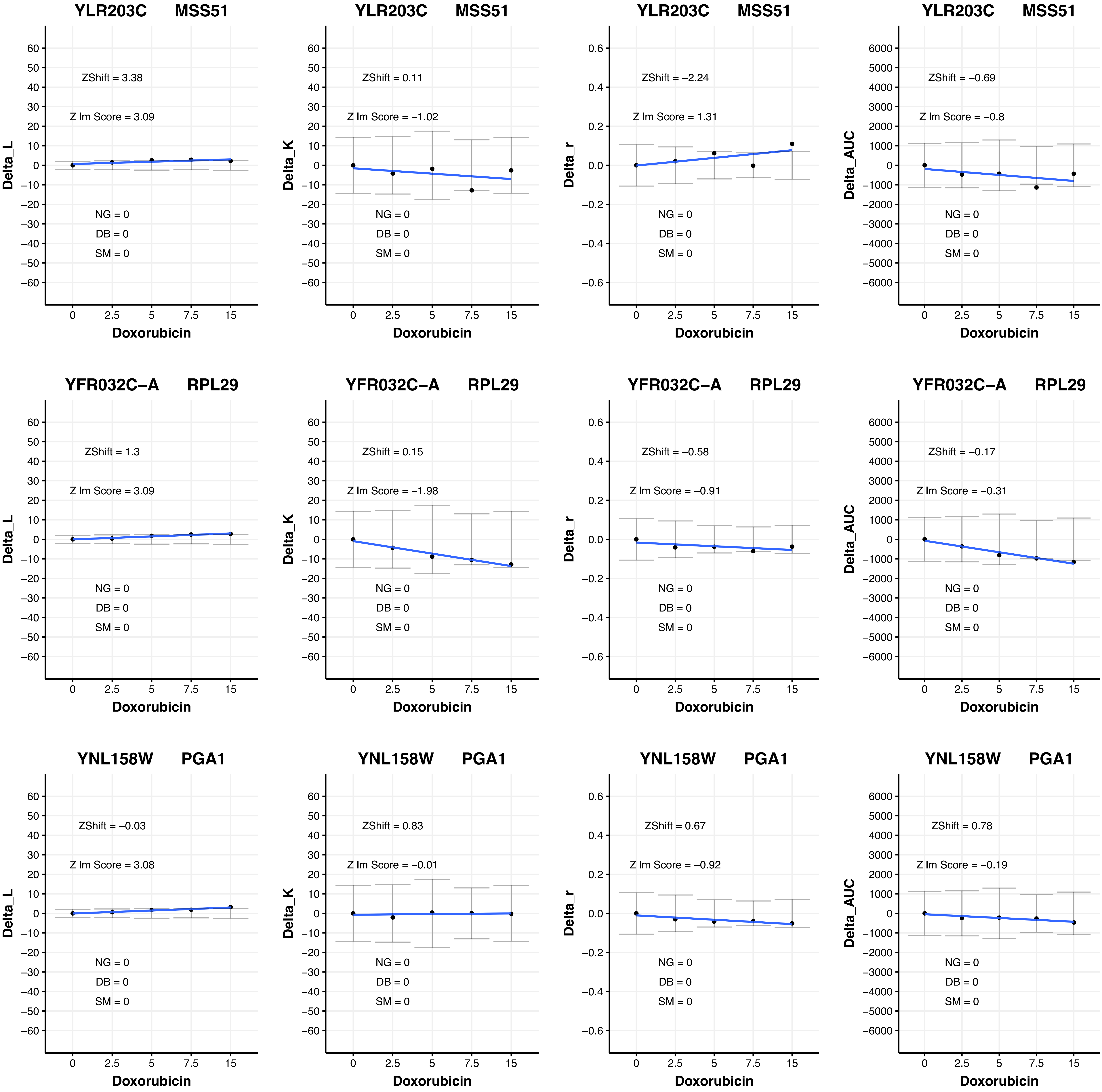
YLR203C MSS51
ZShift = 3.38
Z lm Score = 3.09
NG = 0
DB = 0
SM = 0
Delta_L
Doxorubicin
YLR203C MSS51
ZShift = 0.11
Z lm Score = −1.02
NG = 0
DB = 0
SM = 0
Delta_K
Doxorubicin
YLR203C MSS51
ZShift = −2.24
Z lm Score = 1.31
NG = 0
DB = 0
SM = 0
Delta_r
Doxorubicin
YLR203C MSS51
ZShift = −0.69
Z lm Score = −0.8
NG = 0
DB = 0
SM = 0
Delta_AUC
Doxorubicin
YFR032C−A RPL29
ZShift = 1.3
Z lm Score = 3.09
NG = 0
DB = 0
SM = 0
Delta_L
Doxorubicin
YFR032C−A RPL29
ZShift = 0.15
Z lm Score = −1.98
NG = 0
DB = 0
SM = 0
Delta_K
Doxorubicin
YFR032C−A RPL29
ZShift = −0.58
Z lm Score = −0.91
NG = 0
DB = 0
SM = 0
Delta_r
Doxorubicin
YFR032C−A RPL29
ZShift = −0.17
Z lm Score = −0.31
NG = 0
DB = 0
SM = 0
Delta_AUC
Doxorubicin
YNL158W PGA1
ZShift = −0.03
Z lm Score = 3.08
NG = 0
DB = 0
SM = 0
Delta_L
Doxorubicin
YNL158W PGA1
ZShift = 0.83
Z lm Score = −0.01
NG = 0
DB = 0
SM = 0
Delta_K
Doxorubicin
YNL158W PGA1
ZShift = 0.67
Z lm Score = −0.92
NG = 0
DB = 0
SM = 0
Delta_r
Doxorubicin
YNL158W PGA1
ZShift = 0.78
Z lm Score = −0.19
NG = 0
DB = 0
SM = 0
Delta_AUC
Doxorubicin

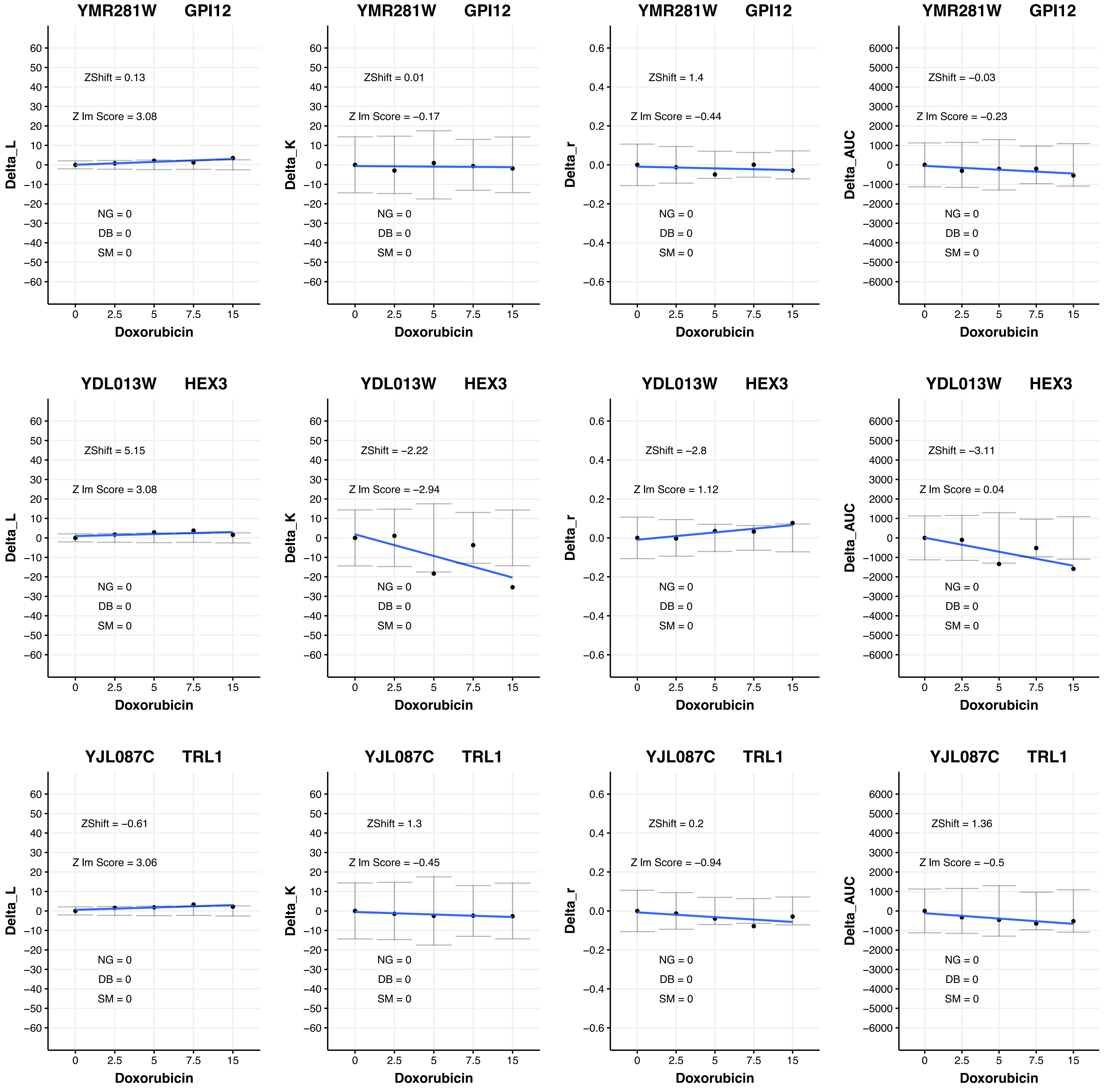

YMR281W GPI12
ZShift = 0.13
Z lm Score = 3.08
NG = 0
DB = 0
SM = 0
Delta_L
Doxorubicin
YMR281W GPI12
ZShift = 0.01
Z lm Score = −0.17
NG = 0
DB = 0
SM = 0
Delta_K
Doxorubicin
YMR281W GPI12
ZShift = 1.4
Z lm Score = −0.44
NG = 0
DB = 0
SM = 0
Delta_r
Doxorubicin
YMR281W GPI12
ZShift = −0.03
Z lm Score = −0.23
NG = 0
DB = 0
SM = 0
Delta_AUC
Doxorubicin
YDL013W HEX3
ZShift = 5.15
Z lm Score = 3.08
NG = 0
DB = 0
SM = 0
Delta_L
Doxorubicin
YDL013W HEX3
ZShift = −2.22
Z lm Score = −2.94
NG = 0
DB = 0
SM = 0
Delta_K
Doxorubicin
YDL013W HEX3
ZShift = −2.8
Z lm Score = 1.12
NG = 0
DB = 0
SM = 0
Delta_r
Doxorubicin
YDL013W HEX3
ZShift = −3.11
Z lm Score = 0.04
NG = 0
DB = 0
SM = 0
Delta_AUC
Doxorubicin
YJL087C TRL1
ZShift = −0.61
Z lm Score = 3.06
NG = 0
DB = 0
SM = 0
Delta_L
Doxorubicin
YJL087C TRL1
ZShift = 1.3
Z lm Score = −0.45
NG = 0
DB = 0
SM = 0
Delta_K
Doxorubicin
YJL087C TRL1
ZShift = 0.2
Z lm Score = −0.94
NG = 0
DB = 0
SM = 0
Delta_r
Doxorubicin
YJL087C TRL1
ZShift = 1.36
Z lm Score = −0.5
NG = 0
DB = 0
SM = 0
Delta_AUC
Doxorubicin

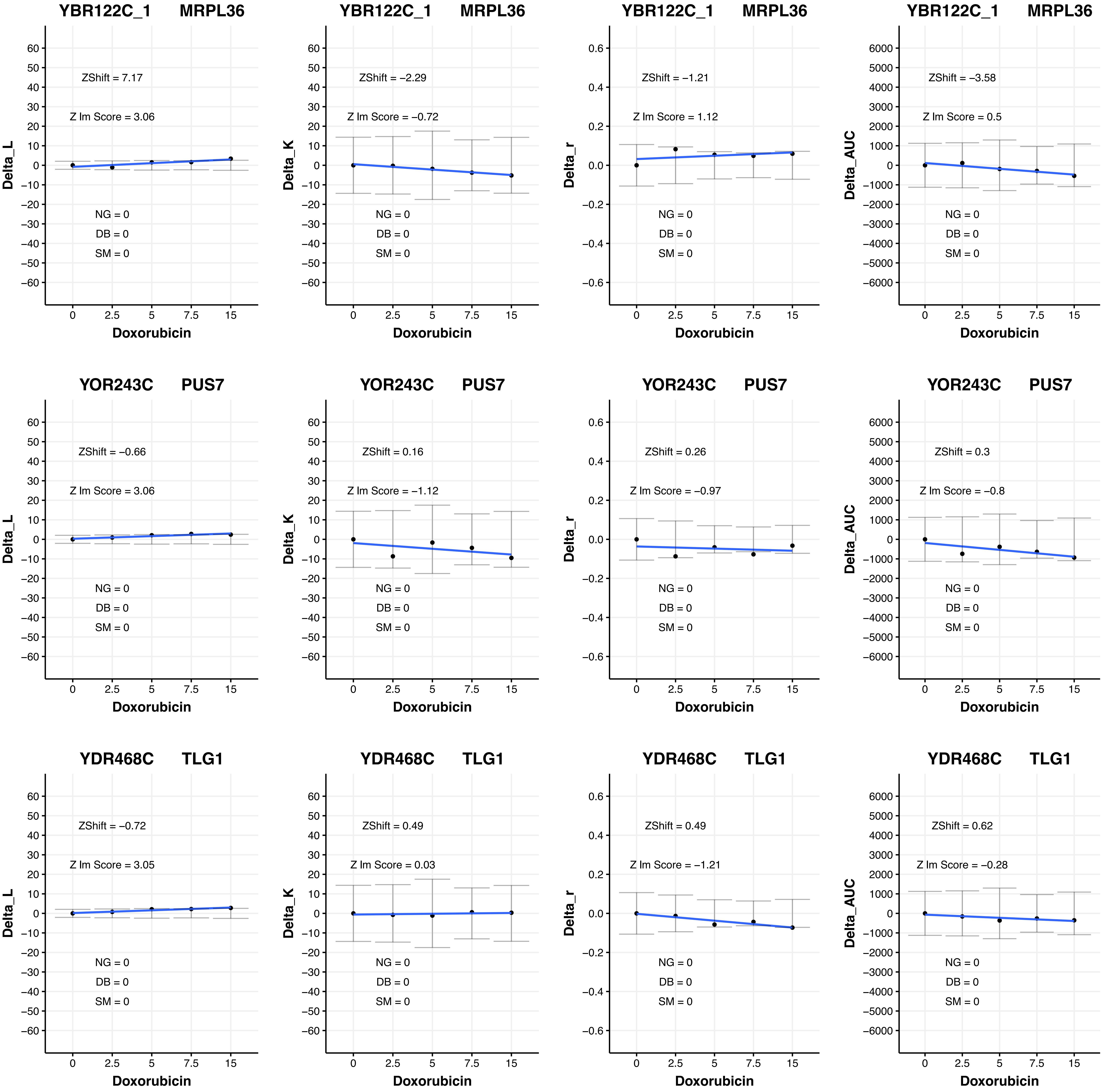
YBR122C_1 MRPL36
ZShift = 7.17
Z lm Score = 3.06
NG = 0
DB = 0
SM = 0
Delta_L
Doxorubicin
YBR122C_1 MRPL36
ZShift = −2.29
Z lm Score = −0.72
NG = 0
DB = 0
SM = 0
Delta_K
Doxorubicin
YBR122C_1 MRPL36
ZShift = −1.21
Z lm Score = 1.12
NG = 0
DB = 0
SM = 0
Delta_r
Doxorubicin
YBR122C_1 MRPL36
ZShift = −3.58
Z lm Score = 0.5
NG = 0
DB = 0
SM = 0
Delta_AUC
Doxorubicin
YOR243C PUS7
ZShift = −0.66
Z lm Score = 3.06
NG = 0
DB = 0
SM = 0
Delta_L
Doxorubicin
YOR243C PUS7
ZShift = 0.16
Z lm Score = −1.12
NG = 0
DB = 0
SM = 0
Delta_K
Doxorubicin
YOR243C PUS7
ZShift = 0.26
Z lm Score = −0.97
NG = 0
DB = 0
SM = 0
Delta_r
Doxorubicin
YOR243C PUS7
ZShift = 0.3
Z lm Score = −0.8
NG = 0
DB = 0
SM = 0
Delta_AUC
Doxorubicin
YDR468C TLG1
ZShift = −0.72
Z lm Score = 3.05
NG = 0
DB = 0
SM = 0
Delta_L
Doxorubicin
YDR468C TLG1
ZShift = 0.49
Z lm Score = 0.03
NG = 0
DB = 0
SM = 0
Delta_K
Doxorubicin
YDR468C TLG1
ZShift = 0.49
Z lm Score = −1.21
NG = 0
DB = 0
SM = 0
Delta_r
Doxorubicin
YDR468C TLG1
ZShift = 0.62
Z lm Score = −0.28
NG = 0
DB = 0
SM = 0
Delta_AUC
Doxorubicin

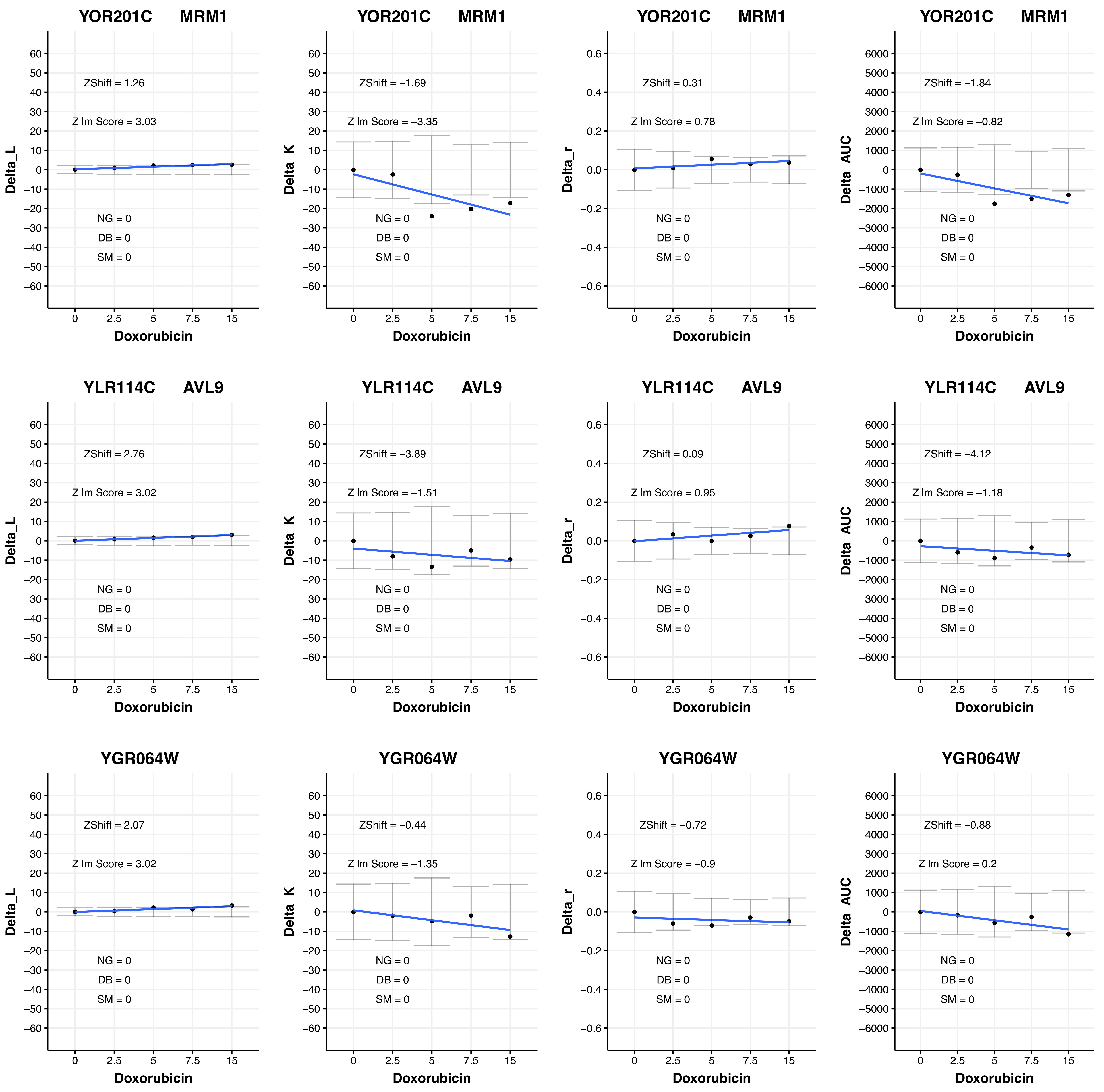
YOR201C MRM1
ZShift = 1.26
Z lm Score = 3.03
NG = 0
DB = 0
SM = 0
Delta_L
Doxorubicin
YOR201C MRM1
ZShift = −1.69
Z lm Score = −3.35
NG = 0
DB = 0
SM = 0
Delta_K
Doxorubicin
YOR201C MRM1
ZShift = 0.31
Z lm Score = 0.78
NG = 0
DB = 0
SM = 0
Delta_r
Doxorubicin
YOR201C MRM1
ZShift = −1.84
Z lm Score = −0.82
NG = 0
DB = 0
SM = 0
Delta_AUC
Doxorubicin
YLR114C AVL9
ZShift = 2.76
Z lm Score = 3.02
NG = 0
DB = 0
SM = 0
Delta_L
Doxorubicin
YLR114C AVL9
ZShift = −3.89
Z lm Score = −1.51
NG = 0
DB = 0
SM = 0
Delta_K
Doxorubicin
YLR114C AVL9
ZShift = 0.09
Z lm Score = 0.95
NG = 0
DB = 0
SM = 0
Delta_r
Doxorubicin
YLR114C AVL9
ZShift = −4.12
Z lm Score = −1.18
NG = 0
DB = 0
SM = 0
Delta_AUC
Doxorubicin
YGR064W
ZShift = 2.07
Z lm Score = 3.02
NG = 0
DB = 0
SM = 0
Delta_L
Doxorubicin
YGR064W
ZShift = −0.44
Z lm Score = −1.35
NG = 0
DB = 0
SM = 0
Delta_K
Doxorubicin
YGR064W
ZShift = −0.72
Z lm Score = −0.9
NG = 0
DB = 0
SM = 0
Delta_r
Doxorubicin
YGR064W
ZShift = −0.88
Z lm Score = 0.2
NG = 0
DB = 0
SM = 0
Delta_AUC
Doxorubicin

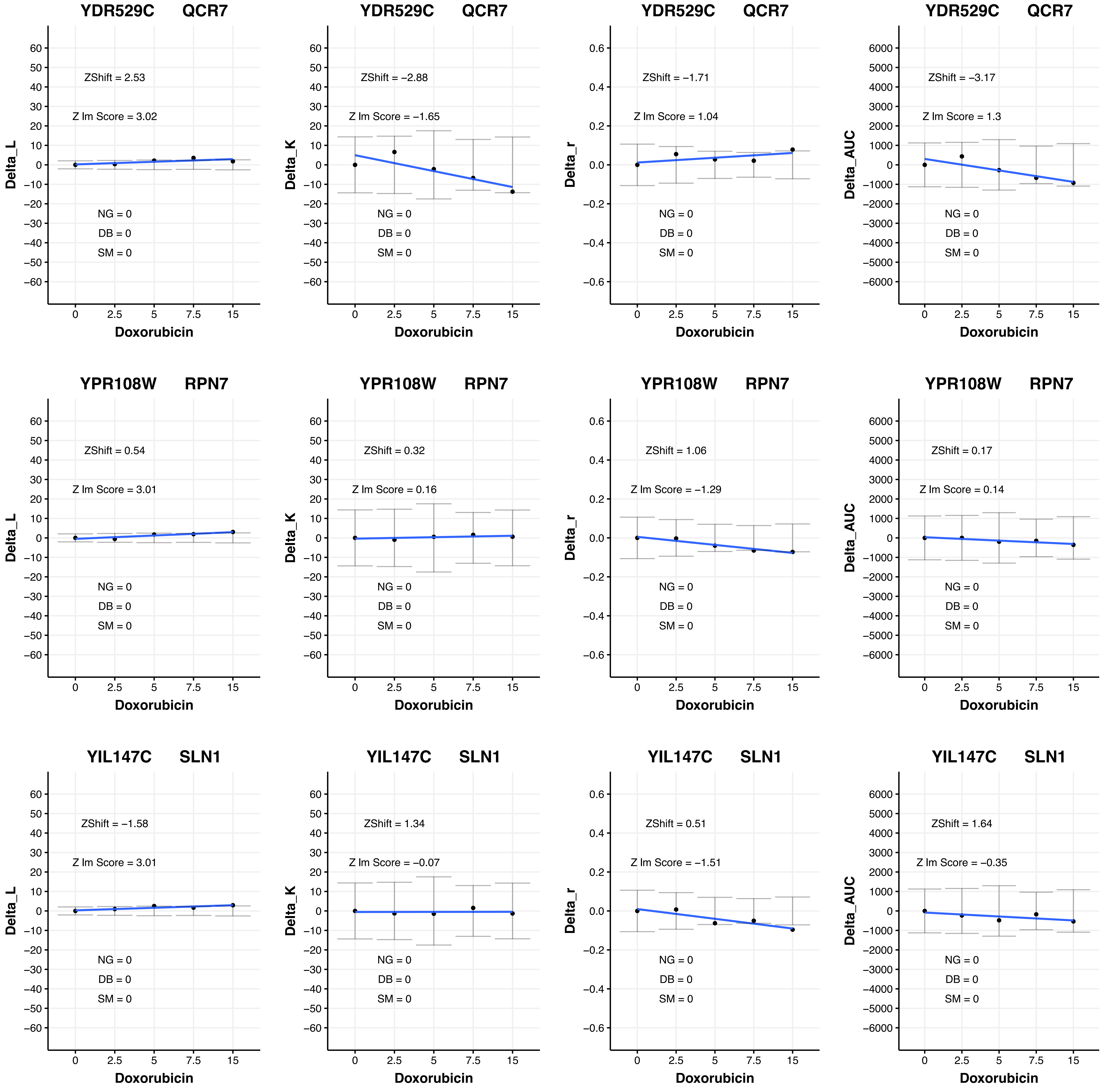
YDR529C QCR7
ZShift = 2.53
Z lm Score = 3.02
NG = 0
DB = 0
SM = 0
Delta_L
Doxorubicin
YDR529C QCR7
ZShift = −2.88
Z lm Score = −1.65
NG = 0
DB = 0
SM = 0
Delta_K
Doxorubicin
YDR529C QCR7
ZShift = −1.71
Z lm Score = 1.04
NG = 0
DB = 0
SM = 0
Delta_r
Doxorubicin
YDR529C QCR7
ZShift = −3.17
Z lm Score = 1.3
NG = 0
DB = 0
SM = 0
Delta_AUC
Doxorubicin
YPR108W RPN7
ZShift = 0.54
Z lm Score = 3.01
NG = 0
DB = 0
SM = 0
Delta_L
Doxorubicin
YPR108W RPN7
ZShift = 0.32
Z lm Score = 0.16
NG = 0
DB = 0
SM = 0
Delta_K
Doxorubicin
YPR108W RPN7
ZShift = 1.06
Z lm Score = −1.29
NG = 0
DB = 0
SM = 0
Delta_r
Doxorubicin
YPR108W RPN7
ZShift = 0.17
Z lm Score = 0.14
NG = 0
DB = 0
SM = 0
Delta_AUC
Doxorubicin
YIL147C SLN1
ZShift = −1.58
Z lm Score = 3.01
NG = 0
DB = 0
SM = 0
Delta_L
Doxorubicin
YIL147C SLN1
ZShift = 1.34
Z lm Score = −0.07
NG = 0
DB = 0
SM = 0
Delta_K
Doxorubicin
YIL147C SLN1
ZShift = 0.51
Z lm Score = −1.51
NG = 0
DB = 0
SM = 0
Delta_r
Doxorubicin
YIL147C SLN1
ZShift = 1.64
Z lm Score = −0.35
NG = 0
DB = 0
SM = 0
Delta_AUC
Doxorubicin

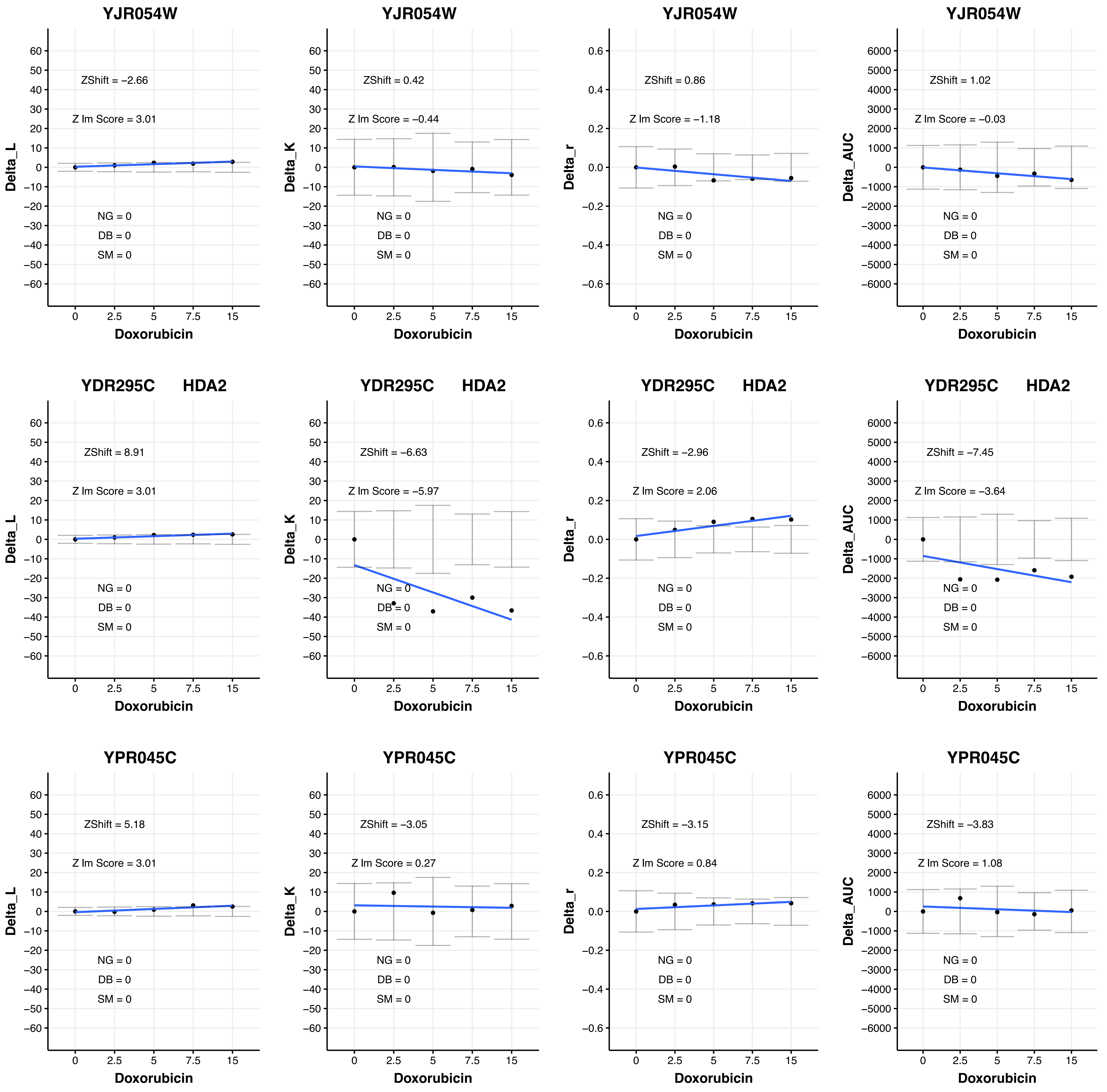

YJR054W
ZShift = −2.66
Z lm Score = 3.01
NG = 0
DB = 0
SM = 0
Delta_L
Doxorubicin
YJR054W
ZShift = 0.42
Z lm Score = −0.44
NG = 0
DB = 0
SM = 0
Delta_K
Doxorubicin
YJR054W
ZShift = 0.86
Z lm Score = −1.18
NG = 0
DB = 0
SM = 0
Delta_r
Doxorubicin
YJR054W
ZShift = 1.02
Z lm Score = −0.03
NG = 0
DB = 0
SM = 0
Delta_AUC
Doxorubicin
YDR295C HDA2
ZShift = 8.91
Z lm Score = 3.01
NG = 0
DB = 0
SM = 0
Delta_L
Doxorubicin
YDR295C HDA2
ZShift = −6.63
Z lm Score = −5.97
NG = 0
DB = 0
SM = 0
Delta_K
Doxorubicin
YDR295C HDA2
ZShift = −2.96
Z lm Score = 2.06
NG = 0
DB = 0
SM = 0
Delta_r
Doxorubicin
YDR295C HDA2
ZShift = −7.45
Z lm Score = −3.64
NG = 0
DB = 0
SM = 0
Delta_AUC
Doxorubicin
YPR045C
ZShift = 5.18
Z lm Score = 3.01
NG = 0
DB = 0
SM = 0
Delta_L
Doxorubicin
YPR045C
ZShift = −3.05
Z lm Score = 0.27
NG = 0
DB = 0
SM = 0
Delta_K
Doxorubicin
YPR045C
ZShift = −3.15
Z lm Score = 0.84
NG = 0
DB = 0
SM = 0
Delta_r
Doxorubicin
YPR045C
ZShift = −3.83
Z lm Score = 1.08
NG = 0
DB = 0
SM = 0
Delta_AUC
Doxorubicin

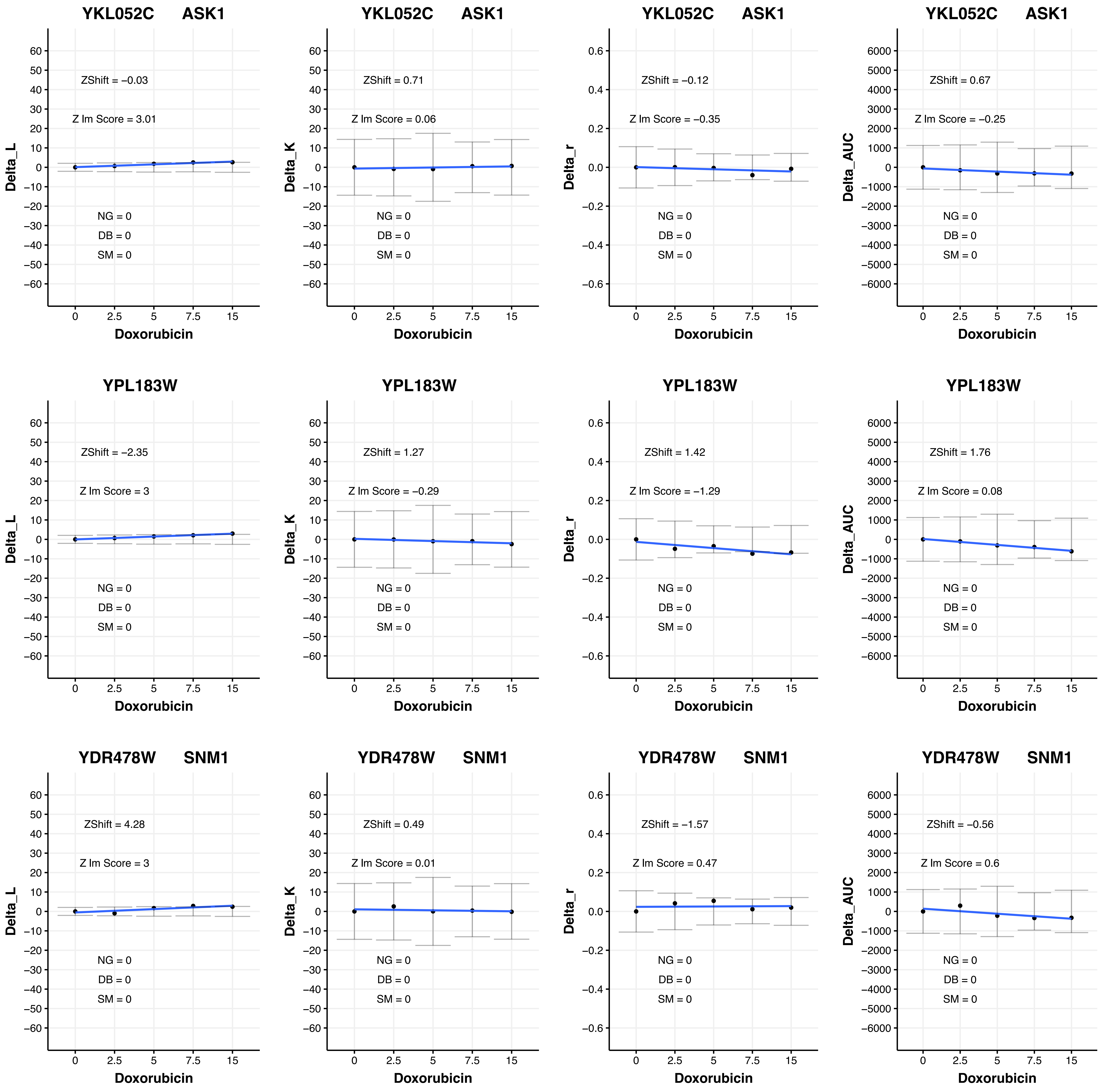

YKL052C ASK1
ZShift = −0.03
Z lm Score = 3.01
NG = 0
DB = 0
SM = 0
Delta_L
Doxorubicin
YKL052C ASK1
ZShift = 0.71
Z lm Score = 0.06
NG = 0
DB = 0
SM = 0
Delta_K
Doxorubicin
YKL052C ASK1
ZShift = −0.12
Z lm Score = −0.35
NG = 0
DB = 0
SM = 0
Delta_r
Doxorubicin
YKL052C ASK1
ZShift = 0.67
Z lm Score = −0.25
NG = 0
DB = 0
SM = 0
Delta_AUC
Doxorubicin
YPL183W
ZShift = −2.35
Z lm Score = 3
NG = 0
DB = 0
SM = 0
Delta_L
Doxorubicin
YPL183W
ZShift = 1.27
Z lm Score = −0.29
NG = 0
DB = 0
SM = 0
Delta_K
Doxorubicin
YPL183W
ZShift = 1.42
Z lm Score = −1.29
NG = 0
DB = 0
SM = 0
Delta_r
Doxorubicin
YPL183W
ZShift = 1.76
Z lm Score = 0.08
NG = 0
DB = 0
SM = 0
Delta_AUC
Doxorubicin
YDR478W SNM1
ZShift = 4.28
Z lm Score = 3
NG = 0
DB = 0
SM = 0
Delta_L
Doxorubicin
YDR478W SNM1
ZShift = 0.49
Z lm Score = 0.01
NG = 0
DB = 0
SM = 0
Delta_K
Doxorubicin
YDR478W SNM1
ZShift = −1.57
Z lm Score = 0.47
NG = 0
DB = 0
SM = 0
Delta_r
Doxorubicin
YDR478W SNM1
ZShift = −0.56
Z lm Score = 0.6
NG = 0
DB = 0
SM = 0
Delta_AUC
Doxorubicin

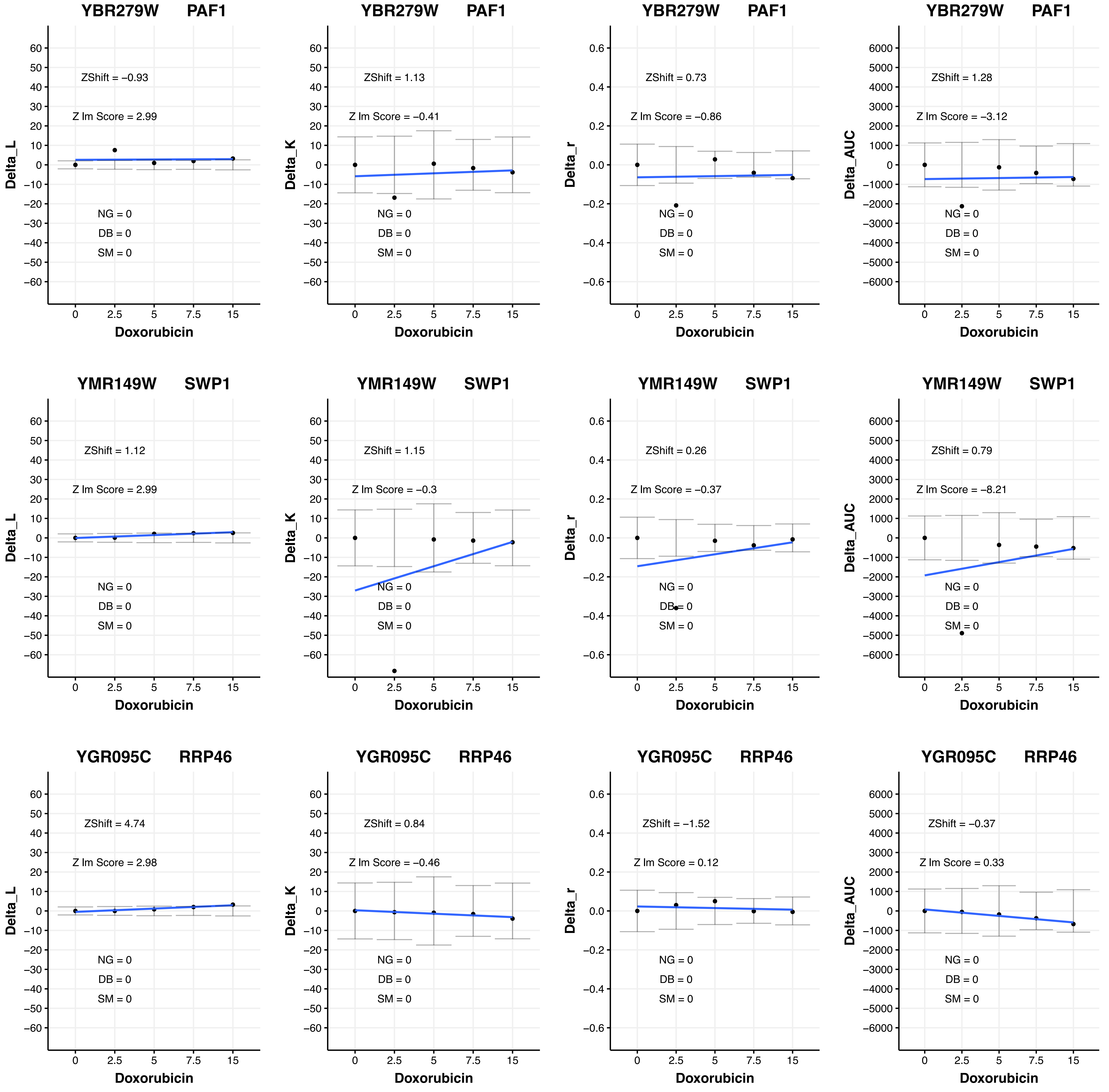
YBR279W PAF1
ZShift = −0.93
Z lm Score = 2.99
NG = 0
DB = 0
SM = 0
Delta_L
Doxorubicin
YBR279W PAF1
ZShift = 1.13
Z lm Score = −0.41
NG = 0
DB = 0
SM = 0
Delta_K
Doxorubicin
YBR279W PAF1
ZShift = 0.73
Z lm Score = −0.86
NG = 0
DB = 0
SM = 0
Delta_r
Doxorubicin
YBR279W PAF1
ZShift = 1.28
Z lm Score = −3.12
NG = 0
DB = 0
SM = 0
Delta_AUC
Doxorubicin
YMR149W SWP1
ZShift = 1.12
Z lm Score = 2.99
NG = 0
DB = 0
SM = 0
Delta_L
Doxorubicin
YMR149W SWP1
ZShift = 1.15
Z lm Score = −0.3
NG = 0
DB = 0
SM = 0
Delta_K
Doxorubicin
YMR149W SWP1
ZShift = 0.26
Z lm Score = −0.37
NG = 0
DB = 0
SM = 0
Delta_r
Doxorubicin
YMR149W SWP1
ZShift = 0.79
Z lm Score = −8.21
NG = 0
DB = 0
SM = 0
Delta_AUC
Doxorubicin
YGR095C RRP46
ZShift = 4.74
Z lm Score = 2.98
NG = 0
DB = 0
SM = 0
Delta_L
Doxorubicin
YGR095C RRP46
ZShift = 0.84
Z lm Score = −0.46
NG = 0
DB = 0
SM = 0
Delta_K
Doxorubicin
YGR095C RRP46
ZShift = −1.52
Z lm Score = 0.12
NG = 0
DB = 0
SM = 0
Delta_r
Doxorubicin
YGR095C RRP46
ZShift = −0.37
Z lm Score = 0.33
NG = 0
DB = 0
SM = 0
Delta_AUC
Doxorubicin

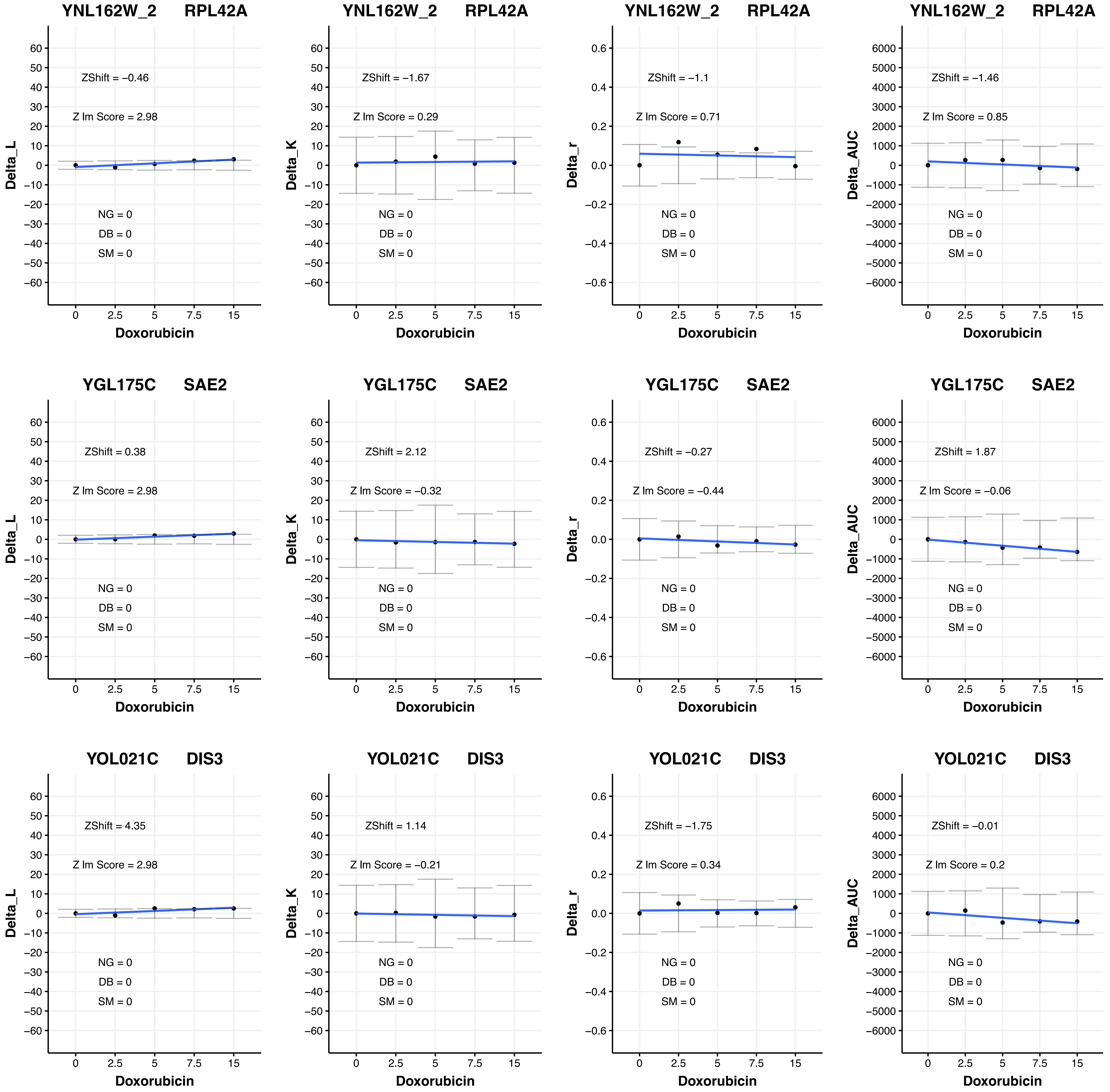

YNL162W_2 RPL42A
ZShift = −0.46
Z lm Score = 2.98
Delta_L
NG = 0
DB = 0
SM = 0
Doxorubicin
YNL162W_2 RPL42A
ZShift = −1.67
Z lm Score = 0.29
Delta_K
NG = 0
DB = 0
SM = 0
Doxorubicin
YNL162W_2 RPL42A
ZShift = −1.1
Z lm Score = 0.71
Delta_r
NG = 0
DB = 0
SM = 0
Doxorubicin
YNL162W_2 RPL42A
ZShift = −1.46
Z lm Score = 0.85
Delta_AUC
NG = 0
DB = 0
SM = 0
Doxorubicin
YGL175C SAE2
ZShift = 0.38
Z lm Score = 2.98
Delta_L
NG = 0
DB = 0
SM = 0
Doxorubicin
YGL175C SAE2
ZShift = 2.12
Z lm Score = −0.32
Delta_K
NG = 0
DB = 0
SM = 0
Doxorubicin
YGL175C SAE2
ZShift = −0.27
Z lm Score = −0.44
Delta_r
NG = 0
DB = 0
SM = 0
Doxorubicin
YGL175C SAE2
ZShift = 1.87
Z lm Score = −0.06
Delta_AUC
NG = 0
DB = 0
SM = 0
Doxorubicin
YOL021C DIS3
ZShift = 4.35
Z lm Score = 2.98
Delta_L
NG = 0
DB = 0
SM = 0
Doxorubicin
YOL021C DIS3
ZShift = 1.14
Z lm Score = −0.21
Delta_K
NG = 0
DB = 0
SM = 0
Doxorubicin
YOL021C DIS3
ZShift = −1.75
Z lm Score = 0.34
Delta_r
NG = 0
DB = 0
SM = 0
Doxorubicin
YOL021C DIS3
ZShift = −0.01
Z lm Score = 0.2
Delta_AUC
NG = 0
DB = 0
SM = 0
Doxorubicin

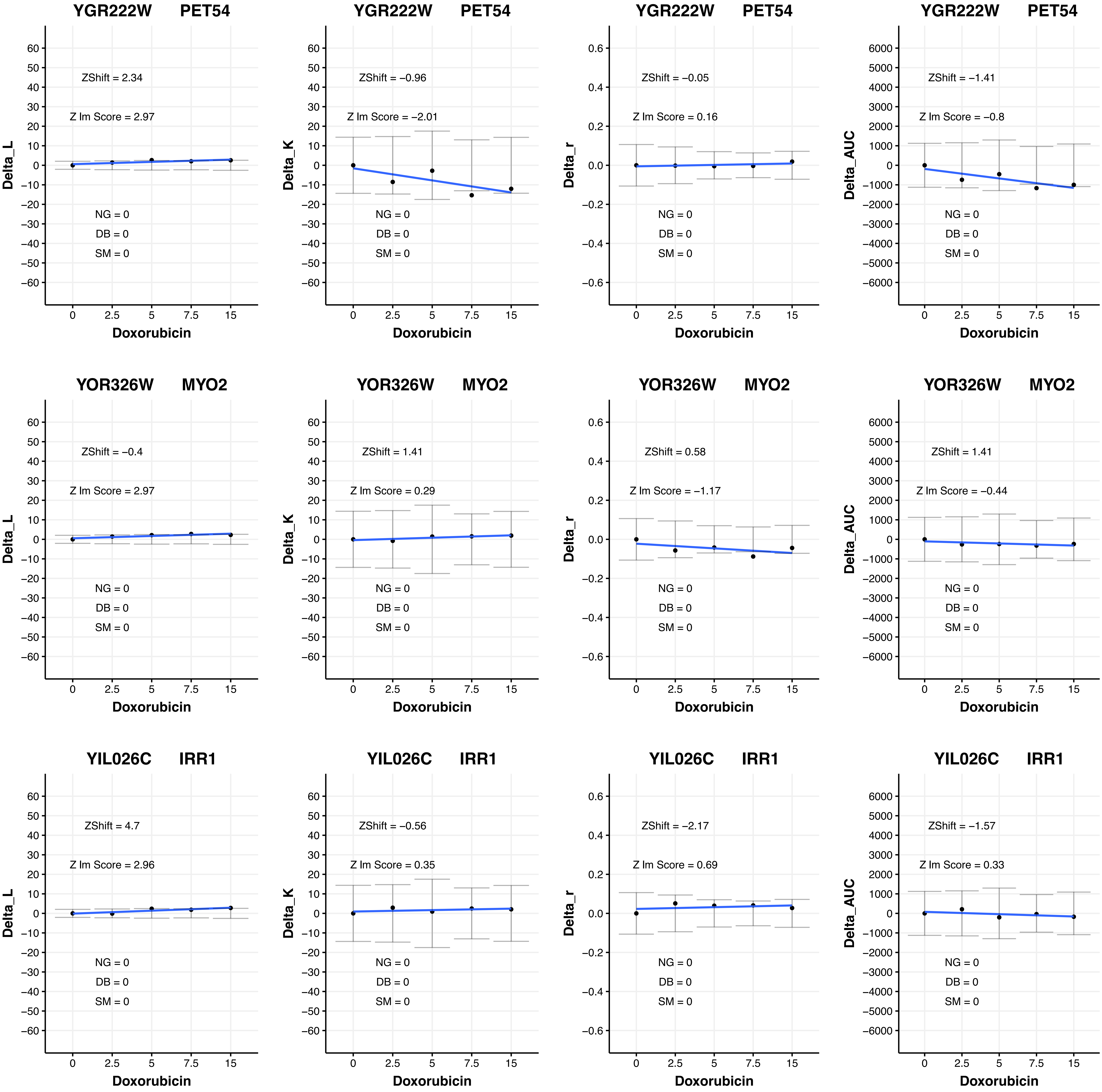
YGR222W PET54
ZShift = 2.34
Z lm Score = 2.97
NG = 0
DB = 0
SM = 0
Delta_L
Doxorubicin
YGR222W PET54
ZShift = −0.96
Z lm Score = −2.01
NG = 0
DB = 0
SM = 0
Delta_K
Doxorubicin
YGR222W PET54
ZShift = −0.05
Z lm Score = 0.16
NG = 0
DB = 0
SM = 0
Delta_r
Doxorubicin
YGR222W PET54
ZShift = −1.41
Z lm Score = −0.8
NG = 0
DB = 0
SM = 0
Delta_AUC
Doxorubicin
YOR326W MYO2
ZShift = −0.4
Z lm Score = 2.97
NG = 0
DB = 0
SM = 0
Delta_L
Doxorubicin
YOR326W MYO2
ZShift = 1.41
Z lm Score = 0.29
NG = 0
DB = 0
SM = 0
Delta_K
Doxorubicin
YOR326W MYO2
ZShift = 0.58
Z lm Score = −1.17
NG = 0
DB = 0
SM = 0
Delta_r
Doxorubicin
YOR326W MYO2
ZShift = 1.41
Z lm Score = −0.44
NG = 0
DB = 0
SM = 0
Delta_AUC
Doxorubicin
YIL026C IRR1
ZShift = 4.7
Z lm Score = 2.96
NG = 0
DB = 0
SM = 0
Delta_L
Doxorubicin
YIL026C IRR1
ZShift = −0.56
Z lm Score = 0.35
NG = 0
DB = 0
SM = 0
Delta_K
Doxorubicin
YIL026C IRR1
ZShift = −2.17
Z lm Score = 0.69
NG = 0
DB = 0
SM = 0
Delta_r
Doxorubicin
YIL026C IRR1
ZShift = −1.57
Z lm Score = 0.33
NG = 0
DB = 0
SM = 0
Delta_AUC
Doxorubicin

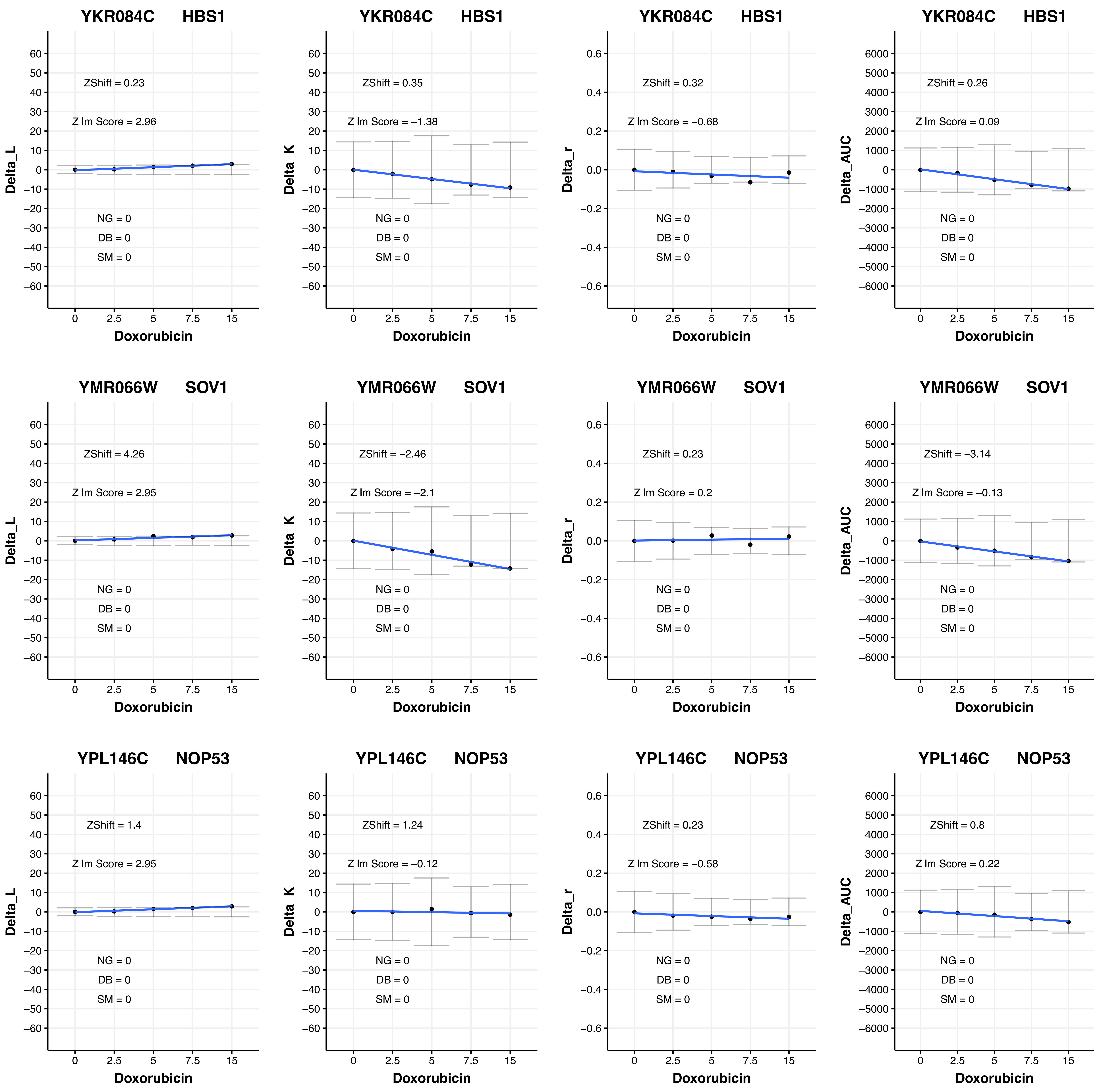

YKR084C HBS1
ZShift = 0.23
Z lm Score = 2.96
NG = 0
DB = 0
SM = 0
Delta_L
Doxorubicin
YKR084C HBS1
ZShift = 0.35
Z lm Score = −1.38
NG = 0
DB = 0
SM = 0
Delta_K
Doxorubicin
YKR084C HBS1
ZShift = 0.32
Z lm Score = −0.68
NG = 0
DB = 0
SM = 0
Delta_r
Doxorubicin
YKR084C HBS1
ZShift = 0.26
Z lm Score = 0.09
NG = 0
DB = 0
SM = 0
Delta_AUC
Doxorubicin
YMR066W SOV1
ZShift = 4.26
Z lm Score = 2.95
NG = 0
DB = 0
SM = 0
Delta_L
Doxorubicin
YMR066W SOV1
ZShift = −2.46
Z lm Score = −2.1
NG = 0
DB = 0
SM = 0
Delta_K
Doxorubicin
YMR066W SOV1
ZShift = 0.23
Z lm Score = 0.2
NG = 0
DB = 0
SM = 0
Delta_r
Doxorubicin
YMR066W SOV1
ZShift = −3.14
Z lm Score = −0.13
NG = 0
DB = 0
SM = 0
Delta_AUC
Doxorubicin
YPL146C NOP53
ZShift = 1.4
Z lm Score = 2.95
NG = 0
DB = 0
SM = 0
Delta_L
Doxorubicin
YPL146C NOP53
ZShift = 1.24
Z lm Score = −0.12
NG = 0
DB = 0
SM = 0
Delta_K
Doxorubicin
YPL146C NOP53
ZShift = 0.23
Z lm Score = −0.58
NG = 0
DB = 0
SM = 0
Delta_r
Doxorubicin
YPL146C NOP53
ZShift = 0.8
Z lm Score = 0.22
NG = 0
DB = 0
SM = 0
Delta_AUC
Doxorubicin

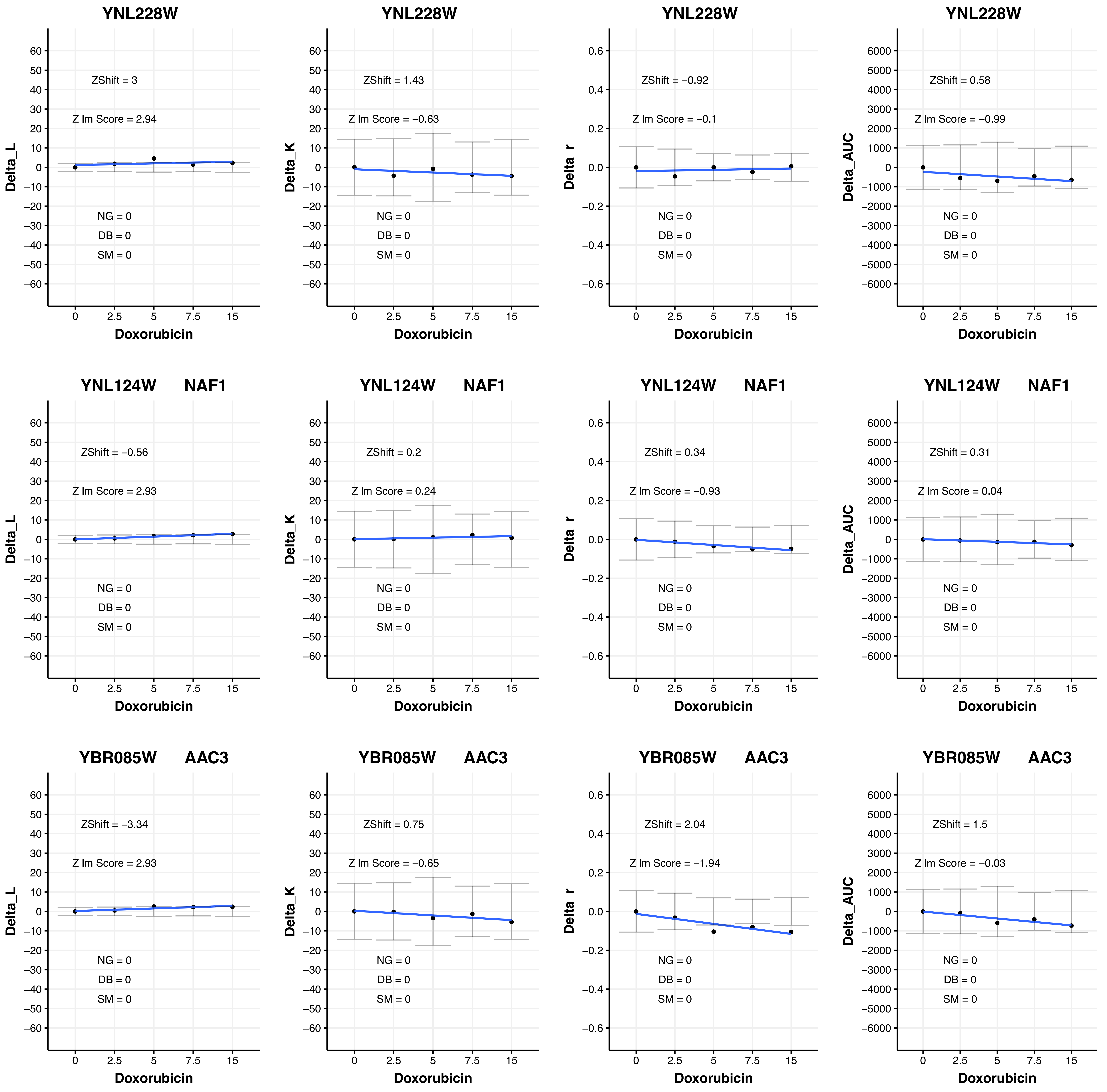
YNL228W
ZShift = 3
Z lm Score = 2.94
NG = 0
DB = 0
SM = 0
Delta_L
Doxorubicin
YNL228W
ZShift = 1.43
Z lm Score = −0.63
NG = 0
DB = 0
SM = 0
Delta_K
Doxorubicin
YNL228W
ZShift = −0.92
Z lm Score = −0.1
NG = 0
DB = 0
SM = 0
Delta_r
Doxorubicin
YNL228W
ZShift = 0.58
Z lm Score = −0.99
NG = 0
DB = 0
SM = 0
Delta_AUC
Doxorubicin
YNL124W NAF1
ZShift = −0.56
Z lm Score = 2.93
NG = 0
DB = 0
SM = 0
Delta_L
Doxorubicin
YNL124W NAF1
ZShift = 0.2
Z lm Score = 0.24
NG = 0
DB = 0
SM = 0
Delta_K
Doxorubicin
YNL124W NAF1
ZShift = 0.34
Z lm Score = −0.93
NG = 0
DB = 0
SM = 0
Delta_r
Doxorubicin
YNL124W NAF1
ZShift = 0.31
Z lm Score = 0.04
NG = 0
DB = 0
SM = 0
Delta_AUC
Doxorubicin
YBR085W AAC3
ZShift = −3.34
Z lm Score = 2.93
NG = 0
DB = 0
SM = 0
Delta_L
Doxorubicin
YBR085W AAC3
ZShift = 0.75
Z lm Score = −0.65
NG = 0
DB = 0
SM = 0
Delta_K
Doxorubicin
YBR085W AAC3
ZShift = 2.04
Z lm Score = −1.94
NG = 0
DB = 0
SM = 0
Delta_r
Doxorubicin
YBR085W AAC3
ZShift = 1.5
Z lm Score = −0.03
NG = 0
DB = 0
SM = 0
Delta_AUC
Doxorubicin

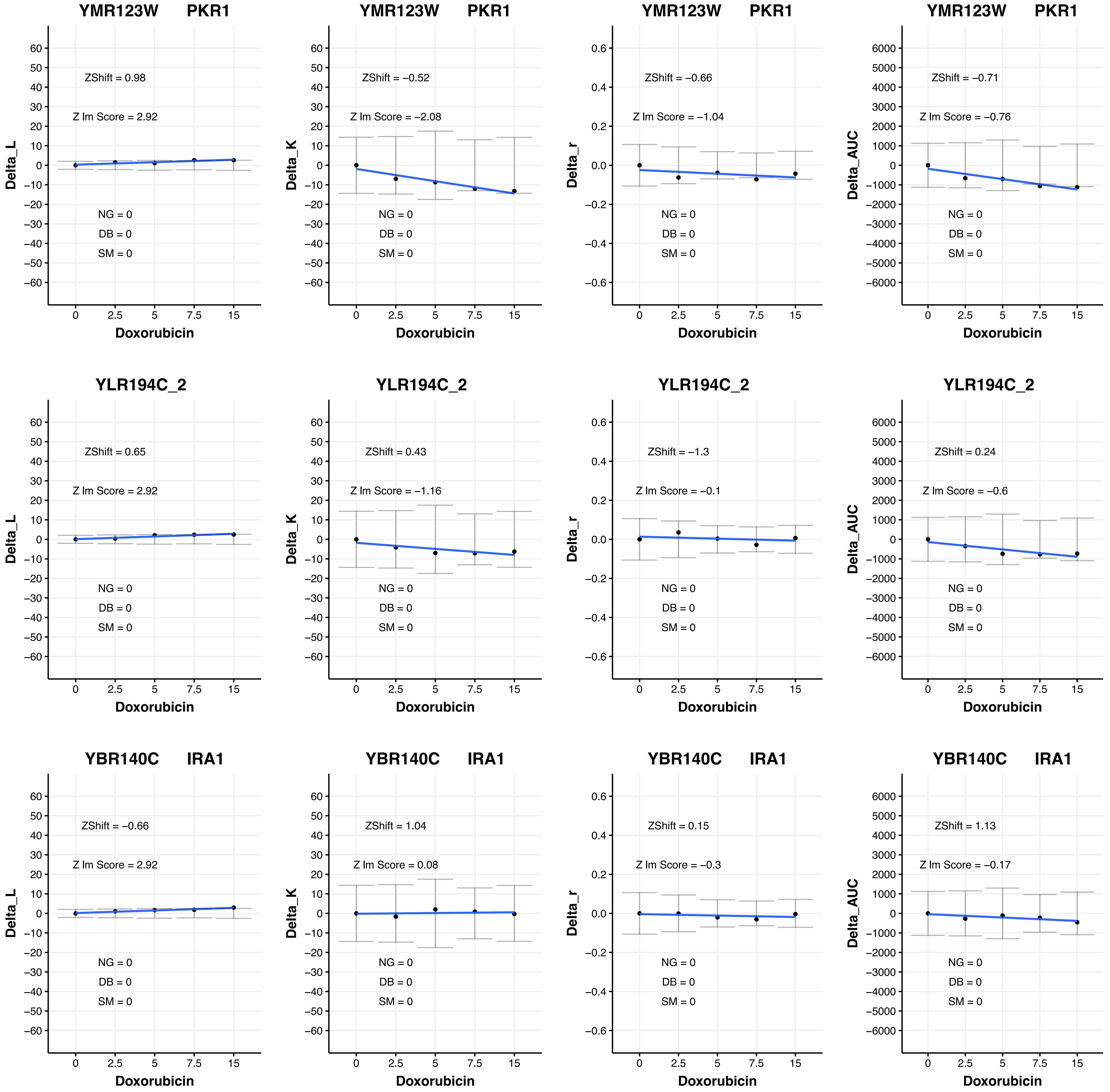
YMR123W PKR1
ZShift = 0.98
Z lm Score = 2.92
NG = 0
DB = 0
SM = 0
Delta_L
Doxorubicin
YMR123W PKR1
ZShift = −0.52
Z lm Score = −2.08
NG = 0
DB = 0
SM = 0
Delta_K
Doxorubicin
YMR123W PKR1
ZShift = −0.66
Z lm Score = −1.04
NG = 0
DB = 0
SM = 0
Delta_r
Doxorubicin
YMR123W PKR1
ZShift = −0.71
Z lm Score = −0.76
NG = 0
DB = 0
SM = 0
Delta_AUC
Doxorubicin
YLR194C_2
ZShift = 0.65
Z lm Score = 2.92
NG = 0
DB = 0
SM = 0
Delta_L
Doxorubicin
YLR194C_2
ZShift = 0.43
Z lm Score = −1.16
NG = 0
DB = 0
SM = 0
Delta_K
Doxorubicin
YLR194C_2
ZShift = −1.3
Z lm Score = −0.1
NG = 0
DB = 0
SM = 0
Delta_r
Doxorubicin
YLR194C_2
ZShift = 0.24
Z lm Score = −0.6
NG = 0
DB = 0
SM = 0
Delta_AUC
Doxorubicin
YBR140C IRA1
ZShift = −0.66
Z lm Score = 2.92
NG = 0
DB = 0
SM = 0
Delta_L
Doxorubicin
YBR140C IRA1
ZShift = 1.04
Z lm Score = 0.08
NG = 0
DB = 0
SM = 0
Delta_K
Doxorubicin
YBR140C IRA1
ZShift = 0.15
Z lm Score = −0.3
NG = 0
DB = 0
SM = 0
Delta_r
Doxorubicin
YBR140C IRA1
ZShift = 1.13
Z lm Score = −0.17
NG = 0
DB = 0
SM = 0
Delta_AUC
Doxorubicin

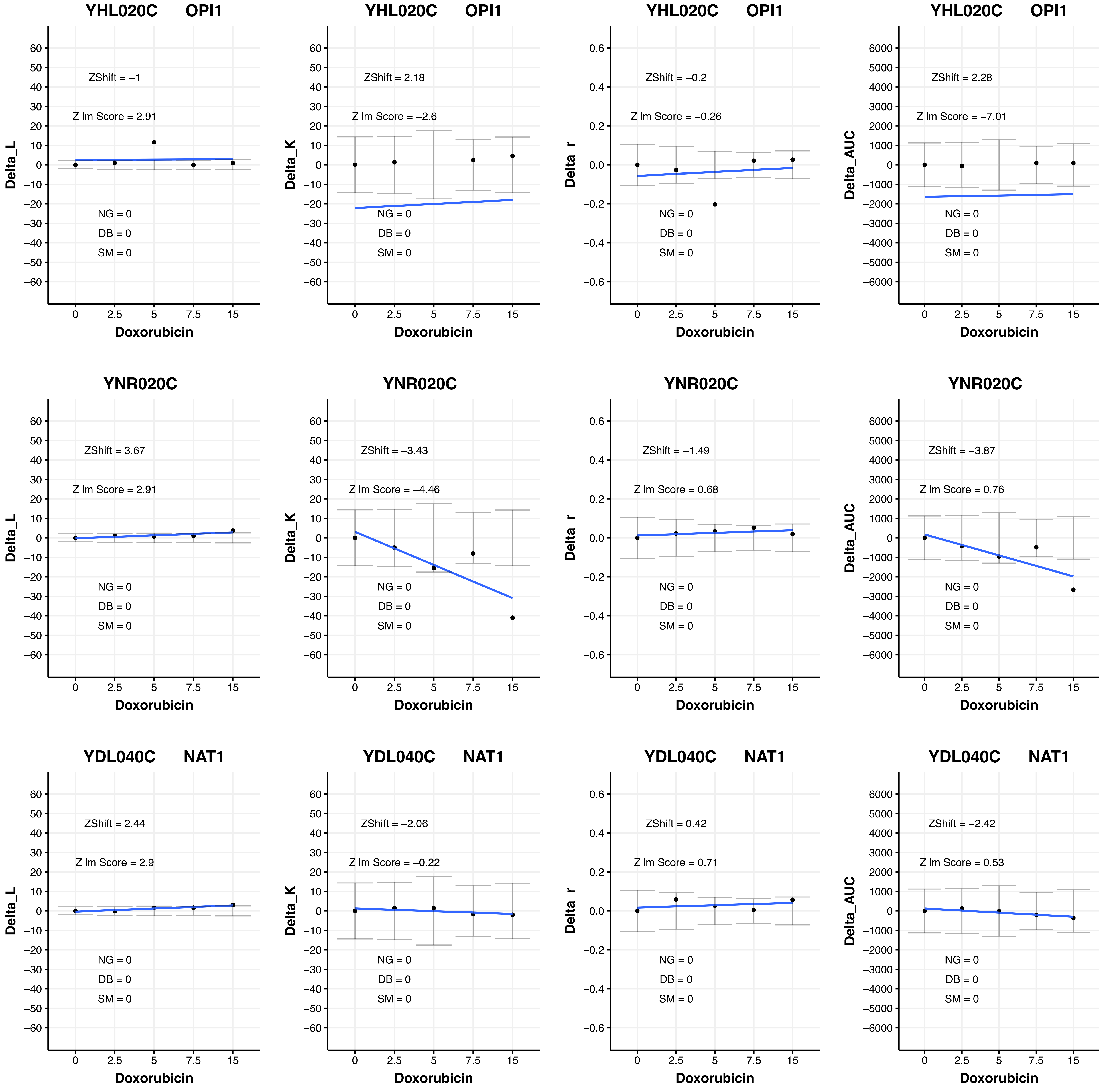

YHL020C OPI1
ZShift = −1
Z lm Score = 2.91
NG = 0
DB = 0
SM = 0
Delta_L
Doxorubicin
YHL020C OPI1
ZShift = 2.18
Z lm Score = −2.6
NG = 0
DB = 0
SM = 0
Delta_K
Doxorubicin
YHL020C OPI1
ZShift = −0.2
Z lm Score = −0.26
NG = 0
DB = 0
SM = 0
Delta_r
Doxorubicin
YHL020C OPI1
ZShift = 2.28
Z lm Score = −7.01
NG = 0
DB = 0
SM = 0
Delta_AUC
Doxorubicin
YNR020C
ZShift = 3.67
Z lm Score = 2.91
NG = 0
DB = 0
SM = 0
Delta_L
Doxorubicin
YNR020C
ZShift = −3.43
Z lm Score = −4.46
NG = 0
DB = 0
SM = 0
Delta_K
Doxorubicin
YNR020C
ZShift = −1.49
Z lm Score = 0.68
NG = 0
DB = 0
SM = 0
Delta_r
Doxorubicin
YNR020C
ZShift = −3.87
Z lm Score = 0.76
NG = 0
DB = 0
SM = 0
Delta_AUC
Doxorubicin
YDL040C NAT1
ZShift = 2.44
Z lm Score = 2.9
NG = 0
DB = 0
SM = 0
Delta_L
Doxorubicin
YDL040C NAT1
ZShift = −2.06
Z lm Score = −0.22
NG = 0
DB = 0
SM = 0
Delta_K
Doxorubicin
YDL040C NAT1
ZShift = 0.42
Z lm Score = 0.71
NG = 0
DB = 0
SM = 0
Delta_r
Doxorubicin
YDL040C NAT1
ZShift = −2.42
Z lm Score = 0.53
NG = 0
DB = 0
SM = 0
Delta_AUC
Doxorubicin

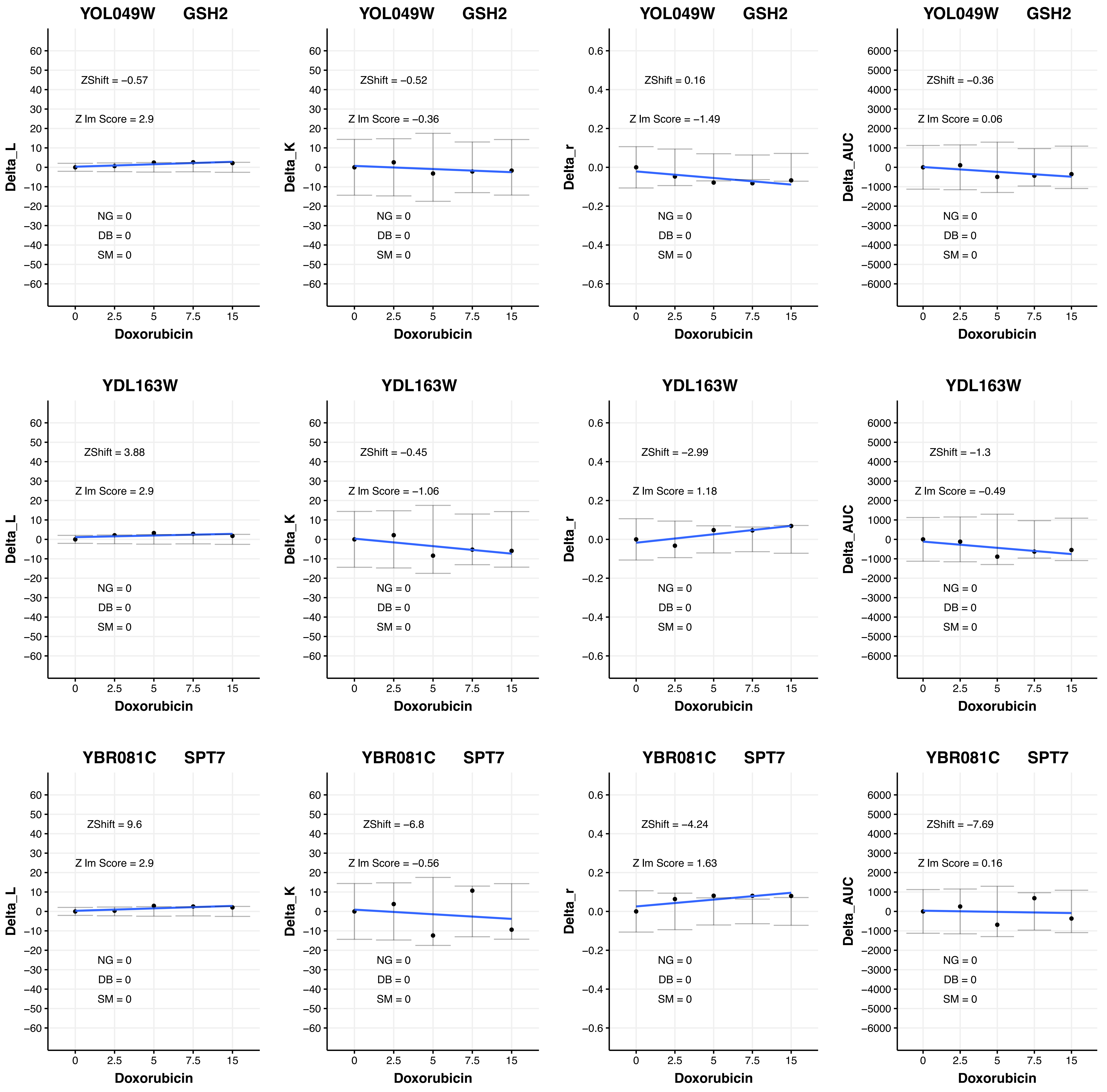
YOL049W GSH2
ZShift = −0.57
Z lm Score = 2.9
NG = 0
DB = 0
SM = 0
Delta_L
Doxorubicin
YOL049W GSH2
ZShift = −0.52
Z lm Score = −0.36
Delta_K
YOL049W GSH2
ZShift = 0.16
Z lm Score = −1.49
Delta_r
YOL049W GSH2
ZShift = −0.36
Z lm Score = 0.06
Delta_AUC
YDL163W
ZShift = 3.88
Z lm Score = 2.9
YDL163W
ZShift = −0.45
Z lm Score = −1.06
YDL163W
ZShift = −2.99
Z lm Score = 1.18
YDL163W
ZShift = −1.3
Z lm Score = −0.49
YBR081C SPT7
ZShift = 9.6
Z lm Score = 2.9
YBR081C SPT7
ZShift = −6.8
Z lm Score = −0.56
YBR081C SPT7
ZShift = −4.24
Z lm Score = 1.63
YBR081C SPT7
ZShift = −7.69
Z lm Score = 0.16

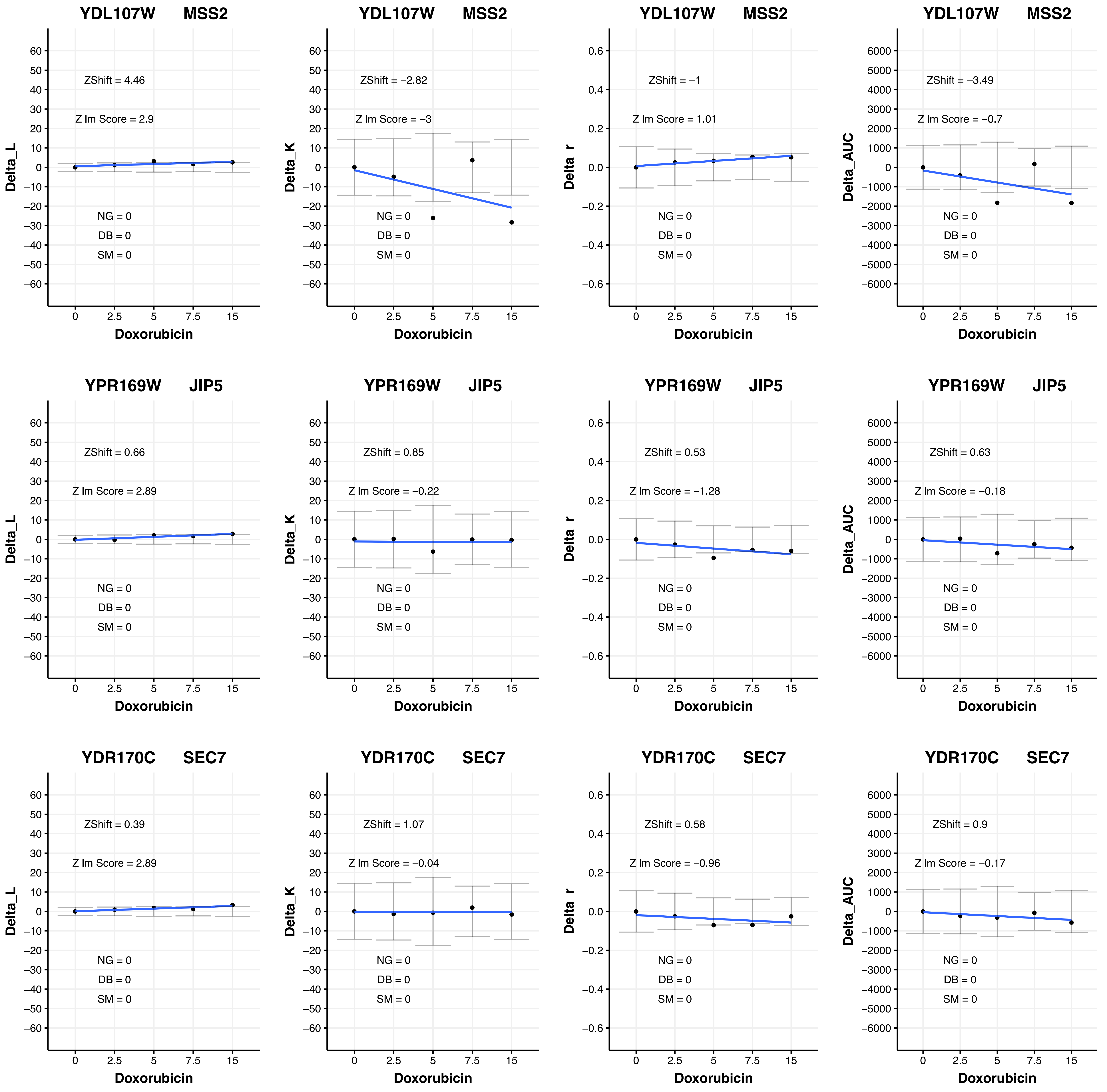

YDL107W MSS2
ZShift = 4.46
Z lm Score = 2.9
NG = 0
DB = 0
SM = 0
Delta_L
Doxorubicin
YDL107W MSS2
ZShift = −2.82
Z lm Score = −3
NG = 0
DB = 0
SM = 0
Delta_K
Doxorubicin
YDL107W MSS2
ZShift = −1
Z lm Score = 1.01
NG = 0
DB = 0
SM = 0
Delta_r
Doxorubicin
YDL107W MSS2
ZShift = −3.49
Z lm Score = −0.7
NG = 0
DB = 0
SM = 0
Delta_AUC
Doxorubicin
YPR169W JIP5
ZShift = 0.66
Z lm Score = 2.89
NG = 0
DB = 0
SM = 0
Delta_L
Doxorubicin
YPR169W JIP5
ZShift = 0.85
Z lm Score = −0.22
NG = 0
DB = 0
SM = 0
Delta_K
Doxorubicin
YPR169W JIP5
ZShift = 0.53
Z lm Score = −1.28
NG = 0
DB = 0
SM = 0
Delta_r
Doxorubicin
YPR169W JIP5
ZShift = 0.63
Z lm Score = −0.18
NG = 0
DB = 0
SM = 0
Delta_AUC
Doxorubicin
YDR170C SEC7
ZShift = 0.39
Z lm Score = 2.89
NG = 0
DB = 0
SM = 0
Delta_L
Doxorubicin
YDR170C SEC7
ZShift = 1.07
Z lm Score = −0.04
NG = 0
DB = 0
SM = 0
Delta_K
Doxorubicin
YDR170C SEC7
ZShift = 0.58
Z lm Score = −0.96
NG = 0
DB = 0
SM = 0
Delta_r
Doxorubicin
YDR170C SEC7
ZShift = 0.9
Z lm Score = −0.17
NG = 0
DB = 0
SM = 0
Delta_AUC
Doxorubicin

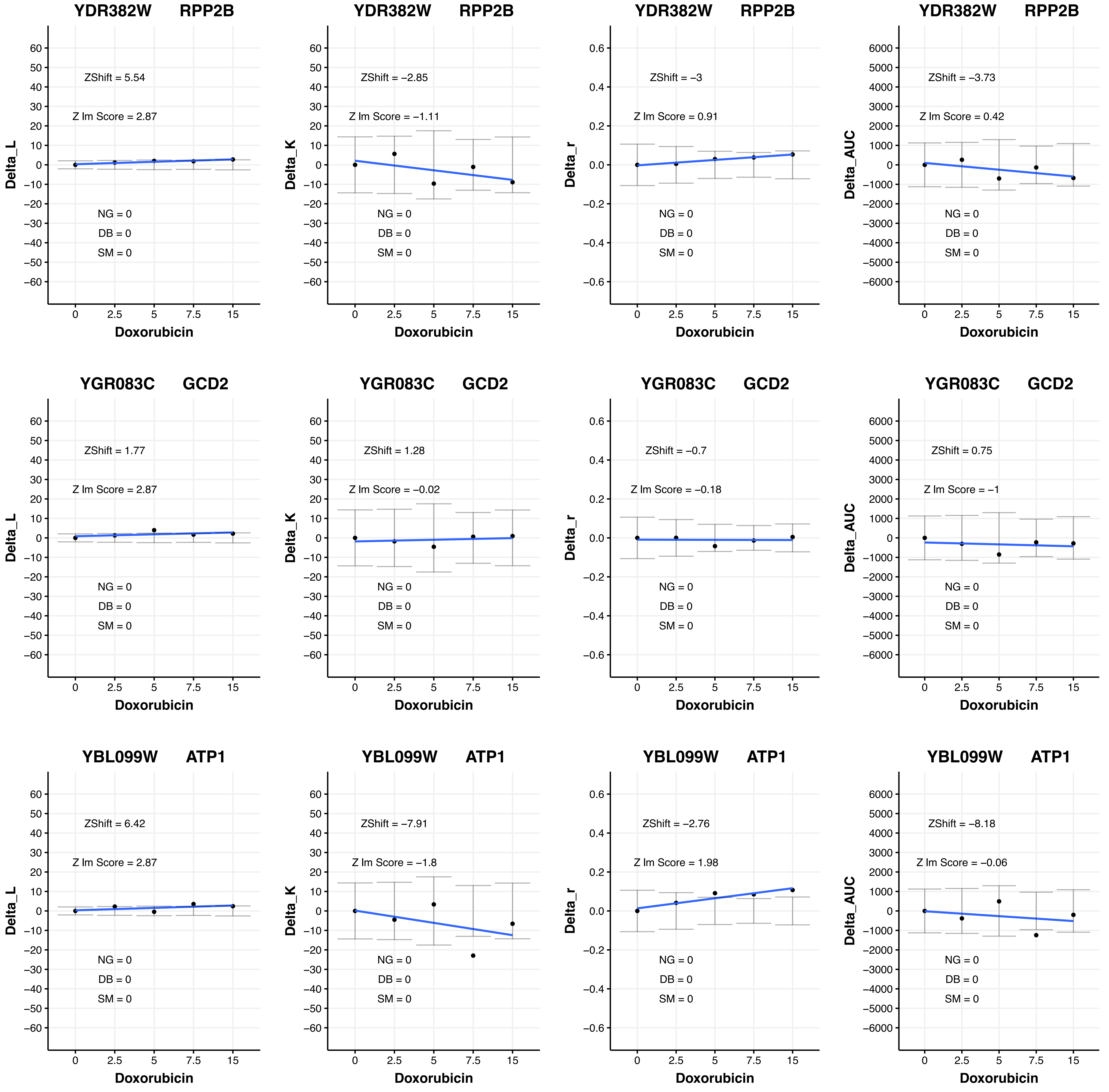
YDR382W RPP2B
ZShift = 5.54
Z lm Score = 2.87
NG = 0
DB = 0
SM = 0
Delta_L
Doxorubicin
ZShift = −2.85
Z lm Score = −1.11
Delta_K
ZShift = −3
Z lm Score = 0.91
Delta_r
ZShift = −3.73
Z lm Score = 0.42
Delta_AUC
YGR083C GCD2
ZShift = 1.77
Z lm Score = 2.87
ZShift = 1.28
Z lm Score = −0.02
ZShift = −0.7
Z lm Score = −0.18
ZShift = 0.75
Z lm Score = −1
YBL099W ATP1
ZShift = 6.42
Z lm Score = 2.87
ZShift = −7.91
Z lm Score = −1.8
ZShift = −2.76
Z lm Score = 1.98
ZShift = −8.18
Z lm Score = −0.06

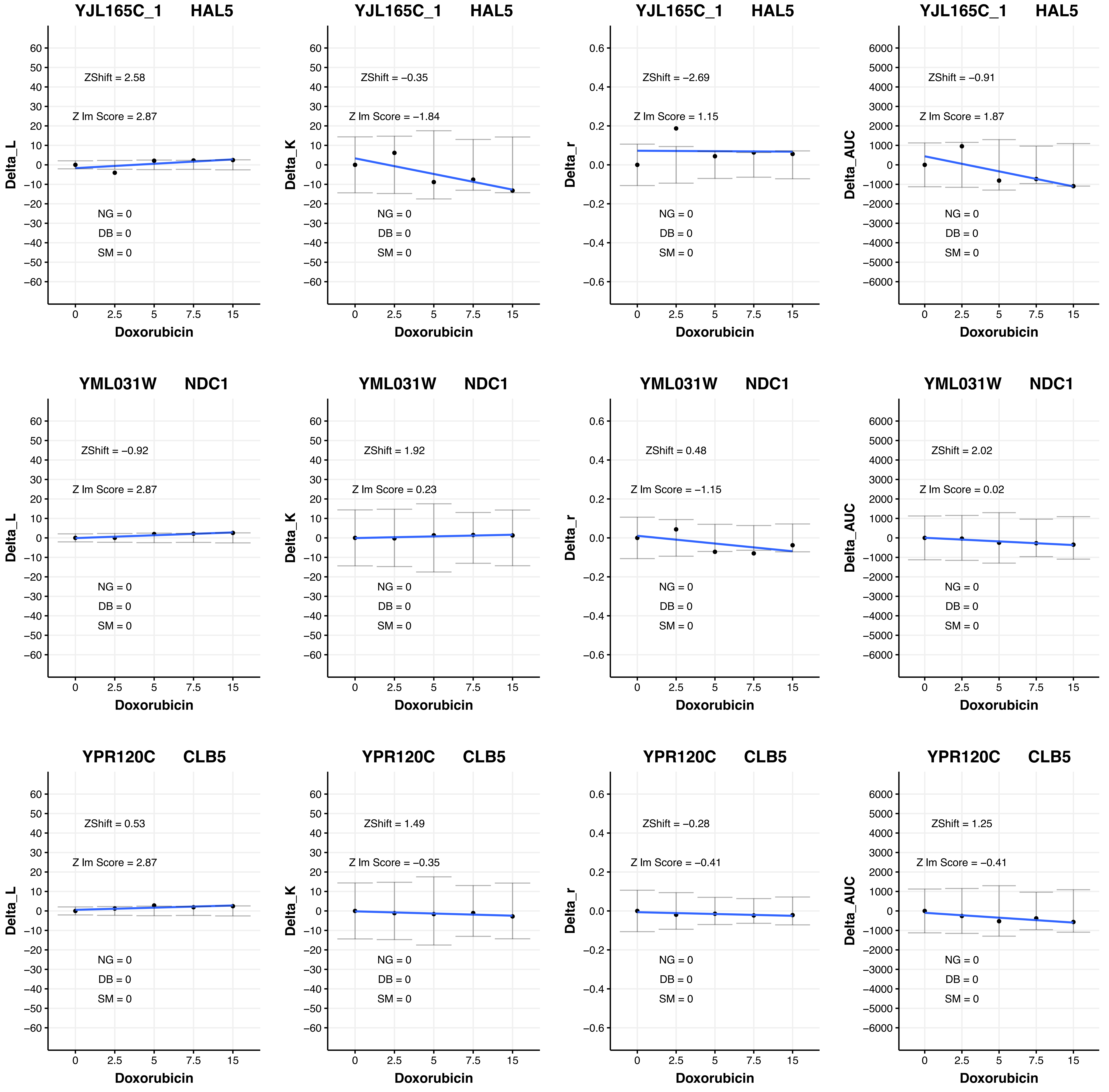
YJL165C_1 HAL5
ZShift = 2.58
Z lm Score = 2.87
NG = 0
DB = 0
SM = 0
Delta_L
Doxorubicin
YJL165C_1 HAL5
ZShift = −0.35
Z lm Score = −1.84
NG = 0
DB = 0
SM = 0
Delta_K
Doxorubicin
YJL165C_1 HAL5
ZShift = −2.69
Z lm Score = 1.15
NG = 0
DB = 0
SM = 0
Delta_r
Doxorubicin
YJL165C_1 HAL5
ZShift = −0.91
Z lm Score = 1.87
NG = 0
DB = 0
SM = 0
Delta_AUC
Doxorubicin
YML031W NDC1
ZShift = −0.92
Z lm Score = 2.87
NG = 0
DB = 0
SM = 0
Delta_L
Doxorubicin
YML031W NDC1
ZShift = 1.92
Z lm Score = 0.23
NG = 0
DB = 0
SM = 0
Delta_K
Doxorubicin
YML031W NDC1
ZShift = 0.48
Z lm Score = −1.15
NG = 0
DB = 0
SM = 0
Delta_r
Doxorubicin
YML031W NDC1
ZShift = 2.02
Z lm Score = 0.02
NG = 0
DB = 0
SM = 0
Delta_AUC
Doxorubicin
YPR120C CLB5
ZShift = 0.53
Z lm Score = 2.87
NG = 0
DB = 0
SM = 0
Delta_L
Doxorubicin
YPR120C CLB5
ZShift = 1.49
Z lm Score = −0.35
NG = 0
DB = 0
SM = 0
Delta_K
Doxorubicin
YPR120C CLB5
ZShift = −0.28
Z lm Score = −0.41
NG = 0
DB = 0
SM = 0
Delta_r
Doxorubicin
YPR120C CLB5
ZShift = 1.25
Z lm Score = −0.41
NG = 0
DB = 0
SM = 0
Delta_AUC
Doxorubicin

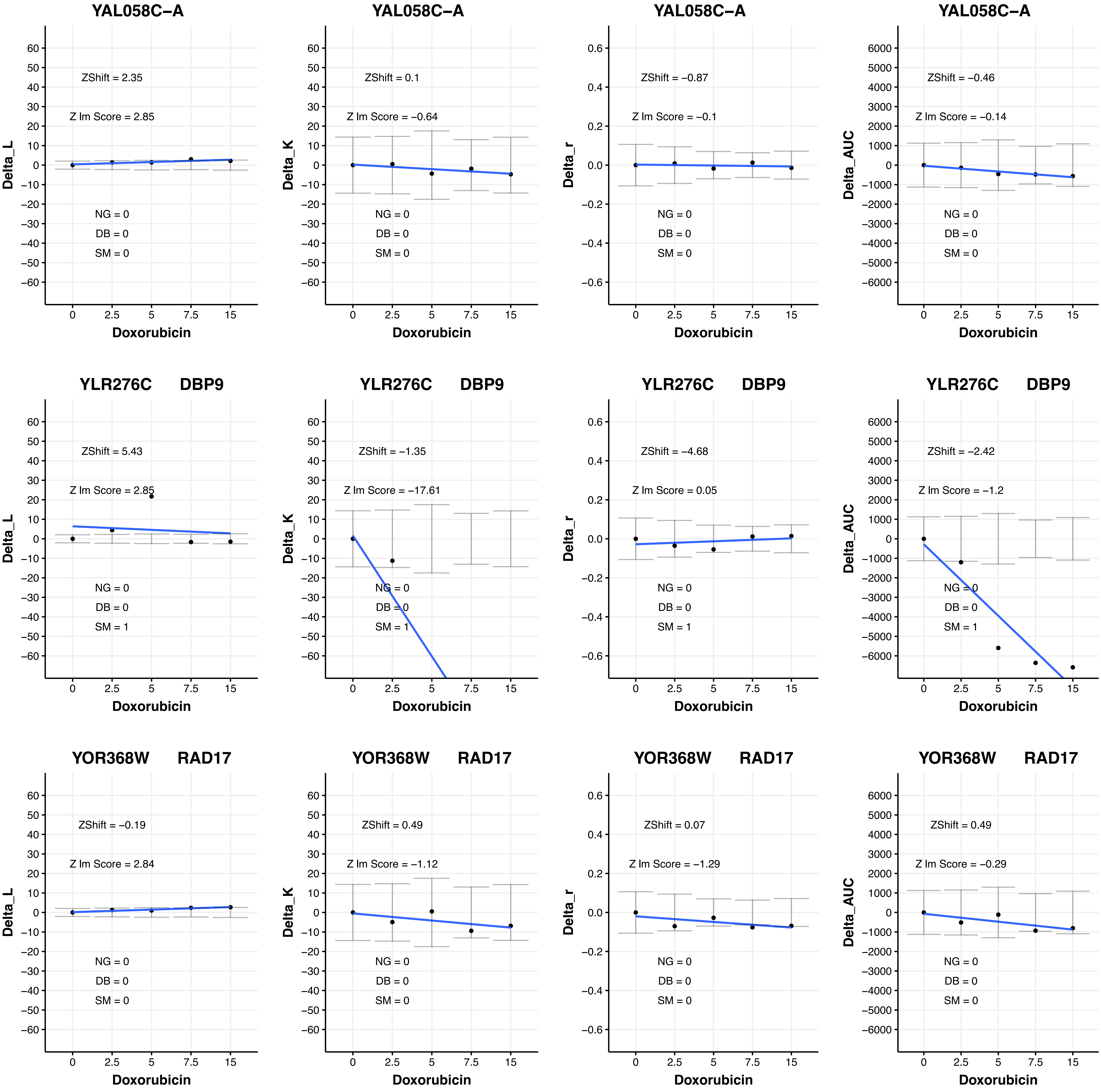
YAL058C−A
Delta_L
ZShift = 2.35
Z lm Score = 2.85
NG = 0
DB = 0
SM = 0
Doxorubicin
YAL058C−A
Delta_K
ZShift = 0.1
Z lm Score = −0.64
NG = 0
DB = 0
SM = 0
Doxorubicin
YAL058C−A
Delta_r
ZShift = −0.87
Z lm Score = −0.1
NG = 0
DB = 0
SM = 0
Doxorubicin
YAL058C−A
Delta_AUC
ZShift = −0.46
Z lm Score = −0.14
NG = 0
DB = 0
SM = 0
Doxorubicin
YLR276C DBP9
Delta_L
ZShift = 5.43
Z lm Score = 2.85
NG = 0
DB = 0
SM = 1
Doxorubicin
YLR276C DBP9
Delta_K
ZShift = −1.35
Z lm Score = −17.61
NG = 0
DB = 0
SM = 1
Doxorubicin
YLR276C DBP9
Delta_r
ZShift = −4.68
Z lm Score = 0.05
NG = 0
DB = 0
SM = 1
Doxorubicin
YLR276C DBP9
Delta_AUC
ZShift = −2.42
Z lm Score = −1.2
NG = 0
DB = 0
SM = 1
Doxorubicin
YOR368W RAD17
Delta_L
ZShift = −0.19
Z lm Score = 2.84
NG = 0
DB = 0
SM = 0
Doxorubicin
YOR368W RAD17
Delta_K
ZShift = 0.49
Z lm Score = −1.12
NG = 0
DB = 0
SM = 0
Doxorubicin
YOR368W RAD17
Delta_r
ZShift = 0.07
Z lm Score = −1.29
NG = 0
DB = 0
SM = 0
Doxorubicin
YOR368W RAD17
Delta_AUC
ZShift = 0.49
Z lm Score = −0.29
NG = 0
DB = 0
SM = 0
Doxorubicin

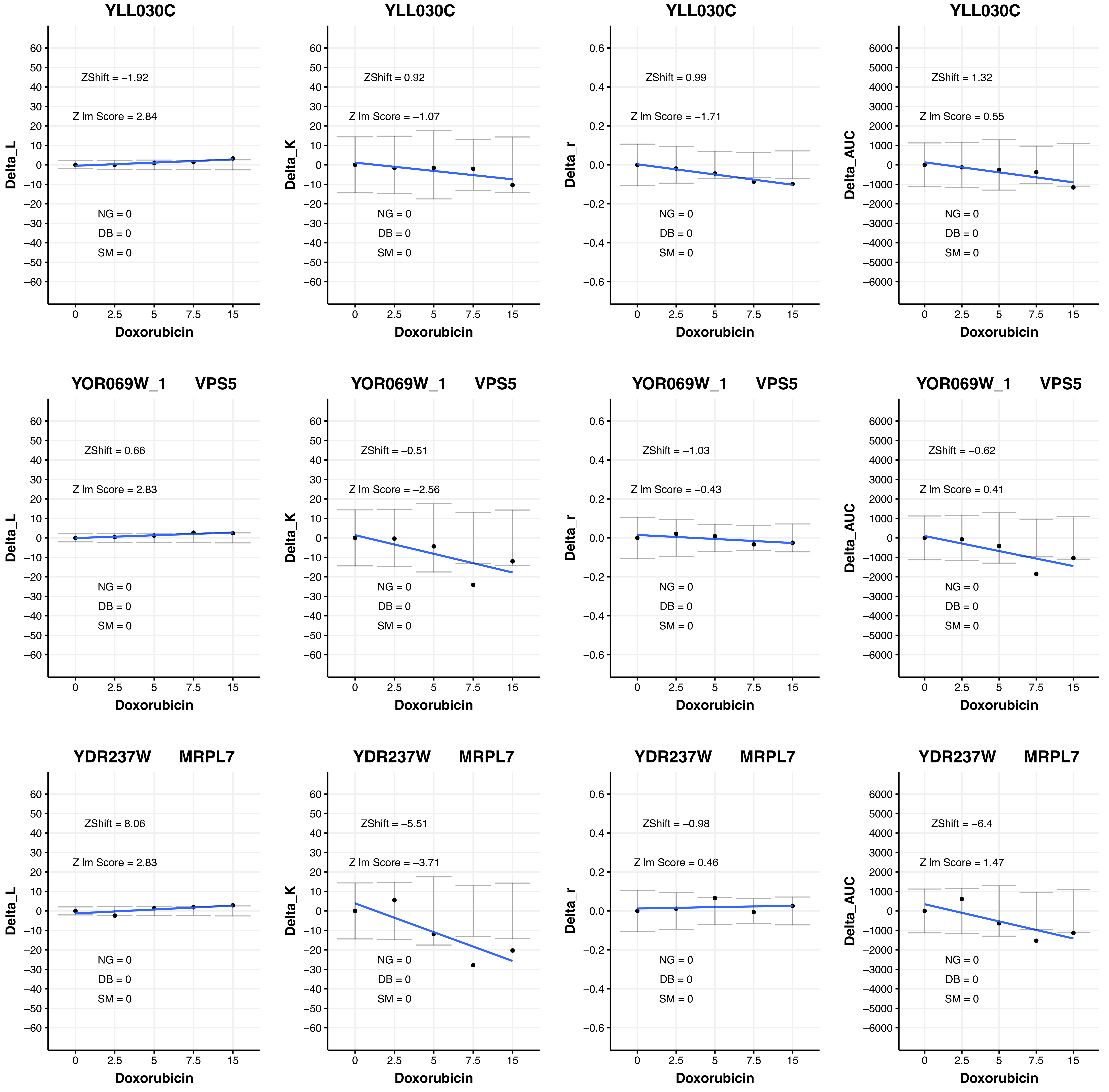

YLL030C
ZShift = −1.92
Z lm Score = 2.84
NG = 0
DB = 0
SM = 0
Delta_L
Doxorubicin
YLL030C
ZShift = 0.92
Z lm Score = −1.07
NG = 0
DB = 0
SM = 0
Delta_K
Doxorubicin
YLL030C
ZShift = 0.99
Z lm Score = −1.71
NG = 0
DB = 0
SM = 0
Delta_r
Doxorubicin
YLL030C
ZShift = 1.32
Z lm Score = 0.55
NG = 0
DB = 0
SM = 0
Delta_AUC
Doxorubicin
YOR069W_1 VPS5
ZShift = 0.66
Z lm Score = 2.83
NG = 0
DB = 0
SM = 0
Delta_L
Doxorubicin
YOR069W_1 VPS5
ZShift = −0.51
Z lm Score = −2.56
NG = 0
DB = 0
SM = 0
Delta_K
Doxorubicin
YOR069W_1 VPS5
ZShift = −1.03
Z lm Score = −0.43
NG = 0
DB = 0
SM = 0
Delta_r
Doxorubicin
YOR069W_1 VPS5
ZShift = −0.62
Z lm Score = 0.41
NG = 0
DB = 0
SM = 0
Delta_AUC
Doxorubicin
YDR237W MRPL7
ZShift = 8.06
Z lm Score = 2.83
NG = 0
DB = 0
SM = 0
Delta_L
Doxorubicin
YDR237W MRPL7
ZShift = −5.51
Z lm Score = −3.71
NG = 0
DB = 0
SM = 0
Delta_K
Doxorubicin
YDR237W MRPL7
ZShift = −0.98
Z lm Score = 0.46
NG = 0
DB = 0
SM = 0
Delta_r
Doxorubicin
YDR237W MRPL7
ZShift = −6.4
Z lm Score = 1.47
NG = 0
DB = 0
SM = 0
Delta_AUC
Doxorubicin

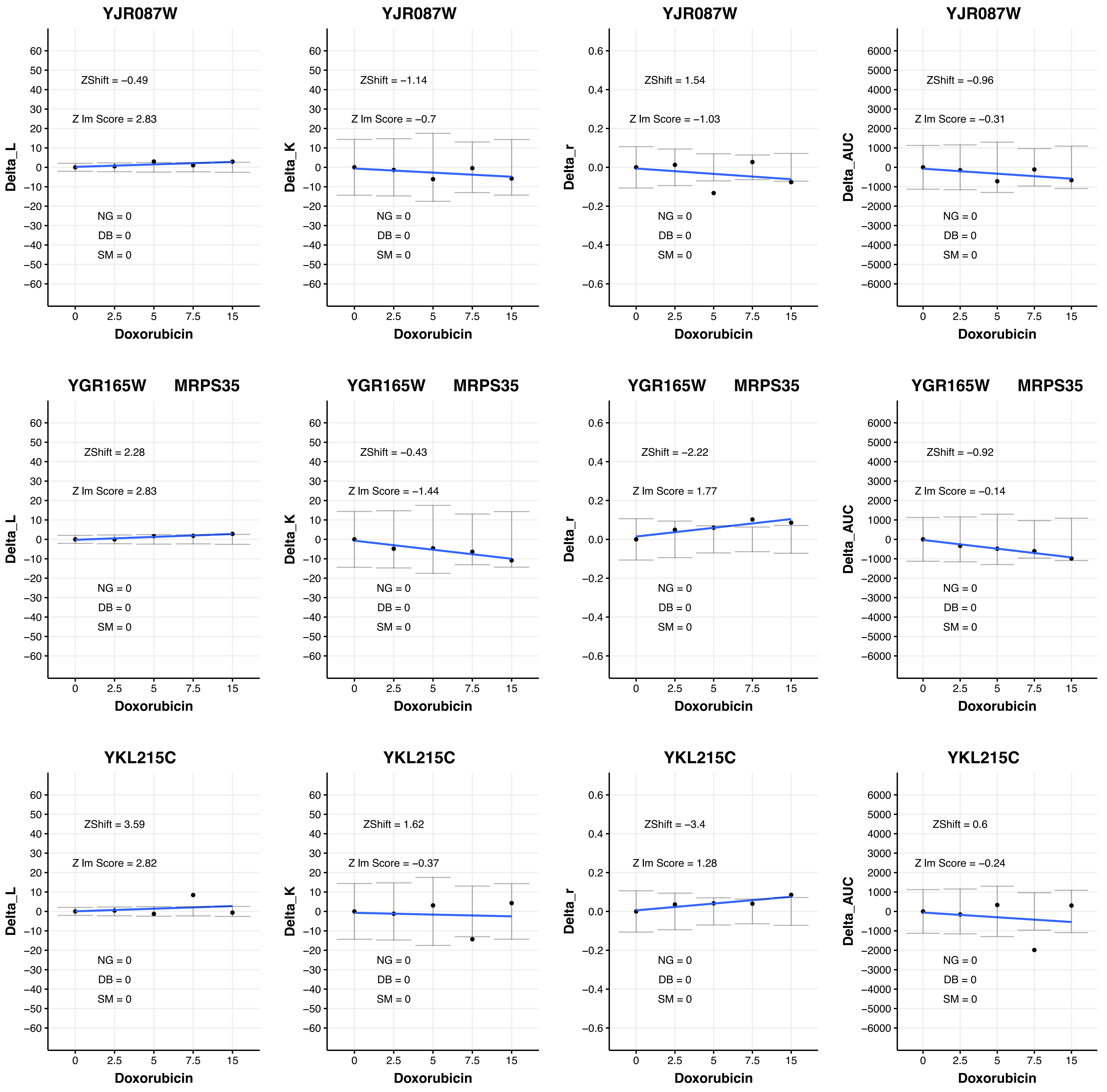

YJR087W
ZShift = −0.49
Z lm Score = 2.83
NG = 0
DB = 0
SM = 0
Delta_L
Doxorubicin
YJR087W
ZShift = −1.14
Z lm Score = −0.7
NG = 0
DB = 0
SM = 0
Delta_K
Doxorubicin
YJR087W
ZShift = 1.54
Z lm Score = −1.03
NG = 0
DB = 0
SM = 0
Delta_r
Doxorubicin
YJR087W
ZShift = −0.96
Z lm Score = −0.31
NG = 0
DB = 0
SM = 0
Delta_AUC
Doxorubicin
YGR165W MRPS35
ZShift = 2.28
Z lm Score = 2.83
NG = 0
DB = 0
SM = 0
Delta_L
Doxorubicin
YGR165W MRPS35
ZShift = −0.43
Z lm Score = −1.44
NG = 0
DB = 0
SM = 0
Delta_K
Doxorubicin
YGR165W MRPS35
ZShift = −2.22
Z lm Score = 1.77
NG = 0
DB = 0
SM = 0
Delta_r
Doxorubicin
YGR165W MRPS35
ZShift = −0.92
Z lm Score = −0.14
NG = 0
DB = 0
SM = 0
Delta_AUC
Doxorubicin
YKL215C
ZShift = 3.59
Z lm Score = 2.82
NG = 0
DB = 0
SM = 0
Delta_L
Doxorubicin
YKL215C
ZShift = 1.62
Z lm Score = −0.37
NG = 0
DB = 0
SM = 0
Delta_K
Doxorubicin
YKL215C
ZShift = −3.4
Z lm Score = 1.28
NG = 0
DB = 0
SM = 0
Delta_r
Doxorubicin
YKL215C
ZShift = 0.6
Z lm Score = −0.24
NG = 0
DB = 0
SM = 0
Delta_AUC
Doxorubicin

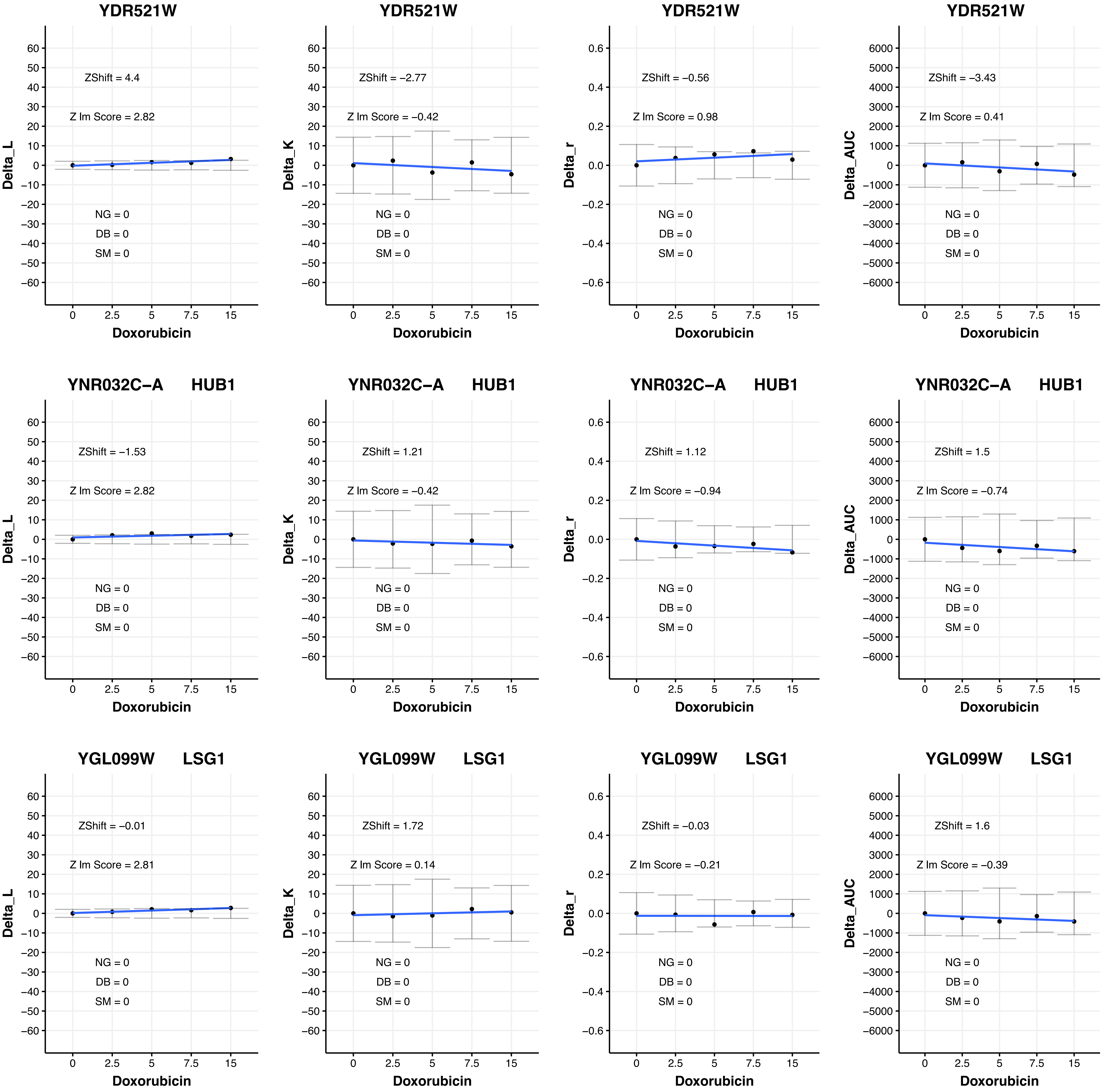

YDR521W
ZShift = 4.4
Z lm Score = 2.82
NG = 0
DB = 0
SM = 0
Delta_L
Doxorubicin
YDR521W
ZShift = −2.77
Z lm Score = −0.42
NG = 0
DB = 0
SM = 0
Delta_K
Doxorubicin
YDR521W
ZShift = −0.56
Z lm Score = 0.98
NG = 0
DB = 0
SM = 0
Delta_r
Doxorubicin
YDR521W
ZShift = −3.43
Z lm Score = 0.41
NG = 0
DB = 0
SM = 0
Delta_AUC
Doxorubicin
YNR032C−A HUB1
ZShift = −1.53
Z lm Score = 2.82
NG = 0
DB = 0
SM = 0
Delta_L
Doxorubicin
YNR032C−A HUB1
ZShift = 1.21
Z lm Score = −0.42
NG = 0
DB = 0
SM = 0
Delta_K
Doxorubicin
YNR032C−A HUB1
ZShift = 1.12
Z lm Score = −0.94
NG = 0
DB = 0
SM = 0
Delta_r
Doxorubicin
YNR032C−A HUB1
ZShift = 1.5
Z lm Score = −0.74
NG = 0
DB = 0
SM = 0
Delta_AUC
Doxorubicin
YGL099W LSG1
ZShift = −0.01
Z lm Score = 2.81
NG = 0
DB = 0
SM = 0
Delta_L
Doxorubicin
YGL099W LSG1
ZShift = 1.72
Z lm Score = 0.14
NG = 0
DB = 0
SM = 0
Delta_K
Doxorubicin
YGL099W LSG1
ZShift = −0.03
Z lm Score = −0.21
NG = 0
DB = 0
SM = 0
Delta_r
Doxorubicin
YGL099W LSG1
ZShift = 1.6
Z lm Score = −0.39
NG = 0
DB = 0
SM = 0
Delta_AUC
Doxorubicin

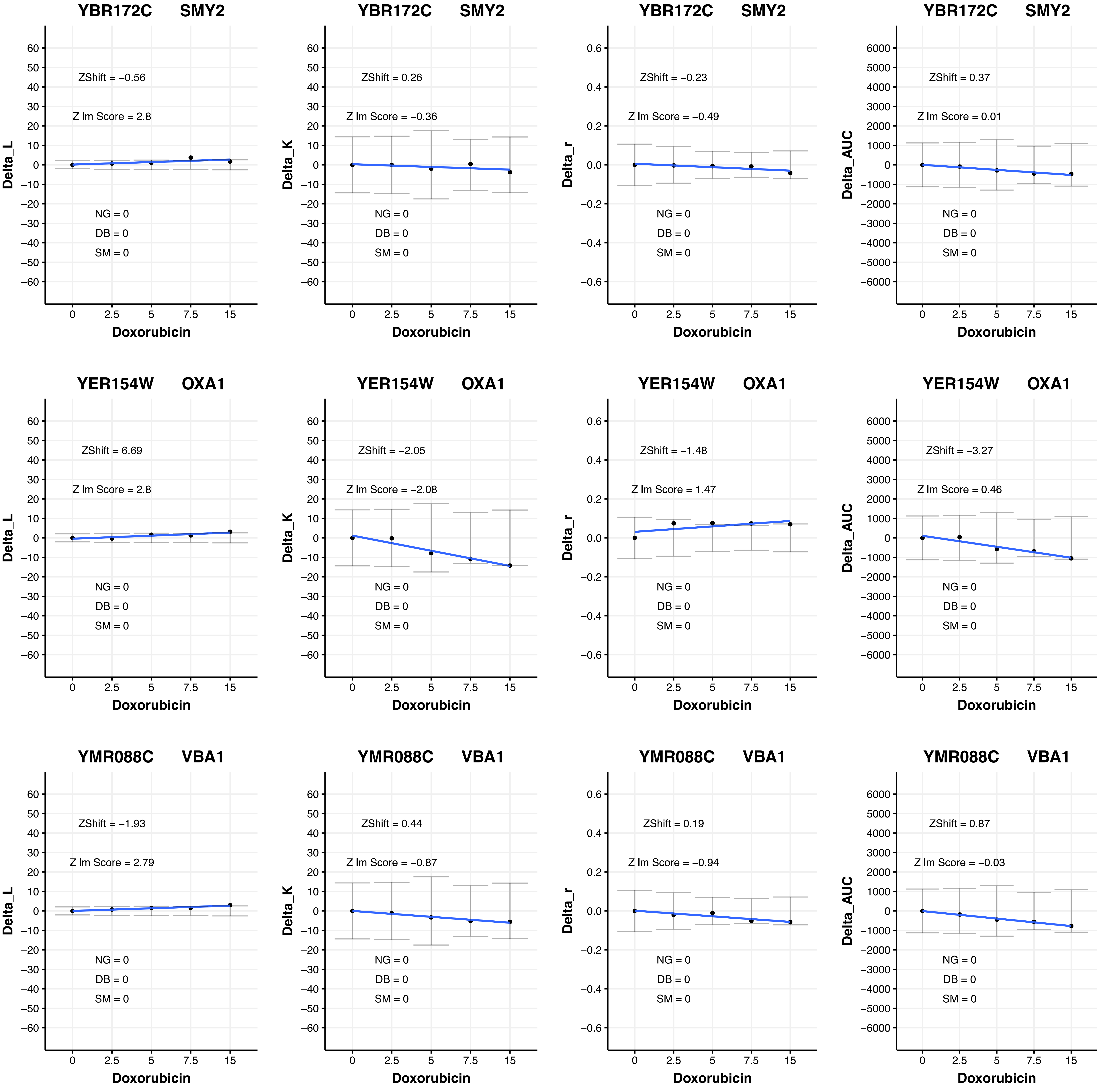
YBR172C SMY2
ZShift = −0.56
Z lm Score = 2.8
NG = 0
DB = 0
SM = 0
Delta_L
Doxorubicin
YBR172C SMY2
ZShift = 0.26
Z lm Score = −0.36
NG = 0
DB = 0
SM = 0
Delta_K
Doxorubicin
YBR172C SMY2
ZShift = −0.23
Z lm Score = −0.49
NG = 0
DB = 0
SM = 0
Delta_r
Doxorubicin
YBR172C SMY2
ZShift = 0.37
Z lm Score = 0.01
NG = 0
DB = 0
SM = 0
Delta_AUC
Doxorubicin
YER154W OXA1
ZShift = 6.69
Z lm Score = 2.8
NG = 0
DB = 0
SM = 0
Delta_L
Doxorubicin
YER154W OXA1
ZShift = −2.05
Z lm Score = −2.08
NG = 0
DB = 0
SM = 0
Delta_K
Doxorubicin
YER154W OXA1
ZShift = −1.48
Z lm Score = 1.47
NG = 0
DB = 0
SM = 0
Delta_r
Doxorubicin
YER154W OXA1
ZShift = −3.27
Z lm Score = 0.46
NG = 0
DB = 0
SM = 0
Delta_AUC
Doxorubicin
YMR088C VBA1
ZShift = −1.93
Z lm Score = 2.79
NG = 0
DB = 0
SM = 0
Delta_L
Doxorubicin
YMR088C VBA1
ZShift = 0.44
Z lm Score = −0.87
NG = 0
DB = 0
SM = 0
Delta_K
Doxorubicin
YMR088C VBA1
ZShift = 0.19
Z lm Score = −0.94
NG = 0
DB = 0
SM = 0
Delta_r
Doxorubicin
YMR088C VBA1
ZShift = 0.87
Z lm Score = −0.03
NG = 0
DB = 0
SM = 0
Delta_AUC
Doxorubicin

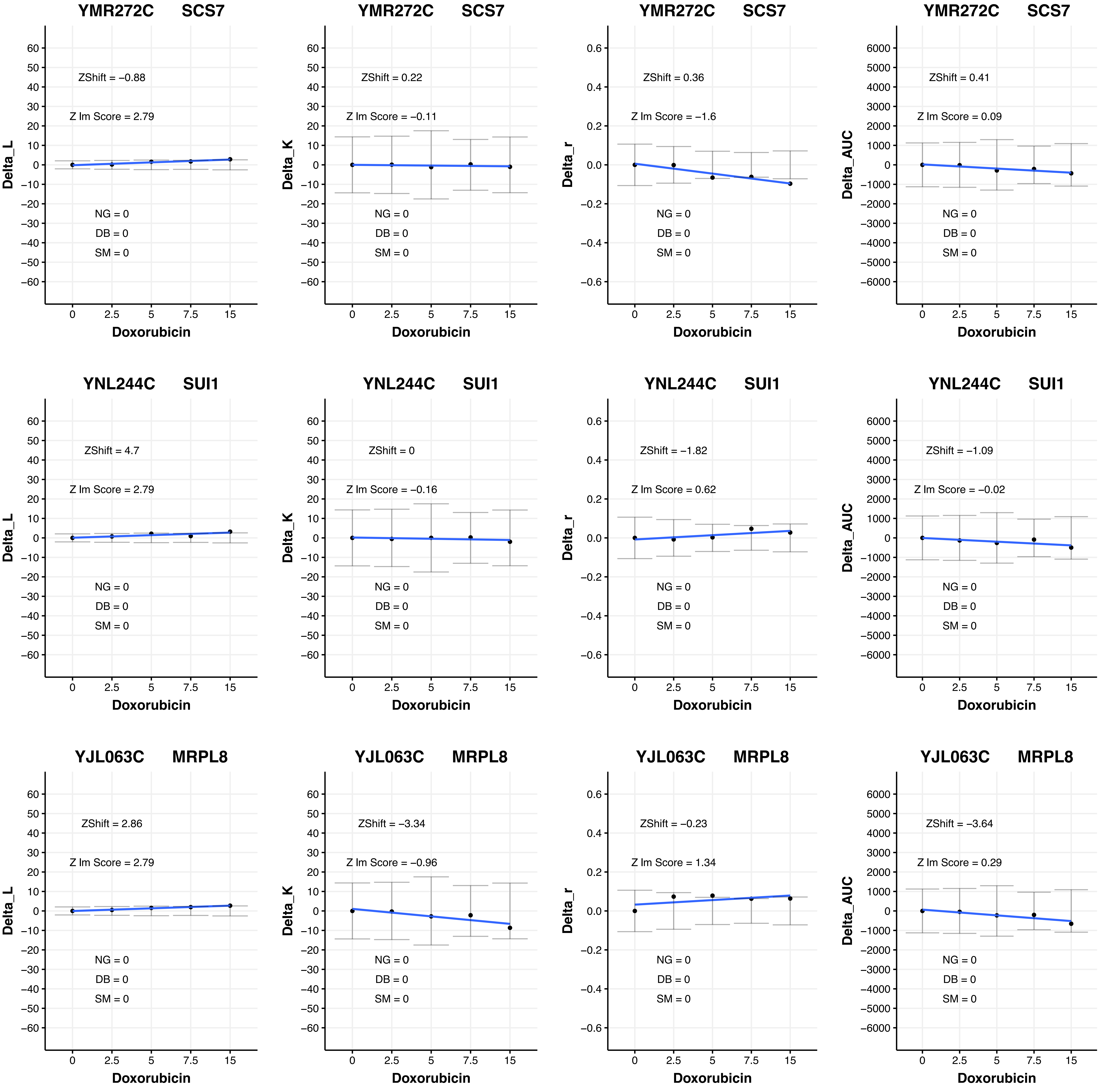

YMR272C SCS7
ZShift = −0.88
Z lm Score = 2.79
NG = 0
DB = 0
SM = 0
Delta_L
Doxorubicin
YMR272C SCS7
ZShift = 0.22
Z lm Score = −0.11
NG = 0
DB = 0
SM = 0
Delta_K
Doxorubicin
YMR272C SCS7
ZShift = 0.36
Z lm Score = −1.6
NG = 0
DB = 0
SM = 0
Delta_r
Doxorubicin
YMR272C SCS7
ZShift = 0.41
Z lm Score = 0.09
NG = 0
DB = 0
SM = 0
Delta_AUC
Doxorubicin
YNL244C SUI1
ZShift = 4.7
Z lm Score = 2.79
NG = 0
DB = 0
SM = 0
Delta_L
Doxorubicin
YNL244C SUI1
ZShift = 0
Z lm Score = −0.16
NG = 0
DB = 0
SM = 0
Delta_K
Doxorubicin
YNL244C SUI1
ZShift = −1.82
Z lm Score = 0.62
NG = 0
DB = 0
SM = 0
Delta_r
Doxorubicin
YNL244C SUI1
ZShift = −1.09
Z lm Score = −0.02
NG = 0
DB = 0
SM = 0
Delta_AUC
Doxorubicin
YJL063C MRPL8
ZShift = 2.86
Z lm Score = 2.79
NG = 0
DB = 0
SM = 0
Delta_L
Doxorubicin
YJL063C MRPL8
ZShift = −3.34
Z lm Score = −0.96
NG = 0
DB = 0
SM = 0
Delta_K
Doxorubicin
YJL063C MRPL8
ZShift = −0.23
Z lm Score = 1.34
NG = 0
DB = 0
SM = 0
Delta_r
Doxorubicin
YJL063C MRPL8
ZShift = −3.64
Z lm Score = 0.29
NG = 0
DB = 0
SM = 0
Delta_AUC
Doxorubicin

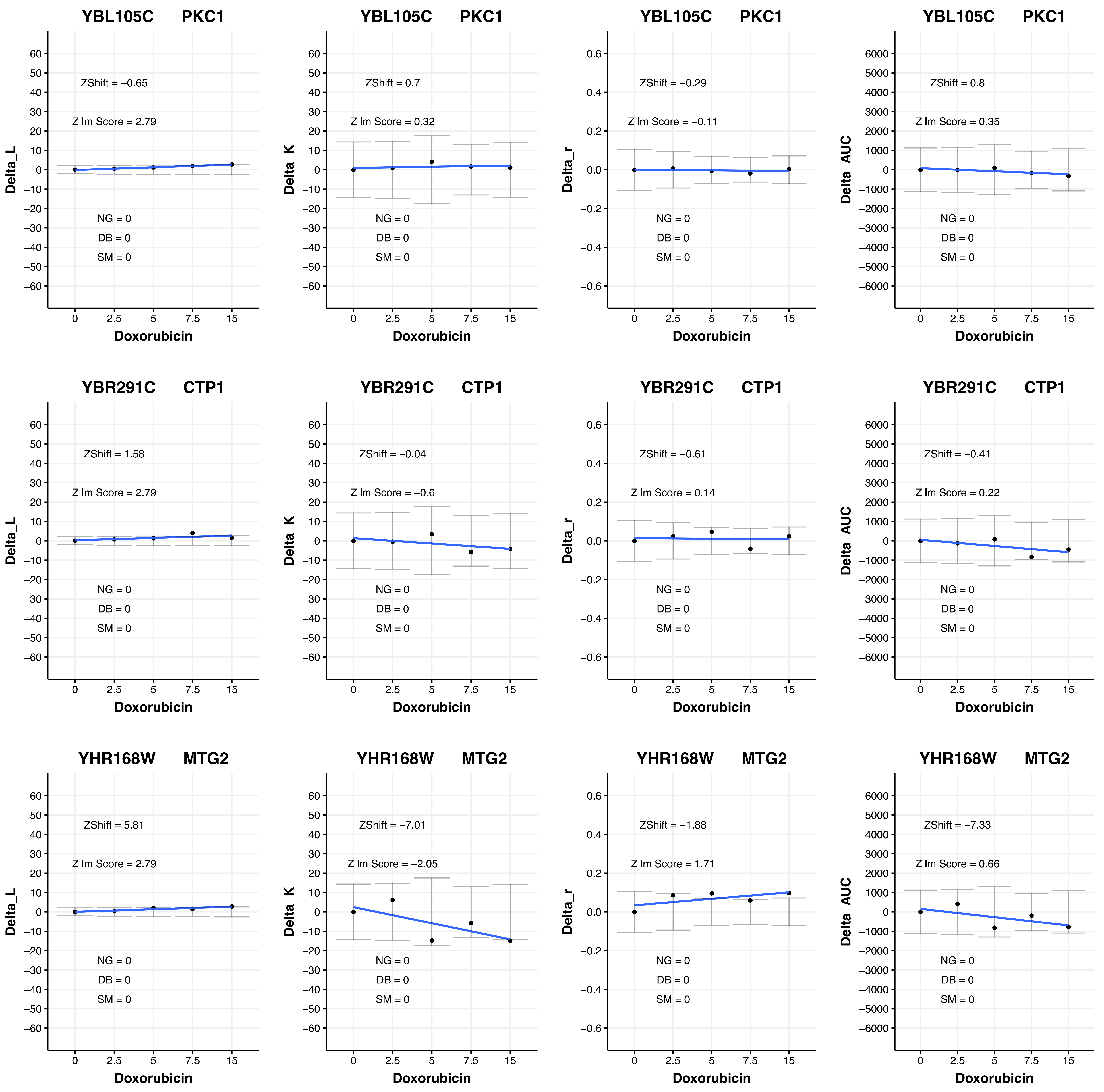
YBL105C PKC1
ZShift = −0.65
Z lm Score = 2.79
NG = 0
DB = 0
SM = 0
Delta_L
Doxorubicin
YBL105C PKC1
ZShift = 0.7
Z lm Score = 0.32
NG = 0
DB = 0
SM = 0
Delta_K
Doxorubicin
YBL105C PKC1
ZShift = −0.29
Z lm Score = −0.11
NG = 0
DB = 0
SM = 0
Delta_r
Doxorubicin
YBL105C PKC1
ZShift = 0.8
Z lm Score = 0.35
NG = 0
DB = 0
SM = 0
Delta_AUC
Doxorubicin
YBR291C CTP1
ZShift = 1.58
Z lm Score = 2.79
NG = 0
DB = 0
SM = 0
Delta_L
Doxorubicin
YBR291C CTP1
ZShift = −0.04
Z lm Score = −0.6
NG = 0
DB = 0
SM = 0
Delta_K
Doxorubicin
YBR291C CTP1
ZShift = −0.61
Z lm Score = 0.14
NG = 0
DB = 0
SM = 0
Delta_r
Doxorubicin
YBR291C CTP1
ZShift = −0.41
Z lm Score = 0.22
NG = 0
DB = 0
SM = 0
Delta_AUC
Doxorubicin
YHR168W MTG2
ZShift = 5.81
Z lm Score = 2.79
NG = 0
DB = 0
SM = 0
Delta_L
Doxorubicin
YHR168W MTG2
ZShift = −7.01
Z lm Score = −2.05
NG = 0
DB = 0
SM = 0
Delta_K
Doxorubicin
YHR168W MTG2
ZShift = −1.88
Z lm Score = 1.71
NG = 0
DB = 0
SM = 0
Delta_r
Doxorubicin
YHR168W MTG2
ZShift = −7.33
Z lm Score = 0.66
NG = 0
DB = 0
SM = 0
Delta_AUC
Doxorubicin

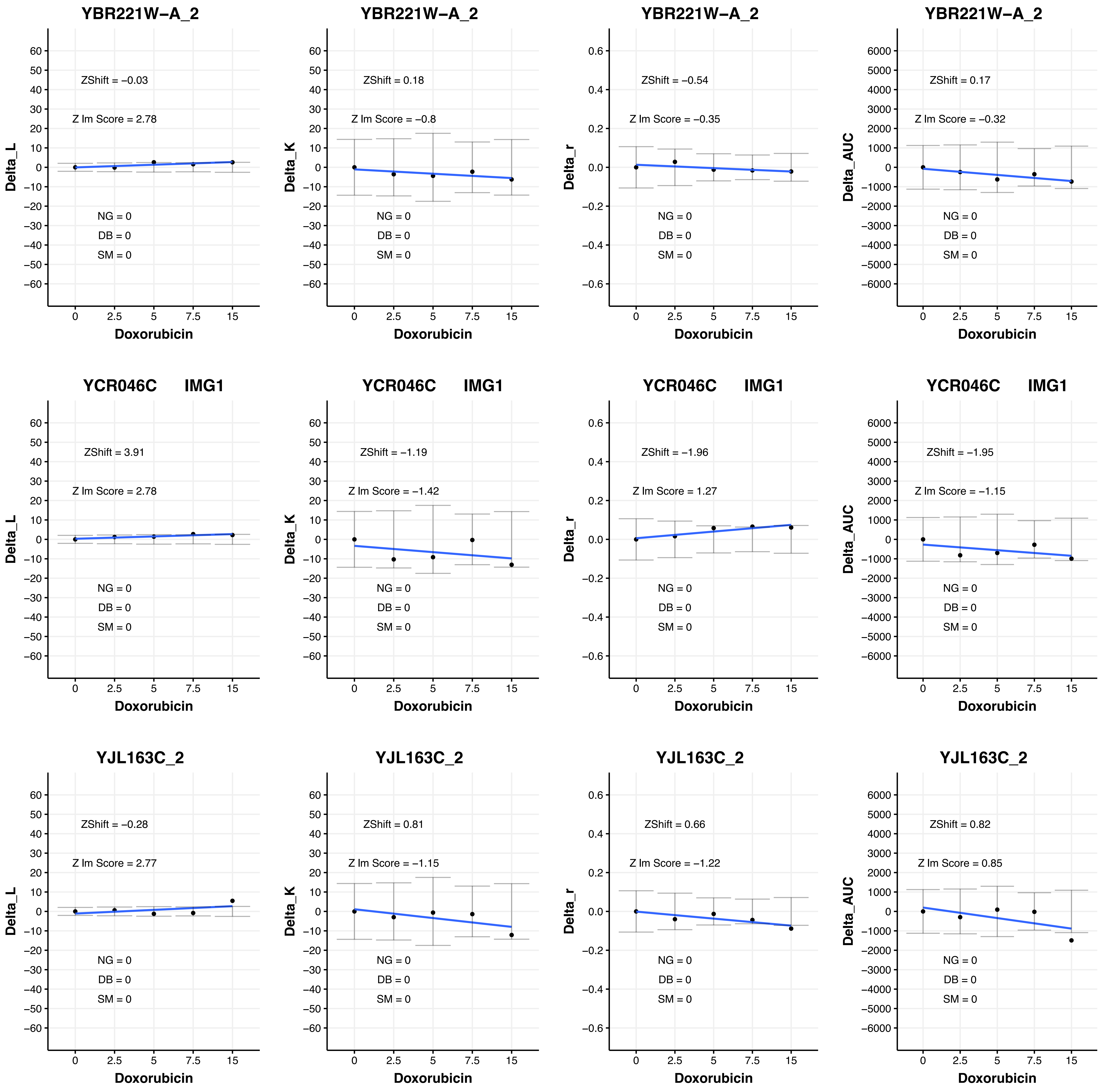
YBR221W−A_2
ZShift = −0.03
Z lm Score = 2.78
NG = 0
DB = 0
SM = 0
Delta_L
Doxorubicin
YBR221W−A_2
ZShift = 0.18
Z lm Score = −0.8
NG = 0
DB = 0
SM = 0
Delta_K
Doxorubicin
YBR221W−A_2
ZShift = −0.54
Z lm Score = −0.35
NG = 0
DB = 0
SM = 0
Delta_r
Doxorubicin
YBR221W−A_2
ZShift = 0.17
Z lm Score = −0.32
NG = 0
DB = 0
SM = 0
Delta_AUC
Doxorubicin
YCR046C IMG1
ZShift = 3.91
Z lm Score = 2.78
NG = 0
DB = 0
SM = 0
Delta_L
Doxorubicin
YCR046C IMG1
ZShift = −1.19
Z lm Score = −1.42
NG = 0
DB = 0
SM = 0
Delta_K
Doxorubicin
YCR046C IMG1
ZShift = −1.96
Z lm Score = 1.27
NG = 0
DB = 0
SM = 0
Delta_r
Doxorubicin
YCR046C IMG1
ZShift = −1.95
Z lm Score = −1.15
NG = 0
DB = 0
SM = 0
Delta_AUC
Doxorubicin
YJL163C_2
ZShift = −0.28
Z lm Score = 2.77
NG = 0
DB = 0
SM = 0
Delta_L
Doxorubicin
YJL163C_2
ZShift = 0.81
Z lm Score = −1.15
NG = 0
DB = 0
SM = 0
Delta_K
Doxorubicin
YJL163C_2
ZShift = 0.66
Z lm Score = −1.22
NG = 0
DB = 0
SM = 0
Delta_r
Doxorubicin
YJL163C_2
ZShift = 0.82
Z lm Score = 0.85
NG = 0
DB = 0
SM = 0
Delta_AUC
Doxorubicin

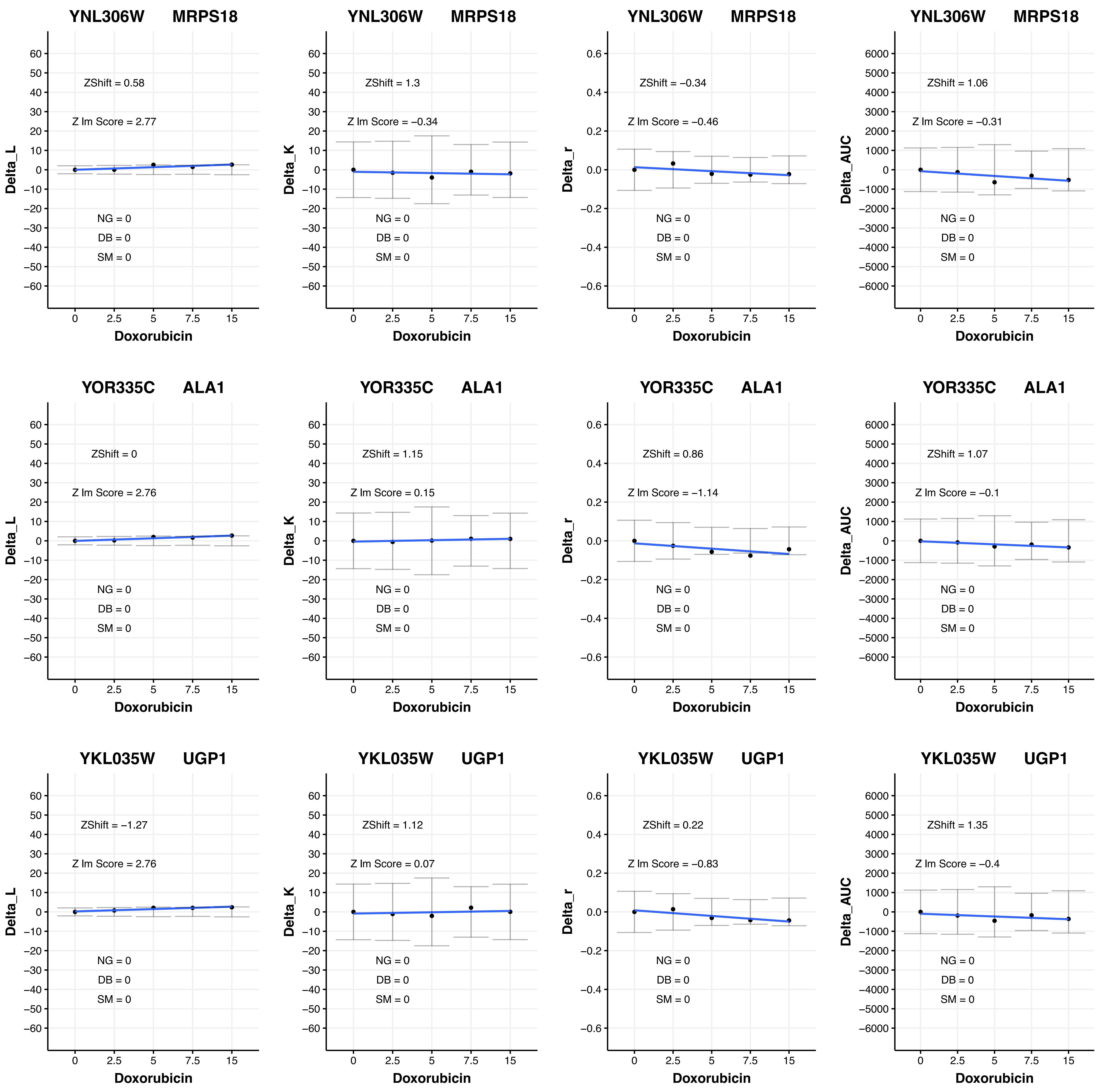
YNL306W MRPS18
ZShift = 0.58
Z lm Score = 2.77
NG = 0
DB = 0
SM = 0
Delta_L
Doxorubicin
YNL306W MRPS18
ZShift = 1.3
Z lm Score = −0.34
NG = 0
DB = 0
SM = 0
Delta_K
Doxorubicin
YNL306W MRPS18
ZShift = −0.34
Z lm Score = −0.46
NG = 0
DB = 0
SM = 0
Delta_r
Doxorubicin
YNL306W MRPS18
ZShift = 1.06
Z lm Score = −0.31
NG = 0
DB = 0
SM = 0
Delta_AUC
Doxorubicin
YOR335C ALA1
ZShift = 0
Z lm Score = 2.76
NG = 0
DB = 0
SM = 0
Delta_L
Doxorubicin
YOR335C ALA1
ZShift = 1.15
Z lm Score = 0.15
NG = 0
DB = 0
SM = 0
Delta_K
Doxorubicin
YOR335C ALA1
ZShift = 0.86
Z lm Score = −1.14
NG = 0
DB = 0
SM = 0
Delta_r
Doxorubicin
YOR335C ALA1
ZShift = 1.07
Z lm Score = −0.1
NG = 0
DB = 0
SM = 0
Delta_AUC
Doxorubicin
YKL035W UGP1
ZShift = −1.27
Z lm Score = 2.76
NG = 0
DB = 0
SM = 0
Delta_L
Doxorubicin
YKL035W UGP1
ZShift = 1.12
Z lm Score = 0.07
NG = 0
DB = 0
SM = 0
Delta_K
Doxorubicin
YKL035W UGP1
ZShift = 0.22
Z lm Score = −0.83
NG = 0
DB = 0
SM = 0
Delta_r
Doxorubicin
YKL035W UGP1
ZShift = 1.35
Z lm Score = −0.4
NG = 0
DB = 0
SM = 0
Delta_AUC
Doxorubicin

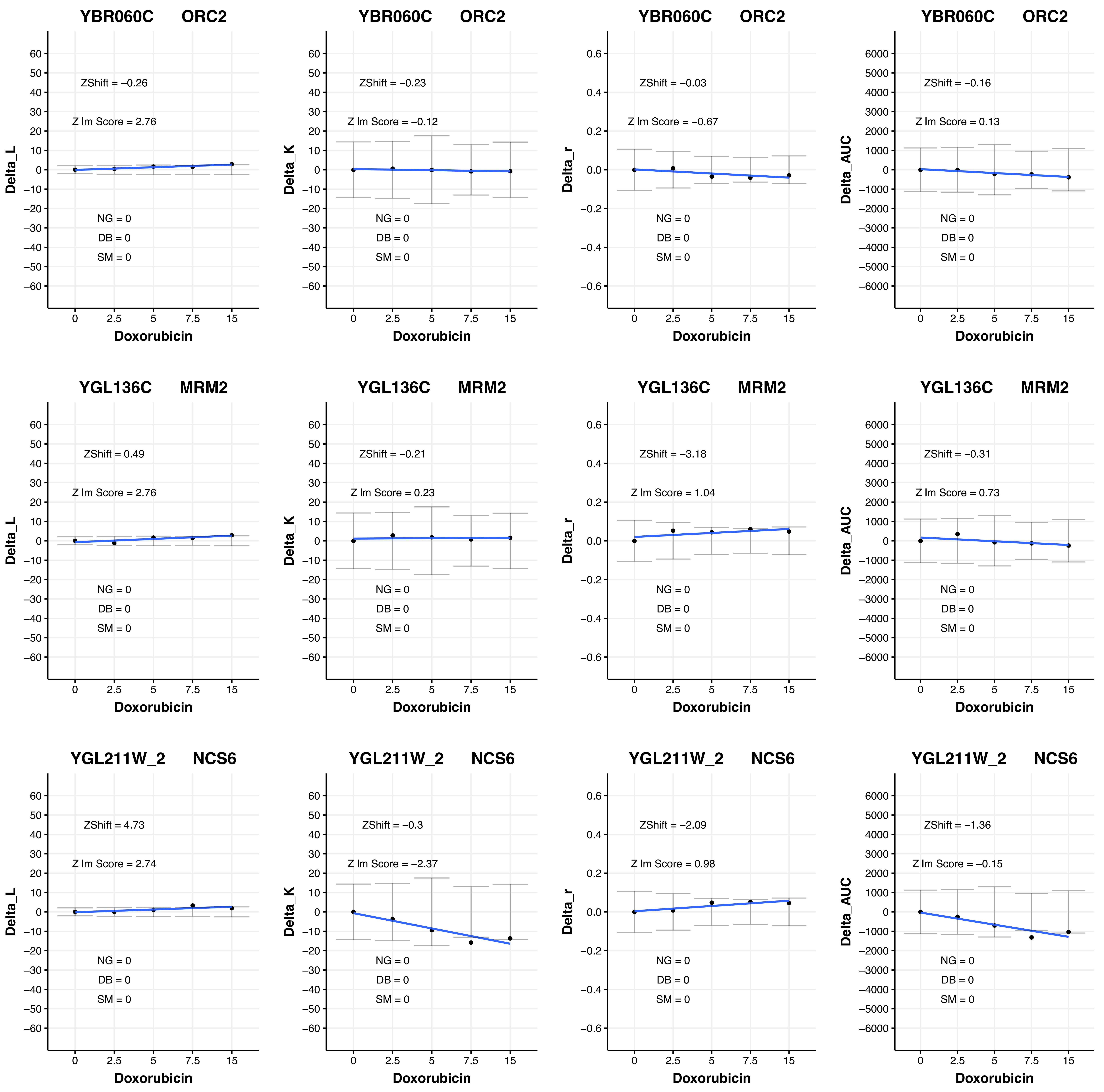
YBR060C ORC2
ZShift = −0.26
Z lm Score = 2.76
NG = 0
DB = 0
SM = 0
Delta_L
Doxorubicin
YBR060C ORC2
ZShift = −0.23
Z lm Score = −0.12
NG = 0
DB = 0
SM = 0
Delta_K
Doxorubicin
YBR060C ORC2
ZShift = −0.03
Z lm Score = −0.67
NG = 0
DB = 0
SM = 0
Delta_r
Doxorubicin
YBR060C ORC2
ZShift = −0.16
Z lm Score = 0.13
NG = 0
DB = 0
SM = 0
Delta_AUC
Doxorubicin
YGL136C MRM2
ZShift = 0.49
Z lm Score = 2.76
NG = 0
DB = 0
SM = 0
Delta_L
Doxorubicin
YGL136C MRM2
ZShift = −0.21
Z lm Score = 0.23
NG = 0
DB = 0
SM = 0
Delta_K
Doxorubicin
YGL136C MRM2
ZShift = −3.18
Z lm Score = 1.04
NG = 0
DB = 0
SM = 0
Delta_r
Doxorubicin
YGL136C MRM2
ZShift = −0.31
Z lm Score = 0.73
NG = 0
DB = 0
SM = 0
Delta_AUC
Doxorubicin
YGL211W_2 NCS6
ZShift = 4.73
Z lm Score = 2.74
NG = 0
DB = 0
SM = 0
Delta_L
Doxorubicin
YGL211W_2 NCS6
ZShift = −0.3
Z lm Score = −2.37
NG = 0
DB = 0
SM = 0
Delta_K
Doxorubicin
YGL211W_2 NCS6
ZShift = −2.09
Z lm Score = 0.98
NG = 0
DB = 0
SM = 0
Delta_r
Doxorubicin
YGL211W_2 NCS6
ZShift = −1.36
Z lm Score = −0.15
NG = 0
DB = 0
SM = 0
Delta_AUC
Doxorubicin

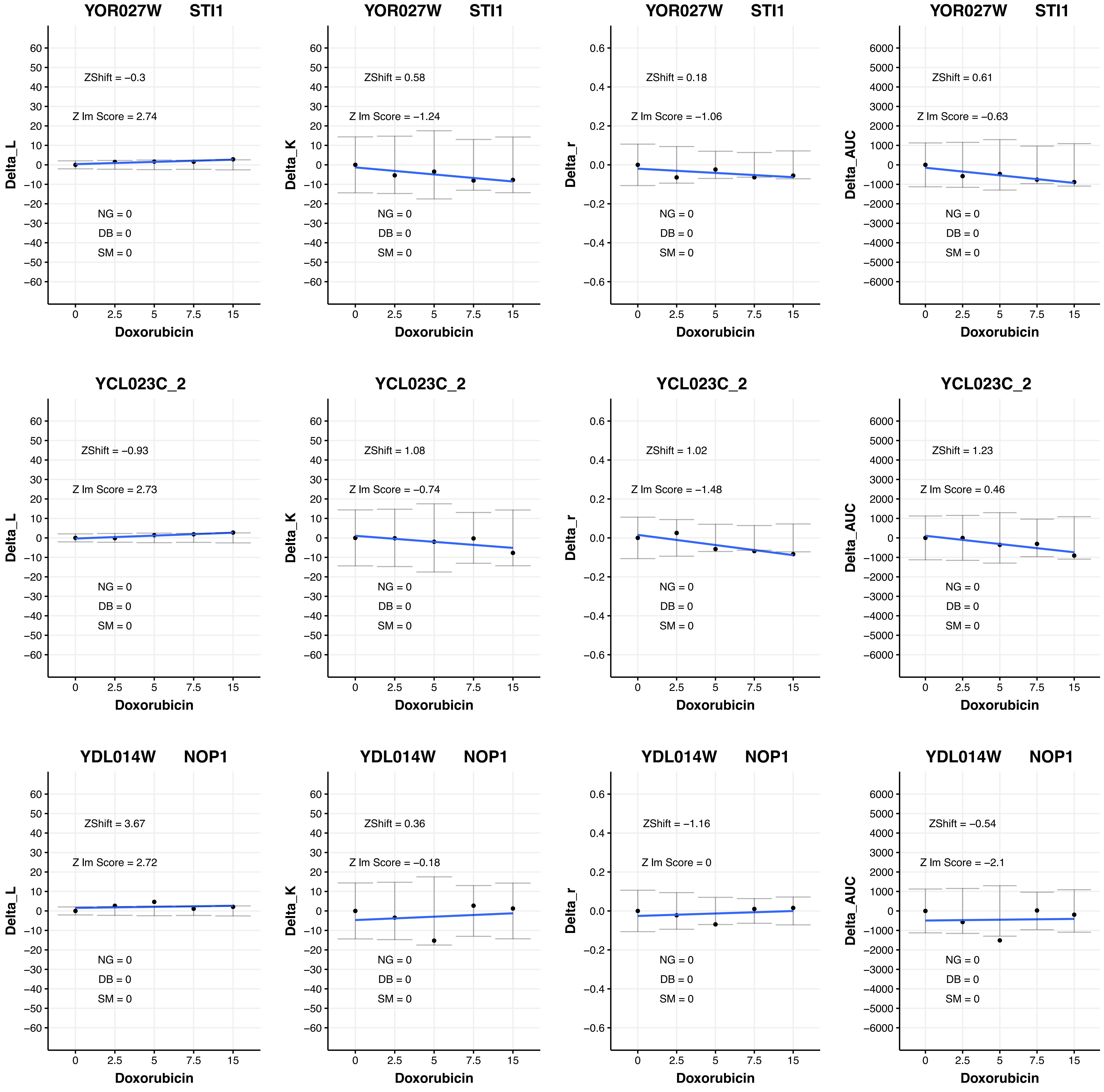
YOR027W STI1
ZShift = −0.3
Z lm Score = 2.74
NG = 0
DB = 0
SM = 0
Delta_L
Doxorubicin
YOR027W STI1
ZShift = 0.58
Z lm Score = −1.24
NG = 0
DB = 0
SM = 0
Delta_K
Doxorubicin
YOR027W STI1
ZShift = 0.18
Z lm Score = −1.06
NG = 0
DB = 0
SM = 0
Delta_r
Doxorubicin
YOR027W STI1
ZShift = 0.61
Z lm Score = −0.63
NG = 0
DB = 0
SM = 0
Delta_AUC
Doxorubicin
YCL023C_2
ZShift = −0.93
Z lm Score = 2.73
NG = 0
DB = 0
SM = 0
Delta_L
Doxorubicin
YCL023C_2
ZShift = 1.08
Z lm Score = −0.74
NG = 0
DB = 0
SM = 0
Delta_K
Doxorubicin
YCL023C_2
ZShift = 1.02
Z lm Score = −1.48
NG = 0
DB = 0
SM = 0
Delta_r
Doxorubicin
YCL023C_2
ZShift = 1.23
Z lm Score = 0.46
NG = 0
DB = 0
SM = 0
Delta_AUC
Doxorubicin
YDL014W NOP1
ZShift = 3.67
Z lm Score = 2.72
NG = 0
DB = 0
SM = 0
Delta_L
Doxorubicin
YDL014W NOP1
ZShift = 0.36
Z lm Score = −0.18
NG = 0
DB = 0
SM = 0
Delta_K
Doxorubicin
YDL014W NOP1
ZShift = −1.16
Z lm Score = 0
NG = 0
DB = 0
SM = 0
Delta_r
Doxorubicin
YDL014W NOP1
ZShift = −0.54
Z lm Score = −2.1
NG = 0
DB = 0
SM = 0
Delta_AUC
Doxorubicin

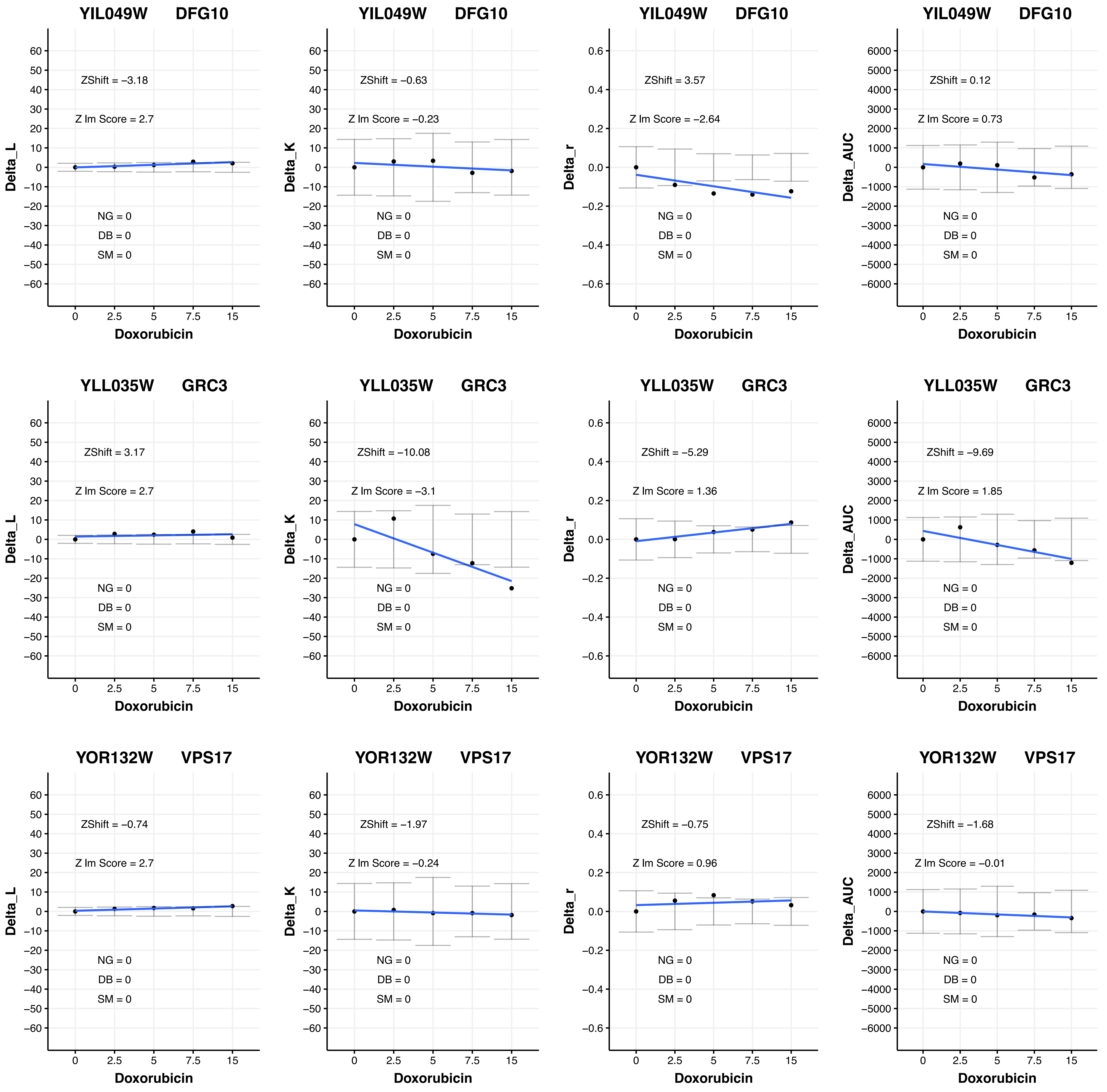

YIL049W DFG10
ZShift = −3.18
Z lm Score = 2.7
NG = 0
DB = 0
SM = 0
Delta_L
Doxorubicin
YIL049W DFG10
ZShift = −0.63
Z lm Score = −0.23
NG = 0
DB = 0
SM = 0
Delta_K
Doxorubicin
YIL049W DFG10
ZShift = 3.57
Z lm Score = −2.64
NG = 0
DB = 0
SM = 0
Delta_r
Doxorubicin
YIL049W DFG10
ZShift = 0.12
Z lm Score = 0.73
NG = 0
DB = 0
SM = 0
Delta_AUC
Doxorubicin
YLL035W GRC3
ZShift = 3.17
Z lm Score = 2.7
NG = 0
DB = 0
SM = 0
Delta_L
Doxorubicin
YLL035W GRC3
ZShift = −10.08
Z lm Score = −3.1
NG = 0
DB = 0
SM = 0
Delta_K
Doxorubicin
YLL035W GRC3
ZShift = −5.29
Z lm Score = 1.36
NG = 0
DB = 0
SM = 0
Delta_r
Doxorubicin
YLL035W GRC3
ZShift = −9.69
Z lm Score = 1.85
NG = 0
DB = 0
SM = 0
Delta_AUC
Doxorubicin
YOR132W VPS17
ZShift = −0.74
Z lm Score = 2.7
NG = 0
DB = 0
SM = 0
Delta_L
Doxorubicin
YOR132W VPS17
ZShift = −1.97
Z lm Score = −0.24
NG = 0
DB = 0
SM = 0
Delta_K
Doxorubicin
YOR132W VPS17
ZShift = −0.75
Z lm Score = 0.96
NG = 0
DB = 0
SM = 0
Delta_r
Doxorubicin
YOR132W VPS17
ZShift = −1.68
Z lm Score = −0.01
NG = 0
DB = 0
SM = 0
Delta_AUC
Doxorubicin

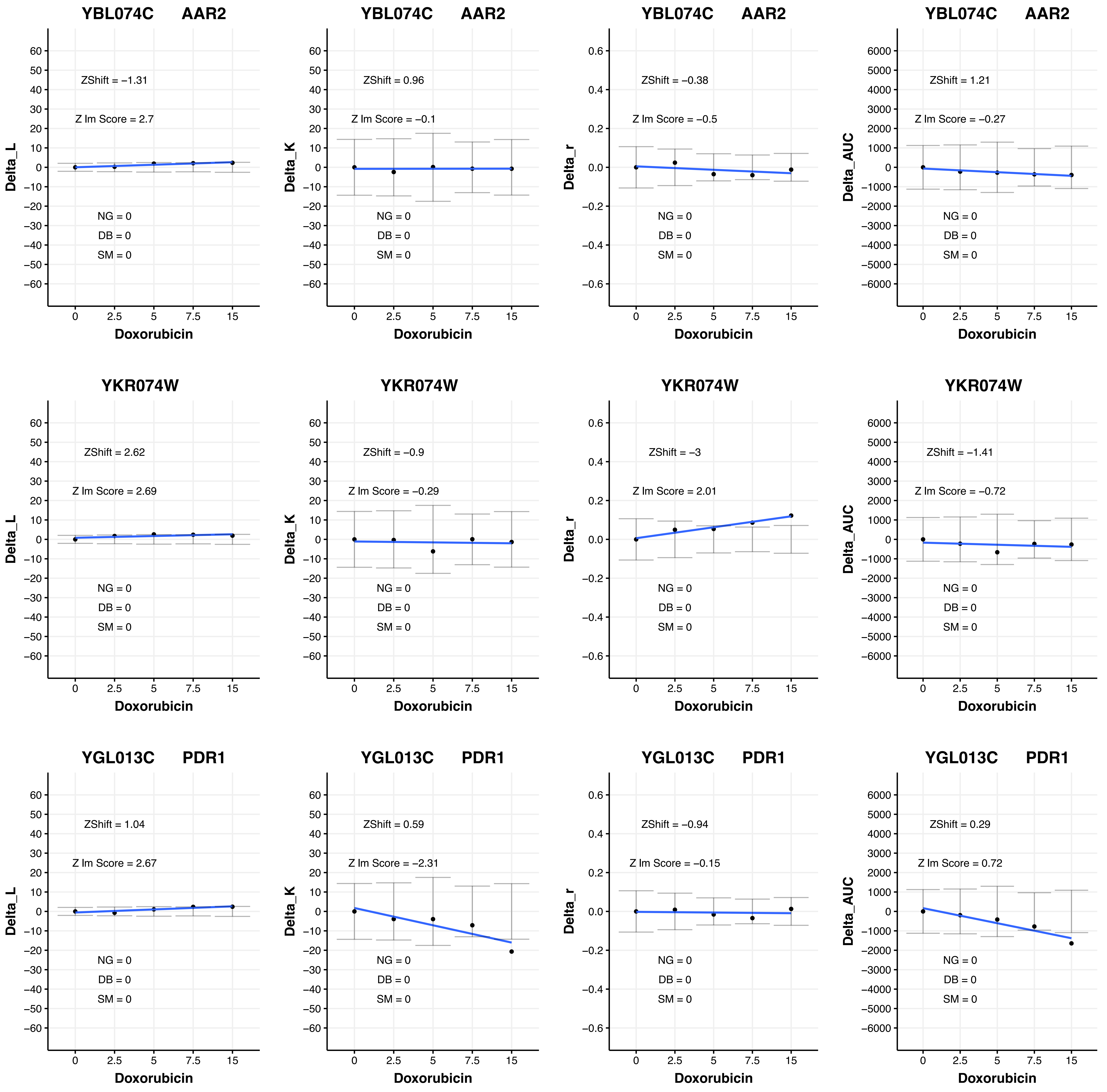

YBL074C AAR2
ZShift = −1.31
Z lm Score = 2.7
Delta_L
NG = 0
DB = 0
SM = 0
Doxorubicin
YBL074C AAR2
ZShift = 0.96
Z lm Score = −0.1
Delta_K
NG = 0
DB = 0
SM = 0
Doxorubicin
YBL074C AAR2
ZShift = −0.38
Z lm Score = −0.5
Delta_r
NG = 0
DB = 0
SM = 0
Doxorubicin
YBL074C AAR2
ZShift = 1.21
Z lm Score = −0.27
Delta_AUC
NG = 0
DB = 0
SM = 0
Doxorubicin
YKR074W
ZShift = 2.62
Z lm Score = 2.69
Delta_L
NG = 0
DB = 0
SM = 0
Doxorubicin
YKR074W
ZShift = −0.9
Z lm Score = −0.29
Delta_K
NG = 0
DB = 0
SM = 0
Doxorubicin
YKR074W
ZShift = −3
Z lm Score = 2.01
Delta_r
NG = 0
DB = 0
SM = 0
Doxorubicin
YKR074W
ZShift = −1.41
Z lm Score = −0.72
Delta_AUC
NG = 0
DB = 0
SM = 0
Doxorubicin
YGL013C PDR1
ZShift = 1.04
Z lm Score = 2.67
Delta_L
NG = 0
DB = 0
SM = 0
Doxorubicin
YGL013C PDR1
ZShift = 0.59
Z lm Score = −2.31
Delta_K
NG = 0
DB = 0
SM = 0
Doxorubicin
YGL013C PDR1
ZShift = −0.94
Z lm Score = −0.15
Delta_r
NG = 0
DB = 0
SM = 0
Doxorubicin
YGL013C PDR1
ZShift = 0.29
Z lm Score = 0.72
Delta_AUC
NG = 0
DB = 0
SM = 0
Doxorubicin

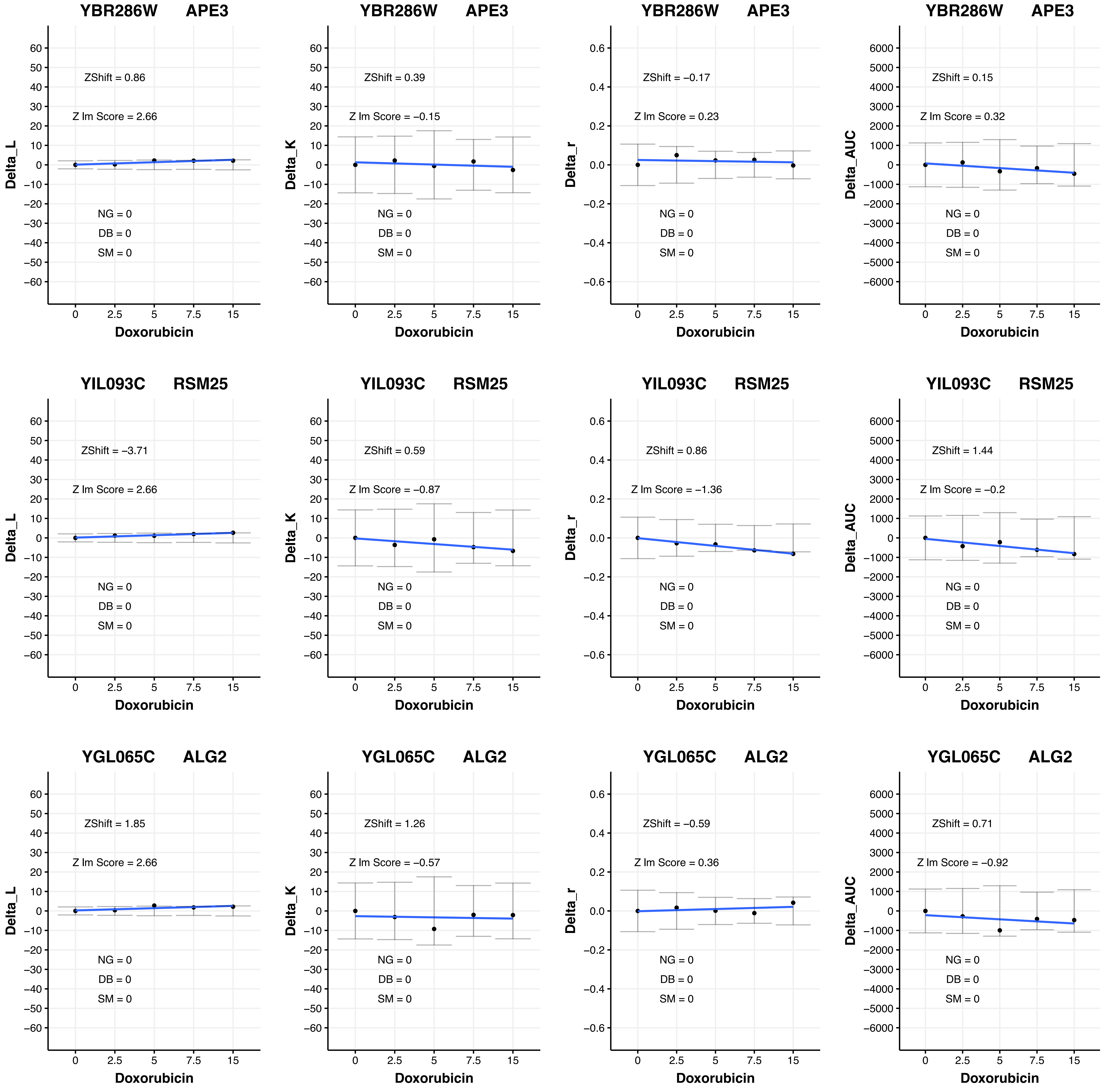
YBR286W APE3
ZShift = 0.86
Z lm Score = 2.66
NG = 0
DB = 0
SM = 0
Delta_L
Doxorubicin
YBR286W APE3
ZShift = 0.39
Z lm Score = −0.15
NG = 0
DB = 0
SM = 0
Delta_K
Doxorubicin
YBR286W APE3
ZShift = −0.17
Z lm Score = 0.23
NG = 0
DB = 0
SM = 0
Delta_r
Doxorubicin
YBR286W APE3
ZShift = 0.15
Z lm Score = 0.32
NG = 0
DB = 0
SM = 0
Delta_AUC
Doxorubicin
YIL093C RSM25
ZShift = −3.71
Z lm Score = 2.66
NG = 0
DB = 0
SM = 0
Delta_L
Doxorubicin
YIL093C RSM25
ZShift = 0.59
Z lm Score = −0.87
NG = 0
DB = 0
SM = 0
Delta_K
Doxorubicin
YIL093C RSM25
ZShift = 0.86
Z lm Score = −1.36
NG = 0
DB = 0
SM = 0
Delta_r
Doxorubicin
YIL093C RSM25
ZShift = 1.44
Z lm Score = −0.2
NG = 0
DB = 0
SM = 0
Delta_AUC
Doxorubicin
YGL065C ALG2
ZShift = 1.85
Z lm Score = 2.66
NG = 0
DB = 0
SM = 0
Delta_L
Doxorubicin
YGL065C ALG2
ZShift = 1.26
Z lm Score = −0.57
NG = 0
DB = 0
SM = 0
Delta_K
Doxorubicin
YGL065C ALG2
ZShift = −0.59
Z lm Score = 0.36
NG = 0
DB = 0
SM = 0
Delta_r
Doxorubicin
YGL065C ALG2
ZShift = 0.71
Z lm Score = −0.92
NG = 0
DB = 0
SM = 0
Delta_AUC
Doxorubicin

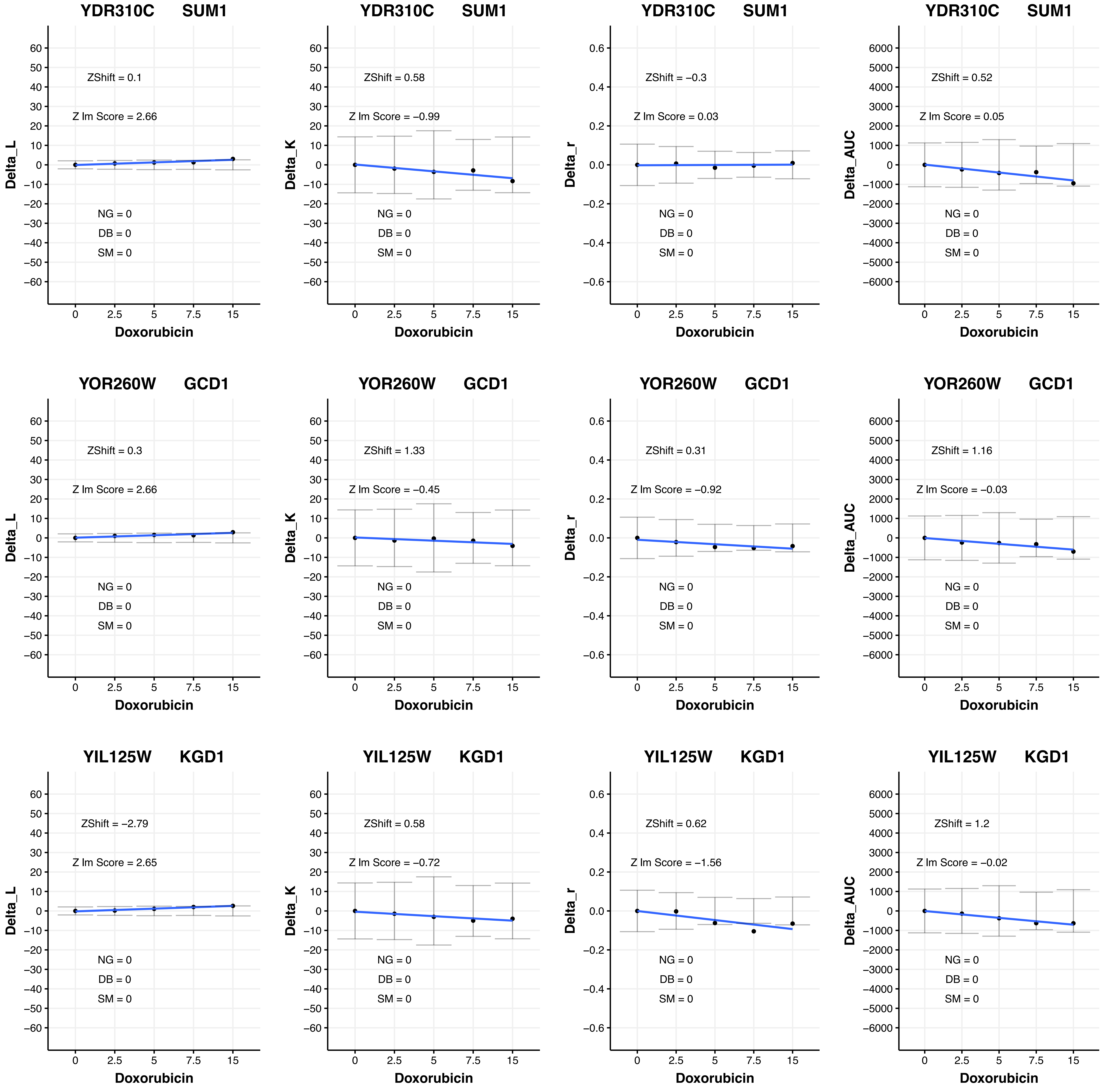
YDR310C SUM1
ZShift = 0.1
Z lm Score = 2.66
NG = 0
DB = 0
SM = 0
Delta_L
Doxorubicin
YDR310C SUM1
ZShift = 0.58
Z lm Score = −0.99
NG = 0
DB = 0
SM = 0
Delta_K
Doxorubicin
YDR310C SUM1
ZShift = −0.3
Z lm Score = 0.03
NG = 0
DB = 0
SM = 0
Delta_r
Doxorubicin
YDR310C SUM1
ZShift = 0.52
Z lm Score = 0.05
NG = 0
DB = 0
SM = 0
Delta_AUC
Doxorubicin
YOR260W GCD1
ZShift = 0.3
Z lm Score = 2.66
NG = 0
DB = 0
SM = 0
Delta_L
Doxorubicin
YOR260W GCD1
ZShift = 1.33
Z lm Score = −0.45
NG = 0
DB = 0
SM = 0
Delta_K
Doxorubicin
YOR260W GCD1
ZShift = 0.31
Z lm Score = −0.92
NG = 0
DB = 0
SM = 0
Delta_r
Doxorubicin
YOR260W GCD1
ZShift = 1.16
Z lm Score = −0.03
NG = 0
DB = 0
SM = 0
Delta_AUC
Doxorubicin
YIL125W KGD1
ZShift = −2.79
Z lm Score = 2.65
NG = 0
DB = 0
SM = 0
Delta_L
Doxorubicin
YIL125W KGD1
ZShift = 0.58
Z lm Score = −0.72
NG = 0
DB = 0
SM = 0
Delta_K
Doxorubicin
YIL125W KGD1
ZShift = 0.62
Z lm Score = −1.56
NG = 0
DB = 0
SM = 0
Delta_r
Doxorubicin
YIL125W KGD1
ZShift = 1.2
Z lm Score = −0.02
NG = 0
DB = 0
SM = 0
Delta_AUC
Doxorubicin

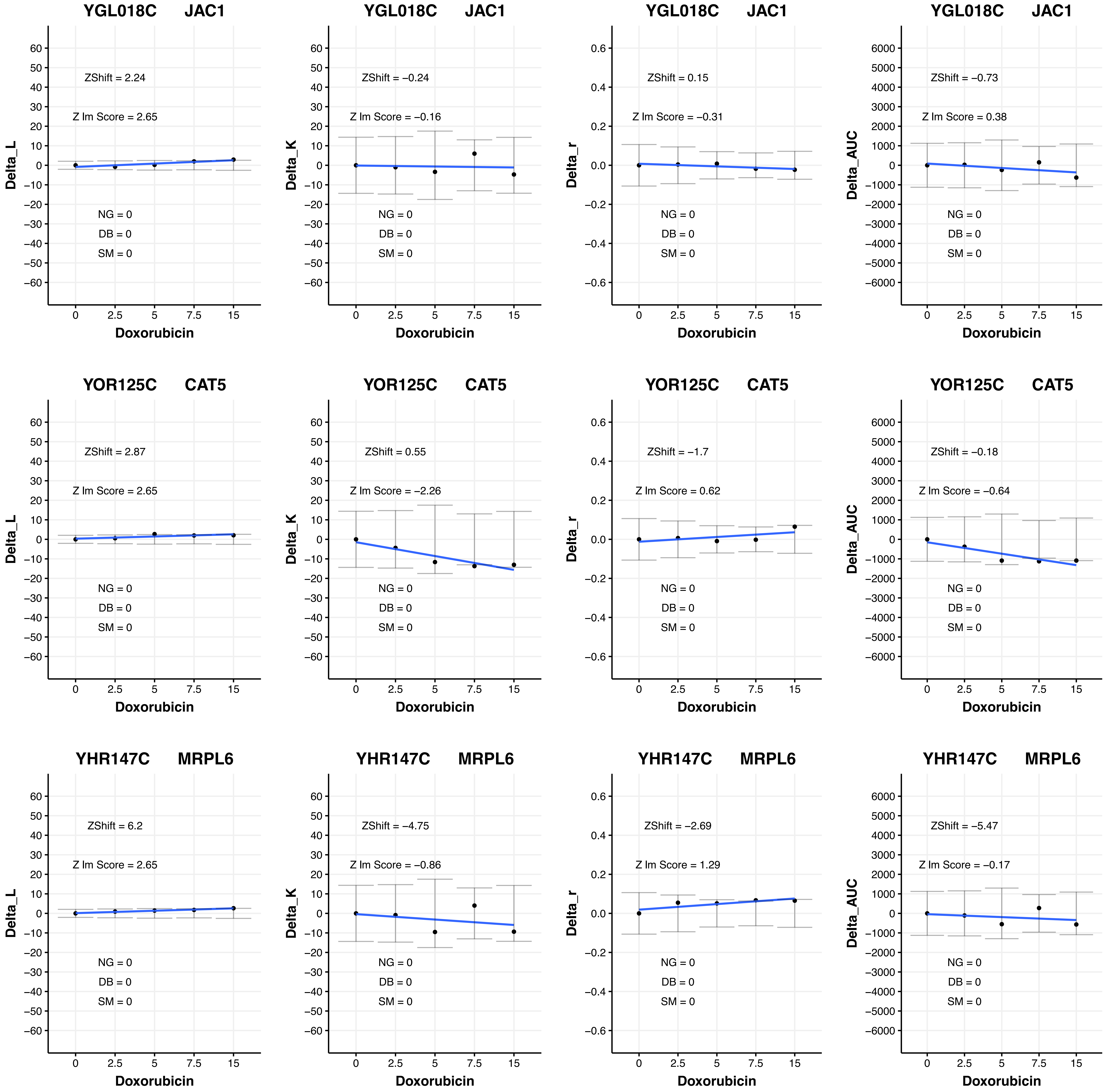
YGL018C JAC1
ZShift = 2.24
Z lm Score = 2.65
NG = 0
DB = 0
SM = 0
Delta_L
Doxorubicin
YGL018C JAC1
ZShift = −0.24
Z lm Score = −0.16
NG = 0
DB = 0
SM = 0
Delta_K
Doxorubicin
YGL018C JAC1
ZShift = 0.15
Z lm Score = −0.31
NG = 0
DB = 0
SM = 0
Delta_r
Doxorubicin
YGL018C JAC1
ZShift = −0.73
Z lm Score = 0.38
NG = 0
DB = 0
SM = 0
Delta_AUC
Doxorubicin
YOR125C CAT5
ZShift = 2.87
Z lm Score = 2.65
NG = 0
DB = 0
SM = 0
Delta_L
Doxorubicin
YOR125C CAT5
ZShift = 0.55
Z lm Score = −2.26
NG = 0
DB = 0
SM = 0
Delta_K
Doxorubicin
YOR125C CAT5
ZShift = −1.7
Z lm Score = 0.62
NG = 0
DB = 0
SM = 0
Delta_r
Doxorubicin
YOR125C CAT5
ZShift = −0.18
Z lm Score = −0.64
NG = 0
DB = 0
SM = 0
Delta_AUC
Doxorubicin
YHR147C MRPL6
ZShift = 6.2
Z lm Score = 2.65
NG = 0
DB = 0
SM = 0
Delta_L
Doxorubicin
YHR147C MRPL6
ZShift = −4.75
Z lm Score = −0.86
NG = 0
DB = 0
SM = 0
Delta_K
Doxorubicin
YHR147C MRPL6
ZShift = −2.69
Z lm Score = 1.29
NG = 0
DB = 0
SM = 0
Delta_r
Doxorubicin
YHR147C MRPL6
ZShift = −5.47
Z lm Score = −0.17
NG = 0
DB = 0
SM = 0
Delta_AUC
Doxorubicin

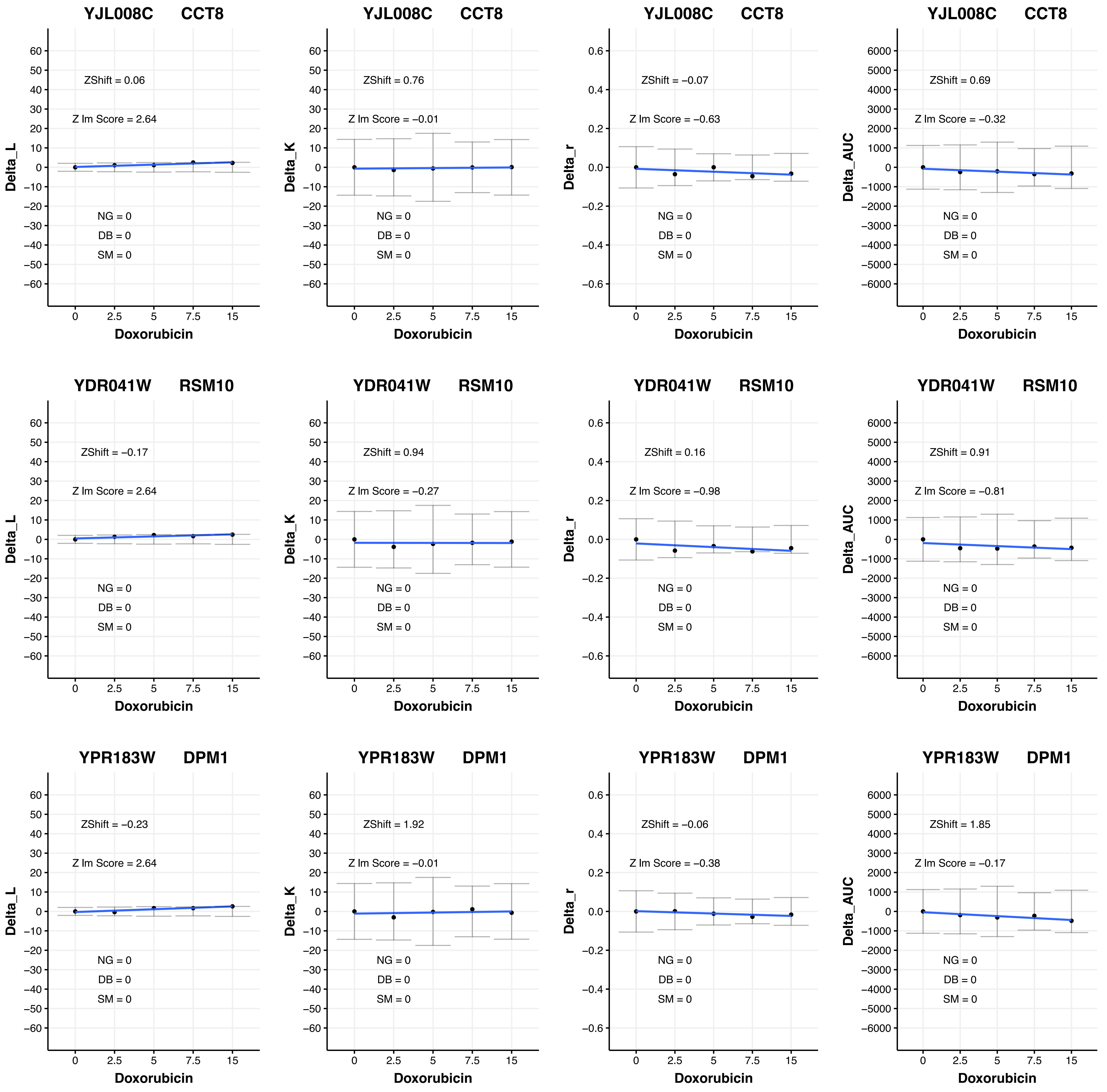
YJL008C CCT8
ZShift = 0.06
Z lm Score = 2.64
NG = 0
DB = 0
SM = 0
Delta_L
Doxorubicin
YJL008C CCT8
ZShift = 0.76
Z lm Score = −0.01
NG = 0
DB = 0
SM = 0
Delta_K
Doxorubicin
YJL008C CCT8
ZShift = −0.07
Z lm Score = −0.63
NG = 0
DB = 0
SM = 0
Delta_r
Doxorubicin
YJL008C CCT8
ZShift = 0.69
Z lm Score = −0.32
NG = 0
DB = 0
SM = 0
Delta_AUC
Doxorubicin
YDR041W RSM10
ZShift = −0.17
Z lm Score = 2.64
NG = 0
DB = 0
SM = 0
Delta_L
Doxorubicin
YDR041W RSM10
ZShift = 0.94
Z lm Score = −0.27
NG = 0
DB = 0
SM = 0
Delta_K
Doxorubicin
YDR041W RSM10
ZShift = 0.16
Z lm Score = −0.98
NG = 0
DB = 0
SM = 0
Delta_r
Doxorubicin
YDR041W RSM10
ZShift = 0.91
Z lm Score = −0.81
NG = 0
DB = 0
SM = 0
Delta_AUC
Doxorubicin
YPR183W DPM1
ZShift = −0.23
Z lm Score = 2.64
NG = 0
DB = 0
SM = 0
Delta_L
Doxorubicin
YPR183W DPM1
ZShift = 1.92
Z lm Score = −0.01
NG = 0
DB = 0
SM = 0
Delta_K
Doxorubicin
YPR183W DPM1
ZShift = −0.06
Z lm Score = −0.38
NG = 0
DB = 0
SM = 0
Delta_r
Doxorubicin
YPR183W DPM1
ZShift = 1.85
Z lm Score = −0.17
NG = 0
DB = 0
SM = 0
Delta_AUC
Doxorubicin

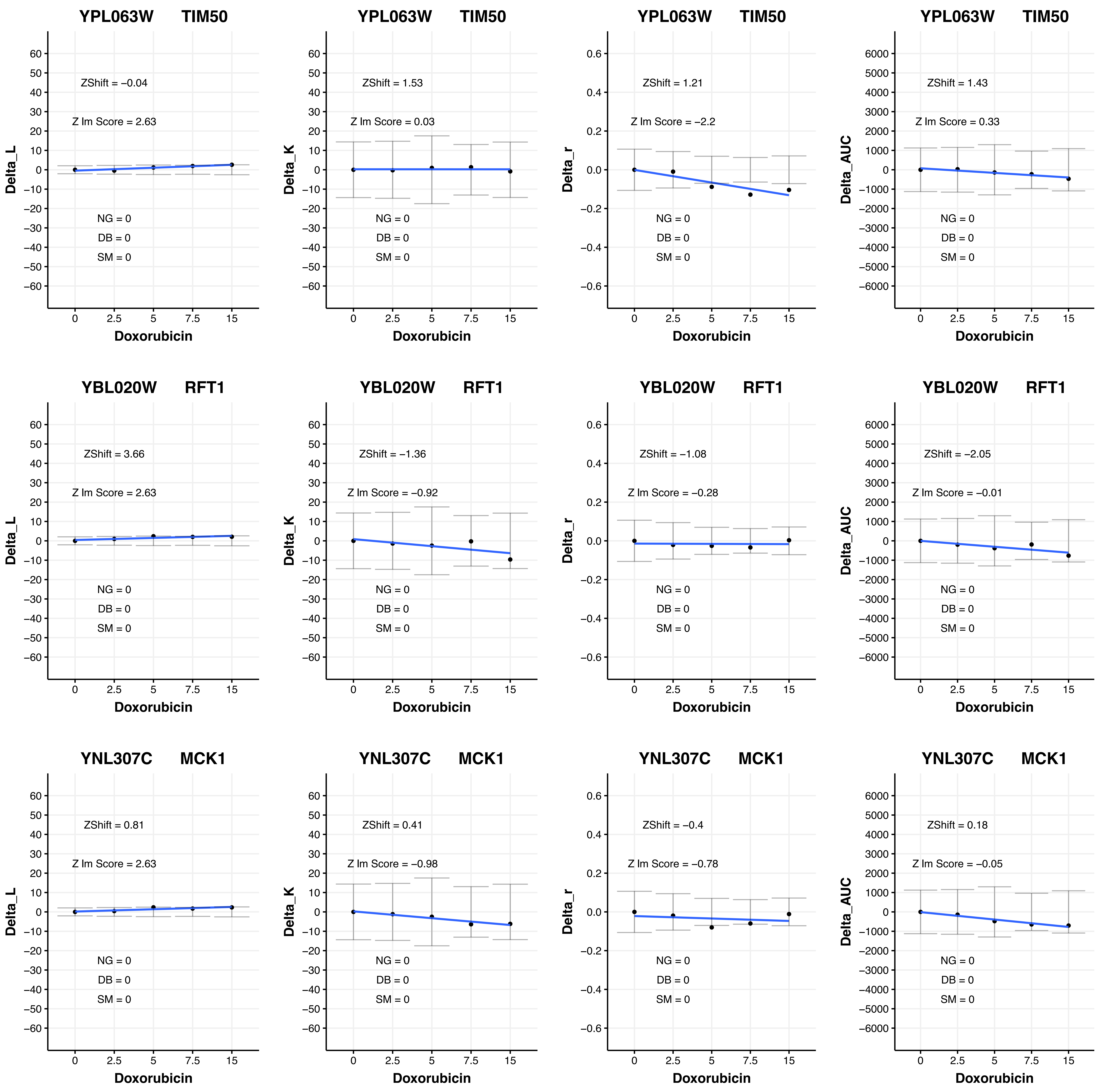
YPL063W TIM50
ZShift = −0.04
Z lm Score = 2.63
NG = 0
DB = 0
SM = 0
Delta_L
Doxorubicin
YPL063W TIM50
ZShift = 1.53
Z lm Score = 0.03
NG = 0
DB = 0
SM = 0
Delta_K
Doxorubicin
YPL063W TIM50
ZShift = 1.21
Z lm Score = −2.2
NG = 0
DB = 0
SM = 0
Delta_r
Doxorubicin
YPL063W TIM50
ZShift = 1.43
Z lm Score = 0.33
NG = 0
DB = 0
SM = 0
Delta_AUC
Doxorubicin
YBL020W RFT1
ZShift = 3.66
Z lm Score = 2.63
NG = 0
DB = 0
SM = 0
Delta_L
Doxorubicin
YBL020W RFT1
ZShift = −1.36
Z lm Score = −0.92
NG = 0
DB = 0
SM = 0
Delta_K
Doxorubicin
YBL020W RFT1
ZShift = −1.08
Z lm Score = −0.28
NG = 0
DB = 0
SM = 0
Delta_r
Doxorubicin
YBL020W RFT1
ZShift = −2.05
Z lm Score = −0.01
NG = 0
DB = 0
SM = 0
Delta_AUC
Doxorubicin
YNL307C MCK1
ZShift = 0.81
Z lm Score = 2.63
NG = 0
DB = 0
SM = 0
Delta_L
Doxorubicin
YNL307C MCK1
ZShift = 0.41
Z lm Score = −0.98
NG = 0
DB = 0
SM = 0
Delta_K
Doxorubicin
YNL307C MCK1
ZShift = −0.4
Z lm Score = −0.78
NG = 0
DB = 0
SM = 0
Delta_r
Doxorubicin
YNL307C MCK1
ZShift = 0.18
Z lm Score = −0.05
NG = 0
DB = 0
SM = 0
Delta_AUC
Doxorubicin

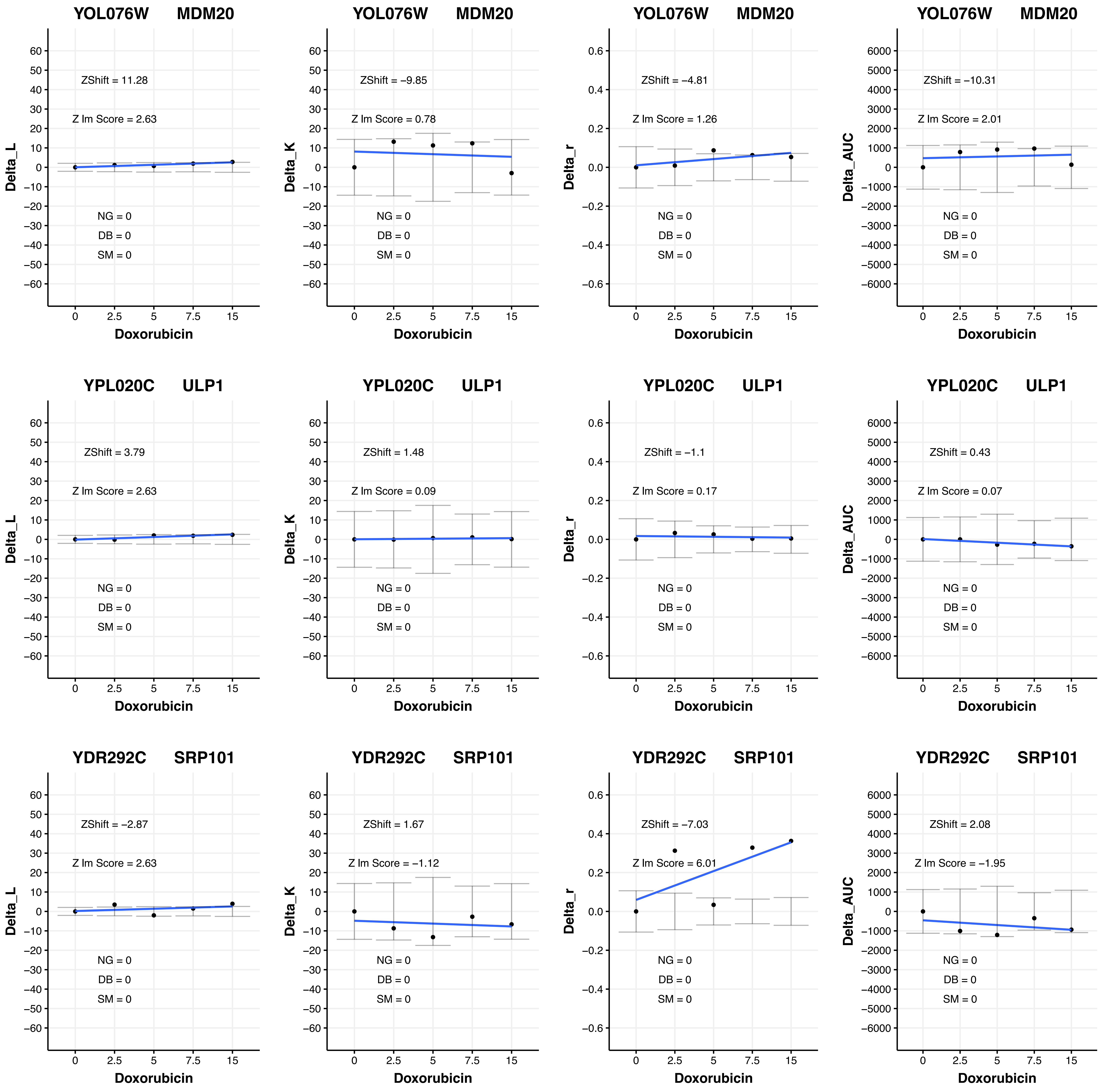

YOL076W MDM20
ZShift = 11.28
Z lm Score = 2.63
NG = 0
DB = 0
SM = 0
Delta_L
Doxorubicin
YOL076W MDM20
ZShift = −9.85
Z lm Score = 0.78
NG = 0
DB = 0
SM = 0
Delta_K
Doxorubicin
YOL076W MDM20
ZShift = −4.81
Z lm Score = 1.26
NG = 0
DB = 0
SM = 0
Delta_r
Doxorubicin
YOL076W MDM20
ZShift = −10.31
Z lm Score = 2.01
NG = 0
DB = 0
SM = 0
Delta_AUC
Doxorubicin
YPL020C ULP1
ZShift = 3.79
Z lm Score = 2.63
NG = 0
DB = 0
SM = 0
Delta_L
Doxorubicin
YPL020C ULP1
ZShift = 1.48
Z lm Score = 0.09
NG = 0
DB = 0
SM = 0
Delta_K
Doxorubicin
YPL020C ULP1
ZShift = −1.1
Z lm Score = 0.17
NG = 0
DB = 0
SM = 0
Delta_r
Doxorubicin
YPL020C ULP1
ZShift = 0.43
Z lm Score = 0.07
NG = 0
DB = 0
SM = 0
Delta_AUC
Doxorubicin
YDR292C SRP101
ZShift = −2.87
Z lm Score = 2.63
NG = 0
DB = 0
SM = 0
Delta_L
Doxorubicin
YDR292C SRP101
ZShift = 1.67
Z lm Score = −1.12
NG = 0
DB = 0
SM = 0
Delta_K
Doxorubicin
YDR292C SRP101
ZShift = −7.03
Z lm Score = 6.01
NG = 0
DB = 0
SM = 0
Delta_r
Doxorubicin
YDR292C SRP101
ZShift = 2.08
Z lm Score = −1.95
NG = 0
DB = 0
SM = 0
Delta_AUC
Doxorubicin

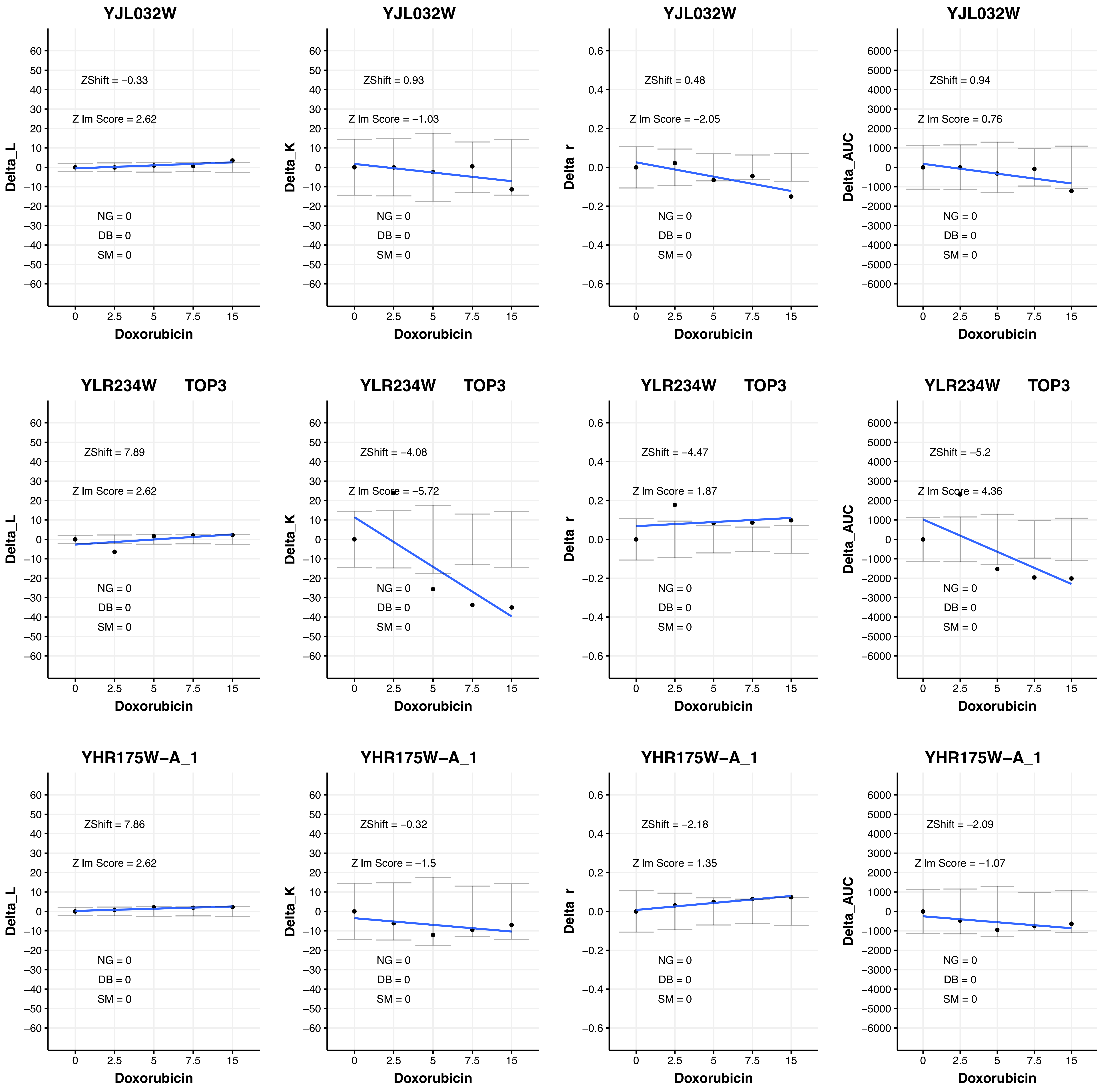

YJL032W
ZShift = −0.33
Z lm Score = 2.62
NG = 0
DB = 0
SM = 0
Delta_L
Doxorubicin
YJL032W
ZShift = 0.93
Z lm Score = −1.03
NG = 0
DB = 0
SM = 0
Delta_K
Doxorubicin
YJL032W
ZShift = 0.48
Z lm Score = −2.05
NG = 0
DB = 0
SM = 0
Delta_r
Doxorubicin
YJL032W
ZShift = 0.94
Z lm Score = 0.76
NG = 0
DB = 0
SM = 0
Delta_AUC
Doxorubicin
YLR234W TOP3
ZShift = 7.89
Z lm Score = 2.62
NG = 0
DB = 0
SM = 0
Delta_L
Doxorubicin
YLR234W TOP3
ZShift = −4.08
Z lm Score = −5.72
NG = 0
DB = 0
SM = 0
Delta_K
Doxorubicin
YLR234W TOP3
ZShift = −4.47
Z lm Score = 1.87
NG = 0
DB = 0
SM = 0
Delta_r
Doxorubicin
YLR234W TOP3
ZShift = −5.2
Z lm Score = 4.36
NG = 0
DB = 0
SM = 0
Delta_AUC
Doxorubicin
YHR175W−A_1
ZShift = 7.86
Z lm Score = 2.62
NG = 0
DB = 0
SM = 0
Delta_L
Doxorubicin
YHR175W−A_1
ZShift = −0.32
Z lm Score = −1.5
NG = 0
DB = 0
SM = 0
Delta_K
Doxorubicin
YHR175W−A_1
ZShift = −2.18
Z lm Score = 1.35
NG = 0
DB = 0
SM = 0
Delta_r
Doxorubicin
YHR175W−A_1
ZShift = −2.09
Z lm Score = −1.07
NG = 0
DB = 0
SM = 0
Delta_AUC
Doxorubicin

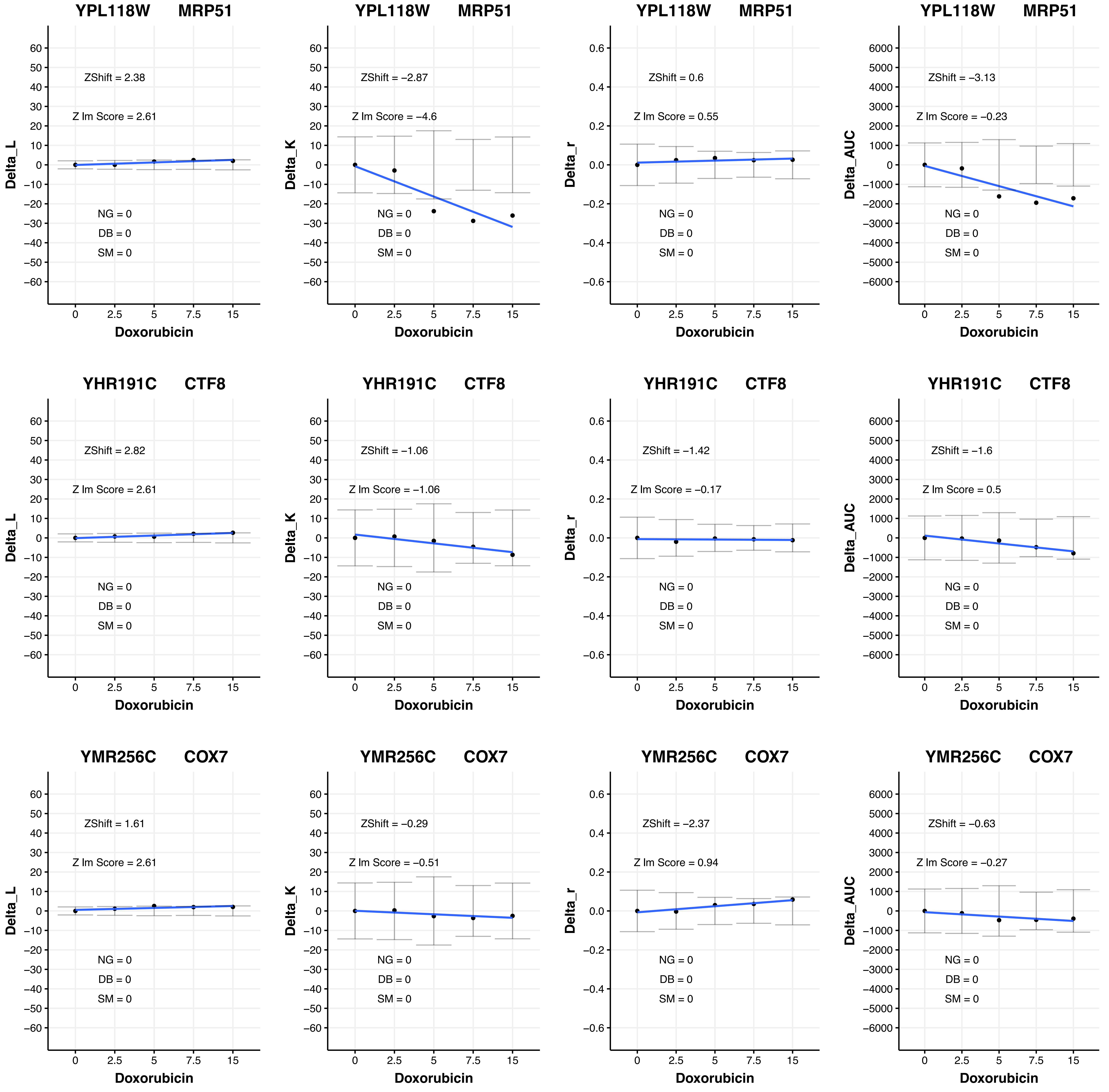
YPL118W MRP51
ZShift = 2.38
Z lm Score = 2.61
NG = 0
DB = 0
SM = 0
Delta_L
Doxorubicin
YPL118W MRP51
ZShift = −2.87
Z lm Score = −4.6
NG = 0
DB = 0
SM = 0
Delta_K
Doxorubicin
YPL118W MRP51
ZShift = 0.6
Z lm Score = 0.55
NG = 0
DB = 0
SM = 0
Delta_r
Doxorubicin
YPL118W MRP51
ZShift = −3.13
Z lm Score = −0.23
NG = 0
DB = 0
SM = 0
Delta_AUC
Doxorubicin
YHR191C CTF8
ZShift = 2.82
Z lm Score = 2.61
NG = 0
DB = 0
SM = 0
Delta_L
Doxorubicin
YHR191C CTF8
ZShift = −1.06
Z lm Score = −1.06
NG = 0
DB = 0
SM = 0
Delta_K
Doxorubicin
YHR191C CTF8
ZShift = −1.42
Z lm Score = −0.17
NG = 0
DB = 0
SM = 0
Delta_r
Doxorubicin
YHR191C CTF8
ZShift = −1.6
Z lm Score = 0.5
NG = 0
DB = 0
SM = 0
Delta_AUC
Doxorubicin
YMR256C COX7
ZShift = 1.61
Z lm Score = 2.61
NG = 0
DB = 0
SM = 0
Delta_L
Doxorubicin
YMR256C COX7
ZShift = −0.29
Z lm Score = −0.51
NG = 0
DB = 0
SM = 0
Delta_K
Doxorubicin
YMR256C COX7
ZShift = −2.37
Z lm Score = 0.94
NG = 0
DB = 0
SM = 0
Delta_r
Doxorubicin
YMR256C COX7
ZShift = −0.63
Z lm Score = −0.27
NG = 0
DB = 0
SM = 0
Delta_AUC
Doxorubicin

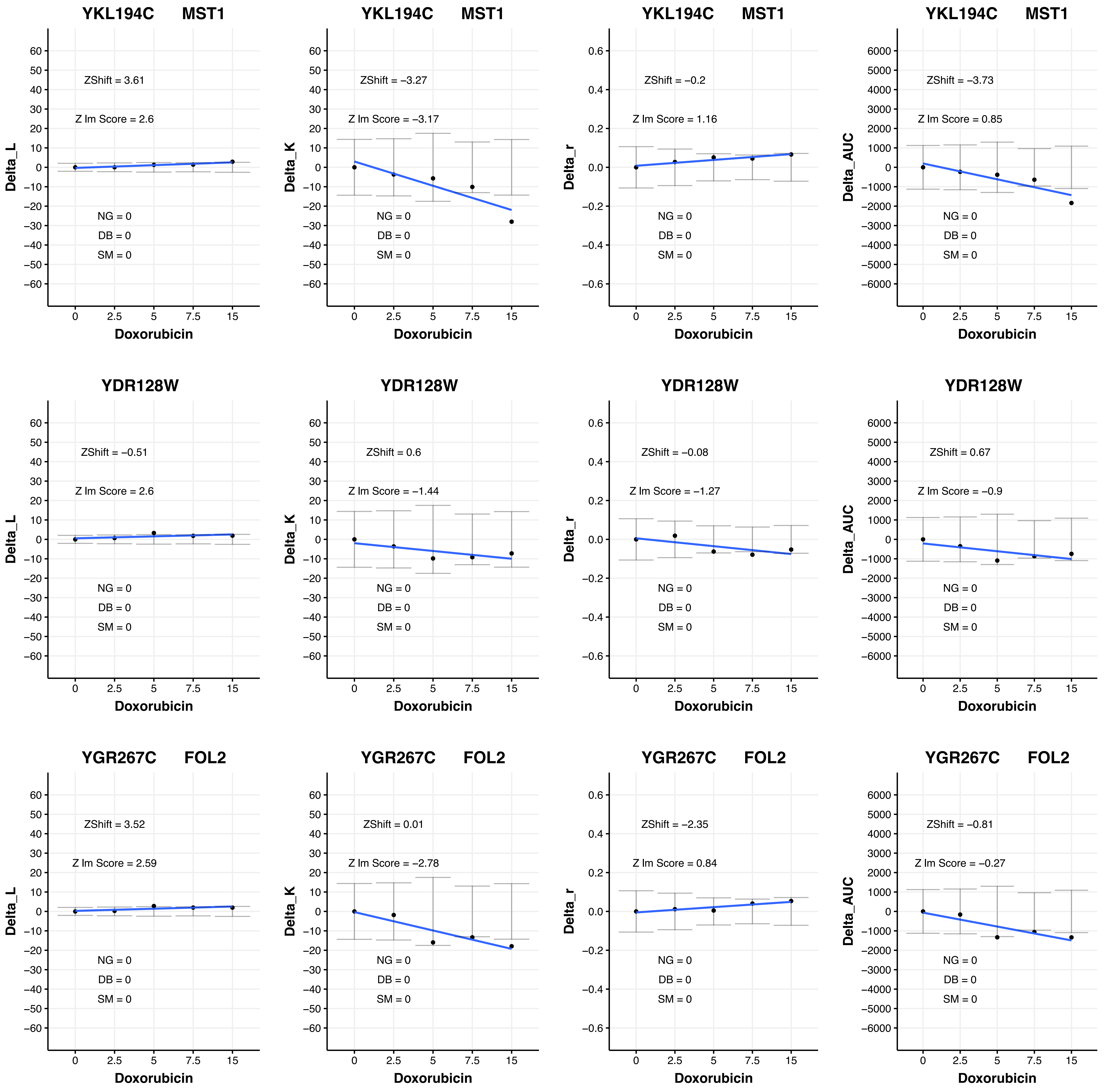

YKL194C MST1
ZShift = 3.61
Z lm Score = 2.6
NG = 0
DB = 0
SM = 0
Delta_L
Doxorubicin
YKL194C MST1
ZShift = −3.27
Z lm Score = −3.17
NG = 0
DB = 0
SM = 0
Delta_K
Doxorubicin
YKL194C MST1
ZShift = −0.2
Z lm Score = 1.16
NG = 0
DB = 0
SM = 0
Delta_r
Doxorubicin
YKL194C MST1
ZShift = −3.73
Z lm Score = 0.85
NG = 0
DB = 0
SM = 0
Delta_AUC
Doxorubicin
YDR128W
ZShift = −0.51
Z lm Score = 2.6
NG = 0
DB = 0
SM = 0
Delta_L
Doxorubicin
YDR128W
ZShift = 0.6
Z lm Score = −1.44
NG = 0
DB = 0
SM = 0
Delta_K
Doxorubicin
YDR128W
ZShift = −0.08
Z lm Score = −1.27
NG = 0
DB = 0
SM = 0
Delta_r
Doxorubicin
YDR128W
ZShift = 0.67
Z lm Score = −0.9
NG = 0
DB = 0
SM = 0
Delta_AUC
Doxorubicin
YGR267C FOL2
ZShift = 3.52
Z lm Score = 2.59
NG = 0
DB = 0
SM = 0
Delta_L
Doxorubicin
YGR267C FOL2
ZShift = 0.01
Z lm Score = −2.78
NG = 0
DB = 0
SM = 0
Delta_K
Doxorubicin
YGR267C FOL2
ZShift = −2.35
Z lm Score = 0.84
NG = 0
DB = 0
SM = 0
Delta_r
Doxorubicin
YGR267C FOL2
ZShift = −0.81
Z lm Score = −0.27
NG = 0
DB = 0
SM = 0
Delta_AUC
Doxorubicin

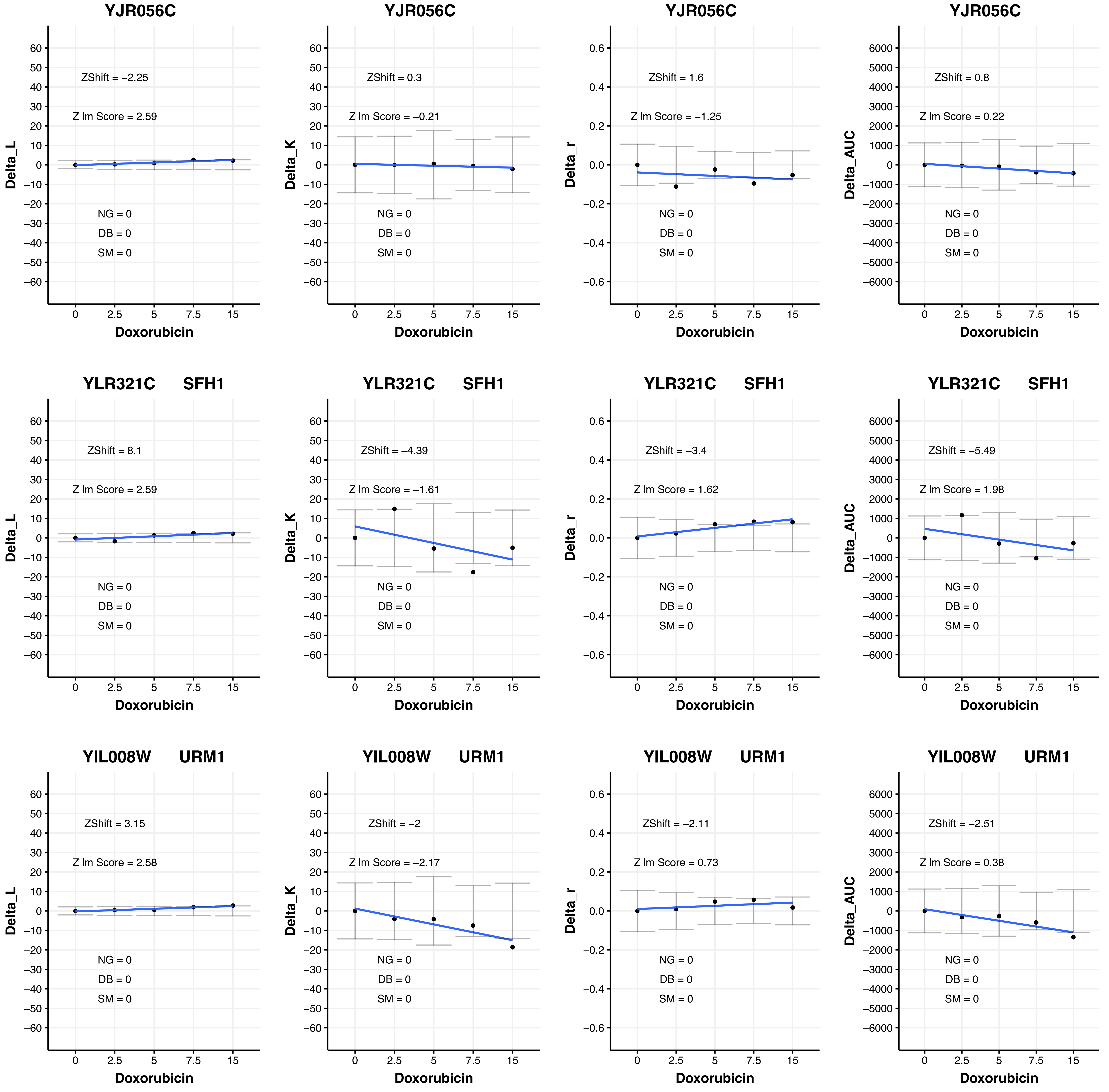

YJR056C
ZShift = −2.25
Z lm Score = 2.59
NG = 0
DB = 0
SM = 0
Delta_L
Doxorubicin
YJR056C
ZShift = 0.3
Z lm Score = −0.21
NG = 0
DB = 0
SM = 0
Delta_K
Doxorubicin
YJR056C
ZShift = 1.6
Z lm Score = −1.25
NG = 0
DB = 0
SM = 0
Delta_r
Doxorubicin
YJR056C
ZShift = 0.8
Z lm Score = 0.22
NG = 0
DB = 0
SM = 0
Delta_AUC
Doxorubicin
YLR321C SFH1
ZShift = 8.1
Z lm Score = 2.59
NG = 0
DB = 0
SM = 0
Delta_L
Doxorubicin
YLR321C SFH1
ZShift = −4.39
Z lm Score = −1.61
NG = 0
DB = 0
SM = 0
Delta_K
Doxorubicin
YLR321C SFH1
ZShift = −3.4
Z lm Score = 1.62
NG = 0
DB = 0
SM = 0
Delta_r
Doxorubicin
YLR321C SFH1
ZShift = −5.49
Z lm Score = 1.98
NG = 0
DB = 0
SM = 0
Delta_AUC
Doxorubicin
YIL008W URM1
ZShift = 3.15
Z lm Score = 2.58
NG = 0
DB = 0
SM = 0
Delta_L
Doxorubicin
YIL008W URM1
ZShift = −2
Z lm Score = −2.17
NG = 0
DB = 0
SM = 0
Delta_K
Doxorubicin
YIL008W URM1
ZShift = −2.11
Z lm Score = 0.73
NG = 0
DB = 0
SM = 0
Delta_r
Doxorubicin
YIL008W URM1
ZShift = −2.51
Z lm Score = 0.38
NG = 0
DB = 0
SM = 0
Delta_AUC
Doxorubicin

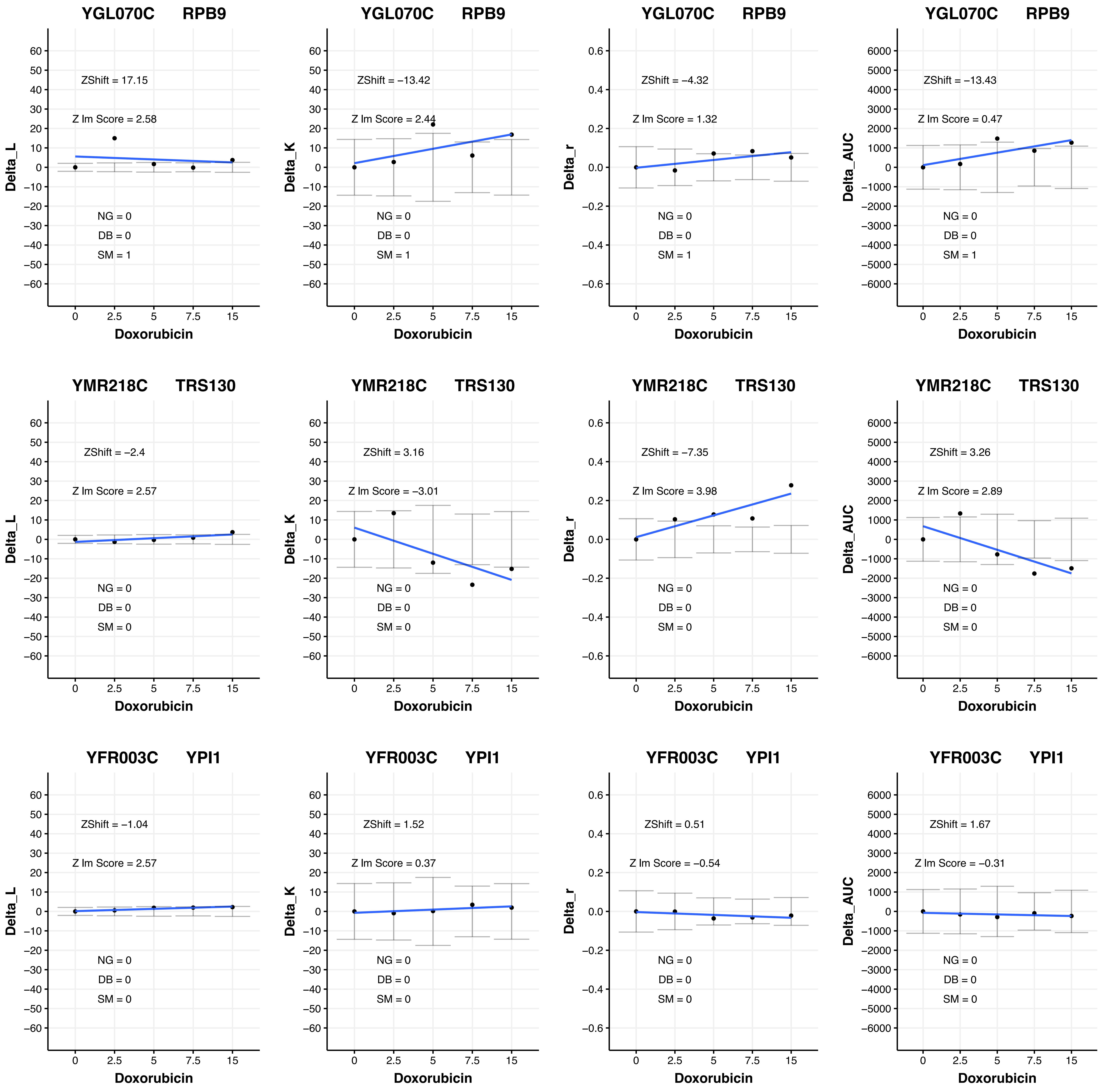
YGL070C RPB9
ZShift = 17.15
Z lm Score = 2.58
NG = 0
DB = 0
SM = 1
Delta_L
Doxorubicin
YGL070C RPB9
ZShift = −13.42
Z lm Score = 2.44
NG = 0
DB = 0
SM = 1
Delta_K
Doxorubicin
YGL070C RPB9
ZShift = −4.32
Z lm Score = 1.32
NG = 0
DB = 0
SM = 1
Delta_r
Doxorubicin
YGL070C RPB9
ZShift = −13.43
Z lm Score = 0.47
NG = 0
DB = 0
SM = 1
Delta_AUC
Doxorubicin
YMR218C TRS130
ZShift = −2.4
Z lm Score = 2.57
NG = 0
DB = 0
SM = 0
Delta_L
Doxorubicin
YMR218C TRS130
ZShift = 3.16
Z lm Score = −3.01
NG = 0
DB = 0
SM = 0
Delta_K
Doxorubicin
YMR218C TRS130
ZShift = −7.35
Z lm Score = 3.98
NG = 0
DB = 0
SM = 0
Delta_r
Doxorubicin
YMR218C TRS130
ZShift = 3.26
Z lm Score = 2.89
NG = 0
DB = 0
SM = 0
Delta_AUC
Doxorubicin
YFR003C YPI1
ZShift = −1.04
Z lm Score = 2.57
NG = 0
DB = 0
SM = 0
Delta_L
Doxorubicin
YFR003C YPI1
ZShift = 1.52
Z lm Score = 0.37
NG = 0
DB = 0
SM = 0
Delta_K
Doxorubicin
YFR003C YPI1
ZShift = 0.51
Z lm Score = −0.54
NG = 0
DB = 0
SM = 0
Delta_r
Doxorubicin
YFR003C YPI1
ZShift = 1.67
Z lm Score = −0.31
NG = 0
DB = 0
SM = 0
Delta_AUC
Doxorubicin

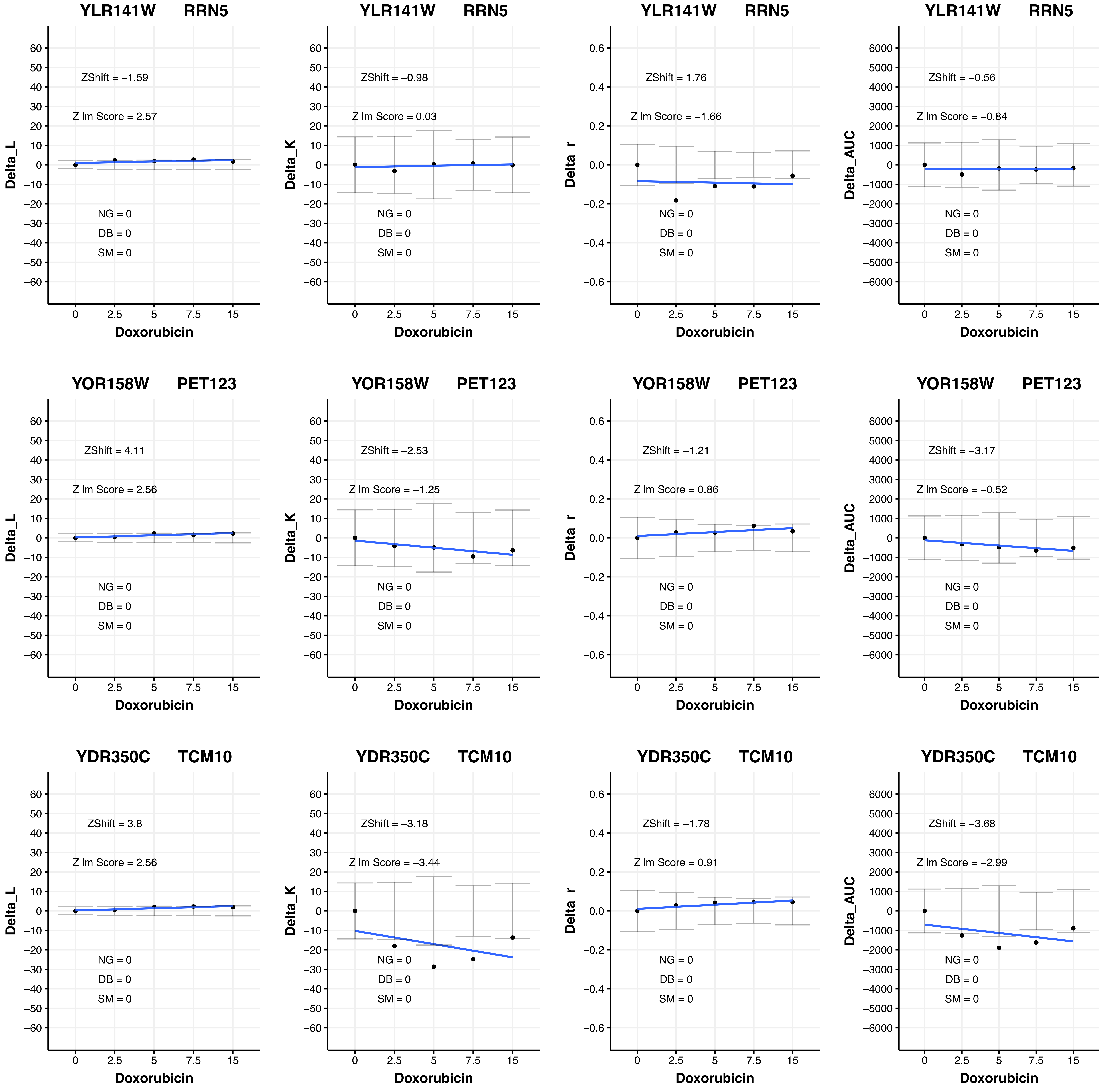
YLR141W RRN5
ZShift = −1.59
Z lm Score = 2.57
NG = 0
DB = 0
SM = 0
Delta_L
Doxorubicin
YLR141W RRN5
ZShift = −0.98
Z lm Score = 0.03
NG = 0
DB = 0
SM = 0
Delta_K
Doxorubicin
YLR141W RRN5
ZShift = 1.76
Z lm Score = −1.66
NG = 0
DB = 0
SM = 0
Delta_r
Doxorubicin
YLR141W RRN5
ZShift = −0.56
Z lm Score = −0.84
NG = 0
DB = 0
SM = 0
Delta_AUC
Doxorubicin
YOR158W PET123
ZShift = 4.11
Z lm Score = 2.56
NG = 0
DB = 0
SM = 0
Delta_L
Doxorubicin
YOR158W PET123
ZShift = −2.53
Z lm Score = −1.25
NG = 0
DB = 0
SM = 0
Delta_K
Doxorubicin
YOR158W PET123
ZShift = −1.21
Z lm Score = 0.86
NG = 0
DB = 0
SM = 0
Delta_r
Doxorubicin
YOR158W PET123
ZShift = −3.17
Z lm Score = −0.52
NG = 0
DB = 0
SM = 0
Delta_AUC
Doxorubicin
YDR350C TCM10
ZShift = 3.8
Z lm Score = 2.56
NG = 0
DB = 0
SM = 0
Delta_L
Doxorubicin
YDR350C TCM10
ZShift = −3.18
Z lm Score = −3.44
NG = 0
DB = 0
SM = 0
Delta_K
Doxorubicin
YDR350C TCM10
ZShift = −1.78
Z lm Score = 0.91
NG = 0
DB = 0
SM = 0
Delta_r
Doxorubicin
YDR350C TCM10
ZShift = −3.68
Z lm Score = −2.99
NG = 0
DB = 0
SM = 0
Delta_AUC
Doxorubicin

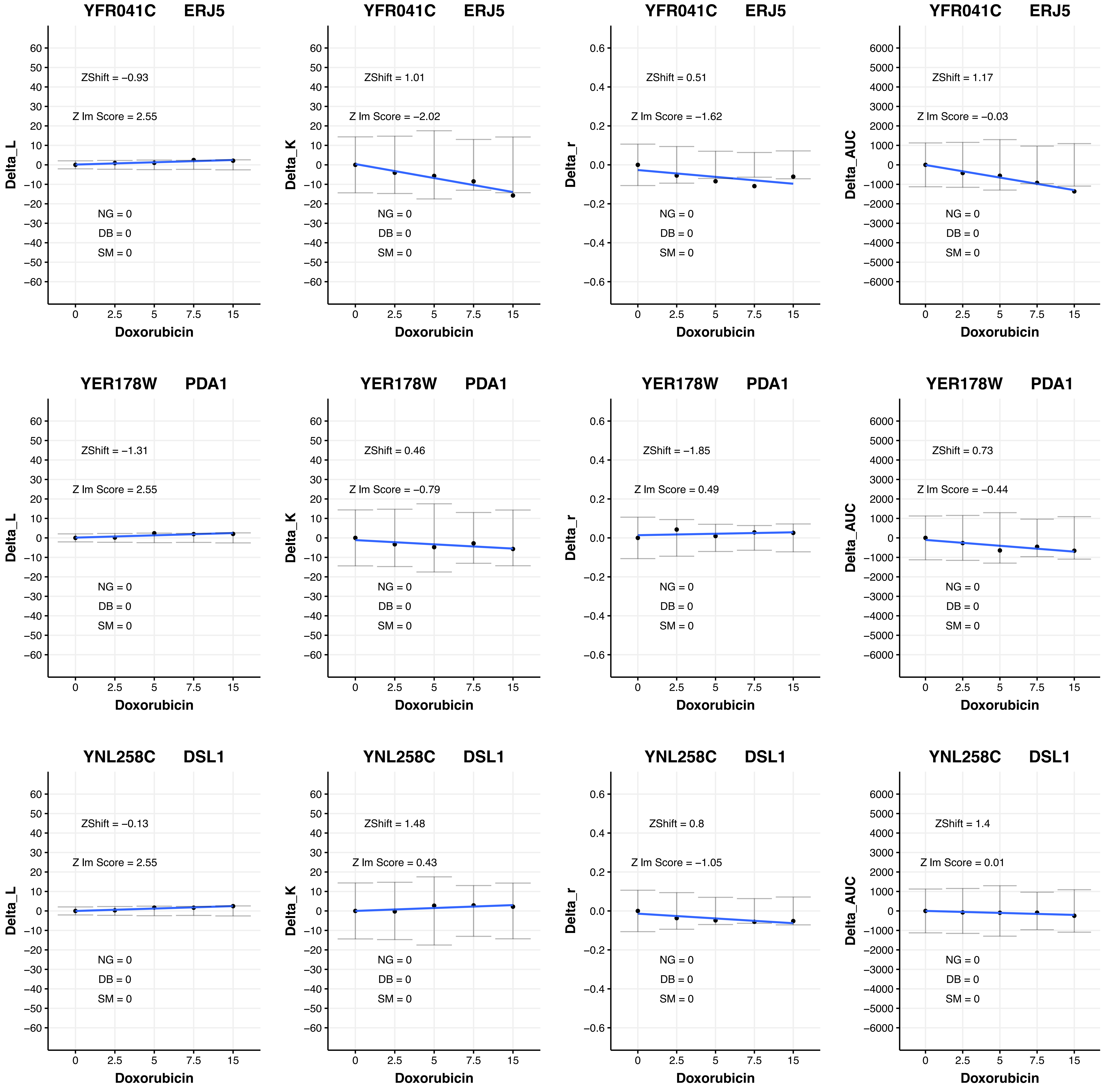
YFR041C ERJ5
ZShift = −0.93
Z lm Score = 2.55
NG = 0
DB = 0
SM = 0
Delta_L
Doxorubicin
YFR041C ERJ5
ZShift = 1.01
Z lm Score = −2.02
NG = 0
DB = 0
SM = 0
Delta_K
Doxorubicin
YFR041C ERJ5
ZShift = 0.51
Z lm Score = −1.62
NG = 0
DB = 0
SM = 0
Delta_r
Doxorubicin
YFR041C ERJ5
ZShift = 1.17
Z lm Score = −0.03
NG = 0
DB = 0
SM = 0
Delta_AUC
Doxorubicin
YER178W PDA1
ZShift = −1.31
Z lm Score = 2.55
NG = 0
DB = 0
SM = 0
Delta_L
Doxorubicin
YER178W PDA1
ZShift = 0.46
Z lm Score = −0.79
NG = 0
DB = 0
SM = 0
Delta_K
Doxorubicin
YER178W PDA1
ZShift = −1.85
Z lm Score = 0.49
NG = 0
DB = 0
SM = 0
Delta_r
Doxorubicin
YER178W PDA1
ZShift = 0.73
Z lm Score = −0.44
NG = 0
DB = 0
SM = 0
Delta_AUC
Doxorubicin
YNL258C DSL1
ZShift = −0.13
Z lm Score = 2.55
NG = 0
DB = 0
SM = 0
Delta_L
Doxorubicin
YNL258C DSL1
ZShift = 1.48
Z lm Score = 0.43
NG = 0
DB = 0
SM = 0
Delta_K
Doxorubicin
YNL258C DSL1
ZShift = 0.8
Z lm Score = −1.05
NG = 0
DB = 0
SM = 0
Delta_r
Doxorubicin
YNL258C DSL1
ZShift = 1.4
Z lm Score = 0.01
NG = 0
DB = 0
SM = 0
Delta_AUC
Doxorubicin

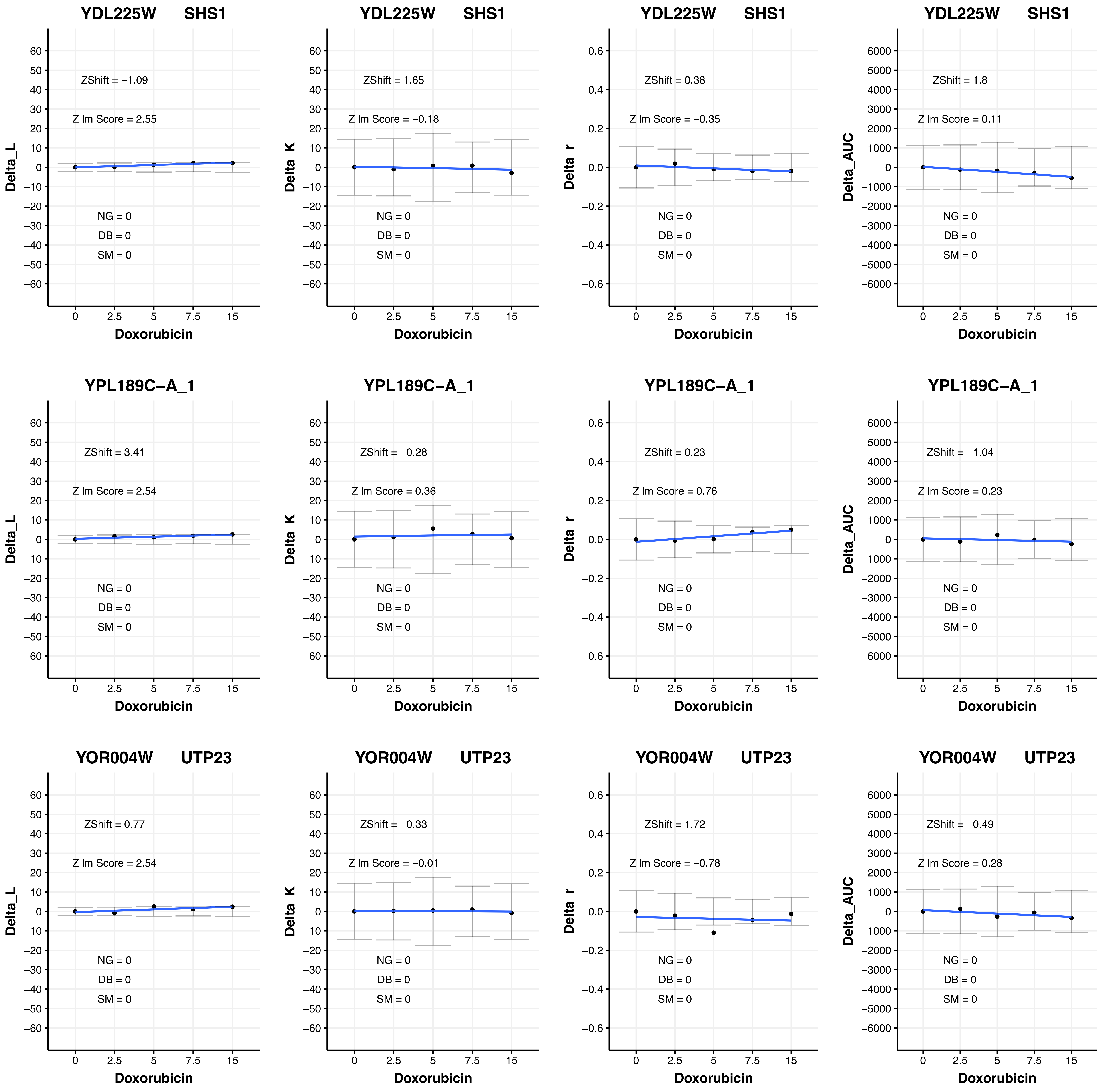
YDL225W SHS1
Delta_L
ZShift = −1.09
Z lm Score = 2.55
NG = 0
DB = 0
SM = 0
Doxorubicin
YDL225W SHS1
Delta_K
ZShift = 1.65
Z lm Score = −0.18
NG = 0
DB = 0
SM = 0
Doxorubicin
YDL225W SHS1
Delta_r
ZShift = 0.38
Z lm Score = −0.35
NG = 0
DB = 0
SM = 0
Doxorubicin
YDL225W SHS1
Delta_AUC
ZShift = 1.8
Z lm Score = 0.11
NG = 0
DB = 0
SM = 0
Doxorubicin
YPL189C−A_1
Delta_L
ZShift = 3.41
Z lm Score = 2.54
NG = 0
DB = 0
SM = 0
Doxorubicin
YPL189C−A_1
Delta_K
ZShift = −0.28
Z lm Score = 0.36
NG = 0
DB = 0
SM = 0
Doxorubicin
YPL189C−A_1
Delta_r
ZShift = 0.23
Z lm Score = 0.76
NG = 0
DB = 0
SM = 0
Doxorubicin
YPL189C−A_1
Delta_AUC
ZShift = −1.04
Z lm Score = 0.23
NG = 0
DB = 0
SM = 0
Doxorubicin
YOR004W UTP23
Delta_L
ZShift = 0.77
Z lm Score = 2.54
NG = 0
DB = 0
SM = 0
Doxorubicin
YOR004W UTP23
Delta_K
ZShift = −0.33
Z lm Score = −0.01
NG = 0
DB = 0
SM = 0
Doxorubicin
YOR004W UTP23
Delta_r
ZShift = 1.72
Z lm Score = −0.78
NG = 0
DB = 0
SM = 0
Doxorubicin
YOR004W UTP23
Delta_AUC
ZShift = −0.49
Z lm Score = 0.28
NG = 0
DB = 0
SM = 0
Doxorubicin

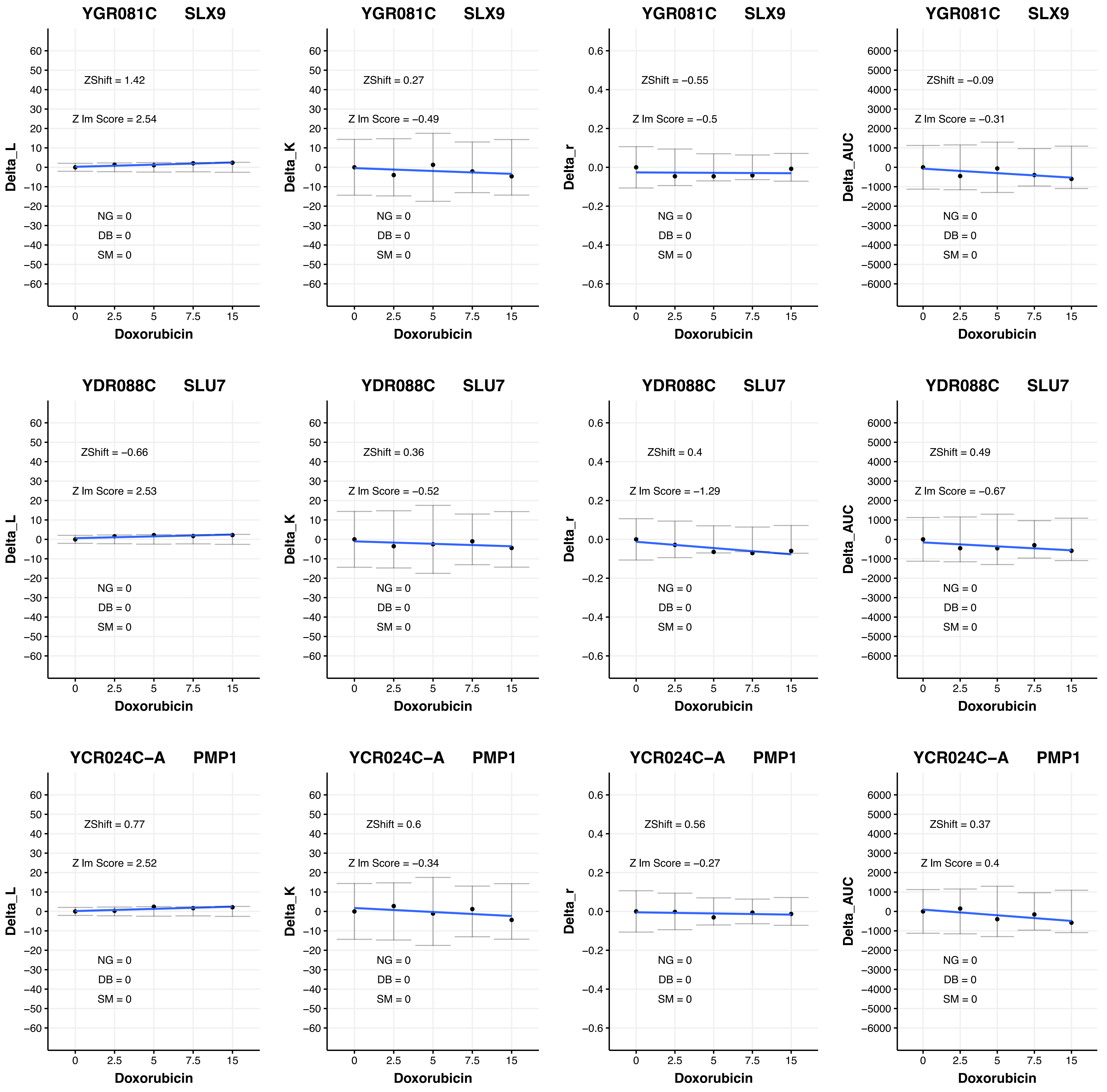

YGR081C SLX9
ZShift = 1.42
Z lm Score = 2.54
Delta_L
NG = 0
DB = 0
SM = 0
Doxorubicin
YGR081C SLX9
ZShift = 0.27
Z lm Score = −0.49
Delta_K
NG = 0
DB = 0
SM = 0
Doxorubicin
YGR081C SLX9
ZShift = −0.55
Z lm Score = −0.5
Delta_r
NG = 0
DB = 0
SM = 0
Doxorubicin
YGR081C SLX9
ZShift = −0.09
Z lm Score = −0.31
Delta_AUC
NG = 0
DB = 0
SM = 0
Doxorubicin
YDR088C SLU7
ZShift = −0.66
Z lm Score = 2.53
Delta_L
NG = 0
DB = 0
SM = 0
Doxorubicin
YDR088C SLU7
ZShift = 0.36
Z lm Score = −0.52
Delta_K
NG = 0
DB = 0
SM = 0
Doxorubicin
YDR088C SLU7
ZShift = 0.4
Z lm Score = −1.29
Delta_r
NG = 0
DB = 0
SM = 0
Doxorubicin
YDR088C SLU7
ZShift = 0.49
Z lm Score = −0.67
Delta_AUC
NG = 0
DB = 0
SM = 0
Doxorubicin
YCR024C−A PMP1
ZShift = 0.77
Z lm Score = 2.52
Delta_L
NG = 0
DB = 0
SM = 0
Doxorubicin
YCR024C−A PMP1
ZShift = 0.6
Z lm Score = −0.34
Delta_K
NG = 0
DB = 0
SM = 0
Doxorubicin
YCR024C−A PMP1
ZShift = 0.56
Z lm Score = −0.27
Delta_r
NG = 0
DB = 0
SM = 0
Doxorubicin
YCR024C−A PMP1
ZShift = 0.37
Z lm Score = 0.4
Delta_AUC
NG = 0
DB = 0
SM = 0
Doxorubicin

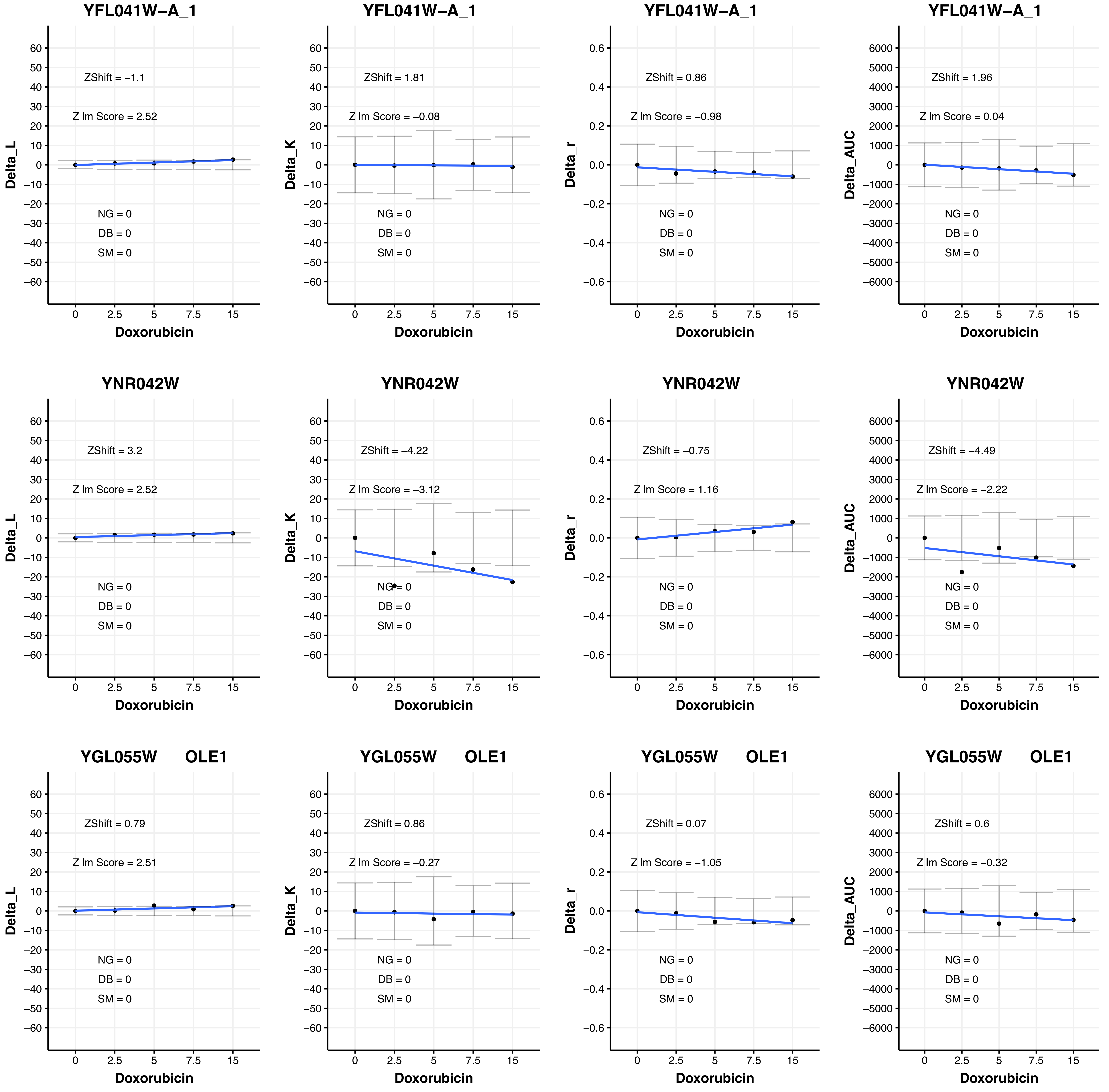
YFL041W−A_1
ZShift = −1.1
Z lm Score = 2.52
NG = 0
DB = 0
SM = 0
Delta_L
Doxorubicin
YFL041W−A_1
ZShift = 1.81
Z lm Score = −0.08
NG = 0
DB = 0
SM = 0
Delta_K
Doxorubicin
YFL041W−A_1
ZShift = 0.86
Z lm Score = −0.98
NG = 0
DB = 0
SM = 0
Delta_r
Doxorubicin
YFL041W−A_1
ZShift = 1.96
Z lm Score = 0.04
NG = 0
DB = 0
SM = 0
Delta_AUC
Doxorubicin
YNR042W
ZShift = 3.2
Z lm Score = 2.52
NG = 0
DB = 0
SM = 0
Delta_L
Doxorubicin
YNR042W
ZShift = −4.22
Z lm Score = −3.12
NG = 0
DB = 0
SM = 0
Delta_K
Doxorubicin
YNR042W
ZShift = −0.75
Z lm Score = 1.16
NG = 0
DB = 0
SM = 0
Delta_r
Doxorubicin
YNR042W
ZShift = −4.49
Z lm Score = −2.22
NG = 0
DB = 0
SM = 0
Delta_AUC
Doxorubicin
YGL055W OLE1
ZShift = 0.79
Z lm Score = 2.51
NG = 0
DB = 0
SM = 0
Delta_L
Doxorubicin
YGL055W OLE1
ZShift = 0.86
Z lm Score = −0.27
NG = 0
DB = 0
SM = 0
Delta_K
Doxorubicin
YGL055W OLE1
ZShift = 0.07
Z lm Score = −1.05
NG = 0
DB = 0
SM = 0
Delta_r
Doxorubicin
YGL055W OLE1
ZShift = 0.6
Z lm Score = −0.32
NG = 0
DB = 0
SM = 0
Delta_AUC
Doxorubicin

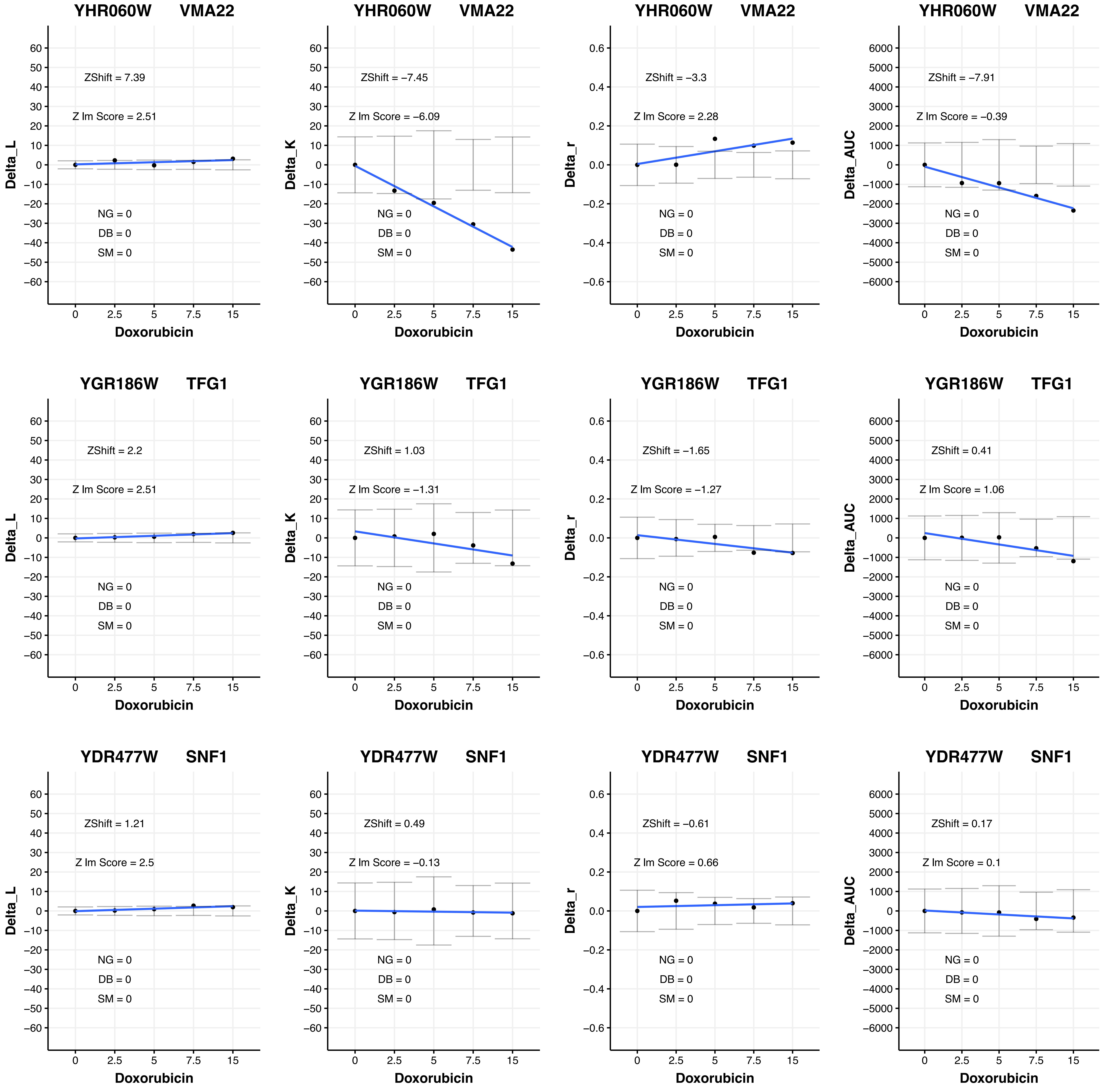
YHR060W VMA22
ZShift = 7.39
Z lm Score = 2.51
NG = 0
DB = 0
SM = 0
Delta_L
Doxorubicin
YHR060W VMA22
ZShift = −7.45
Z lm Score = −6.09
NG = 0
DB = 0
SM = 0
Delta_K
Doxorubicin
YHR060W VMA22
ZShift = −3.3
Z lm Score = 2.28
NG = 0
DB = 0
SM = 0
Delta_r
Doxorubicin
YHR060W VMA22
ZShift = −7.91
Z lm Score = −0.39
NG = 0
DB = 0
SM = 0
Delta_AUC
Doxorubicin
YGR186W TFG1
ZShift = 2.2
Z lm Score = 2.51
NG = 0
DB = 0
SM = 0
Delta_L
Doxorubicin
YGR186W TFG1
ZShift = 1.03
Z lm Score = −1.31
NG = 0
DB = 0
SM = 0
Delta_K
Doxorubicin
YGR186W TFG1
ZShift = −1.65
Z lm Score = −1.27
NG = 0
DB = 0
SM = 0
Delta_r
Doxorubicin
YGR186W TFG1
ZShift = 0.41
Z lm Score = 1.06
NG = 0
DB = 0
SM = 0
Delta_AUC
Doxorubicin
YDR477W SNF1
ZShift = 1.21
Z lm Score = 2.5
NG = 0
DB = 0
SM = 0
Delta_L
Doxorubicin
YDR477W SNF1
ZShift = 0.49
Z lm Score = −0.13
NG = 0
DB = 0
SM = 0
Delta_K
Doxorubicin
YDR477W SNF1
ZShift = −0.61
Z lm Score = 0.66
NG = 0
DB = 0
SM = 0
Delta_r
Doxorubicin
YDR477W SNF1
ZShift = 0.17
Z lm Score = 0.1
NG = 0
DB = 0
SM = 0
Delta_AUC
Doxorubicin

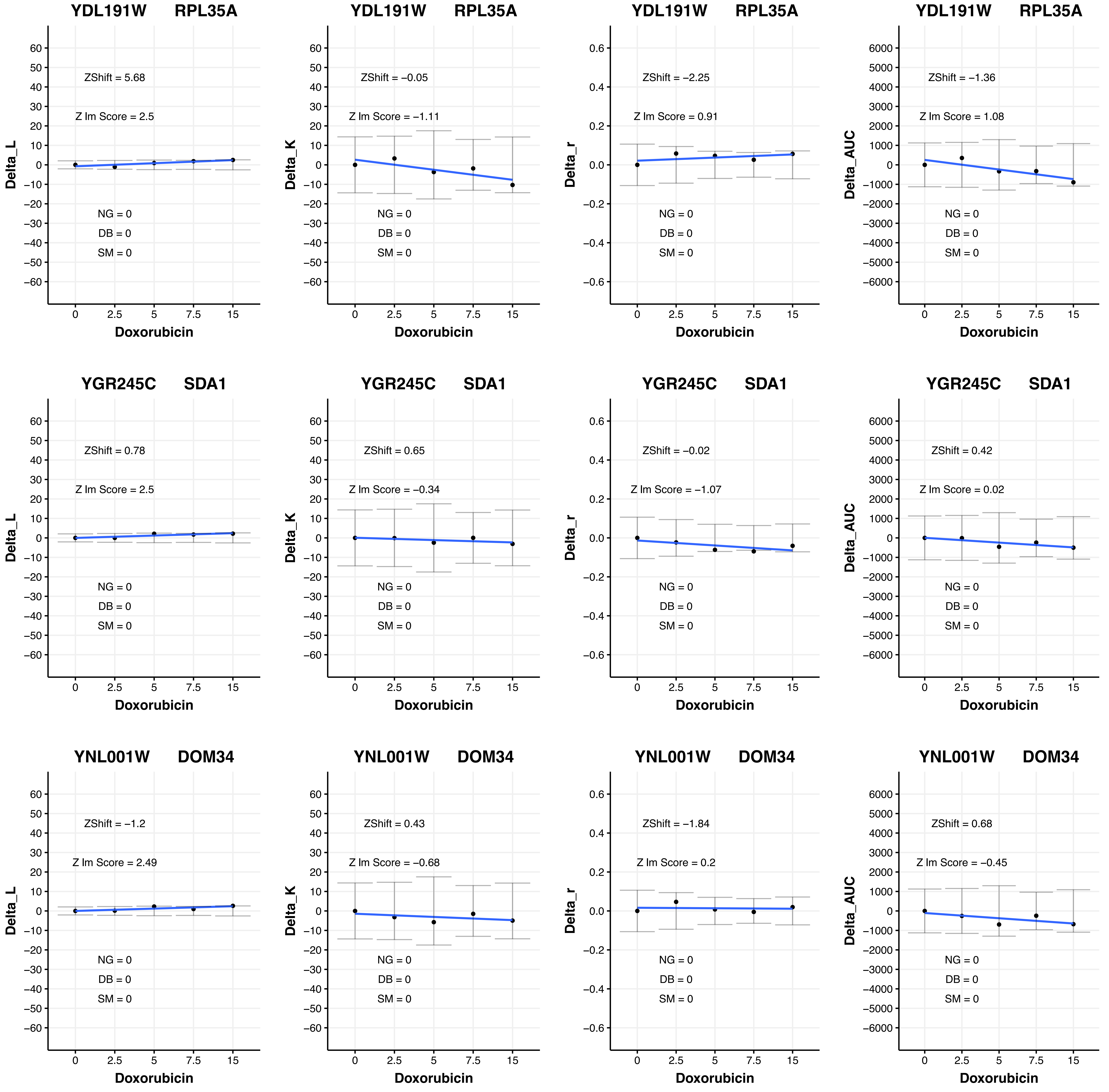
YDL191W RPL35A
ZShift = 5.68
Z lm Score = 2.5
NG = 0
DB = 0
SM = 0
Delta_L
Doxorubicin
YDL191W RPL35A
ZShift = −0.05
Z lm Score = −1.11
NG = 0
DB = 0
SM = 0
Delta_K
Doxorubicin
YDL191W RPL35A
ZShift = −2.25
Z lm Score = 0.91
NG = 0
DB = 0
SM = 0
Delta_r
Doxorubicin
YDL191W RPL35A
ZShift = −1.36
Z lm Score = 1.08
NG = 0
DB = 0
SM = 0
Delta_AUC
Doxorubicin
YGR245C SDA1
ZShift = 0.78
Z lm Score = 2.5
NG = 0
DB = 0
SM = 0
Delta_L
Doxorubicin
YGR245C SDA1
ZShift = 0.65
Z lm Score = −0.34
NG = 0
DB = 0
SM = 0
Delta_K
Doxorubicin
YGR245C SDA1
ZShift = −0.02
Z lm Score = −1.07
NG = 0
DB = 0
SM = 0
Delta_r
Doxorubicin
YGR245C SDA1
ZShift = 0.42
Z lm Score = 0.02
NG = 0
DB = 0
SM = 0
Delta_AUC
Doxorubicin
YNL001W DOM34
ZShift = −1.2
Z lm Score = 2.49
NG = 0
DB = 0
SM = 0
Delta_L
Doxorubicin
YNL001W DOM34
ZShift = 0.43
Z lm Score = −0.68
NG = 0
DB = 0
SM = 0
Delta_K
Doxorubicin
YNL001W DOM34
ZShift = −1.84
Z lm Score = 0.2
NG = 0
DB = 0
SM = 0
Delta_r
Doxorubicin
YNL001W DOM34
ZShift = 0.68
Z lm Score = −0.45
NG = 0
DB = 0
SM = 0
Delta_AUC
Doxorubicin

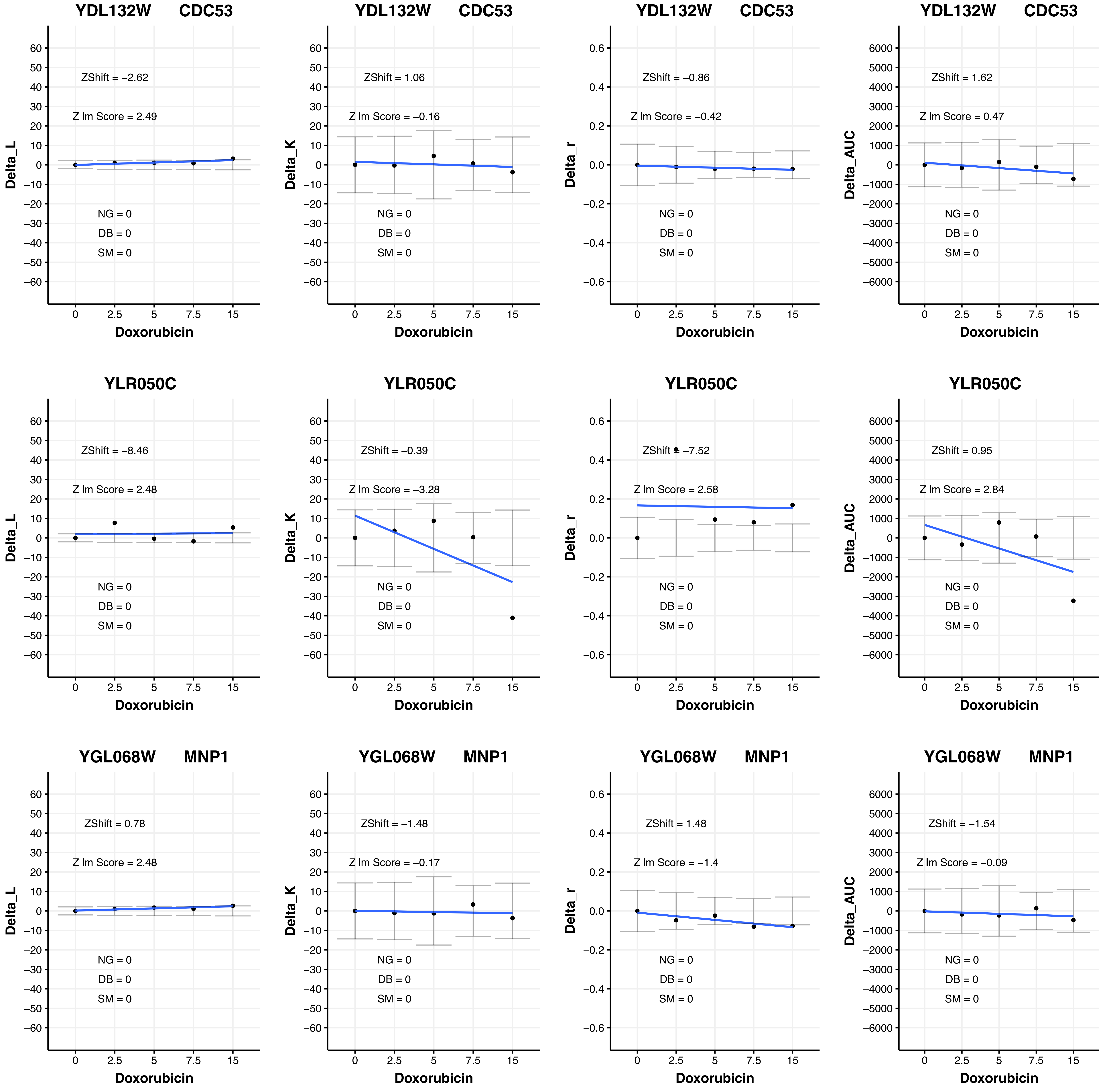
YDL132W CDC53
ZShift = −2.62
Z lm Score = 2.49
NG = 0
DB = 0
SM = 0
Delta_L
Doxorubicin
YDL132W CDC53
ZShift = 1.06
Z lm Score = −0.16
NG = 0
DB = 0
SM = 0
Delta_K
Doxorubicin
YDL132W CDC53
ZShift = −0.86
Z lm Score = −0.42
NG = 0
DB = 0
SM = 0
Delta_r
Doxorubicin
YDL132W CDC53
ZShift = 1.62
Z lm Score = 0.47
NG = 0
DB = 0
SM = 0
Delta_AUC
Doxorubicin
YLR050C
ZShift = −8.46
Z lm Score = 2.48
NG = 0
DB = 0
SM = 0
Delta_L
Doxorubicin
YLR050C
ZShift = −0.39
Z lm Score = −3.28
NG = 0
DB = 0
SM = 0
Delta_K
Doxorubicin
YLR050C
ZShift = −7.52
Z lm Score = 2.58
NG = 0
DB = 0
SM = 0
Delta_r
Doxorubicin
YLR050C
ZShift = 0.95
Z lm Score = 2.84
NG = 0
DB = 0
SM = 0
Delta_AUC
Doxorubicin
YGL068W MNP1
ZShift = 0.78
Z lm Score = 2.48
NG = 0
DB = 0
SM = 0
Delta_L
Doxorubicin
YGL068W MNP1
ZShift = −1.48
Z lm Score = −0.17
NG = 0
DB = 0
SM = 0
Delta_K
Doxorubicin
YGL068W MNP1
ZShift = 1.48
Z lm Score = −1.4
NG = 0
DB = 0
SM = 0
Delta_r
Doxorubicin
YGL068W MNP1
ZShift = −1.54
Z lm Score = −0.09
NG = 0
DB = 0
SM = 0
Delta_AUC
Doxorubicin

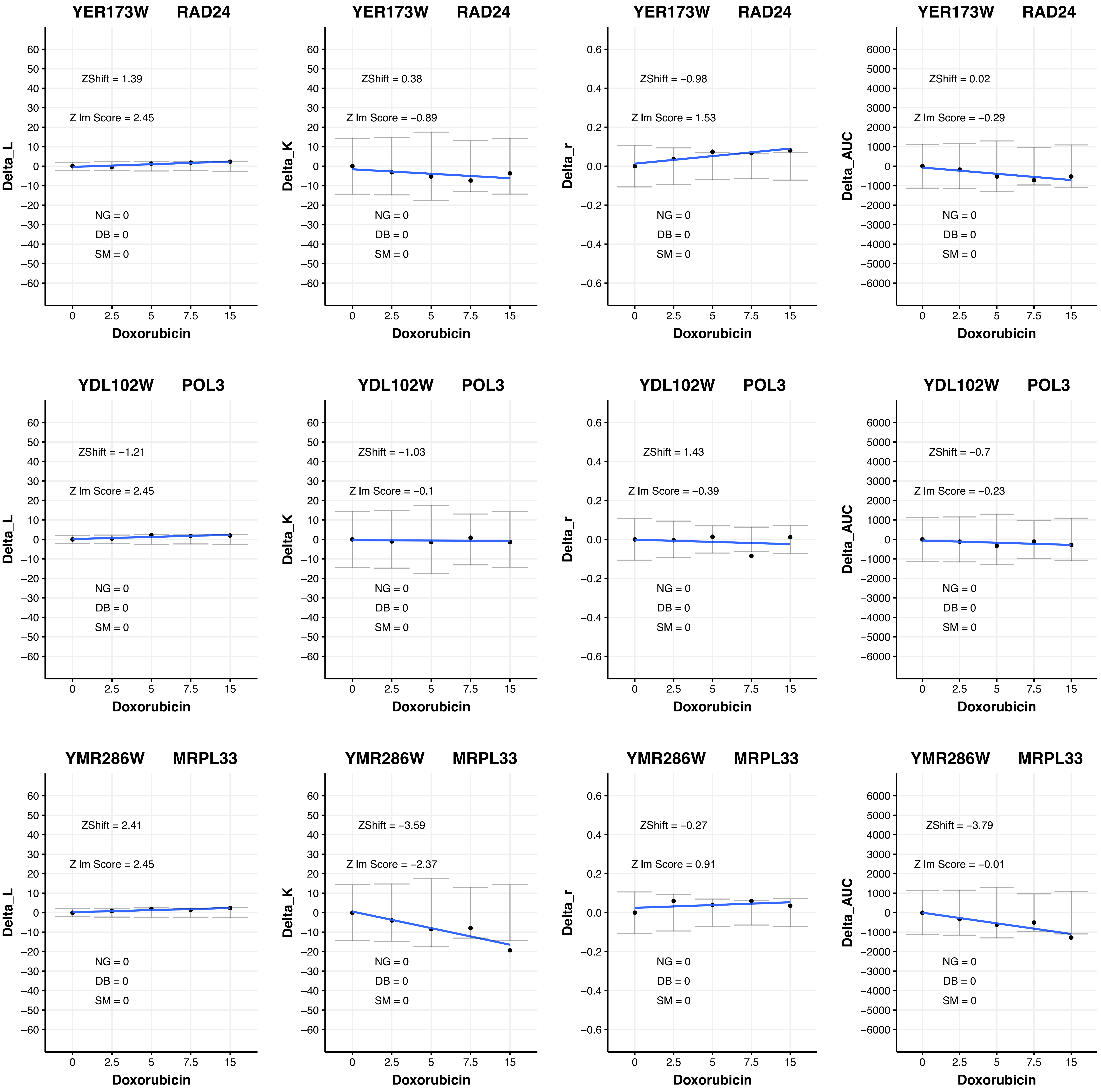

YER173W RAD24
ZShift = 1.39
Z lm Score = 2.45
Delta_L
NG = 0
DB = 0
SM = 0
Doxorubicin
YER173W RAD24
ZShift = 0.38
Z lm Score = −0.89
Delta_K
NG = 0
DB = 0
SM = 0
Doxorubicin
YER173W RAD24
ZShift = −0.98
Z lm Score = 1.53
Delta_r
NG = 0
DB = 0
SM = 0
Doxorubicin
YER173W RAD24
ZShift = 0.02
Z lm Score = −0.29
Delta_AUC
NG = 0
DB = 0
SM = 0
Doxorubicin
YDL102W POL3
ZShift = −1.21
Z lm Score = 2.45
Delta_L
NG = 0
DB = 0
SM = 0
Doxorubicin
YDL102W POL3
ZShift = −1.03
Z lm Score = −0.1
Delta_K
NG = 0
DB = 0
SM = 0
Doxorubicin
YDL102W POL3
ZShift = 1.43
Z lm Score = −0.39
Delta_r
NG = 0
DB = 0
SM = 0
Doxorubicin
YDL102W POL3
ZShift = −0.7
Z lm Score = −0.23
Delta_AUC
NG = 0
DB = 0
SM = 0
Doxorubicin
YMR286W MRPL33
ZShift = 2.41
Z lm Score = 2.45
Delta_L
NG = 0
DB = 0
SM = 0
Doxorubicin
YMR286W MRPL33
ZShift = −3.59
Z lm Score = −2.37
Delta_K
NG = 0
DB = 0
SM = 0
Doxorubicin
YMR286W MRPL33
ZShift = −0.27
Z lm Score = 0.91
Delta_r
NG = 0
DB = 0
SM = 0
Doxorubicin
YMR286W MRPL33
ZShift = −3.79
Z lm Score = −0.01
Delta_AUC
NG = 0
DB = 0
SM = 0
Doxorubicin

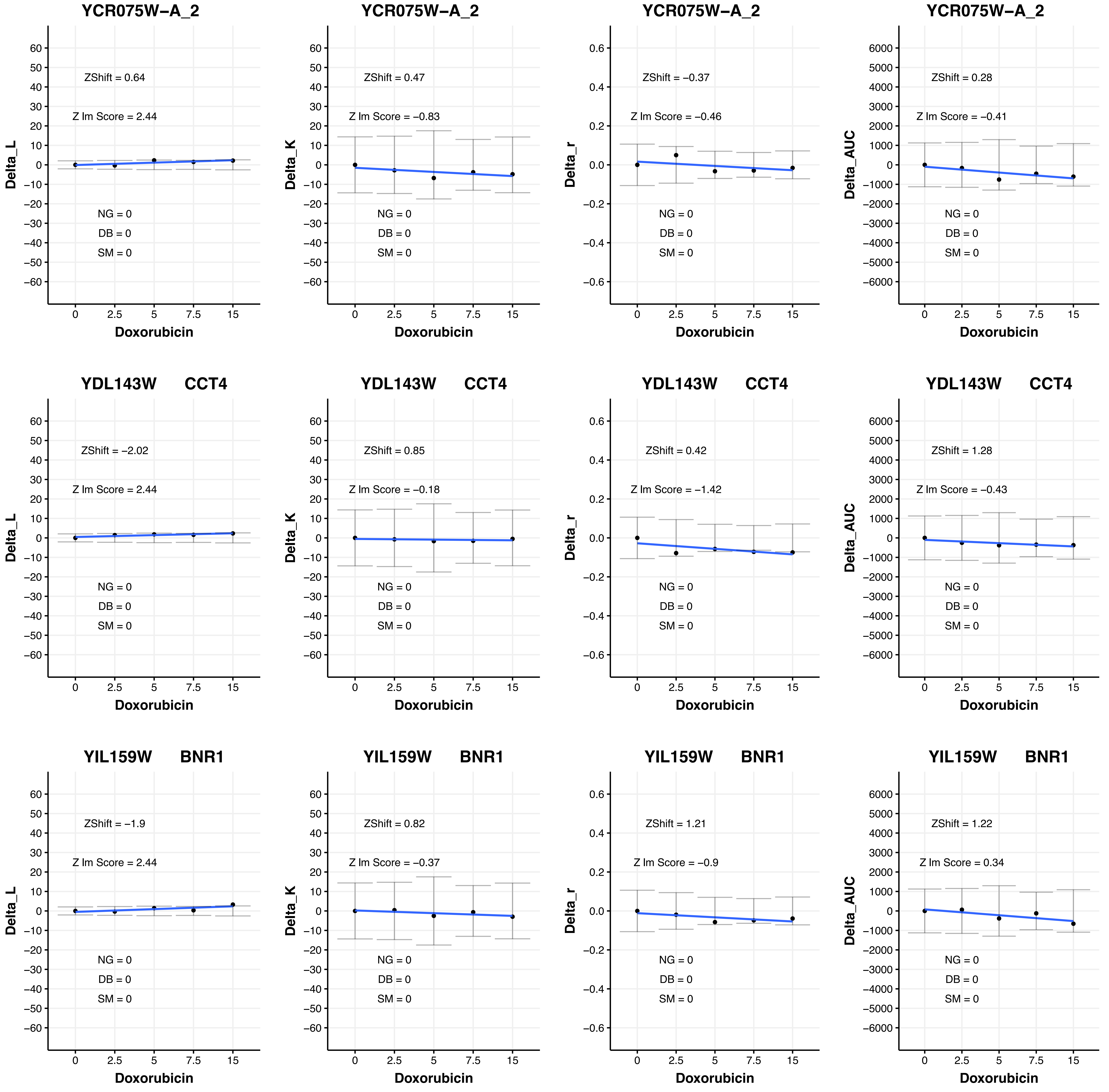
YCR075W−A_2
ZShift = 0.64
Z lm Score = 2.44
NG = 0
DB = 0
SM = 0
Delta_L
Doxorubicin
YCR075W−A_2
ZShift = 0.47
Z lm Score = −0.83
NG = 0
DB = 0
SM = 0
Delta_K
Doxorubicin
YCR075W−A_2
ZShift = −0.37
Z lm Score = −0.46
NG = 0
DB = 0
SM = 0
Delta_r
Doxorubicin
YCR075W−A_2
ZShift = 0.28
Z lm Score = −0.41
NG = 0
DB = 0
SM = 0
Delta_AUC
Doxorubicin
YDL143W CCT4
ZShift = −2.02
Z lm Score = 2.44
NG = 0
DB = 0
SM = 0
Delta_L
Doxorubicin
YDL143W CCT4
ZShift = 0.85
Z lm Score = −0.18
NG = 0
DB = 0
SM = 0
Delta_K
Doxorubicin
YDL143W CCT4
ZShift = 0.42
Z lm Score = −1.42
NG = 0
DB = 0
SM = 0
Delta_r
Doxorubicin
YDL143W CCT4
ZShift = 1.28
Z lm Score = −0.43
NG = 0
DB = 0
SM = 0
Delta_AUC
Doxorubicin
YIL159W BNR1
ZShift = −1.9
Z lm Score = 2.44
NG = 0
DB = 0
SM = 0
Delta_L
Doxorubicin
YIL159W BNR1
ZShift = 0.82
Z lm Score = −0.37
NG = 0
DB = 0
SM = 0
Delta_K
Doxorubicin
YIL159W BNR1
ZShift = 1.21
Z lm Score = −0.9
NG = 0
DB = 0
SM = 0
Delta_r
Doxorubicin
YIL159W BNR1
ZShift = 1.22
Z lm Score = 0.34
NG = 0
DB = 0
SM = 0
Delta_AUC
Doxorubicin

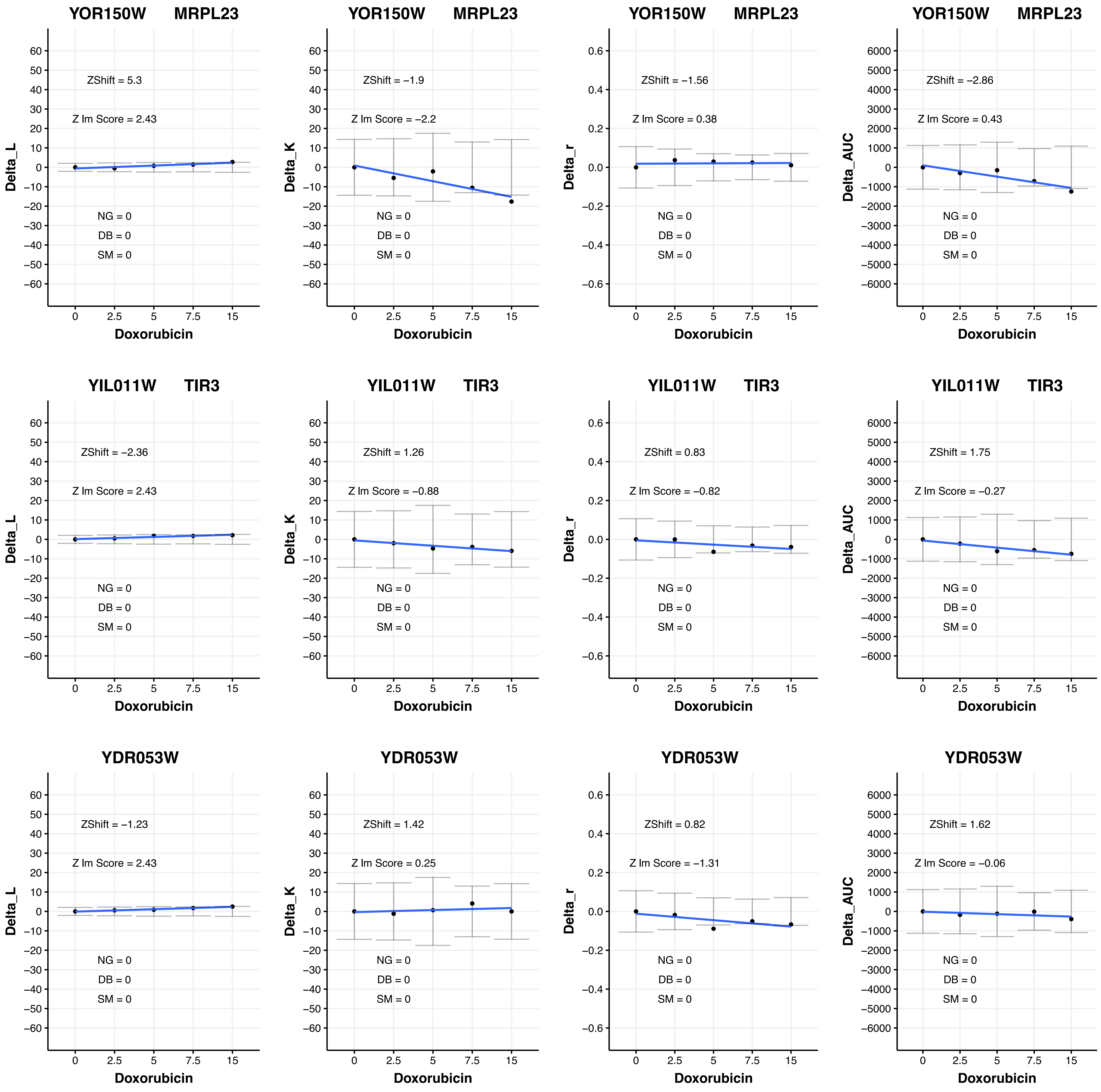
YOR150W MRPL23
ZShift = 5.3
Z lm Score = 2.43
NG = 0
DB = 0
SM = 0
Delta_L
Doxorubicin
YOR150W MRPL23
ZShift = −1.9
Z lm Score = −2.2
NG = 0
DB = 0
SM = 0
Delta_K
Doxorubicin
YOR150W MRPL23
ZShift = −1.56
Z lm Score = 0.38
NG = 0
DB = 0
SM = 0
Delta_r
Doxorubicin
YOR150W MRPL23
ZShift = −2.86
Z lm Score = 0.43
NG = 0
DB = 0
SM = 0
Delta_AUC
Doxorubicin
YIL011W TIR3
ZShift = −2.36
Z lm Score = 2.43
NG = 0
DB = 0
SM = 0
Delta_L
Doxorubicin
YIL011W TIR3
ZShift = 1.26
Z lm Score = −0.88
NG = 0
DB = 0
SM = 0
Delta_K
Doxorubicin
YIL011W TIR3
ZShift = 0.83
Z lm Score = −0.82
NG = 0
DB = 0
SM = 0
Delta_r
Doxorubicin
YIL011W TIR3
ZShift = 1.75
Z lm Score = −0.27
NG = 0
DB = 0
SM = 0
Delta_AUC
Doxorubicin
YDR053W
ZShift = −1.23
Z lm Score = 2.43
NG = 0
DB = 0
SM = 0
Delta_L
Doxorubicin
YDR053W
ZShift = 1.42
Z lm Score = 0.25
NG = 0
DB = 0
SM = 0
Delta_K
Doxorubicin
YDR053W
ZShift = 0.82
Z lm Score = −1.31
NG = 0
DB = 0
SM = 0
Delta_r
Doxorubicin
YDR053W
ZShift = 1.62
Z lm Score = −0.06
NG = 0
DB = 0
SM = 0
Delta_AUC
Doxorubicin

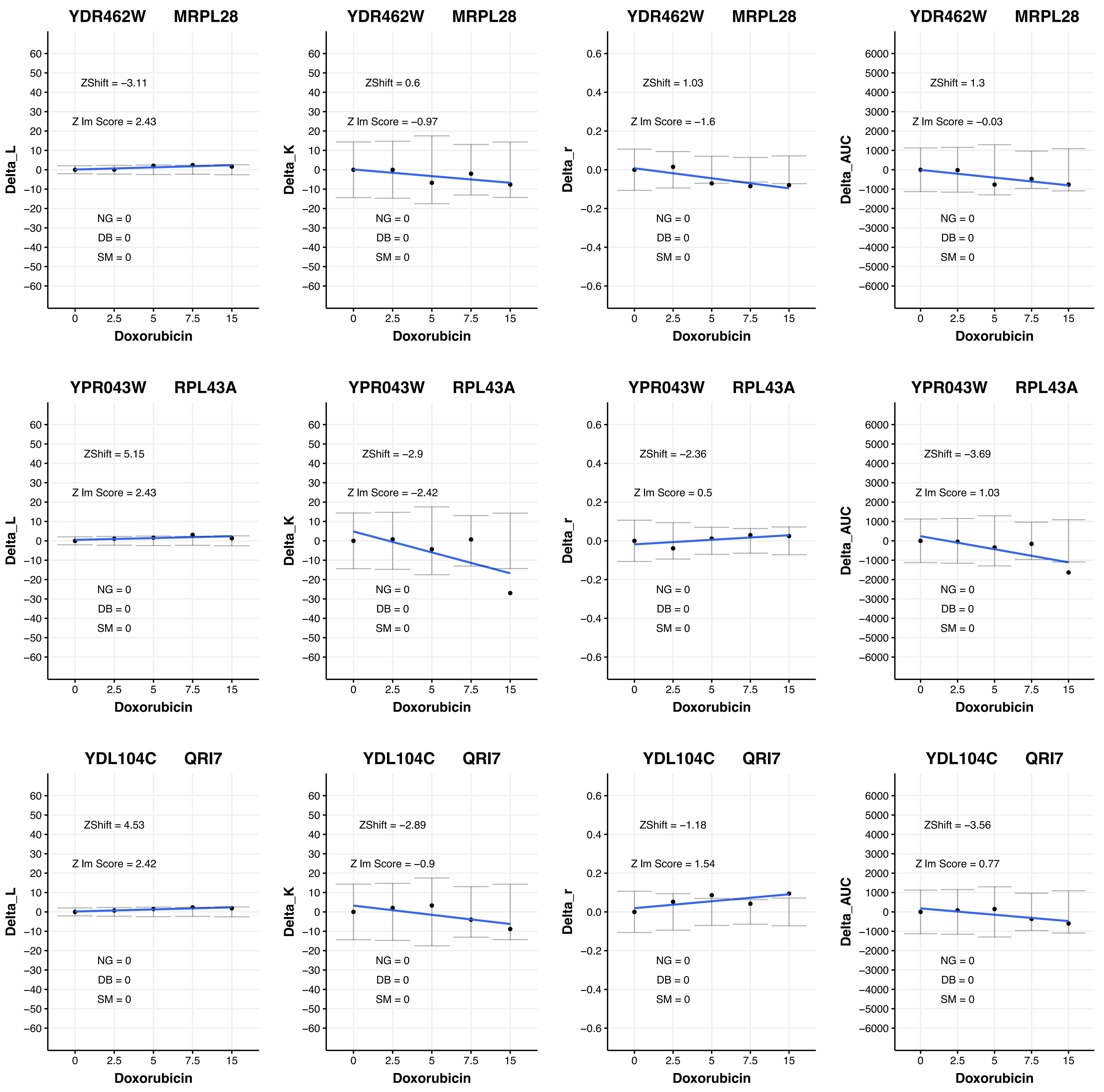
YDR462W MRPL28
ZShift = −3.11
Z lm Score = 2.43
NG = 0
DB = 0
SM = 0
Delta_L
Doxorubicin
YDR462W MRPL28
ZShift = 0.6
Z lm Score = −0.97
NG = 0
DB = 0
SM = 0
Delta_K
Doxorubicin
YDR462W MRPL28
ZShift = 1.03
Z lm Score = −1.6
NG = 0
DB = 0
SM = 0
Delta_r
Doxorubicin
YDR462W MRPL28
ZShift = 1.3
Z lm Score = −0.03
NG = 0
DB = 0
SM = 0
Delta_AUC
Doxorubicin
YPR043W RPL43A
ZShift = 5.15
Z lm Score = 2.43
NG = 0
DB = 0
SM = 0
Delta_L
Doxorubicin
YPR043W RPL43A
ZShift = −2.9
Z lm Score = −2.42
NG = 0
DB = 0
SM = 0
Delta_K
Doxorubicin
YPR043W RPL43A
ZShift = −2.36
Z lm Score = 0.5
NG = 0
DB = 0
SM = 0
Delta_r
Doxorubicin
YPR043W RPL43A
ZShift = −3.69
Z lm Score = 1.03
NG = 0
DB = 0
SM = 0
Delta_AUC
Doxorubicin
YDL104C QRI7
ZShift = 4.53
Z lm Score = 2.42
NG = 0
DB = 0
SM = 0
Delta_L
Doxorubicin
YDL104C QRI7
ZShift = −2.89
Z lm Score = −0.9
NG = 0
DB = 0
SM = 0
Delta_K
Doxorubicin
YDL104C QRI7
ZShift = −1.18
Z lm Score = 1.54
NG = 0
DB = 0
SM = 0
Delta_r
Doxorubicin
YDL104C QRI7
ZShift = −3.56
Z lm Score = 0.77
NG = 0
DB = 0
SM = 0
Delta_AUC
Doxorubicin

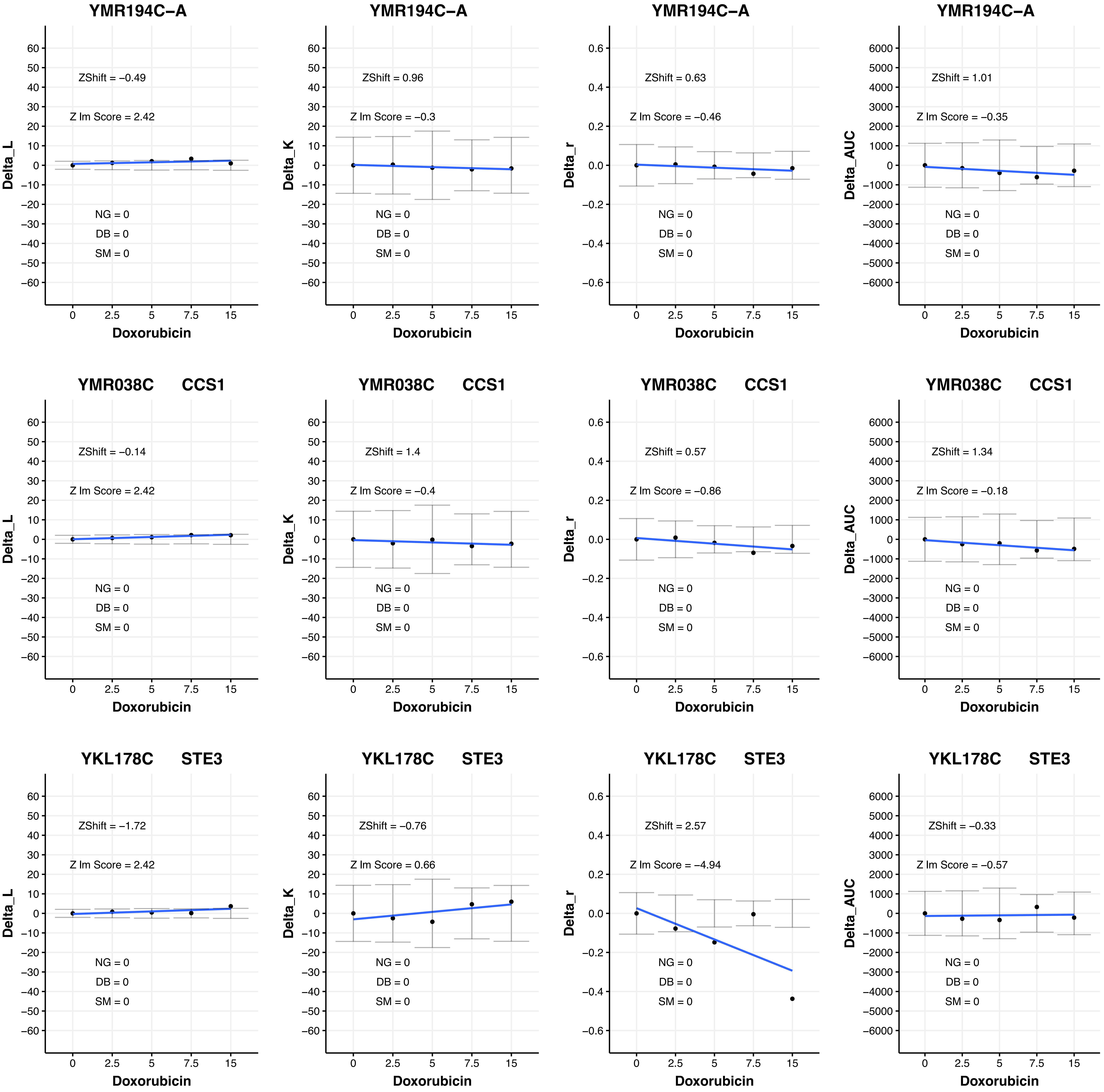
YMR194C−A
Delta_L
Doxorubicin
ZShift = −0.49
Z lm Score = 2.42
NG = 0
DB = 0
SM = 0
YMR194C−A
Delta_K
Doxorubicin
ZShift = 0.96
Z lm Score = −0.3
NG = 0
DB = 0
SM = 0
YMR194C−A
Delta_r
Doxorubicin
ZShift = 0.63
Z lm Score = −0.46
NG = 0
DB = 0
SM = 0
YMR194C−A
Delta_AUC
Doxorubicin
ZShift = 1.01
Z lm Score = −0.35
NG = 0
DB = 0
SM = 0
YMR038C CCS1
Delta_L
Doxorubicin
ZShift = −0.14
Z lm Score = 2.42
NG = 0
DB = 0
SM = 0
YMR038C CCS1
Delta_K
Doxorubicin
ZShift = 1.4
Z lm Score = −0.4
NG = 0
DB = 0
SM = 0
YMR038C CCS1
Delta_r
Doxorubicin
ZShift = 0.57
Z lm Score = −0.86
NG = 0
DB = 0
SM = 0
YMR038C CCS1
Delta_AUC
Doxorubicin
ZShift = 1.34
Z lm Score = −0.18
NG = 0
DB = 0
SM = 0
YKL178C STE3
Delta_L
Doxorubicin
ZShift = −1.72
Z lm Score = 2.42
NG = 0
DB = 0
SM = 0
YKL178C STE3
Delta_K
Doxorubicin
ZShift = −0.76
Z lm Score = 0.66
NG = 0
DB = 0
SM = 0
YKL178C STE3
Delta_r
Doxorubicin
ZShift = 2.57
Z lm Score = −4.94
NG = 0
DB = 0
SM = 0
YKL178C STE3
Delta_AUC
Doxorubicin
ZShift = −0.33
Z lm Score = −0.57
NG = 0
DB = 0
SM = 0

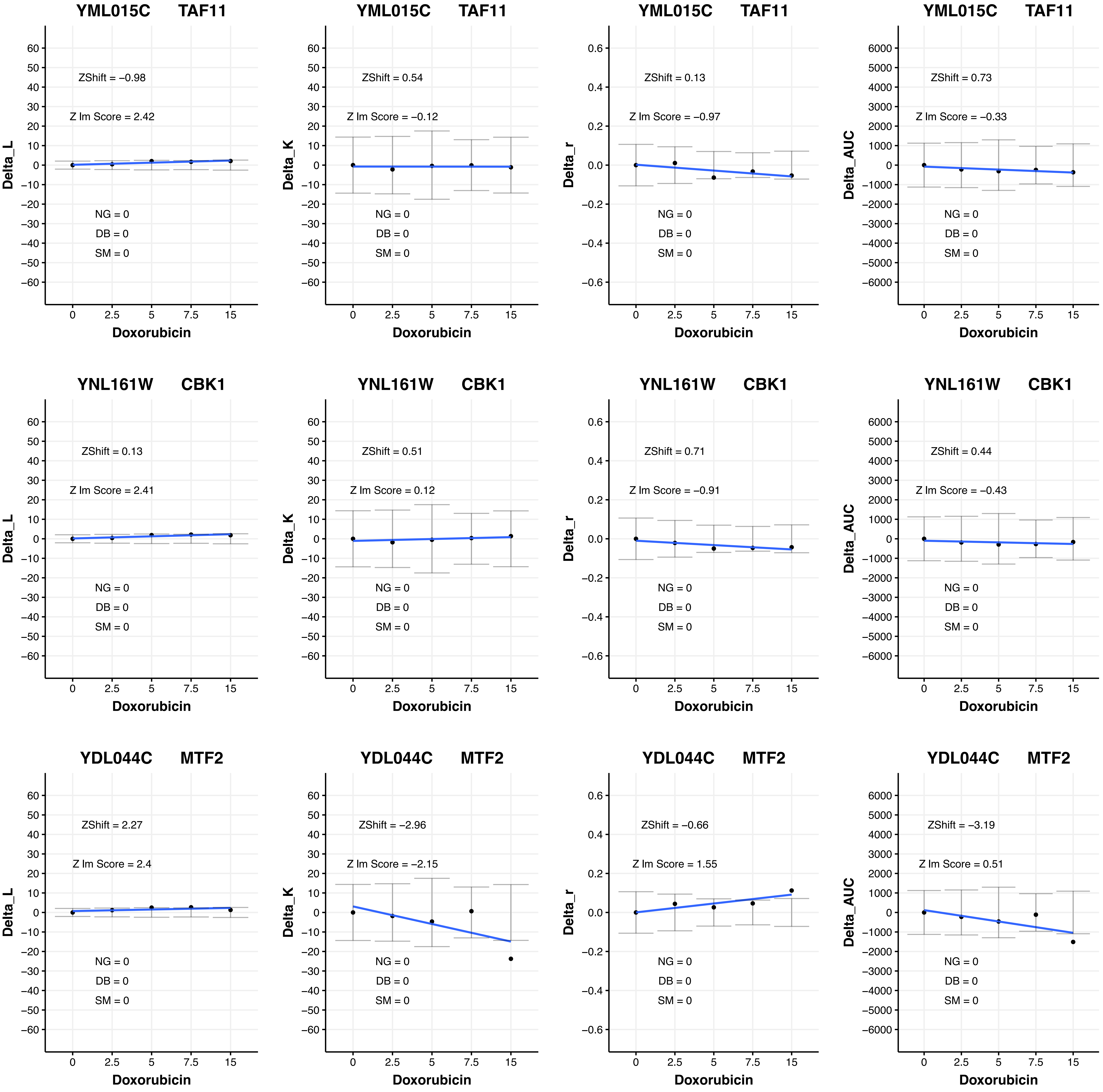
YML015C TAF11
ZShift = −0.98
Z lm Score = 2.42
NG = 0
DB = 0
SM = 0
Delta_L
Doxorubicin
YML015C TAF11
ZShift = 0.54
Z lm Score = −0.12
NG = 0
DB = 0
SM = 0
Delta_K
Doxorubicin
YML015C TAF11
ZShift = 0.13
Z lm Score = −0.97
NG = 0
DB = 0
SM = 0
Delta_r
Doxorubicin
YML015C TAF11
ZShift = 0.73
Z lm Score = −0.33
NG = 0
DB = 0
SM = 0
Delta_AUC
Doxorubicin
YNL161W CBK1
ZShift = 0.13
Z lm Score = 2.41
NG = 0
DB = 0
SM = 0
Delta_L
Doxorubicin
YNL161W CBK1
ZShift = 0.51
Z lm Score = 0.12
NG = 0
DB = 0
SM = 0
Delta_K
Doxorubicin
YNL161W CBK1
ZShift = 0.71
Z lm Score = −0.91
NG = 0
DB = 0
SM = 0
Delta_r
Doxorubicin
YNL161W CBK1
ZShift = 0.44
Z lm Score = −0.43
NG = 0
DB = 0
SM = 0
Delta_AUC
Doxorubicin
YDL044C MTF2
ZShift = 2.27
Z lm Score = 2.4
NG = 0
DB = 0
SM = 0
Delta_L
Doxorubicin
YDL044C MTF2
ZShift = −2.96
Z lm Score = −2.15
NG = 0
DB = 0
SM = 0
Delta_K
Doxorubicin
YDL044C MTF2
ZShift = −0.66
Z lm Score = 1.55
NG = 0
DB = 0
SM = 0
Delta_r
Doxorubicin
YDL044C MTF2
ZShift = −3.19
Z lm Score = 0.51
NG = 0
DB = 0
SM = 0
Delta_AUC
Doxorubicin

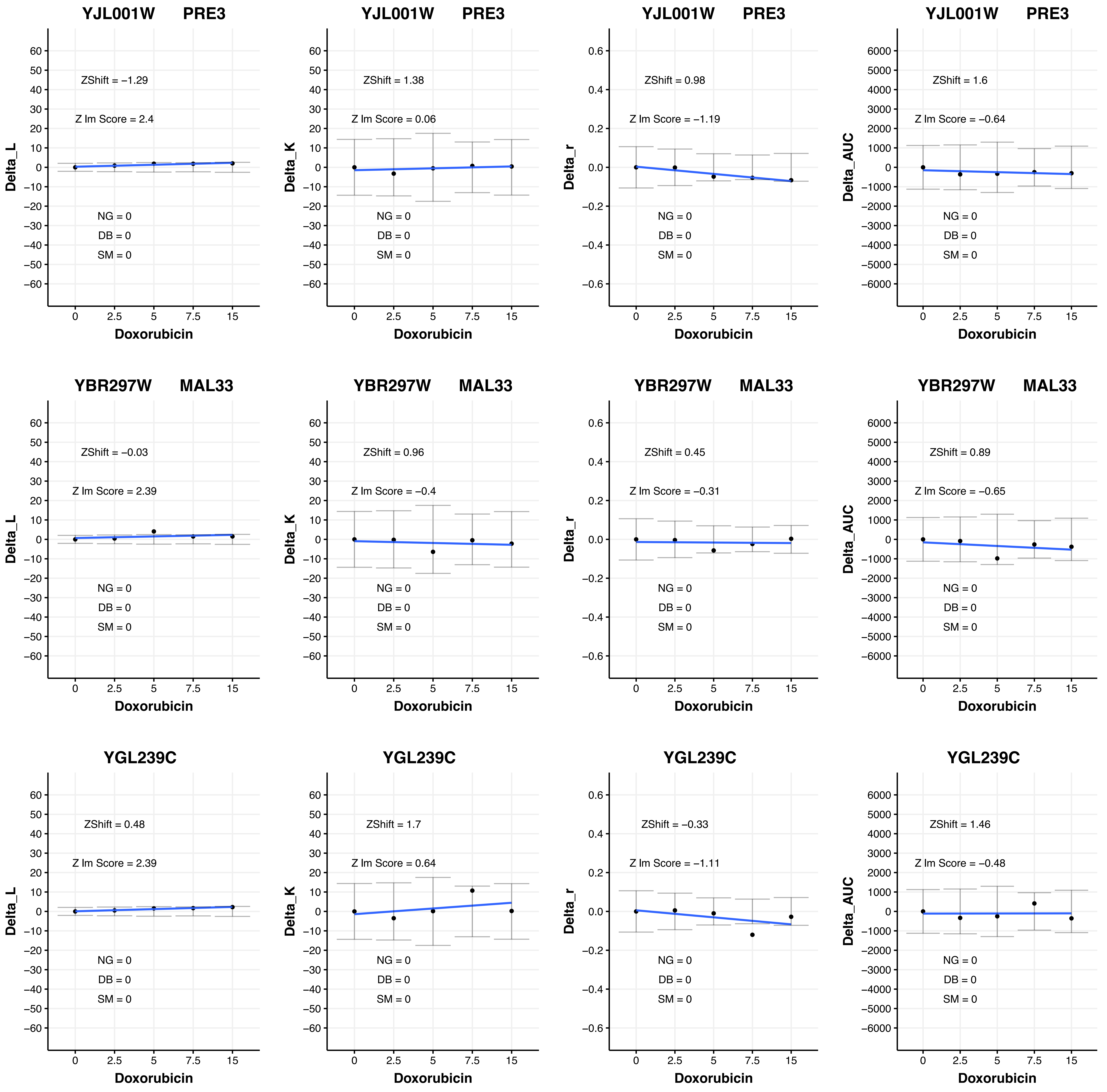
YJL001W PRE3
ZShift = −1.29
Z lm Score = 2.4
NG = 0
DB = 0
SM = 0
Delta_L
Doxorubicin
YJL001W PRE3
ZShift = 1.38
Z lm Score = 0.06
NG = 0
DB = 0
SM = 0
Delta_K
Doxorubicin
YJL001W PRE3
ZShift = 0.98
Z lm Score = −1.19
NG = 0
DB = 0
SM = 0
Delta_r
Doxorubicin
YJL001W PRE3
ZShift = 1.6
Z lm Score = −0.64
NG = 0
DB = 0
SM = 0
Delta_AUC
Doxorubicin
YBR297W MAL33
ZShift = −0.03
Z lm Score = 2.39
NG = 0
DB = 0
SM = 0
Delta_L
Doxorubicin
YBR297W MAL33
ZShift = 0.96
Z lm Score = −0.4
NG = 0
DB = 0
SM = 0
Delta_K
Doxorubicin
YBR297W MAL33
ZShift = 0.45
Z lm Score = −0.31
NG = 0
DB = 0
SM = 0
Delta_r
Doxorubicin
YBR297W MAL33
ZShift = 0.89
Z lm Score = −0.65
NG = 0
DB = 0
SM = 0
Delta_AUC
Doxorubicin
YGL239C
ZShift = 0.48
Z lm Score = 2.39
NG = 0
DB = 0
SM = 0
Delta_L
Doxorubicin
YGL239C
ZShift = 1.7
Z lm Score = 0.64
NG = 0
DB = 0
SM = 0
Delta_K
Doxorubicin
YGL239C
ZShift = −0.33
Z lm Score = −1.11
NG = 0
DB = 0
SM = 0
Delta_r
Doxorubicin
YGL239C
ZShift = 1.46
Z lm Score = −0.48
NG = 0
DB = 0
SM = 0
Delta_AUC
Doxorubicin

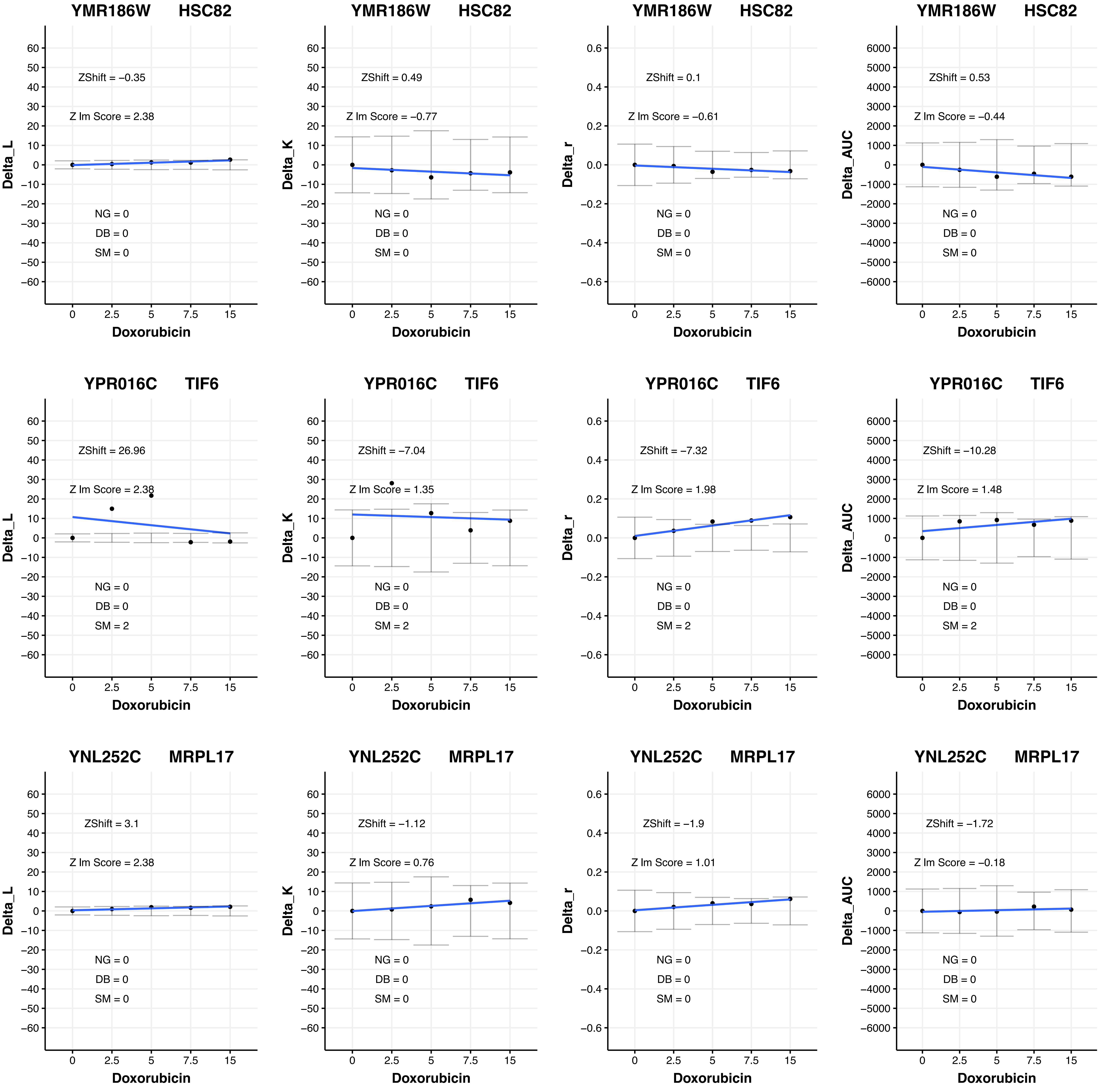
YMR186W HSC82
ZShift = −0.35
Z lm Score = 2.38
NG = 0
DB = 0
SM = 0
Delta_L
Doxorubicin
YMR186W HSC82
ZShift = 0.49
Z lm Score = −0.77
NG = 0
DB = 0
SM = 0
Delta_K
Doxorubicin
YMR186W HSC82
ZShift = 0.1
Z lm Score = −0.61
NG = 0
DB = 0
SM = 0
Delta_r
Doxorubicin
YMR186W HSC82
ZShift = 0.53
Z lm Score = −0.44
NG = 0
DB = 0
SM = 0
Delta_AUC
Doxorubicin
YPR016C TIF6
ZShift = 26.96
Z lm Score = 2.38
NG = 0
DB = 0
SM = 2
Delta_L
Doxorubicin
YPR016C TIF6
ZShift = −7.04
Z lm Score = 1.35
NG = 0
DB = 0
SM = 2
Delta_K
Doxorubicin
YPR016C TIF6
ZShift = −7.32
Z lm Score = 1.98
NG = 0
DB = 0
SM = 2
Delta_r
Doxorubicin
YPR016C TIF6
ZShift = −10.28
Z lm Score = 1.48
NG = 0
DB = 0
SM = 2
Delta_AUC
Doxorubicin
YNL252C MRPL17
ZShift = 3.1
Z lm Score = 2.38
NG = 0
DB = 0
SM = 0
Delta_L
Doxorubicin
YNL252C MRPL17
ZShift = −1.12
Z lm Score = 0.76
NG = 0
DB = 0
SM = 0
Delta_K
Doxorubicin
YNL252C MRPL17
ZShift = −1.9
Z lm Score = 1.01
NG = 0
DB = 0
SM = 0
Delta_r
Doxorubicin
YNL252C MRPL17
ZShift = −1.72
Z lm Score = −0.18
NG = 0
DB = 0
SM = 0
Delta_AUC
Doxorubicin

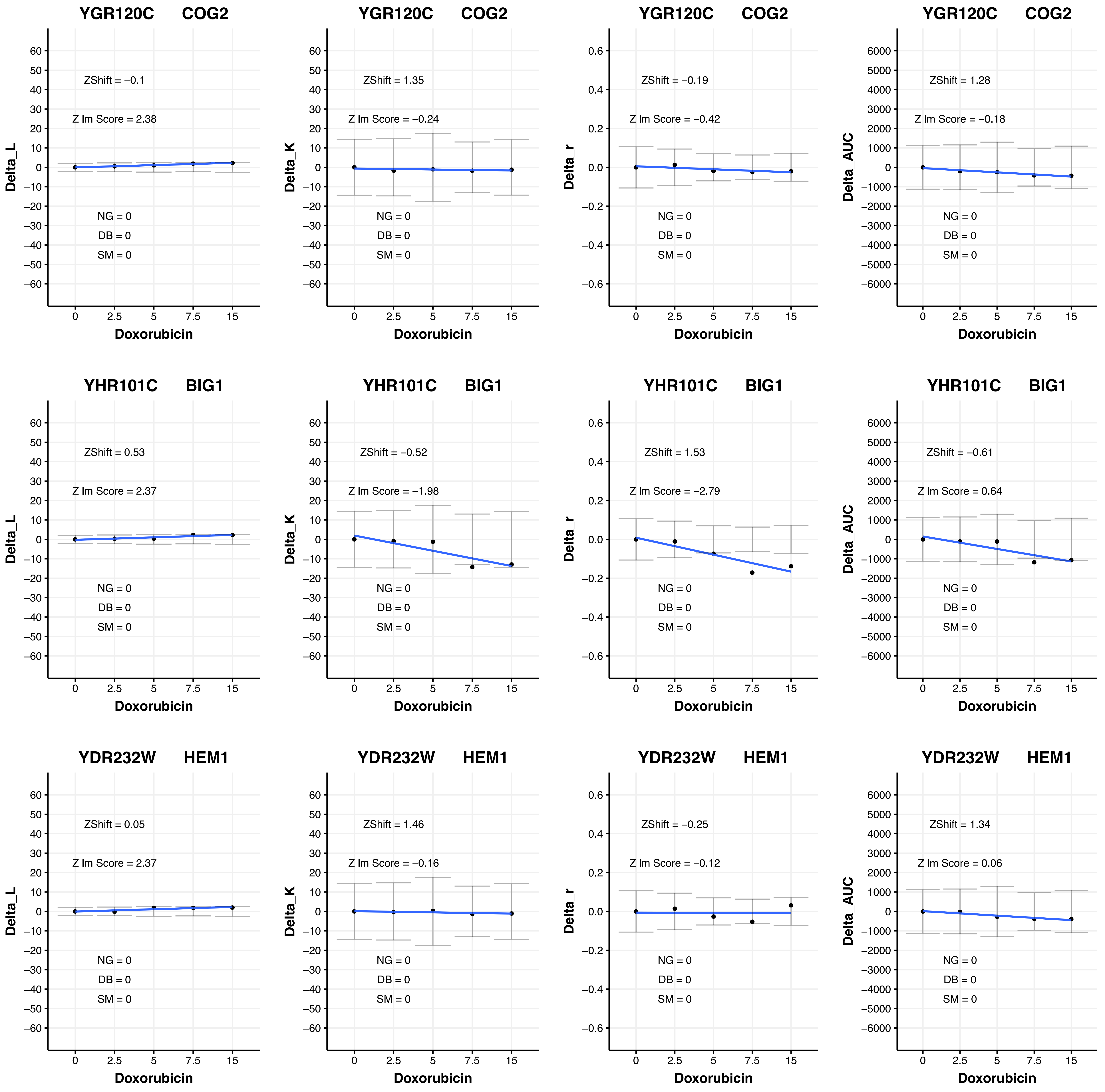
YGR120C COG2
ZShift = −0.1
Z lm Score = 2.38
NG = 0
DB = 0
SM = 0
Delta_L
Doxorubicin
YGR120C COG2
ZShift = 1.35
Z lm Score = −0.24
NG = 0
DB = 0
SM = 0
Delta_K
Doxorubicin
YGR120C COG2
ZShift = −0.19
Z lm Score = −0.42
NG = 0
DB = 0
SM = 0
Delta_r
Doxorubicin
YGR120C COG2
ZShift = 1.28
Z lm Score = −0.18
NG = 0
DB = 0
SM = 0
Delta_AUC
Doxorubicin
YHR101C BIG1
ZShift = 0.53
Z lm Score = 2.37
NG = 0
DB = 0
SM = 0
Delta_L
Doxorubicin
YHR101C BIG1
ZShift = −0.52
Z lm Score = −1.98
NG = 0
DB = 0
SM = 0
Delta_K
Doxorubicin
YHR101C BIG1
ZShift = 1.53
Z lm Score = −2.79
NG = 0
DB = 0
SM = 0
Delta_r
Doxorubicin
YHR101C BIG1
ZShift = −0.61
Z lm Score = 0.64
NG = 0
DB = 0
SM = 0
Delta_AUC
Doxorubicin
YDR232W HEM1
ZShift = 0.05
Z lm Score = 2.37
NG = 0
DB = 0
SM = 0
Delta_L
Doxorubicin
YDR232W HEM1
ZShift = 1.46
Z lm Score = −0.16
NG = 0
DB = 0
SM = 0
Delta_K
Doxorubicin
YDR232W HEM1
ZShift = −0.25
Z lm Score = −0.12
NG = 0
DB = 0
SM = 0
Delta_r
Doxorubicin
YDR232W HEM1
ZShift = 1.34
Z lm Score = 0.06
NG = 0
DB = 0
SM = 0
Delta_AUC
Doxorubicin

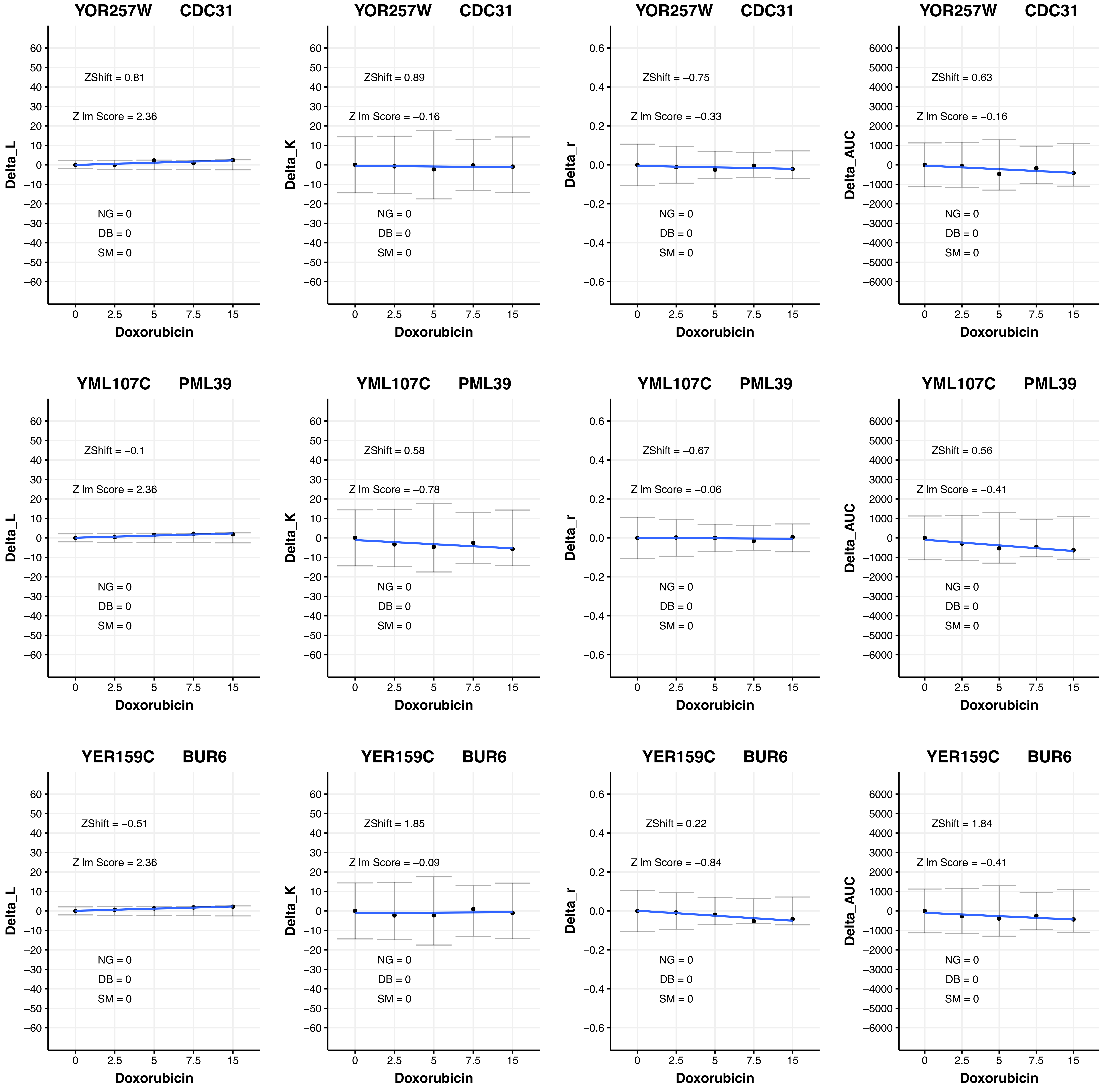

YOR257W CDC31
ZShift = 0.81
Z lm Score = 2.36
NG = 0
DB = 0
SM = 0
Delta_L
Doxorubicin
YOR257W CDC31
ZShift = 0.89
Z lm Score = −0.16
NG = 0
DB = 0
SM = 0
Delta_K
Doxorubicin
YOR257W CDC31
ZShift = −0.75
Z lm Score = −0.33
NG = 0
DB = 0
SM = 0
Delta_r
Doxorubicin
YOR257W CDC31
ZShift = 0.63
Z lm Score = −0.16
NG = 0
DB = 0
SM = 0
Delta_AUC
Doxorubicin
YML107C PML39
ZShift = −0.1
Z lm Score = 2.36
NG = 0
DB = 0
SM = 0
Delta_L
Doxorubicin
YML107C PML39
ZShift = 0.58
Z lm Score = −0.78
NG = 0
DB = 0
SM = 0
Delta_K
Doxorubicin
YML107C PML39
ZShift = −0.67
Z lm Score = −0.06
NG = 0
DB = 0
SM = 0
Delta_r
Doxorubicin
YML107C PML39
ZShift = 0.56
Z lm Score = −0.41
NG = 0
DB = 0
SM = 0
Delta_AUC
Doxorubicin
YER159C BUR6
ZShift = −0.51
Z lm Score = 2.36
NG = 0
DB = 0
SM = 0
Delta_L
Doxorubicin
YER159C BUR6
ZShift = 1.85
Z lm Score = −0.09
NG = 0
DB = 0
SM = 0
Delta_K
Doxorubicin
YER159C BUR6
ZShift = 0.22
Z lm Score = −0.84
NG = 0
DB = 0
SM = 0
Delta_r
Doxorubicin
YER159C BUR6
ZShift = 1.84
Z lm Score = −0.41
NG = 0
DB = 0
SM = 0
Delta_AUC
Doxorubicin

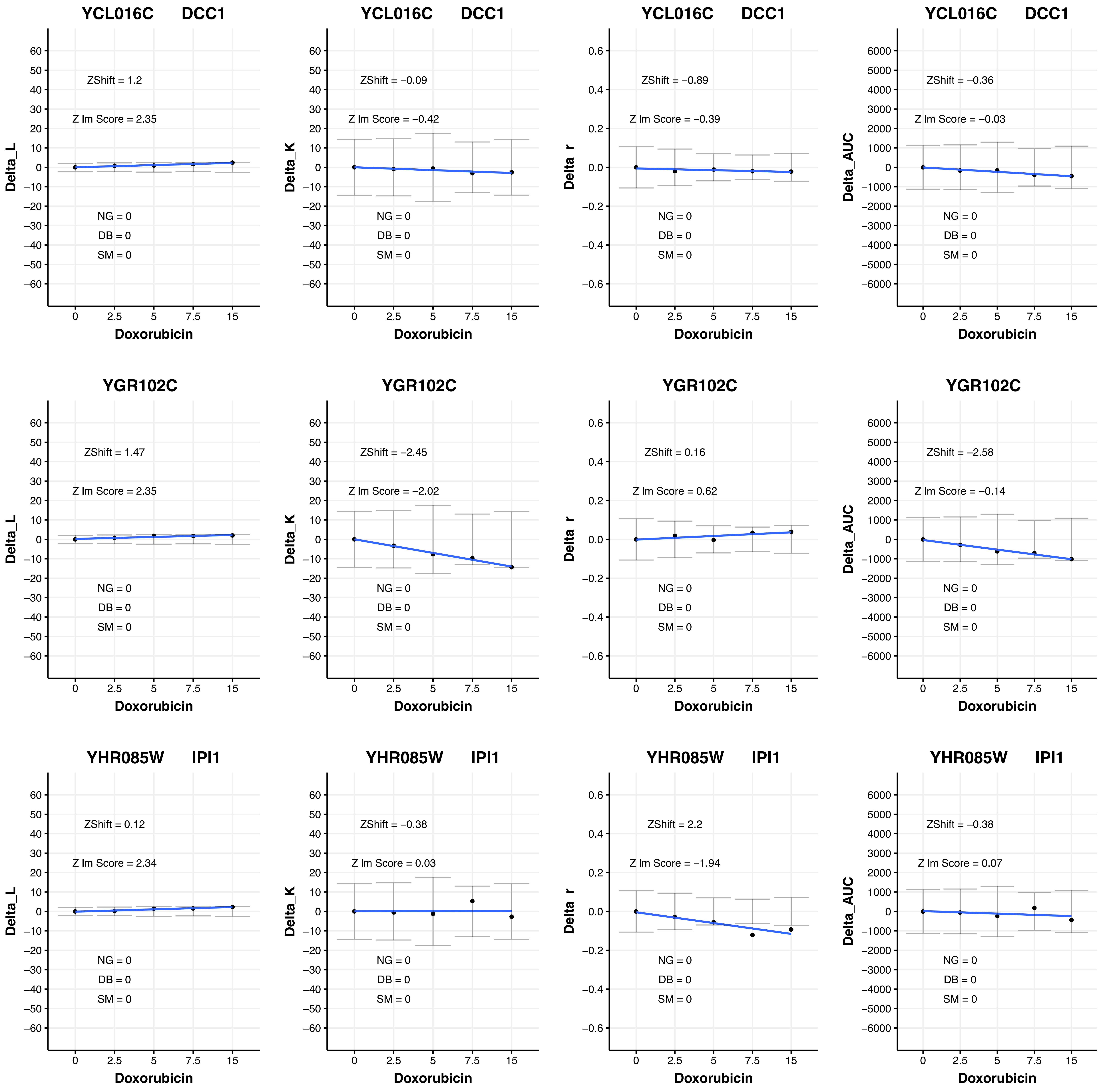

YCL016C DCC1
ZShift = 1.2
Z lm Score = 2.35
NG = 0
DB = 0
SM = 0
Delta_L
Doxorubicin
YCL016C DCC1
ZShift = −0.09
Z lm Score = −0.42
NG = 0
DB = 0
SM = 0
Delta_K
Doxorubicin
YCL016C DCC1
ZShift = −0.89
Z lm Score = −0.39
NG = 0
DB = 0
SM = 0
Delta_r
Doxorubicin
YCL016C DCC1
ZShift = −0.36
Z lm Score = −0.03
NG = 0
DB = 0
SM = 0
Delta_AUC
Doxorubicin
YGR102C
ZShift = 1.47
Z lm Score = 2.35
NG = 0
DB = 0
SM = 0
Delta_L
Doxorubicin
YGR102C
ZShift = −2.45
Z lm Score = −2.02
NG = 0
DB = 0
SM = 0
Delta_K
Doxorubicin
YGR102C
ZShift = 0.16
Z lm Score = 0.62
NG = 0
DB = 0
SM = 0
Delta_r
Doxorubicin
YGR102C
ZShift = −2.58
Z lm Score = −0.14
NG = 0
DB = 0
SM = 0
Delta_AUC
Doxorubicin
YHR085W IPI1
ZShift = 0.12
Z lm Score = 2.34
NG = 0
DB = 0
SM = 0
Delta_L
Doxorubicin
YHR085W IPI1
ZShift = −0.38
Z lm Score = 0.03
NG = 0
DB = 0
SM = 0
Delta_K
Doxorubicin
YHR085W IPI1
ZShift = 2.2
Z lm Score = −1.94
NG = 0
DB = 0
SM = 0
Delta_r
Doxorubicin
YHR085W IPI1
ZShift = −0.38
Z lm Score = 0.07
NG = 0
DB = 0
SM = 0
Delta_AUC
Doxorubicin

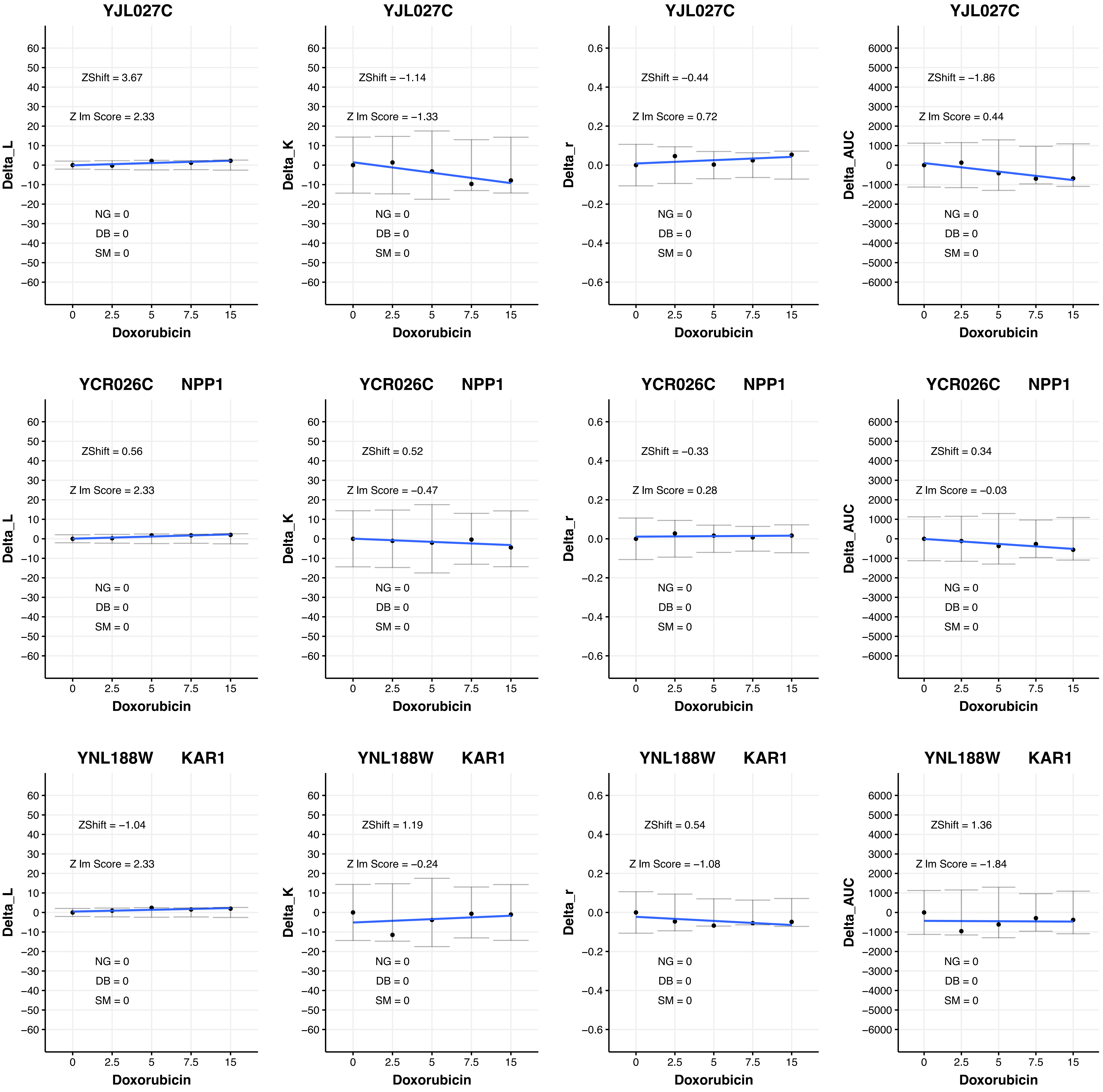

YJL027C
ZShift = 3.67
Z lm Score = 2.33
NG = 0
DB = 0
SM = 0
Delta_L
Doxorubicin
YJL027C
ZShift = −1.14
Z lm Score = −1.33
NG = 0
DB = 0
SM = 0
Delta_K
Doxorubicin
YJL027C
ZShift = −0.44
Z lm Score = 0.72
NG = 0
DB = 0
SM = 0
Delta_r
Doxorubicin
YJL027C
ZShift = −1.86
Z lm Score = 0.44
NG = 0
DB = 0
SM = 0
Delta_AUC
Doxorubicin
YCR026C NPP1
ZShift = 0.56
Z lm Score = 2.33
NG = 0
DB = 0
SM = 0
Delta_L
Doxorubicin
YCR026C NPP1
ZShift = 0.52
Z lm Score = −0.47
NG = 0
DB = 0
SM = 0
Delta_K
Doxorubicin
YCR026C NPP1
ZShift = −0.33
Z lm Score = 0.28
NG = 0
DB = 0
SM = 0
Delta_r
Doxorubicin
YCR026C NPP1
ZShift = 0.34
Z lm Score = −0.03
NG = 0
DB = 0
SM = 0
Delta_AUC
Doxorubicin
YNL188W KAR1
ZShift = −1.04
Z lm Score = 2.33
NG = 0
DB = 0
SM = 0
Delta_L
Doxorubicin
YNL188W KAR1
ZShift = 1.19
Z lm Score = −0.24
NG = 0
DB = 0
SM = 0
Delta_K
Doxorubicin
YNL188W KAR1
ZShift = 0.54
Z lm Score = −1.08
NG = 0
DB = 0
SM = 0
Delta_r
Doxorubicin
YNL188W KAR1
ZShift = 1.36
Z lm Score = −1.84
NG = 0
DB = 0
SM = 0
Delta_AUC
Doxorubicin

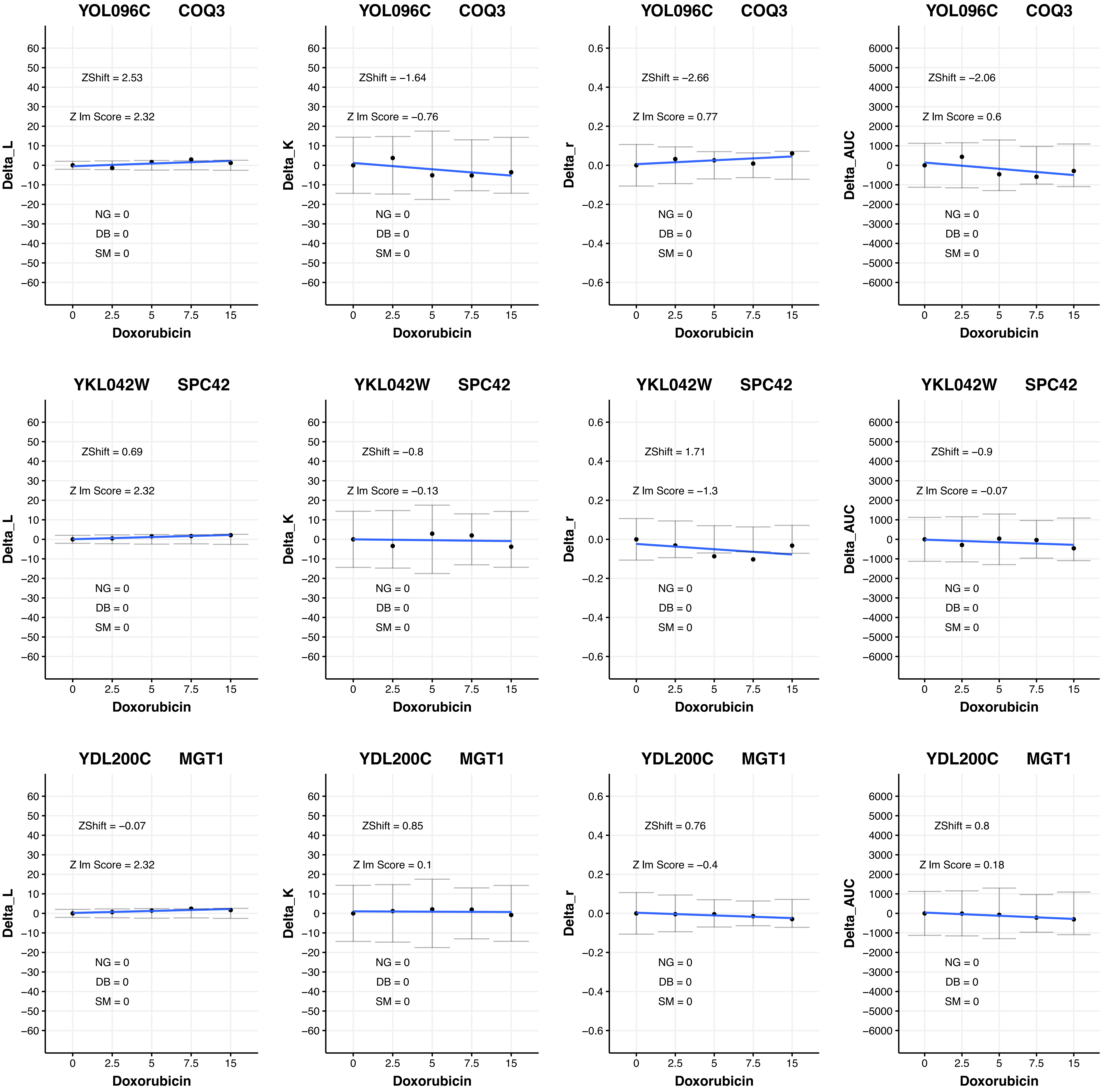
YOL096C COQ3
ZShift = 2.53
Z lm Score = 2.32
NG = 0
DB = 0
SM = 0
Delta_L
Doxorubicin
YOL096C COQ3
ZShift = −1.64
Z lm Score = −0.76
NG = 0
DB = 0
SM = 0
Delta_K
Doxorubicin
YOL096C COQ3
ZShift = −2.66
Z lm Score = 0.77
NG = 0
DB = 0
SM = 0
Delta_r
Doxorubicin
YOL096C COQ3
ZShift = −2.06
Z lm Score = 0.6
NG = 0
DB = 0
SM = 0
Delta_AUC
Doxorubicin
YKL042W SPC42
ZShift = 0.69
Z lm Score = 2.32
NG = 0
DB = 0
SM = 0
Delta_L
Doxorubicin
YKL042W SPC42
ZShift = −0.8
Z lm Score = −0.13
NG = 0
DB = 0
SM = 0
Delta_K
Doxorubicin
YKL042W SPC42
ZShift = 1.71
Z lm Score = −1.3
NG = 0
DB = 0
SM = 0
Delta_r
Doxorubicin
YKL042W SPC42
ZShift = −0.9
Z lm Score = −0.07
NG = 0
DB = 0
SM = 0
Delta_AUC
Doxorubicin
YDL200C MGT1
ZShift = −0.07
Z lm Score = 2.32
NG = 0
DB = 0
SM = 0
Delta_L
Doxorubicin
YDL200C MGT1
ZShift = 0.85
Z lm Score = 0.1
NG = 0
DB = 0
SM = 0
Delta_K
Doxorubicin
YDL200C MGT1
ZShift = 0.76
Z lm Score = −0.4
NG = 0
DB = 0
SM = 0
Delta_r
Doxorubicin
YDL200C MGT1
ZShift = 0.8
Z lm Score = 0.18
NG = 0
DB = 0
SM = 0
Delta_AUC
Doxorubicin

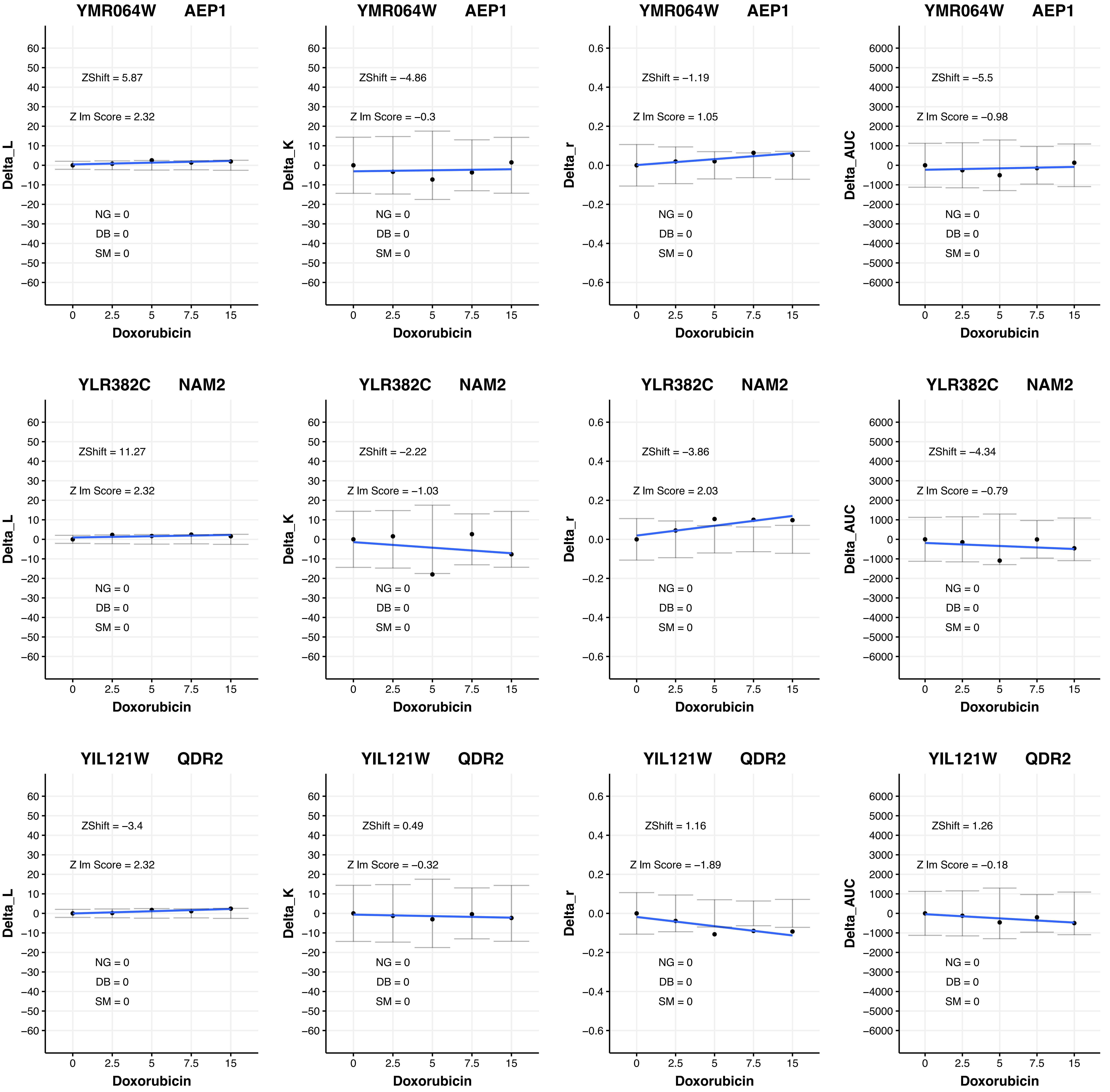
YMR064W AEP1
ZShift = 5.87
Z lm Score = 2.32
NG = 0
DB = 0
SM = 0
Delta_L
Doxorubicin
YMR064W AEP1
ZShift = −4.86
Z lm Score = −0.3
NG = 0
DB = 0
SM = 0
Delta_K
Doxorubicin
YMR064W AEP1
ZShift = −1.19
Z lm Score = 1.05
NG = 0
DB = 0
SM = 0
Delta_r
Doxorubicin
YMR064W AEP1
ZShift = −5.5
Z lm Score = −0.98
NG = 0
DB = 0
SM = 0
Delta_AUC
Doxorubicin
YLR382C NAM2
ZShift = 11.27
Z lm Score = 2.32
NG = 0
DB = 0
SM = 0
Delta_L
Doxorubicin
YLR382C NAM2
ZShift = −2.22
Z lm Score = −1.03
NG = 0
DB = 0
SM = 0
Delta_K
Doxorubicin
YLR382C NAM2
ZShift = −3.86
Z lm Score = 2.03
NG = 0
DB = 0
SM = 0
Delta_r
Doxorubicin
YLR382C NAM2
ZShift = −4.34
Z lm Score = −0.79
NG = 0
DB = 0
SM = 0
Delta_AUC
Doxorubicin
YIL121W QDR2
ZShift = −3.4
Z lm Score = 2.32
NG = 0
DB = 0
SM = 0
Delta_L
Doxorubicin
YIL121W QDR2
ZShift = 0.49
Z lm Score = −0.32
NG = 0
DB = 0
SM = 0
Delta_K
Doxorubicin
YIL121W QDR2
ZShift = 1.16
Z lm Score = −1.89
NG = 0
DB = 0
SM = 0
Delta_r
Doxorubicin
YIL121W QDR2
ZShift = 1.26
Z lm Score = −0.18
NG = 0
DB = 0
SM = 0
Delta_AUC
Doxorubicin

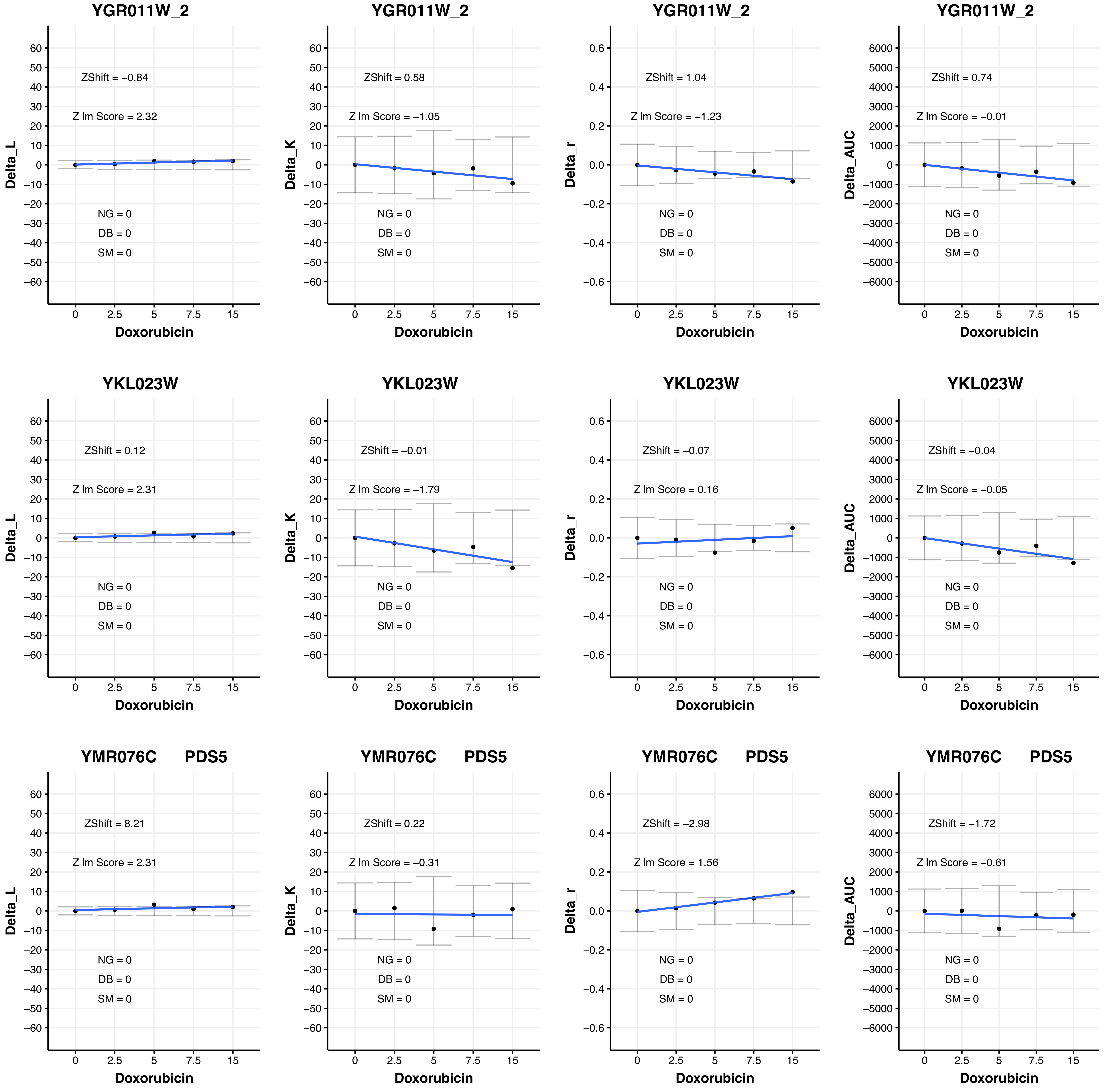

YGR011W_2
ZShift = −0.84
Z lm Score = 2.32
NG = 0
DB = 0
SM = 0
Delta_L
Doxorubicin
YGR011W_2
ZShift = 0.58
Z lm Score = −1.05
NG = 0
DB = 0
SM = 0
Delta_K
Doxorubicin
YGR011W_2
ZShift = 1.04
Z lm Score = −1.23
NG = 0
DB = 0
SM = 0
Delta_r
Doxorubicin
YGR011W_2
ZShift = 0.74
Z lm Score = −0.01
NG = 0
DB = 0
SM = 0
Delta_AUC
Doxorubicin
YKL023W
ZShift = 0.12
Z lm Score = 2.31
NG = 0
DB = 0
SM = 0
Delta_L
Doxorubicin
YKL023W
ZShift = −0.01
Z lm Score = −1.79
NG = 0
DB = 0
SM = 0
Delta_K
Doxorubicin
YKL023W
ZShift = −0.07
Z lm Score = 0.16
NG = 0
DB = 0
SM = 0
Delta_r
Doxorubicin
YKL023W
ZShift = −0.04
Z lm Score = −0.05
NG = 0
DB = 0
SM = 0
Delta_AUC
Doxorubicin
YMR076C PDS5
ZShift = 8.21
Z lm Score = 2.31
NG = 0
DB = 0
SM = 0
Delta_L
Doxorubicin
YMR076C PDS5
ZShift = 0.22
Z lm Score = −0.31
NG = 0
DB = 0
SM = 0
Delta_K
Doxorubicin
YMR076C PDS5
ZShift = −2.98
Z lm Score = 1.56
NG = 0
DB = 0
SM = 0
Delta_r
Doxorubicin
YMR076C PDS5
ZShift = −1.72
Z lm Score = −0.61
NG = 0
DB = 0
SM = 0
Delta_AUC
Doxorubicin

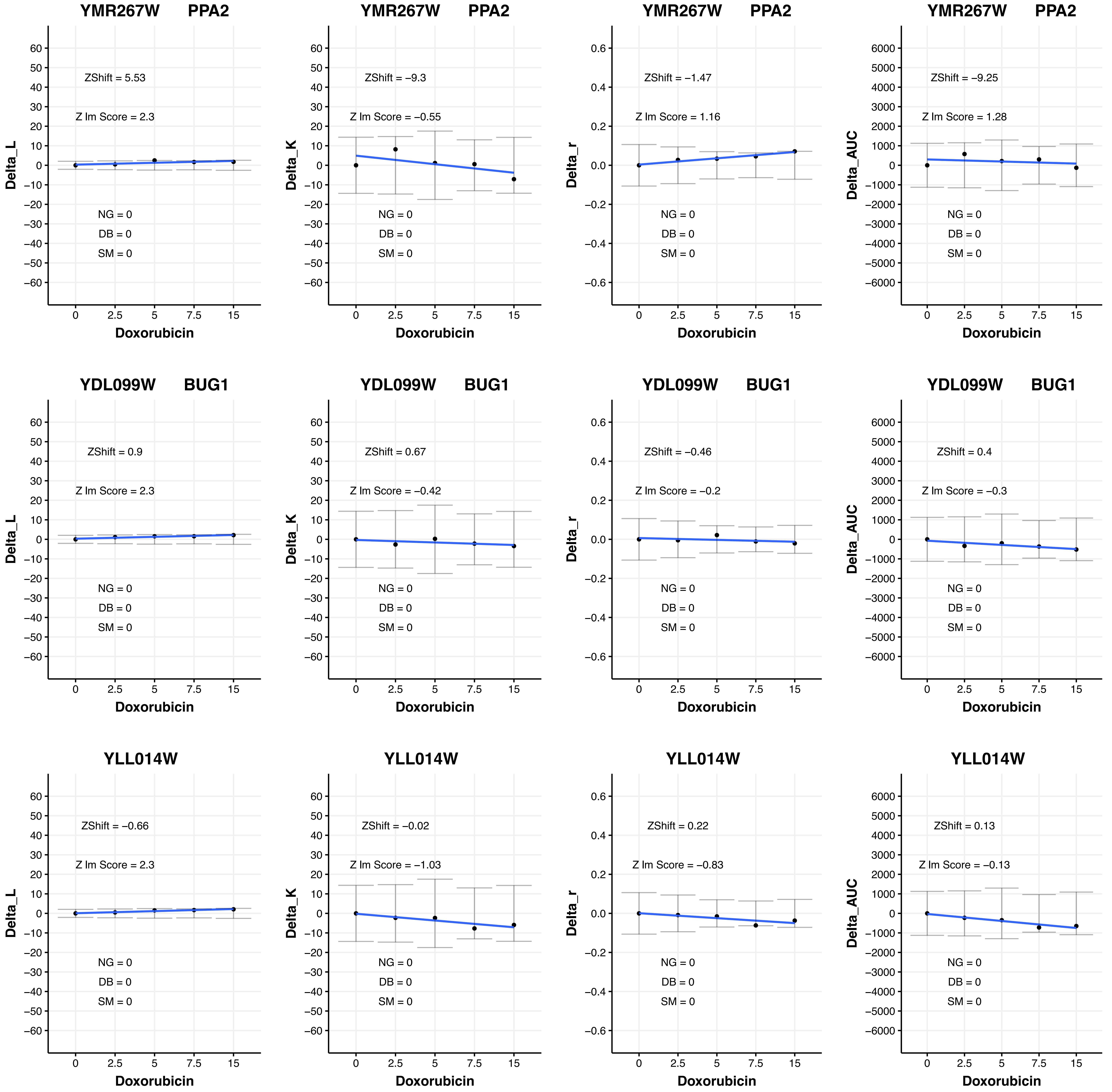

YMR267W PPA2
ZShift = 5.53
Z lm Score = 2.3
NG = 0
DB = 0
SM = 0
Delta_L
Doxorubicin
YMR267W PPA2
ZShift = −9.3
Z lm Score = −0.55
NG = 0
DB = 0
SM = 0
Delta_K
Doxorubicin
YMR267W PPA2
ZShift = −1.47
Z lm Score = 1.16
NG = 0
DB = 0
SM = 0
Delta_r
Doxorubicin
YMR267W PPA2
ZShift = −9.25
Z lm Score = 1.28
NG = 0
DB = 0
SM = 0
Delta_AUC
Doxorubicin
YDL099W BUG1
ZShift = 0.9
Z lm Score = 2.3
NG = 0
DB = 0
SM = 0
Delta_L
Doxorubicin
YDL099W BUG1
ZShift = 0.67
Z lm Score = −0.42
NG = 0
DB = 0
SM = 0
Delta_K
Doxorubicin
YDL099W BUG1
ZShift = −0.46
Z lm Score = −0.2
NG = 0
DB = 0
SM = 0
Delta_r
Doxorubicin
YDL099W BUG1
ZShift = 0.4
Z lm Score = −0.3
NG = 0
DB = 0
SM = 0
Delta_AUC
Doxorubicin
YLL014W
ZShift = −0.66
Z lm Score = 2.3
NG = 0
DB = 0
SM = 0
Delta_L
Doxorubicin
YLL014W
ZShift = −0.02
Z lm Score = −1.03
NG = 0
DB = 0
SM = 0
Delta_K
Doxorubicin
YLL014W
ZShift = 0.22
Z lm Score = −0.83
NG = 0
DB = 0
SM = 0
Delta_r
Doxorubicin
YLL014W
ZShift = 0.13
Z lm Score = −0.13
NG = 0
DB = 0
SM = 0
Delta_AUC
Doxorubicin

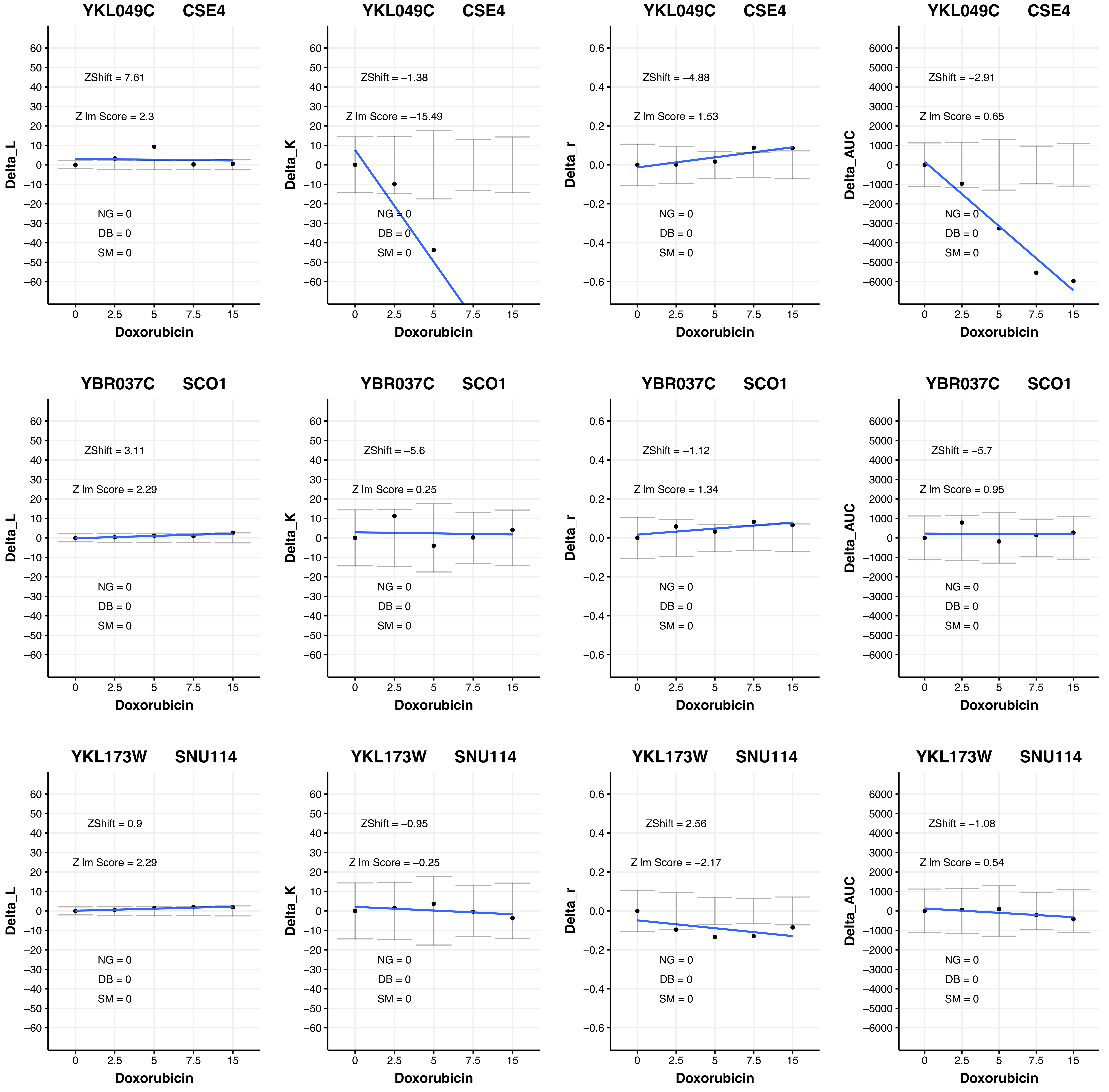
YKL049C CSE4
ZShift = 7.61
Z lm Score = 2.3
NG = 0
DB = 0
SM = 0
Delta_L
Doxorubicin
YKL049C CSE4
ZShift = −1.38
Z lm Score = −15.49
NG = 0
DB = 0
SM = 0
Delta_K
Doxorubicin
YKL049C CSE4
ZShift = −4.88
Z lm Score = 1.53
NG = 0
DB = 0
SM = 0
Delta_r
Doxorubicin
YKL049C CSE4
ZShift = −2.91
Z lm Score = 0.65
NG = 0
DB = 0
SM = 0
Delta_AUC
Doxorubicin
YBR037C SCO1
ZShift = 3.11
Z lm Score = 2.29
NG = 0
DB = 0
SM = 0
Delta_L
Doxorubicin
YBR037C SCO1
ZShift = −5.6
Z lm Score = 0.25
NG = 0
DB = 0
SM = 0
Delta_K
Doxorubicin
YBR037C SCO1
ZShift = −1.12
Z lm Score = 1.34
NG = 0
DB = 0
SM = 0
Delta_r
Doxorubicin
YBR037C SCO1
ZShift = −5.7
Z lm Score = 0.95
NG = 0
DB = 0
SM = 0
Delta_AUC
Doxorubicin
YKL173W SNU114
ZShift = 0.9
Z lm Score = 2.29
NG = 0
DB = 0
SM = 0
Delta_L
Doxorubicin
YKL173W SNU114
ZShift = −0.95
Z lm Score = −0.25
NG = 0
DB = 0
SM = 0
Delta_K
Doxorubicin
YKL173W SNU114
ZShift = 2.56
Z lm Score = −2.17
NG = 0
DB = 0
SM = 0
Delta_r
Doxorubicin
YKL173W SNU114
ZShift = −1.08
Z lm Score = 0.54
NG = 0
DB = 0
SM = 0
Delta_AUC
Doxorubicin

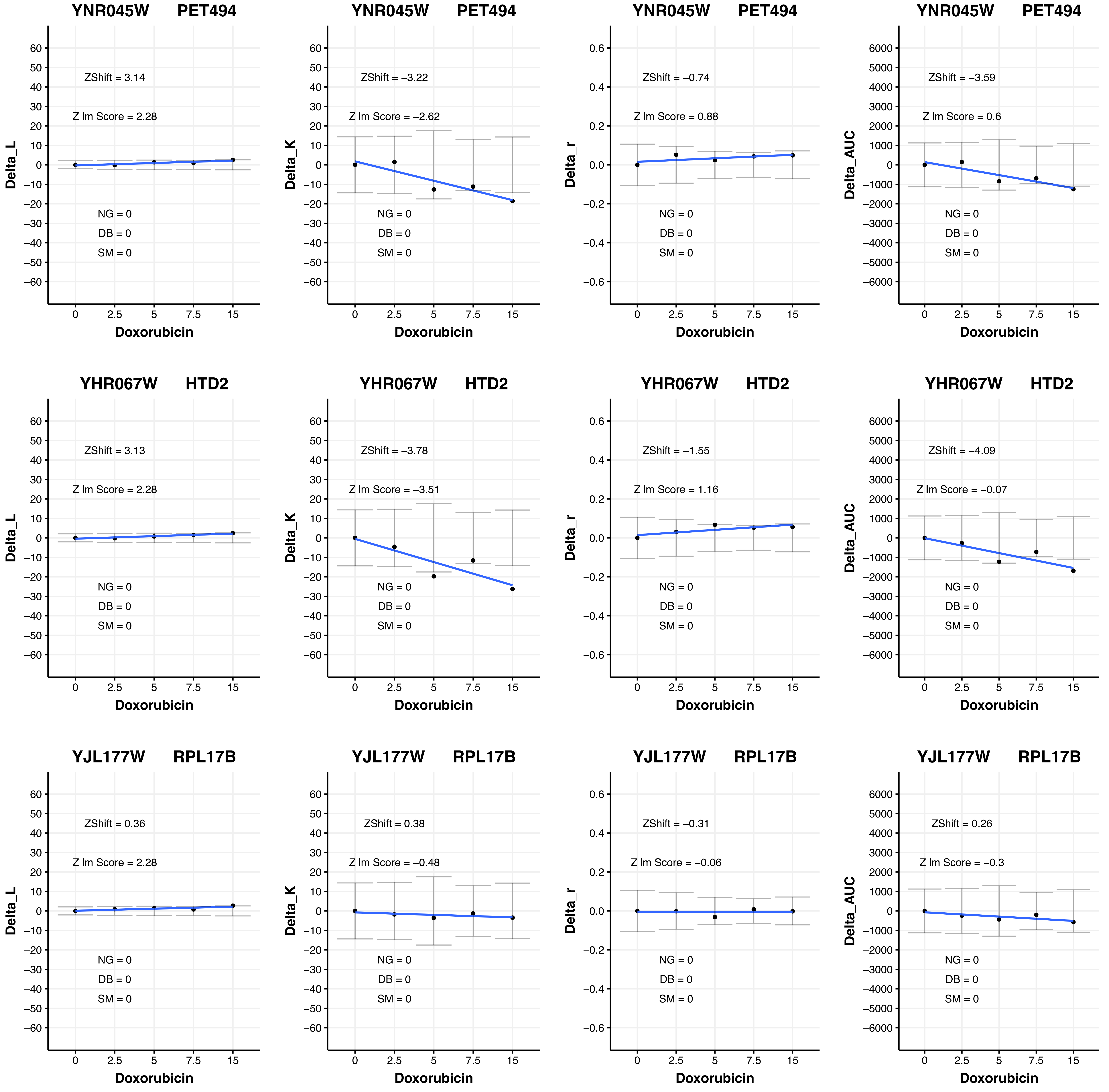

YNR045W PET494
ZShift = 3.14
Z lm Score = 2.28
NG = 0
DB = 0
SM = 0
Delta_L
Doxorubicin
YNR045W PET494
ZShift = −3.22
Z lm Score = −2.62
NG = 0
DB = 0
SM = 0
Delta_K
Doxorubicin
YNR045W PET494
ZShift = −0.74
Z lm Score = 0.88
NG = 0
DB = 0
SM = 0
Delta_r
Doxorubicin
YNR045W PET494
ZShift = −3.59
Z lm Score = 0.6
NG = 0
DB = 0
SM = 0
Delta_AUC
Doxorubicin
YHR067W HTD2
ZShift = 3.13
Z lm Score = 2.28
NG = 0
DB = 0
SM = 0
Delta_L
Doxorubicin
YHR067W HTD2
ZShift = −3.78
Z lm Score = −3.51
NG = 0
DB = 0
SM = 0
Delta_K
Doxorubicin
YHR067W HTD2
ZShift = −1.55
Z lm Score = 1.16
NG = 0
DB = 0
SM = 0
Delta_r
Doxorubicin
YHR067W HTD2
ZShift = −4.09
Z lm Score = −0.07
NG = 0
DB = 0
SM = 0
Delta_AUC
Doxorubicin
YJL177W RPL17B
ZShift = 0.36
Z lm Score = 2.28
NG = 0
DB = 0
SM = 0
Delta_L
Doxorubicin
YJL177W RPL17B
ZShift = 0.38
Z lm Score = −0.48
NG = 0
DB = 0
SM = 0
Delta_K
Doxorubicin
YJL177W RPL17B
ZShift = −0.31
Z lm Score = −0.06
NG = 0
DB = 0
SM = 0
Delta_r
Doxorubicin
YJL177W RPL17B
ZShift = 0.26
Z lm Score = −0.3
NG = 0
DB = 0
SM = 0
Delta_AUC
Doxorubicin

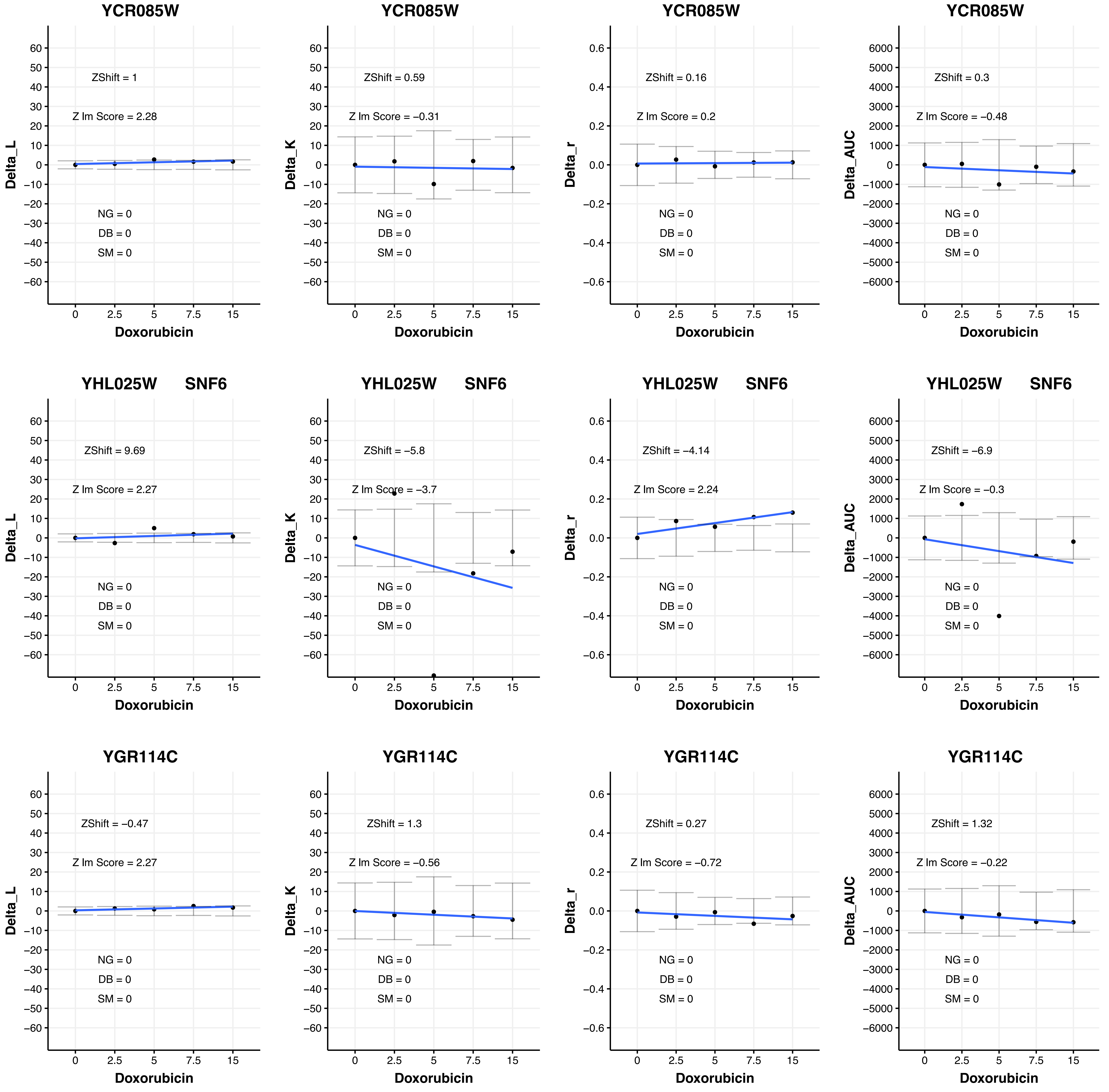

YCR085W
ZShift = 1
Z lm Score = 2.28
NG = 0
DB = 0
SM = 0
Delta_L
Doxorubicin
YCR085W
ZShift = 0.59
Z lm Score = −0.31
NG = 0
DB = 0
SM = 0
Delta_K
Doxorubicin
YCR085W
ZShift = 0.16
Z lm Score = 0.2
NG = 0
DB = 0
SM = 0
Delta_r
Doxorubicin
YCR085W
ZShift = 0.3
Z lm Score = −0.48
NG = 0
DB = 0
SM = 0
Delta_AUC
Doxorubicin
YHL025W SNF6
ZShift = 9.69
Z lm Score = 2.27
NG = 0
DB = 0
SM = 0
Delta_L
Doxorubicin
YHL025W SNF6
ZShift = −5.8
Z lm Score = −3.7
NG = 0
DB = 0
SM = 0
Delta_K
Doxorubicin
YHL025W SNF6
ZShift = −4.14
Z lm Score = 2.24
NG = 0
DB = 0
SM = 0
Delta_r
Doxorubicin
YHL025W SNF6
ZShift = −6.9
Z lm Score = −0.3
NG = 0
DB = 0
SM = 0
Delta_AUC
Doxorubicin
YGR114C
ZShift = −0.47
Z lm Score = 2.27
NG = 0
DB = 0
SM = 0
Delta_L
Doxorubicin
YGR114C
ZShift = 1.3
Z lm Score = −0.56
NG = 0
DB = 0
SM = 0
Delta_K
Doxorubicin
YGR114C
ZShift = 0.27
Z lm Score = −0.72
NG = 0
DB = 0
SM = 0
Delta_r
Doxorubicin
YGR114C
ZShift = 1.32
Z lm Score = −0.22
NG = 0
DB = 0
SM = 0
Delta_AUC
Doxorubicin

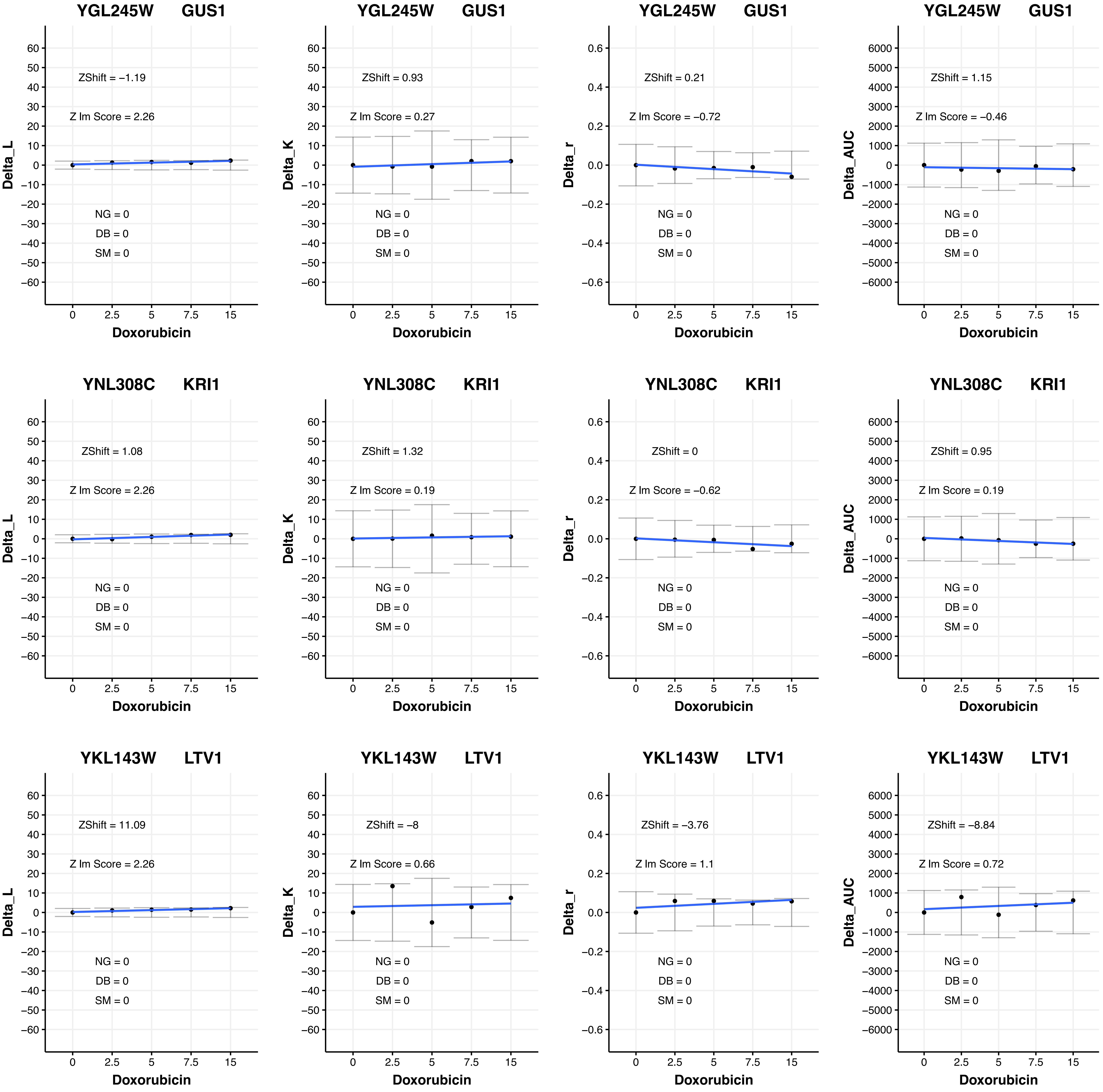
YGL245W GUS1
ZShift = −1.19
Z lm Score = 2.26
NG = 0
DB = 0
SM = 0
Delta_L
Doxorubicin
YGL245W GUS1
ZShift = 0.93
Z lm Score = 0.27
NG = 0
DB = 0
SM = 0
Delta_K
Doxorubicin
YGL245W GUS1
ZShift = 0.21
Z lm Score = −0.72
NG = 0
DB = 0
SM = 0
Delta_r
Doxorubicin
YGL245W GUS1
ZShift = 1.15
Z lm Score = −0.46
NG = 0
DB = 0
SM = 0
Delta_AUC
Doxorubicin
YNL308C KRI1
ZShift = 1.08
Z lm Score = 2.26
NG = 0
DB = 0
SM = 0
Delta_L
Doxorubicin
YNL308C KRI1
ZShift = 1.32
Z lm Score = 0.19
NG = 0
DB = 0
SM = 0
Delta_K
Doxorubicin
YNL308C KRI1
ZShift = 0
Z lm Score = −0.62
NG = 0
DB = 0
SM = 0
Delta_r
Doxorubicin
YNL308C KRI1
ZShift = 0.95
Z lm Score = 0.19
NG = 0
DB = 0
SM = 0
Delta_AUC
Doxorubicin
YKL143W LTV1
ZShift = 11.09
Z lm Score = 2.26
NG = 0
DB = 0
SM = 0
Delta_L
Doxorubicin
YKL143W LTV1
ZShift = −8
Z lm Score = 0.66
NG = 0
DB = 0
SM = 0
Delta_K
Doxorubicin
YKL143W LTV1
ZShift = −3.76
Z lm Score = 1.1
NG = 0
DB = 0
SM = 0
Delta_r
Doxorubicin
YKL143W LTV1
ZShift = −8.84
Z lm Score = 0.72
NG = 0
DB = 0
SM = 0
Delta_AUC
Doxorubicin

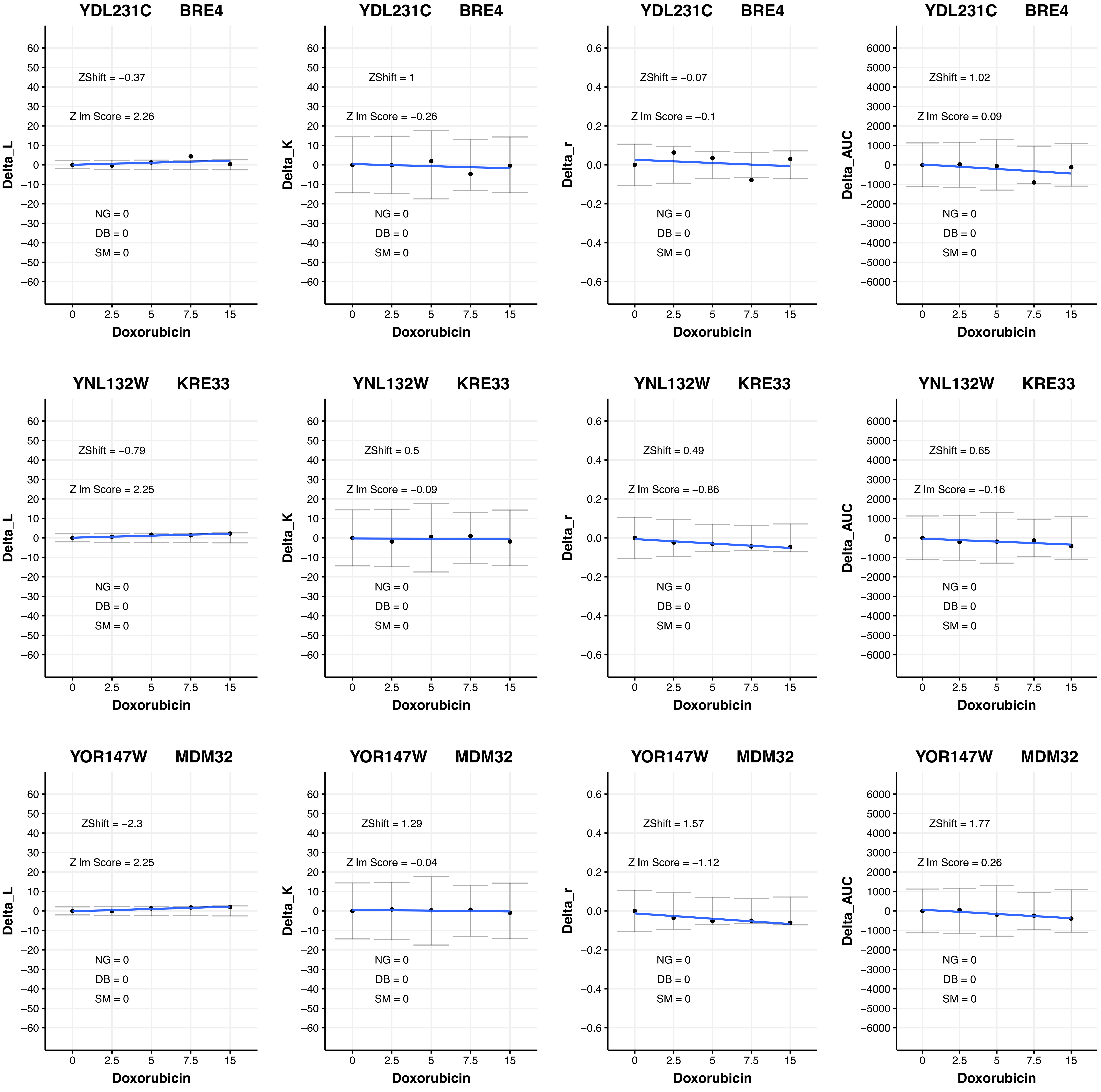

YDL231C BRE4
ZShift = −0.37
Z lm Score = 2.26
NG = 0
DB = 0
SM = 0
Delta_L
Doxorubicin
YDL231C BRE4
ZShift = 1
Z lm Score = −0.26
NG = 0
DB = 0
SM = 0
Delta_K
Doxorubicin
YDL231C BRE4
ZShift = −0.07
Z lm Score = −0.1
NG = 0
DB = 0
SM = 0
Delta_r
Doxorubicin
YDL231C BRE4
ZShift = 1.02
Z lm Score = 0.09
NG = 0
DB = 0
SM = 0
Delta_AUC
Doxorubicin
YNL132W KRE33
ZShift = −0.79
Z lm Score = 2.25
NG = 0
DB = 0
SM = 0
Delta_L
Doxorubicin
YNL132W KRE33
ZShift = 0.5
Z lm Score = −0.09
NG = 0
DB = 0
SM = 0
Delta_K
Doxorubicin
YNL132W KRE33
ZShift = 0.49
Z lm Score = −0.86
NG = 0
DB = 0
SM = 0
Delta_r
Doxorubicin
YNL132W KRE33
ZShift = 0.65
Z lm Score = −0.16
NG = 0
DB = 0
SM = 0
Delta_AUC
Doxorubicin
YOR147W MDM32
ZShift = −2.3
Z lm Score = 2.25
NG = 0
DB = 0
SM = 0
Delta_L
Doxorubicin
YOR147W MDM32
ZShift = 1.29
Z lm Score = −0.04
NG = 0
DB = 0
SM = 0
Delta_K
Doxorubicin
YOR147W MDM32
ZShift = 1.57
Z lm Score = −1.12
NG = 0
DB = 0
SM = 0
Delta_r
Doxorubicin
YOR147W MDM32
ZShift = 1.77
Z lm Score = 0.26
NG = 0
DB = 0
SM = 0
Delta_AUC
Doxorubicin

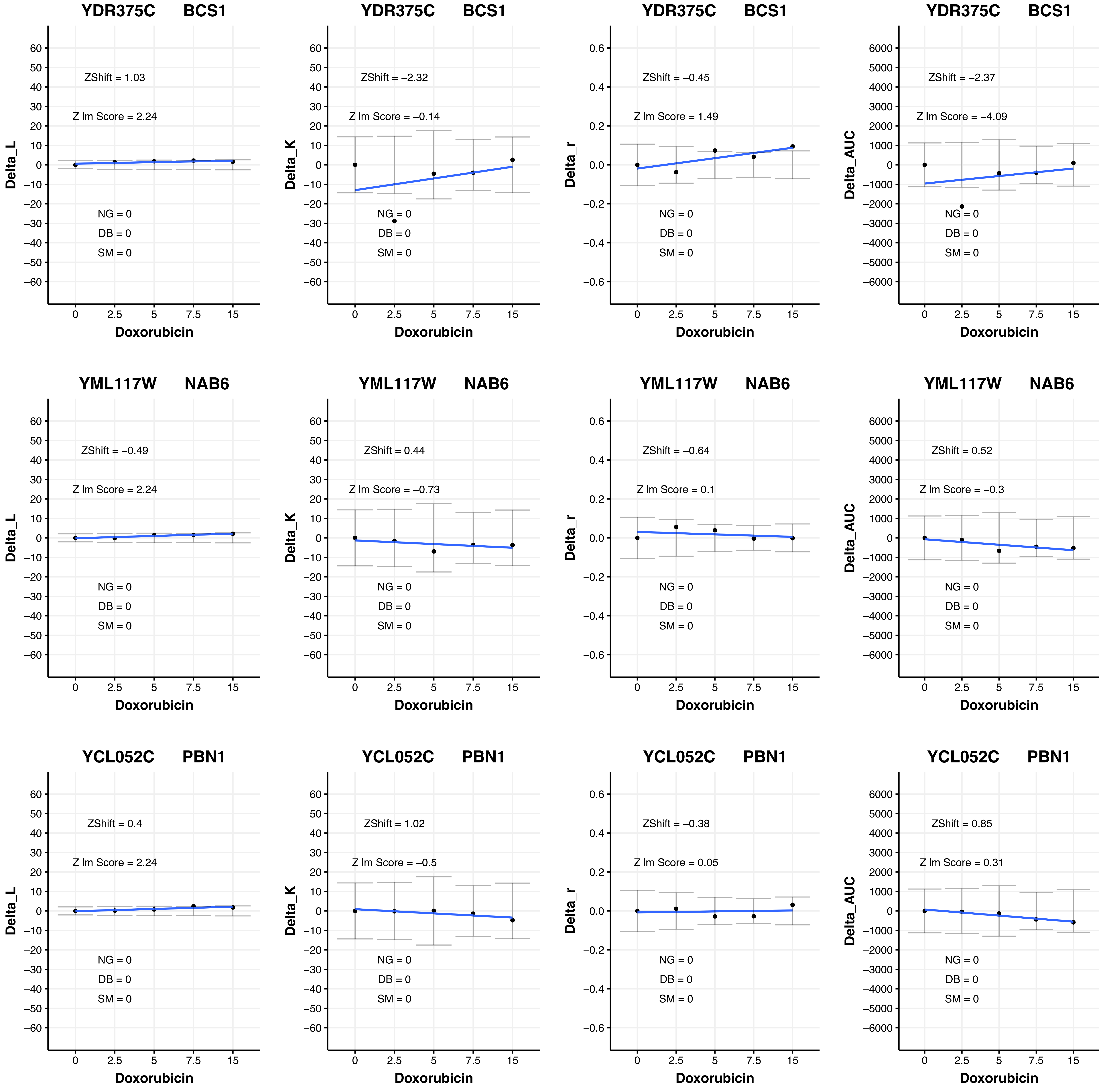
YDR375C BCS1
ZShift = 1.03
Z lm Score = 2.24
NG = 0
DB = 0
SM = 0
Delta_L
Doxorubicin
YDR375C BCS1
ZShift = −2.32
Z lm Score = −0.14
NG = 0
DB = 0
SM = 0
Delta_K
Doxorubicin
YDR375C BCS1
ZShift = −0.45
Z lm Score = 1.49
NG = 0
DB = 0
SM = 0
Delta_r
Doxorubicin
YDR375C BCS1
ZShift = −2.37
Z lm Score = −4.09
NG = 0
DB = 0
SM = 0
Delta_AUC
Doxorubicin
YML117W NAB6
ZShift = −0.49
Z lm Score = 2.24
NG = 0
DB = 0
SM = 0
Delta_L
Doxorubicin
YML117W NAB6
ZShift = 0.44
Z lm Score = −0.73
NG = 0
DB = 0
SM = 0
Delta_K
Doxorubicin
YML117W NAB6
ZShift = −0.64
Z lm Score = 0.1
NG = 0
DB = 0
SM = 0
Delta_r
Doxorubicin
YML117W NAB6
ZShift = 0.52
Z lm Score = −0.3
NG = 0
DB = 0
SM = 0
Delta_AUC
Doxorubicin
YCL052C PBN1
ZShift = 0.4
Z lm Score = 2.24
NG = 0
DB = 0
SM = 0
Delta_L
Doxorubicin
YCL052C PBN1
ZShift = 1.02
Z lm Score = −0.5
NG = 0
DB = 0
SM = 0
Delta_K
Doxorubicin
YCL052C PBN1
ZShift = −0.38
Z lm Score = 0.05
NG = 0
DB = 0
SM = 0
Delta_r
Doxorubicin
YCL052C PBN1
ZShift = 0.85
Z lm Score = 0.31
NG = 0
DB = 0
SM = 0
Delta_AUC
Doxorubicin

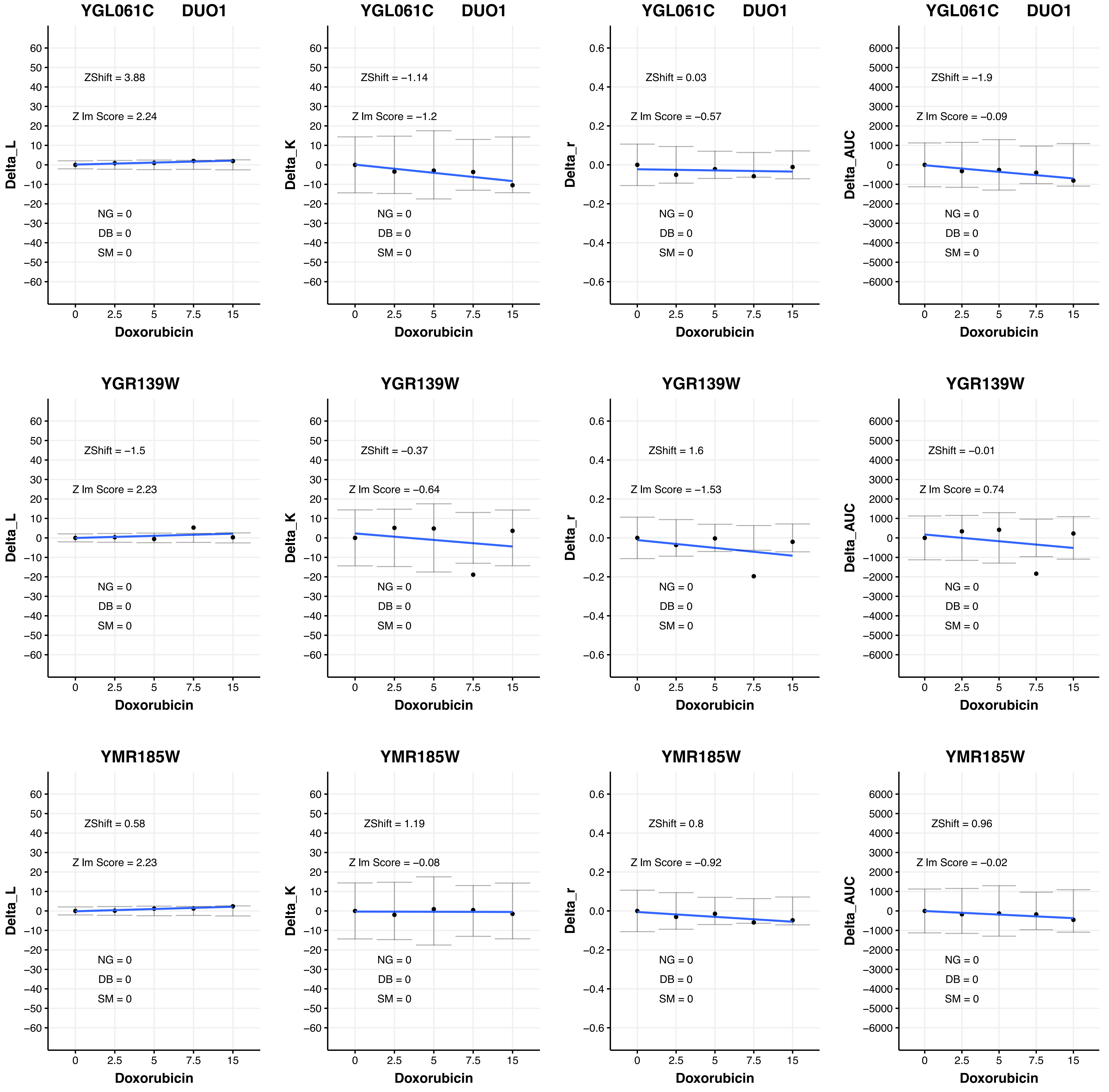

YGL061C DUO1
Delta_L
ZShift = 3.88
Z lm Score = 2.24
NG = 0
DB = 0
SM = 0
Doxorubicin
YGL061C DUO1
Delta_K
ZShift = −1.14
Z lm Score = −1.2
NG = 0
DB = 0
SM = 0
Doxorubicin
YGL061C DUO1
Delta_r
ZShift = 0.03
Z lm Score = −0.57
NG = 0
DB = 0
SM = 0
Doxorubicin
YGL061C DUO1
Delta_AUC
ZShift = −1.9
Z lm Score = −0.09
NG = 0
DB = 0
SM = 0
Doxorubicin
YGR139W
Delta_L
ZShift = −1.5
Z lm Score = 2.23
NG = 0
DB = 0
SM = 0
Doxorubicin
YGR139W
Delta_K
ZShift = −0.37
Z lm Score = −0.64
NG = 0
DB = 0
SM = 0
Doxorubicin
YGR139W
Delta_r
ZShift = 1.6
Z lm Score = −1.53
NG = 0
DB = 0
SM = 0
Doxorubicin
YGR139W
Delta_AUC
ZShift = −0.01
Z lm Score = 0.74
NG = 0
DB = 0
SM = 0
Doxorubicin
YMR185W
Delta_L
ZShift = 0.58
Z lm Score = 2.23
NG = 0
DB = 0
SM = 0
Doxorubicin
YMR185W
Delta_K
ZShift = 1.19
Z lm Score = −0.08
NG = 0
DB = 0
SM = 0
Doxorubicin
YMR185W
Delta_r
ZShift = 0.8
Z lm Score = −0.92
NG = 0
DB = 0
SM = 0
Doxorubicin
YMR185W
Delta_AUC
ZShift = 0.96
Z lm Score = −0.02
NG = 0
DB = 0
SM = 0
Doxorubicin

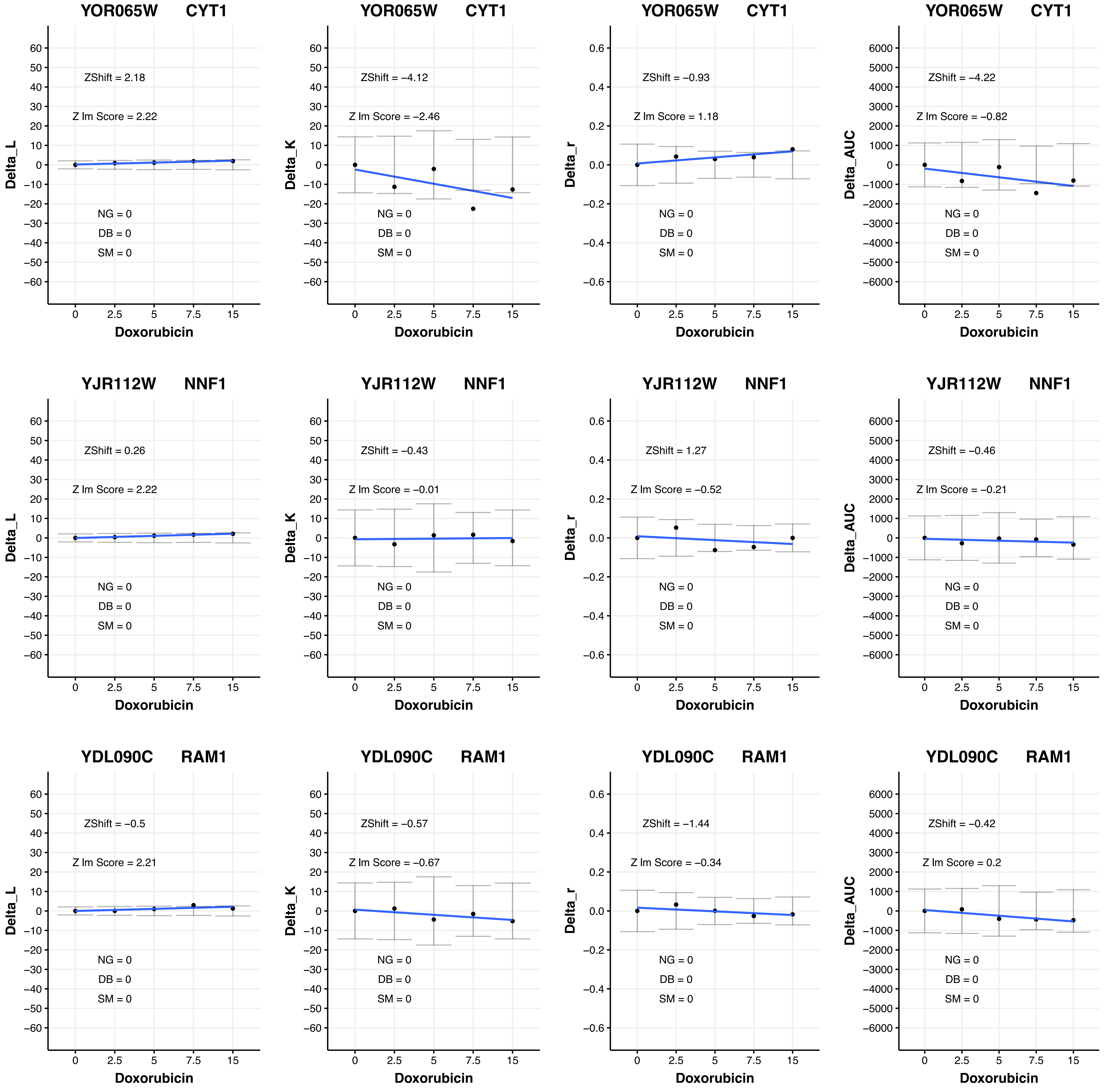
YOR065W CYT1
ZShift = 2.18
Z lm Score = 2.22
NG = 0
DB = 0
SM = 0
Delta_L
Doxorubicin
YOR065W CYT1
ZShift = −4.12
Z lm Score = −2.46
NG = 0
DB = 0
SM = 0
Delta_K
Doxorubicin
YOR065W CYT1
ZShift = −0.93
Z lm Score = 1.18
NG = 0
DB = 0
SM = 0
Delta_r
Doxorubicin
YOR065W CYT1
ZShift = −4.22
Z lm Score = −0.82
NG = 0
DB = 0
SM = 0
Delta_AUC
Doxorubicin
YJR112W NNF1
ZShift = 0.26
Z lm Score = 2.22
NG = 0
DB = 0
SM = 0
Delta_L
Doxorubicin
YJR112W NNF1
ZShift = −0.43
Z lm Score = −0.01
NG = 0
DB = 0
SM = 0
Delta_K
Doxorubicin
YJR112W NNF1
ZShift = 1.27
Z lm Score = −0.52
NG = 0
DB = 0
SM = 0
Delta_r
Doxorubicin
YJR112W NNF1
ZShift = −0.46
Z lm Score = −0.21
NG = 0
DB = 0
SM = 0
Delta_AUC
Doxorubicin
YDL090C RAM1
ZShift = −0.5
Z lm Score = 2.21
NG = 0
DB = 0
SM = 0
Delta_L
Doxorubicin
YDL090C RAM1
ZShift = −0.57
Z lm Score = −0.67
NG = 0
DB = 0
SM = 0
Delta_K
Doxorubicin
YDL090C RAM1
ZShift = −1.44
Z lm Score = −0.34
NG = 0
DB = 0
SM = 0
Delta_r
Doxorubicin
YDL090C RAM1
ZShift = −0.42
Z lm Score = 0.2
NG = 0
DB = 0
SM = 0
Delta_AUC
Doxorubicin

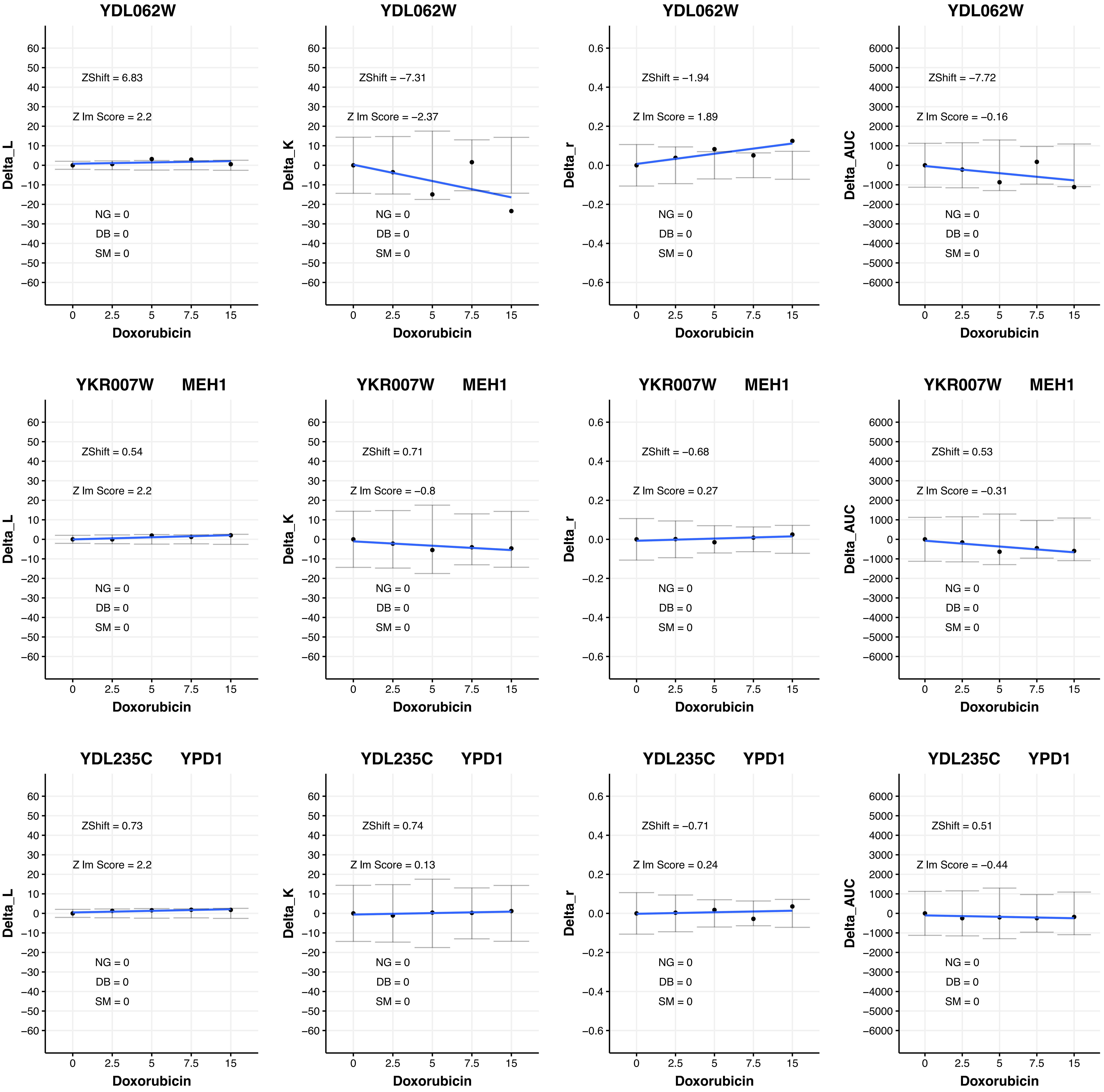
YDL062W
ZShift = 6.83
Z lm Score = 2.2
NG = 0
DB = 0
SM = 0
Delta_L
Doxorubicin
YDL062W
ZShift = −7.31
Z lm Score = −2.37
NG = 0
DB = 0
SM = 0
Delta_K
Doxorubicin
YDL062W
ZShift = −1.94
Z lm Score = 1.89
NG = 0
DB = 0
SM = 0
Delta_r
Doxorubicin
YDL062W
ZShift = −7.72
Z lm Score = −0.16
NG = 0
DB = 0
SM = 0
Delta_AUC
Doxorubicin
YKR007W MEH1
ZShift = 0.54
Z lm Score = 2.2
NG = 0
DB = 0
SM = 0
Delta_L
Doxorubicin
YKR007W MEH1
ZShift = 0.71
Z lm Score = −0.8
NG = 0
DB = 0
SM = 0
Delta_K
Doxorubicin
YKR007W MEH1
ZShift = −0.68
Z lm Score = 0.27
NG = 0
DB = 0
SM = 0
Delta_r
Doxorubicin
YKR007W MEH1
ZShift = 0.53
Z lm Score = −0.31
NG = 0
DB = 0
SM = 0
Delta_AUC
Doxorubicin
YDL235C YPD1
ZShift = 0.73
Z lm Score = 2.2
NG = 0
DB = 0
SM = 0
Delta_L
Doxorubicin
YDL235C YPD1
ZShift = 0.74
Z lm Score = 0.13
NG = 0
DB = 0
SM = 0
Delta_K
Doxorubicin
YDL235C YPD1
ZShift = −0.71
Z lm Score = 0.24
NG = 0
DB = 0
SM = 0
Delta_r
Doxorubicin
YDL235C YPD1
ZShift = 0.51
Z lm Score = −0.44
NG = 0
DB = 0
SM = 0
Delta_AUC
Doxorubicin

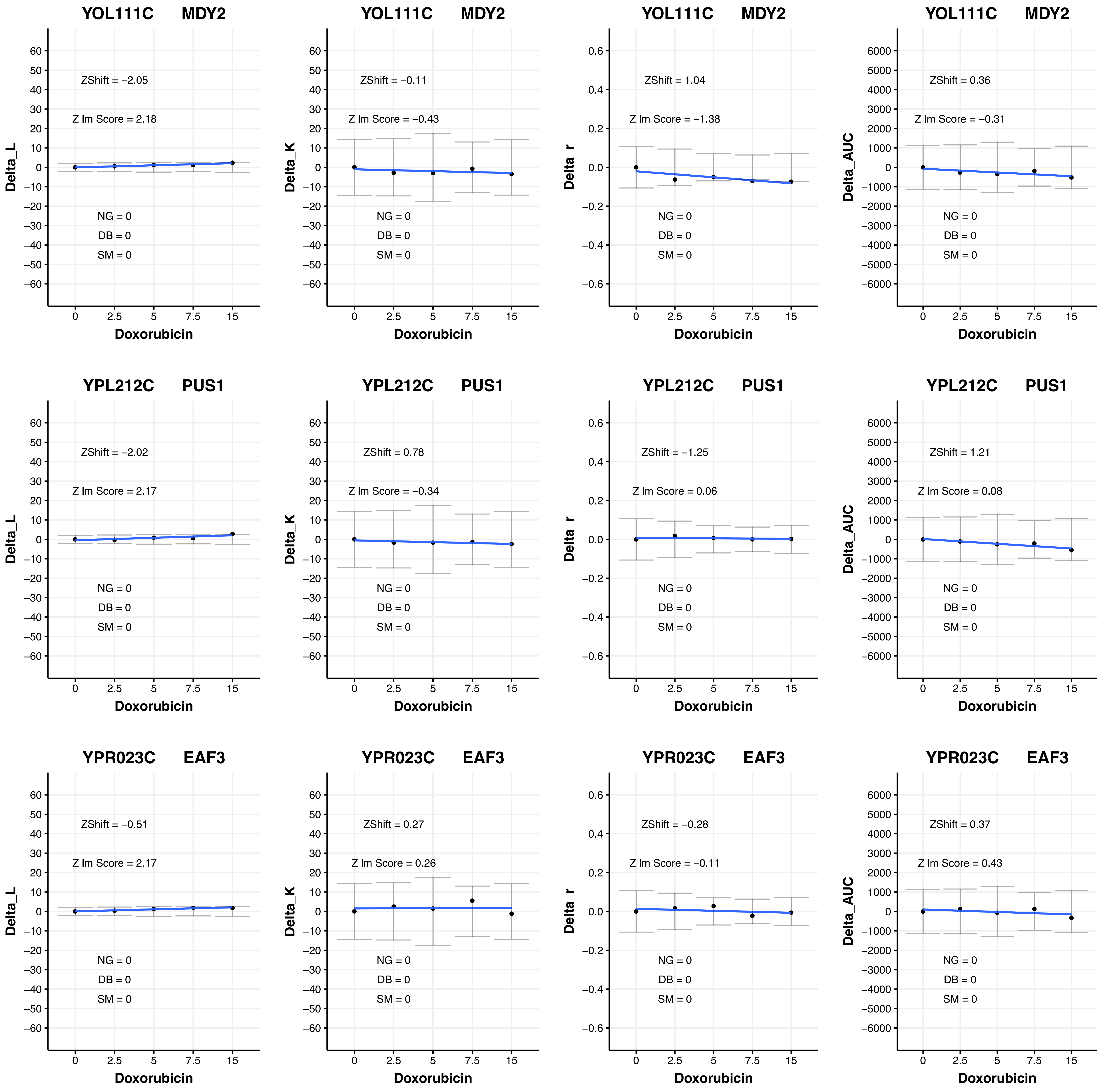
YOL111C MDY2
ZShift = −2.05
Z lm Score = 2.18
Delta_L
NG = 0
DB = 0
SM = 0
Doxorubicin
YOL111C MDY2
ZShift = −0.11
Z lm Score = −0.43
Delta_K
NG = 0
DB = 0
SM = 0
Doxorubicin
YOL111C MDY2
ZShift = 1.04
Z lm Score = −1.38
Delta_r
NG = 0
DB = 0
SM = 0
Doxorubicin
YOL111C MDY2
ZShift = 0.36
Z lm Score = −0.31
Delta_AUC
NG = 0
DB = 0
SM = 0
Doxorubicin
YPL212C PUS1
ZShift = −2.02
Z lm Score = 2.17
Delta_L
NG = 0
DB = 0
SM = 0
Doxorubicin
YPL212C PUS1
ZShift = 0.78
Z lm Score = −0.34
Delta_K
NG = 0
DB = 0
SM = 0
Doxorubicin
YPL212C PUS1
ZShift = −1.25
Z lm Score = 0.06
Delta_r
NG = 0
DB = 0
SM = 0
Doxorubicin
YPL212C PUS1
ZShift = 1.21
Z lm Score = 0.08
Delta_AUC
NG = 0
DB = 0
SM = 0
Doxorubicin
YPR023C EAF3
ZShift = −0.51
Z lm Score = 2.17
Delta_L
NG = 0
DB = 0
SM = 0
Doxorubicin
YPR023C EAF3
ZShift = 0.27
Z lm Score = 0.26
Delta_K
NG = 0
DB = 0
SM = 0
Doxorubicin
YPR023C EAF3
ZShift = −0.28
Z lm Score = −0.11
Delta_r
NG = 0
DB = 0
SM = 0
Doxorubicin
YPR023C EAF3
ZShift = 0.37
Z lm Score = 0.43
Delta_AUC
NG = 0
DB = 0
SM = 0
Doxorubicin

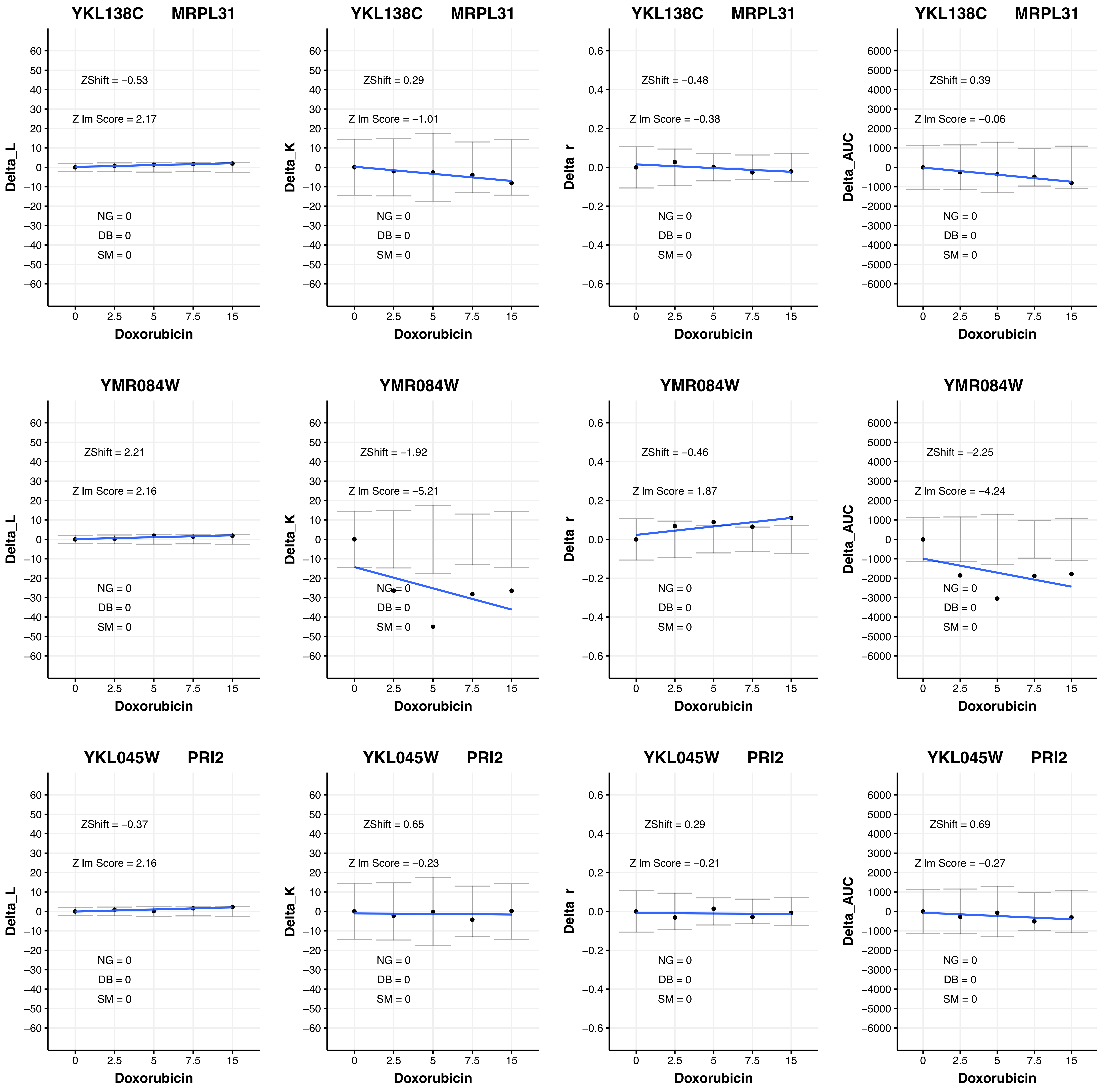
YKL138C MRPL31
ZShift = −0.53
Z lm Score = 2.17
NG = 0
DB = 0
SM = 0
Delta_L
Doxorubicin
YKL138C MRPL31
ZShift = 0.29
Z lm Score = −1.01
NG = 0
DB = 0
SM = 0
Delta_K
Doxorubicin
YKL138C MRPL31
ZShift = −0.48
Z lm Score = −0.38
NG = 0
DB = 0
SM = 0
Delta_r
Doxorubicin
YKL138C MRPL31
ZShift = 0.39
Z lm Score = −0.06
NG = 0
DB = 0
SM = 0
Delta_AUC
Doxorubicin
YMR084W
ZShift = 2.21
Z lm Score = 2.16
NG = 0
DB = 0
SM = 0
Delta_L
Doxorubicin
YMR084W
ZShift = −1.92
Z lm Score = −5.21
NG = 0
DB = 0
SM = 0
Delta_K
Doxorubicin
YMR084W
ZShift = −0.46
Z lm Score = 1.87
NG = 0
DB = 0
SM = 0
Delta_r
Doxorubicin
YMR084W
ZShift = −2.25
Z lm Score = −4.24
NG = 0
DB = 0
SM = 0
Delta_AUC
Doxorubicin
YKL045W PRI2
ZShift = −0.37
Z lm Score = 2.16
NG = 0
DB = 0
SM = 0
Delta_L
Doxorubicin
YKL045W PRI2
ZShift = 0.65
Z lm Score = −0.23
NG = 0
DB = 0
SM = 0
Delta_K
Doxorubicin
YKL045W PRI2
ZShift = 0.29
Z lm Score = −0.21
NG = 0
DB = 0
SM = 0
Delta_r
Doxorubicin
YKL045W PRI2
ZShift = 0.69
Z lm Score = −0.27
NG = 0
DB = 0
SM = 0
Delta_AUC
Doxorubicin

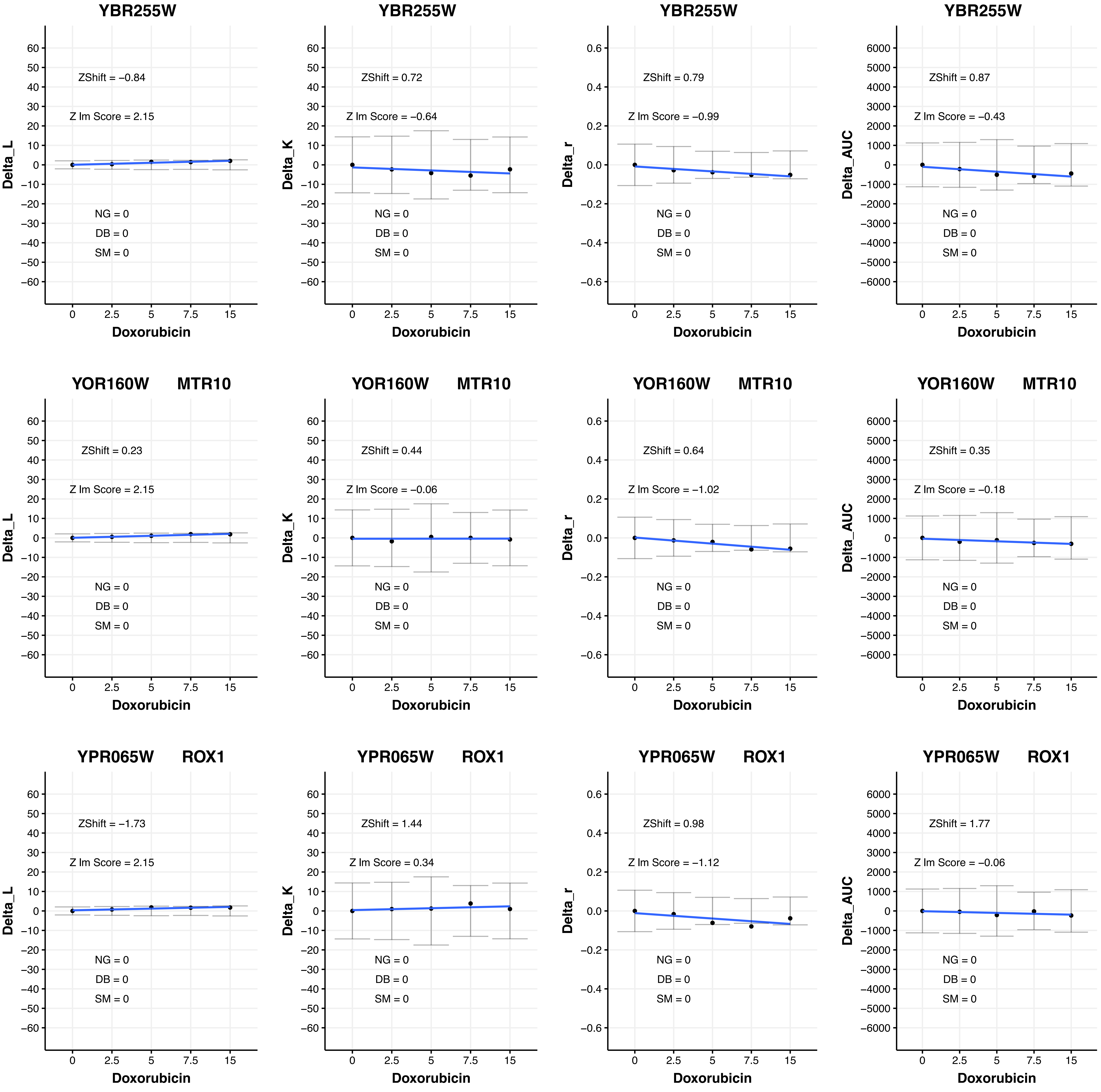
YBR255W
ZShift = −0.84
Z lm Score = 2.15
NG = 0
DB = 0
SM = 0
Delta_L
Doxorubicin
YBR255W
ZShift = 0.72
Z lm Score = −0.64
NG = 0
DB = 0
SM = 0
Delta_K
Doxorubicin
YBR255W
ZShift = 0.79
Z lm Score = −0.99
NG = 0
DB = 0
SM = 0
Delta_r
Doxorubicin
YBR255W
ZShift = 0.87
Z lm Score = −0.43
NG = 0
DB = 0
SM = 0
Delta_AUC
Doxorubicin
YOR160W MTR10
ZShift = 0.23
Z lm Score = 2.15
NG = 0
DB = 0
SM = 0
Delta_L
Doxorubicin
YOR160W MTR10
ZShift = 0.44
Z lm Score = −0.06
NG = 0
DB = 0
SM = 0
Delta_K
Doxorubicin
YOR160W MTR10
ZShift = 0.64
Z lm Score = −1.02
NG = 0
DB = 0
SM = 0
Delta_r
Doxorubicin
YOR160W MTR10
ZShift = 0.35
Z lm Score = −0.18
NG = 0
DB = 0
SM = 0
Delta_AUC
Doxorubicin
YPR065W ROX1
ZShift = −1.73
Z lm Score = 2.15
NG = 0
DB = 0
SM = 0
Delta_L
Doxorubicin
YPR065W ROX1
ZShift = 1.44
Z lm Score = 0.34
NG = 0
DB = 0
SM = 0
Delta_K
Doxorubicin
YPR065W ROX1
ZShift = 0.98
Z lm Score = −1.12
NG = 0
DB = 0
SM = 0
Delta_r
Doxorubicin
YPR065W ROX1
ZShift = 1.77
Z lm Score = −0.06
NG = 0
DB = 0
SM = 0
Delta_AUC
Doxorubicin

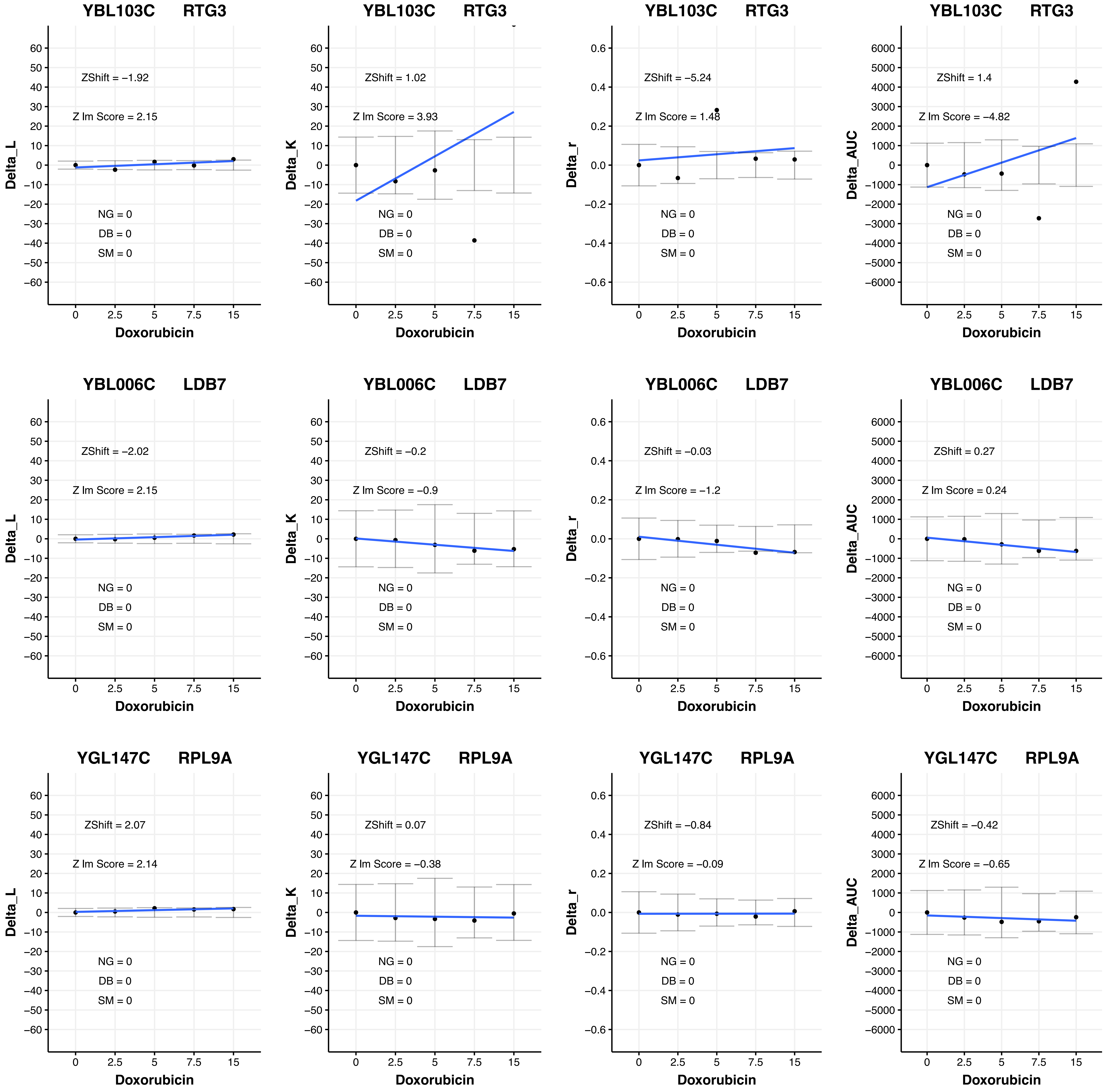
YBL103C RTG3
ZShift = −1.92
Z lm Score = 2.15
NG = 0
DB = 0
SM = 0
Delta_L
Doxorubicin
YBL103C RTG3
ZShift = 1.02
Z lm Score = 3.93
NG = 0
DB = 0
SM = 0
Delta_K
Doxorubicin
YBL103C RTG3
ZShift = −5.24
Z lm Score = 1.48
NG = 0
DB = 0
SM = 0
Delta_r
Doxorubicin
YBL103C RTG3
ZShift = 1.4
Z lm Score = −4.82
NG = 0
DB = 0
SM = 0
Delta_AUC
Doxorubicin
YBL006C LDB7
ZShift = −2.02
Z lm Score = 2.15
NG = 0
DB = 0
SM = 0
Delta_L
Doxorubicin
YBL006C LDB7
ZShift = −0.2
Z lm Score = −0.9
NG = 0
DB = 0
SM = 0
Delta_K
Doxorubicin
YBL006C LDB7
ZShift = −0.03
Z lm Score = −1.2
NG = 0
DB = 0
SM = 0
Delta_r
Doxorubicin
YBL006C LDB7
ZShift = 0.27
Z lm Score = 0.24
NG = 0
DB = 0
SM = 0
Delta_AUC
Doxorubicin
YGL147C RPL9A
ZShift = 2.07
Z lm Score = 2.14
NG = 0
DB = 0
SM = 0
Delta_L
Doxorubicin
YGL147C RPL9A
ZShift = 0.07
Z lm Score = −0.38
NG = 0
DB = 0
SM = 0
Delta_K
Doxorubicin
YGL147C RPL9A
ZShift = −0.84
Z lm Score = −0.09
NG = 0
DB = 0
SM = 0
Delta_r
Doxorubicin
YGL147C RPL9A
ZShift = −0.42
Z lm Score = −0.65
NG = 0
DB = 0
SM = 0
Delta_AUC
Doxorubicin

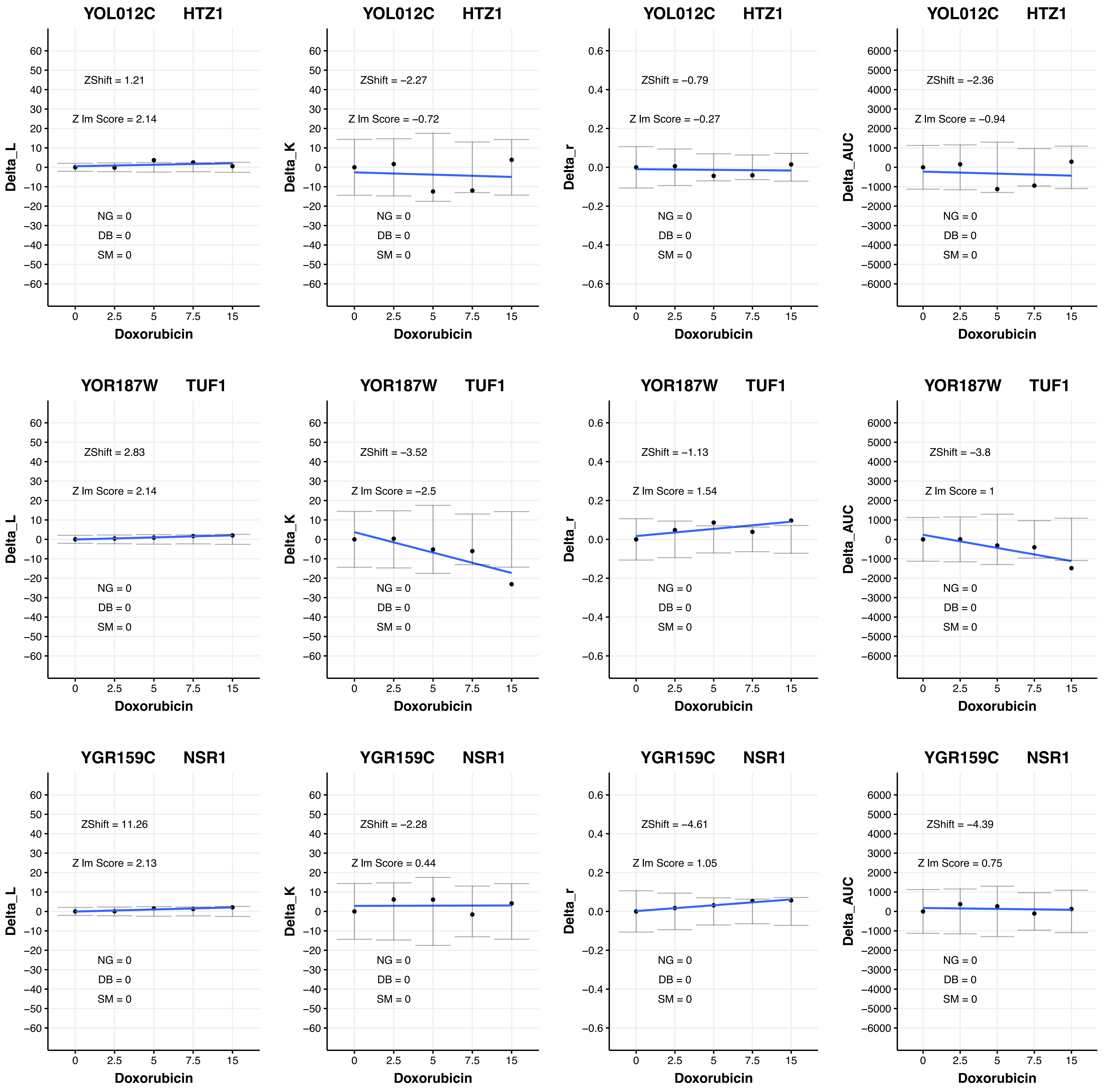

YOL012C HTZ1
ZShift = 1.21
Z lm Score = 2.14
NG = 0
DB = 0
SM = 0
Delta_L
Doxorubicin
YOL012C HTZ1
ZShift = −2.27
Z lm Score = −0.72
NG = 0
DB = 0
SM = 0
Delta_K
Doxorubicin
YOL012C HTZ1
ZShift = −0.79
Z lm Score = −0.27
NG = 0
DB = 0
SM = 0
Delta_r
Doxorubicin
YOL012C HTZ1
ZShift = −2.36
Z lm Score = −0.94
NG = 0
DB = 0
SM = 0
Delta_AUC
Doxorubicin
YOR187W TUF1
ZShift = 2.83
Z lm Score = 2.14
NG = 0
DB = 0
SM = 0
Delta_L
Doxorubicin
YOR187W TUF1
ZShift = −3.52
Z lm Score = −2.5
NG = 0
DB = 0
SM = 0
Delta_K
Doxorubicin
YOR187W TUF1
ZShift = −1.13
Z lm Score = 1.54
NG = 0
DB = 0
SM = 0
Delta_r
Doxorubicin
YOR187W TUF1
ZShift = −3.8
Z lm Score = 1
NG = 0
DB = 0
SM = 0
Delta_AUC
Doxorubicin
YGR159C NSR1
ZShift = 11.26
Z lm Score = 2.13
NG = 0
DB = 0
SM = 0
Delta_L
Doxorubicin
YGR159C NSR1
ZShift = −2.28
Z lm Score = 0.44
NG = 0
DB = 0
SM = 0
Delta_K
Doxorubicin
YGR159C NSR1
ZShift = −4.61
Z lm Score = 1.05
NG = 0
DB = 0
SM = 0
Delta_r
Doxorubicin
YGR159C NSR1
ZShift = −4.39
Z lm Score = 0.75
NG = 0
DB = 0
SM = 0
Delta_AUC
Doxorubicin

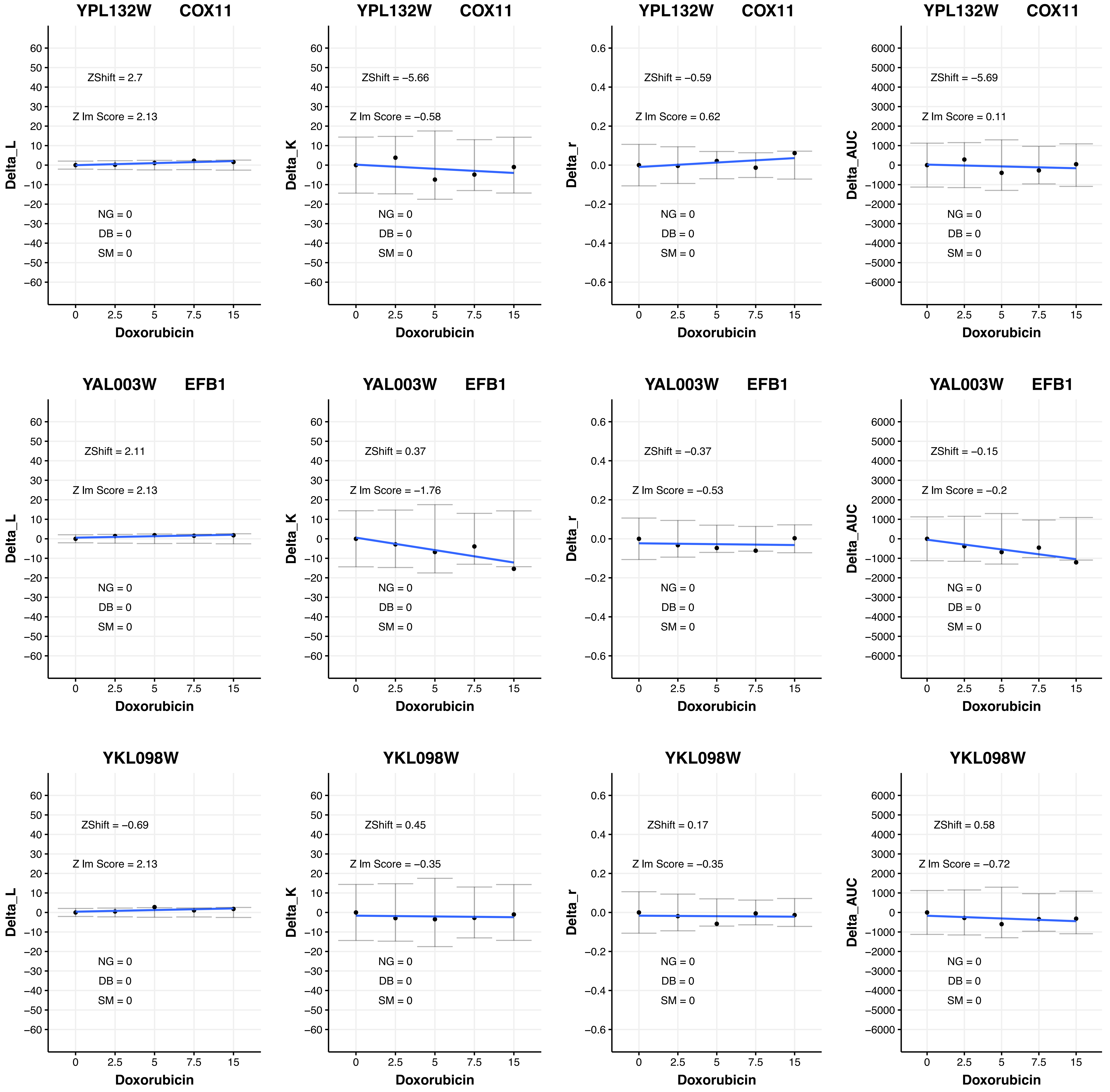
YPL132W COX11
ZShift = 2.7
Z lm Score = 2.13
NG = 0
DB = 0
SM = 0
Delta_L
Doxorubicin
YPL132W COX11
ZShift = −5.66
Z lm Score = −0.58
NG = 0
DB = 0
SM = 0
Delta_K
Doxorubicin
YPL132W COX11
ZShift = −0.59
Z lm Score = 0.62
NG = 0
DB = 0
SM = 0
Delta_r
Doxorubicin
YPL132W COX11
ZShift = −5.69
Z lm Score = 0.11
NG = 0
DB = 0
SM = 0
Delta_AUC
Doxorubicin
YAL003W EFB1
ZShift = 2.11
Z lm Score = 2.13
NG = 0
DB = 0
SM = 0
Delta_L
Doxorubicin
YAL003W EFB1
ZShift = 0.37
Z lm Score = −1.76
NG = 0
DB = 0
SM = 0
Delta_K
Doxorubicin
YAL003W EFB1
ZShift = −0.37
Z lm Score = −0.53
NG = 0
DB = 0
SM = 0
Delta_r
Doxorubicin
YAL003W EFB1
ZShift = −0.15
Z lm Score = −0.2
NG = 0
DB = 0
SM = 0
Delta_AUC
Doxorubicin
YKL098W
ZShift = −0.69
Z lm Score = 2.13
NG = 0
DB = 0
SM = 0
Delta_L
Doxorubicin
YKL098W
ZShift = 0.45
Z lm Score = −0.35
NG = 0
DB = 0
SM = 0
Delta_K
Doxorubicin
YKL098W
ZShift = 0.17
Z lm Score = −0.35
NG = 0
DB = 0
SM = 0
Delta_r
Doxorubicin
YKL098W
ZShift = 0.58
Z lm Score = −0.72
NG = 0
DB = 0
SM = 0
Delta_AUC
Doxorubicin

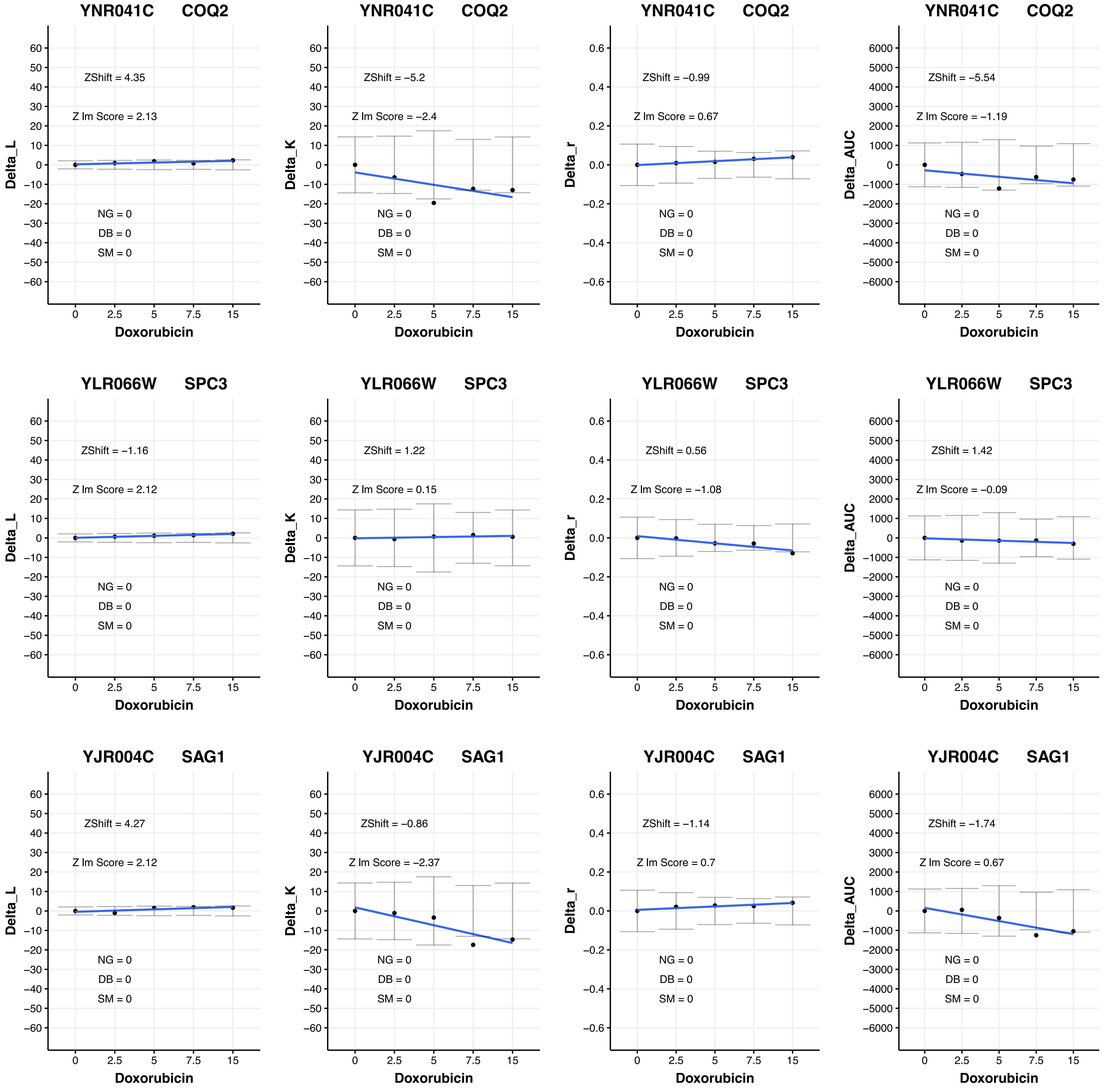

YNR041C COQ2
ZShift = 4.35
Z lm Score = 2.13
NG = 0
DB = 0
SM = 0
Delta_L
Doxorubicin
YNR041C COQ2
ZShift = −5.2
Z lm Score = −2.4
NG = 0
DB = 0
SM = 0
Delta_K
Doxorubicin
YNR041C COQ2
ZShift = −0.99
Z lm Score = 0.67
NG = 0
DB = 0
SM = 0
Delta_r
Doxorubicin
YNR041C COQ2
ZShift = −5.54
Z lm Score = −1.19
NG = 0
DB = 0
SM = 0
Delta_AUC
Doxorubicin
YLR066W SPC3
ZShift = −1.16
Z lm Score = 2.12
NG = 0
DB = 0
SM = 0
Delta_L
Doxorubicin
YLR066W SPC3
ZShift = 1.22
Z lm Score = 0.15
NG = 0
DB = 0
SM = 0
Delta_K
Doxorubicin
YLR066W SPC3
ZShift = 0.56
Z lm Score = −1.08
NG = 0
DB = 0
SM = 0
Delta_r
Doxorubicin
YLR066W SPC3
ZShift = 1.42
Z lm Score = −0.09
NG = 0
DB = 0
SM = 0
Delta_AUC
Doxorubicin
YJR004C SAG1
ZShift = 4.27
Z lm Score = 2.12
NG = 0
DB = 0
SM = 0
Delta_L
Doxorubicin
YJR004C SAG1
ZShift = −0.86
Z lm Score = −2.37
NG = 0
DB = 0
SM = 0
Delta_K
Doxorubicin
YJR004C SAG1
ZShift = −1.14
Z lm Score = 0.7
NG = 0
DB = 0
SM = 0
Delta_r
Doxorubicin
YJR004C SAG1
ZShift = −1.74
Z lm Score = 0.67
NG = 0
DB = 0
SM = 0
Delta_AUC
Doxorubicin

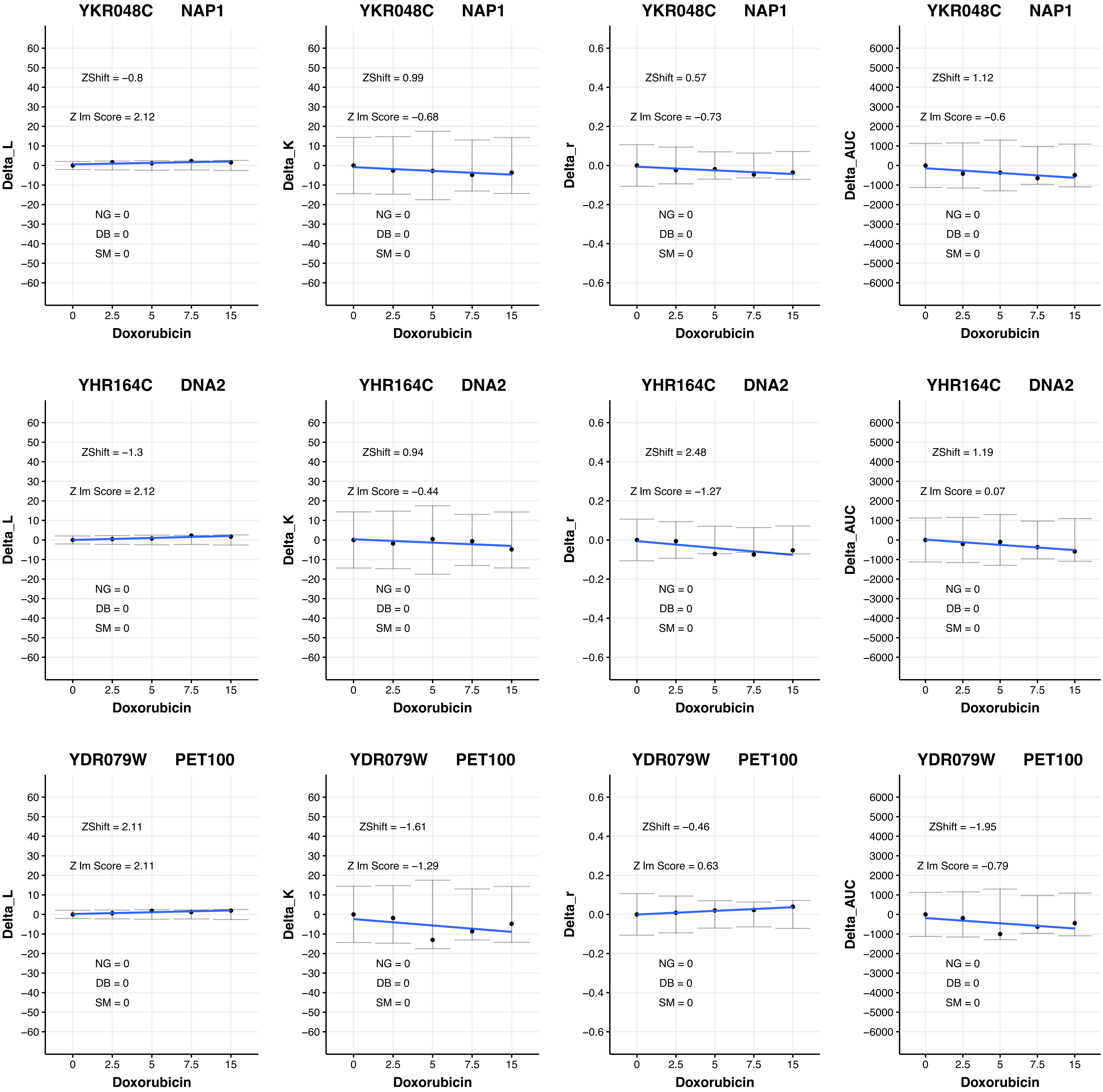
YKR048C NAP1
ZShift = −0.8
Z lm Score = 2.12
NG = 0
DB = 0
SM = 0
Delta_L
Doxorubicin
YKR048C NAP1
ZShift = 0.99
Z lm Score = −0.68
NG = 0
DB = 0
SM = 0
Delta_K
Doxorubicin
YKR048C NAP1
ZShift = 0.57
Z lm Score = −0.73
NG = 0
DB = 0
SM = 0
Delta_r
Doxorubicin
YKR048C NAP1
ZShift = 1.12
Z lm Score = −0.6
NG = 0
DB = 0
SM = 0
Delta_AUC
Doxorubicin
YHR164C DNA2
ZShift = −1.3
Z lm Score = 2.12
NG = 0
DB = 0
SM = 0
Delta_L
Doxorubicin
YHR164C DNA2
ZShift = 0.94
Z lm Score = −0.44
NG = 0
DB = 0
SM = 0
Delta_K
Doxorubicin
YHR164C DNA2
ZShift = 2.48
Z lm Score = −1.27
NG = 0
DB = 0
SM = 0
Delta_r
Doxorubicin
YHR164C DNA2
ZShift = 1.19
Z lm Score = 0.07
NG = 0
DB = 0
SM = 0
Delta_AUC
Doxorubicin
YDR079W PET100
ZShift = 2.11
Z lm Score = 2.11
NG = 0
DB = 0
SM = 0
Delta_L
Doxorubicin
YDR079W PET100
ZShift = −1.61
Z lm Score = −1.29
NG = 0
DB = 0
SM = 0
Delta_K
Doxorubicin
YDR079W PET100
ZShift = −0.46
Z lm Score = 0.63
NG = 0
DB = 0
SM = 0
Delta_r
Doxorubicin
YDR079W PET100
ZShift = −1.95
Z lm Score = −0.79
NG = 0
DB = 0
SM = 0
Delta_AUC
Doxorubicin

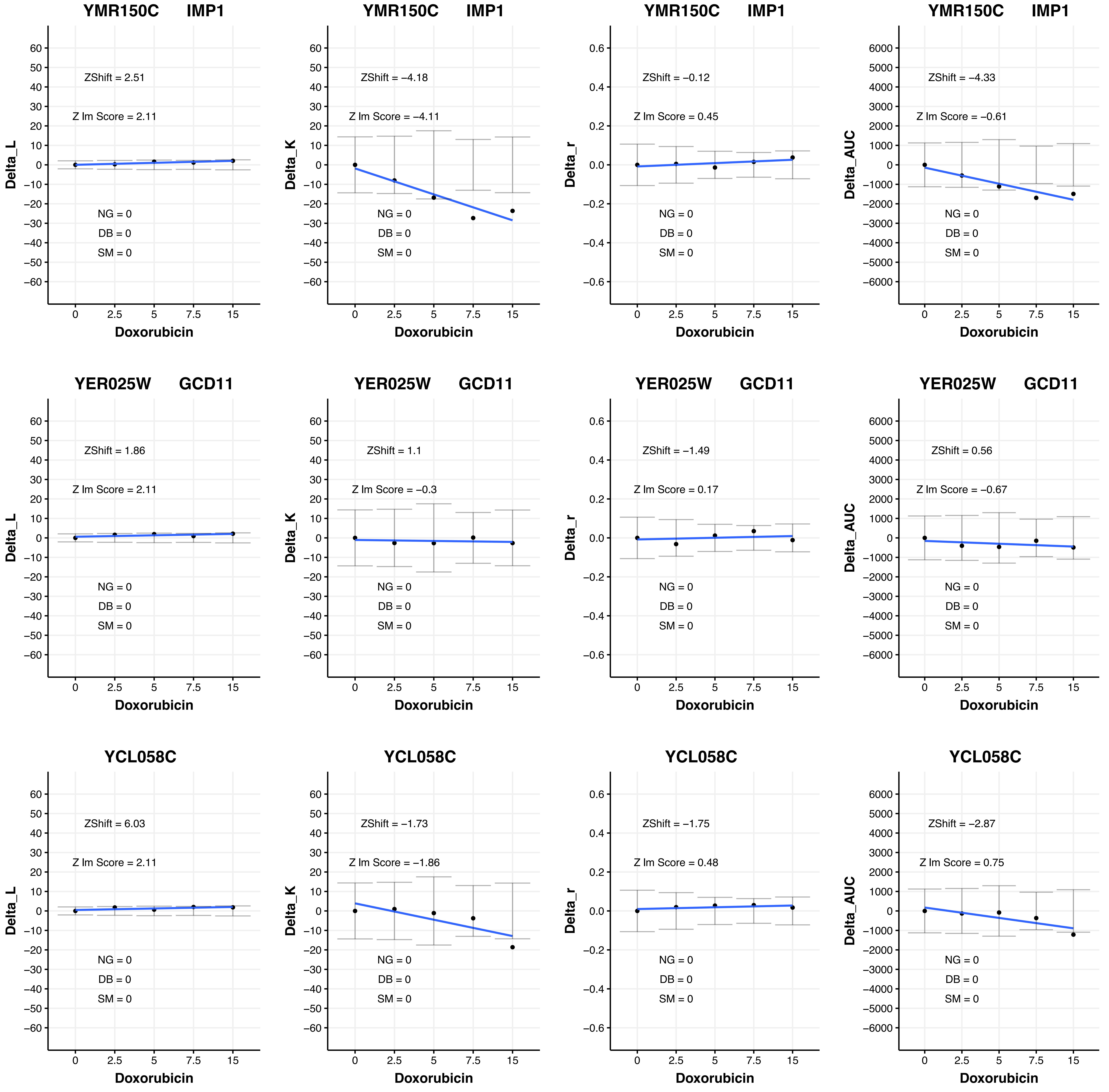
YMR150C IMP1
ZShift = 2.51
Z lm Score = 2.11
NG = 0
DB = 0
SM = 0
Delta_L
Doxorubicin
YMR150C IMP1
ZShift = −4.18
Z lm Score = −4.11
NG = 0
DB = 0
SM = 0
Delta_K
Doxorubicin
YMR150C IMP1
ZShift = −0.12
Z lm Score = 0.45
NG = 0
DB = 0
SM = 0
Delta_r
Doxorubicin
YMR150C IMP1
ZShift = −4.33
Z lm Score = −0.61
NG = 0
DB = 0
SM = 0
Delta_AUC
Doxorubicin
YER025W GCD11
ZShift = 1.86
Z lm Score = 2.11
NG = 0
DB = 0
SM = 0
Delta_L
Doxorubicin
YER025W GCD11
ZShift = 1.1
Z lm Score = −0.3
NG = 0
DB = 0
SM = 0
Delta_K
Doxorubicin
YER025W GCD11
ZShift = −1.49
Z lm Score = 0.17
NG = 0
DB = 0
SM = 0
Delta_r
Doxorubicin
YER025W GCD11
ZShift = 0.56
Z lm Score = −0.67
NG = 0
DB = 0
SM = 0
Delta_AUC
Doxorubicin
YCL058C
ZShift = 6.03
Z lm Score = 2.11
NG = 0
DB = 0
SM = 0
Delta_L
Doxorubicin
YCL058C
ZShift = −1.73
Z lm Score = −1.86
NG = 0
DB = 0
SM = 0
Delta_K
Doxorubicin
YCL058C
ZShift = −1.75
Z lm Score = 0.48
NG = 0
DB = 0
SM = 0
Delta_r
Doxorubicin
YCL058C
ZShift = −2.87
Z lm Score = 0.75
NG = 0
DB = 0
SM = 0
Delta_AUC
Doxorubicin

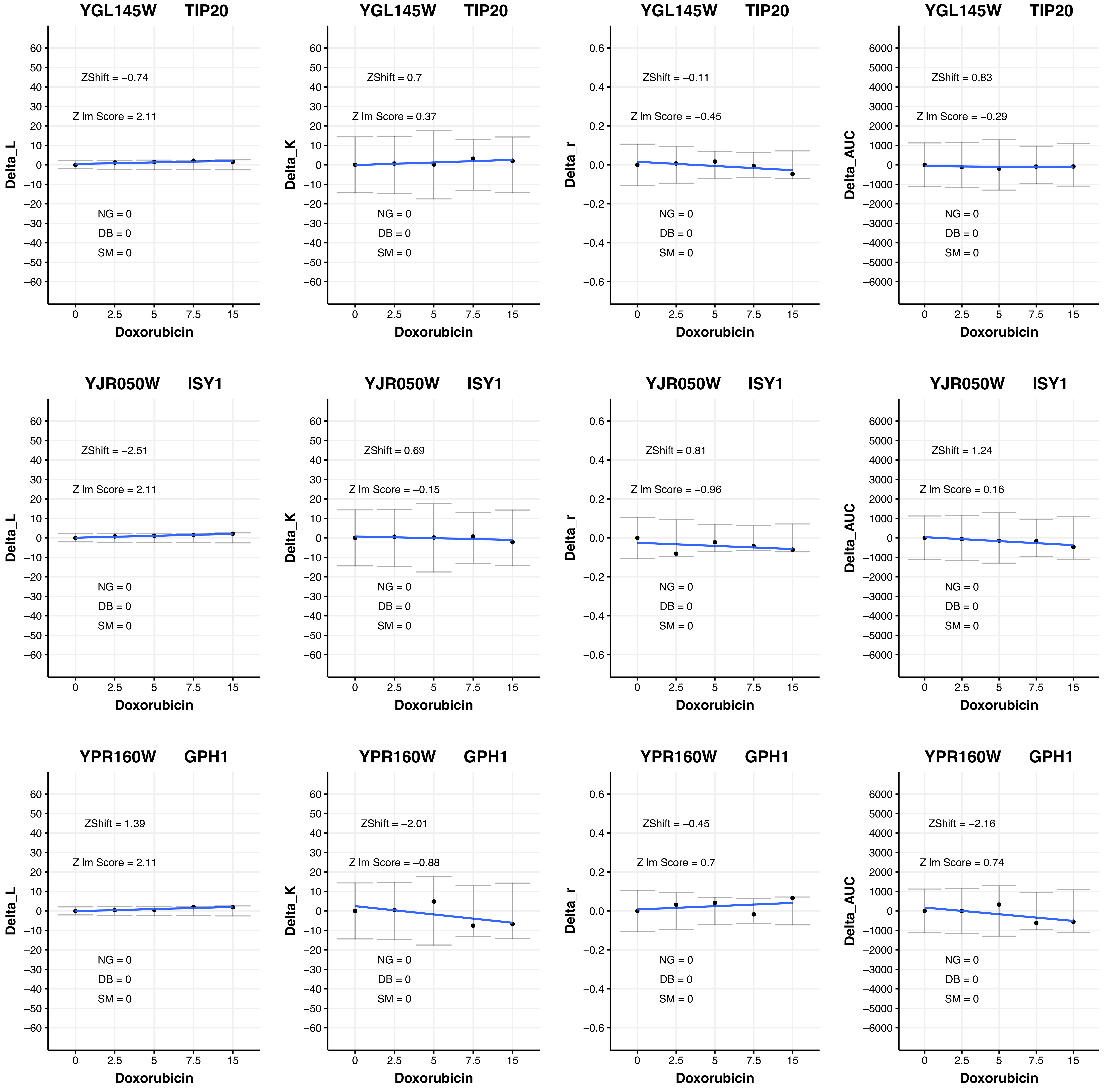
YGL145W TIP20
ZShift = −0.74
Z lm Score = 2.11
NG = 0
DB = 0
SM = 0
Delta_L
Doxorubicin
YGL145W TIP20
ZShift = 0.7
Z lm Score = 0.37
NG = 0
DB = 0
SM = 0
Delta_K
Doxorubicin
YGL145W TIP20
ZShift = −0.11
Z lm Score = −0.45
NG = 0
DB = 0
SM = 0
Delta_r
Doxorubicin
YGL145W TIP20
ZShift = 0.83
Z lm Score = −0.29
NG = 0
DB = 0
SM = 0
Delta_AUC
Doxorubicin
YJR050W ISY1
ZShift = −2.51
Z lm Score = 2.11
NG = 0
DB = 0
SM = 0
Delta_L
Doxorubicin
YJR050W ISY1
ZShift = 0.69
Z lm Score = −0.15
NG = 0
DB = 0
SM = 0
Delta_K
Doxorubicin
YJR050W ISY1
ZShift = 0.81
Z lm Score = −0.96
NG = 0
DB = 0
SM = 0
Delta_r
Doxorubicin
YJR050W ISY1
ZShift = 1.24
Z lm Score = 0.16
NG = 0
DB = 0
SM = 0
Delta_AUC
Doxorubicin
YPR160W GPH1
ZShift = 1.39
Z lm Score = 2.11
NG = 0
DB = 0
SM = 0
Delta_L
Doxorubicin
YPR160W GPH1
ZShift = −2.01
Z lm Score = −0.88
NG = 0
DB = 0
SM = 0
Delta_K
Doxorubicin
YPR160W GPH1
ZShift = −0.45
Z lm Score = 0.7
NG = 0
DB = 0
SM = 0
Delta_r
Doxorubicin
YPR160W GPH1
ZShift = −2.16
Z lm Score = 0.74
NG = 0
DB = 0
SM = 0
Delta_AUC
Doxorubicin

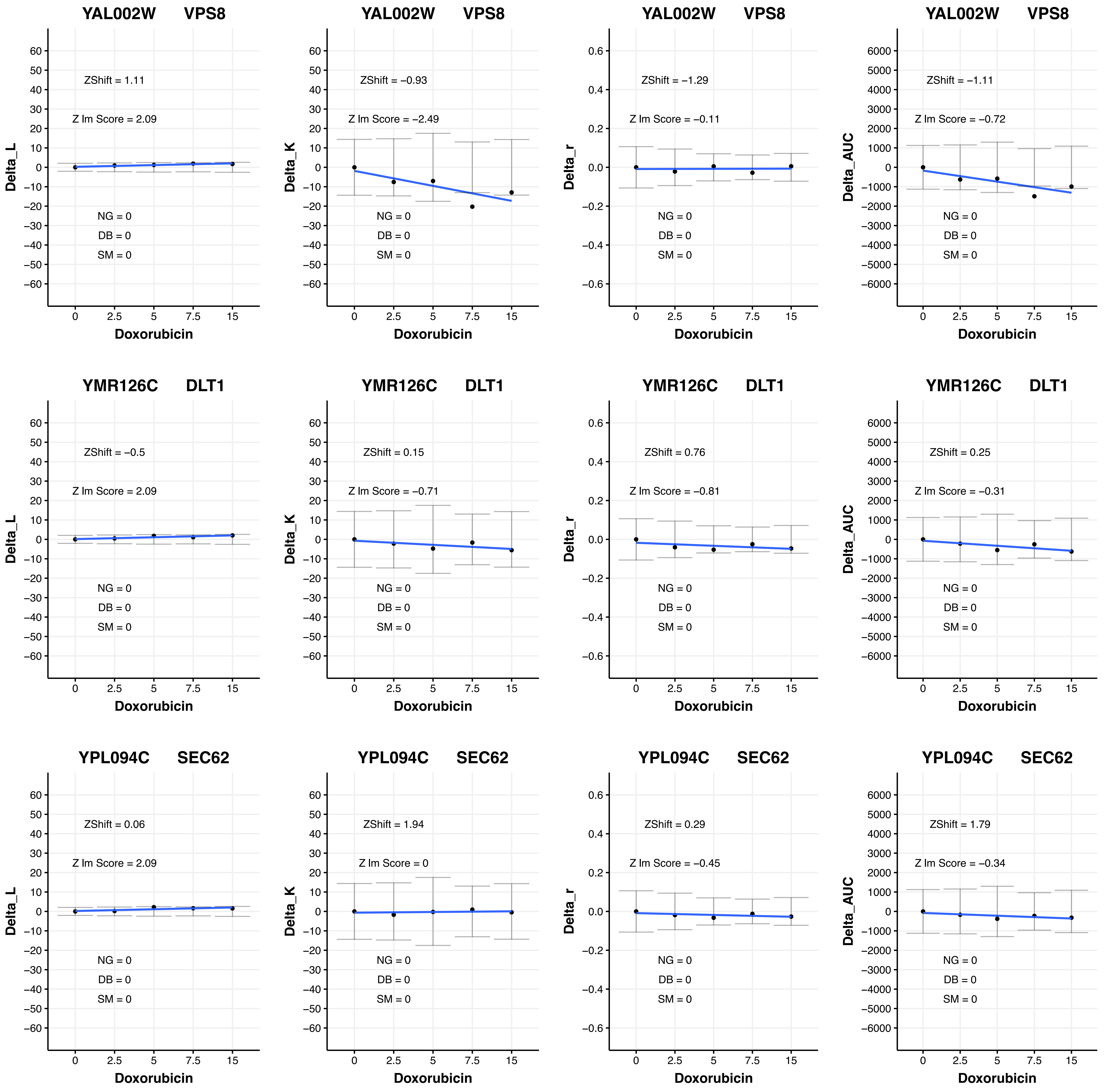
YAL002W VPS8
ZShift = 1.11
Z lm Score = 2.09
NG = 0
DB = 0
SM = 0
Delta_L
Doxorubicin
YAL002W VPS8
ZShift = −0.93
Z lm Score = −2.49
NG = 0
DB = 0
SM = 0
Delta_K
Doxorubicin
YAL002W VPS8
ZShift = −1.29
Z lm Score = −0.11
NG = 0
DB = 0
SM = 0
Delta_r
Doxorubicin
YAL002W VPS8
ZShift = −1.11
Z lm Score = −0.72
NG = 0
DB = 0
SM = 0
Delta_AUC
Doxorubicin
YMR126C DLT1
ZShift = −0.5
Z lm Score = 2.09
NG = 0
DB = 0
SM = 0
Delta_L
Doxorubicin
YMR126C DLT1
ZShift = 0.15
Z lm Score = −0.71
NG = 0
DB = 0
SM = 0
Delta_K
Doxorubicin
YMR126C DLT1
ZShift = 0.76
Z lm Score = −0.81
NG = 0
DB = 0
SM = 0
Delta_r
Doxorubicin
YMR126C DLT1
ZShift = 0.25
Z lm Score = −0.31
NG = 0
DB = 0
SM = 0
Delta_AUC
Doxorubicin
YPL094C SEC62
ZShift = 0.06
Z lm Score = 2.09
NG = 0
DB = 0
SM = 0
Delta_L
Doxorubicin
YPL094C SEC62
ZShift = 1.94
Z lm Score = 0
NG = 0
DB = 0
SM = 0
Delta_K
Doxorubicin
YPL094C SEC62
ZShift = 0.29
Z lm Score = −0.45
NG = 0
DB = 0
SM = 0
Delta_r
Doxorubicin
YPL094C SEC62
ZShift = 1.79
Z lm Score = −0.34
NG = 0
DB = 0
SM = 0
Delta_AUC
Doxorubicin

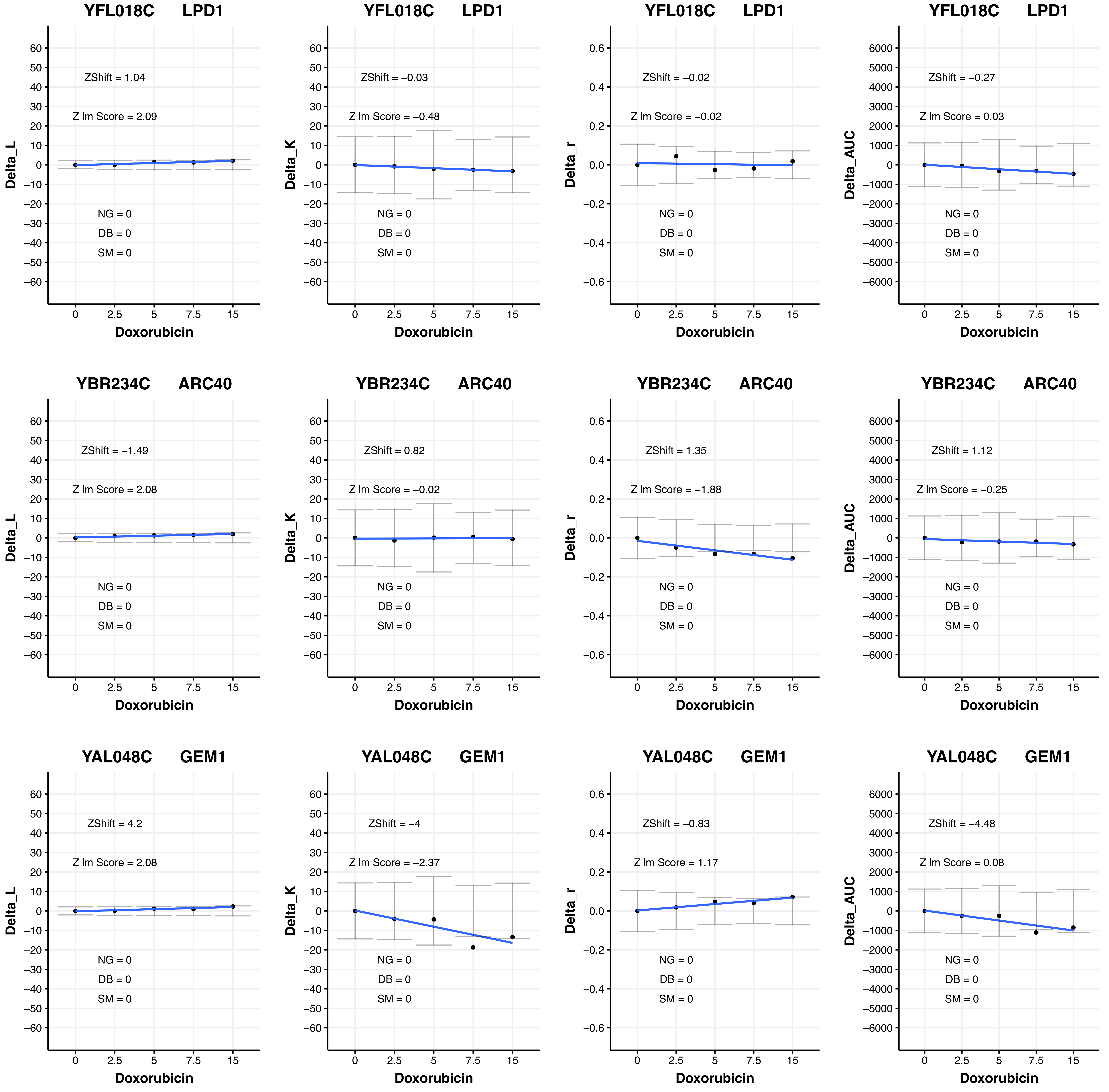
YFL018C LPD1
ZShift = 1.04
Z lm Score = 2.09
NG = 0
DB = 0
SM = 0
Delta_L
Doxorubicin
YFL018C LPD1
ZShift = −0.03
Z lm Score = −0.48
NG = 0
DB = 0
SM = 0
Delta_K
Doxorubicin
YFL018C LPD1
ZShift = −0.02
Z lm Score = −0.02
NG = 0
DB = 0
SM = 0
Delta_r
Doxorubicin
YFL018C LPD1
ZShift = −0.27
Z lm Score = 0.03
NG = 0
DB = 0
SM = 0
Delta_AUC
Doxorubicin
YBR234C ARC40
ZShift = −1.49
Z lm Score = 2.08
NG = 0
DB = 0
SM = 0
Delta_L
Doxorubicin
YBR234C ARC40
ZShift = 0.82
Z lm Score = −0.02
NG = 0
DB = 0
SM = 0
Delta_K
Doxorubicin
YBR234C ARC40
ZShift = 1.35
Z lm Score = −1.88
NG = 0
DB = 0
SM = 0
Delta_r
Doxorubicin
YBR234C ARC40
ZShift = 1.12
Z lm Score = −0.25
NG = 0
DB = 0
SM = 0
Delta_AUC
Doxorubicin
YAL048C GEM1
ZShift = 4.2
Z lm Score = 2.08
NG = 0
DB = 0
SM = 0
Delta_L
Doxorubicin
YAL048C GEM1
ZShift = −4
Z lm Score = −2.37
NG = 0
DB = 0
SM = 0
Delta_K
Doxorubicin
YAL048C GEM1
ZShift = −0.83
Z lm Score = 1.17
NG = 0
DB = 0
SM = 0
Delta_r
Doxorubicin
YAL048C GEM1
ZShift = −4.48
Z lm Score = 0.08
NG = 0
DB = 0
SM = 0
Delta_AUC
Doxorubicin

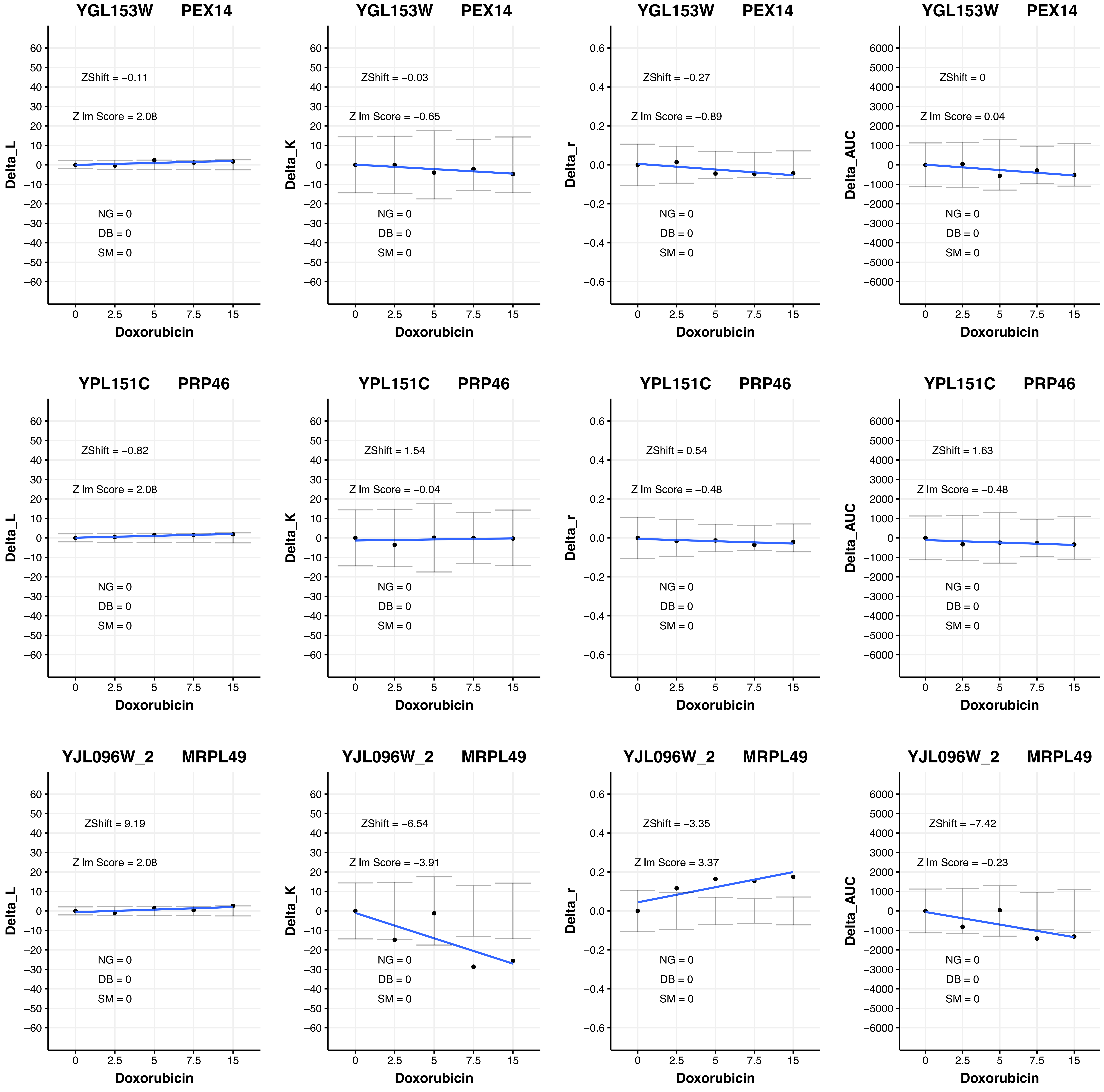
YGL153W PEX14
ZShift = −0.11
Z lm Score = 2.08
NG = 0
DB = 0
SM = 0
Delta_L
Doxorubicin
YGL153W PEX14
ZShift = −0.03
Z lm Score = −0.65
NG = 0
DB = 0
SM = 0
Delta_K
Doxorubicin
YGL153W PEX14
ZShift = −0.27
Z lm Score = −0.89
NG = 0
DB = 0
SM = 0
Delta_r
Doxorubicin
YGL153W PEX14
ZShift = 0
Z lm Score = 0.04
NG = 0
DB = 0
SM = 0
Delta_AUC
Doxorubicin
YPL151C PRP46
ZShift = −0.82
Z lm Score = 2.08
NG = 0
DB = 0
SM = 0
Delta_L
Doxorubicin
YPL151C PRP46
ZShift = 1.54
Z lm Score = −0.04
NG = 0
DB = 0
SM = 0
Delta_K
Doxorubicin
YPL151C PRP46
ZShift = 0.54
Z lm Score = −0.48
NG = 0
DB = 0
SM = 0
Delta_r
Doxorubicin
YPL151C PRP46
ZShift = 1.63
Z lm Score = −0.48
NG = 0
DB = 0
SM = 0
Delta_AUC
Doxorubicin
YJL096W_2 MRPL49
ZShift = 9.19
Z lm Score = 2.08
NG = 0
DB = 0
SM = 0
Delta_L
Doxorubicin
YJL096W_2 MRPL49
ZShift = −6.54
Z lm Score = −3.91
NG = 0
DB = 0
SM = 0
Delta_K
Doxorubicin
YJL096W_2 MRPL49
ZShift = −3.35
Z lm Score = 3.37
NG = 0
DB = 0
SM = 0
Delta_r
Doxorubicin
YJL096W_2 MRPL49
ZShift = −7.42
Z lm Score = −0.23
NG = 0
DB = 0
SM = 0
Delta_AUC
Doxorubicin

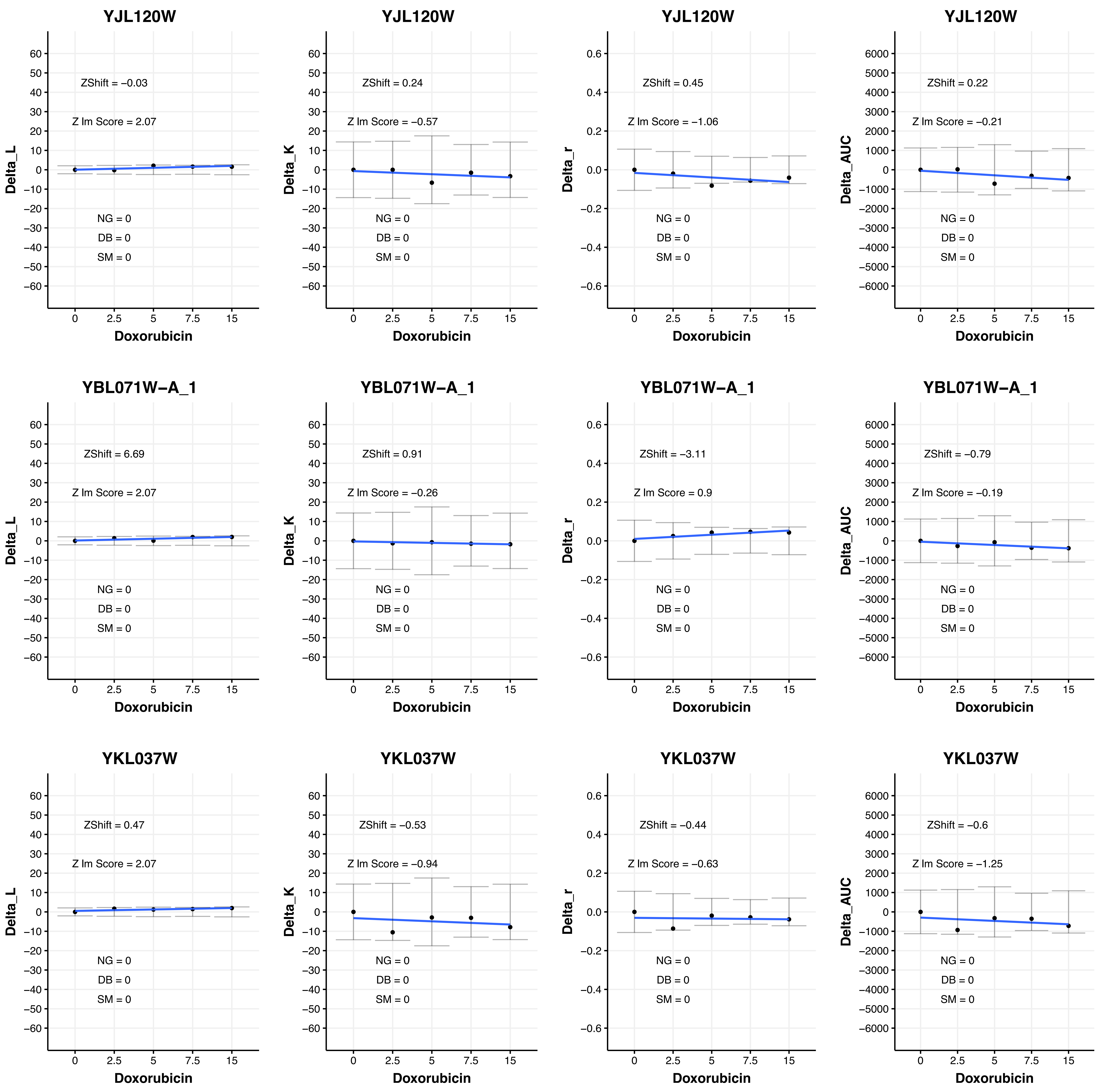
YJL120W
ZShift = −0.03
Z lm Score = 2.07
NG = 0
DB = 0
SM = 0
Delta_L
Doxorubicin
YJL120W
ZShift = 0.24
Z lm Score = −0.57
NG = 0
DB = 0
SM = 0
Delta_K
Doxorubicin
YJL120W
ZShift = 0.45
Z lm Score = −1.06
NG = 0
DB = 0
SM = 0
Delta_r
Doxorubicin
YJL120W
ZShift = 0.22
Z lm Score = −0.21
NG = 0
DB = 0
SM = 0
Delta_AUC
Doxorubicin
YBL071W−A_1
ZShift = 6.69
Z lm Score = 2.07
NG = 0
DB = 0
SM = 0
Delta_L
Doxorubicin
YBL071W−A_1
ZShift = 0.91
Z lm Score = −0.26
NG = 0
DB = 0
SM = 0
Delta_K
Doxorubicin
YBL071W−A_1
ZShift = −3.11
Z lm Score = 0.9
NG = 0
DB = 0
SM = 0
Delta_r
Doxorubicin
YBL071W−A_1
ZShift = −0.79
Z lm Score = −0.19
NG = 0
DB = 0
SM = 0
Delta_AUC
Doxorubicin
YKL037W
ZShift = 0.47
Z lm Score = 2.07
NG = 0
DB = 0
SM = 0
Delta_L
Doxorubicin
YKL037W
ZShift = −0.53
Z lm Score = −0.94
NG = 0
DB = 0
SM = 0
Delta_K
Doxorubicin
YKL037W
ZShift = −0.44
Z lm Score = −0.63
NG = 0
DB = 0
SM = 0
Delta_r
Doxorubicin
YKL037W
ZShift = −0.6
Z lm Score = −1.25
NG = 0
DB = 0
SM = 0
Delta_AUC
Doxorubicin

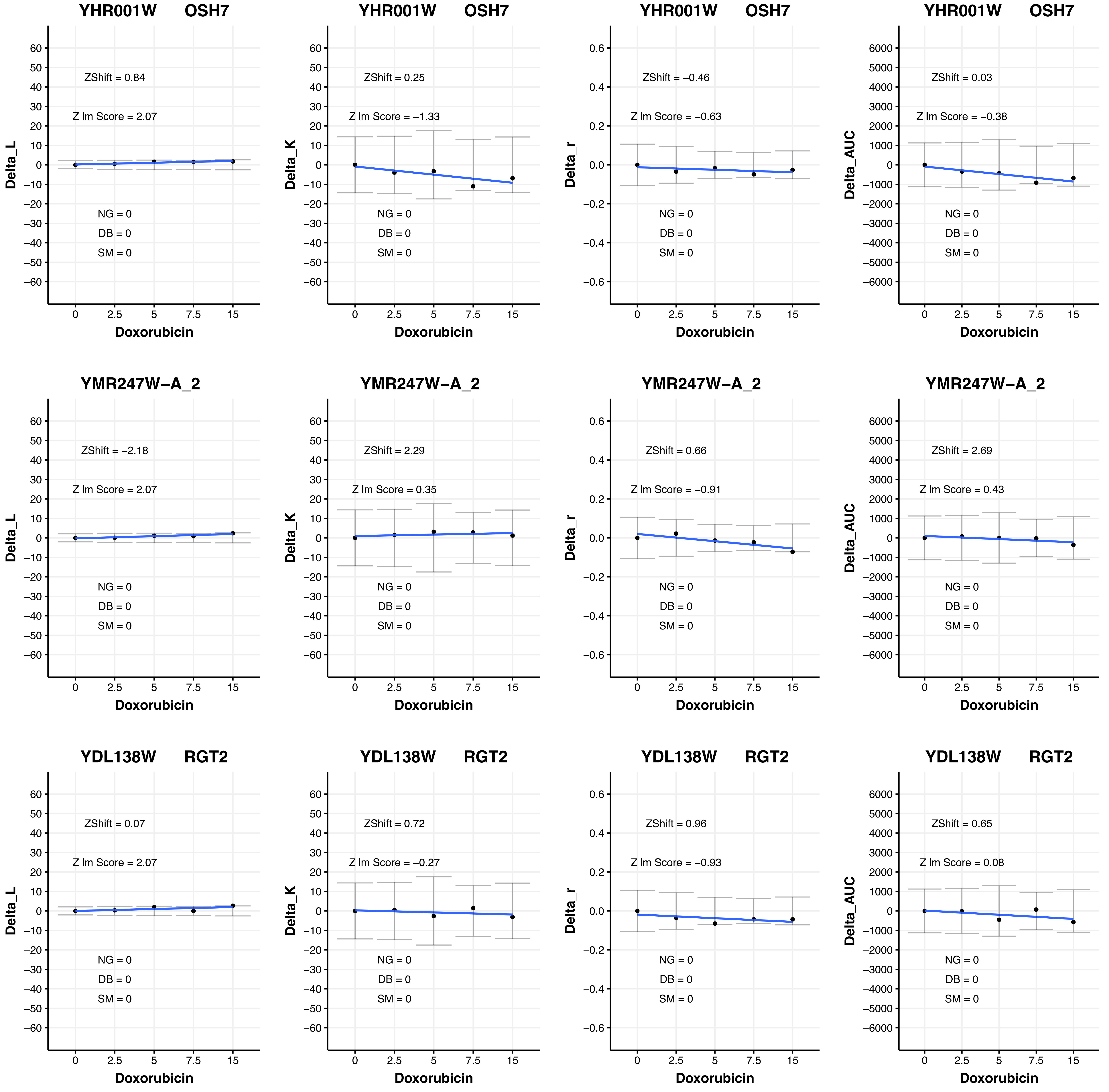
YHR001W OSH7
ZShift = 0.84
Z lm Score = 2.07
NG = 0
DB = 0
SM = 0
Delta_L
Doxorubicin
YHR001W OSH7
ZShift = 0.25
Z lm Score = −1.33
NG = 0
DB = 0
SM = 0
Delta_K
Doxorubicin
YHR001W OSH7
ZShift = −0.46
Z lm Score = −0.63
NG = 0
DB = 0
SM = 0
Delta_r
Doxorubicin
YHR001W OSH7
ZShift = 0.03
Z lm Score = −0.38
NG = 0
DB = 0
SM = 0
Delta_AUC
Doxorubicin
YMR247W−A_2
ZShift = −2.18
Z lm Score = 2.07
NG = 0
DB = 0
SM = 0
Delta_L
Doxorubicin
YMR247W−A_2
ZShift = 2.29
Z lm Score = 0.35
NG = 0
DB = 0
SM = 0
Delta_K
Doxorubicin
YMR247W−A_2
ZShift = 0.66
Z lm Score = −0.91
NG = 0
DB = 0
SM = 0
Delta_r
Doxorubicin
YMR247W−A_2
ZShift = 2.69
Z lm Score = 0.43
NG = 0
DB = 0
SM = 0
Delta_AUC
Doxorubicin
YDL138W RGT2
ZShift = 0.07
Z lm Score = 2.07
NG = 0
DB = 0
SM = 0
Delta_L
Doxorubicin
YDL138W RGT2
ZShift = 0.72
Z lm Score = −0.27
NG = 0
DB = 0
SM = 0
Delta_K
Doxorubicin
YDL138W RGT2
ZShift = 0.96
Z lm Score = −0.93
NG = 0
DB = 0
SM = 0
Delta_r
Doxorubicin
YDL138W RGT2
ZShift = 0.65
Z lm Score = 0.08
NG = 0
DB = 0
SM = 0
Delta_AUC
Doxorubicin

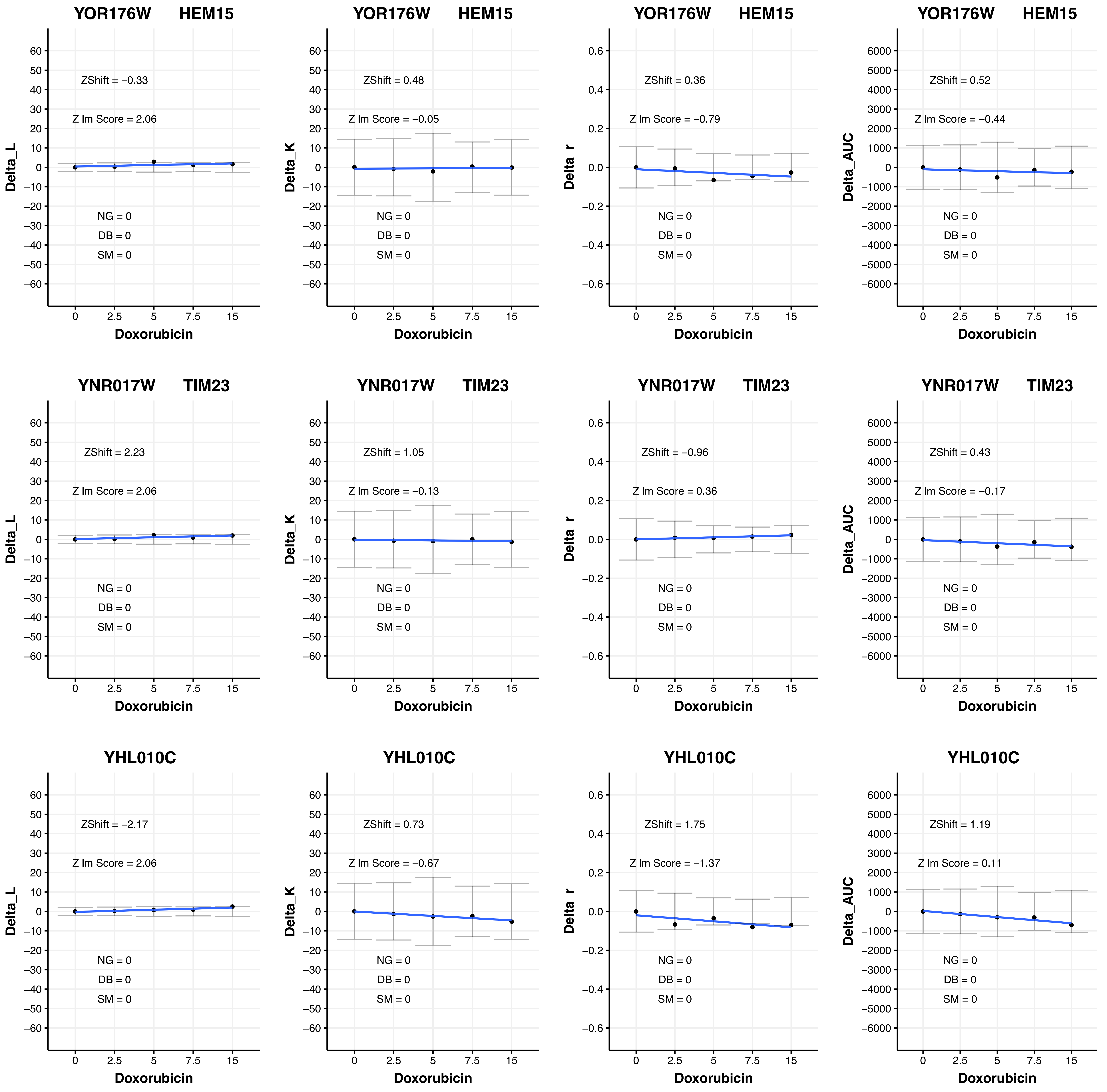

YOR176W HEM15
ZShift = −0.33
Z lm Score = 2.06
Delta_L
NG = 0
DB = 0
SM = 0
Doxorubicin
YOR176W HEM15
ZShift = 0.48
Z lm Score = −0.05
Delta_K
NG = 0
DB = 0
SM = 0
Doxorubicin
YOR176W HEM15
ZShift = 0.36
Z lm Score = −0.79
Delta_r
NG = 0
DB = 0
SM = 0
Doxorubicin
YOR176W HEM15
ZShift = 0.52
Z lm Score = −0.44
Delta_AUC
NG = 0
DB = 0
SM = 0
Doxorubicin
YNR017W TIM23
ZShift = 2.23
Z lm Score = 2.06
Delta_L
NG = 0
DB = 0
SM = 0
Doxorubicin
YNR017W TIM23
ZShift = 1.05
Z lm Score = −0.13
Delta_K
NG = 0
DB = 0
SM = 0
Doxorubicin
YNR017W TIM23
ZShift = −0.96
Z lm Score = 0.36
Delta_r
NG = 0
DB = 0
SM = 0
Doxorubicin
YNR017W TIM23
ZShift = 0.43
Z lm Score = −0.17
Delta_AUC
NG = 0
DB = 0
SM = 0
Doxorubicin
YHL010C
ZShift = −2.17
Z lm Score = 2.06
Delta_L
NG = 0
DB = 0
SM = 0
Doxorubicin
YHL010C
ZShift = 0.73
Z lm Score = −0.67
Delta_K
NG = 0
DB = 0
SM = 0
Doxorubicin
YHL010C
ZShift = 1.75
Z lm Score = −1.37
Delta_r
NG = 0
DB = 0
SM = 0
Doxorubicin
YHL010C
ZShift = 1.19
Z lm Score = 0.11
Delta_AUC
NG = 0
DB = 0
SM = 0
Doxorubicin

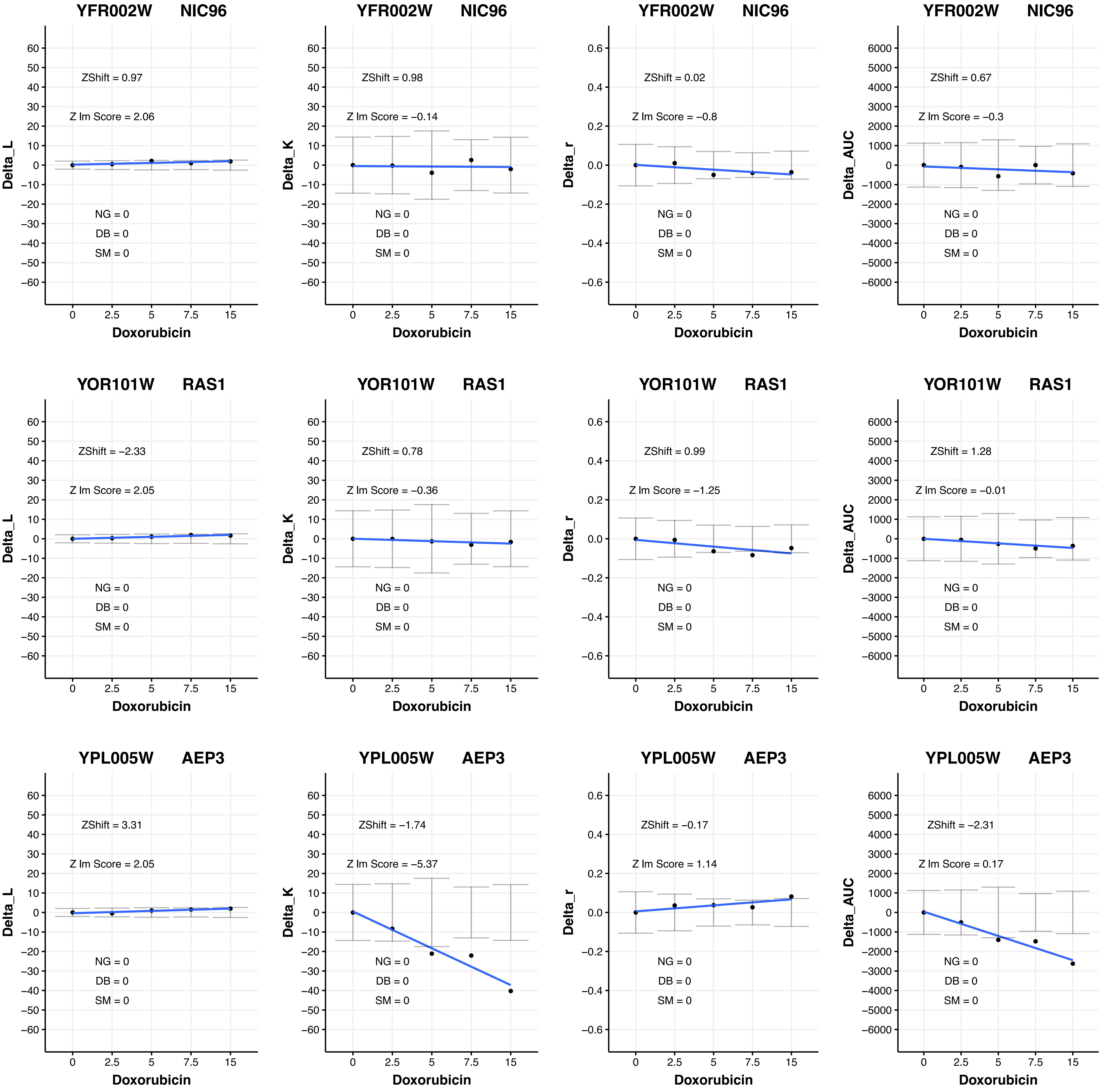
YFR002W NIC96
ZShift = 0.97
Z lm Score = 2.06
Delta_L
NG = 0
DB = 0
SM = 0
Doxorubicin
YFR002W NIC96
ZShift = 0.98
Z lm Score = −0.14
Delta_K
NG = 0
DB = 0
SM = 0
Doxorubicin
YFR002W NIC96
ZShift = 0.02
Z lm Score = −0.8
Delta_r
NG = 0
DB = 0
SM = 0
Doxorubicin
YFR002W NIC96
ZShift = 0.67
Z lm Score = −0.3
Delta_AUC
NG = 0
DB = 0
SM = 0
Doxorubicin
YOR101W RAS1
ZShift = −2.33
Z lm Score = 2.05
Delta_L
NG = 0
DB = 0
SM = 0
Doxorubicin
YOR101W RAS1
ZShift = 0.78
Z lm Score = −0.36
Delta_K
NG = 0
DB = 0
SM = 0
Doxorubicin
YOR101W RAS1
ZShift = 0.99
Z lm Score = −1.25
Delta_r
NG = 0
DB = 0
SM = 0
Doxorubicin
YOR101W RAS1
ZShift = 1.28
Z lm Score = −0.01
Delta_AUC
NG = 0
DB = 0
SM = 0
Doxorubicin
YPL005W AEP3
ZShift = 3.31
Z lm Score = 2.05
Delta_L
NG = 0
DB = 0
SM = 0
Doxorubicin
YPL005W AEP3
ZShift = −1.74
Z lm Score = −5.37
Delta_K
NG = 0
DB = 0
SM = 0
Doxorubicin
YPL005W AEP3
ZShift = −0.17
Z lm Score = 1.14
Delta_r
NG = 0
DB = 0
SM = 0
Doxorubicin
YPL005W AEP3
ZShift = −2.31
Z lm Score = 0.17
Delta_AUC
NG = 0
DB = 0
SM = 0
Doxorubicin

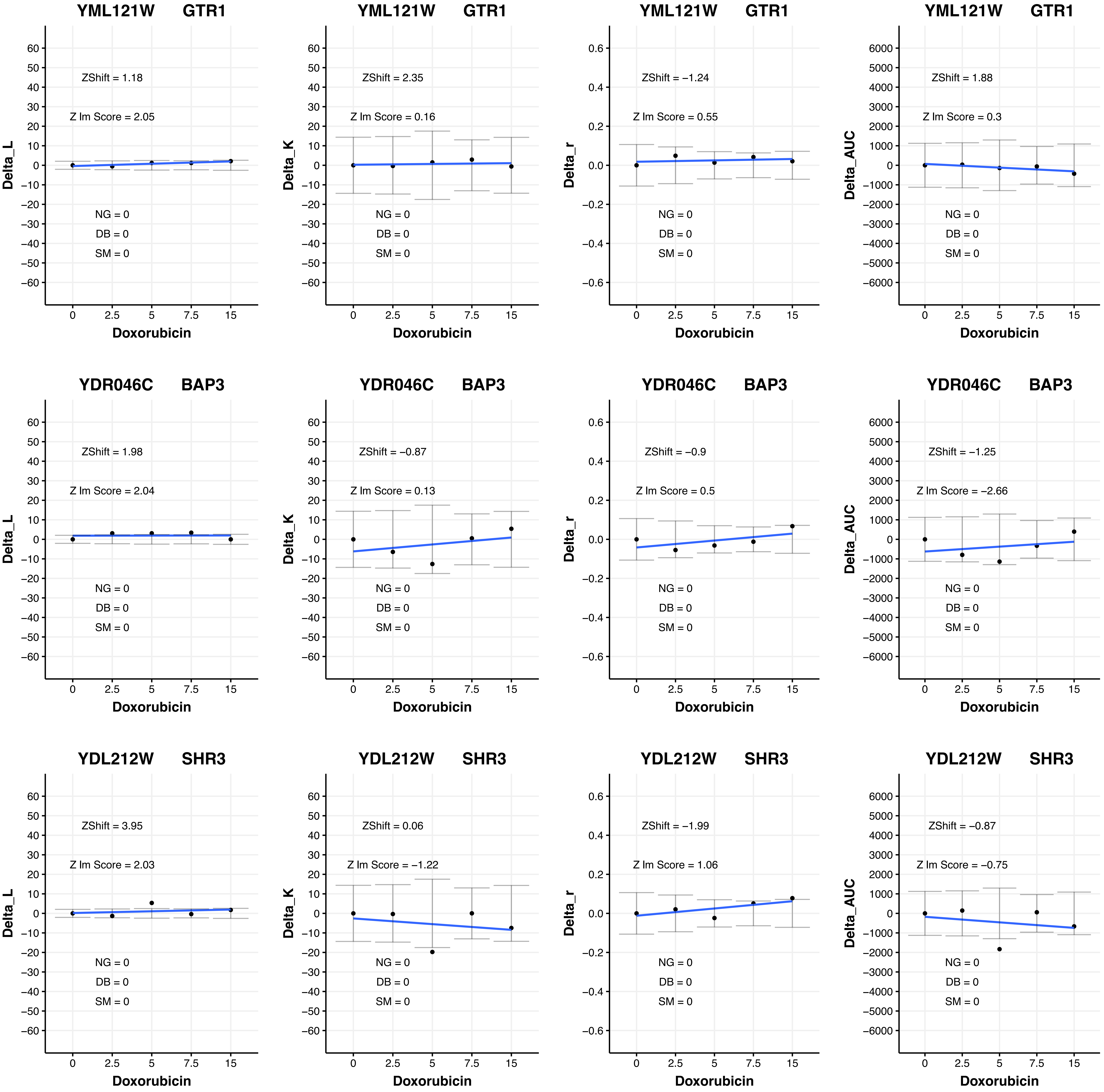

YML121W GTR1
ZShift = 1.18
Z lm Score = 2.05
Delta_L
NG = 0
DB = 0
SM = 0
Doxorubicin
YML121W GTR1
ZShift = 2.35
Z lm Score = 0.16
Delta_K
NG = 0
DB = 0
SM = 0
Doxorubicin
YML121W GTR1
ZShift = −1.24
Z lm Score = 0.55
Delta_r
NG = 0
DB = 0
SM = 0
Doxorubicin
YML121W GTR1
ZShift = 1.88
Z lm Score = 0.3
Delta_AUC
NG = 0
DB = 0
SM = 0
Doxorubicin
YDR046C BAP3
ZShift = 1.98
Z lm Score = 2.04
Delta_L
NG = 0
DB = 0
SM = 0
Doxorubicin
YDR046C BAP3
ZShift = −0.87
Z lm Score = 0.13
Delta_K
NG = 0
DB = 0
SM = 0
Doxorubicin
YDR046C BAP3
ZShift = −0.9
Z lm Score = 0.5
Delta_r
NG = 0
DB = 0
SM = 0
Doxorubicin
YDR046C BAP3
ZShift = −1.25
Z lm Score = −2.66
Delta_AUC
NG = 0
DB = 0
SM = 0
Doxorubicin
YDL212W SHR3
ZShift = 3.95
Z lm Score = 2.03
Delta_L
NG = 0
DB = 0
SM = 0
Doxorubicin
YDL212W SHR3
ZShift = 0.06
Z lm Score = −1.22
Delta_K
NG = 0
DB = 0
SM = 0
Doxorubicin
YDL212W SHR3
ZShift = −1.99
Z lm Score = 1.06
Delta_r
NG = 0
DB = 0
SM = 0
Doxorubicin
YDL212W SHR3
ZShift = −0.87
Z lm Score = −0.75
Delta_AUC
NG = 0
DB = 0
SM = 0
Doxorubicin

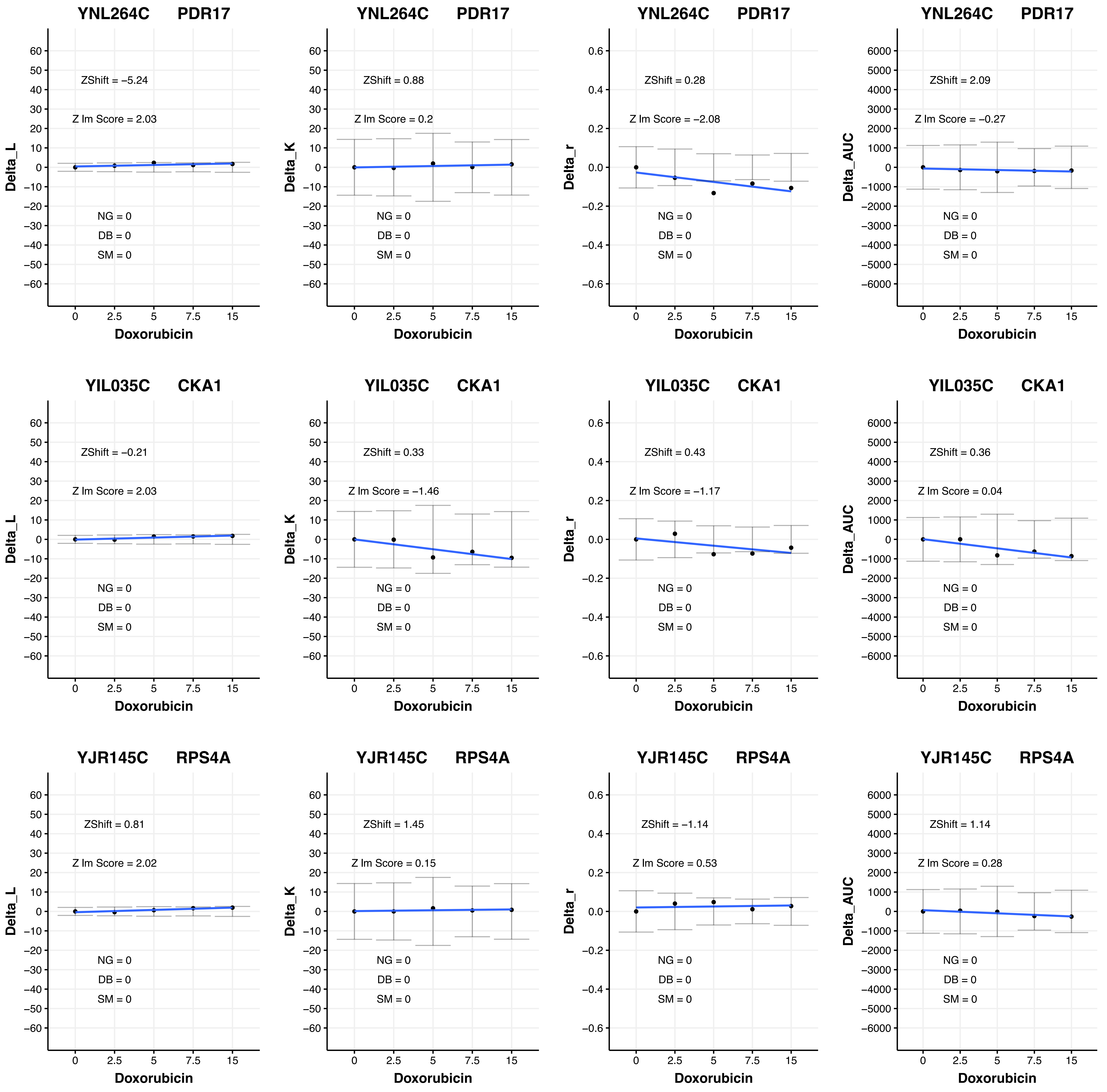

YNL264C PDR17
ZShift = −5.24
Z lm Score = 2.03
NG = 0
DB = 0
SM = 0
Delta_L
Doxorubicin
YNL264C PDR17
ZShift = 0.88
Z lm Score = 0.2
NG = 0
DB = 0
SM = 0
Delta_K
Doxorubicin
YNL264C PDR17
ZShift = 0.28
Z lm Score = −2.08
NG = 0
DB = 0
SM = 0
Delta_r
Doxorubicin
YNL264C PDR17
ZShift = 2.09
Z lm Score = −0.27
NG = 0
DB = 0
SM = 0
Delta_AUC
Doxorubicin
YIL035C CKA1
ZShift = −0.21
Z lm Score = 2.03
NG = 0
DB = 0
SM = 0
Delta_L
Doxorubicin
YIL035C CKA1
ZShift = 0.33
Z lm Score = −1.46
NG = 0
DB = 0
SM = 0
Delta_K
Doxorubicin
YIL035C CKA1
ZShift = 0.43
Z lm Score = −1.17
NG = 0
DB = 0
SM = 0
Delta_r
Doxorubicin
YIL035C CKA1
ZShift = 0.36
Z lm Score = 0.04
NG = 0
DB = 0
SM = 0
Delta_AUC
Doxorubicin
YJR145C RPS4A
ZShift = 0.81
Z lm Score = 2.02
NG = 0
DB = 0
SM = 0
Delta_L
Doxorubicin
YJR145C RPS4A
ZShift = 1.45
Z lm Score = 0.15
NG = 0
DB = 0
SM = 0
Delta_K
Doxorubicin
YJR145C RPS4A
ZShift = −1.14
Z lm Score = 0.53
NG = 0
DB = 0
SM = 0
Delta_r
Doxorubicin
YJR145C RPS4A
ZShift = 1.14
Z lm Score = 0.28
NG = 0
DB = 0
SM = 0
Delta_AUC
Doxorubicin

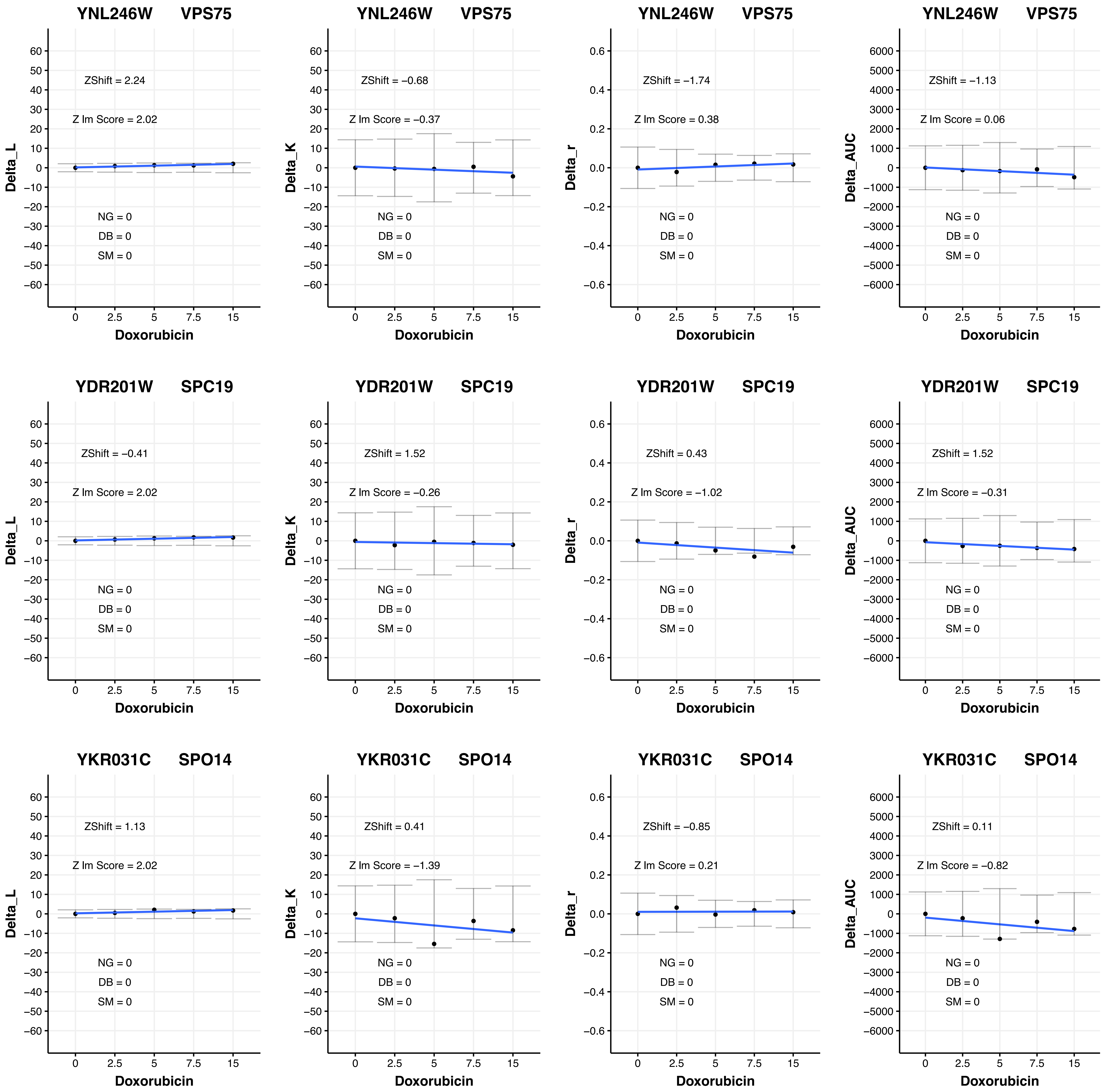
YNL246W VPS75
ZShift = 2.24
Z lm Score = 2.02
NG = 0
DB = 0
SM = 0
Delta_L
Doxorubicin
YNL246W VPS75
ZShift = −0.68
Z lm Score = −0.37
NG = 0
DB = 0
SM = 0
Delta_K
Doxorubicin
YNL246W VPS75
ZShift = −1.74
Z lm Score = 0.38
NG = 0
DB = 0
SM = 0
Delta_r
Doxorubicin
YNL246W VPS75
ZShift = −1.13
Z lm Score = 0.06
NG = 0
DB = 0
SM = 0
Delta_AUC
Doxorubicin
YDR201W SPC19
ZShift = −0.41
Z lm Score = 2.02
NG = 0
DB = 0
SM = 0
Delta_L
Doxorubicin
YDR201W SPC19
ZShift = 1.52
Z lm Score = −0.26
NG = 0
DB = 0
SM = 0
Delta_K
Doxorubicin
YDR201W SPC19
ZShift = 0.43
Z lm Score = −1.02
NG = 0
DB = 0
SM = 0
Delta_r
Doxorubicin
YDR201W SPC19
ZShift = 1.52
Z lm Score = −0.31
NG = 0
DB = 0
SM = 0
Delta_AUC
Doxorubicin
YKR031C SPO14
ZShift = 1.13
Z lm Score = 2.02
NG = 0
DB = 0
SM = 0
Delta_L
Doxorubicin
YKR031C SPO14
ZShift = 0.41
Z lm Score = −1.39
NG = 0
DB = 0
SM = 0
Delta_K
Doxorubicin
YKR031C SPO14
ZShift = −0.85
Z lm Score = 0.21
NG = 0
DB = 0
SM = 0
Delta_r
Doxorubicin
YKR031C SPO14
ZShift = 0.11
Z lm Score = −0.82
NG = 0
DB = 0
SM = 0
Delta_AUC
Doxorubicin

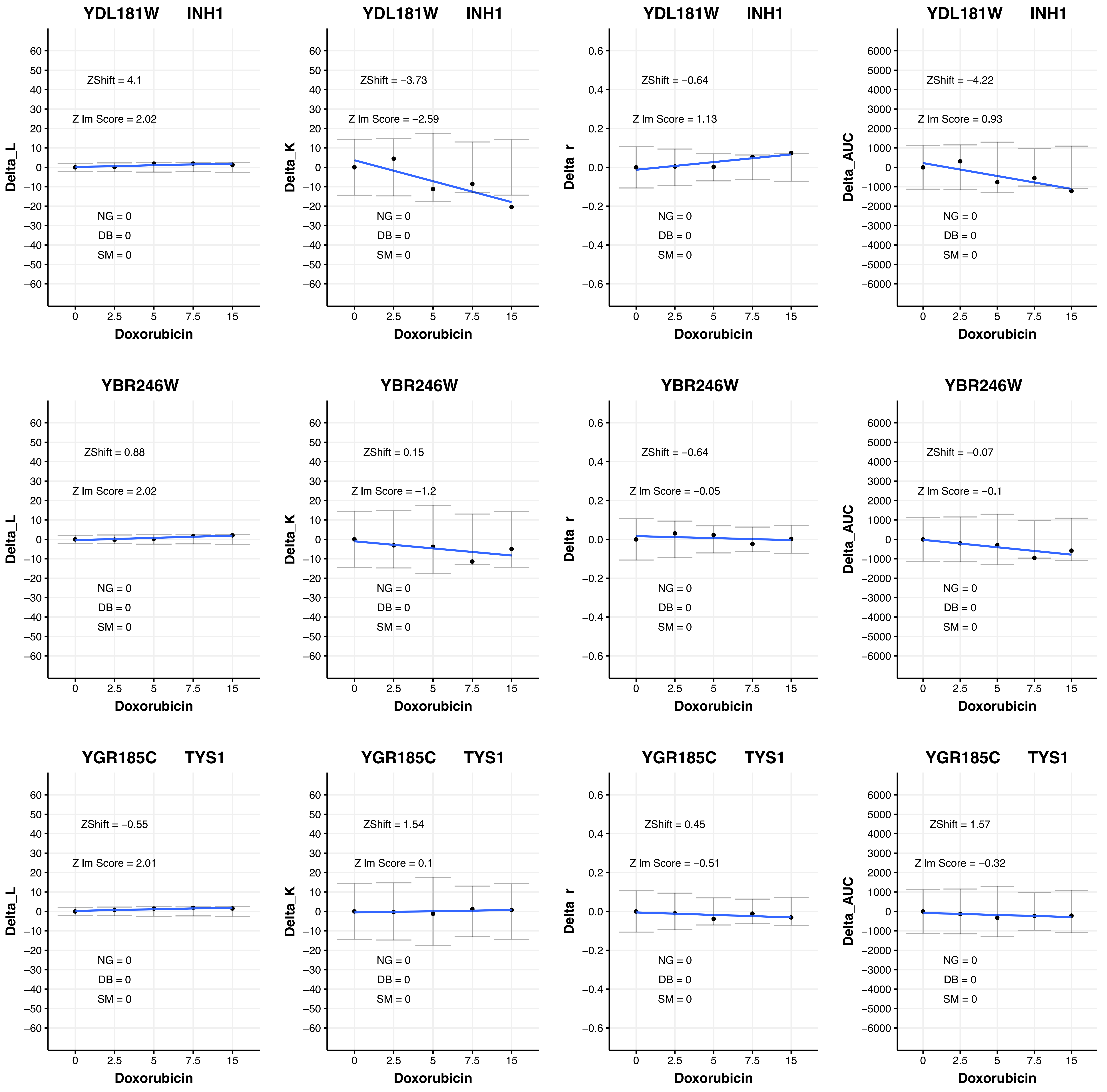
YDL181W INH1
ZShift = 4.1
Z lm Score = 2.02
NG = 0
DB = 0
SM = 0
Delta_L
Doxorubicin
YDL181W INH1
ZShift = −3.73
Z lm Score = −2.59
NG = 0
DB = 0
SM = 0
Delta_K
Doxorubicin
YDL181W INH1
ZShift = −0.64
Z lm Score = 1.13
NG = 0
DB = 0
SM = 0
Delta_r
Doxorubicin
YDL181W INH1
ZShift = −4.22
Z lm Score = 0.93
NG = 0
DB = 0
SM = 0
Delta_AUC
Doxorubicin
YBR246W
ZShift = 0.88
Z lm Score = 2.02
NG = 0
DB = 0
SM = 0
Delta_L
Doxorubicin
YBR246W
ZShift = 0.15
Z lm Score = −1.2
NG = 0
DB = 0
SM = 0
Delta_K
Doxorubicin
YBR246W
ZShift = −0.64
Z lm Score = −0.05
NG = 0
DB = 0
SM = 0
Delta_r
Doxorubicin
YBR246W
ZShift = −0.07
Z lm Score = −0.1
NG = 0
DB = 0
SM = 0
Delta_AUC
Doxorubicin
YGR185C TYS1
ZShift = −0.55
Z lm Score = 2.01
NG = 0
DB = 0
SM = 0
Delta_L
Doxorubicin
YGR185C TYS1
ZShift = 1.54
Z lm Score = 0.1
NG = 0
DB = 0
SM = 0
Delta_K
Doxorubicin
YGR185C TYS1
ZShift = 0.45
Z lm Score = −0.51
NG = 0
DB = 0
SM = 0
Delta_r
Doxorubicin
YGR185C TYS1
ZShift = 1.57
Z lm Score = −0.32
NG = 0
DB = 0
SM = 0
Delta_AUC
Doxorubicin

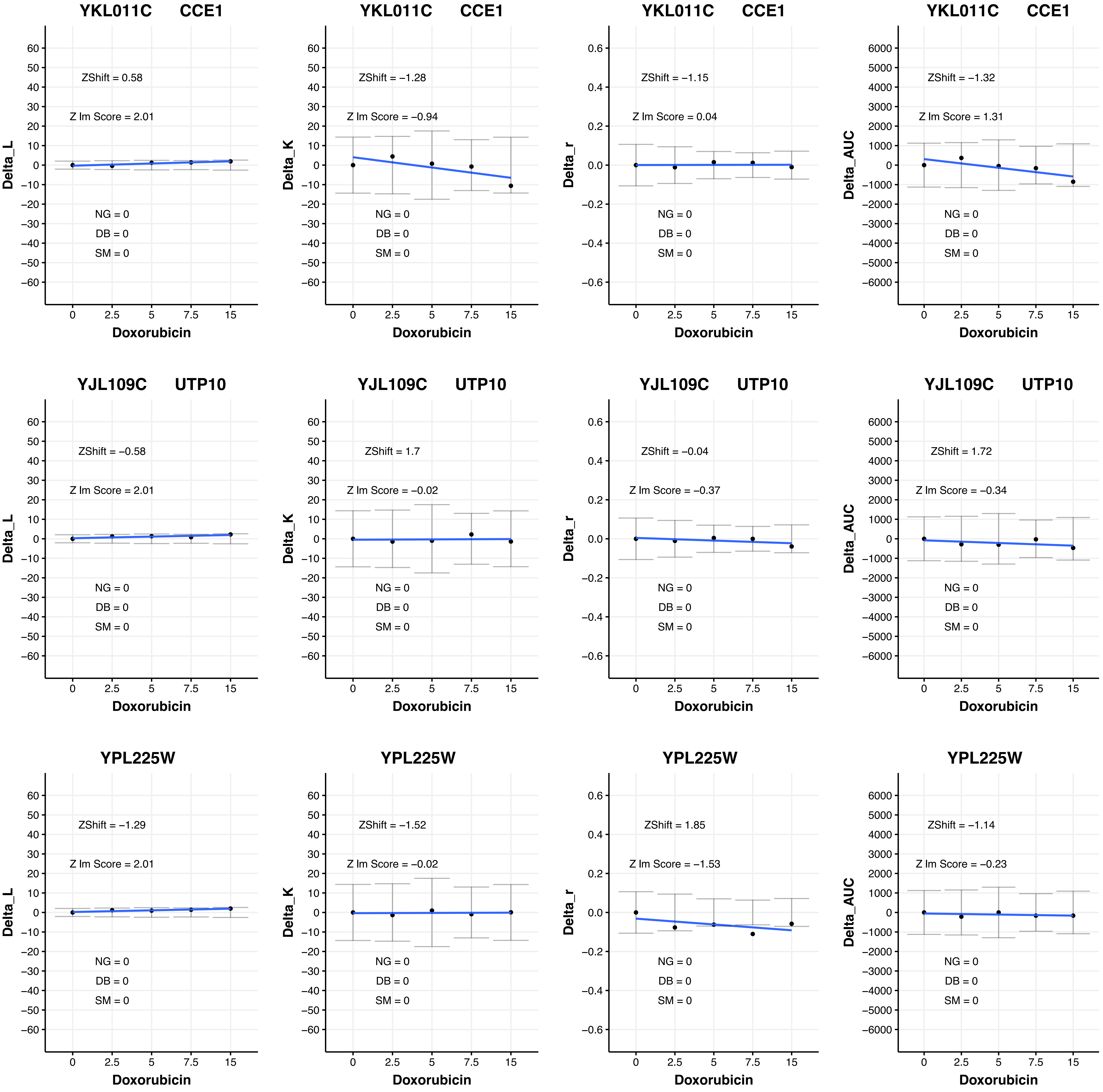
YKL011C CCE1
ZShift = 0.58
Z lm Score = 2.01
NG = 0
DB = 0
SM = 0
Delta_L
Doxorubicin
YKL011C CCE1
ZShift = −1.28
Z lm Score = −0.94
NG = 0
DB = 0
SM = 0
Delta_K
Doxorubicin
YKL011C CCE1
ZShift = −1.15
Z lm Score = 0.04
NG = 0
DB = 0
SM = 0
Delta_r
Doxorubicin
YKL011C CCE1
ZShift = −1.32
Z lm Score = 1.31
NG = 0
DB = 0
SM = 0
Delta_AUC
Doxorubicin
YJL109C UTP10
ZShift = −0.58
Z lm Score = 2.01
NG = 0
DB = 0
SM = 0
Delta_L
Doxorubicin
YJL109C UTP10
ZShift = 1.7
Z lm Score = −0.02
NG = 0
DB = 0
SM = 0
Delta_K
Doxorubicin
YJL109C UTP10
ZShift = −0.04
Z lm Score = −0.37
NG = 0
DB = 0
SM = 0
Delta_r
Doxorubicin
YJL109C UTP10
ZShift = 1.72
Z lm Score = −0.34
NG = 0
DB = 0
SM = 0
Delta_AUC
Doxorubicin
YPL225W
ZShift = −1.29
Z lm Score = 2.01
NG = 0
DB = 0
SM = 0
Delta_L
Doxorubicin
YPL225W
ZShift = −1.52
Z lm Score = −0.02
NG = 0
DB = 0
SM = 0
Delta_K
Doxorubicin
YPL225W
ZShift = 1.85
Z lm Score = −1.53
NG = 0
DB = 0
SM = 0
Delta_r
Doxorubicin
YPL225W
ZShift = −1.14
Z lm Score = −0.23
NG = 0
DB = 0
SM = 0
Delta_AUC
Doxorubicin

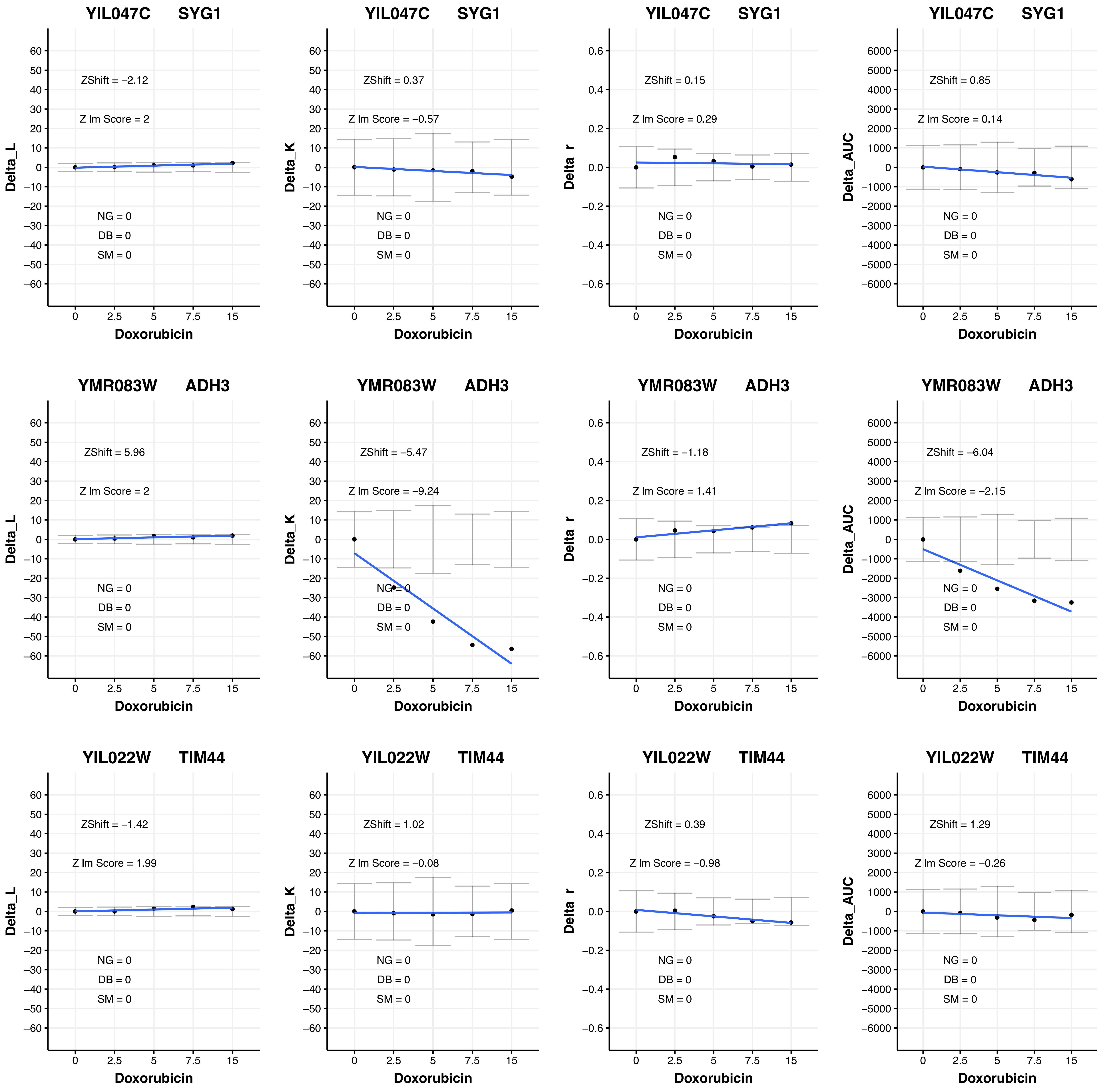

YIL047C SYG1
ZShift = −2.12
Z lm Score = 2
NG = 0
DB = 0
SM = 0
Delta_L
Doxorubicin
YIL047C SYG1
ZShift = 0.37
Z lm Score = −0.57
NG = 0
DB = 0
SM = 0
Delta_K
Doxorubicin
YIL047C SYG1
ZShift = 0.15
Z lm Score = 0.29
NG = 0
DB = 0
SM = 0
Delta_r
Doxorubicin
YIL047C SYG1
ZShift = 0.85
Z lm Score = 0.14
NG = 0
DB = 0
SM = 0
Delta_AUC
Doxorubicin
YMR083W ADH3
ZShift = 5.96
Z lm Score = 2
NG = 0
DB = 0
SM = 0
Delta_L
Doxorubicin
YMR083W ADH3
ZShift = −5.47
Z lm Score = −9.24
NG = 0
DB = 0
SM = 0
Delta_K
Doxorubicin
YMR083W ADH3
ZShift = −1.18
Z lm Score = 1.41
NG = 0
DB = 0
SM = 0
Delta_r
Doxorubicin
YMR083W ADH3
ZShift = −6.04
Z lm Score = −2.15
NG = 0
DB = 0
SM = 0
Delta_AUC
Doxorubicin
YIL022W TIM44
ZShift = −1.42
Z lm Score = 1.99
NG = 0
DB = 0
SM = 0
Delta_L
Doxorubicin
YIL022W TIM44
ZShift = 1.02
Z lm Score = −0.08
NG = 0
DB = 0
SM = 0
Delta_K
Doxorubicin
YIL022W TIM44
ZShift = 0.39
Z lm Score = −0.98
NG = 0
DB = 0
SM = 0
Delta_r
Doxorubicin
YIL022W TIM44
ZShift = 1.29
Z lm Score = −0.26
NG = 0
DB = 0
SM = 0
Delta_AUC
Doxorubicin

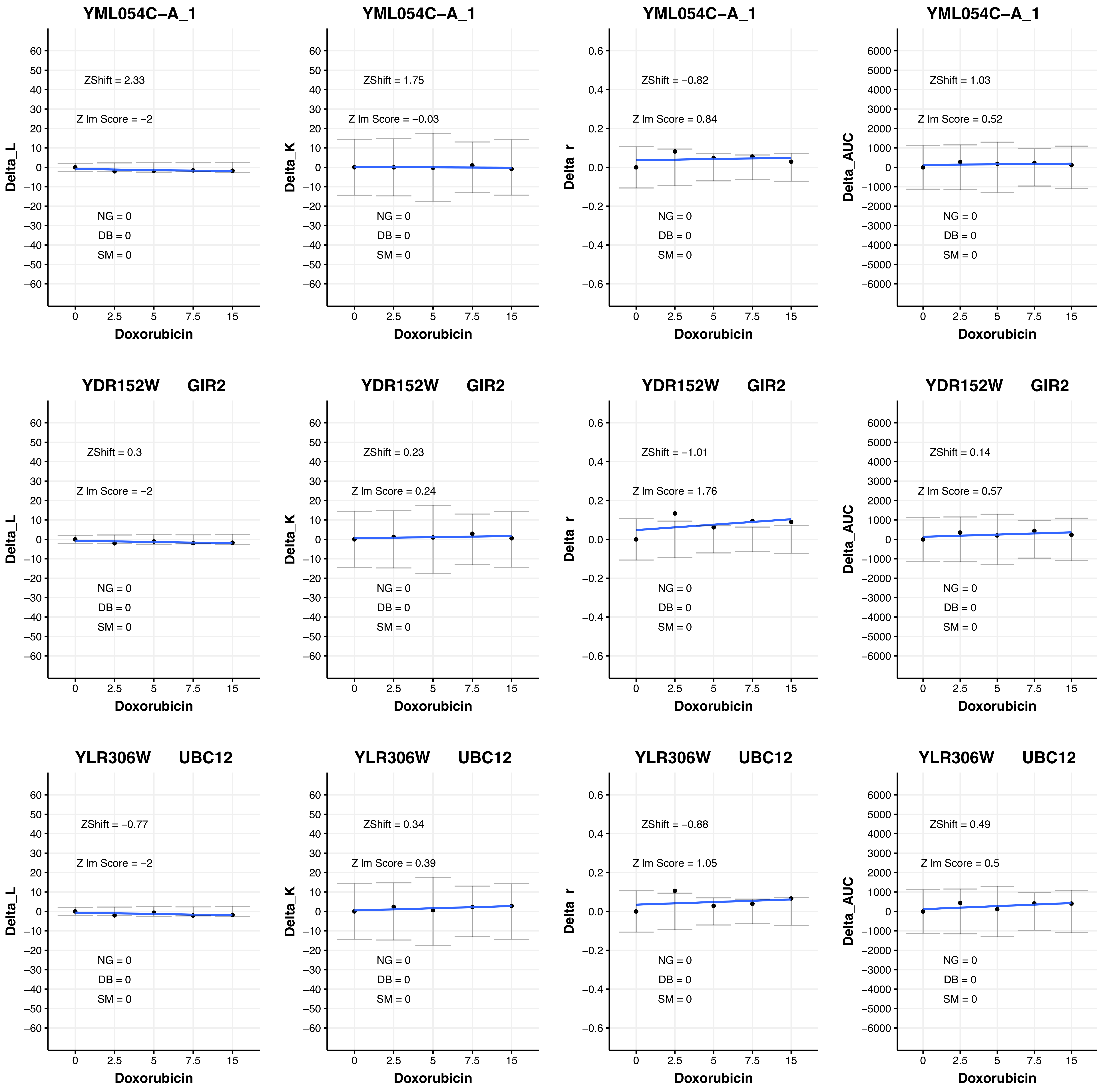
YML054C−A_1
ZShift = 2.33
Z lm Score = −2
NG = 0
DB = 0
SM = 0
Delta_L
Doxorubicin
YML054C−A_1
ZShift = 1.75
Z lm Score = −0.03
NG = 0
DB = 0
SM = 0
Delta_K
Doxorubicin
YML054C−A_1
ZShift = −0.82
Z lm Score = 0.84
NG = 0
DB = 0
SM = 0
Delta_r
Doxorubicin
YML054C−A_1
ZShift = 1.03
Z lm Score = 0.52
NG = 0
DB = 0
SM = 0
Delta_AUC
Doxorubicin
YDR152W GIR2
ZShift = 0.3
Z lm Score = −2
NG = 0
DB = 0
SM = 0
Delta_L
Doxorubicin
YDR152W GIR2
ZShift = 0.23
Z lm Score = 0.24
NG = 0
DB = 0
SM = 0
Delta_K
Doxorubicin
YDR152W GIR2
ZShift = −1.01
Z lm Score = 1.76
NG = 0
DB = 0
SM = 0
Delta_r
Doxorubicin
YDR152W GIR2
ZShift = 0.14
Z lm Score = 0.57
NG = 0
DB = 0
SM = 0
Delta_AUC
Doxorubicin
YLR306W UBC12
ZShift = −0.77
Z lm Score = −2
NG = 0
DB = 0
SM = 0
Delta_L
Doxorubicin
YLR306W UBC12
ZShift = 0.34
Z lm Score = 0.39
NG = 0
DB = 0
SM = 0
Delta_K
Doxorubicin
YLR306W UBC12
ZShift = −0.88
Z lm Score = 1.05
NG = 0
DB = 0
SM = 0
Delta_r
Doxorubicin
YLR306W UBC12
ZShift = 0.49
Z lm Score = 0.5
NG = 0
DB = 0
SM = 0
Delta_AUC
Doxorubicin

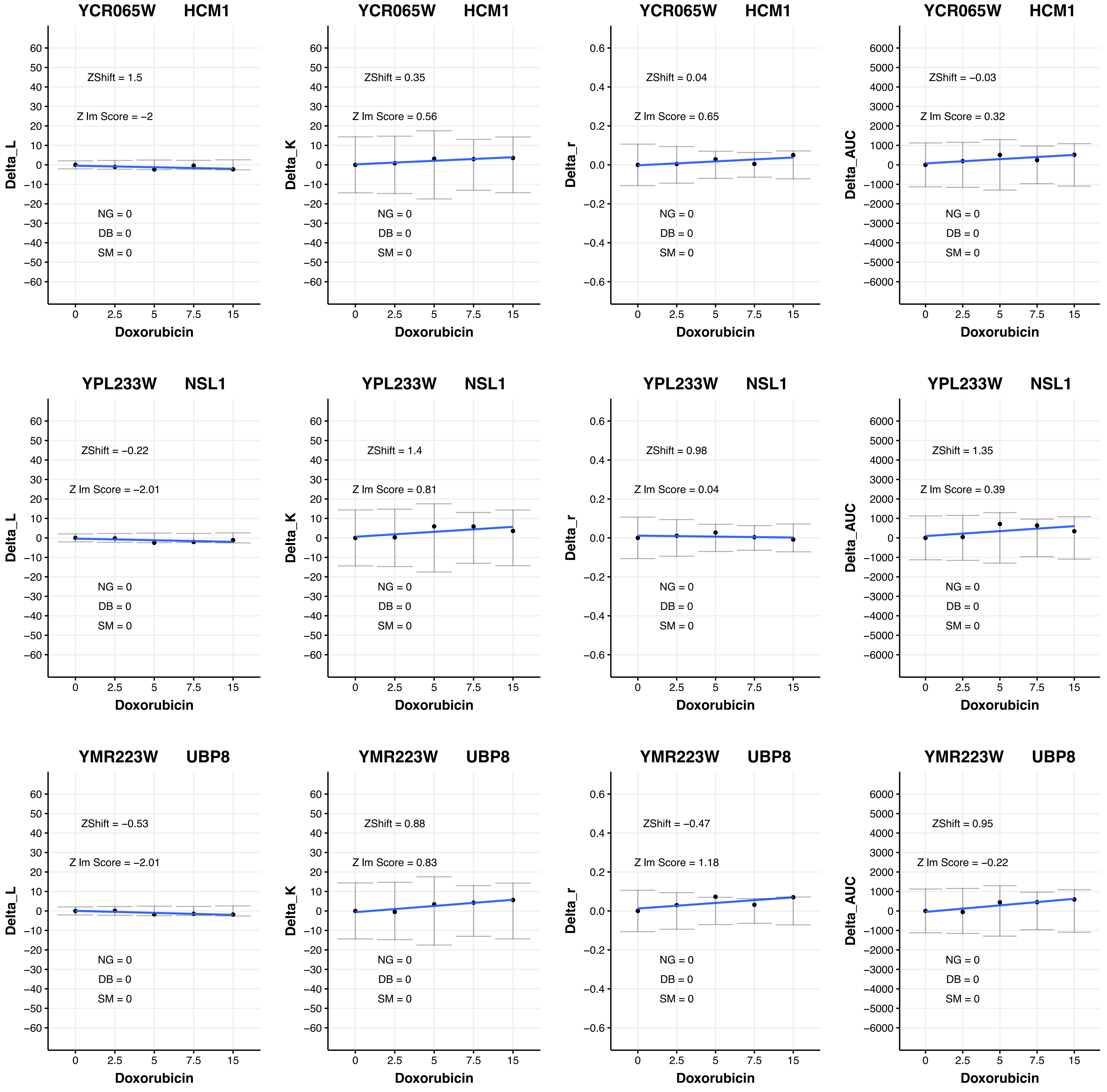
YCR065W HCM1
Delta_L
ZShift = 1.5
Z lm Score = −2
NG = 0
DB = 0
SM = 0
Doxorubicin
YCR065W HCM1
Delta_K
ZShift = 0.35
Z lm Score = 0.56
NG = 0
DB = 0
SM = 0
Doxorubicin
YCR065W HCM1
Delta_r
ZShift = 0.04
Z lm Score = 0.65
NG = 0
DB = 0
SM = 0
Doxorubicin
YCR065W HCM1
Delta_AUC
ZShift = −0.03
Z lm Score = 0.32
NG = 0
DB = 0
SM = 0
Doxorubicin
YPL233W NSL1
Delta_L
ZShift = −0.22
Z lm Score = −2.01
NG = 0
DB = 0
SM = 0
Doxorubicin
YPL233W NSL1
Delta_K
ZShift = 1.4
Z lm Score = 0.81
NG = 0
DB = 0
SM = 0
Doxorubicin
YPL233W NSL1
Delta_r
ZShift = 0.98
Z lm Score = 0.04
NG = 0
DB = 0
SM = 0
Doxorubicin
YPL233W NSL1
Delta_AUC
ZShift = 1.35
Z lm Score = 0.39
NG = 0
DB = 0
SM = 0
Doxorubicin
YMR223W UBP8
Delta_L
ZShift = −0.53
Z lm Score = −2.01
NG = 0
DB = 0
SM = 0
Doxorubicin
YMR223W UBP8
Delta_K
ZShift = 0.88
Z lm Score = 0.83
NG = 0
DB = 0
SM = 0
Doxorubicin
YMR223W UBP8
Delta_r
ZShift = −0.47
Z lm Score = 1.18
NG = 0
DB = 0
SM = 0
Doxorubicin
YMR223W UBP8
Delta_AUC
ZShift = 0.95
Z lm Score = −0.22
NG = 0
DB = 0
SM = 0
Doxorubicin

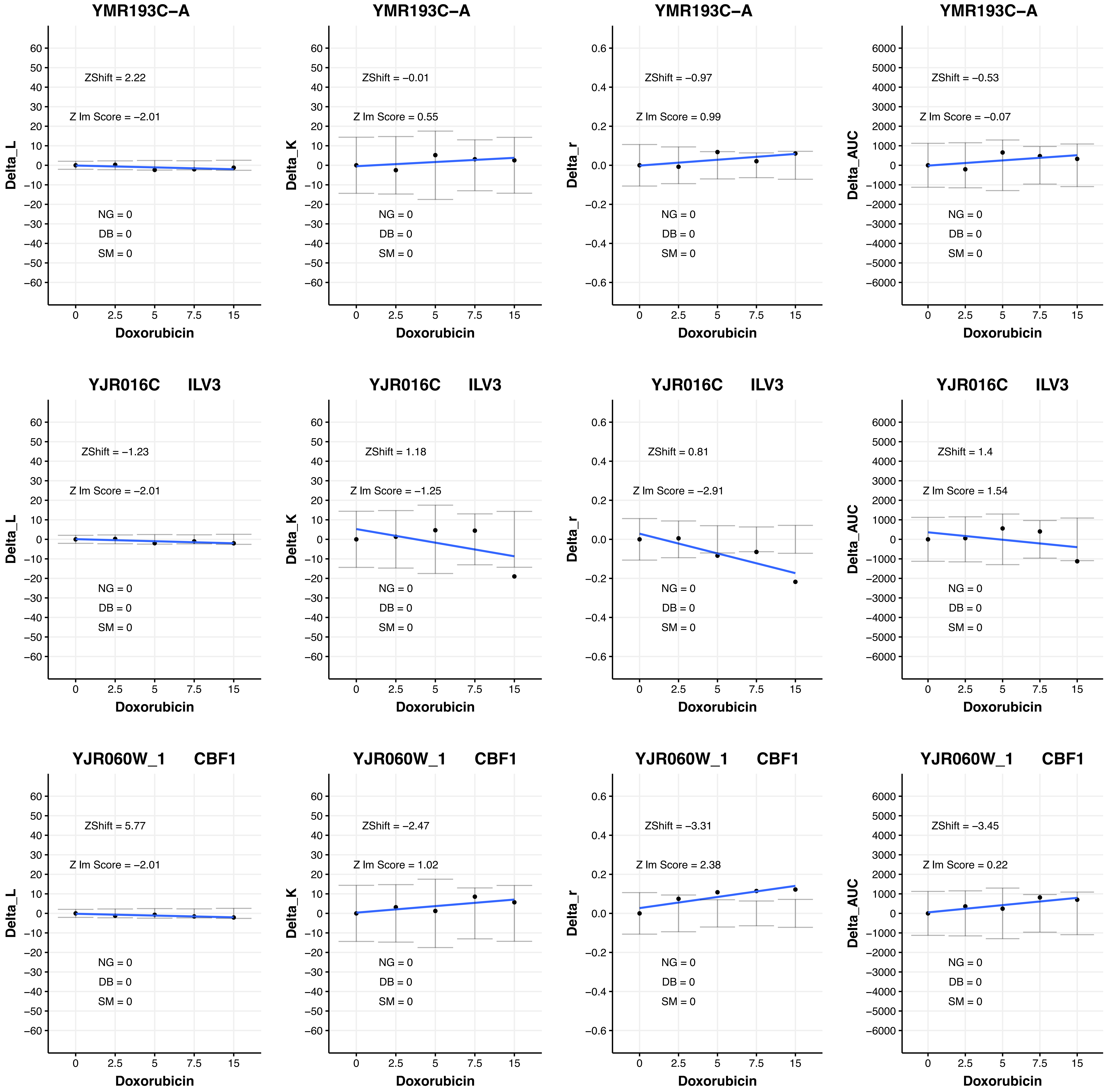
YMR193C−A
ZShift = 2.22
Z lm Score = −2.01
NG = 0
DB = 0
SM = 0
Delta_L
Doxorubicin
YMR193C−A
ZShift = −0.01
Z lm Score = 0.55
NG = 0
DB = 0
SM = 0
Delta_K
Doxorubicin
YMR193C−A
ZShift = −0.97
Z lm Score = 0.99
NG = 0
DB = 0
SM = 0
Delta_r
Doxorubicin
YMR193C−A
ZShift = −0.53
Z lm Score = −0.07
NG = 0
DB = 0
SM = 0
Delta_AUC
Doxorubicin
YJR016C ILV3
ZShift = −1.23
Z lm Score = −2.01
NG = 0
DB = 0
SM = 0
Delta_L
Doxorubicin
YJR016C ILV3
ZShift = 1.18
Z lm Score = −1.25
NG = 0
DB = 0
SM = 0
Delta_K
Doxorubicin
YJR016C ILV3
ZShift = 0.81
Z lm Score = −2.91
NG = 0
DB = 0
SM = 0
Delta_r
Doxorubicin
YJR016C ILV3
ZShift = 1.4
Z lm Score = 1.54
NG = 0
DB = 0
SM = 0
Delta_AUC
Doxorubicin
YJR060W_1 CBF1
ZShift = 5.77
Z lm Score = −2.01
NG = 0
DB = 0
SM = 0
Delta_L
Doxorubicin
YJR060W_1 CBF1
ZShift = −2.47
Z lm Score = 1.02
NG = 0
DB = 0
SM = 0
Delta_K
Doxorubicin
YJR060W_1 CBF1
ZShift = −3.31
Z lm Score = 2.38
NG = 0
DB = 0
SM = 0
Delta_r
Doxorubicin
YJR060W_1 CBF1
ZShift = −3.45
Z lm Score = 0.22
NG = 0
DB = 0
SM = 0
Delta_AUC
Doxorubicin

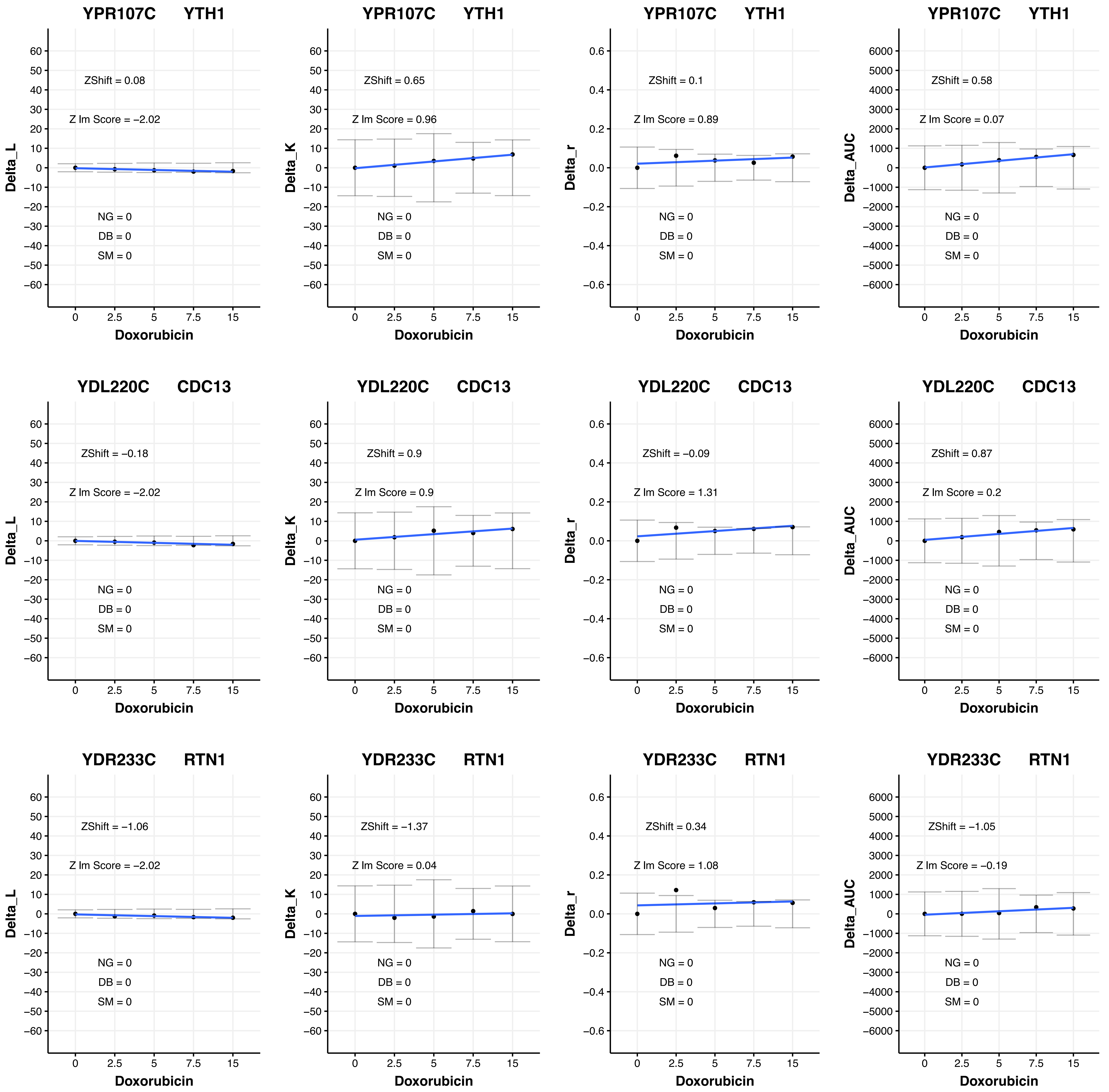
YPR107C YTH1
ZShift = 0.08
Z lm Score = −2.02
NG = 0
DB = 0
SM = 0
Delta_L
Doxorubicin
YPR107C YTH1
ZShift = 0.65
Z lm Score = 0.96
NG = 0
DB = 0
SM = 0
Delta_K
Doxorubicin
YPR107C YTH1
ZShift = 0.1
Z lm Score = 0.89
NG = 0
DB = 0
SM = 0
Delta_r
Doxorubicin
YPR107C YTH1
ZShift = 0.58
Z lm Score = 0.07
NG = 0
DB = 0
SM = 0
Delta_AUC
Doxorubicin
YDL220C CDC13
ZShift = −0.18
Z lm Score = −2.02
NG = 0
DB = 0
SM = 0
Delta_L
Doxorubicin
YDL220C CDC13
ZShift = 0.9
Z lm Score = 0.9
NG = 0
DB = 0
SM = 0
Delta_K
Doxorubicin
YDL220C CDC13
ZShift = −0.09
Z lm Score = 1.31
NG = 0
DB = 0
SM = 0
Delta_r
Doxorubicin
YDL220C CDC13
ZShift = 0.87
Z lm Score = 0.2
NG = 0
DB = 0
SM = 0
Delta_AUC
Doxorubicin
YDR233C RTN1
ZShift = −1.06
Z lm Score = −2.02
NG = 0
DB = 0
SM = 0
Delta_L
Doxorubicin
YDR233C RTN1
ZShift = −1.37
Z lm Score = 0.04
NG = 0
DB = 0
SM = 0
Delta_K
Doxorubicin
YDR233C RTN1
ZShift = 0.34
Z lm Score = 1.08
NG = 0
DB = 0
SM = 0
Delta_r
Doxorubicin
YDR233C RTN1
ZShift = −1.05
Z lm Score = −0.19
NG = 0
DB = 0
SM = 0
Delta_AUC
Doxorubicin

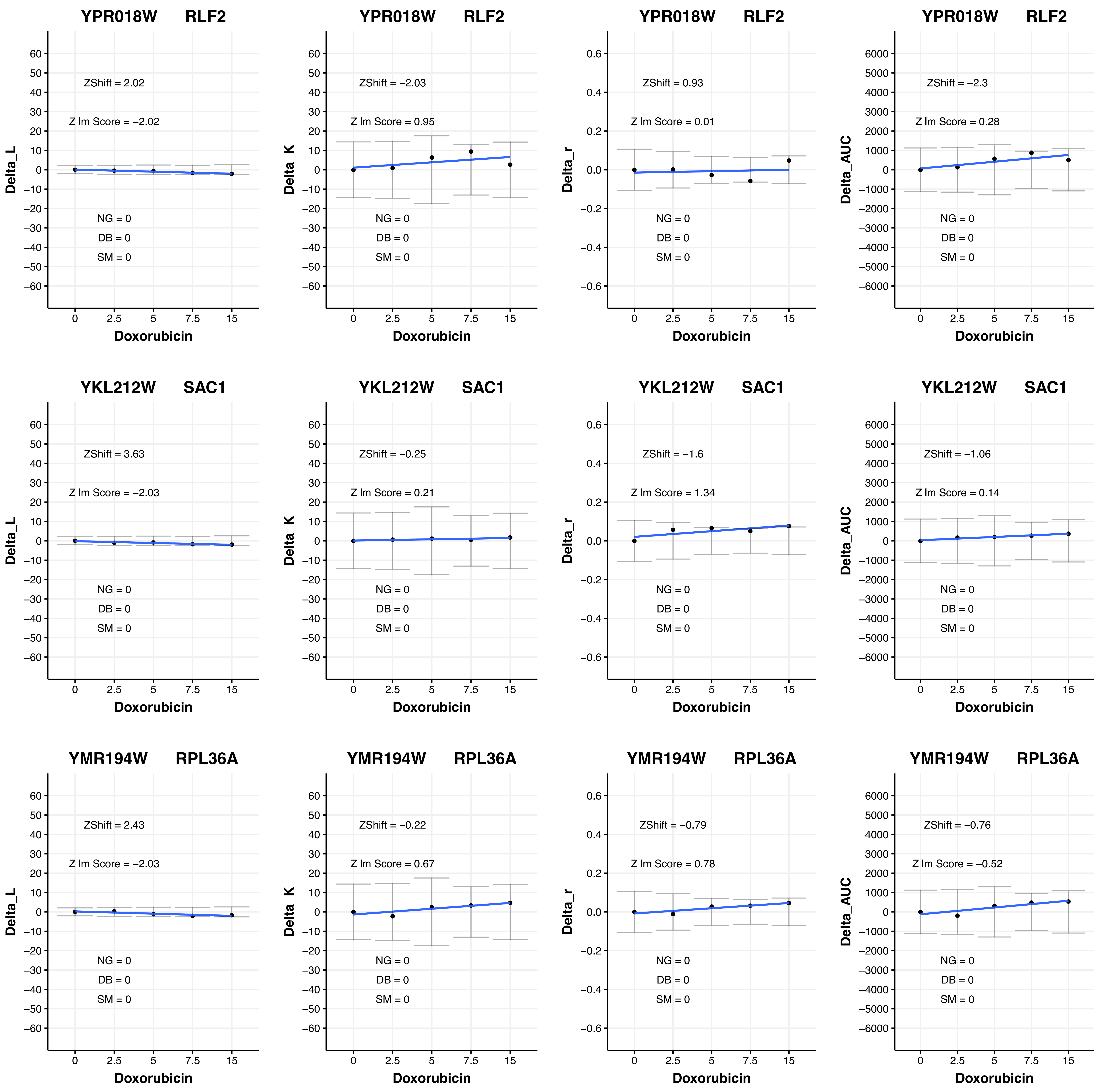

YPR018W RLF2
ZShift = 2.02
Z lm Score = −2.02
NG = 0
DB = 0
SM = 0
Delta_L
Doxorubicin
YPR018W RLF2
ZShift = −2.03
Z lm Score = 0.95
NG = 0
DB = 0
SM = 0
Delta_K
Doxorubicin
YPR018W RLF2
ZShift = 0.93
Z lm Score = 0.01
NG = 0
DB = 0
SM = 0
Delta_r
Doxorubicin
YPR018W RLF2
ZShift = −2.3
Z lm Score = 0.28
NG = 0
DB = 0
SM = 0
Delta_AUC
Doxorubicin
YKL212W SAC1
ZShift = 3.63
Z lm Score = −2.03
NG = 0
DB = 0
SM = 0
Delta_L
Doxorubicin
YKL212W SAC1
ZShift = −0.25
Z lm Score = 0.21
NG = 0
DB = 0
SM = 0
Delta_K
Doxorubicin
YKL212W SAC1
ZShift = −1.6
Z lm Score = 1.34
NG = 0
DB = 0
SM = 0
Delta_r
Doxorubicin
YKL212W SAC1
ZShift = −1.06
Z lm Score = 0.14
NG = 0
DB = 0
SM = 0
Delta_AUC
Doxorubicin
YMR194W RPL36A
ZShift = 2.43
Z lm Score = −2.03
NG = 0
DB = 0
SM = 0
Delta_L
Doxorubicin
YMR194W RPL36A
ZShift = −0.22
Z lm Score = 0.67
NG = 0
DB = 0
SM = 0
Delta_K
Doxorubicin
YMR194W RPL36A
ZShift = −0.79
Z lm Score = 0.78
NG = 0
DB = 0
SM = 0
Delta_r
Doxorubicin
YMR194W RPL36A
ZShift = −0.76
Z lm Score = −0.52
NG = 0
DB = 0
SM = 0
Delta_AUC
Doxorubicin

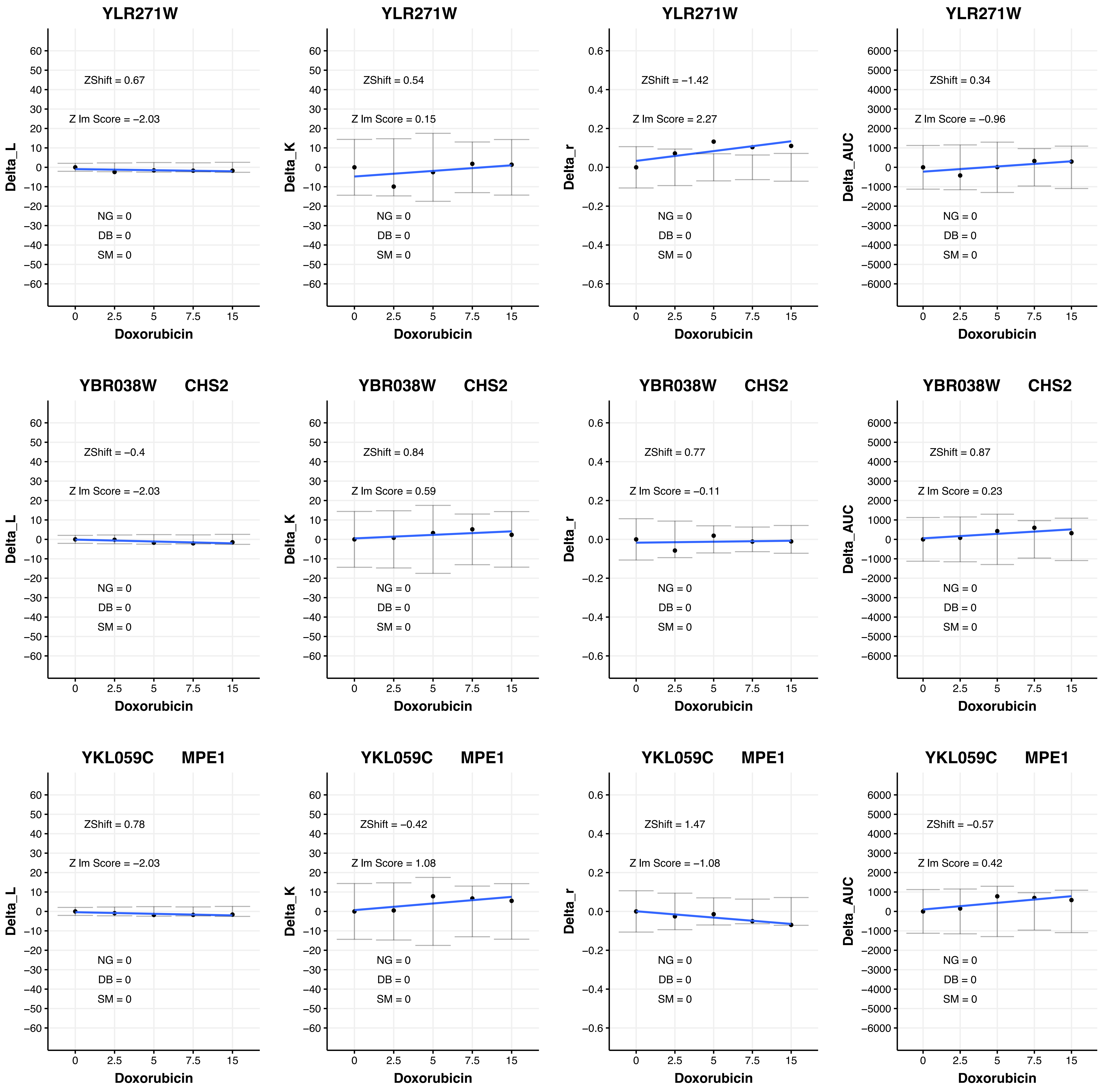
YLR271W
ZShift = 0.67
Z lm Score = −2.03
NG = 0
DB = 0
SM = 0
Delta_L
Doxorubicin
YLR271W
ZShift = 0.54
Z lm Score = 0.15
NG = 0
DB = 0
SM = 0
Delta_K
Doxorubicin
YLR271W
ZShift = −1.42
Z lm Score = 2.27
NG = 0
DB = 0
SM = 0
Delta_r
Doxorubicin
YLR271W
ZShift = 0.34
Z lm Score = −0.96
NG = 0
DB = 0
SM = 0
Delta_AUC
Doxorubicin
YBR038W CHS2
ZShift = −0.4
Z lm Score = −2.03
NG = 0
DB = 0
SM = 0
Delta_L
Doxorubicin
YBR038W CHS2
ZShift = 0.84
Z lm Score = 0.59
NG = 0
DB = 0
SM = 0
Delta_K
Doxorubicin
YBR038W CHS2
ZShift = 0.77
Z lm Score = −0.11
NG = 0
DB = 0
SM = 0
Delta_r
Doxorubicin
YBR038W CHS2
ZShift = 0.87
Z lm Score = 0.23
NG = 0
DB = 0
SM = 0
Delta_AUC
Doxorubicin
YKL059C MPE1
ZShift = 0.78
Z lm Score = −2.03
NG = 0
DB = 0
SM = 0
Delta_L
Doxorubicin
YKL059C MPE1
ZShift = −0.42
Z lm Score = 1.08
NG = 0
DB = 0
SM = 0
Delta_K
Doxorubicin
YKL059C MPE1
ZShift = 1.47
Z lm Score = −1.08
NG = 0
DB = 0
SM = 0
Delta_r
Doxorubicin
YKL059C MPE1
ZShift = −0.57
Z lm Score = 0.42
NG = 0
DB = 0
SM = 0
Delta_AUC
Doxorubicin

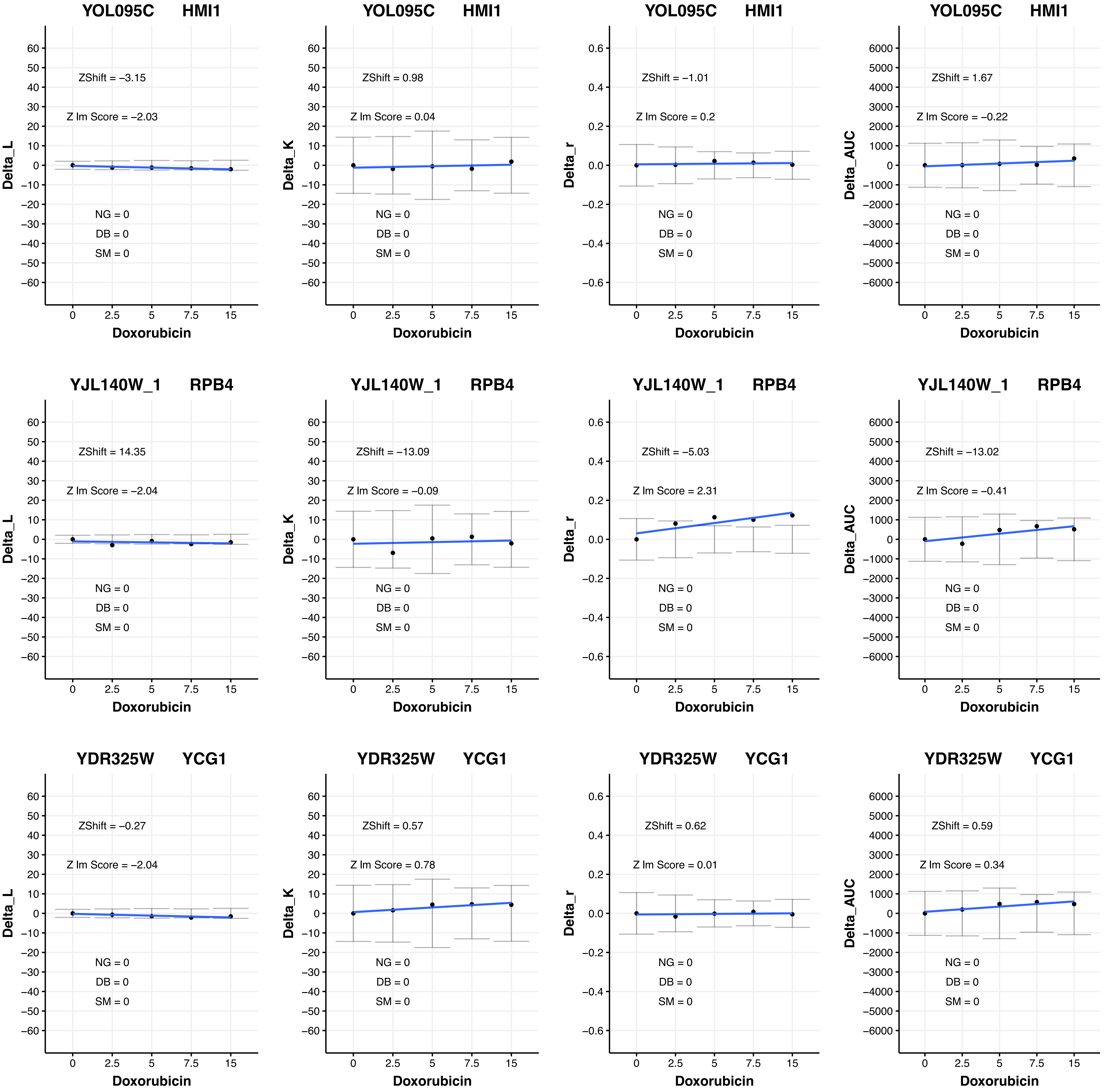
YOL095C HMI1
ZShift = −3.15
Z lm Score = −2.03
NG = 0
DB = 0
SM = 0
Delta_L
Doxorubicin
YOL095C HMI1
ZShift = 0.98
Z lm Score = 0.04
NG = 0
DB = 0
SM = 0
Delta_K
Doxorubicin
YOL095C HMI1
ZShift = −1.01
Z lm Score = 0.2
NG = 0
DB = 0
SM = 0
Delta_r
Doxorubicin
YOL095C HMI1
ZShift = 1.67
Z lm Score = −0.22
NG = 0
DB = 0
SM = 0
Delta_AUC
Doxorubicin
YJL140W_1 RPB4
ZShift = 14.35
Z lm Score = −2.04
NG = 0
DB = 0
SM = 0
Delta_L
Doxorubicin
YJL140W_1 RPB4
ZShift = −13.09
Z lm Score = −0.09
NG = 0
DB = 0
SM = 0
Delta_K
Doxorubicin
YJL140W_1 RPB4
ZShift = −5.03
Z lm Score = 2.31
NG = 0
DB = 0
SM = 0
Delta_r
Doxorubicin
YJL140W_1 RPB4
ZShift = −13.02
Z lm Score = −0.41
NG = 0
DB = 0
SM = 0
Delta_AUC
Doxorubicin
YDR325W YCG1
ZShift = −0.27
Z lm Score = −2.04
NG = 0
DB = 0
SM = 0
Delta_L
Doxorubicin
YDR325W YCG1
ZShift = 0.57
Z lm Score = 0.78
NG = 0
DB = 0
SM = 0
Delta_K
Doxorubicin
YDR325W YCG1
ZShift = 0.62
Z lm Score = 0.01
NG = 0
DB = 0
SM = 0
Delta_r
Doxorubicin
YDR325W YCG1
ZShift = 0.59
Z lm Score = 0.34
NG = 0
DB = 0
SM = 0
Delta_AUC
Doxorubicin

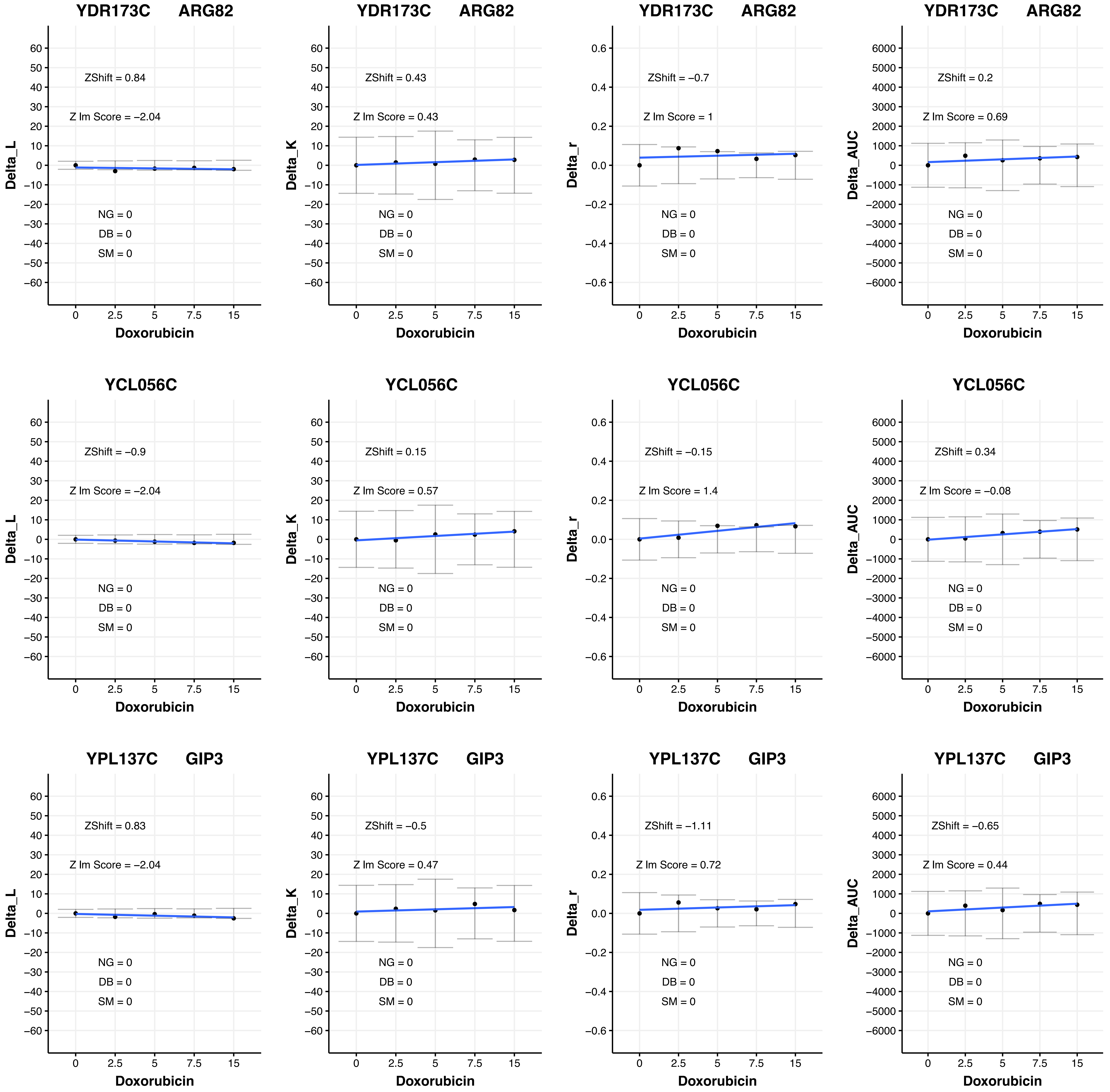
YDR173C ARG82
ZShift = 0.84
Z lm Score = −2.04
NG = 0
DB = 0
SM = 0
Delta_L
Doxorubicin
YDR173C ARG82
ZShift = 0.43
Z lm Score = 0.43
NG = 0
DB = 0
SM = 0
Delta_K
Doxorubicin
YDR173C ARG82
ZShift = −0.7
Z lm Score = 1
NG = 0
DB = 0
SM = 0
Delta_r
Doxorubicin
YDR173C ARG82
ZShift = 0.2
Z lm Score = 0.69
NG = 0
DB = 0
SM = 0
Delta_AUC
Doxorubicin
YCL056C
ZShift = −0.9
Z lm Score = −2.04
NG = 0
DB = 0
SM = 0
Delta_L
Doxorubicin
YCL056C
ZShift = 0.15
Z lm Score = 0.57
NG = 0
DB = 0
SM = 0
Delta_K
Doxorubicin
YCL056C
ZShift = −0.15
Z lm Score = 1.4
NG = 0
DB = 0
SM = 0
Delta_r
Doxorubicin
YCL056C
ZShift = 0.34
Z lm Score = −0.08
NG = 0
DB = 0
SM = 0
Delta_AUC
Doxorubicin
YPL137C GIP3
ZShift = 0.83
Z lm Score = −2.04
NG = 0
DB = 0
SM = 0
Delta_L
Doxorubicin
YPL137C GIP3
ZShift = −0.5
Z lm Score = 0.47
NG = 0
DB = 0
SM = 0
Delta_K
Doxorubicin
YPL137C GIP3
ZShift = −1.11
Z lm Score = 0.72
NG = 0
DB = 0
SM = 0
Delta_r
Doxorubicin
YPL137C GIP3
ZShift = −0.65
Z lm Score = 0.44
NG = 0
DB = 0
SM = 0
Delta_AUC
Doxorubicin

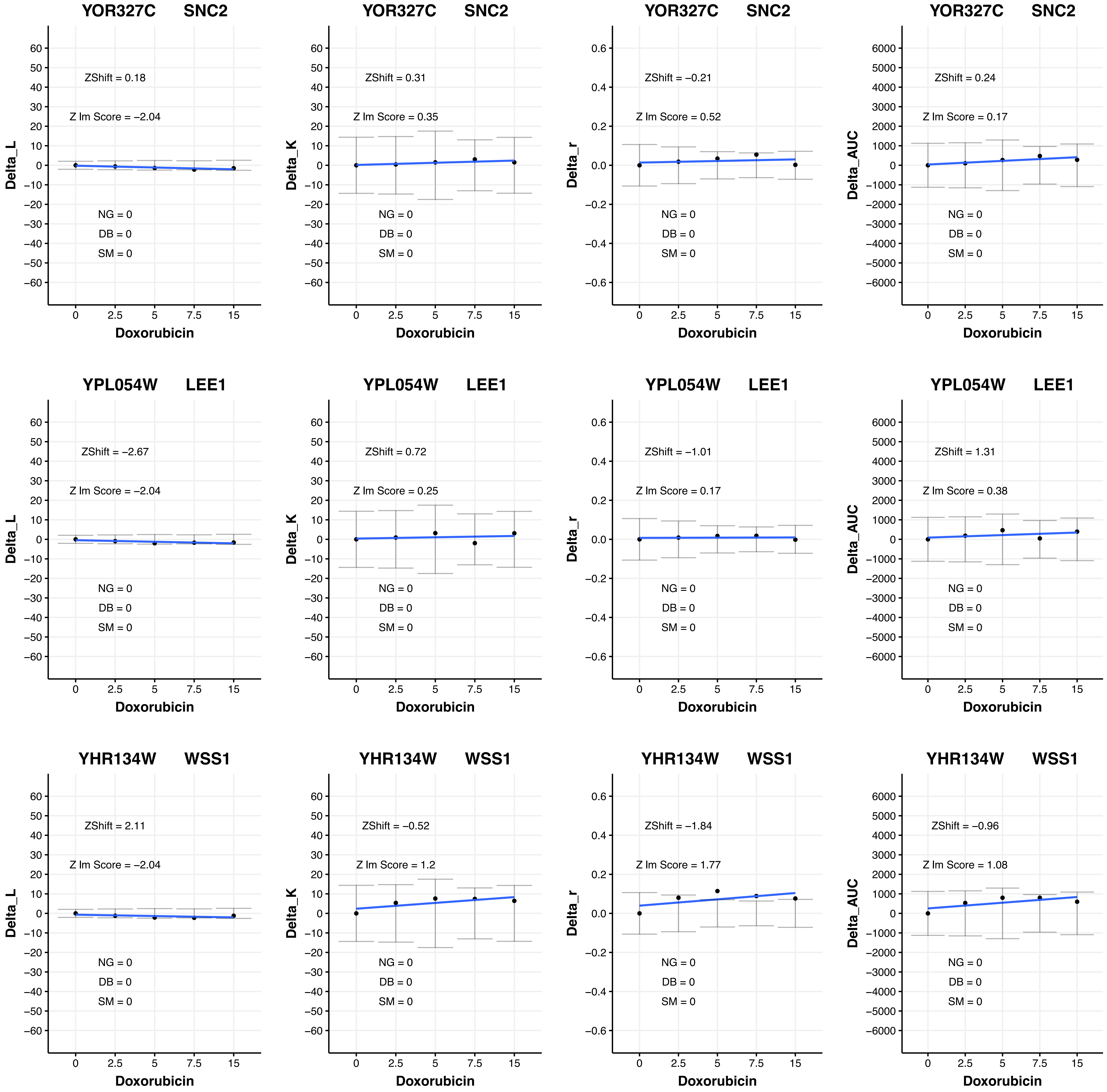
YOR327C SNC2
ZShift = 0.18
Z lm Score = −2.04
NG = 0
DB = 0
SM = 0
Delta_L
Doxorubicin
YOR327C SNC2
ZShift = 0.31
Z lm Score = 0.35
NG = 0
DB = 0
SM = 0
Delta_K
Doxorubicin
YOR327C SNC2
ZShift = −0.21
Z lm Score = 0.52
NG = 0
DB = 0
SM = 0
Delta_r
Doxorubicin
YOR327C SNC2
ZShift = 0.24
Z lm Score = 0.17
NG = 0
DB = 0
SM = 0
Delta_AUC
Doxorubicin
YPL054W LEE1
ZShift = −2.67
Z lm Score = −2.04
NG = 0
DB = 0
SM = 0
Delta_L
Doxorubicin
YPL054W LEE1
ZShift = 0.72
Z lm Score = 0.25
NG = 0
DB = 0
SM = 0
Delta_K
Doxorubicin
YPL054W LEE1
ZShift = −1.01
Z lm Score = 0.17
NG = 0
DB = 0
SM = 0
Delta_r
Doxorubicin
YPL054W LEE1
ZShift = 1.31
Z lm Score = 0.38
NG = 0
DB = 0
SM = 0
Delta_AUC
Doxorubicin
YHR134W WSS1
ZShift = 2.11
Z lm Score = −2.04
NG = 0
DB = 0
SM = 0
Delta_L
Doxorubicin
YHR134W WSS1
ZShift = −0.52
Z lm Score = 1.2
NG = 0
DB = 0
SM = 0
Delta_K
Doxorubicin
YHR134W WSS1
ZShift = −1.84
Z lm Score = 1.77
NG = 0
DB = 0
SM = 0
Delta_r
Doxorubicin
YHR134W WSS1
ZShift = −0.96
Z lm Score = 1.08
NG = 0
DB = 0
SM = 0
Delta_AUC
Doxorubicin

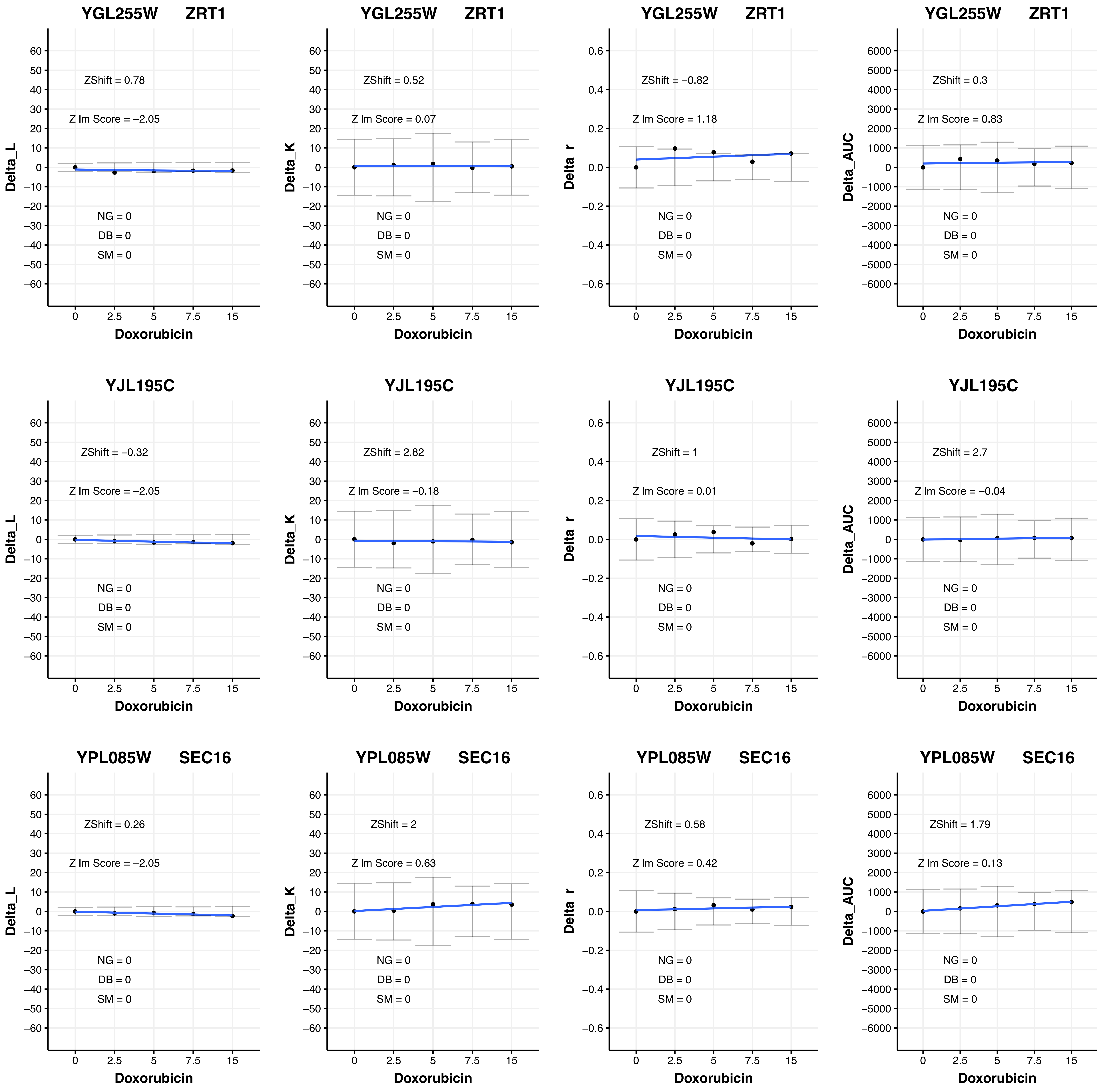

YGL255W ZRT1
ZShift = 0.78
Z lm Score = −2.05
NG = 0
DB = 0
SM = 0
Delta_L
Doxorubicin
YGL255W ZRT1
ZShift = 0.52
Z lm Score = 0.07
NG = 0
DB = 0
SM = 0
Delta_K
Doxorubicin
YGL255W ZRT1
ZShift = −0.82
Z lm Score = 1.18
NG = 0
DB = 0
SM = 0
Delta_r
Doxorubicin
YGL255W ZRT1
ZShift = 0.3
Z lm Score = 0.83
NG = 0
DB = 0
SM = 0
Delta_AUC
Doxorubicin
YJL195C
ZShift = −0.32
Z lm Score = −2.05
NG = 0
DB = 0
SM = 0
Delta_L
Doxorubicin
YJL195C
ZShift = 2.82
Z lm Score = −0.18
NG = 0
DB = 0
SM = 0
Delta_K
Doxorubicin
YJL195C
ZShift = 1
Z lm Score = 0.01
NG = 0
DB = 0
SM = 0
Delta_r
Doxorubicin
YJL195C
ZShift = 2.7
Z lm Score = −0.04
NG = 0
DB = 0
SM = 0
Delta_AUC
Doxorubicin
YPL085W SEC16
ZShift = 0.26
Z lm Score = −2.05
NG = 0
DB = 0
SM = 0
Delta_L
Doxorubicin
YPL085W SEC16
ZShift = 2
Z lm Score = 0.63
NG = 0
DB = 0
SM = 0
Delta_K
Doxorubicin
YPL085W SEC16
ZShift = 0.58
Z lm Score = 0.42
NG = 0
DB = 0
SM = 0
Delta_r
Doxorubicin
YPL085W SEC16
ZShift = 1.79
Z lm Score = 0.13
NG = 0
DB = 0
SM = 0
Delta_AUC
Doxorubicin

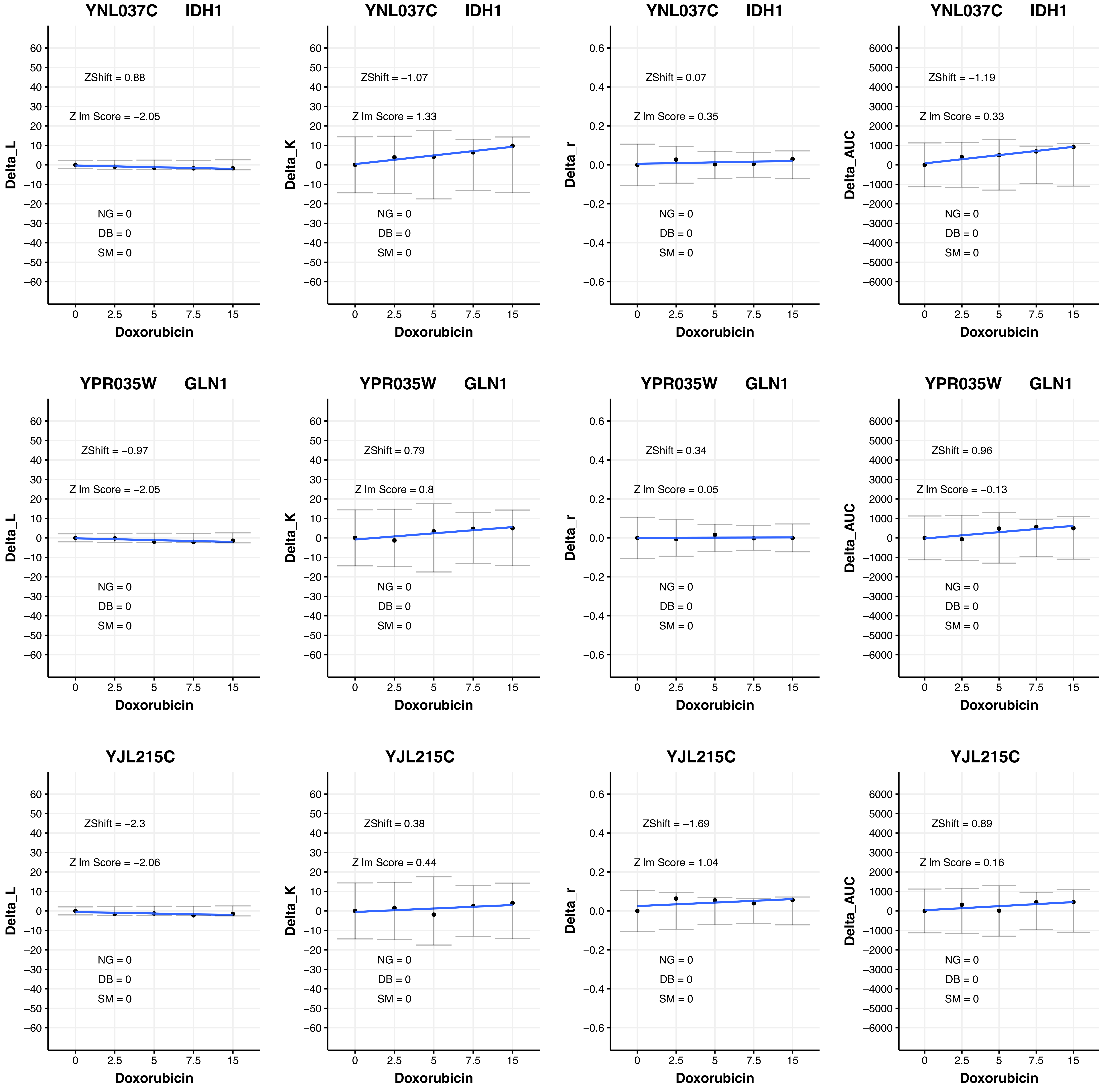
YNL037C IDH1
ZShift = 0.88
Z lm Score = −2.05
NG = 0
DB = 0
SM = 0
Delta_L
Doxorubicin
YNL037C IDH1
ZShift = −1.07
Z lm Score = 1.33
NG = 0
DB = 0
SM = 0
Delta_K
Doxorubicin
YNL037C IDH1
ZShift = 0.07
Z lm Score = 0.35
NG = 0
DB = 0
SM = 0
Delta_r
Doxorubicin
YNL037C IDH1
ZShift = −1.19
Z lm Score = 0.33
NG = 0
DB = 0
SM = 0
Delta_AUC
Doxorubicin
YPR035W GLN1
ZShift = −0.97
Z lm Score = −2.05
NG = 0
DB = 0
SM = 0
Delta_L
Doxorubicin
YPR035W GLN1
ZShift = 0.79
Z lm Score = 0.8
NG = 0
DB = 0
SM = 0
Delta_K
Doxorubicin
YPR035W GLN1
ZShift = 0.34
Z lm Score = 0.05
NG = 0
DB = 0
SM = 0
Delta_r
Doxorubicin
YPR035W GLN1
ZShift = 0.96
Z lm Score = −0.13
NG = 0
DB = 0
SM = 0
Delta_AUC
Doxorubicin
YJL215C
ZShift = −2.3
Z lm Score = −2.06
NG = 0
DB = 0
SM = 0
Delta_L
Doxorubicin
YJL215C
ZShift = 0.38
Z lm Score = 0.44
NG = 0
DB = 0
SM = 0
Delta_K
Doxorubicin
YJL215C
ZShift = −1.69
Z lm Score = 1.04
NG = 0
DB = 0
SM = 0
Delta_r
Doxorubicin
YJL215C
ZShift = 0.89
Z lm Score = 0.16
NG = 0
DB = 0
SM = 0
Delta_AUC
Doxorubicin

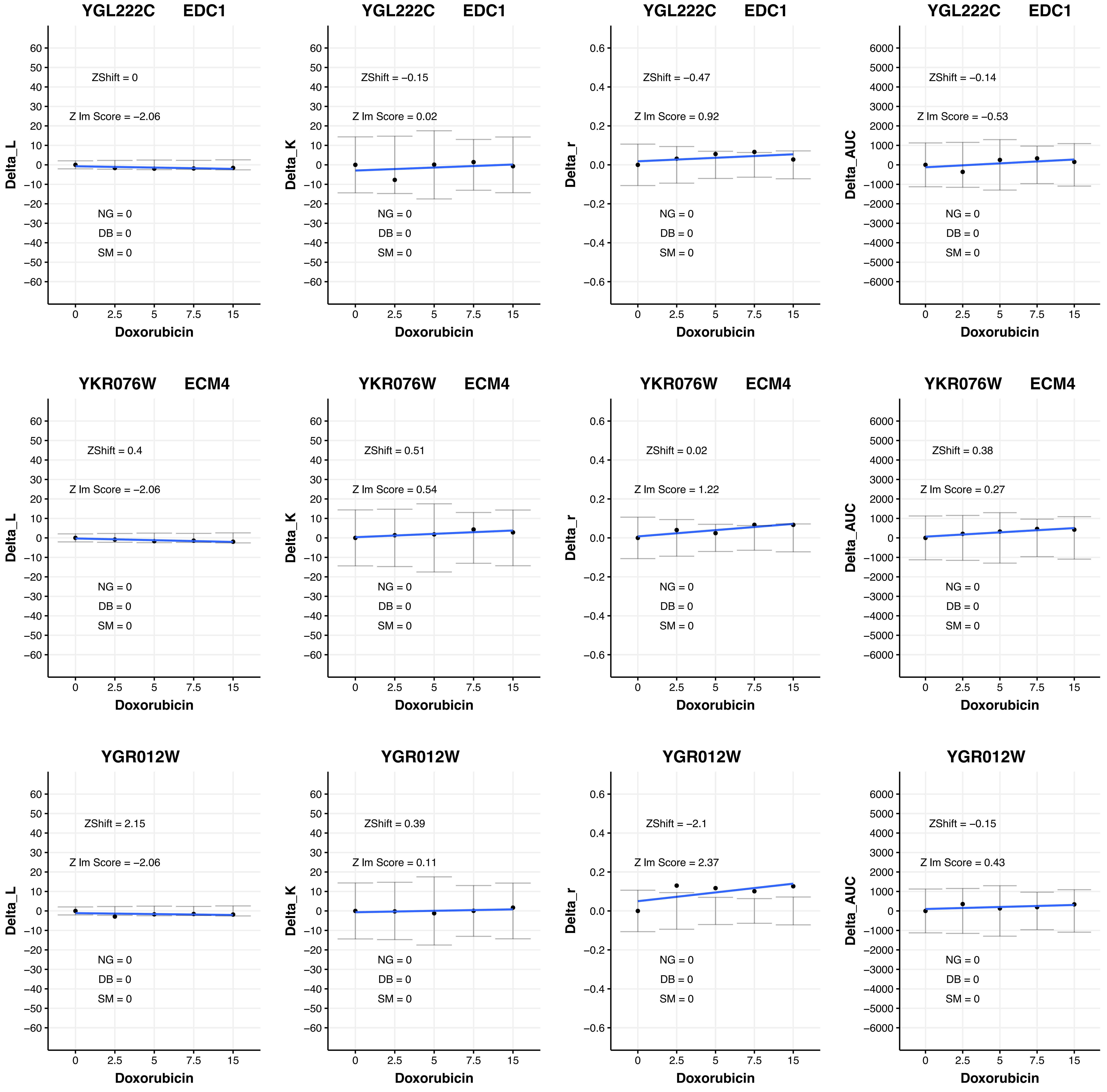
YGL222C EDC1
ZShift = 0
Z lm Score = −2.06
NG = 0
DB = 0
SM = 0
Delta_L
Doxorubicin
YGL222C EDC1
ZShift = −0.15
Z lm Score = 0.02
NG = 0
DB = 0
SM = 0
Delta_K
Doxorubicin
YGL222C EDC1
ZShift = −0.47
Z lm Score = 0.92
NG = 0
DB = 0
SM = 0
Delta_r
Doxorubicin
YGL222C EDC1
ZShift = −0.14
Z lm Score = −0.53
NG = 0
DB = 0
SM = 0
Delta_AUC
Doxorubicin
YKR076W ECM4
ZShift = 0.4
Z lm Score = −2.06
NG = 0
DB = 0
SM = 0
Delta_L
Doxorubicin
YKR076W ECM4
ZShift = 0.51
Z lm Score = 0.54
NG = 0
DB = 0
SM = 0
Delta_K
Doxorubicin
YKR076W ECM4
ZShift = 0.02
Z lm Score = 1.22
NG = 0
DB = 0
SM = 0
Delta_r
Doxorubicin
YKR076W ECM4
ZShift = 0.38
Z lm Score = 0.27
NG = 0
DB = 0
SM = 0
Delta_AUC
Doxorubicin
YGR012W
ZShift = 2.15
Z lm Score = −2.06
NG = 0
DB = 0
SM = 0
Delta_L
Doxorubicin
YGR012W
ZShift = 0.39
Z lm Score = 0.11
NG = 0
DB = 0
SM = 0
Delta_K
Doxorubicin
YGR012W
ZShift = −2.1
Z lm Score = 2.37
NG = 0
DB = 0
SM = 0
Delta_r
Doxorubicin
YGR012W
ZShift = −0.15
Z lm Score = 0.43
NG = 0
DB = 0
SM = 0
Delta_AUC
Doxorubicin

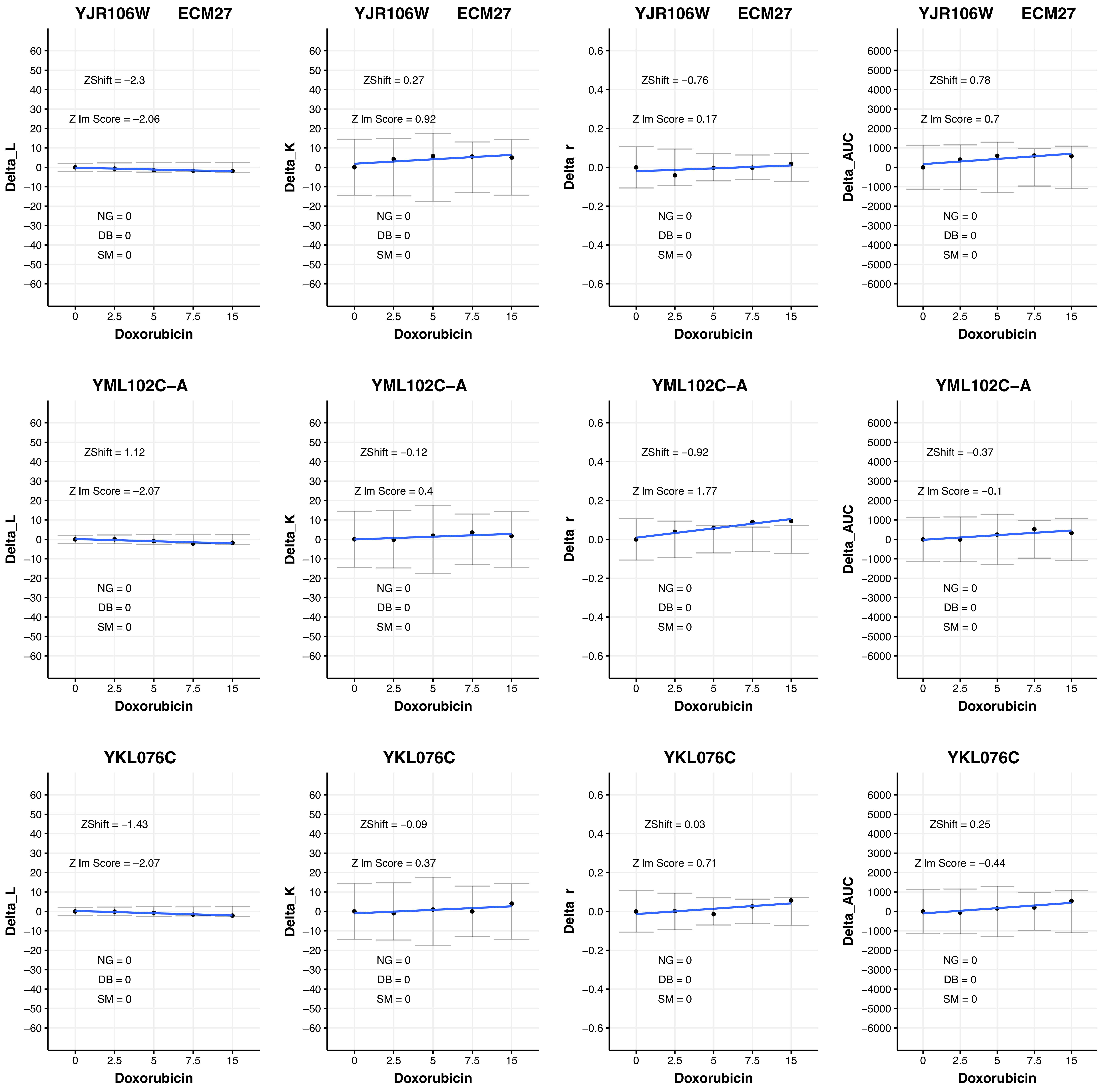
YJR106W ECM27
ZShift = −2.3
Z lm Score = −2.06
NG = 0
DB = 0
SM = 0
Delta_L
Doxorubicin
YJR106W ECM27
ZShift = 0.27
Z lm Score = 0.92
NG = 0
DB = 0
SM = 0
Delta_K
Doxorubicin
YJR106W ECM27
ZShift = −0.76
Z lm Score = 0.17
NG = 0
DB = 0
SM = 0
Delta_r
Doxorubicin
YJR106W ECM27
ZShift = 0.78
Z lm Score = 0.7
NG = 0
DB = 0
SM = 0
Delta_AUC
Doxorubicin
YML102C−A
ZShift = 1.12
Z lm Score = −2.07
NG = 0
DB = 0
SM = 0
Delta_L
Doxorubicin
YML102C−A
ZShift = −0.12
Z lm Score = 0.4
NG = 0
DB = 0
SM = 0
Delta_K
Doxorubicin
YML102C−A
ZShift = −0.92
Z lm Score = 1.77
NG = 0
DB = 0
SM = 0
Delta_r
Doxorubicin
YML102C−A
ZShift = −0.37
Z lm Score = −0.1
NG = 0
DB = 0
SM = 0
Delta_AUC
Doxorubicin
YKL076C
ZShift = −1.43
Z lm Score = −2.07
NG = 0
DB = 0
SM = 0
Delta_L
Doxorubicin
YKL076C
ZShift = −0.09
Z lm Score = 0.37
NG = 0
DB = 0
SM = 0
Delta_K
Doxorubicin
YKL076C
ZShift = 0.03
Z lm Score = 0.71
NG = 0
DB = 0
SM = 0
Delta_r
Doxorubicin
YKL076C
ZShift = 0.25
Z lm Score = −0.44
NG = 0
DB = 0
SM = 0
Delta_AUC
Doxorubicin

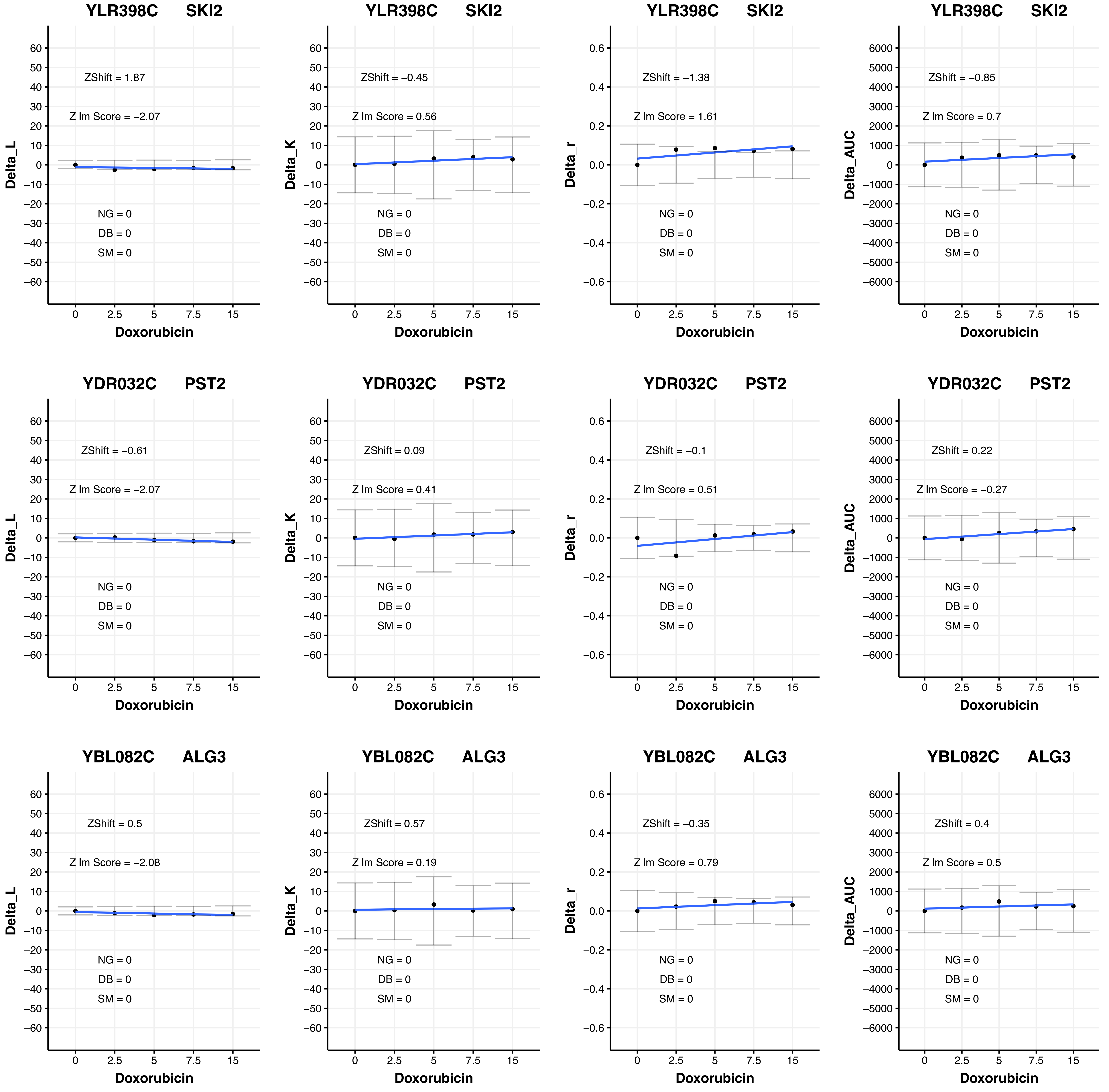

YLR398C SKI2
ZShift = 1.87
Z lm Score = −2.07
NG = 0
DB = 0
SM = 0
Delta_L
Doxorubicin
YLR398C SKI2
ZShift = −0.45
Z lm Score = 0.56
NG = 0
DB = 0
SM = 0
Delta_K
Doxorubicin
YLR398C SKI2
ZShift = −1.38
Z lm Score = 1.61
NG = 0
DB = 0
SM = 0
Delta_r
Doxorubicin
YLR398C SKI2
ZShift = −0.85
Z lm Score = 0.7
NG = 0
DB = 0
SM = 0
Delta_AUC
Doxorubicin
YDR032C PST2
ZShift = −0.61
Z lm Score = −2.07
NG = 0
DB = 0
SM = 0
Delta_L
Doxorubicin
YDR032C PST2
ZShift = 0.09
Z lm Score = 0.41
NG = 0
DB = 0
SM = 0
Delta_K
Doxorubicin
YDR032C PST2
ZShift = −0.1
Z lm Score = 0.51
NG = 0
DB = 0
SM = 0
Delta_r
Doxorubicin
YDR032C PST2
ZShift = 0.22
Z lm Score = −0.27
NG = 0
DB = 0
SM = 0
Delta_AUC
Doxorubicin
YBL082C ALG3
ZShift = 0.5
Z lm Score = −2.08
NG = 0
DB = 0
SM = 0
Delta_L
Doxorubicin
YBL082C ALG3
ZShift = 0.57
Z lm Score = 0.19
NG = 0
DB = 0
SM = 0
Delta_K
Doxorubicin
YBL082C ALG3
ZShift = −0.35
Z lm Score = 0.79
NG = 0
DB = 0
SM = 0
Delta_r
Doxorubicin
YBL082C ALG3
ZShift = 0.4
Z lm Score = 0.5
NG = 0
DB = 0
SM = 0
Delta_AUC
Doxorubicin

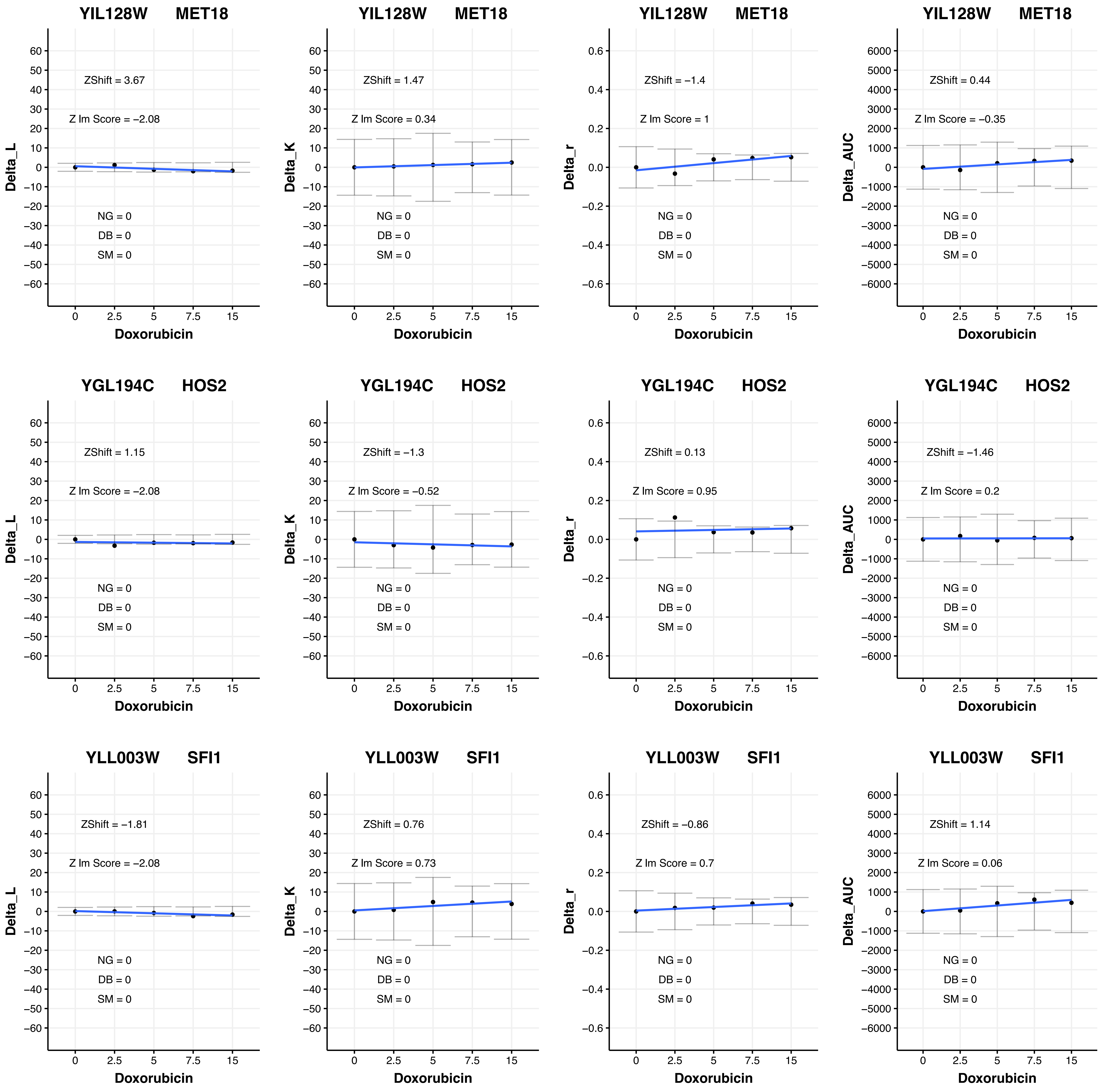
YIL128W MET18
ZShift = 3.67
Z lm Score = −2.08
Delta_L
YIL128W MET18
ZShift = 1.47
Z lm Score = 0.34
Delta_K
YIL128W MET18
ZShift = −1.4
Z lm Score = 1
Delta_r
YIL128W MET18
ZShift = 0.44
Z lm Score = −0.35
Delta_AUC
YGL194C HOS2
ZShift = 1.15
Z lm Score = −2.08
Delta_L
YGL194C HOS2
ZShift = −1.3
Z lm Score = −0.52
Delta_K
YGL194C HOS2
ZShift = 0.13
Z lm Score = 0.95
Delta_r
YGL194C HOS2
ZShift = −1.46
Z lm Score = 0.2
Delta_AUC
YLL003W SFI1
ZShift = −1.81
Z lm Score = −2.08
Delta_L
YLL003W SFI1
ZShift = 0.76
Z lm Score = 0.73
Delta_K
YLL003W SFI1
ZShift = −0.86
Z lm Score = 0.7
Delta_r
YLL003W SFI1
ZShift = 1.14
Z lm Score = 0.06
Delta_AUC
NG = 0
DB = 0
SM = 0
Doxorubicin

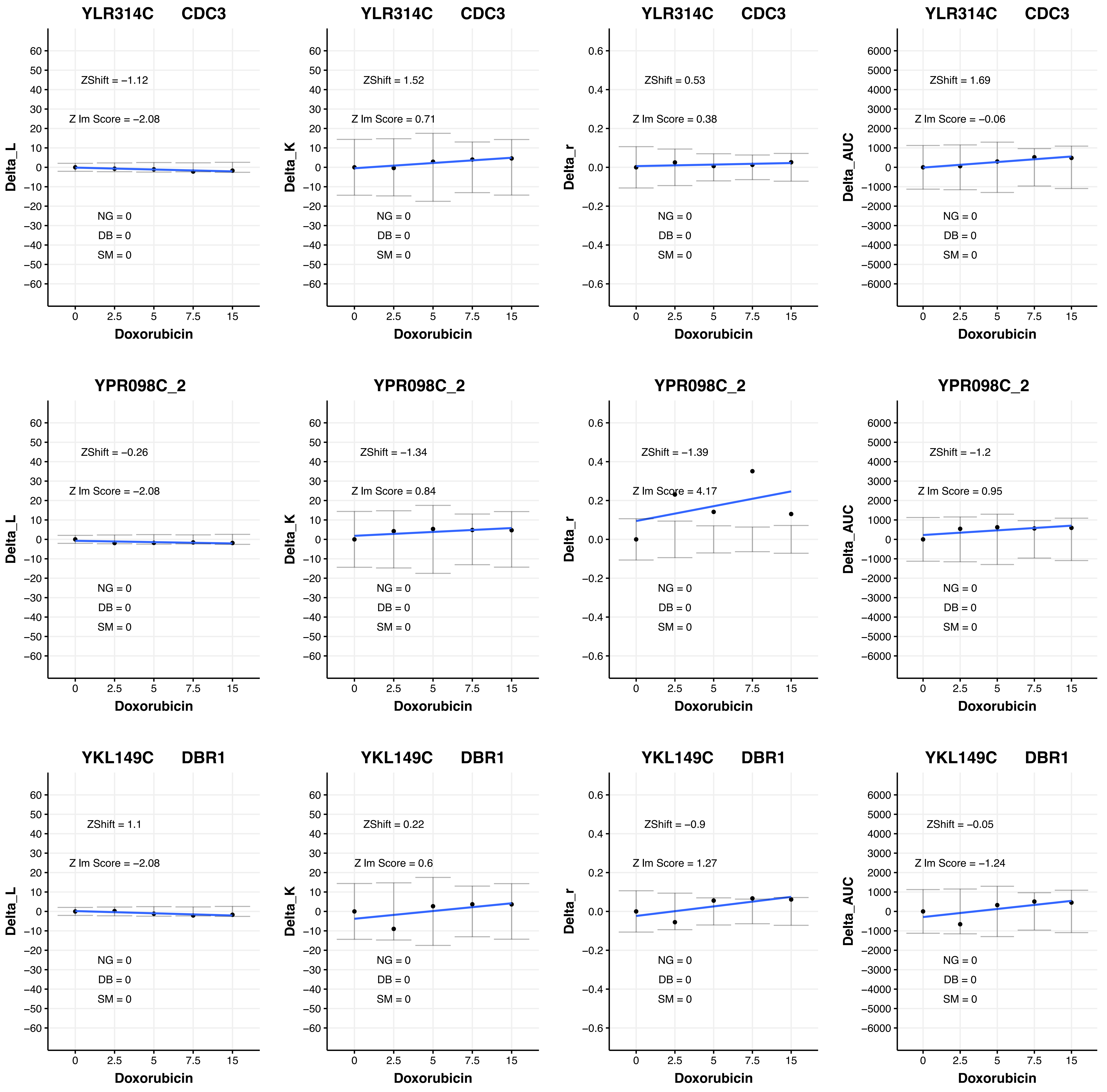
YLR314C CDC3
ZShift = −1.12
Z lm Score = −2.08
NG = 0
DB = 0
SM = 0
Delta_L
Doxorubicin
YLR314C CDC3
ZShift = 1.52
Z lm Score = 0.71
NG = 0
DB = 0
SM = 0
Delta_K
Doxorubicin
YLR314C CDC3
ZShift = 0.53
Z lm Score = 0.38
NG = 0
DB = 0
SM = 0
Delta_r
Doxorubicin
YLR314C CDC3
ZShift = 1.69
Z lm Score = −0.06
NG = 0
DB = 0
SM = 0
Delta_AUC
Doxorubicin
YPR098C_2
ZShift = −0.26
Z lm Score = −2.08
NG = 0
DB = 0
SM = 0
Delta_L
Doxorubicin
YPR098C_2
ZShift = −1.34
Z lm Score = 0.84
NG = 0
DB = 0
SM = 0
Delta_K
Doxorubicin
YPR098C_2
ZShift = −1.39
Z lm Score = 4.17
NG = 0
DB = 0
SM = 0
Delta_r
Doxorubicin
YPR098C_2
ZShift = −1.2
Z lm Score = 0.95
NG = 0
DB = 0
SM = 0
Delta_AUC
Doxorubicin
YKL149C DBR1
ZShift = 1.1
Z lm Score = −2.08
NG = 0
DB = 0
SM = 0
Delta_L
Doxorubicin
YKL149C DBR1
ZShift = 0.22
Z lm Score = 0.6
NG = 0
DB = 0
SM = 0
Delta_K
Doxorubicin
YKL149C DBR1
ZShift = −0.9
Z lm Score = 1.27
NG = 0
DB = 0
SM = 0
Delta_r
Doxorubicin
YKL149C DBR1
ZShift = −0.05
Z lm Score = −1.24
NG = 0
DB = 0
SM = 0
Delta_AUC
Doxorubicin

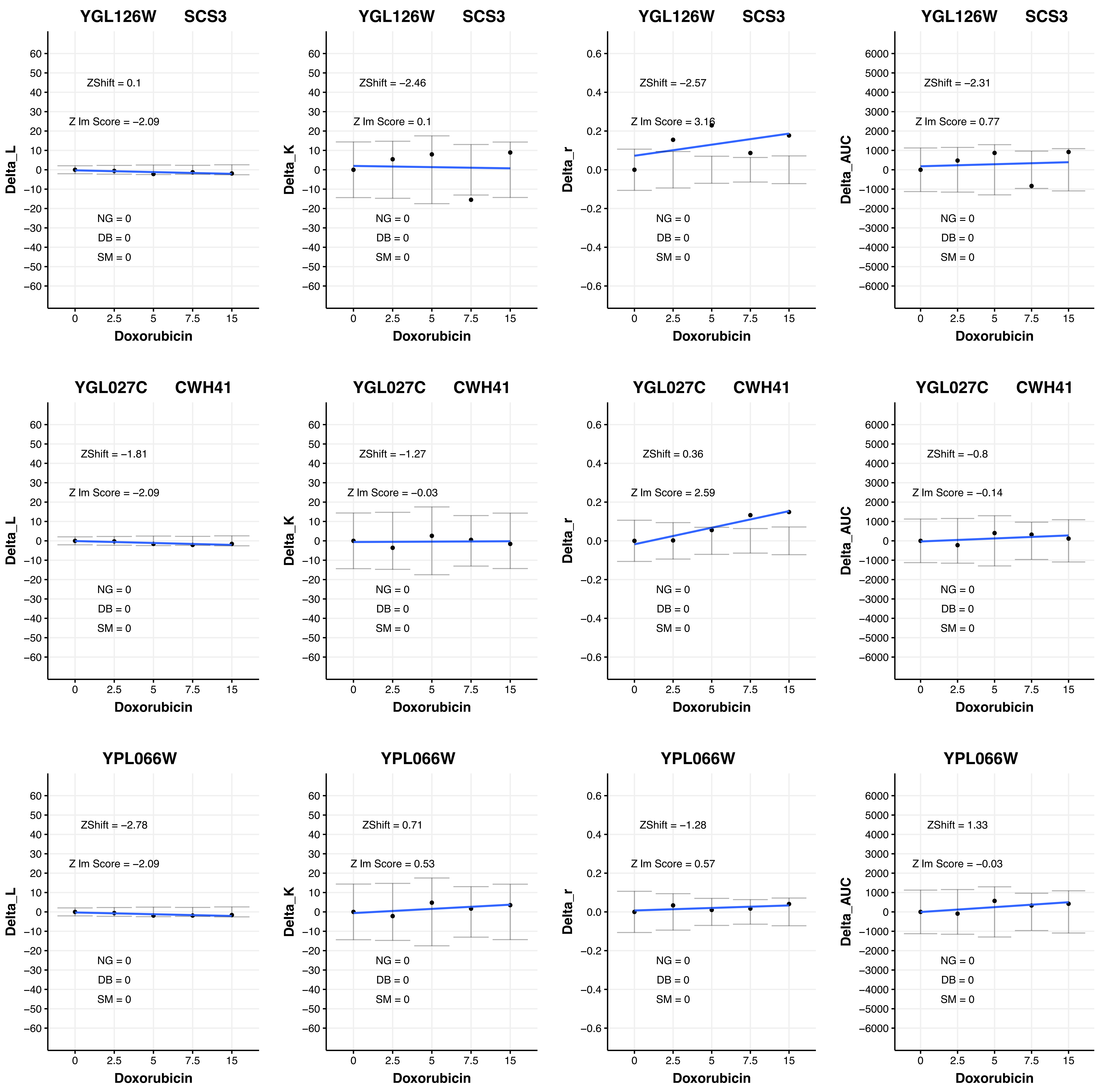
YGL126W SCS3
ZShift = 0.1
Z lm Score = −2.09
NG = 0
DB = 0
SM = 0
Delta_L
Doxorubicin
YGL126W SCS3
ZShift = −2.46
Z lm Score = 0.1
NG = 0
DB = 0
SM = 0
Delta_K
Doxorubicin
YGL126W SCS3
ZShift = −2.57
Z lm Score = 3.16
NG = 0
DB = 0
SM = 0
Delta_r
Doxorubicin
YGL126W SCS3
ZShift = −2.31
Z lm Score = 0.77
NG = 0
DB = 0
SM = 0
Delta_AUC
Doxorubicin
YGL027C CWH41
ZShift = −1.81
Z lm Score = −2.09
NG = 0
DB = 0
SM = 0
Delta_L
Doxorubicin
YGL027C CWH41
ZShift = −1.27
Z lm Score = −0.03
NG = 0
DB = 0
SM = 0
Delta_K
Doxorubicin
YGL027C CWH41
ZShift = 0.36
Z lm Score = 2.59
NG = 0
DB = 0
SM = 0
Delta_r
Doxorubicin
YGL027C CWH41
ZShift = −0.8
Z lm Score = −0.14
NG = 0
DB = 0
SM = 0
Delta_AUC
Doxorubicin
YPL066W
ZShift = −2.78
Z lm Score = −2.09
NG = 0
DB = 0
SM = 0
Delta_L
Doxorubicin
YPL066W
ZShift = 0.71
Z lm Score = 0.53
NG = 0
DB = 0
SM = 0
Delta_K
Doxorubicin
YPL066W
ZShift = −1.28
Z lm Score = 0.57
NG = 0
DB = 0
SM = 0
Delta_r
Doxorubicin
YPL066W
ZShift = 1.33
Z lm Score = −0.03
NG = 0
DB = 0
SM = 0
Delta_AUC
Doxorubicin

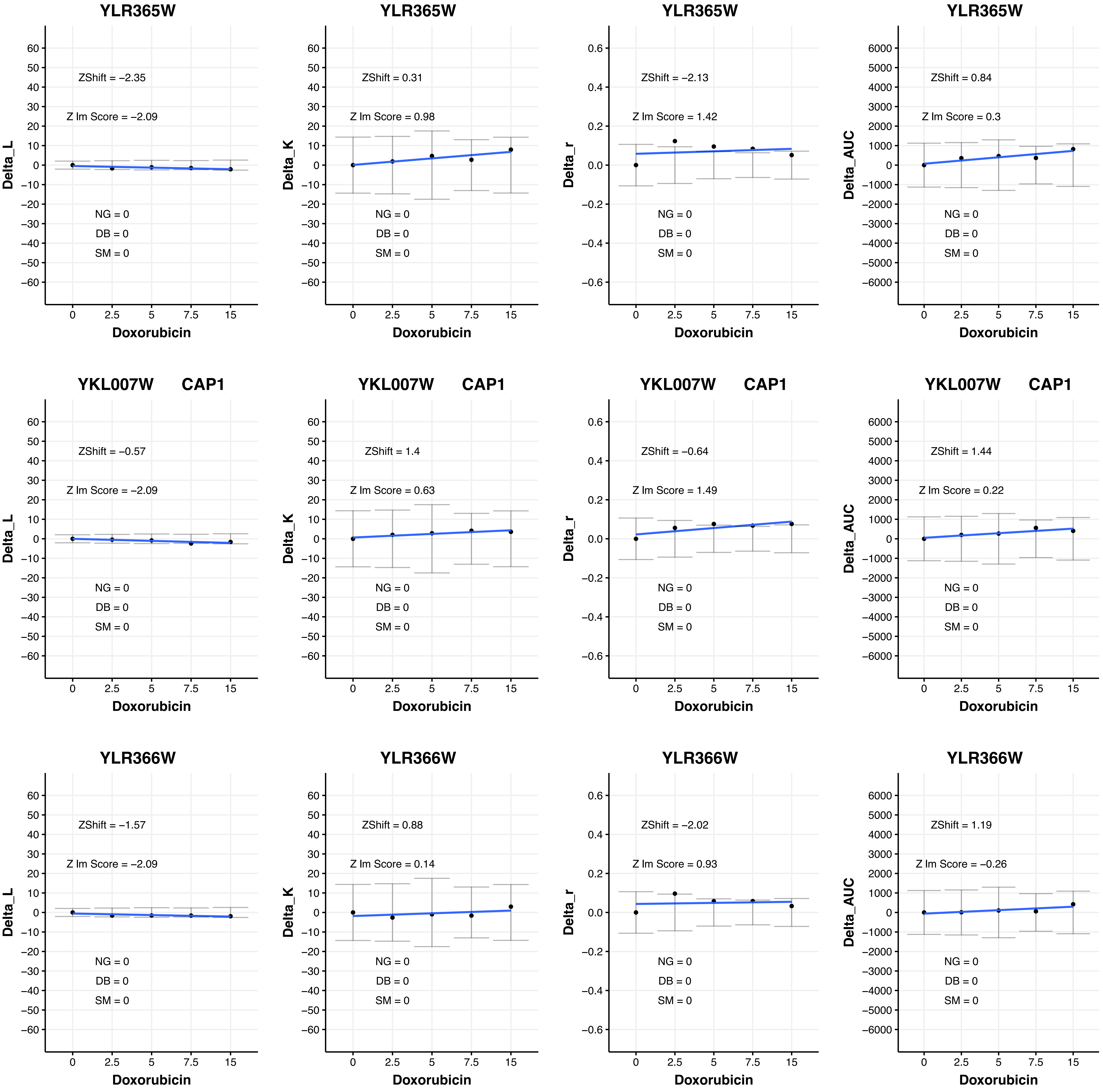
YLR365W
ZShift = −2.35
Z lm Score = −2.09
NG = 0
DB = 0
SM = 0
Delta_L
Doxorubicin
YLR365W
ZShift = 0.31
Z lm Score = 0.98
NG = 0
DB = 0
SM = 0
Delta_K
Doxorubicin
YLR365W
ZShift = −2.13
Z lm Score = 1.42
NG = 0
DB = 0
SM = 0
Delta_r
Doxorubicin
YLR365W
ZShift = 0.84
Z lm Score = 0.3
NG = 0
DB = 0
SM = 0
Delta_AUC
Doxorubicin
YKL007W CAP1
ZShift = −0.57
Z lm Score = −2.09
NG = 0
DB = 0
SM = 0
Delta_L
Doxorubicin
YKL007W CAP1
ZShift = 1.4
Z lm Score = 0.63
NG = 0
DB = 0
SM = 0
Delta_K
Doxorubicin
YKL007W CAP1
ZShift = −0.64
Z lm Score = 1.49
NG = 0
DB = 0
SM = 0
Delta_r
Doxorubicin
YKL007W CAP1
ZShift = 1.44
Z lm Score = 0.22
NG = 0
DB = 0
SM = 0
Delta_AUC
Doxorubicin
YLR366W
ZShift = −1.57
Z lm Score = −2.09
NG = 0
DB = 0
SM = 0
Delta_L
Doxorubicin
YLR366W
ZShift = 0.88
Z lm Score = 0.14
NG = 0
DB = 0
SM = 0
Delta_K
Doxorubicin
YLR366W
ZShift = −2.02
Z lm Score = 0.93
NG = 0
DB = 0
SM = 0
Delta_r
Doxorubicin
YLR366W
ZShift = 1.19
Z lm Score = −0.26
NG = 0
DB = 0
SM = 0
Delta_AUC
Doxorubicin

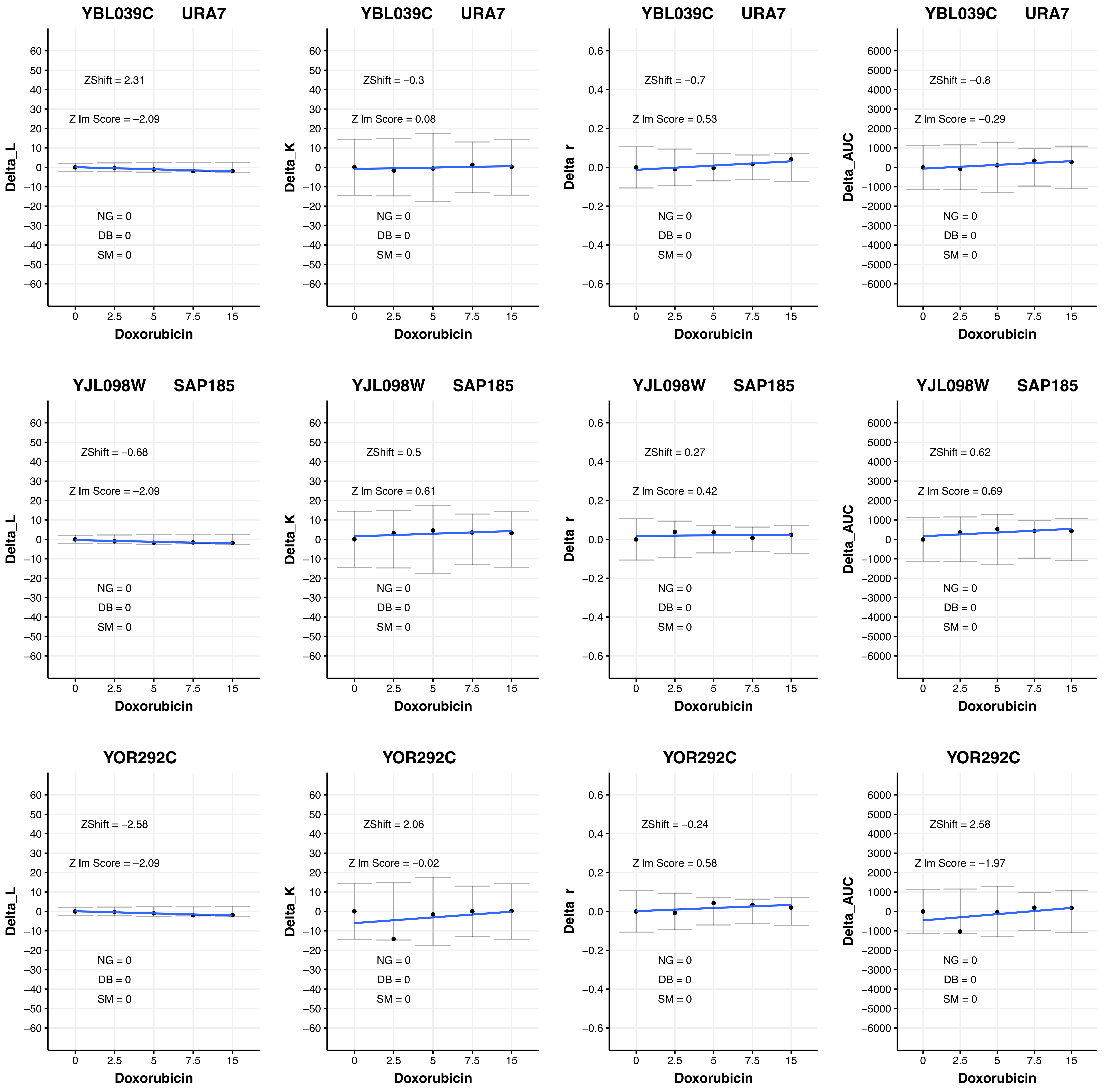

YBL039C URA7
ZShift = 2.31
Z lm Score = −2.09
NG = 0
DB = 0
SM = 0
Delta_L
Doxorubicin
YBL039C URA7
ZShift = −0.3
Z lm Score = 0.08
NG = 0
DB = 0
SM = 0
Delta_K
Doxorubicin
YBL039C URA7
ZShift = −0.7
Z lm Score = 0.53
NG = 0
DB = 0
SM = 0
Delta_r
Doxorubicin
YBL039C URA7
ZShift = −0.8
Z lm Score = −0.29
NG = 0
DB = 0
SM = 0
Delta_AUC
Doxorubicin
YJL098W SAP185
ZShift = −0.68
Z lm Score = −2.09
NG = 0
DB = 0
SM = 0
Delta_L
Doxorubicin
YJL098W SAP185
ZShift = 0.5
Z lm Score = 0.61
NG = 0
DB = 0
SM = 0
Delta_K
Doxorubicin
YJL098W SAP185
ZShift = 0.27
Z lm Score = 0.42
NG = 0
DB = 0
SM = 0
Delta_r
Doxorubicin
YJL098W SAP185
ZShift = 0.62
Z lm Score = 0.69
NG = 0
DB = 0
SM = 0
Delta_AUC
Doxorubicin
YOR292C
ZShift = −2.58
Z lm Score = −2.09
NG = 0
DB = 0
SM = 0
Delta_L
Doxorubicin
YOR292C
ZShift = 2.06
Z lm Score = −0.02
NG = 0
DB = 0
SM = 0
Delta_K
Doxorubicin
YOR292C
ZShift = −0.24
Z lm Score = 0.58
NG = 0
DB = 0
SM = 0
Delta_r
Doxorubicin
YOR292C
ZShift = 2.58
Z lm Score = −1.97
NG = 0
DB = 0
SM = 0
Delta_AUC
Doxorubicin

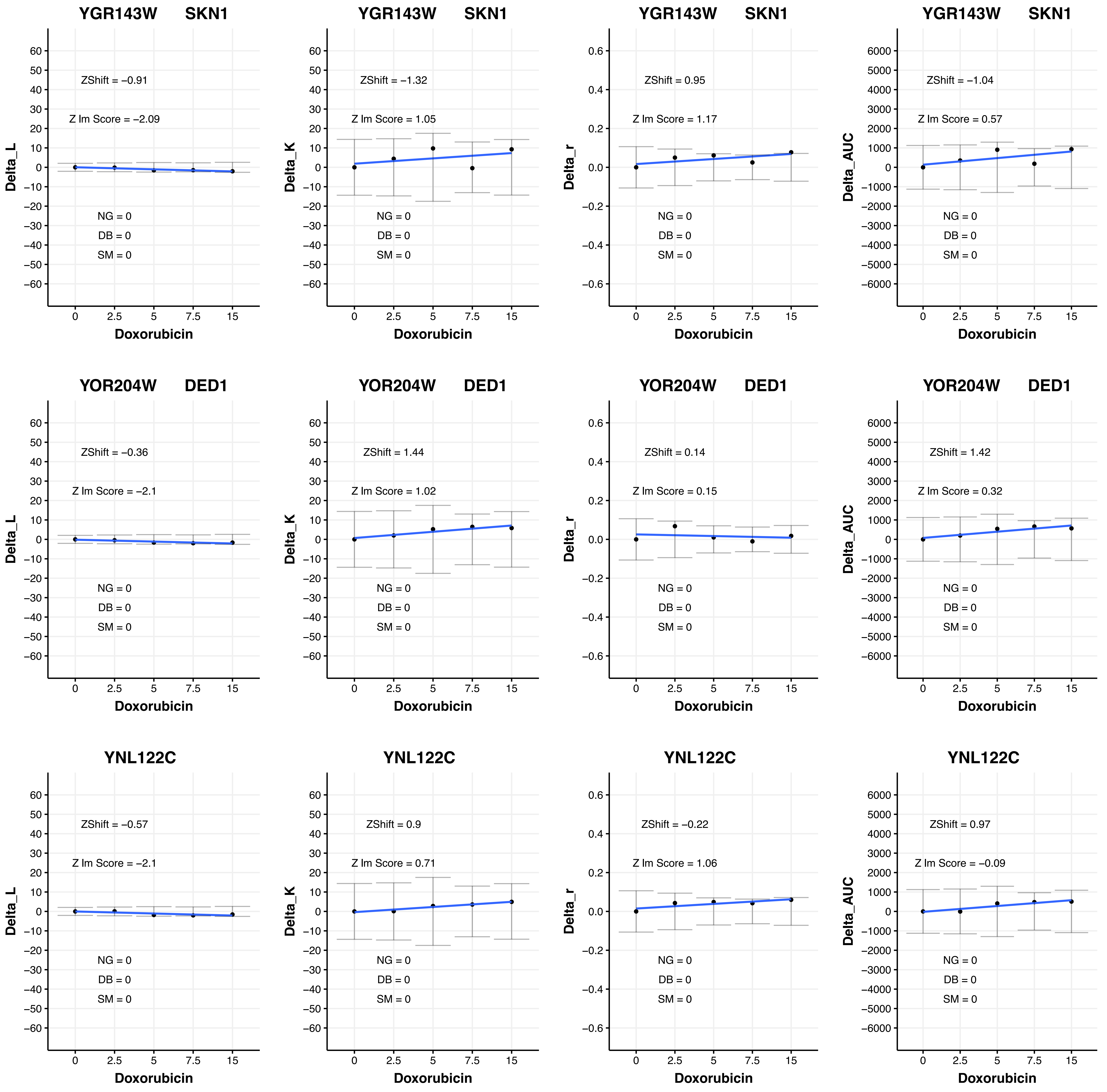

YGR143W SKN1
ZShift = −0.91
Z lm Score = −2.09
NG = 0
DB = 0
SM = 0
Delta_L
Doxorubicin
YGR143W SKN1
ZShift = −1.32
Z lm Score = 1.05
NG = 0
DB = 0
SM = 0
Delta_K
Doxorubicin
YGR143W SKN1
ZShift = 0.95
Z lm Score = 1.17
NG = 0
DB = 0
SM = 0
Delta_r
Doxorubicin
YGR143W SKN1
ZShift = −1.04
Z lm Score = 0.57
NG = 0
DB = 0
SM = 0
Delta_AUC
Doxorubicin
YOR204W DED1
ZShift = −0.36
Z lm Score = −2.1
NG = 0
DB = 0
SM = 0
Delta_L
Doxorubicin
YOR204W DED1
ZShift = 1.44
Z lm Score = 1.02
NG = 0
DB = 0
SM = 0
Delta_K
Doxorubicin
YOR204W DED1
ZShift = 0.14
Z lm Score = 0.15
NG = 0
DB = 0
SM = 0
Delta_r
Doxorubicin
YOR204W DED1
ZShift = 1.42
Z lm Score = 0.32
NG = 0
DB = 0
SM = 0
Delta_AUC
Doxorubicin
YNL122C
ZShift = −0.57
Z lm Score = −2.1
NG = 0
DB = 0
SM = 0
Delta_L
Doxorubicin
YNL122C
ZShift = 0.9
Z lm Score = 0.71
NG = 0
DB = 0
SM = 0
Delta_K
Doxorubicin
YNL122C
ZShift = −0.22
Z lm Score = 1.06
NG = 0
DB = 0
SM = 0
Delta_r
Doxorubicin
YNL122C
ZShift = 0.97
Z lm Score = −0.09
NG = 0
DB = 0
SM = 0
Delta_AUC
Doxorubicin

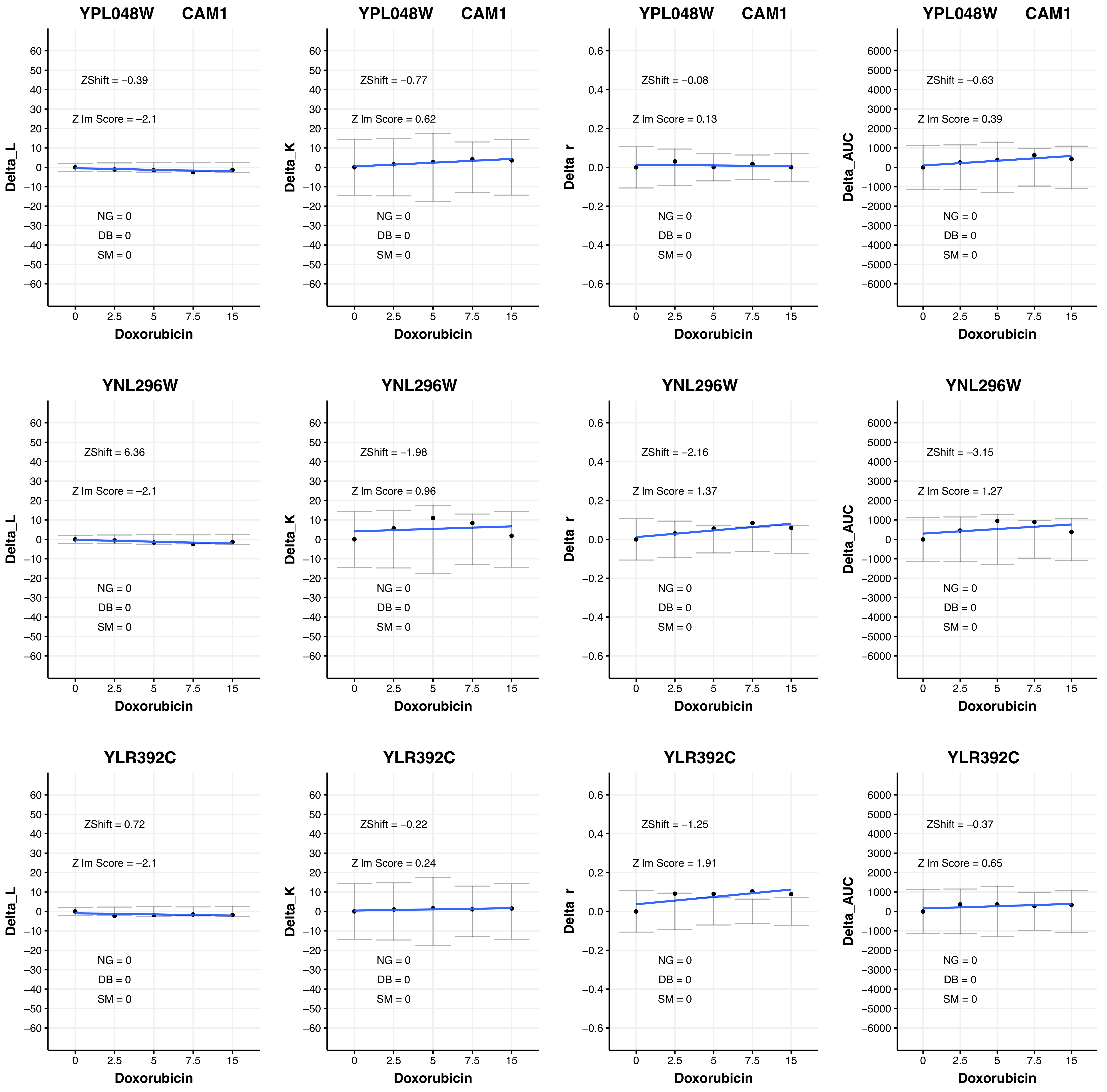
YPL048W CAM1
ZShift = −0.39
Z lm Score = −2.1
NG = 0
DB = 0
SM = 0
Delta_L
Doxorubicin
YPL048W CAM1
ZShift = −0.77
Z lm Score = 0.62
NG = 0
DB = 0
SM = 0
Delta_K
Doxorubicin
YPL048W CAM1
ZShift = −0.08
Z lm Score = 0.13
NG = 0
DB = 0
SM = 0
Delta_r
Doxorubicin
YPL048W CAM1
ZShift = −0.63
Z lm Score = 0.39
NG = 0
DB = 0
SM = 0
Delta_AUC
Doxorubicin
YNL296W
ZShift = 6.36
Z lm Score = −2.1
NG = 0
DB = 0
SM = 0
Delta_L
Doxorubicin
YNL296W
ZShift = −1.98
Z lm Score = 0.96
NG = 0
DB = 0
SM = 0
Delta_K
Doxorubicin
YNL296W
ZShift = −2.16
Z lm Score = 1.37
NG = 0
DB = 0
SM = 0
Delta_r
Doxorubicin
YNL296W
ZShift = −3.15
Z lm Score = 1.27
NG = 0
DB = 0
SM = 0
Delta_AUC
Doxorubicin
YLR392C
ZShift = 0.72
Z lm Score = −2.1
NG = 0
DB = 0
SM = 0
Delta_L
Doxorubicin
YLR392C
ZShift = −0.22
Z lm Score = 0.24
NG = 0
DB = 0
SM = 0
Delta_K
Doxorubicin
YLR392C
ZShift = −1.25
Z lm Score = 1.91
NG = 0
DB = 0
SM = 0
Delta_r
Doxorubicin
YLR392C
ZShift = −0.37
Z lm Score = 0.65
NG = 0
DB = 0
SM = 0
Delta_AUC
Doxorubicin

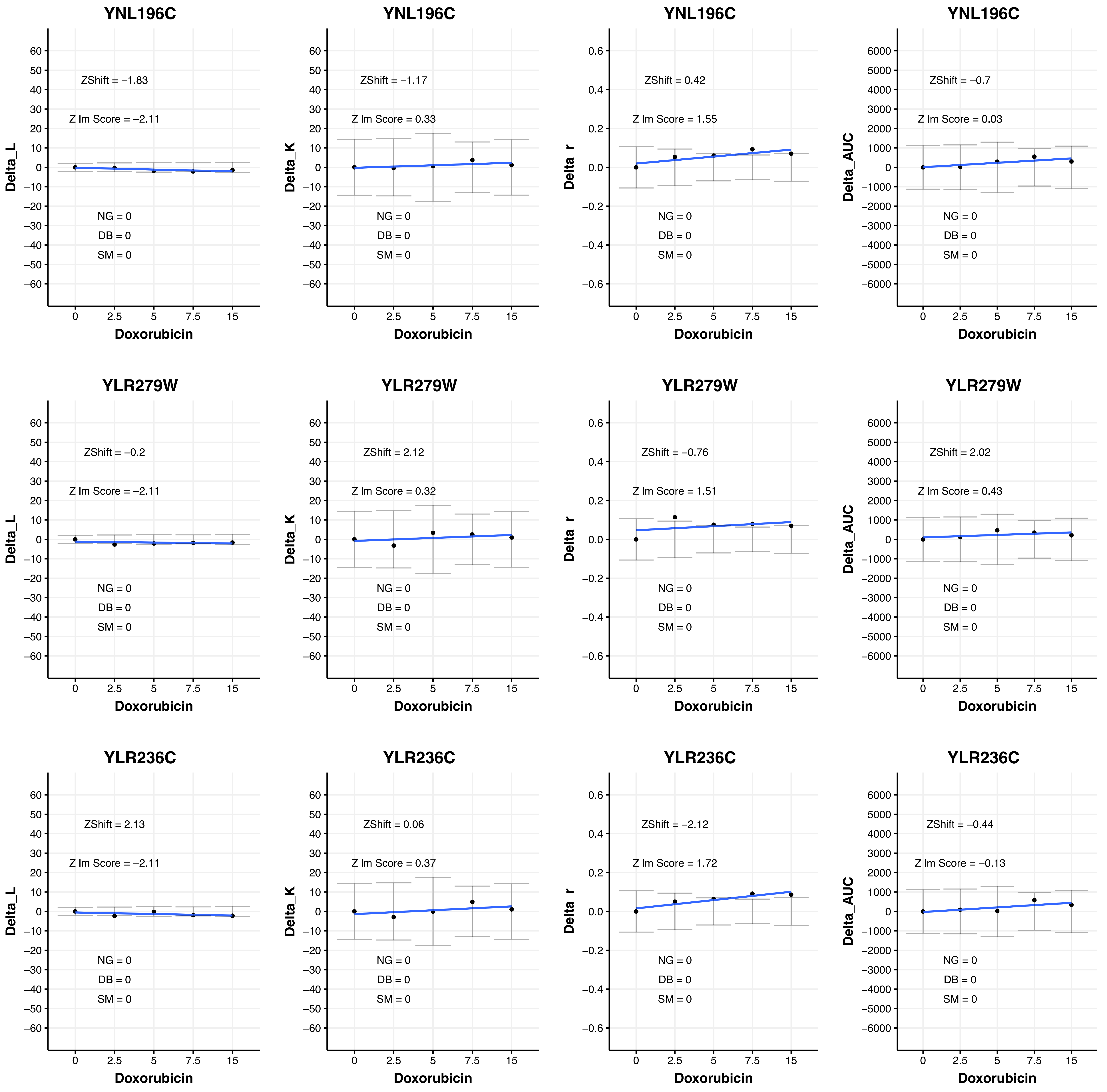
YNL196C
ZShift = −1.83
Z lm Score = −2.11
NG = 0
DB = 0
SM = 0
Delta_L
Doxorubicin
YNL196C
ZShift = −1.17
Z lm Score = 0.33
NG = 0
DB = 0
SM = 0
Delta_K
Doxorubicin
YNL196C
ZShift = 0.42
Z lm Score = 1.55
NG = 0
DB = 0
SM = 0
Delta_r
Doxorubicin
YNL196C
ZShift = −0.7
Z lm Score = 0.03
NG = 0
DB = 0
SM = 0
Delta_AUC
Doxorubicin
YLR279W
ZShift = −0.2
Z lm Score = −2.11
NG = 0
DB = 0
SM = 0
Delta_L
Doxorubicin
YLR279W
ZShift = 2.12
Z lm Score = 0.32
NG = 0
DB = 0
SM = 0
Delta_K
Doxorubicin
YLR279W
ZShift = −0.76
Z lm Score = 1.51
NG = 0
DB = 0
SM = 0
Delta_r
Doxorubicin
YLR279W
ZShift = 2.02
Z lm Score = 0.43
NG = 0
DB = 0
SM = 0
Delta_AUC
Doxorubicin
YLR236C
ZShift = 2.13
Z lm Score = −2.11
NG = 0
DB = 0
SM = 0
Delta_L
Doxorubicin
YLR236C
ZShift = 0.06
Z lm Score = 0.37
NG = 0
DB = 0
SM = 0
Delta_K
Doxorubicin
YLR236C
ZShift = −2.12
Z lm Score = 1.72
NG = 0
DB = 0
SM = 0
Delta_r
Doxorubicin
YLR236C
ZShift = −0.44
Z lm Score = −0.13
NG = 0
DB = 0
SM = 0
Delta_AUC
Doxorubicin

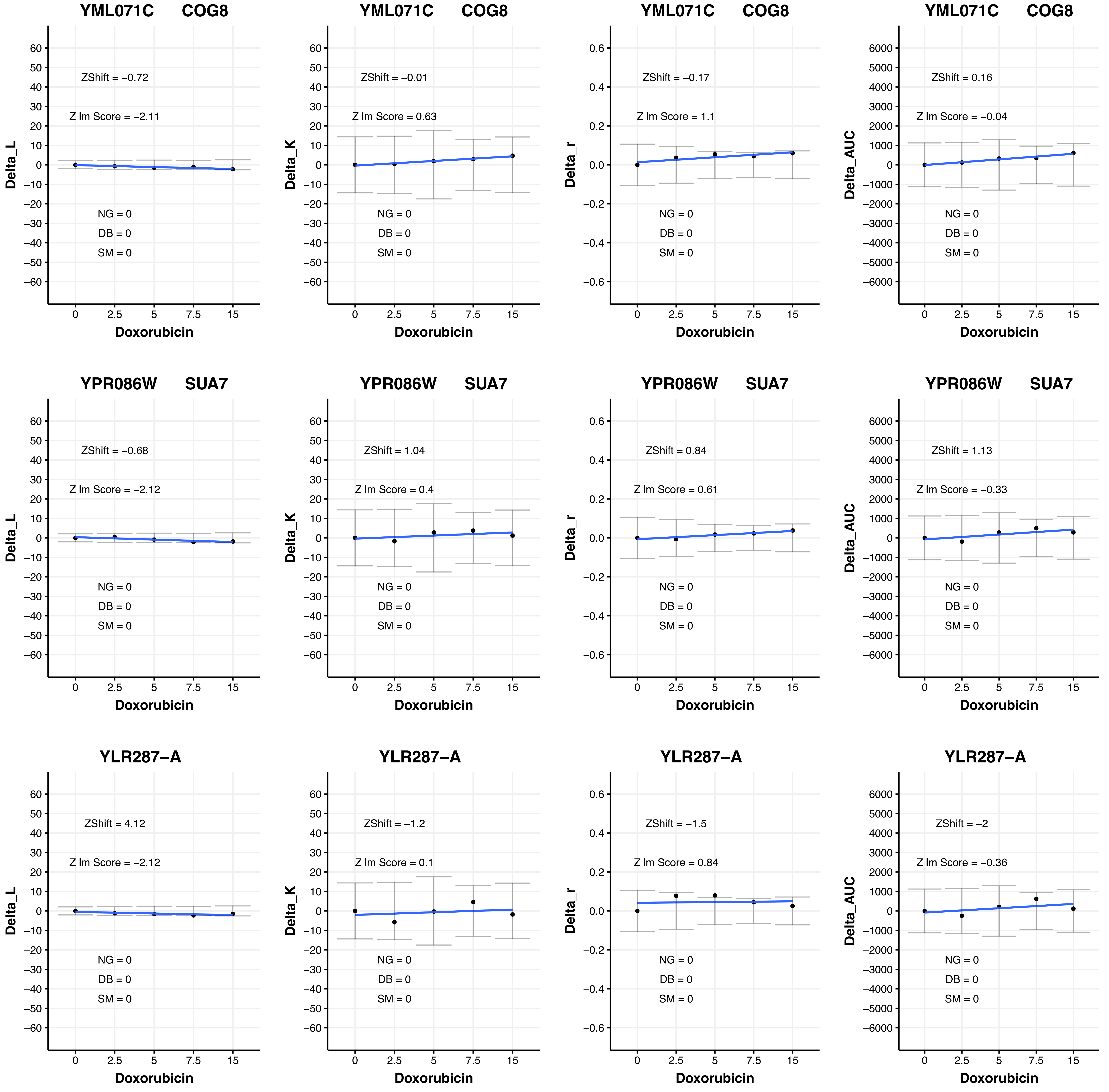

YML071C COG8
ZShift = −0.72
Z lm Score = −2.11
NG = 0
DB = 0
SM = 0
Delta_L
Doxorubicin
YML071C COG8
ZShift = −0.01
Z lm Score = 0.63
NG = 0
DB = 0
SM = 0
Delta_K
Doxorubicin
YML071C COG8
ZShift = −0.17
Z lm Score = 1.1
NG = 0
DB = 0
SM = 0
Delta_r
Doxorubicin
YML071C COG8
ZShift = 0.16
Z lm Score = −0.04
NG = 0
DB = 0
SM = 0
Delta_AUC
Doxorubicin
YPR086W SUA7
ZShift = −0.68
Z lm Score = −2.12
NG = 0
DB = 0
SM = 0
Delta_L
Doxorubicin
YPR086W SUA7
ZShift = 1.04
Z lm Score = 0.4
NG = 0
DB = 0
SM = 0
Delta_K
Doxorubicin
YPR086W SUA7
ZShift = 0.84
Z lm Score = 0.61
NG = 0
DB = 0
SM = 0
Delta_r
Doxorubicin
YPR086W SUA7
ZShift = 1.13
Z lm Score = −0.33
NG = 0
DB = 0
SM = 0
Delta_AUC
Doxorubicin
YLR287−A
ZShift = 4.12
Z lm Score = −2.12
NG = 0
DB = 0
SM = 0
Delta_L
Doxorubicin
YLR287−A
ZShift = −1.2
Z lm Score = 0.1
NG = 0
DB = 0
SM = 0
Delta_K
Doxorubicin
YLR287−A
ZShift = −1.5
Z lm Score = 0.84
NG = 0
DB = 0
SM = 0
Delta_r
Doxorubicin
YLR287−A
ZShift = −2
Z lm Score = −0.36
NG = 0
DB = 0
SM = 0
Delta_AUC
Doxorubicin

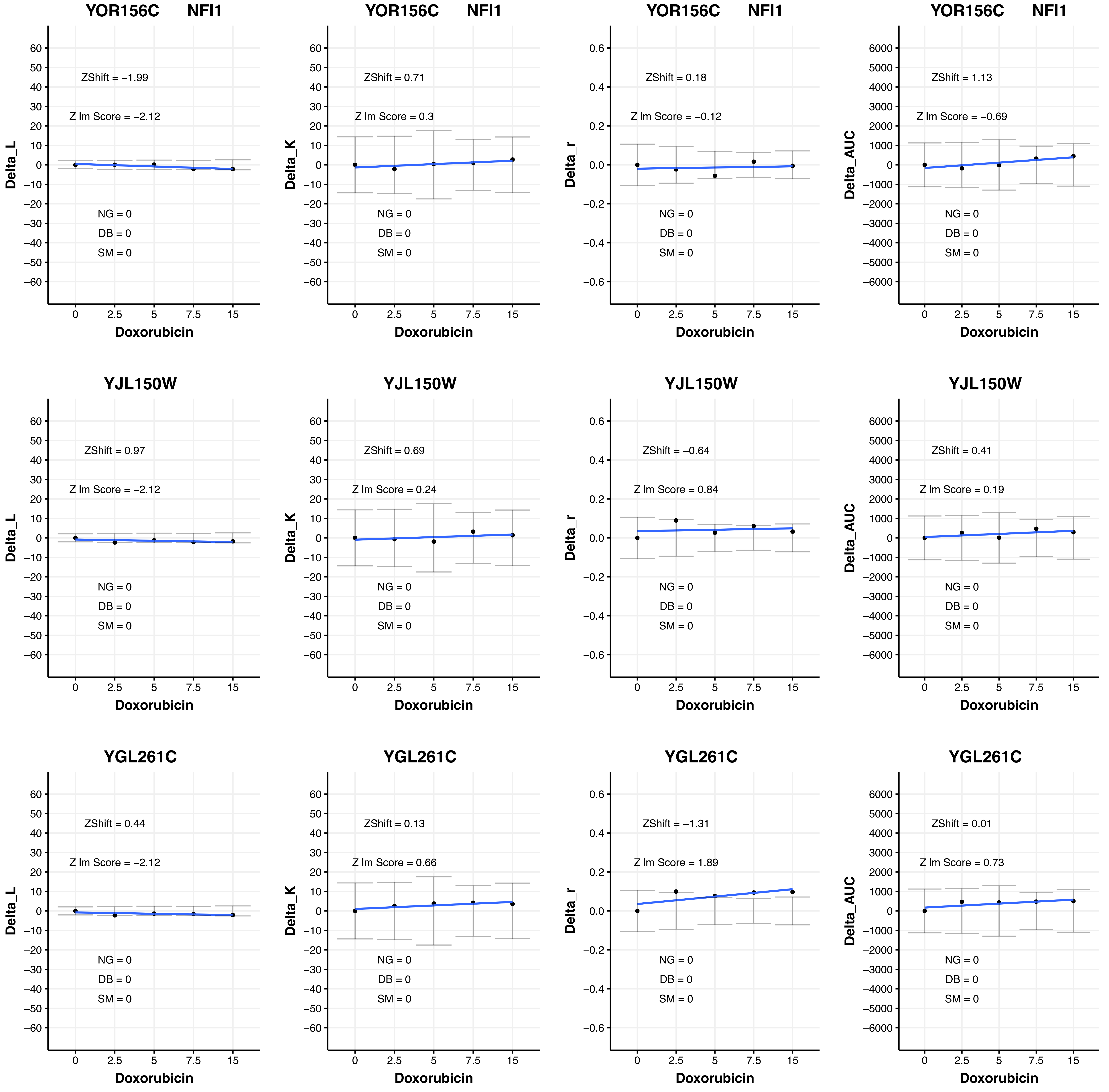
YOR156C NFI1
ZShift = −1.99
Z lm Score = −2.12
NG = 0
DB = 0
SM = 0
Delta_L
Doxorubicin
YOR156C NFI1
ZShift = 0.71
Z lm Score = 0.3
NG = 0
DB = 0
SM = 0
Delta_K
Doxorubicin
YOR156C NFI1
ZShift = 0.18
Z lm Score = −0.12
NG = 0
DB = 0
SM = 0
Delta_r
Doxorubicin
YOR156C NFI1
ZShift = 1.13
Z lm Score = −0.69
NG = 0
DB = 0
SM = 0
Delta_AUC
Doxorubicin
YJL150W
ZShift = 0.97
Z lm Score = −2.12
NG = 0
DB = 0
SM = 0
Delta_L
Doxorubicin
YJL150W
ZShift = 0.69
Z lm Score = 0.24
NG = 0
DB = 0
SM = 0
Delta_K
Doxorubicin
YJL150W
ZShift = −0.64
Z lm Score = 0.84
NG = 0
DB = 0
SM = 0
Delta_r
Doxorubicin
YJL150W
ZShift = 0.41
Z lm Score = 0.19
NG = 0
DB = 0
SM = 0
Delta_AUC
Doxorubicin
YGL261C
ZShift = 0.44
Z lm Score = −2.12
NG = 0
DB = 0
SM = 0
Delta_L
Doxorubicin
YGL261C
ZShift = 0.13
Z lm Score = 0.66
NG = 0
DB = 0
SM = 0
Delta_K
Doxorubicin
YGL261C
ZShift = −1.31
Z lm Score = 1.89
NG = 0
DB = 0
SM = 0
Delta_r
Doxorubicin
YGL261C
ZShift = 0.01
Z lm Score = 0.73
NG = 0
DB = 0
SM = 0
Delta_AUC
Doxorubicin

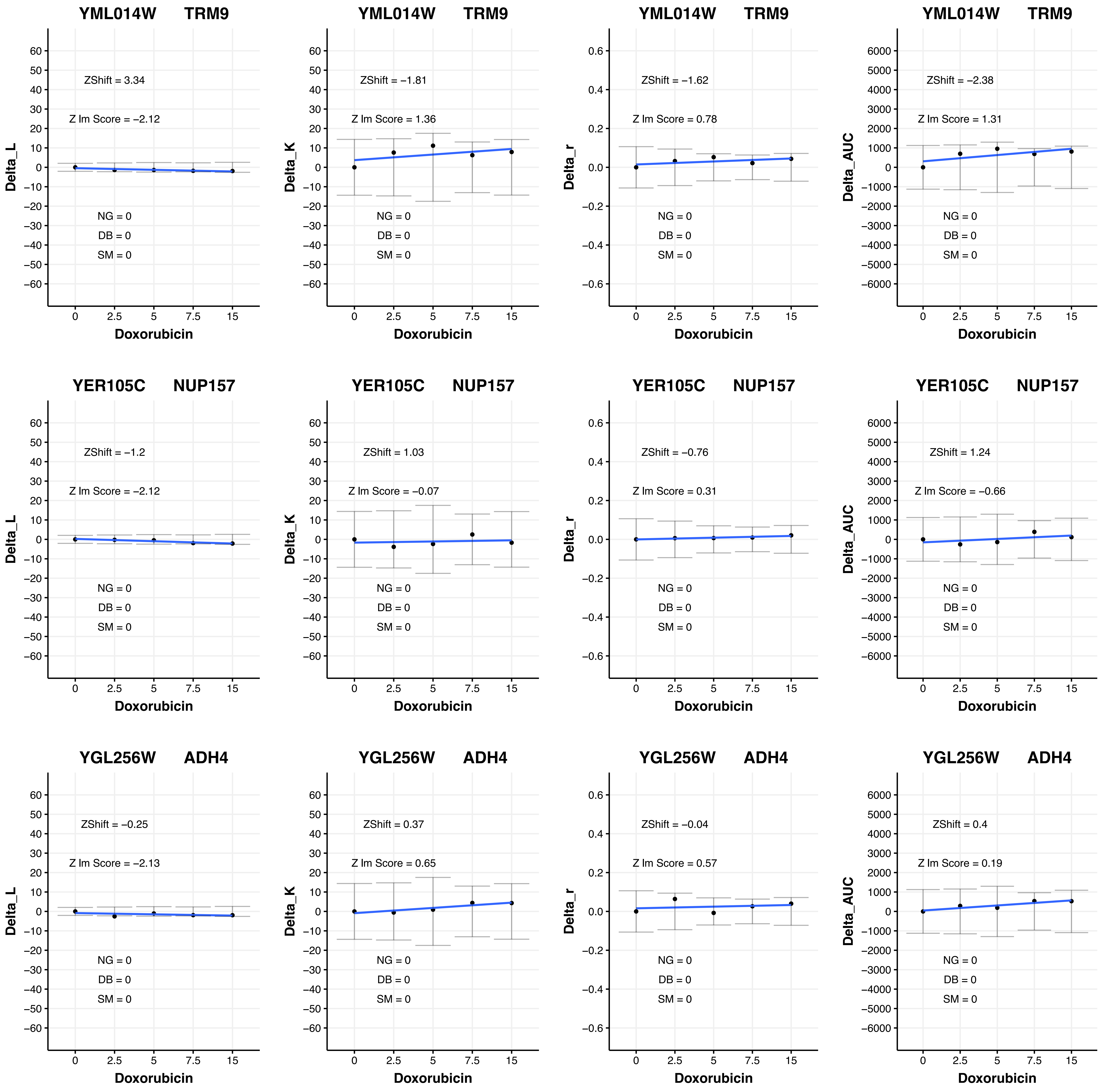

YML014W TRM9
ZShift = 3.34
Z lm Score = −2.12
NG = 0
DB = 0
SM = 0
Delta_L
Doxorubicin
YML014W TRM9
ZShift = −1.81
Z lm Score = 1.36
NG = 0
DB = 0
SM = 0
Delta_K
Doxorubicin
YML014W TRM9
ZShift = −1.62
Z lm Score = 0.78
NG = 0
DB = 0
SM = 0
Delta_r
Doxorubicin
YML014W TRM9
ZShift = −2.38
Z lm Score = 1.31
NG = 0
DB = 0
SM = 0
Delta_AUC
Doxorubicin
YER105C NUP157
ZShift = −1.2
Z lm Score = −2.12
NG = 0
DB = 0
SM = 0
Delta_L
Doxorubicin
YER105C NUP157
ZShift = 1.03
Z lm Score = −0.07
NG = 0
DB = 0
SM = 0
Delta_K
Doxorubicin
YER105C NUP157
ZShift = −0.76
Z lm Score = 0.31
NG = 0
DB = 0
SM = 0
Delta_r
Doxorubicin
YER105C NUP157
ZShift = 1.24
Z lm Score = −0.66
NG = 0
DB = 0
SM = 0
Delta_AUC
Doxorubicin
YGL256W ADH4
ZShift = −0.25
Z lm Score = −2.13
NG = 0
DB = 0
SM = 0
Delta_L
Doxorubicin
YGL256W ADH4
ZShift = 0.37
Z lm Score = 0.65
NG = 0
DB = 0
SM = 0
Delta_K
Doxorubicin
YGL256W ADH4
ZShift = −0.04
Z lm Score = 0.57
NG = 0
DB = 0
SM = 0
Delta_r
Doxorubicin
YGL256W ADH4
ZShift = 0.4
Z lm Score = 0.19
NG = 0
DB = 0
SM = 0
Delta_AUC
Doxorubicin

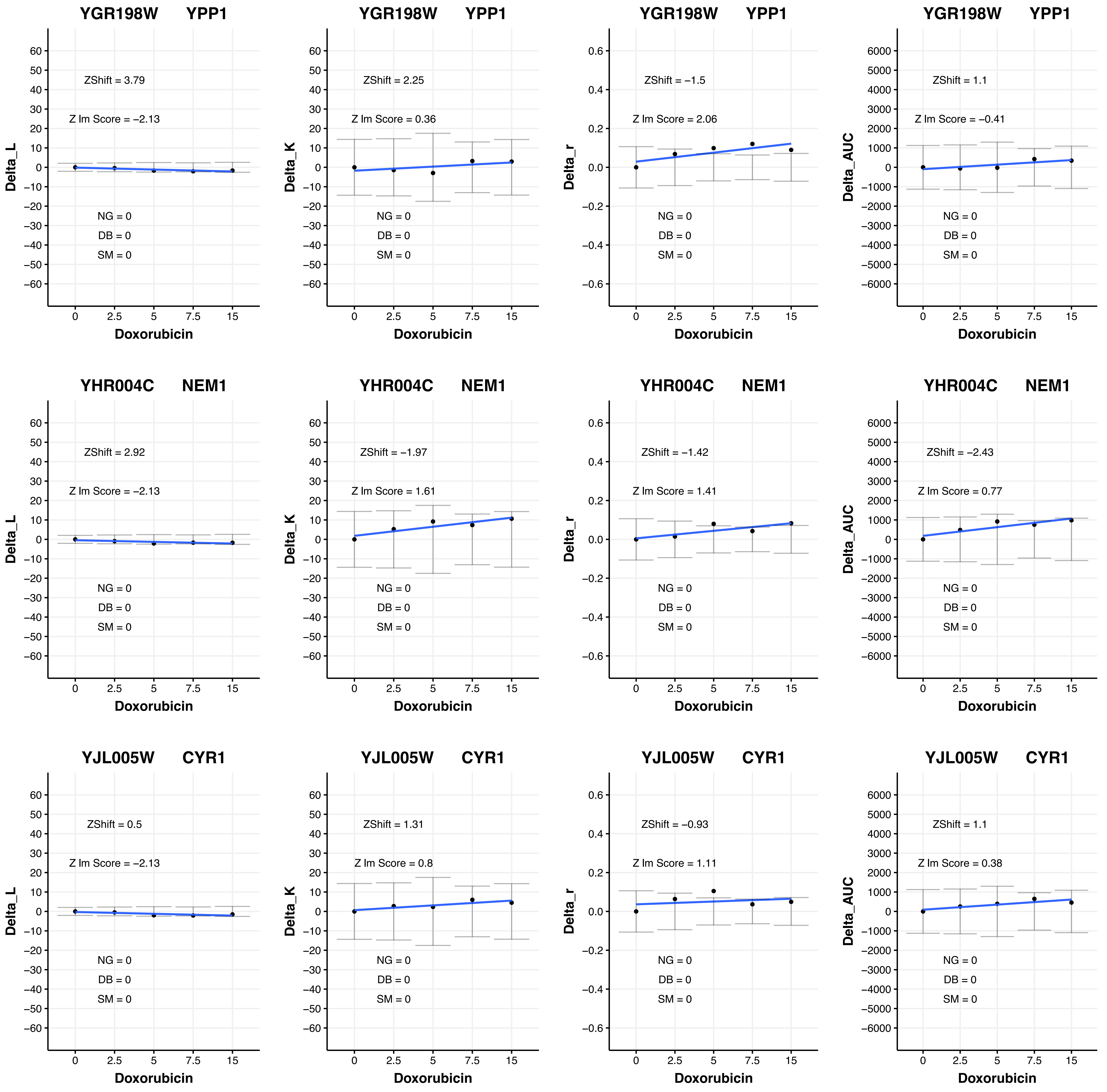
YGR198W YPP1
ZShift = 3.79
Z lm Score = −2.13
NG = 0
DB = 0
SM = 0
Delta_L
Doxorubicin
YGR198W YPP1
ZShift = 2.25
Z lm Score = 0.36
NG = 0
DB = 0
SM = 0
Delta_K
Doxorubicin
YGR198W YPP1
ZShift = −1.5
Z lm Score = 2.06
NG = 0
DB = 0
SM = 0
Delta_r
Doxorubicin
YGR198W YPP1
ZShift = 1.1
Z lm Score = −0.41
NG = 0
DB = 0
SM = 0
Delta_AUC
Doxorubicin
YHR004C NEM1
ZShift = 2.92
Z lm Score = −2.13
NG = 0
DB = 0
SM = 0
Delta_L
Doxorubicin
YHR004C NEM1
ZShift = −1.97
Z lm Score = 1.61
NG = 0
DB = 0
SM = 0
Delta_K
Doxorubicin
YHR004C NEM1
ZShift = −1.42
Z lm Score = 1.41
NG = 0
DB = 0
SM = 0
Delta_r
Doxorubicin
YHR004C NEM1
ZShift = −2.43
Z lm Score = 0.77
NG = 0
DB = 0
SM = 0
Delta_AUC
Doxorubicin
YJL005W CYR1
ZShift = 0.5
Z lm Score = −2.13
NG = 0
DB = 0
SM = 0
Delta_L
Doxorubicin
YJL005W CYR1
ZShift = 1.31
Z lm Score = 0.8
NG = 0
DB = 0
SM = 0
Delta_K
Doxorubicin
YJL005W CYR1
ZShift = −0.93
Z lm Score = 1.11
NG = 0
DB = 0
SM = 0
Delta_r
Doxorubicin
YJL005W CYR1
ZShift = 1.1
Z lm Score = 0.38
NG = 0
DB = 0
SM = 0
Delta_AUC
Doxorubicin

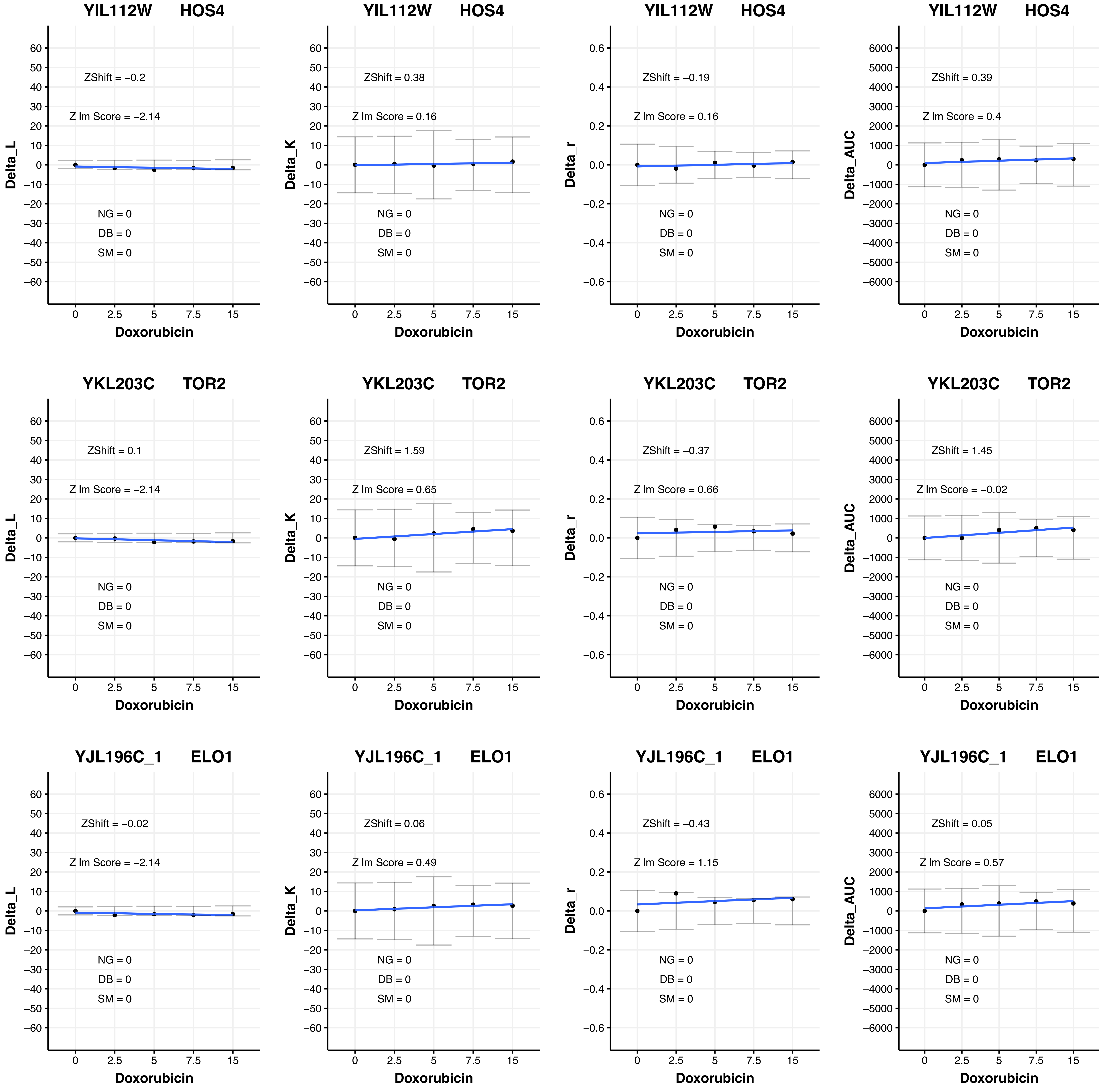
YIL112W HOS4
Delta_L
Doxorubicin
ZShift = −0.2
Z lm Score = −2.14
NG = 0
DB = 0
SM = 0
YIL112W HOS4
Delta_K
Doxorubicin
ZShift = 0.38
Z lm Score = 0.16
NG = 0
DB = 0
SM = 0
YIL112W HOS4
Delta_r
Doxorubicin
ZShift = −0.19
Z lm Score = 0.16
NG = 0
DB = 0
SM = 0
YIL112W HOS4
Delta_AUC
Doxorubicin
ZShift = 0.39
Z lm Score = 0.4
NG = 0
DB = 0
SM = 0
YKL203C TOR2
Delta_L
Doxorubicin
ZShift = 0.1
Z lm Score = −2.14
NG = 0
DB = 0
SM = 0
YKL203C TOR2
Delta_K
Doxorubicin
ZShift = 1.59
Z lm Score = 0.65
NG = 0
DB = 0
SM = 0
YKL203C TOR2
Delta_r
Doxorubicin
ZShift = −0.37
Z lm Score = 0.66
NG = 0
DB = 0
SM = 0
YKL203C TOR2
Delta_AUC
Doxorubicin
ZShift = 1.45
Z lm Score = −0.02
NG = 0
DB = 0
SM = 0
YJL196C_1 ELO1
Delta_L
Doxorubicin
ZShift = −0.02
Z lm Score = −2.14
NG = 0
DB = 0
SM = 0
YJL196C_1 ELO1
Delta_K
Doxorubicin
ZShift = 0.06
Z lm Score = 0.49
NG = 0
DB = 0
SM = 0
YJL196C_1 ELO1
Delta_r
Doxorubicin
ZShift = −0.43
Z lm Score = 1.15
NG = 0
DB = 0
SM = 0
YJL196C_1 ELO1
Delta_AUC
Doxorubicin
ZShift = 0.05
Z lm Score = 0.57
NG = 0
DB = 0
SM = 0

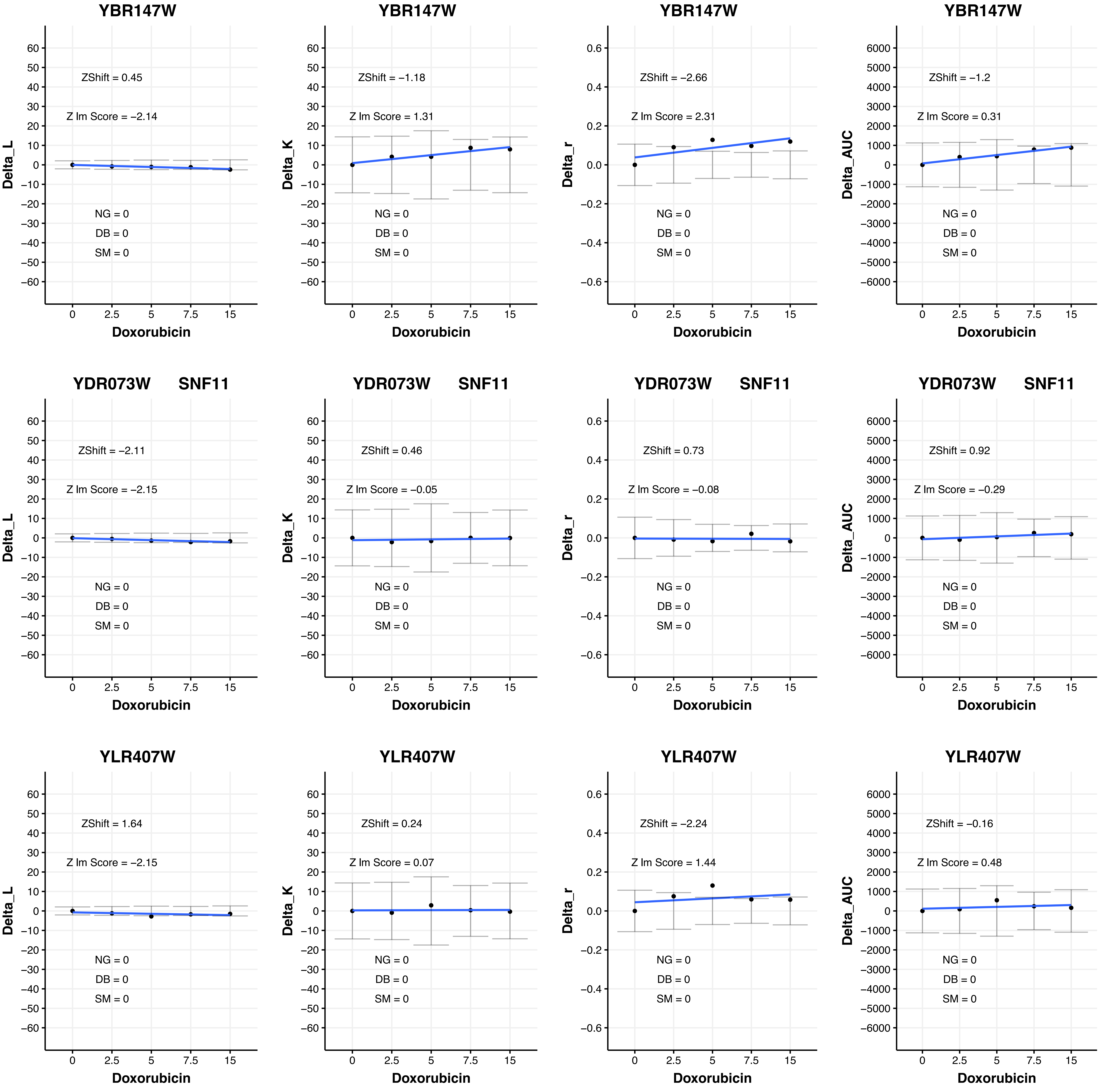
YBR147W
ZShift = 0.45
Z lm Score = −2.14
NG = 0
DB = 0
SM = 0
Delta_L
YBR147W
ZShift = −1.18
Z lm Score = 1.31
NG = 0
DB = 0
SM = 0
Delta_K
YBR147W
ZShift = −2.66
Z lm Score = 2.31
NG = 0
DB = 0
SM = 0
Delta_r
YBR147W
ZShift = −1.2
Z lm Score = 0.31
NG = 0
DB = 0
SM = 0
Delta_AUC
YDR073W SNF11
ZShift = −2.11
Z lm Score = −2.15
NG = 0
DB = 0
SM = 0
Delta_L
YDR073W SNF11
ZShift = 0.46
Z lm Score = −0.05
NG = 0
DB = 0
SM = 0
Delta_K
YDR073W SNF11
ZShift = 0.73
Z lm Score = −0.08
NG = 0
DB = 0
SM = 0
Delta_r
YDR073W SNF11
ZShift = 0.92
Z lm Score = −0.29
NG = 0
DB = 0
SM = 0
Delta_AUC
YLR407W
ZShift = 1.64
Z lm Score = −2.15
NG = 0
DB = 0
SM = 0
Delta_L
YLR407W
ZShift = 0.24
Z lm Score = 0.07
NG = 0
DB = 0
SM = 0
Delta_K
YLR407W
ZShift = −2.24
Z lm Score = 1.44
NG = 0
DB = 0
SM = 0
Delta_r
YLR407W
ZShift = −0.16
Z lm Score = 0.48
NG = 0
DB = 0
SM = 0
Delta_AUC
Doxorubicin
0
2.5
5
7.5
15

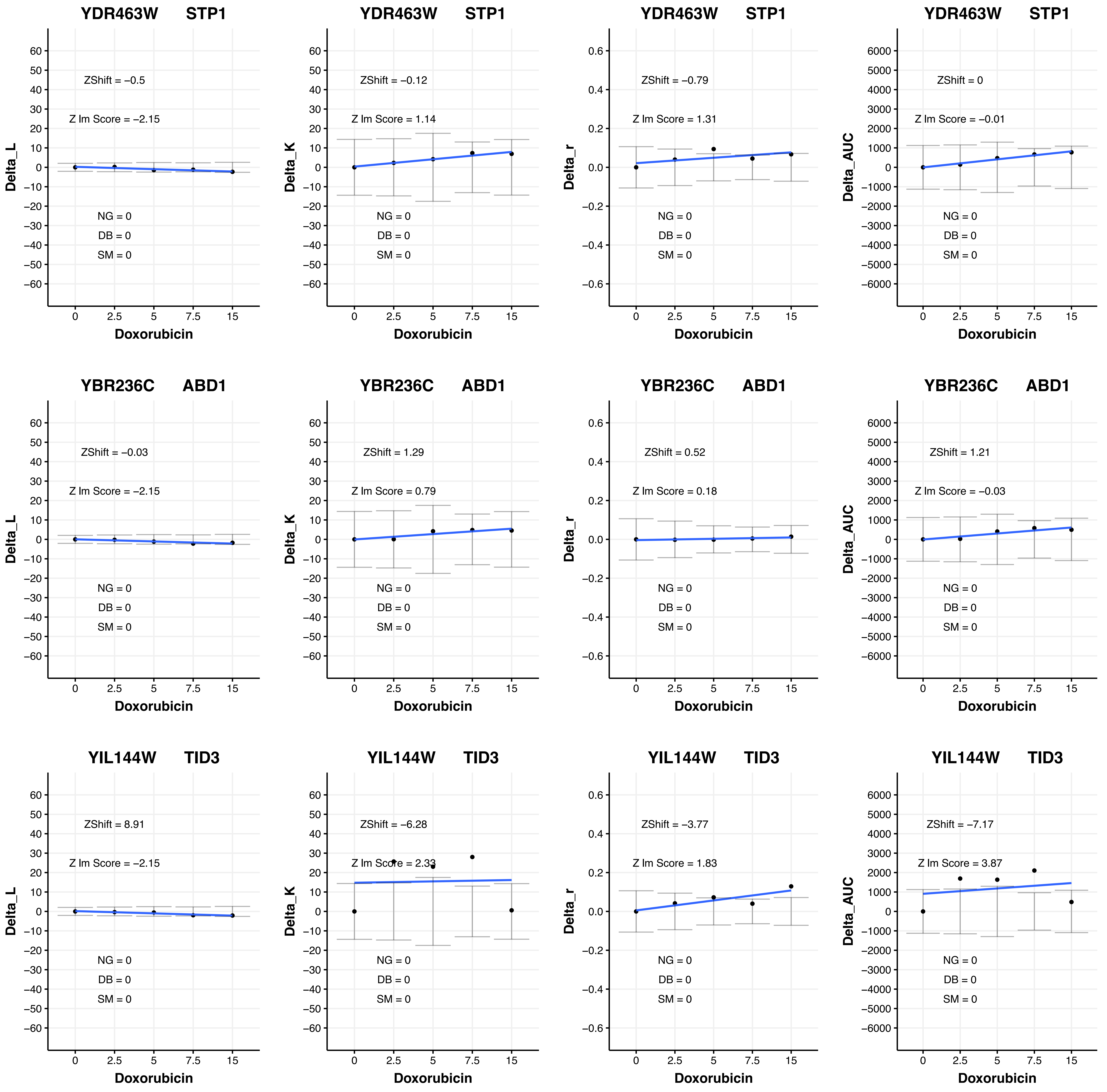

YDR463W STP1
ZShift = −0.5
Z lm Score = −2.15
NG = 0
DB = 0
SM = 0
Delta_L
Doxorubicin
YDR463W STP1
ZShift = −0.12
Z lm Score = 1.14
NG = 0
DB = 0
SM = 0
Delta_K
Doxorubicin
YDR463W STP1
ZShift = −0.79
Z lm Score = 1.31
NG = 0
DB = 0
SM = 0
Delta_r
Doxorubicin
YDR463W STP1
ZShift = 0
Z lm Score = −0.01
NG = 0
DB = 0
SM = 0
Delta_AUC
Doxorubicin
YBR236C ABD1
ZShift = −0.03
Z lm Score = −2.15
NG = 0
DB = 0
SM = 0
Delta_L
Doxorubicin
YBR236C ABD1
ZShift = 1.29
Z lm Score = 0.79
NG = 0
DB = 0
SM = 0
Delta_K
Doxorubicin
YBR236C ABD1
ZShift = 0.52
Z lm Score = 0.18
NG = 0
DB = 0
SM = 0
Delta_r
Doxorubicin
YBR236C ABD1
ZShift = 1.21
Z lm Score = −0.03
NG = 0
DB = 0
SM = 0
Delta_AUC
Doxorubicin
YIL144W TID3
ZShift = 8.91
Z lm Score = −2.15
NG = 0
DB = 0
SM = 0
Delta_L
Doxorubicin
YIL144W TID3
ZShift = −6.28
Z lm Score = 2.33
NG = 0
DB = 0
SM = 0
Delta_K
Doxorubicin
YIL144W TID3
ZShift = −3.77
Z lm Score = 1.83
NG = 0
DB = 0
SM = 0
Delta_r
Doxorubicin
YIL144W TID3
ZShift = −7.17
Z lm Score = 3.87
NG = 0
DB = 0
SM = 0
Delta_AUC
Doxorubicin

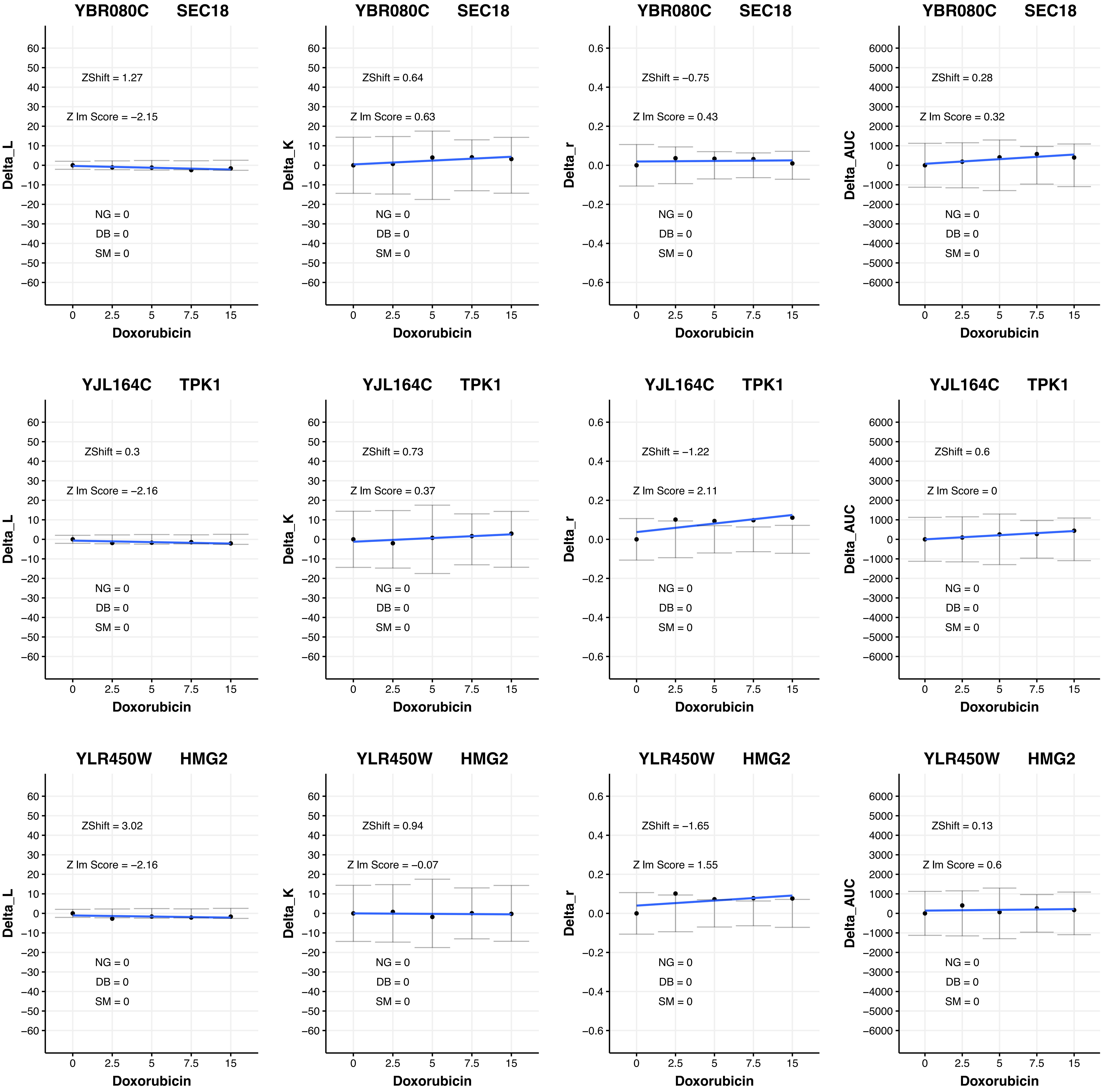

YBR080C SEC18
ZShift = 1.27
Z lm Score = −2.15
NG = 0
DB = 0
SM = 0
Delta_L
Doxorubicin
YBR080C SEC18
ZShift = 0.64
Z lm Score = 0.63
NG = 0
DB = 0
SM = 0
Delta_K
Doxorubicin
YBR080C SEC18
ZShift = −0.75
Z lm Score = 0.43
NG = 0
DB = 0
SM = 0
Delta_r
Doxorubicin
YBR080C SEC18
ZShift = 0.28
Z lm Score = 0.32
NG = 0
DB = 0
SM = 0
Delta_AUC
Doxorubicin
YJL164C TPK1
ZShift = 0.3
Z lm Score = −2.16
NG = 0
DB = 0
SM = 0
Delta_L
Doxorubicin
YJL164C TPK1
ZShift = 0.73
Z lm Score = 0.37
NG = 0
DB = 0
SM = 0
Delta_K
Doxorubicin
YJL164C TPK1
ZShift = −1.22
Z lm Score = 2.11
NG = 0
DB = 0
SM = 0
Delta_r
Doxorubicin
YJL164C TPK1
ZShift = 0.6
Z lm Score = 0
NG = 0
DB = 0
SM = 0
Delta_AUC
Doxorubicin
YLR450W HMG2
ZShift = 3.02
Z lm Score = −2.16
NG = 0
DB = 0
SM = 0
Delta_L
Doxorubicin
YLR450W HMG2
ZShift = 0.94
Z lm Score = −0.07
NG = 0
DB = 0
SM = 0
Delta_K
Doxorubicin
YLR450W HMG2
ZShift = −1.65
Z lm Score = 1.55
NG = 0
DB = 0
SM = 0
Delta_r
Doxorubicin
YLR450W HMG2
ZShift = 0.13
Z lm Score = 0.6
NG = 0
DB = 0
SM = 0
Delta_AUC
Doxorubicin

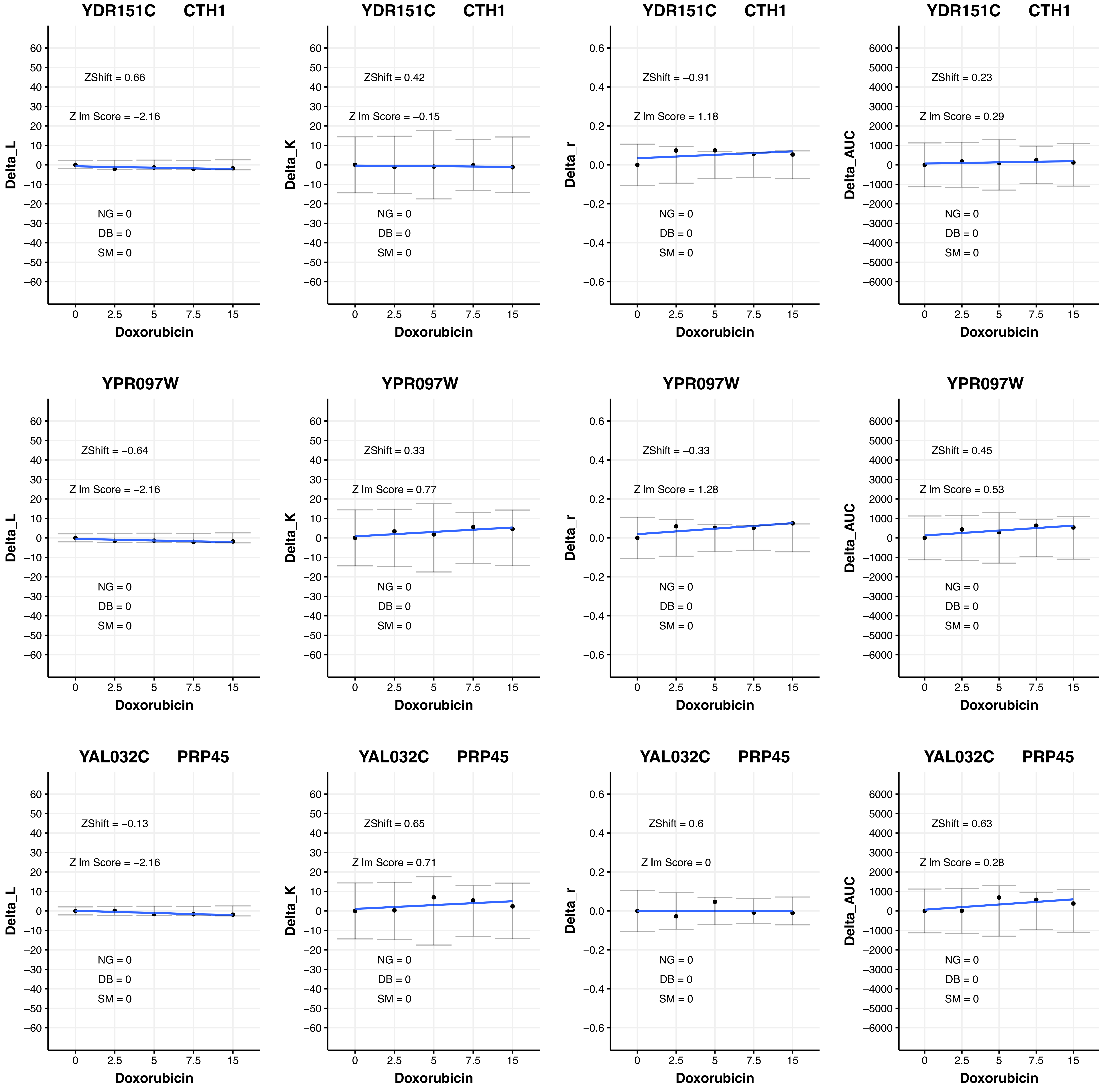
YDR151C CTH1
ZShift = 0.66
Z lm Score = −2.16
NG = 0
DB = 0
SM = 0
Delta_L
Doxorubicin
YDR151C CTH1
ZShift = 0.42
Z lm Score = −0.15
NG = 0
DB = 0
SM = 0
Delta_K
Doxorubicin
YDR151C CTH1
ZShift = −0.91
Z lm Score = 1.18
NG = 0
DB = 0
SM = 0
Delta_r
Doxorubicin
YDR151C CTH1
ZShift = 0.23
Z lm Score = 0.29
NG = 0
DB = 0
SM = 0
Delta_AUC
Doxorubicin
YPR097W
ZShift = −0.64
Z lm Score = −2.16
NG = 0
DB = 0
SM = 0
Delta_L
Doxorubicin
YPR097W
ZShift = 0.33
Z lm Score = 0.77
NG = 0
DB = 0
SM = 0
Delta_K
Doxorubicin
YPR097W
ZShift = −0.33
Z lm Score = 1.28
NG = 0
DB = 0
SM = 0
Delta_r
Doxorubicin
YPR097W
ZShift = 0.45
Z lm Score = 0.53
NG = 0
DB = 0
SM = 0
Delta_AUC
Doxorubicin
YAL032C PRP45
ZShift = −0.13
Z lm Score = −2.16
NG = 0
DB = 0
SM = 0
Delta_L
Doxorubicin
YAL032C PRP45
ZShift = 0.65
Z lm Score = 0.71
NG = 0
DB = 0
SM = 0
Delta_K
Doxorubicin
YAL032C PRP45
ZShift = 0.6
Z lm Score = 0
NG = 0
DB = 0
SM = 0
Delta_r
Doxorubicin
YAL032C PRP45
ZShift = 0.63
Z lm Score = 0.28
NG = 0
DB = 0
SM = 0
Delta_AUC
Doxorubicin

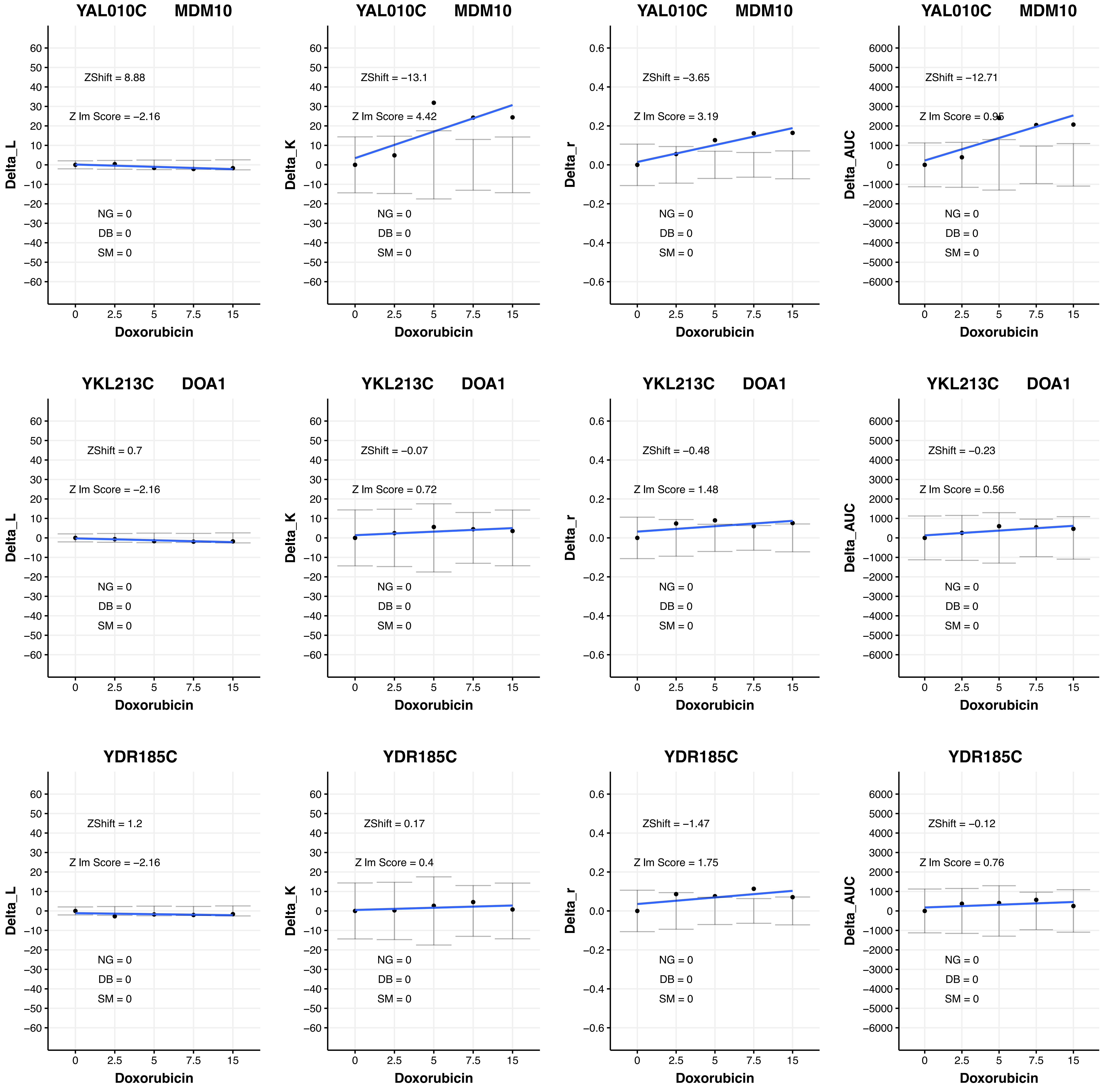

YAL010C MDM10
ZShift = 8.88
Z lm Score = −2.16
NG = 0
DB = 0
SM = 0
Delta_L
Doxorubicin
YAL010C MDM10
ZShift = −13.1
Z lm Score = 4.42
NG = 0
DB = 0
SM = 0
Delta_K
Doxorubicin
YAL010C MDM10
ZShift = −3.65
Z lm Score = 3.19
NG = 0
DB = 0
SM = 0
Delta_r
Doxorubicin
YAL010C MDM10
ZShift = −12.71
Z lm Score = 0.95
NG = 0
DB = 0
SM = 0
Delta_AUC
Doxorubicin
YKL213C DOA1
ZShift = 0.7
Z lm Score = −2.16
NG = 0
DB = 0
SM = 0
Delta_L
Doxorubicin
YKL213C DOA1
ZShift = −0.07
Z lm Score = 0.72
NG = 0
DB = 0
SM = 0
Delta_K
Doxorubicin
YKL213C DOA1
ZShift = −0.48
Z lm Score = 1.48
NG = 0
DB = 0
SM = 0
Delta_r
Doxorubicin
YKL213C DOA1
ZShift = −0.23
Z lm Score = 0.56
NG = 0
DB = 0
SM = 0
Delta_AUC
Doxorubicin
YDR185C
ZShift = 1.2
Z lm Score = −2.16
NG = 0
DB = 0
SM = 0
Delta_L
Doxorubicin
YDR185C
ZShift = 0.17
Z lm Score = 0.4
NG = 0
DB = 0
SM = 0
Delta_K
Doxorubicin
YDR185C
ZShift = −1.47
Z lm Score = 1.75
NG = 0
DB = 0
SM = 0
Delta_r
Doxorubicin
YDR185C
ZShift = −0.12
Z lm Score = 0.76
NG = 0
DB = 0
SM = 0
Delta_AUC
Doxorubicin

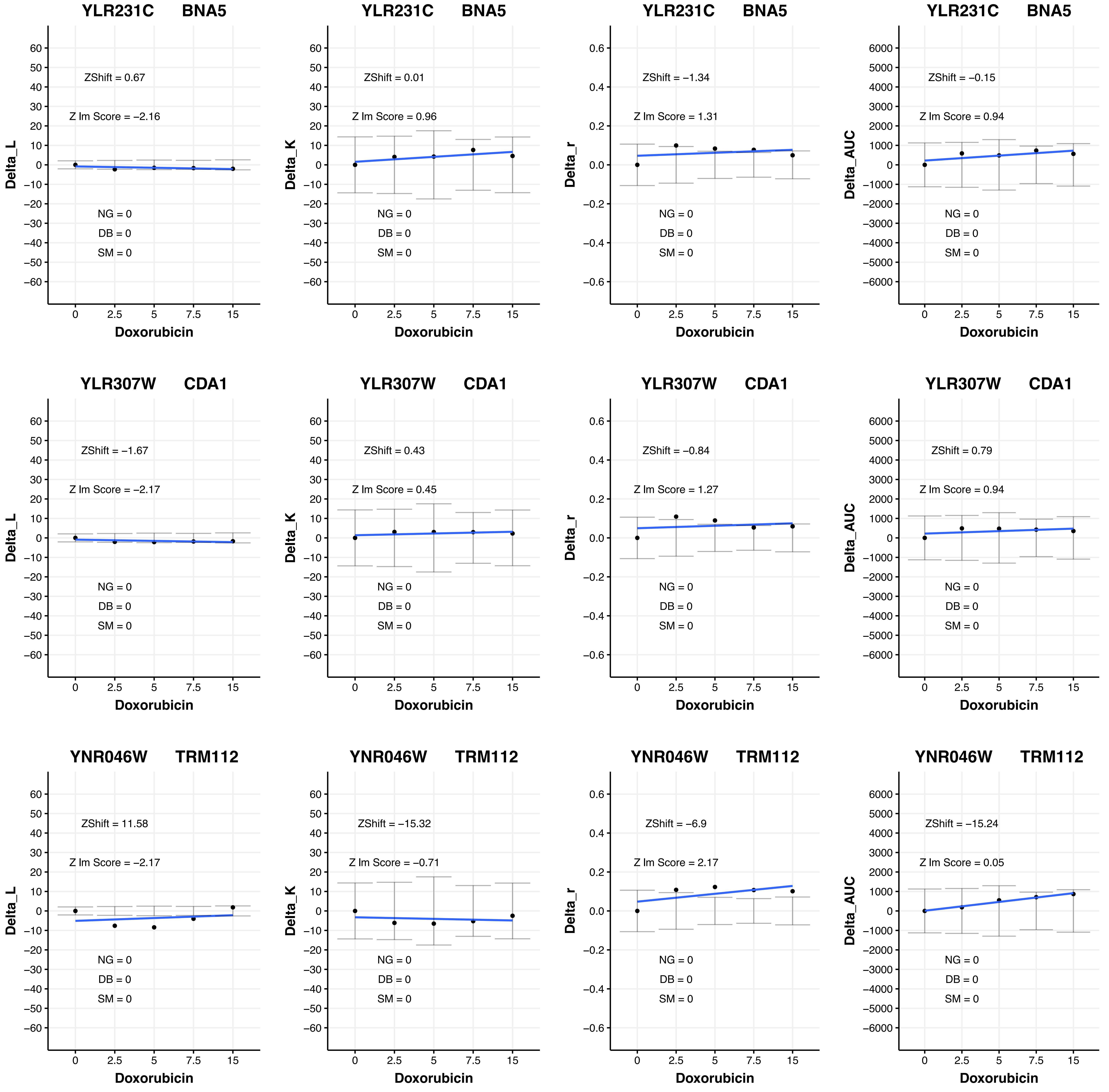

YLR231C BNA5
ZShift = 0.67
Z lm Score = −2.16
NG = 0
DB = 0
SM = 0
Delta_L
Doxorubicin
YLR231C BNA5
ZShift = 0.01
Z lm Score = 0.96
NG = 0
DB = 0
SM = 0
Delta_K
Doxorubicin
YLR231C BNA5
ZShift = −1.34
Z lm Score = 1.31
NG = 0
DB = 0
SM = 0
Delta_r
Doxorubicin
YLR231C BNA5
ZShift = −0.15
Z lm Score = 0.94
NG = 0
DB = 0
SM = 0
Delta_AUC
Doxorubicin
YLR307W CDA1
ZShift = −1.67
Z lm Score = −2.17
NG = 0
DB = 0
SM = 0
Delta_L
Doxorubicin
YLR307W CDA1
ZShift = 0.43
Z lm Score = 0.45
NG = 0
DB = 0
SM = 0
Delta_K
Doxorubicin
YLR307W CDA1
ZShift = −0.84
Z lm Score = 1.27
NG = 0
DB = 0
SM = 0
Delta_r
Doxorubicin
YLR307W CDA1
ZShift = 0.79
Z lm Score = 0.94
NG = 0
DB = 0
SM = 0
Delta_AUC
Doxorubicin
YNR046W TRM112
ZShift = 11.58
Z lm Score = −2.17
NG = 0
DB = 0
SM = 0
Delta_L
Doxorubicin
YNR046W TRM112
ZShift = −15.32
Z lm Score = −0.71
NG = 0
DB = 0
SM = 0
Delta_K
Doxorubicin
YNR046W TRM112
ZShift = −6.9
Z lm Score = 2.17
NG = 0
DB = 0
SM = 0
Delta_r
Doxorubicin
YNR046W TRM112
ZShift = −15.24
Z lm Score = 0.05
NG = 0
DB = 0
SM = 0
Delta_AUC
Doxorubicin

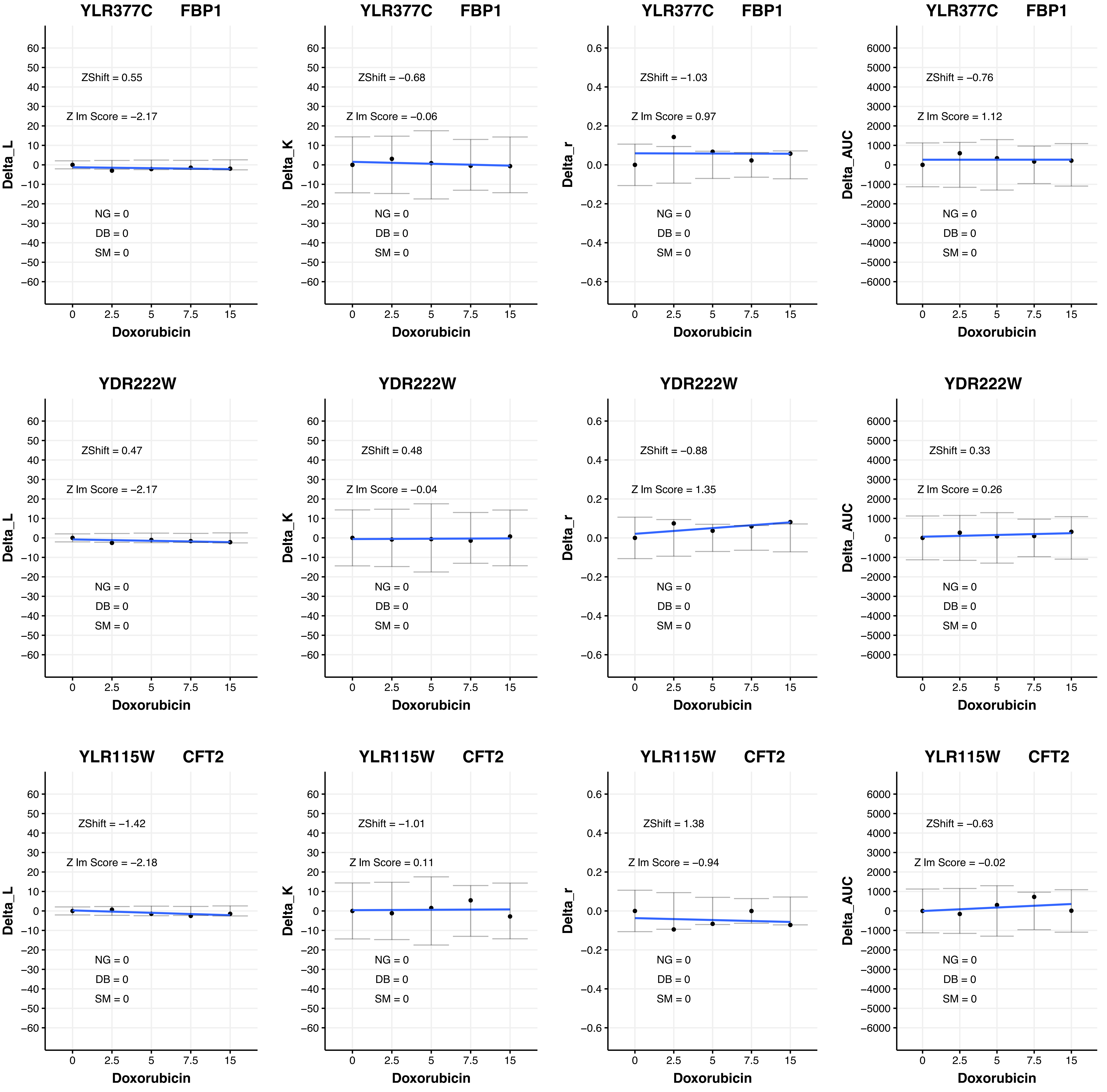

YLR377C FBP1
ZShift = 0.55
Z lm Score = −2.17
NG = 0
DB = 0
SM = 0
Delta_L
Doxorubicin
YLR377C FBP1
ZShift = −0.68
Z lm Score = −0.06
NG = 0
DB = 0
SM = 0
Delta_K
Doxorubicin
YLR377C FBP1
ZShift = −1.03
Z lm Score = 0.97
NG = 0
DB = 0
SM = 0
Delta_r
Doxorubicin
YLR377C FBP1
ZShift = −0.76
Z lm Score = 1.12
NG = 0
DB = 0
SM = 0
Delta_AUC
Doxorubicin
YDR222W
ZShift = 0.47
Z lm Score = −2.17
NG = 0
DB = 0
SM = 0
Delta_L
Doxorubicin
YDR222W
ZShift = 0.48
Z lm Score = −0.04
NG = 0
DB = 0
SM = 0
Delta_K
Doxorubicin
YDR222W
ZShift = −0.88
Z lm Score = 1.35
NG = 0
DB = 0
SM = 0
Delta_r
Doxorubicin
YDR222W
ZShift = 0.33
Z lm Score = 0.26
NG = 0
DB = 0
SM = 0
Delta_AUC
Doxorubicin
YLR115W CFT2
ZShift = −1.42
Z lm Score = −2.18
NG = 0
DB = 0
SM = 0
Delta_L
Doxorubicin
YLR115W CFT2
ZShift = −1.01
Z lm Score = 0.11
NG = 0
DB = 0
SM = 0
Delta_K
Doxorubicin
YLR115W CFT2
ZShift = 1.38
Z lm Score = −0.94
NG = 0
DB = 0
SM = 0
Delta_r
Doxorubicin
YLR115W CFT2
ZShift = −0.63
Z lm Score = −0.02
NG = 0
DB = 0
SM = 0
Delta_AUC
Doxorubicin

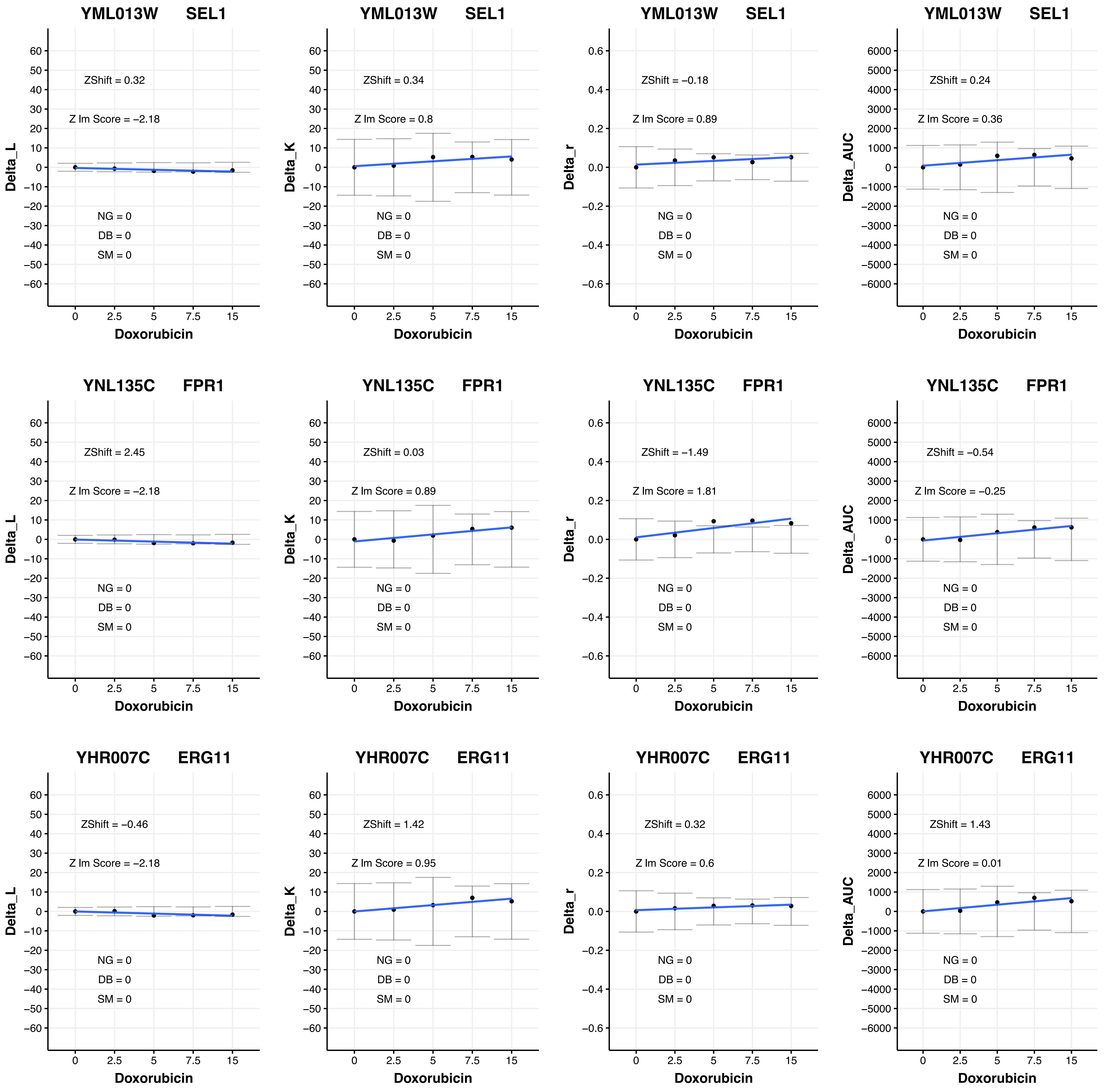

YML013W SEL1
ZShift = 0.32
Z lm Score = −2.18
NG = 0
DB = 0
SM = 0
Delta_L
Doxorubicin
YML013W SEL1
ZShift = 0.34
Z lm Score = 0.8
NG = 0
DB = 0
SM = 0
Delta_K
Doxorubicin
YML013W SEL1
ZShift = −0.18
Z lm Score = 0.89
NG = 0
DB = 0
SM = 0
Delta_r
Doxorubicin
YML013W SEL1
ZShift = 0.24
Z lm Score = 0.36
NG = 0
DB = 0
SM = 0
Delta_AUC
Doxorubicin
YNL135C FPR1
ZShift = 2.45
Z lm Score = −2.18
NG = 0
DB = 0
SM = 0
Delta_L
Doxorubicin
YNL135C FPR1
ZShift = 0.03
Z lm Score = 0.89
NG = 0
DB = 0
SM = 0
Delta_K
Doxorubicin
YNL135C FPR1
ZShift = −1.49
Z lm Score = 1.81
NG = 0
DB = 0
SM = 0
Delta_r
Doxorubicin
YNL135C FPR1
ZShift = −0.54
Z lm Score = −0.25
NG = 0
DB = 0
SM = 0
Delta_AUC
Doxorubicin
YHR007C ERG11
ZShift = −0.46
Z lm Score = −2.18
NG = 0
DB = 0
SM = 0
Delta_L
Doxorubicin
YHR007C ERG11
ZShift = 1.42
Z lm Score = 0.95
NG = 0
DB = 0
SM = 0
Delta_K
Doxorubicin
YHR007C ERG11
ZShift = 0.32
Z lm Score = 0.6
NG = 0
DB = 0
SM = 0
Delta_r
Doxorubicin
YHR007C ERG11
ZShift = 1.43
Z lm Score = 0.01
NG = 0
DB = 0
SM = 0
Delta_AUC
Doxorubicin

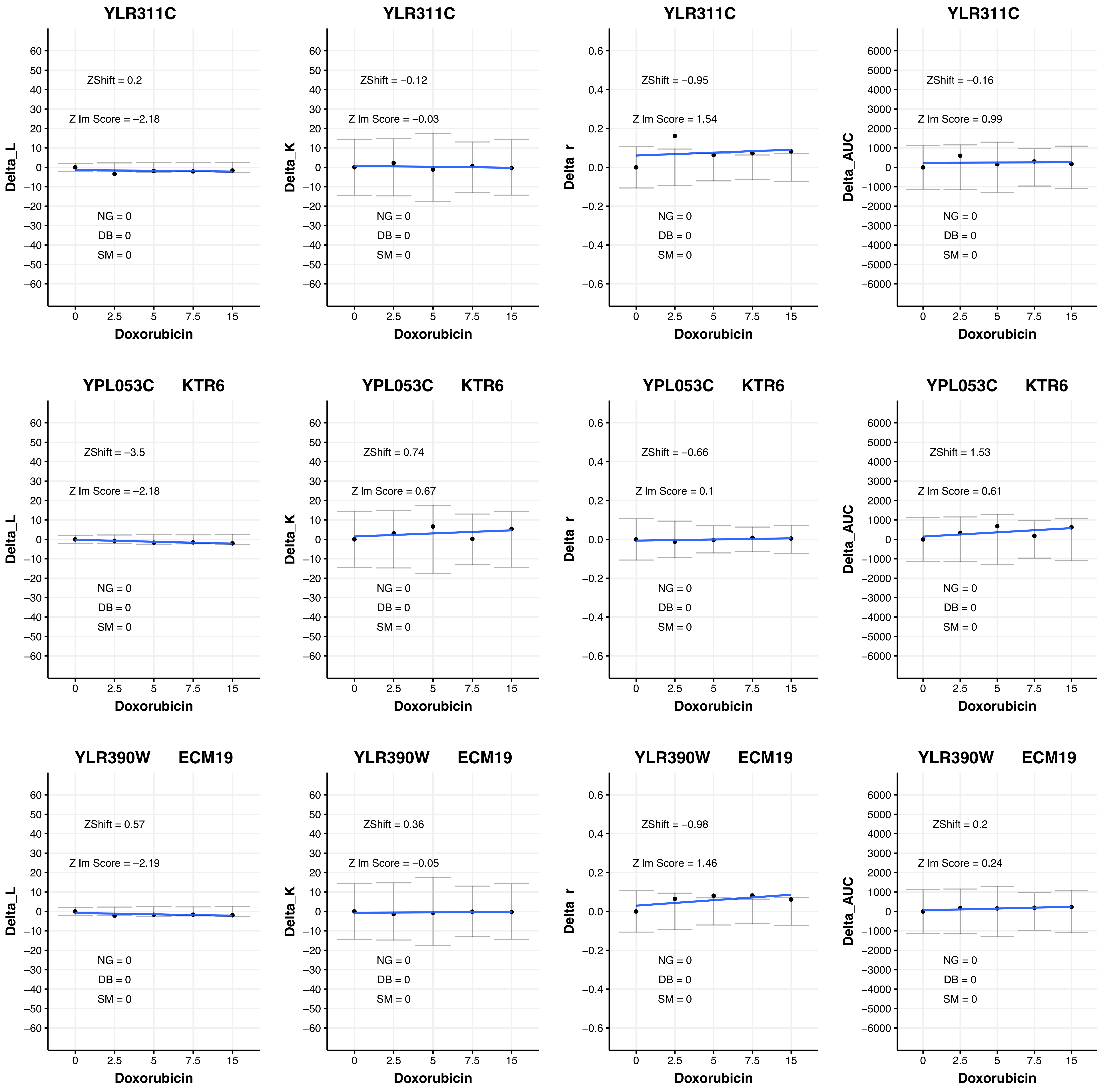
YLR311C
ZShift = 0.2
Z lm Score = −2.18
NG = 0
DB = 0
SM = 0
Delta_L
Doxorubicin
YLR311C
ZShift = −0.12
Z lm Score = −0.03
NG = 0
DB = 0
SM = 0
Delta_K
Doxorubicin
YLR311C
ZShift = −0.95
Z lm Score = 1.54
NG = 0
DB = 0
SM = 0
Delta_r
Doxorubicin
YLR311C
ZShift = −0.16
Z lm Score = 0.99
NG = 0
DB = 0
SM = 0
Delta_AUC
Doxorubicin
YPL053C KTR6
ZShift = −3.5
Z lm Score = −2.18
NG = 0
DB = 0
SM = 0
Delta_L
Doxorubicin
YPL053C KTR6
ZShift = 0.74
Z lm Score = 0.67
NG = 0
DB = 0
SM = 0
Delta_K
Doxorubicin
YPL053C KTR6
ZShift = −0.66
Z lm Score = 0.1
NG = 0
DB = 0
SM = 0
Delta_r
Doxorubicin
YPL053C KTR6
ZShift = 1.53
Z lm Score = 0.61
NG = 0
DB = 0
SM = 0
Delta_AUC
Doxorubicin
YLR390W ECM19
ZShift = 0.57
Z lm Score = −2.19
NG = 0
DB = 0
SM = 0
Delta_L
Doxorubicin
YLR390W ECM19
ZShift = 0.36
Z lm Score = −0.05
NG = 0
DB = 0
SM = 0
Delta_K
Doxorubicin
YLR390W ECM19
ZShift = −0.98
Z lm Score = 1.46
NG = 0
DB = 0
SM = 0
Delta_r
Doxorubicin
YLR390W ECM19
ZShift = 0.2
Z lm Score = 0.24
NG = 0
DB = 0
SM = 0
Delta_AUC
Doxorubicin

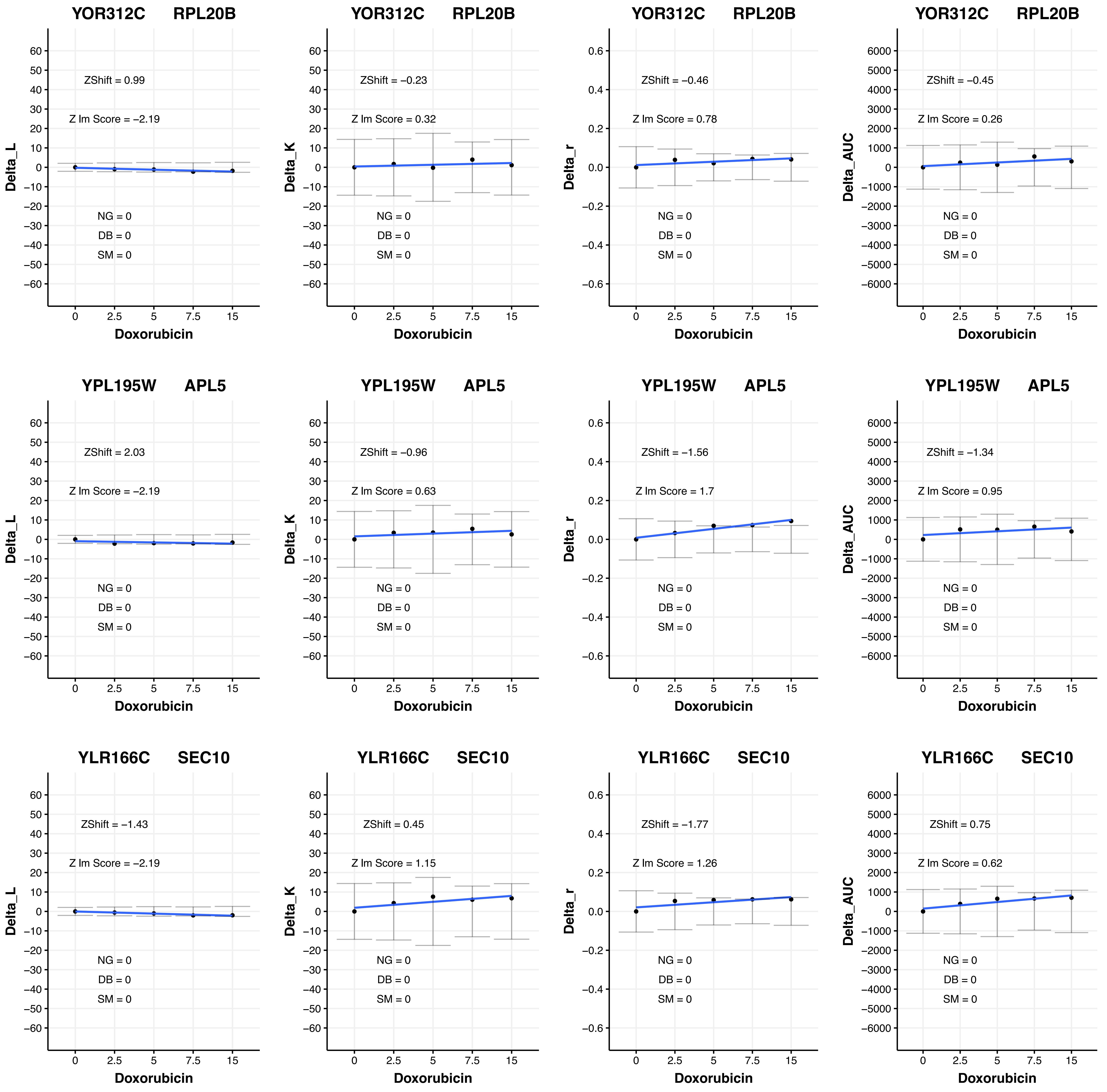

YOR312C RPL20B
ZShift = 0.99
Z lm Score = −2.19
Delta_L
NG = 0
DB = 0
SM = 0
Doxorubicin
YOR312C RPL20B
ZShift = −0.23
Z lm Score = 0.32
Delta_K
NG = 0
DB = 0
SM = 0
Doxorubicin
YOR312C RPL20B
ZShift = −0.46
Z lm Score = 0.78
Delta_r
NG = 0
DB = 0
SM = 0
Doxorubicin
YOR312C RPL20B
ZShift = −0.45
Z lm Score = 0.26
Delta_AUC
NG = 0
DB = 0
SM = 0
Doxorubicin
YPL195W APL5
ZShift = 2.03
Z lm Score = −2.19
Delta_L
NG = 0
DB = 0
SM = 0
Doxorubicin
YPL195W APL5
ZShift = −0.96
Z lm Score = 0.63
Delta_K
NG = 0
DB = 0
SM = 0
Doxorubicin
YPL195W APL5
ZShift = −1.56
Z lm Score = 1.7
Delta_r
NG = 0
DB = 0
SM = 0
Doxorubicin
YPL195W APL5
ZShift = −1.34
Z lm Score = 0.95
Delta_AUC
NG = 0
DB = 0
SM = 0
Doxorubicin
YLR166C SEC10
ZShift = −1.43
Z lm Score = −2.19
Delta_L
NG = 0
DB = 0
SM = 0
Doxorubicin
YLR166C SEC10
ZShift = 0.45
Z lm Score = 1.15
Delta_K
NG = 0
DB = 0
SM = 0
Doxorubicin
YLR166C SEC10
ZShift = −1.77
Z lm Score = 1.26
Delta_r
NG = 0
DB = 0
SM = 0
Doxorubicin
YLR166C SEC10
ZShift = 0.75
Z lm Score = 0.62
Delta_AUC
NG = 0
DB = 0
SM = 0
Doxorubicin

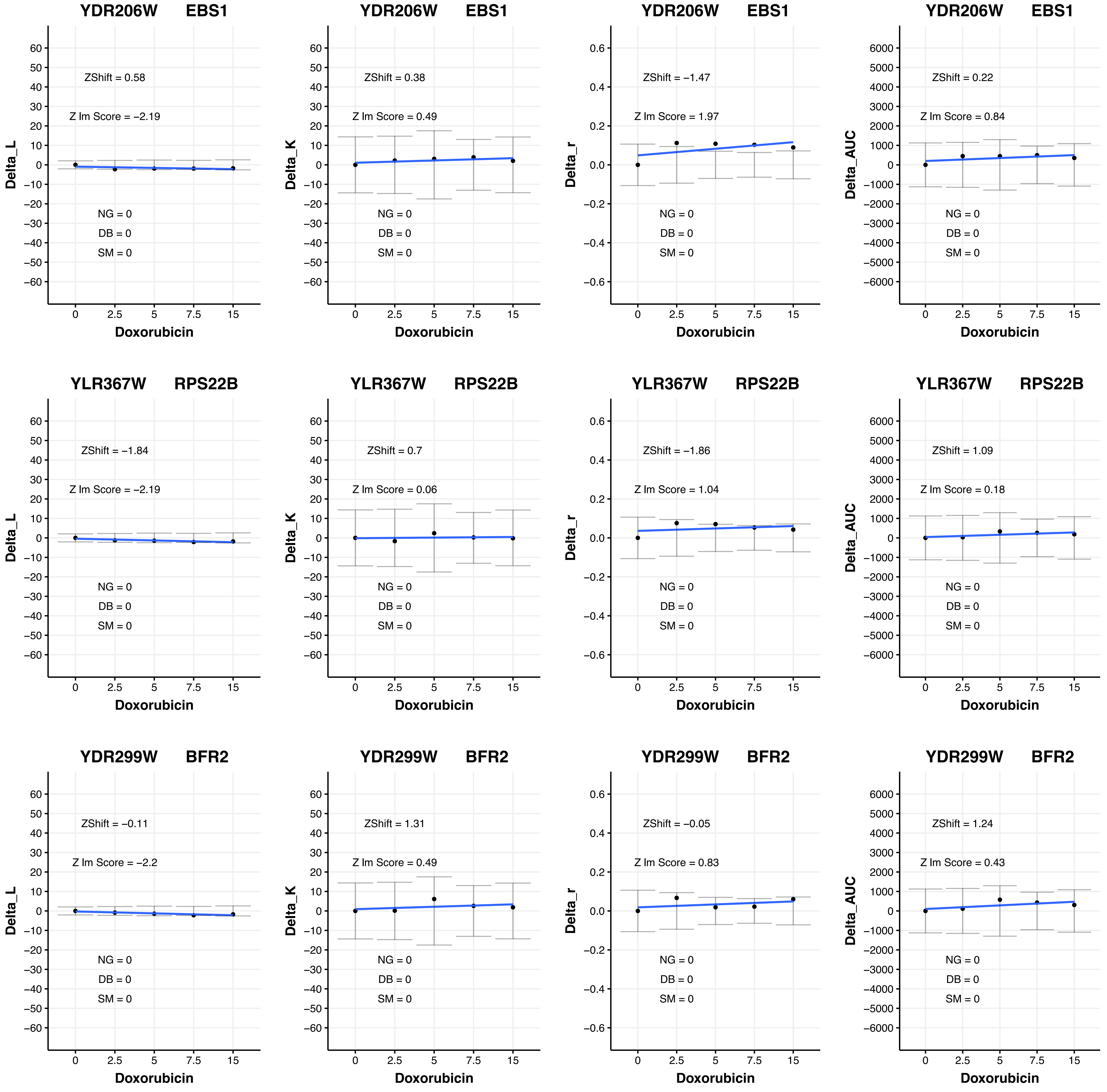
YDR206W EBS1
ZShift = 0.58
Z lm Score = −2.19
NG = 0
DB = 0
SM = 0
Delta_L
Doxorubicin
YDR206W EBS1
ZShift = 0.38
Z lm Score = 0.49
NG = 0
DB = 0
SM = 0
Delta_K
Doxorubicin
YDR206W EBS1
ZShift = −1.47
Z lm Score = 1.97
NG = 0
DB = 0
SM = 0
Delta_r
Doxorubicin
YDR206W EBS1
ZShift = 0.22
Z lm Score = 0.84
NG = 0
DB = 0
SM = 0
Delta_AUC
Doxorubicin
YLR367W RPS22B
ZShift = −1.84
Z lm Score = −2.19
NG = 0
DB = 0
SM = 0
Delta_L
Doxorubicin
YLR367W RPS22B
ZShift = 0.7
Z lm Score = 0.06
NG = 0
DB = 0
SM = 0
Delta_K
Doxorubicin
YLR367W RPS22B
ZShift = −1.86
Z lm Score = 1.04
NG = 0
DB = 0
SM = 0
Delta_r
Doxorubicin
YLR367W RPS22B
ZShift = 1.09
Z lm Score = 0.18
NG = 0
DB = 0
SM = 0
Delta_AUC
Doxorubicin
YDR299W BFR2
ZShift = −0.11
Z lm Score = −2.2
NG = 0
DB = 0
SM = 0
Delta_L
Doxorubicin
YDR299W BFR2
ZShift = 1.31
Z lm Score = 0.49
NG = 0
DB = 0
SM = 0
Delta_K
Doxorubicin
YDR299W BFR2
ZShift = −0.05
Z lm Score = 0.83
NG = 0
DB = 0
SM = 0
Delta_r
Doxorubicin
YDR299W BFR2
ZShift = 1.24
Z lm Score = 0.43
NG = 0
DB = 0
SM = 0
Delta_AUC
Doxorubicin

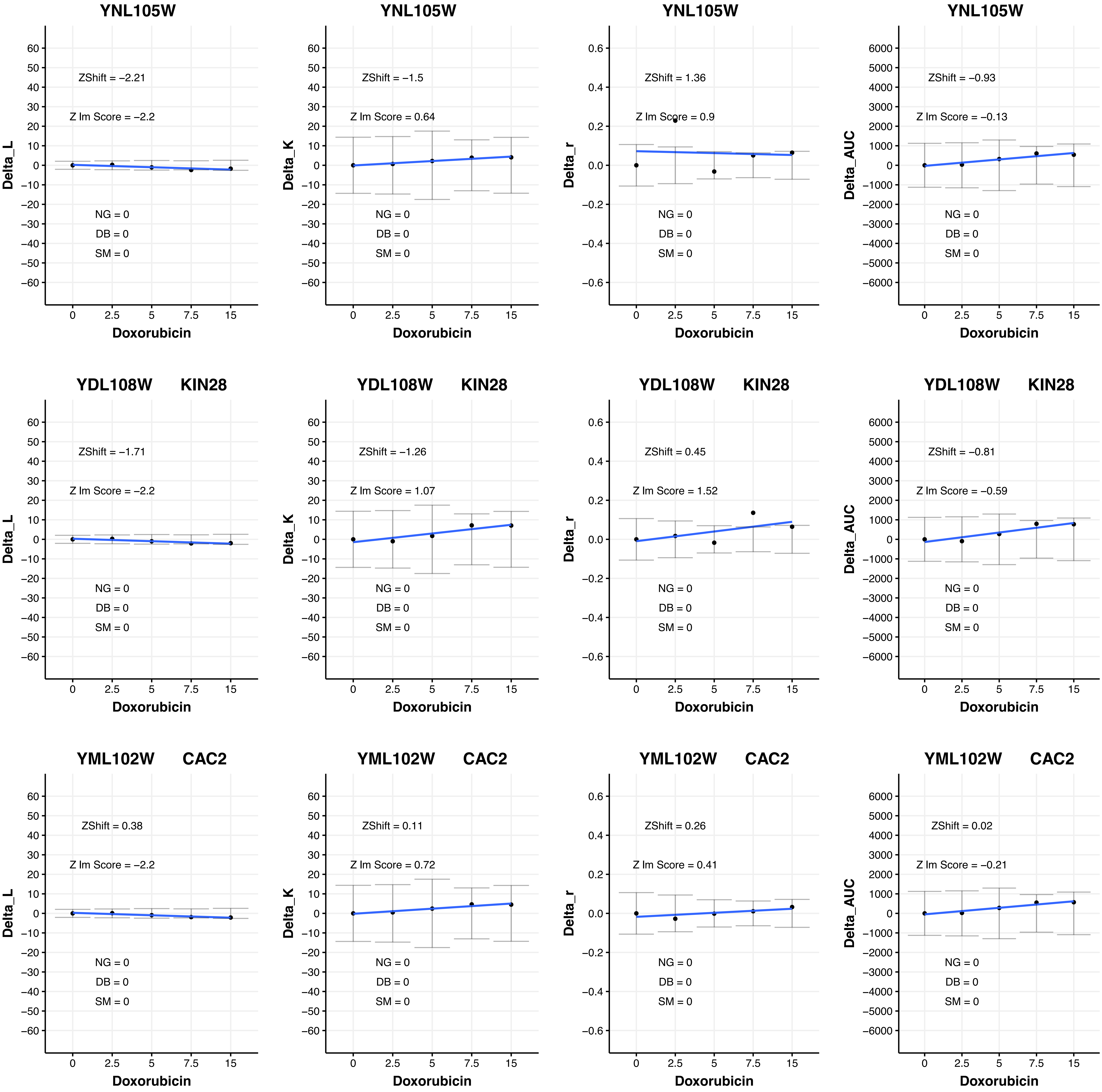
YNL105W
ZShift = −2.21
Z lm Score = −2.2
NG = 0
DB = 0
SM = 0
Delta_L
Doxorubicin
YNL105W
ZShift = −1.5
Z lm Score = 0.64
NG = 0
DB = 0
SM = 0
Delta_K
Doxorubicin
YNL105W
ZShift = 1.36
Z lm Score = 0.9
NG = 0
DB = 0
SM = 0
Delta_r
Doxorubicin
YNL105W
ZShift = −0.93
Z lm Score = −0.13
NG = 0
DB = 0
SM = 0
Delta_AUC
Doxorubicin
YDL108W KIN28
ZShift = −1.71
Z lm Score = −2.2
NG = 0
DB = 0
SM = 0
Delta_L
Doxorubicin
YDL108W KIN28
ZShift = −1.26
Z lm Score = 1.07
NG = 0
DB = 0
SM = 0
Delta_K
Doxorubicin
YDL108W KIN28
ZShift = 0.45
Z lm Score = 1.52
NG = 0
DB = 0
SM = 0
Delta_r
Doxorubicin
YDL108W KIN28
ZShift = −0.81
Z lm Score = −0.59
NG = 0
DB = 0
SM = 0
Delta_AUC
Doxorubicin
YML102W CAC2
ZShift = 0.38
Z lm Score = −2.2
NG = 0
DB = 0
SM = 0
Delta_L
Doxorubicin
YML102W CAC2
ZShift = 0.11
Z lm Score = 0.72
NG = 0
DB = 0
SM = 0
Delta_K
Doxorubicin
YML102W CAC2
ZShift = 0.26
Z lm Score = 0.41
NG = 0
DB = 0
SM = 0
Delta_r
Doxorubicin
YML102W CAC2
ZShift = 0.02
Z lm Score = −0.21
NG = 0
DB = 0
SM = 0
Delta_AUC
Doxorubicin

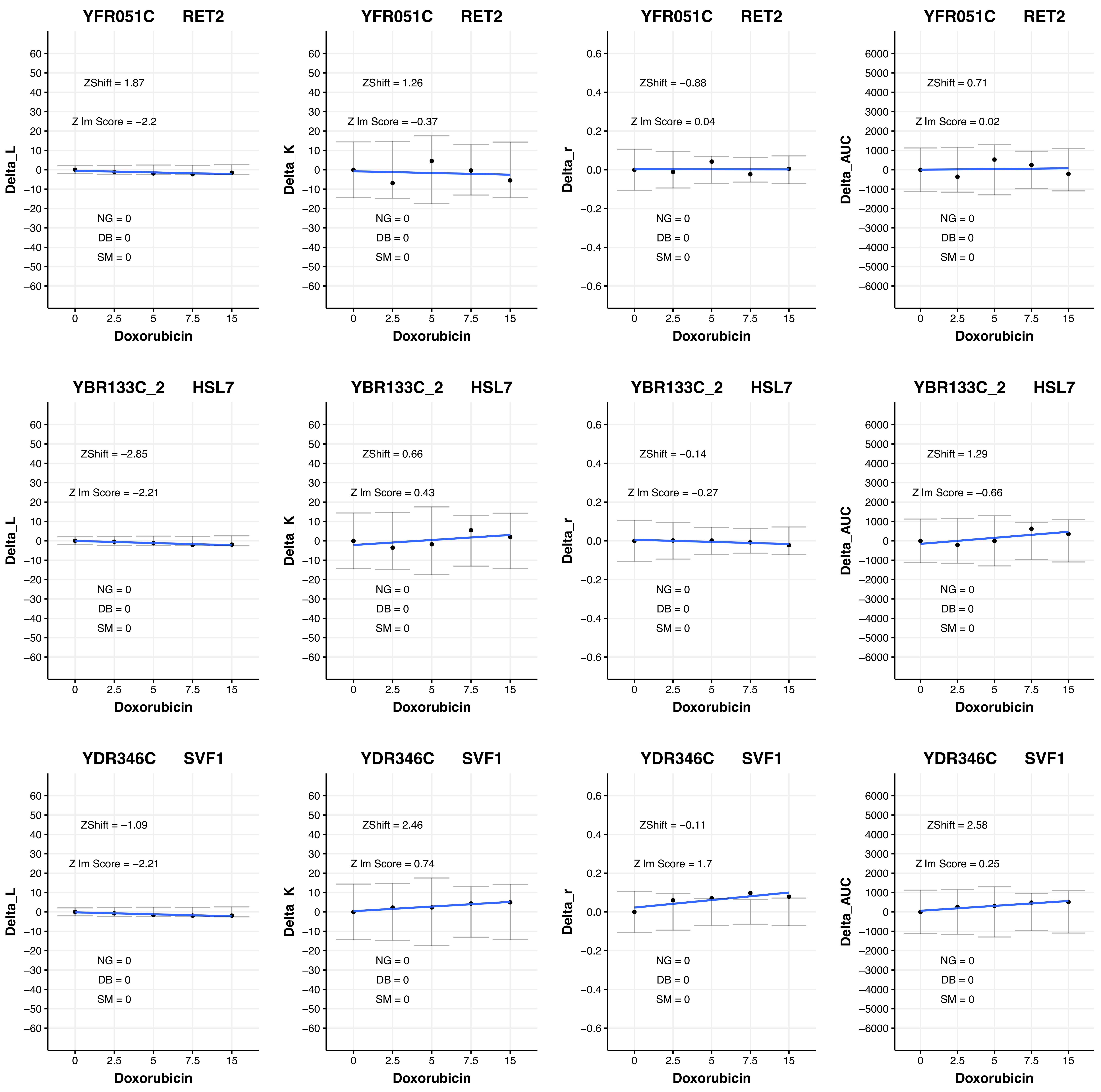
YFR051C RET2
ZShift = 1.87
Z lm Score = −2.2
NG = 0
DB = 0
SM = 0
Delta_L
Doxorubicin
YFR051C RET2
ZShift = 1.26
Z lm Score = −0.37
NG = 0
DB = 0
SM = 0
Delta_K
Doxorubicin
YFR051C RET2
ZShift = −0.88
Z lm Score = 0.04
NG = 0
DB = 0
SM = 0
Delta_r
Doxorubicin
YFR051C RET2
ZShift = 0.71
Z lm Score = 0.02
NG = 0
DB = 0
SM = 0
Delta_AUC
Doxorubicin
YBR133C_2 HSL7
ZShift = −2.85
Z lm Score = −2.21
NG = 0
DB = 0
SM = 0
Delta_L
Doxorubicin
YBR133C_2 HSL7
ZShift = 0.66
Z lm Score = 0.43
NG = 0
DB = 0
SM = 0
Delta_K
Doxorubicin
YBR133C_2 HSL7
ZShift = −0.14
Z lm Score = −0.27
NG = 0
DB = 0
SM = 0
Delta_r
Doxorubicin
YBR133C_2 HSL7
ZShift = 1.29
Z lm Score = −0.66
NG = 0
DB = 0
SM = 0
Delta_AUC
Doxorubicin
YDR346C SVF1
ZShift = −1.09
Z lm Score = −2.21
NG = 0
DB = 0
SM = 0
Delta_L
Doxorubicin
YDR346C SVF1
ZShift = 2.46
Z lm Score = 0.74
NG = 0
DB = 0
SM = 0
Delta_K
Doxorubicin
YDR346C SVF1
ZShift = −0.11
Z lm Score = 1.7
NG = 0
DB = 0
SM = 0
Delta_r
Doxorubicin
YDR346C SVF1
ZShift = 2.58
Z lm Score = 0.25
NG = 0
DB = 0
SM = 0
Delta_AUC
Doxorubicin

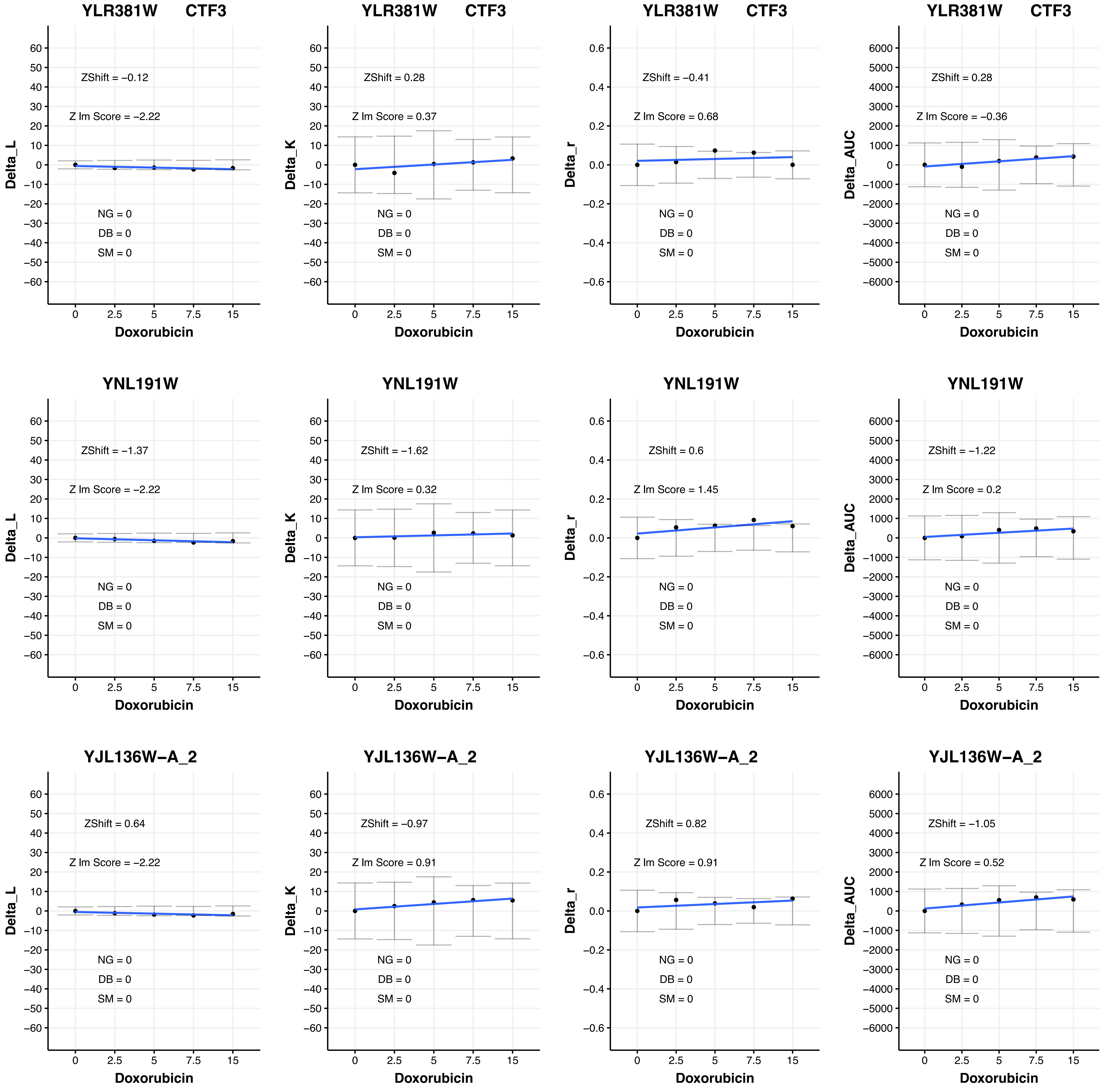
YLR381W CTF3
ZShift = −0.12
Z lm Score = −2.22
NG = 0
DB = 0
SM = 0
Delta_L
Doxorubicin
YLR381W CTF3
ZShift = 0.28
Z lm Score = 0.37
NG = 0
DB = 0
SM = 0
Delta_K
Doxorubicin
YLR381W CTF3
ZShift = −0.41
Z lm Score = 0.68
NG = 0
DB = 0
SM = 0
Delta_r
Doxorubicin
YLR381W CTF3
ZShift = 0.28
Z lm Score = −0.36
NG = 0
DB = 0
SM = 0
Delta_AUC
Doxorubicin
YNL191W
ZShift = −1.37
Z lm Score = −2.22
NG = 0
DB = 0
SM = 0
Delta_L
Doxorubicin
YNL191W
ZShift = −1.62
Z lm Score = 0.32
NG = 0
DB = 0
SM = 0
Delta_K
Doxorubicin
YNL191W
ZShift = 0.6
Z lm Score = 1.45
NG = 0
DB = 0
SM = 0
Delta_r
Doxorubicin
YNL191W
ZShift = −1.22
Z lm Score = 0.2
NG = 0
DB = 0
SM = 0
Delta_AUC
Doxorubicin
YJL136W−A_2
ZShift = 0.64
Z lm Score = −2.22
NG = 0
DB = 0
SM = 0
Delta_L
Doxorubicin
YJL136W−A_2
ZShift = −0.97
Z lm Score = 0.91
NG = 0
DB = 0
SM = 0
Delta_K
Doxorubicin
YJL136W−A_2
ZShift = 0.82
Z lm Score = 0.91
NG = 0
DB = 0
SM = 0
Delta_r
Doxorubicin
YJL136W−A_2
ZShift = −1.05
Z lm Score = 0.52
NG = 0
DB = 0
SM = 0
Delta_AUC
Doxorubicin

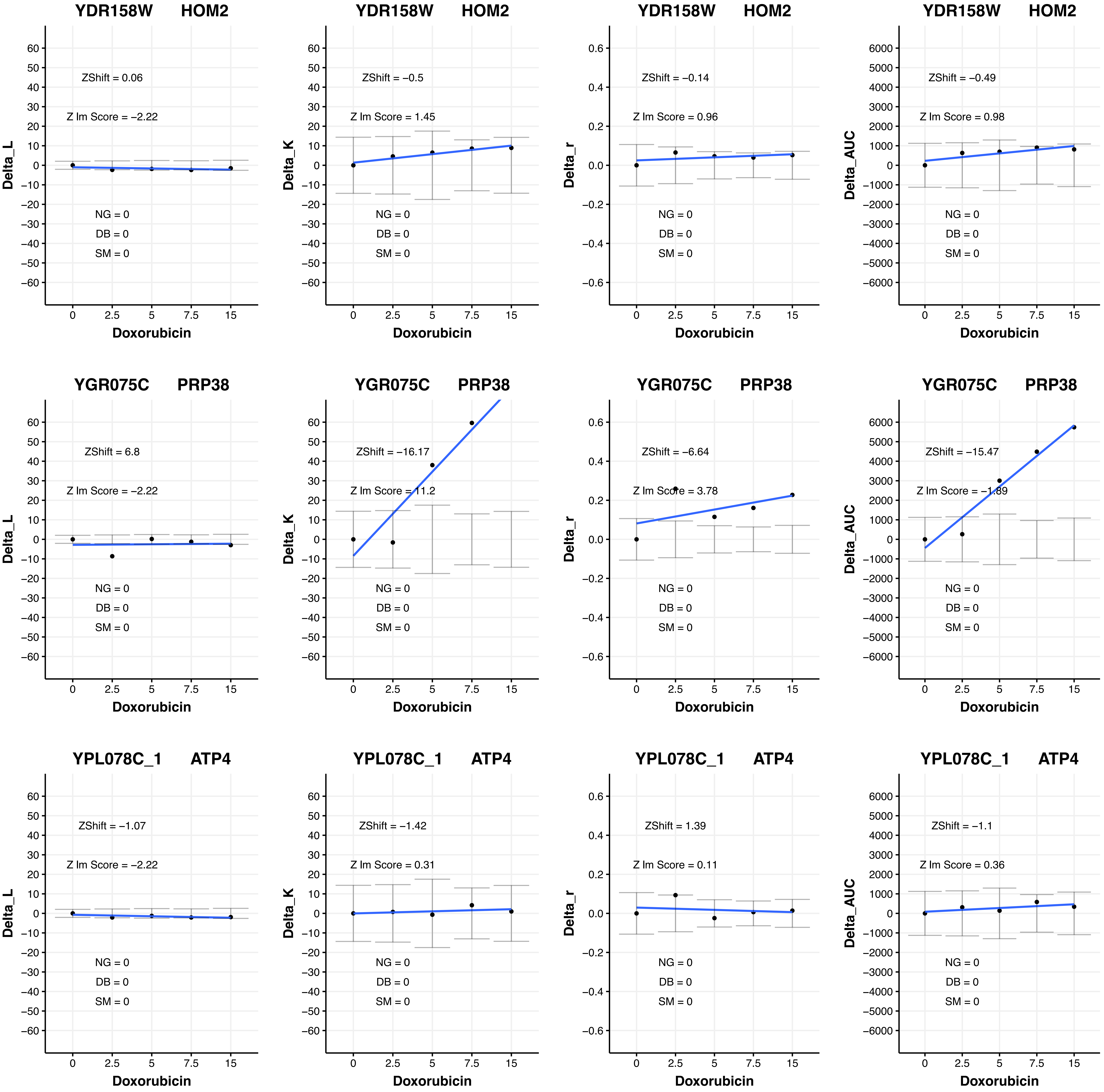
YDR158W HOM2
ZShift = 0.06
Z lm Score = −2.22
NG = 0
DB = 0
SM = 0
Delta_L
Doxorubicin
YDR158W HOM2
ZShift = −0.5
Z lm Score = 1.45
NG = 0
DB = 0
SM = 0
Delta_K
Doxorubicin
YDR158W HOM2
ZShift = −0.14
Z lm Score = 0.96
NG = 0
DB = 0
SM = 0
Delta_r
Doxorubicin
YDR158W HOM2
ZShift = −0.49
Z lm Score = 0.98
NG = 0
DB = 0
SM = 0
Delta_AUC
Doxorubicin
YGR075C PRP38
ZShift = 6.8
Z lm Score = −2.22
NG = 0
DB = 0
SM = 0
Delta_L
Doxorubicin
YGR075C PRP38
ZShift = −16.17
Z lm Score = 11.2
NG = 0
DB = 0
SM = 0
Delta_K
Doxorubicin
YGR075C PRP38
ZShift = −6.64
Z lm Score = 3.78
NG = 0
DB = 0
SM = 0
Delta_r
Doxorubicin
YGR075C PRP38
ZShift = −15.47
Z lm Score = −1.89
NG = 0
DB = 0
SM = 0
Delta_AUC
Doxorubicin
YPL078C_1 ATP4
ZShift = −1.07
Z lm Score = −2.22
NG = 0
DB = 0
SM = 0
Delta_L
Doxorubicin
YPL078C_1 ATP4
ZShift = −1.42
Z lm Score = 0.31
NG = 0
DB = 0
SM = 0
Delta_K
Doxorubicin
YPL078C_1 ATP4
ZShift = 1.39
Z lm Score = 0.11
NG = 0
DB = 0
SM = 0
Delta_r
Doxorubicin
YPL078C_1 ATP4
ZShift = −1.1
Z lm Score = 0.36
NG = 0
DB = 0
SM = 0
Delta_AUC
Doxorubicin

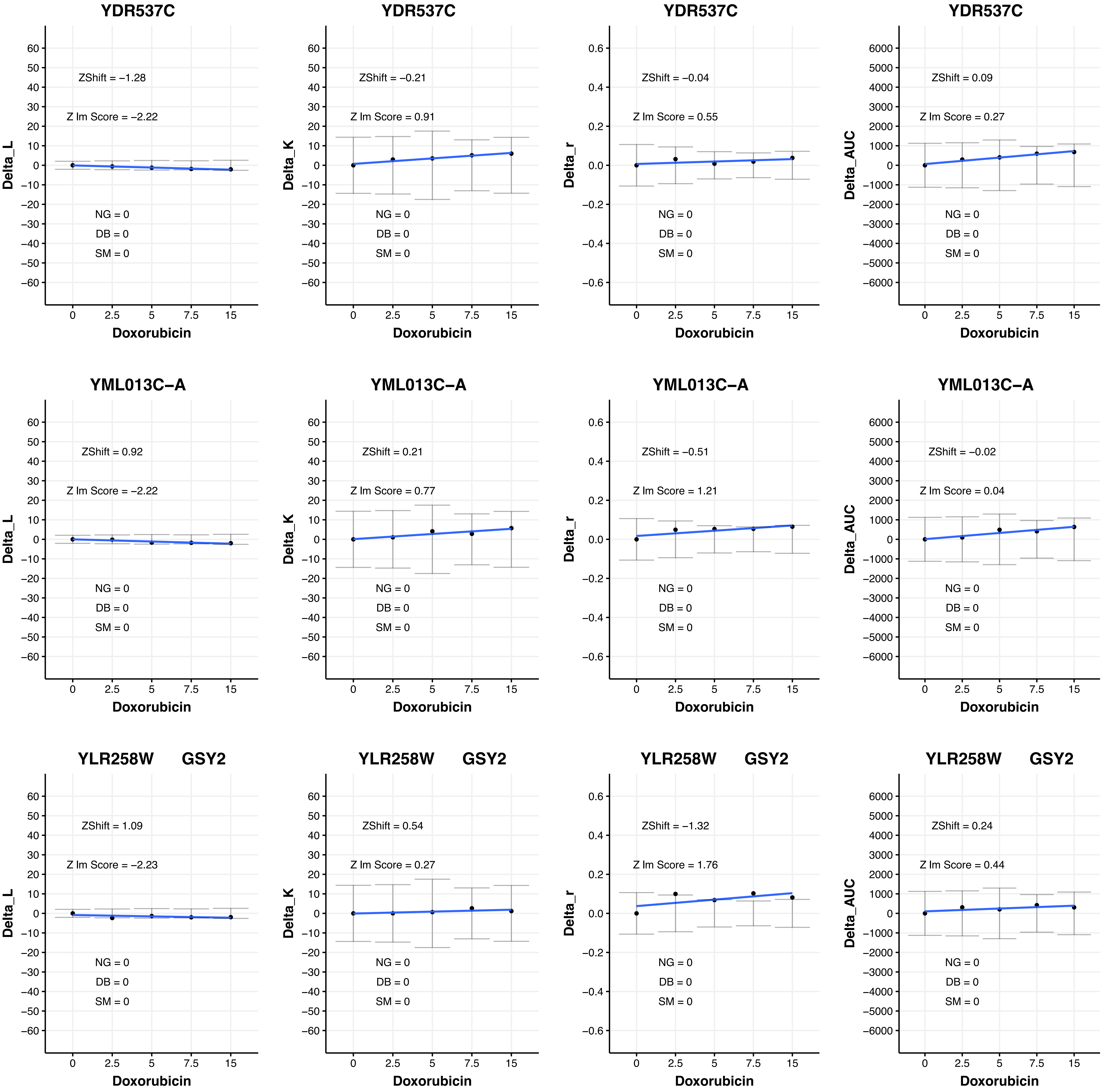

YDR537C
ZShift = −1.28
Z lm Score = −2.22
NG = 0
DB = 0
SM = 0
Delta_L
Doxorubicin
YDR537C
ZShift = −0.21
Z lm Score = 0.91
NG = 0
DB = 0
SM = 0
Delta_K
Doxorubicin
YDR537C
ZShift = −0.04
Z lm Score = 0.55
NG = 0
DB = 0
SM = 0
Delta_r
Doxorubicin
YDR537C
ZShift = 0.09
Z lm Score = 0.27
NG = 0
DB = 0
SM = 0
Delta_AUC
Doxorubicin
YML013C−A
ZShift = 0.92
Z lm Score = −2.22
NG = 0
DB = 0
SM = 0
Delta_L
Doxorubicin
YML013C−A
ZShift = 0.21
Z lm Score = 0.77
NG = 0
DB = 0
SM = 0
Delta_K
Doxorubicin
YML013C−A
ZShift = −0.51
Z lm Score = 1.21
NG = 0
DB = 0
SM = 0
Delta_r
Doxorubicin
YML013C−A
ZShift = −0.02
Z lm Score = 0.04
NG = 0
DB = 0
SM = 0
Delta_AUC
Doxorubicin
YLR258W GSY2
ZShift = 1.09
Z lm Score = −2.23
NG = 0
DB = 0
SM = 0
Delta_L
Doxorubicin
YLR258W GSY2
ZShift = 0.54
Z lm Score = 0.27
NG = 0
DB = 0
SM = 0
Delta_K
Doxorubicin
YLR258W GSY2
ZShift = −1.32
Z lm Score = 1.76
NG = 0
DB = 0
SM = 0
Delta_r
Doxorubicin
YLR258W GSY2
ZShift = 0.24
Z lm Score = 0.44
NG = 0
DB = 0
SM = 0
Delta_AUC
Doxorubicin

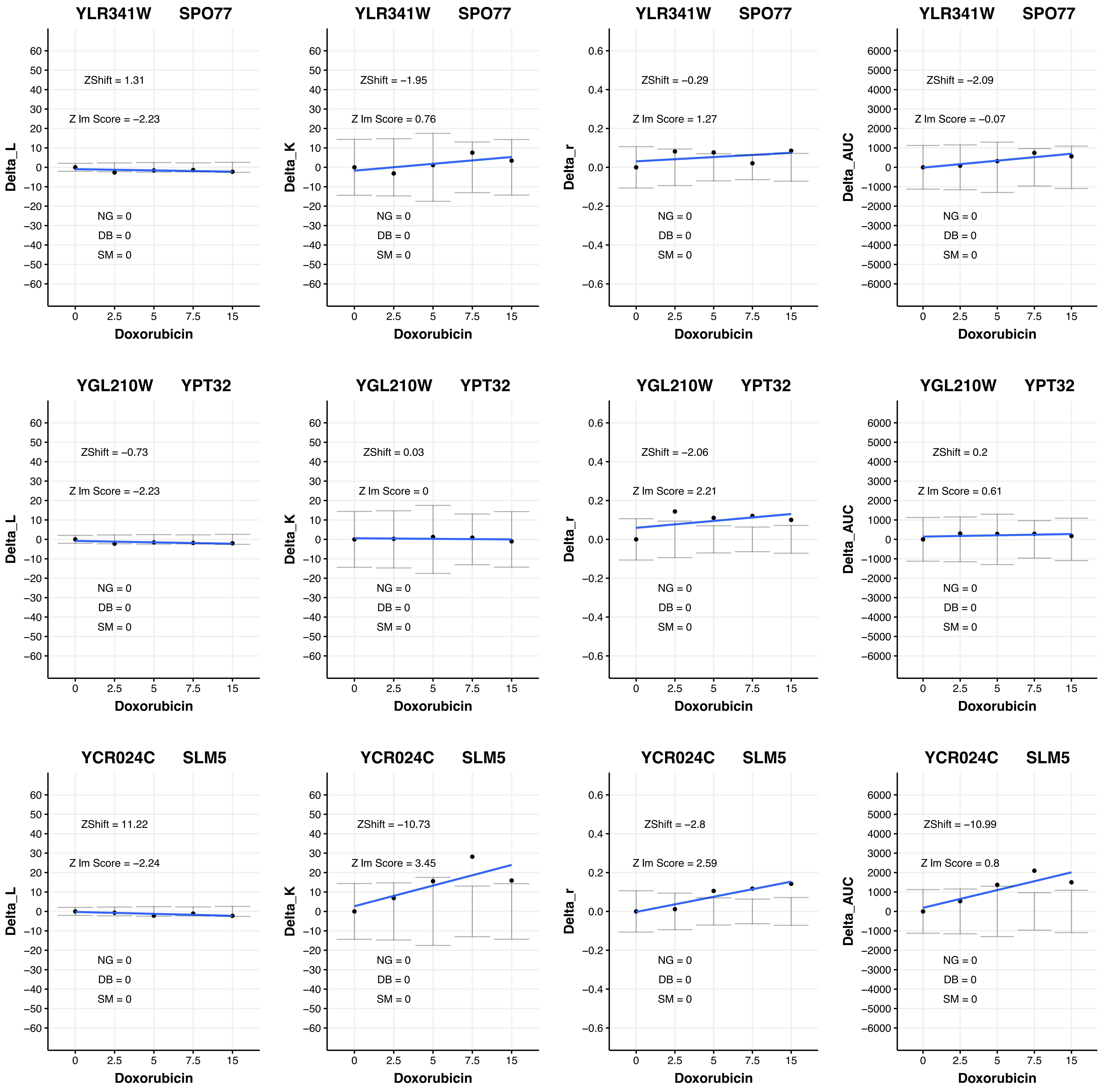

YLR341W SPO77
ZShift = 1.31
Z lm Score = −2.23
NG = 0
DB = 0
SM = 0
Delta_L
Doxorubicin
YLR341W SPO77
ZShift = −1.95
Z lm Score = 0.76
NG = 0
DB = 0
SM = 0
Delta_K
Doxorubicin
YLR341W SPO77
ZShift = −0.29
Z lm Score = 1.27
NG = 0
DB = 0
SM = 0
Delta_r
Doxorubicin
YLR341W SPO77
ZShift = −2.09
Z lm Score = −0.07
NG = 0
DB = 0
SM = 0
Delta_AUC
Doxorubicin
YGL210W YPT32
ZShift = −0.73
Z lm Score = −2.23
NG = 0
DB = 0
SM = 0
Delta_L
Doxorubicin
YGL210W YPT32
ZShift = 0.03
Z lm Score = 0
NG = 0
DB = 0
SM = 0
Delta_K
Doxorubicin
YGL210W YPT32
ZShift = −2.06
Z lm Score = 2.21
NG = 0
DB = 0
SM = 0
Delta_r
Doxorubicin
YGL210W YPT32
ZShift = 0.2
Z lm Score = 0.61
NG = 0
DB = 0
SM = 0
Delta_AUC
Doxorubicin
YCR024C SLM5
ZShift = 11.22
Z lm Score = −2.24
NG = 0
DB = 0
SM = 0
Delta_L
Doxorubicin
YCR024C SLM5
ZShift = −10.73
Z lm Score = 3.45
NG = 0
DB = 0
SM = 0
Delta_K
Doxorubicin
YCR024C SLM5
ZShift = −2.8
Z lm Score = 2.59
NG = 0
DB = 0
SM = 0
Delta_r
Doxorubicin
YCR024C SLM5
ZShift = −10.99
Z lm Score = 0.8
NG = 0
DB = 0
SM = 0
Delta_AUC
Doxorubicin

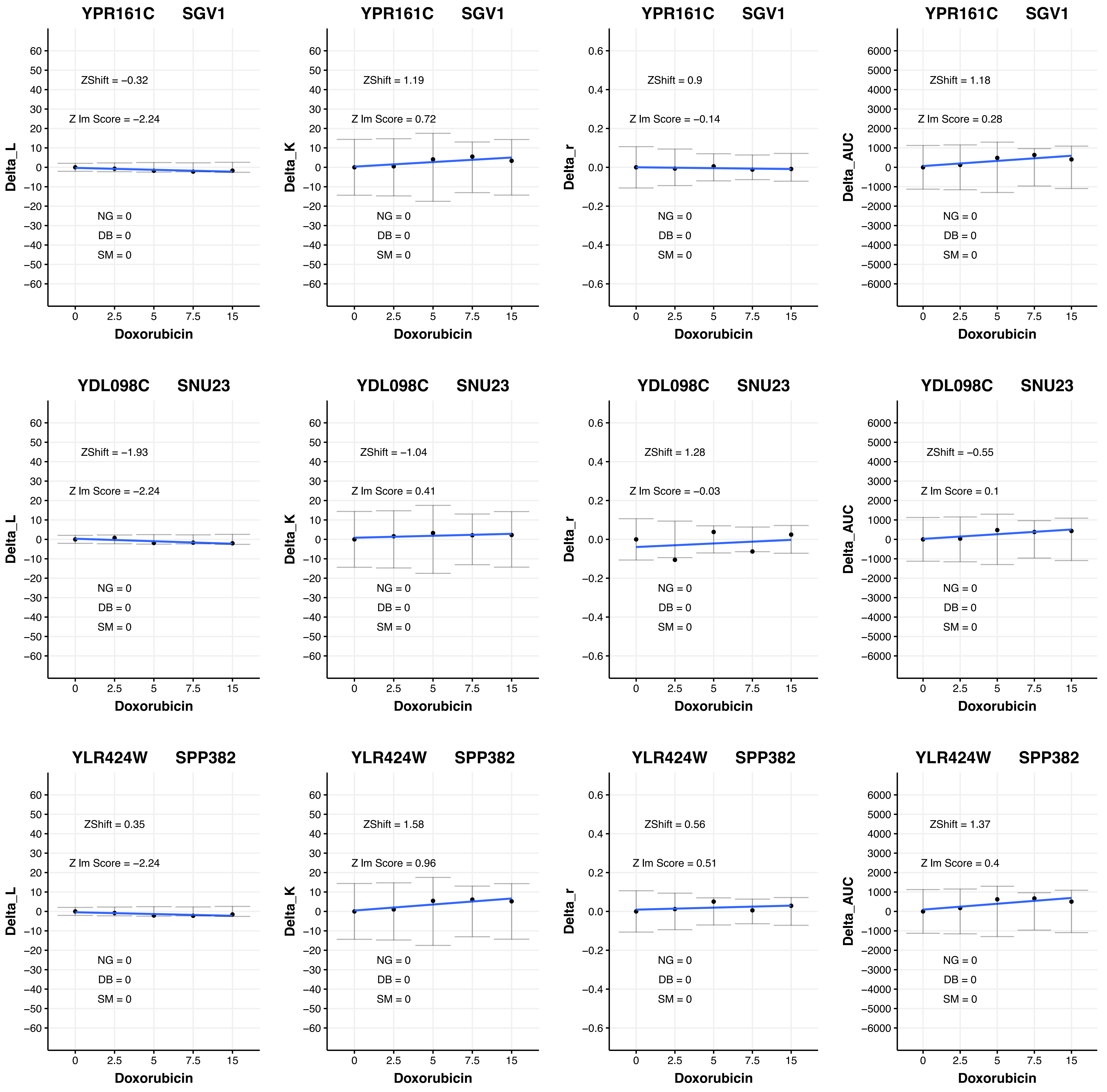
YPR161C SGV1
ZShift = −0.32
Z lm Score = −2.24
NG = 0
DB = 0
SM = 0
Delta_L
Doxorubicin
YPR161C SGV1
ZShift = 1.19
Z lm Score = 0.72
NG = 0
DB = 0
SM = 0
Delta_K
Doxorubicin
YPR161C SGV1
ZShift = 0.9
Z lm Score = −0.14
NG = 0
DB = 0
SM = 0
Delta_r
Doxorubicin
YPR161C SGV1
ZShift = 1.18
Z lm Score = 0.28
NG = 0
DB = 0
SM = 0
Delta_AUC
Doxorubicin
YDL098C SNU23
ZShift = −1.93
Z lm Score = −2.24
NG = 0
DB = 0
SM = 0
Delta_L
Doxorubicin
YDL098C SNU23
ZShift = −1.04
Z lm Score = 0.41
NG = 0
DB = 0
SM = 0
Delta_K
Doxorubicin
YDL098C SNU23
ZShift = 1.28
Z lm Score = −0.03
NG = 0
DB = 0
SM = 0
Delta_r
Doxorubicin
YDL098C SNU23
ZShift = −0.55
Z lm Score = 0.1
NG = 0
DB = 0
SM = 0
Delta_AUC
Doxorubicin
YLR424W SPP382
ZShift = 0.35
Z lm Score = −2.24
NG = 0
DB = 0
SM = 0
Delta_L
Doxorubicin
YLR424W SPP382
ZShift = 1.58
Z lm Score = 0.96
NG = 0
DB = 0
SM = 0
Delta_K
Doxorubicin
YLR424W SPP382
ZShift = 0.56
Z lm Score = 0.51
NG = 0
DB = 0
SM = 0
Delta_r
Doxorubicin
YLR424W SPP382
ZShift = 1.37
Z lm Score = 0.4
NG = 0
DB = 0
SM = 0
Delta_AUC
Doxorubicin

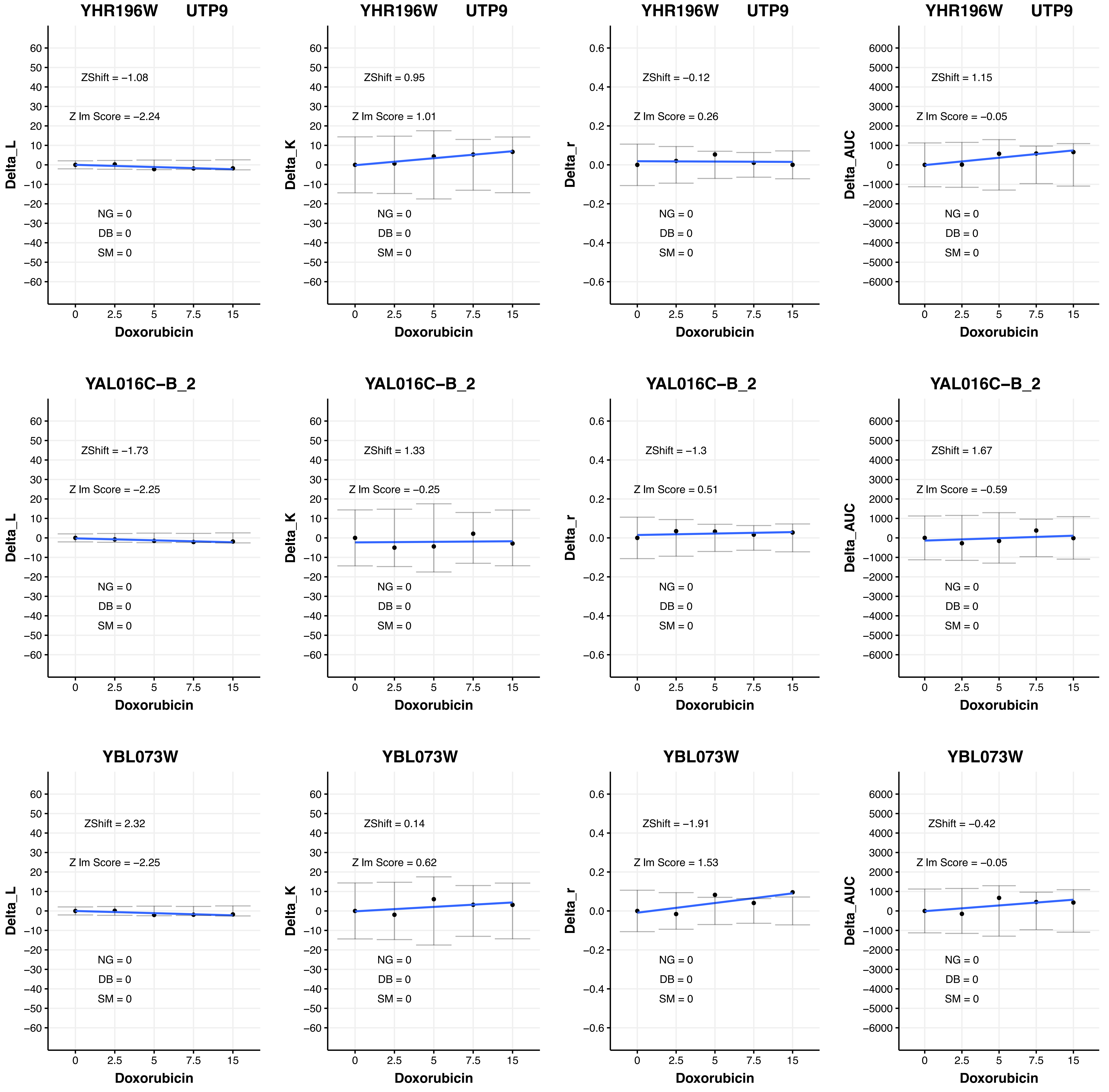
YHR196W UTP9
ZShift = −1.08
Z lm Score = −2.24
NG = 0
DB = 0
SM = 0
Delta_L
Doxorubicin
YHR196W UTP9
ZShift = 0.95
Z lm Score = 1.01
NG = 0
DB = 0
SM = 0
Delta_K
Doxorubicin
YHR196W UTP9
ZShift = −0.12
Z lm Score = 0.26
NG = 0
DB = 0
SM = 0
Delta_r
Doxorubicin
YHR196W UTP9
ZShift = 1.15
Z lm Score = −0.05
NG = 0
DB = 0
SM = 0
Delta_AUC
Doxorubicin
YAL016C−B_2
ZShift = −1.73
Z lm Score = −2.25
NG = 0
DB = 0
SM = 0
Delta_L
Doxorubicin
YAL016C−B_2
ZShift = 1.33
Z lm Score = −0.25
NG = 0
DB = 0
SM = 0
Delta_K
Doxorubicin
YAL016C−B_2
ZShift = −1.3
Z lm Score = 0.51
NG = 0
DB = 0
SM = 0
Delta_r
Doxorubicin
YAL016C−B_2
ZShift = 1.67
Z lm Score = −0.59
NG = 0
DB = 0
SM = 0
Delta_AUC
Doxorubicin
YBL073W
ZShift = 2.32
Z lm Score = −2.25
NG = 0
DB = 0
SM = 0
Delta_L
Doxorubicin
YBL073W
ZShift = 0.14
Z lm Score = 0.62
NG = 0
DB = 0
SM = 0
Delta_K
Doxorubicin
YBL073W
ZShift = −1.91
Z lm Score = 1.53
NG = 0
DB = 0
SM = 0
Delta_r
Doxorubicin
YBL073W
ZShift = −0.42
Z lm Score = −0.05
NG = 0
DB = 0
SM = 0
Delta_AUC
Doxorubicin

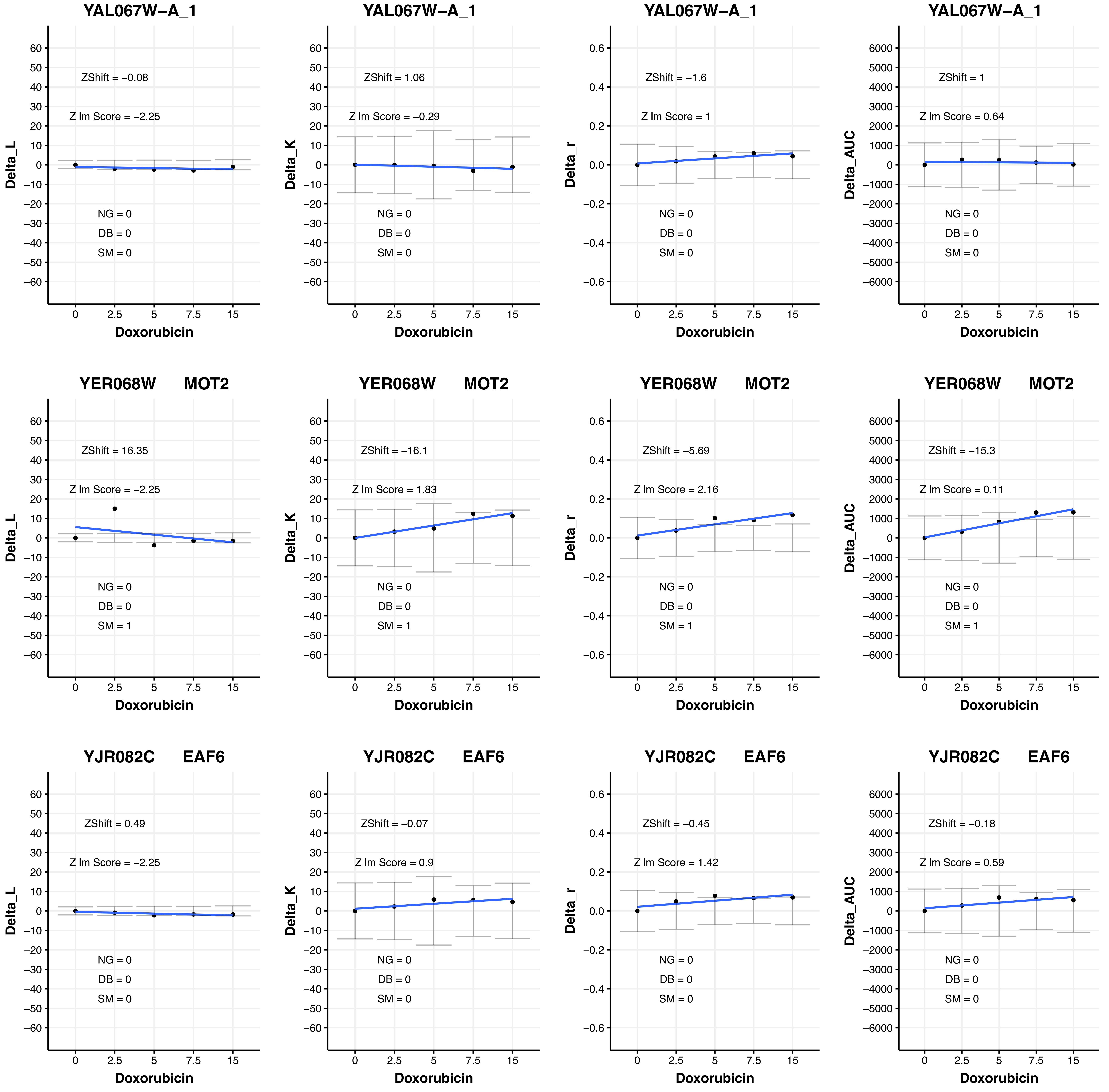

YAL067W−A_1
ZShift = −0.08
Z lm Score = −2.25
NG = 0
DB = 0
SM = 0
Delta_L
Doxorubicin
YAL067W−A_1
ZShift = 1.06
Z lm Score = −0.29
NG = 0
DB = 0
SM = 0
Delta_K
Doxorubicin
YAL067W−A_1
ZShift = −1.6
Z lm Score = 1
NG = 0
DB = 0
SM = 0
Delta_r
Doxorubicin
YAL067W−A_1
ZShift = 1
Z lm Score = 0.64
NG = 0
DB = 0
SM = 0
Delta_AUC
Doxorubicin
YER068W MOT2
ZShift = 16.35
Z lm Score = −2.25
NG = 0
DB = 0
SM = 1
Delta_L
Doxorubicin
YER068W MOT2
ZShift = −16.1
Z lm Score = 1.83
NG = 0
DB = 0
SM = 1
Delta_K
Doxorubicin
YER068W MOT2
ZShift = −5.69
Z lm Score = 2.16
NG = 0
DB = 0
SM = 1
Delta_r
Doxorubicin
YER068W MOT2
ZShift = −15.3
Z lm Score = 0.11
NG = 0
DB = 0
SM = 1
Delta_AUC
Doxorubicin
YJR082C EAF6
ZShift = 0.49
Z lm Score = −2.25
NG = 0
DB = 0
SM = 0
Delta_L
Doxorubicin
YJR082C EAF6
ZShift = −0.07
Z lm Score = 0.9
NG = 0
DB = 0
SM = 0
Delta_K
Doxorubicin
YJR082C EAF6
ZShift = −0.45
Z lm Score = 1.42
NG = 0
DB = 0
SM = 0
Delta_r
Doxorubicin
YJR082C EAF6
ZShift = −0.18
Z lm Score = 0.59
NG = 0
DB = 0
SM = 0
Delta_AUC
Doxorubicin

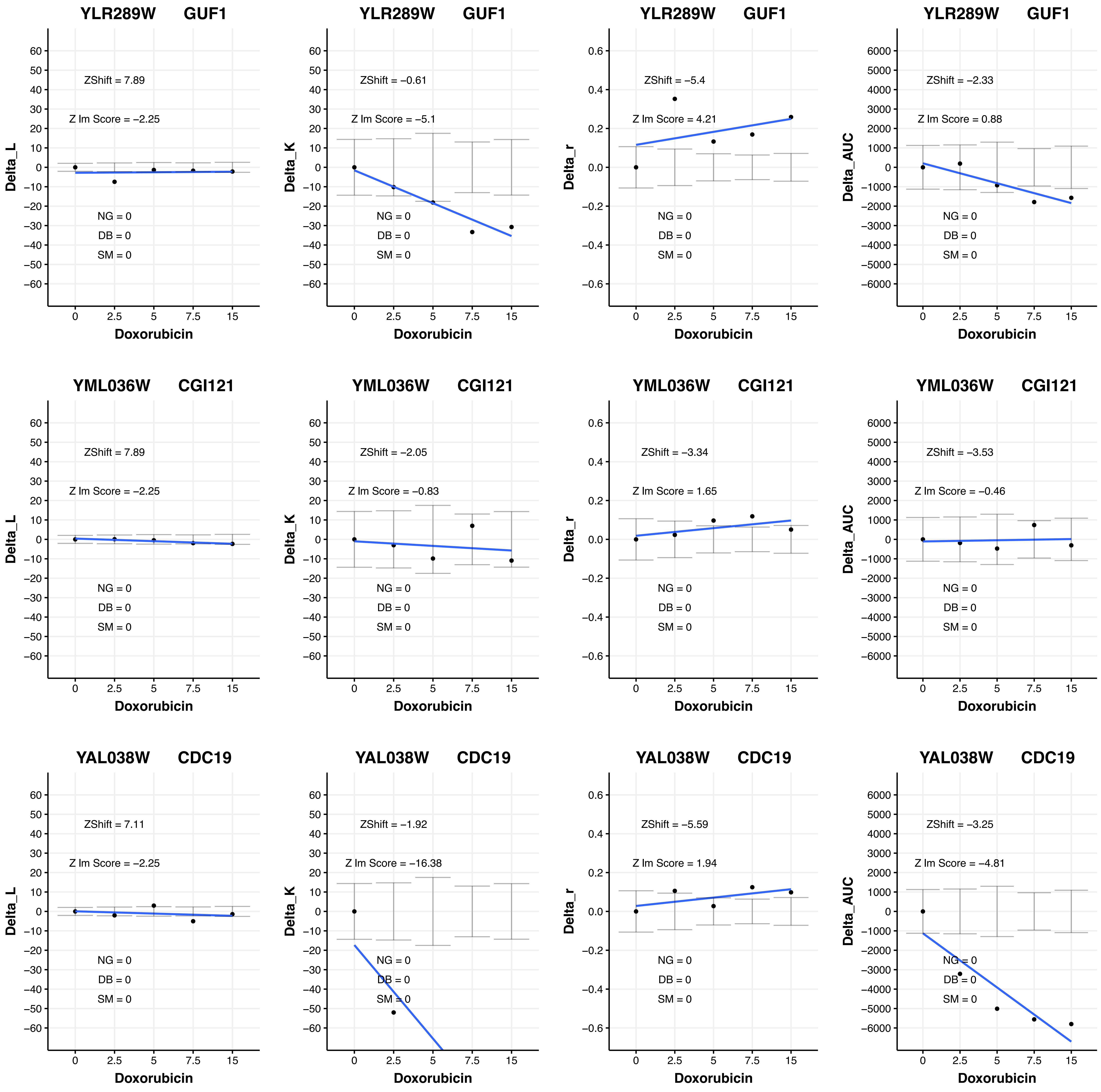
YLR289W GUF1
ZShift = 7.89
Z lm Score = −2.25
NG = 0
DB = 0
SM = 0
Delta_L
Doxorubicin
YLR289W GUF1
ZShift = −0.61
Z lm Score = −5.1
NG = 0
DB = 0
SM = 0
Delta_K
Doxorubicin
YLR289W GUF1
ZShift = −5.4
Z lm Score = 4.21
NG = 0
DB = 0
SM = 0
Delta_r
Doxorubicin
YLR289W GUF1
ZShift = −2.33
Z lm Score = 0.88
NG = 0
DB = 0
SM = 0
Delta_AUC
Doxorubicin
YML036W CGI121
ZShift = 7.89
Z lm Score = −2.25
NG = 0
DB = 0
SM = 0
Delta_L
Doxorubicin
YML036W CGI121
ZShift = −2.05
Z lm Score = −0.83
NG = 0
DB = 0
SM = 0
Delta_K
Doxorubicin
YML036W CGI121
ZShift = −3.34
Z lm Score = 1.65
NG = 0
DB = 0
SM = 0
Delta_r
Doxorubicin
YML036W CGI121
ZShift = −3.53
Z lm Score = −0.46
NG = 0
DB = 0
SM = 0
Delta_AUC
Doxorubicin
YAL038W CDC19
ZShift = 7.11
Z lm Score = −2.25
NG = 0
DB = 0
SM = 0
Delta_L
Doxorubicin
YAL038W CDC19
ZShift = −1.92
Z lm Score = −16.38
NG = 0
DB = 0
SM = 0
Delta_K
Doxorubicin
YAL038W CDC19
ZShift = −5.59
Z lm Score = 1.94
NG = 0
DB = 0
SM = 0
Delta_r
Doxorubicin
YAL038W CDC19
ZShift = −3.25
Z lm Score = −4.81
NG = 0
DB = 0
SM = 0
Delta_AUC
Doxorubicin

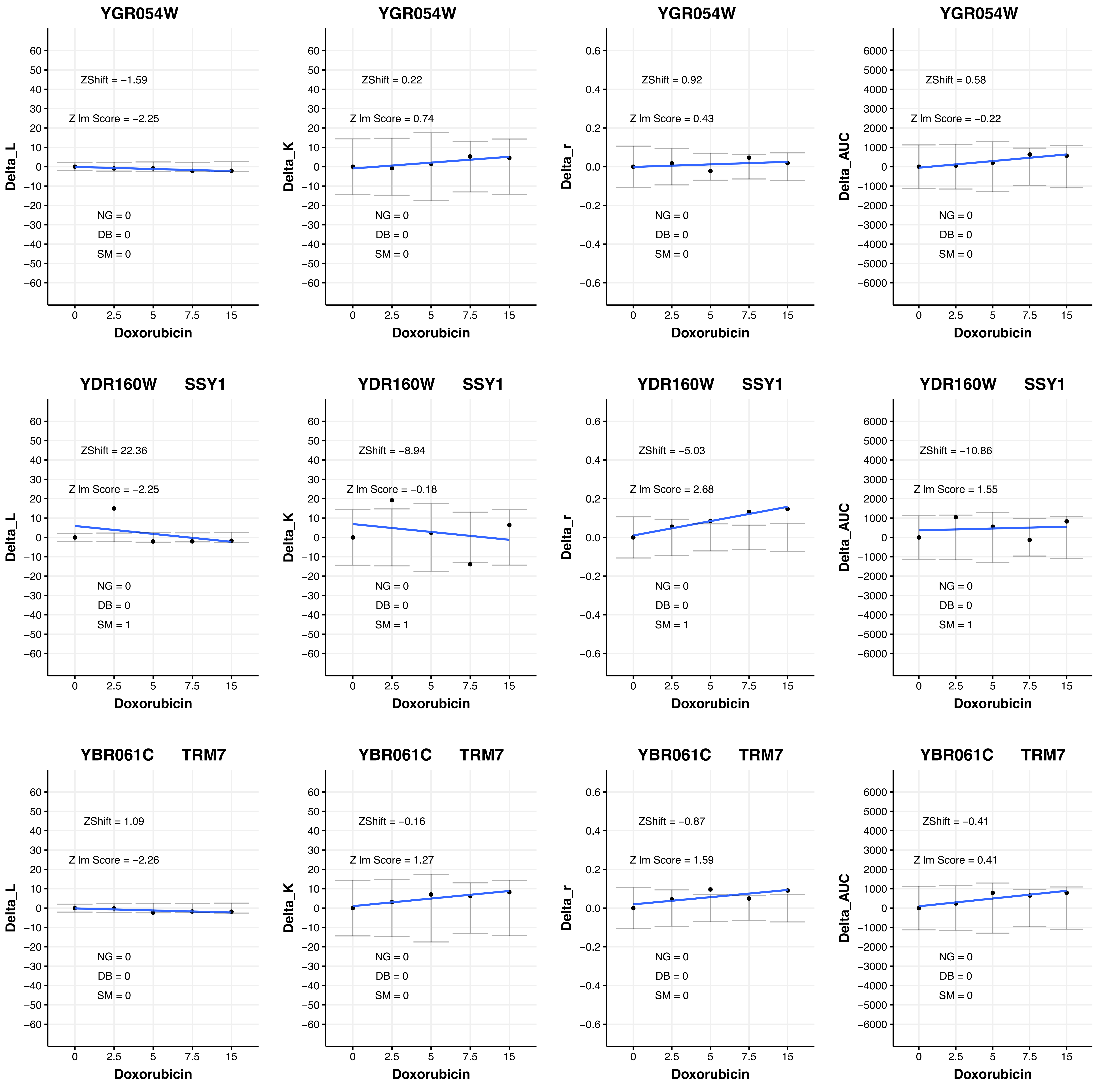
YGR054W
ZShift = −1.59
Z lm Score = −2.25
NG = 0
DB = 0
SM = 0
Delta_L
Doxorubicin
YGR054W
ZShift = 0.22
Z lm Score = 0.74
NG = 0
DB = 0
SM = 0
Delta_K
Doxorubicin
YGR054W
ZShift = 0.92
Z lm Score = 0.43
NG = 0
DB = 0
SM = 0
Delta_r
Doxorubicin
YGR054W
ZShift = 0.58
Z lm Score = −0.22
NG = 0
DB = 0
SM = 0
Delta_AUC
Doxorubicin
YDR160W SSY1
ZShift = 22.36
Z lm Score = −2.25
NG = 0
DB = 0
SM = 1
Delta_L
Doxorubicin
YDR160W SSY1
ZShift = −8.94
Z lm Score = −0.18
NG = 0
DB = 0
SM = 1
Delta_K
Doxorubicin
YDR160W SSY1
ZShift = −5.03
Z lm Score = 2.68
NG = 0
DB = 0
SM = 1
Delta_r
Doxorubicin
YDR160W SSY1
ZShift = −10.86
Z lm Score = 1.55
NG = 0
DB = 0
SM = 1
Delta_AUC
Doxorubicin
YBR061C TRM7
ZShift = 1.09
Z lm Score = −2.26
NG = 0
DB = 0
SM = 0
Delta_L
Doxorubicin
YBR061C TRM7
ZShift = −0.16
Z lm Score = 1.27
NG = 0
DB = 0
SM = 0
Delta_K
Doxorubicin
YBR061C TRM7
ZShift = −0.87
Z lm Score = 1.59
NG = 0
DB = 0
SM = 0
Delta_r
Doxorubicin
YBR061C TRM7
ZShift = −0.41
Z lm Score = 0.41
NG = 0
DB = 0
SM = 0
Delta_AUC
Doxorubicin

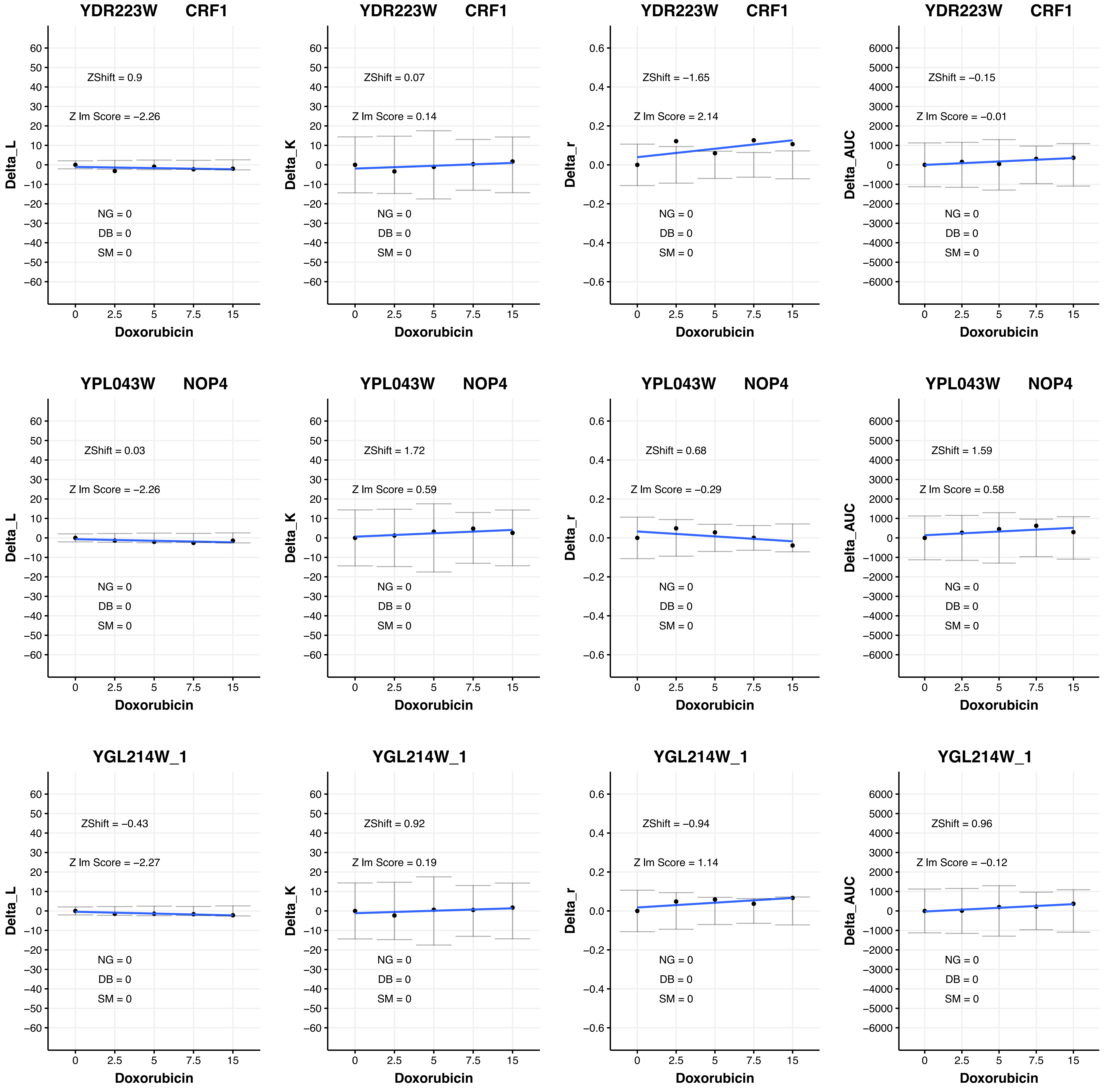
YDR223W CRF1
ZShift = 0.9
Z lm Score = −2.26
NG = 0
DB = 0
SM = 0
Delta_L
Doxorubicin
YDR223W CRF1
ZShift = 0.07
Z lm Score = 0.14
NG = 0
DB = 0
SM = 0
Delta_K
Doxorubicin
YDR223W CRF1
ZShift = −1.65
Z lm Score = 2.14
NG = 0
DB = 0
SM = 0
Delta_r
Doxorubicin
YDR223W CRF1
ZShift = −0.15
Z lm Score = −0.01
NG = 0
DB = 0
SM = 0
Delta_AUC
Doxorubicin
YPL043W NOP4
ZShift = 0.03
Z lm Score = −2.26
NG = 0
DB = 0
SM = 0
Delta_L
Doxorubicin
YPL043W NOP4
ZShift = 1.72
Z lm Score = 0.59
NG = 0
DB = 0
SM = 0
Delta_K
Doxorubicin
YPL043W NOP4
ZShift = 0.68
Z lm Score = −0.29
NG = 0
DB = 0
SM = 0
Delta_r
Doxorubicin
YPL043W NOP4
ZShift = 1.59
Z lm Score = 0.58
NG = 0
DB = 0
SM = 0
Delta_AUC
Doxorubicin
YGL214W_1
ZShift = −0.43
Z lm Score = −2.27
NG = 0
DB = 0
SM = 0
Delta_L
Doxorubicin
YGL214W_1
ZShift = 0.92
Z lm Score = 0.19
NG = 0
DB = 0
SM = 0
Delta_K
Doxorubicin
YGL214W_1
ZShift = −0.94
Z lm Score = 1.14
NG = 0
DB = 0
SM = 0
Delta_r
Doxorubicin
YGL214W_1
ZShift = 0.96
Z lm Score = −0.12
NG = 0
DB = 0
SM = 0
Delta_AUC
Doxorubicin

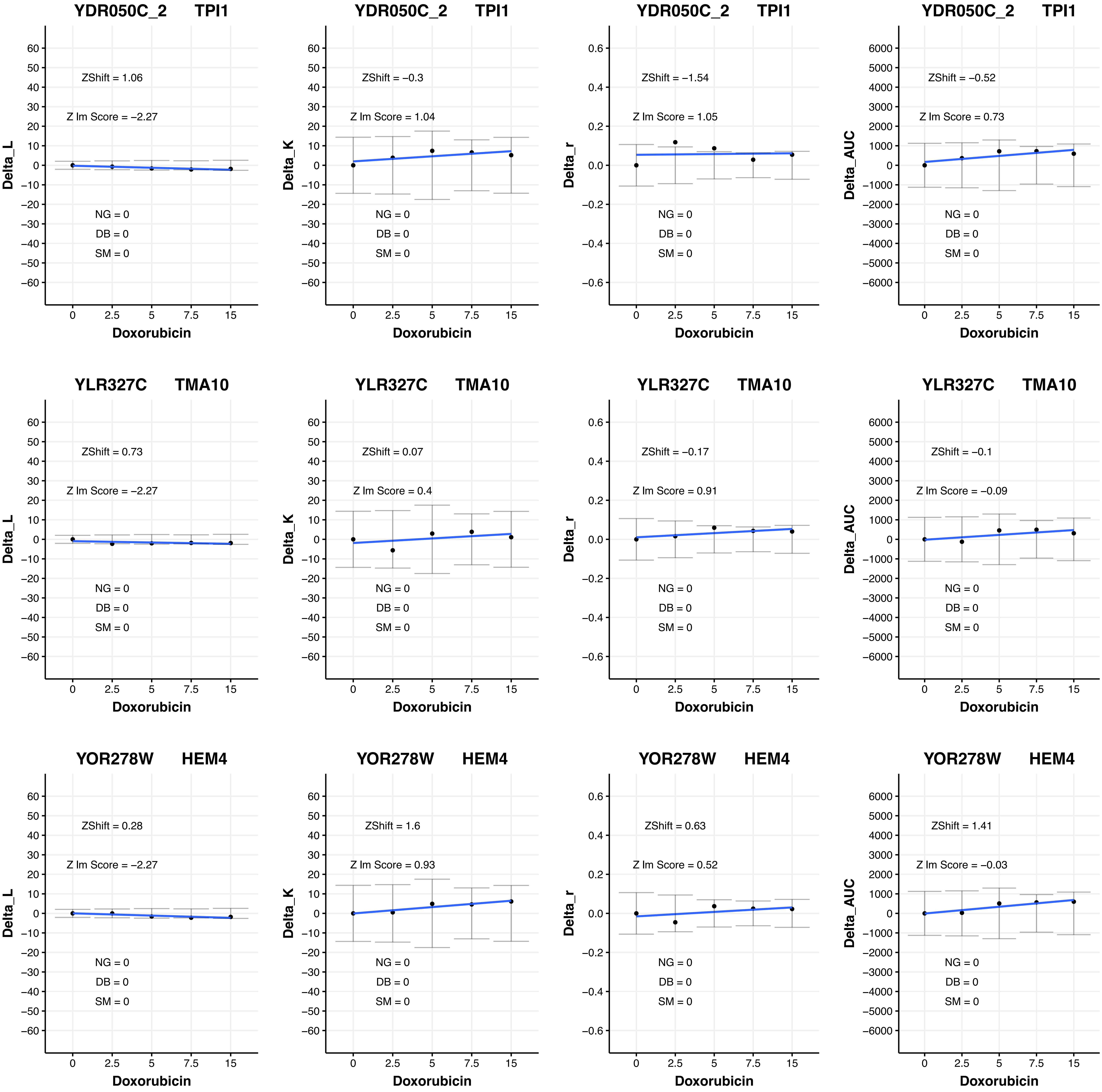
YDR050C_2 TPI1
ZShift = 1.06
Z lm Score = −2.27
NG = 0
DB = 0
SM = 0
Delta_L
Doxorubicin
YDR050C_2 TPI1
ZShift = −0.3
Z lm Score = 1.04
NG = 0
DB = 0
SM = 0
Delta_K
Doxorubicin
YDR050C_2 TPI1
ZShift = −1.54
Z lm Score = 1.05
NG = 0
DB = 0
SM = 0
Delta_r
Doxorubicin
YDR050C_2 TPI1
ZShift = −0.52
Z lm Score = 0.73
NG = 0
DB = 0
SM = 0
Delta_AUC
Doxorubicin
YLR327C TMA10
ZShift = 0.73
Z lm Score = −2.27
NG = 0
DB = 0
SM = 0
Delta_L
Doxorubicin
YLR327C TMA10
ZShift = 0.07
Z lm Score = 0.4
NG = 0
DB = 0
SM = 0
Delta_K
Doxorubicin
YLR327C TMA10
ZShift = −0.17
Z lm Score = 0.91
NG = 0
DB = 0
SM = 0
Delta_r
Doxorubicin
YLR327C TMA10
ZShift = −0.1
Z lm Score = −0.09
NG = 0
DB = 0
SM = 0
Delta_AUC
Doxorubicin
YOR278W HEM4
ZShift = 0.28
Z lm Score = −2.27
NG = 0
DB = 0
SM = 0
Delta_L
Doxorubicin
YOR278W HEM4
ZShift = 1.6
Z lm Score = 0.93
NG = 0
DB = 0
SM = 0
Delta_K
Doxorubicin
YOR278W HEM4
ZShift = 0.63
Z lm Score = 0.52
NG = 0
DB = 0
SM = 0
Delta_r
Doxorubicin
YOR278W HEM4
ZShift = 1.41
Z lm Score = −0.03
NG = 0
DB = 0
SM = 0
Delta_AUC
Doxorubicin

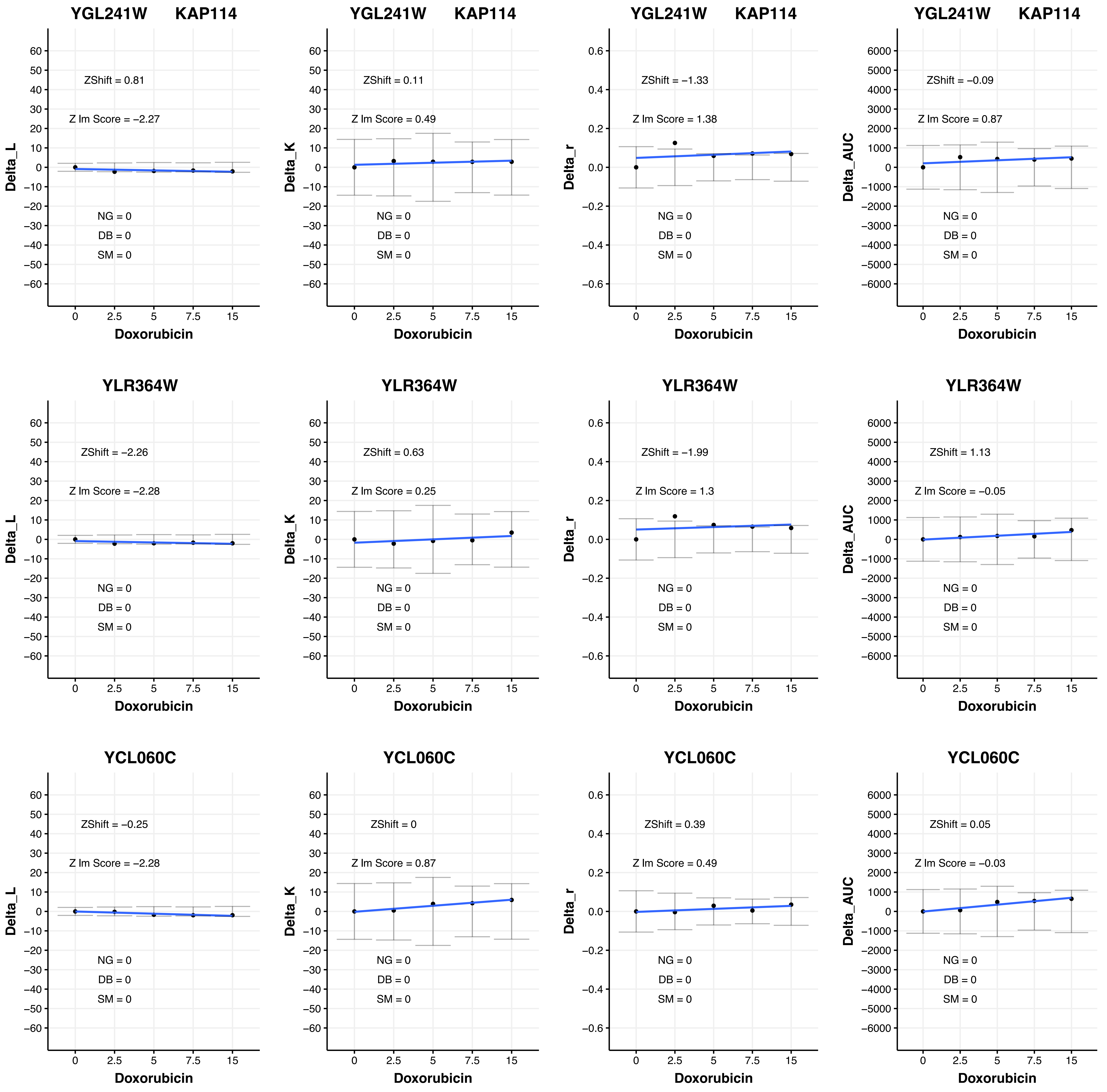

YGL241W KAP114
ZShift = 0.81
Z lm Score = −2.27
NG = 0
DB = 0
SM = 0
Delta_L
Doxorubicin
YGL241W KAP114
ZShift = 0.11
Z lm Score = 0.49
NG = 0
DB = 0
SM = 0
Delta_K
Doxorubicin
YGL241W KAP114
ZShift = −1.33
Z lm Score = 1.38
NG = 0
DB = 0
SM = 0
Delta_r
Doxorubicin
YGL241W KAP114
ZShift = −0.09
Z lm Score = 0.87
NG = 0
DB = 0
SM = 0
Delta_AUC
Doxorubicin
YLR364W
ZShift = −2.26
Z lm Score = −2.28
NG = 0
DB = 0
SM = 0
Delta_L
Doxorubicin
YLR364W
ZShift = 0.63
Z lm Score = 0.25
NG = 0
DB = 0
SM = 0
Delta_K
Doxorubicin
YLR364W
ZShift = −1.99
Z lm Score = 1.3
NG = 0
DB = 0
SM = 0
Delta_r
Doxorubicin
YLR364W
ZShift = 1.13
Z lm Score = −0.05
NG = 0
DB = 0
SM = 0
Delta_AUC
Doxorubicin
YCL060C
ZShift = −0.25
Z lm Score = −2.28
NG = 0
DB = 0
SM = 0
Delta_L
Doxorubicin
YCL060C
ZShift = 0
Z lm Score = 0.87
NG = 0
DB = 0
SM = 0
Delta_K
Doxorubicin
YCL060C
ZShift = 0.39
Z lm Score = 0.49
NG = 0
DB = 0
SM = 0
Delta_r
Doxorubicin
YCL060C
ZShift = 0.05
Z lm Score = −0.03
NG = 0
DB = 0
SM = 0
Delta_AUC
Doxorubicin

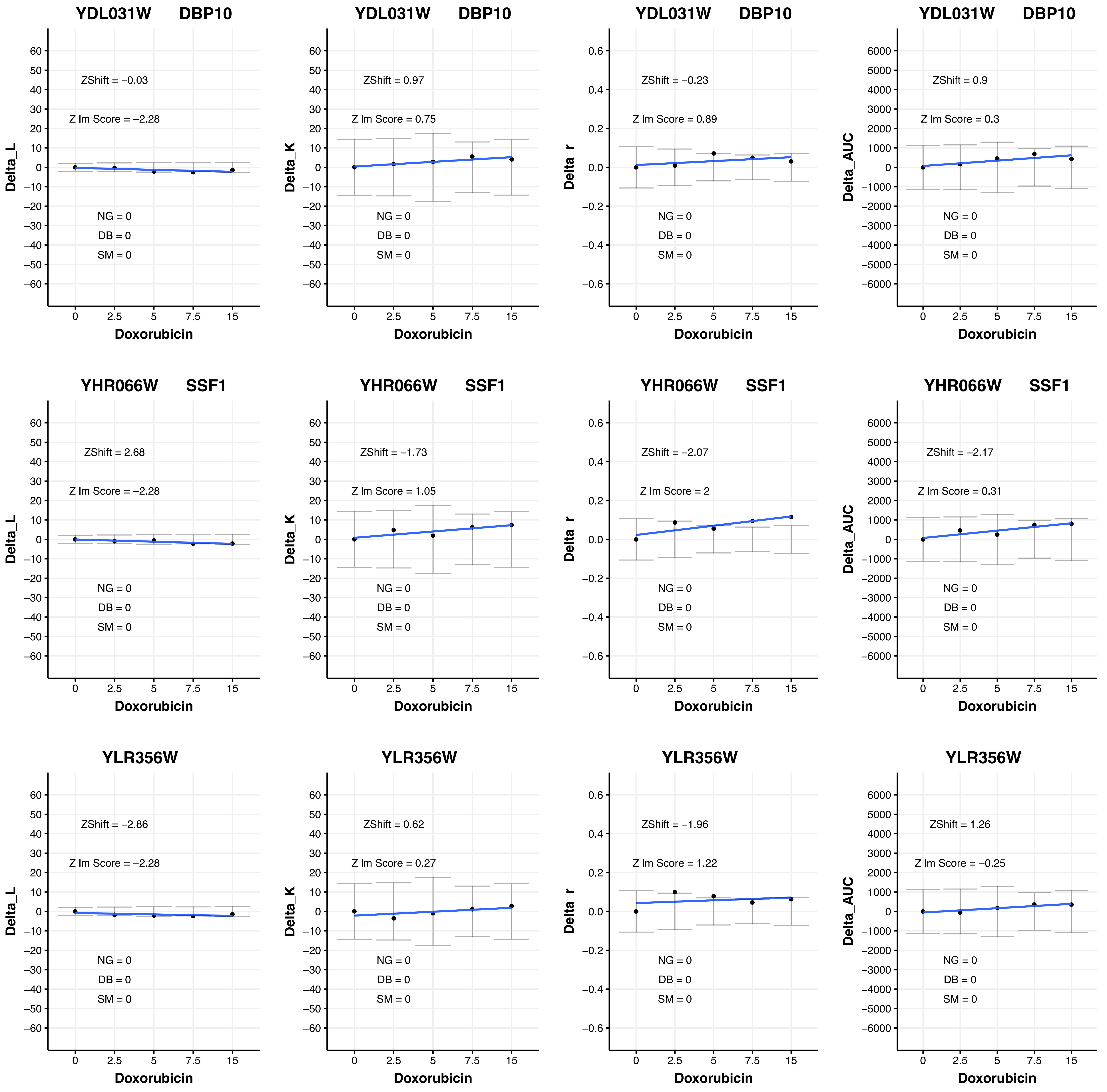
YDL031W DBP10
ZShift = −0.03
Z lm Score = −2.28
NG = 0
DB = 0
SM = 0
Delta_L
Doxorubicin
YDL031W DBP10
ZShift = 0.97
Z lm Score = 0.75
NG = 0
DB = 0
SM = 0
Delta_K
Doxorubicin
YDL031W DBP10
ZShift = −0.23
Z lm Score = 0.89
NG = 0
DB = 0
SM = 0
Delta_r
Doxorubicin
YDL031W DBP10
ZShift = 0.9
Z lm Score = 0.3
NG = 0
DB = 0
SM = 0
Delta_AUC
Doxorubicin
YHR066W SSF1
ZShift = 2.68
Z lm Score = −2.28
NG = 0
DB = 0
SM = 0
Delta_L
Doxorubicin
YHR066W SSF1
ZShift = −1.73
Z lm Score = 1.05
NG = 0
DB = 0
SM = 0
Delta_K
Doxorubicin
YHR066W SSF1
ZShift = −2.07
Z lm Score = 2
NG = 0
DB = 0
SM = 0
Delta_r
Doxorubicin
YHR066W SSF1
ZShift = −2.17
Z lm Score = 0.31
NG = 0
DB = 0
SM = 0
Delta_AUC
Doxorubicin
YLR356W
ZShift = −2.86
Z lm Score = −2.28
NG = 0
DB = 0
SM = 0
Delta_L
Doxorubicin
YLR356W
ZShift = 0.62
Z lm Score = 0.27
NG = 0
DB = 0
SM = 0
Delta_K
Doxorubicin
YLR356W
ZShift = −1.96
Z lm Score = 1.22
NG = 0
DB = 0
SM = 0
Delta_r
Doxorubicin
YLR356W
ZShift = 1.26
Z lm Score = −0.25
NG = 0
DB = 0
SM = 0
Delta_AUC
Doxorubicin

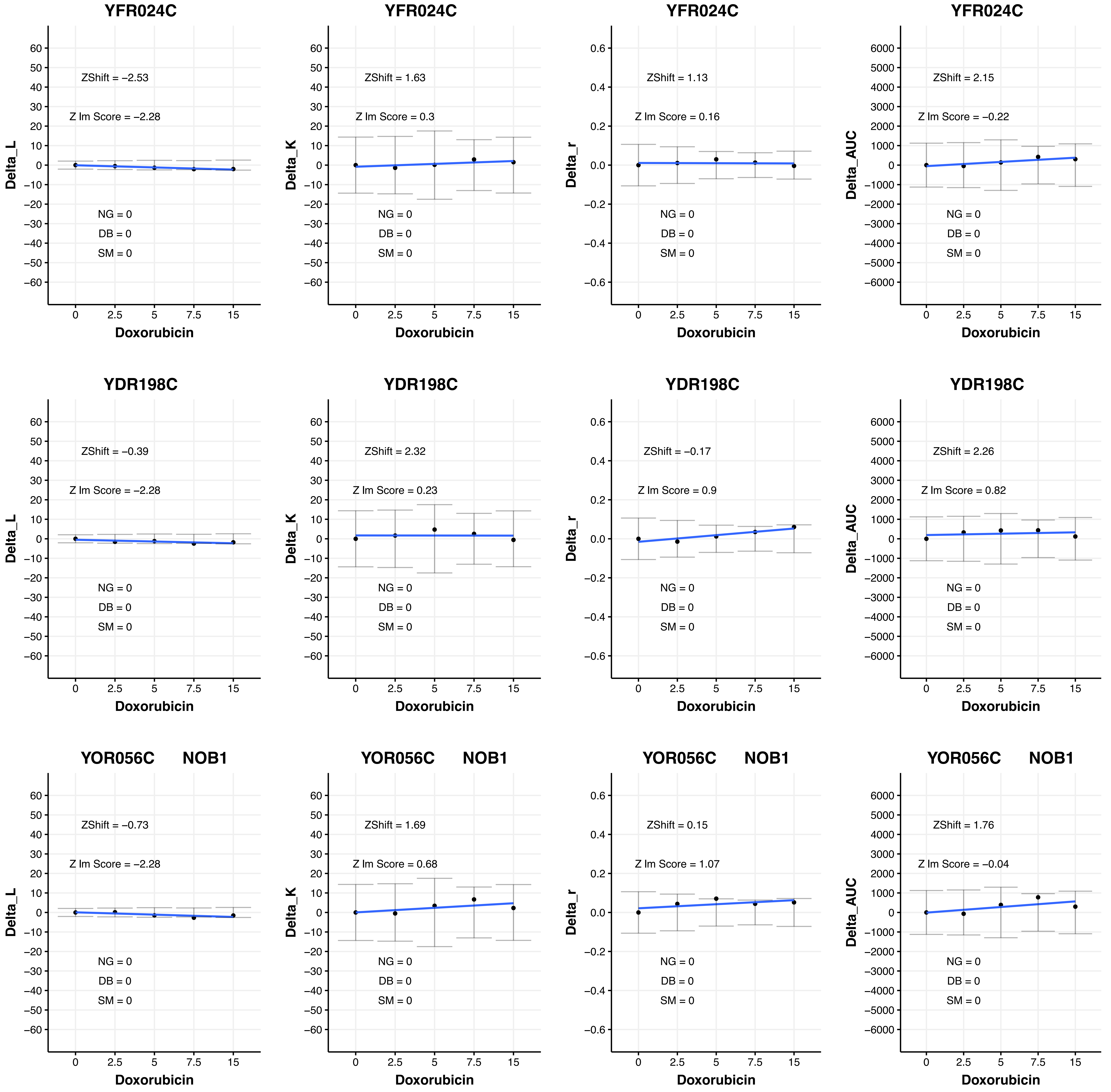
YFR024C
ZShift = −2.53
Z lm Score = −2.28
NG = 0
DB = 0
SM = 0
Delta_L
Doxorubicin
YFR024C
ZShift = 1.63
Z lm Score = 0.3
NG = 0
DB = 0
SM = 0
Delta_K
Doxorubicin
YFR024C
ZShift = 1.13
Z lm Score = 0.16
NG = 0
DB = 0
SM = 0
Delta_r
Doxorubicin
YFR024C
ZShift = 2.15
Z lm Score = −0.22
NG = 0
DB = 0
SM = 0
Delta_AUC
Doxorubicin
YDR198C
ZShift = −0.39
Z lm Score = −2.28
NG = 0
DB = 0
SM = 0
Delta_L
Doxorubicin
YDR198C
ZShift = 2.32
Z lm Score = 0.23
NG = 0
DB = 0
SM = 0
Delta_K
Doxorubicin
YDR198C
ZShift = −0.17
Z lm Score = 0.9
NG = 0
DB = 0
SM = 0
Delta_r
Doxorubicin
YDR198C
ZShift = 2.26
Z lm Score = 0.82
NG = 0
DB = 0
SM = 0
Delta_AUC
Doxorubicin
YOR056C NOB1
ZShift = −0.73
Z lm Score = −2.28
NG = 0
DB = 0
SM = 0
Delta_L
Doxorubicin
YOR056C NOB1
ZShift = 1.69
Z lm Score = 0.68
NG = 0
DB = 0
SM = 0
Delta_K
Doxorubicin
YOR056C NOB1
ZShift = 0.15
Z lm Score = 1.07
NG = 0
DB = 0
SM = 0
Delta_r
Doxorubicin
YOR056C NOB1
ZShift = 1.76
Z lm Score = −0.04
NG = 0
DB = 0
SM = 0
Delta_AUC
Doxorubicin

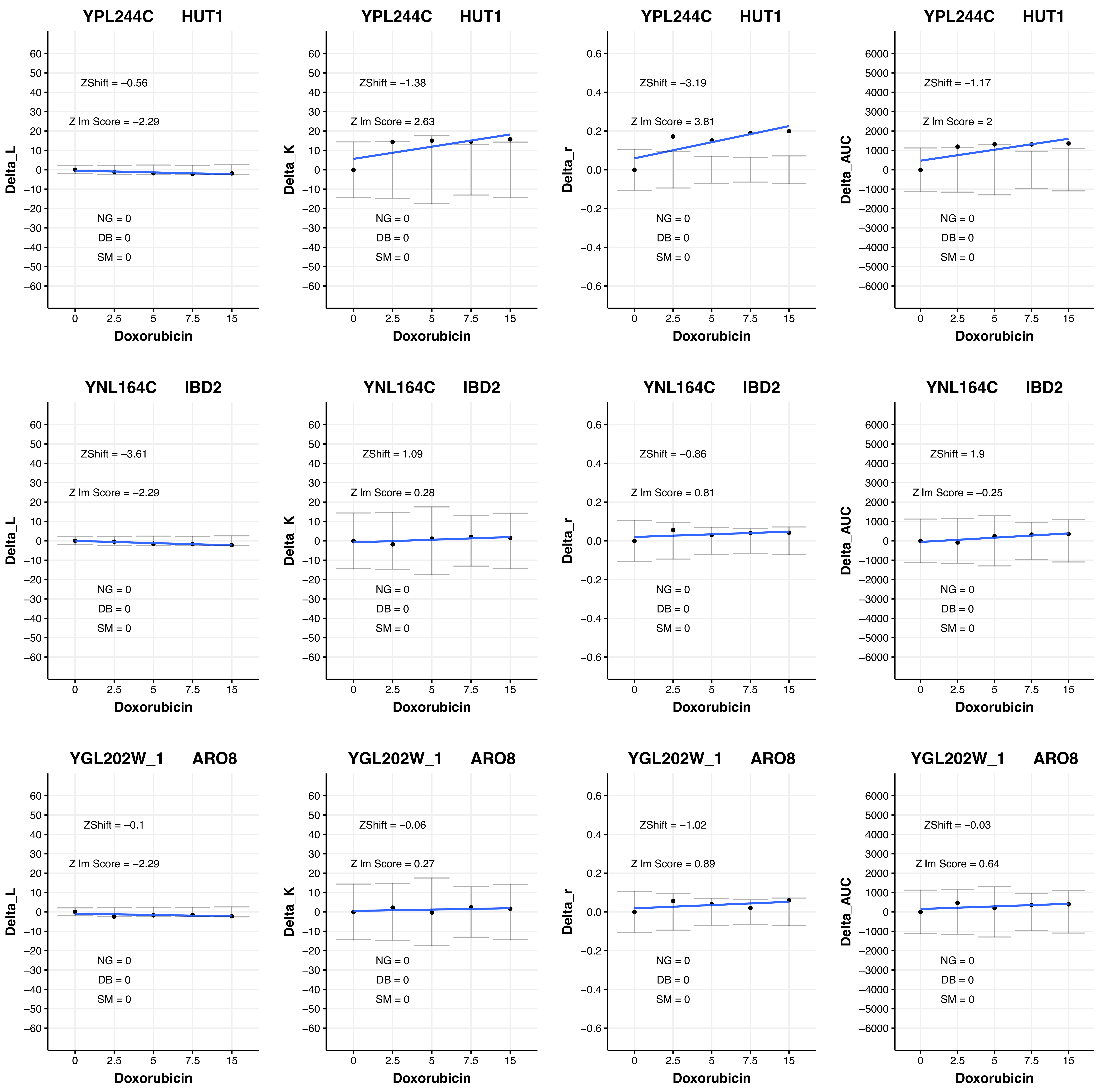
YPL244C HUT1
ZShift = −0.56
Z lm Score = −2.29
NG = 0
DB = 0
SM = 0
Delta_L
Doxorubicin
YPL244C HUT1
ZShift = −1.38
Z lm Score = 2.63
NG = 0
DB = 0
SM = 0
Delta_K
Doxorubicin
YPL244C HUT1
ZShift = −3.19
Z lm Score = 3.81
NG = 0
DB = 0
SM = 0
Delta_r
Doxorubicin
YPL244C HUT1
ZShift = −1.17
Z lm Score = 2
NG = 0
DB = 0
SM = 0
Delta_AUC
Doxorubicin
YNL164C IBD2
ZShift = −3.61
Z lm Score = −2.29
NG = 0
DB = 0
SM = 0
Delta_L
Doxorubicin
YNL164C IBD2
ZShift = 1.09
Z lm Score = 0.28
NG = 0
DB = 0
SM = 0
Delta_K
Doxorubicin
YNL164C IBD2
ZShift = −0.86
Z lm Score = 0.81
NG = 0
DB = 0
SM = 0
Delta_r
Doxorubicin
YNL164C IBD2
ZShift = 1.9
Z lm Score = −0.25
NG = 0
DB = 0
SM = 0
Delta_AUC
Doxorubicin
YGL202W_1 ARO8
ZShift = −0.1
Z lm Score = −2.29
NG = 0
DB = 0
SM = 0
Delta_L
Doxorubicin
YGL202W_1 ARO8
ZShift = −0.06
Z lm Score = 0.27
NG = 0
DB = 0
SM = 0
Delta_K
Doxorubicin
YGL202W_1 ARO8
ZShift = −1.02
Z lm Score = 0.89
NG = 0
DB = 0
SM = 0
Delta_r
Doxorubicin
YGL202W_1 ARO8
ZShift = −0.03
Z lm Score = 0.64
NG = 0
DB = 0
SM = 0
Delta_AUC
Doxorubicin

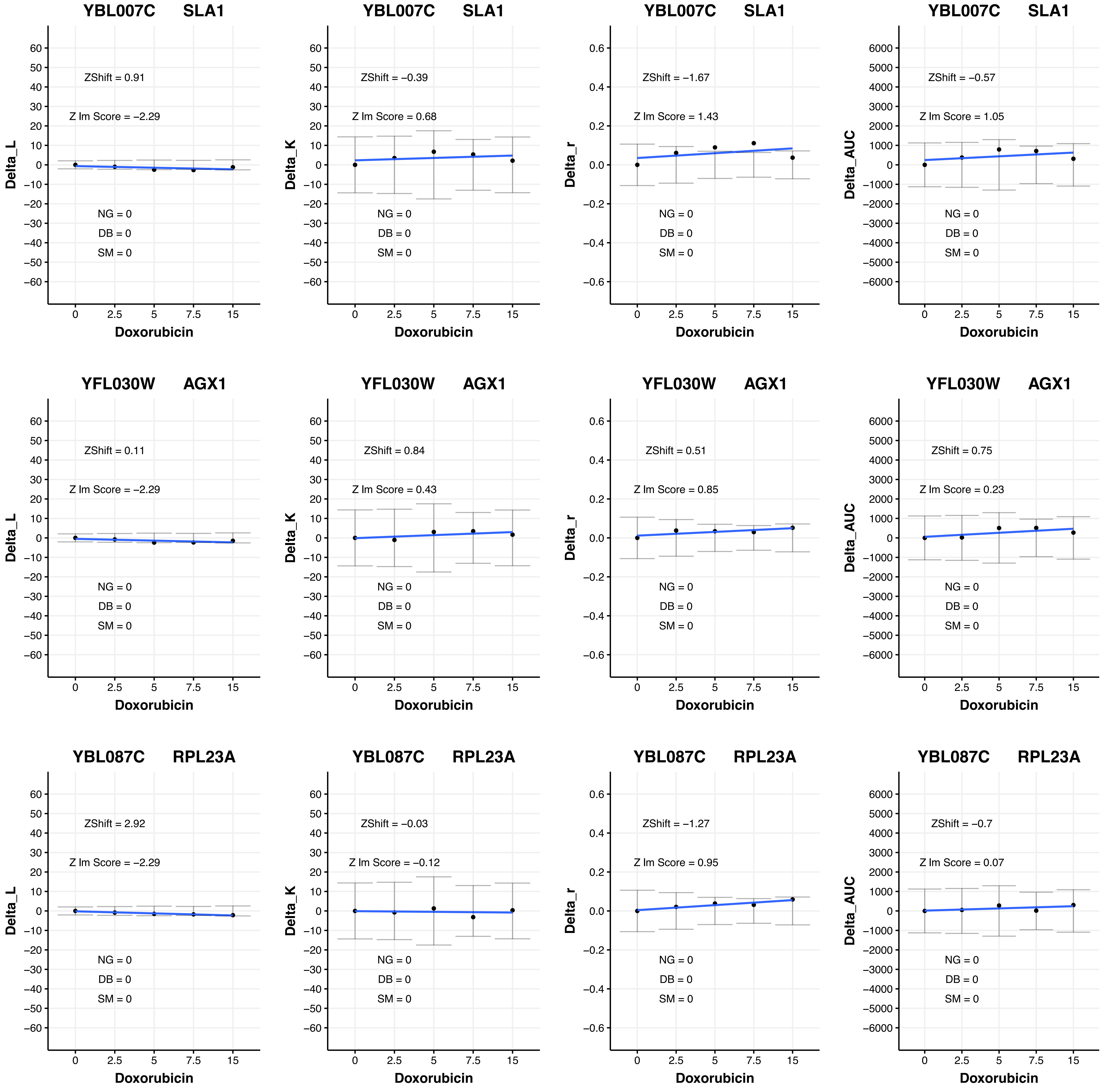

YBL007C SLA1
Delta_L
ZShift = 0.91
Z lm Score = −2.29
NG = 0
DB = 0
SM = 0
Doxorubicin
YBL007C SLA1
Delta_K
ZShift = −0.39
Z lm Score = 0.68
NG = 0
DB = 0
SM = 0
Doxorubicin
YBL007C SLA1
Delta_r
ZShift = −1.67
Z lm Score = 1.43
NG = 0
DB = 0
SM = 0
Doxorubicin
YBL007C SLA1
Delta_AUC
ZShift = −0.57
Z lm Score = 1.05
NG = 0
DB = 0
SM = 0
Doxorubicin
YFL030W AGX1
Delta_L
ZShift = 0.11
Z lm Score = −2.29
NG = 0
DB = 0
SM = 0
Doxorubicin
YFL030W AGX1
Delta_K
ZShift = 0.84
Z lm Score = 0.43
NG = 0
DB = 0
SM = 0
Doxorubicin
YFL030W AGX1
Delta_r
ZShift = 0.51
Z lm Score = 0.85
NG = 0
DB = 0
SM = 0
Doxorubicin
YFL030W AGX1
Delta_AUC
ZShift = 0.75
Z lm Score = 0.23
NG = 0
DB = 0
SM = 0
Doxorubicin
YBL087C RPL23A
Delta_L
ZShift = 2.92
Z lm Score = −2.29
NG = 0
DB = 0
SM = 0
Doxorubicin
YBL087C RPL23A
Delta_K
ZShift = −0.03
Z lm Score = −0.12
NG = 0
DB = 0
SM = 0
Doxorubicin
YBL087C RPL23A
Delta_r
ZShift = −1.27
Z lm Score = 0.95
NG = 0
DB = 0
SM = 0
Doxorubicin
YBL087C RPL23A
Delta_AUC
ZShift = −0.7
Z lm Score = 0.07
NG = 0
DB = 0
SM = 0
Doxorubicin

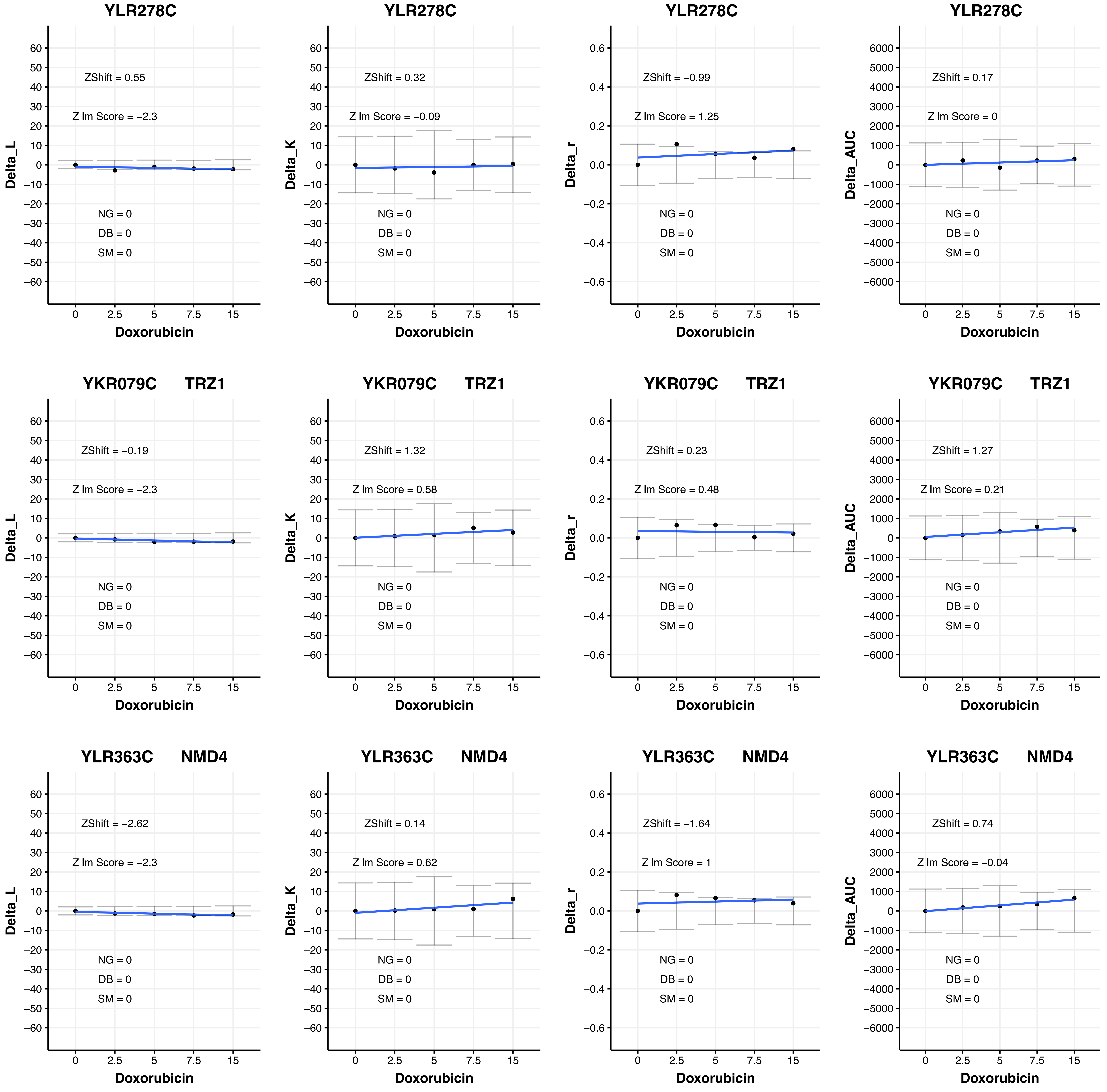

YLR278C
ZShift = 0.55
Z lm Score = −2.3
NG = 0
DB = 0
SM = 0
Delta_L
Doxorubicin
YLR278C
ZShift = 0.32
Z lm Score = −0.09
NG = 0
DB = 0
SM = 0
Delta_K
Doxorubicin
YLR278C
ZShift = −0.99
Z lm Score = 1.25
NG = 0
DB = 0
SM = 0
Delta_r
Doxorubicin
YLR278C
ZShift = 0.17
Z lm Score = 0
NG = 0
DB = 0
SM = 0
Delta_AUC
Doxorubicin
YKR079C TRZ1
ZShift = −0.19
Z lm Score = −2.3
NG = 0
DB = 0
SM = 0
Delta_L
Doxorubicin
YKR079C TRZ1
ZShift = 1.32
Z lm Score = 0.58
NG = 0
DB = 0
SM = 0
Delta_K
Doxorubicin
YKR079C TRZ1
ZShift = 0.23
Z lm Score = 0.48
NG = 0
DB = 0
SM = 0
Delta_r
Doxorubicin
YKR079C TRZ1
ZShift = 1.27
Z lm Score = 0.21
NG = 0
DB = 0
SM = 0
Delta_AUC
Doxorubicin
YLR363C NMD4
ZShift = −2.62
Z lm Score = −2.3
NG = 0
DB = 0
SM = 0
Delta_L
Doxorubicin
YLR363C NMD4
ZShift = 0.14
Z lm Score = 0.62
NG = 0
DB = 0
SM = 0
Delta_K
Doxorubicin
YLR363C NMD4
ZShift = −1.64
Z lm Score = 1
NG = 0
DB = 0
SM = 0
Delta_r
Doxorubicin
YLR363C NMD4
ZShift = 0.74
Z lm Score = −0.04
NG = 0
DB = 0
SM = 0
Delta_AUC
Doxorubicin

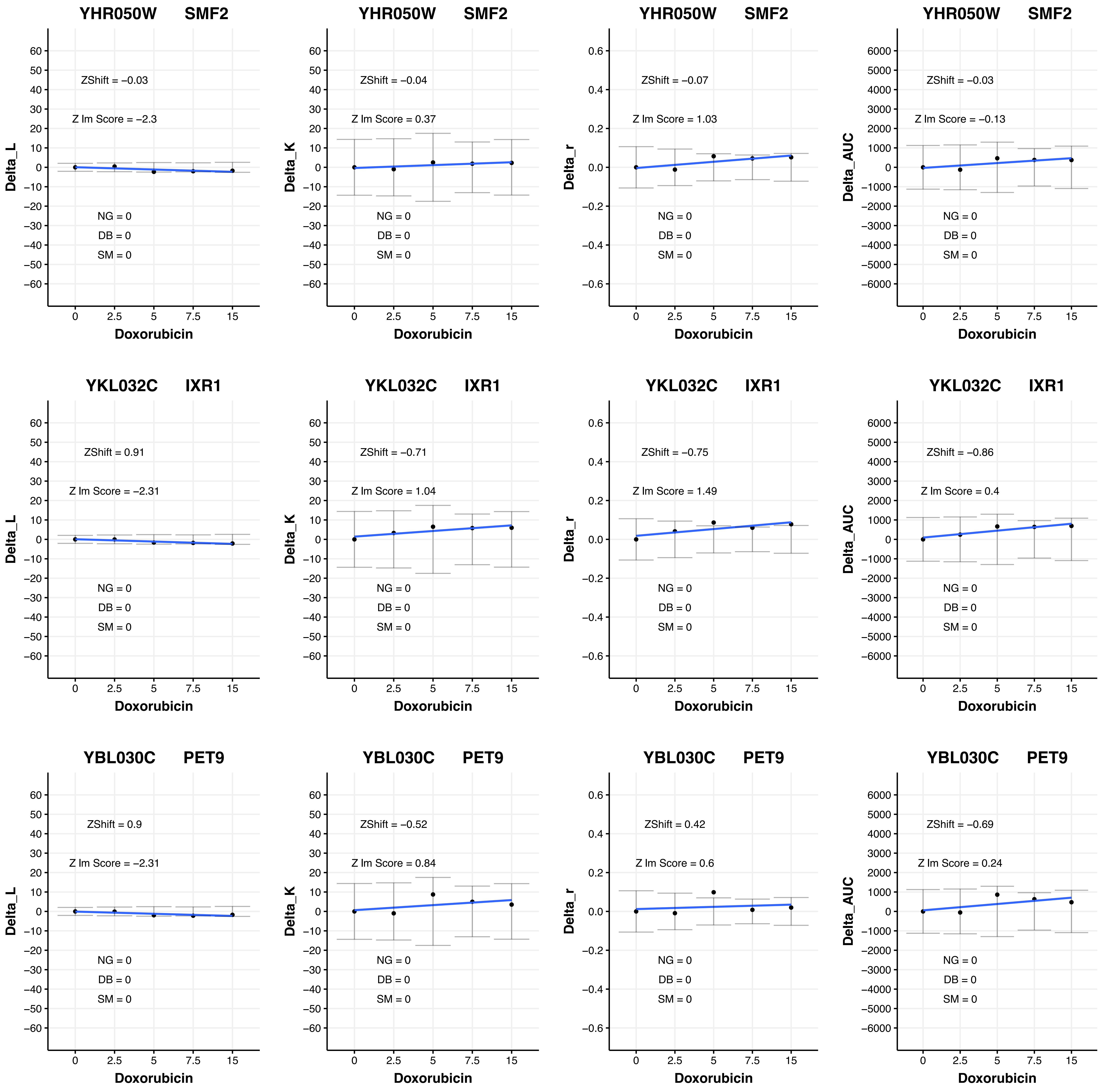
YHR050W SMF2
ZShift = −0.03
Z lm Score = −2.3
NG = 0
DB = 0
SM = 0
Delta_L
Doxorubicin
YHR050W SMF2
ZShift = −0.04
Z lm Score = 0.37
NG = 0
DB = 0
SM = 0
Delta_K
Doxorubicin
YHR050W SMF2
ZShift = −0.07
Z lm Score = 1.03
NG = 0
DB = 0
SM = 0
Delta_r
Doxorubicin
YHR050W SMF2
ZShift = −0.03
Z lm Score = −0.13
NG = 0
DB = 0
SM = 0
Delta_AUC
Doxorubicin
YKL032C IXR1
ZShift = 0.91
Z lm Score = −2.31
NG = 0
DB = 0
SM = 0
Delta_L
Doxorubicin
YKL032C IXR1
ZShift = −0.71
Z lm Score = 1.04
NG = 0
DB = 0
SM = 0
Delta_K
Doxorubicin
YKL032C IXR1
ZShift = −0.75
Z lm Score = 1.49
NG = 0
DB = 0
SM = 0
Delta_r
Doxorubicin
YKL032C IXR1
ZShift = −0.86
Z lm Score = 0.4
NG = 0
DB = 0
SM = 0
Delta_AUC
Doxorubicin
YBL030C PET9
ZShift = 0.9
Z lm Score = −2.31
NG = 0
DB = 0
SM = 0
Delta_L
Doxorubicin
YBL030C PET9
ZShift = −0.52
Z lm Score = 0.84
NG = 0
DB = 0
SM = 0
Delta_K
Doxorubicin
YBL030C PET9
ZShift = 0.42
Z lm Score = 0.6
NG = 0
DB = 0
SM = 0
Delta_r
Doxorubicin
YBL030C PET9
ZShift = −0.69
Z lm Score = 0.24
NG = 0
DB = 0
SM = 0
Delta_AUC
Doxorubicin

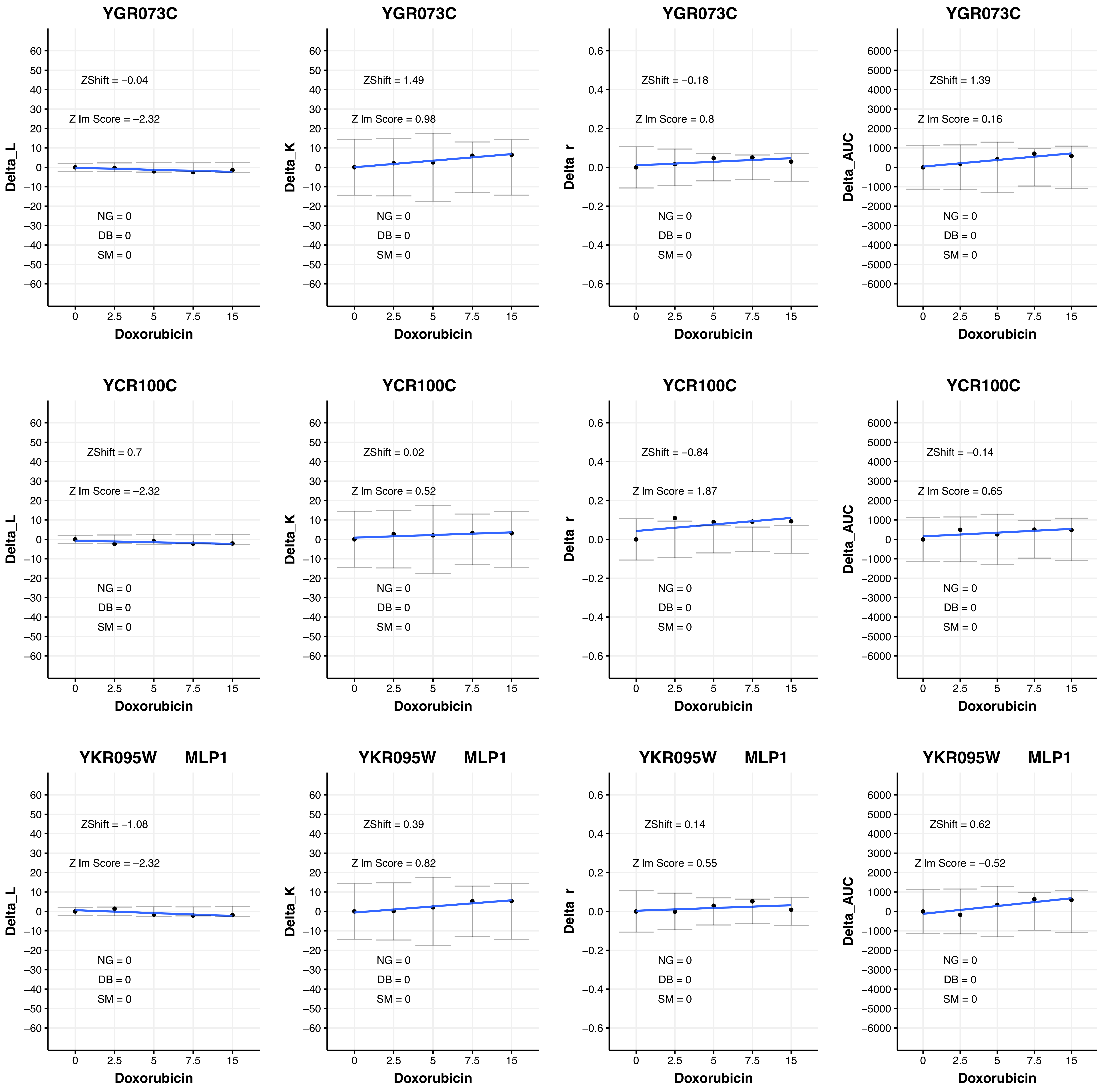

YGR073C
ZShift = −0.04
Z lm Score = −2.32
NG = 0
DB = 0
SM = 0
Delta_L
Doxorubicin
YGR073C
ZShift = 1.49
Z lm Score = 0.98
NG = 0
DB = 0
SM = 0
Delta_K
Doxorubicin
YGR073C
ZShift = −0.18
Z lm Score = 0.8
NG = 0
DB = 0
SM = 0
Delta_r
Doxorubicin
YGR073C
ZShift = 1.39
Z lm Score = 0.16
NG = 0
DB = 0
SM = 0
Delta_AUC
Doxorubicin
YCR100C
ZShift = 0.7
Z lm Score = −2.32
NG = 0
DB = 0
SM = 0
Delta_L
Doxorubicin
YCR100C
ZShift = 0.02
Z lm Score = 0.52
NG = 0
DB = 0
SM = 0
Delta_K
Doxorubicin
YCR100C
ZShift = −0.84
Z lm Score = 1.87
NG = 0
DB = 0
SM = 0
Delta_r
Doxorubicin
YCR100C
ZShift = −0.14
Z lm Score = 0.65
NG = 0
DB = 0
SM = 0
Delta_AUC
Doxorubicin
YKR095W MLP1
ZShift = −1.08
Z lm Score = −2.32
NG = 0
DB = 0
SM = 0
Delta_L
Doxorubicin
YKR095W MLP1
ZShift = 0.39
Z lm Score = 0.82
NG = 0
DB = 0
SM = 0
Delta_K
Doxorubicin
YKR095W MLP1
ZShift = 0.14
Z lm Score = 0.55
NG = 0
DB = 0
SM = 0
Delta_r
Doxorubicin
YKR095W MLP1
ZShift = 0.62
Z lm Score = −0.52
NG = 0
DB = 0
SM = 0
Delta_AUC
Doxorubicin

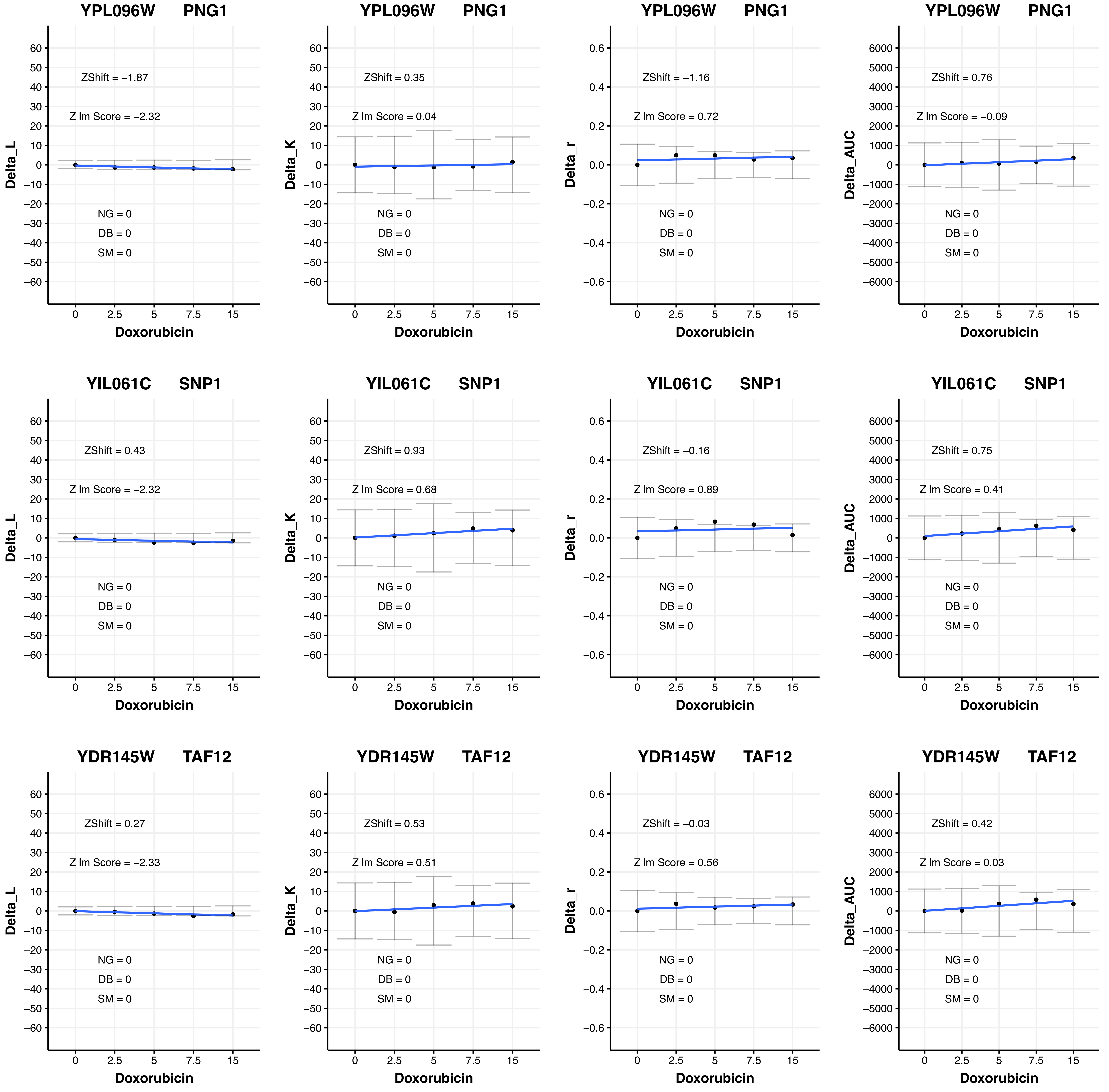
YPL096W PNG1
ZShift = −1.87
Z lm Score = −2.32
NG = 0
DB = 0
SM = 0
Delta_L
Doxorubicin
YPL096W PNG1
ZShift = 0.35
Z lm Score = 0.04
NG = 0
DB = 0
SM = 0
Delta_K
Doxorubicin
YPL096W PNG1
ZShift = −1.16
Z lm Score = 0.72
NG = 0
DB = 0
SM = 0
Delta_r
Doxorubicin
YPL096W PNG1
ZShift = 0.76
Z lm Score = −0.09
NG = 0
DB = 0
SM = 0
Delta_AUC
Doxorubicin
YIL061C SNP1
ZShift = 0.43
Z lm Score = −2.32
NG = 0
DB = 0
SM = 0
Delta_L
Doxorubicin
YIL061C SNP1
ZShift = 0.93
Z lm Score = 0.68
NG = 0
DB = 0
SM = 0
Delta_K
Doxorubicin
YIL061C SNP1
ZShift = −0.16
Z lm Score = 0.89
NG = 0
DB = 0
SM = 0
Delta_r
Doxorubicin
YIL061C SNP1
ZShift = 0.75
Z lm Score = 0.41
NG = 0
DB = 0
SM = 0
Delta_AUC
Doxorubicin
YDR145W TAF12
ZShift = 0.27
Z lm Score = −2.33
NG = 0
DB = 0
SM = 0
Delta_L
Doxorubicin
YDR145W TAF12
ZShift = 0.53
Z lm Score = 0.51
NG = 0
DB = 0
SM = 0
Delta_K
Doxorubicin
YDR145W TAF12
ZShift = −0.03
Z lm Score = 0.56
NG = 0
DB = 0
SM = 0
Delta_r
Doxorubicin
YDR145W TAF12
ZShift = 0.42
Z lm Score = 0.03
NG = 0
DB = 0
SM = 0
Delta_AUC
Doxorubicin

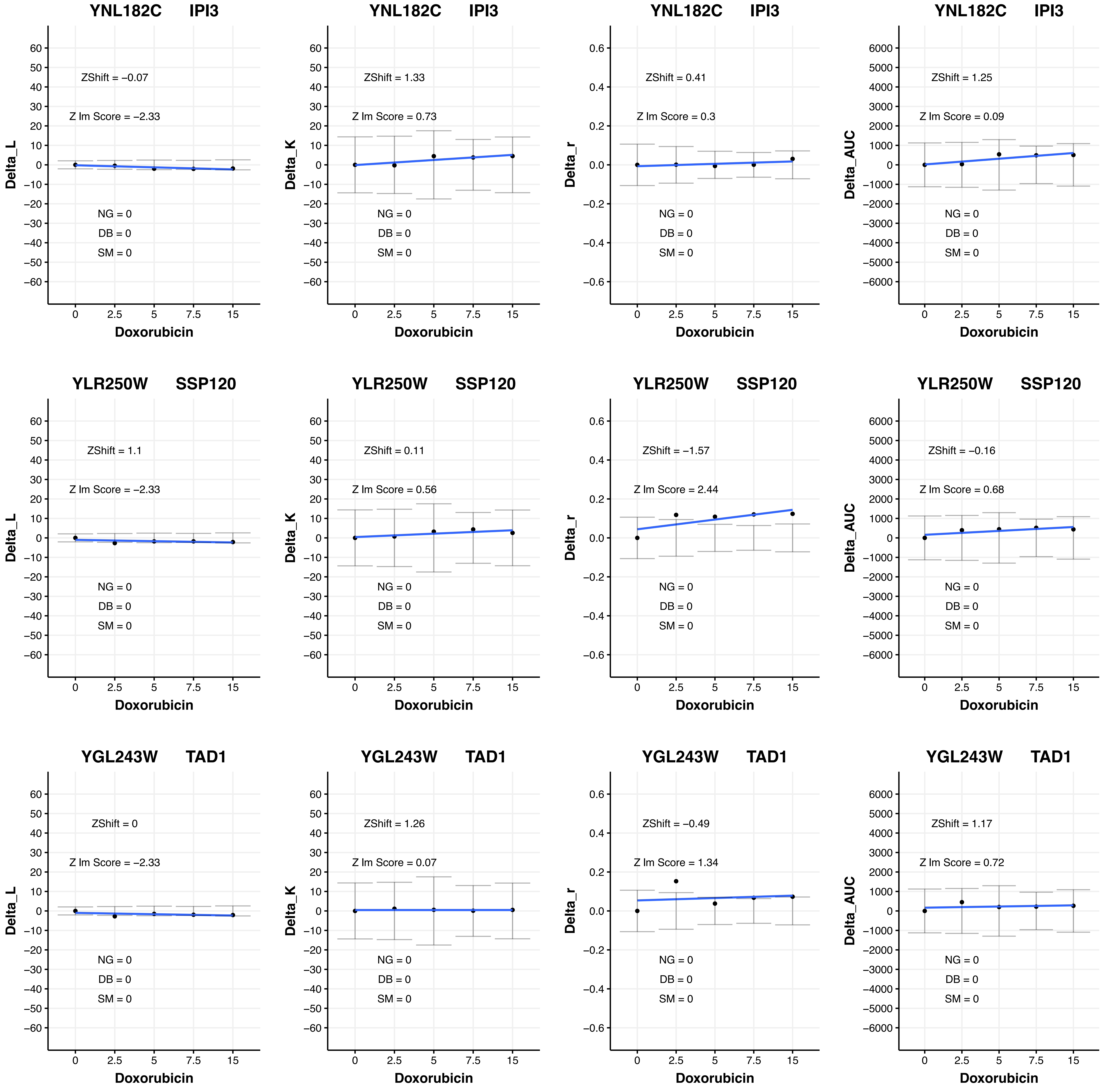
YNL182C IPI3
ZShift = −0.07
Z lm Score = −2.33
Delta_L
NG = 0
DB = 0
SM = 0
Doxorubicin
YNL182C IPI3
ZShift = 1.33
Z lm Score = 0.73
Delta_K
NG = 0
DB = 0
SM = 0
Doxorubicin
YNL182C IPI3
ZShift = 0.41
Z lm Score = 0.3
Delta_r
NG = 0
DB = 0
SM = 0
Doxorubicin
YNL182C IPI3
ZShift = 1.25
Z lm Score = 0.09
Delta_AUC
NG = 0
DB = 0
SM = 0
Doxorubicin
YLR250W SSP120
ZShift = 1.1
Z lm Score = −2.33
Delta_L
NG = 0
DB = 0
SM = 0
Doxorubicin
YLR250W SSP120
ZShift = 0.11
Z lm Score = 0.56
Delta_K
NG = 0
DB = 0
SM = 0
Doxorubicin
YLR250W SSP120
ZShift = −1.57
Z lm Score = 2.44
Delta_r
NG = 0
DB = 0
SM = 0
Doxorubicin
YLR250W SSP120
ZShift = −0.16
Z lm Score = 0.68
Delta_AUC
NG = 0
DB = 0
SM = 0
Doxorubicin
YGL243W TAD1
ZShift = 0
Z lm Score = −2.33
Delta_L
NG = 0
DB = 0
SM = 0
Doxorubicin
YGL243W TAD1
ZShift = 1.26
Z lm Score = 0.07
Delta_K
NG = 0
DB = 0
SM = 0
Doxorubicin
YGL243W TAD1
ZShift = −0.49
Z lm Score = 1.34
Delta_r
NG = 0
DB = 0
SM = 0
Doxorubicin
YGL243W TAD1
ZShift = 1.17
Z lm Score = 0.72
Delta_AUC
NG = 0
DB = 0
SM = 0
Doxorubicin

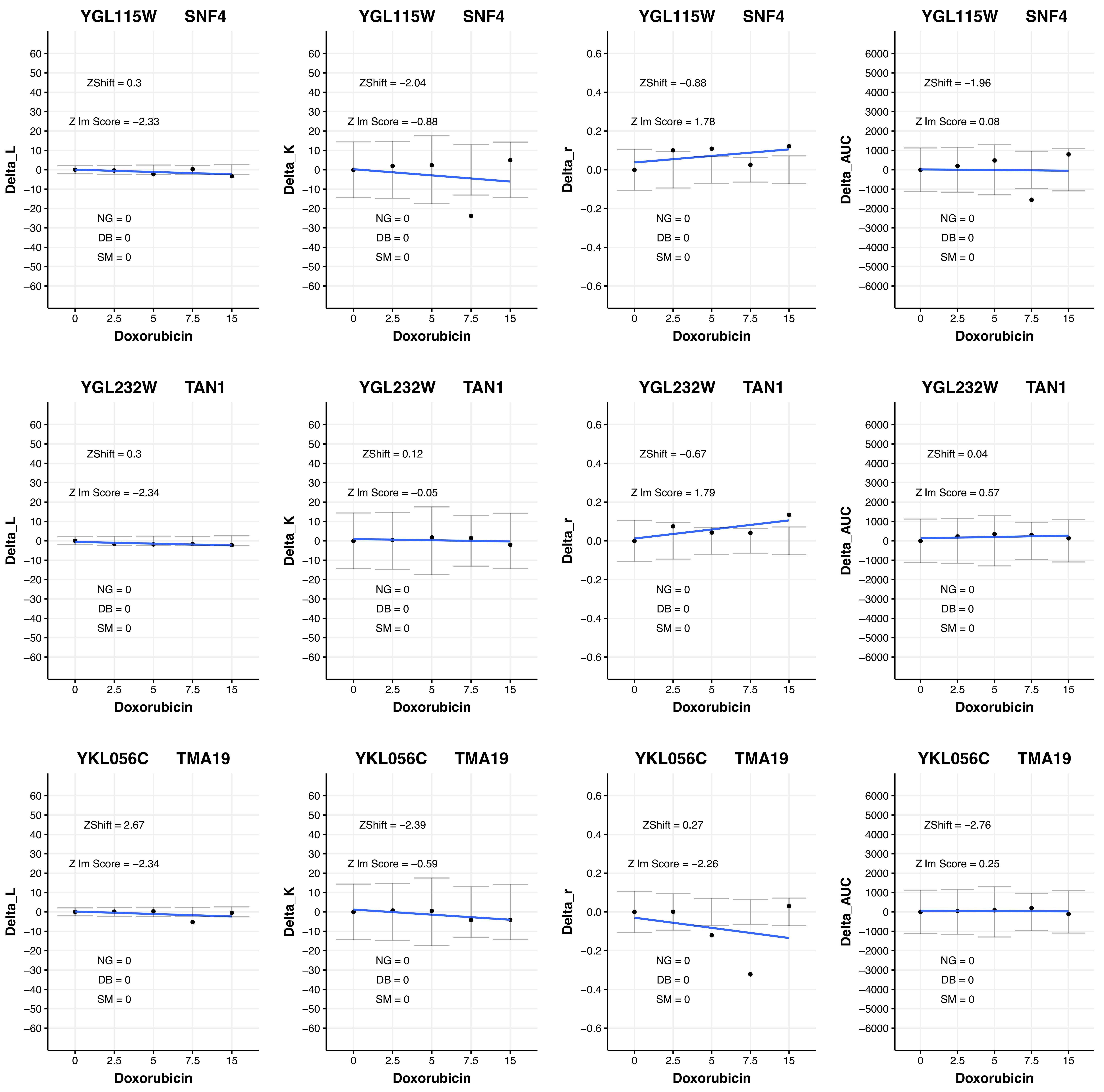

YGL115W SNF4
ZShift = 0.3
Z lm Score = −2.33
NG = 0
DB = 0
SM = 0
Delta_L
Doxorubicin
YGL115W SNF4
ZShift = −2.04
Z lm Score = −0.88
NG = 0
DB = 0
SM = 0
Delta_K
Doxorubicin
YGL115W SNF4
ZShift = −0.88
Z lm Score = 1.78
NG = 0
DB = 0
SM = 0
Delta_r
Doxorubicin
YGL115W SNF4
ZShift = −1.96
Z lm Score = 0.08
NG = 0
DB = 0
SM = 0
Delta_AUC
Doxorubicin
YGL232W TAN1
ZShift = 0.3
Z lm Score = −2.34
NG = 0
DB = 0
SM = 0
Delta_L
Doxorubicin
YGL232W TAN1
ZShift = 0.12
Z lm Score = −0.05
NG = 0
DB = 0
SM = 0
Delta_K
Doxorubicin
YGL232W TAN1
ZShift = −0.67
Z lm Score = 1.79
NG = 0
DB = 0
SM = 0
Delta_r
Doxorubicin
YGL232W TAN1
ZShift = 0.04
Z lm Score = 0.57
NG = 0
DB = 0
SM = 0
Delta_AUC
Doxorubicin
YKL056C TMA19
ZShift = 2.67
Z lm Score = −2.34
NG = 0
DB = 0
SM = 0
Delta_L
Doxorubicin
YKL056C TMA19
ZShift = −2.39
Z lm Score = −0.59
NG = 0
DB = 0
SM = 0
Delta_K
Doxorubicin
YKL056C TMA19
ZShift = 0.27
Z lm Score = −2.26
NG = 0
DB = 0
SM = 0
Delta_r
Doxorubicin
YKL056C TMA19
ZShift = −2.76
Z lm Score = 0.25
NG = 0
DB = 0
SM = 0
Delta_AUC
Doxorubicin

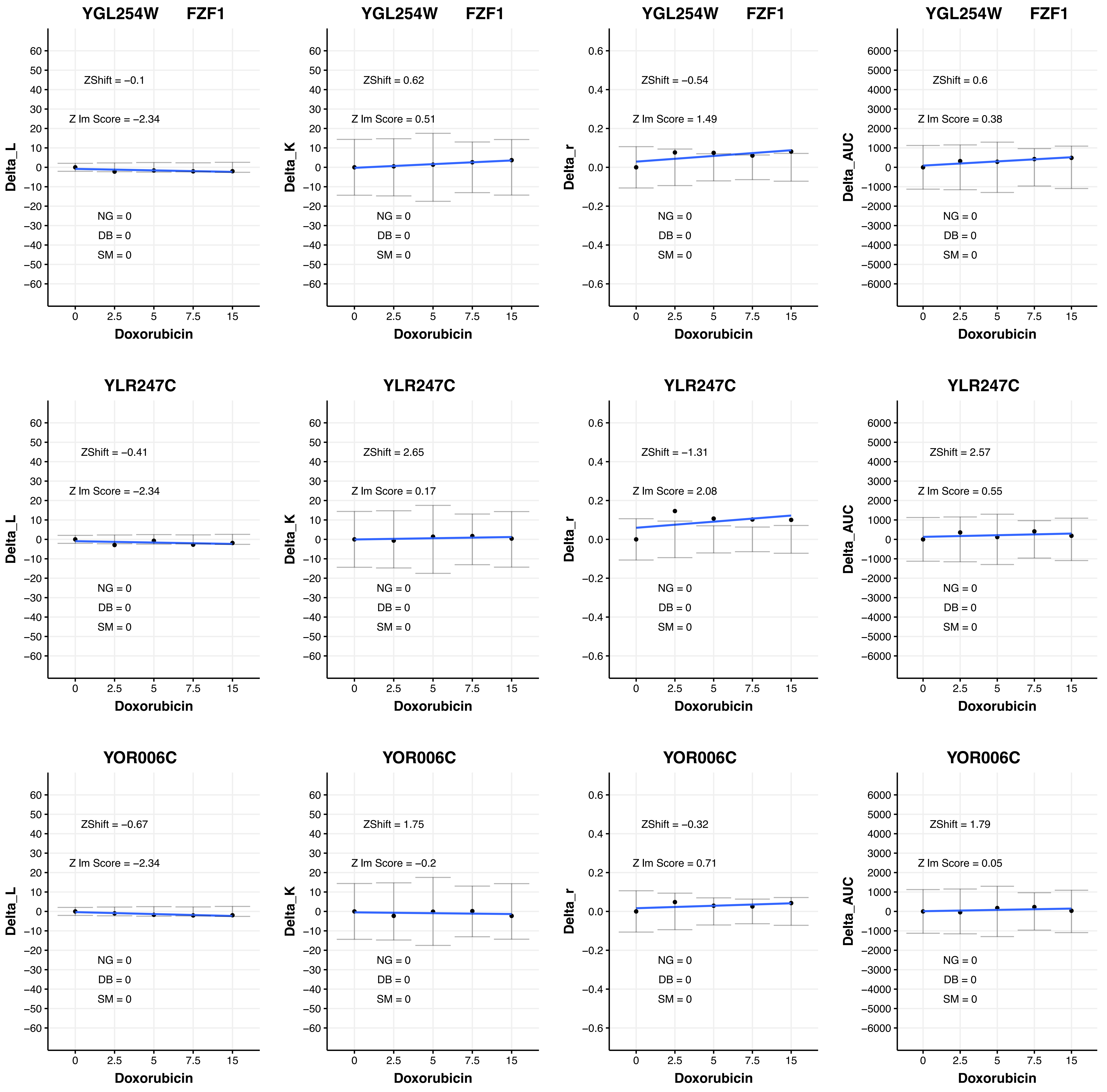
YGL254W FZF1
ZShift = −0.1
Z lm Score = −2.34
NG = 0
DB = 0
SM = 0
Delta_L
Doxorubicin
YGL254W FZF1
ZShift = 0.62
Z lm Score = 0.51
NG = 0
DB = 0
SM = 0
Delta_K
Doxorubicin
YGL254W FZF1
ZShift = −0.54
Z lm Score = 1.49
NG = 0
DB = 0
SM = 0
Delta_r
Doxorubicin
YGL254W FZF1
ZShift = 0.6
Z lm Score = 0.38
NG = 0
DB = 0
SM = 0
Delta_AUC
Doxorubicin
YLR247C
ZShift = −0.41
Z lm Score = −2.34
NG = 0
DB = 0
SM = 0
Delta_L
Doxorubicin
YLR247C
ZShift = 2.65
Z lm Score = 0.17
NG = 0
DB = 0
SM = 0
Delta_K
Doxorubicin
YLR247C
ZShift = −1.31
Z lm Score = 2.08
NG = 0
DB = 0
SM = 0
Delta_r
Doxorubicin
YLR247C
ZShift = 2.57
Z lm Score = 0.55
NG = 0
DB = 0
SM = 0
Delta_AUC
Doxorubicin
YOR006C
ZShift = −0.67
Z lm Score = −2.34
NG = 0
DB = 0
SM = 0
Delta_L
Doxorubicin
YOR006C
ZShift = 1.75
Z lm Score = −0.2
NG = 0
DB = 0
SM = 0
Delta_K
Doxorubicin
YOR006C
ZShift = −0.32
Z lm Score = 0.71
NG = 0
DB = 0
SM = 0
Delta_r
Doxorubicin
YOR006C
ZShift = 1.79
Z lm Score = 0.05
NG = 0
DB = 0
SM = 0
Delta_AUC
Doxorubicin

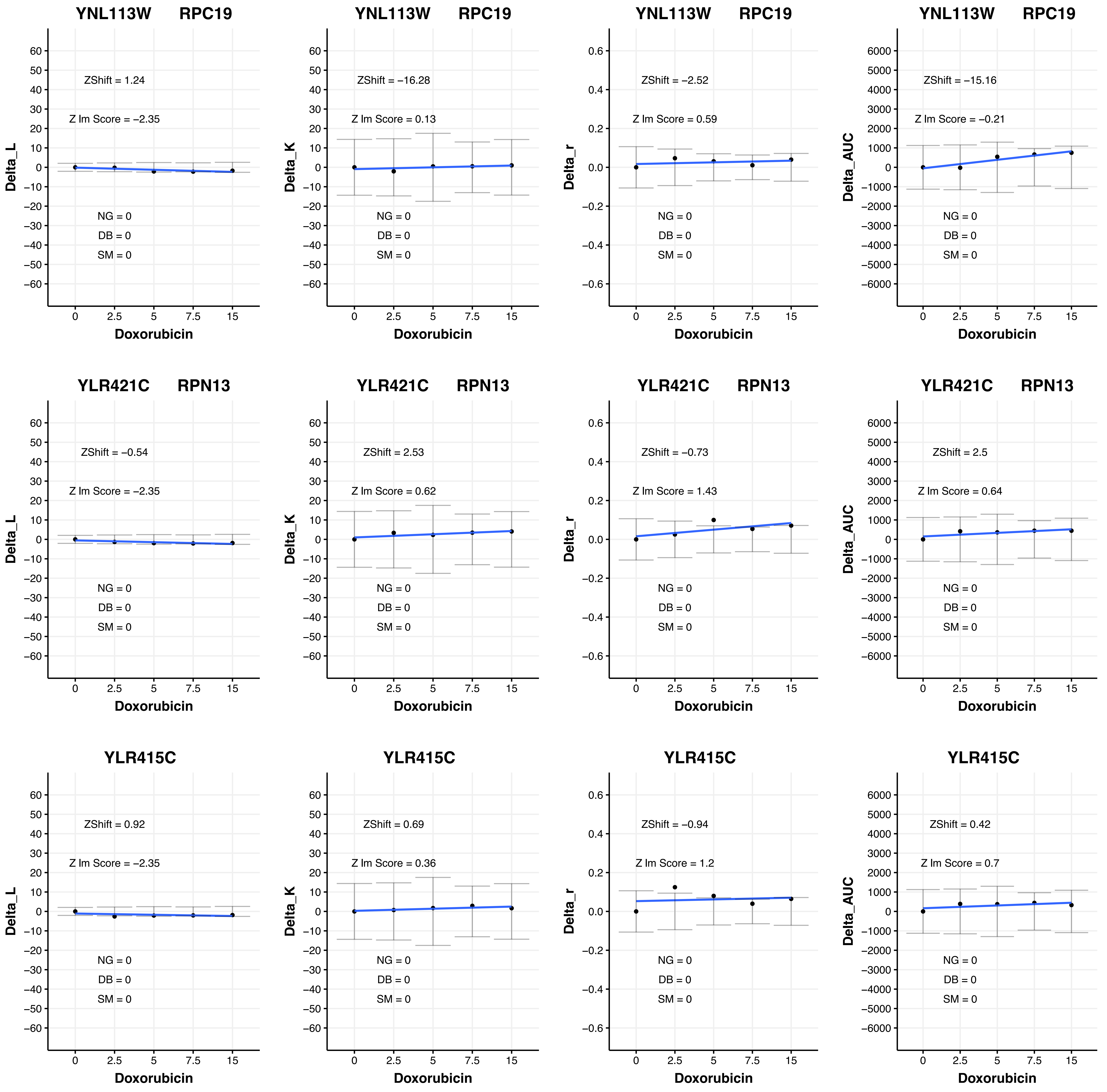

YNL113W RPC19
ZShift = 1.24
Z lm Score = −2.35
NG = 0
DB = 0
SM = 0
Delta_L
Doxorubicin
YNL113W RPC19
ZShift = −16.28
Z lm Score = 0.13
NG = 0
DB = 0
SM = 0
Delta_K
Doxorubicin
YNL113W RPC19
ZShift = −2.52
Z lm Score = 0.59
NG = 0
DB = 0
SM = 0
Delta_r
Doxorubicin
YNL113W RPC19
ZShift = −15.16
Z lm Score = −0.21
NG = 0
DB = 0
SM = 0
Delta_AUC
Doxorubicin
YLR421C RPN13
ZShift = −0.54
Z lm Score = −2.35
NG = 0
DB = 0
SM = 0
Delta_L
Doxorubicin
YLR421C RPN13
ZShift = 2.53
Z lm Score = 0.62
NG = 0
DB = 0
SM = 0
Delta_K
Doxorubicin
YLR421C RPN13
ZShift = −0.73
Z lm Score = 1.43
NG = 0
DB = 0
SM = 0
Delta_r
Doxorubicin
YLR421C RPN13
ZShift = 2.5
Z lm Score = 0.64
NG = 0
DB = 0
SM = 0
Delta_AUC
Doxorubicin
YLR415C
ZShift = 0.92
Z lm Score = −2.35
NG = 0
DB = 0
SM = 0
Delta_L
Doxorubicin
YLR415C
ZShift = 0.69
Z lm Score = 0.36
NG = 0
DB = 0
SM = 0
Delta_K
Doxorubicin
YLR415C
ZShift = −0.94
Z lm Score = 1.2
NG = 0
DB = 0
SM = 0
Delta_r
Doxorubicin
YLR415C
ZShift = 0.42
Z lm Score = 0.7
NG = 0
DB = 0
SM = 0
Delta_AUC
Doxorubicin

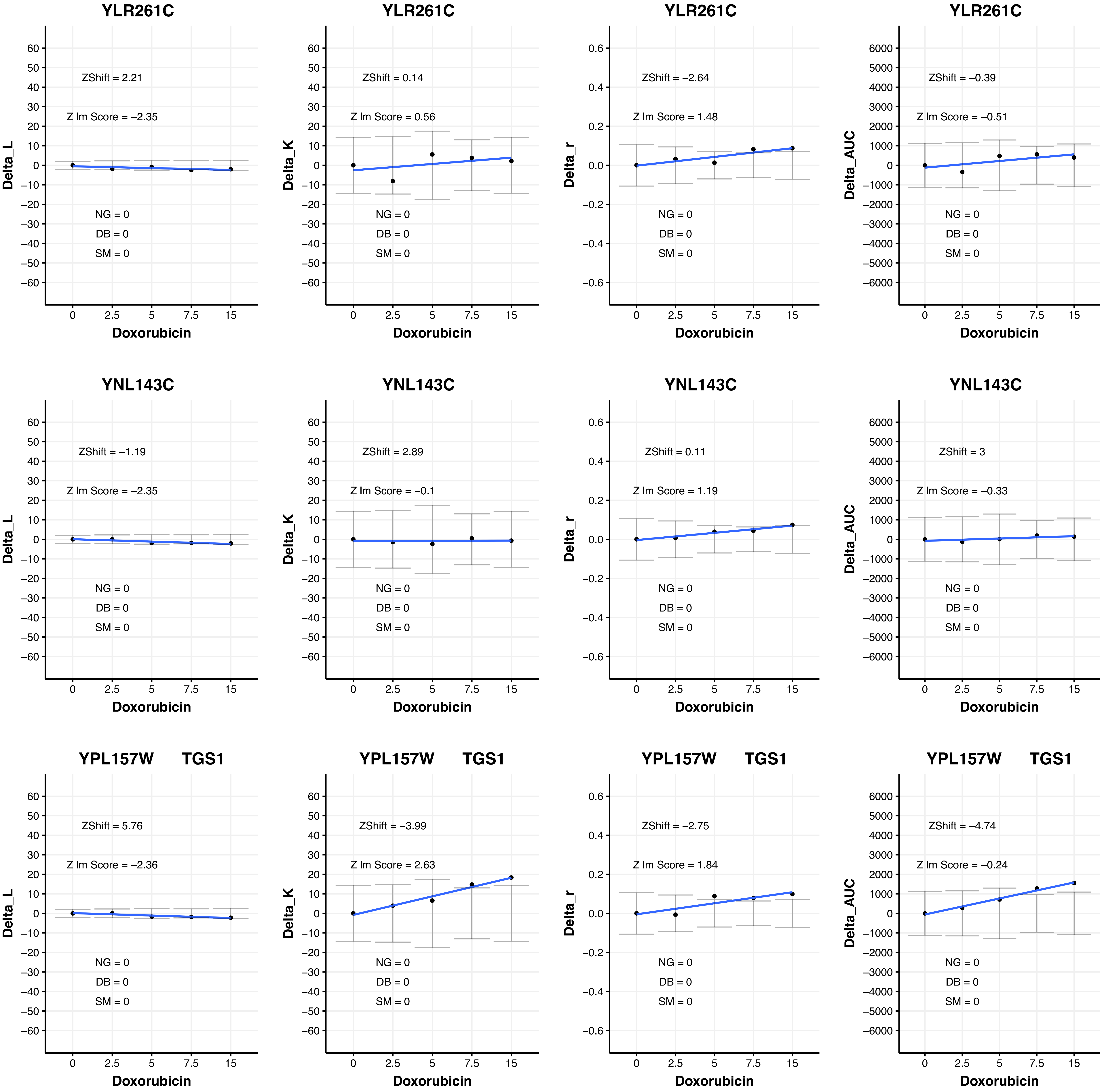
YLR261C
ZShift = 2.21
Z lm Score = −2.35
NG = 0
DB = 0
SM = 0
Delta_L
Doxorubicin
YLR261C
ZShift = 0.14
Z lm Score = 0.56
NG = 0
DB = 0
SM = 0
Delta_K
Doxorubicin
YLR261C
ZShift = −2.64
Z lm Score = 1.48
NG = 0
DB = 0
SM = 0
Delta_r
Doxorubicin
YLR261C
ZShift = −0.39
Z lm Score = −0.51
NG = 0
DB = 0
SM = 0
Delta_AUC
Doxorubicin
YNL143C
ZShift = −1.19
Z lm Score = −2.35
NG = 0
DB = 0
SM = 0
Delta_L
Doxorubicin
YNL143C
ZShift = 2.89
Z lm Score = −0.1
NG = 0
DB = 0
SM = 0
Delta_K
Doxorubicin
YNL143C
ZShift = 0.11
Z lm Score = 1.19
NG = 0
DB = 0
SM = 0
Delta_r
Doxorubicin
YNL143C
ZShift = 3
Z lm Score = −0.33
NG = 0
DB = 0
SM = 0
Delta_AUC
Doxorubicin
YPL157W TGS1
ZShift = 5.76
Z lm Score = −2.36
NG = 0
DB = 0
SM = 0
Delta_L
Doxorubicin
YPL157W TGS1
ZShift = −3.99
Z lm Score = 2.63
NG = 0
DB = 0
SM = 0
Delta_K
Doxorubicin
YPL157W TGS1
ZShift = −2.75
Z lm Score = 1.84
NG = 0
DB = 0
SM = 0
Delta_r
Doxorubicin
YPL157W TGS1
ZShift = −4.74
Z lm Score = −0.24
NG = 0
DB = 0
SM = 0
Delta_AUC
Doxorubicin

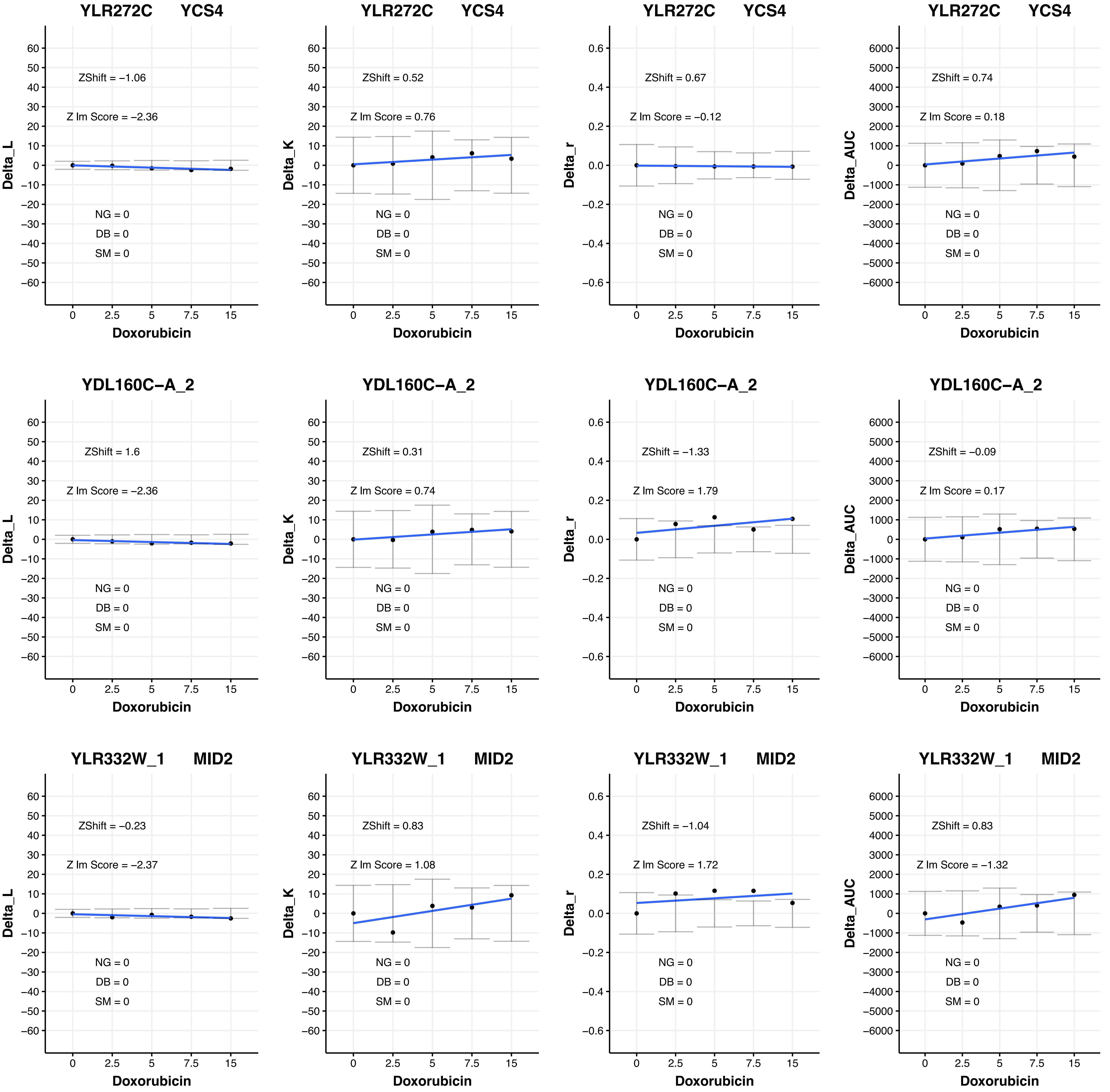

YLR272C YCS4
ZShift = −1.06
Z lm Score = −2.36
NG = 0
DB = 0
SM = 0
Delta_L
Doxorubicin
YLR272C YCS4
ZShift = 0.52
Z lm Score = 0.76
NG = 0
DB = 0
SM = 0
Delta_K
Doxorubicin
YLR272C YCS4
ZShift = 0.67
Z lm Score = −0.12
NG = 0
DB = 0
SM = 0
Delta_r
Doxorubicin
YLR272C YCS4
ZShift = 0.74
Z lm Score = 0.18
NG = 0
DB = 0
SM = 0
Delta_AUC
Doxorubicin
YDL160C−A_2
ZShift = 1.6
Z lm Score = −2.36
NG = 0
DB = 0
SM = 0
Delta_L
Doxorubicin
YDL160C−A_2
ZShift = 0.31
Z lm Score = 0.74
NG = 0
DB = 0
SM = 0
Delta_K
Doxorubicin
YDL160C−A_2
ZShift = −1.33
Z lm Score = 1.79
NG = 0
DB = 0
SM = 0
Delta_r
Doxorubicin
YDL160C−A_2
ZShift = −0.09
Z lm Score = 0.17
NG = 0
DB = 0
SM = 0
Delta_AUC
Doxorubicin
YLR332W_1 MID2
ZShift = −0.23
Z lm Score = −2.37
NG = 0
DB = 0
SM = 0
Delta_L
Doxorubicin
YLR332W_1 MID2
ZShift = 0.83
Z lm Score = 1.08
NG = 0
DB = 0
SM = 0
Delta_K
Doxorubicin
YLR332W_1 MID2
ZShift = −1.04
Z lm Score = 1.72
NG = 0
DB = 0
SM = 0
Delta_r
Doxorubicin
YLR332W_1 MID2
ZShift = 0.83
Z lm Score = −1.32
NG = 0
DB = 0
SM = 0
Delta_AUC
Doxorubicin

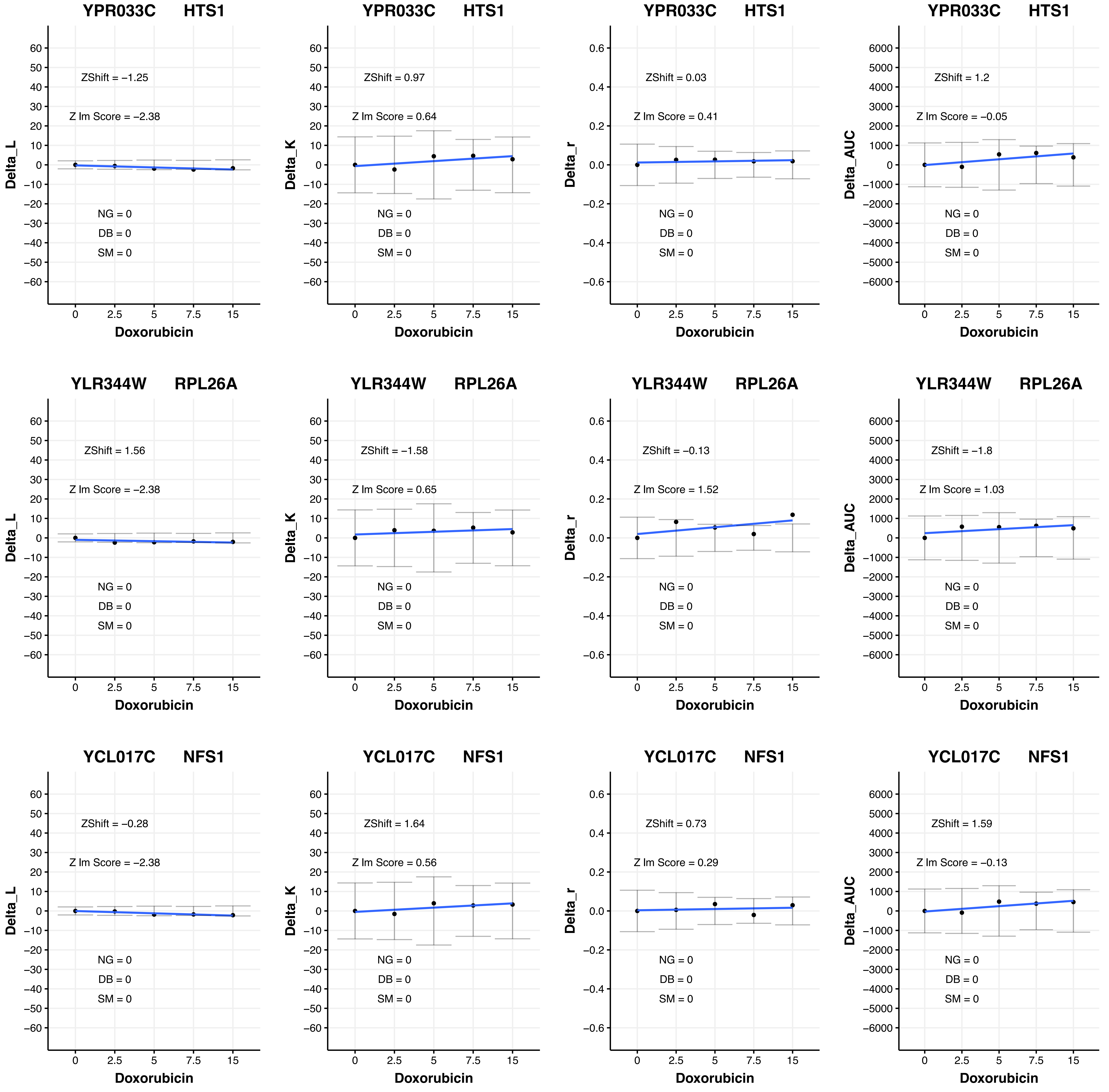
YPR033C HTS1
ZShift = −1.25
Z lm Score = −2.38
NG = 0
DB = 0
SM = 0
Delta_L
Doxorubicin
YPR033C HTS1
ZShift = 0.97
Z lm Score = 0.64
NG = 0
DB = 0
SM = 0
Delta_K
Doxorubicin
YPR033C HTS1
ZShift = 0.03
Z lm Score = 0.41
NG = 0
DB = 0
SM = 0
Delta_r
Doxorubicin
YPR033C HTS1
ZShift = 1.2
Z lm Score = −0.05
NG = 0
DB = 0
SM = 0
Delta_AUC
Doxorubicin
YLR344W RPL26A
ZShift = 1.56
Z lm Score = −2.38
NG = 0
DB = 0
SM = 0
Delta_L
Doxorubicin
YLR344W RPL26A
ZShift = −1.58
Z lm Score = 0.65
NG = 0
DB = 0
SM = 0
Delta_K
Doxorubicin
YLR344W RPL26A
ZShift = −0.13
Z lm Score = 1.52
NG = 0
DB = 0
SM = 0
Delta_r
Doxorubicin
YLR344W RPL26A
ZShift = −1.8
Z lm Score = 1.03
NG = 0
DB = 0
SM = 0
Delta_AUC
Doxorubicin
YCL017C NFS1
ZShift = −0.28
Z lm Score = −2.38
NG = 0
DB = 0
SM = 0
Delta_L
Doxorubicin
YCL017C NFS1
ZShift = 1.64
Z lm Score = 0.56
NG = 0
DB = 0
SM = 0
Delta_K
Doxorubicin
YCL017C NFS1
ZShift = 0.73
Z lm Score = 0.29
NG = 0
DB = 0
SM = 0
Delta_r
Doxorubicin
YCL017C NFS1
ZShift = 1.59
Z lm Score = −0.13
NG = 0
DB = 0
SM = 0
Delta_AUC
Doxorubicin

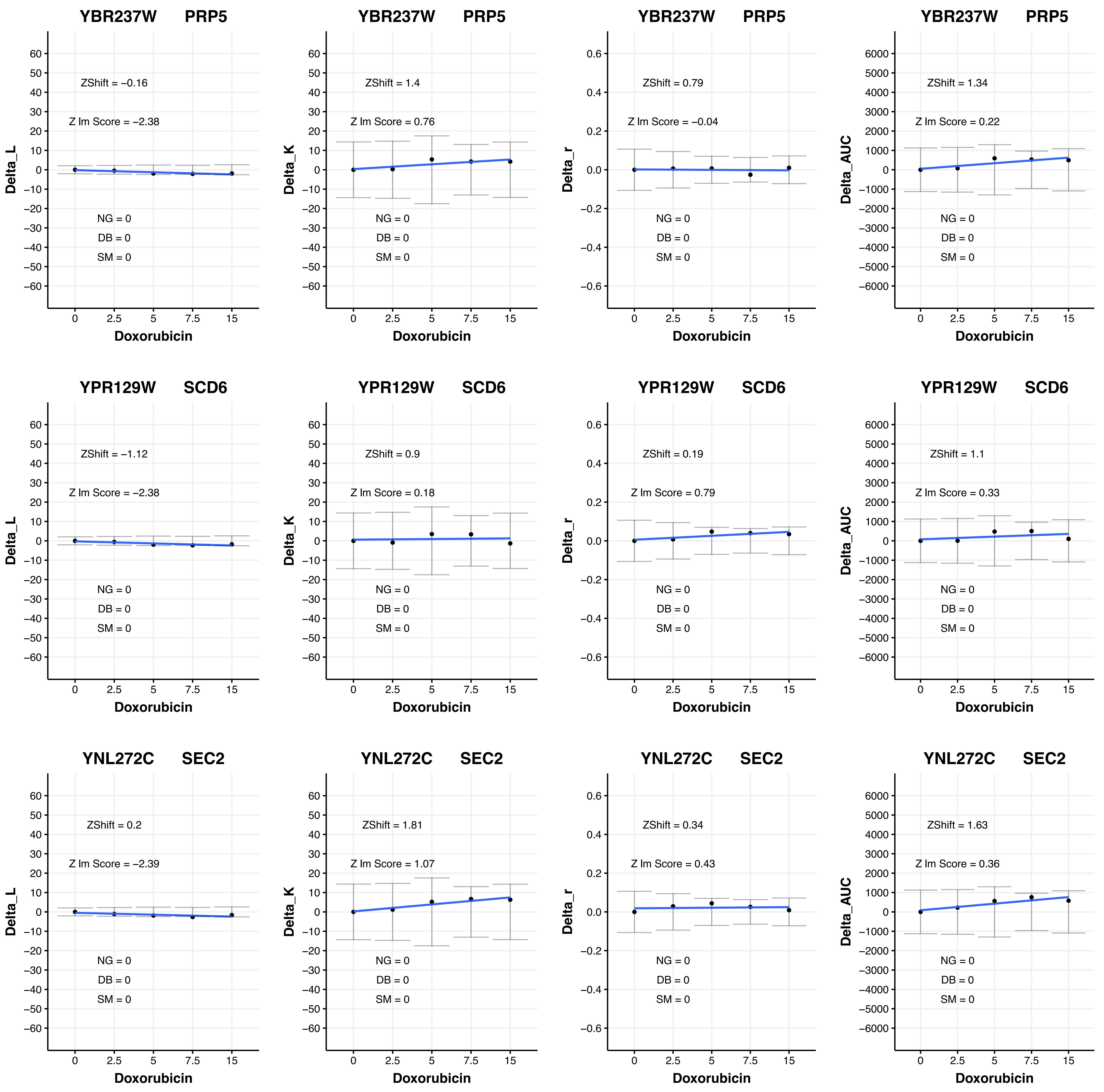
YBR237W PRP5
ZShift = −0.16
Z lm Score = −2.38
NG = 0
DB = 0
SM = 0
Delta_L
Doxorubicin
YBR237W PRP5
ZShift = 1.4
Z lm Score = 0.76
NG = 0
DB = 0
SM = 0
Delta_K
Doxorubicin
YBR237W PRP5
ZShift = 0.79
Z lm Score = −0.04
NG = 0
DB = 0
SM = 0
Delta_r
Doxorubicin
YBR237W PRP5
ZShift = 1.34
Z lm Score = 0.22
NG = 0
DB = 0
SM = 0
Delta_AUC
Doxorubicin
YPR129W SCD6
ZShift = −1.12
Z lm Score = −2.38
NG = 0
DB = 0
SM = 0
Delta_L
Doxorubicin
YPR129W SCD6
ZShift = 0.9
Z lm Score = 0.18
NG = 0
DB = 0
SM = 0
Delta_K
Doxorubicin
YPR129W SCD6
ZShift = 0.19
Z lm Score = 0.79
NG = 0
DB = 0
SM = 0
Delta_r
Doxorubicin
YPR129W SCD6
ZShift = 1.1
Z lm Score = 0.33
NG = 0
DB = 0
SM = 0
Delta_AUC
Doxorubicin
YNL272C SEC2
ZShift = 0.2
Z lm Score = −2.39
NG = 0
DB = 0
SM = 0
Delta_L
Doxorubicin
YNL272C SEC2
ZShift = 1.81
Z lm Score = 1.07
NG = 0
DB = 0
SM = 0
Delta_K
Doxorubicin
YNL272C SEC2
ZShift = 0.34
Z lm Score = 0.43
NG = 0
DB = 0
SM = 0
Delta_r
Doxorubicin
YNL272C SEC2
ZShift = 1.63
Z lm Score = 0.36
NG = 0
DB = 0
SM = 0
Delta_AUC
Doxorubicin

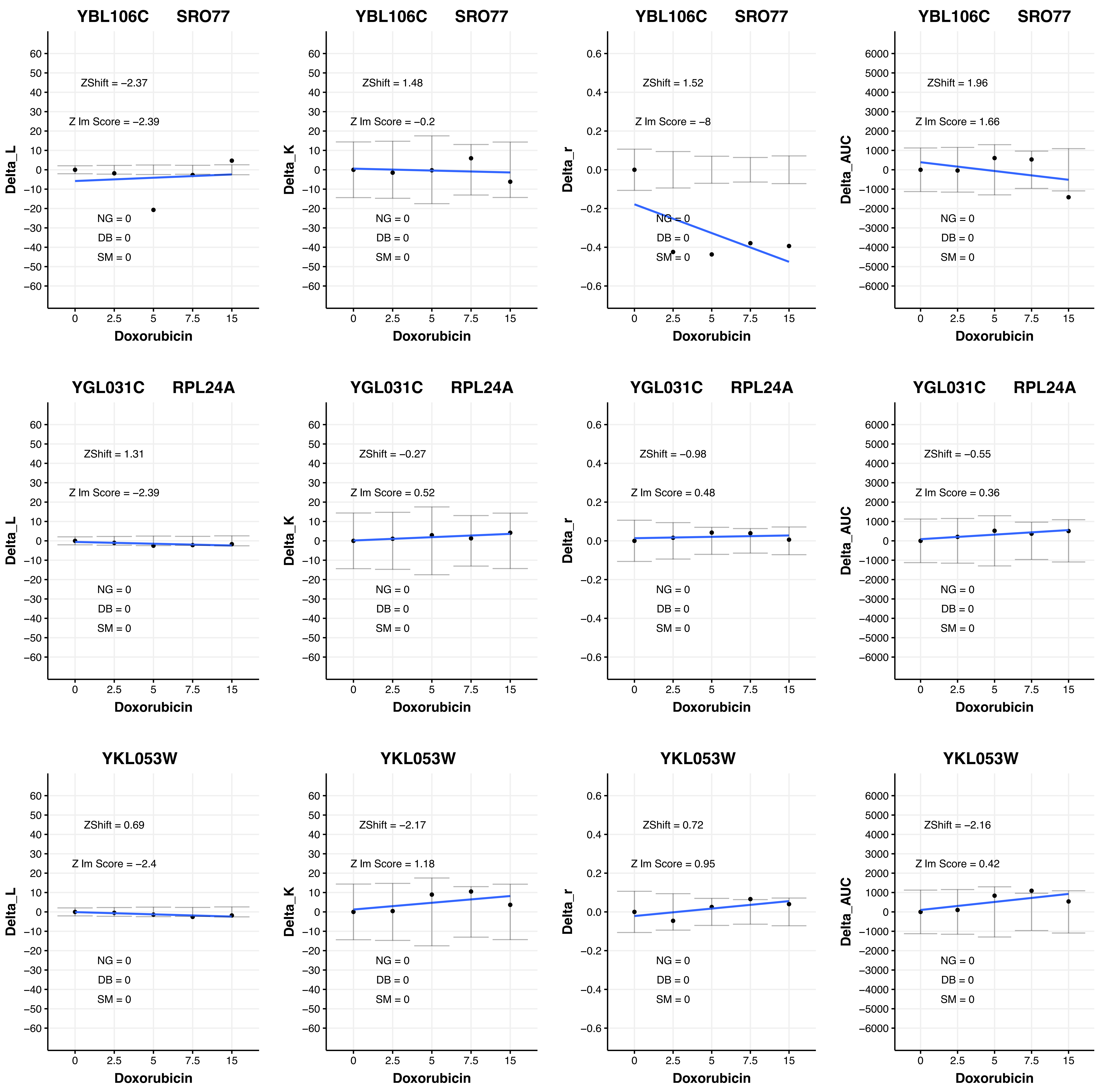

YBL106C SRO77
ZShift = −2.37
Z lm Score = −2.39
NG = 0
DB = 0
SM = 0
Delta_L
Doxorubicin
ZShift = 1.48
Z lm Score = −0.2
Delta_K
ZShift = 1.52
Z lm Score = −8
Delta_r
ZShift = 1.96
Z lm Score = 1.66
Delta_AUC
YGL031C RPL24A
ZShift = 1.31
Z lm Score = −2.39
ZShift = −0.27
Z lm Score = 0.52
ZShift = −0.98
Z lm Score = 0.48
ZShift = −0.55
Z lm Score = 0.36
YKL053W
ZShift = 0.69
Z lm Score = −2.4
ZShift = −2.17
Z lm Score = 1.18
ZShift = 0.72
Z lm Score = 0.95
ZShift = −2.16
Z lm Score = 0.42

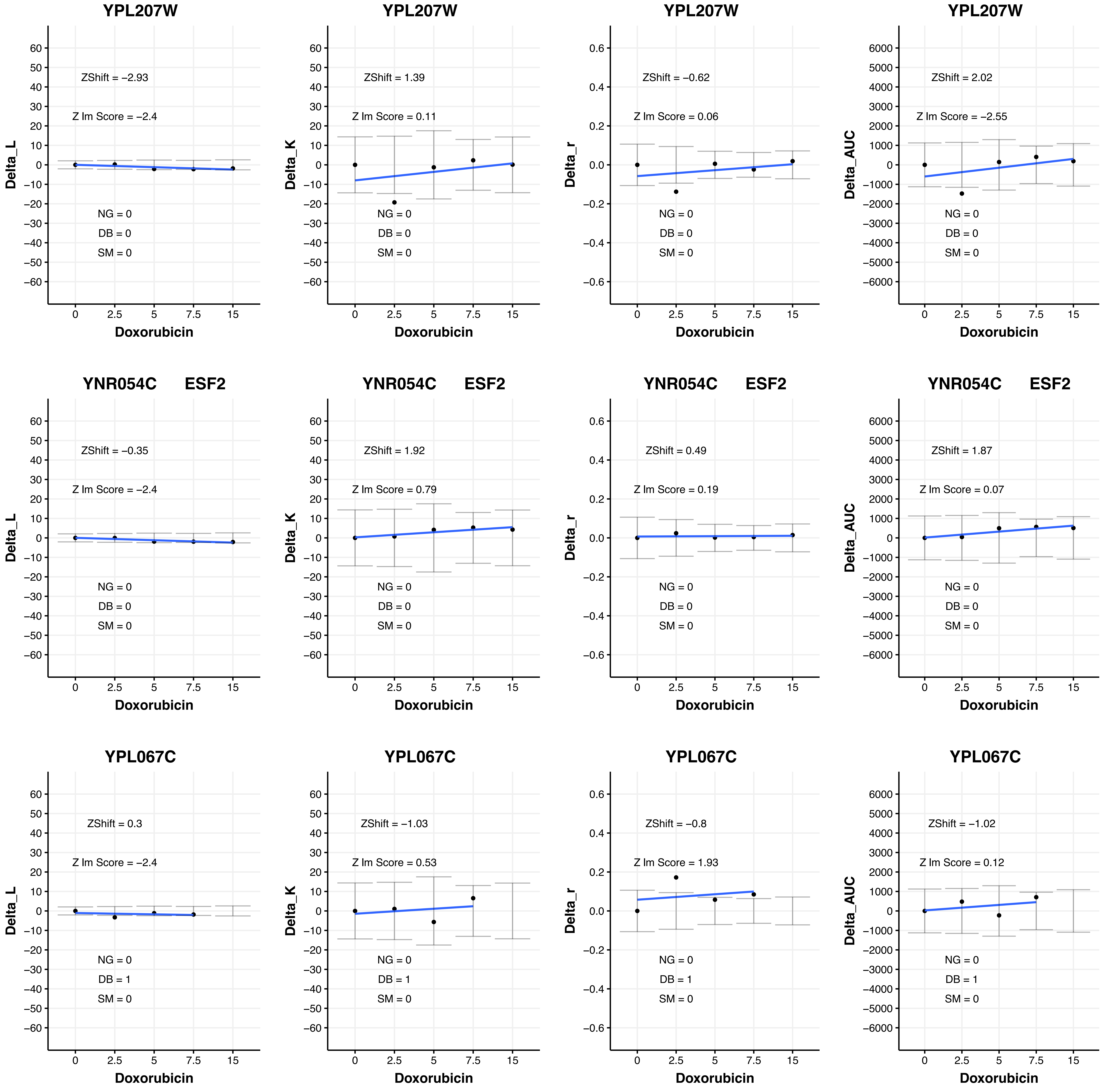
YPL207W
ZShift = −2.93
Z lm Score = −2.4
NG = 0
DB = 0
SM = 0
Delta_L
Doxorubicin
YPL207W
ZShift = 1.39
Z lm Score = 0.11
NG = 0
DB = 0
SM = 0
Delta_K
Doxorubicin
YPL207W
ZShift = −0.62
Z lm Score = 0.06
NG = 0
DB = 0
SM = 0
Delta_r
Doxorubicin
YPL207W
ZShift = 2.02
Z lm Score = −2.55
NG = 0
DB = 0
SM = 0
Delta_AUC
Doxorubicin
YNR054C ESF2
ZShift = −0.35
Z lm Score = −2.4
NG = 0
DB = 0
SM = 0
Delta_L
Doxorubicin
YNR054C ESF2
ZShift = 1.92
Z lm Score = 0.79
NG = 0
DB = 0
SM = 0
Delta_K
Doxorubicin
YNR054C ESF2
ZShift = 0.49
Z lm Score = 0.19
NG = 0
DB = 0
SM = 0
Delta_r
Doxorubicin
YNR054C ESF2
ZShift = 1.87
Z lm Score = 0.07
NG = 0
DB = 0
SM = 0
Delta_AUC
Doxorubicin
YPL067C
ZShift = 0.3
Z lm Score = −2.4
NG = 0
DB = 1
SM = 0
Delta_L
Doxorubicin
YPL067C
ZShift = −1.03
Z lm Score = 0.53
NG = 0
DB = 1
SM = 0
Delta_K
Doxorubicin
YPL067C
ZShift = −0.8
Z lm Score = 1.93
NG = 0
DB = 1
SM = 0
Delta_r
Doxorubicin
YPL067C
ZShift = −1.02
Z lm Score = 0.12
NG = 0
DB = 1
SM = 0
Delta_AUC
Doxorubicin

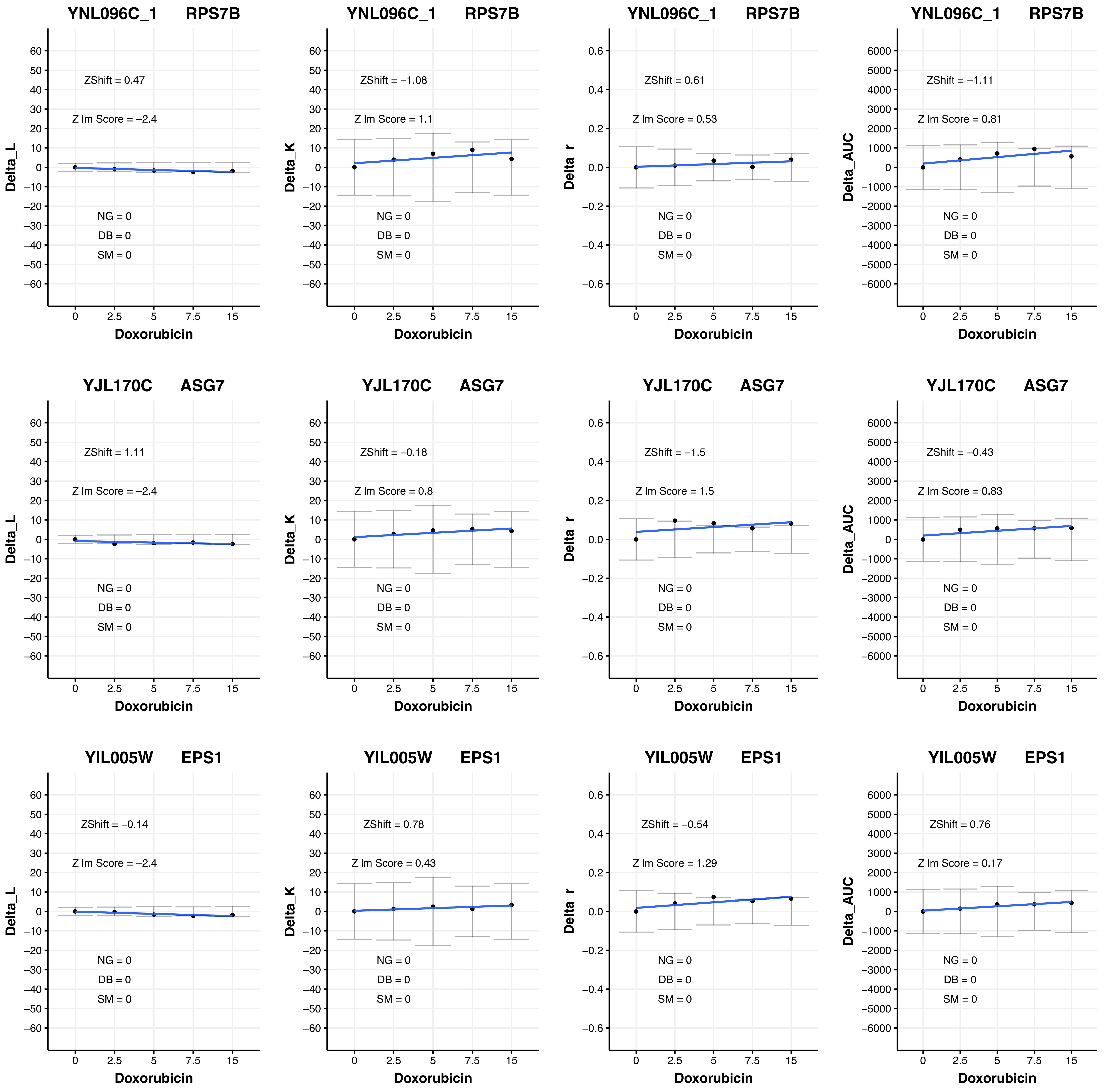
YNL096C_1 RPS7B
ZShift = 0.47
Z lm Score = −2.4
NG = 0
DB = 0
SM = 0
Delta_L
Doxorubicin
YNL096C_1 RPS7B
ZShift = −1.08
Z lm Score = 1.1
NG = 0
DB = 0
SM = 0
Delta_K
Doxorubicin
YNL096C_1 RPS7B
ZShift = 0.61
Z lm Score = 0.53
NG = 0
DB = 0
SM = 0
Delta_r
Doxorubicin
YNL096C_1 RPS7B
ZShift = −1.11
Z lm Score = 0.81
NG = 0
DB = 0
SM = 0
Delta_AUC
Doxorubicin
YJL170C ASG7
ZShift = 1.11
Z lm Score = −2.4
NG = 0
DB = 0
SM = 0
Delta_L
Doxorubicin
YJL170C ASG7
ZShift = −0.18
Z lm Score = 0.8
NG = 0
DB = 0
SM = 0
Delta_K
Doxorubicin
YJL170C ASG7
ZShift = −1.5
Z lm Score = 1.5
NG = 0
DB = 0
SM = 0
Delta_r
Doxorubicin
YJL170C ASG7
ZShift = −0.43
Z lm Score = 0.83
NG = 0
DB = 0
SM = 0
Delta_AUC
Doxorubicin
YIL005W EPS1
ZShift = −0.14
Z lm Score = −2.4
NG = 0
DB = 0
SM = 0
Delta_L
Doxorubicin
YIL005W EPS1
ZShift = 0.78
Z lm Score = 0.43
NG = 0
DB = 0
SM = 0
Delta_K
Doxorubicin
YIL005W EPS1
ZShift = −0.54
Z lm Score = 1.29
NG = 0
DB = 0
SM = 0
Delta_r
Doxorubicin
YIL005W EPS1
ZShift = 0.76
Z lm Score = 0.17
NG = 0
DB = 0
SM = 0
Delta_AUC
Doxorubicin

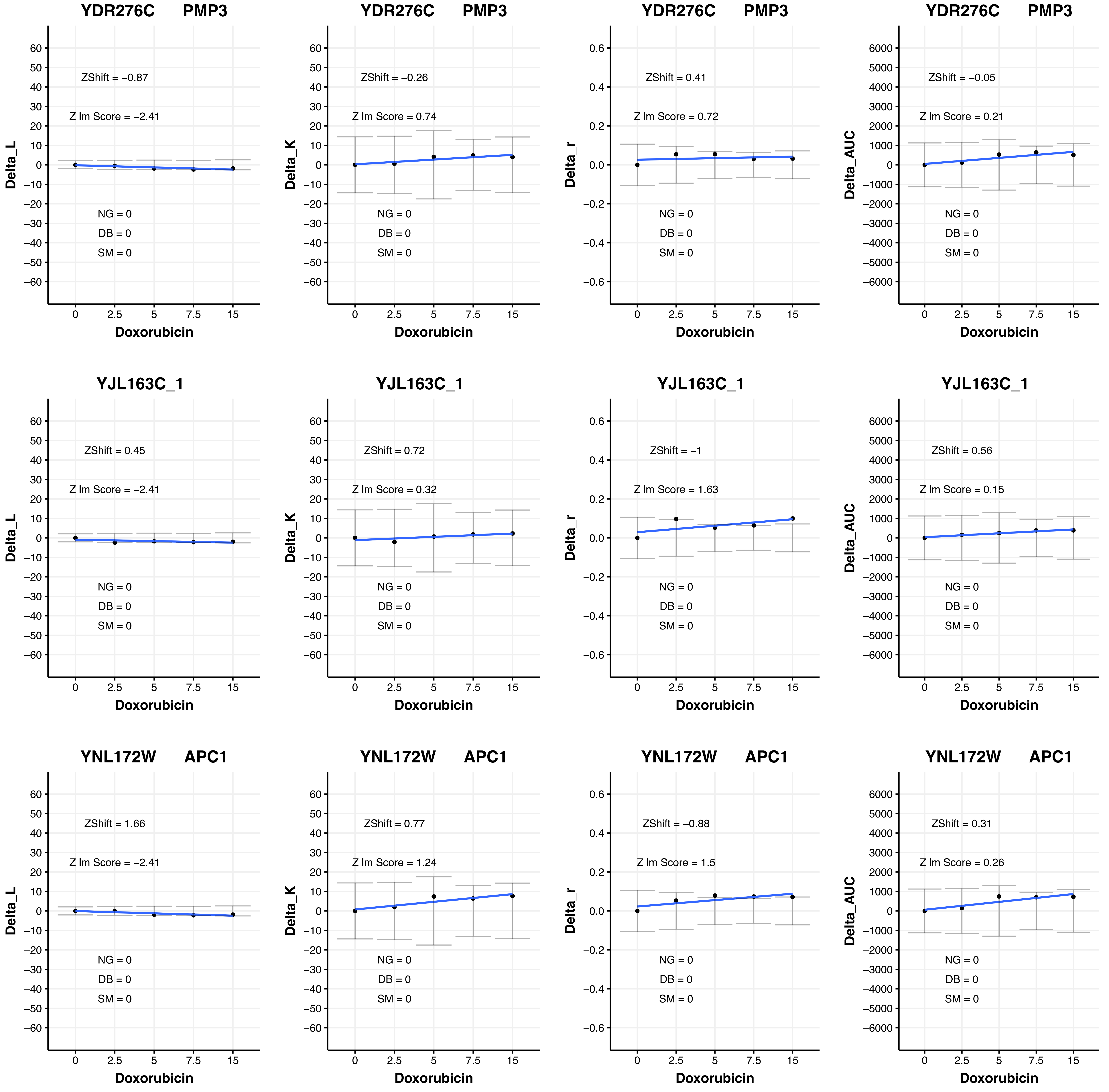
YDR276C PMP3
ZShift = −0.87
Z lm Score = −2.41
NG = 0
DB = 0
SM = 0
Delta_L
Doxorubicin
YDR276C PMP3
ZShift = −0.26
Z lm Score = 0.74
NG = 0
DB = 0
SM = 0
Delta_K
Doxorubicin
YDR276C PMP3
ZShift = 0.41
Z lm Score = 0.72
NG = 0
DB = 0
SM = 0
Delta_r
Doxorubicin
YDR276C PMP3
ZShift = −0.05
Z lm Score = 0.21
NG = 0
DB = 0
SM = 0
Delta_AUC
Doxorubicin
YJL163C_1
ZShift = 0.45
Z lm Score = −2.41
NG = 0
DB = 0
SM = 0
Delta_L
Doxorubicin
YJL163C_1
ZShift = 0.72
Z lm Score = 0.32
NG = 0
DB = 0
SM = 0
Delta_K
Doxorubicin
YJL163C_1
ZShift = −1
Z lm Score = 1.63
NG = 0
DB = 0
SM = 0
Delta_r
Doxorubicin
YJL163C_1
ZShift = 0.56
Z lm Score = 0.15
NG = 0
DB = 0
SM = 0
Delta_AUC
Doxorubicin
YNL172W APC1
ZShift = 1.66
Z lm Score = −2.41
NG = 0
DB = 0
SM = 0
Delta_L
Doxorubicin
YNL172W APC1
ZShift = 0.77
Z lm Score = 1.24
NG = 0
DB = 0
SM = 0
Delta_K
Doxorubicin
YNL172W APC1
ZShift = −0.88
Z lm Score = 1.5
NG = 0
DB = 0
SM = 0
Delta_r
Doxorubicin
YNL172W APC1
ZShift = 0.31
Z lm Score = 0.26
NG = 0
DB = 0
SM = 0
Delta_AUC
Doxorubicin

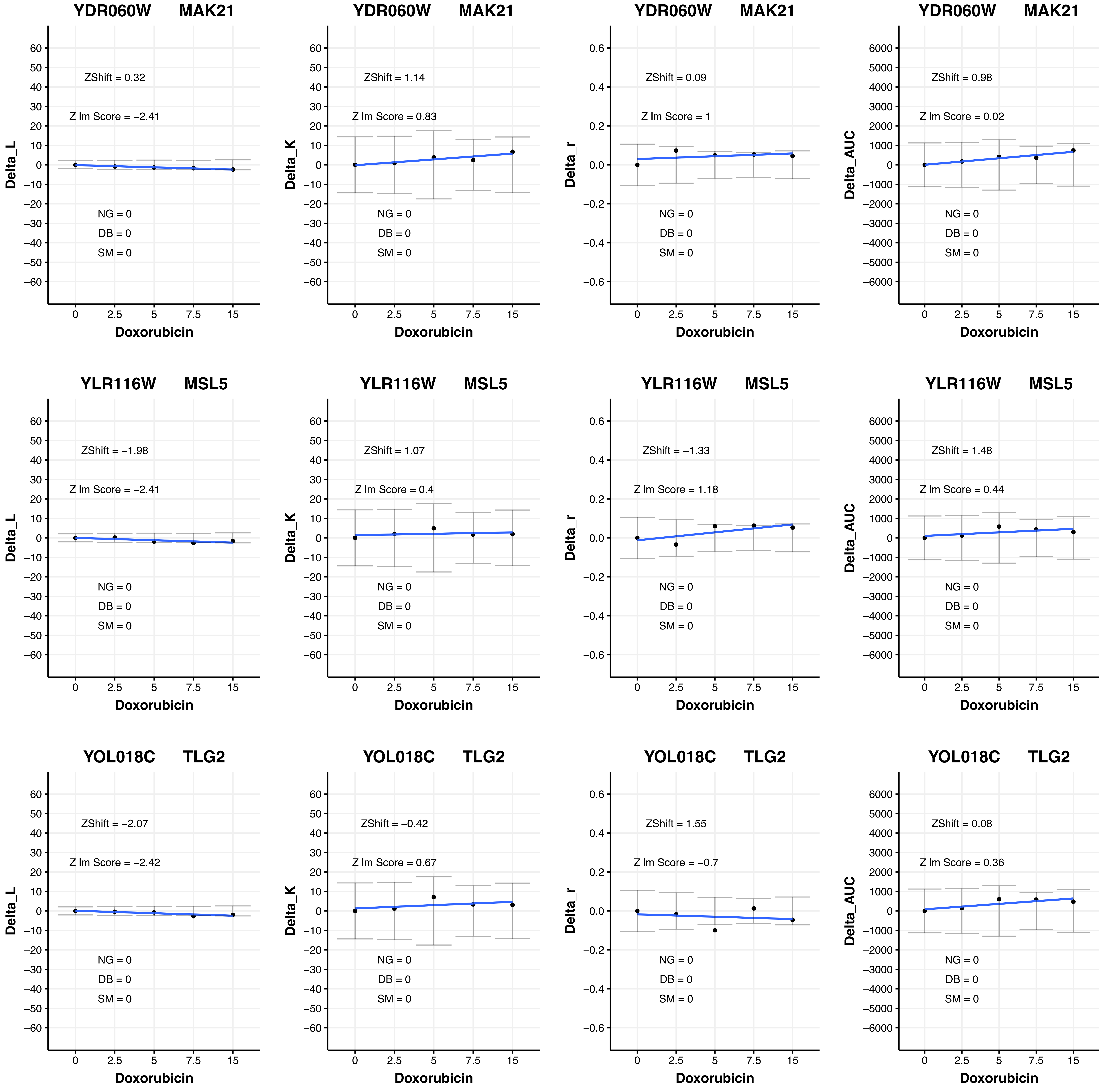
YDR060W MAK21
ZShift = 0.32
Z lm Score = −2.41
NG = 0
DB = 0
SM = 0
Delta_L
Doxorubicin
YDR060W MAK21
ZShift = 1.14
Z lm Score = 0.83
NG = 0
DB = 0
SM = 0
Delta_K
Doxorubicin
YDR060W MAK21
ZShift = 0.09
Z lm Score = 1
NG = 0
DB = 0
SM = 0
Delta_r
Doxorubicin
YDR060W MAK21
ZShift = 0.98
Z lm Score = 0.02
NG = 0
DB = 0
SM = 0
Delta_AUC
Doxorubicin
YLR116W MSL5
ZShift = −1.98
Z lm Score = −2.41
NG = 0
DB = 0
SM = 0
Delta_L
Doxorubicin
YLR116W MSL5
ZShift = 1.07
Z lm Score = 0.4
NG = 0
DB = 0
SM = 0
Delta_K
Doxorubicin
YLR116W MSL5
ZShift = −1.33
Z lm Score = 1.18
NG = 0
DB = 0
SM = 0
Delta_r
Doxorubicin
YLR116W MSL5
ZShift = 1.48
Z lm Score = 0.44
NG = 0
DB = 0
SM = 0
Delta_AUC
Doxorubicin
YOL018C TLG2
ZShift = −2.07
Z lm Score = −2.42
NG = 0
DB = 0
SM = 0
Delta_L
Doxorubicin
YOL018C TLG2
ZShift = −0.42
Z lm Score = 0.67
NG = 0
DB = 0
SM = 0
Delta_K
Doxorubicin
YOL018C TLG2
ZShift = 1.55
Z lm Score = −0.7
NG = 0
DB = 0
SM = 0
Delta_r
Doxorubicin
YOL018C TLG2
ZShift = 0.08
Z lm Score = 0.36
NG = 0
DB = 0
SM = 0
Delta_AUC
Doxorubicin

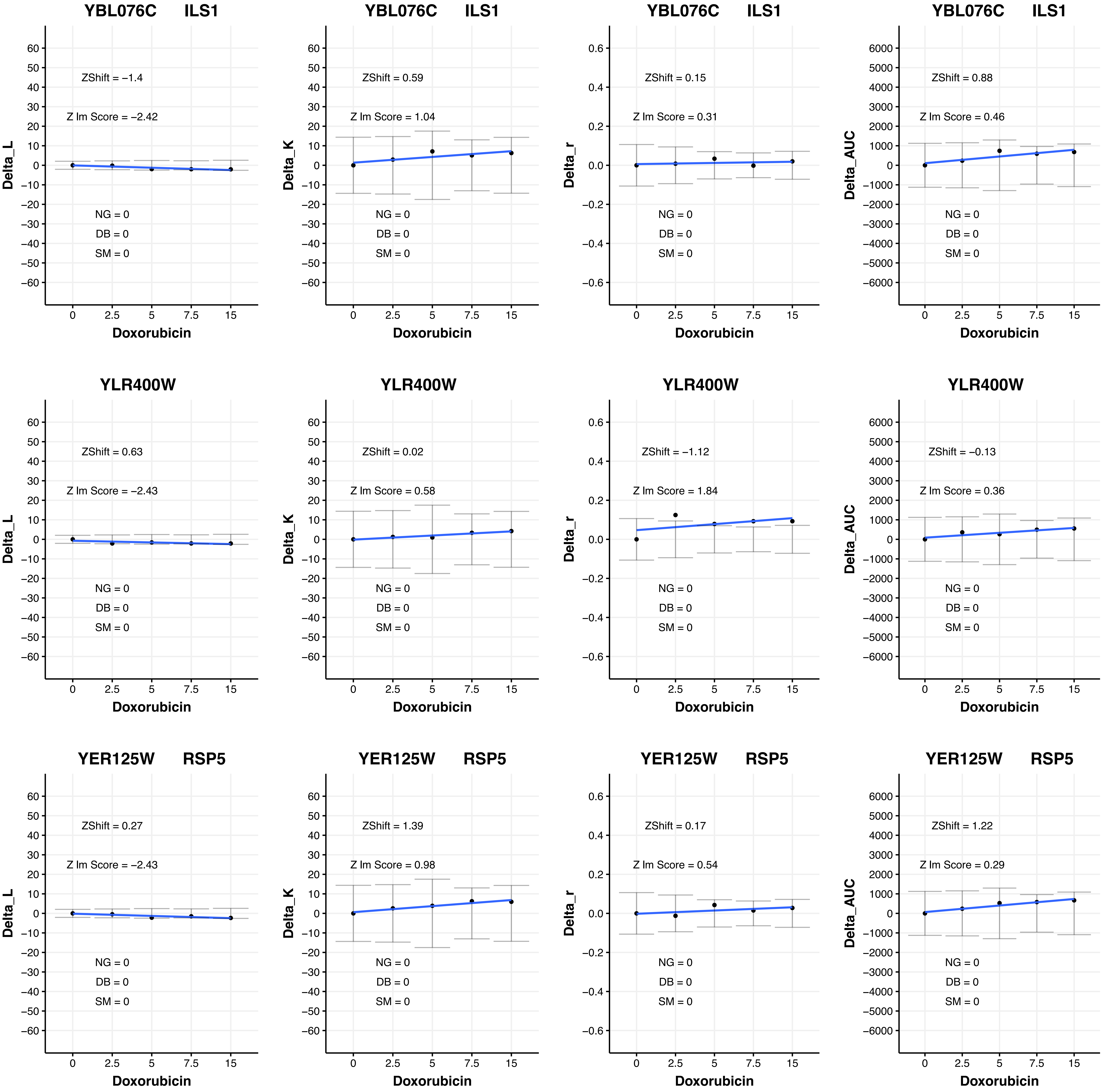
YBL076C ILS1
ZShift = −1.4
Z lm Score = −2.42
NG = 0
DB = 0
SM = 0
Delta_L
YBL076C ILS1
ZShift = 0.59
Z lm Score = 1.04
NG = 0
DB = 0
SM = 0
Delta_K
YBL076C ILS1
ZShift = 0.15
Z lm Score = 0.31
NG = 0
DB = 0
SM = 0
Delta_r
YBL076C ILS1
ZShift = 0.88
Z lm Score = 0.46
NG = 0
DB = 0
SM = 0
Delta_AUC
YLR400W
ZShift = 0.63
Z lm Score = −2.43
NG = 0
DB = 0
SM = 0
Delta_L
YLR400W
ZShift = 0.02
Z lm Score = 0.58
NG = 0
DB = 0
SM = 0
Delta_K
YLR400W
ZShift = −1.12
Z lm Score = 1.84
NG = 0
DB = 0
SM = 0
Delta_r
YLR400W
ZShift = −0.13
Z lm Score = 0.36
NG = 0
DB = 0
SM = 0
Delta_AUC
YER125W RSP5
ZShift = 0.27
Z lm Score = −2.43
NG = 0
DB = 0
SM = 0
Delta_L
YER125W RSP5
ZShift = 1.39
Z lm Score = 0.98
NG = 0
DB = 0
SM = 0
Delta_K
YER125W RSP5
ZShift = 0.17
Z lm Score = 0.54
NG = 0
DB = 0
SM = 0
Delta_r
YER125W RSP5
ZShift = 1.22
Z lm Score = 0.29
NG = 0
DB = 0
SM = 0
Delta_AUC
Doxorubicin
0
2.5
5
7.5
15

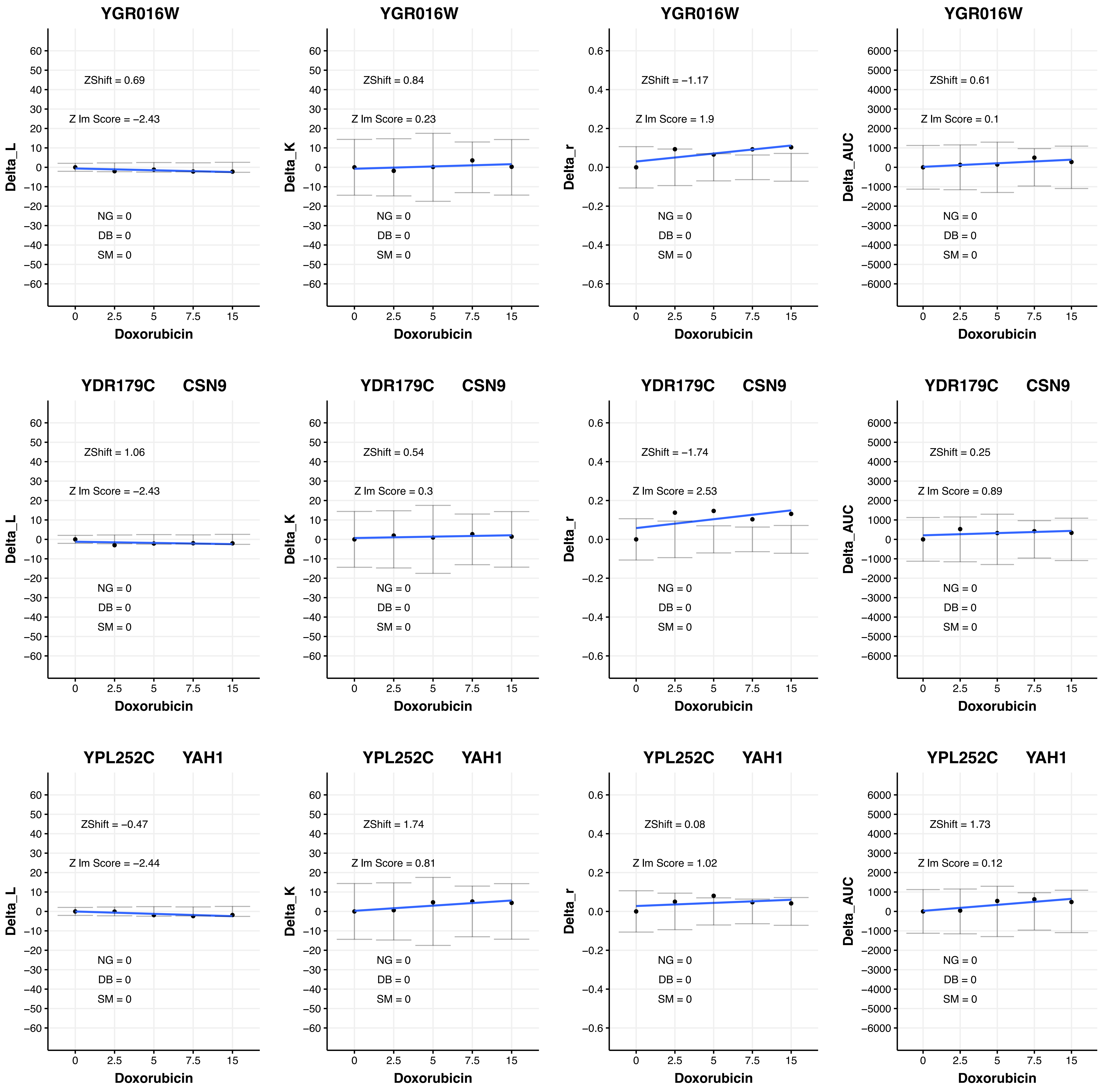
YGR016W
ZShift = 0.69
Z lm Score = −2.43
NG = 0
DB = 0
SM = 0
Delta_L
Doxorubicin
YGR016W
ZShift = 0.84
Z lm Score = 0.23
NG = 0
DB = 0
SM = 0
Delta_K
Doxorubicin
YGR016W
ZShift = −1.17
Z lm Score = 1.9
NG = 0
DB = 0
SM = 0
Delta_r
Doxorubicin
YGR016W
ZShift = 0.61
Z lm Score = 0.1
NG = 0
DB = 0
SM = 0
Delta_AUC
Doxorubicin
YDR179C CSN9
ZShift = 1.06
Z lm Score = −2.43
NG = 0
DB = 0
SM = 0
Delta_L
Doxorubicin
YDR179C CSN9
ZShift = 0.54
Z lm Score = 0.3
NG = 0
DB = 0
SM = 0
Delta_K
Doxorubicin
YDR179C CSN9
ZShift = −1.74
Z lm Score = 2.53
NG = 0
DB = 0
SM = 0
Delta_r
Doxorubicin
YDR179C CSN9
ZShift = 0.25
Z lm Score = 0.89
NG = 0
DB = 0
SM = 0
Delta_AUC
Doxorubicin
YPL252C YAH1
ZShift = −0.47
Z lm Score = −2.44
NG = 0
DB = 0
SM = 0
Delta_L
Doxorubicin
YPL252C YAH1
ZShift = 1.74
Z lm Score = 0.81
NG = 0
DB = 0
SM = 0
Delta_K
Doxorubicin
YPL252C YAH1
ZShift = 0.08
Z lm Score = 1.02
NG = 0
DB = 0
SM = 0
Delta_r
Doxorubicin
YPL252C YAH1
ZShift = 1.73
Z lm Score = 0.12
NG = 0
DB = 0
SM = 0
Delta_AUC
Doxorubicin

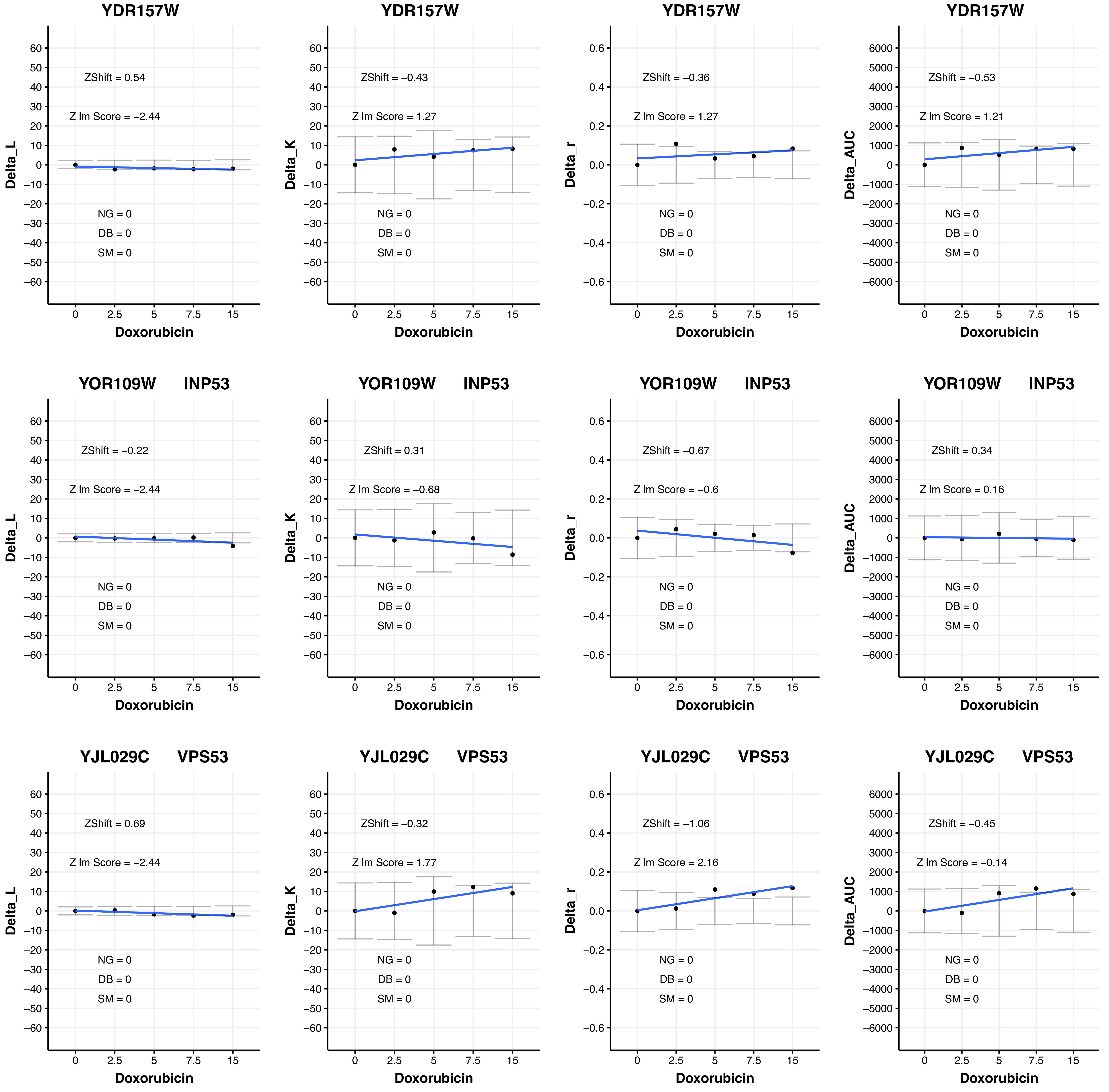

YDR157W
ZShift = 0.54
Z lm Score = −2.44
NG = 0
DB = 0
SM = 0
Delta_L
Doxorubicin
YDR157W
ZShift = −0.43
Z lm Score = 1.27
NG = 0
DB = 0
SM = 0
Delta_K
Doxorubicin
YDR157W
ZShift = −0.36
Z lm Score = 1.27
NG = 0
DB = 0
SM = 0
Delta_r
Doxorubicin
YDR157W
ZShift = −0.53
Z lm Score = 1.21
NG = 0
DB = 0
SM = 0
Delta_AUC
Doxorubicin
YOR109W INP53
ZShift = −0.22
Z lm Score = −2.44
NG = 0
DB = 0
SM = 0
Delta_L
Doxorubicin
YOR109W INP53
ZShift = 0.31
Z lm Score = −0.68
NG = 0
DB = 0
SM = 0
Delta_K
Doxorubicin
YOR109W INP53
ZShift = −0.67
Z lm Score = −0.6
NG = 0
DB = 0
SM = 0
Delta_r
Doxorubicin
YOR109W INP53
ZShift = 0.34
Z lm Score = 0.16
NG = 0
DB = 0
SM = 0
Delta_AUC
Doxorubicin
YJL029C VPS53
ZShift = 0.69
Z lm Score = −2.44
NG = 0
DB = 0
SM = 0
Delta_L
Doxorubicin
YJL029C VPS53
ZShift = −0.32
Z lm Score = 1.77
NG = 0
DB = 0
SM = 0
Delta_K
Doxorubicin
YJL029C VPS53
ZShift = −1.06
Z lm Score = 2.16
NG = 0
DB = 0
SM = 0
Delta_r
Doxorubicin
YJL029C VPS53
ZShift = −0.45
Z lm Score = −0.14
NG = 0
DB = 0
SM = 0
Delta_AUC
Doxorubicin

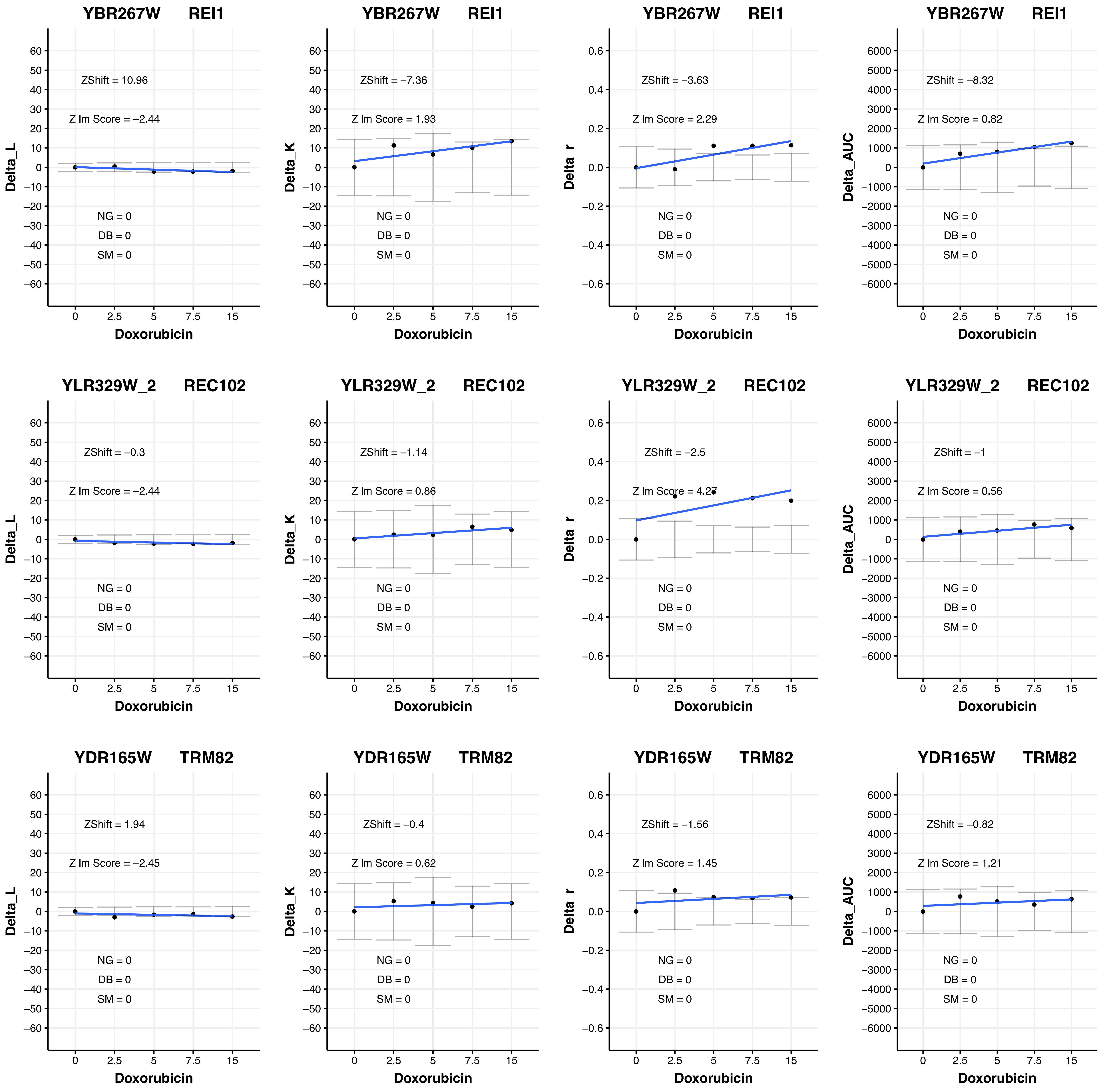
YBR267W REI1
ZShift = 10.96
Z lm Score = −2.44
NG = 0
DB = 0
SM = 0
Delta_L
Doxorubicin
YBR267W REI1
ZShift = −7.36
Z lm Score = 1.93
NG = 0
DB = 0
SM = 0
Delta_K
Doxorubicin
YBR267W REI1
ZShift = −3.63
Z lm Score = 2.29
NG = 0
DB = 0
SM = 0
Delta_r
Doxorubicin
YBR267W REI1
ZShift = −8.32
Z lm Score = 0.82
NG = 0
DB = 0
SM = 0
Delta_AUC
Doxorubicin
YLR329W_2 REC102
ZShift = −0.3
Z lm Score = −2.44
NG = 0
DB = 0
SM = 0
Delta_L
Doxorubicin
YLR329W_2 REC102
ZShift = −1.14
Z lm Score = 0.86
NG = 0
DB = 0
SM = 0
Delta_K
Doxorubicin
YLR329W_2 REC102
ZShift = −2.5
Z lm Score = 4.27
NG = 0
DB = 0
SM = 0
Delta_r
Doxorubicin
YLR329W_2 REC102
ZShift = −1
Z lm Score = 0.56
NG = 0
DB = 0
SM = 0
Delta_AUC
Doxorubicin
YDR165W TRM82
ZShift = 1.94
Z lm Score = −2.45
NG = 0
DB = 0
SM = 0
Delta_L
Doxorubicin
YDR165W TRM82
ZShift = −0.4
Z lm Score = 0.62
NG = 0
DB = 0
SM = 0
Delta_K
Doxorubicin
YDR165W TRM82
ZShift = −1.56
Z lm Score = 1.45
NG = 0
DB = 0
SM = 0
Delta_r
Doxorubicin
YDR165W TRM82
ZShift = −0.82
Z lm Score = 1.21
NG = 0
DB = 0
SM = 0
Delta_AUC
Doxorubicin

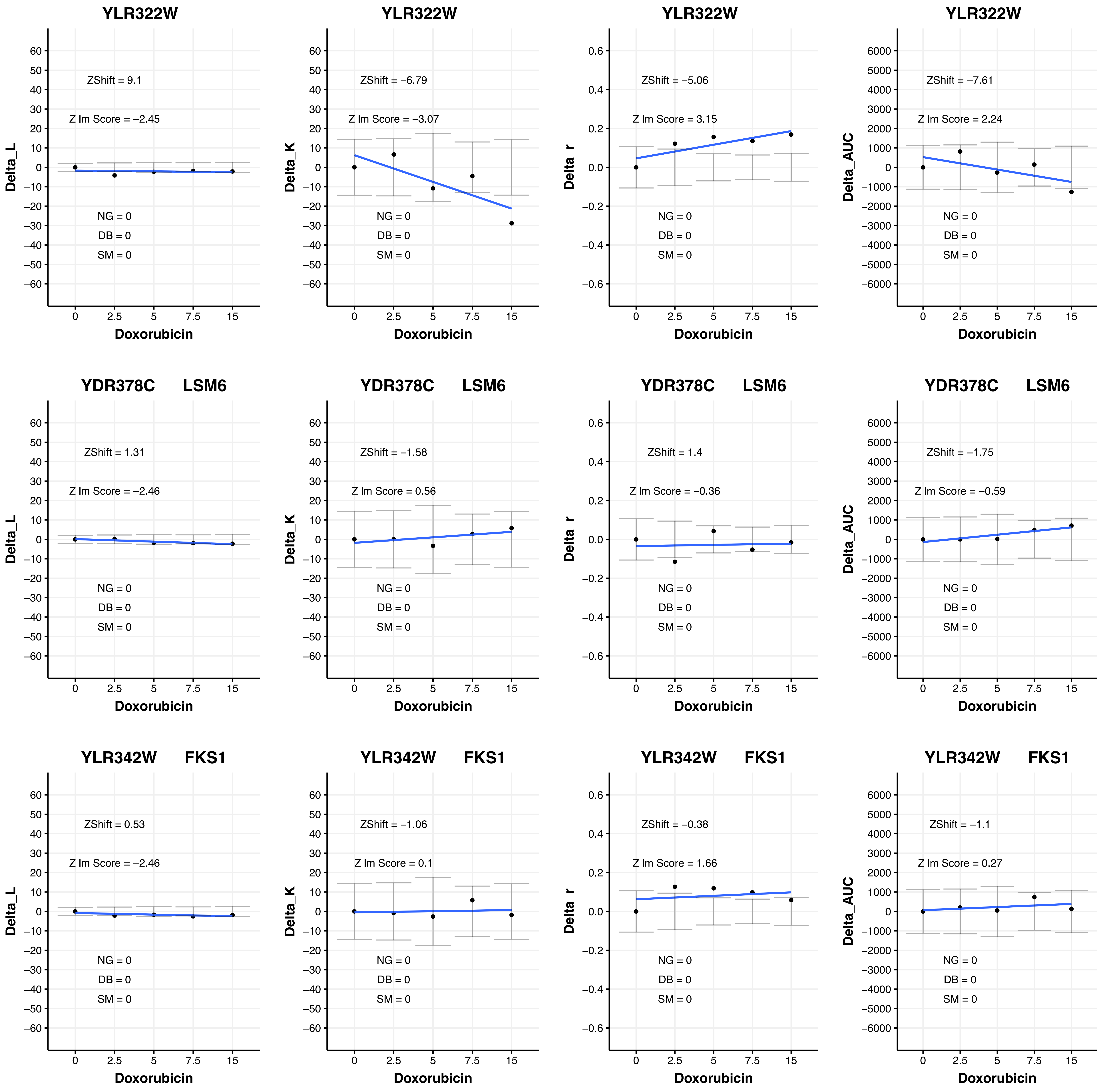

YLR322W
ZShift = 9.1
Z lm Score = −2.45
NG = 0
DB = 0
SM = 0
Delta_L
Doxorubicin
YLR322W
ZShift = −6.79
Z lm Score = −3.07
NG = 0
DB = 0
SM = 0
Delta_K
Doxorubicin
YLR322W
ZShift = −5.06
Z lm Score = 3.15
NG = 0
DB = 0
SM = 0
Delta_r
Doxorubicin
YLR322W
ZShift = −7.61
Z lm Score = 2.24
NG = 0
DB = 0
SM = 0
Delta_AUC
Doxorubicin
YDR378C LSM6
ZShift = 1.31
Z lm Score = −2.46
NG = 0
DB = 0
SM = 0
Delta_L
Doxorubicin
YDR378C LSM6
ZShift = −1.58
Z lm Score = 0.56
NG = 0
DB = 0
SM = 0
Delta_K
Doxorubicin
YDR378C LSM6
ZShift = 1.4
Z lm Score = −0.36
NG = 0
DB = 0
SM = 0
Delta_r
Doxorubicin
YDR378C LSM6
ZShift = −1.75
Z lm Score = −0.59
NG = 0
DB = 0
SM = 0
Delta_AUC
Doxorubicin
YLR342W FKS1
ZShift = 0.53
Z lm Score = −2.46
NG = 0
DB = 0
SM = 0
Delta_L
Doxorubicin
YLR342W FKS1
ZShift = −1.06
Z lm Score = 0.1
NG = 0
DB = 0
SM = 0
Delta_K
Doxorubicin
YLR342W FKS1
ZShift = −0.38
Z lm Score = 1.66
NG = 0
DB = 0
SM = 0
Delta_r
Doxorubicin
YLR342W FKS1
ZShift = −1.1
Z lm Score = 0.27
NG = 0
DB = 0
SM = 0
Delta_AUC
Doxorubicin

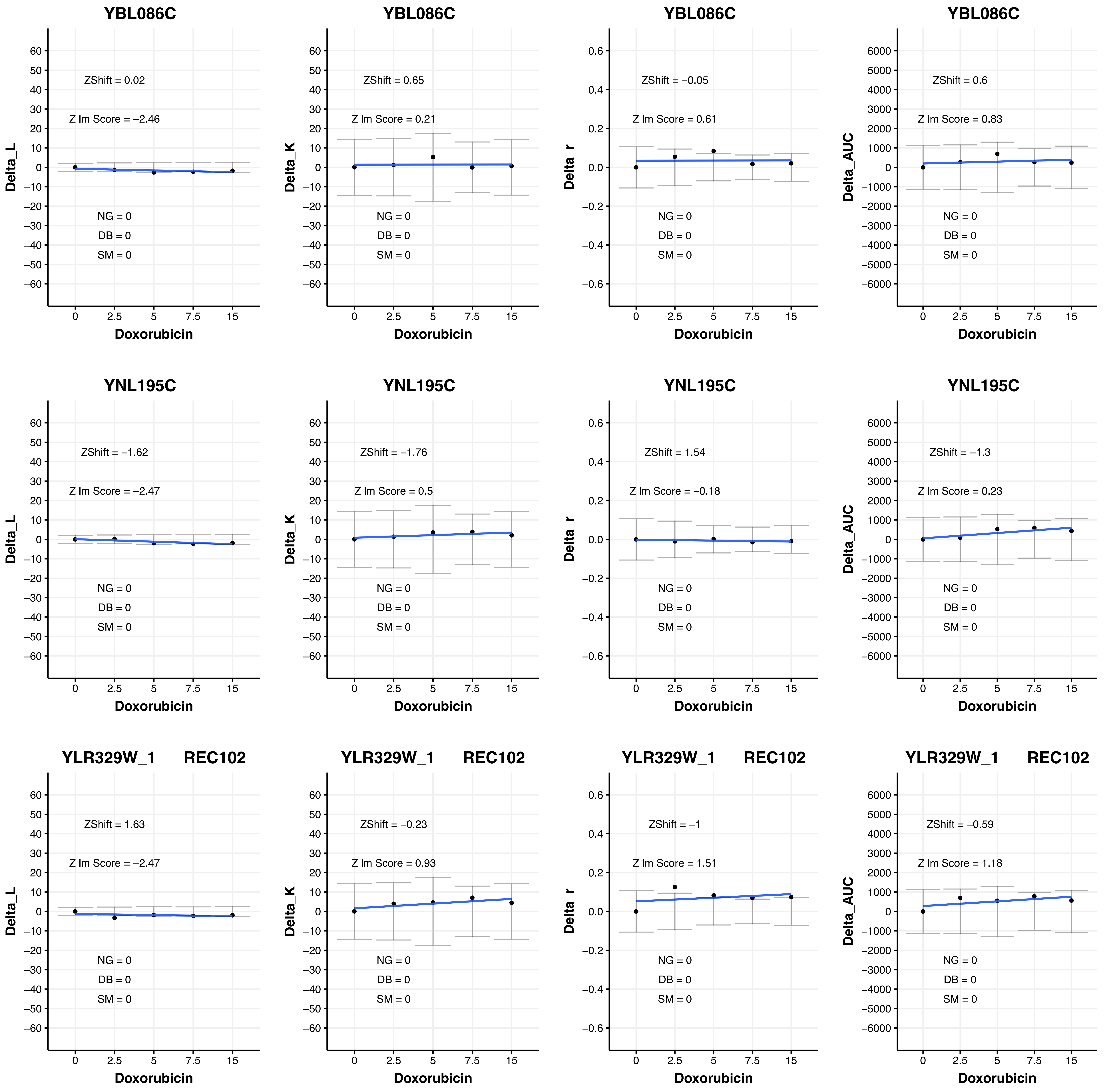

YBL086C
ZShift = 0.02
Z lm Score = −2.46
NG = 0
DB = 0
SM = 0
Delta_L
Doxorubicin
YBL086C
ZShift = 0.65
Z lm Score = 0.21
NG = 0
DB = 0
SM = 0
Delta_K
Doxorubicin
YBL086C
ZShift = −0.05
Z lm Score = 0.61
NG = 0
DB = 0
SM = 0
Delta_r
Doxorubicin
YBL086C
ZShift = 0.6
Z lm Score = 0.83
NG = 0
DB = 0
SM = 0
Delta_AUC
Doxorubicin
YNL195C
ZShift = −1.62
Z lm Score = −2.47
NG = 0
DB = 0
SM = 0
Delta_L
Doxorubicin
YNL195C
ZShift = −1.76
Z lm Score = 0.5
NG = 0
DB = 0
SM = 0
Delta_K
Doxorubicin
YNL195C
ZShift = 1.54
Z lm Score = −0.18
NG = 0
DB = 0
SM = 0
Delta_r
Doxorubicin
YNL195C
ZShift = −1.3
Z lm Score = 0.23
NG = 0
DB = 0
SM = 0
Delta_AUC
Doxorubicin
YLR329W_1 REC102
ZShift = 1.63
Z lm Score = −2.47
NG = 0
DB = 0
SM = 0
Delta_L
Doxorubicin
YLR329W_1 REC102
ZShift = −0.23
Z lm Score = 0.93
NG = 0
DB = 0
SM = 0
Delta_K
Doxorubicin
YLR329W_1 REC102
ZShift = −1
Z lm Score = 1.51
NG = 0
DB = 0
SM = 0
Delta_r
Doxorubicin
YLR329W_1 REC102
ZShift = −0.59
Z lm Score = 1.18
NG = 0
DB = 0
SM = 0
Delta_AUC
Doxorubicin

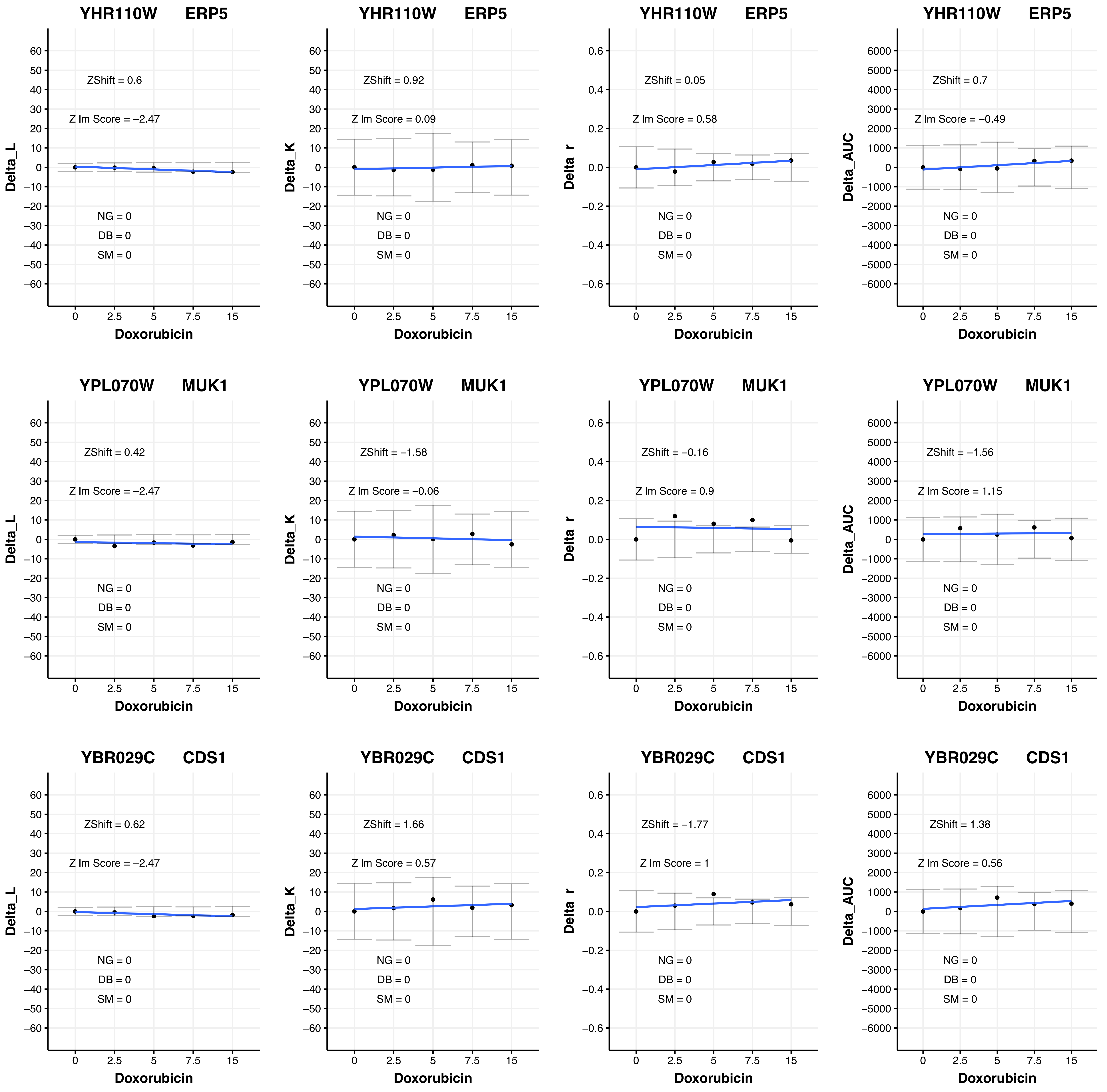

YHR110W ERP5
ZShift = 0.6
Z lm Score = −2.47
NG = 0
DB = 0
SM = 0
Delta_L
Doxorubicin
YHR110W ERP5
ZShift = 0.92
Z lm Score = 0.09
NG = 0
DB = 0
SM = 0
Delta_K
Doxorubicin
YHR110W ERP5
ZShift = 0.05
Z lm Score = 0.58
NG = 0
DB = 0
SM = 0
Delta_r
Doxorubicin
YHR110W ERP5
ZShift = 0.7
Z lm Score = −0.49
NG = 0
DB = 0
SM = 0
Delta_AUC
Doxorubicin
YPL070W MUK1
ZShift = 0.42
Z lm Score = −2.47
NG = 0
DB = 0
SM = 0
Delta_L
Doxorubicin
YPL070W MUK1
ZShift = −1.58
Z lm Score = −0.06
NG = 0
DB = 0
SM = 0
Delta_K
Doxorubicin
YPL070W MUK1
ZShift = −0.16
Z lm Score = 0.9
NG = 0
DB = 0
SM = 0
Delta_r
Doxorubicin
YPL070W MUK1
ZShift = −1.56
Z lm Score = 1.15
NG = 0
DB = 0
SM = 0
Delta_AUC
Doxorubicin
YBR029C CDS1
ZShift = 0.62
Z lm Score = −2.47
NG = 0
DB = 0
SM = 0
Delta_L
Doxorubicin
YBR029C CDS1
ZShift = 1.66
Z lm Score = 0.57
NG = 0
DB = 0
SM = 0
Delta_K
Doxorubicin
YBR029C CDS1
ZShift = −1.77
Z lm Score = 1
NG = 0
DB = 0
SM = 0
Delta_r
Doxorubicin
YBR029C CDS1
ZShift = 1.38
Z lm Score = 0.56
NG = 0
DB = 0
SM = 0
Delta_AUC
Doxorubicin

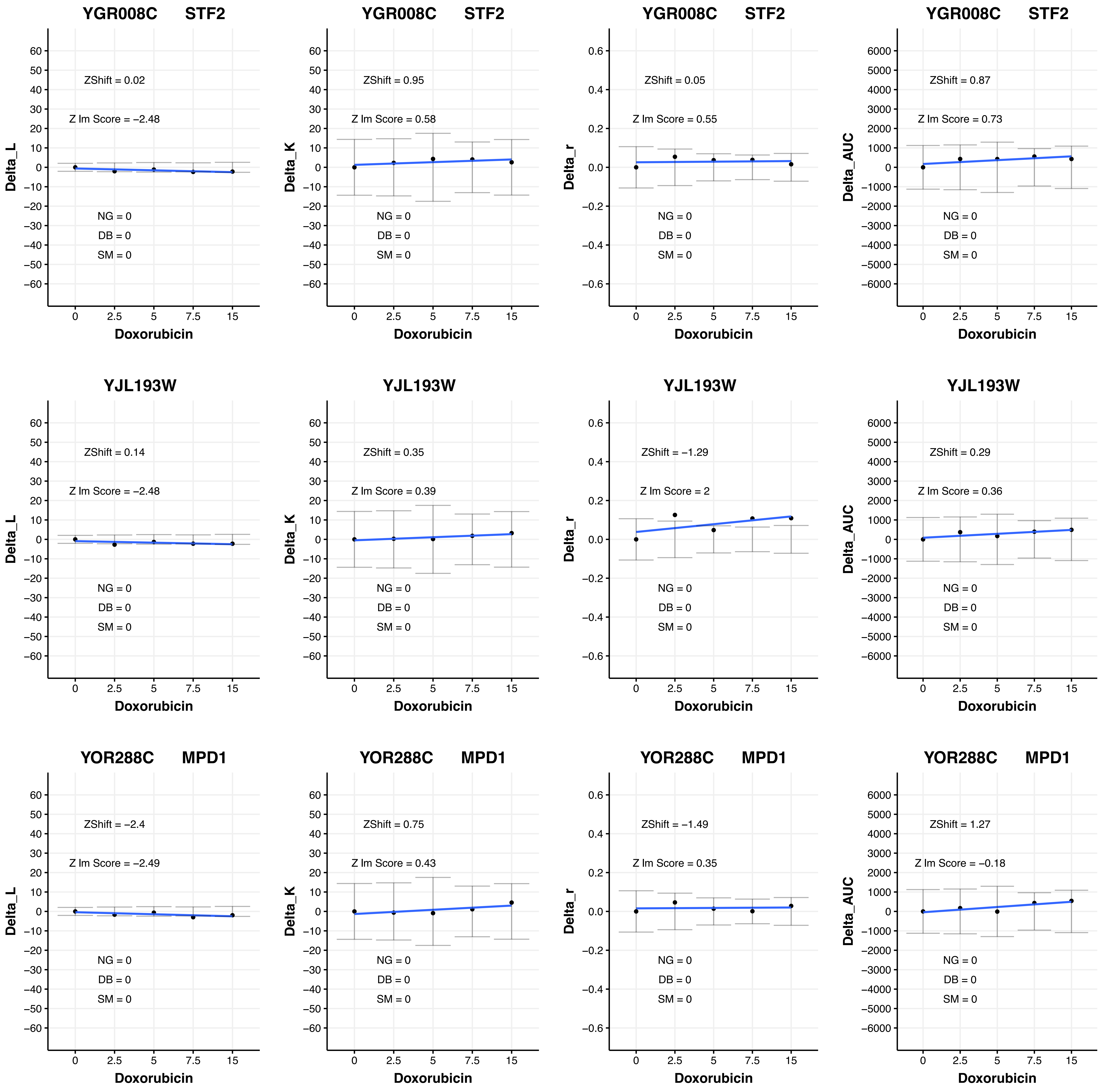
YGR008C STF2
ZShift = 0.02
Z lm Score = −2.48
NG = 0
DB = 0
SM = 0
Delta_L
Doxorubicin
YGR008C STF2
ZShift = 0.95
Z lm Score = 0.58
NG = 0
DB = 0
SM = 0
Delta_K
Doxorubicin
YGR008C STF2
ZShift = 0.05
Z lm Score = 0.55
NG = 0
DB = 0
SM = 0
Delta_r
Doxorubicin
YGR008C STF2
ZShift = 0.87
Z lm Score = 0.73
NG = 0
DB = 0
SM = 0
Delta_AUC
Doxorubicin
YJL193W
ZShift = 0.14
Z lm Score = −2.48
NG = 0
DB = 0
SM = 0
Delta_L
Doxorubicin
YJL193W
ZShift = 0.35
Z lm Score = 0.39
NG = 0
DB = 0
SM = 0
Delta_K
Doxorubicin
YJL193W
ZShift = −1.29
Z lm Score = 2
NG = 0
DB = 0
SM = 0
Delta_r
Doxorubicin
YJL193W
ZShift = 0.29
Z lm Score = 0.36
NG = 0
DB = 0
SM = 0
Delta_AUC
Doxorubicin
YOR288C MPD1
ZShift = −2.4
Z lm Score = −2.49
NG = 0
DB = 0
SM = 0
Delta_L
Doxorubicin
YOR288C MPD1
ZShift = 0.75
Z lm Score = 0.43
NG = 0
DB = 0
SM = 0
Delta_K
Doxorubicin
YOR288C MPD1
ZShift = −1.49
Z lm Score = 0.35
NG = 0
DB = 0
SM = 0
Delta_r
Doxorubicin
YOR288C MPD1
ZShift = 1.27
Z lm Score = −0.18
NG = 0
DB = 0
SM = 0
Delta_AUC
Doxorubicin

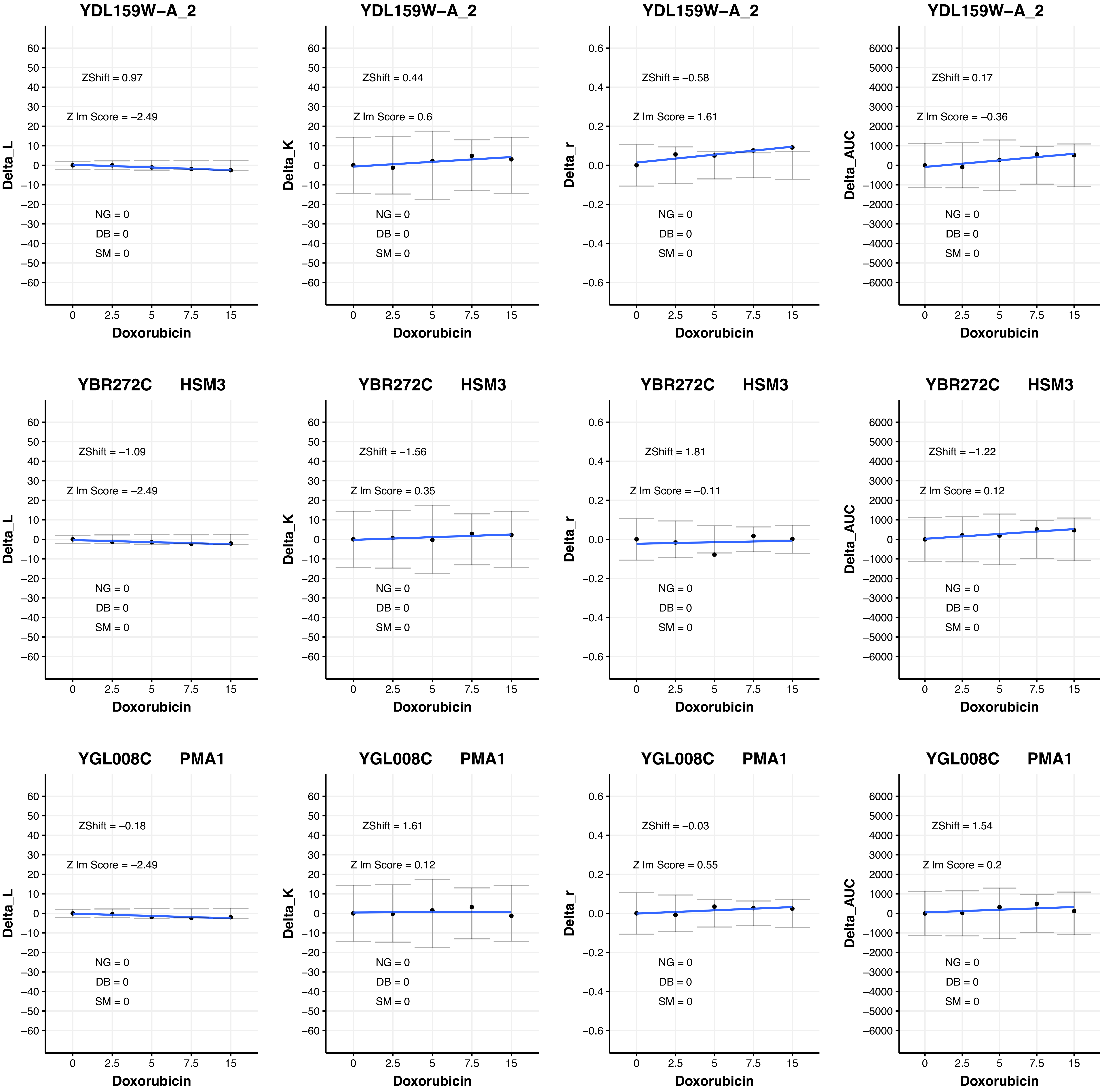
YDL159W−A_2
ZShift = 0.97
Z lm Score = −2.49
NG = 0
DB = 0
SM = 0
Delta_L
Doxorubicin
YDL159W−A_2
ZShift = 0.44
Z lm Score = 0.6
NG = 0
DB = 0
SM = 0
Delta_K
Doxorubicin
YDL159W−A_2
ZShift = −0.58
Z lm Score = 1.61
NG = 0
DB = 0
SM = 0
Delta_r
Doxorubicin
YDL159W−A_2
ZShift = 0.17
Z lm Score = −0.36
NG = 0
DB = 0
SM = 0
Delta_AUC
Doxorubicin
YBR272C HSM3
ZShift = −1.09
Z lm Score = −2.49
NG = 0
DB = 0
SM = 0
Delta_L
Doxorubicin
YBR272C HSM3
ZShift = −1.56
Z lm Score = 0.35
NG = 0
DB = 0
SM = 0
Delta_K
Doxorubicin
YBR272C HSM3
ZShift = 1.81
Z lm Score = −0.11
NG = 0
DB = 0
SM = 0
Delta_r
Doxorubicin
YBR272C HSM3
ZShift = −1.22
Z lm Score = 0.12
NG = 0
DB = 0
SM = 0
Delta_AUC
Doxorubicin
YGL008C PMA1
ZShift = −0.18
Z lm Score = −2.49
NG = 0
DB = 0
SM = 0
Delta_L
Doxorubicin
YGL008C PMA1
ZShift = 1.61
Z lm Score = 0.12
NG = 0
DB = 0
SM = 0
Delta_K
Doxorubicin
YGL008C PMA1
ZShift = −0.03
Z lm Score = 0.55
NG = 0
DB = 0
SM = 0
Delta_r
Doxorubicin
YGL008C PMA1
ZShift = 1.54
Z lm Score = 0.2
NG = 0
DB = 0
SM = 0
Delta_AUC
Doxorubicin

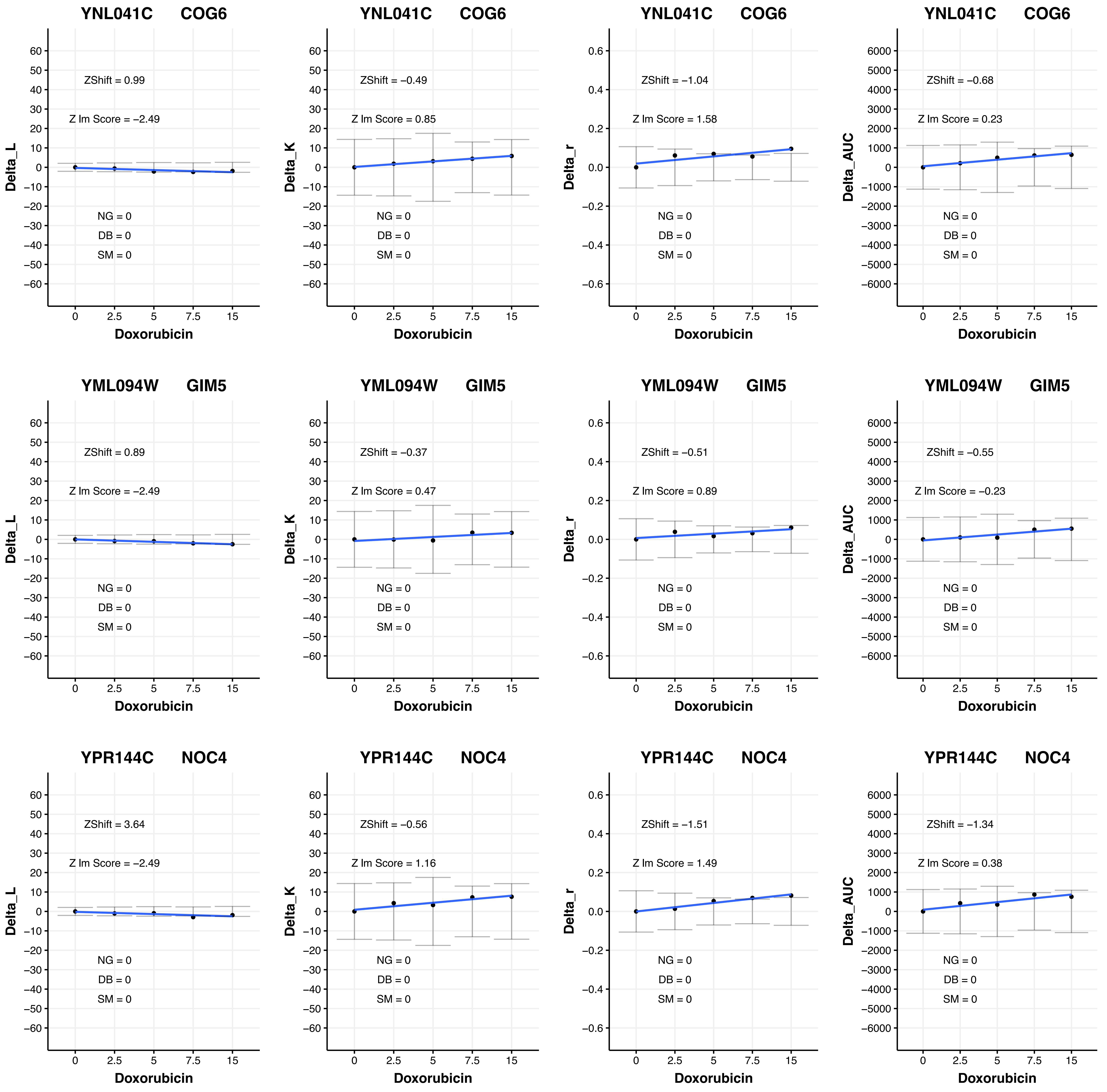

YNL041C COG6
ZShift = 0.99
Z lm Score = −2.49
NG = 0
DB = 0
SM = 0
Delta_L
YNL041C COG6
ZShift = −0.49
Z lm Score = 0.85
NG = 0
DB = 0
SM = 0
Delta_K
YNL041C COG6
ZShift = −1.04
Z lm Score = 1.58
NG = 0
DB = 0
SM = 0
Delta_r
YNL041C COG6
ZShift = −0.68
Z lm Score = 0.23
NG = 0
DB = 0
SM = 0
Delta_AUC
YML094W GIM5
ZShift = 0.89
Z lm Score = −2.49
NG = 0
DB = 0
SM = 0
Delta_L
YML094W GIM5
ZShift = −0.37
Z lm Score = 0.47
NG = 0
DB = 0
SM = 0
Delta_K
YML094W GIM5
ZShift = −0.51
Z lm Score = 0.89
NG = 0
DB = 0
SM = 0
Delta_r
YML094W GIM5
ZShift = −0.55
Z lm Score = −0.23
NG = 0
DB = 0
SM = 0
Delta_AUC
YPR144C NOC4
ZShift = 3.64
Z lm Score = −2.49
NG = 0
DB = 0
SM = 0
Delta_L
YPR144C NOC4
ZShift = −0.56
Z lm Score = 1.16
NG = 0
DB = 0
SM = 0
Delta_K
YPR144C NOC4
ZShift = −1.51
Z lm Score = 1.49
NG = 0
DB = 0
SM = 0
Delta_r
YPR144C NOC4
ZShift = −1.34
Z lm Score = 0.38
NG = 0
DB = 0
SM = 0
Delta_AUC
Doxorubicin
0
2.5
5
7.5
15

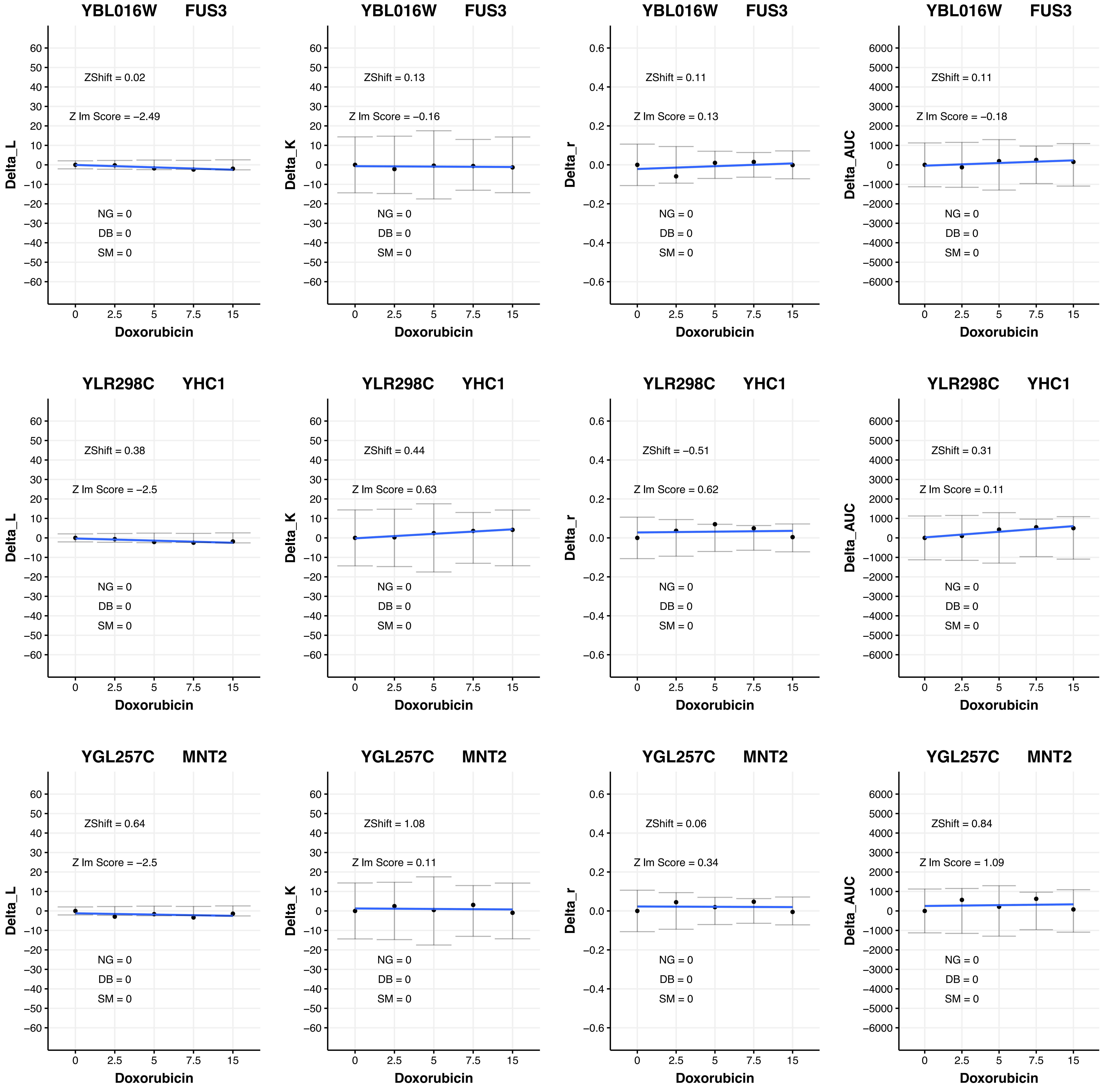
YBL016W FUS3
ZShift = 0.02
Z lm Score = −2.49
NG = 0
DB = 0
SM = 0
Delta_L
Doxorubicin
YBL016W FUS3
ZShift = 0.13
Z lm Score = −0.16
NG = 0
DB = 0
SM = 0
Delta_K
Doxorubicin
YBL016W FUS3
ZShift = 0.11
Z lm Score = 0.13
NG = 0
DB = 0
SM = 0
Delta_r
Doxorubicin
YBL016W FUS3
ZShift = 0.11
Z lm Score = −0.18
NG = 0
DB = 0
SM = 0
Delta_AUC
Doxorubicin
YLR298C YHC1
ZShift = 0.38
Z lm Score = −2.5
NG = 0
DB = 0
SM = 0
Delta_L
Doxorubicin
YLR298C YHC1
ZShift = 0.44
Z lm Score = 0.63
NG = 0
DB = 0
SM = 0
Delta_K
Doxorubicin
YLR298C YHC1
ZShift = −0.51
Z lm Score = 0.62
NG = 0
DB = 0
SM = 0
Delta_r
Doxorubicin
YLR298C YHC1
ZShift = 0.31
Z lm Score = 0.11
NG = 0
DB = 0
SM = 0
Delta_AUC
Doxorubicin
YGL257C MNT2
ZShift = 0.64
Z lm Score = −2.5
NG = 0
DB = 0
SM = 0
Delta_L
Doxorubicin
YGL257C MNT2
ZShift = 1.08
Z lm Score = 0.11
NG = 0
DB = 0
SM = 0
Delta_K
Doxorubicin
YGL257C MNT2
ZShift = 0.06
Z lm Score = 0.34
NG = 0
DB = 0
SM = 0
Delta_r
Doxorubicin
YGL257C MNT2
ZShift = 0.84
Z lm Score = 1.09
NG = 0
DB = 0
SM = 0
Delta_AUC
Doxorubicin

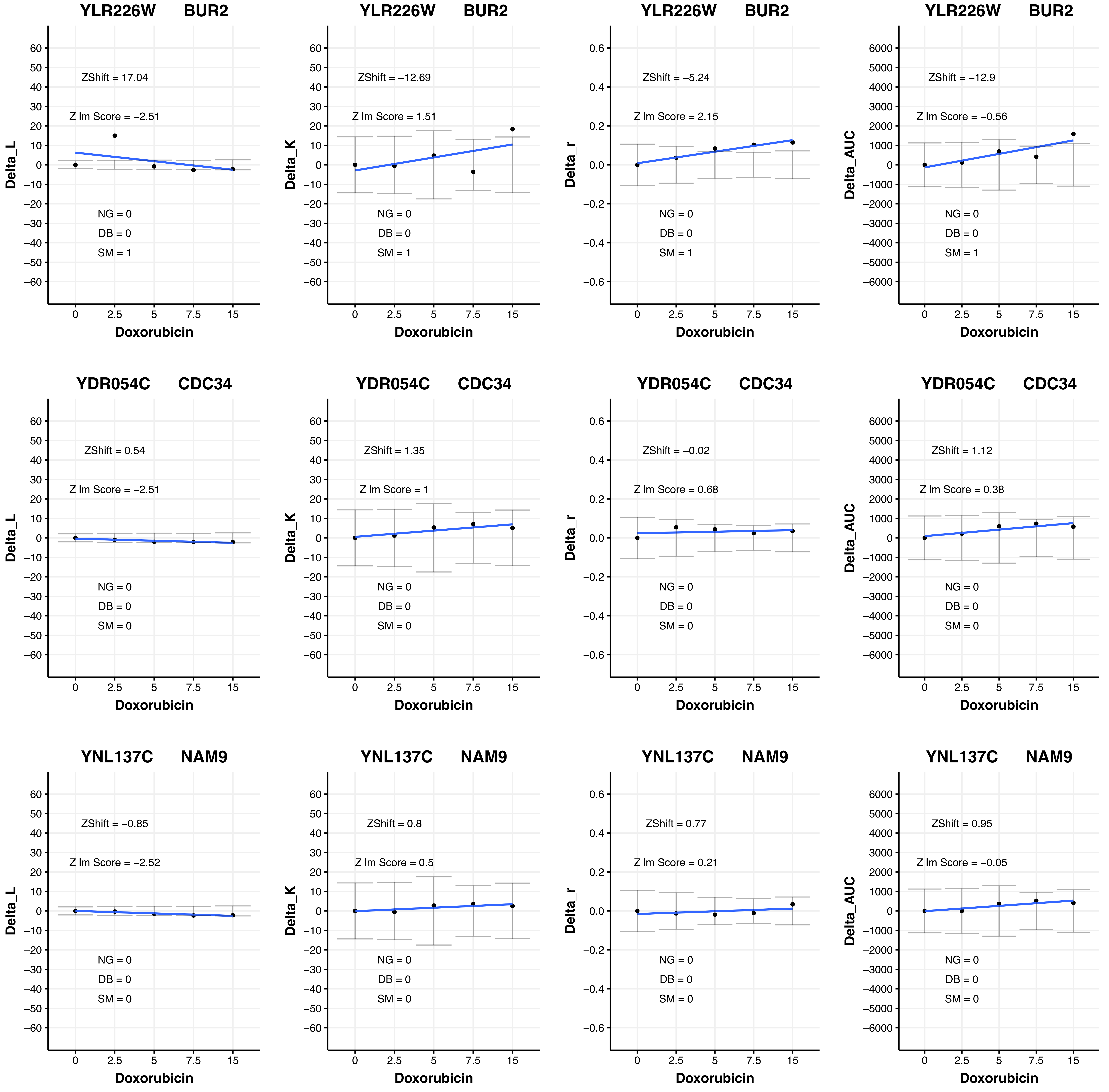

YLR226W BUR2
ZShift = 17.04
Z lm Score = −2.51
NG = 0
DB = 0
SM = 1
Delta_L
Doxorubicin
YLR226W BUR2
ZShift = −12.69
Z lm Score = 1.51
NG = 0
DB = 0
SM = 1
Delta_K
Doxorubicin
YLR226W BUR2
ZShift = −5.24
Z lm Score = 2.15
NG = 0
DB = 0
SM = 1
Delta_r
Doxorubicin
YLR226W BUR2
ZShift = −12.9
Z lm Score = −0.56
NG = 0
DB = 0
SM = 1
Delta_AUC
Doxorubicin
YDR054C CDC34
ZShift = 0.54
Z lm Score = −2.51
NG = 0
DB = 0
SM = 0
Delta_L
Doxorubicin
YDR054C CDC34
ZShift = 1.35
Z lm Score = 1
NG = 0
DB = 0
SM = 0
Delta_K
Doxorubicin
YDR054C CDC34
ZShift = −0.02
Z lm Score = 0.68
NG = 0
DB = 0
SM = 0
Delta_r
Doxorubicin
YDR054C CDC34
ZShift = 1.12
Z lm Score = 0.38
NG = 0
DB = 0
SM = 0
Delta_AUC
Doxorubicin
YNL137C NAM9
ZShift = −0.85
Z lm Score = −2.52
NG = 0
DB = 0
SM = 0
Delta_L
Doxorubicin
YNL137C NAM9
ZShift = 0.8
Z lm Score = 0.5
NG = 0
DB = 0
SM = 0
Delta_K
Doxorubicin
YNL137C NAM9
ZShift = 0.77
Z lm Score = 0.21
NG = 0
DB = 0
SM = 0
Delta_r
Doxorubicin
YNL137C NAM9
ZShift = 0.95
Z lm Score = −0.05
NG = 0
DB = 0
SM = 0
Delta_AUC
Doxorubicin

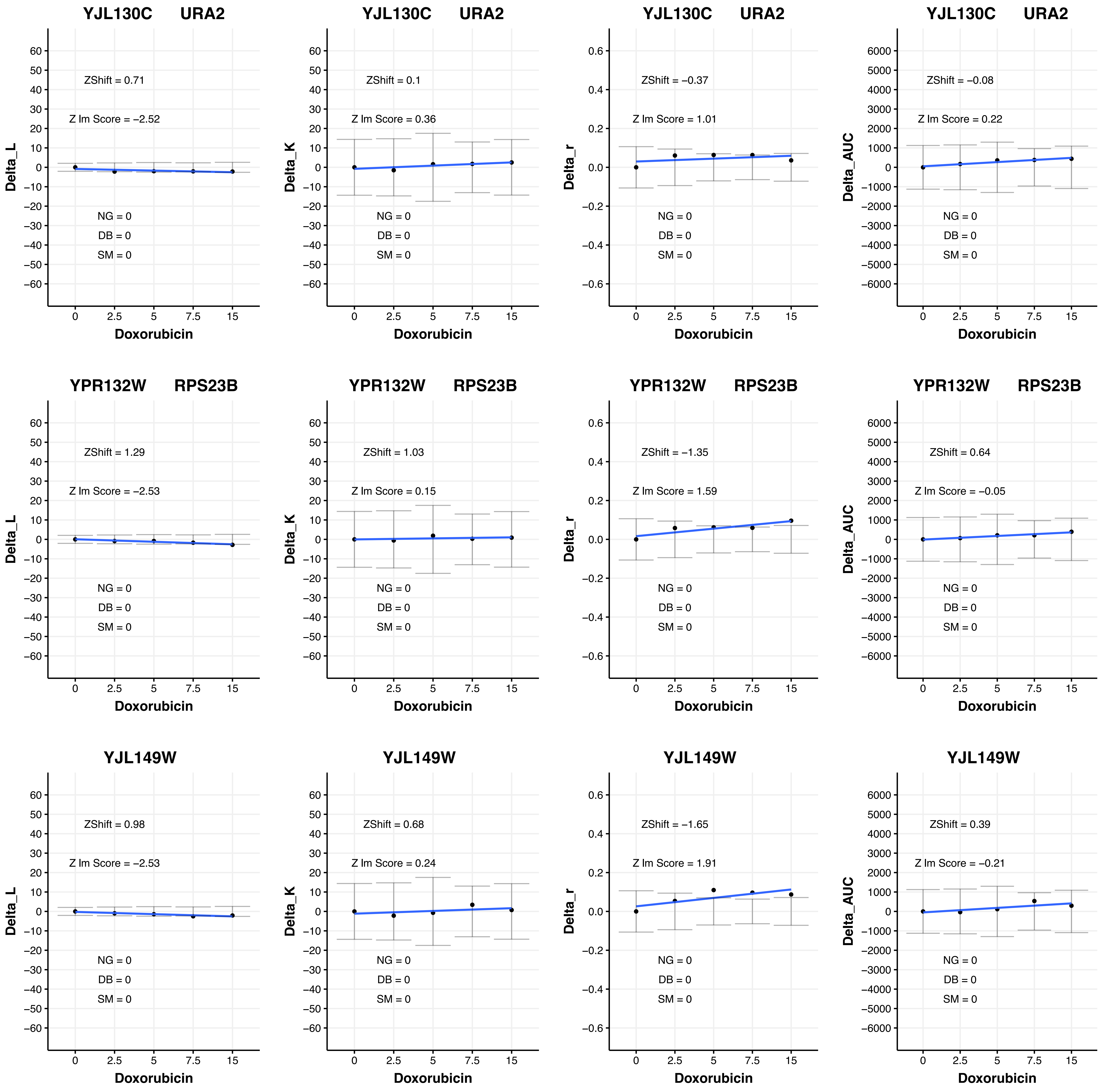

YJL130C URA2
ZShift = 0.71
Z lm Score = −2.52
NG = 0
DB = 0
SM = 0
Delta_L
Doxorubicin
YJL130C URA2
ZShift = 0.1
Z lm Score = 0.36
NG = 0
DB = 0
SM = 0
Delta_K
Doxorubicin
YJL130C URA2
ZShift = −0.37
Z lm Score = 1.01
NG = 0
DB = 0
SM = 0
Delta_r
Doxorubicin
YJL130C URA2
ZShift = −0.08
Z lm Score = 0.22
NG = 0
DB = 0
SM = 0
Delta_AUC
Doxorubicin
YPR132W RPS23B
ZShift = 1.29
Z lm Score = −2.53
NG = 0
DB = 0
SM = 0
Delta_L
Doxorubicin
YPR132W RPS23B
ZShift = 1.03
Z lm Score = 0.15
NG = 0
DB = 0
SM = 0
Delta_K
Doxorubicin
YPR132W RPS23B
ZShift = −1.35
Z lm Score = 1.59
NG = 0
DB = 0
SM = 0
Delta_r
Doxorubicin
YPR132W RPS23B
ZShift = 0.64
Z lm Score = −0.05
NG = 0
DB = 0
SM = 0
Delta_AUC
Doxorubicin
YJL149W
ZShift = 0.98
Z lm Score = −2.53
NG = 0
DB = 0
SM = 0
Delta_L
Doxorubicin
YJL149W
ZShift = 0.68
Z lm Score = 0.24
NG = 0
DB = 0
SM = 0
Delta_K
Doxorubicin
YJL149W
ZShift = −1.65
Z lm Score = 1.91
NG = 0
DB = 0
SM = 0
Delta_r
Doxorubicin
YJL149W
ZShift = 0.39
Z lm Score = −0.21
NG = 0
DB = 0
SM = 0
Delta_AUC
Doxorubicin

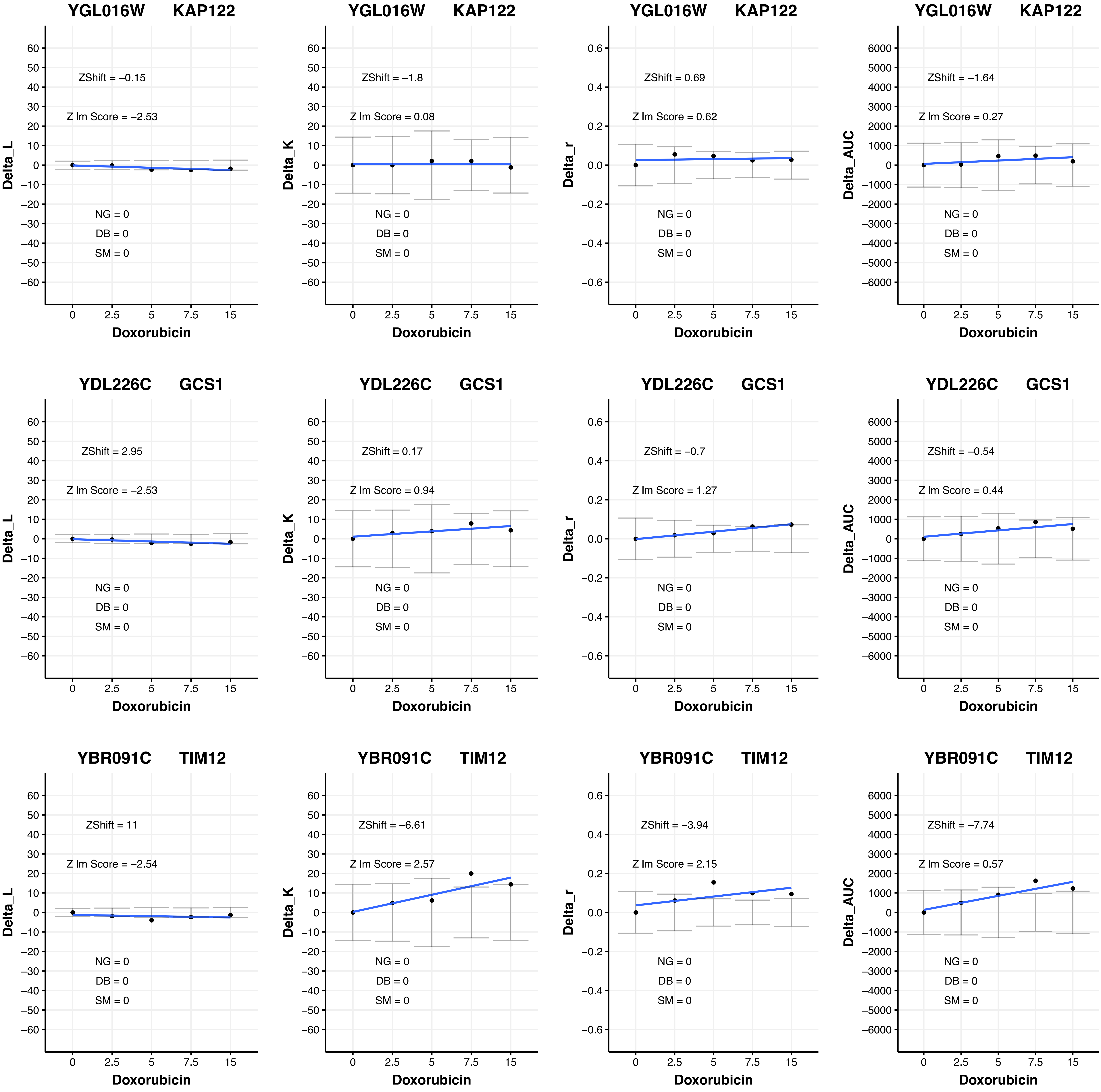
YGL016W KAP122
ZShift = −0.15
Z lm Score = −2.53
NG = 0
DB = 0
SM = 0
Delta_L
Doxorubicin
YGL016W KAP122
ZShift = −1.8
Z lm Score = 0.08
NG = 0
DB = 0
SM = 0
Delta_K
Doxorubicin
YGL016W KAP122
ZShift = 0.69
Z lm Score = 0.62
NG = 0
DB = 0
SM = 0
Delta_r
Doxorubicin
YGL016W KAP122
ZShift = −1.64
Z lm Score = 0.27
NG = 0
DB = 0
SM = 0
Delta_AUC
Doxorubicin
YDL226C GCS1
ZShift = 2.95
Z lm Score = −2.53
NG = 0
DB = 0
SM = 0
Delta_L
Doxorubicin
YDL226C GCS1
ZShift = 0.17
Z lm Score = 0.94
NG = 0
DB = 0
SM = 0
Delta_K
Doxorubicin
YDL226C GCS1
ZShift = −0.7
Z lm Score = 1.27
NG = 0
DB = 0
SM = 0
Delta_r
Doxorubicin
YDL226C GCS1
ZShift = −0.54
Z lm Score = 0.44
NG = 0
DB = 0
SM = 0
Delta_AUC
Doxorubicin
YBR091C TIM12
ZShift = 11
Z lm Score = −2.54
NG = 0
DB = 0
SM = 0
Delta_L
Doxorubicin
YBR091C TIM12
ZShift = −6.61
Z lm Score = 2.57
NG = 0
DB = 0
SM = 0
Delta_K
Doxorubicin
YBR091C TIM12
ZShift = −3.94
Z lm Score = 2.15
NG = 0
DB = 0
SM = 0
Delta_r
Doxorubicin
YBR091C TIM12
ZShift = −7.74
Z lm Score = 0.57
NG = 0
DB = 0
SM = 0
Delta_AUC
Doxorubicin

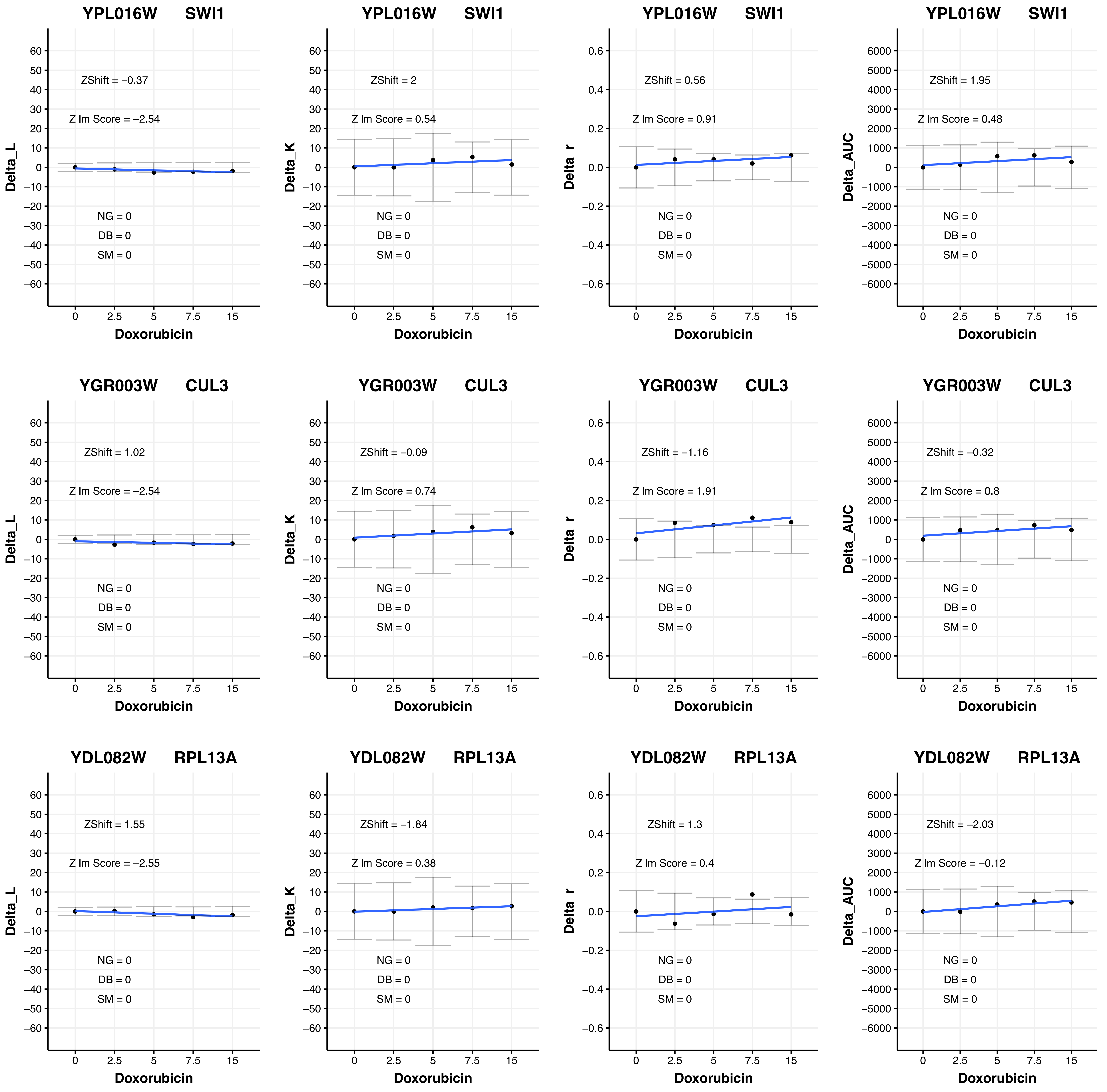
YPL016W SWI1
ZShift = −0.37
Z lm Score = −2.54
NG = 0
DB = 0
SM = 0
Delta_L
Doxorubicin
YPL016W SWI1
ZShift = 2
Z lm Score = 0.54
NG = 0
DB = 0
SM = 0
Delta_K
Doxorubicin
YPL016W SWI1
ZShift = 0.56
Z lm Score = 0.91
NG = 0
DB = 0
SM = 0
Delta_r
Doxorubicin
YPL016W SWI1
ZShift = 1.95
Z lm Score = 0.48
NG = 0
DB = 0
SM = 0
Delta_AUC
Doxorubicin
YGR003W CUL3
ZShift = 1.02
Z lm Score = −2.54
NG = 0
DB = 0
SM = 0
Delta_L
Doxorubicin
YGR003W CUL3
ZShift = −0.09
Z lm Score = 0.74
NG = 0
DB = 0
SM = 0
Delta_K
Doxorubicin
YGR003W CUL3
ZShift = −1.16
Z lm Score = 1.91
NG = 0
DB = 0
SM = 0
Delta_r
Doxorubicin
YGR003W CUL3
ZShift = −0.32
Z lm Score = 0.8
NG = 0
DB = 0
SM = 0
Delta_AUC
Doxorubicin
YDL082W RPL13A
ZShift = 1.55
Z lm Score = −2.55
NG = 0
DB = 0
SM = 0
Delta_L
Doxorubicin
YDL082W RPL13A
ZShift = −1.84
Z lm Score = 0.38
NG = 0
DB = 0
SM = 0
Delta_K
Doxorubicin
YDL082W RPL13A
ZShift = 1.3
Z lm Score = 0.4
NG = 0
DB = 0
SM = 0
Delta_r
Doxorubicin
YDL082W RPL13A
ZShift = −2.03
Z lm Score = −0.12
NG = 0
DB = 0
SM = 0
Delta_AUC
Doxorubicin

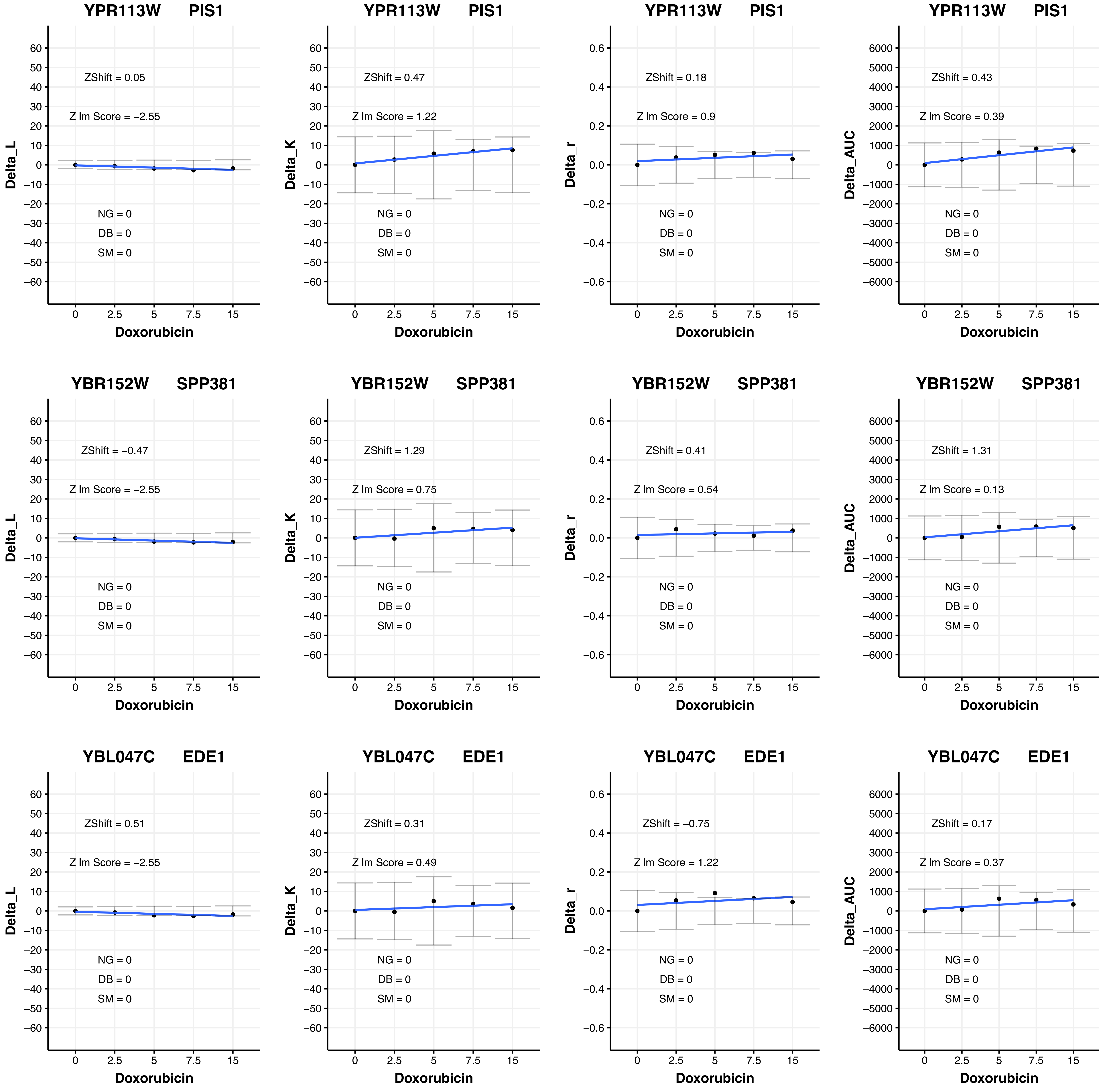

YPR113W PIS1
ZShift = 0.05
Z lm Score = −2.55
NG = 0
DB = 0
SM = 0
Delta_L
Doxorubicin
YPR113W PIS1
ZShift = 0.47
Z lm Score = 1.22
NG = 0
DB = 0
SM = 0
Delta_K
Doxorubicin
YPR113W PIS1
ZShift = 0.18
Z lm Score = 0.9
NG = 0
DB = 0
SM = 0
Delta_r
Doxorubicin
YPR113W PIS1
ZShift = 0.43
Z lm Score = 0.39
NG = 0
DB = 0
SM = 0
Delta_AUC
Doxorubicin
YBR152W SPP381
ZShift = −0.47
Z lm Score = −2.55
NG = 0
DB = 0
SM = 0
Delta_L
Doxorubicin
YBR152W SPP381
ZShift = 1.29
Z lm Score = 0.75
NG = 0
DB = 0
SM = 0
Delta_K
Doxorubicin
YBR152W SPP381
ZShift = 0.41
Z lm Score = 0.54
NG = 0
DB = 0
SM = 0
Delta_r
Doxorubicin
YBR152W SPP381
ZShift = 1.31
Z lm Score = 0.13
NG = 0
DB = 0
SM = 0
Delta_AUC
Doxorubicin
YBL047C EDE1
ZShift = 0.51
Z lm Score = −2.55
NG = 0
DB = 0
SM = 0
Delta_L
Doxorubicin
YBL047C EDE1
ZShift = 0.31
Z lm Score = 0.49
NG = 0
DB = 0
SM = 0
Delta_K
Doxorubicin
YBL047C EDE1
ZShift = −0.75
Z lm Score = 1.22
NG = 0
DB = 0
SM = 0
Delta_r
Doxorubicin
YBL047C EDE1
ZShift = 0.17
Z lm Score = 0.37
NG = 0
DB = 0
SM = 0
Delta_AUC
Doxorubicin

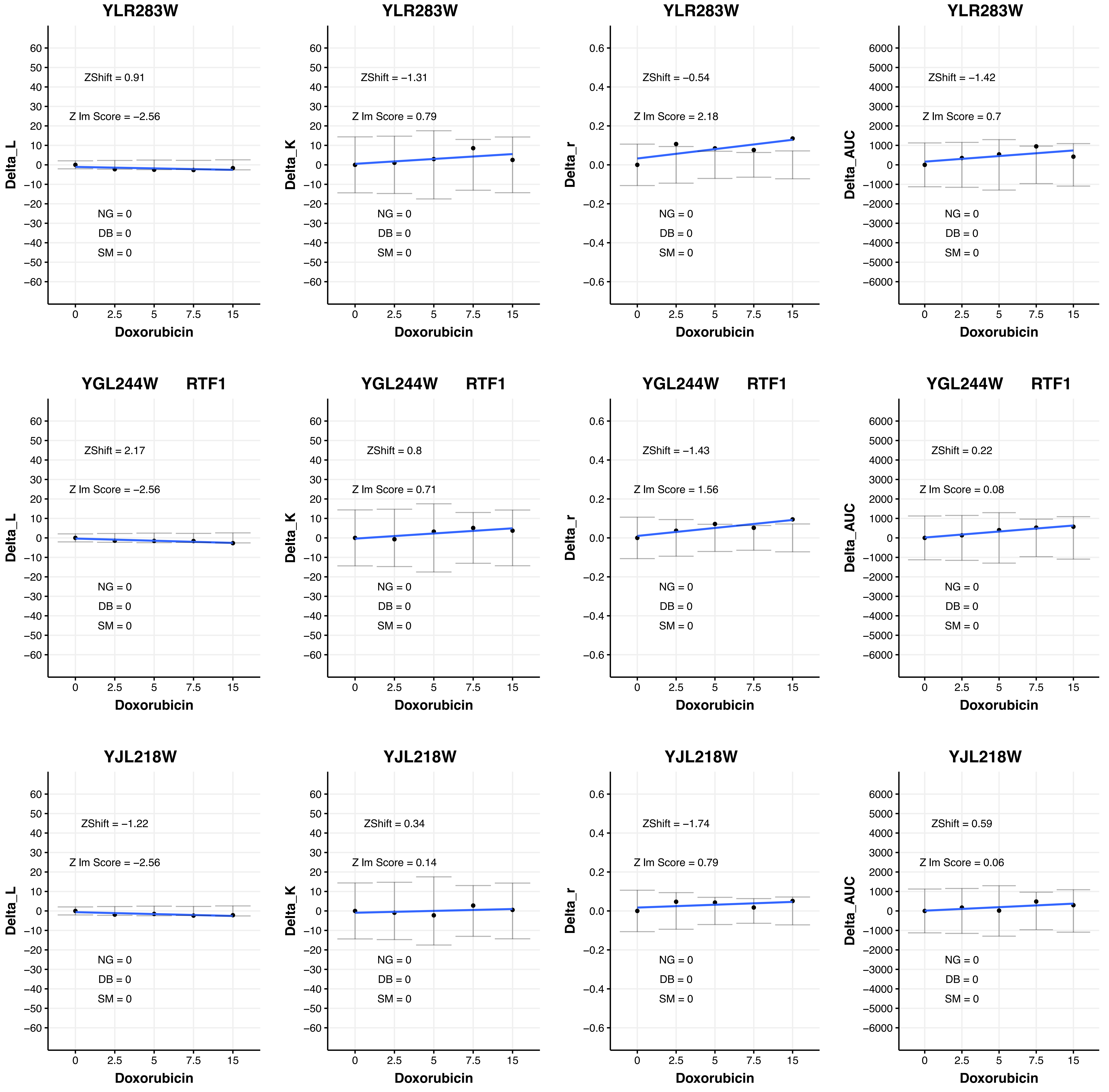

YLR283W
ZShift = 0.91
Z lm Score = −2.56
NG = 0
DB = 0
SM = 0
Delta_L
Doxorubicin
YLR283W
ZShift = −1.31
Z lm Score = 0.79
NG = 0
DB = 0
SM = 0
Delta_K
Doxorubicin
YLR283W
ZShift = −0.54
Z lm Score = 2.18
NG = 0
DB = 0
SM = 0
Delta_r
Doxorubicin
YLR283W
ZShift = −1.42
Z lm Score = 0.7
NG = 0
DB = 0
SM = 0
Delta_AUC
Doxorubicin
YGL244W RTF1
ZShift = 2.17
Z lm Score = −2.56
NG = 0
DB = 0
SM = 0
Delta_L
Doxorubicin
YGL244W RTF1
ZShift = 0.8
Z lm Score = 0.71
NG = 0
DB = 0
SM = 0
Delta_K
Doxorubicin
YGL244W RTF1
ZShift = −1.43
Z lm Score = 1.56
NG = 0
DB = 0
SM = 0
Delta_r
Doxorubicin
YGL244W RTF1
ZShift = 0.22
Z lm Score = 0.08
NG = 0
DB = 0
SM = 0
Delta_AUC
Doxorubicin
YJL218W
ZShift = −1.22
Z lm Score = −2.56
NG = 0
DB = 0
SM = 0
Delta_L
Doxorubicin
YJL218W
ZShift = 0.34
Z lm Score = 0.14
NG = 0
DB = 0
SM = 0
Delta_K
Doxorubicin
YJL218W
ZShift = −1.74
Z lm Score = 0.79
NG = 0
DB = 0
SM = 0
Delta_r
Doxorubicin
YJL218W
ZShift = 0.59
Z lm Score = 0.06
NG = 0
DB = 0
SM = 0
Delta_AUC
Doxorubicin

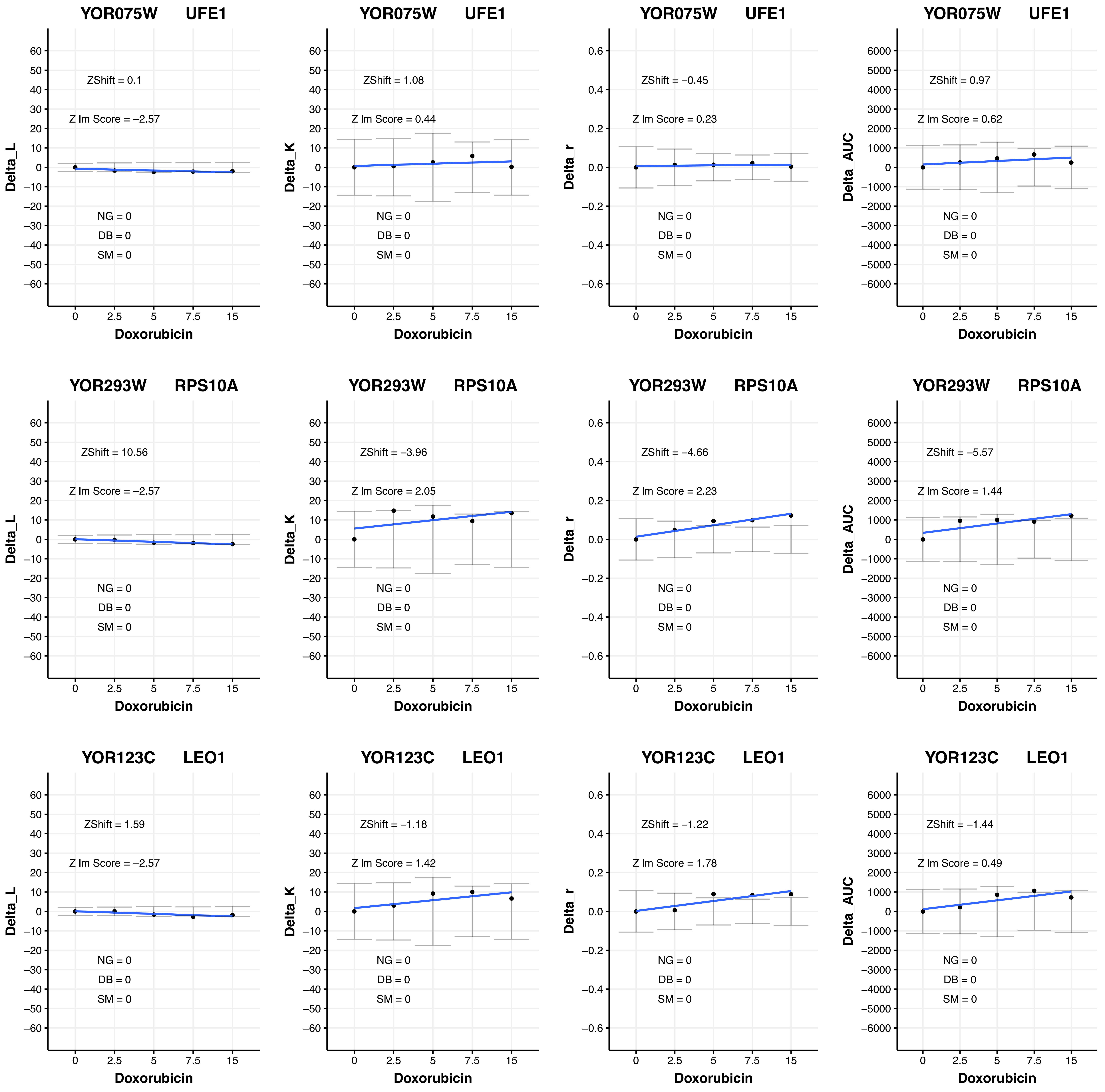
YOR075W UFE1
ZShift = 0.1
Z lm Score = −2.57
NG = 0
DB = 0
SM = 0
Delta_L
Doxorubicin
YOR075W UFE1
ZShift = 1.08
Z lm Score = 0.44
NG = 0
DB = 0
SM = 0
Delta_K
Doxorubicin
YOR075W UFE1
ZShift = −0.45
Z lm Score = 0.23
NG = 0
DB = 0
SM = 0
Delta_r
Doxorubicin
YOR075W UFE1
ZShift = 0.97
Z lm Score = 0.62
NG = 0
DB = 0
SM = 0
Delta_AUC
Doxorubicin
YOR293W RPS10A
ZShift = 10.56
Z lm Score = −2.57
NG = 0
DB = 0
SM = 0
Delta_L
Doxorubicin
YOR293W RPS10A
ZShift = −3.96
Z lm Score = 2.05
NG = 0
DB = 0
SM = 0
Delta_K
Doxorubicin
YOR293W RPS10A
ZShift = −4.66
Z lm Score = 2.23
NG = 0
DB = 0
SM = 0
Delta_r
Doxorubicin
YOR293W RPS10A
ZShift = −5.57
Z lm Score = 1.44
NG = 0
DB = 0
SM = 0
Delta_AUC
Doxorubicin
YOR123C LEO1
ZShift = 1.59
Z lm Score = −2.57
NG = 0
DB = 0
SM = 0
Delta_L
Doxorubicin
YOR123C LEO1
ZShift = −1.18
Z lm Score = 1.42
NG = 0
DB = 0
SM = 0
Delta_K
Doxorubicin
YOR123C LEO1
ZShift = −1.22
Z lm Score = 1.78
NG = 0
DB = 0
SM = 0
Delta_r
Doxorubicin
YOR123C LEO1
ZShift = −1.44
Z lm Score = 0.49
NG = 0
DB = 0
SM = 0
Delta_AUC
Doxorubicin

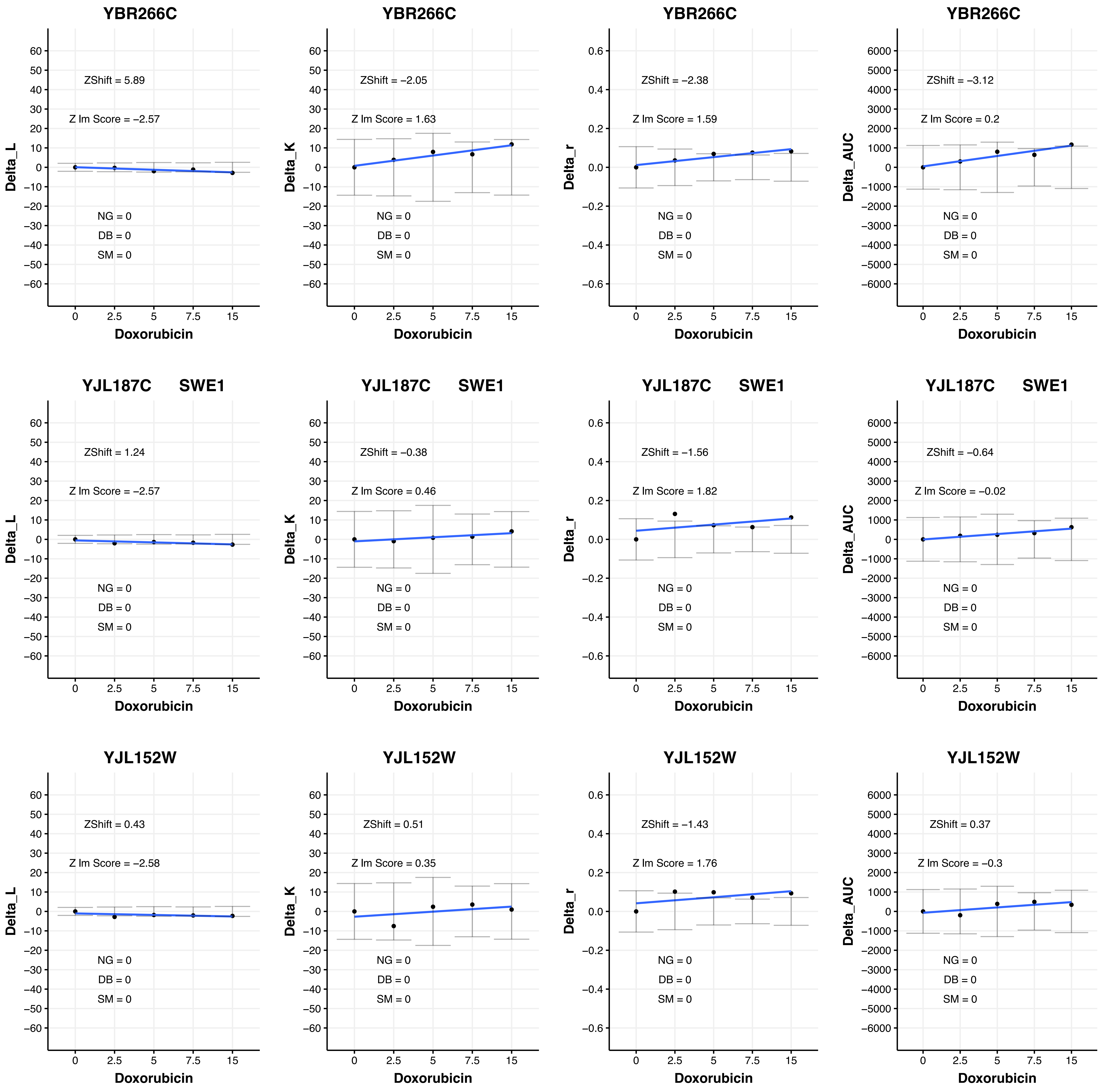
YBR266C
ZShift = 5.89
Z lm Score = −2.57
NG = 0
DB = 0
SM = 0
Delta_L
Doxorubicin
YBR266C
ZShift = −2.05
Z lm Score = 1.63
NG = 0
DB = 0
SM = 0
Delta_K
Doxorubicin
YBR266C
ZShift = −2.38
Z lm Score = 1.59
NG = 0
DB = 0
SM = 0
Delta_r
Doxorubicin
YBR266C
ZShift = −3.12
Z lm Score = 0.2
NG = 0
DB = 0
SM = 0
Delta_AUC
Doxorubicin
YJL187C SWE1
ZShift = 1.24
Z lm Score = −2.57
NG = 0
DB = 0
SM = 0
Delta_L
Doxorubicin
YJL187C SWE1
ZShift = −0.38
Z lm Score = 0.46
NG = 0
DB = 0
SM = 0
Delta_K
Doxorubicin
YJL187C SWE1
ZShift = −1.56
Z lm Score = 1.82
NG = 0
DB = 0
SM = 0
Delta_r
Doxorubicin
YJL187C SWE1
ZShift = −0.64
Z lm Score = −0.02
NG = 0
DB = 0
SM = 0
Delta_AUC
Doxorubicin
YJL152W
ZShift = 0.43
Z lm Score = −2.58
NG = 0
DB = 0
SM = 0
Delta_L
Doxorubicin
YJL152W
ZShift = 0.51
Z lm Score = 0.35
NG = 0
DB = 0
SM = 0
Delta_K
Doxorubicin
YJL152W
ZShift = −1.43
Z lm Score = 1.76
NG = 0
DB = 0
SM = 0
Delta_r
Doxorubicin
YJL152W
ZShift = 0.37
Z lm Score = −0.3
NG = 0
DB = 0
SM = 0
Delta_AUC
Doxorubicin

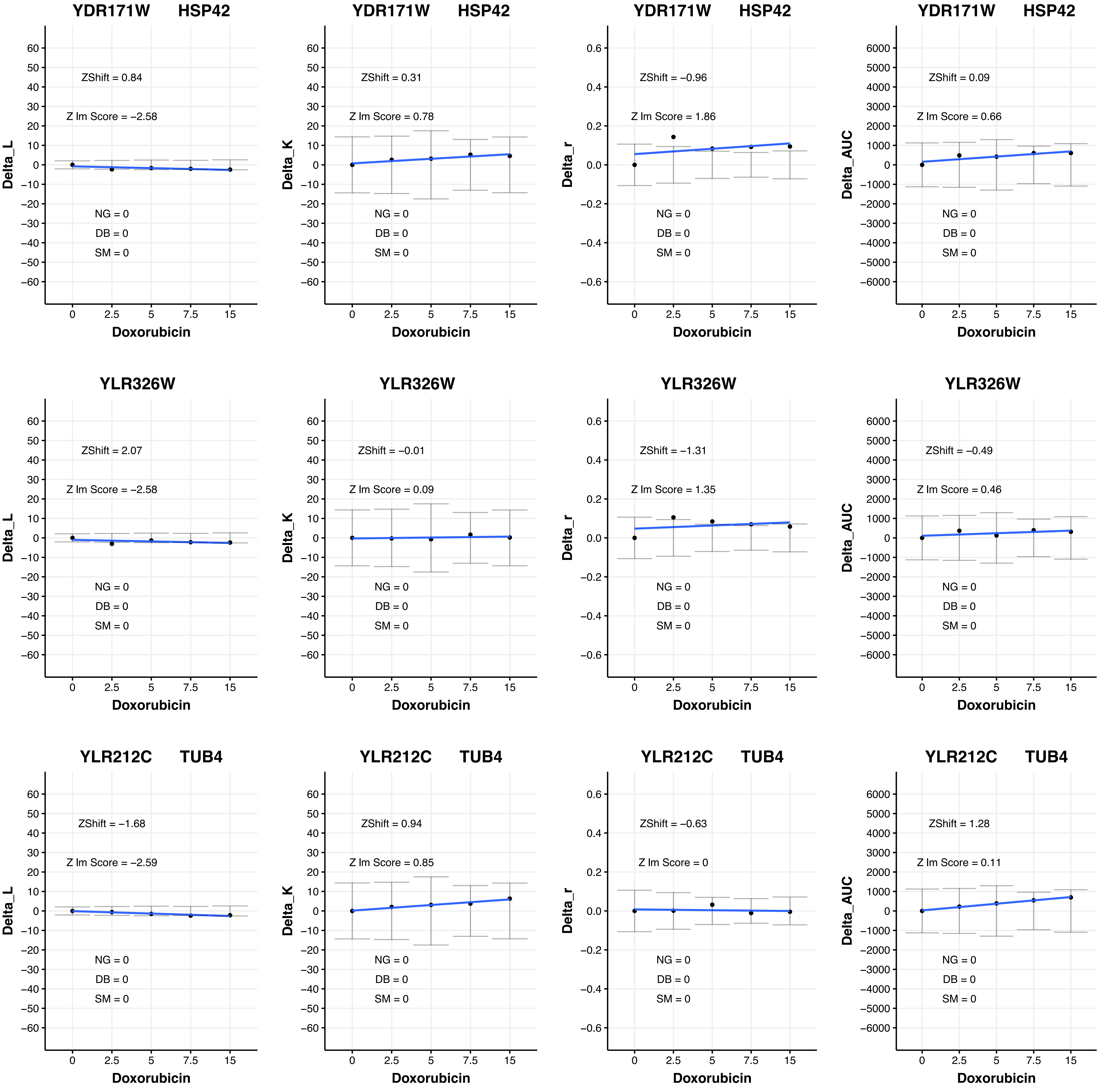
YDR171W HSP42
ZShift = 0.84
Z lm Score = −2.58
NG = 0
DB = 0
SM = 0
Delta_L
Doxorubicin
YDR171W HSP42
ZShift = 0.31
Z lm Score = 0.78
NG = 0
DB = 0
SM = 0
Delta_K
Doxorubicin
YDR171W HSP42
ZShift = −0.96
Z lm Score = 1.86
NG = 0
DB = 0
SM = 0
Delta_r
Doxorubicin
YDR171W HSP42
ZShift = 0.09
Z lm Score = 0.66
NG = 0
DB = 0
SM = 0
Delta_AUC
Doxorubicin
YLR326W
ZShift = 2.07
Z lm Score = −2.58
NG = 0
DB = 0
SM = 0
Delta_L
Doxorubicin
YLR326W
ZShift = −0.01
Z lm Score = 0.09
NG = 0
DB = 0
SM = 0
Delta_K
Doxorubicin
YLR326W
ZShift = −1.31
Z lm Score = 1.35
NG = 0
DB = 0
SM = 0
Delta_r
Doxorubicin
YLR326W
ZShift = −0.49
Z lm Score = 0.46
NG = 0
DB = 0
SM = 0
Delta_AUC
Doxorubicin
YLR212C TUB4
ZShift = −1.68
Z lm Score = −2.59
NG = 0
DB = 0
SM = 0
Delta_L
Doxorubicin
YLR212C TUB4
ZShift = 0.94
Z lm Score = 0.85
NG = 0
DB = 0
SM = 0
Delta_K
Doxorubicin
YLR212C TUB4
ZShift = −0.63
Z lm Score = 0
NG = 0
DB = 0
SM = 0
Delta_r
Doxorubicin
YLR212C TUB4
ZShift = 1.28
Z lm Score = 0.11
NG = 0
DB = 0
SM = 0
Delta_AUC
Doxorubicin

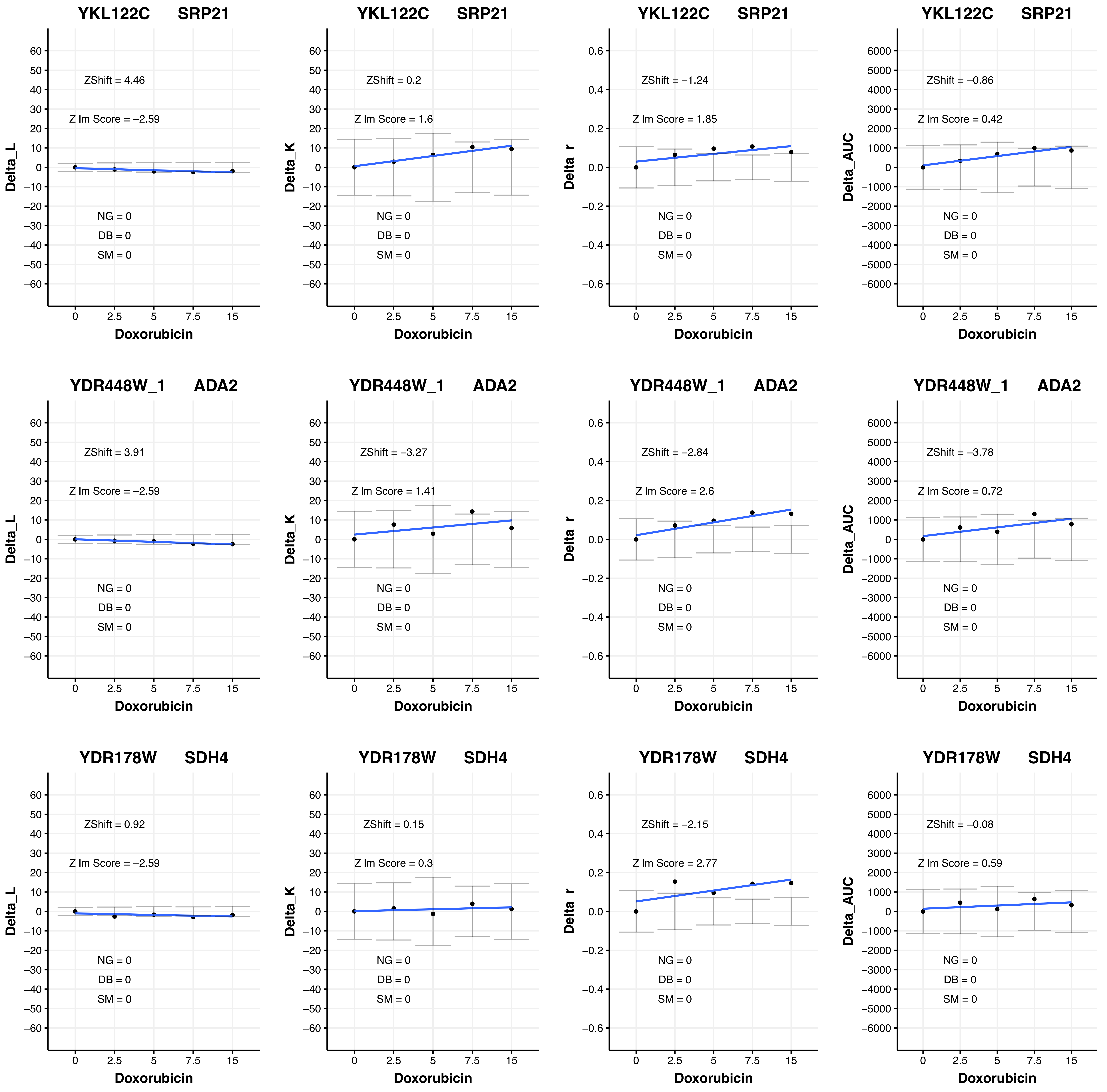

YKL122C SRP21
Delta_L
ZShift = 4.46
Z lm Score = −2.59
NG = 0
DB = 0
SM = 0
Doxorubicin
YKL122C SRP21
Delta_K
ZShift = 0.2
Z lm Score = 1.6
NG = 0
DB = 0
SM = 0
Doxorubicin
YKL122C SRP21
Delta_r
ZShift = −1.24
Z lm Score = 1.85
NG = 0
DB = 0
SM = 0
Doxorubicin
YKL122C SRP21
Delta_AUC
ZShift = −0.86
Z lm Score = 0.42
NG = 0
DB = 0
SM = 0
Doxorubicin
YDR448W_1 ADA2
Delta_L
ZShift = 3.91
Z lm Score = −2.59
NG = 0
DB = 0
SM = 0
Doxorubicin
YDR448W_1 ADA2
Delta_K
ZShift = −3.27
Z lm Score = 1.41
NG = 0
DB = 0
SM = 0
Doxorubicin
YDR448W_1 ADA2
Delta_r
ZShift = −2.84
Z lm Score = 2.6
NG = 0
DB = 0
SM = 0
Doxorubicin
YDR448W_1 ADA2
Delta_AUC
ZShift = −3.78
Z lm Score = 0.72
NG = 0
DB = 0
SM = 0
Doxorubicin
YDR178W SDH4
Delta_L
ZShift = 0.92
Z lm Score = −2.59
NG = 0
DB = 0
SM = 0
Doxorubicin
YDR178W SDH4
Delta_K
ZShift = 0.15
Z lm Score = 0.3
NG = 0
DB = 0
SM = 0
Doxorubicin
YDR178W SDH4
Delta_r
ZShift = −2.15
Z lm Score = 2.77
NG = 0
DB = 0
SM = 0
Doxorubicin
YDR178W SDH4
Delta_AUC
ZShift = −0.08
Z lm Score = 0.59
NG = 0
DB = 0
SM = 0
Doxorubicin

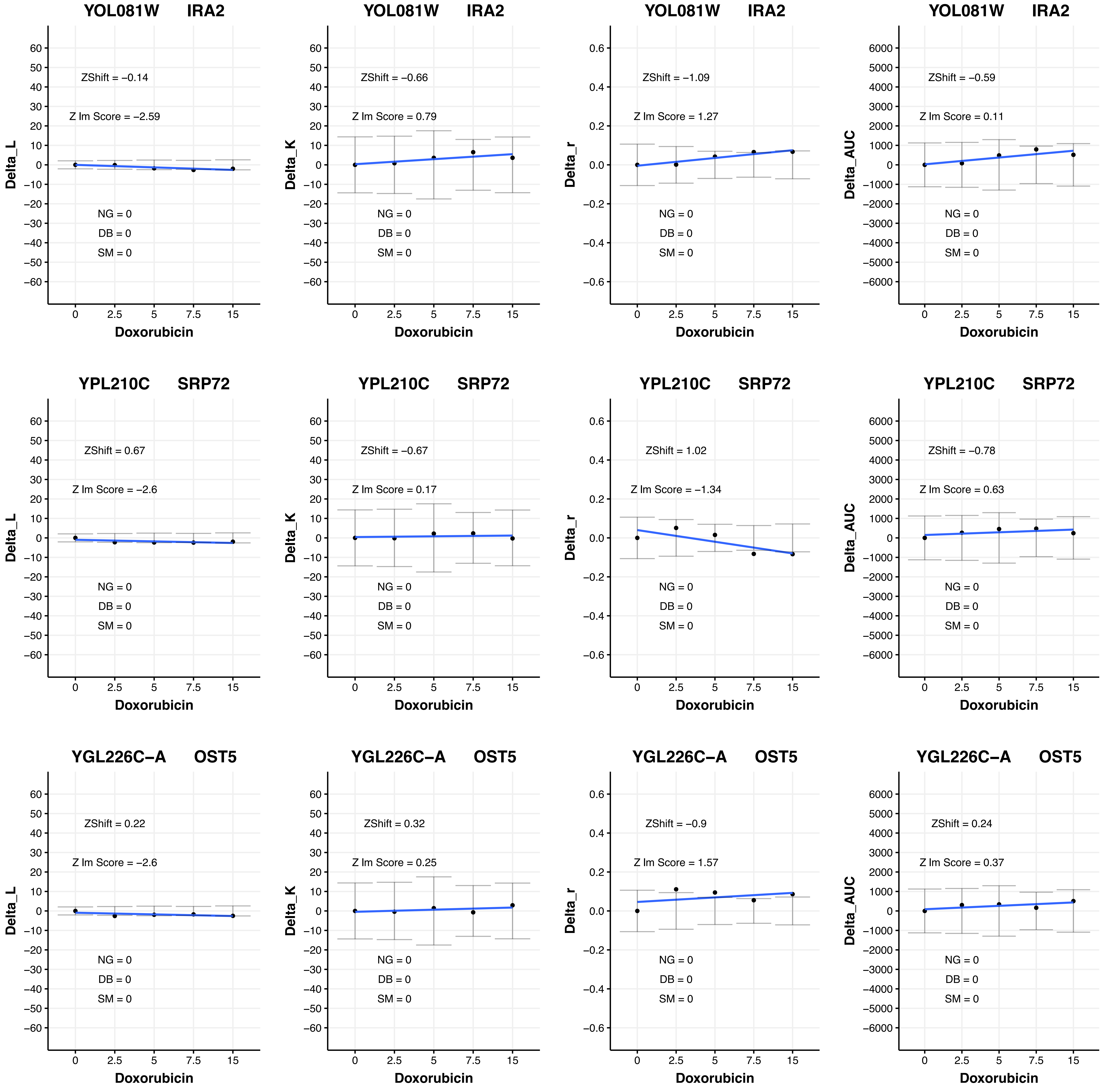

YOL081W IRA2
ZShift = −0.14
Z lm Score = −2.59
NG = 0
DB = 0
SM = 0
Delta_L
Doxorubicin
YOL081W IRA2
ZShift = −0.66
Z lm Score = 0.79
NG = 0
DB = 0
SM = 0
Delta_K
Doxorubicin
YOL081W IRA2
ZShift = −1.09
Z lm Score = 1.27
NG = 0
DB = 0
SM = 0
Delta_r
Doxorubicin
YOL081W IRA2
ZShift = −0.59
Z lm Score = 0.11
NG = 0
DB = 0
SM = 0
Delta_AUC
Doxorubicin
YPL210C SRP72
ZShift = 0.67
Z lm Score = −2.6
NG = 0
DB = 0
SM = 0
Delta_L
Doxorubicin
YPL210C SRP72
ZShift = −0.67
Z lm Score = 0.17
NG = 0
DB = 0
SM = 0
Delta_K
Doxorubicin
YPL210C SRP72
ZShift = 1.02
Z lm Score = −1.34
NG = 0
DB = 0
SM = 0
Delta_r
Doxorubicin
YPL210C SRP72
ZShift = −0.78
Z lm Score = 0.63
NG = 0
DB = 0
SM = 0
Delta_AUC
Doxorubicin
YGL226C−A OST5
ZShift = 0.22
Z lm Score = −2.6
NG = 0
DB = 0
SM = 0
Delta_L
Doxorubicin
YGL226C−A OST5
ZShift = 0.32
Z lm Score = 0.25
NG = 0
DB = 0
SM = 0
Delta_K
Doxorubicin
YGL226C−A OST5
ZShift = −0.9
Z lm Score = 1.57
NG = 0
DB = 0
SM = 0
Delta_r
Doxorubicin
YGL226C−A OST5
ZShift = 0.24
Z lm Score = 0.37
NG = 0
DB = 0
SM = 0
Delta_AUC
Doxorubicin

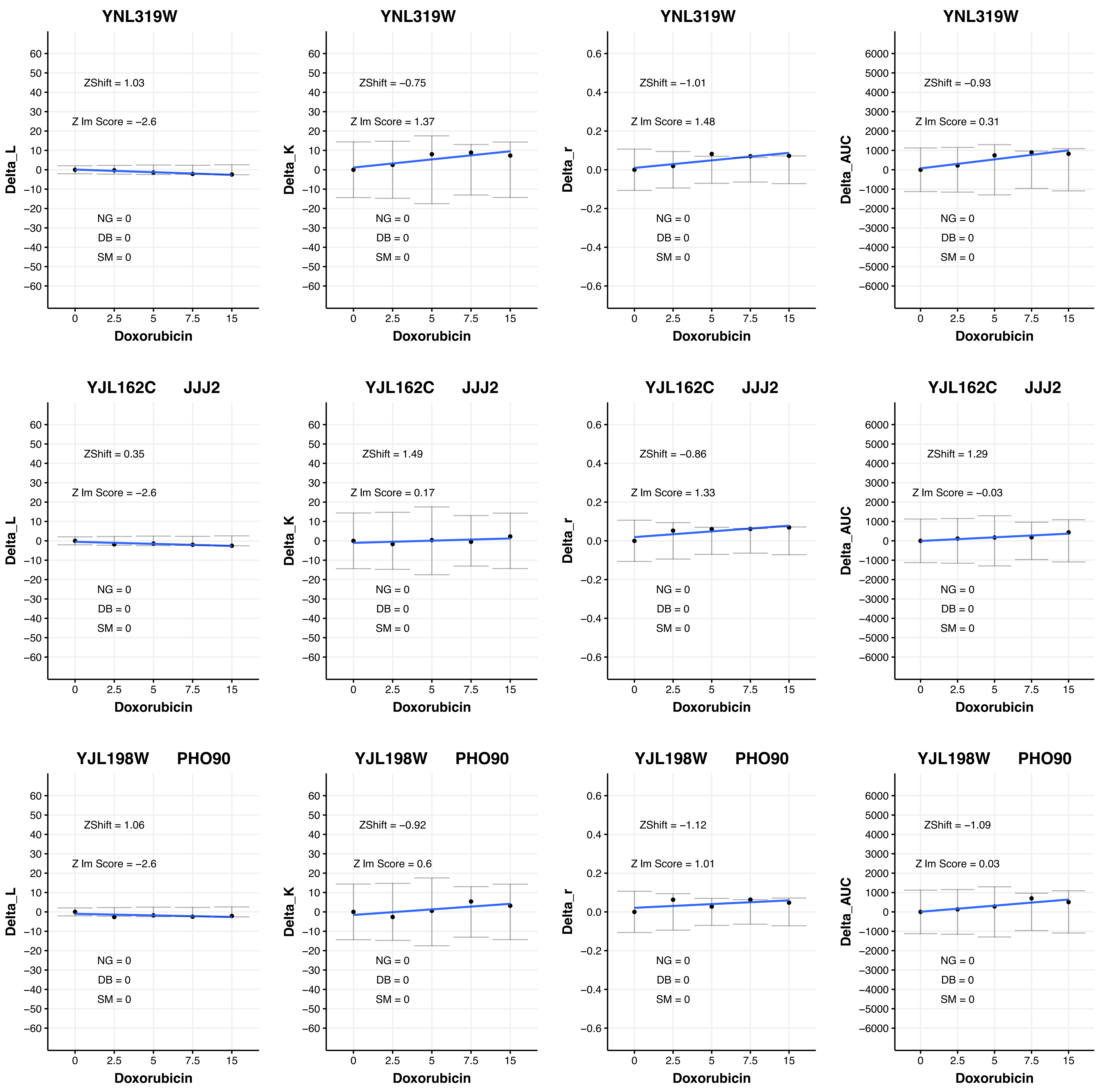

YNL319W
ZShift = 1.03
Z lm Score = −2.6
NG = 0
DB = 0
SM = 0
Delta_L
Doxorubicin
YNL319W
ZShift = −0.75
Z lm Score = 1.37
NG = 0
DB = 0
SM = 0
Delta_K
Doxorubicin
YNL319W
ZShift = −1.01
Z lm Score = 1.48
NG = 0
DB = 0
SM = 0
Delta_r
Doxorubicin
YNL319W
ZShift = −0.93
Z lm Score = 0.31
NG = 0
DB = 0
SM = 0
Delta_AUC
Doxorubicin
YJL162C JJJ2
ZShift = 0.35
Z lm Score = −2.6
NG = 0
DB = 0
SM = 0
Delta_L
Doxorubicin
YJL162C JJJ2
ZShift = 1.49
Z lm Score = 0.17
NG = 0
DB = 0
SM = 0
Delta_K
Doxorubicin
YJL162C JJJ2
ZShift = −0.86
Z lm Score = 1.33
NG = 0
DB = 0
SM = 0
Delta_r
Doxorubicin
YJL162C JJJ2
ZShift = 1.29
Z lm Score = −0.03
NG = 0
DB = 0
SM = 0
Delta_AUC
Doxorubicin
YJL198W PHO90
ZShift = 1.06
Z lm Score = −2.6
NG = 0
DB = 0
SM = 0
Delta_L
Doxorubicin
YJL198W PHO90
ZShift = −0.92
Z lm Score = 0.6
NG = 0
DB = 0
SM = 0
Delta_K
Doxorubicin
YJL198W PHO90
ZShift = −1.12
Z lm Score = 1.01
NG = 0
DB = 0
SM = 0
Delta_r
Doxorubicin
YJL198W PHO90
ZShift = −1.09
Z lm Score = 0.03
NG = 0
DB = 0
SM = 0
Delta_AUC
Doxorubicin

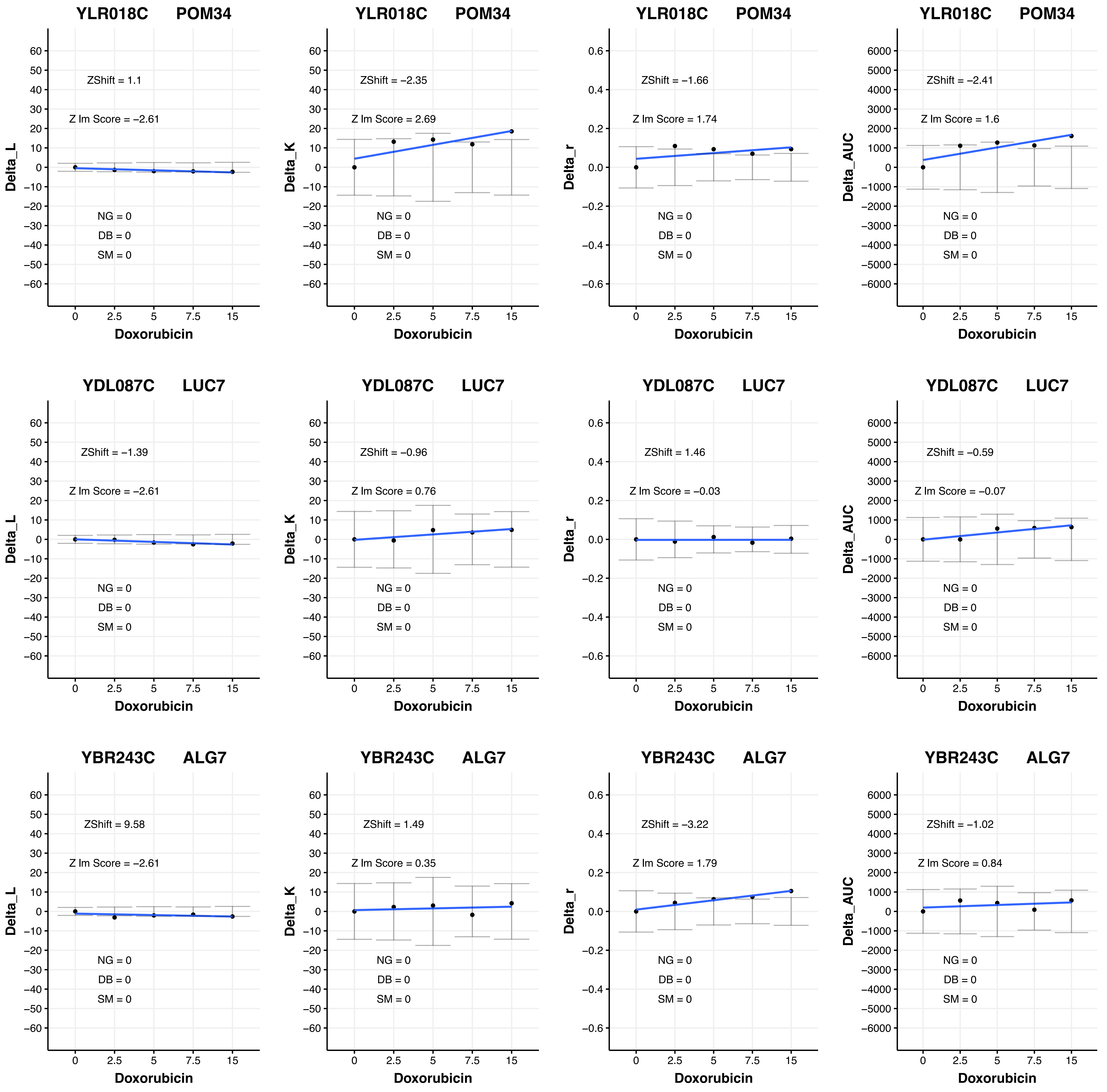

YLR018C POM34
ZShift = 1.1
Z lm Score = −2.61
NG = 0
DB = 0
SM = 0
Delta_L
Doxorubicin
YLR018C POM34
ZShift = −2.35
Z lm Score = 2.69
NG = 0
DB = 0
SM = 0
Delta_K
Doxorubicin
YLR018C POM34
ZShift = −1.66
Z lm Score = 1.74
NG = 0
DB = 0
SM = 0
Delta_r
Doxorubicin
YLR018C POM34
ZShift = −2.41
Z lm Score = 1.6
NG = 0
DB = 0
SM = 0
Delta_AUC
Doxorubicin
YDL087C LUC7
ZShift = −1.39
Z lm Score = −2.61
NG = 0
DB = 0
SM = 0
Delta_L
Doxorubicin
YDL087C LUC7
ZShift = −0.96
Z lm Score = 0.76
NG = 0
DB = 0
SM = 0
Delta_K
Doxorubicin
YDL087C LUC7
ZShift = 1.46
Z lm Score = −0.03
NG = 0
DB = 0
SM = 0
Delta_r
Doxorubicin
YDL087C LUC7
ZShift = −0.59
Z lm Score = −0.07
NG = 0
DB = 0
SM = 0
Delta_AUC
Doxorubicin
YBR243C ALG7
ZShift = 9.58
Z lm Score = −2.61
NG = 0
DB = 0
SM = 0
Delta_L
Doxorubicin
YBR243C ALG7
ZShift = 1.49
Z lm Score = 0.35
NG = 0
DB = 0
SM = 0
Delta_K
Doxorubicin
YBR243C ALG7
ZShift = −3.22
Z lm Score = 1.79
NG = 0
DB = 0
SM = 0
Delta_r
Doxorubicin
YBR243C ALG7
ZShift = −1.02
Z lm Score = 0.84
NG = 0
DB = 0
SM = 0
Delta_AUC
Doxorubicin

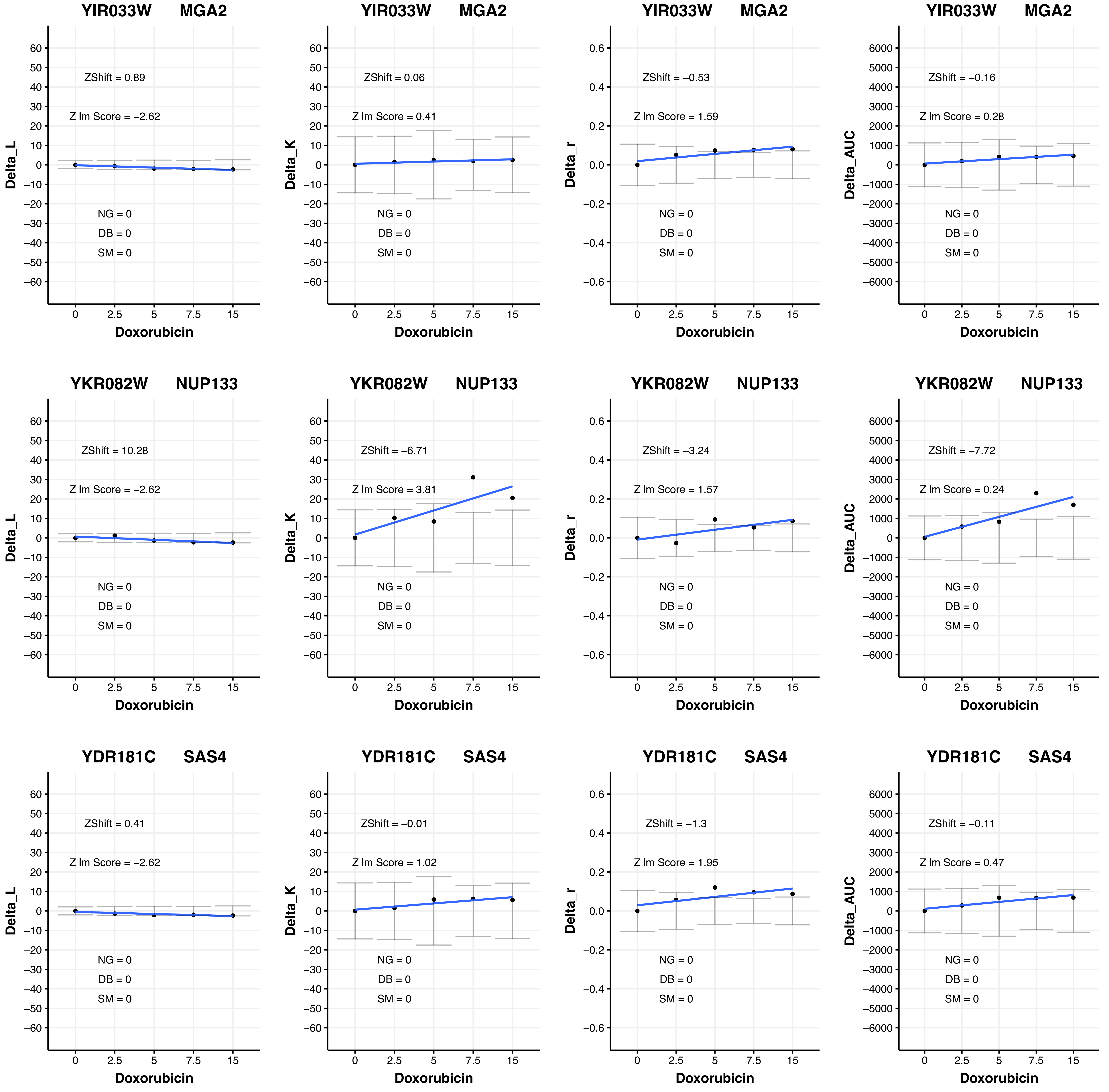
YIR033W MGA2
ZShift = 0.89
Z lm Score = −2.62
NG = 0
DB = 0
SM = 0
Delta_L
Doxorubicin
YIR033W MGA2
ZShift = 0.06
Z lm Score = 0.41
NG = 0
DB = 0
SM = 0
Delta_K
Doxorubicin
YIR033W MGA2
ZShift = −0.53
Z lm Score = 1.59
NG = 0
DB = 0
SM = 0
Delta_r
Doxorubicin
YIR033W MGA2
ZShift = −0.16
Z lm Score = 0.28
NG = 0
DB = 0
SM = 0
Delta_AUC
Doxorubicin
YKR082W NUP133
ZShift = 10.28
Z lm Score = −2.62
NG = 0
DB = 0
SM = 0
Delta_L
Doxorubicin
YKR082W NUP133
ZShift = −6.71
Z lm Score = 3.81
NG = 0
DB = 0
SM = 0
Delta_K
Doxorubicin
YKR082W NUP133
ZShift = −3.24
Z lm Score = 1.57
NG = 0
DB = 0
SM = 0
Delta_r
Doxorubicin
YKR082W NUP133
ZShift = −7.72
Z lm Score = 0.24
NG = 0
DB = 0
SM = 0
Delta_AUC
Doxorubicin
YDR181C SAS4
ZShift = 0.41
Z lm Score = −2.62
NG = 0
DB = 0
SM = 0
Delta_L
Doxorubicin
YDR181C SAS4
ZShift = −0.01
Z lm Score = 1.02
NG = 0
DB = 0
SM = 0
Delta_K
Doxorubicin
YDR181C SAS4
ZShift = −1.3
Z lm Score = 1.95
NG = 0
DB = 0
SM = 0
Delta_r
Doxorubicin
YDR181C SAS4
ZShift = −0.11
Z lm Score = 0.47
NG = 0
DB = 0
SM = 0
Delta_AUC
Doxorubicin

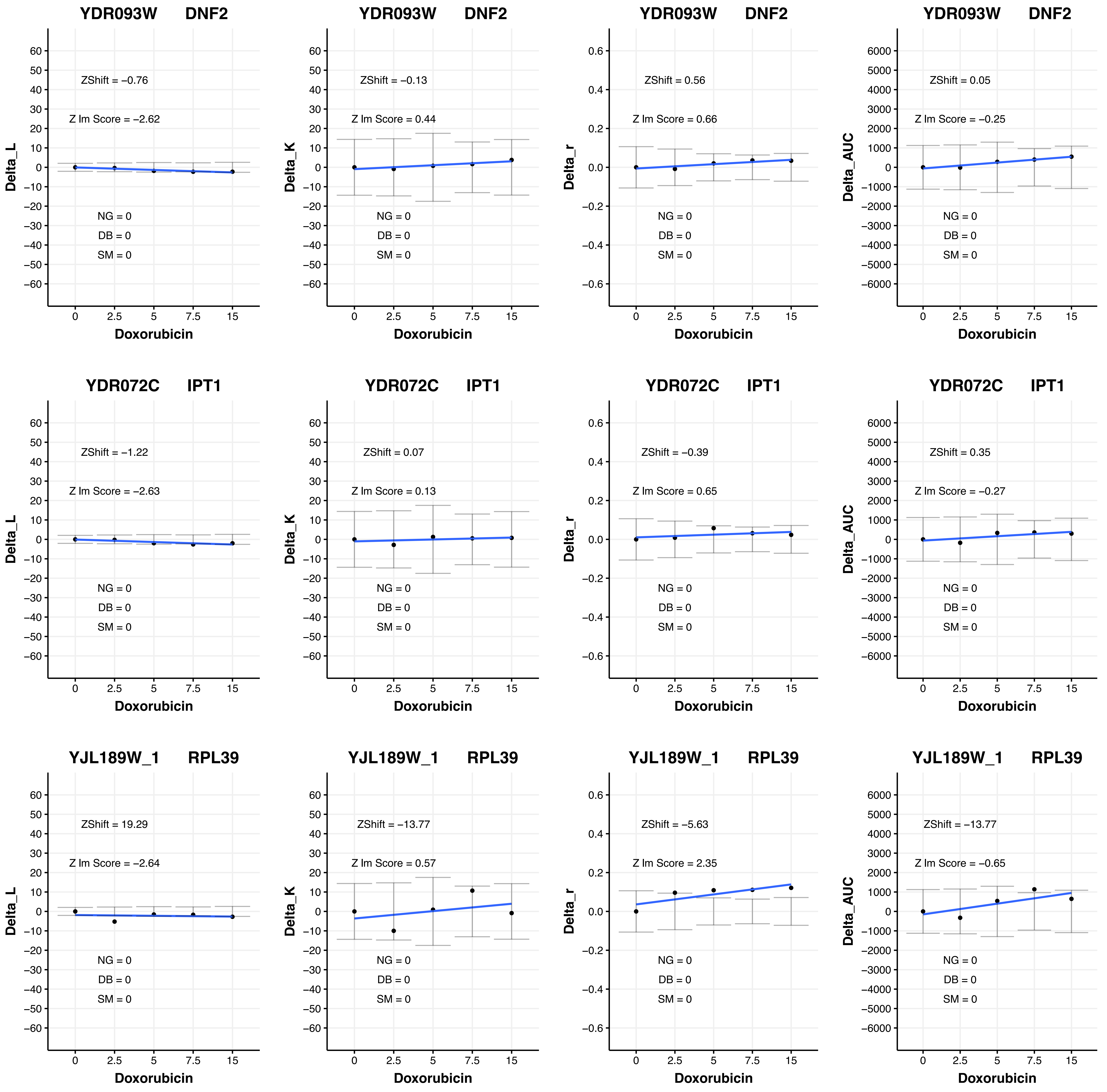
YDR093W DNF2
ZShift = −0.76
Z lm Score = −2.62
NG = 0
DB = 0
SM = 0
Delta_L
Doxorubicin
YDR093W DNF2
ZShift = −0.13
Z lm Score = 0.44
NG = 0
DB = 0
SM = 0
Delta_K
Doxorubicin
YDR093W DNF2
ZShift = 0.56
Z lm Score = 0.66
NG = 0
DB = 0
SM = 0
Delta_r
Doxorubicin
YDR093W DNF2
ZShift = 0.05
Z lm Score = −0.25
NG = 0
DB = 0
SM = 0
Delta_AUC
Doxorubicin
YDR072C IPT1
ZShift = −1.22
Z lm Score = −2.63
NG = 0
DB = 0
SM = 0
Delta_L
Doxorubicin
YDR072C IPT1
ZShift = 0.07
Z lm Score = 0.13
NG = 0
DB = 0
SM = 0
Delta_K
Doxorubicin
YDR072C IPT1
ZShift = −0.39
Z lm Score = 0.65
NG = 0
DB = 0
SM = 0
Delta_r
Doxorubicin
YDR072C IPT1
ZShift = 0.35
Z lm Score = −0.27
NG = 0
DB = 0
SM = 0
Delta_AUC
Doxorubicin
YJL189W_1 RPL39
ZShift = 19.29
Z lm Score = −2.64
NG = 0
DB = 0
SM = 0
Delta_L
Doxorubicin
YJL189W_1 RPL39
ZShift = −13.77
Z lm Score = 0.57
NG = 0
DB = 0
SM = 0
Delta_K
Doxorubicin
YJL189W_1 RPL39
ZShift = −5.63
Z lm Score = 2.35
NG = 0
DB = 0
SM = 0
Delta_r
Doxorubicin
YJL189W_1 RPL39
ZShift = −13.77
Z lm Score = −0.65
NG = 0
DB = 0
SM = 0
Delta_AUC
Doxorubicin

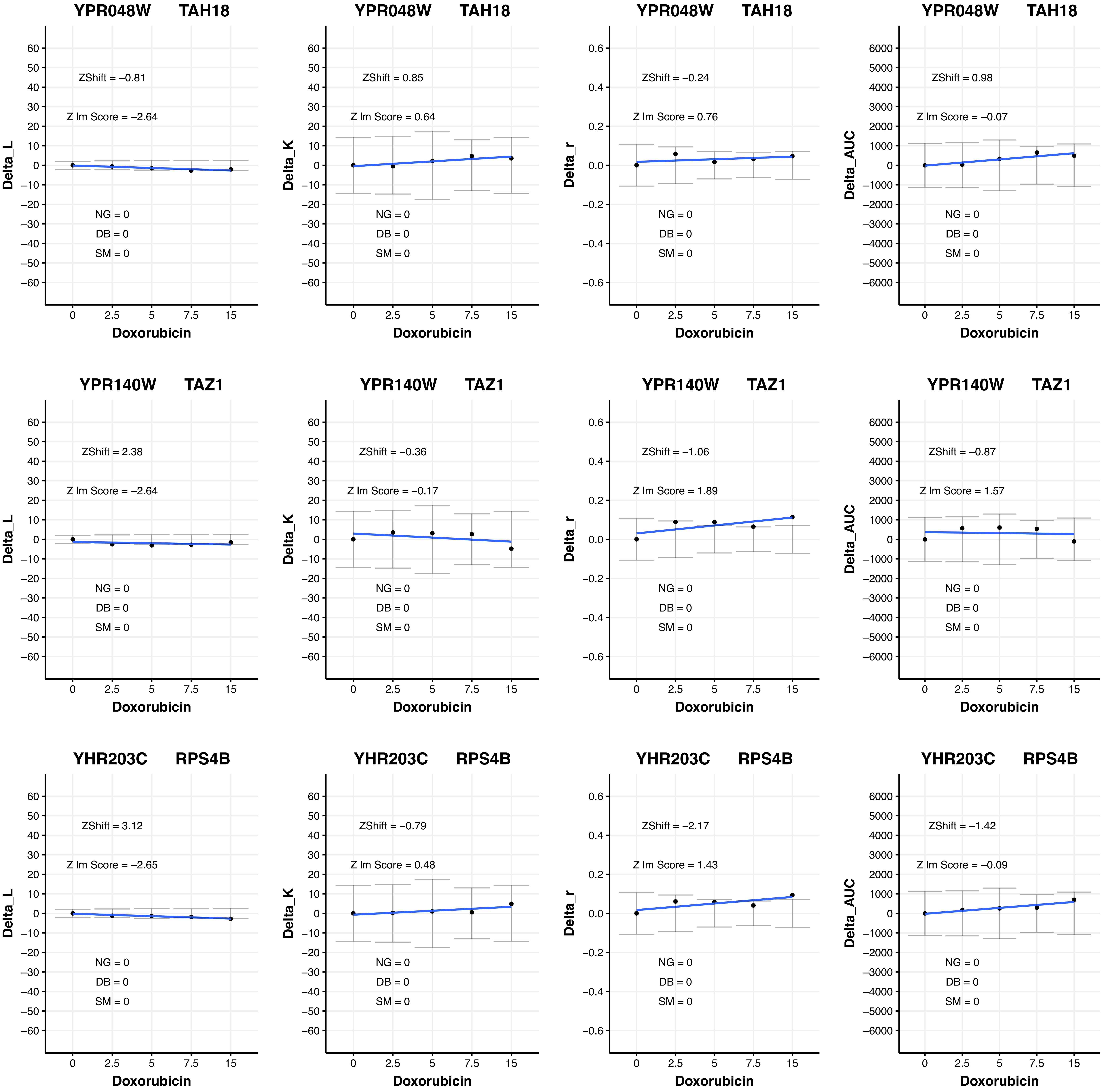
YPR048W TAH18
ZShift = −0.81
Z lm Score = −2.64
NG = 0
DB = 0
SM = 0
Delta_L
Doxorubicin
YPR048W TAH18
ZShift = 0.85
Z lm Score = 0.64
NG = 0
DB = 0
SM = 0
Delta_K
Doxorubicin
YPR048W TAH18
ZShift = −0.24
Z lm Score = 0.76
NG = 0
DB = 0
SM = 0
Delta_r
Doxorubicin
YPR048W TAH18
ZShift = 0.98
Z lm Score = −0.07
NG = 0
DB = 0
SM = 0
Delta_AUC
Doxorubicin
YPR140W TAZ1
ZShift = 2.38
Z lm Score = −2.64
NG = 0
DB = 0
SM = 0
Delta_L
Doxorubicin
YPR140W TAZ1
ZShift = −0.36
Z lm Score = −0.17
NG = 0
DB = 0
SM = 0
Delta_K
Doxorubicin
YPR140W TAZ1
ZShift = −1.06
Z lm Score = 1.89
NG = 0
DB = 0
SM = 0
Delta_r
Doxorubicin
YPR140W TAZ1
ZShift = −0.87
Z lm Score = 1.57
NG = 0
DB = 0
SM = 0
Delta_AUC
Doxorubicin
YHR203C RPS4B
ZShift = 3.12
Z lm Score = −2.65
NG = 0
DB = 0
SM = 0
Delta_L
Doxorubicin
YHR203C RPS4B
ZShift = −0.79
Z lm Score = 0.48
NG = 0
DB = 0
SM = 0
Delta_K
Doxorubicin
YHR203C RPS4B
ZShift = −2.17
Z lm Score = 1.43
NG = 0
DB = 0
SM = 0
Delta_r
Doxorubicin
YHR203C RPS4B
ZShift = −1.42
Z lm Score = −0.09
NG = 0
DB = 0
SM = 0
Delta_AUC
Doxorubicin

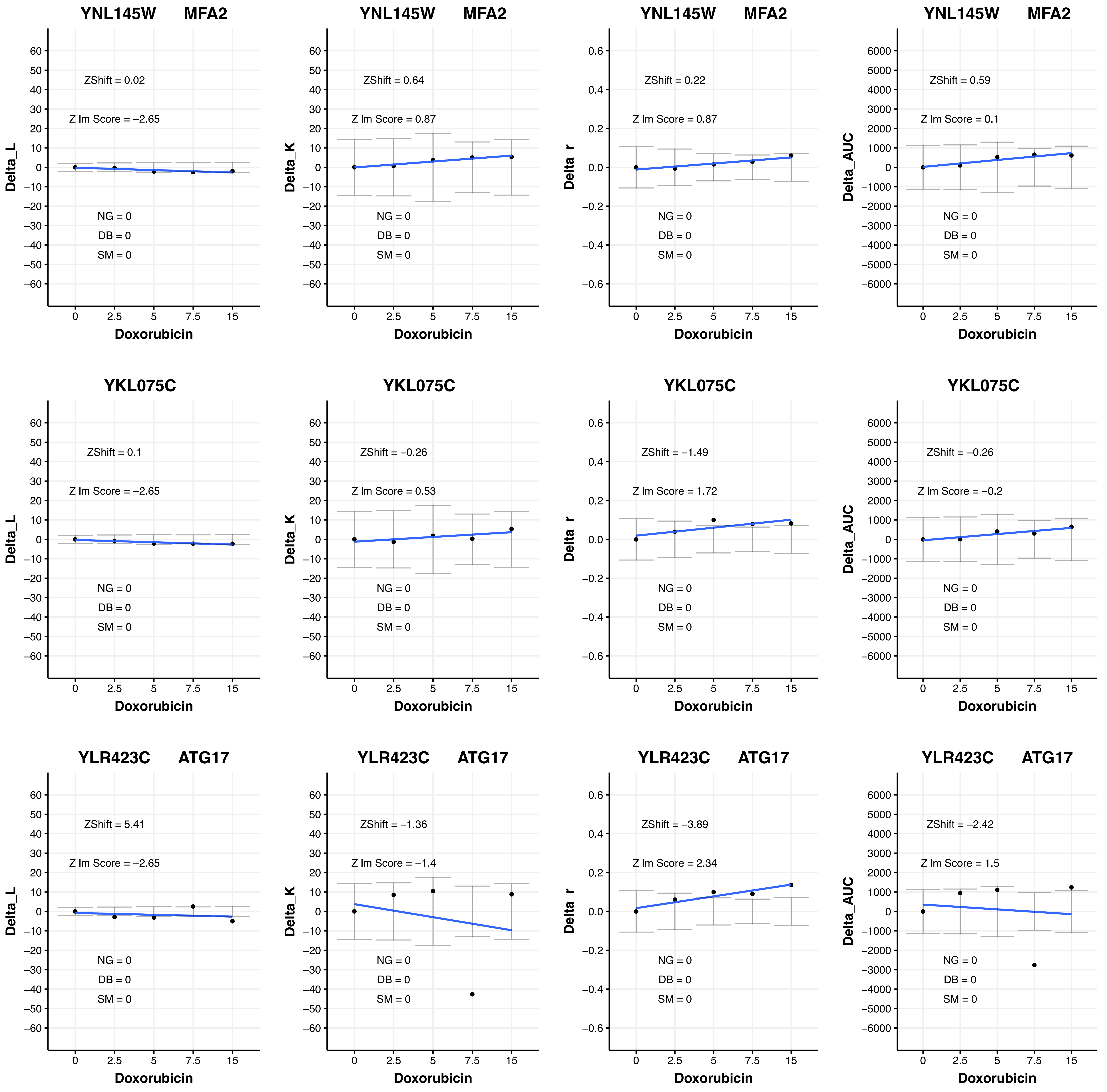
YNL145W MFA2
ZShift = 0.02
Z lm Score = −2.65
NG = 0
DB = 0
SM = 0
Delta_L
Doxorubicin
YNL145W MFA2
ZShift = 0.64
Z lm Score = 0.87
NG = 0
DB = 0
SM = 0
Delta_K
Doxorubicin
YNL145W MFA2
ZShift = 0.22
Z lm Score = 0.87
NG = 0
DB = 0
SM = 0
Delta_r
Doxorubicin
YNL145W MFA2
ZShift = 0.59
Z lm Score = 0.1
NG = 0
DB = 0
SM = 0
Delta_AUC
Doxorubicin
YKL075C
ZShift = 0.1
Z lm Score = −2.65
NG = 0
DB = 0
SM = 0
Delta_L
Doxorubicin
YKL075C
ZShift = −0.26
Z lm Score = 0.53
NG = 0
DB = 0
SM = 0
Delta_K
Doxorubicin
YKL075C
ZShift = −1.49
Z lm Score = 1.72
NG = 0
DB = 0
SM = 0
Delta_r
Doxorubicin
YKL075C
ZShift = −0.26
Z lm Score = −0.2
NG = 0
DB = 0
SM = 0
Delta_AUC
Doxorubicin
YLR423C ATG17
ZShift = 5.41
Z lm Score = −2.65
NG = 0
DB = 0
SM = 0
Delta_L
Doxorubicin
YLR423C ATG17
ZShift = −1.36
Z lm Score = −1.4
NG = 0
DB = 0
SM = 0
Delta_K
Doxorubicin
YLR423C ATG17
ZShift = −3.89
Z lm Score = 2.34
NG = 0
DB = 0
SM = 0
Delta_r
Doxorubicin
YLR423C ATG17
ZShift = −2.42
Z lm Score = 1.5
NG = 0
DB = 0
SM = 0
Delta_AUC
Doxorubicin

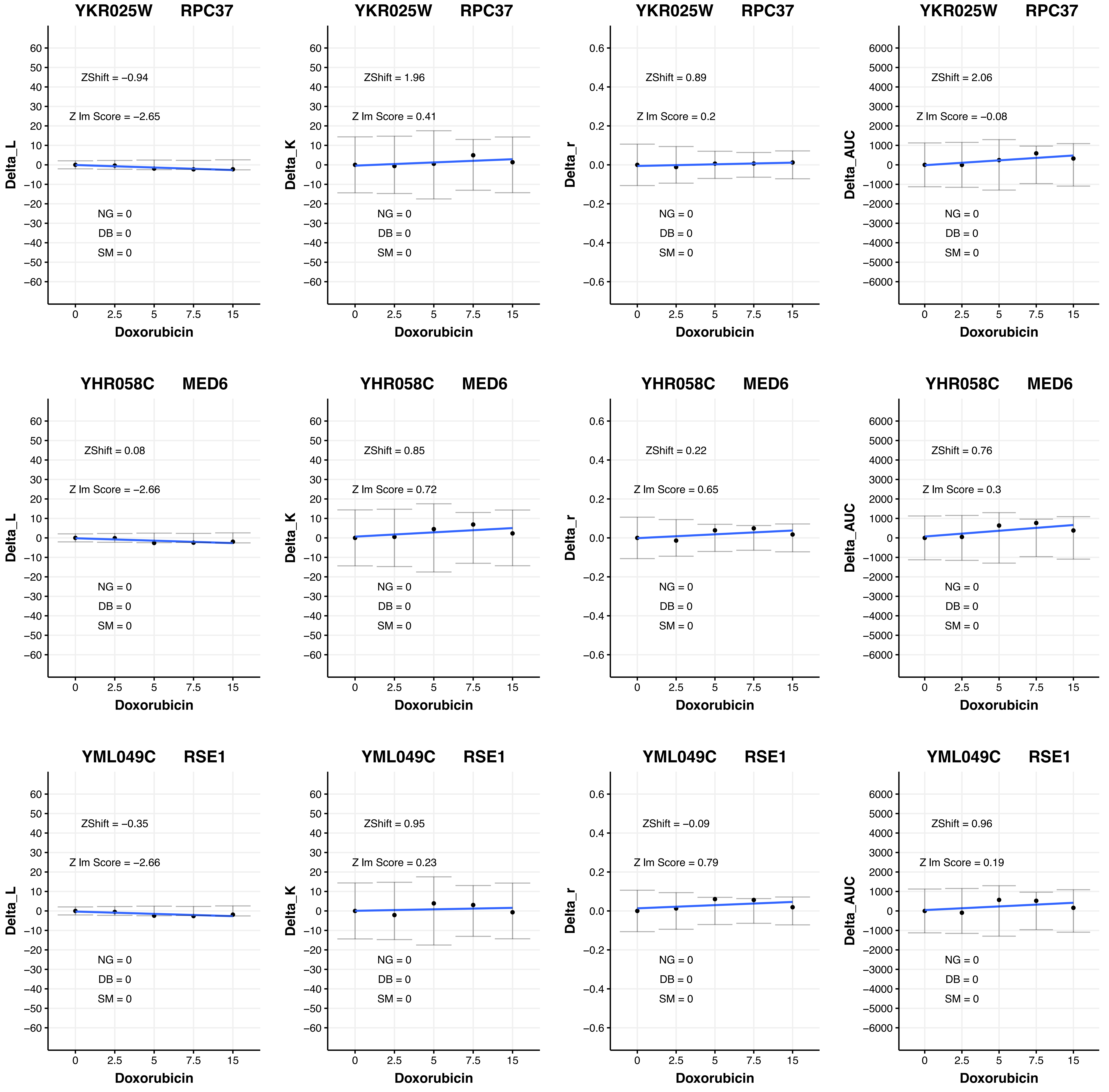
YKR025W RPC37
ZShift = −0.94
Z lm Score = −2.65
NG = 0
DB = 0
SM = 0
Delta_L
Doxorubicin
YKR025W RPC37
ZShift = 1.96
Z lm Score = 0.41
NG = 0
DB = 0
SM = 0
Delta_K
Doxorubicin
YKR025W RPC37
ZShift = 0.89
Z lm Score = 0.2
NG = 0
DB = 0
SM = 0
Delta_r
Doxorubicin
YKR025W RPC37
ZShift = 2.06
Z lm Score = −0.08
NG = 0
DB = 0
SM = 0
Delta_AUC
Doxorubicin
YHR058C MED6
ZShift = 0.08
Z lm Score = −2.66
NG = 0
DB = 0
SM = 0
Delta_L
Doxorubicin
YHR058C MED6
ZShift = 0.85
Z lm Score = 0.72
NG = 0
DB = 0
SM = 0
Delta_K
Doxorubicin
YHR058C MED6
ZShift = 0.22
Z lm Score = 0.65
NG = 0
DB = 0
SM = 0
Delta_r
Doxorubicin
YHR058C MED6
ZShift = 0.76
Z lm Score = 0.3
NG = 0
DB = 0
SM = 0
Delta_AUC
Doxorubicin
YML049C RSE1
ZShift = −0.35
Z lm Score = −2.66
NG = 0
DB = 0
SM = 0
Delta_L
Doxorubicin
YML049C RSE1
ZShift = 0.95
Z lm Score = 0.23
NG = 0
DB = 0
SM = 0
Delta_K
Doxorubicin
YML049C RSE1
ZShift = −0.09
Z lm Score = 0.79
NG = 0
DB = 0
SM = 0
Delta_r
Doxorubicin
YML049C RSE1
ZShift = 0.96
Z lm Score = 0.19
NG = 0
DB = 0
SM = 0
Delta_AUC
Doxorubicin

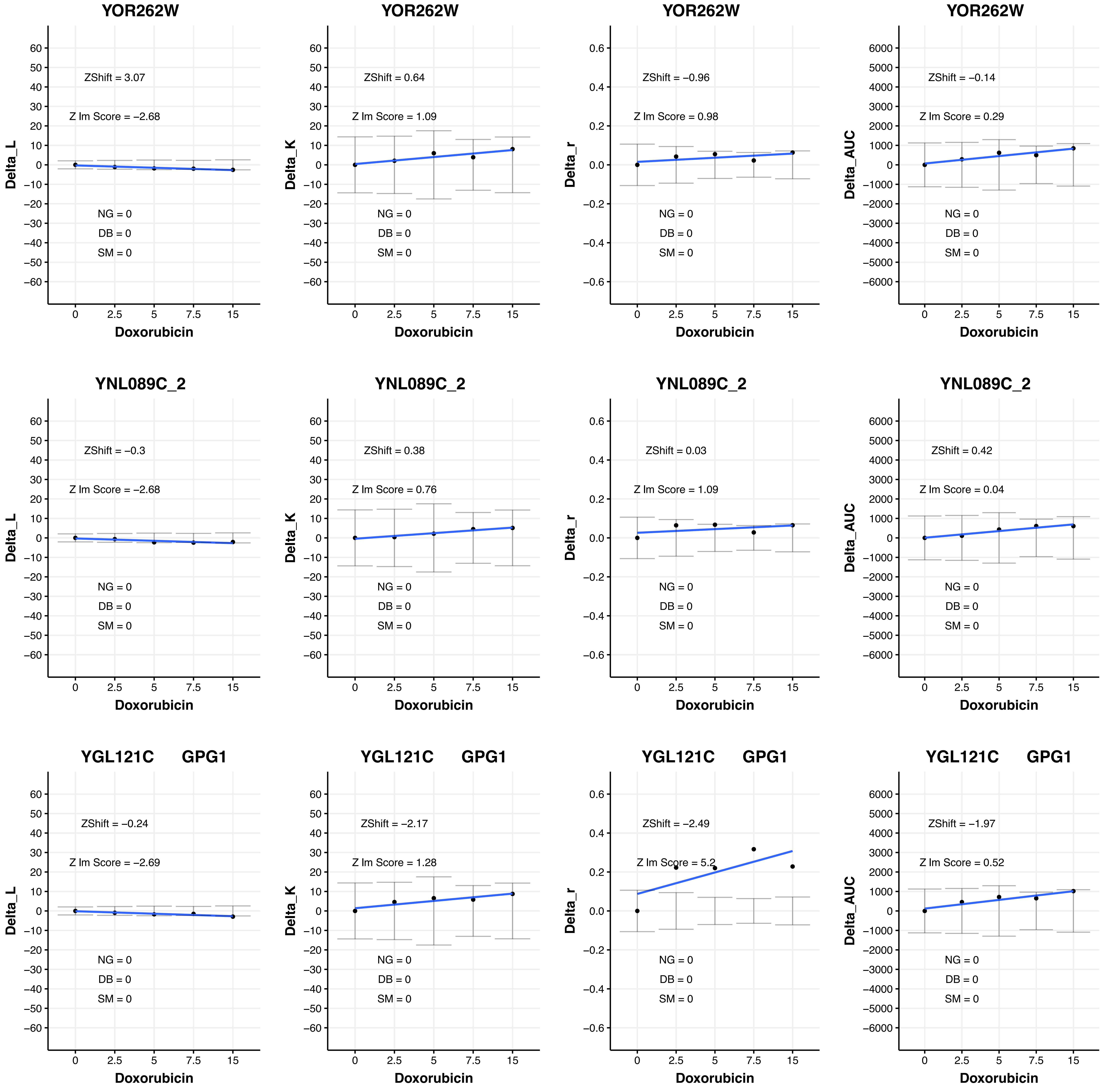
YOR262W
ZShift = 3.07
Z lm Score = −2.68
NG = 0
DB = 0
SM = 0
Delta_L
Doxorubicin
YOR262W
ZShift = 0.64
Z lm Score = 1.09
NG = 0
DB = 0
SM = 0
Delta_K
Doxorubicin
YOR262W
ZShift = −0.96
Z lm Score = 0.98
NG = 0
DB = 0
SM = 0
Delta_r
Doxorubicin
YOR262W
ZShift = −0.14
Z lm Score = 0.29
NG = 0
DB = 0
SM = 0
Delta_AUC
Doxorubicin
YNL089C_2
ZShift = −0.3
Z lm Score = −2.68
NG = 0
DB = 0
SM = 0
Delta_L
Doxorubicin
YNL089C_2
ZShift = 0.38
Z lm Score = 0.76
NG = 0
DB = 0
SM = 0
Delta_K
Doxorubicin
YNL089C_2
ZShift = 0.03
Z lm Score = 1.09
NG = 0
DB = 0
SM = 0
Delta_r
Doxorubicin
YNL089C_2
ZShift = 0.42
Z lm Score = 0.04
NG = 0
DB = 0
SM = 0
Delta_AUC
Doxorubicin
YGL121C GPG1
ZShift = −0.24
Z lm Score = −2.69
NG = 0
DB = 0
SM = 0
Delta_L
Doxorubicin
YGL121C GPG1
ZShift = −2.17
Z lm Score = 1.28
NG = 0
DB = 0
SM = 0
Delta_K
Doxorubicin
YGL121C GPG1
ZShift = −2.49
Z lm Score = 5.2
NG = 0
DB = 0
SM = 0
Delta_r
Doxorubicin
YGL121C GPG1
ZShift = −1.97
Z lm Score = 0.52
NG = 0
DB = 0
SM = 0
Delta_AUC
Doxorubicin

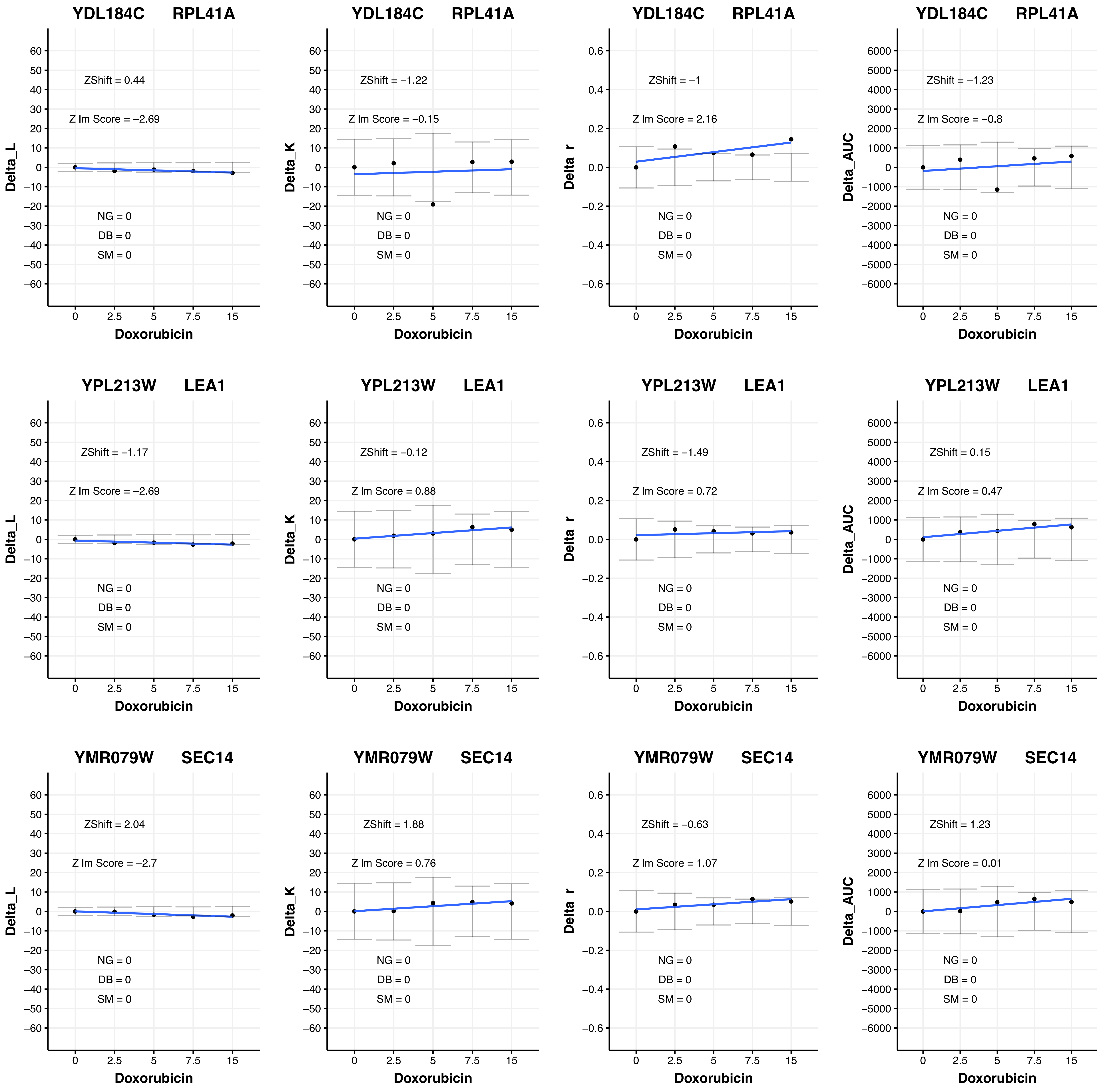
YDL184C RPL41A
ZShift = 0.44
Z lm Score = −2.69
NG = 0
DB = 0
SM = 0
Delta_L
Doxorubicin
YDL184C RPL41A
ZShift = −1.22
Z lm Score = −0.15
NG = 0
DB = 0
SM = 0
Delta_K
Doxorubicin
YDL184C RPL41A
ZShift = −1
Z lm Score = 2.16
NG = 0
DB = 0
SM = 0
Delta_r
Doxorubicin
YDL184C RPL41A
ZShift = −1.23
Z lm Score = −0.8
NG = 0
DB = 0
SM = 0
Delta_AUC
Doxorubicin
YPL213W LEA1
ZShift = −1.17
Z lm Score = −2.69
NG = 0
DB = 0
SM = 0
Delta_L
Doxorubicin
YPL213W LEA1
ZShift = −0.12
Z lm Score = 0.88
NG = 0
DB = 0
SM = 0
Delta_K
Doxorubicin
YPL213W LEA1
ZShift = −1.49
Z lm Score = 0.72
NG = 0
DB = 0
SM = 0
Delta_r
Doxorubicin
YPL213W LEA1
ZShift = 0.15
Z lm Score = 0.47
NG = 0
DB = 0
SM = 0
Delta_AUC
Doxorubicin
YMR079W SEC14
ZShift = 2.04
Z lm Score = −2.7
NG = 0
DB = 0
SM = 0
Delta_L
Doxorubicin
YMR079W SEC14
ZShift = 1.88
Z lm Score = 0.76
NG = 0
DB = 0
SM = 0
Delta_K
Doxorubicin
YMR079W SEC14
ZShift = −0.63
Z lm Score = 1.07
NG = 0
DB = 0
SM = 0
Delta_r
Doxorubicin
YMR079W SEC14
ZShift = 1.23
Z lm Score = 0.01
NG = 0
DB = 0
SM = 0
Delta_AUC
Doxorubicin

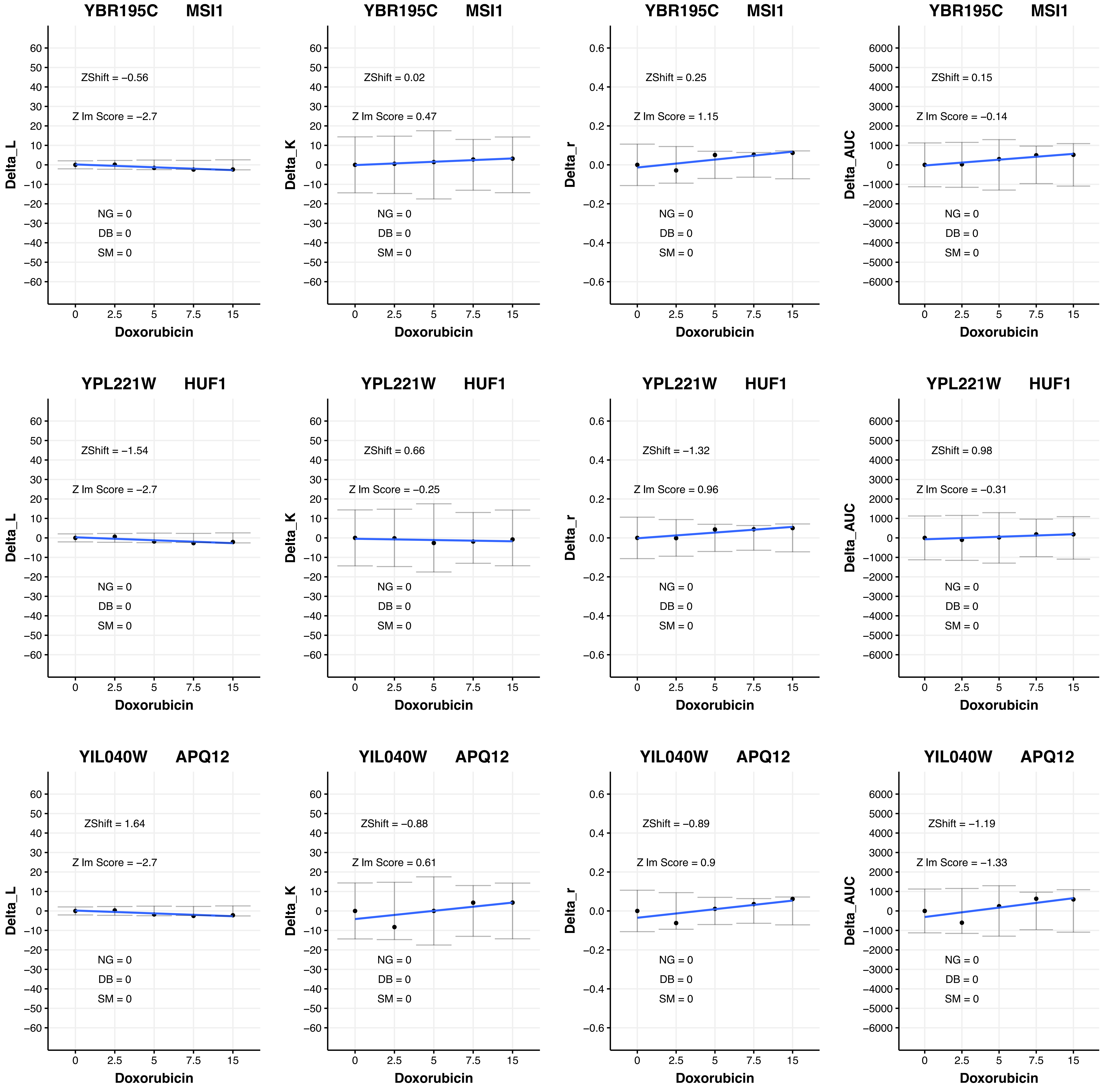
YBR195C MSI1
ZShift = −0.56
Z lm Score = −2.7
NG = 0
DB = 0
SM = 0
Delta_L
Doxorubicin
YBR195C MSI1
ZShift = 0.02
Z lm Score = 0.47
NG = 0
DB = 0
SM = 0
Delta_K
Doxorubicin
YBR195C MSI1
ZShift = 0.25
Z lm Score = 1.15
NG = 0
DB = 0
SM = 0
Delta_r
Doxorubicin
YBR195C MSI1
ZShift = 0.15
Z lm Score = −0.14
NG = 0
DB = 0
SM = 0
Delta_AUC
Doxorubicin
YPL221W HUF1
ZShift = −1.54
Z lm Score = −2.7
NG = 0
DB = 0
SM = 0
Delta_L
Doxorubicin
YPL221W HUF1
ZShift = 0.66
Z lm Score = −0.25
NG = 0
DB = 0
SM = 0
Delta_K
Doxorubicin
YPL221W HUF1
ZShift = −1.32
Z lm Score = 0.96
NG = 0
DB = 0
SM = 0
Delta_r
Doxorubicin
YPL221W HUF1
ZShift = 0.98
Z lm Score = −0.31
NG = 0
DB = 0
SM = 0
Delta_AUC
Doxorubicin
YIL040W APQ12
ZShift = 1.64
Z lm Score = −2.7
NG = 0
DB = 0
SM = 0
Delta_L
Doxorubicin
YIL040W APQ12
ZShift = −0.88
Z lm Score = 0.61
NG = 0
DB = 0
SM = 0
Delta_K
Doxorubicin
YIL040W APQ12
ZShift = −0.89
Z lm Score = 0.9
NG = 0
DB = 0
SM = 0
Delta_r
Doxorubicin
YIL040W APQ12
ZShift = −1.19
Z lm Score = −1.33
NG = 0
DB = 0
SM = 0
Delta_AUC
Doxorubicin

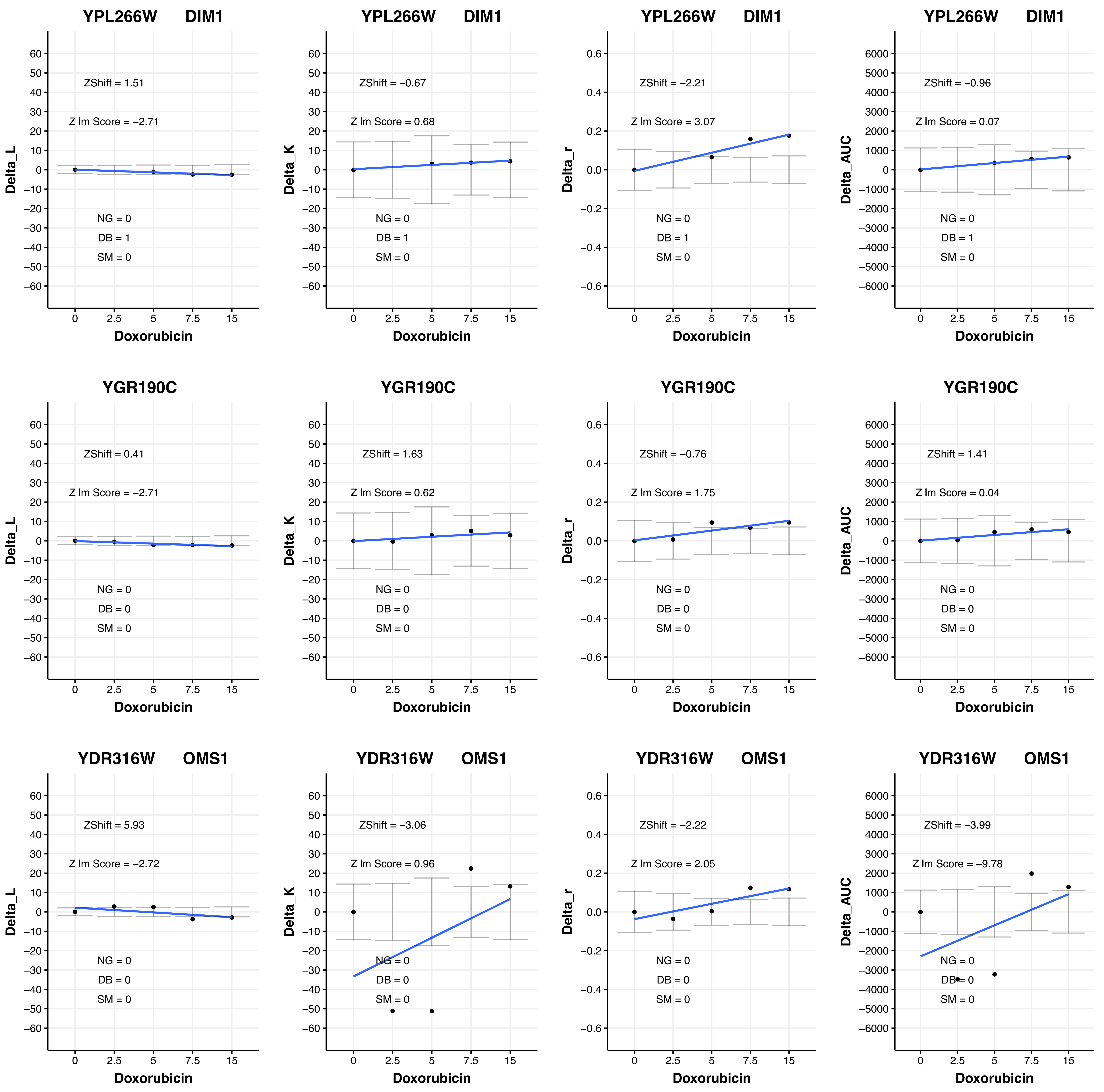
YPL266W DIM1
ZShift = 1.51
Z lm Score = −2.71
NG = 0
DB = 1
SM = 0
Delta_L
Doxorubicin
YPL266W DIM1
ZShift = −0.67
Z lm Score = 0.68
NG = 0
DB = 1
SM = 0
Delta_K
Doxorubicin
YPL266W DIM1
ZShift = −2.21
Z lm Score = 3.07
NG = 0
DB = 1
SM = 0
Delta_r
Doxorubicin
YPL266W DIM1
ZShift = −0.96
Z lm Score = 0.07
NG = 0
DB = 1
SM = 0
Delta_AUC
Doxorubicin
YGR190C
ZShift = 0.41
Z lm Score = −2.71
NG = 0
DB = 0
SM = 0
Delta_L
Doxorubicin
YGR190C
ZShift = 1.63
Z lm Score = 0.62
NG = 0
DB = 0
SM = 0
Delta_K
Doxorubicin
YGR190C
ZShift = −0.76
Z lm Score = 1.75
NG = 0
DB = 0
SM = 0
Delta_r
Doxorubicin
YGR190C
ZShift = 1.41
Z lm Score = 0.04
NG = 0
DB = 0
SM = 0
Delta_AUC
Doxorubicin
YDR316W OMS1
ZShift = 5.93
Z lm Score = −2.72
NG = 0
DB = 0
SM = 0
Delta_L
Doxorubicin
YDR316W OMS1
ZShift = −3.06
Z lm Score = 0.96
NG = 0
DB = 0
SM = 0
Delta_K
Doxorubicin
YDR316W OMS1
ZShift = −2.22
Z lm Score = 2.05
NG = 0
DB = 0
SM = 0
Delta_r
Doxorubicin
YDR316W OMS1
ZShift = −3.99
Z lm Score = −9.78
NG = 0
DB = 0
SM = 0
Delta_AUC
Doxorubicin

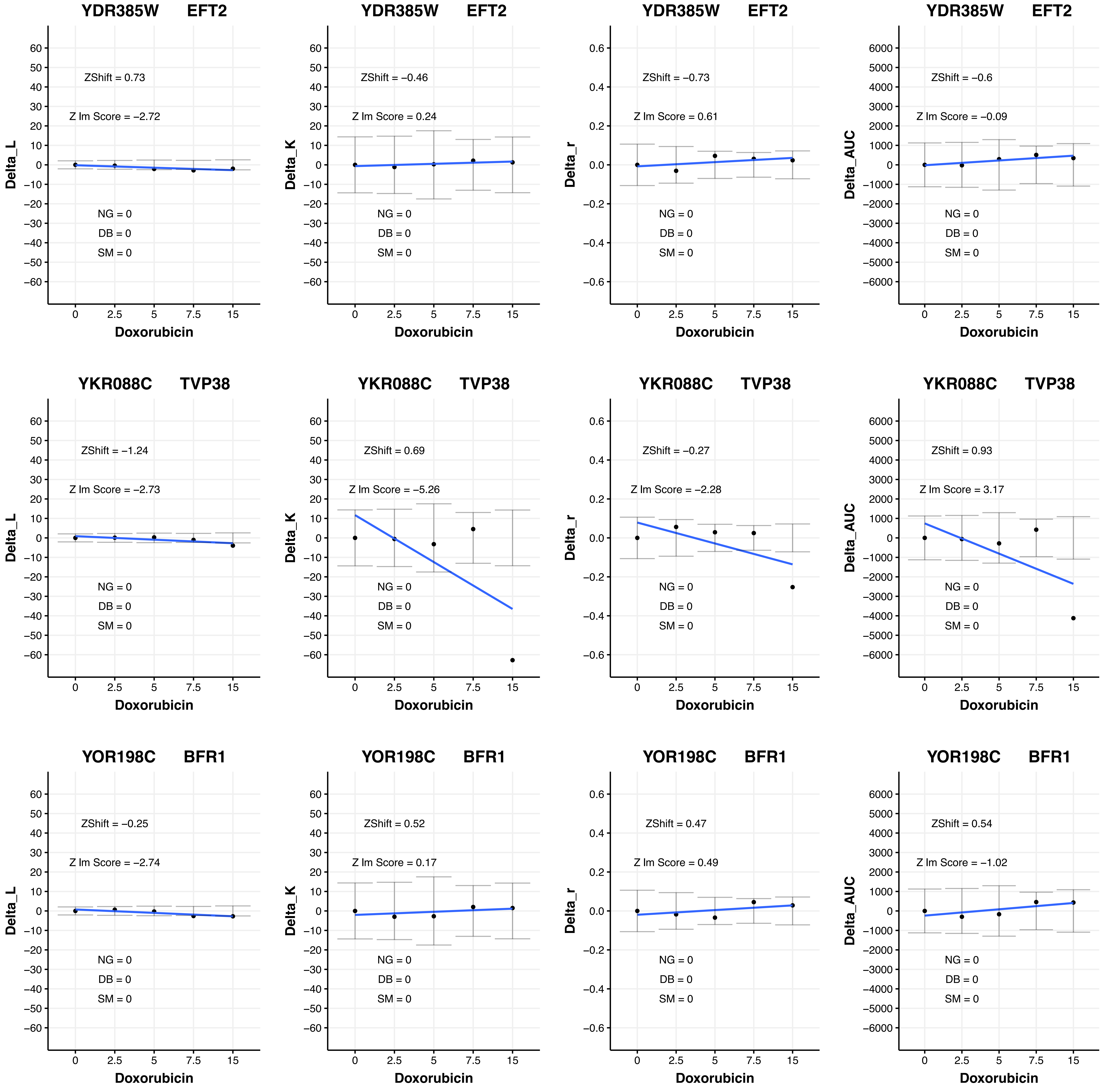
YDR385W EFT2
Delta_L
ZShift = 0.73
Z lm Score = −2.72
NG = 0
DB = 0
SM = 0
Doxorubicin
YDR385W EFT2
Delta_K
ZShift = −0.46
Z lm Score = 0.24
NG = 0
DB = 0
SM = 0
Doxorubicin
YDR385W EFT2
Delta_r
ZShift = −0.73
Z lm Score = 0.61
NG = 0
DB = 0
SM = 0
Doxorubicin
YDR385W EFT2
Delta_AUC
ZShift = −0.6
Z lm Score = −0.09
NG = 0
DB = 0
SM = 0
Doxorubicin
YKR088C TVP38
Delta_L
ZShift = −1.24
Z lm Score = −2.73
NG = 0
DB = 0
SM = 0
Doxorubicin
YKR088C TVP38
Delta_K
ZShift = 0.69
Z lm Score = −5.26
NG = 0
DB = 0
SM = 0
Doxorubicin
YKR088C TVP38
Delta_r
ZShift = −0.27
Z lm Score = −2.28
NG = 0
DB = 0
SM = 0
Doxorubicin
YKR088C TVP38
Delta_AUC
ZShift = 0.93
Z lm Score = 3.17
NG = 0
DB = 0
SM = 0
Doxorubicin
YOR198C BFR1
Delta_L
ZShift = −0.25
Z lm Score = −2.74
NG = 0
DB = 0
SM = 0
Doxorubicin
YOR198C BFR1
Delta_K
ZShift = 0.52
Z lm Score = 0.17
NG = 0
DB = 0
SM = 0
Doxorubicin
YOR198C BFR1
Delta_r
ZShift = 0.47
Z lm Score = 0.49
NG = 0
DB = 0
SM = 0
Doxorubicin
YOR198C BFR1
Delta_AUC
ZShift = 0.54
Z lm Score = −1.02
NG = 0
DB = 0
SM = 0
Doxorubicin

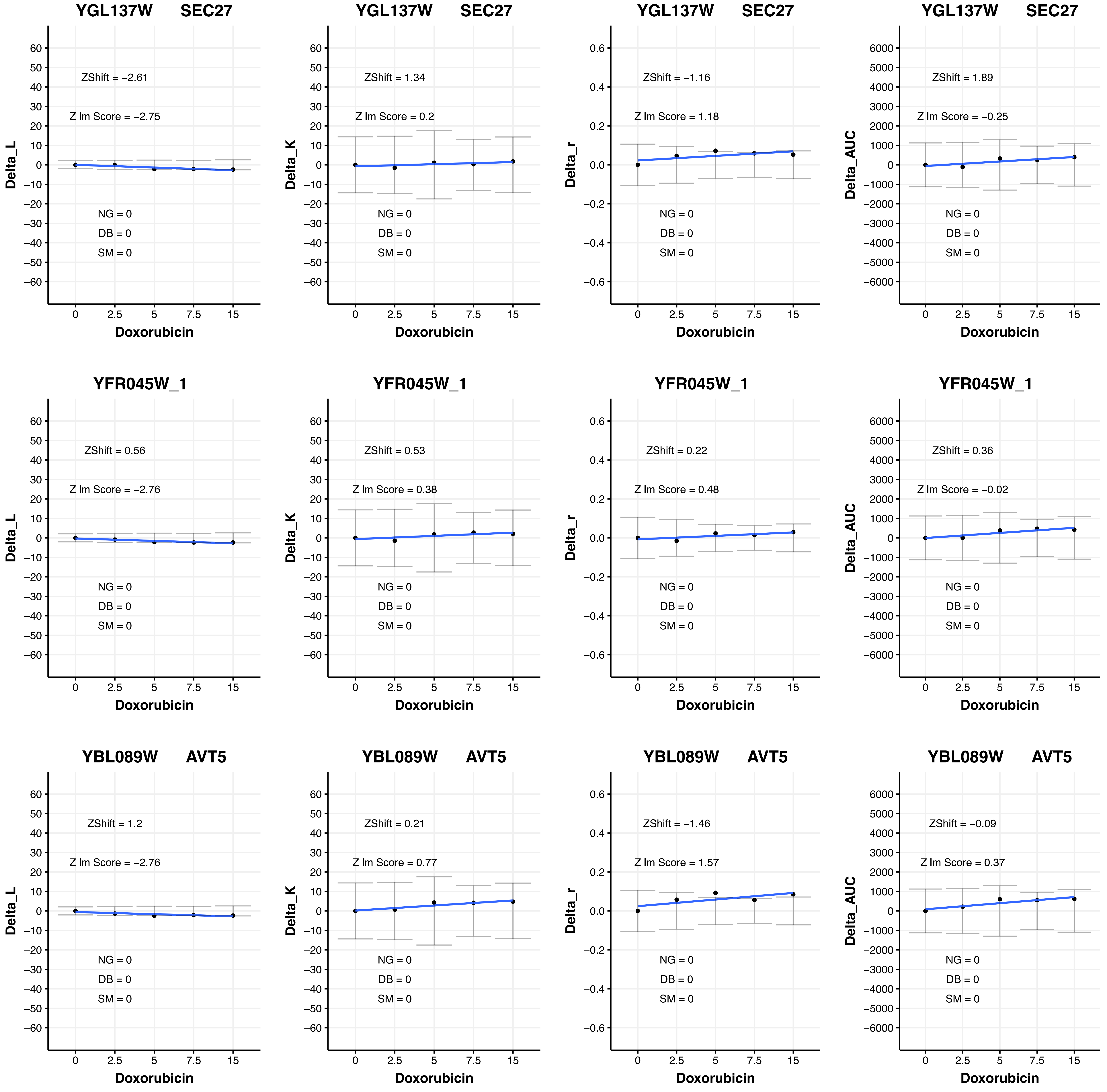
YGL137W SEC27
ZShift = −2.61
Z lm Score = −2.75
NG = 0
DB = 0
SM = 0
Delta_L
Doxorubicin
YGL137W SEC27
ZShift = 1.34
Z lm Score = 0.2
NG = 0
DB = 0
SM = 0
Delta_K
Doxorubicin
YGL137W SEC27
ZShift = −1.16
Z lm Score = 1.18
NG = 0
DB = 0
SM = 0
Delta_r
Doxorubicin
YGL137W SEC27
ZShift = 1.89
Z lm Score = −0.25
NG = 0
DB = 0
SM = 0
Delta_AUC
Doxorubicin
YFR045W_1
ZShift = 0.56
Z lm Score = −2.76
NG = 0
DB = 0
SM = 0
Delta_L
Doxorubicin
YFR045W_1
ZShift = 0.53
Z lm Score = 0.38
NG = 0
DB = 0
SM = 0
Delta_K
Doxorubicin
YFR045W_1
ZShift = 0.22
Z lm Score = 0.48
NG = 0
DB = 0
SM = 0
Delta_r
Doxorubicin
YFR045W_1
ZShift = 0.36
Z lm Score = −0.02
NG = 0
DB = 0
SM = 0
Delta_AUC
Doxorubicin
YBL089W AVT5
ZShift = 1.2
Z lm Score = −2.76
NG = 0
DB = 0
SM = 0
Delta_L
Doxorubicin
YBL089W AVT5
ZShift = 0.21
Z lm Score = 0.77
NG = 0
DB = 0
SM = 0
Delta_K
Doxorubicin
YBL089W AVT5
ZShift = −1.46
Z lm Score = 1.57
NG = 0
DB = 0
SM = 0
Delta_r
Doxorubicin
YBL089W AVT5
ZShift = −0.09
Z lm Score = 0.37
NG = 0
DB = 0
SM = 0
Delta_AUC
Doxorubicin

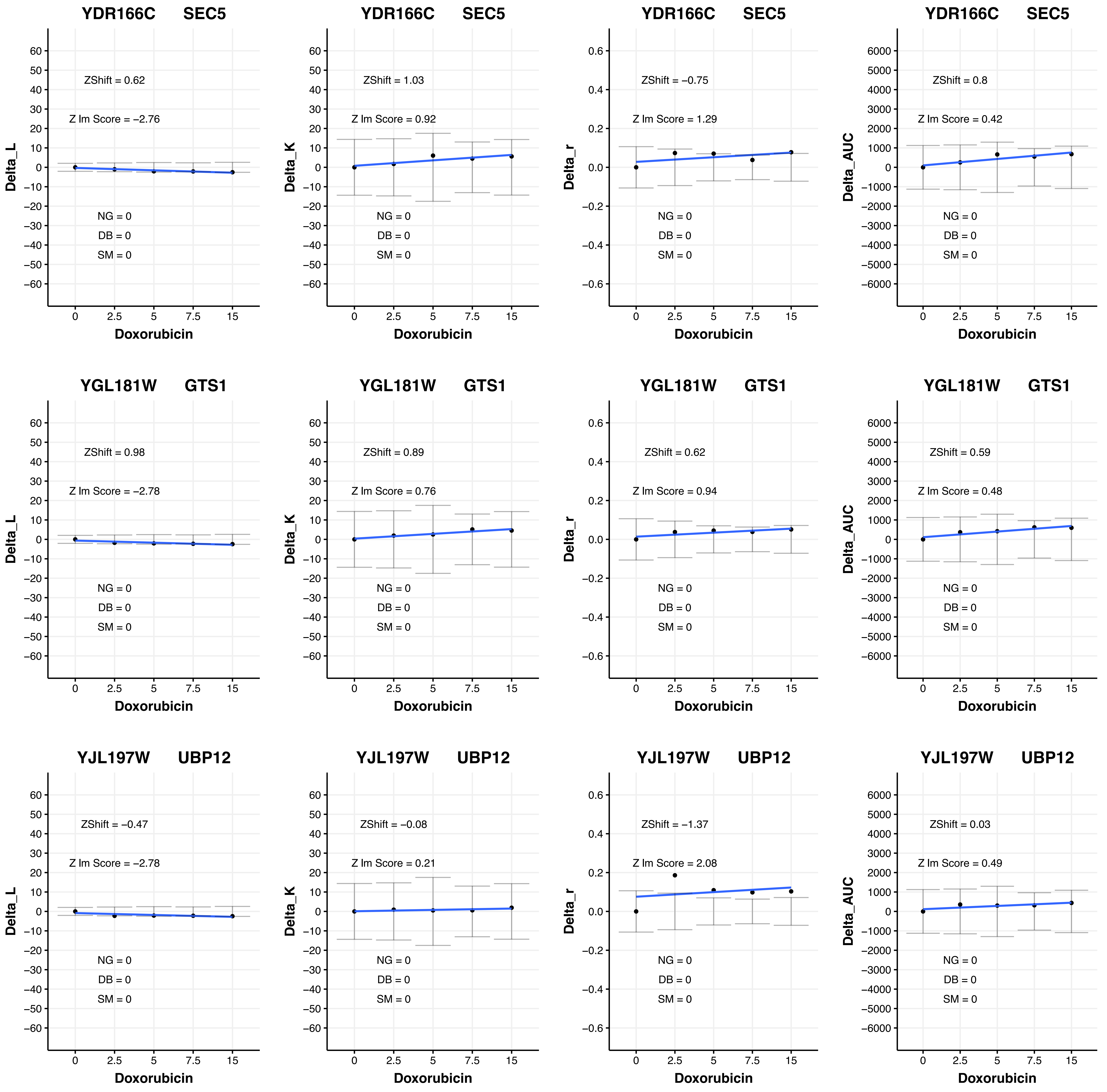
YDR166C SEC5
ZShift = 0.62
Z lm Score = −2.76
NG = 0
DB = 0
SM = 0
Delta_L
Doxorubicin
YDR166C SEC5
ZShift = 1.03
Z lm Score = 0.92
NG = 0
DB = 0
SM = 0
Delta_K
Doxorubicin
YDR166C SEC5
ZShift = −0.75
Z lm Score = 1.29
NG = 0
DB = 0
SM = 0
Delta_r
Doxorubicin
YDR166C SEC5
ZShift = 0.8
Z lm Score = 0.42
NG = 0
DB = 0
SM = 0
Delta_AUC
Doxorubicin
YGL181W GTS1
ZShift = 0.98
Z lm Score = −2.78
NG = 0
DB = 0
SM = 0
Delta_L
Doxorubicin
YGL181W GTS1
ZShift = 0.89
Z lm Score = 0.76
NG = 0
DB = 0
SM = 0
Delta_K
Doxorubicin
YGL181W GTS1
ZShift = 0.62
Z lm Score = 0.94
NG = 0
DB = 0
SM = 0
Delta_r
Doxorubicin
YGL181W GTS1
ZShift = 0.59
Z lm Score = 0.48
NG = 0
DB = 0
SM = 0
Delta_AUC
Doxorubicin
YJL197W UBP12
ZShift = −0.47
Z lm Score = −2.78
NG = 0
DB = 0
SM = 0
Delta_L
Doxorubicin
YJL197W UBP12
ZShift = −0.08
Z lm Score = 0.21
NG = 0
DB = 0
SM = 0
Delta_K
Doxorubicin
YJL197W UBP12
ZShift = −1.37
Z lm Score = 2.08
NG = 0
DB = 0
SM = 0
Delta_r
Doxorubicin
YJL197W UBP12
ZShift = 0.03
Z lm Score = 0.49
NG = 0
DB = 0
SM = 0
Delta_AUC
Doxorubicin

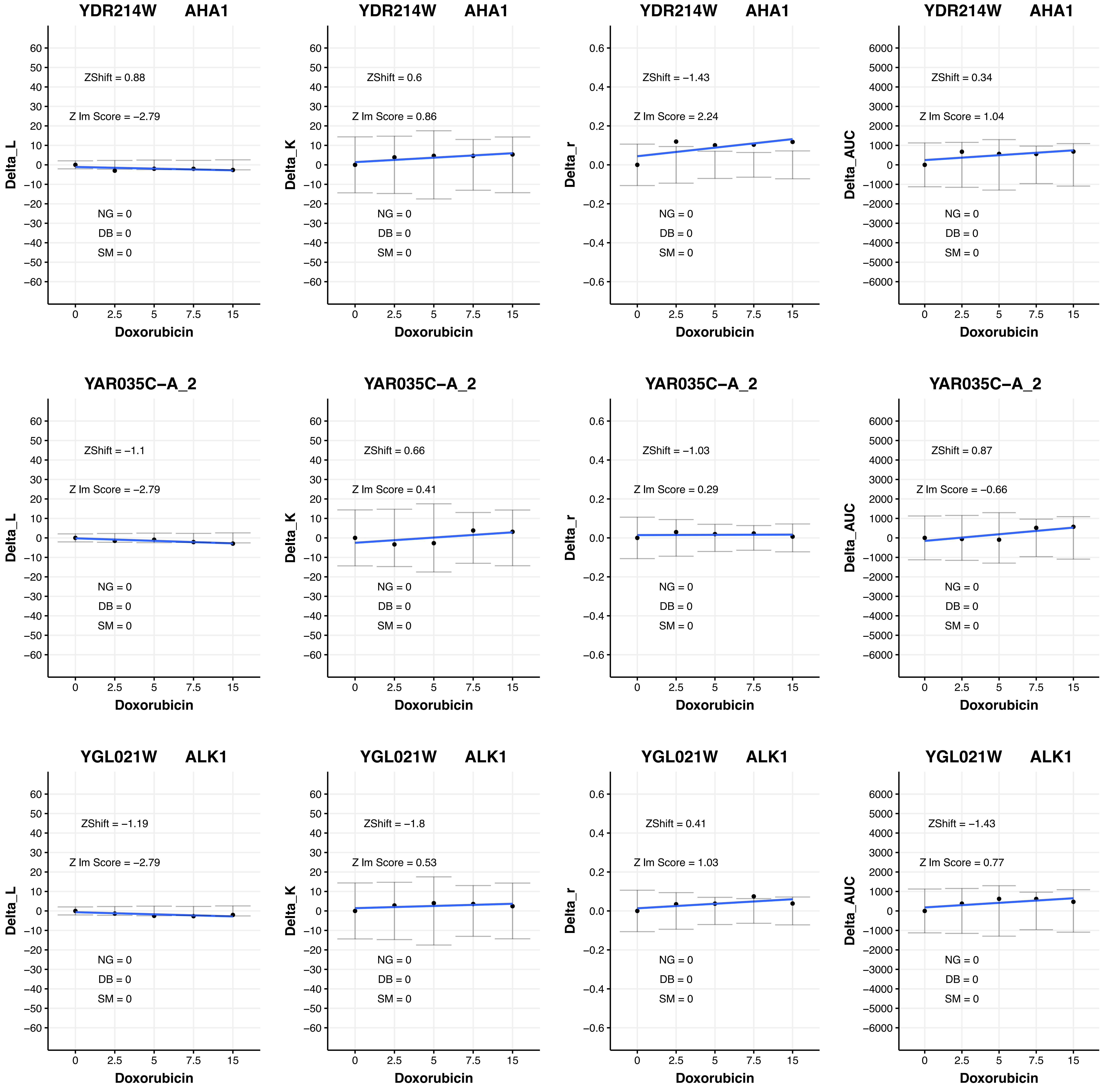

YDR214W AHA1
ZShift = 0.88
Z lm Score = −2.79
NG = 0
DB = 0
SM = 0
Delta_L
Doxorubicin
YDR214W AHA1
ZShift = 0.6
Z lm Score = 0.86
NG = 0
DB = 0
SM = 0
Delta_K
Doxorubicin
YDR214W AHA1
ZShift = −1.43
Z lm Score = 2.24
NG = 0
DB = 0
SM = 0
Delta_r
Doxorubicin
YDR214W AHA1
ZShift = 0.34
Z lm Score = 1.04
NG = 0
DB = 0
SM = 0
Delta_AUC
Doxorubicin
YAR035C−A_2
ZShift = −1.1
Z lm Score = −2.79
NG = 0
DB = 0
SM = 0
Delta_L
Doxorubicin
YAR035C−A_2
ZShift = 0.66
Z lm Score = 0.41
NG = 0
DB = 0
SM = 0
Delta_K
Doxorubicin
YAR035C−A_2
ZShift = −1.03
Z lm Score = 0.29
NG = 0
DB = 0
SM = 0
Delta_r
Doxorubicin
YAR035C−A_2
ZShift = 0.87
Z lm Score = −0.66
NG = 0
DB = 0
SM = 0
Delta_AUC
Doxorubicin
YGL021W ALK1
ZShift = −1.19
Z lm Score = −2.79
NG = 0
DB = 0
SM = 0
Delta_L
Doxorubicin
YGL021W ALK1
ZShift = −1.8
Z lm Score = 0.53
NG = 0
DB = 0
SM = 0
Delta_K
Doxorubicin
YGL021W ALK1
ZShift = 0.41
Z lm Score = 1.03
NG = 0
DB = 0
SM = 0
Delta_r
Doxorubicin
YGL021W ALK1
ZShift = −1.43
Z lm Score = 0.77
NG = 0
DB = 0
SM = 0
Delta_AUC
Doxorubicin

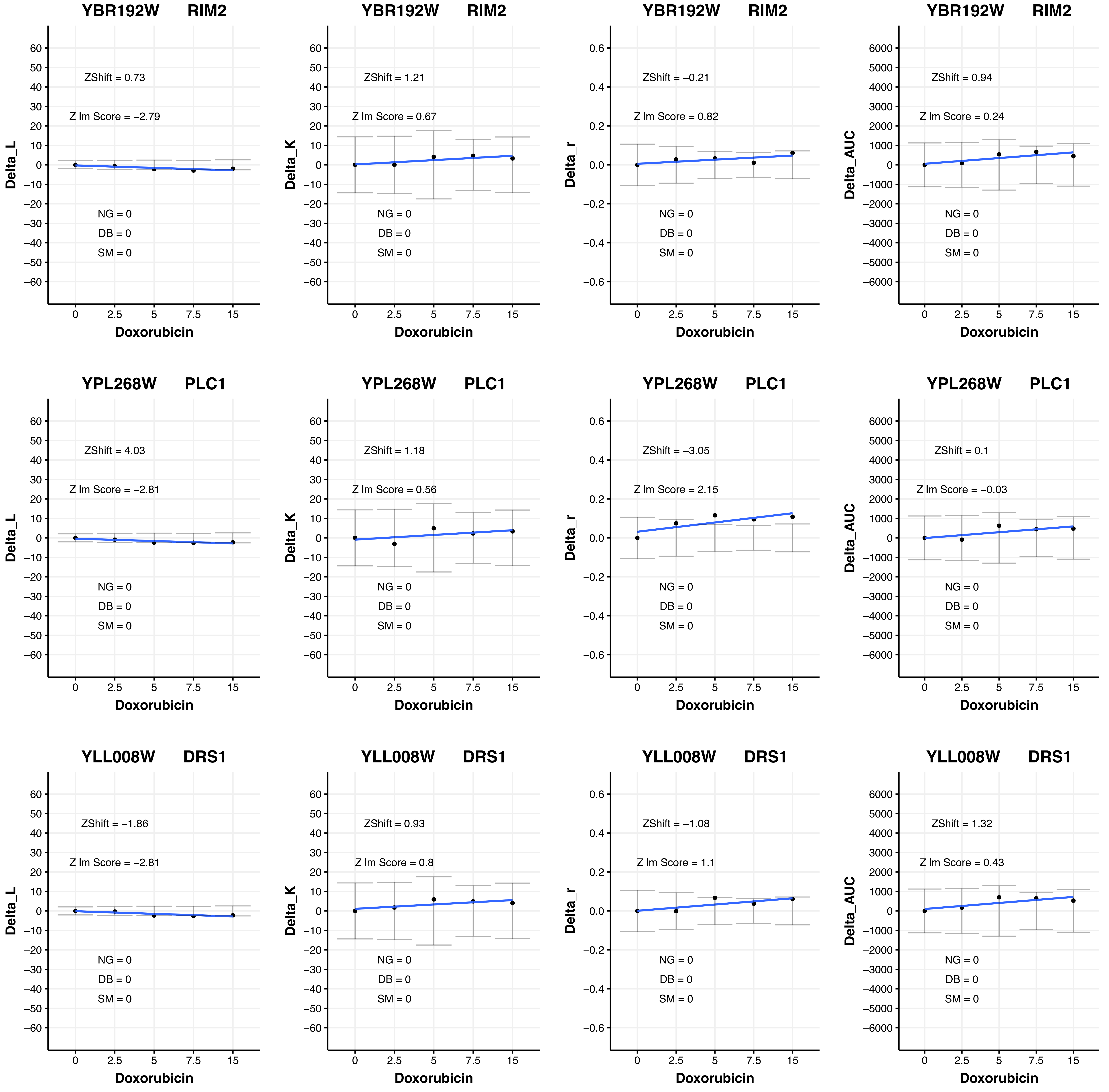

YBR192W RIM2
ZShift = 0.73
Z lm Score = −2.79
NG = 0
DB = 0
SM = 0
Delta_L
Doxorubicin
YBR192W RIM2
ZShift = 1.21
Z lm Score = 0.67
NG = 0
DB = 0
SM = 0
Delta_K
Doxorubicin
YBR192W RIM2
ZShift = −0.21
Z lm Score = 0.82
NG = 0
DB = 0
SM = 0
Delta_r
Doxorubicin
YBR192W RIM2
ZShift = 0.94
Z lm Score = 0.24
NG = 0
DB = 0
SM = 0
Delta_AUC
Doxorubicin
YPL268W PLC1
ZShift = 4.03
Z lm Score = −2.81
NG = 0
DB = 0
SM = 0
Delta_L
Doxorubicin
YPL268W PLC1
ZShift = 1.18
Z lm Score = 0.56
NG = 0
DB = 0
SM = 0
Delta_K
Doxorubicin
YPL268W PLC1
ZShift = −3.05
Z lm Score = 2.15
NG = 0
DB = 0
SM = 0
Delta_r
Doxorubicin
YPL268W PLC1
ZShift = 0.1
Z lm Score = −0.03
NG = 0
DB = 0
SM = 0
Delta_AUC
Doxorubicin
YLL008W DRS1
ZShift = −1.86
Z lm Score = −2.81
NG = 0
DB = 0
SM = 0
Delta_L
Doxorubicin
YLL008W DRS1
ZShift = 0.93
Z lm Score = 0.8
NG = 0
DB = 0
SM = 0
Delta_K
Doxorubicin
YLL008W DRS1
ZShift = −1.08
Z lm Score = 1.1
NG = 0
DB = 0
SM = 0
Delta_r
Doxorubicin
YLL008W DRS1
ZShift = 1.32
Z lm Score = 0.43
NG = 0
DB = 0
SM = 0
Delta_AUC
Doxorubicin

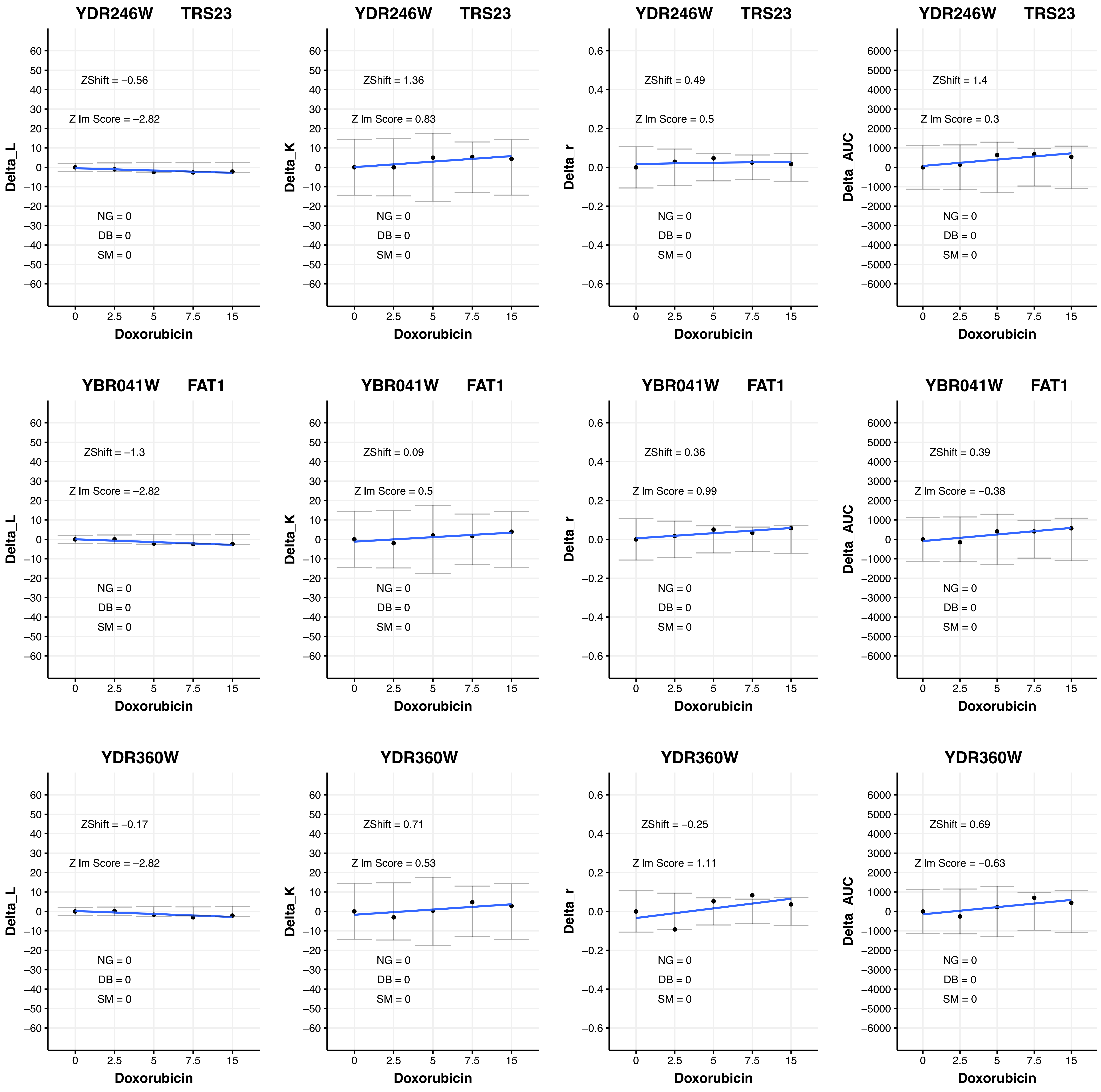
YDR246W TRS23
ZShift = −0.56
Z lm Score = −2.82
NG = 0
DB = 0
SM = 0
Delta_L
Doxorubicin
YDR246W TRS23
ZShift = 1.36
Z lm Score = 0.83
NG = 0
DB = 0
SM = 0
Delta_K
Doxorubicin
YDR246W TRS23
ZShift = 0.49
Z lm Score = 0.5
NG = 0
DB = 0
SM = 0
Delta_r
Doxorubicin
YDR246W TRS23
ZShift = 1.4
Z lm Score = 0.3
NG = 0
DB = 0
SM = 0
Delta_AUC
Doxorubicin
YBR041W FAT1
ZShift = −1.3
Z lm Score = −2.82
NG = 0
DB = 0
SM = 0
Delta_L
Doxorubicin
YBR041W FAT1
ZShift = 0.09
Z lm Score = 0.5
NG = 0
DB = 0
SM = 0
Delta_K
Doxorubicin
YBR041W FAT1
ZShift = 0.36
Z lm Score = 0.99
NG = 0
DB = 0
SM = 0
Delta_r
Doxorubicin
YBR041W FAT1
ZShift = 0.39
Z lm Score = −0.38
NG = 0
DB = 0
SM = 0
Delta_AUC
Doxorubicin
YDR360W
ZShift = −0.17
Z lm Score = −2.82
NG = 0
DB = 0
SM = 0
Delta_L
Doxorubicin
YDR360W
ZShift = 0.71
Z lm Score = 0.53
NG = 0
DB = 0
SM = 0
Delta_K
Doxorubicin
YDR360W
ZShift = −0.25
Z lm Score = 1.11
NG = 0
DB = 0
SM = 0
Delta_r
Doxorubicin
YDR360W
ZShift = 0.69
Z lm Score = −0.63
NG = 0
DB = 0
SM = 0
Delta_AUC
Doxorubicin

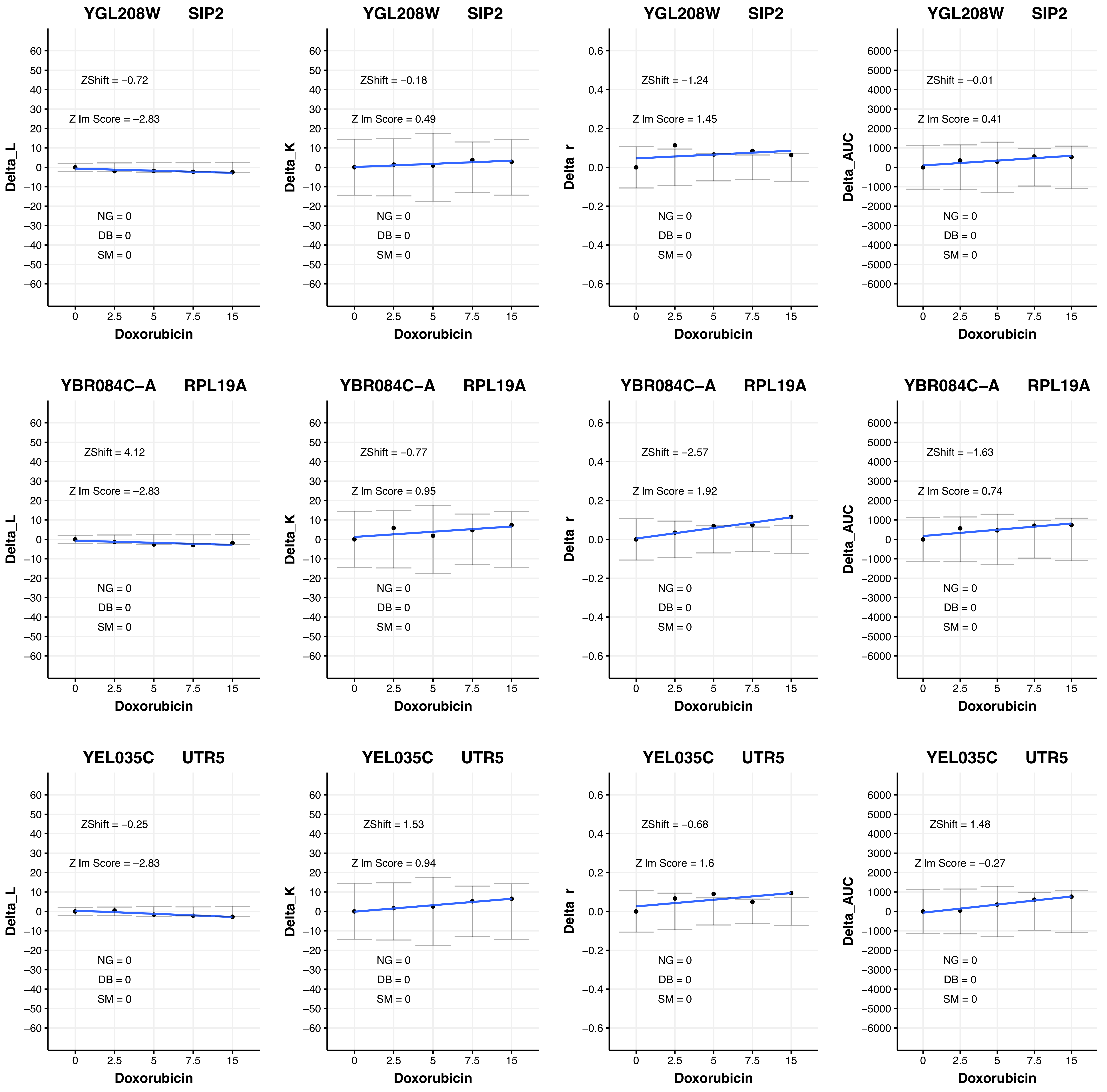

YGL208W SIP2
ZShift = −0.72
Z lm Score = −2.83
NG = 0
DB = 0
SM = 0
Delta_L
Doxorubicin
YGL208W SIP2
ZShift = −0.18
Z lm Score = 0.49
NG = 0
DB = 0
SM = 0
Delta_K
Doxorubicin
YGL208W SIP2
ZShift = −1.24
Z lm Score = 1.45
NG = 0
DB = 0
SM = 0
Delta_r
Doxorubicin
YGL208W SIP2
ZShift = −0.01
Z lm Score = 0.41
NG = 0
DB = 0
SM = 0
Delta_AUC
Doxorubicin
YBR084C−A RPL19A
ZShift = 4.12
Z lm Score = −2.83
NG = 0
DB = 0
SM = 0
Delta_L
Doxorubicin
YBR084C−A RPL19A
ZShift = −0.77
Z lm Score = 0.95
NG = 0
DB = 0
SM = 0
Delta_K
Doxorubicin
YBR084C−A RPL19A
ZShift = −2.57
Z lm Score = 1.92
NG = 0
DB = 0
SM = 0
Delta_r
Doxorubicin
YBR084C−A RPL19A
ZShift = −1.63
Z lm Score = 0.74
NG = 0
DB = 0
SM = 0
Delta_AUC
Doxorubicin
YEL035C UTR5
ZShift = −0.25
Z lm Score = −2.83
NG = 0
DB = 0
SM = 0
Delta_L
Doxorubicin
YEL035C UTR5
ZShift = 1.53
Z lm Score = 0.94
NG = 0
DB = 0
SM = 0
Delta_K
Doxorubicin
YEL035C UTR5
ZShift = −0.68
Z lm Score = 1.6
NG = 0
DB = 0
SM = 0
Delta_r
Doxorubicin
YEL035C UTR5
ZShift = 1.48
Z lm Score = −0.27
NG = 0
DB = 0
SM = 0
Delta_AUC
Doxorubicin

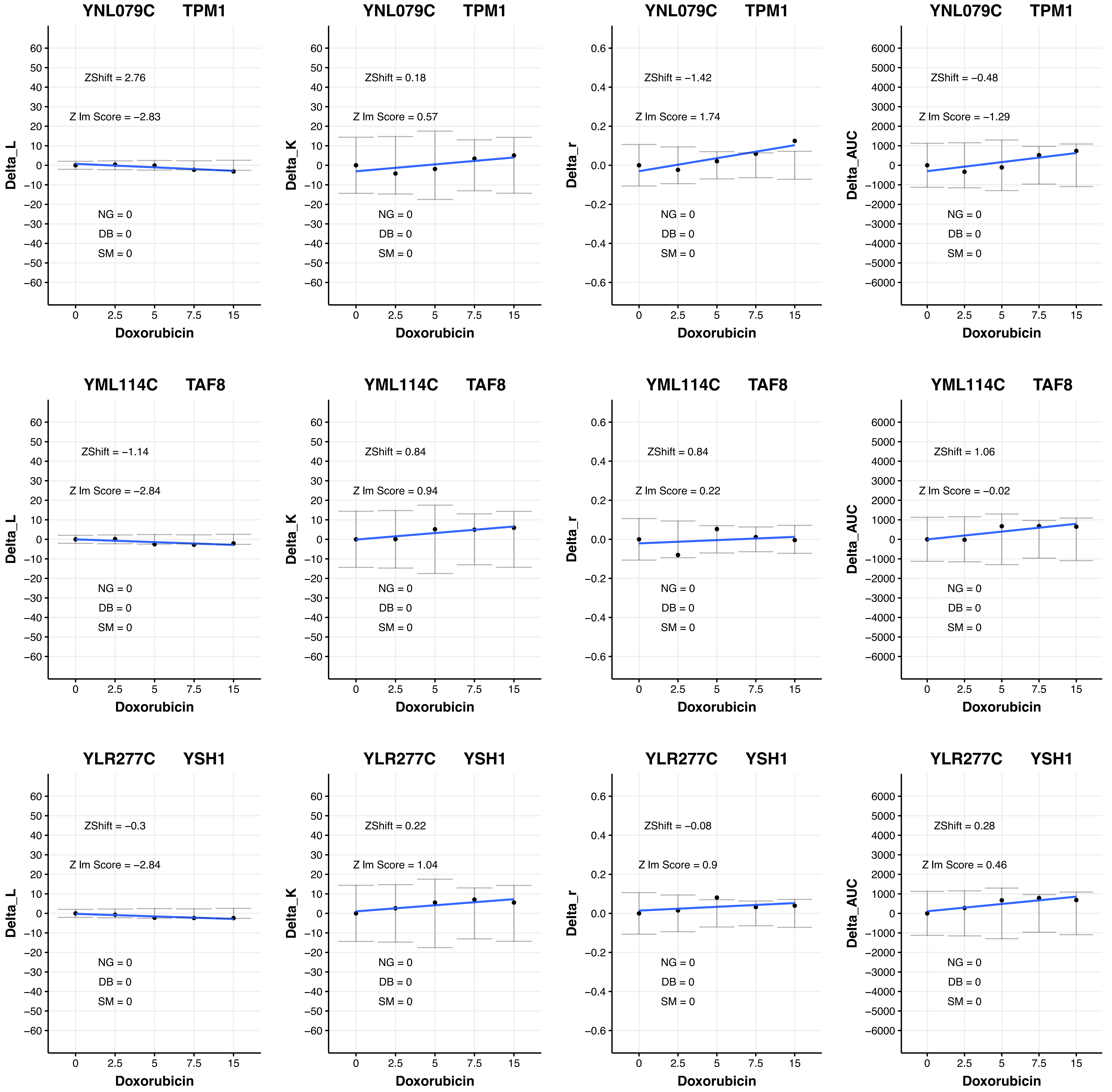
YNL079C TPM1
ZShift = 2.76
Z lm Score = −2.83
NG = 0
DB = 0
SM = 0
Delta_L
Doxorubicin
YNL079C TPM1
ZShift = 0.18
Z lm Score = 0.57
NG = 0
DB = 0
SM = 0
Delta_K
Doxorubicin
YNL079C TPM1
ZShift = −1.42
Z lm Score = 1.74
NG = 0
DB = 0
SM = 0
Delta_r
Doxorubicin
YNL079C TPM1
ZShift = −0.48
Z lm Score = −1.29
NG = 0
DB = 0
SM = 0
Delta_AUC
Doxorubicin
YML114C TAF8
ZShift = −1.14
Z lm Score = −2.84
NG = 0
DB = 0
SM = 0
Delta_L
Doxorubicin
YML114C TAF8
ZShift = 0.84
Z lm Score = 0.94
NG = 0
DB = 0
SM = 0
Delta_K
Doxorubicin
YML114C TAF8
ZShift = 0.84
Z lm Score = 0.22
NG = 0
DB = 0
SM = 0
Delta_r
Doxorubicin
YML114C TAF8
ZShift = 1.06
Z lm Score = −0.02
NG = 0
DB = 0
SM = 0
Delta_AUC
Doxorubicin
YLR277C YSH1
ZShift = −0.3
Z lm Score = −2.84
NG = 0
DB = 0
SM = 0
Delta_L
Doxorubicin
YLR277C YSH1
ZShift = 0.22
Z lm Score = 1.04
NG = 0
DB = 0
SM = 0
Delta_K
Doxorubicin
YLR277C YSH1
ZShift = −0.08
Z lm Score = 0.9
NG = 0
DB = 0
SM = 0
Delta_r
Doxorubicin
YLR277C YSH1
ZShift = 0.28
Z lm Score = 0.46
NG = 0
DB = 0
SM = 0
Delta_AUC
Doxorubicin

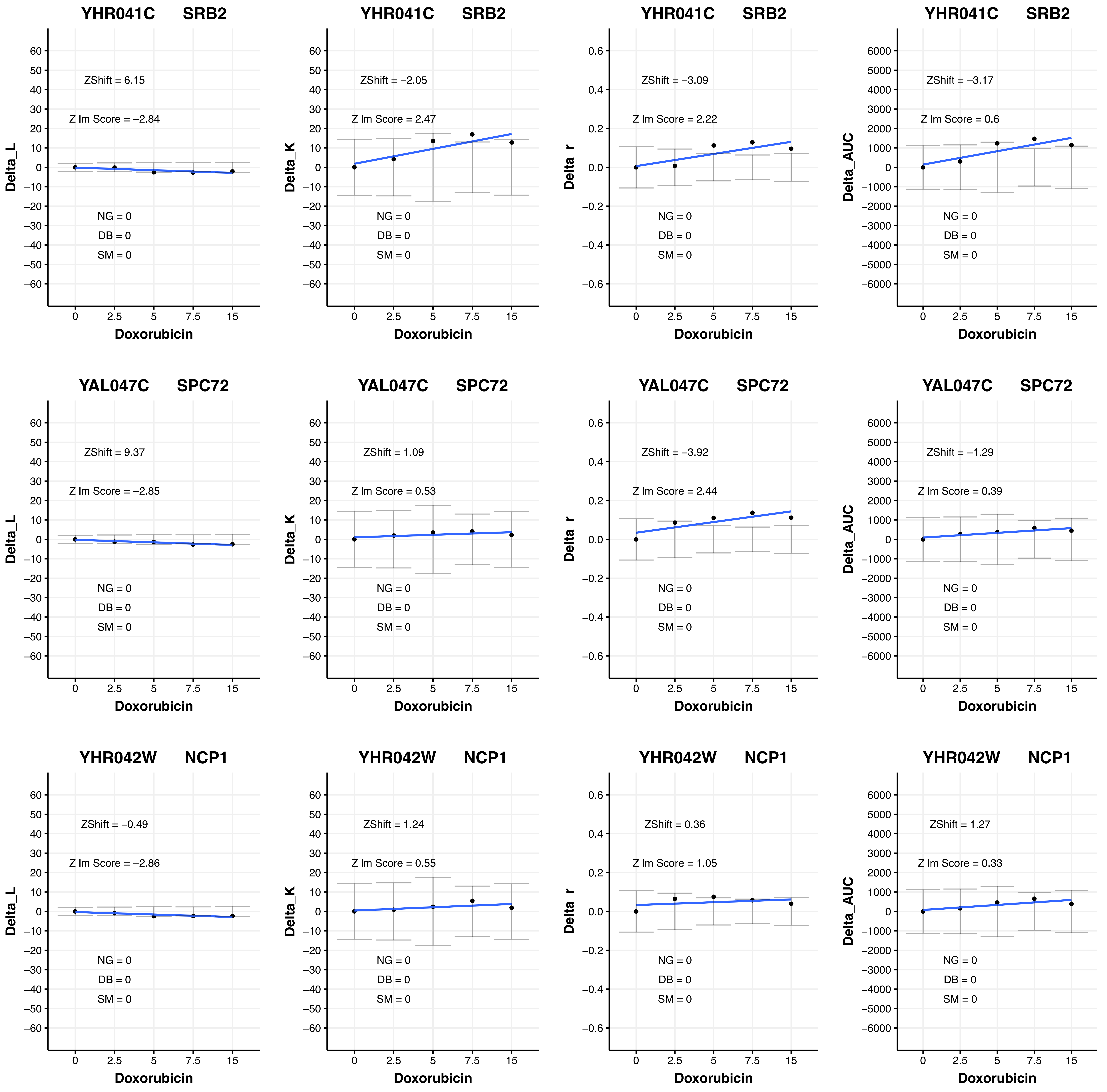
YHR041C SRB2
ZShift = 6.15
Z lm Score = −2.84
NG = 0
DB = 0
SM = 0
Delta_L
Doxorubicin
YHR041C SRB2
ZShift = −2.05
Z lm Score = 2.47
NG = 0
DB = 0
SM = 0
Delta_K
Doxorubicin
YHR041C SRB2
ZShift = −3.09
Z lm Score = 2.22
NG = 0
DB = 0
SM = 0
Delta_r
Doxorubicin
YHR041C SRB2
ZShift = −3.17
Z lm Score = 0.6
NG = 0
DB = 0
SM = 0
Delta_AUC
Doxorubicin
YAL047C SPC72
ZShift = 9.37
Z lm Score = −2.85
NG = 0
DB = 0
SM = 0
Delta_L
Doxorubicin
YAL047C SPC72
ZShift = 1.09
Z lm Score = 0.53
NG = 0
DB = 0
SM = 0
Delta_K
Doxorubicin
YAL047C SPC72
ZShift = −3.92
Z lm Score = 2.44
NG = 0
DB = 0
SM = 0
Delta_r
Doxorubicin
YAL047C SPC72
ZShift = −1.29
Z lm Score = 0.39
NG = 0
DB = 0
SM = 0
Delta_AUC
Doxorubicin
YHR042W NCP1
ZShift = −0.49
Z lm Score = −2.86
NG = 0
DB = 0
SM = 0
Delta_L
Doxorubicin
YHR042W NCP1
ZShift = 1.24
Z lm Score = 0.55
NG = 0
DB = 0
SM = 0
Delta_K
Doxorubicin
YHR042W NCP1
ZShift = 0.36
Z lm Score = 1.05
NG = 0
DB = 0
SM = 0
Delta_r
Doxorubicin
YHR042W NCP1
ZShift = 1.27
Z lm Score = 0.33
NG = 0
DB = 0
SM = 0
Delta_AUC
Doxorubicin

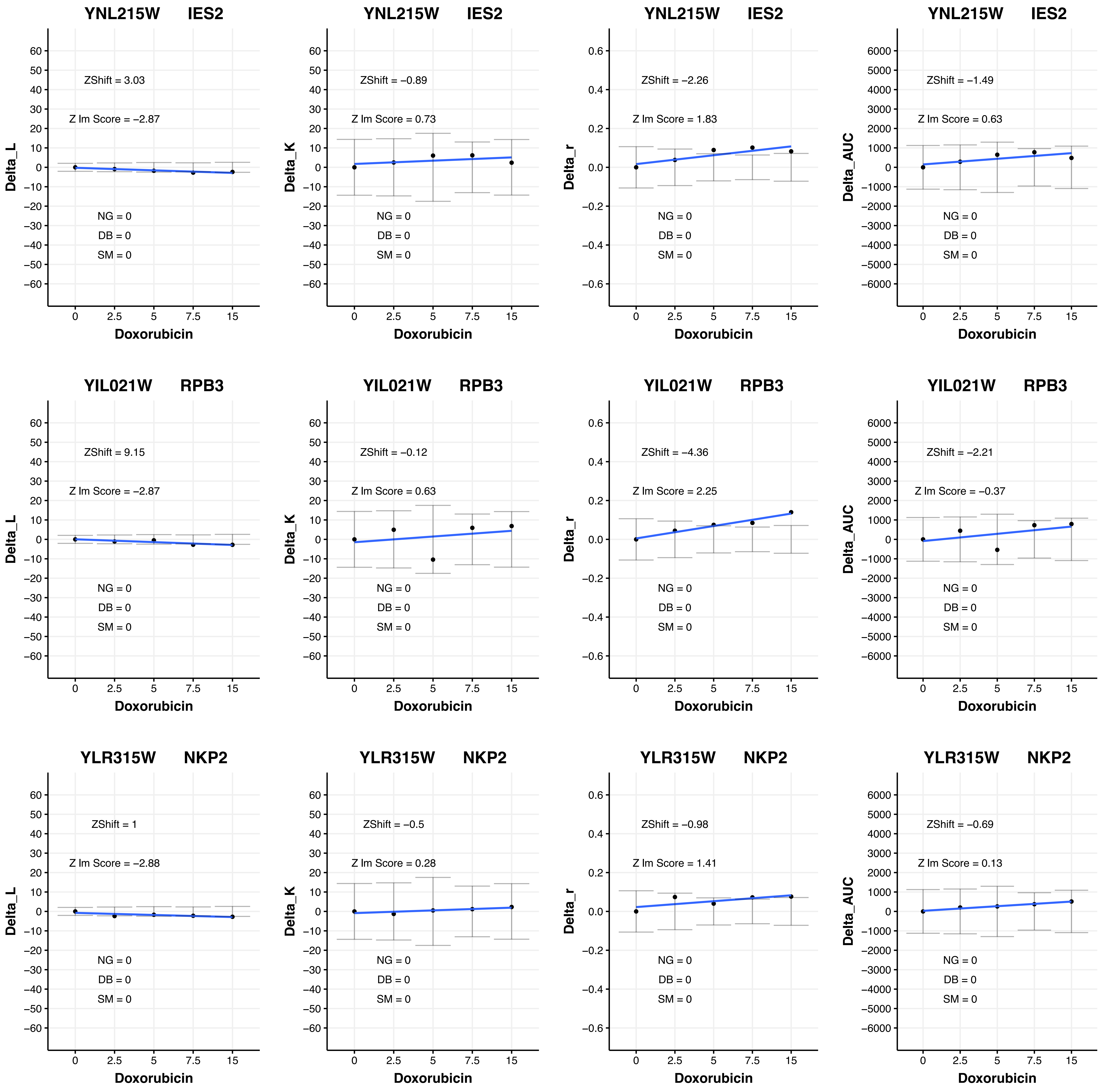
YNL215W IES2
ZShift = 3.03
Z lm Score = −2.87
NG = 0
DB = 0
SM = 0
Delta_L
Doxorubicin
YNL215W IES2
ZShift = −0.89
Z lm Score = 0.73
NG = 0
DB = 0
SM = 0
Delta_K
Doxorubicin
YNL215W IES2
ZShift = −2.26
Z lm Score = 1.83
NG = 0
DB = 0
SM = 0
Delta_r
Doxorubicin
YNL215W IES2
ZShift = −1.49
Z lm Score = 0.63
NG = 0
DB = 0
SM = 0
Delta_AUC
Doxorubicin
YIL021W RPB3
ZShift = 9.15
Z lm Score = −2.87
NG = 0
DB = 0
SM = 0
Delta_L
Doxorubicin
YIL021W RPB3
ZShift = −0.12
Z lm Score = 0.63
NG = 0
DB = 0
SM = 0
Delta_K
Doxorubicin
YIL021W RPB3
ZShift = −4.36
Z lm Score = 2.25
NG = 0
DB = 0
SM = 0
Delta_r
Doxorubicin
YIL021W RPB3
ZShift = −2.21
Z lm Score = −0.37
NG = 0
DB = 0
SM = 0
Delta_AUC
Doxorubicin
YLR315W NKP2
ZShift = 1
Z lm Score = −2.88
NG = 0
DB = 0
SM = 0
Delta_L
Doxorubicin
YLR315W NKP2
ZShift = −0.5
Z lm Score = 0.28
NG = 0
DB = 0
SM = 0
Delta_K
Doxorubicin
YLR315W NKP2
ZShift = −0.98
Z lm Score = 1.41
NG = 0
DB = 0
SM = 0
Delta_r
Doxorubicin
YLR315W NKP2
ZShift = −0.69
Z lm Score = 0.13
NG = 0
DB = 0
SM = 0
Delta_AUC
Doxorubicin

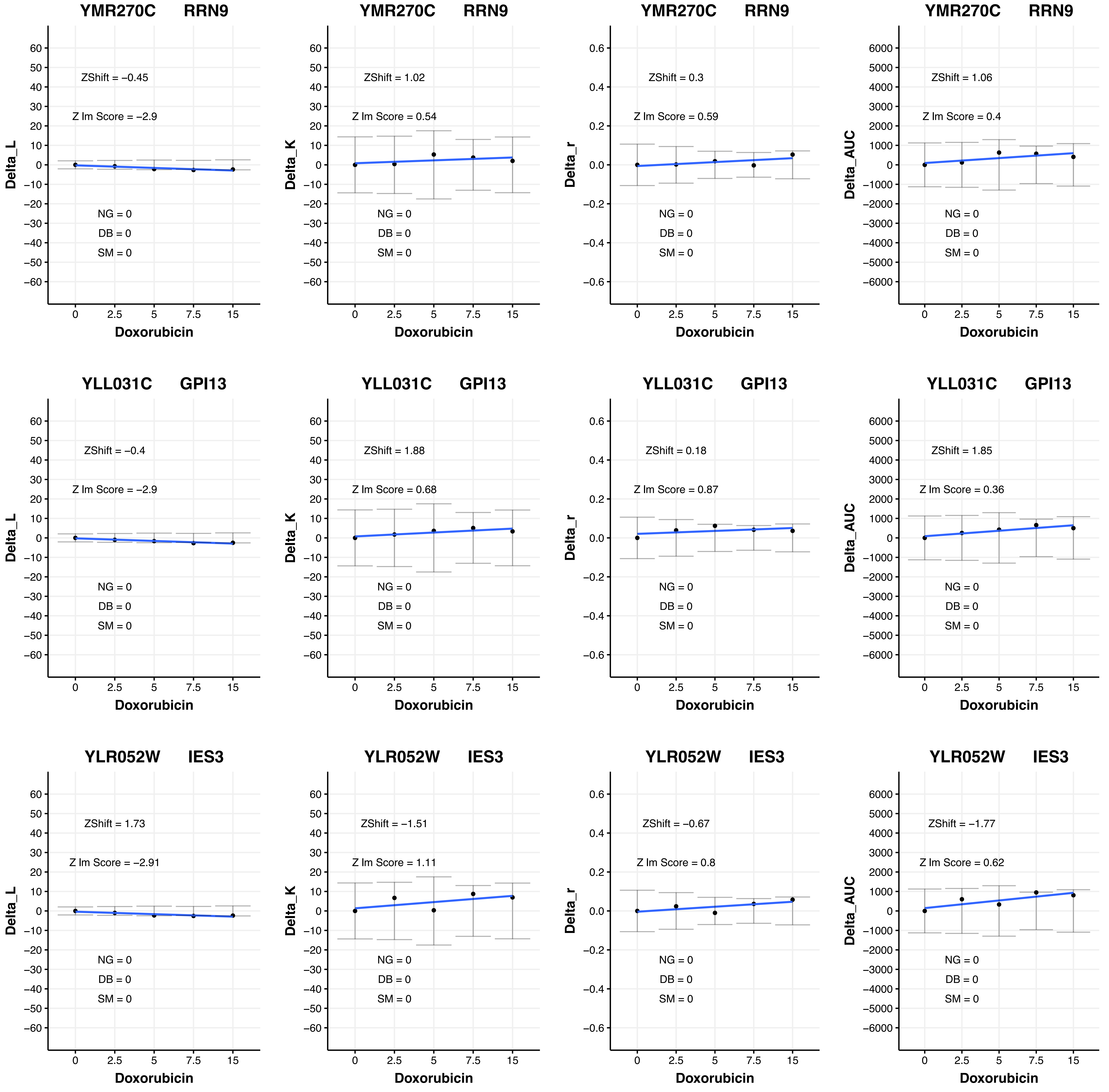
YMR270C RRN9
ZShift = −0.45
Z lm Score = −2.9
NG = 0
DB = 0
SM = 0
Delta_L
Doxorubicin
YMR270C RRN9
ZShift = 1.02
Z lm Score = 0.54
NG = 0
DB = 0
SM = 0
Delta_K
Doxorubicin
YMR270C RRN9
ZShift = 0.3
Z lm Score = 0.59
NG = 0
DB = 0
SM = 0
Delta_r
Doxorubicin
YMR270C RRN9
ZShift = 1.06
Z lm Score = 0.4
NG = 0
DB = 0
SM = 0
Delta_AUC
Doxorubicin
YLL031C GPI13
ZShift = −0.4
Z lm Score = −2.9
NG = 0
DB = 0
SM = 0
Delta_L
Doxorubicin
YLL031C GPI13
ZShift = 1.88
Z lm Score = 0.68
NG = 0
DB = 0
SM = 0
Delta_K
Doxorubicin
YLL031C GPI13
ZShift = 0.18
Z lm Score = 0.87
NG = 0
DB = 0
SM = 0
Delta_r
Doxorubicin
YLL031C GPI13
ZShift = 1.85
Z lm Score = 0.36
NG = 0
DB = 0
SM = 0
Delta_AUC
Doxorubicin
YLR052W IES3
ZShift = 1.73
Z lm Score = −2.91
NG = 0
DB = 0
SM = 0
Delta_L
Doxorubicin
YLR052W IES3
ZShift = −1.51
Z lm Score = 1.11
NG = 0
DB = 0
SM = 0
Delta_K
Doxorubicin
YLR052W IES3
ZShift = −0.67
Z lm Score = 0.8
NG = 0
DB = 0
SM = 0
Delta_r
Doxorubicin
YLR052W IES3
ZShift = −1.77
Z lm Score = 0.62
NG = 0
DB = 0
SM = 0
Delta_AUC
Doxorubicin

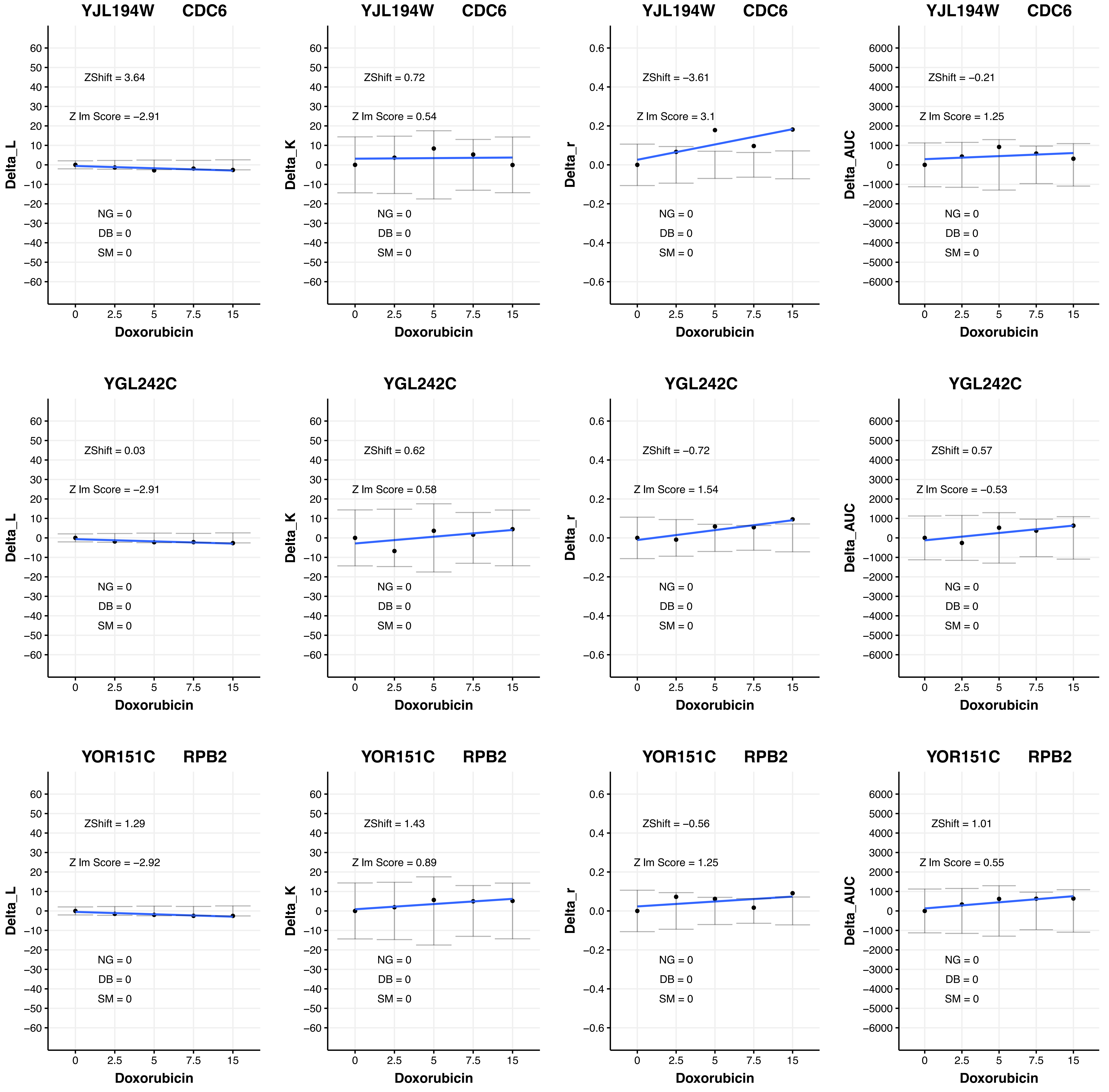

YJL194W CDC6
ZShift = 3.64
Z lm Score = −2.91
NG = 0
DB = 0
SM = 0
Delta_L
Doxorubicin
YJL194W CDC6
ZShift = 0.72
Z lm Score = 0.54
NG = 0
DB = 0
SM = 0
Delta_K
Doxorubicin
YJL194W CDC6
ZShift = −3.61
Z lm Score = 3.1
NG = 0
DB = 0
SM = 0
Delta_r
Doxorubicin
YJL194W CDC6
ZShift = −0.21
Z lm Score = 1.25
NG = 0
DB = 0
SM = 0
Delta_AUC
Doxorubicin
YGL242C
ZShift = 0.03
Z lm Score = −2.91
NG = 0
DB = 0
SM = 0
Delta_L
Doxorubicin
YGL242C
ZShift = 0.62
Z lm Score = 0.58
NG = 0
DB = 0
SM = 0
Delta_K
Doxorubicin
YGL242C
ZShift = −0.72
Z lm Score = 1.54
NG = 0
DB = 0
SM = 0
Delta_r
Doxorubicin
YGL242C
ZShift = 0.57
Z lm Score = −0.53
NG = 0
DB = 0
SM = 0
Delta_AUC
Doxorubicin
YOR151C RPB2
ZShift = 1.29
Z lm Score = −2.92
NG = 0
DB = 0
SM = 0
Delta_L
Doxorubicin
YOR151C RPB2
ZShift = 1.43
Z lm Score = 0.89
NG = 0
DB = 0
SM = 0
Delta_K
Doxorubicin
YOR151C RPB2
ZShift = −0.56
Z lm Score = 1.25
NG = 0
DB = 0
SM = 0
Delta_r
Doxorubicin
YOR151C RPB2
ZShift = 1.01
Z lm Score = 0.55
NG = 0
DB = 0
SM = 0
Delta_AUC
Doxorubicin

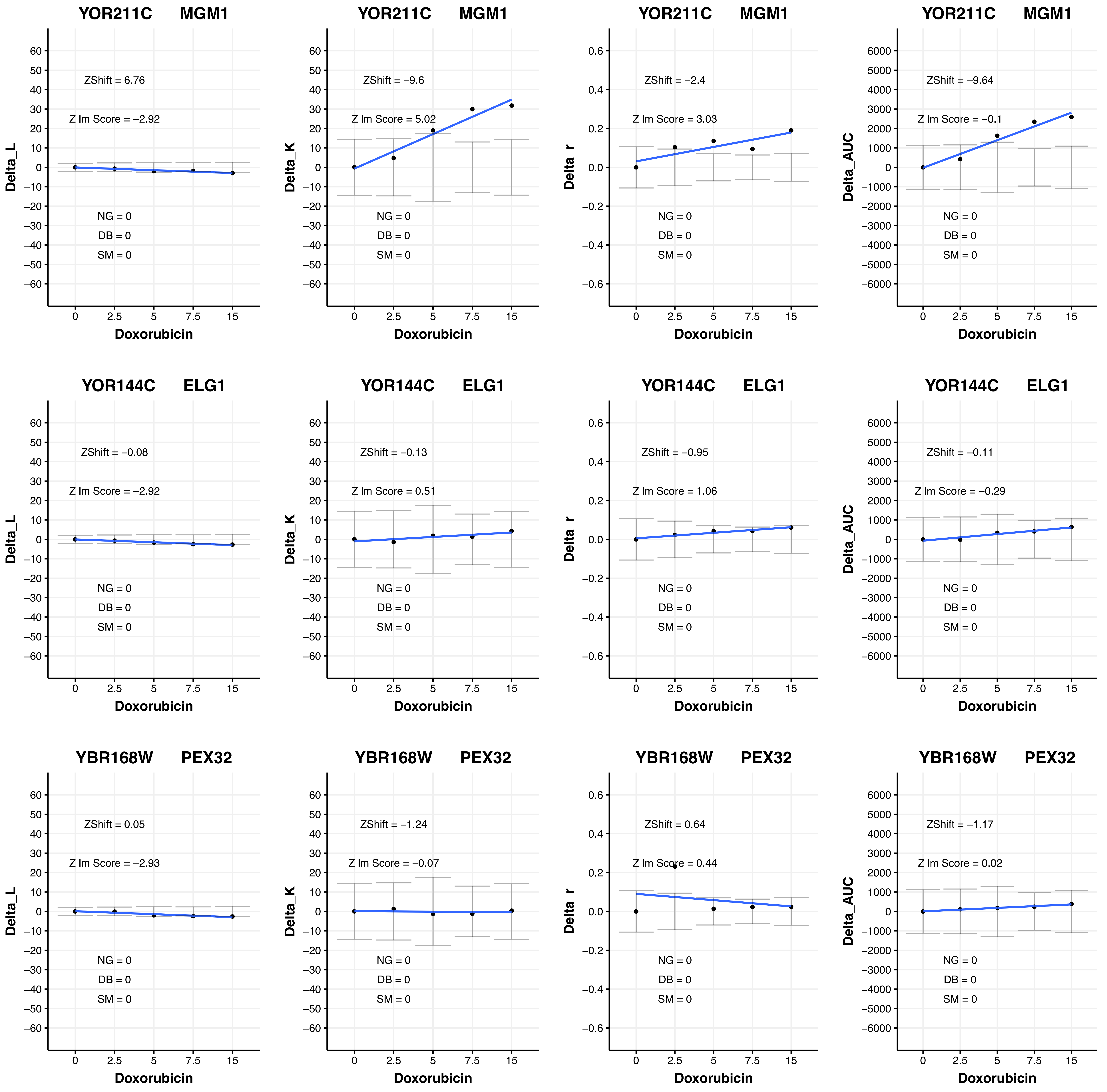
YOR211C MGM1
ZShift = 6.76
Z lm Score = −2.92
NG = 0
DB = 0
SM = 0
Delta_L
Doxorubicin
YOR211C MGM1
ZShift = −9.6
Z lm Score = 5.02
NG = 0
DB = 0
SM = 0
Delta_K
Doxorubicin
YOR211C MGM1
ZShift = −2.4
Z lm Score = 3.03
NG = 0
DB = 0
SM = 0
Delta_r
Doxorubicin
YOR211C MGM1
ZShift = −9.64
Z lm Score = −0.1
NG = 0
DB = 0
SM = 0
Delta_AUC
Doxorubicin
YOR144C ELG1
ZShift = −0.08
Z lm Score = −2.92
NG = 0
DB = 0
SM = 0
Delta_L
Doxorubicin
YOR144C ELG1
ZShift = −0.13
Z lm Score = 0.51
NG = 0
DB = 0
SM = 0
Delta_K
Doxorubicin
YOR144C ELG1
ZShift = −0.95
Z lm Score = 1.06
NG = 0
DB = 0
SM = 0
Delta_r
Doxorubicin
YOR144C ELG1
ZShift = −0.11
Z lm Score = −0.29
NG = 0
DB = 0
SM = 0
Delta_AUC
Doxorubicin
YBR168W PEX32
ZShift = 0.05
Z lm Score = −2.93
NG = 0
DB = 0
SM = 0
Delta_L
Doxorubicin
YBR168W PEX32
ZShift = −1.24
Z lm Score = −0.07
NG = 0
DB = 0
SM = 0
Delta_K
Doxorubicin
YBR168W PEX32
ZShift = 0.64
Z lm Score = 0.44
NG = 0
DB = 0
SM = 0
Delta_r
Doxorubicin
YBR168W PEX32
ZShift = −1.17
Z lm Score = 0.02
NG = 0
DB = 0
SM = 0
Delta_AUC
Doxorubicin

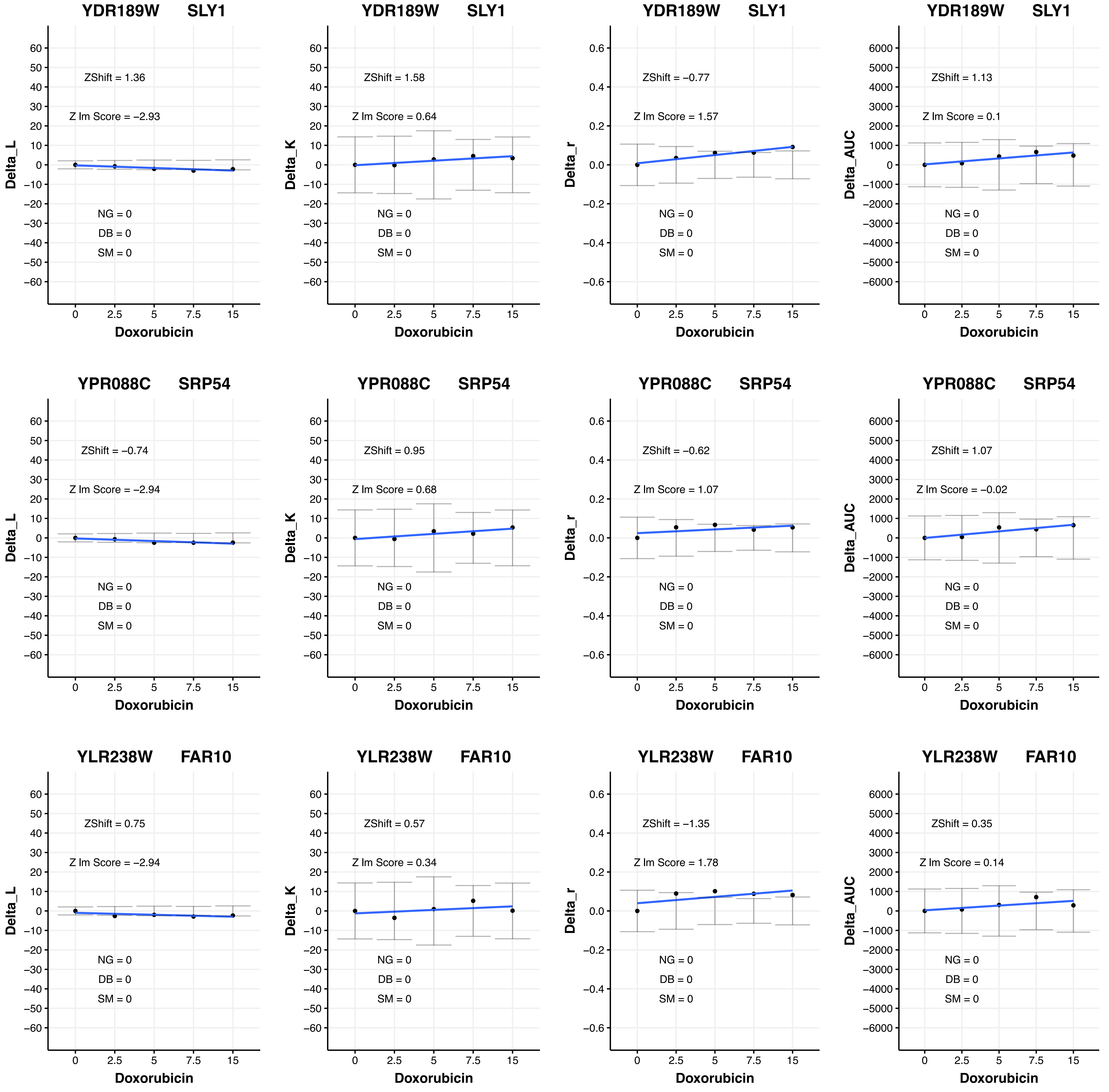
YDR189W SLY1
ZShift = 1.36
Z lm Score = −2.93
NG = 0
DB = 0
SM = 0
Delta_L
Doxorubicin
YDR189W SLY1
ZShift = 1.58
Z lm Score = 0.64
NG = 0
DB = 0
SM = 0
Delta_K
Doxorubicin
YDR189W SLY1
ZShift = −0.77
Z lm Score = 1.57
NG = 0
DB = 0
SM = 0
Delta_r
Doxorubicin
YDR189W SLY1
ZShift = 1.13
Z lm Score = 0.1
NG = 0
DB = 0
SM = 0
Delta_AUC
Doxorubicin
YPR088C SRP54
ZShift = −0.74
Z lm Score = −2.94
NG = 0
DB = 0
SM = 0
Delta_L
Doxorubicin
YPR088C SRP54
ZShift = 0.95
Z lm Score = 0.68
NG = 0
DB = 0
SM = 0
Delta_K
Doxorubicin
YPR088C SRP54
ZShift = −0.62
Z lm Score = 1.07
NG = 0
DB = 0
SM = 0
Delta_r
Doxorubicin
YPR088C SRP54
ZShift = 1.07
Z lm Score = −0.02
NG = 0
DB = 0
SM = 0
Delta_AUC
Doxorubicin
YLR238W FAR10
ZShift = 0.75
Z lm Score = −2.94
NG = 0
DB = 0
SM = 0
Delta_L
Doxorubicin
YLR238W FAR10
ZShift = 0.57
Z lm Score = 0.34
NG = 0
DB = 0
SM = 0
Delta_K
Doxorubicin
YLR238W FAR10
ZShift = −1.35
Z lm Score = 1.78
NG = 0
DB = 0
SM = 0
Delta_r
Doxorubicin
YLR238W FAR10
ZShift = 0.35
Z lm Score = 0.14
NG = 0
DB = 0
SM = 0
Delta_AUC
Doxorubicin

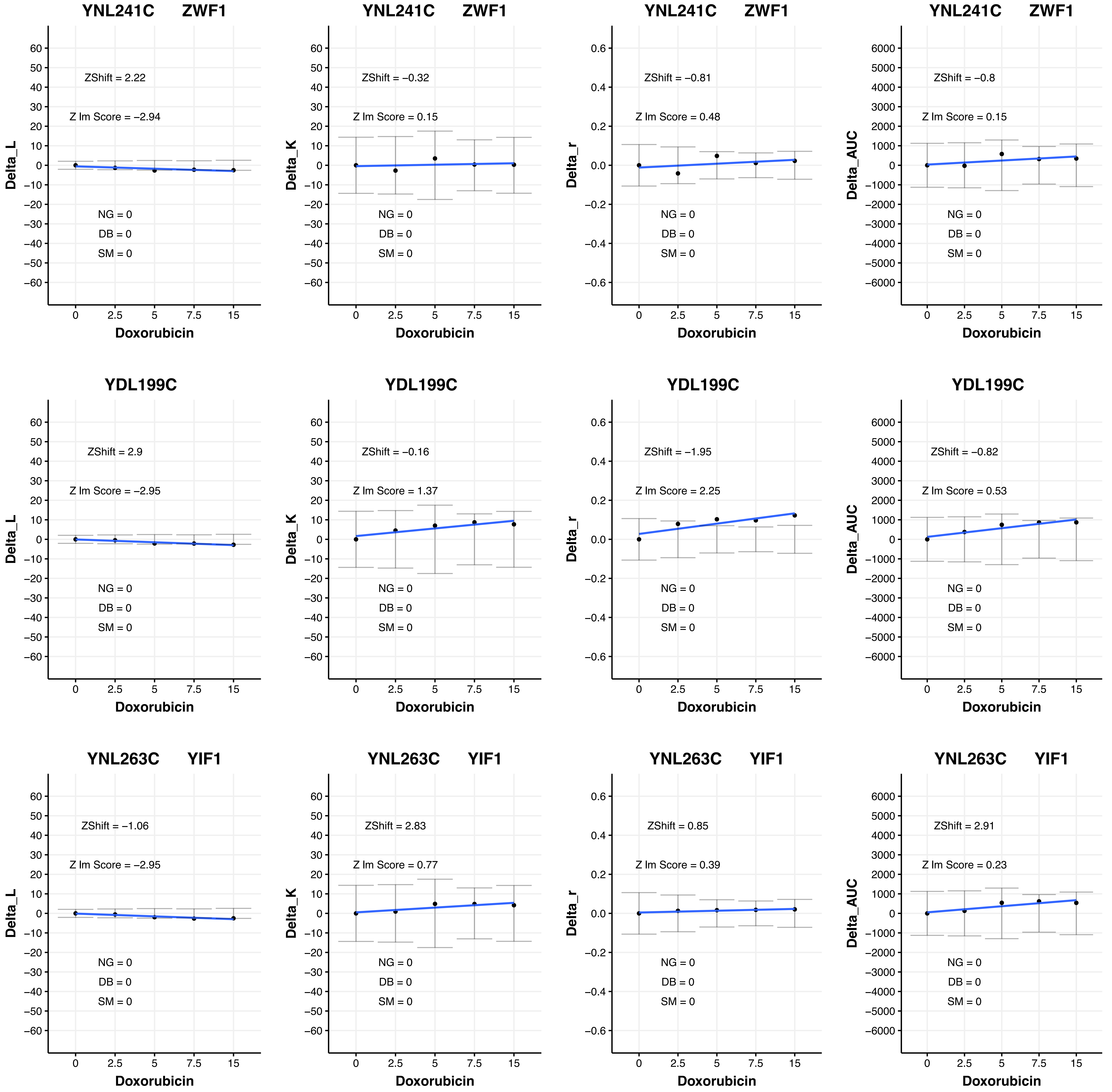

YNL241C ZWF1
ZShift = 2.22
Z lm Score = −2.94
NG = 0
DB = 0
SM = 0
Delta_L
Doxorubicin
YNL241C ZWF1
ZShift = −0.32
Z lm Score = 0.15
NG = 0
DB = 0
SM = 0
Delta_K
Doxorubicin
YNL241C ZWF1
ZShift = −0.81
Z lm Score = 0.48
NG = 0
DB = 0
SM = 0
Delta_r
Doxorubicin
YNL241C ZWF1
ZShift = −0.8
Z lm Score = 0.15
NG = 0
DB = 0
SM = 0
Delta_AUC
Doxorubicin
YDL199C
ZShift = 2.9
Z lm Score = −2.95
NG = 0
DB = 0
SM = 0
Delta_L
Doxorubicin
YDL199C
ZShift = −0.16
Z lm Score = 1.37
NG = 0
DB = 0
SM = 0
Delta_K
Doxorubicin
YDL199C
ZShift = −1.95
Z lm Score = 2.25
NG = 0
DB = 0
SM = 0
Delta_r
Doxorubicin
YDL199C
ZShift = −0.82
Z lm Score = 0.53
NG = 0
DB = 0
SM = 0
Delta_AUC
Doxorubicin
YNL263C YIF1
ZShift = −1.06
Z lm Score = −2.95
NG = 0
DB = 0
SM = 0
Delta_L
Doxorubicin
YNL263C YIF1
ZShift = 2.83
Z lm Score = 0.77
NG = 0
DB = 0
SM = 0
Delta_K
Doxorubicin
YNL263C YIF1
ZShift = 0.85
Z lm Score = 0.39
NG = 0
DB = 0
SM = 0
Delta_r
Doxorubicin
YNL263C YIF1
ZShift = 2.91
Z lm Score = 0.23
NG = 0
DB = 0
SM = 0
Delta_AUC
Doxorubicin

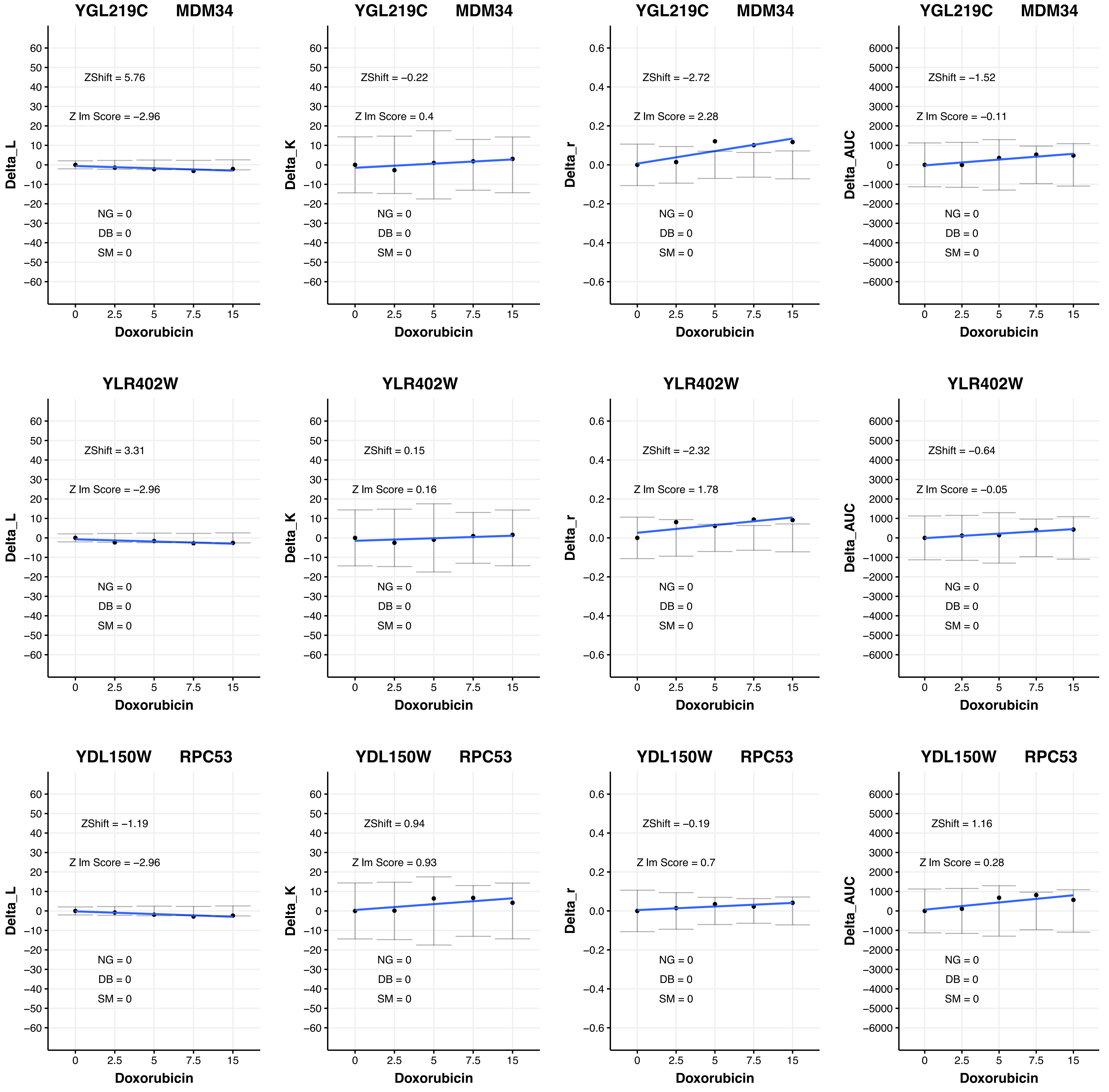

YGL219C MDM34
ZShift = 5.76
Z lm Score = −2.96
NG = 0
DB = 0
SM = 0
Delta_L
Doxorubicin
YGL219C MDM34
ZShift = −0.22
Z lm Score = 0.4
NG = 0
DB = 0
SM = 0
Delta_K
Doxorubicin
YGL219C MDM34
ZShift = −2.72
Z lm Score = 2.28
NG = 0
DB = 0
SM = 0
Delta_r
Doxorubicin
YGL219C MDM34
ZShift = −1.52
Z lm Score = −0.11
NG = 0
DB = 0
SM = 0
Delta_AUC
Doxorubicin
YLR402W
ZShift = 3.31
Z lm Score = −2.96
NG = 0
DB = 0
SM = 0
Delta_L
Doxorubicin
YLR402W
ZShift = 0.15
Z lm Score = 0.16
NG = 0
DB = 0
SM = 0
Delta_K
Doxorubicin
YLR402W
ZShift = −2.32
Z lm Score = 1.78
NG = 0
DB = 0
SM = 0
Delta_r
Doxorubicin
YLR402W
ZShift = −0.64
Z lm Score = −0.05
NG = 0
DB = 0
SM = 0
Delta_AUC
Doxorubicin
YDL150W RPC53
ZShift = −1.19
Z lm Score = −2.96
NG = 0
DB = 0
SM = 0
Delta_L
Doxorubicin
YDL150W RPC53
ZShift = 0.94
Z lm Score = 0.93
NG = 0
DB = 0
SM = 0
Delta_K
Doxorubicin
YDL150W RPC53
ZShift = −0.19
Z lm Score = 0.7
NG = 0
DB = 0
SM = 0
Delta_r
Doxorubicin
YDL150W RPC53
ZShift = 1.16
Z lm Score = 0.28
NG = 0
DB = 0
SM = 0
Delta_AUC
Doxorubicin

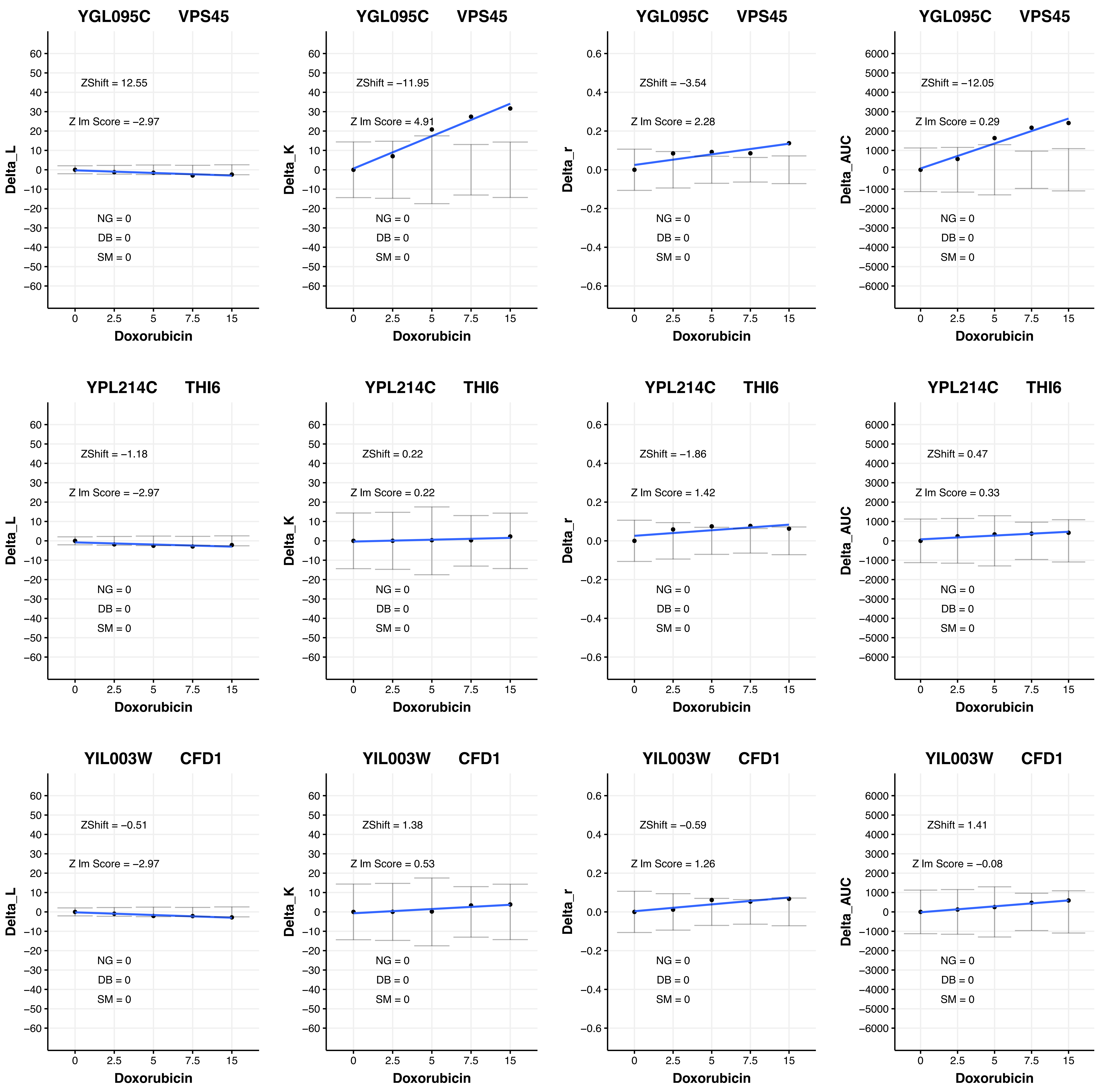
YGL095C VPS45
ZShift = 12.55
Z lm Score = −2.97
NG = 0
DB = 0
SM = 0
Delta_L
Doxorubicin
YGL095C VPS45
ZShift = −11.95
Z lm Score = 4.91
NG = 0
DB = 0
SM = 0
Delta_K
Doxorubicin
YGL095C VPS45
ZShift = −3.54
Z lm Score = 2.28
NG = 0
DB = 0
SM = 0
Delta_r
Doxorubicin
YGL095C VPS45
ZShift = −12.05
Z lm Score = 0.29
NG = 0
DB = 0
SM = 0
Delta_AUC
Doxorubicin
YPL214C THI6
ZShift = −1.18
Z lm Score = −2.97
NG = 0
DB = 0
SM = 0
Delta_L
Doxorubicin
YPL214C THI6
ZShift = 0.22
Z lm Score = 0.22
NG = 0
DB = 0
SM = 0
Delta_K
Doxorubicin
YPL214C THI6
ZShift = −1.86
Z lm Score = 1.42
NG = 0
DB = 0
SM = 0
Delta_r
Doxorubicin
YPL214C THI6
ZShift = 0.47
Z lm Score = 0.33
NG = 0
DB = 0
SM = 0
Delta_AUC
Doxorubicin
YIL003W CFD1
ZShift = −0.51
Z lm Score = −2.97
NG = 0
DB = 0
SM = 0
Delta_L
Doxorubicin
YIL003W CFD1
ZShift = 1.38
Z lm Score = 0.53
NG = 0
DB = 0
SM = 0
Delta_K
Doxorubicin
YIL003W CFD1
ZShift = −0.59
Z lm Score = 1.26
NG = 0
DB = 0
SM = 0
Delta_r
Doxorubicin
YIL003W CFD1
ZShift = 1.41
Z lm Score = −0.08
NG = 0
DB = 0
SM = 0
Delta_AUC
Doxorubicin

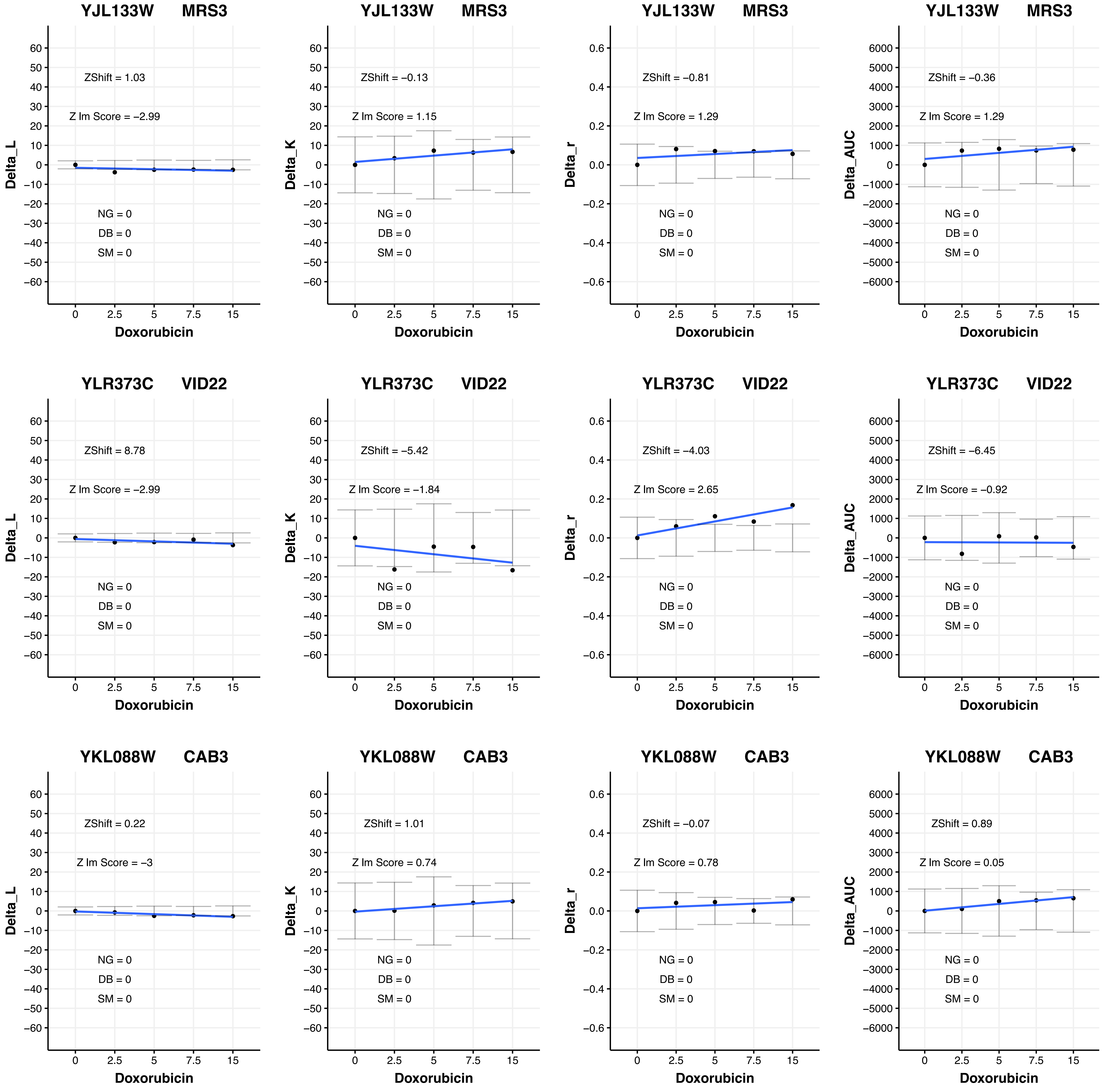

YJL133W MRS3
ZShift = 1.03
Z lm Score = −2.99
NG = 0
DB = 0
SM = 0
Delta_L
Doxorubicin
YJL133W MRS3
ZShift = −0.13
Z lm Score = 1.15
NG = 0
DB = 0
SM = 0
Delta_K
Doxorubicin
YJL133W MRS3
ZShift = −0.81
Z lm Score = 1.29
NG = 0
DB = 0
SM = 0
Delta_r
Doxorubicin
YJL133W MRS3
ZShift = −0.36
Z lm Score = 1.29
NG = 0
DB = 0
SM = 0
Delta_AUC
Doxorubicin
YLR373C VID22
ZShift = 8.78
Z lm Score = −2.99
NG = 0
DB = 0
SM = 0
Delta_L
Doxorubicin
YLR373C VID22
ZShift = −5.42
Z lm Score = −1.84
NG = 0
DB = 0
SM = 0
Delta_K
Doxorubicin
YLR373C VID22
ZShift = −4.03
Z lm Score = 2.65
NG = 0
DB = 0
SM = 0
Delta_r
Doxorubicin
YLR373C VID22
ZShift = −6.45
Z lm Score = −0.92
NG = 0
DB = 0
SM = 0
Delta_AUC
Doxorubicin
YKL088W CAB3
ZShift = 0.22
Z lm Score = −3
NG = 0
DB = 0
SM = 0
Delta_L
Doxorubicin
YKL088W CAB3
ZShift = 1.01
Z lm Score = 0.74
NG = 0
DB = 0
SM = 0
Delta_K
Doxorubicin
YKL088W CAB3
ZShift = −0.07
Z lm Score = 0.78
NG = 0
DB = 0
SM = 0
Delta_r
Doxorubicin
YKL088W CAB3
ZShift = 0.89
Z lm Score = 0.05
NG = 0
DB = 0
SM = 0
Delta_AUC
Doxorubicin

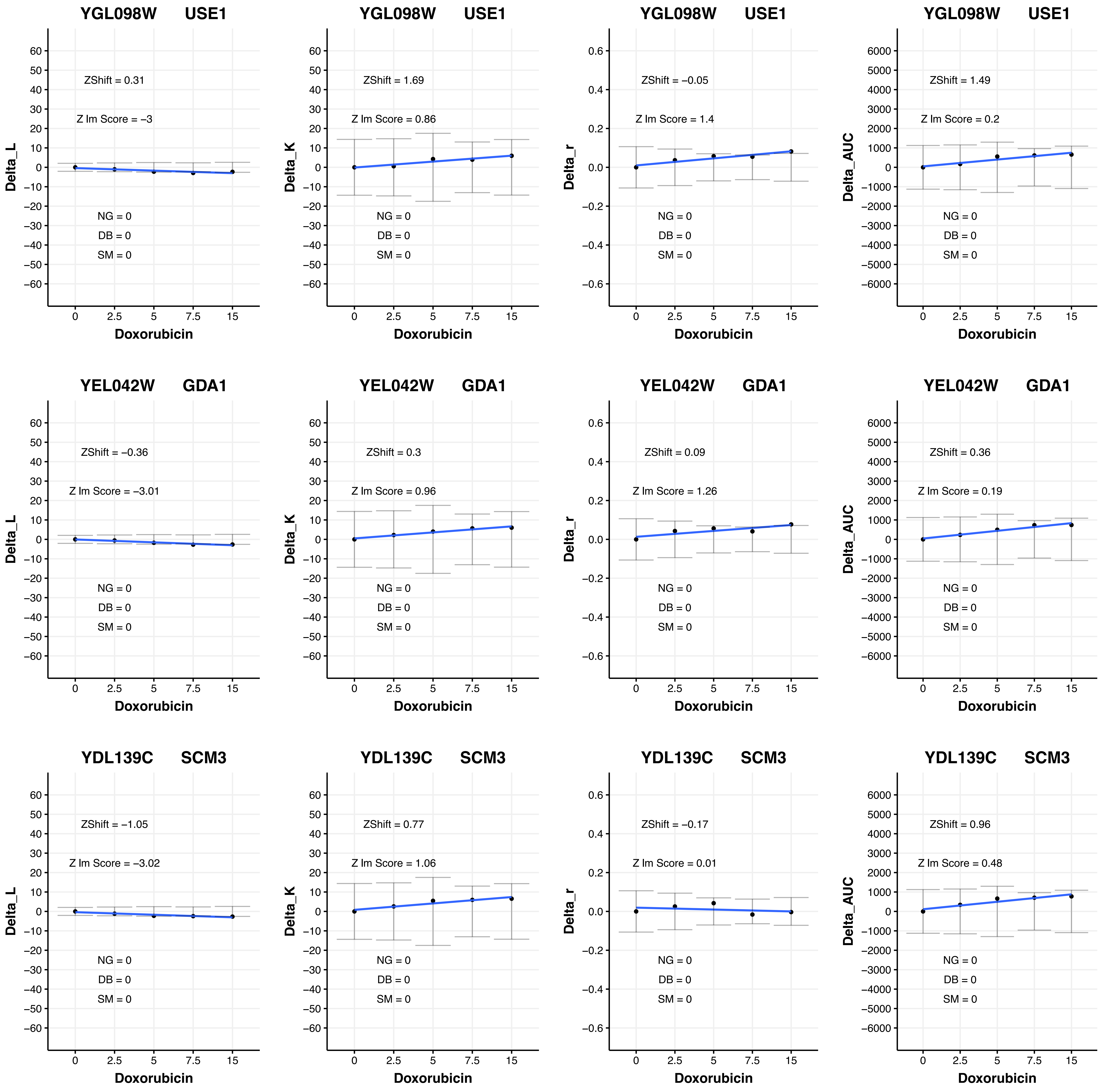
YGL098W USE1
ZShift = 0.31
Z lm Score = −3
NG = 0
DB = 0
SM = 0
Delta_L
Doxorubicin
YGL098W USE1
ZShift = 1.69
Z lm Score = 0.86
NG = 0
DB = 0
SM = 0
Delta_K
Doxorubicin
YGL098W USE1
ZShift = −0.05
Z lm Score = 1.4
NG = 0
DB = 0
SM = 0
Delta_r
Doxorubicin
YGL098W USE1
ZShift = 1.49
Z lm Score = 0.2
NG = 0
DB = 0
SM = 0
Delta_AUC
Doxorubicin
YEL042W GDA1
ZShift = −0.36
Z lm Score = −3.01
NG = 0
DB = 0
SM = 0
Delta_L
Doxorubicin
YEL042W GDA1
ZShift = 0.3
Z lm Score = 0.96
NG = 0
DB = 0
SM = 0
Delta_K
Doxorubicin
YEL042W GDA1
ZShift = 0.09
Z lm Score = 1.26
NG = 0
DB = 0
SM = 0
Delta_r
Doxorubicin
YEL042W GDA1
ZShift = 0.36
Z lm Score = 0.19
NG = 0
DB = 0
SM = 0
Delta_AUC
Doxorubicin
YDL139C SCM3
ZShift = −1.05
Z lm Score = −3.02
NG = 0
DB = 0
SM = 0
Delta_L
Doxorubicin
YDL139C SCM3
ZShift = 0.77
Z lm Score = 1.06
NG = 0
DB = 0
SM = 0
Delta_K
Doxorubicin
YDL139C SCM3
ZShift = −0.17
Z lm Score = 0.01
NG = 0
DB = 0
SM = 0
Delta_r
Doxorubicin
YDL139C SCM3
ZShift = 0.96
Z lm Score = 0.48
NG = 0
DB = 0
SM = 0
Delta_AUC
Doxorubicin

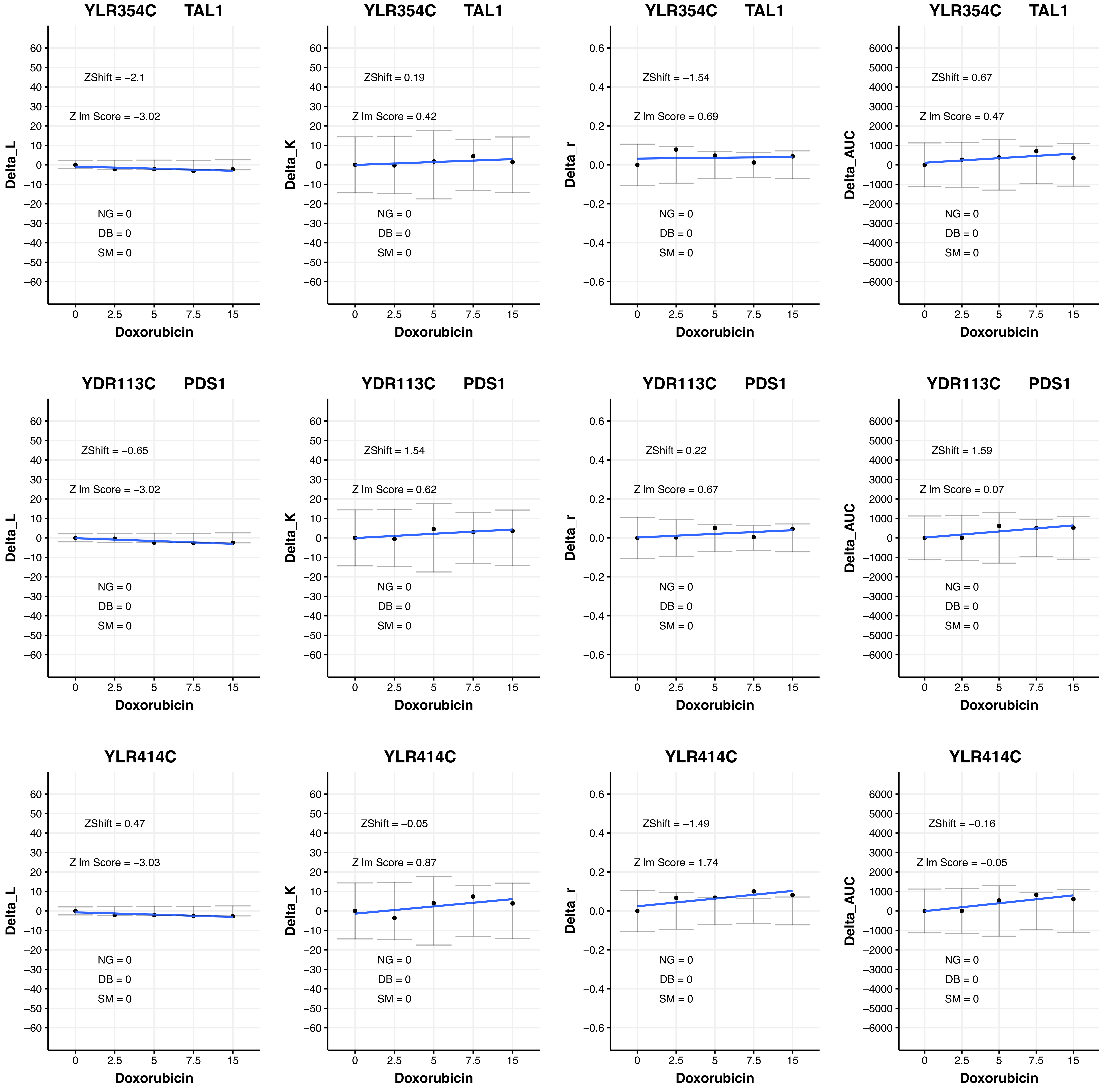
YLR354C TAL1
ZShift = −2.1
Z lm Score = −3.02
NG = 0
DB = 0
SM = 0
Delta_L
Doxorubicin
YLR354C TAL1
ZShift = 0.19
Z lm Score = 0.42
NG = 0
DB = 0
SM = 0
Delta_K
Doxorubicin
YLR354C TAL1
ZShift = −1.54
Z lm Score = 0.69
NG = 0
DB = 0
SM = 0
Delta_r
Doxorubicin
YLR354C TAL1
ZShift = 0.67
Z lm Score = 0.47
NG = 0
DB = 0
SM = 0
Delta_AUC
Doxorubicin
YDR113C PDS1
ZShift = −0.65
Z lm Score = −3.02
NG = 0
DB = 0
SM = 0
Delta_L
Doxorubicin
YDR113C PDS1
ZShift = 1.54
Z lm Score = 0.62
NG = 0
DB = 0
SM = 0
Delta_K
Doxorubicin
YDR113C PDS1
ZShift = 0.22
Z lm Score = 0.67
NG = 0
DB = 0
SM = 0
Delta_r
Doxorubicin
YDR113C PDS1
ZShift = 1.59
Z lm Score = 0.07
NG = 0
DB = 0
SM = 0
Delta_AUC
Doxorubicin
YLR414C
ZShift = 0.47
Z lm Score = −3.03
NG = 0
DB = 0
SM = 0
Delta_L
Doxorubicin
YLR414C
ZShift = −0.05
Z lm Score = 0.87
NG = 0
DB = 0
SM = 0
Delta_K
Doxorubicin
YLR414C
ZShift = −1.49
Z lm Score = 1.74
NG = 0
DB = 0
SM = 0
Delta_r
Doxorubicin
YLR414C
ZShift = −0.16
Z lm Score = −0.05
NG = 0
DB = 0
SM = 0
Delta_AUC
Doxorubicin

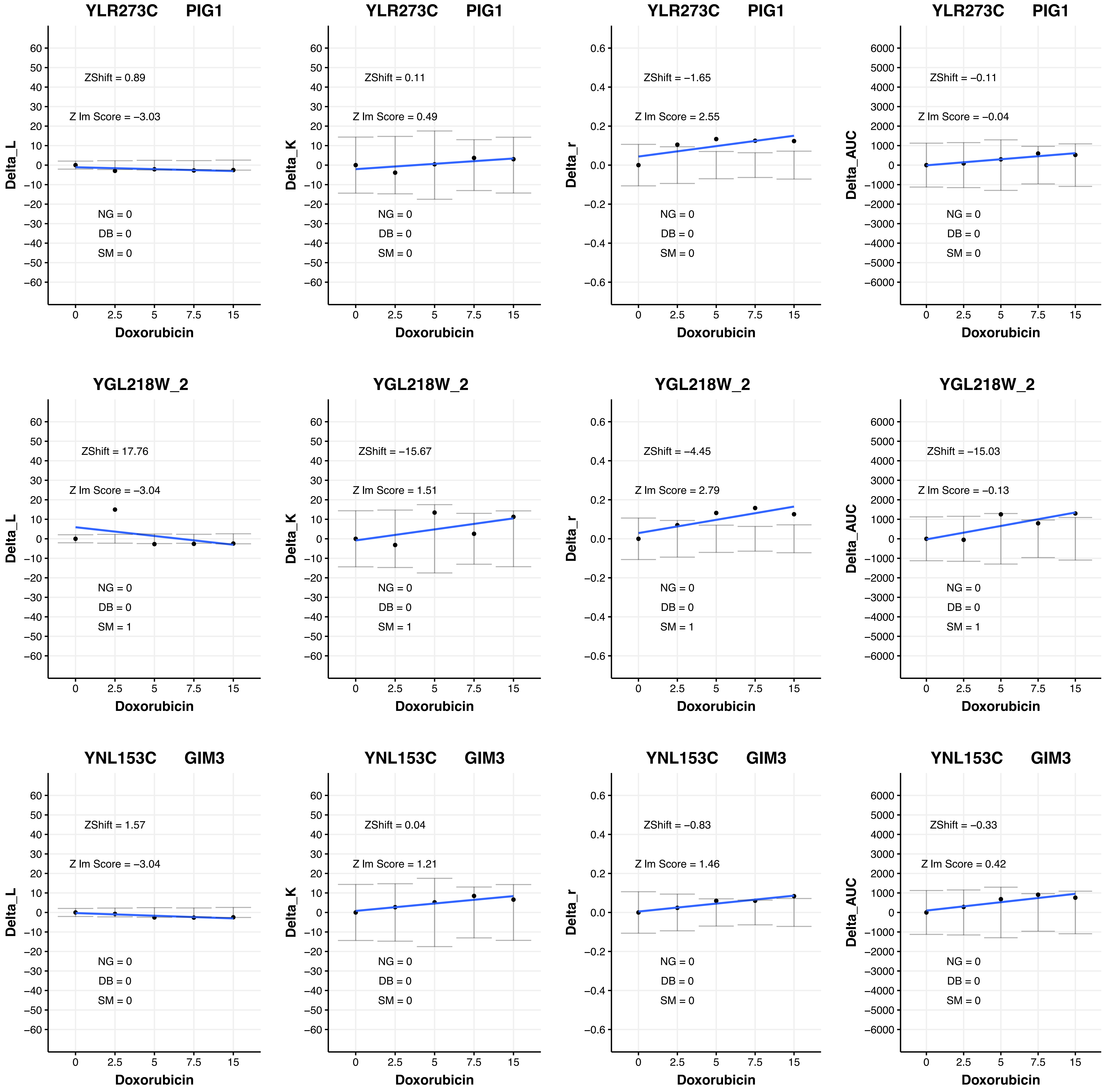
YLR273C PIG1
ZShift = 0.89
Z lm Score = −3.03
NG = 0
DB = 0
SM = 0
Delta_L
Doxorubicin
YLR273C PIG1
ZShift = 0.11
Z lm Score = 0.49
NG = 0
DB = 0
SM = 0
Delta_K
Doxorubicin
YLR273C PIG1
ZShift = −1.65
Z lm Score = 2.55
NG = 0
DB = 0
SM = 0
Delta_r
Doxorubicin
YLR273C PIG1
ZShift = −0.11
Z lm Score = −0.04
NG = 0
DB = 0
SM = 0
Delta_AUC
Doxorubicin
YGL218W_2
ZShift = 17.76
Z lm Score = −3.04
NG = 0
DB = 0
SM = 1
Delta_L
Doxorubicin
YGL218W_2
ZShift = −15.67
Z lm Score = 1.51
NG = 0
DB = 0
SM = 1
Delta_K
Doxorubicin
YGL218W_2
ZShift = −4.45
Z lm Score = 2.79
NG = 0
DB = 0
SM = 1
Delta_r
Doxorubicin
YGL218W_2
ZShift = −15.03
Z lm Score = −0.13
NG = 0
DB = 0
SM = 1
Delta_AUC
Doxorubicin
YNL153C GIM3
ZShift = 1.57
Z lm Score = −3.04
NG = 0
DB = 0
SM = 0
Delta_L
Doxorubicin
YNL153C GIM3
ZShift = 0.04
Z lm Score = 1.21
NG = 0
DB = 0
SM = 0
Delta_K
Doxorubicin
YNL153C GIM3
ZShift = −0.83
Z lm Score = 1.46
NG = 0
DB = 0
SM = 0
Delta_r
Doxorubicin
YNL153C GIM3
ZShift = −0.33
Z lm Score = 0.42
NG = 0
DB = 0
SM = 0
Delta_AUC
Doxorubicin

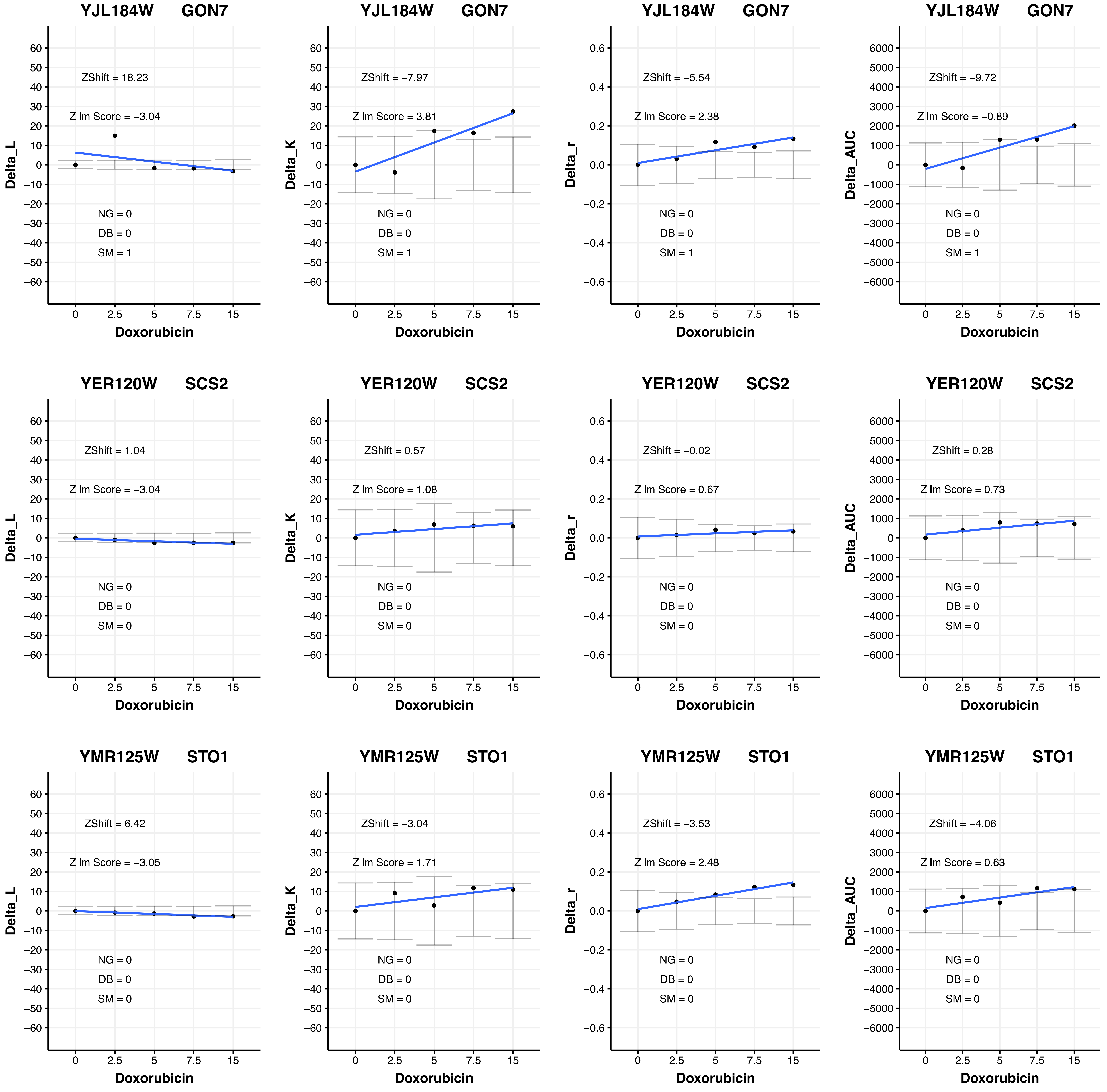
YJL184W GON7
ZShift = 18.23
Z lm Score = −3.04
NG = 0
DB = 0
SM = 1
Delta_L
Doxorubicin
YJL184W GON7
ZShift = −7.97
Z lm Score = 3.81
NG = 0
DB = 0
SM = 1
Delta_K
Doxorubicin
YJL184W GON7
ZShift = −5.54
Z lm Score = 2.38
NG = 0
DB = 0
SM = 1
Delta_r
Doxorubicin
YJL184W GON7
ZShift = −9.72
Z lm Score = −0.89
NG = 0
DB = 0
SM = 1
Delta_AUC
Doxorubicin
YER120W SCS2
ZShift = 1.04
Z lm Score = −3.04
NG = 0
DB = 0
SM = 0
Delta_L
Doxorubicin
YER120W SCS2
ZShift = 0.57
Z lm Score = 1.08
NG = 0
DB = 0
SM = 0
Delta_K
Doxorubicin
YER120W SCS2
ZShift = −0.02
Z lm Score = 0.67
NG = 0
DB = 0
SM = 0
Delta_r
Doxorubicin
YER120W SCS2
ZShift = 0.28
Z lm Score = 0.73
NG = 0
DB = 0
SM = 0
Delta_AUC
Doxorubicin
YMR125W STO1
ZShift = 6.42
Z lm Score = −3.05
NG = 0
DB = 0
SM = 0
Delta_L
Doxorubicin
YMR125W STO1
ZShift = −3.04
Z lm Score = 1.71
NG = 0
DB = 0
SM = 0
Delta_K
Doxorubicin
YMR125W STO1
ZShift = −3.53
Z lm Score = 2.48
NG = 0
DB = 0
SM = 0
Delta_r
Doxorubicin
YMR125W STO1
ZShift = −4.06
Z lm Score = 0.63
NG = 0
DB = 0
SM = 0
Delta_AUC
Doxorubicin

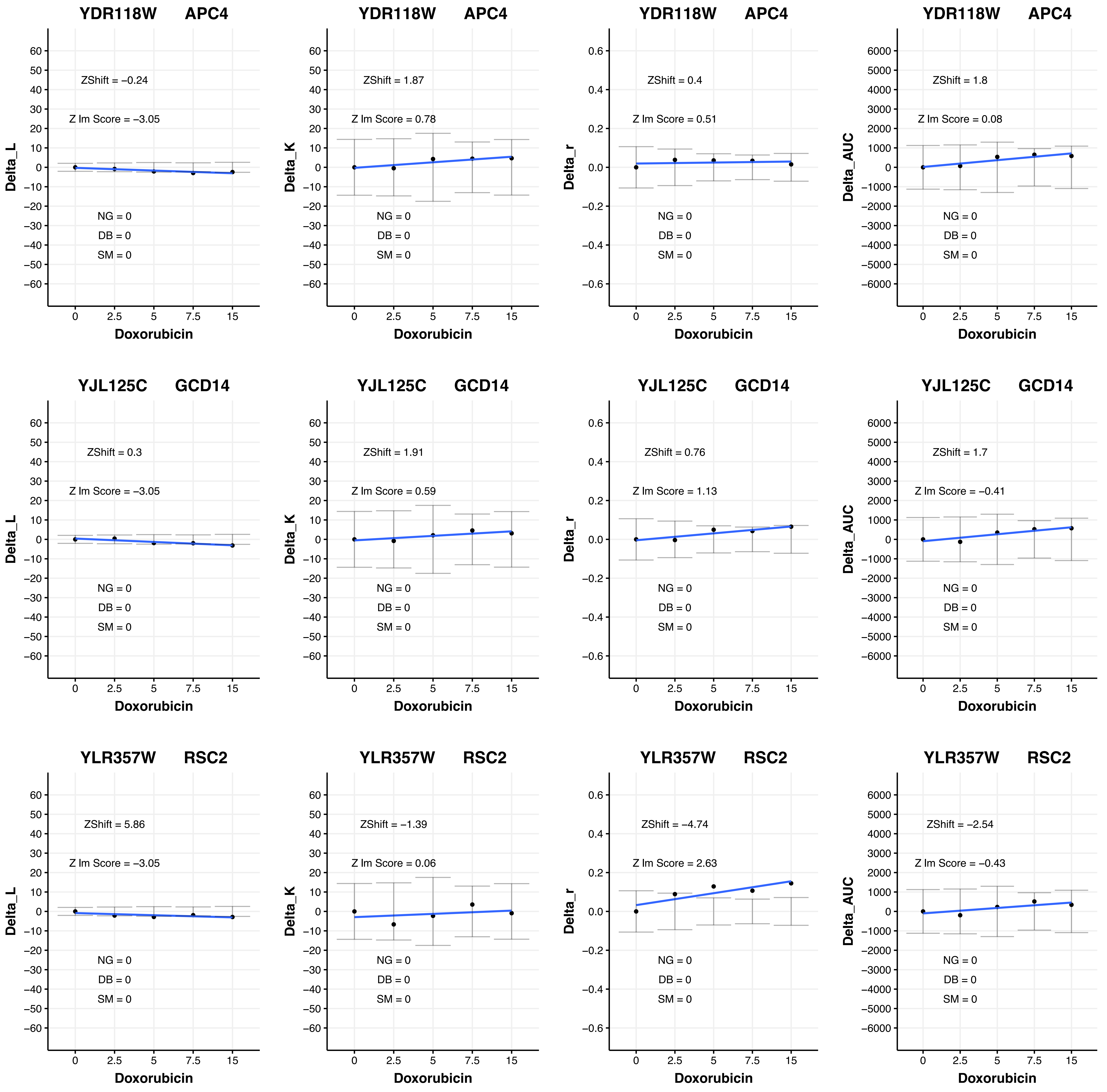

YDR118W APC4
ZShift = −0.24
Z lm Score = −3.05
NG = 0
DB = 0
SM = 0
Delta_L
Doxorubicin
YDR118W APC4
ZShift = 1.87
Z lm Score = 0.78
NG = 0
DB = 0
SM = 0
Delta_K
Doxorubicin
YDR118W APC4
ZShift = 0.4
Z lm Score = 0.51
NG = 0
DB = 0
SM = 0
Delta_r
Doxorubicin
YDR118W APC4
ZShift = 1.8
Z lm Score = 0.08
NG = 0
DB = 0
SM = 0
Delta_AUC
Doxorubicin
YJL125C GCD14
ZShift = 0.3
Z lm Score = −3.05
NG = 0
DB = 0
SM = 0
Delta_L
Doxorubicin
YJL125C GCD14
ZShift = 1.91
Z lm Score = 0.59
NG = 0
DB = 0
SM = 0
Delta_K
Doxorubicin
YJL125C GCD14
ZShift = 0.76
Z lm Score = 1.13
NG = 0
DB = 0
SM = 0
Delta_r
Doxorubicin
YJL125C GCD14
ZShift = 1.7
Z lm Score = −0.41
NG = 0
DB = 0
SM = 0
Delta_AUC
Doxorubicin
YLR357W RSC2
ZShift = 5.86
Z lm Score = −3.05
NG = 0
DB = 0
SM = 0
Delta_L
Doxorubicin
YLR357W RSC2
ZShift = −1.39
Z lm Score = 0.06
NG = 0
DB = 0
SM = 0
Delta_K
Doxorubicin
YLR357W RSC2
ZShift = −4.74
Z lm Score = 2.63
NG = 0
DB = 0
SM = 0
Delta_r
Doxorubicin
YLR357W RSC2
ZShift = −2.54
Z lm Score = −0.43
NG = 0
DB = 0
SM = 0
Delta_AUC
Doxorubicin

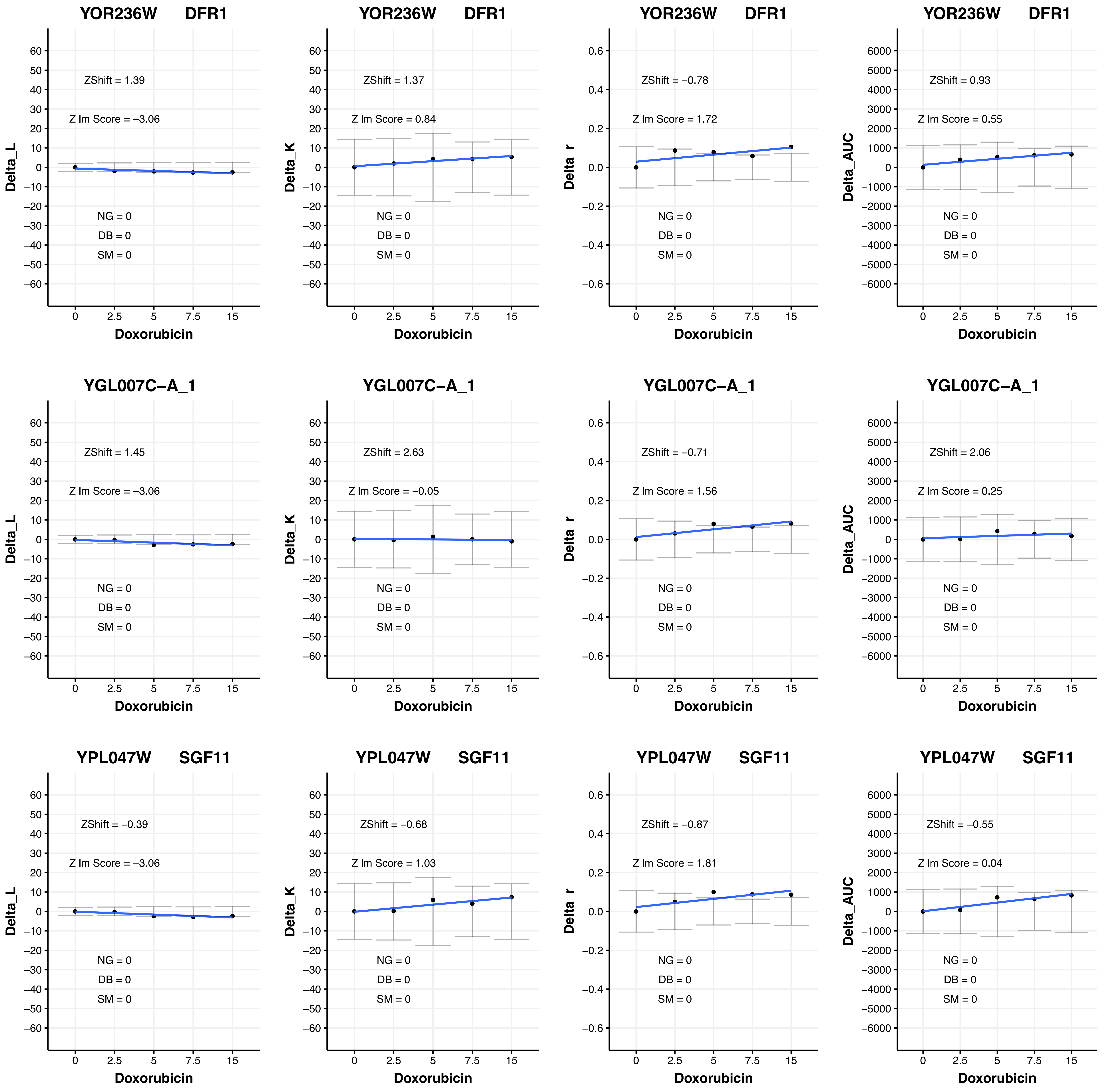
YOR236W DFR1
ZShift = 1.39
Z lm Score = −3.06
NG = 0
DB = 0
SM = 0
Delta_L
Doxorubicin
YOR236W DFR1
ZShift = 1.37
Z lm Score = 0.84
NG = 0
DB = 0
SM = 0
Delta_K
Doxorubicin
YOR236W DFR1
ZShift = −0.78
Z lm Score = 1.72
NG = 0
DB = 0
SM = 0
Delta_r
Doxorubicin
YOR236W DFR1
ZShift = 0.93
Z lm Score = 0.55
NG = 0
DB = 0
SM = 0
Delta_AUC
Doxorubicin
YGL007C−A_1
ZShift = 1.45
Z lm Score = −3.06
NG = 0
DB = 0
SM = 0
Delta_L
Doxorubicin
YGL007C−A_1
ZShift = 2.63
Z lm Score = −0.05
NG = 0
DB = 0
SM = 0
Delta_K
Doxorubicin
YGL007C−A_1
ZShift = −0.71
Z lm Score = 1.56
NG = 0
DB = 0
SM = 0
Delta_r
Doxorubicin
YGL007C−A_1
ZShift = 2.06
Z lm Score = 0.25
NG = 0
DB = 0
SM = 0
Delta_AUC
Doxorubicin
YPL047W SGF11
ZShift = −0.39
Z lm Score = −3.06
NG = 0
DB = 0
SM = 0
Delta_L
Doxorubicin
YPL047W SGF11
ZShift = −0.68
Z lm Score = 1.03
NG = 0
DB = 0
SM = 0
Delta_K
Doxorubicin
YPL047W SGF11
ZShift = −0.87
Z lm Score = 1.81
NG = 0
DB = 0
SM = 0
Delta_r
Doxorubicin
YPL047W SGF11
ZShift = −0.55
Z lm Score = 0.04
NG = 0
DB = 0
SM = 0
Delta_AUC
Doxorubicin

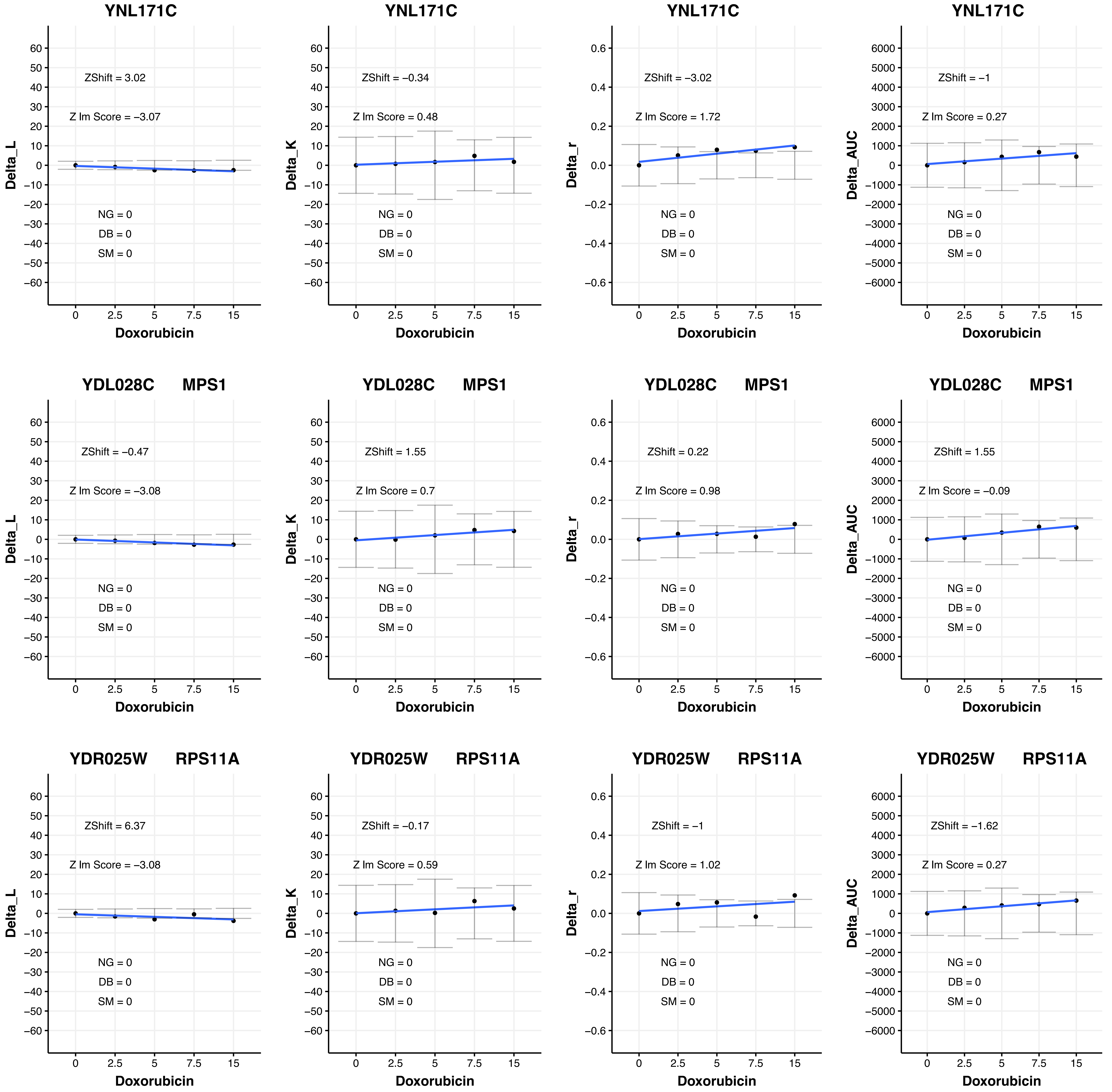
YNL171C
ZShift = 3.02
Z lm Score = −3.07
NG = 0
DB = 0
SM = 0
Delta_L
Doxorubicin
YNL171C
ZShift = −0.34
Z lm Score = 0.48
NG = 0
DB = 0
SM = 0
Delta_K
Doxorubicin
YNL171C
ZShift = −3.02
Z lm Score = 1.72
NG = 0
DB = 0
SM = 0
Delta_r
Doxorubicin
YNL171C
ZShift = −1
Z lm Score = 0.27
NG = 0
DB = 0
SM = 0
Delta_AUC
Doxorubicin
YDL028C MPS1
ZShift = −0.47
Z lm Score = −3.08
NG = 0
DB = 0
SM = 0
Delta_L
Doxorubicin
YDL028C MPS1
ZShift = 1.55
Z lm Score = 0.7
NG = 0
DB = 0
SM = 0
Delta_K
Doxorubicin
YDL028C MPS1
ZShift = 0.22
Z lm Score = 0.98
NG = 0
DB = 0
SM = 0
Delta_r
Doxorubicin
YDL028C MPS1
ZShift = 1.55
Z lm Score = −0.09
NG = 0
DB = 0
SM = 0
Delta_AUC
Doxorubicin
YDR025W RPS11A
ZShift = 6.37
Z lm Score = −3.08
NG = 0
DB = 0
SM = 0
Delta_L
Doxorubicin
YDR025W RPS11A
ZShift = −0.17
Z lm Score = 0.59
NG = 0
DB = 0
SM = 0
Delta_K
Doxorubicin
YDR025W RPS11A
ZShift = −1
Z lm Score = 1.02
NG = 0
DB = 0
SM = 0
Delta_r
Doxorubicin
YDR025W RPS11A
ZShift = −1.62
Z lm Score = 0.27
NG = 0
DB = 0
SM = 0
Delta_AUC
Doxorubicin

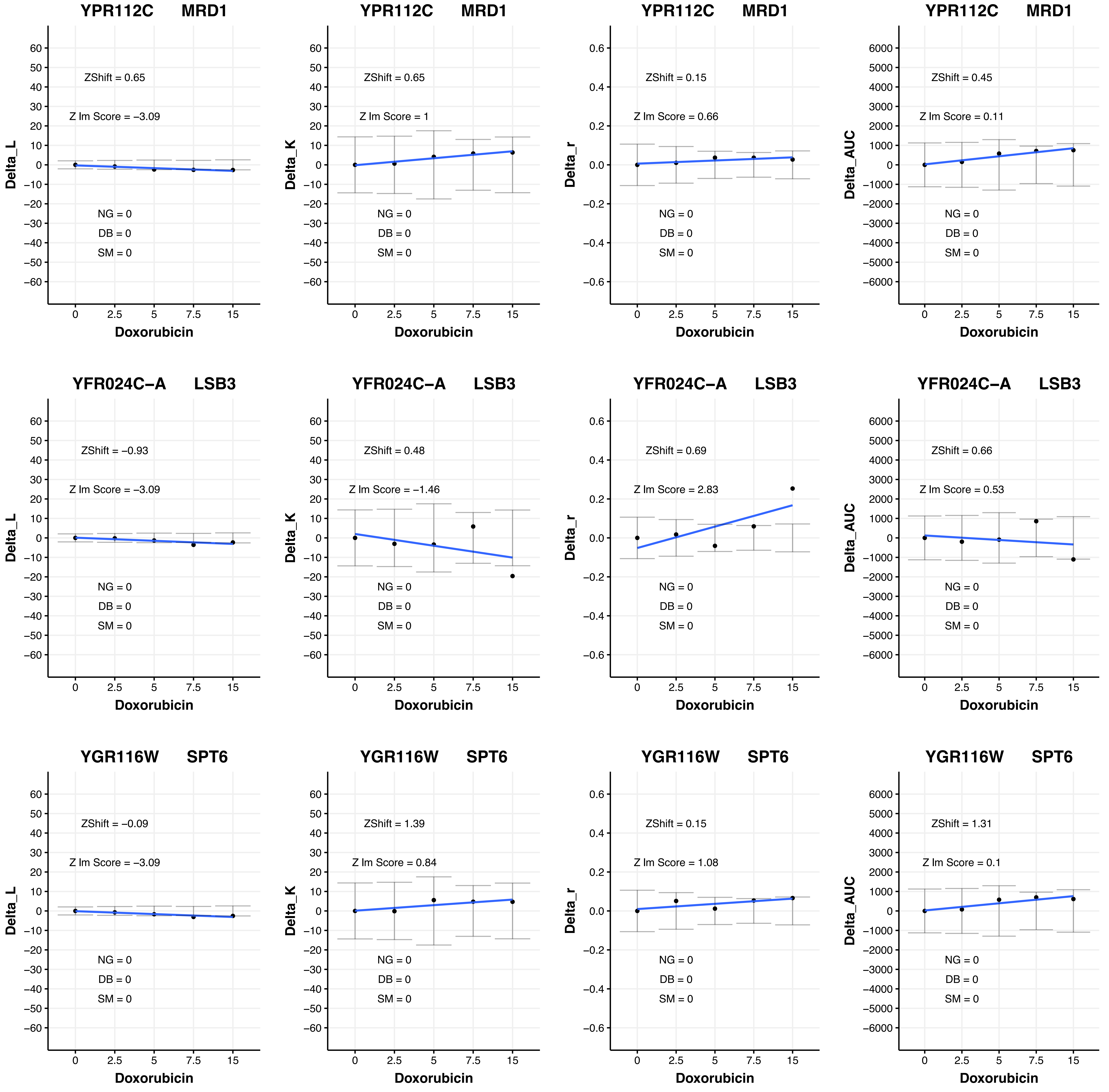
YPR112C MRD1
ZShift = 0.65
Z Im Score = −3.09
NG = 0
DB = 0
SM = 0
Delta_L
Doxorubicin
YPR112C MRD1
ZShift = 0.65
Z Im Score = 1
NG = 0
DB = 0
SM = 0
Delta_K
Doxorubicin
YPR112C MRD1
ZShift = 0.15
Z Im Score = 0.66
NG = 0
DB = 0
SM = 0
Delta_r
Doxorubicin
YPR112C MRD1
ZShift = 0.45
Z Im Score = 0.11
NG = 0
DB = 0
SM = 0
Delta_AUC
Doxorubicin
YFR024C−A LSB3
ZShift = −0.93
Z Im Score = −3.09
NG = 0
DB = 0
SM = 0
Delta_L
Doxorubicin
YFR024C−A LSB3
ZShift = 0.48
Z Im Score = −1.46
NG = 0
DB = 0
SM = 0
Delta_K
Doxorubicin
YFR024C−A LSB3
ZShift = 0.69
Z Im Score = 2.83
NG = 0
DB = 0
SM = 0
Delta_r
Doxorubicin
YFR024C−A LSB3
ZShift = 0.66
Z Im Score = 0.53
NG = 0
DB = 0
SM = 0
Delta_AUC
Doxorubicin
YGR116W SPT6
ZShift = −0.09
Z Im Score = −3.09
NG = 0
DB = 0
SM = 0
Delta_L
Doxorubicin
YGR116W SPT6
ZShift = 1.39
Z Im Score = 0.84
NG = 0
DB = 0
SM = 0
Delta_K
Doxorubicin
YGR116W SPT6
ZShift = 0.15
Z Im Score = 1.08
NG = 0
DB = 0
SM = 0
Delta_r
Doxorubicin
YGR116W SPT6
ZShift = 1.31
Z Im Score = 0.1
NG = 0
DB = 0
SM = 0
Delta_AUC
Doxorubicin

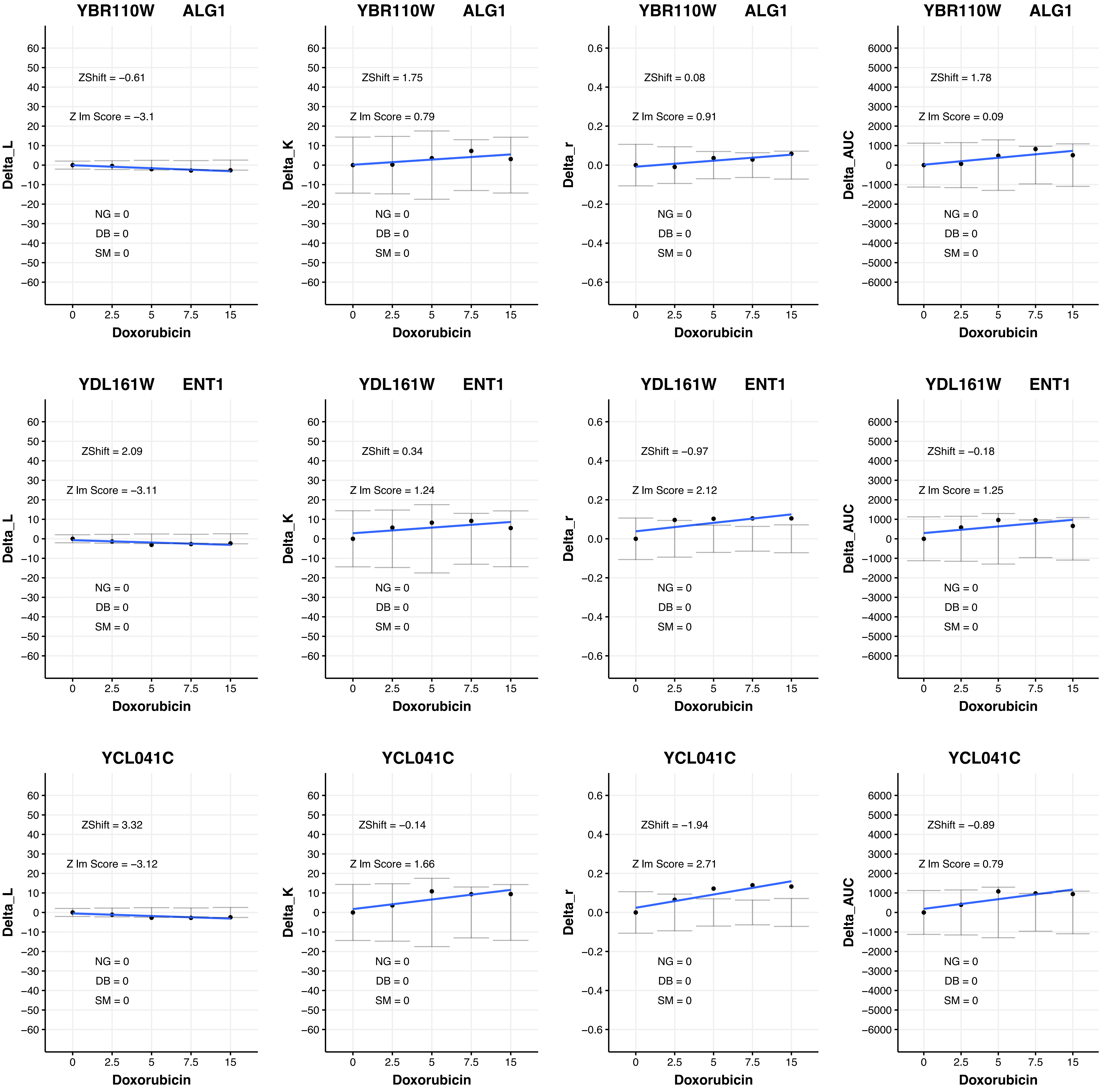

YBR110W ALG1
ZShift = −0.61
Z lm Score = −3.1
NG = 0
DB = 0
SM = 0
Delta_L
Doxorubicin
YBR110W ALG1
ZShift = 1.75
Z lm Score = 0.79
NG = 0
DB = 0
SM = 0
Delta_K
Doxorubicin
YBR110W ALG1
ZShift = 0.08
Z lm Score = 0.91
NG = 0
DB = 0
SM = 0
Delta_r
Doxorubicin
YBR110W ALG1
ZShift = 1.78
Z lm Score = 0.09
NG = 0
DB = 0
SM = 0
Delta_AUC
Doxorubicin
YDL161W ENT1
ZShift = 2.09
Z lm Score = −3.11
NG = 0
DB = 0
SM = 0
Delta_L
Doxorubicin
YDL161W ENT1
ZShift = 0.34
Z lm Score = 1.24
NG = 0
DB = 0
SM = 0
Delta_K
Doxorubicin
YDL161W ENT1
ZShift = −0.97
Z lm Score = 2.12
NG = 0
DB = 0
SM = 0
Delta_r
Doxorubicin
YDL161W ENT1
ZShift = −0.18
Z lm Score = 1.25
NG = 0
DB = 0
SM = 0
Delta_AUC
Doxorubicin
YCL041C
ZShift = 3.32
Z lm Score = −3.12
NG = 0
DB = 0
SM = 0
Delta_L
Doxorubicin
YCL041C
ZShift = −0.14
Z lm Score = 1.66
NG = 0
DB = 0
SM = 0
Delta_K
Doxorubicin
YCL041C
ZShift = −1.94
Z lm Score = 2.71
NG = 0
DB = 0
SM = 0
Delta_r
Doxorubicin
YCL041C
ZShift = −0.89
Z lm Score = 0.79
NG = 0
DB = 0
SM = 0
Delta_AUC
Doxorubicin

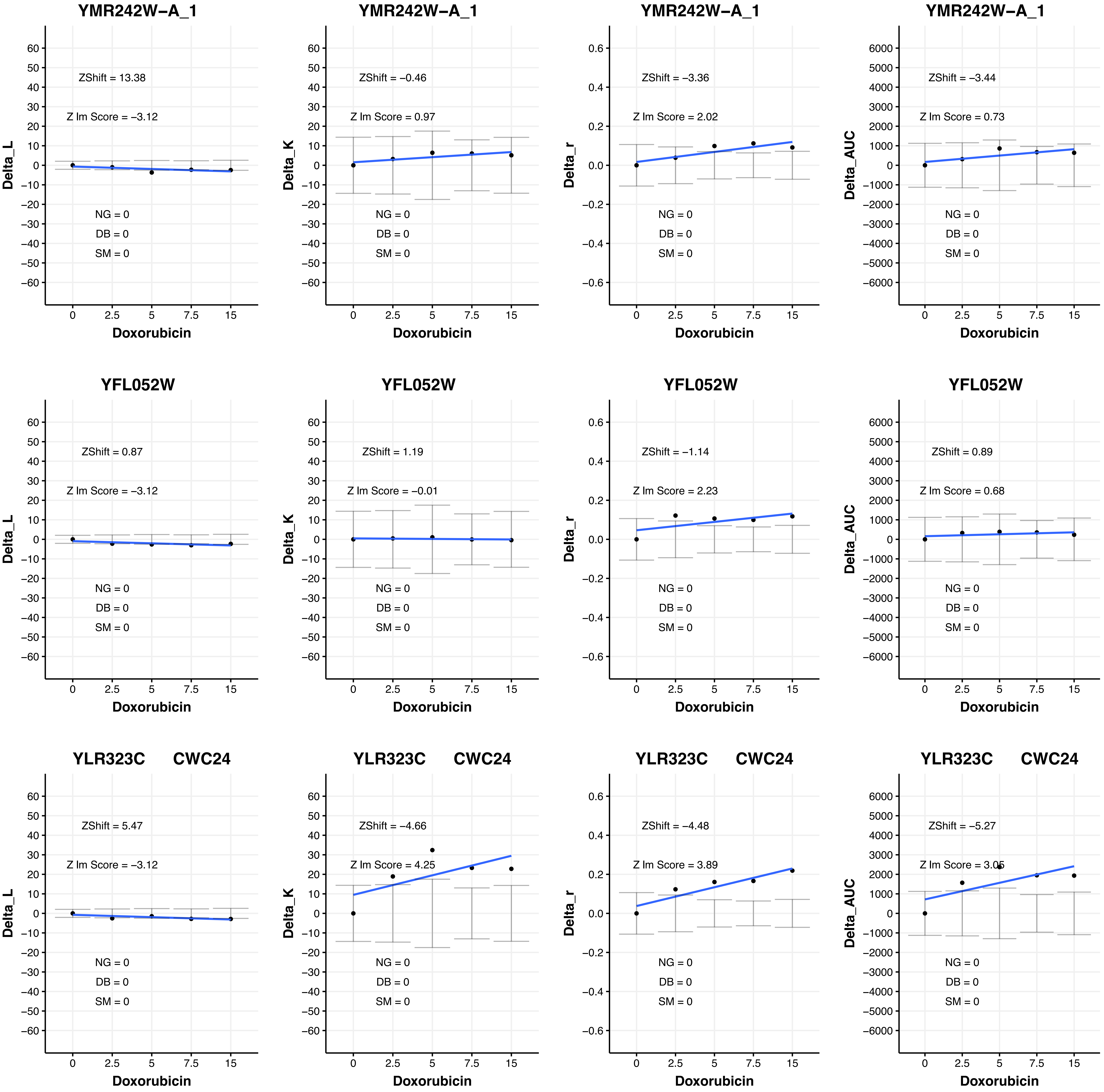

YMR242W-A_1
ZShift = 13.38
Z lm Score = −3.12
NG = 0
DB = 0
SM = 0
Delta_L
Doxorubicin
YMR242W-A_1
ZShift = −0.46
Z lm Score = 0.97
NG = 0
DB = 0
SM = 0
Delta_K
Doxorubicin
YMR242W-A_1
ZShift = −3.36
Z lm Score = 2.02
NG = 0
DB = 0
SM = 0
Delta_r
Doxorubicin
YMR242W-A_1
ZShift = −3.44
Z lm Score = 0.73
NG = 0
DB = 0
SM = 0
Delta_AUC
Doxorubicin
YFL052W
ZShift = 0.87
Z lm Score = −3.12
NG = 0
DB = 0
SM = 0
Delta_L
Doxorubicin
YFL052W
ZShift = 1.19
Z lm Score = −0.01
NG = 0
DB = 0
SM = 0
Delta_K
Doxorubicin
YFL052W
ZShift = −1.14
Z lm Score = 2.23
NG = 0
DB = 0
SM = 0
Delta_r
Doxorubicin
YFL052W
ZShift = 0.89
Z lm Score = 0.68
NG = 0
DB = 0
SM = 0
Delta_AUC
Doxorubicin
YLR323C CWC24
ZShift = 5.47
Z lm Score = −3.12
NG = 0
DB = 0
SM = 0
Delta_L
Doxorubicin
YLR323C CWC24
ZShift = −4.66
Z lm Score = 4.25
NG = 0
DB = 0
SM = 0
Delta_K
Doxorubicin
YLR323C CWC24
ZShift = −4.48
Z lm Score = 3.89
NG = 0
DB = 0
SM = 0
Delta_r
Doxorubicin
YLR323C CWC24
ZShift = −5.27
Z lm Score = 3.05
NG = 0
DB = 0
SM = 0
Delta_AUC
Doxorubicin

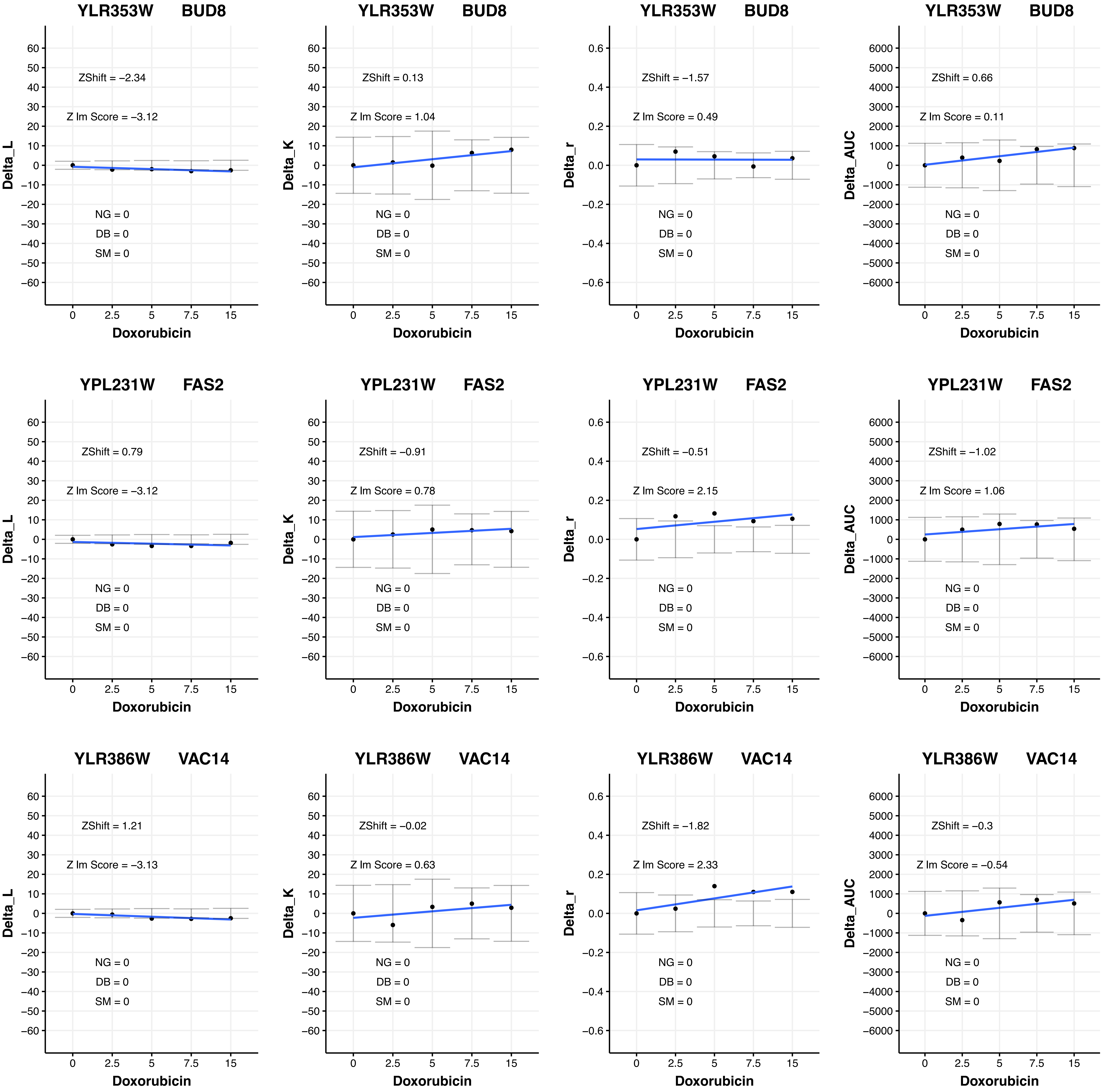

YLR353W BUD8
ZShift = −2.34
Z lm Score = −3.12
NG = 0
DB = 0
SM = 0
Delta_L
Doxorubicin
YLR353W BUD8
ZShift = 0.13
Z lm Score = 1.04
NG = 0
DB = 0
SM = 0
Delta_K
Doxorubicin
YLR353W BUD8
ZShift = −1.57
Z lm Score = 0.49
NG = 0
DB = 0
SM = 0
Delta_r
Doxorubicin
YLR353W BUD8
ZShift = 0.66
Z lm Score = 0.11
NG = 0
DB = 0
SM = 0
Delta_AUC
Doxorubicin
YPL231W FAS2
ZShift = 0.79
Z lm Score = −3.12
NG = 0
DB = 0
SM = 0
Delta_L
Doxorubicin
YPL231W FAS2
ZShift = −0.91
Z lm Score = 0.78
NG = 0
DB = 0
SM = 0
Delta_K
Doxorubicin
YPL231W FAS2
ZShift = −0.51
Z lm Score = 2.15
NG = 0
DB = 0
SM = 0
Delta_r
Doxorubicin
YPL231W FAS2
ZShift = −1.02
Z lm Score = 1.06
NG = 0
DB = 0
SM = 0
Delta_AUC
Doxorubicin
YLR386W VAC14
ZShift = 1.21
Z lm Score = −3.13
NG = 0
DB = 0
SM = 0
Delta_L
Doxorubicin
YLR386W VAC14
ZShift = −0.02
Z lm Score = 0.63
NG = 0
DB = 0
SM = 0
Delta_K
Doxorubicin
YLR386W VAC14
ZShift = −1.82
Z lm Score = 2.33
NG = 0
DB = 0
SM = 0
Delta_r
Doxorubicin
YLR386W VAC14
ZShift = −0.3
Z lm Score = −0.54
NG = 0
DB = 0
SM = 0
Delta_AUC
Doxorubicin

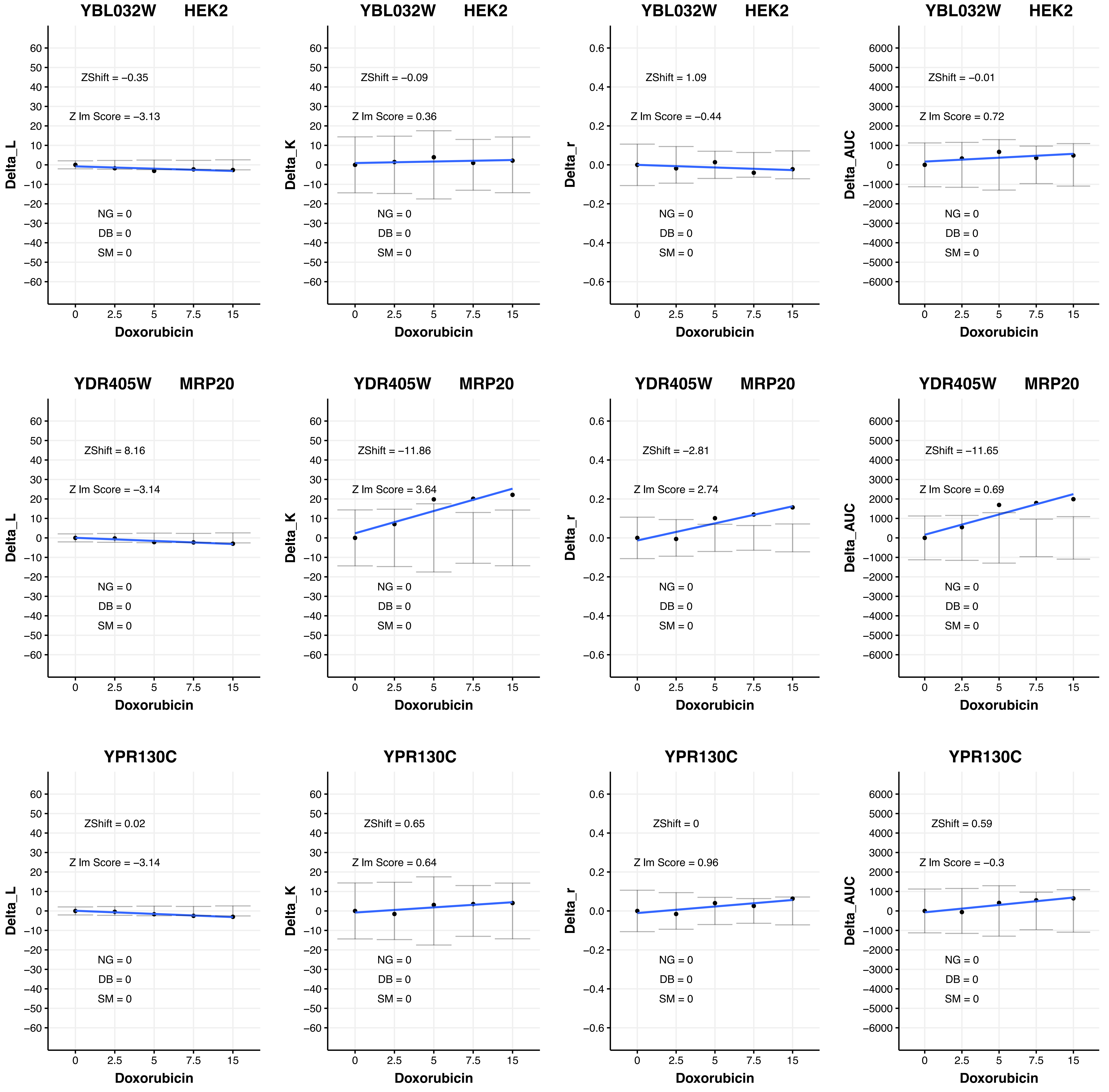
YBL032W HEK2
ZShift = −0.35
Z lm Score = −3.13
NG = 0
DB = 0
SM = 0
Delta_L
Doxorubicin
YBL032W HEK2
ZShift = −0.09
Z lm Score = 0.36
NG = 0
DB = 0
SM = 0
Delta_K
Doxorubicin
YBL032W HEK2
ZShift = 1.09
Z lm Score = −0.44
NG = 0
DB = 0
SM = 0
Delta_r
Doxorubicin
YBL032W HEK2
ZShift = −0.01
Z lm Score = 0.72
NG = 0
DB = 0
SM = 0
Delta_AUC
Doxorubicin
YDR405W MRP20
ZShift = 8.16
Z lm Score = −3.14
NG = 0
DB = 0
SM = 0
Delta_L
Doxorubicin
YDR405W MRP20
ZShift = −11.86
Z lm Score = 3.64
NG = 0
DB = 0
SM = 0
Delta_K
Doxorubicin
YDR405W MRP20
ZShift = −2.81
Z lm Score = 2.74
NG = 0
DB = 0
SM = 0
Delta_r
Doxorubicin
YDR405W MRP20
ZShift = −11.65
Z lm Score = 0.69
NG = 0
DB = 0
SM = 0
Delta_AUC
Doxorubicin
YPR130C
ZShift = 0.02
Z lm Score = −3.14
NG = 0
DB = 0
SM = 0
Delta_L
Doxorubicin
YPR130C
ZShift = 0.65
Z lm Score = 0.64
NG = 0
DB = 0
SM = 0
Delta_K
Doxorubicin
YPR130C
ZShift = 0
Z lm Score = 0.96
NG = 0
DB = 0
SM = 0
Delta_r
Doxorubicin
YPR130C
ZShift = 0.59
Z lm Score = −0.3
NG = 0
DB = 0
SM = 0
Delta_AUC
Doxorubicin

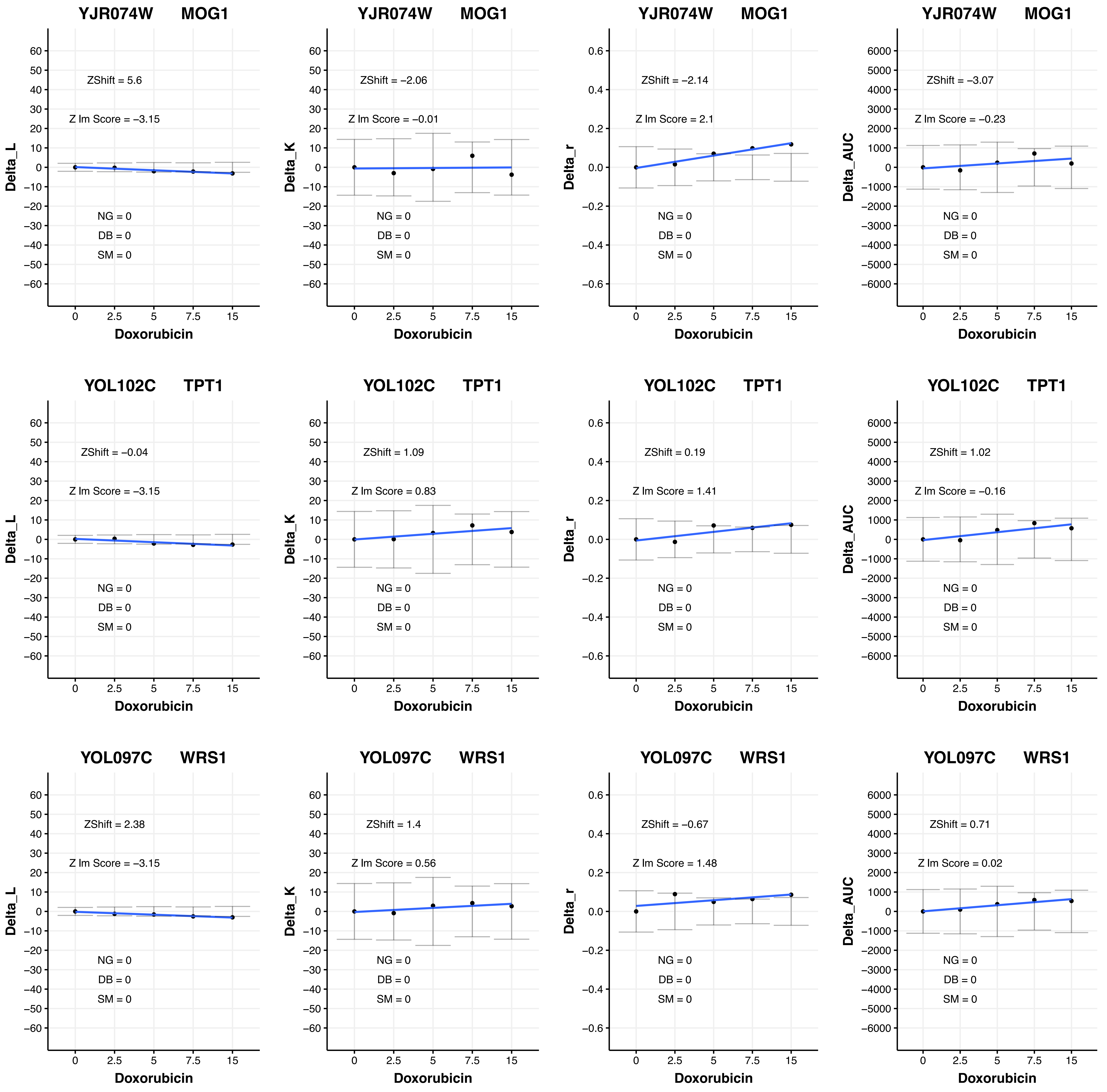

YJR074W MOG1
Delta_L
ZShift = 5.6
Z lm Score = −3.15
NG = 0
DB = 0
SM = 0
Doxorubicin
YJR074W MOG1
Delta_K
ZShift = −2.06
Z lm Score = −0.01
NG = 0
DB = 0
SM = 0
Doxorubicin
YJR074W MOG1
Delta_r
ZShift = −2.14
Z lm Score = 2.1
NG = 0
DB = 0
SM = 0
Doxorubicin
YJR074W MOG1
Delta_AUC
ZShift = −3.07
Z lm Score = −0.23
NG = 0
DB = 0
SM = 0
Doxorubicin
YOL102C TPT1
Delta_L
ZShift = −0.04
Z lm Score = −3.15
NG = 0
DB = 0
SM = 0
Doxorubicin
YOL102C TPT1
Delta_K
ZShift = 1.09
Z lm Score = 0.83
NG = 0
DB = 0
SM = 0
Doxorubicin
YOL102C TPT1
Delta_r
ZShift = 0.19
Z lm Score = 1.41
NG = 0
DB = 0
SM = 0
Doxorubicin
YOL102C TPT1
Delta_AUC
ZShift = 1.02
Z lm Score = −0.16
NG = 0
DB = 0
SM = 0
Doxorubicin
YOL097C WRS1
Delta_L
ZShift = 2.38
Z lm Score = −3.15
NG = 0
DB = 0
SM = 0
Doxorubicin
YOL097C WRS1
Delta_K
ZShift = 1.4
Z lm Score = 0.56
NG = 0
DB = 0
SM = 0
Doxorubicin
YOL097C WRS1
Delta_r
ZShift = −0.67
Z lm Score = 1.48
NG = 0
DB = 0
SM = 0
Doxorubicin
YOL097C WRS1
Delta_AUC
ZShift = 0.71
Z lm Score = 0.02
NG = 0
DB = 0
SM = 0
Doxorubicin

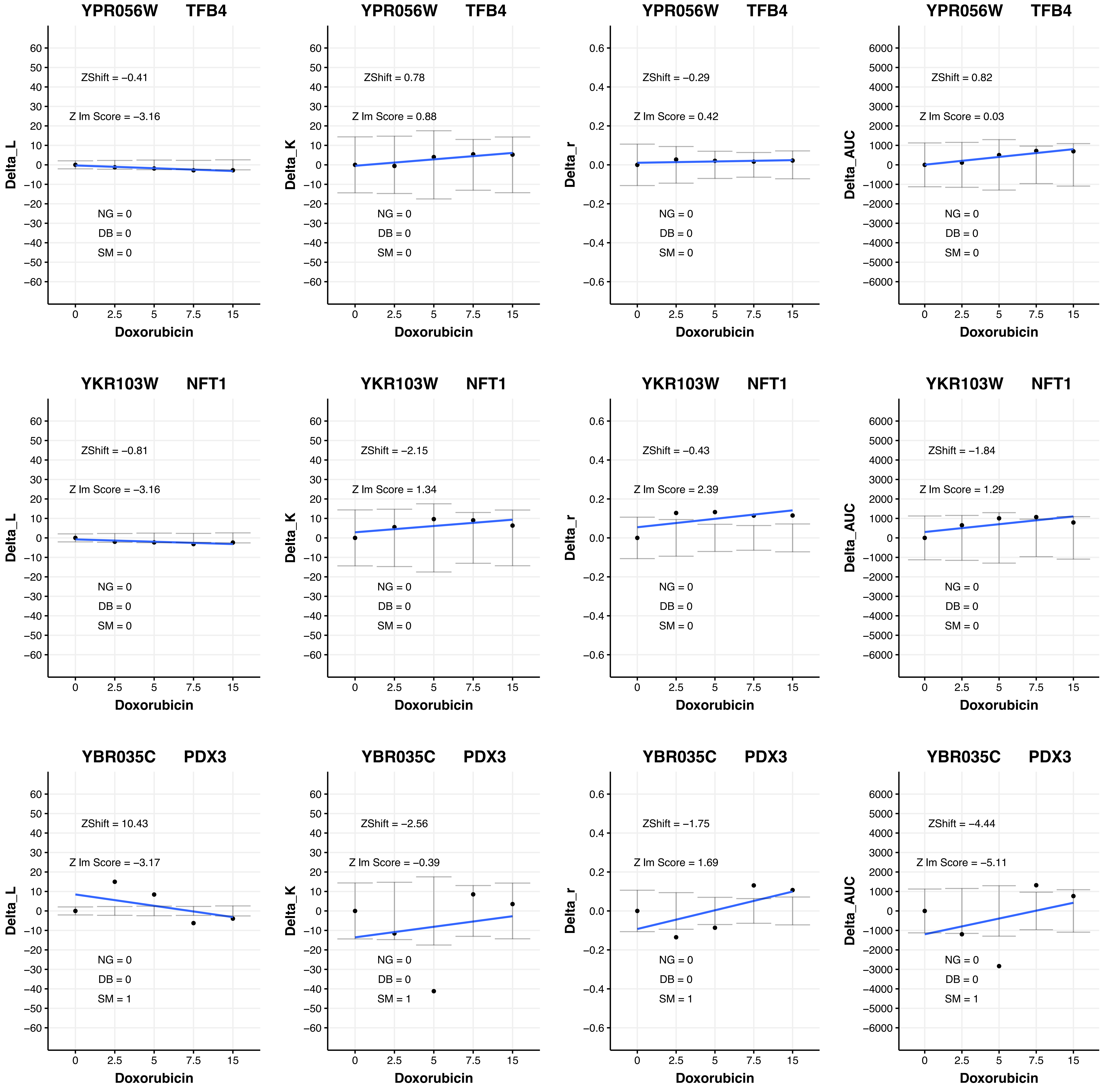
YPR056W TFB4
ZShift = −0.41
Z lm Score = −3.16
NG = 0
DB = 0
SM = 0
Delta_L
Doxorubicin
YPR056W TFB4
ZShift = 0.78
Z lm Score = 0.88
NG = 0
DB = 0
SM = 0
Delta_K
Doxorubicin
YPR056W TFB4
ZShift = −0.29
Z lm Score = 0.42
NG = 0
DB = 0
SM = 0
Delta_r
Doxorubicin
YPR056W TFB4
ZShift = 0.82
Z lm Score = 0.03
NG = 0
DB = 0
SM = 0
Delta_AUC
Doxorubicin
YKR103W NFT1
ZShift = −0.81
Z lm Score = −3.16
NG = 0
DB = 0
SM = 0
Delta_L
Doxorubicin
YKR103W NFT1
ZShift = −2.15
Z lm Score = 1.34
NG = 0
DB = 0
SM = 0
Delta_K
Doxorubicin
YKR103W NFT1
ZShift = −0.43
Z lm Score = 2.39
NG = 0
DB = 0
SM = 0
Delta_r
Doxorubicin
YKR103W NFT1
ZShift = −1.84
Z lm Score = 1.29
NG = 0
DB = 0
SM = 0
Delta_AUC
Doxorubicin
YBR035C PDX3
ZShift = 10.43
Z lm Score = −3.17
NG = 0
DB = 0
SM = 1
Delta_L
Doxorubicin
YBR035C PDX3
ZShift = −2.56
Z lm Score = −0.39
NG = 0
DB = 0
SM = 1
Delta_K
Doxorubicin
YBR035C PDX3
ZShift = −1.75
Z lm Score = 1.69
NG = 0
DB = 0
SM = 1
Delta_r
Doxorubicin
YBR035C PDX3
ZShift = −4.44
Z lm Score = −5.11
NG = 0
DB = 0
SM = 1
Delta_AUC
Doxorubicin

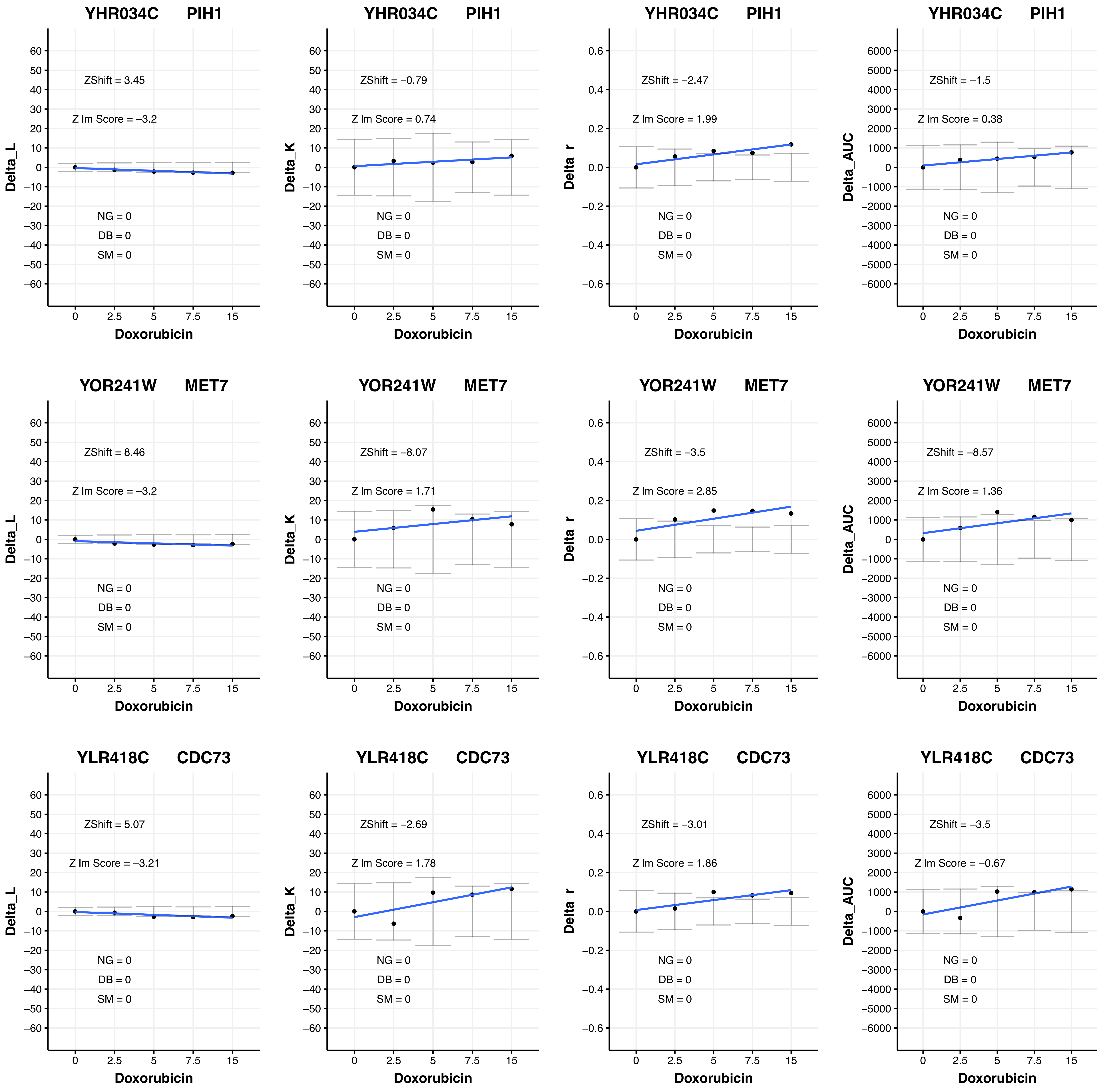

YHR034C PIH1
ZShift = 3.45
Z lm Score = −3.2
NG = 0
DB = 0
SM = 0
Delta_L
Doxorubicin
YHR034C PIH1
ZShift = −0.79
Z lm Score = 0.74
NG = 0
DB = 0
SM = 0
Delta_K
Doxorubicin
YHR034C PIH1
ZShift = −2.47
Z lm Score = 1.99
NG = 0
DB = 0
SM = 0
Delta_r
Doxorubicin
YHR034C PIH1
ZShift = −1.5
Z lm Score = 0.38
NG = 0
DB = 0
SM = 0
Delta_AUC
Doxorubicin
YOR241W MET7
ZShift = 8.46
Z lm Score = −3.2
NG = 0
DB = 0
SM = 0
Delta_L
Doxorubicin
YOR241W MET7
ZShift = −8.07
Z lm Score = 1.71
NG = 0
DB = 0
SM = 0
Delta_K
Doxorubicin
YOR241W MET7
ZShift = −3.5
Z lm Score = 2.85
NG = 0
DB = 0
SM = 0
Delta_r
Doxorubicin
YOR241W MET7
ZShift = −8.57
Z lm Score = 1.36
NG = 0
DB = 0
SM = 0
Delta_AUC
Doxorubicin
YLR418C CDC73
ZShift = 5.07
Z lm Score = −3.21
NG = 0
DB = 0
SM = 0
Delta_L
Doxorubicin
YLR418C CDC73
ZShift = −2.69
Z lm Score = 1.78
NG = 0
DB = 0
SM = 0
Delta_K
Doxorubicin
YLR418C CDC73
ZShift = −3.01
Z lm Score = 1.86
NG = 0
DB = 0
SM = 0
Delta_r
Doxorubicin
YLR418C CDC73
ZShift = −3.5
Z lm Score = −0.67
NG = 0
DB = 0
SM = 0
Delta_AUC
Doxorubicin

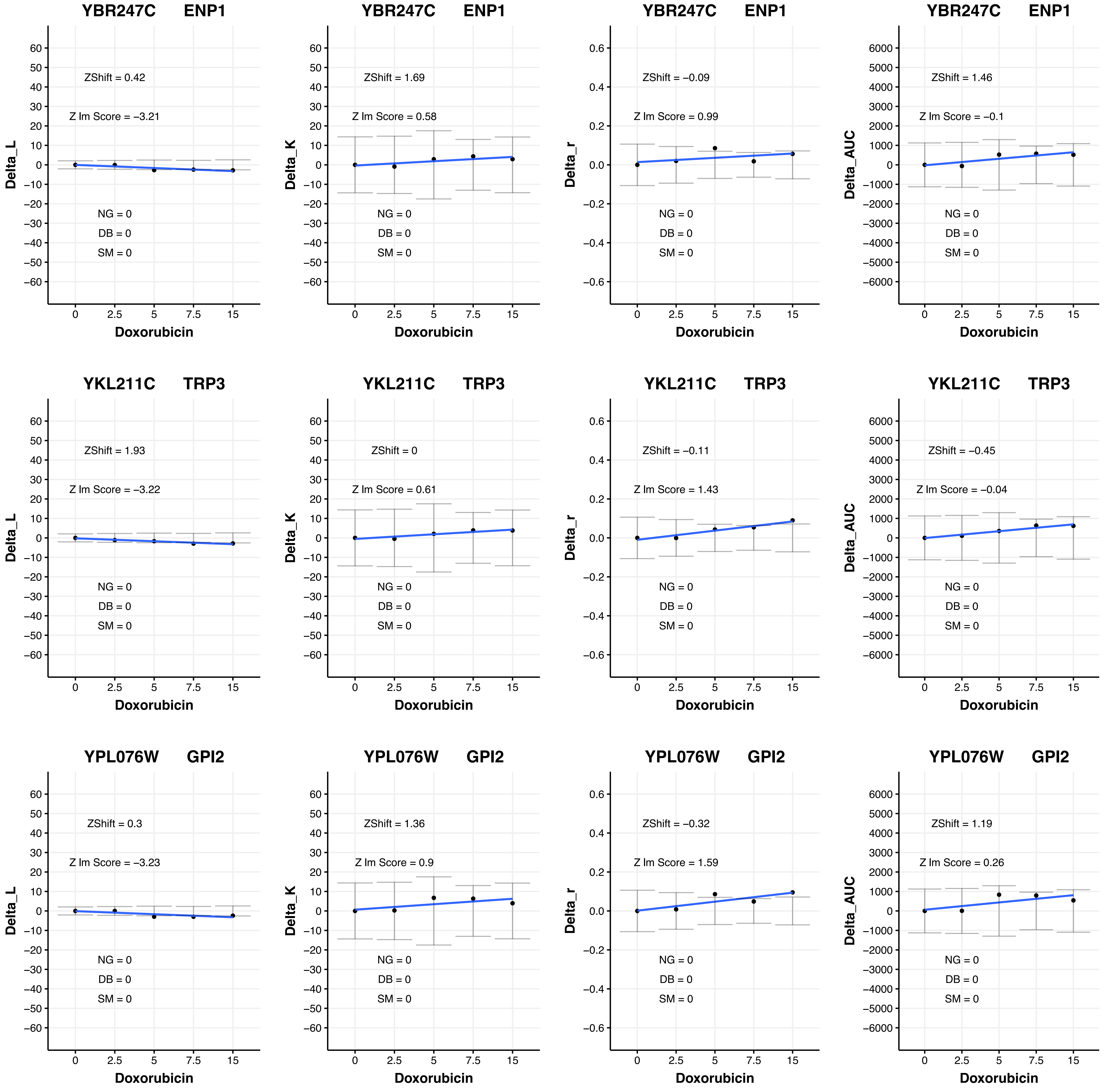

YBR247C ENP1
ZShift = 0.42
Z lm Score = −3.21
NG = 0
DB = 0
SM = 0
Delta_L
Doxorubicin
YBR247C ENP1
ZShift = 1.69
Z lm Score = 0.58
NG = 0
DB = 0
SM = 0
Delta_K
Doxorubicin
YBR247C ENP1
ZShift = −0.09
Z lm Score = 0.99
NG = 0
DB = 0
SM = 0
Delta_r
Doxorubicin
YBR247C ENP1
ZShift = 1.46
Z lm Score = −0.1
NG = 0
DB = 0
SM = 0
Delta_AUC
Doxorubicin
YKL211C TRP3
ZShift = 1.93
Z lm Score = −3.22
NG = 0
DB = 0
SM = 0
Delta_L
Doxorubicin
YKL211C TRP3
ZShift = 0
Z lm Score = 0.61
NG = 0
DB = 0
SM = 0
Delta_K
Doxorubicin
YKL211C TRP3
ZShift = −0.11
Z lm Score = 1.43
NG = 0
DB = 0
SM = 0
Delta_r
Doxorubicin
YKL211C TRP3
ZShift = −0.45
Z lm Score = −0.04
NG = 0
DB = 0
SM = 0
Delta_AUC
Doxorubicin
YPL076W GPI2
ZShift = 0.3
Z lm Score = −3.23
NG = 0
DB = 0
SM = 0
Delta_L
Doxorubicin
YPL076W GPI2
ZShift = 1.36
Z lm Score = 0.9
NG = 0
DB = 0
SM = 0
Delta_K
Doxorubicin
YPL076W GPI2
ZShift = −0.32
Z lm Score = 1.59
NG = 0
DB = 0
SM = 0
Delta_r
Doxorubicin
YPL076W GPI2
ZShift = 1.19
Z lm Score = 0.26
NG = 0
DB = 0
SM = 0
Delta_AUC
Doxorubicin

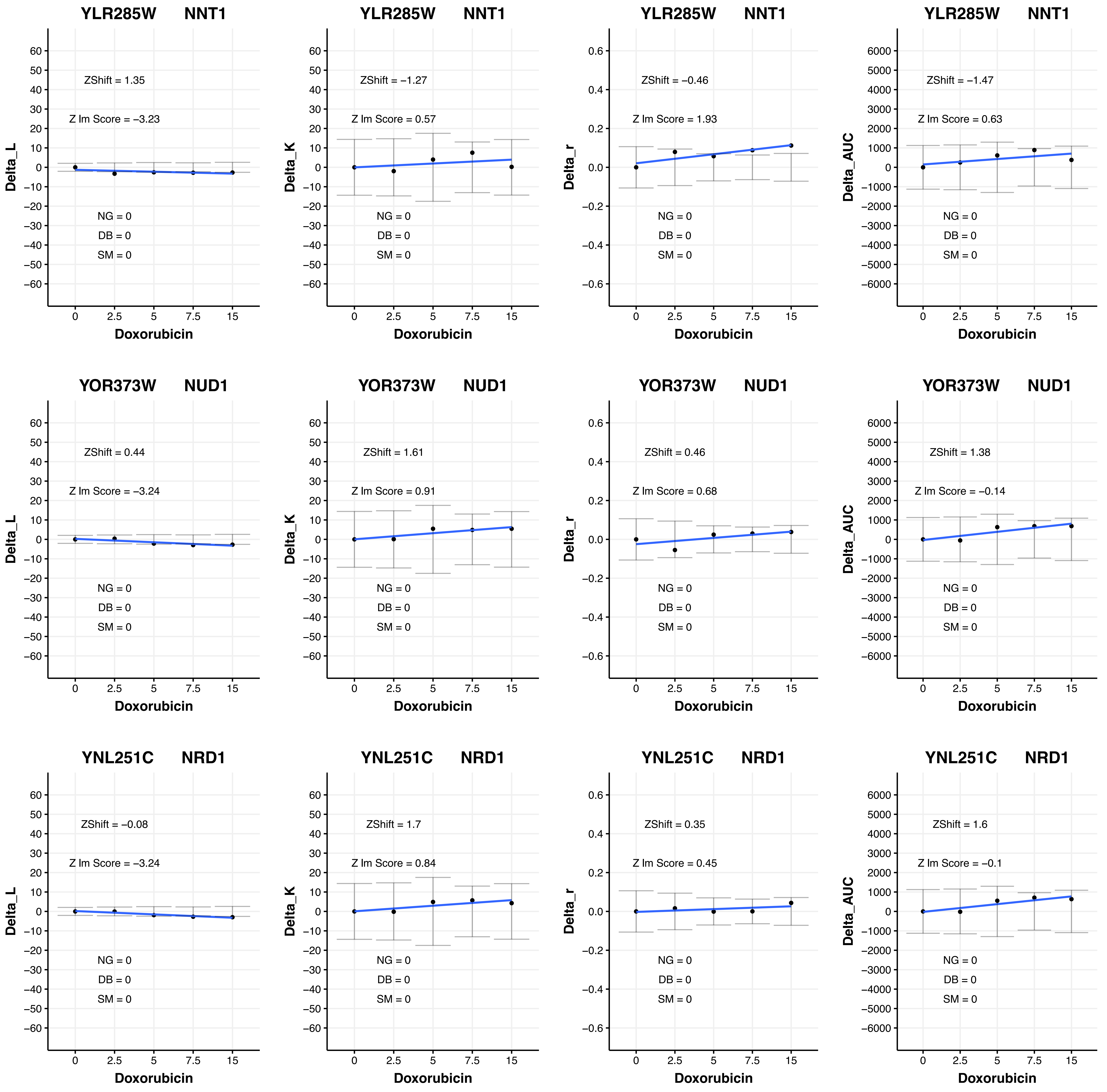

YLR285W NNT1
ZShift = 1.35
Z lm Score = −3.23
NG = 0
DB = 0
SM = 0
Delta_L
Doxorubicin
YLR285W NNT1
ZShift = −1.27
Z lm Score = 0.57
NG = 0
DB = 0
SM = 0
Delta_K
Doxorubicin
YLR285W NNT1
ZShift = −0.46
Z lm Score = 1.93
NG = 0
DB = 0
SM = 0
Delta_r
Doxorubicin
YLR285W NNT1
ZShift = −1.47
Z lm Score = 0.63
NG = 0
DB = 0
SM = 0
Delta_AUC
Doxorubicin
YOR373W NUD1
ZShift = 0.44
Z lm Score = −3.24
NG = 0
DB = 0
SM = 0
Delta_L
Doxorubicin
YOR373W NUD1
ZShift = 1.61
Z lm Score = 0.91
NG = 0
DB = 0
SM = 0
Delta_K
Doxorubicin
YOR373W NUD1
ZShift = 0.46
Z lm Score = 0.68
NG = 0
DB = 0
SM = 0
Delta_r
Doxorubicin
YOR373W NUD1
ZShift = 1.38
Z lm Score = −0.14
NG = 0
DB = 0
SM = 0
Delta_AUC
Doxorubicin
YNL251C NRD1
ZShift = −0.08
Z lm Score = −3.24
NG = 0
DB = 0
SM = 0
Delta_L
Doxorubicin
YNL251C NRD1
ZShift = 1.7
Z lm Score = 0.84
NG = 0
DB = 0
SM = 0
Delta_K
Doxorubicin
YNL251C NRD1
ZShift = 0.35
Z lm Score = 0.45
NG = 0
DB = 0
SM = 0
Delta_r
Doxorubicin
YNL251C NRD1
ZShift = 1.6
Z lm Score = −0.1
NG = 0
DB = 0
SM = 0
Delta_AUC
Doxorubicin

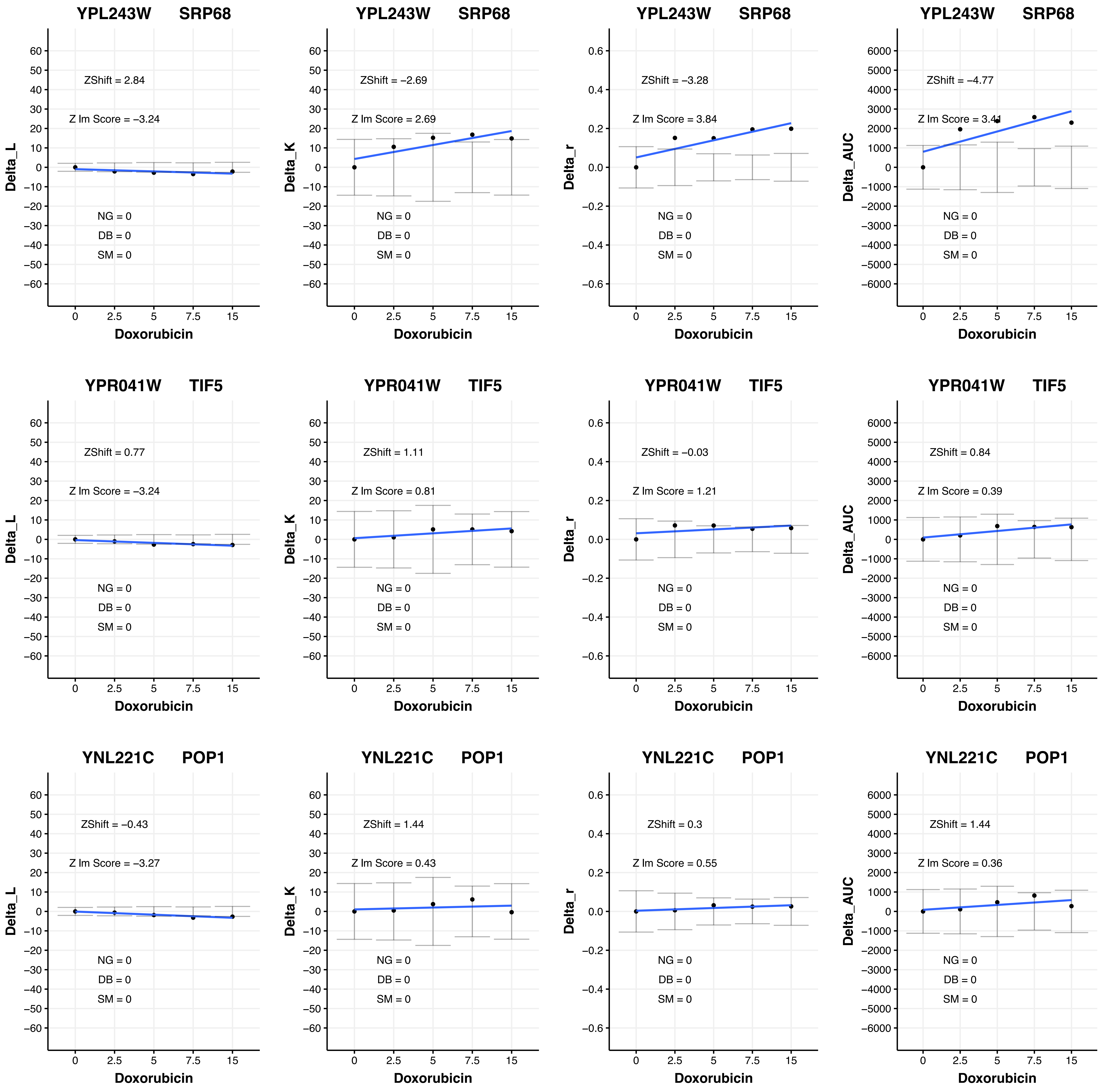

YPL243W SRP68
ZShift = 2.84
Z lm Score = −3.24
NG = 0
DB = 0
SM = 0
Delta_L
Doxorubicin
YPL243W SRP68
ZShift = −2.69
Z lm Score = 2.69
NG = 0
DB = 0
SM = 0
Delta_K
Doxorubicin
YPL243W SRP68
ZShift = −3.28
Z lm Score = 3.84
NG = 0
DB = 0
SM = 0
Delta_r
Doxorubicin
YPL243W SRP68
ZShift = −4.77
Z lm Score = 3.41
NG = 0
DB = 0
SM = 0
Delta_AUC
Doxorubicin
YPR041W TIF5
ZShift = 0.77
Z lm Score = −3.24
NG = 0
DB = 0
SM = 0
Delta_L
Doxorubicin
YPR041W TIF5
ZShift = 1.11
Z lm Score = 0.81
NG = 0
DB = 0
SM = 0
Delta_K
Doxorubicin
YPR041W TIF5
ZShift = −0.03
Z lm Score = 1.21
NG = 0
DB = 0
SM = 0
Delta_r
Doxorubicin
YPR041W TIF5
ZShift = 0.84
Z lm Score = 0.39
NG = 0
DB = 0
SM = 0
Delta_AUC
Doxorubicin
YNL221C POP1
ZShift = −0.43
Z lm Score = −3.27
NG = 0
DB = 0
SM = 0
Delta_L
Doxorubicin
YNL221C POP1
ZShift = 1.44
Z lm Score = 0.43
NG = 0
DB = 0
SM = 0
Delta_K
Doxorubicin
YNL221C POP1
ZShift = 0.3
Z lm Score = 0.55
NG = 0
DB = 0
SM = 0
Delta_r
Doxorubicin
YNL221C POP1
ZShift = 1.44
Z lm Score = 0.36
NG = 0
DB = 0
SM = 0
Delta_AUC
Doxorubicin

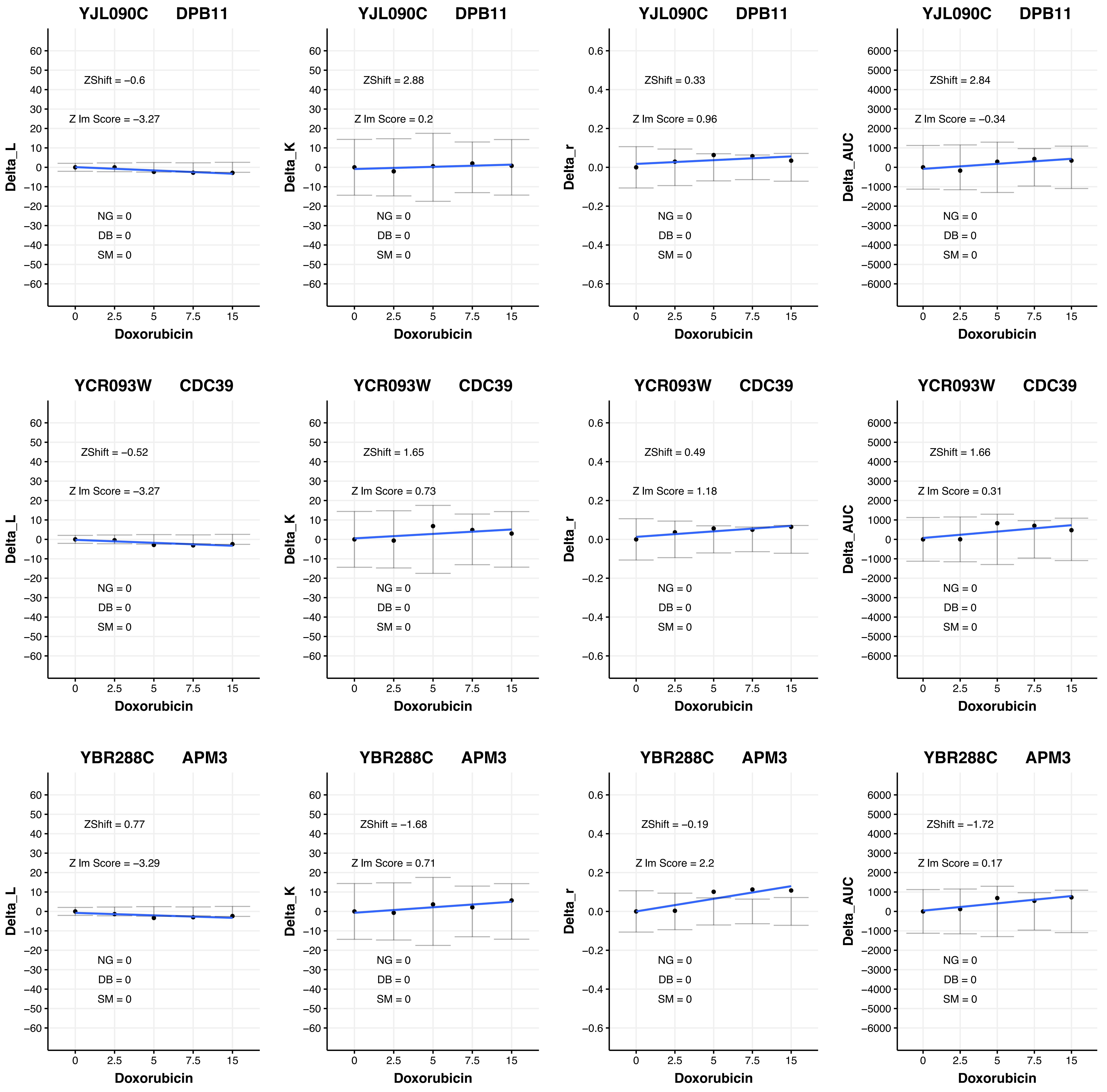
YJL090C DPB11
ZShift = −0.6
Z lm Score = −3.27
NG = 0
DB = 0
SM = 0
Delta_L
Doxorubicin
YJL090C DPB11
ZShift = 2.88
Z lm Score = 0.2
NG = 0
DB = 0
SM = 0
Delta_K
Doxorubicin
YJL090C DPB11
ZShift = 0.33
Z lm Score = 0.96
NG = 0
DB = 0
SM = 0
Delta_r
Doxorubicin
YJL090C DPB11
ZShift = 2.84
Z lm Score = −0.34
NG = 0
DB = 0
SM = 0
Delta_AUC
Doxorubicin
YCR093W CDC39
ZShift = −0.52
Z lm Score = −3.27
NG = 0
DB = 0
SM = 0
Delta_L
Doxorubicin
YCR093W CDC39
ZShift = 1.65
Z lm Score = 0.73
NG = 0
DB = 0
SM = 0
Delta_K
Doxorubicin
YCR093W CDC39
ZShift = 0.49
Z lm Score = 1.18
NG = 0
DB = 0
SM = 0
Delta_r
Doxorubicin
YCR093W CDC39
ZShift = 1.66
Z lm Score = 0.31
NG = 0
DB = 0
SM = 0
Delta_AUC
Doxorubicin
YBR288C APM3
ZShift = 0.77
Z lm Score = −3.29
NG = 0
DB = 0
SM = 0
Delta_L
Doxorubicin
YBR288C APM3
ZShift = −1.68
Z lm Score = 0.71
NG = 0
DB = 0
SM = 0
Delta_K
Doxorubicin
YBR288C APM3
ZShift = −0.19
Z lm Score = 2.2
NG = 0
DB = 0
SM = 0
Delta_r
Doxorubicin
YBR288C APM3
ZShift = −1.72
Z lm Score = 0.17
NG = 0
DB = 0
SM = 0
Delta_AUC
Doxorubicin

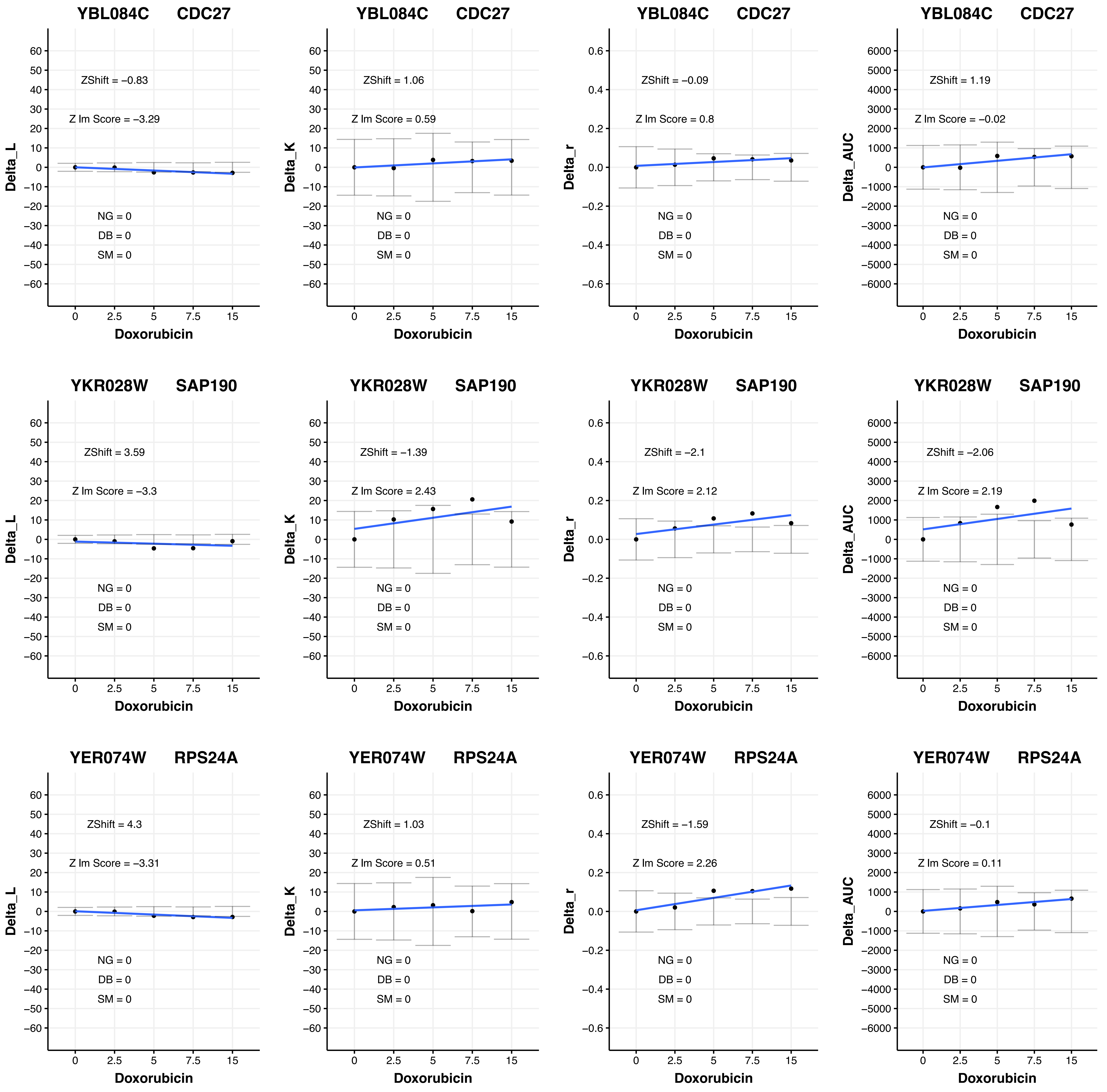

YBL084C CDC27
ZShift = −0.83
Z lm Score = −3.29
Delta_L
NG = 0
DB = 0
SM = 0
Doxorubicin
YBL084C CDC27
ZShift = 1.06
Z lm Score = 0.59
Delta_K
NG = 0
DB = 0
SM = 0
Doxorubicin
YBL084C CDC27
ZShift = −0.09
Z lm Score = 0.8
Delta_r
NG = 0
DB = 0
SM = 0
Doxorubicin
YBL084C CDC27
ZShift = 1.19
Z lm Score = −0.02
Delta_AUC
NG = 0
DB = 0
SM = 0
Doxorubicin
YKR028W SAP190
ZShift = 3.59
Z lm Score = −3.3
Delta_L
NG = 0
DB = 0
SM = 0
Doxorubicin
YKR028W SAP190
ZShift = −1.39
Z lm Score = 2.43
Delta_K
NG = 0
DB = 0
SM = 0
Doxorubicin
YKR028W SAP190
ZShift = −2.1
Z lm Score = 2.12
Delta_r
NG = 0
DB = 0
SM = 0
Doxorubicin
YKR028W SAP190
ZShift = −2.06
Z lm Score = 2.19
Delta_AUC
NG = 0
DB = 0
SM = 0
Doxorubicin
YER074W RPS24A
ZShift = 4.3
Z lm Score = −3.31
Delta_L
NG = 0
DB = 0
SM = 0
Doxorubicin
YER074W RPS24A
ZShift = 1.03
Z lm Score = 0.51
Delta_K
NG = 0
DB = 0
SM = 0
Doxorubicin
YER074W RPS24A
ZShift = −1.59
Z lm Score = 2.26
Delta_r
NG = 0
DB = 0
SM = 0
Doxorubicin
YER074W RPS24A
ZShift = −0.1
Z lm Score = 0.11
Delta_AUC
NG = 0
DB = 0
SM = 0
Doxorubicin

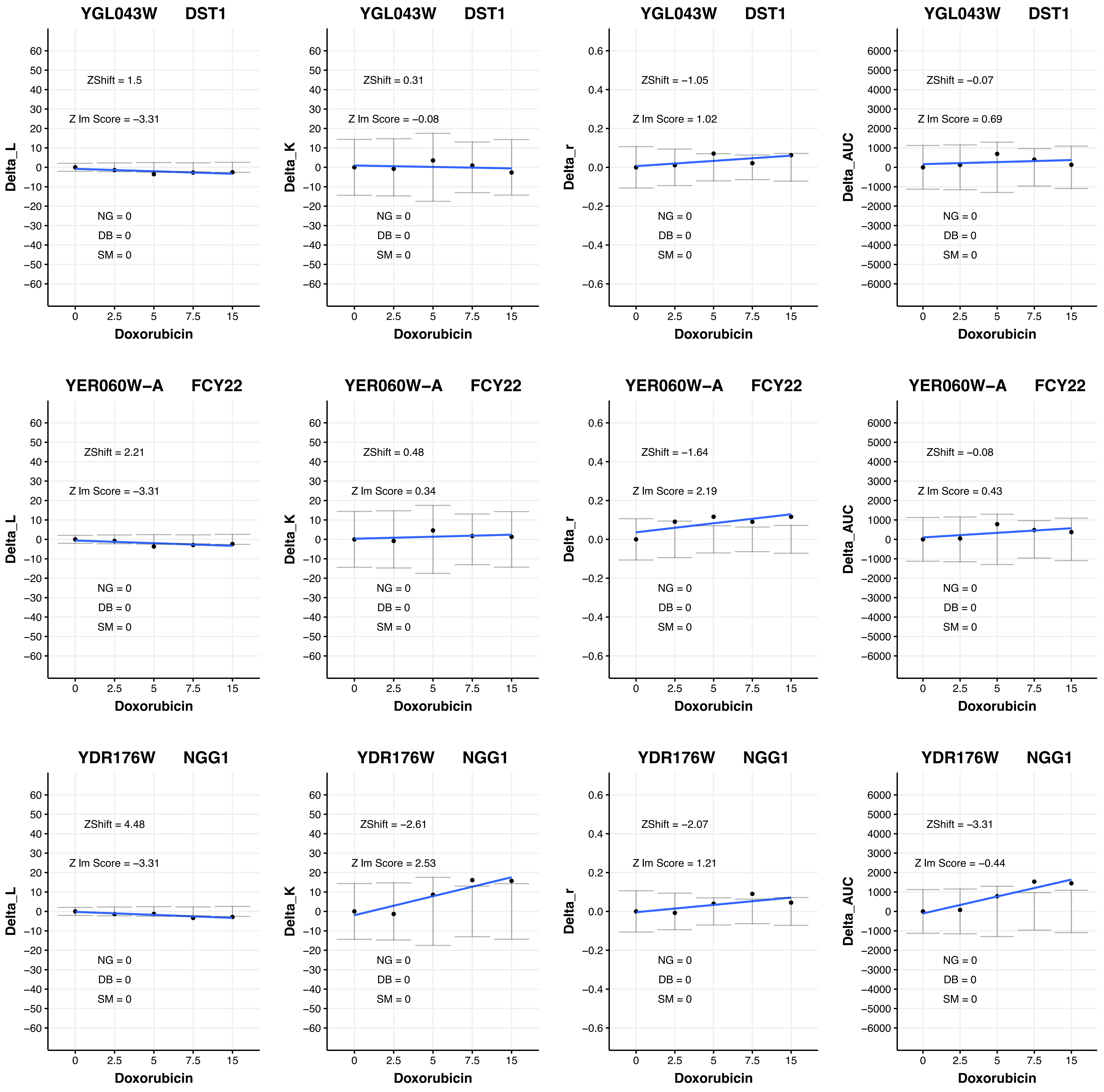

YGL043W DST1
ZShift = 1.5
Z lm Score = −3.31
NG = 0
DB = 0
SM = 0
Delta_L
Doxorubicin
YGL043W DST1
ZShift = 0.31
Z lm Score = −0.08
NG = 0
DB = 0
SM = 0
Delta_K
Doxorubicin
YGL043W DST1
ZShift = −1.05
Z lm Score = 1.02
NG = 0
DB = 0
SM = 0
Delta_r
Doxorubicin
YGL043W DST1
ZShift = −0.07
Z lm Score = 0.69
NG = 0
DB = 0
SM = 0
Delta_AUC
Doxorubicin
YER060W−A FCY22
ZShift = 2.21
Z lm Score = −3.31
NG = 0
DB = 0
SM = 0
Delta_L
Doxorubicin
YER060W−A FCY22
ZShift = 0.48
Z lm Score = 0.34
NG = 0
DB = 0
SM = 0
Delta_K
Doxorubicin
YER060W−A FCY22
ZShift = −1.64
Z lm Score = 2.19
NG = 0
DB = 0
SM = 0
Delta_r
Doxorubicin
YER060W−A FCY22
ZShift = −0.08
Z lm Score = 0.43
NG = 0
DB = 0
SM = 0
Delta_AUC
Doxorubicin
YDR176W NGG1
ZShift = 4.48
Z lm Score = −3.31
NG = 0
DB = 0
SM = 0
Delta_L
Doxorubicin
YDR176W NGG1
ZShift = −2.61
Z lm Score = 2.53
NG = 0
DB = 0
SM = 0
Delta_K
Doxorubicin
YDR176W NGG1
ZShift = −2.07
Z lm Score = 1.21
NG = 0
DB = 0
SM = 0
Delta_r
Doxorubicin
YDR176W NGG1
ZShift = −3.31
Z lm Score = −0.44
NG = 0
DB = 0
SM = 0
Delta_AUC
Doxorubicin

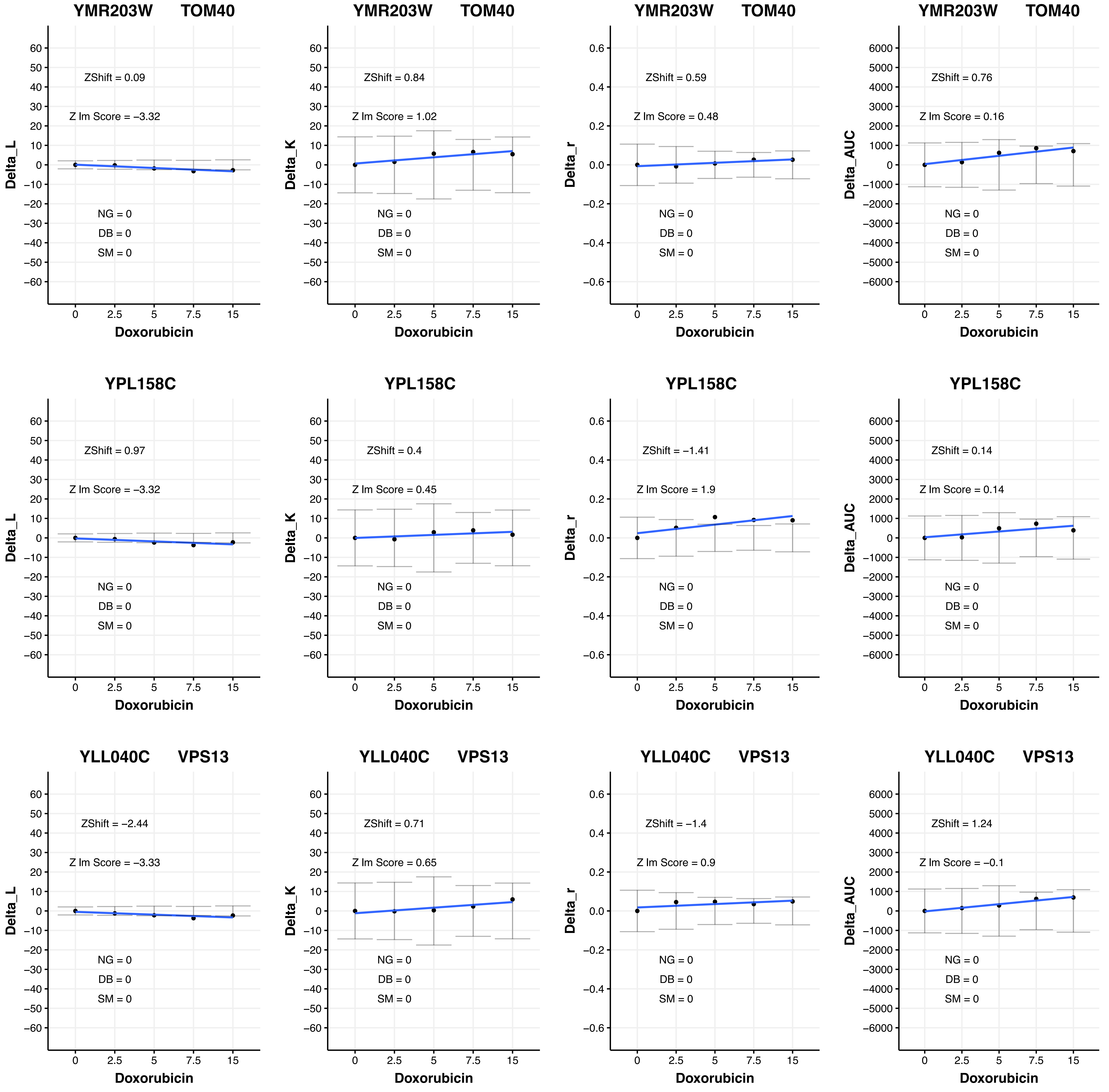

YMR203W TOM40
ZShift = 0.09
Z lm Score = −3.32
NG = 0
DB = 0
SM = 0
Delta_L
Doxorubicin
YMR203W TOM40
ZShift = 0.84
Z lm Score = 1.02
NG = 0
DB = 0
SM = 0
Delta_K
Doxorubicin
YMR203W TOM40
ZShift = 0.59
Z lm Score = 0.48
NG = 0
DB = 0
SM = 0
Delta_r
Doxorubicin
YMR203W TOM40
ZShift = 0.76
Z lm Score = 0.16
NG = 0
DB = 0
SM = 0
Delta_AUC
Doxorubicin
YPL158C
ZShift = 0.97
Z lm Score = −3.32
NG = 0
DB = 0
SM = 0
Delta_L
Doxorubicin
YPL158C
ZShift = 0.4
Z lm Score = 0.45
NG = 0
DB = 0
SM = 0
Delta_K
Doxorubicin
YPL158C
ZShift = −1.41
Z lm Score = 1.9
NG = 0
DB = 0
SM = 0
Delta_r
Doxorubicin
YPL158C
ZShift = 0.14
Z lm Score = 0.14
NG = 0
DB = 0
SM = 0
Delta_AUC
Doxorubicin
YLL040C VPS13
ZShift = −2.44
Z lm Score = −3.33
NG = 0
DB = 0
SM = 0
Delta_L
Doxorubicin
YLL040C VPS13
ZShift = 0.71
Z lm Score = 0.65
NG = 0
DB = 0
SM = 0
Delta_K
Doxorubicin
YLL040C VPS13
ZShift = −1.4
Z lm Score = 0.9
NG = 0
DB = 0
SM = 0
Delta_r
Doxorubicin
YLL040C VPS13
ZShift = 1.24
Z lm Score = −0.1
NG = 0
DB = 0
SM = 0
Delta_AUC
Doxorubicin

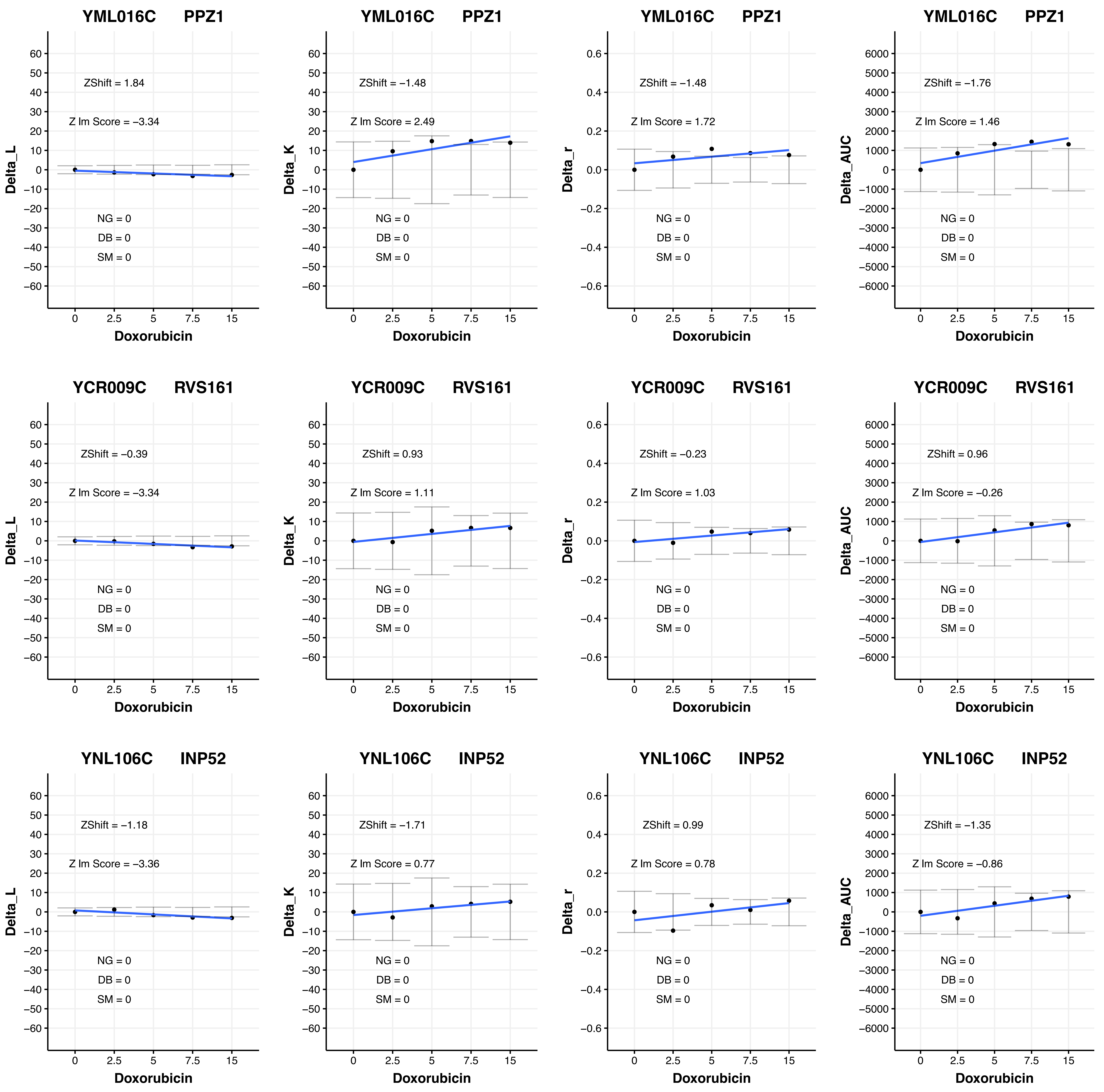

YML016C PPZ1
ZShift = 1.84
Z lm Score = −3.34
NG = 0
DB = 0
SM = 0
Delta_L
Doxorubicin
YML016C PPZ1
ZShift = −1.48
Z lm Score = 2.49
NG = 0
DB = 0
SM = 0
Delta_K
Doxorubicin
YML016C PPZ1
ZShift = −1.48
Z lm Score = 1.72
NG = 0
DB = 0
SM = 0
Delta_r
Doxorubicin
YML016C PPZ1
ZShift = −1.76
Z lm Score = 1.46
NG = 0
DB = 0
SM = 0
Delta_AUC
Doxorubicin
YCR009C RVS161
ZShift = −0.39
Z lm Score = −3.34
NG = 0
DB = 0
SM = 0
Delta_L
Doxorubicin
YCR009C RVS161
ZShift = 0.93
Z lm Score = 1.11
NG = 0
DB = 0
SM = 0
Delta_K
Doxorubicin
YCR009C RVS161
ZShift = −0.23
Z lm Score = 1.03
NG = 0
DB = 0
SM = 0
Delta_r
Doxorubicin
YCR009C RVS161
ZShift = 0.96
Z lm Score = −0.26
NG = 0
DB = 0
SM = 0
Delta_AUC
Doxorubicin
YNL106C INP52
ZShift = −1.18
Z lm Score = −3.36
NG = 0
DB = 0
SM = 0
Delta_L
Doxorubicin
YNL106C INP52
ZShift = −1.71
Z lm Score = 0.77
NG = 0
DB = 0
SM = 0
Delta_K
Doxorubicin
YNL106C INP52
ZShift = 0.99
Z lm Score = 0.78
NG = 0
DB = 0
SM = 0
Delta_r
Doxorubicin
YNL106C INP52
ZShift = −1.35
Z lm Score = −0.86
NG = 0
DB = 0
SM = 0
Delta_AUC
Doxorubicin

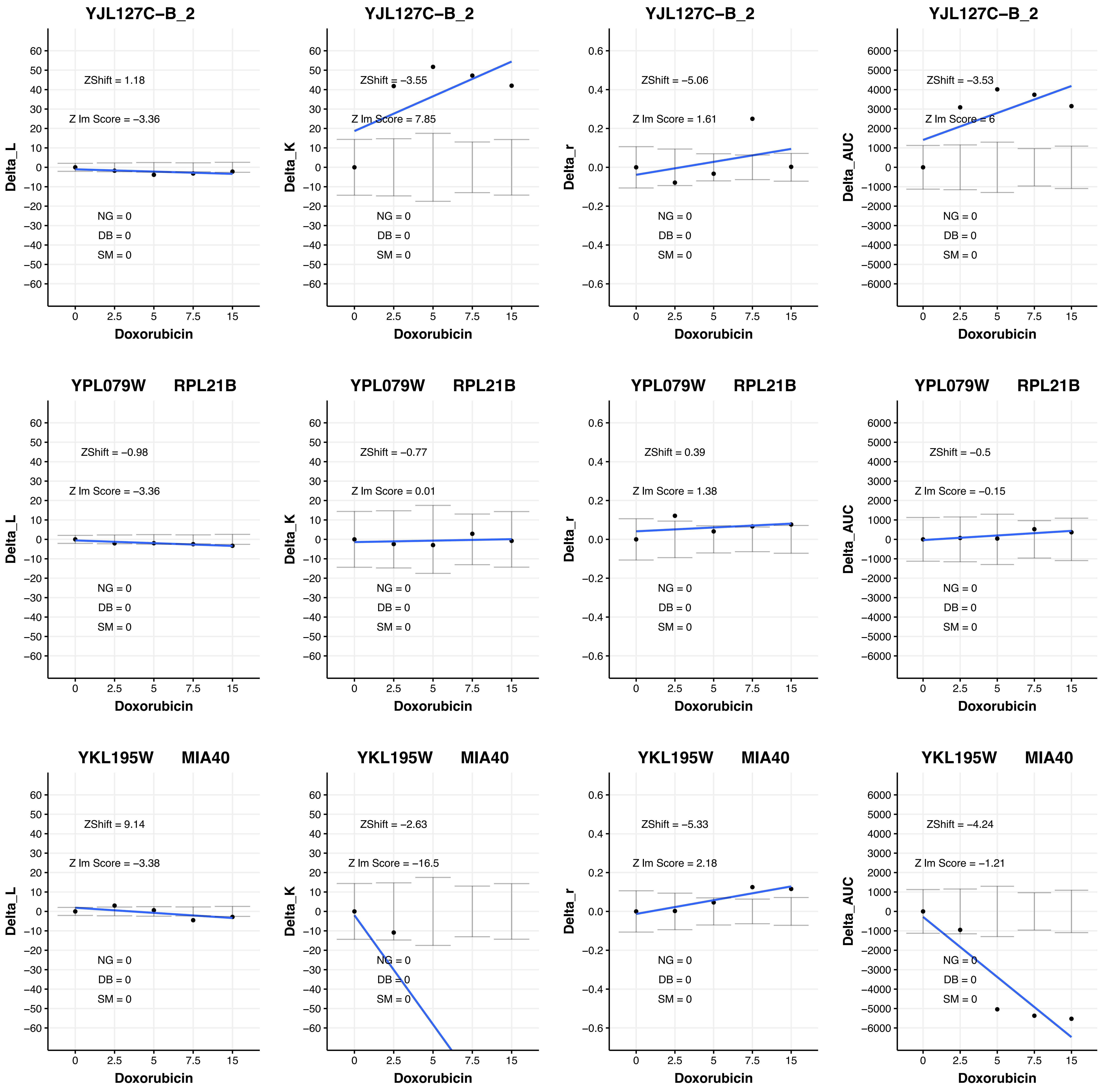

YJL127C−B_2
ZShift = 1.18
Z lm Score = −3.36
NG = 0
DB = 0
SM = 0
Delta_L
Doxorubicin
YJL127C−B_2
ZShift = −3.55
Z lm Score = 7.85
NG = 0
DB = 0
SM = 0
Delta_K
Doxorubicin
YJL127C−B_2
ZShift = −5.06
Z lm Score = 1.61
NG = 0
DB = 0
SM = 0
Delta_r
Doxorubicin
YJL127C−B_2
ZShift = −3.53
Z lm Score = 6
NG = 0
DB = 0
SM = 0
Delta_AUC
Doxorubicin
YPL079W RPL21B
ZShift = −0.98
Z lm Score = −3.36
NG = 0
DB = 0
SM = 0
Delta_L
Doxorubicin
YPL079W RPL21B
ZShift = −0.77
Z lm Score = 0.01
NG = 0
DB = 0
SM = 0
Delta_K
Doxorubicin
YPL079W RPL21B
ZShift = 0.39
Z lm Score = 1.38
NG = 0
DB = 0
SM = 0
Delta_r
Doxorubicin
YPL079W RPL21B
ZShift = −0.5
Z lm Score = −0.15
NG = 0
DB = 0
SM = 0
Delta_AUC
Doxorubicin
YKL195W MIA40
ZShift = 9.14
Z lm Score = −3.38
NG = 0
DB = 0
SM = 0
Delta_L
Doxorubicin
YKL195W MIA40
ZShift = −2.63
Z lm Score = −16.5
NG = 0
DB = 0
SM = 0
Delta_K
Doxorubicin
YKL195W MIA40
ZShift = −5.33
Z lm Score = 2.18
NG = 0
DB = 0
SM = 0
Delta_r
Doxorubicin
YKL195W MIA40
ZShift = −4.24
Z lm Score = −1.21
NG = 0
DB = 0
SM = 0
Delta_AUC
Doxorubicin

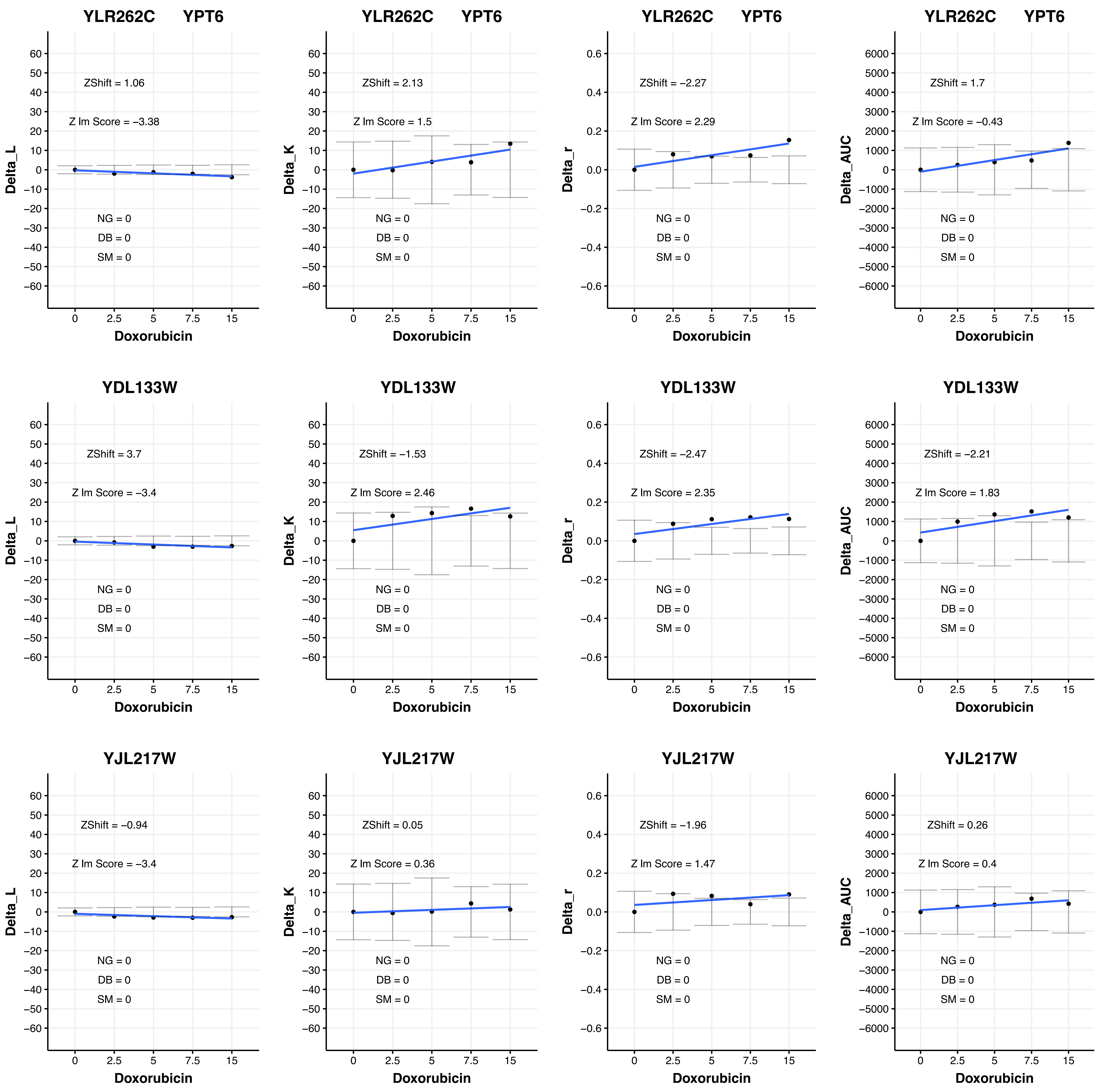
YLR262C YPT6
ZShift = 1.06
Z lm Score = −3.38
NG = 0
DB = 0
SM = 0
Delta_L
Doxorubicin
YLR262C YPT6
ZShift = 2.13
Z lm Score = 1.5
NG = 0
DB = 0
SM = 0
Delta_K
Doxorubicin
YLR262C YPT6
ZShift = −2.27
Z lm Score = 2.29
NG = 0
DB = 0
SM = 0
Delta_r
Doxorubicin
YLR262C YPT6
ZShift = 1.7
Z lm Score = −0.43
NG = 0
DB = 0
SM = 0
Delta_AUC
Doxorubicin
YDL133W
ZShift = 3.7
Z lm Score = −3.4
NG = 0
DB = 0
SM = 0
Delta_L
Doxorubicin
YDL133W
ZShift = −1.53
Z lm Score = 2.46
NG = 0
DB = 0
SM = 0
Delta_K
Doxorubicin
YDL133W
ZShift = −2.47
Z lm Score = 2.35
NG = 0
DB = 0
SM = 0
Delta_r
Doxorubicin
YDL133W
ZShift = −2.21
Z lm Score = 1.83
NG = 0
DB = 0
SM = 0
Delta_AUC
Doxorubicin
YJL217W
ZShift = −0.94
Z lm Score = −3.4
NG = 0
DB = 0
SM = 0
Delta_L
Doxorubicin
YJL217W
ZShift = 0.05
Z lm Score = 0.36
NG = 0
DB = 0
SM = 0
Delta_K
Doxorubicin
YJL217W
ZShift = −1.96
Z lm Score = 1.47
NG = 0
DB = 0
SM = 0
Delta_r
Doxorubicin
YJL217W
ZShift = 0.26
Z lm Score = 0.4
NG = 0
DB = 0
SM = 0
Delta_AUC
Doxorubicin

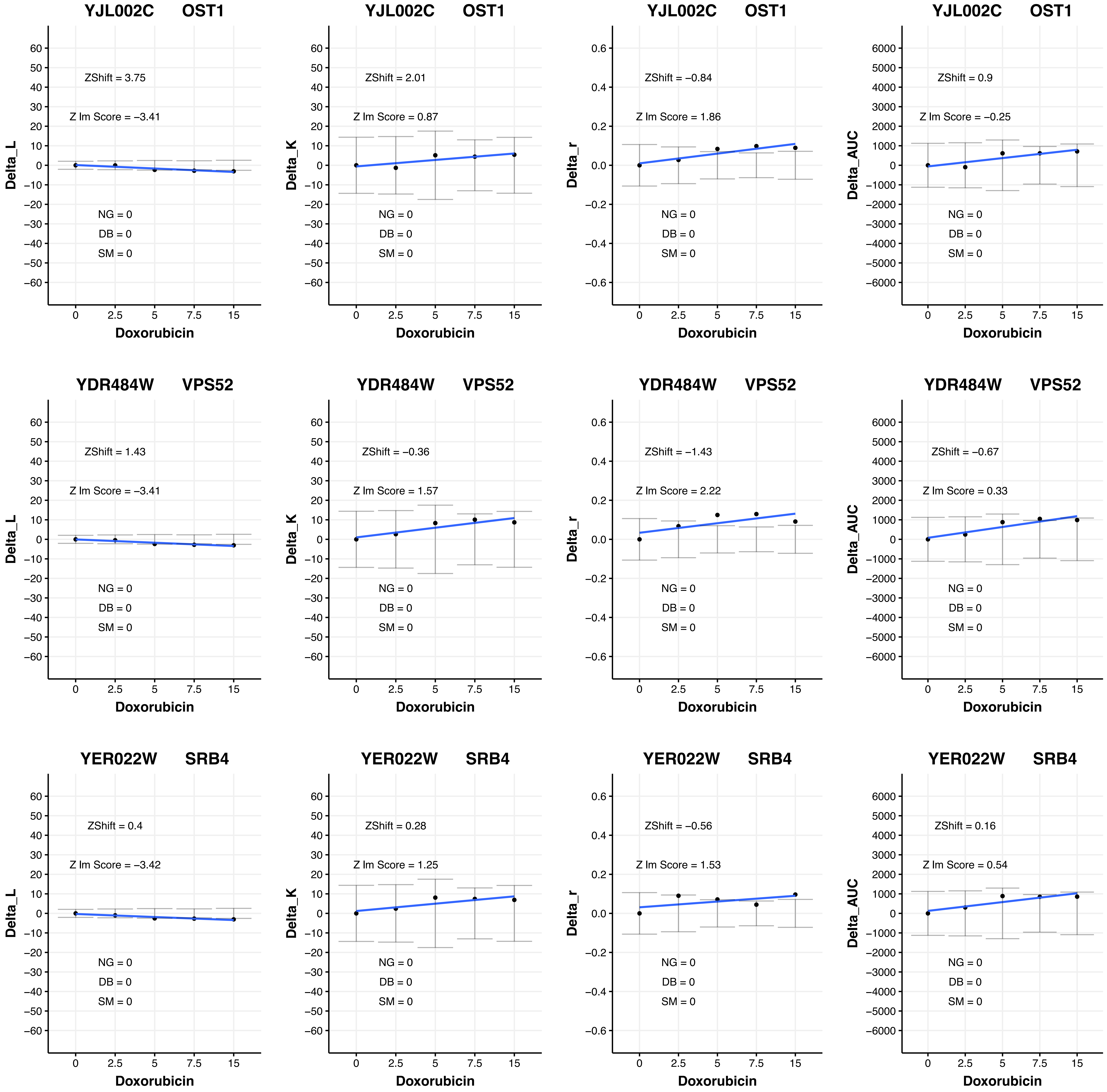
YJL002C OST1
ZShift = 3.75
Z lm Score = −3.41
NG = 0
DB = 0
SM = 0
Delta_L
Doxorubicin
YJL002C OST1
ZShift = 2.01
Z lm Score = 0.87
NG = 0
DB = 0
SM = 0
Delta_K
Doxorubicin
YJL002C OST1
ZShift = −0.84
Z lm Score = 1.86
NG = 0
DB = 0
SM = 0
Delta_r
Doxorubicin
YJL002C OST1
ZShift = 0.9
Z lm Score = −0.25
NG = 0
DB = 0
SM = 0
Delta_AUC
Doxorubicin
YDR484W VPS52
ZShift = 1.43
Z lm Score = −3.41
NG = 0
DB = 0
SM = 0
Delta_L
Doxorubicin
YDR484W VPS52
ZShift = −0.36
Z lm Score = 1.57
NG = 0
DB = 0
SM = 0
Delta_K
Doxorubicin
YDR484W VPS52
ZShift = −1.43
Z lm Score = 2.22
NG = 0
DB = 0
SM = 0
Delta_r
Doxorubicin
YDR484W VPS52
ZShift = −0.67
Z lm Score = 0.33
NG = 0
DB = 0
SM = 0
Delta_AUC
Doxorubicin
YER022W SRB4
ZShift = 0.4
Z lm Score = −3.42
NG = 0
DB = 0
SM = 0
Delta_L
Doxorubicin
YER022W SRB4
ZShift = 0.28
Z lm Score = 1.25
NG = 0
DB = 0
SM = 0
Delta_K
Doxorubicin
YER022W SRB4
ZShift = −0.56
Z lm Score = 1.53
NG = 0
DB = 0
SM = 0
Delta_r
Doxorubicin
YER022W SRB4
ZShift = 0.16
Z lm Score = 0.54
NG = 0
DB = 0
SM = 0
Delta_AUC
Doxorubicin

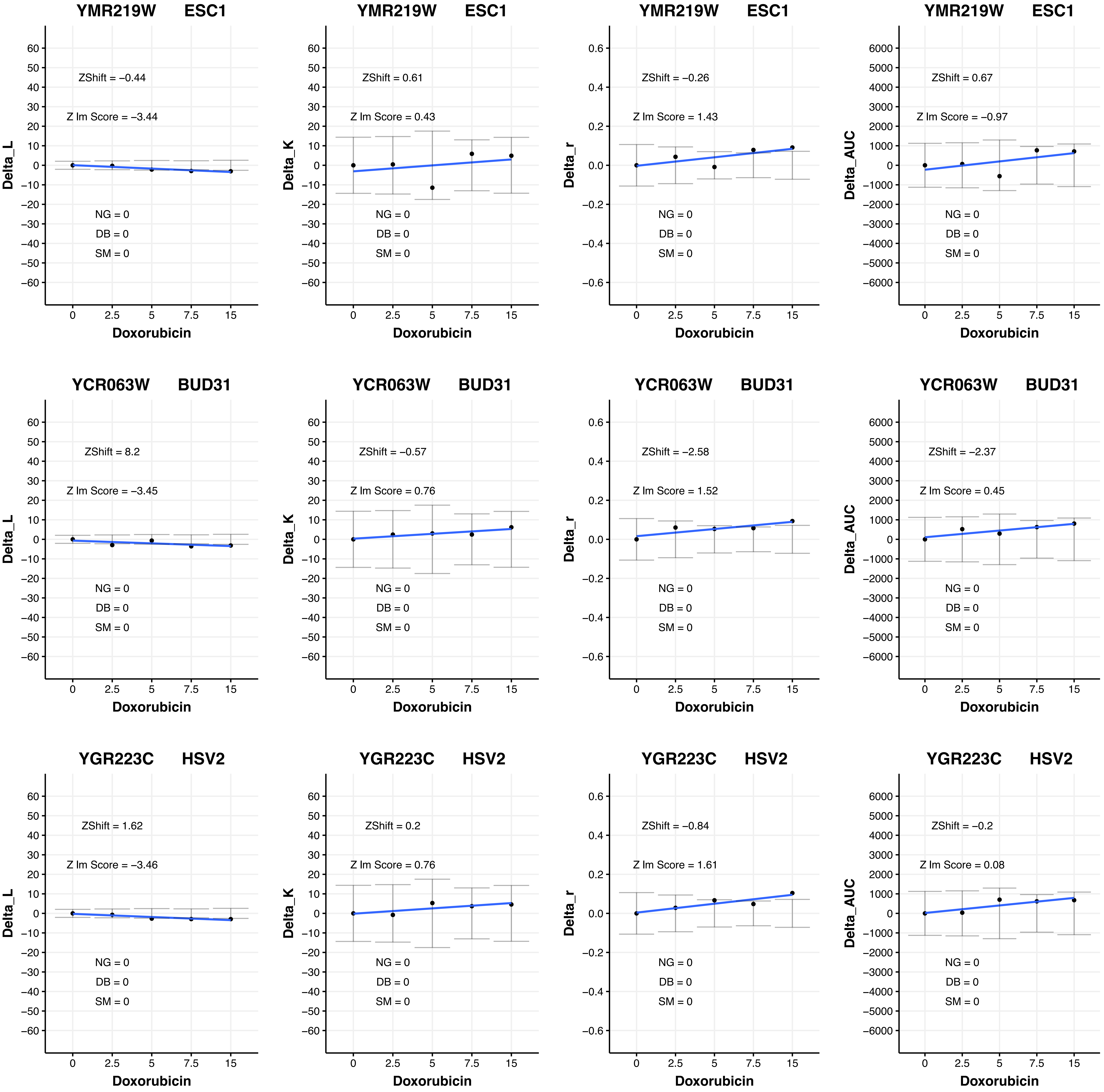

YMR219W ESC1
ZShift = −0.44
Z lm Score = −3.44
NG = 0
DB = 0
SM = 0
Delta_L
Doxorubicin
YMR219W ESC1
ZShift = 0.61
Z lm Score = 0.43
NG = 0
DB = 0
SM = 0
Delta_K
Doxorubicin
YMR219W ESC1
ZShift = −0.26
Z lm Score = 1.43
NG = 0
DB = 0
SM = 0
Delta_r
Doxorubicin
YMR219W ESC1
ZShift = 0.67
Z lm Score = −0.97
NG = 0
DB = 0
SM = 0
Delta_AUC
Doxorubicin
YCR063W BUD31
ZShift = 8.2
Z lm Score = −3.45
NG = 0
DB = 0
SM = 0
Delta_L
Doxorubicin
YCR063W BUD31
ZShift = −0.57
Z lm Score = 0.76
NG = 0
DB = 0
SM = 0
Delta_K
Doxorubicin
YCR063W BUD31
ZShift = −2.58
Z lm Score = 1.52
NG = 0
DB = 0
SM = 0
Delta_r
Doxorubicin
YCR063W BUD31
ZShift = −2.37
Z lm Score = 0.45
NG = 0
DB = 0
SM = 0
Delta_AUC
Doxorubicin
YGR223C HSV2
ZShift = 1.62
Z lm Score = −3.46
NG = 0
DB = 0
SM = 0
Delta_L
Doxorubicin
YGR223C HSV2
ZShift = 0.2
Z lm Score = 0.76
NG = 0
DB = 0
SM = 0
Delta_K
Doxorubicin
YGR223C HSV2
ZShift = −0.84
Z lm Score = 1.61
NG = 0
DB = 0
SM = 0
Delta_r
Doxorubicin
YGR223C HSV2
ZShift = −0.2
Z lm Score = 0.08
NG = 0
DB = 0
SM = 0
Delta_AUC
Doxorubicin

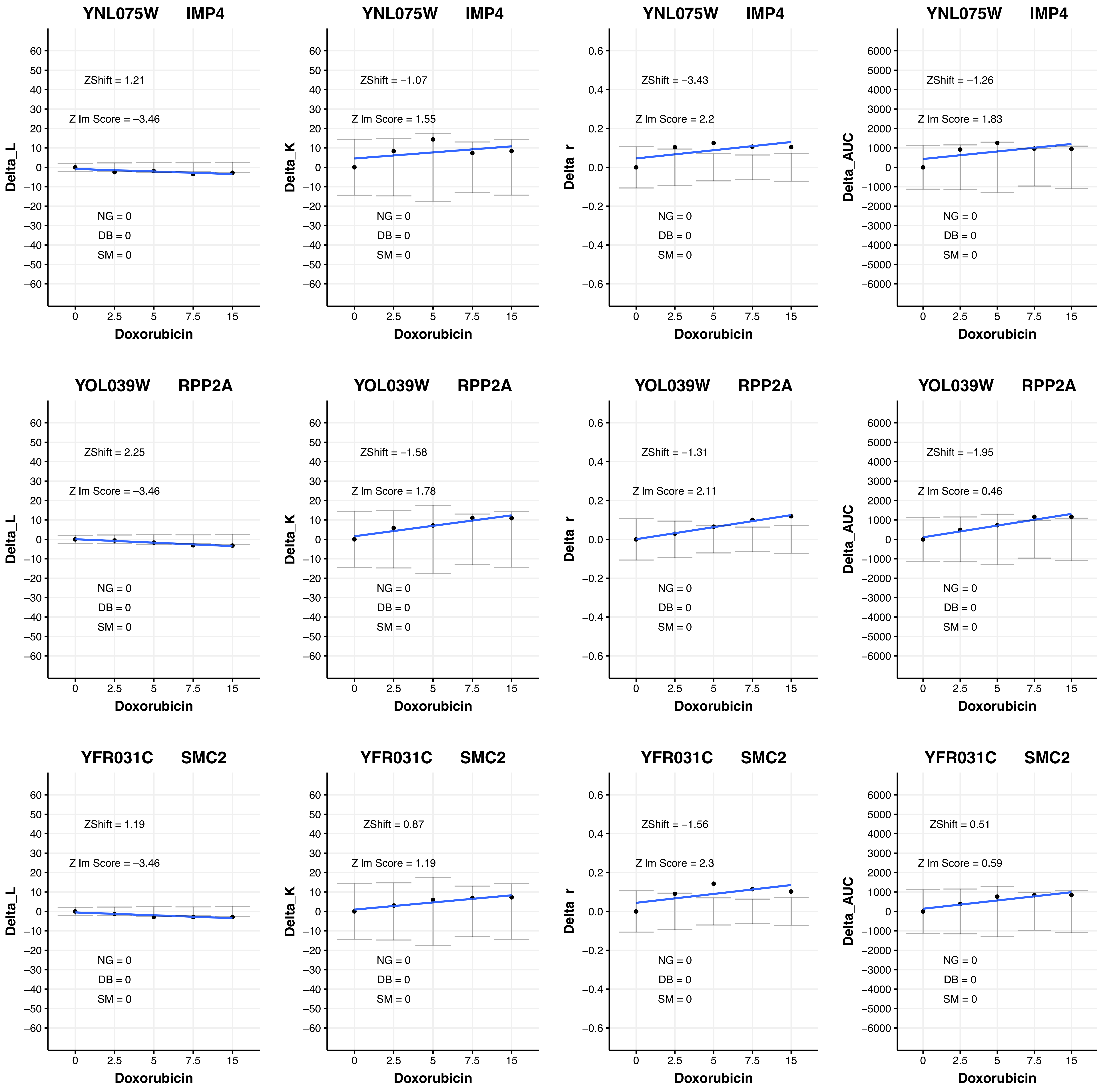
YNL075W IMP4
ZShift = 1.21
Z lm Score = −3.46
NG = 0
DB = 0
SM = 0
Delta_L
Doxorubicin
YNL075W IMP4
ZShift = −1.07
Z lm Score = 1.55
NG = 0
DB = 0
SM = 0
Delta_K
Doxorubicin
YNL075W IMP4
ZShift = −3.43
Z lm Score = 2.2
NG = 0
DB = 0
SM = 0
Delta_r
Doxorubicin
YNL075W IMP4
ZShift = −1.26
Z lm Score = 1.83
NG = 0
DB = 0
SM = 0
Delta_AUC
Doxorubicin
YOL039W RPP2A
ZShift = 2.25
Z lm Score = −3.46
NG = 0
DB = 0
SM = 0
Delta_L
Doxorubicin
YOL039W RPP2A
ZShift = −1.58
Z lm Score = 1.78
NG = 0
DB = 0
SM = 0
Delta_K
Doxorubicin
YOL039W RPP2A
ZShift = −1.31
Z lm Score = 2.11
NG = 0
DB = 0
SM = 0
Delta_r
Doxorubicin
YOL039W RPP2A
ZShift = −1.95
Z lm Score = 0.46
NG = 0
DB = 0
SM = 0
Delta_AUC
Doxorubicin
YFR031C SMC2
ZShift = 1.19
Z lm Score = −3.46
NG = 0
DB = 0
SM = 0
Delta_L
Doxorubicin
YFR031C SMC2
ZShift = 0.87
Z lm Score = 1.19
NG = 0
DB = 0
SM = 0
Delta_K
Doxorubicin
YFR031C SMC2
ZShift = −1.56
Z lm Score = 2.3
NG = 0
DB = 0
SM = 0
Delta_r
Doxorubicin
YFR031C SMC2
ZShift = 0.51
Z lm Score = 0.59
NG = 0
DB = 0
SM = 0
Delta_AUC
Doxorubicin

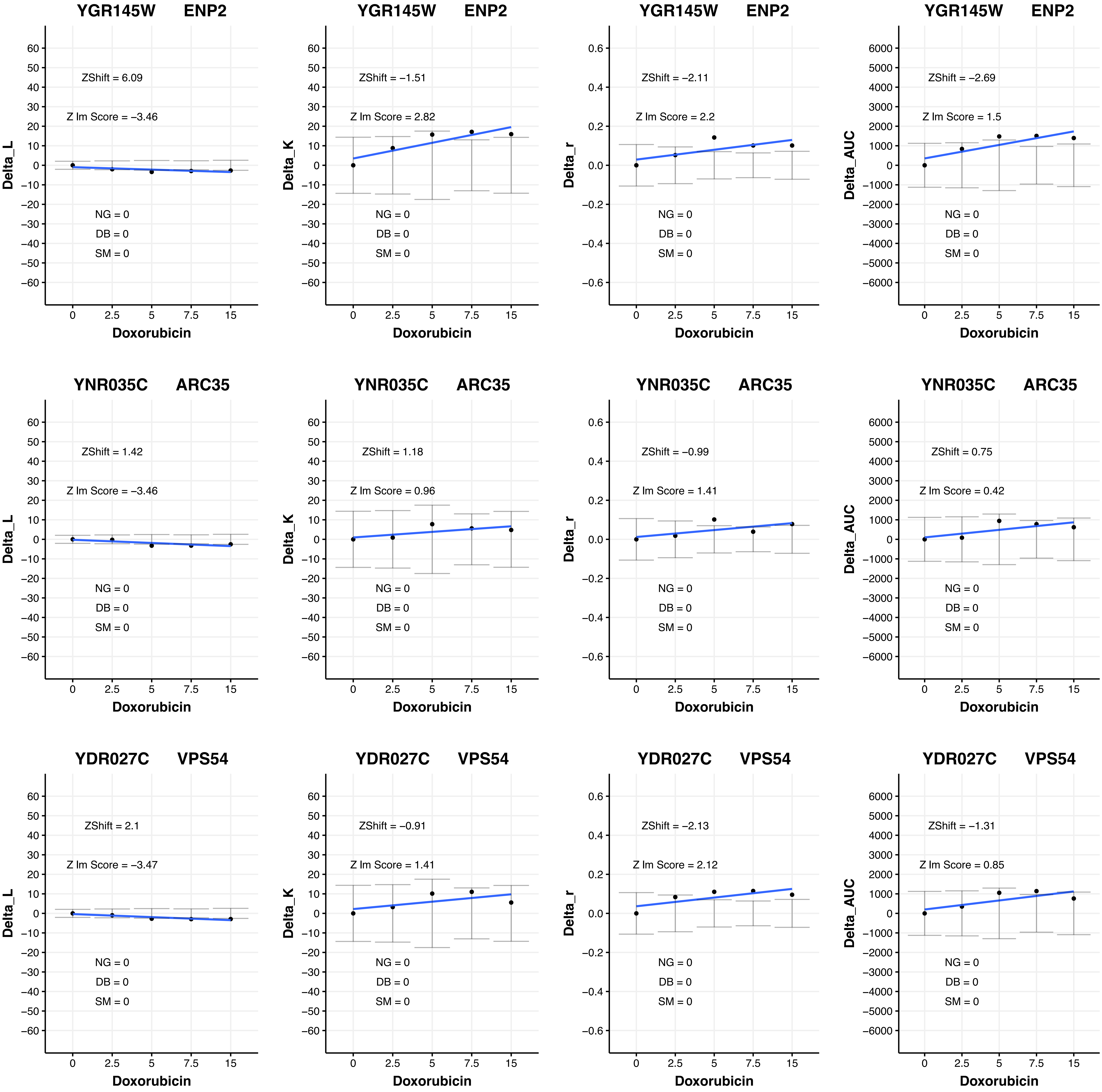

YGR145W ENP2
ZShift = 6.09
Z lm Score = −3.46
NG = 0
DB = 0
SM = 0
Delta_L
Doxorubicin
YGR145W ENP2
ZShift = −1.51
Z lm Score = 2.82
NG = 0
DB = 0
SM = 0
Delta_K
Doxorubicin
YGR145W ENP2
ZShift = −2.11
Z lm Score = 2.2
NG = 0
DB = 0
SM = 0
Delta_r
Doxorubicin
YGR145W ENP2
ZShift = −2.69
Z lm Score = 1.5
NG = 0
DB = 0
SM = 0
Delta_AUC
Doxorubicin
YNR035C ARC35
ZShift = 1.42
Z lm Score = −3.46
NG = 0
DB = 0
SM = 0
Delta_L
Doxorubicin
YNR035C ARC35
ZShift = 1.18
Z lm Score = 0.96
NG = 0
DB = 0
SM = 0
Delta_K
Doxorubicin
YNR035C ARC35
ZShift = −0.99
Z lm Score = 1.41
NG = 0
DB = 0
SM = 0
Delta_r
Doxorubicin
YNR035C ARC35
ZShift = 0.75
Z lm Score = 0.42
NG = 0
DB = 0
SM = 0
Delta_AUC
Doxorubicin
YDR027C VPS54
ZShift = 2.1
Z lm Score = −3.47
NG = 0
DB = 0
SM = 0
Delta_L
Doxorubicin
YDR027C VPS54
ZShift = −0.91
Z lm Score = 1.41
NG = 0
DB = 0
SM = 0
Delta_K
Doxorubicin
YDR027C VPS54
ZShift = −2.13
Z lm Score = 2.12
NG = 0
DB = 0
SM = 0
Delta_r
Doxorubicin
YDR027C VPS54
ZShift = −1.31
Z lm Score = 0.85
NG = 0
DB = 0
SM = 0
Delta_AUC
Doxorubicin

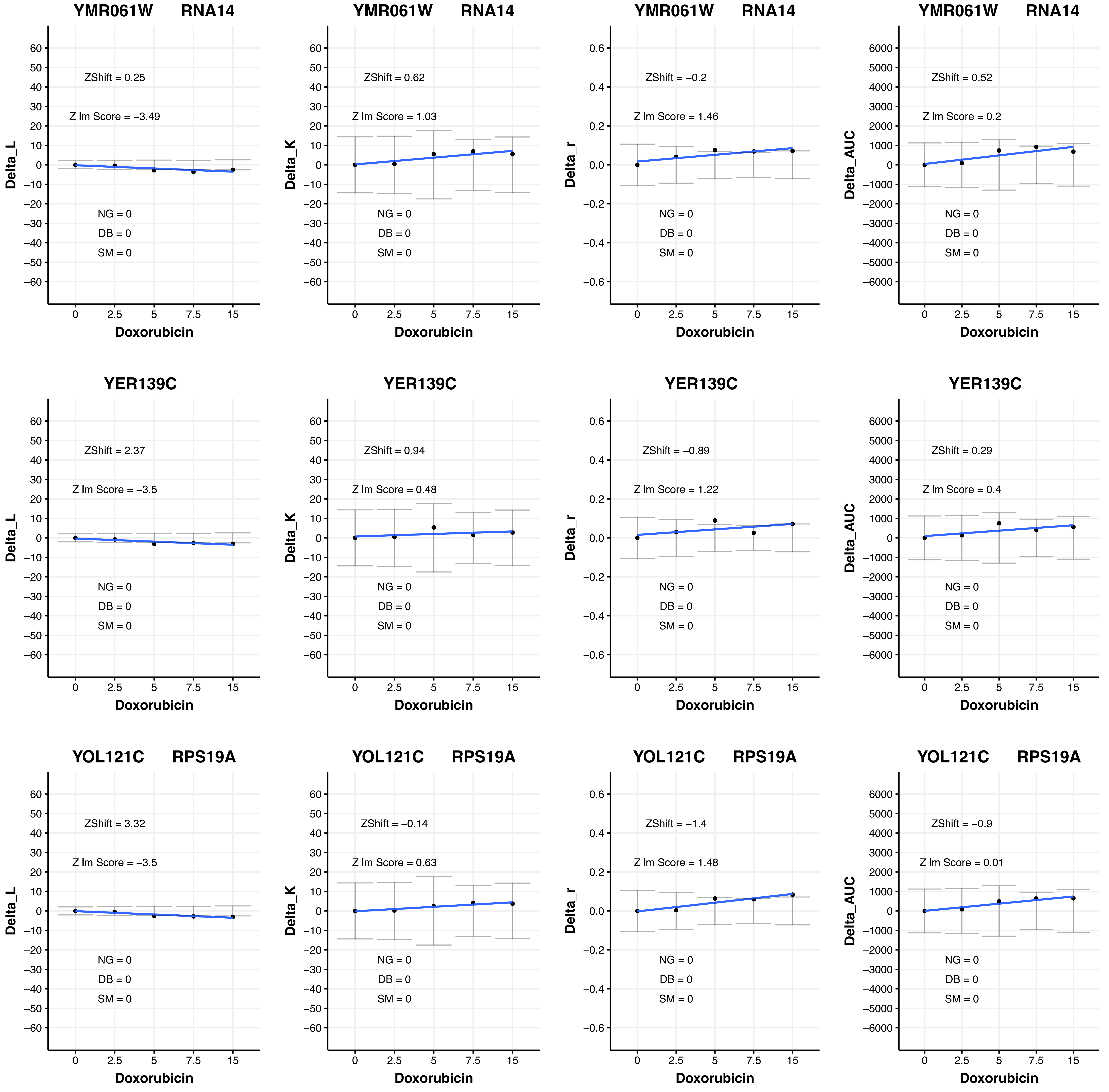
YMR061W RNA14
ZShift = 0.25
Z lm Score = −3.49
NG = 0
DB = 0
SM = 0
Delta_L
Doxorubicin
YMR061W RNA14
ZShift = 0.62
Z lm Score = 1.03
NG = 0
DB = 0
SM = 0
Delta_K
Doxorubicin
YMR061W RNA14
ZShift = −0.2
Z lm Score = 1.46
NG = 0
DB = 0
SM = 0
Delta_r
Doxorubicin
YMR061W RNA14
ZShift = 0.52
Z lm Score = 0.2
NG = 0
DB = 0
SM = 0
Delta_AUC
Doxorubicin
YER139C
ZShift = 2.37
Z lm Score = −3.5
NG = 0
DB = 0
SM = 0
Delta_L
Doxorubicin
YER139C
ZShift = 0.94
Z lm Score = 0.48
NG = 0
DB = 0
SM = 0
Delta_K
Doxorubicin
YER139C
ZShift = −0.89
Z lm Score = 1.22
NG = 0
DB = 0
SM = 0
Delta_r
Doxorubicin
YER139C
ZShift = 0.29
Z lm Score = 0.4
NG = 0
DB = 0
SM = 0
Delta_AUC
Doxorubicin
YOL121C RPS19A
ZShift = 3.32
Z lm Score = −3.5
NG = 0
DB = 0
SM = 0
Delta_L
Doxorubicin
YOL121C RPS19A
ZShift = −0.14
Z lm Score = 0.63
NG = 0
DB = 0
SM = 0
Delta_K
Doxorubicin
YOL121C RPS19A
ZShift = −1.4
Z lm Score = 1.48
NG = 0
DB = 0
SM = 0
Delta_r
Doxorubicin
YOL121C RPS19A
ZShift = −0.9
Z lm Score = 0.01
NG = 0
DB = 0
SM = 0
Delta_AUC
Doxorubicin

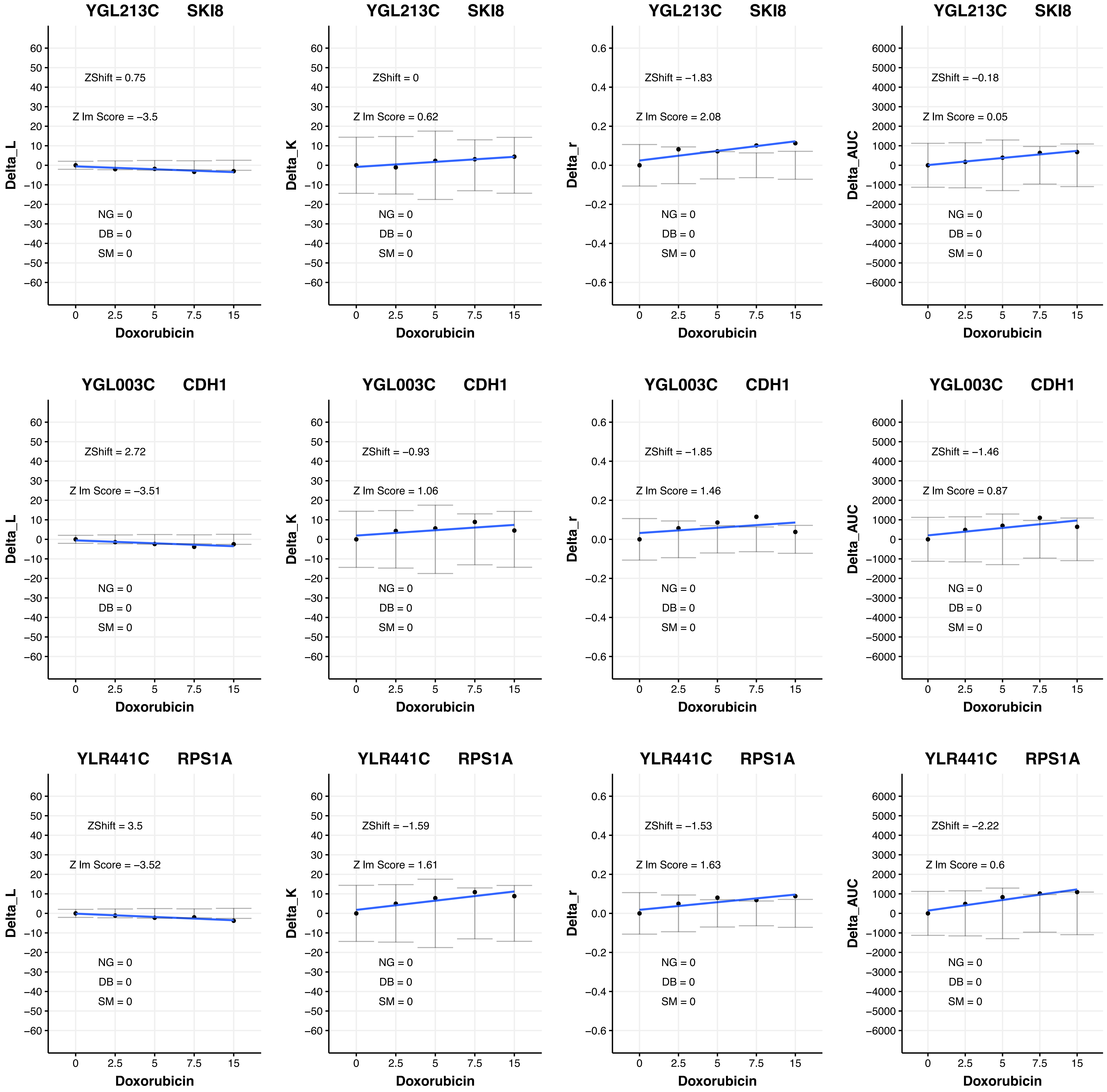

YGL213C SKI8
ZShift = 0.75
Z lm Score = −3.5
NG = 0
DB = 0
SM = 0
Delta_L
Doxorubicin
YGL213C SKI8
ZShift = 0
Z lm Score = 0.62
NG = 0
DB = 0
SM = 0
Delta_K
Doxorubicin
YGL213C SKI8
ZShift = −1.83
Z lm Score = 2.08
NG = 0
DB = 0
SM = 0
Delta_r
Doxorubicin
YGL213C SKI8
ZShift = −0.18
Z lm Score = 0.05
NG = 0
DB = 0
SM = 0
Delta_AUC
Doxorubicin
YGL003C CDH1
ZShift = 2.72
Z lm Score = −3.51
NG = 0
DB = 0
SM = 0
Delta_L
Doxorubicin
YGL003C CDH1
ZShift = −0.93
Z lm Score = 1.06
NG = 0
DB = 0
SM = 0
Delta_K
Doxorubicin
YGL003C CDH1
ZShift = −1.85
Z lm Score = 1.46
NG = 0
DB = 0
SM = 0
Delta_r
Doxorubicin
YGL003C CDH1
ZShift = −1.46
Z lm Score = 0.87
NG = 0
DB = 0
SM = 0
Delta_AUC
Doxorubicin
YLR441C RPS1A
ZShift = 3.5
Z lm Score = −3.52
NG = 0
DB = 0
SM = 0
Delta_L
Doxorubicin
YLR441C RPS1A
ZShift = −1.59
Z lm Score = 1.61
NG = 0
DB = 0
SM = 0
Delta_K
Doxorubicin
YLR441C RPS1A
ZShift = −1.53
Z lm Score = 1.63
NG = 0
DB = 0
SM = 0
Delta_r
Doxorubicin
YLR441C RPS1A
ZShift = −2.22
Z lm Score = 0.6
NG = 0
DB = 0
SM = 0
Delta_AUC
Doxorubicin

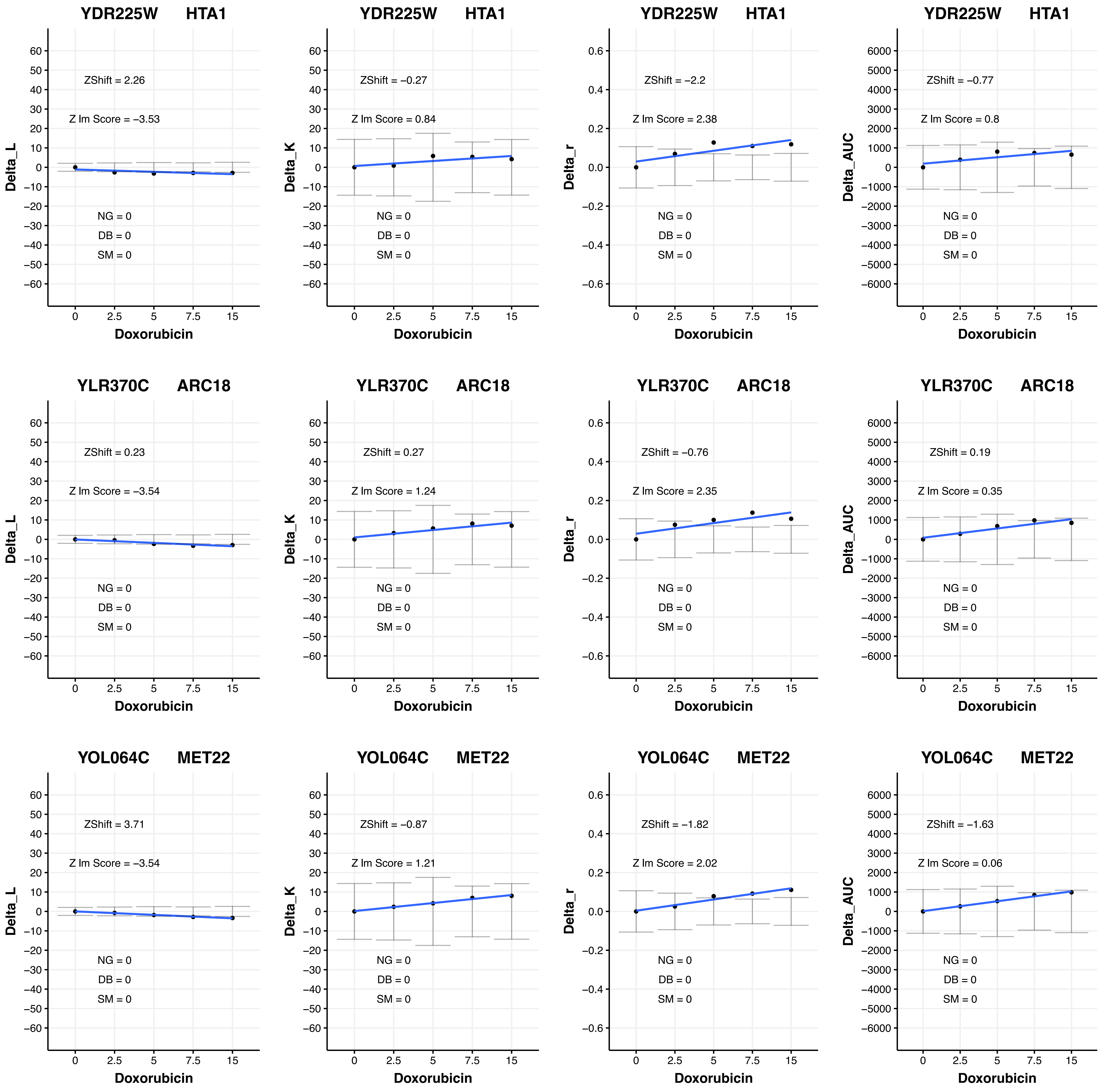

YDR225W HTA1
ZShift = 2.26
Z lm Score = −3.53
NG = 0
DB = 0
SM = 0
Delta_L
Doxorubicin
YDR225W HTA1
ZShift = −0.27
Z lm Score = 0.84
NG = 0
DB = 0
SM = 0
Delta_K
Doxorubicin
YDR225W HTA1
ZShift = −2.2
Z lm Score = 2.38
NG = 0
DB = 0
SM = 0
Delta_r
Doxorubicin
YDR225W HTA1
ZShift = −0.77
Z lm Score = 0.8
NG = 0
DB = 0
SM = 0
Delta_AUC
Doxorubicin
YLR370C ARC18
ZShift = 0.23
Z lm Score = −3.54
NG = 0
DB = 0
SM = 0
Delta_L
Doxorubicin
YLR370C ARC18
ZShift = 0.27
Z lm Score = 1.24
NG = 0
DB = 0
SM = 0
Delta_K
Doxorubicin
YLR370C ARC18
ZShift = −0.76
Z lm Score = 2.35
NG = 0
DB = 0
SM = 0
Delta_r
Doxorubicin
YLR370C ARC18
ZShift = 0.19
Z lm Score = 0.35
NG = 0
DB = 0
SM = 0
Delta_AUC
Doxorubicin
YOL064C MET22
ZShift = 3.71
Z lm Score = −3.54
NG = 0
DB = 0
SM = 0
Delta_L
Doxorubicin
YOL064C MET22
ZShift = −0.87
Z lm Score = 1.21
NG = 0
DB = 0
SM = 0
Delta_K
Doxorubicin
YOL064C MET22
ZShift = −1.82
Z lm Score = 2.02
NG = 0
DB = 0
SM = 0
Delta_r
Doxorubicin
YOL064C MET22
ZShift = −1.63
Z lm Score = 0.06
NG = 0
DB = 0
SM = 0
Delta_AUC
Doxorubicin

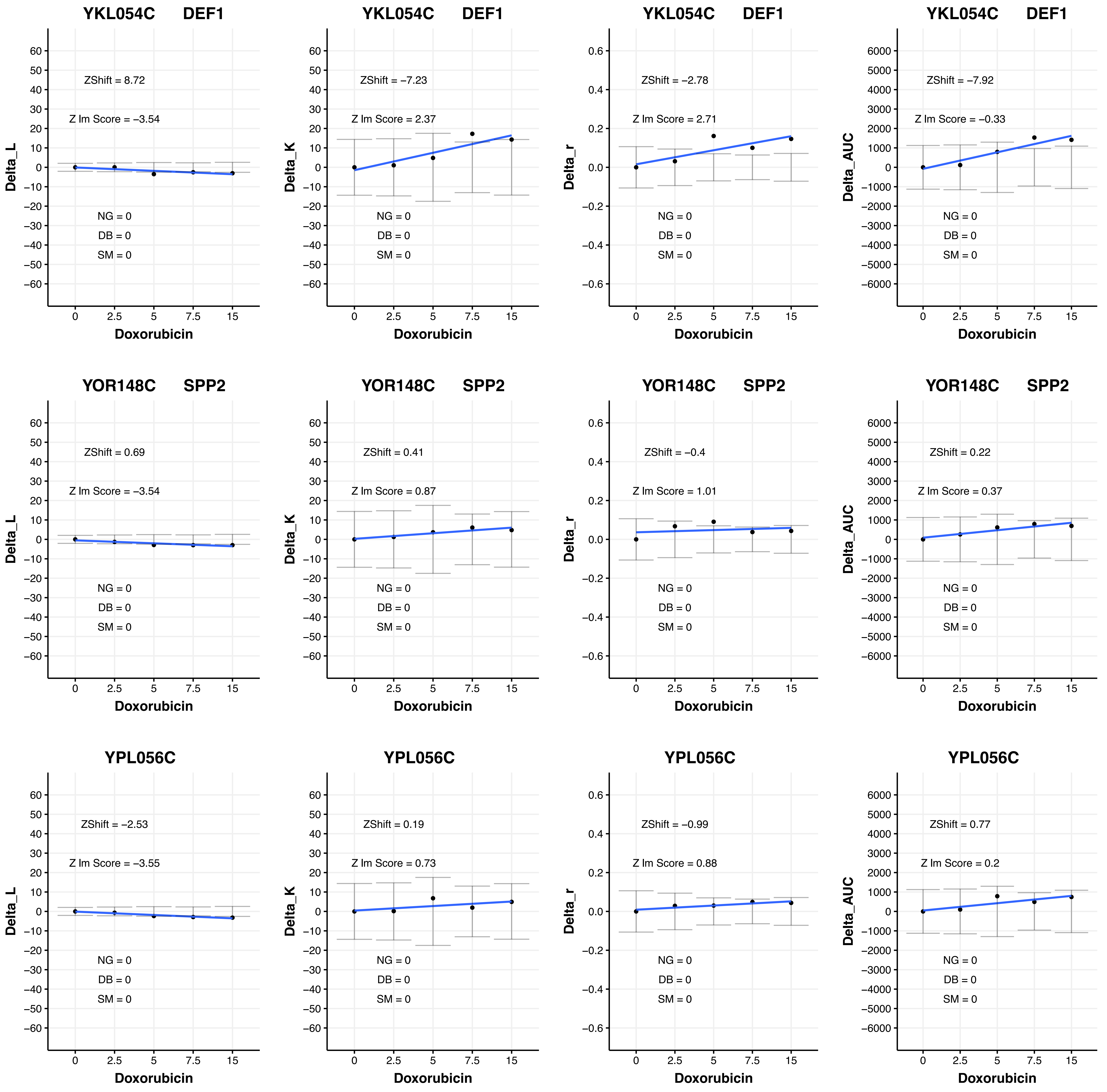

YKL054C DEF1
ZShift = 8.72
Z lm Score = −3.54
NG = 0
DB = 0
SM = 0
Delta_L
Doxorubicin
YKL054C DEF1
ZShift = −7.23
Z lm Score = 2.37
NG = 0
DB = 0
SM = 0
Delta_K
Doxorubicin
YKL054C DEF1
ZShift = −2.78
Z lm Score = 2.71
NG = 0
DB = 0
SM = 0
Delta_r
Doxorubicin
YKL054C DEF1
ZShift = −7.92
Z lm Score = −0.33
NG = 0
DB = 0
SM = 0
Delta_AUC
Doxorubicin
YOR148C SPP2
ZShift = 0.69
Z lm Score = −3.54
NG = 0
DB = 0
SM = 0
Delta_L
Doxorubicin
YOR148C SPP2
ZShift = 0.41
Z lm Score = 0.87
NG = 0
DB = 0
SM = 0
Delta_K
Doxorubicin
YOR148C SPP2
ZShift = −0.4
Z lm Score = 1.01
NG = 0
DB = 0
SM = 0
Delta_r
Doxorubicin
YOR148C SPP2
ZShift = 0.22
Z lm Score = 0.37
NG = 0
DB = 0
SM = 0
Delta_AUC
Doxorubicin
YPL056C
ZShift = −2.53
Z lm Score = −3.55
NG = 0
DB = 0
SM = 0
Delta_L
Doxorubicin
YPL056C
ZShift = 0.19
Z lm Score = 0.73
NG = 0
DB = 0
SM = 0
Delta_K
Doxorubicin
YPL056C
ZShift = −0.99
Z lm Score = 0.88
NG = 0
DB = 0
SM = 0
Delta_r
Doxorubicin
YPL056C
ZShift = 0.77
Z lm Score = 0.2
NG = 0
DB = 0
SM = 0
Delta_AUC
Doxorubicin

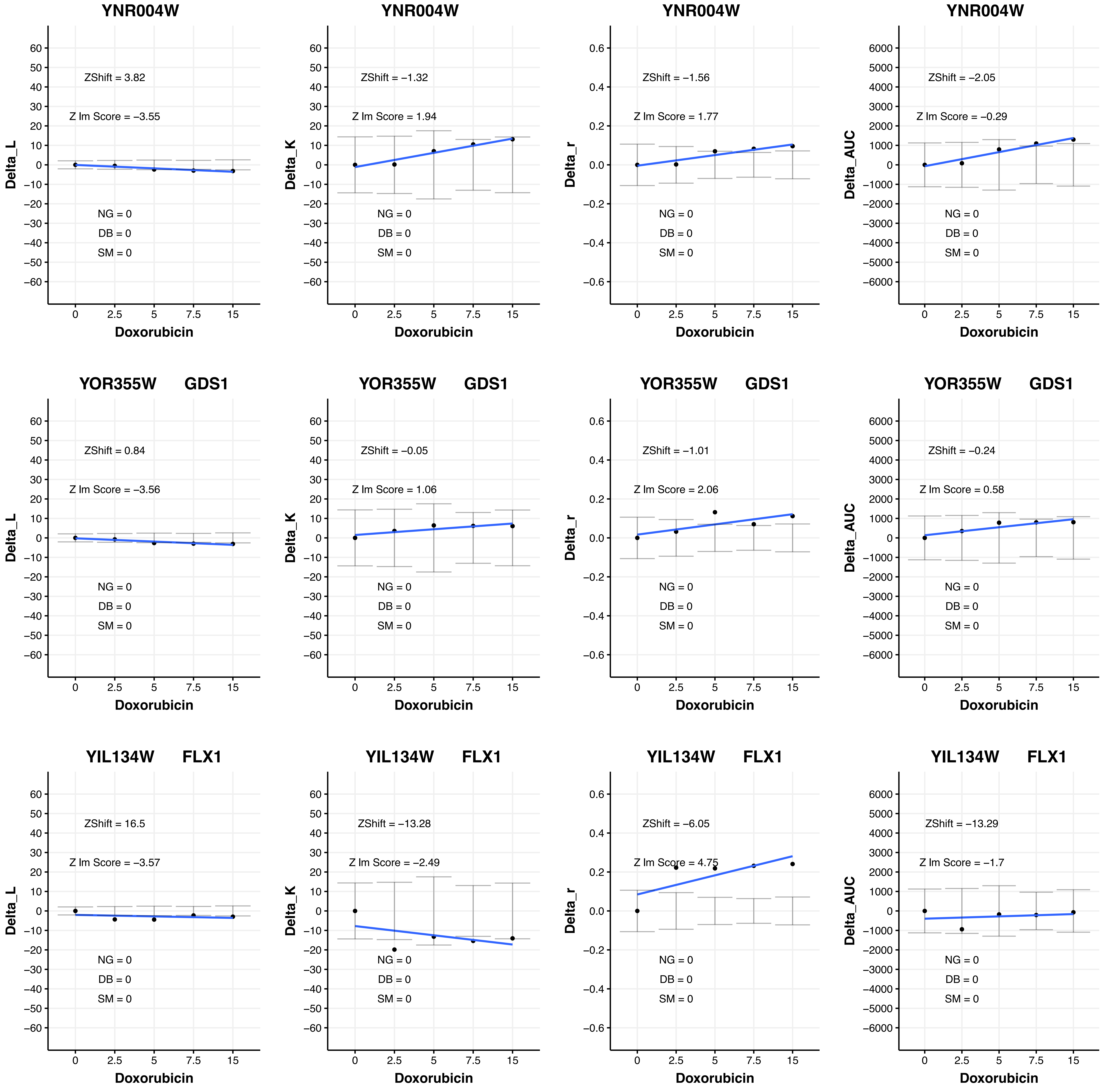
YNR004W
ZShift = 3.82
Z lm Score = −3.55
NG = 0
DB = 0
SM = 0
Delta_L
Doxorubicin
YNR004W
ZShift = −1.32
Z lm Score = 1.94
NG = 0
DB = 0
SM = 0
Delta_K
Doxorubicin
YNR004W
ZShift = −1.56
Z lm Score = 1.77
NG = 0
DB = 0
SM = 0
Delta_r
Doxorubicin
YNR004W
ZShift = −2.05
Z lm Score = −0.29
NG = 0
DB = 0
SM = 0
Delta_AUC
Doxorubicin
YOR355W GDS1
ZShift = 0.84
Z lm Score = −3.56
NG = 0
DB = 0
SM = 0
Delta_L
Doxorubicin
YOR355W GDS1
ZShift = −0.05
Z lm Score = 1.06
NG = 0
DB = 0
SM = 0
Delta_K
Doxorubicin
YOR355W GDS1
ZShift = −1.01
Z lm Score = 2.06
NG = 0
DB = 0
SM = 0
Delta_r
Doxorubicin
YOR355W GDS1
ZShift = −0.24
Z lm Score = 0.58
NG = 0
DB = 0
SM = 0
Delta_AUC
Doxorubicin
YIL134W FLX1
ZShift = 16.5
Z lm Score = −3.57
NG = 0
DB = 0
SM = 0
Delta_L
Doxorubicin
YIL134W FLX1
ZShift = −13.28
Z lm Score = −2.49
NG = 0
DB = 0
SM = 0
Delta_K
Doxorubicin
YIL134W FLX1
ZShift = −6.05
Z lm Score = 4.75
NG = 0
DB = 0
SM = 0
Delta_r
Doxorubicin
YIL134W FLX1
ZShift = −13.29
Z lm Score = −1.7
NG = 0
DB = 0
SM = 0
Delta_AUC
Doxorubicin

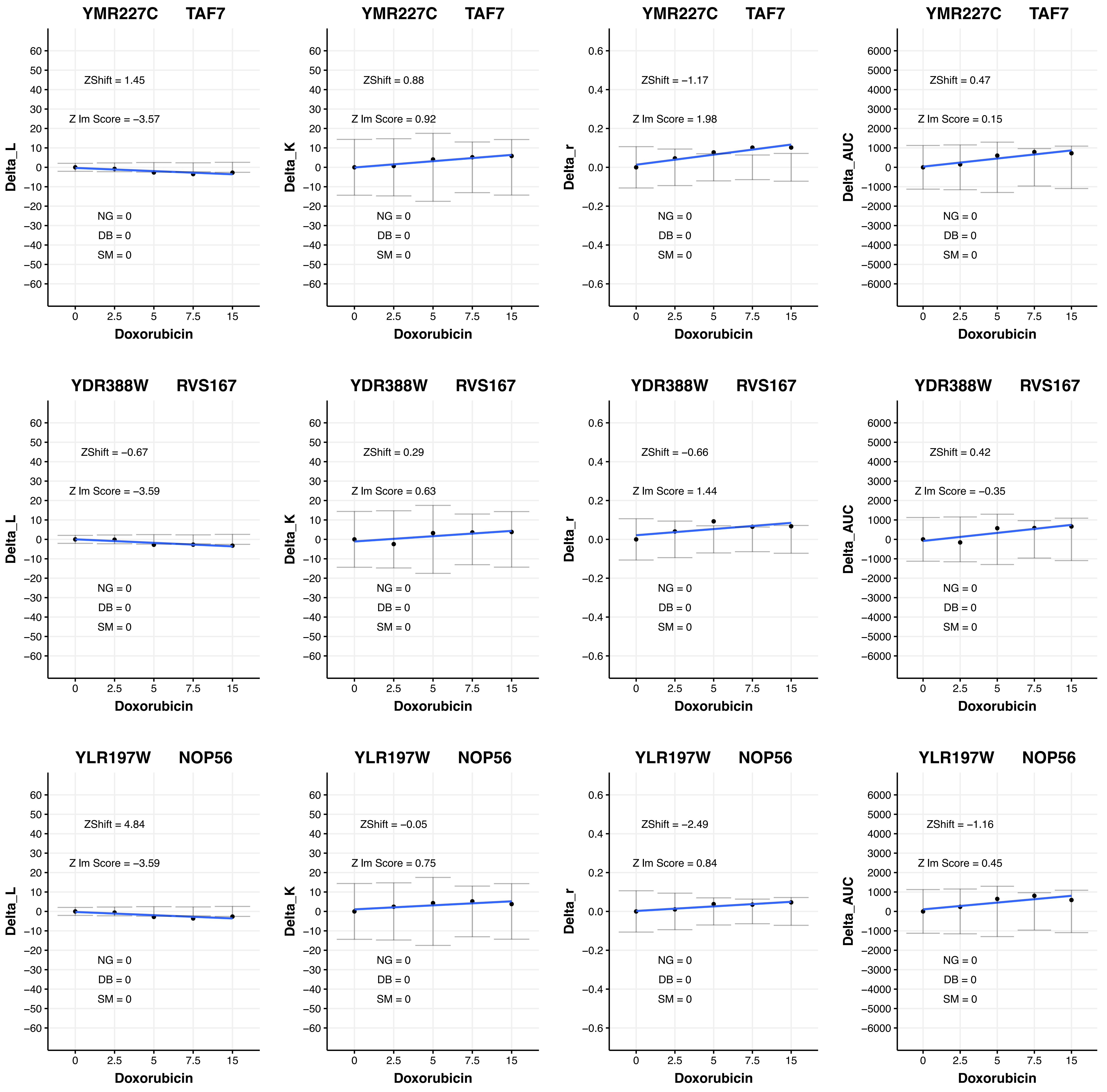

YMR227C TAF7
ZShift = 1.45
Z lm Score = −3.57
NG = 0
DB = 0
SM = 0
Delta_L
Doxorubicin
YMR227C TAF7
ZShift = 0.88
Z lm Score = 0.92
NG = 0
DB = 0
SM = 0
Delta_K
Doxorubicin
YMR227C TAF7
ZShift = −1.17
Z lm Score = 1.98
NG = 0
DB = 0
SM = 0
Delta_r
Doxorubicin
YMR227C TAF7
ZShift = 0.47
Z lm Score = 0.15
NG = 0
DB = 0
SM = 0
Delta_AUC
Doxorubicin
YDR388W RVS167
ZShift = −0.67
Z lm Score = −3.59
NG = 0
DB = 0
SM = 0
Delta_L
Doxorubicin
YDR388W RVS167
ZShift = 0.29
Z lm Score = 0.63
NG = 0
DB = 0
SM = 0
Delta_K
Doxorubicin
YDR388W RVS167
ZShift = −0.66
Z lm Score = 1.44
NG = 0
DB = 0
SM = 0
Delta_r
Doxorubicin
YDR388W RVS167
ZShift = 0.42
Z lm Score = −0.35
NG = 0
DB = 0
SM = 0
Delta_AUC
Doxorubicin
YLR197W NOP56
ZShift = 4.84
Z lm Score = −3.59
NG = 0
DB = 0
SM = 0
Delta_L
Doxorubicin
YLR197W NOP56
ZShift = −0.05
Z lm Score = 0.75
NG = 0
DB = 0
SM = 0
Delta_K
Doxorubicin
YLR197W NOP56
ZShift = −2.49
Z lm Score = 0.84
NG = 0
DB = 0
SM = 0
Delta_r
Doxorubicin
YLR197W NOP56
ZShift = −1.16
Z lm Score = 0.45
NG = 0
DB = 0
SM = 0
Delta_AUC
Doxorubicin

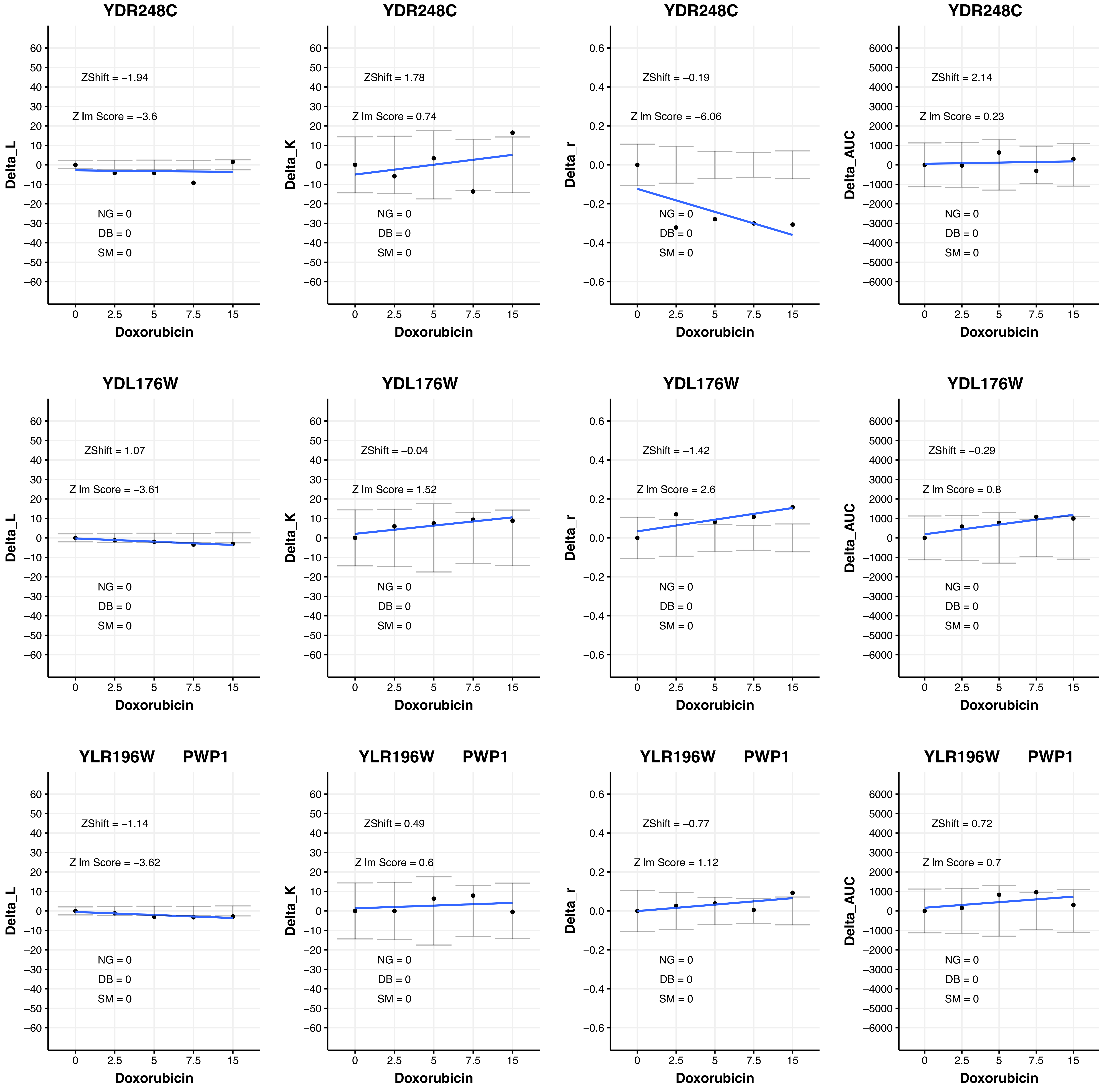

YDR248C
Delta_L
ZShift = −1.94
Z lm Score = −3.6
NG = 0
DB = 0
SM = 0
Doxorubicin
YDR248C
Delta_K
ZShift = 1.78
Z lm Score = 0.74
NG = 0
DB = 0
SM = 0
Doxorubicin
YDR248C
Delta_r
ZShift = −0.19
Z lm Score = −6.06
NG = 0
DB = 0
SM = 0
Doxorubicin
YDR248C
Delta_AUC
ZShift = 2.14
Z lm Score = 0.23
NG = 0
DB = 0
SM = 0
Doxorubicin
YDL176W
Delta_L
ZShift = 1.07
Z lm Score = −3.61
NG = 0
DB = 0
SM = 0
Doxorubicin
YDL176W
Delta_K
ZShift = −0.04
Z lm Score = 1.52
NG = 0
DB = 0
SM = 0
Doxorubicin
YDL176W
Delta_r
ZShift = −1.42
Z lm Score = 2.6
NG = 0
DB = 0
SM = 0
Doxorubicin
YDL176W
Delta_AUC
ZShift = −0.29
Z lm Score = 0.8
NG = 0
DB = 0
SM = 0
Doxorubicin
YLR196W PWP1
Delta_L
ZShift = −1.14
Z lm Score = −3.62
NG = 0
DB = 0
SM = 0
Doxorubicin
YLR196W PWP1
Delta_K
ZShift = 0.49
Z lm Score = 0.6
NG = 0
DB = 0
SM = 0
Doxorubicin
YLR196W PWP1
Delta_r
ZShift = −0.77
Z lm Score = 1.12
NG = 0
DB = 0
SM = 0
Doxorubicin
YLR196W PWP1
Delta_AUC
ZShift = 0.72
Z lm Score = 0.7
NG = 0
DB = 0
SM = 0
Doxorubicin

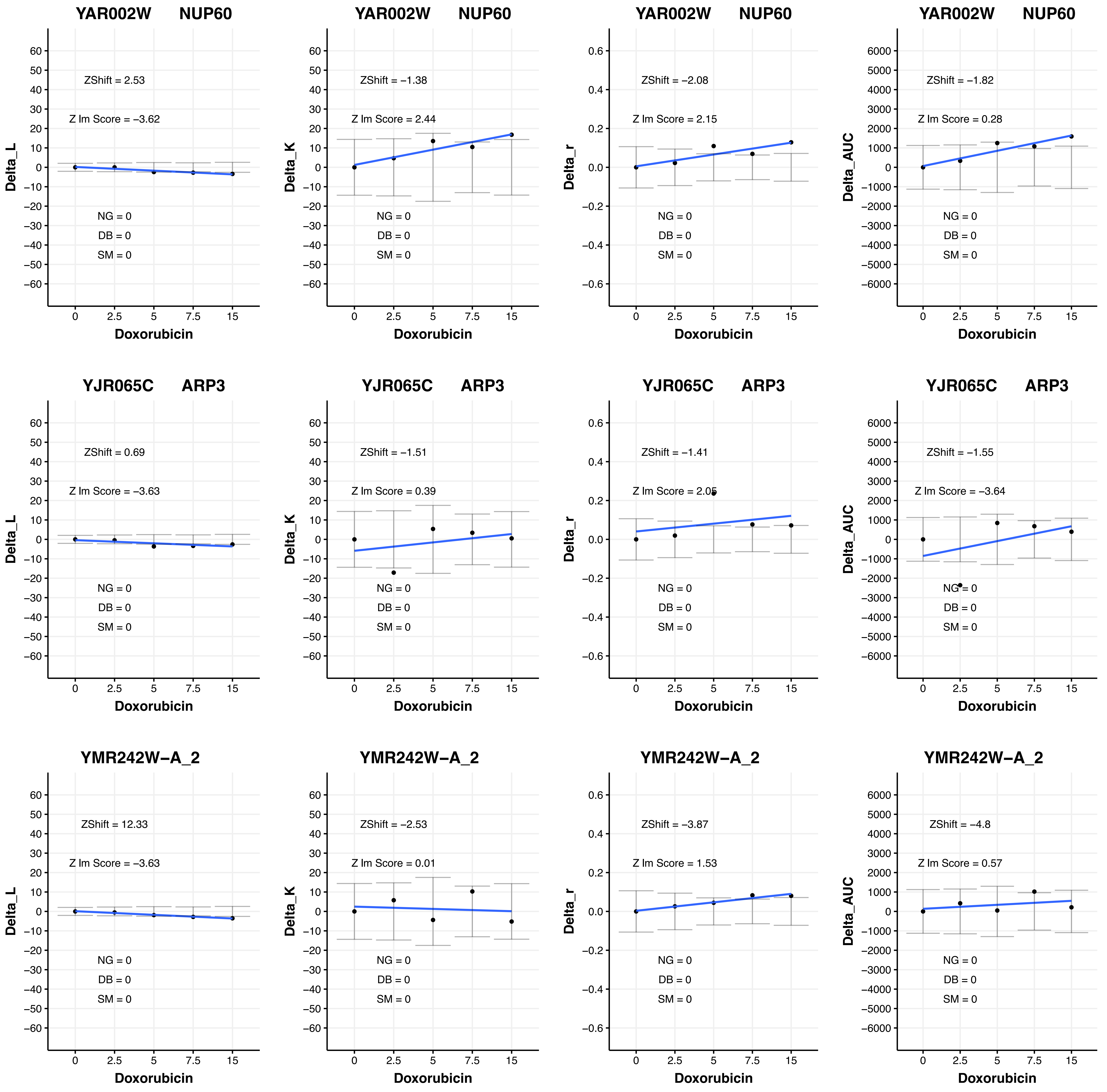

YAR002W NUP60
ZShift = 2.53
Z lm Score = −3.62
NG = 0
DB = 0
SM = 0
Delta_L
Doxorubicin
YAR002W NUP60
ZShift = −1.38
Z lm Score = 2.44
NG = 0
DB = 0
SM = 0
Delta_K
Doxorubicin
YAR002W NUP60
ZShift = −2.08
Z lm Score = 2.15
NG = 0
DB = 0
SM = 0
Delta_r
Doxorubicin
YAR002W NUP60
ZShift = −1.82
Z lm Score = 0.28
NG = 0
DB = 0
SM = 0
Delta_AUC
Doxorubicin
YJR065C ARP3
ZShift = 0.69
Z lm Score = −3.63
NG = 0
DB = 0
SM = 0
Delta_L
Doxorubicin
YJR065C ARP3
ZShift = −1.51
Z lm Score = 0.39
NG = 0
DB = 0
SM = 0
Delta_K
Doxorubicin
YJR065C ARP3
ZShift = −1.41
Z lm Score = 2.05
NG = 0
DB = 0
SM = 0
Delta_r
Doxorubicin
YJR065C ARP3
ZShift = −1.55
Z lm Score = −3.64
NG = 0
DB = 0
SM = 0
Delta_AUC
Doxorubicin
YMR242W−A_2
ZShift = 12.33
Z lm Score = −3.63
NG = 0
DB = 0
SM = 0
Delta_L
Doxorubicin
YMR242W−A_2
ZShift = −2.53
Z lm Score = 0.01
NG = 0
DB = 0
SM = 0
Delta_K
Doxorubicin
YMR242W−A_2
ZShift = −3.87
Z lm Score = 1.53
NG = 0
DB = 0
SM = 0
Delta_r
Doxorubicin
YMR242W−A_2
ZShift = −4.8
Z lm Score = 0.57
NG = 0
DB = 0
SM = 0
Delta_AUC
Doxorubicin

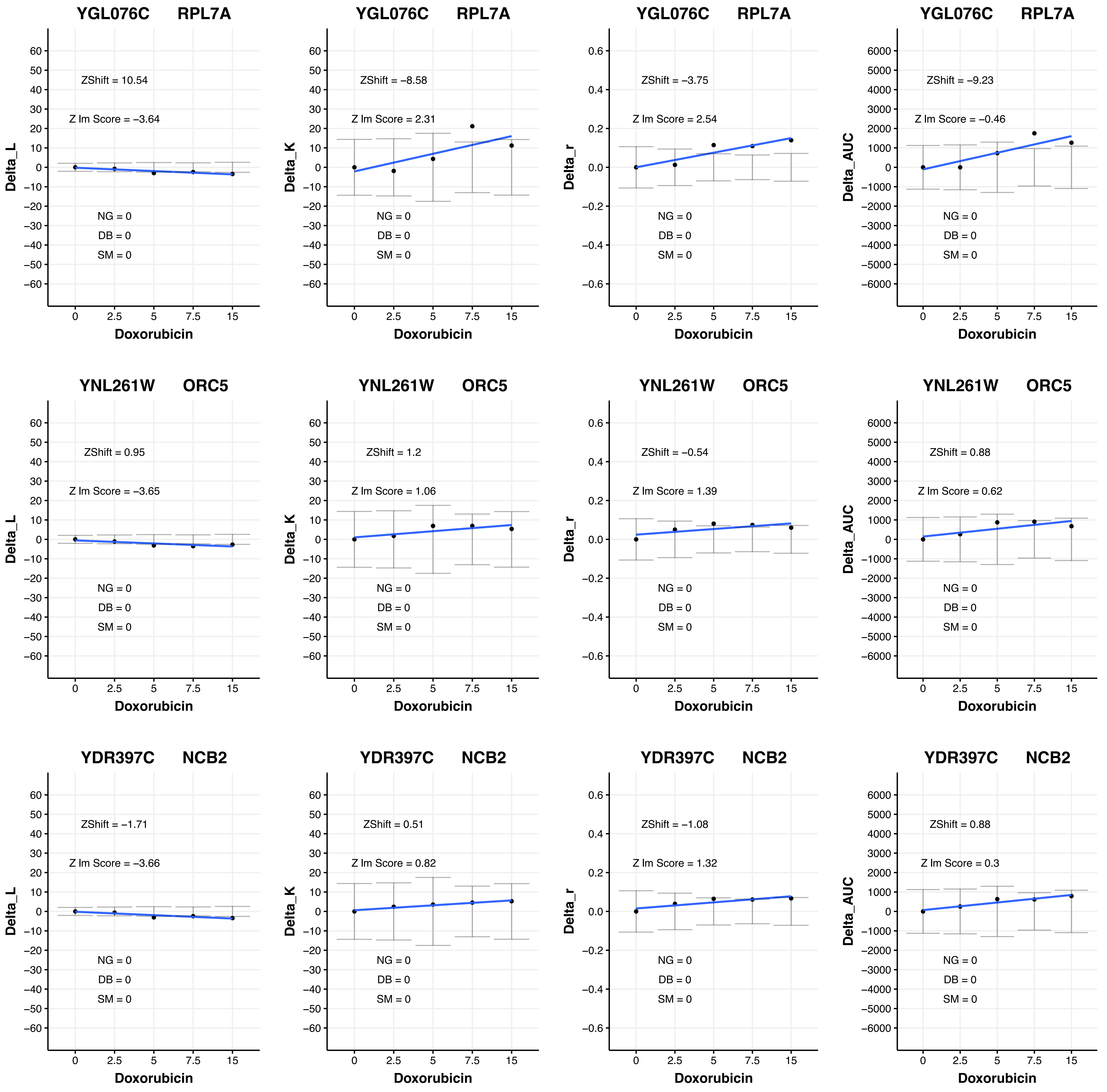
YGL076C RPL7A
ZShift = 10.54
Z lm Score = −3.64
NG = 0
DB = 0
SM = 0
Delta_L
Doxorubicin
YGL076C RPL7A
ZShift = −8.58
Z lm Score = 2.31
NG = 0
DB = 0
SM = 0
Delta_K
Doxorubicin
YGL076C RPL7A
ZShift = −3.75
Z lm Score = 2.54
NG = 0
DB = 0
SM = 0
Delta_r
Doxorubicin
YGL076C RPL7A
ZShift = −9.23
Z lm Score = −0.46
NG = 0
DB = 0
SM = 0
Delta_AUC
Doxorubicin
YNL261W ORC5
ZShift = 0.95
Z lm Score = −3.65
NG = 0
DB = 0
SM = 0
Delta_L
Doxorubicin
YNL261W ORC5
ZShift = 1.2
Z lm Score = 1.06
NG = 0
DB = 0
SM = 0
Delta_K
Doxorubicin
YNL261W ORC5
ZShift = −0.54
Z lm Score = 1.39
NG = 0
DB = 0
SM = 0
Delta_r
Doxorubicin
YNL261W ORC5
ZShift = 0.88
Z lm Score = 0.62
NG = 0
DB = 0
SM = 0
Delta_AUC
Doxorubicin
YDR397C NCB2
ZShift = −1.71
Z lm Score = −3.66
NG = 0
DB = 0
SM = 0
Delta_L
Doxorubicin
YDR397C NCB2
ZShift = 0.51
Z lm Score = 0.82
NG = 0
DB = 0
SM = 0
Delta_K
Doxorubicin
YDR397C NCB2
ZShift = −1.08
Z lm Score = 1.32
NG = 0
DB = 0
SM = 0
Delta_r
Doxorubicin
YDR397C NCB2
ZShift = 0.88
Z lm Score = 0.3
NG = 0
DB = 0
SM = 0
Delta_AUC
Doxorubicin

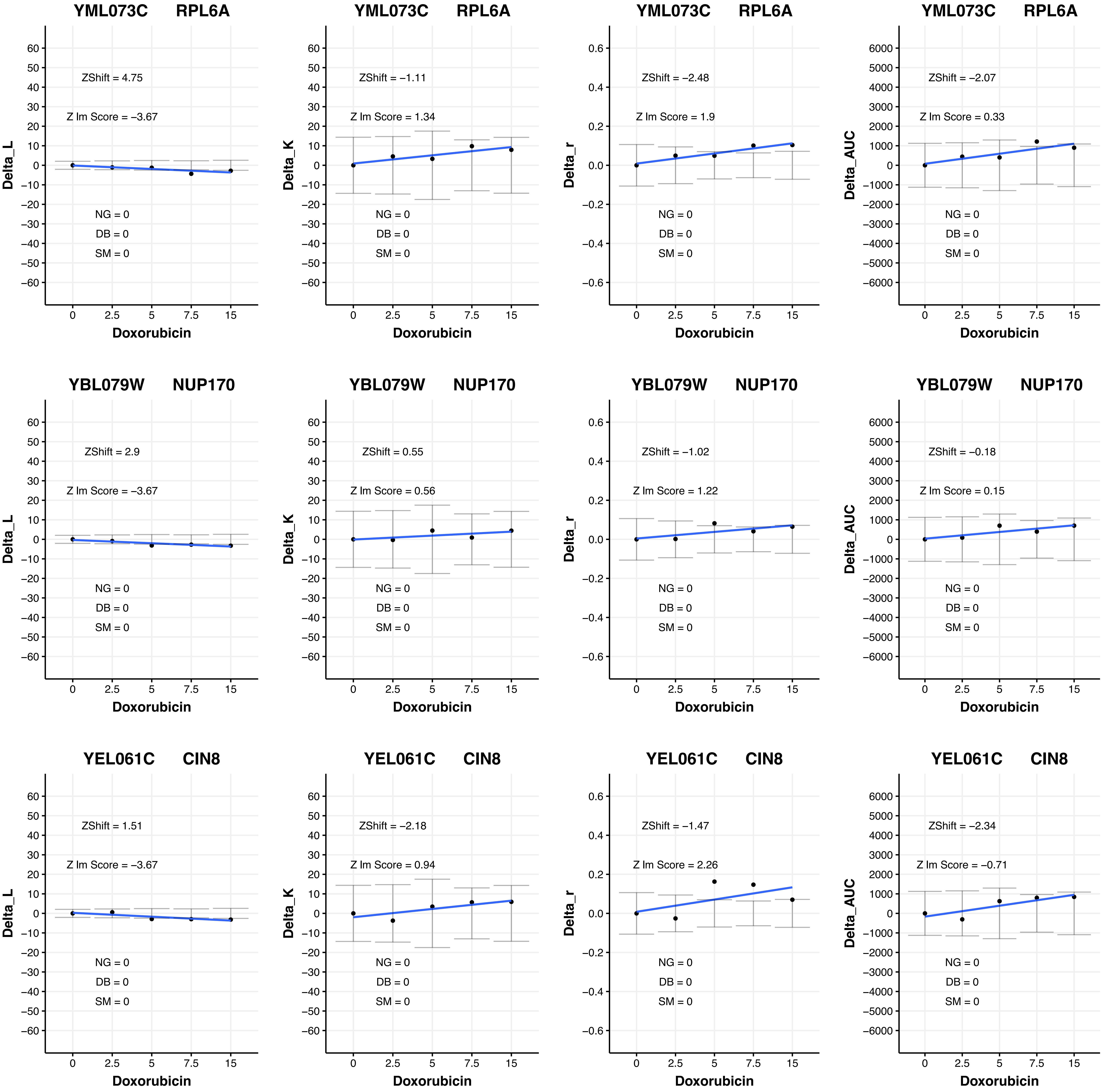

YML073C RPL6A
ZShift = 4.75
Z lm Score = −3.67
NG = 0
DB = 0
SM = 0
Delta_L
Doxorubicin
YML073C RPL6A
ZShift = −1.11
Z lm Score = 1.34
NG = 0
DB = 0
SM = 0
Delta_K
Doxorubicin
YML073C RPL6A
ZShift = −2.48
Z lm Score = 1.9
NG = 0
DB = 0
SM = 0
Delta_r
Doxorubicin
YML073C RPL6A
ZShift = −2.07
Z lm Score = 0.33
NG = 0
DB = 0
SM = 0
Delta_AUC
Doxorubicin
YBL079W NUP170
ZShift = 2.9
Z lm Score = −3.67
NG = 0
DB = 0
SM = 0
Delta_L
Doxorubicin
YBL079W NUP170
ZShift = 0.55
Z lm Score = 0.56
NG = 0
DB = 0
SM = 0
Delta_K
Doxorubicin
YBL079W NUP170
ZShift = −1.02
Z lm Score = 1.22
NG = 0
DB = 0
SM = 0
Delta_r
Doxorubicin
YBL079W NUP170
ZShift = −0.18
Z lm Score = 0.15
NG = 0
DB = 0
SM = 0
Delta_AUC
Doxorubicin
YEL061C CIN8
ZShift = 1.51
Z lm Score = −3.67
NG = 0
DB = 0
SM = 0
Delta_L
Doxorubicin
YEL061C CIN8
ZShift = −2.18
Z lm Score = 0.94
NG = 0
DB = 0
SM = 0
Delta_K
Doxorubicin
YEL061C CIN8
ZShift = −1.47
Z lm Score = 2.26
NG = 0
DB = 0
SM = 0
Delta_r
Doxorubicin
YEL061C CIN8
ZShift = −2.34
Z lm Score = −0.71
NG = 0
DB = 0
SM = 0
Delta_AUC
Doxorubicin

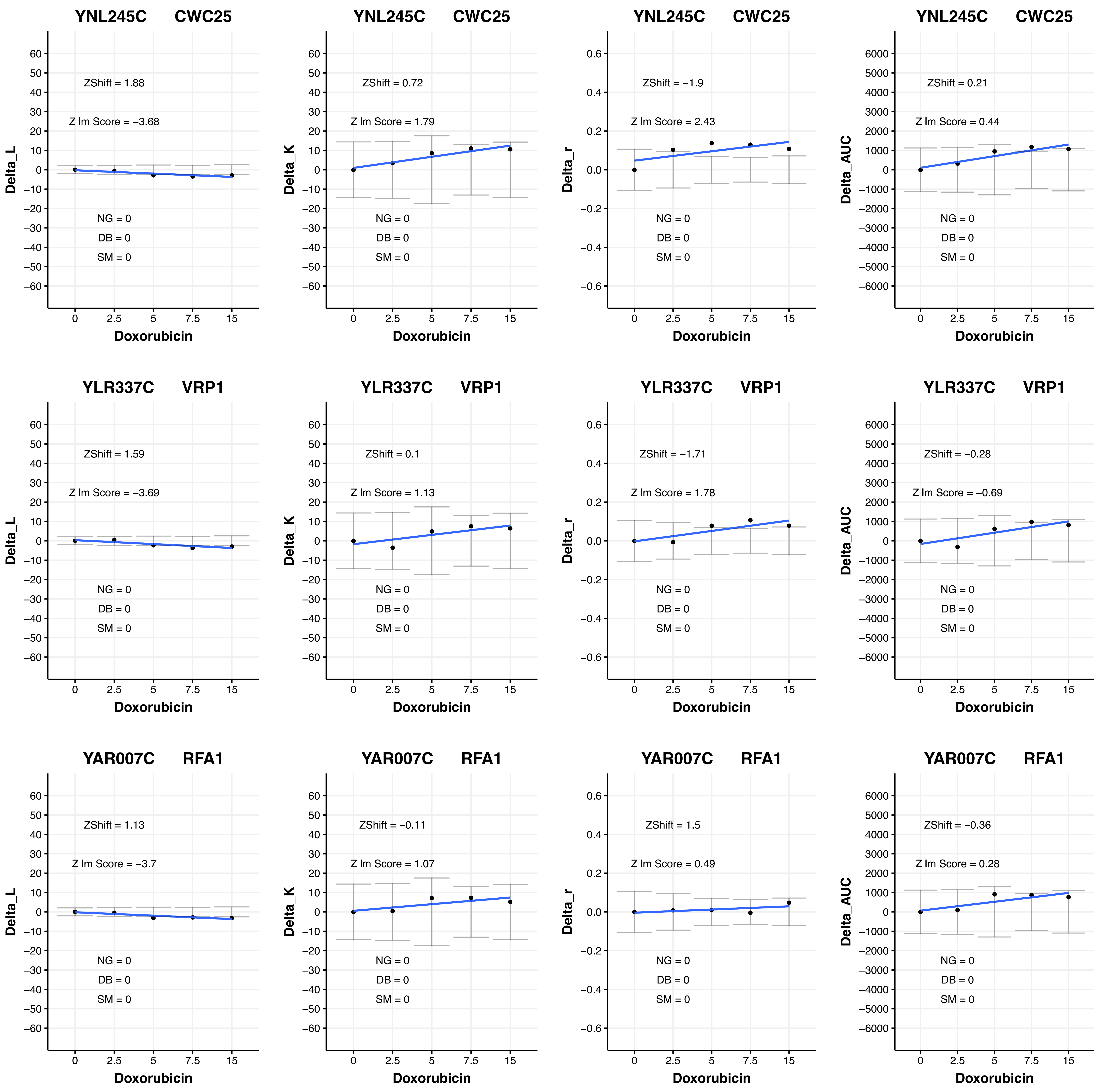
YNL245C CWC25
ZShift = 1.88
Z lm Score = −3.68
NG = 0
DB = 0
SM = 0
Delta_L
Doxorubicin
YNL245C CWC25
ZShift = 0.72
Z lm Score = 1.79
NG = 0
DB = 0
SM = 0
Delta_K
Doxorubicin
YNL245C CWC25
ZShift = −1.9
Z lm Score = 2.43
NG = 0
DB = 0
SM = 0
Delta_r
Doxorubicin
YNL245C CWC25
ZShift = 0.21
Z lm Score = 0.44
NG = 0
DB = 0
SM = 0
Delta_AUC
Doxorubicin
YLR337C VRP1
ZShift = 1.59
Z lm Score = −3.69
NG = 0
DB = 0
SM = 0
Delta_L
Doxorubicin
YLR337C VRP1
ZShift = 0.1
Z lm Score = 1.13
NG = 0
DB = 0
SM = 0
Delta_K
Doxorubicin
YLR337C VRP1
ZShift = −1.71
Z lm Score = 1.78
NG = 0
DB = 0
SM = 0
Delta_r
Doxorubicin
YLR337C VRP1
ZShift = −0.28
Z lm Score = −0.69
NG = 0
DB = 0
SM = 0
Delta_AUC
Doxorubicin
YAR007C RFA1
ZShift = 1.13
Z lm Score = −3.7
NG = 0
DB = 0
SM = 0
Delta_L
Doxorubicin
YAR007C RFA1
ZShift = −0.11
Z lm Score = 1.07
NG = 0
DB = 0
SM = 0
Delta_K
Doxorubicin
YAR007C RFA1
ZShift = 1.5
Z lm Score = 0.49
NG = 0
DB = 0
SM = 0
Delta_r
Doxorubicin
YAR007C RFA1
ZShift = −0.36
Z lm Score = 0.28
NG = 0
DB = 0
SM = 0
Delta_AUC
Doxorubicin

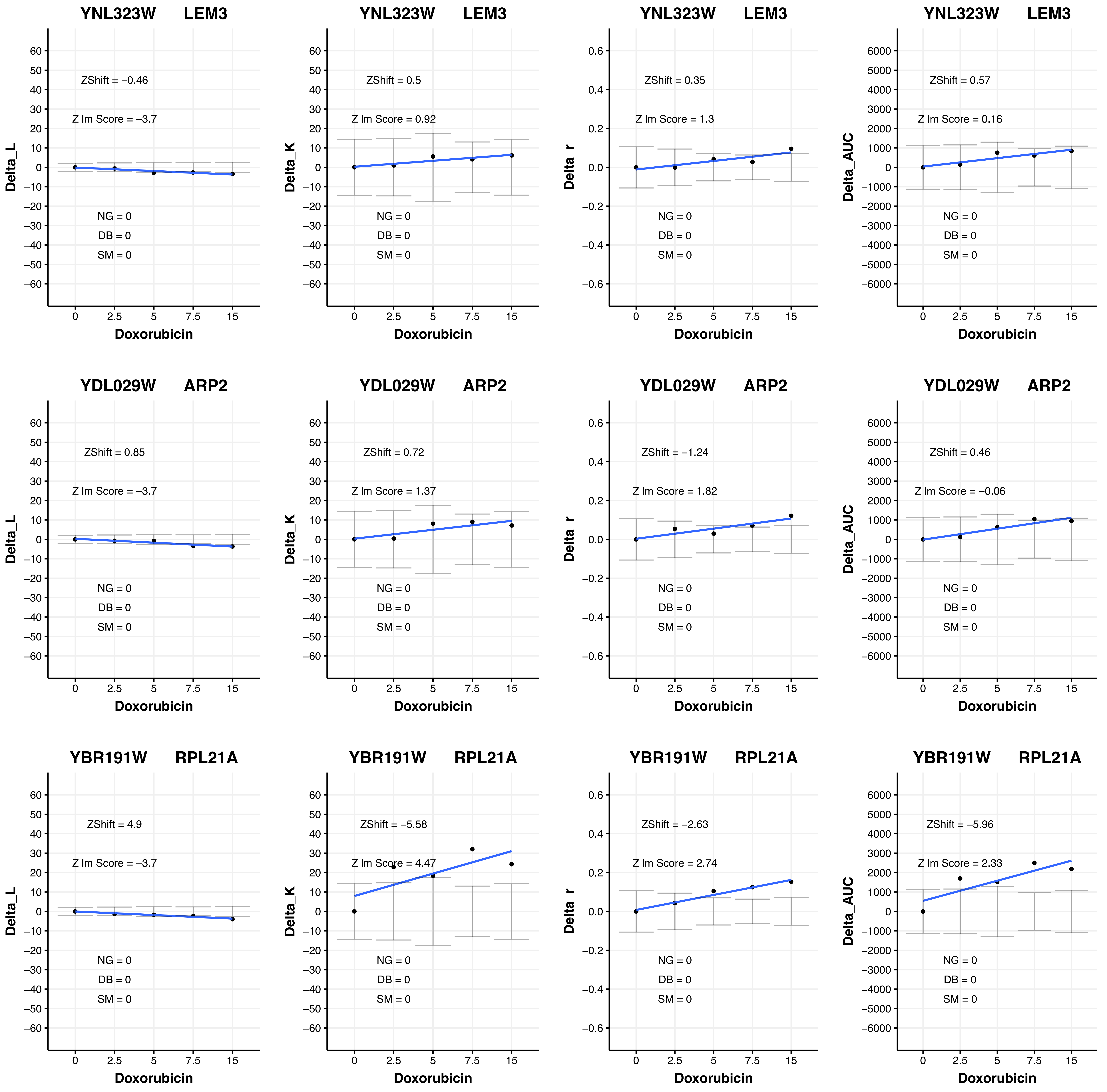

YNL323W LEM3
ZShift = −0.46
Z lm Score = −3.7
NG = 0
DB = 0
SM = 0
Delta_L
Doxorubicin
YNL323W LEM3
ZShift = 0.5
Z lm Score = 0.92
NG = 0
DB = 0
SM = 0
Delta_K
Doxorubicin
YNL323W LEM3
ZShift = 0.35
Z lm Score = 1.3
NG = 0
DB = 0
SM = 0
Delta_r
Doxorubicin
YNL323W LEM3
ZShift = 0.57
Z lm Score = 0.16
NG = 0
DB = 0
SM = 0
Delta_AUC
Doxorubicin
YDL029W ARP2
ZShift = 0.85
Z lm Score = −3.7
NG = 0
DB = 0
SM = 0
Delta_L
Doxorubicin
YDL029W ARP2
ZShift = 0.72
Z lm Score = 1.37
NG = 0
DB = 0
SM = 0
Delta_K
Doxorubicin
YDL029W ARP2
ZShift = −1.24
Z lm Score = 1.82
NG = 0
DB = 0
SM = 0
Delta_r
Doxorubicin
YDL029W ARP2
ZShift = 0.46
Z lm Score = −0.06
NG = 0
DB = 0
SM = 0
Delta_AUC
Doxorubicin
YBR191W RPL21A
ZShift = 4.9
Z lm Score = −3.7
NG = 0
DB = 0
SM = 0
Delta_L
Doxorubicin
YBR191W RPL21A
ZShift = −5.58
Z lm Score = 4.47
NG = 0
DB = 0
SM = 0
Delta_K
Doxorubicin
YBR191W RPL21A
ZShift = −2.63
Z lm Score = 2.74
NG = 0
DB = 0
SM = 0
Delta_r
Doxorubicin
YBR191W RPL21A
ZShift = −5.96
Z lm Score = 2.33
NG = 0
DB = 0
SM = 0
Delta_AUC
Doxorubicin

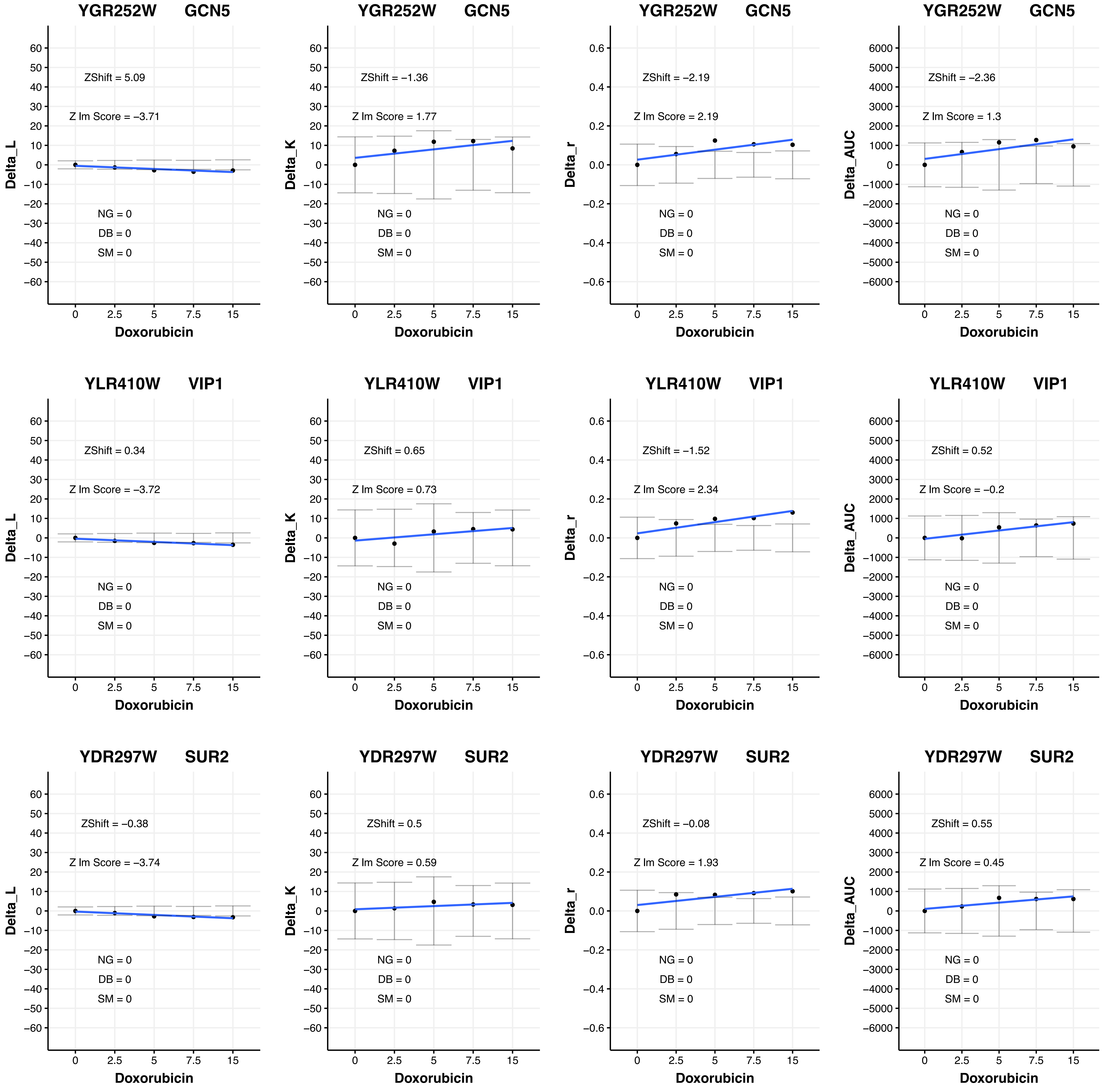
YGR252W GCN5
Delta_L
ZShift = 5.09
Z lm Score = −3.71
NG = 0
DB = 0
SM = 0
Doxorubicin
YGR252W GCN5
Delta_K
ZShift = −1.36
Z lm Score = 1.77
NG = 0
DB = 0
SM = 0
Doxorubicin
YGR252W GCN5
Delta_r
ZShift = −2.19
Z lm Score = 2.19
NG = 0
DB = 0
SM = 0
Doxorubicin
YGR252W GCN5
Delta_AUC
ZShift = −2.36
Z lm Score = 1.3
NG = 0
DB = 0
SM = 0
Doxorubicin
YLR410W VIP1
Delta_L
ZShift = 0.34
Z lm Score = −3.72
NG = 0
DB = 0
SM = 0
Doxorubicin
YLR410W VIP1
Delta_K
ZShift = 0.65
Z lm Score = 0.73
NG = 0
DB = 0
SM = 0
Doxorubicin
YLR410W VIP1
Delta_r
ZShift = −1.52
Z lm Score = 2.34
NG = 0
DB = 0
SM = 0
Doxorubicin
YLR410W VIP1
Delta_AUC
ZShift = 0.52
Z lm Score = −0.2
NG = 0
DB = 0
SM = 0
Doxorubicin
YDR297W SUR2
Delta_L
ZShift = −0.38
Z lm Score = −3.74
NG = 0
DB = 0
SM = 0
Doxorubicin
YDR297W SUR2
Delta_K
ZShift = 0.5
Z lm Score = 0.59
NG = 0
DB = 0
SM = 0
Doxorubicin
YDR297W SUR2
Delta_r
ZShift = −0.08
Z lm Score = 1.93
NG = 0
DB = 0
SM = 0
Doxorubicin
YDR297W SUR2
Delta_AUC
ZShift = 0.55
Z lm Score = 0.45
NG = 0
DB = 0
SM = 0
Doxorubicin

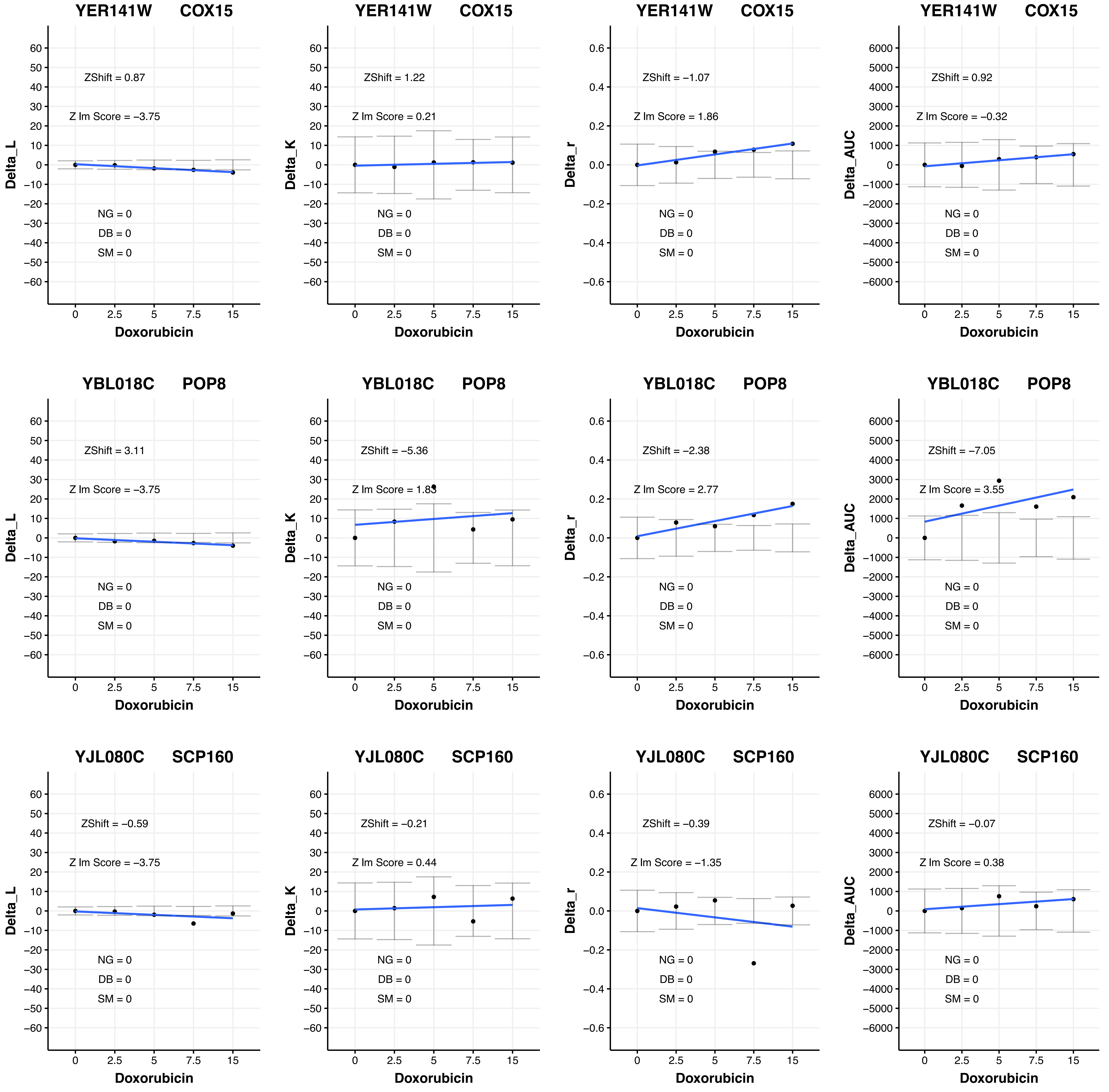

YER141W COX15
ZShift = 0.87
Z lm Score = −3.75
NG = 0
DB = 0
SM = 0
Delta_L
Doxorubicin
YER141W COX15
ZShift = 1.22
Z lm Score = 0.21
NG = 0
DB = 0
SM = 0
Delta_K
Doxorubicin
YER141W COX15
ZShift = −1.07
Z lm Score = 1.86
NG = 0
DB = 0
SM = 0
Delta_r
Doxorubicin
YER141W COX15
ZShift = 0.92
Z lm Score = −0.32
NG = 0
DB = 0
SM = 0
Delta_AUC
Doxorubicin
YBL018C POP8
ZShift = 3.11
Z lm Score = −3.75
NG = 0
DB = 0
SM = 0
Delta_L
Doxorubicin
YBL018C POP8
ZShift = −5.36
Z lm Score = 1.83
NG = 0
DB = 0
SM = 0
Delta_K
Doxorubicin
YBL018C POP8
ZShift = −2.38
Z lm Score = 2.77
NG = 0
DB = 0
SM = 0
Delta_r
Doxorubicin
YBL018C POP8
ZShift = −7.05
Z lm Score = 3.55
NG = 0
DB = 0
SM = 0
Delta_AUC
Doxorubicin
YJL080C SCP160
ZShift = −0.59
Z lm Score = −3.75
NG = 0
DB = 0
SM = 0
Delta_L
Doxorubicin
YJL080C SCP160
ZShift = −0.21
Z lm Score = 0.44
NG = 0
DB = 0
SM = 0
Delta_K
Doxorubicin
YJL080C SCP160
ZShift = −0.39
Z lm Score = −1.35
NG = 0
DB = 0
SM = 0
Delta_r
Doxorubicin
YJL080C SCP160
ZShift = −0.07
Z lm Score = 0.38
NG = 0
DB = 0
SM = 0
Delta_AUC
Doxorubicin

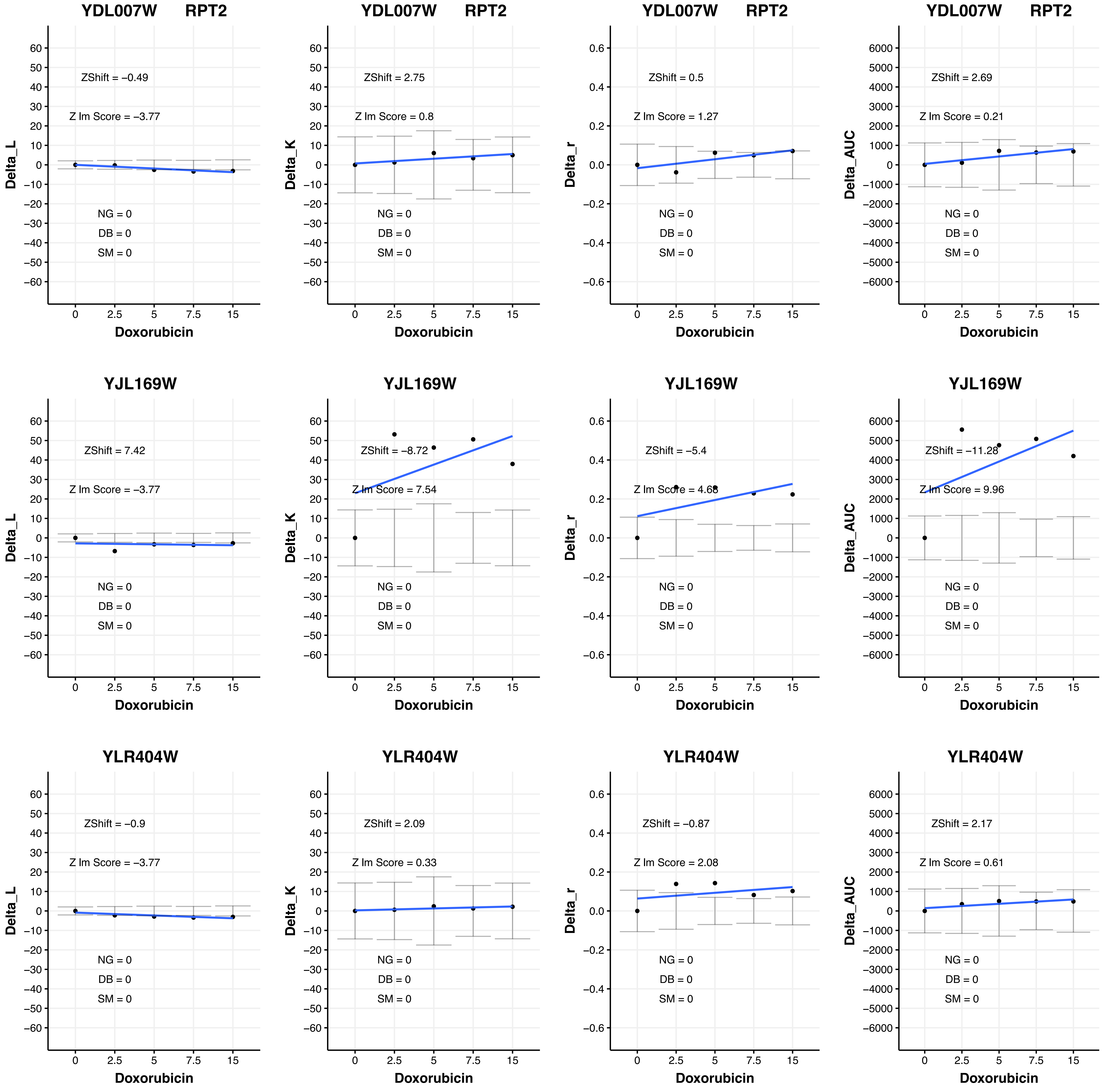
YDL007W RPT2
ZShift = −0.49
Z lm Score = −3.77
NG = 0
DB = 0
SM = 0
Delta_L
Doxorubicin
YDL007W RPT2
ZShift = 2.75
Z lm Score = 0.8
NG = 0
DB = 0
SM = 0
Delta_K
Doxorubicin
YDL007W RPT2
ZShift = 0.5
Z lm Score = 1.27
NG = 0
DB = 0
SM = 0
Delta_r
Doxorubicin
YDL007W RPT2
ZShift = 2.69
Z lm Score = 0.21
NG = 0
DB = 0
SM = 0
Delta_AUC
Doxorubicin
YJL169W
ZShift = 7.42
Z lm Score = −3.77
NG = 0
DB = 0
SM = 0
Delta_L
Doxorubicin
YJL169W
ZShift = −8.72
Z lm Score = 7.54
NG = 0
DB = 0
SM = 0
Delta_K
Doxorubicin
YJL169W
ZShift = −5.4
Z lm Score = 4.68
NG = 0
DB = 0
SM = 0
Delta_r
Doxorubicin
YJL169W
ZShift = −11.28
Z lm Score = 9.96
NG = 0
DB = 0
SM = 0
Delta_AUC
Doxorubicin
YLR404W
ZShift = −0.9
Z lm Score = −3.77
NG = 0
DB = 0
SM = 0
Delta_L
Doxorubicin
YLR404W
ZShift = 2.09
Z lm Score = 0.33
NG = 0
DB = 0
SM = 0
Delta_K
Doxorubicin
YLR404W
ZShift = −0.87
Z lm Score = 2.08
NG = 0
DB = 0
SM = 0
Delta_r
Doxorubicin
YLR404W
ZShift = 2.17
Z lm Score = 0.61
NG = 0
DB = 0
SM = 0
Delta_AUC
Doxorubicin

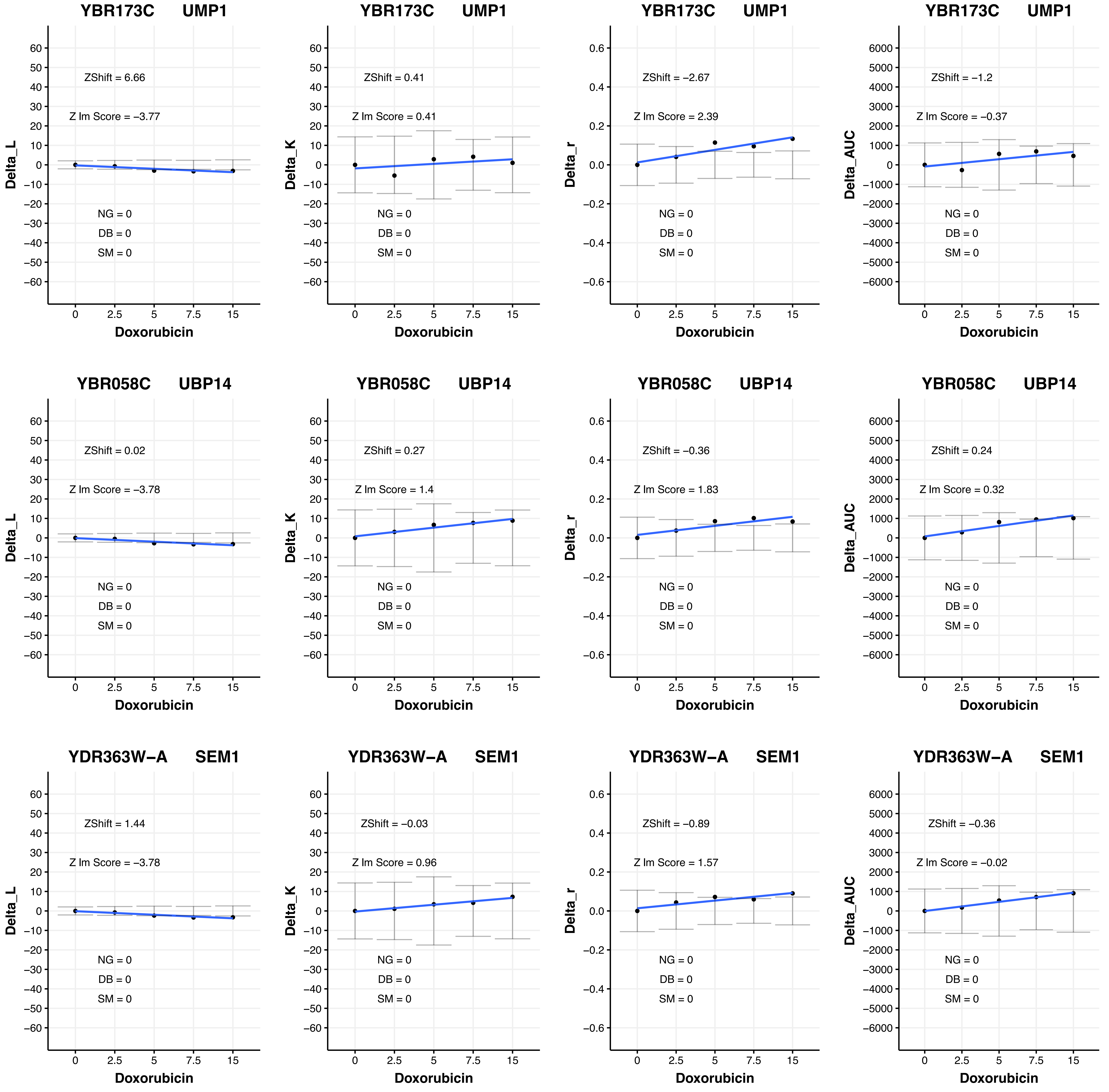

YBR173C UMP1
ZShift = 6.66
Z lm Score = −3.77
NG = 0
DB = 0
SM = 0
Delta_L
Doxorubicin
YBR173C UMP1
ZShift = 0.41
Z lm Score = 0.41
NG = 0
DB = 0
SM = 0
Delta_K
Doxorubicin
YBR173C UMP1
ZShift = −2.67
Z lm Score = 2.39
NG = 0
DB = 0
SM = 0
Delta_r
Doxorubicin
YBR173C UMP1
ZShift = −1.2
Z lm Score = −0.37
NG = 0
DB = 0
SM = 0
Delta_AUC
Doxorubicin
YBR058C UBP14
ZShift = 0.02
Z lm Score = −3.78
NG = 0
DB = 0
SM = 0
Delta_L
Doxorubicin
YBR058C UBP14
ZShift = 0.27
Z lm Score = 1.4
NG = 0
DB = 0
SM = 0
Delta_K
Doxorubicin
YBR058C UBP14
ZShift = −0.36
Z lm Score = 1.83
NG = 0
DB = 0
SM = 0
Delta_r
Doxorubicin
YBR058C UBP14
ZShift = 0.24
Z lm Score = 0.32
NG = 0
DB = 0
SM = 0
Delta_AUC
Doxorubicin
YDR363W−A SEM1
ZShift = 1.44
Z lm Score = −3.78
NG = 0
DB = 0
SM = 0
Delta_L
Doxorubicin
YDR363W−A SEM1
ZShift = −0.03
Z lm Score = 0.96
NG = 0
DB = 0
SM = 0
Delta_K
Doxorubicin
YDR363W−A SEM1
ZShift = −0.89
Z lm Score = 1.57
NG = 0
DB = 0
SM = 0
Delta_r
Doxorubicin
YDR363W−A SEM1
ZShift = −0.36
Z lm Score = −0.02
NG = 0
DB = 0
SM = 0
Delta_AUC
Doxorubicin

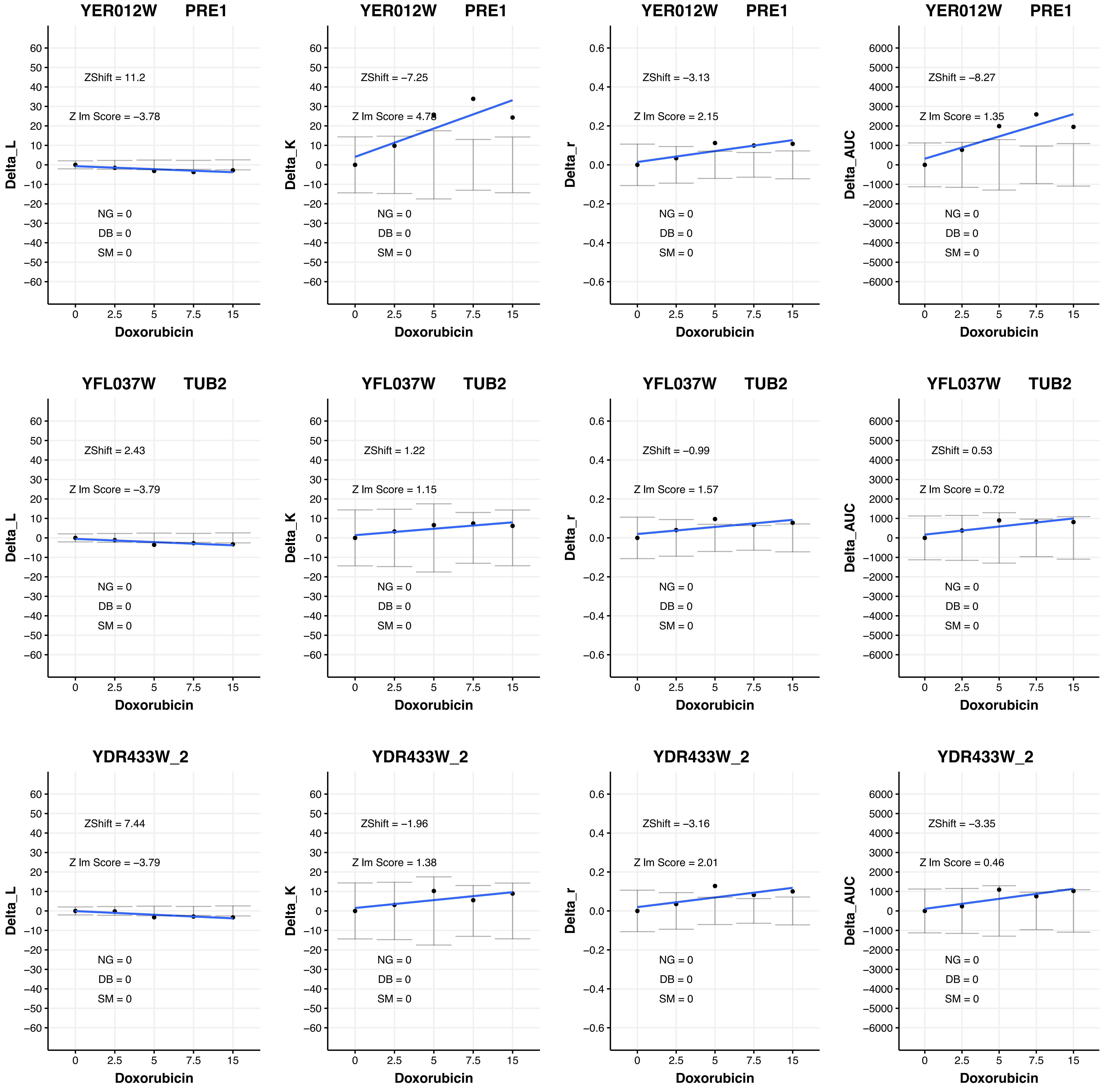
YER012W PRE1
ZShift = 11.2
Z lm Score = −3.78
NG = 0
DB = 0
SM = 0
Delta_L
Doxorubicin
YER012W PRE1
ZShift = −7.25
Z lm Score = 4.78
NG = 0
DB = 0
SM = 0
Delta_K
Doxorubicin
YER012W PRE1
ZShift = −3.13
Z lm Score = 2.15
NG = 0
DB = 0
SM = 0
Delta_r
Doxorubicin
YER012W PRE1
ZShift = −8.27
Z lm Score = 1.35
NG = 0
DB = 0
SM = 0
Delta_AUC
Doxorubicin
YFL037W TUB2
ZShift = 2.43
Z lm Score = −3.79
NG = 0
DB = 0
SM = 0
Delta_L
Doxorubicin
YFL037W TUB2
ZShift = 1.22
Z lm Score = 1.15
NG = 0
DB = 0
SM = 0
Delta_K
Doxorubicin
YFL037W TUB2
ZShift = −0.99
Z lm Score = 1.57
NG = 0
DB = 0
SM = 0
Delta_r
Doxorubicin
YFL037W TUB2
ZShift = 0.53
Z lm Score = 0.72
NG = 0
DB = 0
SM = 0
Delta_AUC
Doxorubicin
YDR433W_2
ZShift = 7.44
Z lm Score = −3.79
NG = 0
DB = 0
SM = 0
Delta_L
Doxorubicin
YDR433W_2
ZShift = −1.96
Z lm Score = 1.38
NG = 0
DB = 0
SM = 0
Delta_K
Doxorubicin
YDR433W_2
ZShift = −3.16
Z lm Score = 2.01
NG = 0
DB = 0
SM = 0
Delta_r
Doxorubicin
YDR433W_2
ZShift = −3.35
Z lm Score = 0.46
NG = 0
DB = 0
SM = 0
Delta_AUC
Doxorubicin

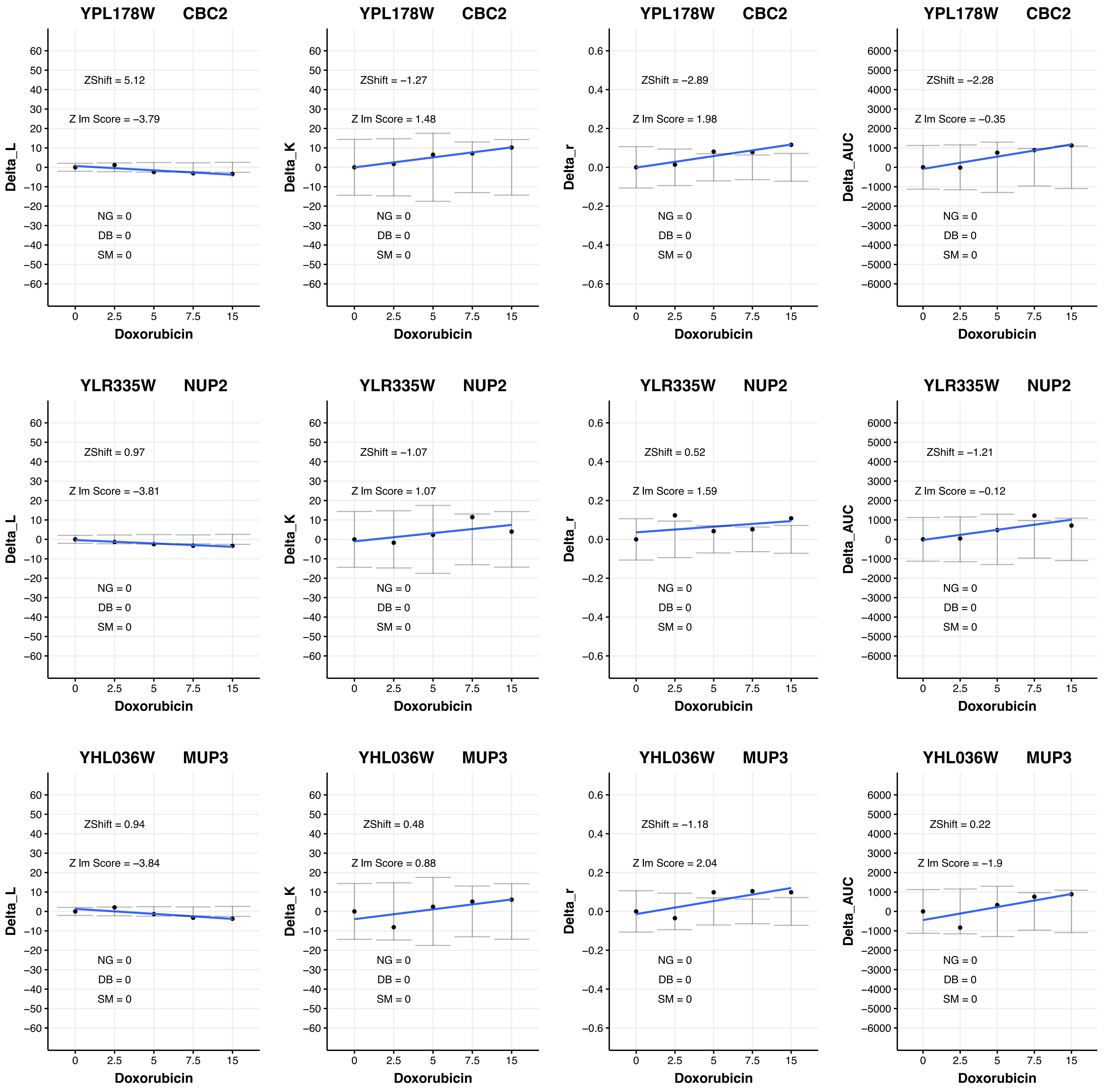
YPL178W CBC2
ZShift = 5.12
Z lm Score = −3.79
NG = 0
DB = 0
SM = 0
Delta_L
Doxorubicin
YPL178W CBC2
ZShift = −1.27
Z lm Score = 1.48
NG = 0
DB = 0
SM = 0
Delta_K
Doxorubicin
YPL178W CBC2
ZShift = −2.89
Z lm Score = 1.98
NG = 0
DB = 0
SM = 0
Delta_r
Doxorubicin
YPL178W CBC2
ZShift = −2.28
Z lm Score = −0.35
NG = 0
DB = 0
SM = 0
Delta_AUC
Doxorubicin
YLR335W NUP2
ZShift = 0.97
Z lm Score = −3.81
NG = 0
DB = 0
SM = 0
Delta_L
Doxorubicin
YLR335W NUP2
ZShift = −1.07
Z lm Score = 1.07
NG = 0
DB = 0
SM = 0
Delta_K
Doxorubicin
YLR335W NUP2
ZShift = 0.52
Z lm Score = 1.59
NG = 0
DB = 0
SM = 0
Delta_r
Doxorubicin
YLR335W NUP2
ZShift = −1.21
Z lm Score = −0.12
NG = 0
DB = 0
SM = 0
Delta_AUC
Doxorubicin
YHL036W MUP3
ZShift = 0.94
Z lm Score = −3.84
NG = 0
DB = 0
SM = 0
Delta_L
Doxorubicin
YHL036W MUP3
ZShift = 0.48
Z lm Score = 0.88
NG = 0
DB = 0
SM = 0
Delta_K
Doxorubicin
YHL036W MUP3
ZShift = −1.18
Z lm Score = 2.04
NG = 0
DB = 0
SM = 0
Delta_r
Doxorubicin
YHL036W MUP3
ZShift = 0.22
Z lm Score = −1.9
NG = 0
DB = 0
SM = 0
Delta_AUC
Doxorubicin

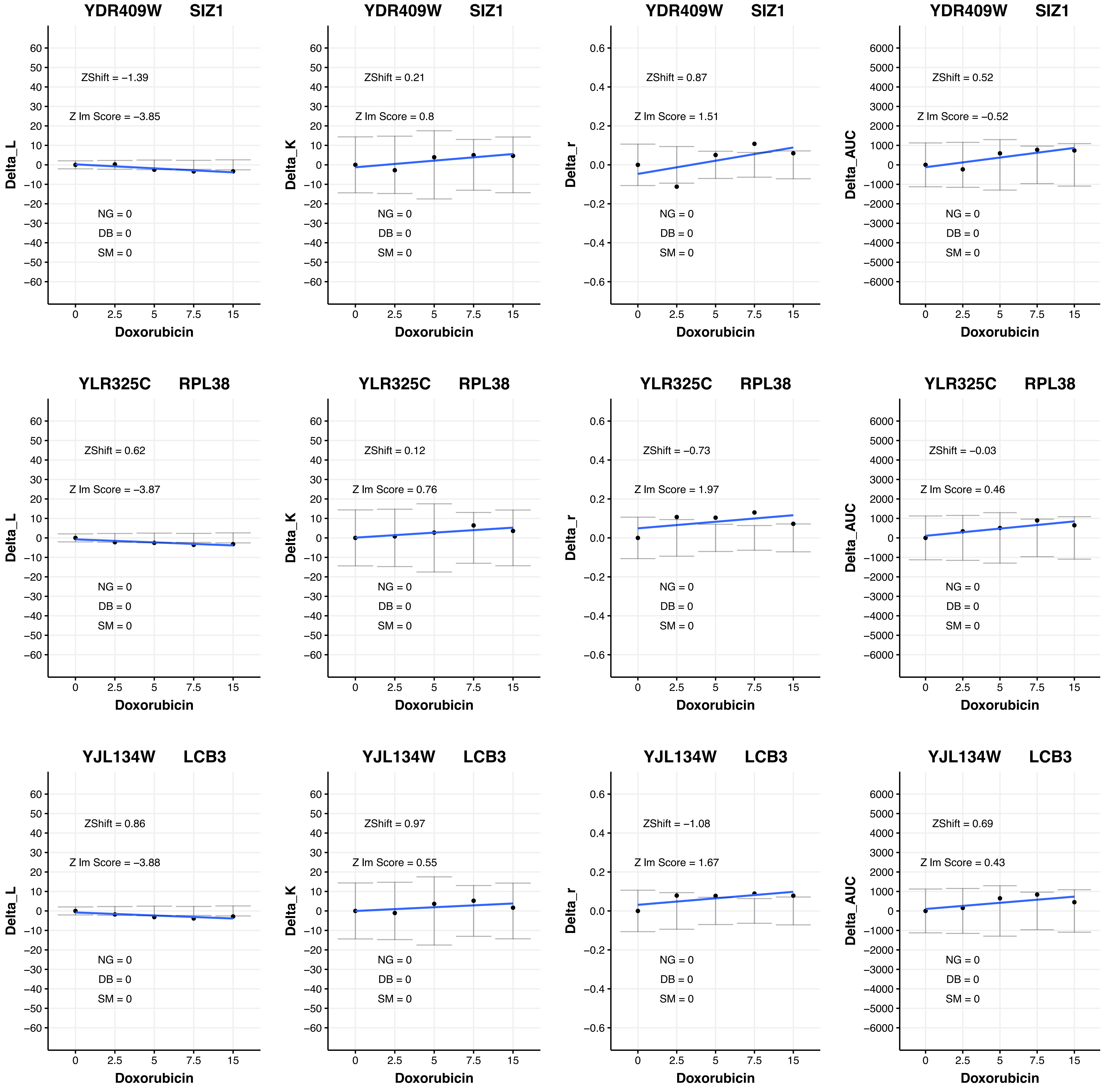
YDR409W SIZ1
ZShift = −1.39
Z lm Score = −3.85
NG = 0
DB = 0
SM = 0
Delta_L
Doxorubicin
YDR409W SIZ1
ZShift = 0.21
Z lm Score = 0.8
NG = 0
DB = 0
SM = 0
Delta_K
Doxorubicin
YDR409W SIZ1
ZShift = 0.87
Z lm Score = 1.51
NG = 0
DB = 0
SM = 0
Delta_r
Doxorubicin
YDR409W SIZ1
ZShift = 0.52
Z lm Score = −0.52
NG = 0
DB = 0
SM = 0
Delta_AUC
Doxorubicin
YLR325C RPL38
ZShift = 0.62
Z lm Score = −3.87
NG = 0
DB = 0
SM = 0
Delta_L
Doxorubicin
YLR325C RPL38
ZShift = 0.12
Z lm Score = 0.76
NG = 0
DB = 0
SM = 0
Delta_K
Doxorubicin
YLR325C RPL38
ZShift = −0.73
Z lm Score = 1.97
NG = 0
DB = 0
SM = 0
Delta_r
Doxorubicin
YLR325C RPL38
ZShift = −0.03
Z lm Score = 0.46
NG = 0
DB = 0
SM = 0
Delta_AUC
Doxorubicin
YJL134W LCB3
ZShift = 0.86
Z lm Score = −3.88
NG = 0
DB = 0
SM = 0
Delta_L
Doxorubicin
YJL134W LCB3
ZShift = 0.97
Z lm Score = 0.55
NG = 0
DB = 0
SM = 0
Delta_K
Doxorubicin
YJL134W LCB3
ZShift = −1.08
Z lm Score = 1.67
NG = 0
DB = 0
SM = 0
Delta_r
Doxorubicin
YJL134W LCB3
ZShift = 0.69
Z lm Score = 0.43
NG = 0
DB = 0
SM = 0
Delta_AUC
Doxorubicin

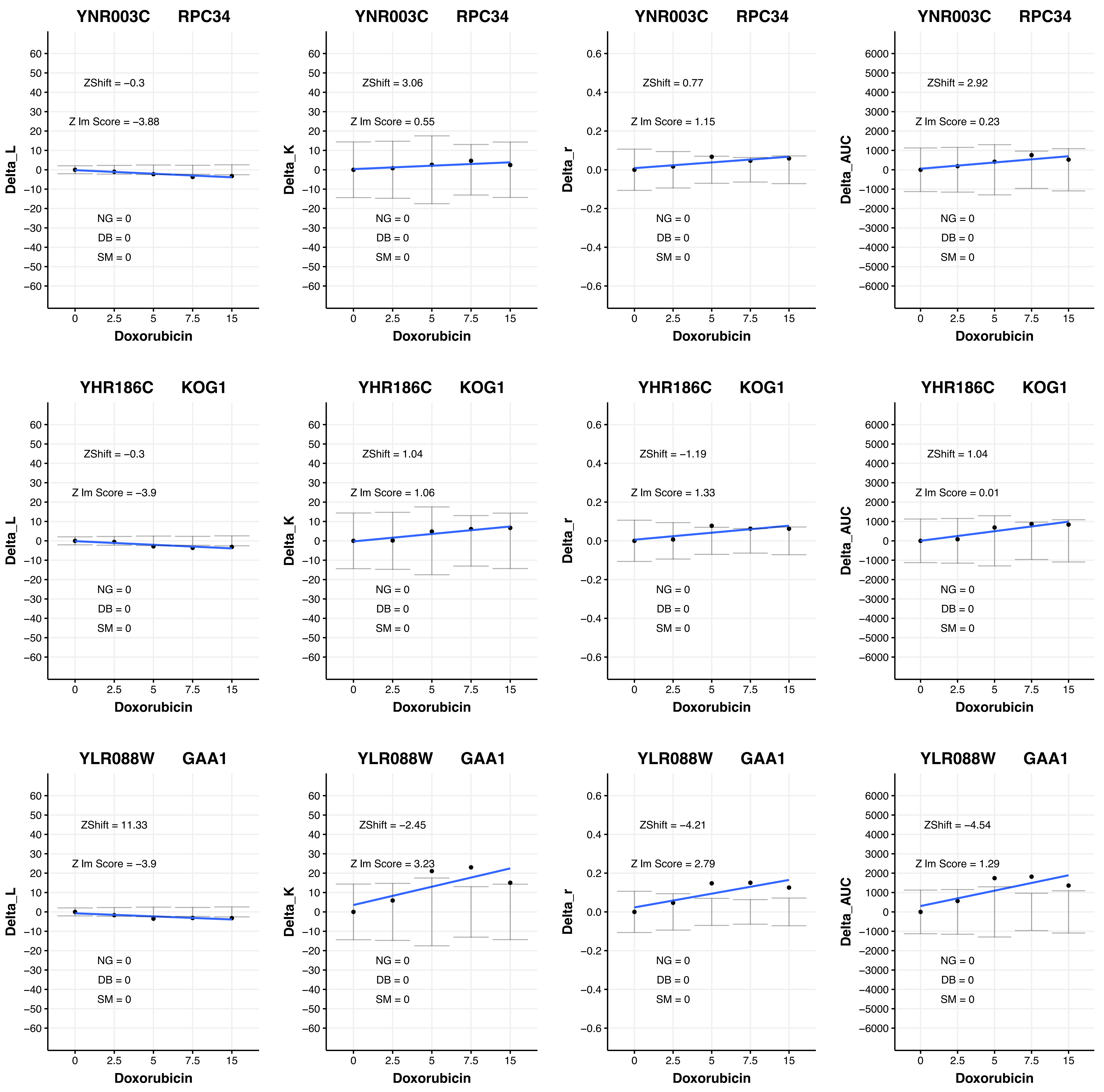
YNR003C RPC34
ZShift = −0.3
Z lm Score = −3.88
NG = 0
DB = 0
SM = 0
Delta_L
Doxorubicin
YNR003C RPC34
ZShift = 3.06
Z lm Score = 0.55
NG = 0
DB = 0
SM = 0
Delta_K
Doxorubicin
YNR003C RPC34
ZShift = 0.77
Z lm Score = 1.15
NG = 0
DB = 0
SM = 0
Delta_r
Doxorubicin
YNR003C RPC34
ZShift = 2.92
Z lm Score = 0.23
NG = 0
DB = 0
SM = 0
Delta_AUC
Doxorubicin
YHR186C KOG1
ZShift = −0.3
Z lm Score = −3.9
NG = 0
DB = 0
SM = 0
Delta_L
Doxorubicin
YHR186C KOG1
ZShift = 1.04
Z lm Score = 1.06
NG = 0
DB = 0
SM = 0
Delta_K
Doxorubicin
YHR186C KOG1
ZShift = −1.19
Z lm Score = 1.33
NG = 0
DB = 0
SM = 0
Delta_r
Doxorubicin
YHR186C KOG1
ZShift = 1.04
Z lm Score = 0.01
NG = 0
DB = 0
SM = 0
Delta_AUC
Doxorubicin
YLR088W GAA1
ZShift = 11.33
Z lm Score = −3.9
NG = 0
DB = 0
SM = 0
Delta_L
Doxorubicin
YLR088W GAA1
ZShift = −2.45
Z lm Score = 3.23
NG = 0
DB = 0
SM = 0
Delta_K
Doxorubicin
YLR088W GAA1
ZShift = −4.21
Z lm Score = 2.79
NG = 0
DB = 0
SM = 0
Delta_r
Doxorubicin
YLR088W GAA1
ZShift = −4.54
Z lm Score = 1.29
NG = 0
DB = 0
SM = 0
Delta_AUC
Doxorubicin

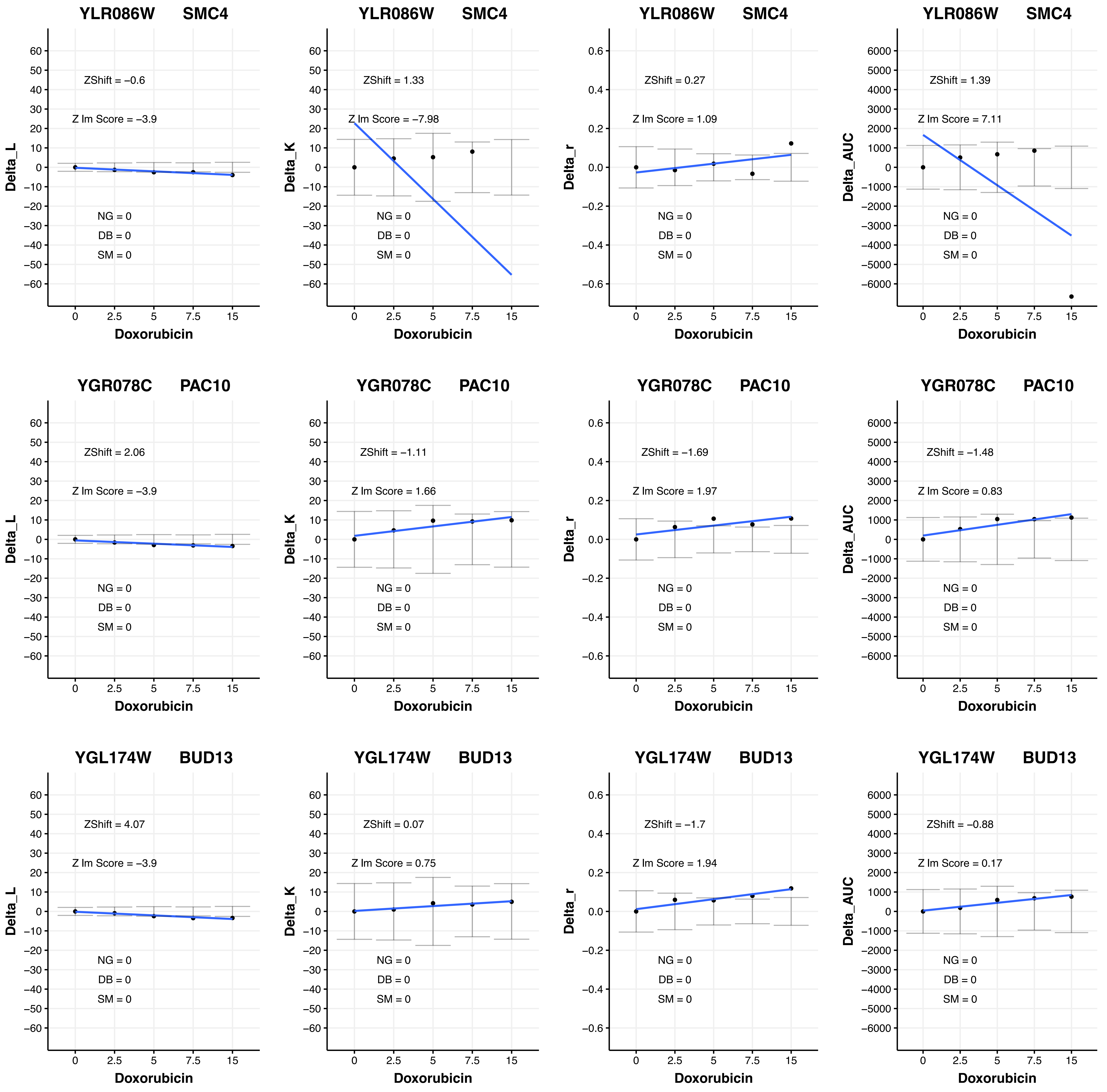
YLR086W SMC4
ZShift = −0.6
Z lm Score = −3.9
NG = 0
DB = 0
SM = 0
Delta_L
Doxorubicin
YLR086W SMC4
ZShift = 1.33
Z lm Score = −7.98
NG = 0
DB = 0
SM = 0
Delta_K
Doxorubicin
YLR086W SMC4
ZShift = 0.27
Z lm Score = 1.09
NG = 0
DB = 0
SM = 0
Delta_r
Doxorubicin
YLR086W SMC4
ZShift = 1.39
Z lm Score = 7.11
NG = 0
DB = 0
SM = 0
Delta_AUC
Doxorubicin
YGR078C PAC10
ZShift = 2.06
Z lm Score = −3.9
NG = 0
DB = 0
SM = 0
Delta_L
Doxorubicin
YGR078C PAC10
ZShift = −1.11
Z lm Score = 1.66
NG = 0
DB = 0
SM = 0
Delta_K
Doxorubicin
YGR078C PAC10
ZShift = −1.69
Z lm Score = 1.97
NG = 0
DB = 0
SM = 0
Delta_r
Doxorubicin
YGR078C PAC10
ZShift = −1.48
Z lm Score = 0.83
NG = 0
DB = 0
SM = 0
Delta_AUC
Doxorubicin
YGL174W BUD13
ZShift = 4.07
Z lm Score = −3.9
NG = 0
DB = 0
SM = 0
Delta_L
Doxorubicin
YGL174W BUD13
ZShift = 0.07
Z lm Score = 0.75
NG = 0
DB = 0
SM = 0
Delta_K
Doxorubicin
YGL174W BUD13
ZShift = −1.7
Z lm Score = 1.94
NG = 0
DB = 0
SM = 0
Delta_r
Doxorubicin
YGL174W BUD13
ZShift = −0.88
Z lm Score = 0.17
NG = 0
DB = 0
SM = 0
Delta_AUC
Doxorubicin

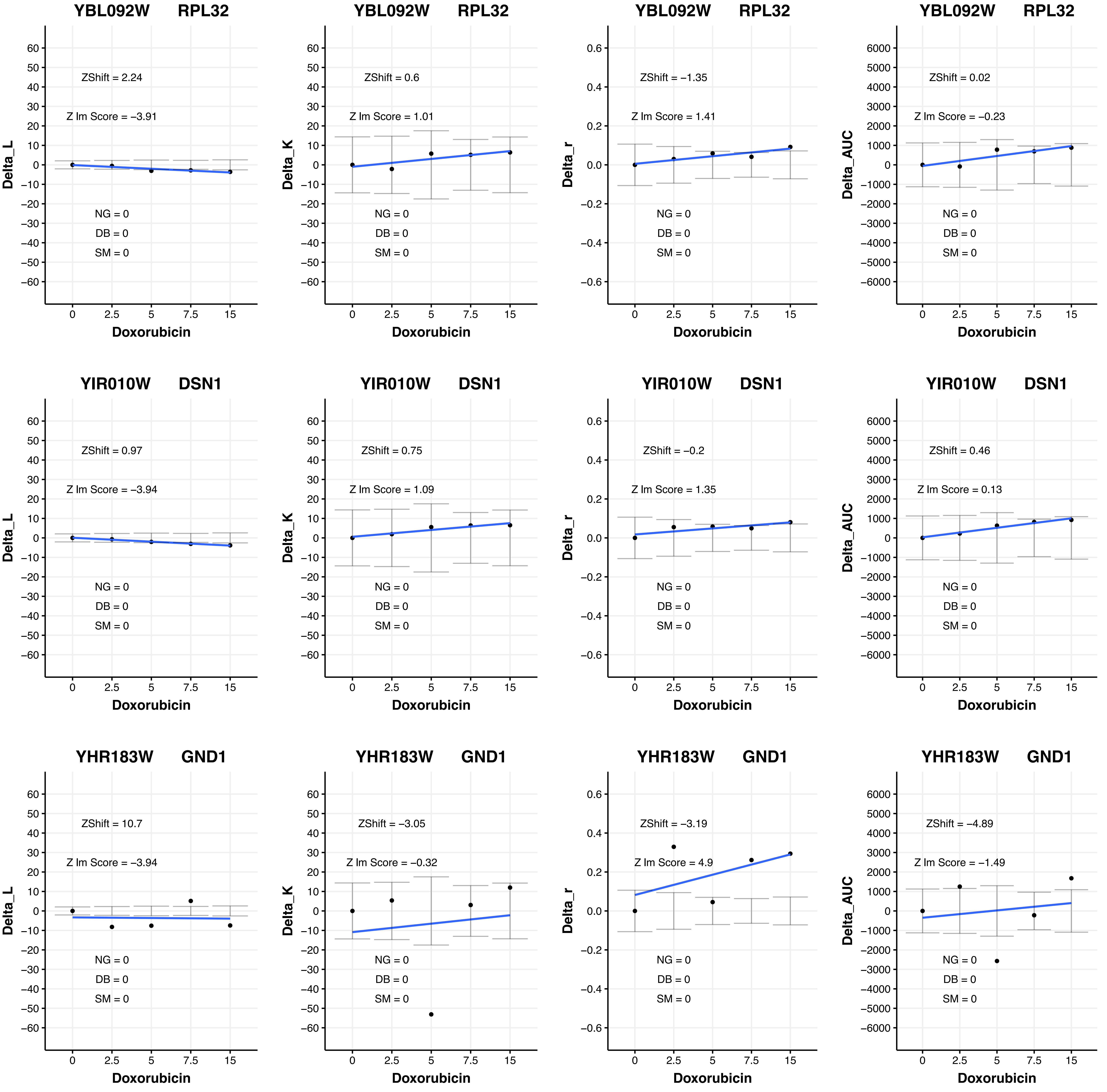
YBL092W RPL32
ZShift = 2.24
Z lm Score = −3.91
NG = 0
DB = 0
SM = 0
Delta_L
Doxorubicin
YBL092W RPL32
ZShift = 0.6
Z lm Score = 1.01
NG = 0
DB = 0
SM = 0
Delta_K
Doxorubicin
YBL092W RPL32
ZShift = −1.35
Z lm Score = 1.41
NG = 0
DB = 0
SM = 0
Delta_r
Doxorubicin
YBL092W RPL32
ZShift = 0.02
Z lm Score = −0.23
NG = 0
DB = 0
SM = 0
Delta_AUC
Doxorubicin
YIR010W DSN1
ZShift = 0.97
Z lm Score = −3.94
NG = 0
DB = 0
SM = 0
Delta_L
Doxorubicin
YIR010W DSN1
ZShift = 0.75
Z lm Score = 1.09
NG = 0
DB = 0
SM = 0
Delta_K
Doxorubicin
YIR010W DSN1
ZShift = −0.2
Z lm Score = 1.35
NG = 0
DB = 0
SM = 0
Delta_r
Doxorubicin
YIR010W DSN1
ZShift = 0.46
Z lm Score = 0.13
NG = 0
DB = 0
SM = 0
Delta_AUC
Doxorubicin
YHR183W GND1
ZShift = 10.7
Z lm Score = −3.94
NG = 0
DB = 0
SM = 0
Delta_L
Doxorubicin
YHR183W GND1
ZShift = −3.05
Z lm Score = −0.32
NG = 0
DB = 0
SM = 0
Delta_K
Doxorubicin
YHR183W GND1
ZShift = −3.19
Z lm Score = 4.9
NG = 0
DB = 0
SM = 0
Delta_r
Doxorubicin
YHR183W GND1
ZShift = −4.89
Z lm Score = −1.49
NG = 0
DB = 0
SM = 0
Delta_AUC
Doxorubicin

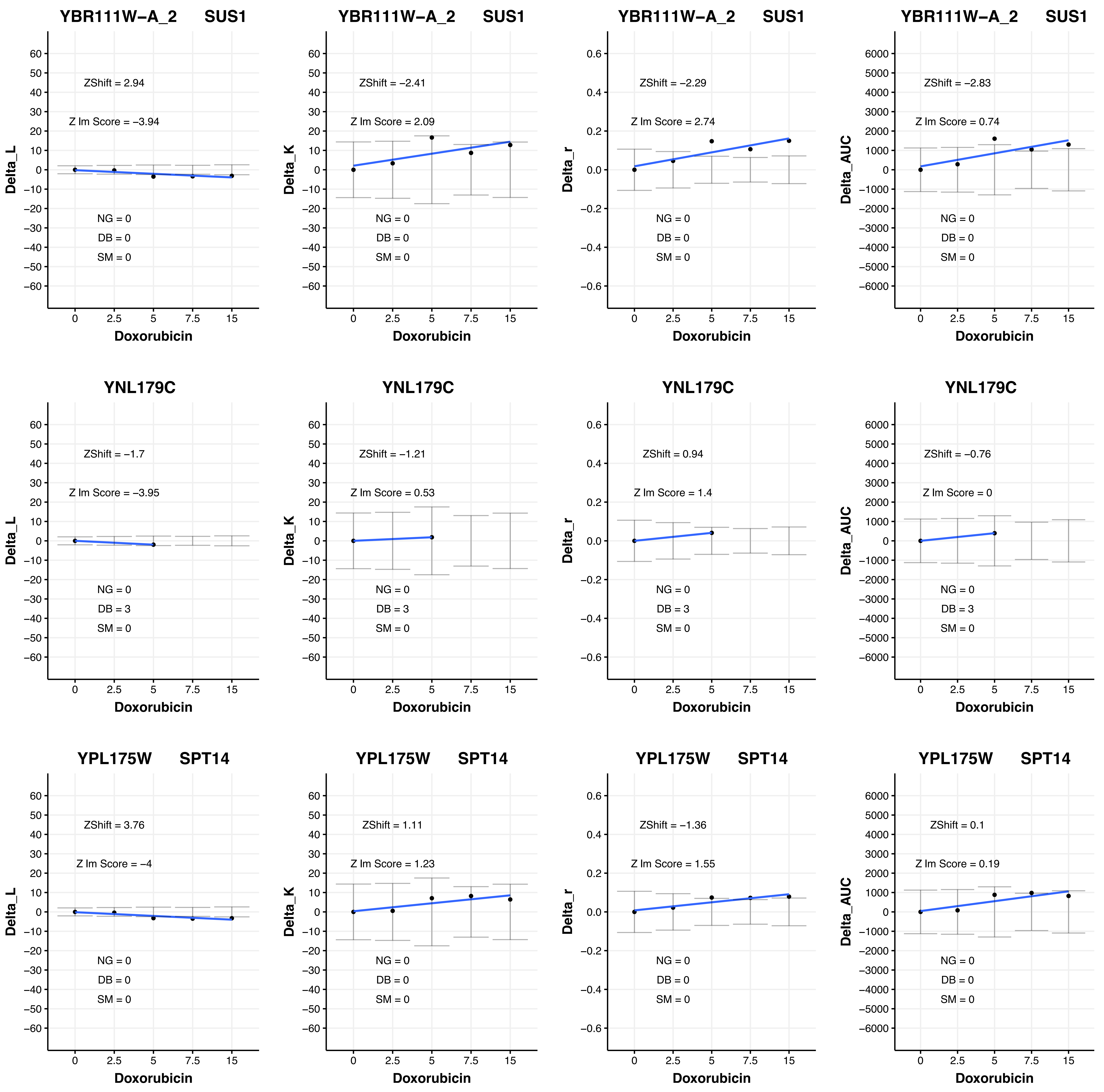

YBR111W-A_2 SUS1
ZShift = 2.94
Z lm Score = −3.94
NG = 0
DB = 0
SM = 0
Delta_L
Doxorubicin
YBR111W-A_2 SUS1
ZShift = −2.41
Z lm Score = 2.09
NG = 0
DB = 0
SM = 0
Delta_K
Doxorubicin
YBR111W-A_2 SUS1
ZShift = −2.29
Z lm Score = 2.74
NG = 0
DB = 0
SM = 0
Delta_r
Doxorubicin
YBR111W-A_2 SUS1
ZShift = −2.83
Z lm Score = 0.74
NG = 0
DB = 0
SM = 0
Delta_AUC
Doxorubicin
YNL179C
ZShift = −1.7
Z lm Score = −3.95
NG = 0
DB = 3
SM = 0
Delta_L
Doxorubicin
YNL179C
ZShift = −1.21
Z lm Score = 0.53
NG = 0
DB = 3
SM = 0
Delta_K
Doxorubicin
YNL179C
ZShift = 0.94
Z lm Score = 1.4
NG = 0
DB = 3
SM = 0
Delta_r
Doxorubicin
YNL179C
ZShift = −0.76
Z lm Score = 0
NG = 0
DB = 3
SM = 0
Delta_AUC
Doxorubicin
YPL175W SPT14
ZShift = 3.76
Z lm Score = −4
NG = 0
DB = 0
SM = 0
Delta_L
Doxorubicin
YPL175W SPT14
ZShift = 1.11
Z lm Score = 1.23
NG = 0
DB = 0
SM = 0
Delta_K
Doxorubicin
YPL175W SPT14
ZShift = −1.36
Z lm Score = 1.55
NG = 0
DB = 0
SM = 0
Delta_r
Doxorubicin
YPL175W SPT14
ZShift = 0.1
Z lm Score = 0.19
NG = 0
DB = 0
SM = 0
Delta_AUC
Doxorubicin

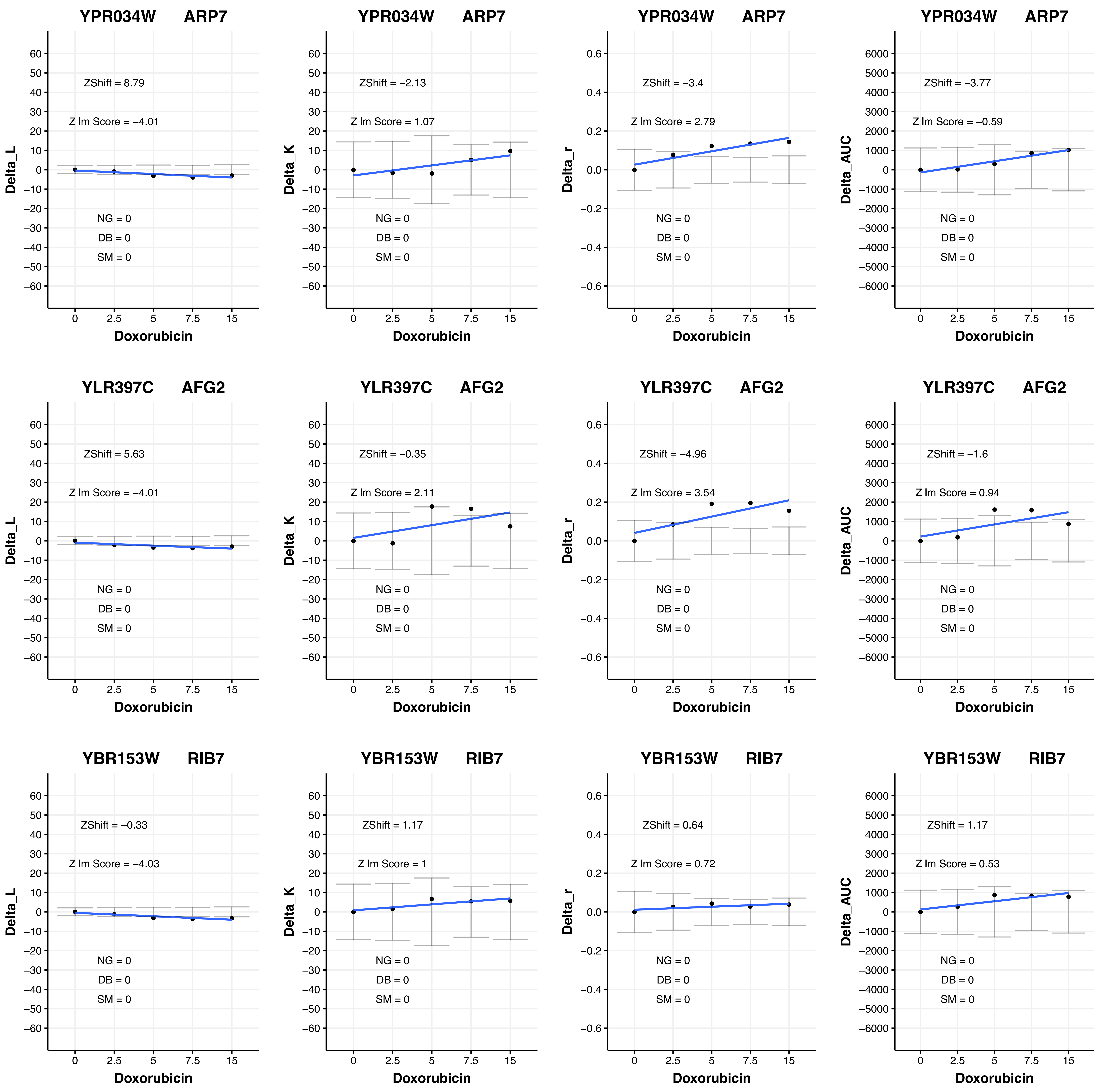

YPR034W ARP7
ZShift = 8.79
Z lm Score = −4.01
NG = 0
DB = 0
SM = 0
Delta_L
Doxorubicin
YPR034W ARP7
ZShift = −2.13
Z lm Score = 1.07
NG = 0
DB = 0
SM = 0
Delta_K
Doxorubicin
YPR034W ARP7
ZShift = −3.4
Z lm Score = 2.79
NG = 0
DB = 0
SM = 0
Delta_r
Doxorubicin
YPR034W ARP7
ZShift = −3.77
Z lm Score = −0.59
NG = 0
DB = 0
SM = 0
Delta_AUC
Doxorubicin
YLR397C AFG2
ZShift = 5.63
Z lm Score = −4.01
NG = 0
DB = 0
SM = 0
Delta_L
Doxorubicin
YLR397C AFG2
ZShift = −0.35
Z lm Score = 2.11
NG = 0
DB = 0
SM = 0
Delta_K
Doxorubicin
YLR397C AFG2
ZShift = −4.96
Z lm Score = 3.54
NG = 0
DB = 0
SM = 0
Delta_r
Doxorubicin
YLR397C AFG2
ZShift = −1.6
Z lm Score = 0.94
NG = 0
DB = 0
SM = 0
Delta_AUC
Doxorubicin
YBR153W RIB7
ZShift = −0.33
Z lm Score = −4.03
NG = 0
DB = 0
SM = 0
Delta_L
Doxorubicin
YBR153W RIB7
ZShift = 1.17
Z lm Score = 1
NG = 0
DB = 0
SM = 0
Delta_K
Doxorubicin
YBR153W RIB7
ZShift = 0.64
Z lm Score = 0.72
NG = 0
DB = 0
SM = 0
Delta_r
Doxorubicin
YBR153W RIB7
ZShift = 1.17
Z lm Score = 0.53
NG = 0
DB = 0
SM = 0
Delta_AUC
Doxorubicin

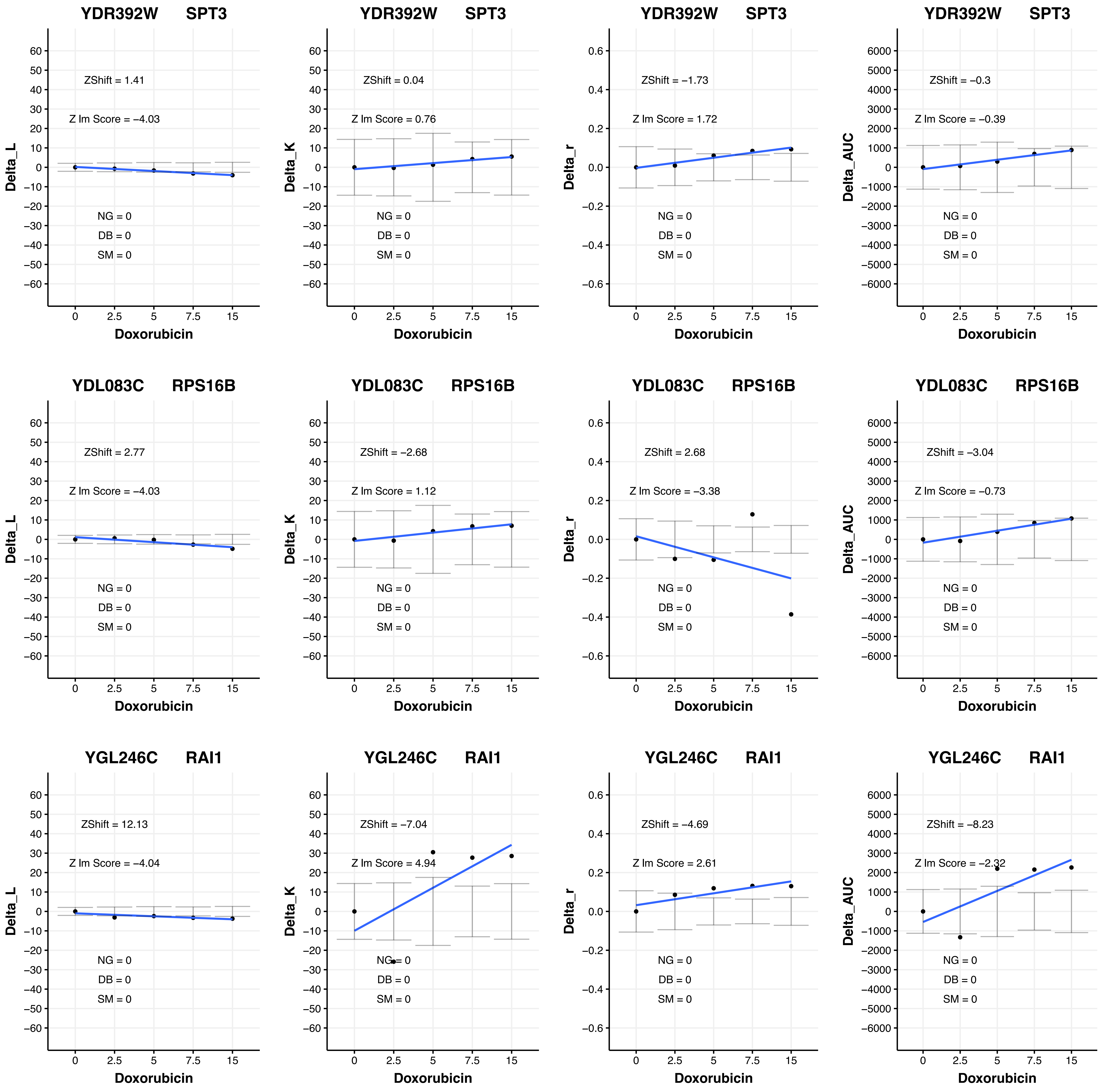

YDR392W SPT3
ZShift = 1.41
Z lm Score = −4.03
NG = 0
DB = 0
SM = 0
Delta_L
Doxorubicin
YDR392W SPT3
ZShift = 0.04
Z lm Score = 0.76
NG = 0
DB = 0
SM = 0
Delta_K
Doxorubicin
YDR392W SPT3
ZShift = −1.73
Z lm Score = 1.72
NG = 0
DB = 0
SM = 0
Delta_r
Doxorubicin
YDR392W SPT3
ZShift = −0.3
Z lm Score = −0.39
NG = 0
DB = 0
SM = 0
Delta_AUC
Doxorubicin
YDL083C RPS16B
ZShift = 2.77
Z lm Score = −4.03
NG = 0
DB = 0
SM = 0
Delta_L
Doxorubicin
YDL083C RPS16B
ZShift = −2.68
Z lm Score = 1.12
NG = 0
DB = 0
SM = 0
Delta_K
Doxorubicin
YDL083C RPS16B
ZShift = 2.68
Z lm Score = −3.38
NG = 0
DB = 0
SM = 0
Delta_r
Doxorubicin
YDL083C RPS16B
ZShift = −3.04
Z lm Score = −0.73
NG = 0
DB = 0
SM = 0
Delta_AUC
Doxorubicin
YGL246C RAI1
ZShift = 12.13
Z lm Score = −4.04
NG = 0
DB = 0
SM = 0
Delta_L
Doxorubicin
YGL246C RAI1
ZShift = −7.04
Z lm Score = 4.94
NG = 0
DB = 0
SM = 0
Delta_K
Doxorubicin
YGL246C RAI1
ZShift = −4.69
Z lm Score = 2.61
NG = 0
DB = 0
SM = 0
Delta_r
Doxorubicin
YGL246C RAI1
ZShift = −8.23
Z lm Score = −2.32
NG = 0
DB = 0
SM = 0
Delta_AUC
Doxorubicin

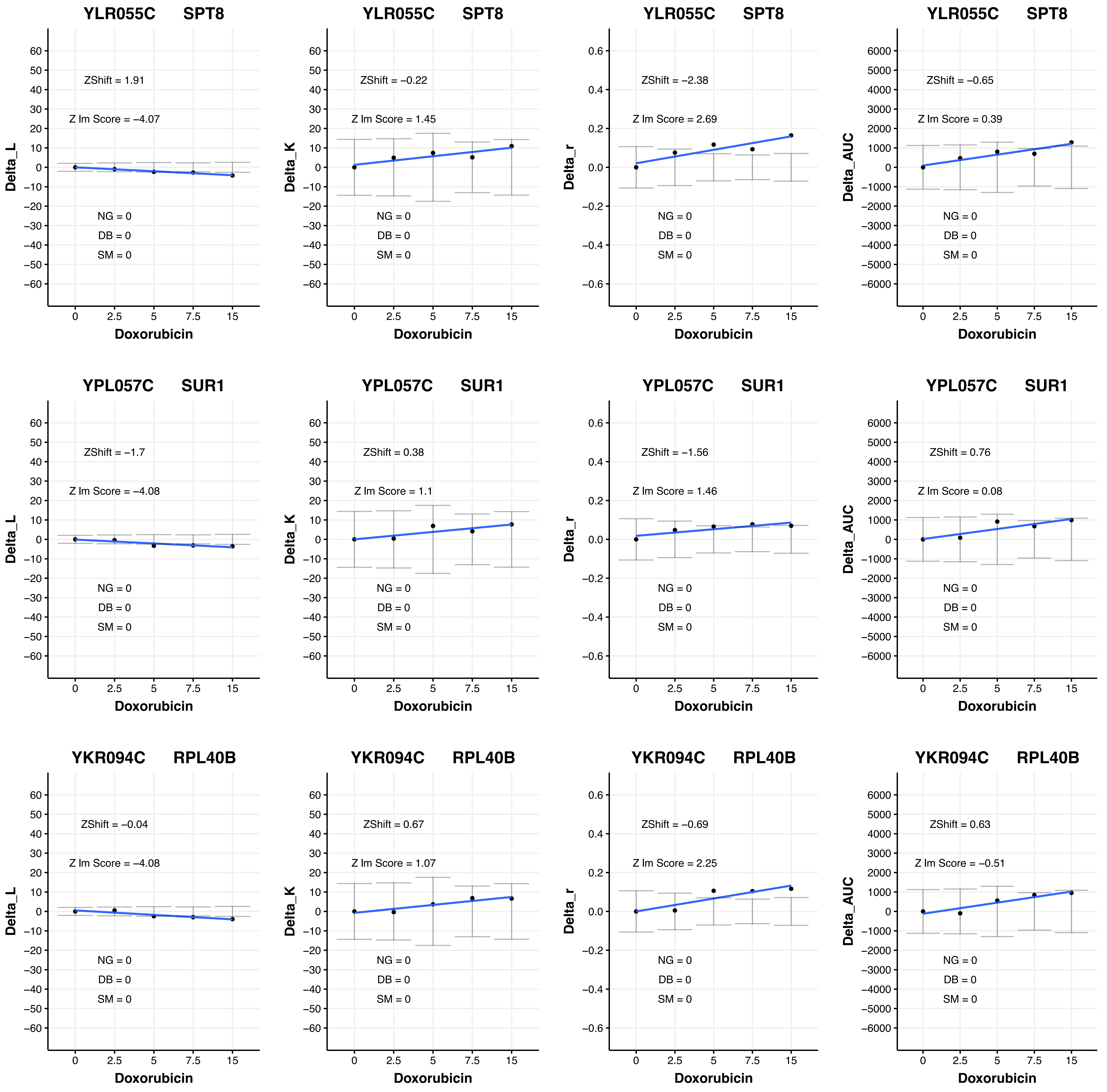

YLR055C SPT8
ZShift = 1.91
Z lm Score = −4.07
Delta_L
ZShift = −0.22
Z lm Score = 1.45
Delta_K
ZShift = −2.38
Z lm Score = 2.69
Delta_r
ZShift = −0.65
Z lm Score = 0.39
Delta_AUC
YPL057C SUR1
ZShift = −1.7
Z lm Score = −4.08
ZShift = 0.38
Z lm Score = 1.1
ZShift = −1.56
Z lm Score = 1.46
ZShift = 0.76
Z lm Score = 0.08
YKR094C RPL40B
ZShift = −0.04
Z lm Score = −4.08
ZShift = 0.67
Z lm Score = 1.07
ZShift = −0.69
Z lm Score = 2.25
ZShift = 0.63
Z lm Score = −0.51
NG = 0
DB = 0
SM = 0
Doxorubicin

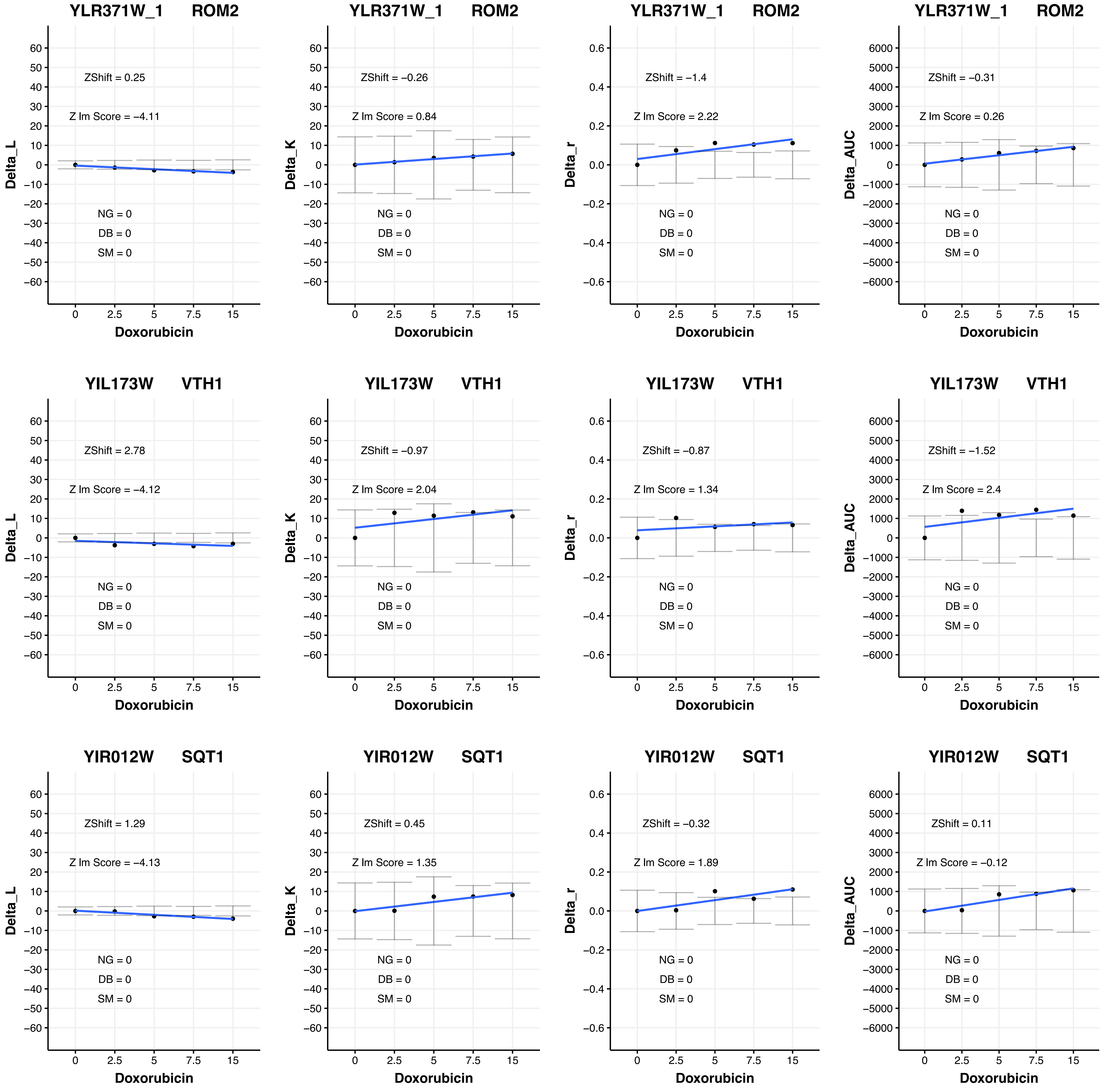
YLR371W_1 ROM2
ZShift = 0.25
Z lm Score = −4.11
NG = 0
DB = 0
SM = 0
Delta_L
Doxorubicin
YLR371W_1 ROM2
ZShift = −0.26
Z lm Score = 0.84
NG = 0
DB = 0
SM = 0
Delta_K
Doxorubicin
YLR371W_1 ROM2
ZShift = −1.4
Z lm Score = 2.22
NG = 0
DB = 0
SM = 0
Delta_r
Doxorubicin
YLR371W_1 ROM2
ZShift = −0.31
Z lm Score = 0.26
NG = 0
DB = 0
SM = 0
Delta_AUC
Doxorubicin
YIL173W VTH1
ZShift = 2.78
Z lm Score = −4.12
NG = 0
DB = 0
SM = 0
Delta_L
Doxorubicin
YIL173W VTH1
ZShift = −0.97
Z lm Score = 2.04
NG = 0
DB = 0
SM = 0
Delta_K
Doxorubicin
YIL173W VTH1
ZShift = −0.87
Z lm Score = 1.34
NG = 0
DB = 0
SM = 0
Delta_r
Doxorubicin
YIL173W VTH1
ZShift = −1.52
Z lm Score = 2.4
NG = 0
DB = 0
SM = 0
Delta_AUC
Doxorubicin
YIR012W SQT1
ZShift = 1.29
Z lm Score = −4.13
NG = 0
DB = 0
SM = 0
Delta_L
Doxorubicin
YIR012W SQT1
ZShift = 0.45
Z lm Score = 1.35
NG = 0
DB = 0
SM = 0
Delta_K
Doxorubicin
YIR012W SQT1
ZShift = −0.32
Z lm Score = 1.89
NG = 0
DB = 0
SM = 0
Delta_r
Doxorubicin
YIR012W SQT1
ZShift = 0.11
Z lm Score = −0.12
NG = 0
DB = 0
SM = 0
Delta_AUC
Doxorubicin

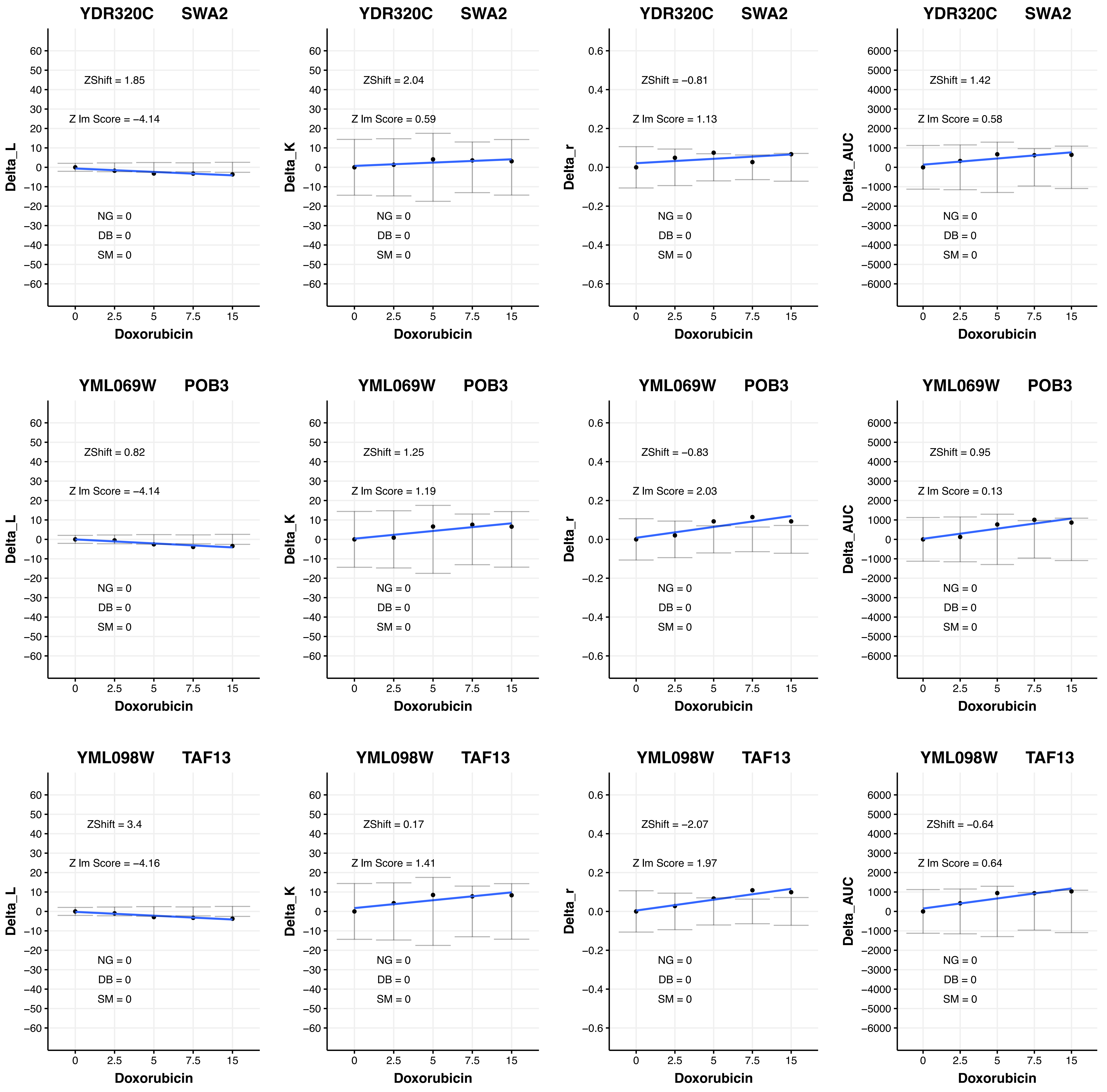
YDR320C SWA2
ZShift = 1.85
Z lm Score = −4.14
NG = 0
DB = 0
SM = 0
Delta_L
Doxorubicin
YDR320C SWA2
ZShift = 2.04
Z lm Score = 0.59
NG = 0
DB = 0
SM = 0
Delta_K
Doxorubicin
YDR320C SWA2
ZShift = −0.81
Z lm Score = 1.13
NG = 0
DB = 0
SM = 0
Delta_r
Doxorubicin
YDR320C SWA2
ZShift = 1.42
Z lm Score = 0.58
NG = 0
DB = 0
SM = 0
Delta_AUC
Doxorubicin
YML069W POB3
ZShift = 0.82
Z lm Score = −4.14
NG = 0
DB = 0
SM = 0
Delta_L
Doxorubicin
YML069W POB3
ZShift = 1.25
Z lm Score = 1.19
NG = 0
DB = 0
SM = 0
Delta_K
Doxorubicin
YML069W POB3
ZShift = −0.83
Z lm Score = 2.03
NG = 0
DB = 0
SM = 0
Delta_r
Doxorubicin
YML069W POB3
ZShift = 0.95
Z lm Score = 0.13
NG = 0
DB = 0
SM = 0
Delta_AUC
Doxorubicin
YML098W TAF13
ZShift = 3.4
Z lm Score = −4.16
NG = 0
DB = 0
SM = 0
Delta_L
Doxorubicin
YML098W TAF13
ZShift = 0.17
Z lm Score = 1.41
NG = 0
DB = 0
SM = 0
Delta_K
Doxorubicin
YML098W TAF13
ZShift = −2.07
Z lm Score = 1.97
NG = 0
DB = 0
SM = 0
Delta_r
Doxorubicin
YML098W TAF13
ZShift = −0.64
Z lm Score = 0.64
NG = 0
DB = 0
SM = 0
Delta_AUC
Doxorubicin

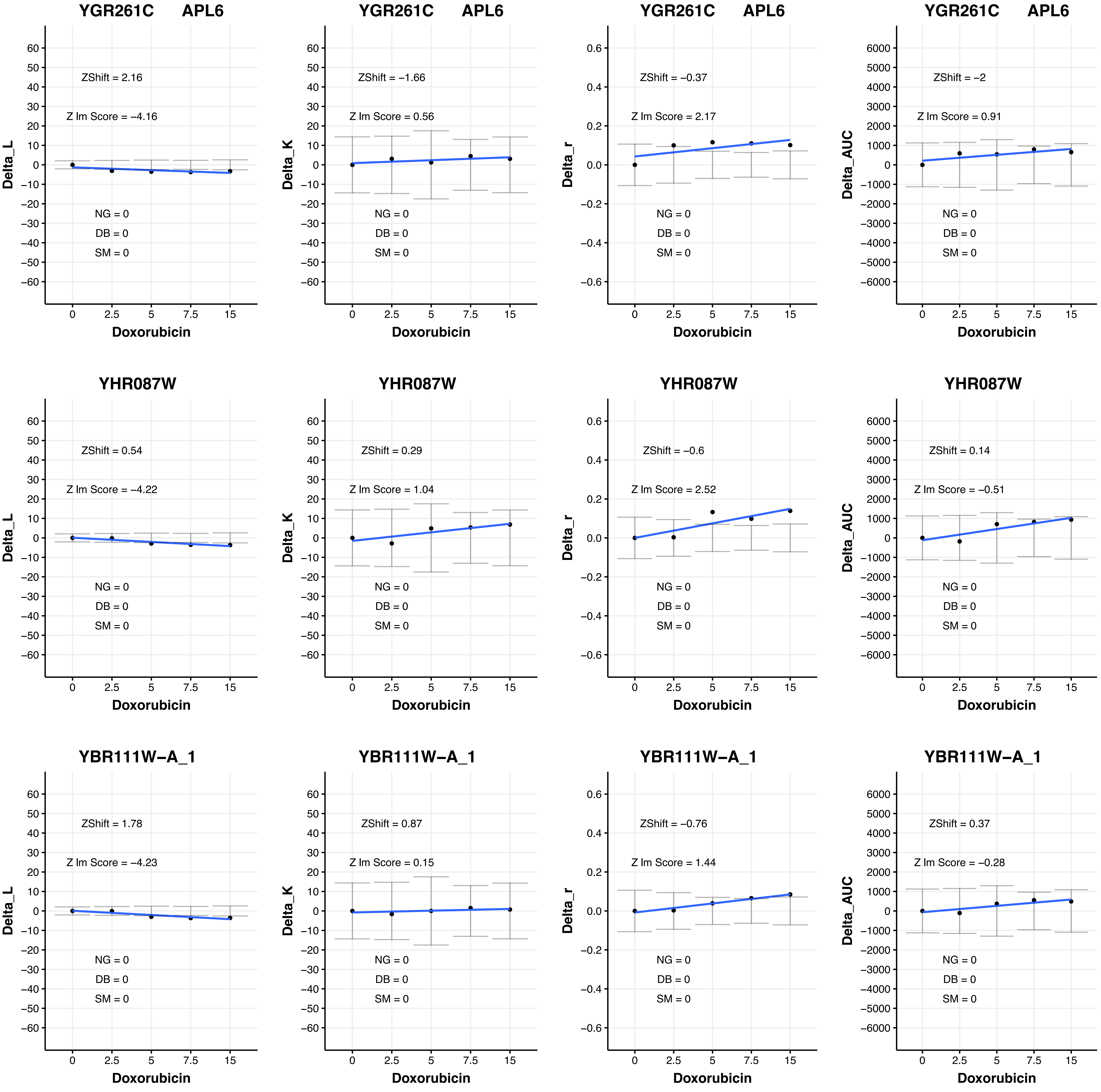
YGR261C APL6
ZShift = 2.16
Z lm Score = −4.16
NG = 0
DB = 0
SM = 0
Delta_L
Doxorubicin
YGR261C APL6
ZShift = −1.66
Z lm Score = 0.56
NG = 0
DB = 0
SM = 0
Delta_K
Doxorubicin
YGR261C APL6
ZShift = −0.37
Z lm Score = 2.17
NG = 0
DB = 0
SM = 0
Delta_r
Doxorubicin
YGR261C APL6
ZShift = −2
Z lm Score = 0.91
NG = 0
DB = 0
SM = 0
Delta_AUC
Doxorubicin
YHR087W
ZShift = 0.54
Z lm Score = −4.22
NG = 0
DB = 0
SM = 0
Delta_L
Doxorubicin
YHR087W
ZShift = 0.29
Z lm Score = 1.04
NG = 0
DB = 0
SM = 0
Delta_K
Doxorubicin
YHR087W
ZShift = −0.6
Z lm Score = 2.52
NG = 0
DB = 0
SM = 0
Delta_r
Doxorubicin
YHR087W
ZShift = 0.14
Z lm Score = −0.51
NG = 0
DB = 0
SM = 0
Delta_AUC
Doxorubicin
YBR111W−A_1
ZShift = 1.78
Z lm Score = −4.23
NG = 0
DB = 0
SM = 0
Delta_L
Doxorubicin
YBR111W−A_1
ZShift = 0.87
Z lm Score = 0.15
NG = 0
DB = 0
SM = 0
Delta_K
Doxorubicin
YBR111W−A_1
ZShift = −0.76
Z lm Score = 1.44
NG = 0
DB = 0
SM = 0
Delta_r
Doxorubicin
YBR111W−A_1
ZShift = 0.37
Z lm Score = −0.28
NG = 0
DB = 0
SM = 0
Delta_AUC
Doxorubicin

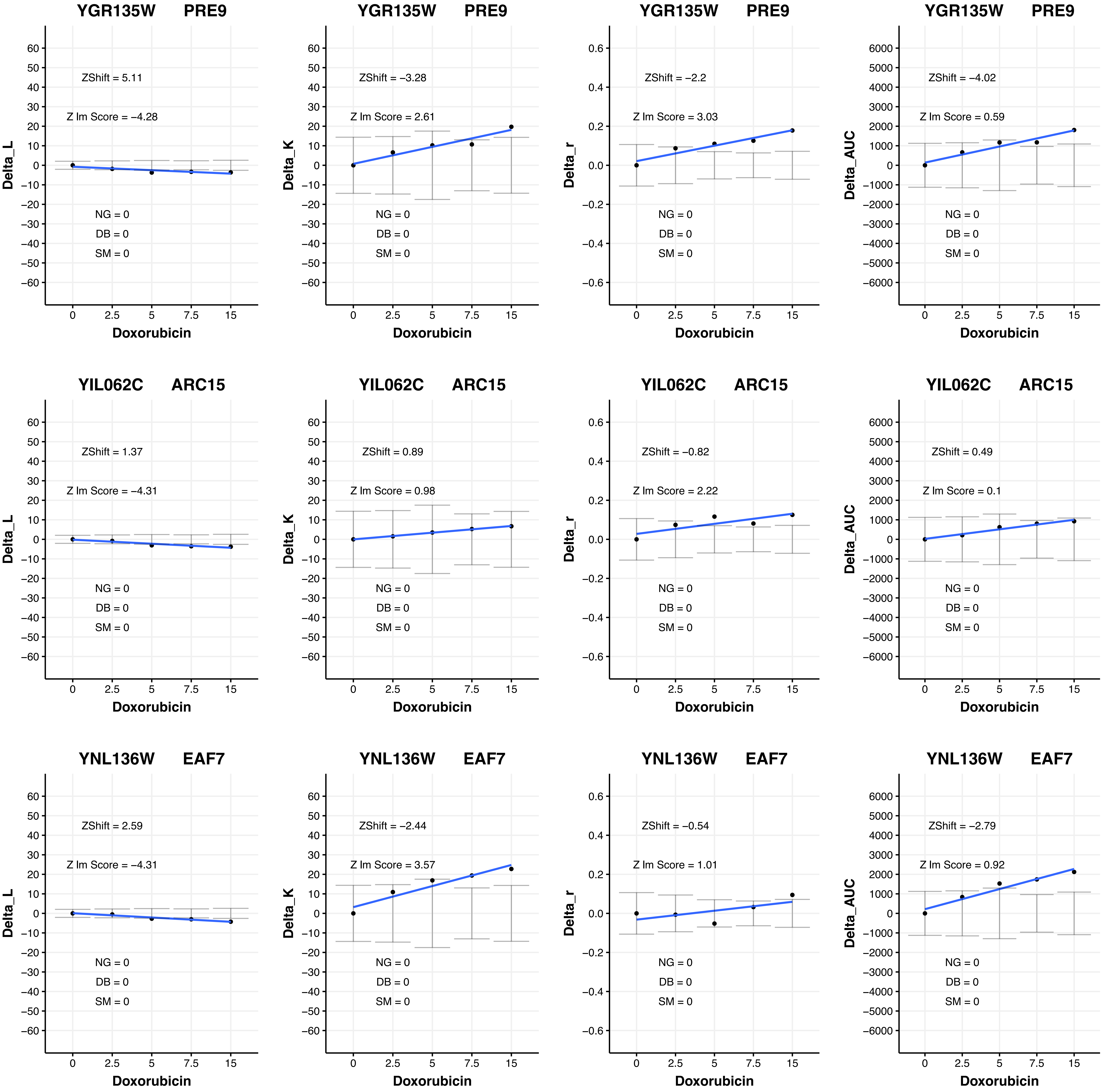
YGR135W PRE9
ZShift = 5.11
Z lm Score = −4.28
NG = 0
DB = 0
SM = 0
Delta_L
Doxorubicin
YGR135W PRE9
ZShift = −3.28
Z lm Score = 2.61
NG = 0
DB = 0
SM = 0
Delta_K
Doxorubicin
YGR135W PRE9
ZShift = −2.2
Z lm Score = 3.03
NG = 0
DB = 0
SM = 0
Delta_r
Doxorubicin
YGR135W PRE9
ZShift = −4.02
Z lm Score = 0.59
NG = 0
DB = 0
SM = 0
Delta_AUC
Doxorubicin
YIL062C ARC15
ZShift = 1.37
Z lm Score = −4.31
NG = 0
DB = 0
SM = 0
Delta_L
Doxorubicin
YIL062C ARC15
ZShift = 0.89
Z lm Score = 0.98
NG = 0
DB = 0
SM = 0
Delta_K
Doxorubicin
YIL062C ARC15
ZShift = −0.82
Z lm Score = 2.22
NG = 0
DB = 0
SM = 0
Delta_r
Doxorubicin
YIL062C ARC15
ZShift = 0.49
Z lm Score = 0.1
NG = 0
DB = 0
SM = 0
Delta_AUC
Doxorubicin
YNL136W EAF7
ZShift = 2.59
Z lm Score = −4.31
NG = 0
DB = 0
SM = 0
Delta_L
Doxorubicin
YNL136W EAF7
ZShift = −2.44
Z lm Score = 3.57
NG = 0
DB = 0
SM = 0
Delta_K
Doxorubicin
YNL136W EAF7
ZShift = −0.54
Z lm Score = 1.01
NG = 0
DB = 0
SM = 0
Delta_r
Doxorubicin
YNL136W EAF7
ZShift = −2.79
Z lm Score = 0.92
NG = 0
DB = 0
SM = 0
Delta_AUC
Doxorubicin

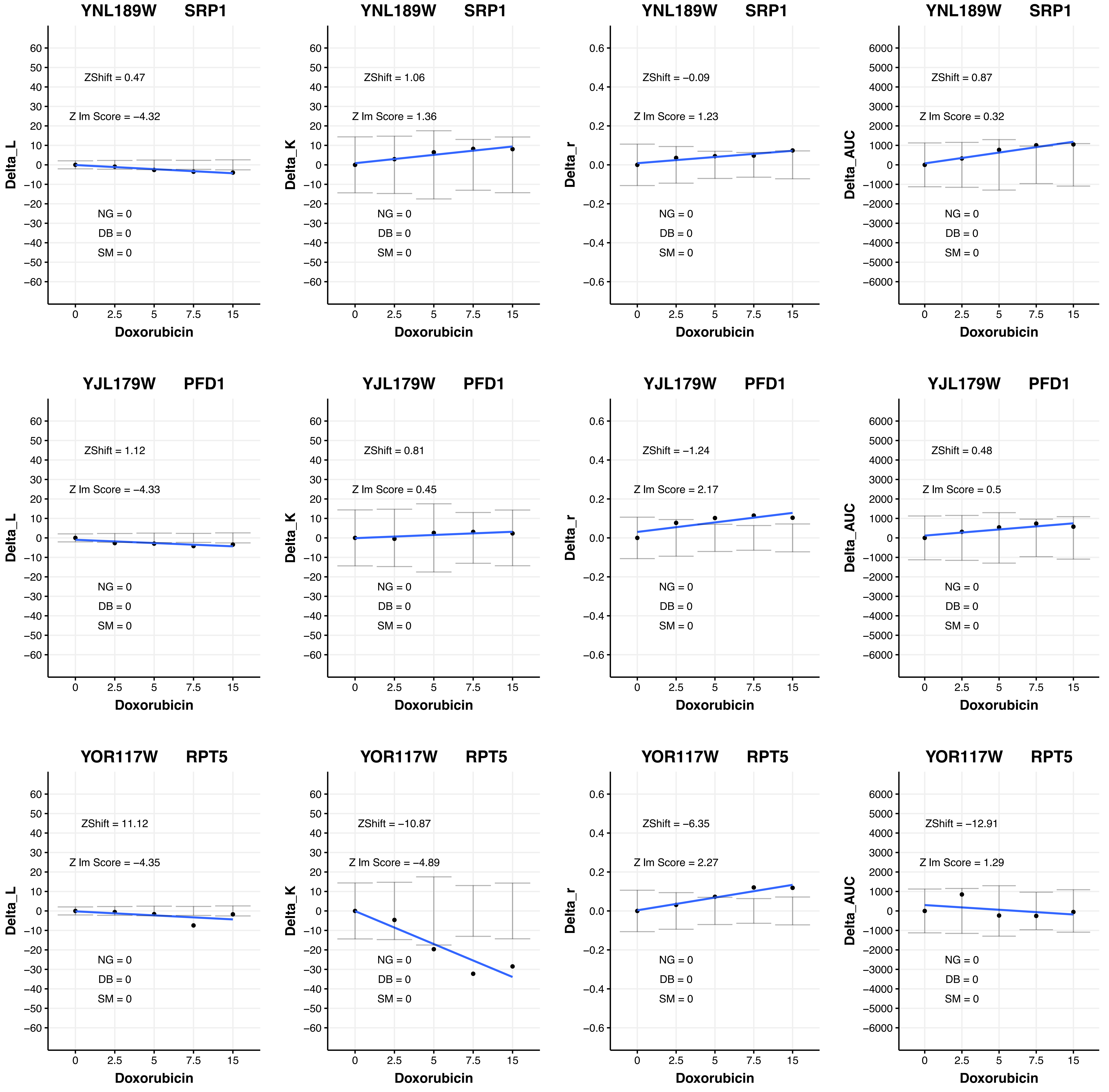
YNL189W SRP1
ZShift = 0.47
Z lm Score = −4.32
NG = 0
DB = 0
SM = 0
Delta_L
Doxorubicin
YNL189W SRP1
ZShift = 1.06
Z lm Score = 1.36
NG = 0
DB = 0
SM = 0
Delta_K
Doxorubicin
YNL189W SRP1
ZShift = −0.09
Z lm Score = 1.23
NG = 0
DB = 0
SM = 0
Delta_r
Doxorubicin
YNL189W SRP1
ZShift = 0.87
Z lm Score = 0.32
NG = 0
DB = 0
SM = 0
Delta_AUC
Doxorubicin
YJL179W PFD1
ZShift = 1.12
Z lm Score = −4.33
NG = 0
DB = 0
SM = 0
Delta_L
Doxorubicin
YJL179W PFD1
ZShift = 0.81
Z lm Score = 0.45
NG = 0
DB = 0
SM = 0
Delta_K
Doxorubicin
YJL179W PFD1
ZShift = −1.24
Z lm Score = 2.17
NG = 0
DB = 0
SM = 0
Delta_r
Doxorubicin
YJL179W PFD1
ZShift = 0.48
Z lm Score = 0.5
NG = 0
DB = 0
SM = 0
Delta_AUC
Doxorubicin
YOR117W RPT5
ZShift = 11.12
Z lm Score = −4.35
NG = 0
DB = 0
SM = 0
Delta_L
Doxorubicin
YOR117W RPT5
ZShift = −10.87
Z lm Score = −4.89
NG = 0
DB = 0
SM = 0
Delta_K
Doxorubicin
YOR117W RPT5
ZShift = −6.35
Z lm Score = 2.27
NG = 0
DB = 0
SM = 0
Delta_r
Doxorubicin
YOR117W RPT5
ZShift = −12.91
Z lm Score = 1.29
NG = 0
DB = 0
SM = 0
Delta_AUC
Doxorubicin

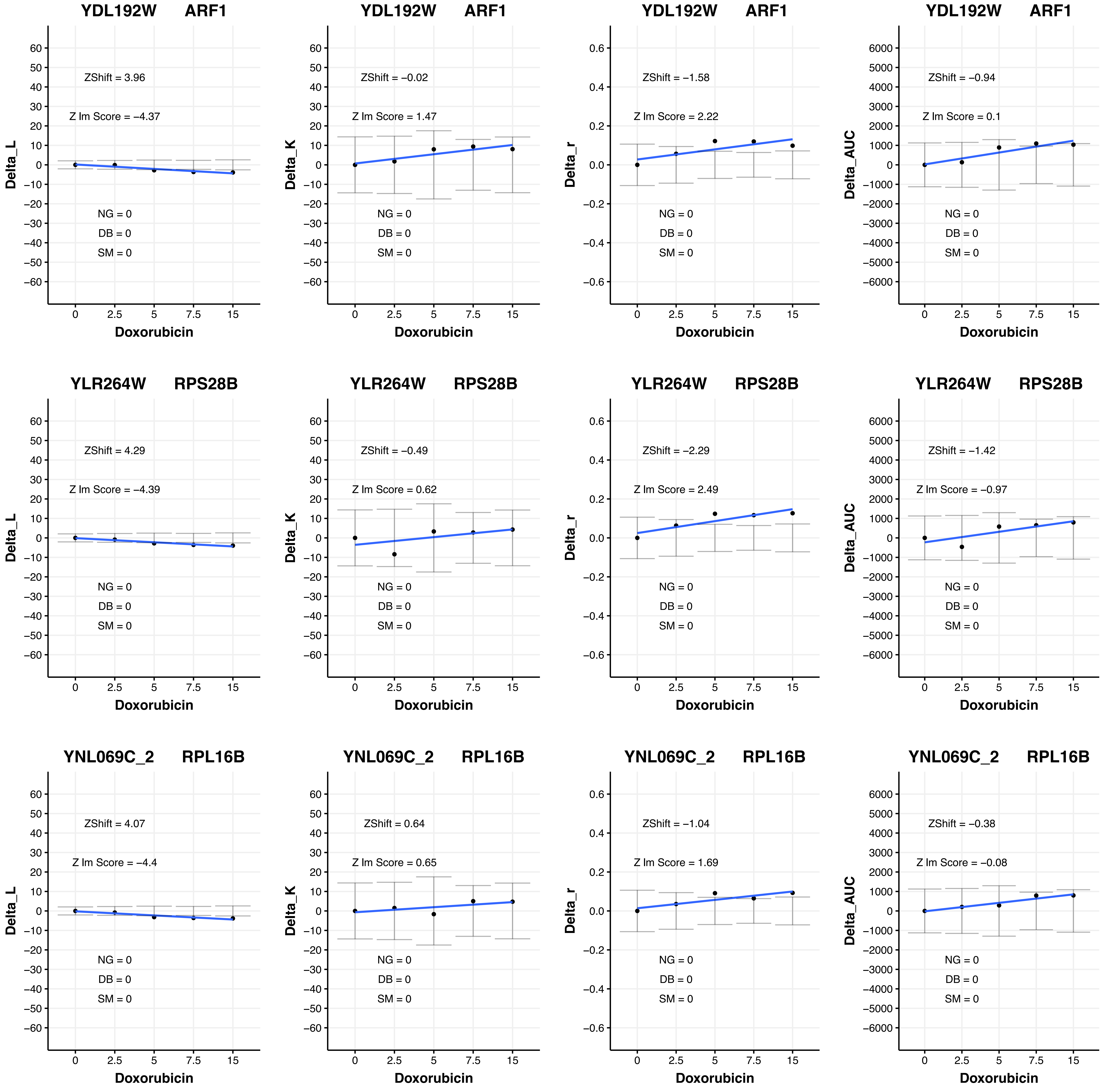

YDL192W ARF1
ZShift = 3.96
Z lm Score = −4.37
Delta_L
NG = 0
DB = 0
SM = 0
Doxorubicin
YDL192W ARF1
ZShift = −0.02
Z lm Score = 1.47
Delta_K
NG = 0
DB = 0
SM = 0
Doxorubicin
YDL192W ARF1
ZShift = −1.58
Z lm Score = 2.22
Delta_r
NG = 0
DB = 0
SM = 0
Doxorubicin
YDL192W ARF1
ZShift = −0.94
Z lm Score = 0.1
Delta_AUC
NG = 0
DB = 0
SM = 0
Doxorubicin
YLR264W RPS28B
ZShift = 4.29
Z lm Score = −4.39
Delta_L
NG = 0
DB = 0
SM = 0
Doxorubicin
YLR264W RPS28B
ZShift = −0.49
Z lm Score = 0.62
Delta_K
NG = 0
DB = 0
SM = 0
Doxorubicin
YLR264W RPS28B
ZShift = −2.29
Z lm Score = 2.49
Delta_r
NG = 0
DB = 0
SM = 0
Doxorubicin
YLR264W RPS28B
ZShift = −1.42
Z lm Score = −0.97
Delta_AUC
NG = 0
DB = 0
SM = 0
Doxorubicin
YNL069C_2 RPL16B
ZShift = 4.07
Z lm Score = −4.4
Delta_L
NG = 0
DB = 0
SM = 0
Doxorubicin
YNL069C_2 RPL16B
ZShift = 0.64
Z lm Score = 0.65
Delta_K
NG = 0
DB = 0
SM = 0
Doxorubicin
YNL069C_2 RPL16B
ZShift = −1.04
Z lm Score = 1.69
Delta_r
NG = 0
DB = 0
SM = 0
Doxorubicin
YNL069C_2 RPL16B
ZShift = −0.38
Z lm Score = −0.08
Delta_AUC
NG = 0
DB = 0
SM = 0
Doxorubicin

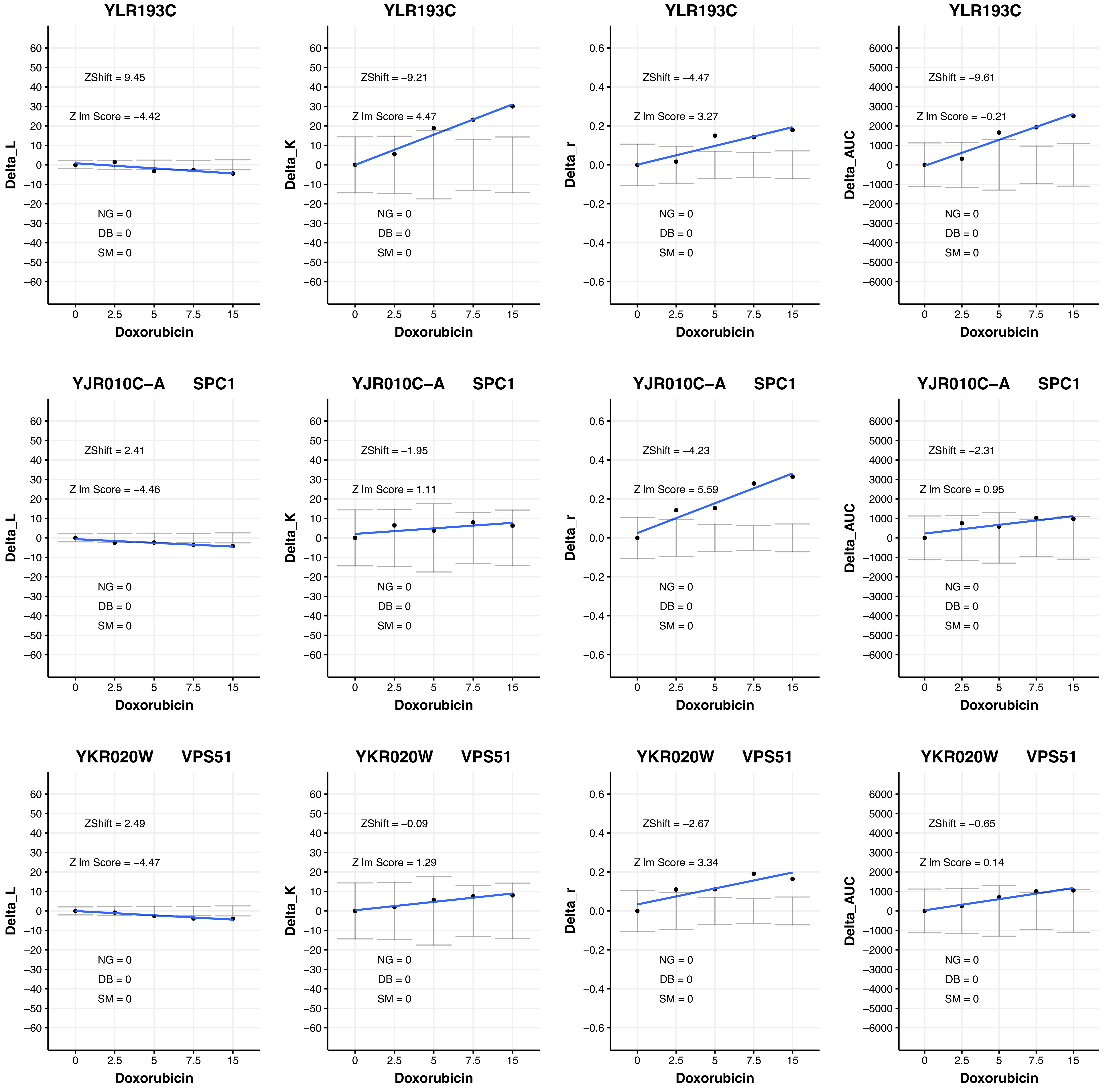
YLR193C
ZShift = 9.45
Z lm Score = −4.42
NG = 0
DB = 0
SM = 0
Delta_L
Doxorubicin
YLR193C
ZShift = −9.21
Z lm Score = 4.47
NG = 0
DB = 0
SM = 0
Delta_K
Doxorubicin
YLR193C
ZShift = −4.47
Z lm Score = 3.27
NG = 0
DB = 0
SM = 0
Delta_r
Doxorubicin
YLR193C
ZShift = −9.61
Z lm Score = −0.21
NG = 0
DB = 0
SM = 0
Delta_AUC
Doxorubicin
YJR010C−A SPC1
ZShift = 2.41
Z lm Score = −4.46
NG = 0
DB = 0
SM = 0
Delta_L
Doxorubicin
YJR010C−A SPC1
ZShift = −1.95
Z lm Score = 1.11
NG = 0
DB = 0
SM = 0
Delta_K
Doxorubicin
YJR010C−A SPC1
ZShift = −4.23
Z lm Score = 5.59
NG = 0
DB = 0
SM = 0
Delta_r
Doxorubicin
YJR010C−A SPC1
ZShift = −2.31
Z lm Score = 0.95
NG = 0
DB = 0
SM = 0
Delta_AUC
Doxorubicin
YKR020W VPS51
ZShift = 2.49
Z lm Score = −4.47
NG = 0
DB = 0
SM = 0
Delta_L
Doxorubicin
YKR020W VPS51
ZShift = −0.09
Z lm Score = 1.29
NG = 0
DB = 0
SM = 0
Delta_K
Doxorubicin
YKR020W VPS51
ZShift = −2.67
Z lm Score = 3.34
NG = 0
DB = 0
SM = 0
Delta_r
Doxorubicin
YKR020W VPS51
ZShift = −0.65
Z lm Score = 0.14
NG = 0
DB = 0
SM = 0
Delta_AUC
Doxorubicin

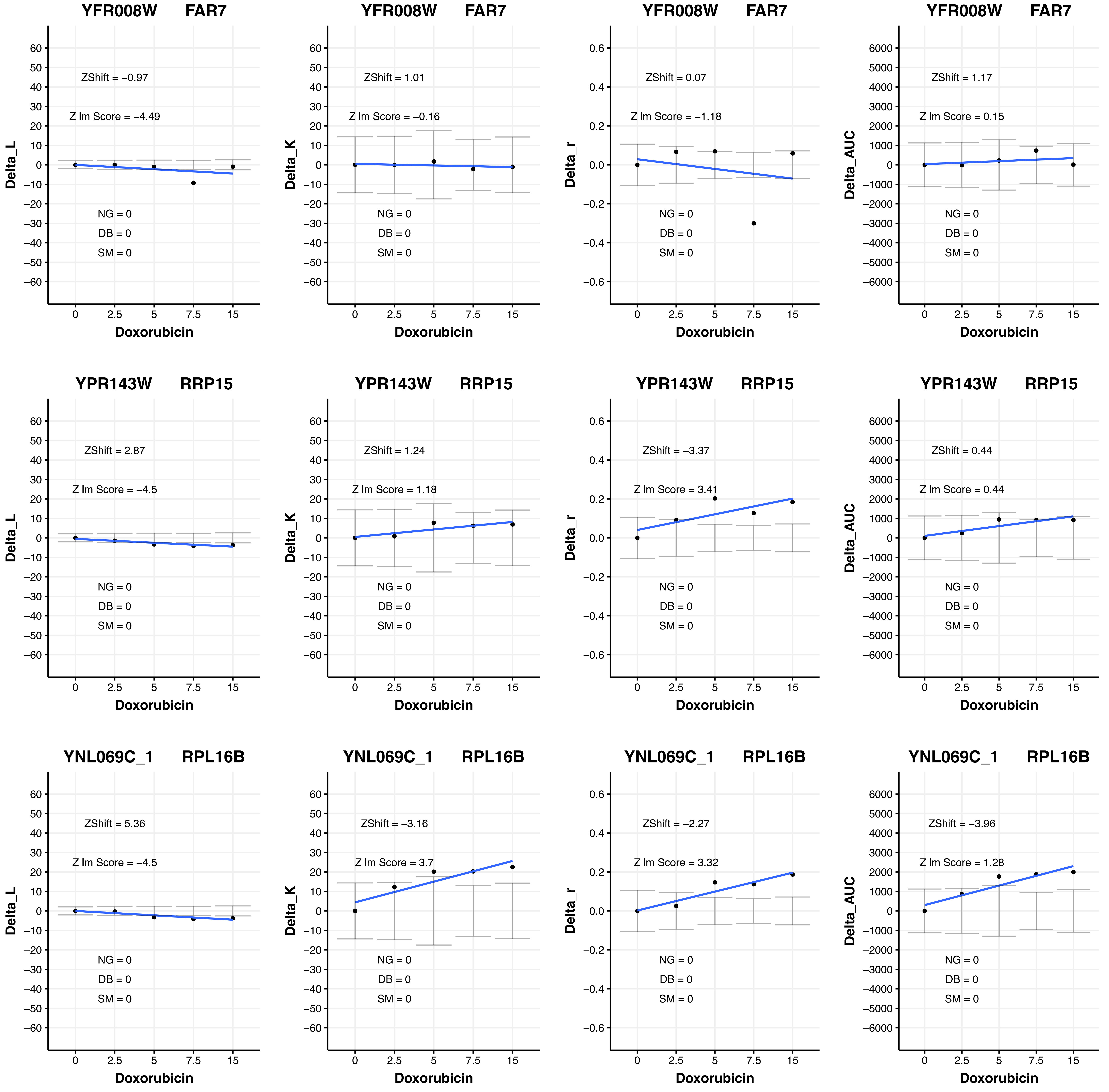
YFR008W FAR7
ZShift = −0.97
Z lm Score = −4.49
NG = 0
DB = 0
SM = 0
Delta_L
Doxorubicin
YFR008W FAR7
ZShift = 1.01
Z lm Score = −0.16
NG = 0
DB = 0
SM = 0
Delta_K
Doxorubicin
YFR008W FAR7
ZShift = 0.07
Z lm Score = −1.18
NG = 0
DB = 0
SM = 0
Delta_r
Doxorubicin
YFR008W FAR7
ZShift = 1.17
Z lm Score = 0.15
NG = 0
DB = 0
SM = 0
Delta_AUC
Doxorubicin
YPR143W RRP15
ZShift = 2.87
Z lm Score = −4.5
NG = 0
DB = 0
SM = 0
Delta_L
Doxorubicin
YPR143W RRP15
ZShift = 1.24
Z lm Score = 1.18
NG = 0
DB = 0
SM = 0
Delta_K
Doxorubicin
YPR143W RRP15
ZShift = −3.37
Z lm Score = 3.41
NG = 0
DB = 0
SM = 0
Delta_r
Doxorubicin
YPR143W RRP15
ZShift = 0.44
Z lm Score = 0.44
NG = 0
DB = 0
SM = 0
Delta_AUC
Doxorubicin
YNL069C_1 RPL16B
ZShift = 5.36
Z lm Score = −4.5
NG = 0
DB = 0
SM = 0
Delta_L
Doxorubicin
YNL069C_1 RPL16B
ZShift = −3.16
Z lm Score = 3.7
NG = 0
DB = 0
SM = 0
Delta_K
Doxorubicin
YNL069C_1 RPL16B
ZShift = −2.27
Z lm Score = 3.32
NG = 0
DB = 0
SM = 0
Delta_r
Doxorubicin
YNL069C_1 RPL16B
ZShift = −3.96
Z lm Score = 1.28
NG = 0
DB = 0
SM = 0
Delta_AUC
Doxorubicin

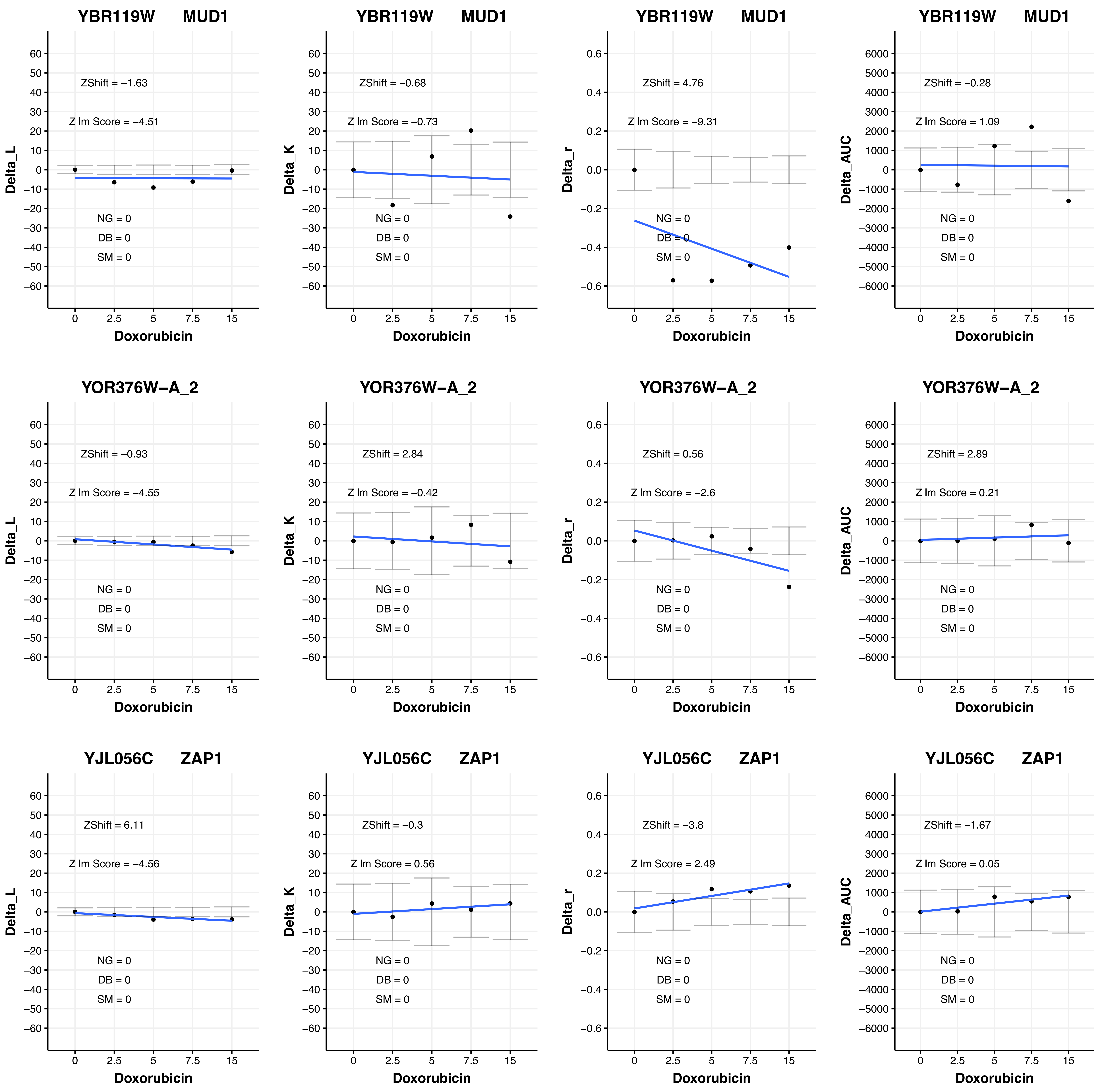
YBR119W MUD1
ZShift = −1.63
Z lm Score = −4.51
NG = 0
DB = 0
SM = 0
Delta_L
Doxorubicin
YBR119W MUD1
ZShift = −0.68
Z lm Score = −0.73
NG = 0
DB = 0
SM = 0
Delta_K
Doxorubicin
YBR119W MUD1
ZShift = 4.76
Z lm Score = −9.31
NG = 0
DB = 0
SM = 0
Delta_r
Doxorubicin
YBR119W MUD1
ZShift = −0.28
Z lm Score = 1.09
NG = 0
DB = 0
SM = 0
Delta_AUC
Doxorubicin
YOR376W−A_2
ZShift = −0.93
Z lm Score = −4.55
NG = 0
DB = 0
SM = 0
Delta_L
Doxorubicin
YOR376W−A_2
ZShift = 2.84
Z lm Score = −0.42
NG = 0
DB = 0
SM = 0
Delta_K
Doxorubicin
YOR376W−A_2
ZShift = 0.56
Z lm Score = −2.6
NG = 0
DB = 0
SM = 0
Delta_r
Doxorubicin
YOR376W−A_2
ZShift = 2.89
Z lm Score = 0.21
NG = 0
DB = 0
SM = 0
Delta_AUC
Doxorubicin
YJL056C ZAP1
ZShift = 6.11
Z lm Score = −4.56
NG = 0
DB = 0
SM = 0
Delta_L
Doxorubicin
YJL056C ZAP1
ZShift = −0.3
Z lm Score = 0.56
NG = 0
DB = 0
SM = 0
Delta_K
Doxorubicin
YJL056C ZAP1
ZShift = −3.8
Z lm Score = 2.49
NG = 0
DB = 0
SM = 0
Delta_r
Doxorubicin
YJL056C ZAP1
ZShift = −1.67
Z lm Score = 0.05
NG = 0
DB = 0
SM = 0
Delta_AUC
Doxorubicin

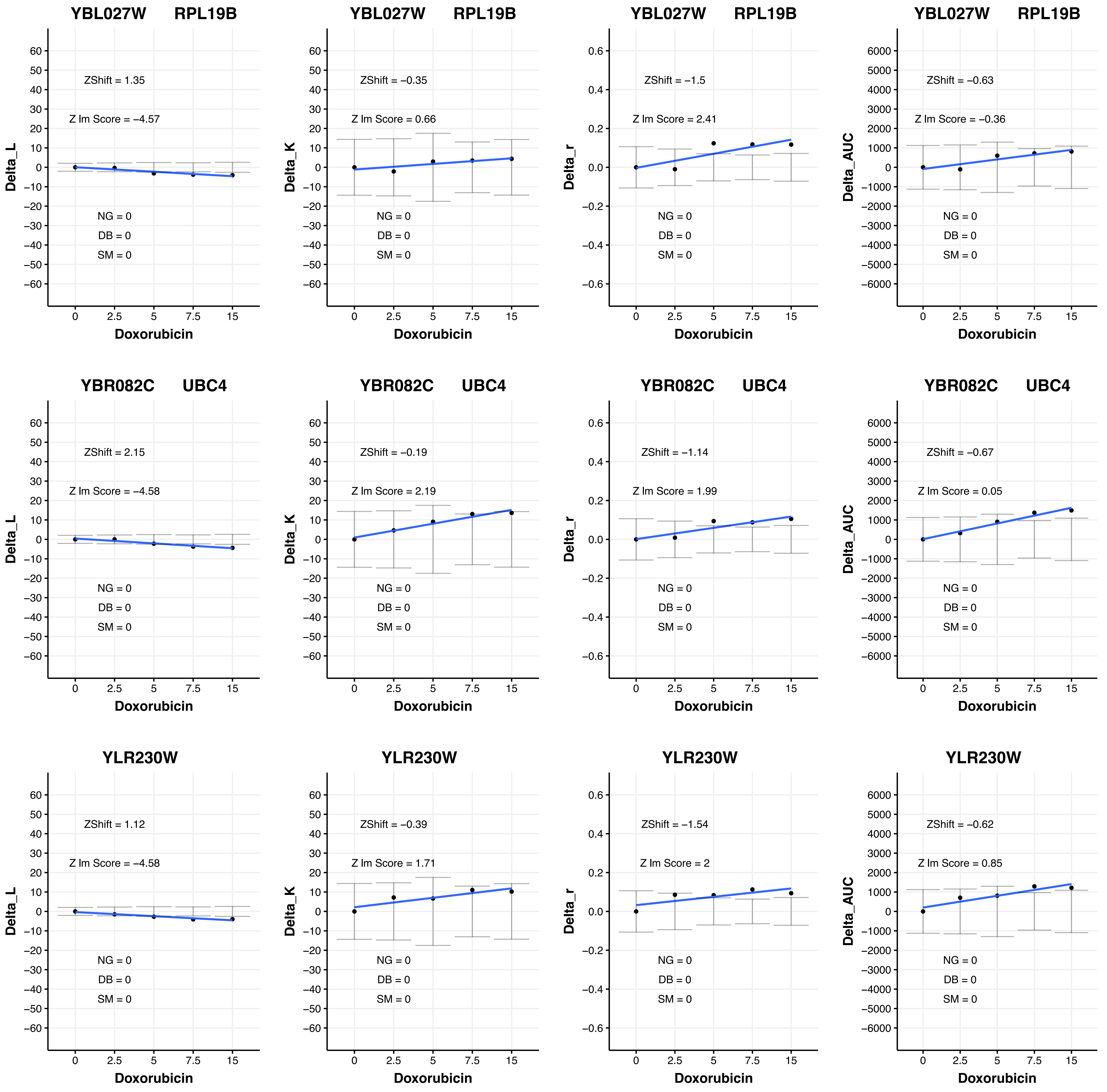
YBL027W RPL19B
ZShift = 1.35
Z lm Score = −4.57
NG = 0
DB = 0
SM = 0
Delta_L
Doxorubicin
YBL027W RPL19B
ZShift = −0.35
Z lm Score = 0.66
NG = 0
DB = 0
SM = 0
Delta_K
Doxorubicin
YBL027W RPL19B
ZShift = −1.5
Z lm Score = 2.41
NG = 0
DB = 0
SM = 0
Delta_r
Doxorubicin
YBL027W RPL19B
ZShift = −0.63
Z lm Score = −0.36
NG = 0
DB = 0
SM = 0
Delta_AUC
Doxorubicin
YBR082C UBC4
ZShift = 2.15
Z lm Score = −4.58
NG = 0
DB = 0
SM = 0
Delta_L
Doxorubicin
YBR082C UBC4
ZShift = −0.19
Z lm Score = 2.19
NG = 0
DB = 0
SM = 0
Delta_K
Doxorubicin
YBR082C UBC4
ZShift = −1.14
Z lm Score = 1.99
NG = 0
DB = 0
SM = 0
Delta_r
Doxorubicin
YBR082C UBC4
ZShift = −0.67
Z lm Score = 0.05
NG = 0
DB = 0
SM = 0
Delta_AUC
Doxorubicin
YLR230W
ZShift = 1.12
Z lm Score = −4.58
NG = 0
DB = 0
SM = 0
Delta_L
Doxorubicin
YLR230W
ZShift = −0.39
Z lm Score = 1.71
NG = 0
DB = 0
SM = 0
Delta_K
Doxorubicin
YLR230W
ZShift = −1.54
Z lm Score = 2
NG = 0
DB = 0
SM = 0
Delta_r
Doxorubicin
YLR230W
ZShift = −0.62
Z lm Score = 0.85
NG = 0
DB = 0
SM = 0
Delta_AUC
Doxorubicin

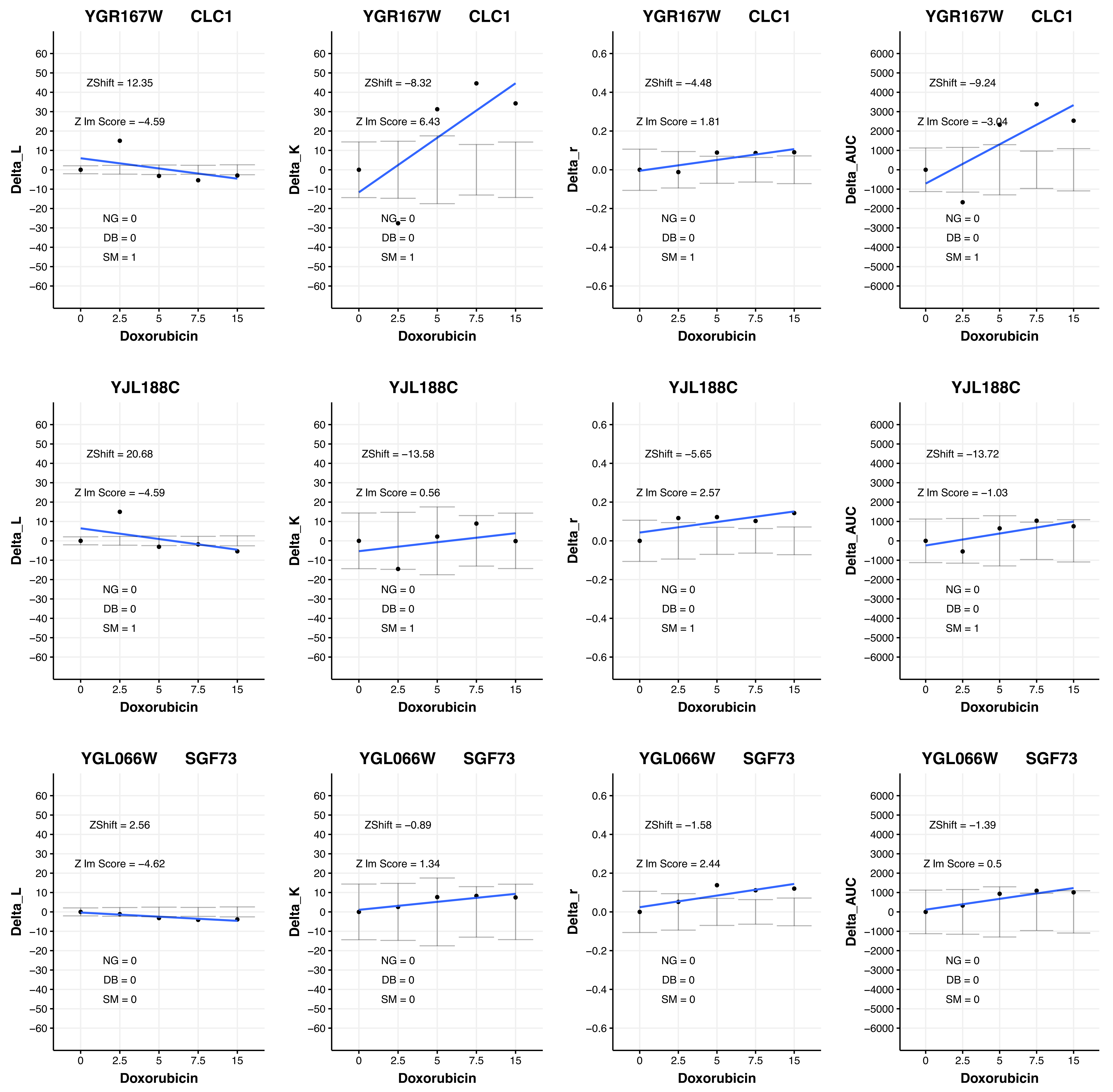
YGR167W CLC1
ZShift = 12.35
Z lm Score = −4.59
NG = 0
DB = 0
SM = 1
Delta_L
Doxorubicin
YGR167W CLC1
ZShift = −8.32
Z lm Score = 6.43
NG = 0
DB = 0
SM = 1
Delta_K
Doxorubicin
YGR167W CLC1
ZShift = −4.48
Z lm Score = 1.81
NG = 0
DB = 0
SM = 1
Delta_r
Doxorubicin
YGR167W CLC1
ZShift = −9.24
Z lm Score = −3.04
NG = 0
DB = 0
SM = 1
Delta_AUC
Doxorubicin
YJL188C
ZShift = 20.68
Z lm Score = −4.59
NG = 0
DB = 0
SM = 1
Delta_L
Doxorubicin
YJL188C
ZShift = −13.58
Z lm Score = 0.56
NG = 0
DB = 0
SM = 1
Delta_K
Doxorubicin
YJL188C
ZShift = −5.65
Z lm Score = 2.57
NG = 0
DB = 0
SM = 1
Delta_r
Doxorubicin
YJL188C
ZShift = −13.72
Z lm Score = −1.03
NG = 0
DB = 0
SM = 1
Delta_AUC
Doxorubicin
YGL066W SGF73
ZShift = 2.56
Z lm Score = −4.62
NG = 0
DB = 0
SM = 0
Delta_L
Doxorubicin
YGL066W SGF73
ZShift = −0.89
Z lm Score = 1.34
NG = 0
DB = 0
SM = 0
Delta_K
Doxorubicin
YGL066W SGF73
ZShift = −1.58
Z lm Score = 2.44
NG = 0
DB = 0
SM = 0
Delta_r
Doxorubicin
YGL066W SGF73
ZShift = −1.39
Z lm Score = 0.5
NG = 0
DB = 0
SM = 0
Delta_AUC
Doxorubicin

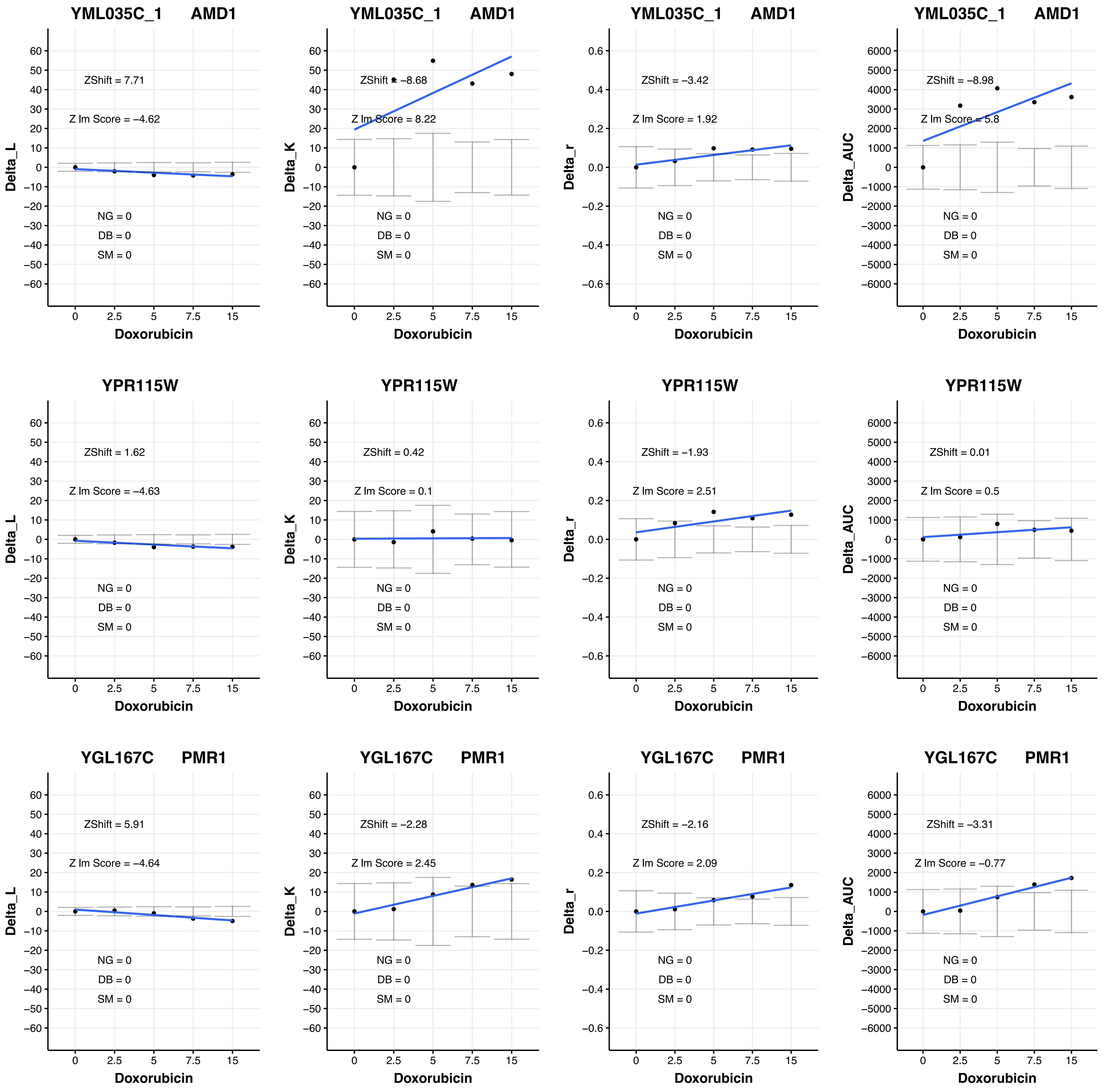

YML035C_1 AMD1
ZShift = 7.71
Z lm Score = −4.62
NG = 0
DB = 0
SM = 0
Delta_L
Doxorubicin
YML035C_1 AMD1
ZShift = −8.68
Z lm Score = 8.22
NG = 0
DB = 0
SM = 0
Delta_K
Doxorubicin
YML035C_1 AMD1
ZShift = −3.42
Z lm Score = 1.92
NG = 0
DB = 0
SM = 0
Delta_r
Doxorubicin
YML035C_1 AMD1
ZShift = −8.98
Z lm Score = 5.8
NG = 0
DB = 0
SM = 0
Delta_AUC
Doxorubicin
YPR115W
ZShift = 1.62
Z lm Score = −4.63
NG = 0
DB = 0
SM = 0
Delta_L
Doxorubicin
YPR115W
ZShift = 0.42
Z lm Score = 0.1
NG = 0
DB = 0
SM = 0
Delta_K
Doxorubicin
YPR115W
ZShift = −1.93
Z lm Score = 2.51
NG = 0
DB = 0
SM = 0
Delta_r
Doxorubicin
YPR115W
ZShift = 0.01
Z lm Score = 0.5
NG = 0
DB = 0
SM = 0
Delta_AUC
Doxorubicin
YGL167C PMR1
ZShift = 5.91
Z lm Score = −4.64
NG = 0
DB = 0
SM = 0
Delta_L
Doxorubicin
YGL167C PMR1
ZShift = −2.28
Z lm Score = 2.45
NG = 0
DB = 0
SM = 0
Delta_K
Doxorubicin
YGL167C PMR1
ZShift = −2.16
Z lm Score = 2.09
NG = 0
DB = 0
SM = 0
Delta_r
Doxorubicin
YGL167C PMR1
ZShift = −3.31
Z lm Score = −0.77
NG = 0
DB = 0
SM = 0
Delta_AUC
Doxorubicin

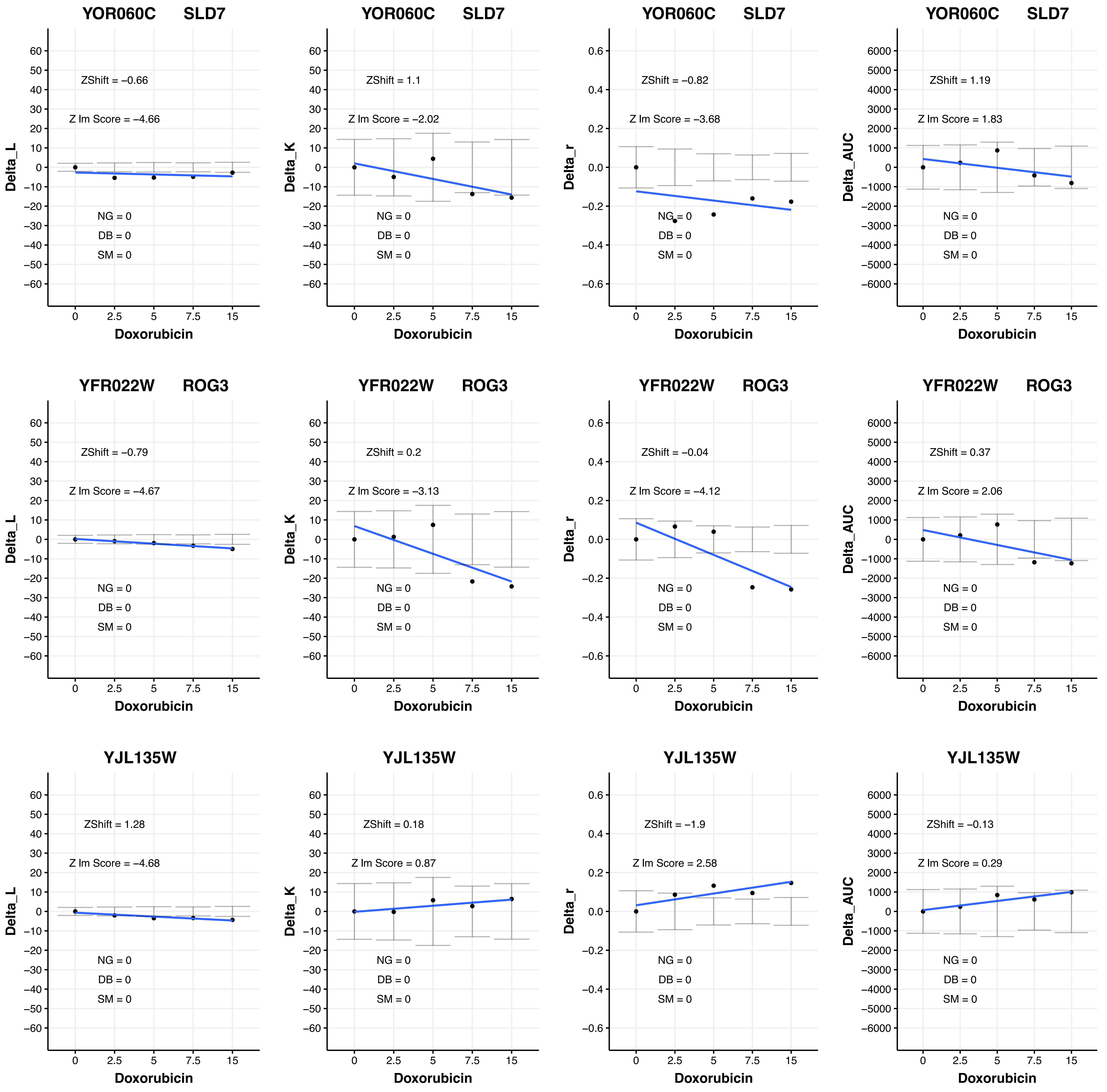
YOR060C SLD7
ZShift = −0.66
Z lm Score = −4.66
NG = 0
DB = 0
SM = 0
Delta_L
Doxorubicin
YOR060C SLD7
ZShift = 1.1
Z lm Score = −2.02
NG = 0
DB = 0
SM = 0
Delta_K
Doxorubicin
YOR060C SLD7
ZShift = −0.82
Z lm Score = −3.68
NG = 0
DB = 0
SM = 0
Delta_r
Doxorubicin
YOR060C SLD7
ZShift = 1.19
Z lm Score = 1.83
NG = 0
DB = 0
SM = 0
Delta_AUC
Doxorubicin
YFR022W ROG3
ZShift = −0.79
Z lm Score = −4.67
NG = 0
DB = 0
SM = 0
Delta_L
Doxorubicin
YFR022W ROG3
ZShift = 0.2
Z lm Score = −3.13
NG = 0
DB = 0
SM = 0
Delta_K
Doxorubicin
YFR022W ROG3
ZShift = −0.04
Z lm Score = −4.12
NG = 0
DB = 0
SM = 0
Delta_r
Doxorubicin
YFR022W ROG3
ZShift = 0.37
Z lm Score = 2.06
NG = 0
DB = 0
SM = 0
Delta_AUC
Doxorubicin
YJL135W
ZShift = 1.28
Z lm Score = −4.68
NG = 0
DB = 0
SM = 0
Delta_L
Doxorubicin
YJL135W
ZShift = 0.18
Z lm Score = 0.87
NG = 0
DB = 0
SM = 0
Delta_K
Doxorubicin
YJL135W
ZShift = −1.9
Z lm Score = 2.58
NG = 0
DB = 0
SM = 0
Delta_r
Doxorubicin
YJL135W
ZShift = −0.13
Z lm Score = 0.29
NG = 0
DB = 0
SM = 0
Delta_AUC
Doxorubicin

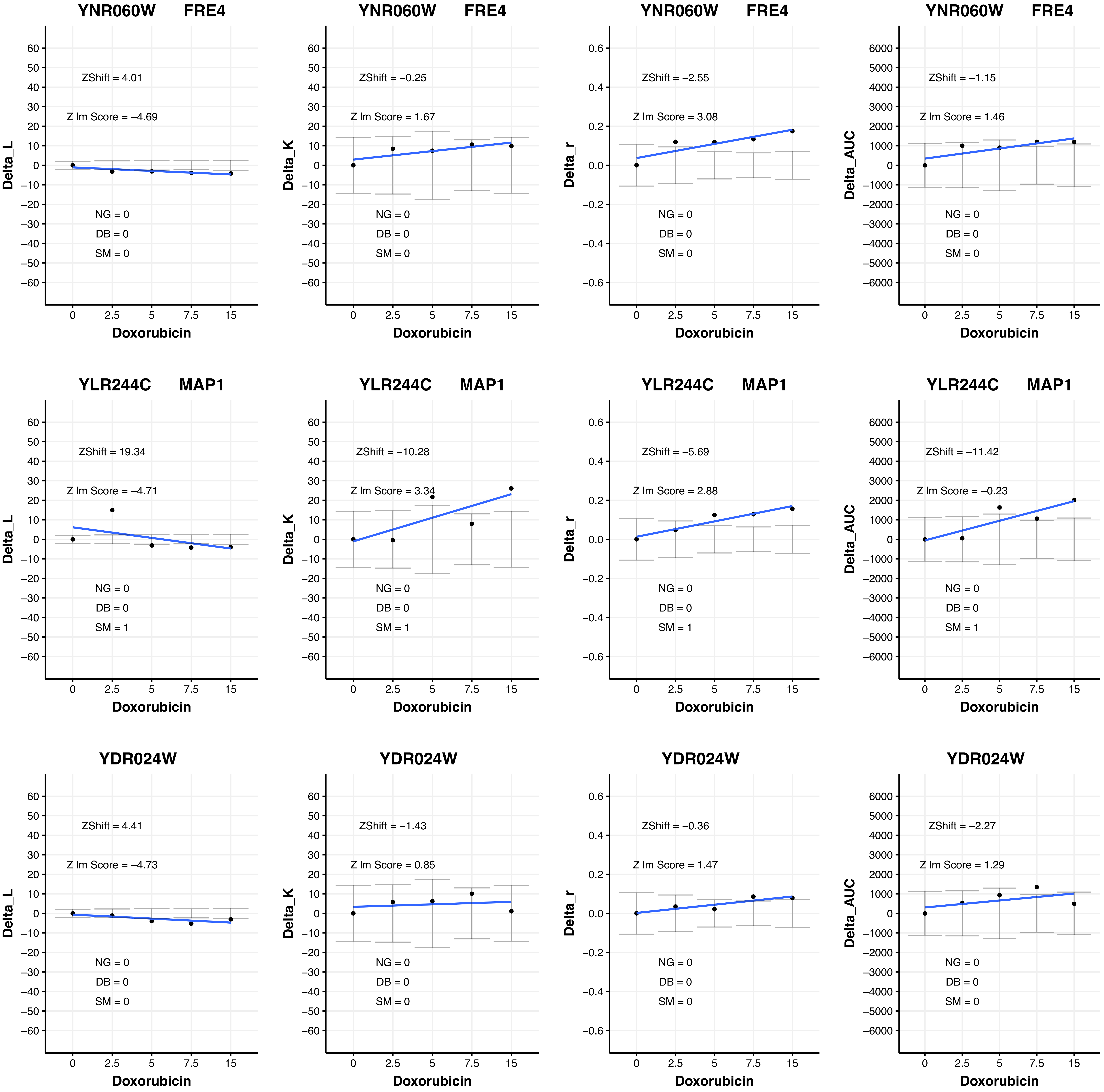
YNR060W FRE4
ZShift = 4.01
Z lm Score = −4.69
NG = 0
DB = 0
SM = 0
Delta_L
Doxorubicin
YNR060W FRE4
ZShift = −0.25
Z lm Score = 1.67
NG = 0
DB = 0
SM = 0
Delta_K
Doxorubicin
YNR060W FRE4
ZShift = −2.55
Z lm Score = 3.08
NG = 0
DB = 0
SM = 0
Delta_r
Doxorubicin
YNR060W FRE4
ZShift = −1.15
Z lm Score = 1.46
NG = 0
DB = 0
SM = 0
Delta_AUC
Doxorubicin
YLR244C MAP1
ZShift = 19.34
Z lm Score = −4.71
NG = 0
DB = 0
SM = 1
Delta_L
Doxorubicin
YLR244C MAP1
ZShift = −10.28
Z lm Score = 3.34
NG = 0
DB = 0
SM = 1
Delta_K
Doxorubicin
YLR244C MAP1
ZShift = −5.69
Z lm Score = 2.88
NG = 0
DB = 0
SM = 1
Delta_r
Doxorubicin
YLR244C MAP1
ZShift = −11.42
Z lm Score = −0.23
NG = 0
DB = 0
SM = 1
Delta_AUC
Doxorubicin
YDR024W
ZShift = 4.41
Z lm Score = −4.73
NG = 0
DB = 0
SM = 0
Delta_L
Doxorubicin
YDR024W
ZShift = −1.43
Z lm Score = 0.85
NG = 0
DB = 0
SM = 0
Delta_K
Doxorubicin
YDR024W
ZShift = −0.36
Z lm Score = 1.47
NG = 0
DB = 0
SM = 0
Delta_r
Doxorubicin
YDR024W
ZShift = −2.27
Z lm Score = 1.29
NG = 0
DB = 0
SM = 0
Delta_AUC
Doxorubicin

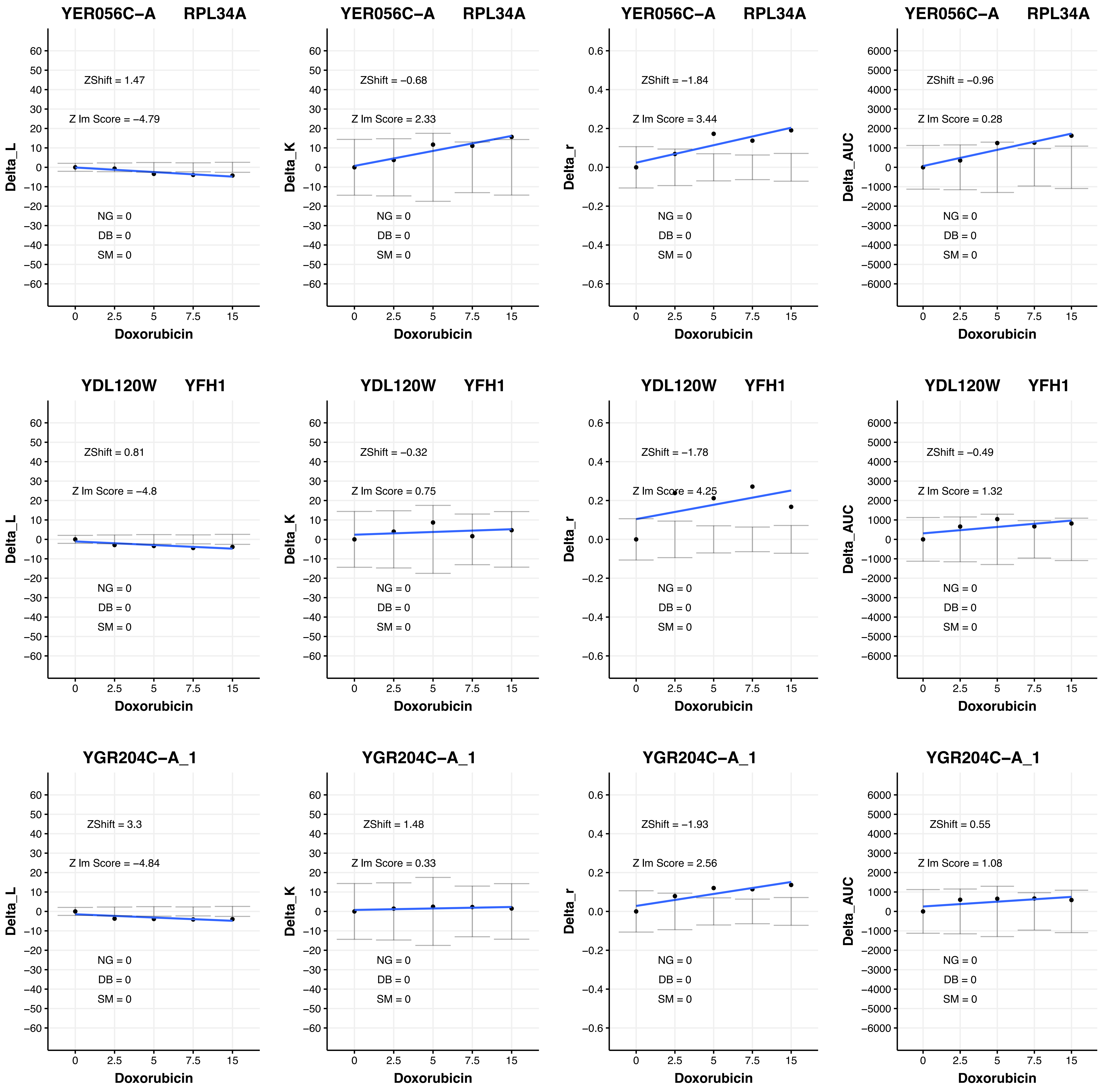

YER056C−A RPL34A
ZShift = 1.47
Z lm Score = −4.79
NG = 0
DB = 0
SM = 0
Delta_L
Doxorubicin
YER056C−A RPL34A
ZShift = −0.68
Z lm Score = 2.33
NG = 0
DB = 0
SM = 0
Delta_K
Doxorubicin
YER056C−A RPL34A
ZShift = −1.84
Z lm Score = 3.44
NG = 0
DB = 0
SM = 0
Delta_r
Doxorubicin
YER056C−A RPL34A
ZShift = −0.96
Z lm Score = 0.28
NG = 0
DB = 0
SM = 0
Delta_AUC
Doxorubicin
YDL120W YFH1
ZShift = 0.81
Z lm Score = −4.8
NG = 0
DB = 0
SM = 0
Delta_L
Doxorubicin
YDL120W YFH1
ZShift = −0.32
Z lm Score = 0.75
NG = 0
DB = 0
SM = 0
Delta_K
Doxorubicin
YDL120W YFH1
ZShift = −1.78
Z lm Score = 4.25
NG = 0
DB = 0
SM = 0
Delta_r
Doxorubicin
YDL120W YFH1
ZShift = −0.49
Z lm Score = 1.32
NG = 0
DB = 0
SM = 0
Delta_AUC
Doxorubicin
YGR204C−A_1
ZShift = 3.3
Z lm Score = −4.84
NG = 0
DB = 0
SM = 0
Delta_L
Doxorubicin
YGR204C−A_1
ZShift = 1.48
Z lm Score = 0.33
NG = 0
DB = 0
SM = 0
Delta_K
Doxorubicin
YGR204C−A_1
ZShift = −1.93
Z lm Score = 2.56
NG = 0
DB = 0
SM = 0
Delta_r
Doxorubicin
YGR204C−A_1
ZShift = 0.55
Z lm Score = 1.08
NG = 0
DB = 0
SM = 0
Delta_AUC
Doxorubicin

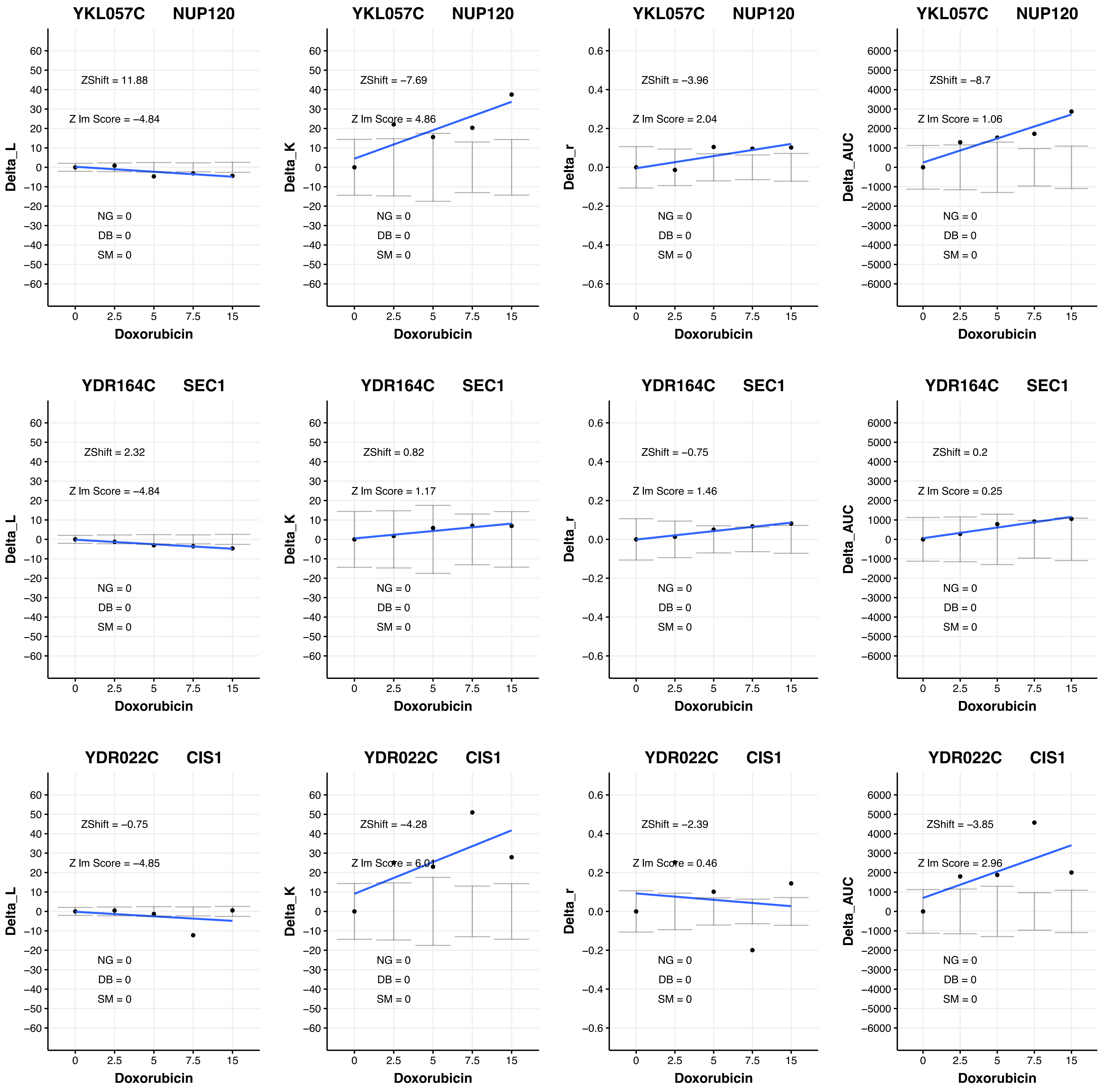

YKL057C NUP120
ZShift = 11.88
Z lm Score = −4.84
NG = 0
DB = 0
SM = 0
Delta_L
Doxorubicin
YKL057C NUP120
ZShift = −7.69
Z lm Score = 4.86
NG = 0
DB = 0
SM = 0
Delta_K
Doxorubicin
YKL057C NUP120
ZShift = −3.96
Z lm Score = 2.04
NG = 0
DB = 0
SM = 0
Delta_r
Doxorubicin
YKL057C NUP120
ZShift = −8.7
Z lm Score = 1.06
NG = 0
DB = 0
SM = 0
Delta_AUC
Doxorubicin
YDR164C SEC1
ZShift = 2.32
Z lm Score = −4.84
NG = 0
DB = 0
SM = 0
Delta_L
Doxorubicin
YDR164C SEC1
ZShift = 0.82
Z lm Score = 1.17
NG = 0
DB = 0
SM = 0
Delta_K
Doxorubicin
YDR164C SEC1
ZShift = −0.75
Z lm Score = 1.46
NG = 0
DB = 0
SM = 0
Delta_r
Doxorubicin
YDR164C SEC1
ZShift = 0.2
Z lm Score = 0.25
NG = 0
DB = 0
SM = 0
Delta_AUC
Doxorubicin
YDR022C CIS1
ZShift = −0.75
Z lm Score = −4.85
NG = 0
DB = 0
SM = 0
Delta_L
Doxorubicin
YDR022C CIS1
ZShift = −4.28
Z lm Score = 6.01
NG = 0
DB = 0
SM = 0
Delta_K
Doxorubicin
YDR022C CIS1
ZShift = −2.39
Z lm Score = 0.46
NG = 0
DB = 0
SM = 0
Delta_r
Doxorubicin
YDR022C CIS1
ZShift = −3.85
Z lm Score = 2.96
NG = 0
DB = 0
SM = 0
Delta_AUC
Doxorubicin

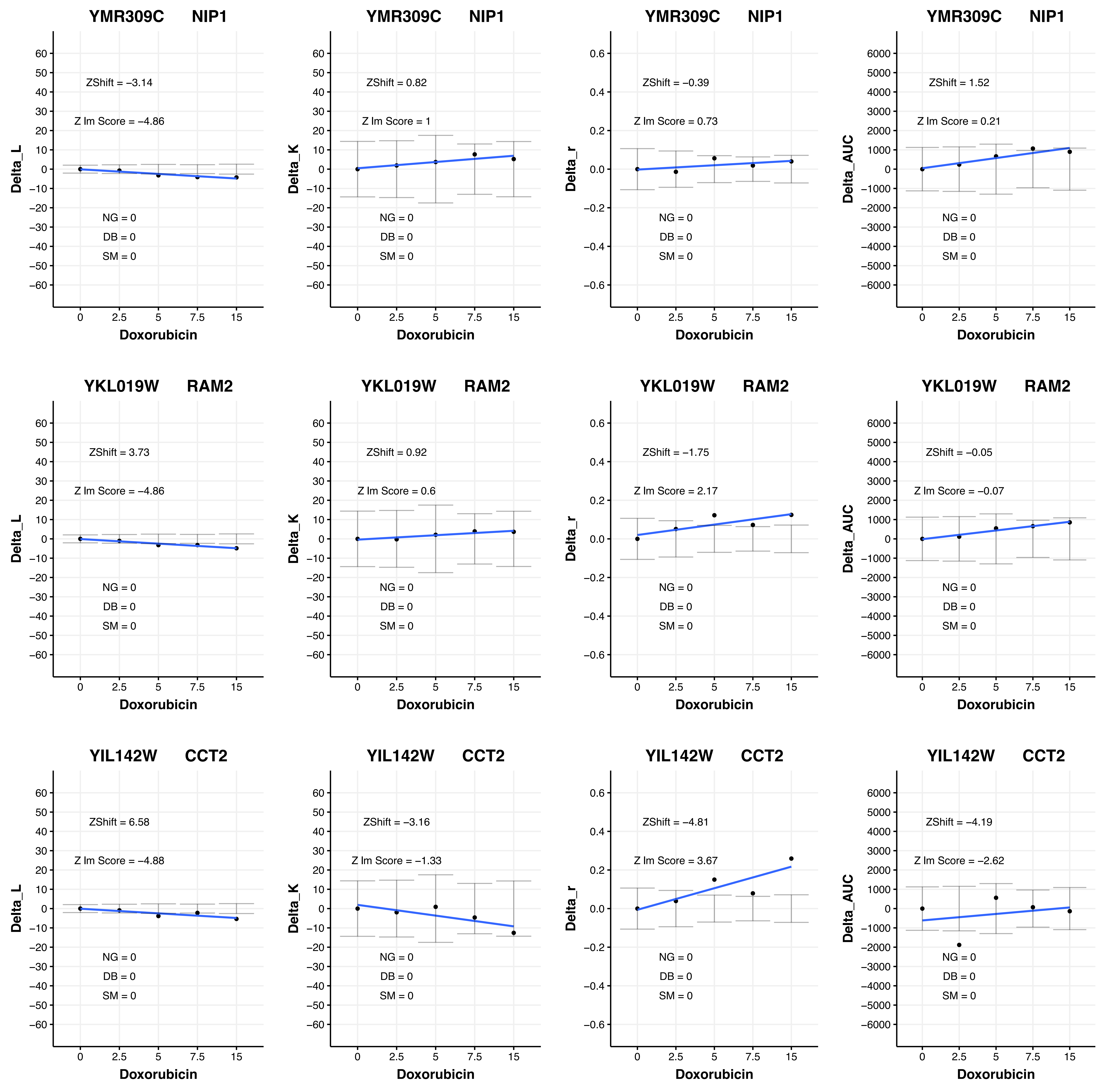
YMR309C NIP1
ZShift = −3.14
Z lm Score = −4.86
NG = 0
DB = 0
SM = 0
Delta_L
Doxorubicin
YMR309C NIP1
ZShift = 0.82
Z lm Score = 1
NG = 0
DB = 0
SM = 0
Delta_K
Doxorubicin
YMR309C NIP1
ZShift = −0.39
Z lm Score = 0.73
NG = 0
DB = 0
SM = 0
Delta_r
Doxorubicin
YMR309C NIP1
ZShift = 1.52
Z lm Score = 0.21
NG = 0
DB = 0
SM = 0
Delta_AUC
Doxorubicin
YKL019W RAM2
ZShift = 3.73
Z lm Score = −4.86
NG = 0
DB = 0
SM = 0
Delta_L
Doxorubicin
YKL019W RAM2
ZShift = 0.92
Z lm Score = 0.6
NG = 0
DB = 0
SM = 0
Delta_K
Doxorubicin
YKL019W RAM2
ZShift = −1.75
Z lm Score = 2.17
NG = 0
DB = 0
SM = 0
Delta_r
Doxorubicin
YKL019W RAM2
ZShift = −0.05
Z lm Score = −0.07
NG = 0
DB = 0
SM = 0
Delta_AUC
Doxorubicin
YIL142W CCT2
ZShift = 6.58
Z lm Score = −4.88
NG = 0
DB = 0
SM = 0
Delta_L
Doxorubicin
YIL142W CCT2
ZShift = −3.16
Z lm Score = −1.33
NG = 0
DB = 0
SM = 0
Delta_K
Doxorubicin
YIL142W CCT2
ZShift = −4.81
Z lm Score = 3.67
NG = 0
DB = 0
SM = 0
Delta_r
Doxorubicin
YIL142W CCT2
ZShift = −4.19
Z lm Score = −2.62
NG = 0
DB = 0
SM = 0
Delta_AUC
Doxorubicin

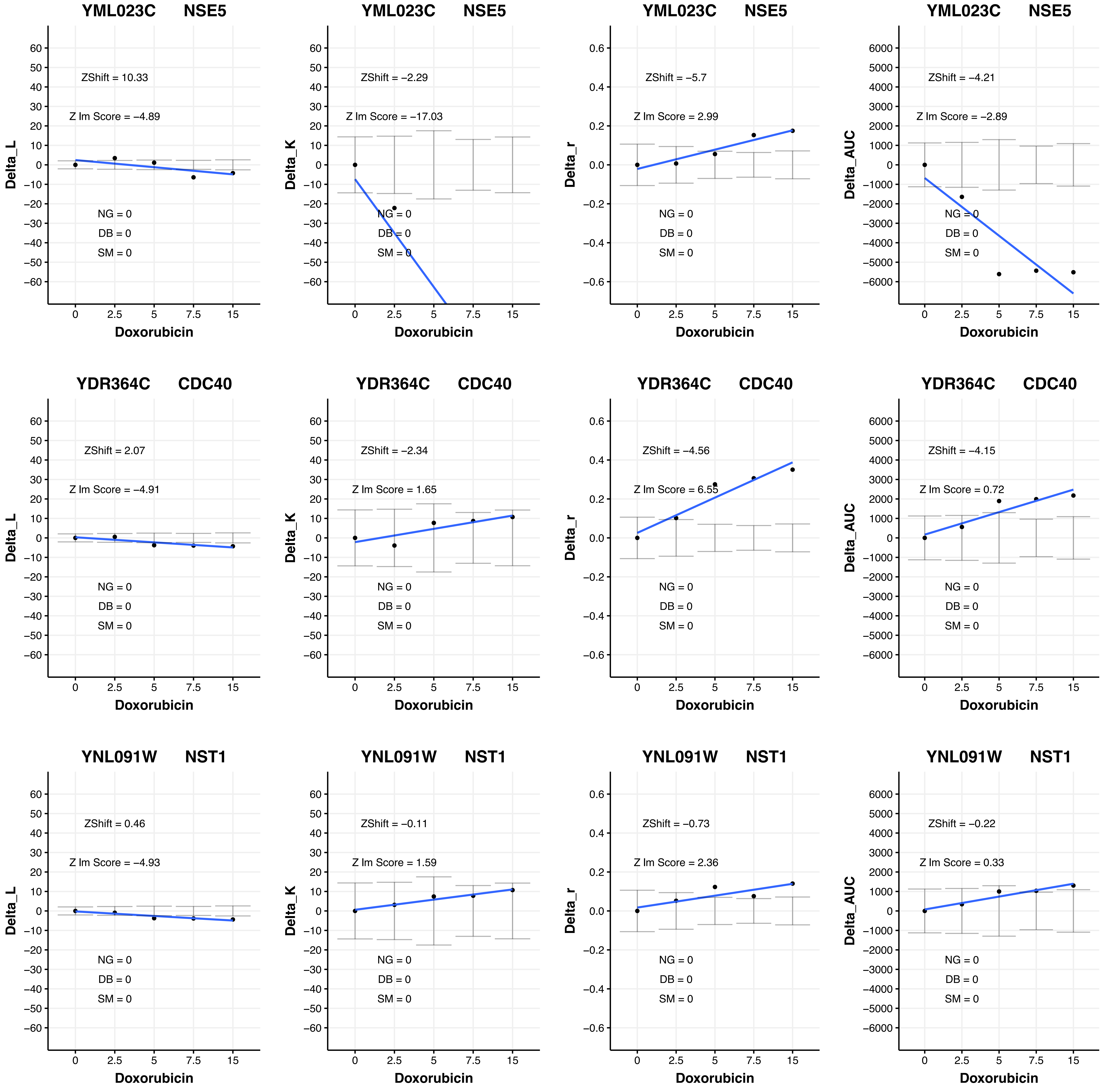
YML023C NSE5
ZShift = 10.33
Z lm Score = −4.89
NG = 0
DB = 0
SM = 0
Delta_L
Doxorubicin
YML023C NSE5
ZShift = −2.29
Z lm Score = −17.03
NG = 0
DB = 0
SM = 0
Delta_K
Doxorubicin
YML023C NSE5
ZShift = −5.7
Z lm Score = 2.99
NG = 0
DB = 0
SM = 0
Delta_r
Doxorubicin
YML023C NSE5
ZShift = −4.21
Z lm Score = −2.89
NG = 0
DB = 0
SM = 0
Delta_AUC
Doxorubicin
YDR364C CDC40
ZShift = 2.07
Z lm Score = −4.91
NG = 0
DB = 0
SM = 0
Delta_L
Doxorubicin
YDR364C CDC40
ZShift = −2.34
Z lm Score = 1.65
NG = 0
DB = 0
SM = 0
Delta_K
Doxorubicin
YDR364C CDC40
ZShift = −4.56
Z lm Score = 6.55
NG = 0
DB = 0
SM = 0
Delta_r
Doxorubicin
YDR364C CDC40
ZShift = −4.15
Z lm Score = 0.72
NG = 0
DB = 0
SM = 0
Delta_AUC
Doxorubicin
YNL091W NST1
ZShift = 0.46
Z lm Score = −4.93
NG = 0
DB = 0
SM = 0
Delta_L
Doxorubicin
YNL091W NST1
ZShift = −0.11
Z lm Score = 1.59
NG = 0
DB = 0
SM = 0
Delta_K
Doxorubicin
YNL091W NST1
ZShift = −0.73
Z lm Score = 2.36
NG = 0
DB = 0
SM = 0
Delta_r
Doxorubicin
YNL091W NST1
ZShift = −0.22
Z lm Score = 0.33
NG = 0
DB = 0
SM = 0
Delta_AUC
Doxorubicin

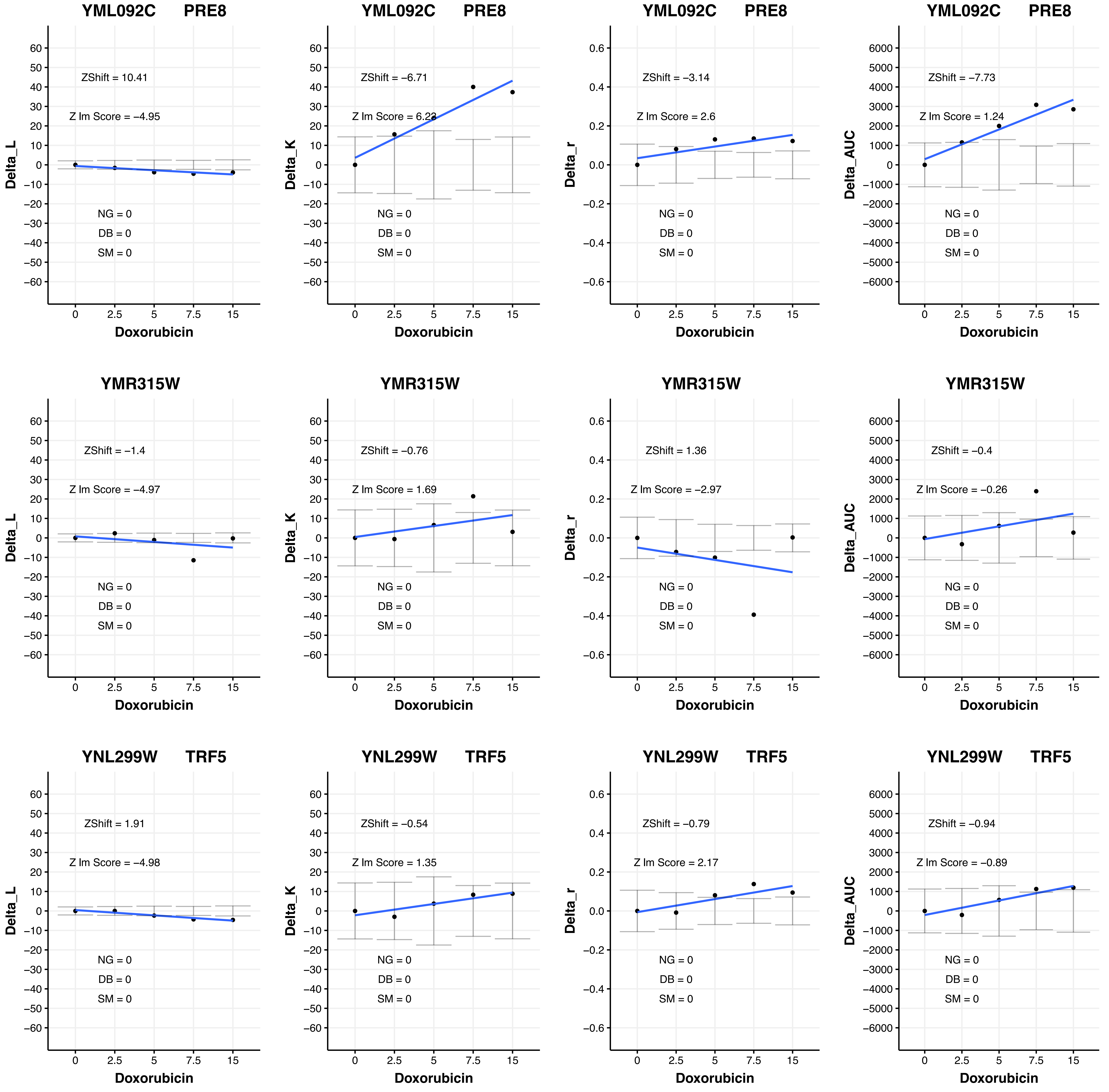
YML092C PRE8
ZShift = 10.41
Z lm Score = −4.95
Delta_L
NG = 0
DB = 0
SM = 0
Doxorubicin
YML092C PRE8
ZShift = −6.71
Z lm Score = 6.22
Delta_K
NG = 0
DB = 0
SM = 0
Doxorubicin
YML092C PRE8
ZShift = −3.14
Z lm Score = 2.6
Delta_r
NG = 0
DB = 0
SM = 0
Doxorubicin
YML092C PRE8
ZShift = −7.73
Z lm Score = 1.24
Delta_AUC
NG = 0
DB = 0
SM = 0
Doxorubicin
YMR315W
ZShift = −1.4
Z lm Score = −4.97
Delta_L
NG = 0
DB = 0
SM = 0
Doxorubicin
YMR315W
ZShift = −0.76
Z lm Score = 1.69
Delta_K
NG = 0
DB = 0
SM = 0
Doxorubicin
YMR315W
ZShift = 1.36
Z lm Score = −2.97
Delta_r
NG = 0
DB = 0
SM = 0
Doxorubicin
YMR315W
ZShift = −0.4
Z lm Score = −0.26
Delta_AUC
NG = 0
DB = 0
SM = 0
Doxorubicin
YNL299W TRF5
ZShift = 1.91
Z lm Score = −4.98
Delta_L
NG = 0
DB = 0
SM = 0
Doxorubicin
YNL299W TRF5
ZShift = −0.54
Z lm Score = 1.35
Delta_K
NG = 0
DB = 0
SM = 0
Doxorubicin
YNL299W TRF5
ZShift = −0.79
Z lm Score = 2.17
Delta_r
NG = 0
DB = 0
SM = 0
Doxorubicin
YNL299W TRF5
ZShift = −0.94
Z lm Score = −0.89
Delta_AUC
NG = 0
DB = 0
SM = 0
Doxorubicin

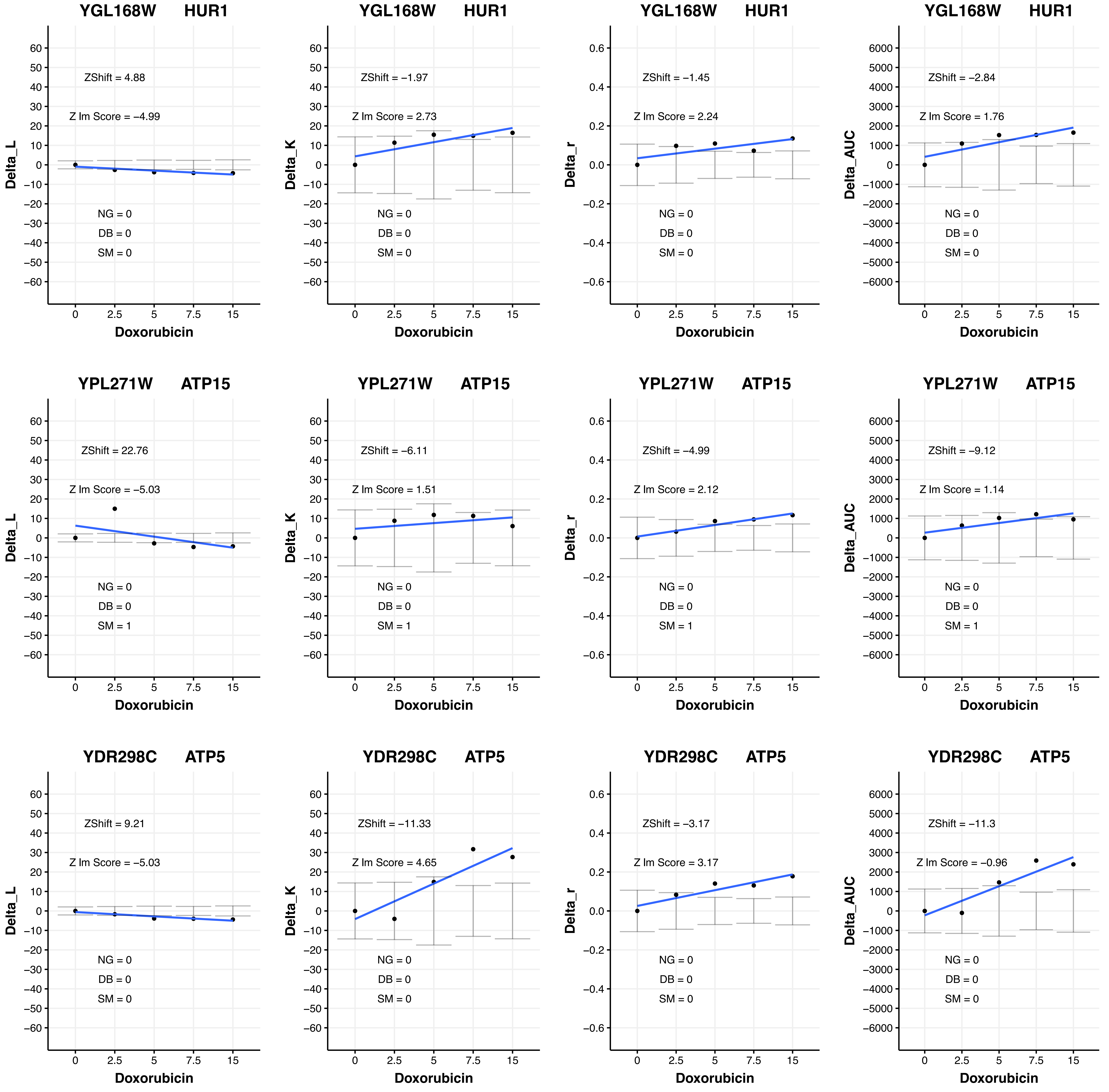

YGL168W HUR1
ZShift = 4.88
Z lm Score = −4.99
NG = 0
DB = 0
SM = 0
Delta_L
Doxorubicin
YGL168W HUR1
ZShift = −1.97
Z lm Score = 2.73
NG = 0
DB = 0
SM = 0
Delta_K
Doxorubicin
YGL168W HUR1
ZShift = −1.45
Z lm Score = 2.24
NG = 0
DB = 0
SM = 0
Delta_r
Doxorubicin
YGL168W HUR1
ZShift = −2.84
Z lm Score = 1.76
NG = 0
DB = 0
SM = 0
Delta_AUC
Doxorubicin
YPL271W ATP15
ZShift = 22.76
Z lm Score = −5.03
NG = 0
DB = 0
SM = 1
Delta_L
Doxorubicin
YPL271W ATP15
ZShift = −6.11
Z lm Score = 1.51
NG = 0
DB = 0
SM = 1
Delta_K
Doxorubicin
YPL271W ATP15
ZShift = −4.99
Z lm Score = 2.12
NG = 0
DB = 0
SM = 1
Delta_r
Doxorubicin
YPL271W ATP15
ZShift = −9.12
Z lm Score = 1.14
NG = 0
DB = 0
SM = 1
Delta_AUC
Doxorubicin
YDR298C ATP5
ZShift = 9.21
Z lm Score = −5.03
NG = 0
DB = 0
SM = 0
Delta_L
Doxorubicin
YDR298C ATP5
ZShift = −11.33
Z lm Score = 4.65
NG = 0
DB = 0
SM = 0
Delta_K
Doxorubicin
YDR298C ATP5
ZShift = −3.17
Z lm Score = 3.17
NG = 0
DB = 0
SM = 0
Delta_r
Doxorubicin
YDR298C ATP5
ZShift = −11.3
Z lm Score = −0.96
NG = 0
DB = 0
SM = 0
Delta_AUC
Doxorubicin

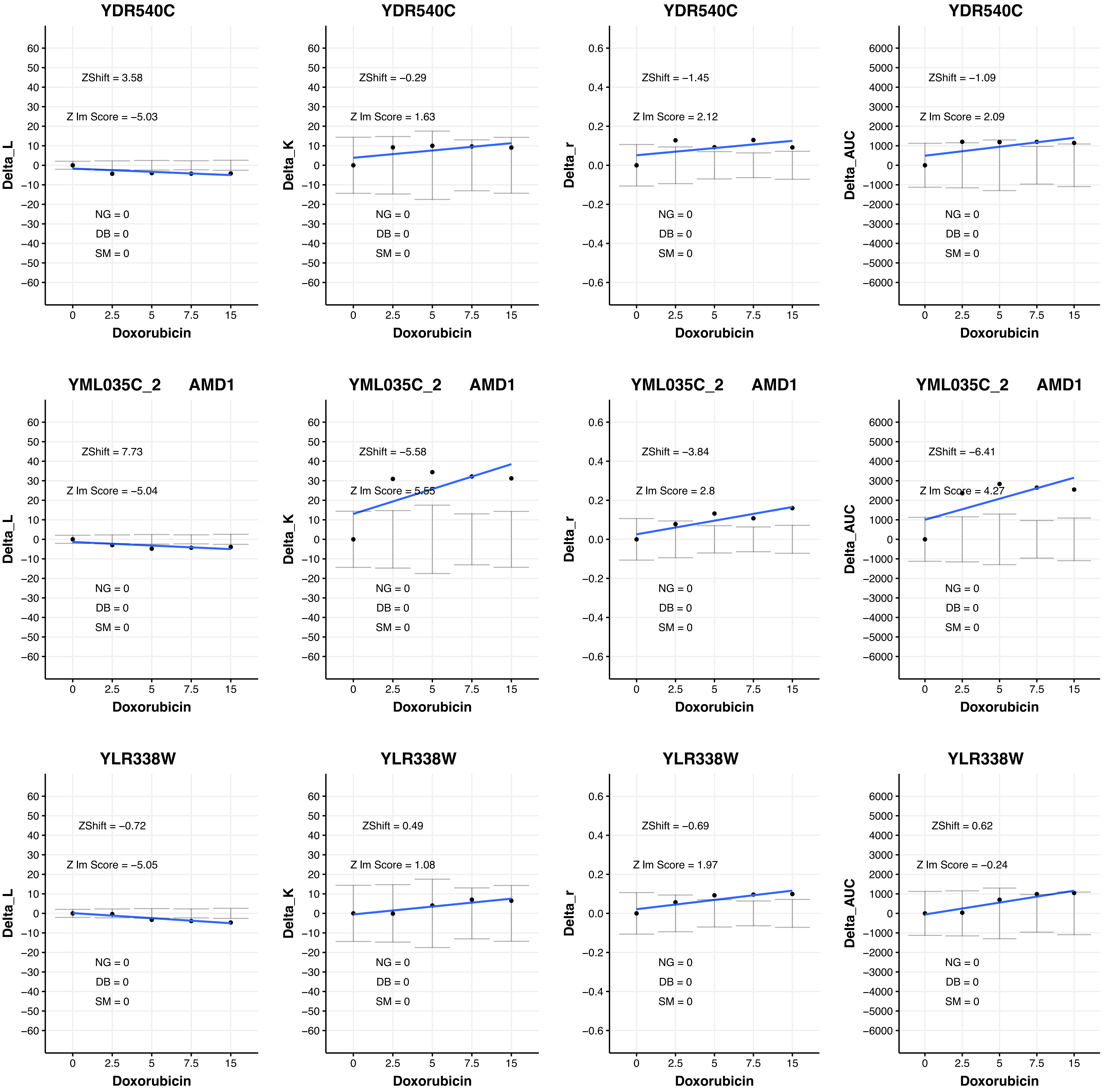

YDR540C
ZShift = 3.58
Z lm Score = −5.03
NG = 0
DB = 0
SM = 0
Delta_L
Doxorubicin
YDR540C
ZShift = −0.29
Z lm Score = 1.63
NG = 0
DB = 0
SM = 0
Delta_K
Doxorubicin
YDR540C
ZShift = −1.45
Z lm Score = 2.12
NG = 0
DB = 0
SM = 0
Delta_r
Doxorubicin
YDR540C
ZShift = −1.09
Z lm Score = 2.09
NG = 0
DB = 0
SM = 0
Delta_AUC
Doxorubicin
YML035C_2 AMD1
ZShift = 7.73
Z lm Score = −5.04
NG = 0
DB = 0
SM = 0
Delta_L
Doxorubicin
YML035C_2 AMD1
ZShift = −5.58
Z lm Score = 5.55
NG = 0
DB = 0
SM = 0
Delta_K
Doxorubicin
YML035C_2 AMD1
ZShift = −3.84
Z lm Score = 2.8
NG = 0
DB = 0
SM = 0
Delta_r
Doxorubicin
YML035C_2 AMD1
ZShift = −6.41
Z lm Score = 4.27
NG = 0
DB = 0
SM = 0
Delta_AUC
Doxorubicin
YLR338W
ZShift = −0.72
Z lm Score = −5.05
NG = 0
DB = 0
SM = 0
Delta_L
Doxorubicin
YLR338W
ZShift = 0.49
Z lm Score = 1.08
NG = 0
DB = 0
SM = 0
Delta_K
Doxorubicin
YLR338W
ZShift = −0.69
Z lm Score = 1.97
NG = 0
DB = 0
SM = 0
Delta_r
Doxorubicin
YLR338W
ZShift = 0.62
Z lm Score = −0.24
NG = 0
DB = 0
SM = 0
Delta_AUC
Doxorubicin
0 2.5 5 7.5 15

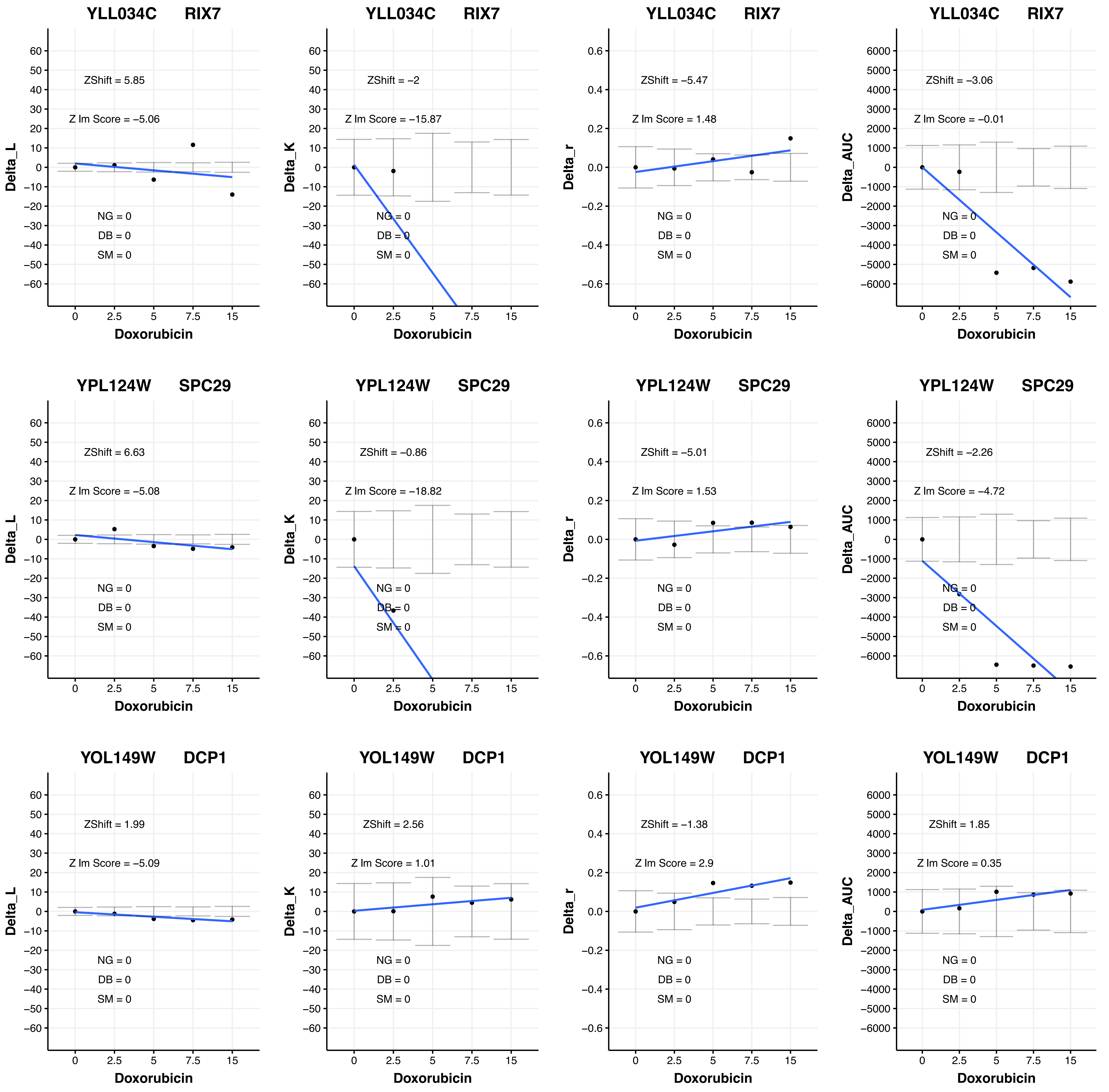

YLL034C RIX7
ZShift = 5.85
Z lm Score = −5.06
NG = 0
DB = 0
SM = 0
Delta_L
Doxorubicin
YLL034C RIX7
ZShift = −2
Z lm Score = −15.87
NG = 0
DB = 0
SM = 0
Delta_K
Doxorubicin
YLL034C RIX7
ZShift = −5.47
Z lm Score = 1.48
NG = 0
DB = 0
SM = 0
Delta_r
Doxorubicin
YLL034C RIX7
ZShift = −3.06
Z lm Score = −0.01
NG = 0
DB = 0
SM = 0
Delta_AUC
Doxorubicin
YPL124W SPC29
ZShift = 6.63
Z lm Score = −5.08
NG = 0
DB = 0
SM = 0
Delta_L
Doxorubicin
YPL124W SPC29
ZShift = −0.86
Z lm Score = −18.82
NG = 0
DB = 0
SM = 0
Delta_K
Doxorubicin
YPL124W SPC29
ZShift = −5.01
Z lm Score = 1.53
NG = 0
DB = 0
SM = 0
Delta_r
Doxorubicin
YPL124W SPC29
ZShift = −2.26
Z lm Score = −4.72
NG = 0
DB = 0
SM = 0
Delta_AUC
Doxorubicin
YOL149W DCP1
ZShift = 1.99
Z lm Score = −5.09
NG = 0
DB = 0
SM = 0
Delta_L
Doxorubicin
YOL149W DCP1
ZShift = 2.56
Z lm Score = 1.01
NG = 0
DB = 0
SM = 0
Delta_K
Doxorubicin
YOL149W DCP1
ZShift = −1.38
Z lm Score = 2.9
NG = 0
DB = 0
SM = 0
Delta_r
Doxorubicin
YOL149W DCP1
ZShift = 1.85
Z lm Score = 0.35
NG = 0
DB = 0
SM = 0
Delta_AUC
Doxorubicin

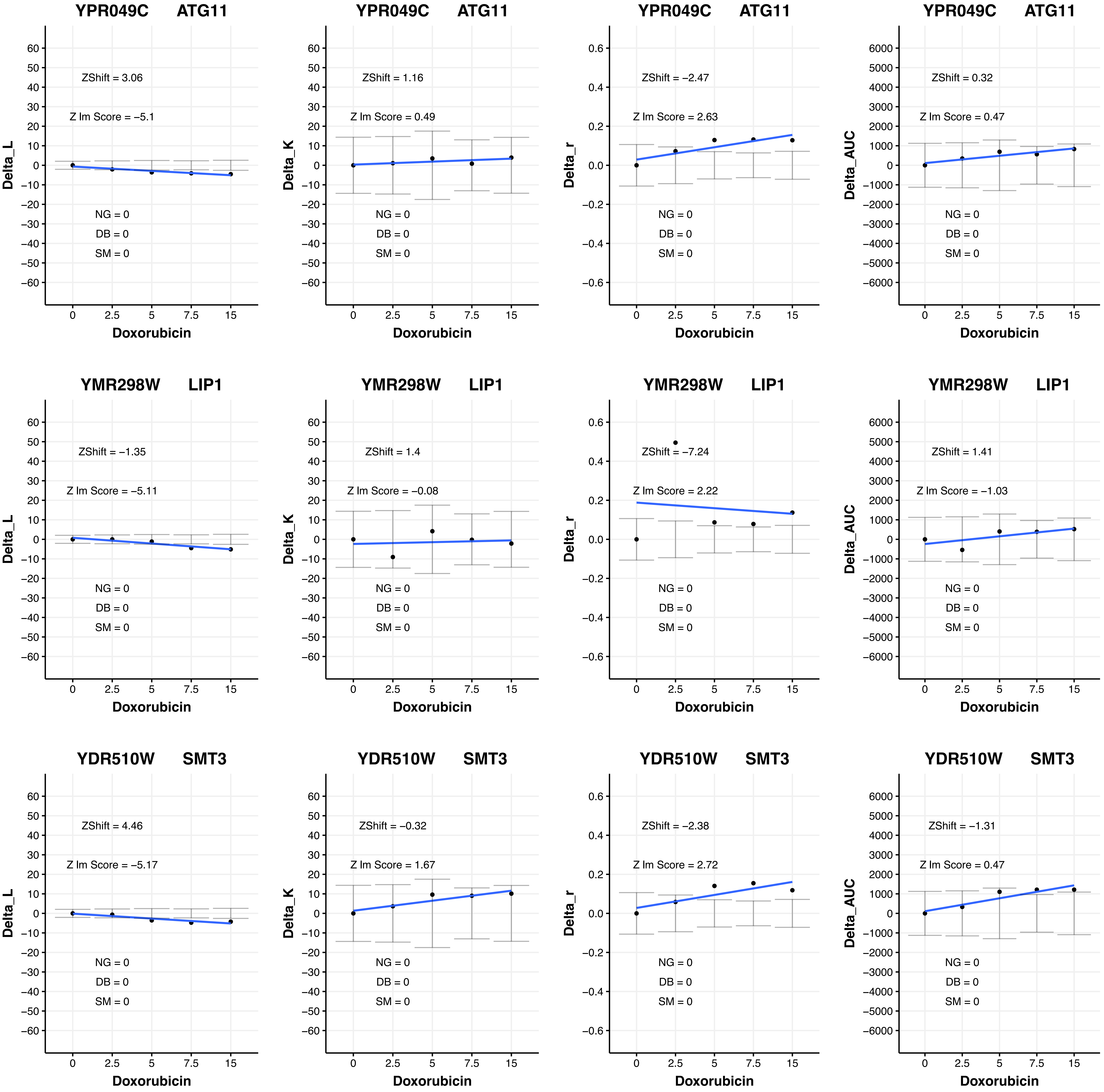
YPR049C ATG11
ZShift = 3.06
Z lm Score = −5.1
NG = 0
DB = 0
SM = 0
Delta_L
Doxorubicin
YPR049C ATG11
ZShift = 1.16
Z lm Score = 0.49
NG = 0
DB = 0
SM = 0
Delta_K
Doxorubicin
YPR049C ATG11
ZShift = −2.47
Z lm Score = 2.63
NG = 0
DB = 0
SM = 0
Delta_r
Doxorubicin
YPR049C ATG11
ZShift = 0.32
Z lm Score = 0.47
NG = 0
DB = 0
SM = 0
Delta_AUC
Doxorubicin
YMR298W LIP1
ZShift = −1.35
Z lm Score = −5.11
NG = 0
DB = 0
SM = 0
Delta_L
Doxorubicin
YMR298W LIP1
ZShift = 1.4
Z lm Score = −0.08
NG = 0
DB = 0
SM = 0
Delta_K
Doxorubicin
YMR298W LIP1
ZShift = −7.24
Z lm Score = 2.22
NG = 0
DB = 0
SM = 0
Delta_r
Doxorubicin
YMR298W LIP1
ZShift = 1.41
Z lm Score = −1.03
NG = 0
DB = 0
SM = 0
Delta_AUC
Doxorubicin
YDR510W SMT3
ZShift = 4.46
Z lm Score = −5.17
NG = 0
DB = 0
SM = 0
Delta_L
Doxorubicin
YDR510W SMT3
ZShift = −0.32
Z lm Score = 1.67
NG = 0
DB = 0
SM = 0
Delta_K
Doxorubicin
YDR510W SMT3
ZShift = −2.38
Z lm Score = 2.72
NG = 0
DB = 0
SM = 0
Delta_r
Doxorubicin
YDR510W SMT3
ZShift = −1.31
Z lm Score = 0.47
NG = 0
DB = 0
SM = 0
Delta_AUC
Doxorubicin

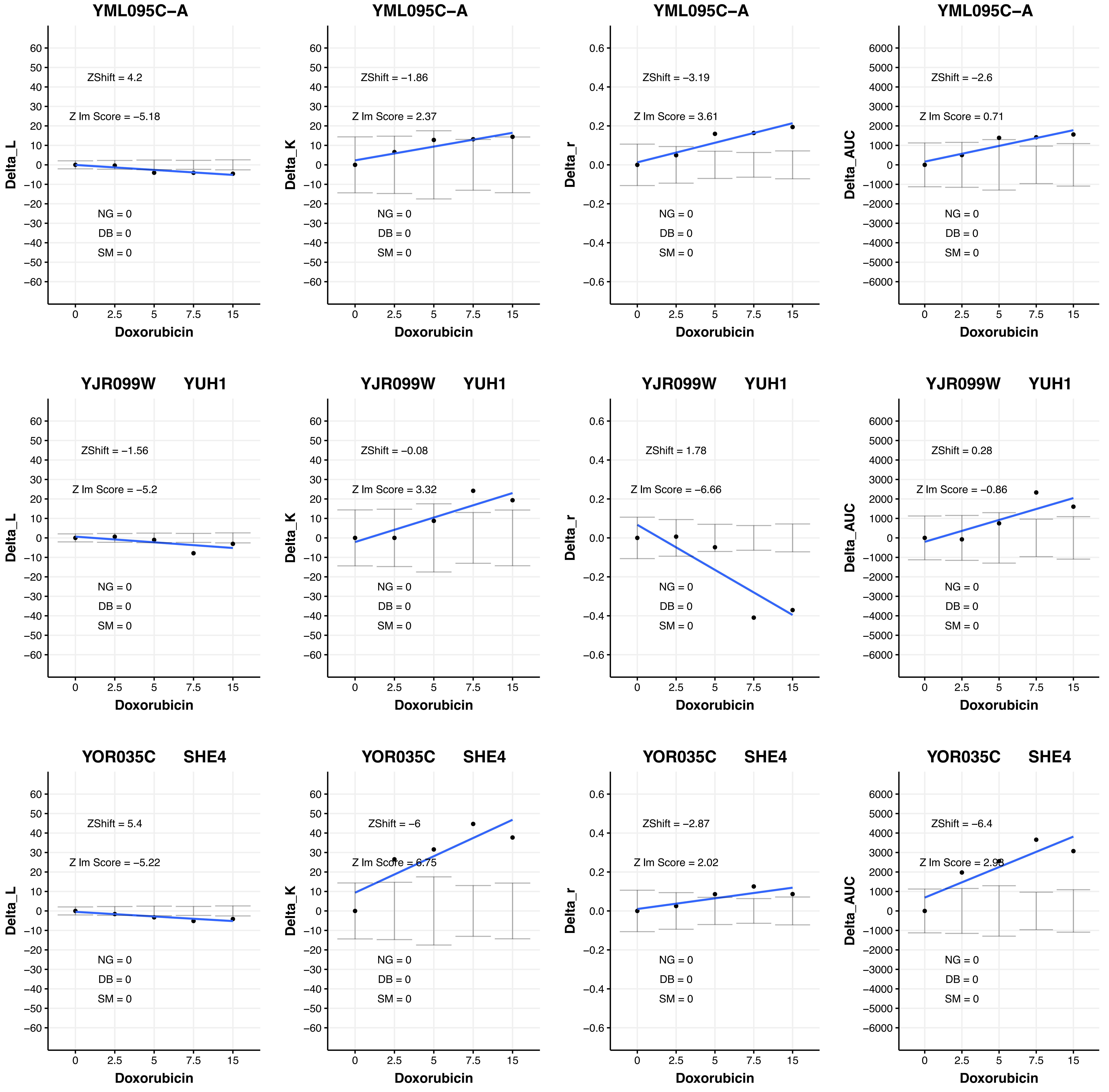
YML095C−A
ZShift = 4.2
Z lm Score = −5.18
NG = 0
DB = 0
SM = 0
Delta_L
Doxorubicin
YML095C−A
ZShift = −1.86
Z lm Score = 2.37
NG = 0
DB = 0
SM = 0
Delta_K
Doxorubicin
YML095C−A
ZShift = −3.19
Z lm Score = 3.61
NG = 0
DB = 0
SM = 0
Delta_r
Doxorubicin
YML095C−A
ZShift = −2.6
Z lm Score = 0.71
NG = 0
DB = 0
SM = 0
Delta_AUC
Doxorubicin
YJR099W YUH1
ZShift = −1.56
Z lm Score = −5.2
NG = 0
DB = 0
SM = 0
Delta_L
Doxorubicin
YJR099W YUH1
ZShift = −0.08
Z lm Score = 3.32
NG = 0
DB = 0
SM = 0
Delta_K
Doxorubicin
YJR099W YUH1
ZShift = 1.78
Z lm Score = −6.66
NG = 0
DB = 0
SM = 0
Delta_r
Doxorubicin
YJR099W YUH1
ZShift = 0.28
Z lm Score = −0.86
NG = 0
DB = 0
SM = 0
Delta_AUC
Doxorubicin
YOR035C SHE4
ZShift = 5.4
Z lm Score = −5.22
NG = 0
DB = 0
SM = 0
Delta_L
Doxorubicin
YOR035C SHE4
ZShift = −6
Z lm Score = 6.75
NG = 0
DB = 0
SM = 0
Delta_K
Doxorubicin
YOR035C SHE4
ZShift = −2.87
Z lm Score = 2.02
NG = 0
DB = 0
SM = 0
Delta_r
Doxorubicin
YOR035C SHE4
ZShift = −6.4
Z lm Score = 2.98
NG = 0
DB = 0
SM = 0
Delta_AUC
Doxorubicin

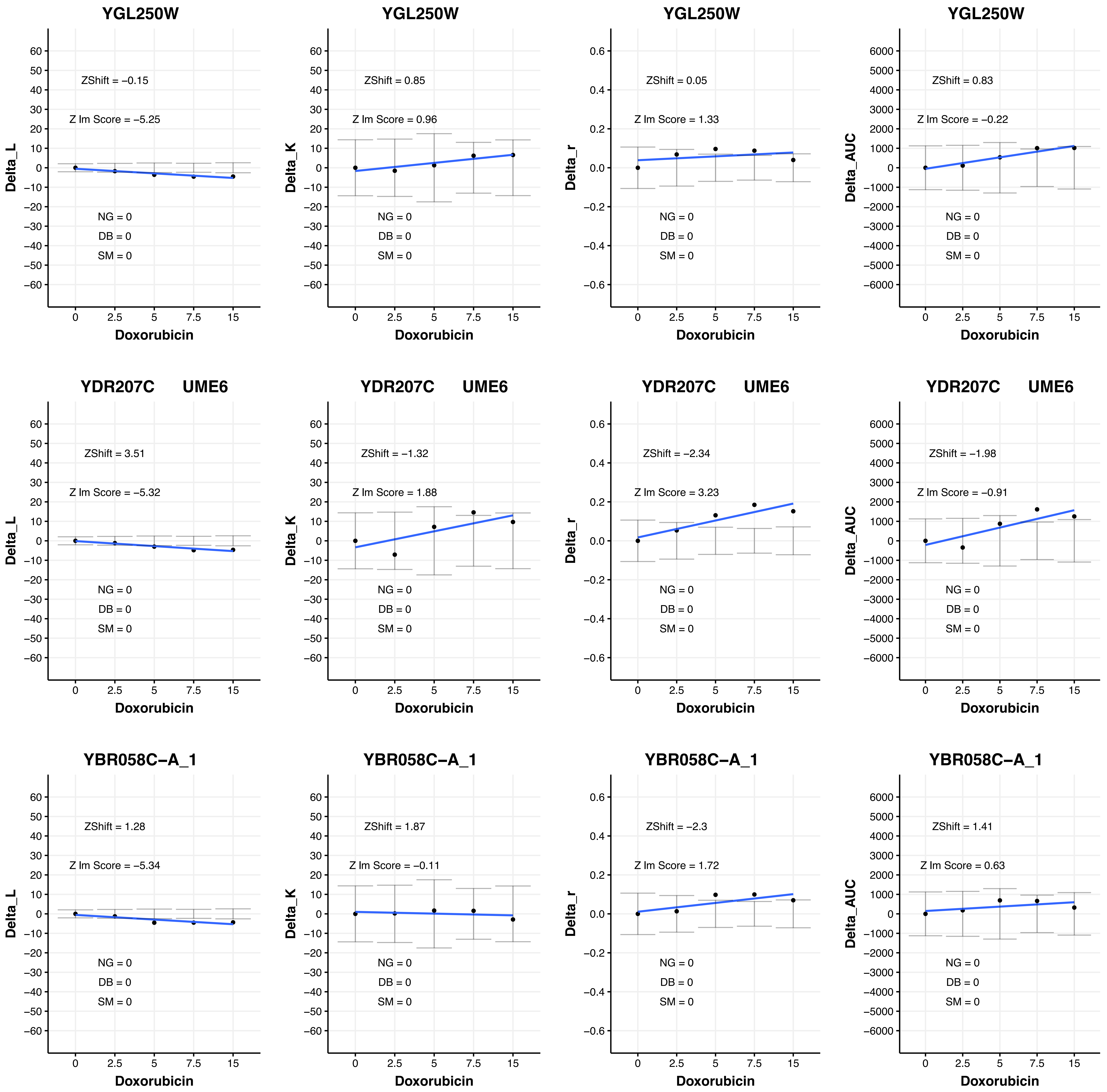
YGL250W
ZShift = −0.15
Z lm Score = −5.25
NG = 0
DB = 0
SM = 0
Delta_L
Doxorubicin
YGL250W
ZShift = 0.85
Z lm Score = 0.96
NG = 0
DB = 0
SM = 0
Delta_K
Doxorubicin
YGL250W
ZShift = 0.05
Z lm Score = 1.33
NG = 0
DB = 0
SM = 0
Delta_r
Doxorubicin
YGL250W
ZShift = 0.83
Z lm Score = −0.22
NG = 0
DB = 0
SM = 0
Delta_AUC
Doxorubicin
YDR207C UME6
ZShift = 3.51
Z lm Score = −5.32
NG = 0
DB = 0
SM = 0
Delta_L
Doxorubicin
YDR207C UME6
ZShift = −1.32
Z lm Score = 1.88
NG = 0
DB = 0
SM = 0
Delta_K
Doxorubicin
YDR207C UME6
ZShift = −2.34
Z lm Score = 3.23
NG = 0
DB = 0
SM = 0
Delta_r
Doxorubicin
YDR207C UME6
ZShift = −1.98
Z lm Score = −0.91
NG = 0
DB = 0
SM = 0
Delta_AUC
Doxorubicin
YBR058C−A_1
ZShift = 1.28
Z lm Score = −5.34
NG = 0
DB = 0
SM = 0
Delta_L
Doxorubicin
YBR058C−A_1
ZShift = 1.87
Z lm Score = −0.11
NG = 0
DB = 0
SM = 0
Delta_K
Doxorubicin
YBR058C−A_1
ZShift = −2.3
Z lm Score = 1.72
NG = 0
DB = 0
SM = 0
Delta_r
Doxorubicin
YBR058C−A_1
ZShift = 1.41
Z lm Score = 0.63
NG = 0
DB = 0
SM = 0
Delta_AUC
Doxorubicin

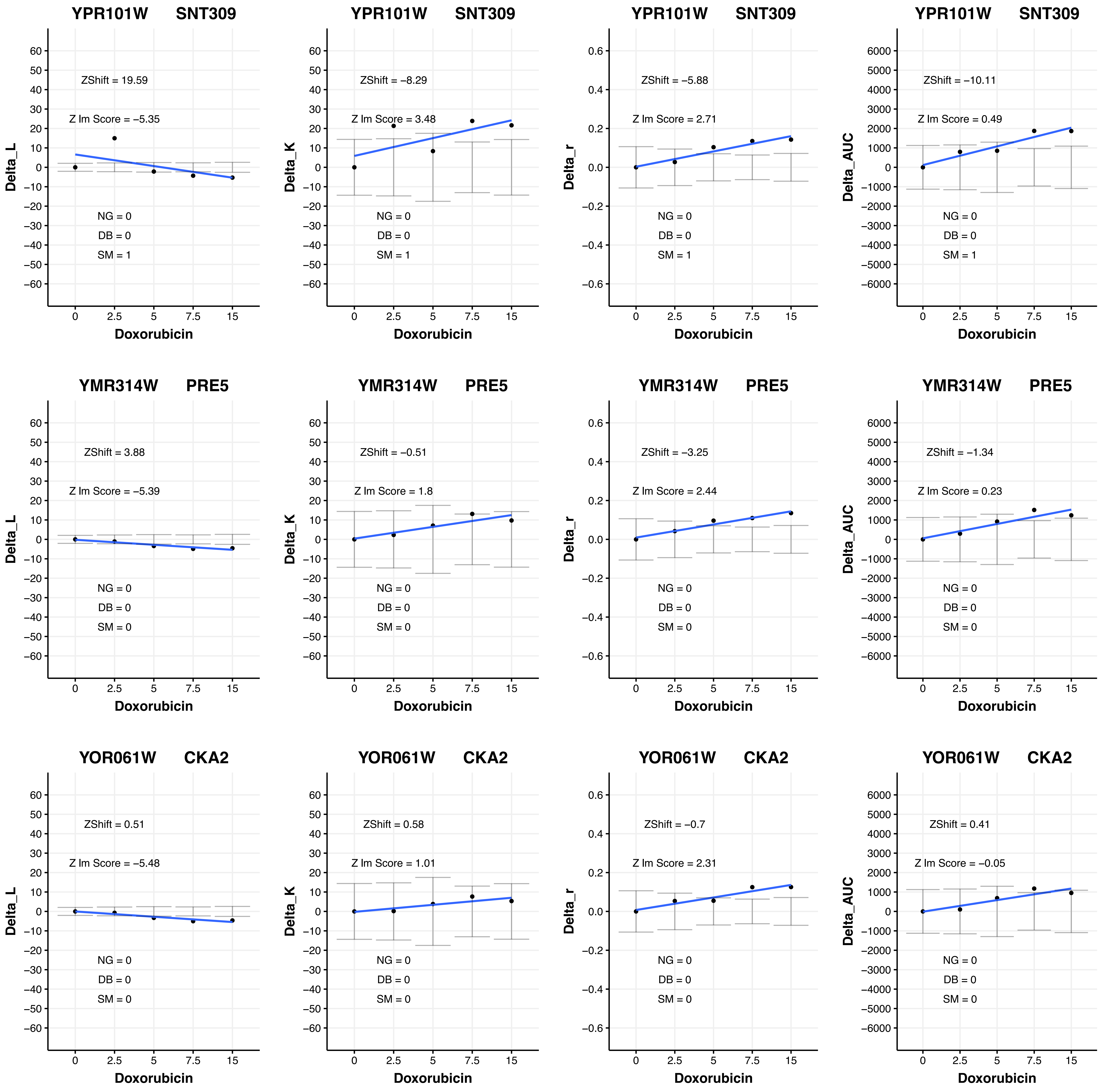

YPR101W SNT309
ZShift = 19.59
Z lm Score = −5.35
NG = 0
DB = 0
SM = 1
Delta_L
Doxorubicin
YPR101W SNT309
ZShift = −8.29
Z lm Score = 3.48
NG = 0
DB = 0
SM = 1
Delta_K
Doxorubicin
YPR101W SNT309
ZShift = −5.88
Z lm Score = 2.71
NG = 0
DB = 0
SM = 1
Delta_r
Doxorubicin
YPR101W SNT309
ZShift = −10.11
Z lm Score = 0.49
NG = 0
DB = 0
SM = 1
Delta_AUC
Doxorubicin
YMR314W PRE5
ZShift = 3.88
Z lm Score = −5.39
NG = 0
DB = 0
SM = 0
Delta_L
Doxorubicin
YMR314W PRE5
ZShift = −0.51
Z lm Score = 1.8
NG = 0
DB = 0
SM = 0
Delta_K
Doxorubicin
YMR314W PRE5
ZShift = −3.25
Z lm Score = 2.44
NG = 0
DB = 0
SM = 0
Delta_r
Doxorubicin
YMR314W PRE5
ZShift = −1.34
Z lm Score = 0.23
NG = 0
DB = 0
SM = 0
Delta_AUC
Doxorubicin
YOR061W CKA2
ZShift = 0.51
Z lm Score = −5.48
NG = 0
DB = 0
SM = 0
Delta_L
Doxorubicin
YOR061W CKA2
ZShift = 0.58
Z lm Score = 1.01
NG = 0
DB = 0
SM = 0
Delta_K
Doxorubicin
YOR061W CKA2
ZShift = −0.7
Z lm Score = 2.31
NG = 0
DB = 0
SM = 0
Delta_r
Doxorubicin
YOR061W CKA2
ZShift = 0.41
Z lm Score = −0.05
NG = 0
DB = 0
SM = 0
Delta_AUC
Doxorubicin

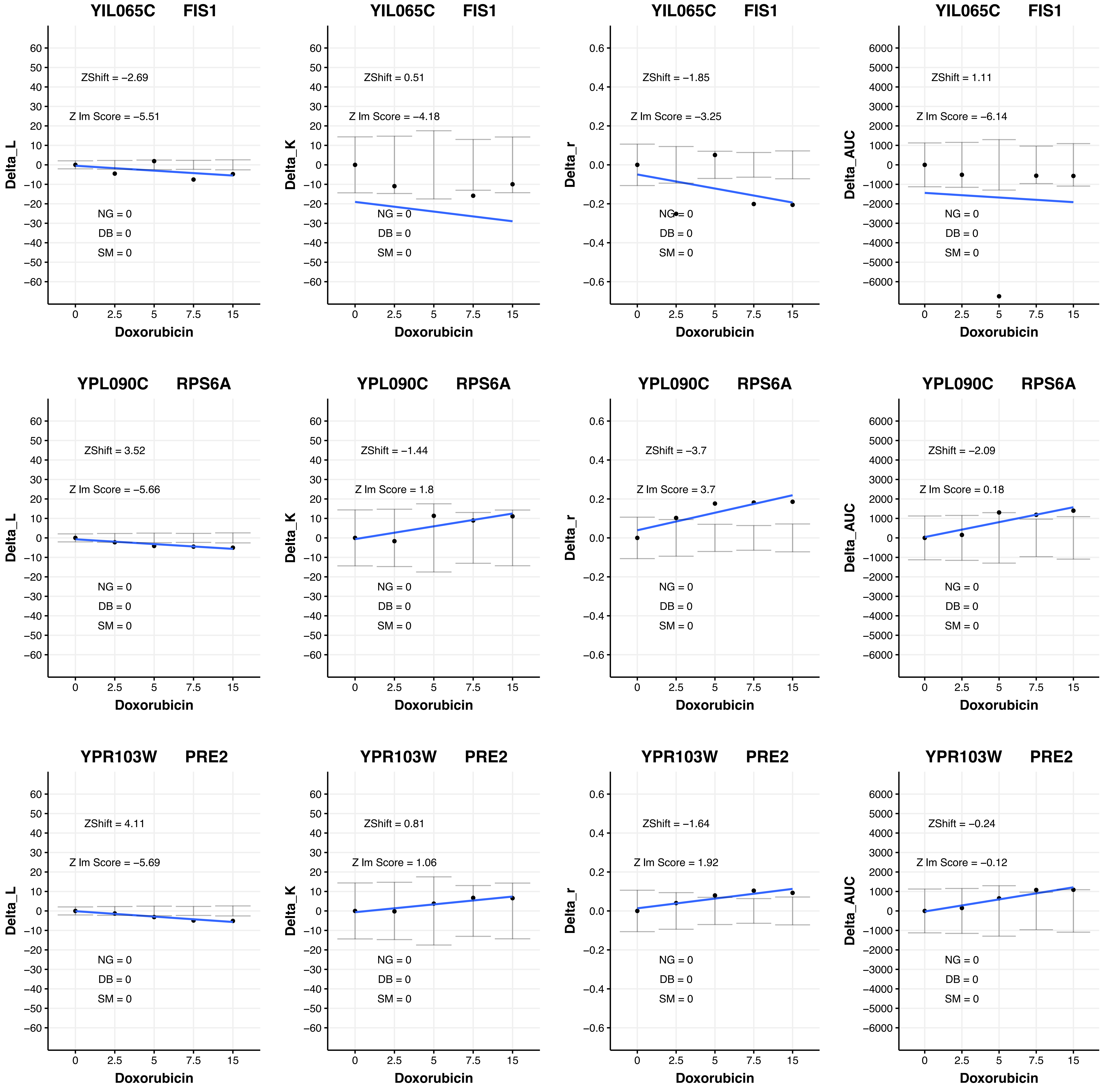

YIL065C FIS1
ZShift = −2.69
Z lm Score = −5.51
NG = 0
DB = 0
SM = 0
Delta_L
Doxorubicin
YIL065C FIS1
ZShift = 0.51
Z lm Score = −4.18
NG = 0
DB = 0
SM = 0
Delta_K
Doxorubicin
YIL065C FIS1
ZShift = −1.85
Z lm Score = −3.25
NG = 0
DB = 0
SM = 0
Delta_r
Doxorubicin
YIL065C FIS1
ZShift = 1.11
Z lm Score = −6.14
NG = 0
DB = 0
SM = 0
Delta_AUC
Doxorubicin
YPL090C RPS6A
ZShift = 3.52
Z lm Score = −5.66
NG = 0
DB = 0
SM = 0
Delta_L
Doxorubicin
YPL090C RPS6A
ZShift = −1.44
Z lm Score = 1.8
NG = 0
DB = 0
SM = 0
Delta_K
Doxorubicin
YPL090C RPS6A
ZShift = −3.7
Z lm Score = 3.7
NG = 0
DB = 0
SM = 0
Delta_r
Doxorubicin
YPL090C RPS6A
ZShift = −2.09
Z lm Score = 0.18
NG = 0
DB = 0
SM = 0
Delta_AUC
Doxorubicin
YPR103W PRE2
ZShift = 4.11
Z lm Score = −5.69
NG = 0
DB = 0
SM = 0
Delta_L
Doxorubicin
YPR103W PRE2
ZShift = 0.81
Z lm Score = 1.06
NG = 0
DB = 0
SM = 0
Delta_K
Doxorubicin
YPR103W PRE2
ZShift = −1.64
Z lm Score = 1.92
NG = 0
DB = 0
SM = 0
Delta_r
Doxorubicin
YPR103W PRE2
ZShift = −0.24
Z lm Score = −0.12
NG = 0
DB = 0
SM = 0
Delta_AUC
Doxorubicin

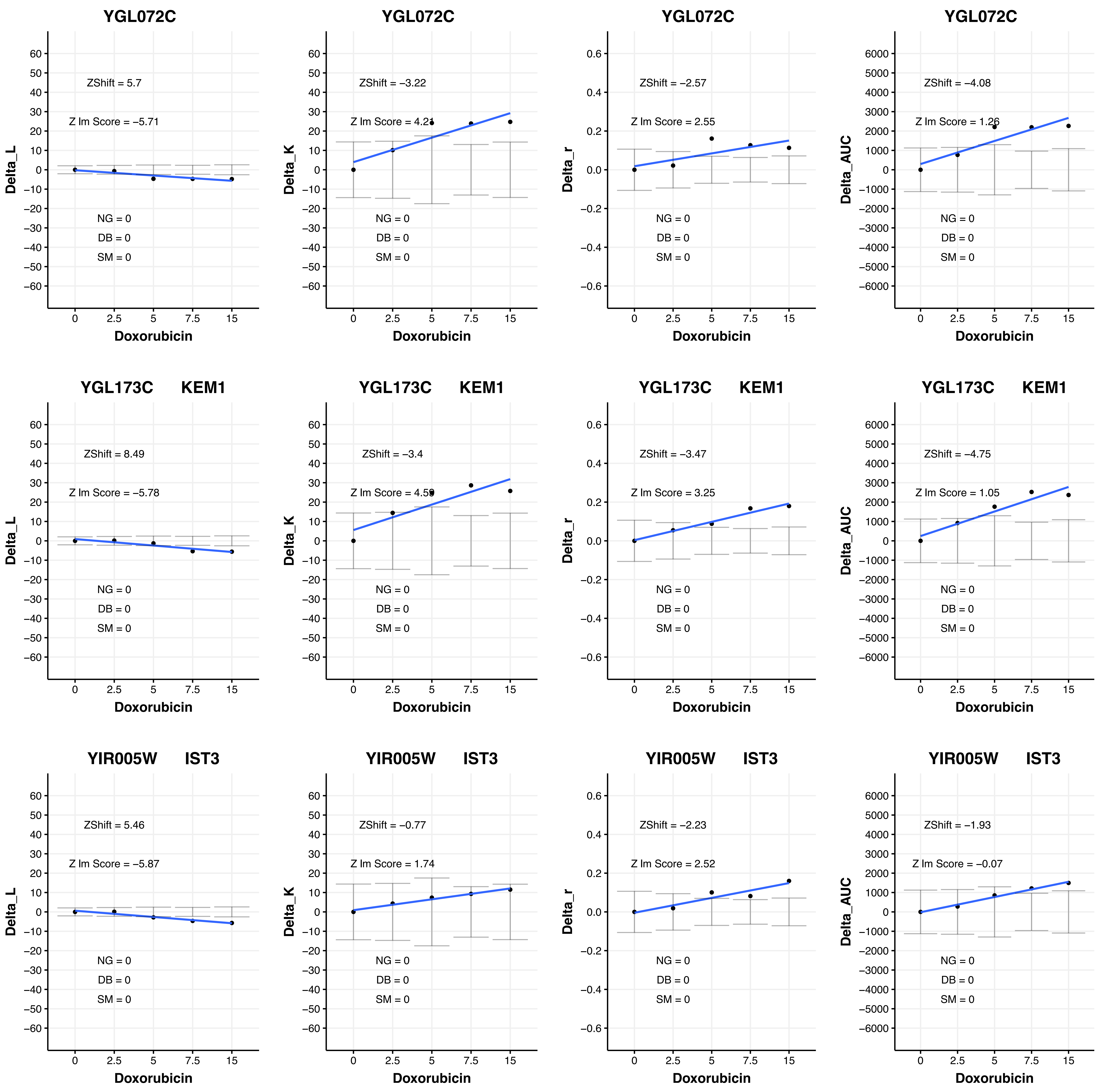

YGL072C
ZShift = 5.7
Z lm Score = −5.71
NG = 0
DB = 0
SM = 0
Delta_L
Doxorubicin
YGL072C
ZShift = −3.22
Z lm Score = 4.21
NG = 0
DB = 0
SM = 0
Delta_K
Doxorubicin
YGL072C
ZShift = −2.57
Z lm Score = 2.55
NG = 0
DB = 0
SM = 0
Delta_r
Doxorubicin
YGL072C
ZShift = −4.08
Z lm Score = 1.26
NG = 0
DB = 0
SM = 0
Delta_AUC
Doxorubicin
YGL173C KEM1
ZShift = 8.49
Z lm Score = −5.78
NG = 0
DB = 0
SM = 0
Delta_L
Doxorubicin
YGL173C KEM1
ZShift = −3.4
Z lm Score = 4.59
NG = 0
DB = 0
SM = 0
Delta_K
Doxorubicin
YGL173C KEM1
ZShift = −3.47
Z lm Score = 3.25
NG = 0
DB = 0
SM = 0
Delta_r
Doxorubicin
YGL173C KEM1
ZShift = −4.75
Z lm Score = 1.05
NG = 0
DB = 0
SM = 0
Delta_AUC
Doxorubicin
YIR005W IST3
ZShift = 5.46
Z lm Score = −5.87
NG = 0
DB = 0
SM = 0
Delta_L
Doxorubicin
YIR005W IST3
ZShift = −0.77
Z lm Score = 1.74
NG = 0
DB = 0
SM = 0
Delta_K
Doxorubicin
YIR005W IST3
ZShift = −2.23
Z lm Score = 2.52
NG = 0
DB = 0
SM = 0
Delta_r
Doxorubicin
YIR005W IST3
ZShift = −1.93
Z lm Score = −0.07
NG = 0
DB = 0
SM = 0
Delta_AUC
Doxorubicin

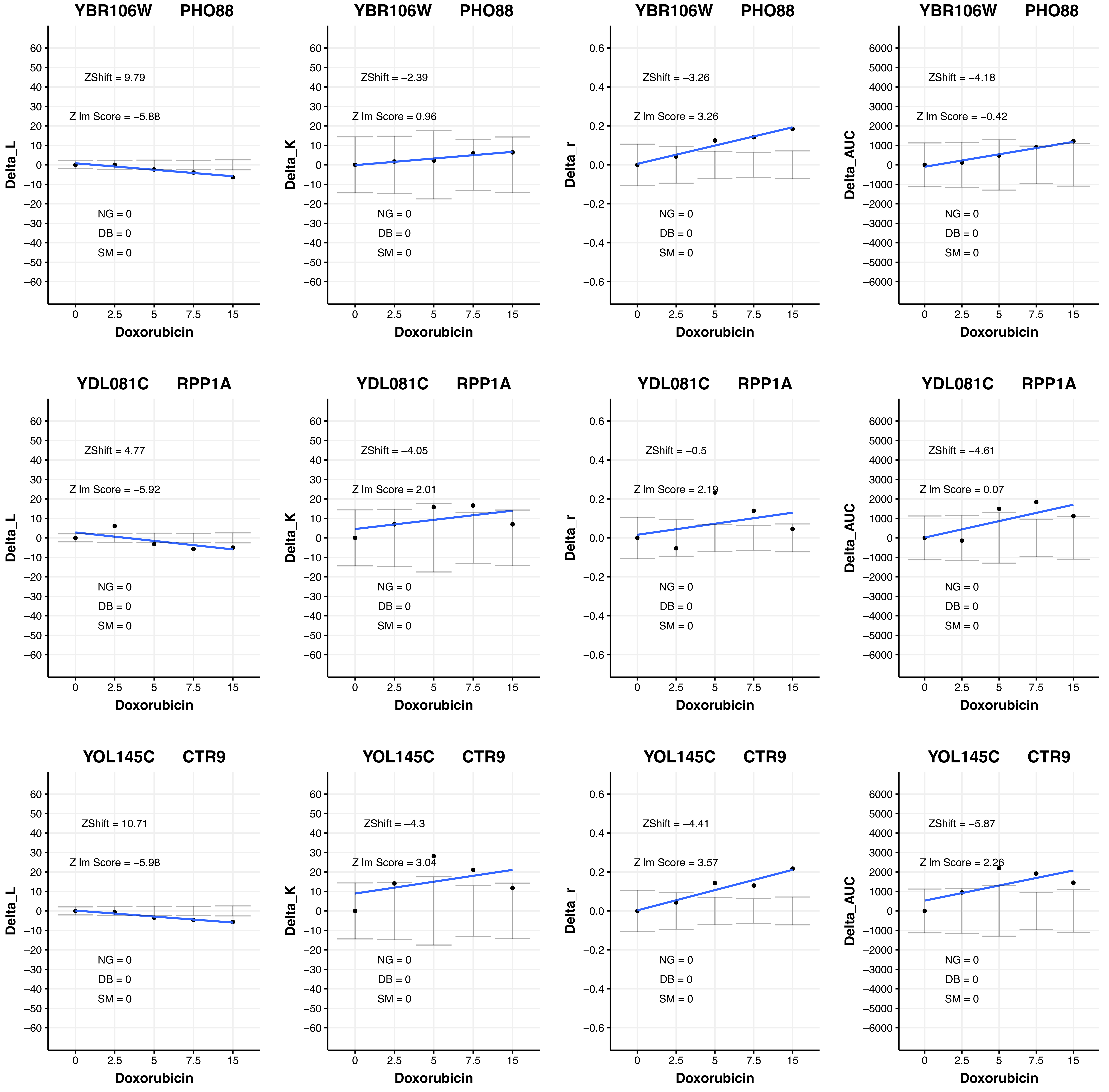
YBR106W PHO88
ZShift = 9.79
Z lm Score = −5.88
NG = 0
DB = 0
SM = 0
Delta_L
Doxorubicin
YBR106W PHO88
ZShift = −2.39
Z lm Score = 0.96
NG = 0
DB = 0
SM = 0
Delta_K
Doxorubicin
YBR106W PHO88
ZShift = −3.26
Z lm Score = 3.26
NG = 0
DB = 0
SM = 0
Delta_r
Doxorubicin
YBR106W PHO88
ZShift = −4.18
Z lm Score = −0.42
NG = 0
DB = 0
SM = 0
Delta_AUC
Doxorubicin
YDL081C RPP1A
ZShift = 4.77
Z lm Score = −5.92
NG = 0
DB = 0
SM = 0
Delta_L
Doxorubicin
YDL081C RPP1A
ZShift = −4.05
Z lm Score = 2.01
NG = 0
DB = 0
SM = 0
Delta_K
Doxorubicin
YDL081C RPP1A
ZShift = −0.5
Z lm Score = 2.19
NG = 0
DB = 0
SM = 0
Delta_r
Doxorubicin
YDL081C RPP1A
ZShift = −4.61
Z lm Score = 0.07
NG = 0
DB = 0
SM = 0
Delta_AUC
Doxorubicin
YOL145C CTR9
ZShift = 10.71
Z lm Score = −5.98
NG = 0
DB = 0
SM = 0
Delta_L
Doxorubicin
YOL145C CTR9
ZShift = −4.3
Z lm Score = 3.04
NG = 0
DB = 0
SM = 0
Delta_K
Doxorubicin
YOL145C CTR9
ZShift = −4.41
Z lm Score = 3.57
NG = 0
DB = 0
SM = 0
Delta_r
Doxorubicin
YOL145C CTR9
ZShift = −5.87
Z lm Score = 2.26
NG = 0
DB = 0
SM = 0
Delta_AUC
Doxorubicin

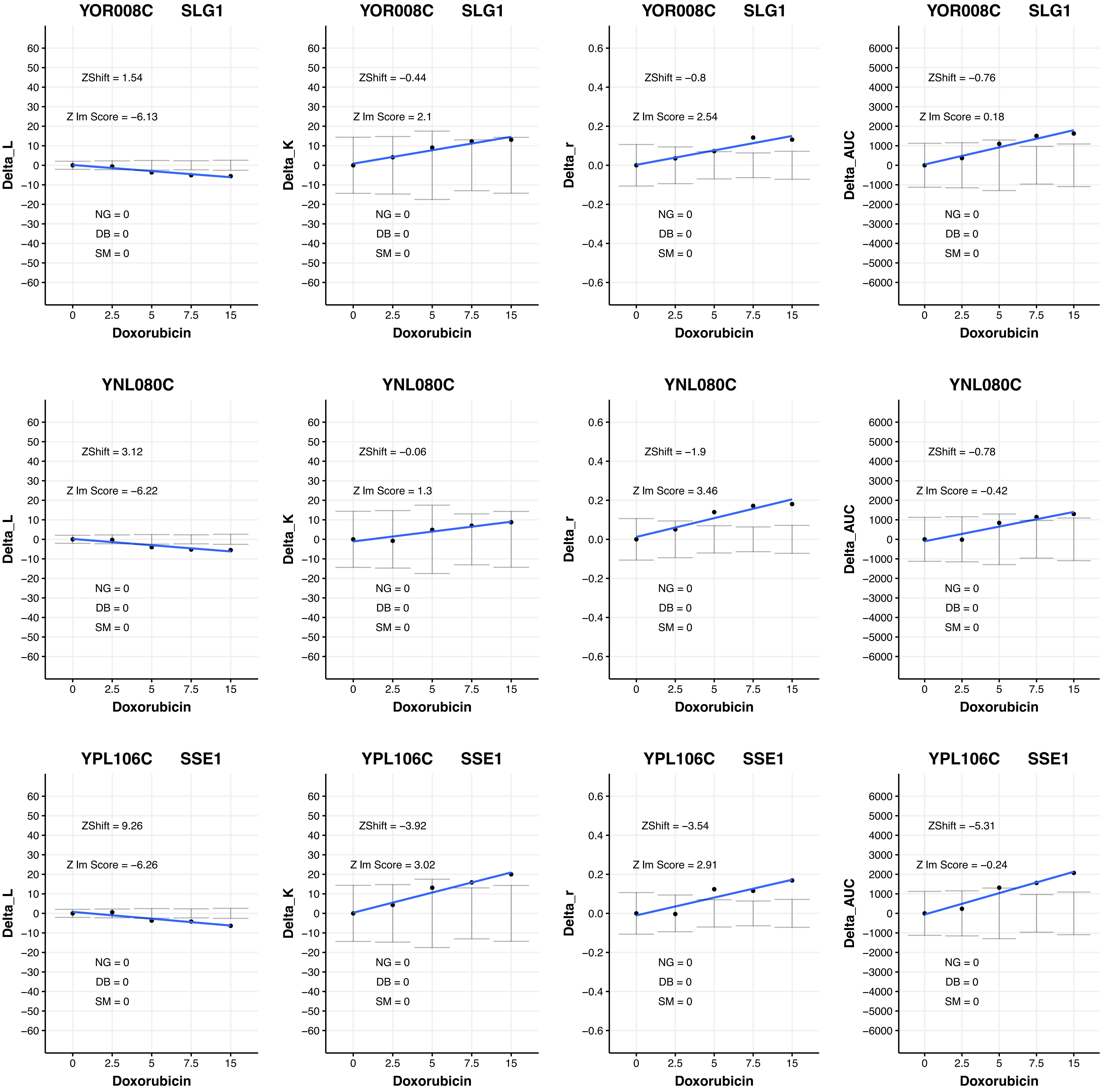

YOR008C SLG1
ZShift = 1.54
Z lm Score = −6.13
NG = 0
DB = 0
SM = 0
Delta_L
Doxorubicin
YOR008C SLG1
ZShift = −0.44
Z lm Score = 2.1
NG = 0
DB = 0
SM = 0
Delta_K
Doxorubicin
YOR008C SLG1
ZShift = −0.8
Z lm Score = 2.54
NG = 0
DB = 0
SM = 0
Delta_r
Doxorubicin
YOR008C SLG1
ZShift = −0.76
Z lm Score = 0.18
NG = 0
DB = 0
SM = 0
Delta_AUC
Doxorubicin
YNL080C
ZShift = 3.12
Z lm Score = −6.22
NG = 0
DB = 0
SM = 0
Delta_L
Doxorubicin
YNL080C
ZShift = −0.06
Z lm Score = 1.3
NG = 0
DB = 0
SM = 0
Delta_K
Doxorubicin
YNL080C
ZShift = −1.9
Z lm Score = 3.46
NG = 0
DB = 0
SM = 0
Delta_r
Doxorubicin
YNL080C
ZShift = −0.78
Z lm Score = −0.42
NG = 0
DB = 0
SM = 0
Delta_AUC
Doxorubicin
YPL106C SSE1
ZShift = 9.26
Z lm Score = −6.26
NG = 0
DB = 0
SM = 0
Delta_L
Doxorubicin
YPL106C SSE1
ZShift = −3.92
Z lm Score = 3.02
NG = 0
DB = 0
SM = 0
Delta_K
Doxorubicin
YPL106C SSE1
ZShift = −3.54
Z lm Score = 2.91
NG = 0
DB = 0
SM = 0
Delta_r
Doxorubicin
YPL106C SSE1
ZShift = −5.31
Z lm Score = −0.24
NG = 0
DB = 0
SM = 0
Delta_AUC
Doxorubicin

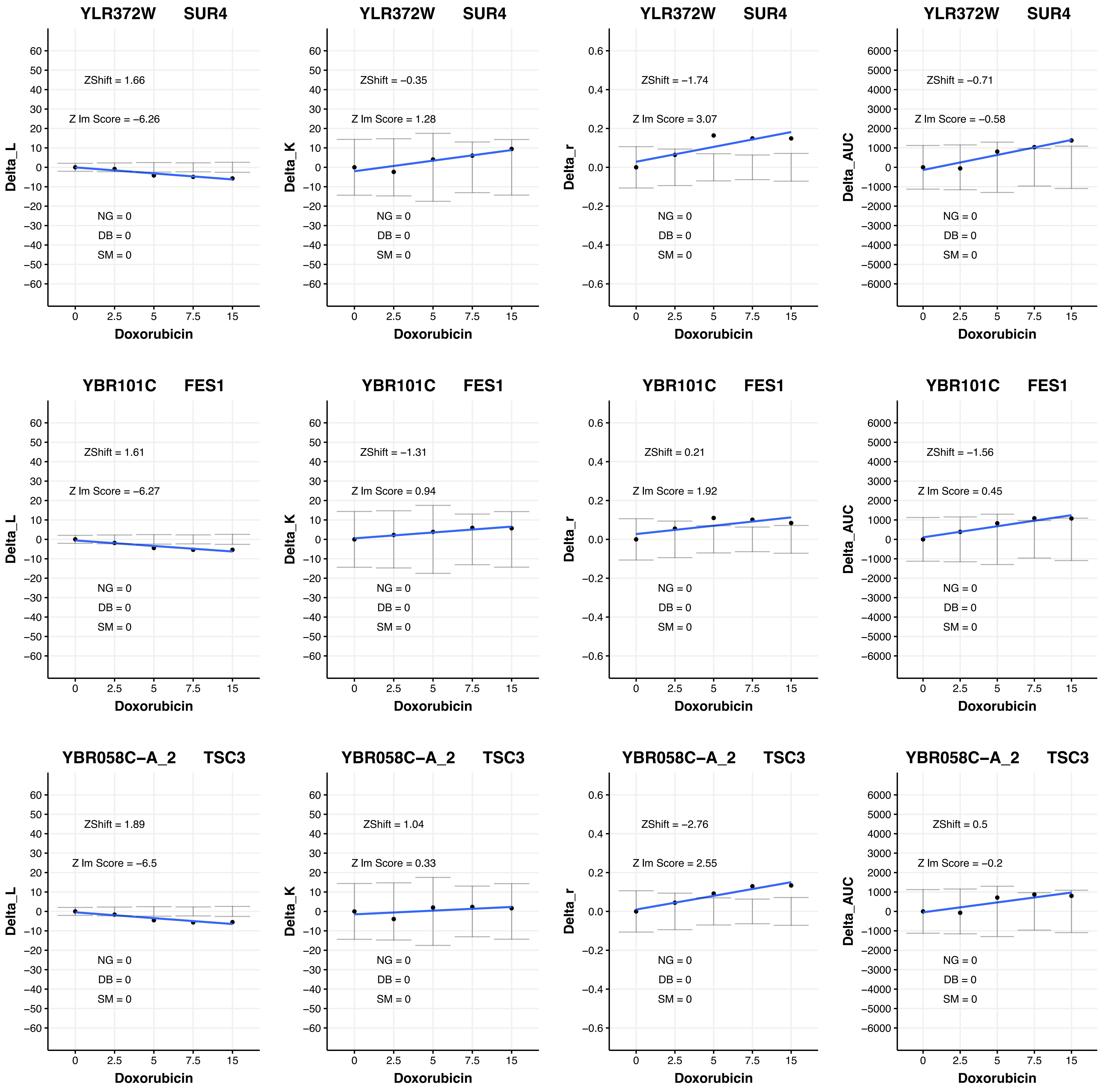
YLR372W SUR4
ZShift = 1.66
Z lm Score = −6.26
NG = 0
DB = 0
SM = 0
Delta_L
Doxorubicin
YLR372W SUR4
ZShift = −0.35
Z lm Score = 1.28
NG = 0
DB = 0
SM = 0
Delta_K
Doxorubicin
YLR372W SUR4
ZShift = −1.74
Z lm Score = 3.07
NG = 0
DB = 0
SM = 0
Delta_r
Doxorubicin
YLR372W SUR4
ZShift = −0.71
Z lm Score = −0.58
NG = 0
DB = 0
SM = 0
Delta_AUC
Doxorubicin
YBR101C FES1
ZShift = 1.61
Z lm Score = −6.27
NG = 0
DB = 0
SM = 0
Delta_L
Doxorubicin
YBR101C FES1
ZShift = −1.31
Z lm Score = 0.94
NG = 0
DB = 0
SM = 0
Delta_K
Doxorubicin
YBR101C FES1
ZShift = 0.21
Z lm Score = 1.92
NG = 0
DB = 0
SM = 0
Delta_r
Doxorubicin
YBR101C FES1
ZShift = −1.56
Z lm Score = 0.45
NG = 0
DB = 0
SM = 0
Delta_AUC
Doxorubicin
YBR058C−A_2 TSC3
ZShift = 1.89
Z lm Score = −6.5
NG = 0
DB = 0
SM = 0
Delta_L
Doxorubicin
YBR058C−A_2 TSC3
ZShift = 1.04
Z lm Score = 0.33
NG = 0
DB = 0
SM = 0
Delta_K
Doxorubicin
YBR058C−A_2 TSC3
ZShift = −2.76
Z lm Score = 2.55
NG = 0
DB = 0
SM = 0
Delta_r
Doxorubicin
YBR058C−A_2 TSC3
ZShift = 0.5
Z lm Score = −0.2
NG = 0
DB = 0
SM = 0
Delta_AUC
Doxorubicin

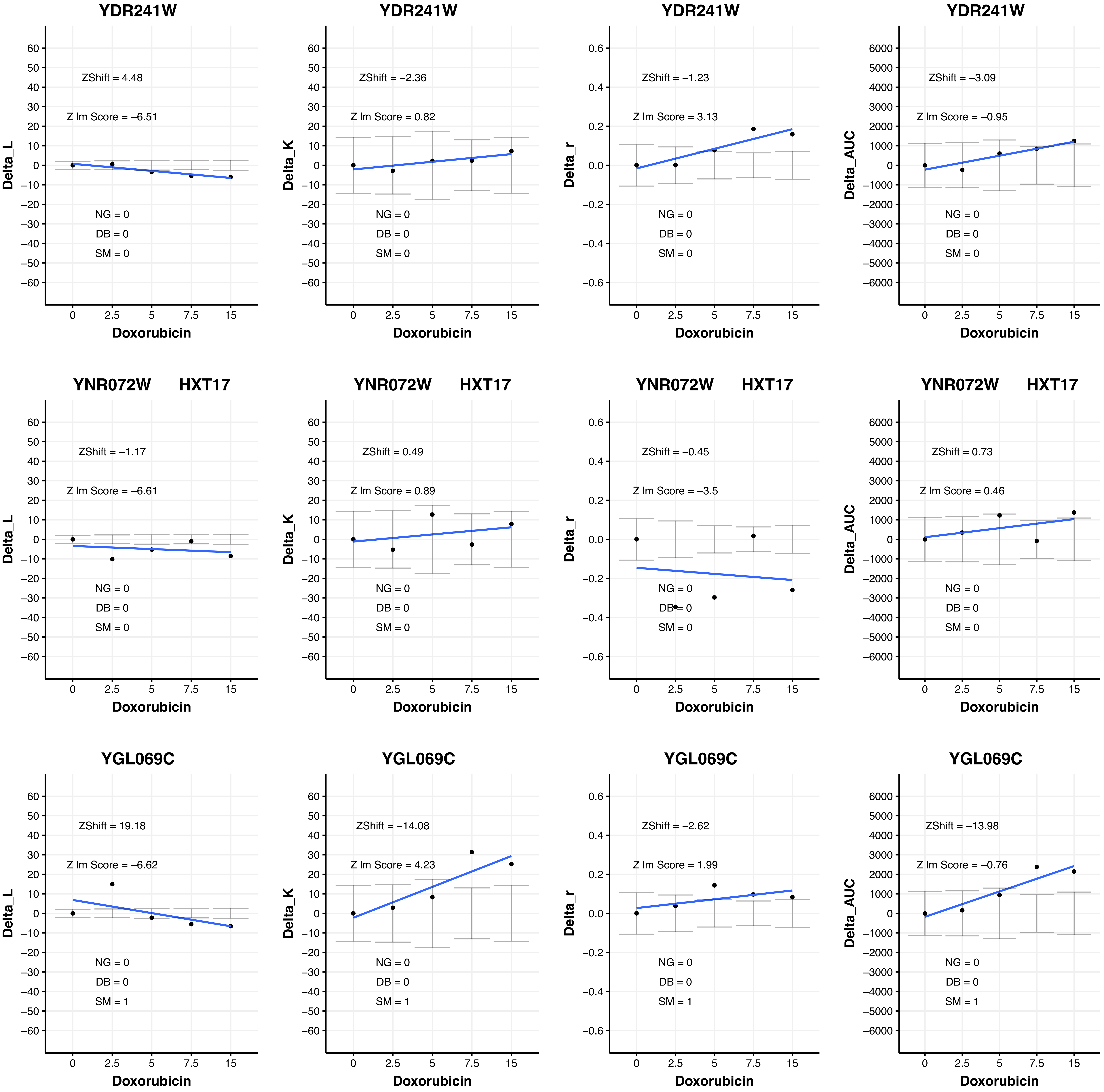

YDR241W
ZShift = 4.48
Z lm Score = −6.51
NG = 0
DB = 0
SM = 0
Delta_L
Doxorubicin
YDR241W
ZShift = −2.36
Z lm Score = 0.82
NG = 0
DB = 0
SM = 0
Delta_K
Doxorubicin
YDR241W
ZShift = −1.23
Z lm Score = 3.13
NG = 0
DB = 0
SM = 0
Delta_r
Doxorubicin
YDR241W
ZShift = −3.09
Z lm Score = −0.95
NG = 0
DB = 0
SM = 0
Delta_AUC
Doxorubicin
YNR072W HXT17
ZShift = −1.17
Z lm Score = −6.61
NG = 0
DB = 0
SM = 0
Delta_L
Doxorubicin
YNR072W HXT17
ZShift = 0.49
Z lm Score = 0.89
NG = 0
DB = 0
SM = 0
Delta_K
Doxorubicin
YNR072W HXT17
ZShift = −0.45
Z lm Score = −3.5
NG = 0
DB = 0
SM = 0
Delta_r
Doxorubicin
YNR072W HXT17
ZShift = 0.73
Z lm Score = 0.46
NG = 0
DB = 0
SM = 0
Delta_AUC
Doxorubicin
YGL069C
ZShift = 19.18
Z lm Score = −6.62
NG = 0
DB = 0
SM = 1
Delta_L
Doxorubicin
YGL069C
ZShift = −14.08
Z lm Score = 4.23
NG = 0
DB = 0
SM = 1
Delta_K
Doxorubicin
YGL069C
ZShift = −2.62
Z lm Score = 1.99
NG = 0
DB = 0
SM = 1
Delta_r
Doxorubicin
YGL069C
ZShift = −13.98
Z lm Score = −0.76
NG = 0
DB = 0
SM = 1
Delta_AUC
Doxorubicin

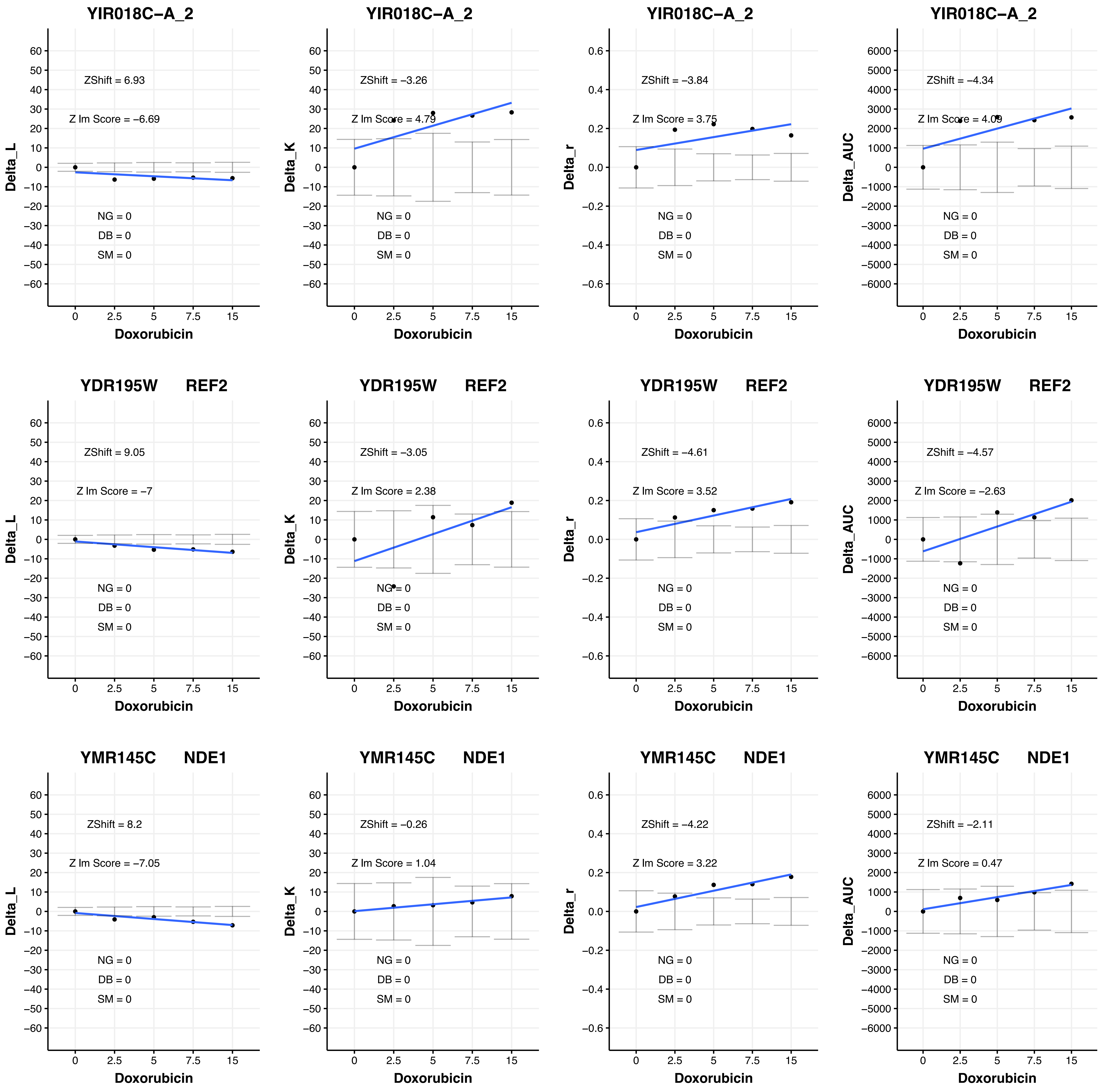

YIR018C-A_2
ZShift = 6.93
Z lm Score = −6.69
NG = 0
DB = 0
SM = 0
Delta_L
Doxorubicin
YIR018C-A_2
ZShift = −3.26
Z lm Score = 4.79
NG = 0
DB = 0
SM = 0
Delta_K
Doxorubicin
YIR018C-A_2
ZShift = −3.84
Z lm Score = 3.75
NG = 0
DB = 0
SM = 0
Delta_r
Doxorubicin
YIR018C-A_2
ZShift = −4.34
Z lm Score = 4.09
NG = 0
DB = 0
SM = 0
Delta_AUC
Doxorubicin
YDR195W REF2
ZShift = 9.05
Z lm Score = −7
NG = 0
DB = 0
SM = 0
Delta_L
Doxorubicin
YDR195W REF2
ZShift = −3.05
Z lm Score = 2.38
NG = 0
DB = 0
SM = 0
Delta_K
Doxorubicin
YDR195W REF2
ZShift = −4.61
Z lm Score = 3.52
NG = 0
DB = 0
SM = 0
Delta_r
Doxorubicin
YDR195W REF2
ZShift = −4.57
Z lm Score = −2.63
NG = 0
DB = 0
SM = 0
Delta_AUC
Doxorubicin
YMR145C NDE1
ZShift = 8.2
Z lm Score = −7.05
NG = 0
DB = 0
SM = 0
Delta_L
Doxorubicin
YMR145C NDE1
ZShift = −0.26
Z lm Score = 1.04
NG = 0
DB = 0
SM = 0
Delta_K
Doxorubicin
YMR145C NDE1
ZShift = −4.22
Z lm Score = 3.22
NG = 0
DB = 0
SM = 0
Delta_r
Doxorubicin
YMR145C NDE1
ZShift = −2.11
Z lm Score = 0.47
NG = 0
DB = 0
SM = 0
Delta_AUC
Doxorubicin

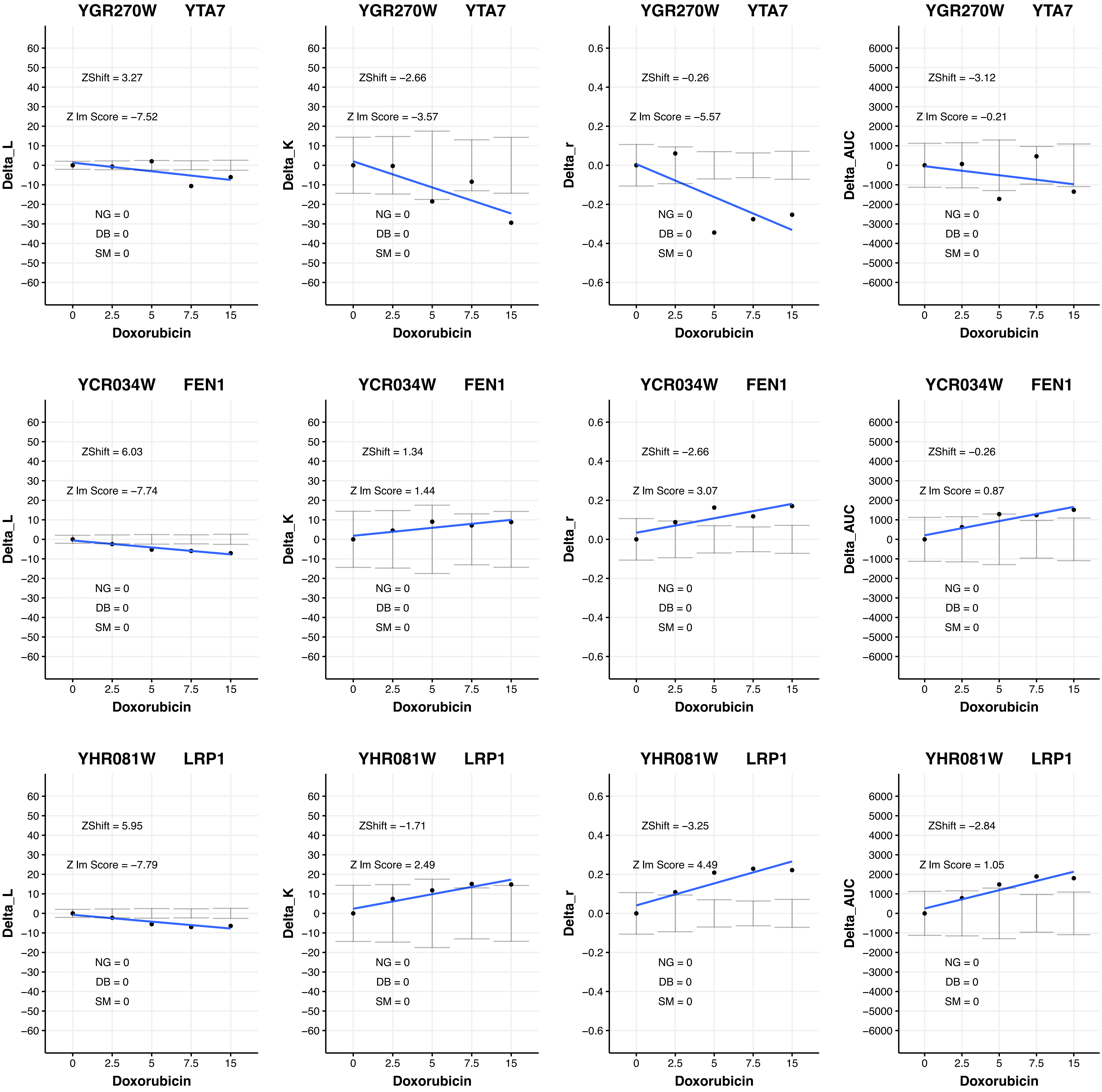

YGR270W YTA7
ZShift = 3.27
Z lm Score = −7.52
NG = 0
DB = 0
SM = 0
Delta_L
Doxorubicin
YGR270W YTA7
ZShift = −2.66
Z lm Score = −3.57
NG = 0
DB = 0
SM = 0
Delta_K
Doxorubicin
YGR270W YTA7
ZShift = −0.26
Z lm Score = −5.57
NG = 0
DB = 0
SM = 0
Delta_r
Doxorubicin
YGR270W YTA7
ZShift = −3.12
Z lm Score = −0.21
NG = 0
DB = 0
SM = 0
Delta_AUC
Doxorubicin
YCR034W FEN1
ZShift = 6.03
Z lm Score = −7.74
NG = 0
DB = 0
SM = 0
Delta_L
Doxorubicin
YCR034W FEN1
ZShift = 1.34
Z lm Score = 1.44
NG = 0
DB = 0
SM = 0
Delta_K
Doxorubicin
YCR034W FEN1
ZShift = −2.66
Z lm Score = 3.07
NG = 0
DB = 0
SM = 0
Delta_r
Doxorubicin
YCR034W FEN1
ZShift = −0.26
Z lm Score = 0.87
NG = 0
DB = 0
SM = 0
Delta_AUC
Doxorubicin
YHR081W LRP1
ZShift = 5.95
Z lm Score = −7.79
NG = 0
DB = 0
SM = 0
Delta_L
Doxorubicin
YHR081W LRP1
ZShift = −1.71
Z lm Score = 2.49
NG = 0
DB = 0
SM = 0
Delta_K
Doxorubicin
YHR081W LRP1
ZShift = −3.25
Z lm Score = 4.49
NG = 0
DB = 0
SM = 0
Delta_r
Doxorubicin
YHR081W LRP1
ZShift = −2.84
Z lm Score = 1.05
NG = 0
DB = 0
SM = 0
Delta_AUC
Doxorubicin

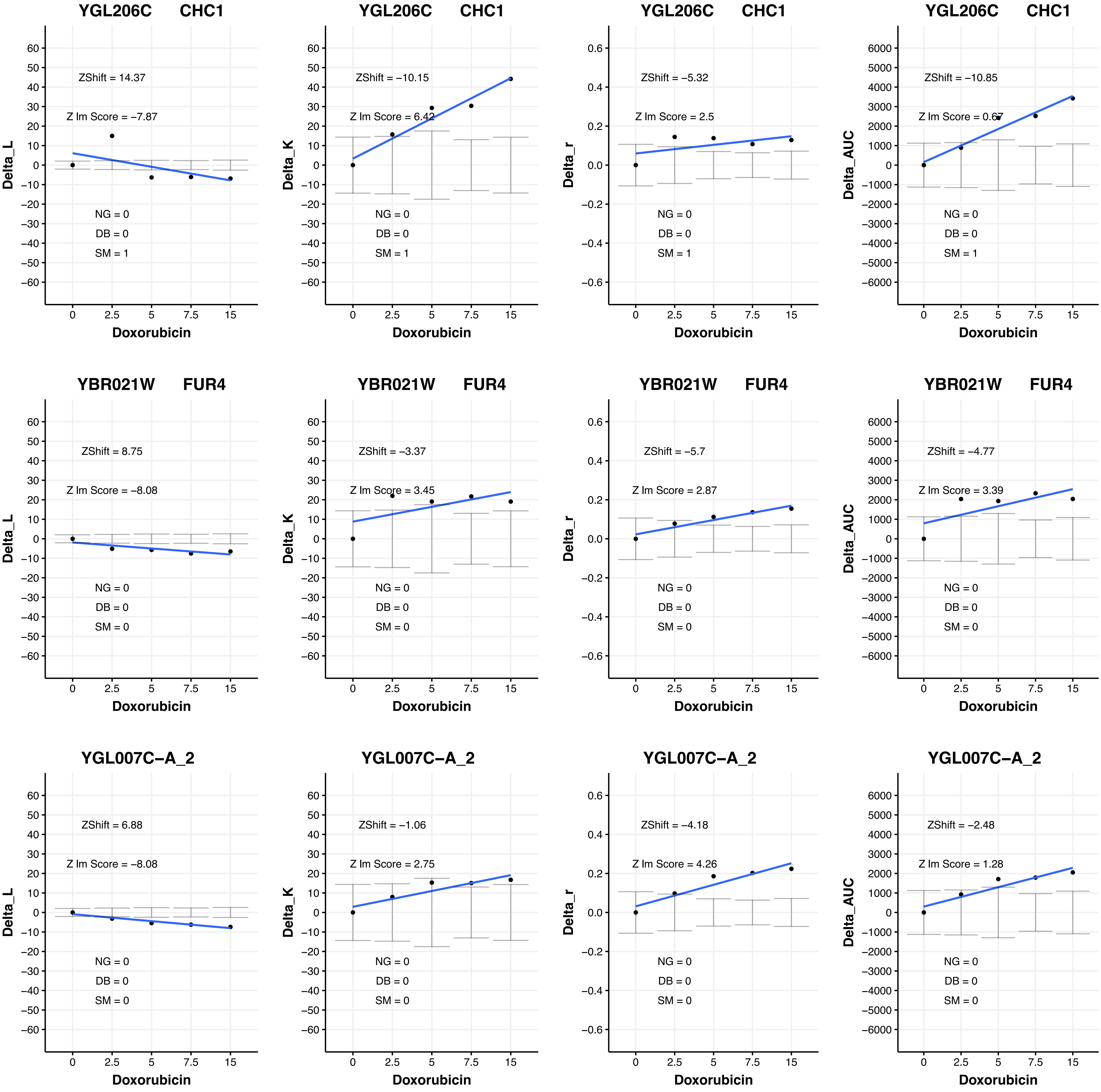
YGL206C CHC1
ZShift = 14.37
Z lm Score = −7.87
NG = 0
DB = 0
SM = 1
Delta_L
Doxorubicin
YGL206C CHC1
ZShift = −10.15
Z lm Score = 6.42
NG = 0
DB = 0
SM = 1
Delta_K
Doxorubicin
YGL206C CHC1
ZShift = −5.32
Z lm Score = 2.5
NG = 0
DB = 0
SM = 1
Delta_r
Doxorubicin
YGL206C CHC1
ZShift = −10.85
Z lm Score = 0.67
NG = 0
DB = 0
SM = 1
Delta_AUC
Doxorubicin
YBR021W FUR4
ZShift = 8.75
Z lm Score = −8.08
NG = 0
DB = 0
SM = 0
Delta_L
Doxorubicin
YBR021W FUR4
ZShift = −3.37
Z lm Score = 3.45
NG = 0
DB = 0
SM = 0
Delta_K
Doxorubicin
YBR021W FUR4
ZShift = −5.7
Z lm Score = 2.87
NG = 0
DB = 0
SM = 0
Delta_r
Doxorubicin
YBR021W FUR4
ZShift = −4.77
Z lm Score = 3.39
NG = 0
DB = 0
SM = 0
Delta_AUC
Doxorubicin
YGL007C−A_2
ZShift = 6.88
Z lm Score = −8.08
NG = 0
DB = 0
SM = 0
Delta_L
Doxorubicin
YGL007C−A_2
ZShift = −1.06
Z lm Score = 2.75
NG = 0
DB = 0
SM = 0
Delta_K
Doxorubicin
YGL007C−A_2
ZShift = −4.18
Z lm Score = 4.26
NG = 0
DB = 0
SM = 0
Delta_r
Doxorubicin
YGL007C−A_2
ZShift = −2.48
Z lm Score = 1.28
NG = 0
DB = 0
SM = 0
Delta_AUC
Doxorubicin

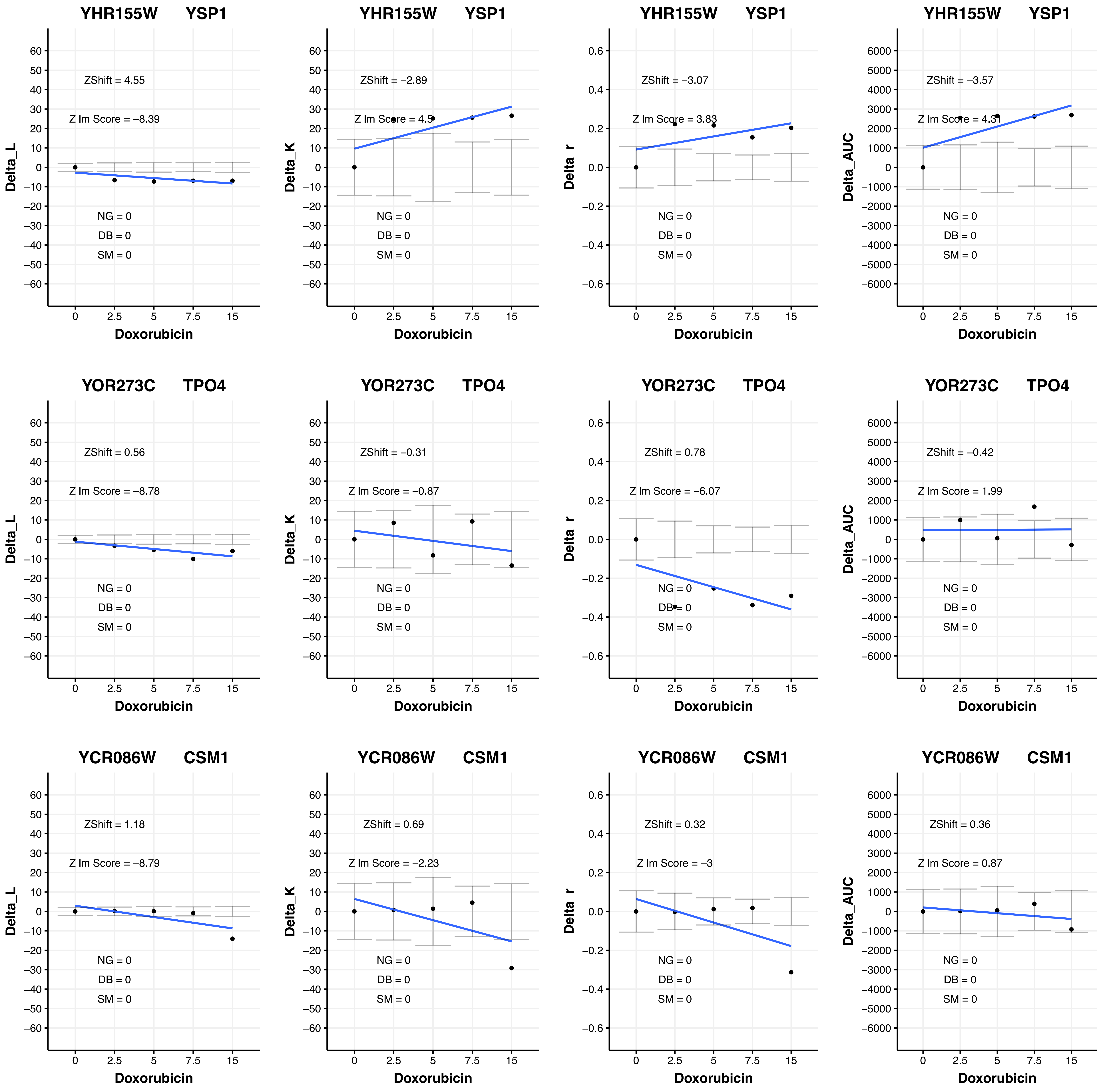

YHR155W YSP1
ZShift = 4.55
Z lm Score = −8.39
NG = 0
DB = 0
SM = 0
Delta_L
Doxorubicin
YHR155W YSP1
ZShift = −2.89
Z lm Score = 4.5
NG = 0
DB = 0
SM = 0
Delta_K
Doxorubicin
YHR155W YSP1
ZShift = −3.07
Z lm Score = 3.83
NG = 0
DB = 0
SM = 0
Delta_r
Doxorubicin
YHR155W YSP1
ZShift = −3.57
Z lm Score = 4.31
NG = 0
DB = 0
SM = 0
Delta_AUC
Doxorubicin
YOR273C TPO4
ZShift = 0.56
Z lm Score = −8.78
NG = 0
DB = 0
SM = 0
Delta_L
Doxorubicin
YOR273C TPO4
ZShift = −0.31
Z lm Score = −0.87
NG = 0
DB = 0
SM = 0
Delta_K
Doxorubicin
YOR273C TPO4
ZShift = 0.78
Z lm Score = −6.07
NG = 0
DB = 0
SM = 0
Delta_r
Doxorubicin
YOR273C TPO4
ZShift = −0.42
Z lm Score = 1.99
NG = 0
DB = 0
SM = 0
Delta_AUC
Doxorubicin
YCR086W CSM1
ZShift = 1.18
Z lm Score = −8.79
NG = 0
DB = 0
SM = 0
Delta_L
Doxorubicin
YCR086W CSM1
ZShift = 0.69
Z lm Score = −2.23
NG = 0
DB = 0
SM = 0
Delta_K
Doxorubicin
YCR086W CSM1
ZShift = 0.32
Z lm Score = −3
NG = 0
DB = 0
SM = 0
Delta_r
Doxorubicin
YCR086W CSM1
ZShift = 0.36
Z lm Score = 0.87
NG = 0
DB = 0
SM = 0
Delta_AUC
Doxorubicin

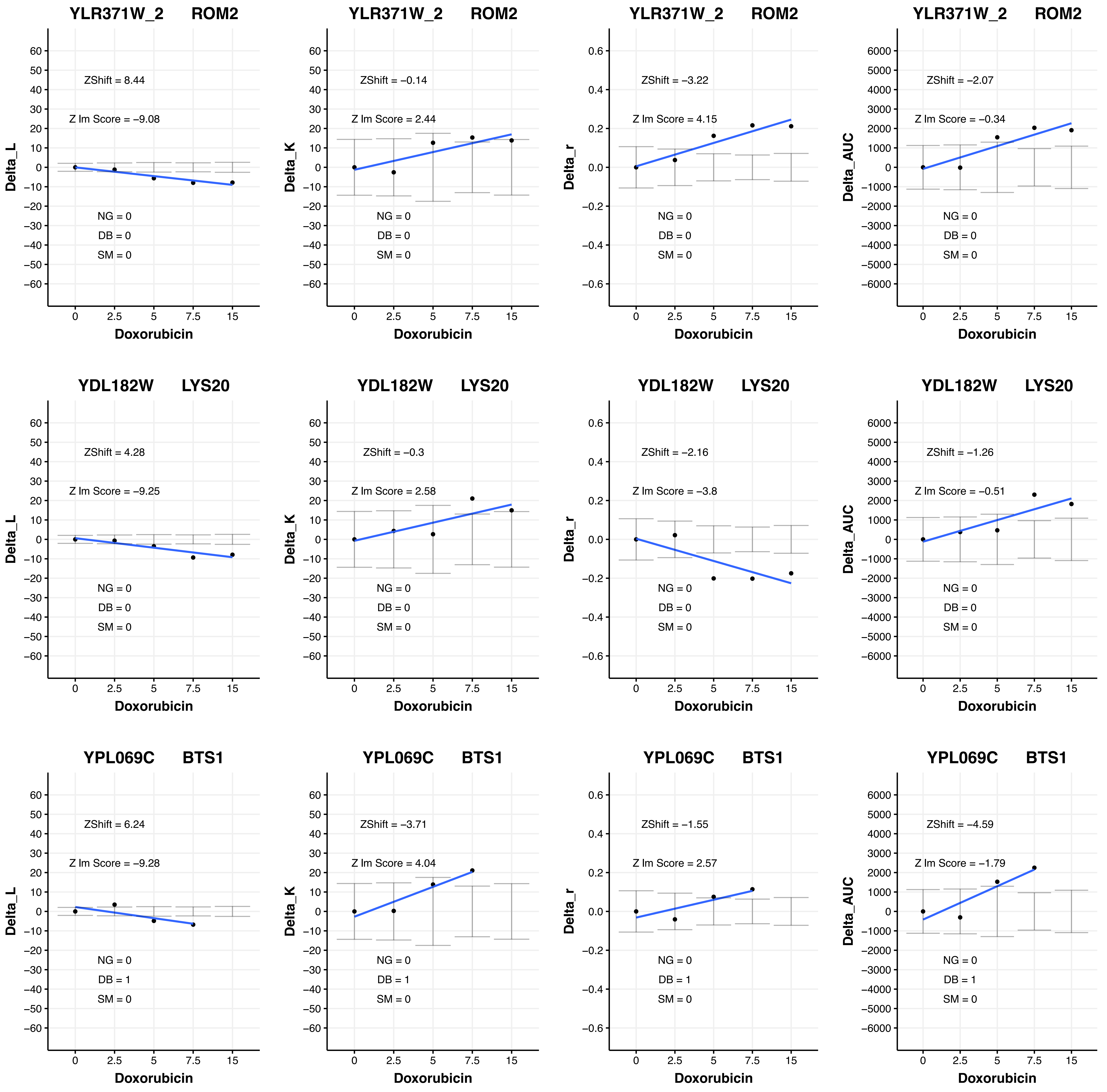

YLR371W_2 ROM2
ZShift = 8.44
Z lm Score = −9.08
NG = 0
DB = 0
SM = 0
Delta_L
Doxorubicin
YLR371W_2 ROM2
ZShift = −0.14
Z lm Score = 2.44
NG = 0
DB = 0
SM = 0
Delta_K
Doxorubicin
YLR371W_2 ROM2
ZShift = −3.22
Z lm Score = 4.15
NG = 0
DB = 0
SM = 0
Delta_r
Doxorubicin
YLR371W_2 ROM2
ZShift = −2.07
Z lm Score = −0.34
NG = 0
DB = 0
SM = 0
Delta_AUC
Doxorubicin
YDL182W LYS20
ZShift = 4.28
Z lm Score = −9.25
NG = 0
DB = 0
SM = 0
Delta_L
Doxorubicin
YDL182W LYS20
ZShift = −0.3
Z lm Score = 2.58
NG = 0
DB = 0
SM = 0
Delta_K
Doxorubicin
YDL182W LYS20
ZShift = −2.16
Z lm Score = −3.8
NG = 0
DB = 0
SM = 0
Delta_r
Doxorubicin
YDL182W LYS20
ZShift = −1.26
Z lm Score = −0.51
NG = 0
DB = 0
SM = 0
Delta_AUC
Doxorubicin
YPL069C BTS1
ZShift = 6.24
Z lm Score = −9.28
NG = 0
DB = 1
SM = 0
Delta_L
Doxorubicin
YPL069C BTS1
ZShift = −3.71
Z lm Score = 4.04
NG = 0
DB = 1
SM = 0
Delta_K
Doxorubicin
YPL069C BTS1
ZShift = −1.55
Z lm Score = 2.57
NG = 0
DB = 1
SM = 0
Delta_r
Doxorubicin
YPL069C BTS1
ZShift = −4.59
Z lm Score = −1.79
NG = 0
DB = 1
SM = 0
Delta_AUC
Doxorubicin

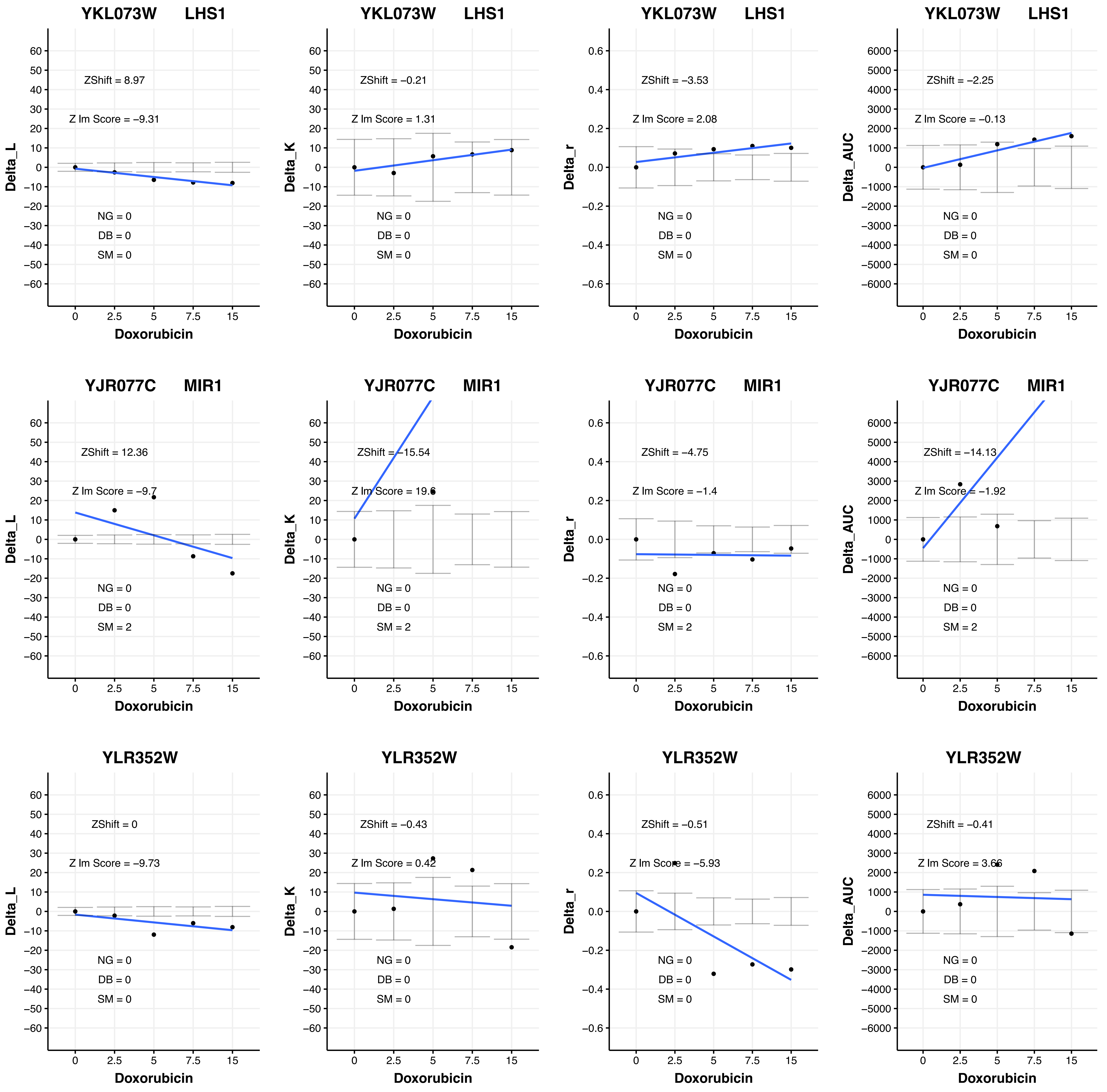
YKL073W LHS1
ZShift = 8.97
Z lm Score = −9.31
NG = 0
DB = 0
SM = 0
Delta_L
Doxorubicin
YKL073W LHS1
ZShift = −0.21
Z lm Score = 1.31
NG = 0
DB = 0
SM = 0
Delta_K
Doxorubicin
YKL073W LHS1
ZShift = −3.53
Z lm Score = 2.08
NG = 0
DB = 0
SM = 0
Delta_r
Doxorubicin
YKL073W LHS1
ZShift = −2.25
Z lm Score = −0.13
NG = 0
DB = 0
SM = 0
Delta_AUC
Doxorubicin
YJR077C MIR1
ZShift = 12.36
Z lm Score = −9.7
NG = 0
DB = 0
SM = 2
Delta_L
Doxorubicin
YJR077C MIR1
ZShift = −15.54
Z lm Score = 19.6
NG = 0
DB = 0
SM = 2
Delta_K
Doxorubicin
YJR077C MIR1
ZShift = −4.75
Z lm Score = −1.4
NG = 0
DB = 0
SM = 2
Delta_r
Doxorubicin
YJR077C MIR1
ZShift = −14.13
Z lm Score = −1.92
NG = 0
DB = 0
SM = 2
Delta_AUC
Doxorubicin
YLR352W
ZShift = 0
Z lm Score = −9.73
NG = 0
DB = 0
SM = 0
Delta_L
Doxorubicin
YLR352W
ZShift = −0.43
Z lm Score = 0.42
NG = 0
DB = 0
SM = 0
Delta_K
Doxorubicin
YLR352W
ZShift = −0.51
Z lm Score = −5.93
NG = 0
DB = 0
SM = 0
Delta_r
Doxorubicin
YLR352W
ZShift = −0.41
Z lm Score = 3.66
NG = 0
DB = 0
SM = 0
Delta_AUC
Doxorubicin

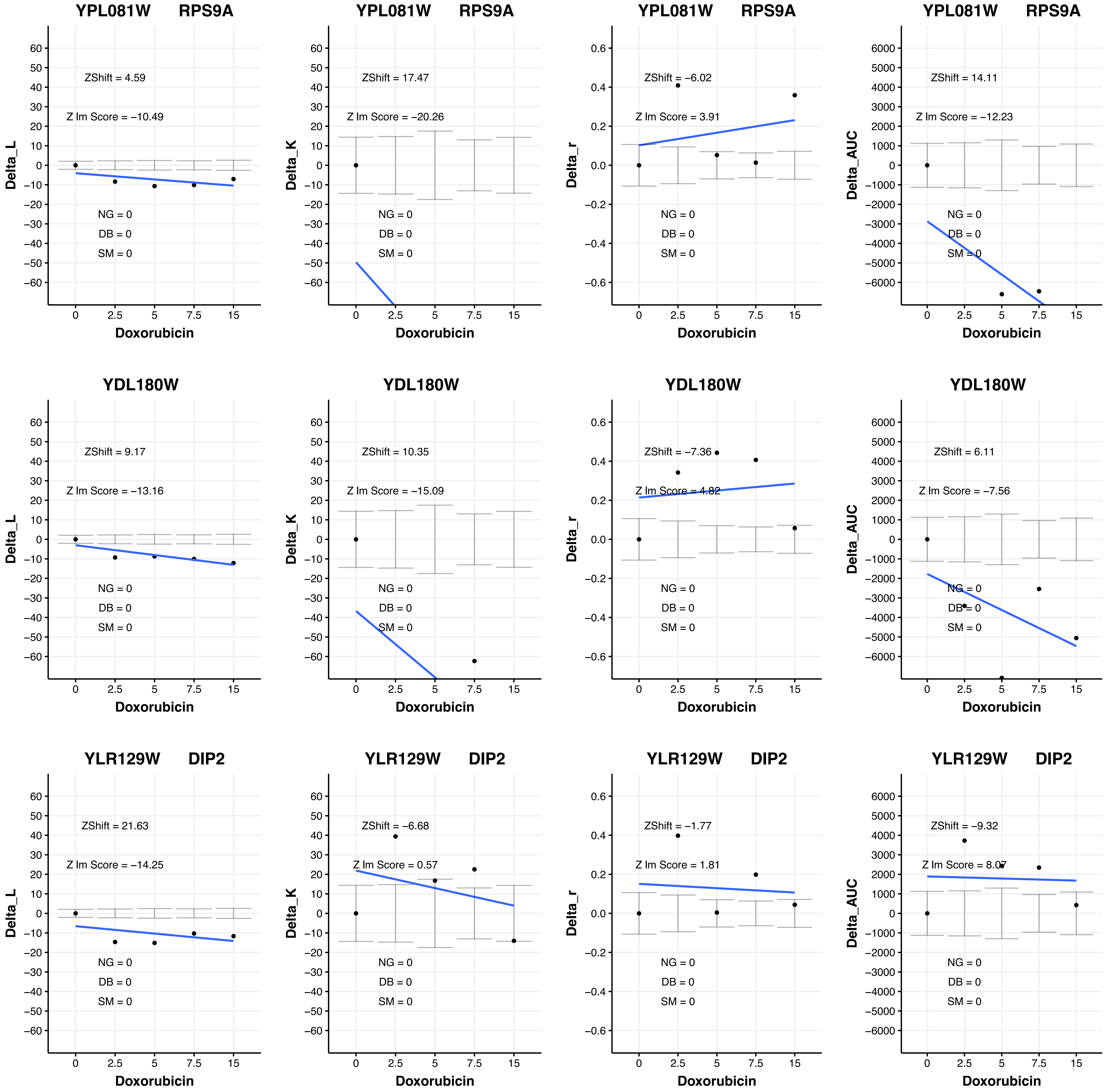
YPL081W RPS9A
ZShift = 4.59
Z lm Score = −10.49
NG = 0
DB = 0
SM = 0
Delta_L
Doxorubicin
YPL081W RPS9A
ZShift = 17.47
Z lm Score = −20.26
NG = 0
DB = 0
SM = 0
Delta_K
Doxorubicin
YPL081W RPS9A
ZShift = −6.02
Z lm Score = 3.91
NG = 0
DB = 0
SM = 0
Delta_r
Doxorubicin
YPL081W RPS9A
ZShift = 14.11
Z lm Score = −12.23
NG = 0
DB = 0
SM = 0
Delta_AUC
Doxorubicin
YDL180W
ZShift = 9.17
Z lm Score = −13.16
NG = 0
DB = 0
SM = 0
Delta_L
Doxorubicin
YDL180W
ZShift = 10.35
Z lm Score = −15.09
NG = 0
DB = 0
SM = 0
Delta_K
Doxorubicin
YDL180W
ZShift = −7.36
Z lm Score = 4.82
NG = 0
DB = 0
SM = 0
Delta_r
Doxorubicin
YDL180W
ZShift = 6.11
Z lm Score = −7.56
NG = 0
DB = 0
SM = 0
Delta_AUC
Doxorubicin
YLR129W DIP2
ZShift = 21.63
Z lm Score = −14.25
NG = 0
DB = 0
SM = 0
Delta_L
Doxorubicin
YLR129W DIP2
ZShift = −6.68
Z lm Score = 0.57
NG = 0
DB = 0
SM = 0
Delta_K
Doxorubicin
YLR129W DIP2
ZShift = −1.77
Z lm Score = 1.81
NG = 0
DB = 0
SM = 0
Delta_r
Doxorubicin
YLR129W DIP2
ZShift = −9.32
Z lm Score = 8.07
NG = 0
DB = 0
SM = 0
Delta_AUC
Doxorubicin

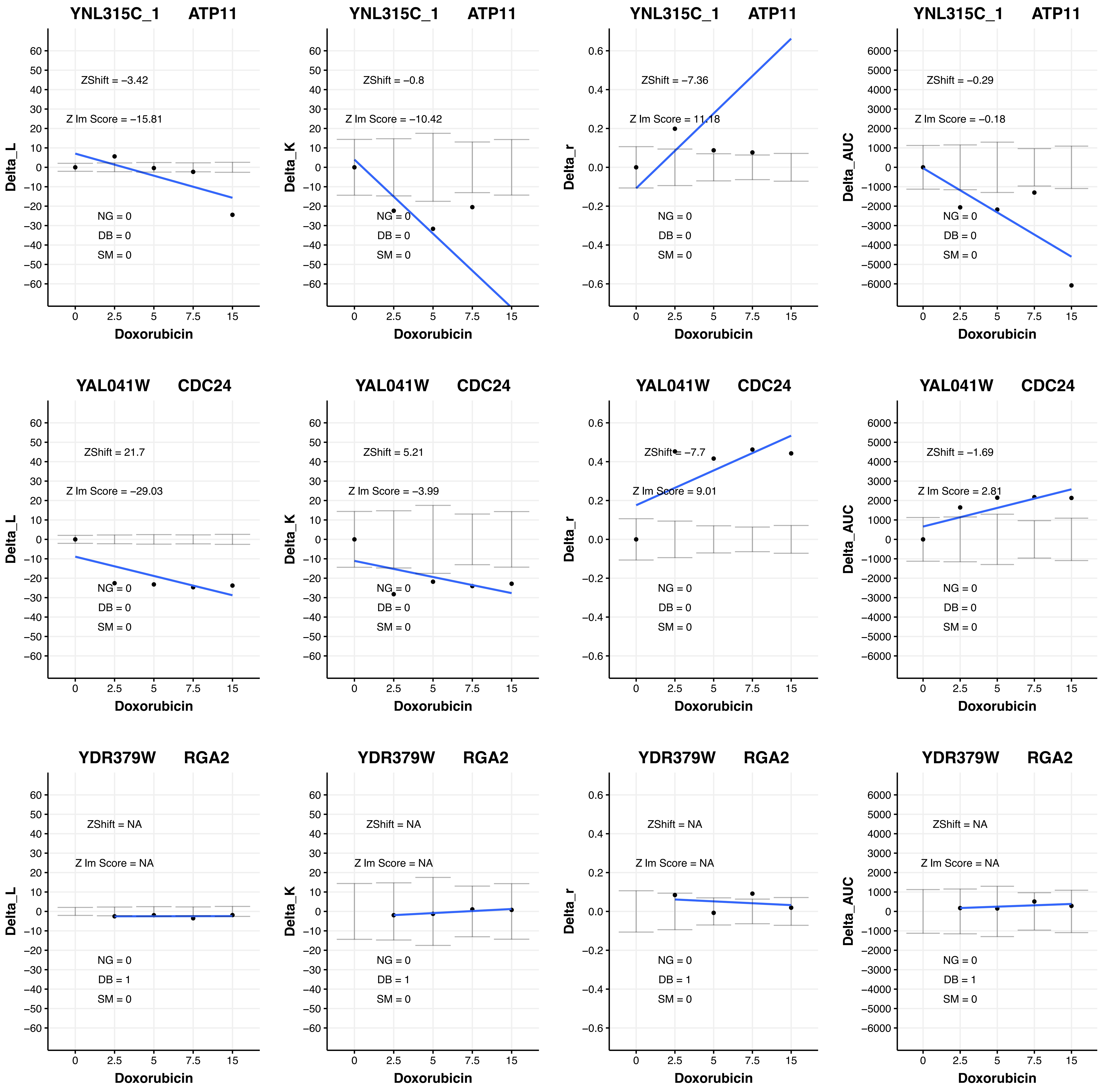

YNL315C_1 ATP11
ZShift = −3.42
Z lm Score = −15.81
NG = 0
DB = 0
SM = 0
Delta_L
Doxorubicin
YNL315C_1 ATP11
ZShift = −0.8
Z lm Score = −10.42
NG = 0
DB = 0
SM = 0
Delta_K
Doxorubicin
YNL315C_1 ATP11
ZShift = −7.36
Z lm Score = 11.18
NG = 0
DB = 0
SM = 0
Delta_r
Doxorubicin
YNL315C_1 ATP11
ZShift = −0.29
Z lm Score = −0.18
NG = 0
DB = 0
SM = 0
Delta_AUC
Doxorubicin
YAL041W CDC24
ZShift = 21.7
Z lm Score = −29.03
NG = 0
DB = 0
SM = 0
Delta_L
Doxorubicin
YAL041W CDC24
ZShift = 5.21
Z lm Score = −3.99
NG = 0
DB = 0
SM = 0
Delta_K
Doxorubicin
YAL041W CDC24
ZShift = −7.7
Z lm Score = 9.01
NG = 0
DB = 0
SM = 0
Delta_r
Doxorubicin
YAL041W CDC24
ZShift = −1.69
Z lm Score = 2.81
NG = 0
DB = 0
SM = 0
Delta_AUC
Doxorubicin
YDR379W RGA2
ZShift = NA
Z lm Score = NA
NG = 0
DB = 1
SM = 0
Delta_L
Doxorubicin
YDR379W RGA2
ZShift = NA
Z lm Score = NA
NG = 0
DB = 1
SM = 0
Delta_K
Doxorubicin
YDR379W RGA2
ZShift = NA
Z lm Score = NA
NG = 0
DB = 1
SM = 0
Delta_r
Doxorubicin
YDR379W RGA2
ZShift = NA
Z lm Score = NA
NG = 0
DB = 1
SM = 0
Delta_AUC
Doxorubicin
