## Additional File 4 for "A yeast phenomic model for the influence of Warburg metabolism on genetic buffering of doxorubicin": B - RF_InteractionPlots_Doxo_HLD.pdf

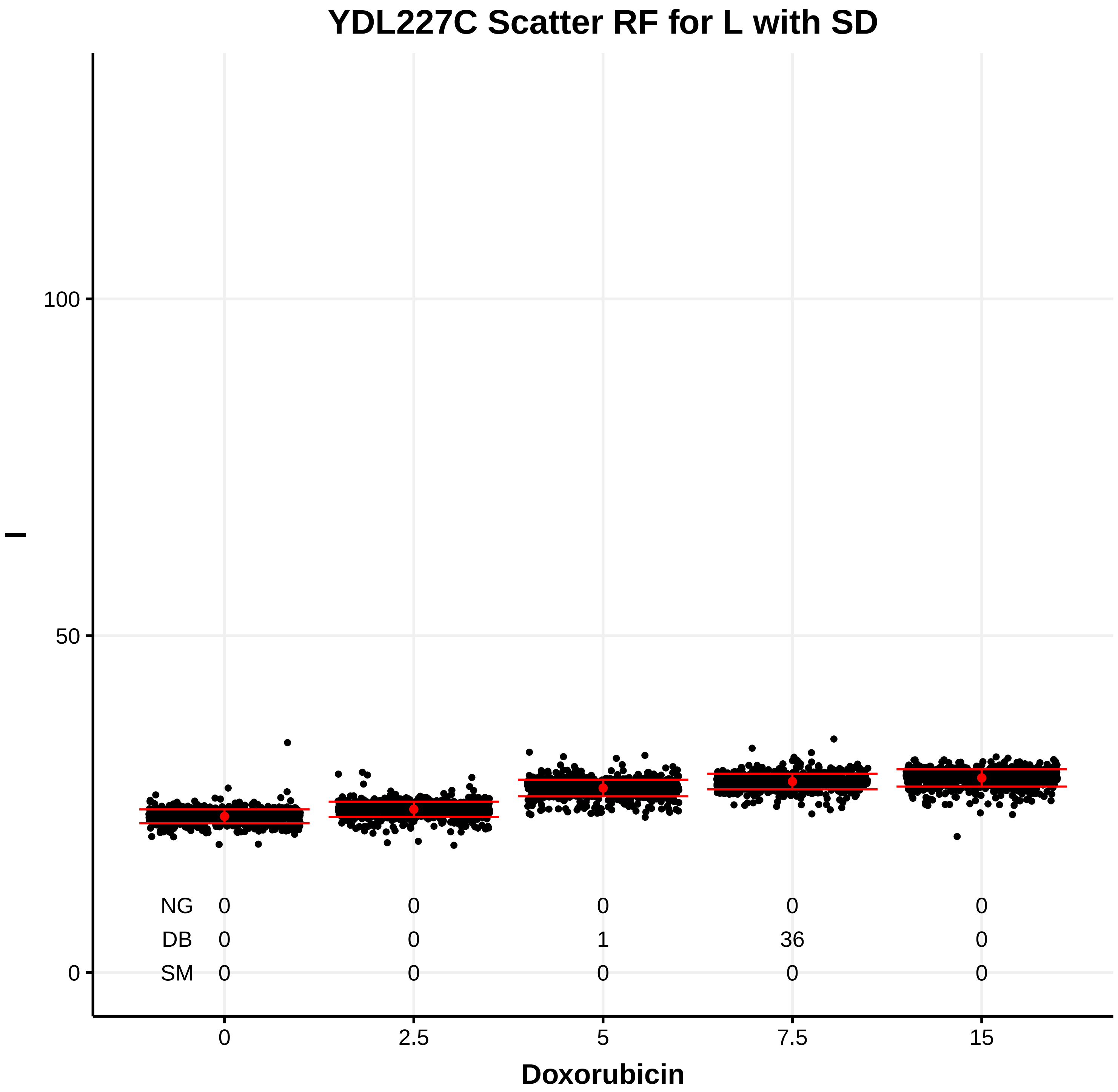
YDL227C Scatter RF for L with SD
l
0
50
100
NG 0 0 0 0 0
DB 0 0 1 36 0
SM 0 0 0 0 0
0
2.5
5
7.5
15
Doxorubicin

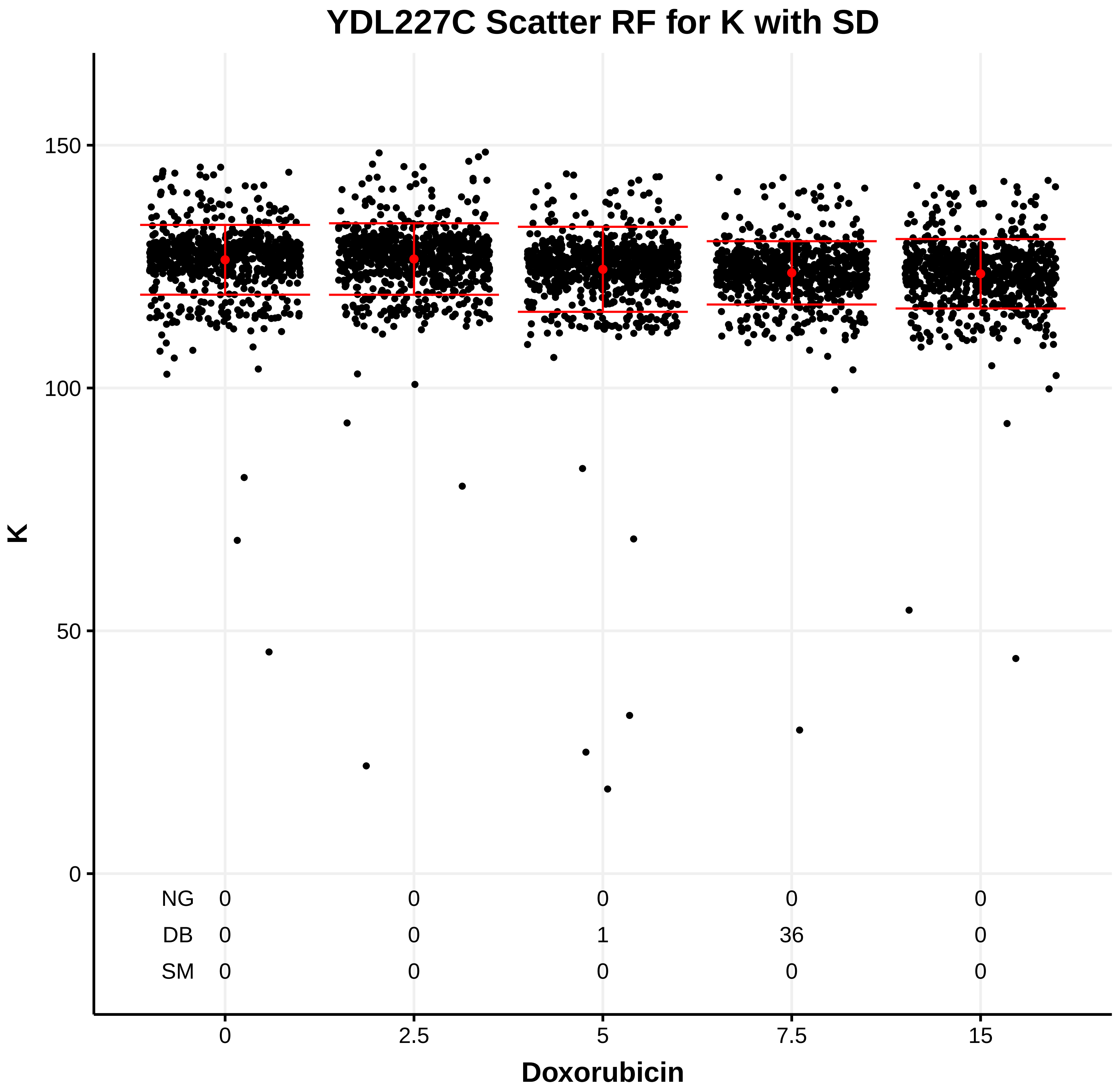
YDL227C Scatter RF for K with SD
K
0
50
100
150
NG 0 0 0 0 0
DB 0 0 1 36 0
SM 0 0 0 0 0
0
2.5
5
7.5
15
Doxorubicin

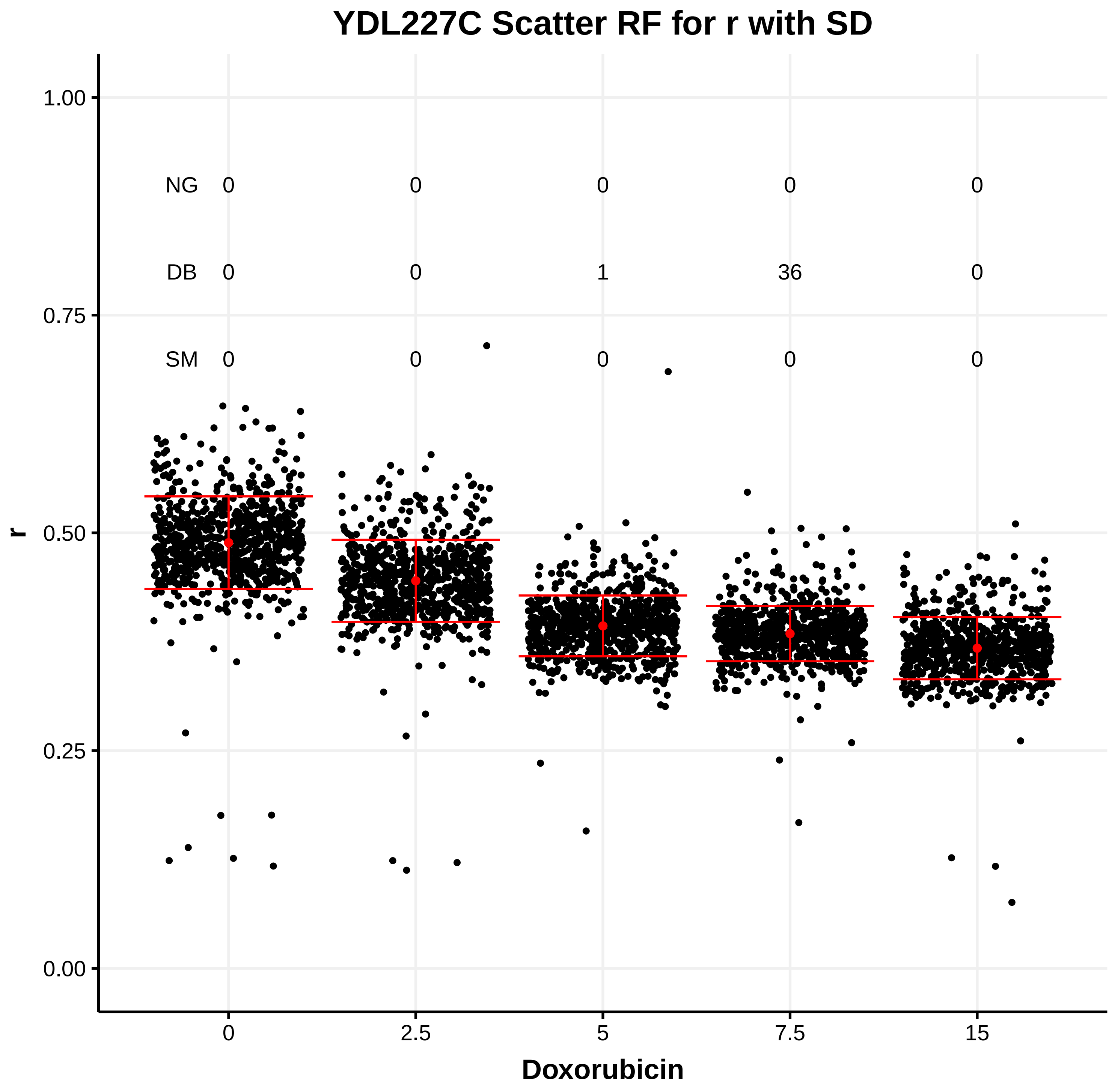
YDL227C Scatter RF for r with SD
r
0.00
0.25
0.50
0.75
1.00
NG 0 0 0 0 0
DB 0 0 1 36 0
SM 0 0 0 0 0
0
2.5
5
7.5
15
Doxorubicin

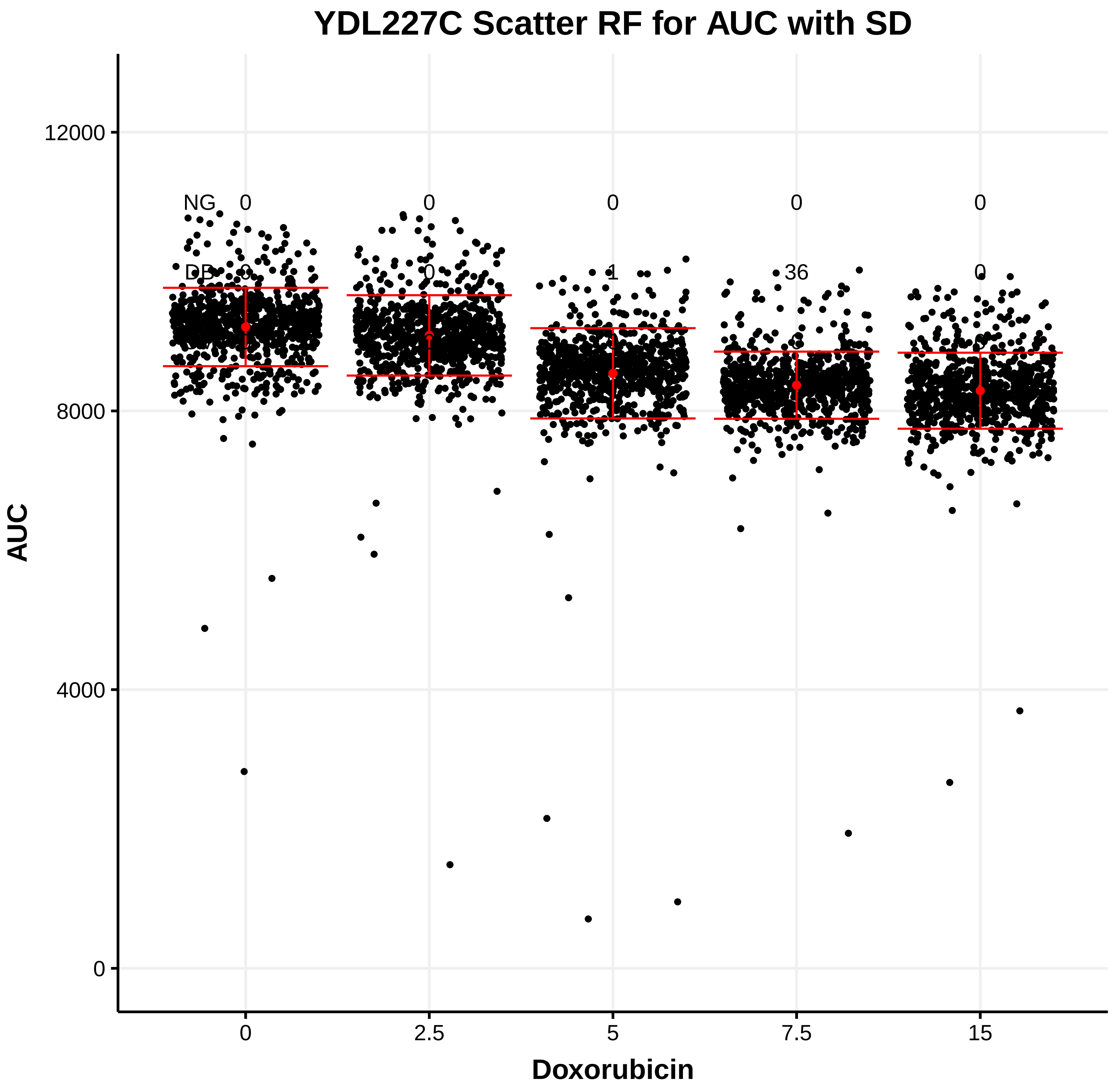
YDL227C Scatter RF for AUC with SD
AUC
0
4000
8000
12000
NG 0 0 0 0 0
DB 0 0 1 36 0
0
2.5
5
7.5
15
Doxorubicin

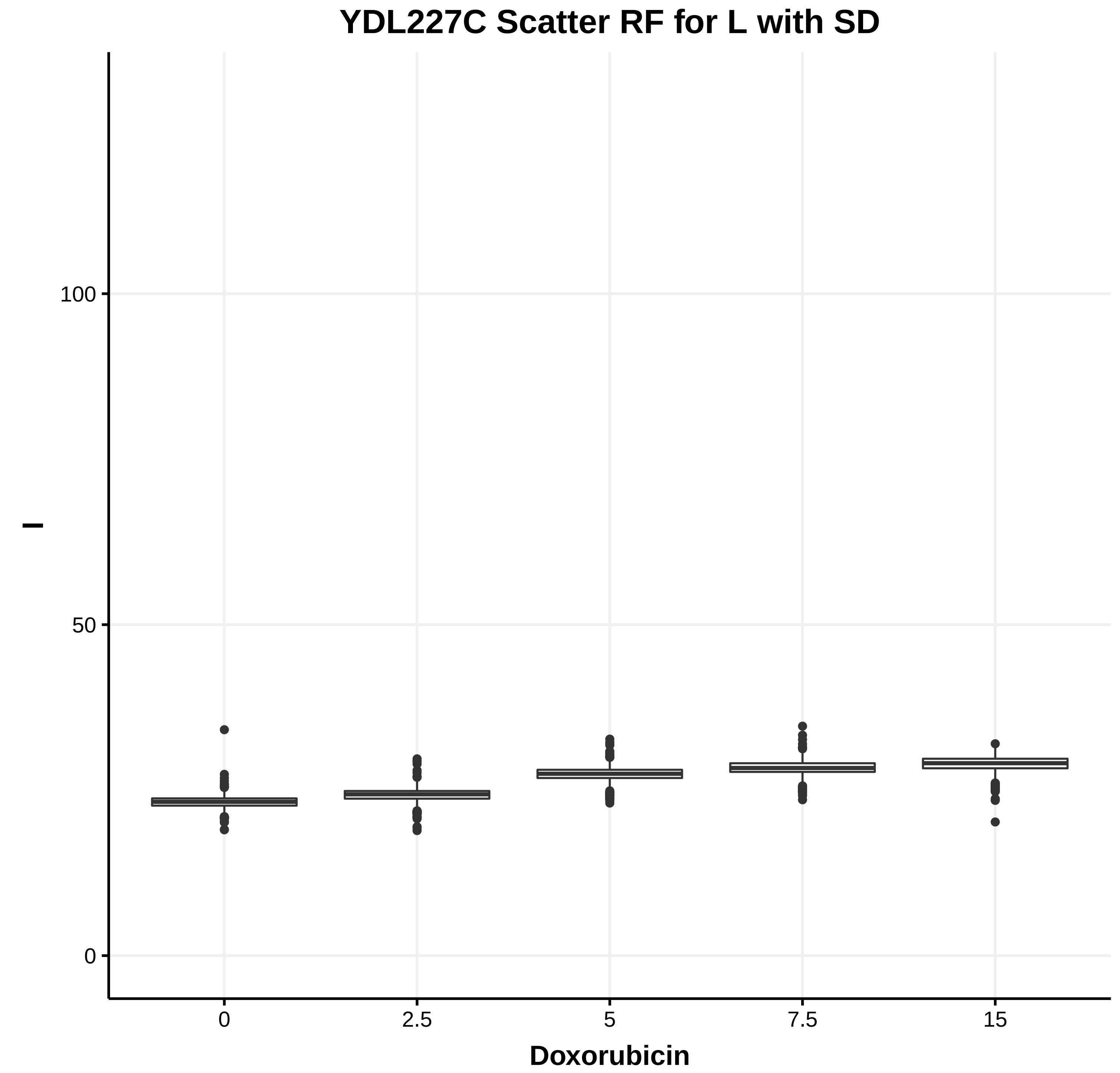
YDL227C Scatter RF for L with SD
l
0
50
100
0
2.5
5
7.5
15
Doxorubicin

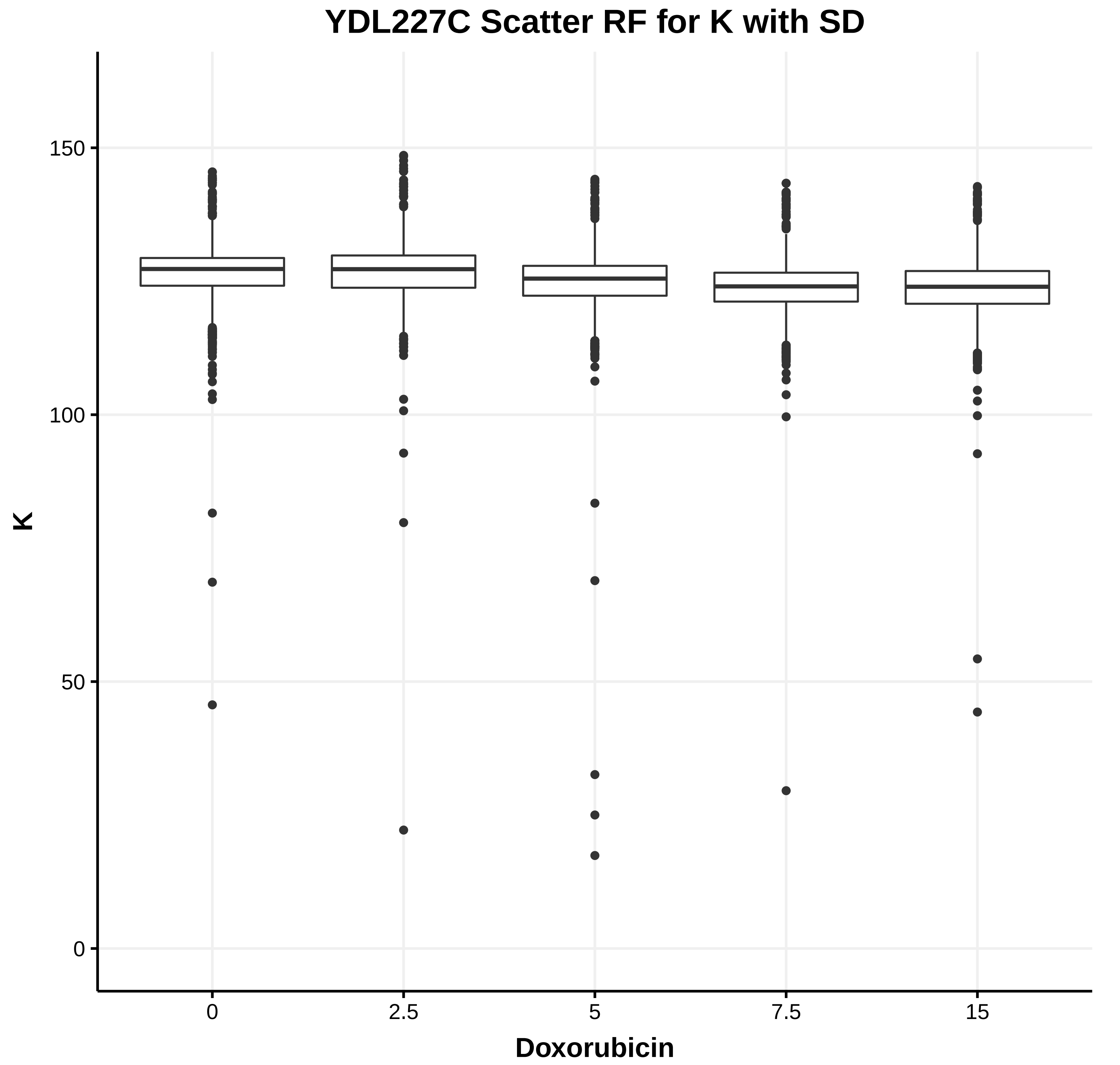
YDL227C Scatter RF for K with SD
K
0
50
100
150
0
2.5
5
7.5
15
Doxorubicin

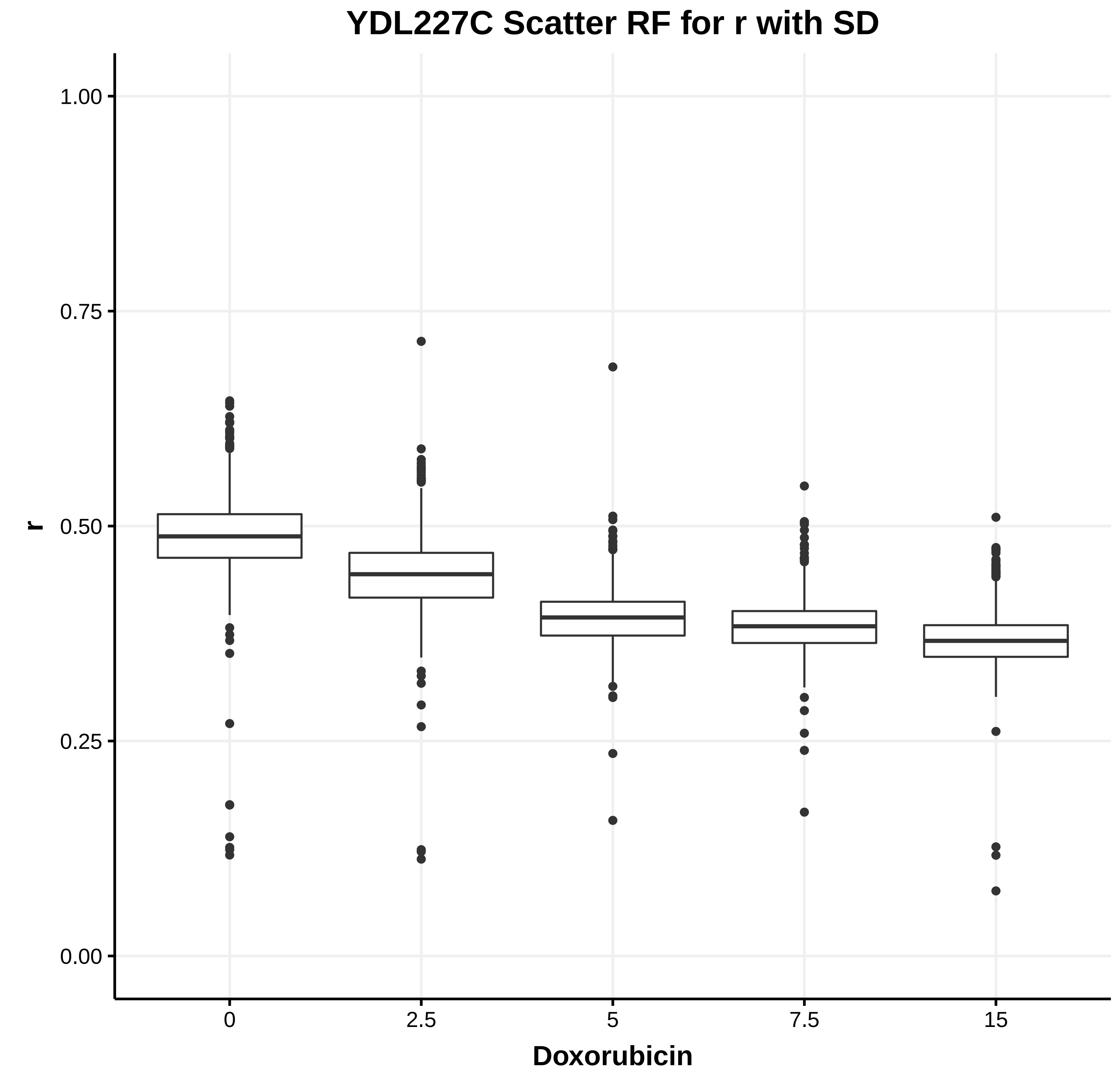
YDL227C Scatter RF for r with SD
r
0.00
0.25
0.50
0.75
1.00
0
2.5
5
7.5
15
Doxorubicin

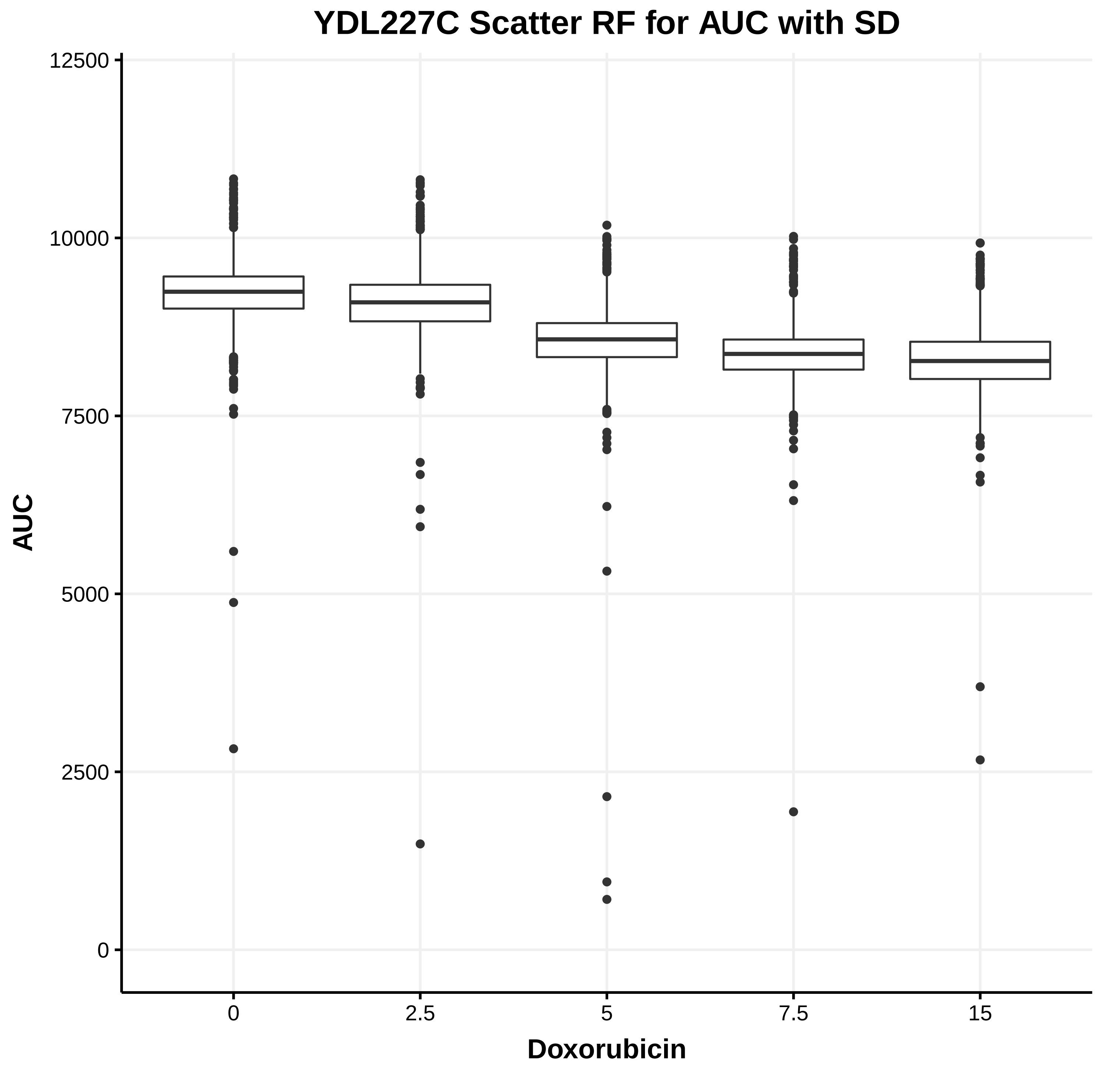
YDL227C Scatter RF for AUC with SD
AUC
0
2500
5000
7500
10000
12500
0
2.5
5
7.5
15
Doxorubicin

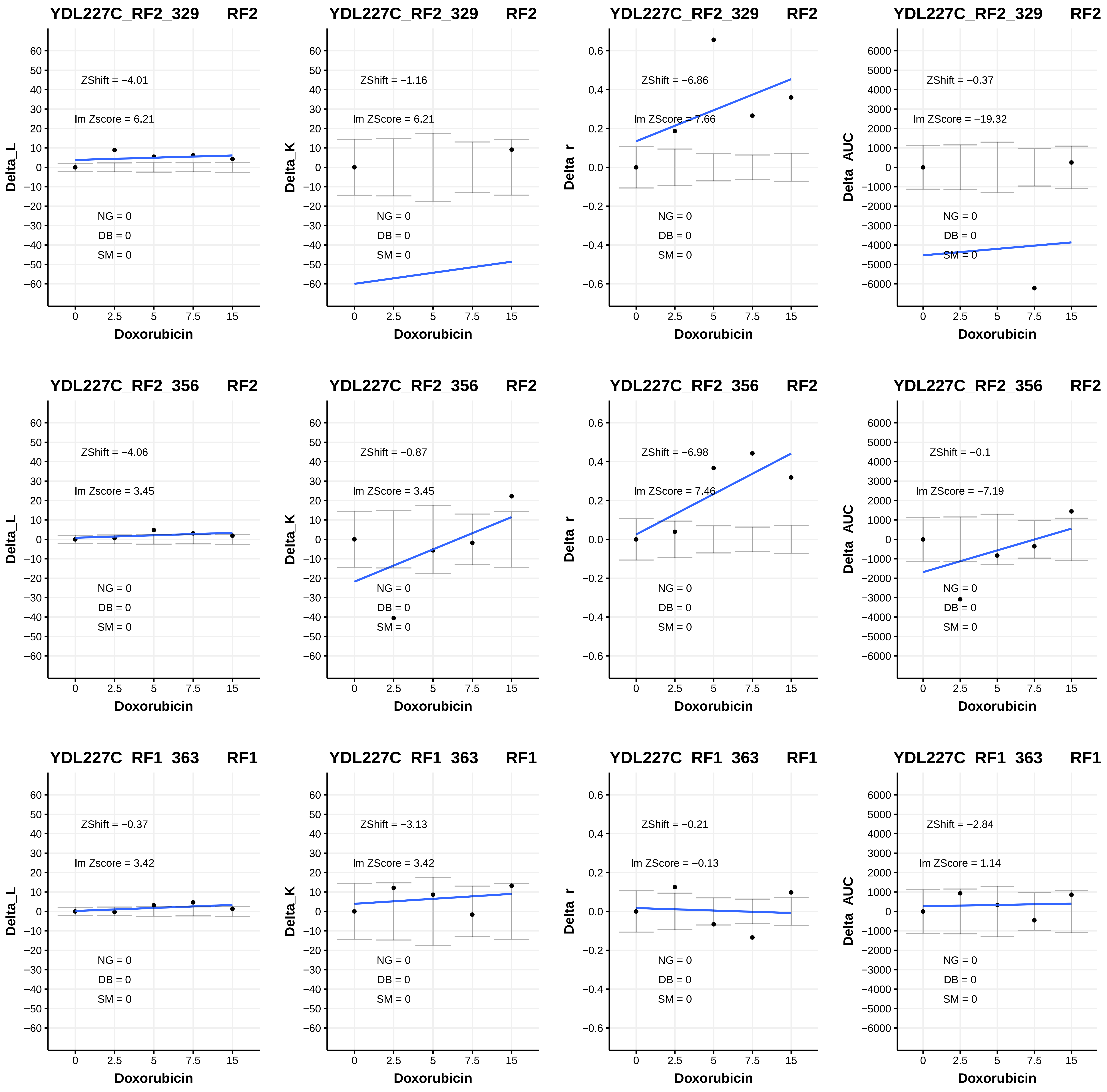
YDL227C_RF2_329 RF2
ZShift = −4.01
lm Zscore = 6.21
NG = 0
DB = 0
SM = 0
Delta_L
Doxorubicin
YDL227C_RF2_329 RF2
ZShift = −1.16
lm ZScore = 6.21
NG = 0
DB = 0
SM = 0
Delta_K
Doxorubicin
YDL227C_RF2_329 RF2
ZShift = −6.86
lm ZScore = 7.66
NG = 0
DB = 0
SM = 0
Delta_r
Doxorubicin
YDL227C_RF2_329 RF2
ZShift = −0.37
lm ZScore = −19.32
NG = 0
DB = 0
SM = 0
Delta_AUC
Doxorubicin
YDL227C_RF2_356 RF2
ZShift = −4.06
lm Zscore = 3.45
NG = 0
DB = 0
SM = 0
Delta_L
Doxorubicin
YDL227C_RF2_356 RF2
ZShift = −0.87
lm ZScore = 3.45
NG = 0
DB = 0
SM = 0
Delta_K
Doxorubicin
YDL227C_RF2_356 RF2
ZShift = −6.98
lm ZScore = 7.46
NG = 0
DB = 0
SM = 0
Delta_r
Doxorubicin
YDL227C_RF2_356 RF2
ZShift = −0.1
lm ZScore = −7.19
NG = 0
DB = 0
SM = 0
Delta_AUC
Doxorubicin
YDL227C_RF1_363 RF1
ZShift = −0.37
lm Zscore = 3.42
NG = 0
DB = 0
SM = 0
Delta_L
Doxorubicin
YDL227C_RF1_363 RF1
ZShift = −3.13
lm ZScore = 3.42
NG = 0
DB = 0
SM = 0
Delta_K
Doxorubicin
YDL227C_RF1_363 RF1
ZShift = −0.21
lm ZScore = −0.13
NG = 0
DB = 0
SM = 0
Delta_r
Doxorubicin
YDL227C_RF1_363 RF1
ZShift = −2.84
lm ZScore = 1.14
NG = 0
DB = 0
SM = 0
Delta_AUC
Doxorubicin

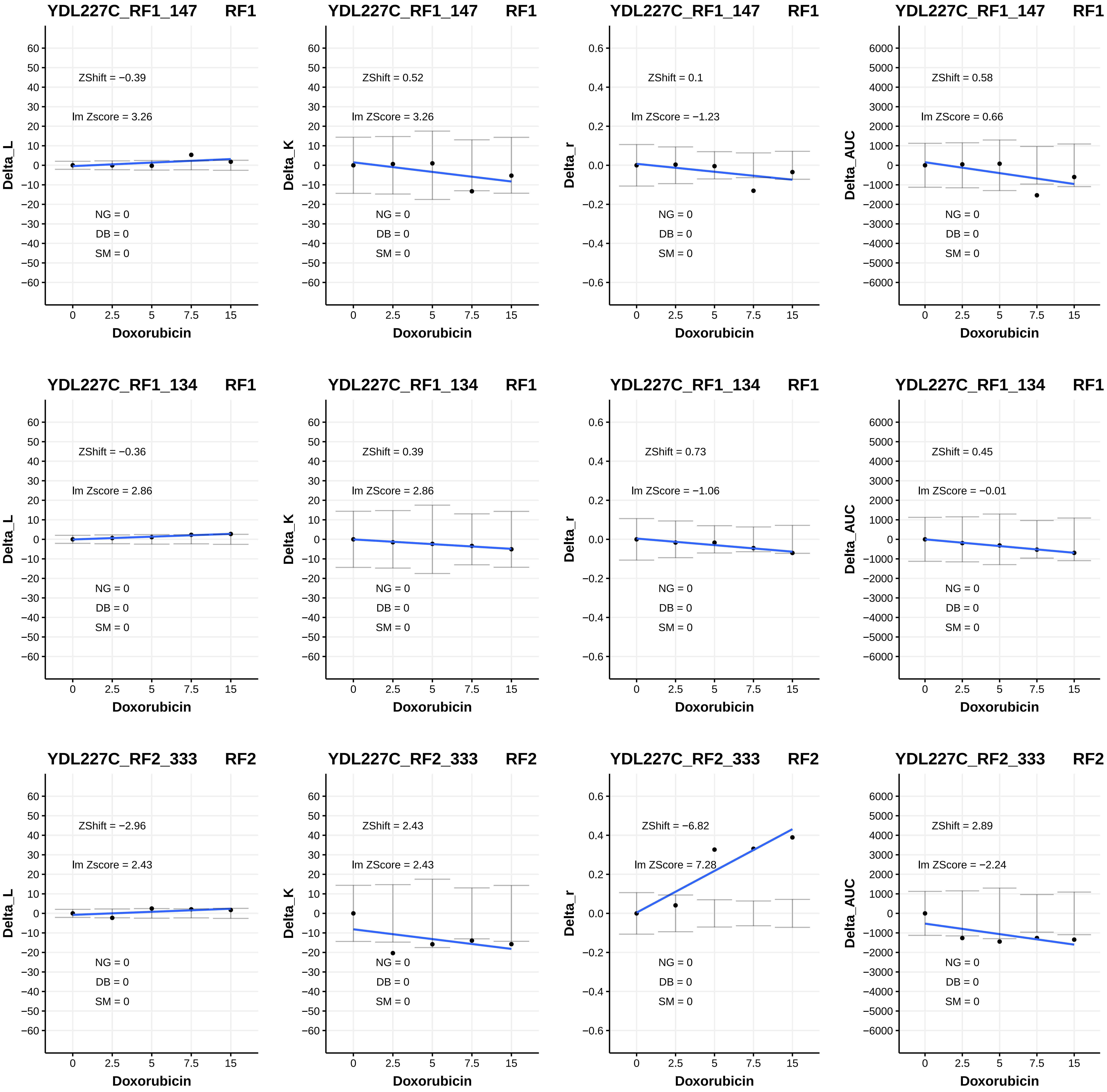

YDL227C_RF1_147 RF1
ZShift = −0.39
lm Zscore = 3.26
NG = 0
DB = 0
SM = 0
Delta_L
Doxorubicin
YDL227C_RF1_147 RF1
ZShift = 0.52
lm ZScore = 3.26
NG = 0
DB = 0
SM = 0
Delta_K
Doxorubicin
YDL227C_RF1_147 RF1
ZShift = 0.1
lm ZScore = −1.23
NG = 0
DB = 0
SM = 0
Delta_r
Doxorubicin
YDL227C_RF1_147 RF1
ZShift = 0.58
lm ZScore = 0.66
NG = 0
DB = 0
SM = 0
Delta_AUC
Doxorubicin
YDL227C_RF1_134 RF1
ZShift = −0.36
lm Zscore = 2.86
NG = 0
DB = 0
SM = 0
Delta_L
Doxorubicin
YDL227C_RF1_134 RF1
ZShift = 0.39
lm ZScore = 2.86
NG = 0
DB = 0
SM = 0
Delta_K
Doxorubicin
YDL227C_RF1_134 RF1
ZShift = 0.73
lm ZScore = −1.06
NG = 0
DB = 0
SM = 0
Delta_r
Doxorubicin
YDL227C_RF1_134 RF1
ZShift = 0.45
lm ZScore = −0.01
NG = 0
DB = 0
SM = 0
Delta_AUC
Doxorubicin
YDL227C_RF2_333 RF2
ZShift = −2.96
lm Zscore = 2.43
NG = 0
DB = 0
SM = 0
Delta_L
Doxorubicin
YDL227C_RF2_333 RF2
ZShift = 2.43
lm ZScore = 2.43
NG = 0
DB = 0
SM = 0
Delta_K
Doxorubicin
YDL227C_RF2_333 RF2
ZShift = −6.82
lm ZScore = 7.28
NG = 0
DB = 0
SM = 0
Delta_r
Doxorubicin
YDL227C_RF2_333 RF2
ZShift = 2.89
lm ZScore = −2.24
NG = 0
DB = 0
SM = 0
Delta_AUC
Doxorubicin

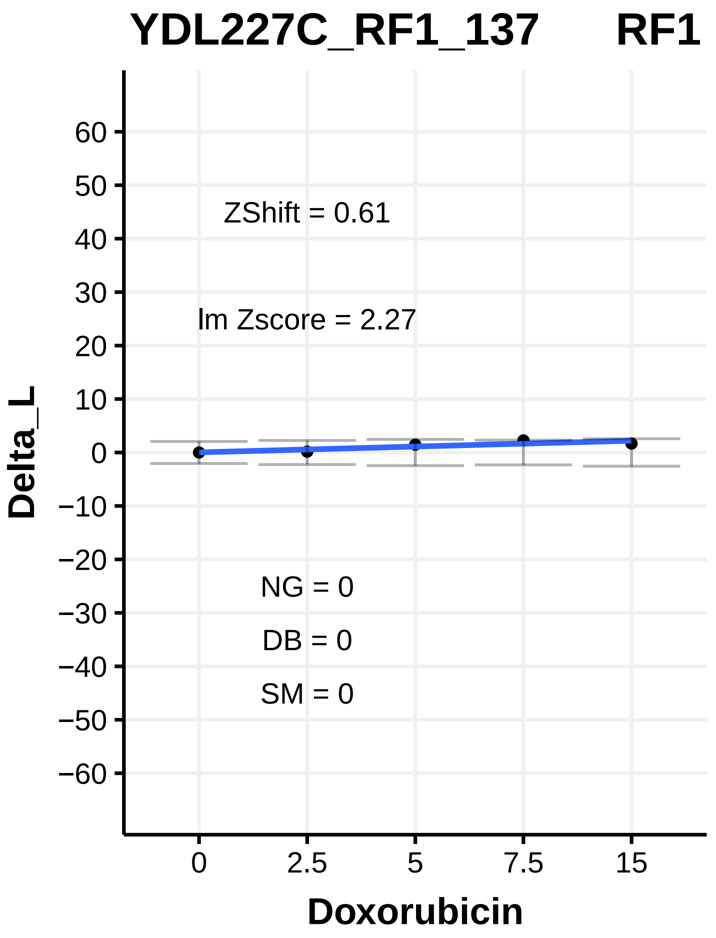
YDL227C_RF1_137 RF1
ZShift = 0.61
lm Zscore = 2.27
NG = 0
DB = 0
SM = 0
Delta_L
Doxorubicin

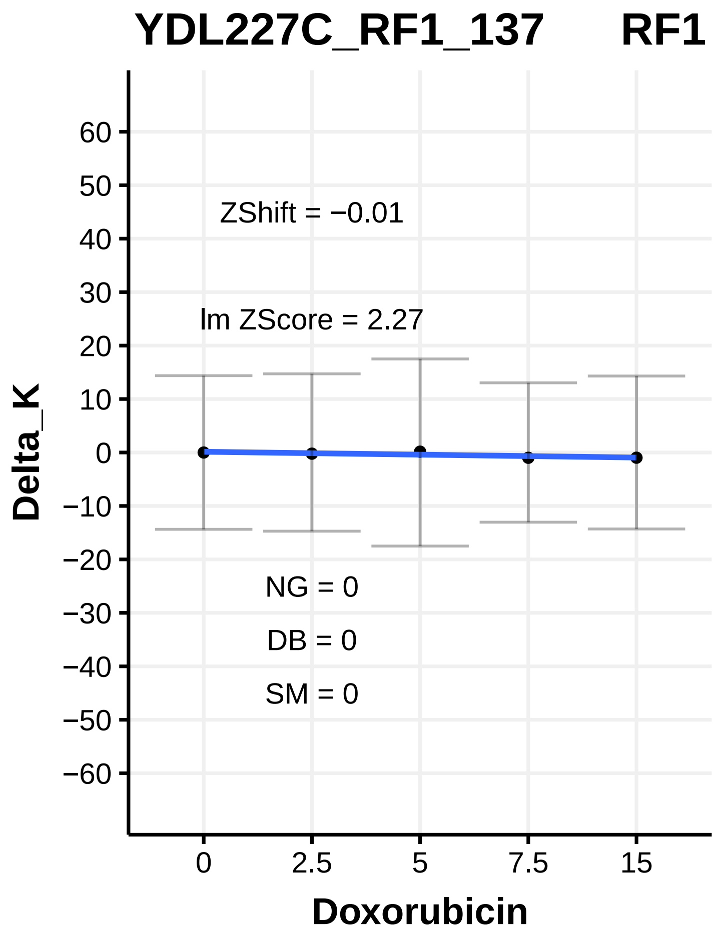
YDL227C_RF1_137 RF1
ZShift = −0.01
lm ZScore = 2.27
NG = 0
DB = 0
SM = 0
Delta_K
Doxorubicin

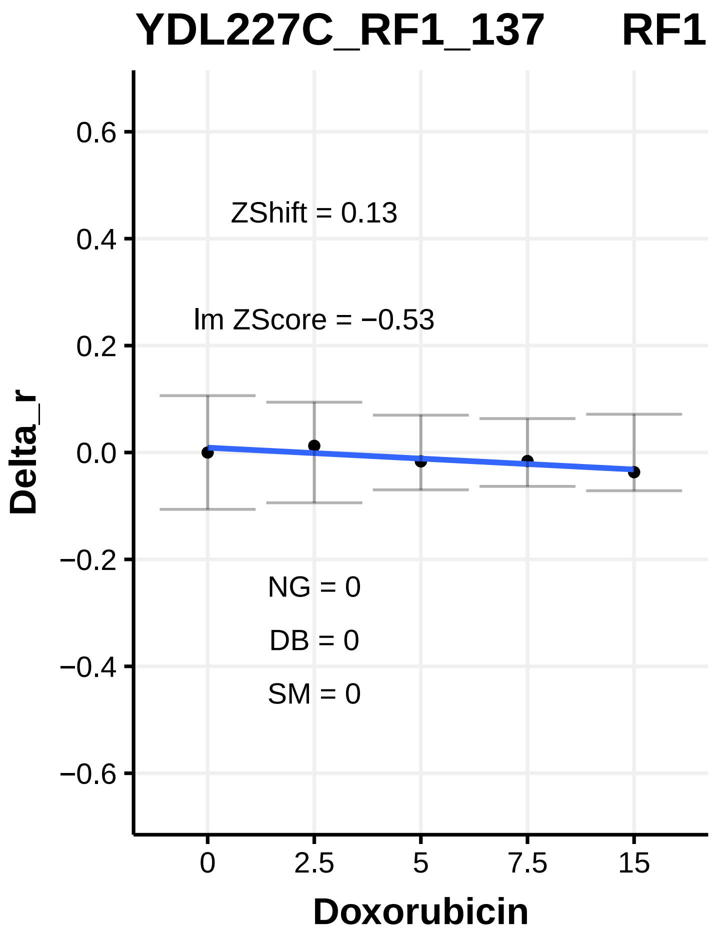
YDL227C_RF1_137 RF1
ZShift = 0.13
lm ZScore = −0.53
NG = 0
DB = 0
SM = 0
Delta_r
Doxorubicin

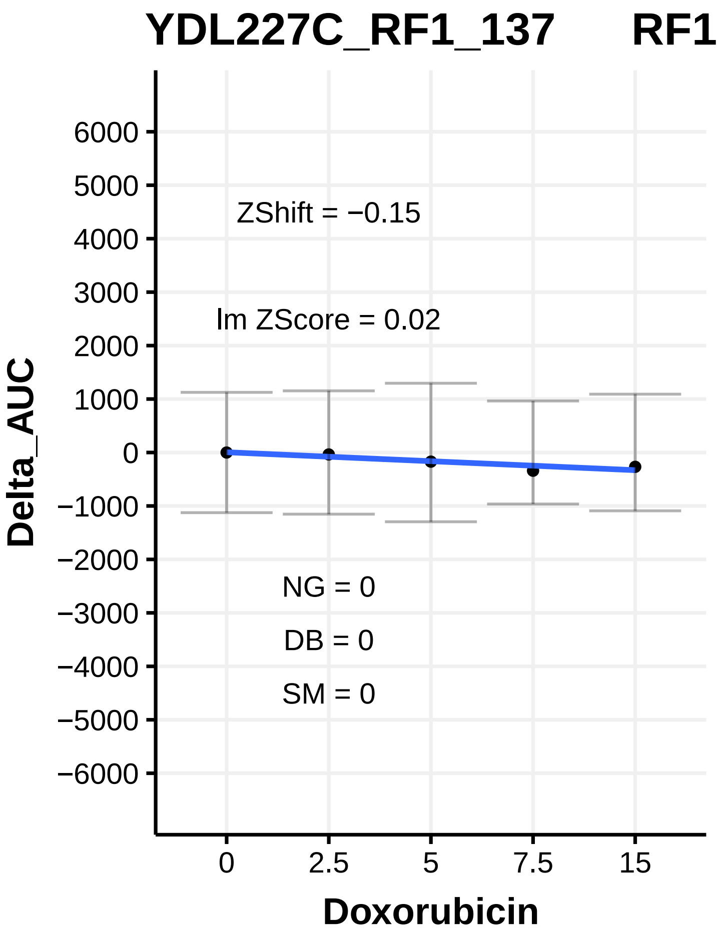
YDL227C_RF1_137 RF1
ZShift = −0.15
lm ZScore = 0.02
NG = 0
DB = 0
SM = 0
Delta_AUC
Doxorubicin

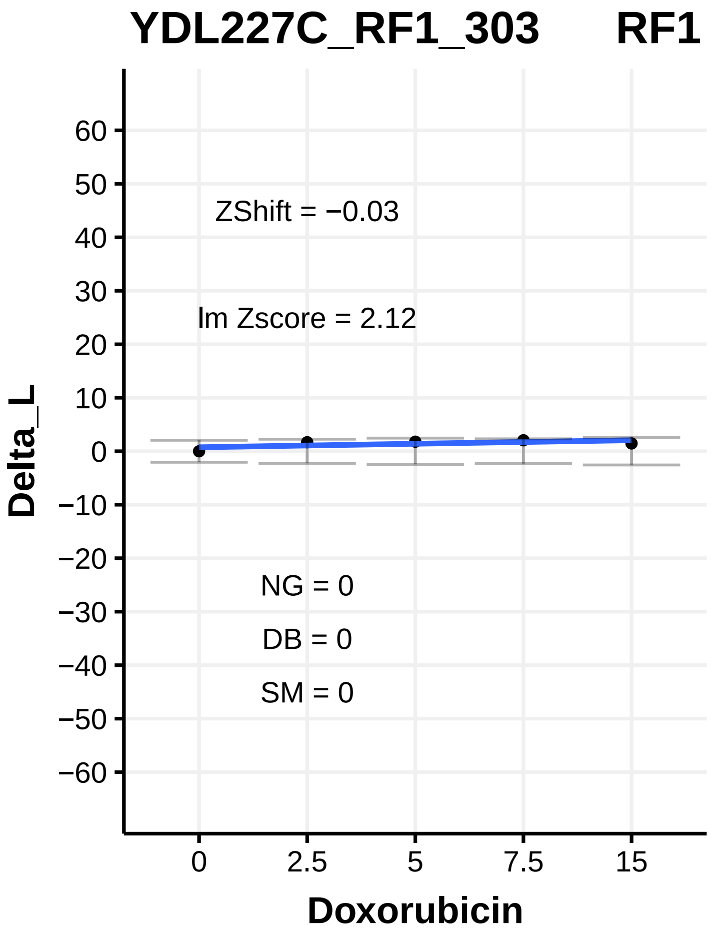
YDL227C_RF1_303 RF1
ZShift = −0.03
lm Zscore = 2.12
NG = 0
DB = 0
SM = 0
Delta_L
Doxorubicin

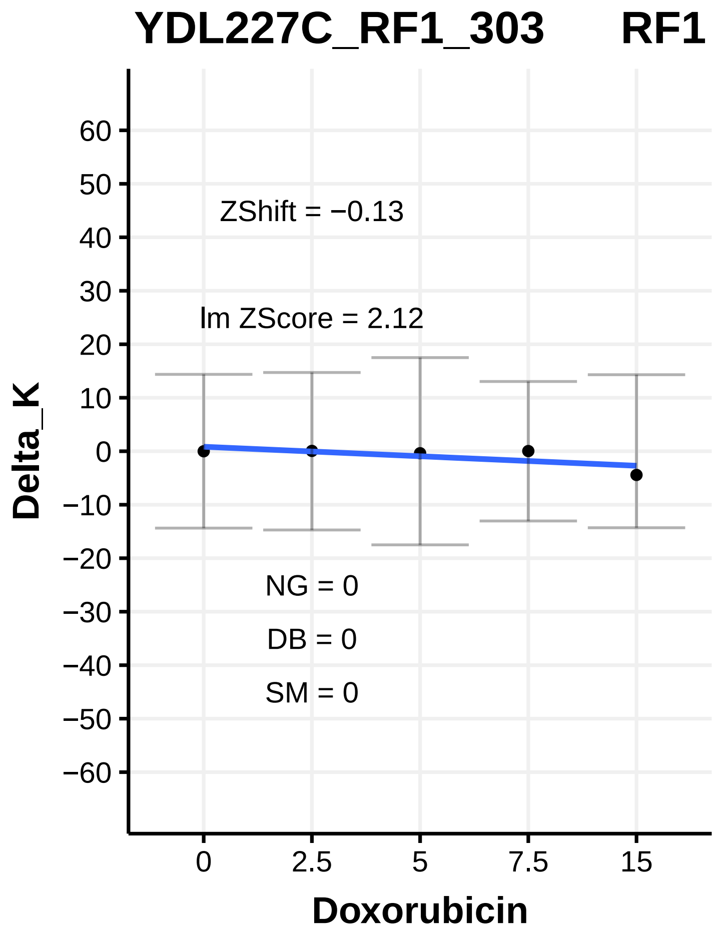
YDL227C_RF1_303 RF1
ZShift = −0.13
lm ZScore = 2.12
NG = 0
DB = 0
SM = 0
Delta_K
Doxorubicin

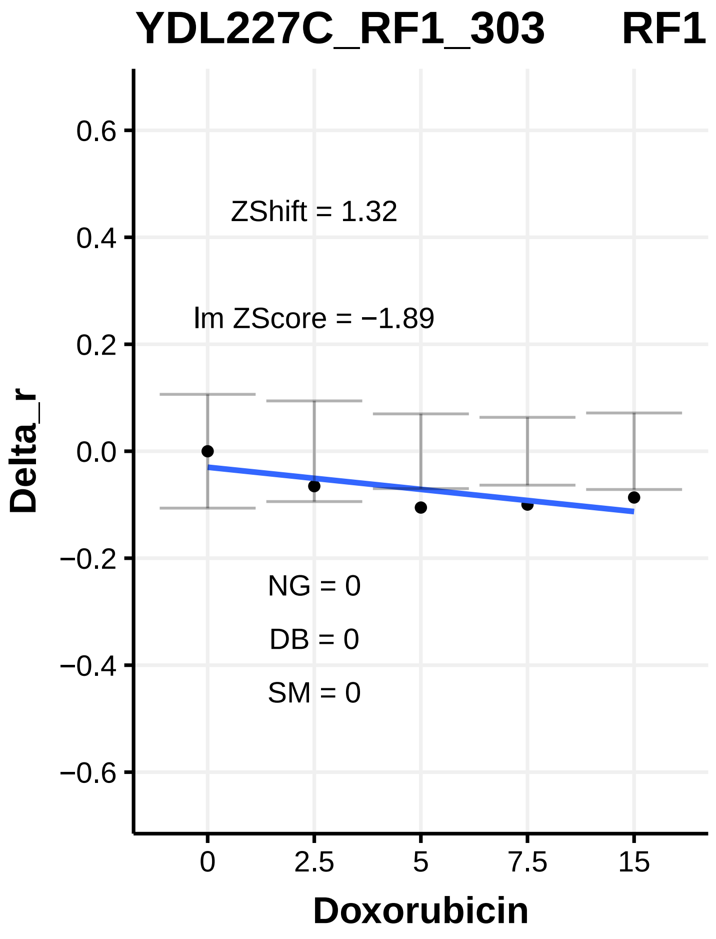
YDL227C_RF1_303 RF1
ZShift = 1.32
lm ZScore = −1.89
NG = 0
DB = 0
SM = 0
Delta_r
Doxorubicin

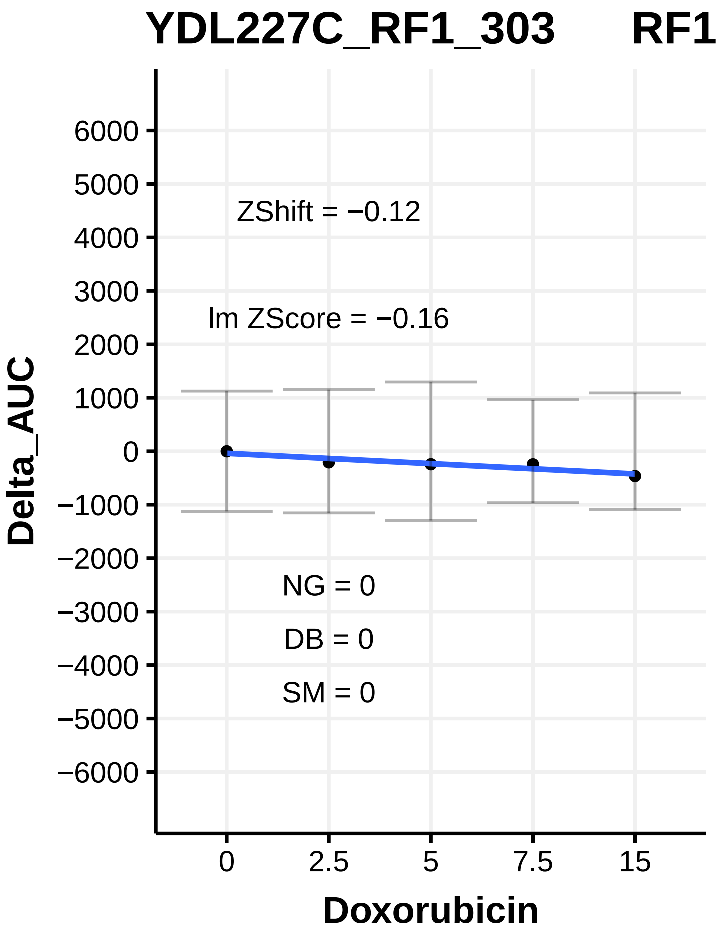
YDL227C_RF1_303 RF1
ZShift = −0.12
lm ZScore = −0.16
NG = 0
DB = 0
SM = 0
Delta_AUC
Doxorubicin

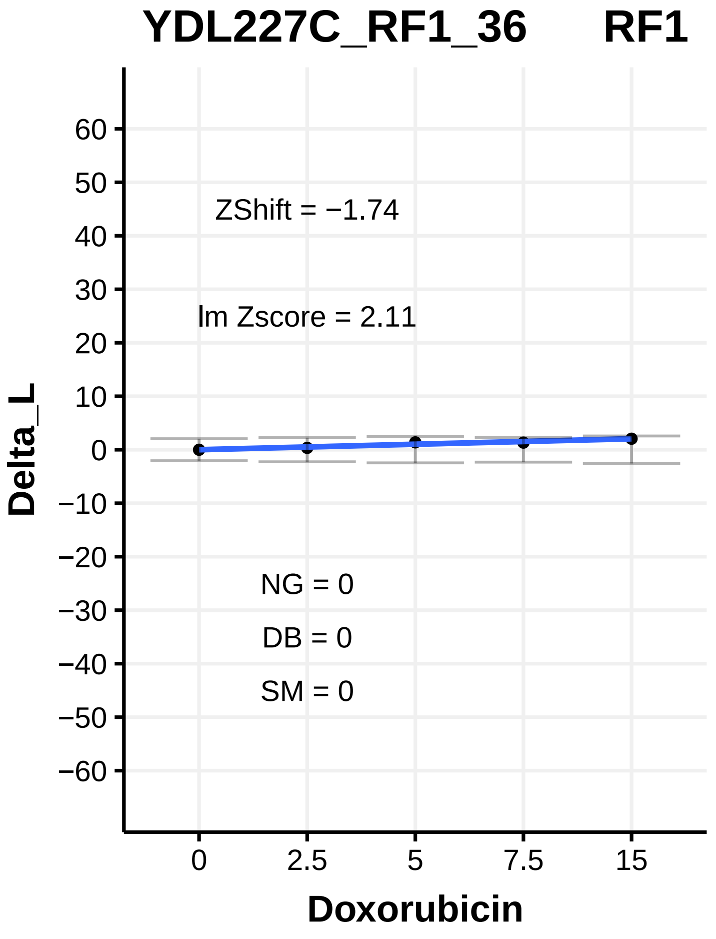
YDL227C_RF1_36 RF1
ZShift = −1.74
lm Zscore = 2.11
NG = 0
DB = 0
SM = 0
Delta_L
Doxorubicin

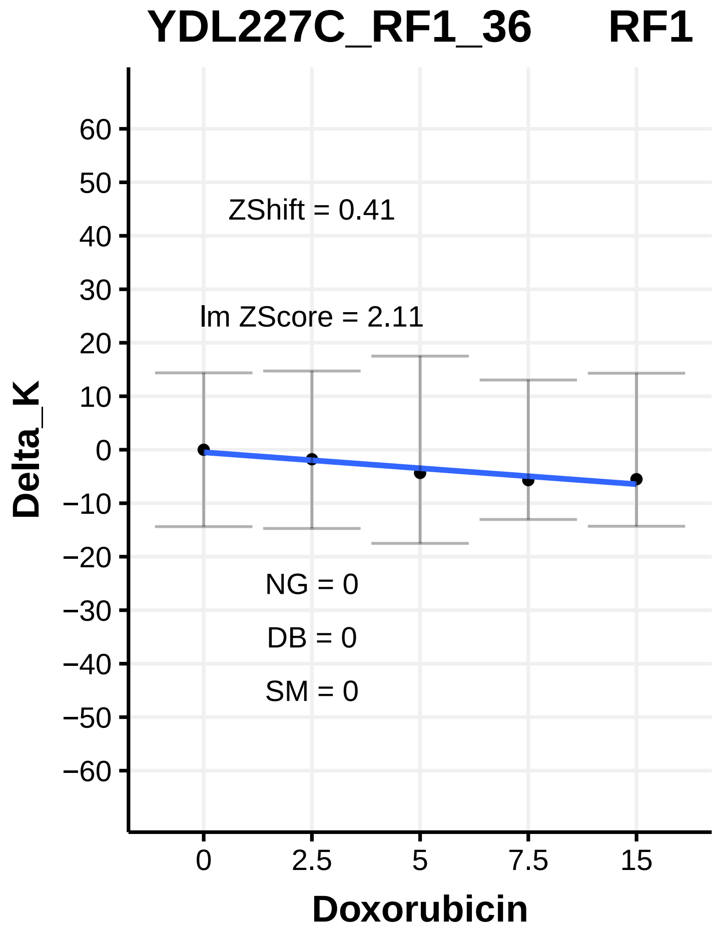
YDL227C_RF1_36 RF1
ZShift = 0.41
lm ZScore = 2.11
NG = 0
DB = 0
SM = 0
Delta_K
Doxorubicin

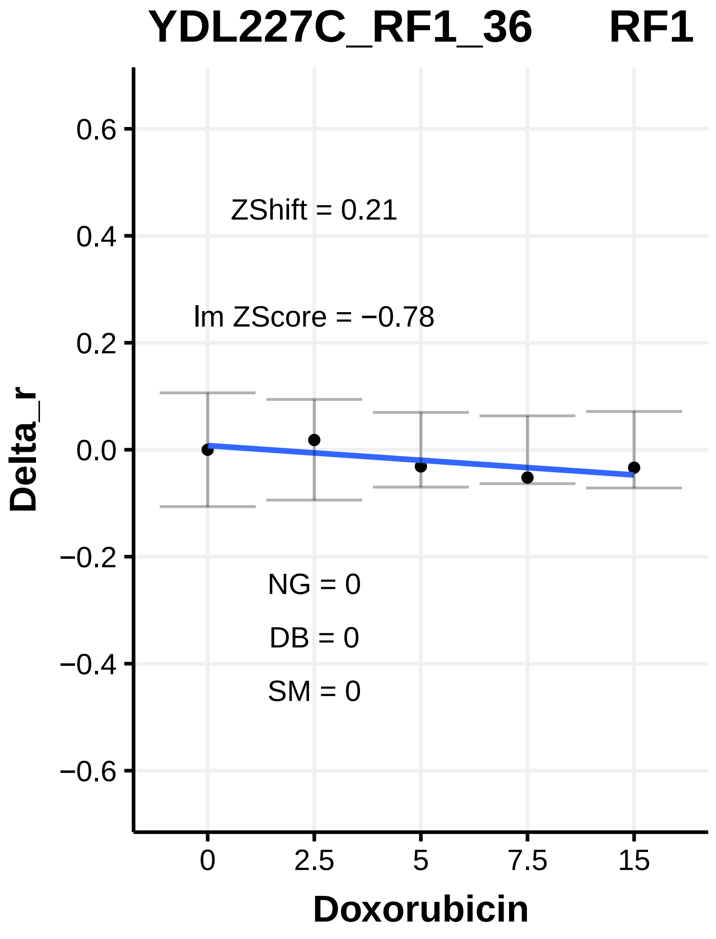
YDL227C_RF1_36 RF1
ZShift = 0.21
lm ZScore = −0.78
NG = 0
DB = 0
SM = 0
Delta_r
Doxorubicin

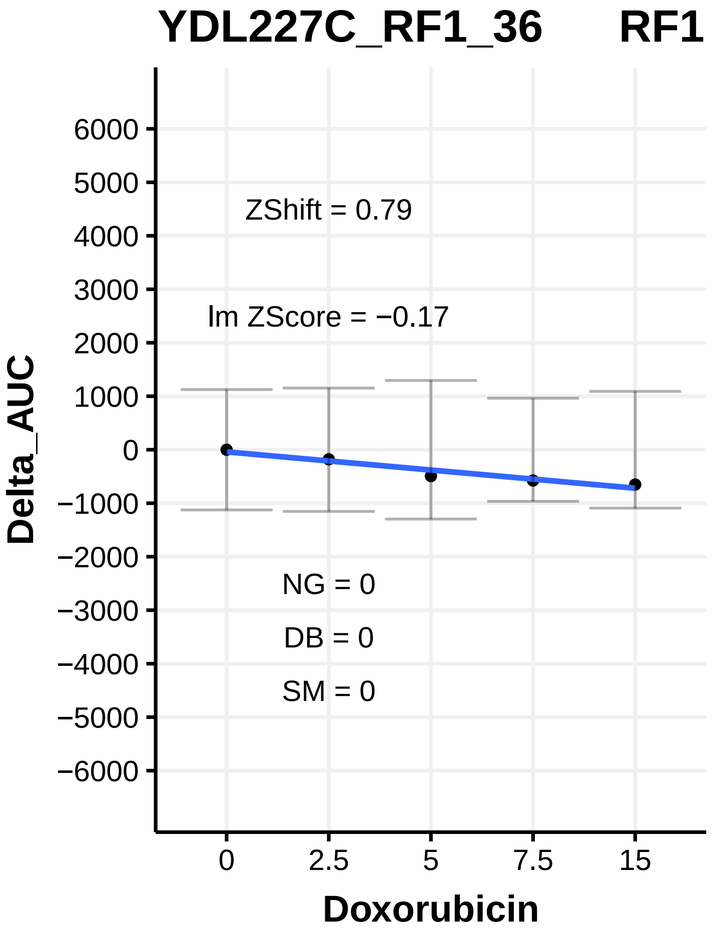
YDL227C_RF1_36 RF1
ZShift = 0.79
lm ZScore = −0.17
NG = 0
DB = 0
SM = 0
Delta_AUC
Doxorubicin

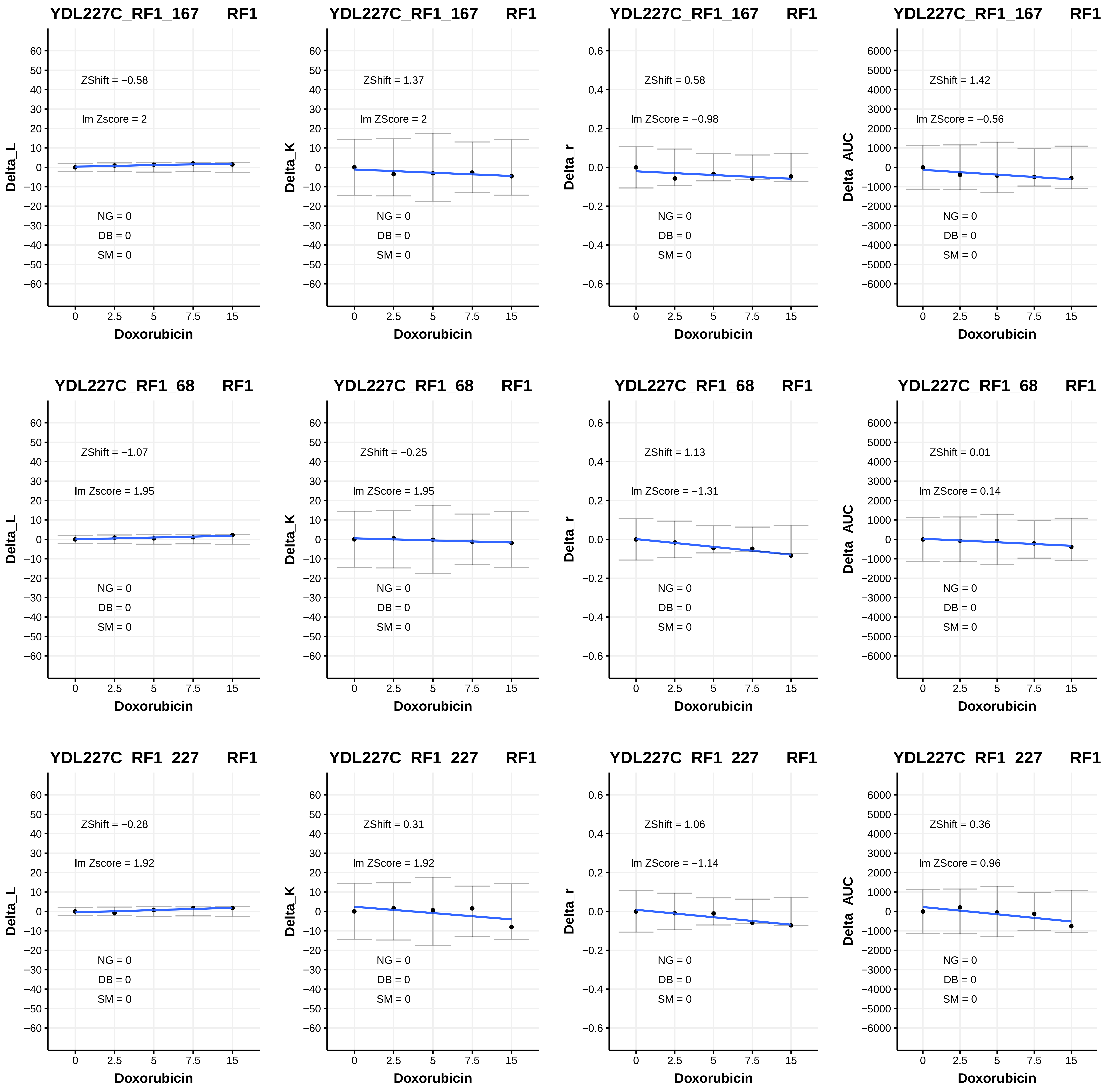

YDL227C_RF1_167 RF1
ZShift = −0.58
lm Zscore = 2
NG = 0
DB = 0
SM = 0
Delta_L
Doxorubicin
YDL227C_RF1_167 RF1
ZShift = 1.37
lm ZScore = 2
NG = 0
DB = 0
SM = 0
Delta_K
Doxorubicin
YDL227C_RF1_167 RF1
ZShift = 0.58
lm ZScore = −0.98
NG = 0
DB = 0
SM = 0
Delta_r
Doxorubicin
YDL227C_RF1_167 RF1
ZShift = 1.42
lm ZScore = −0.56
NG = 0
DB = 0
SM = 0
Delta_AUC
Doxorubicin
YDL227C_RF1_68 RF1
ZShift = −1.07
lm Zscore = 1.95
NG = 0
DB = 0
SM = 0
Delta_L
Doxorubicin
YDL227C_RF1_68 RF1
ZShift = −0.25
lm ZScore = 1.95
NG = 0
DB = 0
SM = 0
Delta_K
Doxorubicin
YDL227C_RF1_68 RF1
ZShift = 1.13
lm ZScore = −1.31
NG = 0
DB = 0
SM = 0
Delta_r
Doxorubicin
YDL227C_RF1_68 RF1
ZShift = 0.01
lm ZScore = 0.14
NG = 0
DB = 0
SM = 0
Delta_AUC
Doxorubicin
YDL227C_RF1_227 RF1
ZShift = −0.28
lm Zscore = 1.92
NG = 0
DB = 0
SM = 0
Delta_L
Doxorubicin
YDL227C_RF1_227 RF1
ZShift = 0.31
lm ZScore = 1.92
NG = 0
DB = 0
SM = 0
Delta_K
Doxorubicin
YDL227C_RF1_227 RF1
ZShift = 1.06
lm ZScore = −1.14
NG = 0
DB = 0
SM = 0
Delta_r
Doxorubicin
YDL227C_RF1_227 RF1
ZShift = 0.36
lm ZScore = 0.96
NG = 0
DB = 0
SM = 0
Delta_AUC
Doxorubicin

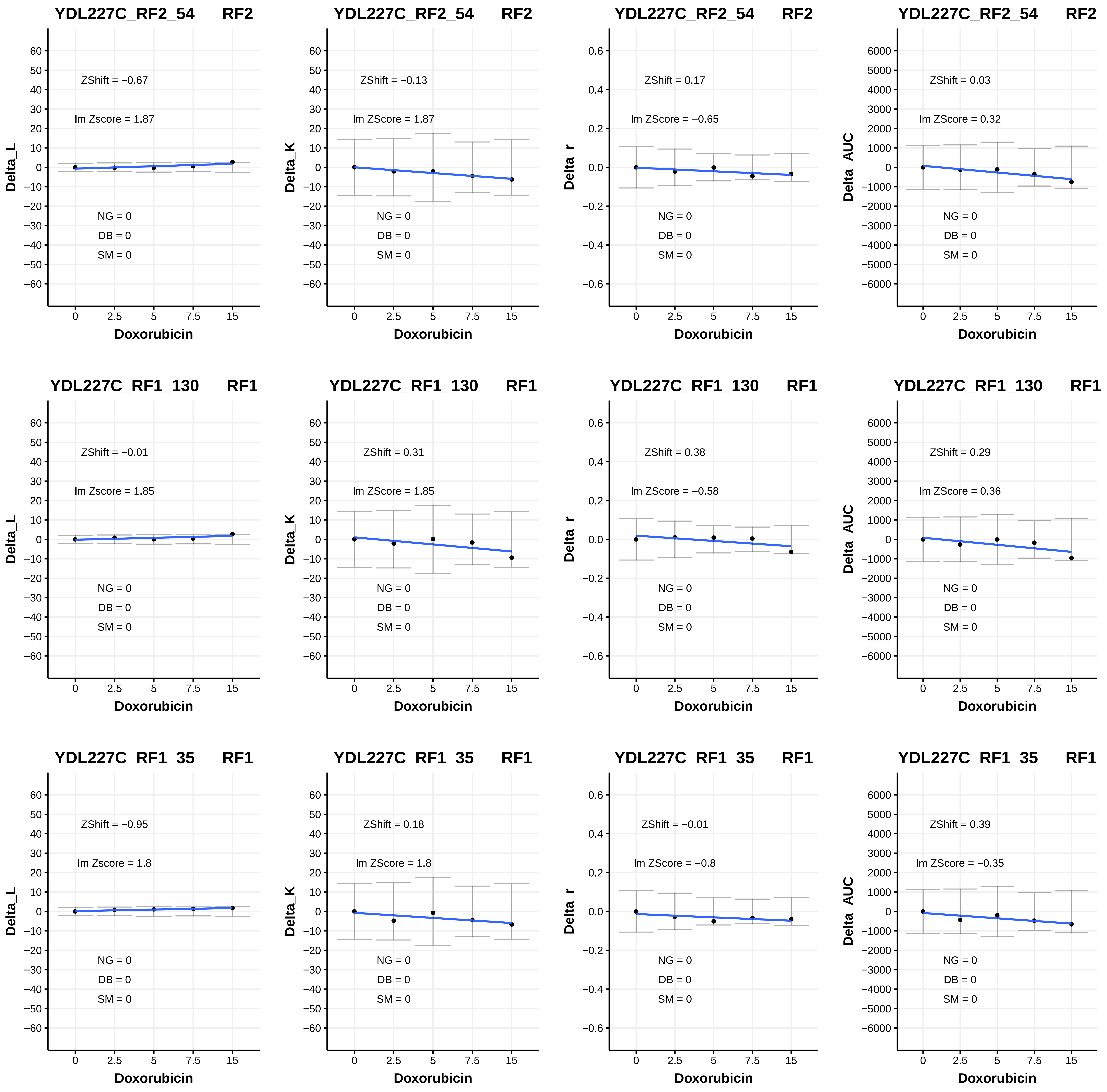
YDL227C_RF2_54 RF2
ZShift = −0.67
lm Zscore = 1.87
NG = 0
DB = 0
SM = 0
Delta_L
Doxorubicin
YDL227C_RF2_54 RF2
ZShift = −0.13
lm ZScore = 1.87
NG = 0
DB = 0
SM = 0
Delta_K
Doxorubicin
YDL227C_RF2_54 RF2
ZShift = 0.17
lm ZScore = −0.65
NG = 0
DB = 0
SM = 0
Delta_r
Doxorubicin
YDL227C_RF2_54 RF2
ZShift = 0.03
lm ZScore = 0.32
NG = 0
DB = 0
SM = 0
Delta_AUC
Doxorubicin
YDL227C_RF1_130 RF1
ZShift = −0.01
lm Zscore = 1.85
NG = 0
DB = 0
SM = 0
Delta_L
Doxorubicin
YDL227C_RF1_130 RF1
ZShift = 0.31
lm ZScore = 1.85
NG = 0
DB = 0
SM = 0
Delta_K
Doxorubicin
YDL227C_RF1_130 RF1
ZShift = 0.38
lm ZScore = −0.58
NG = 0
DB = 0
SM = 0
Delta_r
Doxorubicin
YDL227C_RF1_130 RF1
ZShift = 0.29
lm ZScore = 0.36
NG = 0
DB = 0
SM = 0
Delta_AUC
Doxorubicin
YDL227C_RF1_35 RF1
ZShift = −0.95
lm Zscore = 1.8
NG = 0
DB = 0
SM = 0
Delta_L
Doxorubicin
YDL227C_RF1_35 RF1
ZShift = 0.18
lm ZScore = 1.8
NG = 0
DB = 0
SM = 0
Delta_K
Doxorubicin
YDL227C_RF1_35 RF1
ZShift = −0.01
lm ZScore = −0.8
NG = 0
DB = 0
SM = 0
Delta_r
Doxorubicin
YDL227C_RF1_35 RF1
ZShift = 0.39
lm ZScore = −0.35
NG = 0
DB = 0
SM = 0
Delta_AUC
Doxorubicin

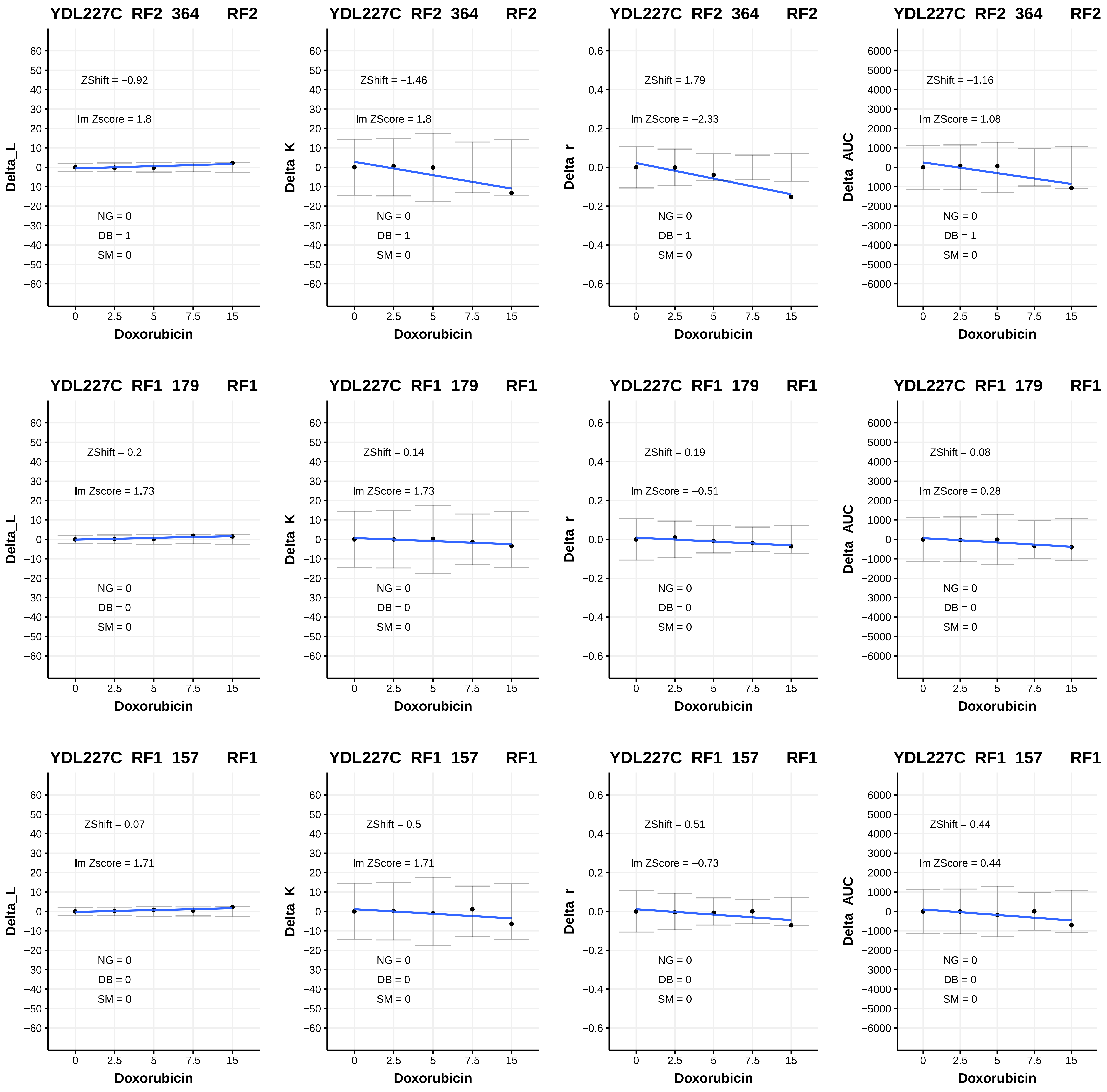

YDL227C_RF2_364 RF2
ZShift = −0.92
lm Zscore = 1.8
NG = 0
DB = 1
SM = 0
Delta_L
Doxorubicin
YDL227C_RF2_364 RF2
ZShift = −1.46
lm ZScore = 1.8
NG = 0
DB = 1
SM = 0
Delta_K
Doxorubicin
YDL227C_RF2_364 RF2
ZShift = 1.79
lm ZScore = −2.33
NG = 0
DB = 1
SM = 0
Delta_r
Doxorubicin
YDL227C_RF2_364 RF2
ZShift = −1.16
lm ZScore = 1.08
NG = 0
DB = 1
SM = 0
Delta_AUC
Doxorubicin
YDL227C_RF1_179 RF1
ZShift = 0.2
lm Zscore = 1.73
NG = 0
DB = 0
SM = 0
Delta_L
Doxorubicin
YDL227C_RF1_179 RF1
ZShift = 0.14
lm ZScore = 1.73
NG = 0
DB = 0
SM = 0
Delta_K
Doxorubicin
YDL227C_RF1_179 RF1
ZShift = 0.19
lm ZScore = −0.51
NG = 0
DB = 0
SM = 0
Delta_r
Doxorubicin
YDL227C_RF1_179 RF1
ZShift = 0.08
lm ZScore = 0.28
NG = 0
DB = 0
SM = 0
Delta_AUC
Doxorubicin
YDL227C_RF1_157 RF1
ZShift = 0.07
lm Zscore = 1.71
NG = 0
DB = 0
SM = 0
Delta_L
Doxorubicin
YDL227C_RF1_157 RF1
ZShift = 0.5
lm ZScore = 1.71
NG = 0
DB = 0
SM = 0
Delta_K
Doxorubicin
YDL227C_RF1_157 RF1
ZShift = 0.51
lm ZScore = −0.73
NG = 0
DB = 0
SM = 0
Delta_r
Doxorubicin
YDL227C_RF1_157 RF1
ZShift = 0.44
lm ZScore = 0.44
NG = 0
DB = 0
SM = 0
Delta_AUC
Doxorubicin

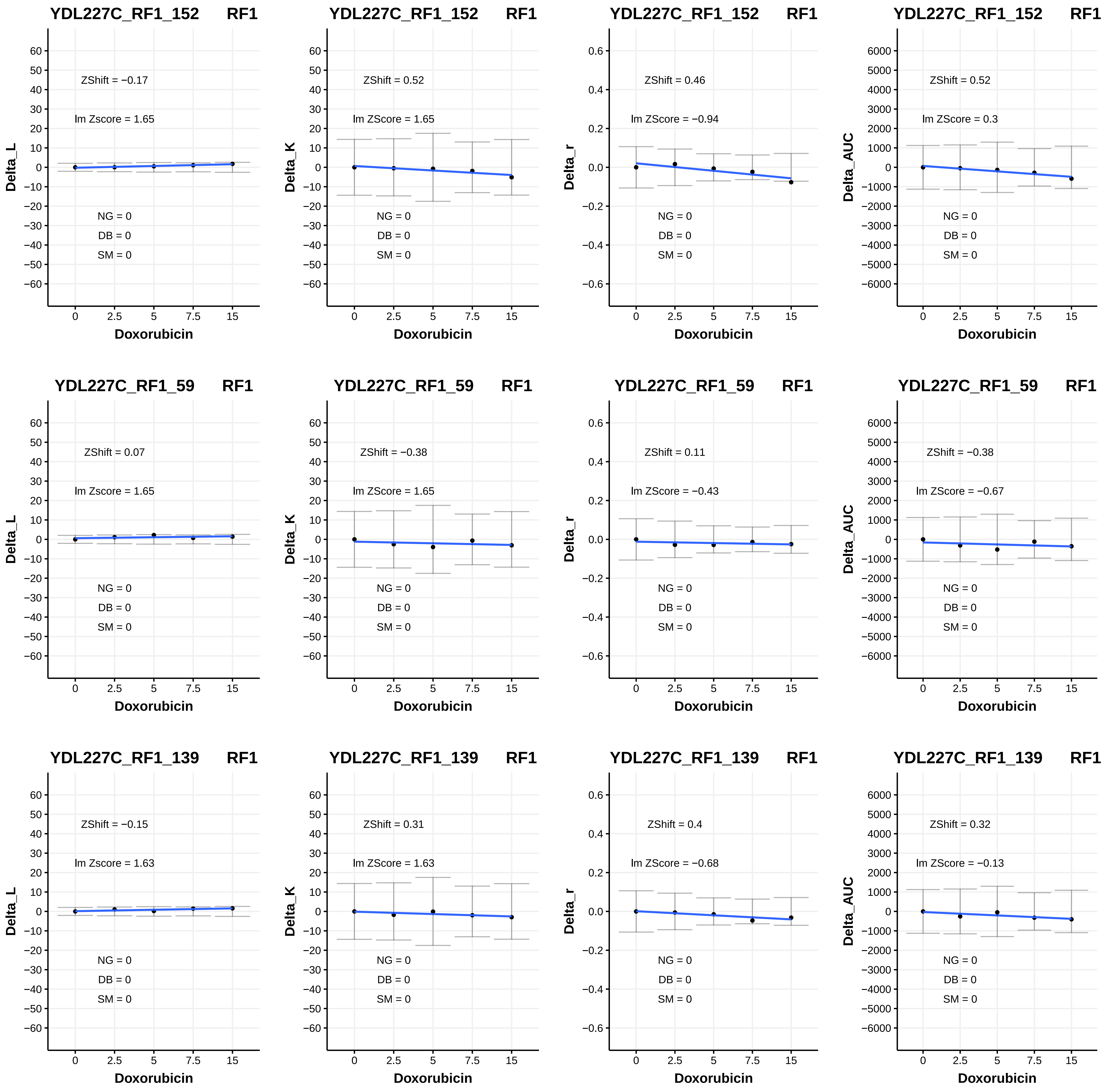
YDL227C_RF1_152 RF1
ZShift = −0.17
lm Zscore = 1.65
NG = 0
DB = 0
SM = 0
Delta_L
Doxorubicin
YDL227C_RF1_152 RF1
ZShift = 0.52
lm ZScore = 1.65
NG = 0
DB = 0
SM = 0
Delta_K
Doxorubicin
YDL227C_RF1_152 RF1
ZShift = 0.46
lm ZScore = −0.94
NG = 0
DB = 0
SM = 0
Delta_r
Doxorubicin
YDL227C_RF1_152 RF1
ZShift = 0.52
lm ZScore = 0.3
NG = 0
DB = 0
SM = 0
Delta_AUC
Doxorubicin
YDL227C_RF1_59 RF1
ZShift = 0.07
lm Zscore = 1.65
NG = 0
DB = 0
SM = 0
Delta_L
Doxorubicin
YDL227C_RF1_59 RF1
ZShift = −0.38
lm ZScore = 1.65
NG = 0
DB = 0
SM = 0
Delta_K
Doxorubicin
YDL227C_RF1_59 RF1
ZShift = 0.11
lm ZScore = −0.43
NG = 0
DB = 0
SM = 0
Delta_r
Doxorubicin
YDL227C_RF1_59 RF1
ZShift = −0.38
lm ZScore = −0.67
NG = 0
DB = 0
SM = 0
Delta_AUC
Doxorubicin
YDL227C_RF1_139 RF1
ZShift = −0.15
lm Zscore = 1.63
NG = 0
DB = 0
SM = 0
Delta_L
Doxorubicin
YDL227C_RF1_139 RF1
ZShift = 0.31
lm ZScore = 1.63
NG = 0
DB = 0
SM = 0
Delta_K
Doxorubicin
YDL227C_RF1_139 RF1
ZShift = 0.4
lm ZScore = −0.68
NG = 0
DB = 0
SM = 0
Delta_r
Doxorubicin
YDL227C_RF1_139 RF1
ZShift = 0.32
lm ZScore = −0.13
NG = 0
DB = 0
SM = 0
Delta_AUC
Doxorubicin

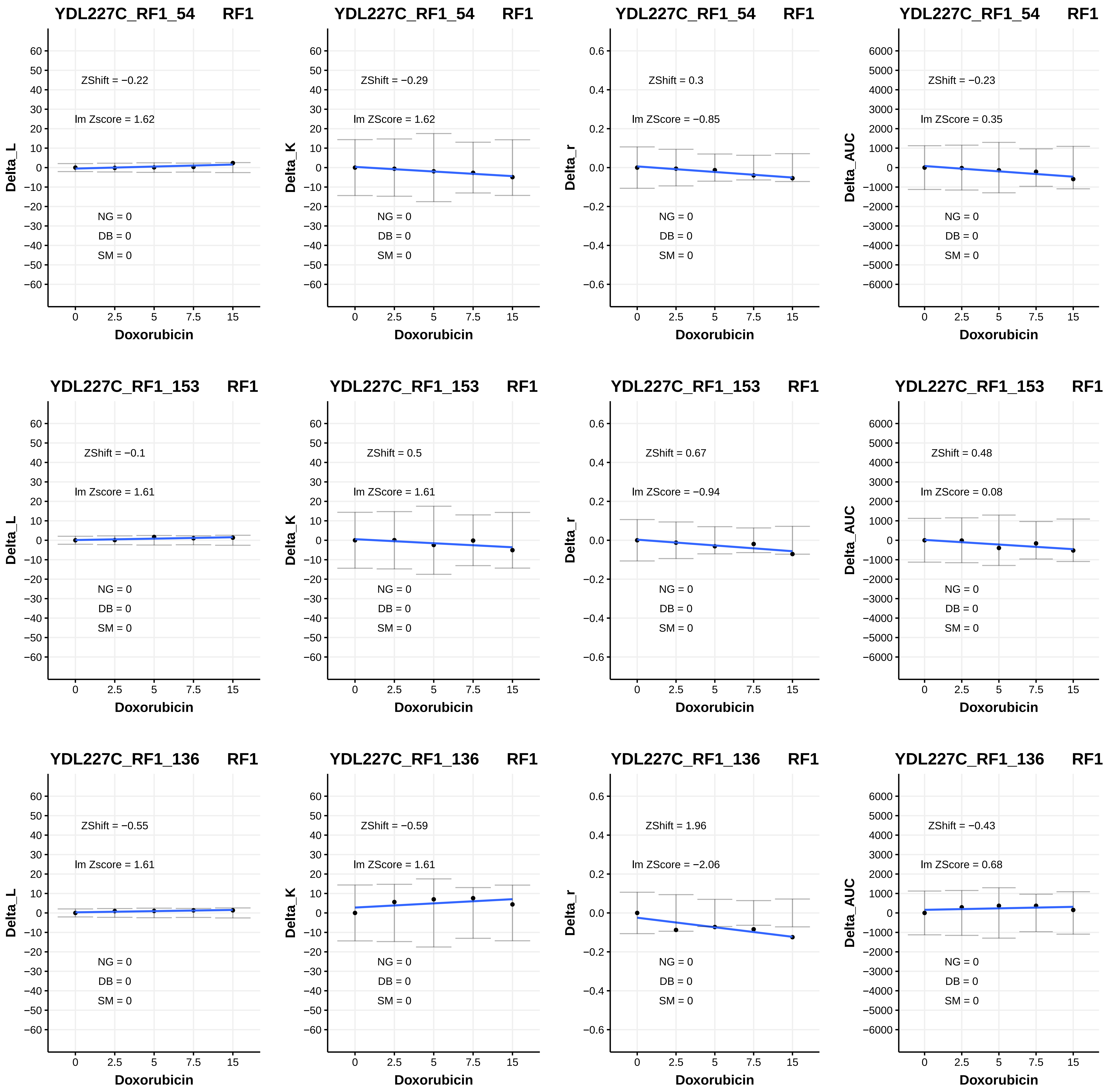
YDL227C_RF1_54 RF1
ZShift = −0.22
lm Zscore = 1.62
NG = 0
DB = 0
SM = 0
Delta_L
Doxorubicin
YDL227C_RF1_54 RF1
ZShift = −0.29
lm ZScore = 1.62
NG = 0
DB = 0
SM = 0
Delta_K
Doxorubicin
YDL227C_RF1_54 RF1
ZShift = 0.3
lm ZScore = −0.85
NG = 0
DB = 0
SM = 0
Delta_r
Doxorubicin
YDL227C_RF1_54 RF1
ZShift = −0.23
lm ZScore = 0.35
NG = 0
DB = 0
SM = 0
Delta_AUC
Doxorubicin
YDL227C_RF1_153 RF1
ZShift = −0.1
lm Zscore = 1.61
NG = 0
DB = 0
SM = 0
Delta_L
Doxorubicin
YDL227C_RF1_153 RF1
ZShift = 0.5
lm ZScore = 1.61
NG = 0
DB = 0
SM = 0
Delta_K
Doxorubicin
YDL227C_RF1_153 RF1
ZShift = 0.67
lm ZScore = −0.94
NG = 0
DB = 0
SM = 0
Delta_r
Doxorubicin
YDL227C_RF1_153 RF1
ZShift = 0.48
lm ZScore = 0.08
NG = 0
DB = 0
SM = 0
Delta_AUC
Doxorubicin
YDL227C_RF1_136 RF1
ZShift = −0.55
lm Zscore = 1.61
NG = 0
DB = 0
SM = 0
Delta_L
Doxorubicin
YDL227C_RF1_136 RF1
ZShift = −0.59
lm ZScore = 1.61
NG = 0
DB = 0
SM = 0
Delta_K
Doxorubicin
YDL227C_RF1_136 RF1
ZShift = 1.96
lm ZScore = −2.06
NG = 0
DB = 0
SM = 0
Delta_r
Doxorubicin
YDL227C_RF1_136 RF1
ZShift = −0.43
lm ZScore = 0.68
NG = 0
DB = 0
SM = 0
Delta_AUC
Doxorubicin

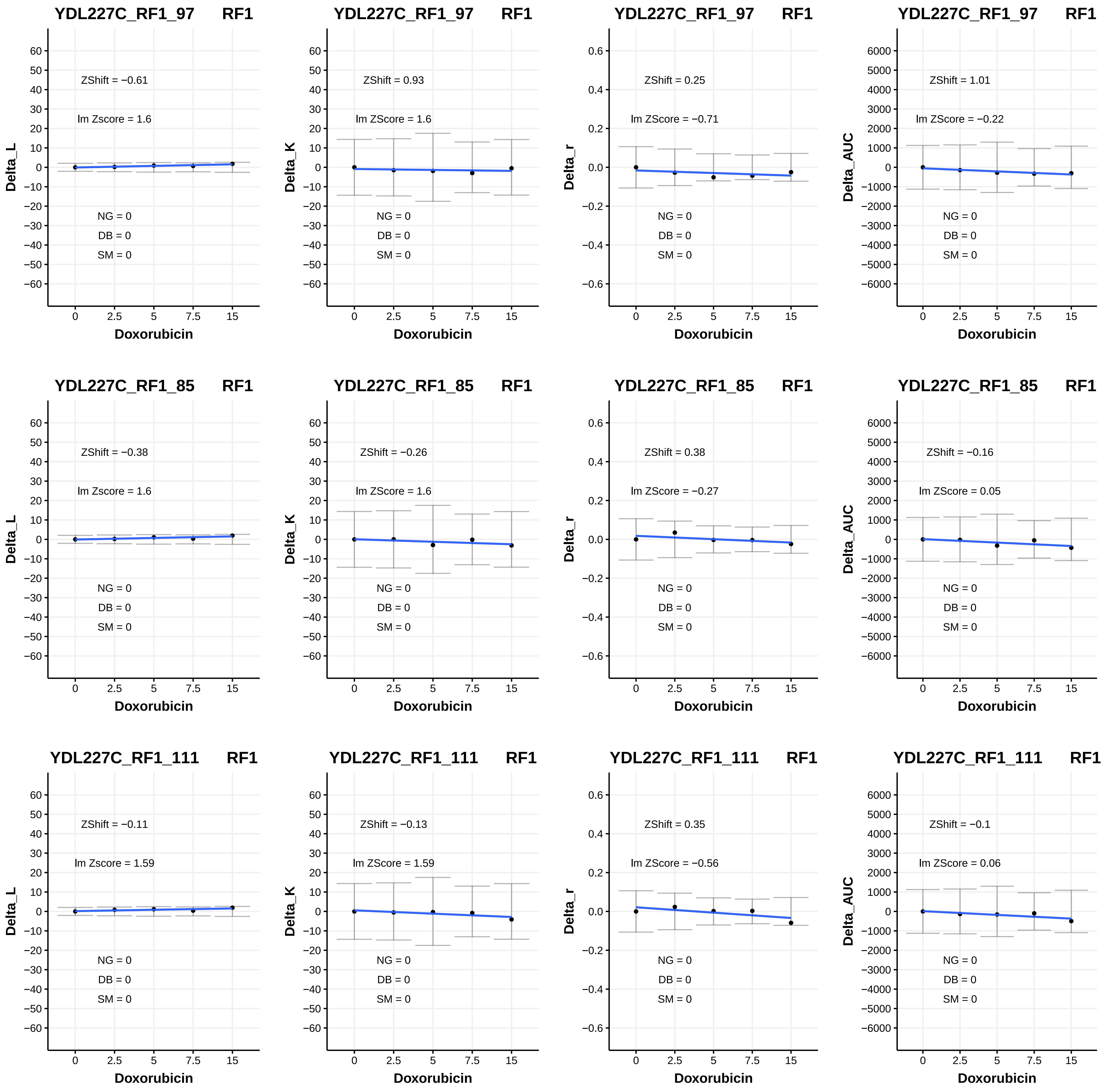

YDL227C_RF1_97 RF1
ZShift = −0.61
lm Zscore = 1.6
NG = 0
DB = 0
SM = 0
Delta_L
Doxorubicin
YDL227C_RF1_97 RF1
ZShift = 0.93
lm ZScore = 1.6
NG = 0
DB = 0
SM = 0
Delta_K
Doxorubicin
YDL227C_RF1_97 RF1
ZShift = 0.25
lm ZScore = −0.71
NG = 0
DB = 0
SM = 0
Delta_r
Doxorubicin
YDL227C_RF1_97 RF1
ZShift = 1.01
lm ZScore = −0.22
NG = 0
DB = 0
SM = 0
Delta_AUC
Doxorubicin
YDL227C_RF1_85 RF1
ZShift = −0.38
lm Zscore = 1.6
NG = 0
DB = 0
SM = 0
Delta_L
Doxorubicin
YDL227C_RF1_85 RF1
ZShift = −0.26
lm ZScore = 1.6
NG = 0
DB = 0
SM = 0
Delta_K
Doxorubicin
YDL227C_RF1_85 RF1
ZShift = 0.38
lm ZScore = −0.27
NG = 0
DB = 0
SM = 0
Delta_r
Doxorubicin
YDL227C_RF1_85 RF1
ZShift = −0.16
lm ZScore = 0.05
NG = 0
DB = 0
SM = 0
Delta_AUC
Doxorubicin
YDL227C_RF1_111 RF1
ZShift = −0.11
lm Zscore = 1.59
NG = 0
DB = 0
SM = 0
Delta_L
Doxorubicin
YDL227C_RF1_111 RF1
ZShift = −0.13
lm ZScore = 1.59
NG = 0
DB = 0
SM = 0
Delta_K
Doxorubicin
YDL227C_RF1_111 RF1
ZShift = 0.35
lm ZScore = −0.56
NG = 0
DB = 0
SM = 0
Delta_r
Doxorubicin
YDL227C_RF1_111 RF1
ZShift = −0.1
lm ZScore = 0.06
NG = 0
DB = 0
SM = 0
Delta_AUC
Doxorubicin

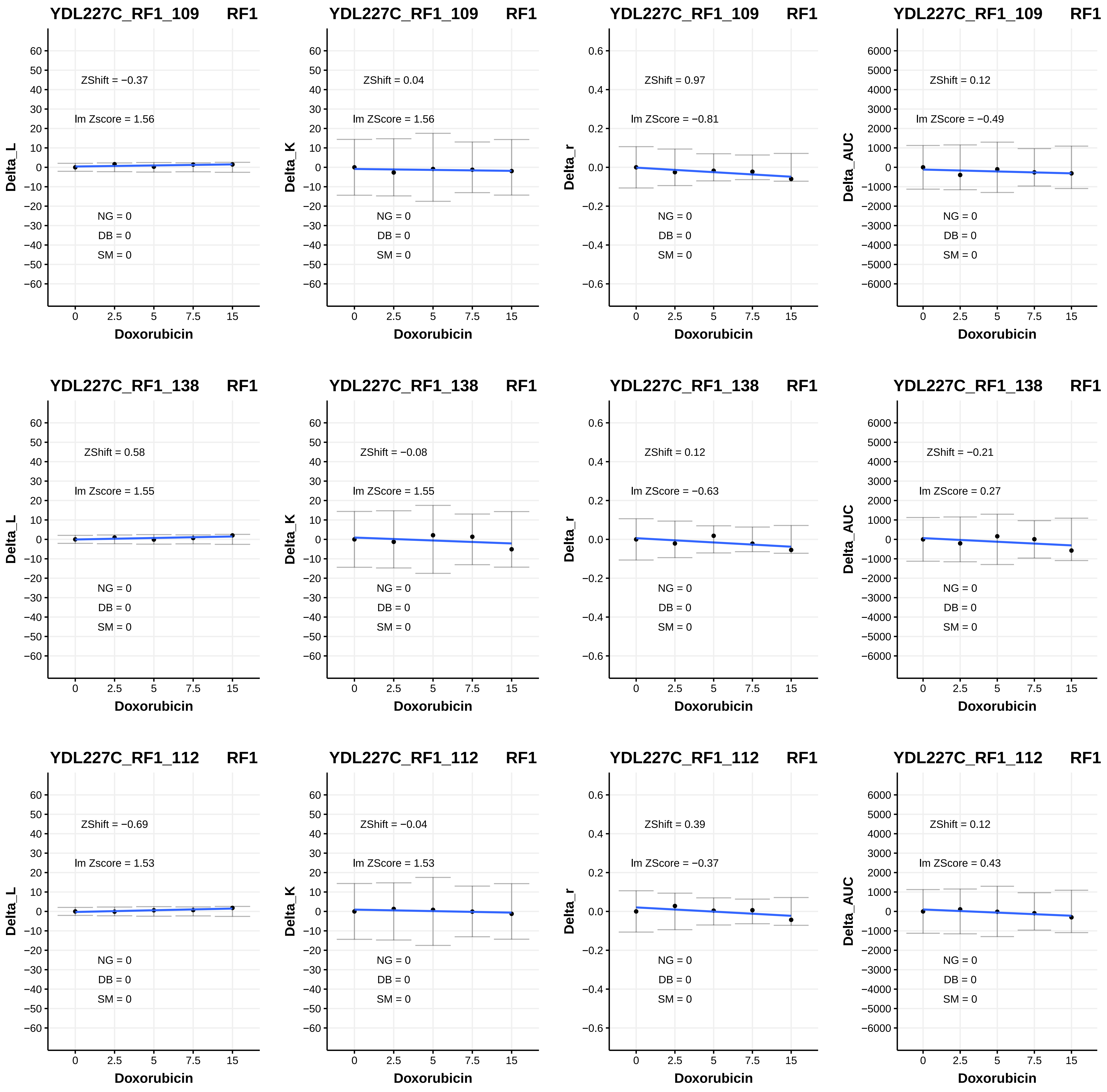
YDL227C_RF1_109 RF1
ZShift = −0.37
lm Zscore = 1.56
NG = 0
DB = 0
SM = 0
Delta_L
Doxorubicin
YDL227C_RF1_109 RF1
ZShift = 0.04
lm ZScore = 1.56
NG = 0
DB = 0
SM = 0
Delta_K
Doxorubicin
YDL227C_RF1_109 RF1
ZShift = 0.97
lm ZScore = −0.81
NG = 0
DB = 0
SM = 0
Delta_r
Doxorubicin
YDL227C_RF1_109 RF1
ZShift = 0.12
lm ZScore = −0.49
NG = 0
DB = 0
SM = 0
Delta_AUC
Doxorubicin
YDL227C_RF1_138 RF1
ZShift = 0.58
lm Zscore = 1.55
NG = 0
DB = 0
SM = 0
Delta_L
Doxorubicin
YDL227C_RF1_138 RF1
ZShift = −0.08
lm ZScore = 1.55
NG = 0
DB = 0
SM = 0
Delta_K
Doxorubicin
YDL227C_RF1_138 RF1
ZShift = 0.12
lm ZScore = −0.63
NG = 0
DB = 0
SM = 0
Delta_r
Doxorubicin
YDL227C_RF1_138 RF1
ZShift = −0.21
lm ZScore = 0.27
NG = 0
DB = 0
SM = 0
Delta_AUC
Doxorubicin
YDL227C_RF1_112 RF1
ZShift = −0.69
lm Zscore = 1.53
NG = 0
DB = 0
SM = 0
Delta_L
Doxorubicin
YDL227C_RF1_112 RF1
ZShift = −0.04
lm ZScore = 1.53
NG = 0
DB = 0
SM = 0
Delta_K
Doxorubicin
YDL227C_RF1_112 RF1
ZShift = 0.39
lm ZScore = −0.37
NG = 0
DB = 0
SM = 0
Delta_r
Doxorubicin
YDL227C_RF1_112 RF1
ZShift = 0.12
lm ZScore = 0.43
NG = 0
DB = 0
SM = 0
Delta_AUC
Doxorubicin

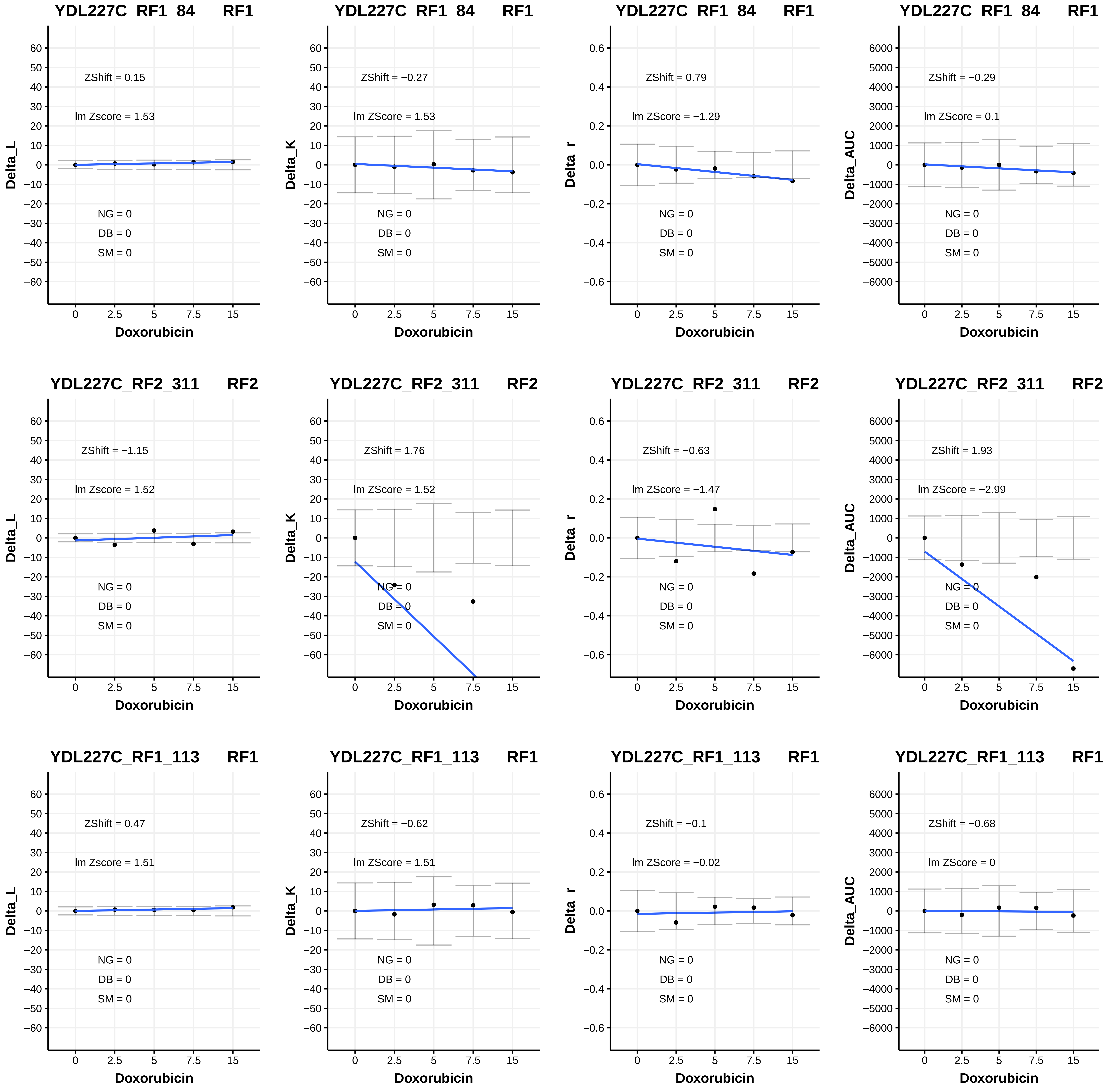
YDL227C_RF1_84 RF1
Delta_L
ZShift = 0.15
lm Zscore = 1.53
NG = 0
DB = 0
SM = 0
Doxorubicin
YDL227C_RF1_84 RF1
Delta_K
ZShift = −0.27
lm ZScore = 1.53
NG = 0
DB = 0
SM = 0
Doxorubicin
YDL227C_RF1_84 RF1
Delta_r
ZShift = 0.79
lm ZScore = −1.29
NG = 0
DB = 0
SM = 0
Doxorubicin
YDL227C_RF1_84 RF1
Delta_AUC
ZShift = −0.29
lm ZScore = 0.1
NG = 0
DB = 0
SM = 0
Doxorubicin
YDL227C_RF2_311 RF2
Delta_L
ZShift = −1.15
lm Zscore = 1.52
NG = 0
DB = 0
SM = 0
Doxorubicin
YDL227C_RF2_311 RF2
Delta_K
ZShift = 1.76
lm ZScore = 1.52
NG = 0
DB = 0
SM = 0
Doxorubicin
YDL227C_RF2_311 RF2
Delta_r
ZShift = −0.63
lm ZScore = −1.47
NG = 0
DB = 0
SM = 0
Doxorubicin
YDL227C_RF2_311 RF2
Delta_AUC
ZShift = 1.93
lm ZScore = −2.99
NG = 0
DB = 0
SM = 0
Doxorubicin
YDL227C_RF1_113 RF1
Delta_L
ZShift = 0.47
lm Zscore = 1.51
NG = 0
DB = 0
SM = 0
Doxorubicin
YDL227C_RF1_113 RF1
Delta_K
ZShift = −0.62
lm ZScore = 1.51
NG = 0
DB = 0
SM = 0
Doxorubicin
YDL227C_RF1_113 RF1
Delta_r
ZShift = −0.1
lm ZScore = −0.02
NG = 0
DB = 0
SM = 0
Doxorubicin
YDL227C_RF1_113 RF1
Delta_AUC
ZShift = −0.68
lm ZScore = 0
NG = 0
DB = 0
SM = 0
Doxorubicin

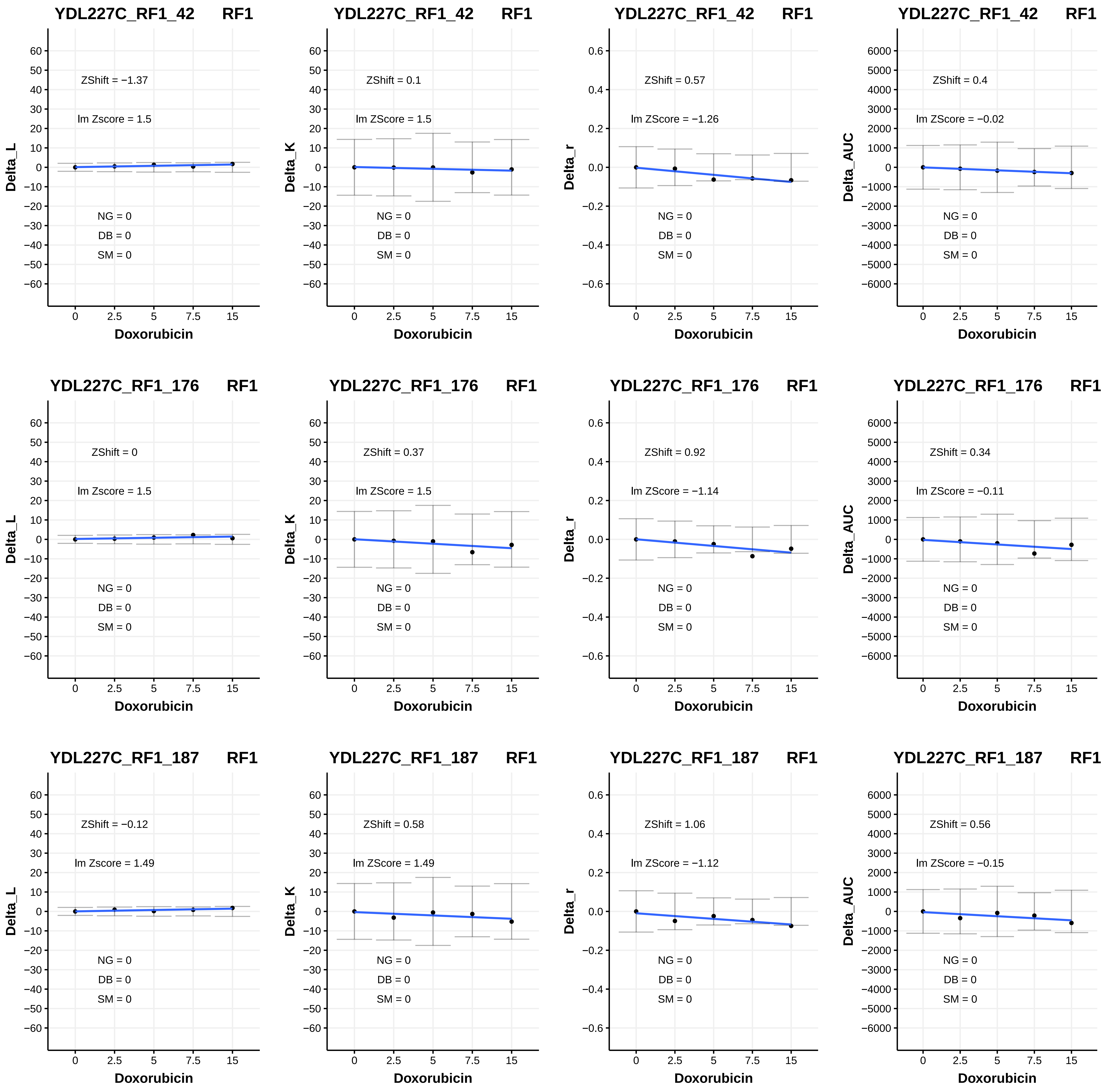

YDL227C_RF1_42 RF1
ZShift = −1.37
lm Zscore = 1.5
NG = 0
DB = 0
SM = 0
Delta_L
Doxorubicin
YDL227C_RF1_42 RF1
ZShift = 0.1
lm ZScore = 1.5
NG = 0
DB = 0
SM = 0
Delta_K
Doxorubicin
YDL227C_RF1_42 RF1
ZShift = 0.57
lm ZScore = −1.26
NG = 0
DB = 0
SM = 0
Delta_r
Doxorubicin
YDL227C_RF1_42 RF1
ZShift = 0.4
lm ZScore = −0.02
NG = 0
DB = 0
SM = 0
Delta_AUC
Doxorubicin
YDL227C_RF1_176 RF1
ZShift = 0
lm Zscore = 1.5
NG = 0
DB = 0
SM = 0
Delta_L
Doxorubicin
YDL227C_RF1_176 RF1
ZShift = 0.37
lm ZScore = 1.5
NG = 0
DB = 0
SM = 0
Delta_K
Doxorubicin
YDL227C_RF1_176 RF1
ZShift = 0.92
lm ZScore = −1.14
NG = 0
DB = 0
SM = 0
Delta_r
Doxorubicin
YDL227C_RF1_176 RF1
ZShift = 0.34
lm ZScore = −0.11
NG = 0
DB = 0
SM = 0
Delta_AUC
Doxorubicin
YDL227C_RF1_187 RF1
ZShift = −0.12
lm Zscore = 1.49
NG = 0
DB = 0
SM = 0
Delta_L
Doxorubicin
YDL227C_RF1_187 RF1
ZShift = 0.58
lm ZScore = 1.49
NG = 0
DB = 0
SM = 0
Delta_K
Doxorubicin
YDL227C_RF1_187 RF1
ZShift = 1.06
lm ZScore = −1.12
NG = 0
DB = 0
SM = 0
Delta_r
Doxorubicin
YDL227C_RF1_187 RF1
ZShift = 0.56
lm ZScore = −0.15
NG = 0
DB = 0
SM = 0
Delta_AUC
Doxorubicin

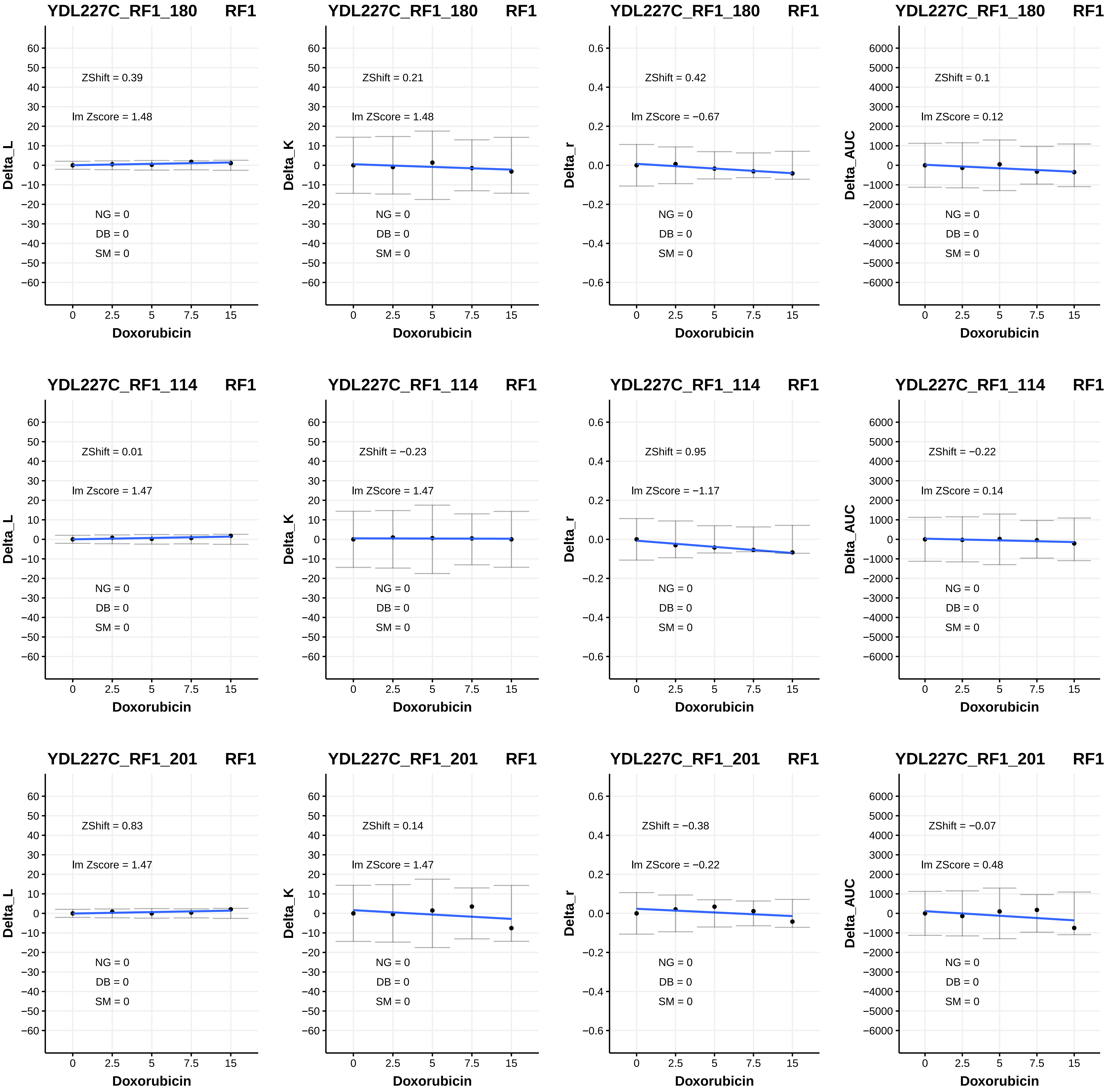

YDL227C_RF1_180 RF1
ZShift = 0.39
lm Zscore = 1.48
NG = 0
DB = 0
SM = 0
Delta_L
Doxorubicin
YDL227C_RF1_180 RF1
ZShift = 0.21
lm ZScore = 1.48
NG = 0
DB = 0
SM = 0
Delta_K
Doxorubicin
YDL227C_RF1_180 RF1
ZShift = 0.42
lm ZScore = −0.67
NG = 0
DB = 0
SM = 0
Delta_r
Doxorubicin
YDL227C_RF1_180 RF1
ZShift = 0.1
lm ZScore = 0.12
NG = 0
DB = 0
SM = 0
Delta_AUC
Doxorubicin
YDL227C_RF1_114 RF1
ZShift = 0.01
lm Zscore = 1.47
NG = 0
DB = 0
SM = 0
Delta_L
Doxorubicin
YDL227C_RF1_114 RF1
ZShift = −0.23
lm ZScore = 1.47
NG = 0
DB = 0
SM = 0
Delta_K
Doxorubicin
YDL227C_RF1_114 RF1
ZShift = 0.95
lm ZScore = −1.17
NG = 0
DB = 0
SM = 0
Delta_r
Doxorubicin
YDL227C_RF1_114 RF1
ZShift = −0.22
lm ZScore = 0.14
NG = 0
DB = 0
SM = 0
Delta_AUC
Doxorubicin
YDL227C_RF1_201 RF1
ZShift = 0.83
lm Zscore = 1.47
NG = 0
DB = 0
SM = 0
Delta_L
Doxorubicin
YDL227C_RF1_201 RF1
ZShift = 0.14
lm ZScore = 1.47
NG = 0
DB = 0
SM = 0
Delta_K
Doxorubicin
YDL227C_RF1_201 RF1
ZShift = −0.38
lm ZScore = −0.22
NG = 0
DB = 0
SM = 0
Delta_r
Doxorubicin
YDL227C_RF1_201 RF1
ZShift = −0.07
lm ZScore = 0.48
NG = 0
DB = 0
SM = 0
Delta_AUC
Doxorubicin

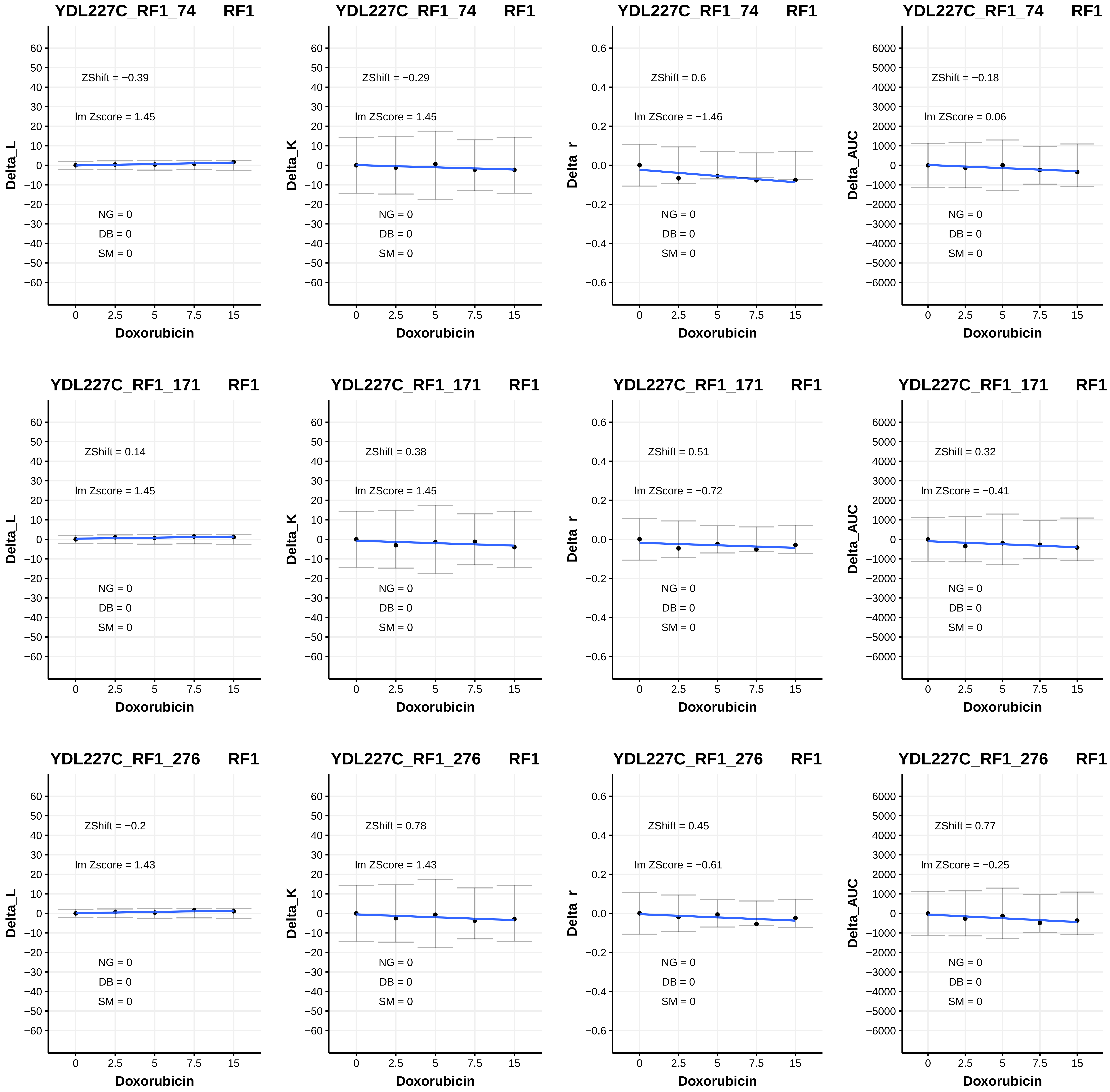
YDL227C_RF1_74 RF1
ZShift = −0.39
lm Zscore = 1.45
NG = 0
DB = 0
SM = 0
Delta_L
Doxorubicin
YDL227C_RF1_74 RF1
ZShift = −0.29
lm ZScore = 1.45
NG = 0
DB = 0
SM = 0
Delta_K
Doxorubicin
YDL227C_RF1_74 RF1
ZShift = 0.6
lm ZScore = −1.46
NG = 0
DB = 0
SM = 0
Delta_r
Doxorubicin
YDL227C_RF1_74 RF1
ZShift = −0.18
lm ZScore = 0.06
NG = 0
DB = 0
SM = 0
Delta_AUC
Doxorubicin
YDL227C_RF1_171 RF1
ZShift = 0.14
lm Zscore = 1.45
NG = 0
DB = 0
SM = 0
Delta_L
Doxorubicin
YDL227C_RF1_171 RF1
ZShift = 0.38
lm ZScore = 1.45
NG = 0
DB = 0
SM = 0
Delta_K
Doxorubicin
YDL227C_RF1_171 RF1
ZShift = 0.51
lm ZScore = −0.72
NG = 0
DB = 0
SM = 0
Delta_r
Doxorubicin
YDL227C_RF1_171 RF1
ZShift = 0.32
lm ZScore = −0.41
NG = 0
DB = 0
SM = 0
Delta_AUC
Doxorubicin
YDL227C_RF1_276 RF1
ZShift = −0.2
lm Zscore = 1.43
NG = 0
DB = 0
SM = 0
Delta_L
Doxorubicin
YDL227C_RF1_276 RF1
ZShift = 0.78
lm ZScore = 1.43
NG = 0
DB = 0
SM = 0
Delta_K
Doxorubicin
YDL227C_RF1_276 RF1
ZShift = 0.45
lm ZScore = −0.61
NG = 0
DB = 0
SM = 0
Delta_r
Doxorubicin
YDL227C_RF1_276 RF1
ZShift = 0.77
lm ZScore = −0.25
NG = 0
DB = 0
SM = 0
Delta_AUC
Doxorubicin

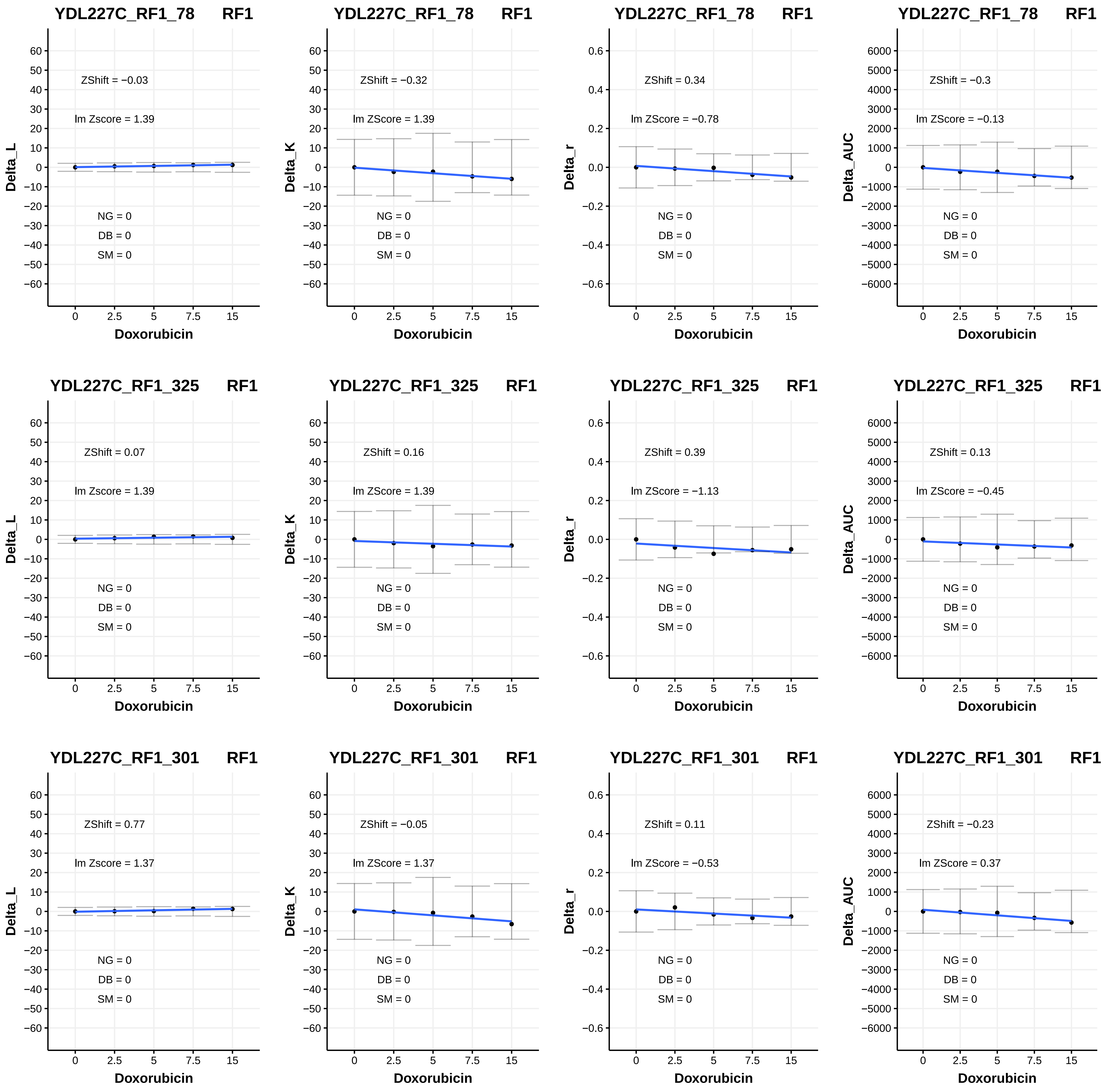
YDL227C_RF1_78 RF1
ZShift = −0.03
lm Zscore = 1.39
NG = 0
DB = 0
SM = 0
Delta_L
Doxorubicin
YDL227C_RF1_78 RF1
ZShift = −0.32
lm ZScore = 1.39
NG = 0
DB = 0
SM = 0
Delta_K
Doxorubicin
YDL227C_RF1_78 RF1
ZShift = 0.34
lm ZScore = −0.78
NG = 0
DB = 0
SM = 0
Delta_r
Doxorubicin
YDL227C_RF1_78 RF1
ZShift = −0.3
lm ZScore = −0.13
NG = 0
DB = 0
SM = 0
Delta_AUC
Doxorubicin
YDL227C_RF1_325 RF1
ZShift = 0.07
lm Zscore = 1.39
NG = 0
DB = 0
SM = 0
Delta_L
Doxorubicin
YDL227C_RF1_325 RF1
ZShift = 0.16
lm ZScore = 1.39
NG = 0
DB = 0
SM = 0
Delta_K
Doxorubicin
YDL227C_RF1_325 RF1
ZShift = 0.39
lm ZScore = −1.13
NG = 0
DB = 0
SM = 0
Delta_r
Doxorubicin
YDL227C_RF1_325 RF1
ZShift = 0.13
lm ZScore = −0.45
NG = 0
DB = 0
SM = 0
Delta_AUC
Doxorubicin
YDL227C_RF1_301 RF1
ZShift = 0.77
lm Zscore = 1.37
NG = 0
DB = 0
SM = 0
Delta_L
Doxorubicin
YDL227C_RF1_301 RF1
ZShift = −0.05
lm ZScore = 1.37
NG = 0
DB = 0
SM = 0
Delta_K
Doxorubicin
YDL227C_RF1_301 RF1
ZShift = 0.11
lm ZScore = −0.53
NG = 0
DB = 0
SM = 0
Delta_r
Doxorubicin
YDL227C_RF1_301 RF1
ZShift = −0.23
lm ZScore = 0.37
NG = 0
DB = 0
SM = 0
Delta_AUC
Doxorubicin

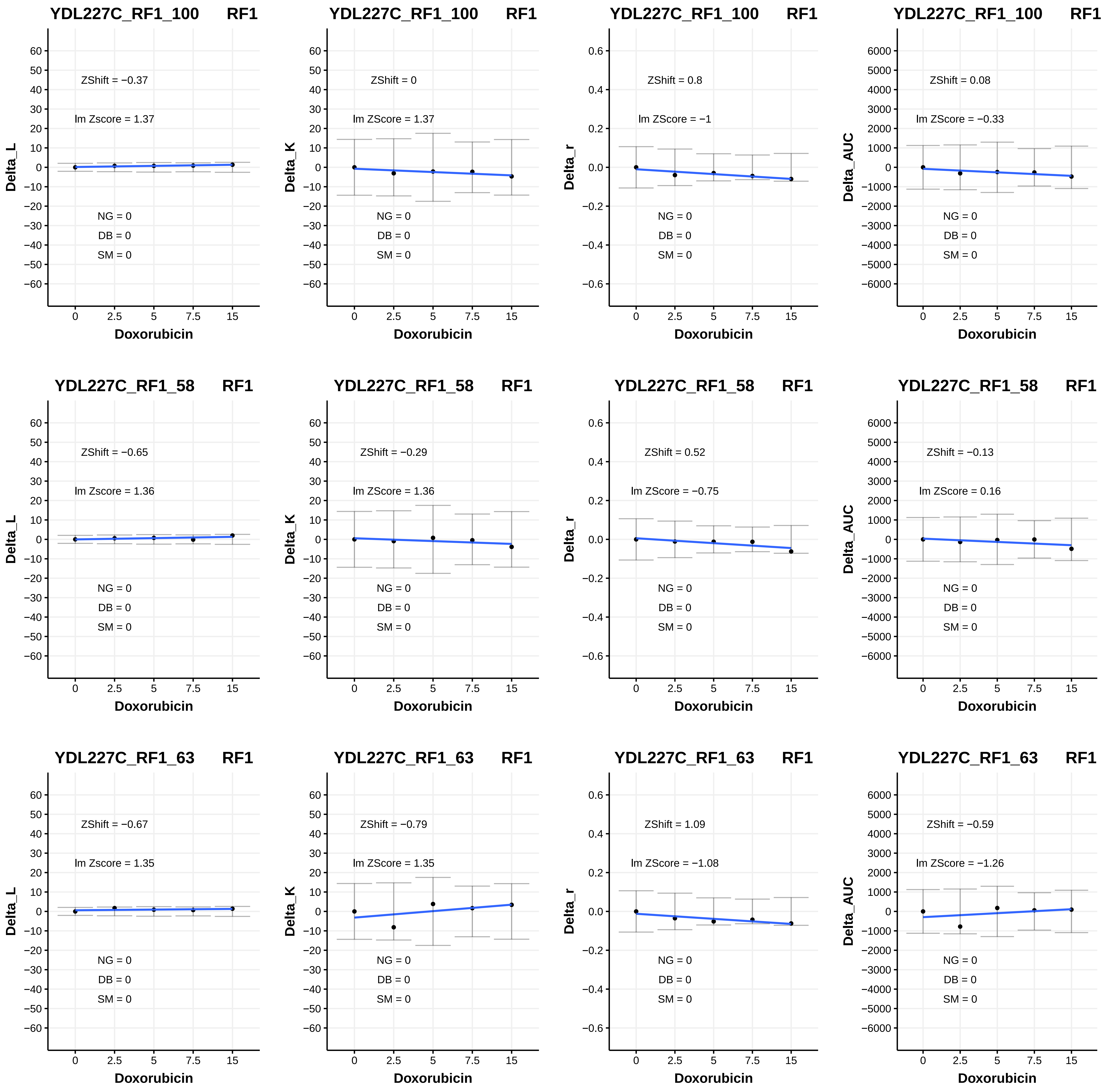
YDL227C_RF1_100 RF1
ZShift = −0.37
lm Zscore = 1.37
NG = 0
DB = 0
SM = 0
Delta_L
Doxorubicin
YDL227C_RF1_100 RF1
ZShift = 0
lm ZScore = 1.37
NG = 0
DB = 0
SM = 0
Delta_K
Doxorubicin
YDL227C_RF1_100 RF1
ZShift = 0.8
lm ZScore = −1
NG = 0
DB = 0
SM = 0
Delta_r
Doxorubicin
YDL227C_RF1_100 RF1
ZShift = 0.08
lm ZScore = −0.33
NG = 0
DB = 0
SM = 0
Delta_AUC
Doxorubicin
YDL227C_RF1_58 RF1
ZShift = −0.65
lm Zscore = 1.36
NG = 0
DB = 0
SM = 0
Delta_L
Doxorubicin
YDL227C_RF1_58 RF1
ZShift = −0.29
lm ZScore = 1.36
NG = 0
DB = 0
SM = 0
Delta_K
Doxorubicin
YDL227C_RF1_58 RF1
ZShift = 0.52
lm ZScore = −0.75
NG = 0
DB = 0
SM = 0
Delta_r
Doxorubicin
YDL227C_RF1_58 RF1
ZShift = −0.13
lm ZScore = 0.16
NG = 0
DB = 0
SM = 0
Delta_AUC
Doxorubicin
YDL227C_RF1_63 RF1
ZShift = −0.67
lm Zscore = 1.35
NG = 0
DB = 0
SM = 0
Delta_L
Doxorubicin
YDL227C_RF1_63 RF1
ZShift = −0.79
lm ZScore = 1.35
NG = 0
DB = 0
SM = 0
Delta_K
Doxorubicin
YDL227C_RF1_63 RF1
ZShift = 1.09
lm ZScore = −1.08
NG = 0
DB = 0
SM = 0
Delta_r
Doxorubicin
YDL227C_RF1_63 RF1
ZShift = −0.59
lm ZScore = −1.26
NG = 0
DB = 0
SM = 0
Delta_AUC
Doxorubicin

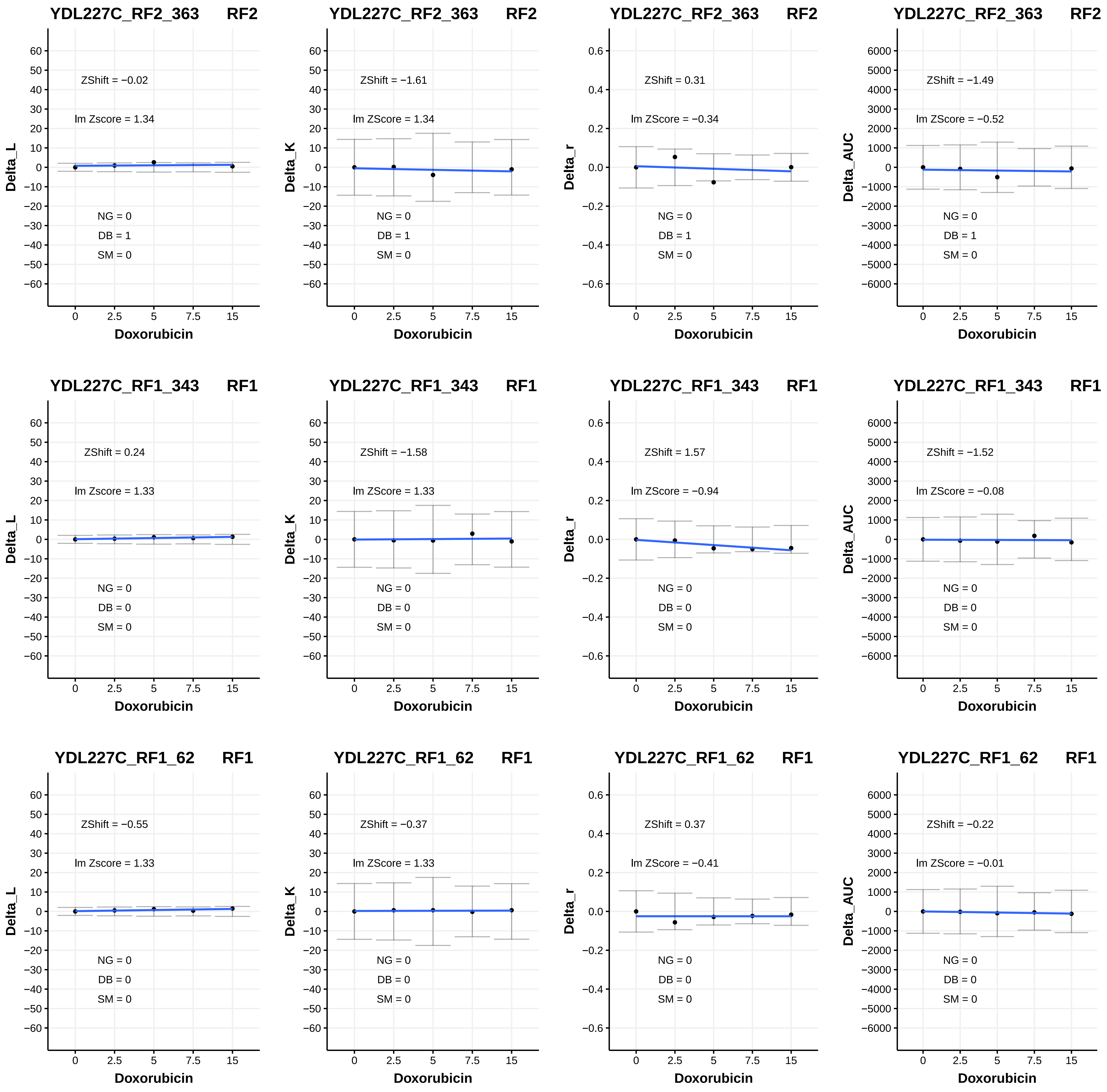
YDL227C_RF2_363 RF2
ZShift = −0.02
lm Zscore = 1.34
NG = 0
DB = 1
SM = 0
Delta_L
Doxorubicin
YDL227C_RF2_363 RF2
ZShift = −1.61
lm ZScore = 1.34
NG = 0
DB = 1
SM = 0
Delta_K
Doxorubicin
YDL227C_RF2_363 RF2
ZShift = 0.31
lm ZScore = −0.34
NG = 0
DB = 1
SM = 0
Delta_r
Doxorubicin
YDL227C_RF2_363 RF2
ZShift = −1.49
lm ZScore = −0.52
NG = 0
DB = 1
SM = 0
Delta_AUC
Doxorubicin
YDL227C_RF1_343 RF1
ZShift = 0.24
lm Zscore = 1.33
NG = 0
DB = 0
SM = 0
Delta_L
Doxorubicin
YDL227C_RF1_343 RF1
ZShift = −1.58
lm ZScore = 1.33
NG = 0
DB = 0
SM = 0
Delta_K
Doxorubicin
YDL227C_RF1_343 RF1
ZShift = 1.57
lm ZScore = −0.94
NG = 0
DB = 0
SM = 0
Delta_r
Doxorubicin
YDL227C_RF1_343 RF1
ZShift = −1.52
lm ZScore = −0.08
NG = 0
DB = 0
SM = 0
Delta_AUC
Doxorubicin
YDL227C_RF1_62 RF1
ZShift = −0.55
lm Zscore = 1.33
NG = 0
DB = 0
SM = 0
Delta_L
Doxorubicin
YDL227C_RF1_62 RF1
ZShift = −0.37
lm ZScore = 1.33
NG = 0
DB = 0
SM = 0
Delta_K
Doxorubicin
YDL227C_RF1_62 RF1
ZShift = 0.37
lm ZScore = −0.41
NG = 0
DB = 0
SM = 0
Delta_r
Doxorubicin
YDL227C_RF1_62 RF1
ZShift = −0.22
lm ZScore = −0.01
NG = 0
DB = 0
SM = 0
Delta_AUC
Doxorubicin

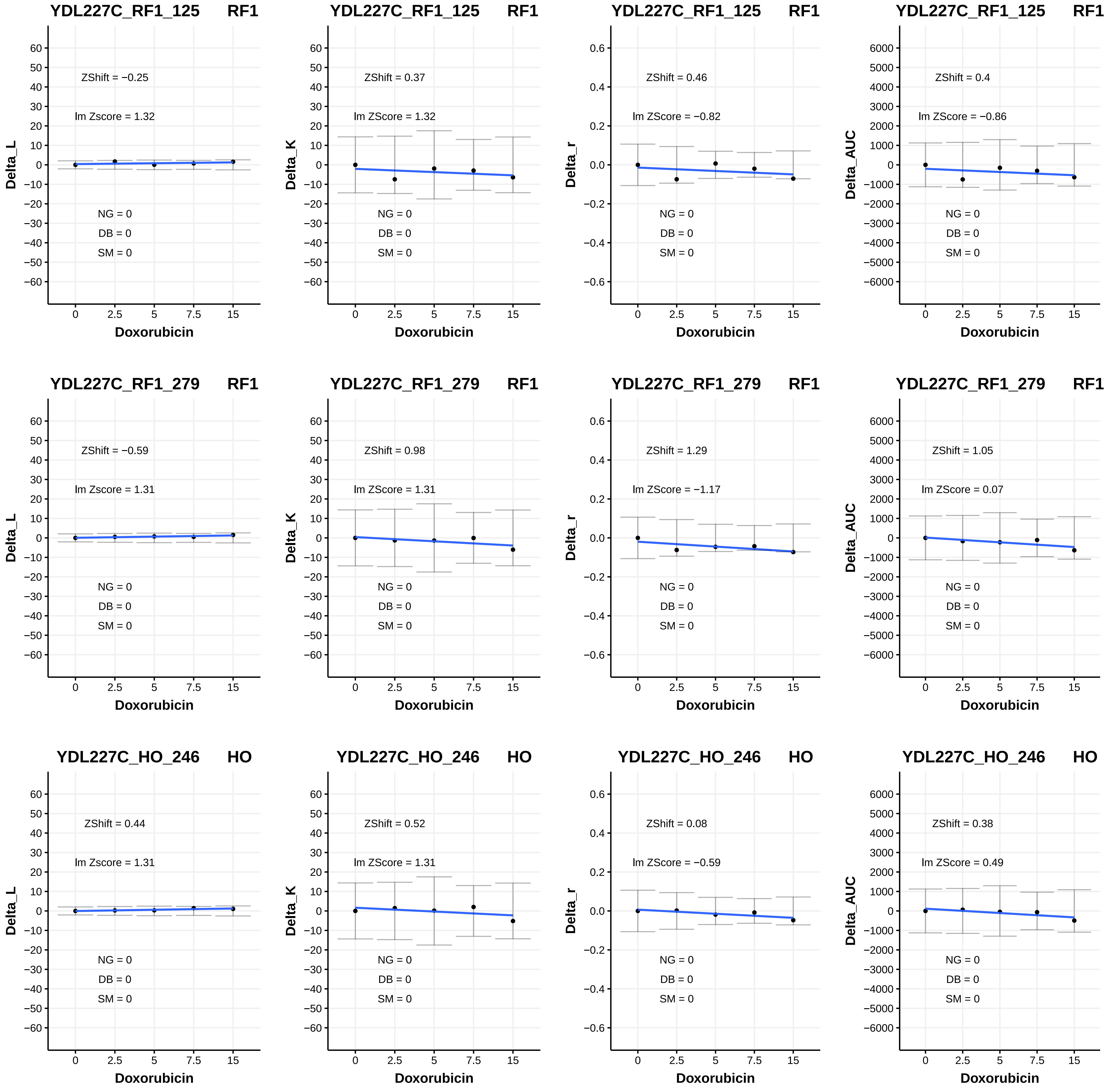
YDL227C_RF1_125 RF1
ZShift = −0.25
lm Zscore = 1.32
NG = 0
DB = 0
SM = 0
Delta_L
Doxorubicin
YDL227C_RF1_125 RF1
ZShift = 0.37
lm ZScore = 1.32
NG = 0
DB = 0
SM = 0
Delta_K
Doxorubicin
YDL227C_RF1_125 RF1
ZShift = 0.46
lm ZScore = −0.82
NG = 0
DB = 0
SM = 0
Delta_r
Doxorubicin
YDL227C_RF1_125 RF1
ZShift = 0.4
lm ZScore = −0.86
NG = 0
DB = 0
SM = 0
Delta_AUC
Doxorubicin
YDL227C_RF1_279 RF1
ZShift = −0.59
lm Zscore = 1.31
NG = 0
DB = 0
SM = 0
Delta_L
Doxorubicin
YDL227C_RF1_279 RF1
ZShift = 0.98
lm ZScore = 1.31
NG = 0
DB = 0
SM = 0
Delta_K
Doxorubicin
YDL227C_RF1_279 RF1
ZShift = 1.29
lm ZScore = −1.17
NG = 0
DB = 0
SM = 0
Delta_r
Doxorubicin
YDL227C_RF1_279 RF1
ZShift = 1.05
lm ZScore = 0.07
NG = 0
DB = 0
SM = 0
Delta_AUC
Doxorubicin
YDL227C_HO_246 HO
ZShift = 0.44
lm Zscore = 1.31
NG = 0
DB = 0
SM = 0
Delta_L
Doxorubicin
YDL227C_HO_246 HO
ZShift = 0.52
lm ZScore = 1.31
NG = 0
DB = 0
SM = 0
Delta_K
Doxorubicin
YDL227C_HO_246 HO
ZShift = 0.08
lm ZScore = −0.59
NG = 0
DB = 0
SM = 0
Delta_r
Doxorubicin
YDL227C_HO_246 HO
ZShift = 0.38
lm ZScore = 0.49
NG = 0
DB = 0
SM = 0
Delta_AUC
Doxorubicin

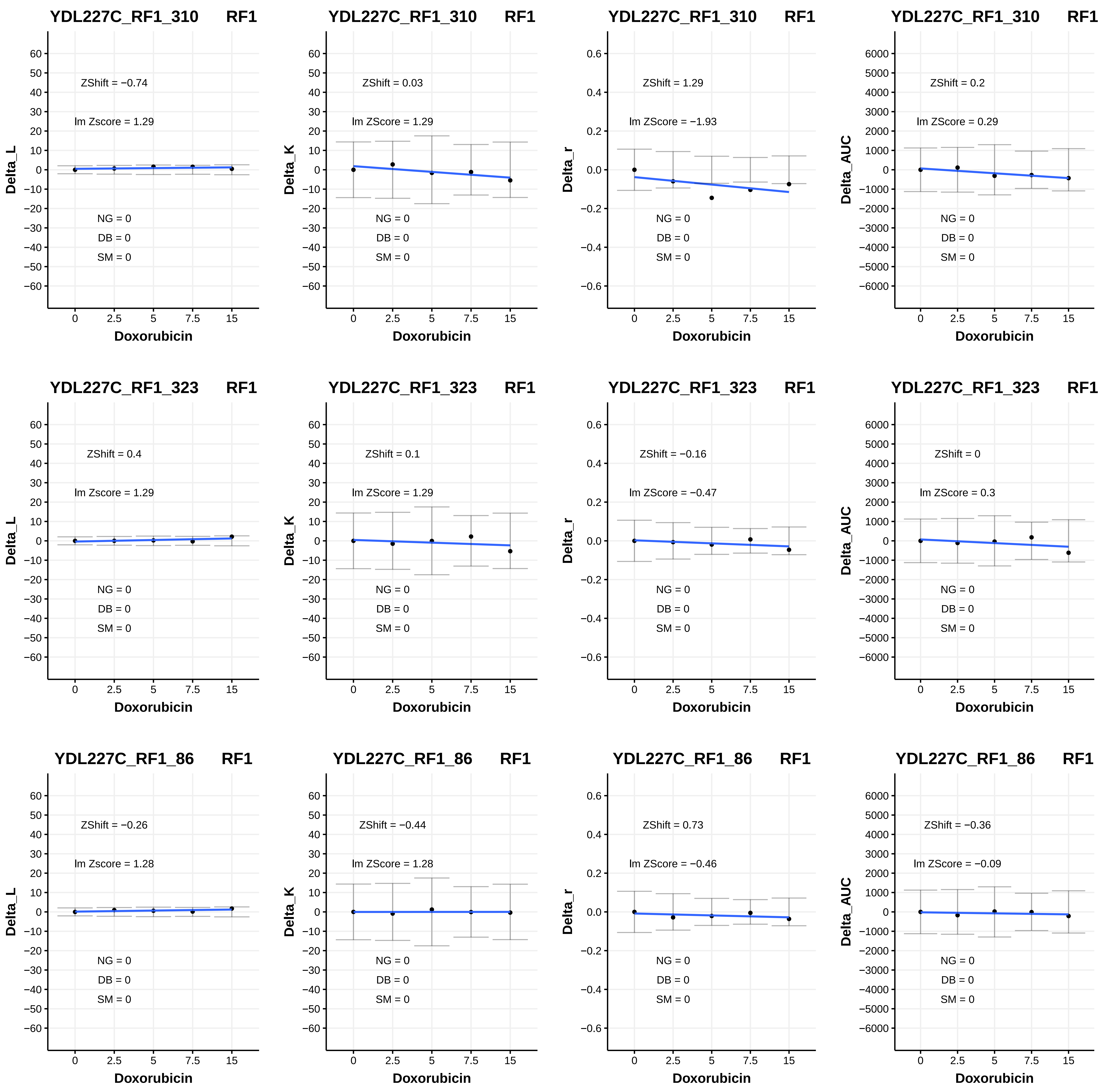

YDL227C_RF1_310 RF1
ZShift = −0.74
lm Zscore = 1.29
NG = 0
DB = 0
SM = 0
Delta_L
Doxorubicin
YDL227C_RF1_310 RF1
ZShift = 0.03
lm ZScore = 1.29
NG = 0
DB = 0
SM = 0
Delta_K
Doxorubicin
YDL227C_RF1_310 RF1
ZShift = 1.29
lm ZScore = −1.93
NG = 0
DB = 0
SM = 0
Delta_r
Doxorubicin
YDL227C_RF1_310 RF1
ZShift = 0.2
lm ZScore = 0.29
NG = 0
DB = 0
SM = 0
Delta_AUC
Doxorubicin
YDL227C_RF1_323 RF1
ZShift = 0.4
lm Zscore = 1.29
NG = 0
DB = 0
SM = 0
Delta_L
Doxorubicin
YDL227C_RF1_323 RF1
ZShift = 0.1
lm ZScore = 1.29
NG = 0
DB = 0
SM = 0
Delta_K
Doxorubicin
YDL227C_RF1_323 RF1
ZShift = −0.16
lm ZScore = −0.47
NG = 0
DB = 0
SM = 0
Delta_r
Doxorubicin
YDL227C_RF1_323 RF1
ZShift = 0
lm ZScore = 0.3
NG = 0
DB = 0
SM = 0
Delta_AUC
Doxorubicin
YDL227C_RF1_86 RF1
ZShift = −0.26
lm Zscore = 1.28
NG = 0
DB = 0
SM = 0
Delta_L
Doxorubicin
YDL227C_RF1_86 RF1
ZShift = −0.44
lm ZScore = 1.28
NG = 0
DB = 0
SM = 0
Delta_K
Doxorubicin
YDL227C_RF1_86 RF1
ZShift = 0.73
lm ZScore = −0.46
NG = 0
DB = 0
SM = 0
Delta_r
Doxorubicin
YDL227C_RF1_86 RF1
ZShift = −0.36
lm ZScore = −0.09
NG = 0
DB = 0
SM = 0
Delta_AUC
Doxorubicin

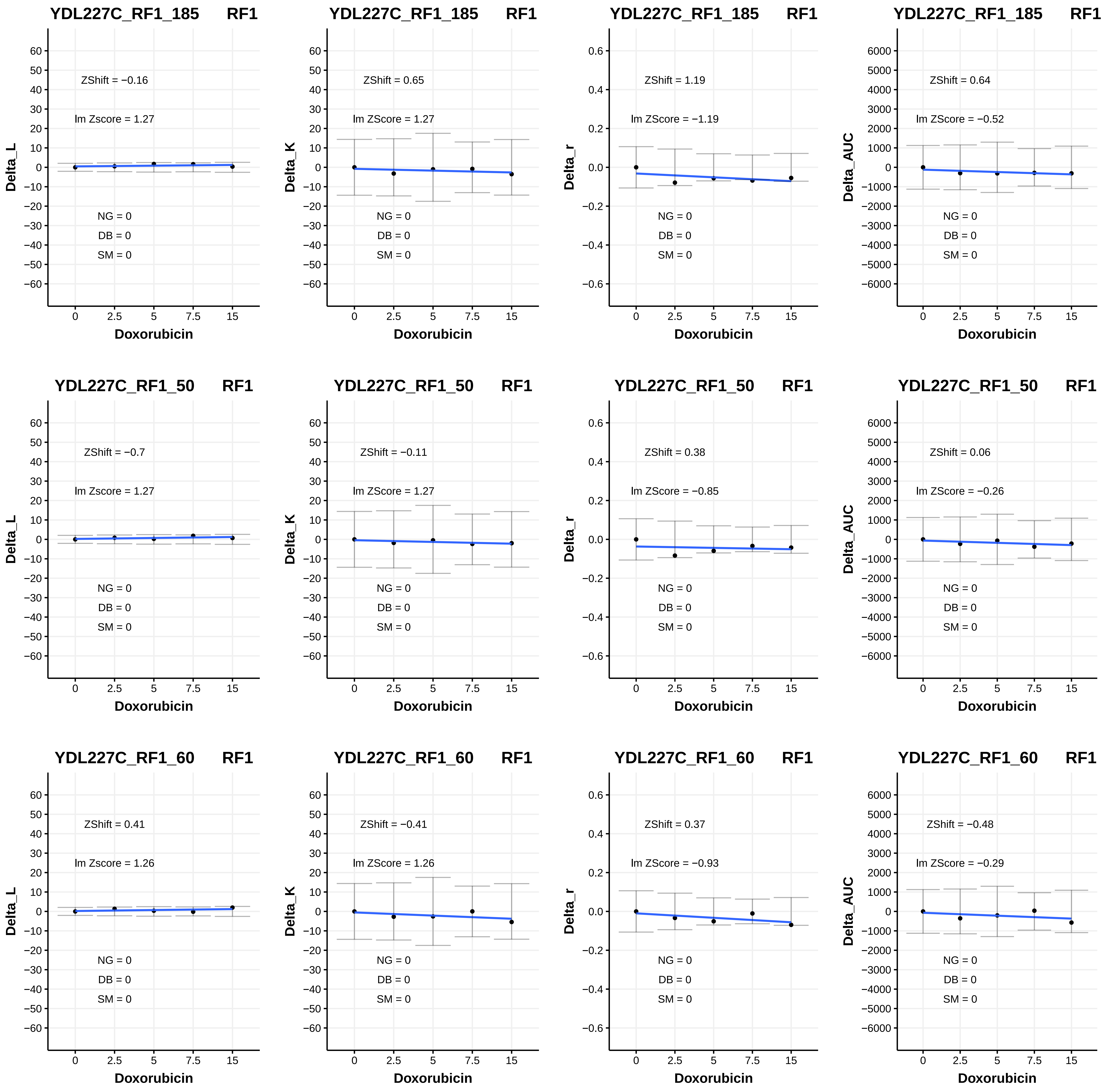
YDL227C_RF1_185 RF1
ZShift = −0.16
lm Zscore = 1.27
NG = 0
DB = 0
SM = 0
Delta_L
Doxorubicin
YDL227C_RF1_185 RF1
ZShift = 0.65
lm ZScore = 1.27
NG = 0
DB = 0
SM = 0
Delta_K
Doxorubicin
YDL227C_RF1_185 RF1
ZShift = 1.19
lm ZScore = −1.19
NG = 0
DB = 0
SM = 0
Delta_r
Doxorubicin
YDL227C_RF1_185 RF1
ZShift = 0.64
lm ZScore = −0.52
NG = 0
DB = 0
SM = 0
Delta_AUC
Doxorubicin
YDL227C_RF1_50 RF1
ZShift = −0.7
lm Zscore = 1.27
NG = 0
DB = 0
SM = 0
Delta_L
Doxorubicin
YDL227C_RF1_50 RF1
ZShift = −0.11
lm ZScore = 1.27
NG = 0
DB = 0
SM = 0
Delta_K
Doxorubicin
YDL227C_RF1_50 RF1
ZShift = 0.38
lm ZScore = −0.85
NG = 0
DB = 0
SM = 0
Delta_r
Doxorubicin
YDL227C_RF1_50 RF1
ZShift = 0.06
lm ZScore = −0.26
NG = 0
DB = 0
SM = 0
Delta_AUC
Doxorubicin
YDL227C_RF1_60 RF1
ZShift = 0.41
lm Zscore = 1.26
NG = 0
DB = 0
SM = 0
Delta_L
Doxorubicin
YDL227C_RF1_60 RF1
ZShift = −0.41
lm ZScore = 1.26
NG = 0
DB = 0
SM = 0
Delta_K
Doxorubicin
YDL227C_RF1_60 RF1
ZShift = 0.37
lm ZScore = −0.93
NG = 0
DB = 0
SM = 0
Delta_r
Doxorubicin
YDL227C_RF1_60 RF1
ZShift = −0.48
lm ZScore = −0.29
NG = 0
DB = 0
SM = 0
Delta_AUC
Doxorubicin

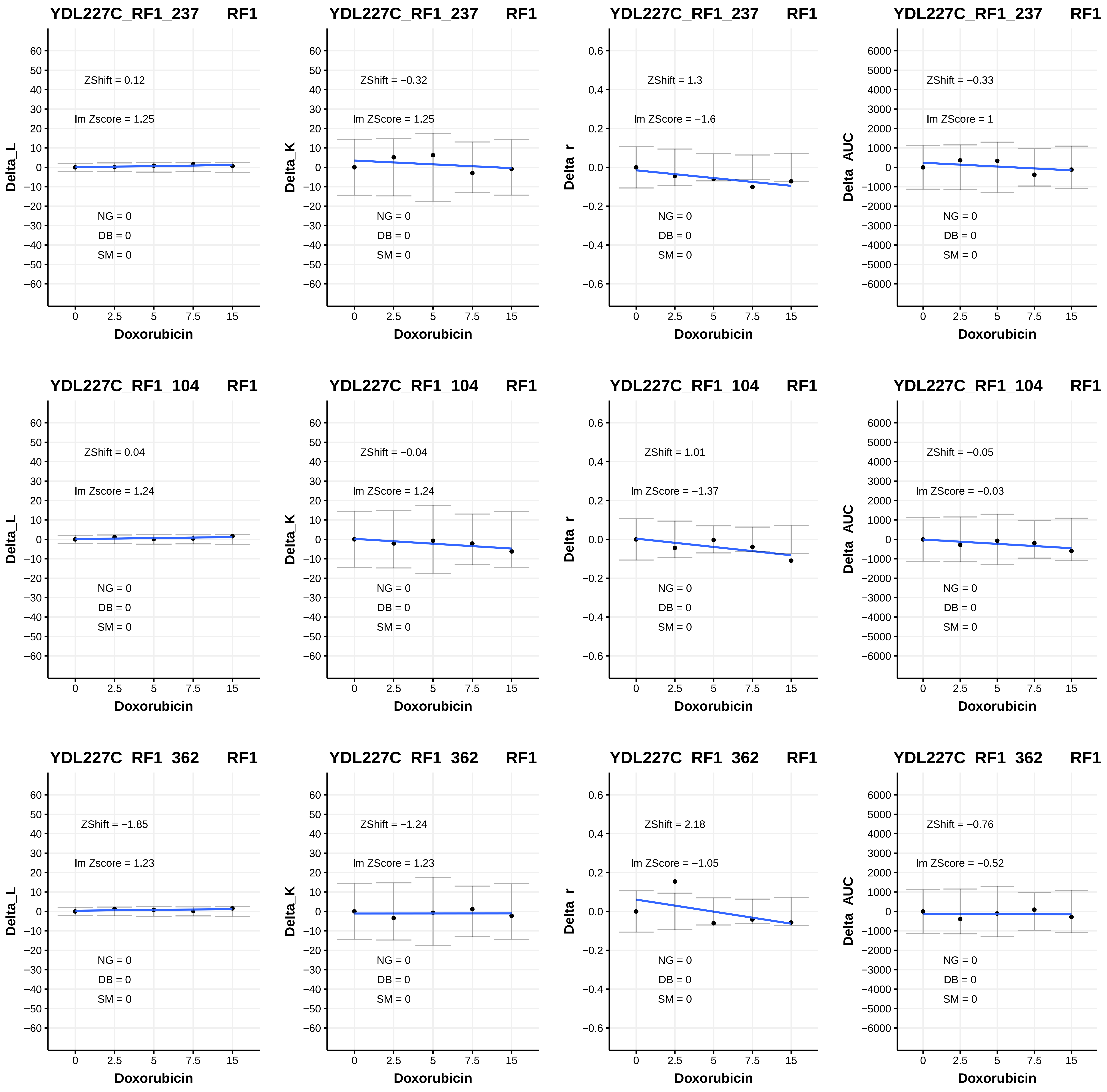
YDL227C_RF1_237 RF1
ZShift = 0.12
lm Zscore = 1.25
NG = 0
DB = 0
SM = 0
Delta_L
Doxorubicin
YDL227C_RF1_237 RF1
ZShift = −0.32
lm ZScore = 1.25
NG = 0
DB = 0
SM = 0
Delta_K
Doxorubicin
YDL227C_RF1_237 RF1
ZShift = 1.3
lm ZScore = −1.6
NG = 0
DB = 0
SM = 0
Delta_r
Doxorubicin
YDL227C_RF1_237 RF1
ZShift = −0.33
lm ZScore = 1
NG = 0
DB = 0
SM = 0
Delta_AUC
Doxorubicin
YDL227C_RF1_104 RF1
ZShift = 0.04
lm Zscore = 1.24
NG = 0
DB = 0
SM = 0
Delta_L
Doxorubicin
YDL227C_RF1_104 RF1
ZShift = −0.04
lm ZScore = 1.24
NG = 0
DB = 0
SM = 0
Delta_K
Doxorubicin
YDL227C_RF1_104 RF1
ZShift = 1.01
lm ZScore = −1.37
NG = 0
DB = 0
SM = 0
Delta_r
Doxorubicin
YDL227C_RF1_104 RF1
ZShift = −0.05
lm ZScore = −0.03
NG = 0
DB = 0
SM = 0
Delta_AUC
Doxorubicin
YDL227C_RF1_362 RF1
ZShift = −1.85
lm Zscore = 1.23
NG = 0
DB = 0
SM = 0
Delta_L
Doxorubicin
YDL227C_RF1_362 RF1
ZShift = −1.24
lm ZScore = 1.23
NG = 0
DB = 0
SM = 0
Delta_K
Doxorubicin
YDL227C_RF1_362 RF1
ZShift = 2.18
lm ZScore = −1.05
NG = 0
DB = 0
SM = 0
Delta_r
Doxorubicin
YDL227C_RF1_362 RF1
ZShift = −0.76
lm ZScore = −0.52
NG = 0
DB = 0
SM = 0
Delta_AUC
Doxorubicin

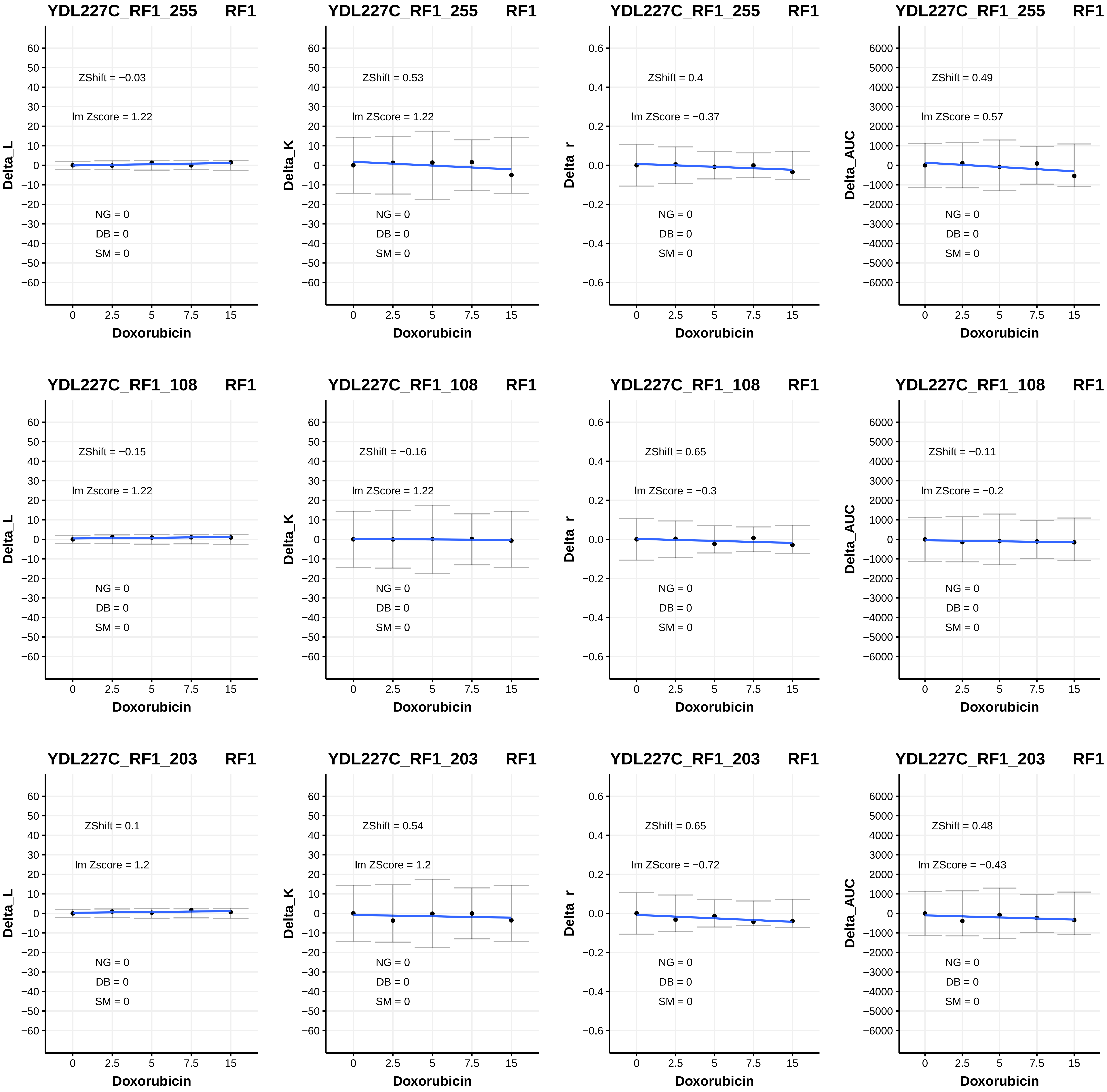
YDL227C_RF1_255 RF1
ZShift = −0.03
lm Zscore = 1.22
NG = 0
DB = 0
SM = 0
Delta_L
Doxorubicin
YDL227C_RF1_255 RF1
ZShift = 0.53
lm ZScore = 1.22
NG = 0
DB = 0
SM = 0
Delta_K
Doxorubicin
YDL227C_RF1_255 RF1
ZShift = 0.4
lm ZScore = −0.37
NG = 0
DB = 0
SM = 0
Delta_r
Doxorubicin
YDL227C_RF1_255 RF1
ZShift = 0.49
lm ZScore = 0.57
NG = 0
DB = 0
SM = 0
Delta_AUC
Doxorubicin
YDL227C_RF1_108 RF1
ZShift = −0.15
lm Zscore = 1.22
NG = 0
DB = 0
SM = 0
Delta_L
Doxorubicin
YDL227C_RF1_108 RF1
ZShift = −0.16
lm ZScore = 1.22
NG = 0
DB = 0
SM = 0
Delta_K
Doxorubicin
YDL227C_RF1_108 RF1
ZShift = 0.65
lm ZScore = −0.3
NG = 0
DB = 0
SM = 0
Delta_r
Doxorubicin
YDL227C_RF1_108 RF1
ZShift = −0.11
lm ZScore = −0.2
NG = 0
DB = 0
SM = 0
Delta_AUC
Doxorubicin
YDL227C_RF1_203 RF1
ZShift = 0.1
lm Zscore = 1.2
NG = 0
DB = 0
SM = 0
Delta_L
Doxorubicin
YDL227C_RF1_203 RF1
ZShift = 0.54
lm ZScore = 1.2
NG = 0
DB = 0
SM = 0
Delta_K
Doxorubicin
YDL227C_RF1_203 RF1
ZShift = 0.65
lm ZScore = −0.72
NG = 0
DB = 0
SM = 0
Delta_r
Doxorubicin
YDL227C_RF1_203 RF1
ZShift = 0.48
lm ZScore = −0.43
NG = 0
DB = 0
SM = 0
Delta_AUC
Doxorubicin

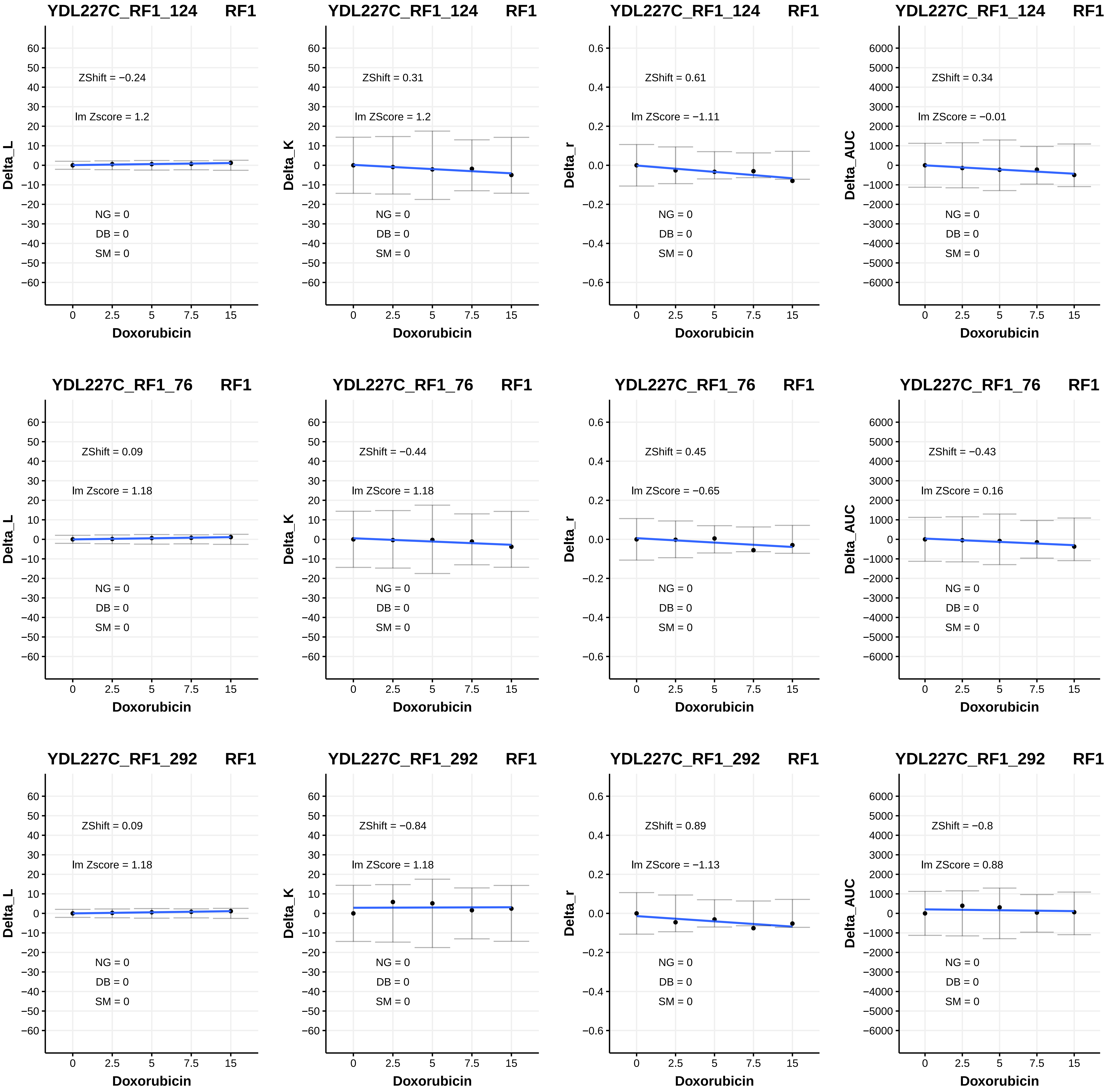
YDL227C_RF1_124 RF1
ZShift = −0.24
lm Zscore = 1.2
NG = 0
DB = 0
SM = 0
Delta_L
Doxorubicin
YDL227C_RF1_124 RF1
ZShift = 0.31
lm ZScore = 1.2
NG = 0
DB = 0
SM = 0
Delta_K
Doxorubicin
YDL227C_RF1_124 RF1
ZShift = 0.61
lm ZScore = −1.11
NG = 0
DB = 0
SM = 0
Delta_r
Doxorubicin
YDL227C_RF1_124 RF1
ZShift = 0.34
lm ZScore = −0.01
NG = 0
DB = 0
SM = 0
Delta_AUC
Doxorubicin
YDL227C_RF1_76 RF1
ZShift = 0.09
lm Zscore = 1.18
NG = 0
DB = 0
SM = 0
Delta_L
Doxorubicin
YDL227C_RF1_76 RF1
ZShift = −0.44
lm ZScore = 1.18
NG = 0
DB = 0
SM = 0
Delta_K
Doxorubicin
YDL227C_RF1_76 RF1
ZShift = 0.45
lm ZScore = −0.65
NG = 0
DB = 0
SM = 0
Delta_r
Doxorubicin
YDL227C_RF1_76 RF1
ZShift = −0.43
lm ZScore = 0.16
NG = 0
DB = 0
SM = 0
Delta_AUC
Doxorubicin
YDL227C_RF1_292 RF1
ZShift = 0.09
lm Zscore = 1.18
NG = 0
DB = 0
SM = 0
Delta_L
Doxorubicin
YDL227C_RF1_292 RF1
ZShift = −0.84
lm ZScore = 1.18
NG = 0
DB = 0
SM = 0
Delta_K
Doxorubicin
YDL227C_RF1_292 RF1
ZShift = 0.89
lm ZScore = −1.13
NG = 0
DB = 0
SM = 0
Delta_r
Doxorubicin
YDL227C_RF1_292 RF1
ZShift = −0.8
lm ZScore = 0.88
NG = 0
DB = 0
SM = 0
Delta_AUC
Doxorubicin

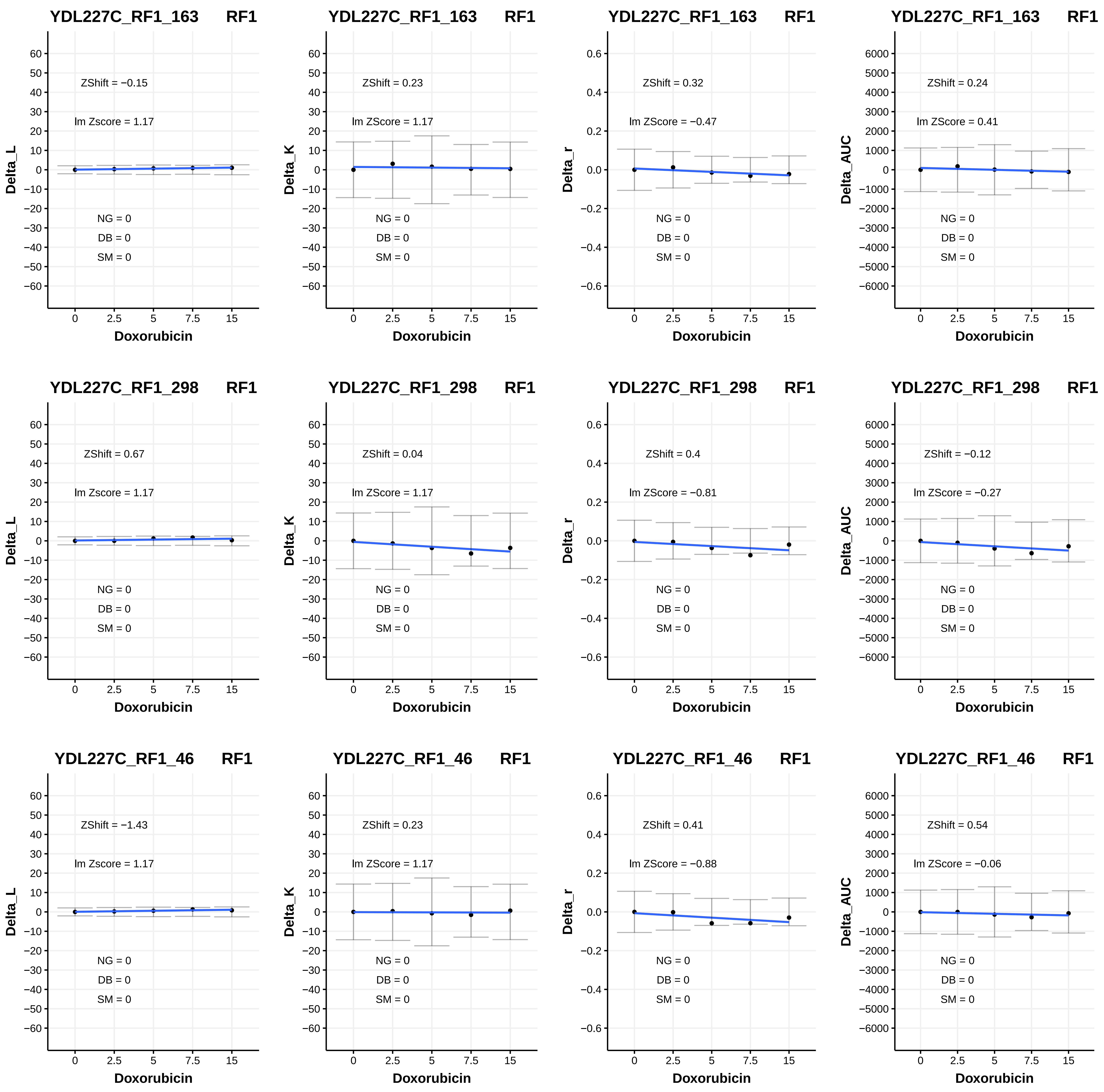
YDL227C_RF1_163 RF1
ZShift = −0.15
lm Zscore = 1.17
NG = 0
DB = 0
SM = 0
Delta_L
Doxorubicin
YDL227C_RF1_163 RF1
ZShift = 0.23
lm ZScore = 1.17
NG = 0
DB = 0
SM = 0
Delta_K
Doxorubicin
YDL227C_RF1_163 RF1
ZShift = 0.32
lm ZScore = −0.47
NG = 0
DB = 0
SM = 0
Delta_r
Doxorubicin
YDL227C_RF1_163 RF1
ZShift = 0.24
lm ZScore = 0.41
NG = 0
DB = 0
SM = 0
Delta_AUC
Doxorubicin
YDL227C_RF1_298 RF1
ZShift = 0.67
lm Zscore = 1.17
NG = 0
DB = 0
SM = 0
Delta_L
Doxorubicin
YDL227C_RF1_298 RF1
ZShift = 0.04
lm ZScore = 1.17
NG = 0
DB = 0
SM = 0
Delta_K
Doxorubicin
YDL227C_RF1_298 RF1
ZShift = 0.4
lm ZScore = −0.81
NG = 0
DB = 0
SM = 0
Delta_r
Doxorubicin
YDL227C_RF1_298 RF1
ZShift = −0.12
lm ZScore = −0.27
NG = 0
DB = 0
SM = 0
Delta_AUC
Doxorubicin
YDL227C_RF1_46 RF1
ZShift = −1.43
lm Zscore = 1.17
NG = 0
DB = 0
SM = 0
Delta_L
Doxorubicin
YDL227C_RF1_46 RF1
ZShift = 0.23
lm ZScore = 1.17
NG = 0
DB = 0
SM = 0
Delta_K
Doxorubicin
YDL227C_RF1_46 RF1
ZShift = 0.41
lm ZScore = −0.88
NG = 0
DB = 0
SM = 0
Delta_r
Doxorubicin
YDL227C_RF1_46 RF1
ZShift = 0.54
lm ZScore = −0.06
NG = 0
DB = 0
SM = 0
Delta_AUC
Doxorubicin

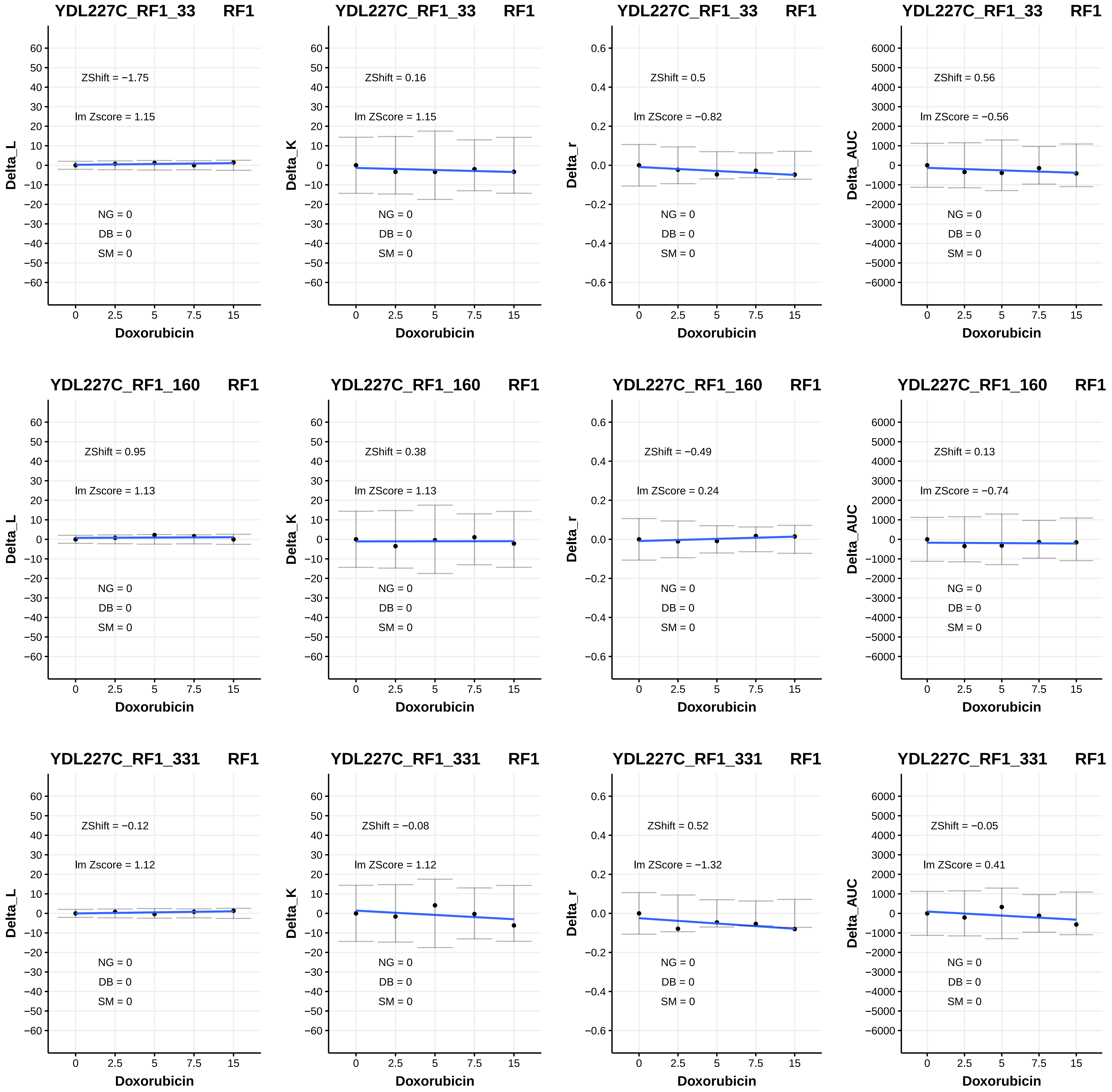
YDL227C_RF1_33 RF1
ZShift = −1.75
lm Zscore = 1.15
NG = 0
DB = 0
SM = 0
Delta_L
Doxorubicin
YDL227C_RF1_33 RF1
ZShift = 0.16
lm ZScore = 1.15
NG = 0
DB = 0
SM = 0
Delta_K
Doxorubicin
YDL227C_RF1_33 RF1
ZShift = 0.5
lm ZScore = −0.82
NG = 0
DB = 0
SM = 0
Delta_r
Doxorubicin
YDL227C_RF1_33 RF1
ZShift = 0.56
lm ZScore = −0.56
NG = 0
DB = 0
SM = 0
Delta_AUC
Doxorubicin
YDL227C_RF1_160 RF1
ZShift = 0.95
lm Zscore = 1.13
NG = 0
DB = 0
SM = 0
Delta_L
Doxorubicin
YDL227C_RF1_160 RF1
ZShift = 0.38
lm ZScore = 1.13
NG = 0
DB = 0
SM = 0
Delta_K
Doxorubicin
YDL227C_RF1_160 RF1
ZShift = −0.49
lm ZScore = 0.24
NG = 0
DB = 0
SM = 0
Delta_r
Doxorubicin
YDL227C_RF1_160 RF1
ZShift = 0.13
lm ZScore = −0.74
NG = 0
DB = 0
SM = 0
Delta_AUC
Doxorubicin
YDL227C_RF1_331 RF1
ZShift = −0.12
lm Zscore = 1.12
NG = 0
DB = 0
SM = 0
Delta_L
Doxorubicin
YDL227C_RF1_331 RF1
ZShift = −0.08
lm ZScore = 1.12
NG = 0
DB = 0
SM = 0
Delta_K
Doxorubicin
YDL227C_RF1_331 RF1
ZShift = 0.52
lm ZScore = −1.32
NG = 0
DB = 0
SM = 0
Delta_r
Doxorubicin
YDL227C_RF1_331 RF1
ZShift = −0.05
lm ZScore = 0.41
NG = 0
DB = 0
SM = 0
Delta_AUC
Doxorubicin
0
2.5
5
7.5
15

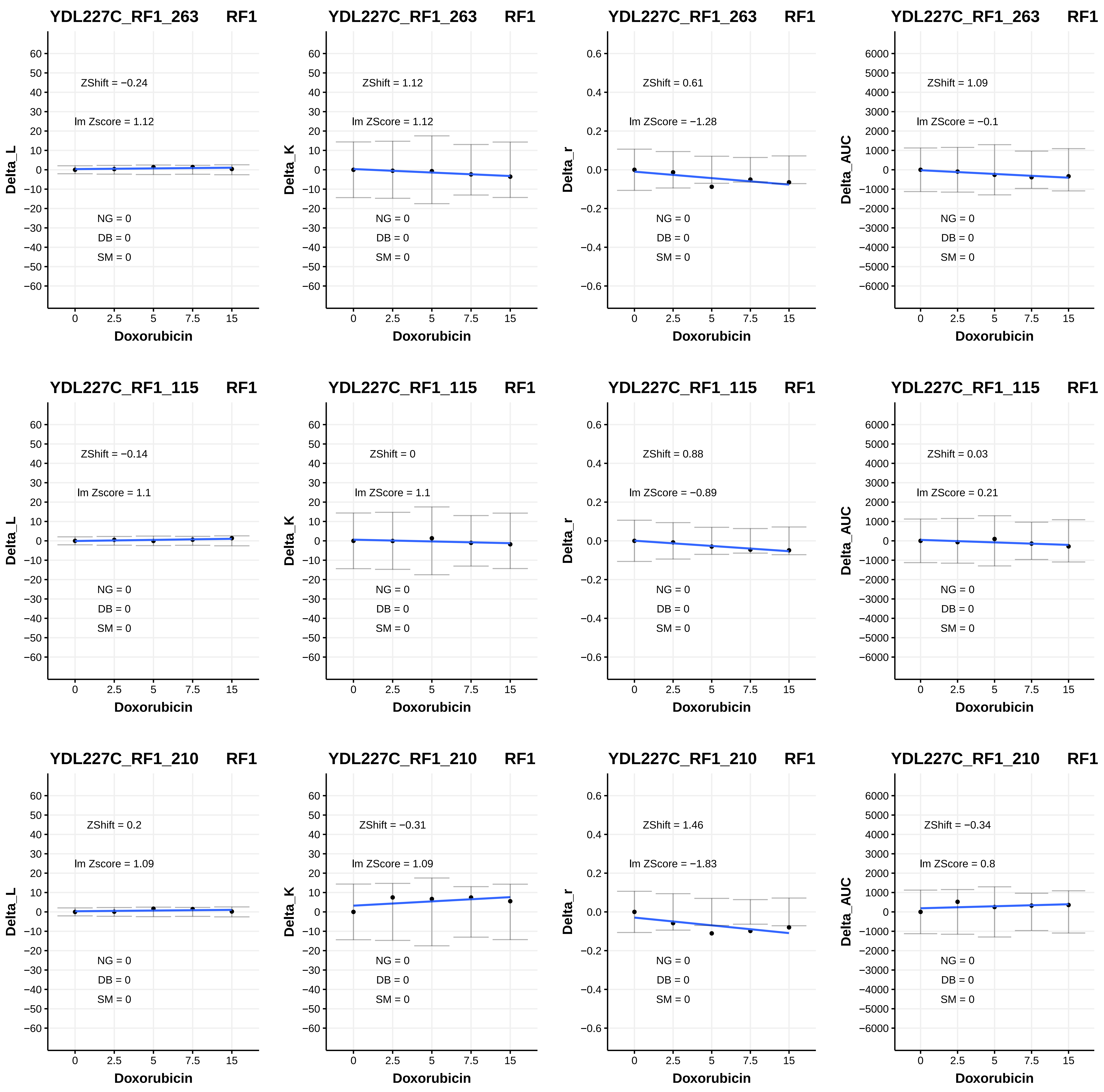

YDL227C_RF1_263 RF1
ZShift = −0.24
lm Zscore = 1.12
NG = 0
DB = 0
SM = 0
Delta_L
Doxorubicin
YDL227C_RF1_263 RF1
ZShift = 1.12
lm ZScore = 1.12
NG = 0
DB = 0
SM = 0
Delta_K
Doxorubicin
YDL227C_RF1_263 RF1
ZShift = 0.61
lm ZScore = −1.28
NG = 0
DB = 0
SM = 0
Delta_r
Doxorubicin
YDL227C_RF1_263 RF1
ZShift = 1.09
lm ZScore = −0.1
NG = 0
DB = 0
SM = 0
Delta_AUC
Doxorubicin
YDL227C_RF1_115 RF1
ZShift = −0.14
lm Zscore = 1.1
NG = 0
DB = 0
SM = 0
Delta_L
Doxorubicin
YDL227C_RF1_115 RF1
ZShift = 0
lm ZScore = 1.1
NG = 0
DB = 0
SM = 0
Delta_K
Doxorubicin
YDL227C_RF1_115 RF1
ZShift = 0.88
lm ZScore = −0.89
NG = 0
DB = 0
SM = 0
Delta_r
Doxorubicin
YDL227C_RF1_115 RF1
ZShift = 0.03
lm ZScore = 0.21
NG = 0
DB = 0
SM = 0
Delta_AUC
Doxorubicin
YDL227C_RF1_210 RF1
ZShift = 0.2
lm Zscore = 1.09
NG = 0
DB = 0
SM = 0
Delta_L
Doxorubicin
YDL227C_RF1_210 RF1
ZShift = −0.31
lm ZScore = 1.09
NG = 0
DB = 0
SM = 0
Delta_K
Doxorubicin
YDL227C_RF1_210 RF1
ZShift = 1.46
lm ZScore = −1.83
NG = 0
DB = 0
SM = 0
Delta_r
Doxorubicin
YDL227C_RF1_210 RF1
ZShift = −0.34
lm ZScore = 0.8
NG = 0
DB = 0
SM = 0
Delta_AUC
Doxorubicin

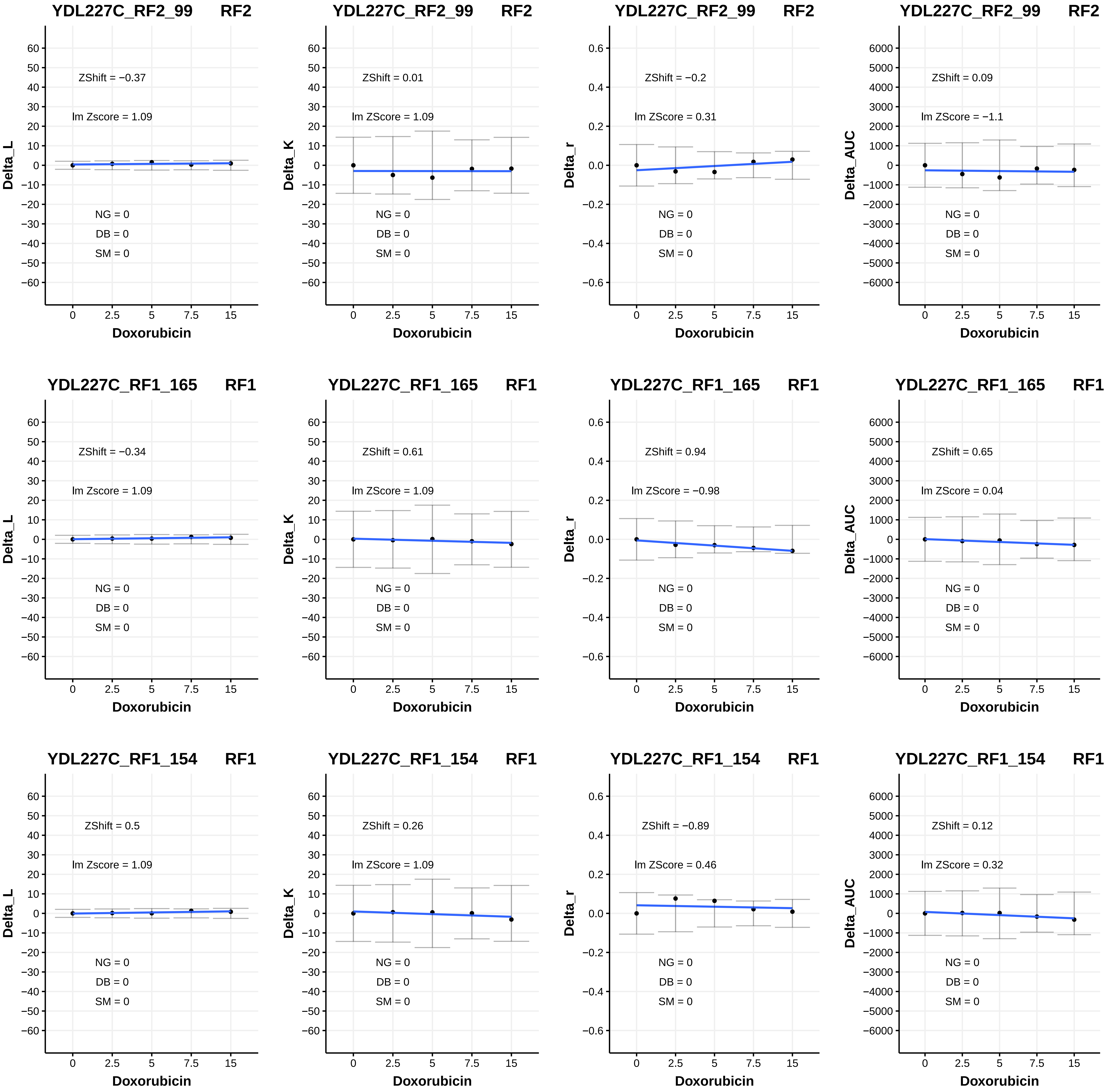

YDL227C_RF2_99 RF2
ZShift = −0.37
lm Zscore = 1.09
NG = 0
DB = 0
SM = 0
Delta_L
Doxorubicin
YDL227C_RF2_99 RF2
ZShift = 0.01
lm ZScore = 1.09
NG = 0
DB = 0
SM = 0
Delta_K
Doxorubicin
YDL227C_RF2_99 RF2
ZShift = −0.2
lm ZScore = 0.31
NG = 0
DB = 0
SM = 0
Delta_r
Doxorubicin
YDL227C_RF2_99 RF2
ZShift = 0.09
lm ZScore = −1.1
NG = 0
DB = 0
SM = 0
Delta_AUC
Doxorubicin
YDL227C_RF1_165 RF1
ZShift = −0.34
lm Zscore = 1.09
NG = 0
DB = 0
SM = 0
Delta_L
Doxorubicin
YDL227C_RF1_165 RF1
ZShift = 0.61
lm ZScore = 1.09
NG = 0
DB = 0
SM = 0
Delta_K
Doxorubicin
YDL227C_RF1_165 RF1
ZShift = 0.94
lm ZScore = −0.98
NG = 0
DB = 0
SM = 0
Delta_r
Doxorubicin
YDL227C_RF1_165 RF1
ZShift = 0.65
lm ZScore = 0.04
NG = 0
DB = 0
SM = 0
Delta_AUC
Doxorubicin
YDL227C_RF1_154 RF1
ZShift = 0.5
lm Zscore = 1.09
NG = 0
DB = 0
SM = 0
Delta_L
Doxorubicin
YDL227C_RF1_154 RF1
ZShift = 0.26
lm ZScore = 1.09
NG = 0
DB = 0
SM = 0
Delta_K
Doxorubicin
YDL227C_RF1_154 RF1
ZShift = −0.89
lm ZScore = 0.46
NG = 0
DB = 0
SM = 0
Delta_r
Doxorubicin
YDL227C_RF1_154 RF1
ZShift = 0.12
lm ZScore = 0.32
NG = 0
DB = 0
SM = 0
Delta_AUC
Doxorubicin

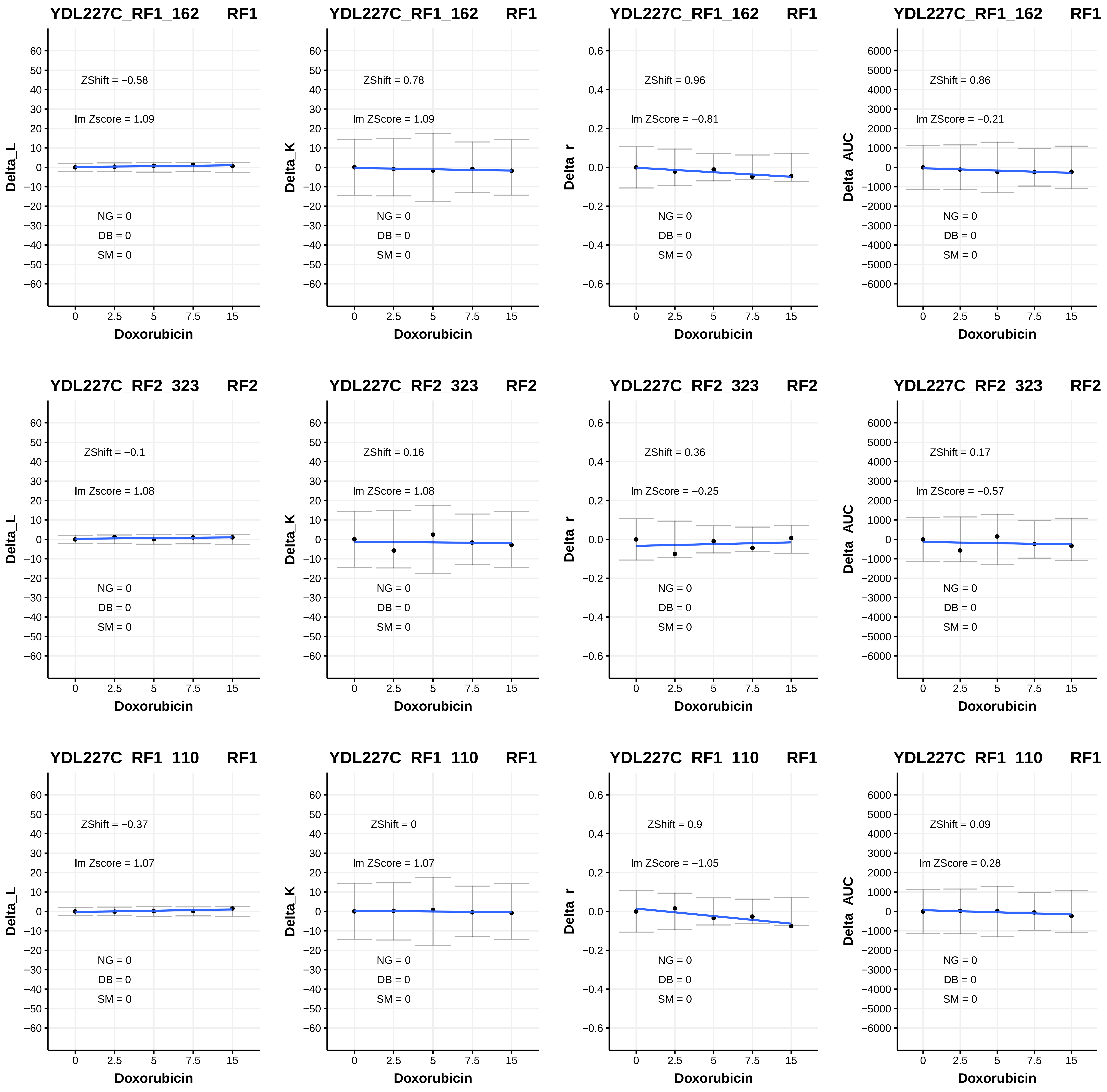
YDL227C_RF1_162 RF1
ZShift = −0.58
lm Zscore = 1.09
NG = 0
DB = 0
SM = 0
Delta_L
Doxorubicin
YDL227C_RF1_162 RF1
ZShift = 0.78
lm ZScore = 1.09
NG = 0
DB = 0
SM = 0
Delta_K
Doxorubicin
YDL227C_RF1_162 RF1
ZShift = 0.96
lm ZScore = −0.81
NG = 0
DB = 0
SM = 0
Delta_r
Doxorubicin
YDL227C_RF1_162 RF1
ZShift = 0.86
lm ZScore = −0.21
NG = 0
DB = 0
SM = 0
Delta_AUC
Doxorubicin
YDL227C_RF2_323 RF2
ZShift = −0.1
lm Zscore = 1.08
NG = 0
DB = 0
SM = 0
Delta_L
Doxorubicin
YDL227C_RF2_323 RF2
ZShift = 0.16
lm ZScore = 1.08
NG = 0
DB = 0
SM = 0
Delta_K
Doxorubicin
YDL227C_RF2_323 RF2
ZShift = 0.36
lm ZScore = −0.25
NG = 0
DB = 0
SM = 0
Delta_r
Doxorubicin
YDL227C_RF2_323 RF2
ZShift = 0.17
lm ZScore = −0.57
NG = 0
DB = 0
SM = 0
Delta_AUC
Doxorubicin
YDL227C_RF1_110 RF1
ZShift = −0.37
lm Zscore = 1.07
NG = 0
DB = 0
SM = 0
Delta_L
Doxorubicin
YDL227C_RF1_110 RF1
ZShift = 0
lm ZScore = 1.07
NG = 0
DB = 0
SM = 0
Delta_K
Doxorubicin
YDL227C_RF1_110 RF1
ZShift = 0.9
lm ZScore = −1.05
NG = 0
DB = 0
SM = 0
Delta_r
Doxorubicin
YDL227C_RF1_110 RF1
ZShift = 0.09
lm ZScore = 0.28
NG = 0
DB = 0
SM = 0
Delta_AUC
Doxorubicin

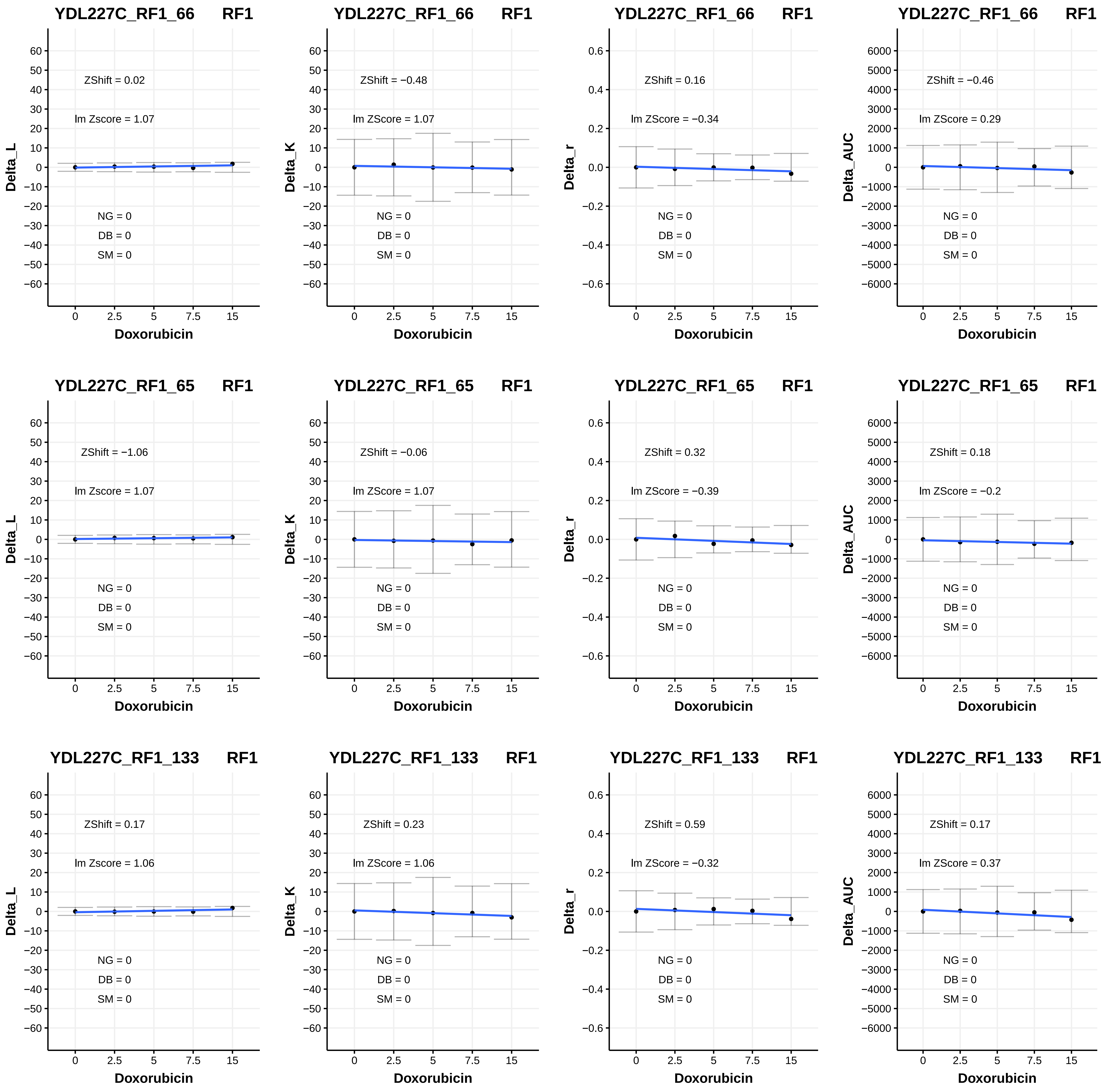

YDL227C_RF1_66 RF1
ZShift = 0.02
lm Zscore = 1.07
NG = 0
DB = 0
SM = 0
Delta_L
Doxorubicin
YDL227C_RF1_66 RF1
ZShift = −0.48
lm ZScore = 1.07
NG = 0
DB = 0
SM = 0
Delta_K
Doxorubicin
YDL227C_RF1_66 RF1
ZShift = 0.16
lm ZScore = −0.34
NG = 0
DB = 0
SM = 0
Delta_r
Doxorubicin
YDL227C_RF1_66 RF1
ZShift = −0.46
lm ZScore = 0.29
NG = 0
DB = 0
SM = 0
Delta_AUC
Doxorubicin
YDL227C_RF1_65 RF1
ZShift = −1.06
lm Zscore = 1.07
NG = 0
DB = 0
SM = 0
Delta_L
Doxorubicin
YDL227C_RF1_65 RF1
ZShift = −0.06
lm ZScore = 1.07
NG = 0
DB = 0
SM = 0
Delta_K
Doxorubicin
YDL227C_RF1_65 RF1
ZShift = 0.32
lm ZScore = −0.39
NG = 0
DB = 0
SM = 0
Delta_r
Doxorubicin
YDL227C_RF1_65 RF1
ZShift = 0.18
lm ZScore = −0.2
NG = 0
DB = 0
SM = 0
Delta_AUC
Doxorubicin
YDL227C_RF1_133 RF1
ZShift = 0.17
lm Zscore = 1.06
NG = 0
DB = 0
SM = 0
Delta_L
Doxorubicin
YDL227C_RF1_133 RF1
ZShift = 0.23
lm ZScore = 1.06
NG = 0
DB = 0
SM = 0
Delta_K
Doxorubicin
YDL227C_RF1_133 RF1
ZShift = 0.59
lm ZScore = −0.32
NG = 0
DB = 0
SM = 0
Delta_r
Doxorubicin
YDL227C_RF1_133 RF1
ZShift = 0.17
lm ZScore = 0.37
NG = 0
DB = 0
SM = 0
Delta_AUC
Doxorubicin

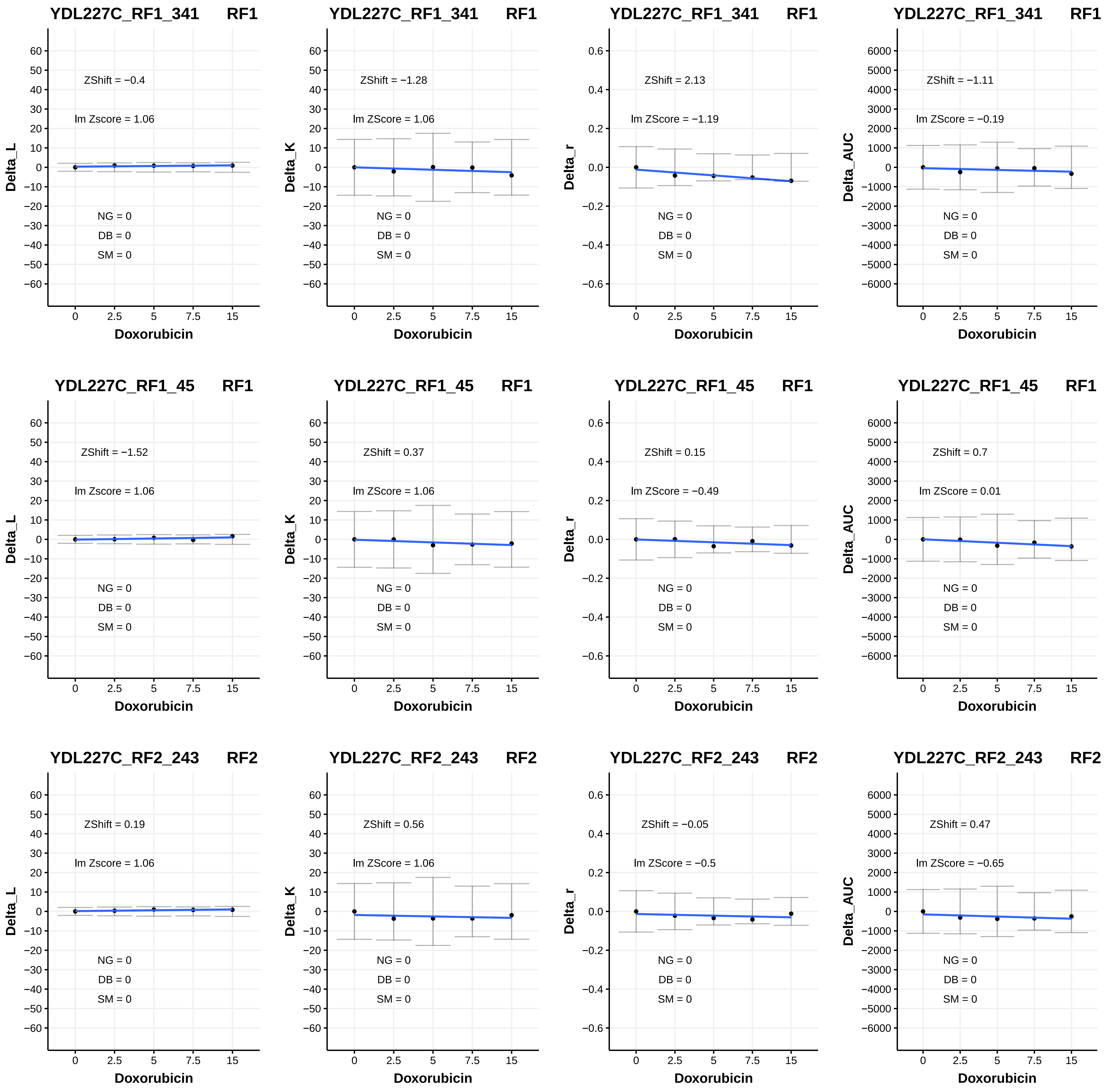
YDL227C_RF1_341 RF1
ZShift = −0.4
lm Zscore = 1.06
NG = 0
DB = 0
SM = 0
Delta_L
Doxorubicin
YDL227C_RF1_341 RF1
ZShift = −1.28
lm ZScore = 1.06
NG = 0
DB = 0
SM = 0
Delta_K
Doxorubicin
YDL227C_RF1_341 RF1
ZShift = 2.13
lm ZScore = −1.19
NG = 0
DB = 0
SM = 0
Delta_r
Doxorubicin
YDL227C_RF1_341 RF1
ZShift = −1.11
lm ZScore = −0.19
NG = 0
DB = 0
SM = 0
Delta_AUC
Doxorubicin
YDL227C_RF1_45 RF1
ZShift = −1.52
lm Zscore = 1.06
NG = 0
DB = 0
SM = 0
Delta_L
Doxorubicin
YDL227C_RF1_45 RF1
ZShift = 0.37
lm ZScore = 1.06
NG = 0
DB = 0
SM = 0
Delta_K
Doxorubicin
YDL227C_RF1_45 RF1
ZShift = 0.15
lm ZScore = −0.49
NG = 0
DB = 0
SM = 0
Delta_r
Doxorubicin
YDL227C_RF1_45 RF1
ZShift = 0.7
lm ZScore = 0.01
NG = 0
DB = 0
SM = 0
Delta_AUC
Doxorubicin
YDL227C_RF2_243 RF2
ZShift = 0.19
lm Zscore = 1.06
NG = 0
DB = 0
SM = 0
Delta_L
Doxorubicin
YDL227C_RF2_243 RF2
ZShift = 0.56
lm ZScore = 1.06
NG = 0
DB = 0
SM = 0
Delta_K
Doxorubicin
YDL227C_RF2_243 RF2
ZShift = −0.05
lm ZScore = −0.5
NG = 0
DB = 0
SM = 0
Delta_r
Doxorubicin
YDL227C_RF2_243 RF2
ZShift = 0.47
lm ZScore = −0.65
NG = 0
DB = 0
SM = 0
Delta_AUC
Doxorubicin

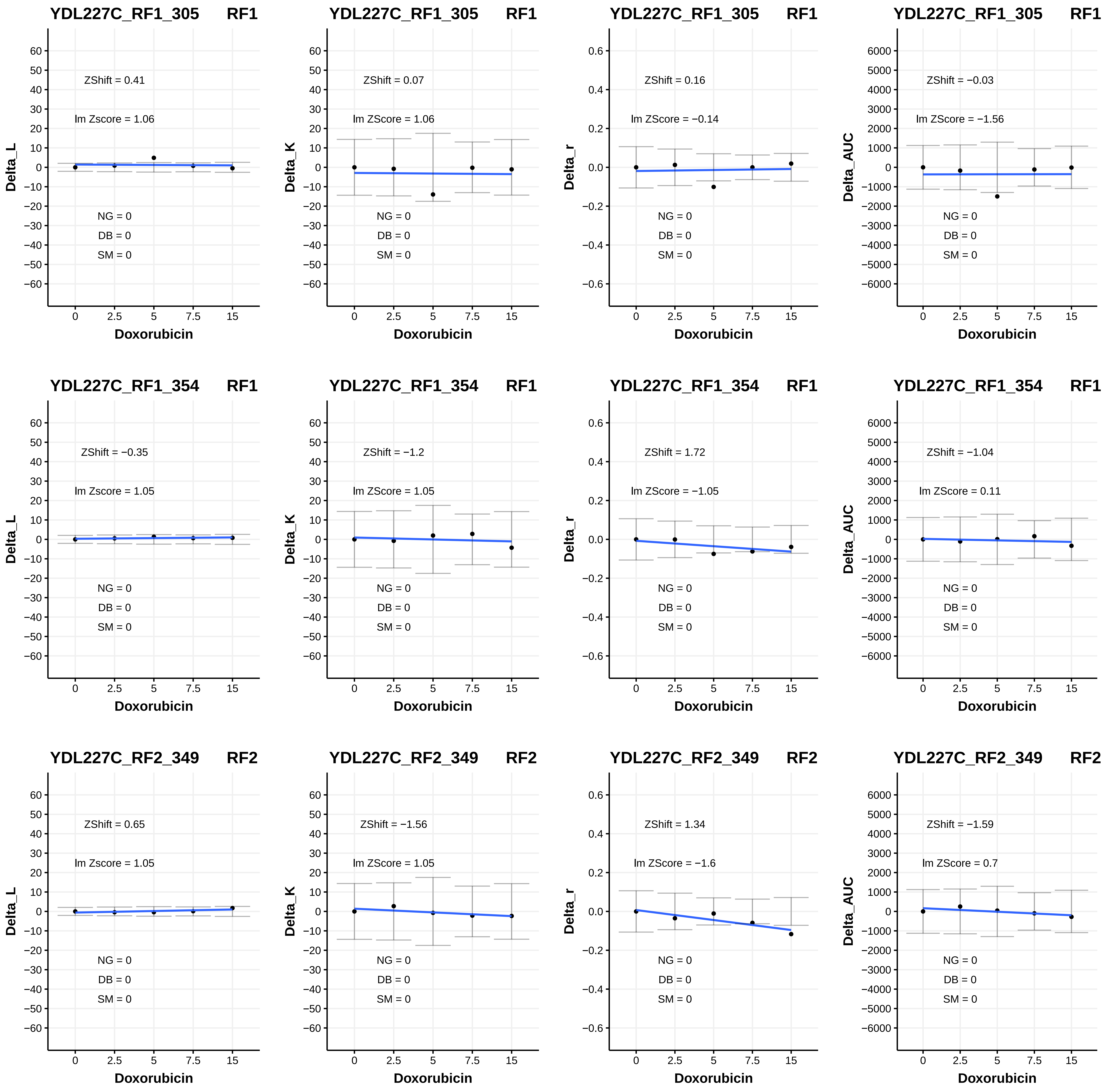

YDL227C_RF1_305 RF1
ZShift = 0.41
lm Zscore = 1.06
NG = 0
DB = 0
SM = 0
Delta_L
Doxorubicin
YDL227C_RF1_305 RF1
ZShift = 0.07
lm ZScore = 1.06
NG = 0
DB = 0
SM = 0
Delta_K
Doxorubicin
YDL227C_RF1_305 RF1
ZShift = 0.16
lm ZScore = −0.14
NG = 0
DB = 0
SM = 0
Delta_r
Doxorubicin
YDL227C_RF1_305 RF1
ZShift = −0.03
lm ZScore = −1.56
NG = 0
DB = 0
SM = 0
Delta_AUC
Doxorubicin
YDL227C_RF1_354 RF1
ZShift = −0.35
lm Zscore = 1.05
NG = 0
DB = 0
SM = 0
Delta_L
Doxorubicin
YDL227C_RF1_354 RF1
ZShift = −1.2
lm ZScore = 1.05
NG = 0
DB = 0
SM = 0
Delta_K
Doxorubicin
YDL227C_RF1_354 RF1
ZShift = 1.72
lm ZScore = −1.05
NG = 0
DB = 0
SM = 0
Delta_r
Doxorubicin
YDL227C_RF1_354 RF1
ZShift = −1.04
lm ZScore = 0.11
NG = 0
DB = 0
SM = 0
Delta_AUC
Doxorubicin
YDL227C_RF2_349 RF2
ZShift = 0.65
lm Zscore = 1.05
NG = 0
DB = 0
SM = 0
Delta_L
Doxorubicin
YDL227C_RF2_349 RF2
ZShift = −1.56
lm ZScore = 1.05
NG = 0
DB = 0
SM = 0
Delta_K
Doxorubicin
YDL227C_RF2_349 RF2
ZShift = 1.34
lm ZScore = −1.6
NG = 0
DB = 0
SM = 0
Delta_r
Doxorubicin
YDL227C_RF2_349 RF2
ZShift = −1.59
lm ZScore = 0.7
NG = 0
DB = 0
SM = 0
Delta_AUC
Doxorubicin

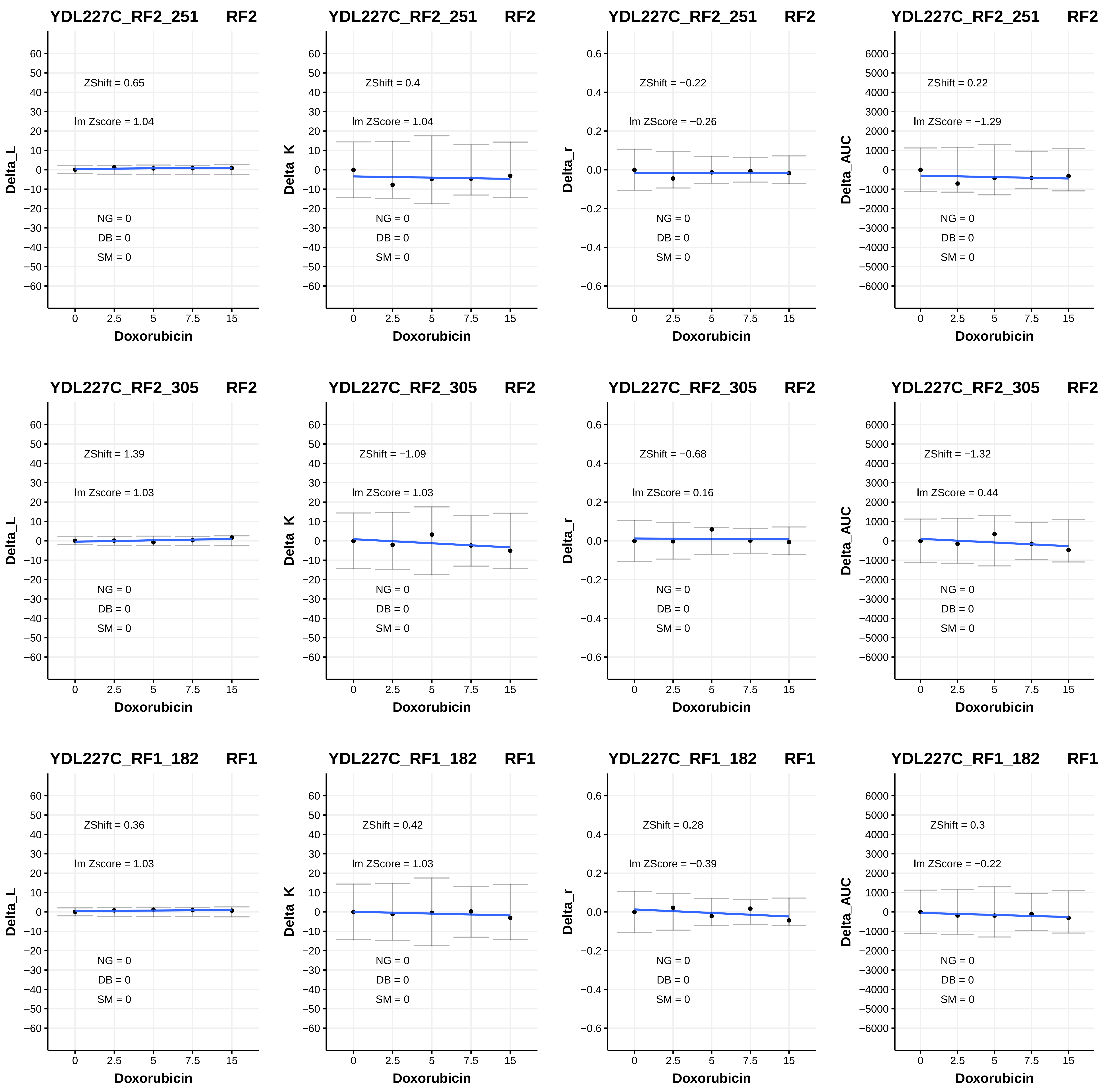
YDL227C_RF2_251 RF2
ZShift = 0.65
lm Zscore = 1.04
NG = 0
DB = 0
SM = 0
Delta_L
Doxorubicin
YDL227C_RF2_251 RF2
ZShift = 0.4
lm ZScore = 1.04
NG = 0
DB = 0
SM = 0
Delta_K
Doxorubicin
YDL227C_RF2_251 RF2
ZShift = −0.22
lm ZScore = −0.26
NG = 0
DB = 0
SM = 0
Delta_r
Doxorubicin
YDL227C_RF2_251 RF2
ZShift = 0.22
lm ZScore = −1.29
NG = 0
DB = 0
SM = 0
Delta_AUC
Doxorubicin
YDL227C_RF2_305 RF2
ZShift = 1.39
lm Zscore = 1.03
NG = 0
DB = 0
SM = 0
Delta_L
Doxorubicin
YDL227C_RF2_305 RF2
ZShift = −1.09
lm ZScore = 1.03
NG = 0
DB = 0
SM = 0
Delta_K
Doxorubicin
YDL227C_RF2_305 RF2
ZShift = −0.68
lm ZScore = 0.16
NG = 0
DB = 0
SM = 0
Delta_r
Doxorubicin
YDL227C_RF2_305 RF2
ZShift = −1.32
lm ZScore = 0.44
NG = 0
DB = 0
SM = 0
Delta_AUC
Doxorubicin
YDL227C_RF1_182 RF1
ZShift = 0.36
lm Zscore = 1.03
NG = 0
DB = 0
SM = 0
Delta_L
Doxorubicin
YDL227C_RF1_182 RF1
ZShift = 0.42
lm ZScore = 1.03
NG = 0
DB = 0
SM = 0
Delta_K
Doxorubicin
YDL227C_RF1_182 RF1
ZShift = 0.28
lm ZScore = −0.39
NG = 0
DB = 0
SM = 0
Delta_r
Doxorubicin
YDL227C_RF1_182 RF1
ZShift = 0.3
lm ZScore = −0.22
NG = 0
DB = 0
SM = 0
Delta_AUC
Doxorubicin

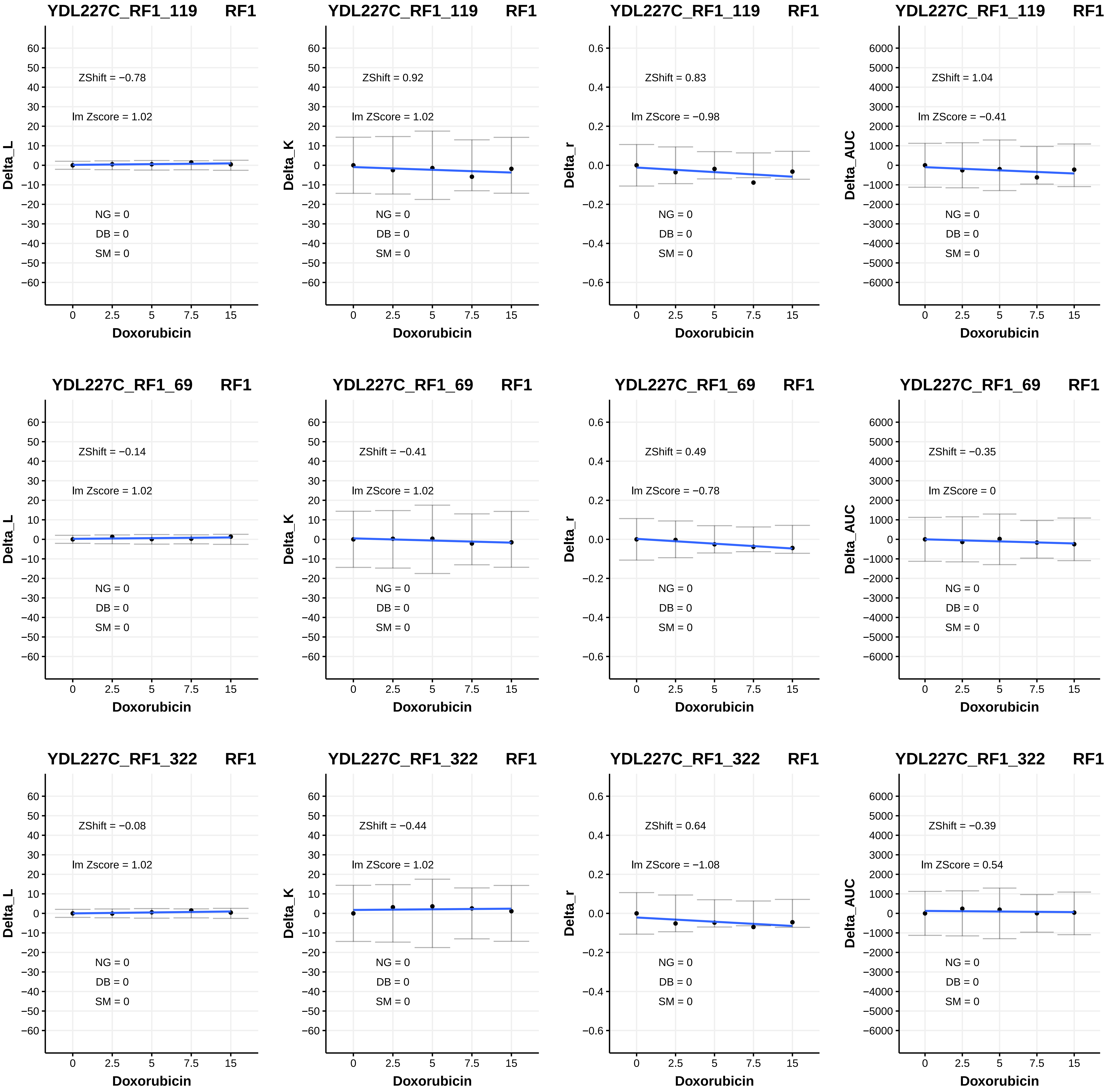
YDL227C_RF1_119 RF1
ZShift = −0.78
lm Zscore = 1.02
NG = 0
DB = 0
SM = 0
Delta_L
Doxorubicin
YDL227C_RF1_119 RF1
ZShift = 0.92
lm ZScore = 1.02
NG = 0
DB = 0
SM = 0
Delta_K
Doxorubicin
YDL227C_RF1_119 RF1
ZShift = 0.83
lm ZScore = −0.98
NG = 0
DB = 0
SM = 0
Delta_r
Doxorubicin
YDL227C_RF1_119 RF1
ZShift = 1.04
lm ZScore = −0.41
NG = 0
DB = 0
SM = 0
Delta_AUC
Doxorubicin
YDL227C_RF1_69 RF1
ZShift = −0.14
lm Zscore = 1.02
NG = 0
DB = 0
SM = 0
Delta_L
Doxorubicin
YDL227C_RF1_69 RF1
ZShift = −0.41
lm ZScore = 1.02
NG = 0
DB = 0
SM = 0
Delta_K
Doxorubicin
YDL227C_RF1_69 RF1
ZShift = 0.49
lm ZScore = −0.78
NG = 0
DB = 0
SM = 0
Delta_r
Doxorubicin
YDL227C_RF1_69 RF1
ZShift = −0.35
lm ZScore = 0
NG = 0
DB = 0
SM = 0
Delta_AUC
Doxorubicin
YDL227C_RF1_322 RF1
ZShift = −0.08
lm Zscore = 1.02
NG = 0
DB = 0
SM = 0
Delta_L
Doxorubicin
YDL227C_RF1_322 RF1
ZShift = −0.44
lm ZScore = 1.02
NG = 0
DB = 0
SM = 0
Delta_K
Doxorubicin
YDL227C_RF1_322 RF1
ZShift = 0.64
lm ZScore = −1.08
NG = 0
DB = 0
SM = 0
Delta_r
Doxorubicin
YDL227C_RF1_322 RF1
ZShift = −0.39
lm ZScore = 0.54
NG = 0
DB = 0
SM = 0
Delta_AUC
Doxorubicin

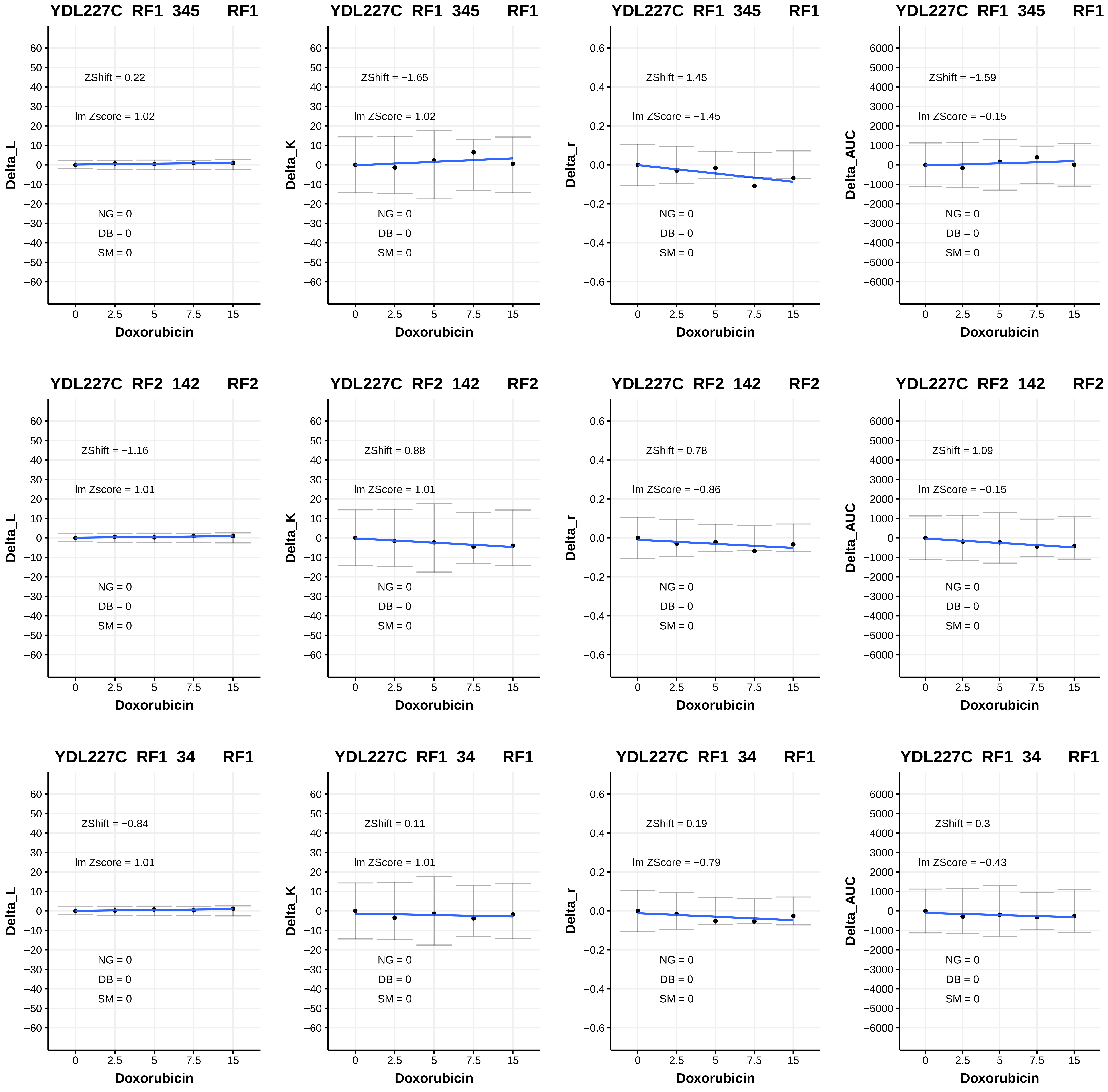
YDL227C_RF1_345 RF1
ZShift = 0.22
lm Zscore = 1.02
NG = 0
DB = 0
SM = 0
Delta_L
Doxorubicin
YDL227C_RF1_345 RF1
ZShift = −1.65
lm ZScore = 1.02
NG = 0
DB = 0
SM = 0
Delta_K
Doxorubicin
YDL227C_RF1_345 RF1
ZShift = 1.45
lm ZScore = −1.45
NG = 0
DB = 0
SM = 0
Delta_r
Doxorubicin
YDL227C_RF1_345 RF1
ZShift = −1.59
lm ZScore = −0.15
NG = 0
DB = 0
SM = 0
Delta_AUC
Doxorubicin
YDL227C_RF2_142 RF2
ZShift = −1.16
lm Zscore = 1.01
NG = 0
DB = 0
SM = 0
Delta_L
Doxorubicin
YDL227C_RF2_142 RF2
ZShift = 0.88
lm ZScore = 1.01
NG = 0
DB = 0
SM = 0
Delta_K
Doxorubicin
YDL227C_RF2_142 RF2
ZShift = 0.78
lm ZScore = −0.86
NG = 0
DB = 0
SM = 0
Delta_r
Doxorubicin
YDL227C_RF2_142 RF2
ZShift = 1.09
lm ZScore = −0.15
NG = 0
DB = 0
SM = 0
Delta_AUC
Doxorubicin
YDL227C_RF1_34 RF1
ZShift = −0.84
lm Zscore = 1.01
NG = 0
DB = 0
SM = 0
Delta_L
Doxorubicin
YDL227C_RF1_34 RF1
ZShift = 0.11
lm ZScore = 1.01
NG = 0
DB = 0
SM = 0
Delta_K
Doxorubicin
YDL227C_RF1_34 RF1
ZShift = 0.19
lm ZScore = −0.79
NG = 0
DB = 0
SM = 0
Delta_r
Doxorubicin
YDL227C_RF1_34 RF1
ZShift = 0.3
lm ZScore = −0.43
NG = 0
DB = 0
SM = 0
Delta_AUC
Doxorubicin

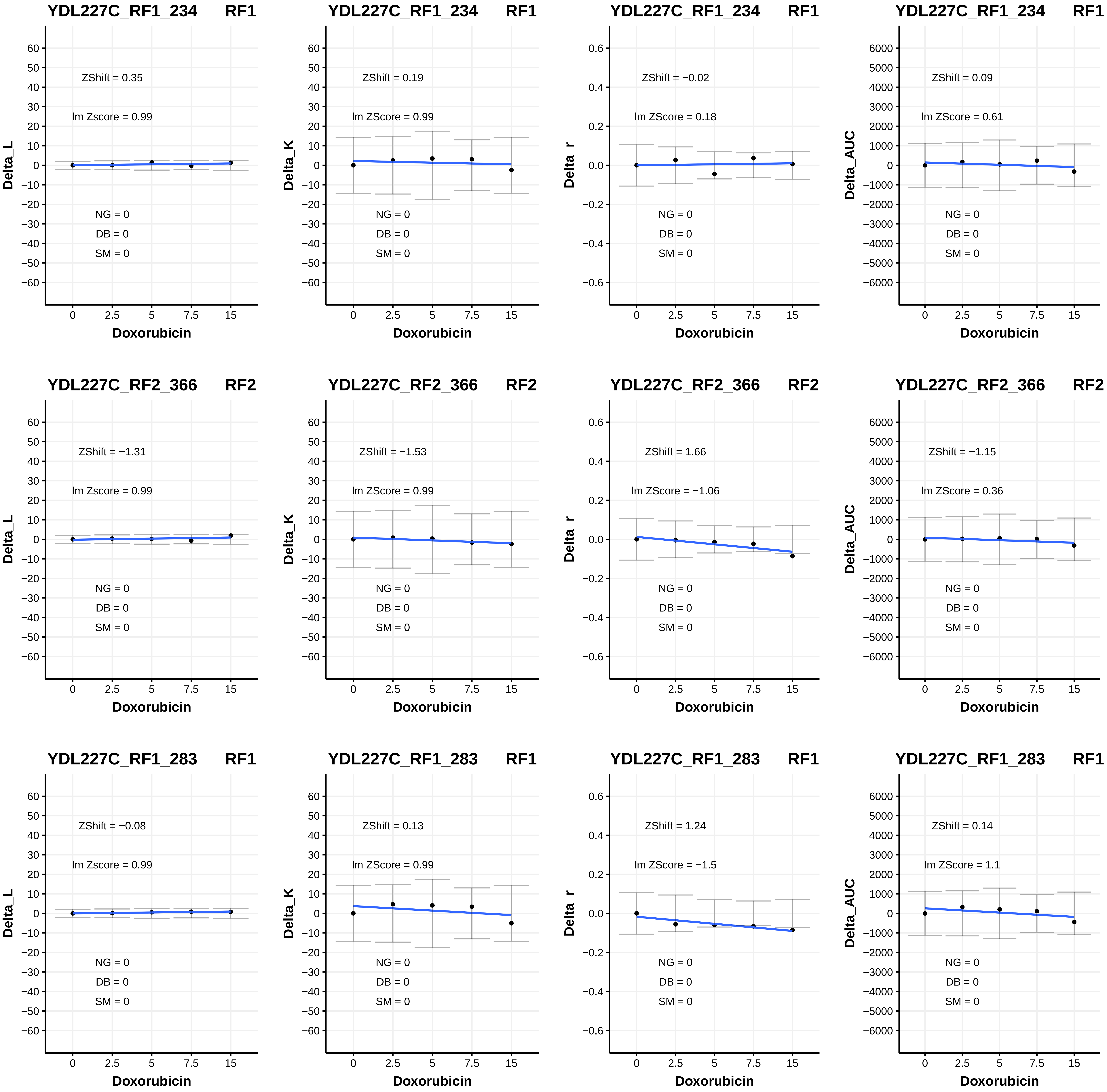

YDL227C_RF1_234 RF1
ZShift = 0.35
lm Zscore = 0.99
NG = 0
DB = 0
SM = 0
Delta_L
Doxorubicin
YDL227C_RF1_234 RF1
ZShift = 0.19
lm ZScore = 0.99
NG = 0
DB = 0
SM = 0
Delta_K
Doxorubicin
YDL227C_RF1_234 RF1
ZShift = −0.02
lm ZScore = 0.18
NG = 0
DB = 0
SM = 0
Delta_r
Doxorubicin
YDL227C_RF1_234 RF1
ZShift = 0.09
lm ZScore = 0.61
NG = 0
DB = 0
SM = 0
Delta_AUC
Doxorubicin
YDL227C_RF2_366 RF2
ZShift = −1.31
lm Zscore = 0.99
NG = 0
DB = 0
SM = 0
Delta_L
Doxorubicin
YDL227C_RF2_366 RF2
ZShift = −1.53
lm ZScore = 0.99
NG = 0
DB = 0
SM = 0
Delta_K
Doxorubicin
YDL227C_RF2_366 RF2
ZShift = 1.66
lm ZScore = −1.06
NG = 0
DB = 0
SM = 0
Delta_r
Doxorubicin
YDL227C_RF2_366 RF2
ZShift = −1.15
lm ZScore = 0.36
NG = 0
DB = 0
SM = 0
Delta_AUC
Doxorubicin
YDL227C_RF1_283 RF1
ZShift = −0.08
lm Zscore = 0.99
NG = 0
DB = 0
SM = 0
Delta_L
Doxorubicin
YDL227C_RF1_283 RF1
ZShift = 0.13
lm ZScore = 0.99
NG = 0
DB = 0
SM = 0
Delta_K
Doxorubicin
YDL227C_RF1_283 RF1
ZShift = 1.24
lm ZScore = −1.5
NG = 0
DB = 0
SM = 0
Delta_r
Doxorubicin
YDL227C_RF1_283 RF1
ZShift = 0.14
lm ZScore = 1.1
NG = 0
DB = 0
SM = 0
Delta_AUC
Doxorubicin

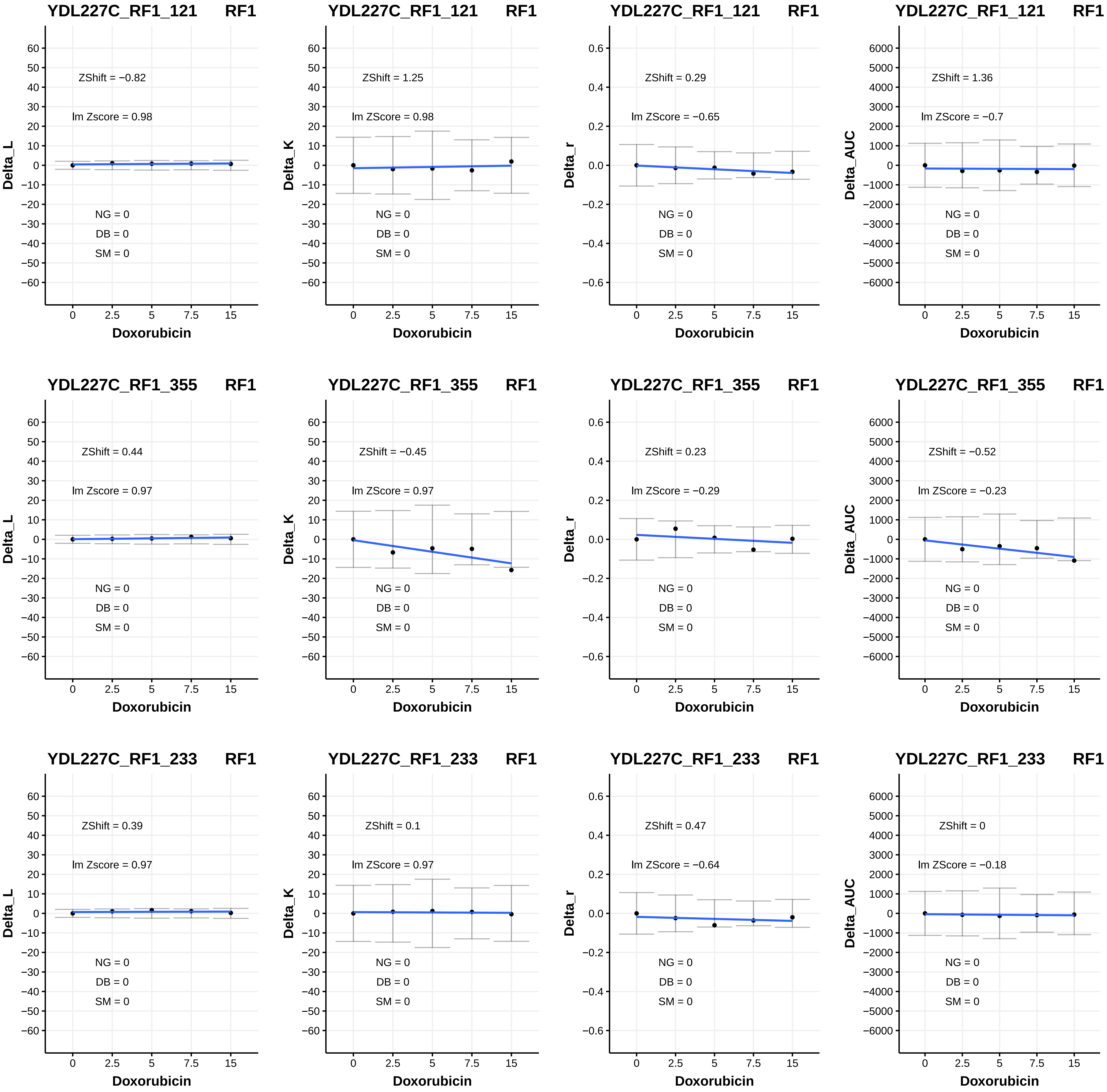

YDL227C_RF1_121 RF1
ZShift = −0.82
lm Zscore = 0.98
NG = 0
DB = 0
SM = 0
Delta_L
Doxorubicin
YDL227C_RF1_121 RF1
ZShift = 1.25
lm ZScore = 0.98
NG = 0
DB = 0
SM = 0
Delta_K
Doxorubicin
YDL227C_RF1_121 RF1
ZShift = 0.29
lm ZScore = −0.65
NG = 0
DB = 0
SM = 0
Delta_r
Doxorubicin
YDL227C_RF1_121 RF1
ZShift = 1.36
lm ZScore = −0.7
NG = 0
DB = 0
SM = 0
Delta_AUC
Doxorubicin
YDL227C_RF1_355 RF1
ZShift = 0.44
lm Zscore = 0.97
NG = 0
DB = 0
SM = 0
Delta_L
Doxorubicin
YDL227C_RF1_355 RF1
ZShift = −0.45
lm ZScore = 0.97
NG = 0
DB = 0
SM = 0
Delta_K
Doxorubicin
YDL227C_RF1_355 RF1
ZShift = 0.23
lm ZScore = −0.29
NG = 0
DB = 0
SM = 0
Delta_r
Doxorubicin
YDL227C_RF1_355 RF1
ZShift = −0.52
lm ZScore = −0.23
NG = 0
DB = 0
SM = 0
Delta_AUC
Doxorubicin
YDL227C_RF1_233 RF1
ZShift = 0.39
lm Zscore = 0.97
NG = 0
DB = 0
SM = 0
Delta_L
Doxorubicin
YDL227C_RF1_233 RF1
ZShift = 0.1
lm ZScore = 0.97
NG = 0
DB = 0
SM = 0
Delta_K
Doxorubicin
YDL227C_RF1_233 RF1
ZShift = 0.47
lm ZScore = −0.64
NG = 0
DB = 0
SM = 0
Delta_r
Doxorubicin
YDL227C_RF1_233 RF1
ZShift = 0
lm ZScore = −0.18
NG = 0
DB = 0
SM = 0
Delta_AUC
Doxorubicin

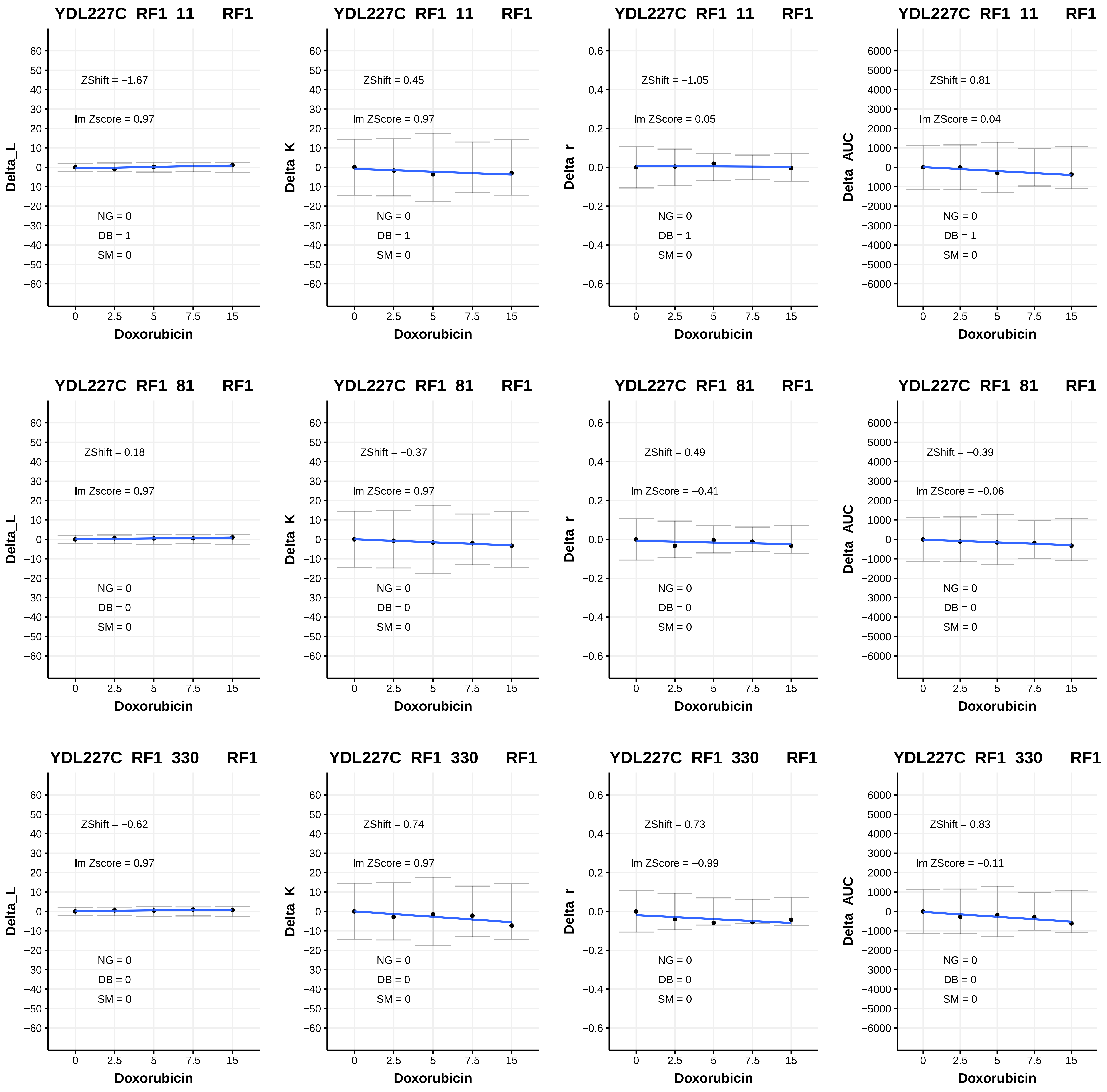

YDL227C_RF1_11 RF1
ZShift = −1.67
lm Zscore = 0.97
NG = 0
DB = 1
SM = 0
Delta_L
Doxorubicin
YDL227C_RF1_11 RF1
ZShift = 0.45
lm ZScore = 0.97
NG = 0
DB = 1
SM = 0
Delta_K
Doxorubicin
YDL227C_RF1_11 RF1
ZShift = −1.05
lm ZScore = 0.05
NG = 0
DB = 1
SM = 0
Delta_r
Doxorubicin
YDL227C_RF1_11 RF1
ZShift = 0.81
lm ZScore = 0.04
NG = 0
DB = 1
SM = 0
Delta_AUC
Doxorubicin
YDL227C_RF1_81 RF1
ZShift = 0.18
lm Zscore = 0.97
NG = 0
DB = 0
SM = 0
Delta_L
Doxorubicin
YDL227C_RF1_81 RF1
ZShift = −0.37
lm ZScore = 0.97
NG = 0
DB = 0
SM = 0
Delta_K
Doxorubicin
YDL227C_RF1_81 RF1
ZShift = 0.49
lm ZScore = −0.41
NG = 0
DB = 0
SM = 0
Delta_r
Doxorubicin
YDL227C_RF1_81 RF1
ZShift = −0.39
lm ZScore = −0.06
NG = 0
DB = 0
SM = 0
Delta_AUC
Doxorubicin
YDL227C_RF1_330 RF1
ZShift = −0.62
lm Zscore = 0.97
NG = 0
DB = 0
SM = 0
Delta_L
Doxorubicin
YDL227C_RF1_330 RF1
ZShift = 0.74
lm ZScore = 0.97
NG = 0
DB = 0
SM = 0
Delta_K
Doxorubicin
YDL227C_RF1_330 RF1
ZShift = 0.73
lm ZScore = −0.99
NG = 0
DB = 0
SM = 0
Delta_r
Doxorubicin
YDL227C_RF1_330 RF1
ZShift = 0.83
lm ZScore = −0.11
NG = 0
DB = 0
SM = 0
Delta_AUC
Doxorubicin

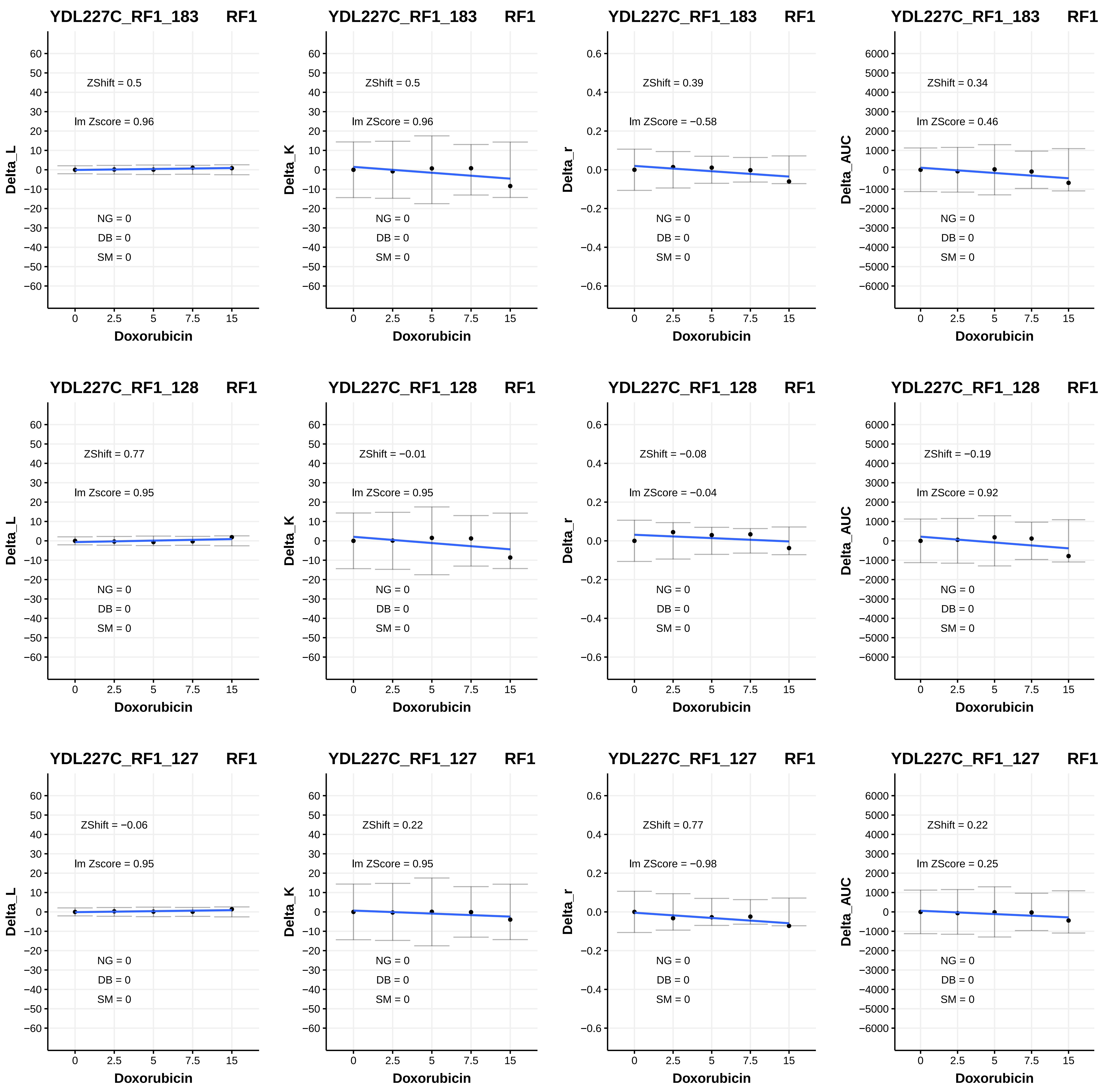
YDL227C_RF1_183 RF1
ZShift = 0.5
lm Zscore = 0.96
NG = 0
DB = 0
SM = 0
Delta_L
Doxorubicin
YDL227C_RF1_183 RF1
ZShift = 0.5
lm ZScore = 0.96
NG = 0
DB = 0
SM = 0
Delta_K
Doxorubicin
YDL227C_RF1_183 RF1
ZShift = 0.39
lm ZScore = −0.58
NG = 0
DB = 0
SM = 0
Delta_r
Doxorubicin
YDL227C_RF1_183 RF1
ZShift = 0.34
lm ZScore = 0.46
NG = 0
DB = 0
SM = 0
Delta_AUC
Doxorubicin
YDL227C_RF1_128 RF1
ZShift = 0.77
lm Zscore = 0.95
NG = 0
DB = 0
SM = 0
Delta_L
Doxorubicin
YDL227C_RF1_128 RF1
ZShift = −0.01
lm ZScore = 0.95
NG = 0
DB = 0
SM = 0
Delta_K
Doxorubicin
YDL227C_RF1_128 RF1
ZShift = −0.08
lm ZScore = −0.04
NG = 0
DB = 0
SM = 0
Delta_r
Doxorubicin
YDL227C_RF1_128 RF1
ZShift = −0.19
lm ZScore = 0.92
NG = 0
DB = 0
SM = 0
Delta_AUC
Doxorubicin
YDL227C_RF1_127 RF1
ZShift = −0.06
lm Zscore = 0.95
NG = 0
DB = 0
SM = 0
Delta_L
Doxorubicin
YDL227C_RF1_127 RF1
ZShift = 0.22
lm ZScore = 0.95
NG = 0
DB = 0
SM = 0
Delta_K
Doxorubicin
YDL227C_RF1_127 RF1
ZShift = 0.77
lm ZScore = −0.98
NG = 0
DB = 0
SM = 0
Delta_r
Doxorubicin
YDL227C_RF1_127 RF1
ZShift = 0.22
lm ZScore = 0.25
NG = 0
DB = 0
SM = 0
Delta_AUC
Doxorubicin

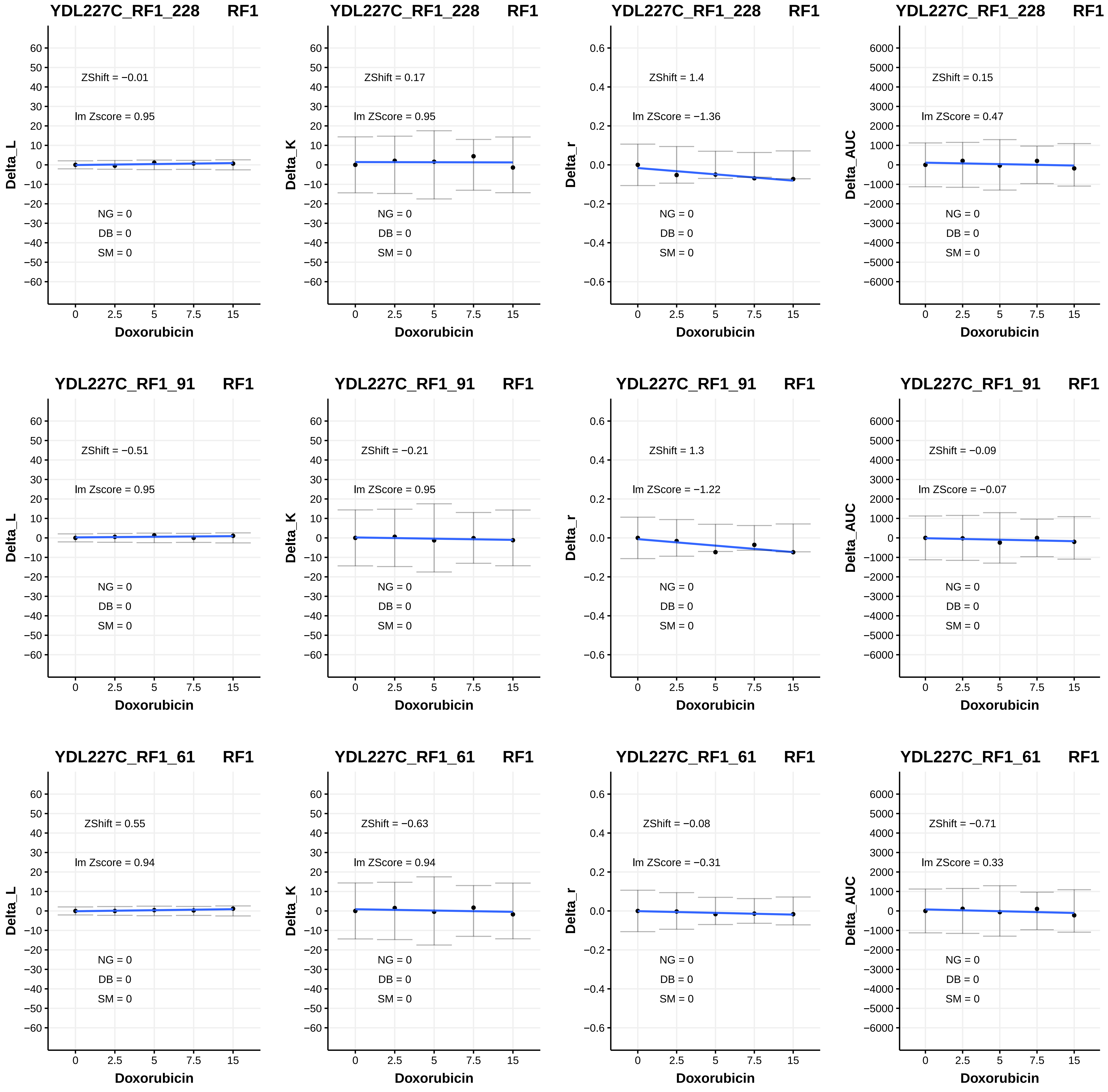
YDL227C_RF1_228 RF1
ZShift = −0.01
lm Zscore = 0.95
NG = 0
DB = 0
SM = 0
Delta_L
Doxorubicin
YDL227C_RF1_228 RF1
ZShift = 0.17
lm ZScore = 0.95
NG = 0
DB = 0
SM = 0
Delta_K
Doxorubicin
YDL227C_RF1_228 RF1
ZShift = 1.4
lm ZScore = −1.36
NG = 0
DB = 0
SM = 0
Delta_r
Doxorubicin
YDL227C_RF1_228 RF1
ZShift = 0.15
lm ZScore = 0.47
NG = 0
DB = 0
SM = 0
Delta_AUC
Doxorubicin
YDL227C_RF1_91 RF1
ZShift = −0.51
lm Zscore = 0.95
NG = 0
DB = 0
SM = 0
Delta_L
Doxorubicin
YDL227C_RF1_91 RF1
ZShift = −0.21
lm ZScore = 0.95
NG = 0
DB = 0
SM = 0
Delta_K
Doxorubicin
YDL227C_RF1_91 RF1
ZShift = 1.3
lm ZScore = −1.22
NG = 0
DB = 0
SM = 0
Delta_r
Doxorubicin
YDL227C_RF1_91 RF1
ZShift = −0.09
lm ZScore = −0.07
NG = 0
DB = 0
SM = 0
Delta_AUC
Doxorubicin
YDL227C_RF1_61 RF1
ZShift = 0.55
lm Zscore = 0.94
NG = 0
DB = 0
SM = 0
Delta_L
Doxorubicin
YDL227C_RF1_61 RF1
ZShift = −0.63
lm ZScore = 0.94
NG = 0
DB = 0
SM = 0
Delta_K
Doxorubicin
YDL227C_RF1_61 RF1
ZShift = −0.08
lm ZScore = −0.31
NG = 0
DB = 0
SM = 0
Delta_r
Doxorubicin
YDL227C_RF1_61 RF1
ZShift = −0.71
lm ZScore = 0.33
NG = 0
DB = 0
SM = 0
Delta_AUC
Doxorubicin

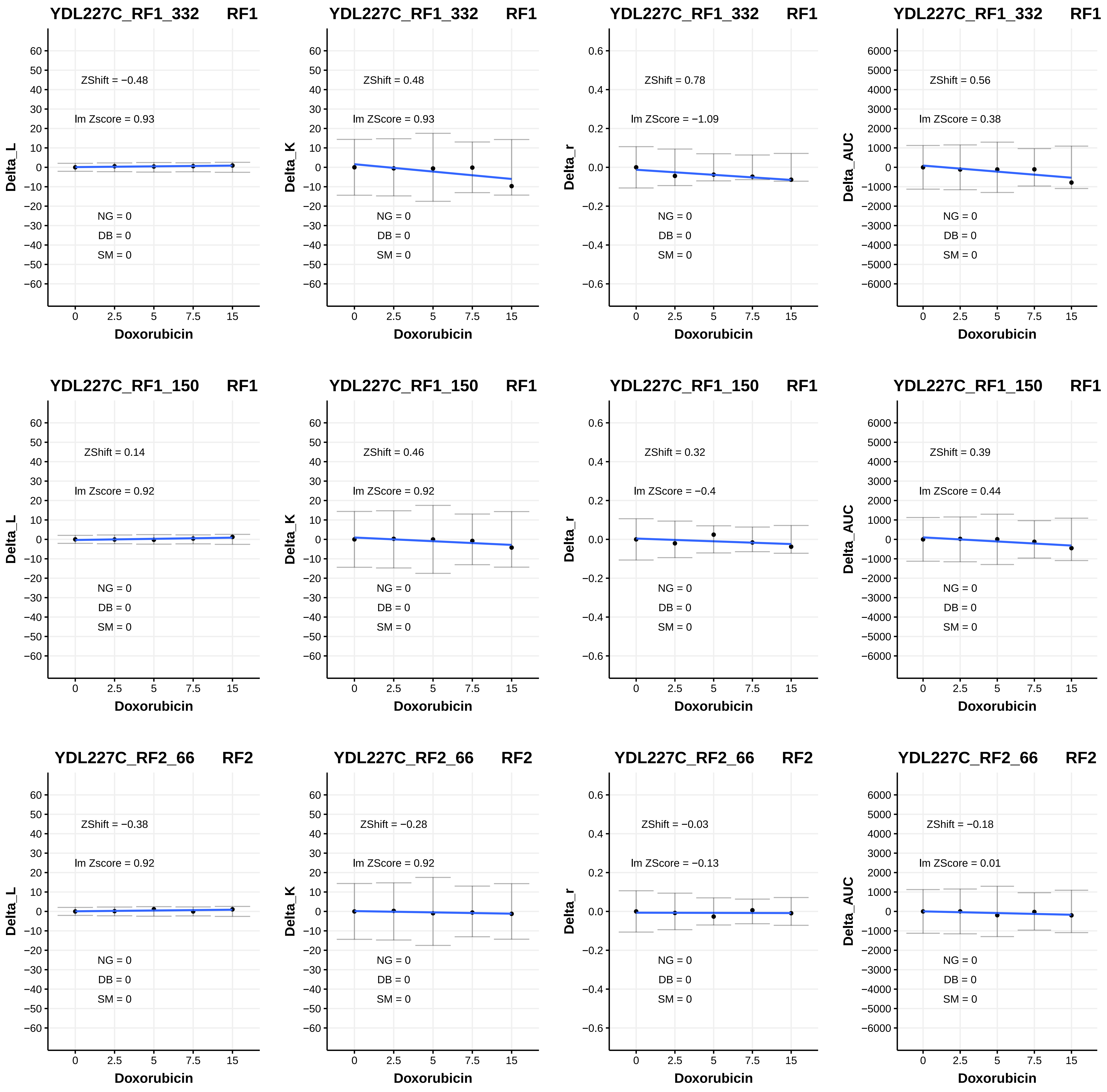

YDL227C_RF1_332 RF1
ZShift = −0.48
lm Zscore = 0.93
NG = 0
DB = 0
SM = 0
Delta_L
Doxorubicin
YDL227C_RF1_332 RF1
ZShift = 0.48
lm ZScore = 0.93
NG = 0
DB = 0
SM = 0
Delta_K
Doxorubicin
YDL227C_RF1_332 RF1
ZShift = 0.78
lm ZScore = −1.09
NG = 0
DB = 0
SM = 0
Delta_r
Doxorubicin
YDL227C_RF1_332 RF1
ZShift = 0.56
lm ZScore = 0.38
NG = 0
DB = 0
SM = 0
Delta_AUC
Doxorubicin
YDL227C_RF1_150 RF1
ZShift = 0.14
lm Zscore = 0.92
NG = 0
DB = 0
SM = 0
Delta_L
Doxorubicin
YDL227C_RF1_150 RF1
ZShift = 0.46
lm ZScore = 0.92
NG = 0
DB = 0
SM = 0
Delta_K
Doxorubicin
YDL227C_RF1_150 RF1
ZShift = 0.32
lm ZScore = −0.4
NG = 0
DB = 0
SM = 0
Delta_r
Doxorubicin
YDL227C_RF1_150 RF1
ZShift = 0.39
lm ZScore = 0.44
NG = 0
DB = 0
SM = 0
Delta_AUC
Doxorubicin
YDL227C_RF2_66 RF2
ZShift = −0.38
lm Zscore = 0.92
NG = 0
DB = 0
SM = 0
Delta_L
Doxorubicin
YDL227C_RF2_66 RF2
ZShift = −0.28
lm ZScore = 0.92
NG = 0
DB = 0
SM = 0
Delta_K
Doxorubicin
YDL227C_RF2_66 RF2
ZShift = −0.03
lm ZScore = −0.13
NG = 0
DB = 0
SM = 0
Delta_r
Doxorubicin
YDL227C_RF2_66 RF2
ZShift = −0.18
lm ZScore = 0.01
NG = 0
DB = 0
SM = 0
Delta_AUC
Doxorubicin

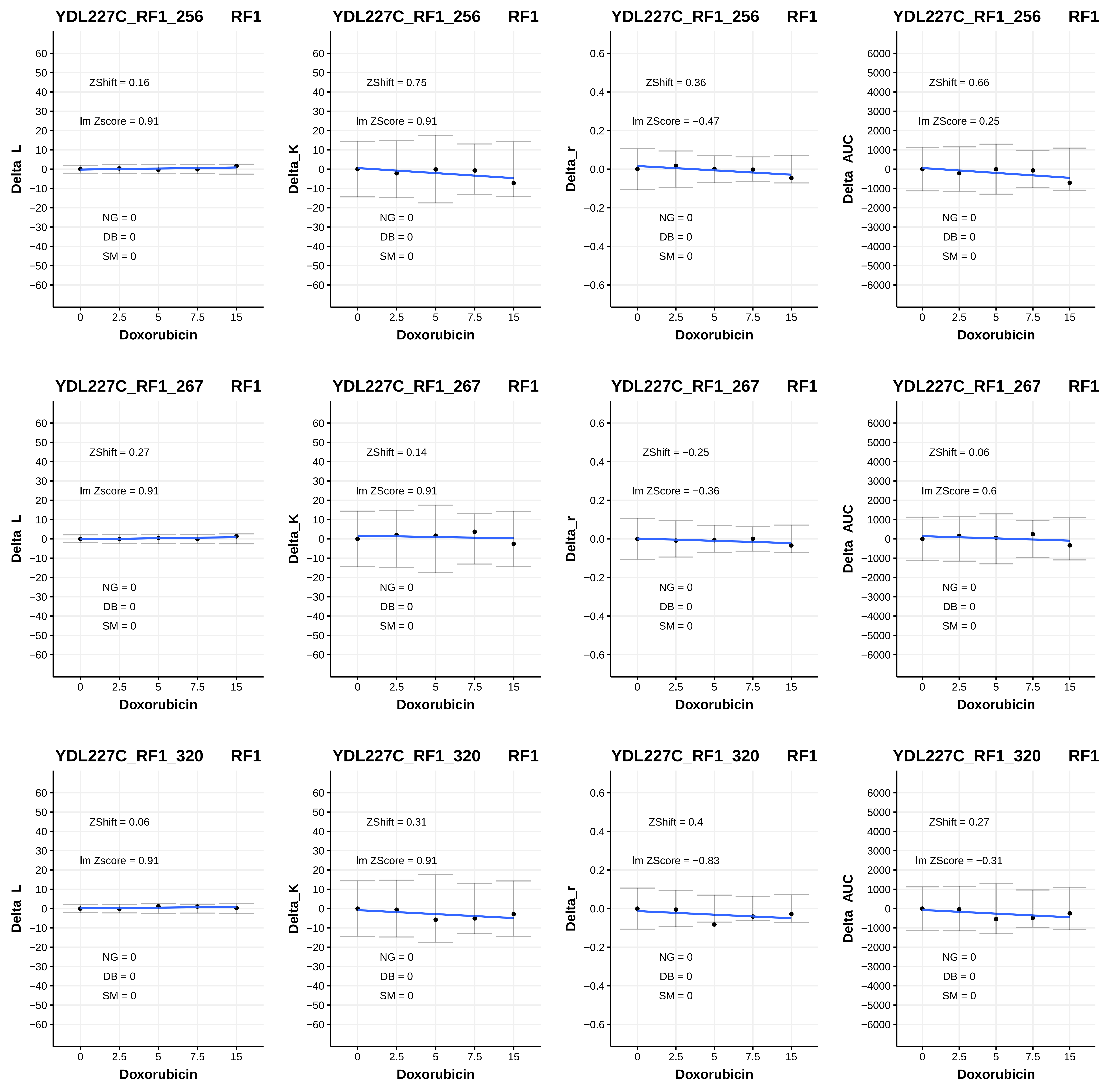
YDL227C_RF1_256 RF1
ZShift = 0.16
lm Zscore = 0.91
NG = 0
DB = 0
SM = 0
Delta_L
Doxorubicin
YDL227C_RF1_256 RF1
ZShift = 0.75
lm ZScore = 0.91
NG = 0
DB = 0
SM = 0
Delta_K
Doxorubicin
YDL227C_RF1_256 RF1
ZShift = 0.36
lm ZScore = −0.47
NG = 0
DB = 0
SM = 0
Delta_r
Doxorubicin
YDL227C_RF1_256 RF1
ZShift = 0.66
lm ZScore = 0.25
NG = 0
DB = 0
SM = 0
Delta_AUC
Doxorubicin
YDL227C_RF1_267 RF1
ZShift = 0.27
lm Zscore = 0.91
NG = 0
DB = 0
SM = 0
Delta_L
Doxorubicin
YDL227C_RF1_267 RF1
ZShift = 0.14
lm ZScore = 0.91
NG = 0
DB = 0
SM = 0
Delta_K
Doxorubicin
YDL227C_RF1_267 RF1
ZShift = −0.25
lm ZScore = −0.36
NG = 0
DB = 0
SM = 0
Delta_r
Doxorubicin
YDL227C_RF1_267 RF1
ZShift = 0.06
lm ZScore = 0.6
NG = 0
DB = 0
SM = 0
Delta_AUC
Doxorubicin
YDL227C_RF1_320 RF1
ZShift = 0.06
lm Zscore = 0.91
NG = 0
DB = 0
SM = 0
Delta_L
Doxorubicin
YDL227C_RF1_320 RF1
ZShift = 0.31
lm ZScore = 0.91
NG = 0
DB = 0
SM = 0
Delta_K
Doxorubicin
YDL227C_RF1_320 RF1
ZShift = 0.4
lm ZScore = −0.83
NG = 0
DB = 0
SM = 0
Delta_r
Doxorubicin
YDL227C_RF1_320 RF1
ZShift = 0.27
lm ZScore = −0.31
NG = 0
DB = 0
SM = 0
Delta_AUC
Doxorubicin

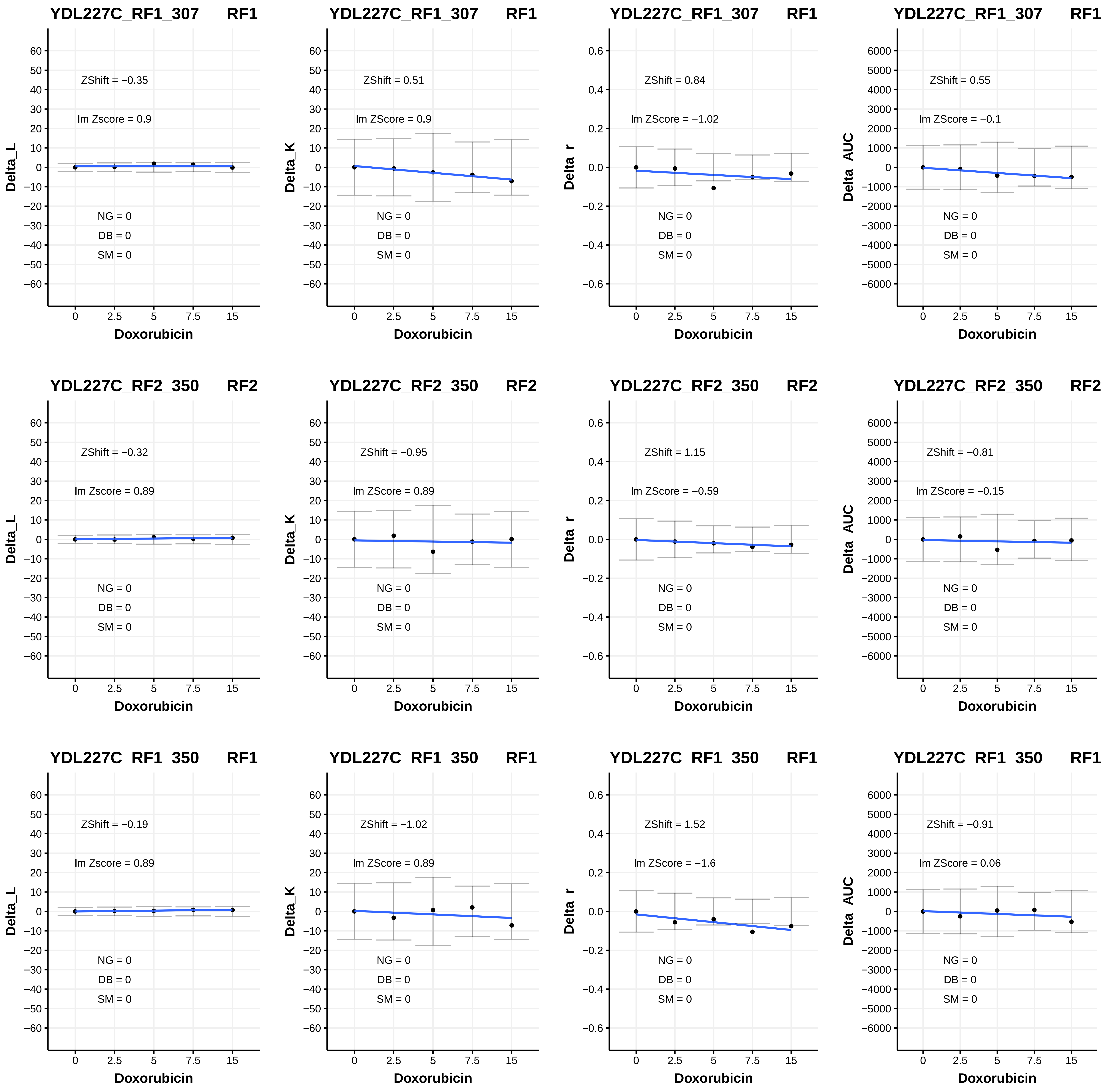

YDL227C_RF1_307 RF1
ZShift = −0.35
lm Zscore = 0.9
NG = 0
DB = 0
SM = 0
Delta_L
Doxorubicin
YDL227C_RF1_307 RF1
ZShift = 0.51
lm ZScore = 0.9
NG = 0
DB = 0
SM = 0
Delta_K
Doxorubicin
YDL227C_RF1_307 RF1
ZShift = 0.84
lm ZScore = −1.02
NG = 0
DB = 0
SM = 0
Delta_r
Doxorubicin
YDL227C_RF1_307 RF1
ZShift = 0.55
lm ZScore = −0.1
NG = 0
DB = 0
SM = 0
Delta_AUC
Doxorubicin
YDL227C_RF2_350 RF2
ZShift = −0.32
lm Zscore = 0.89
NG = 0
DB = 0
SM = 0
Delta_L
Doxorubicin
YDL227C_RF2_350 RF2
ZShift = −0.95
lm ZScore = 0.89
NG = 0
DB = 0
SM = 0
Delta_K
Doxorubicin
YDL227C_RF2_350 RF2
ZShift = 1.15
lm ZScore = −0.59
NG = 0
DB = 0
SM = 0
Delta_r
Doxorubicin
YDL227C_RF2_350 RF2
ZShift = −0.81
lm ZScore = −0.15
NG = 0
DB = 0
SM = 0
Delta_AUC
Doxorubicin
YDL227C_RF1_350 RF1
ZShift = −0.19
lm Zscore = 0.89
NG = 0
DB = 0
SM = 0
Delta_L
Doxorubicin
YDL227C_RF1_350 RF1
ZShift = −1.02
lm ZScore = 0.89
NG = 0
DB = 0
SM = 0
Delta_K
Doxorubicin
YDL227C_RF1_350 RF1
ZShift = 1.52
lm ZScore = −1.6
NG = 0
DB = 0
SM = 0
Delta_r
Doxorubicin
YDL227C_RF1_350 RF1
ZShift = −0.91
lm ZScore = 0.06
NG = 0
DB = 0
SM = 0
Delta_AUC
Doxorubicin

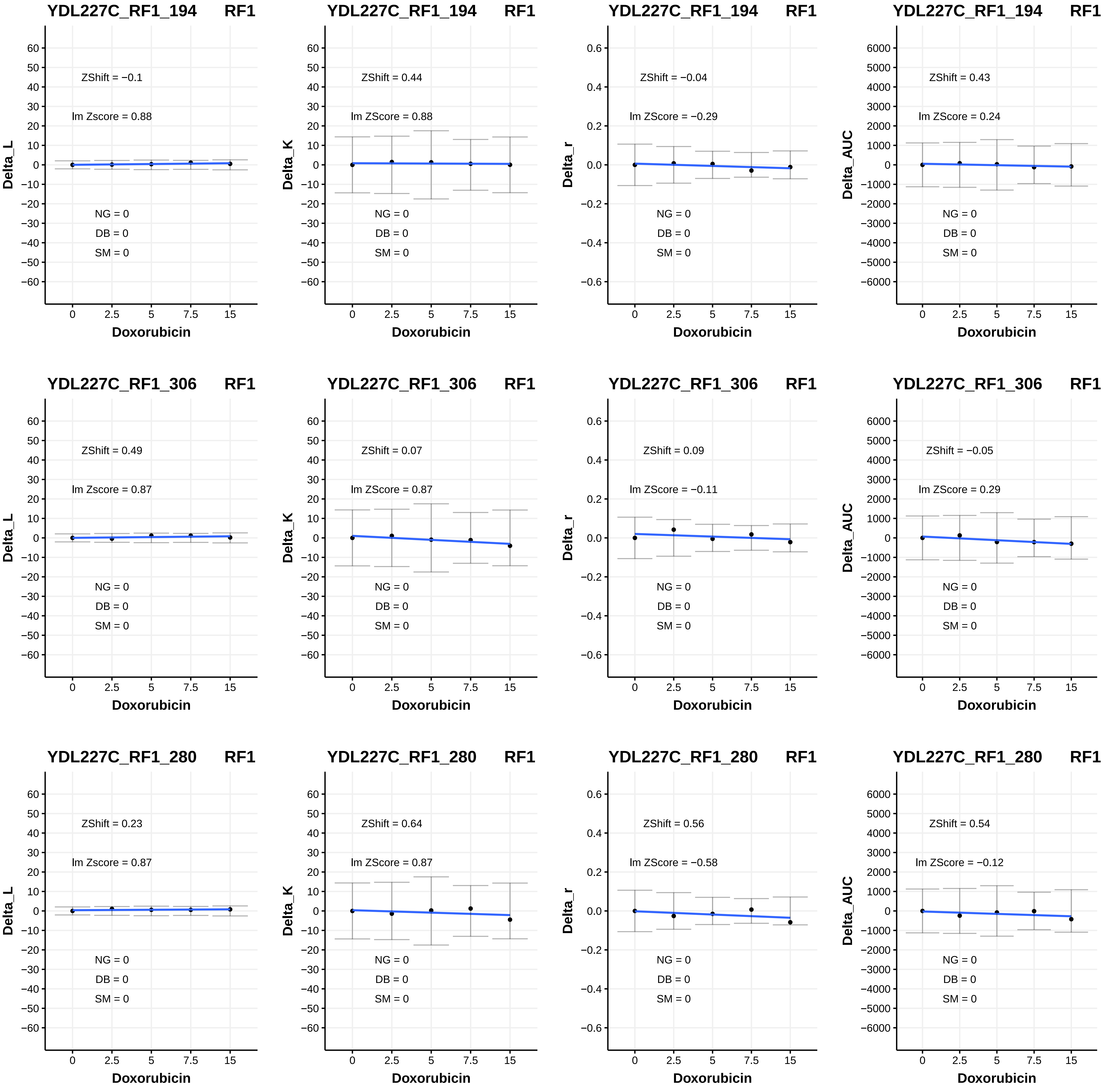

YDL227C_RF1_194 RF1
ZShift = −0.1
lm Zscore = 0.88
NG = 0
DB = 0
SM = 0
Delta_L
Doxorubicin
YDL227C_RF1_194 RF1
ZShift = 0.44
lm ZScore = 0.88
NG = 0
DB = 0
SM = 0
Delta_K
Doxorubicin
YDL227C_RF1_194 RF1
ZShift = −0.04
lm ZScore = −0.29
NG = 0
DB = 0
SM = 0
Delta_r
Doxorubicin
YDL227C_RF1_194 RF1
ZShift = 0.43
lm ZScore = 0.24
NG = 0
DB = 0
SM = 0
Delta_AUC
Doxorubicin
YDL227C_RF1_306 RF1
ZShift = 0.49
lm Zscore = 0.87
NG = 0
DB = 0
SM = 0
Delta_L
Doxorubicin
YDL227C_RF1_306 RF1
ZShift = 0.07
lm ZScore = 0.87
NG = 0
DB = 0
SM = 0
Delta_K
Doxorubicin
YDL227C_RF1_306 RF1
ZShift = 0.09
lm ZScore = −0.11
NG = 0
DB = 0
SM = 0
Delta_r
Doxorubicin
YDL227C_RF1_306 RF1
ZShift = −0.05
lm ZScore = 0.29
NG = 0
DB = 0
SM = 0
Delta_AUC
Doxorubicin
YDL227C_RF1_280 RF1
ZShift = 0.23
lm Zscore = 0.87
NG = 0
DB = 0
SM = 0
Delta_L
Doxorubicin
YDL227C_RF1_280 RF1
ZShift = 0.64
lm ZScore = 0.87
NG = 0
DB = 0
SM = 0
Delta_K
Doxorubicin
YDL227C_RF1_280 RF1
ZShift = 0.56
lm ZScore = −0.58
NG = 0
DB = 0
SM = 0
Delta_r
Doxorubicin
YDL227C_RF1_280 RF1
ZShift = 0.54
lm ZScore = −0.12
NG = 0
DB = 0
SM = 0
Delta_AUC
Doxorubicin

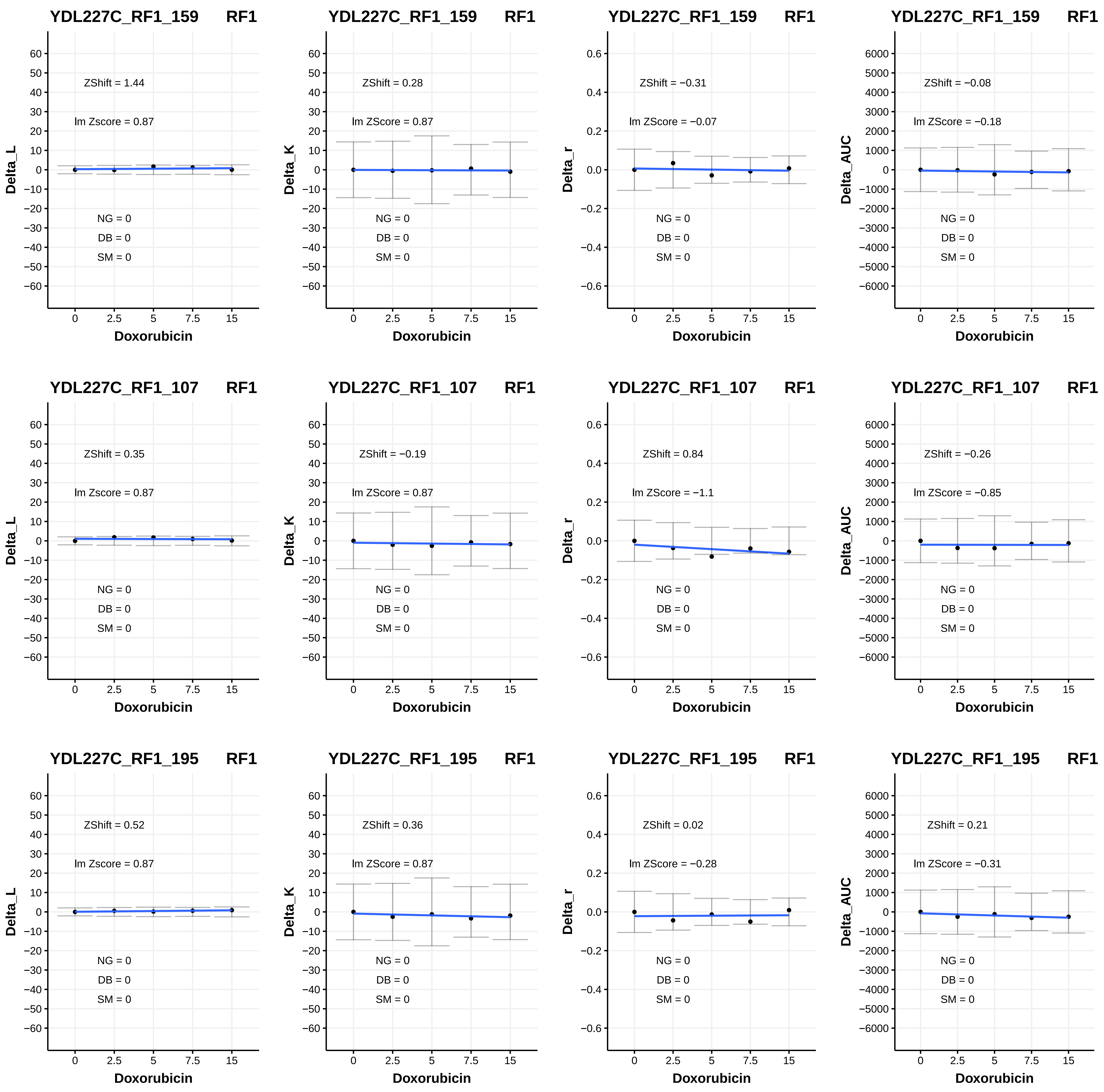

YDL227C_RF1_159 RF1
ZShift = 1.44
lm Zscore = 0.87
NG = 0
DB = 0
SM = 0
Delta_L
Doxorubicin
YDL227C_RF1_159 RF1
ZShift = 0.28
lm ZScore = 0.87
NG = 0
DB = 0
SM = 0
Delta_K
Doxorubicin
YDL227C_RF1_159 RF1
ZShift = −0.31
lm ZScore = −0.07
NG = 0
DB = 0
SM = 0
Delta_r
Doxorubicin
YDL227C_RF1_159 RF1
ZShift = −0.08
lm ZScore = −0.18
NG = 0
DB = 0
SM = 0
Delta_AUC
Doxorubicin
YDL227C_RF1_107 RF1
ZShift = 0.35
lm Zscore = 0.87
NG = 0
DB = 0
SM = 0
Delta_L
Doxorubicin
YDL227C_RF1_107 RF1
ZShift = −0.19
lm ZScore = 0.87
NG = 0
DB = 0
SM = 0
Delta_K
Doxorubicin
YDL227C_RF1_107 RF1
ZShift = 0.84
lm ZScore = −1.1
NG = 0
DB = 0
SM = 0
Delta_r
Doxorubicin
YDL227C_RF1_107 RF1
ZShift = −0.26
lm ZScore = −0.85
NG = 0
DB = 0
SM = 0
Delta_AUC
Doxorubicin
YDL227C_RF1_195 RF1
ZShift = 0.52
lm Zscore = 0.87
NG = 0
DB = 0
SM = 0
Delta_L
Doxorubicin
YDL227C_RF1_195 RF1
ZShift = 0.36
lm ZScore = 0.87
NG = 0
DB = 0
SM = 0
Delta_K
Doxorubicin
YDL227C_RF1_195 RF1
ZShift = 0.02
lm ZScore = −0.28
NG = 0
DB = 0
SM = 0
Delta_r
Doxorubicin
YDL227C_RF1_195 RF1
ZShift = 0.21
lm ZScore = −0.31
NG = 0
DB = 0
SM = 0
Delta_AUC
Doxorubicin

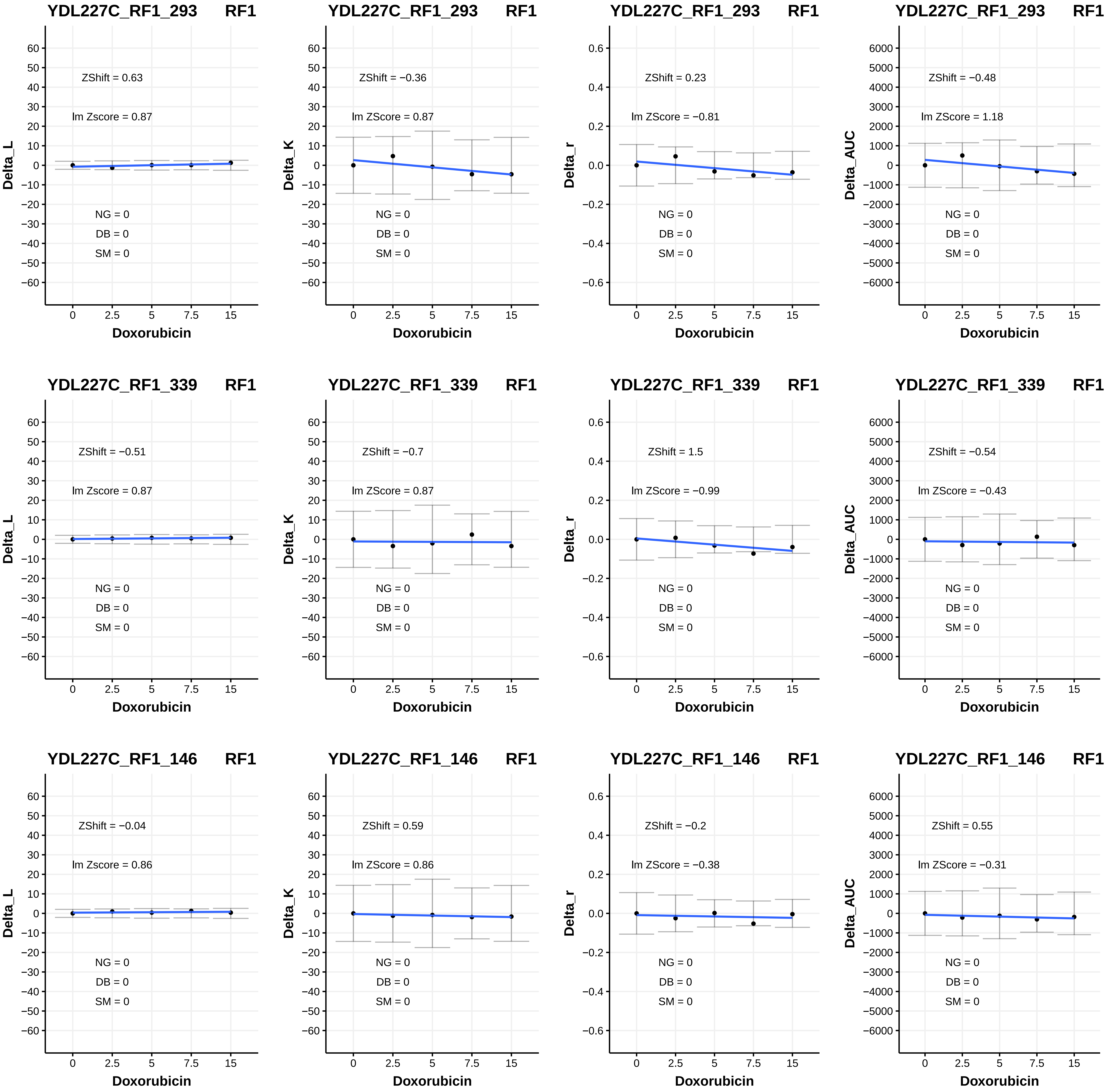
YDL227C_RF1_293 RF1
ZShift = 0.63
lm Zscore = 0.87
NG = 0
DB = 0
SM = 0
Delta_L
Doxorubicin
YDL227C_RF1_293 RF1
ZShift = −0.36
lm ZScore = 0.87
NG = 0
DB = 0
SM = 0
Delta_K
Doxorubicin
YDL227C_RF1_293 RF1
ZShift = 0.23
lm ZScore = −0.81
NG = 0
DB = 0
SM = 0
Delta_r
Doxorubicin
YDL227C_RF1_293 RF1
ZShift = −0.48
lm ZScore = 1.18
NG = 0
DB = 0
SM = 0
Delta_AUC
Doxorubicin
YDL227C_RF1_339 RF1
ZShift = −0.51
lm Zscore = 0.87
NG = 0
DB = 0
SM = 0
Delta_L
Doxorubicin
YDL227C_RF1_339 RF1
ZShift = −0.7
lm ZScore = 0.87
NG = 0
DB = 0
SM = 0
Delta_K
Doxorubicin
YDL227C_RF1_339 RF1
ZShift = 1.5
lm ZScore = −0.99
NG = 0
DB = 0
SM = 0
Delta_r
Doxorubicin
YDL227C_RF1_339 RF1
ZShift = −0.54
lm ZScore = −0.43
NG = 0
DB = 0
SM = 0
Delta_AUC
Doxorubicin
YDL227C_RF1_146 RF1
ZShift = −0.04
lm Zscore = 0.86
NG = 0
DB = 0
SM = 0
Delta_L
Doxorubicin
YDL227C_RF1_146 RF1
ZShift = 0.59
lm ZScore = 0.86
NG = 0
DB = 0
SM = 0
Delta_K
Doxorubicin
YDL227C_RF1_146 RF1
ZShift = −0.2
lm ZScore = −0.38
NG = 0
DB = 0
SM = 0
Delta_r
Doxorubicin
YDL227C_RF1_146 RF1
ZShift = 0.55
lm ZScore = −0.31
NG = 0
DB = 0
SM = 0
Delta_AUC
Doxorubicin

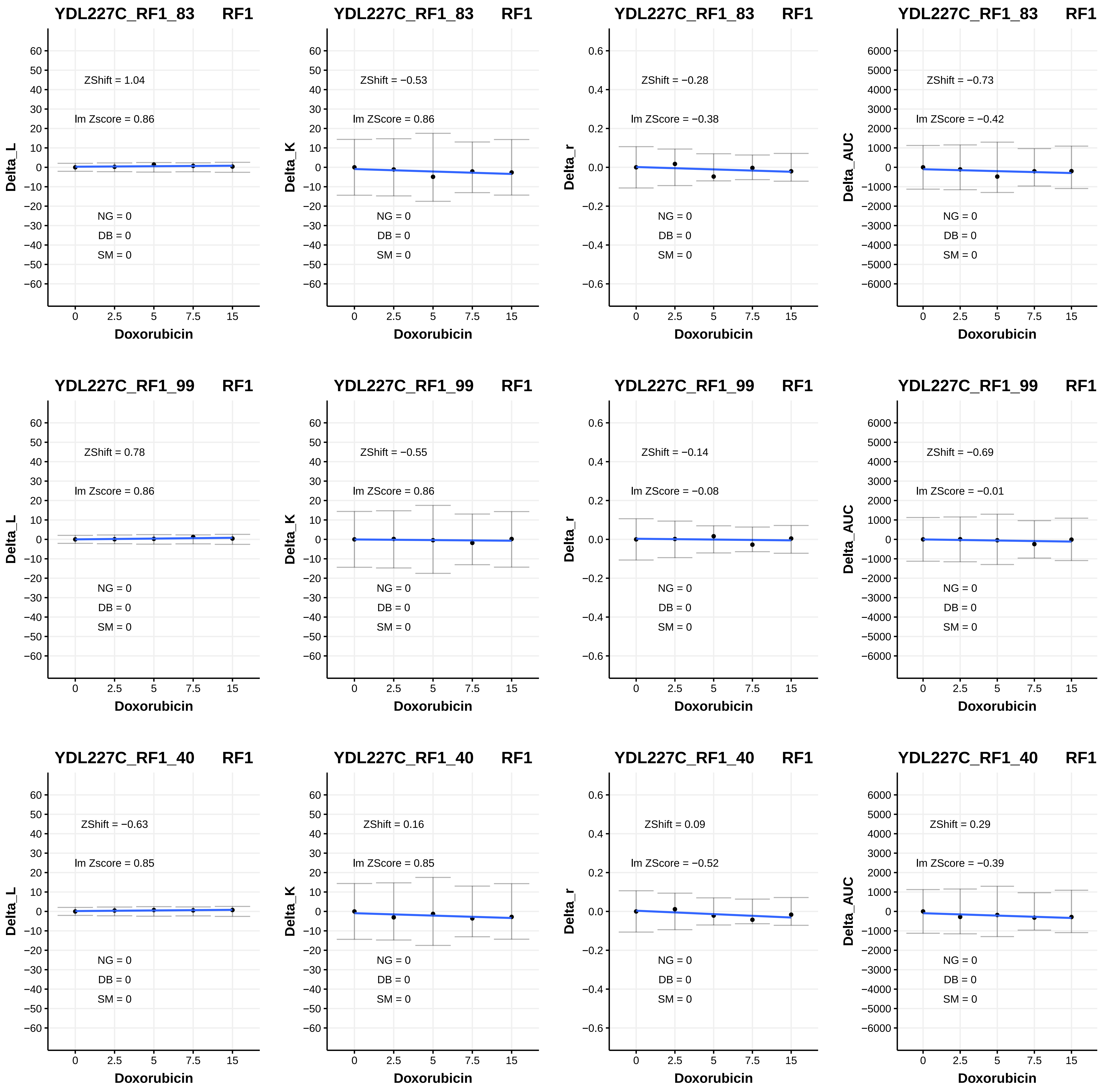
YDL227C_RF1_83 RF1
ZShift = 1.04
lm Zscore = 0.86
NG = 0
DB = 0
SM = 0
Delta_L
Doxorubicin
YDL227C_RF1_83 RF1
ZShift = −0.53
lm ZScore = 0.86
NG = 0
DB = 0
SM = 0
Delta_K
Doxorubicin
YDL227C_RF1_83 RF1
ZShift = −0.28
lm ZScore = −0.38
NG = 0
DB = 0
SM = 0
Delta_r
Doxorubicin
YDL227C_RF1_83 RF1
ZShift = −0.73
lm ZScore = −0.42
NG = 0
DB = 0
SM = 0
Delta_AUC
Doxorubicin
YDL227C_RF1_99 RF1
ZShift = 0.78
lm Zscore = 0.86
NG = 0
DB = 0
SM = 0
Delta_L
Doxorubicin
YDL227C_RF1_99 RF1
ZShift = −0.55
lm ZScore = 0.86
NG = 0
DB = 0
SM = 0
Delta_K
Doxorubicin
YDL227C_RF1_99 RF1
ZShift = −0.14
lm ZScore = −0.08
NG = 0
DB = 0
SM = 0
Delta_r
Doxorubicin
YDL227C_RF1_99 RF1
ZShift = −0.69
lm ZScore = −0.01
NG = 0
DB = 0
SM = 0
Delta_AUC
Doxorubicin
YDL227C_RF1_40 RF1
ZShift = −0.63
lm Zscore = 0.85
NG = 0
DB = 0
SM = 0
Delta_L
Doxorubicin
YDL227C_RF1_40 RF1
ZShift = 0.16
lm ZScore = 0.85
NG = 0
DB = 0
SM = 0
Delta_K
Doxorubicin
YDL227C_RF1_40 RF1
ZShift = 0.09
lm ZScore = −0.52
NG = 0
DB = 0
SM = 0
Delta_r
Doxorubicin
YDL227C_RF1_40 RF1
ZShift = 0.29
lm ZScore = −0.39
NG = 0
DB = 0
SM = 0
Delta_AUC
Doxorubicin

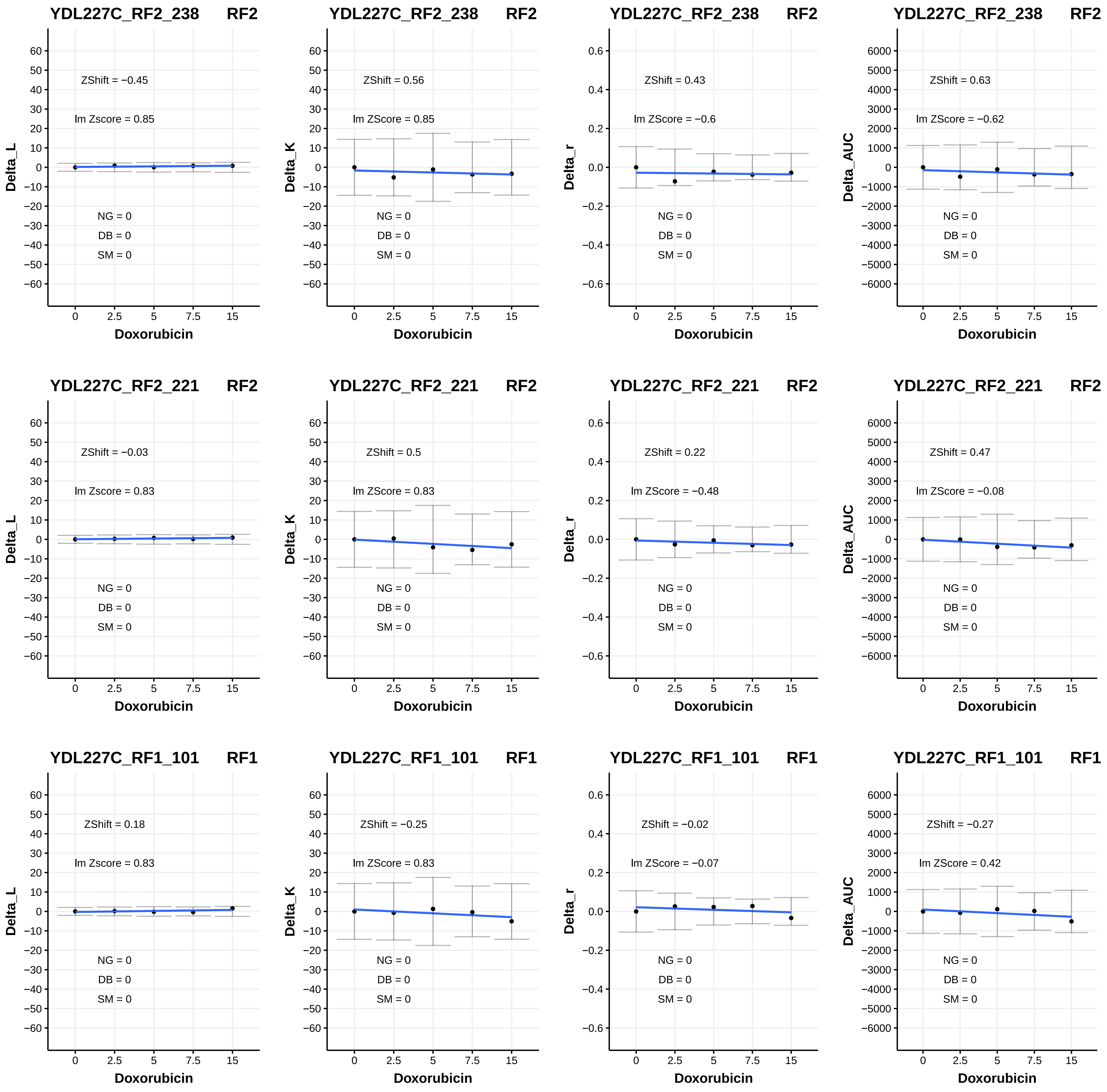
YDL227C_RF2_238 RF2
ZShift = −0.45
lm Zscore = 0.85
NG = 0
DB = 0
SM = 0
Delta_L
Doxorubicin
YDL227C_RF2_238 RF2
ZShift = 0.56
lm ZScore = 0.85
NG = 0
DB = 0
SM = 0
Delta_K
Doxorubicin
YDL227C_RF2_238 RF2
ZShift = 0.43
lm ZScore = −0.6
NG = 0
DB = 0
SM = 0
Delta_r
Doxorubicin
YDL227C_RF2_238 RF2
ZShift = 0.63
lm ZScore = −0.62
NG = 0
DB = 0
SM = 0
Delta_AUC
Doxorubicin
YDL227C_RF2_221 RF2
ZShift = −0.03
lm Zscore = 0.83
NG = 0
DB = 0
SM = 0
Delta_L
Doxorubicin
YDL227C_RF2_221 RF2
ZShift = 0.5
lm ZScore = 0.83
NG = 0
DB = 0
SM = 0
Delta_K
Doxorubicin
YDL227C_RF2_221 RF2
ZShift = 0.22
lm ZScore = −0.48
NG = 0
DB = 0
SM = 0
Delta_r
Doxorubicin
YDL227C_RF2_221 RF2
ZShift = 0.47
lm ZScore = −0.08
NG = 0
DB = 0
SM = 0
Delta_AUC
Doxorubicin
YDL227C_RF1_101 RF1
ZShift = 0.18
lm Zscore = 0.83
NG = 0
DB = 0
SM = 0
Delta_L
Doxorubicin
YDL227C_RF1_101 RF1
ZShift = −0.25
lm ZScore = 0.83
NG = 0
DB = 0
SM = 0
Delta_K
Doxorubicin
YDL227C_RF1_101 RF1
ZShift = −0.02
lm ZScore = −0.07
NG = 0
DB = 0
SM = 0
Delta_r
Doxorubicin
YDL227C_RF1_101 RF1
ZShift = −0.27
lm ZScore = 0.42
NG = 0
DB = 0
SM = 0
Delta_AUC
Doxorubicin

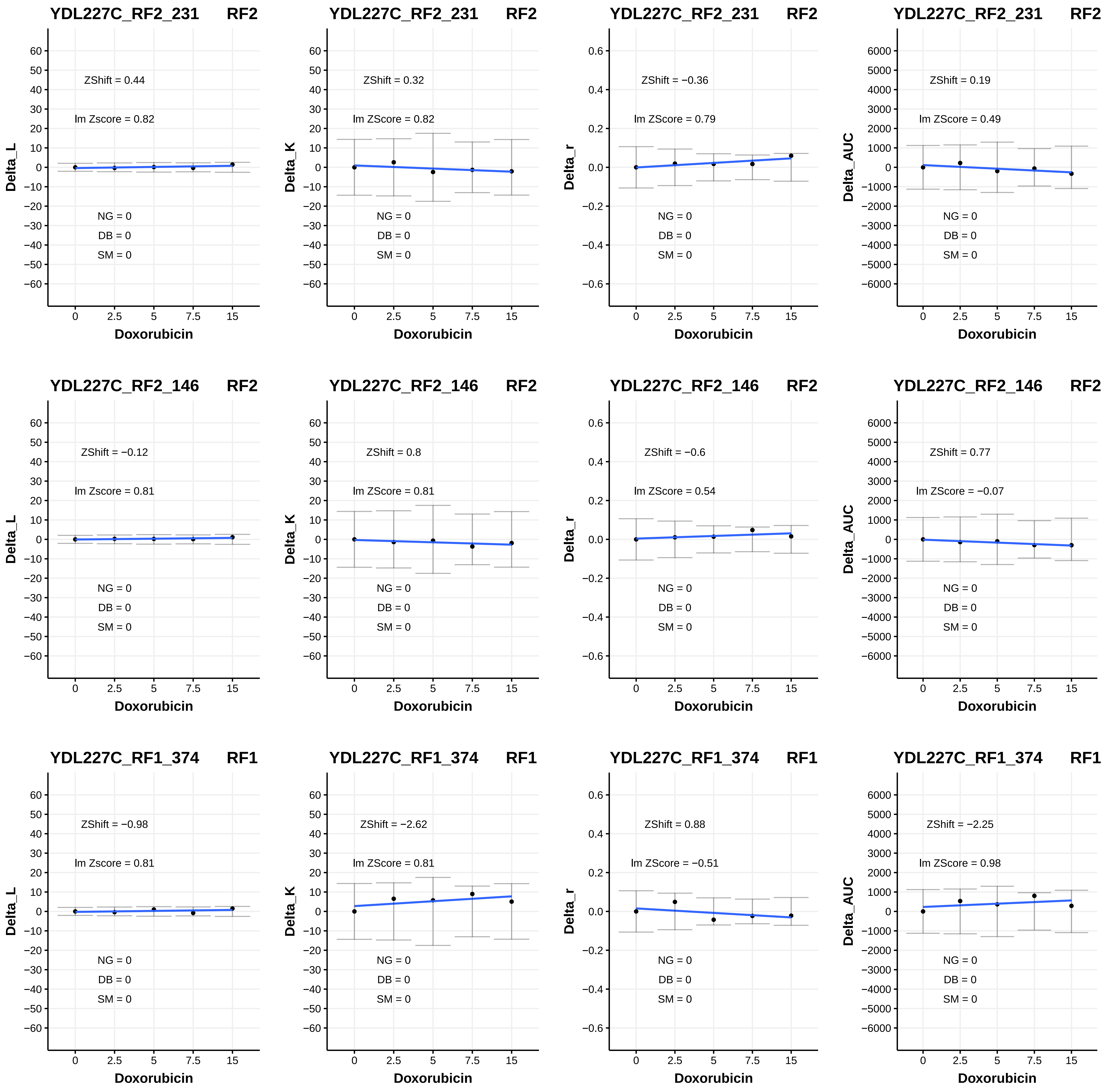
YDL227C_RF2_231 RF2
Delta_L
ZShift = 0.44
lm Zscore = 0.82
NG = 0
DB = 0
SM = 0
Doxorubicin
YDL227C_RF2_231 RF2
Delta_K
ZShift = 0.32
lm ZScore = 0.82
NG = 0
DB = 0
SM = 0
Doxorubicin
YDL227C_RF2_231 RF2
Delta_r
ZShift = −0.36
lm ZScore = 0.79
NG = 0
DB = 0
SM = 0
Doxorubicin
YDL227C_RF2_231 RF2
Delta_AUC
ZShift = 0.19
lm ZScore = 0.49
NG = 0
DB = 0
SM = 0
Doxorubicin
YDL227C_RF2_146 RF2
Delta_L
ZShift = −0.12
lm Zscore = 0.81
NG = 0
DB = 0
SM = 0
Doxorubicin
YDL227C_RF2_146 RF2
Delta_K
ZShift = 0.8
lm ZScore = 0.81
NG = 0
DB = 0
SM = 0
Doxorubicin
YDL227C_RF2_146 RF2
Delta_r
ZShift = −0.6
lm ZScore = 0.54
NG = 0
DB = 0
SM = 0
Doxorubicin
YDL227C_RF2_146 RF2
Delta_AUC
ZShift = 0.77
lm ZScore = −0.07
NG = 0
DB = 0
SM = 0
Doxorubicin
YDL227C_RF1_374 RF1
Delta_L
ZShift = −0.98
lm Zscore = 0.81
NG = 0
DB = 0
SM = 0
Doxorubicin
YDL227C_RF1_374 RF1
Delta_K
ZShift = −2.62
lm ZScore = 0.81
NG = 0
DB = 0
SM = 0
Doxorubicin
YDL227C_RF1_374 RF1
Delta_r
ZShift = 0.88
lm ZScore = −0.51
NG = 0
DB = 0
SM = 0
Doxorubicin
YDL227C_RF1_374 RF1
Delta_AUC
ZShift = −2.25
lm ZScore = 0.98
NG = 0
DB = 0
SM = 0
Doxorubicin

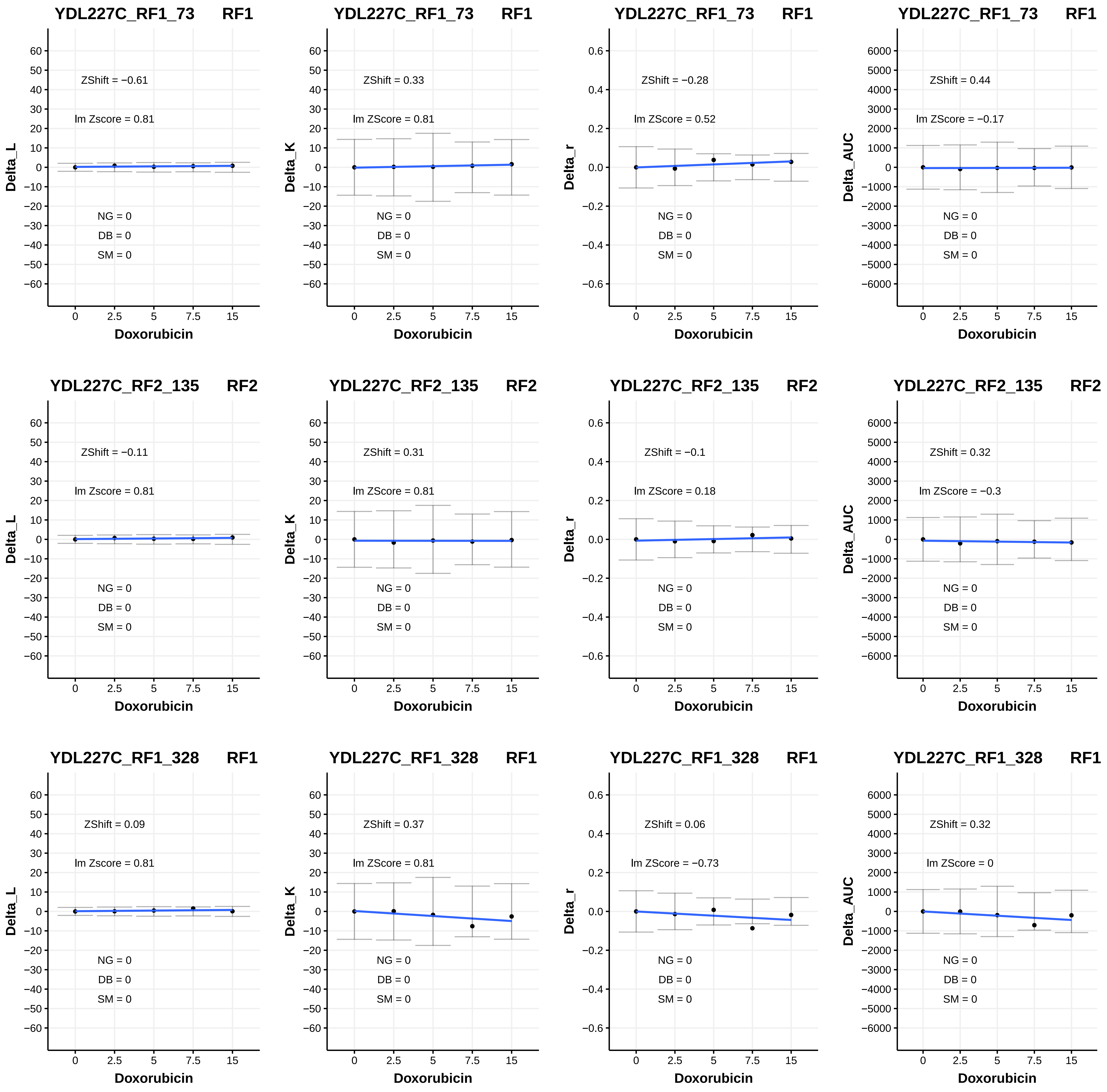

YDL227C_RF1_73 RF1
ZShift = −0.61
lm Zscore = 0.81
NG = 0
DB = 0
SM = 0
Delta_L
Doxorubicin
YDL227C_RF1_73 RF1
ZShift = 0.33
lm ZScore = 0.81
NG = 0
DB = 0
SM = 0
Delta_K
Doxorubicin
YDL227C_RF1_73 RF1
ZShift = −0.28
lm ZScore = 0.52
NG = 0
DB = 0
SM = 0
Delta_r
Doxorubicin
YDL227C_RF1_73 RF1
ZShift = 0.44
lm ZScore = −0.17
NG = 0
DB = 0
SM = 0
Delta_AUC
Doxorubicin
YDL227C_RF2_135 RF2
ZShift = −0.11
lm Zscore = 0.81
NG = 0
DB = 0
SM = 0
Delta_L
Doxorubicin
YDL227C_RF2_135 RF2
ZShift = 0.31
lm ZScore = 0.81
NG = 0
DB = 0
SM = 0
Delta_K
Doxorubicin
YDL227C_RF2_135 RF2
ZShift = −0.1
lm ZScore = 0.18
NG = 0
DB = 0
SM = 0
Delta_r
Doxorubicin
YDL227C_RF2_135 RF2
ZShift = 0.32
lm ZScore = −0.3
NG = 0
DB = 0
SM = 0
Delta_AUC
Doxorubicin
YDL227C_RF1_328 RF1
ZShift = 0.09
lm Zscore = 0.81
NG = 0
DB = 0
SM = 0
Delta_L
Doxorubicin
YDL227C_RF1_328 RF1
ZShift = 0.37
lm ZScore = 0.81
NG = 0
DB = 0
SM = 0
Delta_K
Doxorubicin
YDL227C_RF1_328 RF1
ZShift = 0.06
lm ZScore = −0.73
NG = 0
DB = 0
SM = 0
Delta_r
Doxorubicin
YDL227C_RF1_328 RF1
ZShift = 0.32
lm ZScore = 0
NG = 0
DB = 0
SM = 0
Delta_AUC
Doxorubicin

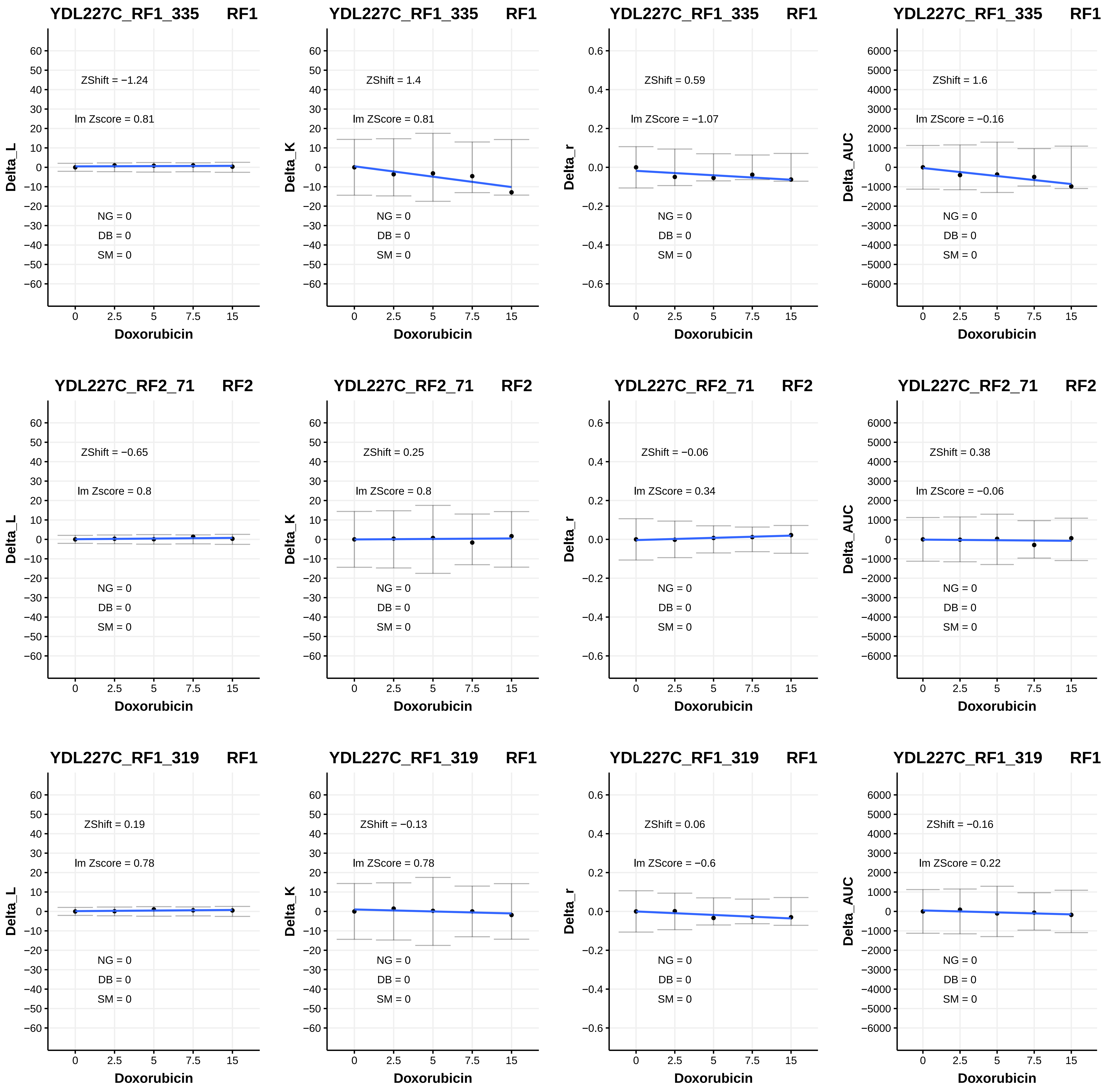

YDL227C_RF1_335 RF1
ZShift = −1.24
lm Zscore = 0.81
NG = 0
DB = 0
SM = 0
Delta_L
Doxorubicin
YDL227C_RF1_335 RF1
ZShift = 1.4
lm ZScore = 0.81
NG = 0
DB = 0
SM = 0
Delta_K
Doxorubicin
YDL227C_RF1_335 RF1
ZShift = 0.59
lm ZScore = −1.07
NG = 0
DB = 0
SM = 0
Delta_r
Doxorubicin
YDL227C_RF1_335 RF1
ZShift = 1.6
lm ZScore = −0.16
NG = 0
DB = 0
SM = 0
Delta_AUC
Doxorubicin
YDL227C_RF2_71 RF2
ZShift = −0.65
lm Zscore = 0.8
NG = 0
DB = 0
SM = 0
Delta_L
Doxorubicin
YDL227C_RF2_71 RF2
ZShift = 0.25
lm ZScore = 0.8
NG = 0
DB = 0
SM = 0
Delta_K
Doxorubicin
YDL227C_RF2_71 RF2
ZShift = −0.06
lm ZScore = 0.34
NG = 0
DB = 0
SM = 0
Delta_r
Doxorubicin
YDL227C_RF2_71 RF2
ZShift = 0.38
lm ZScore = −0.06
NG = 0
DB = 0
SM = 0
Delta_AUC
Doxorubicin
YDL227C_RF1_319 RF1
ZShift = 0.19
lm Zscore = 0.78
NG = 0
DB = 0
SM = 0
Delta_L
Doxorubicin
YDL227C_RF1_319 RF1
ZShift = −0.13
lm ZScore = 0.78
NG = 0
DB = 0
SM = 0
Delta_K
Doxorubicin
YDL227C_RF1_319 RF1
ZShift = 0.06
lm ZScore = −0.6
NG = 0
DB = 0
SM = 0
Delta_r
Doxorubicin
YDL227C_RF1_319 RF1
ZShift = −0.16
lm ZScore = 0.22
NG = 0
DB = 0
SM = 0
Delta_AUC
Doxorubicin

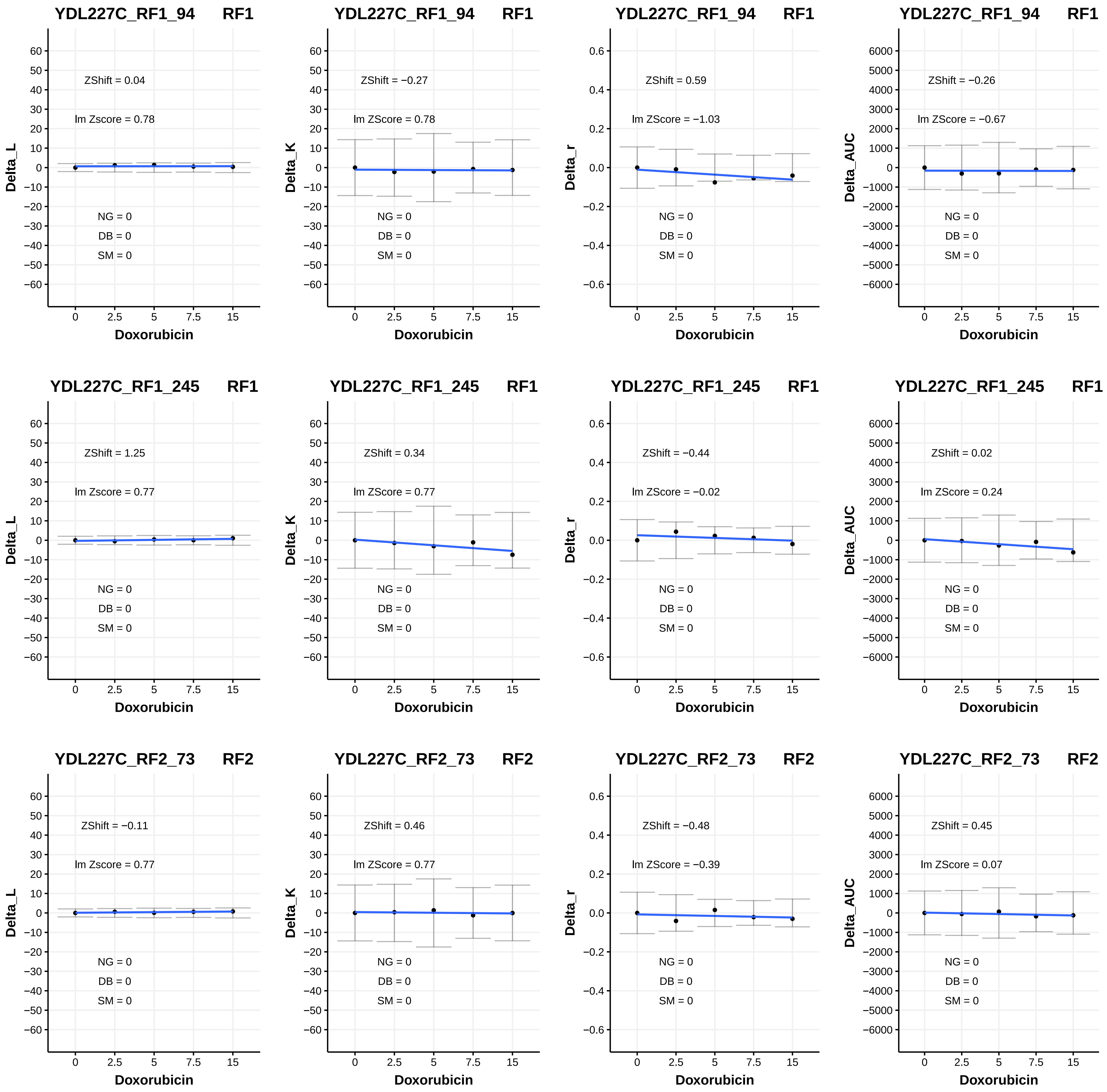
YDL227C_RF1_94 RF1
ZShift = 0.04
lm Zscore = 0.78
NG = 0
DB = 0
SM = 0
Delta_L
Doxorubicin
YDL227C_RF1_94 RF1
ZShift = −0.27
lm ZScore = 0.78
NG = 0
DB = 0
SM = 0
Delta_K
Doxorubicin
YDL227C_RF1_94 RF1
ZShift = 0.59
lm ZScore = −1.03
NG = 0
DB = 0
SM = 0
Delta_r
Doxorubicin
YDL227C_RF1_94 RF1
ZShift = −0.26
lm ZScore = −0.67
NG = 0
DB = 0
SM = 0
Delta_AUC
Doxorubicin
YDL227C_RF1_245 RF1
ZShift = 1.25
lm Zscore = 0.77
NG = 0
DB = 0
SM = 0
Delta_L
Doxorubicin
YDL227C_RF1_245 RF1
ZShift = 0.34
lm ZScore = 0.77
NG = 0
DB = 0
SM = 0
Delta_K
Doxorubicin
YDL227C_RF1_245 RF1
ZShift = −0.44
lm ZScore = −0.02
NG = 0
DB = 0
SM = 0
Delta_r
Doxorubicin
YDL227C_RF1_245 RF1
ZShift = 0.02
lm ZScore = 0.24
NG = 0
DB = 0
SM = 0
Delta_AUC
Doxorubicin
YDL227C_RF2_73 RF2
ZShift = −0.11
lm Zscore = 0.77
NG = 0
DB = 0
SM = 0
Delta_L
Doxorubicin
YDL227C_RF2_73 RF2
ZShift = 0.46
lm ZScore = 0.77
NG = 0
DB = 0
SM = 0
Delta_K
Doxorubicin
YDL227C_RF2_73 RF2
ZShift = −0.48
lm ZScore = −0.39
NG = 0
DB = 0
SM = 0
Delta_r
Doxorubicin
YDL227C_RF2_73 RF2
ZShift = 0.45
lm ZScore = 0.07
NG = 0
DB = 0
SM = 0
Delta_AUC
Doxorubicin

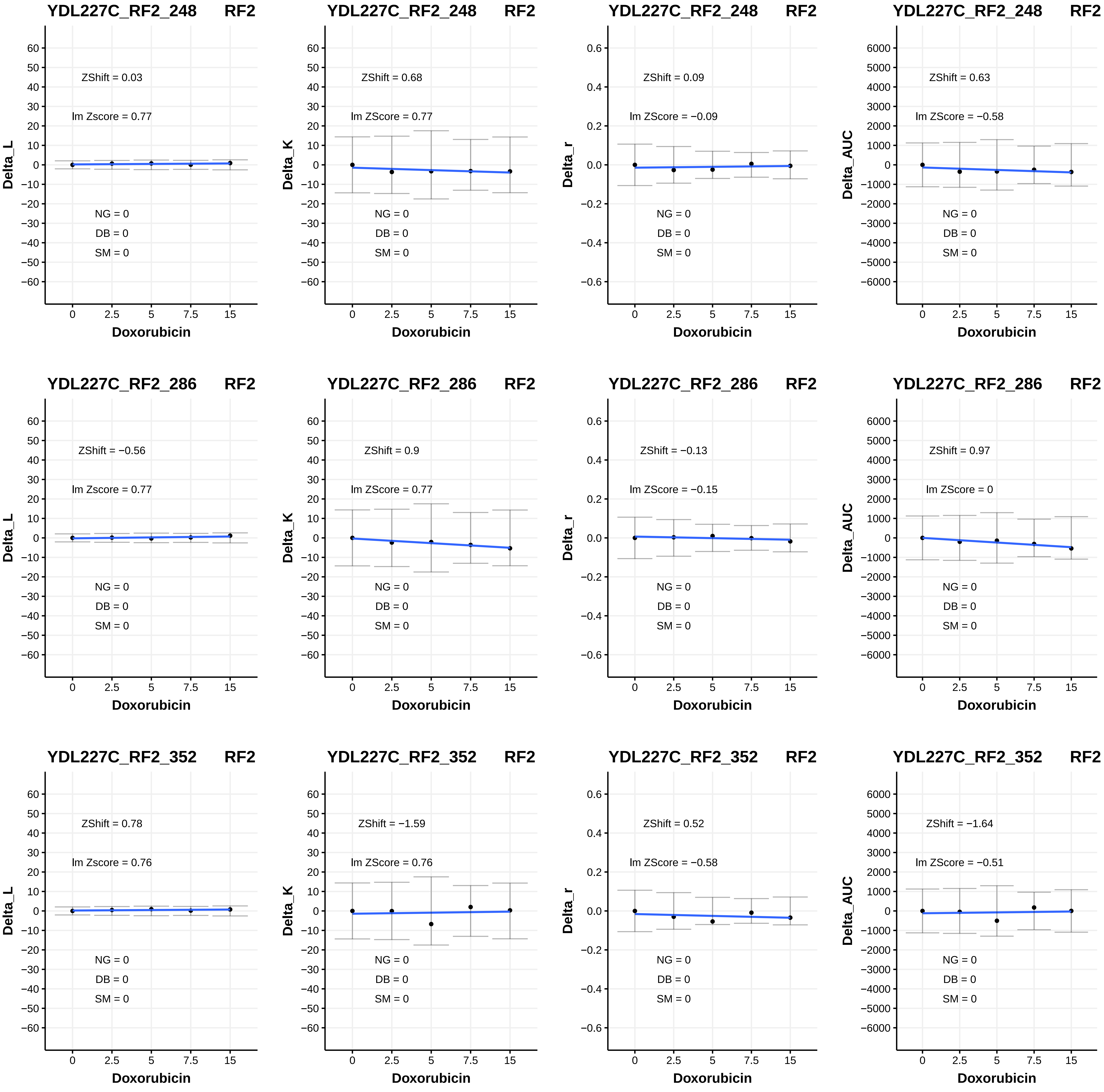

YDL227C_RF2_248 RF2
Delta_L
ZShift = 0.03
lm Zscore = 0.77
NG = 0
DB = 0
SM = 0
Doxorubicin
YDL227C_RF2_248 RF2
Delta_K
ZShift = 0.68
lm ZScore = 0.77
NG = 0
DB = 0
SM = 0
Doxorubicin
YDL227C_RF2_248 RF2
Delta_r
ZShift = 0.09
lm ZScore = −0.09
NG = 0
DB = 0
SM = 0
Doxorubicin
YDL227C_RF2_248 RF2
Delta_AUC
ZShift = 0.63
lm ZScore = −0.58
NG = 0
DB = 0
SM = 0
Doxorubicin
YDL227C_RF2_286 RF2
Delta_L
ZShift = −0.56
lm Zscore = 0.77
NG = 0
DB = 0
SM = 0
Doxorubicin
YDL227C_RF2_286 RF2
Delta_K
ZShift = 0.9
lm ZScore = 0.77
NG = 0
DB = 0
SM = 0
Doxorubicin
YDL227C_RF2_286 RF2
Delta_r
ZShift = −0.13
lm ZScore = −0.15
NG = 0
DB = 0
SM = 0
Doxorubicin
YDL227C_RF2_286 RF2
Delta_AUC
ZShift = 0.97
lm ZScore = 0
NG = 0
DB = 0
SM = 0
Doxorubicin
YDL227C_RF2_352 RF2
Delta_L
ZShift = 0.78
lm Zscore = 0.76
NG = 0
DB = 0
SM = 0
Doxorubicin
YDL227C_RF2_352 RF2
Delta_K
ZShift = −1.59
lm ZScore = 0.76
NG = 0
DB = 0
SM = 0
Doxorubicin
YDL227C_RF2_352 RF2
Delta_r
ZShift = 0.52
lm ZScore = −0.58
NG = 0
DB = 0
SM = 0
Doxorubicin
YDL227C_RF2_352 RF2
Delta_AUC
ZShift = −1.64
lm ZScore = −0.51
NG = 0
DB = 0
SM = 0
Doxorubicin

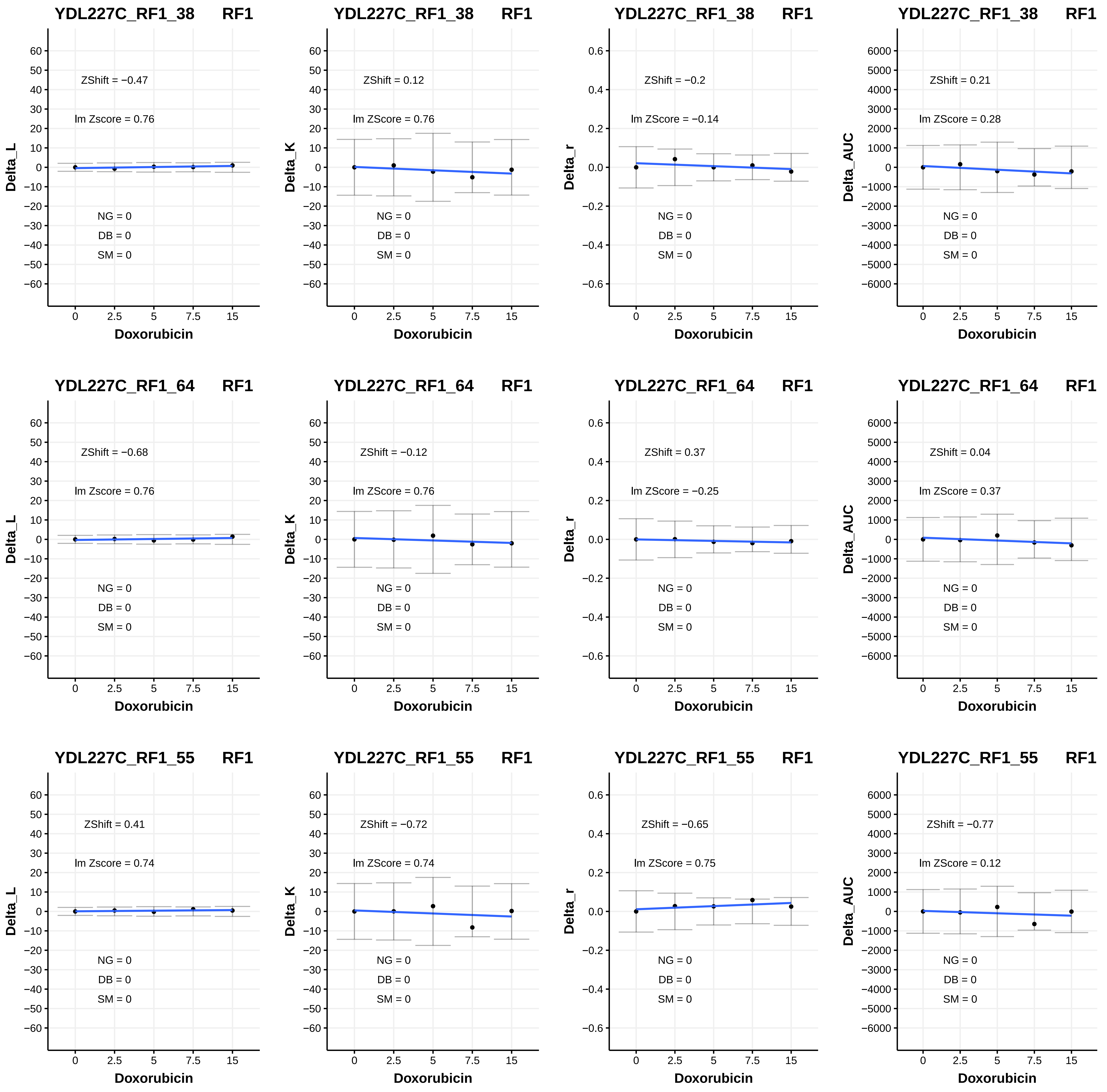

YDL227C_RF1_38 RF1
ZShift = −0.47
lm Zscore = 0.76
NG = 0
DB = 0
SM = 0
Delta_L
Doxorubicin
YDL227C_RF1_38 RF1
ZShift = 0.12
lm ZScore = 0.76
NG = 0
DB = 0
SM = 0
Delta_K
Doxorubicin
YDL227C_RF1_38 RF1
ZShift = −0.2
lm ZScore = −0.14
NG = 0
DB = 0
SM = 0
Delta_r
Doxorubicin
YDL227C_RF1_38 RF1
ZShift = 0.21
lm ZScore = 0.28
NG = 0
DB = 0
SM = 0
Delta_AUC
Doxorubicin
YDL227C_RF1_64 RF1
ZShift = −0.68
lm Zscore = 0.76
NG = 0
DB = 0
SM = 0
Delta_L
Doxorubicin
YDL227C_RF1_64 RF1
ZShift = −0.12
lm ZScore = 0.76
NG = 0
DB = 0
SM = 0
Delta_K
Doxorubicin
YDL227C_RF1_64 RF1
ZShift = 0.37
lm ZScore = −0.25
NG = 0
DB = 0
SM = 0
Delta_r
Doxorubicin
YDL227C_RF1_64 RF1
ZShift = 0.04
lm ZScore = 0.37
NG = 0
DB = 0
SM = 0
Delta_AUC
Doxorubicin
YDL227C_RF1_55 RF1
ZShift = 0.41
lm Zscore = 0.74
NG = 0
DB = 0
SM = 0
Delta_L
Doxorubicin
YDL227C_RF1_55 RF1
ZShift = −0.72
lm ZScore = 0.74
NG = 0
DB = 0
SM = 0
Delta_K
Doxorubicin
YDL227C_RF1_55 RF1
ZShift = −0.65
lm ZScore = 0.75
NG = 0
DB = 0
SM = 0
Delta_r
Doxorubicin
YDL227C_RF1_55 RF1
ZShift = −0.77
lm ZScore = 0.12
NG = 0
DB = 0
SM = 0
Delta_AUC
Doxorubicin

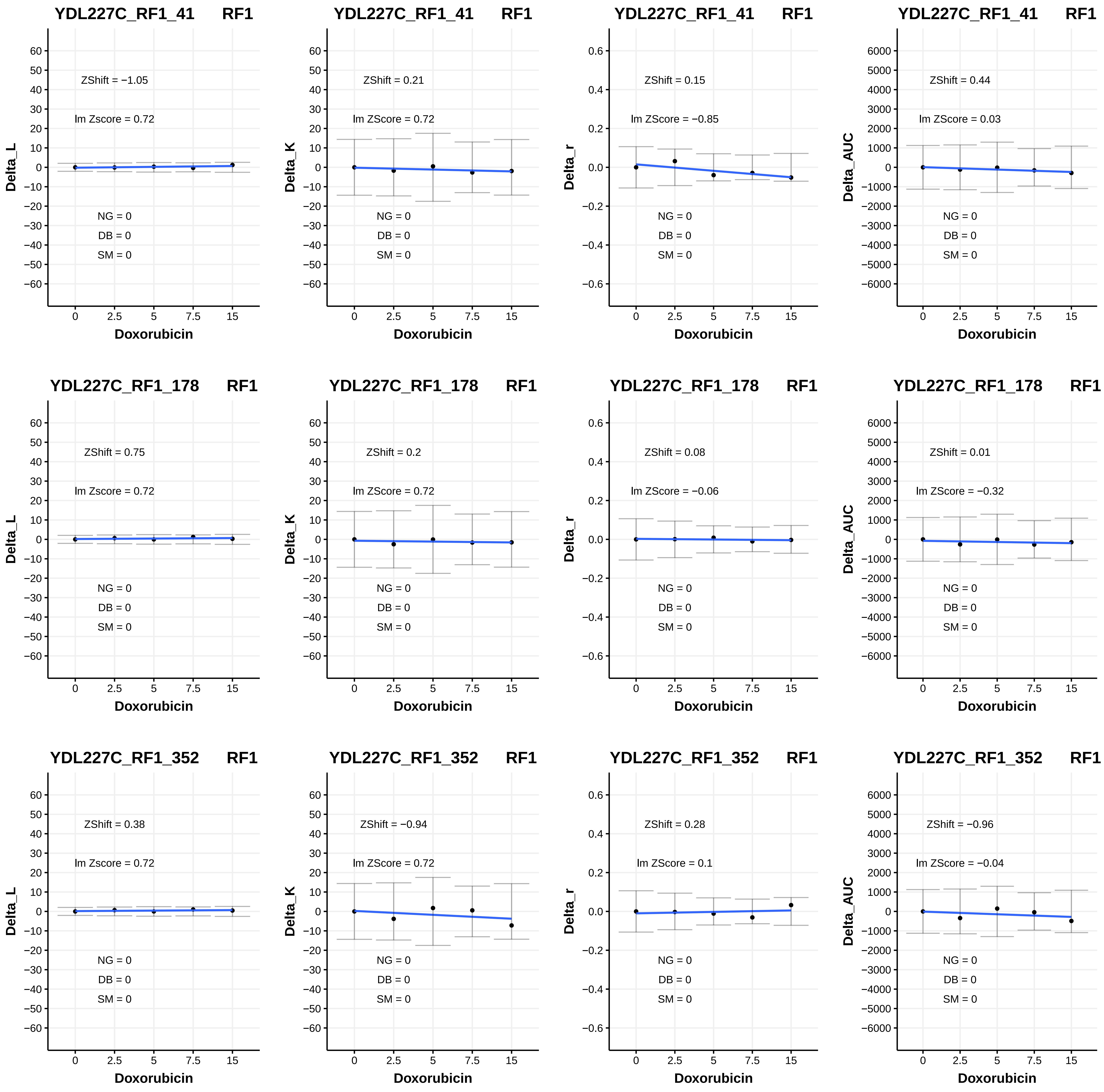
YDL227C_RF1_41 RF1
ZShift = −1.05
lm Zscore = 0.72
NG = 0
DB = 0
SM = 0
Delta_L
Doxorubicin
YDL227C_RF1_41 RF1
ZShift = 0.21
lm ZScore = 0.72
NG = 0
DB = 0
SM = 0
Delta_K
Doxorubicin
YDL227C_RF1_41 RF1
ZShift = 0.15
lm ZScore = −0.85
NG = 0
DB = 0
SM = 0
Delta_r
Doxorubicin
YDL227C_RF1_41 RF1
ZShift = 0.44
lm ZScore = 0.03
NG = 0
DB = 0
SM = 0
Delta_AUC
Doxorubicin
YDL227C_RF1_178 RF1
ZShift = 0.75
lm Zscore = 0.72
NG = 0
DB = 0
SM = 0
Delta_L
Doxorubicin
YDL227C_RF1_178 RF1
ZShift = 0.2
lm ZScore = 0.72
NG = 0
DB = 0
SM = 0
Delta_K
Doxorubicin
YDL227C_RF1_178 RF1
ZShift = 0.08
lm ZScore = −0.06
NG = 0
DB = 0
SM = 0
Delta_r
Doxorubicin
YDL227C_RF1_178 RF1
ZShift = 0.01
lm ZScore = −0.32
NG = 0
DB = 0
SM = 0
Delta_AUC
Doxorubicin
YDL227C_RF1_352 RF1
ZShift = 0.38
lm Zscore = 0.72
NG = 0
DB = 0
SM = 0
Delta_L
Doxorubicin
YDL227C_RF1_352 RF1
ZShift = −0.94
lm ZScore = 0.72
NG = 0
DB = 0
SM = 0
Delta_K
Doxorubicin
YDL227C_RF1_352 RF1
ZShift = 0.28
lm ZScore = 0.1
NG = 0
DB = 0
SM = 0
Delta_r
Doxorubicin
YDL227C_RF1_352 RF1
ZShift = −0.96
lm ZScore = −0.04
NG = 0
DB = 0
SM = 0
Delta_AUC
Doxorubicin

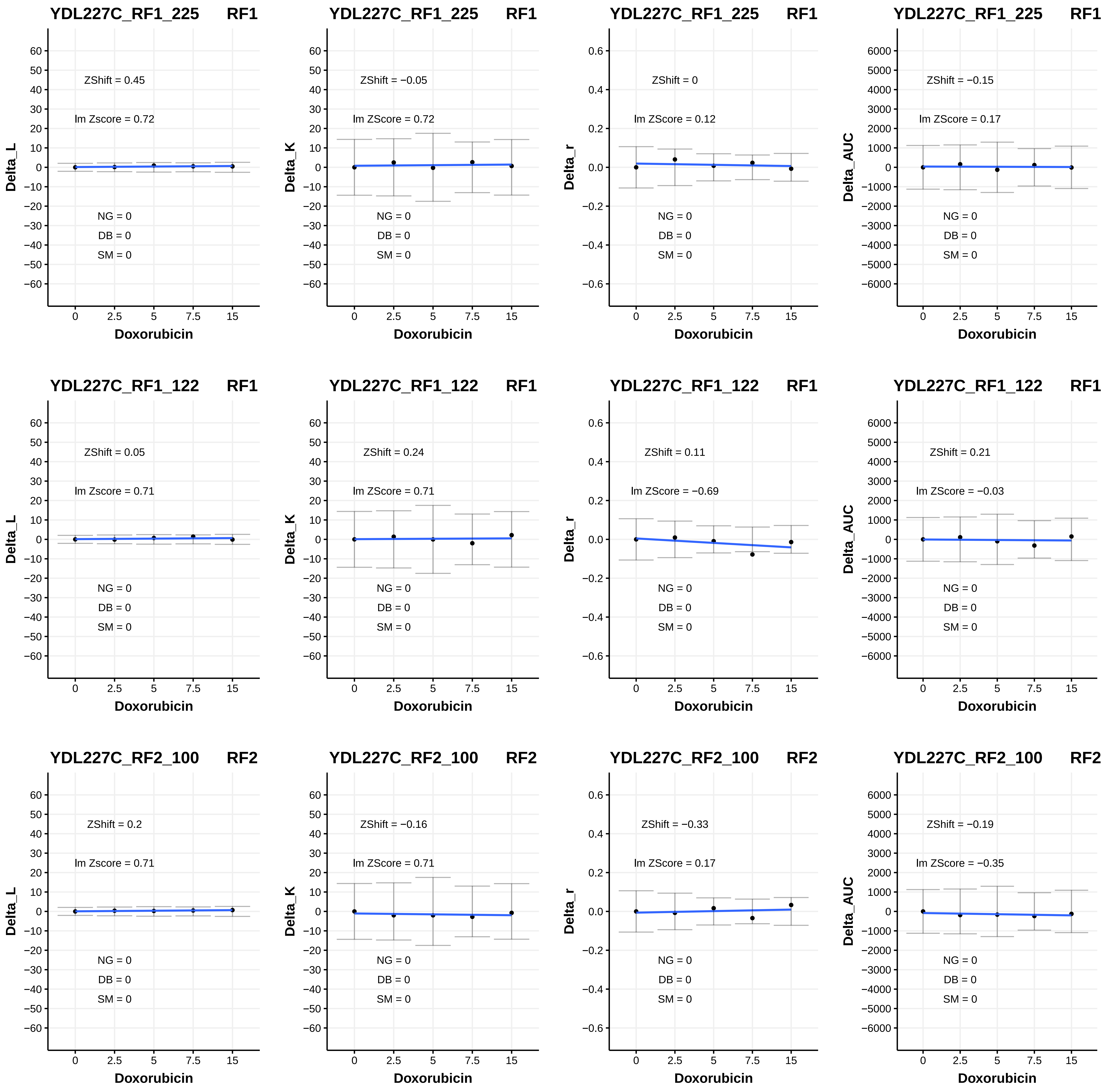
YDL227C_RF1_225 RF1
ZShift = 0.45
lm Zscore = 0.72
NG = 0
DB = 0
SM = 0
Delta_L
Doxorubicin
YDL227C_RF1_225 RF1
ZShift = −0.05
lm ZScore = 0.72
NG = 0
DB = 0
SM = 0
Delta_K
Doxorubicin
YDL227C_RF1_225 RF1
ZShift = 0
lm ZScore = 0.12
NG = 0
DB = 0
SM = 0
Delta_r
Doxorubicin
YDL227C_RF1_225 RF1
ZShift = −0.15
lm ZScore = 0.17
NG = 0
DB = 0
SM = 0
Delta_AUC
Doxorubicin
YDL227C_RF1_122 RF1
ZShift = 0.05
lm Zscore = 0.71
NG = 0
DB = 0
SM = 0
Delta_L
Doxorubicin
YDL227C_RF1_122 RF1
ZShift = 0.24
lm ZScore = 0.71
NG = 0
DB = 0
SM = 0
Delta_K
Doxorubicin
YDL227C_RF1_122 RF1
ZShift = 0.11
lm ZScore = −0.69
NG = 0
DB = 0
SM = 0
Delta_r
Doxorubicin
YDL227C_RF1_122 RF1
ZShift = 0.21
lm ZScore = −0.03
NG = 0
DB = 0
SM = 0
Delta_AUC
Doxorubicin
YDL227C_RF2_100 RF2
ZShift = 0.2
lm Zscore = 0.71
NG = 0
DB = 0
SM = 0
Delta_L
Doxorubicin
YDL227C_RF2_100 RF2
ZShift = −0.16
lm ZScore = 0.71
NG = 0
DB = 0
SM = 0
Delta_K
Doxorubicin
YDL227C_RF2_100 RF2
ZShift = −0.33
lm ZScore = 0.17
NG = 0
DB = 0
SM = 0
Delta_r
Doxorubicin
YDL227C_RF2_100 RF2
ZShift = −0.19
lm ZScore = −0.35
NG = 0
DB = 0
SM = 0
Delta_AUC
Doxorubicin

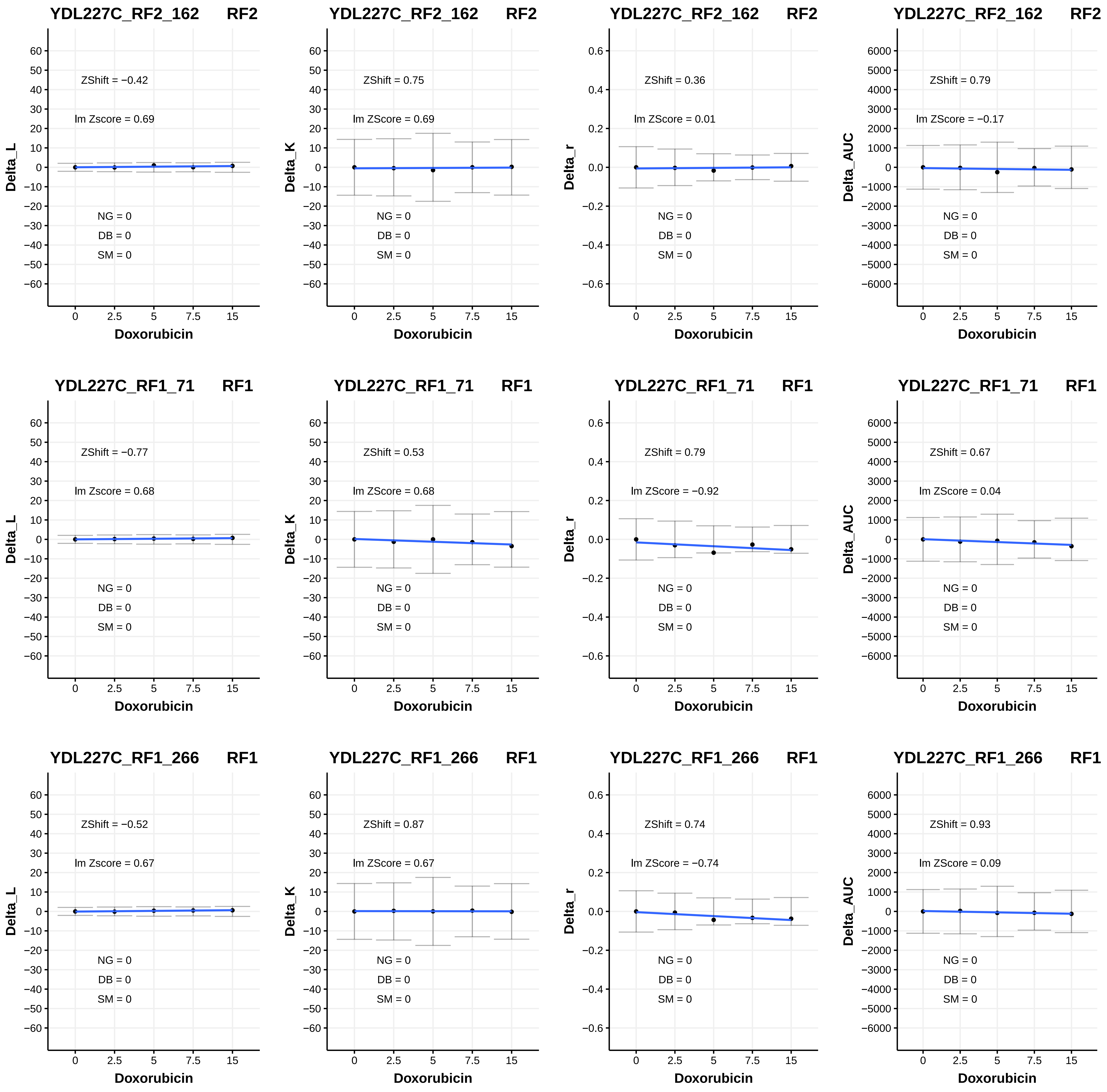

YDL227C_RF2_162 RF2
ZShift = −0.42
lm Zscore = 0.69
NG = 0
DB = 0
SM = 0
Delta_L
Doxorubicin
YDL227C_RF2_162 RF2
ZShift = 0.75
lm ZScore = 0.69
NG = 0
DB = 0
SM = 0
Delta_K
Doxorubicin
YDL227C_RF2_162 RF2
ZShift = 0.36
lm ZScore = 0.01
NG = 0
DB = 0
SM = 0
Delta_r
Doxorubicin
YDL227C_RF2_162 RF2
ZShift = 0.79
lm ZScore = −0.17
NG = 0
DB = 0
SM = 0
Delta_AUC
Doxorubicin
YDL227C_RF1_71 RF1
ZShift = −0.77
lm Zscore = 0.68
NG = 0
DB = 0
SM = 0
Delta_L
Doxorubicin
YDL227C_RF1_71 RF1
ZShift = 0.53
lm ZScore = 0.68
NG = 0
DB = 0
SM = 0
Delta_K
Doxorubicin
YDL227C_RF1_71 RF1
ZShift = 0.79
lm ZScore = −0.92
NG = 0
DB = 0
SM = 0
Delta_r
Doxorubicin
YDL227C_RF1_71 RF1
ZShift = 0.67
lm ZScore = 0.04
NG = 0
DB = 0
SM = 0
Delta_AUC
Doxorubicin
YDL227C_RF1_266 RF1
ZShift = −0.52
lm Zscore = 0.67
NG = 0
DB = 0
SM = 0
Delta_L
Doxorubicin
YDL227C_RF1_266 RF1
ZShift = 0.87
lm ZScore = 0.67
NG = 0
DB = 0
SM = 0
Delta_K
Doxorubicin
YDL227C_RF1_266 RF1
ZShift = 0.74
lm ZScore = −0.74
NG = 0
DB = 0
SM = 0
Delta_r
Doxorubicin
YDL227C_RF1_266 RF1
ZShift = 0.93
lm ZScore = 0.09
NG = 0
DB = 0
SM = 0
Delta_AUC
Doxorubicin

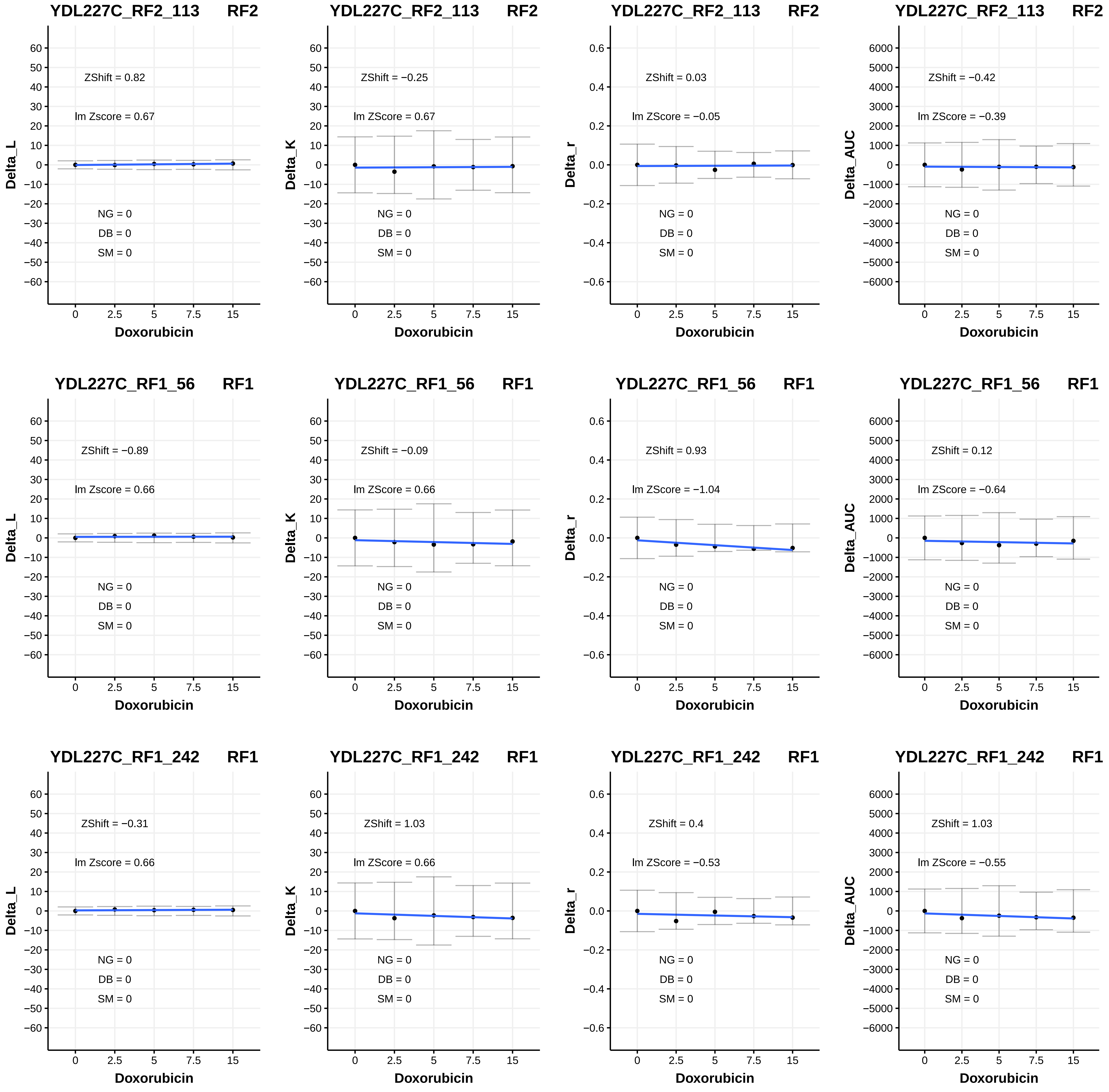

YDL227C_RF2_113 RF2
ZShift = 0.82
lm Zscore = 0.67
NG = 0
DB = 0
SM = 0
Delta_L
Doxorubicin
YDL227C_RF2_113 RF2
ZShift = −0.25
lm ZScore = 0.67
NG = 0
DB = 0
SM = 0
Delta_K
Doxorubicin
YDL227C_RF2_113 RF2
ZShift = 0.03
lm ZScore = −0.05
NG = 0
DB = 0
SM = 0
Delta_r
Doxorubicin
YDL227C_RF2_113 RF2
ZShift = −0.42
lm ZScore = −0.39
NG = 0
DB = 0
SM = 0
Delta_AUC
Doxorubicin
YDL227C_RF1_56 RF1
ZShift = −0.89
lm Zscore = 0.66
NG = 0
DB = 0
SM = 0
Delta_L
Doxorubicin
YDL227C_RF1_56 RF1
ZShift = −0.09
lm ZScore = 0.66
NG = 0
DB = 0
SM = 0
Delta_K
Doxorubicin
YDL227C_RF1_56 RF1
ZShift = 0.93
lm ZScore = −1.04
NG = 0
DB = 0
SM = 0
Delta_r
Doxorubicin
YDL227C_RF1_56 RF1
ZShift = 0.12
lm ZScore = −0.64
NG = 0
DB = 0
SM = 0
Delta_AUC
Doxorubicin
YDL227C_RF1_242 RF1
ZShift = −0.31
lm Zscore = 0.66
NG = 0
DB = 0
SM = 0
Delta_L
Doxorubicin
YDL227C_RF1_242 RF1
ZShift = 1.03
lm ZScore = 0.66
NG = 0
DB = 0
SM = 0
Delta_K
Doxorubicin
YDL227C_RF1_242 RF1
ZShift = 0.4
lm ZScore = −0.53
NG = 0
DB = 0
SM = 0
Delta_r
Doxorubicin
YDL227C_RF1_242 RF1
ZShift = 1.03
lm ZScore = −0.55
NG = 0
DB = 0
SM = 0
Delta_AUC
Doxorubicin

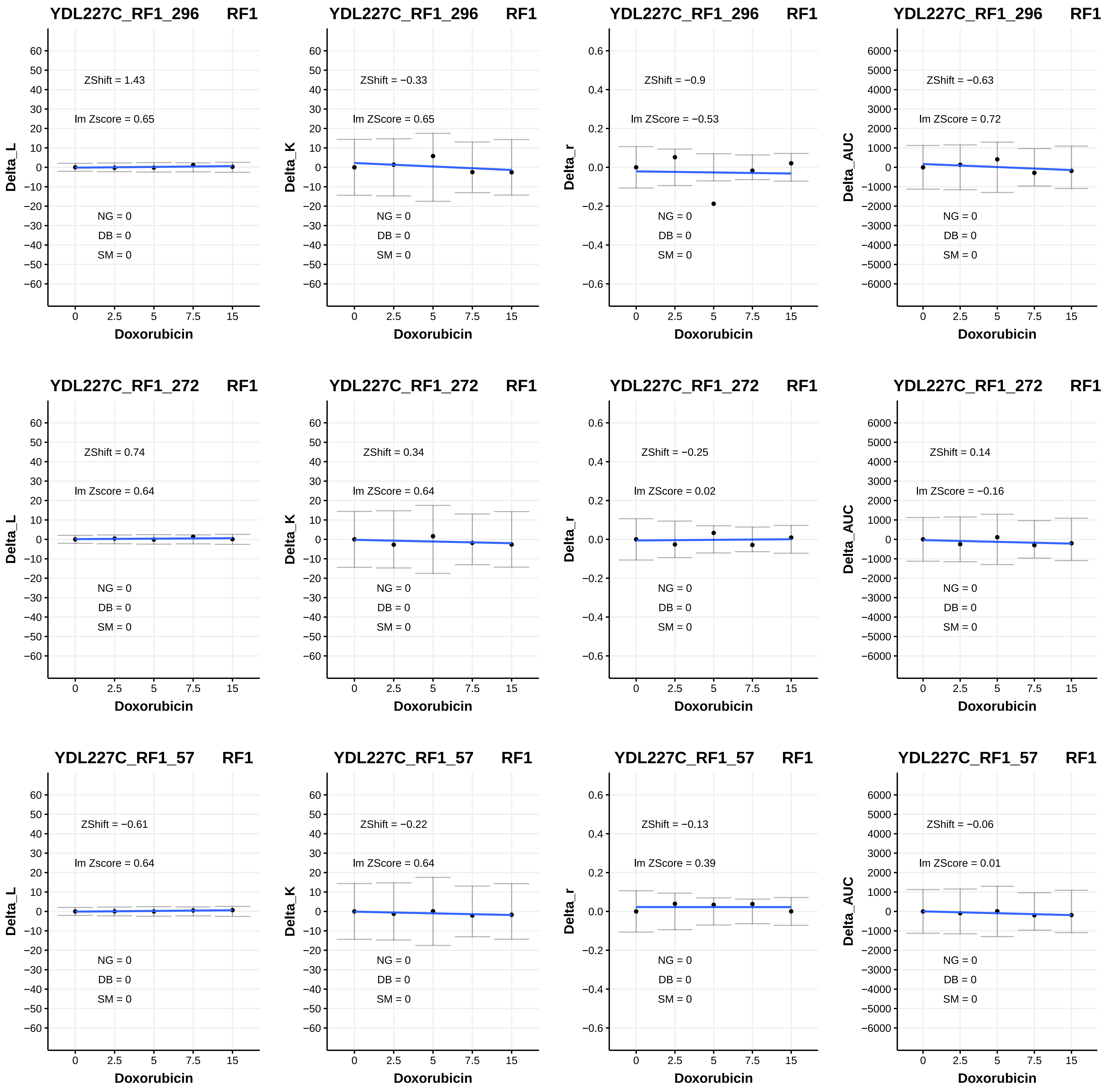
YDL227C_RF1_296 RF1
ZShift = 1.43
lm Zscore = 0.65
NG = 0
DB = 0
SM = 0
Delta_L
Doxorubicin
YDL227C_RF1_296 RF1
ZShift = −0.33
lm ZScore = 0.65
NG = 0
DB = 0
SM = 0
Delta_K
Doxorubicin
YDL227C_RF1_296 RF1
ZShift = −0.9
lm ZScore = −0.53
NG = 0
DB = 0
SM = 0
Delta_r
Doxorubicin
YDL227C_RF1_296 RF1
ZShift = −0.63
lm ZScore = 0.72
NG = 0
DB = 0
SM = 0
Delta_AUC
Doxorubicin
YDL227C_RF1_272 RF1
ZShift = 0.74
lm Zscore = 0.64
NG = 0
DB = 0
SM = 0
Delta_L
Doxorubicin
YDL227C_RF1_272 RF1
ZShift = 0.34
lm ZScore = 0.64
NG = 0
DB = 0
SM = 0
Delta_K
Doxorubicin
YDL227C_RF1_272 RF1
ZShift = −0.25
lm ZScore = 0.02
NG = 0
DB = 0
SM = 0
Delta_r
Doxorubicin
YDL227C_RF1_272 RF1
ZShift = 0.14
lm ZScore = −0.16
NG = 0
DB = 0
SM = 0
Delta_AUC
Doxorubicin
YDL227C_RF1_57 RF1
ZShift = −0.61
lm Zscore = 0.64
NG = 0
DB = 0
SM = 0
Delta_L
Doxorubicin
YDL227C_RF1_57 RF1
ZShift = −0.22
lm ZScore = 0.64
NG = 0
DB = 0
SM = 0
Delta_K
Doxorubicin
YDL227C_RF1_57 RF1
ZShift = −0.13
lm ZScore = 0.39
NG = 0
DB = 0
SM = 0
Delta_r
Doxorubicin
YDL227C_RF1_57 RF1
ZShift = −0.06
lm ZScore = 0.01
NG = 0
DB = 0
SM = 0
Delta_AUC
Doxorubicin

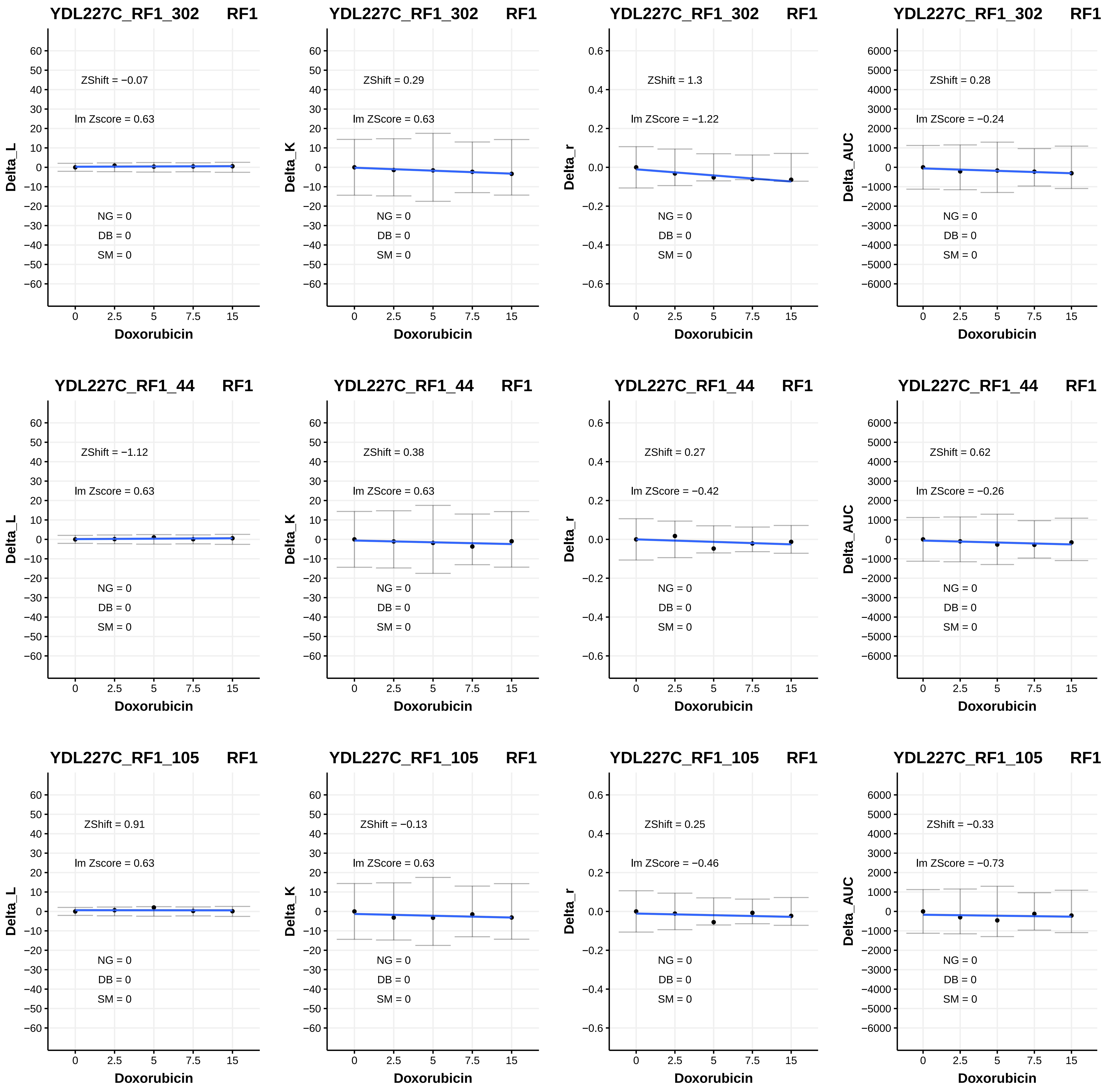
YDL227C_RF1_302 RF1
ZShift = −0.07
lm Zscore = 0.63
NG = 0
DB = 0
SM = 0
Delta_L
Doxorubicin
YDL227C_RF1_302 RF1
ZShift = 0.29
lm ZScore = 0.63
NG = 0
DB = 0
SM = 0
Delta_K
Doxorubicin
YDL227C_RF1_302 RF1
ZShift = 1.3
lm ZScore = −1.22
NG = 0
DB = 0
SM = 0
Delta_r
Doxorubicin
YDL227C_RF1_302 RF1
ZShift = 0.28
lm ZScore = −0.24
NG = 0
DB = 0
SM = 0
Delta_AUC
Doxorubicin
YDL227C_RF1_44 RF1
ZShift = −1.12
lm Zscore = 0.63
NG = 0
DB = 0
SM = 0
Delta_L
Doxorubicin
YDL227C_RF1_44 RF1
ZShift = 0.38
lm ZScore = 0.63
NG = 0
DB = 0
SM = 0
Delta_K
Doxorubicin
YDL227C_RF1_44 RF1
ZShift = 0.27
lm ZScore = −0.42
NG = 0
DB = 0
SM = 0
Delta_r
Doxorubicin
YDL227C_RF1_44 RF1
ZShift = 0.62
lm ZScore = −0.26
NG = 0
DB = 0
SM = 0
Delta_AUC
Doxorubicin
YDL227C_RF1_105 RF1
ZShift = 0.91
lm Zscore = 0.63
NG = 0
DB = 0
SM = 0
Delta_L
Doxorubicin
YDL227C_RF1_105 RF1
ZShift = −0.13
lm ZScore = 0.63
NG = 0
DB = 0
SM = 0
Delta_K
Doxorubicin
YDL227C_RF1_105 RF1
ZShift = 0.25
lm ZScore = −0.46
NG = 0
DB = 0
SM = 0
Delta_r
Doxorubicin
YDL227C_RF1_105 RF1
ZShift = −0.33
lm ZScore = −0.73
NG = 0
DB = 0
SM = 0
Delta_AUC
Doxorubicin

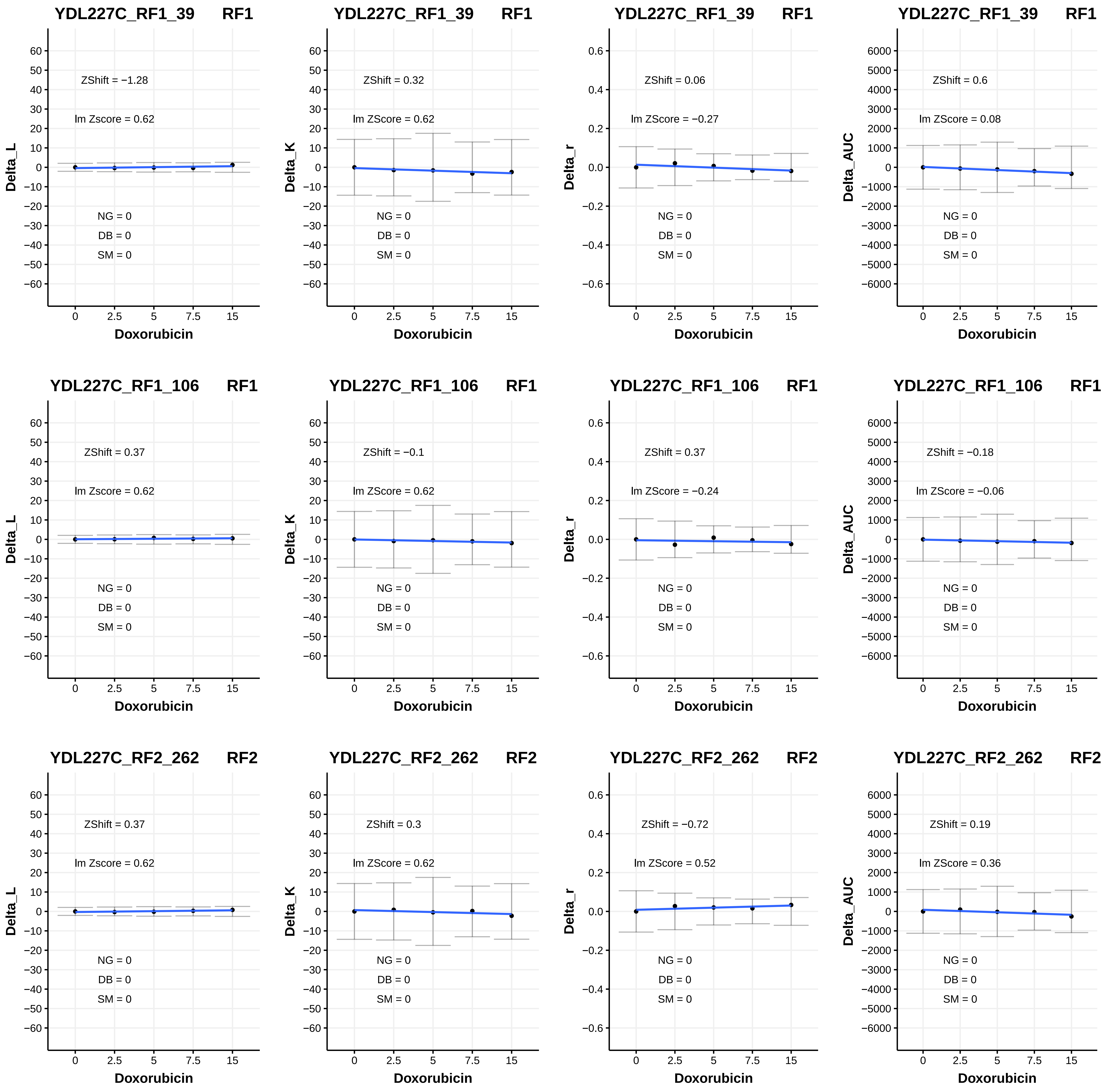

YDL227C_RF1_39 RF1
ZShift = −1.28
lm Zscore = 0.62
NG = 0
DB = 0
SM = 0
Delta_L
Doxorubicin
YDL227C_RF1_39 RF1
ZShift = 0.32
lm ZScore = 0.62
NG = 0
DB = 0
SM = 0
Delta_K
Doxorubicin
YDL227C_RF1_39 RF1
ZShift = 0.06
lm ZScore = −0.27
NG = 0
DB = 0
SM = 0
Delta_r
Doxorubicin
YDL227C_RF1_39 RF1
ZShift = 0.6
lm ZScore = 0.08
NG = 0
DB = 0
SM = 0
Delta_AUC
Doxorubicin
YDL227C_RF1_106 RF1
ZShift = 0.37
lm Zscore = 0.62
NG = 0
DB = 0
SM = 0
Delta_L
Doxorubicin
YDL227C_RF1_106 RF1
ZShift = −0.1
lm ZScore = 0.62
NG = 0
DB = 0
SM = 0
Delta_K
Doxorubicin
YDL227C_RF1_106 RF1
ZShift = 0.37
lm ZScore = −0.24
NG = 0
DB = 0
SM = 0
Delta_r
Doxorubicin
YDL227C_RF1_106 RF1
ZShift = −0.18
lm ZScore = −0.06
NG = 0
DB = 0
SM = 0
Delta_AUC
Doxorubicin
YDL227C_RF2_262 RF2
ZShift = 0.37
lm Zscore = 0.62
NG = 0
DB = 0
SM = 0
Delta_L
Doxorubicin
YDL227C_RF2_262 RF2
ZShift = 0.3
lm ZScore = 0.62
NG = 0
DB = 0
SM = 0
Delta_K
Doxorubicin
YDL227C_RF2_262 RF2
ZShift = −0.72
lm ZScore = 0.52
NG = 0
DB = 0
SM = 0
Delta_r
Doxorubicin
YDL227C_RF2_262 RF2
ZShift = 0.19
lm ZScore = 0.36
NG = 0
DB = 0
SM = 0
Delta_AUC
Doxorubicin

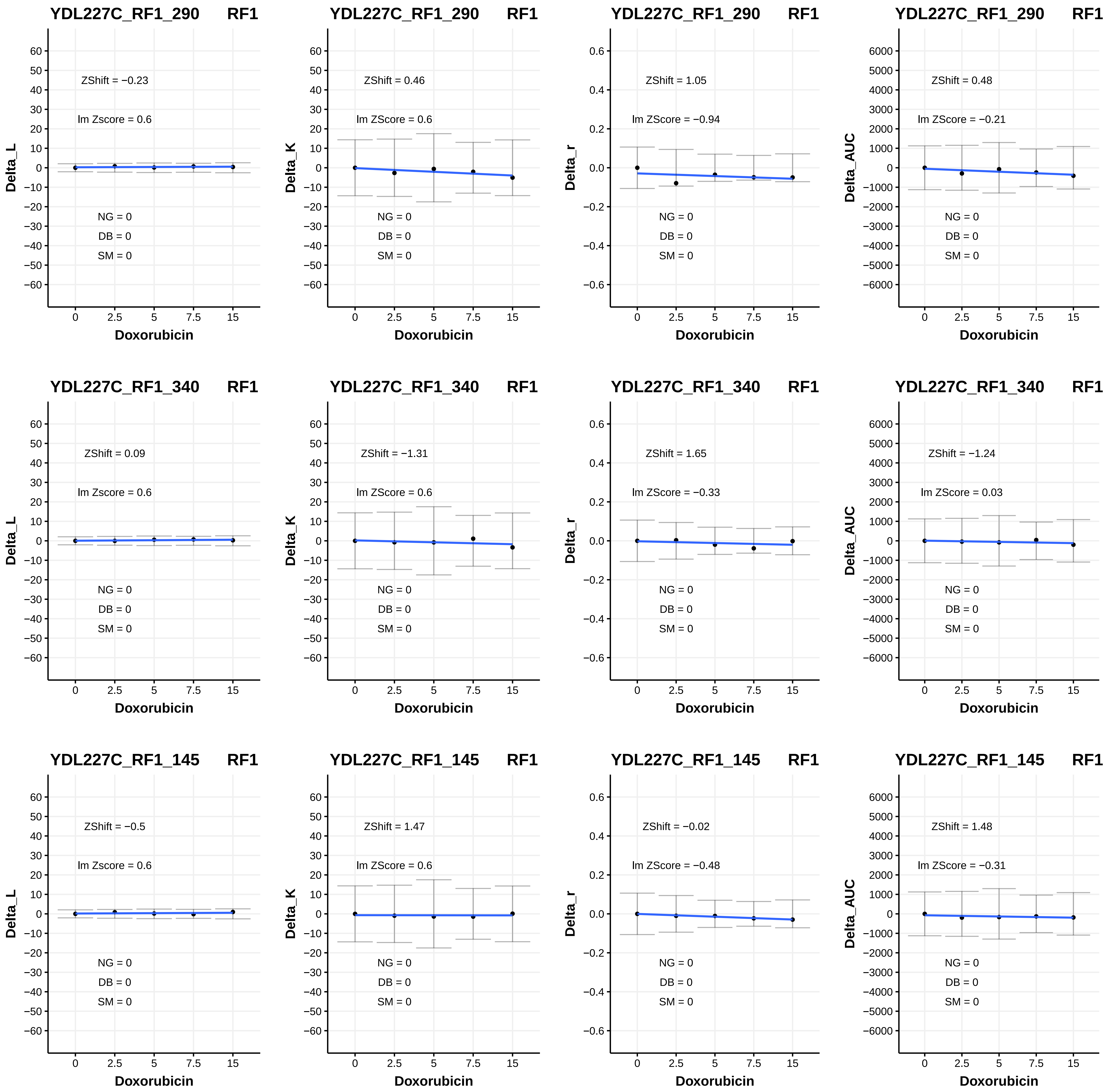
YDL227C_RF1_290 RF1
ZShift = −0.23
lm Zscore = 0.6
NG = 0
DB = 0
SM = 0
Delta_L
Doxorubicin
YDL227C_RF1_290 RF1
ZShift = 0.46
lm ZScore = 0.6
NG = 0
DB = 0
SM = 0
Delta_K
Doxorubicin
YDL227C_RF1_290 RF1
ZShift = 1.05
lm ZScore = −0.94
NG = 0
DB = 0
SM = 0
Delta_r
Doxorubicin
YDL227C_RF1_290 RF1
ZShift = 0.48
lm ZScore = −0.21
NG = 0
DB = 0
SM = 0
Delta_AUC
Doxorubicin
YDL227C_RF1_340 RF1
ZShift = 0.09
lm Zscore = 0.6
NG = 0
DB = 0
SM = 0
Delta_L
Doxorubicin
YDL227C_RF1_340 RF1
ZShift = −1.31
lm ZScore = 0.6
NG = 0
DB = 0
SM = 0
Delta_K
Doxorubicin
YDL227C_RF1_340 RF1
ZShift = 1.65
lm ZScore = −0.33
NG = 0
DB = 0
SM = 0
Delta_r
Doxorubicin
YDL227C_RF1_340 RF1
ZShift = −1.24
lm ZScore = 0.03
NG = 0
DB = 0
SM = 0
Delta_AUC
Doxorubicin
YDL227C_RF1_145 RF1
ZShift = −0.5
lm Zscore = 0.6
NG = 0
DB = 0
SM = 0
Delta_L
Doxorubicin
YDL227C_RF1_145 RF1
ZShift = 1.47
lm ZScore = 0.6
NG = 0
DB = 0
SM = 0
Delta_K
Doxorubicin
YDL227C_RF1_145 RF1
ZShift = −0.02
lm ZScore = −0.48
NG = 0
DB = 0
SM = 0
Delta_r
Doxorubicin
YDL227C_RF1_145 RF1
ZShift = 1.48
lm ZScore = −0.31
NG = 0
DB = 0
SM = 0
Delta_AUC
Doxorubicin

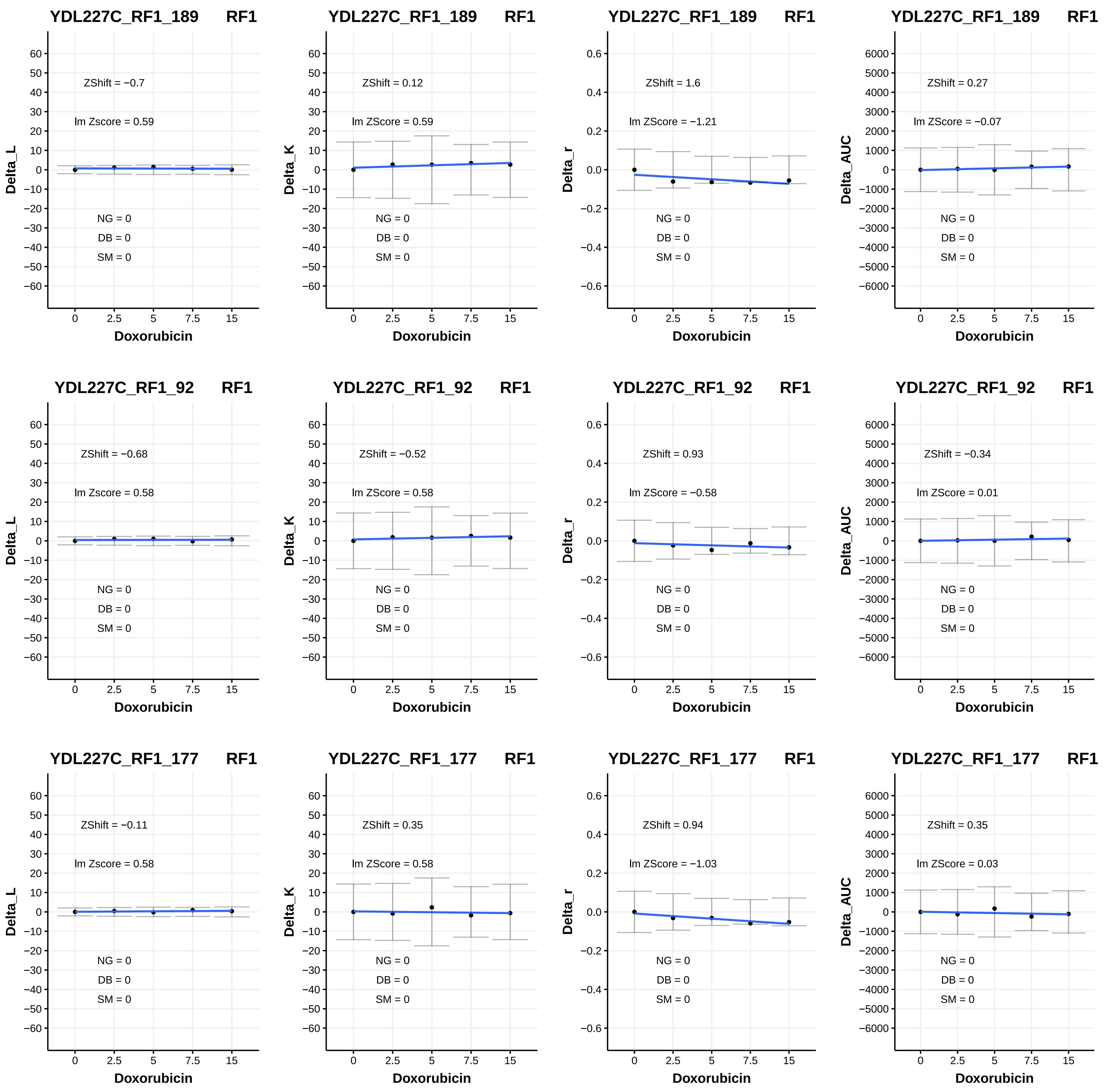
YDL227C_RF1_189 RF1
Delta_L
ZShift = −0.7
lm Zscore = 0.59
NG = 0
DB = 0
SM = 0
Doxorubicin
YDL227C_RF1_189 RF1
Delta_K
ZShift = 0.12
lm ZScore = 0.59
NG = 0
DB = 0
SM = 0
Doxorubicin
YDL227C_RF1_189 RF1
Delta_r
ZShift = 1.6
lm ZScore = −1.21
NG = 0
DB = 0
SM = 0
Doxorubicin
YDL227C_RF1_189 RF1
Delta_AUC
ZShift = 0.27
lm ZScore = −0.07
NG = 0
DB = 0
SM = 0
Doxorubicin
YDL227C_RF1_92 RF1
Delta_L
ZShift = −0.68
lm Zscore = 0.58
NG = 0
DB = 0
SM = 0
Doxorubicin
YDL227C_RF1_92 RF1
Delta_K
ZShift = −0.52
lm ZScore = 0.58
NG = 0
DB = 0
SM = 0
Doxorubicin
YDL227C_RF1_92 RF1
Delta_r
ZShift = 0.93
lm ZScore = −0.58
NG = 0
DB = 0
SM = 0
Doxorubicin
YDL227C_RF1_92 RF1
Delta_AUC
ZShift = −0.34
lm ZScore = 0.01
NG = 0
DB = 0
SM = 0
Doxorubicin
YDL227C_RF1_177 RF1
Delta_L
ZShift = −0.11
lm Zscore = 0.58
NG = 0
DB = 0
SM = 0
Doxorubicin
YDL227C_RF1_177 RF1
Delta_K
ZShift = 0.35
lm ZScore = 0.58
NG = 0
DB = 0
SM = 0
Doxorubicin
YDL227C_RF1_177 RF1
Delta_r
ZShift = 0.94
lm ZScore = −1.03
NG = 0
DB = 0
SM = 0
Doxorubicin
YDL227C_RF1_177 RF1
Delta_AUC
ZShift = 0.35
lm ZScore = 0.03
NG = 0
DB = 0
SM = 0
Doxorubicin

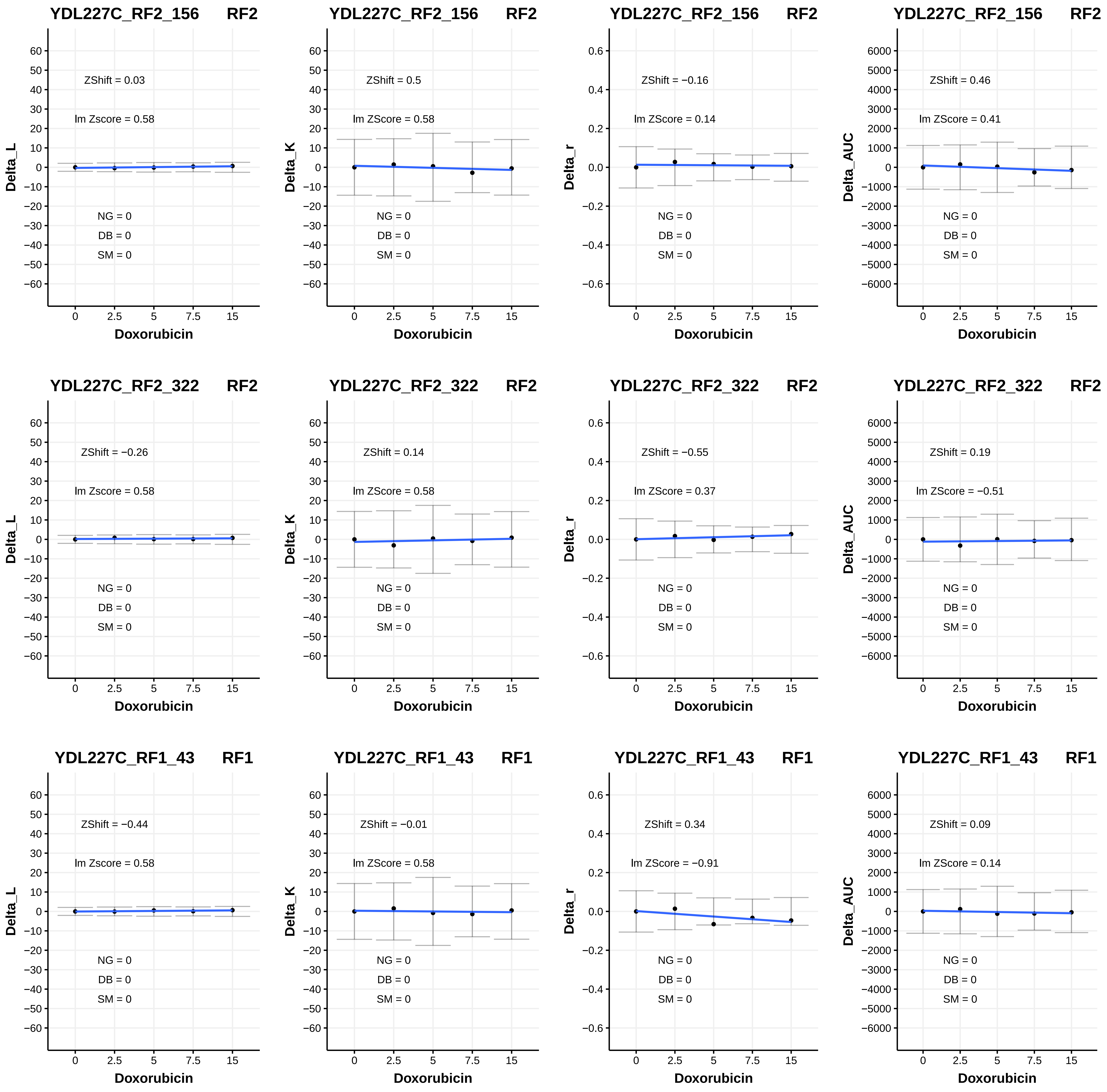
YDL227C_RF2_156 RF2
ZShift = 0.03
lm Zscore = 0.58
NG = 0
DB = 0
SM = 0
Delta_L
Doxorubicin
YDL227C_RF2_156 RF2
ZShift = 0.5
lm ZScore = 0.58
NG = 0
DB = 0
SM = 0
Delta_K
Doxorubicin
YDL227C_RF2_156 RF2
ZShift = −0.16
lm ZScore = 0.14
NG = 0
DB = 0
SM = 0
Delta_r
Doxorubicin
YDL227C_RF2_156 RF2
ZShift = 0.46
lm ZScore = 0.41
NG = 0
DB = 0
SM = 0
Delta_AUC
Doxorubicin
YDL227C_RF2_322 RF2
ZShift = −0.26
lm Zscore = 0.58
NG = 0
DB = 0
SM = 0
Delta_L
Doxorubicin
YDL227C_RF2_322 RF2
ZShift = 0.14
lm ZScore = 0.58
NG = 0
DB = 0
SM = 0
Delta_K
Doxorubicin
YDL227C_RF2_322 RF2
ZShift = −0.55
lm ZScore = 0.37
NG = 0
DB = 0
SM = 0
Delta_r
Doxorubicin
YDL227C_RF2_322 RF2
ZShift = 0.19
lm ZScore = −0.51
NG = 0
DB = 0
SM = 0
Delta_AUC
Doxorubicin
YDL227C_RF1_43 RF1
ZShift = −0.44
lm Zscore = 0.58
NG = 0
DB = 0
SM = 0
Delta_L
Doxorubicin
YDL227C_RF1_43 RF1
ZShift = −0.01
lm ZScore = 0.58
NG = 0
DB = 0
SM = 0
Delta_K
Doxorubicin
YDL227C_RF1_43 RF1
ZShift = 0.34
lm ZScore = −0.91
NG = 0
DB = 0
SM = 0
Delta_r
Doxorubicin
YDL227C_RF1_43 RF1
ZShift = 0.09
lm ZScore = 0.14
NG = 0
DB = 0
SM = 0
Delta_AUC
Doxorubicin

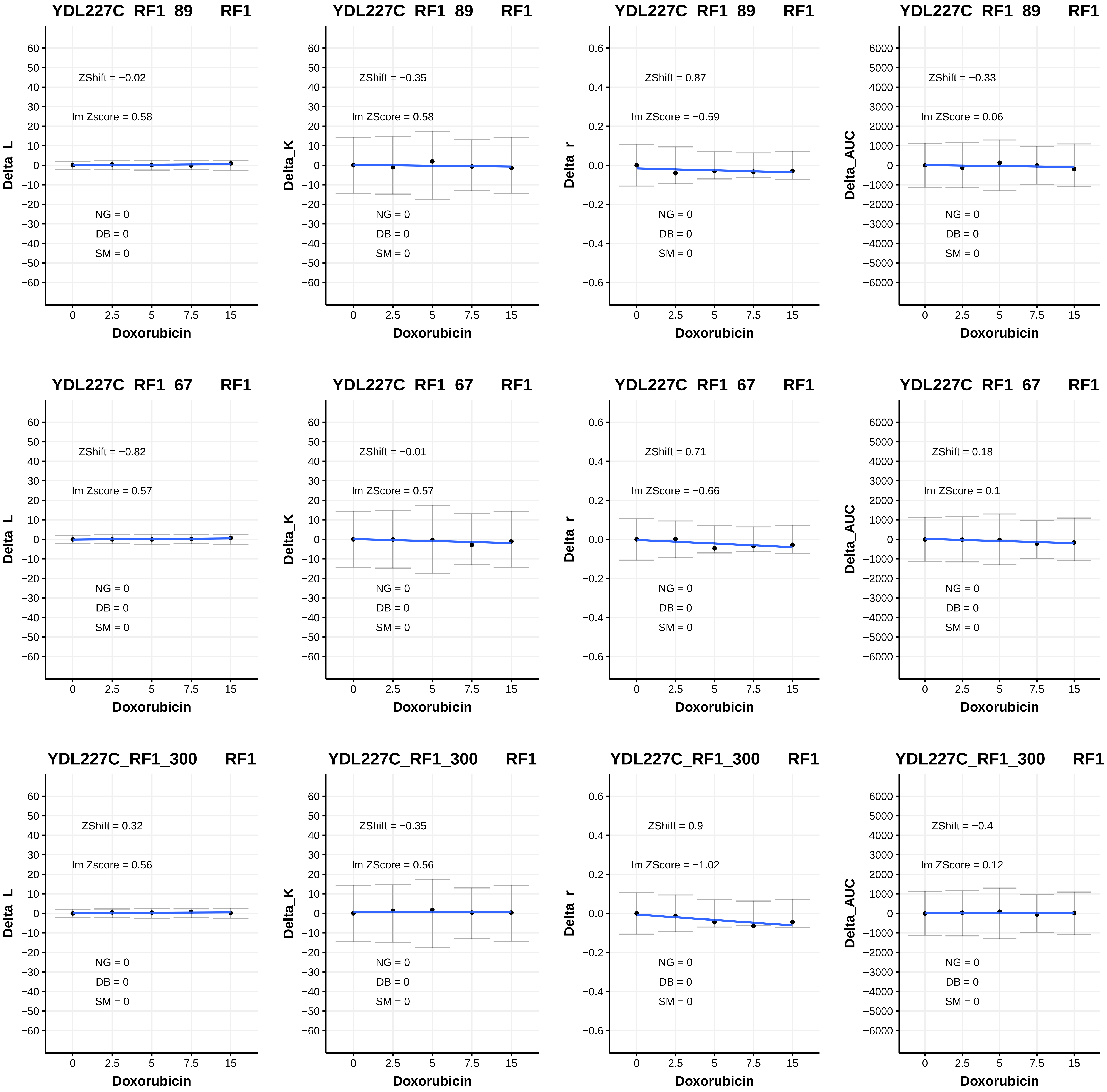

YDL227C_RF1_89 RF1
ZShift = −0.02
lm Zscore = 0.58
NG = 0
DB = 0
SM = 0
Delta_L
Doxorubicin
YDL227C_RF1_89 RF1
ZShift = −0.35
lm ZScore = 0.58
NG = 0
DB = 0
SM = 0
Delta_K
Doxorubicin
YDL227C_RF1_89 RF1
ZShift = 0.87
lm ZScore = −0.59
NG = 0
DB = 0
SM = 0
Delta_r
Doxorubicin
YDL227C_RF1_89 RF1
ZShift = −0.33
lm ZScore = 0.06
NG = 0
DB = 0
SM = 0
Delta_AUC
Doxorubicin
YDL227C_RF1_67 RF1
ZShift = −0.82
lm Zscore = 0.57
NG = 0
DB = 0
SM = 0
Delta_L
Doxorubicin
YDL227C_RF1_67 RF1
ZShift = −0.01
lm ZScore = 0.57
NG = 0
DB = 0
SM = 0
Delta_K
Doxorubicin
YDL227C_RF1_67 RF1
ZShift = 0.71
lm ZScore = −0.66
NG = 0
DB = 0
SM = 0
Delta_r
Doxorubicin
YDL227C_RF1_67 RF1
ZShift = 0.18
lm ZScore = 0.1
NG = 0
DB = 0
SM = 0
Delta_AUC
Doxorubicin
YDL227C_RF1_300 RF1
ZShift = 0.32
lm Zscore = 0.56
NG = 0
DB = 0
SM = 0
Delta_L
Doxorubicin
YDL227C_RF1_300 RF1
ZShift = −0.35
lm ZScore = 0.56
NG = 0
DB = 0
SM = 0
Delta_K
Doxorubicin
YDL227C_RF1_300 RF1
ZShift = 0.9
lm ZScore = −1.02
NG = 0
DB = 0
SM = 0
Delta_r
Doxorubicin
YDL227C_RF1_300 RF1
ZShift = −0.4
lm ZScore = 0.12
NG = 0
DB = 0
SM = 0
Delta_AUC
Doxorubicin

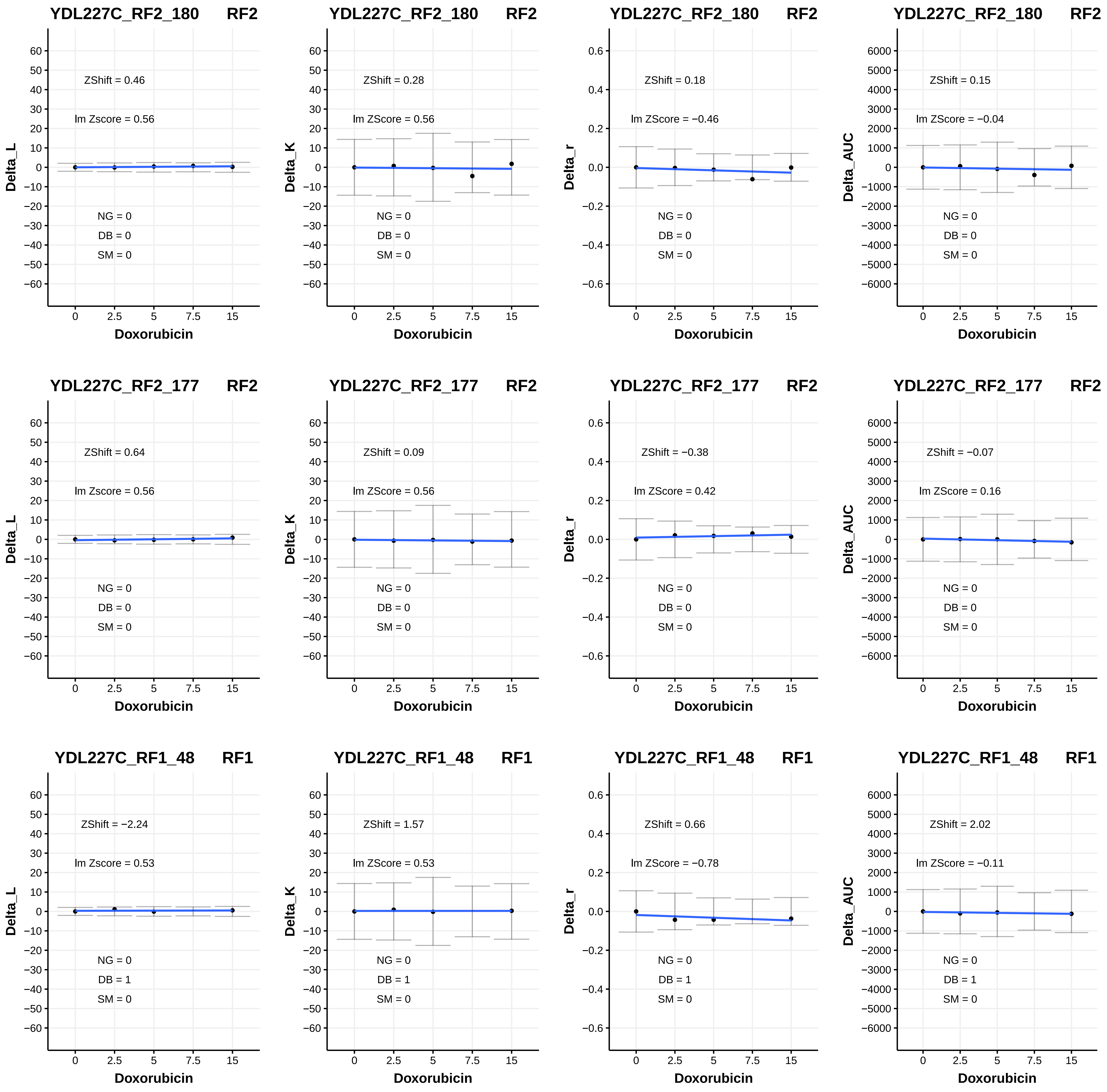

YDL227C_RF2_180 RF2
ZShift = 0.46
lm Zscore = 0.56
NG = 0
DB = 0
SM = 0
Delta_L
Doxorubicin
YDL227C_RF2_180 RF2
ZShift = 0.28
lm ZScore = 0.56
NG = 0
DB = 0
SM = 0
Delta_K
Doxorubicin
YDL227C_RF2_180 RF2
ZShift = 0.18
lm ZScore = −0.46
NG = 0
DB = 0
SM = 0
Delta_r
Doxorubicin
YDL227C_RF2_180 RF2
ZShift = 0.15
lm ZScore = −0.04
NG = 0
DB = 0
SM = 0
Delta_AUC
Doxorubicin
YDL227C_RF2_177 RF2
ZShift = 0.64
lm Zscore = 0.56
NG = 0
DB = 0
SM = 0
Delta_L
Doxorubicin
YDL227C_RF2_177 RF2
ZShift = 0.09
lm ZScore = 0.56
NG = 0
DB = 0
SM = 0
Delta_K
Doxorubicin
YDL227C_RF2_177 RF2
ZShift = −0.38
lm ZScore = 0.42
NG = 0
DB = 0
SM = 0
Delta_r
Doxorubicin
YDL227C_RF2_177 RF2
ZShift = −0.07
lm ZScore = 0.16
NG = 0
DB = 0
SM = 0
Delta_AUC
Doxorubicin
YDL227C_RF1_48 RF1
ZShift = −2.24
lm Zscore = 0.53
NG = 0
DB = 1
SM = 0
Delta_L
Doxorubicin
YDL227C_RF1_48 RF1
ZShift = 1.57
lm ZScore = 0.53
NG = 0
DB = 1
SM = 0
Delta_K
Doxorubicin
YDL227C_RF1_48 RF1
ZShift = 0.66
lm ZScore = −0.78
NG = 0
DB = 1
SM = 0
Delta_r
Doxorubicin
YDL227C_RF1_48 RF1
ZShift = 2.02
lm ZScore = −0.11
NG = 0
DB = 1
SM = 0
Delta_AUC
Doxorubicin

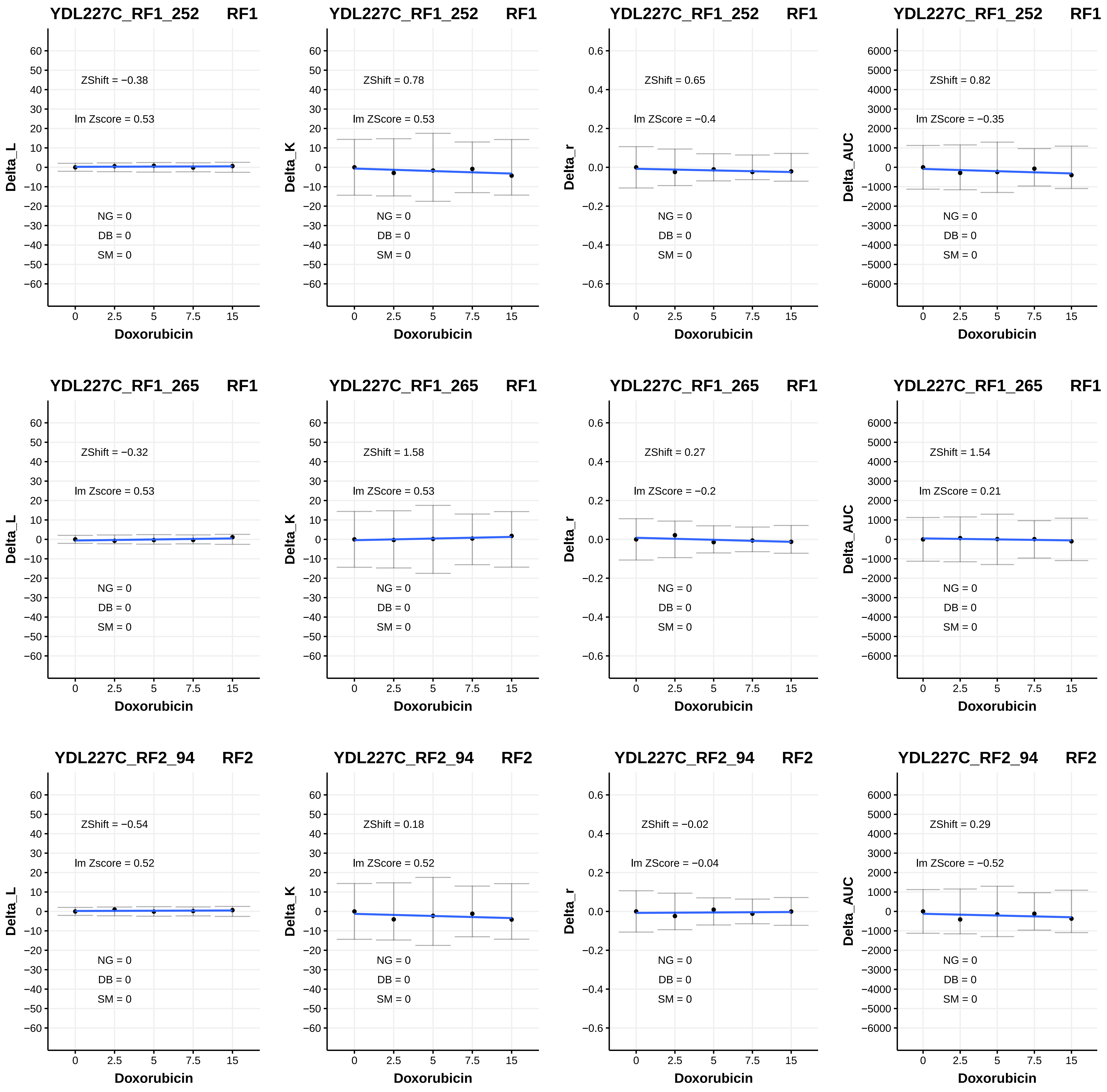

YDL227C_RF1_252 RF1
ZShift = −0.38
lm Zscore = 0.53
NG = 0
DB = 0
SM = 0
Delta_L
Doxorubicin
YDL227C_RF1_252 RF1
ZShift = 0.78
lm ZScore = 0.53
NG = 0
DB = 0
SM = 0
Delta_K
Doxorubicin
YDL227C_RF1_252 RF1
ZShift = 0.65
lm ZScore = −0.4
NG = 0
DB = 0
SM = 0
Delta_r
Doxorubicin
YDL227C_RF1_252 RF1
ZShift = 0.82
lm ZScore = −0.35
NG = 0
DB = 0
SM = 0
Delta_AUC
Doxorubicin
YDL227C_RF1_265 RF1
ZShift = −0.32
lm Zscore = 0.53
NG = 0
DB = 0
SM = 0
Delta_L
Doxorubicin
YDL227C_RF1_265 RF1
ZShift = 1.58
lm ZScore = 0.53
NG = 0
DB = 0
SM = 0
Delta_K
Doxorubicin
YDL227C_RF1_265 RF1
ZShift = 0.27
lm ZScore = −0.2
NG = 0
DB = 0
SM = 0
Delta_r
Doxorubicin
YDL227C_RF1_265 RF1
ZShift = 1.54
lm ZScore = 0.21
NG = 0
DB = 0
SM = 0
Delta_AUC
Doxorubicin
YDL227C_RF2_94 RF2
ZShift = −0.54
lm Zscore = 0.52
NG = 0
DB = 0
SM = 0
Delta_L
Doxorubicin
YDL227C_RF2_94 RF2
ZShift = 0.18
lm ZScore = 0.52
NG = 0
DB = 0
SM = 0
Delta_K
Doxorubicin
YDL227C_RF2_94 RF2
ZShift = −0.02
lm ZScore = −0.04
NG = 0
DB = 0
SM = 0
Delta_r
Doxorubicin
YDL227C_RF2_94 RF2
ZShift = 0.29
lm ZScore = −0.52
NG = 0
DB = 0
SM = 0
Delta_AUC
Doxorubicin

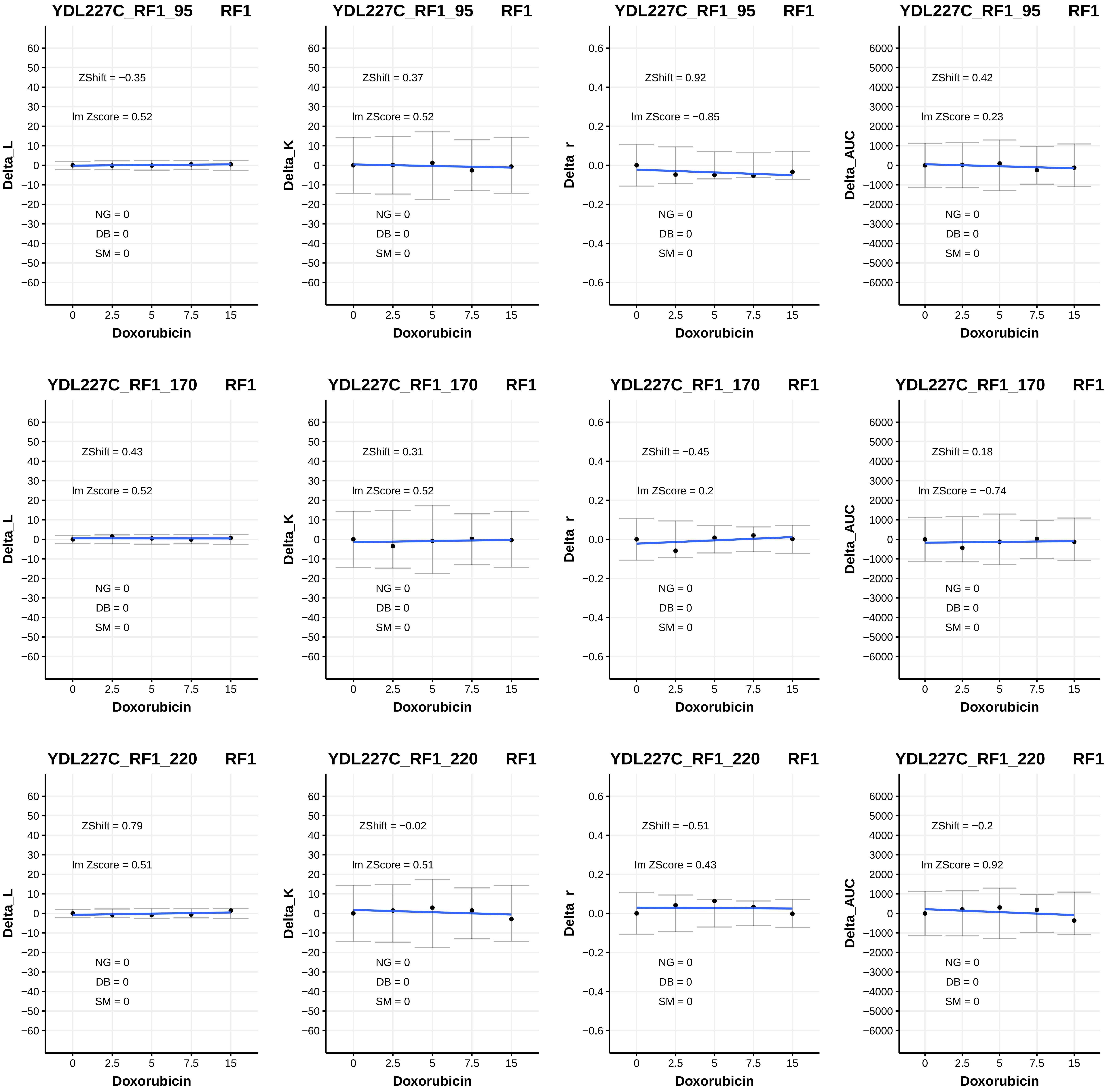

YDL227C_RF1_95 RF1
ZShift = −0.35
lm Zscore = 0.52
NG = 0
DB = 0
SM = 0
Delta_L
Doxorubicin
YDL227C_RF1_95 RF1
ZShift = 0.37
lm ZScore = 0.52
NG = 0
DB = 0
SM = 0
Delta_K
Doxorubicin
YDL227C_RF1_95 RF1
ZShift = 0.92
lm ZScore = −0.85
NG = 0
DB = 0
SM = 0
Delta_r
Doxorubicin
YDL227C_RF1_95 RF1
ZShift = 0.42
lm ZScore = 0.23
NG = 0
DB = 0
SM = 0
Delta_AUC
Doxorubicin
YDL227C_RF1_170 RF1
ZShift = 0.43
lm Zscore = 0.52
NG = 0
DB = 0
SM = 0
Delta_L
Doxorubicin
YDL227C_RF1_170 RF1
ZShift = 0.31
lm ZScore = 0.52
NG = 0
DB = 0
SM = 0
Delta_K
Doxorubicin
YDL227C_RF1_170 RF1
ZShift = −0.45
lm ZScore = 0.2
NG = 0
DB = 0
SM = 0
Delta_r
Doxorubicin
YDL227C_RF1_170 RF1
ZShift = 0.18
lm ZScore = −0.74
NG = 0
DB = 0
SM = 0
Delta_AUC
Doxorubicin
YDL227C_RF1_220 RF1
ZShift = 0.79
lm Zscore = 0.51
NG = 0
DB = 0
SM = 0
Delta_L
Doxorubicin
YDL227C_RF1_220 RF1
ZShift = −0.02
lm ZScore = 0.51
NG = 0
DB = 0
SM = 0
Delta_K
Doxorubicin
YDL227C_RF1_220 RF1
ZShift = −0.51
lm ZScore = 0.43
NG = 0
DB = 0
SM = 0
Delta_r
Doxorubicin
YDL227C_RF1_220 RF1
ZShift = −0.2
lm ZScore = 0.92
NG = 0
DB = 0
SM = 0
Delta_AUC
Doxorubicin

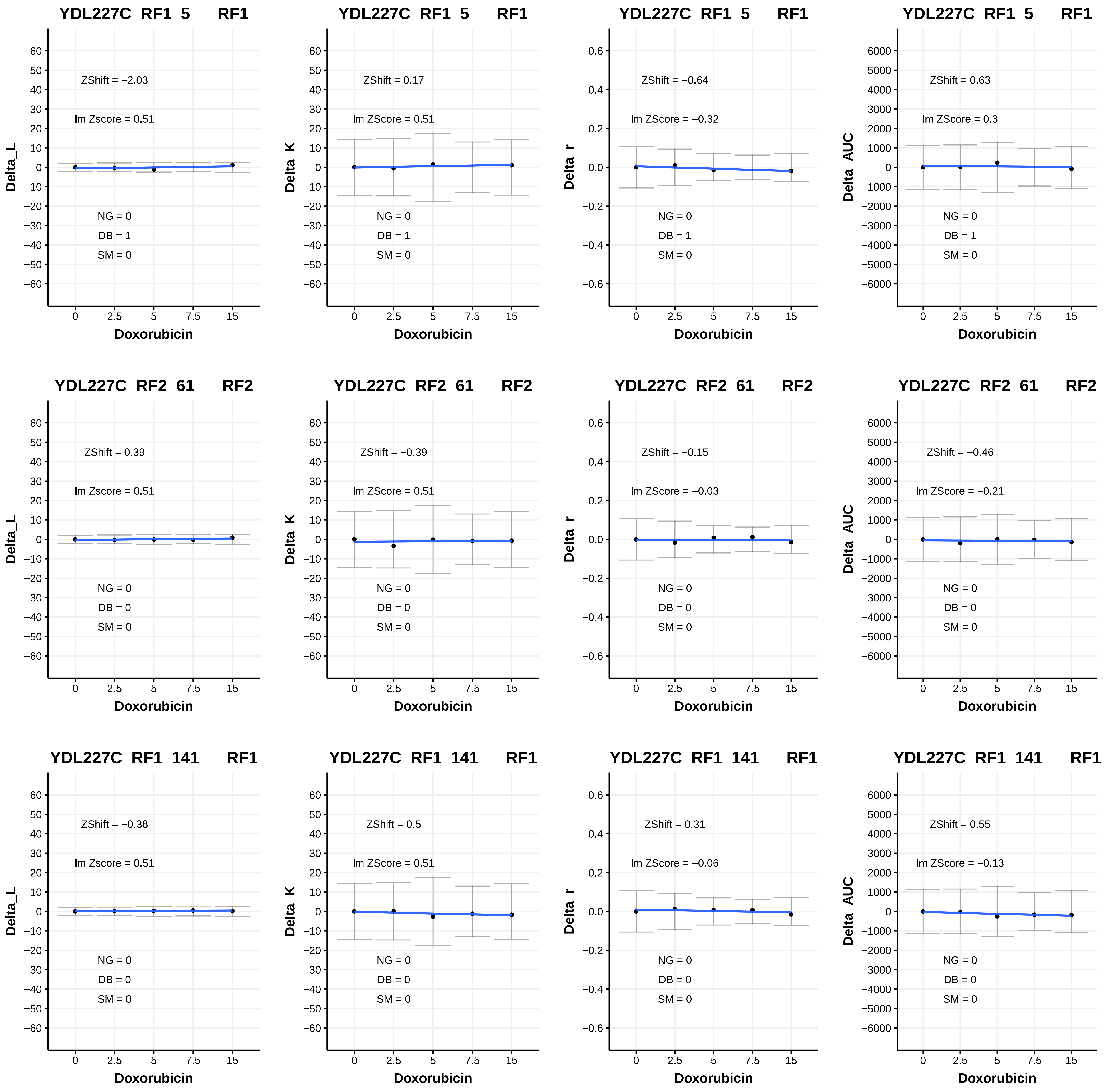
YDL227C_RF1_5 RF1
ZShift = −2.03
lm Zscore = 0.51
NG = 0
DB = 1
SM = 0
Delta_L
Doxorubicin
YDL227C_RF1_5 RF1
ZShift = 0.17
lm ZScore = 0.51
NG = 0
DB = 1
SM = 0
Delta_K
Doxorubicin
YDL227C_RF1_5 RF1
ZShift = −0.64
lm ZScore = −0.32
NG = 0
DB = 1
SM = 0
Delta_r
Doxorubicin
YDL227C_RF1_5 RF1
ZShift = 0.63
lm ZScore = 0.3
NG = 0
DB = 1
SM = 0
Delta_AUC
Doxorubicin
YDL227C_RF2_61 RF2
ZShift = 0.39
lm Zscore = 0.51
NG = 0
DB = 0
SM = 0
Delta_L
Doxorubicin
YDL227C_RF2_61 RF2
ZShift = −0.39
lm ZScore = 0.51
NG = 0
DB = 0
SM = 0
Delta_K
Doxorubicin
YDL227C_RF2_61 RF2
ZShift = −0.15
lm ZScore = −0.03
NG = 0
DB = 0
SM = 0
Delta_r
Doxorubicin
YDL227C_RF2_61 RF2
ZShift = −0.46
lm ZScore = −0.21
NG = 0
DB = 0
SM = 0
Delta_AUC
Doxorubicin
YDL227C_RF1_141 RF1
ZShift = −0.38
lm Zscore = 0.51
NG = 0
DB = 0
SM = 0
Delta_L
Doxorubicin
YDL227C_RF1_141 RF1
ZShift = 0.5
lm ZScore = 0.51
NG = 0
DB = 0
SM = 0
Delta_K
Doxorubicin
YDL227C_RF1_141 RF1
ZShift = 0.31
lm ZScore = −0.06
NG = 0
DB = 0
SM = 0
Delta_r
Doxorubicin
YDL227C_RF1_141 RF1
ZShift = 0.55
lm ZScore = −0.13
NG = 0
DB = 0
SM = 0
Delta_AUC
Doxorubicin

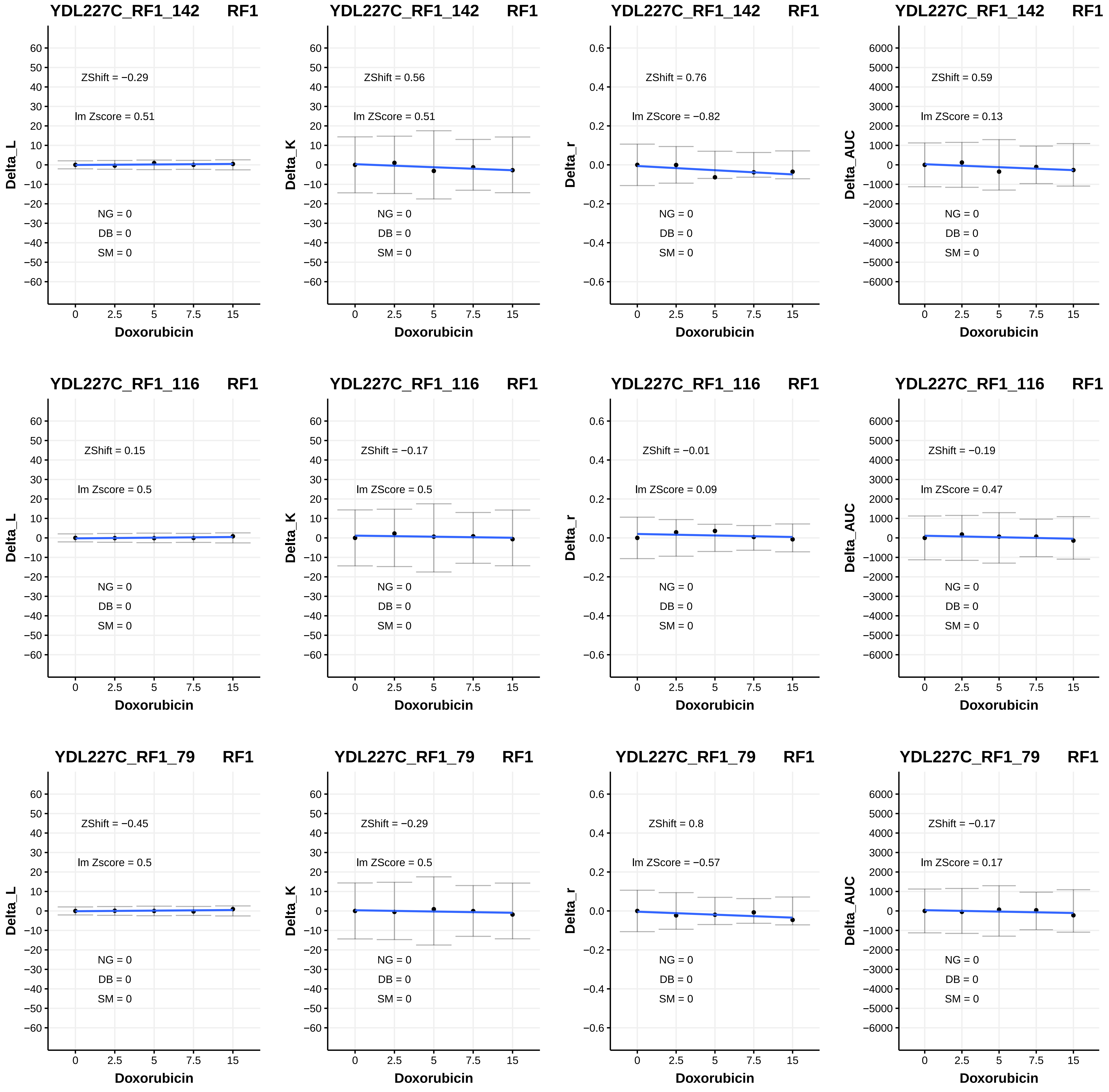
YDL227C_RF1_142 RF1
ZShift = −0.29
lm Zscore = 0.51
NG = 0
DB = 0
SM = 0
Delta_L
Doxorubicin
YDL227C_RF1_142 RF1
ZShift = 0.56
lm ZScore = 0.51
NG = 0
DB = 0
SM = 0
Delta_K
Doxorubicin
YDL227C_RF1_142 RF1
ZShift = 0.76
lm ZScore = −0.82
NG = 0
DB = 0
SM = 0
Delta_r
Doxorubicin
YDL227C_RF1_142 RF1
ZShift = 0.59
lm ZScore = 0.13
NG = 0
DB = 0
SM = 0
Delta_AUC
Doxorubicin
YDL227C_RF1_116 RF1
ZShift = 0.15
lm Zscore = 0.5
NG = 0
DB = 0
SM = 0
Delta_L
Doxorubicin
YDL227C_RF1_116 RF1
ZShift = −0.17
lm ZScore = 0.5
NG = 0
DB = 0
SM = 0
Delta_K
Doxorubicin
YDL227C_RF1_116 RF1
ZShift = −0.01
lm ZScore = 0.09
NG = 0
DB = 0
SM = 0
Delta_r
Doxorubicin
YDL227C_RF1_116 RF1
ZShift = −0.19
lm ZScore = 0.47
NG = 0
DB = 0
SM = 0
Delta_AUC
Doxorubicin
YDL227C_RF1_79 RF1
ZShift = −0.45
lm Zscore = 0.5
NG = 0
DB = 0
SM = 0
Delta_L
Doxorubicin
YDL227C_RF1_79 RF1
ZShift = −0.29
lm ZScore = 0.5
NG = 0
DB = 0
SM = 0
Delta_K
Doxorubicin
YDL227C_RF1_79 RF1
ZShift = 0.8
lm ZScore = −0.57
NG = 0
DB = 0
SM = 0
Delta_r
Doxorubicin
YDL227C_RF1_79 RF1
ZShift = −0.17
lm ZScore = 0.17
NG = 0
DB = 0
SM = 0
Delta_AUC
Doxorubicin

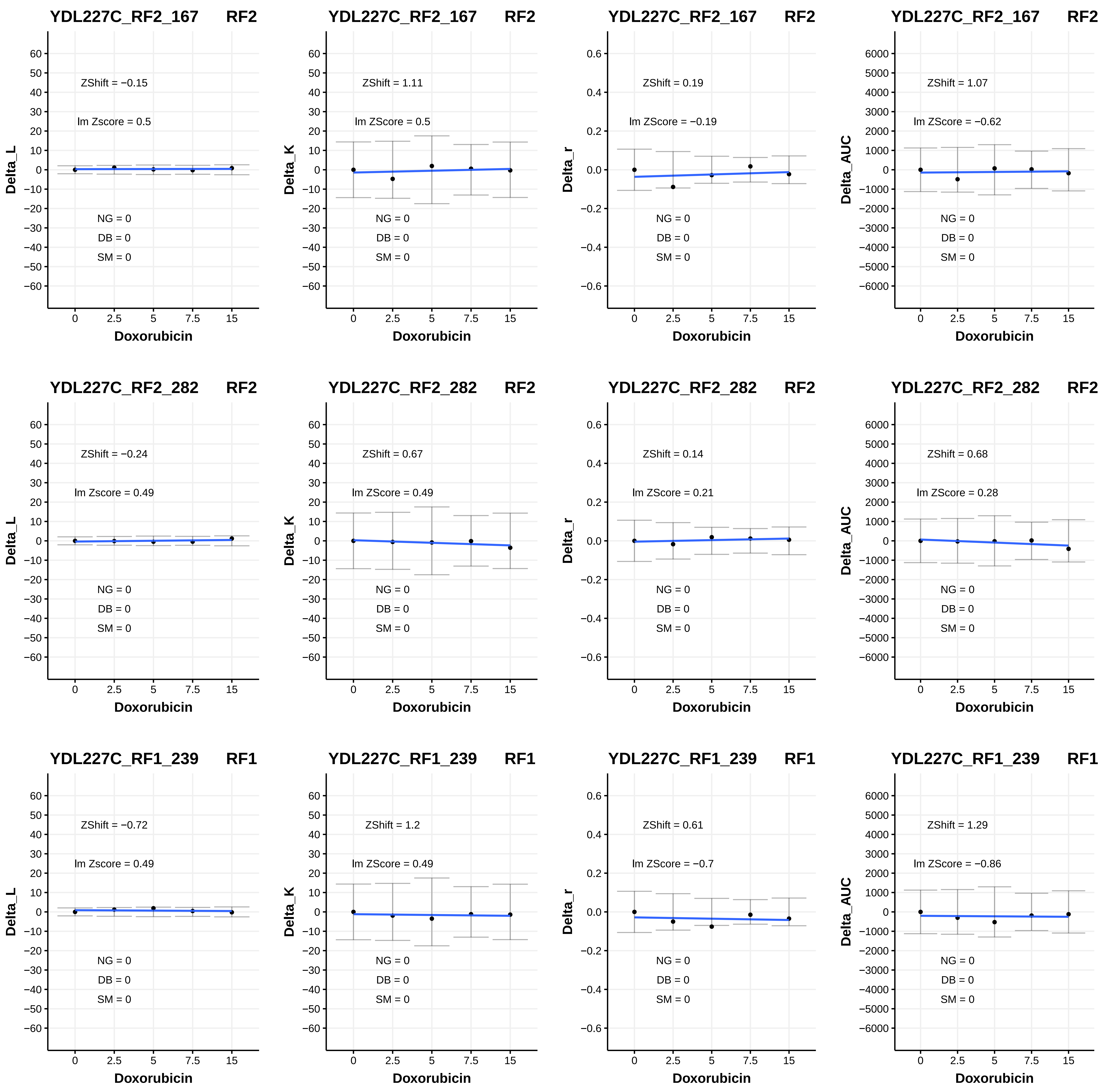

YDL227C_RF2_167 RF2
ZShift = −0.15
lm Zscore = 0.5
NG = 0
DB = 0
SM = 0
Delta_L
Doxorubicin
YDL227C_RF2_167 RF2
ZShift = 1.11
lm ZScore = 0.5
NG = 0
DB = 0
SM = 0
Delta_K
Doxorubicin
YDL227C_RF2_167 RF2
ZShift = 0.19
lm ZScore = −0.19
NG = 0
DB = 0
SM = 0
Delta_r
Doxorubicin
YDL227C_RF2_167 RF2
ZShift = 1.07
lm ZScore = −0.62
NG = 0
DB = 0
SM = 0
Delta_AUC
Doxorubicin
YDL227C_RF2_282 RF2
ZShift = −0.24
lm Zscore = 0.49
NG = 0
DB = 0
SM = 0
Delta_L
Doxorubicin
YDL227C_RF2_282 RF2
ZShift = 0.67
lm ZScore = 0.49
NG = 0
DB = 0
SM = 0
Delta_K
Doxorubicin
YDL227C_RF2_282 RF2
ZShift = 0.14
lm ZScore = 0.21
NG = 0
DB = 0
SM = 0
Delta_r
Doxorubicin
YDL227C_RF2_282 RF2
ZShift = 0.68
lm ZScore = 0.28
NG = 0
DB = 0
SM = 0
Delta_AUC
Doxorubicin
YDL227C_RF1_239 RF1
ZShift = −0.72
lm Zscore = 0.49
NG = 0
DB = 0
SM = 0
Delta_L
Doxorubicin
YDL227C_RF1_239 RF1
ZShift = 1.2
lm ZScore = 0.49
NG = 0
DB = 0
SM = 0
Delta_K
Doxorubicin
YDL227C_RF1_239 RF1
ZShift = 0.61
lm ZScore = −0.7
NG = 0
DB = 0
SM = 0
Delta_r
Doxorubicin
YDL227C_RF1_239 RF1
ZShift = 1.29
lm ZScore = −0.86
NG = 0
DB = 0
SM = 0
Delta_AUC
Doxorubicin

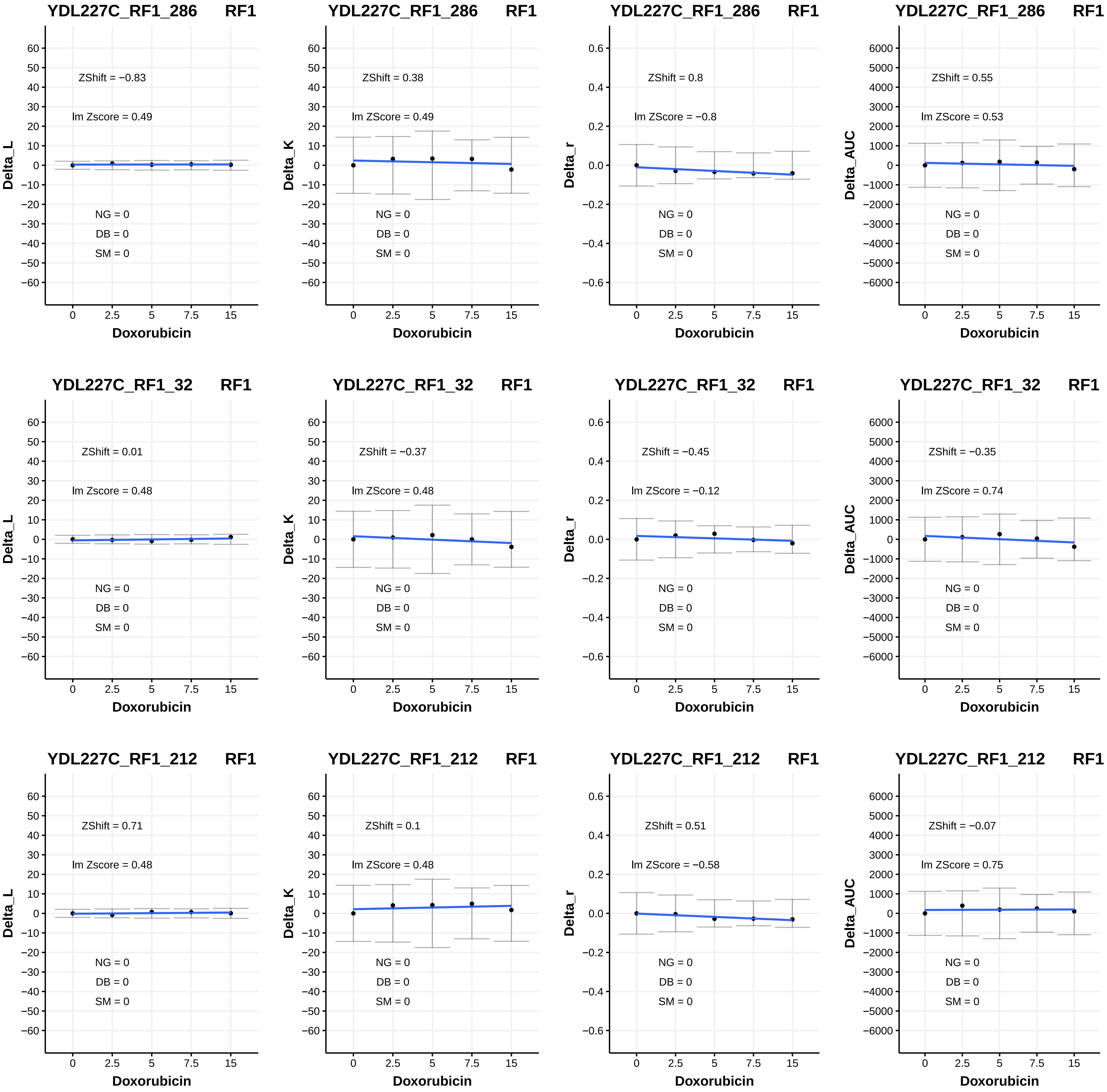
YDL227C_RF1_286 RF1
ZShift = −0.83
lm Zscore = 0.49
NG = 0
DB = 0
SM = 0
Delta_L
Doxorubicin
YDL227C_RF1_286 RF1
ZShift = 0.38
lm ZScore = 0.49
NG = 0
DB = 0
SM = 0
Delta_K
Doxorubicin
YDL227C_RF1_286 RF1
ZShift = 0.8
lm ZScore = −0.8
NG = 0
DB = 0
SM = 0
Delta_r
Doxorubicin
YDL227C_RF1_286 RF1
ZShift = 0.55
lm ZScore = 0.53
NG = 0
DB = 0
SM = 0
Delta_AUC
Doxorubicin
YDL227C_RF1_32 RF1
ZShift = 0.01
lm Zscore = 0.48
NG = 0
DB = 0
SM = 0
Delta_L
Doxorubicin
YDL227C_RF1_32 RF1
ZShift = −0.37
lm ZScore = 0.48
NG = 0
DB = 0
SM = 0
Delta_K
Doxorubicin
YDL227C_RF1_32 RF1
ZShift = −0.45
lm ZScore = −0.12
NG = 0
DB = 0
SM = 0
Delta_r
Doxorubicin
YDL227C_RF1_32 RF1
ZShift = −0.35
lm ZScore = 0.74
NG = 0
DB = 0
SM = 0
Delta_AUC
Doxorubicin
YDL227C_RF1_212 RF1
ZShift = 0.71
lm Zscore = 0.48
NG = 0
DB = 0
SM = 0
Delta_L
Doxorubicin
YDL227C_RF1_212 RF1
ZShift = 0.1
lm ZScore = 0.48
NG = 0
DB = 0
SM = 0
Delta_K
Doxorubicin
YDL227C_RF1_212 RF1
ZShift = 0.51
lm ZScore = −0.58
NG = 0
DB = 0
SM = 0
Delta_r
Doxorubicin
YDL227C_RF1_212 RF1
ZShift = −0.07
lm ZScore = 0.75
NG = 0
DB = 0
SM = 0
Delta_AUC
Doxorubicin

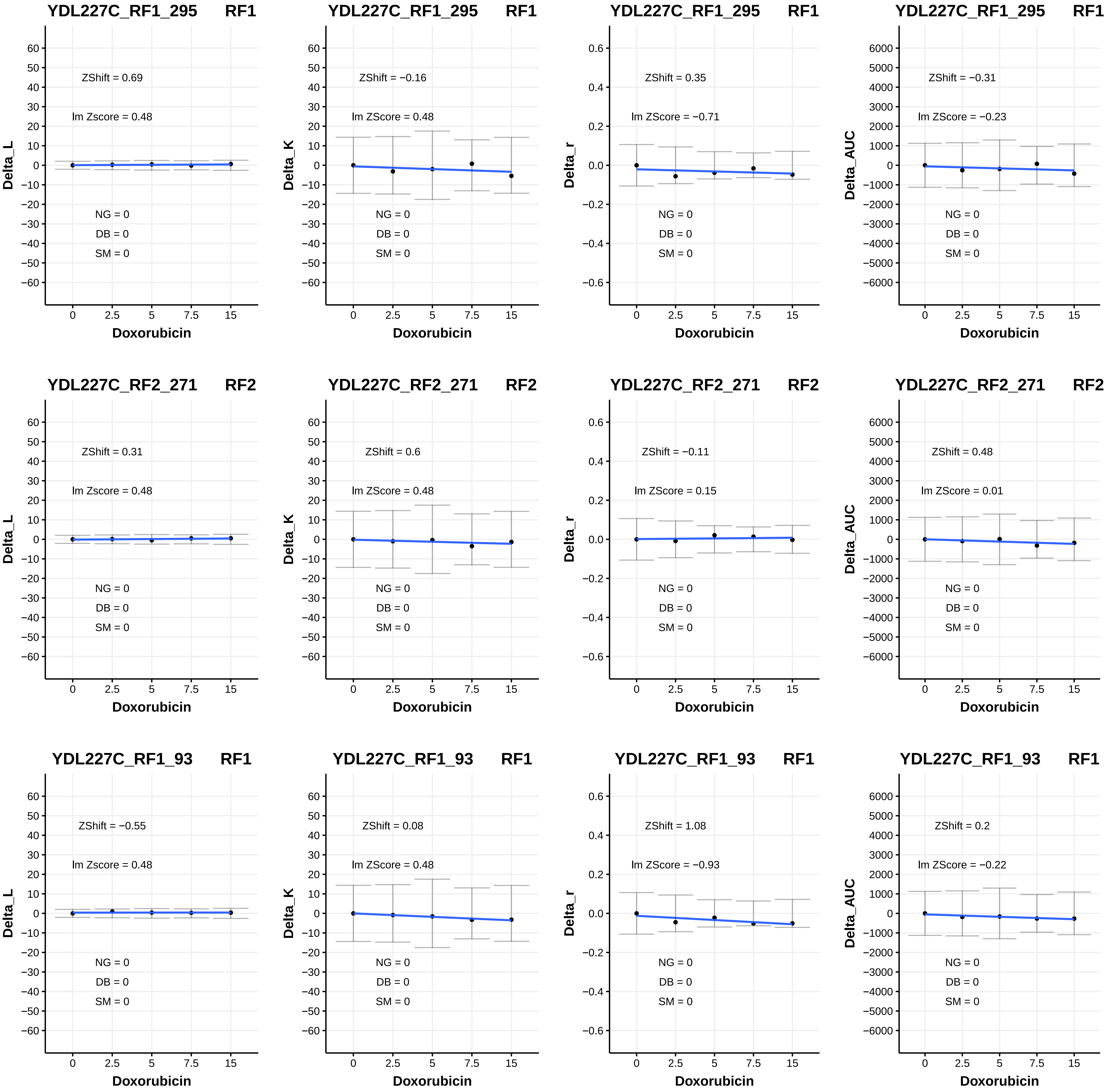

YDL227C_RF1_295 RF1
ZShift = 0.69
lm Zscore = 0.48
NG = 0
DB = 0
SM = 0
Delta_L
Doxorubicin
YDL227C_RF1_295 RF1
ZShift = −0.16
lm ZScore = 0.48
NG = 0
DB = 0
SM = 0
Delta_K
Doxorubicin
YDL227C_RF1_295 RF1
ZShift = 0.35
lm ZScore = −0.71
NG = 0
DB = 0
SM = 0
Delta_r
Doxorubicin
YDL227C_RF1_295 RF1
ZShift = −0.31
lm ZScore = −0.23
NG = 0
DB = 0
SM = 0
Delta_AUC
Doxorubicin
YDL227C_RF2_271 RF2
ZShift = 0.31
lm Zscore = 0.48
NG = 0
DB = 0
SM = 0
Delta_L
Doxorubicin
YDL227C_RF2_271 RF2
ZShift = 0.6
lm ZScore = 0.48
NG = 0
DB = 0
SM = 0
Delta_K
Doxorubicin
YDL227C_RF2_271 RF2
ZShift = −0.11
lm ZScore = 0.15
NG = 0
DB = 0
SM = 0
Delta_r
Doxorubicin
YDL227C_RF2_271 RF2
ZShift = 0.48
lm ZScore = 0.01
NG = 0
DB = 0
SM = 0
Delta_AUC
Doxorubicin
YDL227C_RF1_93 RF1
ZShift = −0.55
lm Zscore = 0.48
NG = 0
DB = 0
SM = 0
Delta_L
Doxorubicin
YDL227C_RF1_93 RF1
ZShift = 0.08
lm ZScore = 0.48
NG = 0
DB = 0
SM = 0
Delta_K
Doxorubicin
YDL227C_RF1_93 RF1
ZShift = 1.08
lm ZScore = −0.93
NG = 0
DB = 0
SM = 0
Delta_r
Doxorubicin
YDL227C_RF1_93 RF1
ZShift = 0.2
lm ZScore = −0.22
NG = 0
DB = 0
SM = 0
Delta_AUC
Doxorubicin

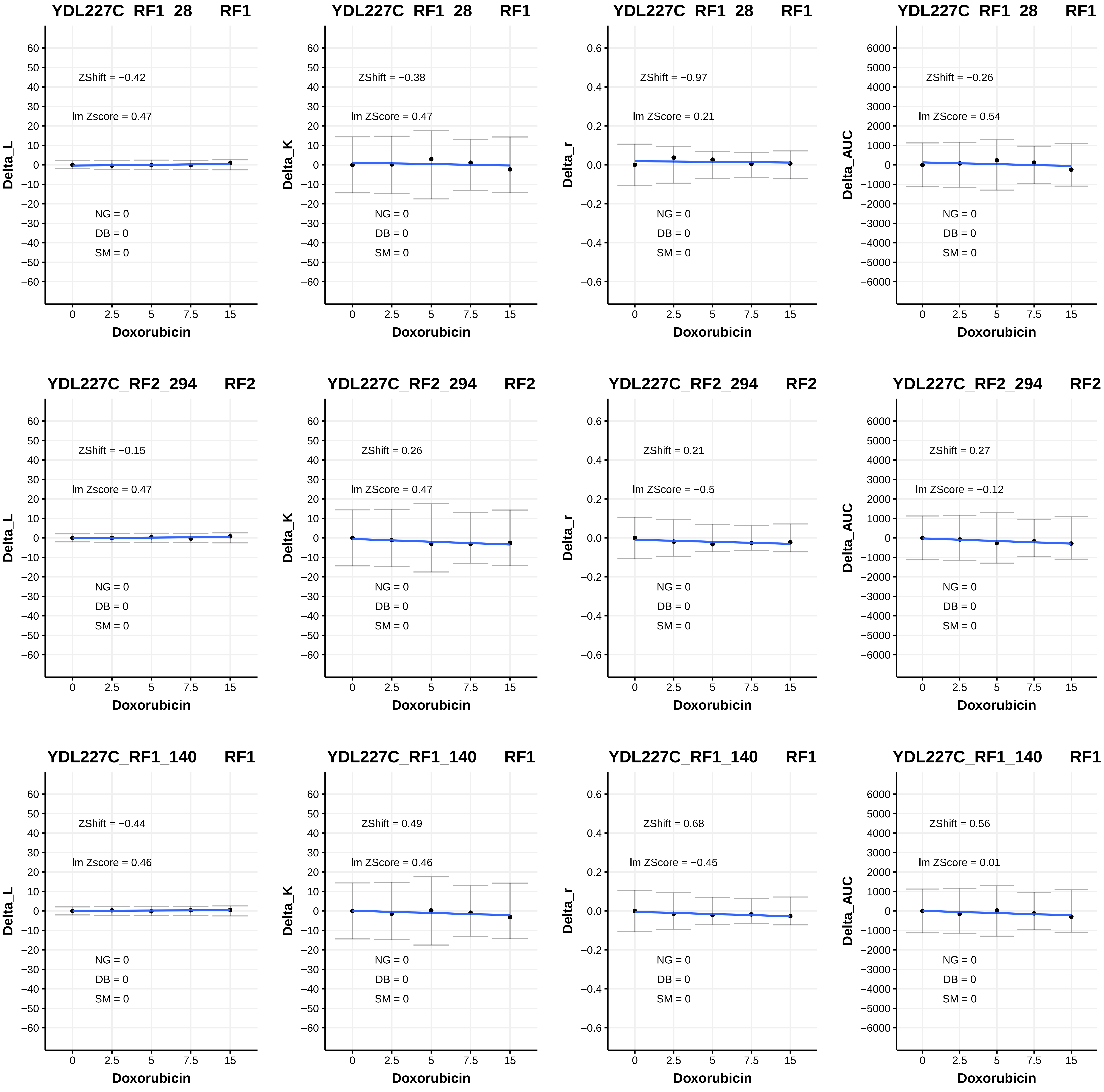

YDL227C_RF1_28 RF1
ZShift = −0.42
lm Zscore = 0.47
NG = 0
DB = 0
SM = 0
Delta_L
Doxorubicin
YDL227C_RF1_28 RF1
ZShift = −0.38
lm ZScore = 0.47
NG = 0
DB = 0
SM = 0
Delta_K
Doxorubicin
YDL227C_RF1_28 RF1
ZShift = −0.97
lm ZScore = 0.21
NG = 0
DB = 0
SM = 0
Delta_r
Doxorubicin
YDL227C_RF1_28 RF1
ZShift = −0.26
lm ZScore = 0.54
NG = 0
DB = 0
SM = 0
Delta_AUC
Doxorubicin
YDL227C_RF2_294 RF2
ZShift = −0.15
lm Zscore = 0.47
NG = 0
DB = 0
SM = 0
Delta_L
Doxorubicin
YDL227C_RF2_294 RF2
ZShift = 0.26
lm ZScore = 0.47
NG = 0
DB = 0
SM = 0
Delta_K
Doxorubicin
YDL227C_RF2_294 RF2
ZShift = 0.21
lm ZScore = −0.5
NG = 0
DB = 0
SM = 0
Delta_r
Doxorubicin
YDL227C_RF2_294 RF2
ZShift = 0.27
lm ZScore = −0.12
NG = 0
DB = 0
SM = 0
Delta_AUC
Doxorubicin
YDL227C_RF1_140 RF1
ZShift = −0.44
lm Zscore = 0.46
NG = 0
DB = 0
SM = 0
Delta_L
Doxorubicin
YDL227C_RF1_140 RF1
ZShift = 0.49
lm ZScore = 0.46
NG = 0
DB = 0
SM = 0
Delta_K
Doxorubicin
YDL227C_RF1_140 RF1
ZShift = 0.68
lm ZScore = −0.45
NG = 0
DB = 0
SM = 0
Delta_r
Doxorubicin
YDL227C_RF1_140 RF1
ZShift = 0.56
lm ZScore = 0.01
NG = 0
DB = 0
SM = 0
Delta_AUC
Doxorubicin

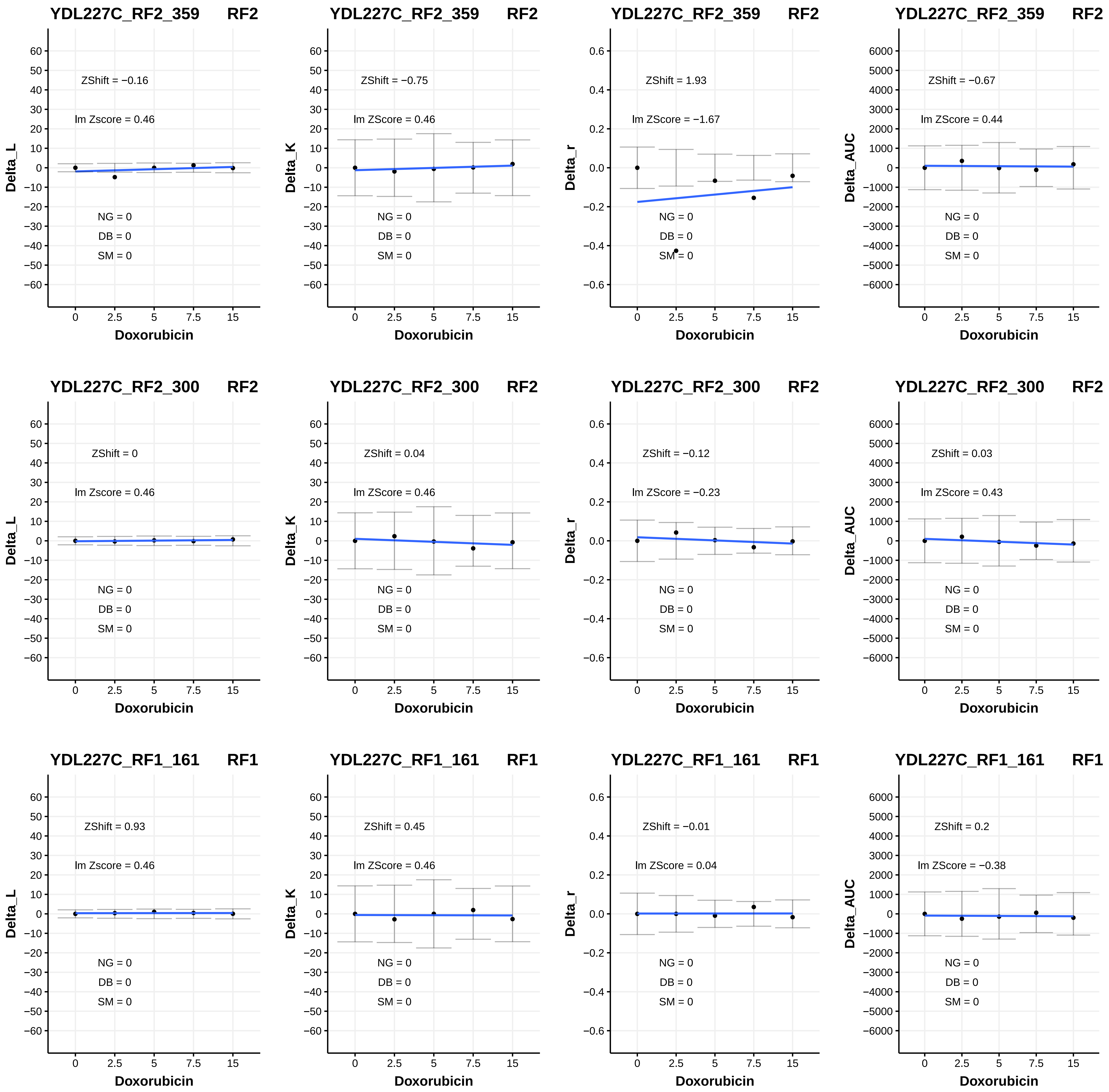

YDL227C_RF2_359 RF2
Delta_L
ZShift = −0.16
lm Zscore = 0.46
NG = 0
DB = 0
SM = 0
Doxorubicin
YDL227C_RF2_359 RF2
Delta_K
ZShift = −0.75
lm ZScore = 0.46
NG = 0
DB = 0
SM = 0
Doxorubicin
YDL227C_RF2_359 RF2
Delta_r
ZShift = 1.93
lm ZScore = −1.67
NG = 0
DB = 0
SM = 0
Doxorubicin
YDL227C_RF2_359 RF2
Delta_AUC
ZShift = −0.67
lm ZScore = 0.44
NG = 0
DB = 0
SM = 0
Doxorubicin
YDL227C_RF2_300 RF2
Delta_L
ZShift = 0
lm Zscore = 0.46
NG = 0
DB = 0
SM = 0
Doxorubicin
YDL227C_RF2_300 RF2
Delta_K
ZShift = 0.04
lm ZScore = 0.46
NG = 0
DB = 0
SM = 0
Doxorubicin
YDL227C_RF2_300 RF2
Delta_r
ZShift = −0.12
lm ZScore = −0.23
NG = 0
DB = 0
SM = 0
Doxorubicin
YDL227C_RF2_300 RF2
Delta_AUC
ZShift = 0.03
lm ZScore = 0.43
NG = 0
DB = 0
SM = 0
Doxorubicin
YDL227C_RF1_161 RF1
Delta_L
ZShift = 0.93
lm Zscore = 0.46
NG = 0
DB = 0
SM = 0
Doxorubicin
YDL227C_RF1_161 RF1
Delta_K
ZShift = 0.45
lm ZScore = 0.46
NG = 0
DB = 0
SM = 0
Doxorubicin
YDL227C_RF1_161 RF1
Delta_r
ZShift = −0.01
lm ZScore = 0.04
NG = 0
DB = 0
SM = 0
Doxorubicin
YDL227C_RF1_161 RF1
Delta_AUC
ZShift = 0.2
lm ZScore = −0.38
NG = 0
DB = 0
SM = 0
Doxorubicin

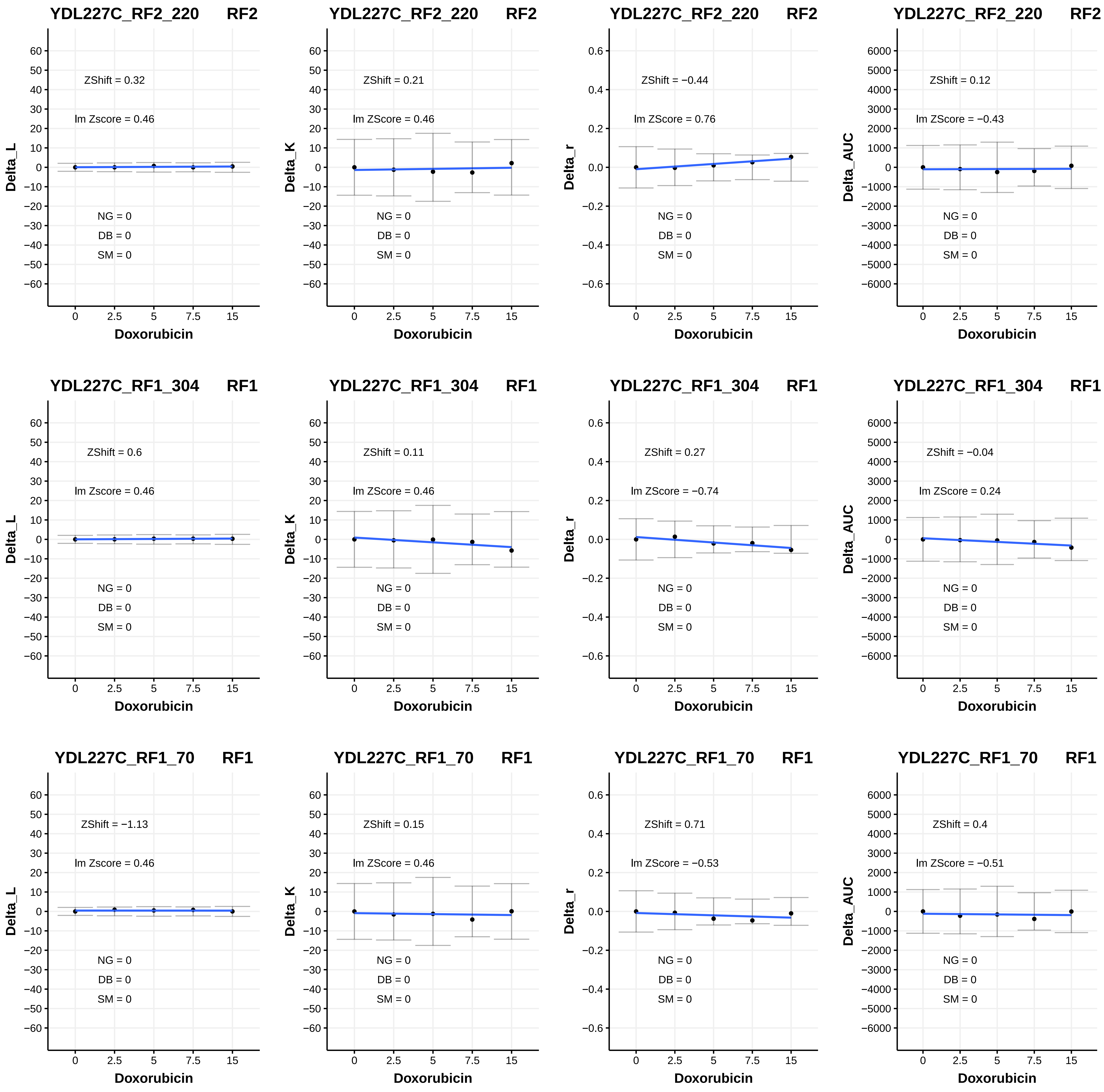
YDL227C_RF2_220 RF2
ZShift = 0.32
lm Zscore = 0.46
NG = 0
DB = 0
SM = 0
Delta_L
Doxorubicin
YDL227C_RF2_220 RF2
ZShift = 0.21
lm ZScore = 0.46
NG = 0
DB = 0
SM = 0
Delta_K
Doxorubicin
YDL227C_RF2_220 RF2
ZShift = −0.44
lm ZScore = 0.76
NG = 0
DB = 0
SM = 0
Delta_r
Doxorubicin
YDL227C_RF2_220 RF2
ZShift = 0.12
lm ZScore = −0.43
NG = 0
DB = 0
SM = 0
Delta_AUC
Doxorubicin
YDL227C_RF1_304 RF1
ZShift = 0.6
lm Zscore = 0.46
NG = 0
DB = 0
SM = 0
Delta_L
Doxorubicin
YDL227C_RF1_304 RF1
ZShift = 0.11
lm ZScore = 0.46
NG = 0
DB = 0
SM = 0
Delta_K
Doxorubicin
YDL227C_RF1_304 RF1
ZShift = 0.27
lm ZScore = −0.74
NG = 0
DB = 0
SM = 0
Delta_r
Doxorubicin
YDL227C_RF1_304 RF1
ZShift = −0.04
lm ZScore = 0.24
NG = 0
DB = 0
SM = 0
Delta_AUC
Doxorubicin
YDL227C_RF1_70 RF1
ZShift = −1.13
lm Zscore = 0.46
NG = 0
DB = 0
SM = 0
Delta_L
Doxorubicin
YDL227C_RF1_70 RF1
ZShift = 0.15
lm ZScore = 0.46
NG = 0
DB = 0
SM = 0
Delta_K
Doxorubicin
YDL227C_RF1_70 RF1
ZShift = 0.71
lm ZScore = −0.53
NG = 0
DB = 0
SM = 0
Delta_r
Doxorubicin
YDL227C_RF1_70 RF1
ZShift = 0.4
lm ZScore = −0.51
NG = 0
DB = 0
SM = 0
Delta_AUC
Doxorubicin

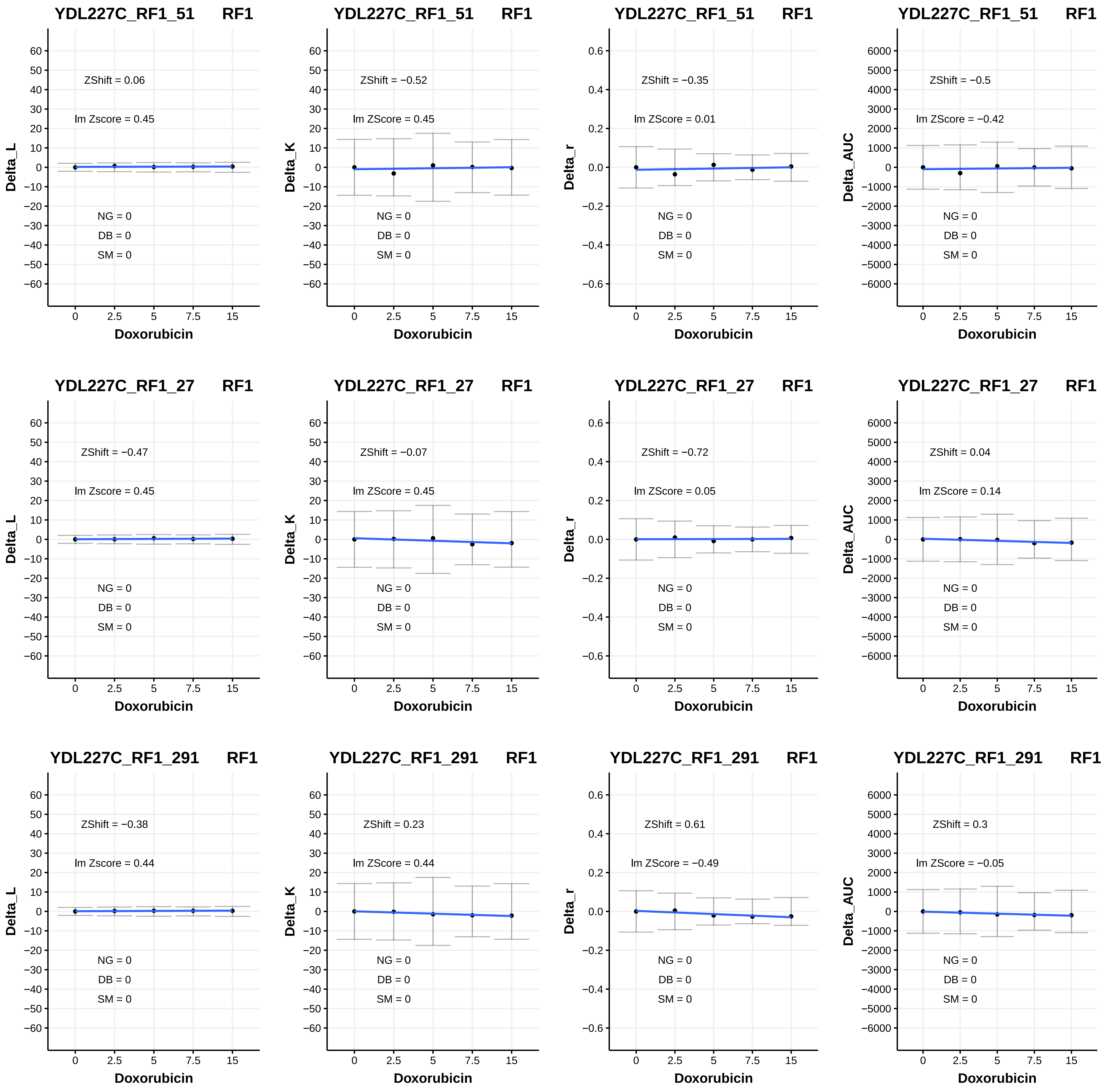
YDL227C_RF1_51 RF1
ZShift = 0.06
lm Zscore = 0.45
NG = 0
DB = 0
SM = 0
Delta_L
Doxorubicin
YDL227C_RF1_51 RF1
ZShift = −0.52
lm ZScore = 0.45
NG = 0
DB = 0
SM = 0
Delta_K
Doxorubicin
YDL227C_RF1_51 RF1
ZShift = −0.35
lm ZScore = 0.01
NG = 0
DB = 0
SM = 0
Delta_r
Doxorubicin
YDL227C_RF1_51 RF1
ZShift = −0.5
lm ZScore = −0.42
NG = 0
DB = 0
SM = 0
Delta_AUC
Doxorubicin
YDL227C_RF1_27 RF1
ZShift = −0.47
lm Zscore = 0.45
NG = 0
DB = 0
SM = 0
Delta_L
Doxorubicin
YDL227C_RF1_27 RF1
ZShift = −0.07
lm ZScore = 0.45
NG = 0
DB = 0
SM = 0
Delta_K
Doxorubicin
YDL227C_RF1_27 RF1
ZShift = −0.72
lm ZScore = 0.05
NG = 0
DB = 0
SM = 0
Delta_r
Doxorubicin
YDL227C_RF1_27 RF1
ZShift = 0.04
lm ZScore = 0.14
NG = 0
DB = 0
SM = 0
Delta_AUC
Doxorubicin
YDL227C_RF1_291 RF1
ZShift = −0.38
lm Zscore = 0.44
NG = 0
DB = 0
SM = 0
Delta_L
Doxorubicin
YDL227C_RF1_291 RF1
ZShift = 0.23
lm ZScore = 0.44
NG = 0
DB = 0
SM = 0
Delta_K
Doxorubicin
YDL227C_RF1_291 RF1
ZShift = 0.61
lm ZScore = −0.49
NG = 0
DB = 0
SM = 0
Delta_r
Doxorubicin
YDL227C_RF1_291 RF1
ZShift = 0.3
lm ZScore = −0.05
NG = 0
DB = 0
SM = 0
Delta_AUC
Doxorubicin

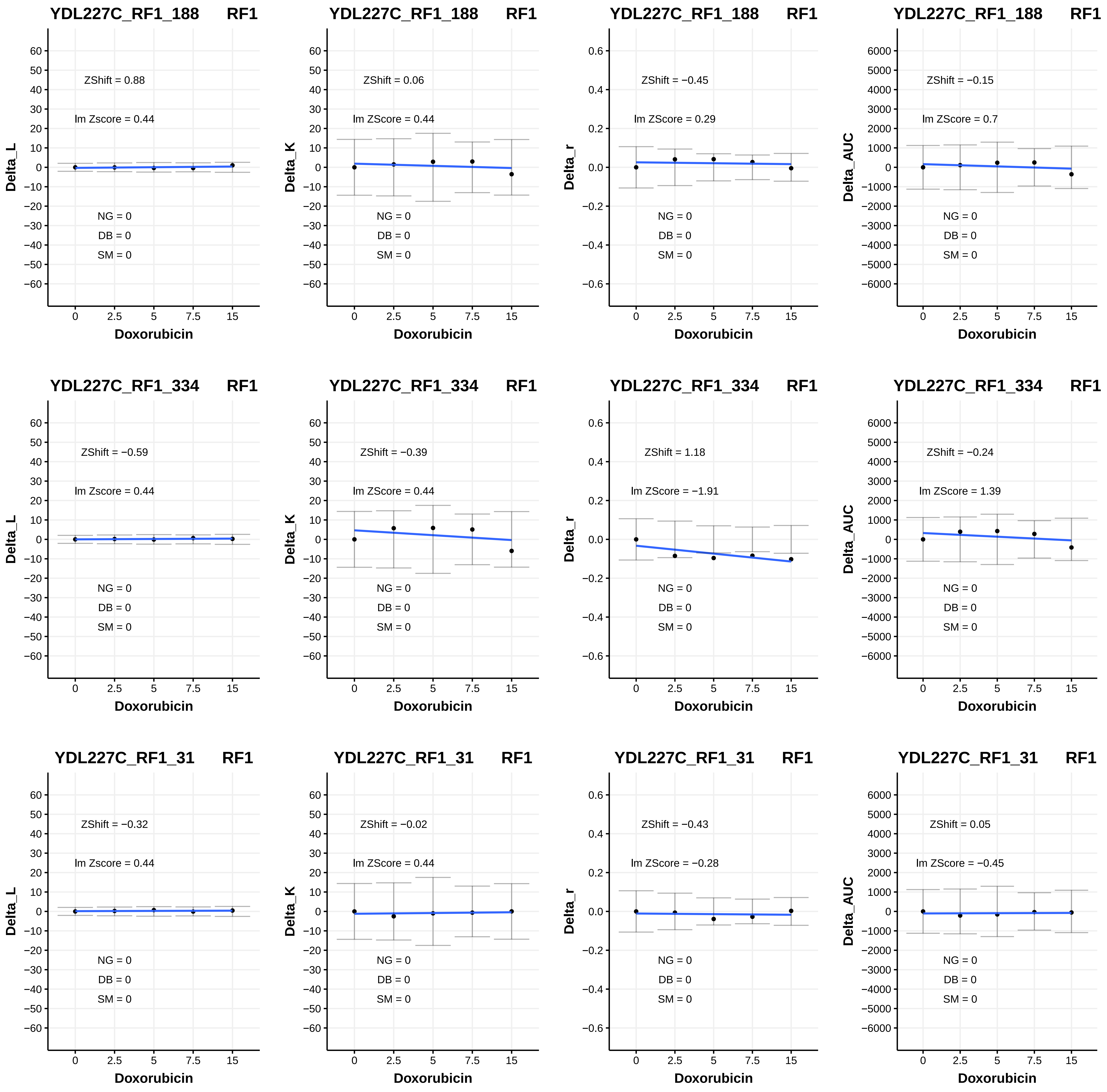
YDL227C_RF1_188 RF1
ZShift = 0.88
lm Zscore = 0.44
NG = 0
DB = 0
SM = 0
Delta_L
Doxorubicin
YDL227C_RF1_188 RF1
ZShift = 0.06
lm ZScore = 0.44
NG = 0
DB = 0
SM = 0
Delta_K
Doxorubicin
YDL227C_RF1_188 RF1
ZShift = −0.45
lm ZScore = 0.29
NG = 0
DB = 0
SM = 0
Delta_r
Doxorubicin
YDL227C_RF1_188 RF1
ZShift = −0.15
lm ZScore = 0.7
NG = 0
DB = 0
SM = 0
Delta_AUC
Doxorubicin
YDL227C_RF1_334 RF1
ZShift = −0.59
lm Zscore = 0.44
NG = 0
DB = 0
SM = 0
Delta_L
Doxorubicin
YDL227C_RF1_334 RF1
ZShift = −0.39
lm ZScore = 0.44
NG = 0
DB = 0
SM = 0
Delta_K
Doxorubicin
YDL227C_RF1_334 RF1
ZShift = 1.18
lm ZScore = −1.91
NG = 0
DB = 0
SM = 0
Delta_r
Doxorubicin
YDL227C_RF1_334 RF1
ZShift = −0.24
lm ZScore = 1.39
NG = 0
DB = 0
SM = 0
Delta_AUC
Doxorubicin
YDL227C_RF1_31 RF1
ZShift = −0.32
lm Zscore = 0.44
NG = 0
DB = 0
SM = 0
Delta_L
Doxorubicin
YDL227C_RF1_31 RF1
ZShift = −0.02
lm ZScore = 0.44
NG = 0
DB = 0
SM = 0
Delta_K
Doxorubicin
YDL227C_RF1_31 RF1
ZShift = −0.43
lm ZScore = −0.28
NG = 0
DB = 0
SM = 0
Delta_r
Doxorubicin
YDL227C_RF1_31 RF1
ZShift = 0.05
lm ZScore = −0.45
NG = 0
DB = 0
SM = 0
Delta_AUC
Doxorubicin

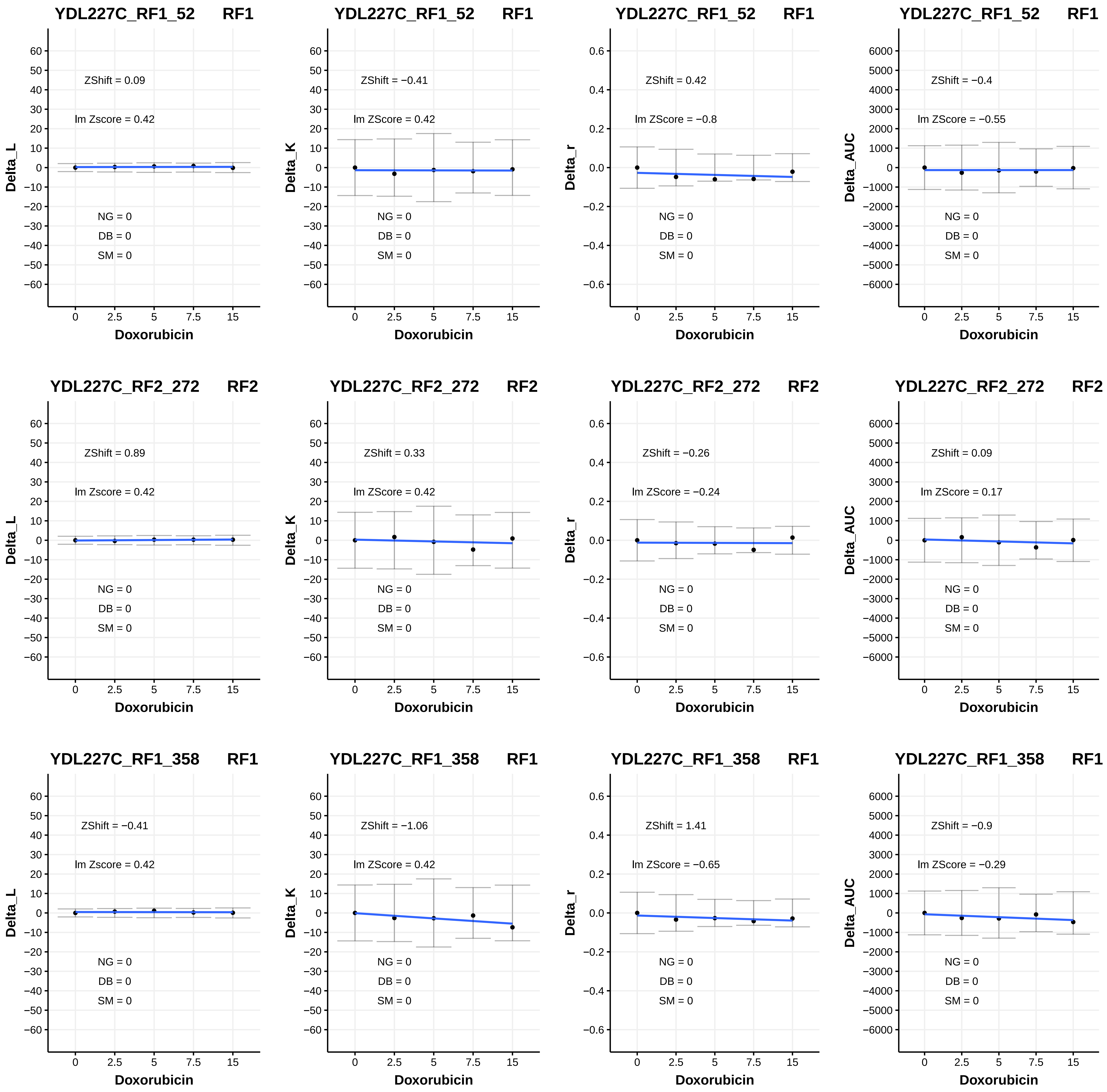
YDL227C_RF1_52 RF1
ZShift = 0.09
lm Zscore = 0.42
NG = 0
DB = 0
SM = 0
Delta_L
Doxorubicin
YDL227C_RF1_52 RF1
ZShift = −0.41
lm ZScore = 0.42
NG = 0
DB = 0
SM = 0
Delta_K
Doxorubicin
YDL227C_RF1_52 RF1
ZShift = 0.42
lm ZScore = −0.8
NG = 0
DB = 0
SM = 0
Delta_r
Doxorubicin
YDL227C_RF1_52 RF1
ZShift = −0.4
lm ZScore = −0.55
NG = 0
DB = 0
SM = 0
Delta_AUC
Doxorubicin
YDL227C_RF2_272 RF2
ZShift = 0.89
lm Zscore = 0.42
NG = 0
DB = 0
SM = 0
Delta_L
Doxorubicin
YDL227C_RF2_272 RF2
ZShift = 0.33
lm ZScore = 0.42
NG = 0
DB = 0
SM = 0
Delta_K
Doxorubicin
YDL227C_RF2_272 RF2
ZShift = −0.26
lm ZScore = −0.24
NG = 0
DB = 0
SM = 0
Delta_r
Doxorubicin
YDL227C_RF2_272 RF2
ZShift = 0.09
lm ZScore = 0.17
NG = 0
DB = 0
SM = 0
Delta_AUC
Doxorubicin
YDL227C_RF1_358 RF1
ZShift = −0.41
lm Zscore = 0.42
NG = 0
DB = 0
SM = 0
Delta_L
Doxorubicin
YDL227C_RF1_358 RF1
ZShift = −1.06
lm ZScore = 0.42
NG = 0
DB = 0
SM = 0
Delta_K
Doxorubicin
YDL227C_RF1_358 RF1
ZShift = 1.41
lm ZScore = −0.65
NG = 0
DB = 0
SM = 0
Delta_r
Doxorubicin
YDL227C_RF1_358 RF1
ZShift = −0.9
lm ZScore = −0.29
NG = 0
DB = 0
SM = 0
Delta_AUC
Doxorubicin

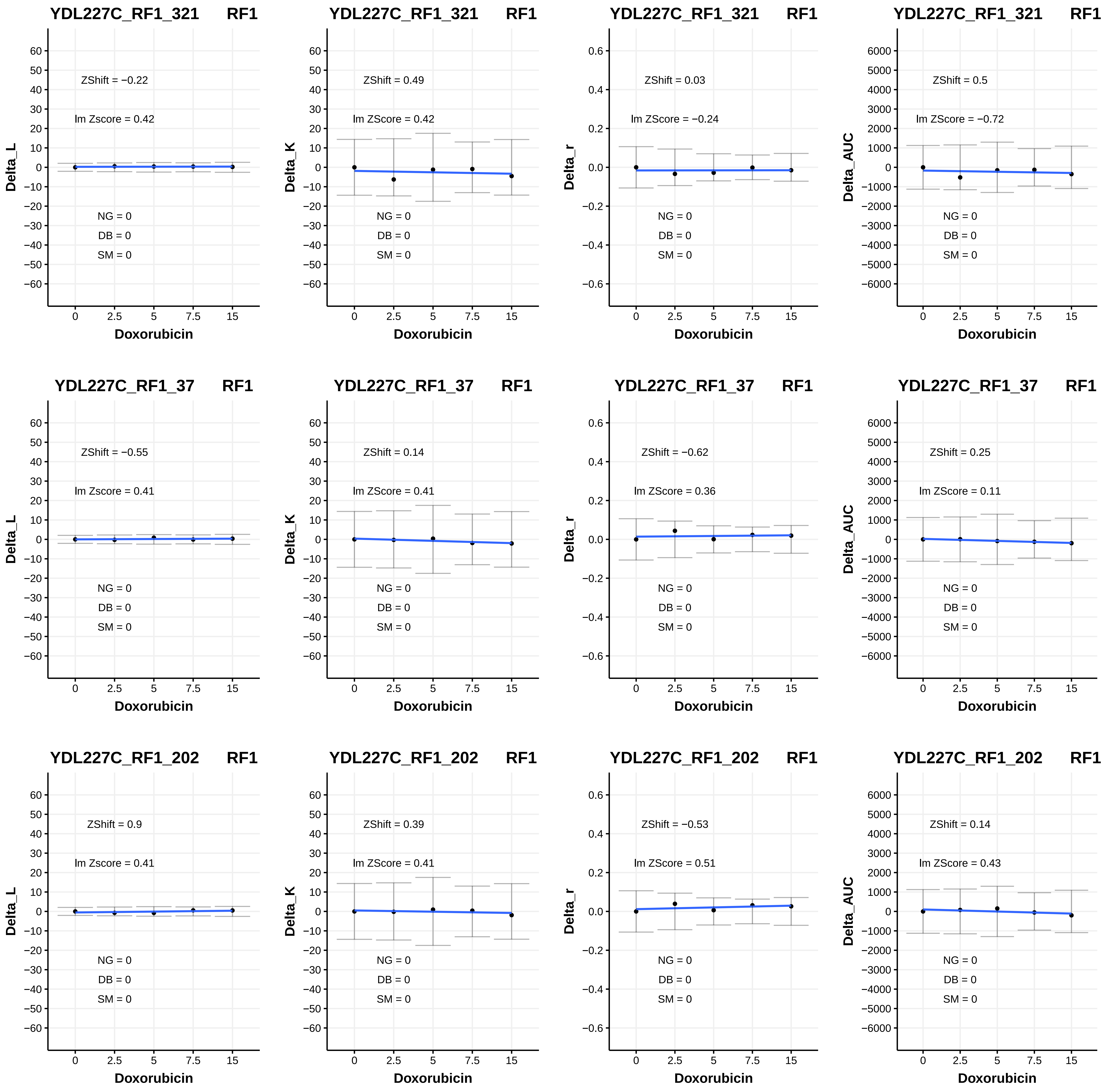
YDL227C_RF1_321 RF1
ZShift = −0.22
lm Zscore = 0.42
NG = 0
DB = 0
SM = 0
Delta_L
Doxorubicin
YDL227C_RF1_321 RF1
ZShift = 0.49
lm ZScore = 0.42
NG = 0
DB = 0
SM = 0
Delta_K
Doxorubicin
YDL227C_RF1_321 RF1
ZShift = 0.03
lm ZScore = −0.24
NG = 0
DB = 0
SM = 0
Delta_r
Doxorubicin
YDL227C_RF1_321 RF1
ZShift = 0.5
lm ZScore = −0.72
NG = 0
DB = 0
SM = 0
Delta_AUC
Doxorubicin
YDL227C_RF1_37 RF1
ZShift = −0.55
lm Zscore = 0.41
NG = 0
DB = 0
SM = 0
Delta_L
Doxorubicin
YDL227C_RF1_37 RF1
ZShift = 0.14
lm ZScore = 0.41
NG = 0
DB = 0
SM = 0
Delta_K
Doxorubicin
YDL227C_RF1_37 RF1
ZShift = −0.62
lm ZScore = 0.36
NG = 0
DB = 0
SM = 0
Delta_r
Doxorubicin
YDL227C_RF1_37 RF1
ZShift = 0.25
lm ZScore = 0.11
NG = 0
DB = 0
SM = 0
Delta_AUC
Doxorubicin
YDL227C_RF1_202 RF1
ZShift = 0.9
lm Zscore = 0.41
NG = 0
DB = 0
SM = 0
Delta_L
Doxorubicin
YDL227C_RF1_202 RF1
ZShift = 0.39
lm ZScore = 0.41
NG = 0
DB = 0
SM = 0
Delta_K
Doxorubicin
YDL227C_RF1_202 RF1
ZShift = −0.53
lm ZScore = 0.51
NG = 0
DB = 0
SM = 0
Delta_r
Doxorubicin
YDL227C_RF1_202 RF1
ZShift = 0.14
lm ZScore = 0.43
NG = 0
DB = 0
SM = 0
Delta_AUC
Doxorubicin

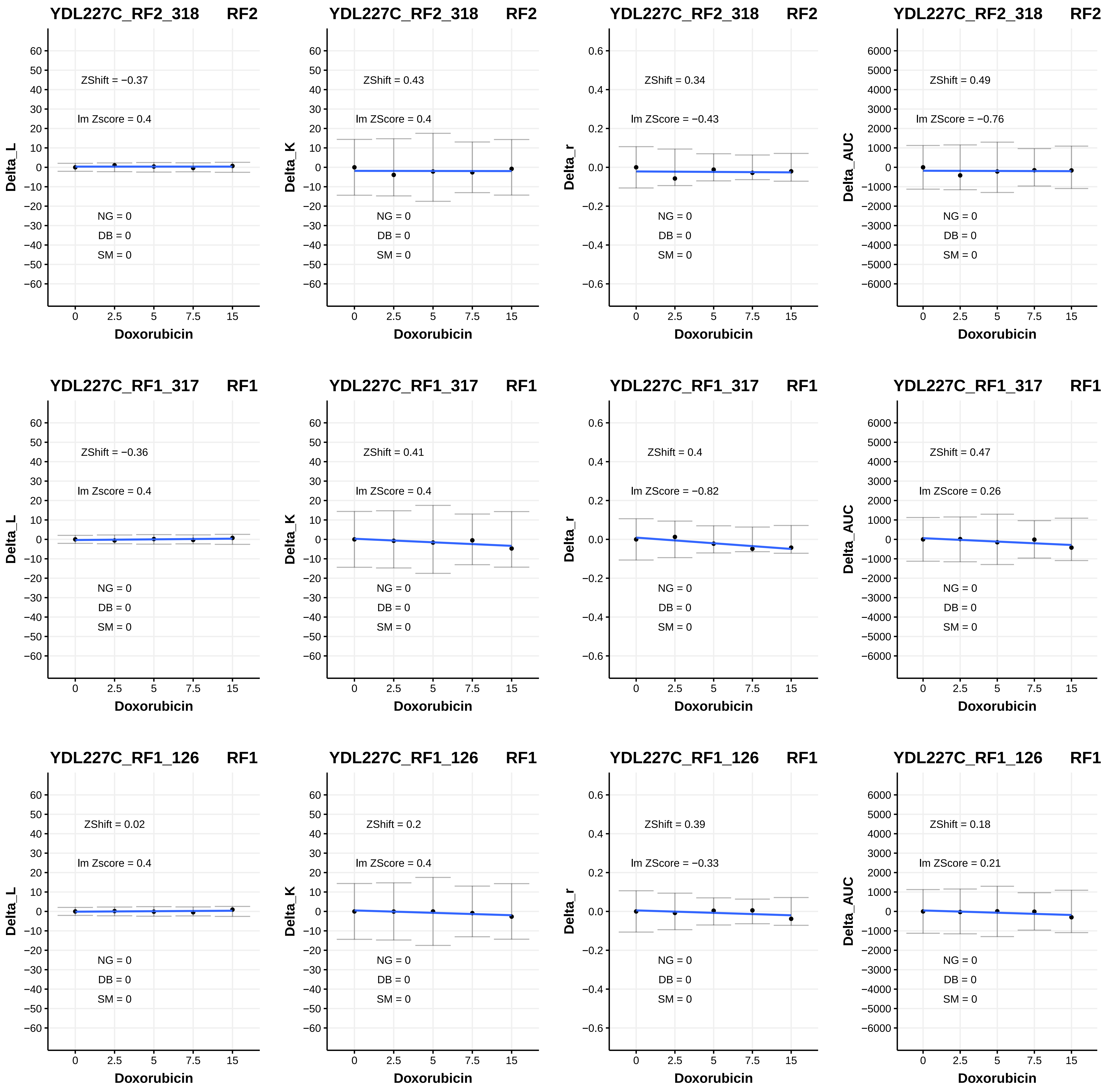

YDL227C_RF2_318 RF2
ZShift = −0.37
lm Zscore = 0.4
NG = 0
DB = 0
SM = 0
Delta_L
Doxorubicin
YDL227C_RF2_318 RF2
ZShift = 0.43
lm ZScore = 0.4
NG = 0
DB = 0
SM = 0
Delta_K
Doxorubicin
YDL227C_RF2_318 RF2
ZShift = 0.34
lm ZScore = −0.43
NG = 0
DB = 0
SM = 0
Delta_r
Doxorubicin
YDL227C_RF2_318 RF2
ZShift = 0.49
lm ZScore = −0.76
NG = 0
DB = 0
SM = 0
Delta_AUC
Doxorubicin
YDL227C_RF1_317 RF1
ZShift = −0.36
lm Zscore = 0.4
NG = 0
DB = 0
SM = 0
Delta_L
Doxorubicin
YDL227C_RF1_317 RF1
ZShift = 0.41
lm ZScore = 0.4
NG = 0
DB = 0
SM = 0
Delta_K
Doxorubicin
YDL227C_RF1_317 RF1
ZShift = 0.4
lm ZScore = −0.82
NG = 0
DB = 0
SM = 0
Delta_r
Doxorubicin
YDL227C_RF1_317 RF1
ZShift = 0.47
lm ZScore = 0.26
NG = 0
DB = 0
SM = 0
Delta_AUC
Doxorubicin
YDL227C_RF1_126 RF1
ZShift = 0.02
lm Zscore = 0.4
NG = 0
DB = 0
SM = 0
Delta_L
Doxorubicin
YDL227C_RF1_126 RF1
ZShift = 0.2
lm ZScore = 0.4
NG = 0
DB = 0
SM = 0
Delta_K
Doxorubicin
YDL227C_RF1_126 RF1
ZShift = 0.39
lm ZScore = −0.33
NG = 0
DB = 0
SM = 0
Delta_r
Doxorubicin
YDL227C_RF1_126 RF1
ZShift = 0.18
lm ZScore = 0.21
NG = 0
DB = 0
SM = 0
Delta_AUC
Doxorubicin

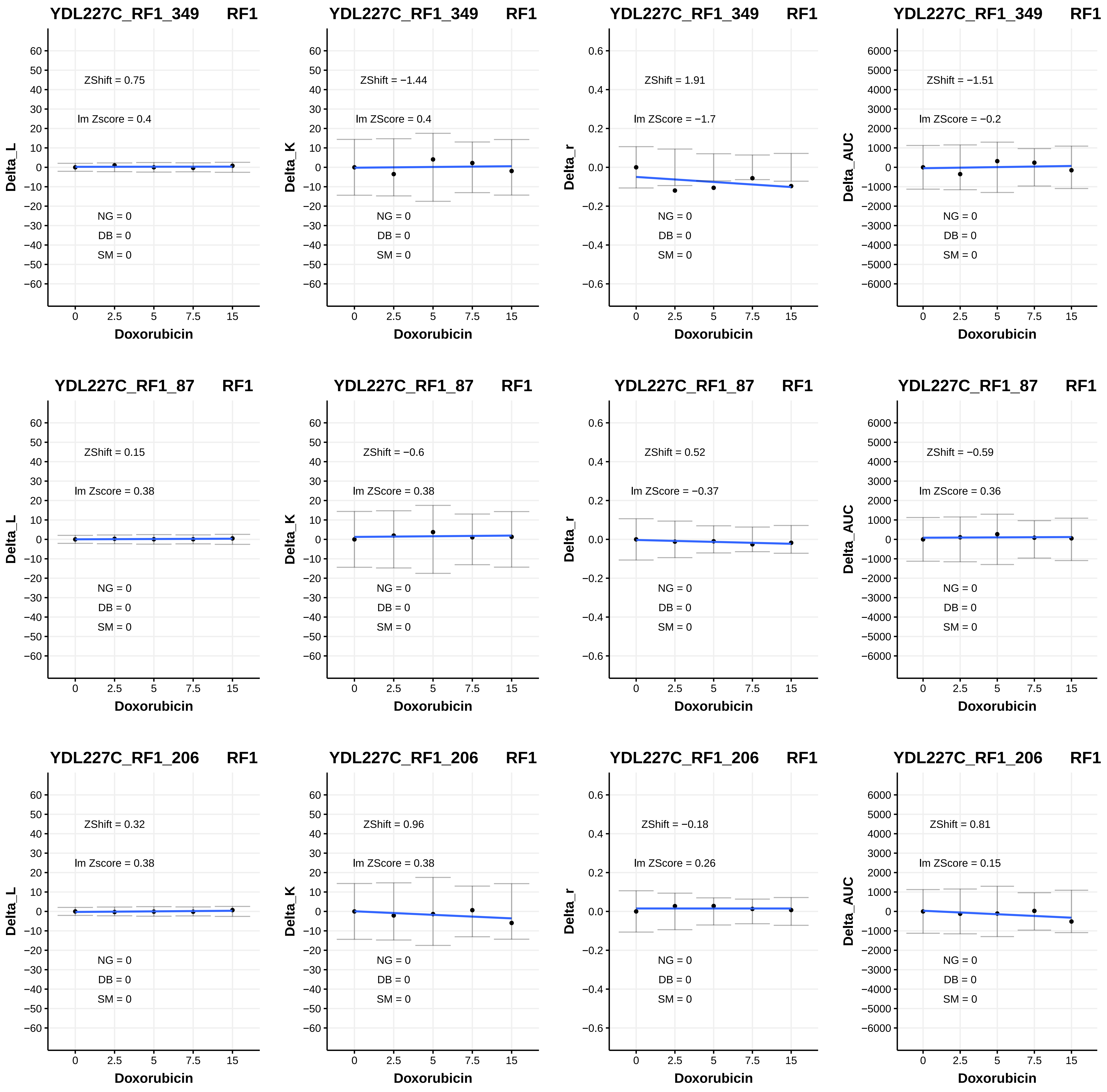
YDL227C_RF1_349 RF1
ZShift = 0.75
lm Zscore = 0.4
NG = 0
DB = 0
SM = 0
Delta_L
Doxorubicin
YDL227C_RF1_349 RF1
ZShift = −1.44
lm ZScore = 0.4
NG = 0
DB = 0
SM = 0
Delta_K
Doxorubicin
YDL227C_RF1_349 RF1
ZShift = 1.91
lm ZScore = −1.7
NG = 0
DB = 0
SM = 0
Delta_r
Doxorubicin
YDL227C_RF1_349 RF1
ZShift = −1.51
lm ZScore = −0.2
NG = 0
DB = 0
SM = 0
Delta_AUC
Doxorubicin
YDL227C_RF1_87 RF1
ZShift = 0.15
lm Zscore = 0.38
NG = 0
DB = 0
SM = 0
Delta_L
Doxorubicin
YDL227C_RF1_87 RF1
ZShift = −0.6
lm ZScore = 0.38
NG = 0
DB = 0
SM = 0
Delta_K
Doxorubicin
YDL227C_RF1_87 RF1
ZShift = 0.52
lm ZScore = −0.37
NG = 0
DB = 0
SM = 0
Delta_r
Doxorubicin
YDL227C_RF1_87 RF1
ZShift = −0.59
lm ZScore = 0.36
NG = 0
DB = 0
SM = 0
Delta_AUC
Doxorubicin
YDL227C_RF1_206 RF1
ZShift = 0.32
lm Zscore = 0.38
NG = 0
DB = 0
SM = 0
Delta_L
Doxorubicin
YDL227C_RF1_206 RF1
ZShift = 0.96
lm ZScore = 0.38
NG = 0
DB = 0
SM = 0
Delta_K
Doxorubicin
YDL227C_RF1_206 RF1
ZShift = −0.18
lm ZScore = 0.26
NG = 0
DB = 0
SM = 0
Delta_r
Doxorubicin
YDL227C_RF1_206 RF1
ZShift = 0.81
lm ZScore = 0.15
NG = 0
DB = 0
SM = 0
Delta_AUC
Doxorubicin

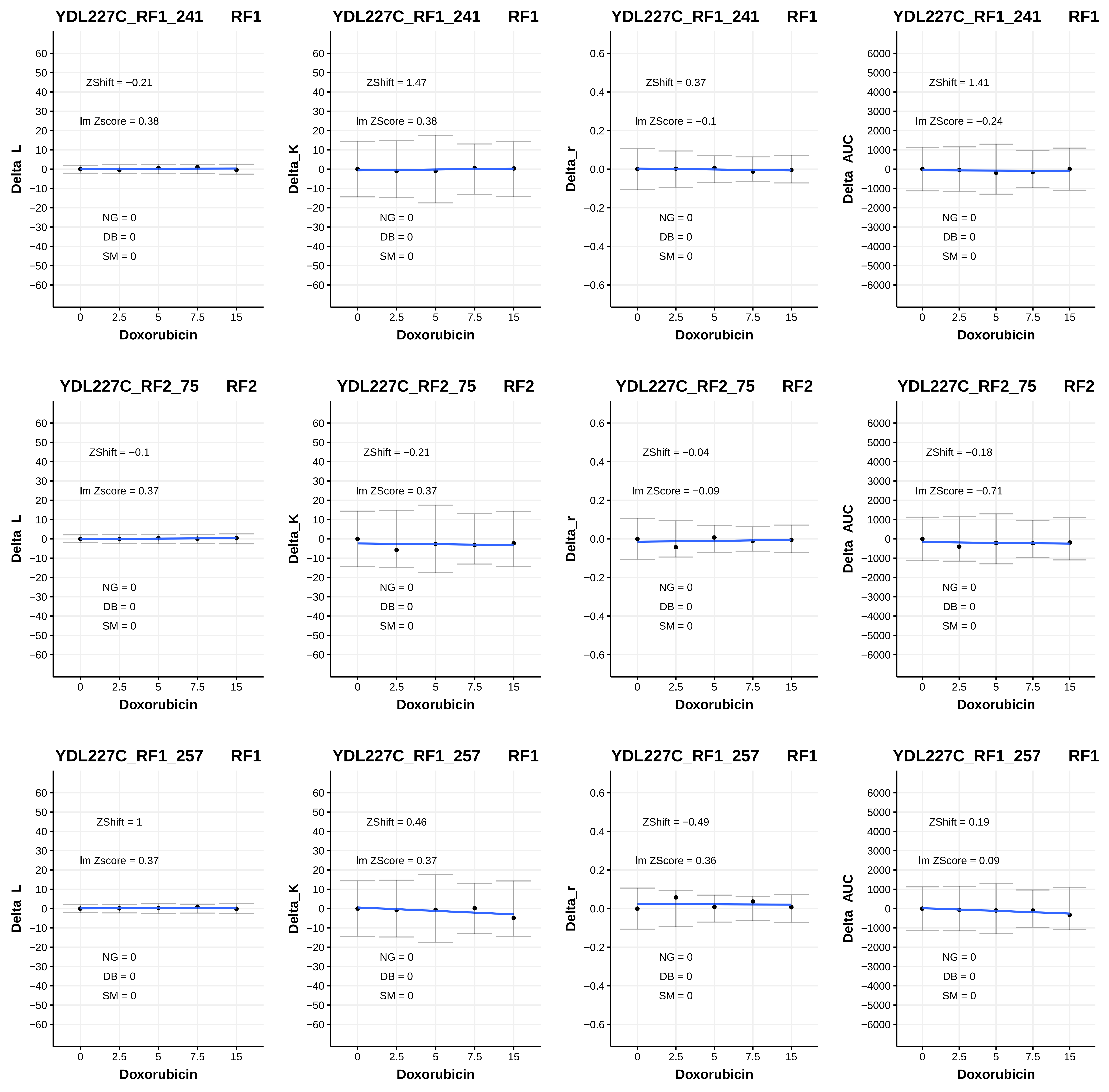

YDL227C_RF1_241 RF1
ZShift = −0.21
lm Zscore = 0.38
NG = 0
DB = 0
SM = 0
Delta_L
Doxorubicin
YDL227C_RF1_241 RF1
ZShift = 1.47
lm ZScore = 0.38
NG = 0
DB = 0
SM = 0
Delta_K
Doxorubicin
YDL227C_RF1_241 RF1
ZShift = 0.37
lm ZScore = −0.1
NG = 0
DB = 0
SM = 0
Delta_r
Doxorubicin
YDL227C_RF1_241 RF1
ZShift = 1.41
lm ZScore = −0.24
NG = 0
DB = 0
SM = 0
Delta_AUC
Doxorubicin
YDL227C_RF2_75 RF2
ZShift = −0.1
lm Zscore = 0.37
NG = 0
DB = 0
SM = 0
Delta_L
Doxorubicin
YDL227C_RF2_75 RF2
ZShift = −0.21
lm ZScore = 0.37
NG = 0
DB = 0
SM = 0
Delta_K
Doxorubicin
YDL227C_RF2_75 RF2
ZShift = −0.04
lm ZScore = −0.09
NG = 0
DB = 0
SM = 0
Delta_r
Doxorubicin
YDL227C_RF2_75 RF2
ZShift = −0.18
lm ZScore = −0.71
NG = 0
DB = 0
SM = 0
Delta_AUC
Doxorubicin
YDL227C_RF1_257 RF1
ZShift = 1
lm Zscore = 0.37
NG = 0
DB = 0
SM = 0
Delta_L
Doxorubicin
YDL227C_RF1_257 RF1
ZShift = 0.46
lm ZScore = 0.37
NG = 0
DB = 0
SM = 0
Delta_K
Doxorubicin
YDL227C_RF1_257 RF1
ZShift = −0.49
lm ZScore = 0.36
NG = 0
DB = 0
SM = 0
Delta_r
Doxorubicin
YDL227C_RF1_257 RF1
ZShift = 0.19
lm ZScore = 0.09
NG = 0
DB = 0
SM = 0
Delta_AUC
Doxorubicin

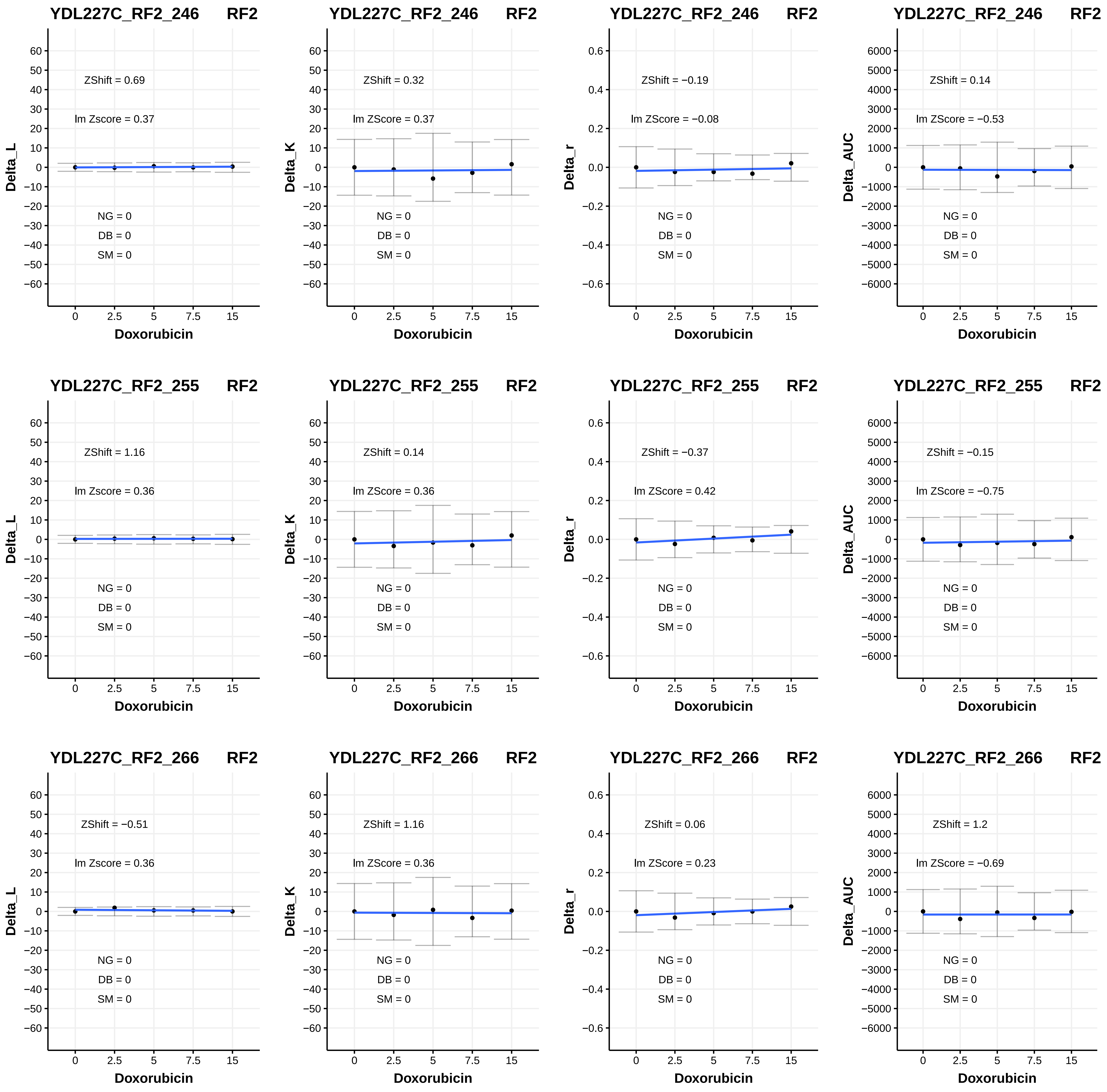

YDL227C_RF2_246 RF2
ZShift = 0.69
lm Zscore = 0.37
NG = 0
DB = 0
SM = 0
Delta_L
Doxorubicin
YDL227C_RF2_246 RF2
ZShift = 0.32
lm ZScore = 0.37
NG = 0
DB = 0
SM = 0
Delta_K
Doxorubicin
YDL227C_RF2_246 RF2
ZShift = −0.19
lm ZScore = −0.08
NG = 0
DB = 0
SM = 0
Delta_r
Doxorubicin
YDL227C_RF2_246 RF2
ZShift = 0.14
lm ZScore = −0.53
NG = 0
DB = 0
SM = 0
Delta_AUC
Doxorubicin
YDL227C_RF2_255 RF2
ZShift = 1.16
lm Zscore = 0.36
NG = 0
DB = 0
SM = 0
Delta_L
Doxorubicin
YDL227C_RF2_255 RF2
ZShift = 0.14
lm ZScore = 0.36
NG = 0
DB = 0
SM = 0
Delta_K
Doxorubicin
YDL227C_RF2_255 RF2
ZShift = −0.37
lm ZScore = 0.42
NG = 0
DB = 0
SM = 0
Delta_r
Doxorubicin
YDL227C_RF2_255 RF2
ZShift = −0.15
lm ZScore = −0.75
NG = 0
DB = 0
SM = 0
Delta_AUC
Doxorubicin
YDL227C_RF2_266 RF2
ZShift = −0.51
lm Zscore = 0.36
NG = 0
DB = 0
SM = 0
Delta_L
Doxorubicin
YDL227C_RF2_266 RF2
ZShift = 1.16
lm ZScore = 0.36
NG = 0
DB = 0
SM = 0
Delta_K
Doxorubicin
YDL227C_RF2_266 RF2
ZShift = 0.06
lm ZScore = 0.23
NG = 0
DB = 0
SM = 0
Delta_r
Doxorubicin
YDL227C_RF2_266 RF2
ZShift = 1.2
lm ZScore = −0.69
NG = 0
DB = 0
SM = 0
Delta_AUC
Doxorubicin

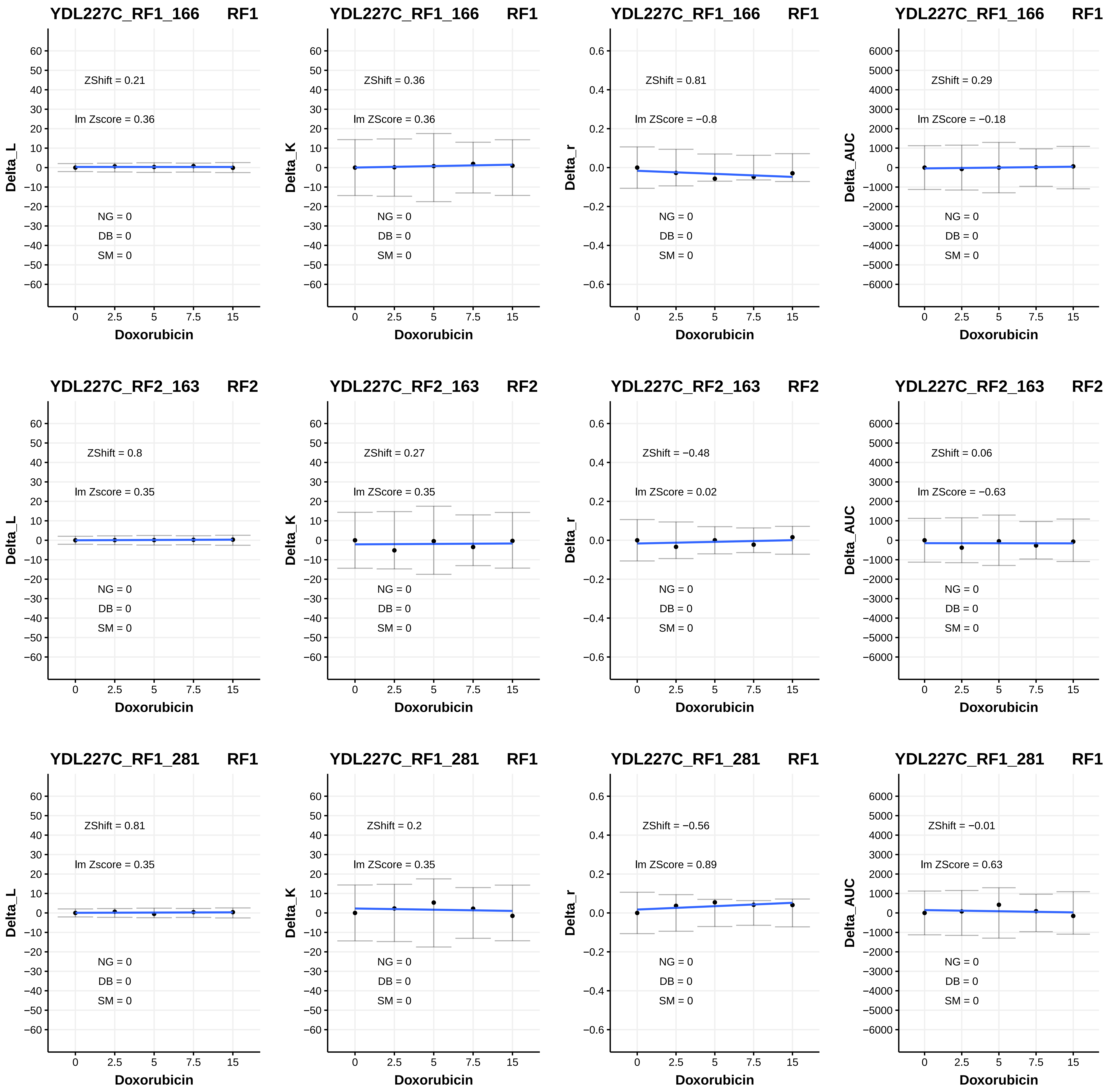

YDL227C_RF1_166 RF1
ZShift = 0.21
lm Zscore = 0.36
NG = 0
DB = 0
SM = 0
Delta_L
Doxorubicin
YDL227C_RF1_166 RF1
ZShift = 0.36
lm ZScore = 0.36
NG = 0
DB = 0
SM = 0
Delta_K
Doxorubicin
YDL227C_RF1_166 RF1
ZShift = 0.81
lm ZScore = −0.8
NG = 0
DB = 0
SM = 0
Delta_r
Doxorubicin
YDL227C_RF1_166 RF1
ZShift = 0.29
lm ZScore = −0.18
NG = 0
DB = 0
SM = 0
Delta_AUC
Doxorubicin
YDL227C_RF2_163 RF2
ZShift = 0.8
lm Zscore = 0.35
NG = 0
DB = 0
SM = 0
Delta_L
Doxorubicin
YDL227C_RF2_163 RF2
ZShift = 0.27
lm ZScore = 0.35
NG = 0
DB = 0
SM = 0
Delta_K
Doxorubicin
YDL227C_RF2_163 RF2
ZShift = −0.48
lm ZScore = 0.02
NG = 0
DB = 0
SM = 0
Delta_r
Doxorubicin
YDL227C_RF2_163 RF2
ZShift = 0.06
lm ZScore = −0.63
NG = 0
DB = 0
SM = 0
Delta_AUC
Doxorubicin
YDL227C_RF1_281 RF1
ZShift = 0.81
lm Zscore = 0.35
NG = 0
DB = 0
SM = 0
Delta_L
Doxorubicin
YDL227C_RF1_281 RF1
ZShift = 0.2
lm ZScore = 0.35
NG = 0
DB = 0
SM = 0
Delta_K
Doxorubicin
YDL227C_RF1_281 RF1
ZShift = −0.56
lm ZScore = 0.89
NG = 0
DB = 0
SM = 0
Delta_r
Doxorubicin
YDL227C_RF1_281 RF1
ZShift = −0.01
lm ZScore = 0.63
NG = 0
DB = 0
SM = 0
Delta_AUC
Doxorubicin

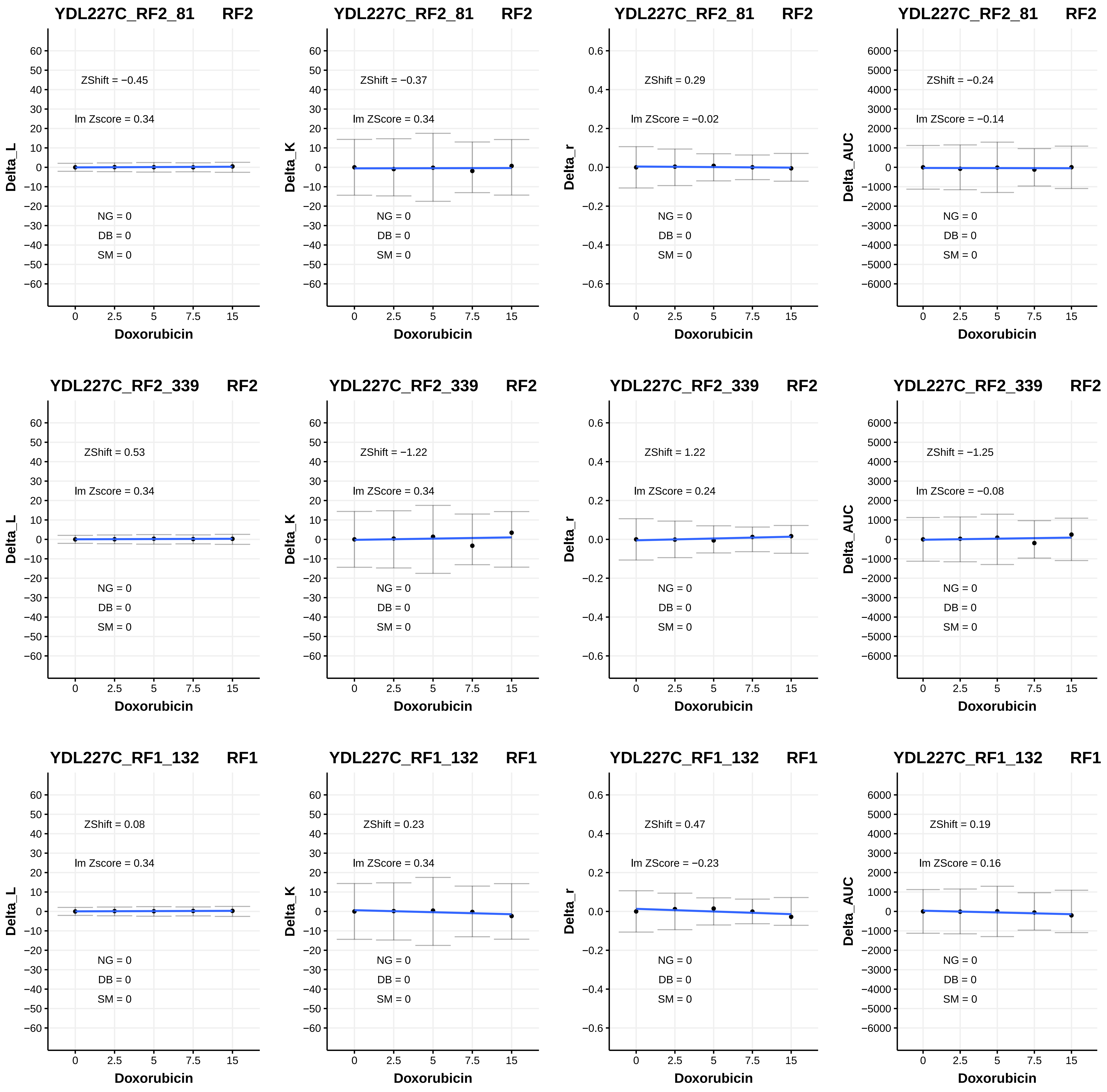

YDL227C_RF2_81 RF2
ZShift = −0.45
lm Zscore = 0.34
NG = 0
DB = 0
SM = 0
Delta_L
Doxorubicin
YDL227C_RF2_81 RF2
ZShift = −0.37
lm ZScore = 0.34
NG = 0
DB = 0
SM = 0
Delta_K
Doxorubicin
YDL227C_RF2_81 RF2
ZShift = 0.29
lm ZScore = −0.02
NG = 0
DB = 0
SM = 0
Delta_r
Doxorubicin
YDL227C_RF2_81 RF2
ZShift = −0.24
lm ZScore = −0.14
NG = 0
DB = 0
SM = 0
Delta_AUC
Doxorubicin
YDL227C_RF2_339 RF2
ZShift = 0.53
lm Zscore = 0.34
NG = 0
DB = 0
SM = 0
Delta_L
Doxorubicin
YDL227C_RF2_339 RF2
ZShift = −1.22
lm ZScore = 0.34
NG = 0
DB = 0
SM = 0
Delta_K
Doxorubicin
YDL227C_RF2_339 RF2
ZShift = 1.22
lm ZScore = 0.24
NG = 0
DB = 0
SM = 0
Delta_r
Doxorubicin
YDL227C_RF2_339 RF2
ZShift = −1.25
lm ZScore = −0.08
NG = 0
DB = 0
SM = 0
Delta_AUC
Doxorubicin
YDL227C_RF1_132 RF1
ZShift = 0.08
lm Zscore = 0.34
NG = 0
DB = 0
SM = 0
Delta_L
Doxorubicin
YDL227C_RF1_132 RF1
ZShift = 0.23
lm ZScore = 0.34
NG = 0
DB = 0
SM = 0
Delta_K
Doxorubicin
YDL227C_RF1_132 RF1
ZShift = 0.47
lm ZScore = −0.23
NG = 0
DB = 0
SM = 0
Delta_r
Doxorubicin
YDL227C_RF1_132 RF1
ZShift = 0.19
lm ZScore = 0.16
NG = 0
DB = 0
SM = 0
Delta_AUC
Doxorubicin

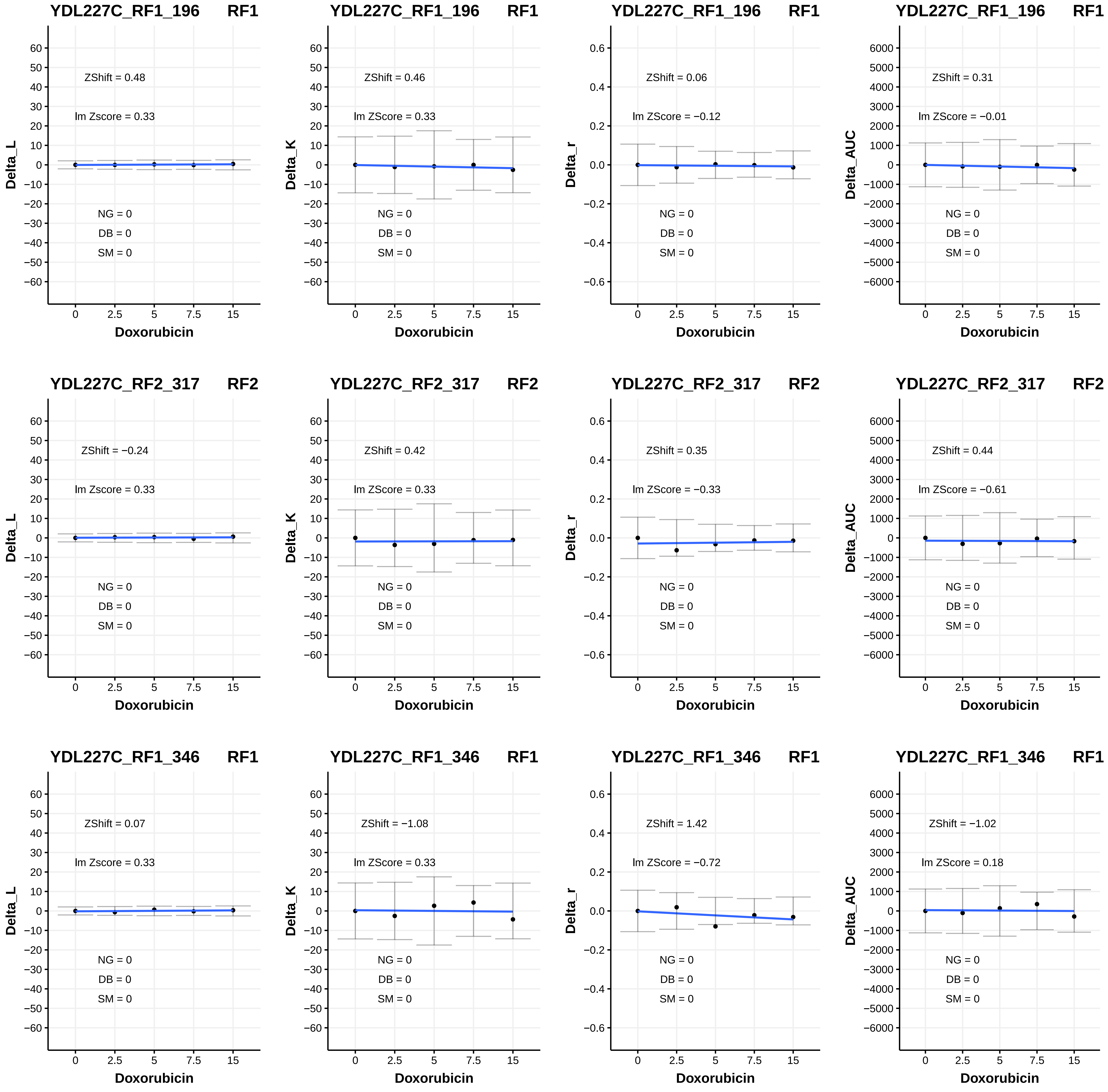

YDL227C_RF1_196 RF1
ZShift = 0.48
lm Zscore = 0.33
NG = 0
DB = 0
SM = 0
Delta_L
Doxorubicin
YDL227C_RF1_196 RF1
ZShift = 0.46
lm ZScore = 0.33
NG = 0
DB = 0
SM = 0
Delta_K
Doxorubicin
YDL227C_RF1_196 RF1
ZShift = 0.06
lm ZScore = −0.12
NG = 0
DB = 0
SM = 0
Delta_r
Doxorubicin
YDL227C_RF1_196 RF1
ZShift = 0.31
lm ZScore = −0.01
NG = 0
DB = 0
SM = 0
Delta_AUC
Doxorubicin
YDL227C_RF2_317 RF2
ZShift = −0.24
lm Zscore = 0.33
NG = 0
DB = 0
SM = 0
Delta_L
Doxorubicin
YDL227C_RF2_317 RF2
ZShift = 0.42
lm ZScore = 0.33
NG = 0
DB = 0
SM = 0
Delta_K
Doxorubicin
YDL227C_RF2_317 RF2
ZShift = 0.35
lm ZScore = −0.33
NG = 0
DB = 0
SM = 0
Delta_r
Doxorubicin
YDL227C_RF2_317 RF2
ZShift = 0.44
lm ZScore = −0.61
NG = 0
DB = 0
SM = 0
Delta_AUC
Doxorubicin
YDL227C_RF1_346 RF1
ZShift = 0.07
lm Zscore = 0.33
NG = 0
DB = 0
SM = 0
Delta_L
Doxorubicin
YDL227C_RF1_346 RF1
ZShift = −1.08
lm ZScore = 0.33
NG = 0
DB = 0
SM = 0
Delta_K
Doxorubicin
YDL227C_RF1_346 RF1
ZShift = 1.42
lm ZScore = −0.72
NG = 0
DB = 0
SM = 0
Delta_r
Doxorubicin
YDL227C_RF1_346 RF1
ZShift = −1.02
lm ZScore = 0.18
NG = 0
DB = 0
SM = 0
Delta_AUC
Doxorubicin

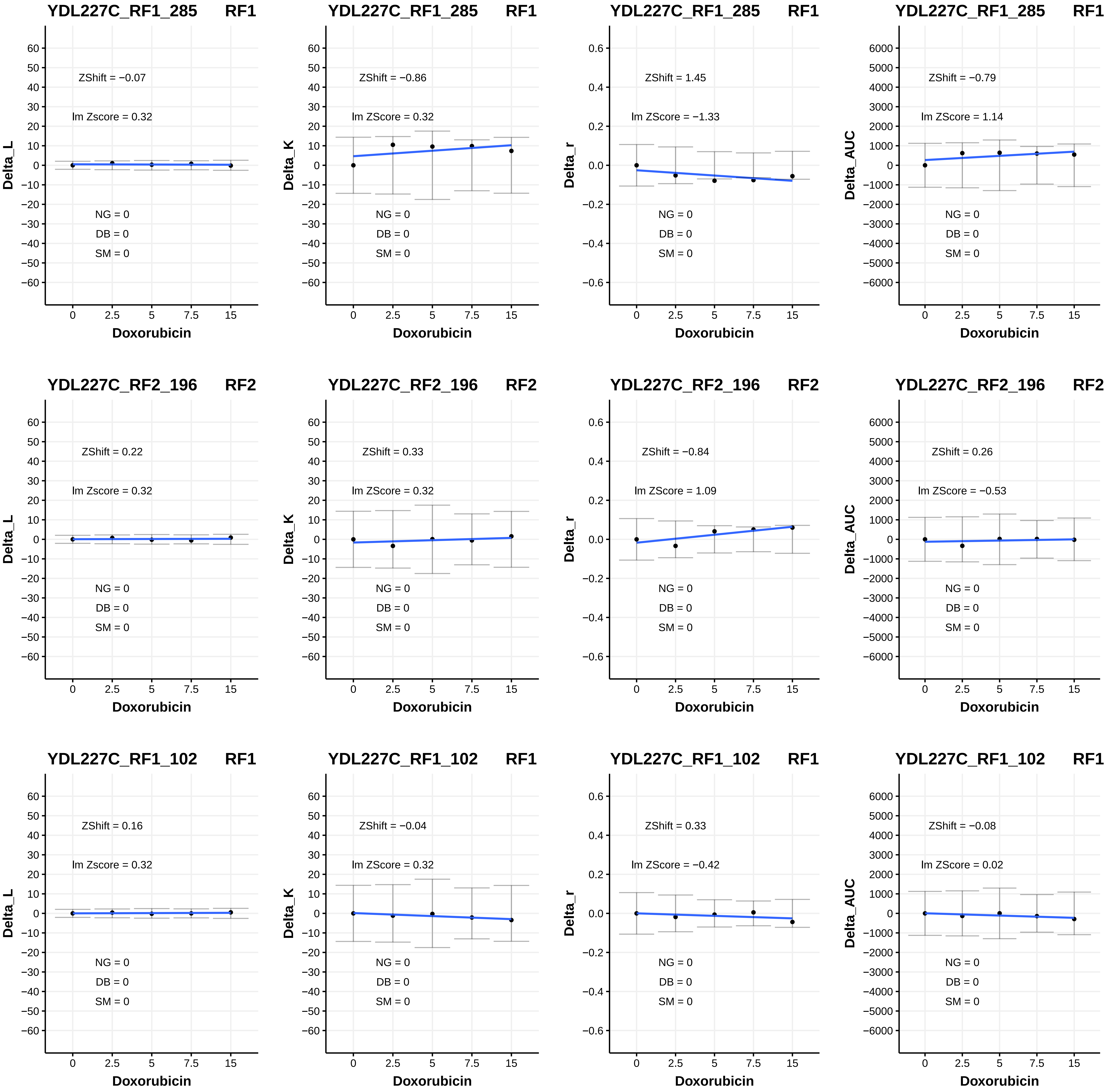

YDL227C_RF1_285 RF1
ZShift = −0.07
lm Zscore = 0.32
NG = 0
DB = 0
SM = 0
Delta_L
Doxorubicin
YDL227C_RF1_285 RF1
ZShift = −0.86
lm ZScore = 0.32
NG = 0
DB = 0
SM = 0
Delta_K
Doxorubicin
YDL227C_RF1_285 RF1
ZShift = 1.45
lm ZScore = −1.33
NG = 0
DB = 0
SM = 0
Delta_r
Doxorubicin
YDL227C_RF1_285 RF1
ZShift = −0.79
lm ZScore = 1.14
NG = 0
DB = 0
SM = 0
Delta_AUC
Doxorubicin
YDL227C_RF2_196 RF2
ZShift = 0.22
lm Zscore = 0.32
NG = 0
DB = 0
SM = 0
Delta_L
Doxorubicin
YDL227C_RF2_196 RF2
ZShift = 0.33
lm ZScore = 0.32
NG = 0
DB = 0
SM = 0
Delta_K
Doxorubicin
YDL227C_RF2_196 RF2
ZShift = −0.84
lm ZScore = 1.09
NG = 0
DB = 0
SM = 0
Delta_r
Doxorubicin
YDL227C_RF2_196 RF2
ZShift = 0.26
lm ZScore = −0.53
NG = 0
DB = 0
SM = 0
Delta_AUC
Doxorubicin
YDL227C_RF1_102 RF1
ZShift = 0.16
lm Zscore = 0.32
NG = 0
DB = 0
SM = 0
Delta_L
Doxorubicin
YDL227C_RF1_102 RF1
ZShift = −0.04
lm ZScore = 0.32
NG = 0
DB = 0
SM = 0
Delta_K
Doxorubicin
YDL227C_RF1_102 RF1
ZShift = 0.33
lm ZScore = −0.42
NG = 0
DB = 0
SM = 0
Delta_r
Doxorubicin
YDL227C_RF1_102 RF1
ZShift = −0.08
lm ZScore = 0.02
NG = 0
DB = 0
SM = 0
Delta_AUC
Doxorubicin

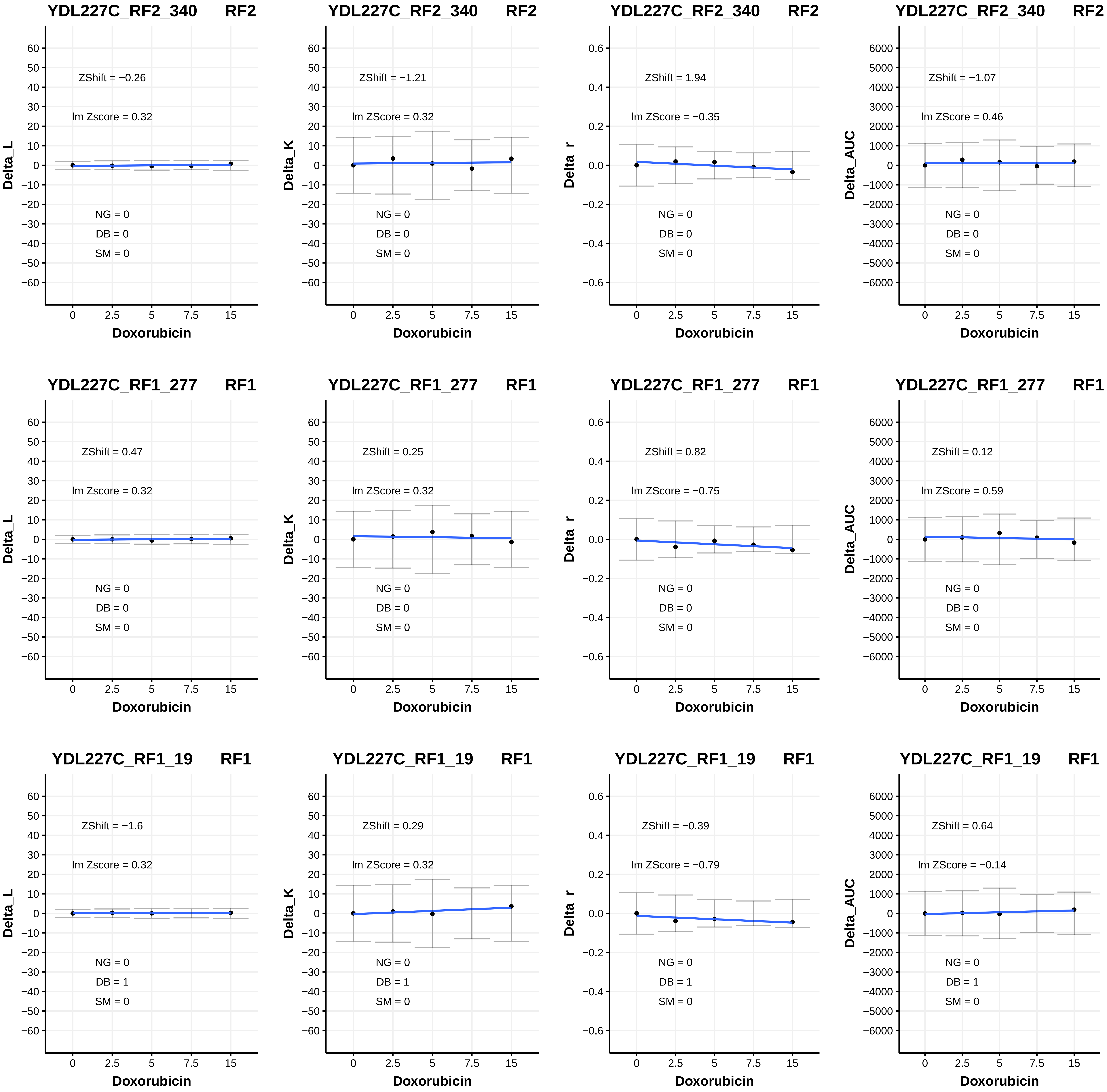
YDL227C_RF2_340 RF2
ZShift = −0.26
lm Zscore = 0.32
NG = 0
DB = 0
SM = 0
Delta_L
YDL227C_RF2_340 RF2
ZShift = −1.21
lm ZScore = 0.32
NG = 0
DB = 0
SM = 0
Delta_K
YDL227C_RF2_340 RF2
ZShift = 1.94
lm ZScore = −0.35
NG = 0
DB = 0
SM = 0
Delta_r
YDL227C_RF2_340 RF2
ZShift = −1.07
lm ZScore = 0.46
NG = 0
DB = 0
SM = 0
Delta_AUC
YDL227C_RF1_277 RF1
ZShift = 0.47
lm Zscore = 0.32
NG = 0
DB = 0
SM = 0
Delta_L
YDL227C_RF1_277 RF1
ZShift = 0.25
lm ZScore = 0.32
NG = 0
DB = 0
SM = 0
Delta_K
YDL227C_RF1_277 RF1
ZShift = 0.82
lm ZScore = −0.75
NG = 0
DB = 0
SM = 0
Delta_r
YDL227C_RF1_277 RF1
ZShift = 0.12
lm ZScore = 0.59
NG = 0
DB = 0
SM = 0
Delta_AUC
YDL227C_RF1_19 RF1
ZShift = −1.6
lm Zscore = 0.32
NG = 0
DB = 1
SM = 0
Delta_L
YDL227C_RF1_19 RF1
ZShift = 0.29
lm ZScore = 0.32
NG = 0
DB = 1
SM = 0
Delta_K
YDL227C_RF1_19 RF1
ZShift = −0.39
lm ZScore = −0.79
NG = 0
DB = 1
SM = 0
Delta_r
YDL227C_RF1_19 RF1
ZShift = 0.64
lm ZScore = −0.14
NG = 0
DB = 1
SM = 0
Delta_AUC
Doxorubicin
0
2.5
5
7.5
15

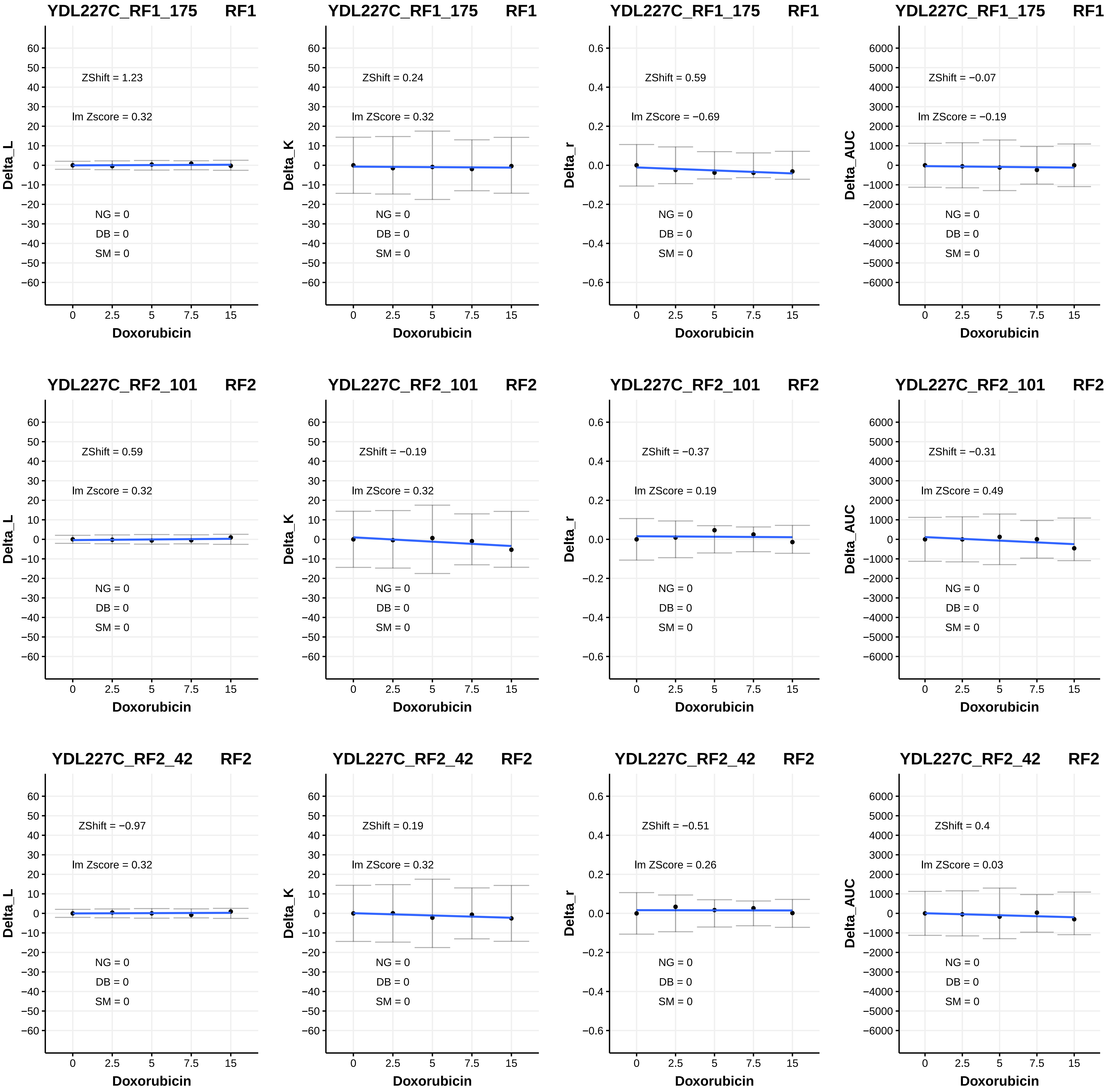
YDL227C_RF1_175 RF1
ZShift = 1.23
lm Zscore = 0.32
NG = 0
DB = 0
SM = 0
Delta_L
Doxorubicin
YDL227C_RF1_175 RF1
ZShift = 0.24
lm ZScore = 0.32
NG = 0
DB = 0
SM = 0
Delta_K
Doxorubicin
YDL227C_RF1_175 RF1
ZShift = 0.59
lm ZScore = −0.69
NG = 0
DB = 0
SM = 0
Delta_r
Doxorubicin
YDL227C_RF1_175 RF1
ZShift = −0.07
lm ZScore = −0.19
NG = 0
DB = 0
SM = 0
Delta_AUC
Doxorubicin
YDL227C_RF2_101 RF2
ZShift = 0.59
lm Zscore = 0.32
NG = 0
DB = 0
SM = 0
Delta_L
Doxorubicin
YDL227C_RF2_101 RF2
ZShift = −0.19
lm ZScore = 0.32
NG = 0
DB = 0
SM = 0
Delta_K
Doxorubicin
YDL227C_RF2_101 RF2
ZShift = −0.37
lm ZScore = 0.19
NG = 0
DB = 0
SM = 0
Delta_r
Doxorubicin
YDL227C_RF2_101 RF2
ZShift = −0.31
lm ZScore = 0.49
NG = 0
DB = 0
SM = 0
Delta_AUC
Doxorubicin
YDL227C_RF2_42 RF2
ZShift = −0.97
lm Zscore = 0.32
NG = 0
DB = 0
SM = 0
Delta_L
Doxorubicin
YDL227C_RF2_42 RF2
ZShift = 0.19
lm ZScore = 0.32
NG = 0
DB = 0
SM = 0
Delta_K
Doxorubicin
YDL227C_RF2_42 RF2
ZShift = −0.51
lm ZScore = 0.26
NG = 0
DB = 0
SM = 0
Delta_r
Doxorubicin
YDL227C_RF2_42 RF2
ZShift = 0.4
lm ZScore = 0.03
NG = 0
DB = 0
SM = 0
Delta_AUC
Doxorubicin

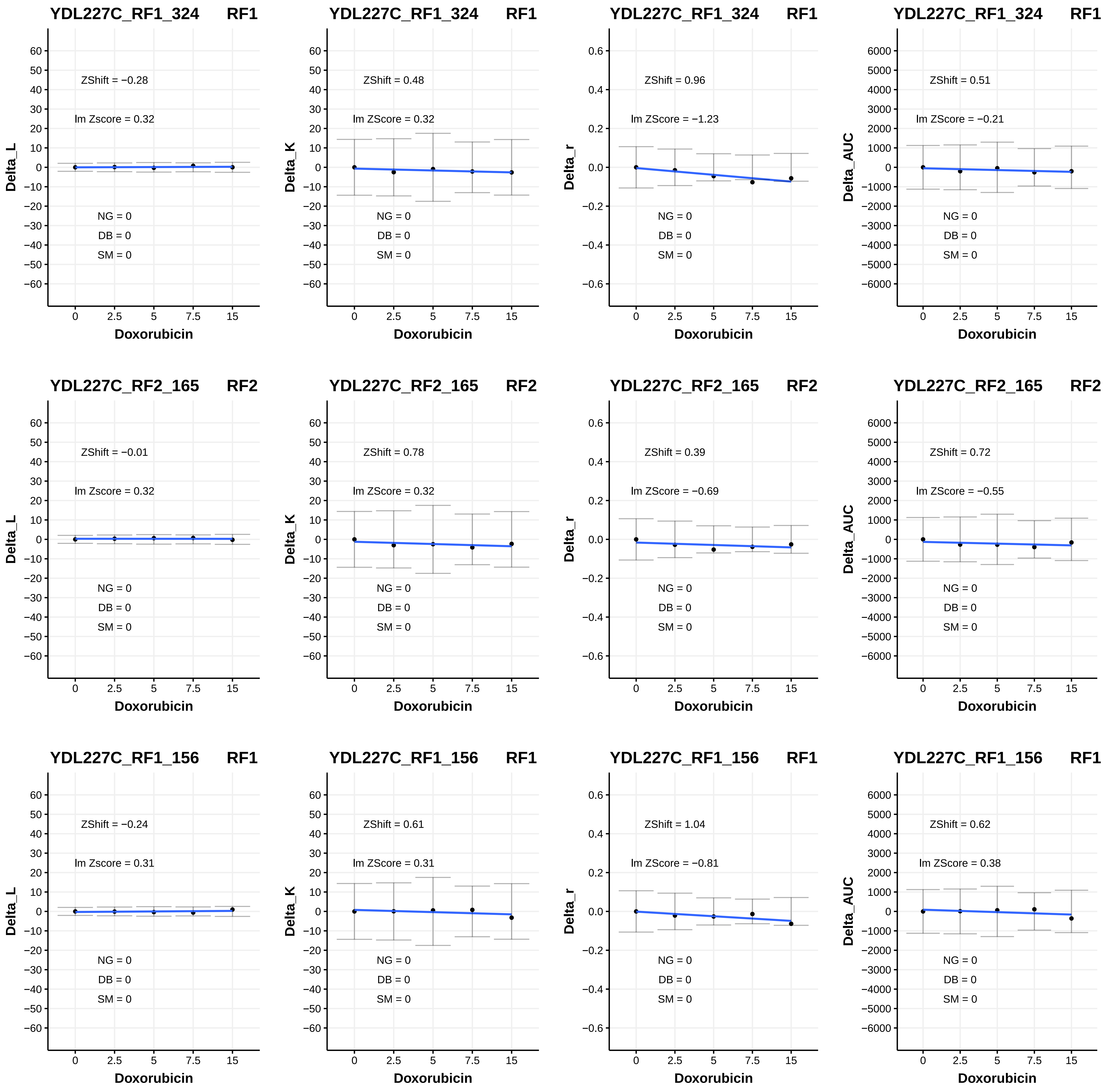

YDL227C_RF1_324 RF1
ZShift = −0.28
lm Zscore = 0.32
NG = 0
DB = 0
SM = 0
Delta_L
Doxorubicin
YDL227C_RF1_324 RF1
ZShift = 0.48
lm ZScore = 0.32
NG = 0
DB = 0
SM = 0
Delta_K
Doxorubicin
YDL227C_RF1_324 RF1
ZShift = 0.96
lm ZScore = −1.23
NG = 0
DB = 0
SM = 0
Delta_r
Doxorubicin
YDL227C_RF1_324 RF1
ZShift = 0.51
lm ZScore = −0.21
NG = 0
DB = 0
SM = 0
Delta_AUC
Doxorubicin
YDL227C_RF2_165 RF2
ZShift = −0.01
lm Zscore = 0.32
NG = 0
DB = 0
SM = 0
Delta_L
Doxorubicin
YDL227C_RF2_165 RF2
ZShift = 0.78
lm ZScore = 0.32
NG = 0
DB = 0
SM = 0
Delta_K
Doxorubicin
YDL227C_RF2_165 RF2
ZShift = 0.39
lm ZScore = −0.69
NG = 0
DB = 0
SM = 0
Delta_r
Doxorubicin
YDL227C_RF2_165 RF2
ZShift = 0.72
lm ZScore = −0.55
NG = 0
DB = 0
SM = 0
Delta_AUC
Doxorubicin
YDL227C_RF1_156 RF1
ZShift = −0.24
lm Zscore = 0.31
NG = 0
DB = 0
SM = 0
Delta_L
Doxorubicin
YDL227C_RF1_156 RF1
ZShift = 0.61
lm ZScore = 0.31
NG = 0
DB = 0
SM = 0
Delta_K
Doxorubicin
YDL227C_RF1_156 RF1
ZShift = 1.04
lm ZScore = −0.81
NG = 0
DB = 0
SM = 0
Delta_r
Doxorubicin
YDL227C_RF1_156 RF1
ZShift = 0.62
lm ZScore = 0.38
NG = 0
DB = 0
SM = 0
Delta_AUC
Doxorubicin

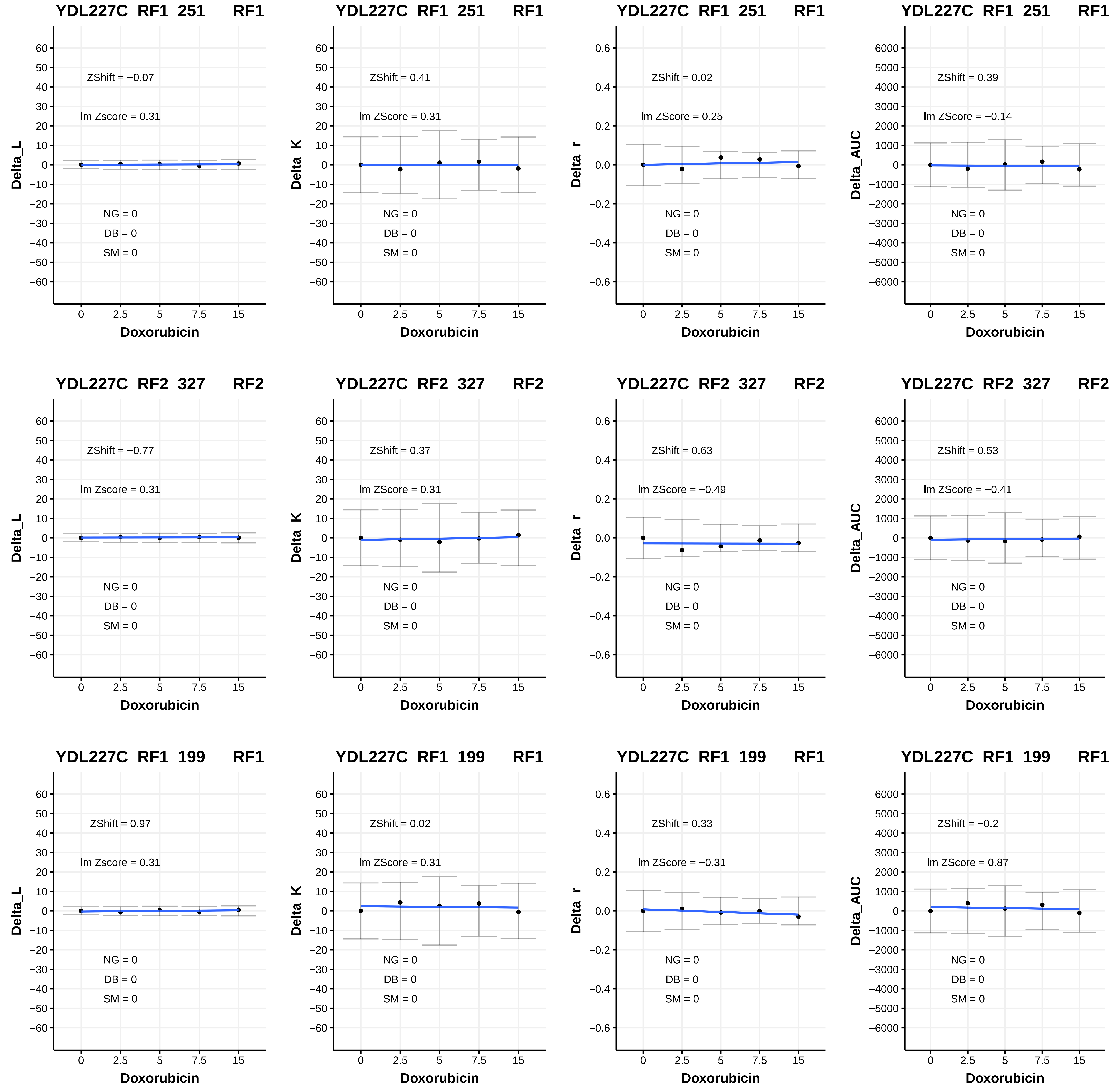
YDL227C_RF1_251 RF1
ZShift = −0.07
lm Zscore = 0.31
NG = 0
DB = 0
SM = 0
Delta_L
Doxorubicin
YDL227C_RF1_251 RF1
ZShift = 0.41
lm ZScore = 0.31
NG = 0
DB = 0
SM = 0
Delta_K
Doxorubicin
YDL227C_RF1_251 RF1
ZShift = 0.02
lm ZScore = 0.25
NG = 0
DB = 0
SM = 0
Delta_r
Doxorubicin
YDL227C_RF1_251 RF1
ZShift = 0.39
lm ZScore = −0.14
NG = 0
DB = 0
SM = 0
Delta_AUC
Doxorubicin
YDL227C_RF2_327 RF2
ZShift = −0.77
lm Zscore = 0.31
NG = 0
DB = 0
SM = 0
Delta_L
Doxorubicin
YDL227C_RF2_327 RF2
ZShift = 0.37
lm ZScore = 0.31
NG = 0
DB = 0
SM = 0
Delta_K
Doxorubicin
YDL227C_RF2_327 RF2
ZShift = 0.63
lm ZScore = −0.49
NG = 0
DB = 0
SM = 0
Delta_r
Doxorubicin
YDL227C_RF2_327 RF2
ZShift = 0.53
lm ZScore = −0.41
NG = 0
DB = 0
SM = 0
Delta_AUC
Doxorubicin
YDL227C_RF1_199 RF1
ZShift = 0.97
lm Zscore = 0.31
NG = 0
DB = 0
SM = 0
Delta_L
Doxorubicin
YDL227C_RF1_199 RF1
ZShift = 0.02
lm ZScore = 0.31
NG = 0
DB = 0
SM = 0
Delta_K
Doxorubicin
YDL227C_RF1_199 RF1
ZShift = 0.33
lm ZScore = −0.31
NG = 0
DB = 0
SM = 0
Delta_r
Doxorubicin
YDL227C_RF1_199 RF1
ZShift = −0.2
lm ZScore = 0.87
NG = 0
DB = 0
SM = 0
Delta_AUC
Doxorubicin

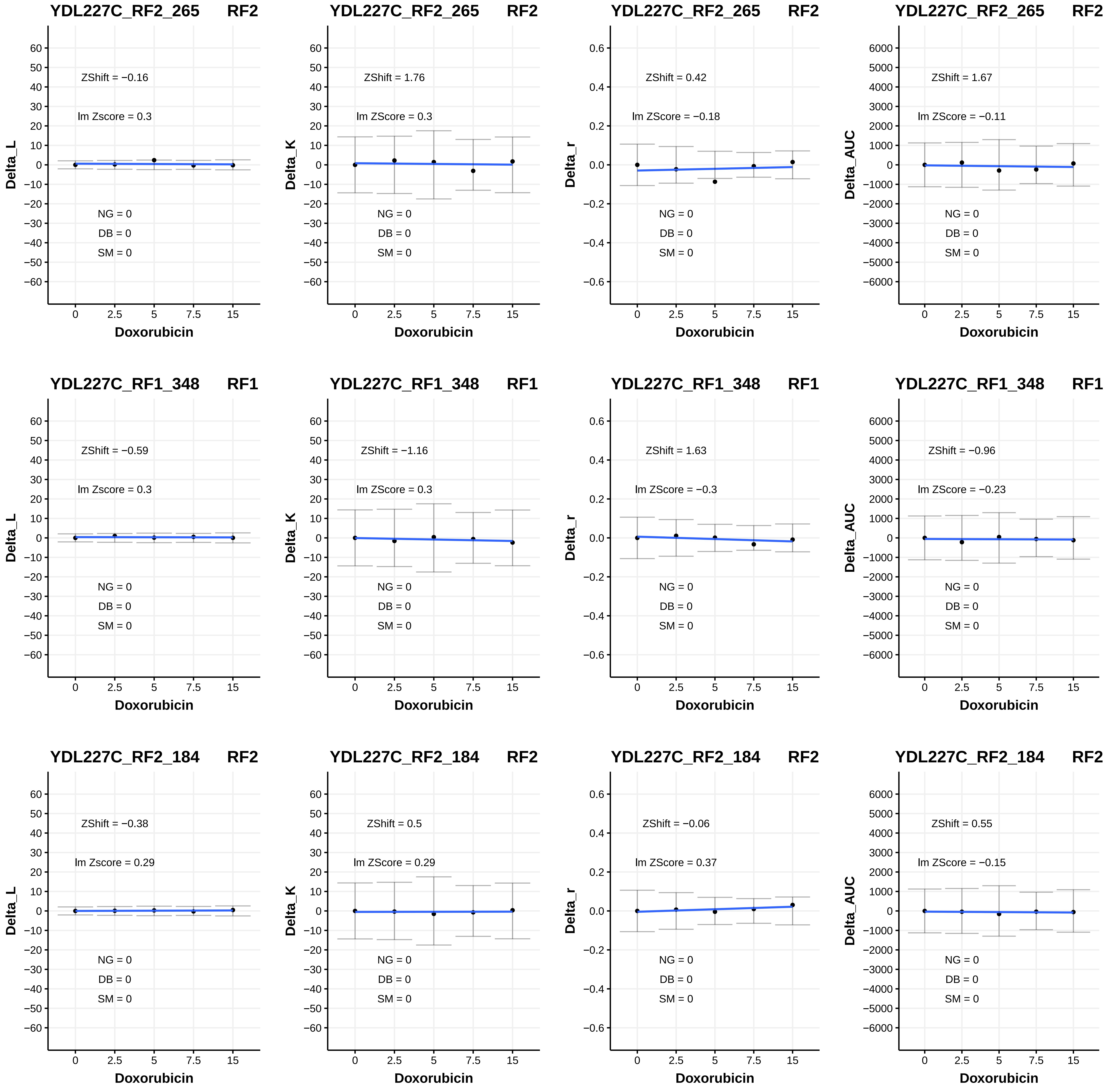
YDL227C_RF2_265 RF2
ZShift = −0.16
lm Zscore = 0.3
NG = 0
DB = 0
SM = 0
Delta_L
Doxorubicin
YDL227C_RF2_265 RF2
ZShift = 1.76
lm ZScore = 0.3
NG = 0
DB = 0
SM = 0
Delta_K
Doxorubicin
YDL227C_RF2_265 RF2
ZShift = 0.42
lm ZScore = −0.18
NG = 0
DB = 0
SM = 0
Delta_r
Doxorubicin
YDL227C_RF2_265 RF2
ZShift = 1.67
lm ZScore = −0.11
NG = 0
DB = 0
SM = 0
Delta_AUC
Doxorubicin
YDL227C_RF1_348 RF1
ZShift = −0.59
lm Zscore = 0.3
NG = 0
DB = 0
SM = 0
Delta_L
Doxorubicin
YDL227C_RF1_348 RF1
ZShift = −1.16
lm ZScore = 0.3
NG = 0
DB = 0
SM = 0
Delta_K
Doxorubicin
YDL227C_RF1_348 RF1
ZShift = 1.63
lm ZScore = −0.3
NG = 0
DB = 0
SM = 0
Delta_r
Doxorubicin
YDL227C_RF1_348 RF1
ZShift = −0.96
lm ZScore = −0.23
NG = 0
DB = 0
SM = 0
Delta_AUC
Doxorubicin
YDL227C_RF2_184 RF2
ZShift = −0.38
lm Zscore = 0.29
NG = 0
DB = 0
SM = 0
Delta_L
Doxorubicin
YDL227C_RF2_184 RF2
ZShift = 0.5
lm ZScore = 0.29
NG = 0
DB = 0
SM = 0
Delta_K
Doxorubicin
YDL227C_RF2_184 RF2
ZShift = −0.06
lm ZScore = 0.37
NG = 0
DB = 0
SM = 0
Delta_r
Doxorubicin
YDL227C_RF2_184 RF2
ZShift = 0.55
lm ZScore = −0.15
NG = 0
DB = 0
SM = 0
Delta_AUC
Doxorubicin

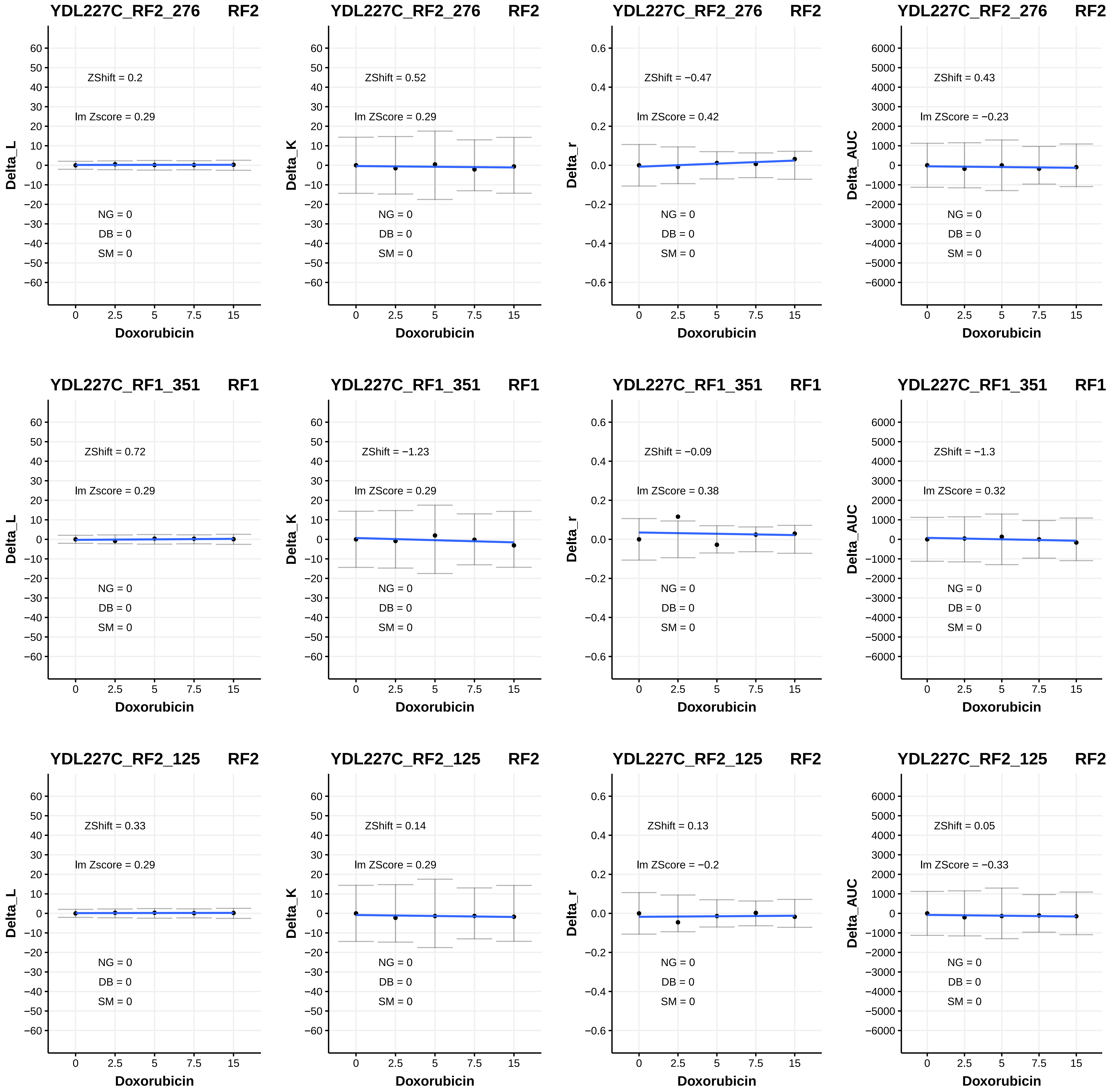
YDL227C_RF2_276 RF2
Delta_L
ZShift = 0.2
lm Zscore = 0.29
NG = 0
DB = 0
SM = 0
Doxorubicin
YDL227C_RF2_276 RF2
Delta_K
ZShift = 0.52
lm ZScore = 0.29
NG = 0
DB = 0
SM = 0
Doxorubicin
YDL227C_RF2_276 RF2
Delta_r
ZShift = −0.47
lm ZScore = 0.42
NG = 0
DB = 0
SM = 0
Doxorubicin
YDL227C_RF2_276 RF2
Delta_AUC
ZShift = 0.43
lm ZScore = −0.23
NG = 0
DB = 0
SM = 0
Doxorubicin
YDL227C_RF1_351 RF1
Delta_L
ZShift = 0.72
lm Zscore = 0.29
NG = 0
DB = 0
SM = 0
Doxorubicin
YDL227C_RF1_351 RF1
Delta_K
ZShift = −1.23
lm ZScore = 0.29
NG = 0
DB = 0
SM = 0
Doxorubicin
YDL227C_RF1_351 RF1
Delta_r
ZShift = −0.09
lm ZScore = 0.38
NG = 0
DB = 0
SM = 0
Doxorubicin
YDL227C_RF1_351 RF1
Delta_AUC
ZShift = −1.3
lm ZScore = 0.32
NG = 0
DB = 0
SM = 0
Doxorubicin
YDL227C_RF2_125 RF2
Delta_L
ZShift = 0.33
lm Zscore = 0.29
NG = 0
DB = 0
SM = 0
Doxorubicin
YDL227C_RF2_125 RF2
Delta_K
ZShift = 0.14
lm ZScore = 0.29
NG = 0
DB = 0
SM = 0
Doxorubicin
YDL227C_RF2_125 RF2
Delta_r
ZShift = 0.13
lm ZScore = −0.2
NG = 0
DB = 0
SM = 0
Doxorubicin
YDL227C_RF2_125 RF2
Delta_AUC
ZShift = 0.05
lm ZScore = −0.33
NG = 0
DB = 0
SM = 0
Doxorubicin

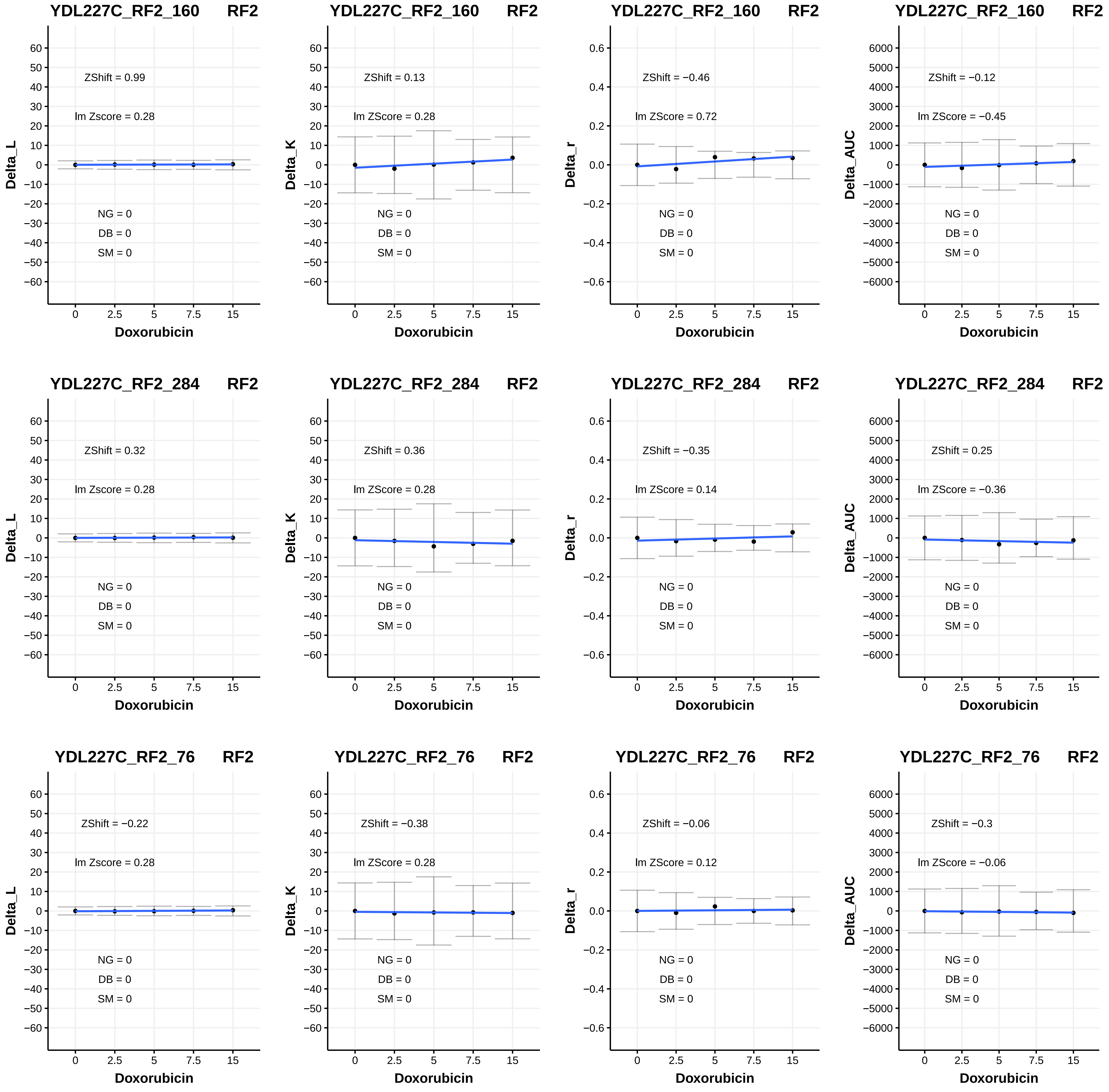

YDL227C_RF2_160 RF2
ZShift = 0.99
lm Zscore = 0.28
NG = 0
DB = 0
SM = 0
Delta_L
Doxorubicin
YDL227C_RF2_160 RF2
ZShift = 0.13
lm ZScore = 0.28
NG = 0
DB = 0
SM = 0
Delta_K
Doxorubicin
YDL227C_RF2_160 RF2
ZShift = −0.46
lm ZScore = 0.72
NG = 0
DB = 0
SM = 0
Delta_r
Doxorubicin
YDL227C_RF2_160 RF2
ZShift = −0.12
lm ZScore = −0.45
NG = 0
DB = 0
SM = 0
Delta_AUC
Doxorubicin
YDL227C_RF2_284 RF2
ZShift = 0.32
lm Zscore = 0.28
NG = 0
DB = 0
SM = 0
Delta_L
Doxorubicin
YDL227C_RF2_284 RF2
ZShift = 0.36
lm ZScore = 0.28
NG = 0
DB = 0
SM = 0
Delta_K
Doxorubicin
YDL227C_RF2_284 RF2
ZShift = −0.35
lm ZScore = 0.14
NG = 0
DB = 0
SM = 0
Delta_r
Doxorubicin
YDL227C_RF2_284 RF2
ZShift = 0.25
lm ZScore = −0.36
NG = 0
DB = 0
SM = 0
Delta_AUC
Doxorubicin
YDL227C_RF2_76 RF2
ZShift = −0.22
lm Zscore = 0.28
NG = 0
DB = 0
SM = 0
Delta_L
Doxorubicin
YDL227C_RF2_76 RF2
ZShift = −0.38
lm ZScore = 0.28
NG = 0
DB = 0
SM = 0
Delta_K
Doxorubicin
YDL227C_RF2_76 RF2
ZShift = −0.06
lm ZScore = 0.12
NG = 0
DB = 0
SM = 0
Delta_r
Doxorubicin
YDL227C_RF2_76 RF2
ZShift = −0.3
lm ZScore = −0.06
NG = 0
DB = 0
SM = 0
Delta_AUC
Doxorubicin

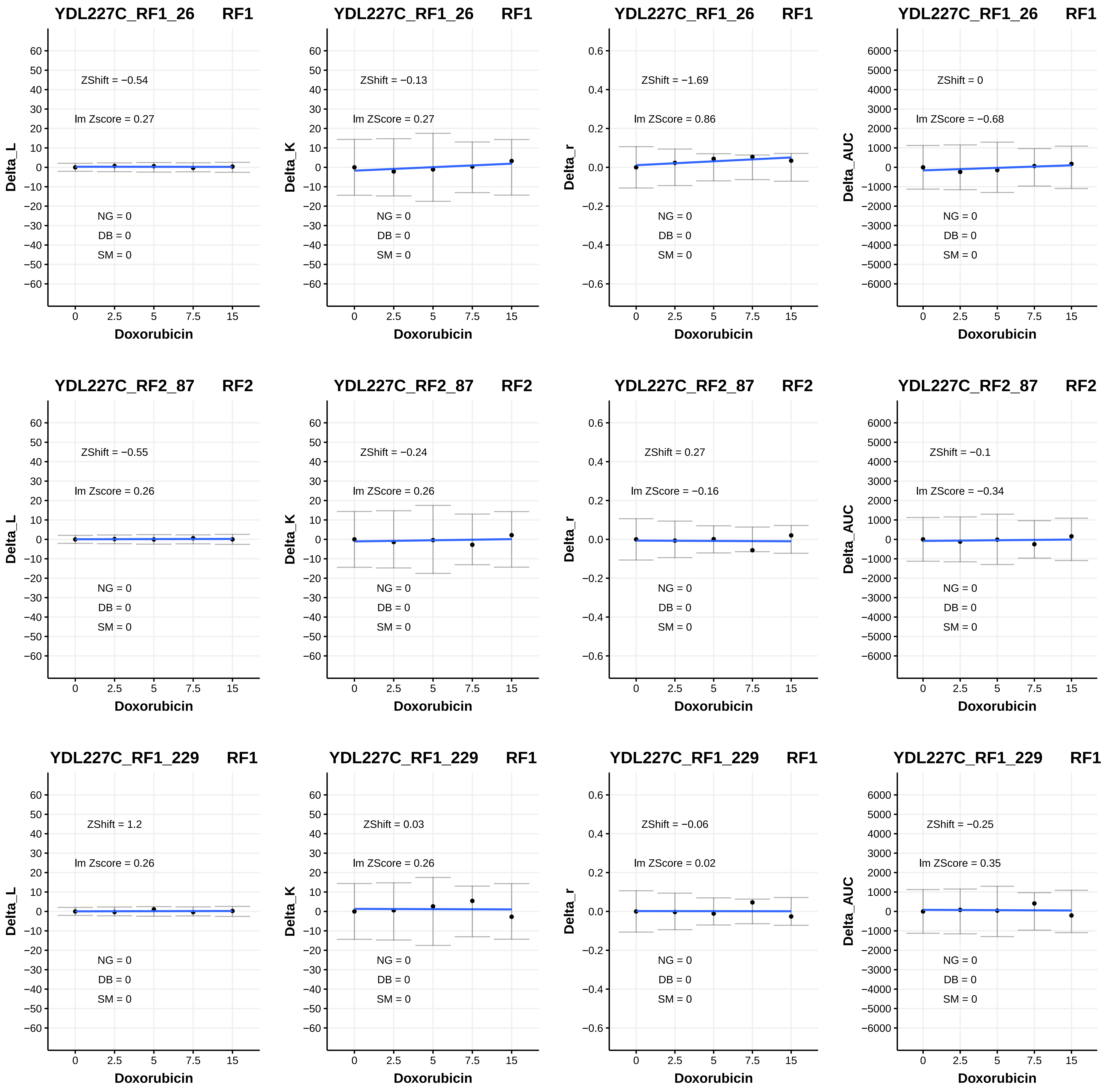
YDL227C_RF1_26 RF1
ZShift = −0.54
lm Zscore = 0.27
NG = 0
DB = 0
SM = 0
Delta_L
Doxorubicin
YDL227C_RF1_26 RF1
ZShift = −0.13
lm ZScore = 0.27
NG = 0
DB = 0
SM = 0
Delta_K
Doxorubicin
YDL227C_RF1_26 RF1
ZShift = −1.69
lm ZScore = 0.86
NG = 0
DB = 0
SM = 0
Delta_r
Doxorubicin
YDL227C_RF1_26 RF1
ZShift = 0
lm ZScore = −0.68
NG = 0
DB = 0
SM = 0
Delta_AUC
Doxorubicin
YDL227C_RF2_87 RF2
ZShift = −0.55
lm Zscore = 0.26
NG = 0
DB = 0
SM = 0
Delta_L
Doxorubicin
YDL227C_RF2_87 RF2
ZShift = −0.24
lm ZScore = 0.26
NG = 0
DB = 0
SM = 0
Delta_K
Doxorubicin
YDL227C_RF2_87 RF2
ZShift = 0.27
lm ZScore = −0.16
NG = 0
DB = 0
SM = 0
Delta_r
Doxorubicin
YDL227C_RF2_87 RF2
ZShift = −0.1
lm ZScore = −0.34
NG = 0
DB = 0
SM = 0
Delta_AUC
Doxorubicin
YDL227C_RF1_229 RF1
ZShift = 1.2
lm Zscore = 0.26
NG = 0
DB = 0
SM = 0
Delta_L
Doxorubicin
YDL227C_RF1_229 RF1
ZShift = 0.03
lm ZScore = 0.26
NG = 0
DB = 0
SM = 0
Delta_K
Doxorubicin
YDL227C_RF1_229 RF1
ZShift = −0.06
lm ZScore = 0.02
NG = 0
DB = 0
SM = 0
Delta_r
Doxorubicin
YDL227C_RF1_229 RF1
ZShift = −0.25
lm ZScore = 0.35
NG = 0
DB = 0
SM = 0
Delta_AUC
Doxorubicin

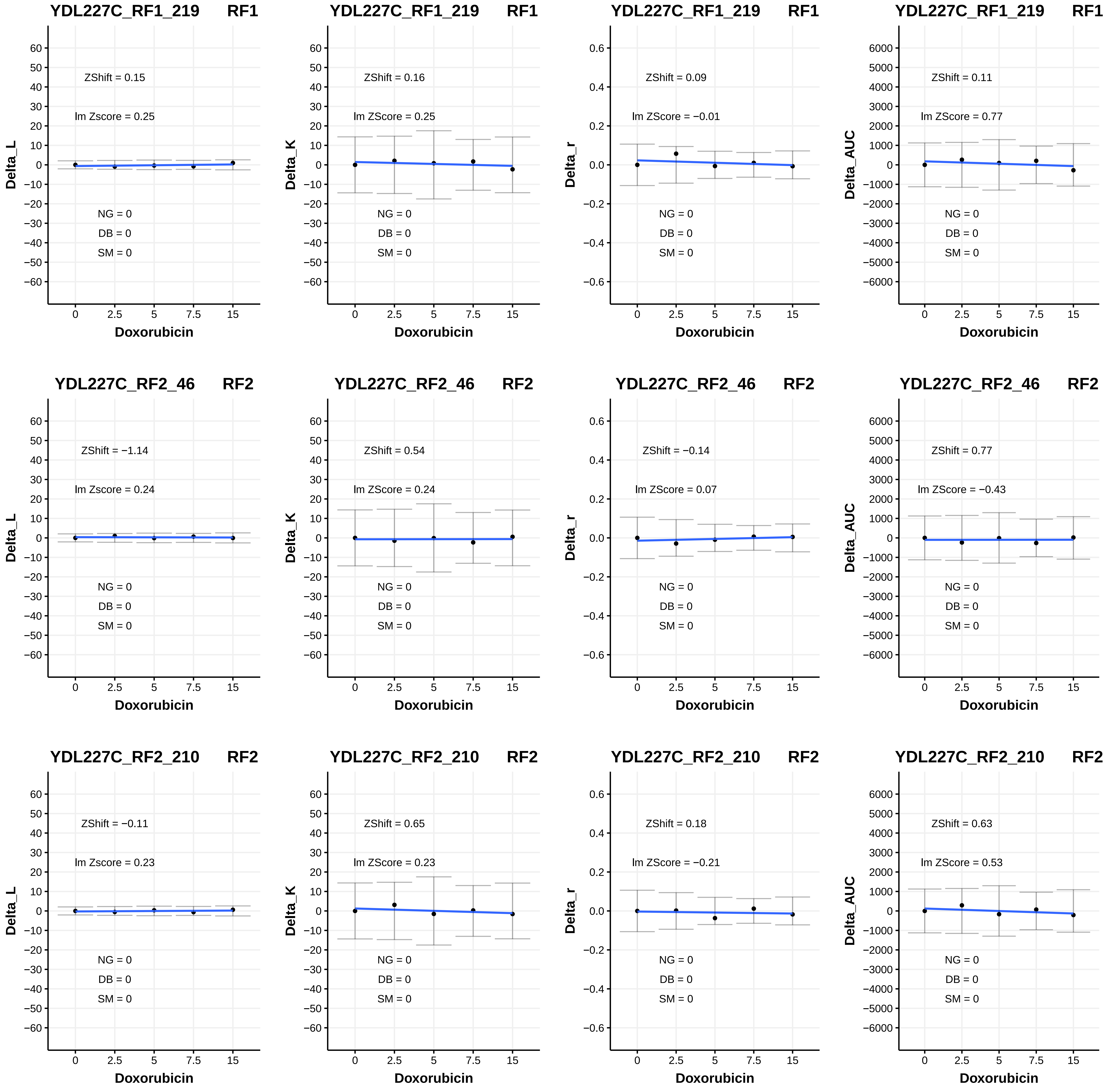

YDL227C_RF1_219 RF1
ZShift = 0.15
lm Zscore = 0.25
NG = 0
DB = 0
SM = 0
Delta_L
Doxorubicin
YDL227C_RF1_219 RF1
ZShift = 0.16
lm ZScore = 0.25
NG = 0
DB = 0
SM = 0
Delta_K
Doxorubicin
YDL227C_RF1_219 RF1
ZShift = 0.09
lm ZScore = −0.01
NG = 0
DB = 0
SM = 0
Delta_r
Doxorubicin
YDL227C_RF1_219 RF1
ZShift = 0.11
lm ZScore = 0.77
NG = 0
DB = 0
SM = 0
Delta_AUC
Doxorubicin
YDL227C_RF2_46 RF2
ZShift = −1.14
lm Zscore = 0.24
NG = 0
DB = 0
SM = 0
Delta_L
Doxorubicin
YDL227C_RF2_46 RF2
ZShift = 0.54
lm ZScore = 0.24
NG = 0
DB = 0
SM = 0
Delta_K
Doxorubicin
YDL227C_RF2_46 RF2
ZShift = −0.14
lm ZScore = 0.07
NG = 0
DB = 0
SM = 0
Delta_r
Doxorubicin
YDL227C_RF2_46 RF2
ZShift = 0.77
lm ZScore = −0.43
NG = 0
DB = 0
SM = 0
Delta_AUC
Doxorubicin
YDL227C_RF2_210 RF2
ZShift = −0.11
lm Zscore = 0.23
NG = 0
DB = 0
SM = 0
Delta_L
Doxorubicin
YDL227C_RF2_210 RF2
ZShift = 0.65
lm ZScore = 0.23
NG = 0
DB = 0
SM = 0
Delta_K
Doxorubicin
YDL227C_RF2_210 RF2
ZShift = 0.18
lm ZScore = −0.21
NG = 0
DB = 0
SM = 0
Delta_r
Doxorubicin
YDL227C_RF2_210 RF2
ZShift = 0.63
lm ZScore = 0.53
NG = 0
DB = 0
SM = 0
Delta_AUC
Doxorubicin

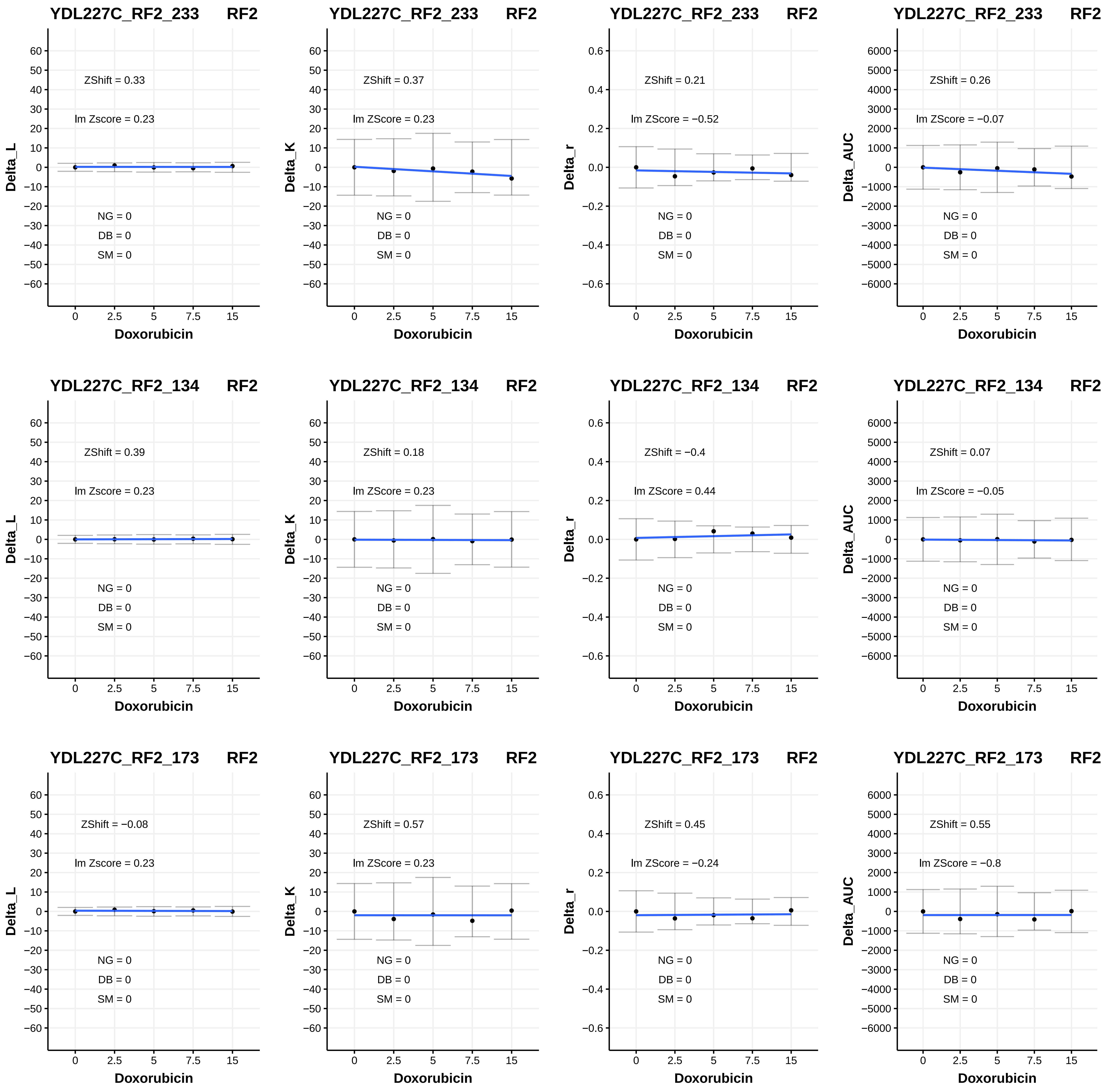

YDL227C_RF2_233 RF2
ZShift = 0.33
lm Zscore = 0.23
NG = 0
DB = 0
SM = 0
Delta_L
Doxorubicin
YDL227C_RF2_233 RF2
ZShift = 0.37
lm ZScore = 0.23
NG = 0
DB = 0
SM = 0
Delta_K
Doxorubicin
YDL227C_RF2_233 RF2
ZShift = 0.21
lm ZScore = −0.52
NG = 0
DB = 0
SM = 0
Delta_r
Doxorubicin
YDL227C_RF2_233 RF2
ZShift = 0.26
lm ZScore = −0.07
NG = 0
DB = 0
SM = 0
Delta_AUC
Doxorubicin
YDL227C_RF2_134 RF2
ZShift = 0.39
lm Zscore = 0.23
NG = 0
DB = 0
SM = 0
Delta_L
Doxorubicin
YDL227C_RF2_134 RF2
ZShift = 0.18
lm ZScore = 0.23
NG = 0
DB = 0
SM = 0
Delta_K
Doxorubicin
YDL227C_RF2_134 RF2
ZShift = −0.4
lm ZScore = 0.44
NG = 0
DB = 0
SM = 0
Delta_r
Doxorubicin
YDL227C_RF2_134 RF2
ZShift = 0.07
lm ZScore = −0.05
NG = 0
DB = 0
SM = 0
Delta_AUC
Doxorubicin
YDL227C_RF2_173 RF2
ZShift = −0.08
lm Zscore = 0.23
NG = 0
DB = 0
SM = 0
Delta_L
Doxorubicin
YDL227C_RF2_173 RF2
ZShift = 0.57
lm ZScore = 0.23
NG = 0
DB = 0
SM = 0
Delta_K
Doxorubicin
YDL227C_RF2_173 RF2
ZShift = 0.45
lm ZScore = −0.24
NG = 0
DB = 0
SM = 0
Delta_r
Doxorubicin
YDL227C_RF2_173 RF2
ZShift = 0.55
lm ZScore = −0.8
NG = 0
DB = 0
SM = 0
Delta_AUC
Doxorubicin

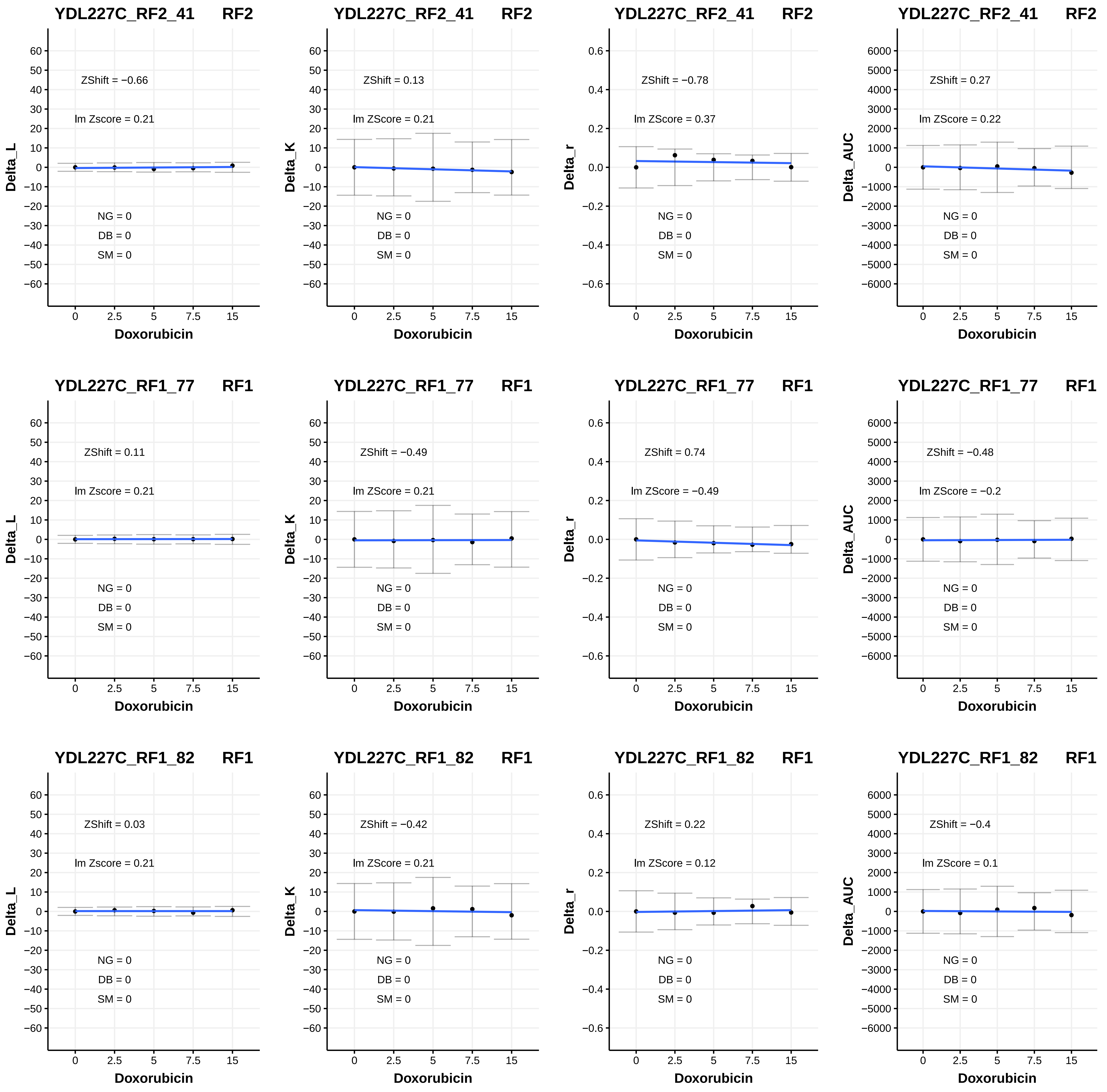
YDL227C_RF2_41 RF2
ZShift = −0.66
lm Zscore = 0.21
NG = 0
DB = 0
SM = 0
Delta_L
Doxorubicin
YDL227C_RF2_41 RF2
ZShift = 0.13
lm ZScore = 0.21
NG = 0
DB = 0
SM = 0
Delta_K
Doxorubicin
YDL227C_RF2_41 RF2
ZShift = −0.78
lm ZScore = 0.37
NG = 0
DB = 0
SM = 0
Delta_r
Doxorubicin
YDL227C_RF2_41 RF2
ZShift = 0.27
lm ZScore = 0.22
NG = 0
DB = 0
SM = 0
Delta_AUC
Doxorubicin
YDL227C_RF1_77 RF1
ZShift = 0.11
lm Zscore = 0.21
NG = 0
DB = 0
SM = 0
Delta_L
Doxorubicin
YDL227C_RF1_77 RF1
ZShift = −0.49
lm ZScore = 0.21
NG = 0
DB = 0
SM = 0
Delta_K
Doxorubicin
YDL227C_RF1_77 RF1
ZShift = 0.74
lm ZScore = −0.49
NG = 0
DB = 0
SM = 0
Delta_r
Doxorubicin
YDL227C_RF1_77 RF1
ZShift = −0.48
lm ZScore = −0.2
NG = 0
DB = 0
SM = 0
Delta_AUC
Doxorubicin
YDL227C_RF1_82 RF1
ZShift = 0.03
lm Zscore = 0.21
NG = 0
DB = 0
SM = 0
Delta_L
Doxorubicin
YDL227C_RF1_82 RF1
ZShift = −0.42
lm ZScore = 0.21
NG = 0
DB = 0
SM = 0
Delta_K
Doxorubicin
YDL227C_RF1_82 RF1
ZShift = 0.22
lm ZScore = 0.12
NG = 0
DB = 0
SM = 0
Delta_r
Doxorubicin
YDL227C_RF1_82 RF1
ZShift = −0.4
lm ZScore = 0.1
NG = 0
DB = 0
SM = 0
Delta_AUC
Doxorubicin

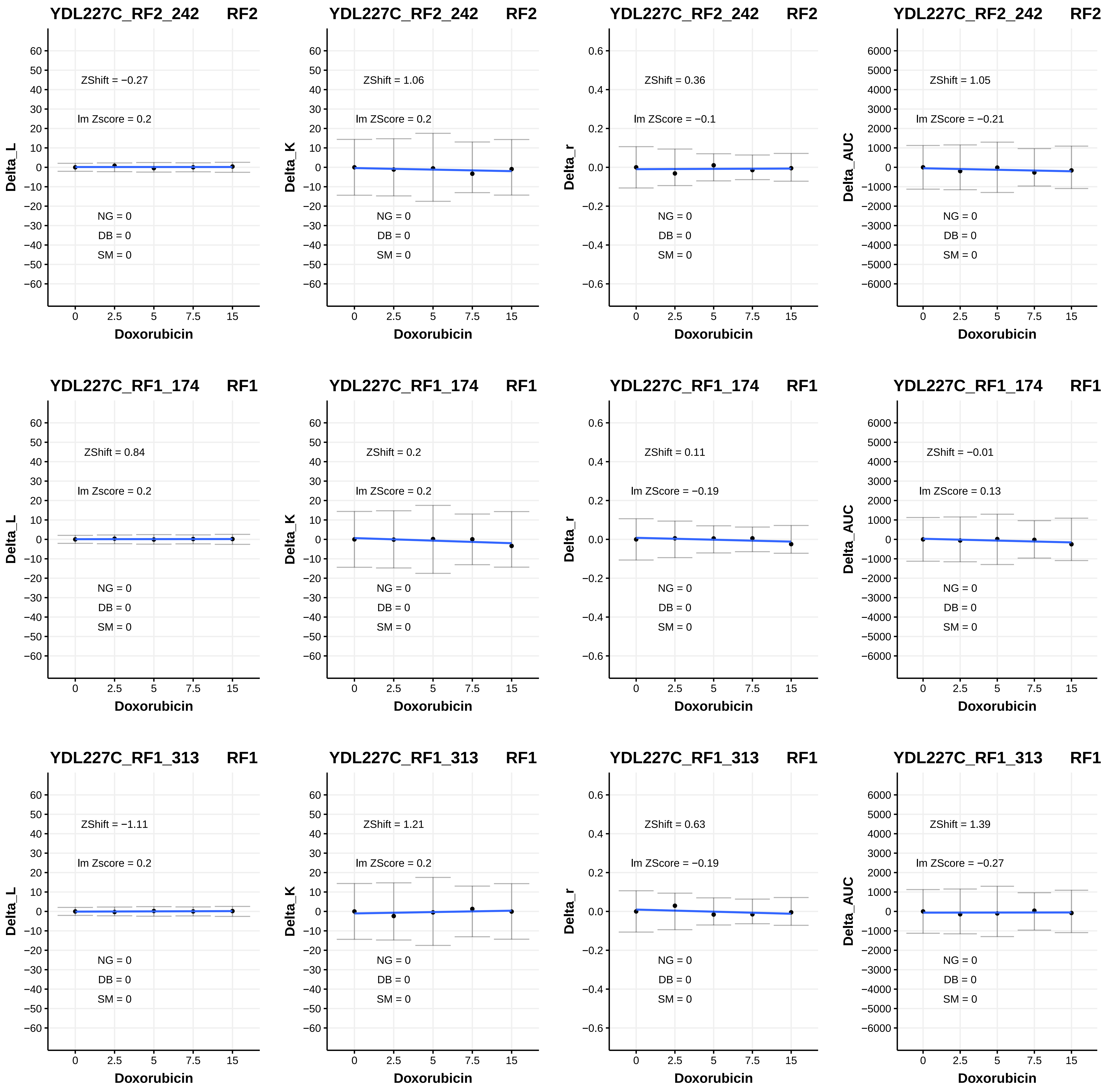
YDL227C_RF2_242 RF2
ZShift = −0.27
lm Zscore = 0.2
NG = 0
DB = 0
SM = 0
Delta_L
Doxorubicin
YDL227C_RF2_242 RF2
ZShift = 1.06
lm ZScore = 0.2
NG = 0
DB = 0
SM = 0
Delta_K
Doxorubicin
YDL227C_RF2_242 RF2
ZShift = 0.36
lm ZScore = −0.1
NG = 0
DB = 0
SM = 0
Delta_r
Doxorubicin
YDL227C_RF2_242 RF2
ZShift = 1.05
lm ZScore = −0.21
NG = 0
DB = 0
SM = 0
Delta_AUC
Doxorubicin
YDL227C_RF1_174 RF1
ZShift = 0.84
lm Zscore = 0.2
NG = 0
DB = 0
SM = 0
Delta_L
Doxorubicin
YDL227C_RF1_174 RF1
ZShift = 0.2
lm ZScore = 0.2
NG = 0
DB = 0
SM = 0
Delta_K
Doxorubicin
YDL227C_RF1_174 RF1
ZShift = 0.11
lm ZScore = −0.19
NG = 0
DB = 0
SM = 0
Delta_r
Doxorubicin
YDL227C_RF1_174 RF1
ZShift = −0.01
lm ZScore = 0.13
NG = 0
DB = 0
SM = 0
Delta_AUC
Doxorubicin
YDL227C_RF1_313 RF1
ZShift = −1.11
lm Zscore = 0.2
NG = 0
DB = 0
SM = 0
Delta_L
Doxorubicin
YDL227C_RF1_313 RF1
ZShift = 1.21
lm ZScore = 0.2
NG = 0
DB = 0
SM = 0
Delta_K
Doxorubicin
YDL227C_RF1_313 RF1
ZShift = 0.63
lm ZScore = −0.19
NG = 0
DB = 0
SM = 0
Delta_r
Doxorubicin
YDL227C_RF1_313 RF1
ZShift = 1.39
lm ZScore = −0.27
NG = 0
DB = 0
SM = 0
Delta_AUC
Doxorubicin

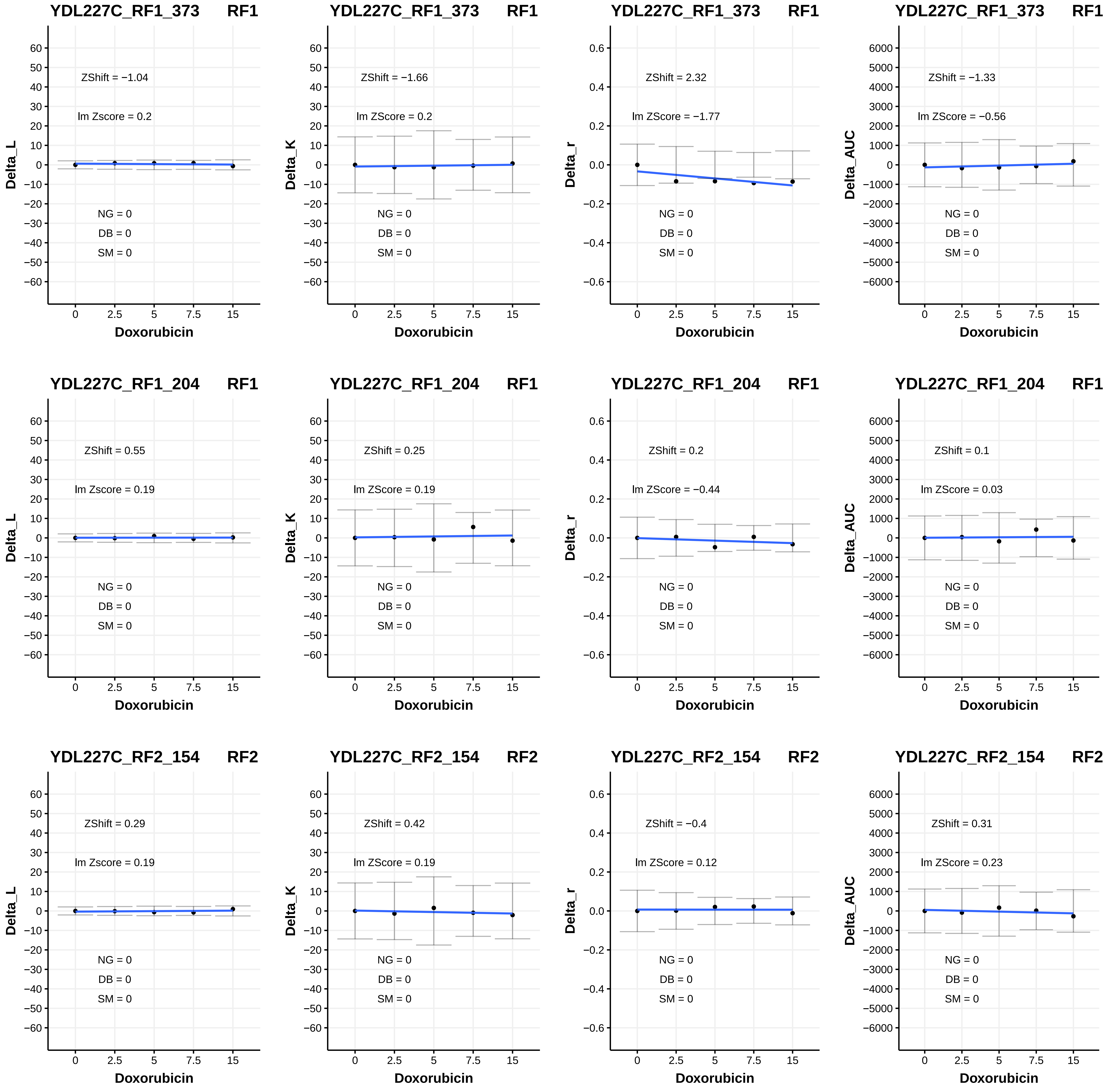
YDL227C_RF1_373 RF1
ZShift = −1.04
lm Zscore = 0.2
NG = 0
DB = 0
SM = 0
Delta_L
Doxorubicin
YDL227C_RF1_373 RF1
ZShift = −1.66
lm ZScore = 0.2
NG = 0
DB = 0
SM = 0
Delta_K
Doxorubicin
YDL227C_RF1_373 RF1
ZShift = 2.32
lm ZScore = −1.77
NG = 0
DB = 0
SM = 0
Delta_r
Doxorubicin
YDL227C_RF1_373 RF1
ZShift = −1.33
lm ZScore = −0.56
NG = 0
DB = 0
SM = 0
Delta_AUC
Doxorubicin
YDL227C_RF1_204 RF1
ZShift = 0.55
lm Zscore = 0.19
NG = 0
DB = 0
SM = 0
Delta_L
Doxorubicin
YDL227C_RF1_204 RF1
ZShift = 0.25
lm ZScore = 0.19
NG = 0
DB = 0
SM = 0
Delta_K
Doxorubicin
YDL227C_RF1_204 RF1
ZShift = 0.2
lm ZScore = −0.44
NG = 0
DB = 0
SM = 0
Delta_r
Doxorubicin
YDL227C_RF1_204 RF1
ZShift = 0.1
lm ZScore = 0.03
NG = 0
DB = 0
SM = 0
Delta_AUC
Doxorubicin
YDL227C_RF2_154 RF2
ZShift = 0.29
lm Zscore = 0.19
NG = 0
DB = 0
SM = 0
Delta_L
Doxorubicin
YDL227C_RF2_154 RF2
ZShift = 0.42
lm ZScore = 0.19
NG = 0
DB = 0
SM = 0
Delta_K
Doxorubicin
YDL227C_RF2_154 RF2
ZShift = −0.4
lm ZScore = 0.12
NG = 0
DB = 0
SM = 0
Delta_r
Doxorubicin
YDL227C_RF2_154 RF2
ZShift = 0.31
lm ZScore = 0.23
NG = 0
DB = 0
SM = 0
Delta_AUC
Doxorubicin

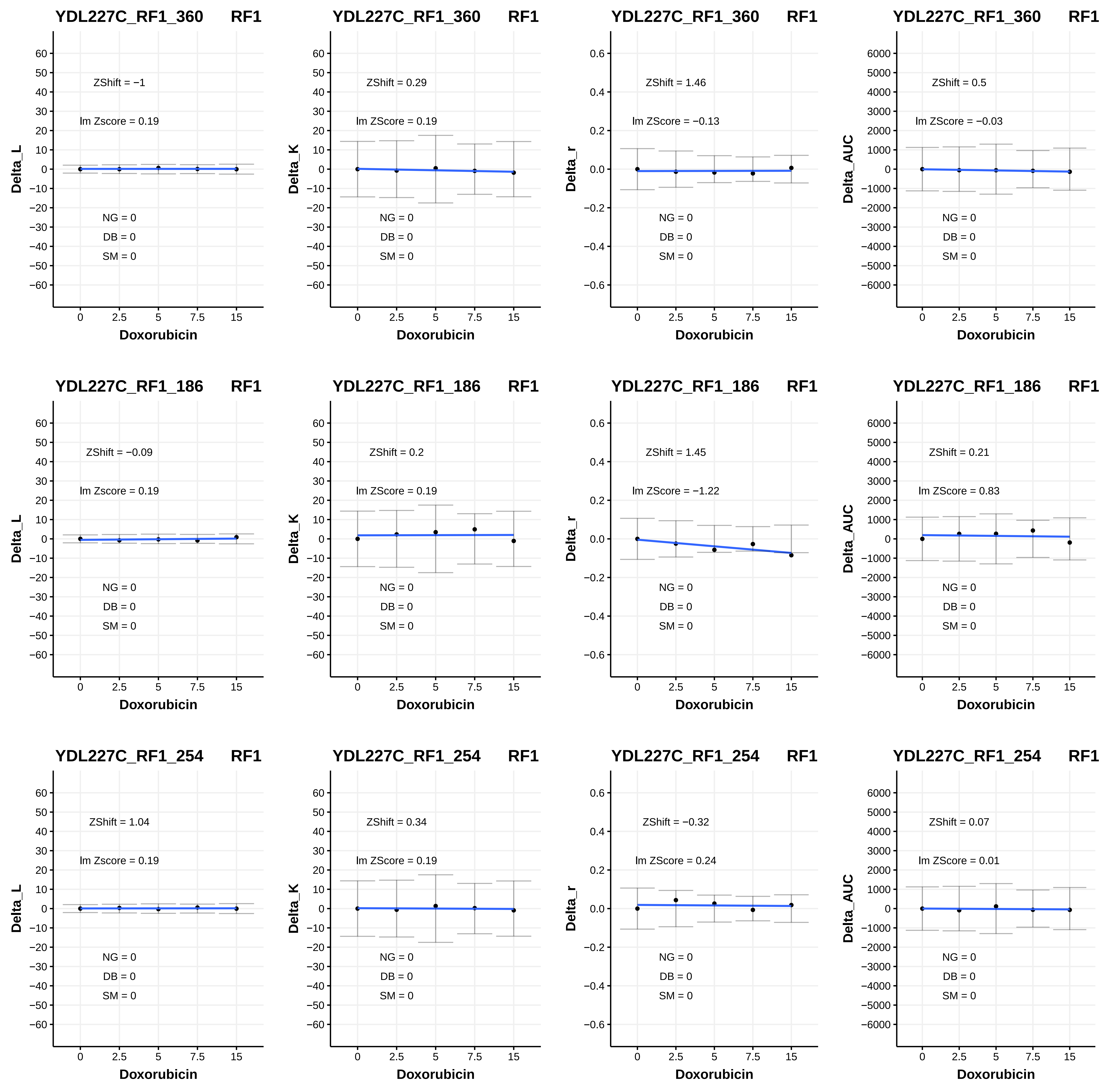
YDL227C_RF1_360 RF1
ZShift = −1
lm Zscore = 0.19
NG = 0
DB = 0
SM = 0
Delta_L
Doxorubicin
YDL227C_RF1_360 RF1
ZShift = 0.29
lm ZScore = 0.19
NG = 0
DB = 0
SM = 0
Delta_K
Doxorubicin
YDL227C_RF1_360 RF1
ZShift = 1.46
lm ZScore = −0.13
NG = 0
DB = 0
SM = 0
Delta_r
Doxorubicin
YDL227C_RF1_360 RF1
ZShift = 0.5
lm ZScore = −0.03
NG = 0
DB = 0
SM = 0
Delta_AUC
Doxorubicin
YDL227C_RF1_186 RF1
ZShift = −0.09
lm Zscore = 0.19
NG = 0
DB = 0
SM = 0
Delta_L
Doxorubicin
YDL227C_RF1_186 RF1
ZShift = 0.2
lm ZScore = 0.19
NG = 0
DB = 0
SM = 0
Delta_K
Doxorubicin
YDL227C_RF1_186 RF1
ZShift = 1.45
lm ZScore = −1.22
NG = 0
DB = 0
SM = 0
Delta_r
Doxorubicin
YDL227C_RF1_186 RF1
ZShift = 0.21
lm ZScore = 0.83
NG = 0
DB = 0
SM = 0
Delta_AUC
Doxorubicin
YDL227C_RF1_254 RF1
ZShift = 1.04
lm Zscore = 0.19
NG = 0
DB = 0
SM = 0
Delta_L
Doxorubicin
YDL227C_RF1_254 RF1
ZShift = 0.34
lm ZScore = 0.19
NG = 0
DB = 0
SM = 0
Delta_K
Doxorubicin
YDL227C_RF1_254 RF1
ZShift = −0.32
lm ZScore = 0.24
NG = 0
DB = 0
SM = 0
Delta_r
Doxorubicin
YDL227C_RF1_254 RF1
ZShift = 0.07
lm ZScore = 0.01
NG = 0
DB = 0
SM = 0
Delta_AUC
Doxorubicin

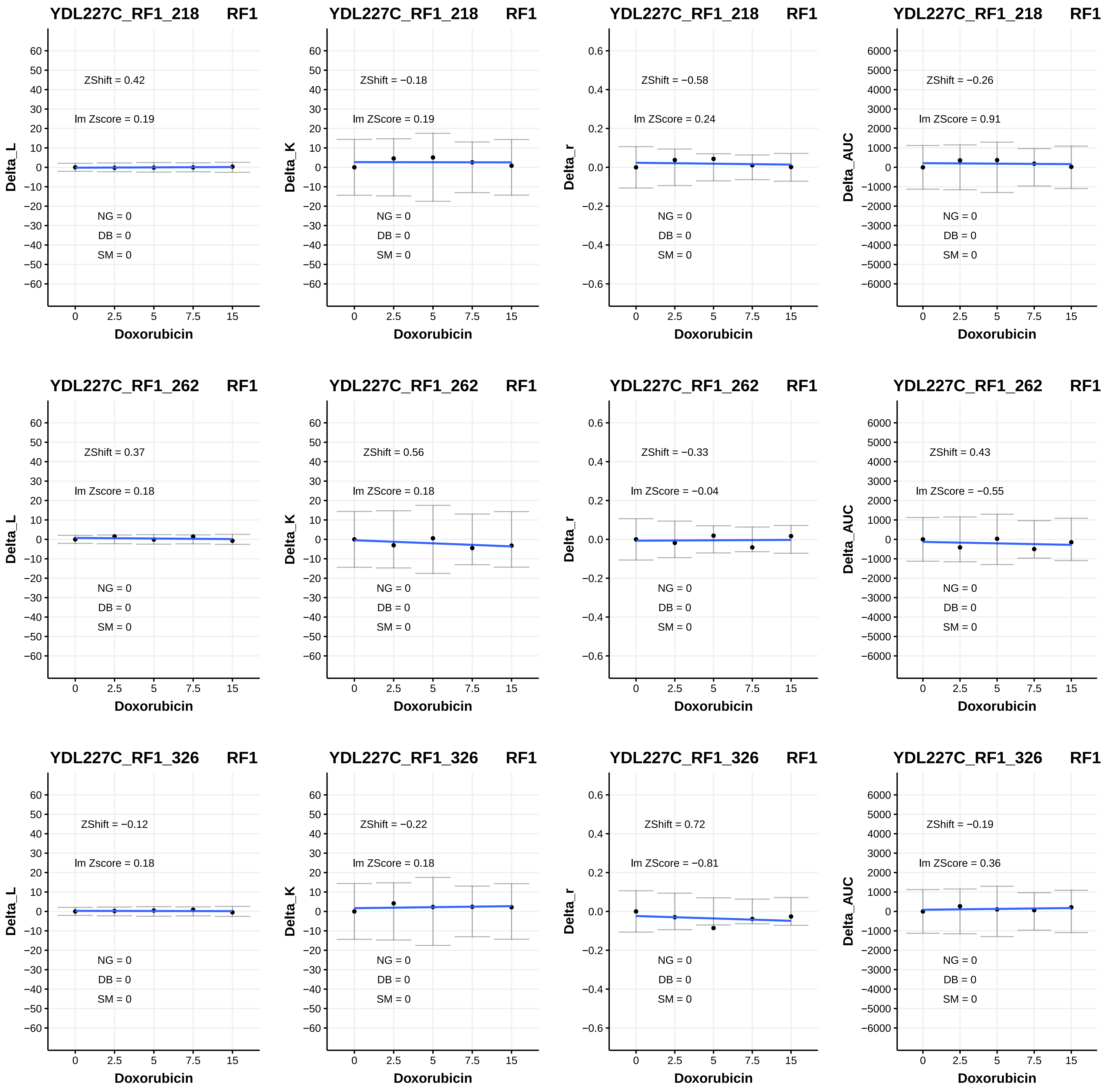

YDL227C_RF1_218 RF1
ZShift = 0.42
lm Zscore = 0.19
NG = 0
DB = 0
SM = 0
Delta_L
Doxorubicin
YDL227C_RF1_218 RF1
ZShift = −0.18
lm ZScore = 0.19
NG = 0
DB = 0
SM = 0
Delta_K
Doxorubicin
YDL227C_RF1_218 RF1
ZShift = −0.58
lm ZScore = 0.24
NG = 0
DB = 0
SM = 0
Delta_r
Doxorubicin
YDL227C_RF1_218 RF1
ZShift = −0.26
lm ZScore = 0.91
NG = 0
DB = 0
SM = 0
Delta_AUC
Doxorubicin
YDL227C_RF1_262 RF1
ZShift = 0.37
lm Zscore = 0.18
NG = 0
DB = 0
SM = 0
Delta_L
Doxorubicin
YDL227C_RF1_262 RF1
ZShift = 0.56
lm ZScore = 0.18
NG = 0
DB = 0
SM = 0
Delta_K
Doxorubicin
YDL227C_RF1_262 RF1
ZShift = −0.33
lm ZScore = −0.04
NG = 0
DB = 0
SM = 0
Delta_r
Doxorubicin
YDL227C_RF1_262 RF1
ZShift = 0.43
lm ZScore = −0.55
NG = 0
DB = 0
SM = 0
Delta_AUC
Doxorubicin
YDL227C_RF1_326 RF1
ZShift = −0.12
lm Zscore = 0.18
NG = 0
DB = 0
SM = 0
Delta_L
Doxorubicin
YDL227C_RF1_326 RF1
ZShift = −0.22
lm ZScore = 0.18
NG = 0
DB = 0
SM = 0
Delta_K
Doxorubicin
YDL227C_RF1_326 RF1
ZShift = 0.72
lm ZScore = −0.81
NG = 0
DB = 0
SM = 0
Delta_r
Doxorubicin
YDL227C_RF1_326 RF1
ZShift = −0.19
lm ZScore = 0.36
NG = 0
DB = 0
SM = 0
Delta_AUC
Doxorubicin

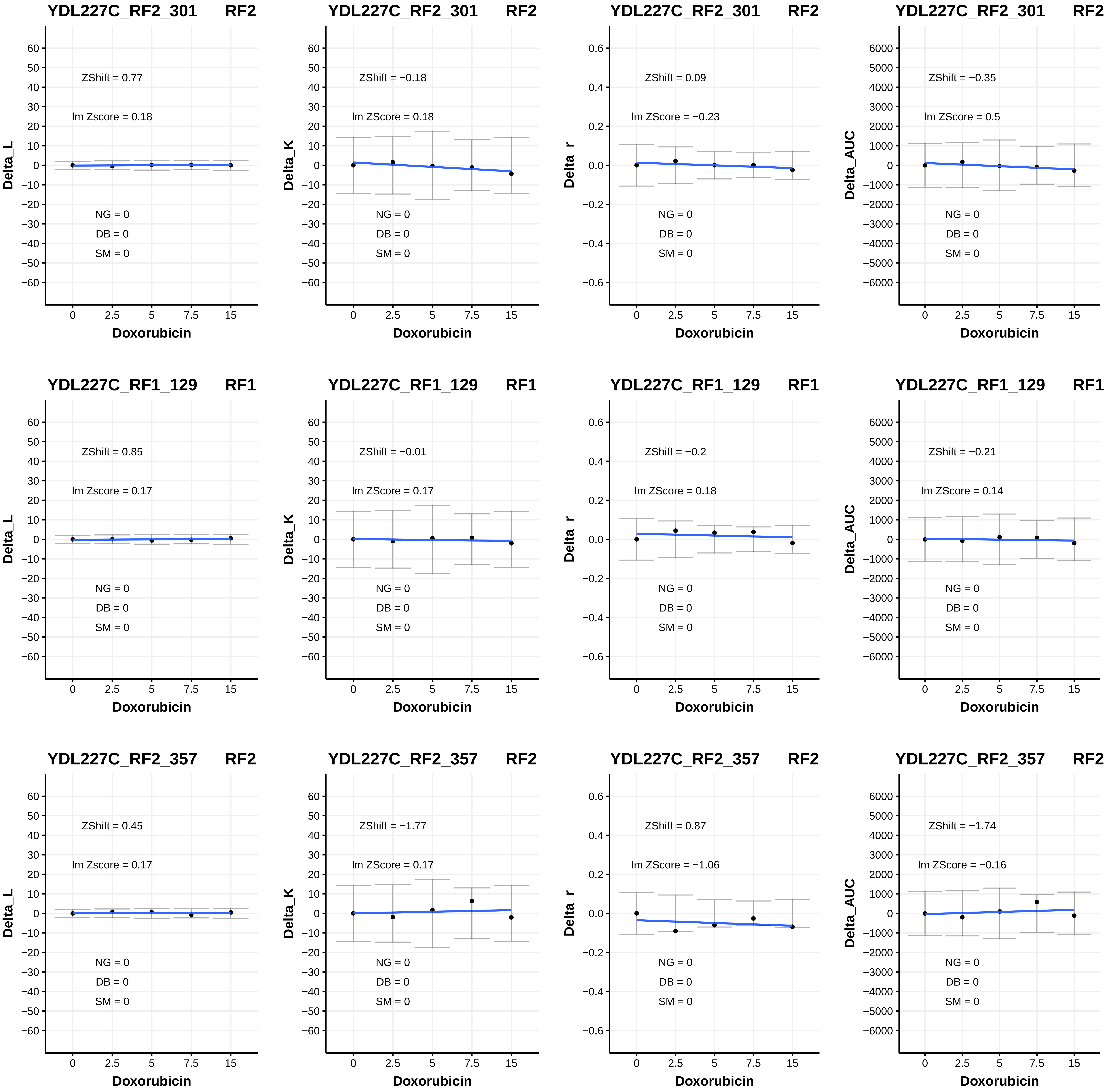
YDL227C_RF2_301 RF2
ZShift = 0.77
lm Zscore = 0.18
NG = 0
DB = 0
SM = 0
Delta_L
Doxorubicin
YDL227C_RF2_301 RF2
ZShift = −0.18
lm ZScore = 0.18
NG = 0
DB = 0
SM = 0
Delta_K
Doxorubicin
YDL227C_RF2_301 RF2
ZShift = 0.09
lm ZScore = −0.23
NG = 0
DB = 0
SM = 0
Delta_r
Doxorubicin
YDL227C_RF2_301 RF2
ZShift = −0.35
lm ZScore = 0.5
NG = 0
DB = 0
SM = 0
Delta_AUC
Doxorubicin
YDL227C_RF1_129 RF1
ZShift = 0.85
lm Zscore = 0.17
NG = 0
DB = 0
SM = 0
Delta_L
Doxorubicin
YDL227C_RF1_129 RF1
ZShift = −0.01
lm ZScore = 0.17
NG = 0
DB = 0
SM = 0
Delta_K
Doxorubicin
YDL227C_RF1_129 RF1
ZShift = −0.2
lm ZScore = 0.18
NG = 0
DB = 0
SM = 0
Delta_r
Doxorubicin
YDL227C_RF1_129 RF1
ZShift = −0.21
lm ZScore = 0.14
NG = 0
DB = 0
SM = 0
Delta_AUC
Doxorubicin
YDL227C_RF2_357 RF2
ZShift = 0.45
lm Zscore = 0.17
NG = 0
DB = 0
SM = 0
Delta_L
Doxorubicin
YDL227C_RF2_357 RF2
ZShift = −1.77
lm ZScore = 0.17
NG = 0
DB = 0
SM = 0
Delta_K
Doxorubicin
YDL227C_RF2_357 RF2
ZShift = 0.87
lm ZScore = −1.06
NG = 0
DB = 0
SM = 0
Delta_r
Doxorubicin
YDL227C_RF2_357 RF2
ZShift = −1.74
lm ZScore = −0.16
NG = 0
DB = 0
SM = 0
Delta_AUC
Doxorubicin

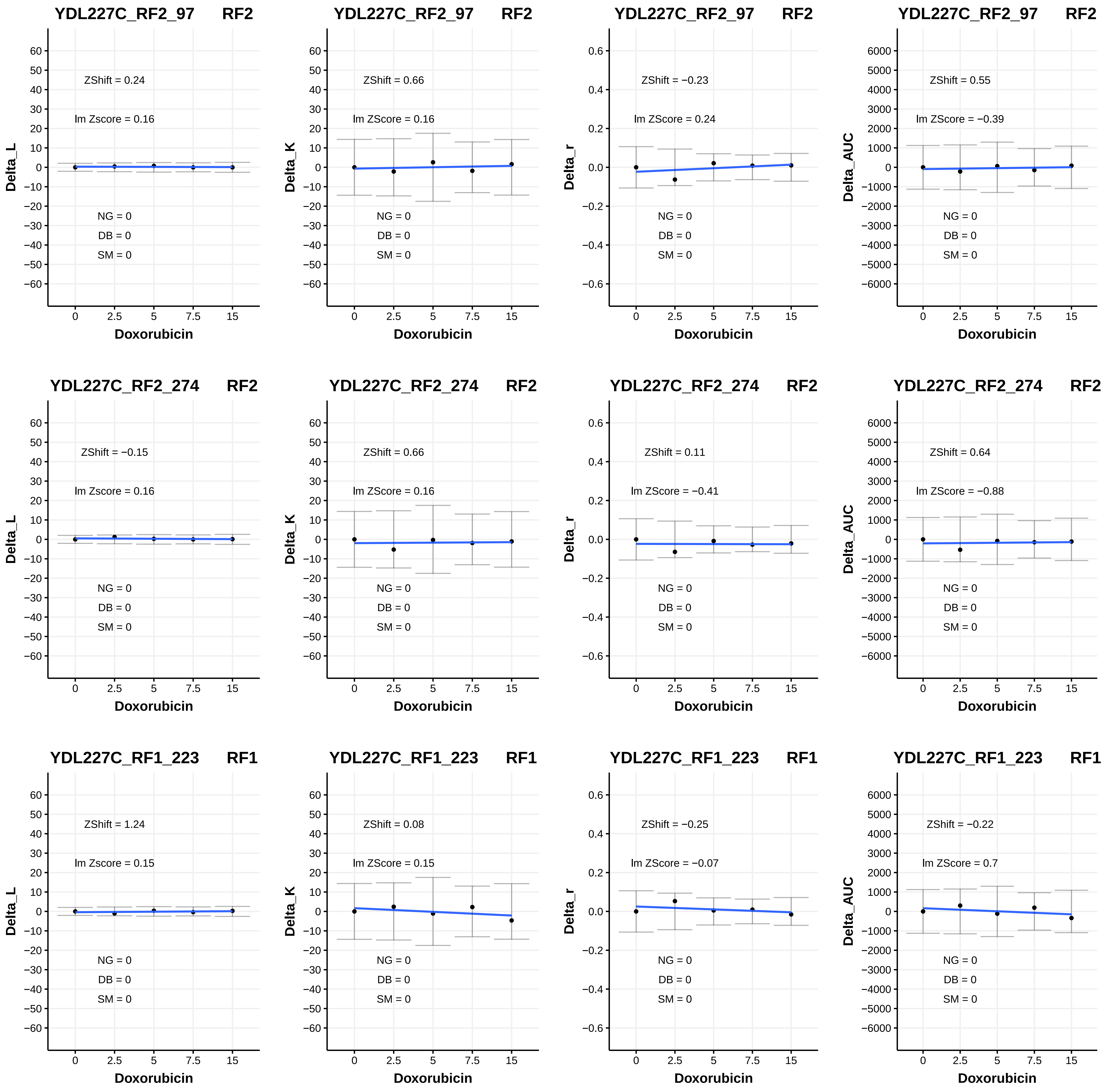

YDL227C_RF2_97 RF2
ZShift = 0.24
lm Zscore = 0.16
NG = 0
DB = 0
SM = 0
Delta_L
Doxorubicin
YDL227C_RF2_97 RF2
ZShift = 0.66
lm ZScore = 0.16
NG = 0
DB = 0
SM = 0
Delta_K
Doxorubicin
YDL227C_RF2_97 RF2
ZShift = −0.23
lm ZScore = 0.24
NG = 0
DB = 0
SM = 0
Delta_r
Doxorubicin
YDL227C_RF2_97 RF2
ZShift = 0.55
lm ZScore = −0.39
NG = 0
DB = 0
SM = 0
Delta_AUC
Doxorubicin
YDL227C_RF2_274 RF2
ZShift = −0.15
lm Zscore = 0.16
NG = 0
DB = 0
SM = 0
Delta_L
Doxorubicin
YDL227C_RF2_274 RF2
ZShift = 0.66
lm ZScore = 0.16
NG = 0
DB = 0
SM = 0
Delta_K
Doxorubicin
YDL227C_RF2_274 RF2
ZShift = 0.11
lm ZScore = −0.41
NG = 0
DB = 0
SM = 0
Delta_r
Doxorubicin
YDL227C_RF2_274 RF2
ZShift = 0.64
lm ZScore = −0.88
NG = 0
DB = 0
SM = 0
Delta_AUC
Doxorubicin
YDL227C_RF1_223 RF1
ZShift = 1.24
lm Zscore = 0.15
NG = 0
DB = 0
SM = 0
Delta_L
Doxorubicin
YDL227C_RF1_223 RF1
ZShift = 0.08
lm ZScore = 0.15
NG = 0
DB = 0
SM = 0
Delta_K
Doxorubicin
YDL227C_RF1_223 RF1
ZShift = −0.25
lm ZScore = −0.07
NG = 0
DB = 0
SM = 0
Delta_r
Doxorubicin
YDL227C_RF1_223 RF1
ZShift = −0.22
lm ZScore = 0.7
NG = 0
DB = 0
SM = 0
Delta_AUC
Doxorubicin

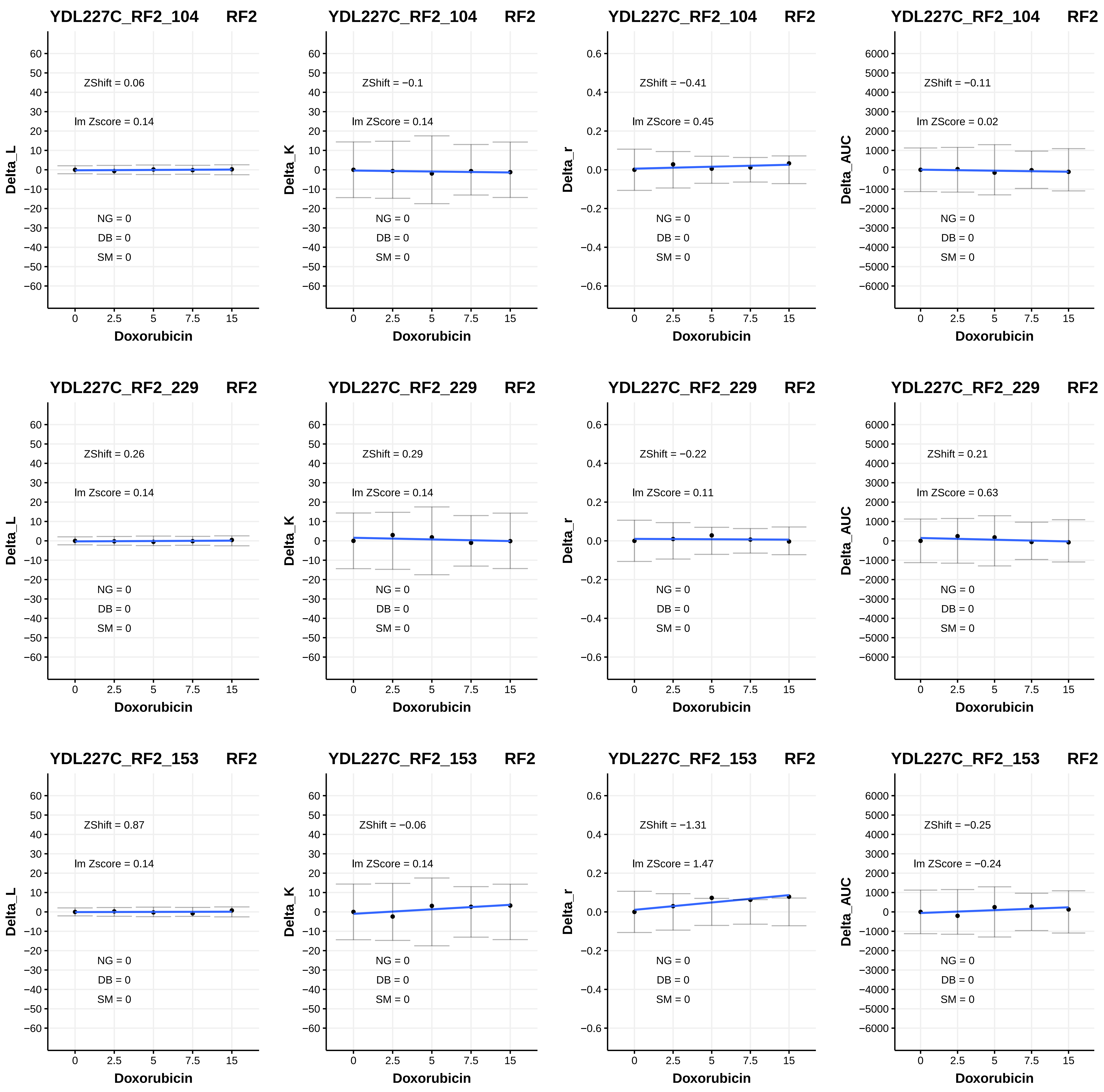
YDL227C_RF2_104 RF2
ZShift = 0.06
lm Zscore = 0.14
NG = 0
DB = 0
SM = 0
Delta_L
Doxorubicin
YDL227C_RF2_104 RF2
ZShift = −0.1
lm ZScore = 0.14
NG = 0
DB = 0
SM = 0
Delta_K
Doxorubicin
YDL227C_RF2_104 RF2
ZShift = −0.41
lm ZScore = 0.45
NG = 0
DB = 0
SM = 0
Delta_r
Doxorubicin
YDL227C_RF2_104 RF2
ZShift = −0.11
lm ZScore = 0.02
NG = 0
DB = 0
SM = 0
Delta_AUC
Doxorubicin
YDL227C_RF2_229 RF2
ZShift = 0.26
lm Zscore = 0.14
NG = 0
DB = 0
SM = 0
Delta_L
Doxorubicin
YDL227C_RF2_229 RF2
ZShift = 0.29
lm ZScore = 0.14
NG = 0
DB = 0
SM = 0
Delta_K
Doxorubicin
YDL227C_RF2_229 RF2
ZShift = −0.22
lm ZScore = 0.11
NG = 0
DB = 0
SM = 0
Delta_r
Doxorubicin
YDL227C_RF2_229 RF2
ZShift = 0.21
lm ZScore = 0.63
NG = 0
DB = 0
SM = 0
Delta_AUC
Doxorubicin
YDL227C_RF2_153 RF2
ZShift = 0.87
lm Zscore = 0.14
NG = 0
DB = 0
SM = 0
Delta_L
Doxorubicin
YDL227C_RF2_153 RF2
ZShift = −0.06
lm ZScore = 0.14
NG = 0
DB = 0
SM = 0
Delta_K
Doxorubicin
YDL227C_RF2_153 RF2
ZShift = −1.31
lm ZScore = 1.47
NG = 0
DB = 0
SM = 0
Delta_r
Doxorubicin
YDL227C_RF2_153 RF2
ZShift = −0.25
lm ZScore = −0.24
NG = 0
DB = 0
SM = 0
Delta_AUC
Doxorubicin

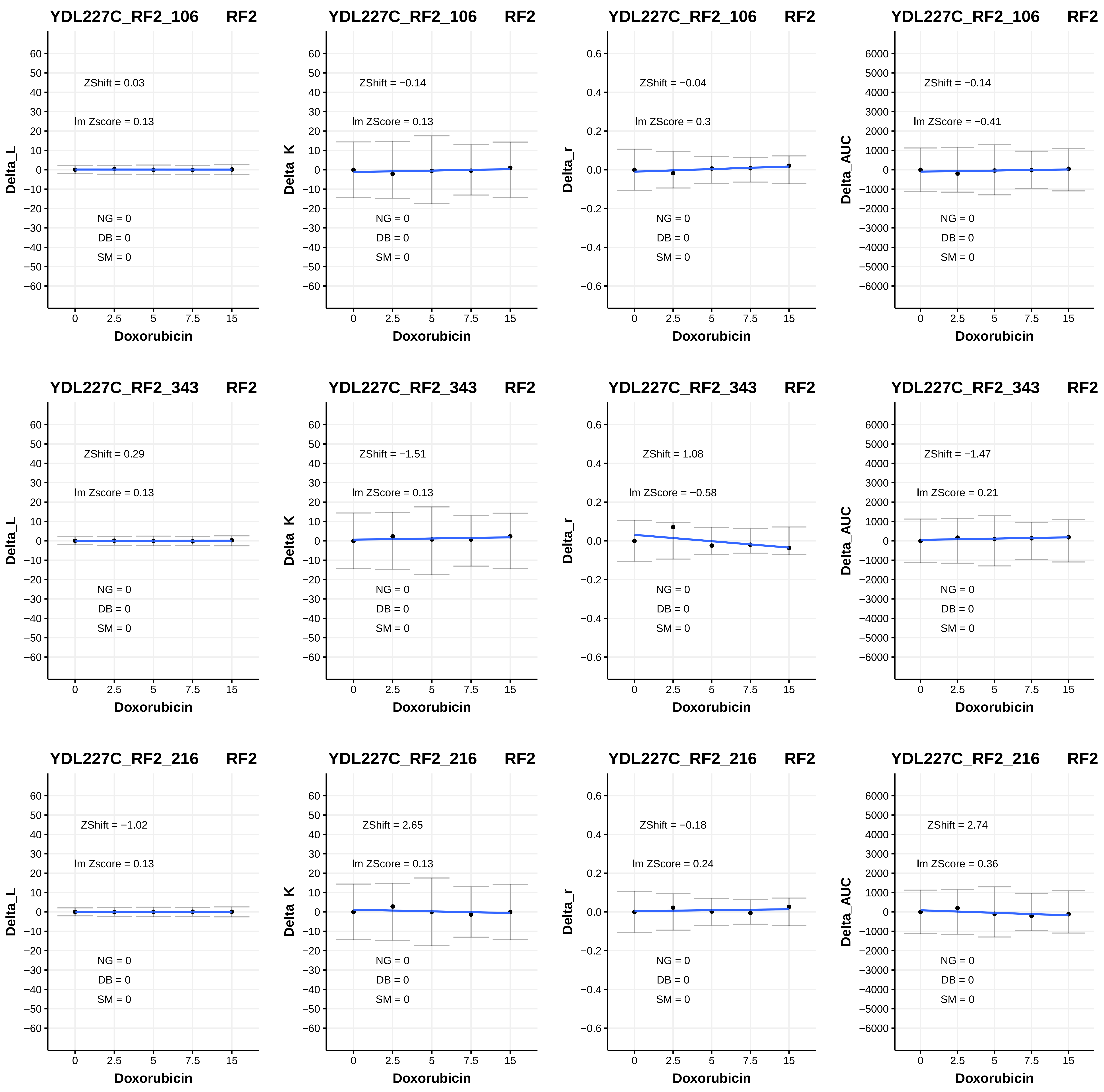

YDL227C_RF2_106 RF2
ZShift = 0.03
lm Zscore = 0.13
NG = 0
DB = 0
SM = 0
Delta_L
Doxorubicin
YDL227C_RF2_106 RF2
ZShift = −0.14
lm ZScore = 0.13
NG = 0
DB = 0
SM = 0
Delta_K
Doxorubicin
YDL227C_RF2_106 RF2
ZShift = −0.04
lm ZScore = 0.3
NG = 0
DB = 0
SM = 0
Delta_r
Doxorubicin
YDL227C_RF2_106 RF2
ZShift = −0.14
lm ZScore = −0.41
NG = 0
DB = 0
SM = 0
Delta_AUC
Doxorubicin
YDL227C_RF2_343 RF2
ZShift = 0.29
lm Zscore = 0.13
NG = 0
DB = 0
SM = 0
Delta_L
Doxorubicin
YDL227C_RF2_343 RF2
ZShift = −1.51
lm ZScore = 0.13
NG = 0
DB = 0
SM = 0
Delta_K
Doxorubicin
YDL227C_RF2_343 RF2
ZShift = 1.08
lm ZScore = −0.58
NG = 0
DB = 0
SM = 0
Delta_r
Doxorubicin
YDL227C_RF2_343 RF2
ZShift = −1.47
lm ZScore = 0.21
NG = 0
DB = 0
SM = 0
Delta_AUC
Doxorubicin
YDL227C_RF2_216 RF2
ZShift = −1.02
lm Zscore = 0.13
NG = 0
DB = 0
SM = 0
Delta_L
Doxorubicin
YDL227C_RF2_216 RF2
ZShift = 2.65
lm ZScore = 0.13
NG = 0
DB = 0
SM = 0
Delta_K
Doxorubicin
YDL227C_RF2_216 RF2
ZShift = −0.18
lm ZScore = 0.24
NG = 0
DB = 0
SM = 0
Delta_r
Doxorubicin
YDL227C_RF2_216 RF2
ZShift = 2.74
lm ZScore = 0.36
NG = 0
DB = 0
SM = 0
Delta_AUC
Doxorubicin

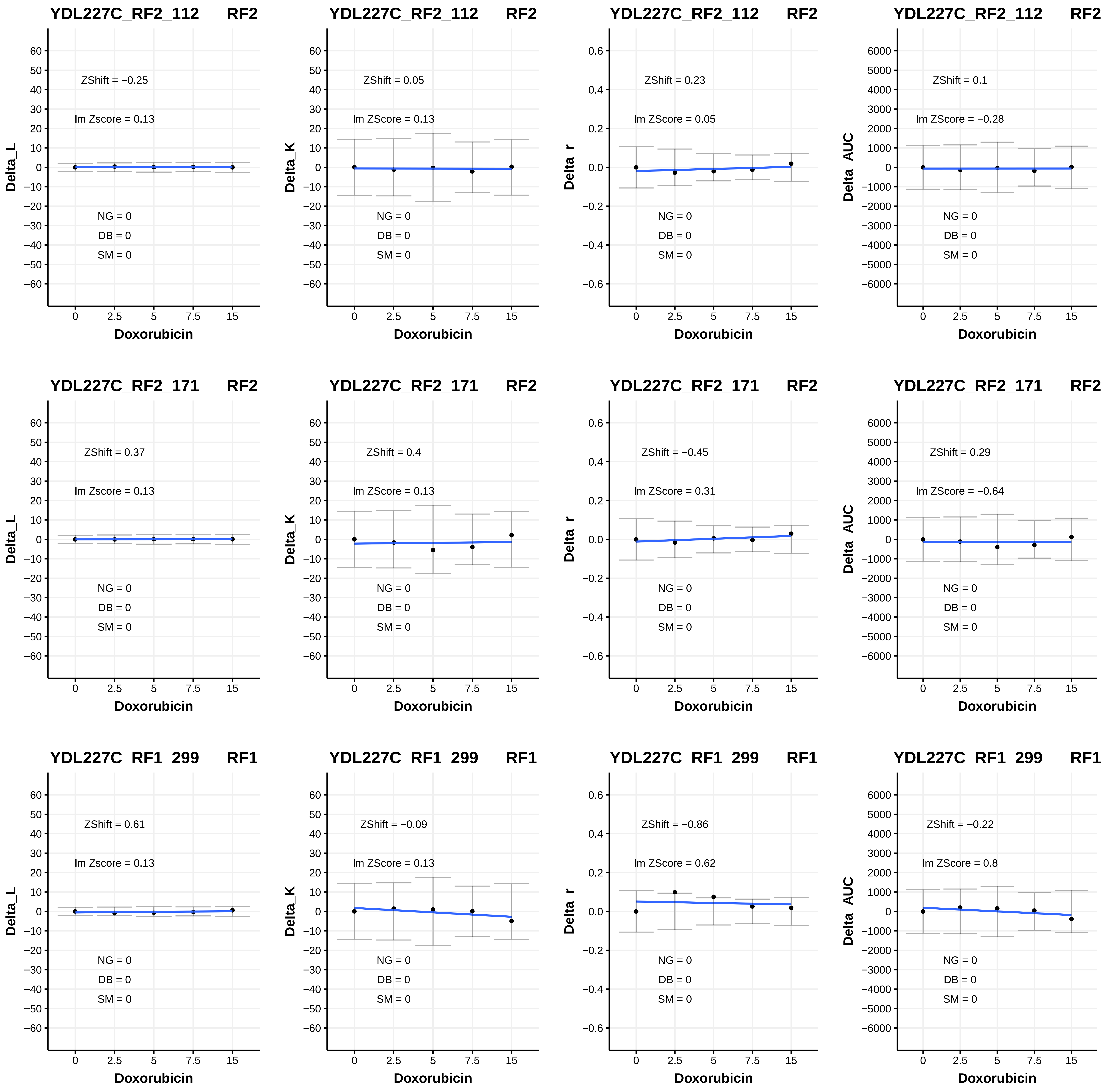
YDL227C_RF2_112 RF2
ZShift = −0.25
lm Zscore = 0.13
NG = 0
DB = 0
SM = 0
Delta_L
Doxorubicin
YDL227C_RF2_112 RF2
ZShift = 0.05
lm ZScore = 0.13
NG = 0
DB = 0
SM = 0
Delta_K
Doxorubicin
YDL227C_RF2_112 RF2
ZShift = 0.23
lm ZScore = 0.05
NG = 0
DB = 0
SM = 0
Delta_r
Doxorubicin
YDL227C_RF2_112 RF2
ZShift = 0.1
lm ZScore = −0.28
NG = 0
DB = 0
SM = 0
Delta_AUC
Doxorubicin
YDL227C_RF2_171 RF2
ZShift = 0.37
lm Zscore = 0.13
NG = 0
DB = 0
SM = 0
Delta_L
Doxorubicin
YDL227C_RF2_171 RF2
ZShift = 0.4
lm ZScore = 0.13
NG = 0
DB = 0
SM = 0
Delta_K
Doxorubicin
YDL227C_RF2_171 RF2
ZShift = −0.45
lm ZScore = 0.31
NG = 0
DB = 0
SM = 0
Delta_r
Doxorubicin
YDL227C_RF2_171 RF2
ZShift = 0.29
lm ZScore = −0.64
NG = 0
DB = 0
SM = 0
Delta_AUC
Doxorubicin
YDL227C_RF1_299 RF1
ZShift = 0.61
lm Zscore = 0.13
NG = 0
DB = 0
SM = 0
Delta_L
Doxorubicin
YDL227C_RF1_299 RF1
ZShift = −0.09
lm ZScore = 0.13
NG = 0
DB = 0
SM = 0
Delta_K
Doxorubicin
YDL227C_RF1_299 RF1
ZShift = −0.86
lm ZScore = 0.62
NG = 0
DB = 0
SM = 0
Delta_r
Doxorubicin
YDL227C_RF1_299 RF1
ZShift = −0.22
lm ZScore = 0.8
NG = 0
DB = 0
SM = 0
Delta_AUC
Doxorubicin

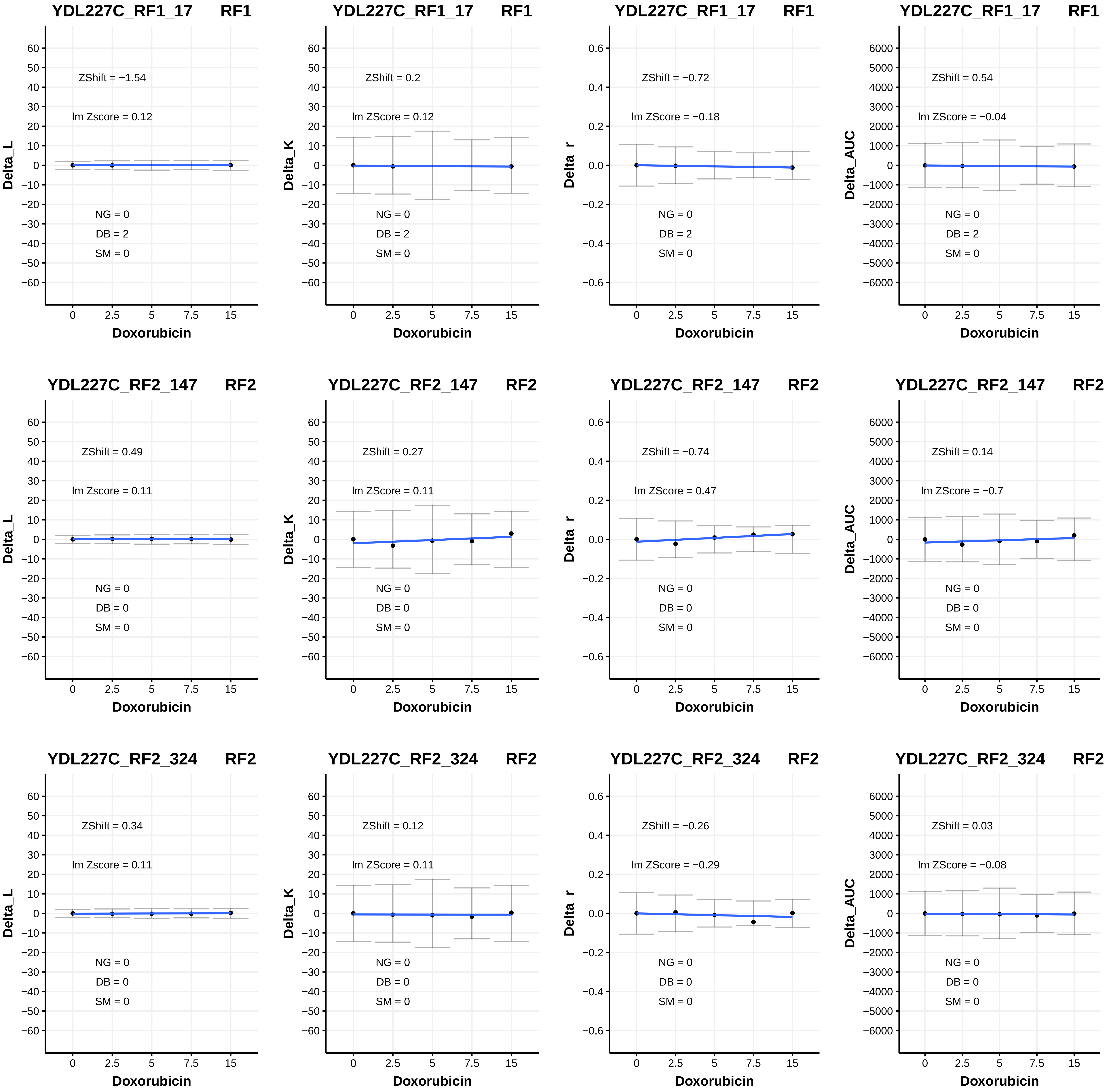
YDL227C_RF1_17 RF1
ZShift = −1.54
lm Zscore = 0.12
NG = 0
DB = 2
SM = 0
Delta_L
Doxorubicin
YDL227C_RF1_17 RF1
ZShift = 0.2
lm ZScore = 0.12
NG = 0
DB = 2
SM = 0
Delta_K
Doxorubicin
YDL227C_RF1_17 RF1
ZShift = −0.72
lm ZScore = −0.18
NG = 0
DB = 2
SM = 0
Delta_r
Doxorubicin
YDL227C_RF1_17 RF1
ZShift = 0.54
lm ZScore = −0.04
NG = 0
DB = 2
SM = 0
Delta_AUC
Doxorubicin
YDL227C_RF2_147 RF2
ZShift = 0.49
lm Zscore = 0.11
NG = 0
DB = 0
SM = 0
Delta_L
Doxorubicin
YDL227C_RF2_147 RF2
ZShift = 0.27
lm ZScore = 0.11
NG = 0
DB = 0
SM = 0
Delta_K
Doxorubicin
YDL227C_RF2_147 RF2
ZShift = −0.74
lm ZScore = 0.47
NG = 0
DB = 0
SM = 0
Delta_r
Doxorubicin
YDL227C_RF2_147 RF2
ZShift = 0.14
lm ZScore = −0.7
NG = 0
DB = 0
SM = 0
Delta_AUC
Doxorubicin
YDL227C_RF2_324 RF2
ZShift = 0.34
lm Zscore = 0.11
NG = 0
DB = 0
SM = 0
Delta_L
Doxorubicin
YDL227C_RF2_324 RF2
ZShift = 0.12
lm ZScore = 0.11
NG = 0
DB = 0
SM = 0
Delta_K
Doxorubicin
YDL227C_RF2_324 RF2
ZShift = −0.26
lm ZScore = −0.29
NG = 0
DB = 0
SM = 0
Delta_r
Doxorubicin
YDL227C_RF2_324 RF2
ZShift = 0.03
lm ZScore = −0.08
NG = 0
DB = 0
SM = 0
Delta_AUC
Doxorubicin

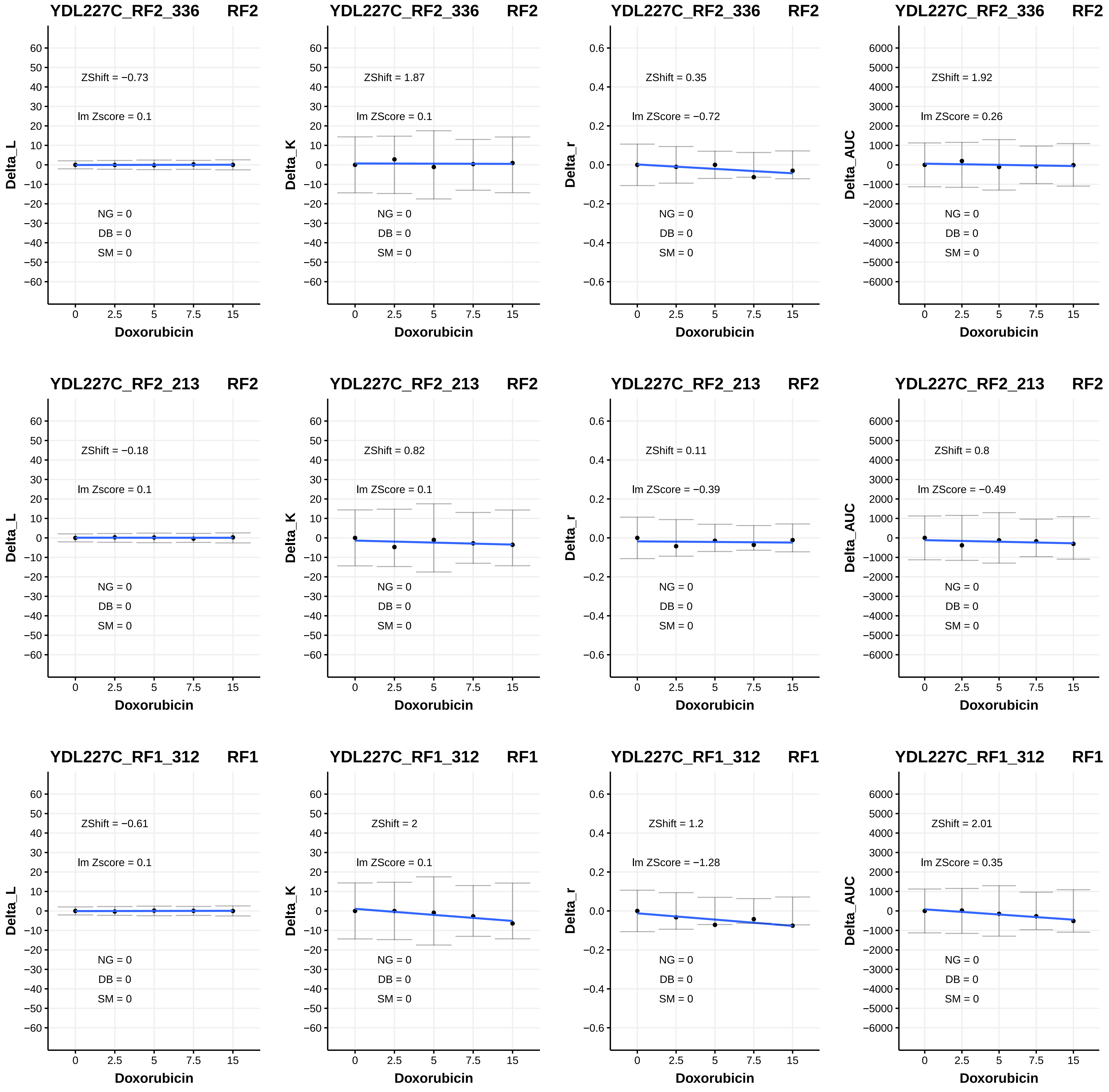
YDL227C_RF2_336 RF2
ZShift = −0.73
lm Zscore = 0.1
NG = 0
DB = 0
SM = 0
Delta_L
Doxorubicin
YDL227C_RF2_336 RF2
ZShift = 1.87
lm ZScore = 0.1
NG = 0
DB = 0
SM = 0
Delta_K
Doxorubicin
YDL227C_RF2_336 RF2
ZShift = 0.35
lm ZScore = −0.72
NG = 0
DB = 0
SM = 0
Delta_r
Doxorubicin
YDL227C_RF2_336 RF2
ZShift = 1.92
lm ZScore = 0.26
NG = 0
DB = 0
SM = 0
Delta_AUC
Doxorubicin
YDL227C_RF2_213 RF2
ZShift = −0.18
lm Zscore = 0.1
NG = 0
DB = 0
SM = 0
Delta_L
Doxorubicin
YDL227C_RF2_213 RF2
ZShift = 0.82
lm ZScore = 0.1
NG = 0
DB = 0
SM = 0
Delta_K
Doxorubicin
YDL227C_RF2_213 RF2
ZShift = 0.11
lm ZScore = −0.39
NG = 0
DB = 0
SM = 0
Delta_r
Doxorubicin
YDL227C_RF2_213 RF2
ZShift = 0.8
lm ZScore = −0.49
NG = 0
DB = 0
SM = 0
Delta_AUC
Doxorubicin
YDL227C_RF1_312 RF1
ZShift = −0.61
lm Zscore = 0.1
NG = 0
DB = 0
SM = 0
Delta_L
Doxorubicin
YDL227C_RF1_312 RF1
ZShift = 2
lm ZScore = 0.1
NG = 0
DB = 0
SM = 0
Delta_K
Doxorubicin
YDL227C_RF1_312 RF1
ZShift = 1.2
lm ZScore = −1.28
NG = 0
DB = 0
SM = 0
Delta_r
Doxorubicin
YDL227C_RF1_312 RF1
ZShift = 2.01
lm ZScore = 0.35
NG = 0
DB = 0
SM = 0
Delta_AUC
Doxorubicin

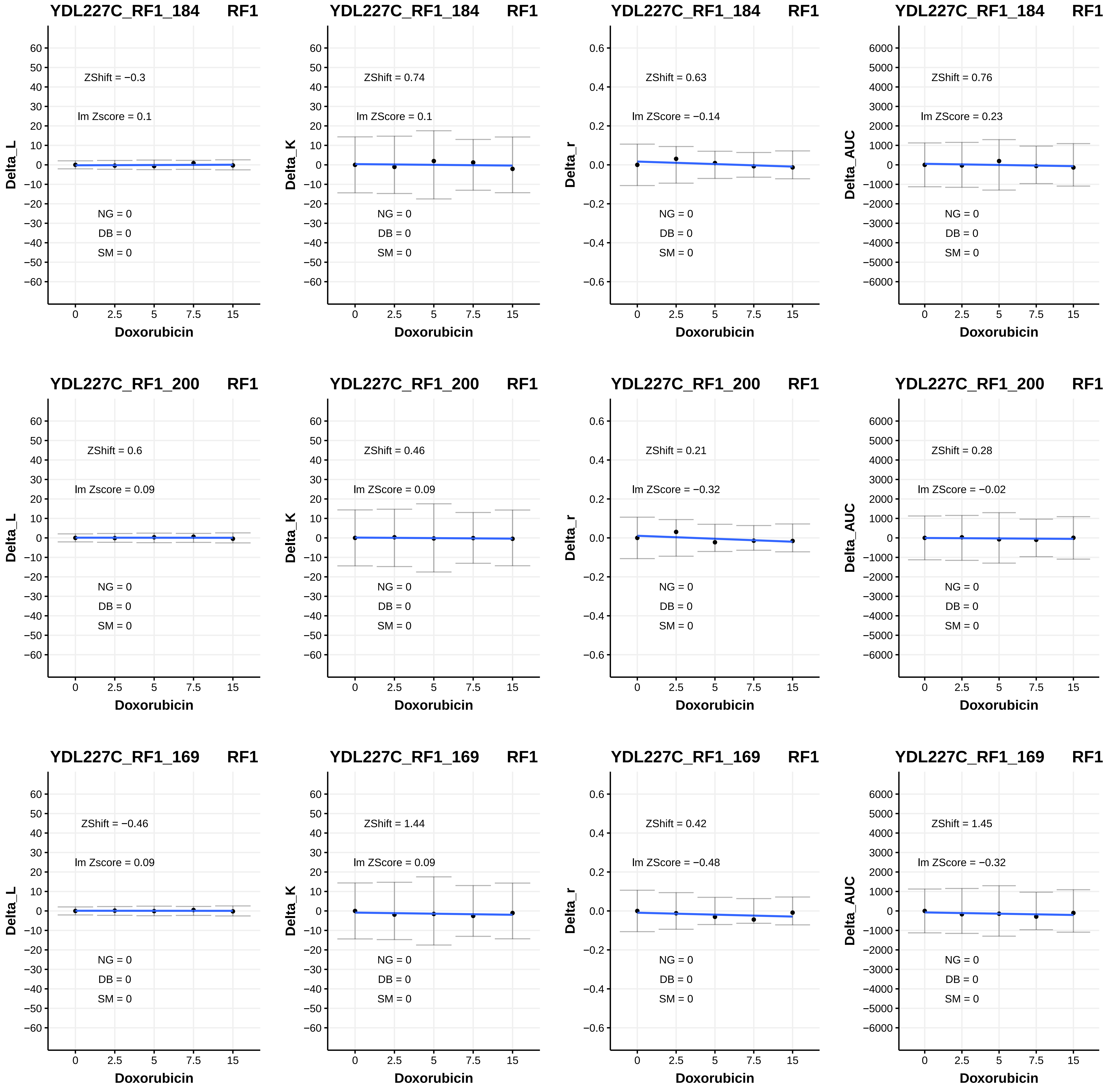

YDL227C_RF1_184 RF1
ZShift = −0.3
lm Zscore = 0.1
NG = 0
DB = 0
SM = 0
Delta_L
Doxorubicin
YDL227C_RF1_184 RF1
ZShift = 0.74
lm ZScore = 0.1
NG = 0
DB = 0
SM = 0
Delta_K
Doxorubicin
YDL227C_RF1_184 RF1
ZShift = 0.63
lm ZScore = −0.14
NG = 0
DB = 0
SM = 0
Delta_r
Doxorubicin
YDL227C_RF1_184 RF1
ZShift = 0.76
lm ZScore = 0.23
NG = 0
DB = 0
SM = 0
Delta_AUC
Doxorubicin
YDL227C_RF1_200 RF1
ZShift = 0.6
lm Zscore = 0.09
NG = 0
DB = 0
SM = 0
Delta_L
Doxorubicin
YDL227C_RF1_200 RF1
ZShift = 0.46
lm ZScore = 0.09
NG = 0
DB = 0
SM = 0
Delta_K
Doxorubicin
YDL227C_RF1_200 RF1
ZShift = 0.21
lm ZScore = −0.32
NG = 0
DB = 0
SM = 0
Delta_r
Doxorubicin
YDL227C_RF1_200 RF1
ZShift = 0.28
lm ZScore = −0.02
NG = 0
DB = 0
SM = 0
Delta_AUC
Doxorubicin
YDL227C_RF1_169 RF1
ZShift = −0.46
lm Zscore = 0.09
NG = 0
DB = 0
SM = 0
Delta_L
Doxorubicin
YDL227C_RF1_169 RF1
ZShift = 1.44
lm ZScore = 0.09
NG = 0
DB = 0
SM = 0
Delta_K
Doxorubicin
YDL227C_RF1_169 RF1
ZShift = 0.42
lm ZScore = −0.48
NG = 0
DB = 0
SM = 0
Delta_r
Doxorubicin
YDL227C_RF1_169 RF1
ZShift = 1.45
lm ZScore = −0.32
NG = 0
DB = 0
SM = 0
Delta_AUC
Doxorubicin

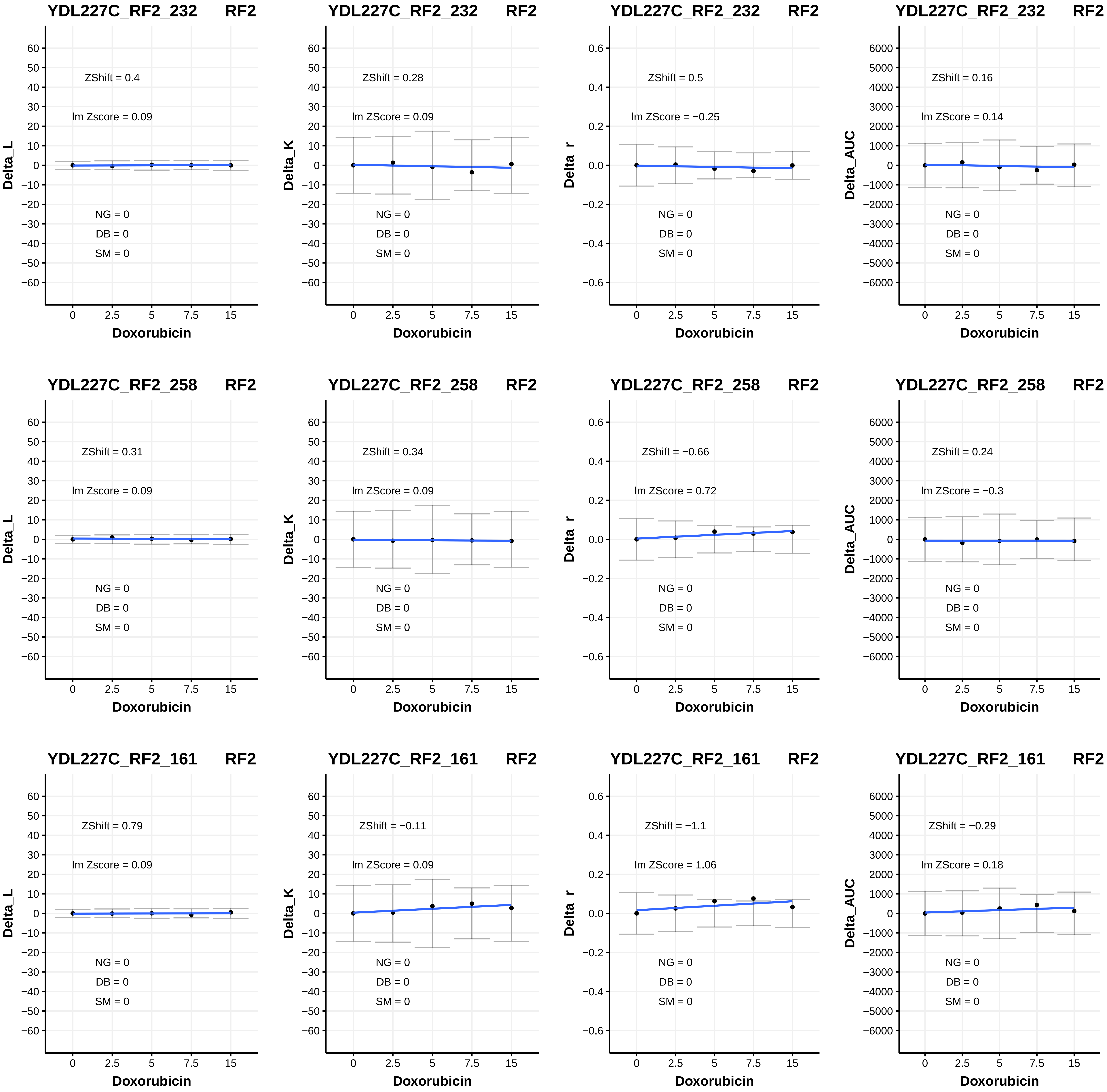
YDL227C_RF2_232 RF2
ZShift = 0.4
lm Zscore = 0.09
NG = 0
DB = 0
SM = 0
Delta_L
Doxorubicin
YDL227C_RF2_232 RF2
ZShift = 0.28
lm ZScore = 0.09
NG = 0
DB = 0
SM = 0
Delta_K
Doxorubicin
YDL227C_RF2_232 RF2
ZShift = 0.5
lm ZScore = −0.25
NG = 0
DB = 0
SM = 0
Delta_r
Doxorubicin
YDL227C_RF2_232 RF2
ZShift = 0.16
lm ZScore = 0.14
NG = 0
DB = 0
SM = 0
Delta_AUC
Doxorubicin
YDL227C_RF2_258 RF2
ZShift = 0.31
lm Zscore = 0.09
NG = 0
DB = 0
SM = 0
Delta_L
Doxorubicin
YDL227C_RF2_258 RF2
ZShift = 0.34
lm ZScore = 0.09
NG = 0
DB = 0
SM = 0
Delta_K
Doxorubicin
YDL227C_RF2_258 RF2
ZShift = −0.66
lm ZScore = 0.72
NG = 0
DB = 0
SM = 0
Delta_r
Doxorubicin
YDL227C_RF2_258 RF2
ZShift = 0.24
lm ZScore = −0.3
NG = 0
DB = 0
SM = 0
Delta_AUC
Doxorubicin
YDL227C_RF2_161 RF2
ZShift = 0.79
lm Zscore = 0.09
NG = 0
DB = 0
SM = 0
Delta_L
Doxorubicin
YDL227C_RF2_161 RF2
ZShift = −0.11
lm ZScore = 0.09
NG = 0
DB = 0
SM = 0
Delta_K
Doxorubicin
YDL227C_RF2_161 RF2
ZShift = −1.1
lm ZScore = 1.06
NG = 0
DB = 0
SM = 0
Delta_r
Doxorubicin
YDL227C_RF2_161 RF2
ZShift = −0.29
lm ZScore = 0.18
NG = 0
DB = 0
SM = 0
Delta_AUC
Doxorubicin

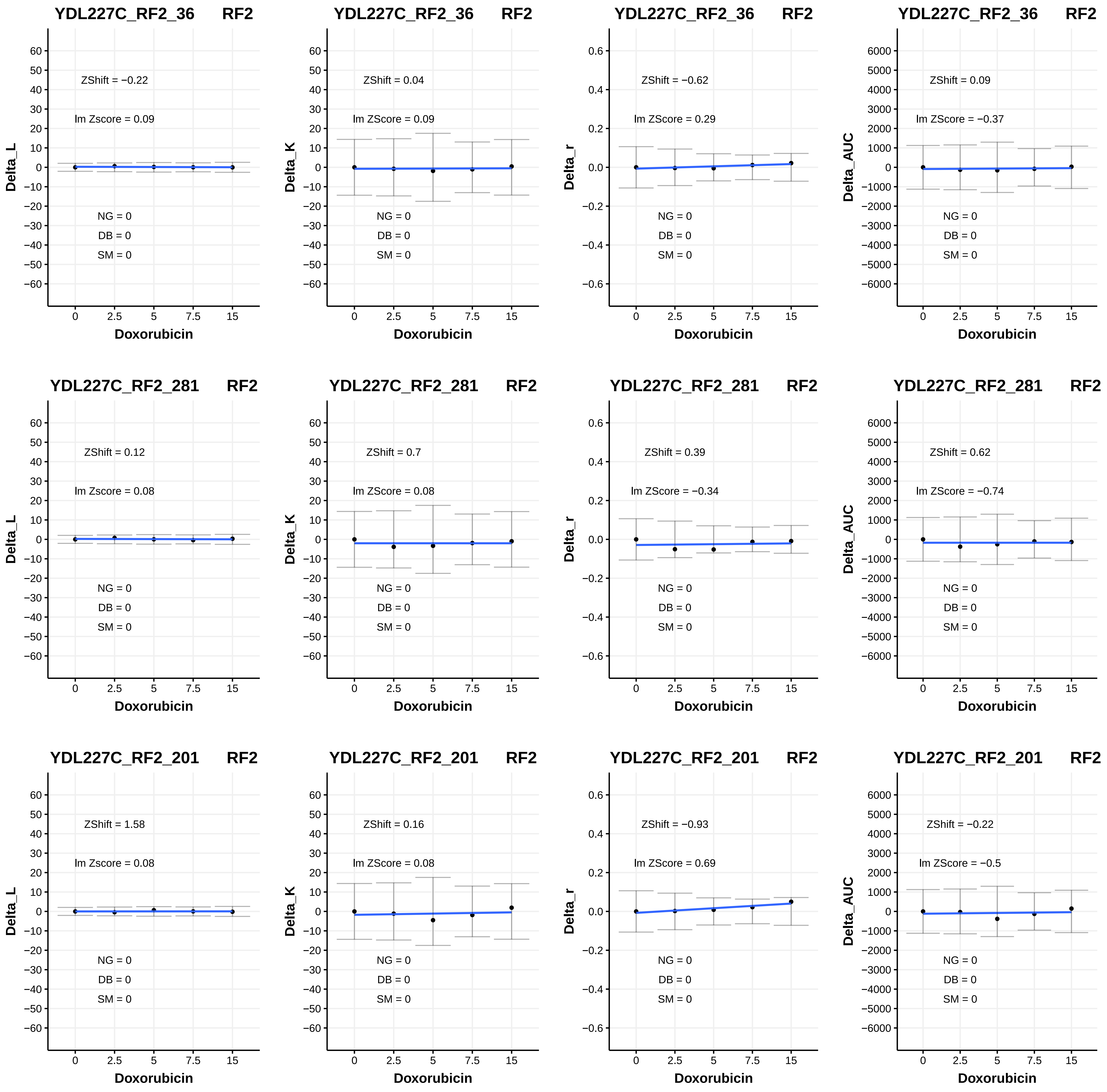
YDL227C_RF2_36 RF2
ZShift = −0.22
lm Zscore = 0.09
NG = 0
DB = 0
SM = 0
Delta_L
Doxorubicin
YDL227C_RF2_36 RF2
ZShift = 0.04
lm ZScore = 0.09
NG = 0
DB = 0
SM = 0
Delta_K
Doxorubicin
YDL227C_RF2_36 RF2
ZShift = −0.62
lm ZScore = 0.29
NG = 0
DB = 0
SM = 0
Delta_r
Doxorubicin
YDL227C_RF2_36 RF2
ZShift = 0.09
lm ZScore = −0.37
NG = 0
DB = 0
SM = 0
Delta_AUC
Doxorubicin
YDL227C_RF2_281 RF2
ZShift = 0.12
lm Zscore = 0.08
NG = 0
DB = 0
SM = 0
Delta_L
Doxorubicin
YDL227C_RF2_281 RF2
ZShift = 0.7
lm ZScore = 0.08
NG = 0
DB = 0
SM = 0
Delta_K
Doxorubicin
YDL227C_RF2_281 RF2
ZShift = 0.39
lm ZScore = −0.34
NG = 0
DB = 0
SM = 0
Delta_r
Doxorubicin
YDL227C_RF2_281 RF2
ZShift = 0.62
lm ZScore = −0.74
NG = 0
DB = 0
SM = 0
Delta_AUC
Doxorubicin
YDL227C_RF2_201 RF2
ZShift = 1.58
lm Zscore = 0.08
NG = 0
DB = 0
SM = 0
Delta_L
Doxorubicin
YDL227C_RF2_201 RF2
ZShift = 0.16
lm ZScore = 0.08
NG = 0
DB = 0
SM = 0
Delta_K
Doxorubicin
YDL227C_RF2_201 RF2
ZShift = −0.93
lm ZScore = 0.69
NG = 0
DB = 0
SM = 0
Delta_r
Doxorubicin
YDL227C_RF2_201 RF2
ZShift = −0.22
lm ZScore = −0.5
NG = 0
DB = 0
SM = 0
Delta_AUC
Doxorubicin

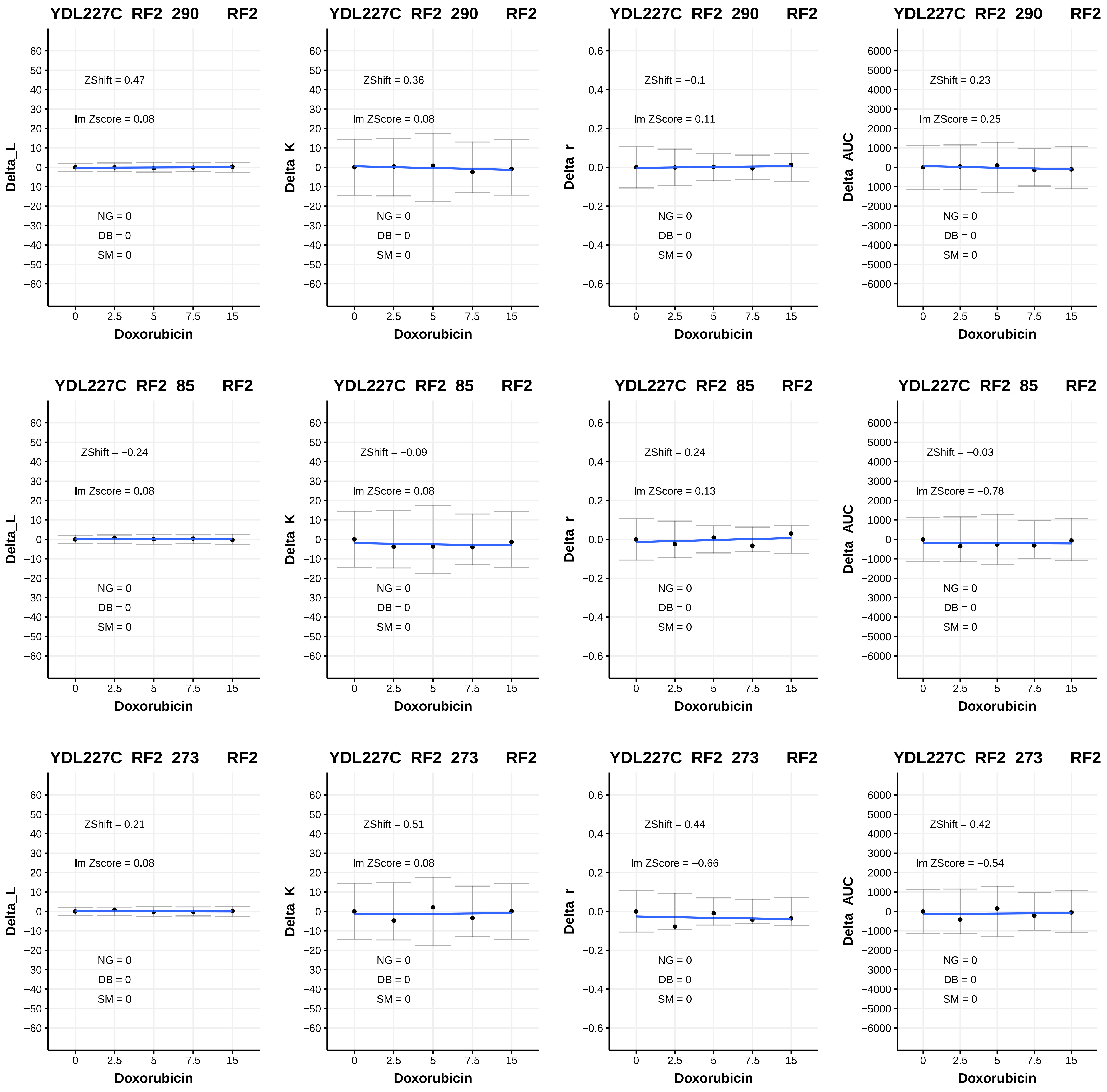

YDL227C_RF2_290 RF2
ZShift = 0.47
lm Zscore = 0.08
NG = 0
DB = 0
SM = 0
Delta_L
Doxorubicin
YDL227C_RF2_290 RF2
ZShift = 0.36
lm ZScore = 0.08
NG = 0
DB = 0
SM = 0
Delta_K
Doxorubicin
YDL227C_RF2_290 RF2
ZShift = −0.1
lm ZScore = 0.11
NG = 0
DB = 0
SM = 0
Delta_r
Doxorubicin
YDL227C_RF2_290 RF2
ZShift = 0.23
lm ZScore = 0.25
NG = 0
DB = 0
SM = 0
Delta_AUC
Doxorubicin
YDL227C_RF2_85 RF2
ZShift = −0.24
lm Zscore = 0.08
NG = 0
DB = 0
SM = 0
Delta_L
Doxorubicin
YDL227C_RF2_85 RF2
ZShift = −0.09
lm ZScore = 0.08
NG = 0
DB = 0
SM = 0
Delta_K
Doxorubicin
YDL227C_RF2_85 RF2
ZShift = 0.24
lm ZScore = 0.13
NG = 0
DB = 0
SM = 0
Delta_r
Doxorubicin
YDL227C_RF2_85 RF2
ZShift = −0.03
lm ZScore = −0.78
NG = 0
DB = 0
SM = 0
Delta_AUC
Doxorubicin
YDL227C_RF2_273 RF2
ZShift = 0.21
lm Zscore = 0.08
NG = 0
DB = 0
SM = 0
Delta_L
Doxorubicin
YDL227C_RF2_273 RF2
ZShift = 0.51
lm ZScore = 0.08
NG = 0
DB = 0
SM = 0
Delta_K
Doxorubicin
YDL227C_RF2_273 RF2
ZShift = 0.44
lm ZScore = −0.66
NG = 0
DB = 0
SM = 0
Delta_r
Doxorubicin
YDL227C_RF2_273 RF2
ZShift = 0.42
lm ZScore = −0.54
NG = 0
DB = 0
SM = 0
Delta_AUC
Doxorubicin

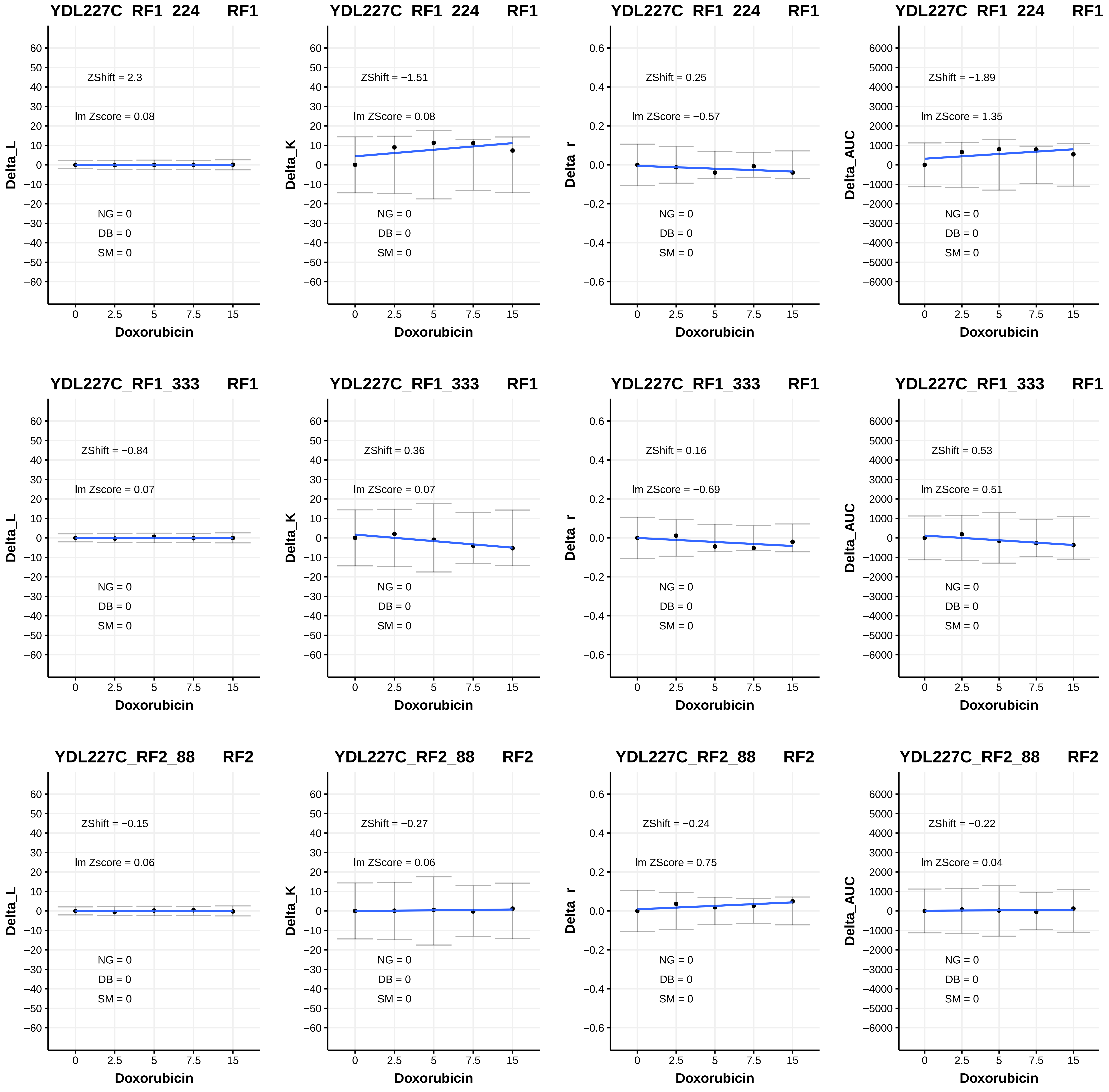

YDL227C_RF1_224 RF1
ZShift = 2.3
lm Zscore = 0.08
NG = 0
DB = 0
SM = 0
Delta_L
Doxorubicin
YDL227C_RF1_224 RF1
ZShift = −1.51
lm ZScore = 0.08
NG = 0
DB = 0
SM = 0
Delta_K
Doxorubicin
YDL227C_RF1_224 RF1
ZShift = 0.25
lm ZScore = −0.57
NG = 0
DB = 0
SM = 0
Delta_r
Doxorubicin
YDL227C_RF1_224 RF1
ZShift = −1.89
lm ZScore = 1.35
NG = 0
DB = 0
SM = 0
Delta_AUC
Doxorubicin
YDL227C_RF1_333 RF1
ZShift = −0.84
lm Zscore = 0.07
NG = 0
DB = 0
SM = 0
Delta_L
Doxorubicin
YDL227C_RF1_333 RF1
ZShift = 0.36
lm ZScore = 0.07
NG = 0
DB = 0
SM = 0
Delta_K
Doxorubicin
YDL227C_RF1_333 RF1
ZShift = 0.16
lm ZScore = −0.69
NG = 0
DB = 0
SM = 0
Delta_r
Doxorubicin
YDL227C_RF1_333 RF1
ZShift = 0.53
lm ZScore = 0.51
NG = 0
DB = 0
SM = 0
Delta_AUC
Doxorubicin
YDL227C_RF2_88 RF2
ZShift = −0.15
lm Zscore = 0.06
NG = 0
DB = 0
SM = 0
Delta_L
Doxorubicin
YDL227C_RF2_88 RF2
ZShift = −0.27
lm ZScore = 0.06
NG = 0
DB = 0
SM = 0
Delta_K
Doxorubicin
YDL227C_RF2_88 RF2
ZShift = −0.24
lm ZScore = 0.75
NG = 0
DB = 0
SM = 0
Delta_r
Doxorubicin
YDL227C_RF2_88 RF2
ZShift = −0.22
lm ZScore = 0.04
NG = 0
DB = 0
SM = 0
Delta_AUC
Doxorubicin

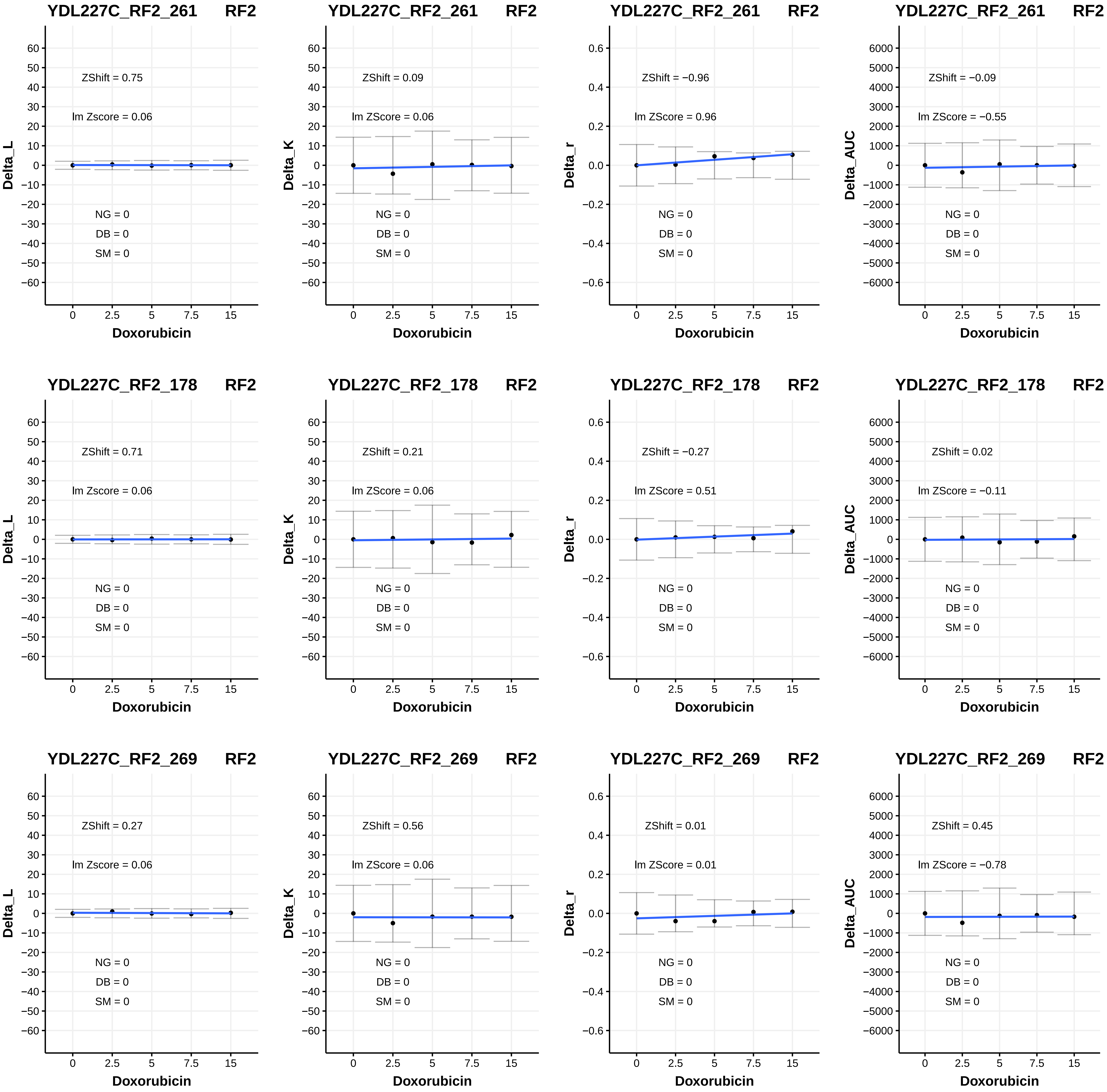

YDL227C_RF2_261 RF2
ZShift = 0.75
lm Zscore = 0.06
NG = 0
DB = 0
SM = 0
Delta_L
Doxorubicin
YDL227C_RF2_261 RF2
ZShift = 0.09
lm ZScore = 0.06
NG = 0
DB = 0
SM = 0
Delta_K
Doxorubicin
YDL227C_RF2_261 RF2
ZShift = −0.96
lm ZScore = 0.96
NG = 0
DB = 0
SM = 0
Delta_r
Doxorubicin
YDL227C_RF2_261 RF2
ZShift = −0.09
lm ZScore = −0.55
NG = 0
DB = 0
SM = 0
Delta_AUC
Doxorubicin
YDL227C_RF2_178 RF2
ZShift = 0.71
lm Zscore = 0.06
NG = 0
DB = 0
SM = 0
Delta_L
Doxorubicin
YDL227C_RF2_178 RF2
ZShift = 0.21
lm ZScore = 0.06
NG = 0
DB = 0
SM = 0
Delta_K
Doxorubicin
YDL227C_RF2_178 RF2
ZShift = −0.27
lm ZScore = 0.51
NG = 0
DB = 0
SM = 0
Delta_r
Doxorubicin
YDL227C_RF2_178 RF2
ZShift = 0.02
lm ZScore = −0.11
NG = 0
DB = 0
SM = 0
Delta_AUC
Doxorubicin
YDL227C_RF2_269 RF2
ZShift = 0.27
lm Zscore = 0.06
NG = 0
DB = 0
SM = 0
Delta_L
Doxorubicin
YDL227C_RF2_269 RF2
ZShift = 0.56
lm ZScore = 0.06
NG = 0
DB = 0
SM = 0
Delta_K
Doxorubicin
YDL227C_RF2_269 RF2
ZShift = 0.01
lm ZScore = 0.01
NG = 0
DB = 0
SM = 0
Delta_r
Doxorubicin
YDL227C_RF2_269 RF2
ZShift = 0.45
lm ZScore = −0.78
NG = 0
DB = 0
SM = 0
Delta_AUC
Doxorubicin

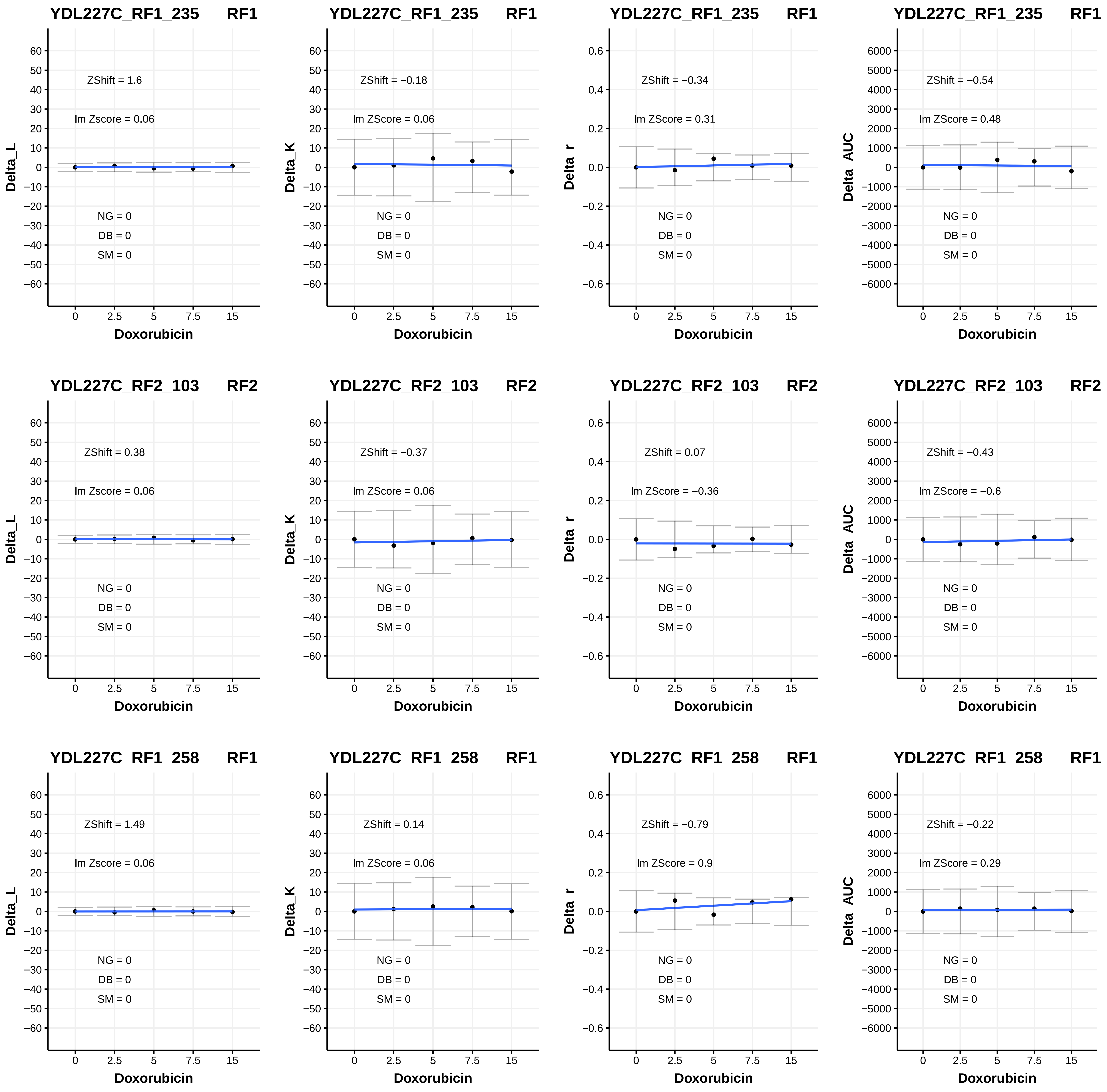
YDL227C_RF1_235 RF1
ZShift = 1.6
lm Zscore = 0.06
NG = 0
DB = 0
SM = 0
Delta_L
Doxorubicin
YDL227C_RF1_235 RF1
ZShift = −0.18
lm ZScore = 0.06
NG = 0
DB = 0
SM = 0
Delta_K
Doxorubicin
YDL227C_RF1_235 RF1
ZShift = −0.34
lm ZScore = 0.31
NG = 0
DB = 0
SM = 0
Delta_r
Doxorubicin
YDL227C_RF1_235 RF1
ZShift = −0.54
lm ZScore = 0.48
NG = 0
DB = 0
SM = 0
Delta_AUC
Doxorubicin
YDL227C_RF2_103 RF2
ZShift = 0.38
lm Zscore = 0.06
NG = 0
DB = 0
SM = 0
Delta_L
Doxorubicin
YDL227C_RF2_103 RF2
ZShift = −0.37
lm ZScore = 0.06
NG = 0
DB = 0
SM = 0
Delta_K
Doxorubicin
YDL227C_RF2_103 RF2
ZShift = 0.07
lm ZScore = −0.36
NG = 0
DB = 0
SM = 0
Delta_r
Doxorubicin
YDL227C_RF2_103 RF2
ZShift = −0.43
lm ZScore = −0.6
NG = 0
DB = 0
SM = 0
Delta_AUC
Doxorubicin
YDL227C_RF1_258 RF1
ZShift = 1.49
lm Zscore = 0.06
NG = 0
DB = 0
SM = 0
Delta_L
Doxorubicin
YDL227C_RF1_258 RF1
ZShift = 0.14
lm ZScore = 0.06
NG = 0
DB = 0
SM = 0
Delta_K
Doxorubicin
YDL227C_RF1_258 RF1
ZShift = −0.79
lm ZScore = 0.9
NG = 0
DB = 0
SM = 0
Delta_r
Doxorubicin
YDL227C_RF1_258 RF1
ZShift = −0.22
lm ZScore = 0.29
NG = 0
DB = 0
SM = 0
Delta_AUC
Doxorubicin

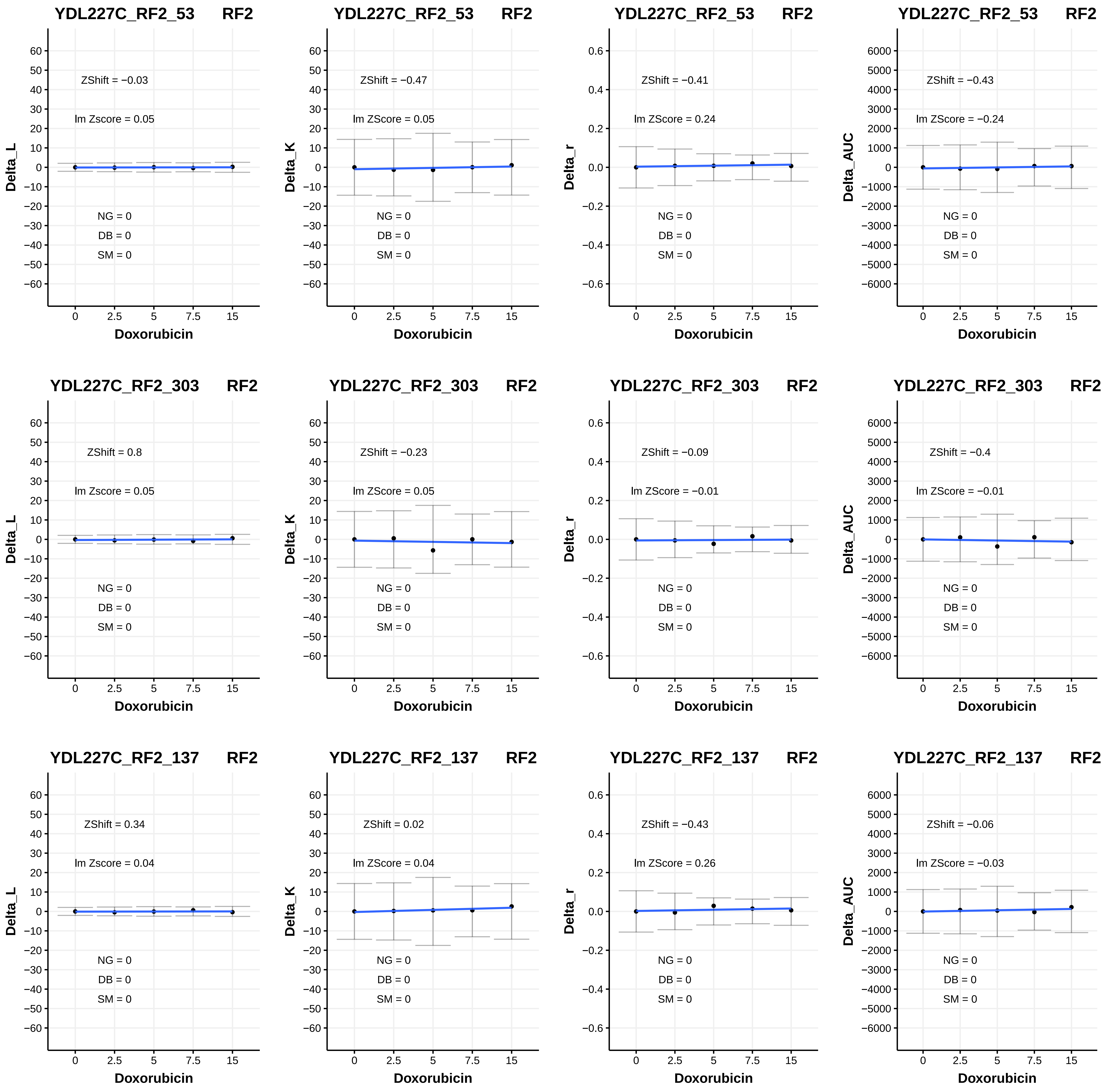
YDL227C_RF2_53 RF2
ZShift = −0.03
lm Zscore = 0.05
NG = 0
DB = 0
SM = 0
Delta_L
Doxorubicin
YDL227C_RF2_53 RF2
ZShift = −0.47
lm ZScore = 0.05
NG = 0
DB = 0
SM = 0
Delta_K
Doxorubicin
YDL227C_RF2_53 RF2
ZShift = −0.41
lm ZScore = 0.24
NG = 0
DB = 0
SM = 0
Delta_r
Doxorubicin
YDL227C_RF2_53 RF2
ZShift = −0.43
lm ZScore = −0.24
NG = 0
DB = 0
SM = 0
Delta_AUC
Doxorubicin
YDL227C_RF2_303 RF2
ZShift = 0.8
lm Zscore = 0.05
NG = 0
DB = 0
SM = 0
Delta_L
Doxorubicin
YDL227C_RF2_303 RF2
ZShift = −0.23
lm ZScore = 0.05
NG = 0
DB = 0
SM = 0
Delta_K
Doxorubicin
YDL227C_RF2_303 RF2
ZShift = −0.09
lm ZScore = −0.01
NG = 0
DB = 0
SM = 0
Delta_r
Doxorubicin
YDL227C_RF2_303 RF2
ZShift = −0.4
lm ZScore = −0.01
NG = 0
DB = 0
SM = 0
Delta_AUC
Doxorubicin
YDL227C_RF2_137 RF2
ZShift = 0.34
lm Zscore = 0.04
NG = 0
DB = 0
SM = 0
Delta_L
Doxorubicin
YDL227C_RF2_137 RF2
ZShift = 0.02
lm ZScore = 0.04
NG = 0
DB = 0
SM = 0
Delta_K
Doxorubicin
YDL227C_RF2_137 RF2
ZShift = −0.43
lm ZScore = 0.26
NG = 0
DB = 0
SM = 0
Delta_r
Doxorubicin
YDL227C_RF2_137 RF2
ZShift = −0.06
lm ZScore = −0.03
NG = 0
DB = 0
SM = 0
Delta_AUC
Doxorubicin

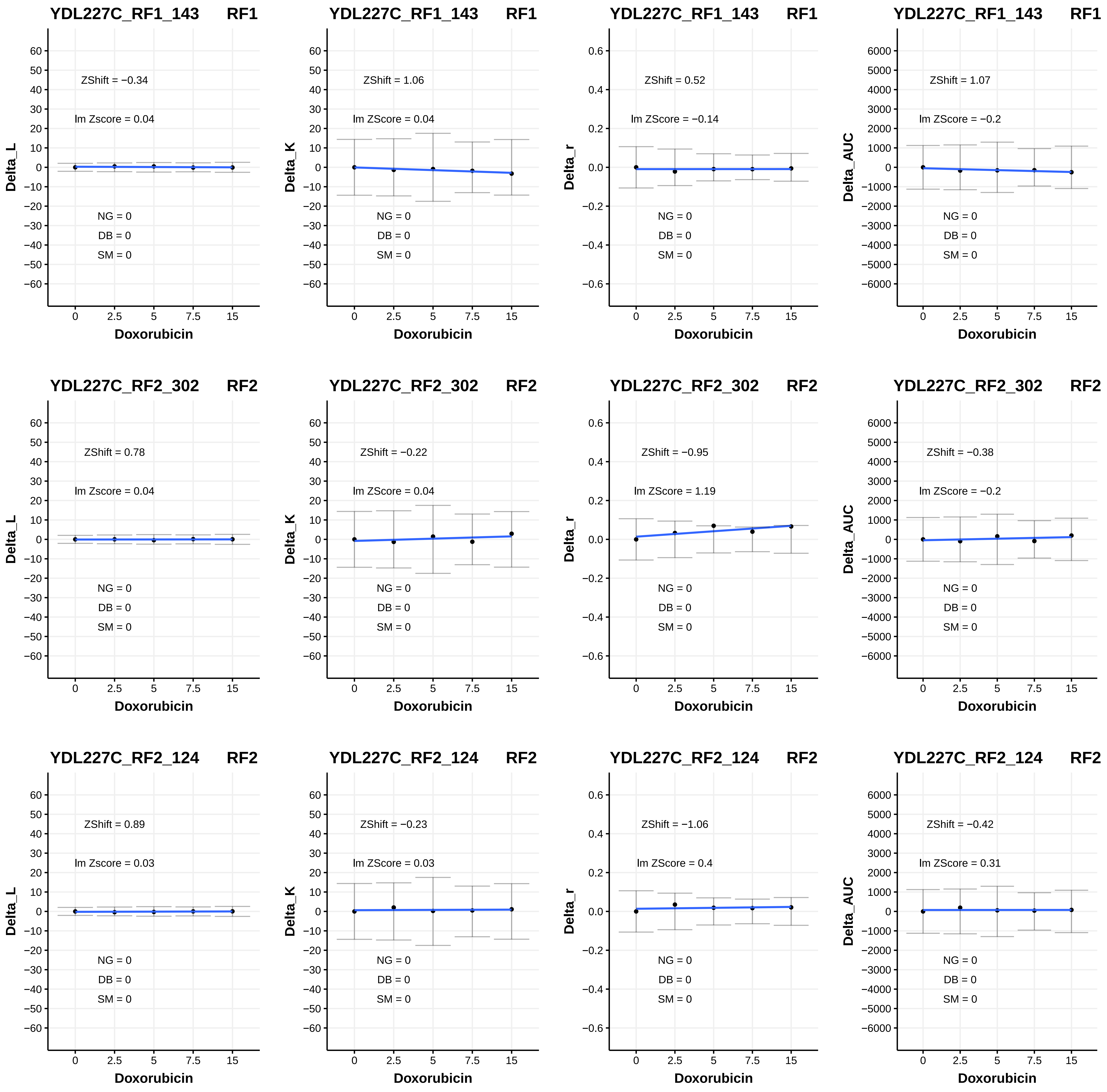
YDL227C_RF1_143 RF1
ZShift = −0.34
lm Zscore = 0.04
NG = 0
DB = 0
SM = 0
Delta_L
Doxorubicin
YDL227C_RF1_143 RF1
ZShift = 1.06
lm ZScore = 0.04
NG = 0
DB = 0
SM = 0
Delta_K
Doxorubicin
YDL227C_RF1_143 RF1
ZShift = 0.52
lm ZScore = −0.14
NG = 0
DB = 0
SM = 0
Delta_r
Doxorubicin
YDL227C_RF1_143 RF1
ZShift = 1.07
lm ZScore = −0.2
NG = 0
DB = 0
SM = 0
Delta_AUC
Doxorubicin
YDL227C_RF2_302 RF2
ZShift = 0.78
lm Zscore = 0.04
NG = 0
DB = 0
SM = 0
Delta_L
Doxorubicin
YDL227C_RF2_302 RF2
ZShift = −0.22
lm ZScore = 0.04
NG = 0
DB = 0
SM = 0
Delta_K
Doxorubicin
YDL227C_RF2_302 RF2
ZShift = −0.95
lm ZScore = 1.19
NG = 0
DB = 0
SM = 0
Delta_r
Doxorubicin
YDL227C_RF2_302 RF2
ZShift = −0.38
lm ZScore = −0.2
NG = 0
DB = 0
SM = 0
Delta_AUC
Doxorubicin
YDL227C_RF2_124 RF2
ZShift = 0.89
lm Zscore = 0.03
NG = 0
DB = 0
SM = 0
Delta_L
Doxorubicin
YDL227C_RF2_124 RF2
ZShift = −0.23
lm ZScore = 0.03
NG = 0
DB = 0
SM = 0
Delta_K
Doxorubicin
YDL227C_RF2_124 RF2
ZShift = −1.06
lm ZScore = 0.4
NG = 0
DB = 0
SM = 0
Delta_r
Doxorubicin
YDL227C_RF2_124 RF2
ZShift = −0.42
lm ZScore = 0.31
NG = 0
DB = 0
SM = 0
Delta_AUC
Doxorubicin

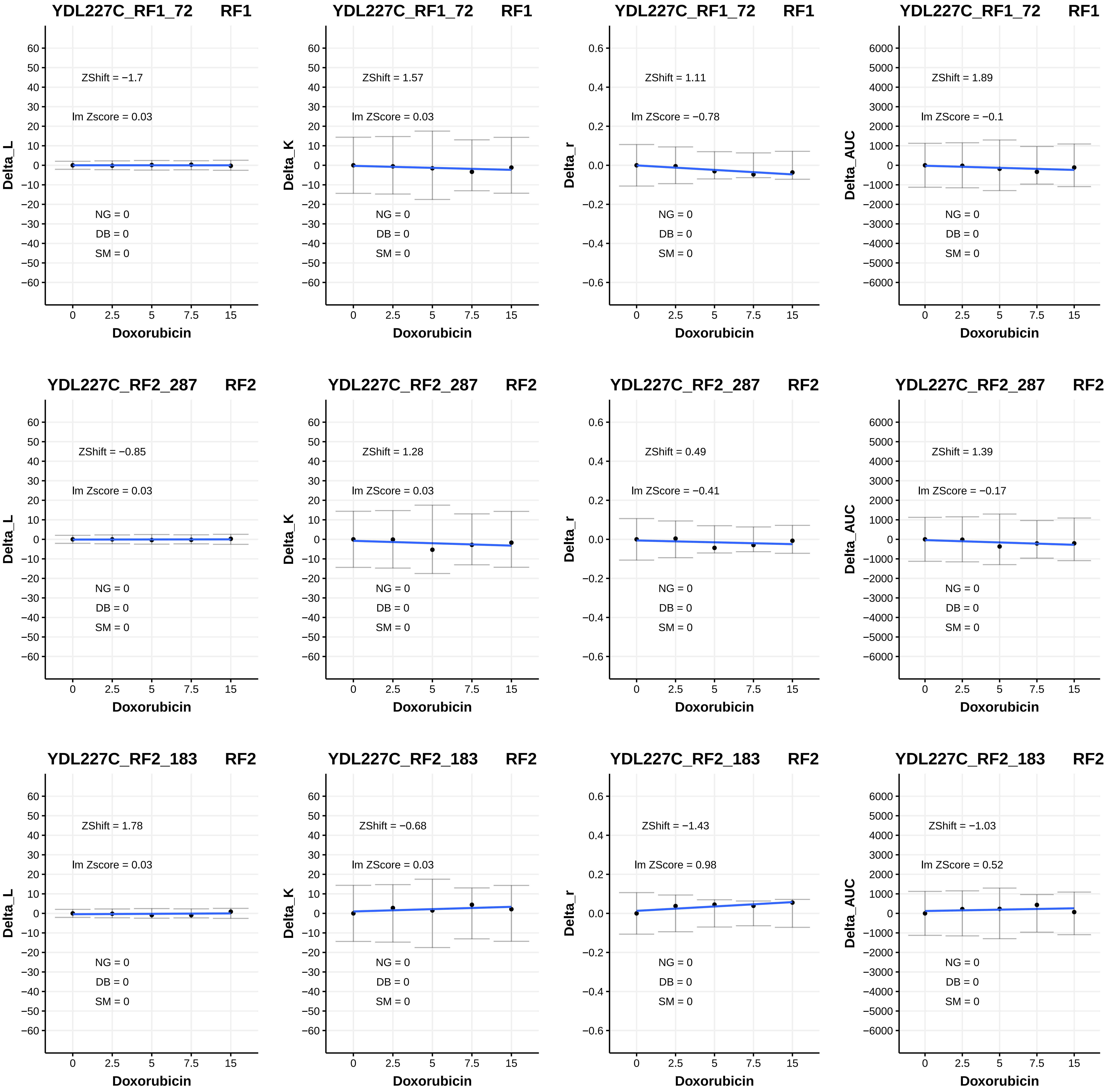

YDL227C_RF1_72 RF1
ZShift = −1.7
lm Zscore = 0.03
NG = 0
DB = 0
SM = 0
Delta_L
Doxorubicin
YDL227C_RF1_72 RF1
ZShift = 1.57
lm ZScore = 0.03
NG = 0
DB = 0
SM = 0
Delta_K
Doxorubicin
YDL227C_RF1_72 RF1
ZShift = 1.11
lm ZScore = −0.78
NG = 0
DB = 0
SM = 0
Delta_r
Doxorubicin
YDL227C_RF1_72 RF1
ZShift = 1.89
lm ZScore = −0.1
NG = 0
DB = 0
SM = 0
Delta_AUC
Doxorubicin
YDL227C_RF2_287 RF2
ZShift = −0.85
lm Zscore = 0.03
NG = 0
DB = 0
SM = 0
Delta_L
Doxorubicin
YDL227C_RF2_287 RF2
ZShift = 1.28
lm ZScore = 0.03
NG = 0
DB = 0
SM = 0
Delta_K
Doxorubicin
YDL227C_RF2_287 RF2
ZShift = 0.49
lm ZScore = −0.41
NG = 0
DB = 0
SM = 0
Delta_r
Doxorubicin
YDL227C_RF2_287 RF2
ZShift = 1.39
lm ZScore = −0.17
NG = 0
DB = 0
SM = 0
Delta_AUC
Doxorubicin
YDL227C_RF2_183 RF2
ZShift = 1.78
lm Zscore = 0.03
NG = 0
DB = 0
SM = 0
Delta_L
Doxorubicin
YDL227C_RF2_183 RF2
ZShift = −0.68
lm ZScore = 0.03
NG = 0
DB = 0
SM = 0
Delta_K
Doxorubicin
YDL227C_RF2_183 RF2
ZShift = −1.43
lm ZScore = 0.98
NG = 0
DB = 0
SM = 0
Delta_r
Doxorubicin
YDL227C_RF2_183 RF2
ZShift = −1.03
lm ZScore = 0.52
NG = 0
DB = 0
SM = 0
Delta_AUC
Doxorubicin

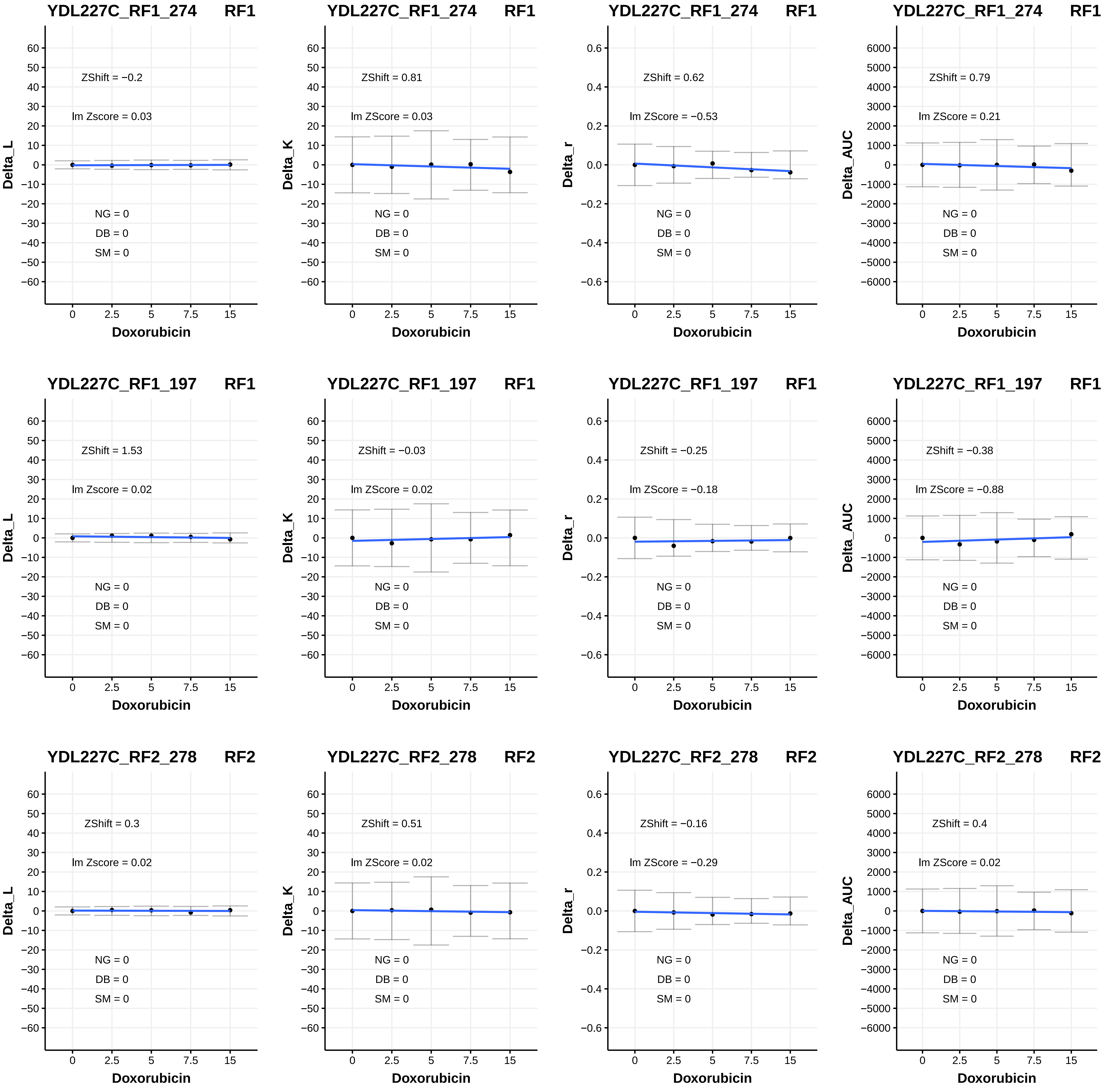
YDL227C_RF1_274 RF1
ZShift = −0.2
lm Zscore = 0.03
NG = 0
DB = 0
SM = 0
Delta_L
Doxorubicin
YDL227C_RF1_274 RF1
ZShift = 0.81
lm ZScore = 0.03
NG = 0
DB = 0
SM = 0
Delta_K
Doxorubicin
YDL227C_RF1_274 RF1
ZShift = 0.62
lm ZScore = −0.53
NG = 0
DB = 0
SM = 0
Delta_r
Doxorubicin
YDL227C_RF1_274 RF1
ZShift = 0.79
lm ZScore = 0.21
NG = 0
DB = 0
SM = 0
Delta_AUC
Doxorubicin
YDL227C_RF1_197 RF1
ZShift = 1.53
lm Zscore = 0.02
NG = 0
DB = 0
SM = 0
Delta_L
Doxorubicin
YDL227C_RF1_197 RF1
ZShift = −0.03
lm ZScore = 0.02
NG = 0
DB = 0
SM = 0
Delta_K
Doxorubicin
YDL227C_RF1_197 RF1
ZShift = −0.25
lm ZScore = −0.18
NG = 0
DB = 0
SM = 0
Delta_r
Doxorubicin
YDL227C_RF1_197 RF1
ZShift = −0.38
lm ZScore = −0.88
NG = 0
DB = 0
SM = 0
Delta_AUC
Doxorubicin
YDL227C_RF2_278 RF2
ZShift = 0.3
lm Zscore = 0.02
NG = 0
DB = 0
SM = 0
Delta_L
Doxorubicin
YDL227C_RF2_278 RF2
ZShift = 0.51
lm ZScore = 0.02
NG = 0
DB = 0
SM = 0
Delta_K
Doxorubicin
YDL227C_RF2_278 RF2
ZShift = −0.16
lm ZScore = −0.29
NG = 0
DB = 0
SM = 0
Delta_r
Doxorubicin
YDL227C_RF2_278 RF2
ZShift = 0.4
lm ZScore = 0.02
NG = 0
DB = 0
SM = 0
Delta_AUC
Doxorubicin

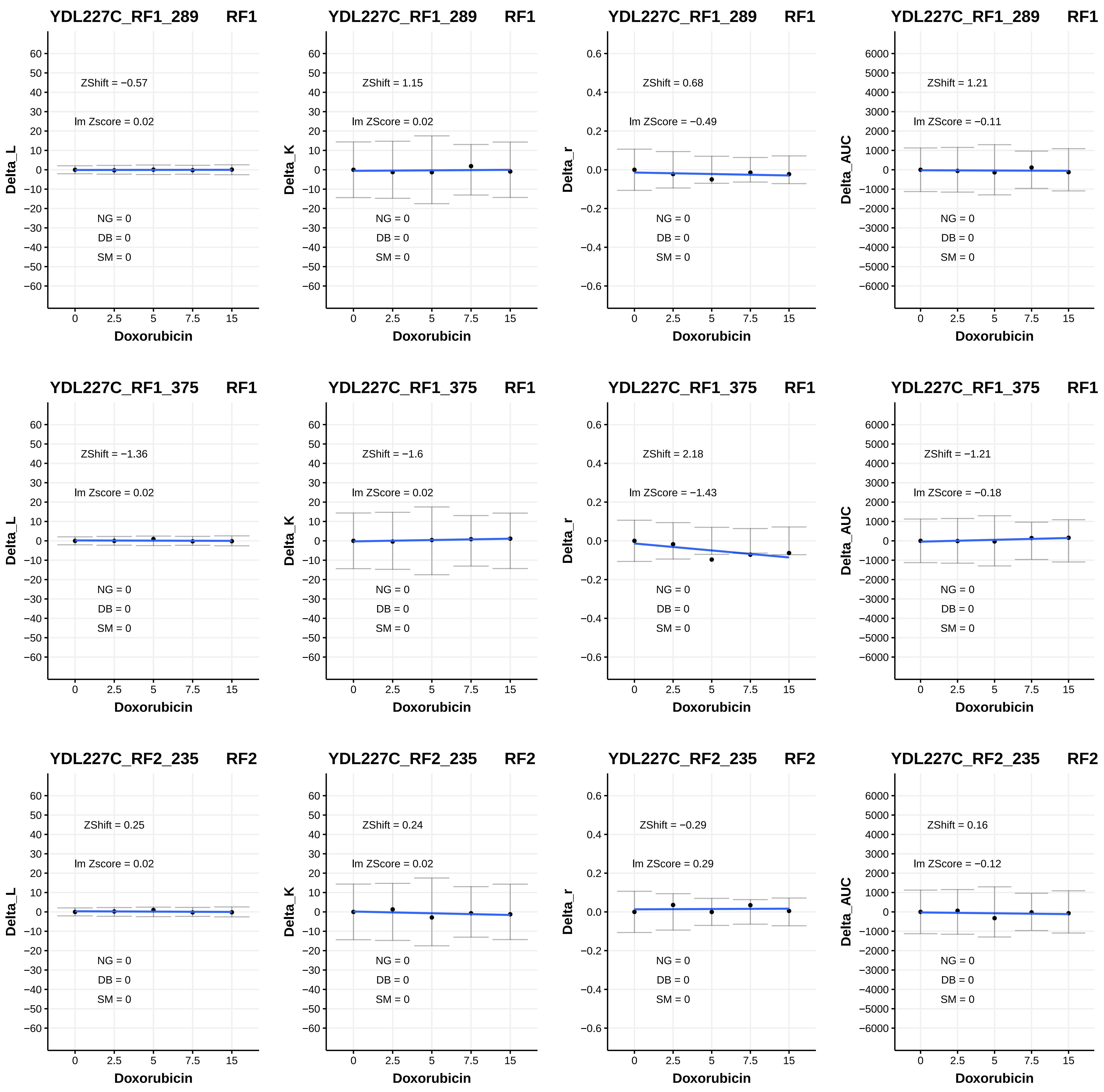
YDL227C_RF1_289 RF1
ZShift = −0.57
lm Zscore = 0.02
NG = 0
DB = 0
SM = 0
Delta_L
Doxorubicin
YDL227C_RF1_289 RF1
ZShift = 1.15
lm ZScore = 0.02
NG = 0
DB = 0
SM = 0
Delta_K
Doxorubicin
YDL227C_RF1_289 RF1
ZShift = 0.68
lm ZScore = −0.49
NG = 0
DB = 0
SM = 0
Delta_r
Doxorubicin
YDL227C_RF1_289 RF1
ZShift = 1.21
lm ZScore = −0.11
NG = 0
DB = 0
SM = 0
Delta_AUC
Doxorubicin
YDL227C_RF1_375 RF1
ZShift = −1.36
lm Zscore = 0.02
NG = 0
DB = 0
SM = 0
Delta_L
Doxorubicin
YDL227C_RF1_375 RF1
ZShift = −1.6
lm ZScore = 0.02
NG = 0
DB = 0
SM = 0
Delta_K
Doxorubicin
YDL227C_RF1_375 RF1
ZShift = 2.18
lm ZScore = −1.43
NG = 0
DB = 0
SM = 0
Delta_r
Doxorubicin
YDL227C_RF1_375 RF1
ZShift = −1.21
lm ZScore = −0.18
NG = 0
DB = 0
SM = 0
Delta_AUC
Doxorubicin
YDL227C_RF2_235 RF2
ZShift = 0.25
lm Zscore = 0.02
NG = 0
DB = 0
SM = 0
Delta_L
Doxorubicin
YDL227C_RF2_235 RF2
ZShift = 0.24
lm ZScore = 0.02
NG = 0
DB = 0
SM = 0
Delta_K
Doxorubicin
YDL227C_RF2_235 RF2
ZShift = −0.29
lm ZScore = 0.29
NG = 0
DB = 0
SM = 0
Delta_r
Doxorubicin
YDL227C_RF2_235 RF2
ZShift = 0.16
lm ZScore = −0.12
NG = 0
DB = 0
SM = 0
Delta_AUC
Doxorubicin

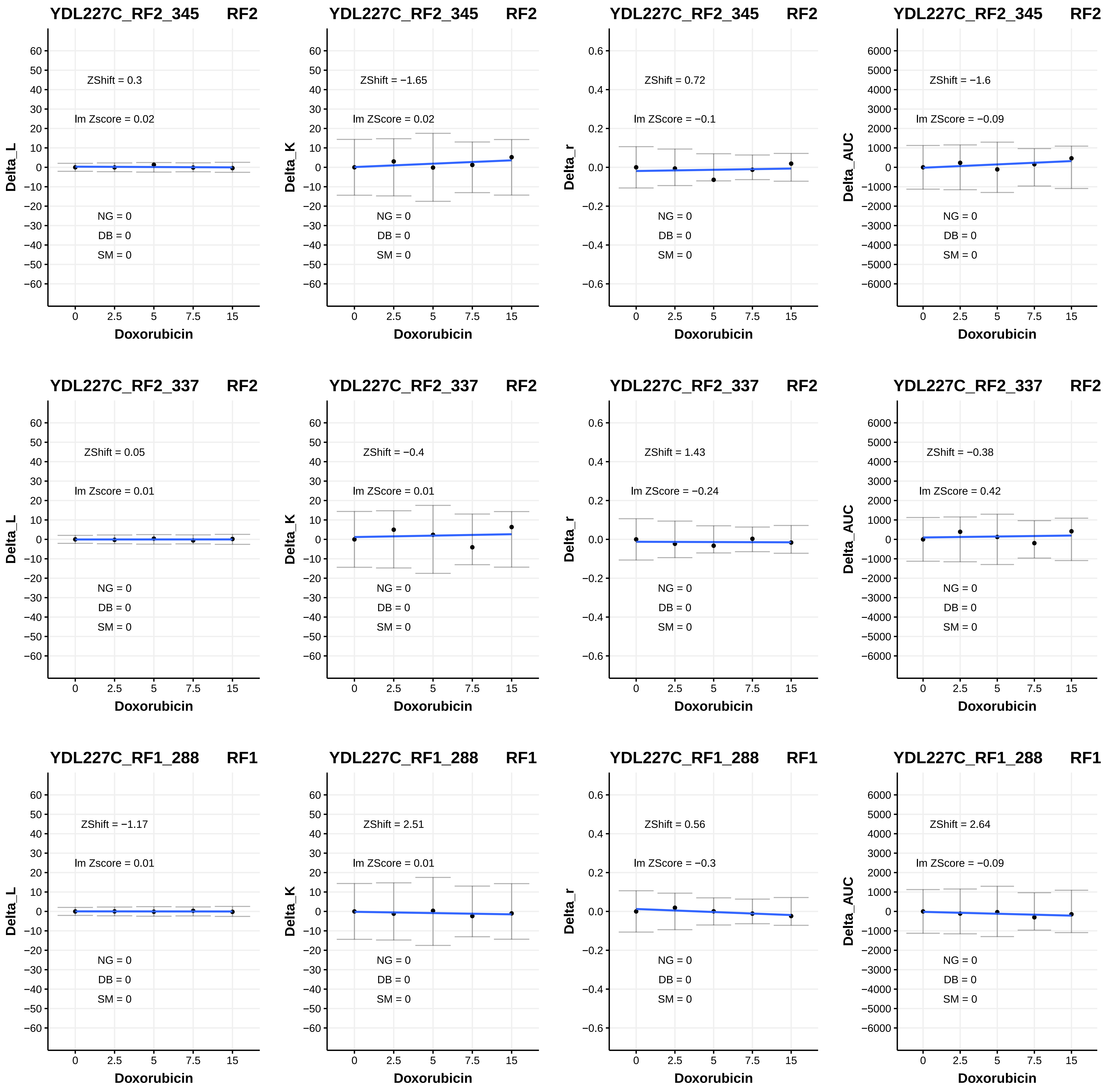
YDL227C_RF2_345 RF2
ZShift = 0.3
lm Zscore = 0.02
NG = 0
DB = 0
SM = 0
Delta_L
Doxorubicin
YDL227C_RF2_345 RF2
ZShift = −1.65
lm ZScore = 0.02
NG = 0
DB = 0
SM = 0
Delta_K
Doxorubicin
YDL227C_RF2_345 RF2
ZShift = 0.72
lm ZScore = −0.1
NG = 0
DB = 0
SM = 0
Delta_r
Doxorubicin
YDL227C_RF2_345 RF2
ZShift = −1.6
lm ZScore = −0.09
NG = 0
DB = 0
SM = 0
Delta_AUC
Doxorubicin
YDL227C_RF2_337 RF2
ZShift = 0.05
lm Zscore = 0.01
NG = 0
DB = 0
SM = 0
Delta_L
Doxorubicin
YDL227C_RF2_337 RF2
ZShift = −0.4
lm ZScore = 0.01
NG = 0
DB = 0
SM = 0
Delta_K
Doxorubicin
YDL227C_RF2_337 RF2
ZShift = 1.43
lm ZScore = −0.24
NG = 0
DB = 0
SM = 0
Delta_r
Doxorubicin
YDL227C_RF2_337 RF2
ZShift = −0.38
lm ZScore = 0.42
NG = 0
DB = 0
SM = 0
Delta_AUC
Doxorubicin
YDL227C_RF1_288 RF1
ZShift = −1.17
lm Zscore = 0.01
NG = 0
DB = 0
SM = 0
Delta_L
Doxorubicin
YDL227C_RF1_288 RF1
ZShift = 2.51
lm ZScore = 0.01
NG = 0
DB = 0
SM = 0
Delta_K
Doxorubicin
YDL227C_RF1_288 RF1
ZShift = 0.56
lm ZScore = −0.3
NG = 0
DB = 0
SM = 0
Delta_r
Doxorubicin
YDL227C_RF1_288 RF1
ZShift = 2.64
lm ZScore = −0.09
NG = 0
DB = 0
SM = 0
Delta_AUC
Doxorubicin

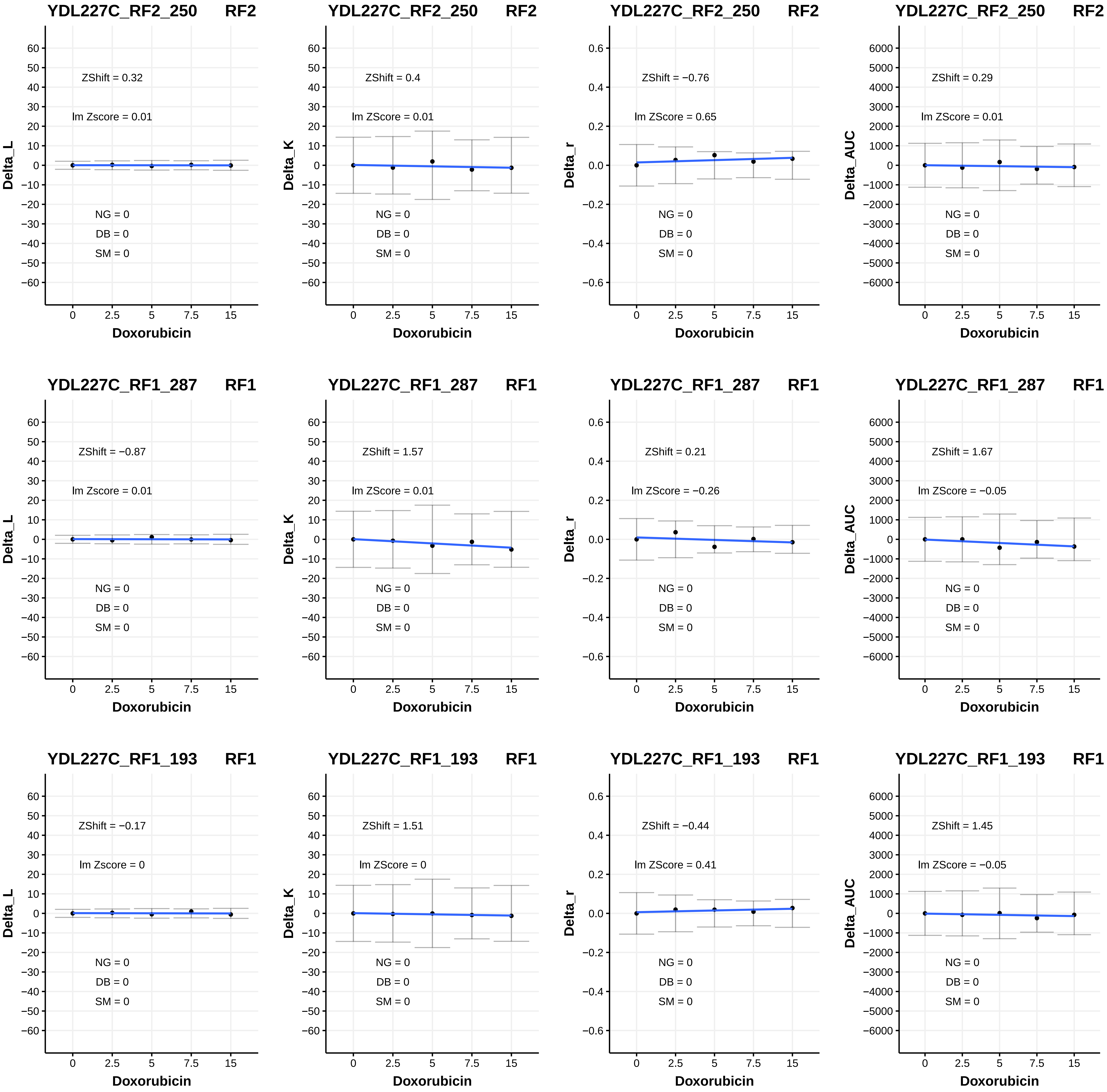

YDL227C_RF2_250 RF2
ZShift = 0.32
lm Zscore = 0.01
NG = 0
DB = 0
SM = 0
Delta_L
Doxorubicin
YDL227C_RF2_250 RF2
ZShift = 0.4
lm ZScore = 0.01
NG = 0
DB = 0
SM = 0
Delta_K
Doxorubicin
YDL227C_RF2_250 RF2
ZShift = −0.76
lm ZScore = 0.65
NG = 0
DB = 0
SM = 0
Delta_r
Doxorubicin
YDL227C_RF2_250 RF2
ZShift = 0.29
lm ZScore = 0.01
NG = 0
DB = 0
SM = 0
Delta_AUC
Doxorubicin
YDL227C_RF1_287 RF1
ZShift = −0.87
lm Zscore = 0.01
NG = 0
DB = 0
SM = 0
Delta_L
Doxorubicin
YDL227C_RF1_287 RF1
ZShift = 1.57
lm ZScore = 0.01
NG = 0
DB = 0
SM = 0
Delta_K
Doxorubicin
YDL227C_RF1_287 RF1
ZShift = 0.21
lm ZScore = −0.26
NG = 0
DB = 0
SM = 0
Delta_r
Doxorubicin
YDL227C_RF1_287 RF1
ZShift = 1.67
lm ZScore = −0.05
NG = 0
DB = 0
SM = 0
Delta_AUC
Doxorubicin
YDL227C_RF1_193 RF1
ZShift = −0.17
lm Zscore = 0
NG = 0
DB = 0
SM = 0
Delta_L
Doxorubicin
YDL227C_RF1_193 RF1
ZShift = 1.51
lm ZScore = 0
NG = 0
DB = 0
SM = 0
Delta_K
Doxorubicin
YDL227C_RF1_193 RF1
ZShift = −0.44
lm ZScore = 0.41
NG = 0
DB = 0
SM = 0
Delta_r
Doxorubicin
YDL227C_RF1_193 RF1
ZShift = 1.45
lm ZScore = −0.05
NG = 0
DB = 0
SM = 0
Delta_AUC
Doxorubicin

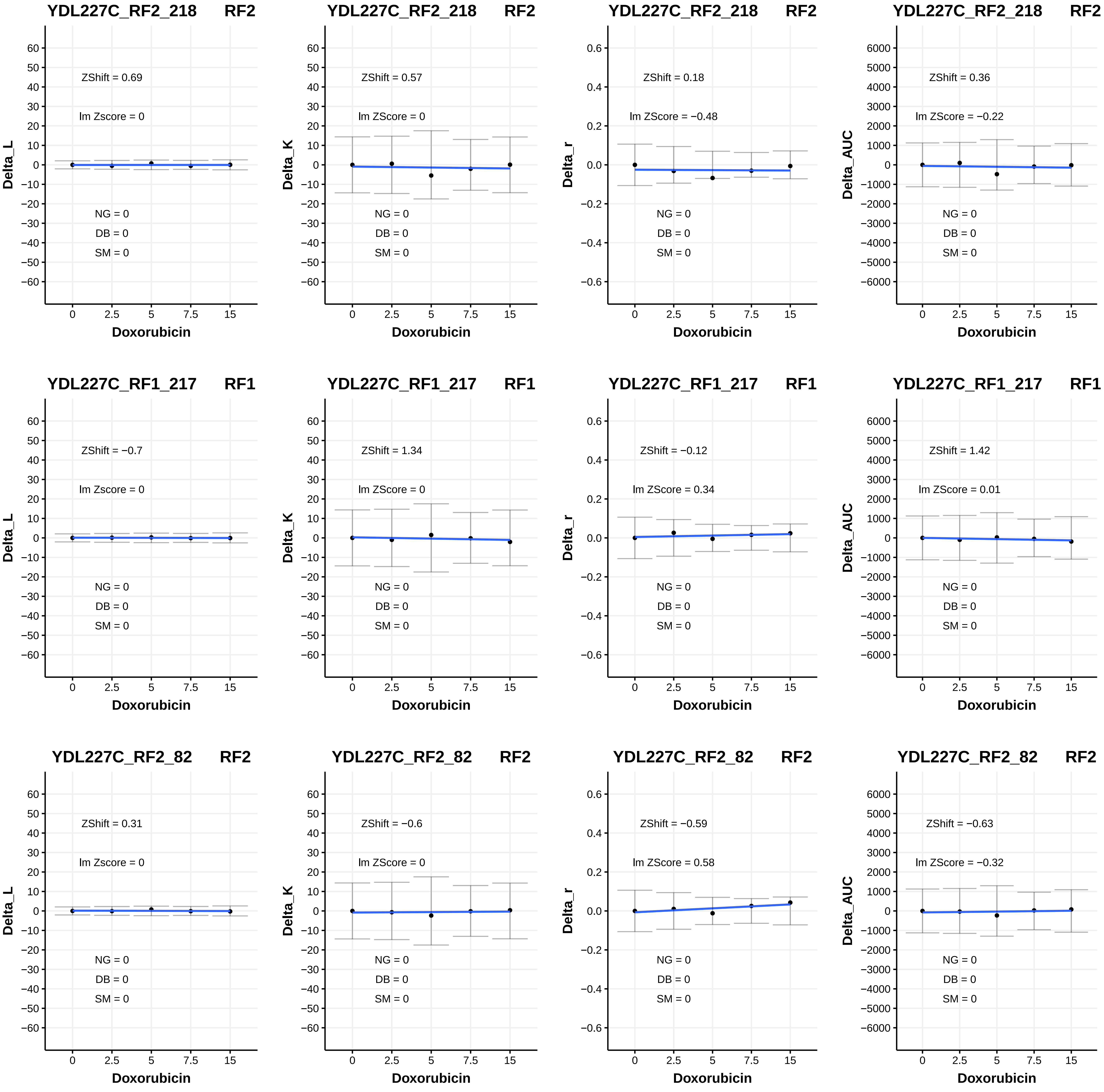

YDL227C_RF2_218 RF2
ZShift = 0.69
lm Zscore = 0
NG = 0
DB = 0
SM = 0
Delta_L
Doxorubicin
YDL227C_RF2_218 RF2
ZShift = 0.57
lm ZScore = 0
NG = 0
DB = 0
SM = 0
Delta_K
Doxorubicin
YDL227C_RF2_218 RF2
ZShift = 0.18
lm ZScore = −0.48
NG = 0
DB = 0
SM = 0
Delta_r
Doxorubicin
YDL227C_RF2_218 RF2
ZShift = 0.36
lm ZScore = −0.22
NG = 0
DB = 0
SM = 0
Delta_AUC
Doxorubicin
YDL227C_RF1_217 RF1
ZShift = −0.7
lm Zscore = 0
NG = 0
DB = 0
SM = 0
Delta_L
Doxorubicin
YDL227C_RF1_217 RF1
ZShift = 1.34
lm ZScore = 0
NG = 0
DB = 0
SM = 0
Delta_K
Doxorubicin
YDL227C_RF1_217 RF1
ZShift = −0.12
lm ZScore = 0.34
NG = 0
DB = 0
SM = 0
Delta_r
Doxorubicin
YDL227C_RF1_217 RF1
ZShift = 1.42
lm ZScore = 0.01
NG = 0
DB = 0
SM = 0
Delta_AUC
Doxorubicin
YDL227C_RF2_82 RF2
ZShift = 0.31
lm Zscore = 0
NG = 0
DB = 0
SM = 0
Delta_L
Doxorubicin
YDL227C_RF2_82 RF2
ZShift = −0.6
lm ZScore = 0
NG = 0
DB = 0
SM = 0
Delta_K
Doxorubicin
YDL227C_RF2_82 RF2
ZShift = −0.59
lm ZScore = 0.58
NG = 0
DB = 0
SM = 0
Delta_r
Doxorubicin
YDL227C_RF2_82 RF2
ZShift = −0.63
lm ZScore = −0.32
NG = 0
DB = 0
SM = 0
Delta_AUC
Doxorubicin

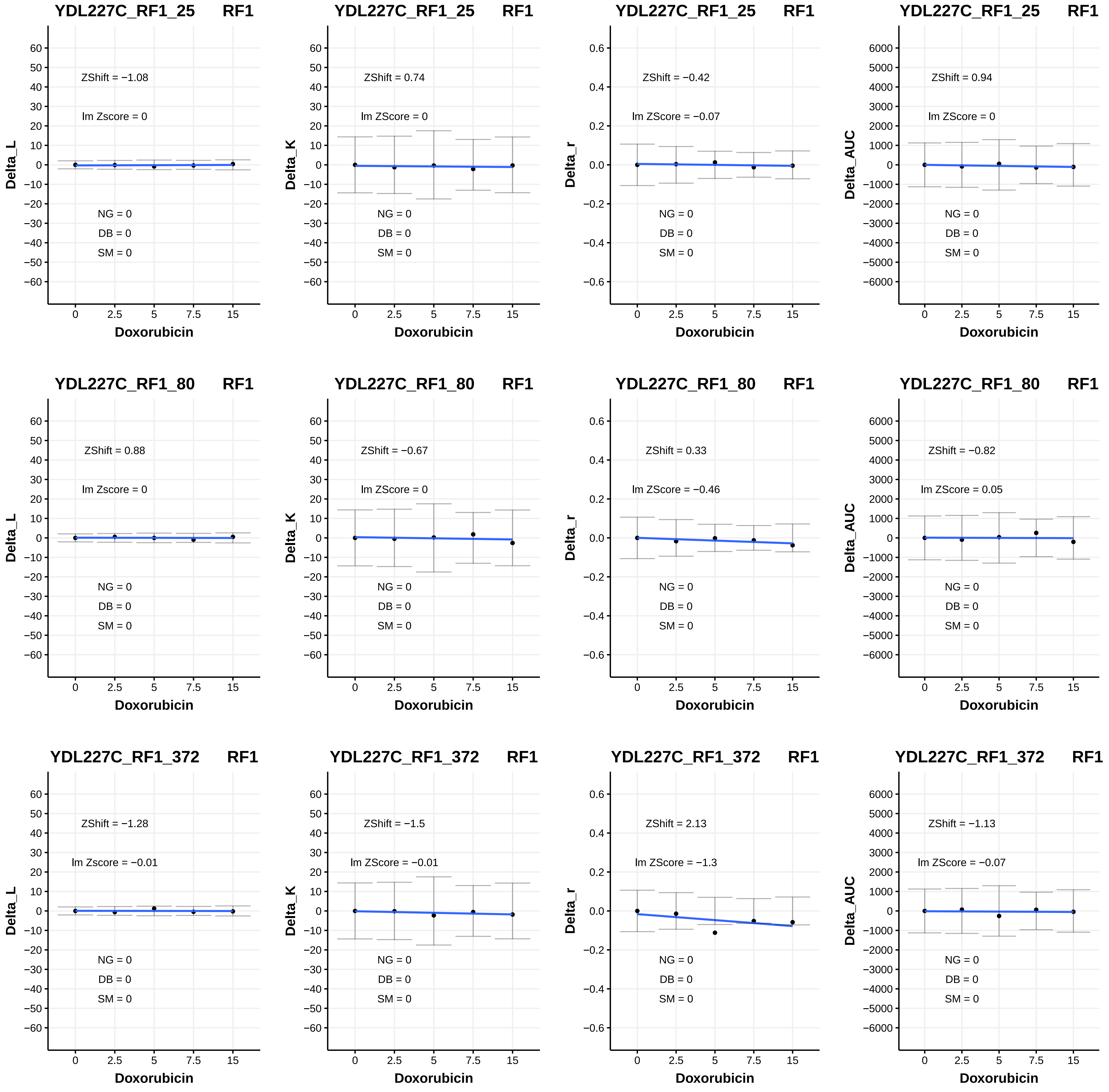
YDL227C_RF1_25 RF1
ZShift = −1.08
lm Zscore = 0
NG = 0
DB = 0
SM = 0
Delta_L
Doxorubicin
YDL227C_RF1_25 RF1
ZShift = 0.74
lm ZScore = 0
NG = 0
DB = 0
SM = 0
Delta_K
Doxorubicin
YDL227C_RF1_25 RF1
ZShift = −0.42
lm ZScore = −0.07
NG = 0
DB = 0
SM = 0
Delta_r
Doxorubicin
YDL227C_RF1_25 RF1
ZShift = 0.94
lm ZScore = 0
NG = 0
DB = 0
SM = 0
Delta_AUC
Doxorubicin
YDL227C_RF1_80 RF1
ZShift = 0.88
lm Zscore = 0
NG = 0
DB = 0
SM = 0
Delta_L
Doxorubicin
YDL227C_RF1_80 RF1
ZShift = −0.67
lm ZScore = 0
NG = 0
DB = 0
SM = 0
Delta_K
Doxorubicin
YDL227C_RF1_80 RF1
ZShift = 0.33
lm ZScore = −0.46
NG = 0
DB = 0
SM = 0
Delta_r
Doxorubicin
YDL227C_RF1_80 RF1
ZShift = −0.82
lm ZScore = 0.05
NG = 0
DB = 0
SM = 0
Delta_AUC
Doxorubicin
YDL227C_RF1_372 RF1
ZShift = −1.28
lm Zscore = −0.01
NG = 0
DB = 0
SM = 0
Delta_L
Doxorubicin
YDL227C_RF1_372 RF1
ZShift = −1.5
lm ZScore = −0.01
NG = 0
DB = 0
SM = 0
Delta_K
Doxorubicin
YDL227C_RF1_372 RF1
ZShift = 2.13
lm ZScore = −1.3
NG = 0
DB = 0
SM = 0
Delta_r
Doxorubicin
YDL227C_RF1_372 RF1
ZShift = −1.13
lm ZScore = −0.07
NG = 0
DB = 0
SM = 0
Delta_AUC
Doxorubicin

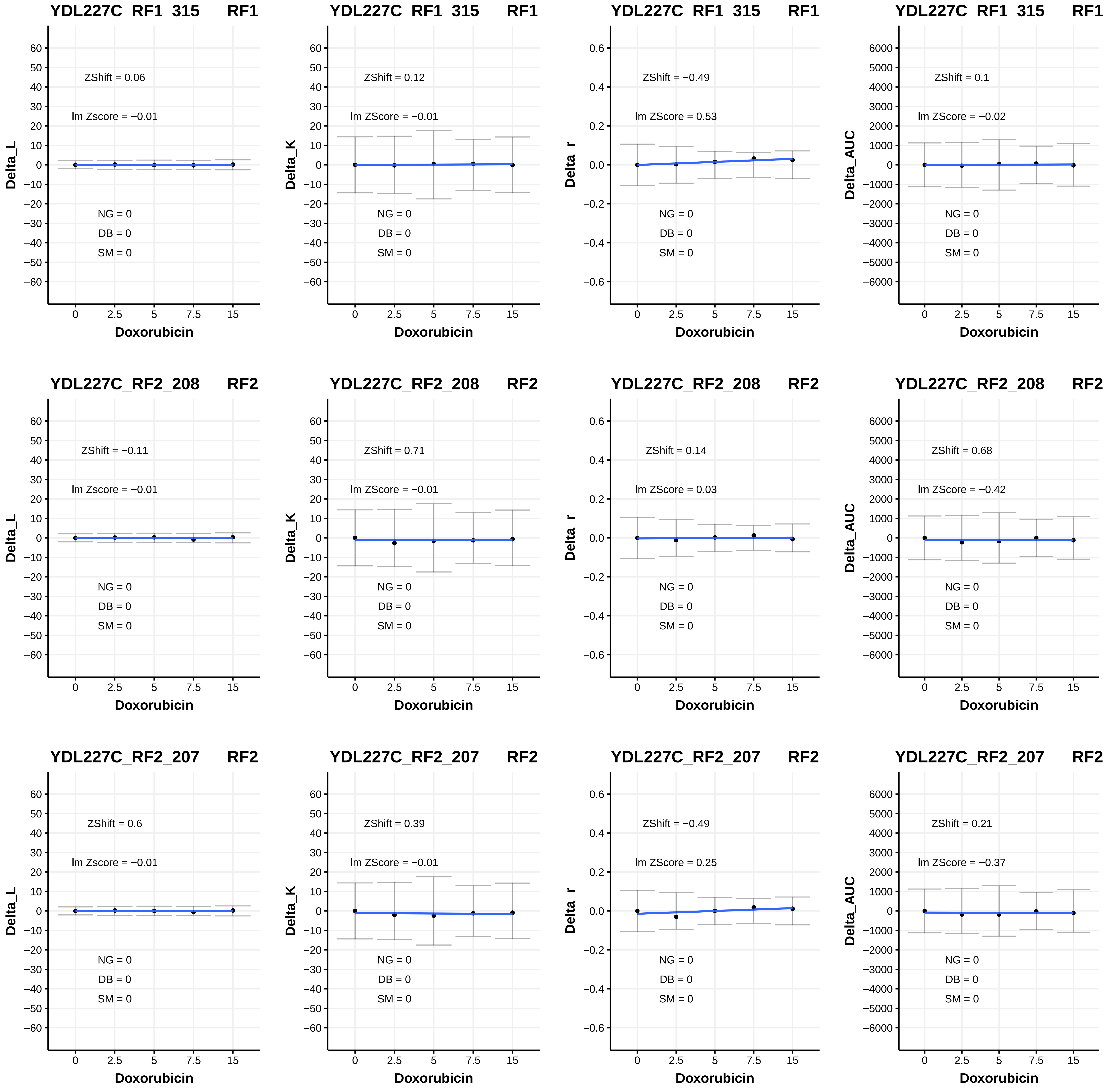
YDL227C_RF1_315 RF1
Delta_L
ZShift = 0.06
lm Zscore = −0.01
NG = 0
DB = 0
SM = 0
Doxorubicin
YDL227C_RF1_315 RF1
Delta_K
ZShift = 0.12
lm ZScore = −0.01
NG = 0
DB = 0
SM = 0
Doxorubicin
YDL227C_RF1_315 RF1
Delta_r
ZShift = −0.49
lm ZScore = 0.53
NG = 0
DB = 0
SM = 0
Doxorubicin
YDL227C_RF1_315 RF1
Delta_AUC
ZShift = 0.1
lm ZScore = −0.02
NG = 0
DB = 0
SM = 0
Doxorubicin
YDL227C_RF2_208 RF2
Delta_L
ZShift = −0.11
lm Zscore = −0.01
NG = 0
DB = 0
SM = 0
Doxorubicin
YDL227C_RF2_208 RF2
Delta_K
ZShift = 0.71
lm ZScore = −0.01
NG = 0
DB = 0
SM = 0
Doxorubicin
YDL227C_RF2_208 RF2
Delta_r
ZShift = 0.14
lm ZScore = 0.03
NG = 0
DB = 0
SM = 0
Doxorubicin
YDL227C_RF2_208 RF2
Delta_AUC
ZShift = 0.68
lm ZScore = −0.42
NG = 0
DB = 0
SM = 0
Doxorubicin
YDL227C_RF2_207 RF2
Delta_L
ZShift = 0.6
lm Zscore = −0.01
NG = 0
DB = 0
SM = 0
Doxorubicin
YDL227C_RF2_207 RF2
Delta_K
ZShift = 0.39
lm ZScore = −0.01
NG = 0
DB = 0
SM = 0
Doxorubicin
YDL227C_RF2_207 RF2
Delta_r
ZShift = −0.49
lm ZScore = 0.25
NG = 0
DB = 0
SM = 0
Doxorubicin
YDL227C_RF2_207 RF2
Delta_AUC
ZShift = 0.21
lm ZScore = −0.37
NG = 0
DB = 0
SM = 0
Doxorubicin

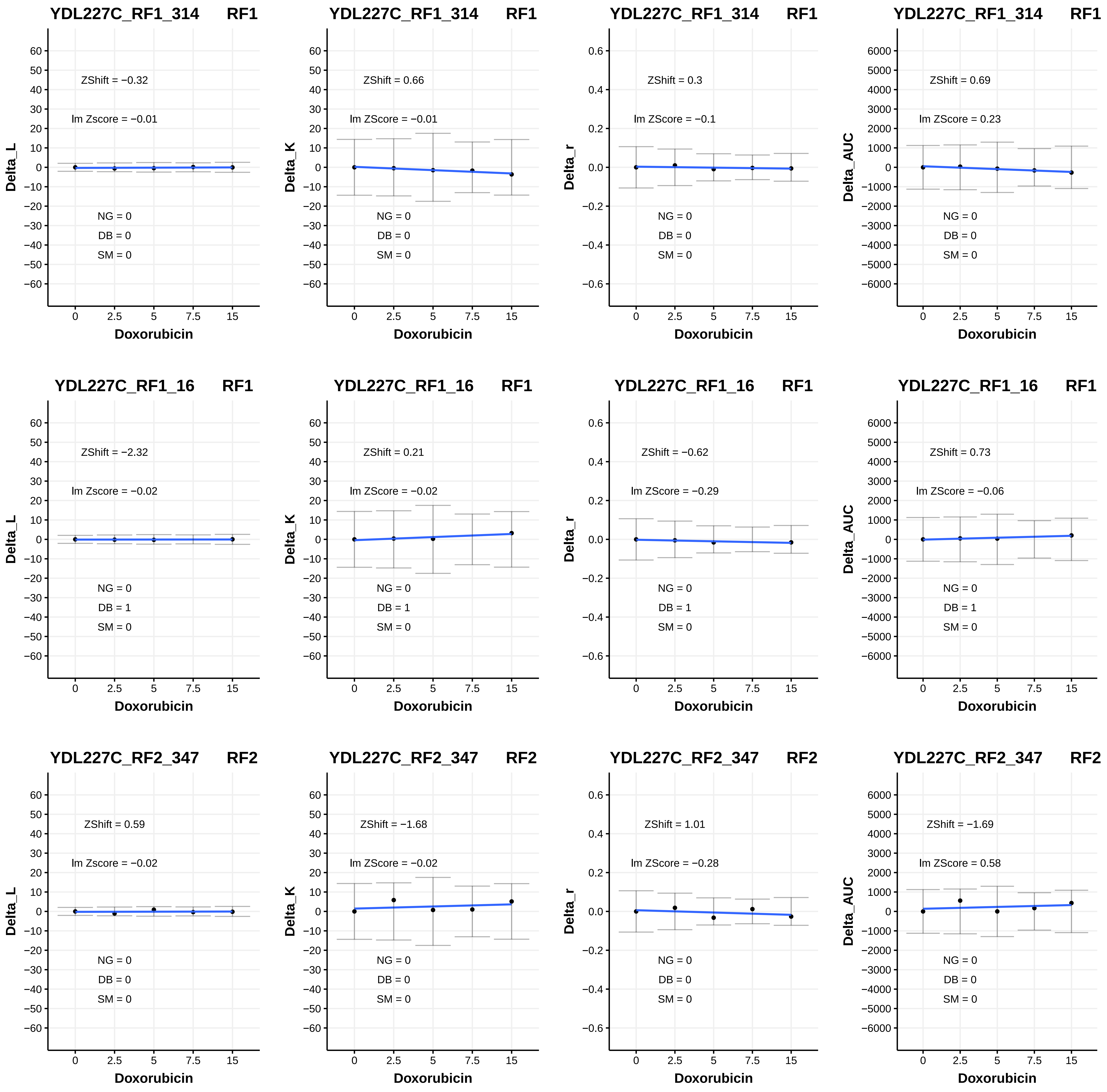

YDL227C_RF1_314 RF1
ZShift = −0.32
lm Zscore = −0.01
NG = 0
DB = 0
SM = 0
Delta_L
Doxorubicin
YDL227C_RF1_314 RF1
ZShift = 0.66
lm ZScore = −0.01
NG = 0
DB = 0
SM = 0
Delta_K
Doxorubicin
YDL227C_RF1_314 RF1
ZShift = 0.3
lm ZScore = −0.1
NG = 0
DB = 0
SM = 0
Delta_r
Doxorubicin
YDL227C_RF1_314 RF1
ZShift = 0.69
lm ZScore = 0.23
NG = 0
DB = 0
SM = 0
Delta_AUC
Doxorubicin
YDL227C_RF1_16 RF1
ZShift = −2.32
lm Zscore = −0.02
NG = 0
DB = 1
SM = 0
Delta_L
Doxorubicin
YDL227C_RF1_16 RF1
ZShift = 0.21
lm ZScore = −0.02
NG = 0
DB = 1
SM = 0
Delta_K
Doxorubicin
YDL227C_RF1_16 RF1
ZShift = −0.62
lm ZScore = −0.29
NG = 0
DB = 1
SM = 0
Delta_r
Doxorubicin
YDL227C_RF1_16 RF1
ZShift = 0.73
lm ZScore = −0.06
NG = 0
DB = 1
SM = 0
Delta_AUC
Doxorubicin
YDL227C_RF2_347 RF2
ZShift = 0.59
lm Zscore = −0.02
NG = 0
DB = 0
SM = 0
Delta_L
Doxorubicin
YDL227C_RF2_347 RF2
ZShift = −1.68
lm ZScore = −0.02
NG = 0
DB = 0
SM = 0
Delta_K
Doxorubicin
YDL227C_RF2_347 RF2
ZShift = 1.01
lm ZScore = −0.28
NG = 0
DB = 0
SM = 0
Delta_r
Doxorubicin
YDL227C_RF2_347 RF2
ZShift = −1.69
lm ZScore = 0.58
NG = 0
DB = 0
SM = 0
Delta_AUC
Doxorubicin

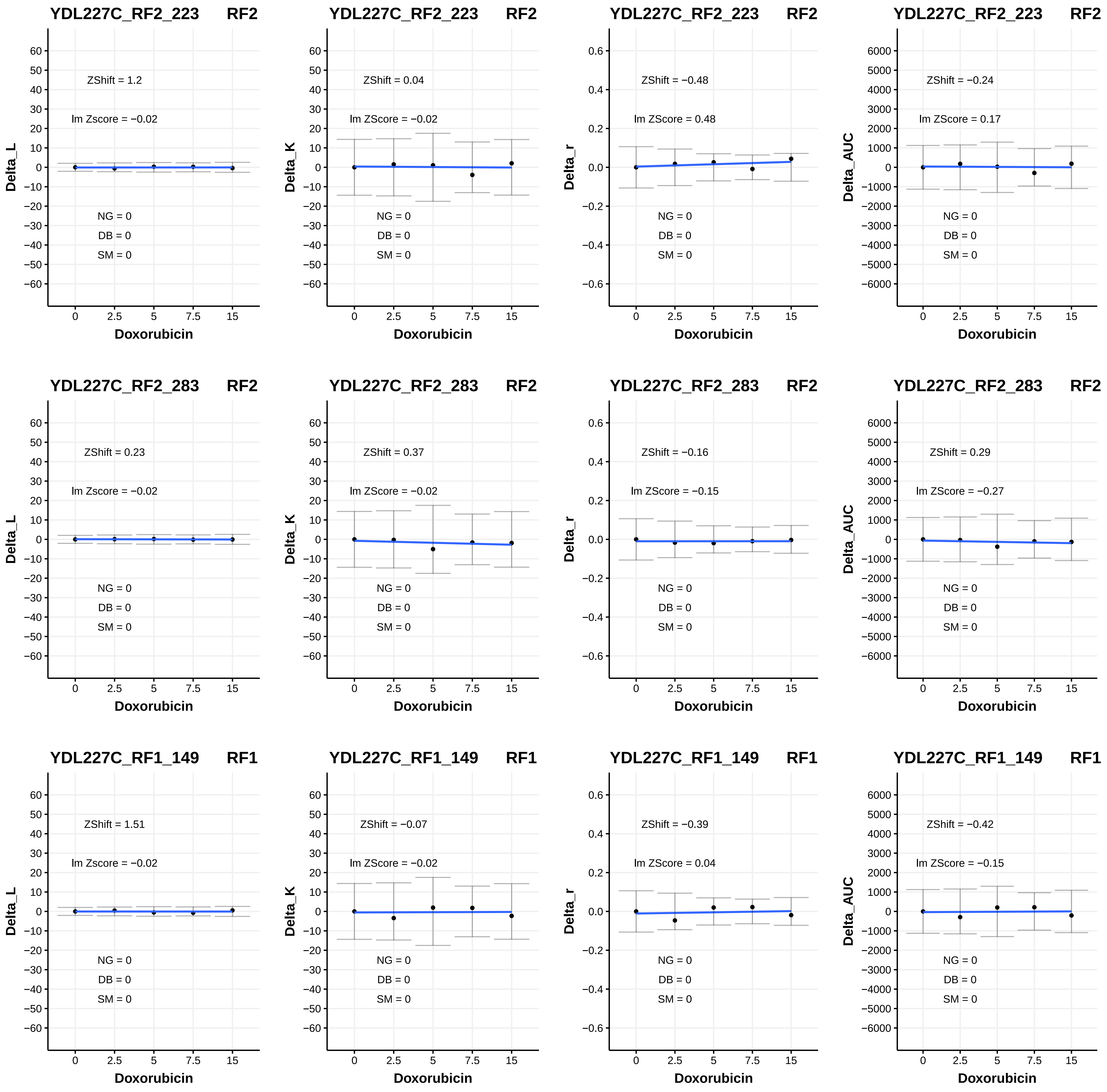
YDL227C_RF2_223 RF2
ZShift = 1.2
lm Zscore = −0.02
NG = 0
DB = 0
SM = 0
Delta_L
Doxorubicin
YDL227C_RF2_223 RF2
ZShift = 0.04
lm ZScore = −0.02
NG = 0
DB = 0
SM = 0
Delta_K
Doxorubicin
YDL227C_RF2_223 RF2
ZShift = −0.48
lm ZScore = 0.48
NG = 0
DB = 0
SM = 0
Delta_r
Doxorubicin
YDL227C_RF2_223 RF2
ZShift = −0.24
lm ZScore = 0.17
NG = 0
DB = 0
SM = 0
Delta_AUC
Doxorubicin
YDL227C_RF2_283 RF2
ZShift = 0.23
lm Zscore = −0.02
NG = 0
DB = 0
SM = 0
Delta_L
Doxorubicin
YDL227C_RF2_283 RF2
ZShift = 0.37
lm ZScore = −0.02
NG = 0
DB = 0
SM = 0
Delta_K
Doxorubicin
YDL227C_RF2_283 RF2
ZShift = −0.16
lm ZScore = −0.15
NG = 0
DB = 0
SM = 0
Delta_r
Doxorubicin
YDL227C_RF2_283 RF2
ZShift = 0.29
lm ZScore = −0.27
NG = 0
DB = 0
SM = 0
Delta_AUC
Doxorubicin
YDL227C_RF1_149 RF1
ZShift = 1.51
lm Zscore = −0.02
NG = 0
DB = 0
SM = 0
Delta_L
Doxorubicin
YDL227C_RF1_149 RF1
ZShift = −0.07
lm ZScore = −0.02
NG = 0
DB = 0
SM = 0
Delta_K
Doxorubicin
YDL227C_RF1_149 RF1
ZShift = −0.39
lm ZScore = 0.04
NG = 0
DB = 0
SM = 0
Delta_r
Doxorubicin
YDL227C_RF1_149 RF1
ZShift = −0.42
lm ZScore = −0.15
NG = 0
DB = 0
SM = 0
Delta_AUC
Doxorubicin

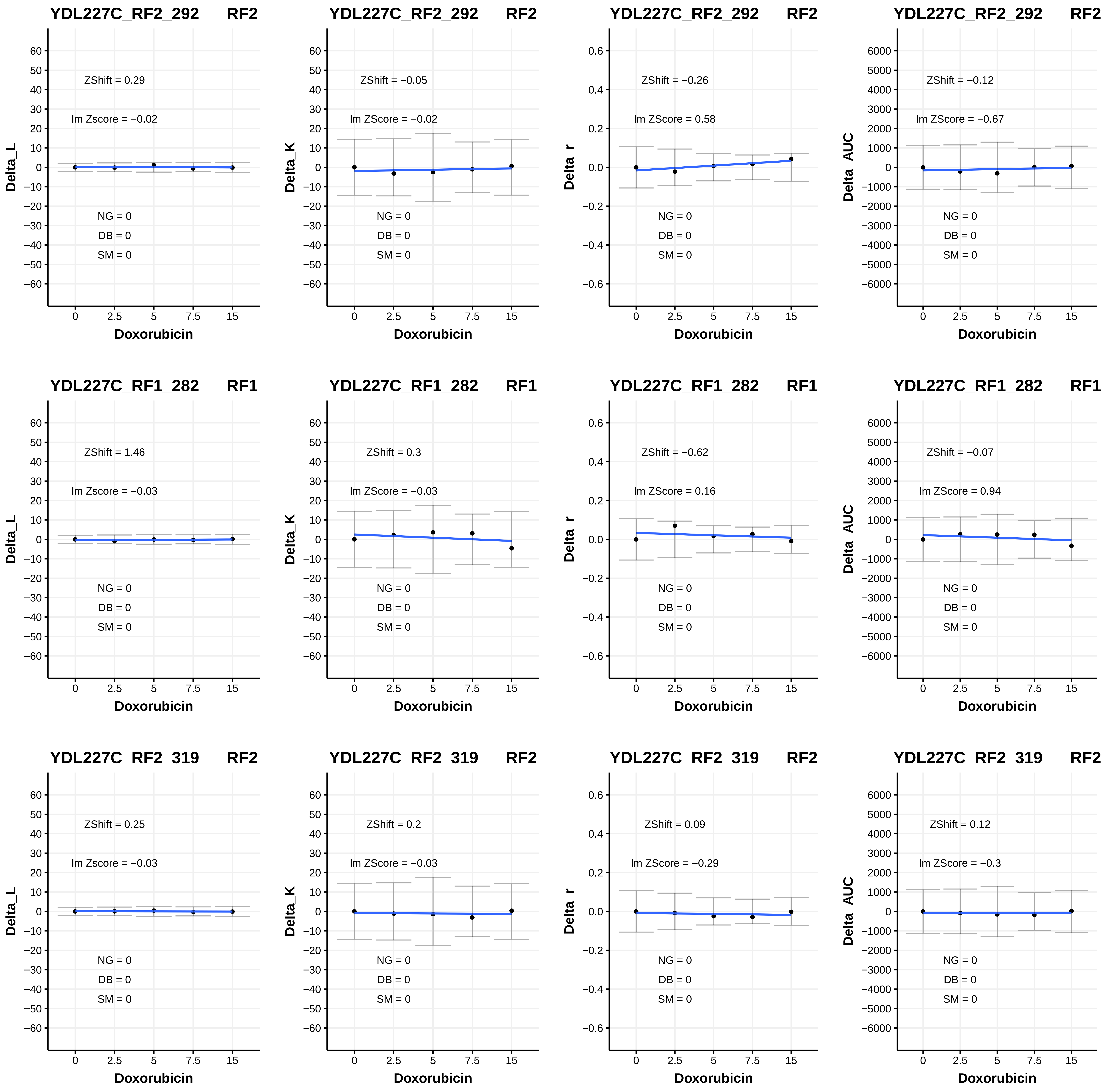
YDL227C_RF2_292 RF2
ZShift = 0.29
lm Zscore = −0.02
NG = 0
DB = 0
SM = 0
Delta_L
Doxorubicin
YDL227C_RF2_292 RF2
ZShift = −0.05
lm ZScore = −0.02
NG = 0
DB = 0
SM = 0
Delta_K
Doxorubicin
YDL227C_RF2_292 RF2
ZShift = −0.26
lm ZScore = 0.58
NG = 0
DB = 0
SM = 0
Delta_r
Doxorubicin
YDL227C_RF2_292 RF2
ZShift = −0.12
lm ZScore = −0.67
NG = 0
DB = 0
SM = 0
Delta_AUC
Doxorubicin
YDL227C_RF1_282 RF1
ZShift = 1.46
lm Zscore = −0.03
NG = 0
DB = 0
SM = 0
Delta_L
Doxorubicin
YDL227C_RF1_282 RF1
ZShift = 0.3
lm ZScore = −0.03
NG = 0
DB = 0
SM = 0
Delta_K
Doxorubicin
YDL227C_RF1_282 RF1
ZShift = −0.62
lm ZScore = 0.16
NG = 0
DB = 0
SM = 0
Delta_r
Doxorubicin
YDL227C_RF1_282 RF1
ZShift = −0.07
lm ZScore = 0.94
NG = 0
DB = 0
SM = 0
Delta_AUC
Doxorubicin
YDL227C_RF2_319 RF2
ZShift = 0.25
lm Zscore = −0.03
NG = 0
DB = 0
SM = 0
Delta_L
Doxorubicin
YDL227C_RF2_319 RF2
ZShift = 0.2
lm ZScore = −0.03
NG = 0
DB = 0
SM = 0
Delta_K
Doxorubicin
YDL227C_RF2_319 RF2
ZShift = 0.09
lm ZScore = −0.29
NG = 0
DB = 0
SM = 0
Delta_r
Doxorubicin
YDL227C_RF2_319 RF2
ZShift = 0.12
lm Zscore = −0.3
NG = 0
DB = 0
SM = 0
Delta_AUC
Doxorubicin

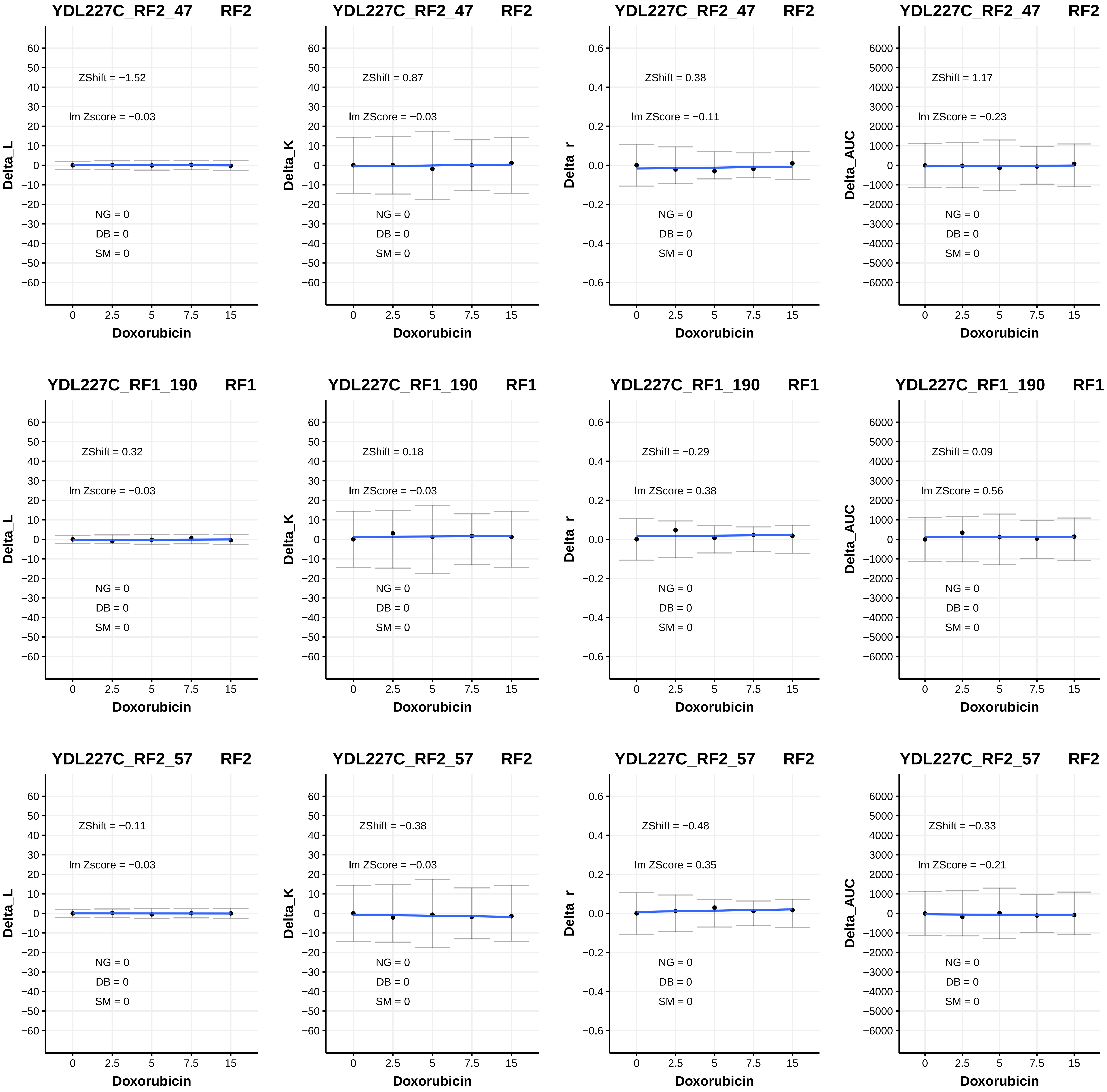

YDL227C_RF2_47 RF2
ZShift = −1.52
lm Zscore = −0.03
NG = 0
DB = 0
SM = 0
Delta_L
Doxorubicin
YDL227C_RF2_47 RF2
ZShift = 0.87
lm ZScore = −0.03
NG = 0
DB = 0
SM = 0
Delta_K
Doxorubicin
YDL227C_RF2_47 RF2
ZShift = 0.38
lm ZScore = −0.11
NG = 0
DB = 0
SM = 0
Delta_r
Doxorubicin
YDL227C_RF2_47 RF2
ZShift = 1.17
lm ZScore = −0.23
NG = 0
DB = 0
SM = 0
Delta_AUC
Doxorubicin
YDL227C_RF1_190 RF1
ZShift = 0.32
lm Zscore = −0.03
NG = 0
DB = 0
SM = 0
Delta_L
Doxorubicin
YDL227C_RF1_190 RF1
ZShift = 0.18
lm ZScore = −0.03
NG = 0
DB = 0
SM = 0
Delta_K
Doxorubicin
YDL227C_RF1_190 RF1
ZShift = −0.29
lm ZScore = 0.38
NG = 0
DB = 0
SM = 0
Delta_r
Doxorubicin
YDL227C_RF1_190 RF1
ZShift = 0.09
lm ZScore = 0.56
NG = 0
DB = 0
SM = 0
Delta_AUC
Doxorubicin
YDL227C_RF2_57 RF2
ZShift = −0.11
lm Zscore = −0.03
NG = 0
DB = 0
SM = 0
Delta_L
Doxorubicin
YDL227C_RF2_57 RF2
ZShift = −0.38
lm ZScore = −0.03
NG = 0
DB = 0
SM = 0
Delta_K
Doxorubicin
YDL227C_RF2_57 RF2
ZShift = −0.48
lm ZScore = 0.35
NG = 0
DB = 0
SM = 0
Delta_r
Doxorubicin
YDL227C_RF2_57 RF2
ZShift = −0.33
lm ZScore = −0.21
NG = 0
DB = 0
SM = 0
Delta_AUC
Doxorubicin

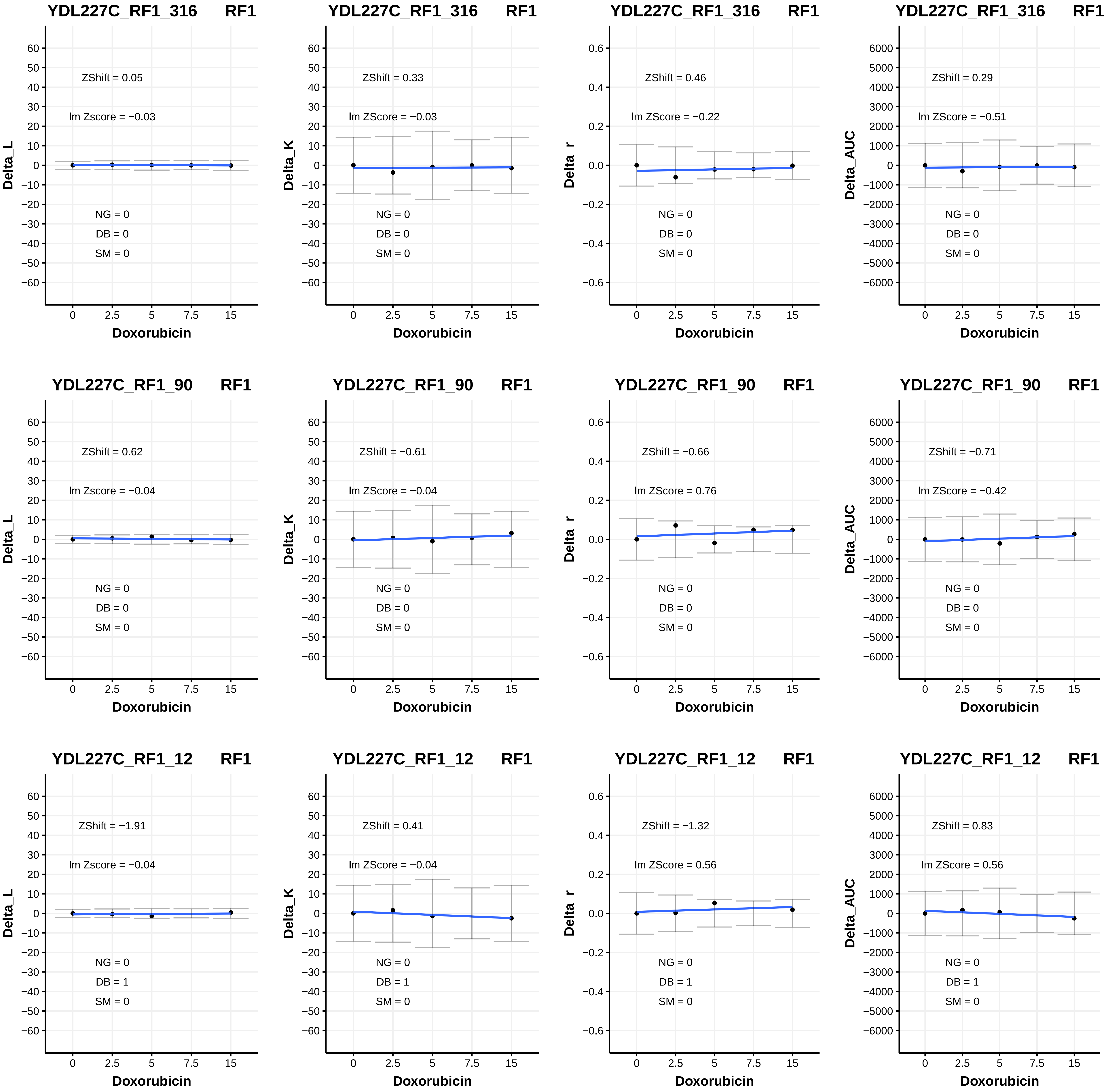

YDL227C_RF1_316 RF1
ZShift = 0.05
lm Zscore = −0.03
NG = 0
DB = 0
SM = 0
Delta_L
Doxorubicin
YDL227C_RF1_316 RF1
ZShift = 0.33
lm ZScore = −0.03
NG = 0
DB = 0
SM = 0
Delta_K
Doxorubicin
YDL227C_RF1_316 RF1
ZShift = 0.46
lm ZScore = −0.22
NG = 0
DB = 0
SM = 0
Delta_r
Doxorubicin
YDL227C_RF1_316 RF1
ZShift = 0.29
lm ZScore = −0.51
NG = 0
DB = 0
SM = 0
Delta_AUC
Doxorubicin
YDL227C_RF1_90 RF1
ZShift = 0.62
lm Zscore = −0.04
NG = 0
DB = 0
SM = 0
Delta_L
Doxorubicin
YDL227C_RF1_90 RF1
ZShift = −0.61
lm ZScore = −0.04
NG = 0
DB = 0
SM = 0
Delta_K
Doxorubicin
YDL227C_RF1_90 RF1
ZShift = −0.66
lm ZScore = 0.76
NG = 0
DB = 0
SM = 0
Delta_r
Doxorubicin
YDL227C_RF1_90 RF1
ZShift = −0.71
lm ZScore = −0.42
NG = 0
DB = 0
SM = 0
Delta_AUC
Doxorubicin
YDL227C_RF1_12 RF1
ZShift = −1.91
lm Zscore = −0.04
NG = 0
DB = 1
SM = 0
Delta_L
Doxorubicin
YDL227C_RF1_12 RF1
ZShift = 0.41
lm ZScore = −0.04
NG = 0
DB = 1
SM = 0
Delta_K
Doxorubicin
YDL227C_RF1_12 RF1
ZShift = −1.32
lm ZScore = 0.56
NG = 0
DB = 1
SM = 0
Delta_r
Doxorubicin
YDL227C_RF1_12 RF1
ZShift = 0.83
lm ZScore = 0.56
NG = 0
DB = 1
SM = 0
Delta_AUC
Doxorubicin

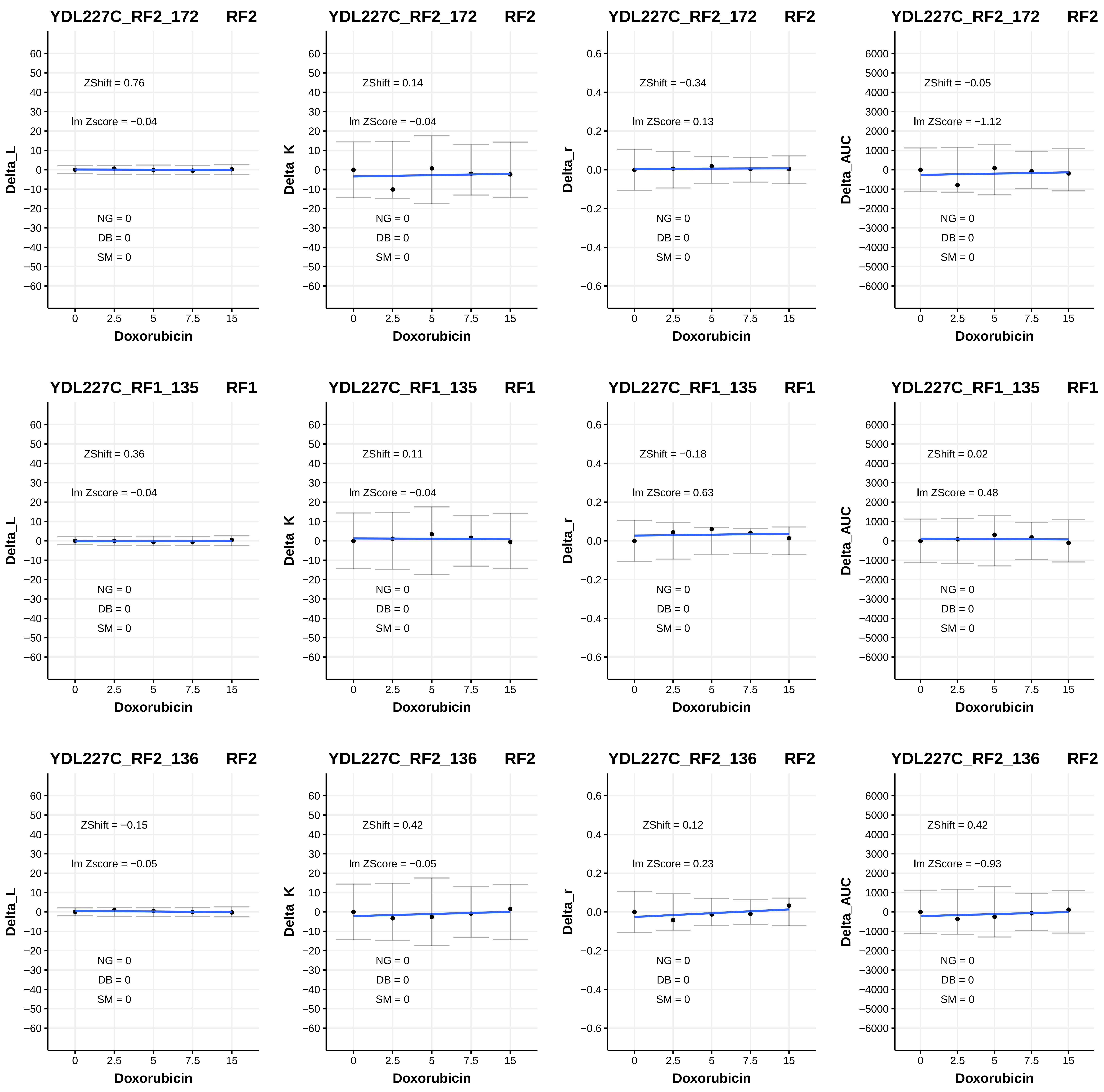

YDL227C_RF2_172 RF2
ZShift = 0.76
lm Zscore = −0.04
NG = 0
DB = 0
SM = 0
Delta_L
Doxorubicin
YDL227C_RF2_172 RF2
ZShift = 0.14
lm ZScore = −0.04
NG = 0
DB = 0
SM = 0
Delta_K
Doxorubicin
YDL227C_RF2_172 RF2
ZShift = −0.34
lm ZScore = 0.13
NG = 0
DB = 0
SM = 0
Delta_r
Doxorubicin
YDL227C_RF2_172 RF2
ZShift = −0.05
lm ZScore = −1.12
NG = 0
DB = 0
SM = 0
Delta_AUC
Doxorubicin
YDL227C_RF1_135 RF1
ZShift = 0.36
lm Zscore = −0.04
NG = 0
DB = 0
SM = 0
Delta_L
Doxorubicin
YDL227C_RF1_135 RF1
ZShift = 0.11
lm ZScore = −0.04
NG = 0
DB = 0
SM = 0
Delta_K
Doxorubicin
YDL227C_RF1_135 RF1
ZShift = −0.18
lm ZScore = 0.63
NG = 0
DB = 0
SM = 0
Delta_r
Doxorubicin
YDL227C_RF1_135 RF1
ZShift = 0.02
lm ZScore = 0.48
NG = 0
DB = 0
SM = 0
Delta_AUC
Doxorubicin
YDL227C_RF2_136 RF2
ZShift = −0.15
lm Zscore = −0.05
NG = 0
DB = 0
SM = 0
Delta_L
Doxorubicin
YDL227C_RF2_136 RF2
ZShift = 0.42
lm ZScore = −0.05
NG = 0
DB = 0
SM = 0
Delta_K
Doxorubicin
YDL227C_RF2_136 RF2
ZShift = 0.12
lm ZScore = 0.23
NG = 0
DB = 0
SM = 0
Delta_r
Doxorubicin
YDL227C_RF2_136 RF2
ZShift = 0.42
lm ZScore = −0.93
NG = 0
DB = 0
SM = 0
Delta_AUC
Doxorubicin

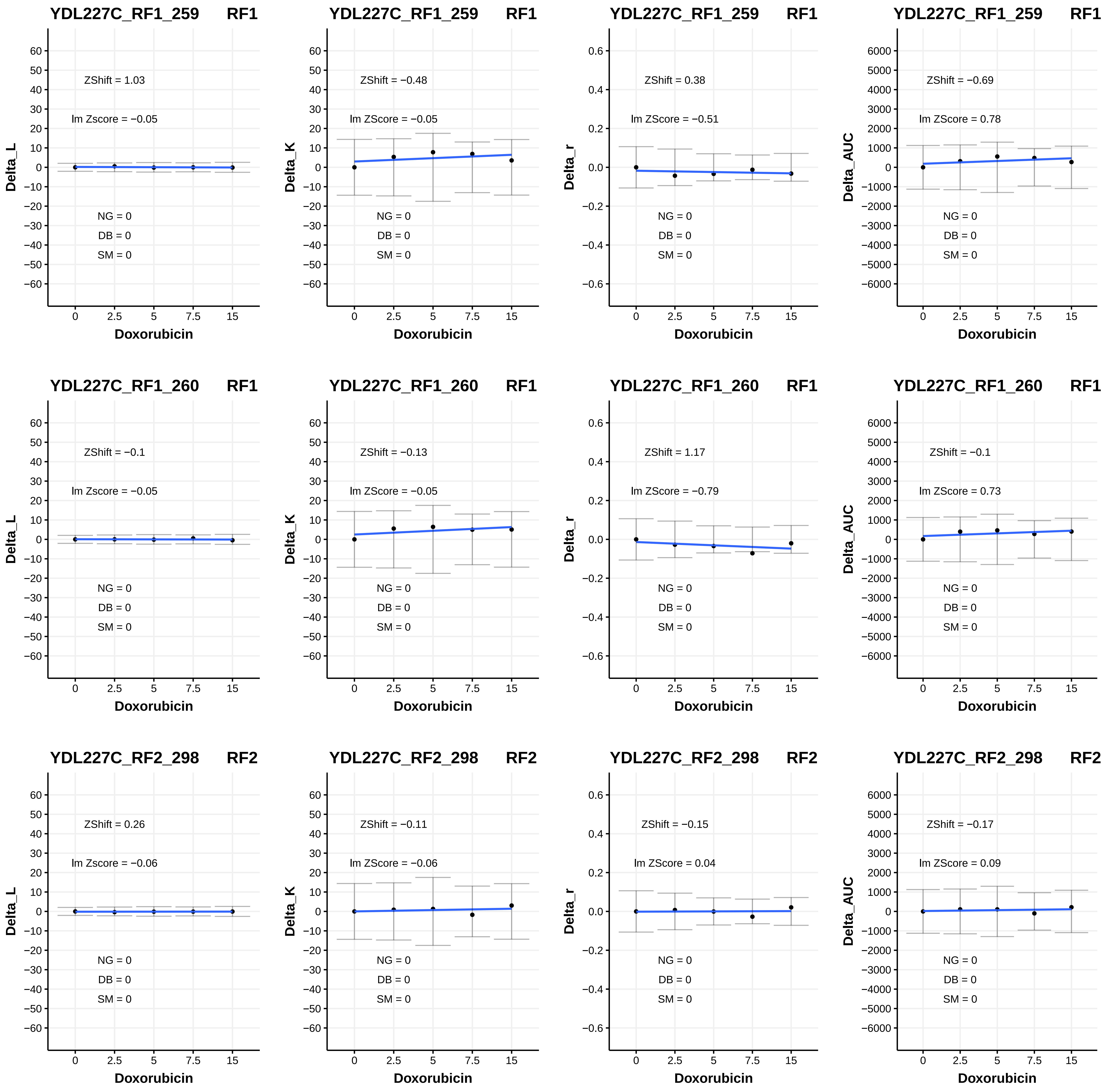

YDL227C_RF1_259 RF1
ZShift = 1.03
lm Zscore = −0.05
NG = 0
DB = 0
SM = 0
Delta_L
Doxorubicin
YDL227C_RF1_259 RF1
ZShift = −0.48
lm ZScore = −0.05
NG = 0
DB = 0
SM = 0
Delta_K
Doxorubicin
YDL227C_RF1_259 RF1
ZShift = 0.38
lm ZScore = −0.51
NG = 0
DB = 0
SM = 0
Delta_r
Doxorubicin
YDL227C_RF1_259 RF1
ZShift = −0.69
lm ZScore = 0.78
NG = 0
DB = 0
SM = 0
Delta_AUC
Doxorubicin
YDL227C_RF1_260 RF1
ZShift = −0.1
lm Zscore = −0.05
NG = 0
DB = 0
SM = 0
Delta_L
Doxorubicin
YDL227C_RF1_260 RF1
ZShift = −0.13
lm ZScore = −0.05
NG = 0
DB = 0
SM = 0
Delta_K
Doxorubicin
YDL227C_RF1_260 RF1
ZShift = 1.17
lm ZScore = −0.79
NG = 0
DB = 0
SM = 0
Delta_r
Doxorubicin
YDL227C_RF1_260 RF1
ZShift = −0.1
lm ZScore = 0.73
NG = 0
DB = 0
SM = 0
Delta_AUC
Doxorubicin
YDL227C_RF2_298 RF2
ZShift = 0.26
lm Zscore = −0.06
NG = 0
DB = 0
SM = 0
Delta_L
Doxorubicin
YDL227C_RF2_298 RF2
ZShift = −0.11
lm ZScore = −0.06
NG = 0
DB = 0
SM = 0
Delta_K
Doxorubicin
YDL227C_RF2_298 RF2
ZShift = −0.15
lm ZScore = 0.04
NG = 0
DB = 0
SM = 0
Delta_r
Doxorubicin
YDL227C_RF2_298 RF2
ZShift = −0.17
lm ZScore = 0.09
NG = 0
DB = 0
SM = 0
Delta_AUC
Doxorubicin

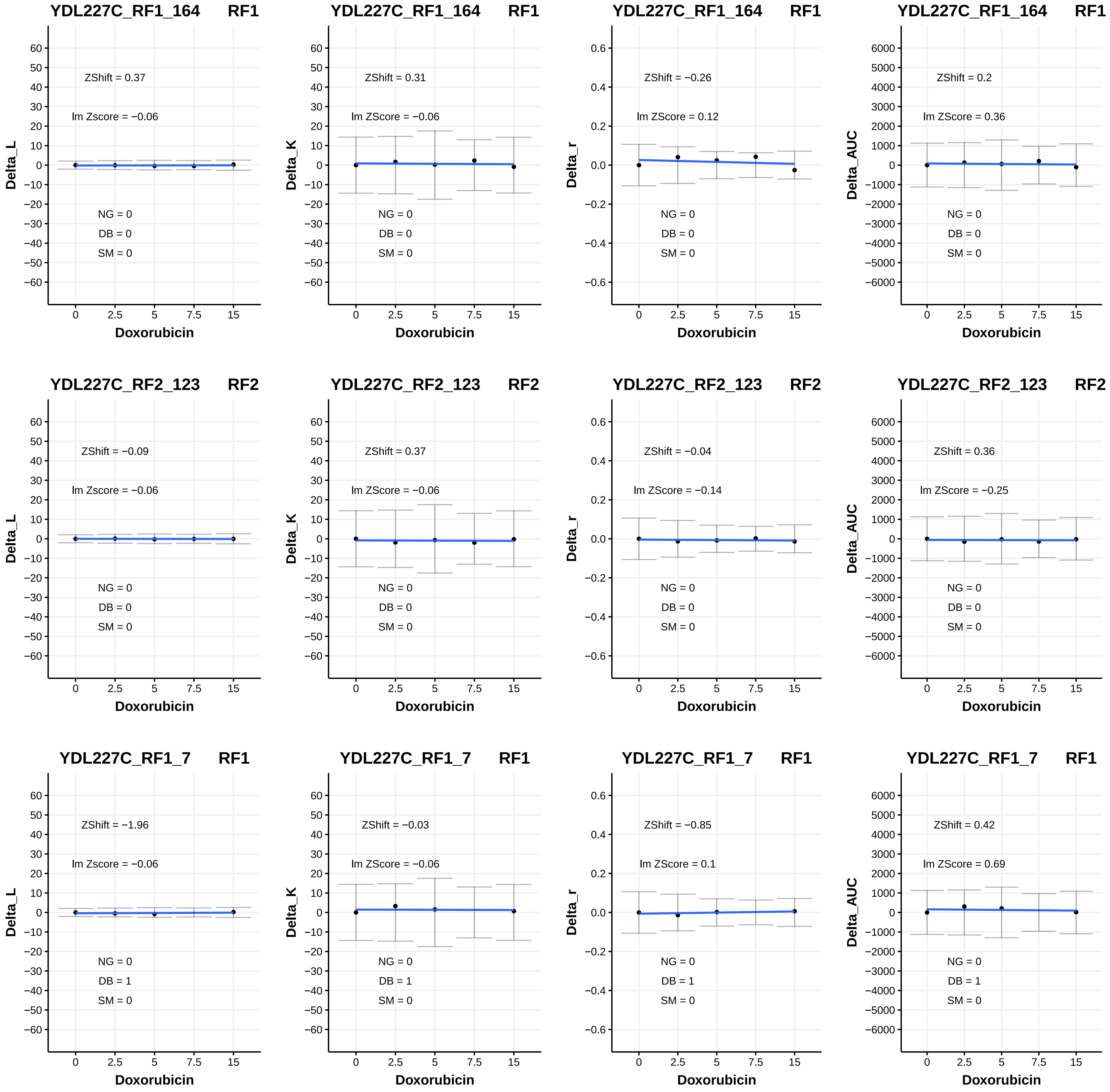

YDL227C_RF1_164 RF1
ZShift = 0.37
lm Zscore = −0.06
NG = 0
DB = 0
SM = 0
Delta_L
Doxorubicin
YDL227C_RF1_164 RF1
ZShift = 0.31
lm ZScore = −0.06
NG = 0
DB = 0
SM = 0
Delta_K
Doxorubicin
YDL227C_RF1_164 RF1
ZShift = −0.26
lm ZScore = 0.12
NG = 0
DB = 0
SM = 0
Delta_r
Doxorubicin
YDL227C_RF1_164 RF1
ZShift = 0.2
lm ZScore = 0.36
NG = 0
DB = 0
SM = 0
Delta_AUC
Doxorubicin
YDL227C_RF2_123 RF2
ZShift = −0.09
lm Zscore = −0.06
NG = 0
DB = 0
SM = 0
Delta_L
Doxorubicin
YDL227C_RF2_123 RF2
ZShift = 0.37
lm ZScore = −0.06
NG = 0
DB = 0
SM = 0
Delta_K
Doxorubicin
YDL227C_RF2_123 RF2
ZShift = −0.04
lm ZScore = −0.14
NG = 0
DB = 0
SM = 0
Delta_r
Doxorubicin
YDL227C_RF2_123 RF2
ZShift = 0.36
lm ZScore = −0.25
NG = 0
DB = 0
SM = 0
Delta_AUC
Doxorubicin
YDL227C_RF1_7 RF1
ZShift = −1.96
lm Zscore = −0.06
NG = 0
DB = 1
SM = 0
Delta_L
Doxorubicin
YDL227C_RF1_7 RF1
ZShift = −0.03
lm ZScore = −0.06
NG = 0
DB = 1
SM = 0
Delta_K
Doxorubicin
YDL227C_RF1_7 RF1
ZShift = −0.85
lm ZScore = 0.1
NG = 0
DB = 1
SM = 0
Delta_r
Doxorubicin
YDL227C_RF1_7 RF1
ZShift = 0.42
lm ZScore = 0.69
NG = 0
DB = 1
SM = 0
Delta_AUC
Doxorubicin

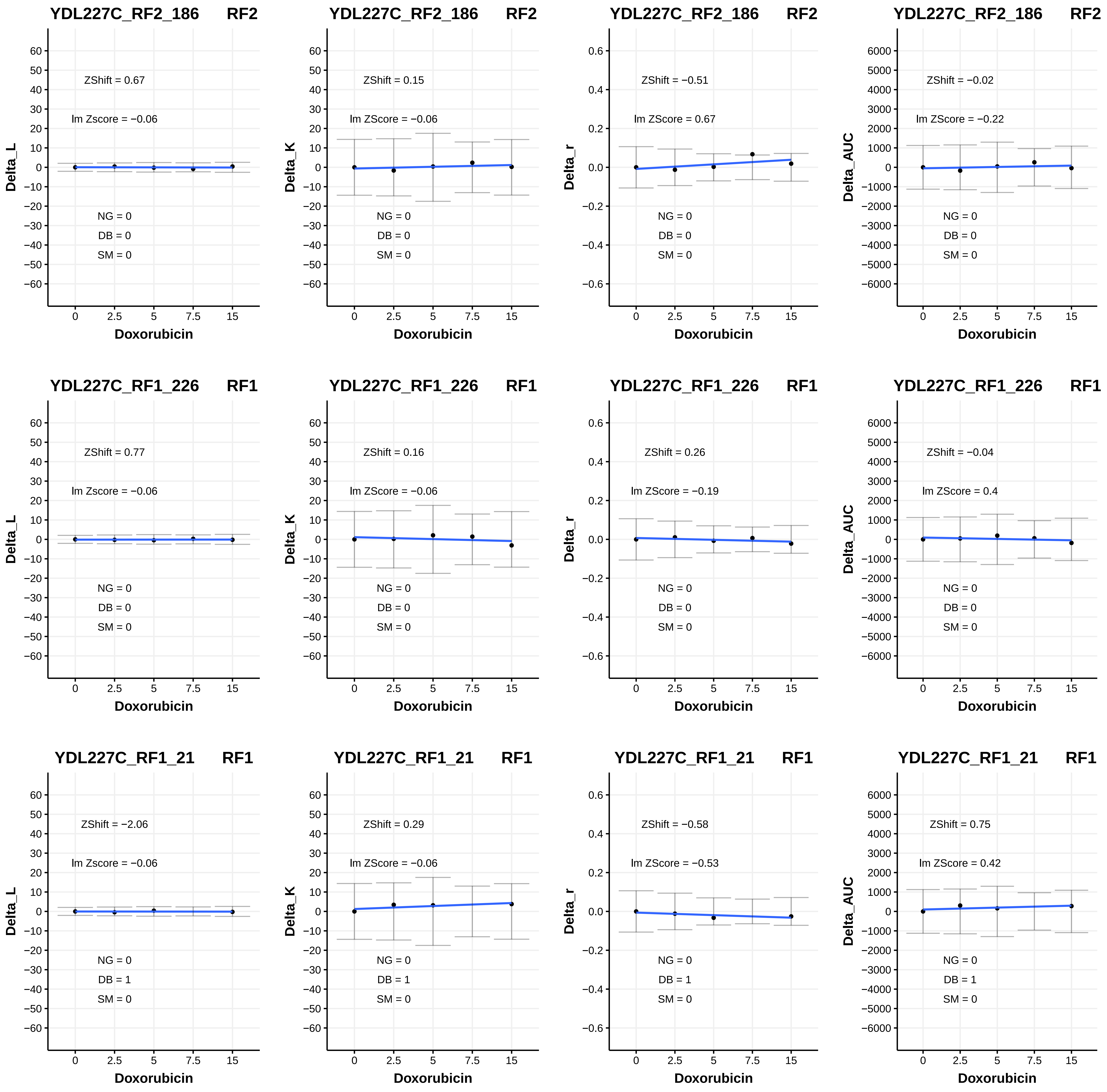
YDL227C_RF2_186 RF2
ZShift = 0.67
lm Zscore = −0.06
NG = 0
DB = 0
SM = 0
Delta_L
Doxorubicin
YDL227C_RF2_186 RF2
ZShift = 0.15
lm ZScore = −0.06
NG = 0
DB = 0
SM = 0
Delta_K
Doxorubicin
YDL227C_RF2_186 RF2
ZShift = −0.51
lm ZScore = 0.67
NG = 0
DB = 0
SM = 0
Delta_r
Doxorubicin
YDL227C_RF2_186 RF2
ZShift = −0.02
lm ZScore = −0.22
NG = 0
DB = 0
SM = 0
Delta_AUC
Doxorubicin
YDL227C_RF1_226 RF1
ZShift = 0.77
lm Zscore = −0.06
NG = 0
DB = 0
SM = 0
Delta_L
Doxorubicin
YDL227C_RF1_226 RF1
ZShift = 0.16
lm ZScore = −0.06
NG = 0
DB = 0
SM = 0
Delta_K
Doxorubicin
YDL227C_RF1_226 RF1
ZShift = 0.26
lm ZScore = −0.19
NG = 0
DB = 0
SM = 0
Delta_r
Doxorubicin
YDL227C_RF1_226 RF1
ZShift = −0.04
lm ZScore = 0.4
NG = 0
DB = 0
SM = 0
Delta_AUC
Doxorubicin
YDL227C_RF1_21 RF1
ZShift = −2.06
lm Zscore = −0.06
NG = 0
DB = 1
SM = 0
Delta_L
Doxorubicin
YDL227C_RF1_21 RF1
ZShift = 0.29
lm ZScore = −0.06
NG = 0
DB = 1
SM = 0
Delta_K
Doxorubicin
YDL227C_RF1_21 RF1
ZShift = −0.58
lm ZScore = −0.53
NG = 0
DB = 1
SM = 0
Delta_r
Doxorubicin
YDL227C_RF1_21 RF1
ZShift = 0.75
lm ZScore = 0.42
NG = 0
DB = 1
SM = 0
Delta_AUC
Doxorubicin

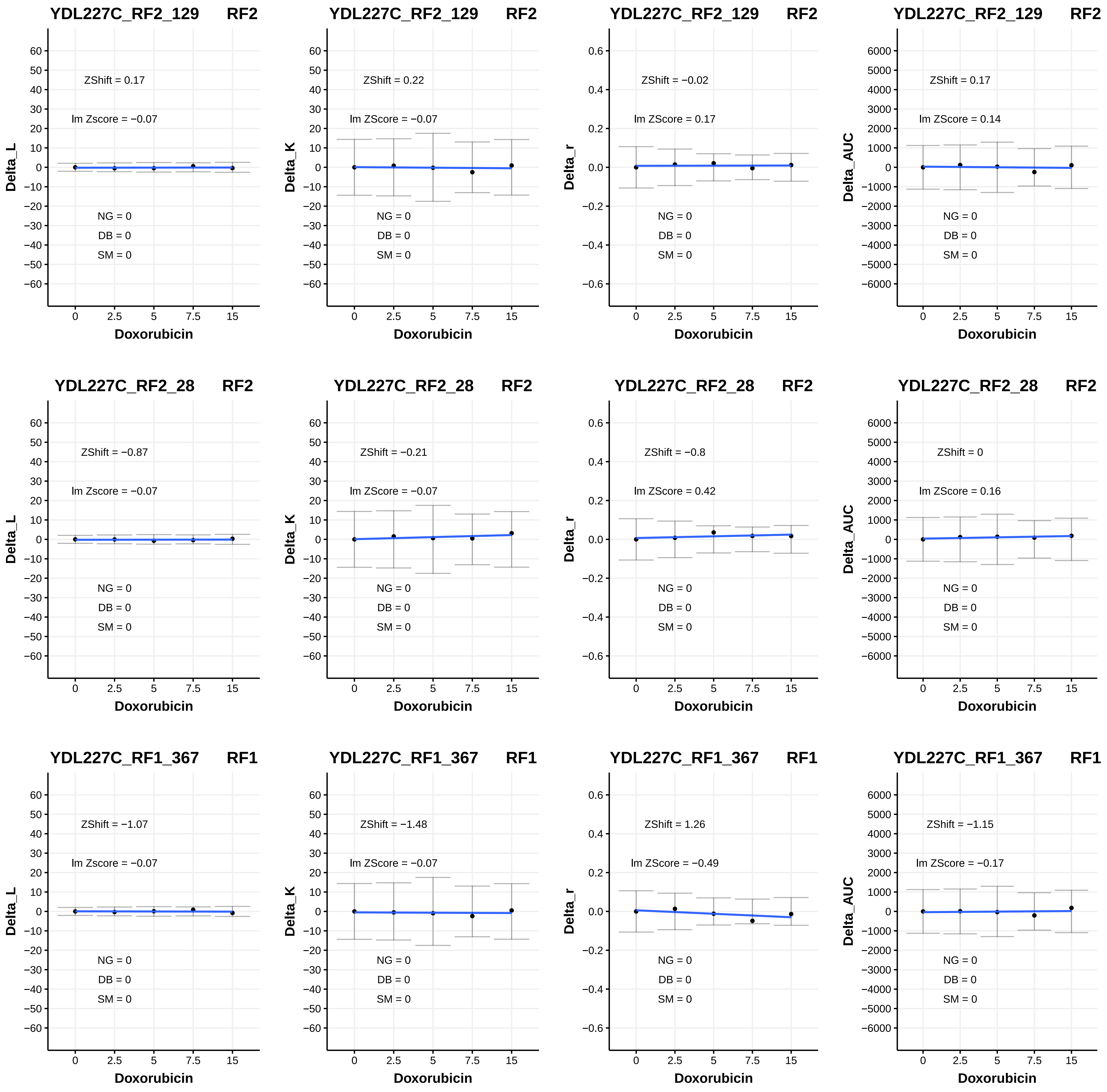
YDL227C_RF2_129 RF2
ZShift = 0.17
lm Zscore = −0.07
NG = 0
DB = 0
SM = 0
Delta_L
Doxorubicin
YDL227C_RF2_129 RF2
ZShift = 0.22
lm ZScore = −0.07
NG = 0
DB = 0
SM = 0
Delta_K
Doxorubicin
YDL227C_RF2_129 RF2
ZShift = −0.02
lm ZScore = 0.17
NG = 0
DB = 0
SM = 0
Delta_r
Doxorubicin
YDL227C_RF2_129 RF2
ZShift = 0.17
lm ZScore = 0.14
NG = 0
DB = 0
SM = 0
Delta_AUC
Doxorubicin
YDL227C_RF2_28 RF2
ZShift = −0.87
lm Zscore = −0.07
NG = 0
DB = 0
SM = 0
Delta_L
Doxorubicin
YDL227C_RF2_28 RF2
ZShift = −0.21
lm ZScore = −0.07
NG = 0
DB = 0
SM = 0
Delta_K
Doxorubicin
YDL227C_RF2_28 RF2
ZShift = −0.8
lm ZScore = 0.42
NG = 0
DB = 0
SM = 0
Delta_r
Doxorubicin
YDL227C_RF2_28 RF2
ZShift = 0
lm ZScore = 0.16
NG = 0
DB = 0
SM = 0
Delta_AUC
Doxorubicin
YDL227C_RF1_367 RF1
ZShift = −1.07
lm Zscore = −0.07
NG = 0
DB = 0
SM = 0
Delta_L
Doxorubicin
YDL227C_RF1_367 RF1
ZShift = −1.48
lm ZScore = −0.07
NG = 0
DB = 0
SM = 0
Delta_K
Doxorubicin
YDL227C_RF1_367 RF1
ZShift = 1.26
lm ZScore = −0.49
NG = 0
DB = 0
SM = 0
Delta_r
Doxorubicin
YDL227C_RF1_367 RF1
ZShift = −1.15
lm ZScore = −0.17
NG = 0
DB = 0
SM = 0
Delta_AUC
Doxorubicin

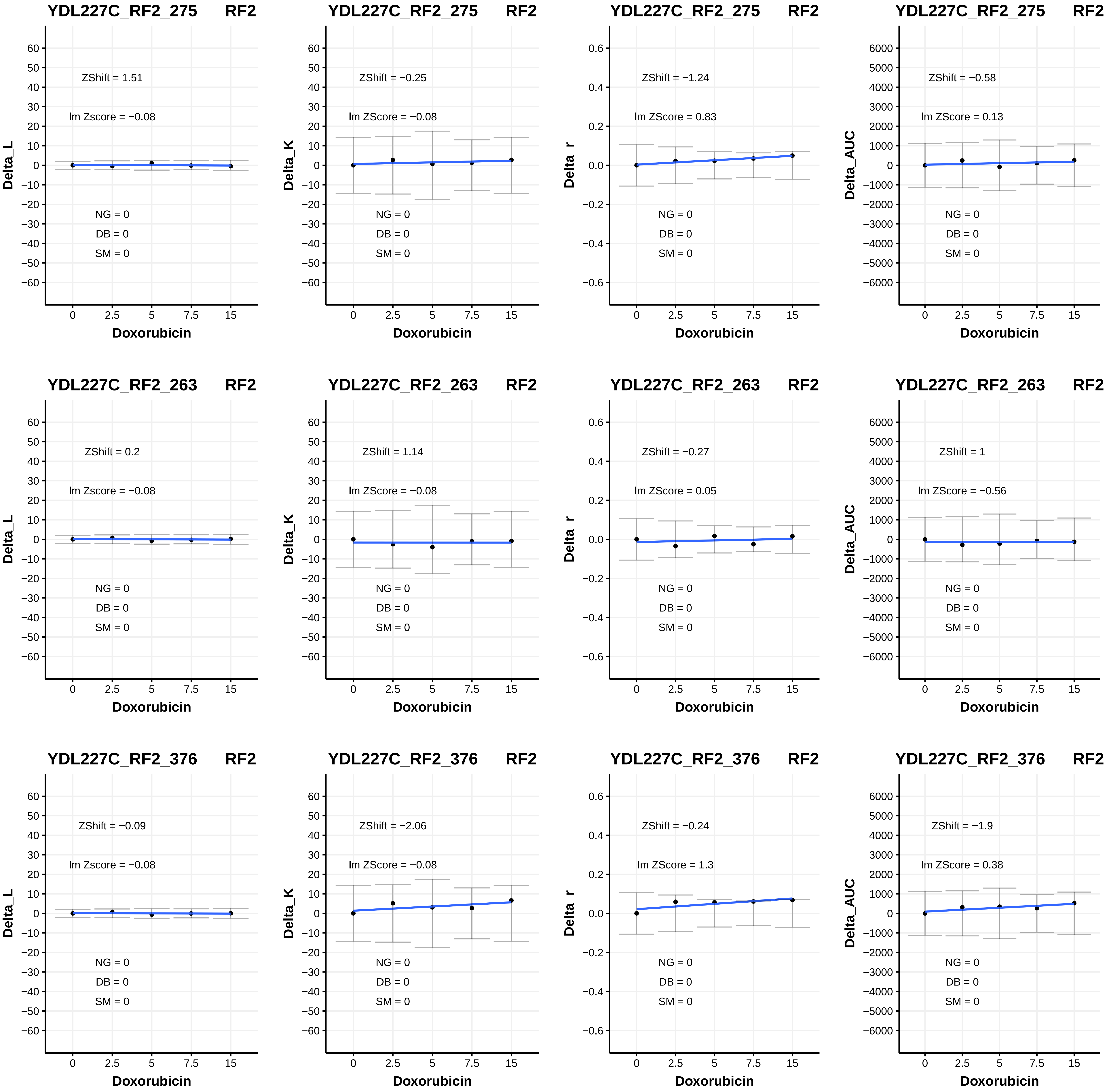
YDL227C_RF2_275 RF2
ZShift = 1.51
lm Zscore = −0.08
NG = 0
DB = 0
SM = 0
Delta_L
Doxorubicin
YDL227C_RF2_275 RF2
ZShift = −0.25
lm ZScore = −0.08
NG = 0
DB = 0
SM = 0
Delta_K
Doxorubicin
YDL227C_RF2_275 RF2
ZShift = −1.24
lm ZScore = 0.83
NG = 0
DB = 0
SM = 0
Delta_r
Doxorubicin
YDL227C_RF2_275 RF2
ZShift = −0.58
lm ZScore = 0.13
NG = 0
DB = 0
SM = 0
Delta_AUC
Doxorubicin
YDL227C_RF2_263 RF2
ZShift = 0.2
lm Zscore = −0.08
NG = 0
DB = 0
SM = 0
Delta_L
Doxorubicin
YDL227C_RF2_263 RF2
ZShift = 1.14
lm ZScore = −0.08
NG = 0
DB = 0
SM = 0
Delta_K
Doxorubicin
YDL227C_RF2_263 RF2
ZShift = −0.27
lm ZScore = 0.05
NG = 0
DB = 0
SM = 0
Delta_r
Doxorubicin
YDL227C_RF2_263 RF2
ZShift = 1
lm ZScore = −0.56
NG = 0
DB = 0
SM = 0
Delta_AUC
Doxorubicin
YDL227C_RF2_376 RF2
ZShift = −0.09
lm Zscore = −0.08
NG = 0
DB = 0
SM = 0
Delta_L
Doxorubicin
YDL227C_RF2_376 RF2
ZShift = −2.06
lm ZScore = −0.08
NG = 0
DB = 0
SM = 0
Delta_K
Doxorubicin
YDL227C_RF2_376 RF2
ZShift = −0.24
lm ZScore = 1.3
NG = 0
DB = 0
SM = 0
Delta_r
Doxorubicin
YDL227C_RF2_376 RF2
ZShift = −1.9
lm ZScore = 0.38
NG = 0
DB = 0
SM = 0
Delta_AUC
Doxorubicin

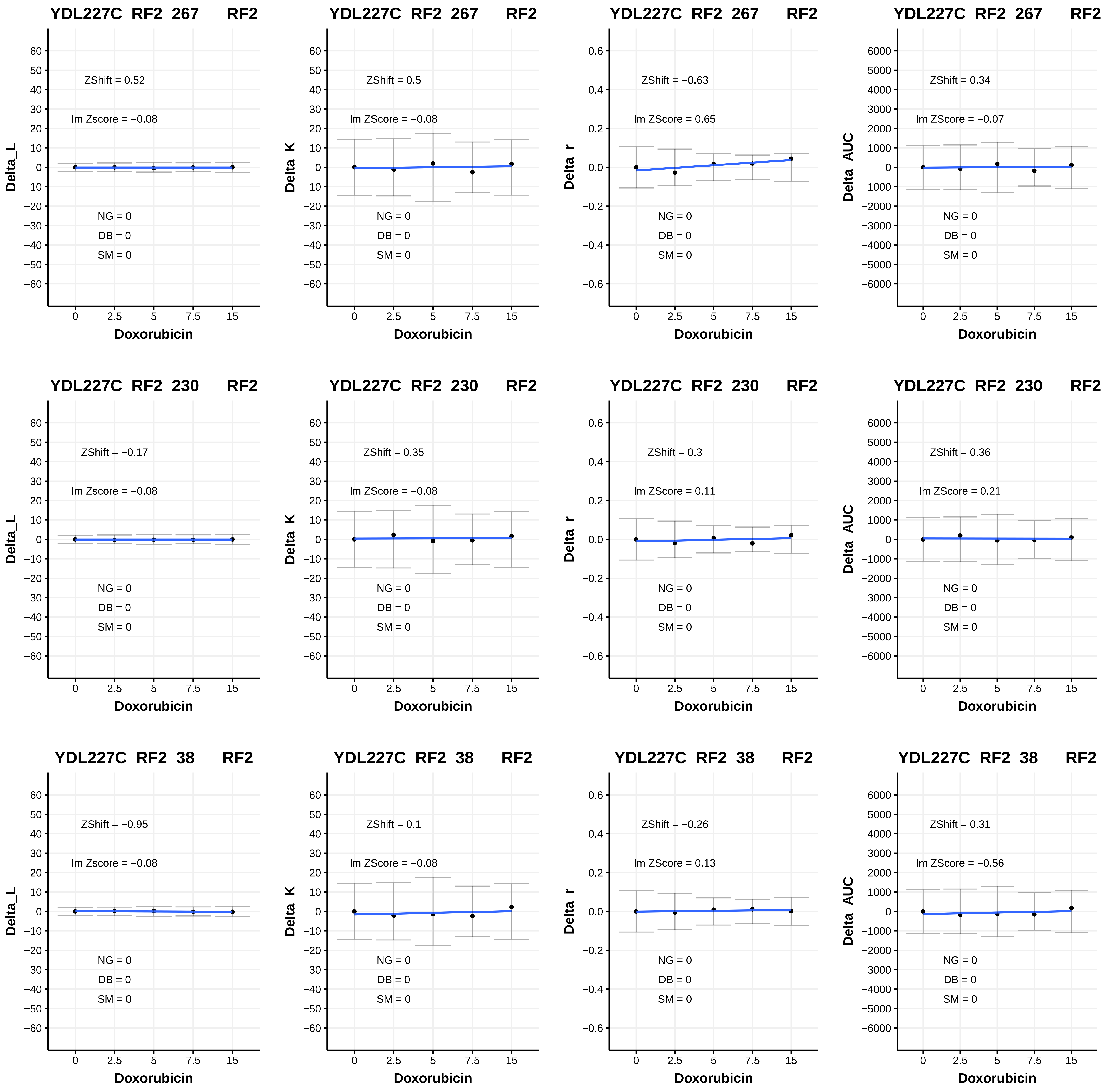
YDL227C_RF2_267 RF2
ZShift = 0.52
lm Zscore = −0.08
NG = 0
DB = 0
SM = 0
Delta_L
Doxorubicin
YDL227C_RF2_267 RF2
ZShift = 0.5
lm ZScore = −0.08
NG = 0
DB = 0
SM = 0
Delta_K
Doxorubicin
YDL227C_RF2_267 RF2
ZShift = −0.63
lm ZScore = 0.65
NG = 0
DB = 0
SM = 0
Delta_r
Doxorubicin
YDL227C_RF2_267 RF2
ZShift = 0.34
lm ZScore = −0.07
NG = 0
DB = 0
SM = 0
Delta_AUC
Doxorubicin
YDL227C_RF2_230 RF2
ZShift = −0.17
lm Zscore = −0.08
NG = 0
DB = 0
SM = 0
Delta_L
Doxorubicin
YDL227C_RF2_230 RF2
ZShift = 0.35
lm ZScore = −0.08
NG = 0
DB = 0
SM = 0
Delta_K
Doxorubicin
YDL227C_RF2_230 RF2
ZShift = 0.3
lm ZScore = 0.11
NG = 0
DB = 0
SM = 0
Delta_r
Doxorubicin
YDL227C_RF2_230 RF2
ZShift = 0.36
lm ZScore = 0.21
NG = 0
DB = 0
SM = 0
Delta_AUC
Doxorubicin
YDL227C_RF2_38 RF2
ZShift = −0.95
lm Zscore = −0.08
NG = 0
DB = 0
SM = 0
Delta_L
Doxorubicin
YDL227C_RF2_38 RF2
ZShift = 0.1
lm ZScore = −0.08
NG = 0
DB = 0
SM = 0
Delta_K
Doxorubicin
YDL227C_RF2_38 RF2
ZShift = −0.26
lm ZScore = 0.13
NG = 0
DB = 0
SM = 0
Delta_r
Doxorubicin
YDL227C_RF2_38 RF2
ZShift = 0.31
lm ZScore = −0.56
NG = 0
DB = 0
SM = 0
Delta_AUC
Doxorubicin

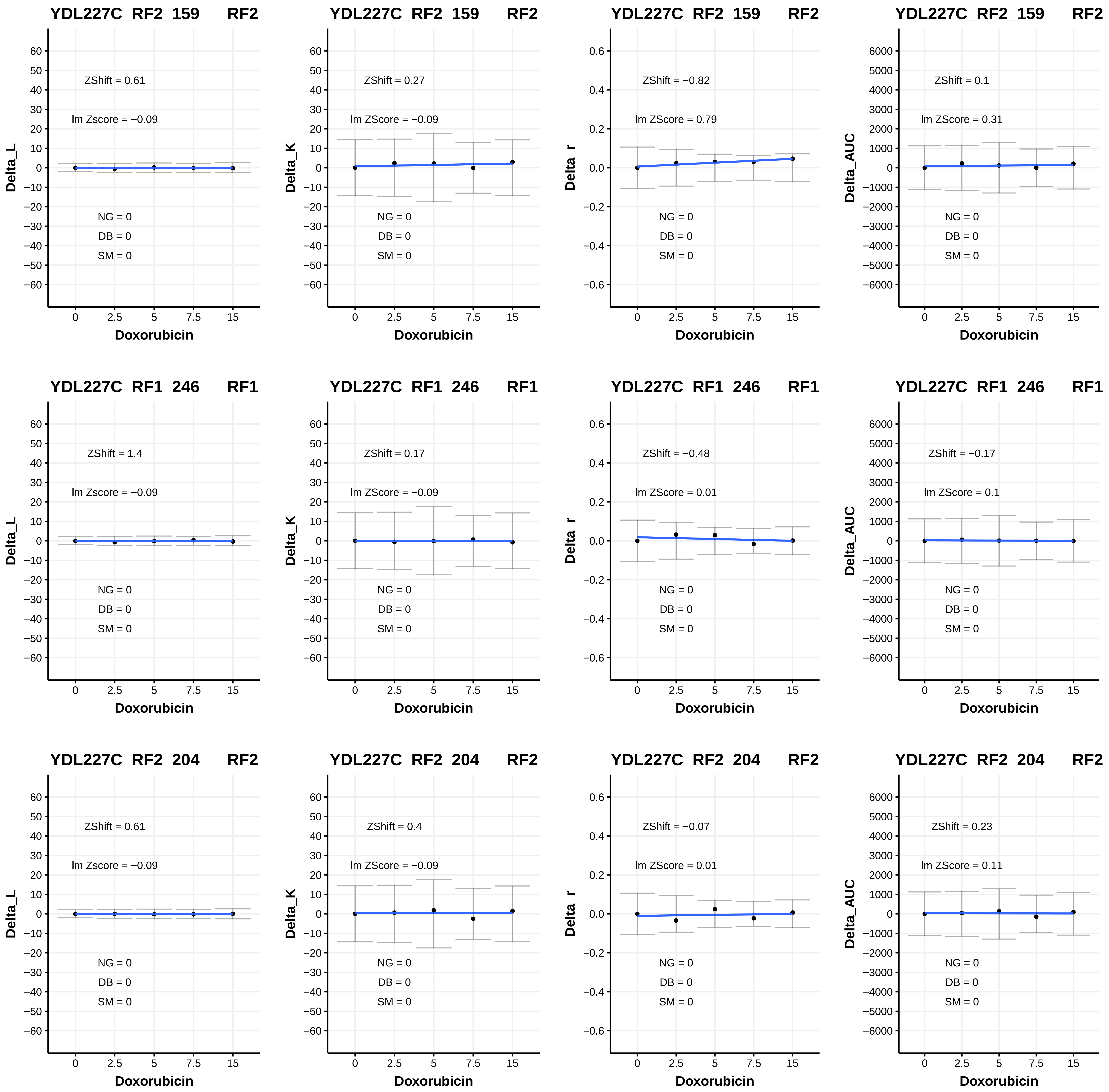
YDL227C_RF2_159 RF2
ZShift = 0.61
lm Zscore = −0.09
NG = 0
DB = 0
SM = 0
Delta_L
Doxorubicin
YDL227C_RF2_159 RF2
ZShift = 0.27
lm ZScore = −0.09
NG = 0
DB = 0
SM = 0
Delta_K
Doxorubicin
YDL227C_RF2_159 RF2
ZShift = −0.82
lm ZScore = 0.79
NG = 0
DB = 0
SM = 0
Delta_r
Doxorubicin
YDL227C_RF2_159 RF2
ZShift = 0.1
lm ZScore = 0.31
NG = 0
DB = 0
SM = 0
Delta_AUC
Doxorubicin
YDL227C_RF1_246 RF1
ZShift = 1.4
lm Zscore = −0.09
NG = 0
DB = 0
SM = 0
Delta_L
Doxorubicin
YDL227C_RF1_246 RF1
ZShift = 0.17
lm ZScore = −0.09
NG = 0
DB = 0
SM = 0
Delta_K
Doxorubicin
YDL227C_RF1_246 RF1
ZShift = −0.48
lm ZScore = 0.01
NG = 0
DB = 0
SM = 0
Delta_r
Doxorubicin
YDL227C_RF1_246 RF1
ZShift = −0.17
lm ZScore = 0.1
NG = 0
DB = 0
SM = 0
Delta_AUC
Doxorubicin
YDL227C_RF2_204 RF2
ZShift = 0.61
lm Zscore = −0.09
NG = 0
DB = 0
SM = 0
Delta_L
Doxorubicin
YDL227C_RF2_204 RF2
ZShift = 0.4
lm ZScore = −0.09
NG = 0
DB = 0
SM = 0
Delta_K
Doxorubicin
YDL227C_RF2_204 RF2
ZShift = −0.07
lm ZScore = 0.01
NG = 0
DB = 0
SM = 0
Delta_r
Doxorubicin
YDL227C_RF2_204 RF2
ZShift = 0.23
lm ZScore = 0.11
NG = 0
DB = 0
SM = 0
Delta_AUC
Doxorubicin

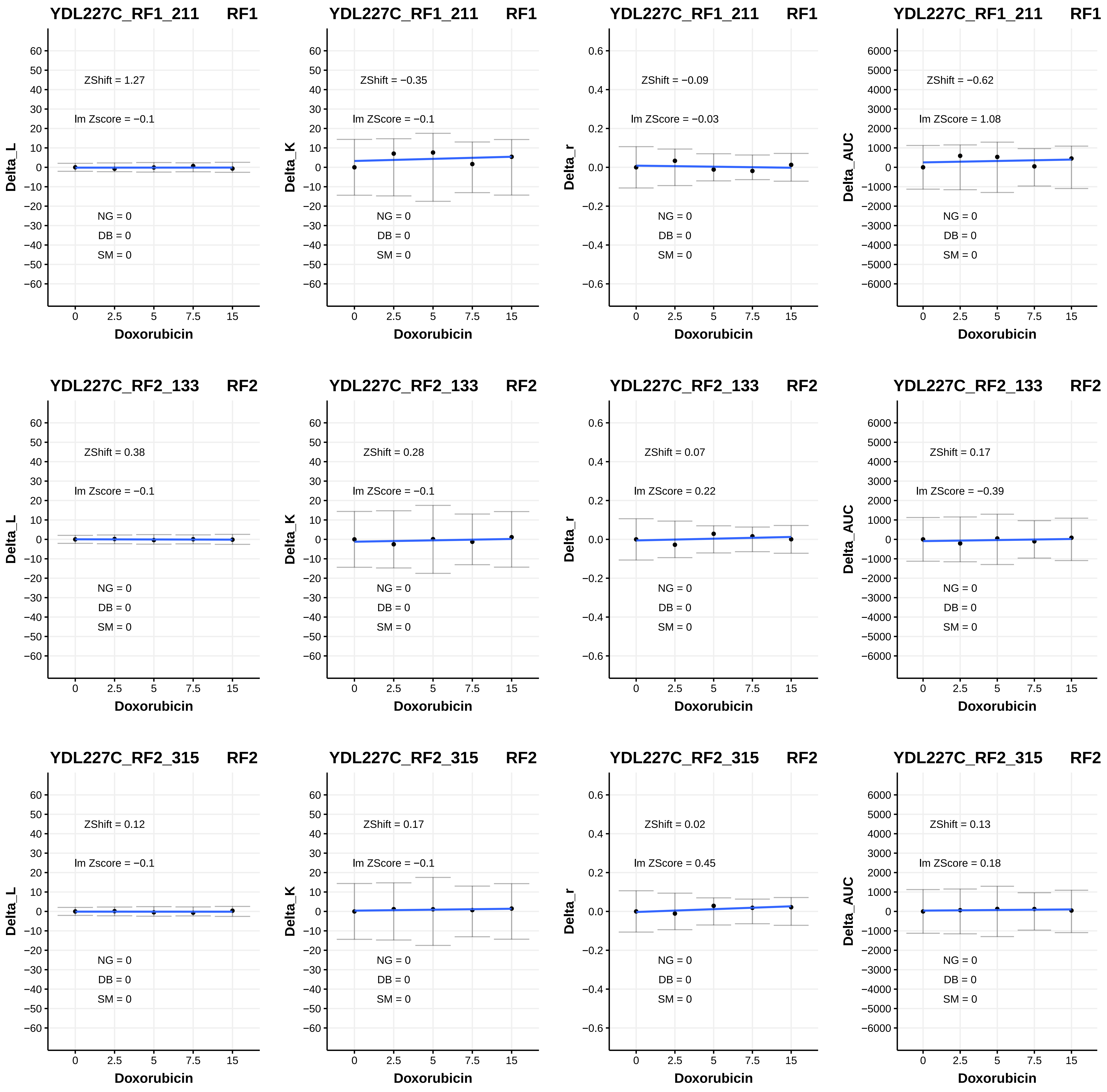
YDL227C_RF1_211 RF1
ZShift = 1.27
lm Zscore = −0.1
NG = 0
DB = 0
SM = 0
Delta_L
Doxorubicin
YDL227C_RF1_211 RF1
ZShift = −0.35
lm ZScore = −0.1
NG = 0
DB = 0
SM = 0
Delta_K
Doxorubicin
YDL227C_RF1_211 RF1
ZShift = −0.09
lm ZScore = −0.03
NG = 0
DB = 0
SM = 0
Delta_r
Doxorubicin
YDL227C_RF1_211 RF1
ZShift = −0.62
lm ZScore = 1.08
NG = 0
DB = 0
SM = 0
Delta_AUC
Doxorubicin
YDL227C_RF2_133 RF2
ZShift = 0.38
lm Zscore = −0.1
NG = 0
DB = 0
SM = 0
Delta_L
Doxorubicin
YDL227C_RF2_133 RF2
ZShift = 0.28
lm ZScore = −0.1
NG = 0
DB = 0
SM = 0
Delta_K
Doxorubicin
YDL227C_RF2_133 RF2
ZShift = 0.07
lm ZScore = 0.22
NG = 0
DB = 0
SM = 0
Delta_r
Doxorubicin
YDL227C_RF2_133 RF2
ZShift = 0.17
lm ZScore = −0.39
NG = 0
DB = 0
SM = 0
Delta_AUC
Doxorubicin
YDL227C_RF2_315 RF2
ZShift = 0.12
lm Zscore = −0.1
NG = 0
DB = 0
SM = 0
Delta_L
Doxorubicin
YDL227C_RF2_315 RF2
ZShift = 0.17
lm ZScore = −0.1
NG = 0
DB = 0
SM = 0
Delta_K
Doxorubicin
YDL227C_RF2_315 RF2
ZShift = 0.02
lm ZScore = 0.45
NG = 0
DB = 0
SM = 0
Delta_r
Doxorubicin
YDL227C_RF2_315 RF2
ZShift = 0.13
lm ZScore = 0.18
NG = 0
DB = 0
SM = 0
Delta_AUC
Doxorubicin

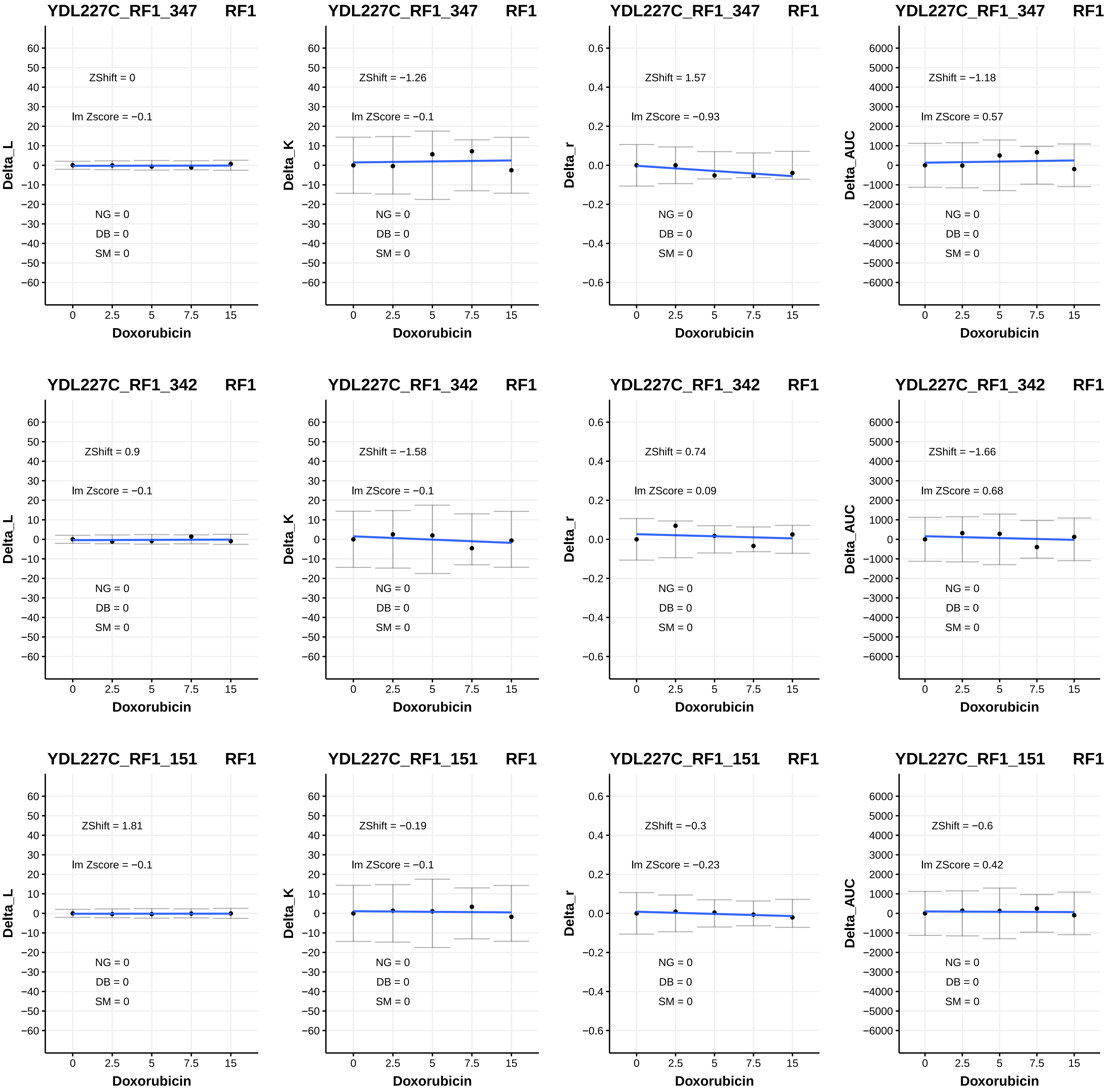

YDL227C_RF1_347 RF1
Delta_L
ZShift = 0
lm Zscore = −0.1
NG = 0
DB = 0
SM = 0
Doxorubicin
YDL227C_RF1_347 RF1
Delta_K
ZShift = −1.26
lm ZScore = −0.1
NG = 0
DB = 0
SM = 0
Doxorubicin
YDL227C_RF1_347 RF1
Delta_r
ZShift = 1.57
lm ZScore = −0.93
NG = 0
DB = 0
SM = 0
Doxorubicin
YDL227C_RF1_347 RF1
Delta_AUC
ZShift = −1.18
lm ZScore = 0.57
NG = 0
DB = 0
SM = 0
Doxorubicin
YDL227C_RF1_342 RF1
Delta_L
ZShift = 0.9
lm Zscore = −0.1
NG = 0
DB = 0
SM = 0
Doxorubicin
YDL227C_RF1_342 RF1
Delta_K
ZShift = −1.58
lm ZScore = −0.1
NG = 0
DB = 0
SM = 0
Doxorubicin
YDL227C_RF1_342 RF1
Delta_r
ZShift = 0.74
lm ZScore = 0.09
NG = 0
DB = 0
SM = 0
Doxorubicin
YDL227C_RF1_342 RF1
Delta_AUC
ZShift = −1.66
lm ZScore = 0.68
NG = 0
DB = 0
SM = 0
Doxorubicin
YDL227C_RF1_151 RF1
Delta_L
ZShift = 1.81
lm Zscore = −0.1
NG = 0
DB = 0
SM = 0
Doxorubicin
YDL227C_RF1_151 RF1
Delta_K
ZShift = −0.19
lm ZScore = −0.1
NG = 0
DB = 0
SM = 0
Doxorubicin
YDL227C_RF1_151 RF1
Delta_r
ZShift = −0.3
lm ZScore = −0.23
NG = 0
DB = 0
SM = 0
Doxorubicin
YDL227C_RF1_151 RF1
Delta_AUC
ZShift = −0.6
lm ZScore = 0.42
NG = 0
DB = 0
SM = 0
Doxorubicin

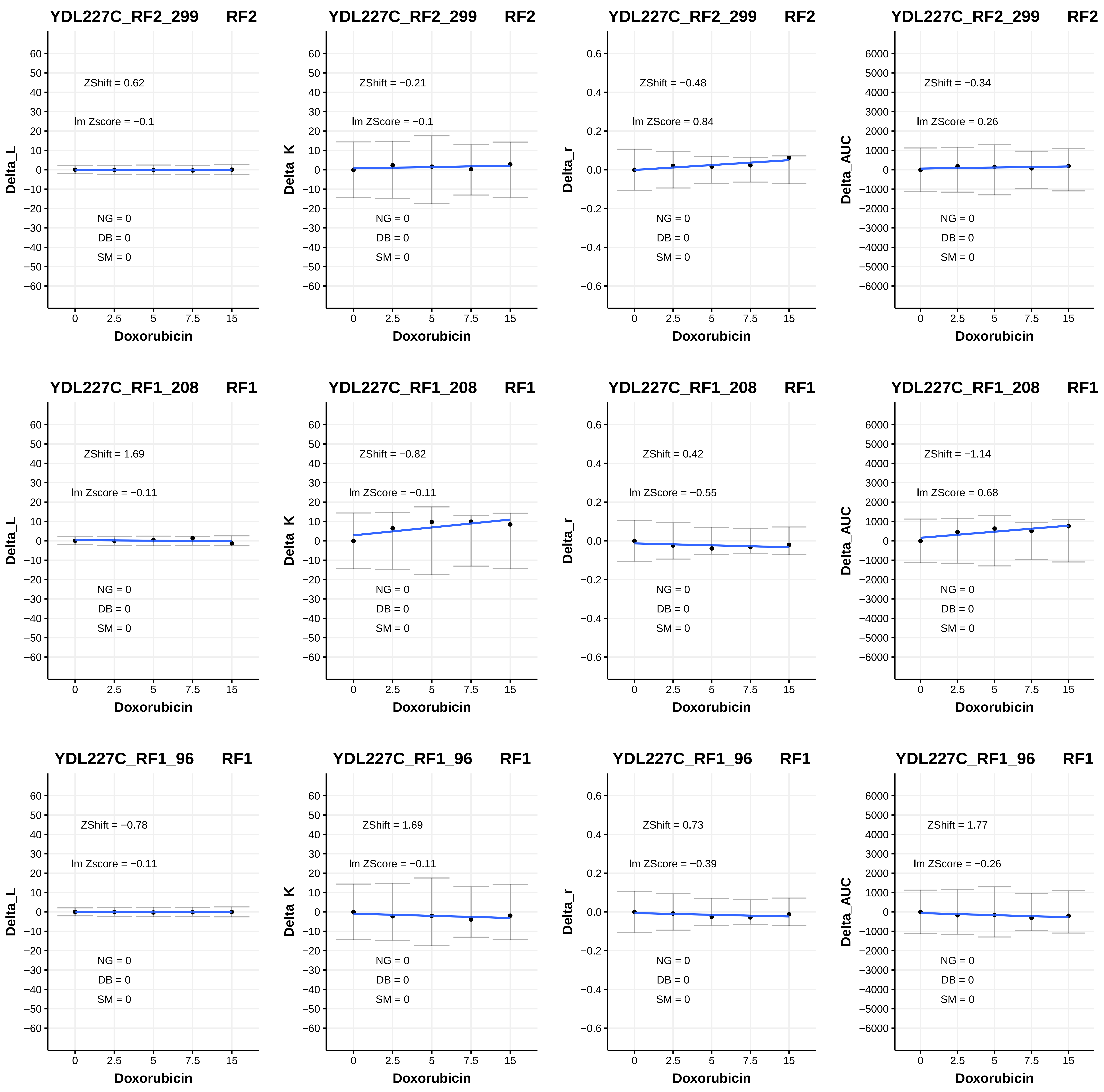
YDL227C_RF2_299 RF2
ZShift = 0.62
lm Zscore = −0.1
Delta_L
NG = 0
DB = 0
SM = 0
Doxorubicin
YDL227C_RF2_299 RF2
ZShift = −0.21
lm ZScore = −0.1
Delta_K
NG = 0
DB = 0
SM = 0
Doxorubicin
YDL227C_RF2_299 RF2
ZShift = −0.48
lm ZScore = 0.84
Delta_r
NG = 0
DB = 0
SM = 0
Doxorubicin
YDL227C_RF2_299 RF2
ZShift = −0.34
lm ZScore = 0.26
Delta_AUC
NG = 0
DB = 0
SM = 0
Doxorubicin
YDL227C_RF1_208 RF1
ZShift = 1.69
lm Zscore = −0.11
Delta_L
NG = 0
DB = 0
SM = 0
Doxorubicin
YDL227C_RF1_208 RF1
ZShift = −0.82
lm ZScore = −0.11
Delta_K
NG = 0
DB = 0
SM = 0
Doxorubicin
YDL227C_RF1_208 RF1
ZShift = 0.42
lm ZScore = −0.55
Delta_r
NG = 0
DB = 0
SM = 0
Doxorubicin
YDL227C_RF1_208 RF1
ZShift = −1.14
lm ZScore = 0.68
Delta_AUC
NG = 0
DB = 0
SM = 0
Doxorubicin
YDL227C_RF1_96 RF1
ZShift = −0.78
lm Zscore = −0.11
Delta_L
NG = 0
DB = 0
SM = 0
Doxorubicin
YDL227C_RF1_96 RF1
ZShift = 1.69
lm ZScore = −0.11
Delta_K
NG = 0
DB = 0
SM = 0
Doxorubicin
YDL227C_RF1_96 RF1
ZShift = 0.73
lm ZScore = −0.39
Delta_r
NG = 0
DB = 0
SM = 0
Doxorubicin
YDL227C_RF1_96 RF1
ZShift = 1.77
lm ZScore = −0.26
Delta_AUC
NG = 0
DB = 0
SM = 0
Doxorubicin

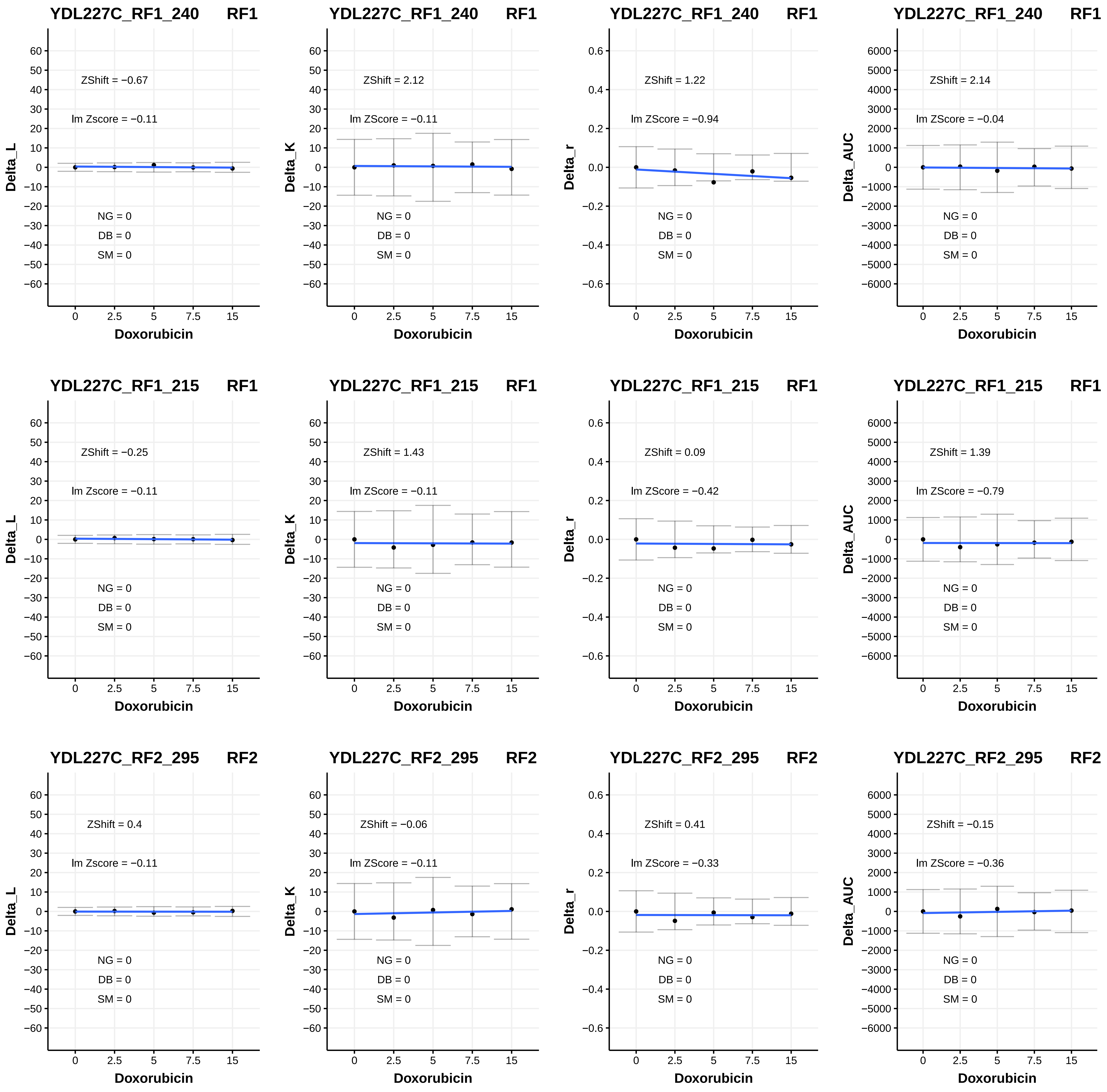
YDL227C_RF1_240 RF1
ZShift = −0.67
lm Zscore = −0.11
NG = 0
DB = 0
SM = 0
Delta_L
Doxorubicin
YDL227C_RF1_240 RF1
ZShift = 2.12
lm ZScore = −0.11
NG = 0
DB = 0
SM = 0
Delta_K
Doxorubicin
YDL227C_RF1_240 RF1
ZShift = 1.22
lm ZScore = −0.94
NG = 0
DB = 0
SM = 0
Delta_r
Doxorubicin
YDL227C_RF1_240 RF1
ZShift = 2.14
lm ZScore = −0.04
NG = 0
DB = 0
SM = 0
Delta_AUC
Doxorubicin
YDL227C_RF1_215 RF1
ZShift = −0.25
lm Zscore = −0.11
NG = 0
DB = 0
SM = 0
Delta_L
Doxorubicin
YDL227C_RF1_215 RF1
ZShift = 1.43
lm ZScore = −0.11
NG = 0
DB = 0
SM = 0
Delta_K
Doxorubicin
YDL227C_RF1_215 RF1
ZShift = 0.09
lm ZScore = −0.42
NG = 0
DB = 0
SM = 0
Delta_r
Doxorubicin
YDL227C_RF1_215 RF1
ZShift = 1.39
lm ZScore = −0.79
NG = 0
DB = 0
SM = 0
Delta_AUC
Doxorubicin
YDL227C_RF2_295 RF2
ZShift = 0.4
lm Zscore = −0.11
NG = 0
DB = 0
SM = 0
Delta_L
Doxorubicin
YDL227C_RF2_295 RF2
ZShift = −0.06
lm ZScore = −0.11
NG = 0
DB = 0
SM = 0
Delta_K
Doxorubicin
YDL227C_RF2_295 RF2
ZShift = 0.41
lm ZScore = −0.33
NG = 0
DB = 0
SM = 0
Delta_r
Doxorubicin
YDL227C_RF2_295 RF2
ZShift = −0.15
lm ZScore = −0.36
NG = 0
DB = 0
SM = 0
Delta_AUC
Doxorubicin

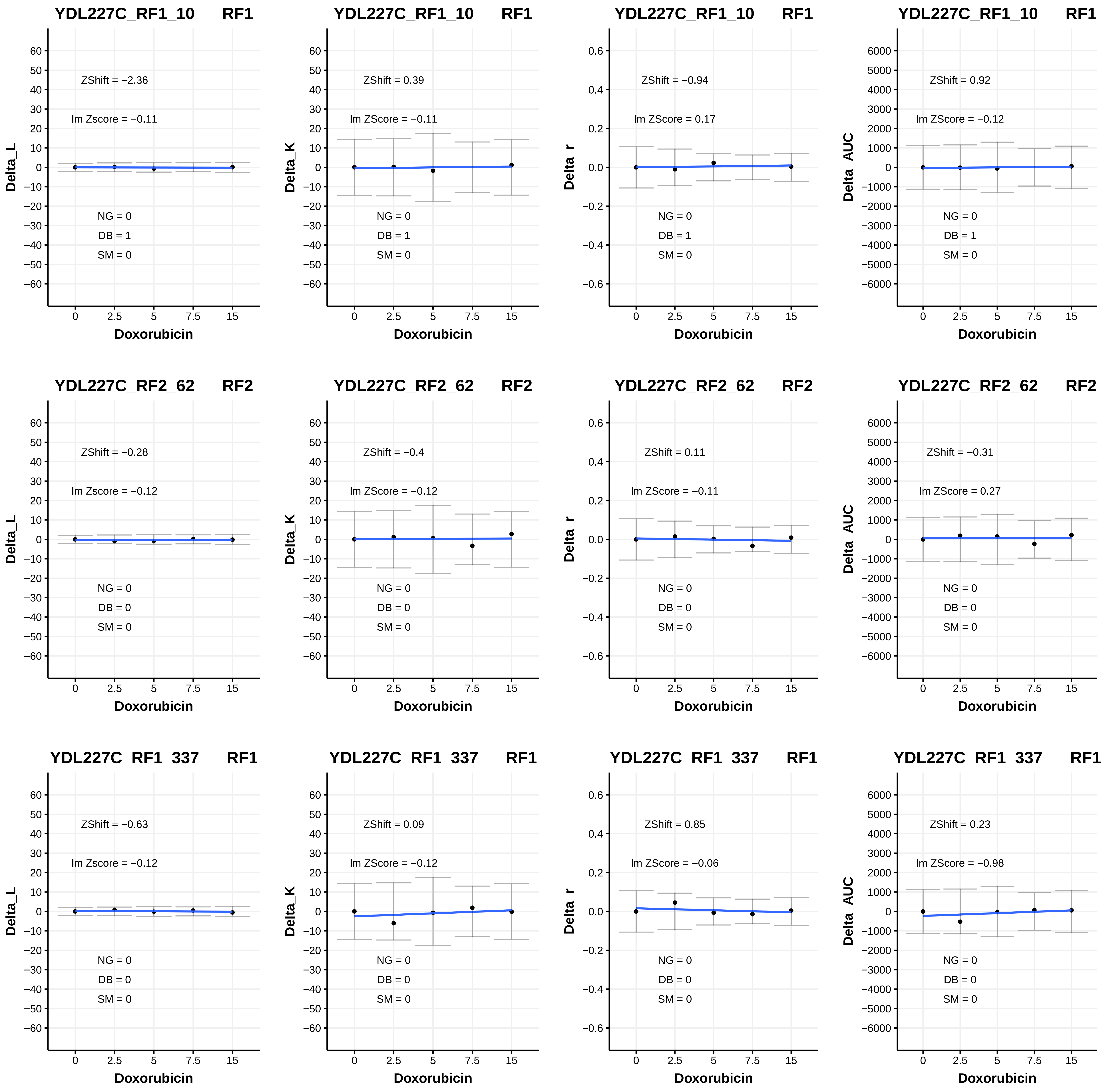

YDL227C_RF1_10 RF1
Delta_L
ZShift = −2.36
lm Zscore = −0.11
NG = 0
DB = 1
SM = 0
Doxorubicin
YDL227C_RF1_10 RF1
Delta_K
ZShift = 0.39
lm ZScore = −0.11
NG = 0
DB = 1
SM = 0
Doxorubicin
YDL227C_RF1_10 RF1
Delta_r
ZShift = −0.94
lm ZScore = 0.17
NG = 0
DB = 1
SM = 0
Doxorubicin
YDL227C_RF1_10 RF1
Delta_AUC
ZShift = 0.92
lm ZScore = −0.12
NG = 0
DB = 1
SM = 0
Doxorubicin
YDL227C_RF2_62 RF2
Delta_L
ZShift = −0.28
lm Zscore = −0.12
NG = 0
DB = 0
SM = 0
Doxorubicin
YDL227C_RF2_62 RF2
Delta_K
ZShift = −0.4
lm ZScore = −0.12
NG = 0
DB = 0
SM = 0
Doxorubicin
YDL227C_RF2_62 RF2
Delta_r
ZShift = 0.11
lm ZScore = −0.11
NG = 0
DB = 0
SM = 0
Doxorubicin
YDL227C_RF2_62 RF2
Delta_AUC
ZShift = −0.31
lm ZScore = 0.27
NG = 0
DB = 0
SM = 0
Doxorubicin
YDL227C_RF1_337 RF1
Delta_L
ZShift = −0.63
lm Zscore = −0.12
NG = 0
DB = 0
SM = 0
Doxorubicin
YDL227C_RF1_337 RF1
Delta_K
ZShift = 0.09
lm ZScore = −0.12
NG = 0
DB = 0
SM = 0
Doxorubicin
YDL227C_RF1_337 RF1
Delta_r
ZShift = 0.85
lm ZScore = −0.06
NG = 0
DB = 0
SM = 0
Doxorubicin
YDL227C_RF1_337 RF1
Delta_AUC
ZShift = 0.23
lm ZScore = −0.98
NG = 0
DB = 0
SM = 0
Doxorubicin

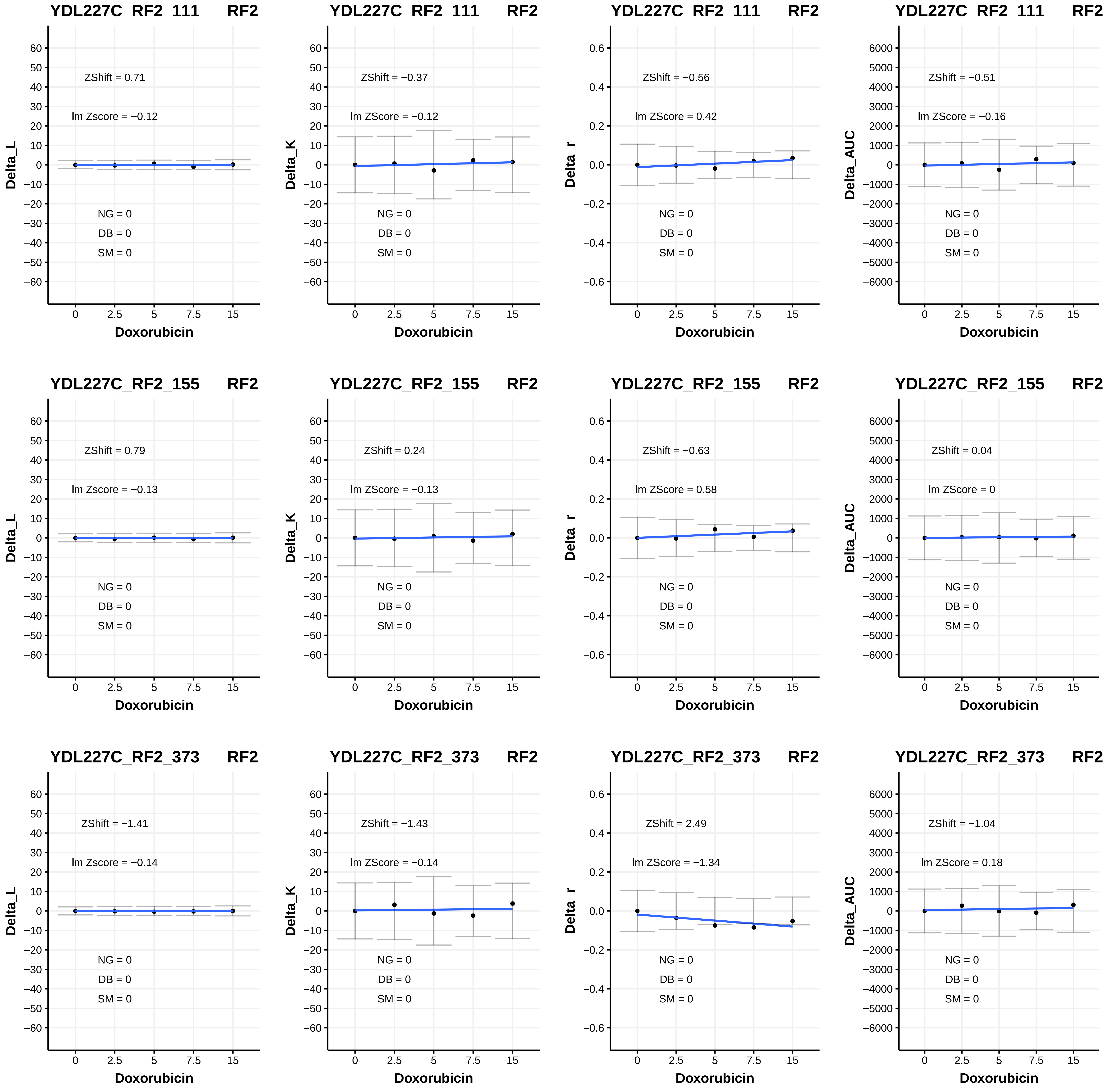
YDL227C_RF2_111 RF2
Delta_L
ZShift = 0.71
lm Zscore = −0.12
NG = 0
DB = 0
SM = 0
Doxorubicin
YDL227C_RF2_111 RF2
Delta_K
ZShift = −0.37
lm ZScore = −0.12
NG = 0
DB = 0
SM = 0
Doxorubicin
YDL227C_RF2_111 RF2
Delta_r
ZShift = −0.56
lm ZScore = 0.42
NG = 0
DB = 0
SM = 0
Doxorubicin
YDL227C_RF2_111 RF2
Delta_AUC
ZShift = −0.51
lm ZScore = −0.16
NG = 0
DB = 0
SM = 0
Doxorubicin
YDL227C_RF2_155 RF2
Delta_L
ZShift = 0.79
lm Zscore = −0.13
NG = 0
DB = 0
SM = 0
Doxorubicin
YDL227C_RF2_155 RF2
Delta_K
ZShift = 0.24
lm ZScore = −0.13
NG = 0
DB = 0
SM = 0
Doxorubicin
YDL227C_RF2_155 RF2
Delta_r
ZShift = −0.63
lm ZScore = 0.58
NG = 0
DB = 0
SM = 0
Doxorubicin
YDL227C_RF2_155 RF2
Delta_AUC
ZShift = 0.04
lm ZScore = 0
NG = 0
DB = 0
SM = 0
Doxorubicin
YDL227C_RF2_373 RF2
Delta_L
ZShift = −1.41
lm Zscore = −0.14
NG = 0
DB = 0
SM = 0
Doxorubicin
YDL227C_RF2_373 RF2
Delta_K
ZShift = −1.43
lm ZScore = −0.14
NG = 0
DB = 0
SM = 0
Doxorubicin
YDL227C_RF2_373 RF2
Delta_r
ZShift = 2.49
lm ZScore = −1.34
NG = 0
DB = 0
SM = 0
Doxorubicin
YDL227C_RF2_373 RF2
Delta_AUC
ZShift = −1.04
lm ZScore = 0.18
NG = 0
DB = 0
SM = 0
Doxorubicin

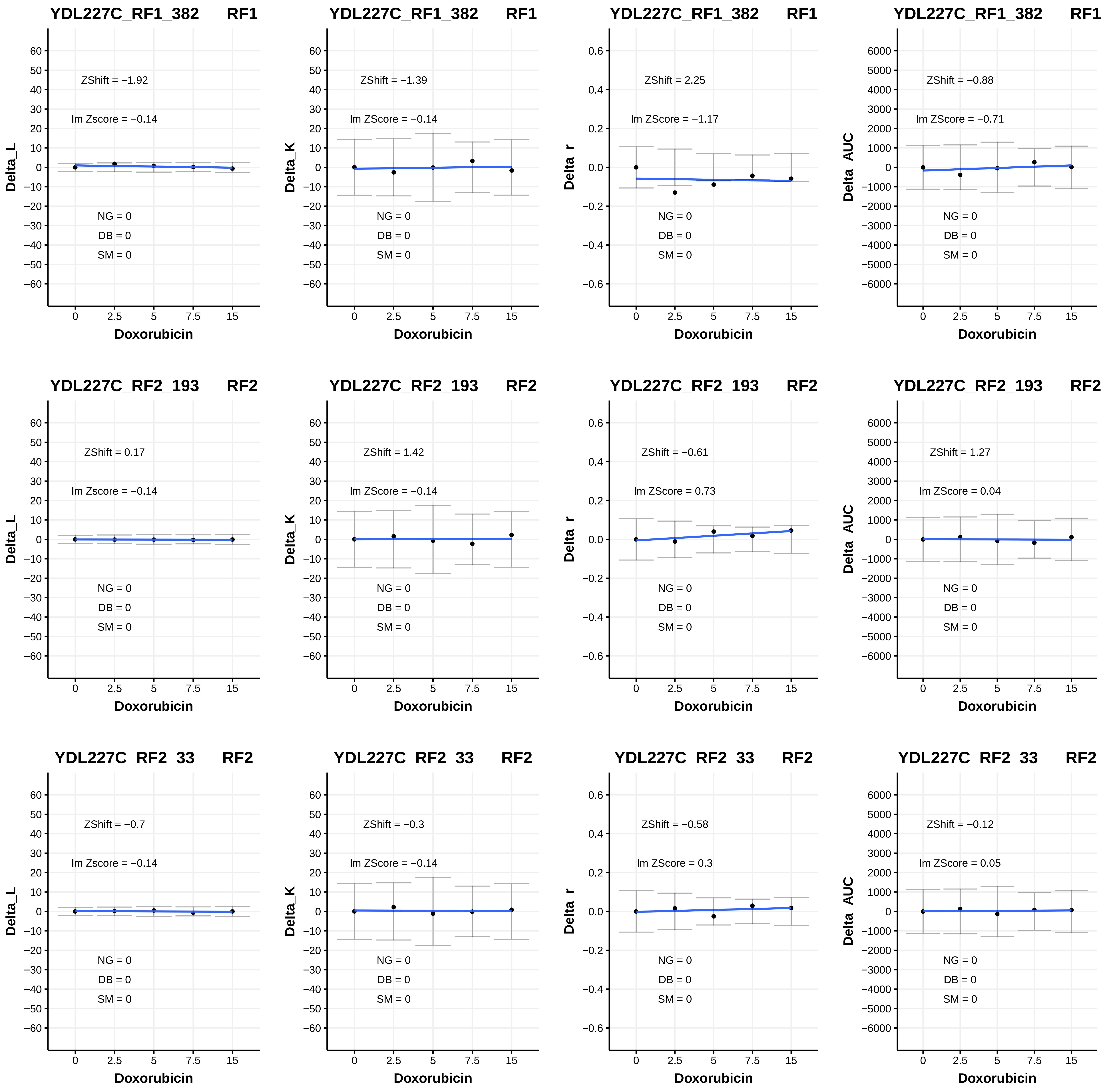
YDL227C_RF1_382 RF1
ZShift = −1.92
lm Zscore = −0.14
NG = 0
DB = 0
SM = 0
Delta_L
Doxorubicin
YDL227C_RF1_382 RF1
ZShift = −1.39
lm ZScore = −0.14
NG = 0
DB = 0
SM = 0
Delta_K
Doxorubicin
YDL227C_RF1_382 RF1
ZShift = 2.25
lm ZScore = −1.17
NG = 0
DB = 0
SM = 0
Delta_r
Doxorubicin
YDL227C_RF1_382 RF1
ZShift = −0.88
lm ZScore = −0.71
NG = 0
DB = 0
SM = 0
Delta_AUC
Doxorubicin
YDL227C_RF2_193 RF2
ZShift = 0.17
lm Zscore = −0.14
NG = 0
DB = 0
SM = 0
Delta_L
Doxorubicin
YDL227C_RF2_193 RF2
ZShift = 1.42
lm ZScore = −0.14
NG = 0
DB = 0
SM = 0
Delta_K
Doxorubicin
YDL227C_RF2_193 RF2
ZShift = −0.61
lm ZScore = 0.73
NG = 0
DB = 0
SM = 0
Delta_r
Doxorubicin
YDL227C_RF2_193 RF2
ZShift = 1.27
lm ZScore = 0.04
NG = 0
DB = 0
SM = 0
Delta_AUC
Doxorubicin
YDL227C_RF2_33 RF2
ZShift = −0.7
lm Zscore = −0.14
NG = 0
DB = 0
SM = 0
Delta_L
Doxorubicin
YDL227C_RF2_33 RF2
ZShift = −0.3
lm ZScore = −0.14
NG = 0
DB = 0
SM = 0
Delta_K
Doxorubicin
YDL227C_RF2_33 RF2
ZShift = −0.58
lm ZScore = 0.3
NG = 0
DB = 0
SM = 0
Delta_r
Doxorubicin
YDL227C_RF2_33 RF2
ZShift = −0.12
lm ZScore = 0.05
NG = 0
DB = 0
SM = 0
Delta_AUC
Doxorubicin

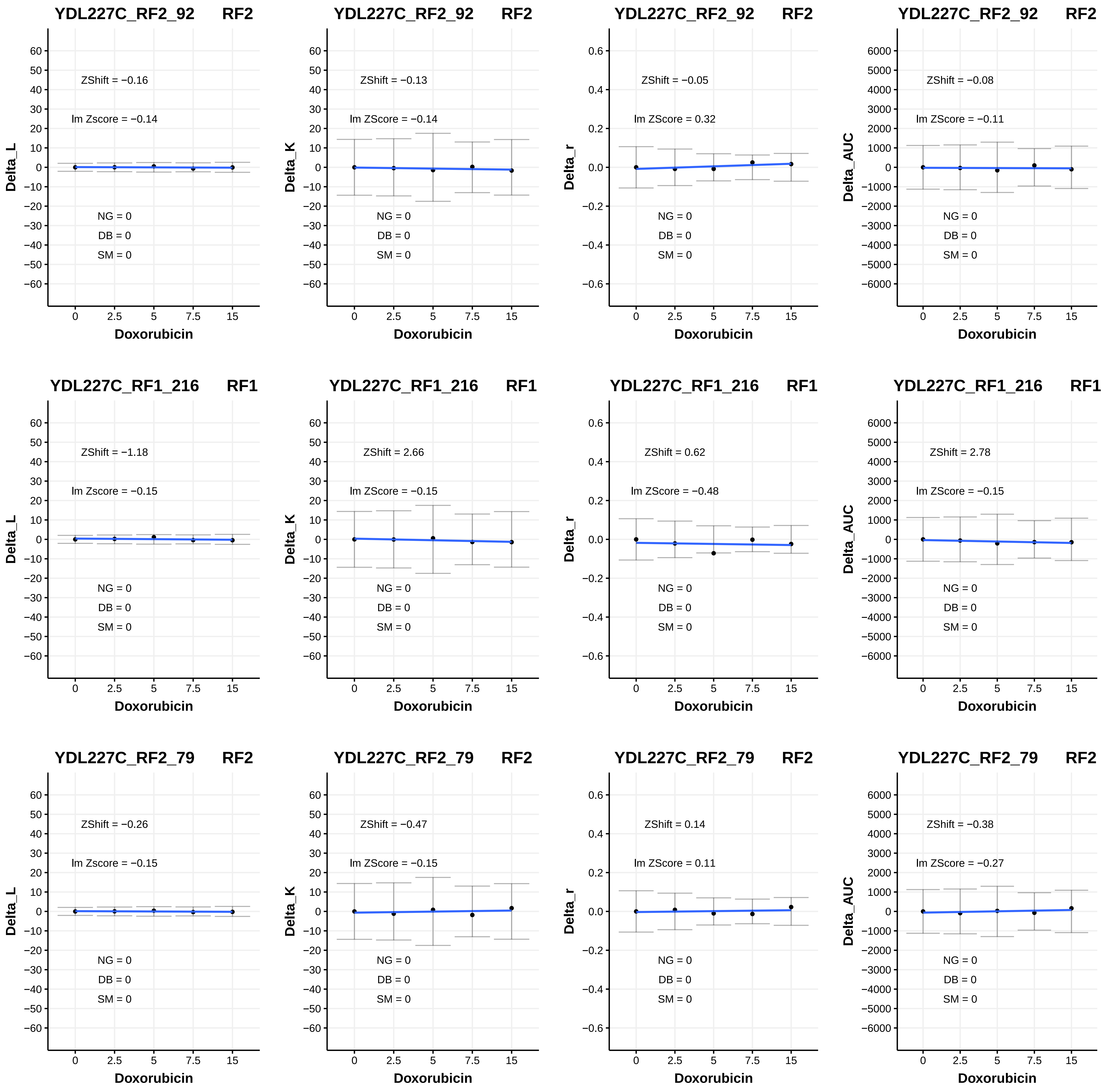
YDL227C_RF2_92 RF2
ZShift = −0.16
lm Zscore = −0.14
NG = 0
DB = 0
SM = 0
Delta_L
Doxorubicin
YDL227C_RF2_92 RF2
ZShift = −0.13
lm ZScore = −0.14
NG = 0
DB = 0
SM = 0
Delta_K
Doxorubicin
YDL227C_RF2_92 RF2
ZShift = −0.05
lm ZScore = 0.32
NG = 0
DB = 0
SM = 0
Delta_r
Doxorubicin
YDL227C_RF2_92 RF2
ZShift = −0.08
lm ZScore = −0.11
NG = 0
DB = 0
SM = 0
Delta_AUC
Doxorubicin
YDL227C_RF1_216 RF1
ZShift = −1.18
lm Zscore = −0.15
NG = 0
DB = 0
SM = 0
Delta_L
Doxorubicin
YDL227C_RF1_216 RF1
ZShift = 2.66
lm ZScore = −0.15
NG = 0
DB = 0
SM = 0
Delta_K
Doxorubicin
YDL227C_RF1_216 RF1
ZShift = 0.62
lm ZScore = −0.48
NG = 0
DB = 0
SM = 0
Delta_r
Doxorubicin
YDL227C_RF1_216 RF1
ZShift = 2.78
lm ZScore = −0.15
NG = 0
DB = 0
SM = 0
Delta_AUC
Doxorubicin
YDL227C_RF2_79 RF2
ZShift = −0.26
lm Zscore = −0.15
NG = 0
DB = 0
SM = 0
Delta_L
Doxorubicin
YDL227C_RF2_79 RF2
ZShift = −0.47
lm ZScore = −0.15
NG = 0
DB = 0
SM = 0
Delta_K
Doxorubicin
YDL227C_RF2_79 RF2
ZShift = 0.14
lm ZScore = 0.11
NG = 0
DB = 0
SM = 0
Delta_r
Doxorubicin
YDL227C_RF2_79 RF2
ZShift = −0.38
lm ZScore = −0.27
NG = 0
DB = 0
SM = 0
Delta_AUC
Doxorubicin

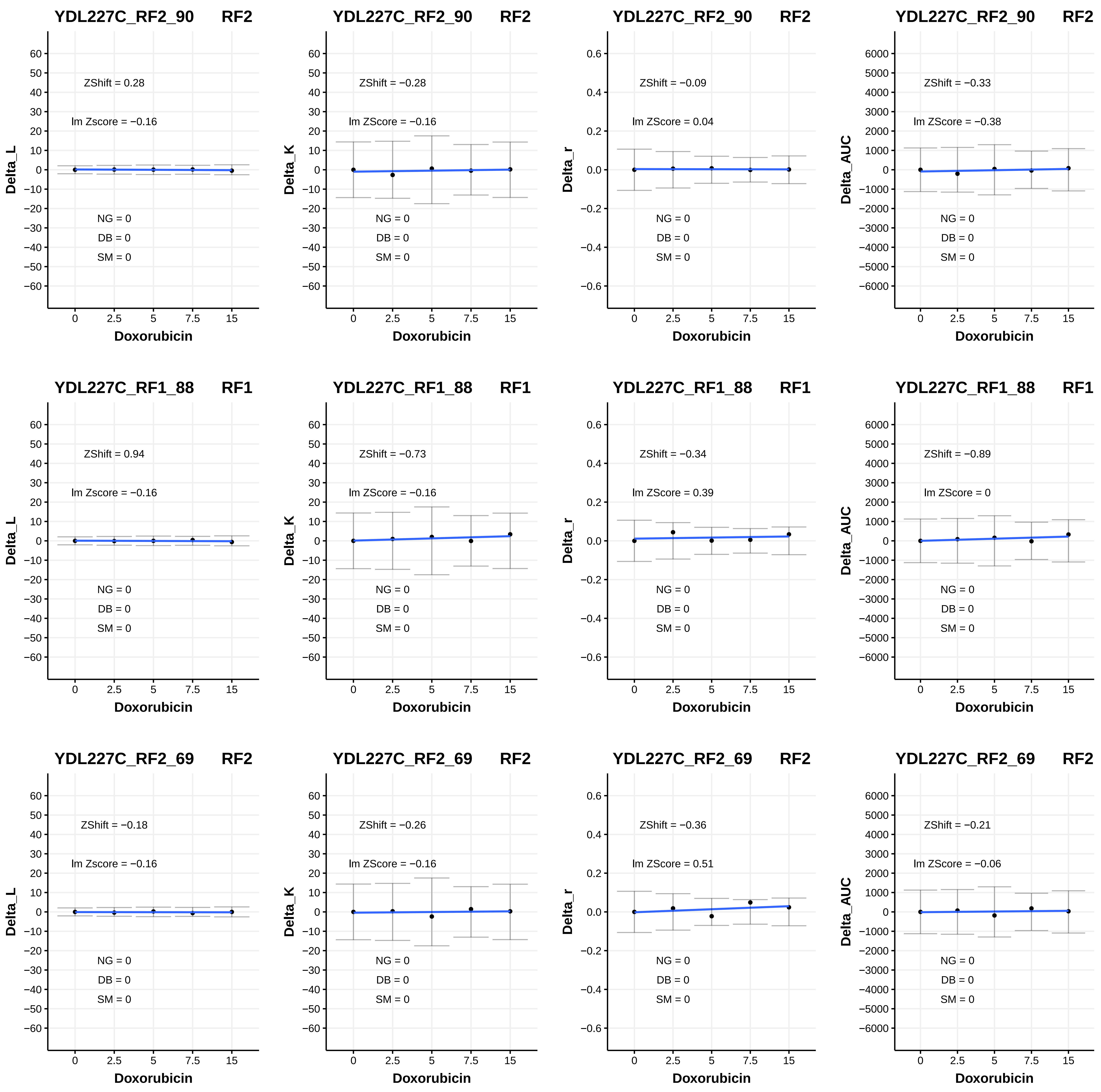
YDL227C_RF2_90 RF2
ZShift = 0.28
lm Zscore = −0.16
NG = 0
DB = 0
SM = 0
Delta_L
Doxorubicin
YDL227C_RF2_90 RF2
ZShift = −0.28
lm ZScore = −0.16
NG = 0
DB = 0
SM = 0
Delta_K
Doxorubicin
YDL227C_RF2_90 RF2
ZShift = −0.09
lm ZScore = 0.04
NG = 0
DB = 0
SM = 0
Delta_r
Doxorubicin
YDL227C_RF2_90 RF2
ZShift = −0.33
lm ZScore = −0.38
NG = 0
DB = 0
SM = 0
Delta_AUC
Doxorubicin
YDL227C_RF1_88 RF1
ZShift = 0.94
lm Zscore = −0.16
NG = 0
DB = 0
SM = 0
Delta_L
Doxorubicin
YDL227C_RF1_88 RF1
ZShift = −0.73
lm ZScore = −0.16
NG = 0
DB = 0
SM = 0
Delta_K
Doxorubicin
YDL227C_RF1_88 RF1
ZShift = −0.34
lm ZScore = 0.39
NG = 0
DB = 0
SM = 0
Delta_r
Doxorubicin
YDL227C_RF1_88 RF1
ZShift = −0.89
lm ZScore = 0
NG = 0
DB = 0
SM = 0
Delta_AUC
Doxorubicin
YDL227C_RF2_69 RF2
ZShift = −0.18
lm Zscore = −0.16
NG = 0
DB = 0
SM = 0
Delta_L
Doxorubicin
YDL227C_RF2_69 RF2
ZShift = −0.26
lm ZScore = −0.16
NG = 0
DB = 0
SM = 0
Delta_K
Doxorubicin
YDL227C_RF2_69 RF2
ZShift = −0.36
lm ZScore = 0.51
NG = 0
DB = 0
SM = 0
Delta_r
Doxorubicin
YDL227C_RF2_69 RF2
ZShift = −0.21
lm ZScore = −0.06
NG = 0
DB = 0
SM = 0
Delta_AUC
Doxorubicin

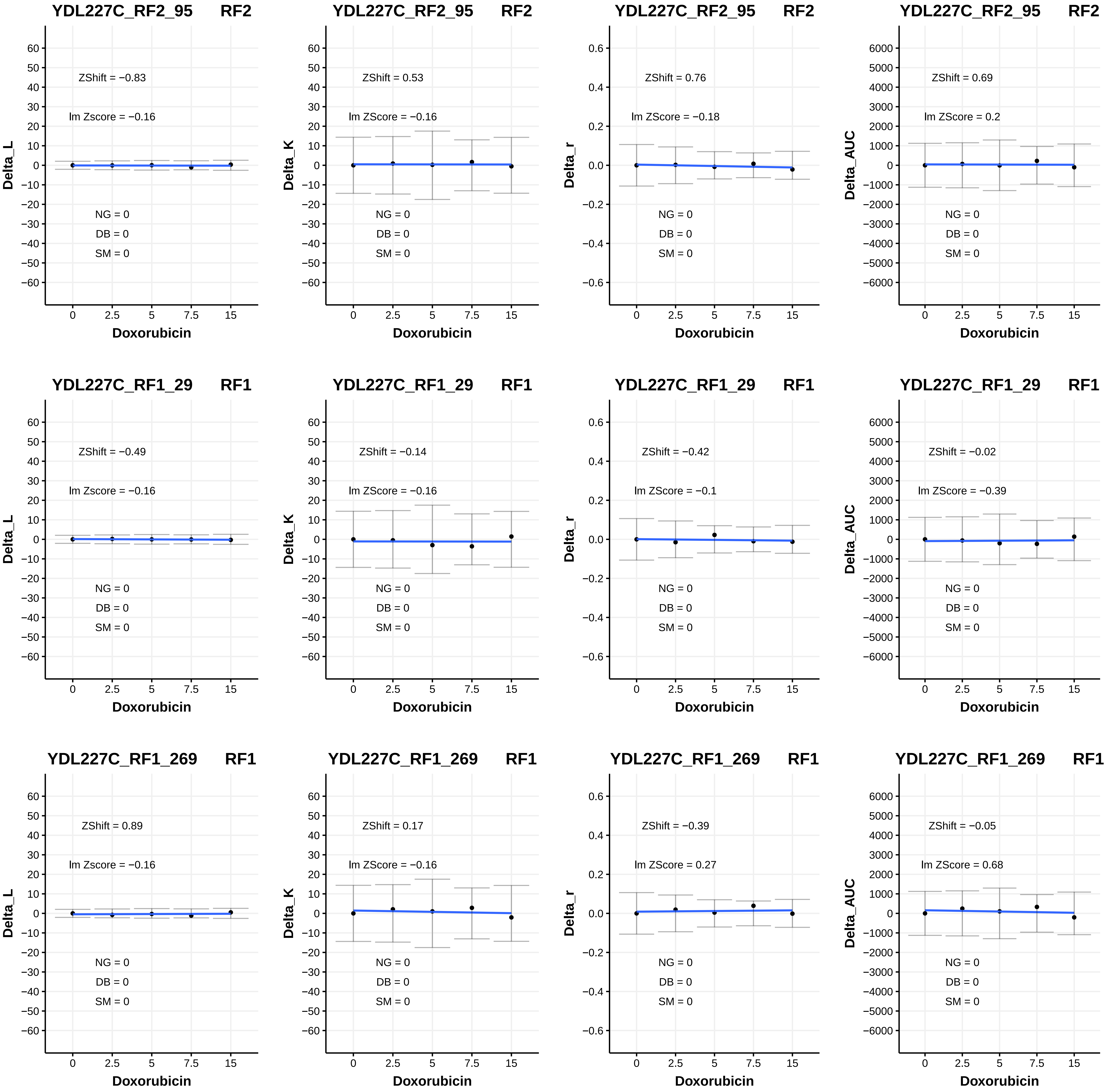
YDL227C_RF2_95 RF2
ZShift = −0.83
lm Zscore = −0.16
NG = 0
DB = 0
SM = 0
Delta_L
Doxorubicin
YDL227C_RF2_95 RF2
ZShift = 0.53
lm ZScore = −0.16
NG = 0
DB = 0
SM = 0
Delta_K
Doxorubicin
YDL227C_RF2_95 RF2
ZShift = 0.76
lm ZScore = −0.18
NG = 0
DB = 0
SM = 0
Delta_r
Doxorubicin
YDL227C_RF2_95 RF2
ZShift = 0.69
lm ZScore = 0.2
NG = 0
DB = 0
SM = 0
Delta_AUC
Doxorubicin
YDL227C_RF1_29 RF1
ZShift = −0.49
lm Zscore = −0.16
NG = 0
DB = 0
SM = 0
Delta_L
Doxorubicin
YDL227C_RF1_29 RF1
ZShift = −0.14
lm ZScore = −0.16
NG = 0
DB = 0
SM = 0
Delta_K
Doxorubicin
YDL227C_RF1_29 RF1
ZShift = −0.42
lm ZScore = −0.1
NG = 0
DB = 0
SM = 0
Delta_r
Doxorubicin
YDL227C_RF1_29 RF1
ZShift = −0.02
lm ZScore = −0.39
NG = 0
DB = 0
SM = 0
Delta_AUC
Doxorubicin
YDL227C_RF1_269 RF1
ZShift = 0.89
lm Zscore = −0.16
NG = 0
DB = 0
SM = 0
Delta_L
Doxorubicin
YDL227C_RF1_269 RF1
ZShift = 0.17
lm ZScore = −0.16
NG = 0
DB = 0
SM = 0
Delta_K
Doxorubicin
YDL227C_RF1_269 RF1
ZShift = −0.39
lm ZScore = 0.27
NG = 0
DB = 0
SM = 0
Delta_r
Doxorubicin
YDL227C_RF1_269 RF1
ZShift = −0.05
lm ZScore = 0.68
NG = 0
DB = 0
SM = 0
Delta_AUC
Doxorubicin

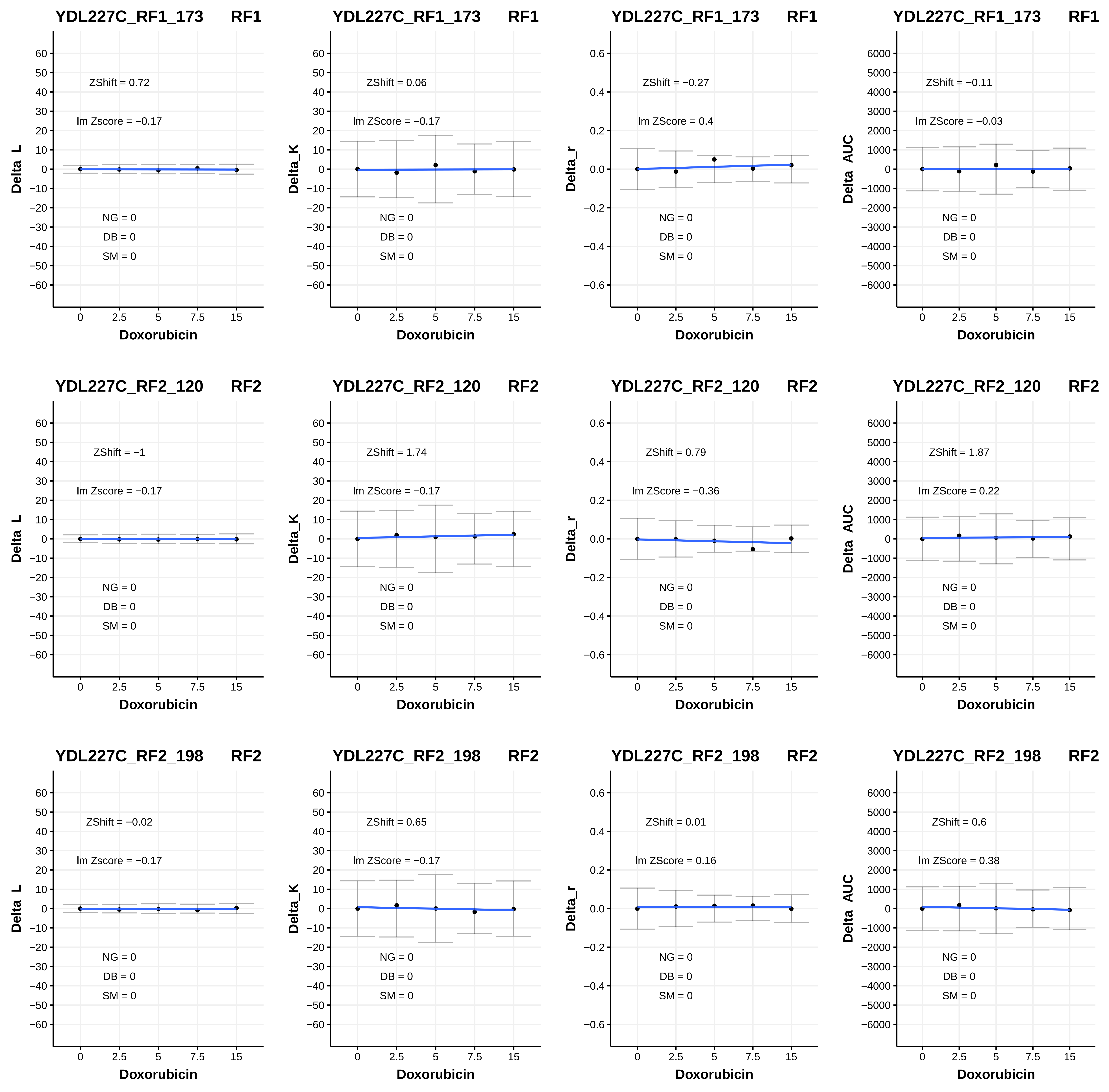

YDL227C_RF1_173 RF1
ZShift = 0.72
lm Zscore = −0.17
NG = 0
DB = 0
SM = 0
Delta_L
Doxorubicin
YDL227C_RF1_173 RF1
ZShift = 0.06
lm ZScore = −0.17
NG = 0
DB = 0
SM = 0
Delta_K
Doxorubicin
YDL227C_RF1_173 RF1
ZShift = −0.27
lm ZScore = 0.4
NG = 0
DB = 0
SM = 0
Delta_r
Doxorubicin
YDL227C_RF1_173 RF1
ZShift = −0.11
lm ZScore = −0.03
NG = 0
DB = 0
SM = 0
Delta_AUC
Doxorubicin
YDL227C_RF2_120 RF2
ZShift = −1
lm Zscore = −0.17
NG = 0
DB = 0
SM = 0
Delta_L
Doxorubicin
YDL227C_RF2_120 RF2
ZShift = 1.74
lm ZScore = −0.17
NG = 0
DB = 0
SM = 0
Delta_K
Doxorubicin
YDL227C_RF2_120 RF2
ZShift = 0.79
lm ZScore = −0.36
NG = 0
DB = 0
SM = 0
Delta_r
Doxorubicin
YDL227C_RF2_120 RF2
ZShift = 1.87
lm ZScore = 0.22
NG = 0
DB = 0
SM = 0
Delta_AUC
Doxorubicin
YDL227C_RF2_198 RF2
ZShift = −0.02
lm Zscore = −0.17
NG = 0
DB = 0
SM = 0
Delta_L
Doxorubicin
YDL227C_RF2_198 RF2
ZShift = 0.65
lm ZScore = −0.17
NG = 0
DB = 0
SM = 0
Delta_K
Doxorubicin
YDL227C_RF2_198 RF2
ZShift = 0.01
lm ZScore = 0.16
NG = 0
DB = 0
SM = 0
Delta_r
Doxorubicin
YDL227C_RF2_198 RF2
ZShift = 0.6
lm ZScore = 0.38
NG = 0
DB = 0
SM = 0
Delta_AUC
Doxorubicin

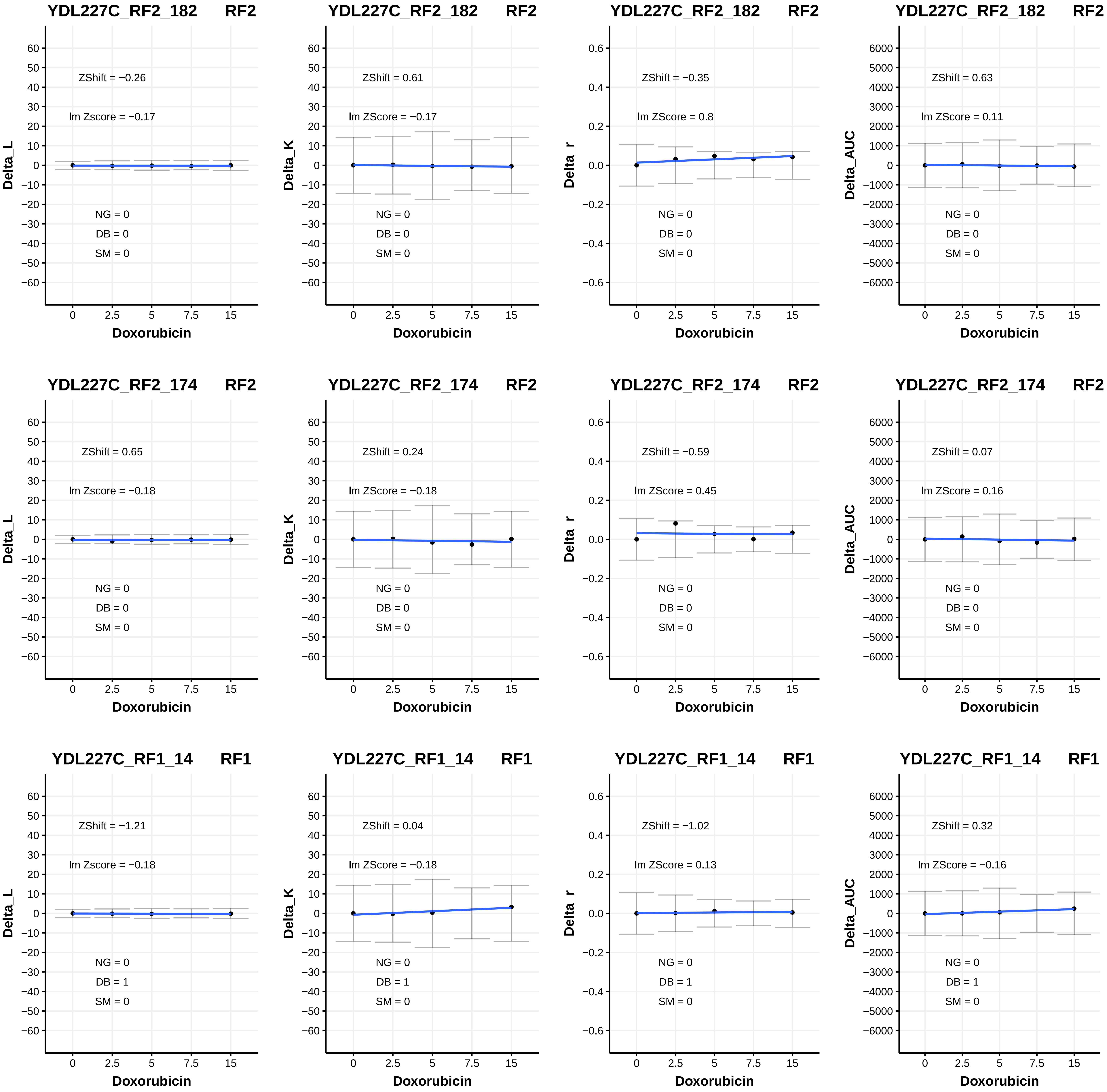

YDL227C_RF2_182 RF2
ZShift = −0.26
lm Zscore = −0.17
NG = 0
DB = 0
SM = 0
Delta_L
Doxorubicin
YDL227C_RF2_182 RF2
ZShift = 0.61
lm ZScore = −0.17
NG = 0
DB = 0
SM = 0
Delta_K
Doxorubicin
YDL227C_RF2_182 RF2
ZShift = −0.35
lm ZScore = 0.8
NG = 0
DB = 0
SM = 0
Delta_r
Doxorubicin
YDL227C_RF2_182 RF2
ZShift = 0.63
lm ZScore = 0.11
NG = 0
DB = 0
SM = 0
Delta_AUC
Doxorubicin
YDL227C_RF2_174 RF2
ZShift = 0.65
lm Zscore = −0.18
NG = 0
DB = 0
SM = 0
Delta_L
Doxorubicin
YDL227C_RF2_174 RF2
ZShift = 0.24
lm ZScore = −0.18
NG = 0
DB = 0
SM = 0
Delta_K
Doxorubicin
YDL227C_RF2_174 RF2
ZShift = −0.59
lm ZScore = 0.45
NG = 0
DB = 0
SM = 0
Delta_r
Doxorubicin
YDL227C_RF2_174 RF2
ZShift = 0.07
lm ZScore = 0.16
NG = 0
DB = 0
SM = 0
Delta_AUC
Doxorubicin
YDL227C_RF1_14 RF1
ZShift = −1.21
lm Zscore = −0.18
NG = 0
DB = 1
SM = 0
Delta_L
Doxorubicin
YDL227C_RF1_14 RF1
ZShift = 0.04
lm ZScore = −0.18
NG = 0
DB = 1
SM = 0
Delta_K
Doxorubicin
YDL227C_RF1_14 RF1
ZShift = −1.02
lm ZScore = 0.13
NG = 0
DB = 1
SM = 0
Delta_r
Doxorubicin
YDL227C_RF1_14 RF1
ZShift = 0.32
lm ZScore = −0.16
NG = 0
DB = 1
SM = 0
Delta_AUC
Doxorubicin

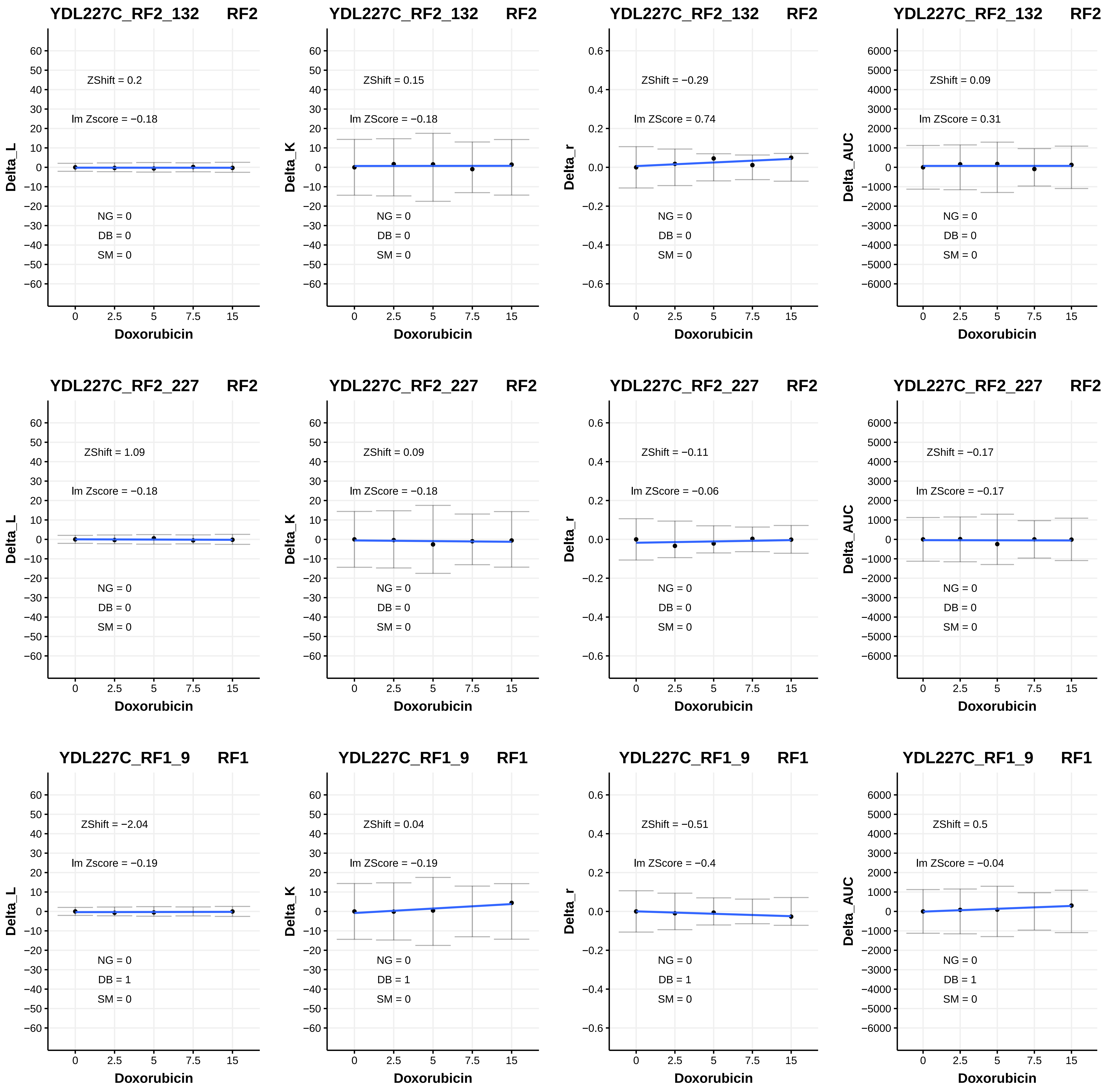
YDL227C_RF2_132 RF2
ZShift = 0.2
lm Zscore = −0.18
NG = 0
DB = 0
SM = 0
Delta_L
Doxorubicin
YDL227C_RF2_132 RF2
ZShift = 0.15
lm ZScore = −0.18
NG = 0
DB = 0
SM = 0
Delta_K
Doxorubicin
YDL227C_RF2_132 RF2
ZShift = −0.29
lm ZScore = 0.74
NG = 0
DB = 0
SM = 0
Delta_r
Doxorubicin
YDL227C_RF2_132 RF2
ZShift = 0.09
lm ZScore = 0.31
NG = 0
DB = 0
SM = 0
Delta_AUC
Doxorubicin
YDL227C_RF2_227 RF2
ZShift = 1.09
lm Zscore = −0.18
NG = 0
DB = 0
SM = 0
Delta_L
Doxorubicin
YDL227C_RF2_227 RF2
ZShift = 0.09
lm ZScore = −0.18
NG = 0
DB = 0
SM = 0
Delta_K
Doxorubicin
YDL227C_RF2_227 RF2
ZShift = −0.11
lm ZScore = −0.06
NG = 0
DB = 0
SM = 0
Delta_r
Doxorubicin
YDL227C_RF2_227 RF2
ZShift = −0.17
lm ZScore = −0.17
NG = 0
DB = 0
SM = 0
Delta_AUC
Doxorubicin
YDL227C_RF1_9 RF1
ZShift = −2.04
lm Zscore = −0.19
NG = 0
DB = 1
SM = 0
Delta_L
Doxorubicin
YDL227C_RF1_9 RF1
ZShift = 0.04
lm ZScore = −0.19
NG = 0
DB = 1
SM = 0
Delta_K
Doxorubicin
YDL227C_RF1_9 RF1
ZShift = −0.51
lm ZScore = −0.4
NG = 0
DB = 1
SM = 0
Delta_r
Doxorubicin
YDL227C_RF1_9 RF1
ZShift = 0.5
lm ZScore = −0.04
NG = 0
DB = 1
SM = 0
Delta_AUC
Doxorubicin

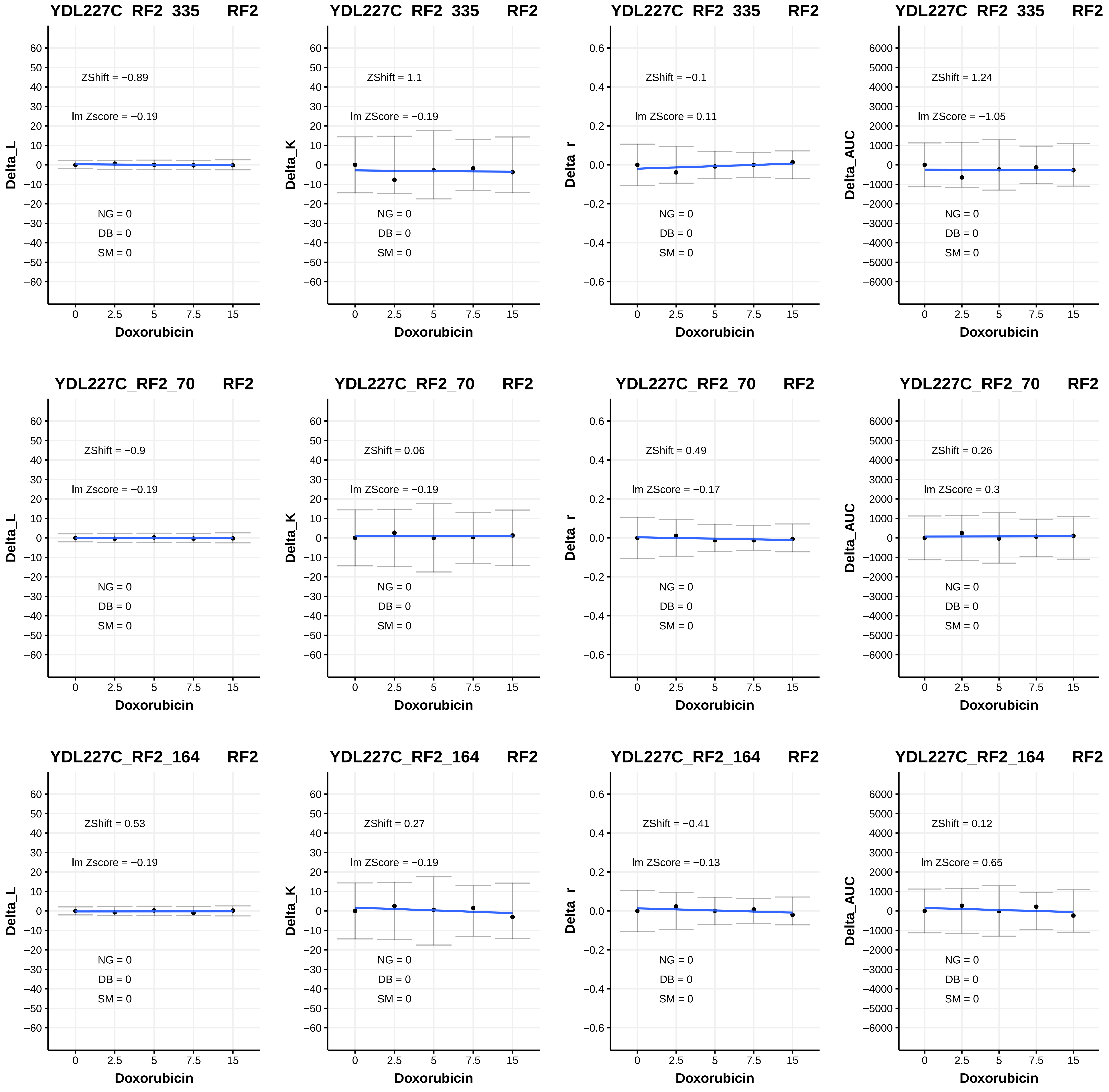

YDL227C_RF2_335 RF2
ZShift = −0.89
lm Zscore = −0.19
NG = 0
DB = 0
SM = 0
Delta_L
Doxorubicin
YDL227C_RF2_335 RF2
ZShift = 1.1
lm ZScore = −0.19
NG = 0
DB = 0
SM = 0
Delta_K
Doxorubicin
YDL227C_RF2_335 RF2
ZShift = −0.1
lm ZScore = 0.11
NG = 0
DB = 0
SM = 0
Delta_r
Doxorubicin
YDL227C_RF2_335 RF2
ZShift = 1.24
lm ZScore = −1.05
NG = 0
DB = 0
SM = 0
Delta_AUC
Doxorubicin
YDL227C_RF2_70 RF2
ZShift = −0.9
lm Zscore = −0.19
NG = 0
DB = 0
SM = 0
Delta_L
Doxorubicin
YDL227C_RF2_70 RF2
ZShift = 0.06
lm ZScore = −0.19
NG = 0
DB = 0
SM = 0
Delta_K
Doxorubicin
YDL227C_RF2_70 RF2
ZShift = 0.49
lm ZScore = −0.17
NG = 0
DB = 0
SM = 0
Delta_r
Doxorubicin
YDL227C_RF2_70 RF2
ZShift = 0.26
lm ZScore = 0.3
NG = 0
DB = 0
SM = 0
Delta_AUC
Doxorubicin
YDL227C_RF2_164 RF2
ZShift = 0.53
lm Zscore = −0.19
NG = 0
DB = 0
SM = 0
Delta_L
Doxorubicin
YDL227C_RF2_164 RF2
ZShift = 0.27
lm ZScore = −0.19
NG = 0
DB = 0
SM = 0
Delta_K
Doxorubicin
YDL227C_RF2_164 RF2
ZShift = −0.41
lm ZScore = −0.13
NG = 0
DB = 0
SM = 0
Delta_r
Doxorubicin
YDL227C_RF2_164 RF2
ZShift = 0.12
lm ZScore = 0.65
NG = 0
DB = 0
SM = 0
Delta_AUC
Doxorubicin

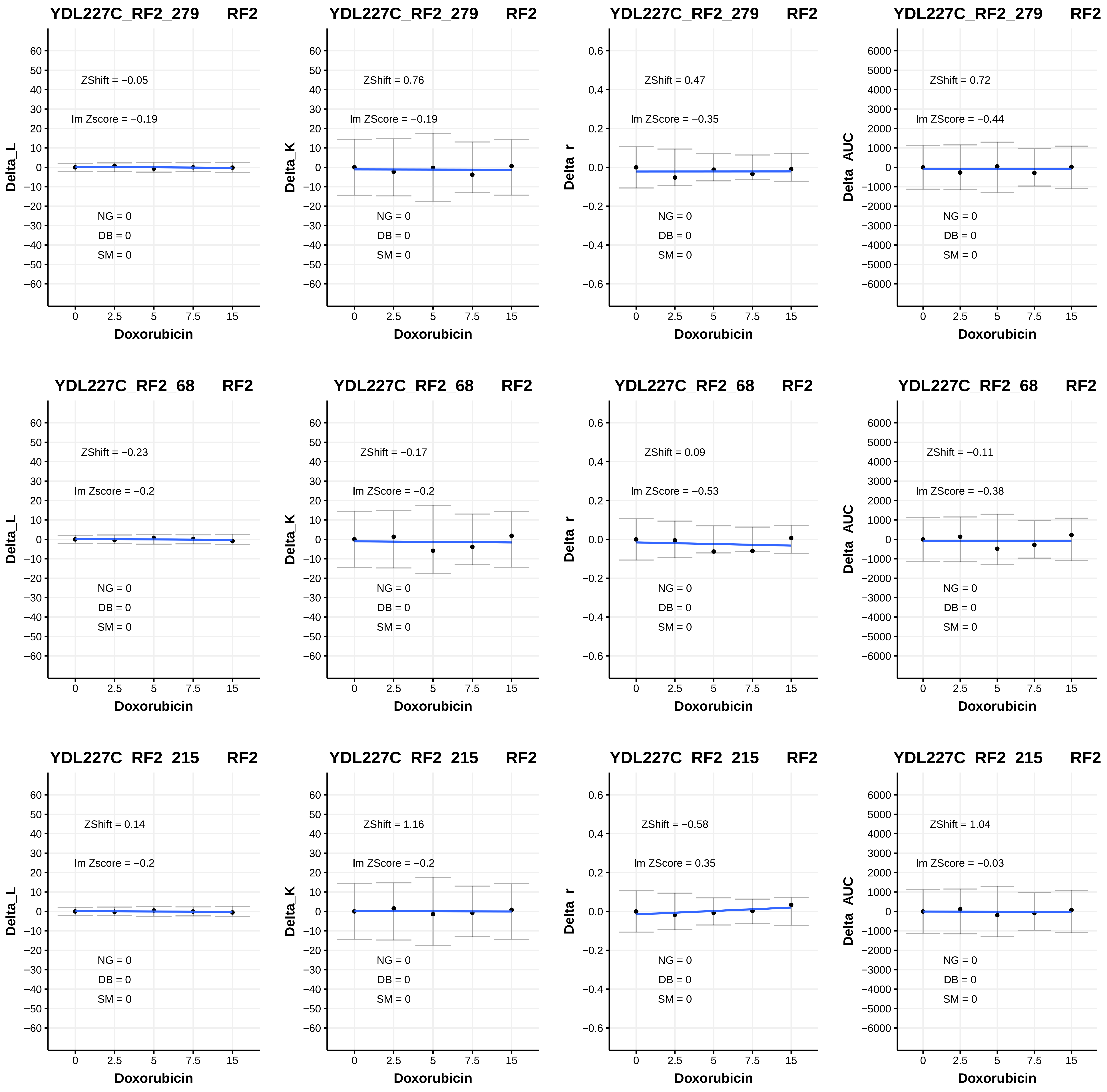
YDL227C_RF2_279 RF2
ZShift = −0.05
lm Zscore = −0.19
NG = 0
DB = 0
SM = 0
Delta_L
Doxorubicin
YDL227C_RF2_279 RF2
ZShift = 0.76
lm ZScore = −0.19
NG = 0
DB = 0
SM = 0
Delta_K
Doxorubicin
YDL227C_RF2_279 RF2
ZShift = 0.47
lm ZScore = −0.35
NG = 0
DB = 0
SM = 0
Delta_r
Doxorubicin
YDL227C_RF2_279 RF2
ZShift = 0.72
lm ZScore = −0.44
NG = 0
DB = 0
SM = 0
Delta_AUC
Doxorubicin
YDL227C_RF2_68 RF2
ZShift = −0.23
lm Zscore = −0.2
NG = 0
DB = 0
SM = 0
Delta_L
Doxorubicin
YDL227C_RF2_68 RF2
ZShift = −0.17
lm ZScore = −0.2
NG = 0
DB = 0
SM = 0
Delta_K
Doxorubicin
YDL227C_RF2_68 RF2
ZShift = 0.09
lm ZScore = −0.53
NG = 0
DB = 0
SM = 0
Delta_r
Doxorubicin
YDL227C_RF2_68 RF2
ZShift = −0.11
lm ZScore = −0.38
NG = 0
DB = 0
SM = 0
Delta_AUC
Doxorubicin
YDL227C_RF2_215 RF2
ZShift = 0.14
lm Zscore = −0.2
NG = 0
DB = 0
SM = 0
Delta_L
Doxorubicin
YDL227C_RF2_215 RF2
ZShift = 1.16
lm ZScore = −0.2
NG = 0
DB = 0
SM = 0
Delta_K
Doxorubicin
YDL227C_RF2_215 RF2
ZShift = −0.58
lm ZScore = 0.35
NG = 0
DB = 0
SM = 0
Delta_r
Doxorubicin
YDL227C_RF2_215 RF2
ZShift = 1.04
lm ZScore = −0.03
NG = 0
DB = 0
SM = 0
Delta_AUC
Doxorubicin

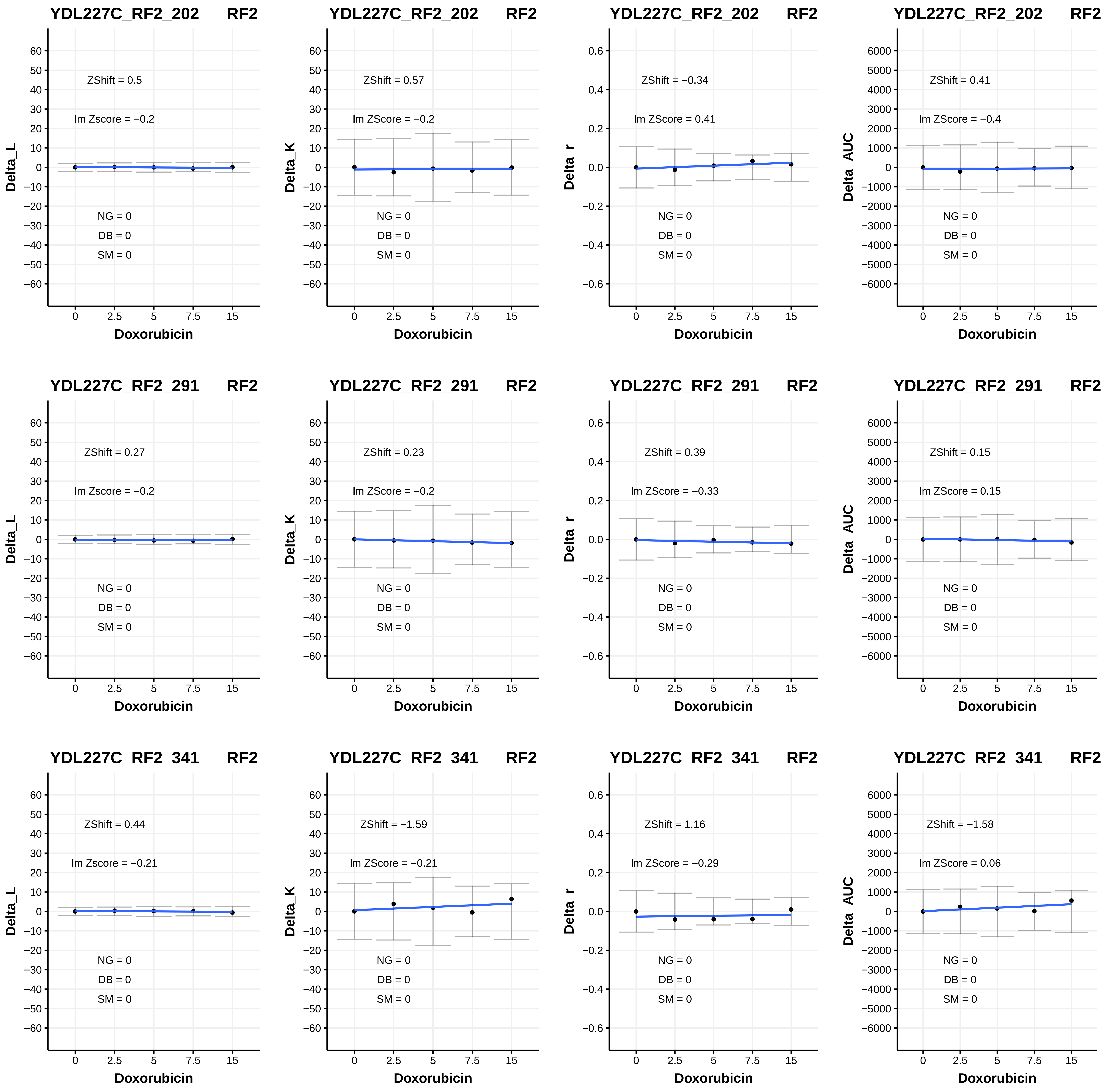
YDL227C_RF2_202 RF2
ZShift = 0.5
lm Zscore = −0.2
NG = 0
DB = 0
SM = 0
Delta_L
Doxorubicin
YDL227C_RF2_202 RF2
ZShift = 0.57
lm ZScore = −0.2
NG = 0
DB = 0
SM = 0
Delta_K
Doxorubicin
YDL227C_RF2_202 RF2
ZShift = −0.34
lm ZScore = 0.41
NG = 0
DB = 0
SM = 0
Delta_r
Doxorubicin
YDL227C_RF2_202 RF2
ZShift = 0.41
lm ZScore = −0.4
NG = 0
DB = 0
SM = 0
Delta_AUC
Doxorubicin
YDL227C_RF2_291 RF2
ZShift = 0.27
lm Zscore = −0.2
NG = 0
DB = 0
SM = 0
Delta_L
Doxorubicin
YDL227C_RF2_291 RF2
ZShift = 0.23
lm ZScore = −0.2
NG = 0
DB = 0
SM = 0
Delta_K
Doxorubicin
YDL227C_RF2_291 RF2
ZShift = 0.39
lm ZScore = −0.33
NG = 0
DB = 0
SM = 0
Delta_r
Doxorubicin
YDL227C_RF2_291 RF2
ZShift = 0.15
lm ZScore = 0.15
NG = 0
DB = 0
SM = 0
Delta_AUC
Doxorubicin
YDL227C_RF2_341 RF2
ZShift = 0.44
lm Zscore = −0.21
NG = 0
DB = 0
SM = 0
Delta_L
Doxorubicin
YDL227C_RF2_341 RF2
ZShift = −1.59
lm ZScore = −0.21
NG = 0
DB = 0
SM = 0
Delta_K
Doxorubicin
YDL227C_RF2_341 RF2
ZShift = 1.16
lm ZScore = −0.29
NG = 0
DB = 0
SM = 0
Delta_r
Doxorubicin
YDL227C_RF2_341 RF2
ZShift = −1.58
lm ZScore = 0.06
NG = 0
DB = 0
SM = 0
Delta_AUC
Doxorubicin

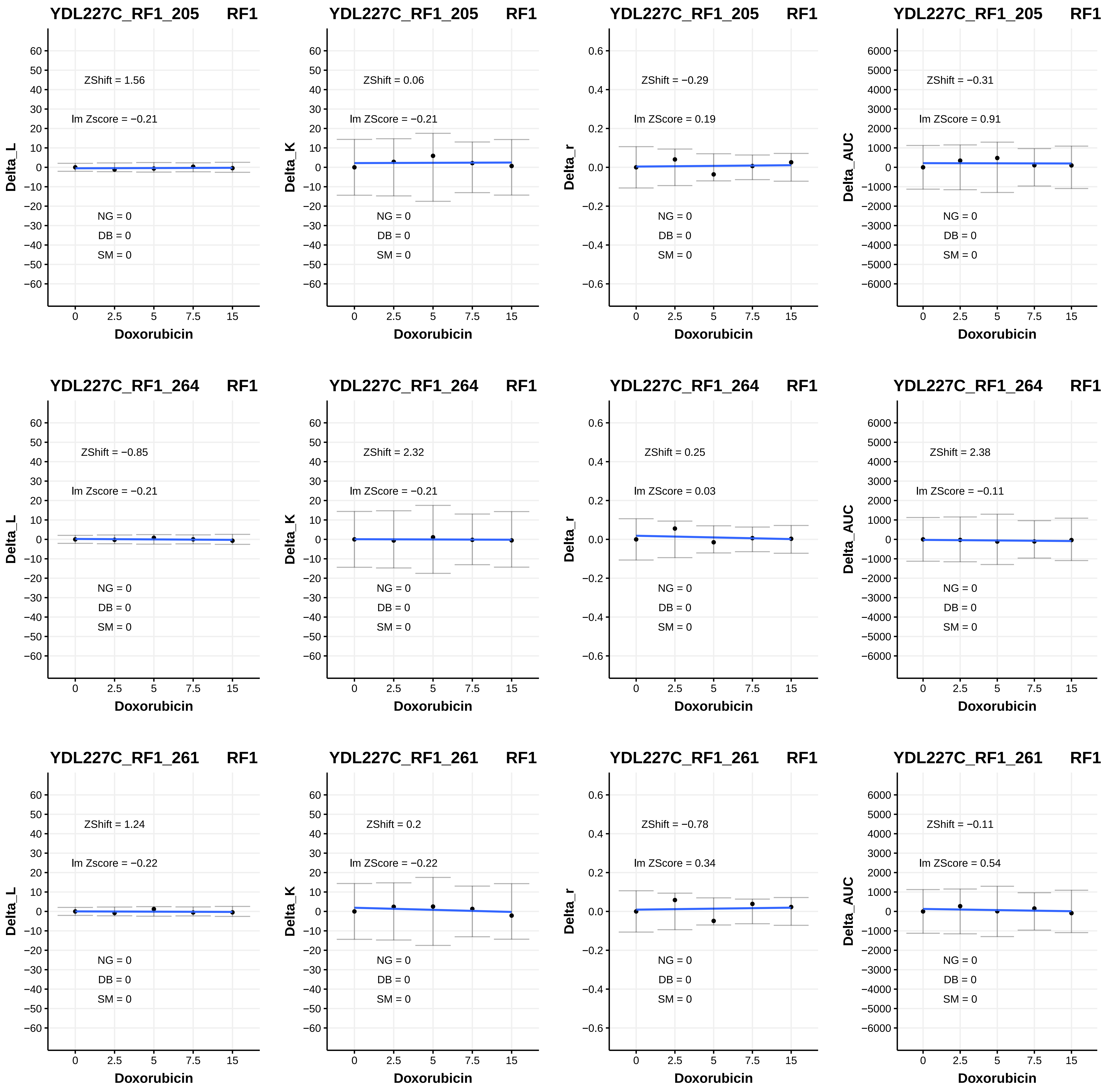
YDL227C_RF1_205 RF1
ZShift = 1.56
lm Zscore = −0.21
NG = 0
DB = 0
SM = 0
Delta_L
Doxorubicin
YDL227C_RF1_205 RF1
ZShift = 0.06
lm ZScore = −0.21
NG = 0
DB = 0
SM = 0
Delta_K
Doxorubicin
YDL227C_RF1_205 RF1
ZShift = −0.29
lm ZScore = 0.19
NG = 0
DB = 0
SM = 0
Delta_r
Doxorubicin
YDL227C_RF1_205 RF1
ZShift = −0.31
lm ZScore = 0.91
NG = 0
DB = 0
SM = 0
Delta_AUC
Doxorubicin
YDL227C_RF1_264 RF1
ZShift = −0.85
lm Zscore = −0.21
NG = 0
DB = 0
SM = 0
Delta_L
Doxorubicin
YDL227C_RF1_264 RF1
ZShift = 2.32
lm ZScore = −0.21
NG = 0
DB = 0
SM = 0
Delta_K
Doxorubicin
YDL227C_RF1_264 RF1
ZShift = 0.25
lm ZScore = 0.03
NG = 0
DB = 0
SM = 0
Delta_r
Doxorubicin
YDL227C_RF1_264 RF1
ZShift = 2.38
lm ZScore = −0.11
NG = 0
DB = 0
SM = 0
Delta_AUC
Doxorubicin
YDL227C_RF1_261 RF1
ZShift = 1.24
lm Zscore = −0.22
NG = 0
DB = 0
SM = 0
Delta_L
Doxorubicin
YDL227C_RF1_261 RF1
ZShift = 0.2
lm ZScore = −0.22
NG = 0
DB = 0
SM = 0
Delta_K
Doxorubicin
YDL227C_RF1_261 RF1
ZShift = −0.78
lm ZScore = 0.34
NG = 0
DB = 0
SM = 0
Delta_r
Doxorubicin
YDL227C_RF1_261 RF1
ZShift = −0.11
lm ZScore = 0.54
NG = 0
DB = 0
SM = 0
Delta_AUC
Doxorubicin

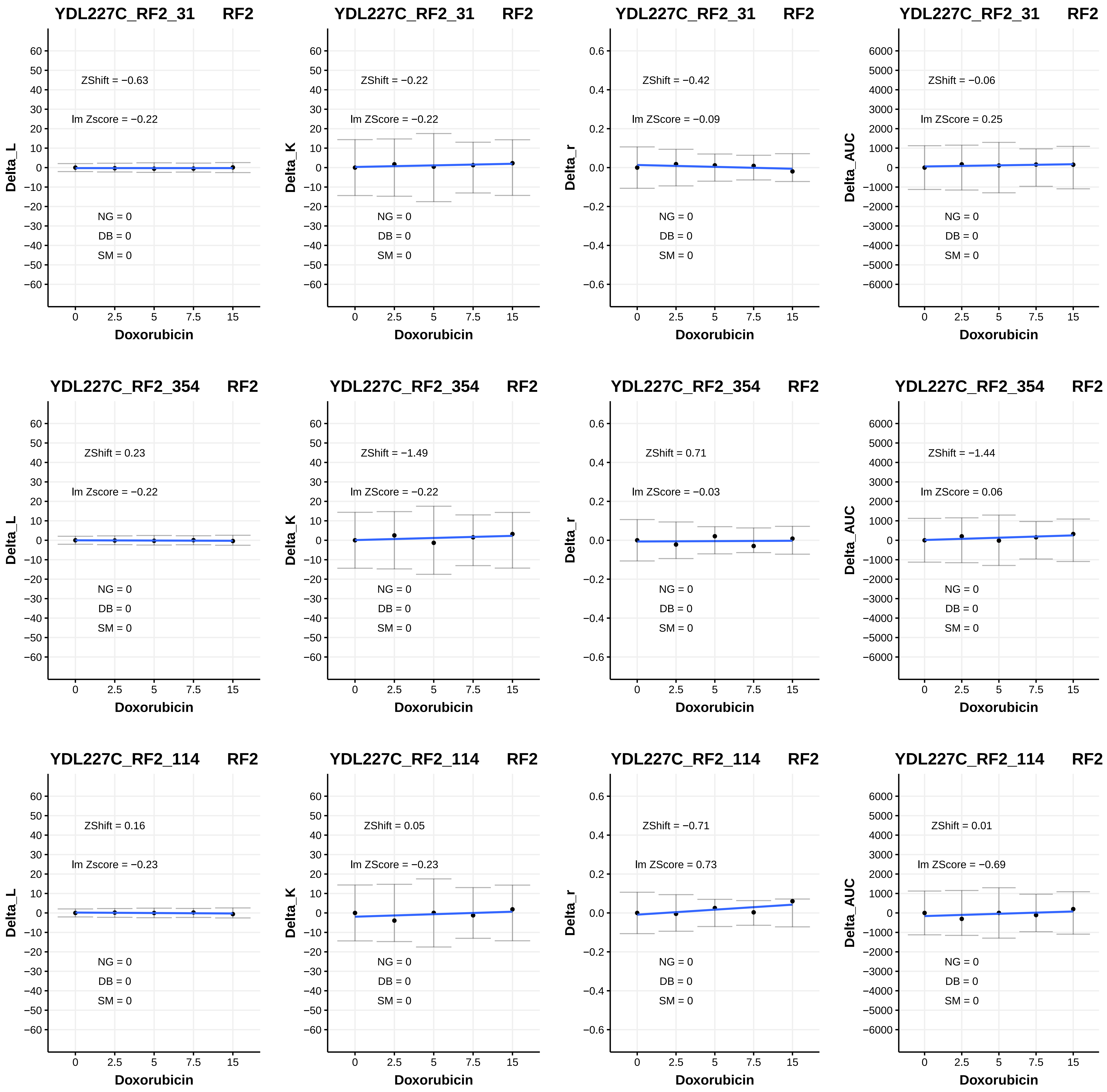

YDL227C_RF2_31 RF2
ZShift = −0.63
lm Zscore = −0.22
NG = 0
DB = 0
SM = 0
Delta_L
Doxorubicin
YDL227C_RF2_31 RF2
ZShift = −0.22
lm ZScore = −0.22
NG = 0
DB = 0
SM = 0
Delta_K
Doxorubicin
YDL227C_RF2_31 RF2
ZShift = −0.42
lm ZScore = −0.09
NG = 0
DB = 0
SM = 0
Delta_r
Doxorubicin
YDL227C_RF2_31 RF2
ZShift = −0.06
lm ZScore = 0.25
NG = 0
DB = 0
SM = 0
Delta_AUC
Doxorubicin
YDL227C_RF2_354 RF2
ZShift = 0.23
lm Zscore = −0.22
NG = 0
DB = 0
SM = 0
Delta_L
Doxorubicin
YDL227C_RF2_354 RF2
ZShift = −1.49
lm ZScore = −0.22
NG = 0
DB = 0
SM = 0
Delta_K
Doxorubicin
YDL227C_RF2_354 RF2
ZShift = 0.71
lm ZScore = −0.03
NG = 0
DB = 0
SM = 0
Delta_r
Doxorubicin
YDL227C_RF2_354 RF2
ZShift = −1.44
lm ZScore = 0.06
NG = 0
DB = 0
SM = 0
Delta_AUC
Doxorubicin
YDL227C_RF2_114 RF2
ZShift = 0.16
lm Zscore = −0.23
NG = 0
DB = 0
SM = 0
Delta_L
Doxorubicin
YDL227C_RF2_114 RF2
ZShift = 0.05
lm ZScore = −0.23
NG = 0
DB = 0
SM = 0
Delta_K
Doxorubicin
YDL227C_RF2_114 RF2
ZShift = −0.71
lm ZScore = 0.73
NG = 0
DB = 0
SM = 0
Delta_r
Doxorubicin
YDL227C_RF2_114 RF2
ZShift = 0.01
lm ZScore = −0.69
NG = 0
DB = 0
SM = 0
Delta_AUC
Doxorubicin

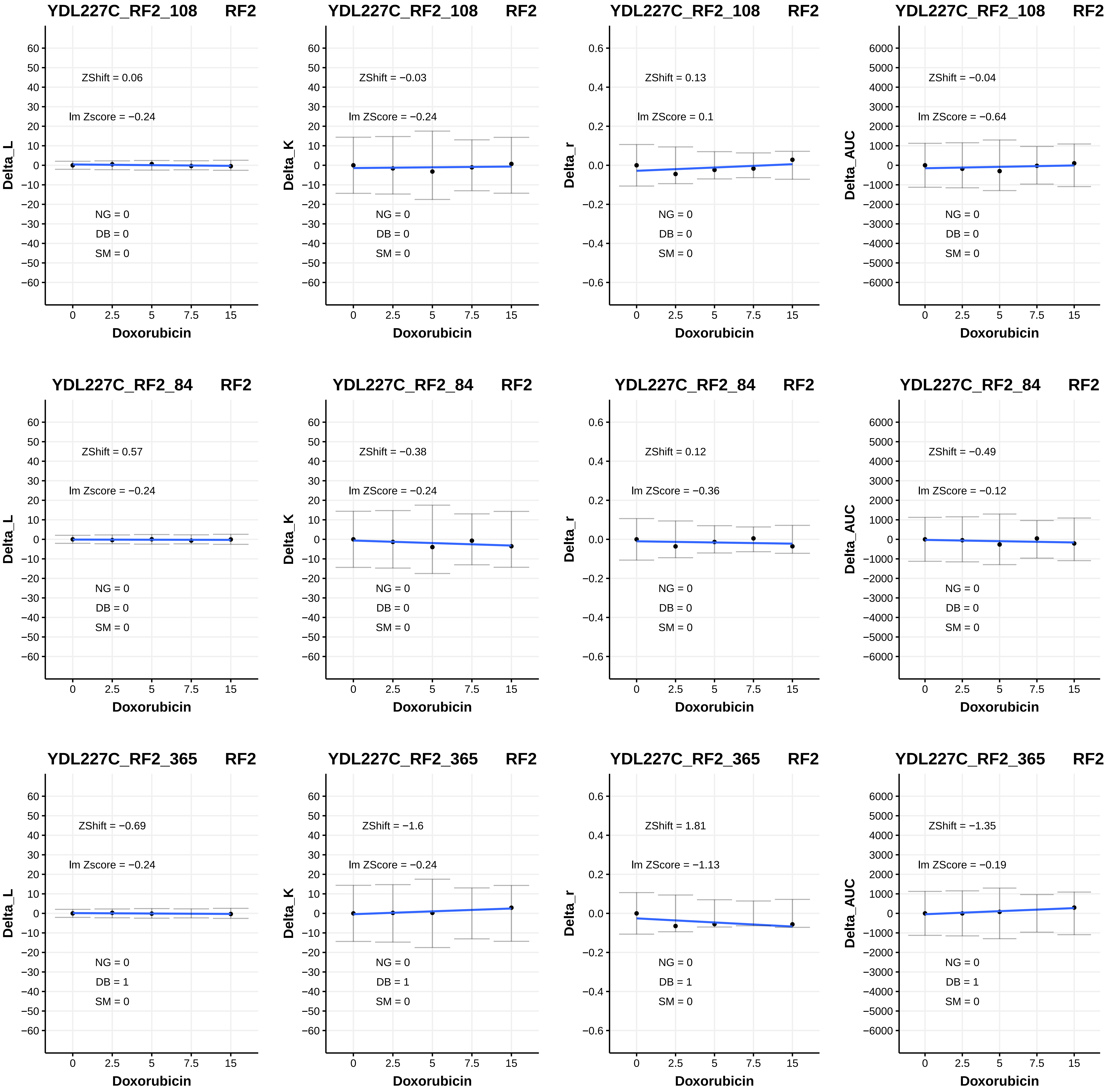
YDL227C_RF2_108 RF2
ZShift = 0.06
lm Zscore = −0.24
NG = 0
DB = 0
SM = 0
Delta_L
Doxorubicin
YDL227C_RF2_108 RF2
ZShift = −0.03
lm ZScore = −0.24
NG = 0
DB = 0
SM = 0
Delta_K
Doxorubicin
YDL227C_RF2_108 RF2
ZShift = 0.13
lm ZScore = 0.1
NG = 0
DB = 0
SM = 0
Delta_r
Doxorubicin
YDL227C_RF2_108 RF2
ZShift = −0.04
lm ZScore = −0.64
NG = 0
DB = 0
SM = 0
Delta_AUC
Doxorubicin
YDL227C_RF2_84 RF2
ZShift = 0.57
lm Zscore = −0.24
NG = 0
DB = 0
SM = 0
Delta_L
Doxorubicin
YDL227C_RF2_84 RF2
ZShift = −0.38
lm ZScore = −0.24
NG = 0
DB = 0
SM = 0
Delta_K
Doxorubicin
YDL227C_RF2_84 RF2
ZShift = 0.12
lm ZScore = −0.36
NG = 0
DB = 0
SM = 0
Delta_r
Doxorubicin
YDL227C_RF2_84 RF2
ZShift = −0.49
lm ZScore = −0.12
NG = 0
DB = 0
SM = 0
Delta_AUC
Doxorubicin
YDL227C_RF2_365 RF2
ZShift = −0.69
lm Zscore = −0.24
NG = 0
DB = 1
SM = 0
Delta_L
Doxorubicin
YDL227C_RF2_365 RF2
ZShift = −1.6
lm ZScore = −0.24
NG = 0
DB = 1
SM = 0
Delta_K
Doxorubicin
YDL227C_RF2_365 RF2
ZShift = 1.81
lm ZScore = −1.13
NG = 0
DB = 1
SM = 0
Delta_r
Doxorubicin
YDL227C_RF2_365 RF2
ZShift = −1.35
lm ZScore = −0.19
NG = 0
DB = 1
SM = 0
Delta_AUC
Doxorubicin

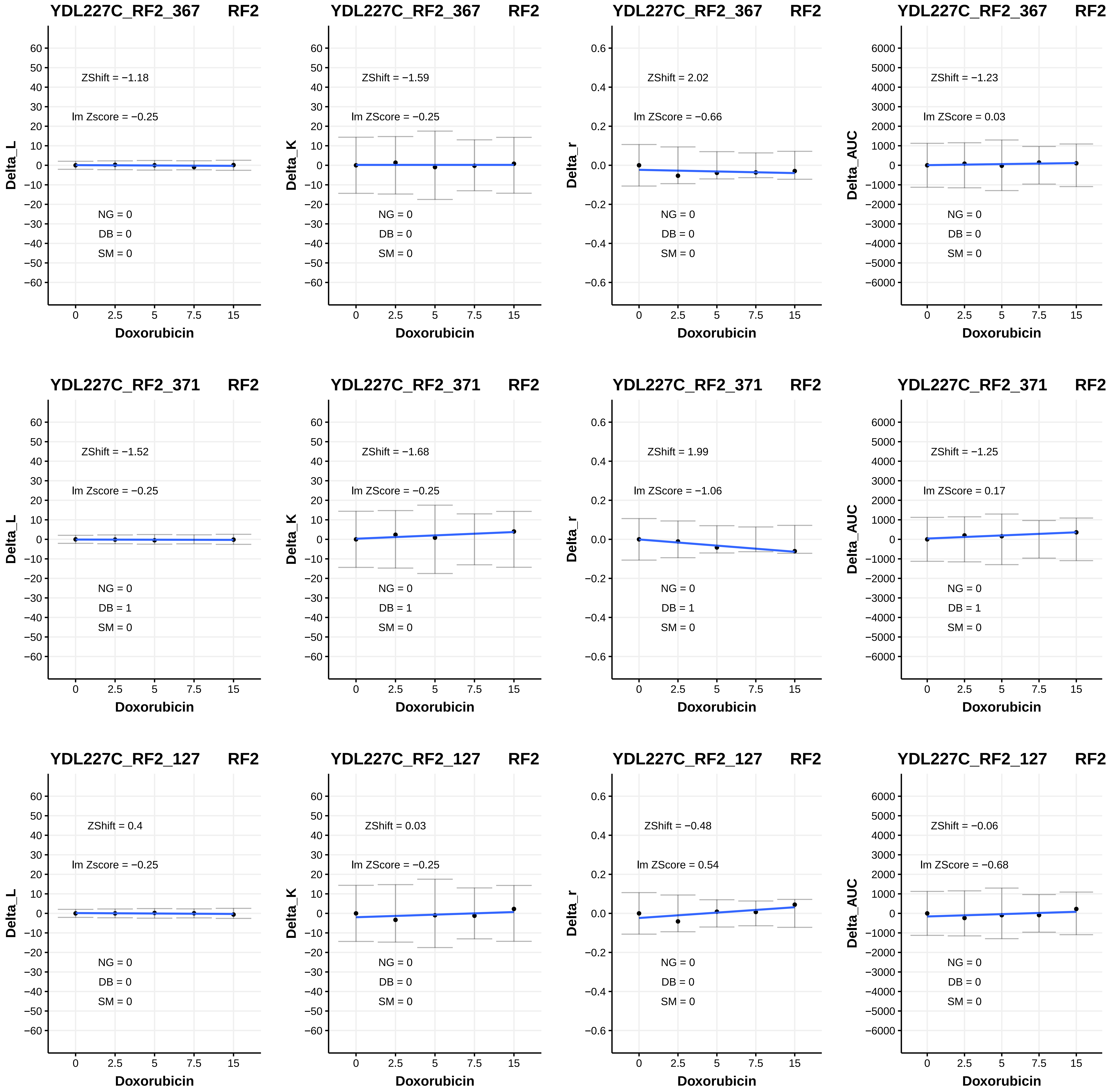
YDL227C_RF2_367 RF2
ZShift = −1.18
lm Zscore = −0.25
NG = 0
DB = 0
SM = 0
Delta_L
Doxorubicin
YDL227C_RF2_367 RF2
ZShift = −1.59
lm ZScore = −0.25
NG = 0
DB = 0
SM = 0
Delta_K
Doxorubicin
YDL227C_RF2_367 RF2
ZShift = 2.02
lm ZScore = −0.66
NG = 0
DB = 0
SM = 0
Delta_r
Doxorubicin
YDL227C_RF2_367 RF2
ZShift = −1.23
lm ZScore = 0.03
NG = 0
DB = 0
SM = 0
Delta_AUC
Doxorubicin
YDL227C_RF2_371 RF2
ZShift = −1.52
lm Zscore = −0.25
NG = 0
DB = 1
SM = 0
Delta_L
Doxorubicin
YDL227C_RF2_371 RF2
ZShift = −1.68
lm ZScore = −0.25
NG = 0
DB = 1
SM = 0
Delta_K
Doxorubicin
YDL227C_RF2_371 RF2
ZShift = 1.99
lm ZScore = −1.06
NG = 0
DB = 1
SM = 0
Delta_r
Doxorubicin
YDL227C_RF2_371 RF2
ZShift = −1.25
lm ZScore = 0.17
NG = 0
DB = 1
SM = 0
Delta_AUC
Doxorubicin
YDL227C_RF2_127 RF2
ZShift = 0.4
lm Zscore = −0.25
NG = 0
DB = 0
SM = 0
Delta_L
Doxorubicin
YDL227C_RF2_127 RF2
ZShift = 0.03
lm ZScore = −0.25
NG = 0
DB = 0
SM = 0
Delta_K
Doxorubicin
YDL227C_RF2_127 RF2
ZShift = −0.48
lm ZScore = 0.54
NG = 0
DB = 0
SM = 0
Delta_r
Doxorubicin
YDL227C_RF2_127 RF2
ZShift = −0.06
lm ZScore = −0.68
NG = 0
DB = 0
SM = 0
Delta_AUC
Doxorubicin

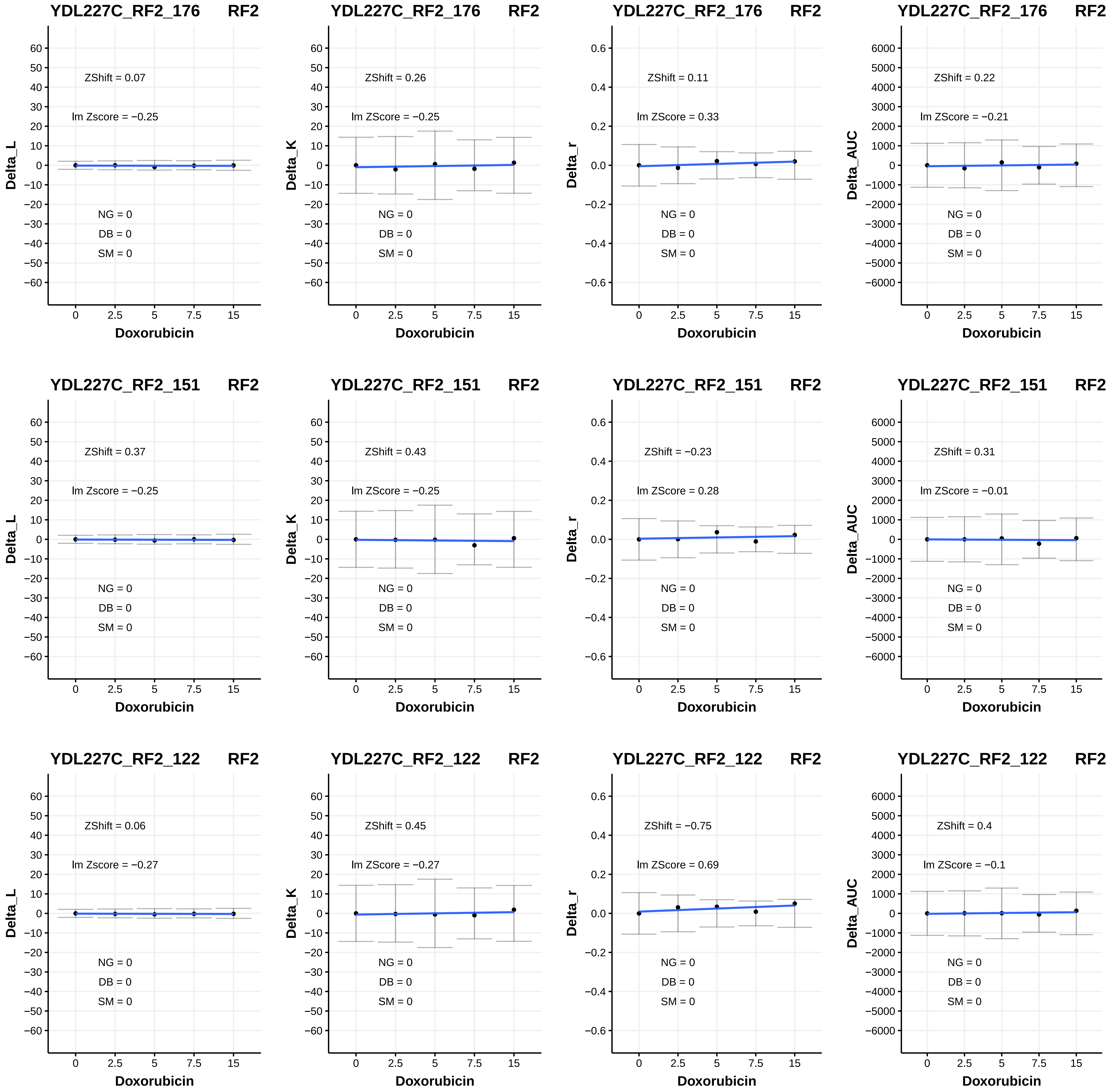
YDL227C_RF2_176 RF2
ZShift = 0.07
lm Zscore = −0.25
NG = 0
DB = 0
SM = 0
Delta_L
Doxorubicin
YDL227C_RF2_176 RF2
ZShift = 0.26
lm ZScore = −0.25
NG = 0
DB = 0
SM = 0
Delta_K
Doxorubicin
YDL227C_RF2_176 RF2
ZShift = 0.11
lm ZScore = 0.33
NG = 0
DB = 0
SM = 0
Delta_r
Doxorubicin
YDL227C_RF2_176 RF2
ZShift = 0.22
lm ZScore = −0.21
NG = 0
DB = 0
SM = 0
Delta_AUC
Doxorubicin
YDL227C_RF2_151 RF2
ZShift = 0.37
lm Zscore = −0.25
NG = 0
DB = 0
SM = 0
Delta_L
Doxorubicin
YDL227C_RF2_151 RF2
ZShift = 0.43
lm ZScore = −0.25
NG = 0
DB = 0
SM = 0
Delta_K
Doxorubicin
YDL227C_RF2_151 RF2
ZShift = −0.23
lm ZScore = 0.28
NG = 0
DB = 0
SM = 0
Delta_r
Doxorubicin
YDL227C_RF2_151 RF2
ZShift = 0.31
lm ZScore = −0.01
NG = 0
DB = 0
SM = 0
Delta_AUC
Doxorubicin
YDL227C_RF2_122 RF2
ZShift = 0.06
lm Zscore = −0.27
NG = 0
DB = 0
SM = 0
Delta_L
Doxorubicin
YDL227C_RF2_122 RF2
ZShift = 0.45
lm ZScore = −0.27
NG = 0
DB = 0
SM = 0
Delta_K
Doxorubicin
YDL227C_RF2_122 RF2
ZShift = −0.75
lm ZScore = 0.69
NG = 0
DB = 0
SM = 0
Delta_r
Doxorubicin
YDL227C_RF2_122 RF2
ZShift = 0.4
lm ZScore = −0.1
NG = 0
DB = 0
SM = 0
Delta_AUC
Doxorubicin

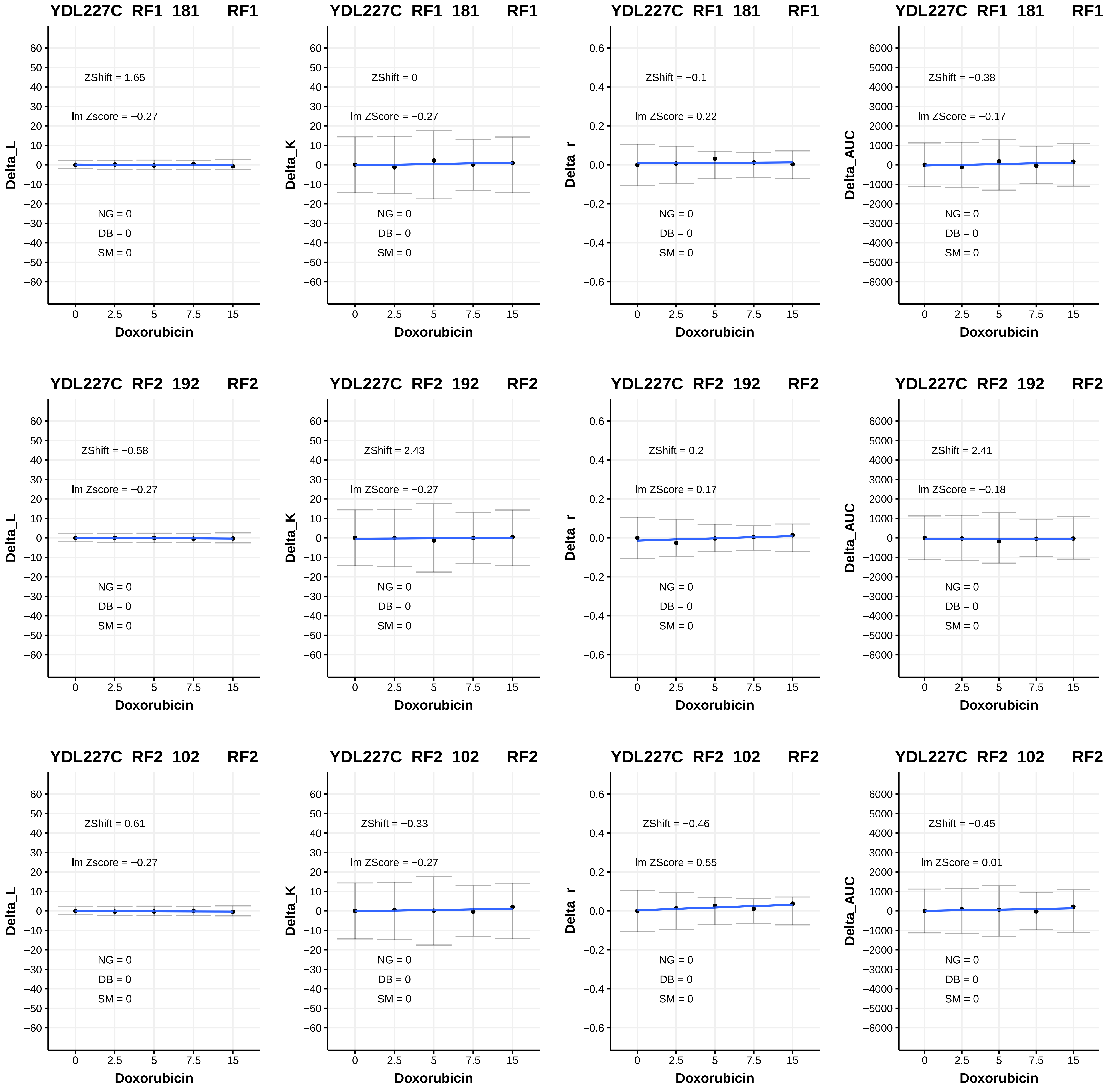

YDL227C_RF1_181 RF1
ZShift = 1.65
lm Zscore = −0.27
NG = 0
DB = 0
SM = 0
Delta_L
Doxorubicin
YDL227C_RF1_181 RF1
ZShift = 0
lm ZScore = −0.27
NG = 0
DB = 0
SM = 0
Delta_K
Doxorubicin
YDL227C_RF1_181 RF1
ZShift = −0.1
lm ZScore = 0.22
NG = 0
DB = 0
SM = 0
Delta_r
Doxorubicin
YDL227C_RF1_181 RF1
ZShift = −0.38
lm ZScore = −0.17
NG = 0
DB = 0
SM = 0
Delta_AUC
Doxorubicin
YDL227C_RF2_192 RF2
ZShift = −0.58
lm Zscore = −0.27
NG = 0
DB = 0
SM = 0
Delta_L
Doxorubicin
YDL227C_RF2_192 RF2
ZShift = 2.43
lm ZScore = −0.27
NG = 0
DB = 0
SM = 0
Delta_K
Doxorubicin
YDL227C_RF2_192 RF2
ZShift = 0.2
lm ZScore = 0.17
NG = 0
DB = 0
SM = 0
Delta_r
Doxorubicin
YDL227C_RF2_192 RF2
ZShift = 2.41
lm ZScore = −0.18
NG = 0
DB = 0
SM = 0
Delta_AUC
Doxorubicin
YDL227C_RF2_102 RF2
ZShift = 0.61
lm Zscore = −0.27
NG = 0
DB = 0
SM = 0
Delta_L
Doxorubicin
YDL227C_RF2_102 RF2
ZShift = −0.33
lm ZScore = −0.27
NG = 0
DB = 0
SM = 0
Delta_K
Doxorubicin
YDL227C_RF2_102 RF2
ZShift = −0.46
lm ZScore = 0.55
NG = 0
DB = 0
SM = 0
Delta_r
Doxorubicin
YDL227C_RF2_102 RF2
ZShift = −0.45
lm ZScore = 0.01
NG = 0
DB = 0
SM = 0
Delta_AUC
Doxorubicin

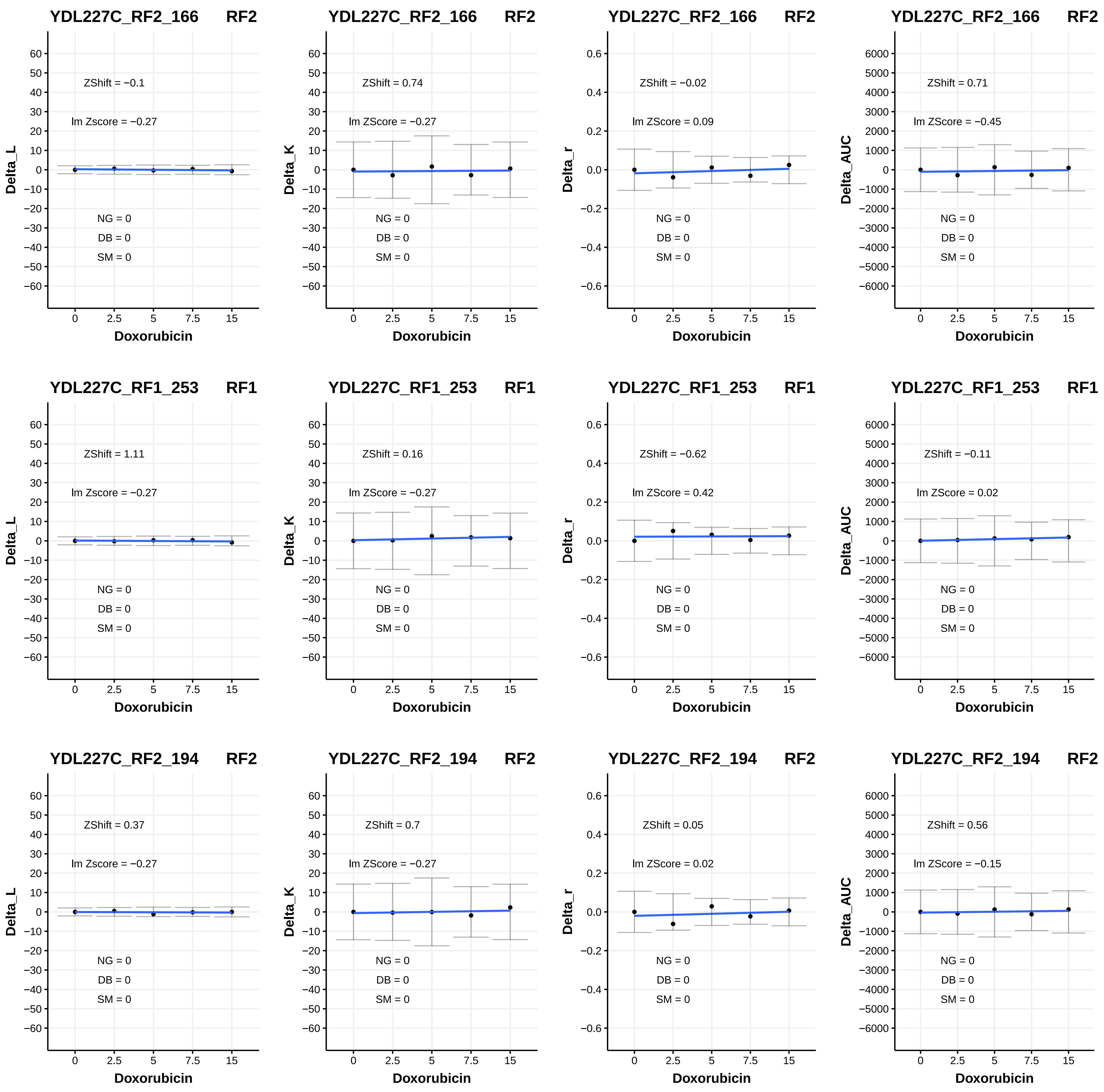

YDL227C_RF2_166 RF2
ZShift = −0.1
lm Zscore = −0.27
NG = 0
DB = 0
SM = 0
Delta_L
Doxorubicin
YDL227C_RF2_166 RF2
ZShift = 0.74
lm ZScore = −0.27
NG = 0
DB = 0
SM = 0
Delta_K
Doxorubicin
YDL227C_RF2_166 RF2
ZShift = −0.02
lm ZScore = 0.09
NG = 0
DB = 0
SM = 0
Delta_r
Doxorubicin
YDL227C_RF2_166 RF2
ZShift = 0.71
lm ZScore = −0.45
NG = 0
DB = 0
SM = 0
Delta_AUC
Doxorubicin
YDL227C_RF1_253 RF1
ZShift = 1.11
lm Zscore = −0.27
NG = 0
DB = 0
SM = 0
Delta_L
Doxorubicin
YDL227C_RF1_253 RF1
ZShift = 0.16
lm ZScore = −0.27
NG = 0
DB = 0
SM = 0
Delta_K
Doxorubicin
YDL227C_RF1_253 RF1
ZShift = −0.62
lm ZScore = 0.42
NG = 0
DB = 0
SM = 0
Delta_r
Doxorubicin
YDL227C_RF1_253 RF1
ZShift = −0.11
lm ZScore = 0.02
NG = 0
DB = 0
SM = 0
Delta_AUC
Doxorubicin
YDL227C_RF2_194 RF2
ZShift = 0.37
lm Zscore = −0.27
NG = 0
DB = 0
SM = 0
Delta_L
Doxorubicin
YDL227C_RF2_194 RF2
ZShift = 0.7
lm ZScore = −0.27
NG = 0
DB = 0
SM = 0
Delta_K
Doxorubicin
YDL227C_RF2_194 RF2
ZShift = 0.05
lm ZScore = 0.02
NG = 0
DB = 0
SM = 0
Delta_r
Doxorubicin
YDL227C_RF2_194 RF2
ZShift = 0.56
lm ZScore = −0.15
NG = 0
DB = 0
SM = 0
Delta_AUC
Doxorubicin

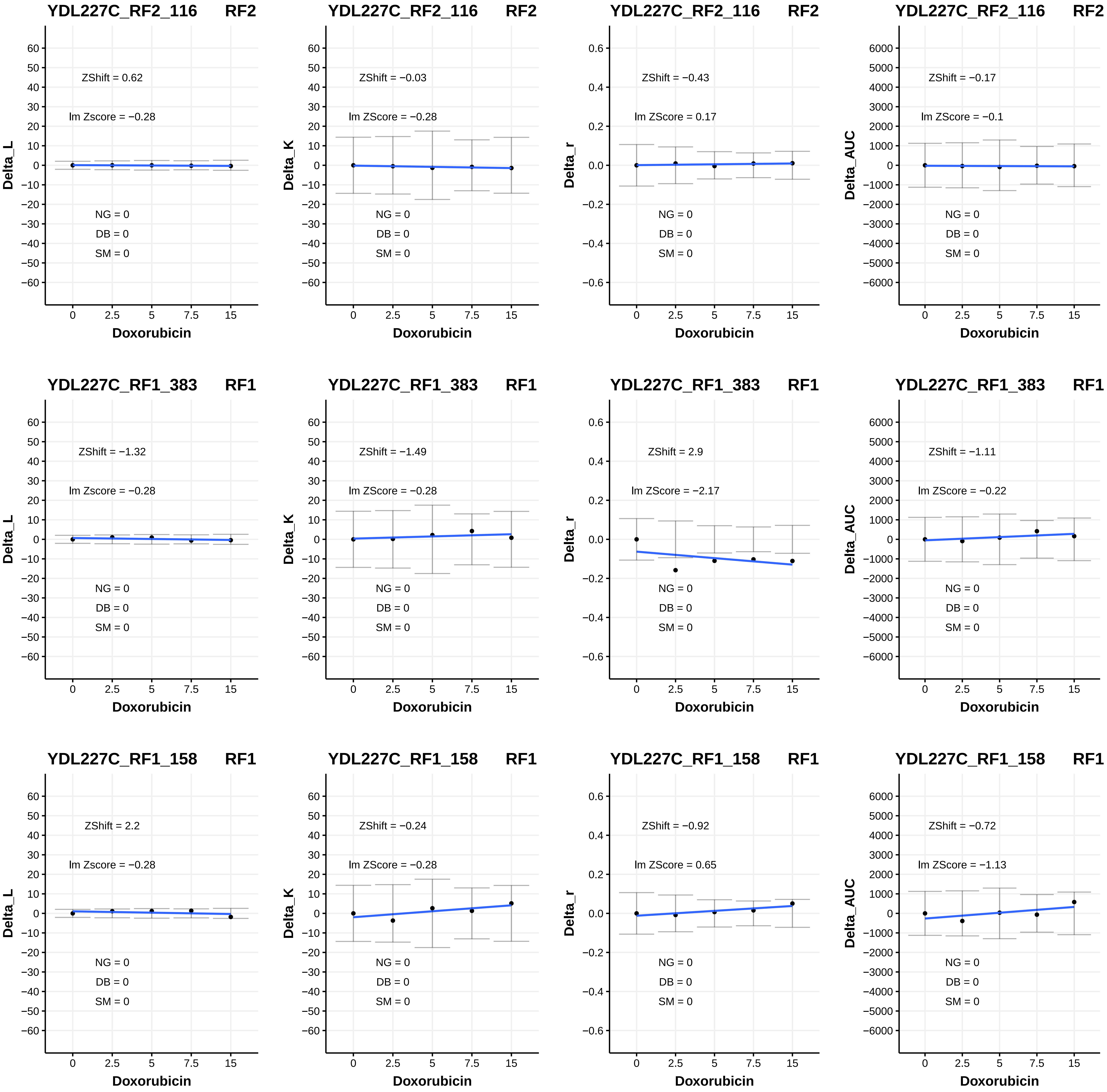
YDL227C_RF2_116 RF2
ZShift = 0.62
lm Zscore = −0.28
NG = 0
DB = 0
SM = 0
Delta_L
Doxorubicin
YDL227C_RF2_116 RF2
ZShift = −0.03
lm ZScore = −0.28
NG = 0
DB = 0
SM = 0
Delta_K
Doxorubicin
YDL227C_RF2_116 RF2
ZShift = −0.43
lm ZScore = 0.17
NG = 0
DB = 0
SM = 0
Delta_r
Doxorubicin
YDL227C_RF2_116 RF2
ZShift = −0.17
lm ZScore = −0.1
NG = 0
DB = 0
SM = 0
Delta_AUC
Doxorubicin
YDL227C_RF1_383 RF1
ZShift = −1.32
lm Zscore = −0.28
NG = 0
DB = 0
SM = 0
Delta_L
Doxorubicin
YDL227C_RF1_383 RF1
ZShift = −1.49
lm ZScore = −0.28
NG = 0
DB = 0
SM = 0
Delta_K
Doxorubicin
YDL227C_RF1_383 RF1
ZShift = 2.9
lm ZScore = −2.17
NG = 0
DB = 0
SM = 0
Delta_r
Doxorubicin
YDL227C_RF1_383 RF1
ZShift = −1.11
lm ZScore = −0.22
NG = 0
DB = 0
SM = 0
Delta_AUC
Doxorubicin
YDL227C_RF1_158 RF1
ZShift = 2.2
lm Zscore = −0.28
NG = 0
DB = 0
SM = 0
Delta_L
Doxorubicin
YDL227C_RF1_158 RF1
ZShift = −0.24
lm ZScore = −0.28
NG = 0
DB = 0
SM = 0
Delta_K
Doxorubicin
YDL227C_RF1_158 RF1
ZShift = −0.92
lm ZScore = 0.65
NG = 0
DB = 0
SM = 0
Delta_r
Doxorubicin
YDL227C_RF1_158 RF1
ZShift = −0.72
lm ZScore = −1.13
NG = 0
DB = 0
SM = 0
Delta_AUC
Doxorubicin

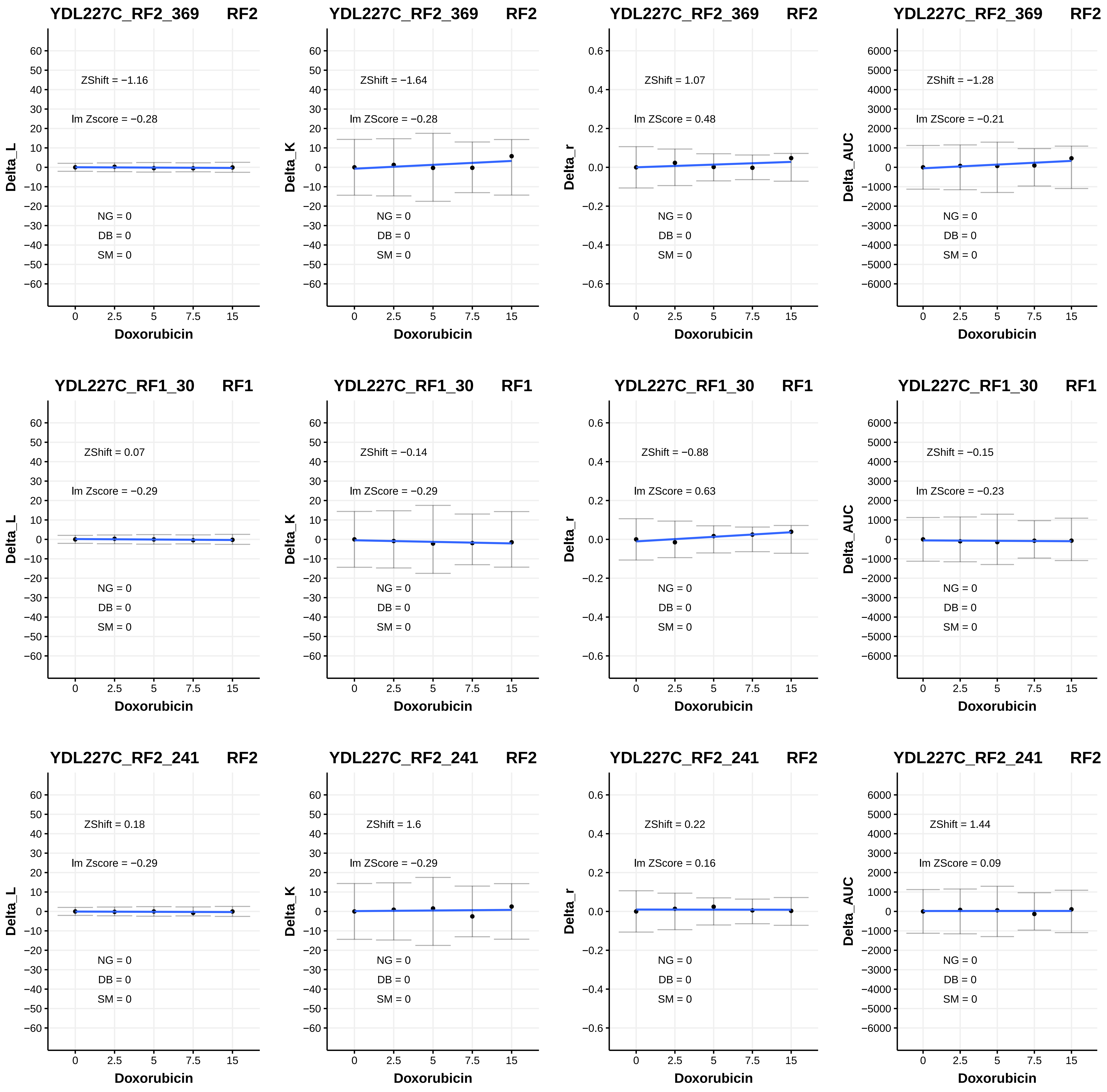

YDL227C_RF2_369 RF2
ZShift = −1.16
lm Zscore = −0.28
NG = 0
DB = 0
SM = 0
Delta_L
Doxorubicin
YDL227C_RF2_369 RF2
ZShift = −1.64
lm ZScore = −0.28
NG = 0
DB = 0
SM = 0
Delta_K
Doxorubicin
YDL227C_RF2_369 RF2
ZShift = 1.07
lm ZScore = 0.48
NG = 0
DB = 0
SM = 0
Delta_r
Doxorubicin
YDL227C_RF2_369 RF2
ZShift = −1.28
lm ZScore = −0.21
NG = 0
DB = 0
SM = 0
Delta_AUC
Doxorubicin
YDL227C_RF1_30 RF1
ZShift = 0.07
lm Zscore = −0.29
NG = 0
DB = 0
SM = 0
Delta_L
Doxorubicin
YDL227C_RF1_30 RF1
ZShift = −0.14
lm ZScore = −0.29
NG = 0
DB = 0
SM = 0
Delta_K
Doxorubicin
YDL227C_RF1_30 RF1
ZShift = −0.88
lm ZScore = 0.63
NG = 0
DB = 0
SM = 0
Delta_r
Doxorubicin
YDL227C_RF1_30 RF1
ZShift = −0.15
lm ZScore = −0.23
NG = 0
DB = 0
SM = 0
Delta_AUC
Doxorubicin
YDL227C_RF2_241 RF2
ZShift = 0.18
lm Zscore = −0.29
NG = 0
DB = 0
SM = 0
Delta_L
Doxorubicin
YDL227C_RF2_241 RF2
ZShift = 1.6
lm ZScore = −0.29
NG = 0
DB = 0
SM = 0
Delta_K
Doxorubicin
YDL227C_RF2_241 RF2
ZShift = 0.22
lm ZScore = 0.16
NG = 0
DB = 0
SM = 0
Delta_r
Doxorubicin
YDL227C_RF2_241 RF2
ZShift = 1.44
lm ZScore = 0.09
NG = 0
DB = 0
SM = 0
Delta_AUC
Doxorubicin

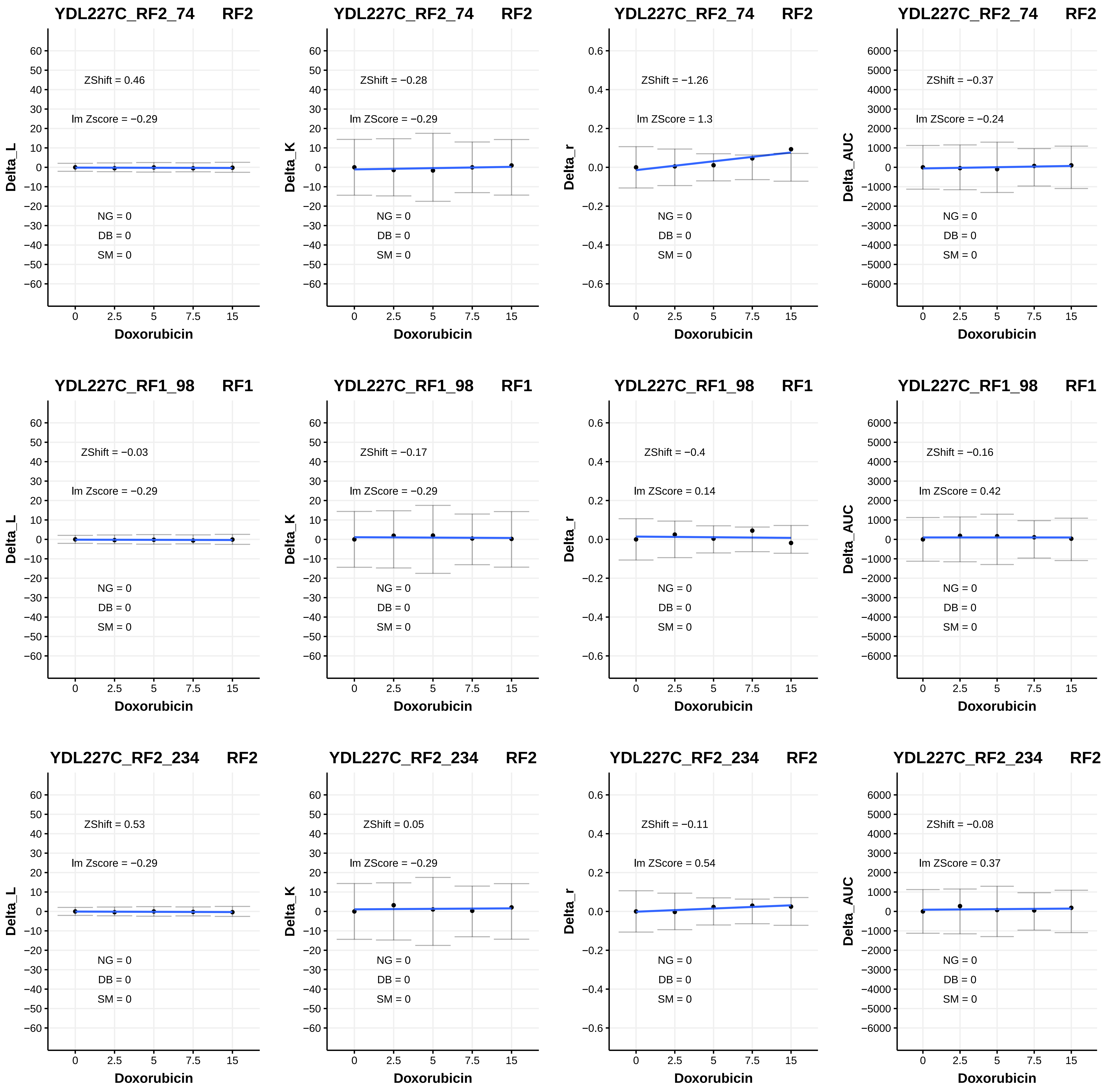
YDL227C_RF2_74 RF2
ZShift = 0.46
lm Zscore = −0.29
NG = 0
DB = 0
SM = 0
Delta_L
Doxorubicin
YDL227C_RF2_74 RF2
ZShift = −0.28
lm ZScore = −0.29
NG = 0
DB = 0
SM = 0
Delta_K
Doxorubicin
YDL227C_RF2_74 RF2
ZShift = −1.26
lm ZScore = 1.3
NG = 0
DB = 0
SM = 0
Delta_r
Doxorubicin
YDL227C_RF2_74 RF2
ZShift = −0.37
lm ZScore = −0.24
NG = 0
DB = 0
SM = 0
Delta_AUC
Doxorubicin
YDL227C_RF1_98 RF1
ZShift = −0.03
lm Zscore = −0.29
NG = 0
DB = 0
SM = 0
Delta_L
Doxorubicin
YDL227C_RF1_98 RF1
ZShift = −0.17
lm ZScore = −0.29
NG = 0
DB = 0
SM = 0
Delta_K
Doxorubicin
YDL227C_RF1_98 RF1
ZShift = −0.4
lm ZScore = 0.14
NG = 0
DB = 0
SM = 0
Delta_r
Doxorubicin
YDL227C_RF1_98 RF1
ZShift = −0.16
lm ZScore = 0.42
NG = 0
DB = 0
SM = 0
Delta_AUC
Doxorubicin
YDL227C_RF2_234 RF2
ZShift = 0.53
lm Zscore = −0.29
NG = 0
DB = 0
SM = 0
Delta_L
Doxorubicin
YDL227C_RF2_234 RF2
ZShift = 0.05
lm ZScore = −0.29
NG = 0
DB = 0
SM = 0
Delta_K
Doxorubicin
YDL227C_RF2_234 RF2
ZShift = −0.11
lm ZScore = 0.54
NG = 0
DB = 0
SM = 0
Delta_r
Doxorubicin
YDL227C_RF2_234 RF2
ZShift = −0.08
lm ZScore = 0.37
NG = 0
DB = 0
SM = 0
Delta_AUC
Doxorubicin

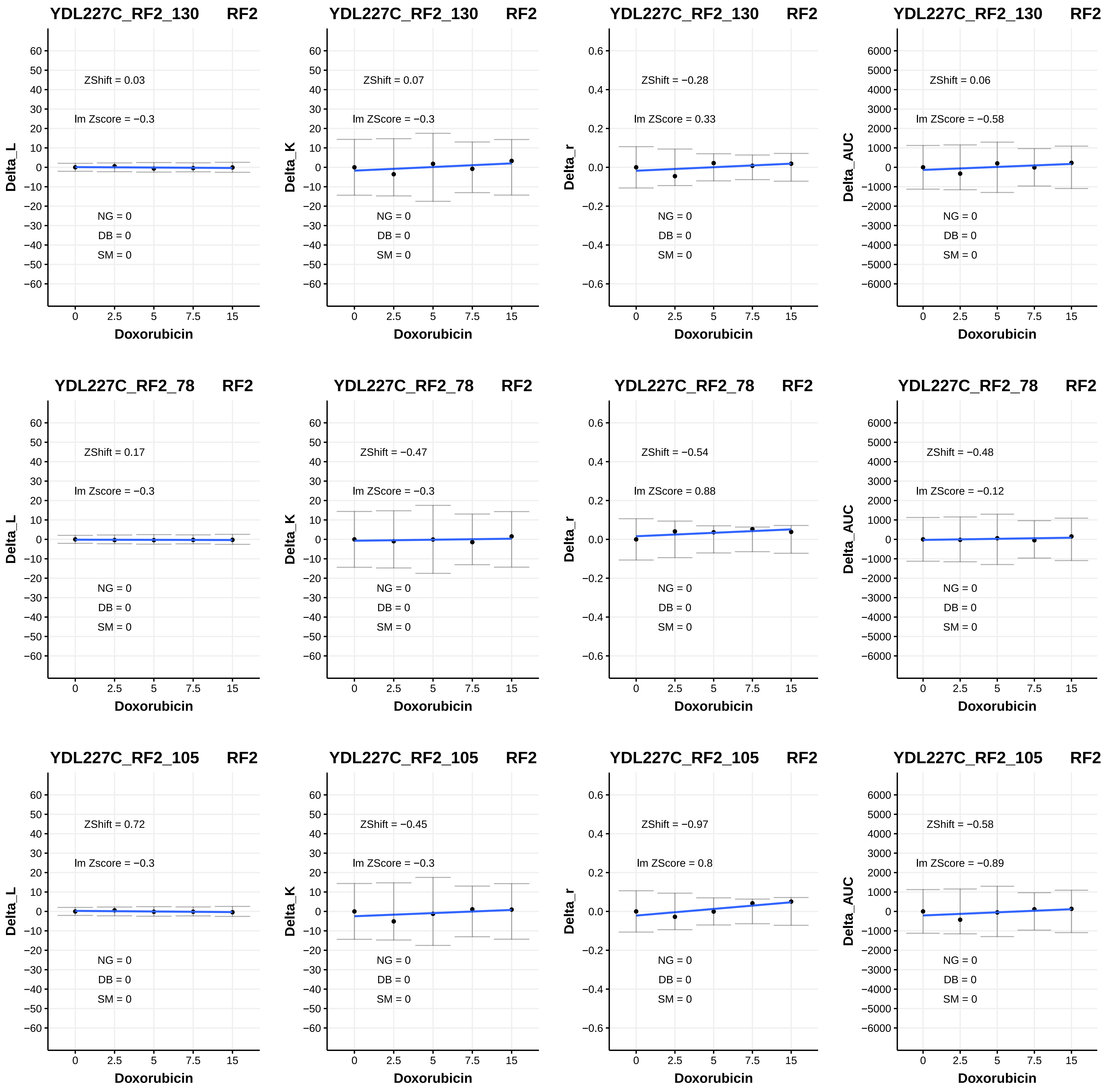
YDL227C_RF2_130 RF2
ZShift = 0.03
lm Zscore = −0.3
NG = 0
DB = 0
SM = 0
Delta_L
Doxorubicin
YDL227C_RF2_130 RF2
ZShift = 0.07
lm ZScore = −0.3
NG = 0
DB = 0
SM = 0
Delta_K
Doxorubicin
YDL227C_RF2_130 RF2
ZShift = −0.28
lm ZScore = 0.33
NG = 0
DB = 0
SM = 0
Delta_r
Doxorubicin
YDL227C_RF2_130 RF2
ZShift = 0.06
lm ZScore = −0.58
NG = 0
DB = 0
SM = 0
Delta_AUC
Doxorubicin
YDL227C_RF2_78 RF2
ZShift = 0.17
lm Zscore = −0.3
NG = 0
DB = 0
SM = 0
Delta_L
Doxorubicin
YDL227C_RF2_78 RF2
ZShift = −0.47
lm ZScore = −0.3
NG = 0
DB = 0
SM = 0
Delta_K
Doxorubicin
YDL227C_RF2_78 RF2
ZShift = −0.54
lm ZScore = 0.88
NG = 0
DB = 0
SM = 0
Delta_r
Doxorubicin
YDL227C_RF2_78 RF2
ZShift = −0.48
lm ZScore = −0.12
NG = 0
DB = 0
SM = 0
Delta_AUC
Doxorubicin
YDL227C_RF2_105 RF2
ZShift = 0.72
lm Zscore = −0.3
NG = 0
DB = 0
SM = 0
Delta_L
Doxorubicin
YDL227C_RF2_105 RF2
ZShift = −0.45
lm ZScore = −0.3
NG = 0
DB = 0
SM = 0
Delta_K
Doxorubicin
YDL227C_RF2_105 RF2
ZShift = −0.97
lm ZScore = 0.8
NG = 0
DB = 0
SM = 0
Delta_r
Doxorubicin
YDL227C_RF2_105 RF2
ZShift = −0.58
lm ZScore = −0.89
NG = 0
DB = 0
SM = 0
Delta_AUC
Doxorubicin

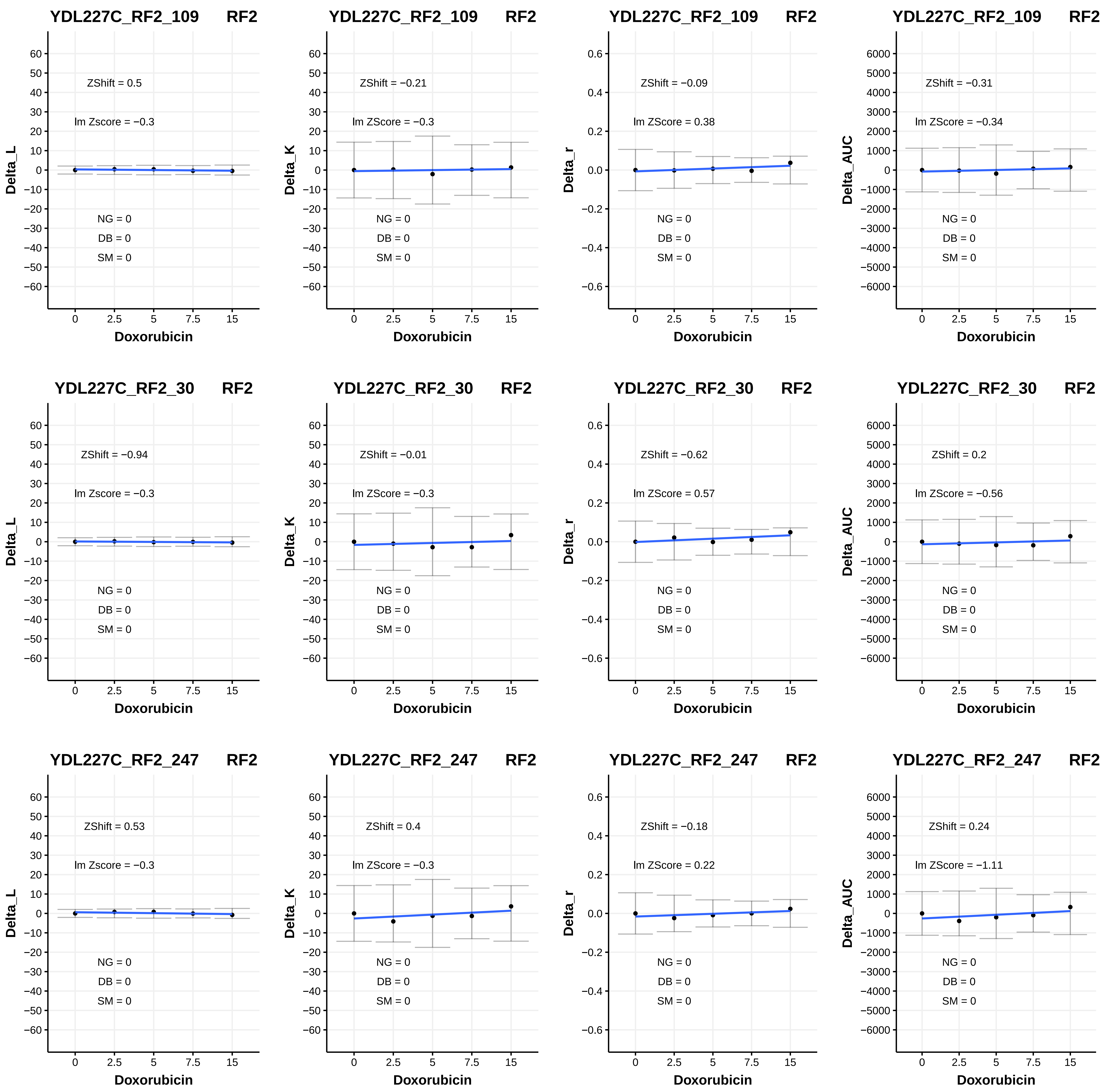

YDL227C_RF2_109 RF2
ZShift = 0.5
lm Zscore = −0.3
NG = 0
DB = 0
SM = 0
Delta_L
Doxorubicin
YDL227C_RF2_109 RF2
ZShift = −0.21
lm ZScore = −0.3
NG = 0
DB = 0
SM = 0
Delta_K
Doxorubicin
YDL227C_RF2_109 RF2
ZShift = −0.09
lm ZScore = 0.38
NG = 0
DB = 0
SM = 0
Delta_r
Doxorubicin
YDL227C_RF2_109 RF2
ZShift = −0.31
lm ZScore = −0.34
NG = 0
DB = 0
SM = 0
Delta_AUC
Doxorubicin
YDL227C_RF2_30 RF2
ZShift = −0.94
lm Zscore = −0.3
NG = 0
DB = 0
SM = 0
Delta_L
Doxorubicin
YDL227C_RF2_30 RF2
ZShift = −0.01
lm ZScore = −0.3
NG = 0
DB = 0
SM = 0
Delta_K
Doxorubicin
YDL227C_RF2_30 RF2
ZShift = −0.62
lm ZScore = 0.57
NG = 0
DB = 0
SM = 0
Delta_r
Doxorubicin
YDL227C_RF2_30 RF2
ZShift = 0.2
lm ZScore = −0.56
NG = 0
DB = 0
SM = 0
Delta_AUC
Doxorubicin
YDL227C_RF2_247 RF2
ZShift = 0.53
lm Zscore = −0.3
NG = 0
DB = 0
SM = 0
Delta_L
Doxorubicin
YDL227C_RF2_247 RF2
ZShift = 0.4
lm ZScore = −0.3
NG = 0
DB = 0
SM = 0
Delta_K
Doxorubicin
YDL227C_RF2_247 RF2
ZShift = −0.18
lm ZScore = 0.22
NG = 0
DB = 0
SM = 0
Delta_r
Doxorubicin
YDL227C_RF2_247 RF2
ZShift = 0.24
lm ZScore = −1.11
NG = 0
DB = 0
SM = 0
Delta_AUC
Doxorubicin

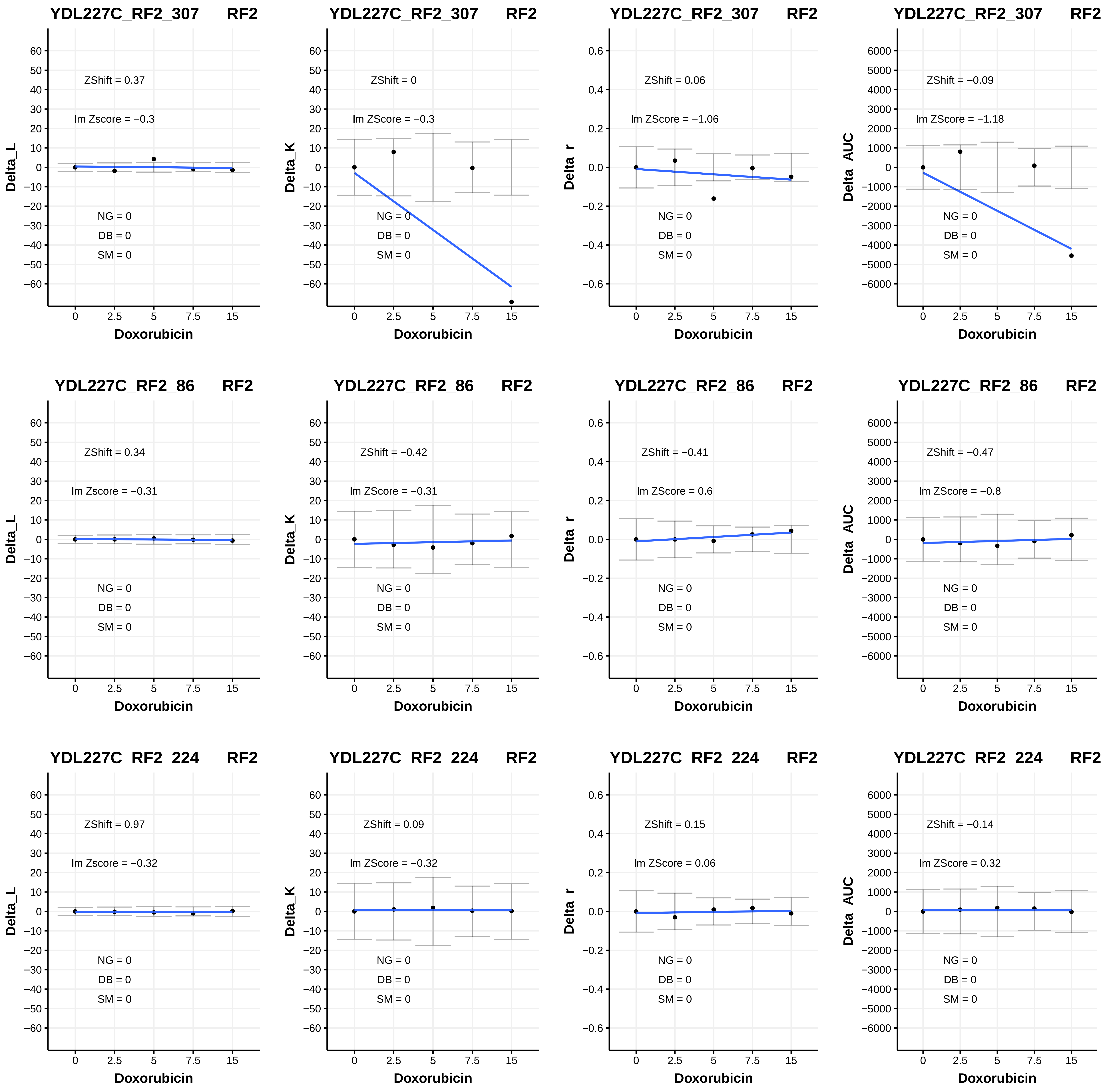
YDL227C_RF2_307 RF2
ZShift = 0.37
lm Zscore = −0.3
NG = 0
DB = 0
SM = 0
Delta_L
Doxorubicin
YDL227C_RF2_307 RF2
ZShift = 0
lm ZScore = −0.3
NG = 0
DB = 0
SM = 0
Delta_K
Doxorubicin
YDL227C_RF2_307 RF2
ZShift = 0.06
lm ZScore = −1.06
NG = 0
DB = 0
SM = 0
Delta_r
Doxorubicin
YDL227C_RF2_307 RF2
ZShift = −0.09
lm ZScore = −1.18
NG = 0
DB = 0
SM = 0
Delta_AUC
Doxorubicin
YDL227C_RF2_86 RF2
ZShift = 0.34
lm Zscore = −0.31
NG = 0
DB = 0
SM = 0
Delta_L
Doxorubicin
YDL227C_RF2_86 RF2
ZShift = −0.42
lm ZScore = −0.31
NG = 0
DB = 0
SM = 0
Delta_K
Doxorubicin
YDL227C_RF2_86 RF2
ZShift = −0.41
lm ZScore = 0.6
NG = 0
DB = 0
SM = 0
Delta_r
Doxorubicin
YDL227C_RF2_86 RF2
ZShift = −0.47
lm ZScore = −0.8
NG = 0
DB = 0
SM = 0
Delta_AUC
Doxorubicin
YDL227C_RF2_224 RF2
ZShift = 0.97
lm Zscore = −0.32
NG = 0
DB = 0
SM = 0
Delta_L
Doxorubicin
YDL227C_RF2_224 RF2
ZShift = 0.09
lm ZScore = −0.32
NG = 0
DB = 0
SM = 0
Delta_K
Doxorubicin
YDL227C_RF2_224 RF2
ZShift = 0.15
lm ZScore = 0.06
NG = 0
DB = 0
SM = 0
Delta_r
Doxorubicin
YDL227C_RF2_224 RF2
ZShift = −0.14
lm ZScore = 0.32
NG = 0
DB = 0
SM = 0
Delta_AUC
Doxorubicin

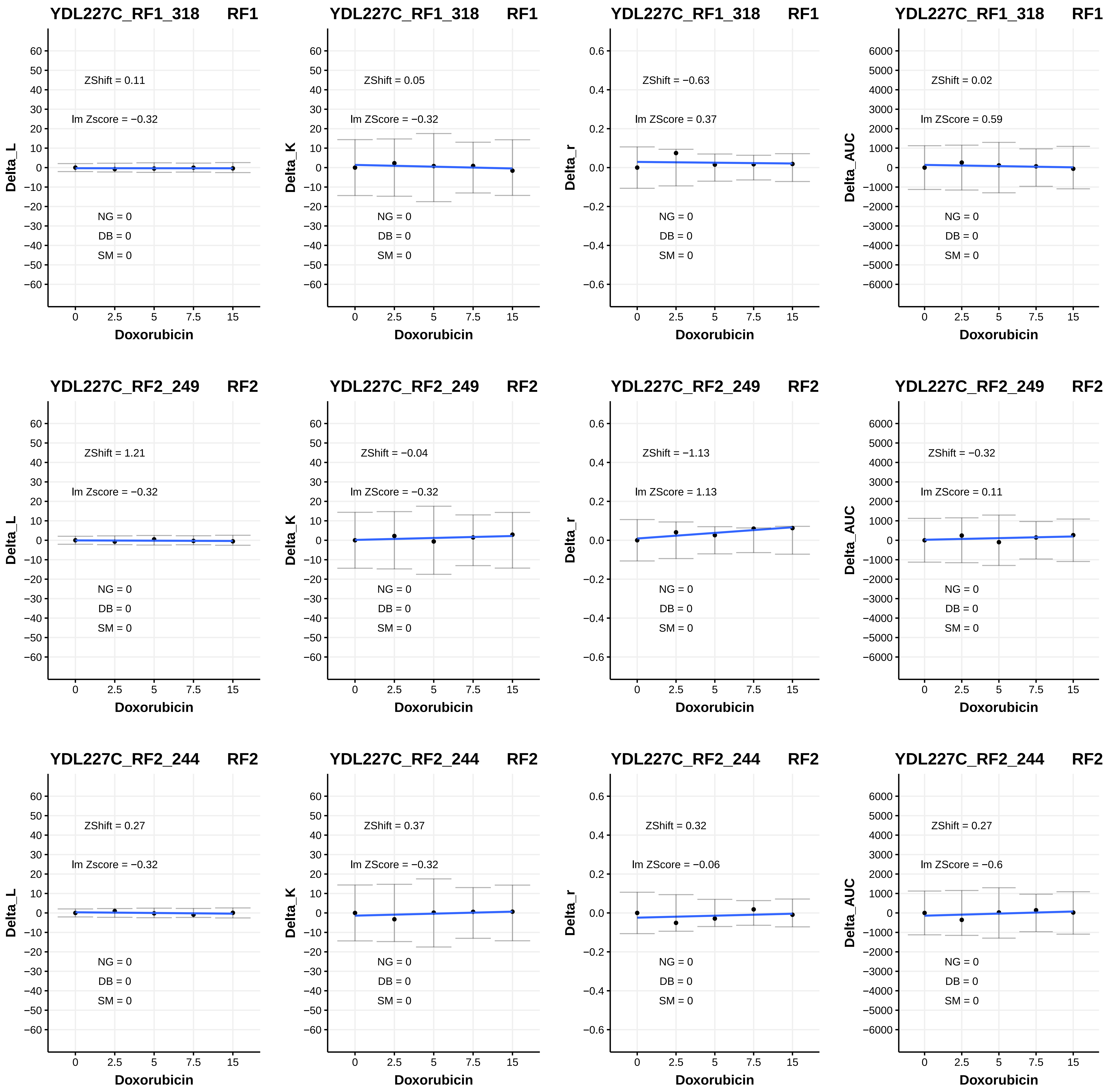
YDL227C_RF1_318 RF1
ZShift = 0.11
lm Zscore = −0.32
NG = 0
DB = 0
SM = 0
Delta_L
Doxorubicin
YDL227C_RF1_318 RF1
ZShift = 0.05
lm ZScore = −0.32
NG = 0
DB = 0
SM = 0
Delta_K
Doxorubicin
YDL227C_RF1_318 RF1
ZShift = −0.63
lm ZScore = 0.37
NG = 0
DB = 0
SM = 0
Delta_r
Doxorubicin
YDL227C_RF1_318 RF1
ZShift = 0.02
lm ZScore = 0.59
NG = 0
DB = 0
SM = 0
Delta_AUC
Doxorubicin
YDL227C_RF2_249 RF2
ZShift = 1.21
lm Zscore = −0.32
NG = 0
DB = 0
SM = 0
Delta_L
Doxorubicin
YDL227C_RF2_249 RF2
ZShift = −0.04
lm ZScore = −0.32
NG = 0
DB = 0
SM = 0
Delta_K
Doxorubicin
YDL227C_RF2_249 RF2
ZShift = −1.13
lm ZScore = 1.13
NG = 0
DB = 0
SM = 0
Delta_r
Doxorubicin
YDL227C_RF2_249 RF2
ZShift = −0.32
lm ZScore = 0.11
NG = 0
DB = 0
SM = 0
Delta_AUC
Doxorubicin
YDL227C_RF2_244 RF2
ZShift = 0.27
lm Zscore = −0.32
NG = 0
DB = 0
SM = 0
Delta_L
Doxorubicin
YDL227C_RF2_244 RF2
ZShift = 0.37
lm ZScore = −0.32
NG = 0
DB = 0
SM = 0
Delta_K
Doxorubicin
YDL227C_RF2_244 RF2
ZShift = 0.32
lm ZScore = −0.06
NG = 0
DB = 0
SM = 0
Delta_r
Doxorubicin
YDL227C_RF2_244 RF2
ZShift = 0.27
lm ZScore = −0.6
NG = 0
DB = 0
SM = 0
Delta_AUC
Doxorubicin

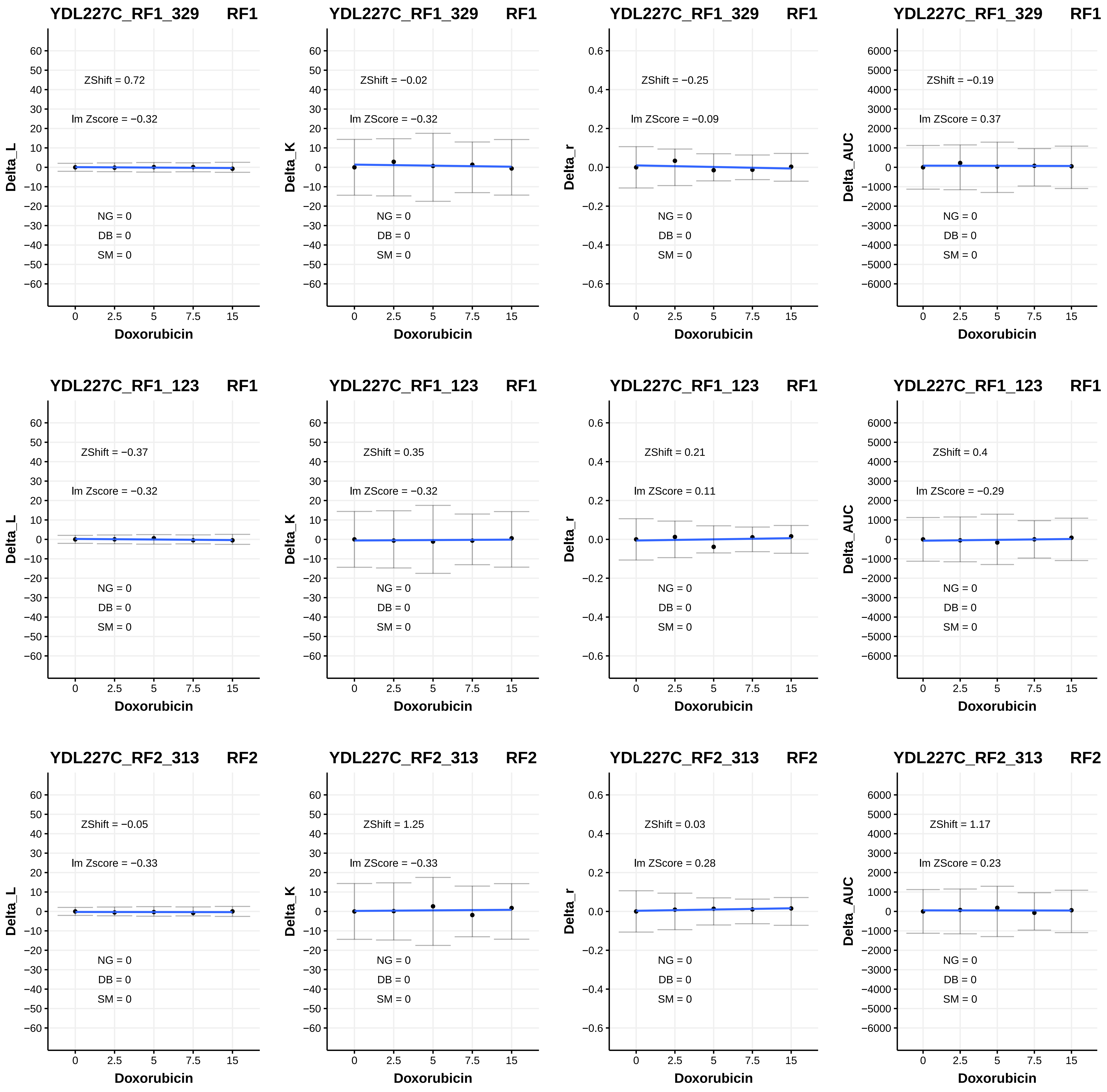

YDL227C_RF1_329 RF1
ZShift = 0.72
lm Zscore = −0.32
NG = 0
DB = 0
SM = 0
Delta_L
Doxorubicin
YDL227C_RF1_329 RF1
ZShift = −0.02
lm ZScore = −0.32
NG = 0
DB = 0
SM = 0
Delta_K
Doxorubicin
YDL227C_RF1_329 RF1
ZShift = −0.25
lm ZScore = −0.09
NG = 0
DB = 0
SM = 0
Delta_r
Doxorubicin
YDL227C_RF1_329 RF1
ZShift = −0.19
lm ZScore = 0.37
NG = 0
DB = 0
SM = 0
Delta_AUC
Doxorubicin
YDL227C_RF1_123 RF1
ZShift = −0.37
lm Zscore = −0.32
NG = 0
DB = 0
SM = 0
Delta_L
Doxorubicin
YDL227C_RF1_123 RF1
ZShift = 0.35
lm ZScore = −0.32
NG = 0
DB = 0
SM = 0
Delta_K
Doxorubicin
YDL227C_RF1_123 RF1
ZShift = 0.21
lm ZScore = 0.11
NG = 0
DB = 0
SM = 0
Delta_r
Doxorubicin
YDL227C_RF1_123 RF1
ZShift = 0.4
lm ZScore = −0.29
NG = 0
DB = 0
SM = 0
Delta_AUC
Doxorubicin
YDL227C_RF2_313 RF2
ZShift = −0.05
lm Zscore = −0.33
NG = 0
DB = 0
SM = 0
Delta_L
Doxorubicin
YDL227C_RF2_313 RF2
ZShift = 1.25
lm ZScore = −0.33
NG = 0
DB = 0
SM = 0
Delta_K
Doxorubicin
YDL227C_RF2_313 RF2
ZShift = 0.03
lm ZScore = 0.28
NG = 0
DB = 0
SM = 0
Delta_r
Doxorubicin
YDL227C_RF2_313 RF2
ZShift = 1.17
lm ZScore = 0.23
NG = 0
DB = 0
SM = 0
Delta_AUC
Doxorubicin

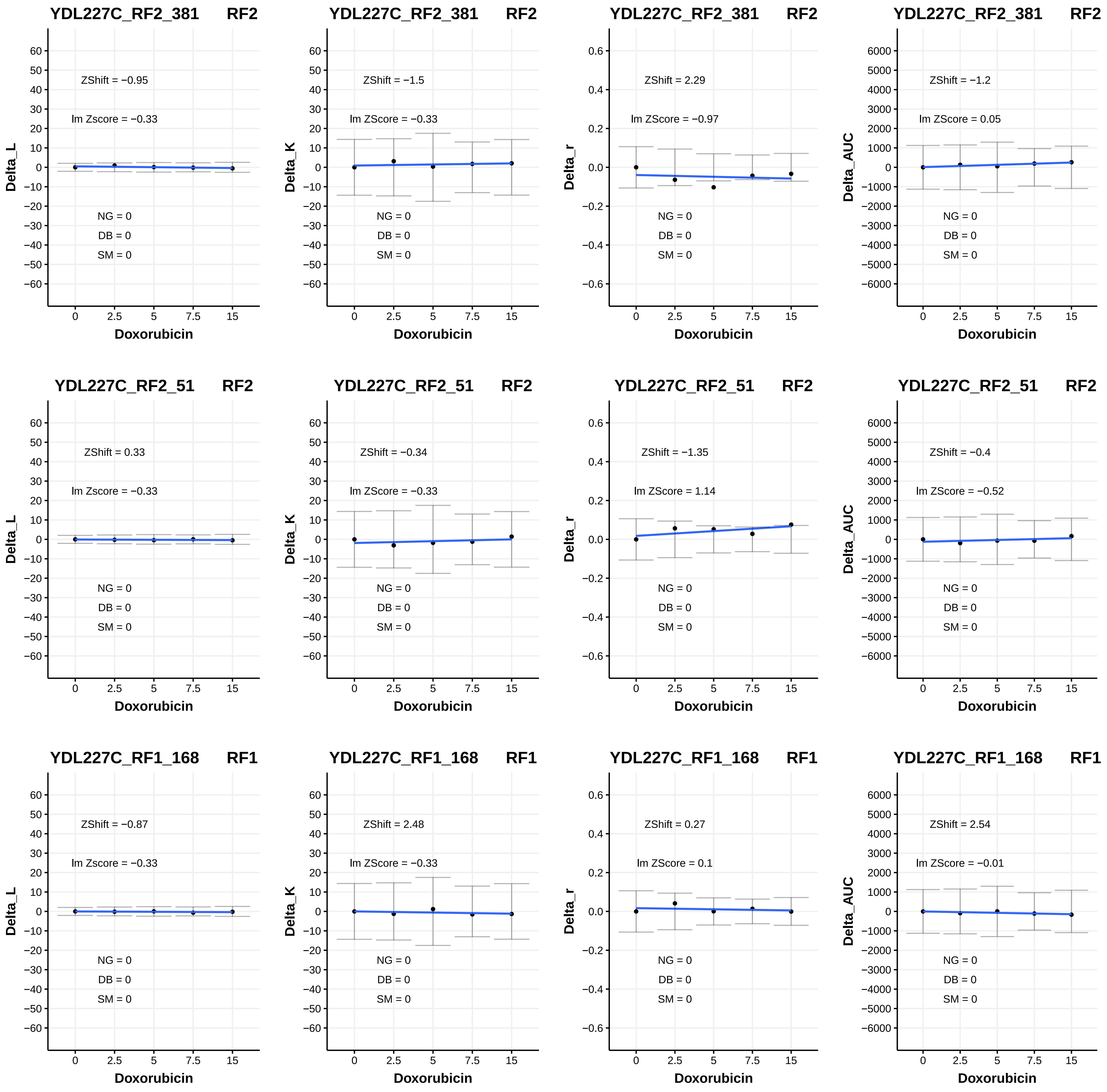
YDL227C_RF2_381 RF2
ZShift = −0.95
lm Zscore = −0.33
NG = 0
DB = 0
SM = 0
Delta_L
Doxorubicin
YDL227C_RF2_381 RF2
ZShift = −1.5
lm ZScore = −0.33
NG = 0
DB = 0
SM = 0
Delta_K
Doxorubicin
YDL227C_RF2_381 RF2
ZShift = 2.29
lm ZScore = −0.97
NG = 0
DB = 0
SM = 0
Delta_r
Doxorubicin
YDL227C_RF2_381 RF2
ZShift = −1.2
lm ZScore = 0.05
NG = 0
DB = 0
SM = 0
Delta_AUC
Doxorubicin
YDL227C_RF2_51 RF2
ZShift = 0.33
lm Zscore = −0.33
NG = 0
DB = 0
SM = 0
Delta_L
Doxorubicin
YDL227C_RF2_51 RF2
ZShift = −0.34
lm ZScore = −0.33
NG = 0
DB = 0
SM = 0
Delta_K
Doxorubicin
YDL227C_RF2_51 RF2
ZShift = −1.35
lm ZScore = 1.14
NG = 0
DB = 0
SM = 0
Delta_r
Doxorubicin
YDL227C_RF2_51 RF2
ZShift = −0.4
lm ZScore = −0.52
NG = 0
DB = 0
SM = 0
Delta_AUC
Doxorubicin
YDL227C_RF1_168 RF1
ZShift = −0.87
lm Zscore = −0.33
NG = 0
DB = 0
SM = 0
Delta_L
Doxorubicin
YDL227C_RF1_168 RF1
ZShift = 2.48
lm ZScore = −0.33
NG = 0
DB = 0
SM = 0
Delta_K
Doxorubicin
YDL227C_RF1_168 RF1
ZShift = 0.27
lm ZScore = 0.1
NG = 0
DB = 0
SM = 0
Delta_r
Doxorubicin
YDL227C_RF1_168 RF1
ZShift = 2.54
lm ZScore = −0.01
NG = 0
DB = 0
SM = 0
Delta_AUC
Doxorubicin

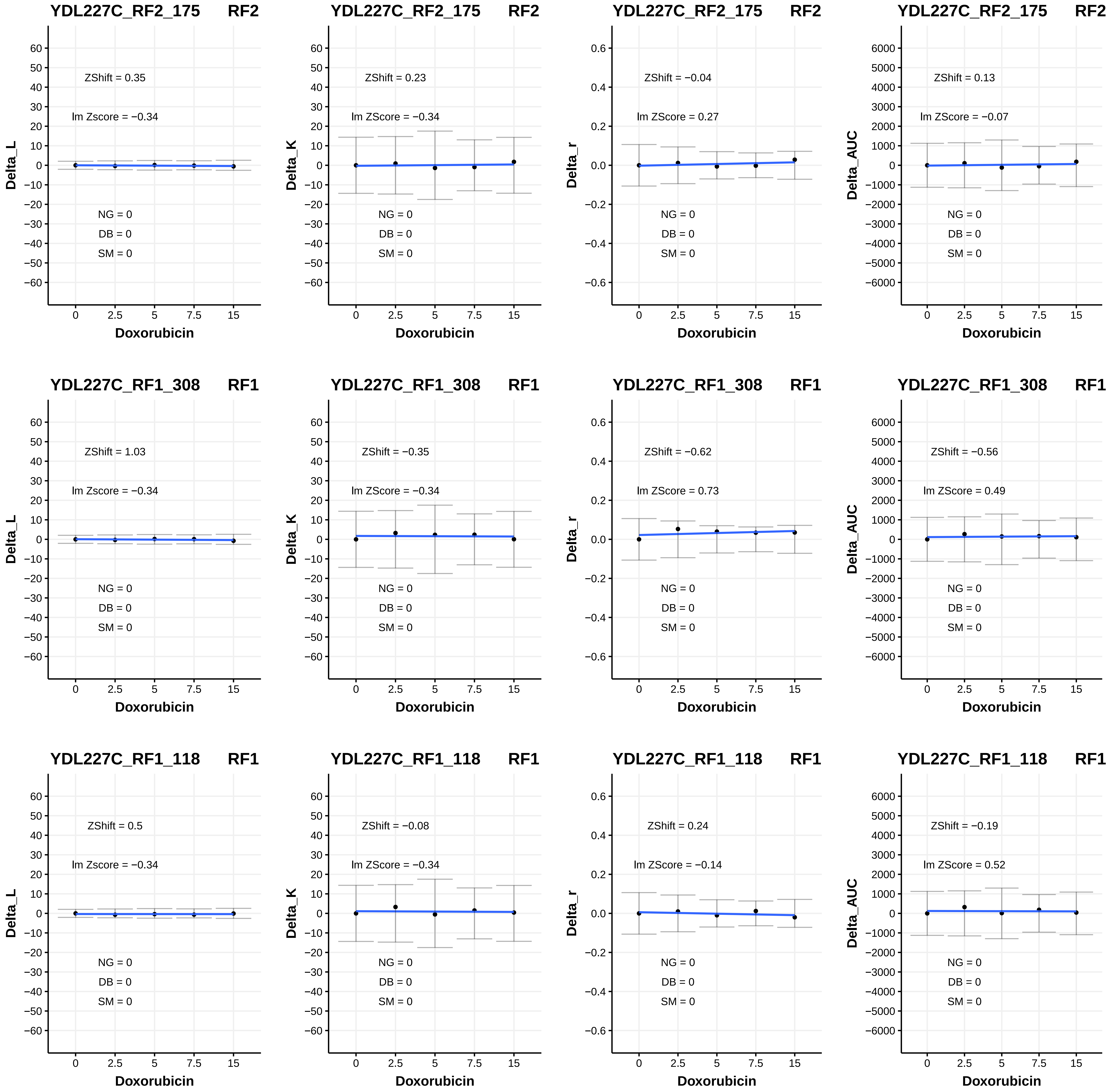
YDL227C_RF2_175 RF2
ZShift = 0.35
lm Zscore = −0.34
NG = 0
DB = 0
SM = 0
Delta_L
Doxorubicin
YDL227C_RF2_175 RF2
ZShift = 0.23
lm ZScore = −0.34
NG = 0
DB = 0
SM = 0
Delta_K
Doxorubicin
YDL227C_RF2_175 RF2
ZShift = −0.04
lm ZScore = 0.27
NG = 0
DB = 0
SM = 0
Delta_r
Doxorubicin
YDL227C_RF2_175 RF2
ZShift = 0.13
lm ZScore = −0.07
NG = 0
DB = 0
SM = 0
Delta_AUC
Doxorubicin
YDL227C_RF1_308 RF1
ZShift = 1.03
lm Zscore = −0.34
NG = 0
DB = 0
SM = 0
Delta_L
Doxorubicin
YDL227C_RF1_308 RF1
ZShift = −0.35
lm ZScore = −0.34
NG = 0
DB = 0
SM = 0
Delta_K
Doxorubicin
YDL227C_RF1_308 RF1
ZShift = −0.62
lm ZScore = 0.73
NG = 0
DB = 0
SM = 0
Delta_r
Doxorubicin
YDL227C_RF1_308 RF1
ZShift = −0.56
lm ZScore = 0.49
NG = 0
DB = 0
SM = 0
Delta_AUC
Doxorubicin
YDL227C_RF1_118 RF1
ZShift = 0.5
lm Zscore = −0.34
NG = 0
DB = 0
SM = 0
Delta_L
Doxorubicin
YDL227C_RF1_118 RF1
ZShift = −0.08
lm ZScore = −0.34
NG = 0
DB = 0
SM = 0
Delta_K
Doxorubicin
YDL227C_RF1_118 RF1
ZShift = 0.24
lm ZScore = −0.14
NG = 0
DB = 0
SM = 0
Delta_r
Doxorubicin
YDL227C_RF1_118 RF1
ZShift = −0.19
lm ZScore = 0.52
NG = 0
DB = 0
SM = 0
Delta_AUC
Doxorubicin

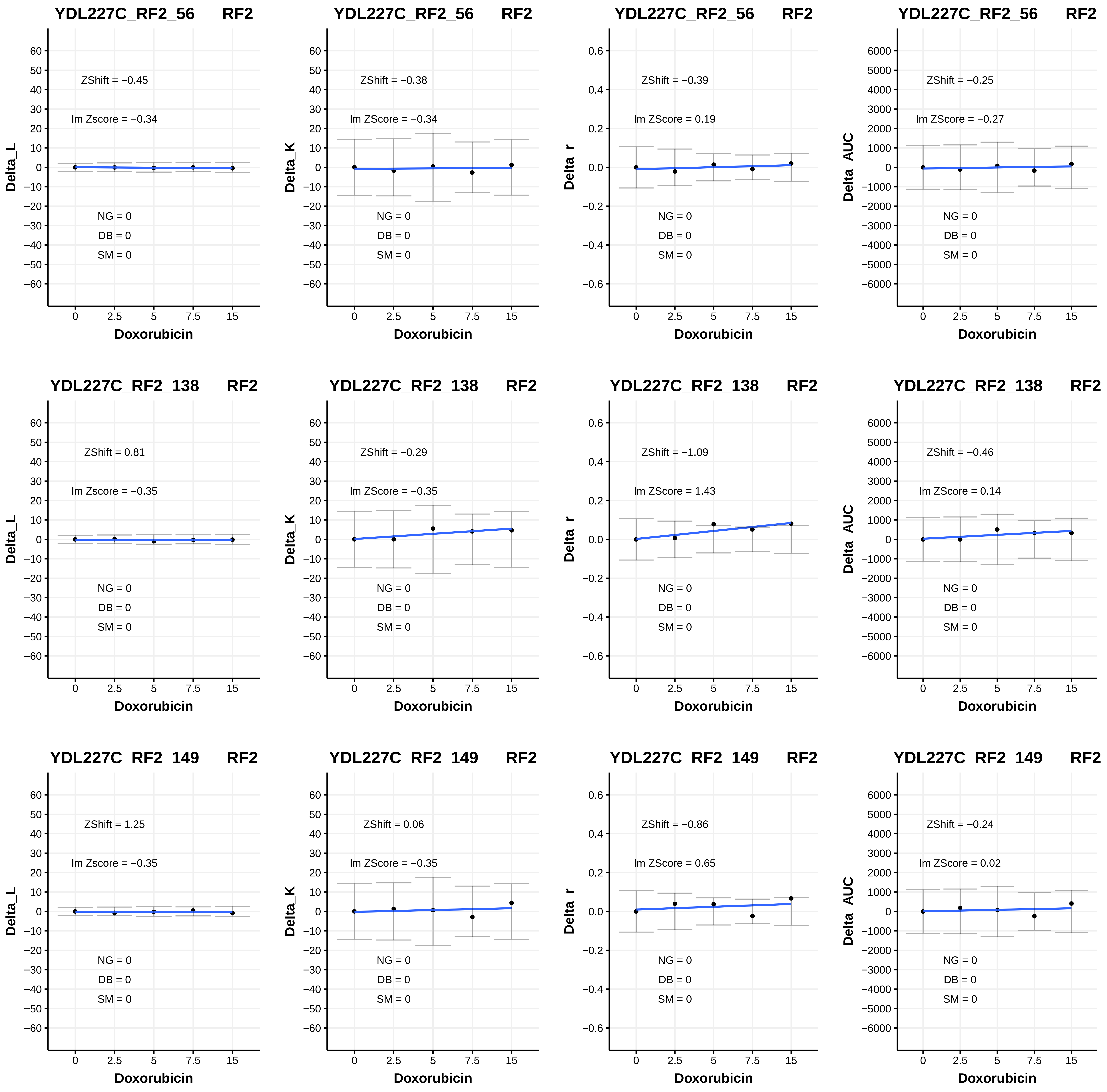
YDL227C_RF2_56 RF2
ZShift = −0.45
lm Zscore = −0.34
NG = 0
DB = 0
SM = 0
Delta_L
Doxorubicin
YDL227C_RF2_56 RF2
ZShift = −0.38
lm ZScore = −0.34
NG = 0
DB = 0
SM = 0
Delta_K
Doxorubicin
YDL227C_RF2_56 RF2
ZShift = −0.39
lm ZScore = 0.19
NG = 0
DB = 0
SM = 0
Delta_r
Doxorubicin
YDL227C_RF2_56 RF2
ZShift = −0.25
lm ZScore = −0.27
NG = 0
DB = 0
SM = 0
Delta_AUC
Doxorubicin
YDL227C_RF2_138 RF2
ZShift = 0.81
lm Zscore = −0.35
NG = 0
DB = 0
SM = 0
Delta_L
Doxorubicin
YDL227C_RF2_138 RF2
ZShift = −0.29
lm ZScore = −0.35
NG = 0
DB = 0
SM = 0
Delta_K
Doxorubicin
YDL227C_RF2_138 RF2
ZShift = −1.09
lm ZScore = 1.43
NG = 0
DB = 0
SM = 0
Delta_r
Doxorubicin
YDL227C_RF2_138 RF2
ZShift = −0.46
lm ZScore = 0.14
NG = 0
DB = 0
SM = 0
Delta_AUC
Doxorubicin
YDL227C_RF2_149 RF2
ZShift = 1.25
lm Zscore = −0.35
NG = 0
DB = 0
SM = 0
Delta_L
Doxorubicin
YDL227C_RF2_149 RF2
ZShift = 0.06
lm ZScore = −0.35
NG = 0
DB = 0
SM = 0
Delta_K
Doxorubicin
YDL227C_RF2_149 RF2
ZShift = −0.86
lm ZScore = 0.65
NG = 0
DB = 0
SM = 0
Delta_r
Doxorubicin
YDL227C_RF2_149 RF2
ZShift = −0.24
lm ZScore = 0.02
NG = 0
DB = 0
SM = 0
Delta_AUC
Doxorubicin

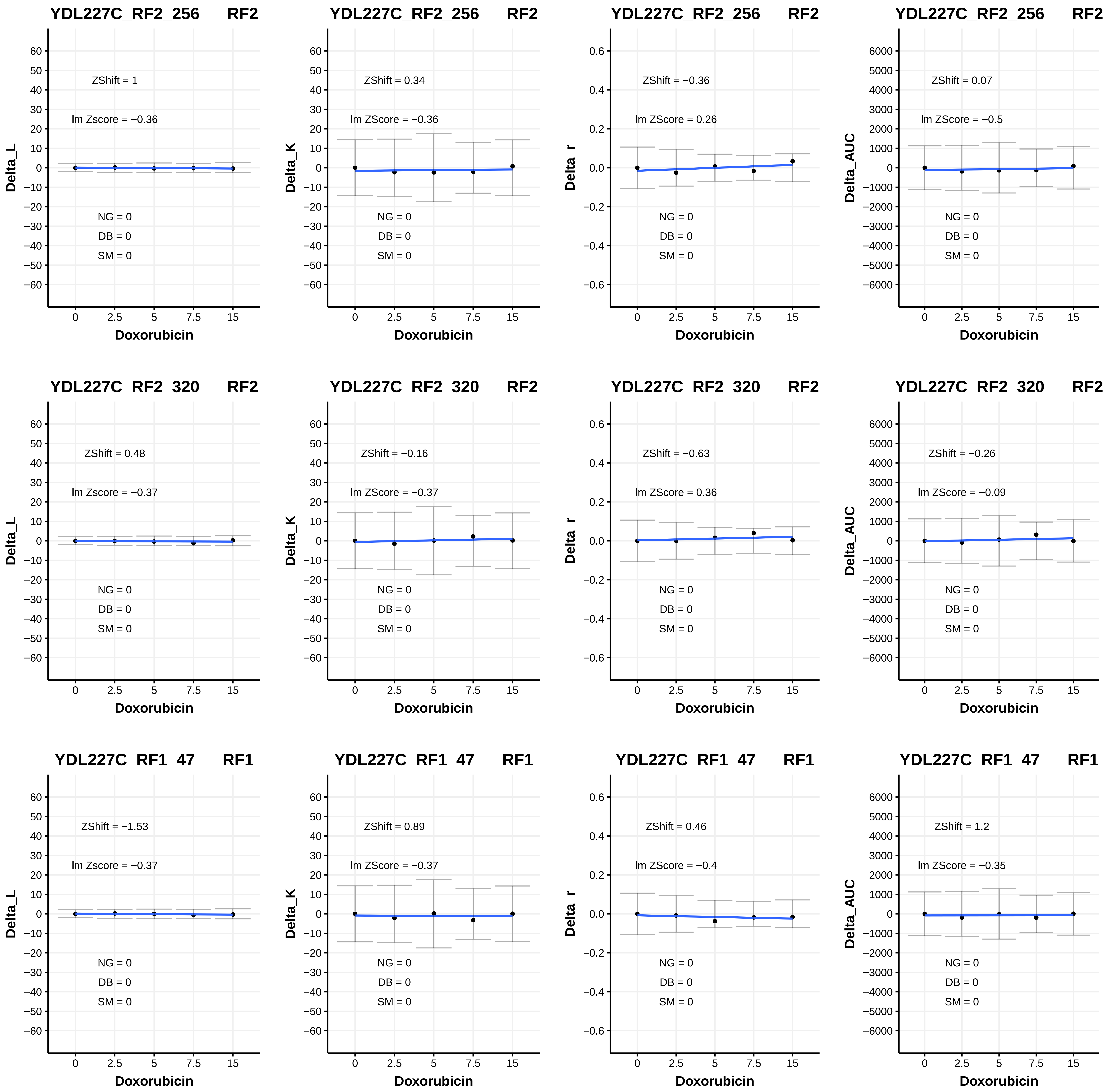

YDL227C_RF2_256 RF2
ZShift = 1
lm Zscore = −0.36
NG = 0
DB = 0
SM = 0
Delta_L
Doxorubicin
YDL227C_RF2_256 RF2
ZShift = 0.34
lm ZScore = −0.36
NG = 0
DB = 0
SM = 0
Delta_K
Doxorubicin
YDL227C_RF2_256 RF2
ZShift = −0.36
lm ZScore = 0.26
NG = 0
DB = 0
SM = 0
Delta_r
Doxorubicin
YDL227C_RF2_256 RF2
ZShift = 0.07
lm ZScore = −0.5
NG = 0
DB = 0
SM = 0
Delta_AUC
Doxorubicin
YDL227C_RF2_320 RF2
ZShift = 0.48
lm Zscore = −0.37
NG = 0
DB = 0
SM = 0
Delta_L
Doxorubicin
YDL227C_RF2_320 RF2
ZShift = −0.16
lm ZScore = −0.37
NG = 0
DB = 0
SM = 0
Delta_K
Doxorubicin
YDL227C_RF2_320 RF2
ZShift = −0.63
lm ZScore = 0.36
NG = 0
DB = 0
SM = 0
Delta_r
Doxorubicin
YDL227C_RF2_320 RF2
ZShift = −0.26
lm ZScore = −0.09
NG = 0
DB = 0
SM = 0
Delta_AUC
Doxorubicin
YDL227C_RF1_47 RF1
ZShift = −1.53
lm Zscore = −0.37
NG = 0
DB = 0
SM = 0
Delta_L
Doxorubicin
YDL227C_RF1_47 RF1
ZShift = 0.89
lm ZScore = −0.37
NG = 0
DB = 0
SM = 0
Delta_K
Doxorubicin
YDL227C_RF1_47 RF1
ZShift = 0.46
lm ZScore = −0.4
NG = 0
DB = 0
SM = 0
Delta_r
Doxorubicin
YDL227C_RF1_47 RF1
ZShift = 1.2
lm ZScore = −0.35
NG = 0
DB = 0
SM = 0
Delta_AUC
Doxorubicin

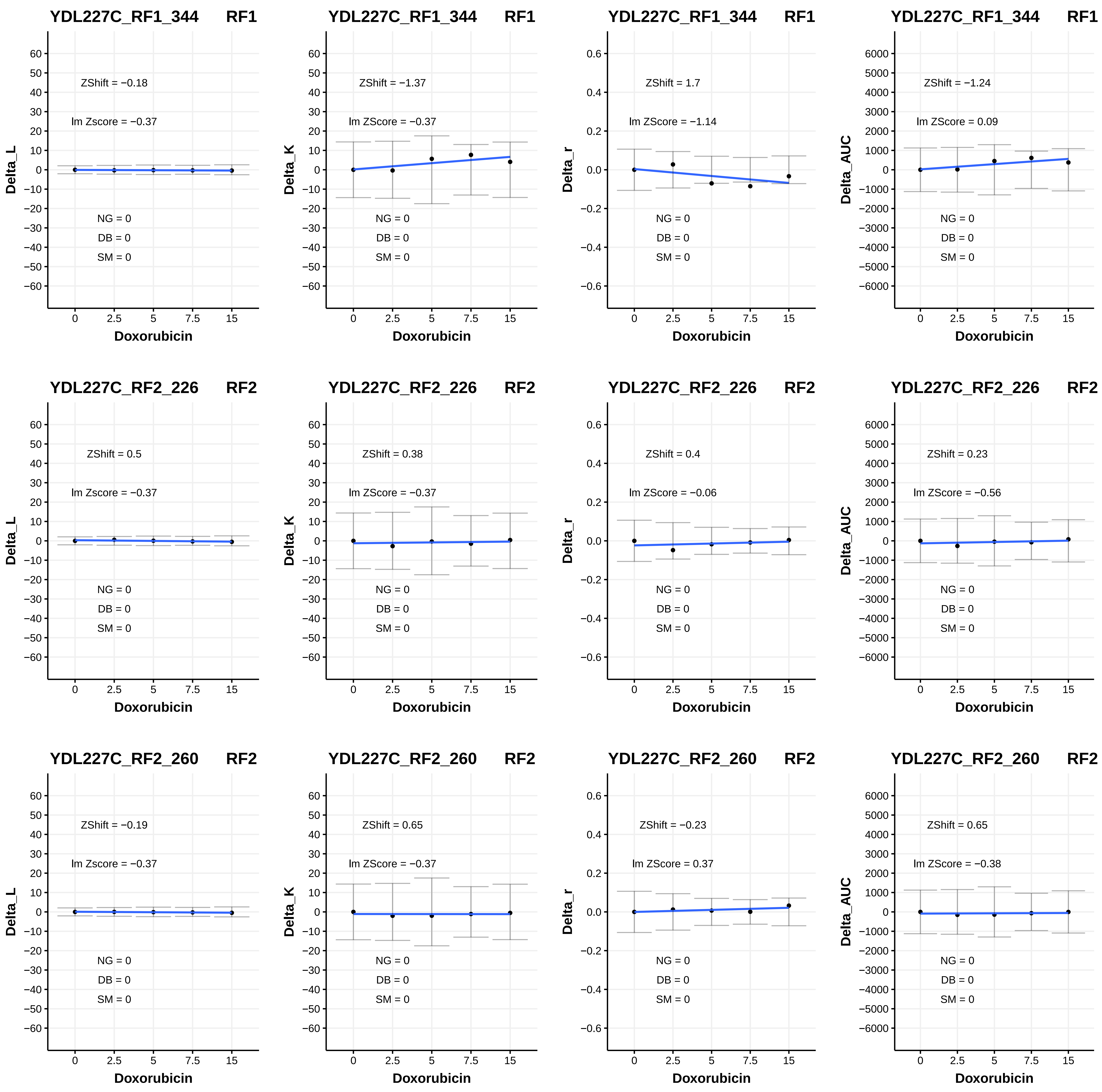
YDL227C_RF1_344 RF1
ZShift = −0.18
lm Zscore = −0.37
NG = 0
DB = 0
SM = 0
Delta_L
Doxorubicin
YDL227C_RF1_344 RF1
ZShift = −1.37
lm ZScore = −0.37
NG = 0
DB = 0
SM = 0
Delta_K
Doxorubicin
YDL227C_RF1_344 RF1
ZShift = 1.7
lm ZScore = −1.14
NG = 0
DB = 0
SM = 0
Delta_r
Doxorubicin
YDL227C_RF1_344 RF1
ZShift = −1.24
lm ZScore = 0.09
NG = 0
DB = 0
SM = 0
Delta_AUC
Doxorubicin
YDL227C_RF2_226 RF2
ZShift = 0.5
lm Zscore = −0.37
NG = 0
DB = 0
SM = 0
Delta_L
Doxorubicin
YDL227C_RF2_226 RF2
ZShift = 0.38
lm ZScore = −0.37
NG = 0
DB = 0
SM = 0
Delta_K
Doxorubicin
YDL227C_RF2_226 RF2
ZShift = 0.4
lm ZScore = −0.06
NG = 0
DB = 0
SM = 0
Delta_r
Doxorubicin
YDL227C_RF2_226 RF2
ZShift = 0.23
lm ZScore = −0.56
NG = 0
DB = 0
SM = 0
Delta_AUC
Doxorubicin
YDL227C_RF2_260 RF2
ZShift = −0.19
lm Zscore = −0.37
NG = 0
DB = 0
SM = 0
Delta_L
Doxorubicin
YDL227C_RF2_260 RF2
ZShift = 0.65
lm ZScore = −0.37
NG = 0
DB = 0
SM = 0
Delta_K
Doxorubicin
YDL227C_RF2_260 RF2
ZShift = −0.23
lm ZScore = 0.37
NG = 0
DB = 0
SM = 0
Delta_r
Doxorubicin
YDL227C_RF2_260 RF2
ZShift = 0.65
lm ZScore = −0.38
NG = 0
DB = 0
SM = 0
Delta_AUC
Doxorubicin

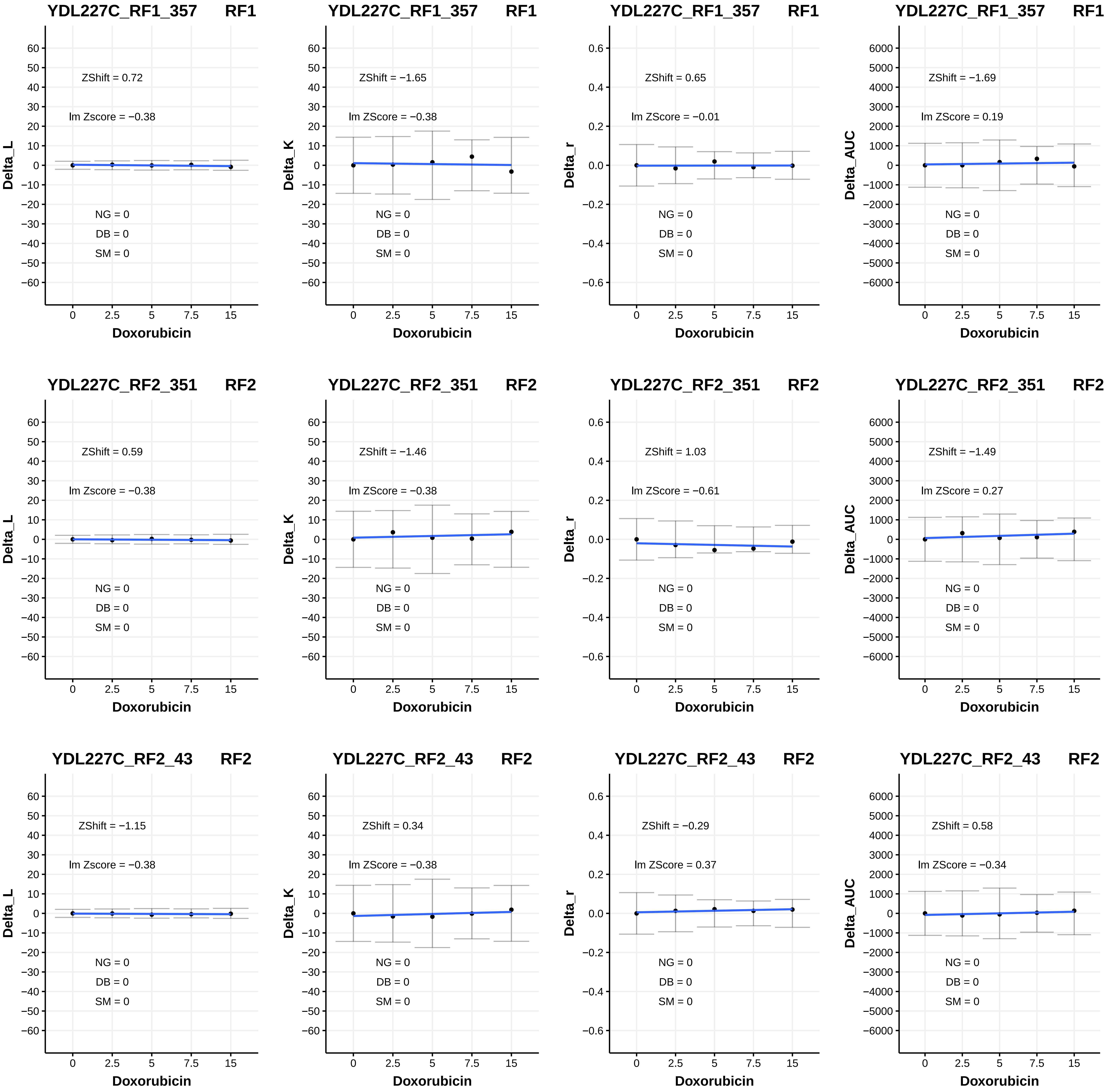
YDL227C_RF1_357 RF1
ZShift = 0.72
lm Zscore = −0.38
NG = 0
DB = 0
SM = 0
Delta_L
Doxorubicin
YDL227C_RF1_357 RF1
ZShift = −1.65
lm ZScore = −0.38
NG = 0
DB = 0
SM = 0
Delta_K
Doxorubicin
YDL227C_RF1_357 RF1
ZShift = 0.65
lm ZScore = −0.01
NG = 0
DB = 0
SM = 0
Delta_r
Doxorubicin
YDL227C_RF1_357 RF1
ZShift = −1.69
lm ZScore = 0.19
NG = 0
DB = 0
SM = 0
Delta_AUC
Doxorubicin
YDL227C_RF2_351 RF2
ZShift = 0.59
lm Zscore = −0.38
NG = 0
DB = 0
SM = 0
Delta_L
Doxorubicin
YDL227C_RF2_351 RF2
ZShift = −1.46
lm ZScore = −0.38
NG = 0
DB = 0
SM = 0
Delta_K
Doxorubicin
YDL227C_RF2_351 RF2
ZShift = 1.03
lm ZScore = −0.61
NG = 0
DB = 0
SM = 0
Delta_r
Doxorubicin
YDL227C_RF2_351 RF2
ZShift = −1.49
lm ZScore = 0.27
NG = 0
DB = 0
SM = 0
Delta_AUC
Doxorubicin
YDL227C_RF2_43 RF2
ZShift = −1.15
lm Zscore = −0.38
NG = 0
DB = 0
SM = 0
Delta_L
Doxorubicin
YDL227C_RF2_43 RF2
ZShift = 0.34
lm ZScore = −0.38
NG = 0
DB = 0
SM = 0
Delta_K
Doxorubicin
YDL227C_RF2_43 RF2
ZShift = −0.29
lm ZScore = 0.37
NG = 0
DB = 0
SM = 0
Delta_r
Doxorubicin
YDL227C_RF2_43 RF2
ZShift = 0.58
lm ZScore = −0.34
NG = 0
DB = 0
SM = 0
Delta_AUC
Doxorubicin

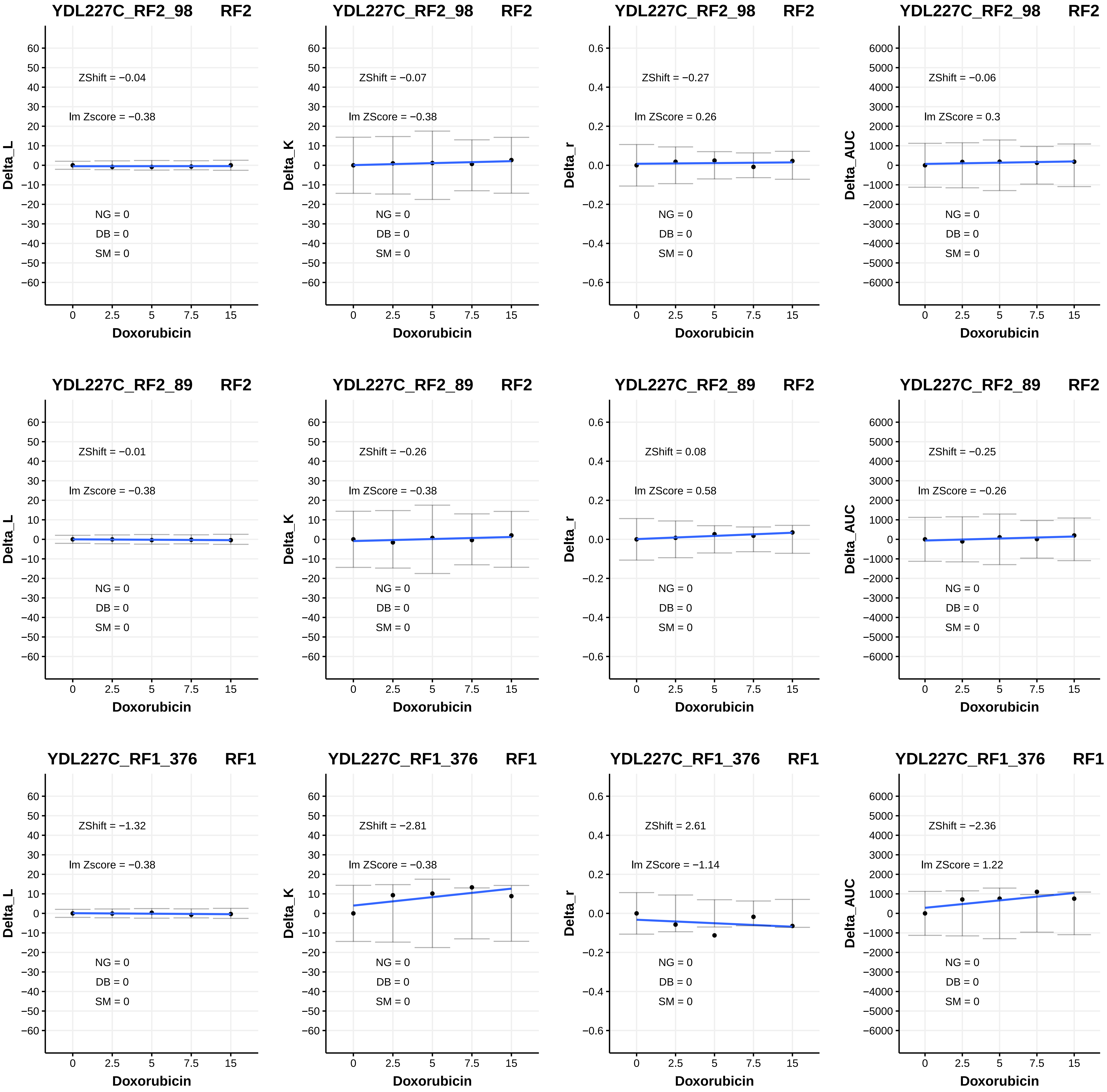
YDL227C_RF2_98 RF2
ZShift = −0.04
lm Zscore = −0.38
NG = 0
DB = 0
SM = 0
Delta_L
Doxorubicin
YDL227C_RF2_98 RF2
ZShift = −0.07
lm ZScore = −0.38
NG = 0
DB = 0
SM = 0
Delta_K
Doxorubicin
YDL227C_RF2_98 RF2
ZShift = −0.27
lm ZScore = 0.26
NG = 0
DB = 0
SM = 0
Delta_r
Doxorubicin
YDL227C_RF2_98 RF2
ZShift = −0.06
lm ZScore = 0.3
NG = 0
DB = 0
SM = 0
Delta_AUC
Doxorubicin
YDL227C_RF2_89 RF2
ZShift = −0.01
lm Zscore = −0.38
NG = 0
DB = 0
SM = 0
Delta_L
Doxorubicin
YDL227C_RF2_89 RF2
ZShift = −0.26
lm ZScore = −0.38
NG = 0
DB = 0
SM = 0
Delta_K
Doxorubicin
YDL227C_RF2_89 RF2
ZShift = 0.08
lm ZScore = 0.58
NG = 0
DB = 0
SM = 0
Delta_r
Doxorubicin
YDL227C_RF2_89 RF2
ZShift = −0.25
lm ZScore = −0.26
NG = 0
DB = 0
SM = 0
Delta_AUC
Doxorubicin
YDL227C_RF1_376 RF1
ZShift = −1.32
lm Zscore = −0.38
NG = 0
DB = 0
SM = 0
Delta_L
Doxorubicin
YDL227C_RF1_376 RF1
ZShift = −2.81
lm ZScore = −0.38
NG = 0
DB = 0
SM = 0
Delta_K
Doxorubicin
YDL227C_RF1_376 RF1
ZShift = 2.61
lm ZScore = −1.14
NG = 0
DB = 0
SM = 0
Delta_r
Doxorubicin
YDL227C_RF1_376 RF1
ZShift = −2.36
lm ZScore = 1.22
NG = 0
DB = 0
SM = 0
Delta_AUC
Doxorubicin

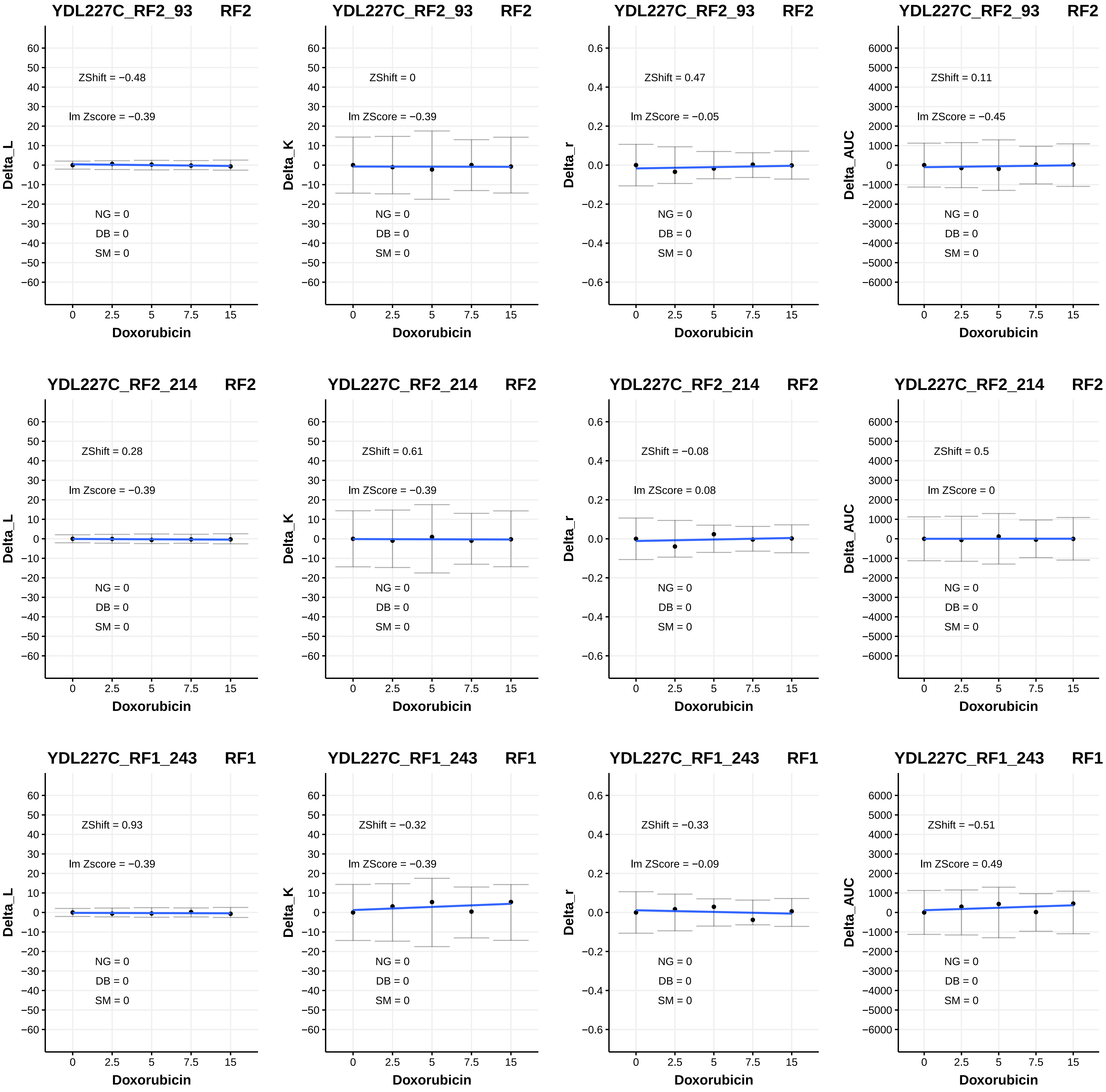
YDL227C_RF2_93 RF2
ZShift = −0.48
lm Zscore = −0.39
NG = 0
DB = 0
SM = 0
Delta_L
Doxorubicin
YDL227C_RF2_93 RF2
ZShift = 0
lm ZScore = −0.39
NG = 0
DB = 0
SM = 0
Delta_K
Doxorubicin
YDL227C_RF2_93 RF2
ZShift = 0.47
lm ZScore = −0.05
NG = 0
DB = 0
SM = 0
Delta_r
Doxorubicin
YDL227C_RF2_93 RF2
ZShift = 0.11
lm ZScore = −0.45
NG = 0
DB = 0
SM = 0
Delta_AUC
Doxorubicin
YDL227C_RF2_214 RF2
ZShift = 0.28
lm Zscore = −0.39
NG = 0
DB = 0
SM = 0
Delta_L
Doxorubicin
YDL227C_RF2_214 RF2
ZShift = 0.61
lm ZScore = −0.39
NG = 0
DB = 0
SM = 0
Delta_K
Doxorubicin
YDL227C_RF2_214 RF2
ZShift = −0.08
lm ZScore = 0.08
NG = 0
DB = 0
SM = 0
Delta_r
Doxorubicin
YDL227C_RF2_214 RF2
ZShift = 0.5
lm ZScore = 0
NG = 0
DB = 0
SM = 0
Delta_AUC
Doxorubicin
YDL227C_RF1_243 RF1
ZShift = 0.93
lm Zscore = −0.39
NG = 0
DB = 0
SM = 0
Delta_L
Doxorubicin
YDL227C_RF1_243 RF1
ZShift = −0.32
lm ZScore = −0.39
NG = 0
DB = 0
SM = 0
Delta_K
Doxorubicin
YDL227C_RF1_243 RF1
ZShift = −0.33
lm ZScore = −0.09
NG = 0
DB = 0
SM = 0
Delta_r
Doxorubicin
YDL227C_RF1_243 RF1
ZShift = −0.51
lm ZScore = 0.49
NG = 0
DB = 0
SM = 0
Delta_AUC
Doxorubicin

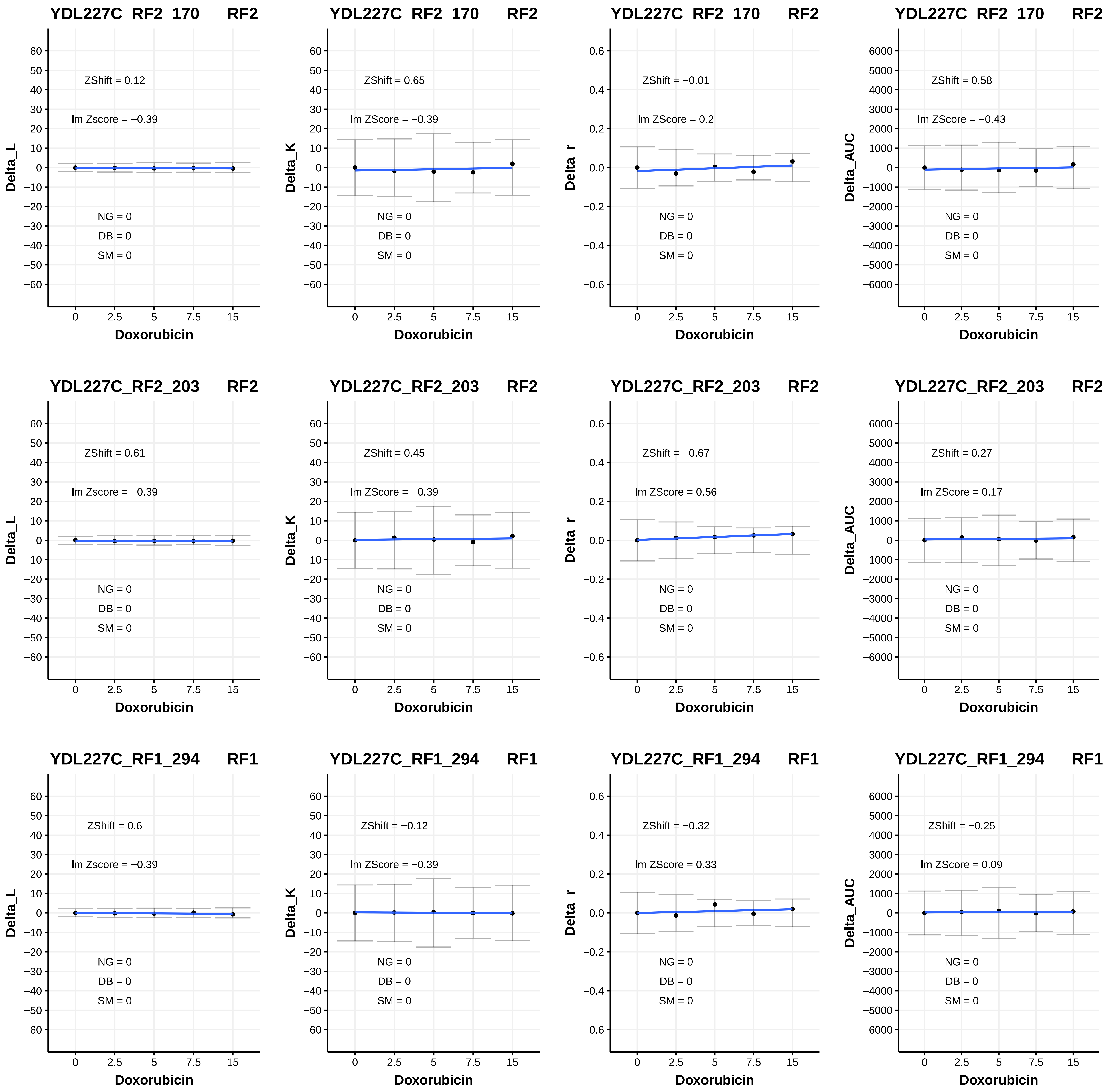

YDL227C_RF2_170 RF2
ZShift = 0.12
lm Zscore = −0.39
NG = 0
DB = 0
SM = 0
Delta_L
Doxorubicin
YDL227C_RF2_170 RF2
ZShift = 0.65
lm ZScore = −0.39
NG = 0
DB = 0
SM = 0
Delta_K
Doxorubicin
YDL227C_RF2_170 RF2
ZShift = −0.01
lm ZScore = 0.2
NG = 0
DB = 0
SM = 0
Delta_r
Doxorubicin
YDL227C_RF2_170 RF2
ZShift = 0.58
lm ZScore = −0.43
NG = 0
DB = 0
SM = 0
Delta_AUC
Doxorubicin
YDL227C_RF2_203 RF2
ZShift = 0.61
lm Zscore = −0.39
NG = 0
DB = 0
SM = 0
Delta_L
Doxorubicin
YDL227C_RF2_203 RF2
ZShift = 0.45
lm ZScore = −0.39
NG = 0
DB = 0
SM = 0
Delta_K
Doxorubicin
YDL227C_RF2_203 RF2
ZShift = −0.67
lm ZScore = 0.56
NG = 0
DB = 0
SM = 0
Delta_r
Doxorubicin
YDL227C_RF2_203 RF2
ZShift = 0.27
lm ZScore = 0.17
NG = 0
DB = 0
SM = 0
Delta_AUC
Doxorubicin
YDL227C_RF1_294 RF1
ZShift = 0.6
lm Zscore = −0.39
NG = 0
DB = 0
SM = 0
Delta_L
Doxorubicin
YDL227C_RF1_294 RF1
ZShift = −0.12
lm ZScore = −0.39
NG = 0
DB = 0
SM = 0
Delta_K
Doxorubicin
YDL227C_RF1_294 RF1
ZShift = −0.32
lm ZScore = 0.33
NG = 0
DB = 0
SM = 0
Delta_r
Doxorubicin
YDL227C_RF1_294 RF1
ZShift = −0.25
lm ZScore = 0.09
NG = 0
DB = 0
SM = 0
Delta_AUC
Doxorubicin

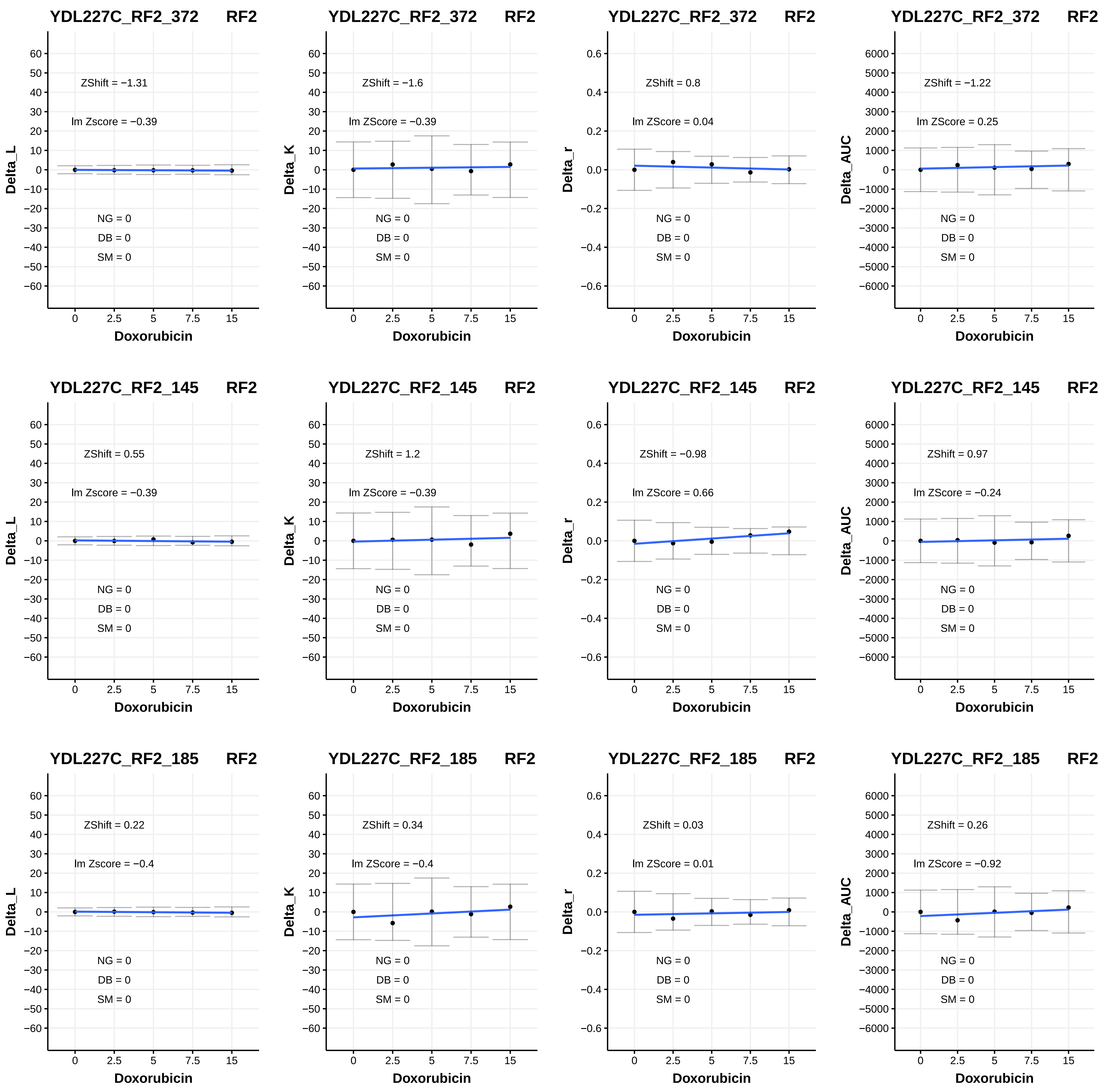
YDL227C_RF2_372 RF2
ZShift = −1.31
lm Zscore = −0.39
NG = 0
DB = 0
SM = 0
Delta_L
Doxorubicin
YDL227C_RF2_372 RF2
ZShift = −1.6
lm ZScore = −0.39
NG = 0
DB = 0
SM = 0
Delta_K
Doxorubicin
YDL227C_RF2_372 RF2
ZShift = 0.8
lm ZScore = 0.04
NG = 0
DB = 0
SM = 0
Delta_r
Doxorubicin
YDL227C_RF2_372 RF2
ZShift = −1.22
lm ZScore = 0.25
NG = 0
DB = 0
SM = 0
Delta_AUC
Doxorubicin
YDL227C_RF2_145 RF2
ZShift = 0.55
lm Zscore = −0.39
NG = 0
DB = 0
SM = 0
Delta_L
Doxorubicin
YDL227C_RF2_145 RF2
ZShift = 1.2
lm ZScore = −0.39
NG = 0
DB = 0
SM = 0
Delta_K
Doxorubicin
YDL227C_RF2_145 RF2
ZShift = −0.98
lm ZScore = 0.66
NG = 0
DB = 0
SM = 0
Delta_r
Doxorubicin
YDL227C_RF2_145 RF2
ZShift = 0.97
lm ZScore = −0.24
NG = 0
DB = 0
SM = 0
Delta_AUC
Doxorubicin
YDL227C_RF2_185 RF2
ZShift = 0.22
lm Zscore = −0.4
NG = 0
DB = 0
SM = 0
Delta_L
Doxorubicin
YDL227C_RF2_185 RF2
ZShift = 0.34
lm ZScore = −0.4
NG = 0
DB = 0
SM = 0
Delta_K
Doxorubicin
YDL227C_RF2_185 RF2
ZShift = 0.03
lm ZScore = 0.01
NG = 0
DB = 0
SM = 0
Delta_r
Doxorubicin
YDL227C_RF2_185 RF2
ZShift = 0.26
lm ZScore = −0.92
NG = 0
DB = 0
SM = 0
Delta_AUC
Doxorubicin

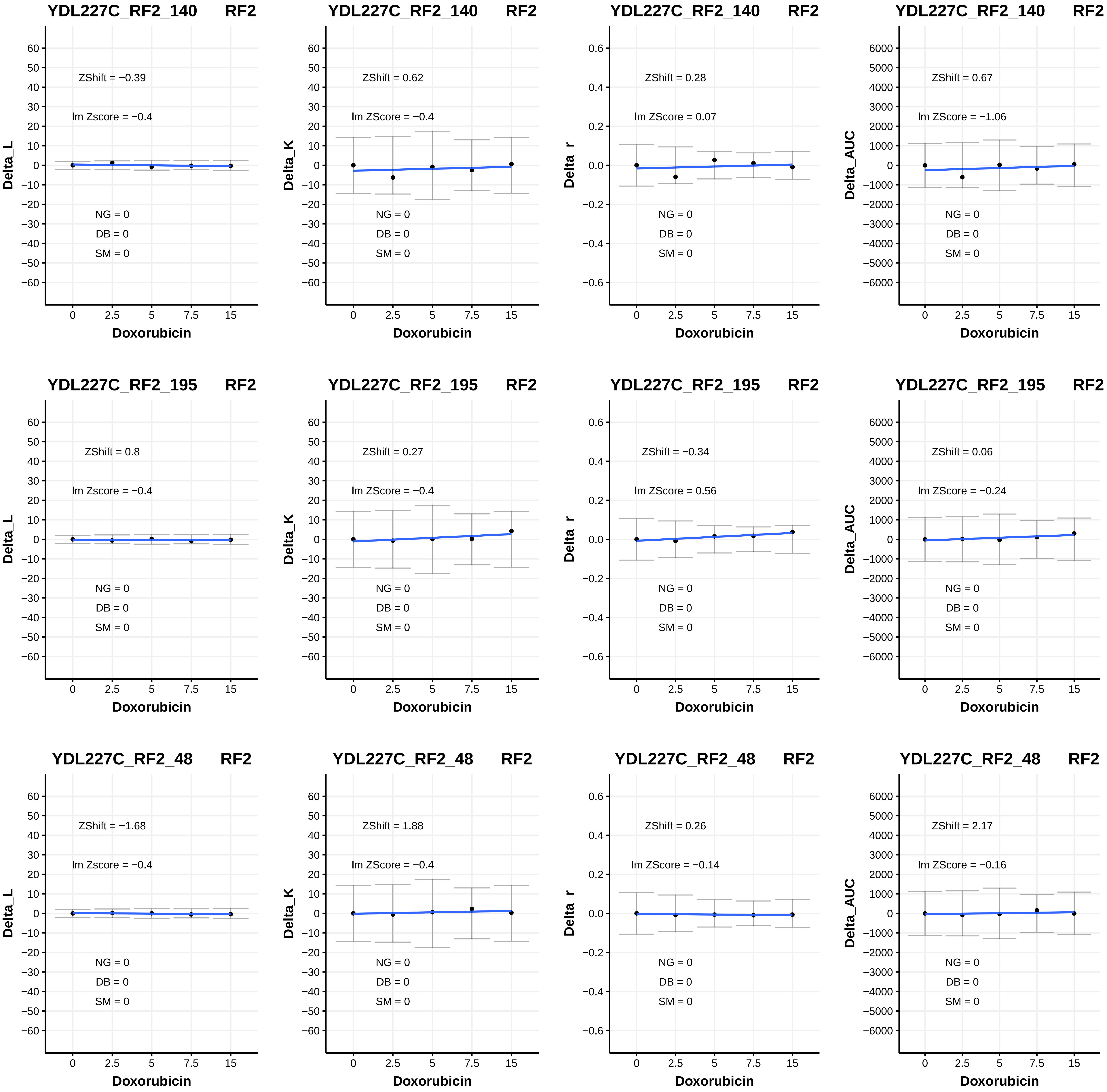

YDL227C_RF2_140 RF2
ZShift = −0.39
lm Zscore = −0.4
NG = 0
DB = 0
SM = 0
Delta_L
Doxorubicin
YDL227C_RF2_140 RF2
ZShift = 0.62
lm ZScore = −0.4
NG = 0
DB = 0
SM = 0
Delta_K
Doxorubicin
YDL227C_RF2_140 RF2
ZShift = 0.28
lm ZScore = 0.07
NG = 0
DB = 0
SM = 0
Delta_r
Doxorubicin
YDL227C_RF2_140 RF2
ZShift = 0.67
lm ZScore = −1.06
NG = 0
DB = 0
SM = 0
Delta_AUC
Doxorubicin
YDL227C_RF2_195 RF2
ZShift = 0.8
lm Zscore = −0.4
NG = 0
DB = 0
SM = 0
Delta_L
Doxorubicin
YDL227C_RF2_195 RF2
ZShift = 0.27
lm ZScore = −0.4
NG = 0
DB = 0
SM = 0
Delta_K
Doxorubicin
YDL227C_RF2_195 RF2
ZShift = −0.34
lm ZScore = 0.56
NG = 0
DB = 0
SM = 0
Delta_r
Doxorubicin
YDL227C_RF2_195 RF2
ZShift = 0.06
lm ZScore = −0.24
NG = 0
DB = 0
SM = 0
Delta_AUC
Doxorubicin
YDL227C_RF2_48 RF2
ZShift = −1.68
lm Zscore = −0.4
NG = 0
DB = 0
SM = 0
Delta_L
Doxorubicin
YDL227C_RF2_48 RF2
ZShift = 1.88
lm ZScore = −0.4
NG = 0
DB = 0
SM = 0
Delta_K
Doxorubicin
YDL227C_RF2_48 RF2
ZShift = 0.26
lm ZScore = −0.14
NG = 0
DB = 0
SM = 0
Delta_r
Doxorubicin
YDL227C_RF2_48 RF2
ZShift = 2.17
lm ZScore = −0.16
NG = 0
DB = 0
SM = 0
Delta_AUC
Doxorubicin

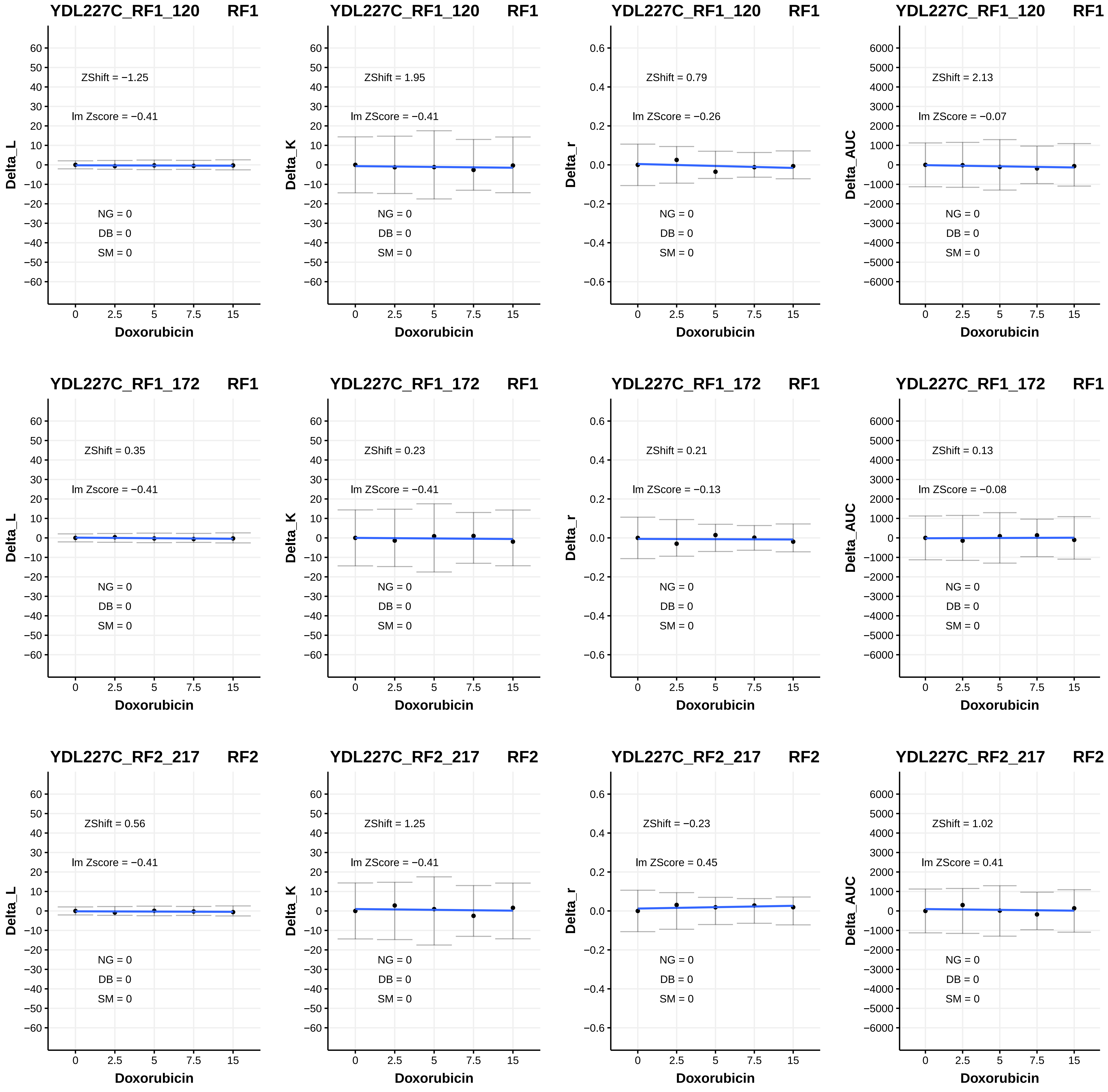

YDL227C_RF1_120 RF1
ZShift = −1.25
lm Zscore = −0.41
NG = 0
DB = 0
SM = 0
Delta_L
Doxorubicin
YDL227C_RF1_120 RF1
ZShift = 1.95
lm ZScore = −0.41
NG = 0
DB = 0
SM = 0
Delta_K
Doxorubicin
YDL227C_RF1_120 RF1
ZShift = 0.79
lm ZScore = −0.26
NG = 0
DB = 0
SM = 0
Delta_r
Doxorubicin
YDL227C_RF1_120 RF1
ZShift = 2.13
lm ZScore = −0.07
NG = 0
DB = 0
SM = 0
Delta_AUC
Doxorubicin
YDL227C_RF1_172 RF1
ZShift = 0.35
lm Zscore = −0.41
NG = 0
DB = 0
SM = 0
Delta_L
Doxorubicin
YDL227C_RF1_172 RF1
ZShift = 0.23
lm ZScore = −0.41
NG = 0
DB = 0
SM = 0
Delta_K
Doxorubicin
YDL227C_RF1_172 RF1
ZShift = 0.21
lm ZScore = −0.13
NG = 0
DB = 0
SM = 0
Delta_r
Doxorubicin
YDL227C_RF1_172 RF1
ZShift = 0.13
lm ZScore = −0.08
NG = 0
DB = 0
SM = 0
Delta_AUC
Doxorubicin
YDL227C_RF2_217 RF2
ZShift = 0.56
lm Zscore = −0.41
NG = 0
DB = 0
SM = 0
Delta_L
Doxorubicin
YDL227C_RF2_217 RF2
ZShift = 1.25
lm ZScore = −0.41
NG = 0
DB = 0
SM = 0
Delta_K
Doxorubicin
YDL227C_RF2_217 RF2
ZShift = −0.23
lm ZScore = 0.45
NG = 0
DB = 0
SM = 0
Delta_r
Doxorubicin
YDL227C_RF2_217 RF2
ZShift = 1.02
lm ZScore = 0.41
NG = 0
DB = 0
SM = 0
Delta_AUC
Doxorubicin

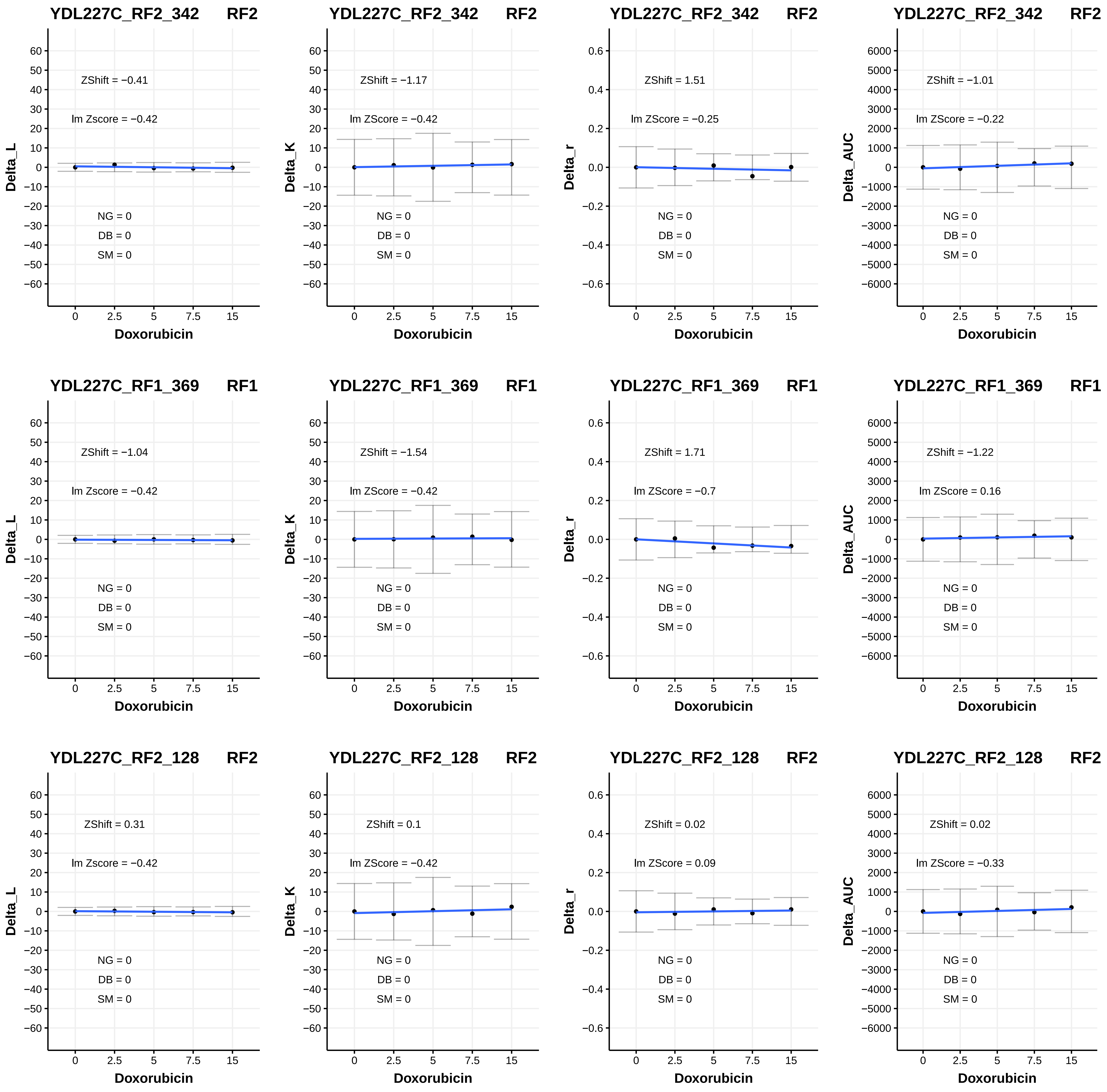

YDL227C_RF2_342 RF2
ZShift = −0.41
lm Zscore = −0.42
NG = 0
DB = 0
SM = 0
Delta_L
Doxorubicin
YDL227C_RF2_342 RF2
ZShift = −1.17
lm ZScore = −0.42
NG = 0
DB = 0
SM = 0
Delta_K
Doxorubicin
YDL227C_RF2_342 RF2
ZShift = 1.51
lm ZScore = −0.25
NG = 0
DB = 0
SM = 0
Delta_r
Doxorubicin
YDL227C_RF2_342 RF2
ZShift = −1.01
lm ZScore = −0.22
NG = 0
DB = 0
SM = 0
Delta_AUC
Doxorubicin
YDL227C_RF1_369 RF1
ZShift = −1.04
lm Zscore = −0.42
NG = 0
DB = 0
SM = 0
Delta_L
Doxorubicin
YDL227C_RF1_369 RF1
ZShift = −1.54
lm ZScore = −0.42
NG = 0
DB = 0
SM = 0
Delta_K
Doxorubicin
YDL227C_RF1_369 RF1
ZShift = 1.71
lm ZScore = −0.7
NG = 0
DB = 0
SM = 0
Delta_r
Doxorubicin
YDL227C_RF1_369 RF1
ZShift = −1.22
lm ZScore = 0.16
NG = 0
DB = 0
SM = 0
Delta_AUC
Doxorubicin
YDL227C_RF2_128 RF2
ZShift = 0.31
lm Zscore = −0.42
NG = 0
DB = 0
SM = 0
Delta_L
Doxorubicin
YDL227C_RF2_128 RF2
ZShift = 0.1
lm ZScore = −0.42
NG = 0
DB = 0
SM = 0
Delta_K
Doxorubicin
YDL227C_RF2_128 RF2
ZShift = 0.02
lm ZScore = 0.09
NG = 0
DB = 0
SM = 0
Delta_r
Doxorubicin
YDL227C_RF2_128 RF2
ZShift = 0.02
lm ZScore = −0.33
NG = 0
DB = 0
SM = 0
Delta_AUC
Doxorubicin

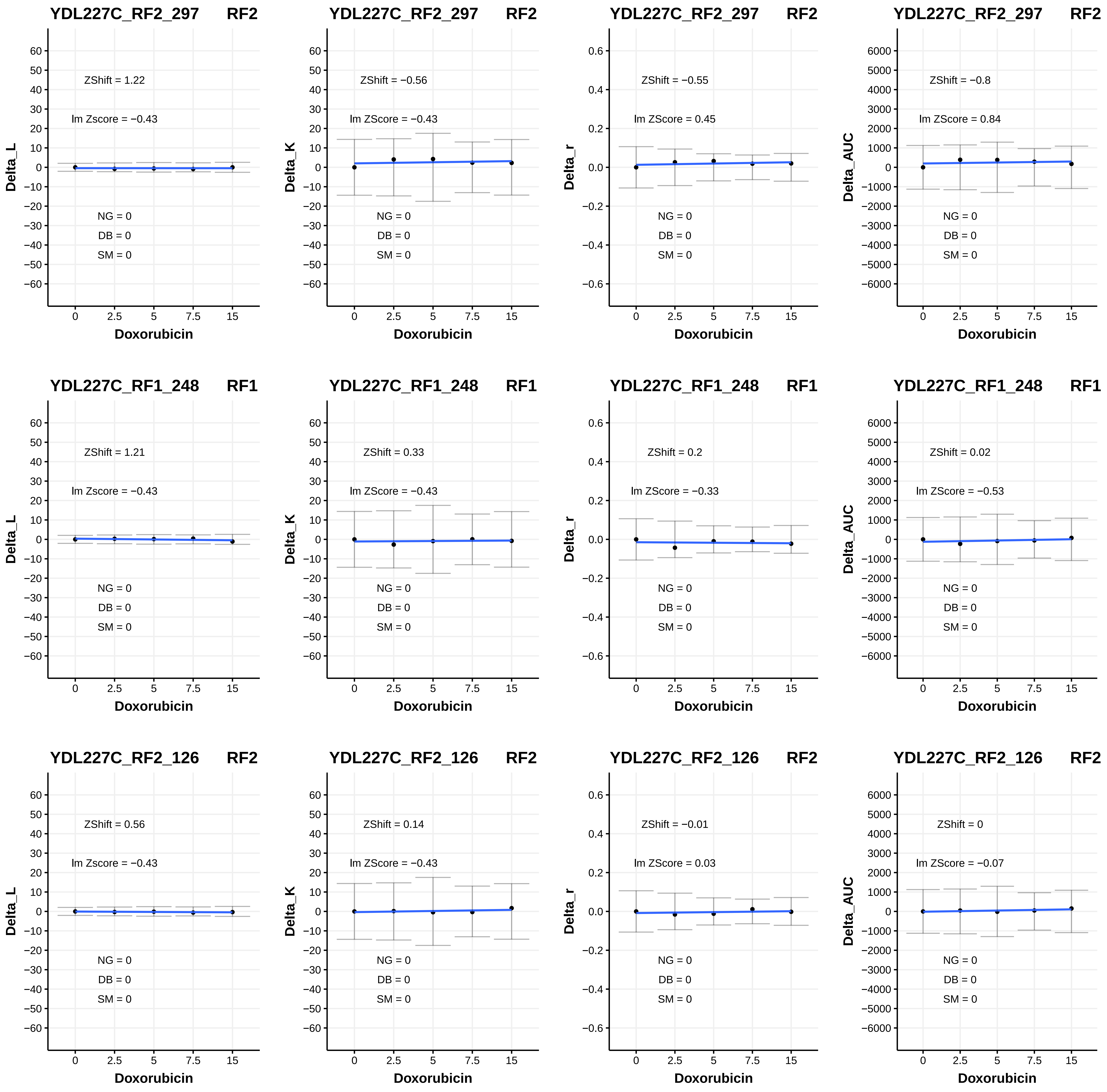

YDL227C_RF2_297 RF2
ZShift = 1.22
lm Zscore = −0.43
NG = 0
DB = 0
SM = 0
Delta_L
Doxorubicin
YDL227C_RF2_297 RF2
ZShift = −0.56
lm ZScore = −0.43
NG = 0
DB = 0
SM = 0
Delta_K
Doxorubicin
YDL227C_RF2_297 RF2
ZShift = −0.55
lm ZScore = 0.45
NG = 0
DB = 0
SM = 0
Delta_r
Doxorubicin
YDL227C_RF2_297 RF2
ZShift = −0.8
lm ZScore = 0.84
NG = 0
DB = 0
SM = 0
Delta_AUC
Doxorubicin
YDL227C_RF1_248 RF1
ZShift = 1.21
lm Zscore = −0.43
NG = 0
DB = 0
SM = 0
Delta_L
Doxorubicin
YDL227C_RF1_248 RF1
ZShift = 0.33
lm ZScore = −0.43
NG = 0
DB = 0
SM = 0
Delta_K
Doxorubicin
YDL227C_RF1_248 RF1
ZShift = 0.2
lm ZScore = −0.33
NG = 0
DB = 0
SM = 0
Delta_r
Doxorubicin
YDL227C_RF1_248 RF1
ZShift = 0.02
lm ZScore = −0.53
NG = 0
DB = 0
SM = 0
Delta_AUC
Doxorubicin
YDL227C_RF2_126 RF2
ZShift = 0.56
lm Zscore = −0.43
NG = 0
DB = 0
SM = 0
Delta_L
Doxorubicin
YDL227C_RF2_126 RF2
ZShift = 0.14
lm ZScore = −0.43
NG = 0
DB = 0
SM = 0
Delta_K
Doxorubicin
YDL227C_RF2_126 RF2
ZShift = −0.01
lm ZScore = 0.03
NG = 0
DB = 0
SM = 0
Delta_r
Doxorubicin
YDL227C_RF2_126 RF2
ZShift = 0
lm ZScore = −0.07
NG = 0
DB = 0
SM = 0
Delta_AUC
Doxorubicin

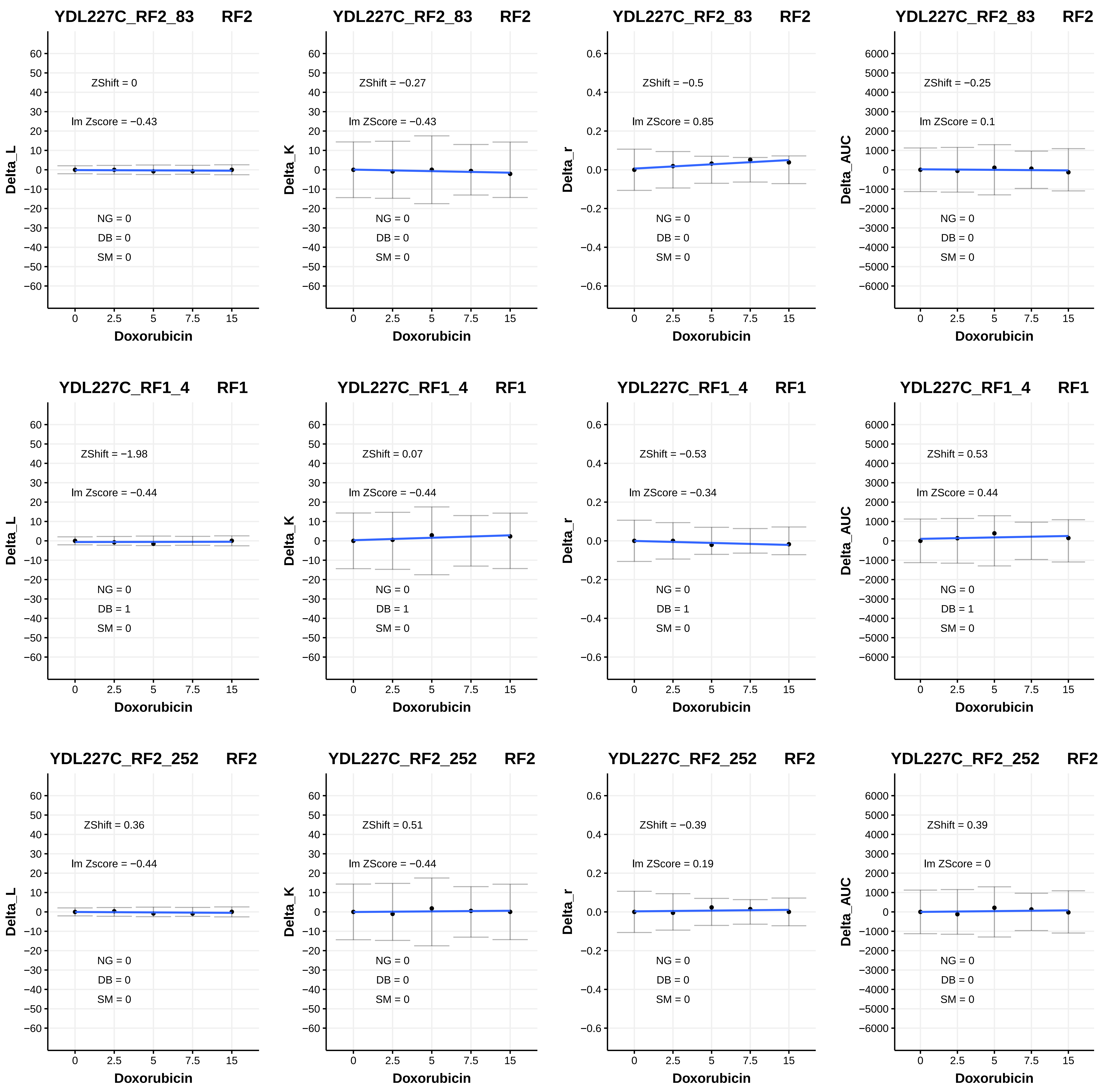
YDL227C_RF2_83 RF2
ZShift = 0
lm Zscore = −0.43
NG = 0
DB = 0
SM = 0
Delta_L
Doxorubicin
YDL227C_RF2_83 RF2
ZShift = −0.27
lm ZScore = −0.43
NG = 0
DB = 0
SM = 0
Delta_K
Doxorubicin
YDL227C_RF2_83 RF2
ZShift = −0.5
lm ZScore = 0.85
NG = 0
DB = 0
SM = 0
Delta_r
Doxorubicin
YDL227C_RF2_83 RF2
ZShift = −0.25
lm ZScore = 0.1
NG = 0
DB = 0
SM = 0
Delta_AUC
Doxorubicin
YDL227C_RF1_4 RF1
ZShift = −1.98
lm Zscore = −0.44
NG = 0
DB = 1
SM = 0
Delta_L
Doxorubicin
YDL227C_RF1_4 RF1
ZShift = 0.07
lm ZScore = −0.44
NG = 0
DB = 1
SM = 0
Delta_K
Doxorubicin
YDL227C_RF1_4 RF1
ZShift = −0.53
lm ZScore = −0.34
NG = 0
DB = 1
SM = 0
Delta_r
Doxorubicin
YDL227C_RF1_4 RF1
ZShift = 0.53
lm ZScore = 0.44
NG = 0
DB = 1
SM = 0
Delta_AUC
Doxorubicin
YDL227C_RF2_252 RF2
ZShift = 0.36
lm Zscore = −0.44
NG = 0
DB = 0
SM = 0
Delta_L
Doxorubicin
YDL227C_RF2_252 RF2
ZShift = 0.51
lm ZScore = −0.44
NG = 0
DB = 0
SM = 0
Delta_K
Doxorubicin
YDL227C_RF2_252 RF2
ZShift = −0.39
lm ZScore = 0.19
NG = 0
DB = 0
SM = 0
Delta_r
Doxorubicin
YDL227C_RF2_252 RF2
ZShift = 0.39
lm ZScore = 0
NG = 0
DB = 0
SM = 0
Delta_AUC
Doxorubicin

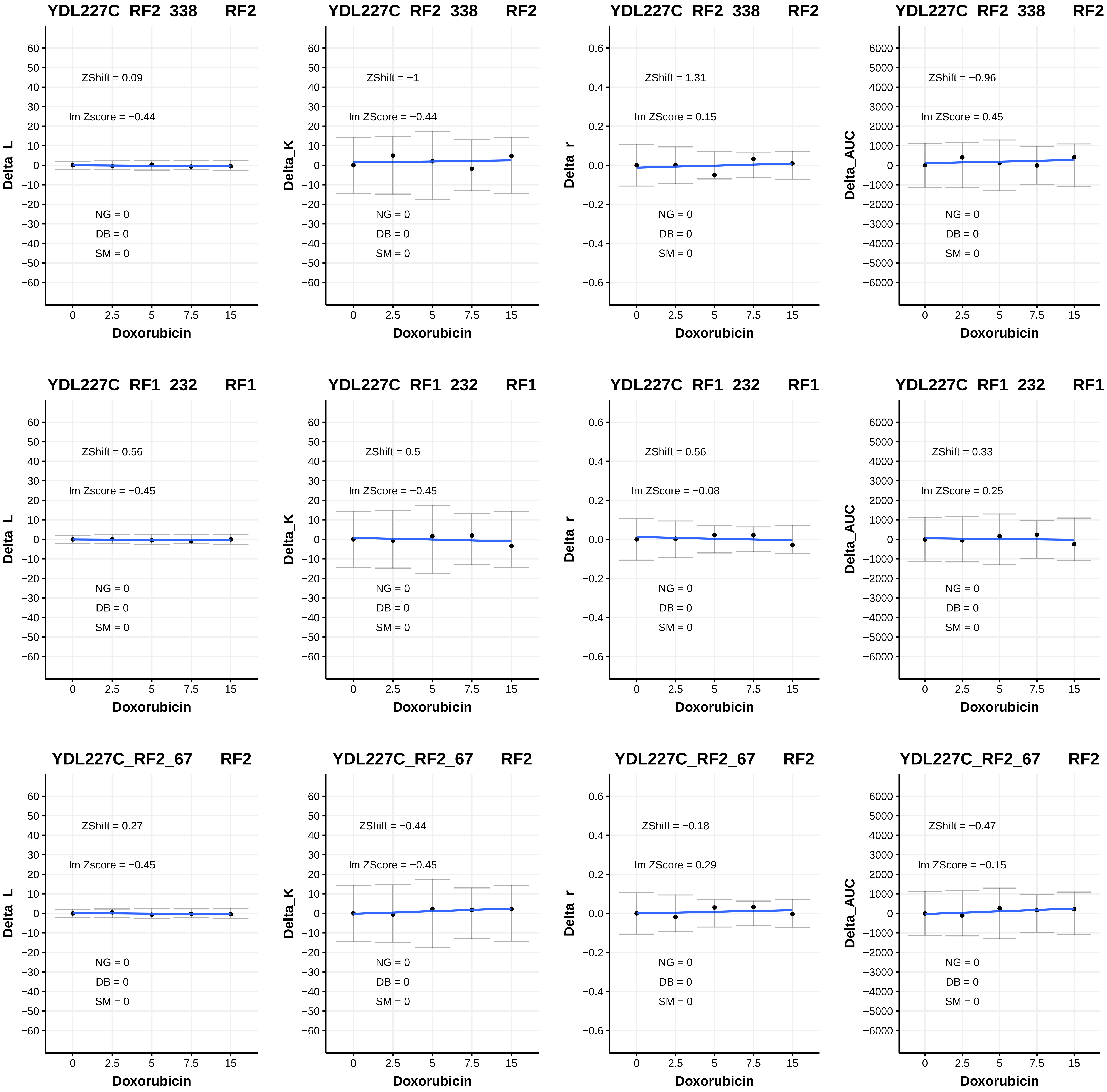
YDL227C_RF2_338 RF2
ZShift = 0.09
lm Zscore = −0.44
NG = 0
DB = 0
SM = 0
Delta_L
Doxorubicin
YDL227C_RF2_338 RF2
ZShift = −1
lm ZScore = −0.44
NG = 0
DB = 0
SM = 0
Delta_K
Doxorubicin
YDL227C_RF2_338 RF2
ZShift = 1.31
lm ZScore = 0.15
NG = 0
DB = 0
SM = 0
Delta_r
Doxorubicin
YDL227C_RF2_338 RF2
ZShift = −0.96
lm ZScore = 0.45
NG = 0
DB = 0
SM = 0
Delta_AUC
Doxorubicin
YDL227C_RF1_232 RF1
ZShift = 0.56
lm Zscore = −0.45
NG = 0
DB = 0
SM = 0
Delta_L
Doxorubicin
YDL227C_RF1_232 RF1
ZShift = 0.5
lm ZScore = −0.45
NG = 0
DB = 0
SM = 0
Delta_K
Doxorubicin
YDL227C_RF1_232 RF1
ZShift = 0.56
lm ZScore = −0.08
NG = 0
DB = 0
SM = 0
Delta_r
Doxorubicin
YDL227C_RF1_232 RF1
ZShift = 0.33
lm ZScore = 0.25
NG = 0
DB = 0
SM = 0
Delta_AUC
Doxorubicin
YDL227C_RF2_67 RF2
ZShift = 0.27
lm Zscore = −0.45
NG = 0
DB = 0
SM = 0
Delta_L
Doxorubicin
YDL227C_RF2_67 RF2
ZShift = −0.44
lm ZScore = −0.45
NG = 0
DB = 0
SM = 0
Delta_K
Doxorubicin
YDL227C_RF2_67 RF2
ZShift = −0.18
lm ZScore = 0.29
NG = 0
DB = 0
SM = 0
Delta_r
Doxorubicin
YDL227C_RF2_67 RF2
ZShift = −0.47
lm ZScore = −0.15
NG = 0
DB = 0
SM = 0
Delta_AUC
Doxorubicin

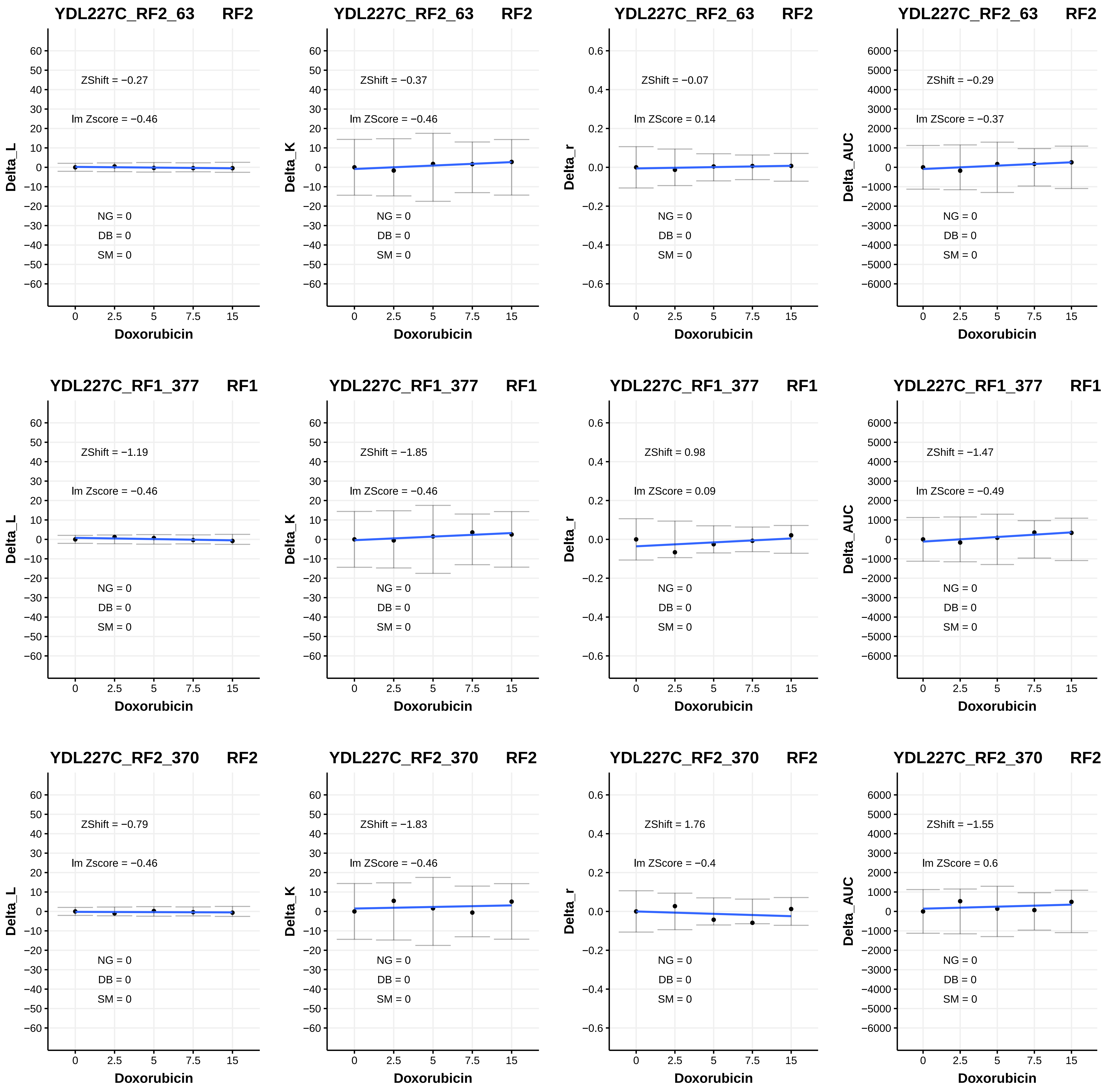
YDL227C_RF2_63 RF2
ZShift = −0.27
lm Zscore = −0.46
NG = 0
DB = 0
SM = 0
Delta_L
Doxorubicin
YDL227C_RF2_63 RF2
ZShift = −0.37
lm ZScore = −0.46
NG = 0
DB = 0
SM = 0
Delta_K
Doxorubicin
YDL227C_RF2_63 RF2
ZShift = −0.07
lm ZScore = 0.14
NG = 0
DB = 0
SM = 0
Delta_r
Doxorubicin
YDL227C_RF2_63 RF2
ZShift = −0.29
lm ZScore = −0.37
NG = 0
DB = 0
SM = 0
Delta_AUC
Doxorubicin
YDL227C_RF1_377 RF1
ZShift = −1.19
lm Zscore = −0.46
NG = 0
DB = 0
SM = 0
Delta_L
Doxorubicin
YDL227C_RF1_377 RF1
ZShift = −1.85
lm ZScore = −0.46
NG = 0
DB = 0
SM = 0
Delta_K
Doxorubicin
YDL227C_RF1_377 RF1
ZShift = 0.98
lm ZScore = 0.09
NG = 0
DB = 0
SM = 0
Delta_r
Doxorubicin
YDL227C_RF1_377 RF1
ZShift = −1.47
lm ZScore = −0.49
NG = 0
DB = 0
SM = 0
Delta_AUC
Doxorubicin
YDL227C_RF2_370 RF2
ZShift = −0.79
lm Zscore = −0.46
NG = 0
DB = 0
SM = 0
Delta_L
Doxorubicin
YDL227C_RF2_370 RF2
ZShift = −1.83
lm ZScore = −0.46
NG = 0
DB = 0
SM = 0
Delta_K
Doxorubicin
YDL227C_RF2_370 RF2
ZShift = 1.76
lm ZScore = −0.4
NG = 0
DB = 0
SM = 0
Delta_r
Doxorubicin
YDL227C_RF2_370 RF2
ZShift = −1.55
lm ZScore = 0.6
NG = 0
DB = 0
SM = 0
Delta_AUC
Doxorubicin

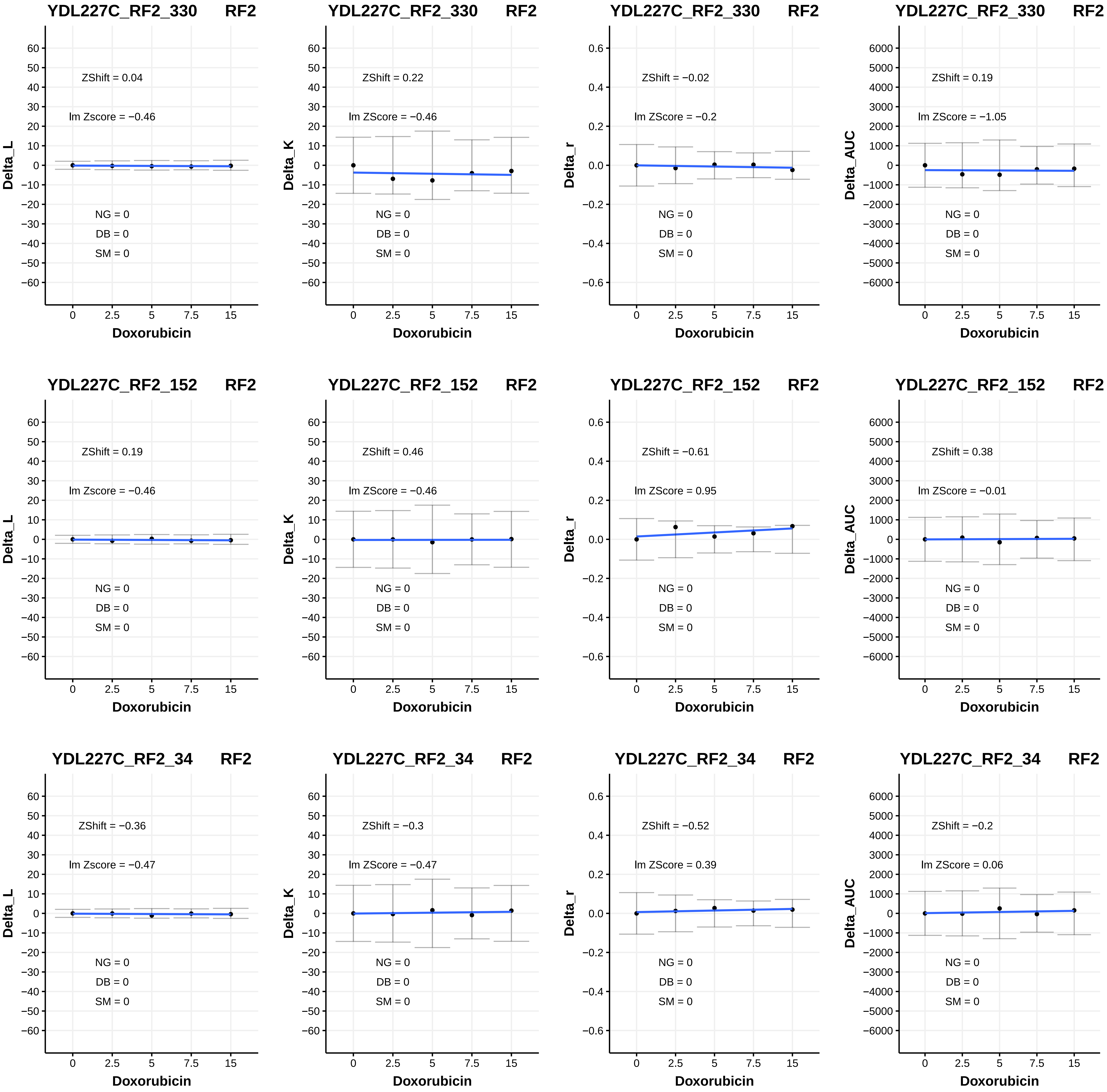
YDL227C_RF2_330 RF2
ZShift = 0.04
lm Zscore = −0.46
NG = 0
DB = 0
SM = 0
Delta_L
Doxorubicin
YDL227C_RF2_330 RF2
ZShift = 0.22
lm ZScore = −0.46
NG = 0
DB = 0
SM = 0
Delta_K
Doxorubicin
YDL227C_RF2_330 RF2
ZShift = −0.02
lm ZScore = −0.2
NG = 0
DB = 0
SM = 0
Delta_r
Doxorubicin
YDL227C_RF2_330 RF2
ZShift = 0.19
lm ZScore = −1.05
NG = 0
DB = 0
SM = 0
Delta_AUC
Doxorubicin
YDL227C_RF2_152 RF2
ZShift = 0.19
lm Zscore = −0.46
NG = 0
DB = 0
SM = 0
Delta_L
Doxorubicin
YDL227C_RF2_152 RF2
ZShift = 0.46
lm ZScore = −0.46
NG = 0
DB = 0
SM = 0
Delta_K
Doxorubicin
YDL227C_RF2_152 RF2
ZShift = −0.61
lm ZScore = 0.95
NG = 0
DB = 0
SM = 0
Delta_r
Doxorubicin
YDL227C_RF2_152 RF2
ZShift = 0.38
lm ZScore = −0.01
NG = 0
DB = 0
SM = 0
Delta_AUC
Doxorubicin
YDL227C_RF2_34 RF2
ZShift = −0.36
lm Zscore = −0.47
NG = 0
DB = 0
SM = 0
Delta_L
Doxorubicin
YDL227C_RF2_34 RF2
ZShift = −0.3
lm ZScore = −0.47
NG = 0
DB = 0
SM = 0
Delta_K
Doxorubicin
YDL227C_RF2_34 RF2
ZShift = −0.52
lm ZScore = 0.39
NG = 0
DB = 0
SM = 0
Delta_r
Doxorubicin
YDL227C_RF2_34 RF2
ZShift = −0.2
lm ZScore = 0.06
NG = 0
DB = 0
SM = 0
Delta_AUC
Doxorubicin

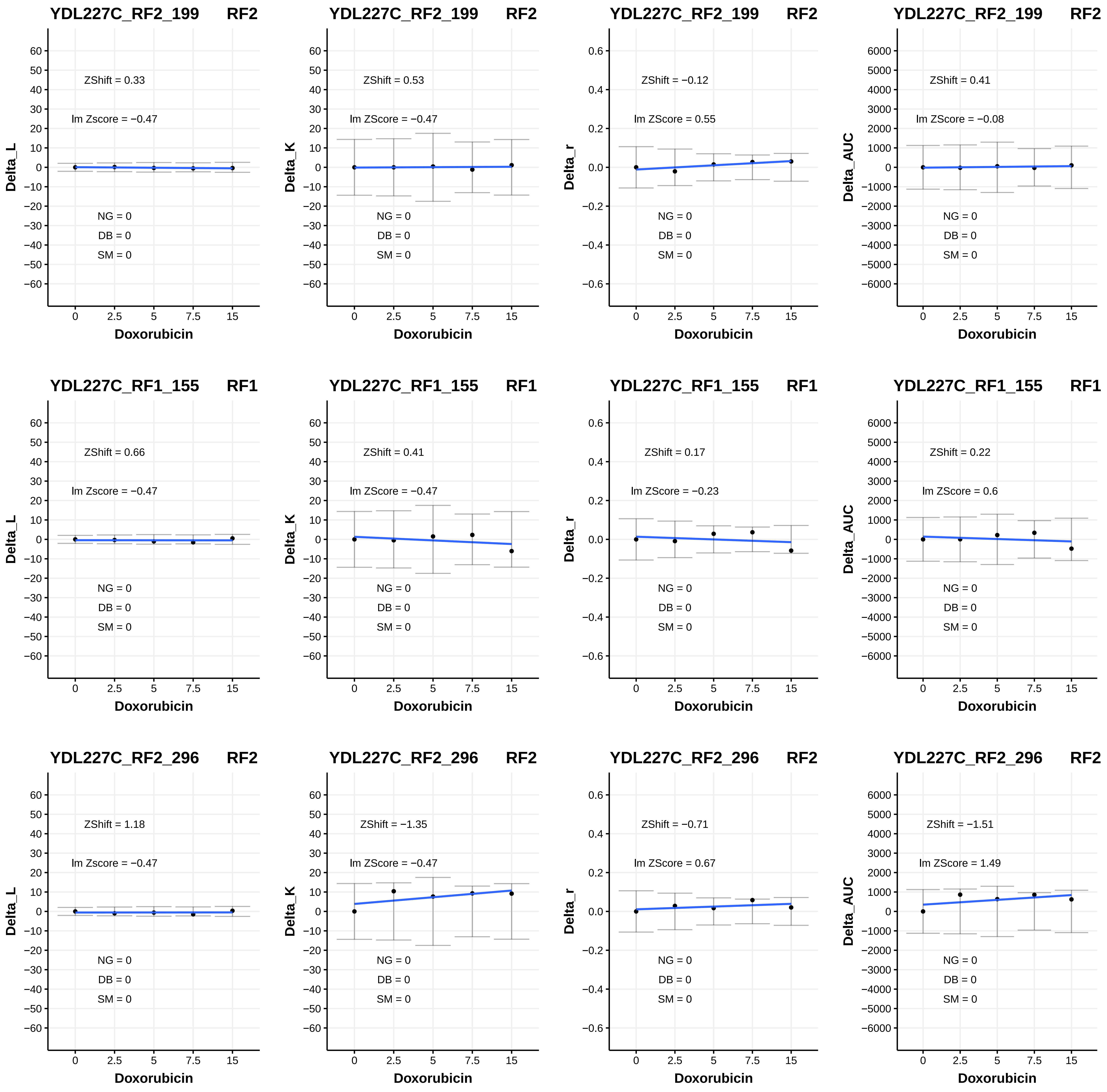

YDL227C_RF2_199 RF2
ZShift = 0.33
lm Zscore = −0.47
NG = 0
DB = 0
SM = 0
Delta_L
Doxorubicin
YDL227C_RF2_199 RF2
ZShift = 0.53
lm ZScore = −0.47
NG = 0
DB = 0
SM = 0
Delta_K
Doxorubicin
YDL227C_RF2_199 RF2
ZShift = −0.12
lm ZScore = 0.55
NG = 0
DB = 0
SM = 0
Delta_r
Doxorubicin
YDL227C_RF2_199 RF2
ZShift = 0.41
lm ZScore = −0.08
NG = 0
DB = 0
SM = 0
Delta_AUC
Doxorubicin
YDL227C_RF1_155 RF1
ZShift = 0.66
lm Zscore = −0.47
NG = 0
DB = 0
SM = 0
Delta_L
Doxorubicin
YDL227C_RF1_155 RF1
ZShift = 0.41
lm ZScore = −0.47
NG = 0
DB = 0
SM = 0
Delta_K
Doxorubicin
YDL227C_RF1_155 RF1
ZShift = 0.17
lm ZScore = −0.23
NG = 0
DB = 0
SM = 0
Delta_r
Doxorubicin
YDL227C_RF1_155 RF1
ZShift = 0.22
lm ZScore = 0.6
NG = 0
DB = 0
SM = 0
Delta_AUC
Doxorubicin
YDL227C_RF2_296 RF2
ZShift = 1.18
lm Zscore = −0.47
NG = 0
DB = 0
SM = 0
Delta_L
Doxorubicin
YDL227C_RF2_296 RF2
ZShift = −1.35
lm ZScore = −0.47
NG = 0
DB = 0
SM = 0
Delta_K
Doxorubicin
YDL227C_RF2_296 RF2
ZShift = −0.71
lm ZScore = 0.67
NG = 0
DB = 0
SM = 0
Delta_r
Doxorubicin
YDL227C_RF2_296 RF2
ZShift = −1.51
lm ZScore = 1.49
NG = 0
DB = 0
SM = 0
Delta_AUC
Doxorubicin

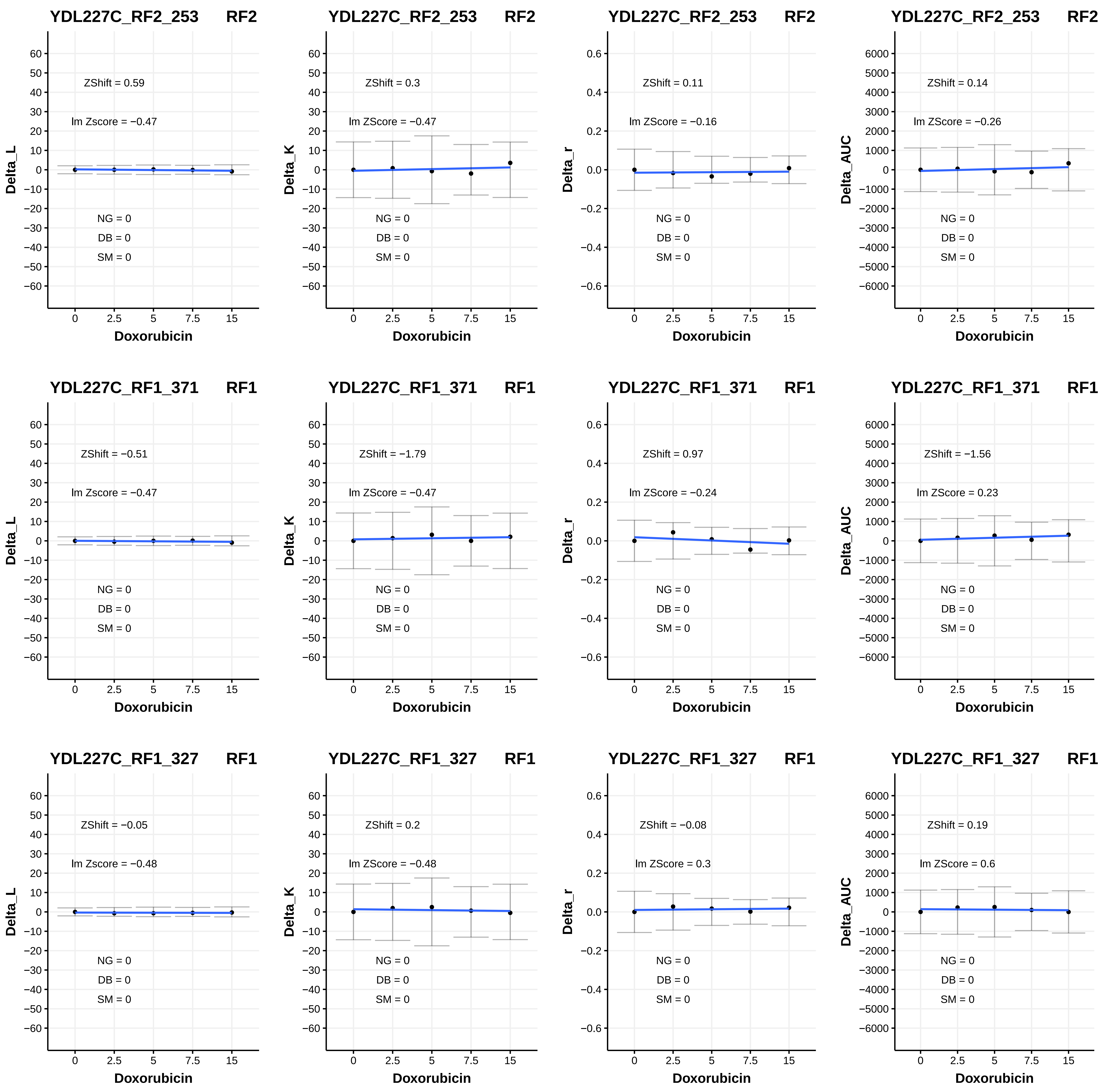
YDL227C_RF2_253 RF2
ZShift = 0.59
lm Zscore = −0.47
NG = 0
DB = 0
SM = 0
Delta_L
Doxorubicin
YDL227C_RF2_253 RF2
ZShift = 0.3
lm ZScore = −0.47
NG = 0
DB = 0
SM = 0
Delta_K
Doxorubicin
YDL227C_RF2_253 RF2
ZShift = 0.11
lm ZScore = −0.16
NG = 0
DB = 0
SM = 0
Delta_r
Doxorubicin
YDL227C_RF2_253 RF2
ZShift = 0.14
lm ZScore = −0.26
NG = 0
DB = 0
SM = 0
Delta_AUC
Doxorubicin
YDL227C_RF1_371 RF1
ZShift = −0.51
lm Zscore = −0.47
NG = 0
DB = 0
SM = 0
Delta_L
Doxorubicin
YDL227C_RF1_371 RF1
ZShift = −1.79
lm ZScore = −0.47
NG = 0
DB = 0
SM = 0
Delta_K
Doxorubicin
YDL227C_RF1_371 RF1
ZShift = 0.97
lm ZScore = −0.24
NG = 0
DB = 0
SM = 0
Delta_r
Doxorubicin
YDL227C_RF1_371 RF1
ZShift = −1.56
lm ZScore = 0.23
NG = 0
DB = 0
SM = 0
Delta_AUC
Doxorubicin
YDL227C_RF1_327 RF1
ZShift = −0.05
lm Zscore = −0.48
NG = 0
DB = 0
SM = 0
Delta_L
Doxorubicin
YDL227C_RF1_327 RF1
ZShift = 0.2
lm ZScore = −0.48
NG = 0
DB = 0
SM = 0
Delta_K
Doxorubicin
YDL227C_RF1_327 RF1
ZShift = −0.08
lm ZScore = 0.3
NG = 0
DB = 0
SM = 0
Delta_r
Doxorubicin
YDL227C_RF1_327 RF1
ZShift = 0.19
lm ZScore = 0.6
NG = 0
DB = 0
SM = 0
Delta_AUC
Doxorubicin

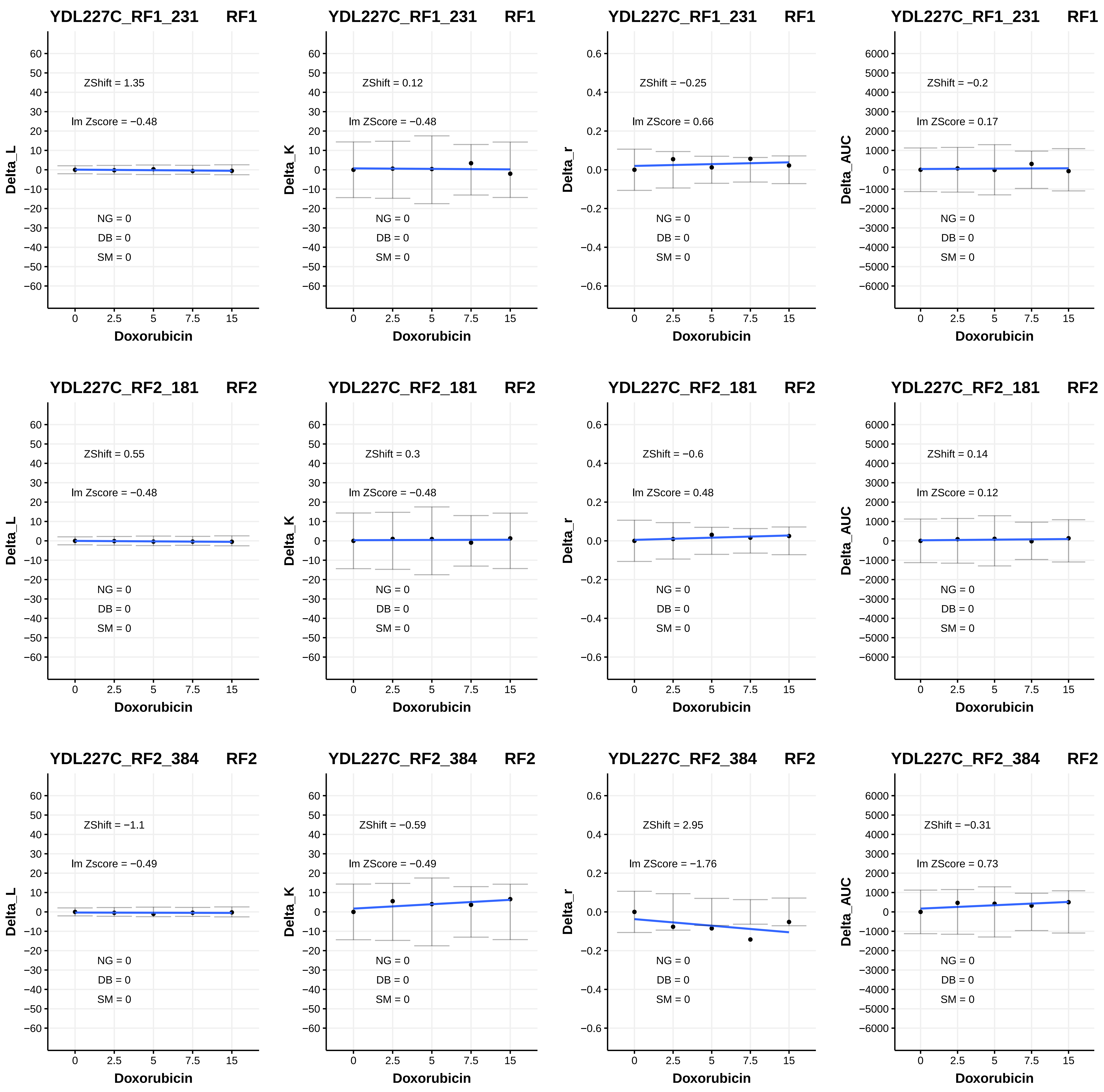

YDL227C_RF1_231 RF1
ZShift = 1.35
lm Zscore = −0.48
NG = 0
DB = 0
SM = 0
Delta_L
Doxorubicin
YDL227C_RF1_231 RF1
ZShift = 0.12
lm ZScore = −0.48
NG = 0
DB = 0
SM = 0
Delta_K
Doxorubicin
YDL227C_RF1_231 RF1
ZShift = −0.25
lm ZScore = 0.66
NG = 0
DB = 0
SM = 0
Delta_r
Doxorubicin
YDL227C_RF1_231 RF1
ZShift = −0.2
lm ZScore = 0.17
NG = 0
DB = 0
SM = 0
Delta_AUC
Doxorubicin
YDL227C_RF2_181 RF2
ZShift = 0.55
lm Zscore = −0.48
NG = 0
DB = 0
SM = 0
Delta_L
Doxorubicin
YDL227C_RF2_181 RF2
ZShift = 0.3
lm ZScore = −0.48
NG = 0
DB = 0
SM = 0
Delta_K
Doxorubicin
YDL227C_RF2_181 RF2
ZShift = −0.6
lm ZScore = 0.48
NG = 0
DB = 0
SM = 0
Delta_r
Doxorubicin
YDL227C_RF2_181 RF2
ZShift = 0.14
lm ZScore = 0.12
NG = 0
DB = 0
SM = 0
Delta_AUC
Doxorubicin
YDL227C_RF2_384 RF2
ZShift = −1.1
lm Zscore = −0.49
NG = 0
DB = 0
SM = 0
Delta_L
Doxorubicin
YDL227C_RF2_384 RF2
ZShift = −0.59
lm ZScore = −0.49
NG = 0
DB = 0
SM = 0
Delta_K
Doxorubicin
YDL227C_RF2_384 RF2
ZShift = 2.95
lm ZScore = −1.76
NG = 0
DB = 0
SM = 0
Delta_r
Doxorubicin
YDL227C_RF2_384 RF2
ZShift = −0.31
lm ZScore = 0.73
NG = 0
DB = 0
SM = 0
Delta_AUC
Doxorubicin

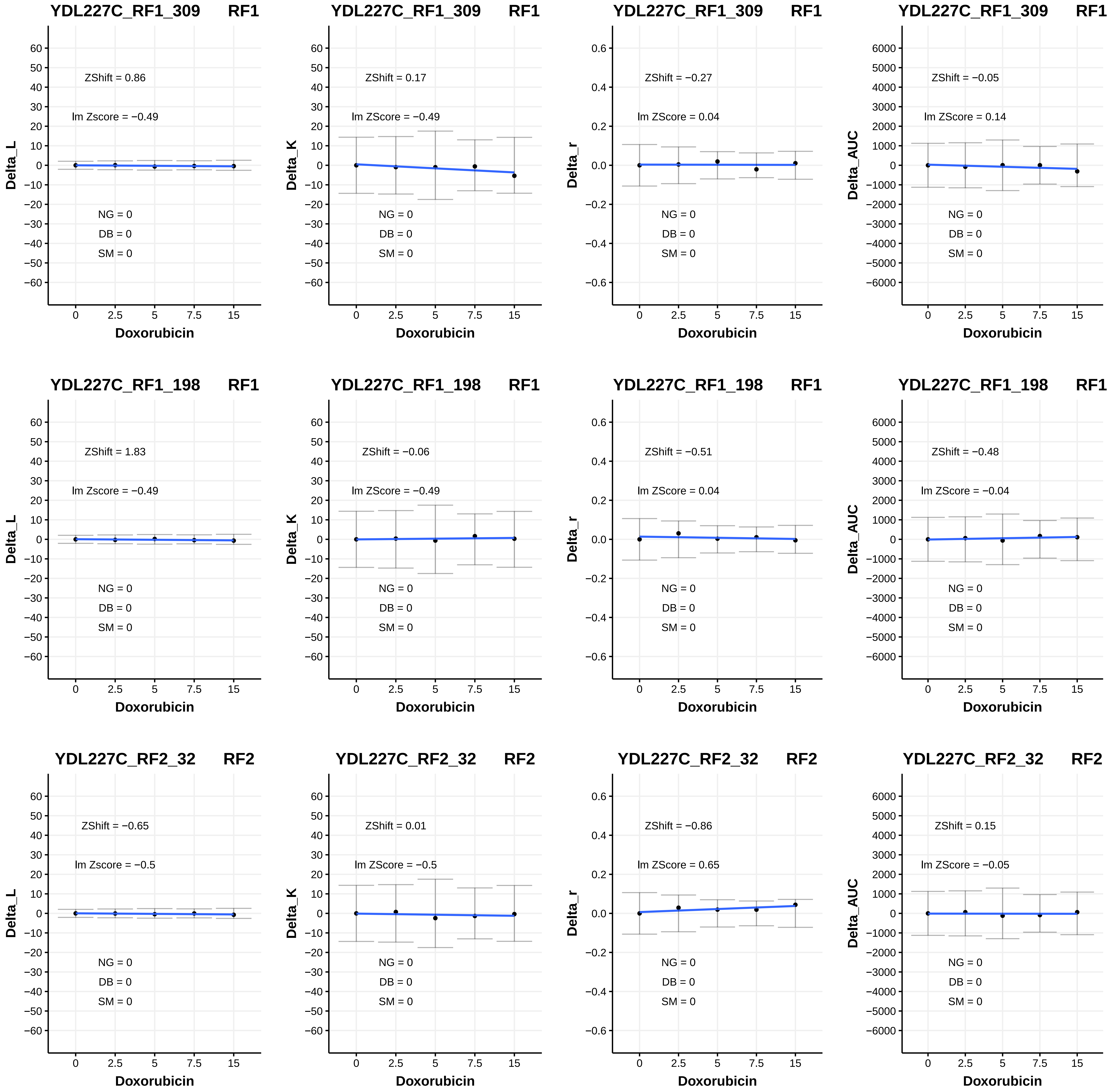
YDL227C_RF1_309 RF1
ZShift = 0.86
lm Zscore = −0.49
NG = 0
DB = 0
SM = 0
Delta_L
Doxorubicin
YDL227C_RF1_309 RF1
ZShift = 0.17
lm ZScore = −0.49
NG = 0
DB = 0
SM = 0
Delta_K
Doxorubicin
YDL227C_RF1_309 RF1
ZShift = −0.27
lm ZScore = 0.04
NG = 0
DB = 0
SM = 0
Delta_r
Doxorubicin
YDL227C_RF1_309 RF1
ZShift = −0.05
lm ZScore = 0.14
NG = 0
DB = 0
SM = 0
Delta_AUC
Doxorubicin
YDL227C_RF1_198 RF1
ZShift = 1.83
lm Zscore = −0.49
NG = 0
DB = 0
SM = 0
Delta_L
Doxorubicin
YDL227C_RF1_198 RF1
ZShift = −0.06
lm ZScore = −0.49
NG = 0
DB = 0
SM = 0
Delta_K
Doxorubicin
YDL227C_RF1_198 RF1
ZShift = −0.51
lm ZScore = 0.04
NG = 0
DB = 0
SM = 0
Delta_r
Doxorubicin
YDL227C_RF1_198 RF1
ZShift = −0.48
lm ZScore = −0.04
NG = 0
DB = 0
SM = 0
Delta_AUC
Doxorubicin
YDL227C_RF2_32 RF2
ZShift = −0.65
lm Zscore = −0.5
NG = 0
DB = 0
SM = 0
Delta_L
Doxorubicin
YDL227C_RF2_32 RF2
ZShift = 0.01
lm ZScore = −0.5
NG = 0
DB = 0
SM = 0
Delta_K
Doxorubicin
YDL227C_RF2_32 RF2
ZShift = −0.86
lm ZScore = 0.65
NG = 0
DB = 0
SM = 0
Delta_r
Doxorubicin
YDL227C_RF2_32 RF2
ZShift = 0.15
lm ZScore = −0.05
NG = 0
DB = 0
SM = 0
Delta_AUC
Doxorubicin

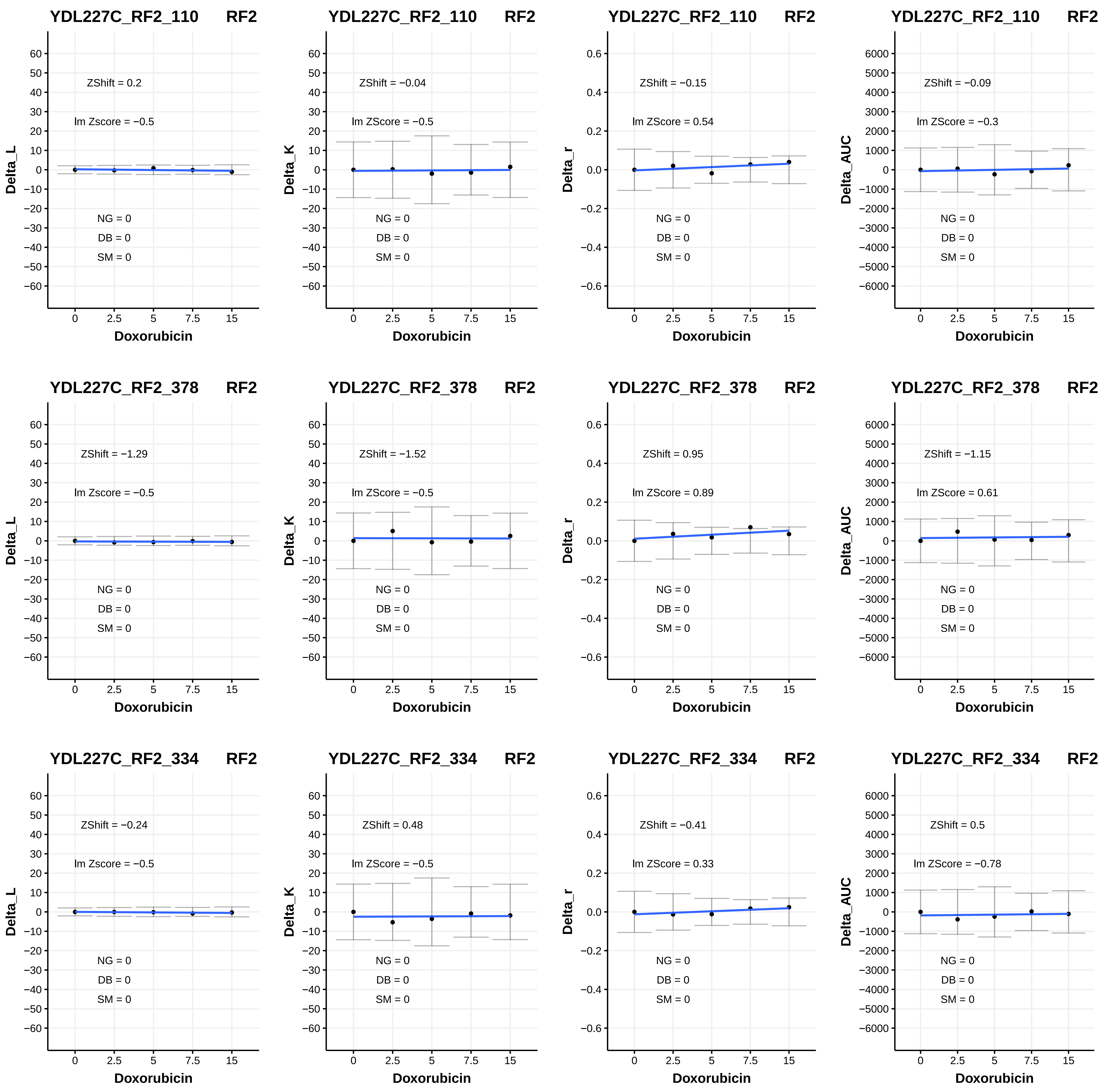
YDL227C_RF2_110 RF2
ZShift = 0.2
lm Zscore = −0.5
NG = 0
DB = 0
SM = 0
Delta_L
Doxorubicin
YDL227C_RF2_110 RF2
ZShift = −0.04
lm ZScore = −0.5
NG = 0
DB = 0
SM = 0
Delta_K
Doxorubicin
YDL227C_RF2_110 RF2
ZShift = −0.15
lm ZScore = 0.54
NG = 0
DB = 0
SM = 0
Delta_r
Doxorubicin
YDL227C_RF2_110 RF2
ZShift = −0.09
lm ZScore = −0.3
NG = 0
DB = 0
SM = 0
Delta_AUC
Doxorubicin
YDL227C_RF2_378 RF2
ZShift = −1.29
lm Zscore = −0.5
NG = 0
DB = 0
SM = 0
Delta_L
Doxorubicin
YDL227C_RF2_378 RF2
ZShift = −1.52
lm ZScore = −0.5
NG = 0
DB = 0
SM = 0
Delta_K
Doxorubicin
YDL227C_RF2_378 RF2
ZShift = 0.95
lm ZScore = 0.89
NG = 0
DB = 0
SM = 0
Delta_r
Doxorubicin
YDL227C_RF2_378 RF2
ZShift = −1.15
lm ZScore = 0.61
NG = 0
DB = 0
SM = 0
Delta_AUC
Doxorubicin
YDL227C_RF2_334 RF2
ZShift = −0.24
lm Zscore = −0.5
NG = 0
DB = 0
SM = 0
Delta_L
Doxorubicin
YDL227C_RF2_334 RF2
ZShift = 0.48
lm ZScore = −0.5
NG = 0
DB = 0
SM = 0
Delta_K
Doxorubicin
YDL227C_RF2_334 RF2
ZShift = −0.41
lm ZScore = 0.33
NG = 0
DB = 0
SM = 0
Delta_r
Doxorubicin
YDL227C_RF2_334 RF2
ZShift = 0.5
lm ZScore = −0.78
NG = 0
DB = 0
SM = 0
Delta_AUC
Doxorubicin

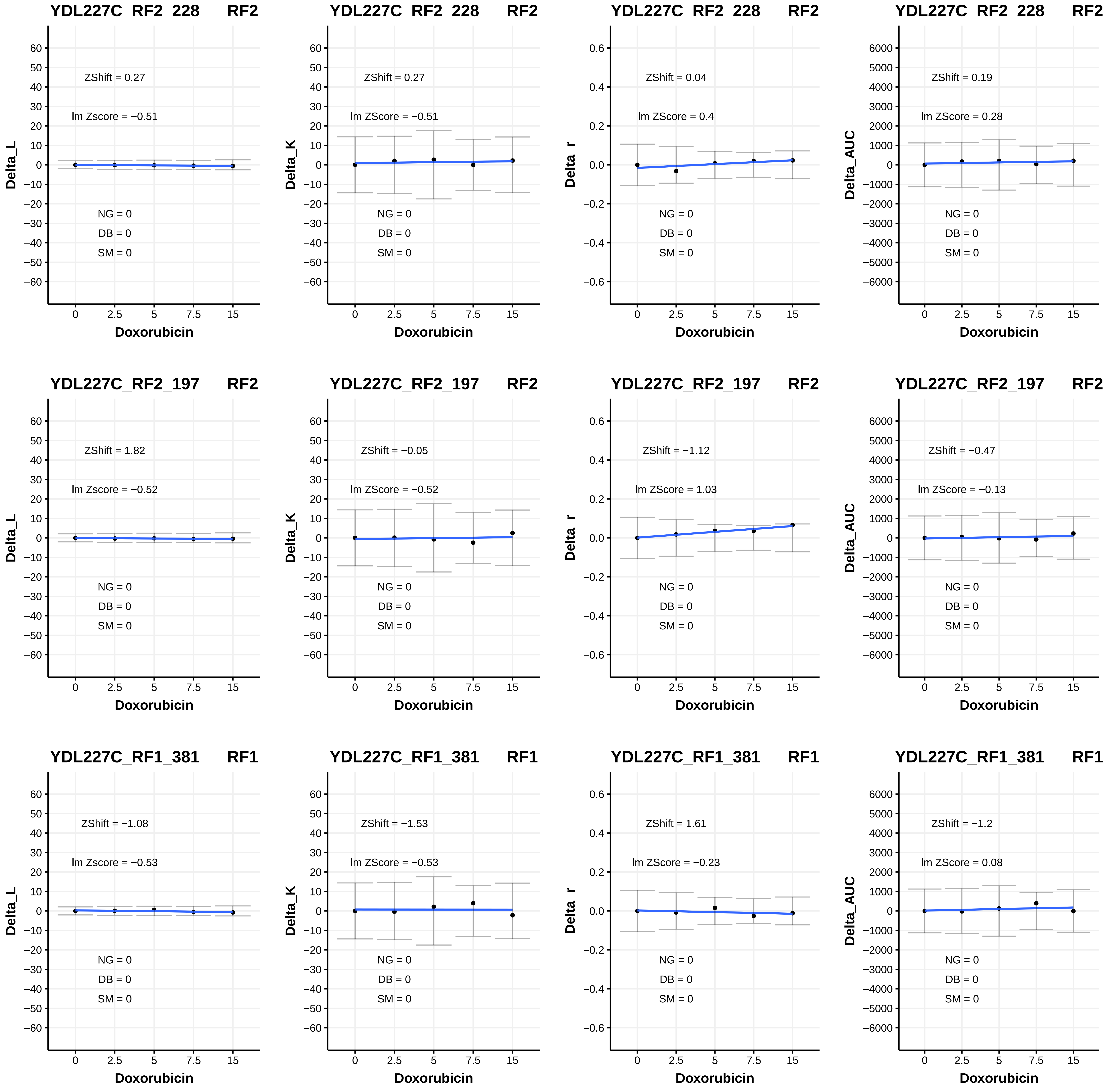
YDL227C_RF2_228 RF2
ZShift = 0.27
lm Zscore = −0.51
NG = 0
DB = 0
SM = 0
Delta_L
Doxorubicin
YDL227C_RF2_228 RF2
ZShift = 0.27
lm ZScore = −0.51
NG = 0
DB = 0
SM = 0
Delta_K
Doxorubicin
YDL227C_RF2_228 RF2
ZShift = 0.04
lm ZScore = 0.4
NG = 0
DB = 0
SM = 0
Delta_r
Doxorubicin
YDL227C_RF2_228 RF2
ZShift = 0.19
lm ZScore = 0.28
NG = 0
DB = 0
SM = 0
Delta_AUC
Doxorubicin
YDL227C_RF2_197 RF2
ZShift = 1.82
lm Zscore = −0.52
NG = 0
DB = 0
SM = 0
Delta_L
Doxorubicin
YDL227C_RF2_197 RF2
ZShift = −0.05
lm ZScore = −0.52
NG = 0
DB = 0
SM = 0
Delta_K
Doxorubicin
YDL227C_RF2_197 RF2
ZShift = −1.12
lm ZScore = 1.03
NG = 0
DB = 0
SM = 0
Delta_r
Doxorubicin
YDL227C_RF2_197 RF2
ZShift = −0.47
lm ZScore = −0.13
NG = 0
DB = 0
SM = 0
Delta_AUC
Doxorubicin
YDL227C_RF1_381 RF1
ZShift = −1.08
lm Zscore = −0.53
NG = 0
DB = 0
SM = 0
Delta_L
Doxorubicin
YDL227C_RF1_381 RF1
ZShift = −1.53
lm ZScore = −0.53
NG = 0
DB = 0
SM = 0
Delta_K
Doxorubicin
YDL227C_RF1_381 RF1
ZShift = 1.61
lm ZScore = −0.23
NG = 0
DB = 0
SM = 0
Delta_r
Doxorubicin
YDL227C_RF1_381 RF1
ZShift = −1.2
lm ZScore = 0.08
NG = 0
DB = 0
SM = 0
Delta_AUC
Doxorubicin

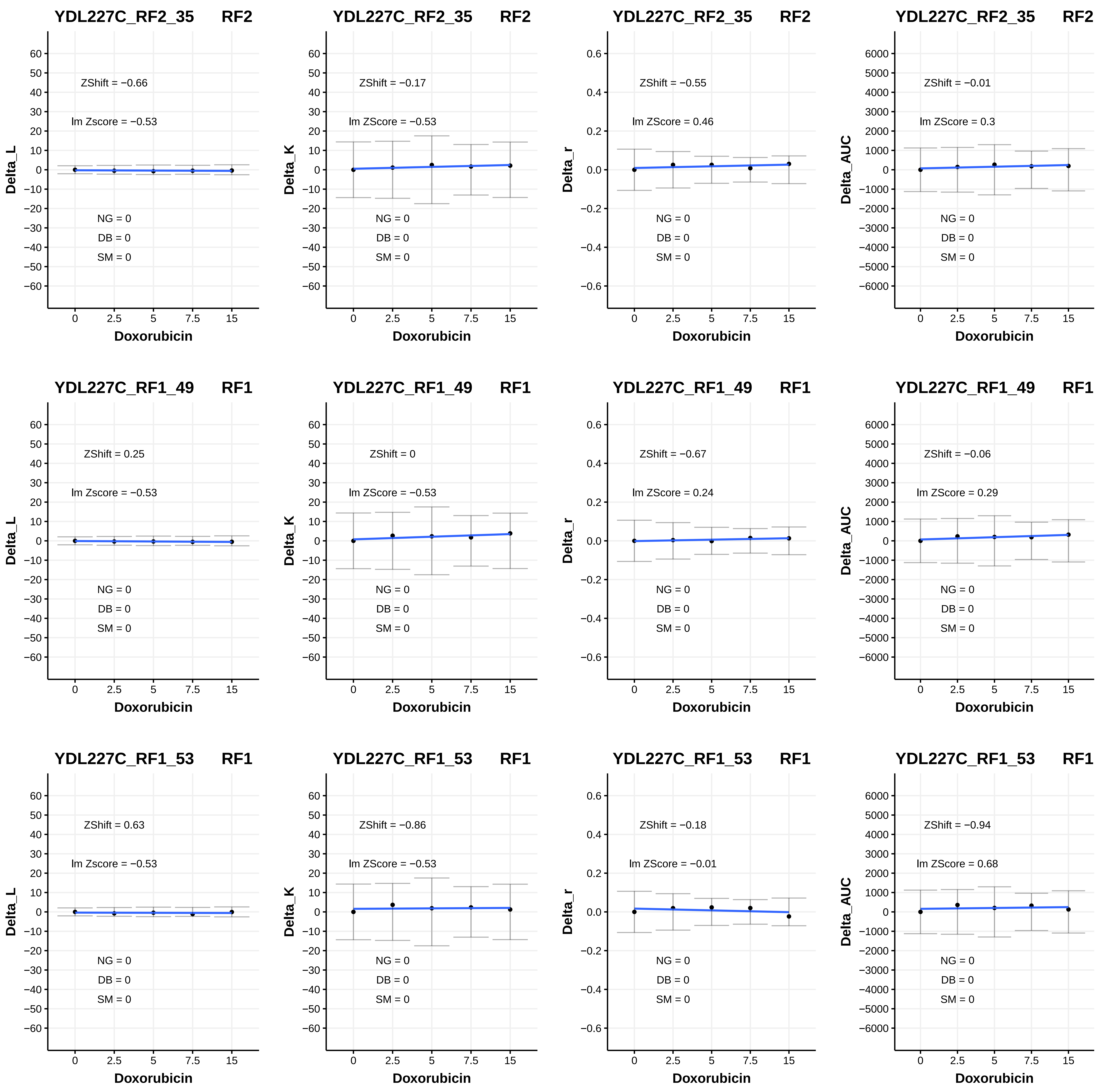
YDL227C_RF2_35 RF2
ZShift = −0.66
lm Zscore = −0.53
NG = 0
DB = 0
SM = 0
Delta_L
Doxorubicin
YDL227C_RF2_35 RF2
ZShift = −0.17
lm ZScore = −0.53
NG = 0
DB = 0
SM = 0
Delta_K
Doxorubicin
YDL227C_RF2_35 RF2
ZShift = −0.55
lm ZScore = 0.46
NG = 0
DB = 0
SM = 0
Delta_r
Doxorubicin
YDL227C_RF2_35 RF2
ZShift = −0.01
lm ZScore = 0.3
NG = 0
DB = 0
SM = 0
Delta_AUC
Doxorubicin
YDL227C_RF1_49 RF1
ZShift = 0.25
lm Zscore = −0.53
NG = 0
DB = 0
SM = 0
Delta_L
Doxorubicin
YDL227C_RF1_49 RF1
ZShift = 0
lm ZScore = −0.53
NG = 0
DB = 0
SM = 0
Delta_K
Doxorubicin
YDL227C_RF1_49 RF1
ZShift = −0.67
lm ZScore = 0.24
NG = 0
DB = 0
SM = 0
Delta_r
Doxorubicin
YDL227C_RF1_49 RF1
ZShift = −0.06
lm ZScore = 0.29
NG = 0
DB = 0
SM = 0
Delta_AUC
Doxorubicin
YDL227C_RF1_53 RF1
ZShift = 0.63
lm Zscore = −0.53
NG = 0
DB = 0
SM = 0
Delta_L
Doxorubicin
YDL227C_RF1_53 RF1
ZShift = −0.86
lm ZScore = −0.53
NG = 0
DB = 0
SM = 0
Delta_K
Doxorubicin
YDL227C_RF1_53 RF1
ZShift = −0.18
lm ZScore = −0.01
NG = 0
DB = 0
SM = 0
Delta_r
Doxorubicin
YDL227C_RF1_53 RF1
ZShift = −0.94
lm ZScore = 0.68
NG = 0
DB = 0
SM = 0
Delta_AUC
Doxorubicin

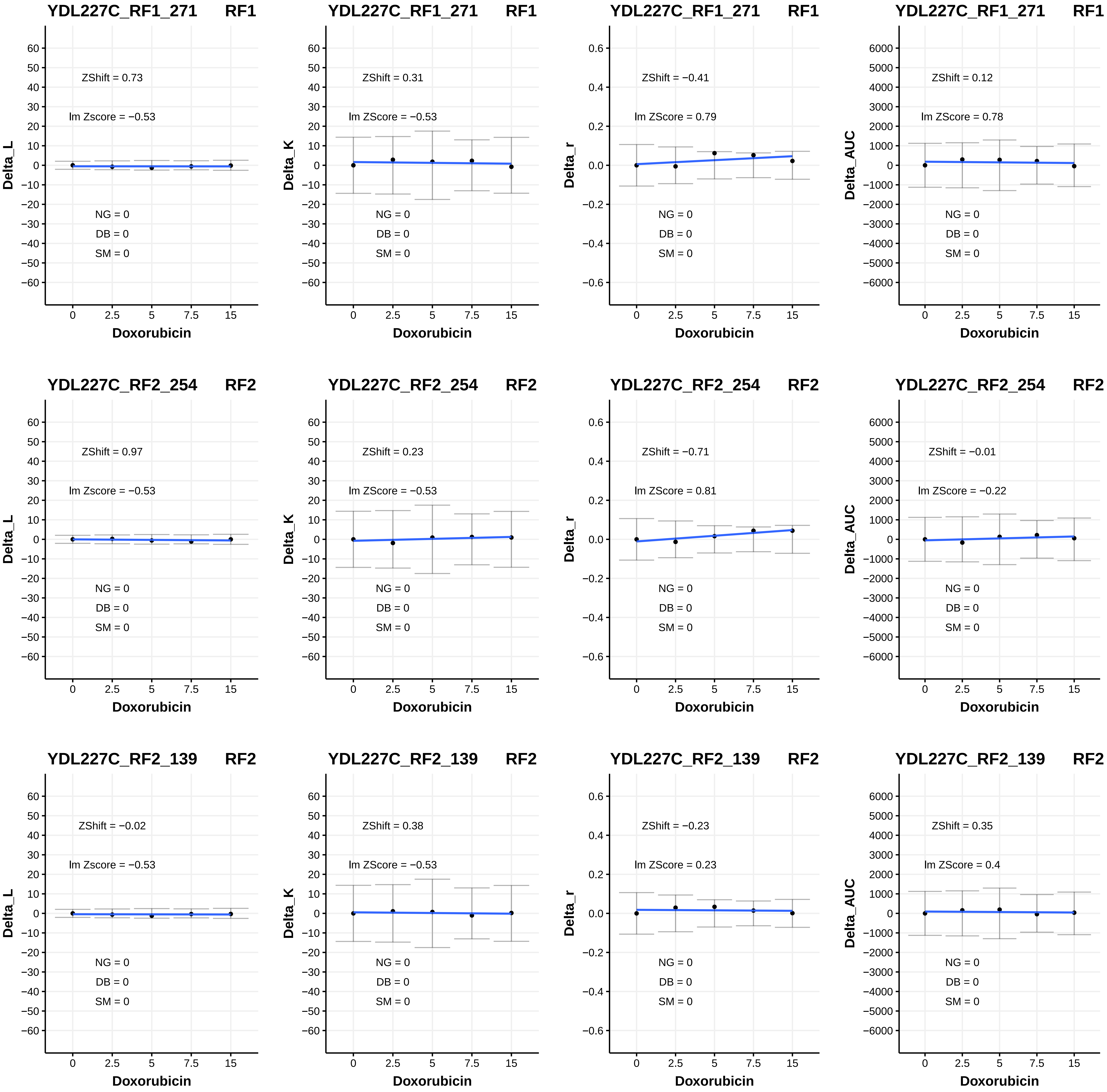

YDL227C_RF1_271 RF1
ZShift = 0.73
lm Zscore = −0.53
NG = 0
DB = 0
SM = 0
Delta_L
Doxorubicin
YDL227C_RF1_271 RF1
ZShift = 0.31
lm ZScore = −0.53
NG = 0
DB = 0
SM = 0
Delta_K
Doxorubicin
YDL227C_RF1_271 RF1
ZShift = −0.41
lm ZScore = 0.79
NG = 0
DB = 0
SM = 0
Delta_r
Doxorubicin
YDL227C_RF1_271 RF1
ZShift = 0.12
lm ZScore = 0.78
NG = 0
DB = 0
SM = 0
Delta_AUC
Doxorubicin
YDL227C_RF2_254 RF2
ZShift = 0.97
lm Zscore = −0.53
NG = 0
DB = 0
SM = 0
Delta_L
Doxorubicin
YDL227C_RF2_254 RF2
ZShift = 0.23
lm ZScore = −0.53
NG = 0
DB = 0
SM = 0
Delta_K
Doxorubicin
YDL227C_RF2_254 RF2
ZShift = −0.71
lm ZScore = 0.81
NG = 0
DB = 0
SM = 0
Delta_r
Doxorubicin
YDL227C_RF2_254 RF2
ZShift = −0.01
lm ZScore = −0.22
NG = 0
DB = 0
SM = 0
Delta_AUC
Doxorubicin
YDL227C_RF2_139 RF2
ZShift = −0.02
lm Zscore = −0.53
NG = 0
DB = 0
SM = 0
Delta_L
Doxorubicin
YDL227C_RF2_139 RF2
ZShift = 0.38
lm ZScore = −0.53
NG = 0
DB = 0
SM = 0
Delta_K
Doxorubicin
YDL227C_RF2_139 RF2
ZShift = −0.23
lm ZScore = 0.23
NG = 0
DB = 0
SM = 0
Delta_r
Doxorubicin
YDL227C_RF2_139 RF2
ZShift = 0.35
lm ZScore = 0.4
NG = 0
DB = 0
SM = 0
Delta_AUC
Doxorubicin

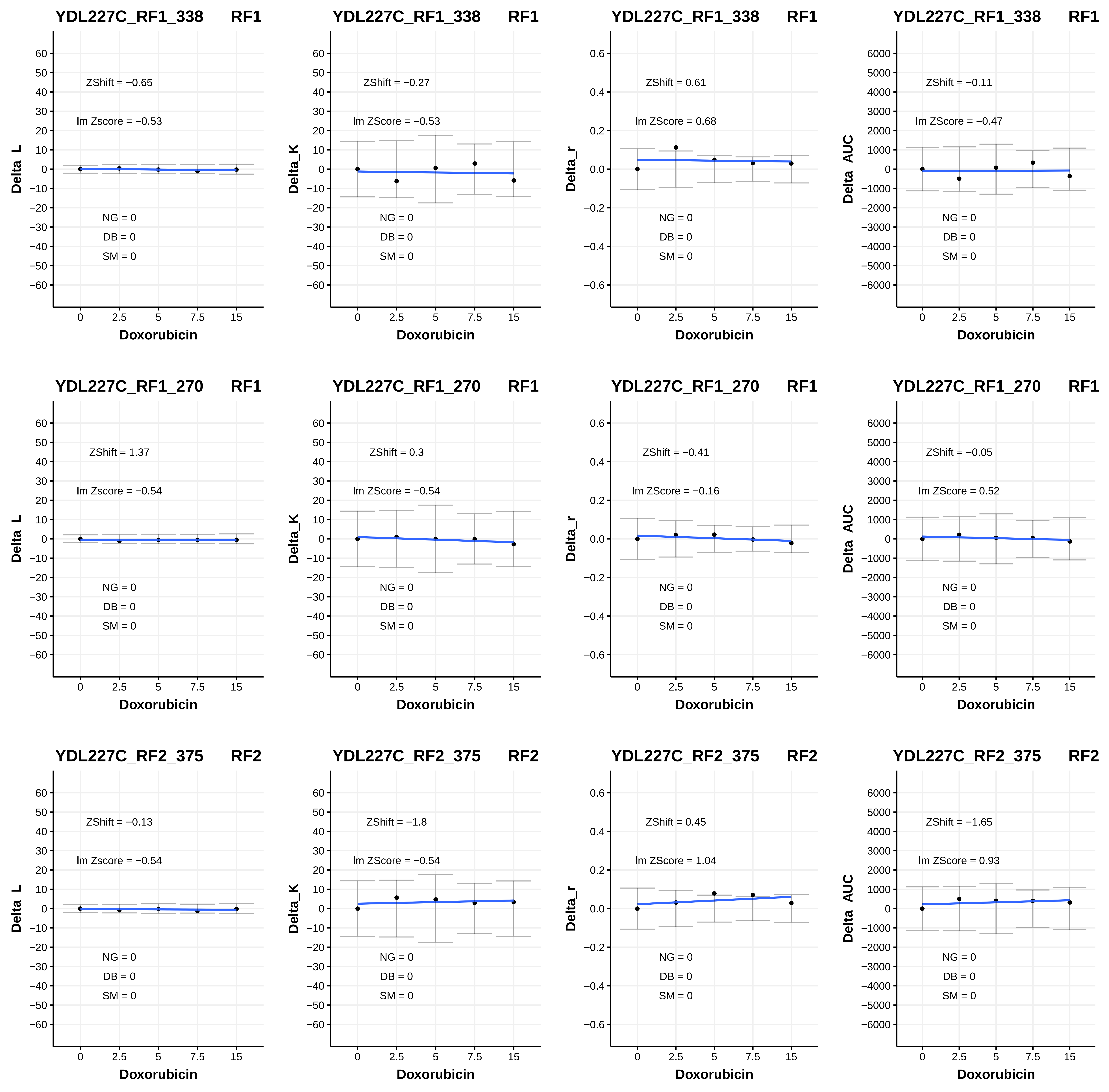

YDL227C_RF1_338 RF1
ZShift = −0.65
lm Zscore = −0.53
NG = 0
DB = 0
SM = 0
Delta_L
Doxorubicin
YDL227C_RF1_338 RF1
ZShift = −0.27
lm ZScore = −0.53
NG = 0
DB = 0
SM = 0
Delta_K
Doxorubicin
YDL227C_RF1_338 RF1
ZShift = 0.61
lm ZScore = 0.68
NG = 0
DB = 0
SM = 0
Delta_r
Doxorubicin
YDL227C_RF1_338 RF1
ZShift = −0.11
lm ZScore = −0.47
NG = 0
DB = 0
SM = 0
Delta_AUC
Doxorubicin
YDL227C_RF1_270 RF1
ZShift = 1.37
lm Zscore = −0.54
NG = 0
DB = 0
SM = 0
Delta_L
Doxorubicin
YDL227C_RF1_270 RF1
ZShift = 0.3
lm ZScore = −0.54
NG = 0
DB = 0
SM = 0
Delta_K
Doxorubicin
YDL227C_RF1_270 RF1
ZShift = −0.41
lm ZScore = −0.16
NG = 0
DB = 0
SM = 0
Delta_r
Doxorubicin
YDL227C_RF1_270 RF1
ZShift = −0.05
lm ZScore = 0.52
NG = 0
DB = 0
SM = 0
Delta_AUC
Doxorubicin
YDL227C_RF2_375 RF2
ZShift = −0.13
lm Zscore = −0.54
NG = 0
DB = 0
SM = 0
Delta_L
Doxorubicin
YDL227C_RF2_375 RF2
ZShift = −1.8
lm ZScore = −0.54
NG = 0
DB = 0
SM = 0
Delta_K
Doxorubicin
YDL227C_RF2_375 RF2
ZShift = 0.45
lm ZScore = 1.04
NG = 0
DB = 0
SM = 0
Delta_r
Doxorubicin
YDL227C_RF2_375 RF2
ZShift = −1.65
lm ZScore = 0.93
NG = 0
DB = 0
SM = 0
Delta_AUC
Doxorubicin

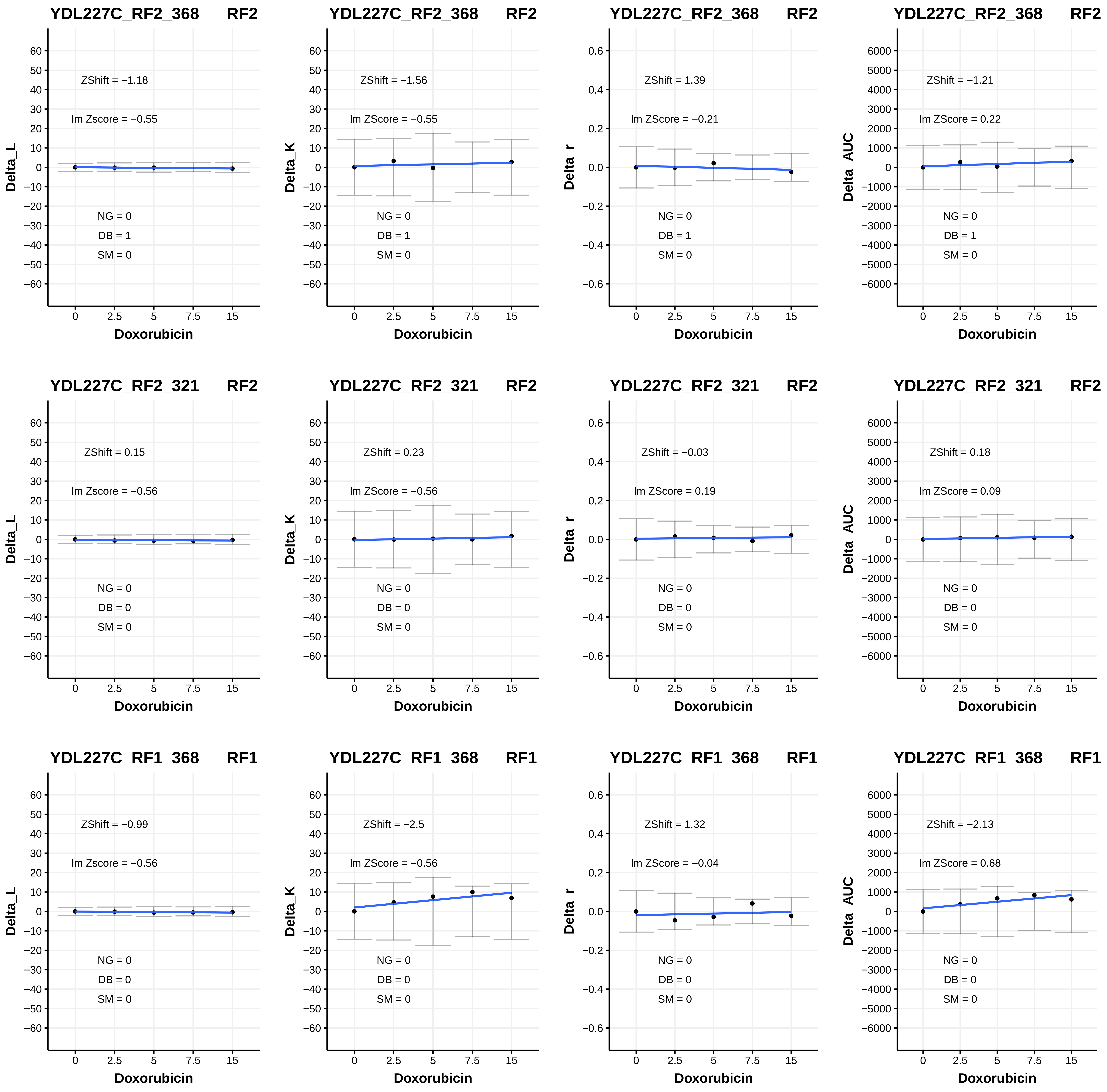
YDL227C_RF2_368 RF2
ZShift = −1.18
lm Zscore = −0.55
NG = 0
DB = 1
SM = 0
Delta_L
Doxorubicin
YDL227C_RF2_368 RF2
ZShift = −1.56
lm ZScore = −0.55
NG = 0
DB = 1
SM = 0
Delta_K
Doxorubicin
YDL227C_RF2_368 RF2
ZShift = 1.39
lm ZScore = −0.21
NG = 0
DB = 1
SM = 0
Delta_r
Doxorubicin
YDL227C_RF2_368 RF2
ZShift = −1.21
lm ZScore = 0.22
NG = 0
DB = 1
SM = 0
Delta_AUC
Doxorubicin
YDL227C_RF2_321 RF2
ZShift = 0.15
lm Zscore = −0.56
NG = 0
DB = 0
SM = 0
Delta_L
Doxorubicin
YDL227C_RF2_321 RF2
ZShift = 0.23
lm ZScore = −0.56
NG = 0
DB = 0
SM = 0
Delta_K
Doxorubicin
YDL227C_RF2_321 RF2
ZShift = −0.03
lm ZScore = 0.19
NG = 0
DB = 0
SM = 0
Delta_r
Doxorubicin
YDL227C_RF2_321 RF2
ZShift = 0.18
lm ZScore = 0.09
NG = 0
DB = 0
SM = 0
Delta_AUC
Doxorubicin
YDL227C_RF1_368 RF1
ZShift = −0.99
lm Zscore = −0.56
NG = 0
DB = 0
SM = 0
Delta_L
Doxorubicin
YDL227C_RF1_368 RF1
ZShift = −2.5
lm ZScore = −0.56
NG = 0
DB = 0
SM = 0
Delta_K
Doxorubicin
YDL227C_RF1_368 RF1
ZShift = 1.32
lm ZScore = −0.04
NG = 0
DB = 0
SM = 0
Delta_r
Doxorubicin
YDL227C_RF1_368 RF1
ZShift = −2.13
lm ZScore = 0.68
NG = 0
DB = 0
SM = 0
Delta_AUC
Doxorubicin

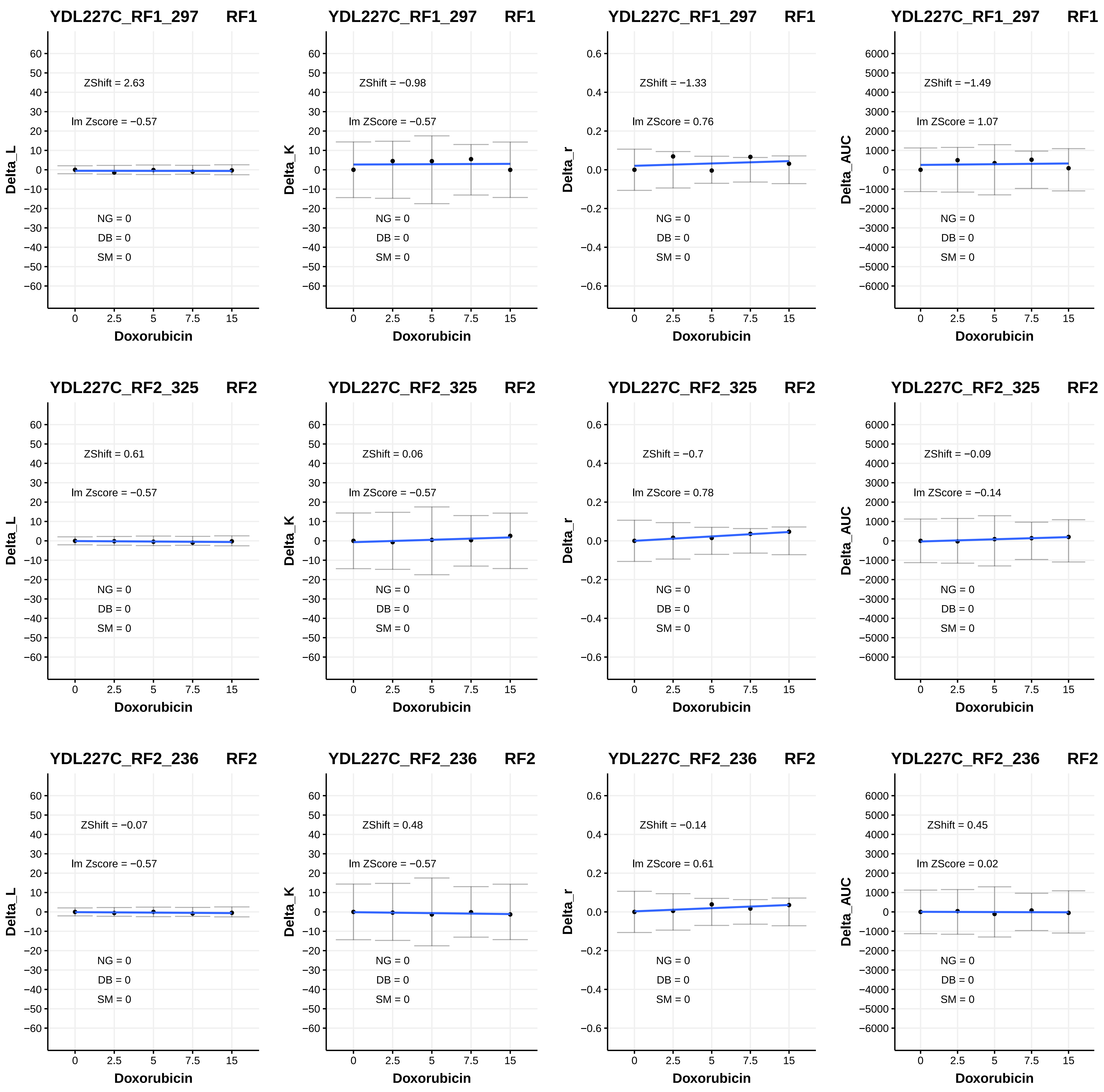
YDL227C_RF1_297 RF1
ZShift = 2.63
lm Zscore = −0.57
NG = 0
DB = 0
SM = 0
Delta_L
Doxorubicin
YDL227C_RF1_297 RF1
ZShift = −0.98
lm ZScore = −0.57
NG = 0
DB = 0
SM = 0
Delta_K
Doxorubicin
YDL227C_RF1_297 RF1
ZShift = −1.33
lm ZScore = 0.76
NG = 0
DB = 0
SM = 0
Delta_r
Doxorubicin
YDL227C_RF1_297 RF1
ZShift = −1.49
lm ZScore = 1.07
NG = 0
DB = 0
SM = 0
Delta_AUC
Doxorubicin
YDL227C_RF2_325 RF2
ZShift = 0.61
lm Zscore = −0.57
NG = 0
DB = 0
SM = 0
Delta_L
Doxorubicin
YDL227C_RF2_325 RF2
ZShift = 0.06
lm ZScore = −0.57
NG = 0
DB = 0
SM = 0
Delta_K
Doxorubicin
YDL227C_RF2_325 RF2
ZShift = −0.7
lm ZScore = 0.78
NG = 0
DB = 0
SM = 0
Delta_r
Doxorubicin
YDL227C_RF2_325 RF2
ZShift = −0.09
lm ZScore = −0.14
NG = 0
DB = 0
SM = 0
Delta_AUC
Doxorubicin
YDL227C_RF2_236 RF2
ZShift = −0.07
lm Zscore = −0.57
NG = 0
DB = 0
SM = 0
Delta_L
Doxorubicin
YDL227C_RF2_236 RF2
ZShift = 0.48
lm ZScore = −0.57
NG = 0
DB = 0
SM = 0
Delta_K
Doxorubicin
YDL227C_RF2_236 RF2
ZShift = −0.14
lm ZScore = 0.61
NG = 0
DB = 0
SM = 0
Delta_r
Doxorubicin
YDL227C_RF2_236 RF2
ZShift = 0.45
lm ZScore = 0.02
NG = 0
DB = 0
SM = 0
Delta_AUC
Doxorubicin

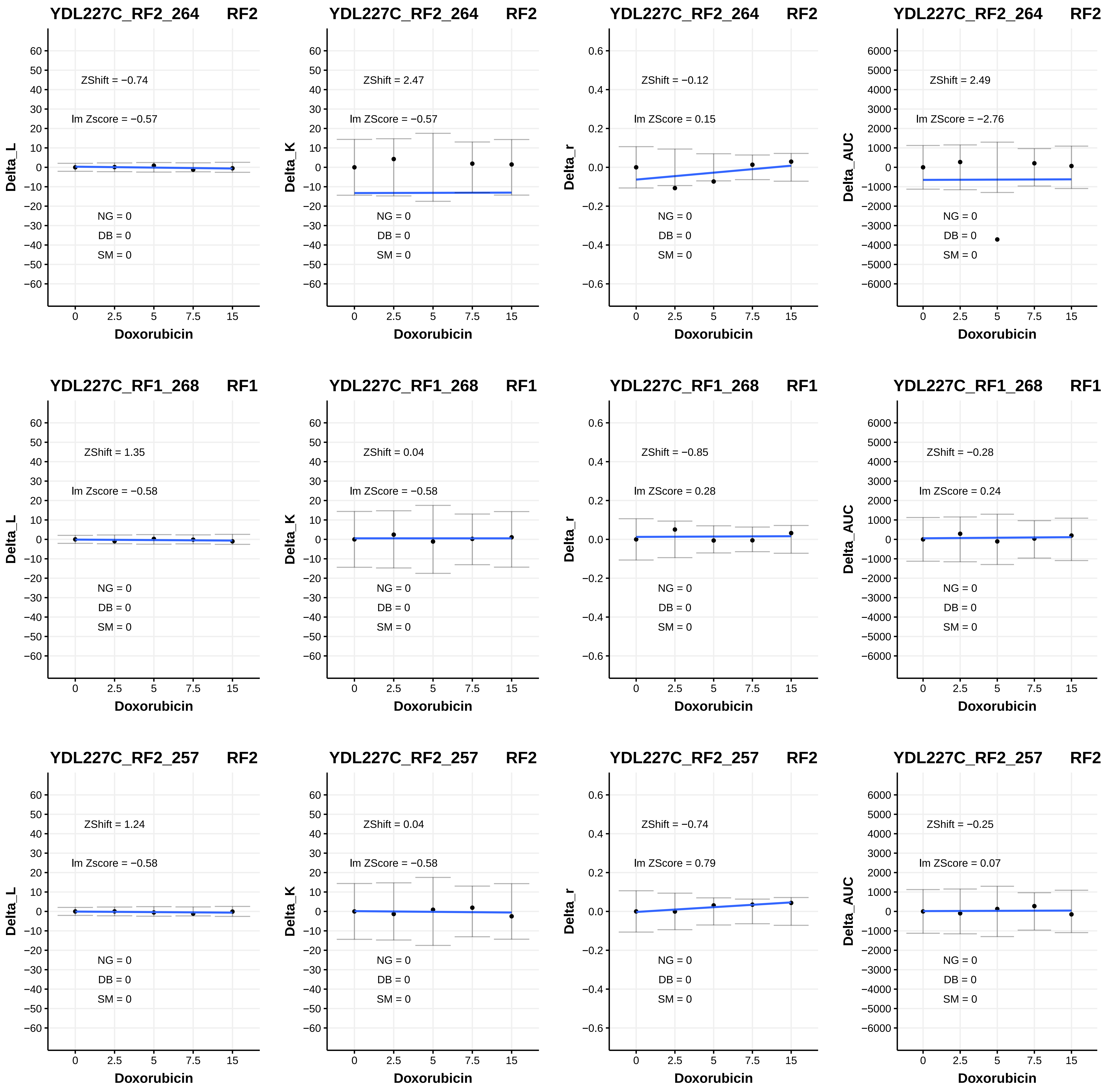

YDL227C_RF2_264 RF2
ZShift = −0.74
lm Zscore = −0.57
NG = 0
DB = 0
SM = 0
Delta_L
Doxorubicin
YDL227C_RF2_264 RF2
ZShift = 2.47
lm ZScore = −0.57
NG = 0
DB = 0
SM = 0
Delta_K
Doxorubicin
YDL227C_RF2_264 RF2
ZShift = −0.12
lm ZScore = 0.15
NG = 0
DB = 0
SM = 0
Delta_r
Doxorubicin
YDL227C_RF2_264 RF2
ZShift = 2.49
lm ZScore = −2.76
NG = 0
DB = 0
SM = 0
Delta_AUC
Doxorubicin
YDL227C_RF1_268 RF1
ZShift = 1.35
lm Zscore = −0.58
NG = 0
DB = 0
SM = 0
Delta_L
Doxorubicin
YDL227C_RF1_268 RF1
ZShift = 0.04
lm ZScore = −0.58
NG = 0
DB = 0
SM = 0
Delta_K
Doxorubicin
YDL227C_RF1_268 RF1
ZShift = −0.85
lm ZScore = 0.28
NG = 0
DB = 0
SM = 0
Delta_r
Doxorubicin
YDL227C_RF1_268 RF1
ZShift = −0.28
lm ZScore = 0.24
NG = 0
DB = 0
SM = 0
Delta_AUC
Doxorubicin
YDL227C_RF2_257 RF2
ZShift = 1.24
lm Zscore = −0.58
NG = 0
DB = 0
SM = 0
Delta_L
Doxorubicin
YDL227C_RF2_257 RF2
ZShift = 0.04
lm ZScore = −0.58
NG = 0
DB = 0
SM = 0
Delta_K
Doxorubicin
YDL227C_RF2_257 RF2
ZShift = −0.74
lm ZScore = 0.79
NG = 0
DB = 0
SM = 0
Delta_r
Doxorubicin
YDL227C_RF2_257 RF2
ZShift = −0.25
lm ZScore = 0.07
NG = 0
DB = 0
SM = 0
Delta_AUC
Doxorubicin

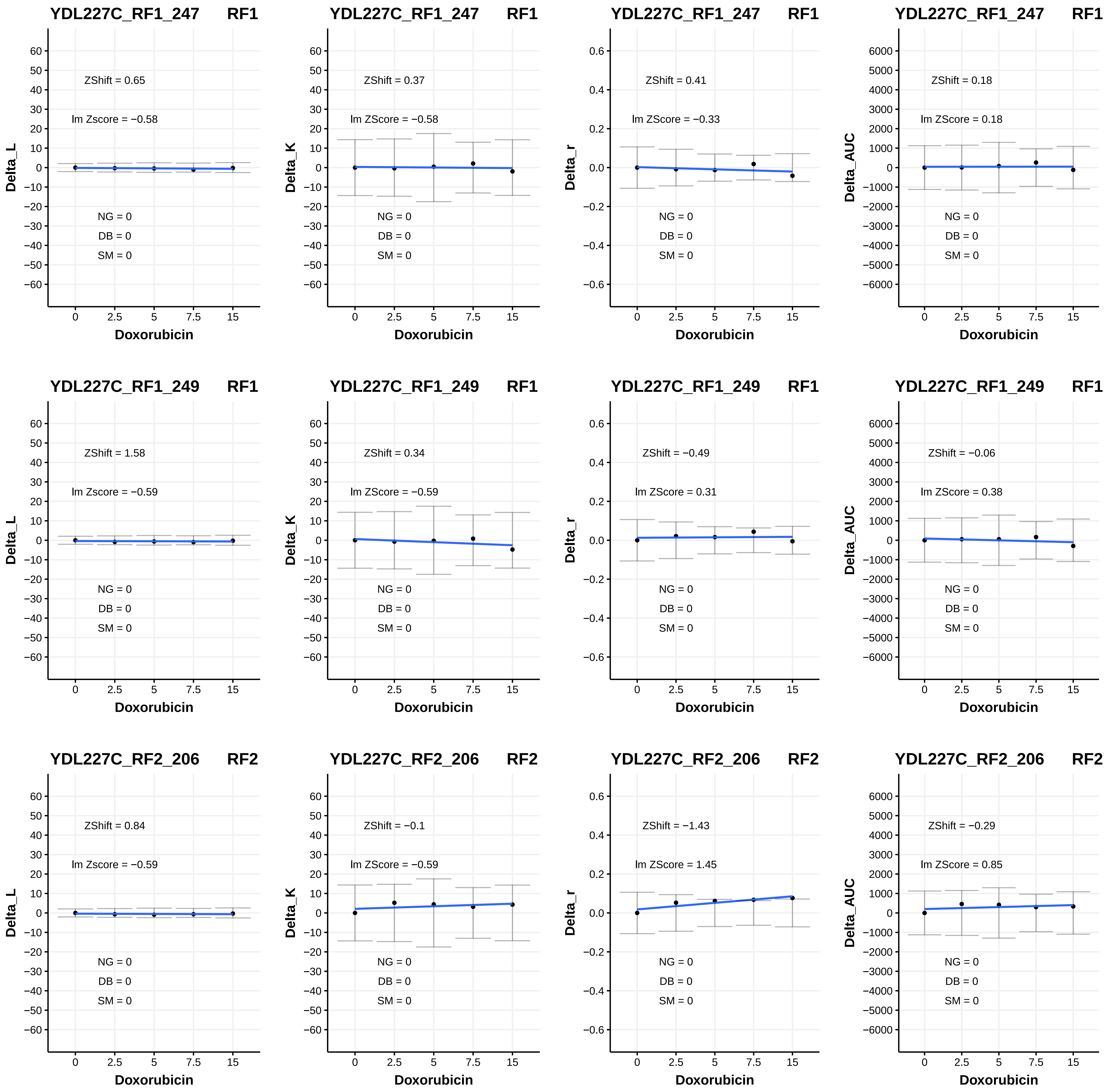
YDL227C_RF1_247 RF1
ZShift = 0.65
lm Zscore = −0.58
NG = 0
DB = 0
SM = 0
Delta_L
Doxorubicin
YDL227C_RF1_247 RF1
ZShift = 0.37
lm ZScore = −0.58
NG = 0
DB = 0
SM = 0
Delta_K
Doxorubicin
YDL227C_RF1_247 RF1
ZShift = 0.41
lm ZScore = −0.33
NG = 0
DB = 0
SM = 0
Delta_r
Doxorubicin
YDL227C_RF1_247 RF1
ZShift = 0.18
lm ZScore = 0.18
NG = 0
DB = 0
SM = 0
Delta_AUC
Doxorubicin
YDL227C_RF1_249 RF1
ZShift = 1.58
lm Zscore = −0.59
NG = 0
DB = 0
SM = 0
Delta_L
Doxorubicin
YDL227C_RF1_249 RF1
ZShift = 0.34
lm ZScore = −0.59
NG = 0
DB = 0
SM = 0
Delta_K
Doxorubicin
YDL227C_RF1_249 RF1
ZShift = −0.49
lm ZScore = 0.31
NG = 0
DB = 0
SM = 0
Delta_r
Doxorubicin
YDL227C_RF1_249 RF1
ZShift = −0.06
lm ZScore = 0.38
NG = 0
DB = 0
SM = 0
Delta_AUC
Doxorubicin
YDL227C_RF2_206 RF2
ZShift = 0.84
lm Zscore = −0.59
NG = 0
DB = 0
SM = 0
Delta_L
Doxorubicin
YDL227C_RF2_206 RF2
ZShift = −0.1
lm ZScore = −0.59
NG = 0
DB = 0
SM = 0
Delta_K
Doxorubicin
YDL227C_RF2_206 RF2
ZShift = −1.43
lm ZScore = 1.45
NG = 0
DB = 0
SM = 0
Delta_r
Doxorubicin
YDL227C_RF2_206 RF2
ZShift = −0.29
lm ZScore = 0.85
NG = 0
DB = 0
SM = 0
Delta_AUC
Doxorubicin

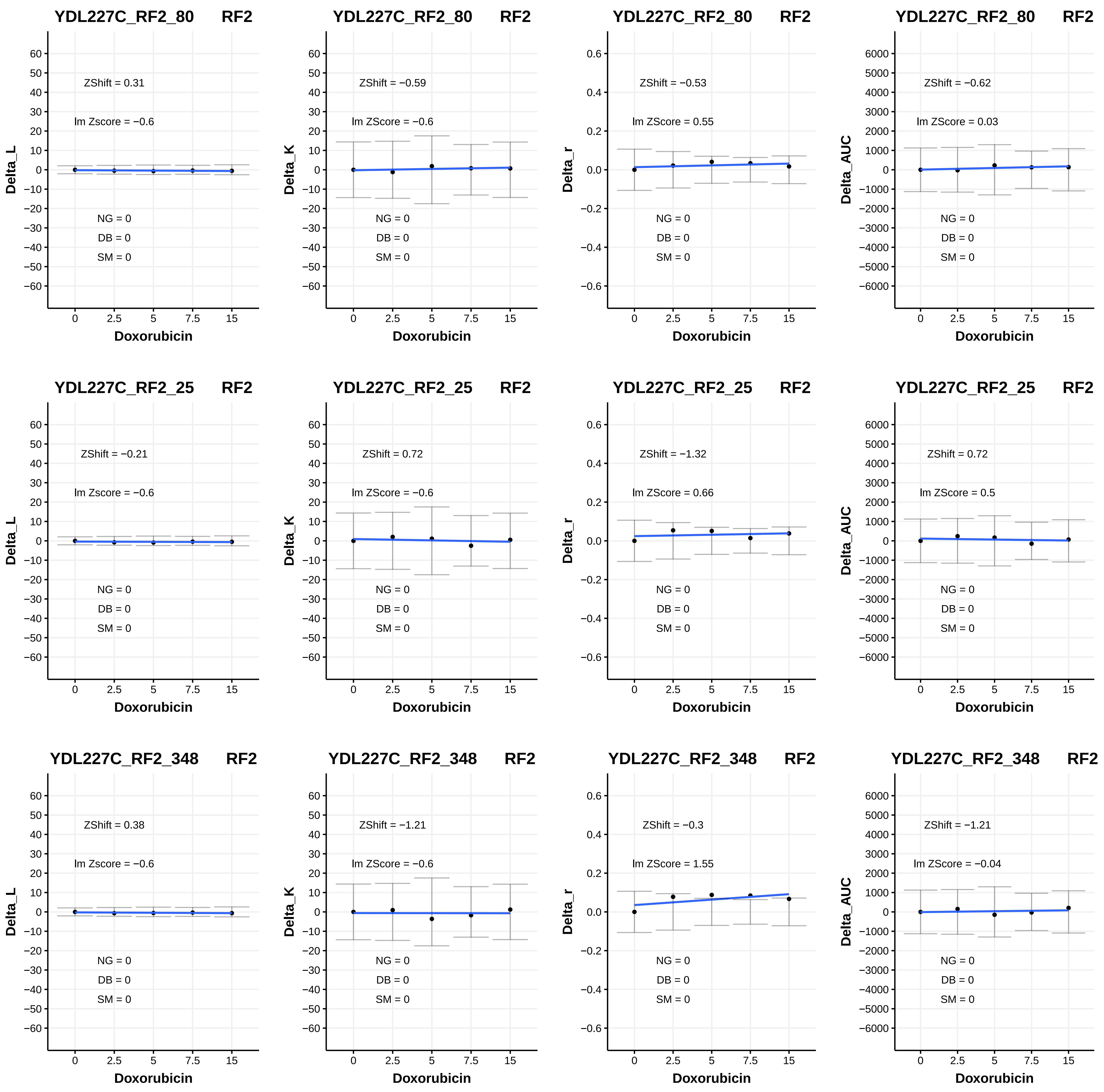

YDL227C_RF2_80 RF2
ZShift = 0.31
lm Zscore = −0.6
NG = 0
DB = 0
SM = 0
Delta_L
Doxorubicin
YDL227C_RF2_80 RF2
ZShift = −0.59
lm ZScore = −0.6
NG = 0
DB = 0
SM = 0
Delta_K
Doxorubicin
YDL227C_RF2_80 RF2
ZShift = −0.53
lm ZScore = 0.55
NG = 0
DB = 0
SM = 0
Delta_r
Doxorubicin
YDL227C_RF2_80 RF2
ZShift = −0.62
lm ZScore = 0.03
NG = 0
DB = 0
SM = 0
Delta_AUC
Doxorubicin
YDL227C_RF2_25 RF2
ZShift = −0.21
lm Zscore = −0.6
NG = 0
DB = 0
SM = 0
Delta_L
Doxorubicin
YDL227C_RF2_25 RF2
ZShift = 0.72
lm ZScore = −0.6
NG = 0
DB = 0
SM = 0
Delta_K
Doxorubicin
YDL227C_RF2_25 RF2
ZShift = −1.32
lm ZScore = 0.66
NG = 0
DB = 0
SM = 0
Delta_r
Doxorubicin
YDL227C_RF2_25 RF2
ZShift = 0.72
lm ZScore = 0.5
NG = 0
DB = 0
SM = 0
Delta_AUC
Doxorubicin
YDL227C_RF2_348 RF2
ZShift = 0.38
lm Zscore = −0.6
NG = 0
DB = 0
SM = 0
Delta_L
Doxorubicin
YDL227C_RF2_348 RF2
ZShift = −1.21
lm ZScore = −0.6
NG = 0
DB = 0
SM = 0
Delta_K
Doxorubicin
YDL227C_RF2_348 RF2
ZShift = −0.3
lm ZScore = 1.55
NG = 0
DB = 0
SM = 0
Delta_r
Doxorubicin
YDL227C_RF2_348 RF2
ZShift = −1.21
lm ZScore = −0.04
NG = 0
DB = 0
SM = 0
Delta_AUC
Doxorubicin

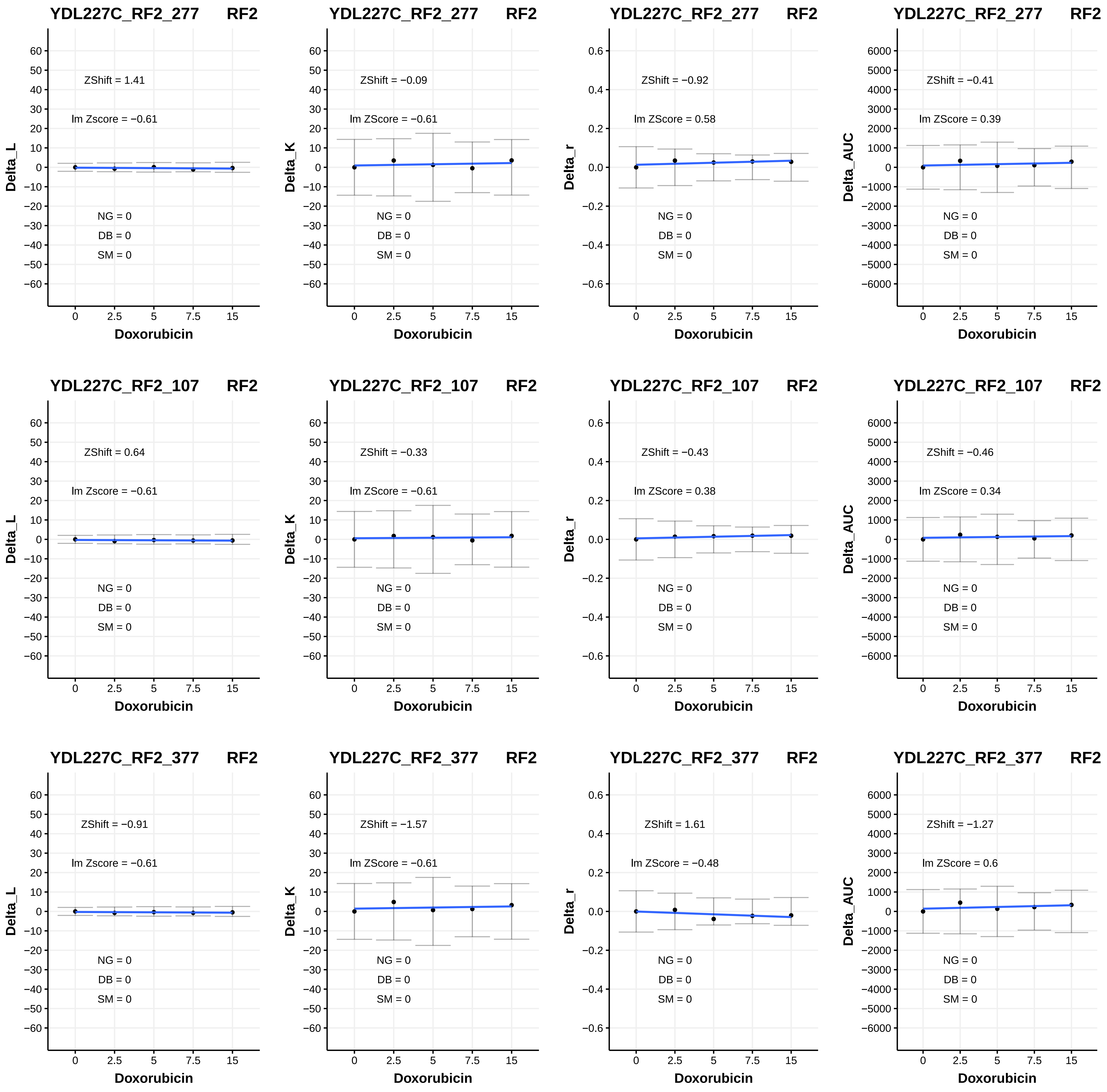
YDL227C_RF2_277 RF2
ZShift = 1.41
lm Zscore = −0.61
NG = 0
DB = 0
SM = 0
Delta_L
Doxorubicin
YDL227C_RF2_277 RF2
ZShift = −0.09
lm ZScore = −0.61
NG = 0
DB = 0
SM = 0
Delta_K
Doxorubicin
YDL227C_RF2_277 RF2
ZShift = −0.92
lm ZScore = 0.58
NG = 0
DB = 0
SM = 0
Delta_r
Doxorubicin
YDL227C_RF2_277 RF2
ZShift = −0.41
lm ZScore = 0.39
NG = 0
DB = 0
SM = 0
Delta_AUC
Doxorubicin
YDL227C_RF2_107 RF2
ZShift = 0.64
lm Zscore = −0.61
NG = 0
DB = 0
SM = 0
Delta_L
Doxorubicin
YDL227C_RF2_107 RF2
ZShift = −0.33
lm ZScore = −0.61
NG = 0
DB = 0
SM = 0
Delta_K
Doxorubicin
YDL227C_RF2_107 RF2
ZShift = −0.43
lm ZScore = 0.38
NG = 0
DB = 0
SM = 0
Delta_r
Doxorubicin
YDL227C_RF2_107 RF2
ZShift = −0.46
lm ZScore = 0.34
NG = 0
DB = 0
SM = 0
Delta_AUC
Doxorubicin
YDL227C_RF2_377 RF2
ZShift = −0.91
lm Zscore = −0.61
NG = 0
DB = 0
SM = 0
Delta_L
Doxorubicin
YDL227C_RF2_377 RF2
ZShift = −1.57
lm ZScore = −0.61
NG = 0
DB = 0
SM = 0
Delta_K
Doxorubicin
YDL227C_RF2_377 RF2
ZShift = 1.61
lm ZScore = −0.48
NG = 0
DB = 0
SM = 0
Delta_r
Doxorubicin
YDL227C_RF2_377 RF2
ZShift = −1.27
lm ZScore = 0.6
NG = 0
DB = 0
SM = 0
Delta_AUC
Doxorubicin

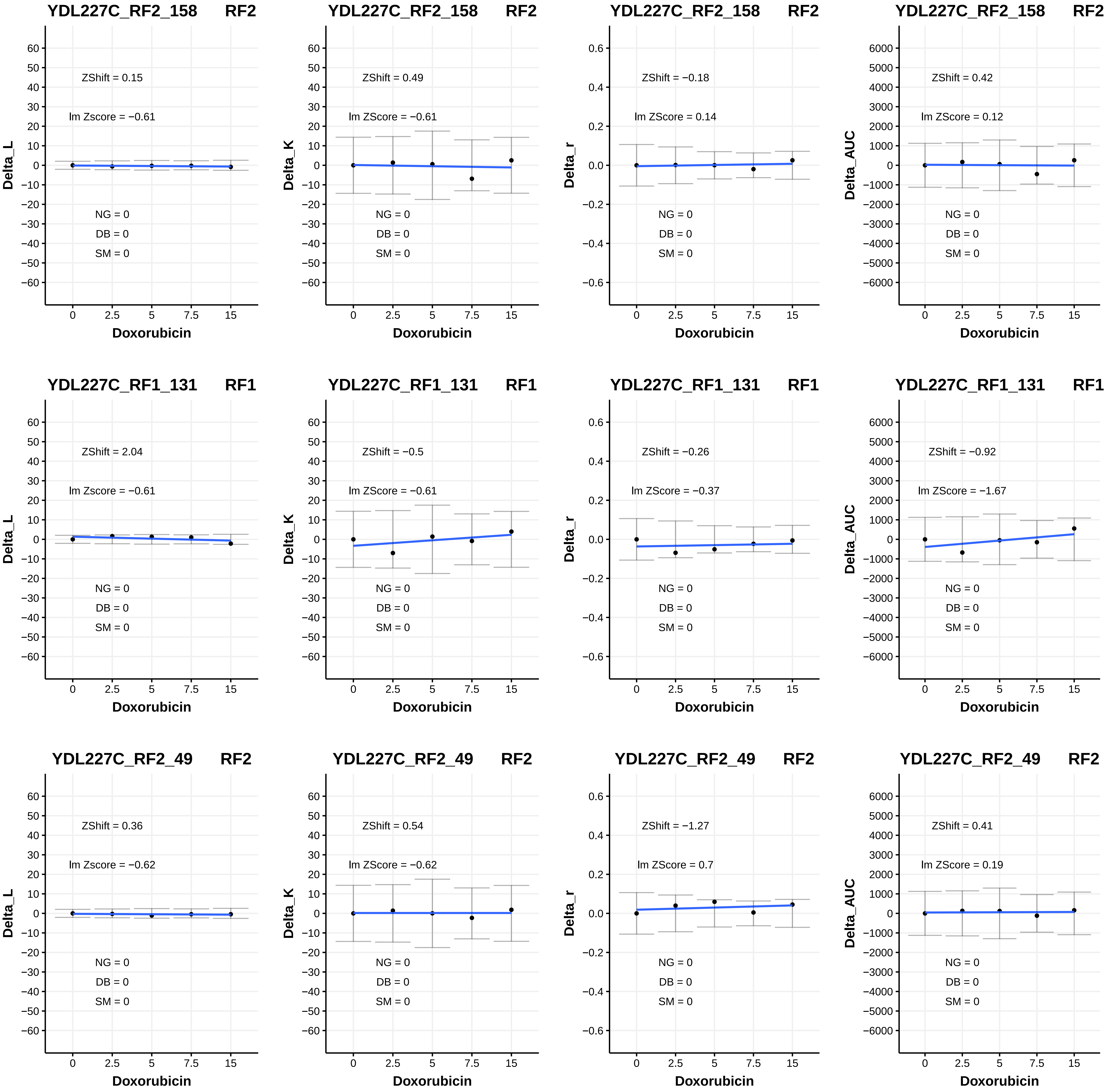
YDL227C_RF2_158 RF2
ZShift = 0.15
lm Zscore = −0.61
NG = 0
DB = 0
SM = 0
Delta_L
Doxorubicin
YDL227C_RF2_158 RF2
ZShift = 0.49
lm ZScore = −0.61
NG = 0
DB = 0
SM = 0
Delta_K
Doxorubicin
YDL227C_RF2_158 RF2
ZShift = −0.18
lm ZScore = 0.14
NG = 0
DB = 0
SM = 0
Delta_r
Doxorubicin
YDL227C_RF2_158 RF2
ZShift = 0.42
lm ZScore = 0.12
NG = 0
DB = 0
SM = 0
Delta_AUC
Doxorubicin
YDL227C_RF1_131 RF1
ZShift = 2.04
lm Zscore = −0.61
NG = 0
DB = 0
SM = 0
Delta_L
Doxorubicin
YDL227C_RF1_131 RF1
ZShift = −0.5
lm ZScore = −0.61
NG = 0
DB = 0
SM = 0
Delta_K
Doxorubicin
YDL227C_RF1_131 RF1
ZShift = −0.26
lm ZScore = −0.37
NG = 0
DB = 0
SM = 0
Delta_r
Doxorubicin
YDL227C_RF1_131 RF1
ZShift = −0.92
lm ZScore = −1.67
NG = 0
DB = 0
SM = 0
Delta_AUC
Doxorubicin
YDL227C_RF2_49 RF2
ZShift = 0.36
lm Zscore = −0.62
NG = 0
DB = 0
SM = 0
Delta_L
Doxorubicin
YDL227C_RF2_49 RF2
ZShift = 0.54
lm ZScore = −0.62
NG = 0
DB = 0
SM = 0
Delta_K
Doxorubicin
YDL227C_RF2_49 RF2
ZShift = −1.27
lm ZScore = 0.7
NG = 0
DB = 0
SM = 0
Delta_r
Doxorubicin
YDL227C_RF2_49 RF2
ZShift = 0.41
lm ZScore = 0.19
NG = 0
DB = 0
SM = 0
Delta_AUC
Doxorubicin

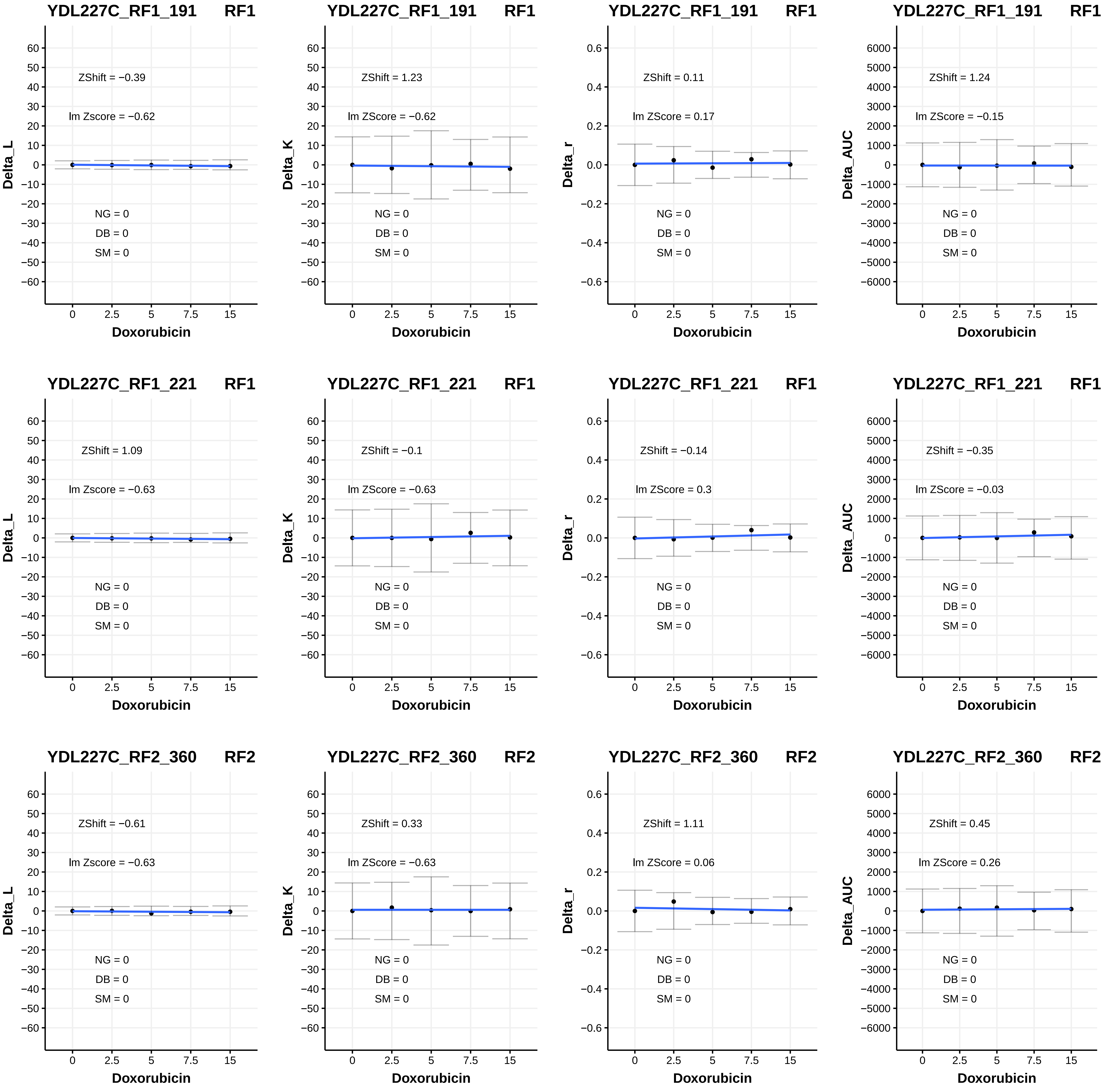
YDL227C_RF1_191 RF1
ZShift = −0.39
lm Zscore = −0.62
NG = 0
DB = 0
SM = 0
Delta_L
Doxorubicin
YDL227C_RF1_191 RF1
ZShift = 1.23
lm ZScore = −0.62
NG = 0
DB = 0
SM = 0
Delta_K
Doxorubicin
YDL227C_RF1_191 RF1
ZShift = 0.11
lm ZScore = 0.17
NG = 0
DB = 0
SM = 0
Delta_r
Doxorubicin
YDL227C_RF1_191 RF1
ZShift = 1.24
lm ZScore = −0.15
NG = 0
DB = 0
SM = 0
Delta_AUC
Doxorubicin
YDL227C_RF1_221 RF1
ZShift = 1.09
lm Zscore = −0.63
NG = 0
DB = 0
SM = 0
Delta_L
Doxorubicin
YDL227C_RF1_221 RF1
ZShift = −0.1
lm ZScore = −0.63
NG = 0
DB = 0
SM = 0
Delta_K
Doxorubicin
YDL227C_RF1_221 RF1
ZShift = −0.14
lm ZScore = 0.3
NG = 0
DB = 0
SM = 0
Delta_r
Doxorubicin
YDL227C_RF1_221 RF1
ZShift = −0.35
lm ZScore = −0.03
NG = 0
DB = 0
SM = 0
Delta_AUC
Doxorubicin
YDL227C_RF2_360 RF2
ZShift = −0.61
lm Zscore = −0.63
NG = 0
DB = 0
SM = 0
Delta_L
Doxorubicin
YDL227C_RF2_360 RF2
ZShift = 0.33
lm ZScore = −0.63
NG = 0
DB = 0
SM = 0
Delta_K
Doxorubicin
YDL227C_RF2_360 RF2
ZShift = 1.11
lm ZScore = 0.06
NG = 0
DB = 0
SM = 0
Delta_r
Doxorubicin
YDL227C_RF2_360 RF2
ZShift = 0.45
lm ZScore = 0.26
NG = 0
DB = 0
SM = 0
Delta_AUC
Doxorubicin

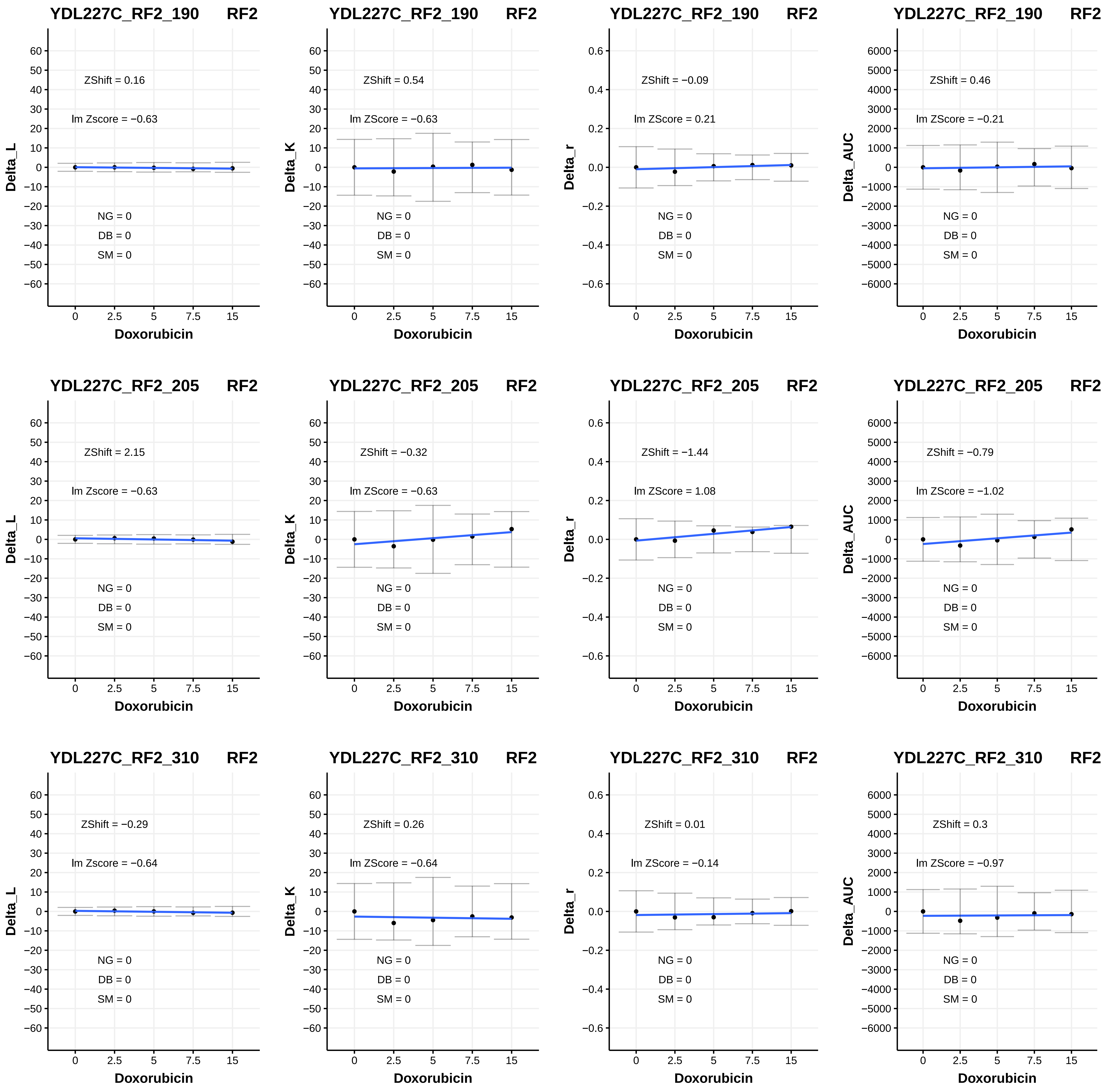

YDL227C_RF2_190 RF2
ZShift = 0.16
lm Zscore = −0.63
NG = 0
DB = 0
SM = 0
Delta_L
Doxorubicin
YDL227C_RF2_190 RF2
ZShift = 0.54
lm ZScore = −0.63
NG = 0
DB = 0
SM = 0
Delta_K
Doxorubicin
YDL227C_RF2_190 RF2
ZShift = −0.09
lm ZScore = 0.21
NG = 0
DB = 0
SM = 0
Delta_r
Doxorubicin
YDL227C_RF2_190 RF2
ZShift = 0.46
lm ZScore = −0.21
NG = 0
DB = 0
SM = 0
Delta_AUC
Doxorubicin
YDL227C_RF2_205 RF2
ZShift = 2.15
lm Zscore = −0.63
NG = 0
DB = 0
SM = 0
Delta_L
Doxorubicin
YDL227C_RF2_205 RF2
ZShift = −0.32
lm ZScore = −0.63
NG = 0
DB = 0
SM = 0
Delta_K
Doxorubicin
YDL227C_RF2_205 RF2
ZShift = −1.44
lm ZScore = 1.08
NG = 0
DB = 0
SM = 0
Delta_r
Doxorubicin
YDL227C_RF2_205 RF2
ZShift = −0.79
lm ZScore = −1.02
NG = 0
DB = 0
SM = 0
Delta_AUC
Doxorubicin
YDL227C_RF2_310 RF2
ZShift = −0.29
lm Zscore = −0.64
NG = 0
DB = 0
SM = 0
Delta_L
Doxorubicin
YDL227C_RF2_310 RF2
ZShift = 0.26
lm ZScore = −0.64
NG = 0
DB = 0
SM = 0
Delta_K
Doxorubicin
YDL227C_RF2_310 RF2
ZShift = 0.01
lm ZScore = −0.14
NG = 0
DB = 0
SM = 0
Delta_r
Doxorubicin
YDL227C_RF2_310 RF2
ZShift = 0.3
lm ZScore = −0.97
NG = 0
DB = 0
SM = 0
Delta_AUC
Doxorubicin

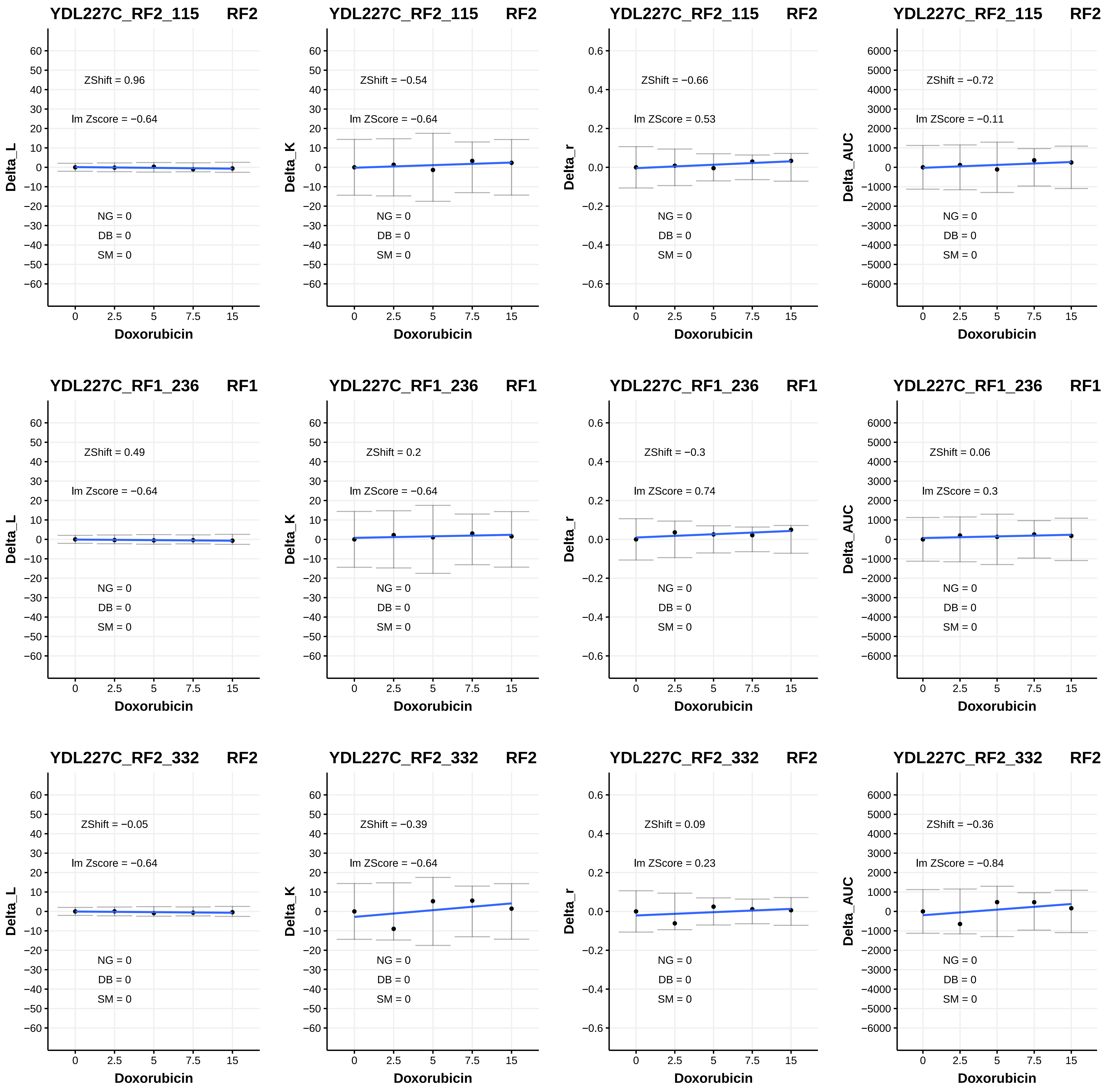

YDL227C_RF2_115 RF2
ZShift = 0.96
lm Zscore = −0.64
NG = 0
DB = 0
SM = 0
Delta_L
Doxorubicin
YDL227C_RF2_115 RF2
ZShift = −0.54
lm ZScore = −0.64
NG = 0
DB = 0
SM = 0
Delta_K
Doxorubicin
YDL227C_RF2_115 RF2
ZShift = −0.66
lm ZScore = 0.53
NG = 0
DB = 0
SM = 0
Delta_r
Doxorubicin
YDL227C_RF2_115 RF2
ZShift = −0.72
lm ZScore = −0.11
NG = 0
DB = 0
SM = 0
Delta_AUC
Doxorubicin
YDL227C_RF1_236 RF1
ZShift = 0.49
lm Zscore = −0.64
NG = 0
DB = 0
SM = 0
Delta_L
Doxorubicin
YDL227C_RF1_236 RF1
ZShift = 0.2
lm ZScore = −0.64
NG = 0
DB = 0
SM = 0
Delta_K
Doxorubicin
YDL227C_RF1_236 RF1
ZShift = −0.3
lm ZScore = 0.74
NG = 0
DB = 0
SM = 0
Delta_r
Doxorubicin
YDL227C_RF1_236 RF1
ZShift = 0.06
lm ZScore = 0.3
NG = 0
DB = 0
SM = 0
Delta_AUC
Doxorubicin
YDL227C_RF2_332 RF2
ZShift = −0.05
lm Zscore = −0.64
NG = 0
DB = 0
SM = 0
Delta_L
Doxorubicin
YDL227C_RF2_332 RF2
ZShift = −0.39
lm ZScore = −0.64
NG = 0
DB = 0
SM = 0
Delta_K
Doxorubicin
YDL227C_RF2_332 RF2
ZShift = 0.09
lm ZScore = 0.23
NG = 0
DB = 0
SM = 0
Delta_r
Doxorubicin
YDL227C_RF2_332 RF2
ZShift = −0.36
lm ZScore = −0.84
NG = 0
DB = 0
SM = 0
Delta_AUC
Doxorubicin

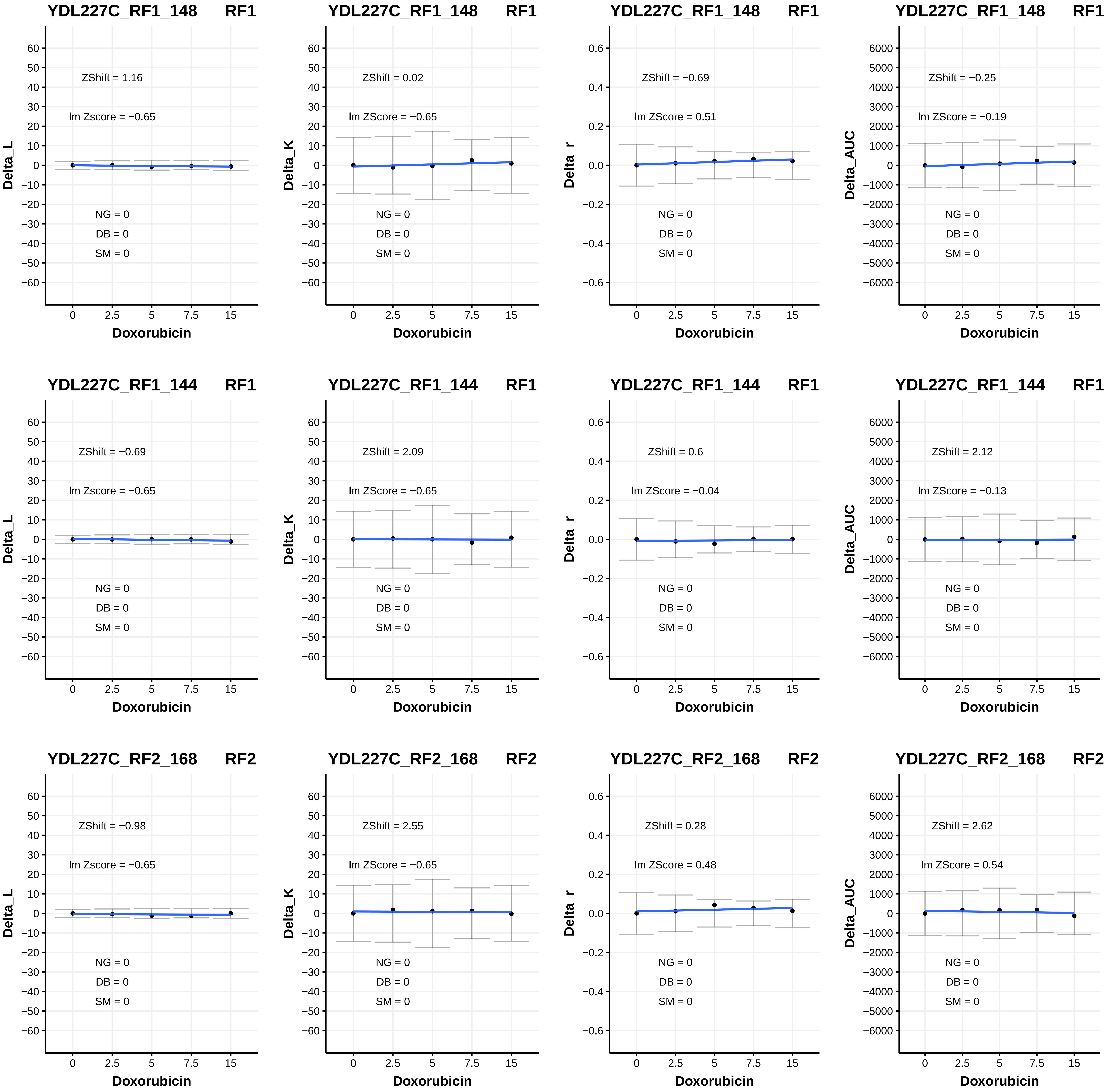

YDL227C_RF1_148 RF1
ZShift = 1.16
lm Zscore = −0.65
NG = 0
DB = 0
SM = 0
Delta_L
Doxorubicin
YDL227C_RF1_148 RF1
ZShift = 0.02
lm ZScore = −0.65
NG = 0
DB = 0
SM = 0
Delta_K
Doxorubicin
YDL227C_RF1_148 RF1
ZShift = −0.69
lm ZScore = 0.51
NG = 0
DB = 0
SM = 0
Delta_r
Doxorubicin
YDL227C_RF1_148 RF1
ZShift = −0.25
lm ZScore = −0.19
NG = 0
DB = 0
SM = 0
Delta_AUC
Doxorubicin
YDL227C_RF1_144 RF1
ZShift = −0.69
lm Zscore = −0.65
NG = 0
DB = 0
SM = 0
Delta_L
Doxorubicin
YDL227C_RF1_144 RF1
ZShift = 2.09
lm ZScore = −0.65
NG = 0
DB = 0
SM = 0
Delta_K
Doxorubicin
YDL227C_RF1_144 RF1
ZShift = 0.6
lm ZScore = −0.04
NG = 0
DB = 0
SM = 0
Delta_r
Doxorubicin
YDL227C_RF1_144 RF1
ZShift = 2.12
lm ZScore = −0.13
NG = 0
DB = 0
SM = 0
Delta_AUC
Doxorubicin
YDL227C_RF2_168 RF2
ZShift = −0.98
lm Zscore = −0.65
NG = 0
DB = 0
SM = 0
Delta_L
Doxorubicin
YDL227C_RF2_168 RF2
ZShift = 2.55
lm ZScore = −0.65
NG = 0
DB = 0
SM = 0
Delta_K
Doxorubicin
YDL227C_RF2_168 RF2
ZShift = 0.28
lm ZScore = 0.48
NG = 0
DB = 0
SM = 0
Delta_r
Doxorubicin
YDL227C_RF2_168 RF2
ZShift = 2.62
lm ZScore = 0.54
NG = 0
DB = 0
SM = 0
Delta_AUC
Doxorubicin

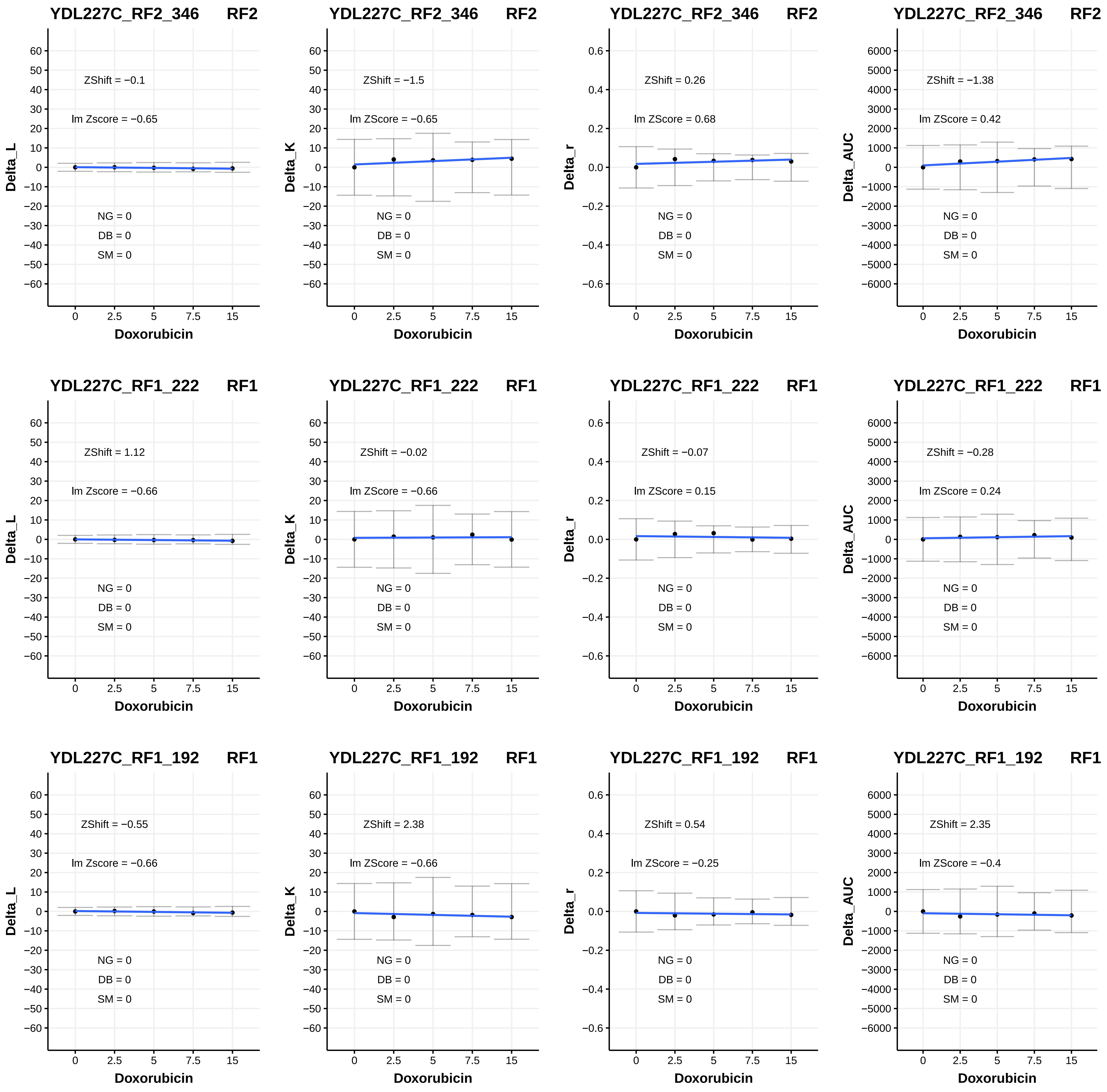
YDL227C_RF2_346 RF2
ZShift = −0.1
lm Zscore = −0.65
NG = 0
DB = 0
SM = 0
Delta_L
Doxorubicin
YDL227C_RF2_346 RF2
ZShift = −1.5
lm ZScore = −0.65
NG = 0
DB = 0
SM = 0
Delta_K
Doxorubicin
YDL227C_RF2_346 RF2
ZShift = 0.26
lm ZScore = 0.68
NG = 0
DB = 0
SM = 0
Delta_r
Doxorubicin
YDL227C_RF2_346 RF2
ZShift = −1.38
lm ZScore = 0.42
NG = 0
DB = 0
SM = 0
Delta_AUC
Doxorubicin
YDL227C_RF1_222 RF1
ZShift = 1.12
lm Zscore = −0.66
NG = 0
DB = 0
SM = 0
Delta_L
Doxorubicin
YDL227C_RF1_222 RF1
ZShift = −0.02
lm ZScore = −0.66
NG = 0
DB = 0
SM = 0
Delta_K
Doxorubicin
YDL227C_RF1_222 RF1
ZShift = −0.07
lm ZScore = 0.15
NG = 0
DB = 0
SM = 0
Delta_r
Doxorubicin
YDL227C_RF1_222 RF1
ZShift = −0.28
lm ZScore = 0.24
NG = 0
DB = 0
SM = 0
Delta_AUC
Doxorubicin
YDL227C_RF1_192 RF1
ZShift = −0.55
lm Zscore = −0.66
NG = 0
DB = 0
SM = 0
Delta_L
Doxorubicin
YDL227C_RF1_192 RF1
ZShift = 2.38
lm ZScore = −0.66
NG = 0
DB = 0
SM = 0
Delta_K
Doxorubicin
YDL227C_RF1_192 RF1
ZShift = 0.54
lm ZScore = −0.25
NG = 0
DB = 0
SM = 0
Delta_r
Doxorubicin
YDL227C_RF1_192 RF1
ZShift = 2.35
lm ZScore = −0.4
NG = 0
DB = 0
SM = 0
Delta_AUC
Doxorubicin

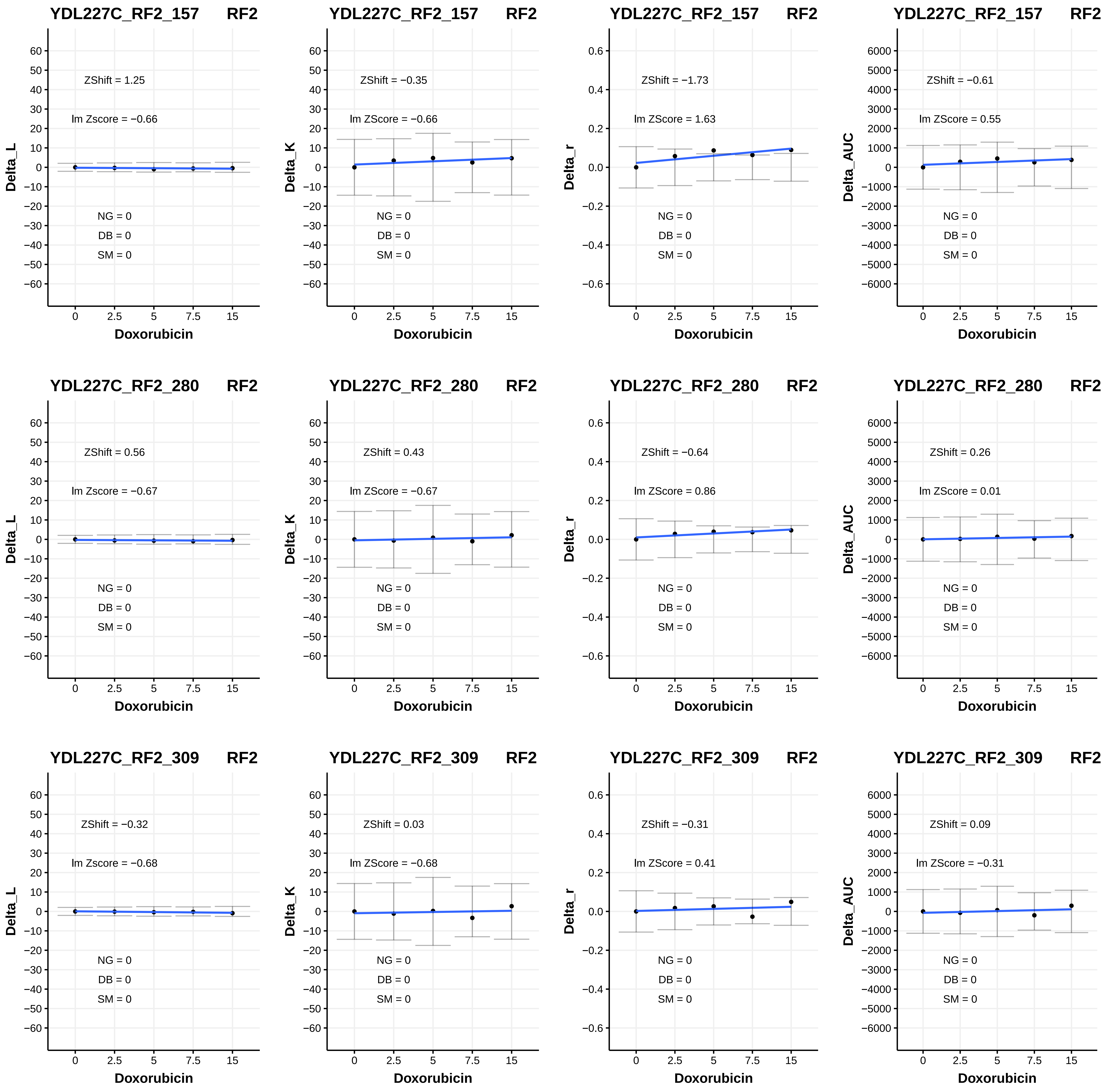
YDL227C_RF2_157 RF2
ZShift = 1.25
lm Zscore = −0.66
NG = 0
DB = 0
SM = 0
Delta_L
Doxorubicin
YDL227C_RF2_157 RF2
ZShift = −0.35
lm ZScore = −0.66
NG = 0
DB = 0
SM = 0
Delta_K
Doxorubicin
YDL227C_RF2_157 RF2
ZShift = −1.73
lm ZScore = 1.63
NG = 0
DB = 0
SM = 0
Delta_r
Doxorubicin
YDL227C_RF2_157 RF2
ZShift = −0.61
lm ZScore = 0.55
NG = 0
DB = 0
SM = 0
Delta_AUC
Doxorubicin
YDL227C_RF2_280 RF2
ZShift = 0.56
lm Zscore = −0.67
NG = 0
DB = 0
SM = 0
Delta_L
Doxorubicin
YDL227C_RF2_280 RF2
ZShift = 0.43
lm ZScore = −0.67
NG = 0
DB = 0
SM = 0
Delta_K
Doxorubicin
YDL227C_RF2_280 RF2
ZShift = −0.64
lm ZScore = 0.86
NG = 0
DB = 0
SM = 0
Delta_r
Doxorubicin
YDL227C_RF2_280 RF2
ZShift = 0.26
lm ZScore = 0.01
NG = 0
DB = 0
SM = 0
Delta_AUC
Doxorubicin
YDL227C_RF2_309 RF2
ZShift = −0.32
lm Zscore = −0.68
NG = 0
DB = 0
SM = 0
Delta_L
Doxorubicin
YDL227C_RF2_309 RF2
ZShift = 0.03
lm ZScore = −0.68
NG = 0
DB = 0
SM = 0
Delta_K
Doxorubicin
YDL227C_RF2_309 RF2
ZShift = −0.31
lm ZScore = 0.41
NG = 0
DB = 0
SM = 0
Delta_r
Doxorubicin
YDL227C_RF2_309 RF2
ZShift = 0.09
lm ZScore = −0.31
NG = 0
DB = 0
SM = 0
Delta_AUC
Doxorubicin

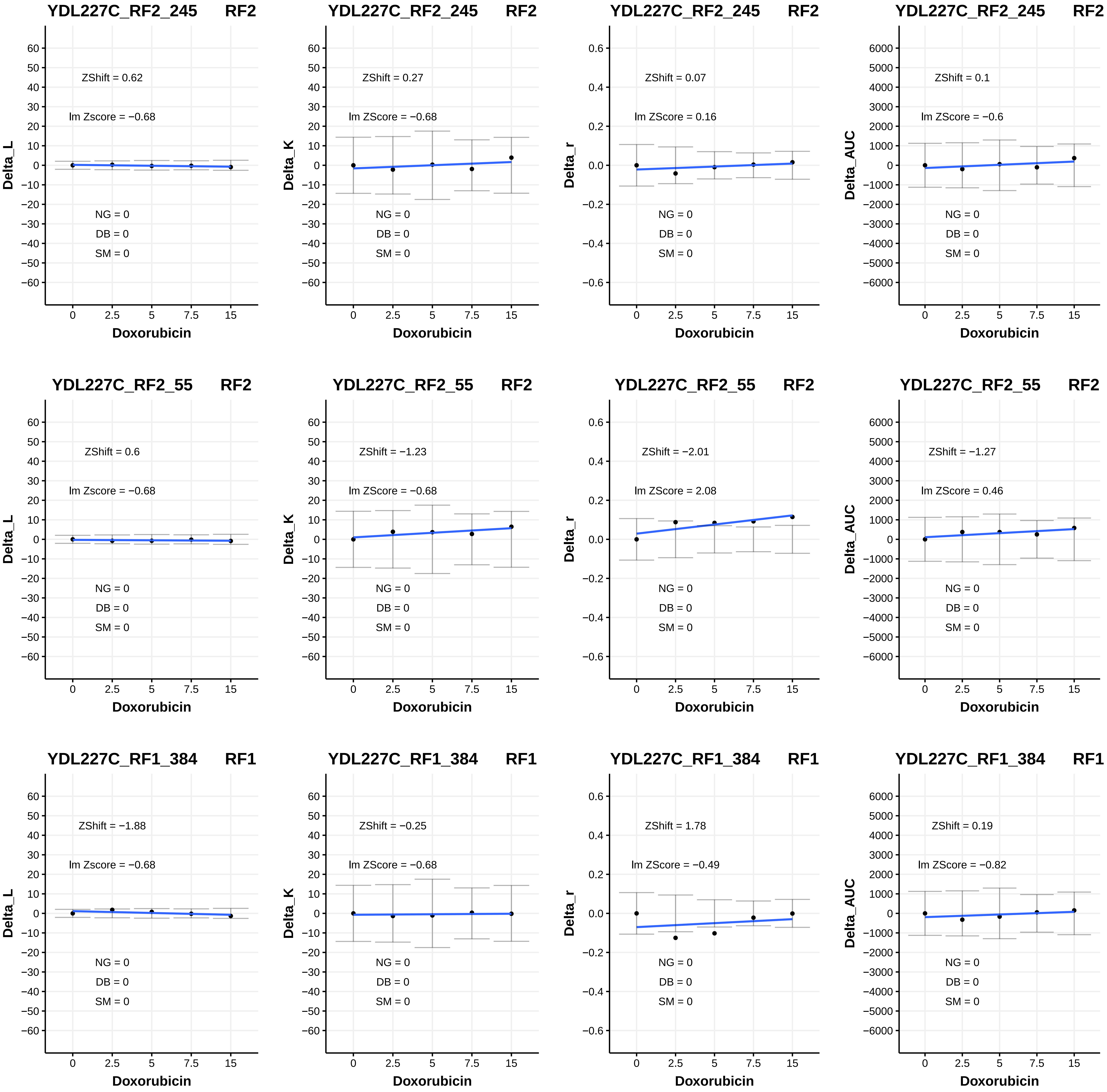
YDL227C_RF2_245 RF2
ZShift = 0.62
lm Zscore = −0.68
NG = 0
DB = 0
SM = 0
Delta_L
Doxorubicin
YDL227C_RF2_245 RF2
ZShift = 0.27
lm ZScore = −0.68
NG = 0
DB = 0
SM = 0
Delta_K
Doxorubicin
YDL227C_RF2_245 RF2
ZShift = 0.07
lm ZScore = 0.16
NG = 0
DB = 0
SM = 0
Delta_r
Doxorubicin
YDL227C_RF2_245 RF2
ZShift = 0.1
lm ZScore = −0.6
NG = 0
DB = 0
SM = 0
Delta_AUC
Doxorubicin
YDL227C_RF2_55 RF2
ZShift = 0.6
lm Zscore = −0.68
NG = 0
DB = 0
SM = 0
Delta_L
Doxorubicin
YDL227C_RF2_55 RF2
ZShift = −1.23
lm ZScore = −0.68
NG = 0
DB = 0
SM = 0
Delta_K
Doxorubicin
YDL227C_RF2_55 RF2
ZShift = −2.01
lm ZScore = 2.08
NG = 0
DB = 0
SM = 0
Delta_r
Doxorubicin
YDL227C_RF2_55 RF2
ZShift = −1.27
lm ZScore = 0.46
NG = 0
DB = 0
SM = 0
Delta_AUC
Doxorubicin
YDL227C_RF1_384 RF1
ZShift = −1.88
lm Zscore = −0.68
NG = 0
DB = 0
SM = 0
Delta_L
Doxorubicin
YDL227C_RF1_384 RF1
ZShift = −0.25
lm ZScore = −0.68
NG = 0
DB = 0
SM = 0
Delta_K
Doxorubicin
YDL227C_RF1_384 RF1
ZShift = 1.78
lm ZScore = −0.49
NG = 0
DB = 0
SM = 0
Delta_r
Doxorubicin
YDL227C_RF1_384 RF1
ZShift = 0.19
lm ZScore = −0.82
NG = 0
DB = 0
SM = 0
Delta_AUC
Doxorubicin

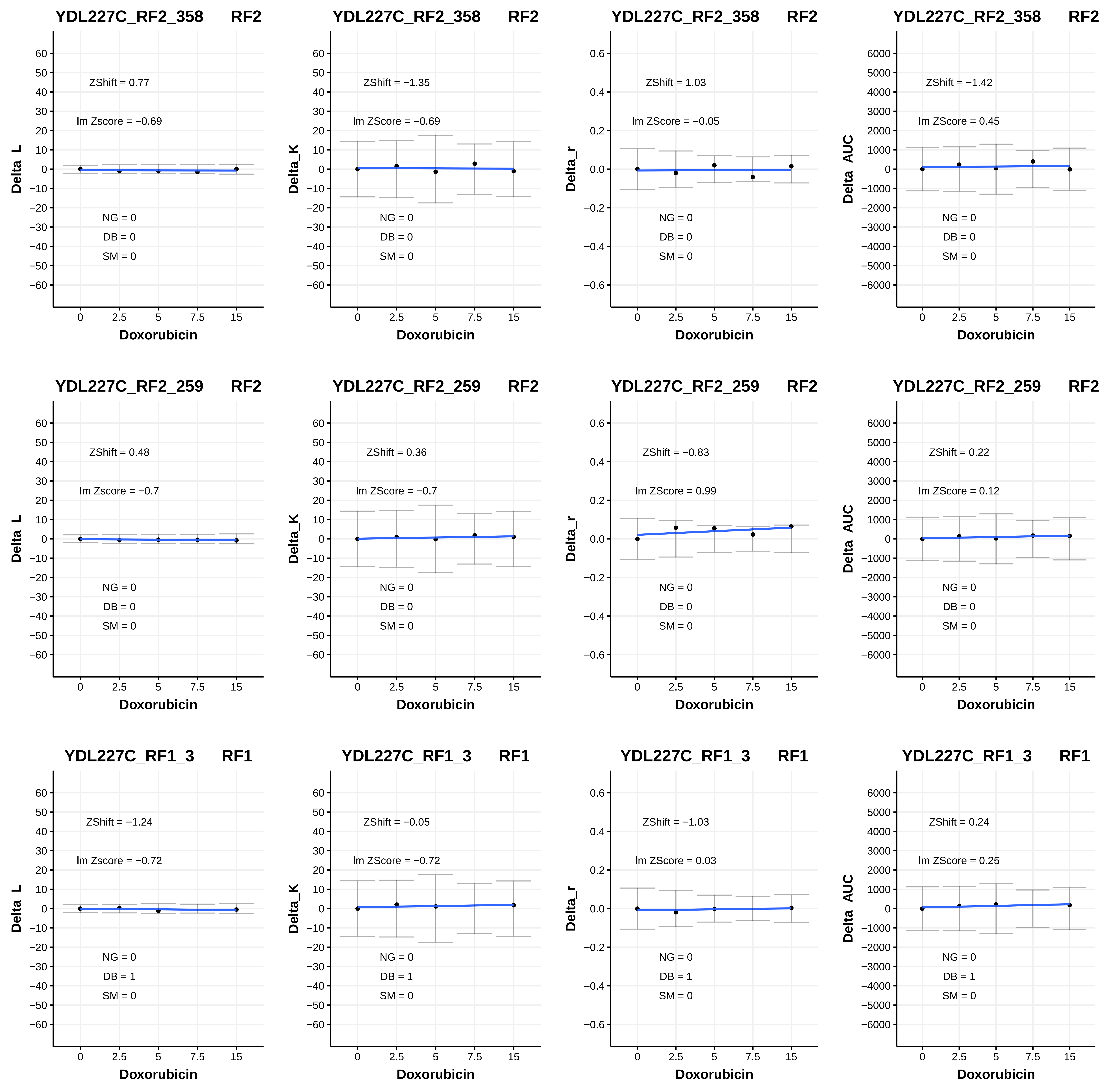

YDL227C_RF2_358 RF2
ZShift = 0.77
lm Zscore = −0.69
NG = 0
DB = 0
SM = 0
Delta_L
Doxorubicin
YDL227C_RF2_358 RF2
ZShift = −1.35
lm ZScore = −0.69
NG = 0
DB = 0
SM = 0
Delta_K
Doxorubicin
YDL227C_RF2_358 RF2
ZShift = 1.03
lm ZScore = −0.05
NG = 0
DB = 0
SM = 0
Delta_r
Doxorubicin
YDL227C_RF2_358 RF2
ZShift = −1.42
lm ZScore = 0.45
NG = 0
DB = 0
SM = 0
Delta_AUC
Doxorubicin
YDL227C_RF2_259 RF2
ZShift = 0.48
lm Zscore = −0.7
NG = 0
DB = 0
SM = 0
Delta_L
Doxorubicin
YDL227C_RF2_259 RF2
ZShift = 0.36
lm ZScore = −0.7
NG = 0
DB = 0
SM = 0
Delta_K
Doxorubicin
YDL227C_RF2_259 RF2
ZShift = −0.83
lm ZScore = 0.99
NG = 0
DB = 0
SM = 0
Delta_r
Doxorubicin
YDL227C_RF2_259 RF2
ZShift = 0.22
lm ZScore = 0.12
NG = 0
DB = 0
SM = 0
Delta_AUC
Doxorubicin
YDL227C_RF1_3 RF1
ZShift = −1.24
lm Zscore = −0.72
NG = 0
DB = 1
SM = 0
Delta_L
Doxorubicin
YDL227C_RF1_3 RF1
ZShift = −0.05
lm ZScore = −0.72
NG = 0
DB = 1
SM = 0
Delta_K
Doxorubicin
YDL227C_RF1_3 RF1
ZShift = −1.03
lm ZScore = 0.03
NG = 0
DB = 1
SM = 0
Delta_r
Doxorubicin
YDL227C_RF1_3 RF1
ZShift = 0.24
lm ZScore = 0.25
NG = 0
DB = 1
SM = 0
Delta_AUC
Doxorubicin

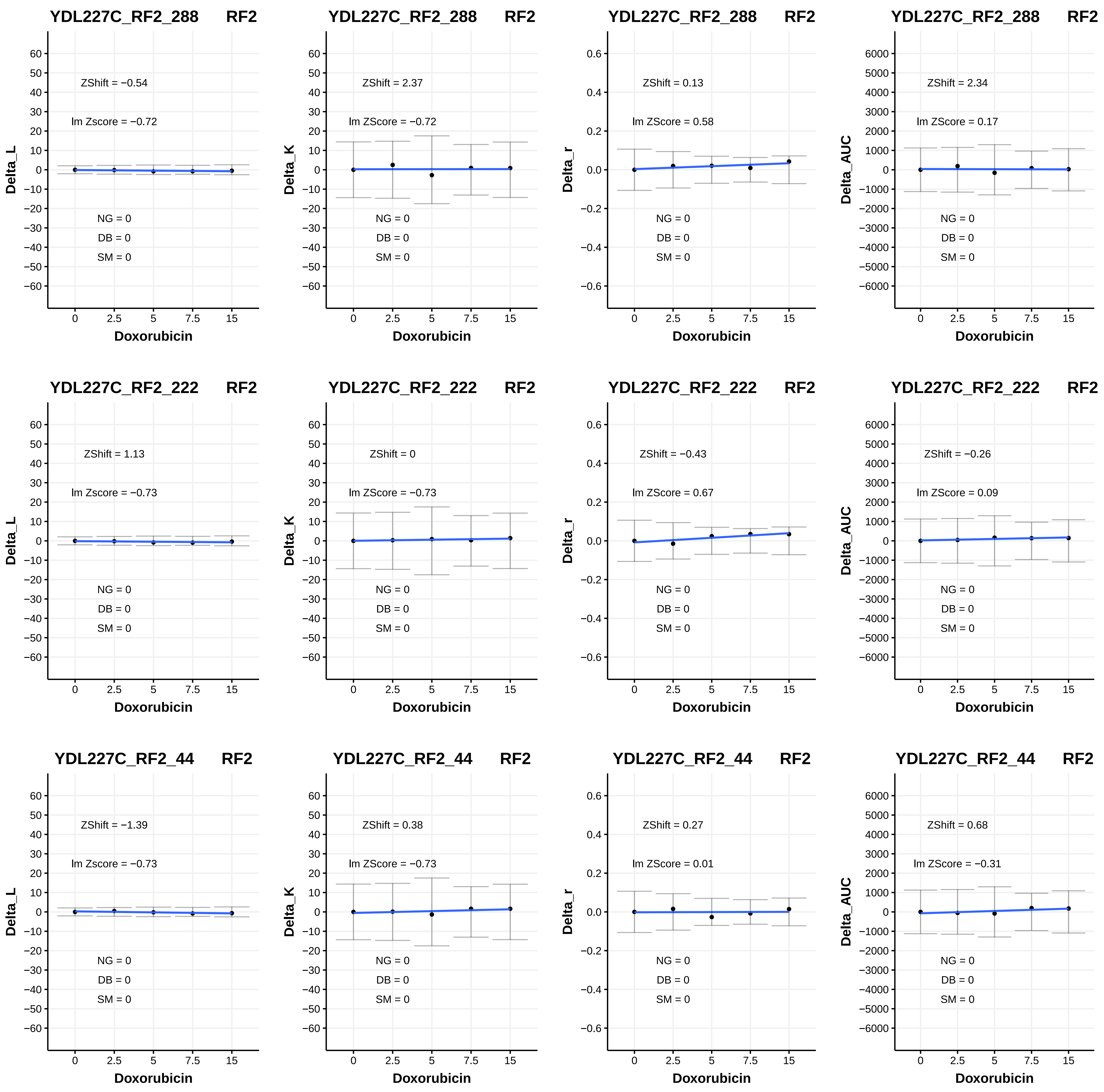

YDL227C_RF2_288 RF2
ZShift = −0.54
lm Zscore = −0.72
NG = 0
DB = 0
SM = 0
Delta_L
Doxorubicin
YDL227C_RF2_288 RF2
ZShift = 2.37
lm ZScore = −0.72
NG = 0
DB = 0
SM = 0
Delta_K
Doxorubicin
YDL227C_RF2_288 RF2
ZShift = 0.13
lm ZScore = 0.58
NG = 0
DB = 0
SM = 0
Delta_r
Doxorubicin
YDL227C_RF2_288 RF2
ZShift = 2.34
lm ZScore = 0.17
NG = 0
DB = 0
SM = 0
Delta_AUC
Doxorubicin
YDL227C_RF2_222 RF2
ZShift = 1.13
lm Zscore = −0.73
NG = 0
DB = 0
SM = 0
Delta_L
Doxorubicin
YDL227C_RF2_222 RF2
ZShift = 0
lm ZScore = −0.73
NG = 0
DB = 0
SM = 0
Delta_K
Doxorubicin
YDL227C_RF2_222 RF2
ZShift = −0.43
lm ZScore = 0.67
NG = 0
DB = 0
SM = 0
Delta_r
Doxorubicin
YDL227C_RF2_222 RF2
ZShift = −0.26
lm ZScore = 0.09
NG = 0
DB = 0
SM = 0
Delta_AUC
Doxorubicin
YDL227C_RF2_44 RF2
ZShift = −1.39
lm Zscore = −0.73
NG = 0
DB = 0
SM = 0
Delta_L
Doxorubicin
YDL227C_RF2_44 RF2
ZShift = 0.38
lm ZScore = −0.73
NG = 0
DB = 0
SM = 0
Delta_K
Doxorubicin
YDL227C_RF2_44 RF2
ZShift = 0.27
lm ZScore = 0.01
NG = 0
DB = 0
SM = 0
Delta_r
Doxorubicin
YDL227C_RF2_44 RF2
ZShift = 0.68
lm ZScore = −0.31
NG = 0
DB = 0
SM = 0
Delta_AUC
Doxorubicin

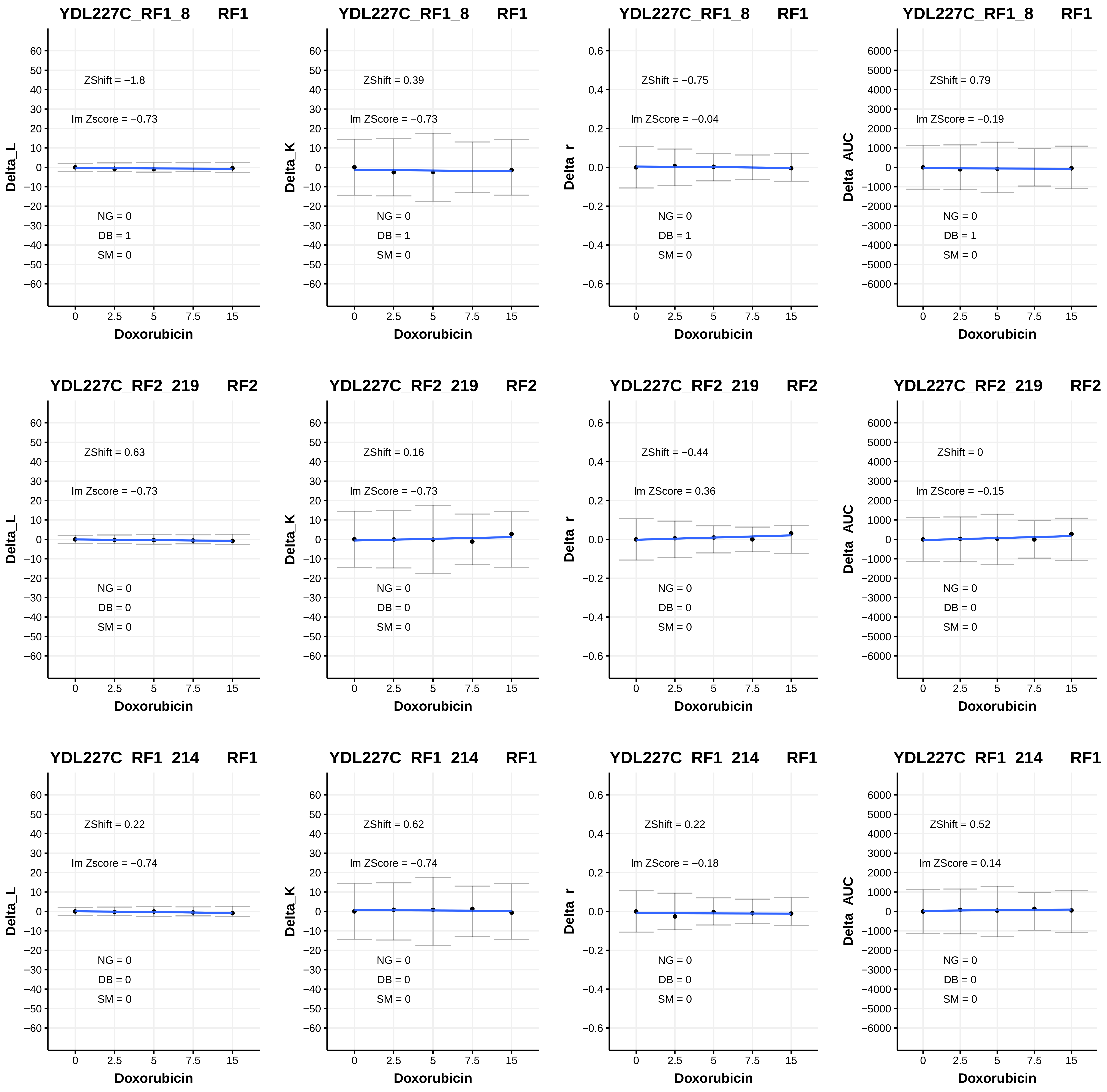
YDL227C_RF1_8 RF1
ZShift = −1.8
lm Zscore = −0.73
NG = 0
DB = 1
SM = 0
Delta_L
Doxorubicin
YDL227C_RF1_8 RF1
ZShift = 0.39
lm ZScore = −0.73
NG = 0
DB = 1
SM = 0
Delta_K
Doxorubicin
YDL227C_RF1_8 RF1
ZShift = −0.75
lm ZScore = −0.04
NG = 0
DB = 1
SM = 0
Delta_r
Doxorubicin
YDL227C_RF1_8 RF1
ZShift = 0.79
lm ZScore = −0.19
NG = 0
DB = 1
SM = 0
Delta_AUC
Doxorubicin
YDL227C_RF2_219 RF2
ZShift = 0.63
lm Zscore = −0.73
NG = 0
DB = 0
SM = 0
Delta_L
Doxorubicin
YDL227C_RF2_219 RF2
ZShift = 0.16
lm ZScore = −0.73
NG = 0
DB = 0
SM = 0
Delta_K
Doxorubicin
YDL227C_RF2_219 RF2
ZShift = −0.44
lm ZScore = 0.36
NG = 0
DB = 0
SM = 0
Delta_r
Doxorubicin
YDL227C_RF2_219 RF2
ZShift = 0
lm ZScore = −0.15
NG = 0
DB = 0
SM = 0
Delta_AUC
Doxorubicin
YDL227C_RF1_214 RF1
ZShift = 0.22
lm Zscore = −0.74
NG = 0
DB = 0
SM = 0
Delta_L
Doxorubicin
YDL227C_RF1_214 RF1
ZShift = 0.62
lm ZScore = −0.74
NG = 0
DB = 0
SM = 0
Delta_K
Doxorubicin
YDL227C_RF1_214 RF1
ZShift = 0.22
lm ZScore = −0.18
NG = 0
DB = 0
SM = 0
Delta_r
Doxorubicin
YDL227C_RF1_214 RF1
ZShift = 0.52
lm ZScore = 0.14
NG = 0
DB = 0
SM = 0
Delta_AUC
Doxorubicin
0 2.5 5 7.5 15

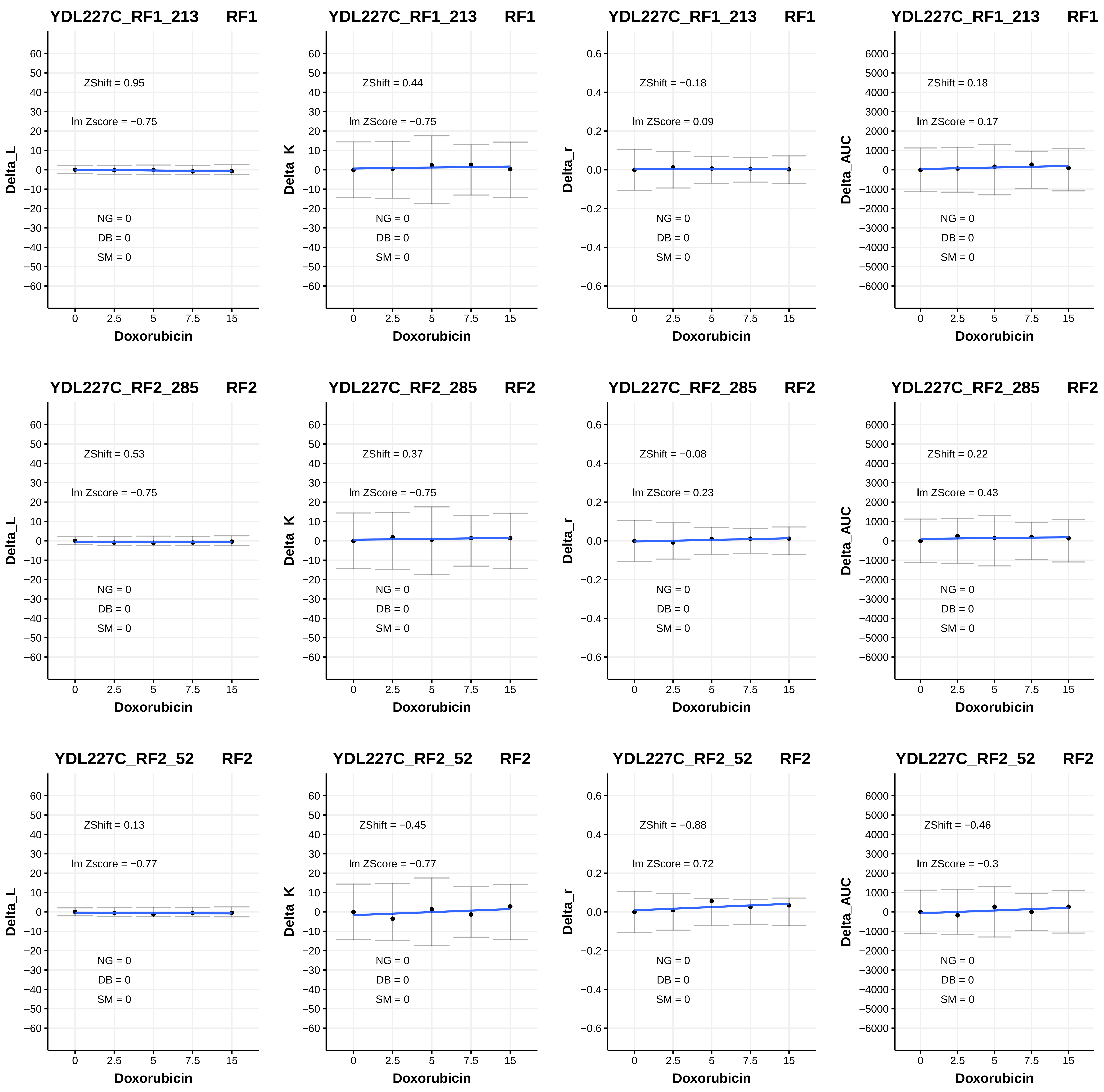
YDL227C_RF1_213 RF1
ZShift = 0.95
lm Zscore = −0.75
NG = 0
DB = 0
SM = 0
Delta_L
Doxorubicin
YDL227C_RF1_213 RF1
ZShift = 0.44
lm ZScore = −0.75
NG = 0
DB = 0
SM = 0
Delta_K
Doxorubicin
YDL227C_RF1_213 RF1
ZShift = −0.18
lm ZScore = 0.09
NG = 0
DB = 0
SM = 0
Delta_r
Doxorubicin
YDL227C_RF1_213 RF1
ZShift = 0.18
lm ZScore = 0.17
NG = 0
DB = 0
SM = 0
Delta_AUC
Doxorubicin
YDL227C_RF2_285 RF2
ZShift = 0.53
lm Zscore = −0.75
NG = 0
DB = 0
SM = 0
Delta_L
Doxorubicin
YDL227C_RF2_285 RF2
ZShift = 0.37
lm ZScore = −0.75
NG = 0
DB = 0
SM = 0
Delta_K
Doxorubicin
YDL227C_RF2_285 RF2
ZShift = −0.08
lm ZScore = 0.23
NG = 0
DB = 0
SM = 0
Delta_r
Doxorubicin
YDL227C_RF2_285 RF2
ZShift = 0.22
lm ZScore = 0.43
NG = 0
DB = 0
SM = 0
Delta_AUC
Doxorubicin
YDL227C_RF2_52 RF2
ZShift = 0.13
lm Zscore = −0.77
NG = 0
DB = 0
SM = 0
Delta_L
Doxorubicin
YDL227C_RF2_52 RF2
ZShift = −0.45
lm ZScore = −0.77
NG = 0
DB = 0
SM = 0
Delta_K
Doxorubicin
YDL227C_RF2_52 RF2
ZShift = −0.88
lm ZScore = 0.72
NG = 0
DB = 0
SM = 0
Delta_r
Doxorubicin
YDL227C_RF2_52 RF2
ZShift = −0.46
lm ZScore = −0.3
NG = 0
DB = 0
SM = 0
Delta_AUC
Doxorubicin

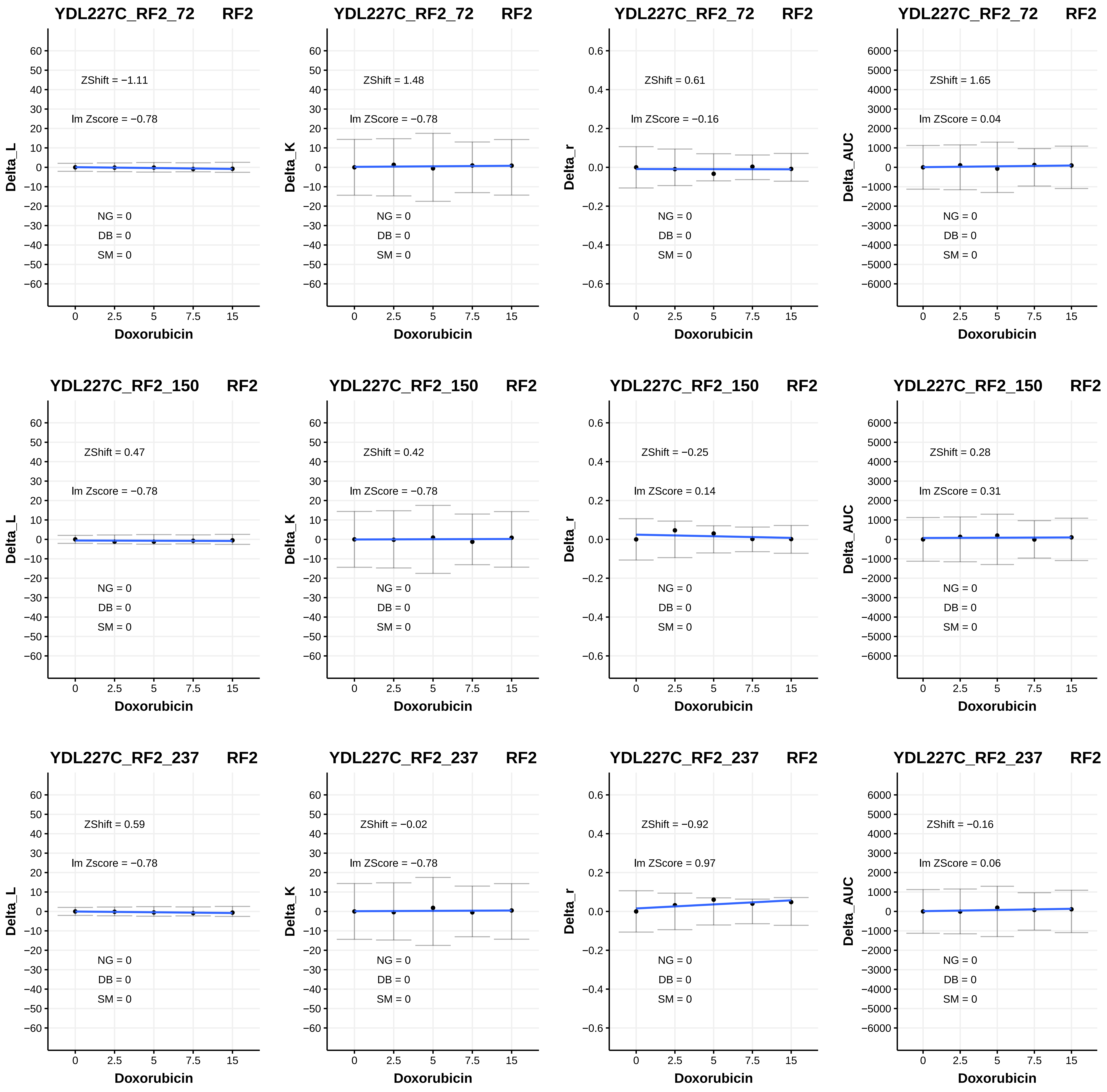
YDL227C_RF2_72 RF2
ZShift = −1.11
lm Zscore = −0.78
NG = 0
DB = 0
SM = 0
Delta_L
Doxorubicin
YDL227C_RF2_72 RF2
ZShift = 1.48
lm ZScore = −0.78
NG = 0
DB = 0
SM = 0
Delta_K
Doxorubicin
YDL227C_RF2_72 RF2
ZShift = 0.61
lm ZScore = −0.16
NG = 0
DB = 0
SM = 0
Delta_r
Doxorubicin
YDL227C_RF2_72 RF2
ZShift = 1.65
lm ZScore = 0.04
NG = 0
DB = 0
SM = 0
Delta_AUC
Doxorubicin
YDL227C_RF2_150 RF2
ZShift = 0.47
lm Zscore = −0.78
NG = 0
DB = 0
SM = 0
Delta_L
Doxorubicin
YDL227C_RF2_150 RF2
ZShift = 0.42
lm ZScore = −0.78
NG = 0
DB = 0
SM = 0
Delta_K
Doxorubicin
YDL227C_RF2_150 RF2
ZShift = −0.25
lm ZScore = 0.14
NG = 0
DB = 0
SM = 0
Delta_r
Doxorubicin
YDL227C_RF2_150 RF2
ZShift = 0.28
lm ZScore = 0.31
NG = 0
DB = 0
SM = 0
Delta_AUC
Doxorubicin
YDL227C_RF2_237 RF2
ZShift = 0.59
lm Zscore = −0.78
NG = 0
DB = 0
SM = 0
Delta_L
Doxorubicin
YDL227C_RF2_237 RF2
ZShift = −0.02
lm ZScore = −0.78
NG = 0
DB = 0
SM = 0
Delta_K
Doxorubicin
YDL227C_RF2_237 RF2
ZShift = −0.92
lm ZScore = 0.97
NG = 0
DB = 0
SM = 0
Delta_r
Doxorubicin
YDL227C_RF2_237 RF2
ZShift = −0.16
lm ZScore = 0.06
NG = 0
DB = 0
SM = 0
Delta_AUC
Doxorubicin

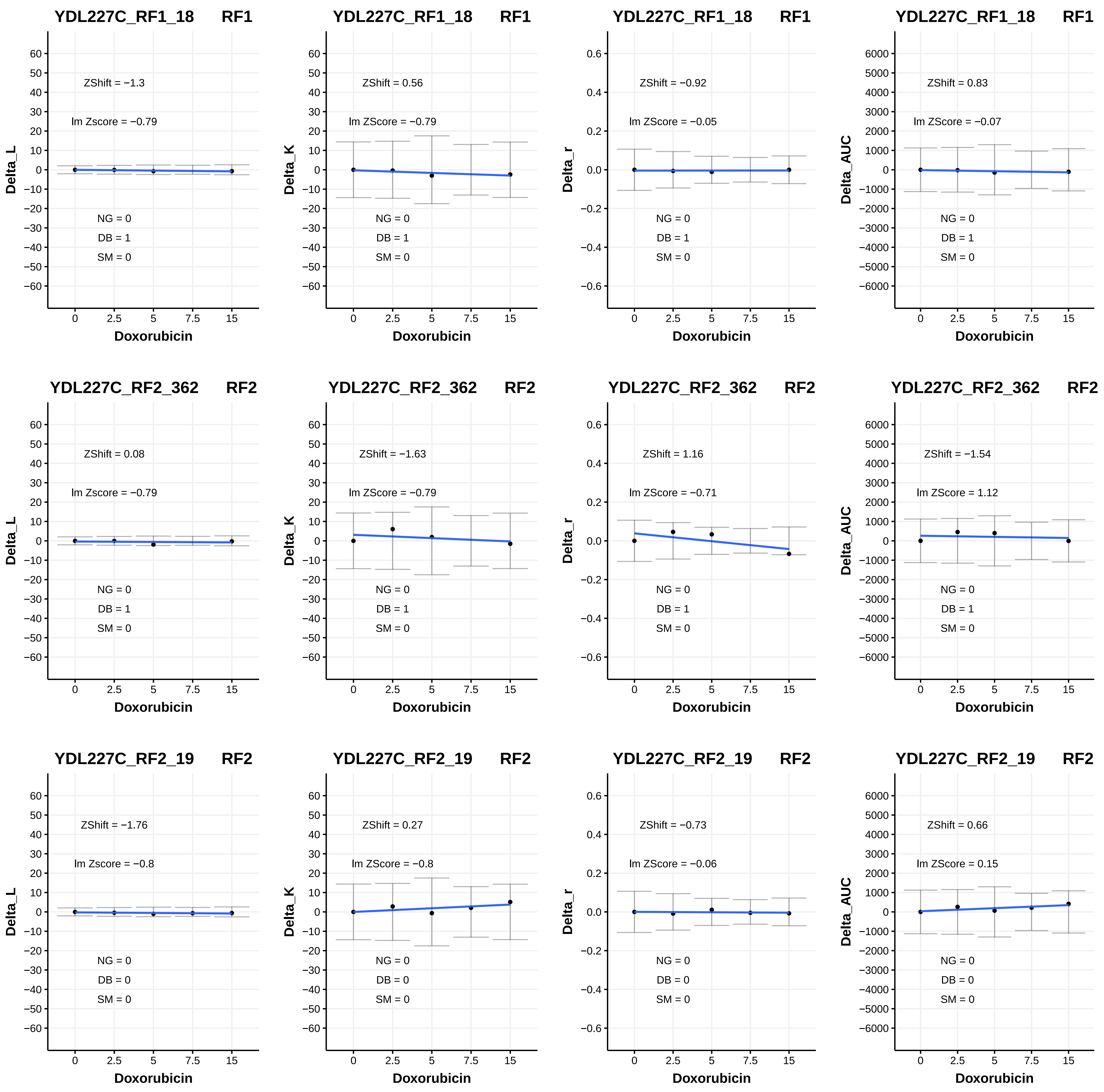

YDL227C_RF1_18 RF1
ZShift = −1.3
lm Zscore = −0.79
NG = 0
DB = 1
SM = 0
Delta_L
Doxorubicin
YDL227C_RF1_18 RF1
ZShift = 0.56
lm ZScore = −0.79
NG = 0
DB = 1
SM = 0
Delta_K
Doxorubicin
YDL227C_RF1_18 RF1
ZShift = −0.92
lm ZScore = −0.05
NG = 0
DB = 1
SM = 0
Delta_r
Doxorubicin
YDL227C_RF1_18 RF1
ZShift = 0.83
lm ZScore = −0.07
NG = 0
DB = 1
SM = 0
Delta_AUC
Doxorubicin
YDL227C_RF2_362 RF2
ZShift = 0.08
lm Zscore = −0.79
NG = 0
DB = 1
SM = 0
Delta_L
Doxorubicin
YDL227C_RF2_362 RF2
ZShift = −1.63
lm ZScore = −0.79
NG = 0
DB = 1
SM = 0
Delta_K
Doxorubicin
YDL227C_RF2_362 RF2
ZShift = 1.16
lm ZScore = −0.71
NG = 0
DB = 1
SM = 0
Delta_r
Doxorubicin
YDL227C_RF2_362 RF2
ZShift = −1.54
lm ZScore = 1.12
NG = 0
DB = 1
SM = 0
Delta_AUC
Doxorubicin
YDL227C_RF2_19 RF2
ZShift = −1.76
lm Zscore = −0.8
NG = 0
DB = 0
SM = 0
Delta_L
Doxorubicin
YDL227C_RF2_19 RF2
ZShift = 0.27
lm ZScore = −0.8
NG = 0
DB = 0
SM = 0
Delta_K
Doxorubicin
YDL227C_RF2_19 RF2
ZShift = −0.73
lm ZScore = −0.06
NG = 0
DB = 0
SM = 0
Delta_r
Doxorubicin
YDL227C_RF2_19 RF2
ZShift = 0.66
lm ZScore = 0.15
NG = 0
DB = 0
SM = 0
Delta_AUC
Doxorubicin

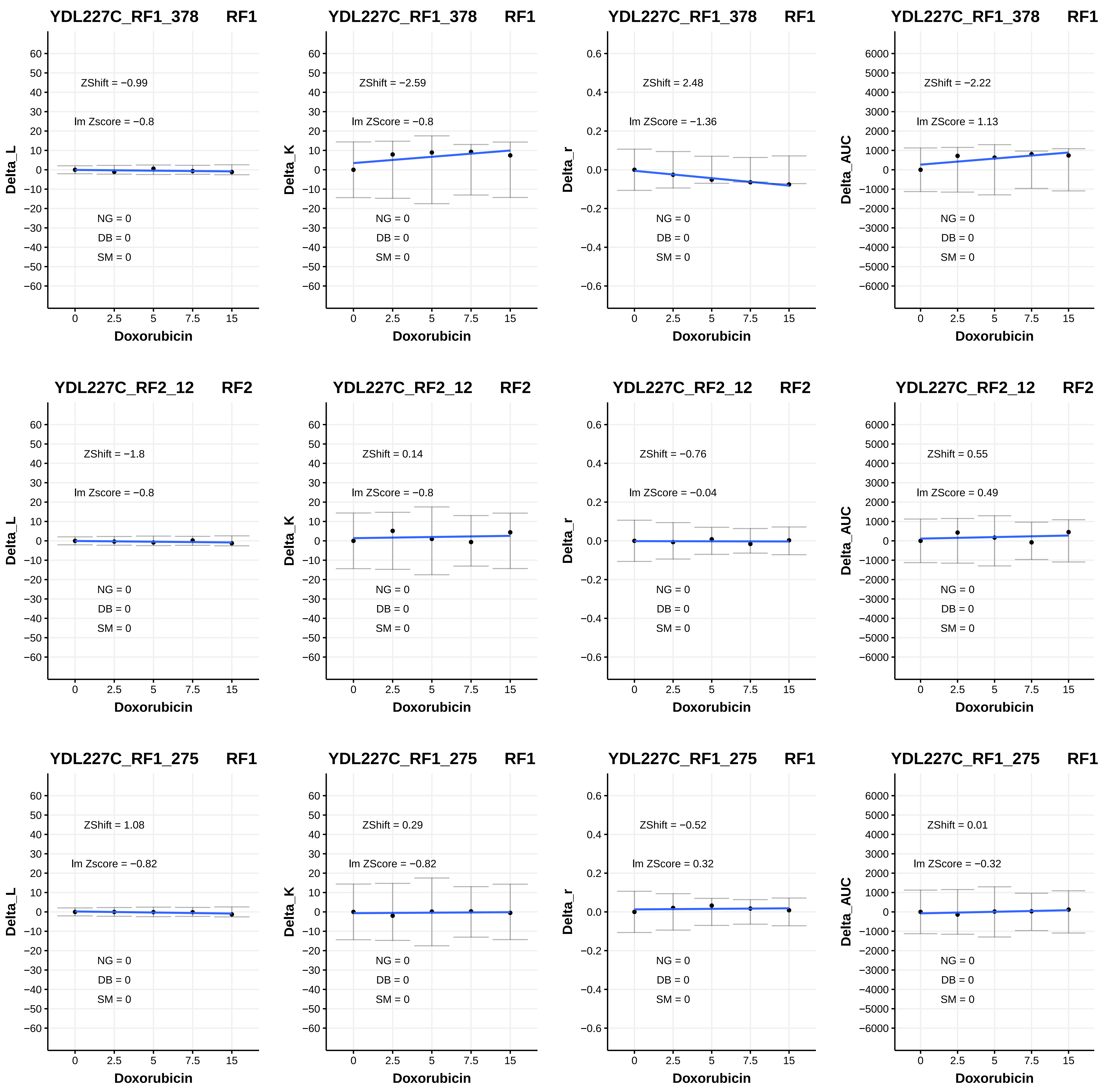
YDL227C_RF1_378 RF1
ZShift = −0.99
lm Zscore = −0.8
NG = 0
DB = 0
SM = 0
Delta_L
Doxorubicin
YDL227C_RF1_378 RF1
ZShift = −2.59
lm ZScore = −0.8
NG = 0
DB = 0
SM = 0
Delta_K
Doxorubicin
YDL227C_RF1_378 RF1
ZShift = 2.48
lm ZScore = −1.36
NG = 0
DB = 0
SM = 0
Delta_r
Doxorubicin
YDL227C_RF1_378 RF1
ZShift = −2.22
lm ZScore = 1.13
NG = 0
DB = 0
SM = 0
Delta_AUC
Doxorubicin
YDL227C_RF2_12 RF2
ZShift = −1.8
lm Zscore = −0.8
NG = 0
DB = 0
SM = 0
Delta_L
Doxorubicin
YDL227C_RF2_12 RF2
ZShift = 0.14
lm ZScore = −0.8
NG = 0
DB = 0
SM = 0
Delta_K
Doxorubicin
YDL227C_RF2_12 RF2
ZShift = −0.76
lm ZScore = −0.04
NG = 0
DB = 0
SM = 0
Delta_r
Doxorubicin
YDL227C_RF2_12 RF2
ZShift = 0.55
lm ZScore = 0.49
NG = 0
DB = 0
SM = 0
Delta_AUC
Doxorubicin
YDL227C_RF1_275 RF1
ZShift = 1.08
lm Zscore = −0.82
NG = 0
DB = 0
SM = 0
Delta_L
Doxorubicin
YDL227C_RF1_275 RF1
ZShift = 0.29
lm ZScore = −0.82
NG = 0
DB = 0
SM = 0
Delta_K
Doxorubicin
YDL227C_RF1_275 RF1
ZShift = −0.52
lm ZScore = 0.32
NG = 0
DB = 0
SM = 0
Delta_r
Doxorubicin
YDL227C_RF1_275 RF1
ZShift = 0.01
lm ZScore = −0.32
NG = 0
DB = 0
SM = 0
Delta_AUC
Doxorubicin

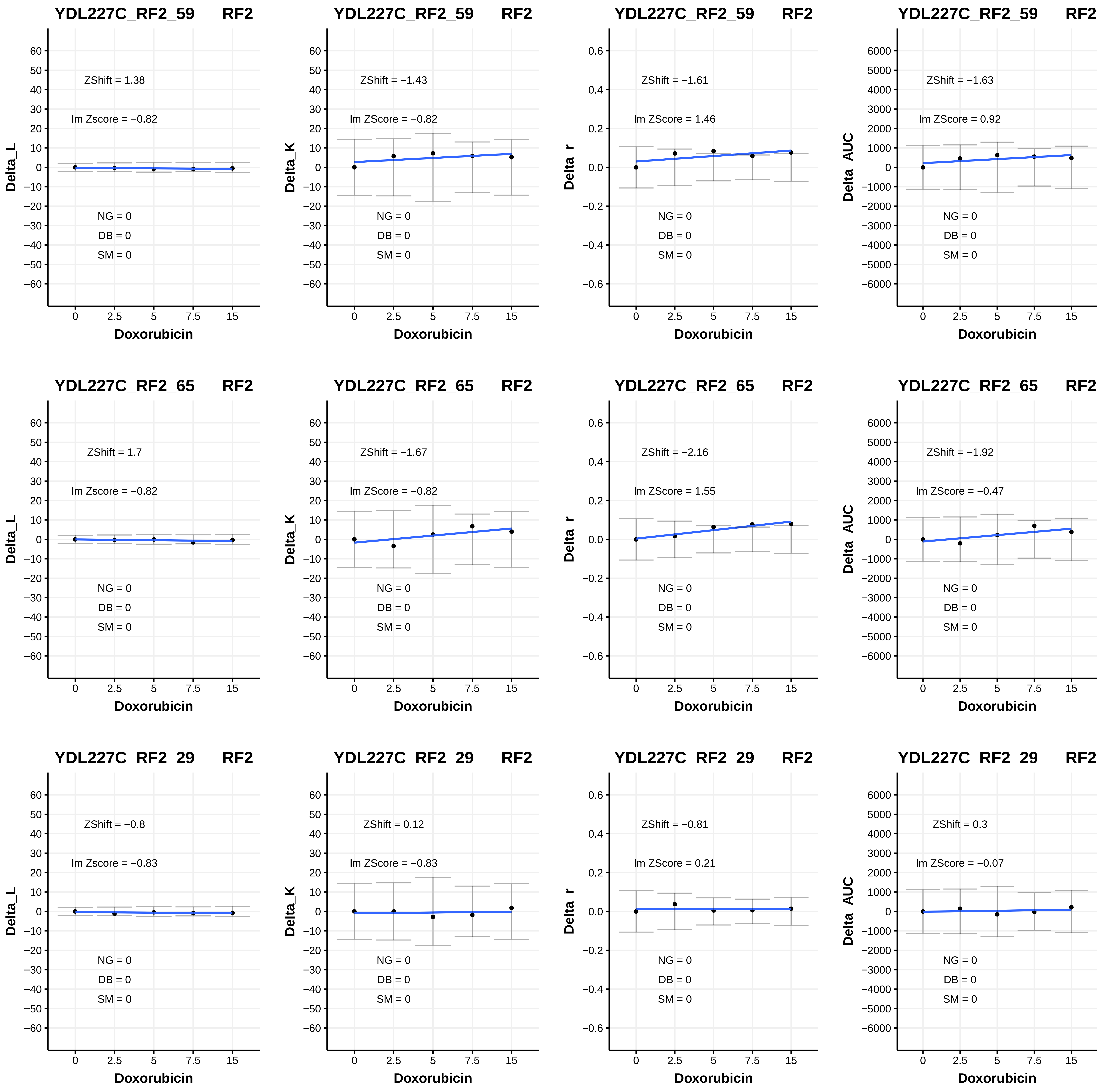
YDL227C_RF2_59 RF2
ZShift = 1.38
lm Zscore = −0.82
NG = 0
DB = 0
SM = 0
Delta_L
Doxorubicin
YDL227C_RF2_59 RF2
ZShift = −1.43
lm ZScore = −0.82
NG = 0
DB = 0
SM = 0
Delta_K
Doxorubicin
YDL227C_RF2_59 RF2
ZShift = −1.61
lm ZScore = 1.46
NG = 0
DB = 0
SM = 0
Delta_r
Doxorubicin
YDL227C_RF2_59 RF2
ZShift = −1.63
lm ZScore = 0.92
NG = 0
DB = 0
SM = 0
Delta_AUC
Doxorubicin
YDL227C_RF2_65 RF2
ZShift = 1.7
lm Zscore = −0.82
NG = 0
DB = 0
SM = 0
Delta_L
Doxorubicin
YDL227C_RF2_65 RF2
ZShift = −1.67
lm ZScore = −0.82
NG = 0
DB = 0
SM = 0
Delta_K
Doxorubicin
YDL227C_RF2_65 RF2
ZShift = −2.16
lm ZScore = 1.55
NG = 0
DB = 0
SM = 0
Delta_r
Doxorubicin
YDL227C_RF2_65 RF2
ZShift = −1.92
lm ZScore = −0.47
NG = 0
DB = 0
SM = 0
Delta_AUC
Doxorubicin
YDL227C_RF2_29 RF2
ZShift = −0.8
lm Zscore = −0.83
NG = 0
DB = 0
SM = 0
Delta_L
Doxorubicin
YDL227C_RF2_29 RF2
ZShift = 0.12
lm ZScore = −0.83
NG = 0
DB = 0
SM = 0
Delta_K
Doxorubicin
YDL227C_RF2_29 RF2
ZShift = −0.81
lm ZScore = 0.21
NG = 0
DB = 0
SM = 0
Delta_r
Doxorubicin
YDL227C_RF2_29 RF2
ZShift = 0.3
lm ZScore = −0.07
NG = 0
DB = 0
SM = 0
Delta_AUC
Doxorubicin

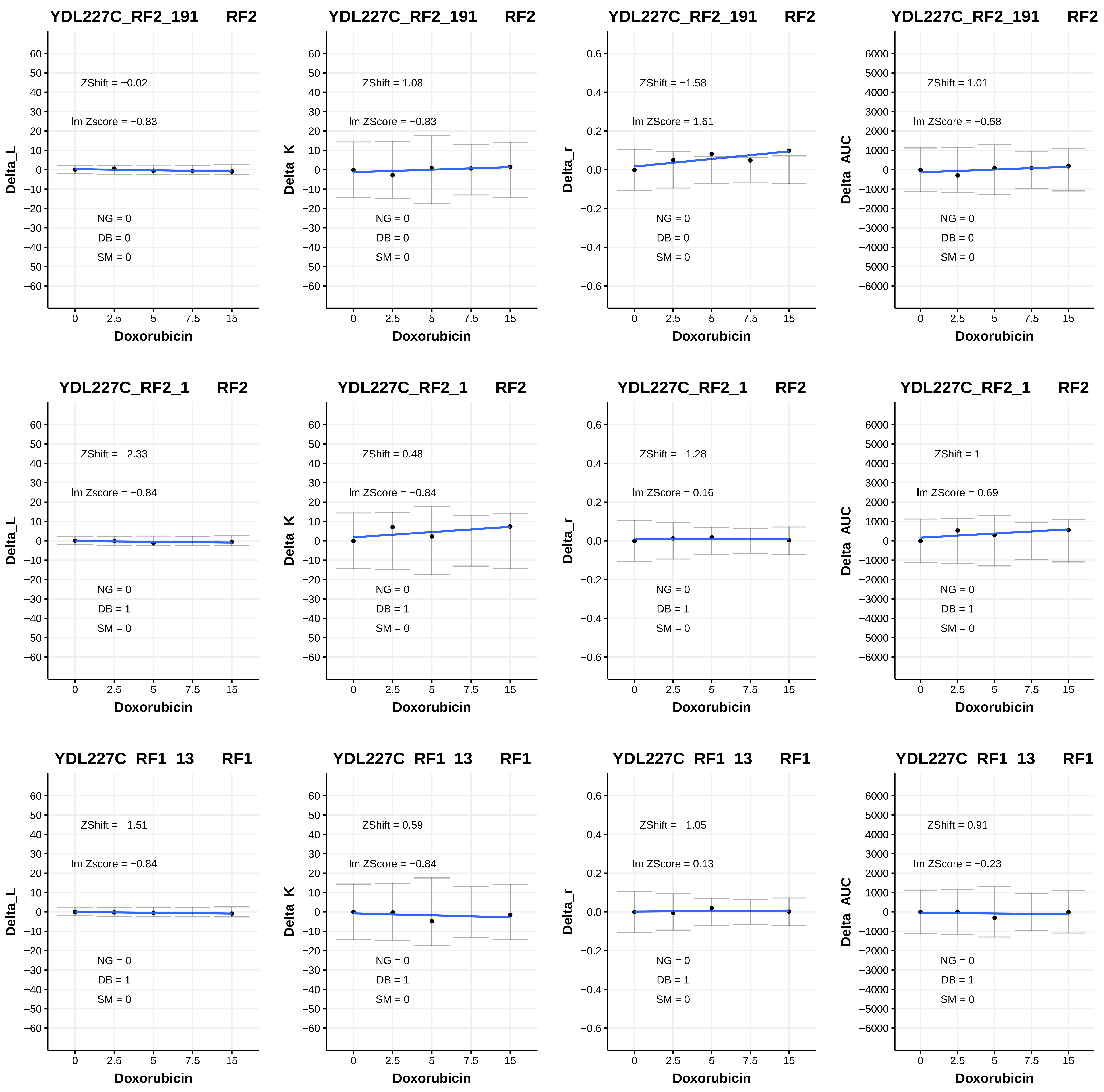
YDL227C_RF2_191 RF2
Delta_L
Doxorubicin
ZShift = −0.02
lm Zscore = −0.83
NG = 0
DB = 0
SM = 0
YDL227C_RF2_191 RF2
Delta_K
Doxorubicin
ZShift = 1.08
lm ZScore = −0.83
NG = 0
DB = 0
SM = 0
YDL227C_RF2_191 RF2
Delta_r
Doxorubicin
ZShift = −1.58
lm ZScore = 1.61
NG = 0
DB = 0
SM = 0
YDL227C_RF2_191 RF2
Delta_AUC
Doxorubicin
ZShift = 1.01
lm ZScore = −0.58
NG = 0
DB = 0
SM = 0
YDL227C_RF2_1 RF2
Delta_L
Doxorubicin
ZShift = −2.33
lm Zscore = −0.84
NG = 0
DB = 1
SM = 0
YDL227C_RF2_1 RF2
Delta_K
Doxorubicin
ZShift = 0.48
lm ZScore = −0.84
NG = 0
DB = 1
SM = 0
YDL227C_RF2_1 RF2
Delta_r
Doxorubicin
ZShift = −1.28
lm ZScore = 0.16
NG = 0
DB = 1
SM = 0
YDL227C_RF2_1 RF2
Delta_AUC
Doxorubicin
ZShift = 1
lm ZScore = 0.69
NG = 0
DB = 1
SM = 0
YDL227C_RF1_13 RF1
Delta_L
Doxorubicin
ZShift = −1.51
lm Zscore = −0.84
NG = 0
DB = 1
SM = 0
YDL227C_RF1_13 RF1
Delta_K
Doxorubicin
ZShift = 0.59
lm ZScore = −0.84
NG = 0
DB = 1
SM = 0
YDL227C_RF1_13 RF1
Delta_r
Doxorubicin
ZShift = −1.05
lm ZScore = 0.13
NG = 0
DB = 1
SM = 0
YDL227C_RF1_13 RF1
Delta_AUC
Doxorubicin
ZShift = 0.91
lm ZScore = −0.23
NG = 0
DB = 1
SM = 0

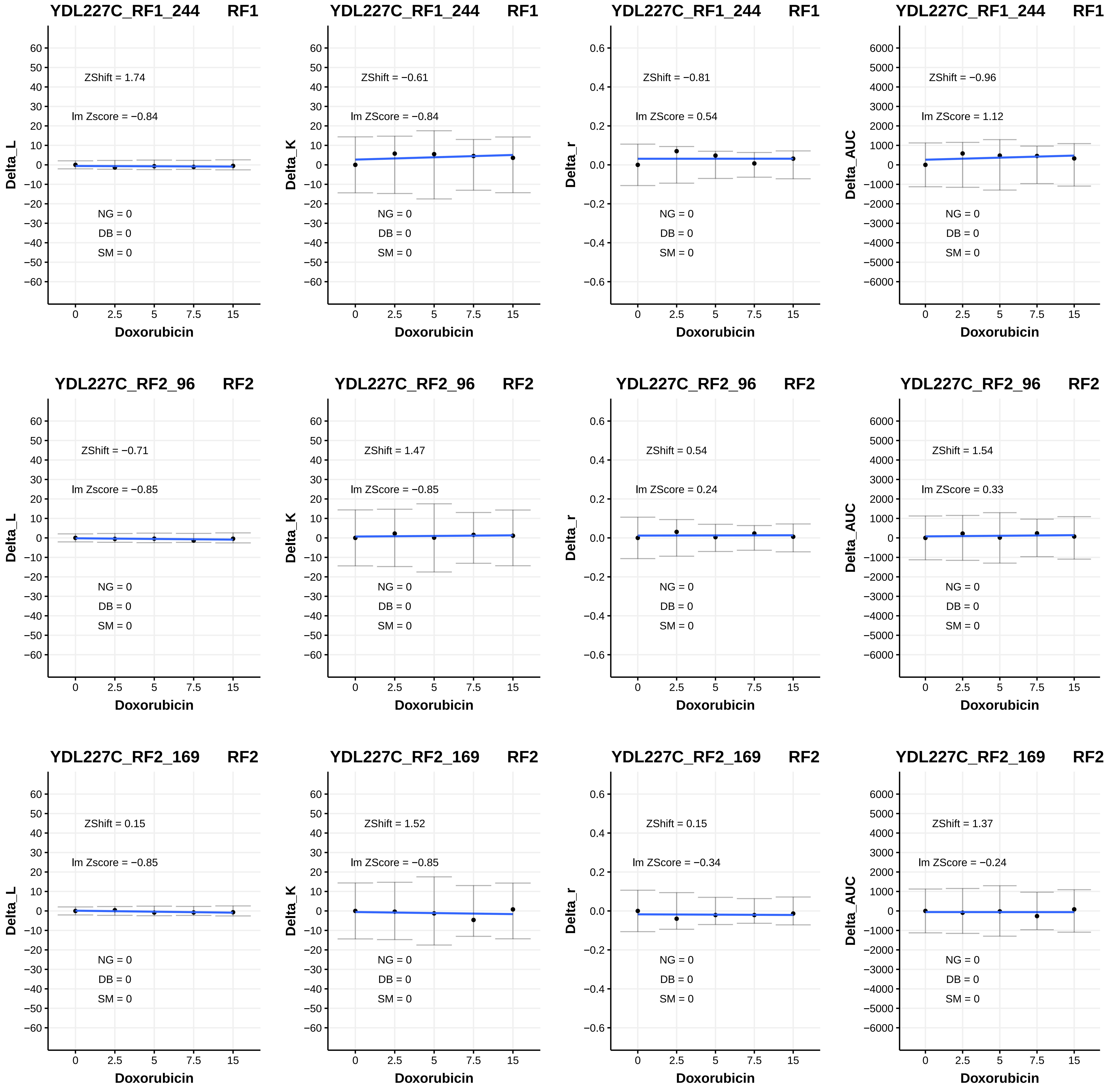
YDL227C_RF1_244 RF1
ZShift = 1.74
lm Zscore = −0.84
NG = 0
DB = 0
SM = 0
Delta_L
Doxorubicin
YDL227C_RF1_244 RF1
ZShift = −0.61
lm ZScore = −0.84
NG = 0
DB = 0
SM = 0
Delta_K
Doxorubicin
YDL227C_RF1_244 RF1
ZShift = −0.81
lm ZScore = 0.54
NG = 0
DB = 0
SM = 0
Delta_r
Doxorubicin
YDL227C_RF1_244 RF1
ZShift = −0.96
lm ZScore = 1.12
NG = 0
DB = 0
SM = 0
Delta_AUC
Doxorubicin
YDL227C_RF2_96 RF2
ZShift = −0.71
lm Zscore = −0.85
NG = 0
DB = 0
SM = 0
Delta_L
Doxorubicin
YDL227C_RF2_96 RF2
ZShift = 1.47
lm ZScore = −0.85
NG = 0
DB = 0
SM = 0
Delta_K
Doxorubicin
YDL227C_RF2_96 RF2
ZShift = 0.54
lm ZScore = 0.24
NG = 0
DB = 0
SM = 0
Delta_r
Doxorubicin
YDL227C_RF2_96 RF2
ZShift = 1.54
lm ZScore = 0.33
NG = 0
DB = 0
SM = 0
Delta_AUC
Doxorubicin
YDL227C_RF2_169 RF2
ZShift = 0.15
lm Zscore = −0.85
NG = 0
DB = 0
SM = 0
Delta_L
Doxorubicin
YDL227C_RF2_169 RF2
ZShift = 1.52
lm ZScore = −0.85
NG = 0
DB = 0
SM = 0
Delta_K
Doxorubicin
YDL227C_RF2_169 RF2
ZShift = 0.15
lm ZScore = −0.34
NG = 0
DB = 0
SM = 0
Delta_r
Doxorubicin
YDL227C_RF2_169 RF2
ZShift = 1.37
lm ZScore = −0.24
NG = 0
DB = 0
SM = 0
Delta_AUC
Doxorubicin

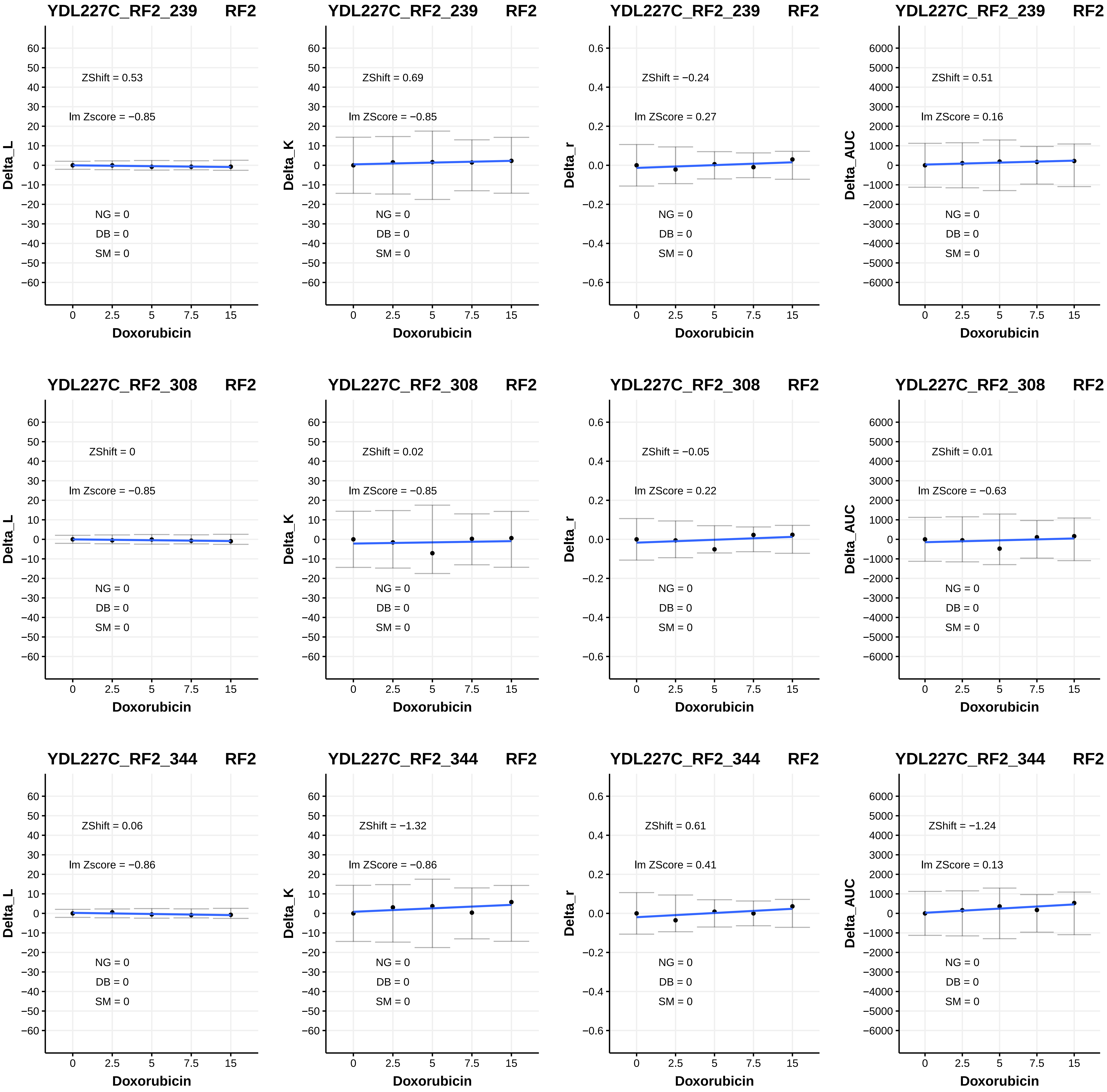

YDL227C_RF2_239 RF2
ZShift = 0.53
lm Zscore = −0.85
NG = 0
DB = 0
SM = 0
Delta_L
Doxorubicin
YDL227C_RF2_239 RF2
ZShift = 0.69
lm ZScore = −0.85
NG = 0
DB = 0
SM = 0
Delta_K
Doxorubicin
YDL227C_RF2_239 RF2
ZShift = −0.24
lm ZScore = 0.27
NG = 0
DB = 0
SM = 0
Delta_r
Doxorubicin
YDL227C_RF2_239 RF2
ZShift = 0.51
lm ZScore = 0.16
NG = 0
DB = 0
SM = 0
Delta_AUC
Doxorubicin
YDL227C_RF2_308 RF2
ZShift = 0
lm Zscore = −0.85
NG = 0
DB = 0
SM = 0
Delta_L
Doxorubicin
YDL227C_RF2_308 RF2
ZShift = 0.02
lm ZScore = −0.85
NG = 0
DB = 0
SM = 0
Delta_K
Doxorubicin
YDL227C_RF2_308 RF2
ZShift = −0.05
lm ZScore = 0.22
NG = 0
DB = 0
SM = 0
Delta_r
Doxorubicin
YDL227C_RF2_308 RF2
ZShift = 0.01
lm ZScore = −0.63
NG = 0
DB = 0
SM = 0
Delta_AUC
Doxorubicin
YDL227C_RF2_344 RF2
ZShift = 0.06
lm Zscore = −0.86
NG = 0
DB = 0
SM = 0
Delta_L
Doxorubicin
YDL227C_RF2_344 RF2
ZShift = −1.32
lm ZScore = −0.86
NG = 0
DB = 0
SM = 0
Delta_K
Doxorubicin
YDL227C_RF2_344 RF2
ZShift = 0.61
lm ZScore = 0.41
NG = 0
DB = 0
SM = 0
Delta_r
Doxorubicin
YDL227C_RF2_344 RF2
ZShift = −1.24
lm ZScore = 0.13
NG = 0
DB = 0
SM = 0
Delta_AUC
Doxorubicin

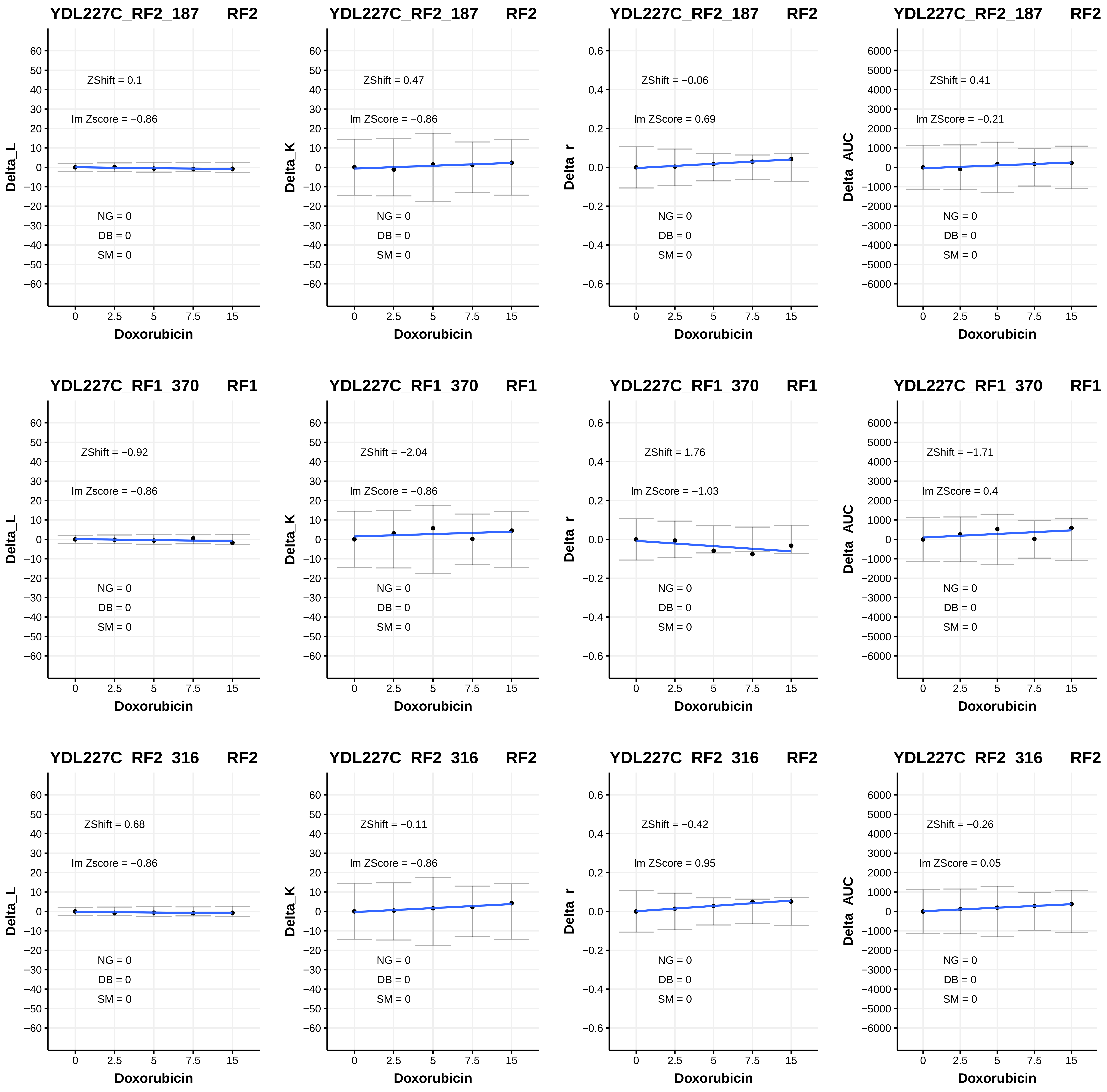

YDL227C_RF2_187 RF2
ZShift = 0.1
lm Zscore = −0.86
NG = 0
DB = 0
SM = 0
Delta_L
Doxorubicin
YDL227C_RF2_187 RF2
ZShift = 0.47
lm ZScore = −0.86
NG = 0
DB = 0
SM = 0
Delta_K
Doxorubicin
YDL227C_RF2_187 RF2
ZShift = −0.06
lm ZScore = 0.69
NG = 0
DB = 0
SM = 0
Delta_r
Doxorubicin
YDL227C_RF2_187 RF2
ZShift = 0.41
lm ZScore = −0.21
NG = 0
DB = 0
SM = 0
Delta_AUC
Doxorubicin
YDL227C_RF1_370 RF1
ZShift = −0.92
lm Zscore = −0.86
NG = 0
DB = 0
SM = 0
Delta_L
Doxorubicin
YDL227C_RF1_370 RF1
ZShift = −2.04
lm ZScore = −0.86
NG = 0
DB = 0
SM = 0
Delta_K
Doxorubicin
YDL227C_RF1_370 RF1
ZShift = 1.76
lm ZScore = −1.03
NG = 0
DB = 0
SM = 0
Delta_r
Doxorubicin
YDL227C_RF1_370 RF1
ZShift = −1.71
lm ZScore = 0.4
NG = 0
DB = 0
SM = 0
Delta_AUC
Doxorubicin
YDL227C_RF2_316 RF2
ZShift = 0.68
lm Zscore = −0.86
NG = 0
DB = 0
SM = 0
Delta_L
Doxorubicin
YDL227C_RF2_316 RF2
ZShift = −0.11
lm ZScore = −0.86
NG = 0
DB = 0
SM = 0
Delta_K
Doxorubicin
YDL227C_RF2_316 RF2
ZShift = −0.42
lm ZScore = 0.95
NG = 0
DB = 0
SM = 0
Delta_r
Doxorubicin
YDL227C_RF2_316 RF2
ZShift = −0.26
lm ZScore = 0.05
NG = 0
DB = 0
SM = 0
Delta_AUC
Doxorubicin

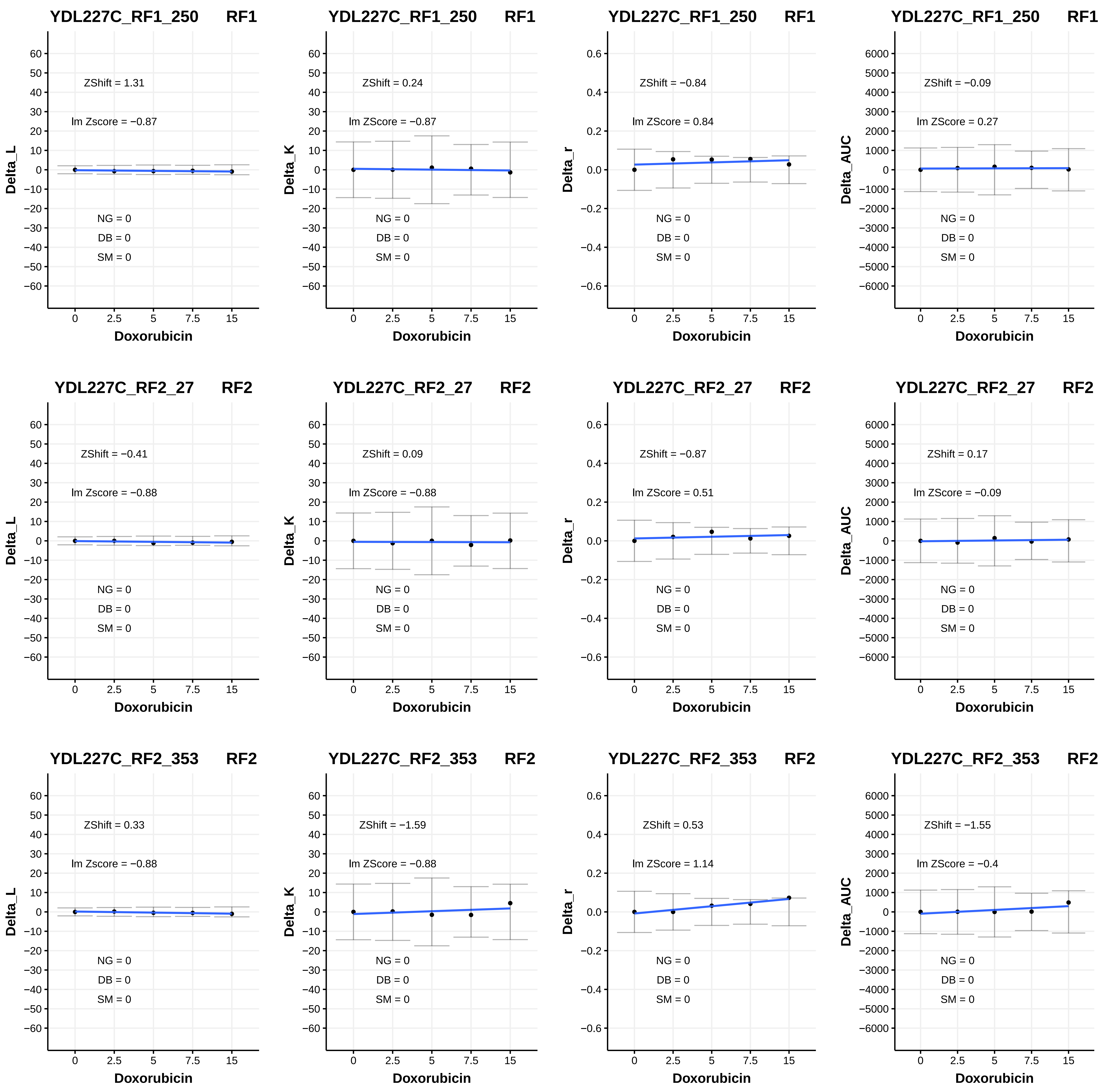

YDL227C_RF1_250 RF1
ZShift = 1.31
lm Zscore = −0.87
NG = 0
DB = 0
SM = 0
Delta_L
Doxorubicin
YDL227C_RF1_250 RF1
ZShift = 0.24
lm ZScore = −0.87
NG = 0
DB = 0
SM = 0
Delta_K
Doxorubicin
YDL227C_RF1_250 RF1
ZShift = −0.84
lm ZScore = 0.84
NG = 0
DB = 0
SM = 0
Delta_r
Doxorubicin
YDL227C_RF1_250 RF1
ZShift = −0.09
lm ZScore = 0.27
NG = 0
DB = 0
SM = 0
Delta_AUC
Doxorubicin
YDL227C_RF2_27 RF2
ZShift = −0.41
lm Zscore = −0.88
NG = 0
DB = 0
SM = 0
Delta_L
Doxorubicin
YDL227C_RF2_27 RF2
ZShift = 0.09
lm ZScore = −0.88
NG = 0
DB = 0
SM = 0
Delta_K
Doxorubicin
YDL227C_RF2_27 RF2
ZShift = −0.87
lm ZScore = 0.51
NG = 0
DB = 0
SM = 0
Delta_r
Doxorubicin
YDL227C_RF2_27 RF2
ZShift = 0.17
lm ZScore = −0.09
NG = 0
DB = 0
SM = 0
Delta_AUC
Doxorubicin
YDL227C_RF2_353 RF2
ZShift = 0.33
lm Zscore = −0.88
NG = 0
DB = 0
SM = 0
Delta_L
Doxorubicin
YDL227C_RF2_353 RF2
ZShift = −1.59
lm ZScore = −0.88
NG = 0
DB = 0
SM = 0
Delta_K
Doxorubicin
YDL227C_RF2_353 RF2
ZShift = 0.53
lm ZScore = 1.14
NG = 0
DB = 0
SM = 0
Delta_r
Doxorubicin
YDL227C_RF2_353 RF2
ZShift = −1.55
lm ZScore = −0.4
NG = 0
DB = 0
SM = 0
Delta_AUC
Doxorubicin

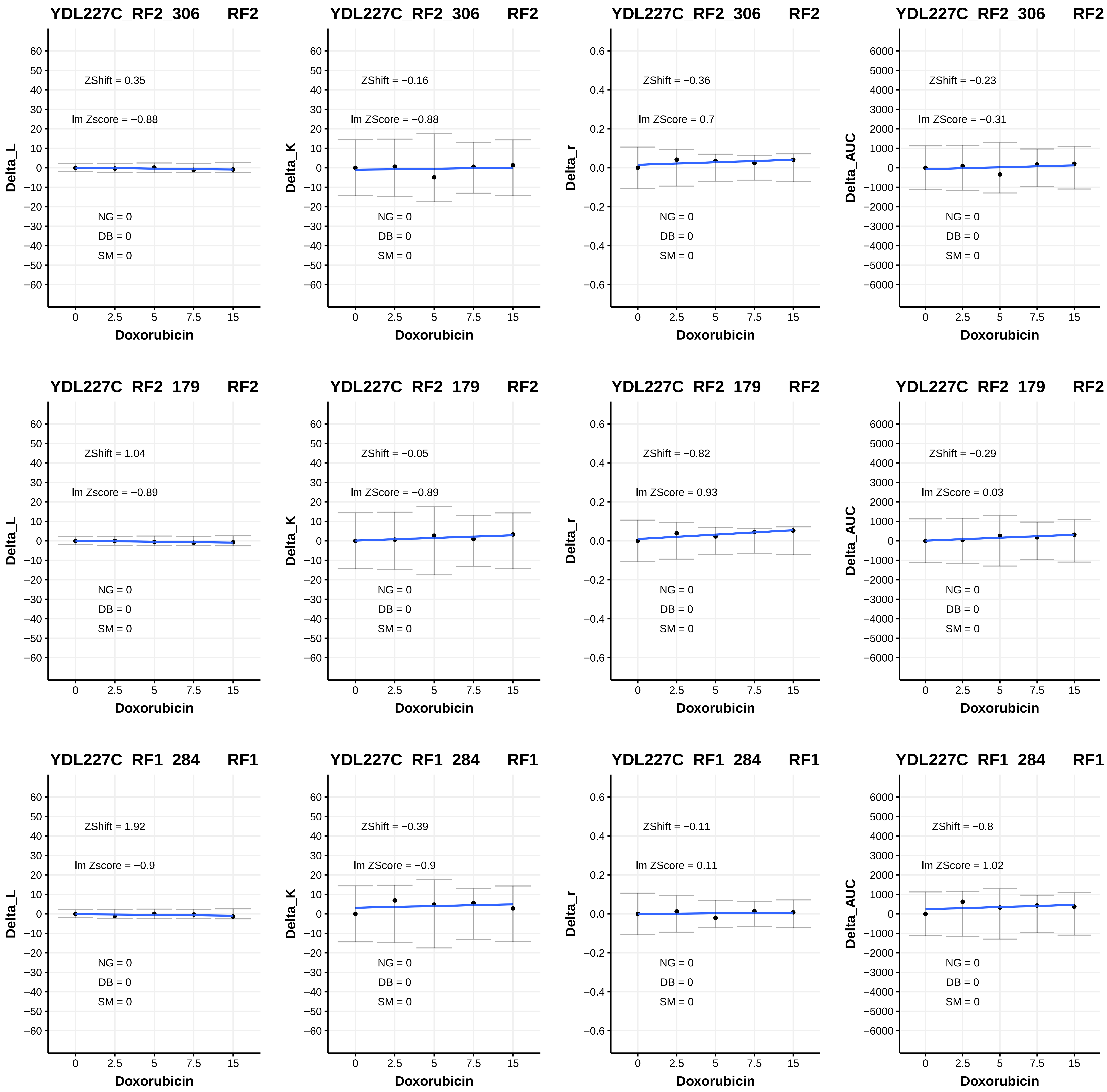

YDL227C_RF2_306 RF2
ZShift = 0.35
lm Zscore = −0.88
NG = 0
DB = 0
SM = 0
Delta_L
Doxorubicin
YDL227C_RF2_306 RF2
ZShift = −0.16
lm ZScore = −0.88
NG = 0
DB = 0
SM = 0
Delta_K
Doxorubicin
YDL227C_RF2_306 RF2
ZShift = −0.36
lm ZScore = 0.7
NG = 0
DB = 0
SM = 0
Delta_r
Doxorubicin
YDL227C_RF2_306 RF2
ZShift = −0.23
lm ZScore = −0.31
NG = 0
DB = 0
SM = 0
Delta_AUC
Doxorubicin
YDL227C_RF2_179 RF2
ZShift = 1.04
lm Zscore = −0.89
NG = 0
DB = 0
SM = 0
Delta_L
Doxorubicin
YDL227C_RF2_179 RF2
ZShift = −0.05
lm ZScore = −0.89
NG = 0
DB = 0
SM = 0
Delta_K
Doxorubicin
YDL227C_RF2_179 RF2
ZShift = −0.82
lm ZScore = 0.93
NG = 0
DB = 0
SM = 0
Delta_r
Doxorubicin
YDL227C_RF2_179 RF2
ZShift = −0.29
lm ZScore = 0.03
NG = 0
DB = 0
SM = 0
Delta_AUC
Doxorubicin
YDL227C_RF1_284 RF1
ZShift = 1.92
lm Zscore = −0.9
NG = 0
DB = 0
SM = 0
Delta_L
Doxorubicin
YDL227C_RF1_284 RF1
ZShift = −0.39
lm ZScore = −0.9
NG = 0
DB = 0
SM = 0
Delta_K
Doxorubicin
YDL227C_RF1_284 RF1
ZShift = −0.11
lm ZScore = 0.11
NG = 0
DB = 0
SM = 0
Delta_r
Doxorubicin
YDL227C_RF1_284 RF1
ZShift = −0.8
lm ZScore = 1.02
NG = 0
DB = 0
SM = 0
Delta_AUC
Doxorubicin

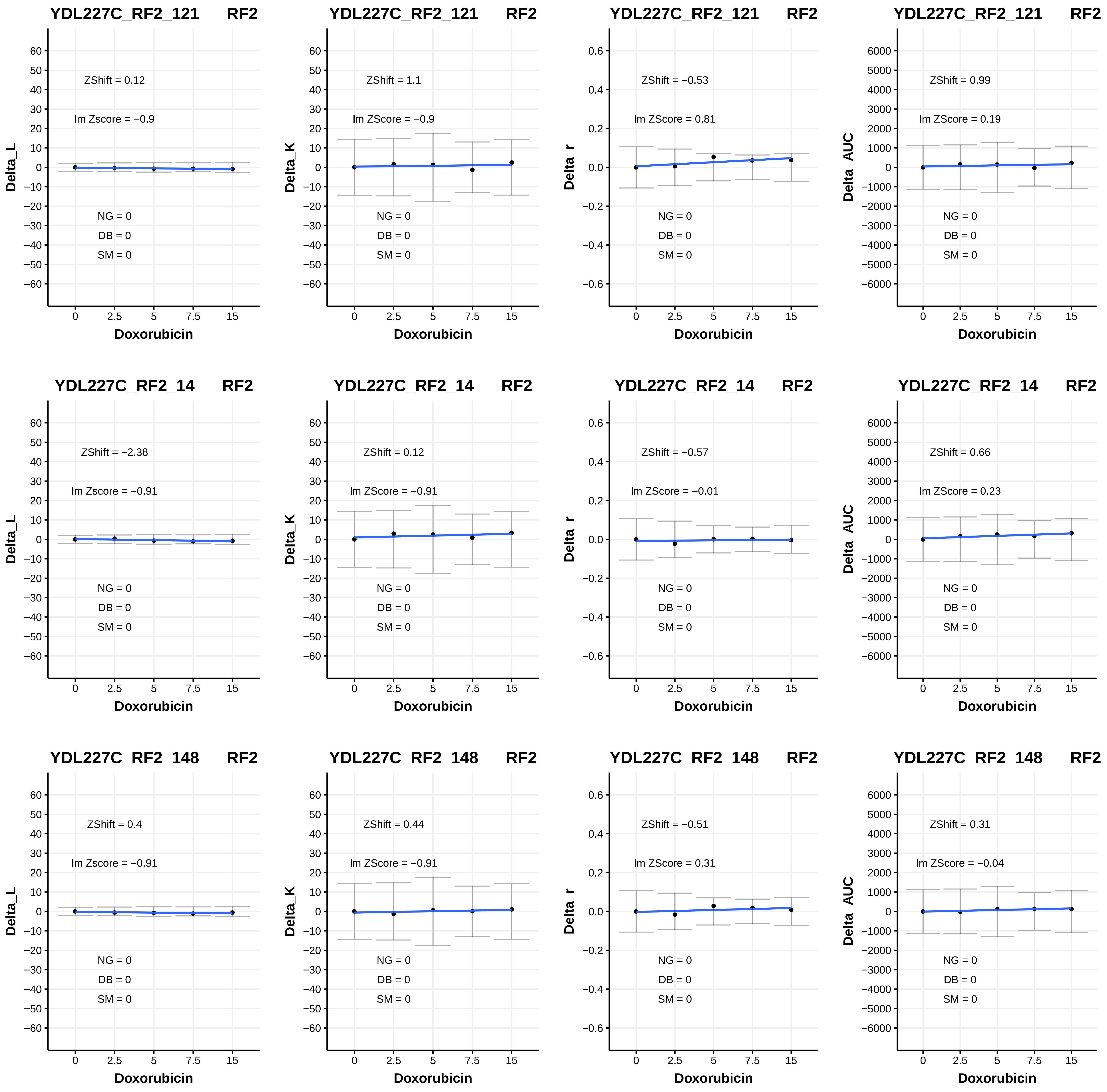
YDL227C_RF2_121 RF2
ZShift = 0.12
lm Zscore = −0.9
NG = 0
DB = 0
SM = 0
Delta_L
Doxorubicin
YDL227C_RF2_121 RF2
ZShift = 1.1
lm ZScore = −0.9
NG = 0
DB = 0
SM = 0
Delta_K
Doxorubicin
YDL227C_RF2_121 RF2
ZShift = −0.53
lm ZScore = 0.81
NG = 0
DB = 0
SM = 0
Delta_r
Doxorubicin
YDL227C_RF2_121 RF2
ZShift = 0.99
lm ZScore = 0.19
NG = 0
DB = 0
SM = 0
Delta_AUC
Doxorubicin
YDL227C_RF2_14 RF2
ZShift = −2.38
lm Zscore = −0.91
NG = 0
DB = 0
SM = 0
Delta_L
Doxorubicin
YDL227C_RF2_14 RF2
ZShift = 0.12
lm ZScore = −0.91
NG = 0
DB = 0
SM = 0
Delta_K
Doxorubicin
YDL227C_RF2_14 RF2
ZShift = −0.57
lm ZScore = −0.01
NG = 0
DB = 0
SM = 0
Delta_r
Doxorubicin
YDL227C_RF2_14 RF2
ZShift = 0.66
lm ZScore = 0.23
NG = 0
DB = 0
SM = 0
Delta_AUC
Doxorubicin
YDL227C_RF2_148 RF2
ZShift = 0.4
lm Zscore = −0.91
NG = 0
DB = 0
SM = 0
Delta_L
Doxorubicin
YDL227C_RF2_148 RF2
ZShift = 0.44
lm ZScore = −0.91
NG = 0
DB = 0
SM = 0
Delta_K
Doxorubicin
YDL227C_RF2_148 RF2
ZShift = −0.51
lm ZScore = 0.31
NG = 0
DB = 0
SM = 0
Delta_r
Doxorubicin
YDL227C_RF2_148 RF2
ZShift = 0.31
lm ZScore = −0.04
NG = 0
DB = 0
SM = 0
Delta_AUC
Doxorubicin

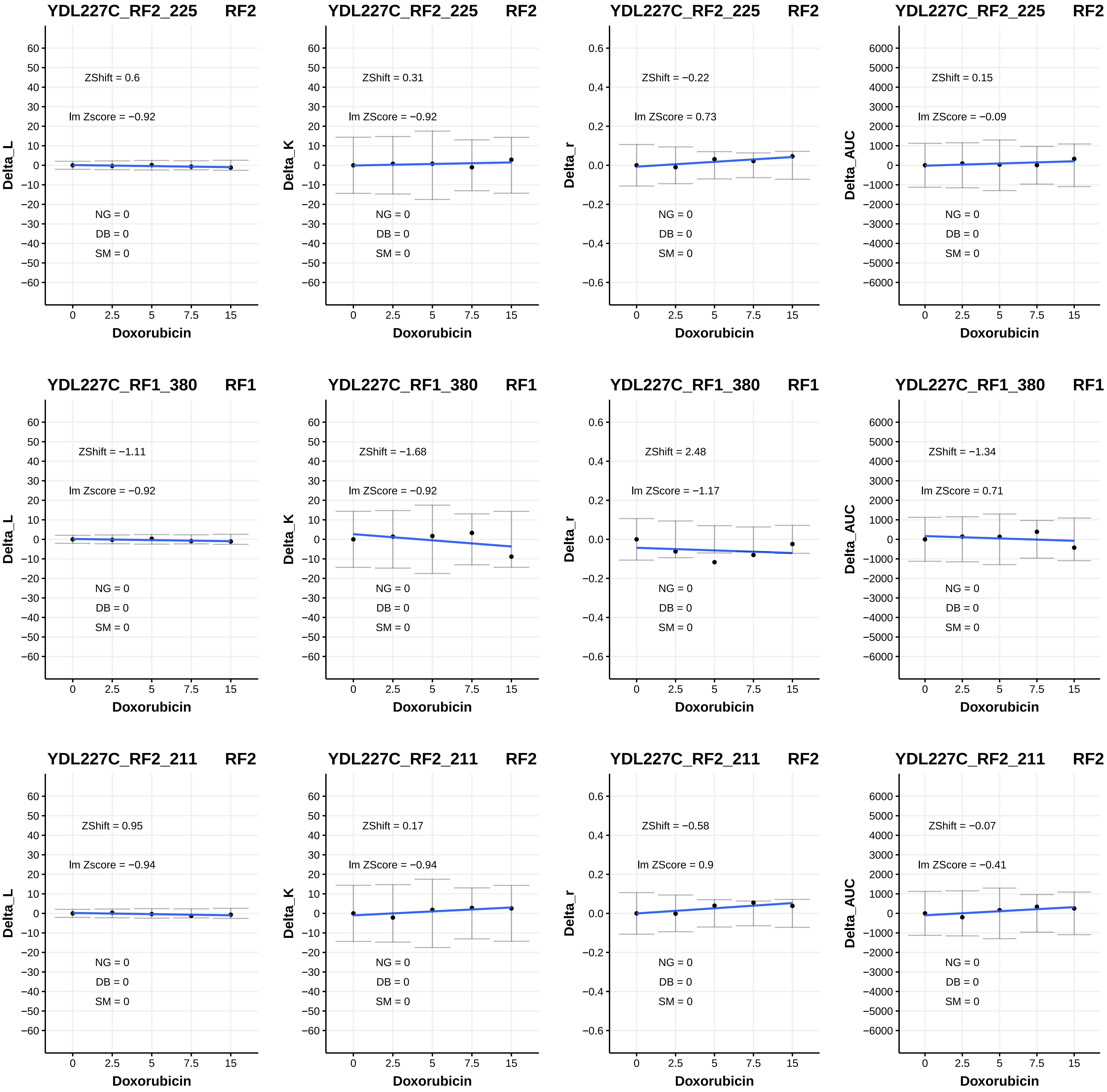

YDL227C_RF2_225 RF2
ZShift = 0.6
lm Zscore = −0.92
NG = 0
DB = 0
SM = 0
Delta_L
Doxorubicin
YDL227C_RF2_225 RF2
ZShift = 0.31
lm ZScore = −0.92
NG = 0
DB = 0
SM = 0
Delta_K
Doxorubicin
YDL227C_RF2_225 RF2
ZShift = −0.22
lm ZScore = 0.73
NG = 0
DB = 0
SM = 0
Delta_r
Doxorubicin
YDL227C_RF2_225 RF2
ZShift = 0.15
lm ZScore = −0.09
NG = 0
DB = 0
SM = 0
Delta_AUC
Doxorubicin
YDL227C_RF1_380 RF1
ZShift = −1.11
lm Zscore = −0.92
NG = 0
DB = 0
SM = 0
Delta_L
Doxorubicin
YDL227C_RF1_380 RF1
ZShift = −1.68
lm ZScore = −0.92
NG = 0
DB = 0
SM = 0
Delta_K
Doxorubicin
YDL227C_RF1_380 RF1
ZShift = 2.48
lm ZScore = −1.17
NG = 0
DB = 0
SM = 0
Delta_r
Doxorubicin
YDL227C_RF1_380 RF1
ZShift = −1.34
lm ZScore = 0.71
NG = 0
DB = 0
SM = 0
Delta_AUC
Doxorubicin
YDL227C_RF2_211 RF2
ZShift = 0.95
lm Zscore = −0.94
NG = 0
DB = 0
SM = 0
Delta_L
Doxorubicin
YDL227C_RF2_211 RF2
ZShift = 0.17
lm ZScore = −0.94
NG = 0
DB = 0
SM = 0
Delta_K
Doxorubicin
YDL227C_RF2_211 RF2
ZShift = −0.58
lm ZScore = 0.9
NG = 0
DB = 0
SM = 0
Delta_r
Doxorubicin
YDL227C_RF2_211 RF2
ZShift = −0.07
lm ZScore = −0.41
NG = 0
DB = 0
SM = 0
Delta_AUC
Doxorubicin

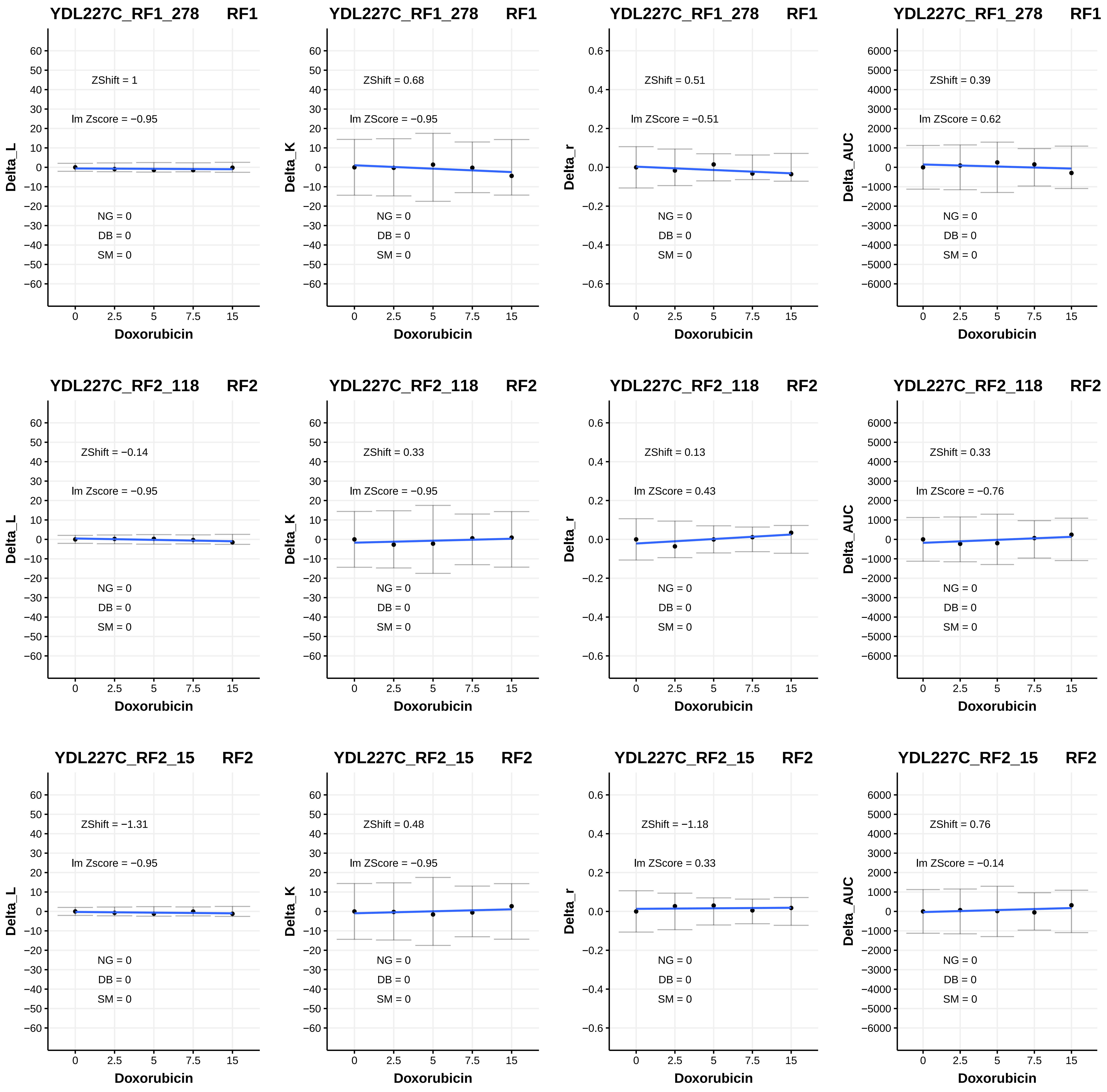

YDL227C_RF1_278 RF1
ZShift = 1
lm Zscore = −0.95
NG = 0
DB = 0
SM = 0
Delta_L
Doxorubicin
YDL227C_RF1_278 RF1
ZShift = 0.68
lm ZScore = −0.95
NG = 0
DB = 0
SM = 0
Delta_K
Doxorubicin
YDL227C_RF1_278 RF1
ZShift = 0.51
lm ZScore = −0.51
NG = 0
DB = 0
SM = 0
Delta_r
Doxorubicin
YDL227C_RF1_278 RF1
ZShift = 0.39
lm ZScore = 0.62
NG = 0
DB = 0
SM = 0
Delta_AUC
Doxorubicin
YDL227C_RF2_118 RF2
ZShift = −0.14
lm Zscore = −0.95
NG = 0
DB = 0
SM = 0
Delta_L
Doxorubicin
YDL227C_RF2_118 RF2
ZShift = 0.33
lm ZScore = −0.95
NG = 0
DB = 0
SM = 0
Delta_K
Doxorubicin
YDL227C_RF2_118 RF2
ZShift = 0.13
lm ZScore = 0.43
NG = 0
DB = 0
SM = 0
Delta_r
Doxorubicin
YDL227C_RF2_118 RF2
ZShift = 0.33
lm ZScore = −0.76
NG = 0
DB = 0
SM = 0
Delta_AUC
Doxorubicin
YDL227C_RF2_15 RF2
ZShift = −1.31
lm Zscore = −0.95
NG = 0
DB = 0
SM = 0
Delta_L
Doxorubicin
YDL227C_RF2_15 RF2
ZShift = 0.48
lm ZScore = −0.95
NG = 0
DB = 0
SM = 0
Delta_K
Doxorubicin
YDL227C_RF2_15 RF2
ZShift = −1.18
lm ZScore = 0.33
NG = 0
DB = 0
SM = 0
Delta_r
Doxorubicin
YDL227C_RF2_15 RF2
ZShift = 0.76
lm ZScore = −0.14
NG = 0
DB = 0
SM = 0
Delta_AUC
Doxorubicin

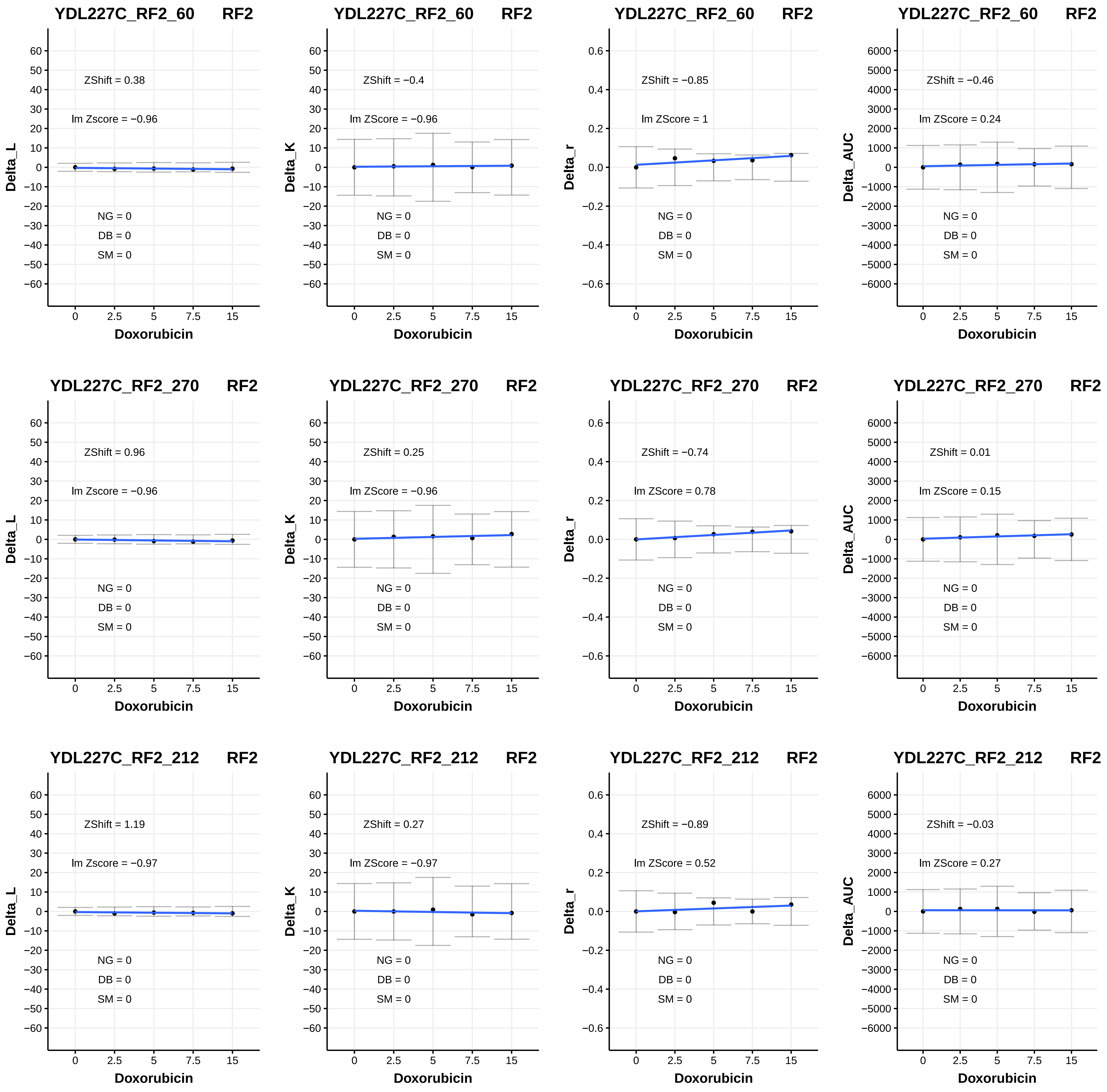
YDL227C_RF2_60 RF2
ZShift = 0.38
lm Zscore = −0.96
NG = 0
DB = 0
SM = 0
Delta_L
Doxorubicin
YDL227C_RF2_60 RF2
ZShift = −0.4
lm ZScore = −0.96
NG = 0
DB = 0
SM = 0
Delta_K
Doxorubicin
YDL227C_RF2_60 RF2
ZShift = −0.85
lm ZScore = 1
NG = 0
DB = 0
SM = 0
Delta_r
Doxorubicin
YDL227C_RF2_60 RF2
ZShift = −0.46
lm ZScore = 0.24
NG = 0
DB = 0
SM = 0
Delta_AUC
Doxorubicin
YDL227C_RF2_270 RF2
ZShift = 0.96
lm Zscore = −0.96
NG = 0
DB = 0
SM = 0
Delta_L
Doxorubicin
YDL227C_RF2_270 RF2
ZShift = 0.25
lm ZScore = −0.96
NG = 0
DB = 0
SM = 0
Delta_K
Doxorubicin
YDL227C_RF2_270 RF2
ZShift = −0.74
lm ZScore = 0.78
NG = 0
DB = 0
SM = 0
Delta_r
Doxorubicin
YDL227C_RF2_270 RF2
ZShift = 0.01
lm ZScore = 0.15
NG = 0
DB = 0
SM = 0
Delta_AUC
Doxorubicin
YDL227C_RF2_212 RF2
ZShift = 1.19
lm Zscore = −0.97
NG = 0
DB = 0
SM = 0
Delta_L
Doxorubicin
YDL227C_RF2_212 RF2
ZShift = 0.27
lm ZScore = −0.97
NG = 0
DB = 0
SM = 0
Delta_K
Doxorubicin
YDL227C_RF2_212 RF2
ZShift = −0.89
lm ZScore = 0.52
NG = 0
DB = 0
SM = 0
Delta_r
Doxorubicin
YDL227C_RF2_212 RF2
ZShift = −0.03
lm ZScore = 0.27
NG = 0
DB = 0
SM = 0
Delta_AUC
Doxorubicin

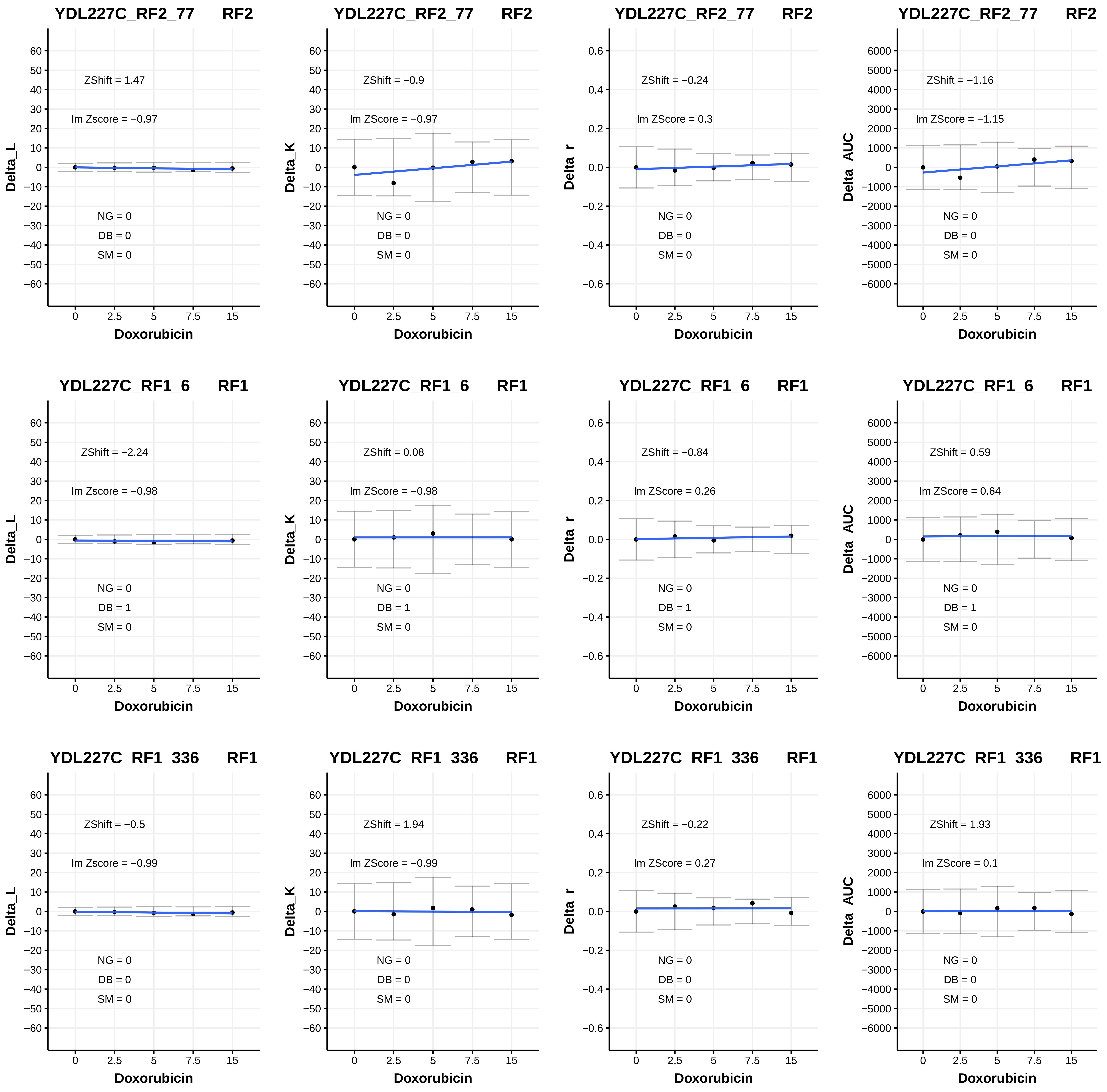

YDL227C_RF2_77 RF2
ZShift = 1.47
lm Zscore = −0.97
NG = 0
DB = 0
SM = 0
Delta_L
Doxorubicin
YDL227C_RF2_77 RF2
ZShift = −0.9
lm ZScore = −0.97
NG = 0
DB = 0
SM = 0
Delta_K
Doxorubicin
YDL227C_RF2_77 RF2
ZShift = −0.24
lm ZScore = 0.3
NG = 0
DB = 0
SM = 0
Delta_r
Doxorubicin
YDL227C_RF2_77 RF2
ZShift = −1.16
lm ZScore = −1.15
NG = 0
DB = 0
SM = 0
Delta_AUC
Doxorubicin
YDL227C_RF1_6 RF1
ZShift = −2.24
lm Zscore = −0.98
NG = 0
DB = 1
SM = 0
Delta_L
Doxorubicin
YDL227C_RF1_6 RF1
ZShift = 0.08
lm ZScore = −0.98
NG = 0
DB = 1
SM = 0
Delta_K
Doxorubicin
YDL227C_RF1_6 RF1
ZShift = −0.84
lm ZScore = 0.26
NG = 0
DB = 1
SM = 0
Delta_r
Doxorubicin
YDL227C_RF1_6 RF1
ZShift = 0.59
lm ZScore = 0.64
NG = 0
DB = 1
SM = 0
Delta_AUC
Doxorubicin
YDL227C_RF1_336 RF1
ZShift = −0.5
lm Zscore = −0.99
NG = 0
DB = 0
SM = 0
Delta_L
Doxorubicin
YDL227C_RF1_336 RF1
ZShift = 1.94
lm ZScore = −0.99
NG = 0
DB = 0
SM = 0
Delta_K
Doxorubicin
YDL227C_RF1_336 RF1
ZShift = −0.22
lm ZScore = 0.27
NG = 0
DB = 0
SM = 0
Delta_r
Doxorubicin
YDL227C_RF1_336 RF1
ZShift = 1.93
lm ZScore = 0.1
NG = 0
DB = 0
SM = 0
Delta_AUC
Doxorubicin

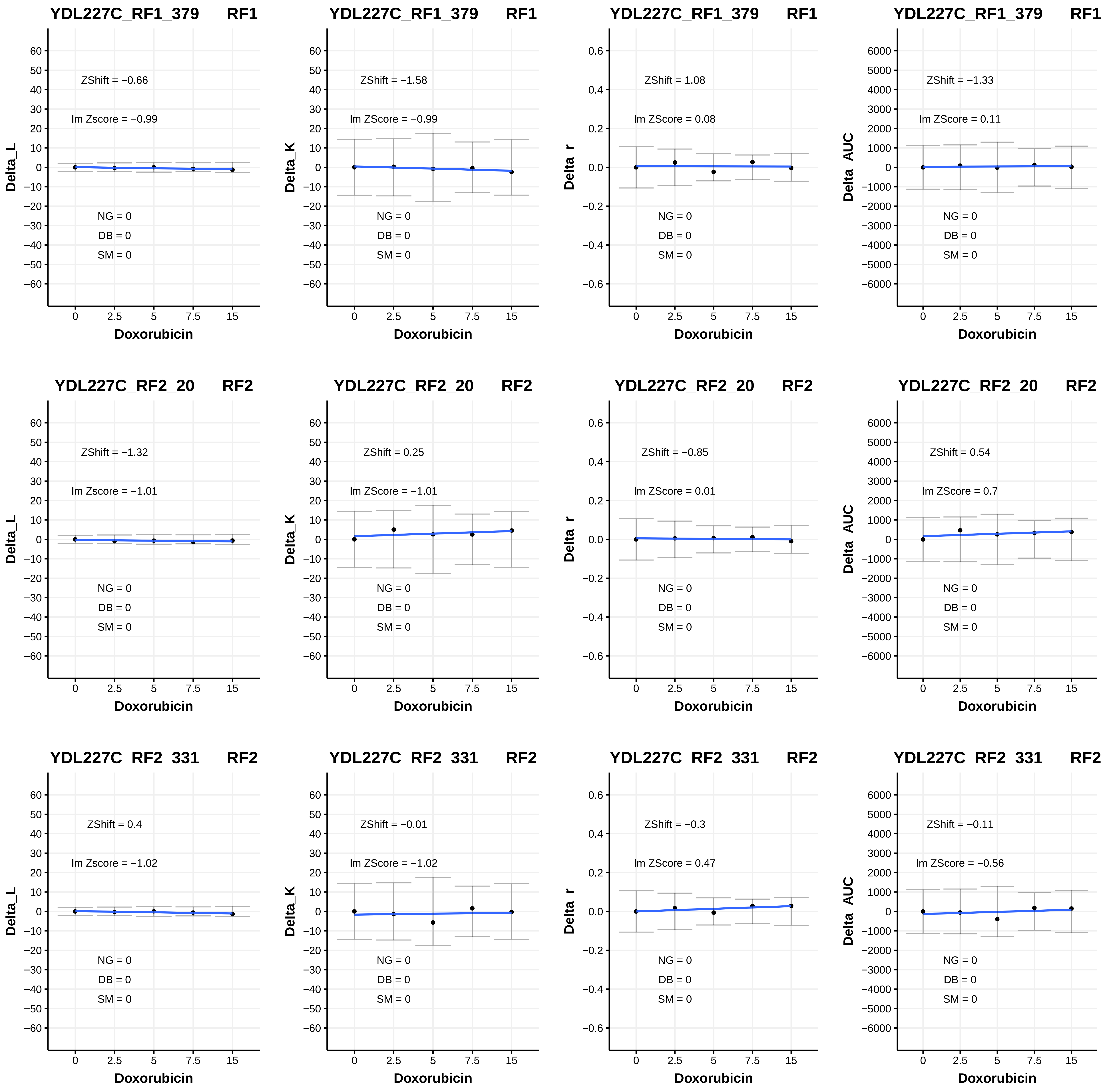

YDL227C_RF1_379 RF1
ZShift = −0.66
lm Zscore = −0.99
NG = 0
DB = 0
SM = 0
Delta_L
Doxorubicin
YDL227C_RF1_379 RF1
ZShift = −1.58
lm ZScore = −0.99
NG = 0
DB = 0
SM = 0
Delta_K
Doxorubicin
YDL227C_RF1_379 RF1
ZShift = 1.08
lm ZScore = 0.08
NG = 0
DB = 0
SM = 0
Delta_r
Doxorubicin
YDL227C_RF1_379 RF1
ZShift = −1.33
lm ZScore = 0.11
NG = 0
DB = 0
SM = 0
Delta_AUC
Doxorubicin
YDL227C_RF2_20 RF2
ZShift = −1.32
lm Zscore = −1.01
NG = 0
DB = 0
SM = 0
Delta_L
Doxorubicin
YDL227C_RF2_20 RF2
ZShift = 0.25
lm ZScore = −1.01
NG = 0
DB = 0
SM = 0
Delta_K
Doxorubicin
YDL227C_RF2_20 RF2
ZShift = −0.85
lm ZScore = 0.01
NG = 0
DB = 0
SM = 0
Delta_r
Doxorubicin
YDL227C_RF2_20 RF2
ZShift = 0.54
lm ZScore = 0.7
NG = 0
DB = 0
SM = 0
Delta_AUC
Doxorubicin
YDL227C_RF2_331 RF2
ZShift = 0.4
lm Zscore = −1.02
NG = 0
DB = 0
SM = 0
Delta_L
Doxorubicin
YDL227C_RF2_331 RF2
ZShift = −0.01
lm ZScore = −1.02
NG = 0
DB = 0
SM = 0
Delta_K
Doxorubicin
YDL227C_RF2_331 RF2
ZShift = −0.3
lm ZScore = 0.47
NG = 0
DB = 0
SM = 0
Delta_r
Doxorubicin
YDL227C_RF2_331 RF2
ZShift = −0.11
lm ZScore = −0.56
NG = 0
DB = 0
SM = 0
Delta_AUC
Doxorubicin

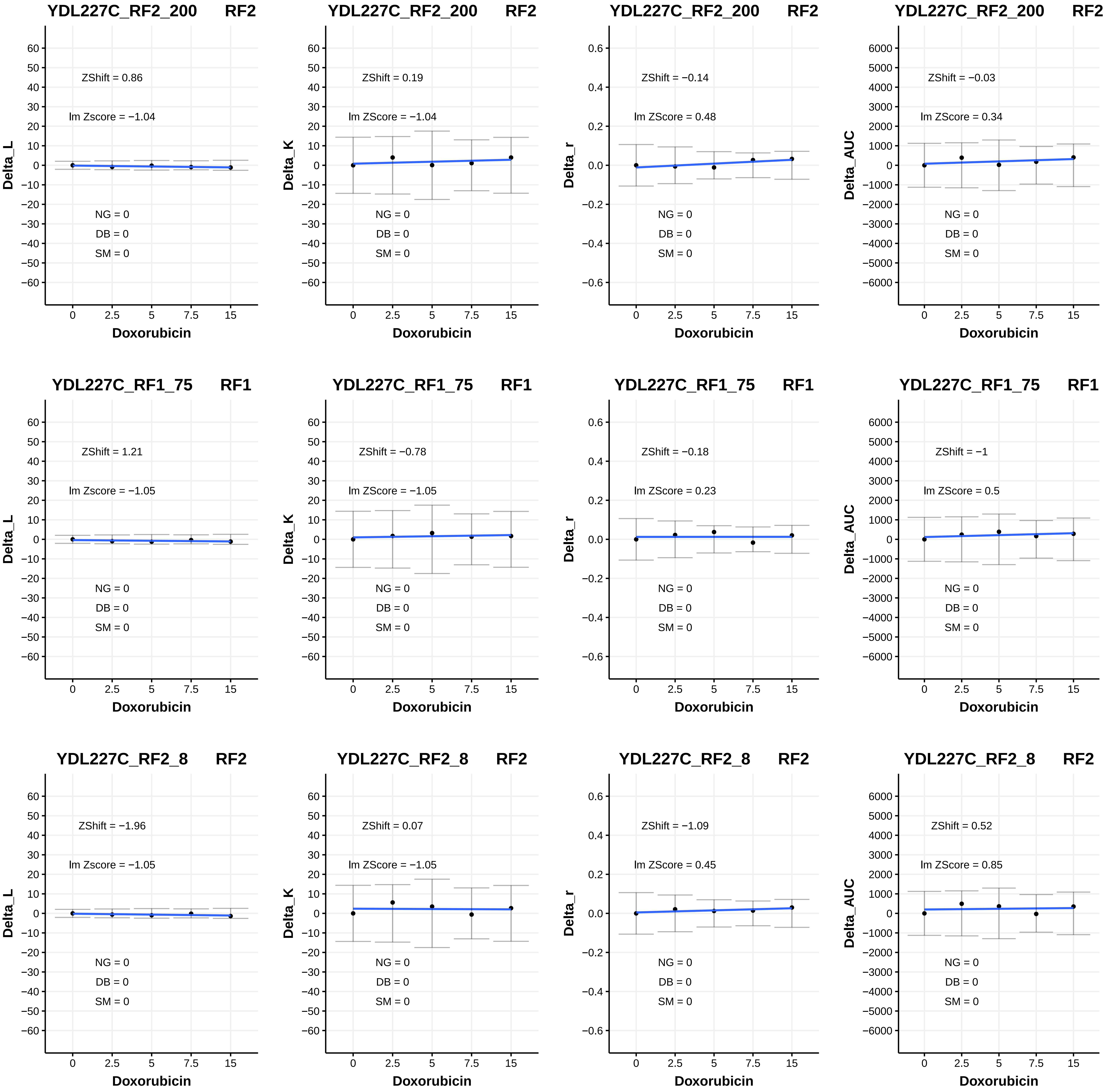
YDL227C_RF2_200 RF2
ZShift = 0.86
lm Zscore = −1.04
NG = 0
DB = 0
SM = 0
Delta_L
Doxorubicin
YDL227C_RF2_200 RF2
ZShift = 0.19
lm ZScore = −1.04
NG = 0
DB = 0
SM = 0
Delta_K
Doxorubicin
YDL227C_RF2_200 RF2
ZShift = −0.14
lm ZScore = 0.48
NG = 0
DB = 0
SM = 0
Delta_r
Doxorubicin
YDL227C_RF2_200 RF2
ZShift = −0.03
lm ZScore = 0.34
NG = 0
DB = 0
SM = 0
Delta_AUC
Doxorubicin
YDL227C_RF1_75 RF1
ZShift = 1.21
lm Zscore = −1.05
NG = 0
DB = 0
SM = 0
Delta_L
Doxorubicin
YDL227C_RF1_75 RF1
ZShift = −0.78
lm ZScore = −1.05
NG = 0
DB = 0
SM = 0
Delta_K
Doxorubicin
YDL227C_RF1_75 RF1
ZShift = −0.18
lm ZScore = 0.23
NG = 0
DB = 0
SM = 0
Delta_r
Doxorubicin
YDL227C_RF1_75 RF1
ZShift = −1
lm ZScore = 0.5
NG = 0
DB = 0
SM = 0
Delta_AUC
Doxorubicin
YDL227C_RF2_8 RF2
ZShift = −1.96
lm Zscore = −1.05
NG = 0
DB = 0
SM = 0
Delta_L
Doxorubicin
YDL227C_RF2_8 RF2
ZShift = 0.07
lm ZScore = −1.05
NG = 0
DB = 0
SM = 0
Delta_K
Doxorubicin
YDL227C_RF2_8 RF2
ZShift = −1.09
lm ZScore = 0.45
NG = 0
DB = 0
SM = 0
Delta_r
Doxorubicin
YDL227C_RF2_8 RF2
ZShift = 0.52
lm ZScore = 0.85
NG = 0
DB = 0
SM = 0
Delta_AUC
Doxorubicin

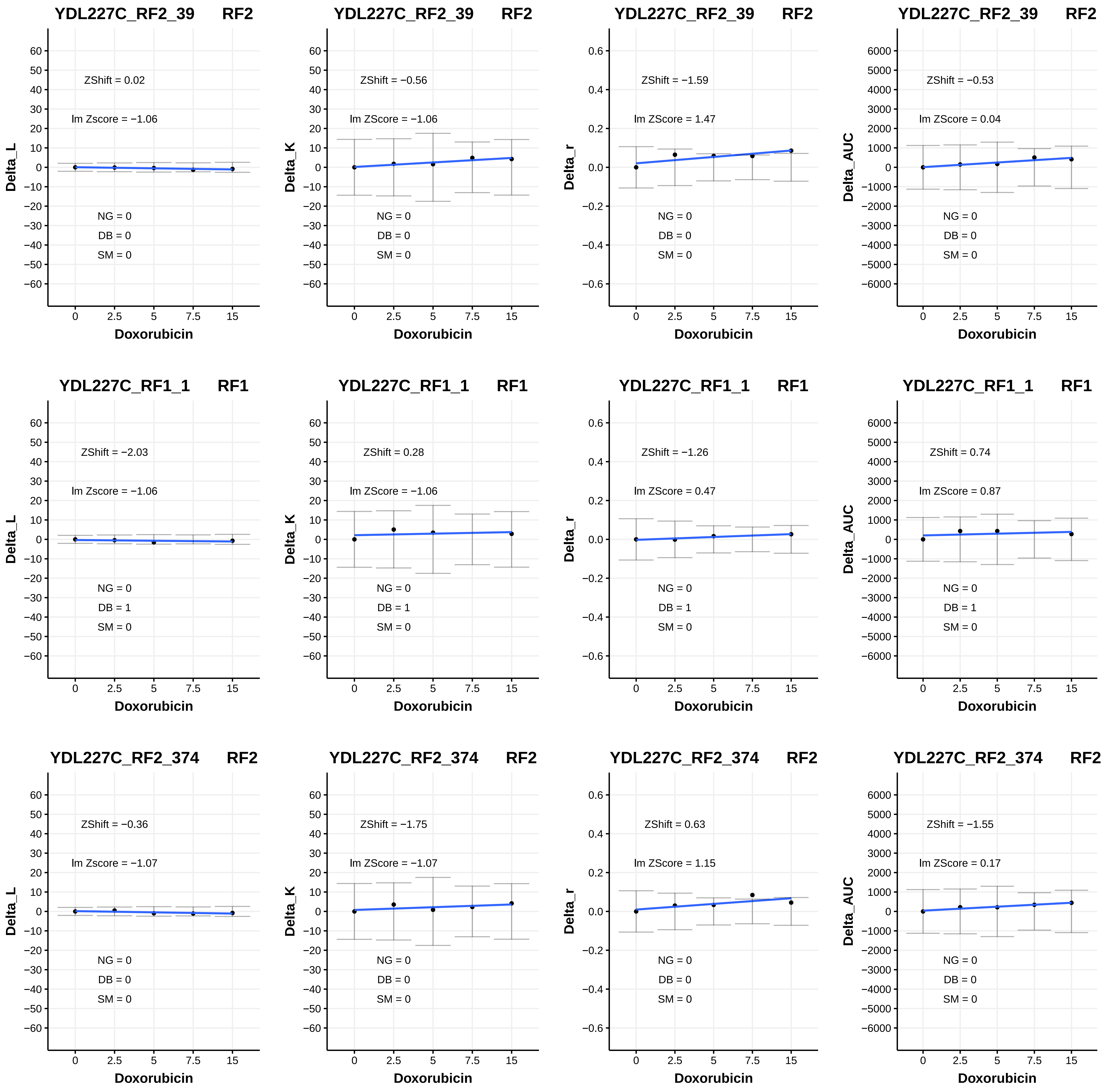
YDL227C_RF2_39 RF2
ZShift = 0.02
lm Zscore = −1.06
NG = 0
DB = 0
SM = 0
Delta_L
Doxorubicin
YDL227C_RF2_39 RF2
ZShift = −0.56
lm ZScore = −1.06
NG = 0
DB = 0
SM = 0
Delta_K
Doxorubicin
YDL227C_RF2_39 RF2
ZShift = −1.59
lm ZScore = 1.47
NG = 0
DB = 0
SM = 0
Delta_r
Doxorubicin
YDL227C_RF2_39 RF2
ZShift = −0.53
lm ZScore = 0.04
NG = 0
DB = 0
SM = 0
Delta_AUC
Doxorubicin
YDL227C_RF1_1 RF1
ZShift = −2.03
lm Zscore = −1.06
NG = 0
DB = 1
SM = 0
Delta_L
Doxorubicin
YDL227C_RF1_1 RF1
ZShift = 0.28
lm ZScore = −1.06
NG = 0
DB = 1
SM = 0
Delta_K
Doxorubicin
YDL227C_RF1_1 RF1
ZShift = −1.26
lm ZScore = 0.47
NG = 0
DB = 1
SM = 0
Delta_r
Doxorubicin
YDL227C_RF1_1 RF1
ZShift = 0.74
lm ZScore = 0.87
NG = 0
DB = 1
SM = 0
Delta_AUC
Doxorubicin
YDL227C_RF2_374 RF2
ZShift = −0.36
lm Zscore = −1.07
NG = 0
DB = 0
SM = 0
Delta_L
Doxorubicin
YDL227C_RF2_374 RF2
ZShift = −1.75
lm ZScore = −1.07
NG = 0
DB = 0
SM = 0
Delta_K
Doxorubicin
YDL227C_RF2_374 RF2
ZShift = 0.63
lm ZScore = 1.15
NG = 0
DB = 0
SM = 0
Delta_r
Doxorubicin
YDL227C_RF2_374 RF2
ZShift = −1.55
lm ZScore = 0.17
NG = 0
DB = 0
SM = 0
Delta_AUC
Doxorubicin

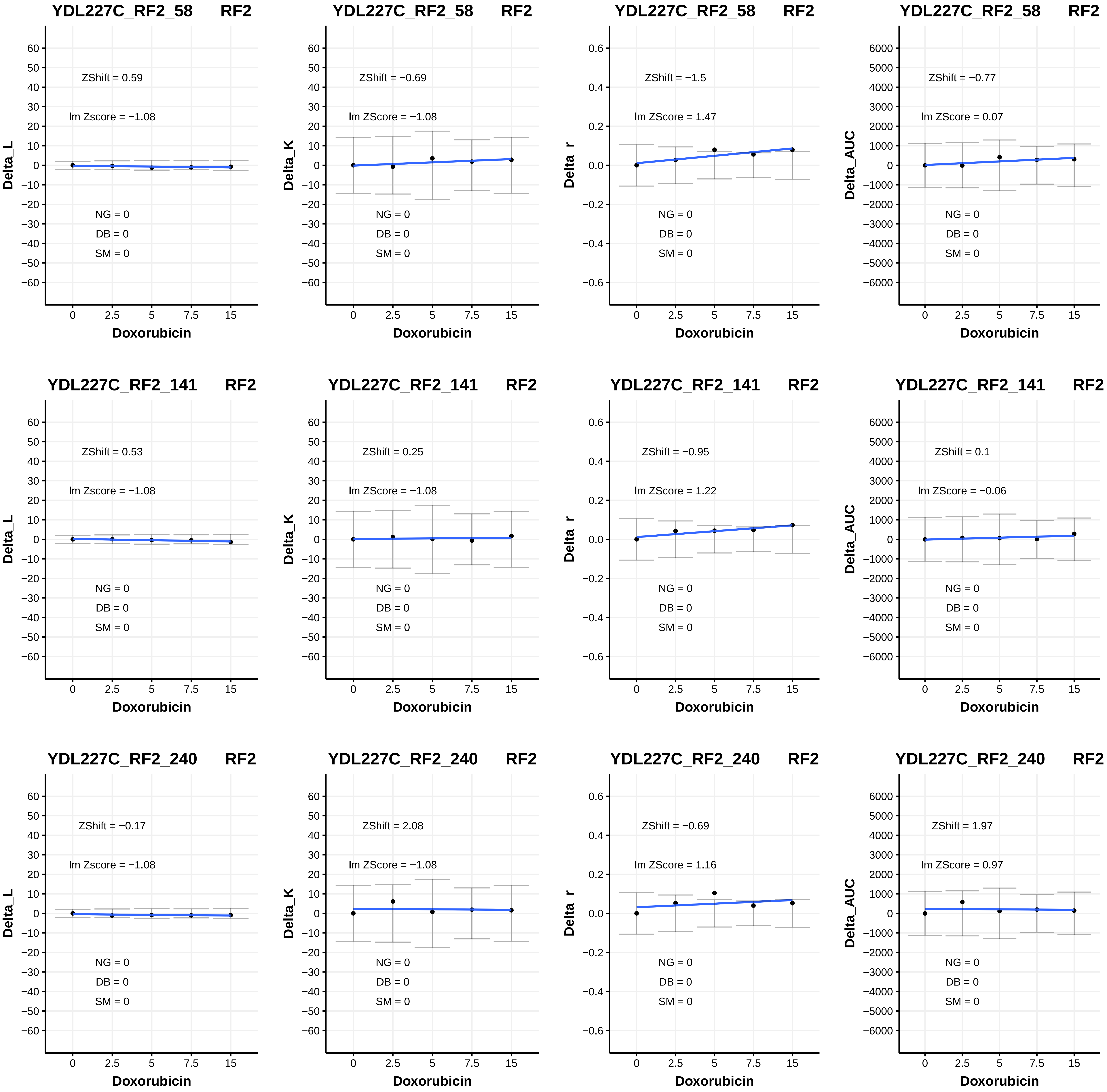
YDL227C_RF2_58 RF2
ZShift = 0.59
lm Zscore = −1.08
NG = 0
DB = 0
SM = 0
Delta_L
Doxorubicin
YDL227C_RF2_58 RF2
ZShift = −0.69
lm ZScore = −1.08
NG = 0
DB = 0
SM = 0
Delta_K
Doxorubicin
YDL227C_RF2_58 RF2
ZShift = −1.5
lm ZScore = 1.47
NG = 0
DB = 0
SM = 0
Delta_r
Doxorubicin
YDL227C_RF2_58 RF2
ZShift = −0.77
lm ZScore = 0.07
NG = 0
DB = 0
SM = 0
Delta_AUC
Doxorubicin
YDL227C_RF2_141 RF2
ZShift = 0.53
lm Zscore = −1.08
NG = 0
DB = 0
SM = 0
Delta_L
Doxorubicin
YDL227C_RF2_141 RF2
ZShift = 0.25
lm ZScore = −1.08
NG = 0
DB = 0
SM = 0
Delta_K
Doxorubicin
YDL227C_RF2_141 RF2
ZShift = −0.95
lm ZScore = 1.22
NG = 0
DB = 0
SM = 0
Delta_r
Doxorubicin
YDL227C_RF2_141 RF2
ZShift = 0.1
lm ZScore = −0.06
NG = 0
DB = 0
SM = 0
Delta_AUC
Doxorubicin
YDL227C_RF2_240 RF2
ZShift = −0.17
lm Zscore = −1.08
NG = 0
DB = 0
SM = 0
Delta_L
Doxorubicin
YDL227C_RF2_240 RF2
ZShift = 2.08
lm ZScore = −1.08
NG = 0
DB = 0
SM = 0
Delta_K
Doxorubicin
YDL227C_RF2_240 RF2
ZShift = −0.69
lm ZScore = 1.16
NG = 0
DB = 0
SM = 0
Delta_r
Doxorubicin
YDL227C_RF2_240 RF2
ZShift = 1.97
lm ZScore = 0.97
NG = 0
DB = 0
SM = 0
Delta_AUC
Doxorubicin

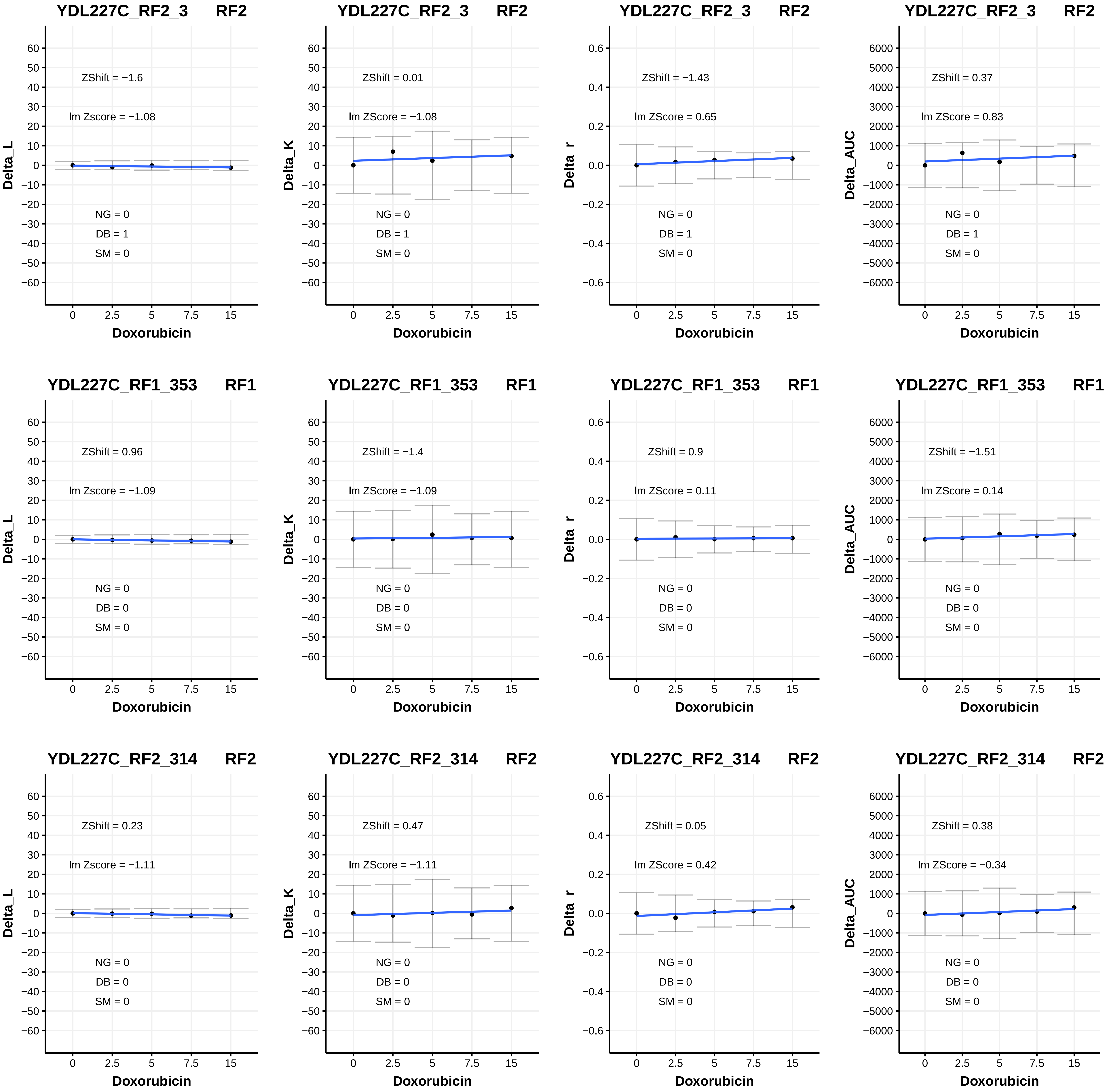

YDL227C_RF2_3 RF2
ZShift = −1.6
lm Zscore = −1.08
NG = 0
DB = 1
SM = 0
Delta_L
Doxorubicin
YDL227C_RF2_3 RF2
ZShift = 0.01
lm ZScore = −1.08
NG = 0
DB = 1
SM = 0
Delta_K
Doxorubicin
YDL227C_RF2_3 RF2
ZShift = −1.43
lm ZScore = 0.65
NG = 0
DB = 1
SM = 0
Delta_r
Doxorubicin
YDL227C_RF2_3 RF2
ZShift = 0.37
lm ZScore = 0.83
NG = 0
DB = 1
SM = 0
Delta_AUC
Doxorubicin
YDL227C_RF1_353 RF1
ZShift = 0.96
lm Zscore = −1.09
NG = 0
DB = 0
SM = 0
Delta_L
Doxorubicin
YDL227C_RF1_353 RF1
ZShift = −1.4
lm ZScore = −1.09
NG = 0
DB = 0
SM = 0
Delta_K
Doxorubicin
YDL227C_RF1_353 RF1
ZShift = 0.9
lm ZScore = 0.11
NG = 0
DB = 0
SM = 0
Delta_r
Doxorubicin
YDL227C_RF1_353 RF1
ZShift = −1.51
lm ZScore = 0.14
NG = 0
DB = 0
SM = 0
Delta_AUC
Doxorubicin
YDL227C_RF2_314 RF2
ZShift = 0.23
lm Zscore = −1.11
NG = 0
DB = 0
SM = 0
Delta_L
Doxorubicin
YDL227C_RF2_314 RF2
ZShift = 0.47
lm ZScore = −1.11
NG = 0
DB = 0
SM = 0
Delta_K
Doxorubicin
YDL227C_RF2_314 RF2
ZShift = 0.05
lm ZScore = 0.42
NG = 0
DB = 0
SM = 0
Delta_r
Doxorubicin
YDL227C_RF2_314 RF2
ZShift = 0.38
lm ZScore = −0.34
NG = 0
DB = 0
SM = 0
Delta_AUC
Doxorubicin

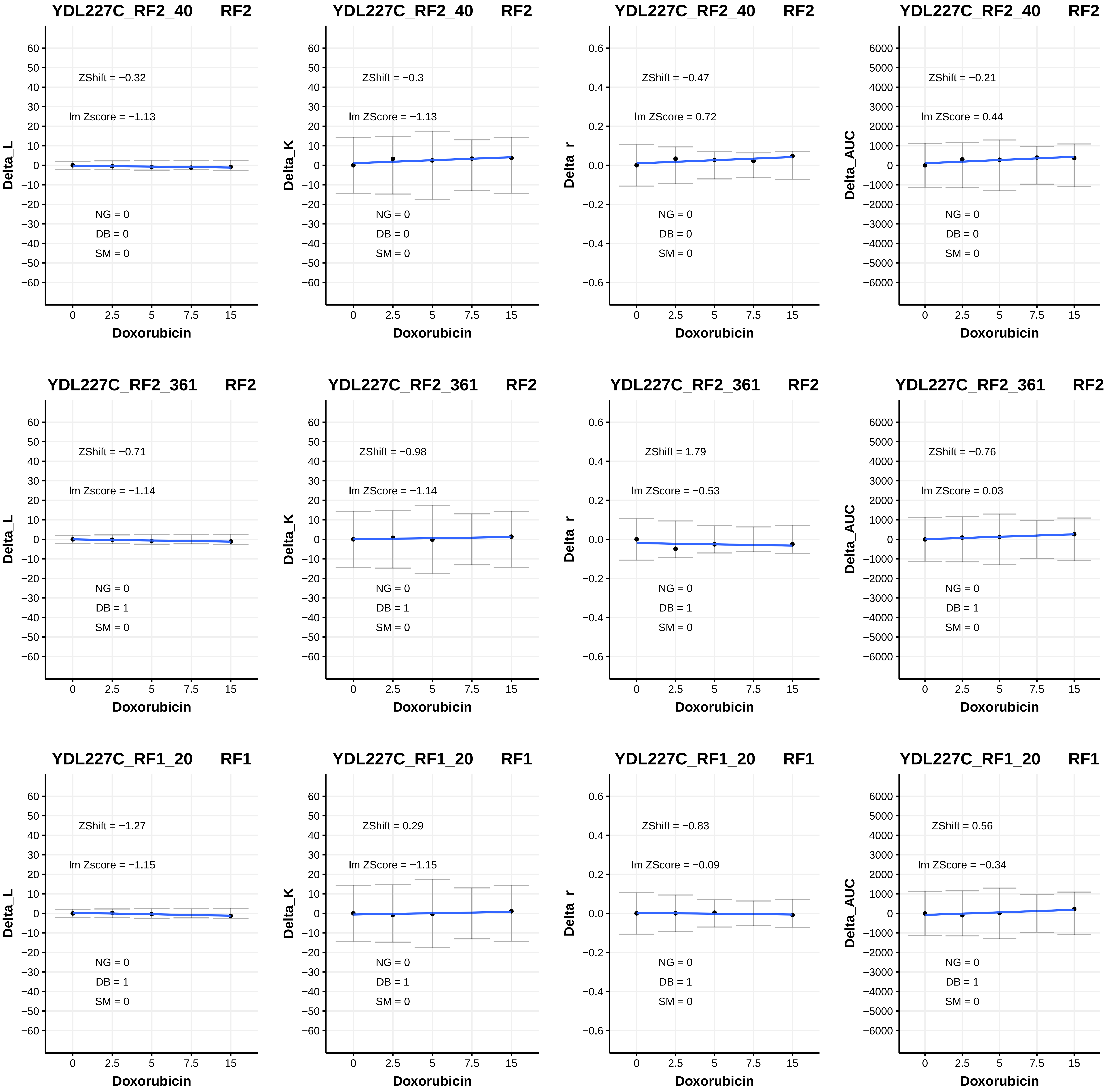

YDL227C_RF2_40 RF2
ZShift = −0.32
lm Zscore = −1.13
NG = 0
DB = 0
SM = 0
Delta_L
Doxorubicin
YDL227C_RF2_40 RF2
ZShift = −0.3
lm ZScore = −1.13
NG = 0
DB = 0
SM = 0
Delta_K
Doxorubicin
YDL227C_RF2_40 RF2
ZShift = −0.47
lm ZScore = 0.72
NG = 0
DB = 0
SM = 0
Delta_r
Doxorubicin
YDL227C_RF2_40 RF2
ZShift = −0.21
lm ZScore = 0.44
NG = 0
DB = 0
SM = 0
Delta_AUC
Doxorubicin
YDL227C_RF2_361 RF2
ZShift = −0.71
lm Zscore = −1.14
NG = 0
DB = 1
SM = 0
Delta_L
Doxorubicin
YDL227C_RF2_361 RF2
ZShift = −0.98
lm ZScore = −1.14
NG = 0
DB = 1
SM = 0
Delta_K
Doxorubicin
YDL227C_RF2_361 RF2
ZShift = 1.79
lm ZScore = −0.53
NG = 0
DB = 1
SM = 0
Delta_r
Doxorubicin
YDL227C_RF2_361 RF2
ZShift = −0.76
lm ZScore = 0.03
NG = 0
DB = 1
SM = 0
Delta_AUC
Doxorubicin
YDL227C_RF1_20 RF1
ZShift = −1.27
lm Zscore = −1.15
NG = 0
DB = 1
SM = 0
Delta_L
Doxorubicin
YDL227C_RF1_20 RF1
ZShift = 0.29
lm ZScore = −1.15
NG = 0
DB = 1
SM = 0
Delta_K
Doxorubicin
YDL227C_RF1_20 RF1
ZShift = −0.83
lm ZScore = −0.09
NG = 0
DB = 1
SM = 0
Delta_r
Doxorubicin
YDL227C_RF1_20 RF1
ZShift = 0.56
lm ZScore = −0.34
NG = 0
DB = 1
SM = 0
Delta_AUC
Doxorubicin

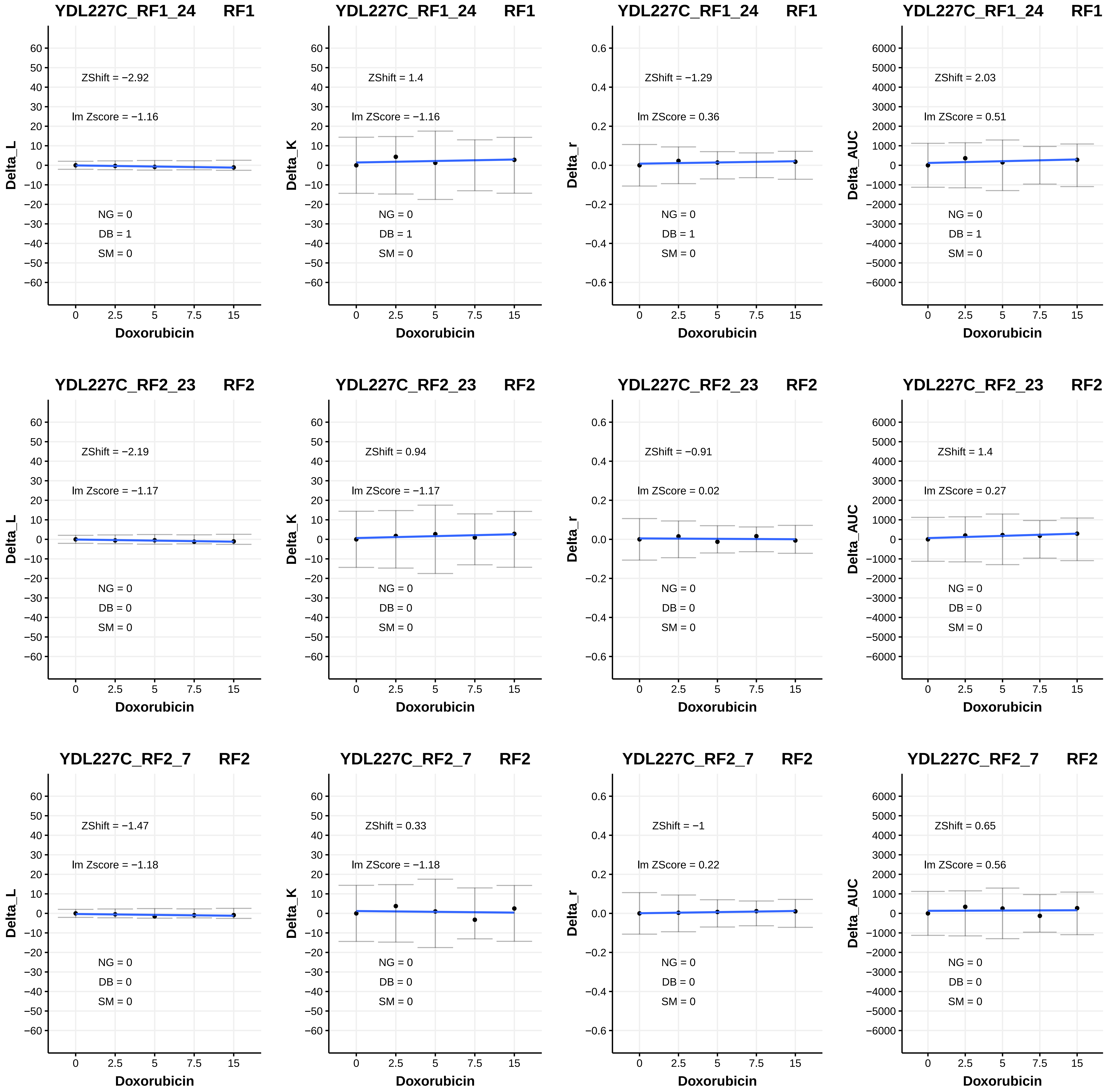

YDL227C_RF1_24 RF1
ZShift = −2.92
lm Zscore = −1.16
NG = 0
DB = 1
SM = 0
Delta_L
Doxorubicin
YDL227C_RF1_24 RF1
ZShift = 1.4
lm ZScore = −1.16
NG = 0
DB = 1
SM = 0
Delta_K
Doxorubicin
YDL227C_RF1_24 RF1
ZShift = −1.29
lm ZScore = 0.36
NG = 0
DB = 1
SM = 0
Delta_r
Doxorubicin
YDL227C_RF1_24 RF1
ZShift = 2.03
lm ZScore = 0.51
NG = 0
DB = 1
SM = 0
Delta_AUC
Doxorubicin
YDL227C_RF2_23 RF2
ZShift = −2.19
lm Zscore = −1.17
NG = 0
DB = 0
SM = 0
Delta_L
Doxorubicin
YDL227C_RF2_23 RF2
ZShift = 0.94
lm ZScore = −1.17
NG = 0
DB = 0
SM = 0
Delta_K
Doxorubicin
YDL227C_RF2_23 RF2
ZShift = −0.91
lm ZScore = 0.02
NG = 0
DB = 0
SM = 0
Delta_r
Doxorubicin
YDL227C_RF2_23 RF2
ZShift = 1.4
lm ZScore = 0.27
NG = 0
DB = 0
SM = 0
Delta_AUC
Doxorubicin
YDL227C_RF2_7 RF2
ZShift = −1.47
lm Zscore = −1.18
NG = 0
DB = 0
SM = 0
Delta_L
Doxorubicin
YDL227C_RF2_7 RF2
ZShift = 0.33
lm ZScore = −1.18
NG = 0
DB = 0
SM = 0
Delta_K
Doxorubicin
YDL227C_RF2_7 RF2
ZShift = −1
lm ZScore = 0.22
NG = 0
DB = 0
SM = 0
Delta_r
Doxorubicin
YDL227C_RF2_7 RF2
ZShift = 0.65
lm ZScore = 0.56
NG = 0
DB = 0
SM = 0
Delta_AUC
Doxorubicin

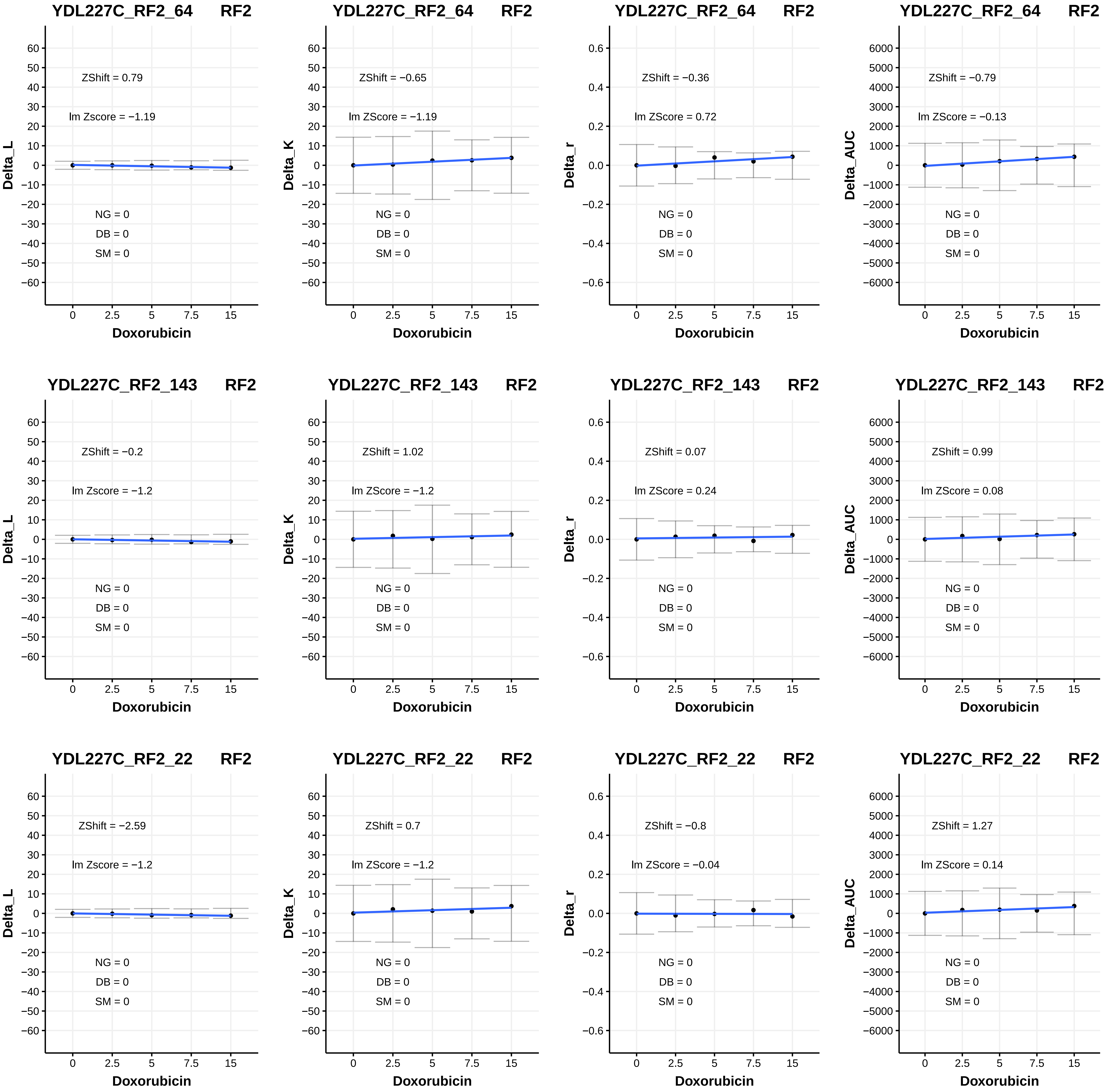

YDL227C_RF2_64 RF2
ZShift = 0.79
lm Zscore = −1.19
NG = 0
DB = 0
SM = 0
Delta_L
Doxorubicin
YDL227C_RF2_64 RF2
ZShift = −0.65
lm ZScore = −1.19
NG = 0
DB = 0
SM = 0
Delta_K
Doxorubicin
YDL227C_RF2_64 RF2
ZShift = −0.36
lm ZScore = 0.72
NG = 0
DB = 0
SM = 0
Delta_r
Doxorubicin
YDL227C_RF2_64 RF2
ZShift = −0.79
lm ZScore = −0.13
NG = 0
DB = 0
SM = 0
Delta_AUC
Doxorubicin
YDL227C_RF2_143 RF2
ZShift = −0.2
lm Zscore = −1.2
NG = 0
DB = 0
SM = 0
Delta_L
Doxorubicin
YDL227C_RF2_143 RF2
ZShift = 1.02
lm ZScore = −1.2
NG = 0
DB = 0
SM = 0
Delta_K
Doxorubicin
YDL227C_RF2_143 RF2
ZShift = 0.07
lm ZScore = 0.24
NG = 0
DB = 0
SM = 0
Delta_r
Doxorubicin
YDL227C_RF2_143 RF2
ZShift = 0.99
lm ZScore = 0.08
NG = 0
DB = 0
SM = 0
Delta_AUC
Doxorubicin
YDL227C_RF2_22 RF2
ZShift = −2.59
lm Zscore = −1.2
NG = 0
DB = 0
SM = 0
Delta_L
Doxorubicin
YDL227C_RF2_22 RF2
ZShift = 0.7
lm ZScore = −1.2
NG = 0
DB = 0
SM = 0
Delta_K
Doxorubicin
YDL227C_RF2_22 RF2
ZShift = −0.8
lm ZScore = −0.04
NG = 0
DB = 0
SM = 0
Delta_r
Doxorubicin
YDL227C_RF2_22 RF2
ZShift = 1.27
lm ZScore = 0.14
NG = 0
DB = 0
SM = 0
Delta_AUC
Doxorubicin

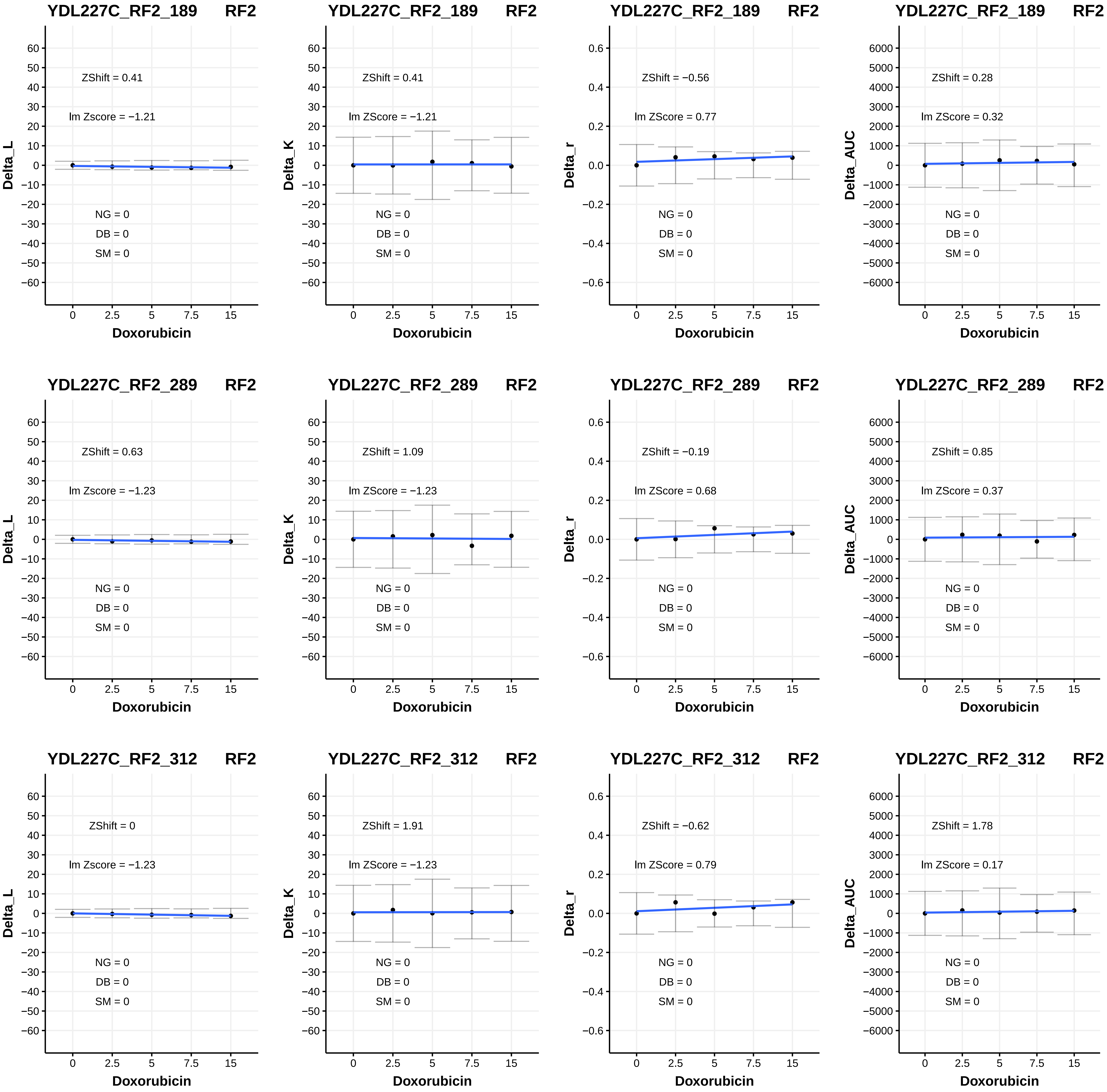
YDL227C_RF2_189 RF2
ZShift = 0.41
lm Zscore = −1.21
NG = 0
DB = 0
SM = 0
Delta_L
Doxorubicin
YDL227C_RF2_189 RF2
ZShift = 0.41
lm ZScore = −1.21
NG = 0
DB = 0
SM = 0
Delta_K
Doxorubicin
YDL227C_RF2_189 RF2
ZShift = −0.56
lm ZScore = 0.77
NG = 0
DB = 0
SM = 0
Delta_r
Doxorubicin
YDL227C_RF2_189 RF2
ZShift = 0.28
lm ZScore = 0.32
NG = 0
DB = 0
SM = 0
Delta_AUC
Doxorubicin
YDL227C_RF2_289 RF2
ZShift = 0.63
lm Zscore = −1.23
NG = 0
DB = 0
SM = 0
Delta_L
Doxorubicin
YDL227C_RF2_289 RF2
ZShift = 1.09
lm ZScore = −1.23
NG = 0
DB = 0
SM = 0
Delta_K
Doxorubicin
YDL227C_RF2_289 RF2
ZShift = −0.19
lm ZScore = 0.68
NG = 0
DB = 0
SM = 0
Delta_r
Doxorubicin
YDL227C_RF2_289 RF2
ZShift = 0.85
lm ZScore = 0.37
NG = 0
DB = 0
SM = 0
Delta_AUC
Doxorubicin
YDL227C_RF2_312 RF2
ZShift = 0
lm Zscore = −1.23
NG = 0
DB = 0
SM = 0
Delta_L
Doxorubicin
YDL227C_RF2_312 RF2
ZShift = 1.91
lm ZScore = −1.23
NG = 0
DB = 0
SM = 0
Delta_K
Doxorubicin
YDL227C_RF2_312 RF2
ZShift = −0.62
lm ZScore = 0.79
NG = 0
DB = 0
SM = 0
Delta_r
Doxorubicin
YDL227C_RF2_312 RF2
ZShift = 1.78
lm ZScore = 0.17
NG = 0
DB = 0
SM = 0
Delta_AUC
Doxorubicin

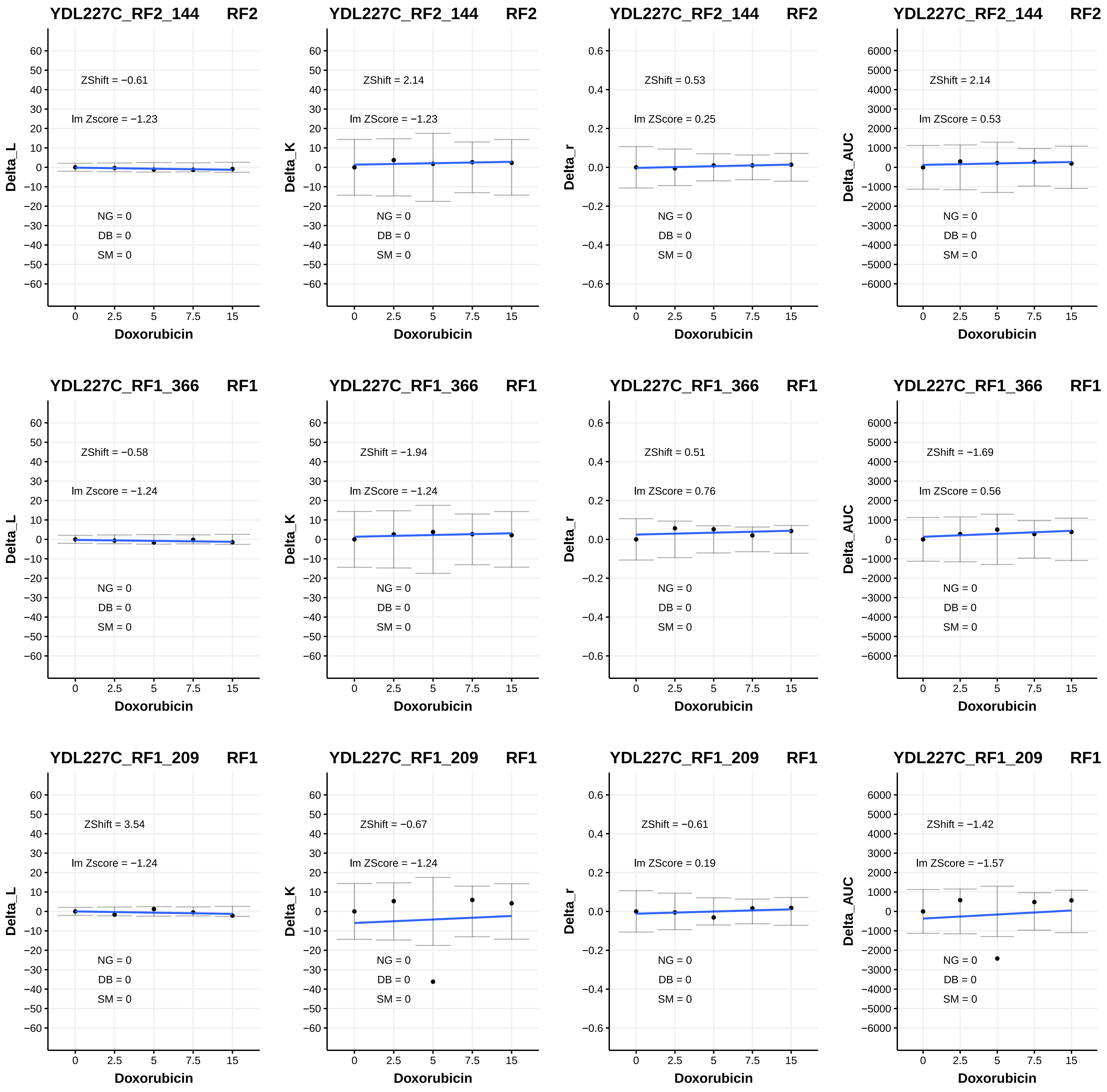

YDL227C_RF2_144 RF2
ZShift = −0.61
lm Zscore = −1.23
NG = 0
DB = 0
SM = 0
Delta_L
Doxorubicin
YDL227C_RF2_144 RF2
ZShift = 2.14
lm ZScore = −1.23
NG = 0
DB = 0
SM = 0
Delta_K
Doxorubicin
YDL227C_RF2_144 RF2
ZShift = 0.53
lm ZScore = 0.25
NG = 0
DB = 0
SM = 0
Delta_r
Doxorubicin
YDL227C_RF2_144 RF2
ZShift = 2.14
lm ZScore = 0.53
NG = 0
DB = 0
SM = 0
Delta_AUC
Doxorubicin
YDL227C_RF1_366 RF1
ZShift = −0.58
lm Zscore = −1.24
NG = 0
DB = 0
SM = 0
Delta_L
Doxorubicin
YDL227C_RF1_366 RF1
ZShift = −1.94
lm ZScore = −1.24
NG = 0
DB = 0
SM = 0
Delta_K
Doxorubicin
YDL227C_RF1_366 RF1
ZShift = 0.51
lm ZScore = 0.76
NG = 0
DB = 0
SM = 0
Delta_r
Doxorubicin
YDL227C_RF1_366 RF1
ZShift = −1.69
lm ZScore = 0.56
NG = 0
DB = 0
SM = 0
Delta_AUC
Doxorubicin
YDL227C_RF1_209 RF1
ZShift = 3.54
lm Zscore = −1.24
NG = 0
DB = 0
SM = 0
Delta_L
Doxorubicin
YDL227C_RF1_209 RF1
ZShift = −0.67
lm ZScore = −1.24
NG = 0
DB = 0
SM = 0
Delta_K
Doxorubicin
YDL227C_RF1_209 RF1
ZShift = −0.61
lm ZScore = 0.19
NG = 0
DB = 0
SM = 0
Delta_r
Doxorubicin
YDL227C_RF1_209 RF1
ZShift = −1.42
lm ZScore = −1.57
NG = 0
DB = 0
SM = 0
Delta_AUC
Doxorubicin

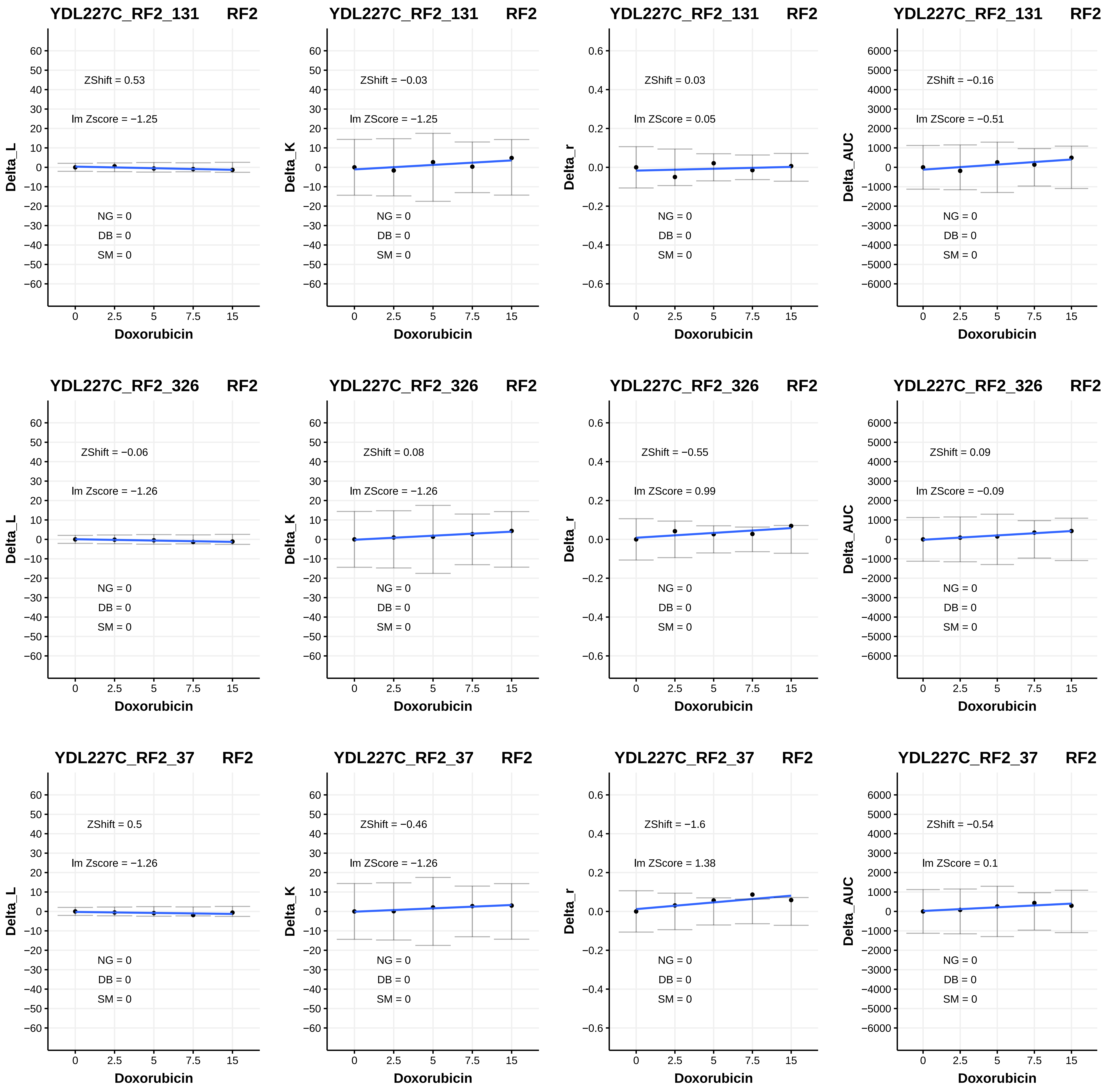
YDL227C_RF2_131 RF2
ZShift = 0.53
lm Zscore = −1.25
NG = 0
DB = 0
SM = 0
Delta_L
Doxorubicin
YDL227C_RF2_131 RF2
ZShift = −0.03
lm ZScore = −1.25
NG = 0
DB = 0
SM = 0
Delta_K
Doxorubicin
YDL227C_RF2_131 RF2
ZShift = 0.03
lm ZScore = 0.05
NG = 0
DB = 0
SM = 0
Delta_r
Doxorubicin
YDL227C_RF2_131 RF2
ZShift = −0.16
lm ZScore = −0.51
NG = 0
DB = 0
SM = 0
Delta_AUC
Doxorubicin
YDL227C_RF2_326 RF2
ZShift = −0.06
lm Zscore = −1.26
NG = 0
DB = 0
SM = 0
Delta_L
Doxorubicin
YDL227C_RF2_326 RF2
ZShift = 0.08
lm ZScore = −1.26
NG = 0
DB = 0
SM = 0
Delta_K
Doxorubicin
YDL227C_RF2_326 RF2
ZShift = −0.55
lm ZScore = 0.99
NG = 0
DB = 0
SM = 0
Delta_r
Doxorubicin
YDL227C_RF2_326 RF2
ZShift = 0.09
lm ZScore = −0.09
NG = 0
DB = 0
SM = 0
Delta_AUC
Doxorubicin
YDL227C_RF2_37 RF2
ZShift = 0.5
lm Zscore = −1.26
NG = 0
DB = 0
SM = 0
Delta_L
Doxorubicin
YDL227C_RF2_37 RF2
ZShift = −0.46
lm ZScore = −1.26
NG = 0
DB = 0
SM = 0
Delta_K
Doxorubicin
YDL227C_RF2_37 RF2
ZShift = −1.6
lm ZScore = 1.38
NG = 0
DB = 0
SM = 0
Delta_r
Doxorubicin
YDL227C_RF2_37 RF2
ZShift = −0.54
lm ZScore = 0.1
NG = 0
DB = 0
SM = 0
Delta_AUC
Doxorubicin

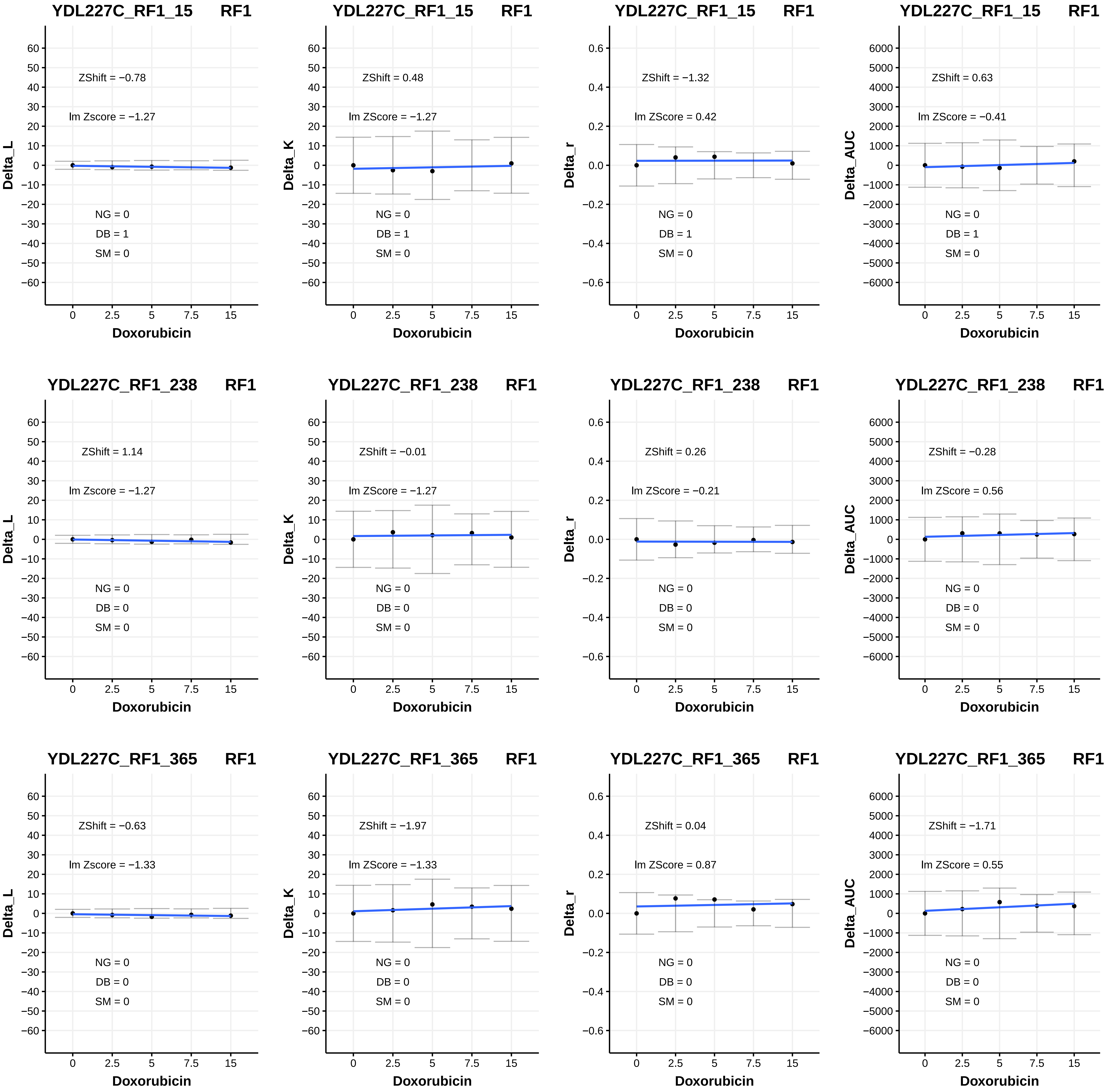

YDL227C_RF1_15 RF1
ZShift = −0.78
lm Zscore = −1.27
NG = 0
DB = 1
SM = 0
Delta_L
Doxorubicin
YDL227C_RF1_15 RF1
ZShift = 0.48
lm ZScore = −1.27
NG = 0
DB = 1
SM = 0
Delta_K
Doxorubicin
YDL227C_RF1_15 RF1
ZShift = −1.32
lm ZScore = 0.42
NG = 0
DB = 1
SM = 0
Delta_r
Doxorubicin
YDL227C_RF1_15 RF1
ZShift = 0.63
lm ZScore = −0.41
NG = 0
DB = 1
SM = 0
Delta_AUC
Doxorubicin
YDL227C_RF1_238 RF1
ZShift = 1.14
lm Zscore = −1.27
NG = 0
DB = 0
SM = 0
Delta_L
Doxorubicin
YDL227C_RF1_238 RF1
ZShift = −0.01
lm ZScore = −1.27
NG = 0
DB = 0
SM = 0
Delta_K
Doxorubicin
YDL227C_RF1_238 RF1
ZShift = 0.26
lm ZScore = −0.21
NG = 0
DB = 0
SM = 0
Delta_r
Doxorubicin
YDL227C_RF1_238 RF1
ZShift = −0.28
lm ZScore = 0.56
NG = 0
DB = 0
SM = 0
Delta_AUC
Doxorubicin
YDL227C_RF1_365 RF1
ZShift = −0.63
lm Zscore = −1.33
NG = 0
DB = 0
SM = 0
Delta_L
Doxorubicin
YDL227C_RF1_365 RF1
ZShift = −1.97
lm ZScore = −1.33
NG = 0
DB = 0
SM = 0
Delta_K
Doxorubicin
YDL227C_RF1_365 RF1
ZShift = 0.04
lm ZScore = 0.87
NG = 0
DB = 0
SM = 0
Delta_r
Doxorubicin
YDL227C_RF1_365 RF1
ZShift = −1.71
lm ZScore = 0.55
NG = 0
DB = 0
SM = 0
Delta_AUC
Doxorubicin

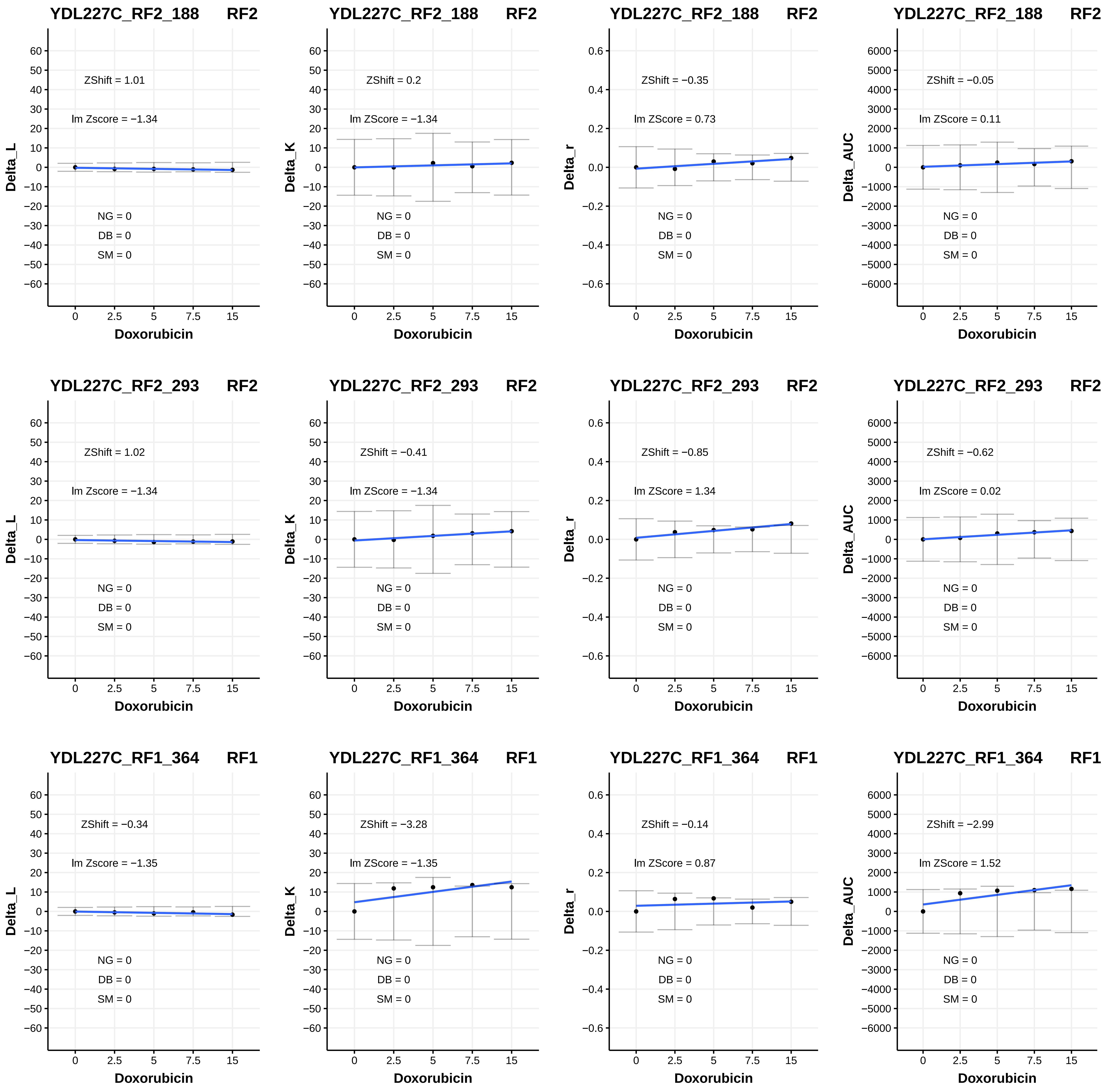
YDL227C_RF2_188 RF2
ZShift = 1.01
lm Zscore = −1.34
NG = 0
DB = 0
SM = 0
Delta_L
Doxorubicin
YDL227C_RF2_188 RF2
ZShift = 0.2
lm ZScore = −1.34
NG = 0
DB = 0
SM = 0
Delta_K
Doxorubicin
YDL227C_RF2_188 RF2
ZShift = −0.35
lm ZScore = 0.73
NG = 0
DB = 0
SM = 0
Delta_r
Doxorubicin
YDL227C_RF2_188 RF2
ZShift = −0.05
lm ZScore = 0.11
NG = 0
DB = 0
SM = 0
Delta_AUC
Doxorubicin
YDL227C_RF2_293 RF2
ZShift = 1.02
lm Zscore = −1.34
NG = 0
DB = 0
SM = 0
Delta_L
Doxorubicin
YDL227C_RF2_293 RF2
ZShift = −0.41
lm ZScore = −1.34
NG = 0
DB = 0
SM = 0
Delta_K
Doxorubicin
YDL227C_RF2_293 RF2
ZShift = −0.85
lm ZScore = 1.34
NG = 0
DB = 0
SM = 0
Delta_r
Doxorubicin
YDL227C_RF2_293 RF2
ZShift = −0.62
lm ZScore = 0.02
NG = 0
DB = 0
SM = 0
Delta_AUC
Doxorubicin
YDL227C_RF1_364 RF1
ZShift = −0.34
lm Zscore = −1.35
NG = 0
DB = 0
SM = 0
Delta_L
Doxorubicin
YDL227C_RF1_364 RF1
ZShift = −3.28
lm ZScore = −1.35
NG = 0
DB = 0
SM = 0
Delta_K
Doxorubicin
YDL227C_RF1_364 RF1
ZShift = −0.14
lm ZScore = 0.87
NG = 0
DB = 0
SM = 0
Delta_r
Doxorubicin
YDL227C_RF1_364 RF1
ZShift = −2.99
lm ZScore = 1.52
NG = 0
DB = 0
SM = 0
Delta_AUC
Doxorubicin

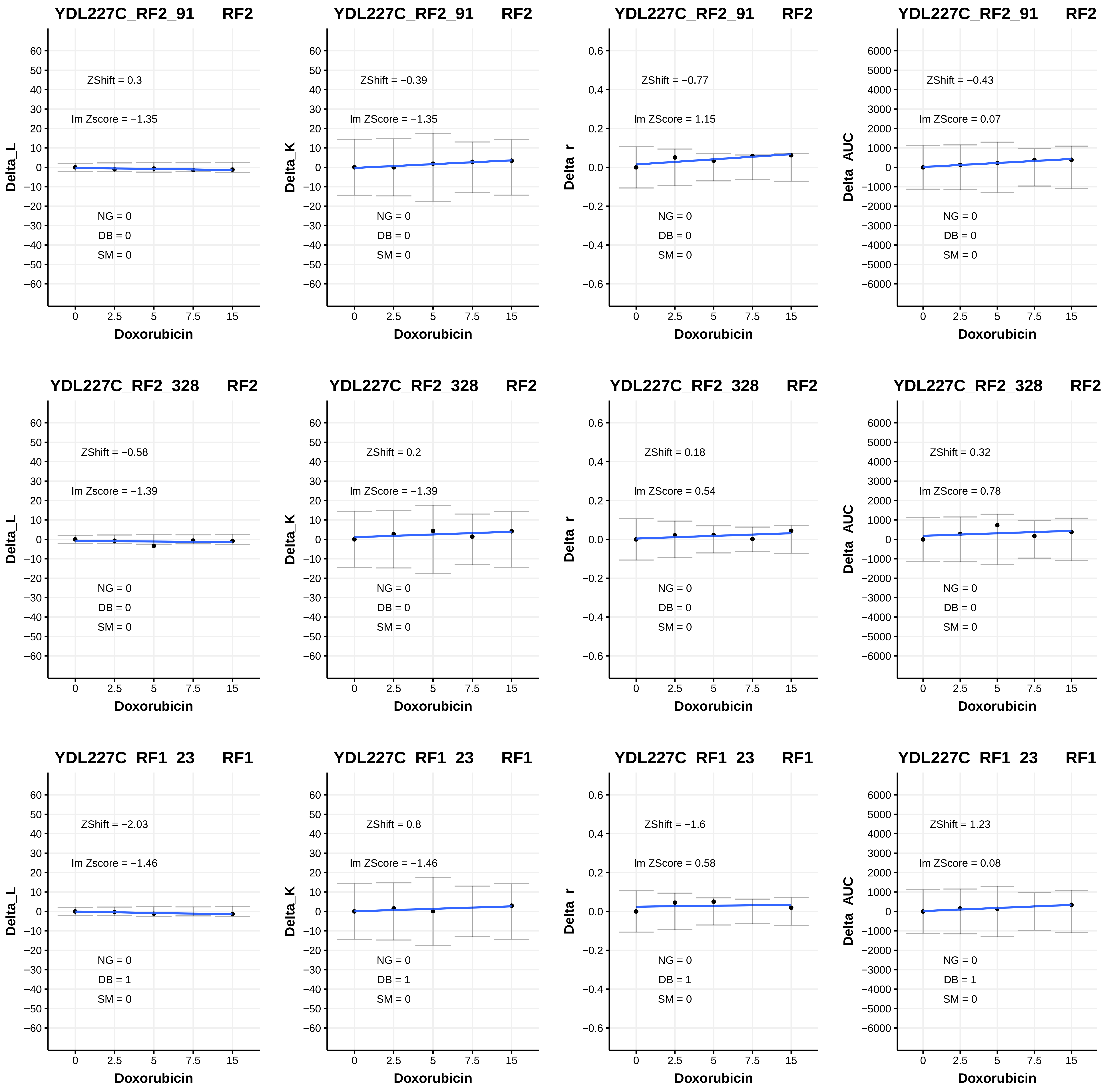

YDL227C_RF2_91 RF2
ZShift = 0.3
lm Zscore = −1.35
NG = 0
DB = 0
SM = 0
Delta_L
Doxorubicin
YDL227C_RF2_91 RF2
ZShift = −0.39
lm ZScore = −1.35
NG = 0
DB = 0
SM = 0
Delta_K
Doxorubicin
YDL227C_RF2_91 RF2
ZShift = −0.77
lm ZScore = 1.15
NG = 0
DB = 0
SM = 0
Delta_r
Doxorubicin
YDL227C_RF2_91 RF2
ZShift = −0.43
lm ZScore = 0.07
NG = 0
DB = 0
SM = 0
Delta_AUC
Doxorubicin
YDL227C_RF2_328 RF2
ZShift = −0.58
lm Zscore = −1.39
NG = 0
DB = 0
SM = 0
Delta_L
Doxorubicin
YDL227C_RF2_328 RF2
ZShift = 0.2
lm ZScore = −1.39
NG = 0
DB = 0
SM = 0
Delta_K
Doxorubicin
YDL227C_RF2_328 RF2
ZShift = 0.18
lm ZScore = 0.54
NG = 0
DB = 0
SM = 0
Delta_r
Doxorubicin
YDL227C_RF2_328 RF2
ZShift = 0.32
lm ZScore = 0.78
NG = 0
DB = 0
SM = 0
Delta_AUC
Doxorubicin
YDL227C_RF1_23 RF1
ZShift = −2.03
lm Zscore = −1.46
NG = 0
DB = 1
SM = 0
Delta_L
Doxorubicin
YDL227C_RF1_23 RF1
ZShift = 0.8
lm ZScore = −1.46
NG = 0
DB = 1
SM = 0
Delta_K
Doxorubicin
YDL227C_RF1_23 RF1
ZShift = −1.6
lm ZScore = 0.58
NG = 0
DB = 1
SM = 0
Delta_r
Doxorubicin
YDL227C_RF1_23 RF1
ZShift = 1.23
lm ZScore = 0.08
NG = 0
DB = 1
SM = 0
Delta_AUC
Doxorubicin

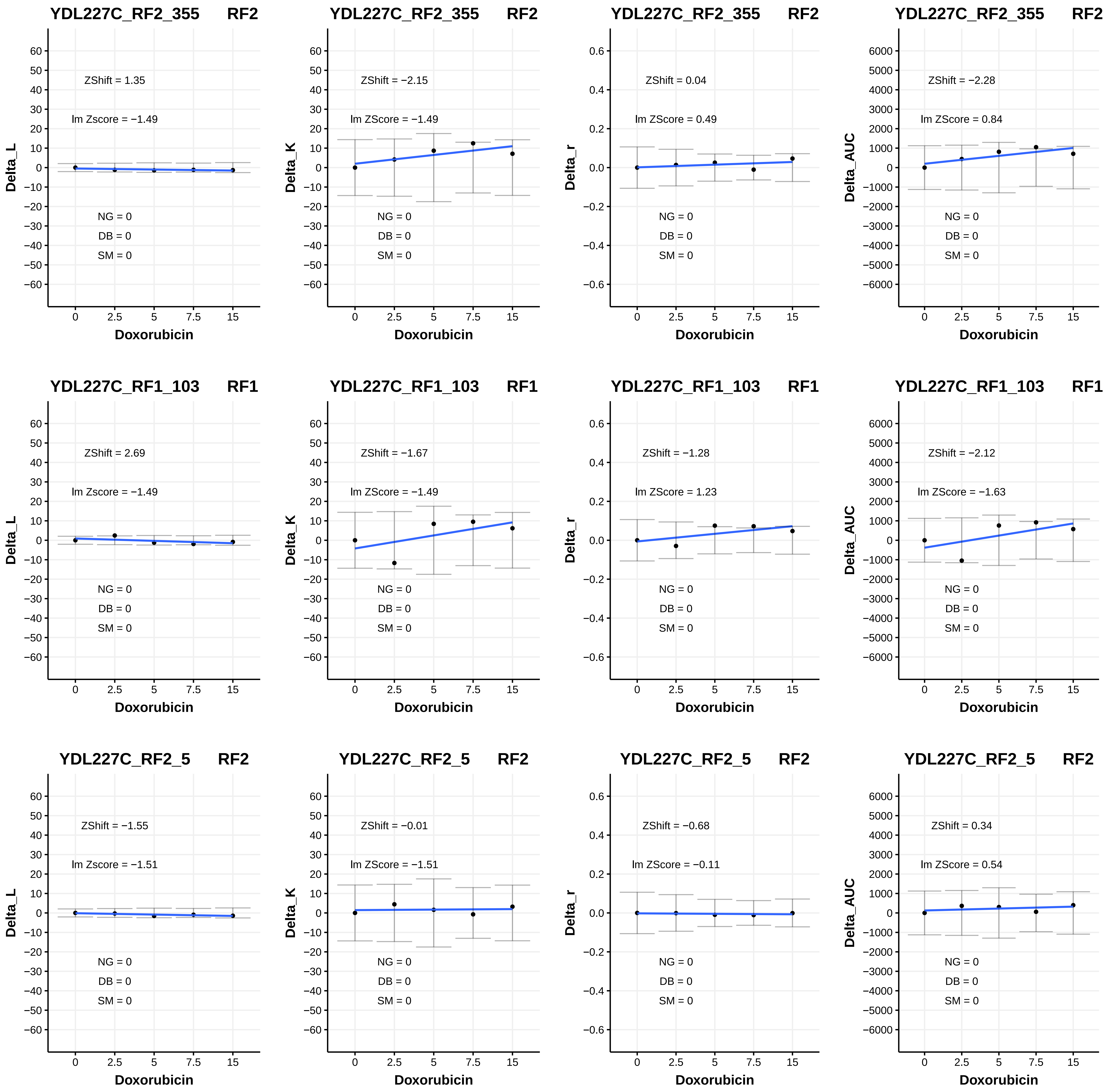
YDL227C_RF2_355 RF2
ZShift = 1.35
lm Zscore = −1.49
NG = 0
DB = 0
SM = 0
Delta_L
Doxorubicin
YDL227C_RF2_355 RF2
ZShift = −2.15
lm ZScore = −1.49
NG = 0
DB = 0
SM = 0
Delta_K
Doxorubicin
YDL227C_RF2_355 RF2
ZShift = 0.04
lm ZScore = 0.49
NG = 0
DB = 0
SM = 0
Delta_r
Doxorubicin
YDL227C_RF2_355 RF2
ZShift = −2.28
lm ZScore = 0.84
NG = 0
DB = 0
SM = 0
Delta_AUC
Doxorubicin
YDL227C_RF1_103 RF1
ZShift = 2.69
lm Zscore = −1.49
NG = 0
DB = 0
SM = 0
Delta_L
Doxorubicin
YDL227C_RF1_103 RF1
ZShift = −1.67
lm ZScore = −1.49
NG = 0
DB = 0
SM = 0
Delta_K
Doxorubicin
YDL227C_RF1_103 RF1
ZShift = −1.28
lm ZScore = 1.23
NG = 0
DB = 0
SM = 0
Delta_r
Doxorubicin
YDL227C_RF1_103 RF1
ZShift = −2.12
lm ZScore = −1.63
NG = 0
DB = 0
SM = 0
Delta_AUC
Doxorubicin
YDL227C_RF2_5 RF2
ZShift = −1.55
lm Zscore = −1.51
NG = 0
DB = 0
SM = 0
Delta_L
Doxorubicin
YDL227C_RF2_5 RF2
ZShift = −0.01
lm ZScore = −1.51
NG = 0
DB = 0
SM = 0
Delta_K
Doxorubicin
YDL227C_RF2_5 RF2
ZShift = −0.68
lm ZScore = −0.11
NG = 0
DB = 0
SM = 0
Delta_r
Doxorubicin
YDL227C_RF2_5 RF2
ZShift = 0.34
lm ZScore = 0.54
NG = 0
DB = 0
SM = 0
Delta_AUC
Doxorubicin

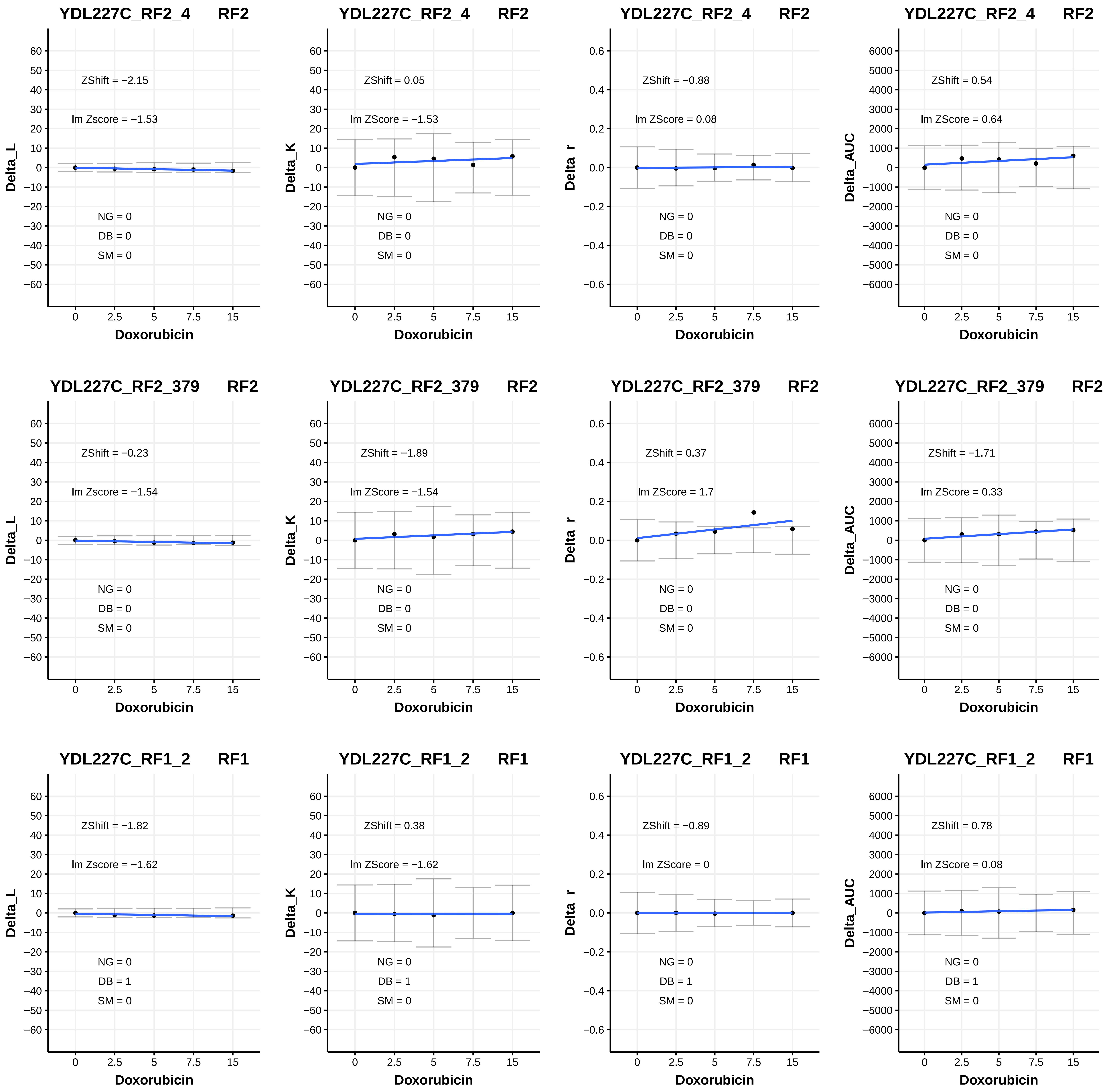

YDL227C_RF2_4 RF2
ZShift = −2.15
lm Zscore = −1.53
NG = 0
DB = 0
SM = 0
Delta_L
Doxorubicin
YDL227C_RF2_4 RF2
ZShift = 0.05
lm ZScore = −1.53
NG = 0
DB = 0
SM = 0
Delta_K
Doxorubicin
YDL227C_RF2_4 RF2
ZShift = −0.88
lm ZScore = 0.08
NG = 0
DB = 0
SM = 0
Delta_r
Doxorubicin
YDL227C_RF2_4 RF2
ZShift = 0.54
lm ZScore = 0.64
NG = 0
DB = 0
SM = 0
Delta_AUC
Doxorubicin
YDL227C_RF2_379 RF2
ZShift = −0.23
lm Zscore = −1.54
NG = 0
DB = 0
SM = 0
Delta_L
Doxorubicin
YDL227C_RF2_379 RF2
ZShift = −1.89
lm ZScore = −1.54
NG = 0
DB = 0
SM = 0
Delta_K
Doxorubicin
YDL227C_RF2_379 RF2
ZShift = 0.37
lm ZScore = 1.7
NG = 0
DB = 0
SM = 0
Delta_r
Doxorubicin
YDL227C_RF2_379 RF2
ZShift = −1.71
lm ZScore = 0.33
NG = 0
DB = 0
SM = 0
Delta_AUC
Doxorubicin
YDL227C_RF1_2 RF1
ZShift = −1.82
lm Zscore = −1.62
NG = 0
DB = 1
SM = 0
Delta_L
Doxorubicin
YDL227C_RF1_2 RF1
ZShift = 0.38
lm ZScore = −1.62
NG = 0
DB = 1
SM = 0
Delta_K
Doxorubicin
YDL227C_RF1_2 RF1
ZShift = −0.89
lm ZScore = 0
NG = 0
DB = 1
SM = 0
Delta_r
Doxorubicin
YDL227C_RF1_2 RF1
ZShift = 0.78
lm ZScore = 0.08
NG = 0
DB = 1
SM = 0
Delta_AUC
Doxorubicin

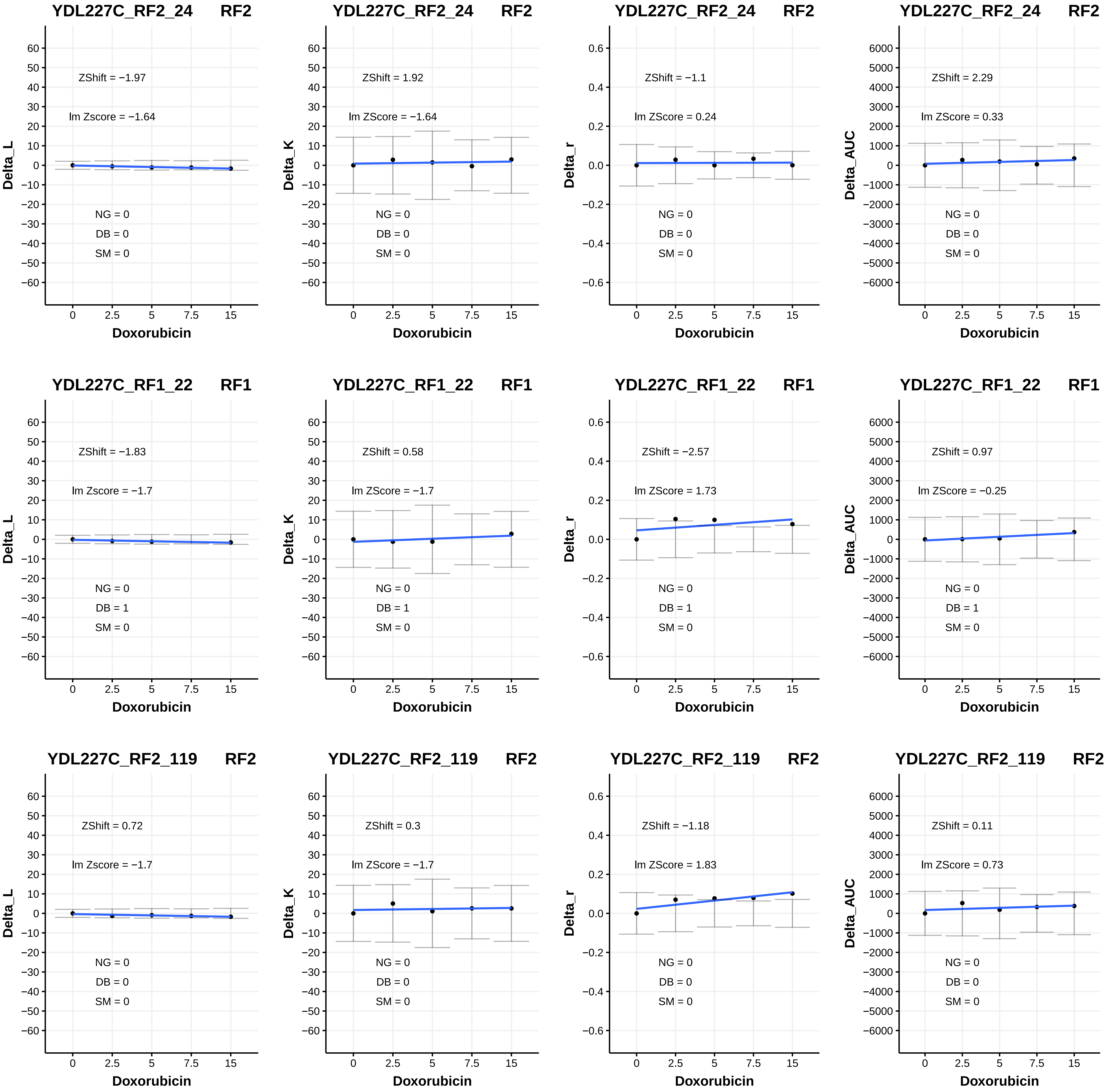

YDL227C_RF2_24 RF2
ZShift = −1.97
lm Zscore = −1.64
NG = 0
DB = 0
SM = 0
Delta_L
Doxorubicin
YDL227C_RF2_24 RF2
ZShift = 1.92
lm ZScore = −1.64
NG = 0
DB = 0
SM = 0
Delta_K
Doxorubicin
YDL227C_RF2_24 RF2
ZShift = −1.1
lm ZScore = 0.24
NG = 0
DB = 0
SM = 0
Delta_r
Doxorubicin
YDL227C_RF2_24 RF2
ZShift = 2.29
lm ZScore = 0.33
NG = 0
DB = 0
SM = 0
Delta_AUC
Doxorubicin
YDL227C_RF1_22 RF1
ZShift = −1.83
lm Zscore = −1.7
NG = 0
DB = 1
SM = 0
Delta_L
Doxorubicin
YDL227C_RF1_22 RF1
ZShift = 0.58
lm ZScore = −1.7
NG = 0
DB = 1
SM = 0
Delta_K
Doxorubicin
YDL227C_RF1_22 RF1
ZShift = −2.57
lm ZScore = 1.73
NG = 0
DB = 1
SM = 0
Delta_r
Doxorubicin
YDL227C_RF1_22 RF1
ZShift = 0.97
lm ZScore = −0.25
NG = 0
DB = 1
SM = 0
Delta_AUC
Doxorubicin
YDL227C_RF2_119 RF2
ZShift = 0.72
lm Zscore = −1.7
NG = 0
DB = 0
SM = 0
Delta_L
Doxorubicin
YDL227C_RF2_119 RF2
ZShift = 0.3
lm ZScore = −1.7
NG = 0
DB = 0
SM = 0
Delta_K
Doxorubicin
YDL227C_RF2_119 RF2
ZShift = −1.18
lm ZScore = 1.83
NG = 0
DB = 0
SM = 0
Delta_r
Doxorubicin
YDL227C_RF2_119 RF2
ZShift = 0.11
lm ZScore = 0.73
NG = 0
DB = 0
SM = 0
Delta_AUC
Doxorubicin

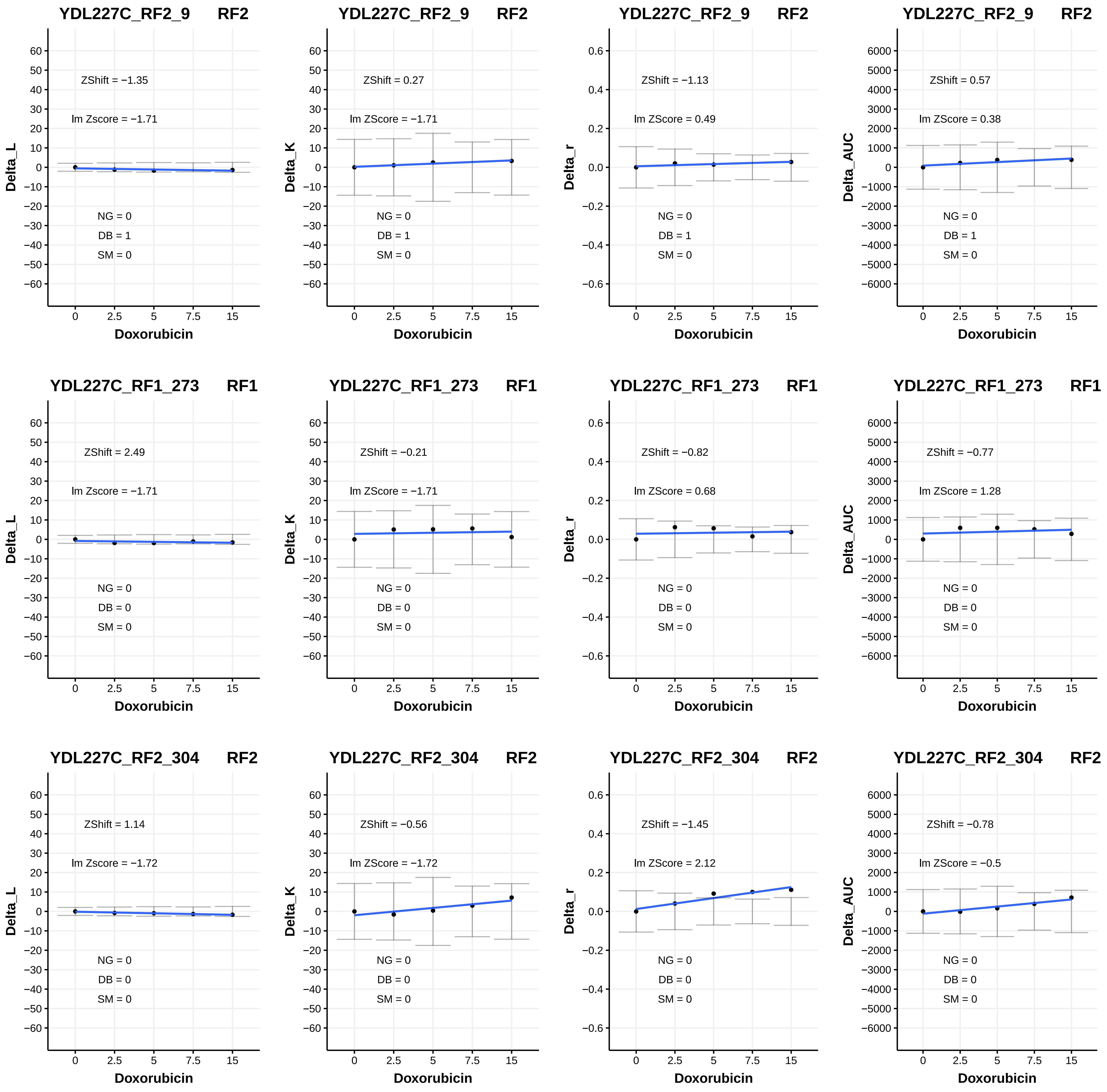

YDL227C_RF2_9 RF2
ZShift = −1.35
lm Zscore = −1.71
NG = 0
DB = 1
SM = 0
Delta_L
Doxorubicin
YDL227C_RF2_9 RF2
ZShift = 0.27
lm ZScore = −1.71
NG = 0
DB = 1
SM = 0
Delta_K
Doxorubicin
YDL227C_RF2_9 RF2
ZShift = −1.13
lm ZScore = 0.49
NG = 0
DB = 1
SM = 0
Delta_r
Doxorubicin
YDL227C_RF2_9 RF2
ZShift = 0.57
lm ZScore = 0.38
NG = 0
DB = 1
SM = 0
Delta_AUC
Doxorubicin
YDL227C_RF1_273 RF1
ZShift = 2.49
lm Zscore = −1.71
NG = 0
DB = 0
SM = 0
Delta_L
Doxorubicin
YDL227C_RF1_273 RF1
ZShift = −0.21
lm ZScore = −1.71
NG = 0
DB = 0
SM = 0
Delta_K
Doxorubicin
YDL227C_RF1_273 RF1
ZShift = −0.82
lm ZScore = 0.68
NG = 0
DB = 0
SM = 0
Delta_r
Doxorubicin
YDL227C_RF1_273 RF1
ZShift = −0.77
lm ZScore = 1.28
NG = 0
DB = 0
SM = 0
Delta_AUC
Doxorubicin
YDL227C_RF2_304 RF2
ZShift = 1.14
lm Zscore = −1.72
NG = 0
DB = 0
SM = 0
Delta_L
Doxorubicin
YDL227C_RF2_304 RF2
ZShift = −0.56
lm ZScore = −1.72
NG = 0
DB = 0
SM = 0
Delta_K
Doxorubicin
YDL227C_RF2_304 RF2
ZShift = −1.45
lm ZScore = 2.12
NG = 0
DB = 0
SM = 0
Delta_r
Doxorubicin
YDL227C_RF2_304 RF2
ZShift = −0.78
lm ZScore = −0.5
NG = 0
DB = 0
SM = 0
Delta_AUC
Doxorubicin

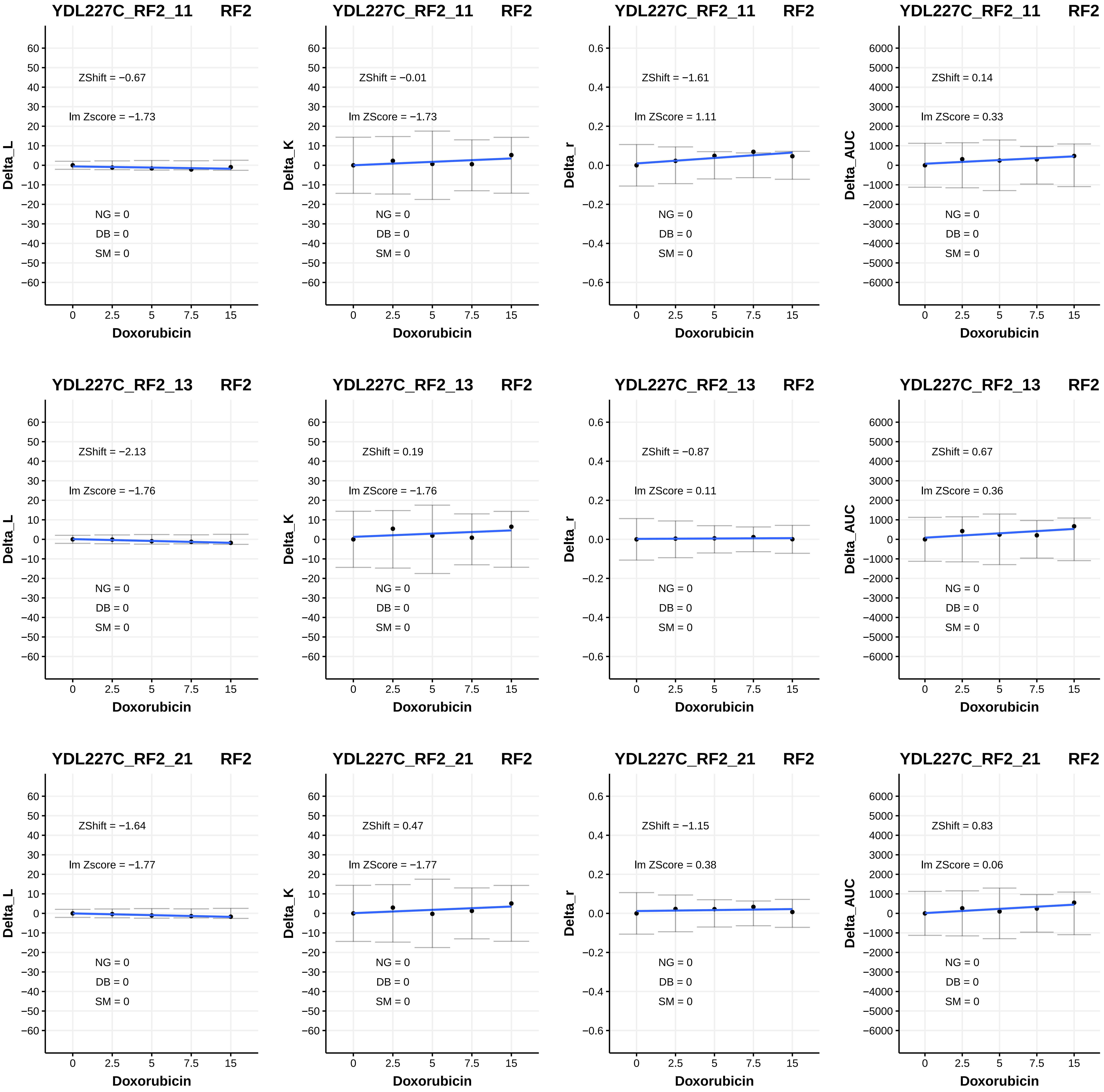

YDL227C_RF2_11 RF2
ZShift = −0.67
lm Zscore = −1.73
NG = 0
DB = 0
SM = 0
Delta_L
Doxorubicin
YDL227C_RF2_11 RF2
ZShift = −0.01
lm ZScore = −1.73
NG = 0
DB = 0
SM = 0
Delta_K
Doxorubicin
YDL227C_RF2_11 RF2
ZShift = −1.61
lm ZScore = 1.11
NG = 0
DB = 0
SM = 0
Delta_r
Doxorubicin
YDL227C_RF2_11 RF2
ZShift = 0.14
lm ZScore = 0.33
NG = 0
DB = 0
SM = 0
Delta_AUC
Doxorubicin
YDL227C_RF2_13 RF2
ZShift = −2.13
lm Zscore = −1.76
NG = 0
DB = 0
SM = 0
Delta_L
Doxorubicin
YDL227C_RF2_13 RF2
ZShift = 0.19
lm ZScore = −1.76
NG = 0
DB = 0
SM = 0
Delta_K
Doxorubicin
YDL227C_RF2_13 RF2
ZShift = −0.87
lm ZScore = 0.11
NG = 0
DB = 0
SM = 0
Delta_r
Doxorubicin
YDL227C_RF2_13 RF2
ZShift = 0.67
lm ZScore = 0.36
NG = 0
DB = 0
SM = 0
Delta_AUC
Doxorubicin
YDL227C_RF2_21 RF2
ZShift = −1.64
lm Zscore = −1.77
NG = 0
DB = 0
SM = 0
Delta_L
Doxorubicin
YDL227C_RF2_21 RF2
ZShift = 0.47
lm ZScore = −1.77
NG = 0
DB = 0
SM = 0
Delta_K
Doxorubicin
YDL227C_RF2_21 RF2
ZShift = −1.15
lm ZScore = 0.38
NG = 0
DB = 0
SM = 0
Delta_r
Doxorubicin
YDL227C_RF2_21 RF2
ZShift = 0.83
lm ZScore = 0.06
NG = 0
DB = 0
SM = 0
Delta_AUC
Doxorubicin

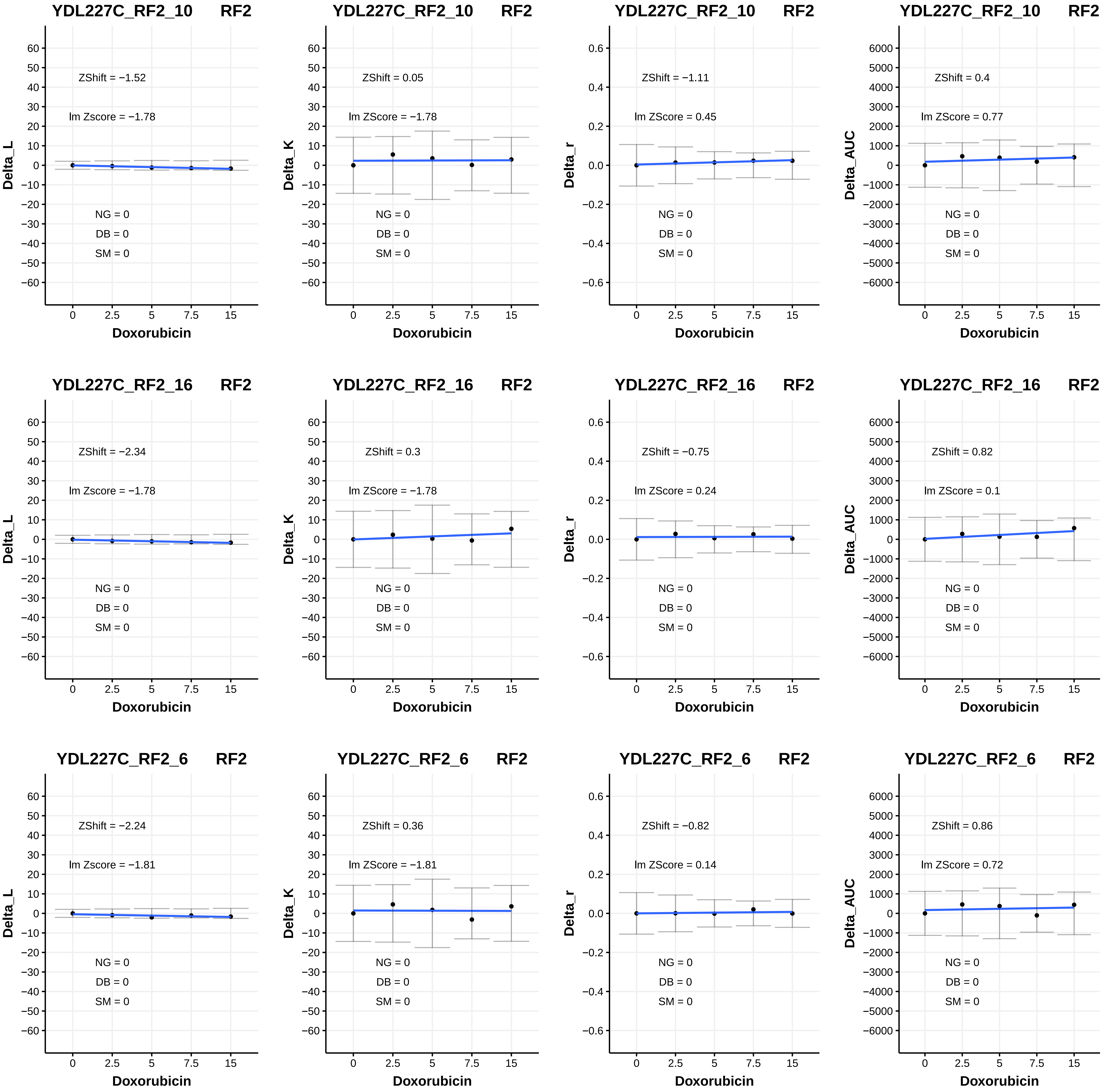
YDL227C_RF2_10 RF2
ZShift = −1.52
lm Zscore = −1.78
NG = 0
DB = 0
SM = 0
Delta_L
Doxorubicin
YDL227C_RF2_10 RF2
ZShift = 0.05
lm ZScore = −1.78
NG = 0
DB = 0
SM = 0
Delta_K
Doxorubicin
YDL227C_RF2_10 RF2
ZShift = −1.11
lm ZScore = 0.45
NG = 0
DB = 0
SM = 0
Delta_r
Doxorubicin
YDL227C_RF2_10 RF2
ZShift = 0.4
lm ZScore = 0.77
NG = 0
DB = 0
SM = 0
Delta_AUC
Doxorubicin
YDL227C_RF2_16 RF2
ZShift = −2.34
lm Zscore = −1.78
NG = 0
DB = 0
SM = 0
Delta_L
Doxorubicin
YDL227C_RF2_16 RF2
ZShift = 0.3
lm ZScore = −1.78
NG = 0
DB = 0
SM = 0
Delta_K
Doxorubicin
YDL227C_RF2_16 RF2
ZShift = −0.75
lm ZScore = 0.24
NG = 0
DB = 0
SM = 0
Delta_r
Doxorubicin
YDL227C_RF2_16 RF2
ZShift = 0.82
lm ZScore = 0.1
NG = 0
DB = 0
SM = 0
Delta_AUC
Doxorubicin
YDL227C_RF2_6 RF2
ZShift = −2.24
lm Zscore = −1.81
NG = 0
DB = 0
SM = 0
Delta_L
Doxorubicin
YDL227C_RF2_6 RF2
ZShift = 0.36
lm ZScore = −1.81
NG = 0
DB = 0
SM = 0
Delta_K
Doxorubicin
YDL227C_RF2_6 RF2
ZShift = −0.82
lm ZScore = 0.14
NG = 0
DB = 0
SM = 0
Delta_r
Doxorubicin
YDL227C_RF2_6 RF2
ZShift = 0.86
lm ZScore = 0.72
NG = 0
DB = 0
SM = 0
Delta_AUC
Doxorubicin

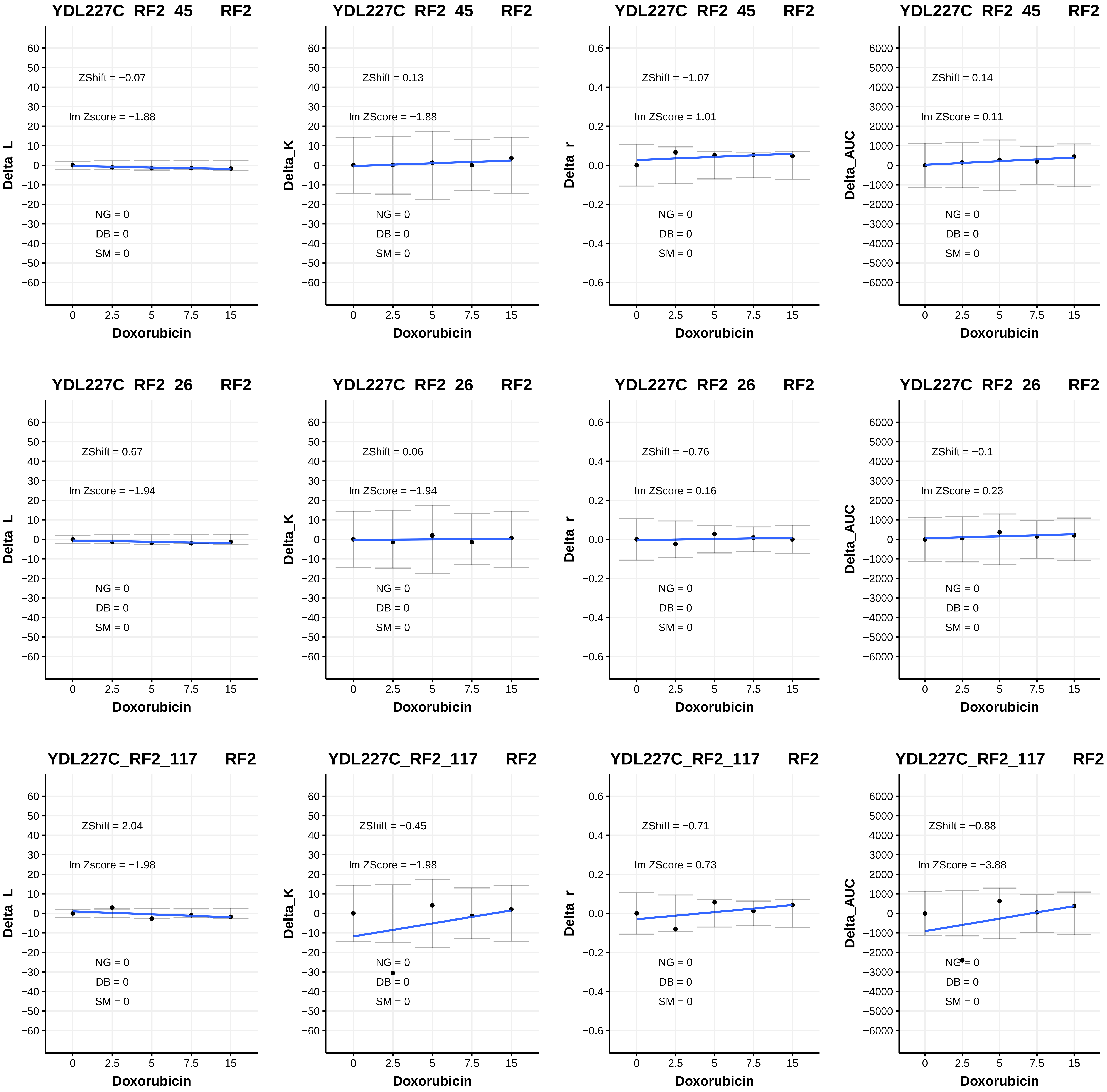

YDL227C_RF2_45 RF2
ZShift = −0.07
lm Zscore = −1.88
NG = 0
DB = 0
SM = 0
Delta_L
Doxorubicin
YDL227C_RF2_45 RF2
ZShift = 0.13
lm ZScore = −1.88
NG = 0
DB = 0
SM = 0
Delta_K
Doxorubicin
YDL227C_RF2_45 RF2
ZShift = −1.07
lm ZScore = 1.01
NG = 0
DB = 0
SM = 0
Delta_r
Doxorubicin
YDL227C_RF2_45 RF2
ZShift = 0.14
lm ZScore = 0.11
NG = 0
DB = 0
SM = 0
Delta_AUC
Doxorubicin
YDL227C_RF2_26 RF2
ZShift = 0.67
lm Zscore = −1.94
NG = 0
DB = 0
SM = 0
Delta_L
Doxorubicin
YDL227C_RF2_26 RF2
ZShift = 0.06
lm ZScore = −1.94
NG = 0
DB = 0
SM = 0
Delta_K
Doxorubicin
YDL227C_RF2_26 RF2
ZShift = −0.76
lm ZScore = 0.16
NG = 0
DB = 0
SM = 0
Delta_r
Doxorubicin
YDL227C_RF2_26 RF2
ZShift = −0.1
lm ZScore = 0.23
NG = 0
DB = 0
SM = 0
Delta_AUC
Doxorubicin
YDL227C_RF2_117 RF2
ZShift = 2.04
lm Zscore = −1.98
NG = 0
DB = 0
SM = 0
Delta_L
Doxorubicin
YDL227C_RF2_117 RF2
ZShift = −0.45
lm ZScore = −1.98
NG = 0
DB = 0
SM = 0
Delta_K
Doxorubicin
YDL227C_RF2_117 RF2
ZShift = −0.71
lm ZScore = 0.73
NG = 0
DB = 0
SM = 0
Delta_r
Doxorubicin
YDL227C_RF2_117 RF2
ZShift = −0.88
lm ZScore = −3.88
NG = 0
DB = 0
SM = 0
Delta_AUC
Doxorubicin

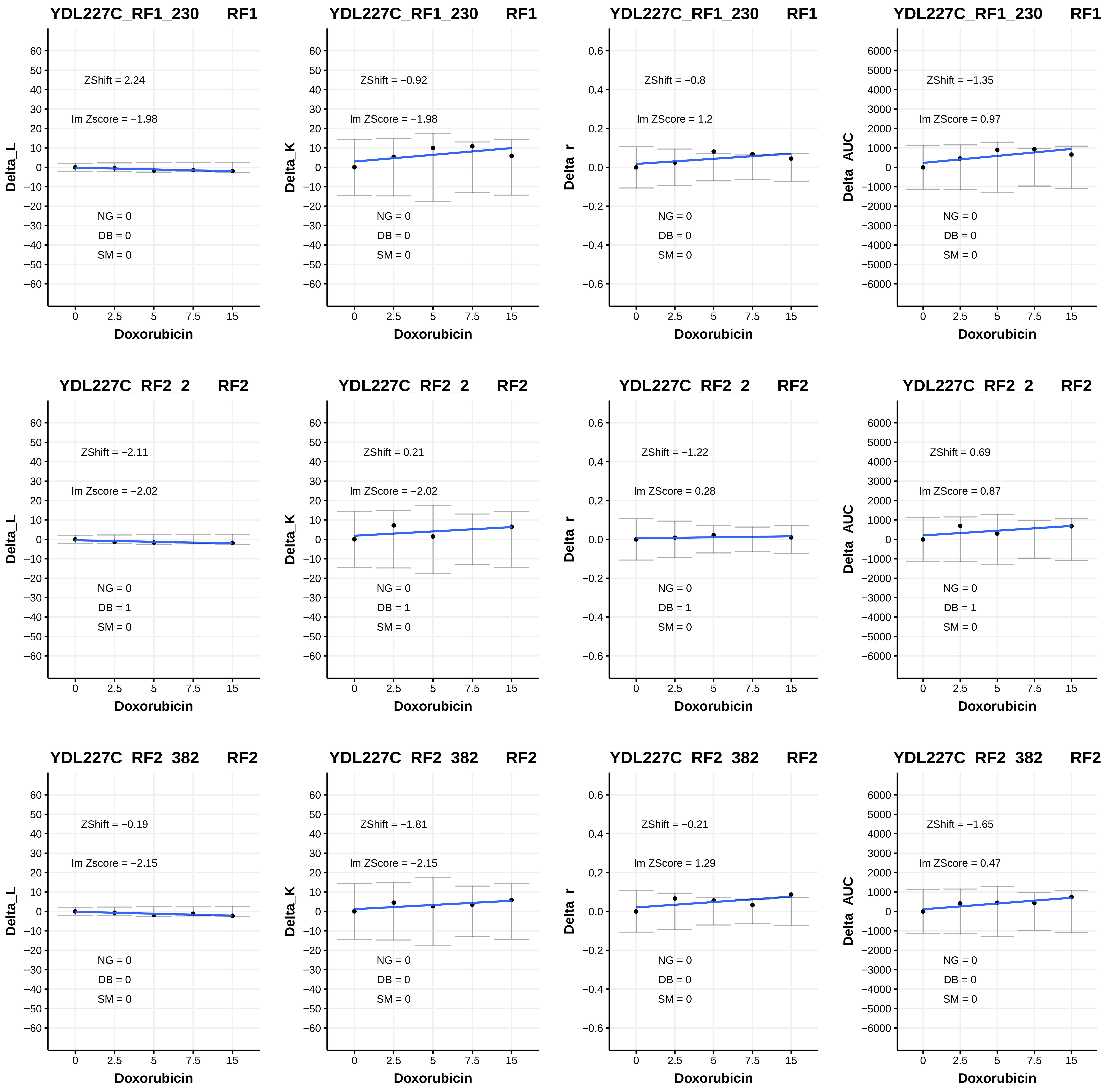
YDL227C_RF1_230 RF1
ZShift = −0.92
lm ZScore = −1.98
Delta_L
ZShift = 2.24
lm Zscore = −1.98
NG = 0
DB = 0
SM = 0
Doxorubicin
YDL227C_RF1_230 RF1
Delta_K
ZShift = −0.92
lm ZScore = −1.98
NG = 0
DB = 0
SM = 0
Doxorubicin
YDL227C_RF1_230 RF1
Delta_r
ZShift = −0.8
lm ZScore = 1.2
NG = 0
DB = 0
SM = 0
Doxorubicin
YDL227C_RF1_230 RF1
Delta_AUC
ZShift = −1.35
lm ZScore = 0.97
NG = 0
DB = 0
SM = 0
Doxorubicin
YDL227C_RF2_2 RF2
Delta_L
ZShift = −2.11
lm Zscore = −2.02
NG = 0
DB = 1
SM = 0
Doxorubicin
YDL227C_RF2_2 RF2
Delta_K
ZShift = 0.21
lm ZScore = −2.02
NG = 0
DB = 1
SM = 0
Doxorubicin
YDL227C_RF2_2 RF2
Delta_r
ZShift = −1.22
lm ZScore = 0.28
NG = 0
DB = 1
SM = 0
Doxorubicin
YDL227C_RF2_2 RF2
Delta_AUC
ZShift = 0.69
lm ZScore = 0.87
NG = 0
DB = 1
SM = 0
Doxorubicin
YDL227C_RF2_382 RF2
Delta_L
ZShift = −0.19
lm Zscore = −2.15
NG = 0
DB = 0
SM = 0
Doxorubicin
YDL227C_RF2_382 RF2
Delta_K
ZShift = −1.81
lm ZScore = −2.15
NG = 0
DB = 0
SM = 0
Doxorubicin
YDL227C_RF2_382 RF2
Delta_r
ZShift = −0.21
lm ZScore = 1.29
NG = 0
DB = 0
SM = 0
Doxorubicin
YDL227C_RF2_382 RF2
Delta_AUC
ZShift = −1.65
lm ZScore = 0.47
NG = 0
DB = 0
SM = 0
Doxorubicin

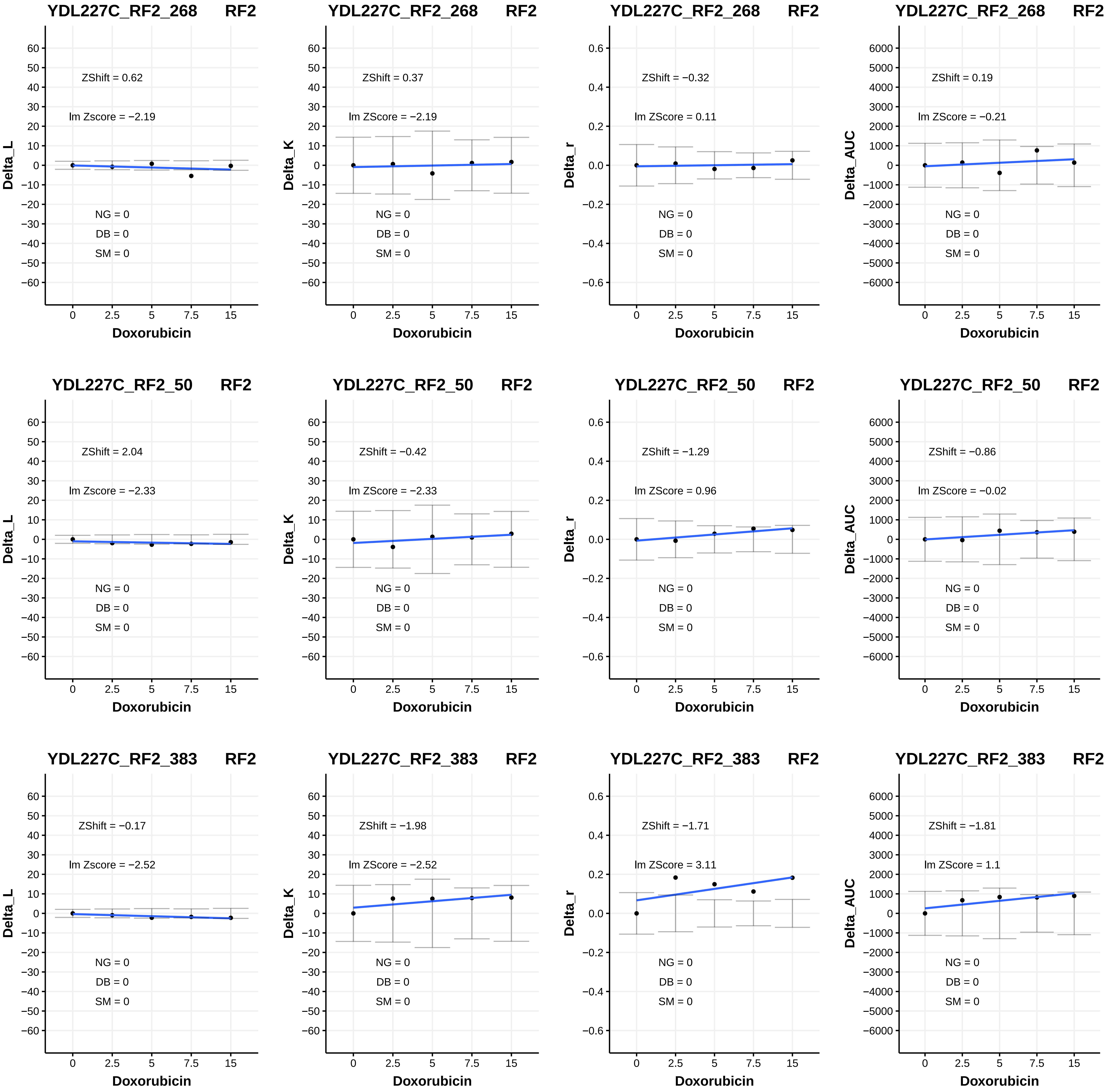

YDL227C_RF2_268 RF2
ZShift = 0.62
lm Zscore = −2.19
NG = 0
DB = 0
SM = 0
Delta_L
Doxorubicin
YDL227C_RF2_268 RF2
ZShift = 0.37
lm ZScore = −2.19
NG = 0
DB = 0
SM = 0
Delta_K
Doxorubicin
YDL227C_RF2_268 RF2
ZShift = −0.32
lm ZScore = 0.11
NG = 0
DB = 0
SM = 0
Delta_r
Doxorubicin
YDL227C_RF2_268 RF2
ZShift = 0.19
lm ZScore = −0.21
NG = 0
DB = 0
SM = 0
Delta_AUC
Doxorubicin
YDL227C_RF2_50 RF2
ZShift = 2.04
lm Zscore = −2.33
NG = 0
DB = 0
SM = 0
Delta_L
Doxorubicin
YDL227C_RF2_50 RF2
ZShift = −0.42
lm ZScore = −2.33
NG = 0
DB = 0
SM = 0
Delta_K
Doxorubicin
YDL227C_RF2_50 RF2
ZShift = −1.29
lm ZScore = 0.96
NG = 0
DB = 0
SM = 0
Delta_r
Doxorubicin
YDL227C_RF2_50 RF2
ZShift = −0.86
lm ZScore = −0.02
NG = 0
DB = 0
SM = 0
Delta_AUC
Doxorubicin
YDL227C_RF2_383 RF2
ZShift = −0.17
lm Zscore = −2.52
NG = 0
DB = 0
SM = 0
Delta_L
Doxorubicin
YDL227C_RF2_383 RF2
ZShift = −1.98
lm ZScore = −2.52
NG = 0
DB = 0
SM = 0
Delta_K
Doxorubicin
YDL227C_RF2_383 RF2
ZShift = −1.71
lm ZScore = 3.11
NG = 0
DB = 0
SM = 0
Delta_r
Doxorubicin
YDL227C_RF2_383 RF2
ZShift = −1.81
lm ZScore = 1.1
NG = 0
DB = 0
SM = 0
Delta_AUC
Doxorubicin

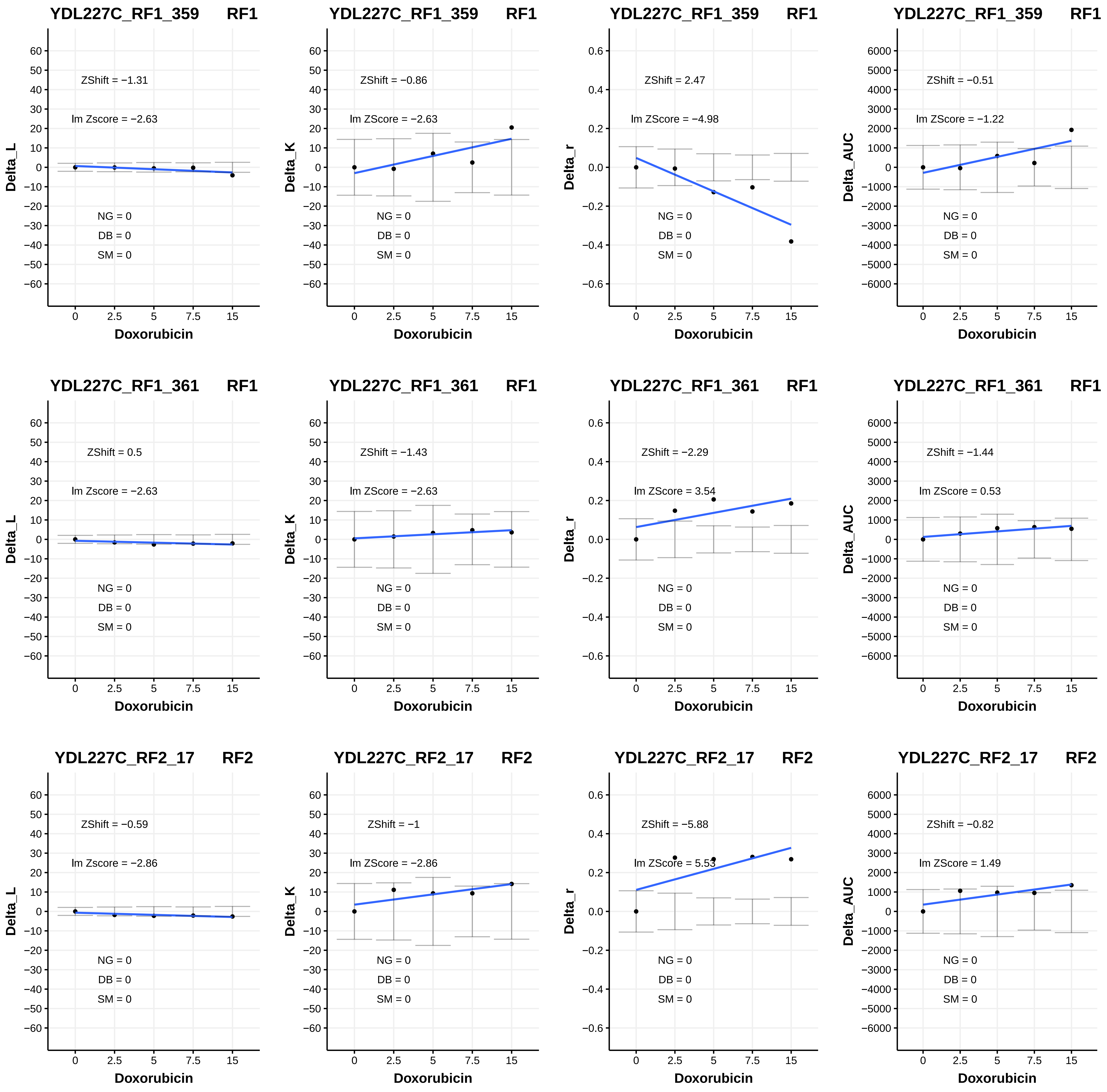
YDL227C_RF1_359 RF1
ZShift = −1.31
lm Zscore = −2.63
NG = 0
DB = 0
SM = 0
Delta_L
Doxorubicin
YDL227C_RF1_359 RF1
ZShift = −0.86
lm ZScore = −2.63
NG = 0
DB = 0
SM = 0
Delta_K
Doxorubicin
YDL227C_RF1_359 RF1
ZShift = 2.47
lm ZScore = −4.98
NG = 0
DB = 0
SM = 0
Delta_r
Doxorubicin
YDL227C_RF1_359 RF1
ZShift = −0.51
lm ZScore = −1.22
NG = 0
DB = 0
SM = 0
Delta_AUC
Doxorubicin
YDL227C_RF1_361 RF1
ZShift = 0.5
lm Zscore = −2.63
NG = 0
DB = 0
SM = 0
Delta_L
Doxorubicin
YDL227C_RF1_361 RF1
ZShift = −1.43
lm ZScore = −2.63
NG = 0
DB = 0
SM = 0
Delta_K
Doxorubicin
YDL227C_RF1_361 RF1
ZShift = −2.29
lm ZScore = 3.54
NG = 0
DB = 0
SM = 0
Delta_r
Doxorubicin
YDL227C_RF1_361 RF1
ZShift = −1.44
lm ZScore = 0.53
NG = 0
DB = 0
SM = 0
Delta_AUC
Doxorubicin
YDL227C_RF2_17 RF2
ZShift = −0.59
lm Zscore = −2.86
NG = 0
DB = 0
SM = 0
Delta_L
Doxorubicin
YDL227C_RF2_17 RF2
ZShift = −1
lm ZScore = −2.86
NG = 0
DB = 0
SM = 0
Delta_K
Doxorubicin
YDL227C_RF2_17 RF2
ZShift = −5.88
lm ZScore = 5.53
NG = 0
DB = 0
SM = 0
Delta_r
Doxorubicin
YDL227C_RF2_17 RF2
ZShift = −0.82
lm ZScore = 1.49
NG = 0
DB = 0
SM = 0
Delta_AUC
Doxorubicin

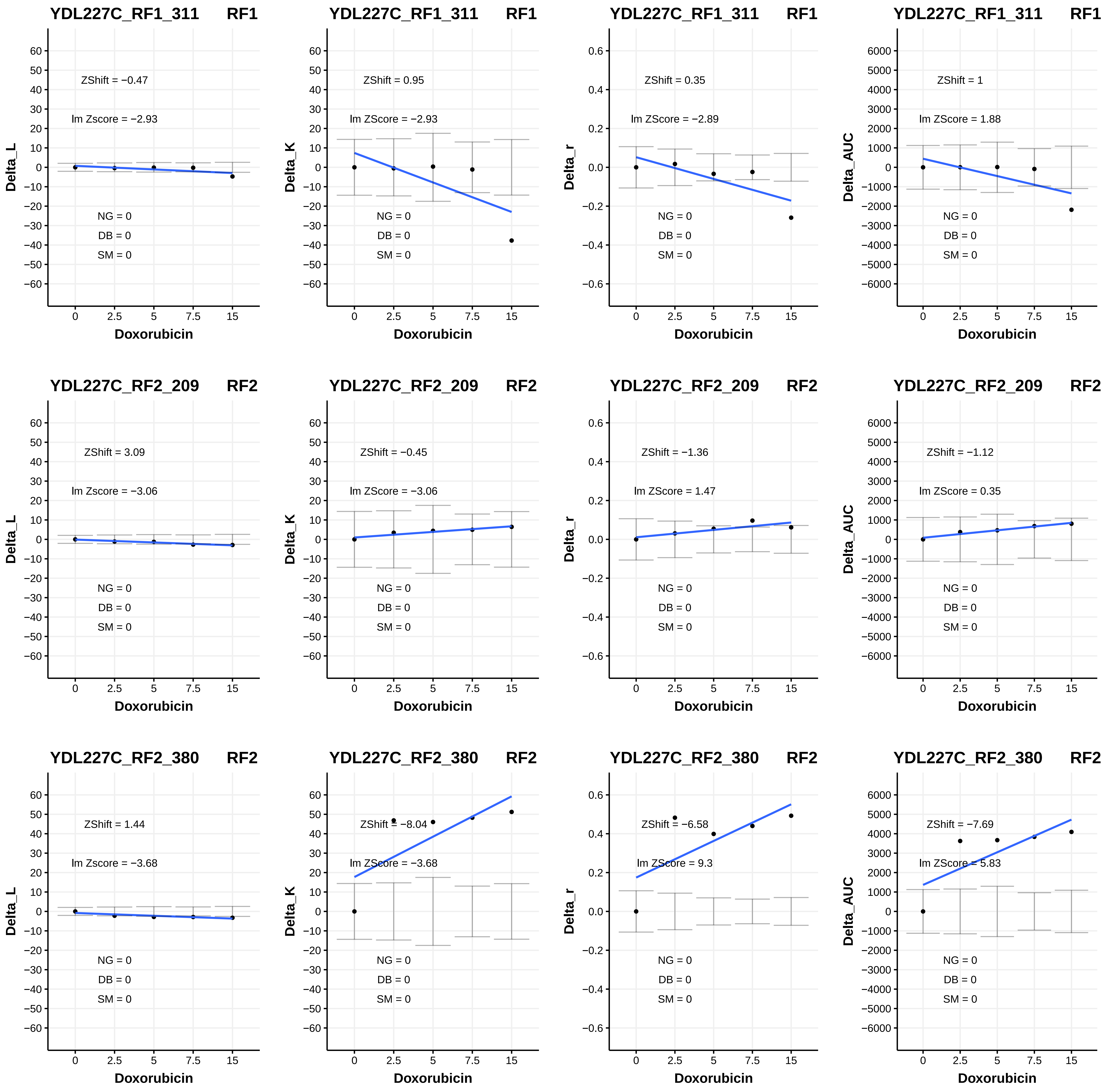
YDL227C_RF1_311 RF1
ZShift = −0.47
lm Zscore = −2.93
NG = 0
DB = 0
SM = 0
Delta_L
Doxorubicin
YDL227C_RF1_311 RF1
ZShift = 0.95
lm ZScore = −2.93
NG = 0
DB = 0
SM = 0
Delta_K
Doxorubicin
YDL227C_RF1_311 RF1
ZShift = 0.35
lm ZScore = −2.89
NG = 0
DB = 0
SM = 0
Delta_r
Doxorubicin
YDL227C_RF1_311 RF1
ZShift = 1
lm ZScore = 1.88
NG = 0
DB = 0
SM = 0
Delta_AUC
Doxorubicin
YDL227C_RF2_209 RF2
ZShift = 3.09
lm Zscore = −3.06
NG = 0
DB = 0
SM = 0
Delta_L
Doxorubicin
YDL227C_RF2_209 RF2
ZShift = −0.45
lm ZScore = −3.06
NG = 0
DB = 0
SM = 0
Delta_K
Doxorubicin
YDL227C_RF2_209 RF2
ZShift = −1.36
lm ZScore = 1.47
NG = 0
DB = 0
SM = 0
Delta_r
Doxorubicin
YDL227C_RF2_209 RF2
ZShift = −1.12
lm ZScore = 0.35
NG = 0
DB = 0
SM = 0
Delta_AUC
Doxorubicin
YDL227C_RF2_380 RF2
ZShift = 1.44
lm Zscore = −3.68
NG = 0
DB = 0
SM = 0
Delta_L
Doxorubicin
YDL227C_RF2_380 RF2
ZShift = −8.04
lm ZScore = −3.68
NG = 0
DB = 0
SM = 0
Delta_K
Doxorubicin
YDL227C_RF2_380 RF2
ZShift = −6.58
lm ZScore = 9.3
NG = 0
DB = 0
SM = 0
Delta_r
Doxorubicin
YDL227C_RF2_380 RF2
ZShift = −7.69
lm ZScore = 5.83
NG = 0
DB = 0
SM = 0
Delta_AUC
Doxorubicin

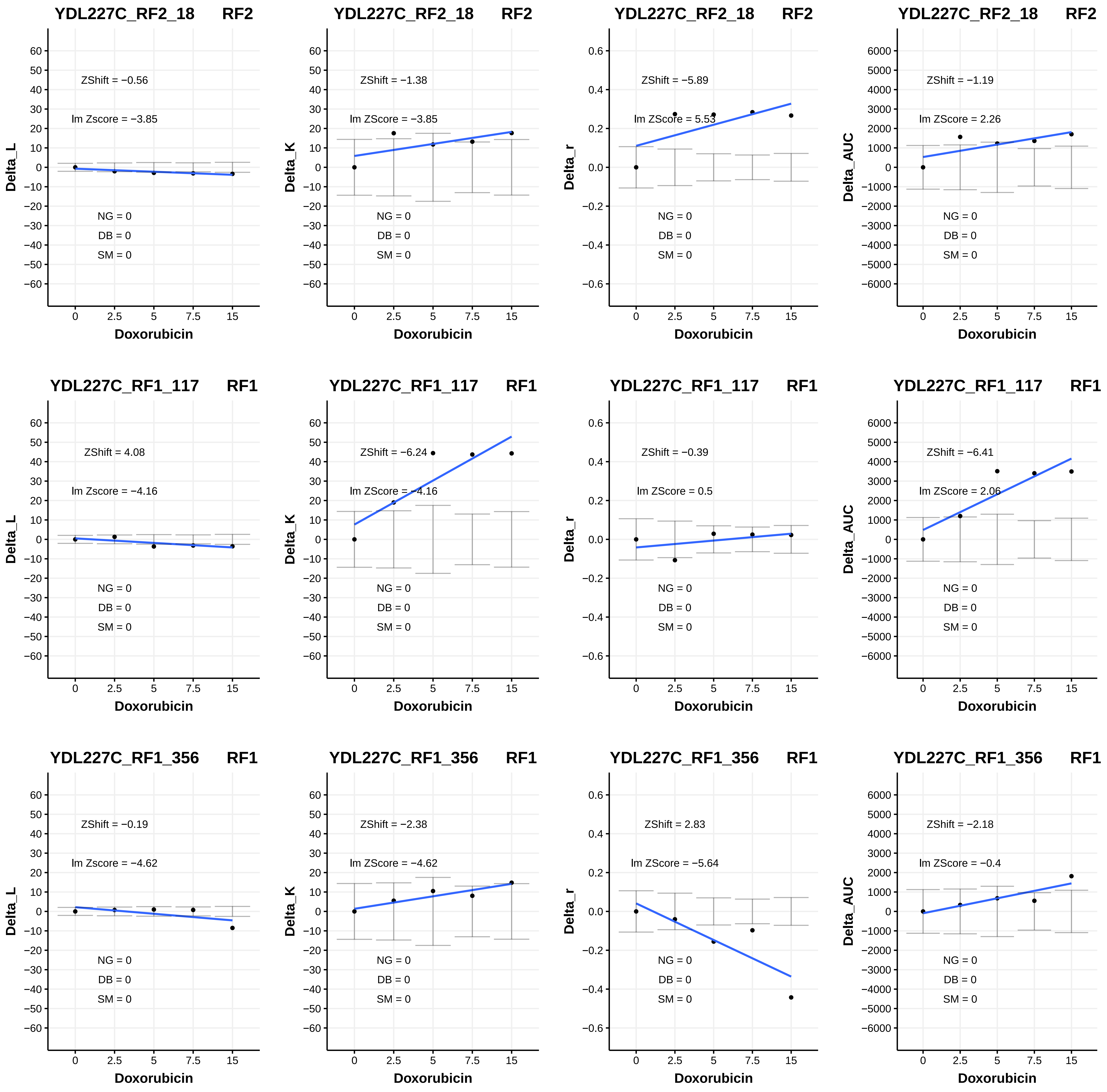

YDL227C_RF2_18 RF2
ZShift = −0.56
lm Zscore = −3.85
NG = 0
DB = 0
SM = 0
Delta_L
Doxorubicin
YDL227C_RF2_18 RF2
ZShift = −1.38
lm ZScore = −3.85
NG = 0
DB = 0
SM = 0
Delta_K
Doxorubicin
YDL227C_RF2_18 RF2
ZShift = −5.89
lm ZScore = 5.53
NG = 0
DB = 0
SM = 0
Delta_r
Doxorubicin
YDL227C_RF2_18 RF2
ZShift = −1.19
lm ZScore = 2.26
NG = 0
DB = 0
SM = 0
Delta_AUC
Doxorubicin
YDL227C_RF1_117 RF1
ZShift = 4.08
lm Zscore = −4.16
NG = 0
DB = 0
SM = 0
Delta_L
Doxorubicin
YDL227C_RF1_117 RF1
ZShift = −6.24
lm ZScore = −4.16
NG = 0
DB = 0
SM = 0
Delta_K
Doxorubicin
YDL227C_RF1_117 RF1
ZShift = −0.39
lm ZScore = 0.5
NG = 0
DB = 0
SM = 0
Delta_r
Doxorubicin
YDL227C_RF1_117 RF1
ZShift = −6.41
lm ZScore = 2.06
NG = 0
DB = 0
SM = 0
Delta_AUC
Doxorubicin
YDL227C_RF1_356 RF1
ZShift = −0.19
lm Zscore = −4.62
NG = 0
DB = 0
SM = 0
Delta_L
Doxorubicin
YDL227C_RF1_356 RF1
ZShift = −2.38
lm ZScore = −4.62
NG = 0
DB = 0
SM = 0
Delta_K
Doxorubicin
YDL227C_RF1_356 RF1
ZShift = 2.83
lm ZScore = −5.64
NG = 0
DB = 0
SM = 0
Delta_r
Doxorubicin
YDL227C_RF1_356 RF1
ZShift = −2.18
lm ZScore = −0.4
NG = 0
DB = 0
SM = 0
Delta_AUC
Doxorubicin

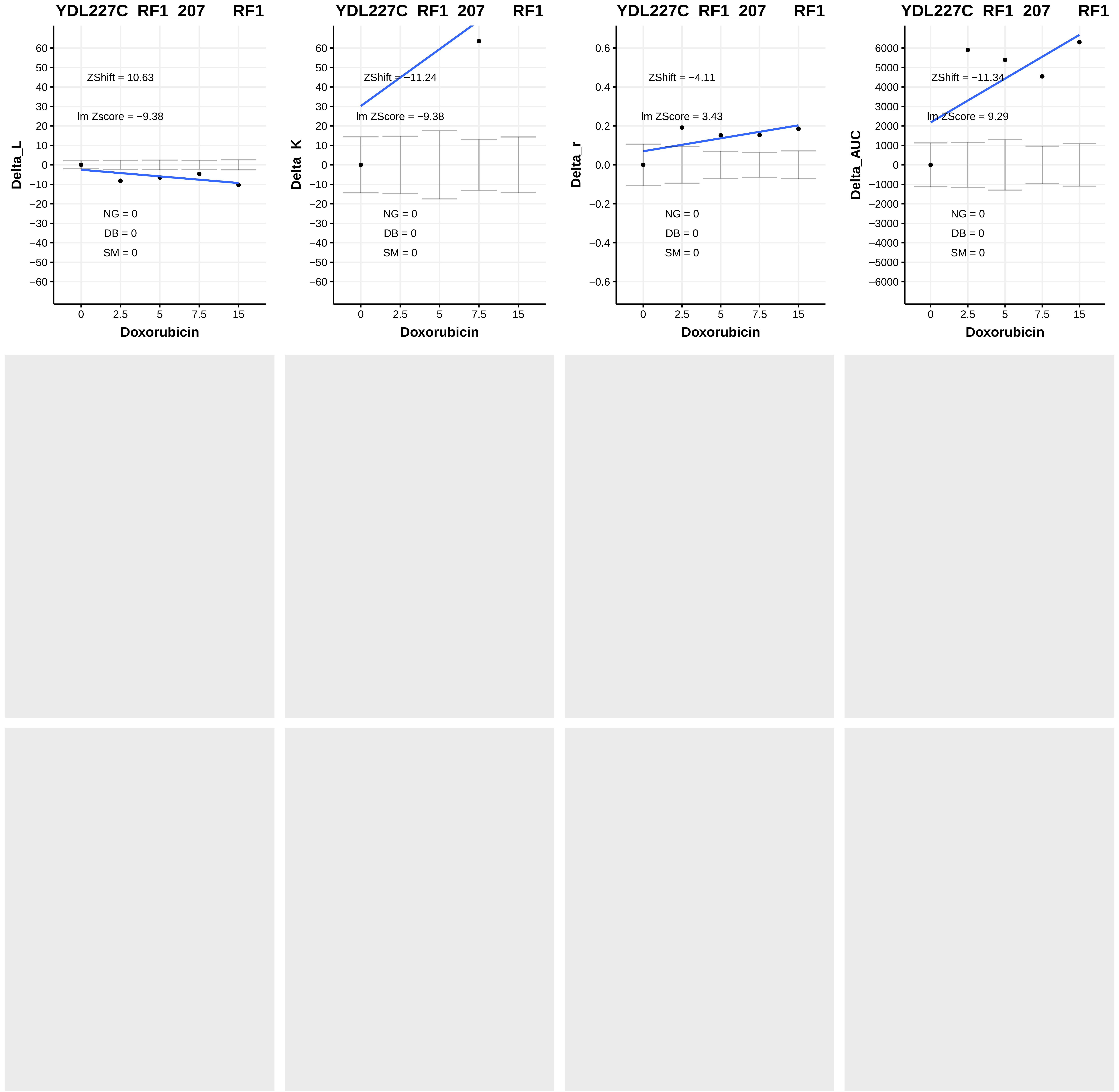
YDL227C_RF1_207 RF1
ZShift = 10.63
lm Zscore = −9.38
NG = 0
DB = 0
SM = 0
Delta_L
Doxorubicin
YDL227C_RF1_207 RF1
ZShift = −11.24
lm ZScore = −9.38
NG = 0
DB = 0
SM = 0
Delta_K
Doxorubicin
YDL227C_RF1_207 RF1
ZShift = −4.11
lm ZScore = 3.43
NG = 0
DB = 0
SM = 0
Delta_r
Doxorubicin
YDL227C_RF1_207 RF1
ZShift = −11.34
lm ZScore = 9.29
NG = 0
DB = 0
SM = 0
Delta_AUC
Doxorubicin
