## Additional File 4 for "A yeast phenomic model for the influence of Warburg metabolism on genetic buffering of doxorubicin": D - RF_InteractionPlots_Doxo_HLD_V.pdf

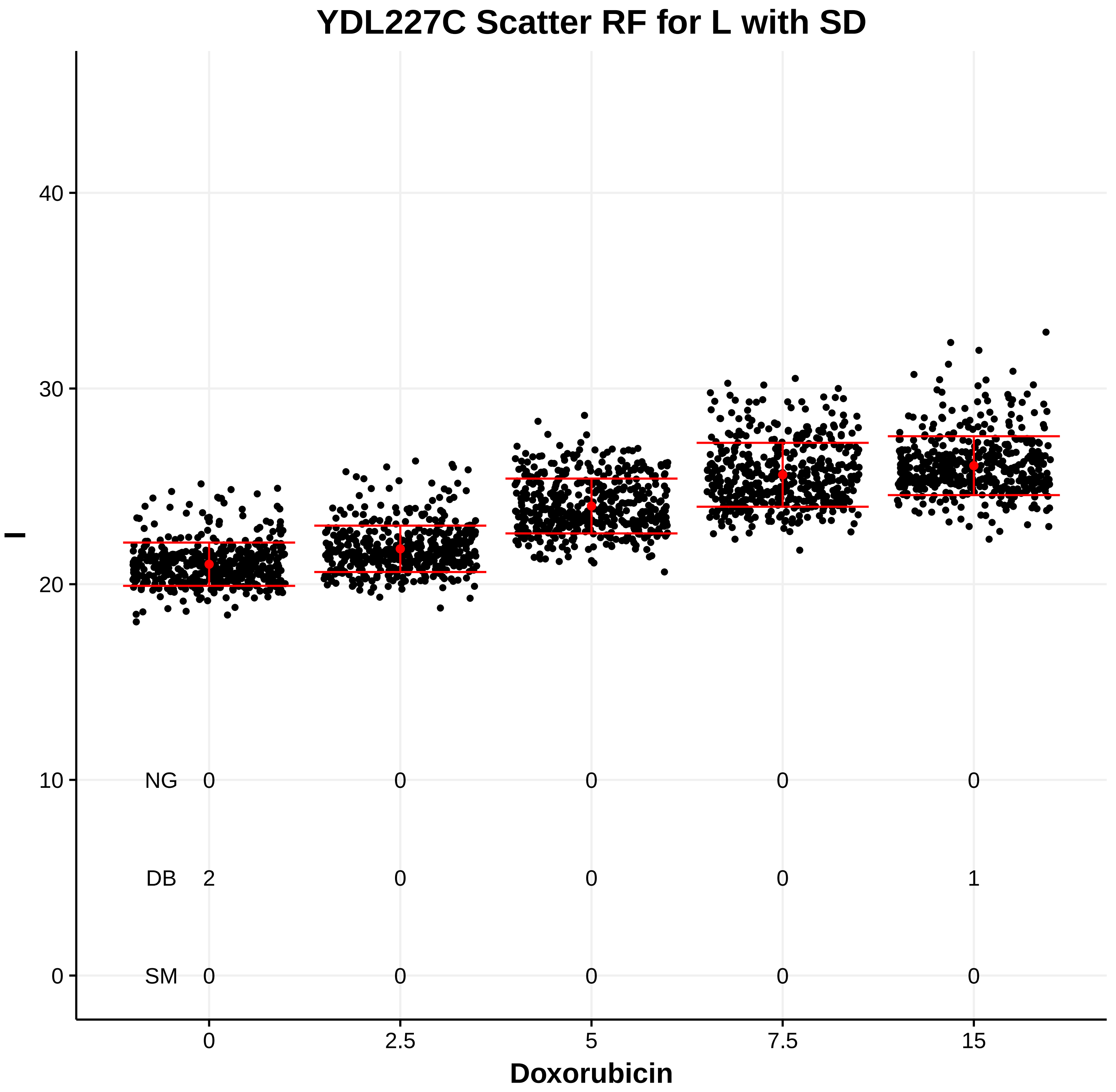

YDL227C Scatter RF for L with SD
l
Doxorubicin
0
2.5
5
7.5
15
NG 0 0 0 0 0
DB 2 0 0 0 1
SM 0 0 0 0 0


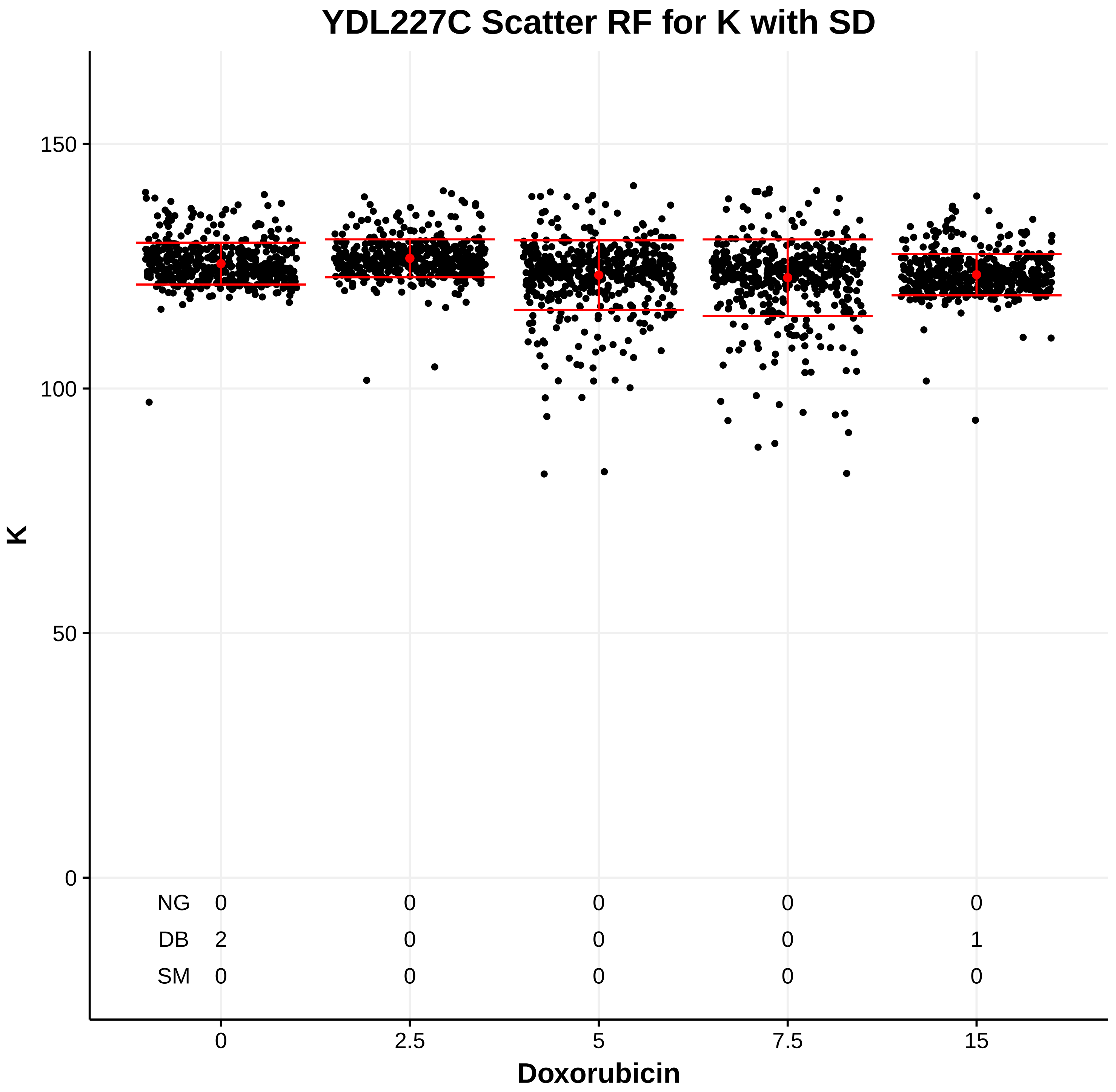

YDL227C Scatter RF for K with SD
K
Doxorubicin
0
2.5
5
7.5
15
NG 0 0 0 0 0
DB 2 0 0 0 1
SM 0 0 0 0 0


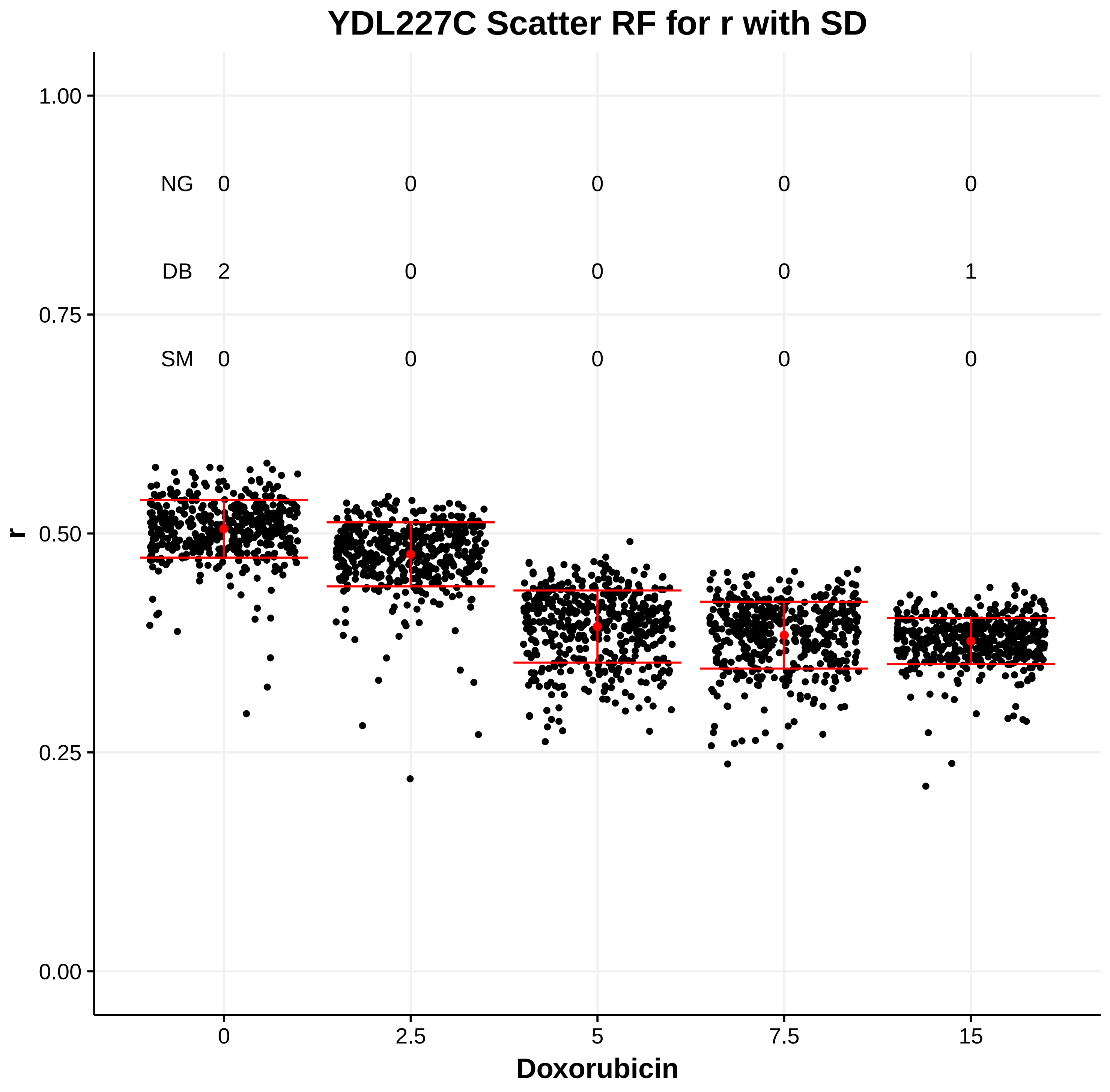

YDL227C Scatter RF for r with SD
r
Doxorubicin
0
2.5
5
7.5
15
NG 0 0 0 0 0
DB 2 0 0 0 1
SM 0 0 0 0 0


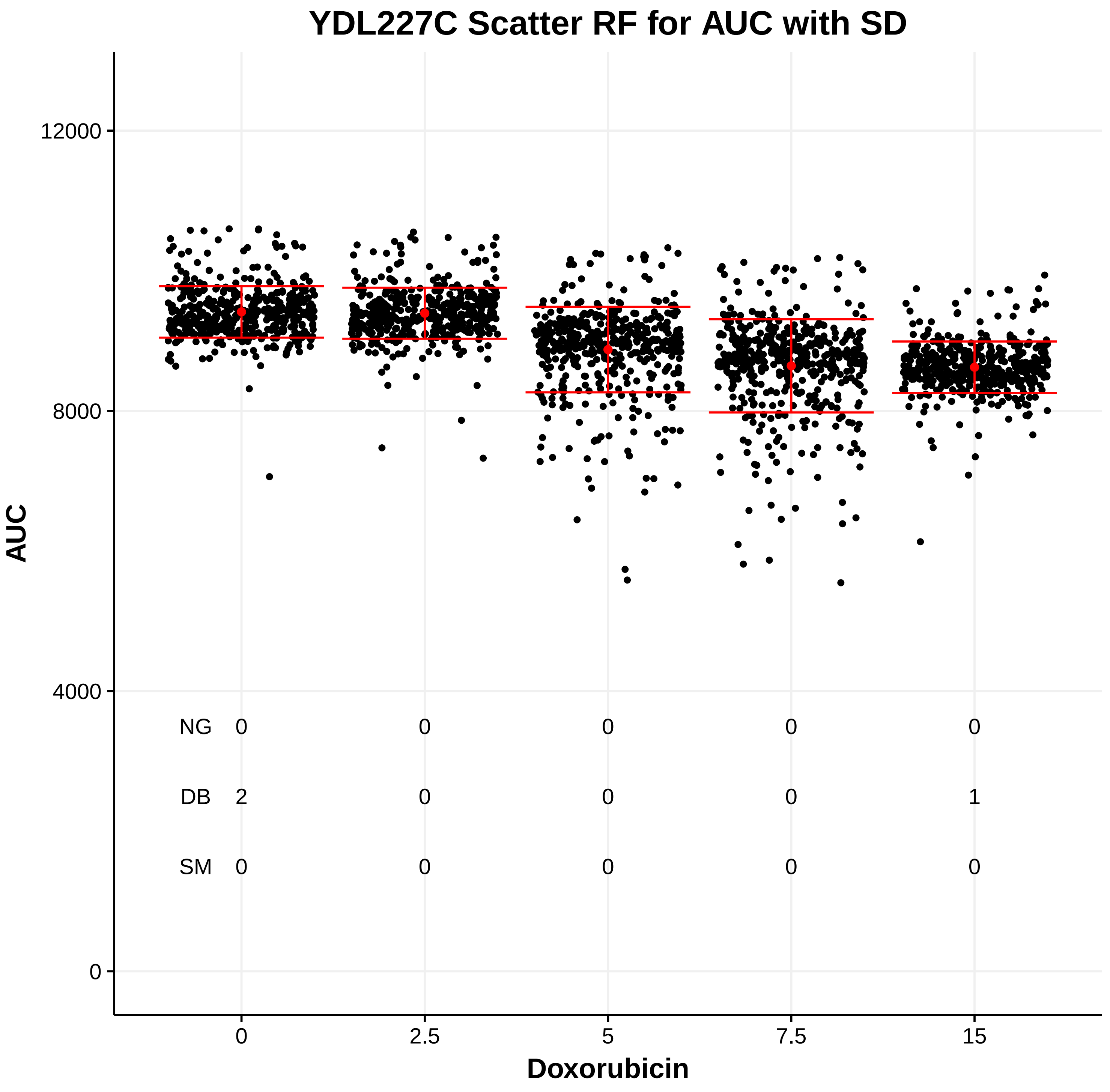

YDL227C Scatter RF for AUC with SD
AUC
Doxorubicin
0
2.5
5
7.5
15
NG 0 0 0 0 0
DB 2 0 0 0 1
SM 0 0 0 0 0

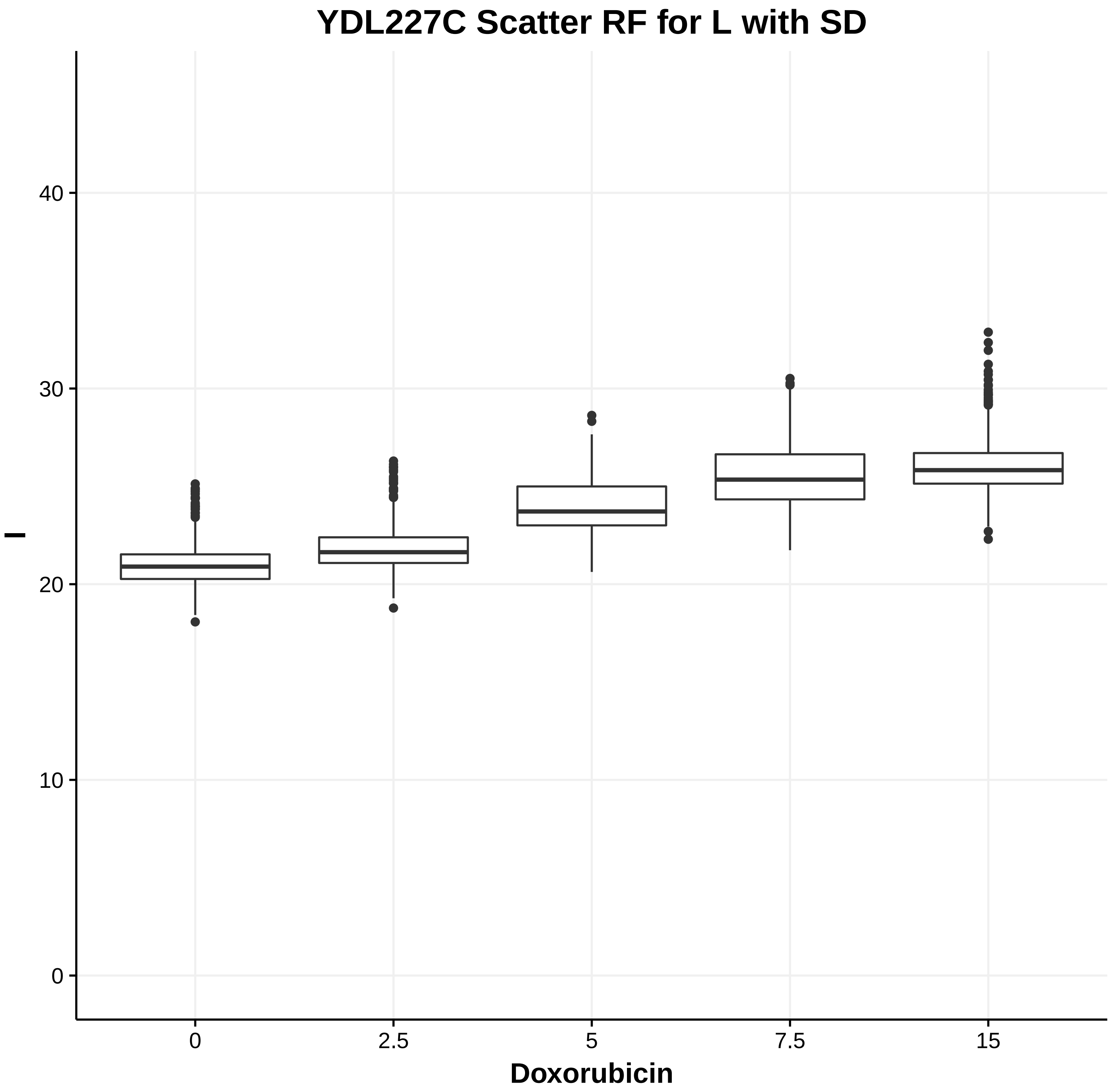
YDL227C Scatter RF for L with SD
l
Doxorubicin
0
10
20
30
40
0
2.5
5
7.5
15

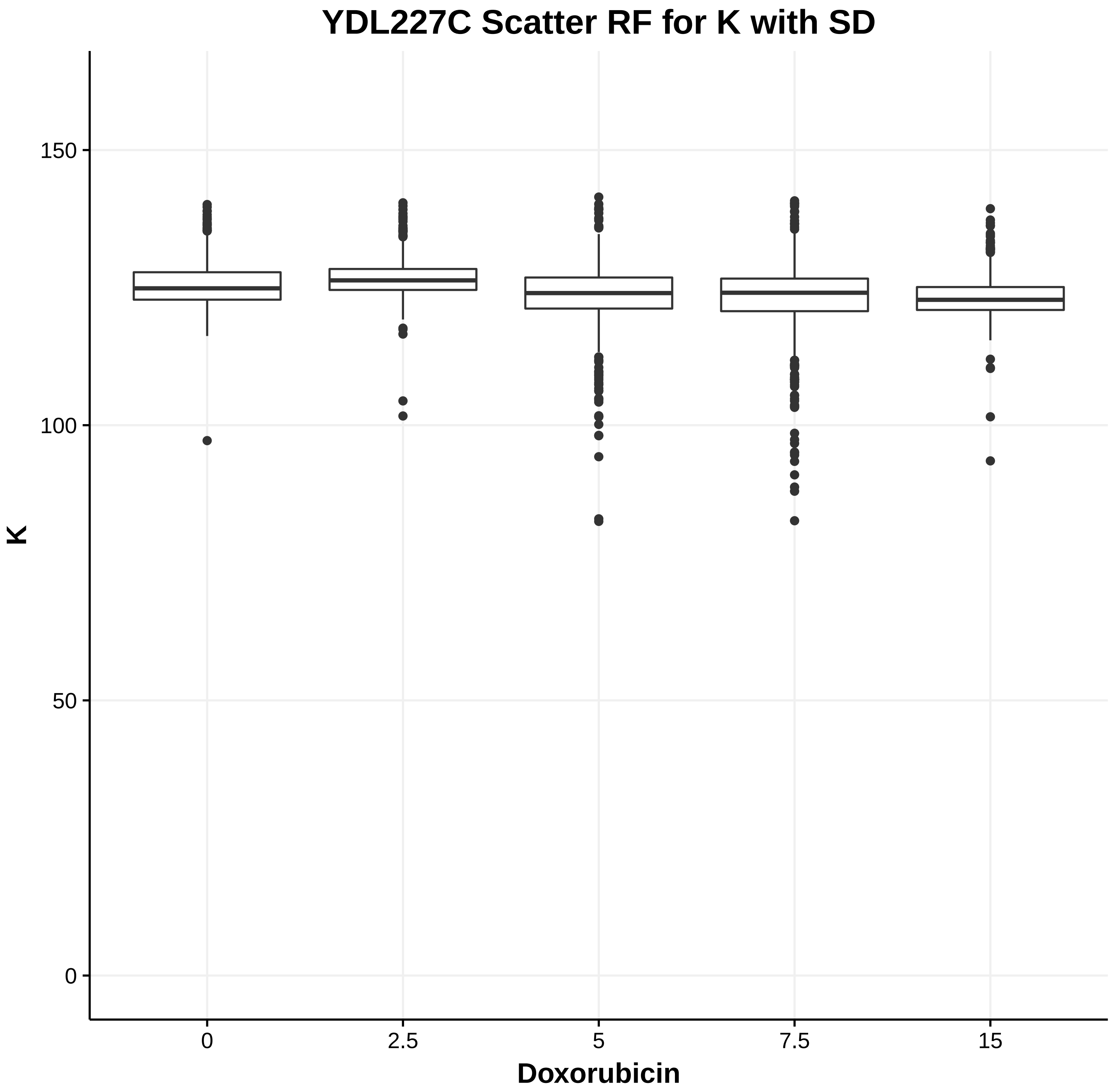
YDL227C Scatter RF for K with SD
K
Doxorubicin
0
50
100
150
0
2.5
5
7.5
15

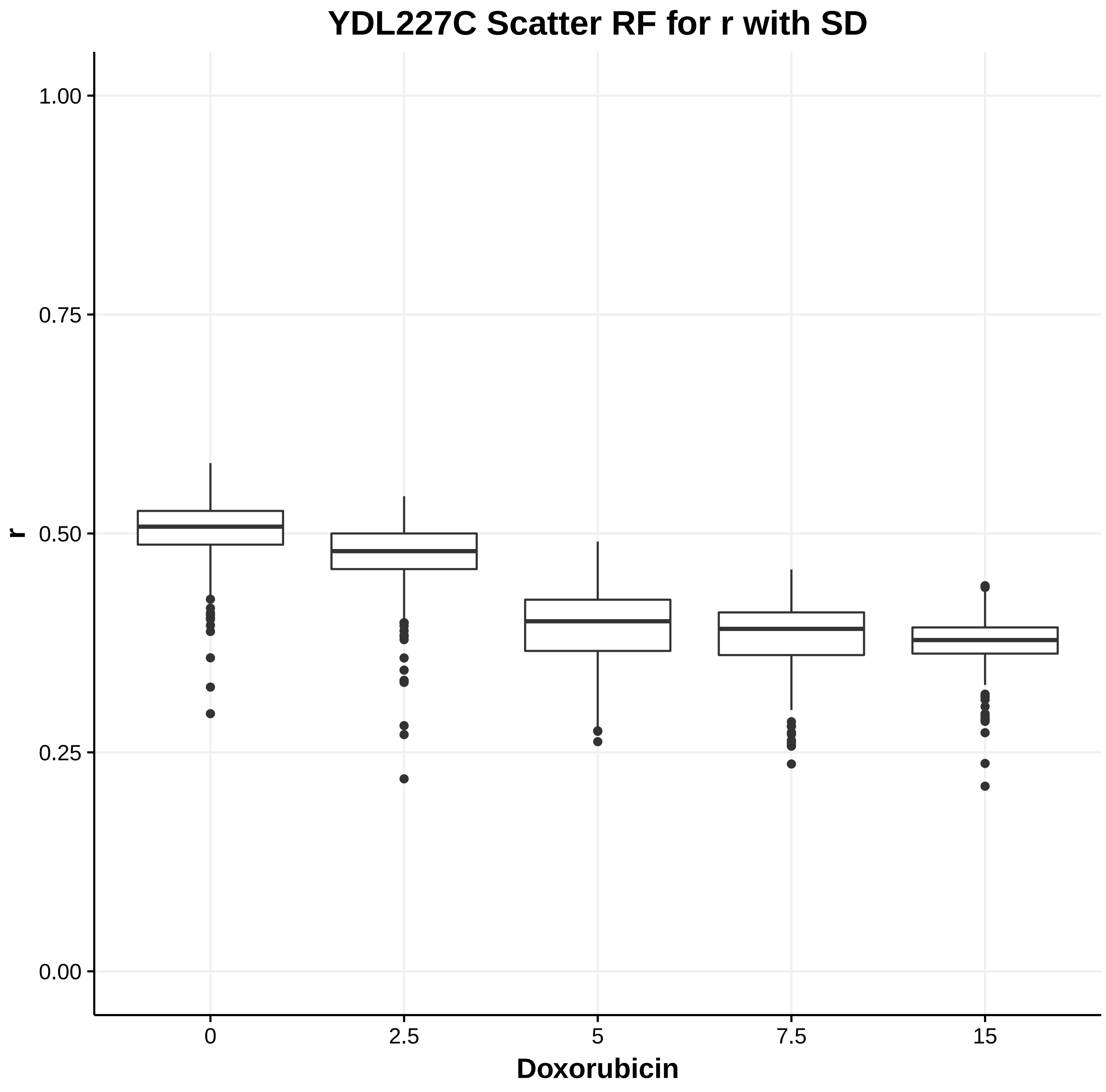
YDL227C Scatter RF for r with SD
r
Doxorubicin
0.00
0.25
0.50
0.75
1.00
0
2.5
5
7.5
15

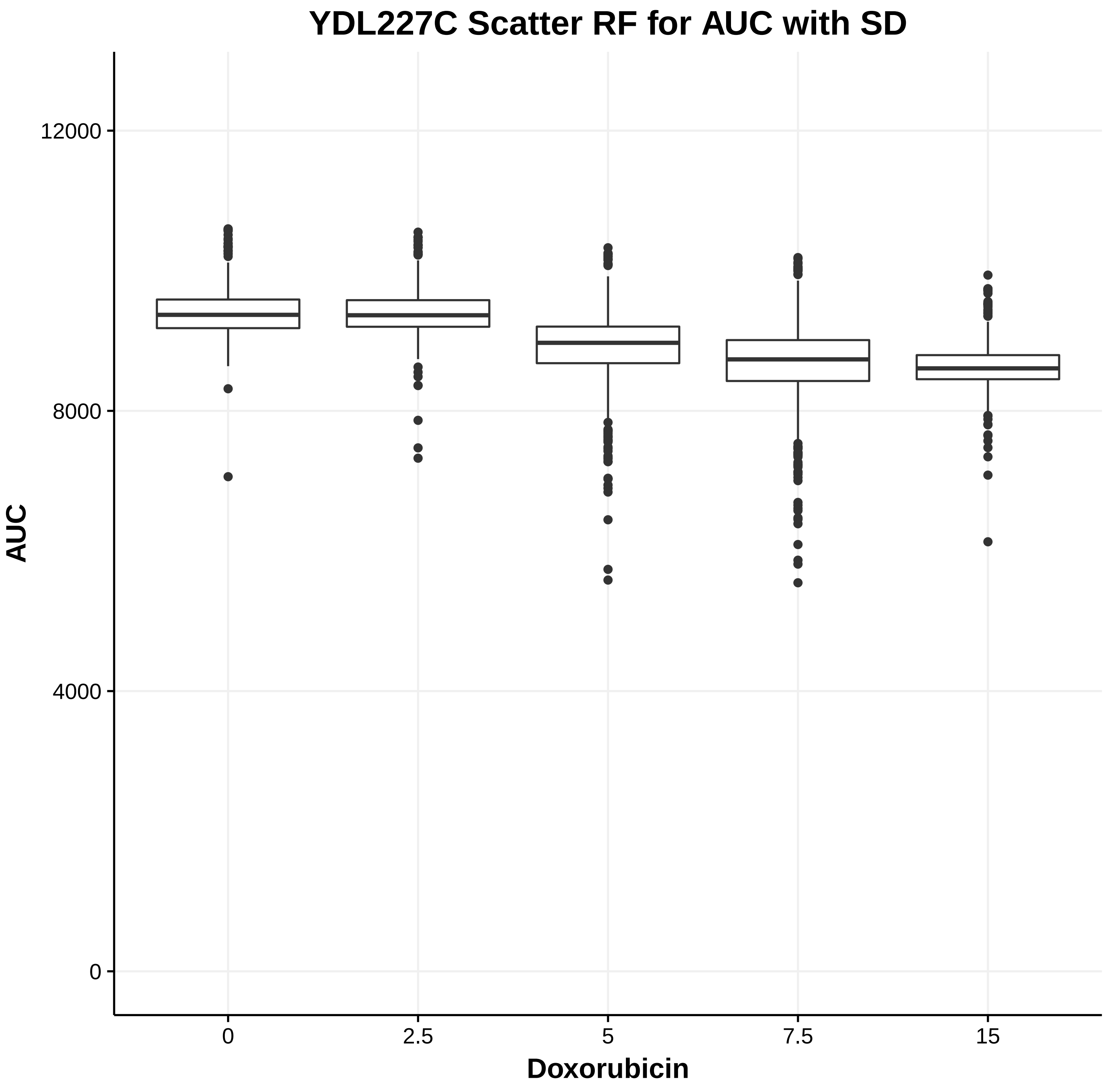
YDL227C Scatter RF for AUC with SD
AUC
Doxorubicin
0
4000
8000
12000
0
2.5
5
7.5
15

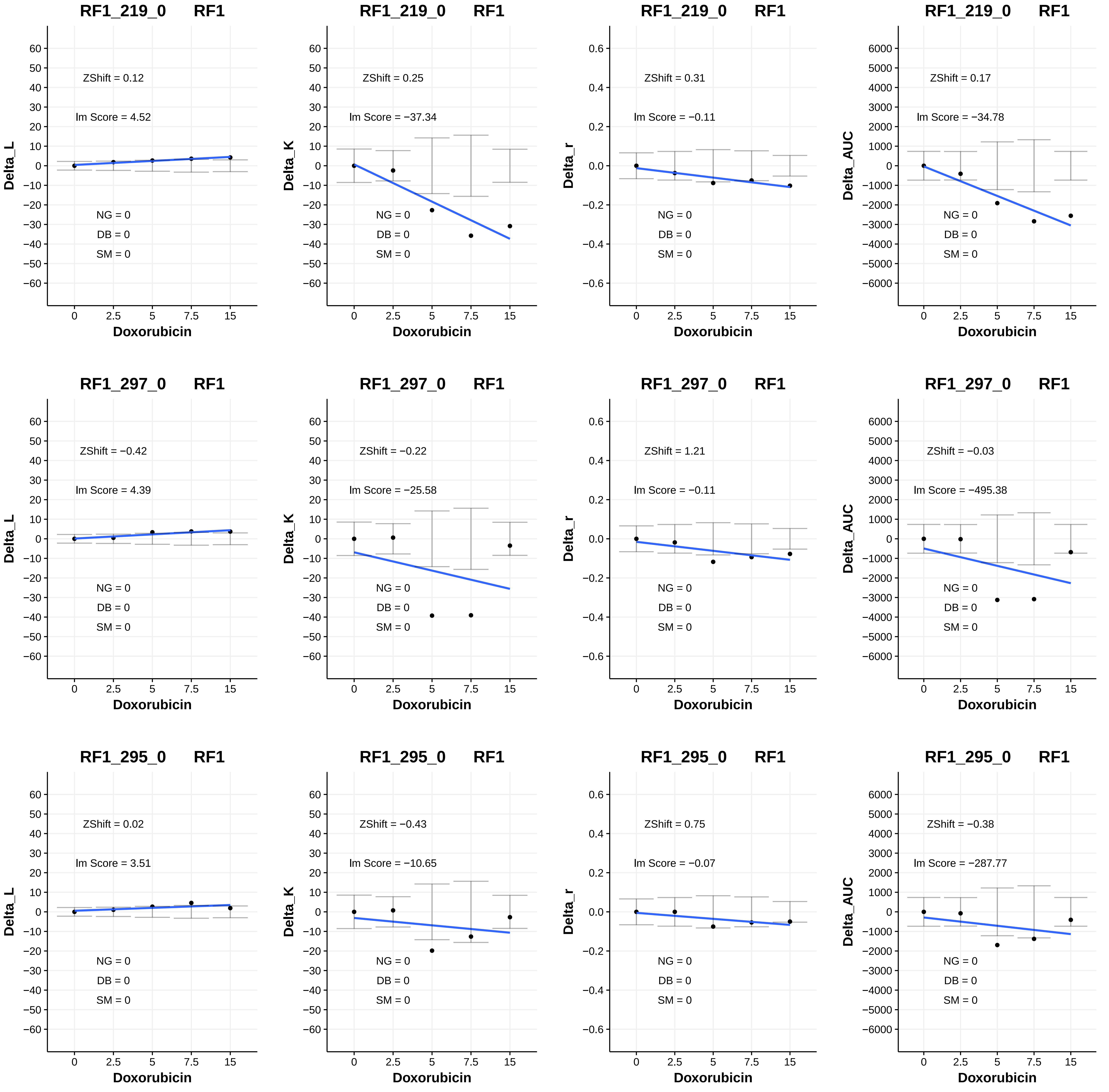
RF1_219_0 RF1
ZShift = 0.12
Im Score = 4.52
NG = 0
DB = 0
SM = 0
Delta_L
Doxorubicin
RF1_219_0 RF1
ZShift = 0.25
Im Score = −37.34
NG = 0
DB = 0
SM = 0
Delta_K
Doxorubicin
RF1_219_0 RF1
ZShift = 0.31
Im Score = −0.11
NG = 0
DB = 0
SM = 0
Delta_r
Doxorubicin
RF1_219_0 RF1
ZShift = 0.17
Im Score = −34.78
NG = 0
DB = 0
SM = 0
Delta_AUC
Doxorubicin
RF1_297_0 RF1
ZShift = −0.42
Im Score = 4.39
NG = 0
DB = 0
SM = 0
Delta_L
Doxorubicin
RF1_297_0 RF1
ZShift = −0.22
Im Score = −25.58
NG = 0
DB = 0
SM = 0
Delta_K
Doxorubicin
RF1_297_0 RF1
ZShift = 1.21
Im Score = −0.11
NG = 0
DB = 0
SM = 0
Delta_r
Doxorubicin
RF1_297_0 RF1
ZShift = −0.03
Im Score = −495.38
NG = 0
DB = 0
SM = 0
Delta_AUC
Doxorubicin
RF1_295_0 RF1
ZShift = 0.02
Im Score = 3.51
NG = 0
DB = 0
SM = 0
Delta_L
Doxorubicin
RF1_295_0 RF1
ZShift = −0.43
Im Score = −10.65
NG = 0
DB = 0
SM = 0
Delta_K
Doxorubicin
RF1_295_0 RF1
ZShift = 0.75
Im Score = −0.07
NG = 0
DB = 0
SM = 0
Delta_r
Doxorubicin
RF1_295_0 RF1
ZShift = −0.38
Im Score = −287.77
NG = 0
DB = 0
SM = 0
Delta_AUC
Doxorubicin

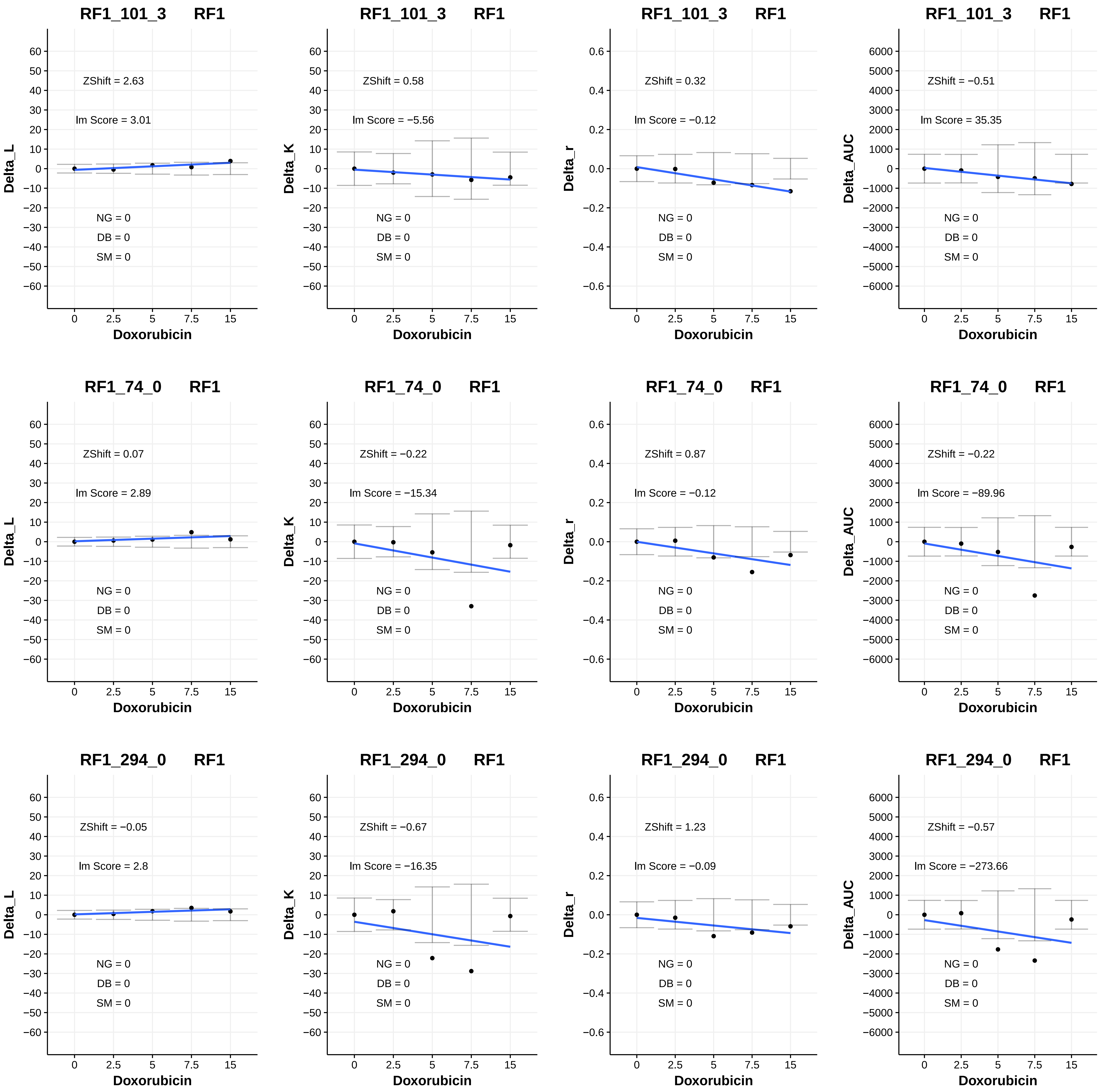
RF1_101_3 RF1
ZShift = 2.63
Im Score = 3.01
Delta_L
NG = 0
DB = 0
SM = 0
Doxorubicin
RF1_101_3 RF1
ZShift = 0.58
Im Score = −5.56
Delta_K
NG = 0
DB = 0
SM = 0
Doxorubicin
RF1_101_3 RF1
ZShift = 0.32
Im Score = −0.12
Delta_r
NG = 0
DB = 0
SM = 0
Doxorubicin
RF1_101_3 RF1
ZShift = −0.51
Im Score = 35.35
Delta_AUC
NG = 0
DB = 0
SM = 0
Doxorubicin
RF1_74_0 RF1
ZShift = 0.07
Im Score = 2.89
Delta_L
NG = 0
DB = 0
SM = 0
Doxorubicin
RF1_74_0 RF1
ZShift = −0.22
Im Score = −15.34
Delta_K
NG = 0
DB = 0
SM = 0
Doxorubicin
RF1_74_0 RF1
ZShift = 0.87
Im Score = −0.12
Delta_r
NG = 0
DB = 0
SM = 0
Doxorubicin
RF1_74_0 RF1
ZShift = −0.22
Im Score = −89.96
Delta_AUC
NG = 0
DB = 0
SM = 0
Doxorubicin
RF1_294_0 RF1
ZShift = −0.05
Im Score = 2.8
Delta_L
NG = 0
DB = 0
SM = 0
Doxorubicin
RF1_294_0 RF1
ZShift = −0.67
Im Score = −16.35
Delta_K
NG = 0
DB = 0
SM = 0
Doxorubicin
RF1_294_0 RF1
ZShift = 1.23
Im Score = −0.09
Delta_r
NG = 0
DB = 0
SM = 0
Doxorubicin
RF1_294_0 RF1
ZShift = −0.57
Im Score = −273.66
Delta_AUC
NG = 0
DB = 0
SM = 0
Doxorubicin

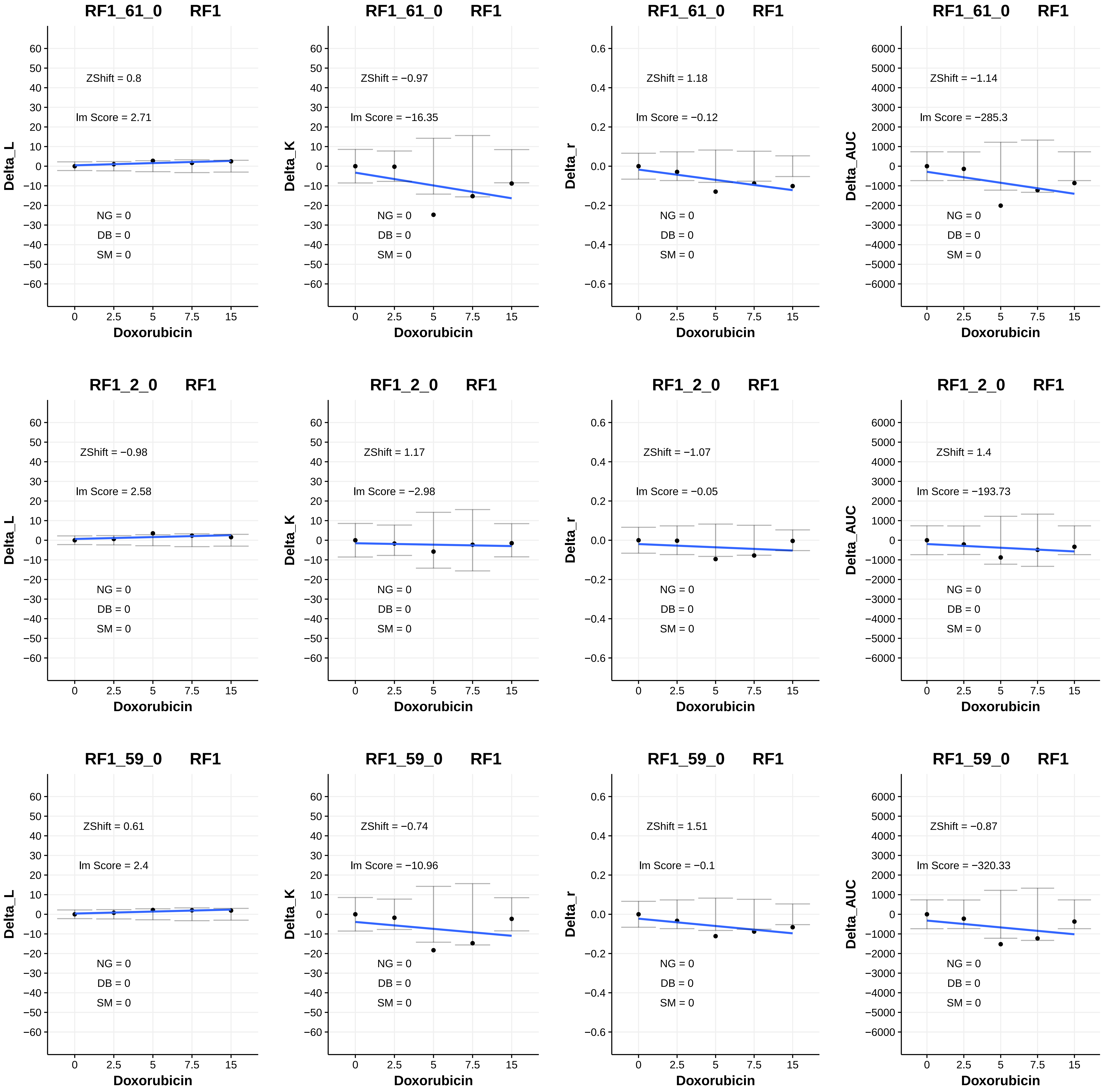
RF1_61_0 RF1
ZShift = 0.8
lm Score = 2.71
NG = 0
DB = 0
SM = 0
Delta_L
Doxorubicin
RF1_61_0 RF1
ZShift = −0.97
lm Score = −16.35
NG = 0
DB = 0
SM = 0
Delta_K
Doxorubicin
RF1_61_0 RF1
ZShift = 1.18
lm Score = −0.12
NG = 0
DB = 0
SM = 0
Delta_r
Doxorubicin
RF1_61_0 RF1
ZShift = −1.14
lm Score = −285.3
NG = 0
DB = 0
SM = 0
Delta_AUC
Doxorubicin
RF1_2_0 RF1
ZShift = −0.98
lm Score = 2.58
NG = 0
DB = 0
SM = 0
Delta_L
Doxorubicin
RF1_2_0 RF1
ZShift = 1.17
lm Score = −2.98
NG = 0
DB = 0
SM = 0
Delta_K
Doxorubicin
RF1_2_0 RF1
ZShift = −1.07
lm Score = −0.05
NG = 0
DB = 0
SM = 0
Delta_r
Doxorubicin
RF1_2_0 RF1
ZShift = 1.4
lm Score = −193.73
NG = 0
DB = 0
SM = 0
Delta_AUC
Doxorubicin
RF1_59_0 RF1
ZShift = 0.61
lm Score = 2.4
NG = 0
DB = 0
SM = 0
Delta_L
Doxorubicin
RF1_59_0 RF1
ZShift = −0.74
lm Score = −10.96
NG = 0
DB = 0
SM = 0
Delta_K
Doxorubicin
RF1_59_0 RF1
ZShift = 1.51
lm Score = −0.1
NG = 0
DB = 0
SM = 0
Delta_r
Doxorubicin
RF1_59_0 RF1
ZShift = −0.87
lm Score = −320.33
NG = 0
DB = 0
SM = 0
Delta_AUC
Doxorubicin

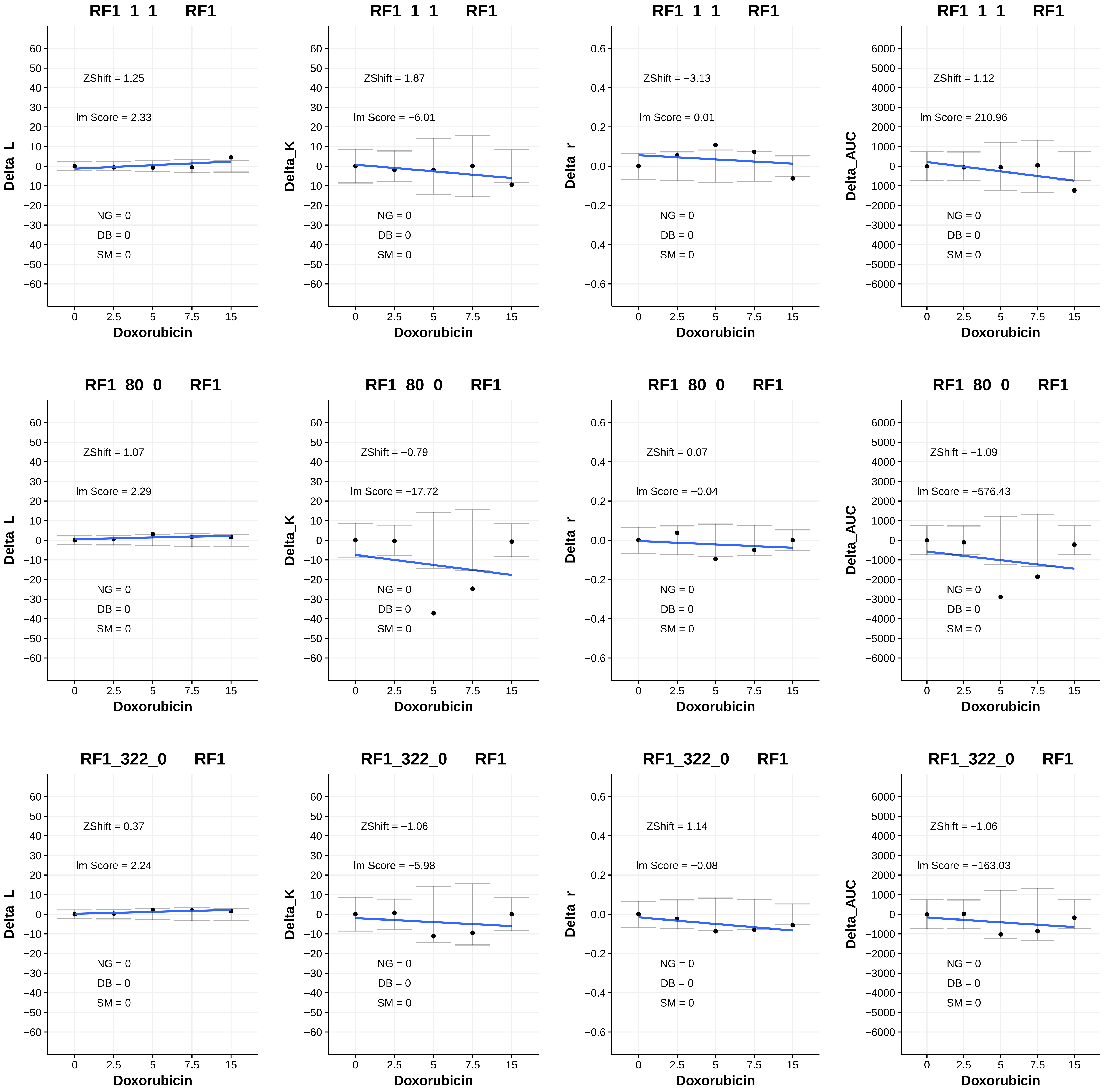
RF1_1_1 RF1
ZShift = 1.25
lm Score = 2.33
NG = 0
DB = 0
SM = 0
Delta_L
Doxorubicin
RF1_1_1 RF1
ZShift = 1.87
lm Score = −6.01
NG = 0
DB = 0
SM = 0
Delta_K
Doxorubicin
RF1_1_1 RF1
ZShift = −3.13
lm Score = 0.01
NG = 0
DB = 0
SM = 0
Delta_r
Doxorubicin
RF1_1_1 RF1
ZShift = 1.12
lm Score = 210.96
NG = 0
DB = 0
SM = 0
Delta_AUC
Doxorubicin
RF1_80_0 RF1
ZShift = 1.07
lm Score = 2.29
NG = 0
DB = 0
SM = 0
Delta_L
Doxorubicin
RF1_80_0 RF1
ZShift = −0.79
lm Score = −17.72
NG = 0
DB = 0
SM = 0
Delta_K
Doxorubicin
RF1_80_0 RF1
ZShift = 0.07
lm Score = −0.04
NG = 0
DB = 0
SM = 0
Delta_r
Doxorubicin
RF1_80_0 RF1
ZShift = −1.09
lm Score = −576.43
NG = 0
DB = 0
SM = 0
Delta_AUC
Doxorubicin
RF1_322_0 RF1
ZShift = 0.37
lm Score = 2.24
NG = 0
DB = 0
SM = 0
Delta_L
Doxorubicin
RF1_322_0 RF1
ZShift = −1.06
lm Score = −5.98
NG = 0
DB = 0
SM = 0
Delta_K
Doxorubicin
RF1_322_0 RF1
ZShift = 1.14
lm Score = −0.08
NG = 0
DB = 0
SM = 0
Delta_r
Doxorubicin
RF1_322_0 RF1
ZShift = −1.06
lm Score = −163.03
NG = 0
DB = 0
SM = 0
Delta_AUC
Doxorubicin

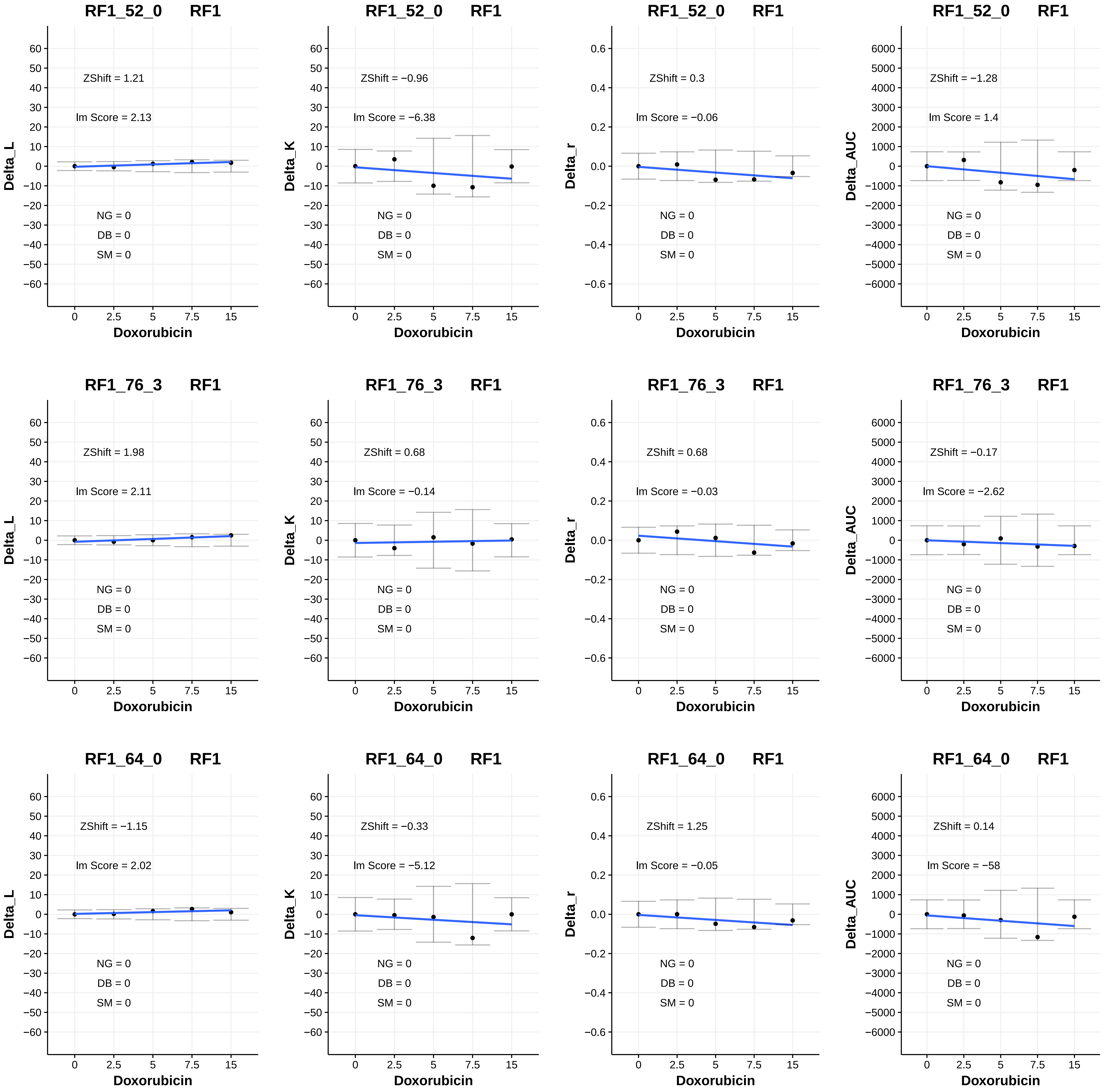
RF1_52_0 RF1
ZShift = 1.21
Im Score = 2.13
NG = 0
DB = 0
SM = 0
Delta_L
Doxorubicin
RF1_52_0 RF1
ZShift = −0.96
Im Score = −6.38
NG = 0
DB = 0
SM = 0
Delta_K
Doxorubicin
RF1_52_0 RF1
ZShift = 0.3
Im Score = −0.06
NG = 0
DB = 0
SM = 0
Delta_r
Doxorubicin
RF1_52_0 RF1
ZShift = −1.28
Im Score = 1.4
NG = 0
DB = 0
SM = 0
Delta_AUC
Doxorubicin
RF1_76_3 RF1
ZShift = 1.98
Im Score = 2.11
NG = 0
DB = 0
SM = 0
Delta_L
Doxorubicin
RF1_76_3 RF1
ZShift = 0.68
Im Score = −0.14
NG = 0
DB = 0
SM = 0
Delta_K
Doxorubicin
RF1_76_3 RF1
ZShift = 0.68
Im Score = −0.03
NG = 0
DB = 0
SM = 0
Delta_r
Doxorubicin
RF1_76_3 RF1
ZShift = −0.17
Im Score = −2.62
NG = 0
DB = 0
SM = 0
Delta_AUC
Doxorubicin
RF1_64_0 RF1
ZShift = −1.15
Im Score = 2.02
NG = 0
DB = 0
SM = 0
Delta_L
Doxorubicin
RF1_64_0 RF1
ZShift = −0.33
Im Score = −5.12
NG = 0
DB = 0
SM = 0
Delta_K
Doxorubicin
RF1_64_0 RF1
ZShift = 1.25
Im Score = −0.05
NG = 0
DB = 0
SM = 0
Delta_r
Doxorubicin
RF1_64_0 RF1
ZShift = 0.14
Im Score = −58
NG = 0
DB = 0
SM = 0
Delta_AUC
Doxorubicin

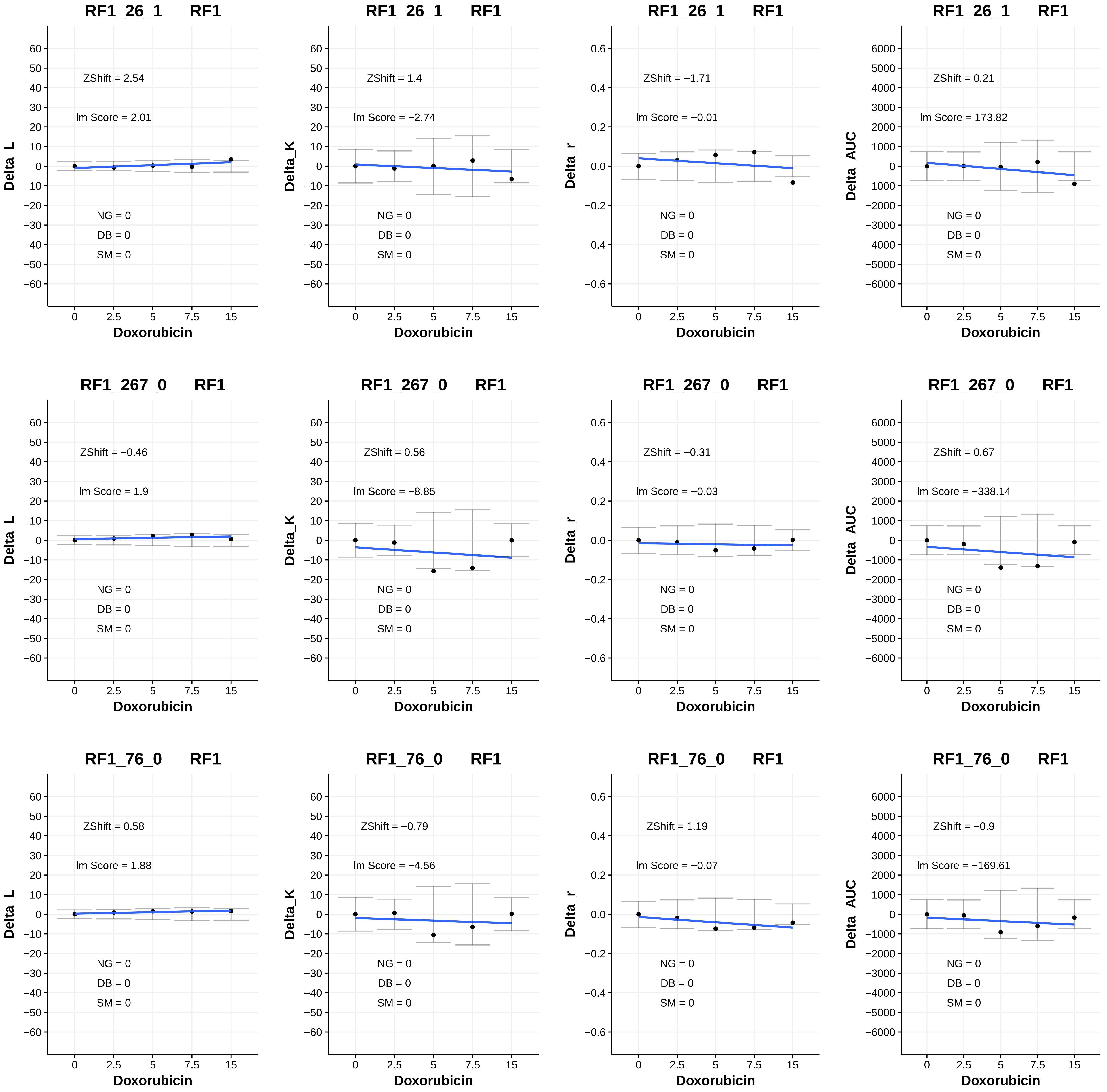
RF1_26_1 RF1
ZShift = 2.54
lm Score = 2.01
NG = 0
DB = 0
SM = 0
Delta_L
Doxorubicin
RF1_26_1 RF1
ZShift = 1.4
lm Score = −2.74
NG = 0
DB = 0
SM = 0
Delta_K
Doxorubicin
RF1_26_1 RF1
ZShift = −1.71
lm Score = −0.01
NG = 0
DB = 0
SM = 0
Delta_r
Doxorubicin
RF1_26_1 RF1
ZShift = 0.21
lm Score = 173.82
NG = 0
DB = 0
SM = 0
Delta_AUC
Doxorubicin
RF1_267_0 RF1
ZShift = −0.46
lm Score = 1.9
NG = 0
DB = 0
SM = 0
Delta_L
Doxorubicin
RF1_267_0 RF1
ZShift = 0.56
lm Score = −8.85
NG = 0
DB = 0
SM = 0
Delta_K
Doxorubicin
RF1_267_0 RF1
ZShift = −0.31
lm Score = −0.03
NG = 0
DB = 0
SM = 0
Delta_r
Doxorubicin
RF1_267_0 RF1
ZShift = 0.67
lm Score = −338.14
NG = 0
DB = 0
SM = 0
Delta_AUC
Doxorubicin
RF1_76_0 RF1
ZShift = 0.58
lm Score = 1.88
NG = 0
DB = 0
SM = 0
Delta_L
Doxorubicin
RF1_76_0 RF1
ZShift = −0.79
lm Score = −4.56
NG = 0
DB = 0
SM = 0
Delta_K
Doxorubicin
RF1_76_0 RF1
ZShift = 1.19
lm Score = −0.07
NG = 0
DB = 0
SM = 0
Delta_r
Doxorubicin
RF1_76_0 RF1
ZShift = −0.9
lm Score = −169.61
NG = 0
DB = 0
SM = 0
Delta_AUC
Doxorubicin

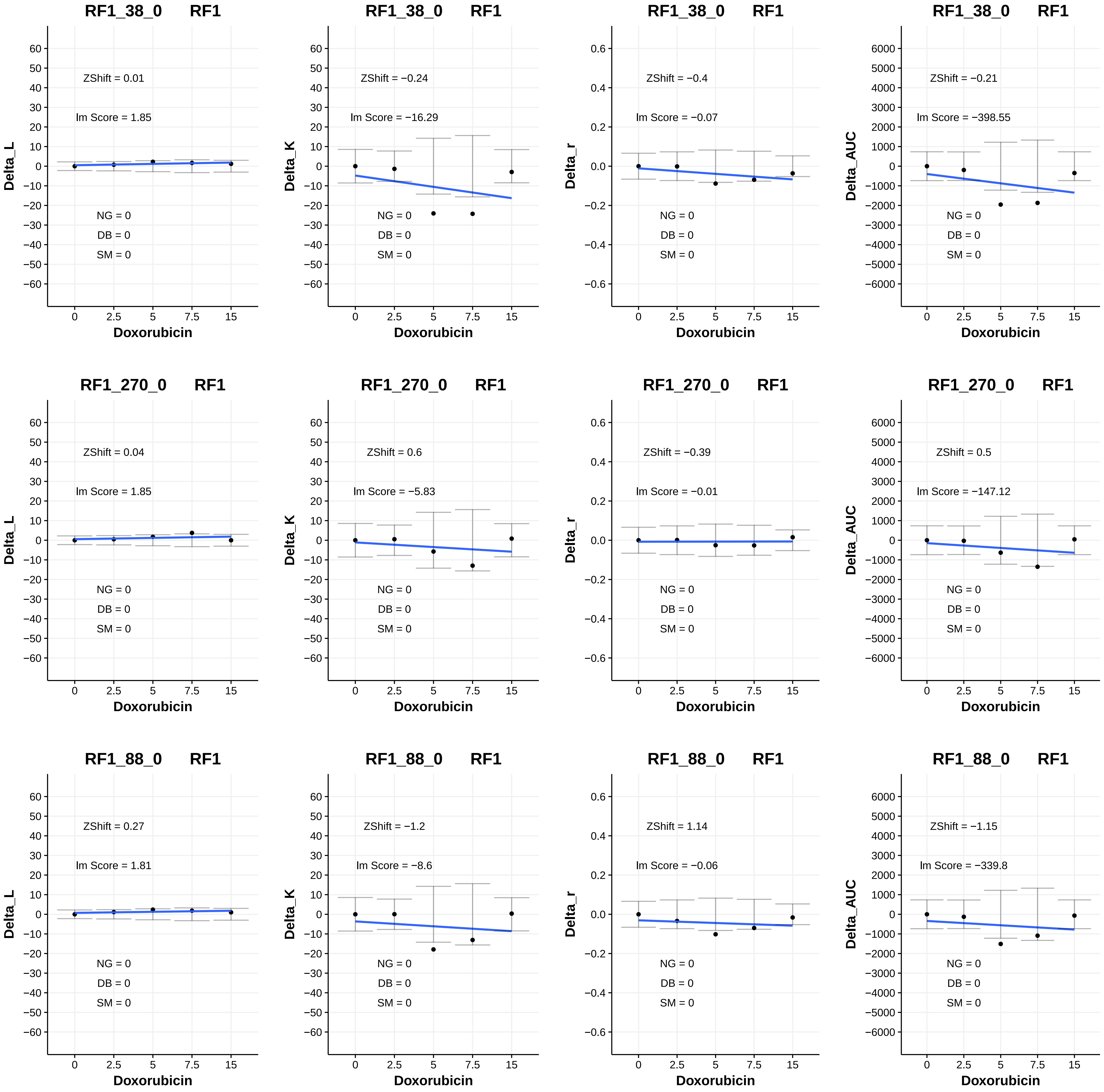
RF1_38_0 RF1
ZShift = 0.01
Im Score = 1.85
NG = 0
DB = 0
SM = 0
Delta_L
Doxorubicin
RF1_38_0 RF1
ZShift = −0.24
Im Score = −16.29
NG = 0
DB = 0
SM = 0
Delta_K
Doxorubicin
RF1_38_0 RF1
ZShift = −0.4
Im Score = −0.07
NG = 0
DB = 0
SM = 0
Delta_r
Doxorubicin
RF1_38_0 RF1
ZShift = −0.21
Im Score = −398.55
NG = 0
DB = 0
SM = 0
Delta_AUC
Doxorubicin
RF1_270_0 RF1
ZShift = 0.04
Im Score = 1.85
NG = 0
DB = 0
SM = 0
Delta_L
Doxorubicin
RF1_270_0 RF1
ZShift = 0.6
Im Score = −5.83
NG = 0
DB = 0
SM = 0
Delta_K
Doxorubicin
RF1_270_0 RF1
ZShift = −0.39
Im Score = −0.01
NG = 0
DB = 0
SM = 0
Delta_r
Doxorubicin
RF1_270_0 RF1
ZShift = 0.5
Im Score = −147.12
NG = 0
DB = 0
SM = 0
Delta_AUC
Doxorubicin
RF1_88_0 RF1
ZShift = 0.27
Im Score = 1.81
NG = 0
DB = 0
SM = 0
Delta_L
Doxorubicin
RF1_88_0 RF1
ZShift = −1.2
Im Score = −8.6
NG = 0
DB = 0
SM = 0
Delta_K
Doxorubicin
RF1_88_0 RF1
ZShift = 1.14
Im Score = −0.06
NG = 0
DB = 0
SM = 0
Delta_r
Doxorubicin
RF1_88_0 RF1
ZShift = −1.15
Im Score = −339.8
NG = 0
DB = 0
SM = 0
Delta_AUC
Doxorubicin

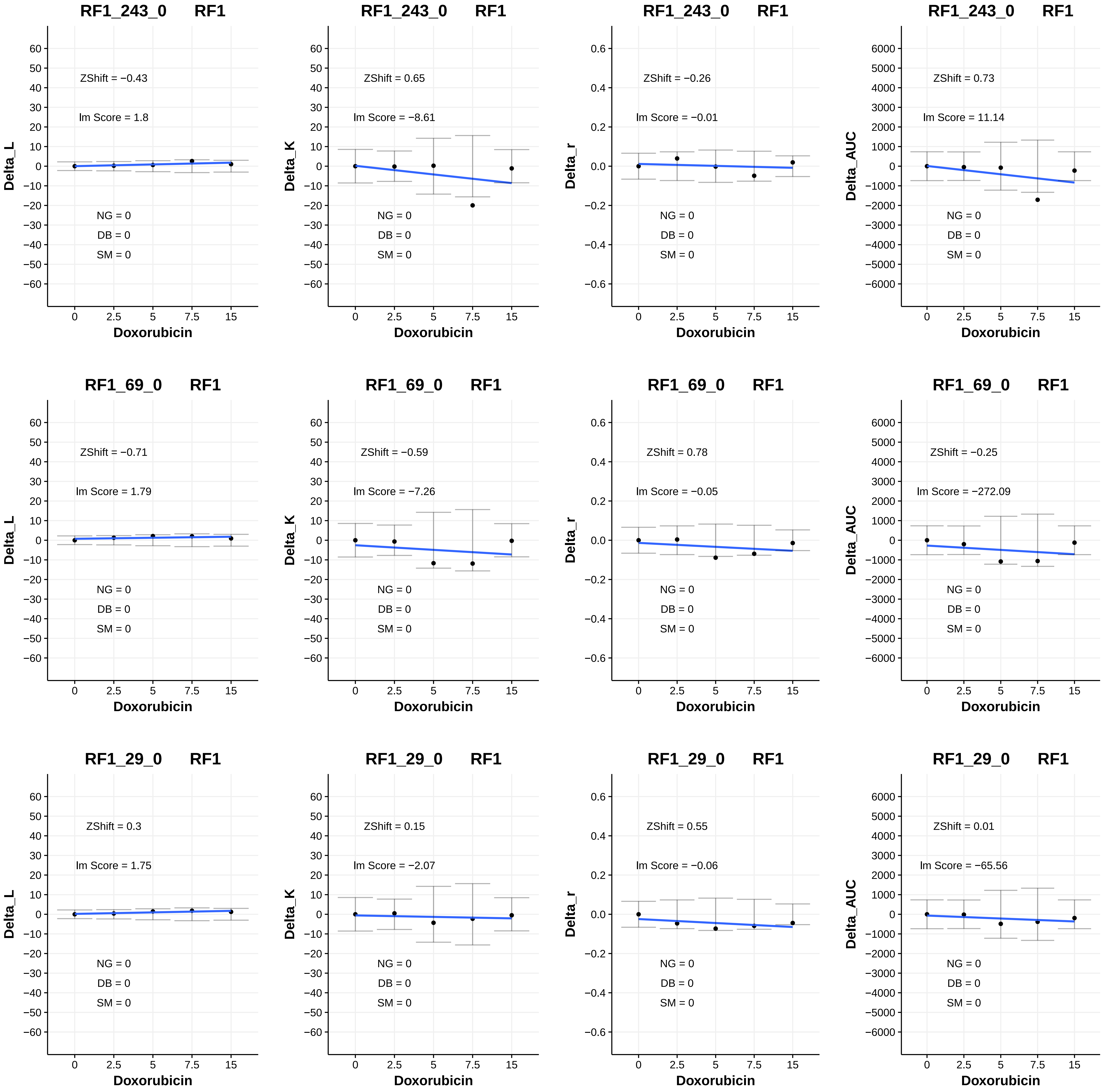

RF1_243_0 RF1
ZShift = −0.43
lm Score = 1.8
NG = 0
DB = 0
SM = 0
Delta_L
Doxorubicin
RF1_243_0 RF1
ZShift = 0.65
lm Score = −8.61
NG = 0
DB = 0
SM = 0
Delta_K
Doxorubicin
RF1_243_0 RF1
ZShift = −0.26
lm Score = −0.01
NG = 0
DB = 0
SM = 0
Delta_r
Doxorubicin
RF1_243_0 RF1
ZShift = 0.73
lm Score = 11.14
NG = 0
DB = 0
SM = 0
Delta_AUC
Doxorubicin
RF1_69_0 RF1
ZShift = −0.71
lm Score = 1.79
NG = 0
DB = 0
SM = 0
Delta_L
Doxorubicin
RF1_69_0 RF1
ZShift = −0.59
lm Score = −7.26
NG = 0
DB = 0
SM = 0
Delta_K
Doxorubicin
RF1_69_0 RF1
ZShift = 0.78
lm Score = −0.05
NG = 0
DB = 0
SM = 0
Delta_r
Doxorubicin
RF1_69_0 RF1
ZShift = −0.25
lm Score = −272.09
NG = 0
DB = 0
SM = 0
Delta_AUC
Doxorubicin
RF1_29_0 RF1
ZShift = 0.3
lm Score = 1.75
NG = 0
DB = 0
SM = 0
Delta_L
Doxorubicin
RF1_29_0 RF1
ZShift = 0.15
lm Score = −2.07
NG = 0
DB = 0
SM = 0
Delta_K
Doxorubicin
RF1_29_0 RF1
ZShift = 0.55
lm Score = −0.06
NG = 0
DB = 0
SM = 0
Delta_r
Doxorubicin
RF1_29_0 RF1
ZShift = 0.01
lm Score = −65.56
NG = 0
DB = 0
SM = 0
Delta_AUC
Doxorubicin

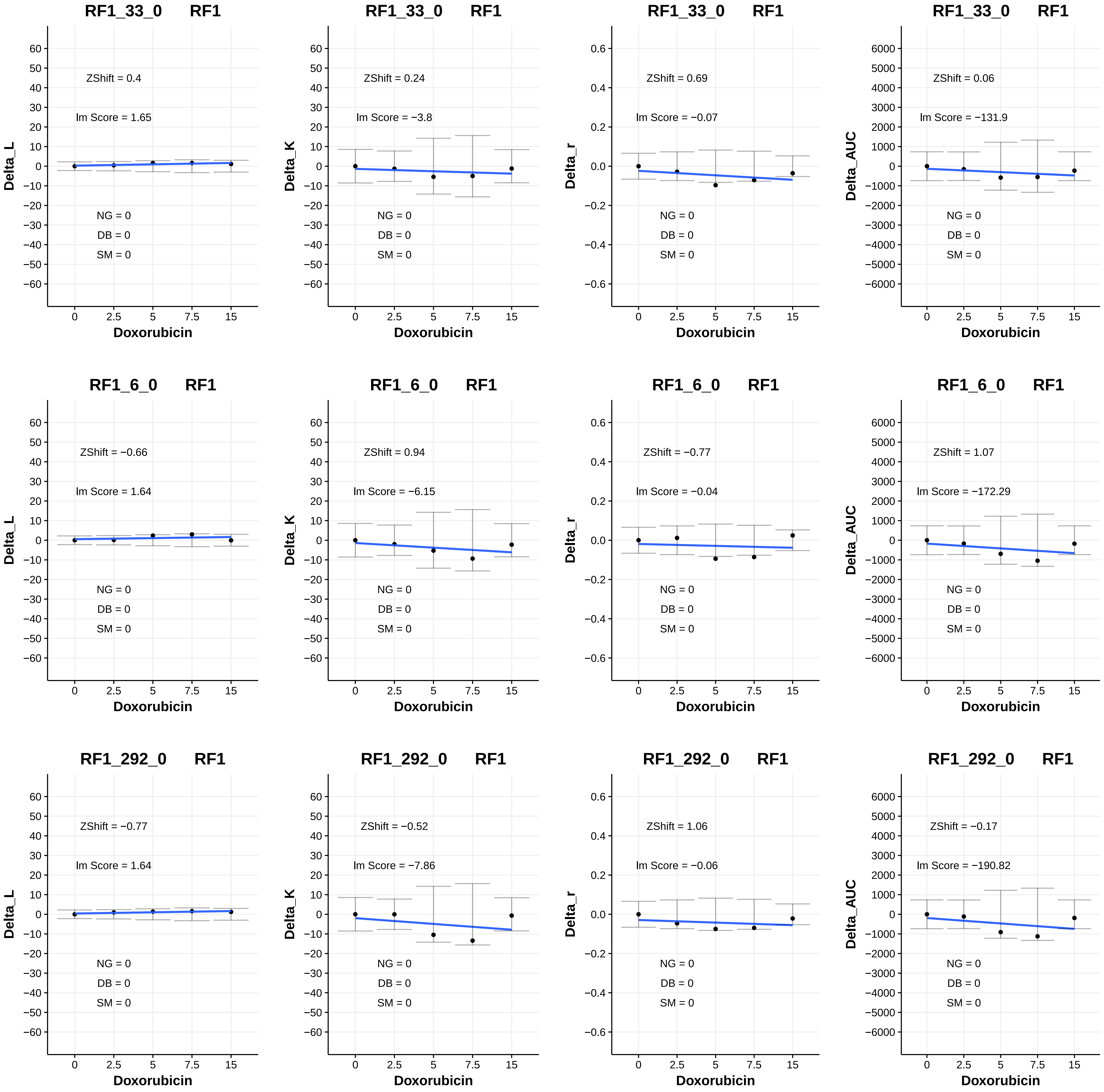

RF1_33_0 RF1
ZShift = 0.4
Im Score = 1.65
NG = 0
DB = 0
SM = 0
Delta_L
Doxorubicin
RF1_33_0 RF1
ZShift = 0.24
Im Score = −3.8
NG = 0
DB = 0
SM = 0
Delta_K
Doxorubicin
RF1_33_0 RF1
ZShift = 0.69
Im Score = −0.07
NG = 0
DB = 0
SM = 0
Delta_r
Doxorubicin
RF1_33_0 RF1
ZShift = 0.06
Im Score = −131.9
NG = 0
DB = 0
SM = 0
Delta_AUC
Doxorubicin
RF1_6_0 RF1
ZShift = −0.66
Im Score = 1.64
NG = 0
DB = 0
SM = 0
Delta_L
Doxorubicin
RF1_6_0 RF1
ZShift = 0.94
Im Score = −6.15
NG = 0
DB = 0
SM = 0
Delta_K
Doxorubicin
RF1_6_0 RF1
ZShift = −0.77
Im Score = −0.04
NG = 0
DB = 0
SM = 0
Delta_r
Doxorubicin
RF1_6_0 RF1
ZShift = 1.07
Im Score = −172.29
NG = 0
DB = 0
SM = 0
Delta_AUC
Doxorubicin
RF1_292_0 RF1
ZShift = −0.77
Im Score = 1.64
NG = 0
DB = 0
SM = 0
Delta_L
Doxorubicin
RF1_292_0 RF1
ZShift = −0.52
Im Score = −7.86
NG = 0
DB = 0
SM = 0
Delta_K
Doxorubicin
RF1_292_0 RF1
ZShift = 1.06
Im Score = −0.06
NG = 0
DB = 0
SM = 0
Delta_r
Doxorubicin
RF1_292_0 RF1
ZShift = −0.17
Im Score = −190.82
NG = 0
DB = 0
SM = 0
Delta_AUC
Doxorubicin

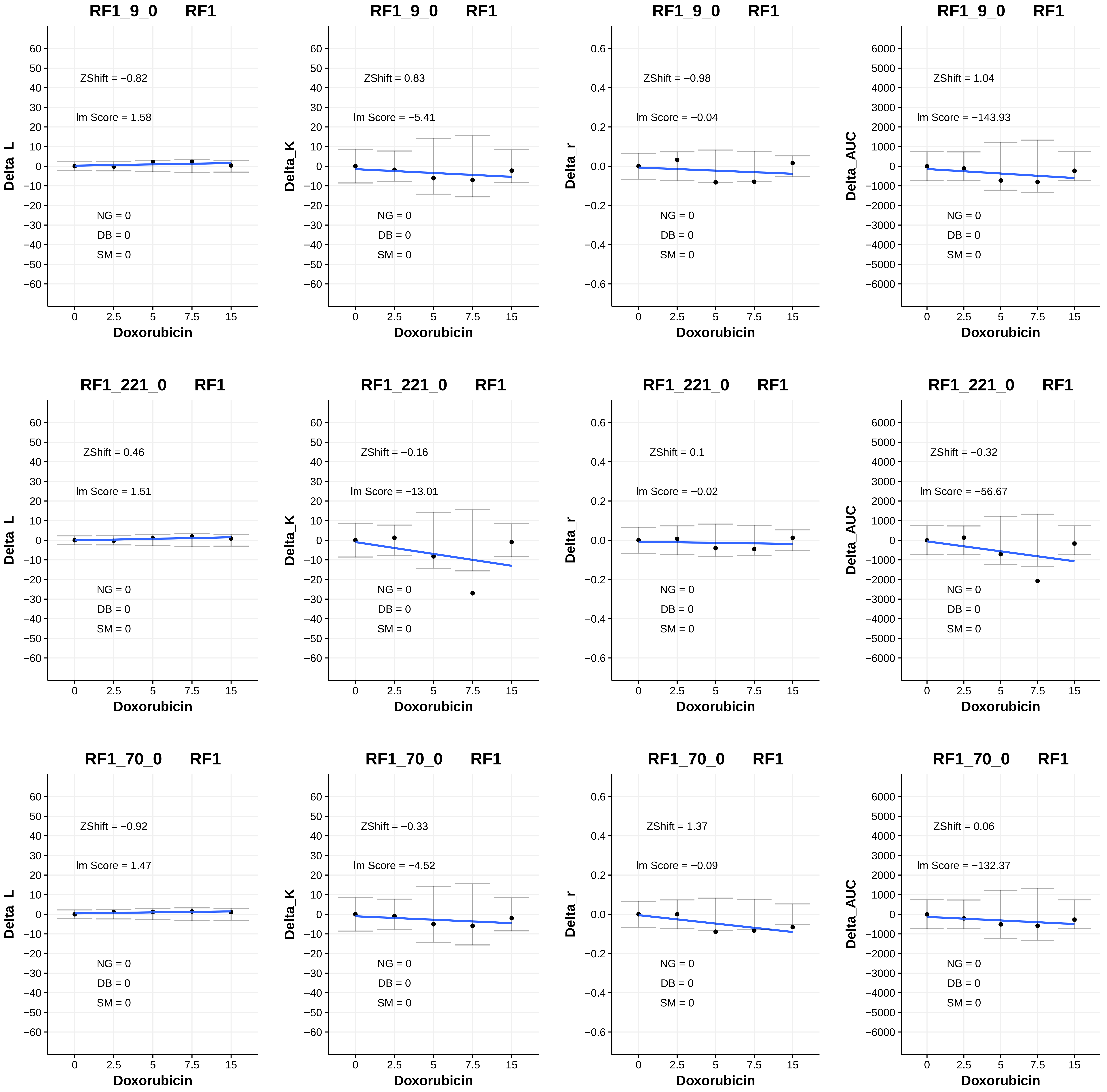
RF1_9_0 RF1
ZShift = −0.82
lm Score = 1.58
NG = 0
DB = 0
SM = 0
Delta_L
Doxorubicin
RF1_9_0 RF1
ZShift = 0.83
lm Score = −5.41
NG = 0
DB = 0
SM = 0
Delta_K
Doxorubicin
RF1_9_0 RF1
ZShift = −0.98
lm Score = −0.04
NG = 0
DB = 0
SM = 0
Delta_r
Doxorubicin
RF1_9_0 RF1
ZShift = 1.04
lm Score = −143.93
NG = 0
DB = 0
SM = 0
Delta_AUC
Doxorubicin
RF1_221_0 RF1
ZShift = 0.46
lm Score = 1.51
NG = 0
DB = 0
SM = 0
Delta_L
Doxorubicin
RF1_221_0 RF1
ZShift = −0.16
lm Score = −13.01
NG = 0
DB = 0
SM = 0
Delta_K
Doxorubicin
RF1_221_0 RF1
ZShift = 0.1
lm Score = −0.02
NG = 0
DB = 0
SM = 0
Delta_r
Doxorubicin
RF1_221_0 RF1
ZShift = −0.32
lm Score = −56.67
NG = 0
DB = 0
SM = 0
Delta_AUC
Doxorubicin
RF1_70_0 RF1
ZShift = −0.92
lm Score = 1.47
NG = 0
DB = 0
SM = 0
Delta_L
Doxorubicin
RF1_70_0 RF1
ZShift = −0.33
lm Score = −4.52
NG = 0
DB = 0
SM = 0
Delta_K
Doxorubicin
RF1_70_0 RF1
ZShift = 1.37
lm Score = −0.09
NG = 0
DB = 0
SM = 0
Delta_r
Doxorubicin
RF1_70_0 RF1
ZShift = 0.06
lm Score = −132.37
NG = 0
DB = 0
SM = 0
Delta_AUC
Doxorubicin

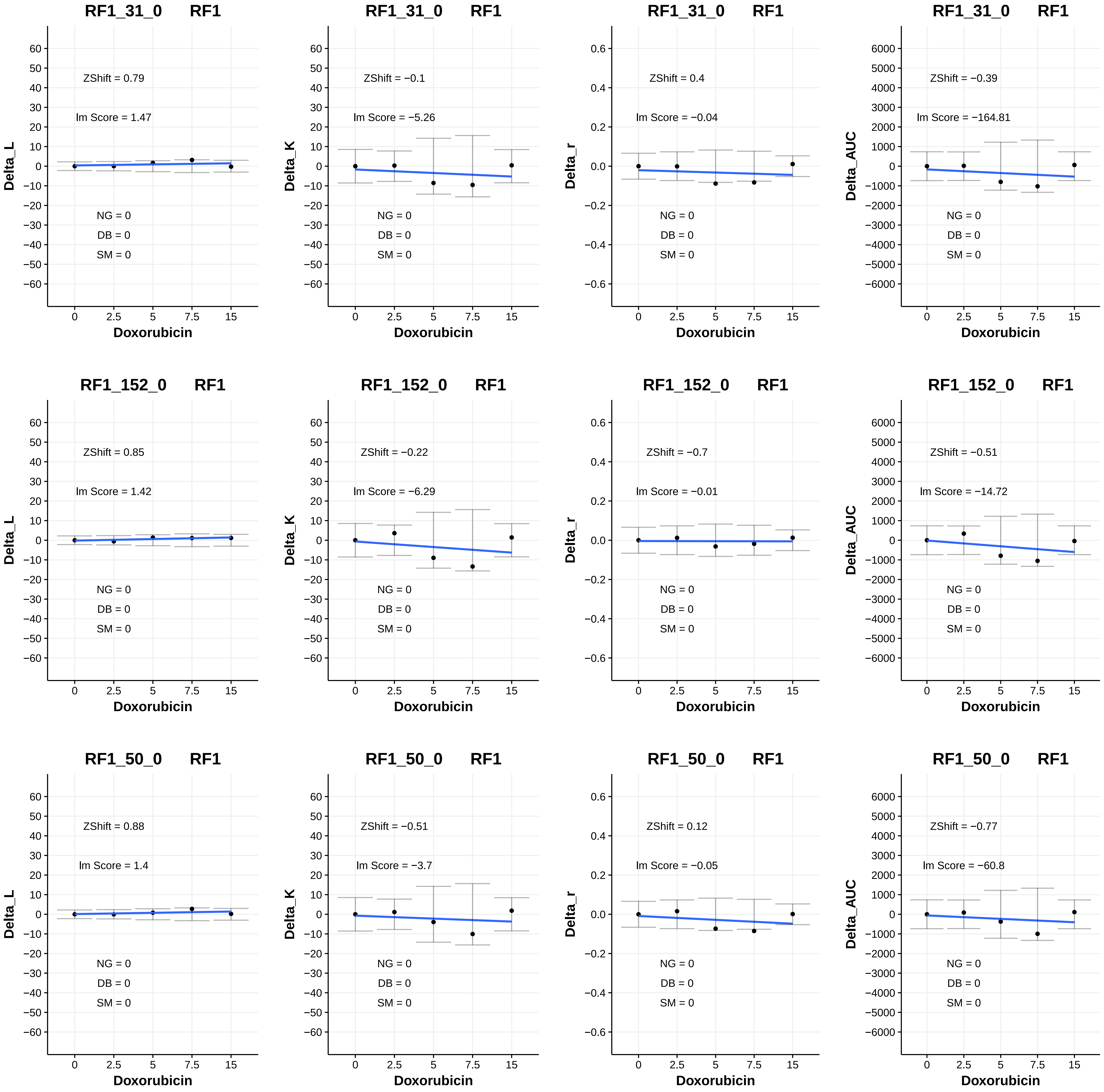
RF1_31_0 RF1
ZShift = 0.79
lm Score = 1.47
NG = 0
DB = 0
SM = 0
Delta_L
Doxorubicin
RF1_31_0 RF1
ZShift = −0.1
lm Score = −5.26
NG = 0
DB = 0
SM = 0
Delta_K
Doxorubicin
RF1_31_0 RF1
ZShift = 0.4
lm Score = −0.04
NG = 0
DB = 0
SM = 0
Delta_r
Doxorubicin
RF1_31_0 RF1
ZShift = −0.39
lm Score = −164.81
NG = 0
DB = 0
SM = 0
Delta_AUC
Doxorubicin
RF1_152_0 RF1
ZShift = 0.85
lm Score = 1.42
NG = 0
DB = 0
SM = 0
Delta_L
Doxorubicin
RF1_152_0 RF1
ZShift = −0.22
lm Score = −6.29
NG = 0
DB = 0
SM = 0
Delta_K
Doxorubicin
RF1_152_0 RF1
ZShift = −0.7
lm Score = −0.01
NG = 0
DB = 0
SM = 0
Delta_r
Doxorubicin
RF1_152_0 RF1
ZShift = −0.51
lm Score = −14.72
NG = 0
DB = 0
SM = 0
Delta_AUC
Doxorubicin
RF1_50_0 RF1
ZShift = 0.88
lm Score = 1.4
NG = 0
DB = 0
SM = 0
Delta_L
Doxorubicin
RF1_50_0 RF1
ZShift = −0.51
lm Score = −3.7
NG = 0
DB = 0
SM = 0
Delta_K
Doxorubicin
RF1_50_0 RF1
ZShift = 0.12
lm Score = −0.05
NG = 0
DB = 0
SM = 0
Delta_r
Doxorubicin
RF1_50_0 RF1
ZShift = −0.77
lm Score = −60.8
NG = 0
DB = 0
SM = 0
Delta_AUC
Doxorubicin

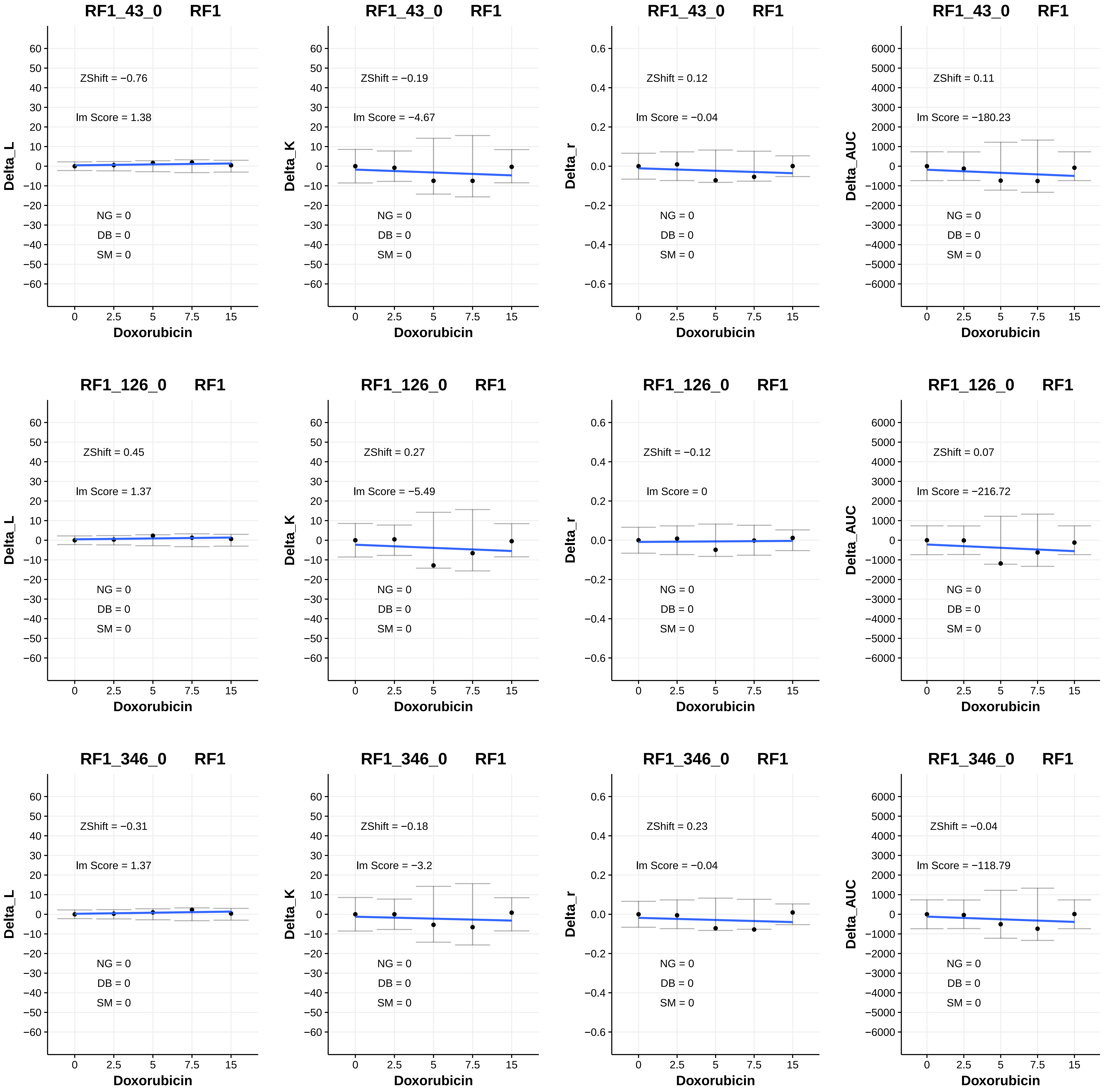
RF1_43_0 RF1
ZShift = −0.76
lm Score = 1.38
NG = 0
DB = 0
SM = 0
Delta_L
Doxorubicin
RF1_43_0 RF1
ZShift = −0.19
lm Score = −4.67
NG = 0
DB = 0
SM = 0
Delta_K
Doxorubicin
RF1_43_0 RF1
ZShift = 0.12
lm Score = −0.04
NG = 0
DB = 0
SM = 0
Delta_r
Doxorubicin
RF1_43_0 RF1
ZShift = 0.11
lm Score = −180.23
NG = 0
DB = 0
SM = 0
Delta_AUC
Doxorubicin
RF1_126_0 RF1
ZShift = 0.45
lm Score = 1.37
NG = 0
DB = 0
SM = 0
Delta_L
Doxorubicin
RF1_126_0 RF1
ZShift = 0.27
lm Score = −5.49
NG = 0
DB = 0
SM = 0
Delta_K
Doxorubicin
RF1_126_0 RF1
ZShift = −0.12
lm Score = 0
NG = 0
DB = 0
SM = 0
Delta_r
Doxorubicin
RF1_126_0 RF1
ZShift = 0.07
lm Score = −216.72
NG = 0
DB = 0
SM = 0
Delta_AUC
Doxorubicin
RF1_346_0 RF1
ZShift = −0.31
lm Score = 1.37
NG = 0
DB = 0
SM = 0
Delta_L
Doxorubicin
RF1_346_0 RF1
ZShift = −0.18
lm Score = −3.2
NG = 0
DB = 0
SM = 0
Delta_K
Doxorubicin
RF1_346_0 RF1
ZShift = 0.23
lm Score = −0.04
NG = 0
DB = 0
SM = 0
Delta_r
Doxorubicin
RF1_346_0 RF1
ZShift = −0.04
lm Score = −118.79
NG = 0
DB = 0
SM = 0
Delta_AUC
Doxorubicin

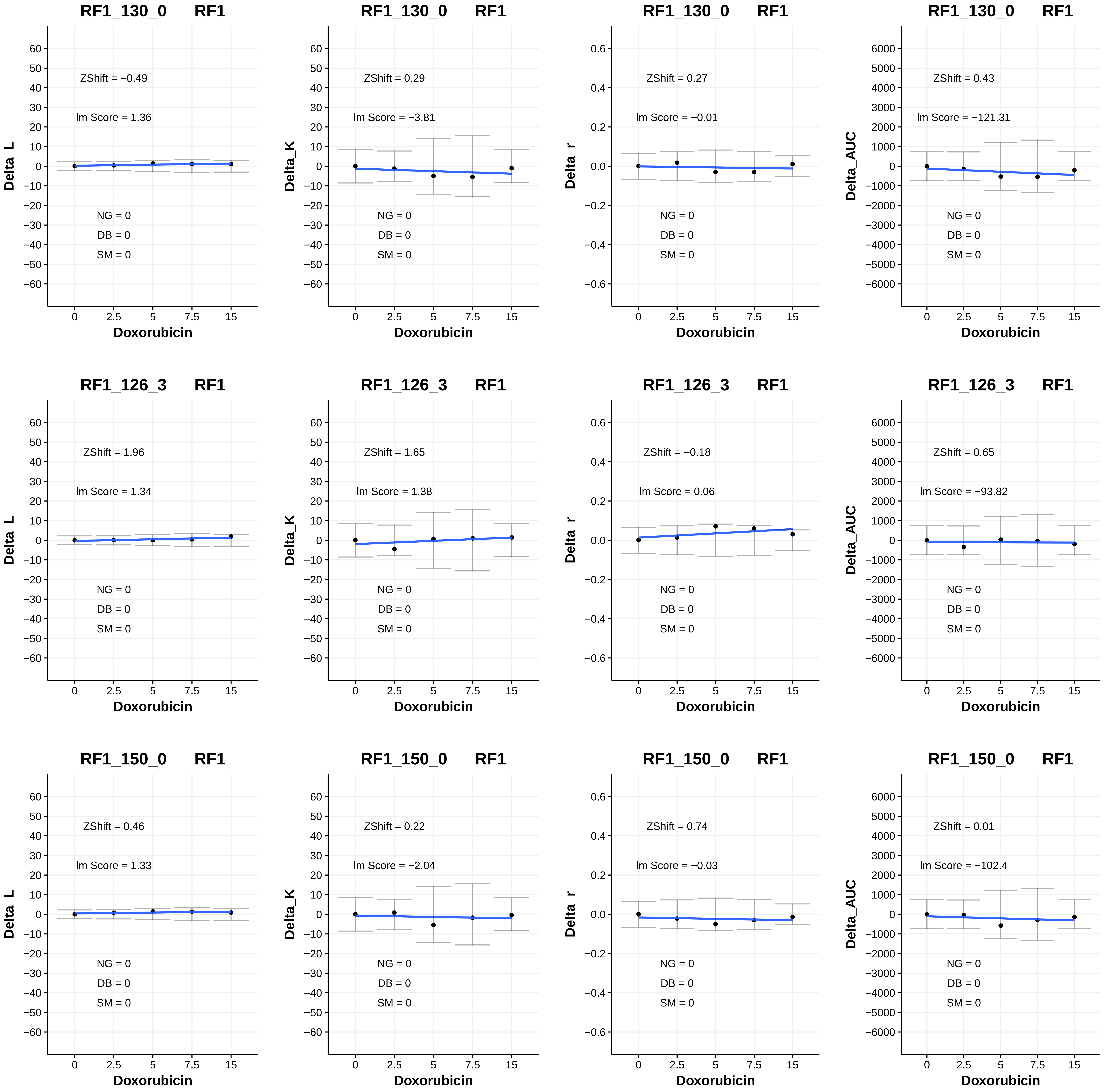
RF1_130_0 RF1
ZShift = −0.49
lm Score = 1.36
NG = 0
DB = 0
SM = 0
Delta_L
Doxorubicin
RF1_130_0 RF1
ZShift = 0.29
lm Score = −3.81
NG = 0
DB = 0
SM = 0
Delta_K
Doxorubicin
RF1_130_0 RF1
ZShift = 0.27
lm Score = −0.01
NG = 0
DB = 0
SM = 0
Delta_r
Doxorubicin
RF1_130_0 RF1
ZShift = 0.43
lm Score = −121.31
NG = 0
DB = 0
SM = 0
Delta_AUC
Doxorubicin
RF1_126_3 RF1
ZShift = 1.96
lm Score = 1.34
NG = 0
DB = 0
SM = 0
Delta_L
Doxorubicin
RF1_126_3 RF1
ZShift = 1.65
lm Score = 1.38
NG = 0
DB = 0
SM = 0
Delta_K
Doxorubicin
RF1_126_3 RF1
ZShift = −0.18
lm Score = 0.06
NG = 0
DB = 0
SM = 0
Delta_r
Doxorubicin
RF1_126_3 RF1
ZShift = 0.65
lm Score = −93.82
NG = 0
DB = 0
SM = 0
Delta_AUC
Doxorubicin
RF1_150_0 RF1
ZShift = 0.46
lm Score = 1.33
NG = 0
DB = 0
SM = 0
Delta_L
Doxorubicin
RF1_150_0 RF1
ZShift = 0.22
lm Score = −2.04
NG = 0
DB = 0
SM = 0
Delta_K
Doxorubicin
RF1_150_0 RF1
ZShift = 0.74
lm Score = −0.03
NG = 0
DB = 0
SM = 0
Delta_r
Doxorubicin
RF1_150_0 RF1
ZShift = 0.01
lm Score = −102.4
NG = 0
DB = 0
SM = 0
Delta_AUC
Doxorubicin

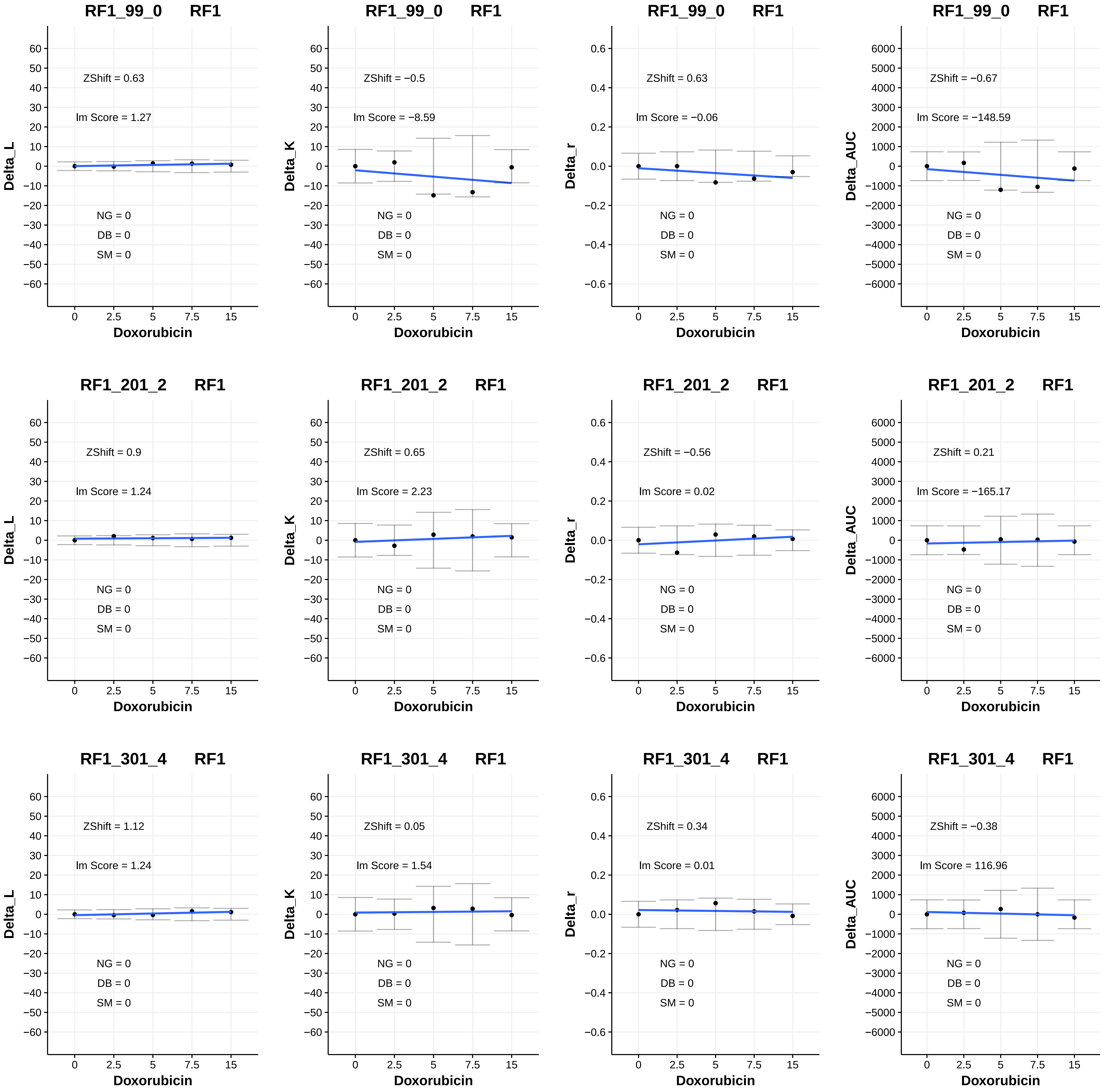
RF1_99_0 RF1
ZShift = 0.63
lm Score = 1.27
NG = 0
DB = 0
SM = 0
Delta_L
Doxorubicin
RF1_99_0 RF1
ZShift = −0.5
lm Score = −8.59
NG = 0
DB = 0
SM = 0
Delta_K
Doxorubicin
RF1_99_0 RF1
ZShift = 0.63
lm Score = −0.06
NG = 0
DB = 0
SM = 0
Delta_r
Doxorubicin
RF1_99_0 RF1
ZShift = −0.67
lm Score = −148.59
NG = 0
DB = 0
SM = 0
Delta_AUC
Doxorubicin
RF1_201_2 RF1
ZShift = 0.9
lm Score = 1.24
NG = 0
DB = 0
SM = 0
Delta_L
Doxorubicin
RF1_201_2 RF1
ZShift = 0.65
lm Score = 2.23
NG = 0
DB = 0
SM = 0
Delta_K
Doxorubicin
RF1_201_2 RF1
ZShift = −0.56
lm Score = 0.02
NG = 0
DB = 0
SM = 0
Delta_r
Doxorubicin
RF1_201_2 RF1
ZShift = 0.21
lm Score = −165.17
NG = 0
DB = 0
SM = 0
Delta_AUC
Doxorubicin
RF1_301_4 RF1
ZShift = 1.12
lm Score = 1.24
NG = 0
DB = 0
SM = 0
Delta_L
Doxorubicin
RF1_301_4 RF1
ZShift = 0.05
lm Score = 1.54
NG = 0
DB = 0
SM = 0
Delta_K
Doxorubicin
RF1_301_4 RF1
ZShift = 0.34
lm Score = 0.01
NG = 0
DB = 0
SM = 0
Delta_r
Doxorubicin
RF1_301_4 RF1
ZShift = −0.38
lm Score = 116.96
NG = 0
DB = 0
SM = 0
Delta_AUC
Doxorubicin

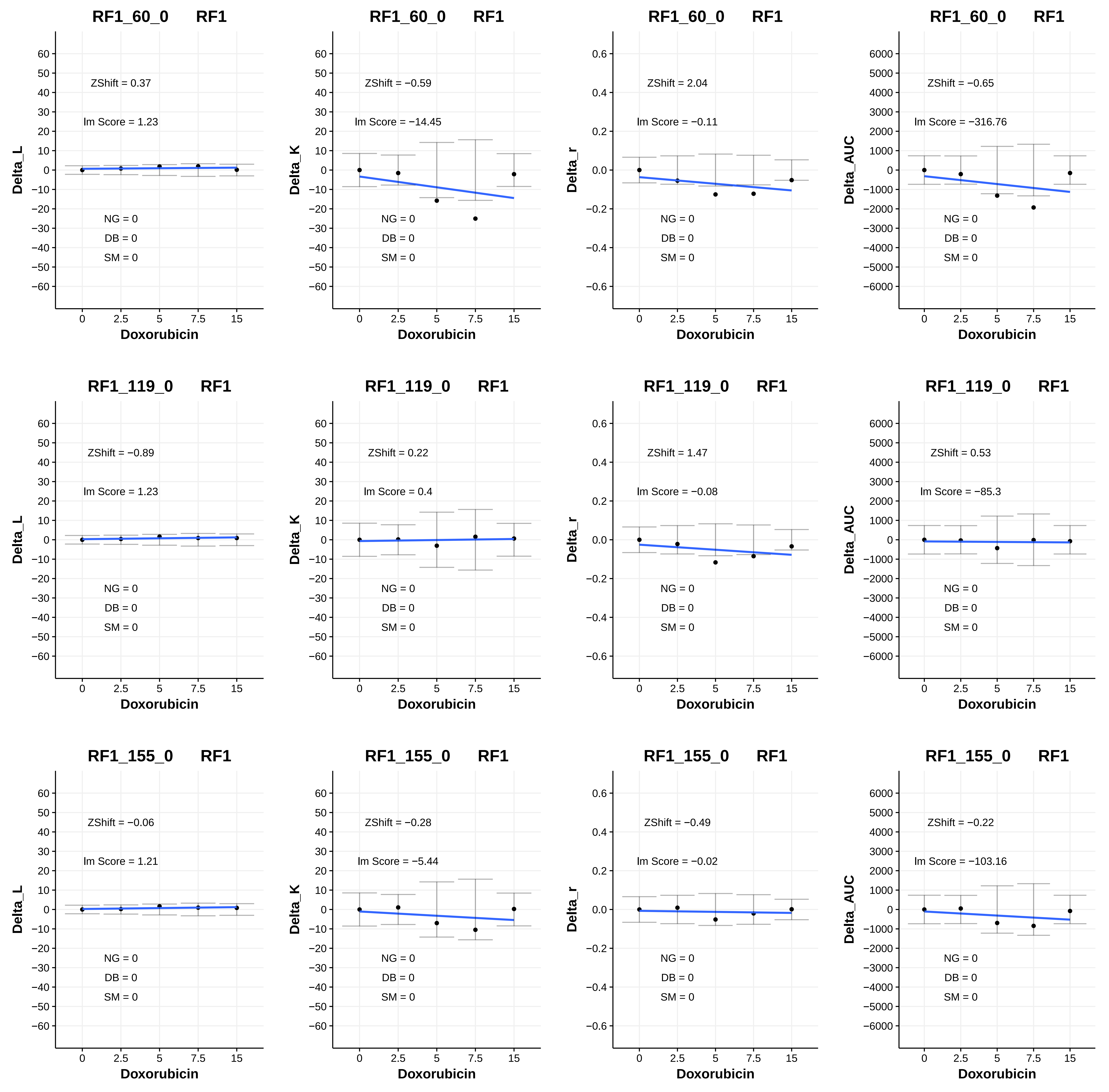
RF1_60_0 RF1
ZShift = 0.37
Im Score = 1.23
NG = 0
DB = 0
SM = 0
Delta_L
Doxorubicin
RF1_60_0 RF1
ZShift = −0.59
Im Score = −14.45
NG = 0
DB = 0
SM = 0
Delta_K
Doxorubicin
RF1_60_0 RF1
ZShift = 2.04
Im Score = −0.11
NG = 0
DB = 0
SM = 0
Delta_r
Doxorubicin
RF1_60_0 RF1
ZShift = −0.65
Im Score = −316.76
NG = 0
DB = 0
SM = 0
Delta_AUC
Doxorubicin
RF1_119_0 RF1
ZShift = −0.89
Im Score = 1.23
NG = 0
DB = 0
SM = 0
Delta_L
Doxorubicin
RF1_119_0 RF1
ZShift = 0.22
Im Score = 0.4
NG = 0
DB = 0
SM = 0
Delta_K
Doxorubicin
RF1_119_0 RF1
ZShift = 1.47
Im Score = −0.08
NG = 0
DB = 0
SM = 0
Delta_r
Doxorubicin
RF1_119_0 RF1
ZShift = 0.53
Im Score = −85.3
NG = 0
DB = 0
SM = 0
Delta_AUC
Doxorubicin
RF1_155_0 RF1
ZShift = −0.06
Im Score = 1.21
NG = 0
DB = 0
SM = 0
Delta_L
Doxorubicin
RF1_155_0 RF1
ZShift = −0.28
Im Score = −5.44
NG = 0
DB = 0
SM = 0
Delta_K
Doxorubicin
RF1_155_0 RF1
ZShift = −0.49
Im Score = −0.02
NG = 0
DB = 0
SM = 0
Delta_r
Doxorubicin
RF1_155_0 RF1
ZShift = −0.22
Im Score = −103.16
NG = 0
DB = 0
SM = 0
Delta_AUC
Doxorubicin

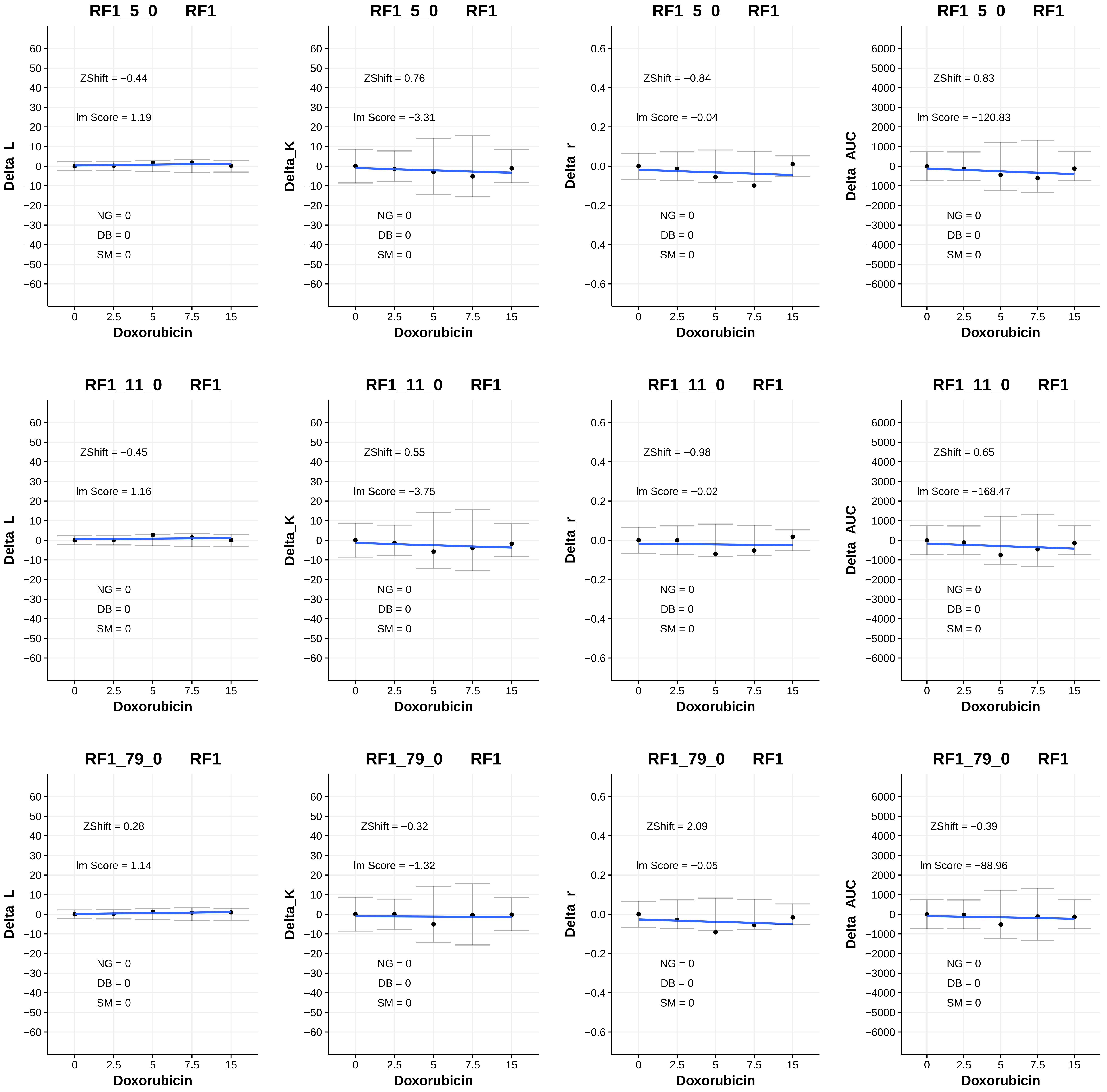

RF1_5_0 RF1
ZShift = −0.44
lm Score = 1.19
NG = 0
DB = 0
SM = 0
Delta_L
Doxorubicin
RF1_5_0 RF1
ZShift = 0.76
lm Score = −3.31
NG = 0
DB = 0
SM = 0
Delta_K
Doxorubicin
RF1_5_0 RF1
ZShift = −0.84
lm Score = −0.04
NG = 0
DB = 0
SM = 0
Delta_r
Doxorubicin
RF1_5_0 RF1
ZShift = 0.83
lm Score = −120.83
NG = 0
DB = 0
SM = 0
Delta_AUC
Doxorubicin
RF1_11_0 RF1
ZShift = −0.45
lm Score = 1.16
NG = 0
DB = 0
SM = 0
Delta_L
Doxorubicin
RF1_11_0 RF1
ZShift = 0.55
lm Score = −3.75
NG = 0
DB = 0
SM = 0
Delta_K
Doxorubicin
RF1_11_0 RF1
ZShift = −0.98
lm Score = −0.02
NG = 0
DB = 0
SM = 0
Delta_r
Doxorubicin
RF1_11_0 RF1
ZShift = 0.65
lm Score = −168.47
NG = 0
DB = 0
SM = 0
Delta_AUC
Doxorubicin
RF1_79_0 RF1
ZShift = 0.28
lm Score = 1.14
NG = 0
DB = 0
SM = 0
Delta_L
Doxorubicin
RF1_79_0 RF1
ZShift = −0.32
lm Score = −1.32
NG = 0
DB = 0
SM = 0
Delta_K
Doxorubicin
RF1_79_0 RF1
ZShift = 2.09
lm Score = −0.05
NG = 0
DB = 0
SM = 0
Delta_r
Doxorubicin
RF1_79_0 RF1
ZShift = −0.39
lm Score = −88.96
NG = 0
DB = 0
SM = 0
Delta_AUC
Doxorubicin

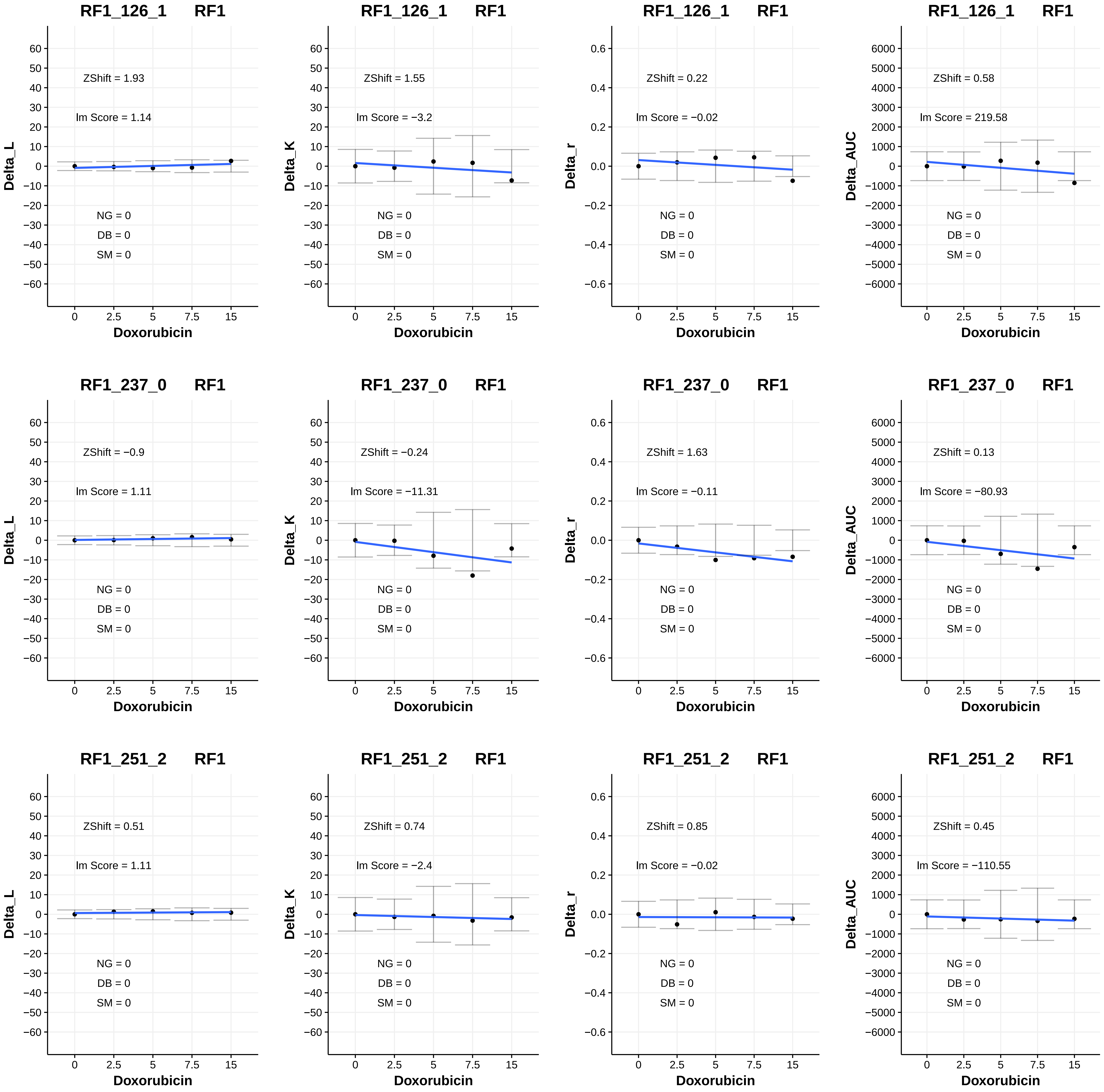
RF1_126_1 RF1
ZShift = 1.93
lm Score = 1.14
NG = 0
DB = 0
SM = 0
Delta_L
Doxorubicin
RF1_126_1 RF1
ZShift = 1.55
lm Score = −3.2
NG = 0
DB = 0
SM = 0
Delta_K
Doxorubicin
RF1_126_1 RF1
ZShift = 0.22
lm Score = −0.02
NG = 0
DB = 0
SM = 0
Delta_r
Doxorubicin
RF1_126_1 RF1
ZShift = 0.58
lm Score = 219.58
NG = 0
DB = 0
SM = 0
Delta_AUC
Doxorubicin
RF1_237_0 RF1
ZShift = −0.9
lm Score = 1.11
NG = 0
DB = 0
SM = 0
Delta_L
Doxorubicin
RF1_237_0 RF1
ZShift = −0.24
lm Score = −11.31
NG = 0
DB = 0
SM = 0
Delta_K
Doxorubicin
RF1_237_0 RF1
ZShift = 1.63
lm Score = −0.11
NG = 0
DB = 0
SM = 0
Delta_r
Doxorubicin
RF1_237_0 RF1
ZShift = 0.13
lm Score = −80.93
NG = 0
DB = 0
SM = 0
Delta_AUC
Doxorubicin
RF1_251_2 RF1
ZShift = 0.51
lm Score = 1.11
NG = 0
DB = 0
SM = 0
Delta_L
Doxorubicin
RF1_251_2 RF1
ZShift = 0.74
lm Score = −2.4
NG = 0
DB = 0
SM = 0
Delta_K
Doxorubicin
RF1_251_2 RF1
ZShift = 0.85
lm Score = −0.02
NG = 0
DB = 0
SM = 0
Delta_r
Doxorubicin
RF1_251_2 RF1
ZShift = 0.45
lm Score = −110.55
NG = 0
DB = 0
SM = 0
Delta_AUC
Doxorubicin

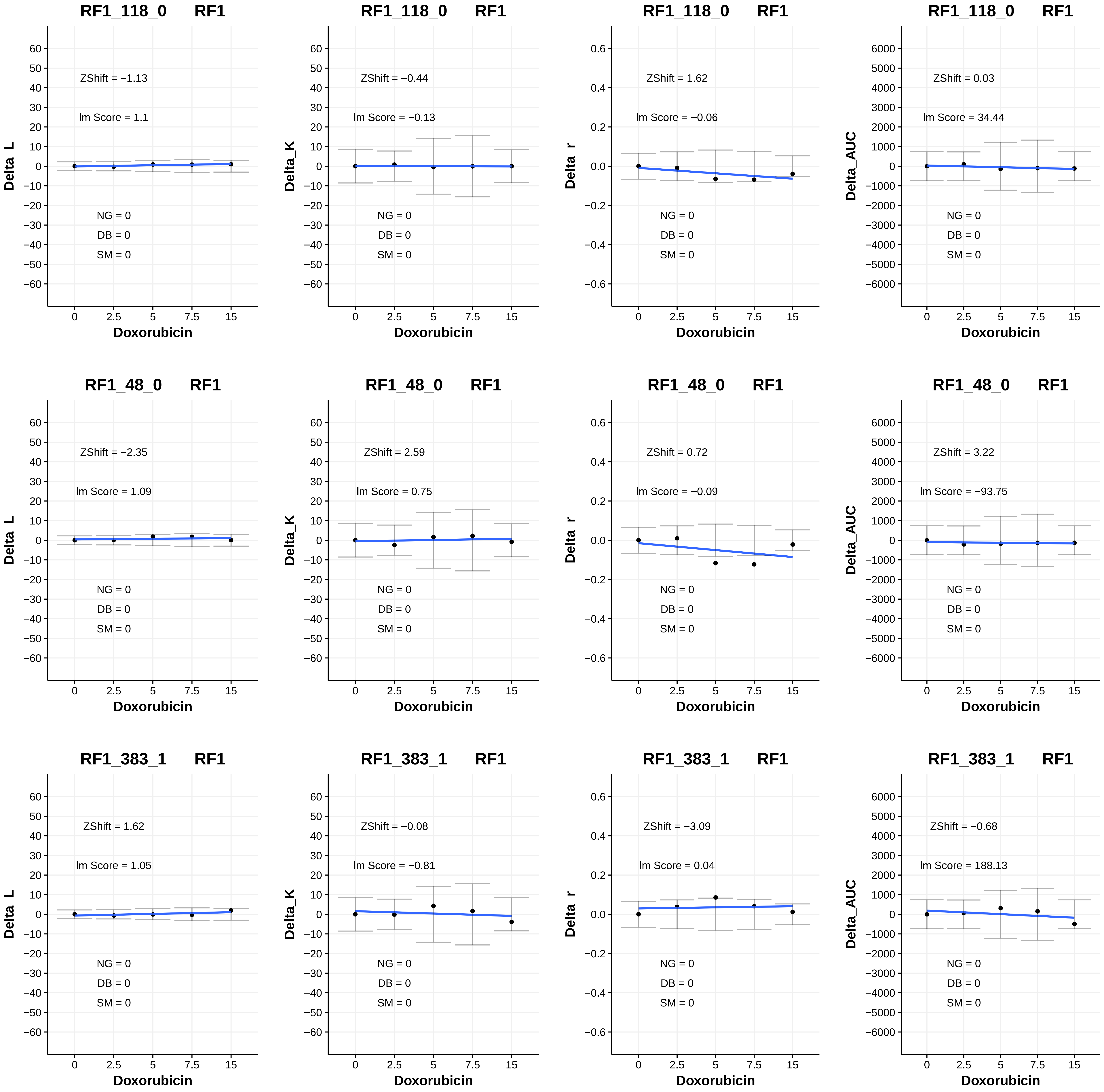

RF1_118_0 RF1
ZShift = −1.13
lm Score = 1.1
NG = 0
DB = 0
SM = 0
Delta_L
Doxorubicin
RF1_118_0 RF1
ZShift = −0.44
lm Score = −0.13
NG = 0
DB = 0
SM = 0
Delta_K
Doxorubicin
RF1_118_0 RF1
ZShift = 1.62
lm Score = −0.06
NG = 0
DB = 0
SM = 0
Delta_r
Doxorubicin
RF1_118_0 RF1
ZShift = 0.03
lm Score = 34.44
NG = 0
DB = 0
SM = 0
Delta_AUC
Doxorubicin
RF1_48_0 RF1
ZShift = −2.35
lm Score = 1.09
NG = 0
DB = 0
SM = 0
Delta_L
Doxorubicin
RF1_48_0 RF1
ZShift = 2.59
lm Score = 0.75
NG = 0
DB = 0
SM = 0
Delta_K
Doxorubicin
RF1_48_0 RF1
ZShift = 0.72
lm Score = −0.09
NG = 0
DB = 0
SM = 0
Delta_r
Doxorubicin
RF1_48_0 RF1
ZShift = 3.22
lm Score = −93.75
NG = 0
DB = 0
SM = 0
Delta_AUC
Doxorubicin
RF1_383_1 RF1
ZShift = 1.62
lm Score = 1.05
NG = 0
DB = 0
SM = 0
Delta_L
Doxorubicin
RF1_383_1 RF1
ZShift = −0.08
lm Score = −0.81
NG = 0
DB = 0
SM = 0
Delta_K
Doxorubicin
RF1_383_1 RF1
ZShift = −3.09
lm Score = 0.04
NG = 0
DB = 0
SM = 0
Delta_r
Doxorubicin
RF1_383_1 RF1
ZShift = −0.68
lm Score = 188.13
NG = 0
DB = 0
SM = 0
Delta_AUC
Doxorubicin

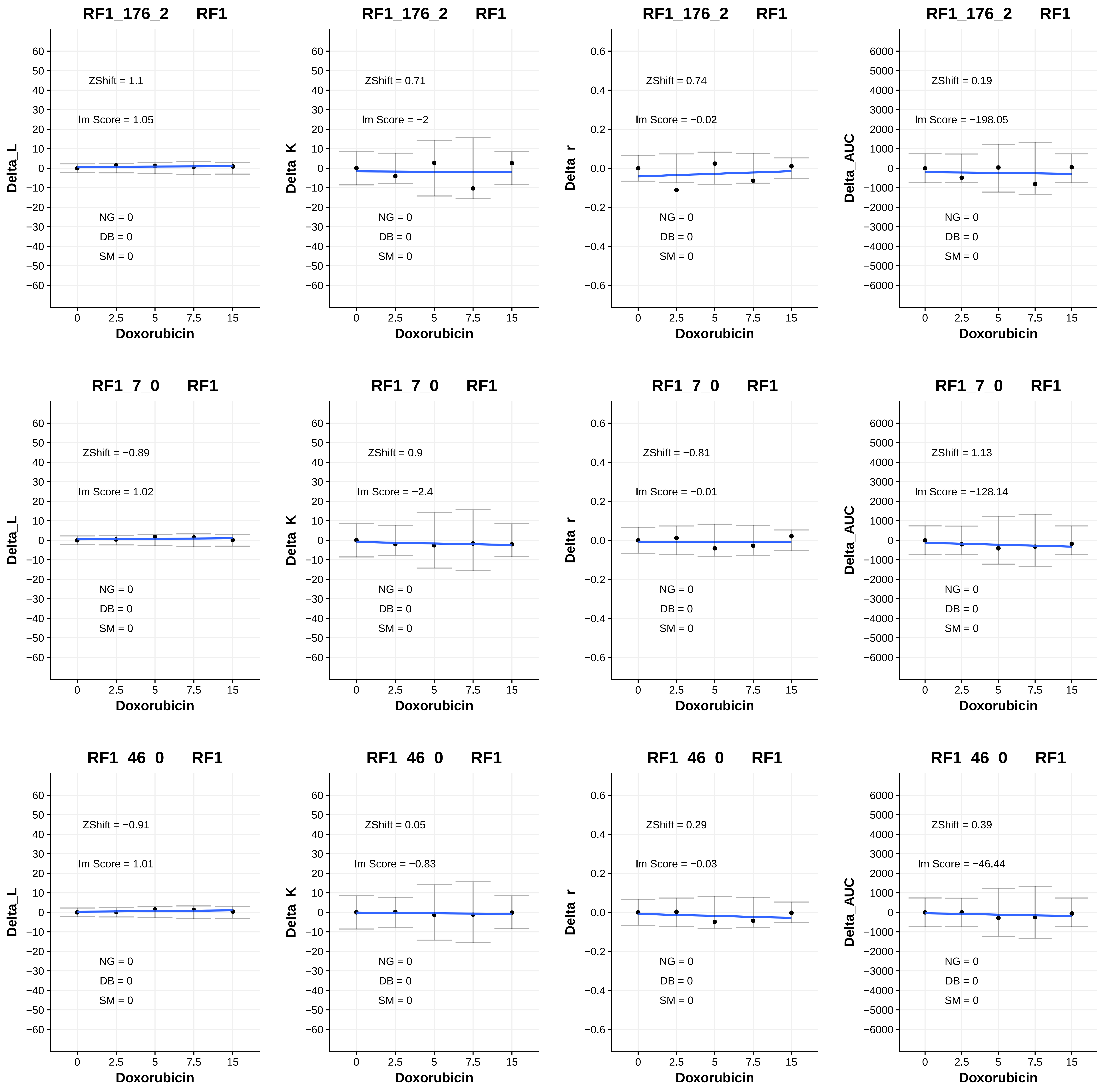
RF1_176_2 RF1
ZShift = 1.1
lm Score = 1.05
NG = 0
DB = 0
SM = 0
Delta_L
Doxorubicin
RF1_176_2 RF1
ZShift = 0.71
lm Score = −2
NG = 0
DB = 0
SM = 0
Delta_K
Doxorubicin
RF1_176_2 RF1
ZShift = 0.74
lm Score = −0.02
NG = 0
DB = 0
SM = 0
Delta_r
Doxorubicin
RF1_176_2 RF1
ZShift = 0.19
lm Score = −198.05
NG = 0
DB = 0
SM = 0
Delta_AUC
Doxorubicin
RF1_7_0 RF1
ZShift = −0.89
lm Score = 1.02
NG = 0
DB = 0
SM = 0
Delta_L
Doxorubicin
RF1_7_0 RF1
ZShift = 0.9
lm Score = −2.4
NG = 0
DB = 0
SM = 0
Delta_K
Doxorubicin
RF1_7_0 RF1
ZShift = −0.81
lm Score = −0.01
NG = 0
DB = 0
SM = 0
Delta_r
Doxorubicin
RF1_7_0 RF1
ZShift = 1.13
lm Score = −128.14
NG = 0
DB = 0
SM = 0
Delta_AUC
Doxorubicin
RF1_46_0 RF1
ZShift = −0.91
lm Score = 1.01
NG = 0
DB = 0
SM = 0
Delta_L
Doxorubicin
RF1_46_0 RF1
ZShift = 0.05
lm Score = −0.83
NG = 0
DB = 0
SM = 0
Delta_K
Doxorubicin
RF1_46_0 RF1
ZShift = 0.29
lm Score = −0.03
NG = 0
DB = 0
SM = 0
Delta_r
Doxorubicin
RF1_46_0 RF1
ZShift = 0.39
lm Score = −46.44
NG = 0
DB = 0
SM = 0
Delta_AUC
Doxorubicin

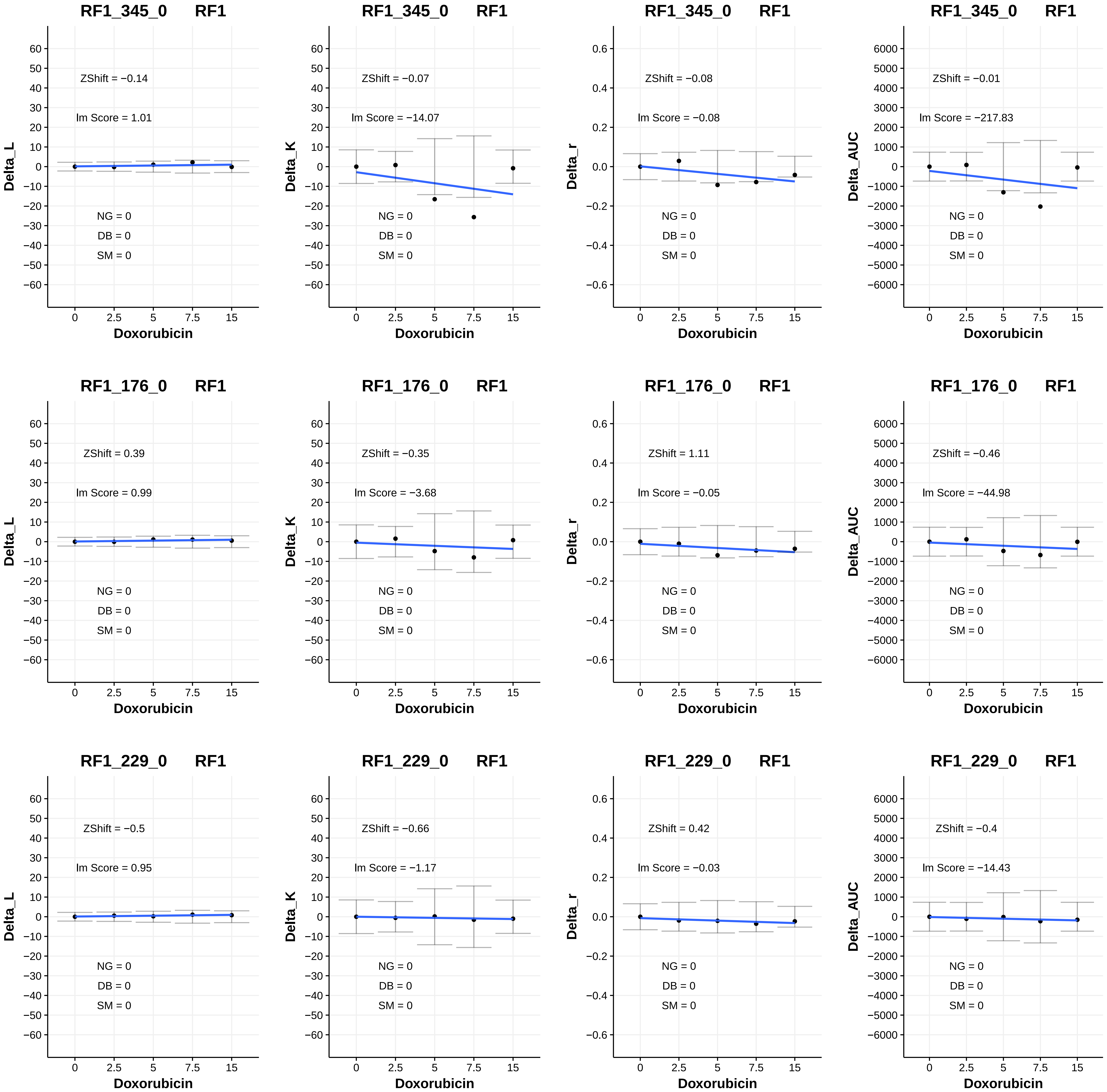
RF1_345_0 RF1
ZShift = −0.14
lm Score = 1.01
NG = 0
DB = 0
SM = 0
Delta_L
Doxorubicin
RF1_345_0 RF1
ZShift = −0.07
lm Score = −14.07
NG = 0
DB = 0
SM = 0
Delta_K
Doxorubicin
RF1_345_0 RF1
ZShift = −0.08
lm Score = −0.08
NG = 0
DB = 0
SM = 0
Delta_r
Doxorubicin
RF1_345_0 RF1
ZShift = −0.01
lm Score = −217.83
NG = 0
DB = 0
SM = 0
Delta_AUC
Doxorubicin
RF1_176_0 RF1
ZShift = 0.39
lm Score = 0.99
NG = 0
DB = 0
SM = 0
Delta_L
Doxorubicin
RF1_176_0 RF1
ZShift = −0.35
lm Score = −3.68
NG = 0
DB = 0
SM = 0
Delta_K
Doxorubicin
RF1_176_0 RF1
ZShift = 1.11
lm Score = −0.05
NG = 0
DB = 0
SM = 0
Delta_r
Doxorubicin
RF1_176_0 RF1
ZShift = −0.46
lm Score = −44.98
NG = 0
DB = 0
SM = 0
Delta_AUC
Doxorubicin
RF1_229_0 RF1
ZShift = −0.5
lm Score = 0.95
NG = 0
DB = 0
SM = 0
Delta_L
Doxorubicin
RF1_229_0 RF1
ZShift = −0.66
lm Score = −1.17
NG = 0
DB = 0
SM = 0
Delta_K
Doxorubicin
RF1_229_0 RF1
ZShift = 0.42
lm Score = −0.03
NG = 0
DB = 0
SM = 0
Delta_r
Doxorubicin
RF1_229_0 RF1
ZShift = −0.4
lm Score = −14.43
NG = 0
DB = 0
SM = 0
Delta_AUC
Doxorubicin

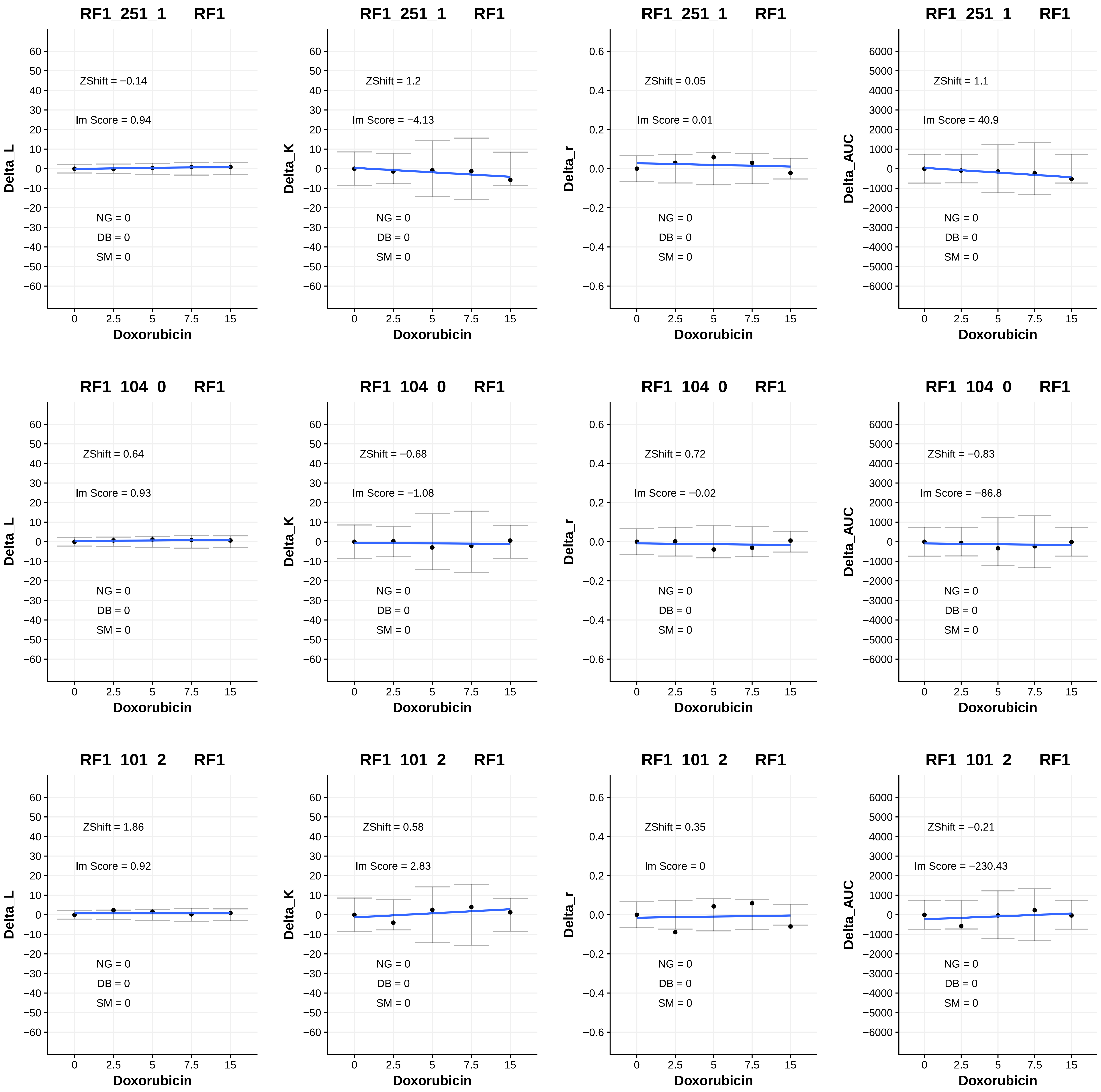
RF1_251_1 RF1
ZShift = −0.14
lm Score = 0.94
NG = 0
DB = 0
SM = 0
Delta_L
Doxorubicin
RF1_251_1 RF1
ZShift = 1.2
lm Score = −4.13
NG = 0
DB = 0
SM = 0
Delta_K
Doxorubicin
RF1_251_1 RF1
ZShift = 0.05
lm Score = 0.01
NG = 0
DB = 0
SM = 0
Delta_r
Doxorubicin
RF1_251_1 RF1
ZShift = 1.1
lm Score = 40.9
NG = 0
DB = 0
SM = 0
Delta_AUC
Doxorubicin
RF1_104_0 RF1
ZShift = 0.64
lm Score = 0.93
NG = 0
DB = 0
SM = 0
Delta_L
Doxorubicin
RF1_104_0 RF1
ZShift = −0.68
lm Score = −1.08
NG = 0
DB = 0
SM = 0
Delta_K
Doxorubicin
RF1_104_0 RF1
ZShift = 0.72
lm Score = −0.02
NG = 0
DB = 0
SM = 0
Delta_r
Doxorubicin
RF1_104_0 RF1
ZShift = −0.83
lm Score = −86.8
NG = 0
DB = 0
SM = 0
Delta_AUC
Doxorubicin
RF1_101_2 RF1
ZShift = 1.86
lm Score = 0.92
NG = 0
DB = 0
SM = 0
Delta_L
Doxorubicin
RF1_101_2 RF1
ZShift = 0.58
lm Score = 2.83
NG = 0
DB = 0
SM = 0
Delta_K
Doxorubicin
RF1_101_2 RF1
ZShift = 0.35
lm Score = 0
NG = 0
DB = 0
SM = 0
Delta_r
Doxorubicin
RF1_101_2 RF1
ZShift = −0.21
lm Score = −230.43
NG = 0
DB = 0
SM = 0
Delta_AUC
Doxorubicin

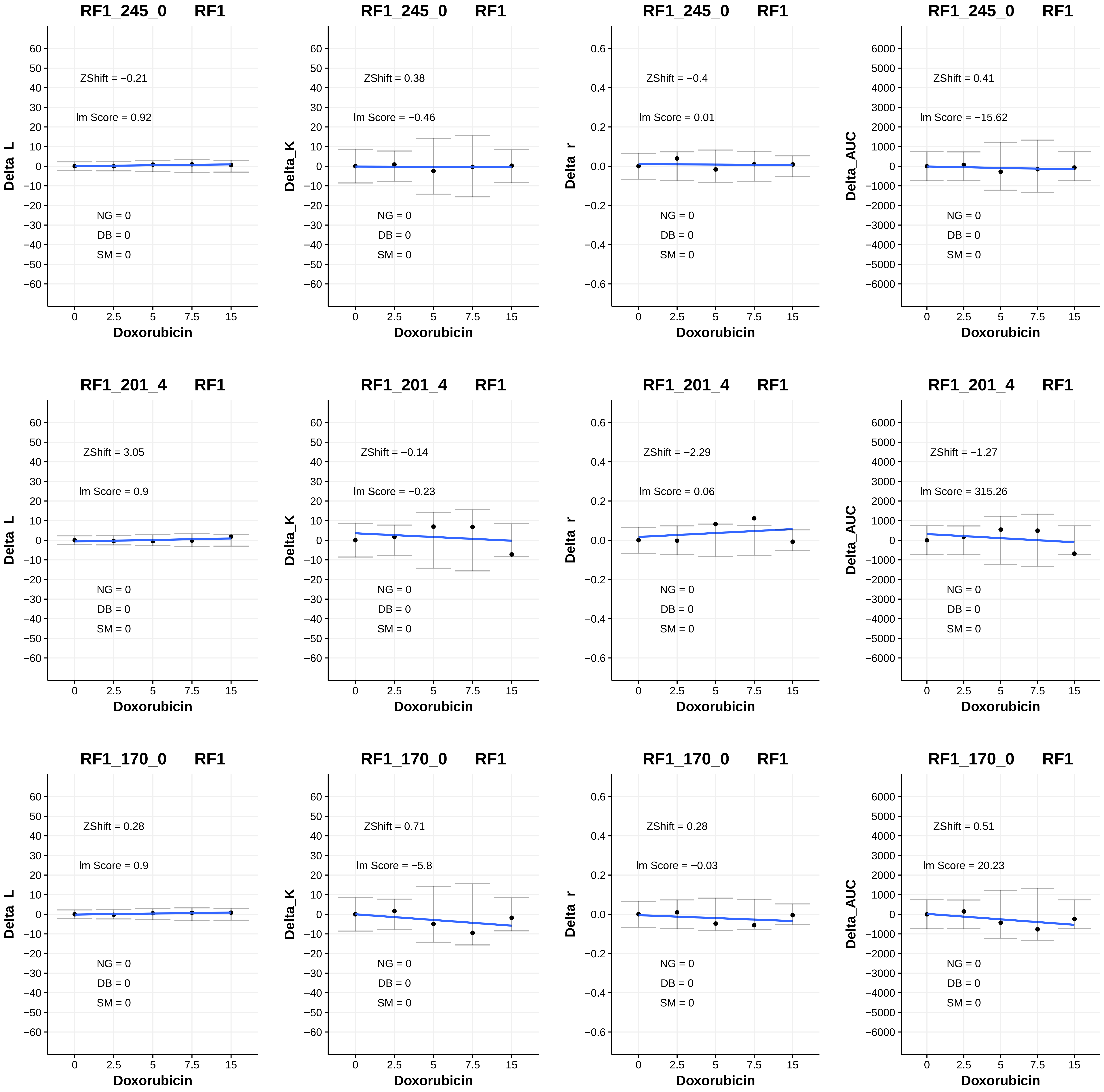
RF1_245_0 RF1
ZShift = −0.21
lm Score = 0.92
NG = 0
DB = 0
SM = 0
Delta_L
Doxorubicin
RF1_245_0 RF1
ZShift = 0.38
lm Score = −0.46
NG = 0
DB = 0
SM = 0
Delta_K
Doxorubicin
RF1_245_0 RF1
ZShift = −0.4
lm Score = 0.01
NG = 0
DB = 0
SM = 0
Delta_r
Doxorubicin
RF1_245_0 RF1
ZShift = 0.41
lm Score = −15.62
NG = 0
DB = 0
SM = 0
Delta_AUC
Doxorubicin
RF1_201_4 RF1
ZShift = 3.05
lm Score = 0.9
NG = 0
DB = 0
SM = 0
Delta_L
Doxorubicin
RF1_201_4 RF1
ZShift = −0.14
lm Score = −0.23
NG = 0
DB = 0
SM = 0
Delta_K
Doxorubicin
RF1_201_4 RF1
ZShift = −2.29
lm Score = 0.06
NG = 0
DB = 0
SM = 0
Delta_r
Doxorubicin
RF1_201_4 RF1
ZShift = −1.27
lm Score = 315.26
NG = 0
DB = 0
SM = 0
Delta_AUC
Doxorubicin
RF1_170_0 RF1
ZShift = 0.28
lm Score = 0.9
NG = 0
DB = 0
SM = 0
Delta_L
Doxorubicin
RF1_170_0 RF1
ZShift = 0.71
lm Score = −5.8
NG = 0
DB = 0
SM = 0
Delta_K
Doxorubicin
RF1_170_0 RF1
ZShift = 0.28
lm Score = −0.03
NG = 0
DB = 0
SM = 0
Delta_r
Doxorubicin
RF1_170_0 RF1
ZShift = 0.51
lm Score = 20.23
NG = 0
DB = 0
SM = 0
Delta_AUC
Doxorubicin

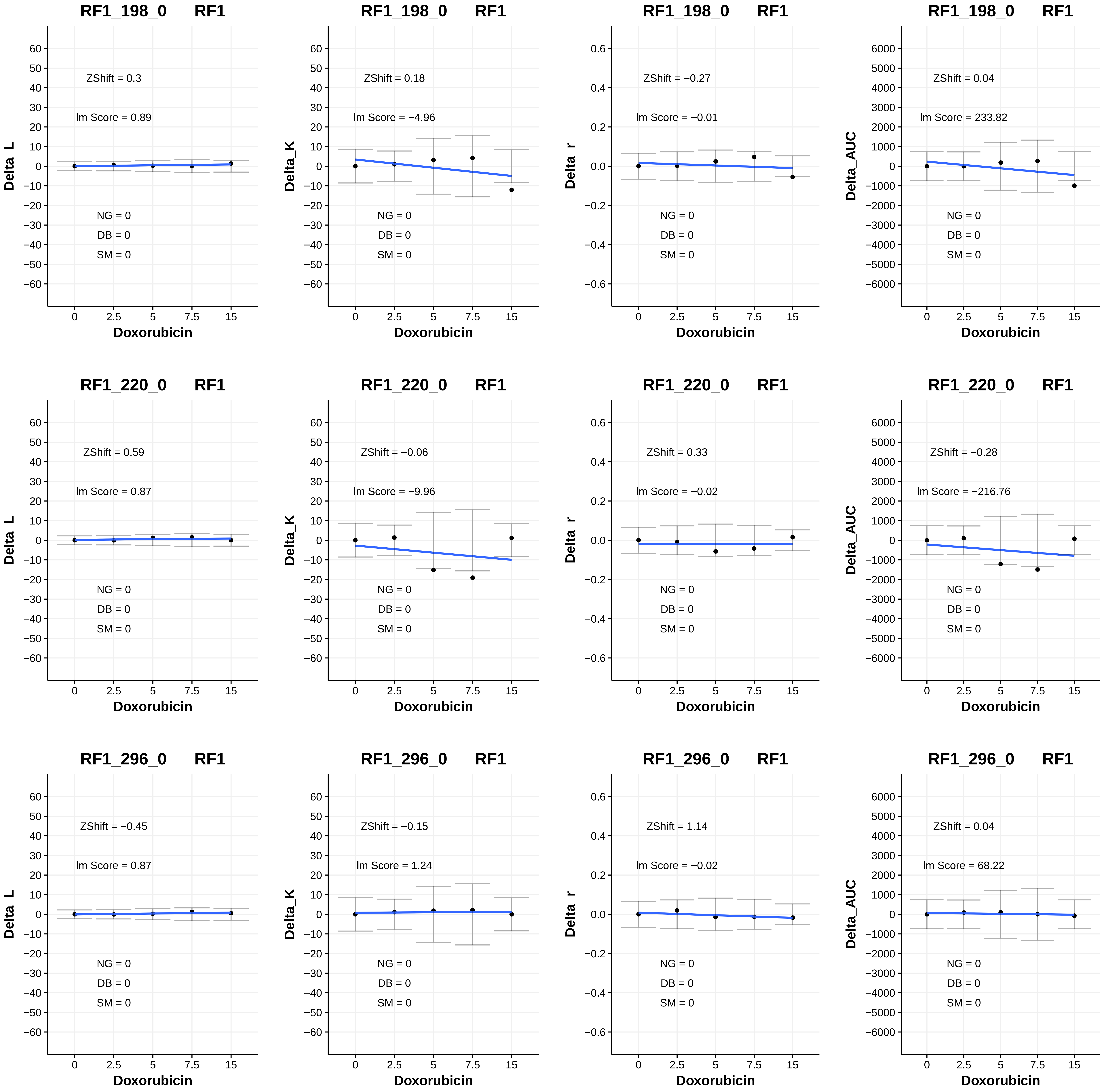
RF1_198_0 RF1
ZShift = 0.3
Im Score = 0.89
NG = 0
DB = 0
SM = 0
Delta_L
Doxorubicin
RF1_198_0 RF1
ZShift = 0.18
Im Score = −4.96
NG = 0
DB = 0
SM = 0
Delta_K
Doxorubicin
RF1_198_0 RF1
ZShift = −0.27
Im Score = −0.01
NG = 0
DB = 0
SM = 0
Delta_r
Doxorubicin
RF1_198_0 RF1
ZShift = 0.04
Im Score = 233.82
NG = 0
DB = 0
SM = 0
Delta_AUC
Doxorubicin
RF1_220_0 RF1
ZShift = 0.59
Im Score = 0.87
NG = 0
DB = 0
SM = 0
Delta_L
Doxorubicin
RF1_220_0 RF1
ZShift = −0.06
Im Score = −9.96
NG = 0
DB = 0
SM = 0
Delta_K
Doxorubicin
RF1_220_0 RF1
ZShift = 0.33
Im Score = −0.02
NG = 0
DB = 0
SM = 0
Delta_r
Doxorubicin
RF1_220_0 RF1
ZShift = −0.28
Im Score = −216.76
NG = 0
DB = 0
SM = 0
Delta_AUC
Doxorubicin
RF1_296_0 RF1
ZShift = −0.45
Im Score = 0.87
NG = 0
DB = 0
SM = 0
Delta_L
Doxorubicin
RF1_296_0 RF1
ZShift = −0.15
Im Score = 1.24
NG = 0
DB = 0
SM = 0
Delta_K
Doxorubicin
RF1_296_0 RF1
ZShift = 1.14
Im Score = −0.02
NG = 0
DB = 0
SM = 0
Delta_r
Doxorubicin
RF1_296_0 RF1
ZShift = 0.04
Im Score = 68.22
NG = 0
DB = 0
SM = 0
Delta_AUC
Doxorubicin

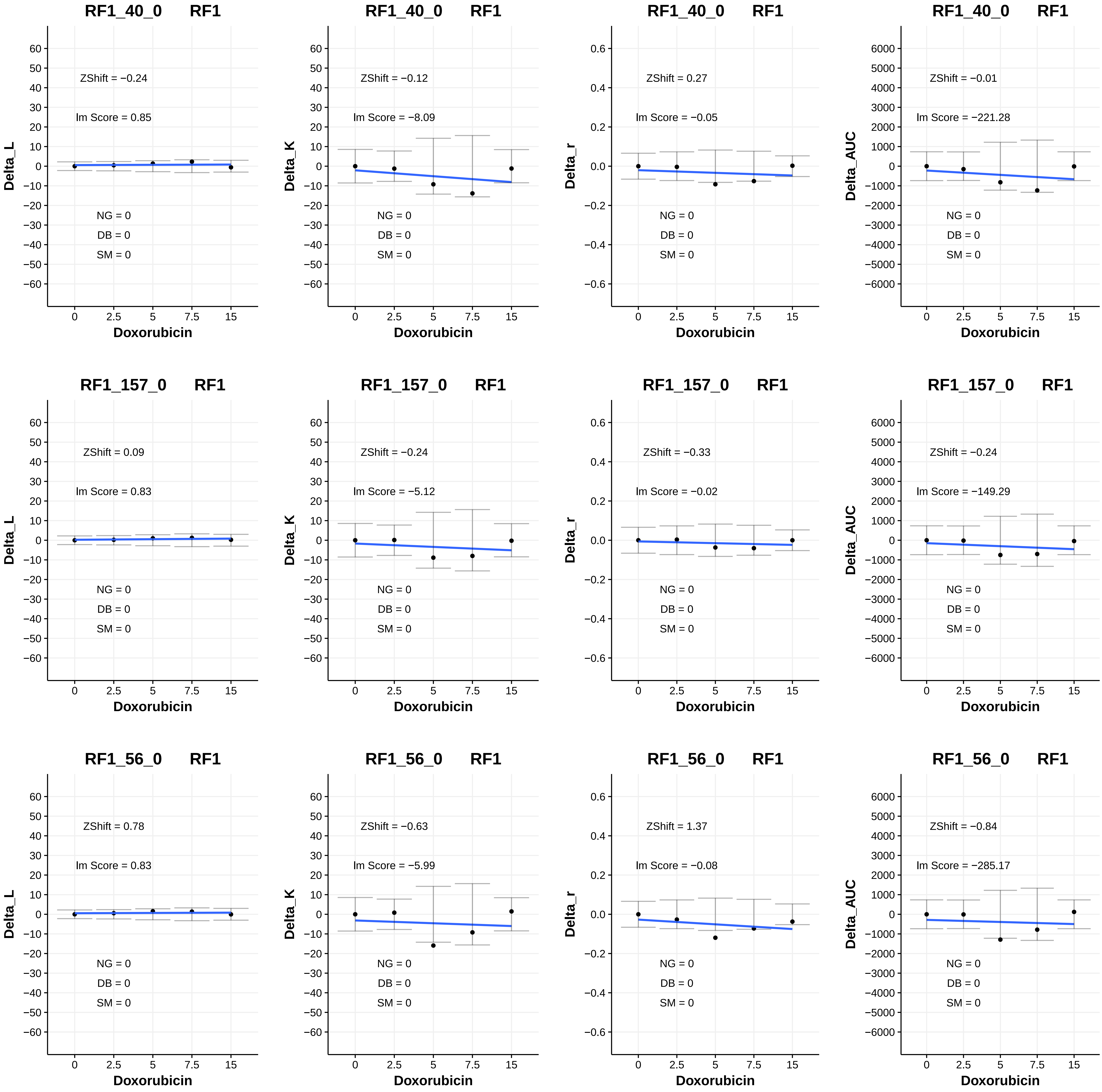
RF1_40_0 RF1
ZShift = −0.24
Im Score = 0.85
NG = 0
DB = 0
SM = 0
Delta_L
Doxorubicin
RF1_40_0 RF1
ZShift = −0.12
Im Score = −8.09
NG = 0
DB = 0
SM = 0
Delta_K
Doxorubicin
RF1_40_0 RF1
ZShift = 0.27
Im Score = −0.05
NG = 0
DB = 0
SM = 0
Delta_r
Doxorubicin
RF1_40_0 RF1
ZShift = −0.01
Im Score = −221.28
NG = 0
DB = 0
SM = 0
Delta_AUC
Doxorubicin
RF1_157_0 RF1
ZShift = 0.09
Im Score = 0.83
NG = 0
DB = 0
SM = 0
Delta_L
Doxorubicin
RF1_157_0 RF1
ZShift = −0.24
Im Score = −5.12
NG = 0
DB = 0
SM = 0
Delta_K
Doxorubicin
RF1_157_0 RF1
ZShift = −0.33
Im Score = −0.02
NG = 0
DB = 0
SM = 0
Delta_r
Doxorubicin
RF1_157_0 RF1
ZShift = −0.24
Im Score = −149.29
NG = 0
DB = 0
SM = 0
Delta_AUC
Doxorubicin
RF1_56_0 RF1
ZShift = 0.78
Im Score = 0.83
NG = 0
DB = 0
SM = 0
Delta_L
Doxorubicin
RF1_56_0 RF1
ZShift = −0.63
Im Score = −5.99
NG = 0
DB = 0
SM = 0
Delta_K
Doxorubicin
RF1_56_0 RF1
ZShift = 1.37
Im Score = −0.08
NG = 0
DB = 0
SM = 0
Delta_r
Doxorubicin
RF1_56_0 RF1
ZShift = −0.84
Im Score = −285.17
NG = 0
DB = 0
SM = 0
Delta_AUC
Doxorubicin

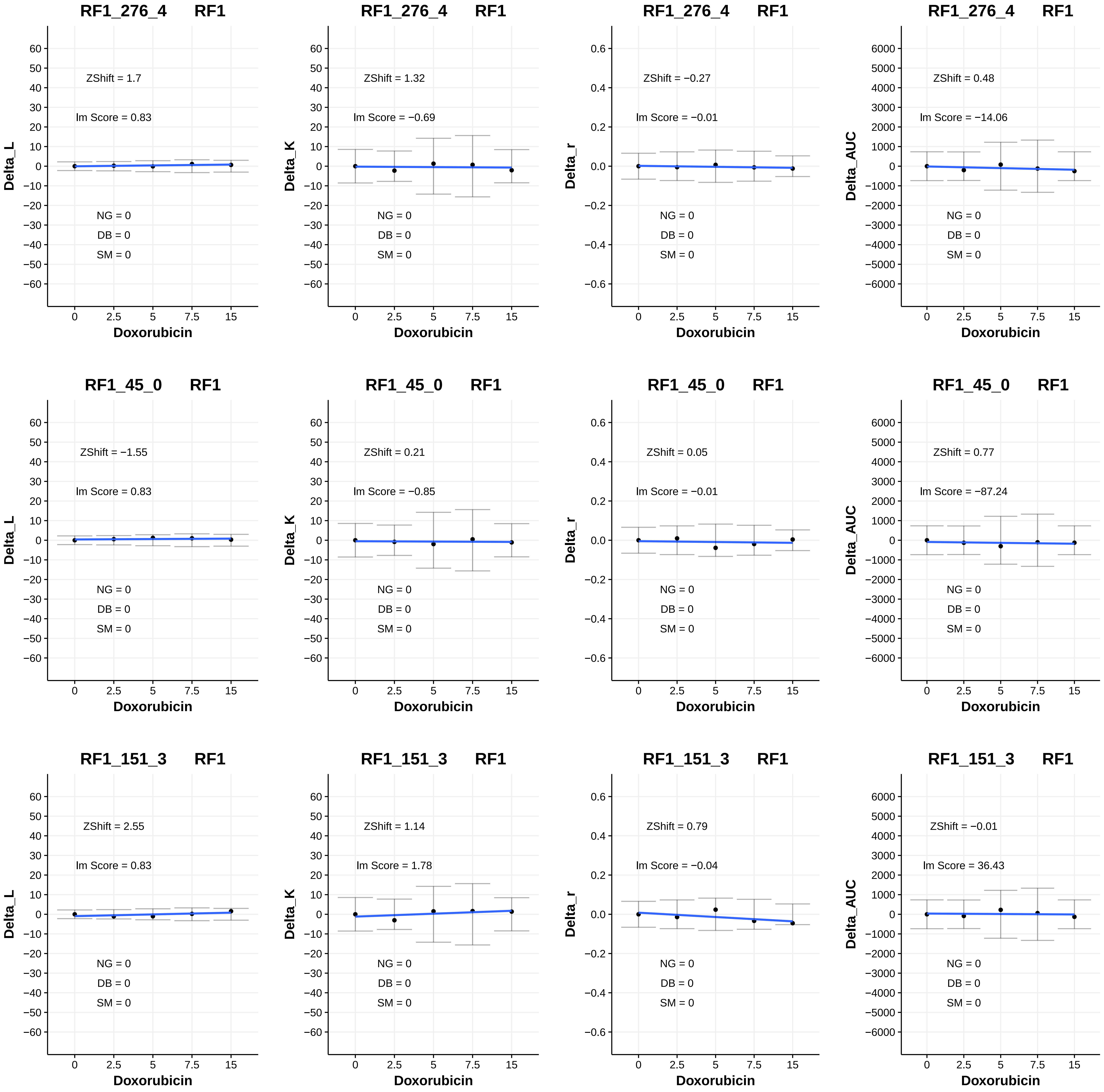
RF1_276_4 RF1
ZShift = 1.7
Im Score = 0.83
NG = 0
DB = 0
SM = 0
Delta_L
Doxorubicin
RF1_276_4 RF1
ZShift = 1.32
Im Score = −0.69
NG = 0
DB = 0
SM = 0
Delta_K
Doxorubicin
RF1_276_4 RF1
ZShift = −0.27
Im Score = −0.01
NG = 0
DB = 0
SM = 0
Delta_r
Doxorubicin
RF1_276_4 RF1
ZShift = 0.48
Im Score = −14.06
NG = 0
DB = 0
SM = 0
Delta_AUC
Doxorubicin
RF1_45_0 RF1
ZShift = −1.55
Im Score = 0.83
NG = 0
DB = 0
SM = 0
Delta_L
Doxorubicin
RF1_45_0 RF1
ZShift = 0.21
Im Score = −0.85
NG = 0
DB = 0
SM = 0
Delta_K
Doxorubicin
RF1_45_0 RF1
ZShift = 0.05
Im Score = −0.01
NG = 0
DB = 0
SM = 0
Delta_r
Doxorubicin
RF1_45_0 RF1
ZShift = 0.77
Im Score = −87.24
NG = 0
DB = 0
SM = 0
Delta_AUC
Doxorubicin
RF1_151_3 RF1
ZShift = 2.55
Im Score = 0.83
NG = 0
DB = 0
SM = 0
Delta_L
Doxorubicin
RF1_151_3 RF1
ZShift = 1.14
Im Score = 1.78
NG = 0
DB = 0
SM = 0
Delta_K
Doxorubicin
RF1_151_3 RF1
ZShift = 0.79
Im Score = −0.04
NG = 0
DB = 0
SM = 0
Delta_r
Doxorubicin
RF1_151_3 RF1
ZShift = −0.01
Im Score = 36.43
NG = 0
DB = 0
SM = 0
Delta_AUC
Doxorubicin

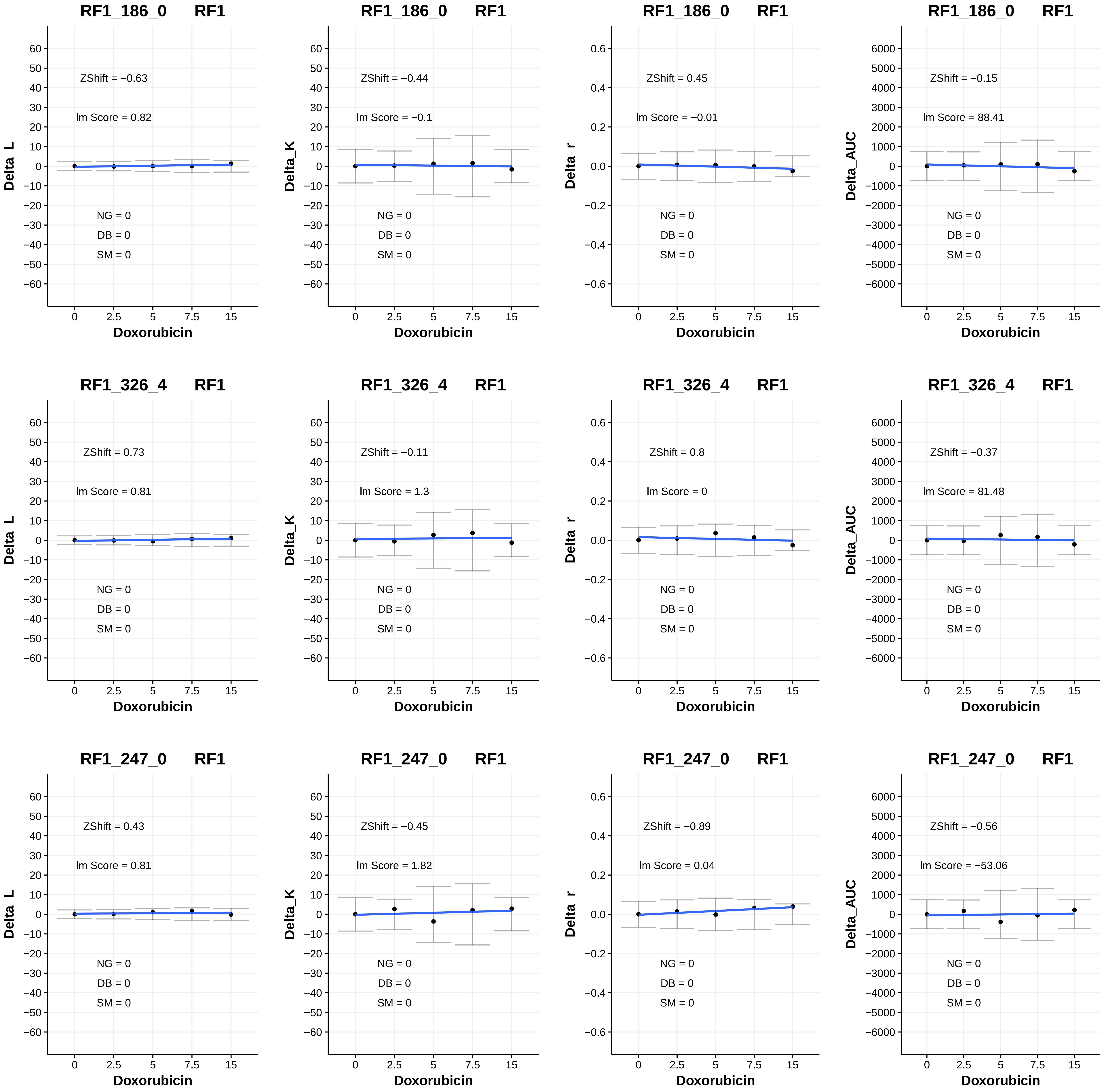
RF1_186_0 RF1
ZShift = −0.63
lm Score = 0.82
NG = 0
DB = 0
SM = 0
Delta_L
Doxorubicin
RF1_186_0 RF1
ZShift = −0.44
lm Score = −0.1
NG = 0
DB = 0
SM = 0
Delta_K
Doxorubicin
RF1_186_0 RF1
ZShift = 0.45
lm Score = −0.01
NG = 0
DB = 0
SM = 0
Delta_r
Doxorubicin
RF1_186_0 RF1
ZShift = −0.15
lm Score = 88.41
NG = 0
DB = 0
SM = 0
Delta_AUC
Doxorubicin
RF1_326_4 RF1
ZShift = 0.73
lm Score = 0.81
NG = 0
DB = 0
SM = 0
Delta_L
Doxorubicin
RF1_326_4 RF1
ZShift = −0.11
lm Score = 1.3
NG = 0
DB = 0
SM = 0
Delta_K
Doxorubicin
RF1_326_4 RF1
ZShift = 0.8
lm Score = 0
NG = 0
DB = 0
SM = 0
Delta_r
Doxorubicin
RF1_326_4 RF1
ZShift = −0.37
lm Score = 81.48
NG = 0
DB = 0
SM = 0
Delta_AUC
Doxorubicin
RF1_247_0 RF1
ZShift = 0.43
lm Score = 0.81
NG = 0
DB = 0
SM = 0
Delta_L
Doxorubicin
RF1_247_0 RF1
ZShift = −0.45
lm Score = 1.82
NG = 0
DB = 0
SM = 0
Delta_K
Doxorubicin
RF1_247_0 RF1
ZShift = −0.89
lm Score = 0.04
NG = 0
DB = 0
SM = 0
Delta_r
Doxorubicin
RF1_247_0 RF1
ZShift = −0.56
lm Score = −53.06
NG = 0
DB = 0
SM = 0
Delta_AUC
Doxorubicin

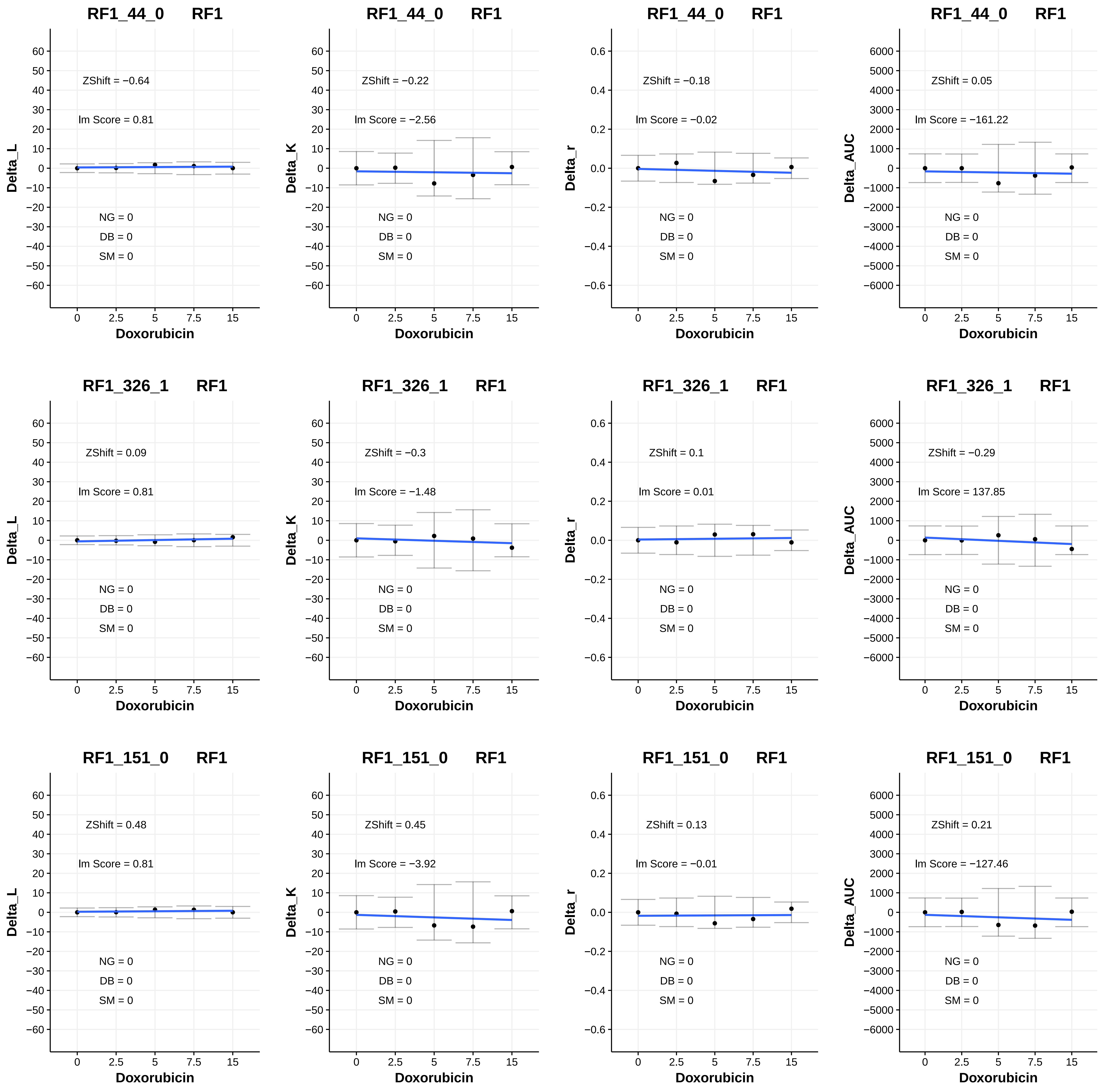

RF1_44_0 RF1
ZShift = −0.64
Im Score = 0.81
NG = 0
DB = 0
SM = 0
Delta_L
Doxorubicin
RF1_44_0 RF1
ZShift = −0.22
Im Score = −2.56
NG = 0
DB = 0
SM = 0
Delta_K
Doxorubicin
RF1_44_0 RF1
ZShift = −0.18
Im Score = −0.02
NG = 0
DB = 0
SM = 0
Delta_r
Doxorubicin
RF1_44_0 RF1
ZShift = 0.05
Im Score = −161.22
NG = 0
DB = 0
SM = 0
Delta_AUC
Doxorubicin
RF1_326_1 RF1
ZShift = 0.09
Im Score = 0.81
NG = 0
DB = 0
SM = 0
Delta_L
Doxorubicin
RF1_326_1 RF1
ZShift = −0.3
Im Score = −1.48
NG = 0
DB = 0
SM = 0
Delta_K
Doxorubicin
RF1_326_1 RF1
ZShift = 0.1
Im Score = 0.01
NG = 0
DB = 0
SM = 0
Delta_r
Doxorubicin
RF1_326_1 RF1
ZShift = −0.29
Im Score = 137.85
NG = 0
DB = 0
SM = 0
Delta_AUC
Doxorubicin
RF1_151_0 RF1
ZShift = 0.48
Im Score = 0.81
NG = 0
DB = 0
SM = 0
Delta_L
Doxorubicin
RF1_151_0 RF1
ZShift = 0.45
Im Score = −3.92
NG = 0
DB = 0
SM = 0
Delta_K
Doxorubicin
RF1_151_0 RF1
ZShift = 0.13
Im Score = −0.01
NG = 0
DB = 0
SM = 0
Delta_r
Doxorubicin
RF1_151_0 RF1
ZShift = 0.21
Im Score = −127.46
NG = 0
DB = 0
SM = 0
Delta_AUC
Doxorubicin

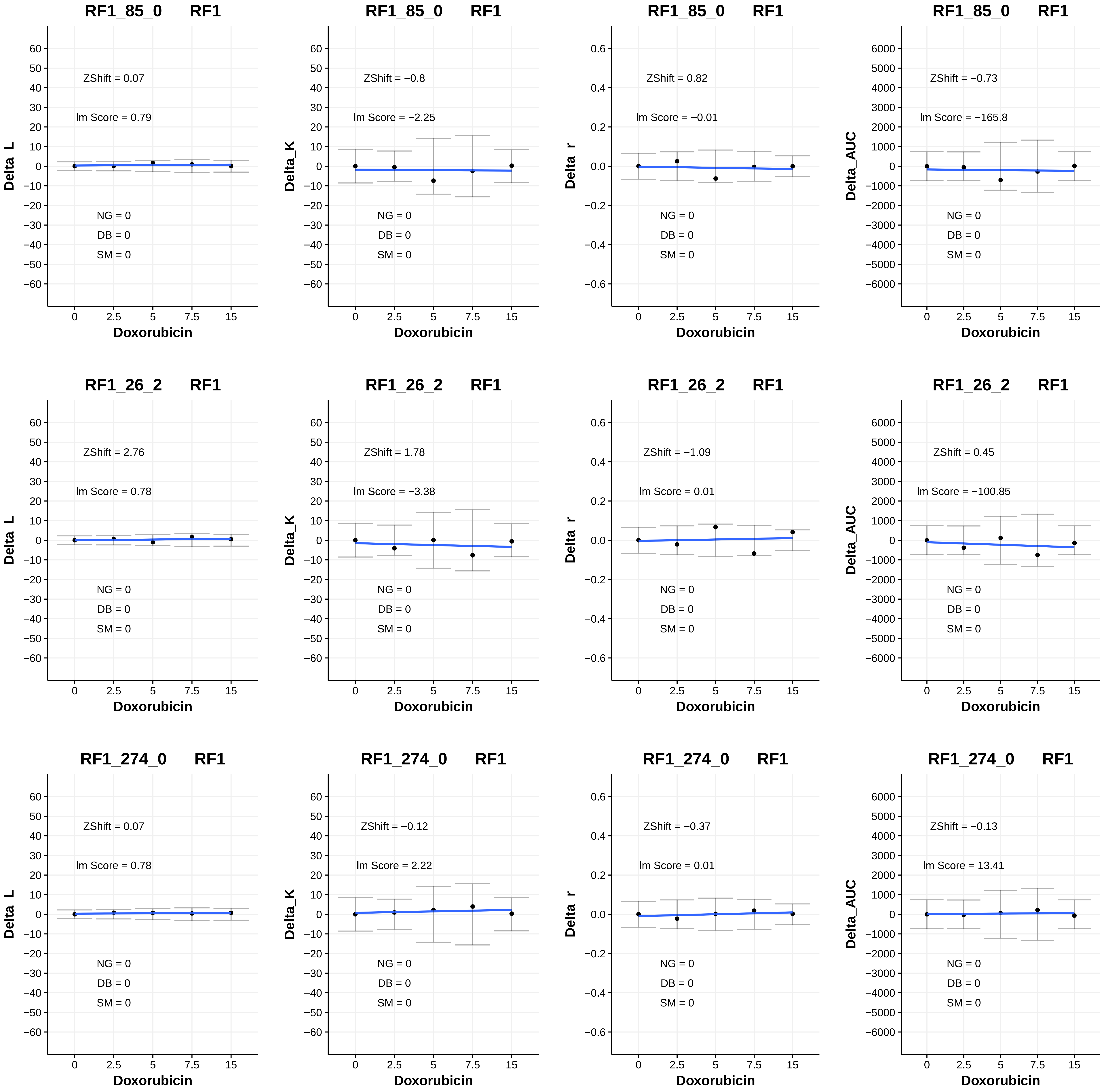
RF1_85_0 RF1
ZShift = 0.07
Im Score = 0.79
NG = 0
DB = 0
SM = 0
Delta_L
Doxorubicin
RF1_85_0 RF1
ZShift = −0.8
Im Score = −2.25
NG = 0
DB = 0
SM = 0
Delta_K
Doxorubicin
RF1_85_0 RF1
ZShift = 0.82
Im Score = −0.01
NG = 0
DB = 0
SM = 0
Delta_r
Doxorubicin
RF1_85_0 RF1
ZShift = −0.73
Im Score = −165.8
NG = 0
DB = 0
SM = 0
Delta_AUC
Doxorubicin
RF1_26_2 RF1
ZShift = 2.76
Im Score = 0.78
NG = 0
DB = 0
SM = 0
Delta_L
Doxorubicin
RF1_26_2 RF1
ZShift = 1.78
Im Score = −3.38
NG = 0
DB = 0
SM = 0
Delta_K
Doxorubicin
RF1_26_2 RF1
ZShift = −1.09
Im Score = 0.01
NG = 0
DB = 0
SM = 0
Delta_r
Doxorubicin
RF1_26_2 RF1
ZShift = 0.45
Im Score = −100.85
NG = 0
DB = 0
SM = 0
Delta_AUC
Doxorubicin
RF1_274_0 RF1
ZShift = 0.07
Im Score = 0.78
NG = 0
DB = 0
SM = 0
Delta_L
Doxorubicin
RF1_274_0 RF1
ZShift = −0.12
Im Score = 2.22
NG = 0
DB = 0
SM = 0
Delta_K
Doxorubicin
RF1_274_0 RF1
ZShift = −0.37
Im Score = 0.01
NG = 0
DB = 0
SM = 0
Delta_r
Doxorubicin
RF1_274_0 RF1
ZShift = −0.13
Im Score = 13.41
NG = 0
DB = 0
SM = 0
Delta_AUC
Doxorubicin

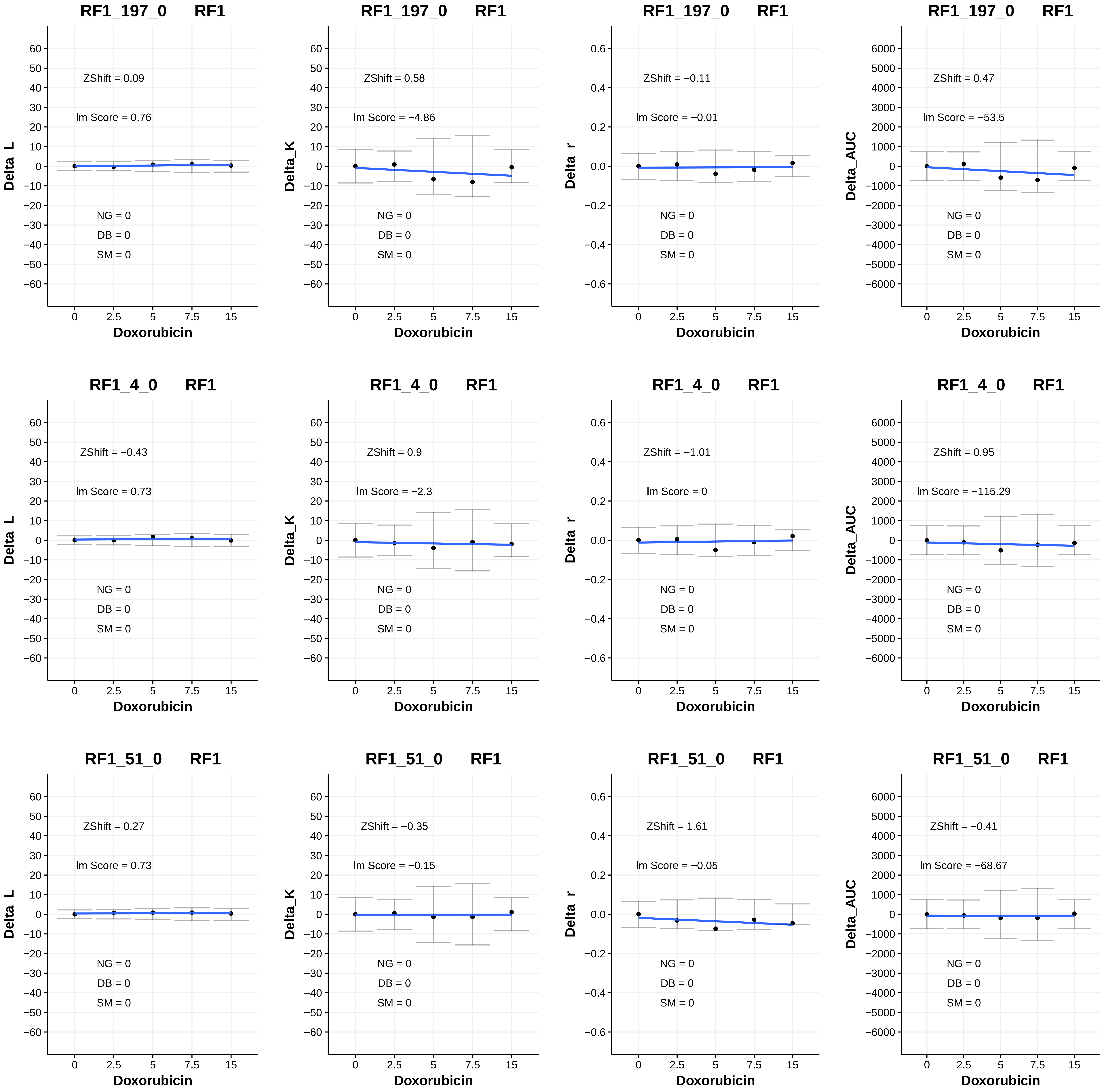
RF1_197_0 RF1
ZShift = 0.09
lm Score = 0.76
NG = 0
DB = 0
SM = 0
Delta_L
Doxorubicin
RF1_197_0 RF1
ZShift = 0.58
lm Score = −4.86
NG = 0
DB = 0
SM = 0
Delta_K
Doxorubicin
RF1_197_0 RF1
ZShift = −0.11
lm Score = −0.01
NG = 0
DB = 0
SM = 0
Delta_r
Doxorubicin
RF1_197_0 RF1
ZShift = 0.47
lm Score = −53.5
NG = 0
DB = 0
SM = 0
Delta_AUC
Doxorubicin
RF1_4_0 RF1
ZShift = −0.43
lm Score = 0.73
NG = 0
DB = 0
SM = 0
Delta_L
Doxorubicin
RF1_4_0 RF1
ZShift = 0.9
lm Score = −2.3
NG = 0
DB = 0
SM = 0
Delta_K
Doxorubicin
RF1_4_0 RF1
ZShift = −1.01
lm Score = 0
NG = 0
DB = 0
SM = 0
Delta_r
Doxorubicin
RF1_4_0 RF1
ZShift = 0.95
lm Score = −115.29
NG = 0
DB = 0
SM = 0
Delta_AUC
Doxorubicin
RF1_51_0 RF1
ZShift = 0.27
lm Score = 0.73
NG = 0
DB = 0
SM = 0
Delta_L
Doxorubicin
RF1_51_0 RF1
ZShift = −0.35
lm Score = −0.15
NG = 0
DB = 0
SM = 0
Delta_K
Doxorubicin
RF1_51_0 RF1
ZShift = 1.61
lm Score = −0.05
NG = 0
DB = 0
SM = 0
Delta_r
Doxorubicin
RF1_51_0 RF1
ZShift = −0.41
lm Score = −68.67
NG = 0
DB = 0
SM = 0
Delta_AUC
Doxorubicin

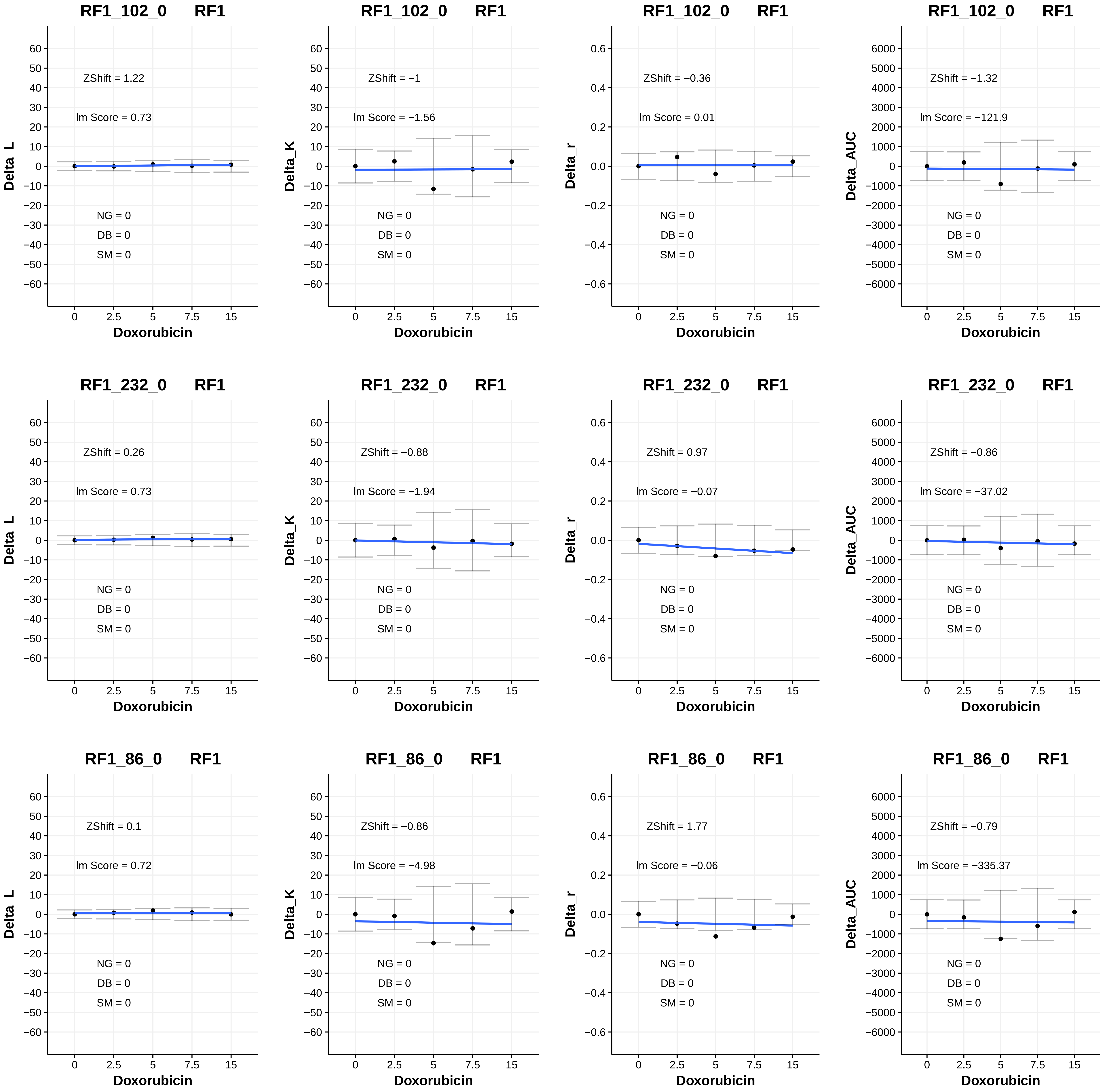
RF1_102_0 RF1
ZShift = 1.22
lm Score = 0.73
NG = 0
DB = 0
SM = 0
Delta_L
Doxorubicin
RF1_102_0 RF1
ZShift = −1
lm Score = −1.56
NG = 0
DB = 0
SM = 0
Delta_K
Doxorubicin
RF1_102_0 RF1
ZShift = −0.36
lm Score = 0.01
NG = 0
DB = 0
SM = 0
Delta_r
Doxorubicin
RF1_102_0 RF1
ZShift = −1.32
lm Score = −121.9
NG = 0
DB = 0
SM = 0
Delta_AUC
Doxorubicin
RF1_232_0 RF1
ZShift = 0.26
lm Score = 0.73
NG = 0
DB = 0
SM = 0
Delta_L
Doxorubicin
RF1_232_0 RF1
ZShift = −0.88
lm Score = −1.94
NG = 0
DB = 0
SM = 0
Delta_K
Doxorubicin
RF1_232_0 RF1
ZShift = 0.97
lm Score = −0.07
NG = 0
DB = 0
SM = 0
Delta_r
Doxorubicin
RF1_232_0 RF1
ZShift = −0.86
lm Score = −37.02
NG = 0
DB = 0
SM = 0
Delta_AUC
Doxorubicin
RF1_86_0 RF1
ZShift = 0.1
lm Score = 0.72
NG = 0
DB = 0
SM = 0
Delta_L
Doxorubicin
RF1_86_0 RF1
ZShift = −0.86
lm Score = −4.98
NG = 0
DB = 0
SM = 0
Delta_K
Doxorubicin
RF1_86_0 RF1
ZShift = 1.77
lm Score = −0.06
NG = 0
DB = 0
SM = 0
Delta_r
Doxorubicin
RF1_86_0 RF1
ZShift = −0.79
lm Score = −335.37
NG = 0
DB = 0
SM = 0
Delta_AUC
Doxorubicin

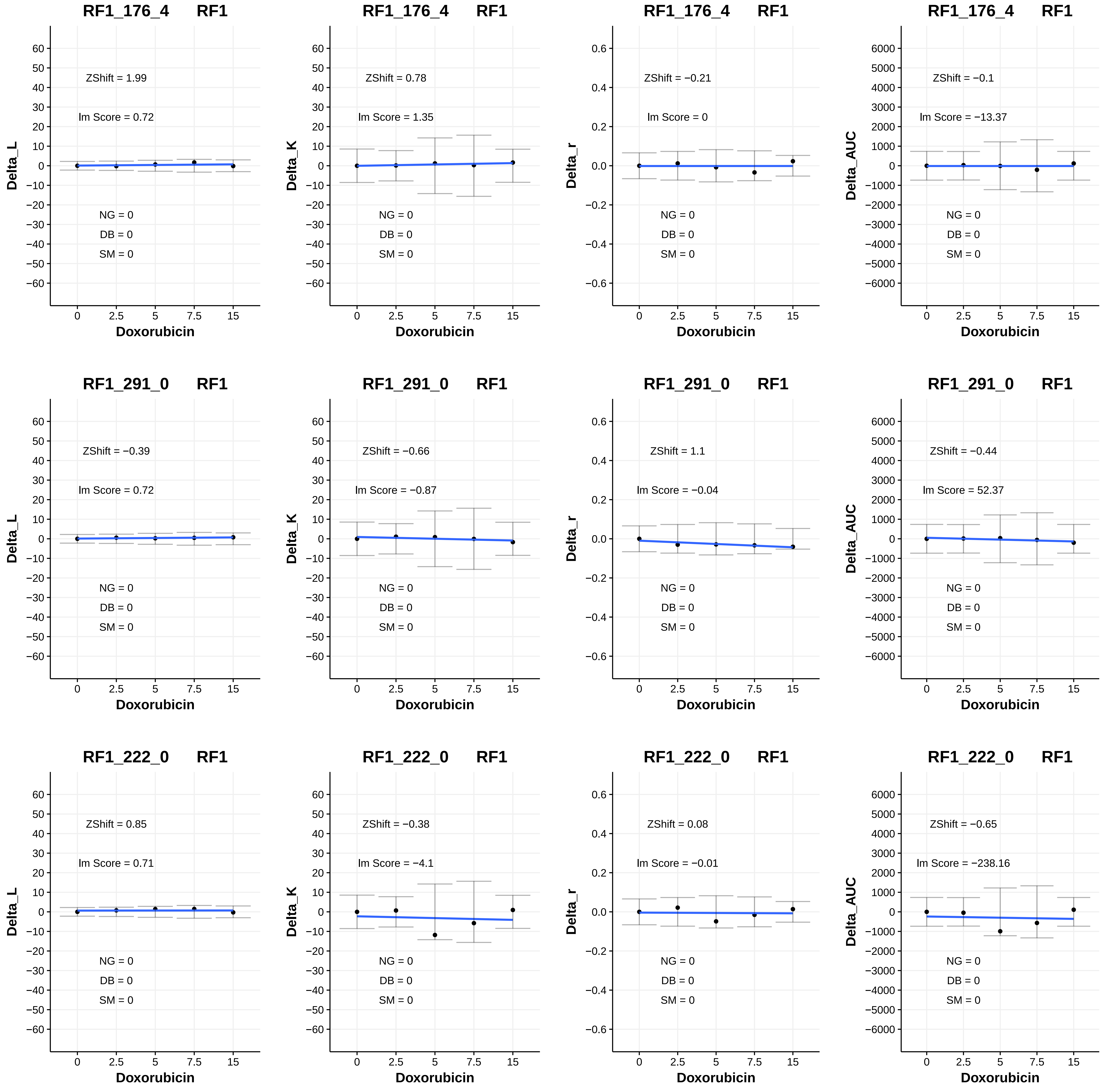

RF1_176_4 RF1
ZShift = 1.99
lm Score = 0.72
NG = 0
DB = 0
SM = 0
Delta_L
Doxorubicin
RF1_176_4 RF1
ZShift = 0.78
lm Score = 1.35
NG = 0
DB = 0
SM = 0
Delta_K
Doxorubicin
RF1_176_4 RF1
ZShift = −0.21
lm Score = 0
NG = 0
DB = 0
SM = 0
Delta_r
Doxorubicin
RF1_176_4 RF1
ZShift = −0.1
lm Score = −13.37
NG = 0
DB = 0
SM = 0
Delta_AUC
Doxorubicin
RF1_291_0 RF1
ZShift = −0.39
lm Score = 0.72
NG = 0
DB = 0
SM = 0
Delta_L
Doxorubicin
RF1_291_0 RF1
ZShift = −0.66
lm Score = −0.87
NG = 0
DB = 0
SM = 0
Delta_K
Doxorubicin
RF1_291_0 RF1
ZShift = 1.1
lm Score = −0.04
NG = 0
DB = 0
SM = 0
Delta_r
Doxorubicin
RF1_291_0 RF1
ZShift = −0.44
lm Score = 52.37
NG = 0
DB = 0
SM = 0
Delta_AUC
Doxorubicin
RF1_222_0 RF1
ZShift = 0.85
lm Score = 0.71
NG = 0
DB = 0
SM = 0
Delta_L
Doxorubicin
RF1_222_0 RF1
ZShift = −0.38
lm Score = −4.1
NG = 0
DB = 0
SM = 0
Delta_K
Doxorubicin
RF1_222_0 RF1
ZShift = 0.08
lm Score = −0.01
NG = 0
DB = 0
SM = 0
Delta_r
Doxorubicin
RF1_222_0 RF1
ZShift = −0.65
lm Score = −238.16
NG = 0
DB = 0
SM = 0
Delta_AUC
Doxorubicin

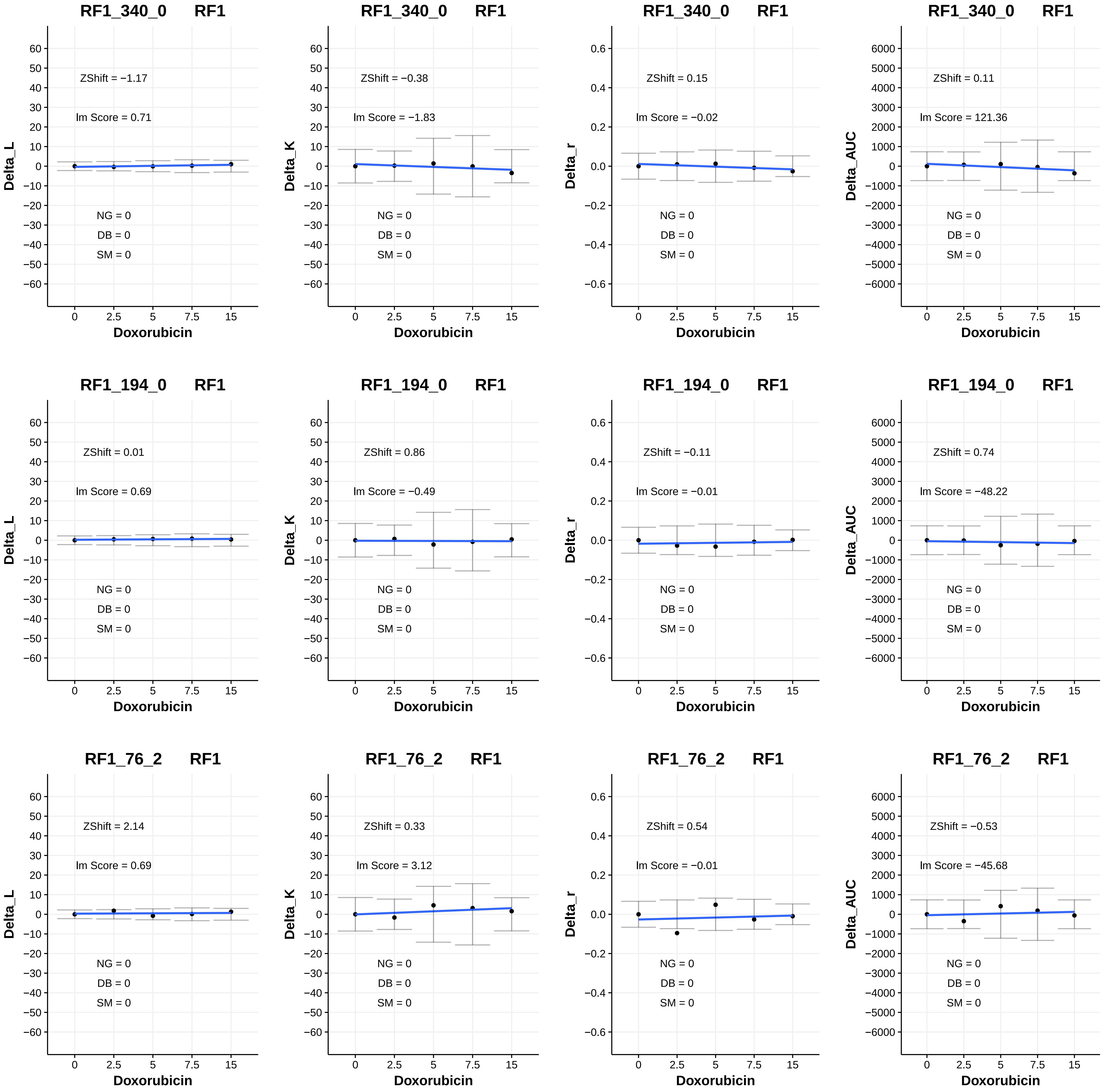

RF1_340_0 RF1
ZShift = −1.17
lm Score = 0.71
NG = 0
DB = 0
SM = 0
Delta_L
Doxorubicin
RF1_340_0 RF1
ZShift = −0.38
lm Score = −1.83
NG = 0
DB = 0
SM = 0
Delta_K
Doxorubicin
RF1_340_0 RF1
ZShift = 0.15
lm Score = −0.02
NG = 0
DB = 0
SM = 0
Delta_r
Doxorubicin
RF1_340_0 RF1
ZShift = 0.11
lm Score = 121.36
NG = 0
DB = 0
SM = 0
Delta_AUC
Doxorubicin
RF1_194_0 RF1
ZShift = 0.01
lm Score = 0.69
NG = 0
DB = 0
SM = 0
Delta_L
Doxorubicin
RF1_194_0 RF1
ZShift = 0.86
lm Score = −0.49
NG = 0
DB = 0
SM = 0
Delta_K
Doxorubicin
RF1_194_0 RF1
ZShift = −0.11
lm Score = −0.01
NG = 0
DB = 0
SM = 0
Delta_r
Doxorubicin
RF1_194_0 RF1
ZShift = 0.74
lm Score = −48.22
NG = 0
DB = 0
SM = 0
Delta_AUC
Doxorubicin
RF1_76_2 RF1
ZShift = 2.14
lm Score = 0.69
NG = 0
DB = 0
SM = 0
Delta_L
Doxorubicin
RF1_76_2 RF1
ZShift = 0.33
lm Score = 3.12
NG = 0
DB = 0
SM = 0
Delta_K
Doxorubicin
RF1_76_2 RF1
ZShift = 0.54
lm Score = −0.01
NG = 0
DB = 0
SM = 0
Delta_r
Doxorubicin
RF1_76_2 RF1
ZShift = −0.53
lm Score = −45.68
NG = 0
DB = 0
SM = 0
Delta_AUC
Doxorubicin

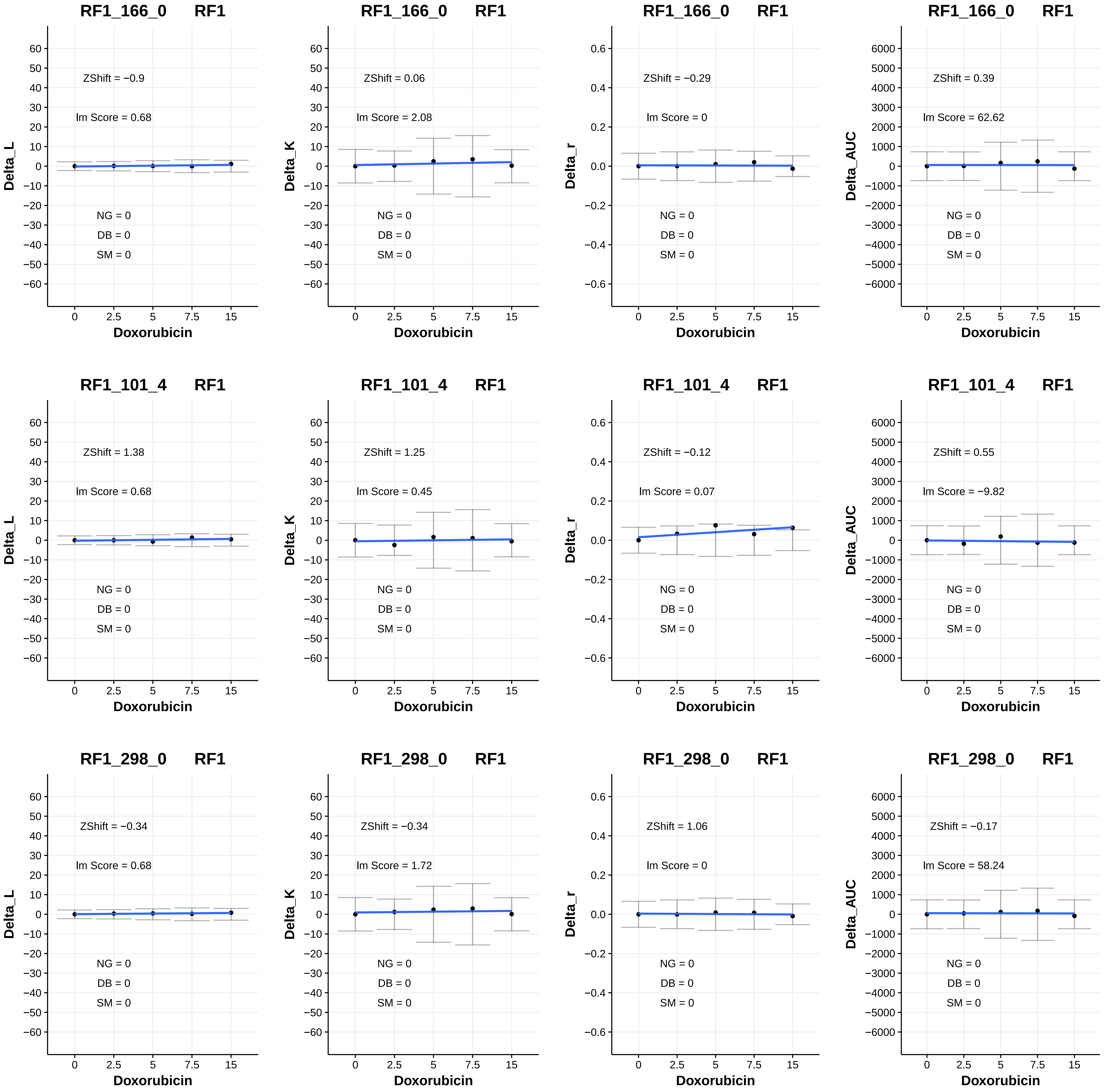

RF1_166_0 RF1
ZShift = −0.9
Im Score = 0.68
NG = 0
DB = 0
SM = 0
Delta_L
Doxorubicin
RF1_166_0 RF1
ZShift = 0.06
Im Score = 2.08
NG = 0
DB = 0
SM = 0
Delta_K
Doxorubicin
RF1_166_0 RF1
ZShift = −0.29
Im Score = 0
NG = 0
DB = 0
SM = 0
Delta_r
Doxorubicin
RF1_166_0 RF1
ZShift = 0.39
Im Score = 62.62
NG = 0
DB = 0
SM = 0
Delta_AUC
Doxorubicin
RF1_101_4 RF1
ZShift = 1.38
Im Score = 0.68
NG = 0
DB = 0
SM = 0
Delta_L
Doxorubicin
RF1_101_4 RF1
ZShift = 1.25
Im Score = 0.45
NG = 0
DB = 0
SM = 0
Delta_K
Doxorubicin
RF1_101_4 RF1
ZShift = −0.12
Im Score = 0.07
NG = 0
DB = 0
SM = 0
Delta_r
Doxorubicin
RF1_101_4 RF1
ZShift = 0.55
Im Score = −9.82
NG = 0
DB = 0
SM = 0
Delta_AUC
Doxorubicin
RF1_298_0 RF1
ZShift = −0.34
Im Score = 0.68
NG = 0
DB = 0
SM = 0
Delta_L
Doxorubicin
RF1_298_0 RF1
ZShift = −0.34
Im Score = 1.72
NG = 0
DB = 0
SM = 0
Delta_K
Doxorubicin
RF1_298_0 RF1
ZShift = 1.06
Im Score = 0
NG = 0
DB = 0
SM = 0
Delta_r
Doxorubicin
RF1_298_0 RF1
ZShift = −0.17
Im Score = 58.24
NG = 0
DB = 0
SM = 0
Delta_AUC
Doxorubicin

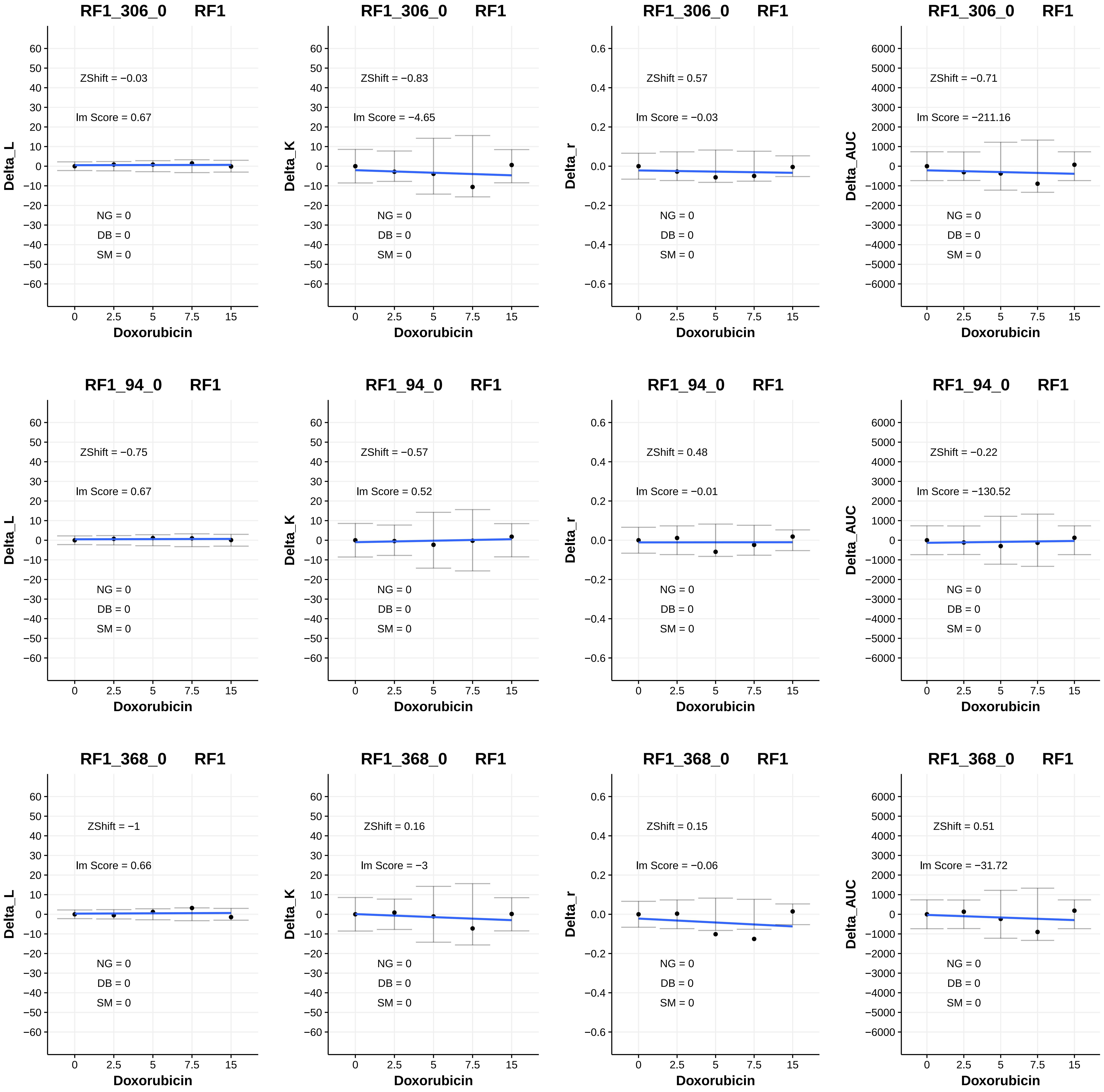
RF1_306_0 RF1
ZShift = −0.03
Im Score = 0.67
NG = 0
DB = 0
SM = 0
Delta_L
Doxorubicin
RF1_306_0 RF1
ZShift = −0.83
Im Score = −4.65
NG = 0
DB = 0
SM = 0
Delta_K
Doxorubicin
RF1_306_0 RF1
ZShift = 0.57
Im Score = −0.03
NG = 0
DB = 0
SM = 0
Delta_r
Doxorubicin
RF1_306_0 RF1
ZShift = −0.71
Im Score = −211.16
NG = 0
DB = 0
SM = 0
Delta_AUC
Doxorubicin
RF1_94_0 RF1
ZShift = −0.75
Im Score = 0.67
NG = 0
DB = 0
SM = 0
Delta_L
Doxorubicin
RF1_94_0 RF1
ZShift = −0.57
Im Score = 0.52
NG = 0
DB = 0
SM = 0
Delta_K
Doxorubicin
RF1_94_0 RF1
ZShift = 0.48
Im Score = −0.01
NG = 0
DB = 0
SM = 0
Delta_r
Doxorubicin
RF1_94_0 RF1
ZShift = −0.22
Im Score = −130.52
NG = 0
DB = 0
SM = 0
Delta_AUC
Doxorubicin
RF1_368_0 RF1
ZShift = −1
Im Score = 0.66
NG = 0
DB = 0
SM = 0
Delta_L
Doxorubicin
RF1_368_0 RF1
ZShift = 0.16
Im Score = −3
NG = 0
DB = 0
SM = 0
Delta_K
Doxorubicin
RF1_368_0 RF1
ZShift = 0.15
Im Score = −0.06
NG = 0
DB = 0
SM = 0
Delta_r
Doxorubicin
RF1_368_0 RF1
ZShift = 0.51
Im Score = −31.72
NG = 0
DB = 0
SM = 0
Delta_AUC
Doxorubicin

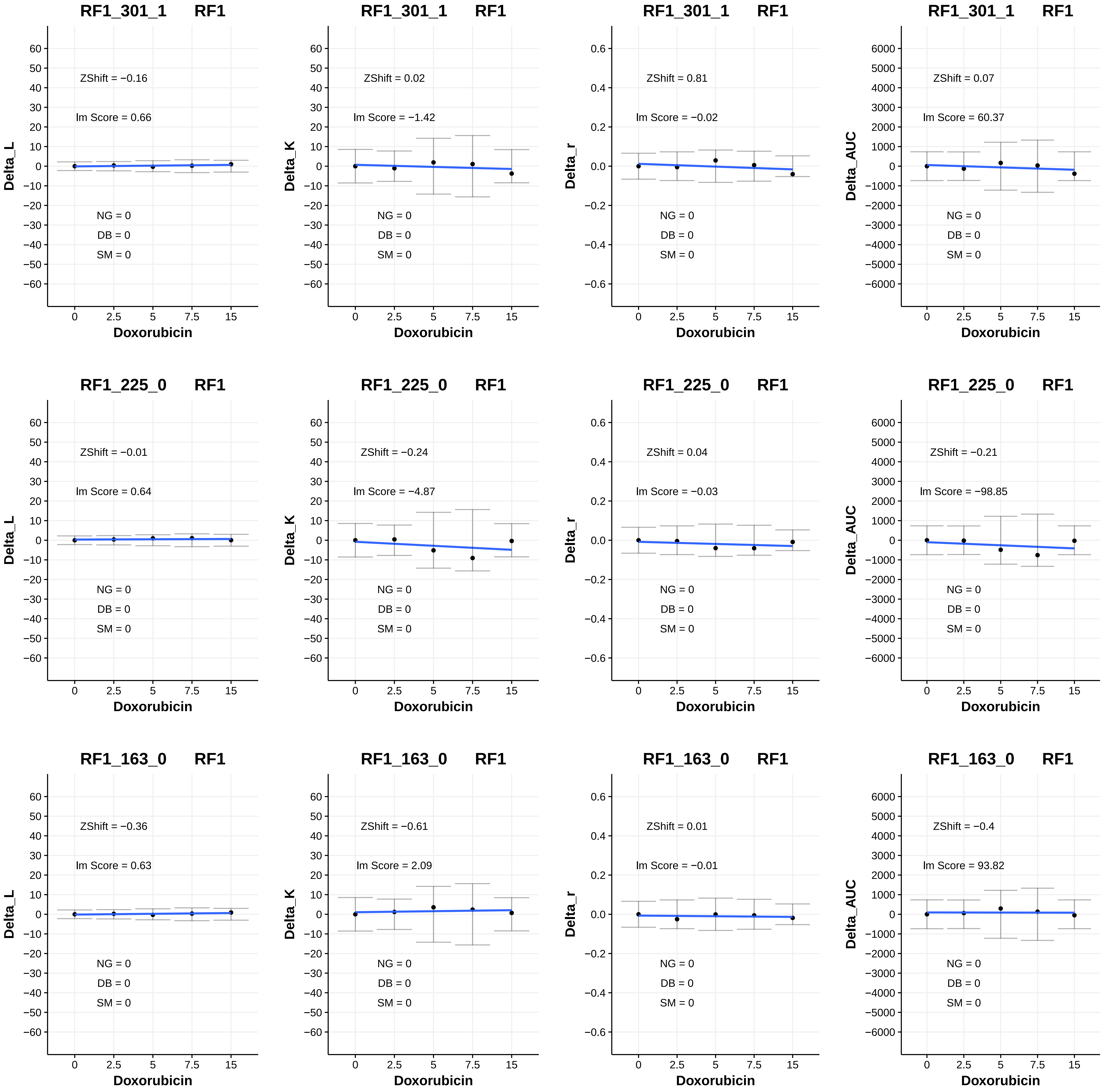
RF1_301_1 RF1
ZShift = −0.16
lm Score = 0.66
NG = 0
DB = 0
SM = 0
Delta_L
Doxorubicin
RF1_301_1 RF1
ZShift = 0.02
lm Score = −1.42
NG = 0
DB = 0
SM = 0
Delta_K
Doxorubicin
RF1_301_1 RF1
ZShift = 0.81
lm Score = −0.02
NG = 0
DB = 0
SM = 0
Delta_r
Doxorubicin
RF1_301_1 RF1
ZShift = 0.07
lm Score = 60.37
NG = 0
DB = 0
SM = 0
Delta_AUC
Doxorubicin
RF1_225_0 RF1
ZShift = −0.01
lm Score = 0.64
NG = 0
DB = 0
SM = 0
Delta_L
Doxorubicin
RF1_225_0 RF1
ZShift = −0.24
lm Score = −4.87
NG = 0
DB = 0
SM = 0
Delta_K
Doxorubicin
RF1_225_0 RF1
ZShift = 0.04
lm Score = −0.03
NG = 0
DB = 0
SM = 0
Delta_r
Doxorubicin
RF1_225_0 RF1
ZShift = −0.21
lm Score = −98.85
NG = 0
DB = 0
SM = 0
Delta_AUC
Doxorubicin
RF1_163_0 RF1
ZShift = −0.36
lm Score = 0.63
NG = 0
DB = 0
SM = 0
Delta_L
Doxorubicin
RF1_163_0 RF1
ZShift = −0.61
lm Score = 2.09
NG = 0
DB = 0
SM = 0
Delta_K
Doxorubicin
RF1_163_0 RF1
ZShift = 0.01
lm Score = −0.01
NG = 0
DB = 0
SM = 0
Delta_r
Doxorubicin
RF1_163_0 RF1
ZShift = −0.4
lm Score = 93.82
NG = 0
DB = 0
SM = 0
Delta_AUC
Doxorubicin

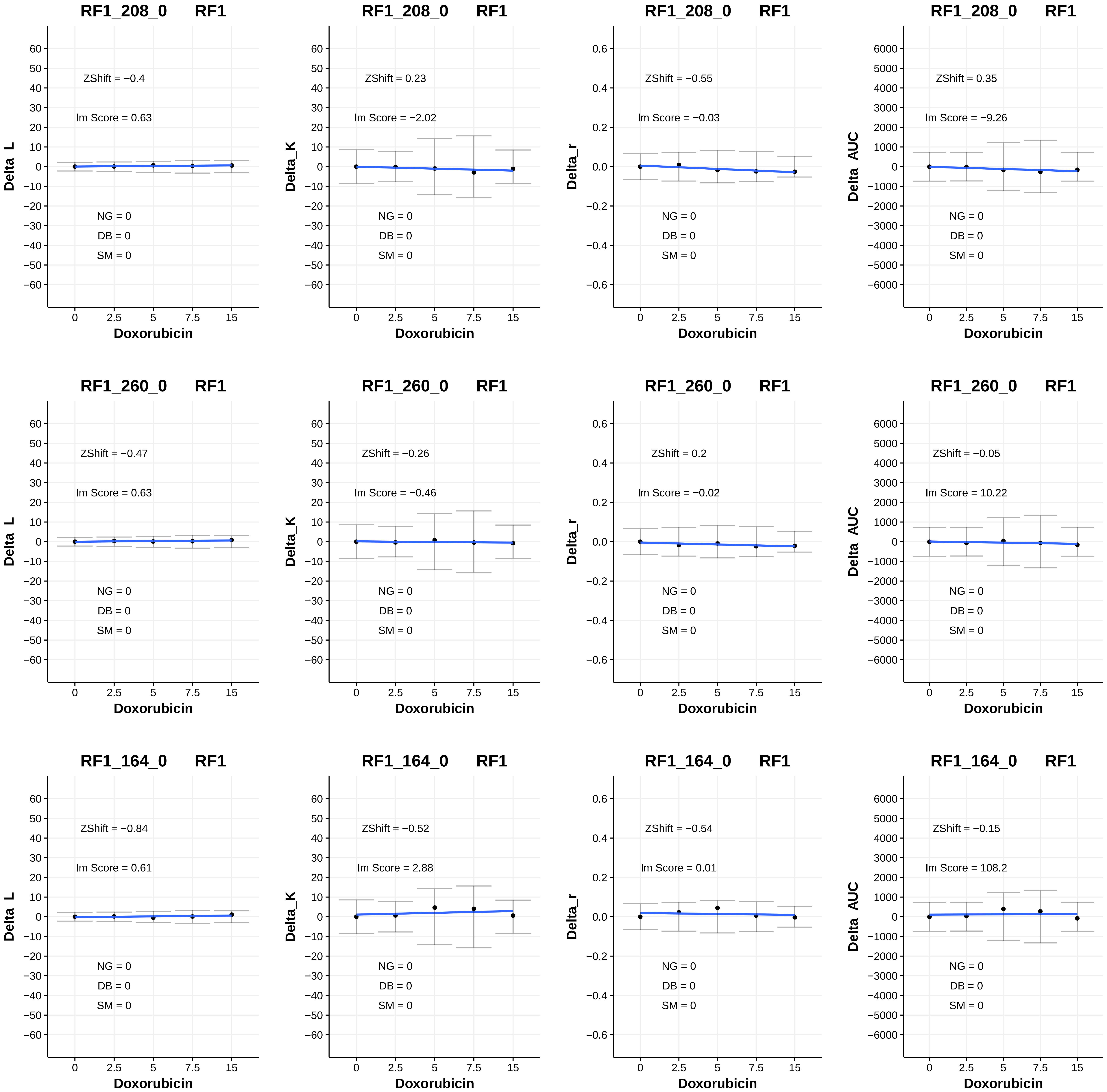
RF1_208_0 RF1
ZShift = −0.4
lm Score = 0.63
NG = 0
DB = 0
SM = 0
Delta_L
Doxorubicin
RF1_208_0 RF1
ZShift = 0.23
lm Score = −2.02
NG = 0
DB = 0
SM = 0
Delta_K
Doxorubicin
RF1_208_0 RF1
ZShift = −0.55
lm Score = −0.03
NG = 0
DB = 0
SM = 0
Delta_r
Doxorubicin
RF1_208_0 RF1
ZShift = 0.35
lm Score = −9.26
NG = 0
DB = 0
SM = 0
Delta_AUC
Doxorubicin
RF1_260_0 RF1
ZShift = −0.47
lm Score = 0.63
NG = 0
DB = 0
SM = 0
Delta_L
Doxorubicin
RF1_260_0 RF1
ZShift = −0.26
lm Score = −0.46
NG = 0
DB = 0
SM = 0
Delta_K
Doxorubicin
RF1_260_0 RF1
ZShift = 0.2
lm Score = −0.02
NG = 0
DB = 0
SM = 0
Delta_r
Doxorubicin
RF1_260_0 RF1
ZShift = −0.05
lm Score = 10.22
NG = 0
DB = 0
SM = 0
Delta_AUC
Doxorubicin
RF1_164_0 RF1
ZShift = −0.84
lm Score = 0.61
NG = 0
DB = 0
SM = 0
Delta_L
Doxorubicin
RF1_164_0 RF1
ZShift = −0.52
lm Score = 2.88
NG = 0
DB = 0
SM = 0
Delta_K
Doxorubicin
RF1_164_0 RF1
ZShift = −0.54
lm Score = 0.01
NG = 0
DB = 0
SM = 0
Delta_r
Doxorubicin
RF1_164_0 RF1
ZShift = −0.15
lm Score = 108.2
NG = 0
DB = 0
SM = 0
Delta_AUC
Doxorubicin

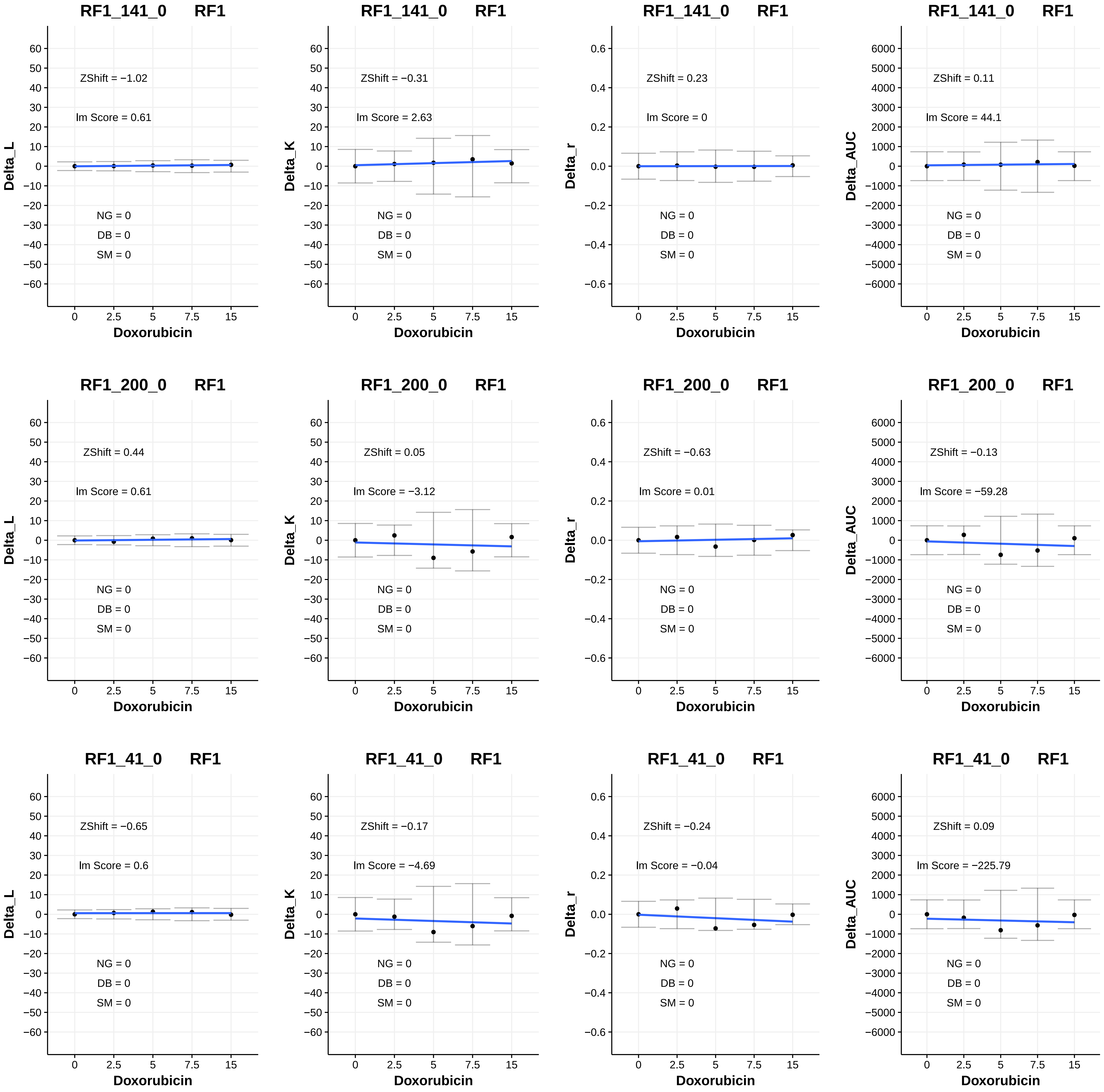
RF1_141_0 RF1
ZShift = −1.02
lm Score = 0.61
NG = 0
DB = 0
SM = 0
Delta_L
Doxorubicin
RF1_141_0 RF1
ZShift = −0.31
lm Score = 2.63
NG = 0
DB = 0
SM = 0
Delta_K
Doxorubicin
RF1_141_0 RF1
ZShift = 0.23
lm Score = 0
NG = 0
DB = 0
SM = 0
Delta_r
Doxorubicin
RF1_141_0 RF1
ZShift = 0.11
lm Score = 44.1
NG = 0
DB = 0
SM = 0
Delta_AUC
Doxorubicin
RF1_200_0 RF1
ZShift = 0.44
lm Score = 0.61
NG = 0
DB = 0
SM = 0
Delta_L
Doxorubicin
RF1_200_0 RF1
ZShift = 0.05
lm Score = −3.12
NG = 0
DB = 0
SM = 0
Delta_K
Doxorubicin
RF1_200_0 RF1
ZShift = −0.63
lm Score = 0.01
NG = 0
DB = 0
SM = 0
Delta_r
Doxorubicin
RF1_200_0 RF1
ZShift = −0.13
lm Score = −59.28
NG = 0
DB = 0
SM = 0
Delta_AUC
Doxorubicin
RF1_41_0 RF1
ZShift = −0.65
lm Score = 0.6
NG = 0
DB = 0
SM = 0
Delta_L
Doxorubicin
RF1_41_0 RF1
ZShift = −0.17
lm Score = −4.69
NG = 0
DB = 0
SM = 0
Delta_K
Doxorubicin
RF1_41_0 RF1
ZShift = −0.24
lm Score = −0.04
NG = 0
DB = 0
SM = 0
Delta_r
Doxorubicin
RF1_41_0 RF1
ZShift = 0.09
lm Score = −225.79
NG = 0
DB = 0
SM = 0
Delta_AUC
Doxorubicin

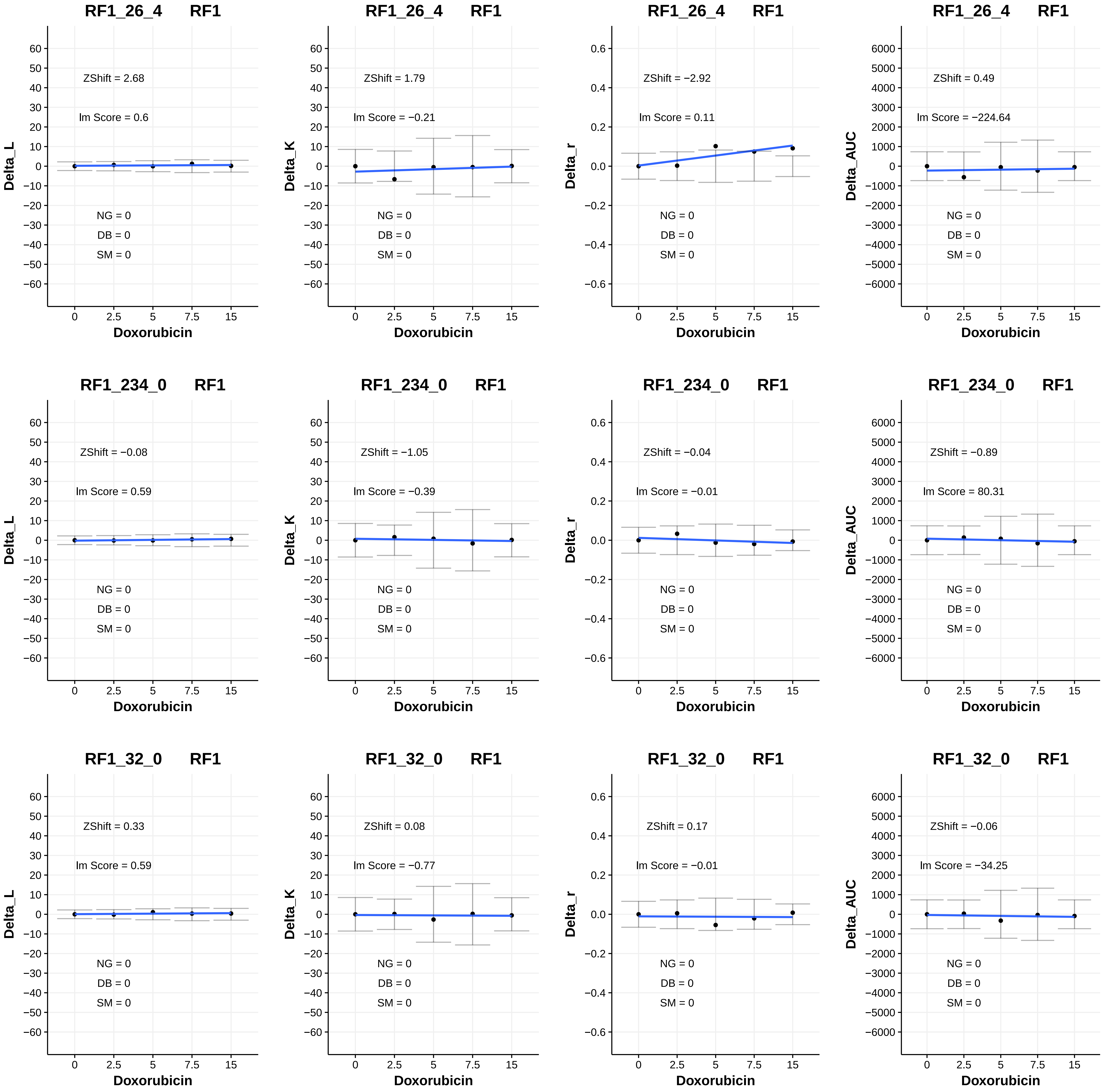

RF1_26_4 RF1
ZShift = 2.68
lm Score = 0.6
NG = 0
DB = 0
SM = 0
Delta_L
Doxorubicin
RF1_26_4 RF1
ZShift = 1.79
lm Score = −0.21
NG = 0
DB = 0
SM = 0
Delta_K
Doxorubicin
RF1_26_4 RF1
ZShift = −2.92
lm Score = 0.11
NG = 0
DB = 0
SM = 0
Delta_r
Doxorubicin
RF1_26_4 RF1
ZShift = 0.49
lm Score = −224.64
NG = 0
DB = 0
SM = 0
Delta_AUC
Doxorubicin
RF1_234_0 RF1
ZShift = −0.08
lm Score = 0.59
NG = 0
DB = 0
SM = 0
Delta_L
Doxorubicin
RF1_234_0 RF1
ZShift = −1.05
lm Score = −0.39
NG = 0
DB = 0
SM = 0
Delta_K
Doxorubicin
RF1_234_0 RF1
ZShift = −0.04
lm Score = −0.01
NG = 0
DB = 0
SM = 0
Delta_r
Doxorubicin
RF1_234_0 RF1
ZShift = −0.89
lm Score = 80.31
NG = 0
DB = 0
SM = 0
Delta_AUC
Doxorubicin
RF1_32_0 RF1
ZShift = 0.33
lm Score = 0.59
NG = 0
DB = 0
SM = 0
Delta_L
Doxorubicin
RF1_32_0 RF1
ZShift = 0.08
lm Score = −0.77
NG = 0
DB = 0
SM = 0
Delta_K
Doxorubicin
RF1_32_0 RF1
ZShift = 0.17
lm Score = −0.01
NG = 0
DB = 0
SM = 0
Delta_r
Doxorubicin
RF1_32_0 RF1
ZShift = −0.06
lm Score = −34.25
NG = 0
DB = 0
SM = 0
Delta_AUC
Doxorubicin

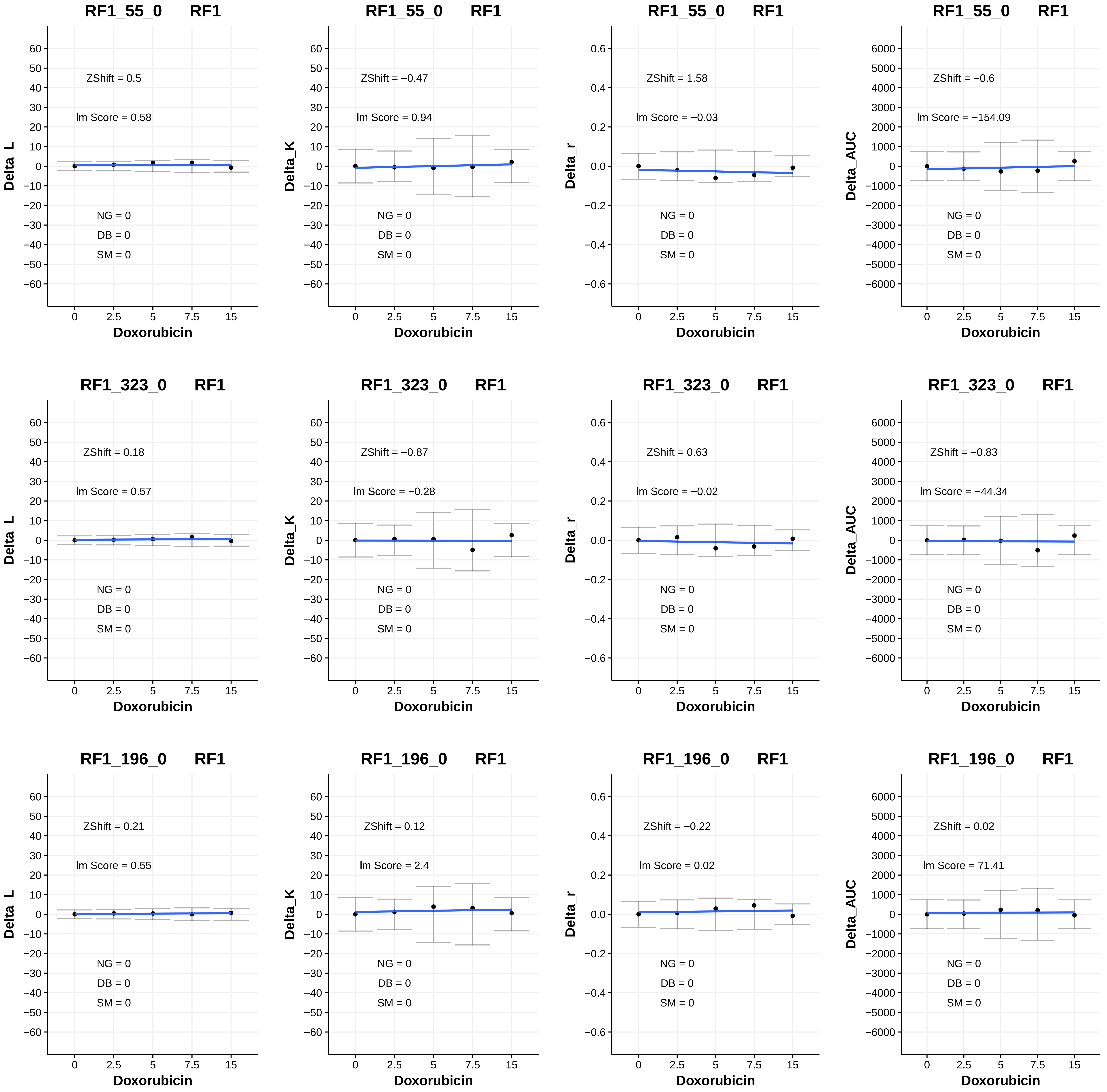
RF1_55_0 RF1
ZShift = 0.5
Im Score = 0.58
Delta_L
NG = 0
DB = 0
SM = 0
Doxorubicin
RF1_55_0 RF1
ZShift = −0.47
Im Score = 0.94
Delta_K
NG = 0
DB = 0
SM = 0
Doxorubicin
RF1_55_0 RF1
ZShift = 1.58
Im Score = −0.03
Delta_r
NG = 0
DB = 0
SM = 0
Doxorubicin
RF1_55_0 RF1
ZShift = −0.6
Im Score = −154.09
Delta_AUC
NG = 0
DB = 0
SM = 0
Doxorubicin
RF1_323_0 RF1
ZShift = 0.18
Im Score = 0.57
Delta_L
NG = 0
DB = 0
SM = 0
Doxorubicin
RF1_323_0 RF1
ZShift = −0.87
Im Score = −0.28
Delta_K
NG = 0
DB = 0
SM = 0
Doxorubicin
RF1_323_0 RF1
ZShift = 0.63
Im Score = −0.02
Delta_r
NG = 0
DB = 0
SM = 0
Doxorubicin
RF1_323_0 RF1
ZShift = −0.83
Im Score = −44.34
Delta_AUC
NG = 0
DB = 0
SM = 0
Doxorubicin
RF1_196_0 RF1
ZShift = 0.21
Im Score = 0.55
Delta_L
NG = 0
DB = 0
SM = 0
Doxorubicin
RF1_196_0 RF1
ZShift = 0.12
Im Score = 2.4
Delta_K
NG = 0
DB = 0
SM = 0
Doxorubicin
RF1_196_0 RF1
ZShift = −0.22
Im Score = 0.02
Delta_r
NG = 0
DB = 0
SM = 0
Doxorubicin
RF1_196_0 RF1
ZShift = 0.02
Im Score = 71.41
Delta_AUC
NG = 0
DB = 0
SM = 0
Doxorubicin

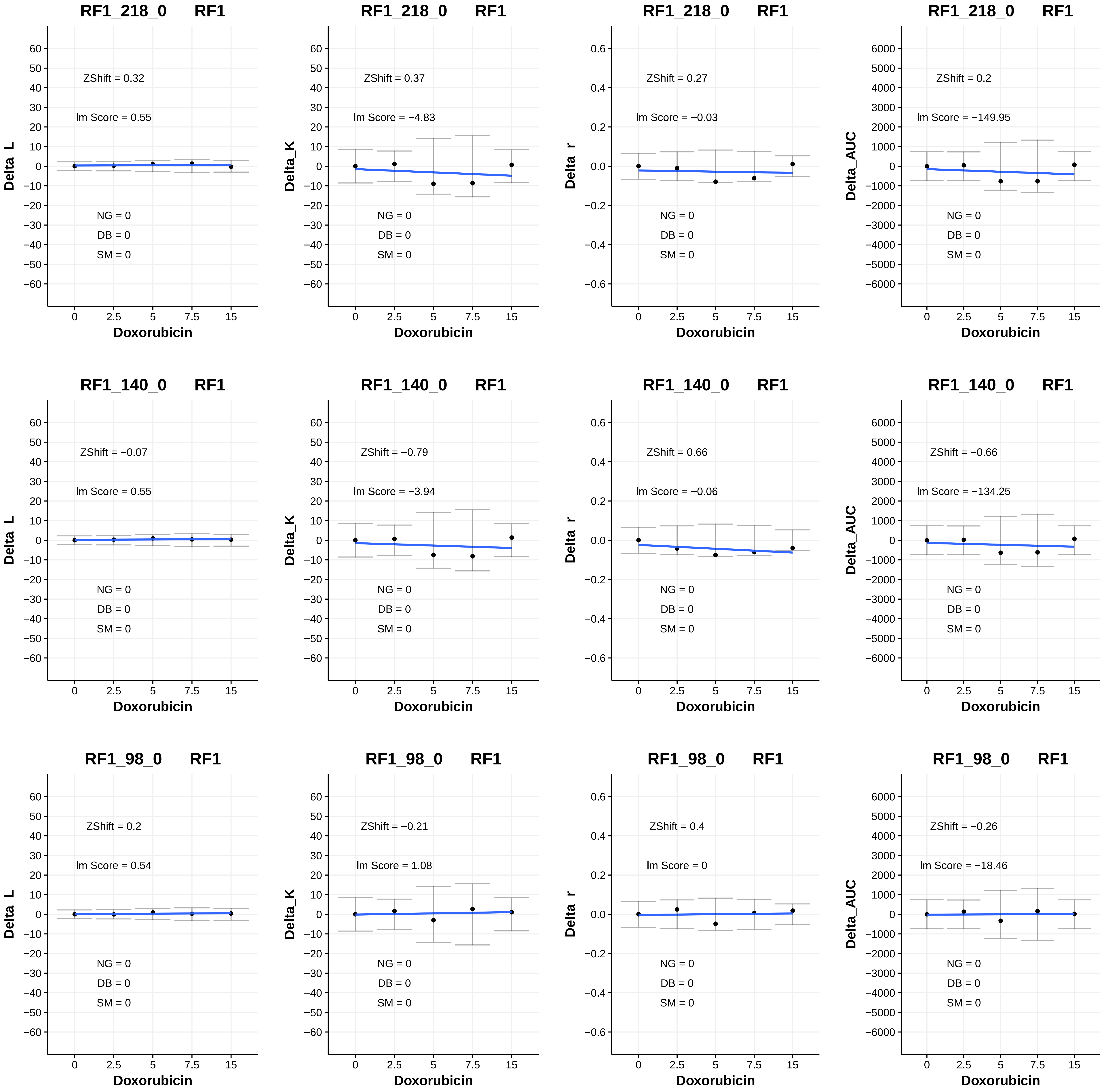
RF1_218_0 RF1
ZShift = 0.32
Im Score = 0.55
NG = 0
DB = 0
SM = 0
Delta_L
Doxorubicin
RF1_218_0 RF1
ZShift = 0.37
Im Score = −4.83
Delta_K
RF1_218_0 RF1
ZShift = 0.27
Im Score = −0.03
Delta_r
RF1_218_0 RF1
ZShift = 0.2
Im Score = −149.95
Delta_AUC
RF1_140_0 RF1
ZShift = −0.07
Im Score = 0.55
RF1_140_0 RF1
ZShift = −0.79
Im Score = −3.94
RF1_140_0 RF1
ZShift = 0.66
Im Score = −0.06
RF1_140_0 RF1
ZShift = −0.66
Im Score = −134.25
RF1_98_0 RF1
ZShift = 0.2
Im Score = 0.54
RF1_98_0 RF1
ZShift = −0.21
Im Score = 1.08
RF1_98_0 RF1
ZShift = 0.4
Im Score = 0
RF1_98_0 RF1
ZShift = −0.26
Im Score = −18.46

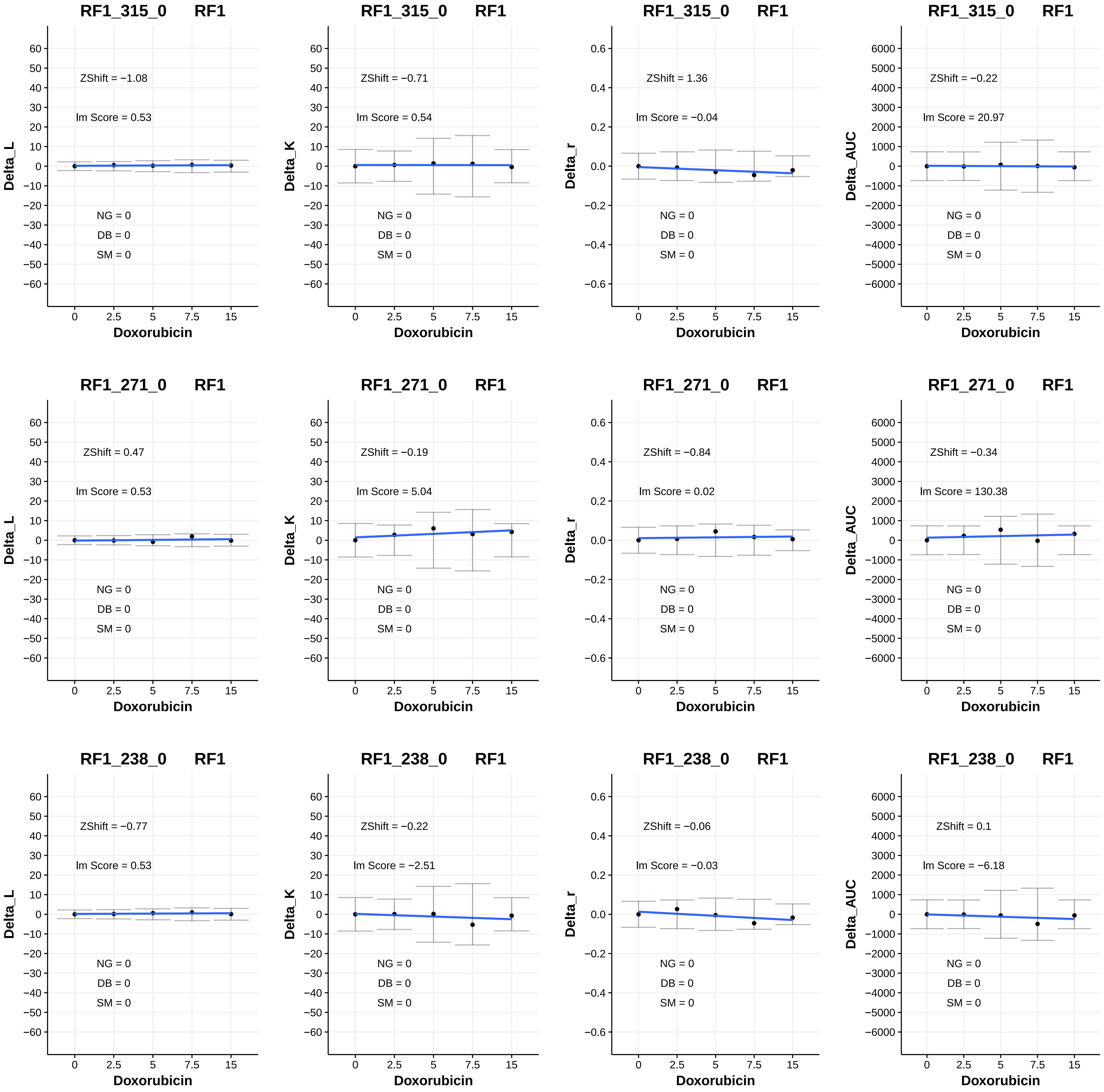
RF1_315_0 RF1
ZShift = −1.08
Im Score = 0.53
NG = 0
DB = 0
SM = 0
Delta_L
Doxorubicin
RF1_315_0 RF1
ZShift = −0.71
Im Score = 0.54
NG = 0
DB = 0
SM = 0
Delta_K
Doxorubicin
RF1_315_0 RF1
ZShift = 1.36
Im Score = −0.04
NG = 0
DB = 0
SM = 0
Delta_r
Doxorubicin
RF1_315_0 RF1
ZShift = −0.22
Im Score = 20.97
NG = 0
DB = 0
SM = 0
Delta_AUC
Doxorubicin
RF1_271_0 RF1
ZShift = 0.47
Im Score = 0.53
NG = 0
DB = 0
SM = 0
Delta_L
Doxorubicin
RF1_271_0 RF1
ZShift = −0.19
Im Score = 5.04
NG = 0
DB = 0
SM = 0
Delta_K
Doxorubicin
RF1_271_0 RF1
ZShift = −0.84
Im Score = 0.02
NG = 0
DB = 0
SM = 0
Delta_r
Doxorubicin
RF1_271_0 RF1
ZShift = −0.34
Im Score = 130.38
NG = 0
DB = 0
SM = 0
Delta_AUC
Doxorubicin
RF1_238_0 RF1
ZShift = −0.77
Im Score = 0.53
NG = 0
DB = 0
SM = 0
Delta_L
Doxorubicin
RF1_238_0 RF1
ZShift = −0.22
Im Score = −2.51
NG = 0
DB = 0
SM = 0
Delta_K
Doxorubicin
RF1_238_0 RF1
ZShift = −0.06
Im Score = −0.03
NG = 0
DB = 0
SM = 0
Delta_r
Doxorubicin
RF1_238_0 RF1
ZShift = 0.1
Im Score = −6.18
NG = 0
DB = 0
SM = 0
Delta_AUC
Doxorubicin

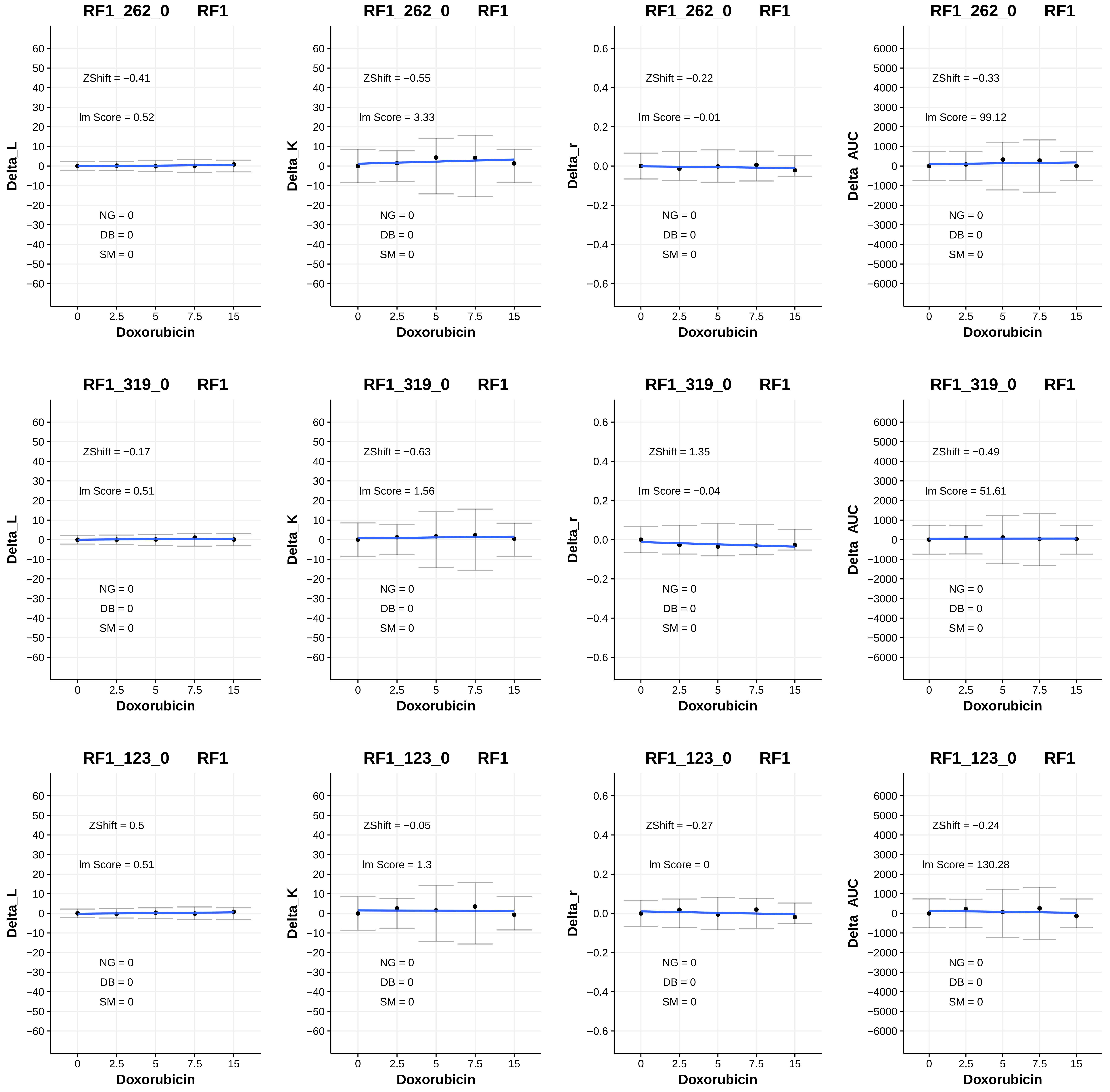
RF1_262_0 RF1
ZShift = −0.41
Im Score = 0.52
NG = 0
DB = 0
SM = 0
Delta_L
Doxorubicin
RF1_262_0 RF1
ZShift = −0.55
Im Score = 3.33
NG = 0
DB = 0
SM = 0
Delta_K
Doxorubicin
RF1_262_0 RF1
ZShift = −0.22
Im Score = −0.01
NG = 0
DB = 0
SM = 0
Delta_r
Doxorubicin
RF1_262_0 RF1
ZShift = −0.33
Im Score = 99.12
NG = 0
DB = 0
SM = 0
Delta_AUC
Doxorubicin
RF1_319_0 RF1
ZShift = −0.17
Im Score = 0.51
NG = 0
DB = 0
SM = 0
Delta_L
Doxorubicin
RF1_319_0 RF1
ZShift = −0.63
Im Score = 1.56
NG = 0
DB = 0
SM = 0
Delta_K
Doxorubicin
RF1_319_0 RF1
ZShift = 1.35
Im Score = −0.04
NG = 0
DB = 0
SM = 0
Delta_r
Doxorubicin
RF1_319_0 RF1
ZShift = −0.49
Im Score = 51.61
NG = 0
DB = 0
SM = 0
Delta_AUC
Doxorubicin
RF1_123_0 RF1
ZShift = 0.5
Im Score = 0.51
NG = 0
DB = 0
SM = 0
Delta_L
Doxorubicin
RF1_123_0 RF1
ZShift = −0.05
Im Score = 1.3
NG = 0
DB = 0
SM = 0
Delta_K
Doxorubicin
RF1_123_0 RF1
ZShift = −0.27
Im Score = 0
NG = 0
DB = 0
SM = 0
Delta_r
Doxorubicin
RF1_123_0 RF1
ZShift = −0.24
Im Score = 130.28
NG = 0
DB = 0
SM = 0
Delta_AUC
Doxorubicin

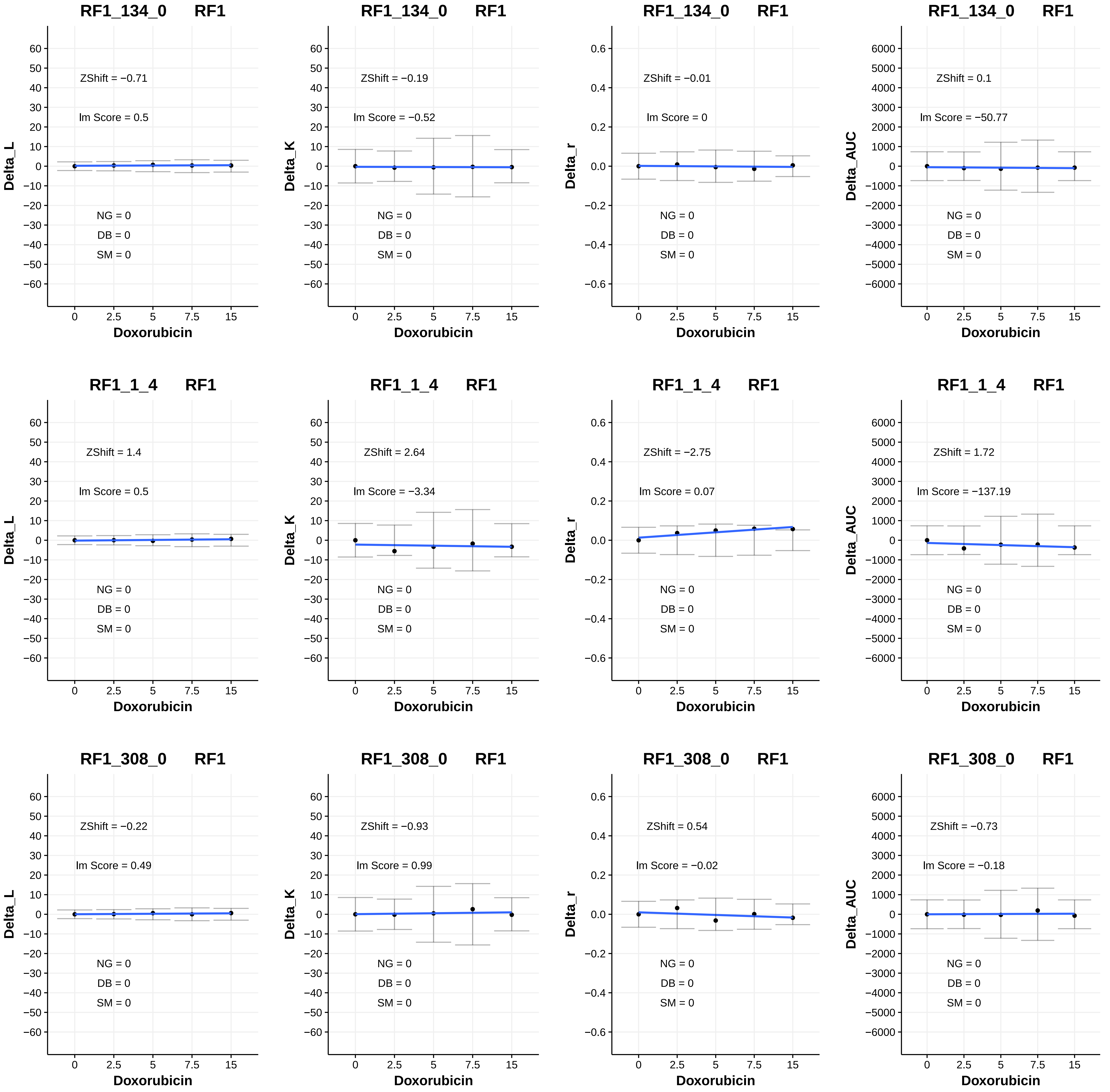
RF1_134_0 RF1
ZShift = −0.71
lm Score = 0.5
NG = 0
DB = 0
SM = 0
Delta_L
Doxorubicin
RF1_134_0 RF1
ZShift = −0.19
lm Score = −0.52
NG = 0
DB = 0
SM = 0
Delta_K
Doxorubicin
RF1_134_0 RF1
ZShift = −0.01
lm Score = 0
NG = 0
DB = 0
SM = 0
Delta_r
Doxorubicin
RF1_134_0 RF1
ZShift = 0.1
lm Score = −50.77
NG = 0
DB = 0
SM = 0
Delta_AUC
Doxorubicin
RF1_1_4 RF1
ZShift = 1.4
lm Score = 0.5
NG = 0
DB = 0
SM = 0
Delta_L
Doxorubicin
RF1_1_4 RF1
ZShift = 2.64
lm Score = −3.34
NG = 0
DB = 0
SM = 0
Delta_K
Doxorubicin
RF1_1_4 RF1
ZShift = −2.75
lm Score = 0.07
NG = 0
DB = 0
SM = 0
Delta_r
Doxorubicin
RF1_1_4 RF1
ZShift = 1.72
lm Score = −137.19
NG = 0
DB = 0
SM = 0
Delta_AUC
Doxorubicin
RF1_308_0 RF1
ZShift = −0.22
lm Score = 0.49
NG = 0
DB = 0
SM = 0
Delta_L
Doxorubicin
RF1_308_0 RF1
ZShift = −0.93
lm Score = 0.99
NG = 0
DB = 0
SM = 0
Delta_K
Doxorubicin
RF1_308_0 RF1
ZShift = 0.54
lm Score = −0.02
NG = 0
DB = 0
SM = 0
Delta_r
Doxorubicin
RF1_308_0 RF1
ZShift = −0.73
lm Score = −0.18
NG = 0
DB = 0
SM = 0
Delta_AUC
Doxorubicin

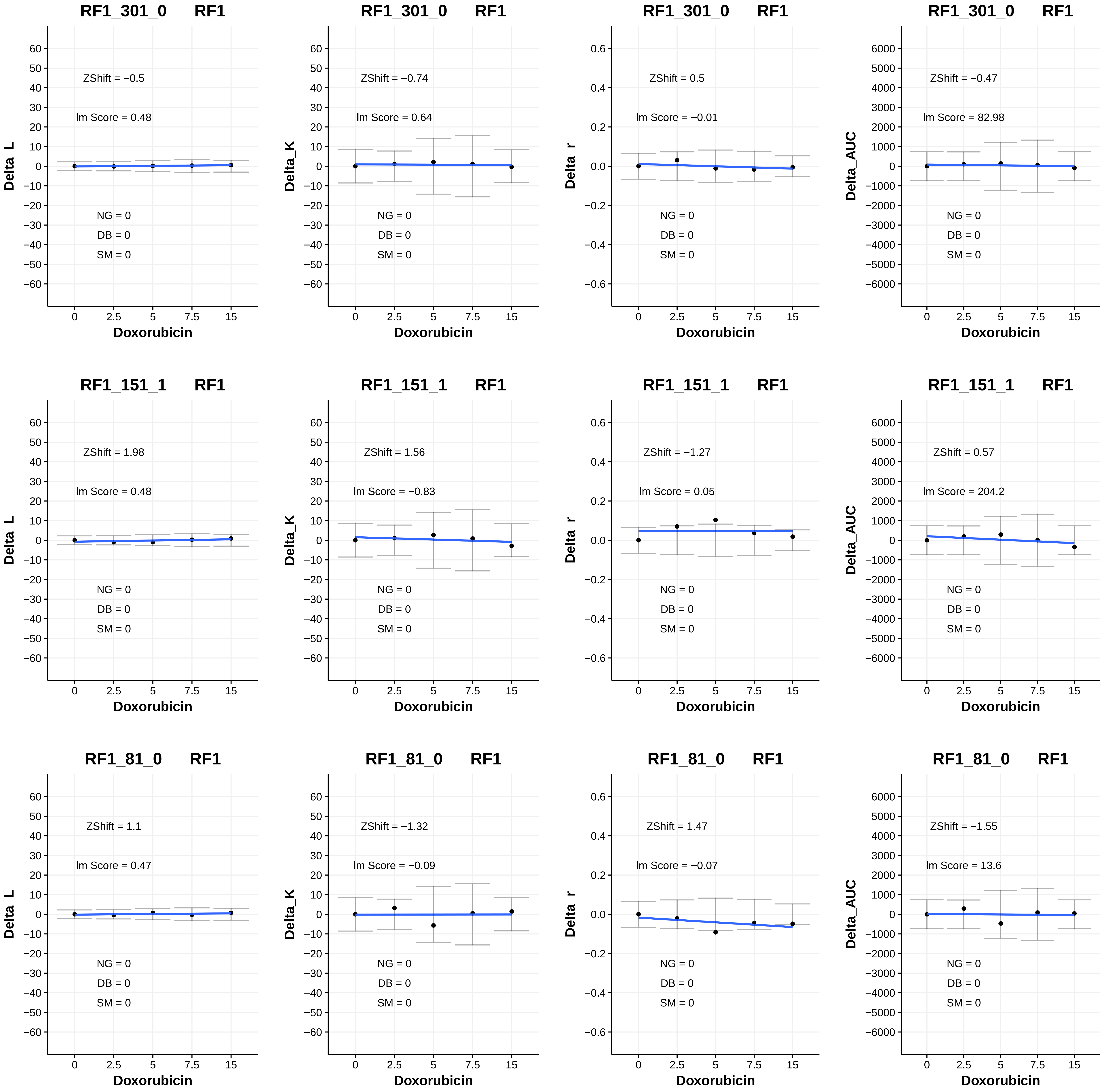
RF1_301_0 RF1
Delta_L
ZShift = −0.5
Im Score = 0.48
NG = 0
DB = 0
SM = 0
Doxorubicin
RF1_301_0 RF1
Delta_K
ZShift = −0.74
Im Score = 0.64
NG = 0
DB = 0
SM = 0
Doxorubicin
RF1_301_0 RF1
Delta_r
ZShift = 0.5
Im Score = −0.01
NG = 0
DB = 0
SM = 0
Doxorubicin
RF1_301_0 RF1
Delta_AUC
ZShift = −0.47
Im Score = 82.98
NG = 0
DB = 0
SM = 0
Doxorubicin
RF1_151_1 RF1
Delta_L
ZShift = 1.98
Im Score = 0.48
NG = 0
DB = 0
SM = 0
Doxorubicin
RF1_151_1 RF1
Delta_K
ZShift = 1.56
Im Score = −0.83
NG = 0
DB = 0
SM = 0
Doxorubicin
RF1_151_1 RF1
Delta_r
ZShift = −1.27
Im Score = 0.05
NG = 0
DB = 0
SM = 0
Doxorubicin
RF1_151_1 RF1
Delta_AUC
ZShift = 0.57
Im Score = 204.2
NG = 0
DB = 0
SM = 0
Doxorubicin
RF1_81_0 RF1
Delta_L
ZShift = 1.1
Im Score = 0.47
NG = 0
DB = 0
SM = 0
Doxorubicin
RF1_81_0 RF1
Delta_K
ZShift = −1.32
Im Score = −0.09
NG = 0
DB = 0
SM = 0
Doxorubicin
RF1_81_0 RF1
Delta_r
ZShift = 1.47
Im Score = −0.07
NG = 0
DB = 0
SM = 0
Doxorubicin
RF1_81_0 RF1
Delta_AUC
ZShift = −1.55
Im Score = 13.6
NG = 0
DB = 0
SM = 0
Doxorubicin

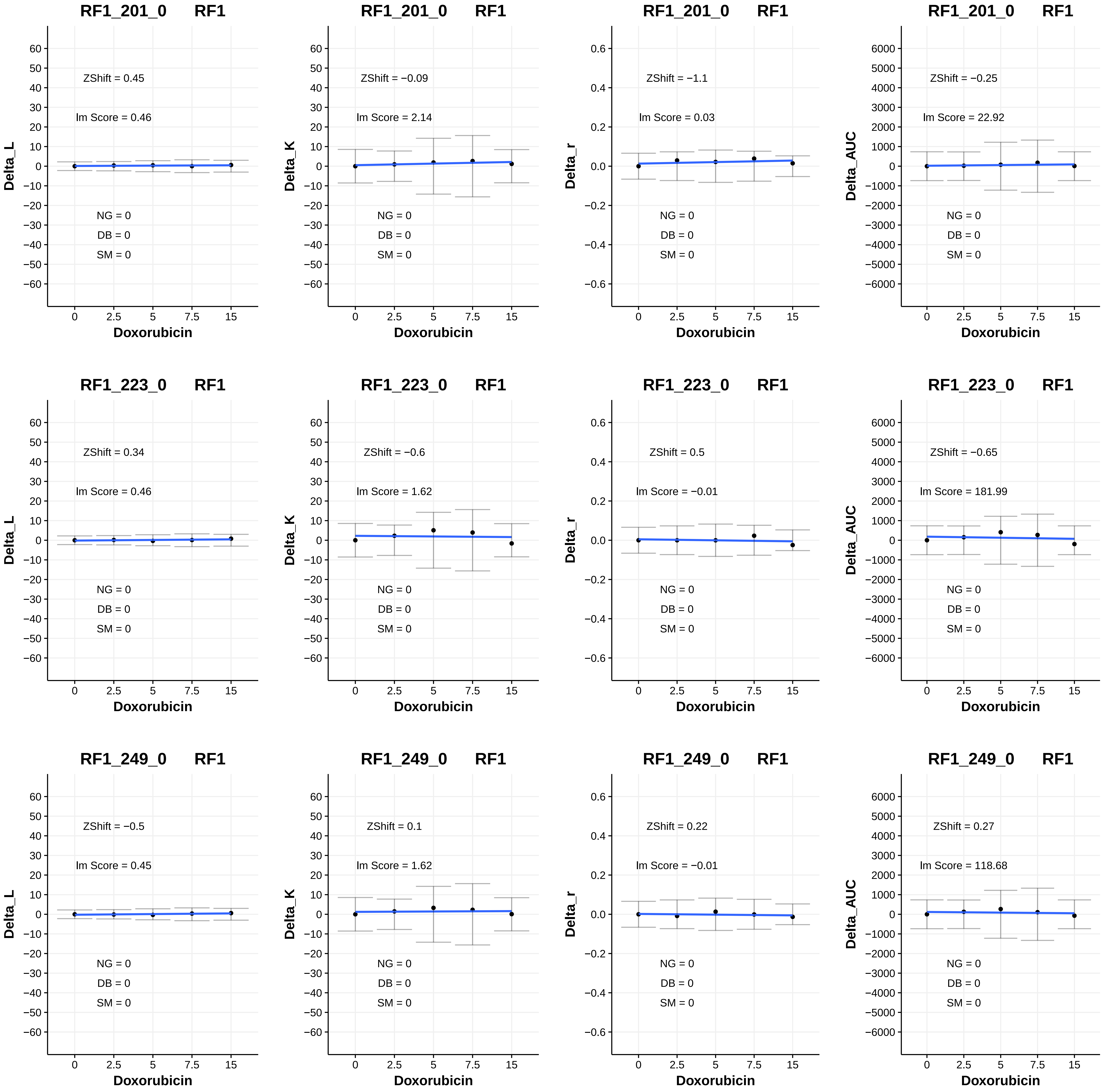
RF1_201_0 RF1
ZShift = 0.45
Im Score = 0.46
NG = 0
DB = 0
SM = 0
Delta_L
Doxorubicin
RF1_201_0 RF1
ZShift = −0.09
Im Score = 2.14
NG = 0
DB = 0
SM = 0
Delta_K
Doxorubicin
RF1_201_0 RF1
ZShift = −1.1
Im Score = 0.03
NG = 0
DB = 0
SM = 0
Delta_r
Doxorubicin
RF1_201_0 RF1
ZShift = −0.25
Im Score = 22.92
NG = 0
DB = 0
SM = 0
Delta_AUC
Doxorubicin
RF1_223_0 RF1
ZShift = 0.34
Im Score = 0.46
NG = 0
DB = 0
SM = 0
Delta_L
Doxorubicin
RF1_223_0 RF1
ZShift = −0.6
Im Score = 1.62
NG = 0
DB = 0
SM = 0
Delta_K
Doxorubicin
RF1_223_0 RF1
ZShift = 0.5
Im Score = −0.01
NG = 0
DB = 0
SM = 0
Delta_r
Doxorubicin
RF1_223_0 RF1
ZShift = −0.65
Im Score = 181.99
NG = 0
DB = 0
SM = 0
Delta_AUC
Doxorubicin
RF1_249_0 RF1
ZShift = −0.5
Im Score = 0.45
NG = 0
DB = 0
SM = 0
Delta_L
Doxorubicin
RF1_249_0 RF1
ZShift = 0.1
Im Score = 1.62
NG = 0
DB = 0
SM = 0
Delta_K
Doxorubicin
RF1_249_0 RF1
ZShift = 0.22
Im Score = −0.01
NG = 0
DB = 0
SM = 0
Delta_r
Doxorubicin
RF1_249_0 RF1
ZShift = 0.27
Im Score = 118.68
NG = 0
DB = 0
SM = 0
Delta_AUC
Doxorubicin
0
2.5
5
7.5
15

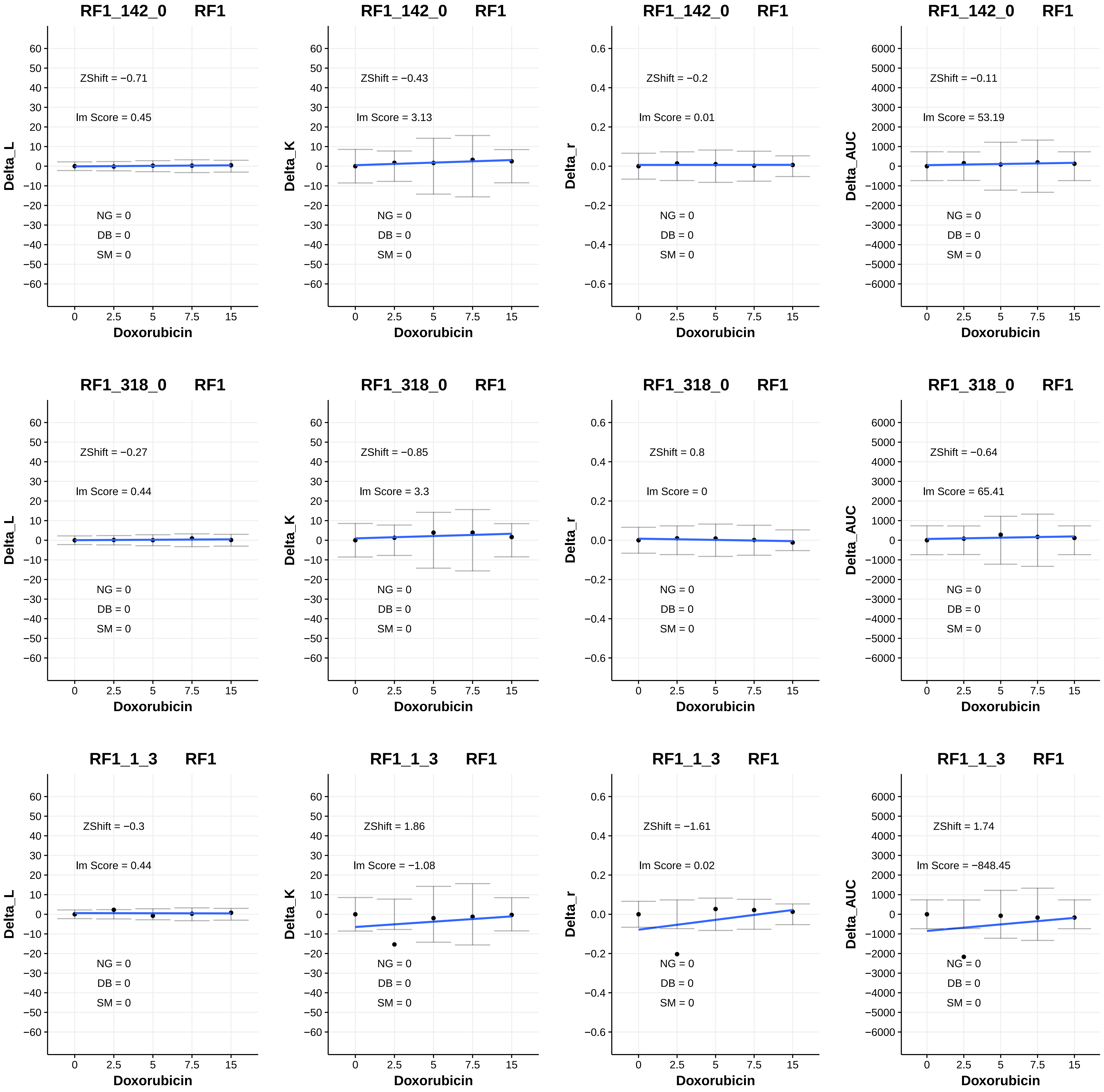
RF1_142_0 RF1
ZShift = −0.71
lm Score = 0.45
NG = 0
DB = 0
SM = 0
Delta_L
Doxorubicin
RF1_142_0 RF1
ZShift = −0.43
lm Score = 3.13
NG = 0
DB = 0
SM = 0
Delta_K
Doxorubicin
RF1_142_0 RF1
ZShift = −0.2
lm Score = 0.01
NG = 0
DB = 0
SM = 0
Delta_r
Doxorubicin
RF1_142_0 RF1
ZShift = −0.11
lm Score = 53.19
NG = 0
DB = 0
SM = 0
Delta_AUC
Doxorubicin
RF1_318_0 RF1
ZShift = −0.27
lm Score = 0.44
NG = 0
DB = 0
SM = 0
Delta_L
Doxorubicin
RF1_318_0 RF1
ZShift = −0.85
lm Score = 3.3
NG = 0
DB = 0
SM = 0
Delta_K
Doxorubicin
RF1_318_0 RF1
ZShift = 0.8
lm Score = 0
NG = 0
DB = 0
SM = 0
Delta_r
Doxorubicin
RF1_318_0 RF1
ZShift = −0.64
lm Score = 65.41
NG = 0
DB = 0
SM = 0
Delta_AUC
Doxorubicin
RF1_1_3 RF1
ZShift = −0.3
lm Score = 0.44
NG = 0
DB = 0
SM = 0
Delta_L
Doxorubicin
RF1_1_3 RF1
ZShift = 1.86
lm Score = −1.08
NG = 0
DB = 0
SM = 0
Delta_K
Doxorubicin
RF1_1_3 RF1
ZShift = −1.61
lm Score = 0.02
NG = 0
DB = 0
SM = 0
Delta_r
Doxorubicin
RF1_1_3 RF1
ZShift = 1.74
lm Score = −848.45
NG = 0
DB = 0
SM = 0
Delta_AUC
Doxorubicin

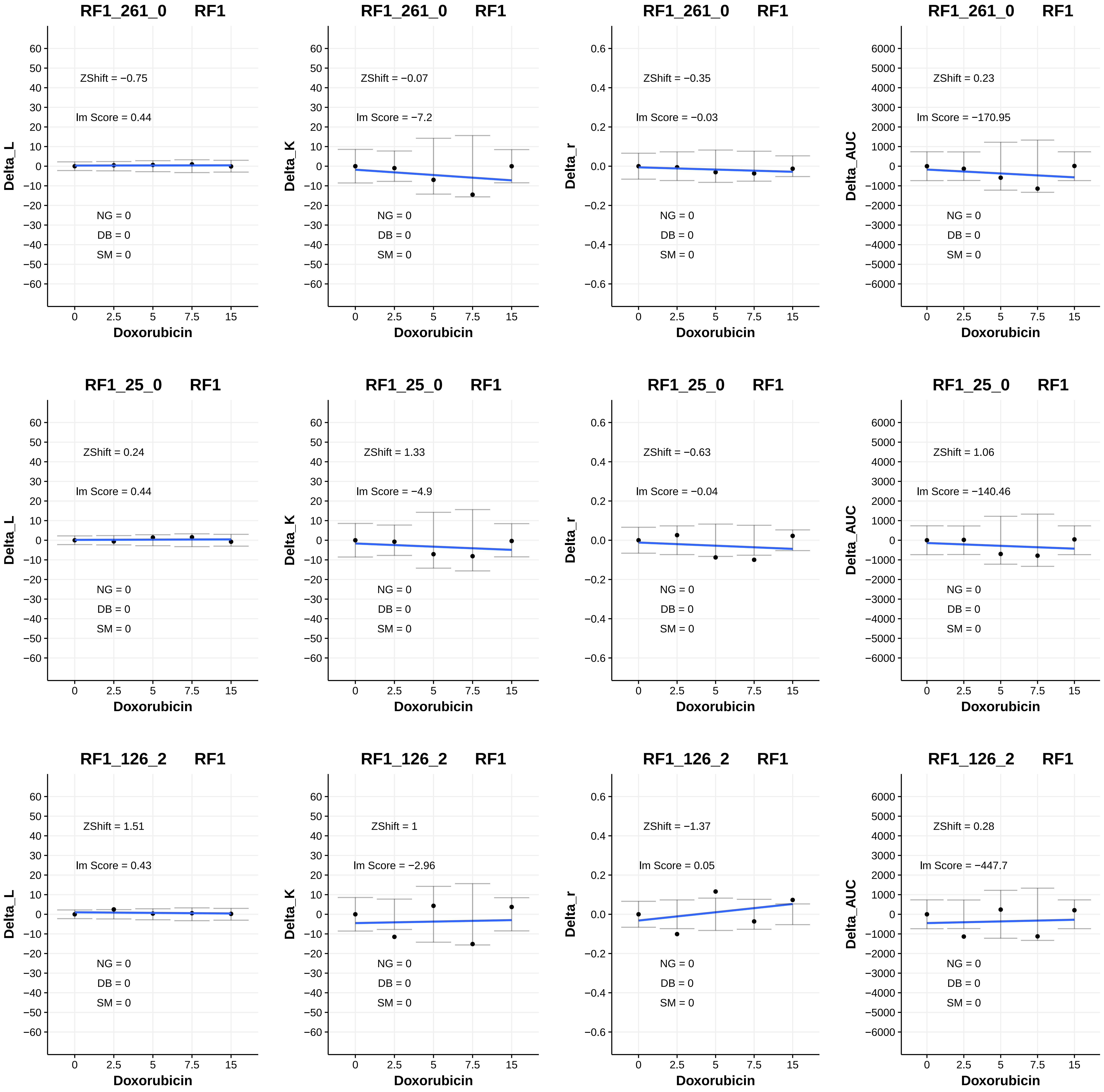
RF1_261_0 RF1
Delta_L
ZShift = −0.75
Im Score = 0.44
NG = 0
DB = 0
SM = 0
Doxorubicin
RF1_261_0 RF1
Delta_K
ZShift = −0.07
Im Score = −7.2
NG = 0
DB = 0
SM = 0
Doxorubicin
RF1_261_0 RF1
Delta_r
ZShift = −0.35
Im Score = −0.03
NG = 0
DB = 0
SM = 0
Doxorubicin
RF1_261_0 RF1
Delta_AUC
ZShift = 0.23
Im Score = −170.95
NG = 0
DB = 0
SM = 0
Doxorubicin
RF1_25_0 RF1
Delta_L
ZShift = 0.24
Im Score = 0.44
NG = 0
DB = 0
SM = 0
Doxorubicin
RF1_25_0 RF1
Delta_K
ZShift = 1.33
Im Score = −4.9
NG = 0
DB = 0
SM = 0
Doxorubicin
RF1_25_0 RF1
Delta_r
ZShift = −0.63
Im Score = −0.04
NG = 0
DB = 0
SM = 0
Doxorubicin
RF1_25_0 RF1
Delta_AUC
ZShift = 1.06
Im Score = −140.46
NG = 0
DB = 0
SM = 0
Doxorubicin
RF1_126_2 RF1
Delta_L
ZShift = 1.51
Im Score = 0.43
NG = 0
DB = 0
SM = 0
Doxorubicin
RF1_126_2 RF1
Delta_K
ZShift = 1
Im Score = −2.96
NG = 0
DB = 0
SM = 0
Doxorubicin
RF1_126_2 RF1
Delta_r
ZShift = −1.37
Im Score = 0.05
NG = 0
DB = 0
SM = 0
Doxorubicin
RF1_126_2 RF1
Delta_AUC
ZShift = 0.28
Im Score = −447.7
NG = 0
DB = 0
SM = 0
Doxorubicin

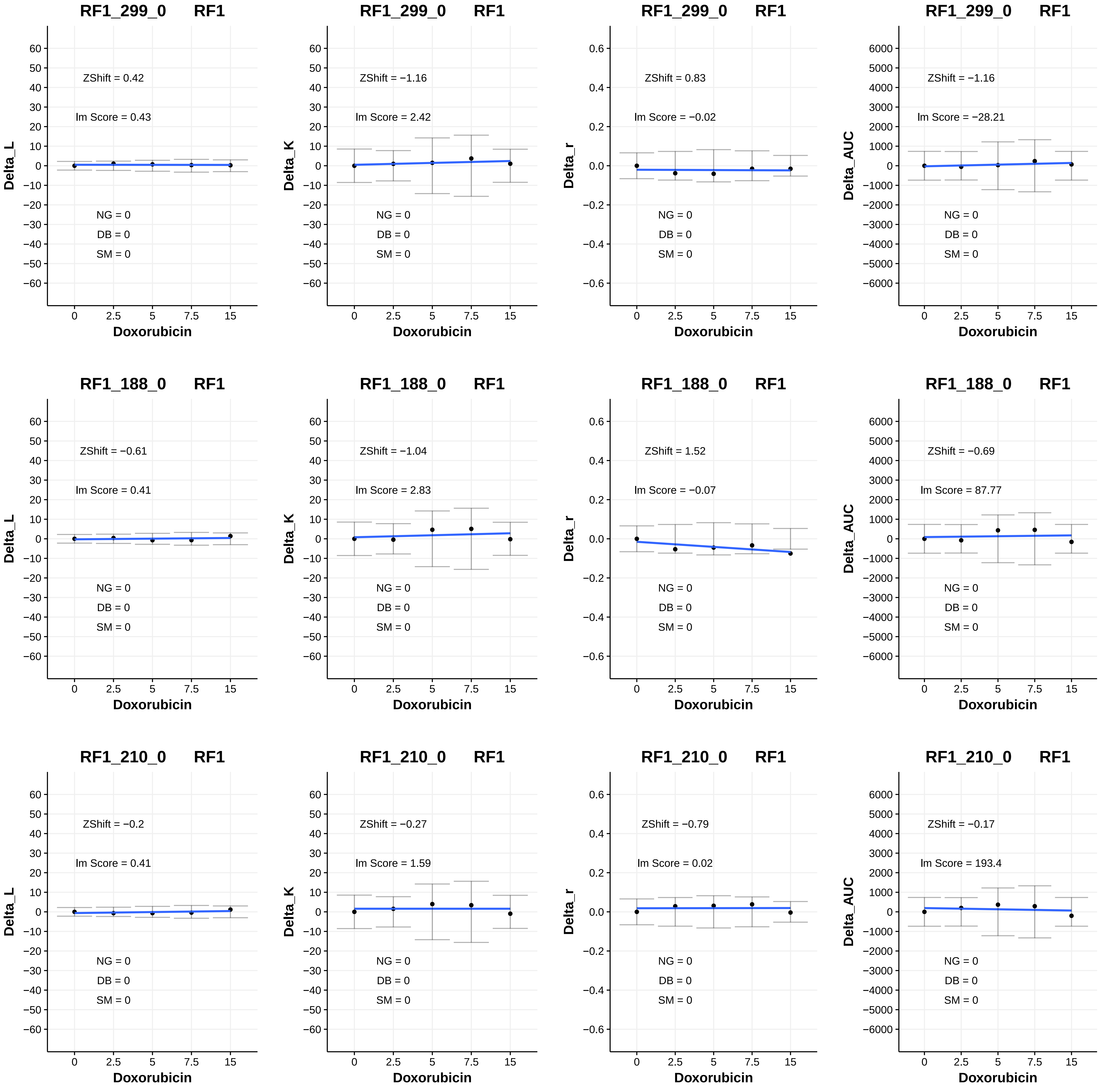
RF1_299_0 RF1
ZShift = 0.42
lm Score = 0.43
NG = 0
DB = 0
SM = 0
Delta_L
Doxorubicin
RF1_299_0 RF1
ZShift = −1.16
lm Score = 2.42
NG = 0
DB = 0
SM = 0
Delta_K
Doxorubicin
RF1_299_0 RF1
ZShift = 0.83
lm Score = −0.02
NG = 0
DB = 0
SM = 0
Delta_r
Doxorubicin
RF1_299_0 RF1
ZShift = −1.16
lm Score = −28.21
NG = 0
DB = 0
SM = 0
Delta_AUC
Doxorubicin
RF1_188_0 RF1
ZShift = −0.61
lm Score = 0.41
NG = 0
DB = 0
SM = 0
Delta_L
Doxorubicin
RF1_188_0 RF1
ZShift = −1.04
lm Score = 2.83
NG = 0
DB = 0
SM = 0
Delta_K
Doxorubicin
RF1_188_0 RF1
ZShift = 1.52
lm Score = −0.07
NG = 0
DB = 0
SM = 0
Delta_r
Doxorubicin
RF1_188_0 RF1
ZShift = −0.69
lm Score = 87.77
NG = 0
DB = 0
SM = 0
Delta_AUC
Doxorubicin
RF1_210_0 RF1
ZShift = −0.2
lm Score = 0.41
NG = 0
DB = 0
SM = 0
Delta_L
Doxorubicin
RF1_210_0 RF1
ZShift = −0.27
lm Score = 1.59
NG = 0
DB = 0
SM = 0
Delta_K
Doxorubicin
RF1_210_0 RF1
ZShift = −0.79
lm Score = 0.02
NG = 0
DB = 0
SM = 0
Delta_r
Doxorubicin
RF1_210_0 RF1
ZShift = −0.17
lm Score = 193.4
NG = 0
DB = 0
SM = 0
Delta_AUC
Doxorubicin

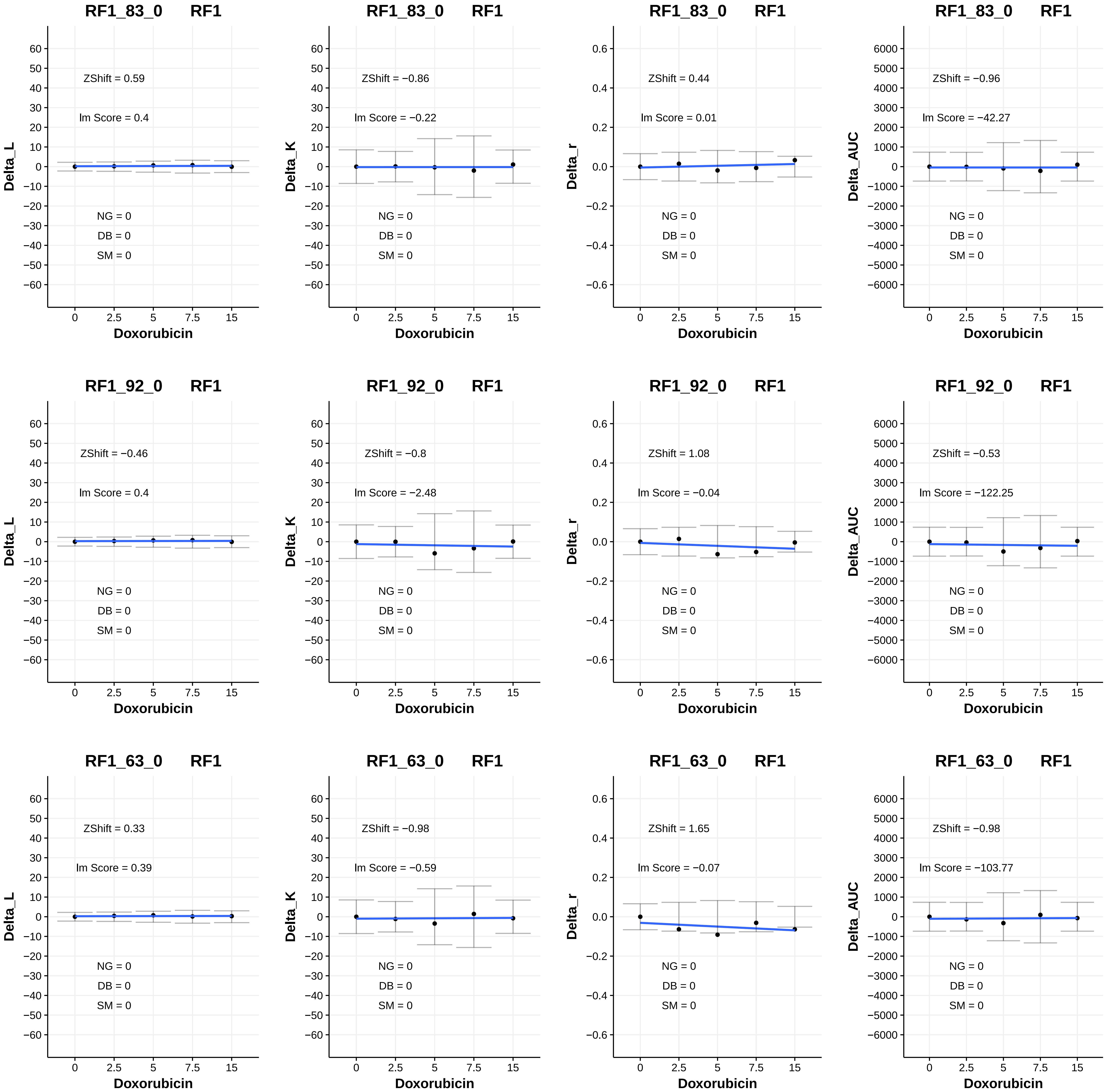

RF1_83_0 RF1
ZShift = 0.59
lm Score = 0.4
NG = 0
DB = 0
SM = 0
Delta_L
Doxorubicin
RF1_83_0 RF1
ZShift = −0.86
lm Score = −0.22
NG = 0
DB = 0
SM = 0
Delta_K
Doxorubicin
RF1_83_0 RF1
ZShift = 0.44
lm Score = 0.01
NG = 0
DB = 0
SM = 0
Delta_r
Doxorubicin
RF1_83_0 RF1
ZShift = −0.96
lm Score = −42.27
NG = 0
DB = 0
SM = 0
Delta_AUC
Doxorubicin
RF1_92_0 RF1
ZShift = −0.46
lm Score = 0.4
NG = 0
DB = 0
SM = 0
Delta_L
Doxorubicin
RF1_92_0 RF1
ZShift = −0.8
lm Score = −2.48
NG = 0
DB = 0
SM = 0
Delta_K
Doxorubicin
RF1_92_0 RF1
ZShift = 1.08
lm Score = −0.04
NG = 0
DB = 0
SM = 0
Delta_r
Doxorubicin
RF1_92_0 RF1
ZShift = −0.53
lm Score = −122.25
NG = 0
DB = 0
SM = 0
Delta_AUC
Doxorubicin
RF1_63_0 RF1
ZShift = 0.33
lm Score = 0.39
NG = 0
DB = 0
SM = 0
Delta_L
Doxorubicin
RF1_63_0 RF1
ZShift = −0.98
lm Score = −0.59
NG = 0
DB = 0
SM = 0
Delta_K
Doxorubicin
RF1_63_0 RF1
ZShift = 1.65
lm Score = −0.07
NG = 0
DB = 0
SM = 0
Delta_r
Doxorubicin
RF1_63_0 RF1
ZShift = −0.98
lm Score = −103.77
NG = 0
DB = 0
SM = 0
Delta_AUC
Doxorubicin

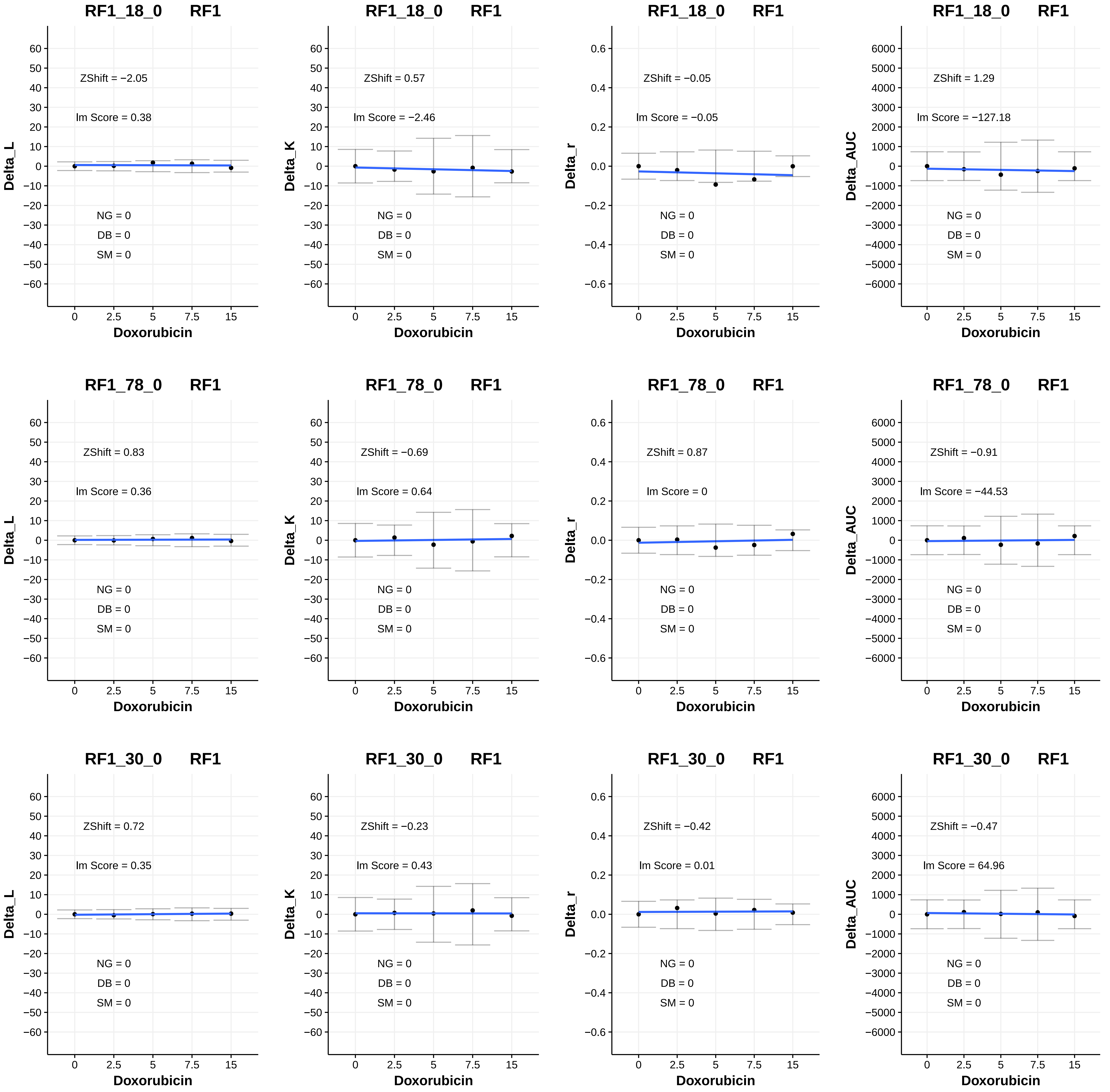
RF1_18_0 RF1
ZShift = −2.05
lm Score = 0.38
NG = 0
DB = 0
SM = 0
Delta_L
Doxorubicin
RF1_18_0 RF1
ZShift = 0.57
lm Score = −2.46
NG = 0
DB = 0
SM = 0
Delta_K
Doxorubicin
RF1_18_0 RF1
ZShift = −0.05
lm Score = −0.05
NG = 0
DB = 0
SM = 0
Delta_r
Doxorubicin
RF1_18_0 RF1
ZShift = 1.29
lm Score = −127.18
NG = 0
DB = 0
SM = 0
Delta_AUC
Doxorubicin
RF1_78_0 RF1
ZShift = 0.83
lm Score = 0.36
NG = 0
DB = 0
SM = 0
Delta_L
Doxorubicin
RF1_78_0 RF1
ZShift = −0.69
lm Score = 0.64
NG = 0
DB = 0
SM = 0
Delta_K
Doxorubicin
RF1_78_0 RF1
ZShift = 0.87
lm Score = 0
NG = 0
DB = 0
SM = 0
Delta_r
Doxorubicin
RF1_78_0 RF1
ZShift = −0.91
lm Score = −44.53
NG = 0
DB = 0
SM = 0
Delta_AUC
Doxorubicin
RF1_30_0 RF1
ZShift = 0.72
lm Score = 0.35
NG = 0
DB = 0
SM = 0
Delta_L
Doxorubicin
RF1_30_0 RF1
ZShift = −0.23
lm Score = 0.43
NG = 0
DB = 0
SM = 0
Delta_K
Doxorubicin
RF1_30_0 RF1
ZShift = −0.42
lm Score = 0.01
NG = 0
DB = 0
SM = 0
Delta_r
Doxorubicin
RF1_30_0 RF1
ZShift = −0.47
lm Score = 64.96
NG = 0
DB = 0
SM = 0
Delta_AUC
Doxorubicin

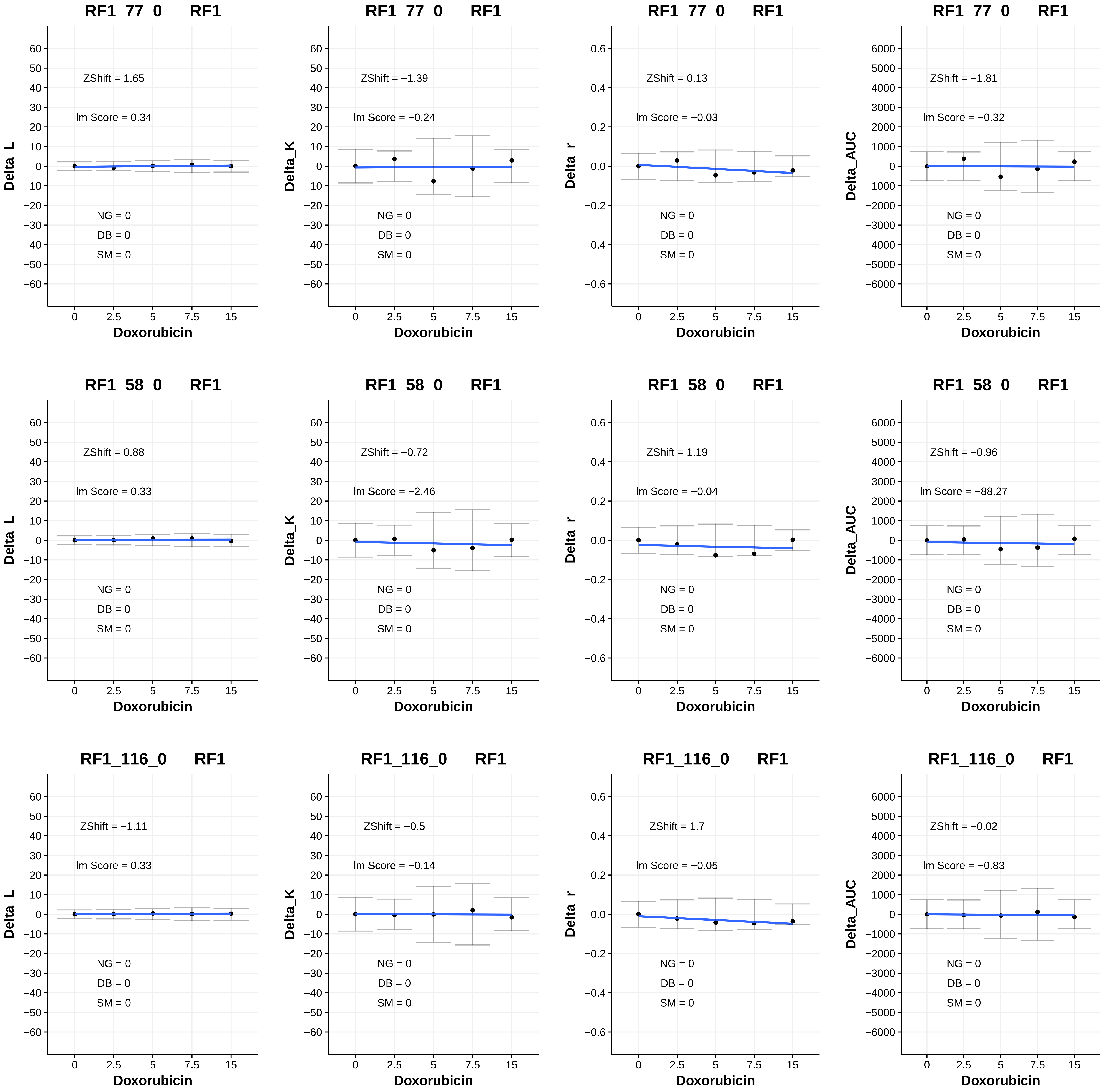
RF1_77_0 RF1
ZShift = 1.65
lm Score = 0.34
NG = 0
DB = 0
SM = 0
Delta_L
Doxorubicin
RF1_77_0 RF1
ZShift = −1.39
lm Score = −0.24
NG = 0
DB = 0
SM = 0
Delta_K
Doxorubicin
RF1_77_0 RF1
ZShift = 0.13
lm Score = −0.03
NG = 0
DB = 0
SM = 0
Delta_r
Doxorubicin
RF1_77_0 RF1
ZShift = −1.81
lm Score = −0.32
NG = 0
DB = 0
SM = 0
Delta_AUC
Doxorubicin
RF1_58_0 RF1
ZShift = 0.88
lm Score = 0.33
NG = 0
DB = 0
SM = 0
Delta_L
Doxorubicin
RF1_58_0 RF1
ZShift = −0.72
lm Score = −2.46
NG = 0
DB = 0
SM = 0
Delta_K
Doxorubicin
RF1_58_0 RF1
ZShift = 1.19
lm Score = −0.04
NG = 0
DB = 0
SM = 0
Delta_r
Doxorubicin
RF1_58_0 RF1
ZShift = −0.96
lm Score = −88.27
NG = 0
DB = 0
SM = 0
Delta_AUC
Doxorubicin
RF1_116_0 RF1
ZShift = −1.11
lm Score = 0.33
NG = 0
DB = 0
SM = 0
Delta_L
Doxorubicin
RF1_116_0 RF1
ZShift = −0.5
lm Score = −0.14
NG = 0
DB = 0
SM = 0
Delta_K
Doxorubicin
RF1_116_0 RF1
ZShift = 1.7
lm Score = −0.05
NG = 0
DB = 0
SM = 0
Delta_r
Doxorubicin
RF1_116_0 RF1
ZShift = −0.02
lm Score = −0.83
NG = 0
DB = 0
SM = 0
Delta_AUC
Doxorubicin

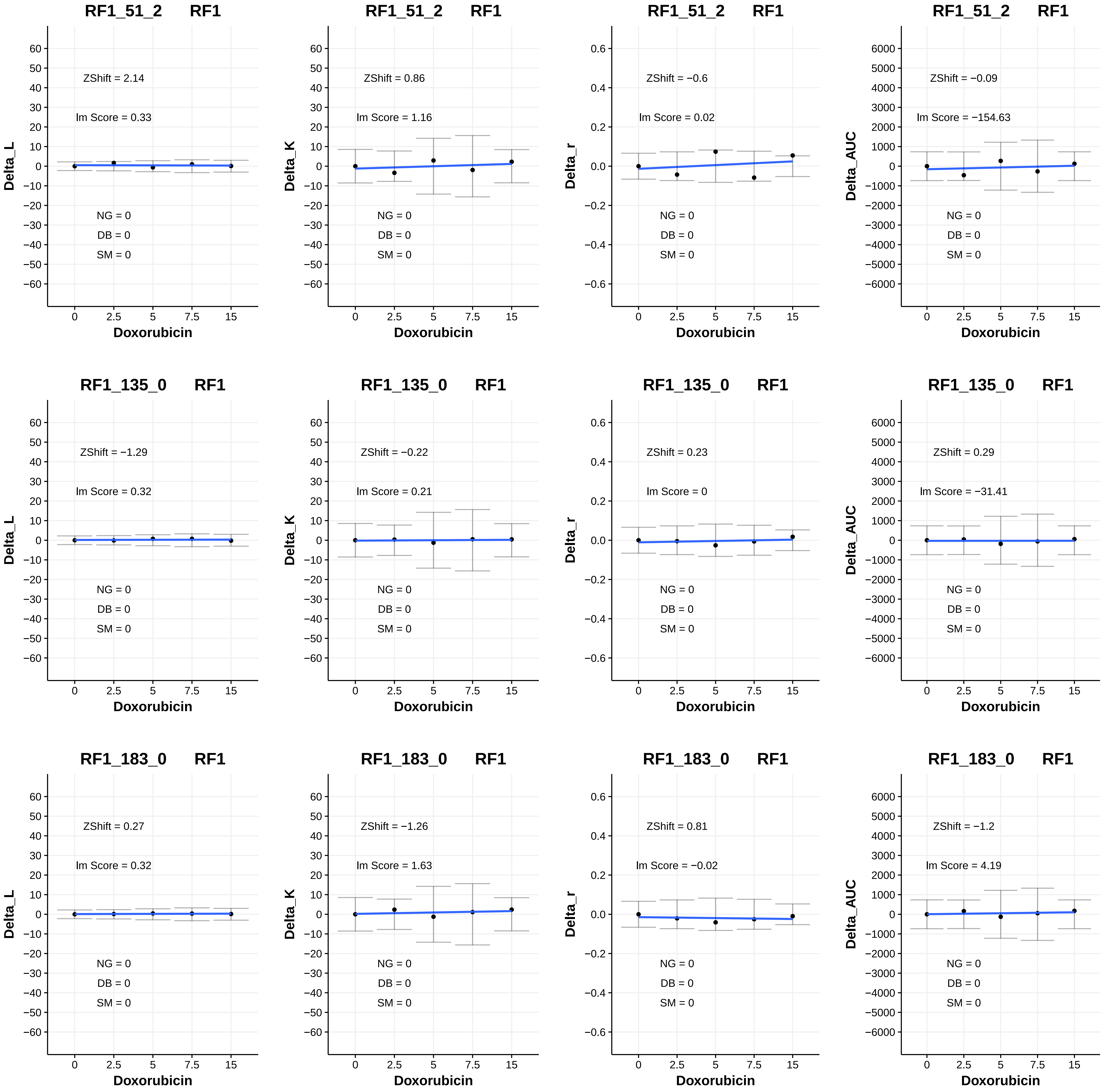

RF1_51_2 RF1
ZShift = 2.14
lm Score = 0.33
NG = 0
DB = 0
SM = 0
Delta_L
Doxorubicin
RF1_51_2 RF1
ZShift = 0.86
lm Score = 1.16
NG = 0
DB = 0
SM = 0
Delta_K
Doxorubicin
RF1_51_2 RF1
ZShift = −0.6
lm Score = 0.02
NG = 0
DB = 0
SM = 0
Delta_r
Doxorubicin
RF1_51_2 RF1
ZShift = −0.09
lm Score = −154.63
NG = 0
DB = 0
SM = 0
Delta_AUC
Doxorubicin
RF1_135_0 RF1
ZShift = −1.29
lm Score = 0.32
NG = 0
DB = 0
SM = 0
Delta_L
Doxorubicin
RF1_135_0 RF1
ZShift = −0.22
lm Score = 0.21
NG = 0
DB = 0
SM = 0
Delta_K
Doxorubicin
RF1_135_0 RF1
ZShift = 0.23
lm Score = 0
NG = 0
DB = 0
SM = 0
Delta_r
Doxorubicin
RF1_135_0 RF1
ZShift = 0.29
lm Score = −31.41
NG = 0
DB = 0
SM = 0
Delta_AUC
Doxorubicin
RF1_183_0 RF1
ZShift = 0.27
lm Score = 0.32
NG = 0
DB = 0
SM = 0
Delta_L
Doxorubicin
RF1_183_0 RF1
ZShift = −1.26
lm Score = 1.63
NG = 0
DB = 0
SM = 0
Delta_K
Doxorubicin
RF1_183_0 RF1
ZShift = 0.81
lm Score = −0.02
NG = 0
DB = 0
SM = 0
Delta_r
Doxorubicin
RF1_183_0 RF1
ZShift = −1.2
lm Score = 4.19
NG = 0
DB = 0
SM = 0
Delta_AUC
Doxorubicin

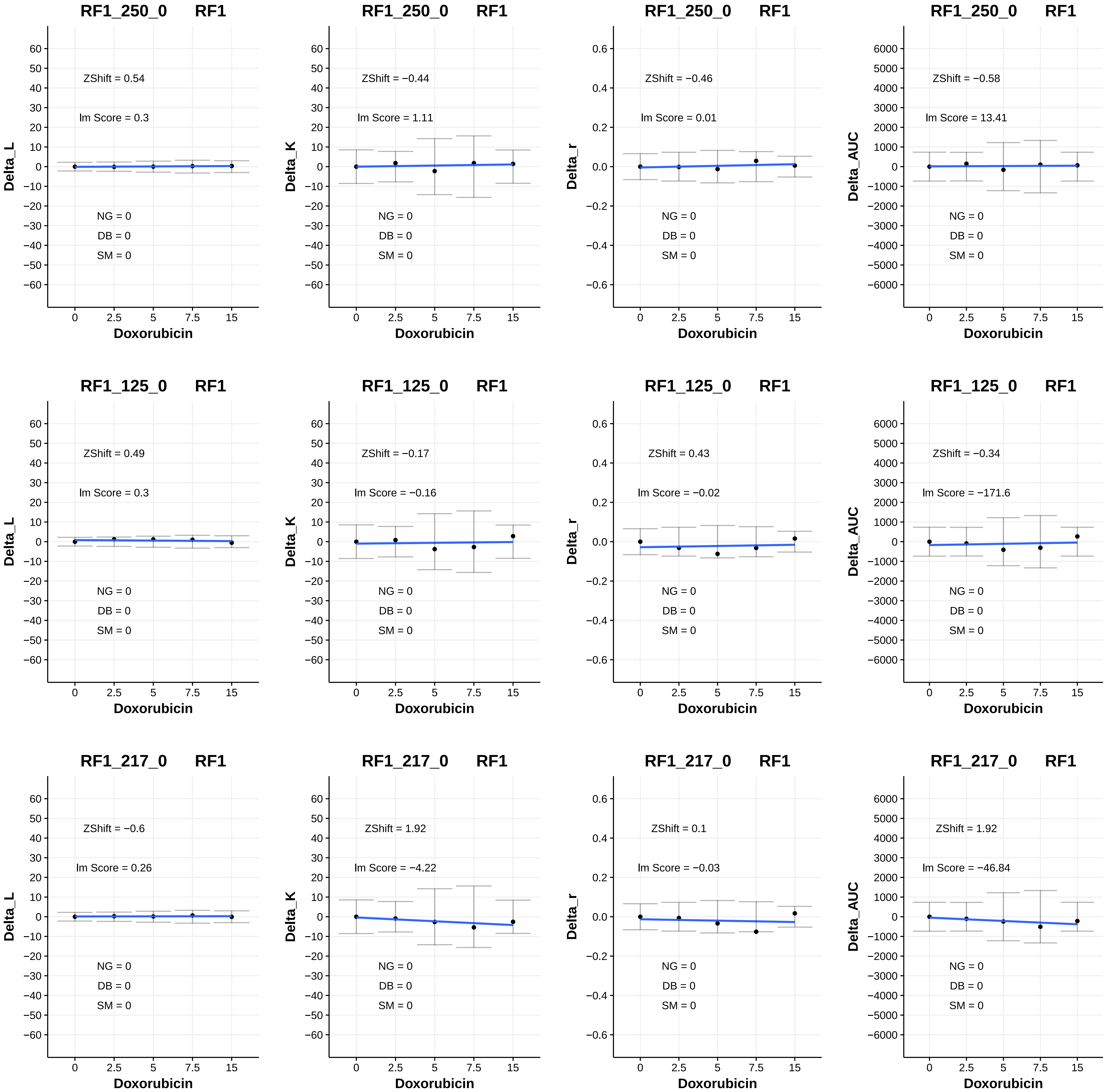

RF1_250_0 RF1
ZShift = 0.54
lm Score = 0.3
NG = 0
DB = 0
SM = 0
Delta_L
Doxorubicin
RF1_250_0 RF1
ZShift = −0.44
lm Score = 1.11
NG = 0
DB = 0
SM = 0
Delta_K
Doxorubicin
RF1_250_0 RF1
ZShift = −0.46
lm Score = 0.01
NG = 0
DB = 0
SM = 0
Delta_r
Doxorubicin
RF1_250_0 RF1
ZShift = −0.58
lm Score = 13.41
NG = 0
DB = 0
SM = 0
Delta_AUC
Doxorubicin
RF1_125_0 RF1
ZShift = 0.49
lm Score = 0.3
NG = 0
DB = 0
SM = 0
Delta_L
Doxorubicin
RF1_125_0 RF1
ZShift = −0.17
lm Score = −0.16
NG = 0
DB = 0
SM = 0
Delta_K
Doxorubicin
RF1_125_0 RF1
ZShift = 0.43
lm Score = −0.02
NG = 0
DB = 0
SM = 0
Delta_r
Doxorubicin
RF1_125_0 RF1
ZShift = −0.34
lm Score = −171.6
NG = 0
DB = 0
SM = 0
Delta_AUC
Doxorubicin
RF1_217_0 RF1
ZShift = −0.6
lm Score = 0.26
NG = 0
DB = 0
SM = 0
Delta_L
Doxorubicin
RF1_217_0 RF1
ZShift = 1.92
lm Score = −4.22
NG = 0
DB = 0
SM = 0
Delta_K
Doxorubicin
RF1_217_0 RF1
ZShift = 0.1
lm Score = −0.03
NG = 0
DB = 0
SM = 0
Delta_r
Doxorubicin
RF1_217_0 RF1
ZShift = 1.92
lm Score = −46.84
NG = 0
DB = 0
SM = 0
Delta_AUC
Doxorubicin

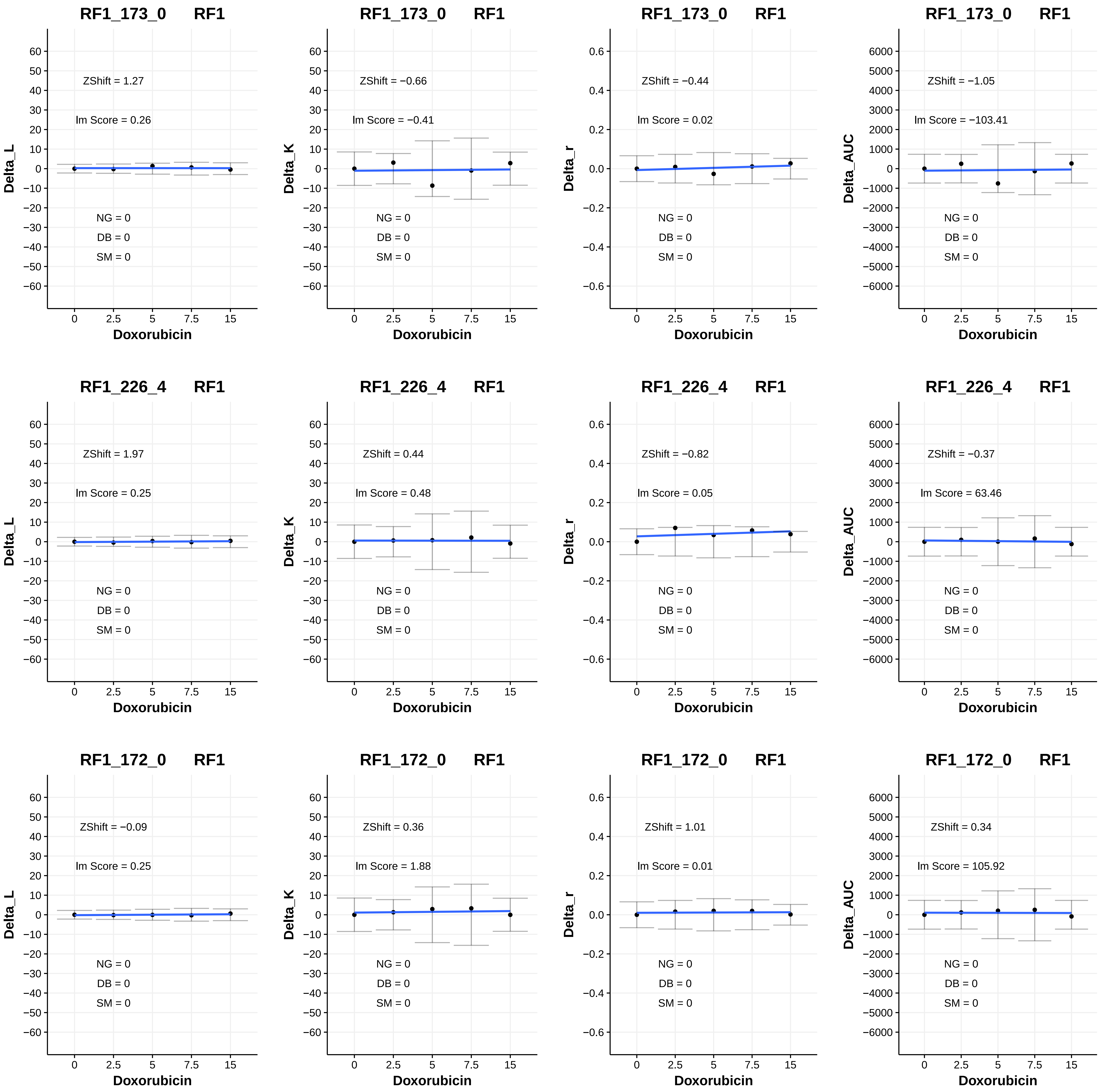

RF1_173_0 RF1
ZShift = 1.27
lm Score = 0.26
NG = 0
DB = 0
SM = 0
Delta_L
Doxorubicin
RF1_173_0 RF1
ZShift = −0.66
lm Score = −0.41
NG = 0
DB = 0
SM = 0
Delta_K
Doxorubicin
RF1_173_0 RF1
ZShift = −0.44
lm Score = 0.02
NG = 0
DB = 0
SM = 0
Delta_r
Doxorubicin
RF1_173_0 RF1
ZShift = −1.05
lm Score = −103.41
NG = 0
DB = 0
SM = 0
Delta_AUC
Doxorubicin
RF1_226_4 RF1
ZShift = 1.97
lm Score = 0.25
NG = 0
DB = 0
SM = 0
Delta_L
Doxorubicin
RF1_226_4 RF1
ZShift = 0.44
lm Score = 0.48
NG = 0
DB = 0
SM = 0
Delta_K
Doxorubicin
RF1_226_4 RF1
ZShift = −0.82
lm Score = 0.05
NG = 0
DB = 0
SM = 0
Delta_r
Doxorubicin
RF1_226_4 RF1
ZShift = −0.37
lm Score = 63.46
NG = 0
DB = 0
SM = 0
Delta_AUC
Doxorubicin
RF1_172_0 RF1
ZShift = −0.09
lm Score = 0.25
NG = 0
DB = 0
SM = 0
Delta_L
Doxorubicin
RF1_172_0 RF1
ZShift = 0.36
lm Score = 1.88
NG = 0
DB = 0
SM = 0
Delta_K
Doxorubicin
RF1_172_0 RF1
ZShift = 1.01
lm Score = 0.01
NG = 0
DB = 0
SM = 0
Delta_r
Doxorubicin
RF1_172_0 RF1
ZShift = 0.34
lm Score = 105.92
NG = 0
DB = 0
SM = 0
Delta_AUC
Doxorubicin

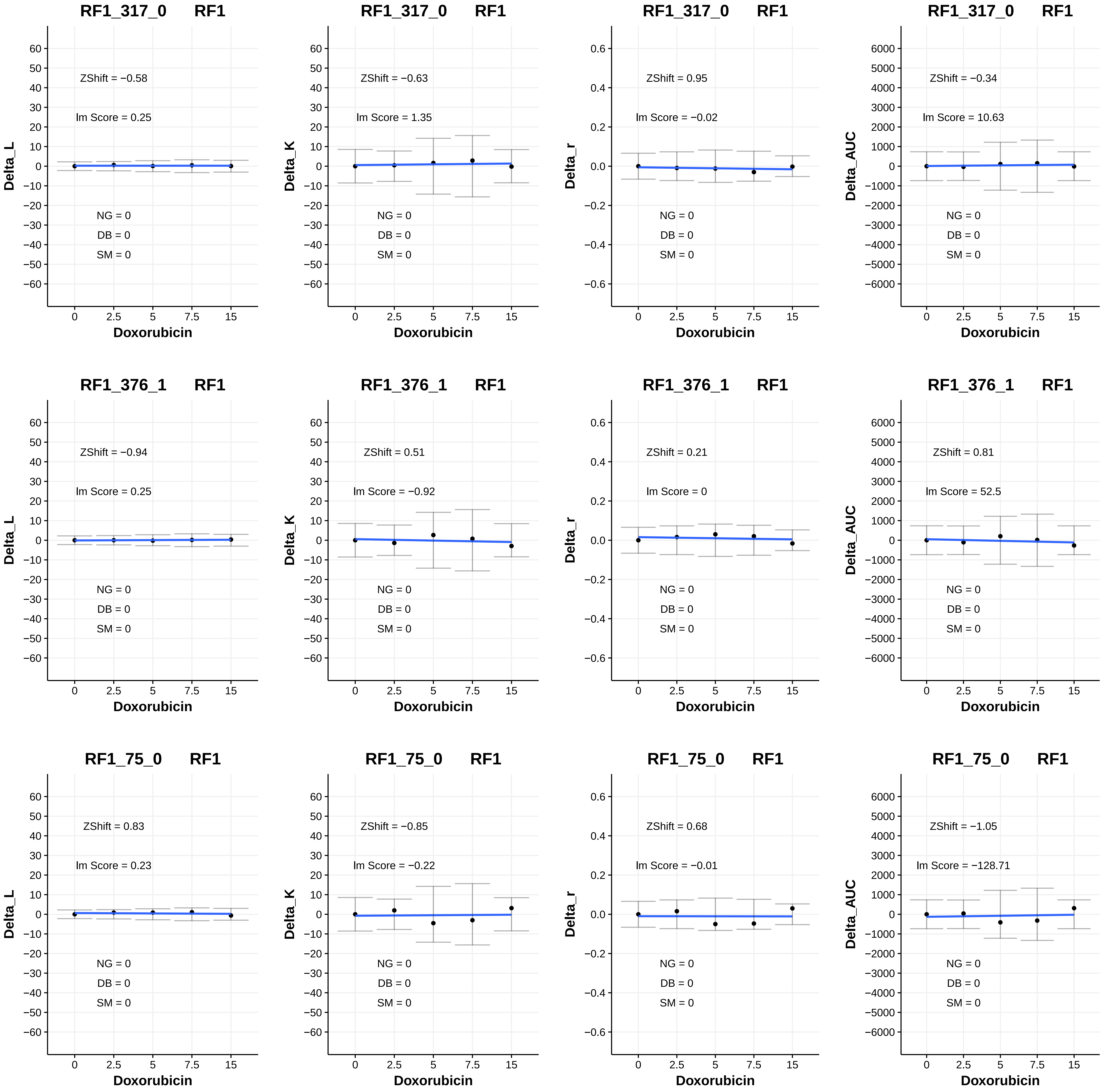

RF1_317_0 RF1
ZShift = −0.58
lm Score = 0.25
NG = 0
DB = 0
SM = 0
Delta_L
Doxorubicin
RF1_317_0 RF1
ZShift = −0.63
lm Score = 1.35
NG = 0
DB = 0
SM = 0
Delta_K
Doxorubicin
RF1_317_0 RF1
ZShift = 0.95
lm Score = −0.02
NG = 0
DB = 0
SM = 0
Delta_r
Doxorubicin
RF1_317_0 RF1
ZShift = −0.34
lm Score = 10.63
NG = 0
DB = 0
SM = 0
Delta_AUC
Doxorubicin
RF1_376_1 RF1
ZShift = −0.94
lm Score = 0.25
NG = 0
DB = 0
SM = 0
Delta_L
Doxorubicin
RF1_376_1 RF1
ZShift = 0.51
lm Score = −0.92
NG = 0
DB = 0
SM = 0
Delta_K
Doxorubicin
RF1_376_1 RF1
ZShift = 0.21
lm Score = 0
NG = 0
DB = 0
SM = 0
Delta_r
Doxorubicin
RF1_376_1 RF1
ZShift = 0.81
lm Score = 52.5
NG = 0
DB = 0
SM = 0
Delta_AUC
Doxorubicin
RF1_75_0 RF1
ZShift = 0.83
lm Score = 0.23
NG = 0
DB = 0
SM = 0
Delta_L
Doxorubicin
RF1_75_0 RF1
ZShift = −0.85
lm Score = −0.22
NG = 0
DB = 0
SM = 0
Delta_K
Doxorubicin
RF1_75_0 RF1
ZShift = 0.68
lm Score = −0.01
NG = 0
DB = 0
SM = 0
Delta_r
Doxorubicin
RF1_75_0 RF1
ZShift = −1.05
lm Score = −128.71
NG = 0
DB = 0
SM = 0
Delta_AUC
Doxorubicin

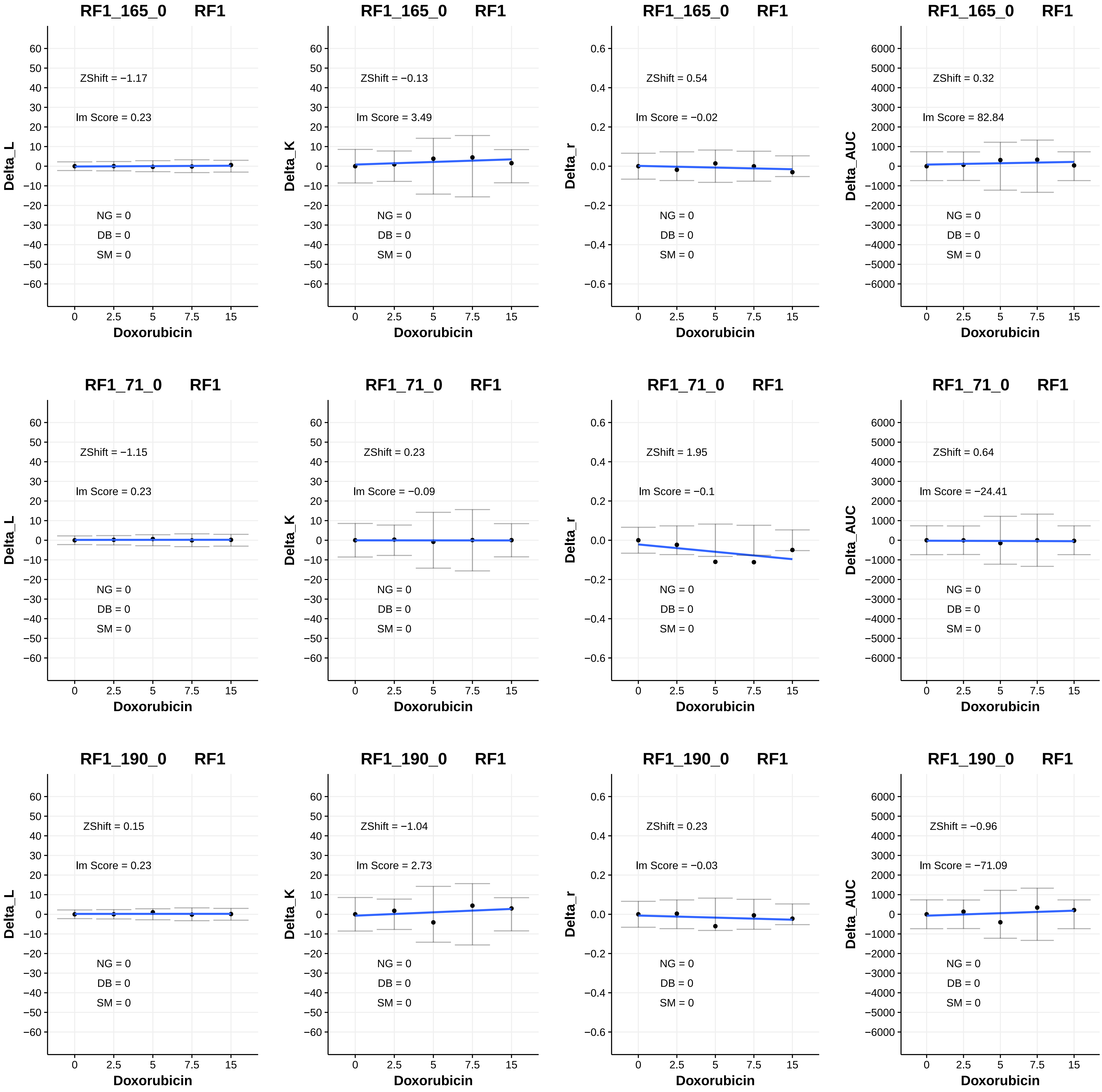
RF1_165_0 RF1
ZShift = −1.17
lm Score = 0.23
NG = 0
DB = 0
SM = 0
Delta_L
Doxorubicin
RF1_165_0 RF1
ZShift = −0.13
lm Score = 3.49
NG = 0
DB = 0
SM = 0
Delta_K
Doxorubicin
RF1_165_0 RF1
ZShift = 0.54
lm Score = −0.02
NG = 0
DB = 0
SM = 0
Delta_r
Doxorubicin
RF1_165_0 RF1
ZShift = 0.32
lm Score = 82.84
NG = 0
DB = 0
SM = 0
Delta_AUC
Doxorubicin
RF1_71_0 RF1
ZShift = −1.15
lm Score = 0.23
NG = 0
DB = 0
SM = 0
Delta_L
Doxorubicin
RF1_71_0 RF1
ZShift = 0.23
lm Score = −0.09
NG = 0
DB = 0
SM = 0
Delta_K
Doxorubicin
RF1_71_0 RF1
ZShift = 1.95
lm Score = −0.1
NG = 0
DB = 0
SM = 0
Delta_r
Doxorubicin
RF1_71_0 RF1
ZShift = 0.64
lm Score = −24.41
NG = 0
DB = 0
SM = 0
Delta_AUC
Doxorubicin
RF1_190_0 RF1
ZShift = 0.15
lm Score = 0.23
NG = 0
DB = 0
SM = 0
Delta_L
Doxorubicin
RF1_190_0 RF1
ZShift = −1.04
lm Score = 2.73
NG = 0
DB = 0
SM = 0
Delta_K
Doxorubicin
RF1_190_0 RF1
ZShift = 0.23
lm Score = −0.03
NG = 0
DB = 0
SM = 0
Delta_r
Doxorubicin
RF1_190_0 RF1
ZShift = −0.96
lm Score = −71.09
NG = 0
DB = 0
SM = 0
Delta_AUC
Doxorubicin

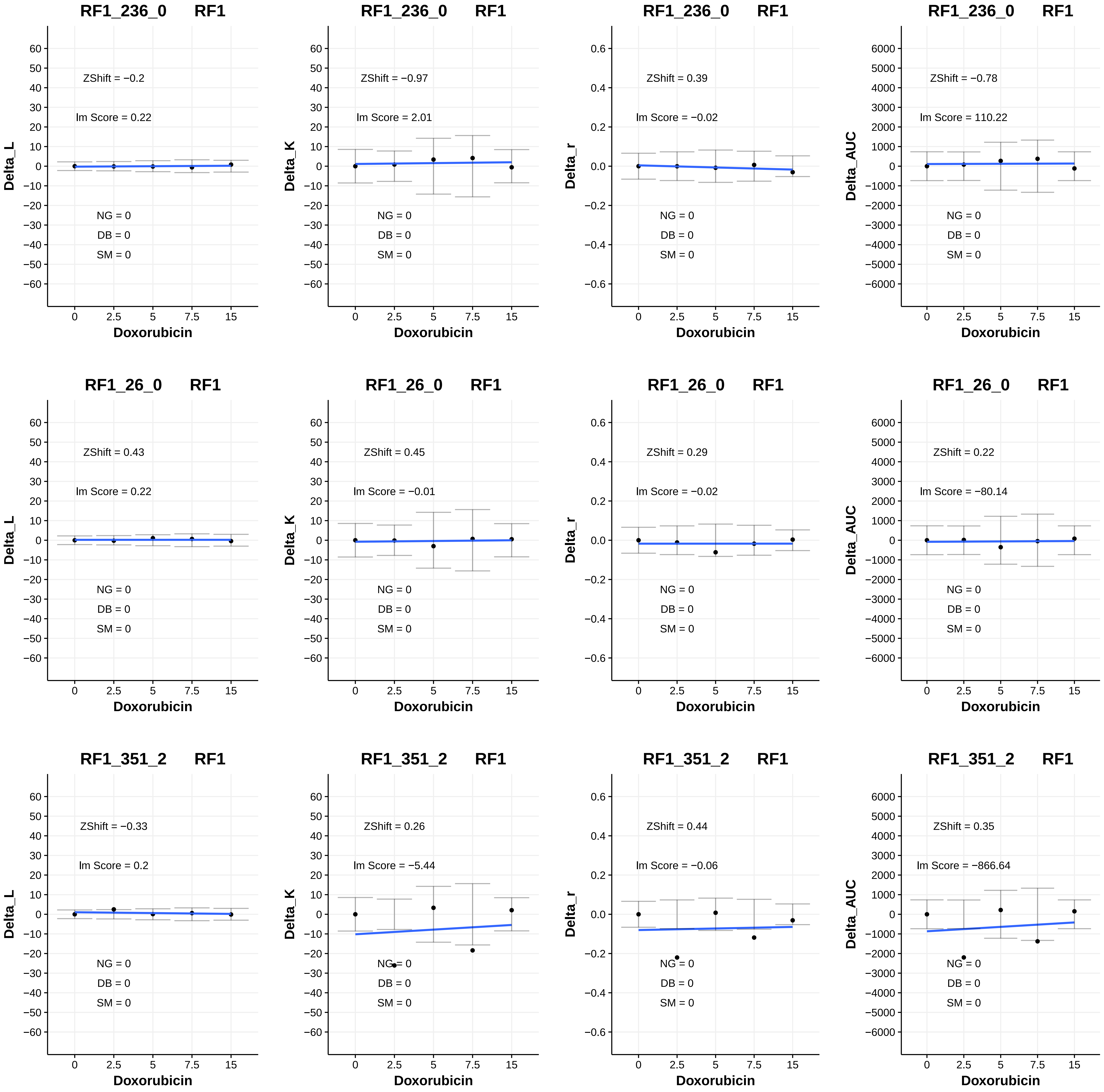

RF1_236_0 RF1
ZShift = −0.2
lm Score = 0.22
NG = 0
DB = 0
SM = 0
Delta_L
Doxorubicin
RF1_236_0 RF1
ZShift = −0.97
lm Score = 2.01
NG = 0
DB = 0
SM = 0
Delta_K
Doxorubicin
RF1_236_0 RF1
ZShift = 0.39
lm Score = −0.02
NG = 0
DB = 0
SM = 0
Delta_r
Doxorubicin
RF1_236_0 RF1
ZShift = −0.78
lm Score = 110.22
NG = 0
DB = 0
SM = 0
Delta_AUC
Doxorubicin
RF1_26_0 RF1
ZShift = 0.43
lm Score = 0.22
NG = 0
DB = 0
SM = 0
Delta_L
Doxorubicin
RF1_26_0 RF1
ZShift = 0.45
lm Score = −0.01
NG = 0
DB = 0
SM = 0
Delta_K
Doxorubicin
RF1_26_0 RF1
ZShift = 0.29
lm Score = −0.02
NG = 0
DB = 0
SM = 0
Delta_r
Doxorubicin
RF1_26_0 RF1
ZShift = 0.22
lm Score = −80.14
NG = 0
DB = 0
SM = 0
Delta_AUC
Doxorubicin
RF1_351_2 RF1
ZShift = −0.33
lm Score = 0.2
NG = 0
DB = 0
SM = 0
Delta_L
Doxorubicin
RF1_351_2 RF1
ZShift = 0.26
lm Score = −5.44
NG = 0
DB = 0
SM = 0
Delta_K
Doxorubicin
RF1_351_2 RF1
ZShift = 0.44
lm Score = −0.06
NG = 0
DB = 0
SM = 0
Delta_r
Doxorubicin
RF1_351_2 RF1
ZShift = 0.35
lm Score = −866.64
NG = 0
DB = 0
SM = 0
Delta_AUC
Doxorubicin

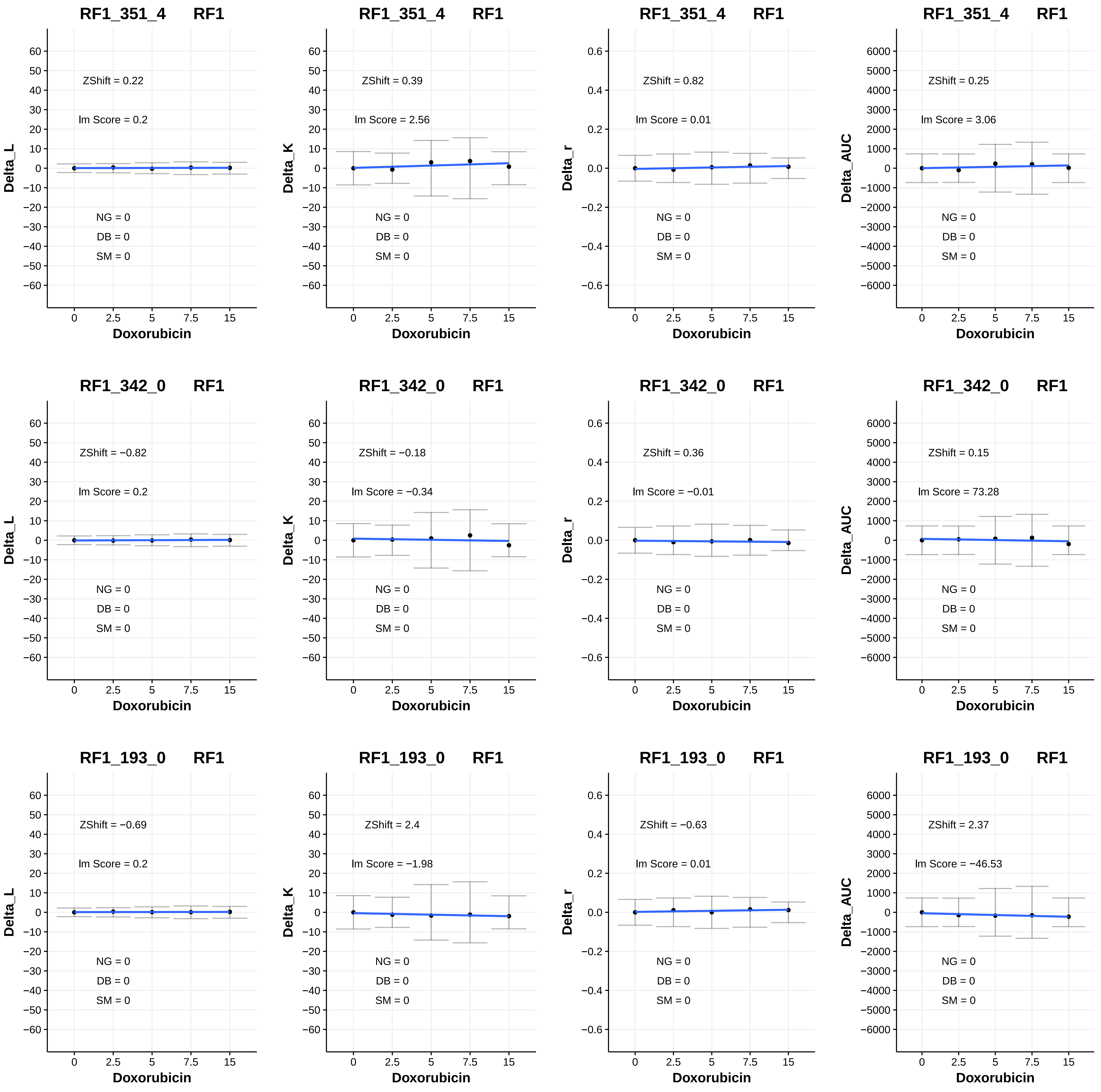
RF1_351_4 RF1
ZShift = 0.22
lm Score = 0.2
NG = 0
DB = 0
SM = 0
Delta_L
Doxorubicin
RF1_351_4 RF1
ZShift = 0.39
lm Score = 2.56
NG = 0
DB = 0
SM = 0
Delta_K
Doxorubicin
RF1_351_4 RF1
ZShift = 0.82
lm Score = 0.01
NG = 0
DB = 0
SM = 0
Delta_r
Doxorubicin
RF1_351_4 RF1
ZShift = 0.25
lm Score = 3.06
NG = 0
DB = 0
SM = 0
Delta_AUC
Doxorubicin
RF1_342_0 RF1
ZShift = −0.82
lm Score = 0.2
NG = 0
DB = 0
SM = 0
Delta_L
Doxorubicin
RF1_342_0 RF1
ZShift = −0.18
lm Score = −0.34
NG = 0
DB = 0
SM = 0
Delta_K
Doxorubicin
RF1_342_0 RF1
ZShift = 0.36
lm Score = −0.01
NG = 0
DB = 0
SM = 0
Delta_r
Doxorubicin
RF1_342_0 RF1
ZShift = 0.15
lm Score = 73.28
NG = 0
DB = 0
SM = 0
Delta_AUC
Doxorubicin
RF1_193_0 RF1
ZShift = −0.69
lm Score = 0.2
NG = 0
DB = 0
SM = 0
Delta_L
Doxorubicin
RF1_193_0 RF1
ZShift = 2.4
lm Score = −1.98
NG = 0
DB = 0
SM = 0
Delta_K
Doxorubicin
RF1_193_0 RF1
ZShift = −0.63
lm Score = 0.01
NG = 0
DB = 0
SM = 0
Delta_r
Doxorubicin
RF1_193_0 RF1
ZShift = 2.37
lm Score = −46.53
NG = 0
DB = 0
SM = 0
Delta_AUC
Doxorubicin

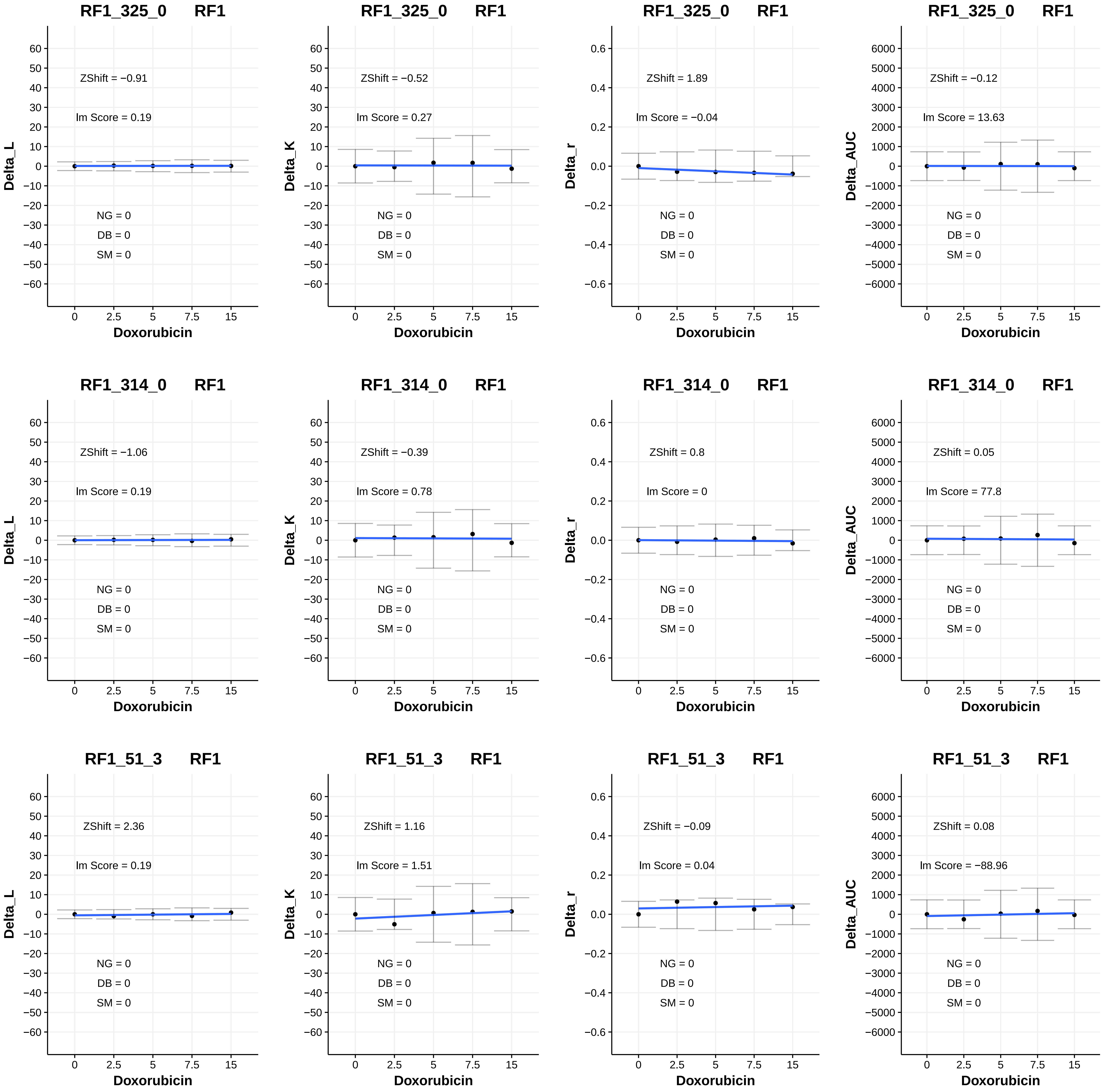
RF1_325_0 RF1
ZShift = −0.91
lm Score = 0.19
NG = 0
DB = 0
SM = 0
Delta_L
Doxorubicin
RF1_325_0 RF1
ZShift = −0.52
lm Score = 0.27
NG = 0
DB = 0
SM = 0
Delta_K
Doxorubicin
RF1_325_0 RF1
ZShift = 1.89
lm Score = −0.04
NG = 0
DB = 0
SM = 0
Delta_r
Doxorubicin
RF1_325_0 RF1
ZShift = −0.12
lm Score = 13.63
NG = 0
DB = 0
SM = 0
Delta_AUC
Doxorubicin
RF1_314_0 RF1
ZShift = −1.06
lm Score = 0.19
NG = 0
DB = 0
SM = 0
Delta_L
Doxorubicin
RF1_314_0 RF1
ZShift = −0.39
lm Score = 0.78
NG = 0
DB = 0
SM = 0
Delta_K
Doxorubicin
RF1_314_0 RF1
ZShift = 0.8
lm Score = 0
NG = 0
DB = 0
SM = 0
Delta_r
Doxorubicin
RF1_314_0 RF1
ZShift = 0.05
lm Score = 77.8
NG = 0
DB = 0
SM = 0
Delta_AUC
Doxorubicin
RF1_51_3 RF1
ZShift = 2.36
lm Score = 0.19
NG = 0
DB = 0
SM = 0
Delta_L
Doxorubicin
RF1_51_3 RF1
ZShift = 1.16
lm Score = 1.51
NG = 0
DB = 0
SM = 0
Delta_K
Doxorubicin
RF1_51_3 RF1
ZShift = −0.09
lm Score = 0.04
NG = 0
DB = 0
SM = 0
Delta_r
Doxorubicin
RF1_51_3 RF1
ZShift = 0.08
lm Score = −88.96
NG = 0
DB = 0
SM = 0
Delta_AUC
Doxorubicin

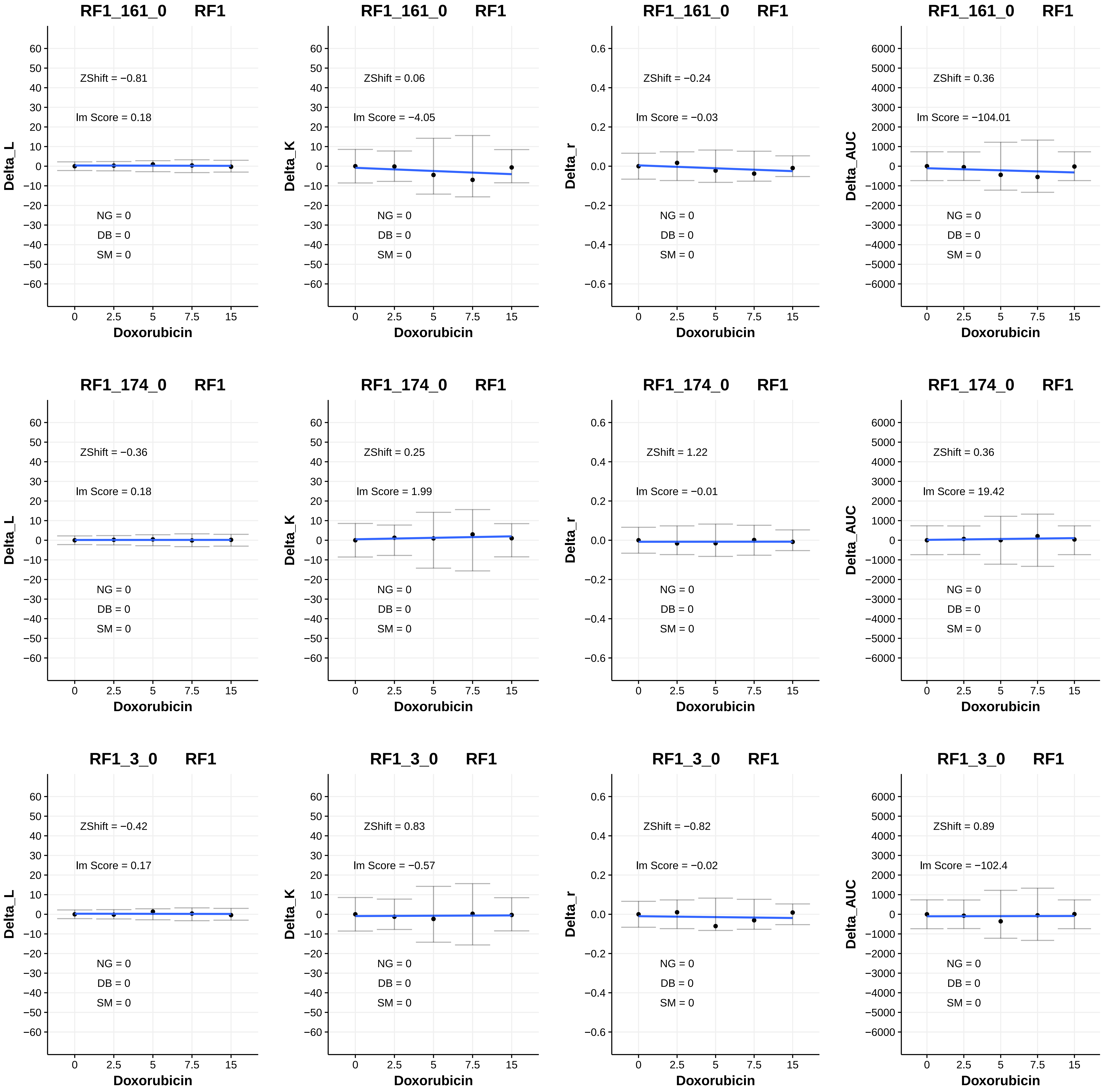

RF1_161_0 RF1
ZShift = −0.81
Im Score = 0.18
NG = 0
DB = 0
SM = 0
Delta_L
Doxorubicin
RF1_161_0 RF1
ZShift = 0.06
Im Score = −4.05
NG = 0
DB = 0
SM = 0
Delta_K
Doxorubicin
RF1_161_0 RF1
ZShift = −0.24
Im Score = −0.03
NG = 0
DB = 0
SM = 0
Delta_r
Doxorubicin
RF1_161_0 RF1
ZShift = 0.36
Im Score = −104.01
NG = 0
DB = 0
SM = 0
Delta_AUC
Doxorubicin
RF1_174_0 RF1
ZShift = −0.36
Im Score = 0.18
NG = 0
DB = 0
SM = 0
Delta_L
Doxorubicin
RF1_174_0 RF1
ZShift = 0.25
Im Score = 1.99
NG = 0
DB = 0
SM = 0
Delta_K
Doxorubicin
RF1_174_0 RF1
ZShift = 1.22
Im Score = −0.01
NG = 0
DB = 0
SM = 0
Delta_r
Doxorubicin
RF1_174_0 RF1
ZShift = 0.36
Im Score = 19.42
NG = 0
DB = 0
SM = 0
Delta_AUC
Doxorubicin
RF1_3_0 RF1
ZShift = −0.42
Im Score = 0.17
NG = 0
DB = 0
SM = 0
Delta_L
Doxorubicin
RF1_3_0 RF1
ZShift = 0.83
Im Score = −0.57
NG = 0
DB = 0
SM = 0
Delta_K
Doxorubicin
RF1_3_0 RF1
ZShift = −0.82
Im Score = −0.02
NG = 0
DB = 0
SM = 0
Delta_r
Doxorubicin
RF1_3_0 RF1
ZShift = 0.89
Im Score = −102.4
NG = 0
DB = 0
SM = 0
Delta_AUC
Doxorubicin

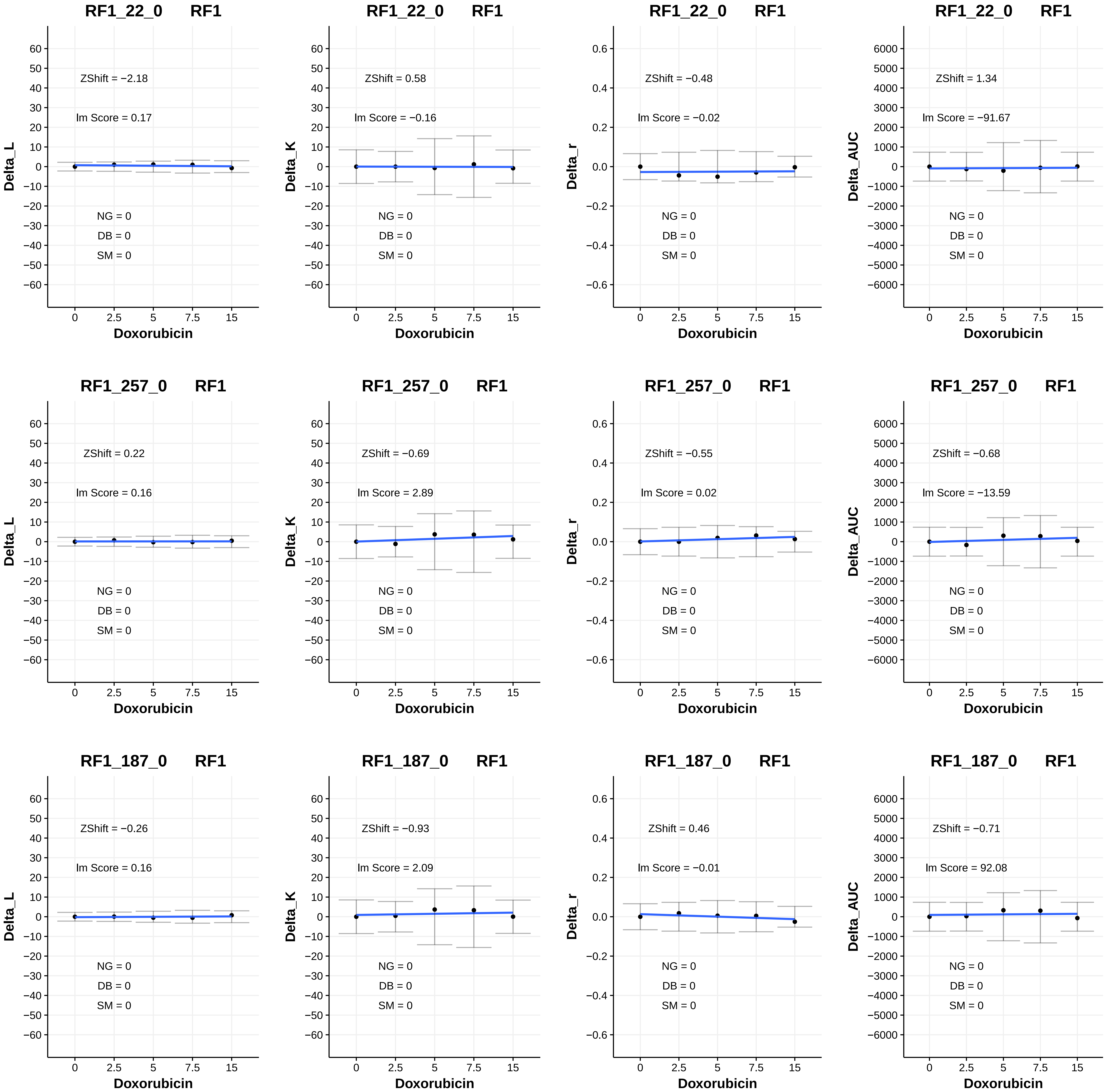
RF1_22_0 RF1
Delta_L
ZShift = −2.18
lm Score = 0.17
NG = 0
DB = 0
SM = 0
Doxorubicin
RF1_22_0 RF1
Delta_K
ZShift = 0.58
lm Score = −0.16
NG = 0
DB = 0
SM = 0
Doxorubicin
RF1_22_0 RF1
Delta_r
ZShift = −0.48
lm Score = −0.02
NG = 0
DB = 0
SM = 0
Doxorubicin
RF1_22_0 RF1
Delta_AUC
ZShift = 1.34
lm Score = −91.67
NG = 0
DB = 0
SM = 0
Doxorubicin
RF1_257_0 RF1
Delta_L
ZShift = 0.22
lm Score = 0.16
NG = 0
DB = 0
SM = 0
Doxorubicin
RF1_257_0 RF1
Delta_K
ZShift = −0.69
lm Score = 2.89
NG = 0
DB = 0
SM = 0
Doxorubicin
RF1_257_0 RF1
Delta_r
ZShift = −0.55
lm Score = 0.02
NG = 0
DB = 0
SM = 0
Doxorubicin
RF1_257_0 RF1
Delta_AUC
ZShift = −0.68
lm Score = −13.59
NG = 0
DB = 0
SM = 0
Doxorubicin
RF1_187_0 RF1
Delta_L
ZShift = −0.26
lm Score = 0.16
NG = 0
DB = 0
SM = 0
Doxorubicin
RF1_187_0 RF1
Delta_K
ZShift = −0.93
lm Score = 2.09
NG = 0
DB = 0
SM = 0
Doxorubicin
RF1_187_0 RF1
Delta_r
ZShift = 0.46
lm Score = −0.01
NG = 0
DB = 0
SM = 0
Doxorubicin
RF1_187_0 RF1
Delta_AUC
ZShift = −0.71
lm Score = 92.08
NG = 0
DB = 0
SM = 0
Doxorubicin

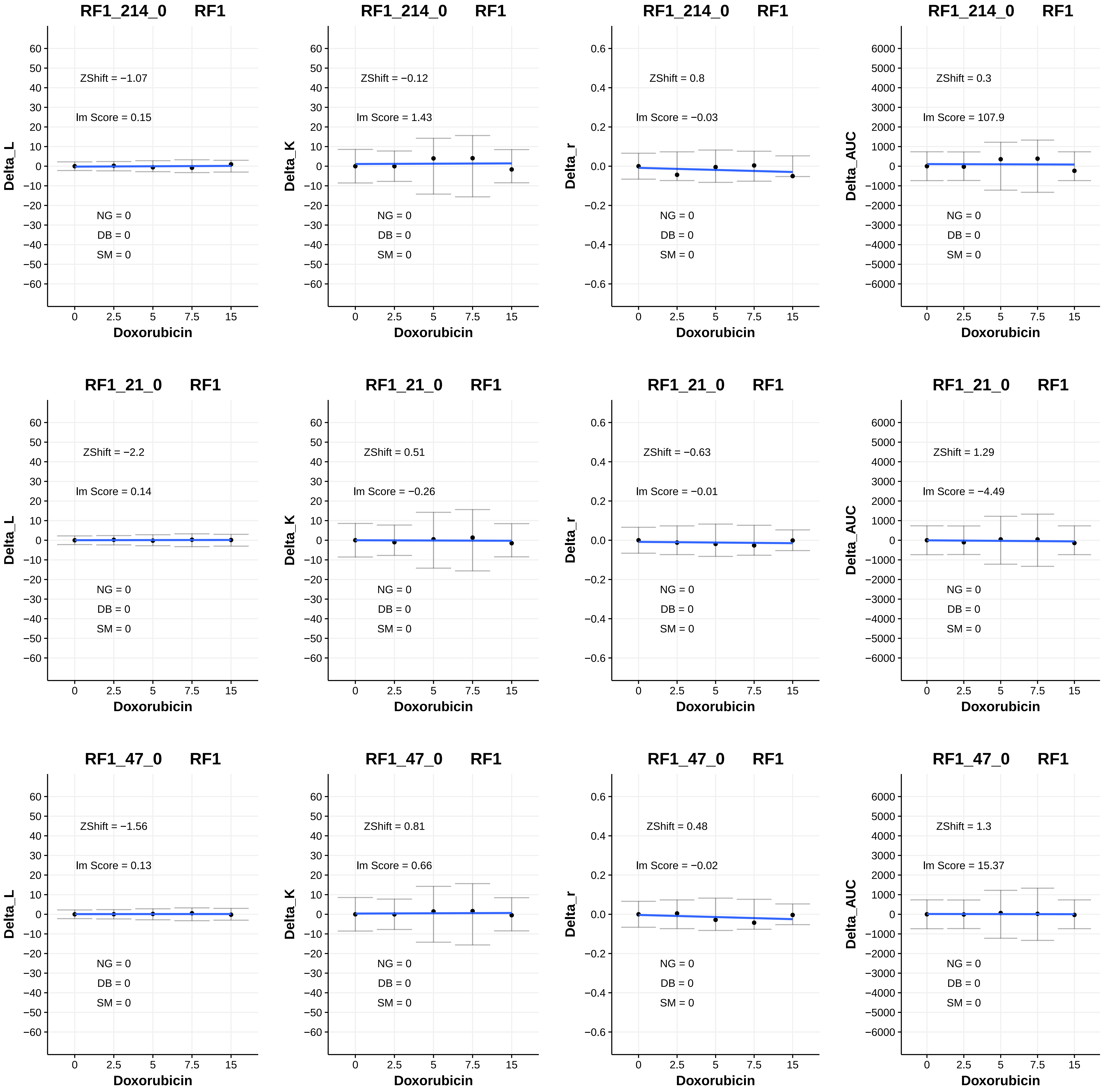

RF1_214_0 RF1
ZShift = −1.07
Im Score = 0.15
NG = 0
DB = 0
SM = 0
Delta_L
Doxorubicin
RF1_214_0 RF1
ZShift = −0.12
Im Score = 1.43
NG = 0
DB = 0
SM = 0
Delta_K
Doxorubicin
RF1_214_0 RF1
ZShift = 0.8
Im Score = −0.03
NG = 0
DB = 0
SM = 0
Delta_r
Doxorubicin
RF1_214_0 RF1
ZShift = 0.3
Im Score = 107.9
NG = 0
DB = 0
SM = 0
Delta_AUC
Doxorubicin
RF1_21_0 RF1
ZShift = −2.2
Im Score = 0.14
NG = 0
DB = 0
SM = 0
Delta_L
Doxorubicin
RF1_21_0 RF1
ZShift = 0.51
Im Score = −0.26
NG = 0
DB = 0
SM = 0
Delta_K
Doxorubicin
RF1_21_0 RF1
ZShift = −0.63
Im Score = −0.01
NG = 0
DB = 0
SM = 0
Delta_r
Doxorubicin
RF1_21_0 RF1
ZShift = 1.29
Im Score = −4.49
NG = 0
DB = 0
SM = 0
Delta_AUC
Doxorubicin
RF1_47_0 RF1
ZShift = −1.56
Im Score = 0.13
NG = 0
DB = 0
SM = 0
Delta_L
Doxorubicin
RF1_47_0 RF1
ZShift = 0.81
Im Score = 0.66
NG = 0
DB = 0
SM = 0
Delta_K
Doxorubicin
RF1_47_0 RF1
ZShift = 0.48
Im Score = −0.02
NG = 0
DB = 0
SM = 0
Delta_r
Doxorubicin
RF1_47_0 RF1
ZShift = 1.3
Im Score = 15.37
NG = 0
DB = 0
SM = 0
Delta_AUC
Doxorubicin

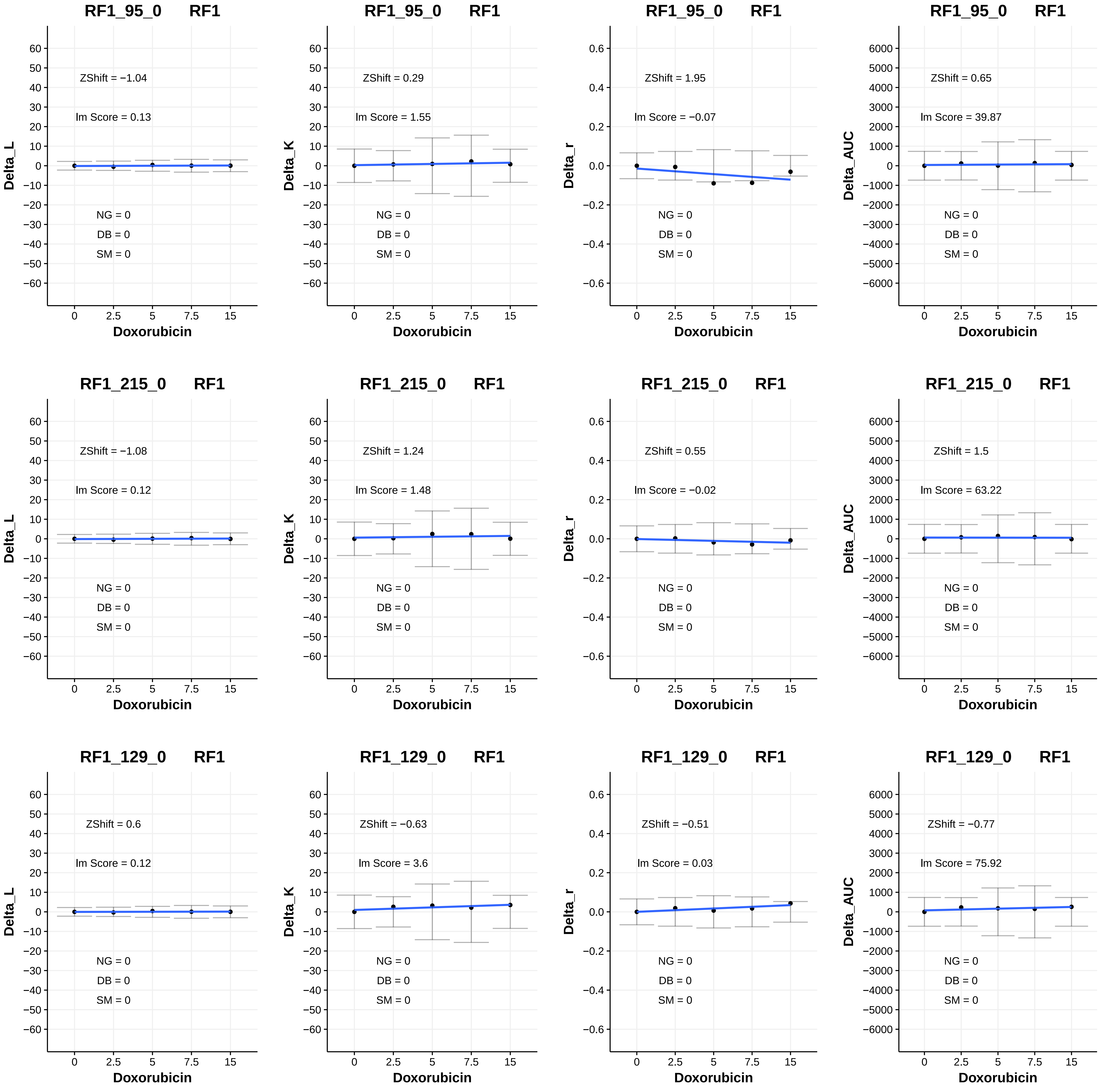

RF1_95_0 RF1
ZShift = −1.04
lm Score = 0.13
NG = 0
DB = 0
SM = 0
Delta_L
Doxorubicin
RF1_95_0 RF1
ZShift = 0.29
lm Score = 1.55
NG = 0
DB = 0
SM = 0
Delta_K
Doxorubicin
RF1_95_0 RF1
ZShift = 1.95
lm Score = −0.07
NG = 0
DB = 0
SM = 0
Delta_r
Doxorubicin
RF1_95_0 RF1
ZShift = 0.65
lm Score = 39.87
NG = 0
DB = 0
SM = 0
Delta_AUC
Doxorubicin
RF1_215_0 RF1
ZShift = −1.08
lm Score = 0.12
NG = 0
DB = 0
SM = 0
Delta_L
Doxorubicin
RF1_215_0 RF1
ZShift = 1.24
lm Score = 1.48
NG = 0
DB = 0
SM = 0
Delta_K
Doxorubicin
RF1_215_0 RF1
ZShift = 0.55
lm Score = −0.02
NG = 0
DB = 0
SM = 0
Delta_r
Doxorubicin
RF1_215_0 RF1
ZShift = 1.5
lm Score = 63.22
NG = 0
DB = 0
SM = 0
Delta_AUC
Doxorubicin
RF1_129_0 RF1
ZShift = 0.6
lm Score = 0.12
NG = 0
DB = 0
SM = 0
Delta_L
Doxorubicin
RF1_129_0 RF1
ZShift = −0.63
lm Score = 3.6
NG = 0
DB = 0
SM = 0
Delta_K
Doxorubicin
RF1_129_0 RF1
ZShift = −0.51
lm Score = 0.03
NG = 0
DB = 0
SM = 0
Delta_r
Doxorubicin
RF1_129_0 RF1
ZShift = −0.77
lm Score = 75.92
NG = 0
DB = 0
SM = 0
Delta_AUC
Doxorubicin

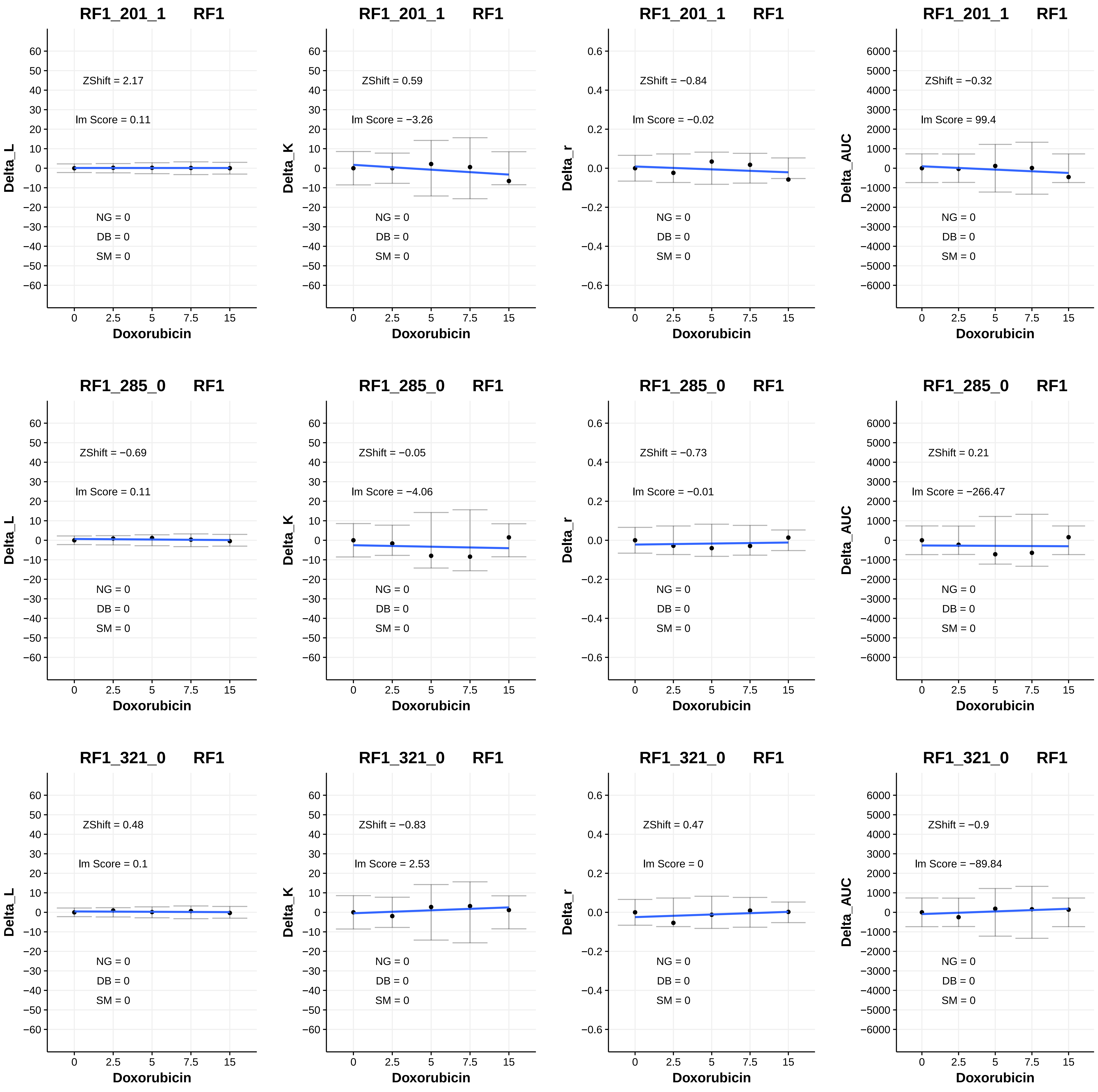
RF1_201_1 RF1
Delta_L
ZShift = 2.17
lm Score = 0.11
NG = 0
DB = 0
SM = 0
Doxorubicin
RF1_201_1 RF1
Delta_K
ZShift = 0.59
lm Score = −3.26
NG = 0
DB = 0
SM = 0
Doxorubicin
RF1_201_1 RF1
Delta_r
ZShift = −0.84
lm Score = −0.02
NG = 0
DB = 0
SM = 0
Doxorubicin
RF1_201_1 RF1
Delta_AUC
ZShift = −0.32
lm Score = 99.4
NG = 0
DB = 0
SM = 0
Doxorubicin
RF1_285_0 RF1
Delta_L
ZShift = −0.69
lm Score = 0.11
NG = 0
DB = 0
SM = 0
Doxorubicin
RF1_285_0 RF1
Delta_K
ZShift = −0.05
lm Score = −4.06
NG = 0
DB = 0
SM = 0
Doxorubicin
RF1_285_0 RF1
Delta_r
ZShift = −0.73
lm Score = −0.01
NG = 0
DB = 0
SM = 0
Doxorubicin
RF1_285_0 RF1
Delta_AUC
ZShift = 0.21
lm Score = −266.47
NG = 0
DB = 0
SM = 0
Doxorubicin
RF1_321_0 RF1
Delta_L
ZShift = 0.48
lm Score = 0.1
NG = 0
DB = 0
SM = 0
Doxorubicin
RF1_321_0 RF1
Delta_K
ZShift = −0.83
lm Score = 2.53
NG = 0
DB = 0
SM = 0
Doxorubicin
RF1_321_0 RF1
Delta_r
ZShift = 0.47
lm Score = 0
NG = 0
DB = 0
SM = 0
Doxorubicin
RF1_321_0 RF1
Delta_AUC
ZShift = −0.9
lm Score = −89.84
NG = 0
DB = 0
SM = 0
Doxorubicin

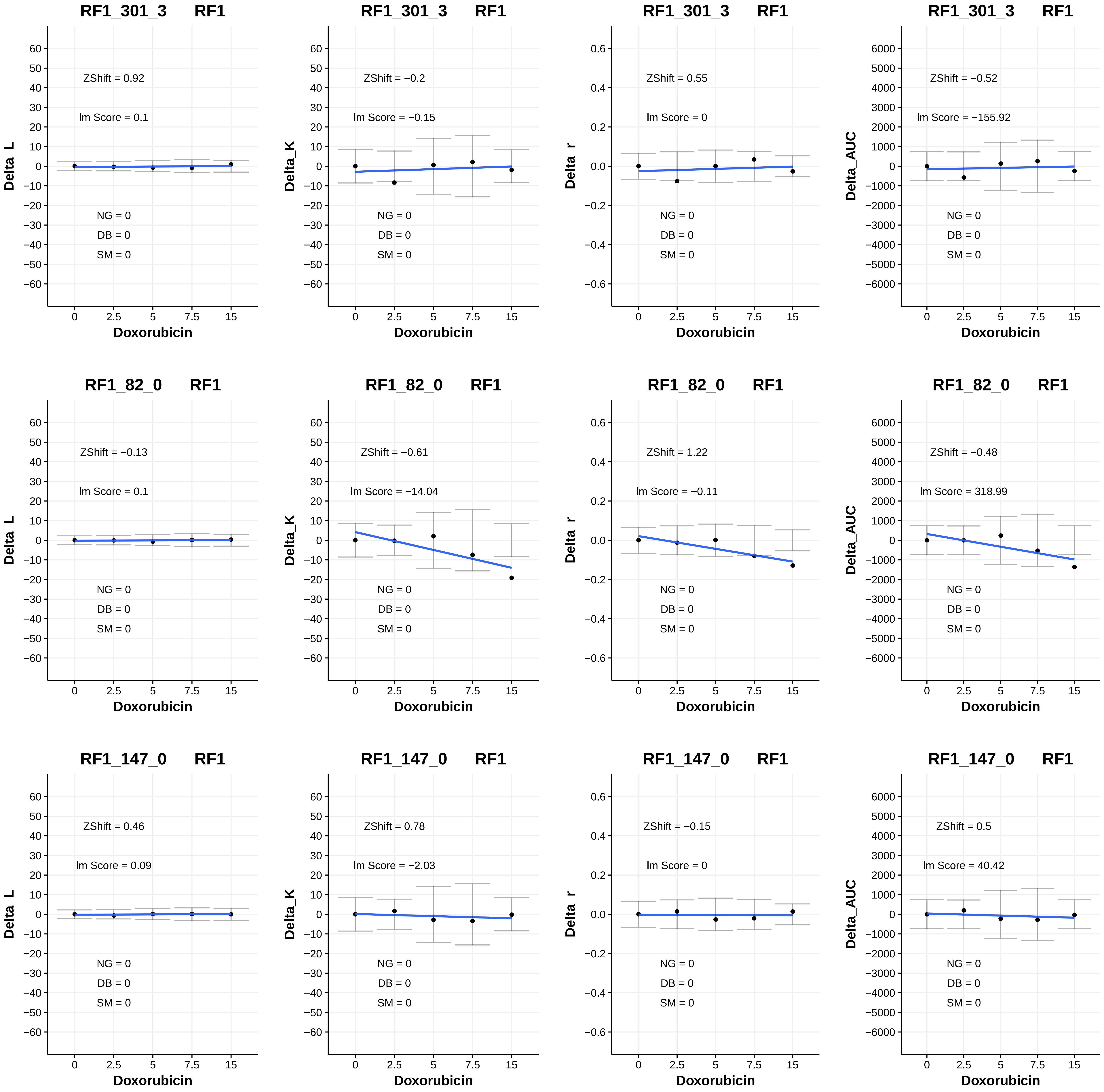

RF1_301_3 RF1
ZShift = 0.92
lm Score = 0.1
NG = 0
DB = 0
SM = 0
Delta_L
Doxorubicin
RF1_301_3 RF1
ZShift = −0.2
lm Score = −0.15
NG = 0
DB = 0
SM = 0
Delta_K
Doxorubicin
RF1_301_3 RF1
ZShift = 0.55
lm Score = 0
NG = 0
DB = 0
SM = 0
Delta_r
Doxorubicin
RF1_301_3 RF1
ZShift = −0.52
lm Score = −155.92
NG = 0
DB = 0
SM = 0
Delta_AUC
Doxorubicin
RF1_82_0 RF1
ZShift = −0.13
lm Score = 0.1
NG = 0
DB = 0
SM = 0
Delta_L
Doxorubicin
RF1_82_0 RF1
ZShift = −0.61
lm Score = −14.04
NG = 0
DB = 0
SM = 0
Delta_K
Doxorubicin
RF1_82_0 RF1
ZShift = 1.22
lm Score = −0.11
NG = 0
DB = 0
SM = 0
Delta_r
Doxorubicin
RF1_82_0 RF1
ZShift = −0.48
lm Score = 318.99
NG = 0
DB = 0
SM = 0
Delta_AUC
Doxorubicin
RF1_147_0 RF1
ZShift = 0.46
lm Score = 0.09
NG = 0
DB = 0
SM = 0
Delta_L
Doxorubicin
RF1_147_0 RF1
ZShift = 0.78
lm Score = −2.03
NG = 0
DB = 0
SM = 0
Delta_K
Doxorubicin
RF1_147_0 RF1
ZShift = −0.15
lm Score = 0
NG = 0
DB = 0
SM = 0
Delta_r
Doxorubicin
RF1_147_0 RF1
ZShift = 0.5
lm Score = 40.42
NG = 0
DB = 0
SM = 0
Delta_AUC
Doxorubicin

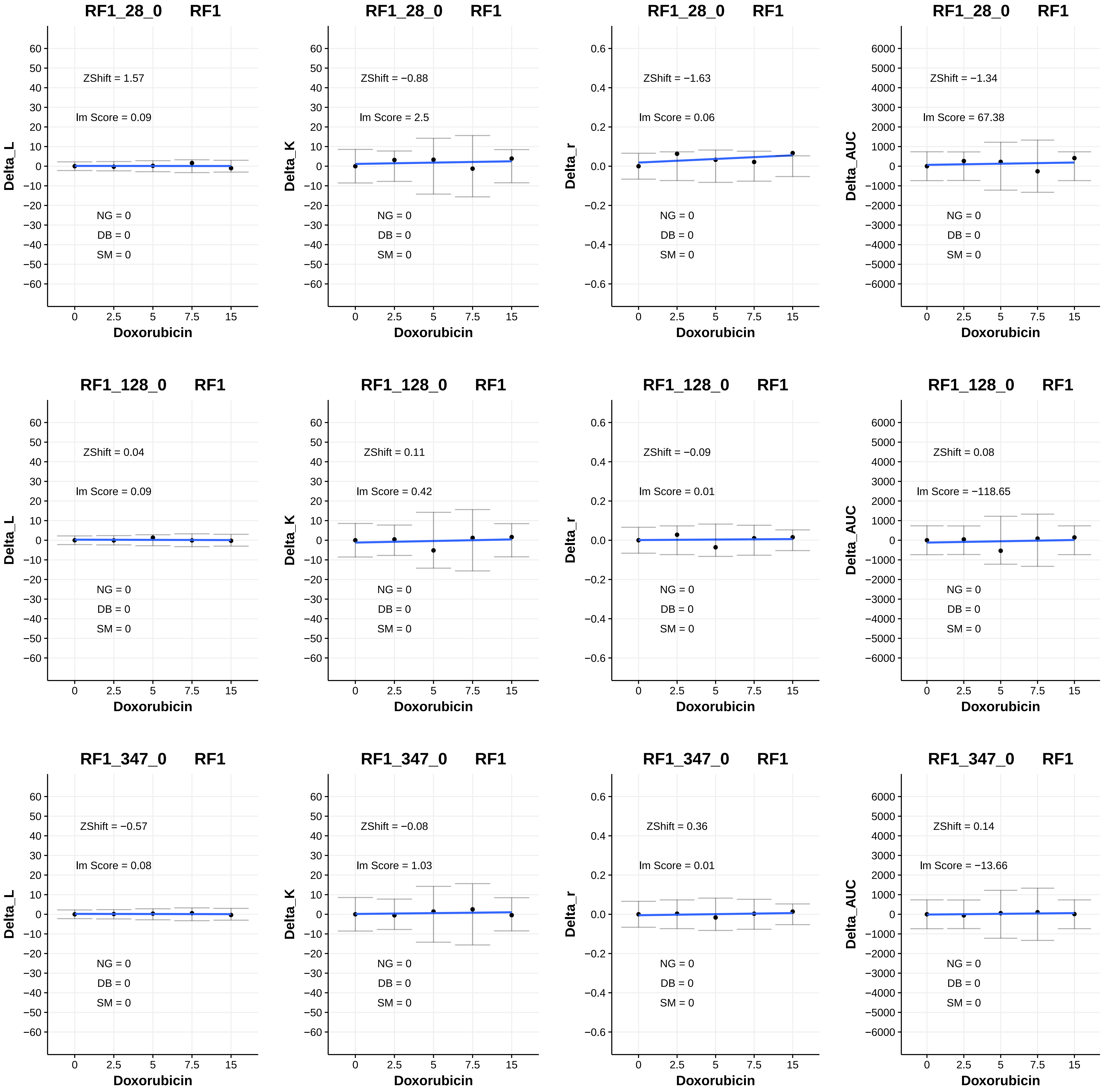

RF1_28_0 RF1
ZShift = 1.57
lm Score = 0.09
NG = 0
DB = 0
SM = 0
Delta_L
Doxorubicin
RF1_28_0 RF1
ZShift = −0.88
lm Score = 2.5
NG = 0
DB = 0
SM = 0
Delta_K
Doxorubicin
RF1_28_0 RF1
ZShift = −1.63
lm Score = 0.06
NG = 0
DB = 0
SM = 0
Delta_r
Doxorubicin
RF1_28_0 RF1
ZShift = −1.34
lm Score = 67.38
NG = 0
DB = 0
SM = 0
Delta_AUC
Doxorubicin
RF1_128_0 RF1
ZShift = 0.04
lm Score = 0.09
NG = 0
DB = 0
SM = 0
Delta_L
Doxorubicin
RF1_128_0 RF1
ZShift = 0.11
lm Score = 0.42
NG = 0
DB = 0
SM = 0
Delta_K
Doxorubicin
RF1_128_0 RF1
ZShift = −0.09
lm Score = 0.01
NG = 0
DB = 0
SM = 0
Delta_r
Doxorubicin
RF1_128_0 RF1
ZShift = 0.08
lm Score = −118.65
NG = 0
DB = 0
SM = 0
Delta_AUC
Doxorubicin
RF1_347_0 RF1
ZShift = −0.57
lm Score = 0.08
NG = 0
DB = 0
SM = 0
Delta_L
Doxorubicin
RF1_347_0 RF1
ZShift = −0.08
lm Score = 1.03
NG = 0
DB = 0
SM = 0
Delta_K
Doxorubicin
RF1_347_0 RF1
ZShift = 0.36
lm Score = 0.01
NG = 0
DB = 0
SM = 0
Delta_r
Doxorubicin
RF1_347_0 RF1
ZShift = 0.14
lm Score = −13.66
NG = 0
DB = 0
SM = 0
Delta_AUC
Doxorubicin

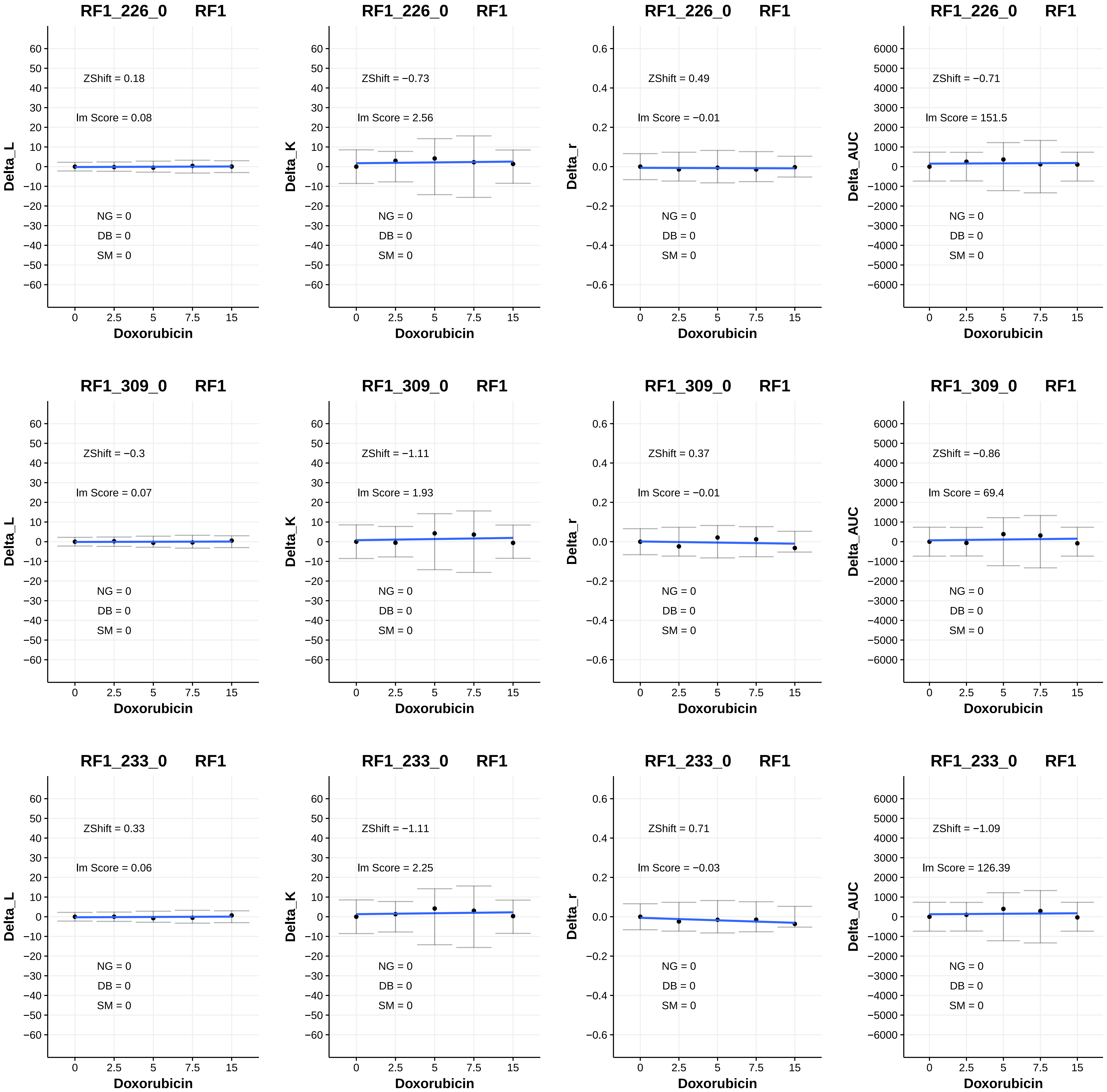

RF1_226_0 RF1
ZShift = 0.18
Im Score = 0.08
NG = 0
DB = 0
SM = 0
Delta_L
Doxorubicin
RF1_226_0 RF1
ZShift = −0.73
Im Score = 2.56
NG = 0
DB = 0
SM = 0
Delta_K
Doxorubicin
RF1_226_0 RF1
ZShift = 0.49
Im Score = −0.01
NG = 0
DB = 0
SM = 0
Delta_r
Doxorubicin
RF1_226_0 RF1
ZShift = −0.71
Im Score = 151.5
NG = 0
DB = 0
SM = 0
Delta_AUC
Doxorubicin
RF1_309_0 RF1
ZShift = −0.3
Im Score = 0.07
NG = 0
DB = 0
SM = 0
Delta_L
Doxorubicin
RF1_309_0 RF1
ZShift = −1.11
Im Score = 1.93
NG = 0
DB = 0
SM = 0
Delta_K
Doxorubicin
RF1_309_0 RF1
ZShift = 0.37
Im Score = −0.01
NG = 0
DB = 0
SM = 0
Delta_r
Doxorubicin
RF1_309_0 RF1
ZShift = −0.86
Im Score = 69.4
NG = 0
DB = 0
SM = 0
Delta_AUC
Doxorubicin
RF1_233_0 RF1
ZShift = 0.33
Im Score = 0.06
NG = 0
DB = 0
SM = 0
Delta_L
Doxorubicin
RF1_233_0 RF1
ZShift = −1.11
Im Score = 2.25
NG = 0
DB = 0
SM = 0
Delta_K
Doxorubicin
RF1_233_0 RF1
ZShift = 0.71
Im Score = −0.03
NG = 0
DB = 0
SM = 0
Delta_r
Doxorubicin
RF1_233_0 RF1
ZShift = −1.09
Im Score = 126.39
NG = 0
DB = 0
SM = 0
Delta_AUC
Doxorubicin

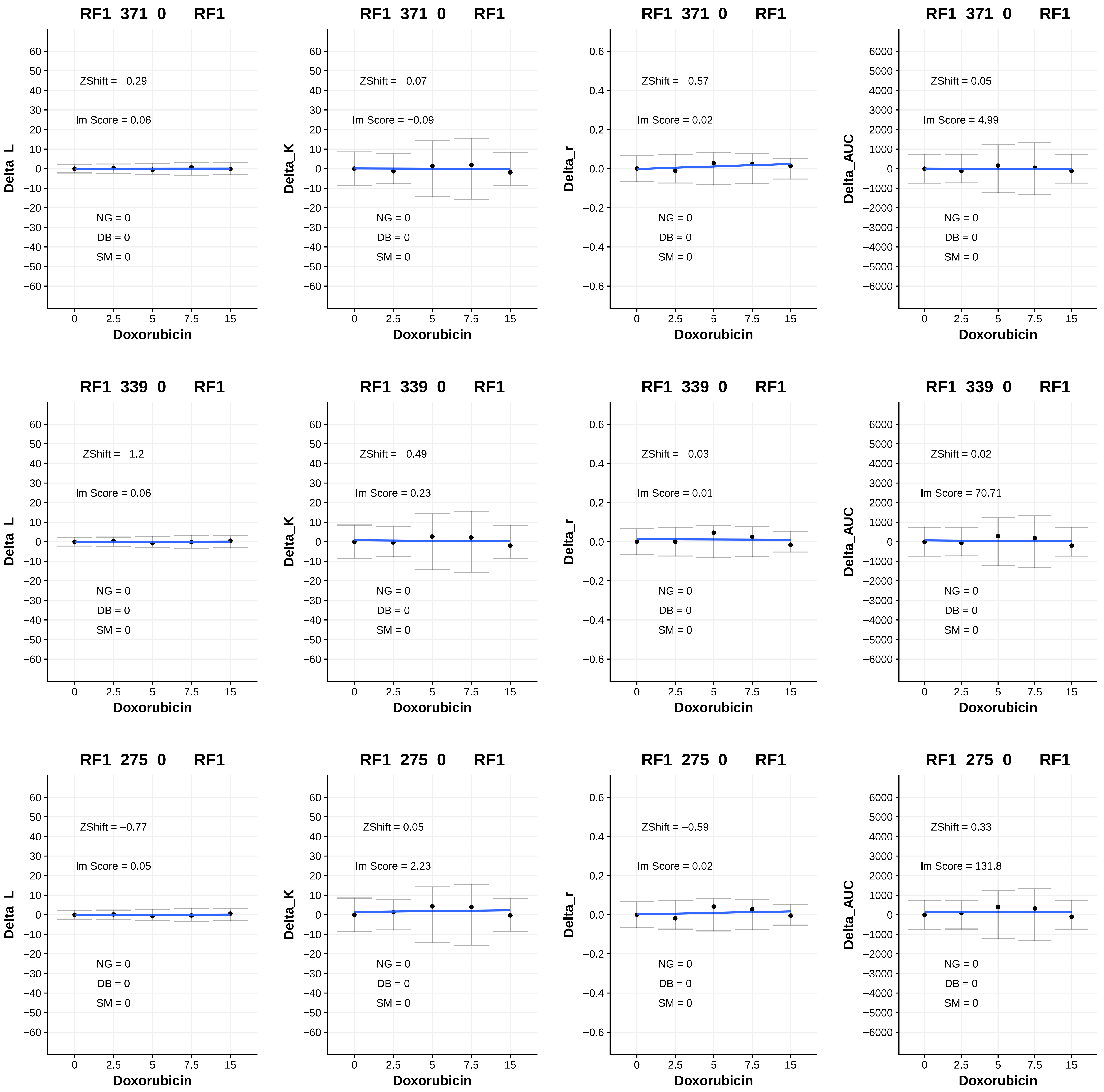
RF1_371_0 RF1
ZShift = −0.29
lm Score = 0.06
NG = 0
DB = 0
SM = 0
Delta_L
Doxorubicin
RF1_371_0 RF1
ZShift = −0.07
lm Score = −0.09
Delta_K
RF1_371_0 RF1
ZShift = −0.57
lm Score = 0.02
Delta_r
RF1_371_0 RF1
ZShift = 0.05
lm Score = 4.99
Delta_AUC
RF1_339_0 RF1
ZShift = −1.2
lm Score = 0.06
RF1_339_0 RF1
ZShift = −0.49
lm Score = 0.23
RF1_339_0 RF1
ZShift = −0.03
lm Score = 0.01
RF1_339_0 RF1
ZShift = 0.02
lm Score = 70.71
RF1_275_0 RF1
ZShift = −0.77
lm Score = 0.05
RF1_275_0 RF1
ZShift = 0.05
lm Score = 2.23
RF1_275_0 RF1
ZShift = −0.59
lm Score = 0.02
RF1_275_0 RF1
ZShift = 0.33
lm Score = 131.8

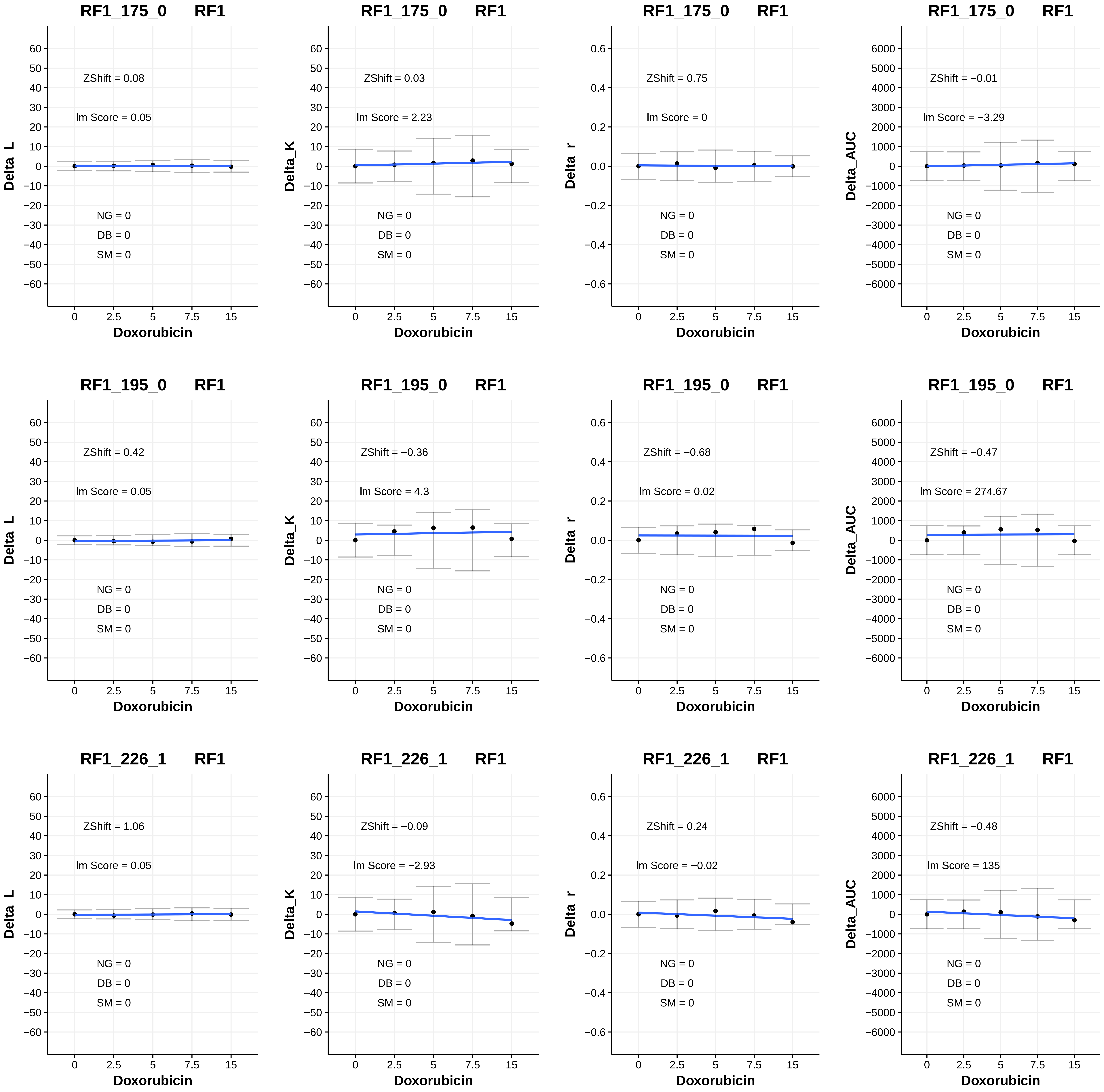
RF1_175_0 RF1
ZShift = 0.08
lm Score = 0.05
NG = 0
DB = 0
SM = 0
Delta_L
Doxorubicin
RF1_175_0 RF1
ZShift = 0.03
lm Score = 2.23
NG = 0
DB = 0
SM = 0
Delta_K
Doxorubicin
RF1_175_0 RF1
ZShift = 0.75
lm Score = 0
NG = 0
DB = 0
SM = 0
Delta_r
Doxorubicin
RF1_175_0 RF1
ZShift = −0.01
lm Score = −3.29
NG = 0
DB = 0
SM = 0
Delta_AUC
Doxorubicin
RF1_195_0 RF1
ZShift = 0.42
lm Score = 0.05
NG = 0
DB = 0
SM = 0
Delta_L
Doxorubicin
RF1_195_0 RF1
ZShift = −0.36
lm Score = 4.3
NG = 0
DB = 0
SM = 0
Delta_K
Doxorubicin
RF1_195_0 RF1
ZShift = −0.68
lm Score = 0.02
NG = 0
DB = 0
SM = 0
Delta_r
Doxorubicin
RF1_195_0 RF1
ZShift = −0.47
lm Score = 274.67
NG = 0
DB = 0
SM = 0
Delta_AUC
Doxorubicin
RF1_226_1 RF1
ZShift = 1.06
lm Score = 0.05
NG = 0
DB = 0
SM = 0
Delta_L
Doxorubicin
RF1_226_1 RF1
ZShift = −0.09
lm Score = −2.93
NG = 0
DB = 0
SM = 0
Delta_K
Doxorubicin
RF1_226_1 RF1
ZShift = 0.24
lm Score = −0.02
NG = 0
DB = 0
SM = 0
Delta_r
Doxorubicin
RF1_226_1 RF1
ZShift = −0.48
lm Score = 135
NG = 0
DB = 0
SM = 0
Delta_AUC
Doxorubicin

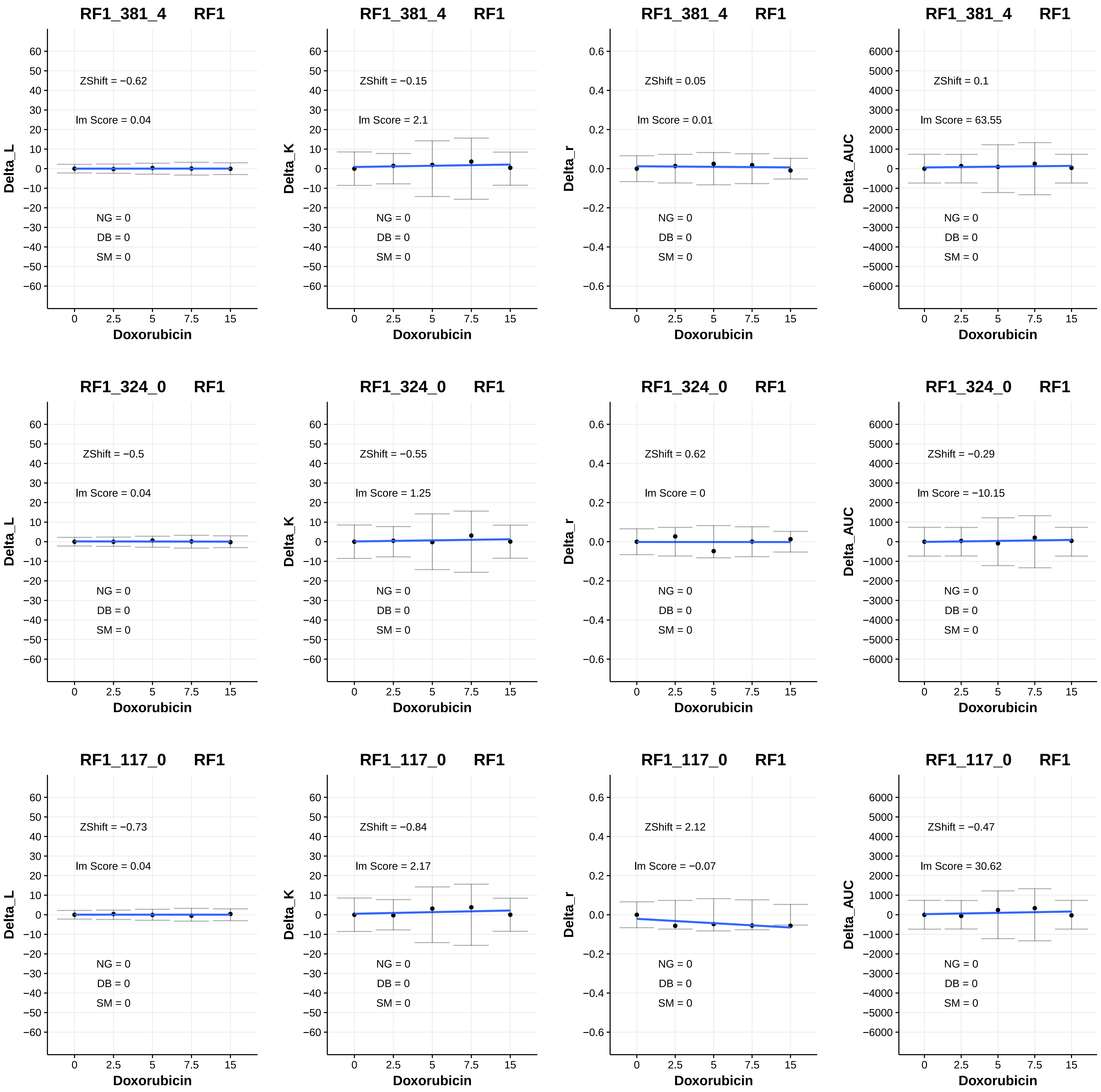

RF1_381_4 RF1
ZShift = −0.62
lm Score = 0.04
NG = 0
DB = 0
SM = 0
Delta_L
Doxorubicin
RF1_381_4 RF1
ZShift = −0.15
lm Score = 2.1
NG = 0
DB = 0
SM = 0
Delta_K
Doxorubicin
RF1_381_4 RF1
ZShift = 0.05
lm Score = 0.01
NG = 0
DB = 0
SM = 0
Delta_r
Doxorubicin
RF1_381_4 RF1
ZShift = 0.1
lm Score = 63.55
NG = 0
DB = 0
SM = 0
Delta_AUC
Doxorubicin
RF1_324_0 RF1
ZShift = −0.5
lm Score = 0.04
NG = 0
DB = 0
SM = 0
Delta_L
Doxorubicin
RF1_324_0 RF1
ZShift = −0.55
lm Score = 1.25
NG = 0
DB = 0
SM = 0
Delta_K
Doxorubicin
RF1_324_0 RF1
ZShift = 0.62
lm Score = 0
NG = 0
DB = 0
SM = 0
Delta_r
Doxorubicin
RF1_324_0 RF1
ZShift = −0.29
lm Score = −10.15
NG = 0
DB = 0
SM = 0
Delta_AUC
Doxorubicin
RF1_117_0 RF1
ZShift = −0.73
lm Score = 0.04
NG = 0
DB = 0
SM = 0
Delta_L
Doxorubicin
RF1_117_0 RF1
ZShift = −0.84
lm Score = 2.17
NG = 0
DB = 0
SM = 0
Delta_K
Doxorubicin
RF1_117_0 RF1
ZShift = 2.12
lm Score = −0.07
NG = 0
DB = 0
SM = 0
Delta_r
Doxorubicin
RF1_117_0 RF1
ZShift = −0.47
lm Score = 30.62
NG = 0
DB = 0
SM = 0
Delta_AUC
Doxorubicin

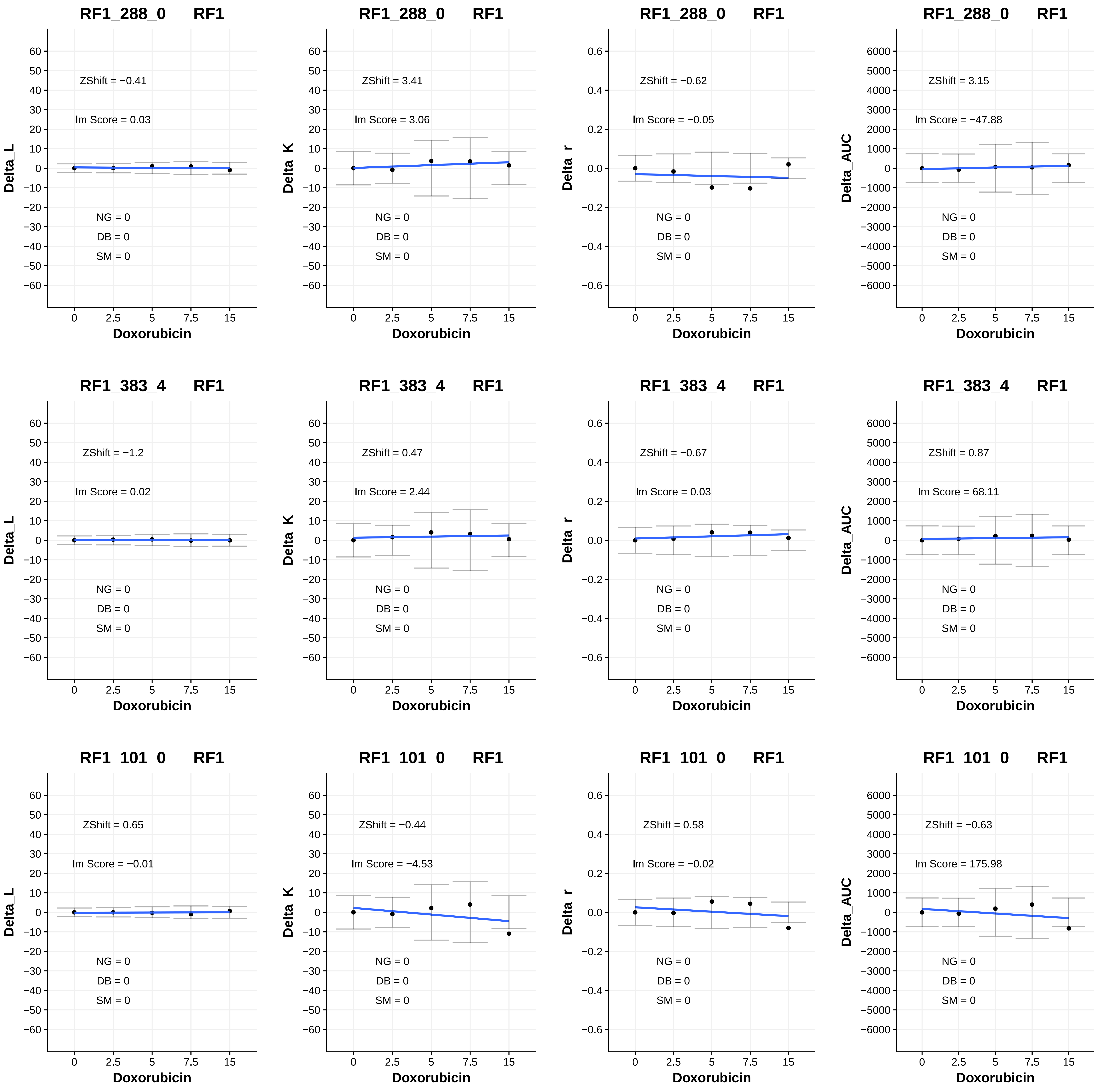
RF1_288_0 RF1
ZShift = −0.41
Im Score = 0.03
NG = 0
DB = 0
SM = 0
Delta_L
Doxorubicin
RF1_288_0 RF1
ZShift = 3.41
Im Score = 3.06
NG = 0
DB = 0
SM = 0
Delta_K
Doxorubicin
RF1_288_0 RF1
ZShift = −0.62
Im Score = −0.05
NG = 0
DB = 0
SM = 0
Delta_r
Doxorubicin
RF1_288_0 RF1
ZShift = 3.15
Im Score = −47.88
NG = 0
DB = 0
SM = 0
Delta_AUC
Doxorubicin
RF1_383_4 RF1
ZShift = −1.2
Im Score = 0.02
NG = 0
DB = 0
SM = 0
Delta_L
Doxorubicin
RF1_383_4 RF1
ZShift = 0.47
Im Score = 2.44
NG = 0
DB = 0
SM = 0
Delta_K
Doxorubicin
RF1_383_4 RF1
ZShift = −0.67
Im Score = 0.03
NG = 0
DB = 0
SM = 0
Delta_r
Doxorubicin
RF1_383_4 RF1
ZShift = 0.87
Im Score = 68.11
NG = 0
DB = 0
SM = 0
Delta_AUC
Doxorubicin
RF1_101_0 RF1
ZShift = 0.65
Im Score = −0.01
NG = 0
DB = 0
SM = 0
Delta_L
Doxorubicin
RF1_101_0 RF1
ZShift = −0.44
Im Score = −4.53
NG = 0
DB = 0
SM = 0
Delta_K
Doxorubicin
RF1_101_0 RF1
ZShift = 0.58
Im Score = −0.02
NG = 0
DB = 0
SM = 0
Delta_r
Doxorubicin
RF1_101_0 RF1
ZShift = −0.63
Im Score = 175.98
NG = 0
DB = 0
SM = 0
Delta_AUC
Doxorubicin

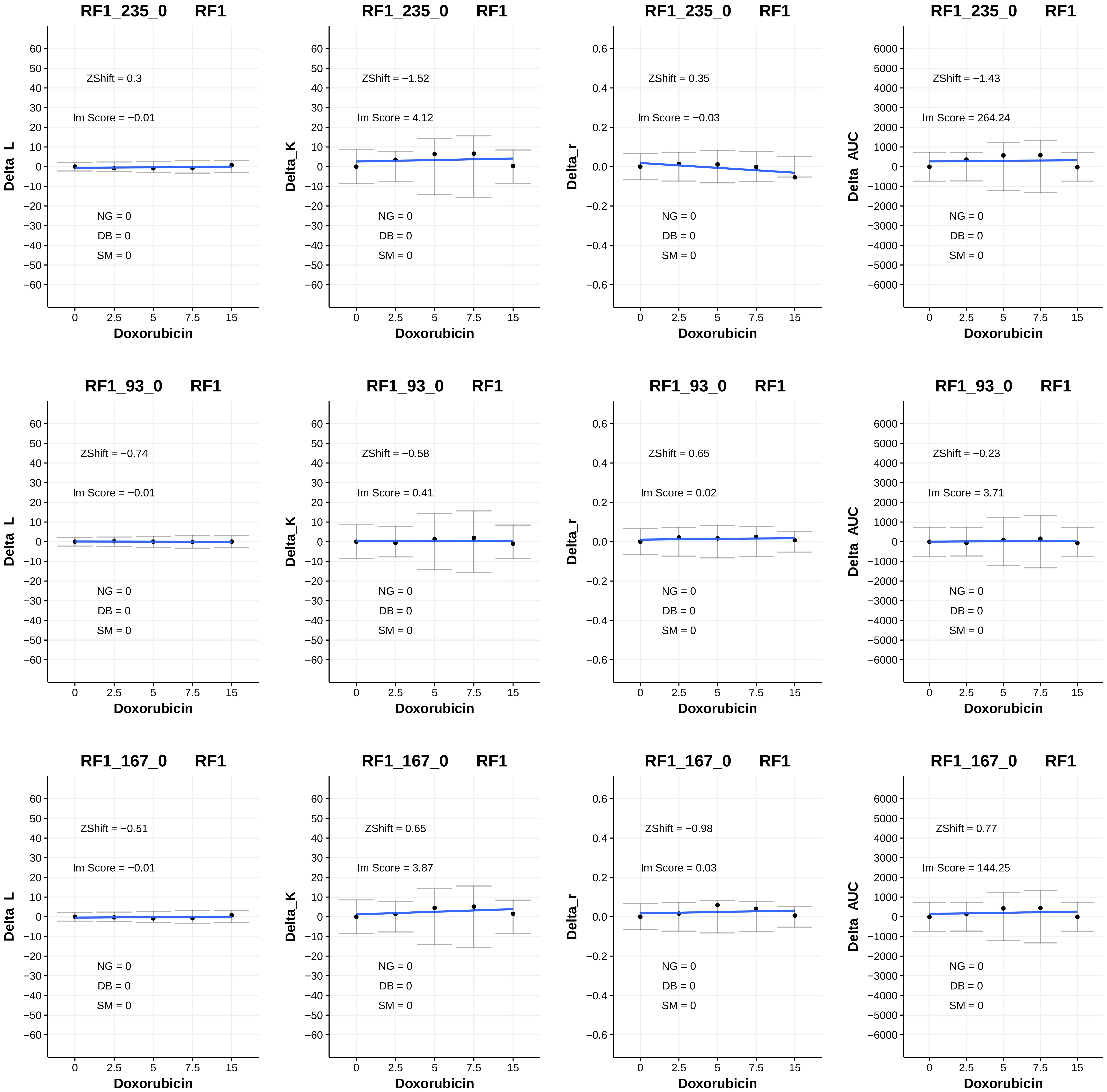
RF1_235_0 RF1
ZShift = 0.3
Im Score = −0.01
NG = 0
DB = 0
SM = 0
Delta_L
Doxorubicin
RF1_235_0 RF1
ZShift = −1.52
Im Score = 4.12
NG = 0
DB = 0
SM = 0
Delta_K
Doxorubicin
RF1_235_0 RF1
ZShift = 0.35
Im Score = −0.03
NG = 0
DB = 0
SM = 0
Delta_r
Doxorubicin
RF1_235_0 RF1
ZShift = −1.43
Im Score = 264.24
NG = 0
DB = 0
SM = 0
Delta_AUC
Doxorubicin
RF1_93_0 RF1
ZShift = −0.74
Im Score = −0.01
NG = 0
DB = 0
SM = 0
Delta_L
Doxorubicin
RF1_93_0 RF1
ZShift = −0.58
Im Score = 0.41
NG = 0
DB = 0
SM = 0
Delta_K
Doxorubicin
RF1_93_0 RF1
ZShift = 0.65
Im Score = 0.02
NG = 0
DB = 0
SM = 0
Delta_r
Doxorubicin
RF1_93_0 RF1
ZShift = −0.23
Im Score = 3.71
NG = 0
DB = 0
SM = 0
Delta_AUC
Doxorubicin
RF1_167_0 RF1
ZShift = −0.51
Im Score = −0.01
NG = 0
DB = 0
SM = 0
Delta_L
Doxorubicin
RF1_167_0 RF1
ZShift = 0.65
Im Score = 3.87
NG = 0
DB = 0
SM = 0
Delta_K
Doxorubicin
RF1_167_0 RF1
ZShift = −0.98
Im Score = 0.03
NG = 0
DB = 0
SM = 0
Delta_r
Doxorubicin
RF1_167_0 RF1
ZShift = 0.77
Im Score = 144.25
NG = 0
DB = 0
SM = 0
Delta_AUC
Doxorubicin

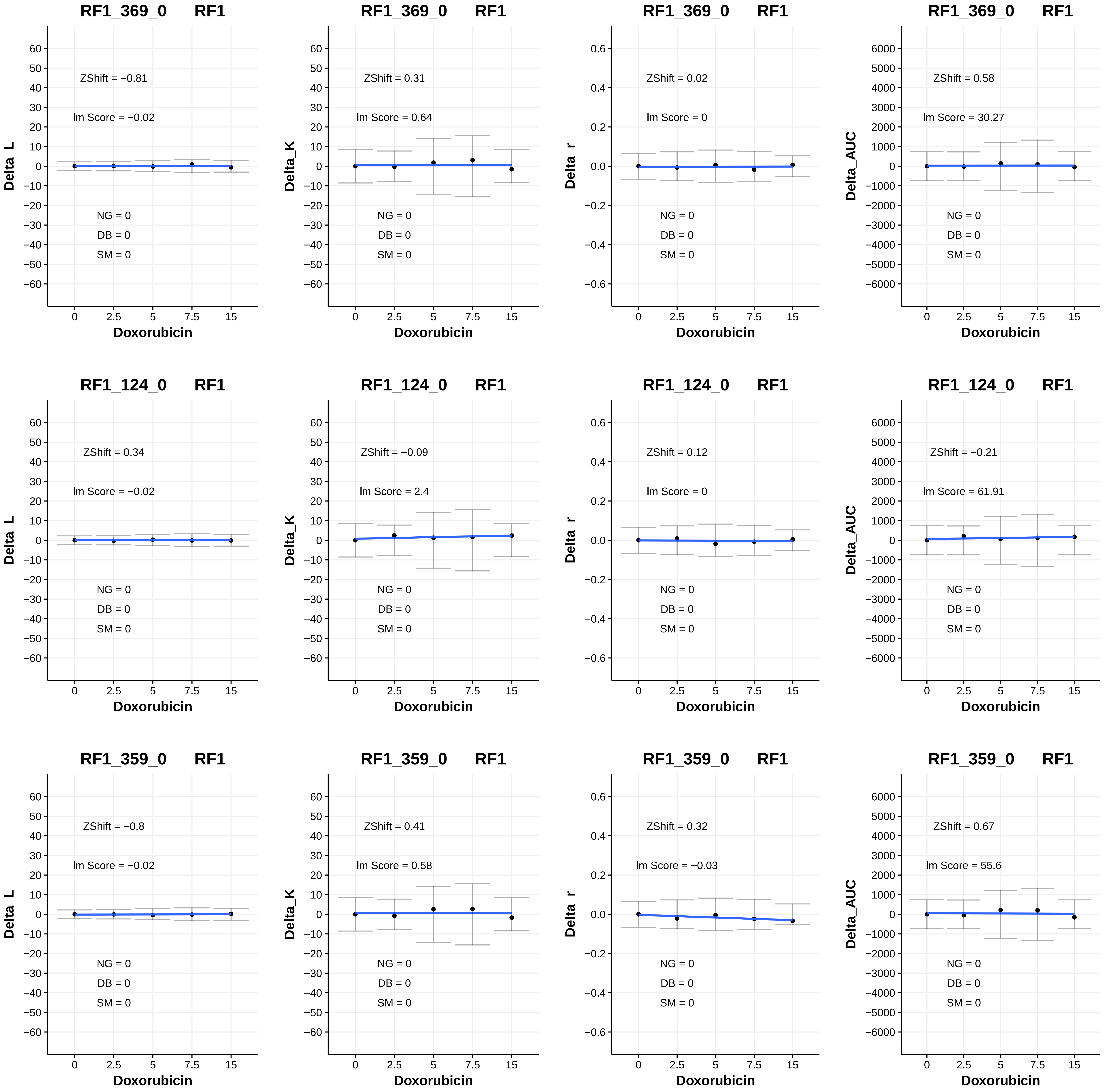
RF1_369_0 RF1
ZShift = −0.81
lm Score = −0.02
NG = 0
DB = 0
SM = 0
Delta_L
Doxorubicin
RF1_369_0 RF1
ZShift = 0.31
lm Score = 0.64
NG = 0
DB = 0
SM = 0
Delta_K
Doxorubicin
RF1_369_0 RF1
ZShift = 0.02
lm Score = 0
NG = 0
DB = 0
SM = 0
Delta_r
Doxorubicin
RF1_369_0 RF1
ZShift = 0.58
lm Score = 30.27
NG = 0
DB = 0
SM = 0
Delta_AUC
Doxorubicin
RF1_124_0 RF1
ZShift = 0.34
lm Score = −0.02
NG = 0
DB = 0
SM = 0
Delta_L
Doxorubicin
RF1_124_0 RF1
ZShift = −0.09
lm Score = 2.4
NG = 0
DB = 0
SM = 0
Delta_K
Doxorubicin
RF1_124_0 RF1
ZShift = 0.12
lm Score = 0
NG = 0
DB = 0
SM = 0
Delta_r
Doxorubicin
RF1_124_0 RF1
ZShift = −0.21
lm Score = 61.91
NG = 0
DB = 0
SM = 0
Delta_AUC
Doxorubicin
RF1_359_0 RF1
ZShift = −0.8
lm Score = −0.02
NG = 0
DB = 0
SM = 0
Delta_L
Doxorubicin
RF1_359_0 RF1
ZShift = 0.41
lm Score = 0.58
NG = 0
DB = 0
SM = 0
Delta_K
Doxorubicin
RF1_359_0 RF1
ZShift = 0.32
lm Score = −0.03
NG = 0
DB = 0
SM = 0
Delta_r
Doxorubicin
RF1_359_0 RF1
ZShift = 0.67
lm Score = 55.6
NG = 0
DB = 0
SM = 0
Delta_AUC
Doxorubicin

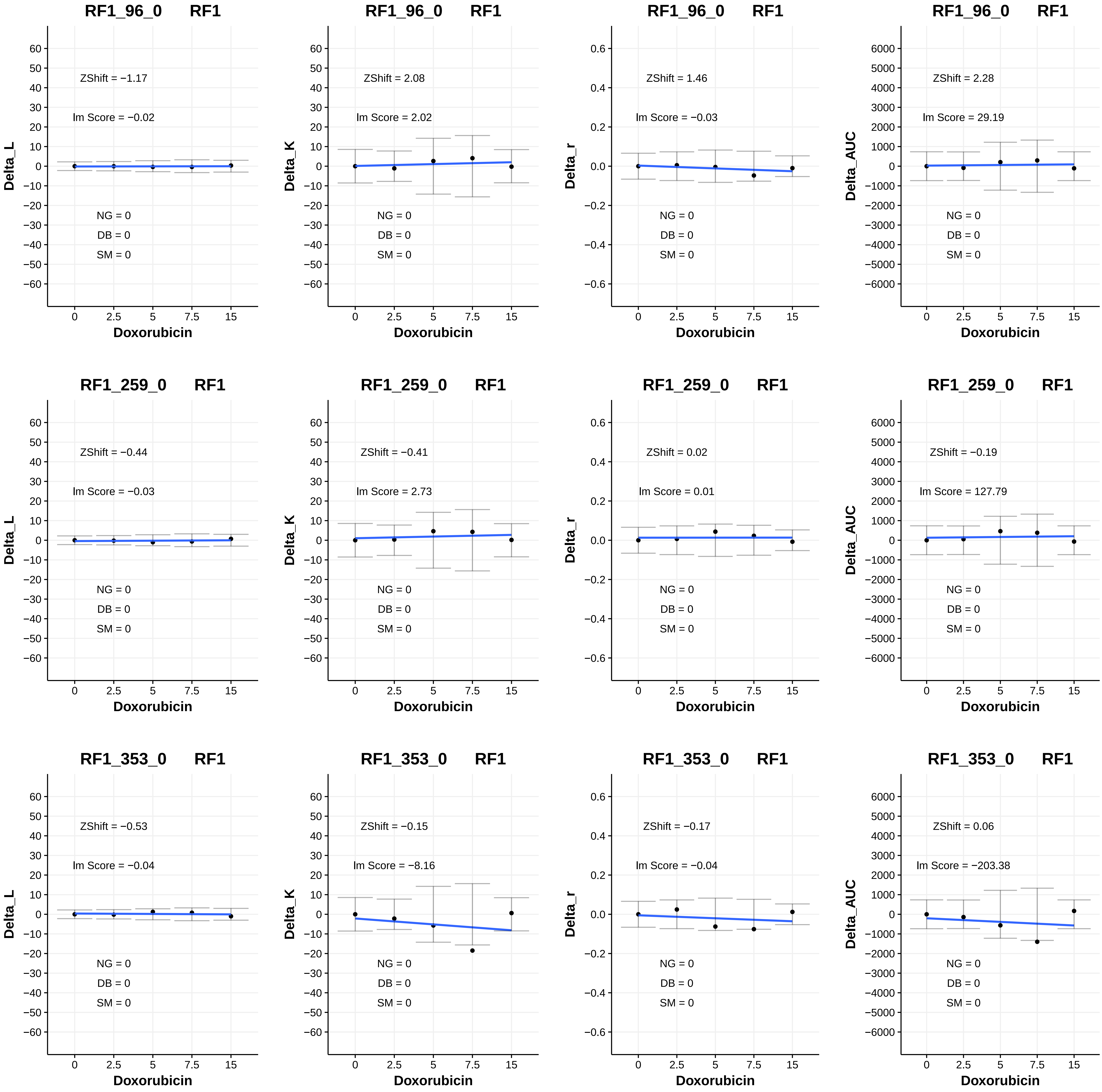

RF1_96_0 RF1
ZShift = −1.17
lm Score = −0.02
NG = 0
DB = 0
SM = 0
Delta_L
RF1_96_0 RF1
ZShift = 2.08
lm Score = 2.02
NG = 0
DB = 0
SM = 0
Delta_K
RF1_96_0 RF1
ZShift = 1.46
lm Score = −0.03
NG = 0
DB = 0
SM = 0
Delta_r
RF1_96_0 RF1
ZShift = 2.28
lm Score = 29.19
NG = 0
DB = 0
SM = 0
Delta_AUC
RF1_259_0 RF1
ZShift = −0.44
lm Score = −0.03
NG = 0
DB = 0
SM = 0
Delta_L
RF1_259_0 RF1
ZShift = −0.41
lm Score = 2.73
NG = 0
DB = 0
SM = 0
Delta_K
RF1_259_0 RF1
ZShift = 0.02
lm Score = 0.01
NG = 0
DB = 0
SM = 0
Delta_r
RF1_259_0 RF1
ZShift = −0.19
lm Score = 127.79
NG = 0
DB = 0
SM = 0
Delta_AUC
RF1_353_0 RF1
ZShift = −0.53
lm Score = −0.04
NG = 0
DB = 0
SM = 0
Delta_L
RF1_353_0 RF1
ZShift = −0.15
lm Score = −8.16
NG = 0
DB = 0
SM = 0
Delta_K
RF1_353_0 RF1
ZShift = −0.17
lm Score = −0.04
NG = 0
DB = 0
SM = 0
Delta_r
RF1_353_0 RF1
ZShift = 0.06
lm Score = −203.38
NG = 0
DB = 0
SM = 0
Delta_AUC
Doxorubicin
0 2.5 5 7.5 15

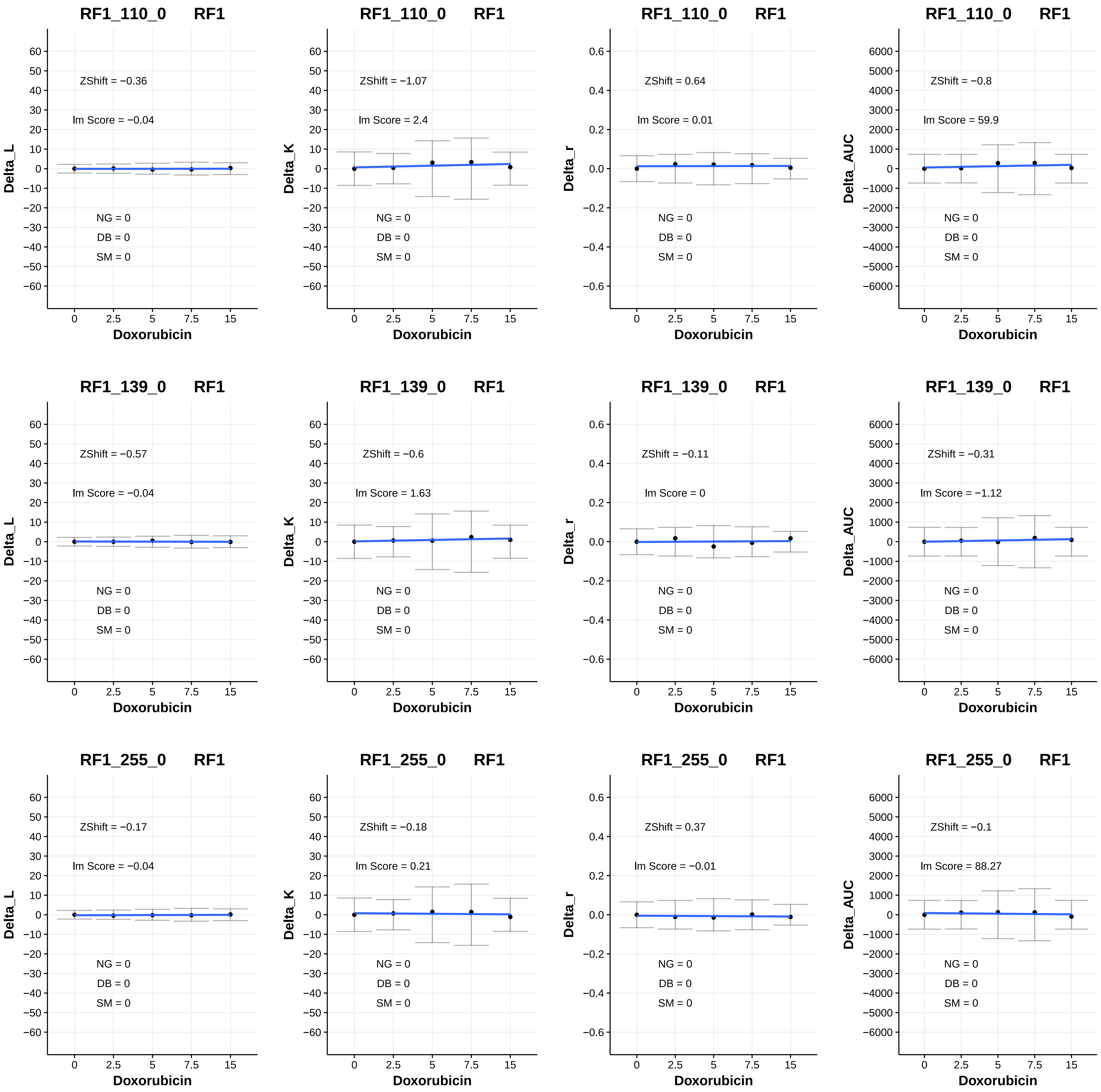
RF1_110_0 RF1
ZShift = −0.36
lm Score = −0.04
NG = 0
DB = 0
SM = 0
Delta_L
Doxorubicin
RF1_110_0 RF1
ZShift = −1.07
lm Score = 2.4
NG = 0
DB = 0
SM = 0
Delta_K
Doxorubicin
RF1_110_0 RF1
ZShift = 0.64
lm Score = 0.01
NG = 0
DB = 0
SM = 0
Delta_r
Doxorubicin
RF1_110_0 RF1
ZShift = −0.8
lm Score = 59.9
NG = 0
DB = 0
SM = 0
Delta_AUC
Doxorubicin
RF1_139_0 RF1
ZShift = −0.57
lm Score = −0.04
NG = 0
DB = 0
SM = 0
Delta_L
Doxorubicin
RF1_139_0 RF1
ZShift = −0.6
lm Score = 1.63
NG = 0
DB = 0
SM = 0
Delta_K
Doxorubicin
RF1_139_0 RF1
ZShift = −0.11
lm Score = 0
NG = 0
DB = 0
SM = 0
Delta_r
Doxorubicin
RF1_139_0 RF1
ZShift = −0.31
lm Score = −1.12
NG = 0
DB = 0
SM = 0
Delta_AUC
Doxorubicin
RF1_255_0 RF1
ZShift = −0.17
lm Score = −0.04
NG = 0
DB = 0
SM = 0
Delta_L
Doxorubicin
RF1_255_0 RF1
ZShift = −0.18
lm Score = 0.21
NG = 0
DB = 0
SM = 0
Delta_K
Doxorubicin
RF1_255_0 RF1
ZShift = 0.37
lm Score = −0.01
NG = 0
DB = 0
SM = 0
Delta_r
Doxorubicin
RF1_255_0 RF1
ZShift = −0.1
lm Score = 88.27
NG = 0
DB = 0
SM = 0
Delta_AUC
Doxorubicin

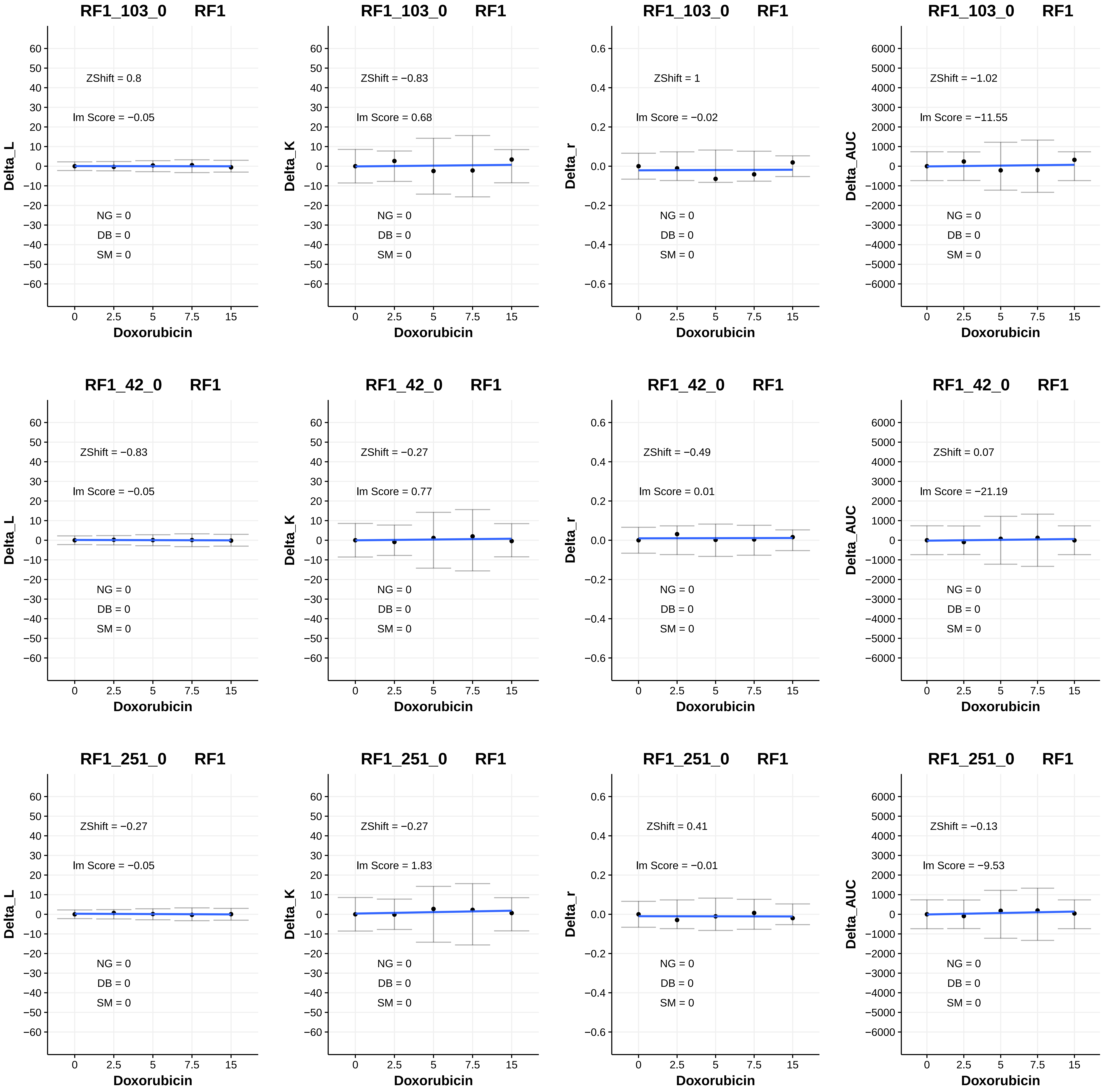
RF1_103_0 RF1
ZShift = 0.8
lm Score = −0.05
NG = 0
DB = 0
SM = 0
Delta_L
Doxorubicin
RF1_103_0 RF1
ZShift = −0.83
lm Score = 0.68
NG = 0
DB = 0
SM = 0
Delta_K
Doxorubicin
RF1_103_0 RF1
ZShift = 1
lm Score = −0.02
NG = 0
DB = 0
SM = 0
Delta_r
Doxorubicin
RF1_103_0 RF1
ZShift = −1.02
lm Score = −11.55
NG = 0
DB = 0
SM = 0
Delta_AUC
Doxorubicin
RF1_42_0 RF1
ZShift = −0.83
lm Score = −0.05
NG = 0
DB = 0
SM = 0
Delta_L
Doxorubicin
RF1_42_0 RF1
ZShift = −0.27
lm Score = 0.77
NG = 0
DB = 0
SM = 0
Delta_K
Doxorubicin
RF1_42_0 RF1
ZShift = −0.49
lm Score = 0.01
NG = 0
DB = 0
SM = 0
Delta_r
Doxorubicin
RF1_42_0 RF1
ZShift = 0.07
lm Score = −21.19
NG = 0
DB = 0
SM = 0
Delta_AUC
Doxorubicin
RF1_251_0 RF1
ZShift = −0.27
lm Score = −0.05
NG = 0
DB = 0
SM = 0
Delta_L
Doxorubicin
RF1_251_0 RF1
ZShift = −0.27
lm Score = 1.83
NG = 0
DB = 0
SM = 0
Delta_K
Doxorubicin
RF1_251_0 RF1
ZShift = 0.41
lm Score = −0.01
NG = 0
DB = 0
SM = 0
Delta_r
Doxorubicin
RF1_251_0 RF1
ZShift = −0.13
lm Score = −9.53
NG = 0
DB = 0
SM = 0
Delta_AUC
Doxorubicin

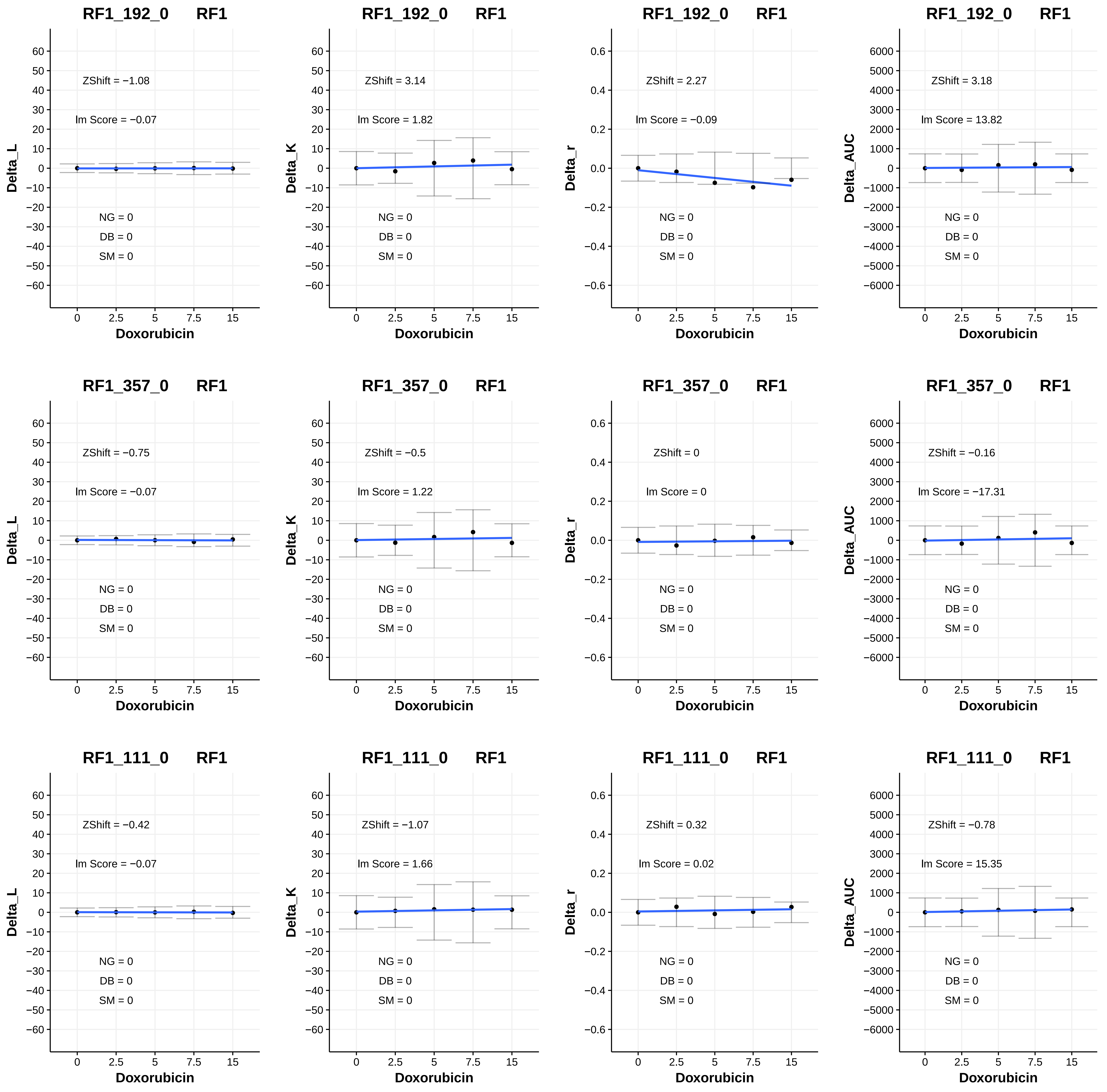
RF1_192_0 RF1
Delta_L
ZShift = −1.08
lm Score = −0.07
NG = 0
DB = 0
SM = 0
Doxorubicin
RF1_192_0 RF1
Delta_K
ZShift = 3.14
lm Score = 1.82
NG = 0
DB = 0
SM = 0
Doxorubicin
RF1_192_0 RF1
Delta_r
ZShift = 2.27
lm Score = −0.09
NG = 0
DB = 0
SM = 0
Doxorubicin
RF1_192_0 RF1
Delta_AUC
ZShift = 3.18
lm Score = 13.82
NG = 0
DB = 0
SM = 0
Doxorubicin
RF1_357_0 RF1
Delta_L
ZShift = −0.75
lm Score = −0.07
NG = 0
DB = 0
SM = 0
Doxorubicin
RF1_357_0 RF1
Delta_K
ZShift = −0.5
lm Score = 1.22
NG = 0
DB = 0
SM = 0
Doxorubicin
RF1_357_0 RF1
Delta_r
ZShift = 0
lm Score = 0
NG = 0
DB = 0
SM = 0
Doxorubicin
RF1_357_0 RF1
Delta_AUC
ZShift = −0.16
lm Score = −17.31
NG = 0
DB = 0
SM = 0
Doxorubicin
RF1_111_0 RF1
Delta_L
ZShift = −0.42
lm Score = −0.07
NG = 0
DB = 0
SM = 0
Doxorubicin
RF1_111_0 RF1
Delta_K
ZShift = −1.07
lm Score = 1.66
NG = 0
DB = 0
SM = 0
Doxorubicin
RF1_111_0 RF1
Delta_r
ZShift = 0.32
lm Score = 0.02
NG = 0
DB = 0
SM = 0
Doxorubicin
RF1_111_0 RF1
Delta_AUC
ZShift = −0.78
lm Score = 15.35
NG = 0
DB = 0
SM = 0
Doxorubicin

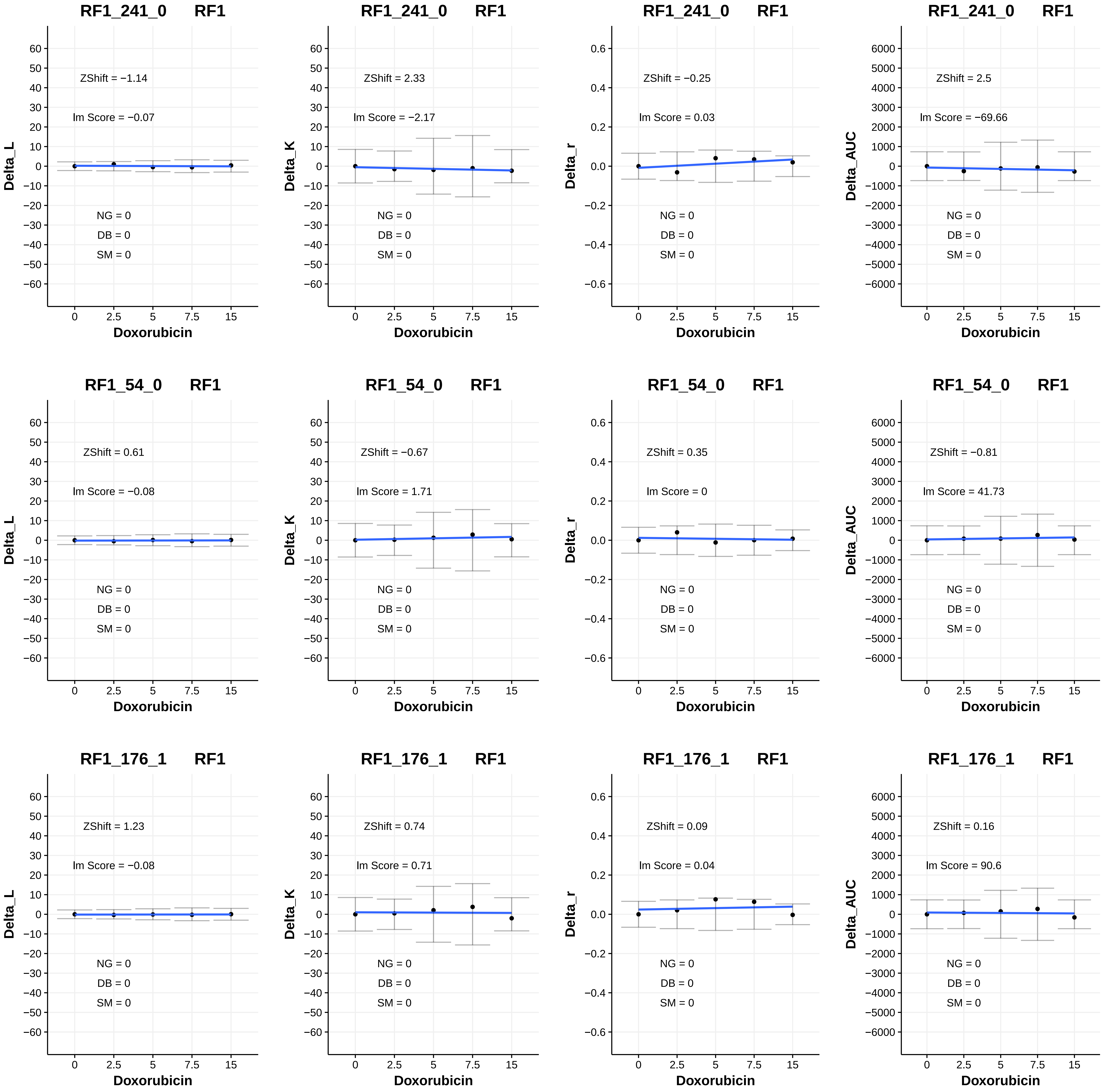

RF1_241_0 RF1
ZShift = −1.14
lm Score = −0.07
NG = 0
DB = 0
SM = 0
Delta_L
Doxorubicin
RF1_241_0 RF1
ZShift = 2.33
lm Score = −2.17
NG = 0
DB = 0
SM = 0
Delta_K
Doxorubicin
RF1_241_0 RF1
ZShift = −0.25
lm Score = 0.03
NG = 0
DB = 0
SM = 0
Delta_r
Doxorubicin
RF1_241_0 RF1
ZShift = 2.5
lm Score = −69.66
NG = 0
DB = 0
SM = 0
Delta_AUC
Doxorubicin
RF1_54_0 RF1
ZShift = 0.61
lm Score = −0.08
NG = 0
DB = 0
SM = 0
Delta_L
Doxorubicin
RF1_54_0 RF1
ZShift = −0.67
lm Score = 1.71
NG = 0
DB = 0
SM = 0
Delta_K
Doxorubicin
RF1_54_0 RF1
ZShift = 0.35
lm Score = 0
NG = 0
DB = 0
SM = 0
Delta_r
Doxorubicin
RF1_54_0 RF1
ZShift = −0.81
lm Score = 41.73
NG = 0
DB = 0
SM = 0
Delta_AUC
Doxorubicin
RF1_176_1 RF1
ZShift = 1.23
lm Score = −0.08
NG = 0
DB = 0
SM = 0
Delta_L
Doxorubicin
RF1_176_1 RF1
ZShift = 0.74
lm Score = 0.71
NG = 0
DB = 0
SM = 0
Delta_K
Doxorubicin
RF1_176_1 RF1
ZShift = 0.09
lm Score = 0.04
NG = 0
DB = 0
SM = 0
Delta_r
Doxorubicin
RF1_176_1 RF1
ZShift = 0.16
lm Score = 90.6
NG = 0
DB = 0
SM = 0
Delta_AUC
Doxorubicin

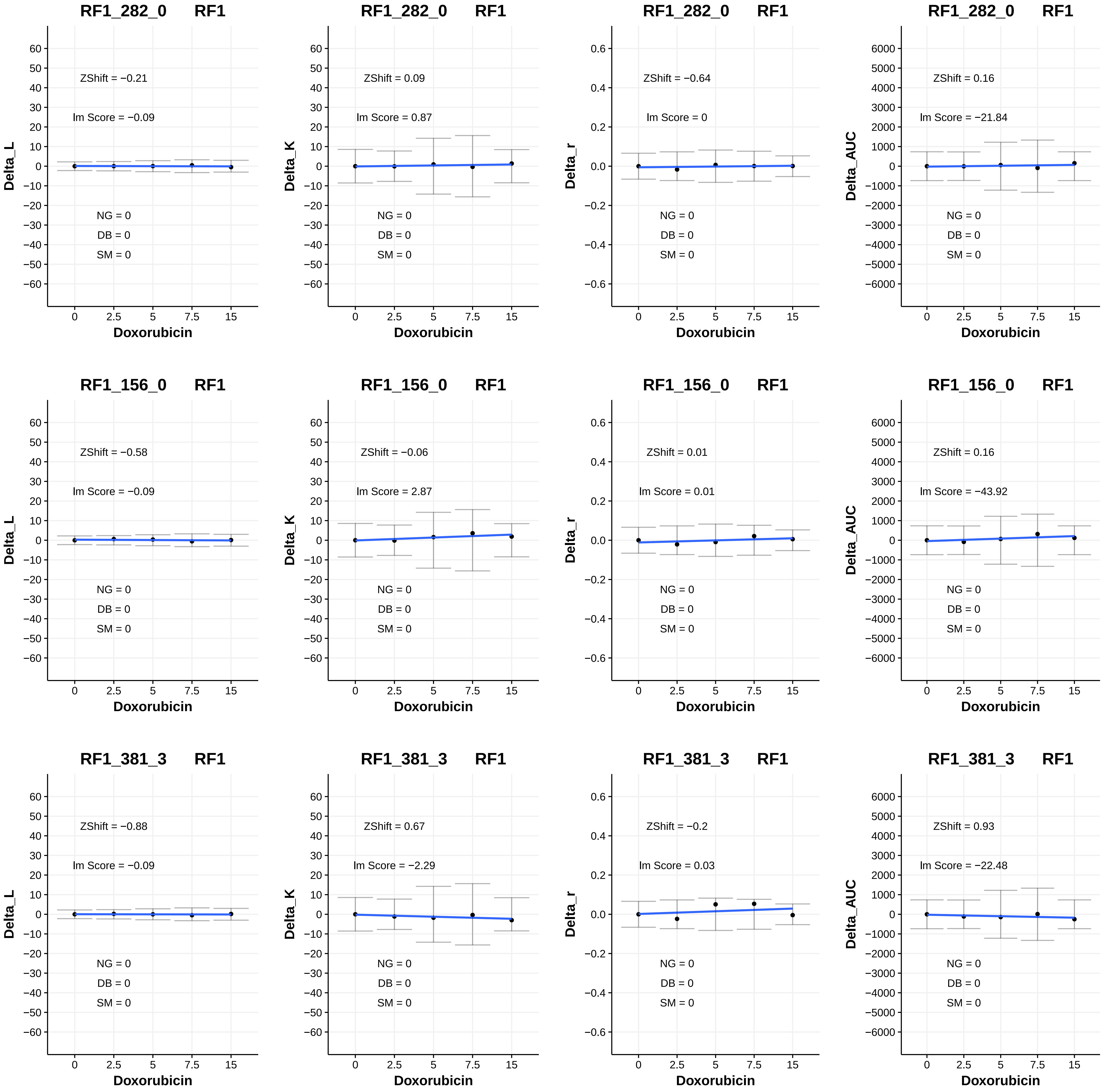
RF1_282_0 RF1
ZShift = −0.21
lm Score = −0.09
NG = 0
DB = 0
SM = 0
Delta_L
Doxorubicin
RF1_282_0 RF1
ZShift = 0.09
lm Score = 0.87
NG = 0
DB = 0
SM = 0
Delta_K
Doxorubicin
RF1_282_0 RF1
ZShift = −0.64
lm Score = 0
NG = 0
DB = 0
SM = 0
Delta_r
Doxorubicin
RF1_282_0 RF1
ZShift = 0.16
lm Score = −21.84
NG = 0
DB = 0
SM = 0
Delta_AUC
Doxorubicin
RF1_156_0 RF1
ZShift = −0.58
lm Score = −0.09
NG = 0
DB = 0
SM = 0
Delta_L
Doxorubicin
RF1_156_0 RF1
ZShift = −0.06
lm Score = 2.87
NG = 0
DB = 0
SM = 0
Delta_K
Doxorubicin
RF1_156_0 RF1
ZShift = 0.01
lm Score = 0.01
NG = 0
DB = 0
SM = 0
Delta_r
Doxorubicin
RF1_156_0 RF1
ZShift = 0.16
lm Score = −43.92
NG = 0
DB = 0
SM = 0
Delta_AUC
Doxorubicin
RF1_381_3 RF1
ZShift = −0.88
lm Score = −0.09
NG = 0
DB = 0
SM = 0
Delta_L
Doxorubicin
RF1_381_3 RF1
ZShift = 0.67
lm Score = −2.29
NG = 0
DB = 0
SM = 0
Delta_K
Doxorubicin
RF1_381_3 RF1
ZShift = −0.2
lm Score = 0.03
NG = 0
DB = 0
SM = 0
Delta_r
Doxorubicin
RF1_381_3 RF1
ZShift = 0.93
lm Score = −22.48
NG = 0
DB = 0
SM = 0
Delta_AUC
Doxorubicin

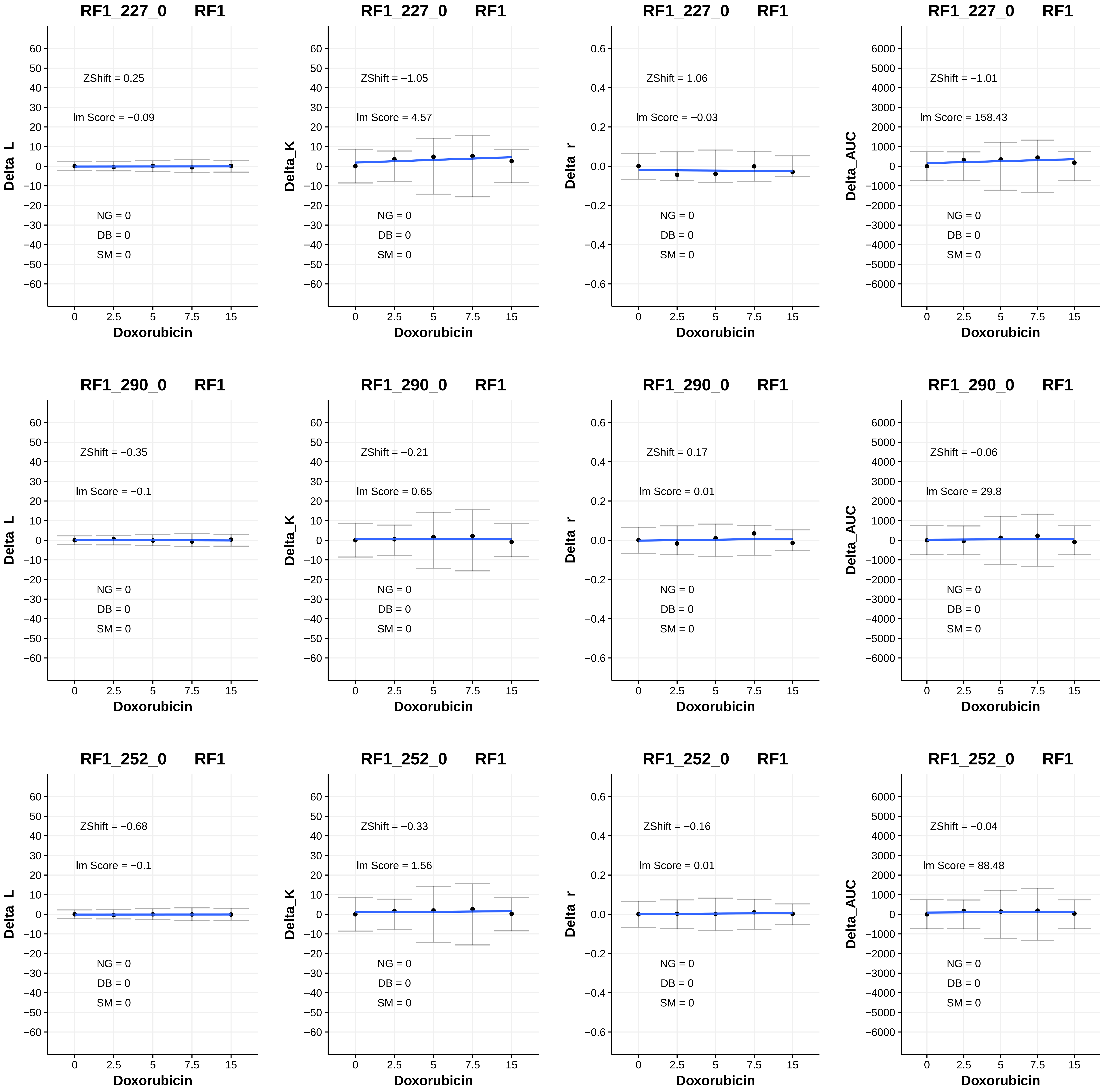

RF1_227_0 RF1
ZShift = 0.25
Im Score = −0.09
NG = 0
DB = 0
SM = 0
Delta_L
Doxorubicin
RF1_227_0 RF1
ZShift = −1.05
Im Score = 4.57
NG = 0
DB = 0
SM = 0
Delta_K
Doxorubicin
RF1_227_0 RF1
ZShift = 1.06
Im Score = −0.03
NG = 0
DB = 0
SM = 0
Delta_r
Doxorubicin
RF1_227_0 RF1
ZShift = −1.01
Im Score = 158.43
NG = 0
DB = 0
SM = 0
Delta_AUC
Doxorubicin
RF1_290_0 RF1
ZShift = −0.35
Im Score = −0.1
NG = 0
DB = 0
SM = 0
Delta_L
Doxorubicin
RF1_290_0 RF1
ZShift = −0.21
Im Score = 0.65
NG = 0
DB = 0
SM = 0
Delta_K
Doxorubicin
RF1_290_0 RF1
ZShift = 0.17
Im Score = 0.01
NG = 0
DB = 0
SM = 0
Delta_r
Doxorubicin
RF1_290_0 RF1
ZShift = −0.06
Im Score = 29.8
NG = 0
DB = 0
SM = 0
Delta_AUC
Doxorubicin
RF1_252_0 RF1
ZShift = −0.68
Im Score = −0.1
NG = 0
DB = 0
SM = 0
Delta_L
Doxorubicin
RF1_252_0 RF1
ZShift = −0.33
Im Score = 1.56
NG = 0
DB = 0
SM = 0
Delta_K
Doxorubicin
RF1_252_0 RF1
ZShift = −0.16
Im Score = 0.01
NG = 0
DB = 0
SM = 0
Delta_r
Doxorubicin
RF1_252_0 RF1
ZShift = −0.04
Im Score = 88.48
NG = 0
DB = 0
SM = 0
Delta_AUC
Doxorubicin

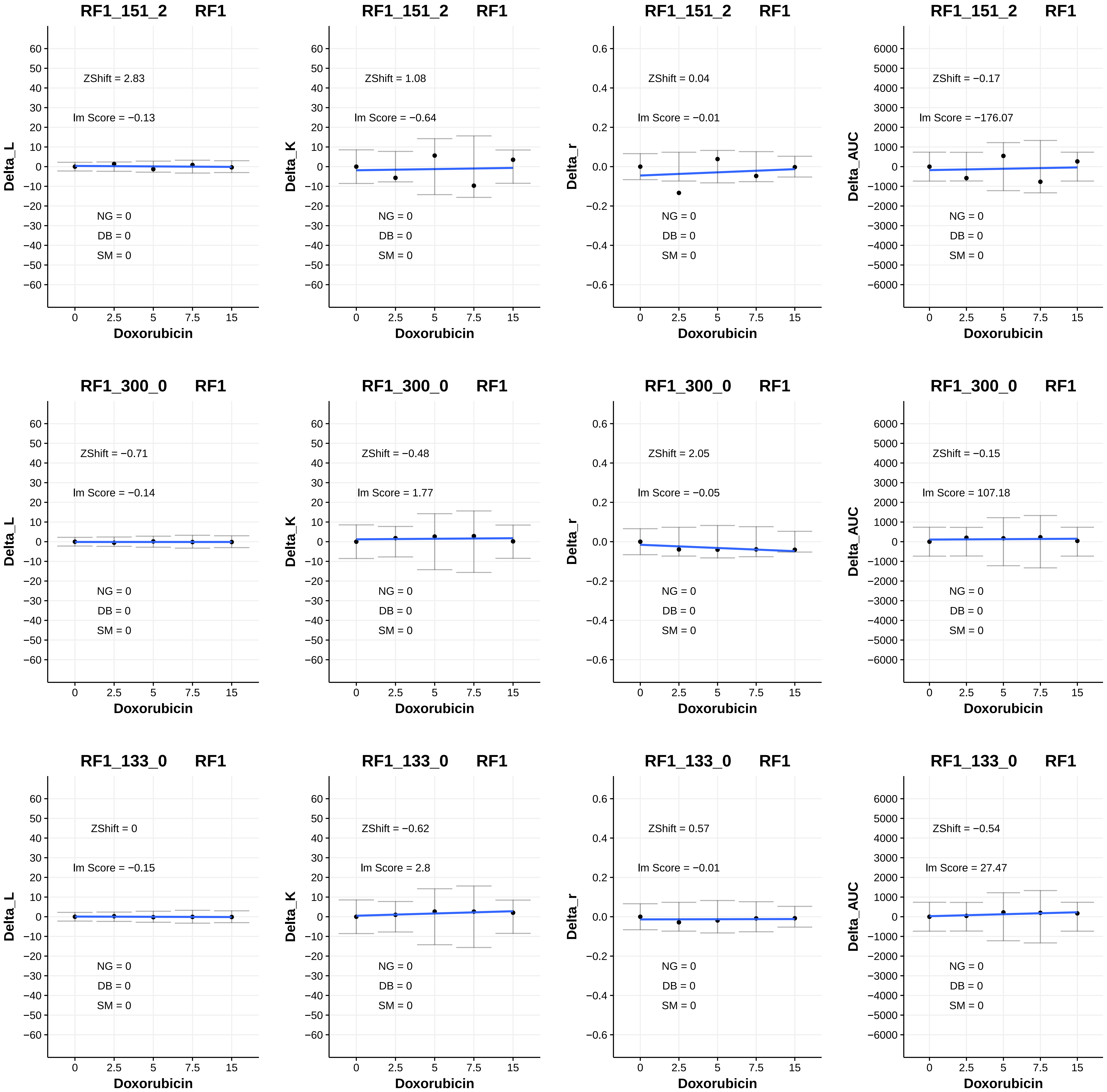
RF1_151_2 RF1
Delta_L
ZShift = 2.83
lm Score = −0.13
NG = 0
DB = 0
SM = 0
Doxorubicin
RF1_151_2 RF1
Delta_K
ZShift = 1.08
lm Score = −0.64
NG = 0
DB = 0
SM = 0
Doxorubicin
RF1_151_2 RF1
Delta_r
ZShift = 0.04
lm Score = −0.01
NG = 0
DB = 0
SM = 0
Doxorubicin
RF1_151_2 RF1
Delta_AUC
ZShift = −0.17
lm Score = −176.07
NG = 0
DB = 0
SM = 0
Doxorubicin
RF1_300_0 RF1
Delta_L
ZShift = −0.71
lm Score = −0.14
NG = 0
DB = 0
SM = 0
Doxorubicin
RF1_300_0 RF1
Delta_K
ZShift = −0.48
lm Score = 1.77
NG = 0
DB = 0
SM = 0
Doxorubicin
RF1_300_0 RF1
Delta_r
ZShift = 2.05
lm Score = −0.05
NG = 0
DB = 0
SM = 0
Doxorubicin
RF1_300_0 RF1
Delta_AUC
ZShift = −0.15
lm Score = 107.18
NG = 0
DB = 0
SM = 0
Doxorubicin
RF1_133_0 RF1
Delta_L
ZShift = 0
lm Score = −0.15
NG = 0
DB = 0
SM = 0
Doxorubicin
RF1_133_0 RF1
Delta_K
ZShift = −0.62
lm Score = 2.8
NG = 0
DB = 0
SM = 0
Doxorubicin
RF1_133_0 RF1
Delta_r
ZShift = 0.57
lm Score = −0.01
NG = 0
DB = 0
SM = 0
Doxorubicin
RF1_133_0 RF1
Delta_AUC
ZShift = −0.54
lm Score = 27.47
NG = 0
DB = 0
SM = 0
Doxorubicin

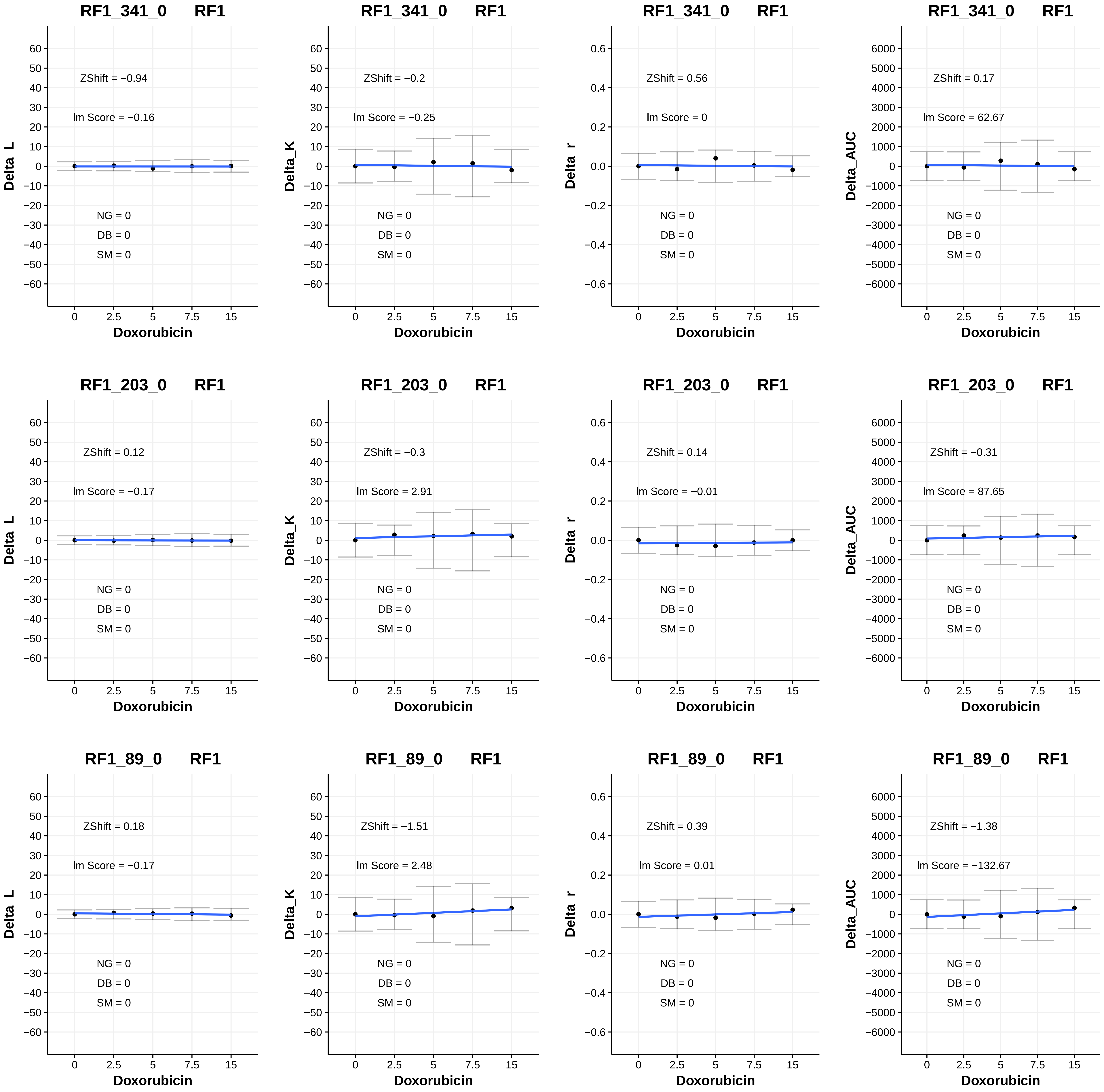
RF1_341_0 RF1
ZShift = −0.94
lm Score = −0.16
NG = 0
DB = 0
SM = 0
Delta_L
Doxorubicin
RF1_341_0 RF1
ZShift = −0.2
lm Score = −0.25
NG = 0
DB = 0
SM = 0
Delta_K
Doxorubicin
RF1_341_0 RF1
ZShift = 0.56
lm Score = 0
NG = 0
DB = 0
SM = 0
Delta_r
Doxorubicin
RF1_341_0 RF1
ZShift = 0.17
lm Score = 62.67
NG = 0
DB = 0
SM = 0
Delta_AUC
Doxorubicin
RF1_203_0 RF1
ZShift = 0.12
lm Score = −0.17
NG = 0
DB = 0
SM = 0
Delta_L
Doxorubicin
RF1_203_0 RF1
ZShift = −0.3
lm Score = 2.91
NG = 0
DB = 0
SM = 0
Delta_K
Doxorubicin
RF1_203_0 RF1
ZShift = 0.14
lm Score = −0.01
NG = 0
DB = 0
SM = 0
Delta_r
Doxorubicin
RF1_203_0 RF1
ZShift = −0.31
lm Score = 87.65
NG = 0
DB = 0
SM = 0
Delta_AUC
Doxorubicin
RF1_89_0 RF1
ZShift = 0.18
lm Score = −0.17
NG = 0
DB = 0
SM = 0
Delta_L
Doxorubicin
RF1_89_0 RF1
ZShift = −1.51
lm Score = 2.48
NG = 0
DB = 0
SM = 0
Delta_K
Doxorubicin
RF1_89_0 RF1
ZShift = 0.39
lm Score = 0.01
NG = 0
DB = 0
SM = 0
Delta_r
Doxorubicin
RF1_89_0 RF1
ZShift = −1.38
lm Score = −132.67
NG = 0
DB = 0
SM = 0
Delta_AUC
Doxorubicin

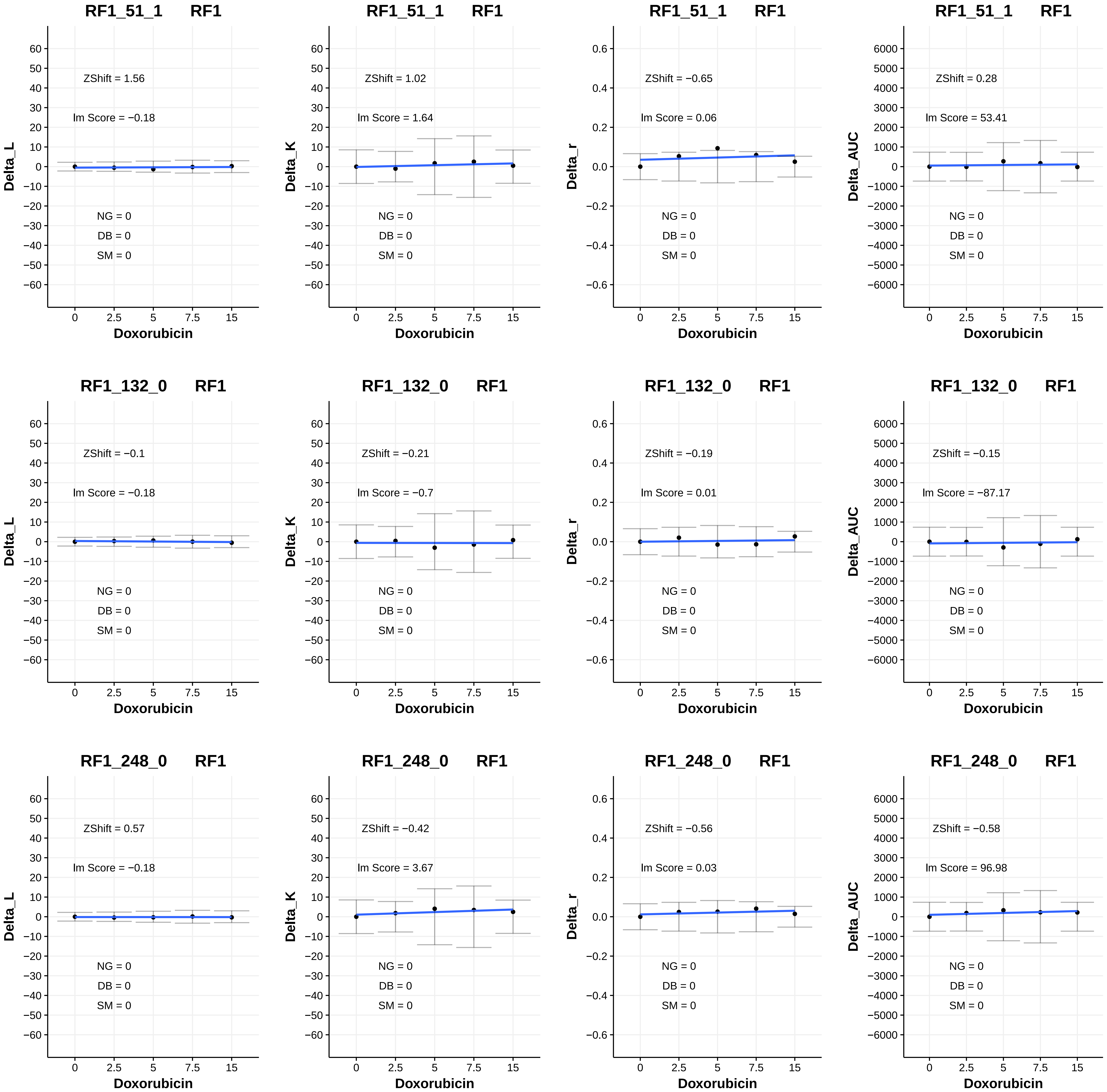
RF1_51_1 RF1
ZShift = 1.56
lm Score = −0.18
NG = 0
DB = 0
SM = 0
Delta_L
Doxorubicin
RF1_51_1 RF1
ZShift = 1.02
lm Score = 1.64
NG = 0
DB = 0
SM = 0
Delta_K
Doxorubicin
RF1_51_1 RF1
ZShift = −0.65
lm Score = 0.06
NG = 0
DB = 0
SM = 0
Delta_r
Doxorubicin
RF1_51_1 RF1
ZShift = 0.28
lm Score = 53.41
NG = 0
DB = 0
SM = 0
Delta_AUC
Doxorubicin
RF1_132_0 RF1
ZShift = −0.1
lm Score = −0.18
NG = 0
DB = 0
SM = 0
Delta_L
Doxorubicin
RF1_132_0 RF1
ZShift = −0.21
lm Score = −0.7
NG = 0
DB = 0
SM = 0
Delta_K
Doxorubicin
RF1_132_0 RF1
ZShift = −0.19
lm Score = 0.01
NG = 0
DB = 0
SM = 0
Delta_r
Doxorubicin
RF1_132_0 RF1
ZShift = −0.15
lm Score = −87.17
NG = 0
DB = 0
SM = 0
Delta_AUC
Doxorubicin
RF1_248_0 RF1
ZShift = 0.57
lm Score = −0.18
NG = 0
DB = 0
SM = 0
Delta_L
Doxorubicin
RF1_248_0 RF1
ZShift = −0.42
lm Score = 3.67
NG = 0
DB = 0
SM = 0
Delta_K
Doxorubicin
RF1_248_0 RF1
ZShift = −0.56
lm Score = 0.03
NG = 0
DB = 0
SM = 0
Delta_r
Doxorubicin
RF1_248_0 RF1
ZShift = −0.58
lm Score = 96.98
NG = 0
DB = 0
SM = 0
Delta_AUC
Doxorubicin

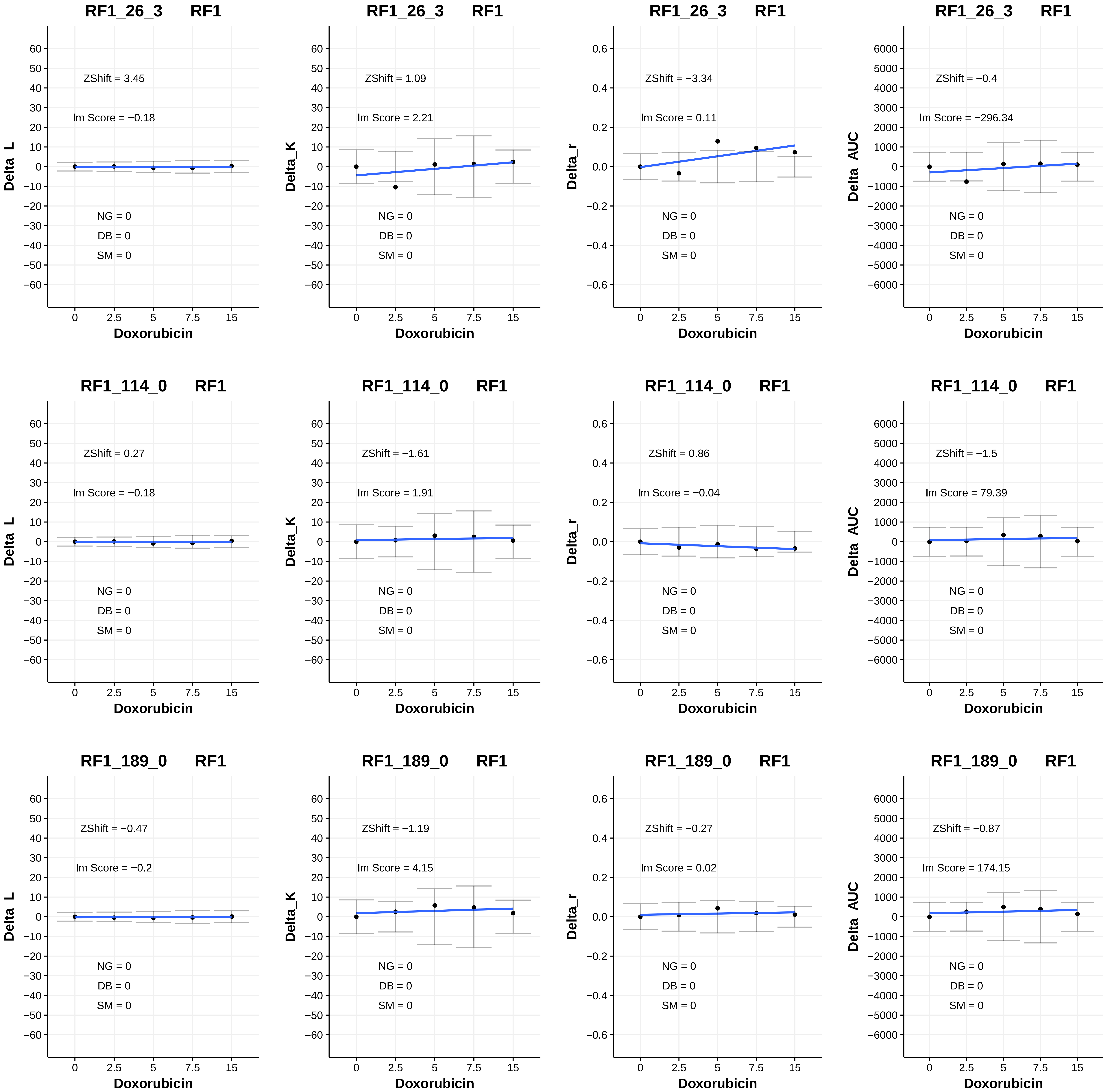
RF1_26_3 RF1
Delta_L
ZShift = 3.45
lm Score = −0.18
NG = 0
DB = 0
SM = 0
Doxorubicin
RF1_26_3 RF1
Delta_K
ZShift = 1.09
lm Score = 2.21
NG = 0
DB = 0
SM = 0
Doxorubicin
RF1_26_3 RF1
Delta_r
ZShift = −3.34
lm Score = 0.11
NG = 0
DB = 0
SM = 0
Doxorubicin
RF1_26_3 RF1
Delta_AUC
ZShift = −0.4
lm Score = −296.34
NG = 0
DB = 0
SM = 0
Doxorubicin
RF1_114_0 RF1
Delta_L
ZShift = 0.27
lm Score = −0.18
NG = 0
DB = 0
SM = 0
Doxorubicin
RF1_114_0 RF1
Delta_K
ZShift = −1.61
lm Score = 1.91
NG = 0
DB = 0
SM = 0
Doxorubicin
RF1_114_0 RF1
Delta_r
ZShift = 0.86
lm Score = −0.04
NG = 0
DB = 0
SM = 0
Doxorubicin
RF1_114_0 RF1
Delta_AUC
ZShift = −1.5
lm Score = 79.39
NG = 0
DB = 0
SM = 0
Doxorubicin
RF1_189_0 RF1
Delta_L
ZShift = −0.47
lm Score = −0.2
NG = 0
DB = 0
SM = 0
Doxorubicin
RF1_189_0 RF1
Delta_K
ZShift = −1.19
lm Score = 4.15
NG = 0
DB = 0
SM = 0
Doxorubicin
RF1_189_0 RF1
Delta_r
ZShift = −0.27
lm Score = 0.02
NG = 0
DB = 0
SM = 0
Doxorubicin
RF1_189_0 RF1
Delta_AUC
ZShift = −0.87
lm Score = 174.15
NG = 0
DB = 0
SM = 0
Doxorubicin

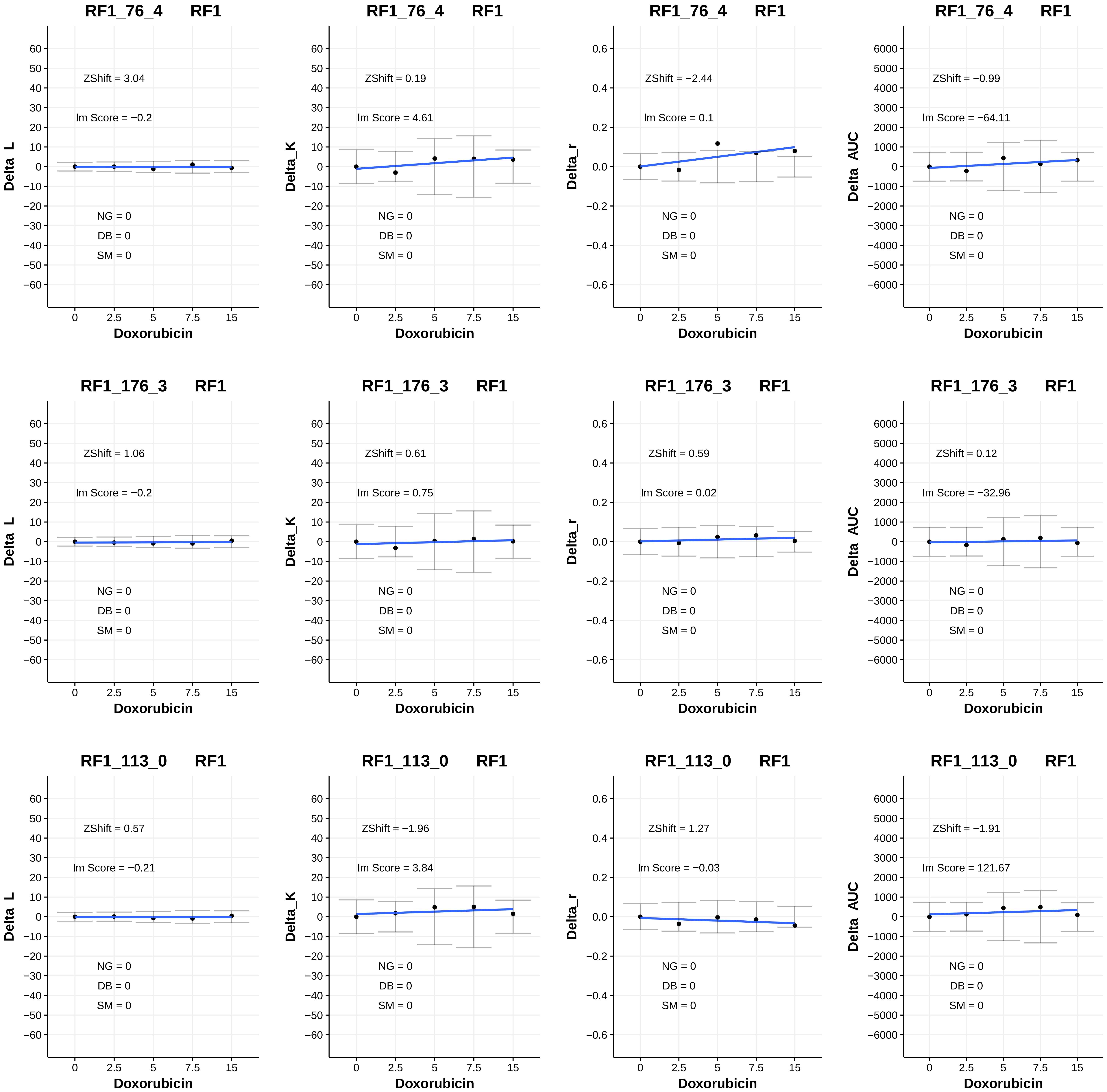

RF1_76_4 RF1
Delta_L
ZShift = 3.04
lm Score = −0.2
NG = 0
DB = 0
SM = 0
Doxorubicin
RF1_76_4 RF1
Delta_K
ZShift = 0.19
lm Score = 4.61
NG = 0
DB = 0
SM = 0
Doxorubicin
RF1_76_4 RF1
Delta_r
ZShift = −2.44
lm Score = 0.1
NG = 0
DB = 0
SM = 0
Doxorubicin
RF1_76_4 RF1
Delta_AUC
ZShift = −0.99
lm Score = −64.11
NG = 0
DB = 0
SM = 0
Doxorubicin
RF1_176_3 RF1
Delta_L
ZShift = 1.06
lm Score = −0.2
NG = 0
DB = 0
SM = 0
Doxorubicin
RF1_176_3 RF1
Delta_K
ZShift = 0.61
lm Score = 0.75
NG = 0
DB = 0
SM = 0
Doxorubicin
RF1_176_3 RF1
Delta_r
ZShift = 0.59
lm Score = 0.02
NG = 0
DB = 0
SM = 0
Doxorubicin
RF1_176_3 RF1
Delta_AUC
ZShift = 0.12
lm Score = −32.96
NG = 0
DB = 0
SM = 0
Doxorubicin
RF1_113_0 RF1
Delta_L
ZShift = 0.57
lm Score = −0.21
NG = 0
DB = 0
SM = 0
Doxorubicin
RF1_113_0 RF1
Delta_K
ZShift = −1.96
lm Score = 3.84
NG = 0
DB = 0
SM = 0
Doxorubicin
RF1_113_0 RF1
Delta_r
ZShift = 1.27
lm Score = −0.03
NG = 0
DB = 0
SM = 0
Doxorubicin
RF1_113_0 RF1
Delta_AUC
ZShift = −1.91
lm Score = 121.67
NG = 0
DB = 0
SM = 0
Doxorubicin

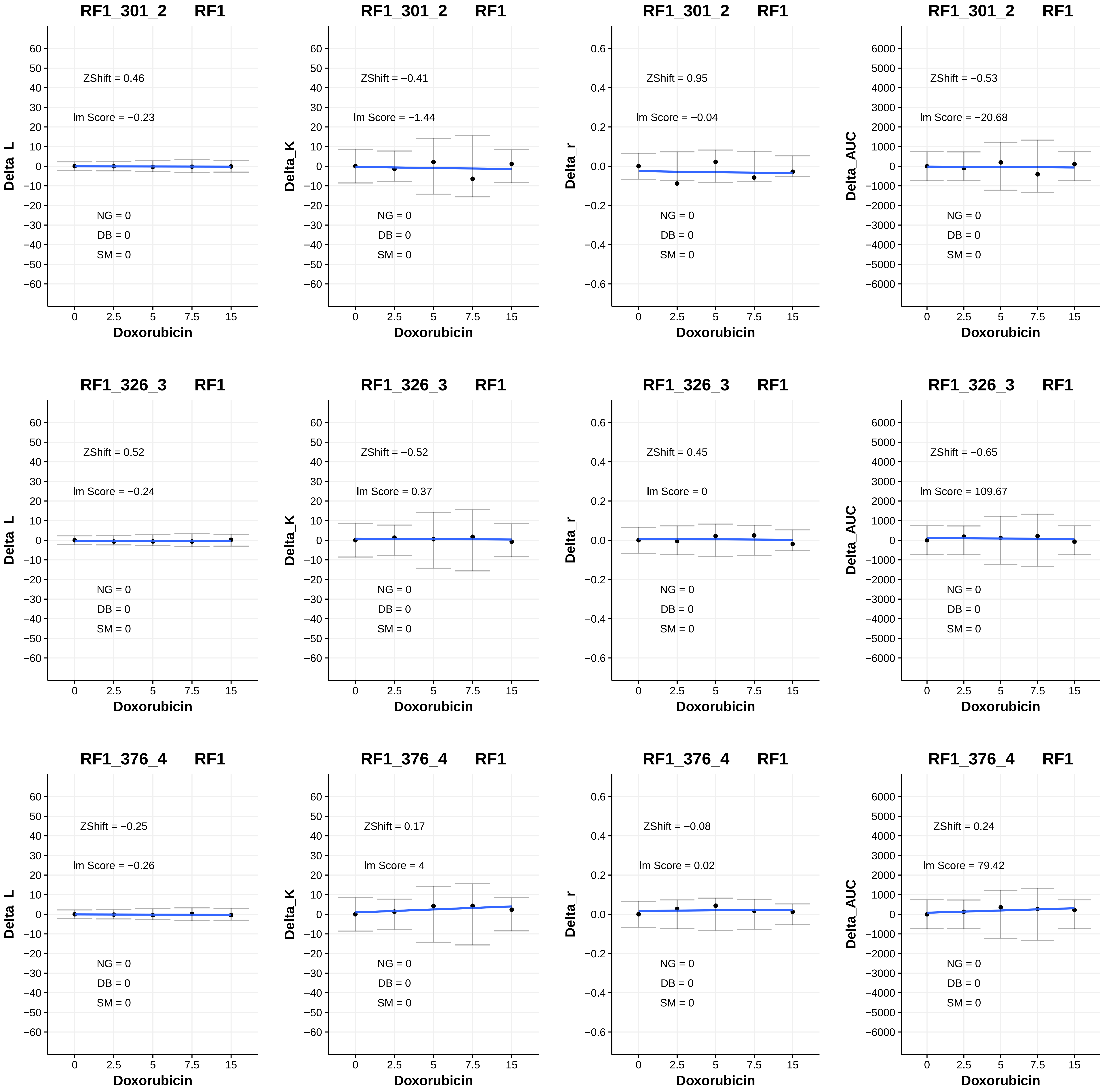

RF1_301_2 RF1
ZShift = 0.46
lm Score = −0.23
NG = 0
DB = 0
SM = 0
Delta_L
Doxorubicin
RF1_301_2 RF1
ZShift = −0.41
lm Score = −1.44
NG = 0
DB = 0
SM = 0
Delta_K
Doxorubicin
RF1_301_2 RF1
ZShift = 0.95
lm Score = −0.04
NG = 0
DB = 0
SM = 0
Delta_r
Doxorubicin
RF1_301_2 RF1
ZShift = −0.53
lm Score = −20.68
NG = 0
DB = 0
SM = 0
Delta_AUC
Doxorubicin
RF1_326_3 RF1
ZShift = 0.52
lm Score = −0.24
NG = 0
DB = 0
SM = 0
Delta_L
Doxorubicin
RF1_326_3 RF1
ZShift = −0.52
lm Score = 0.37
NG = 0
DB = 0
SM = 0
Delta_K
Doxorubicin
RF1_326_3 RF1
ZShift = 0.45
lm Score = 0
NG = 0
DB = 0
SM = 0
Delta_r
Doxorubicin
RF1_326_3 RF1
ZShift = −0.65
lm Score = 109.67
NG = 0
DB = 0
SM = 0
Delta_AUC
Doxorubicin
RF1_376_4 RF1
ZShift = −0.25
lm Score = −0.26
NG = 0
DB = 0
SM = 0
Delta_L
Doxorubicin
RF1_376_4 RF1
ZShift = 0.17
lm Score = 4
NG = 0
DB = 0
SM = 0
Delta_K
Doxorubicin
RF1_376_4 RF1
ZShift = −0.08
lm Score = 0.02
NG = 0
DB = 0
SM = 0
Delta_r
Doxorubicin
RF1_376_4 RF1
ZShift = 0.24
lm Score = 79.42
NG = 0
DB = 0
SM = 0
Delta_AUC
Doxorubicin

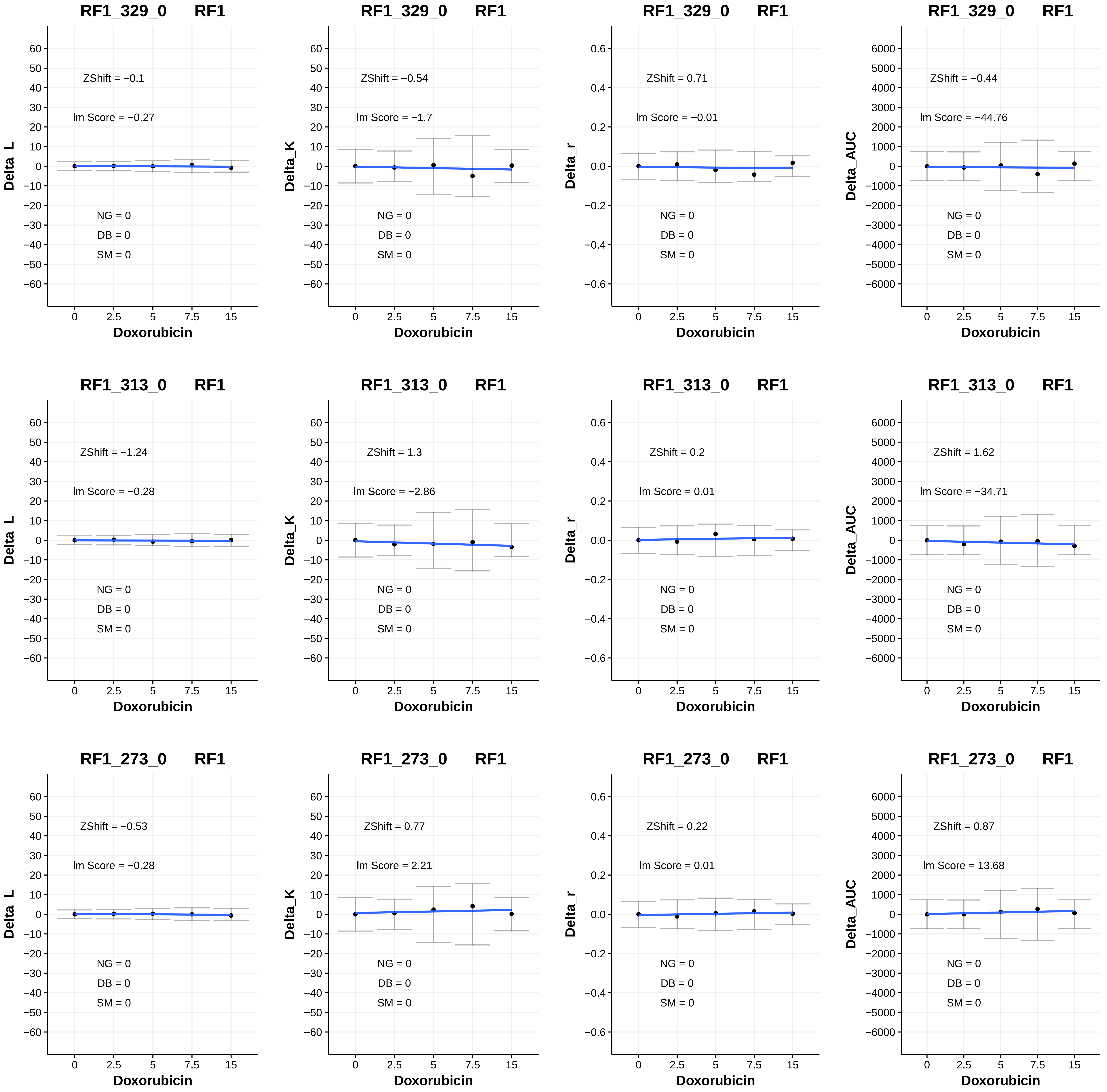
RF1_329_0 RF1
ZShift = −0.1
Im Score = −0.27
NG = 0
DB = 0
SM = 0
Delta_L
Doxorubicin
RF1_329_0 RF1
ZShift = −0.54
Im Score = −1.7
NG = 0
DB = 0
SM = 0
Delta_K
Doxorubicin
RF1_329_0 RF1
ZShift = 0.71
Im Score = −0.01
NG = 0
DB = 0
SM = 0
Delta_r
Doxorubicin
RF1_329_0 RF1
ZShift = −0.44
Im Score = −44.76
NG = 0
DB = 0
SM = 0
Delta_AUC
Doxorubicin
RF1_313_0 RF1
ZShift = −1.24
Im Score = −0.28
NG = 0
DB = 0
SM = 0
Delta_L
Doxorubicin
RF1_313_0 RF1
ZShift = 1.3
Im Score = −2.86
NG = 0
DB = 0
SM = 0
Delta_K
Doxorubicin
RF1_313_0 RF1
ZShift = 0.2
Im Score = 0.01
NG = 0
DB = 0
SM = 0
Delta_r
Doxorubicin
RF1_313_0 RF1
ZShift = 1.62
Im Score = −34.71
NG = 0
DB = 0
SM = 0
Delta_AUC
Doxorubicin
RF1_273_0 RF1
ZShift = −0.53
Im Score = −0.28
NG = 0
DB = 0
SM = 0
Delta_L
Doxorubicin
RF1_273_0 RF1
ZShift = 0.77
Im Score = 2.21
NG = 0
DB = 0
SM = 0
Delta_K
Doxorubicin
RF1_273_0 RF1
ZShift = 0.22
Im Score = 0.01
NG = 0
DB = 0
SM = 0
Delta_r
Doxorubicin
RF1_273_0 RF1
ZShift = 0.87
Im Score = 13.68
NG = 0
DB = 0
SM = 0
Delta_AUC
Doxorubicin

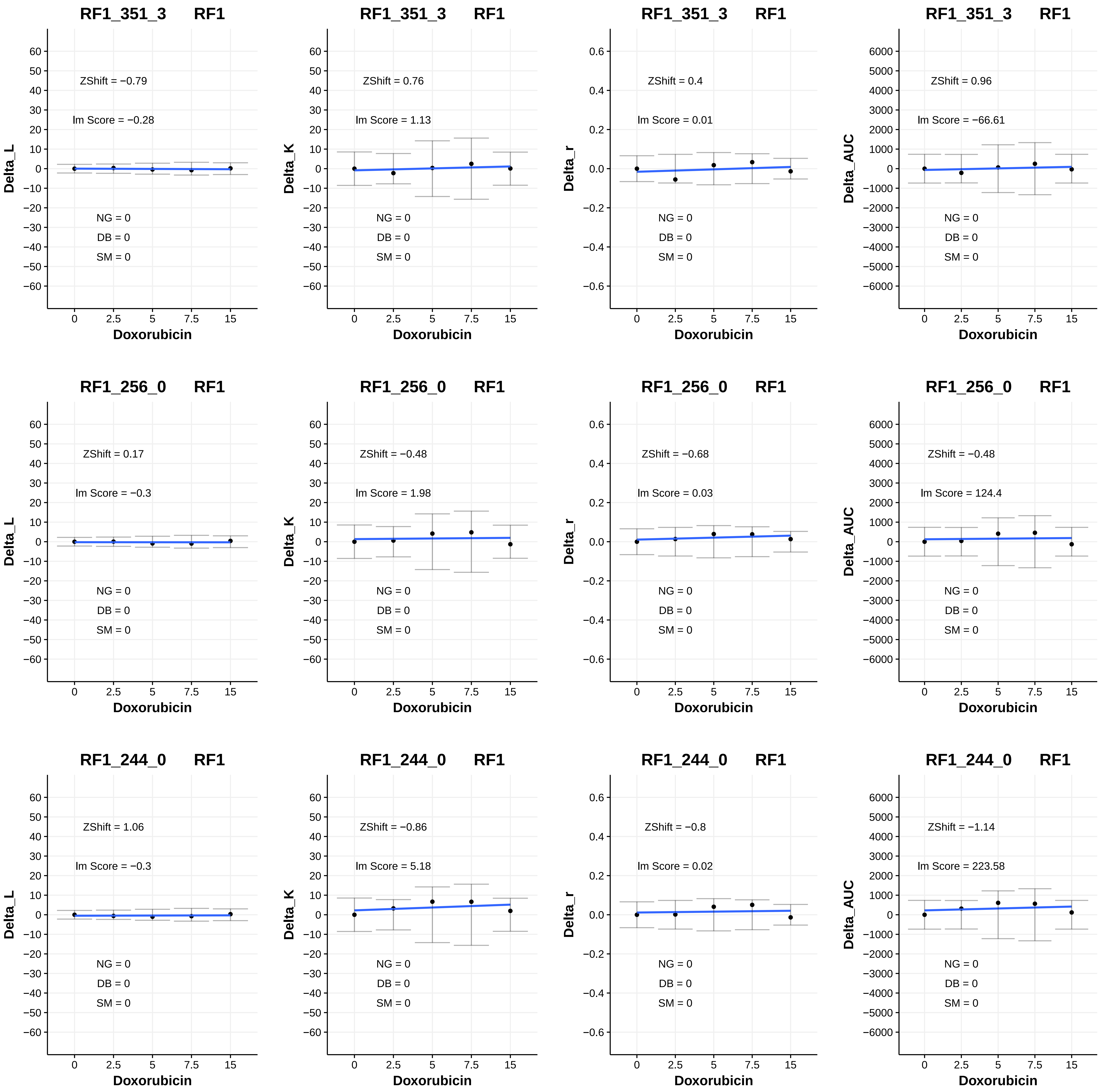
RF1_351_3 RF1
ZShift = −0.79
lm Score = −0.28
NG = 0
DB = 0
SM = 0
Delta_L
Doxorubicin
RF1_351_3 RF1
ZShift = 0.76
lm Score = 1.13
NG = 0
DB = 0
SM = 0
Delta_K
Doxorubicin
RF1_351_3 RF1
ZShift = 0.4
lm Score = 0.01
NG = 0
DB = 0
SM = 0
Delta_r
Doxorubicin
RF1_351_3 RF1
ZShift = 0.96
lm Score = −66.61
NG = 0
DB = 0
SM = 0
Delta_AUC
Doxorubicin
RF1_256_0 RF1
ZShift = 0.17
lm Score = −0.3
NG = 0
DB = 0
SM = 0
Delta_L
Doxorubicin
RF1_256_0 RF1
ZShift = −0.48
lm Score = 1.98
NG = 0
DB = 0
SM = 0
Delta_K
Doxorubicin
RF1_256_0 RF1
ZShift = −0.68
lm Score = 0.03
NG = 0
DB = 0
SM = 0
Delta_r
Doxorubicin
RF1_256_0 RF1
ZShift = −0.48
lm Score = 124.4
NG = 0
DB = 0
SM = 0
Delta_AUC
Doxorubicin
RF1_244_0 RF1
ZShift = 1.06
lm Score = −0.3
NG = 0
DB = 0
SM = 0
Delta_L
Doxorubicin
RF1_244_0 RF1
ZShift = −0.86
lm Score = 5.18
NG = 0
DB = 0
SM = 0
Delta_K
Doxorubicin
RF1_244_0 RF1
ZShift = −0.8
lm Score = 0.02
NG = 0
DB = 0
SM = 0
Delta_r
Doxorubicin
RF1_244_0 RF1
ZShift = −1.14
lm Score = 223.58
NG = 0
DB = 0
SM = 0
Delta_AUC
Doxorubicin

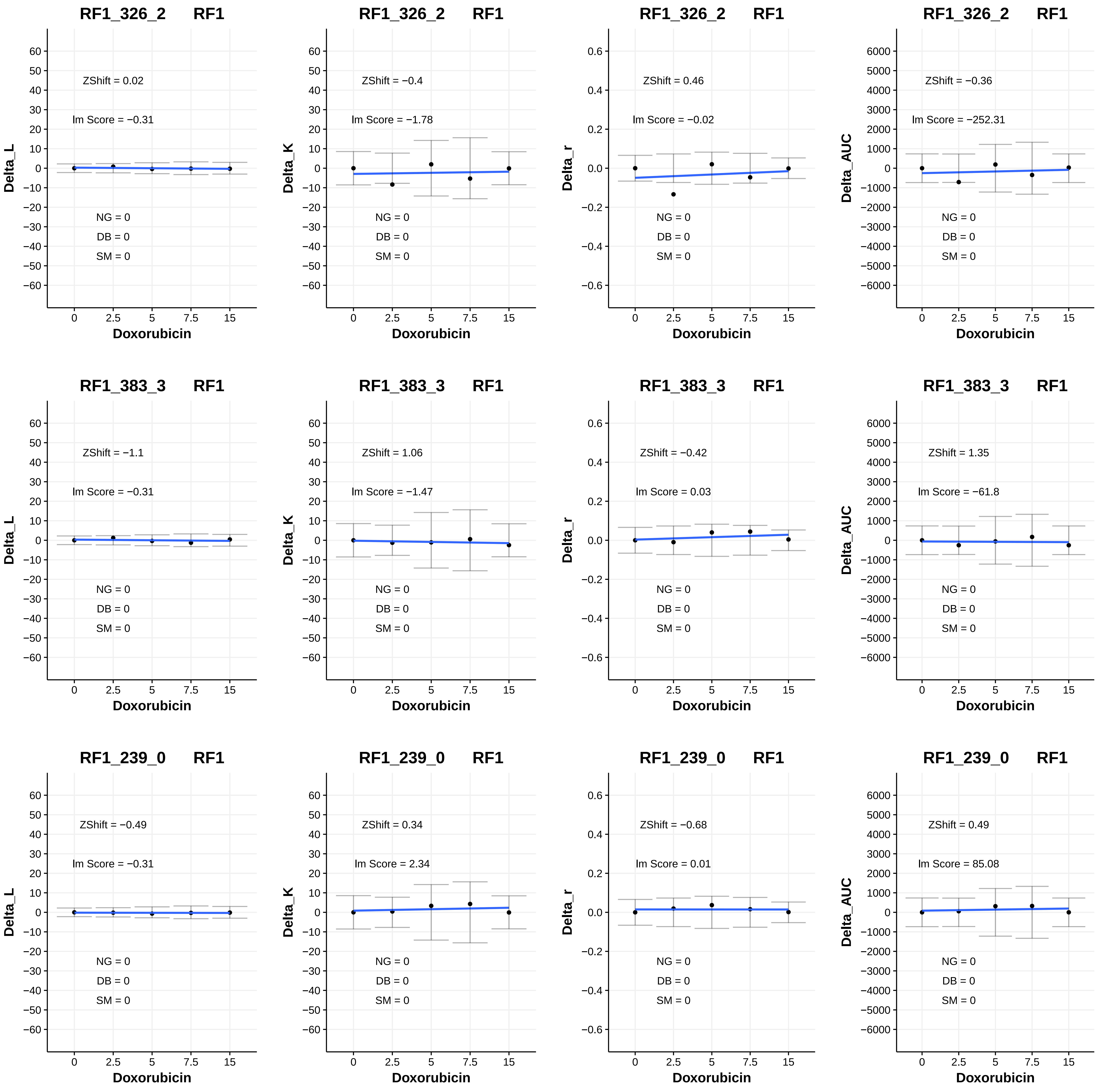
RF1_326_2 RF1
ZShift = 0.02
Im Score = −0.31
NG = 0
DB = 0
SM = 0
Delta_L
Doxorubicin
RF1_326_2 RF1
ZShift = −0.4
Im Score = −1.78
NG = 0
DB = 0
SM = 0
Delta_K
Doxorubicin
RF1_326_2 RF1
ZShift = 0.46
Im Score = −0.02
NG = 0
DB = 0
SM = 0
Delta_r
Doxorubicin
RF1_326_2 RF1
ZShift = −0.36
Im Score = −252.31
NG = 0
DB = 0
SM = 0
Delta_AUC
Doxorubicin
RF1_383_3 RF1
ZShift = −1.1
Im Score = −0.31
NG = 0
DB = 0
SM = 0
Delta_L
Doxorubicin
RF1_383_3 RF1
ZShift = 1.06
Im Score = −1.47
NG = 0
DB = 0
SM = 0
Delta_K
Doxorubicin
RF1_383_3 RF1
ZShift = −0.42
Im Score = 0.03
NG = 0
DB = 0
SM = 0
Delta_r
Doxorubicin
RF1_383_3 RF1
ZShift = 1.35
Im Score = −61.8
NG = 0
DB = 0
SM = 0
Delta_AUC
Doxorubicin
RF1_239_0 RF1
ZShift = −0.49
Im Score = −0.31
NG = 0
DB = 0
SM = 0
Delta_L
Doxorubicin
RF1_239_0 RF1
ZShift = 0.34
Im Score = 2.34
NG = 0
DB = 0
SM = 0
Delta_K
Doxorubicin
RF1_239_0 RF1
ZShift = −0.68
Im Score = 0.01
NG = 0
DB = 0
SM = 0
Delta_r
Doxorubicin
RF1_239_0 RF1
ZShift = 0.49
Im Score = 85.08
NG = 0
DB = 0
SM = 0
Delta_AUC
Doxorubicin

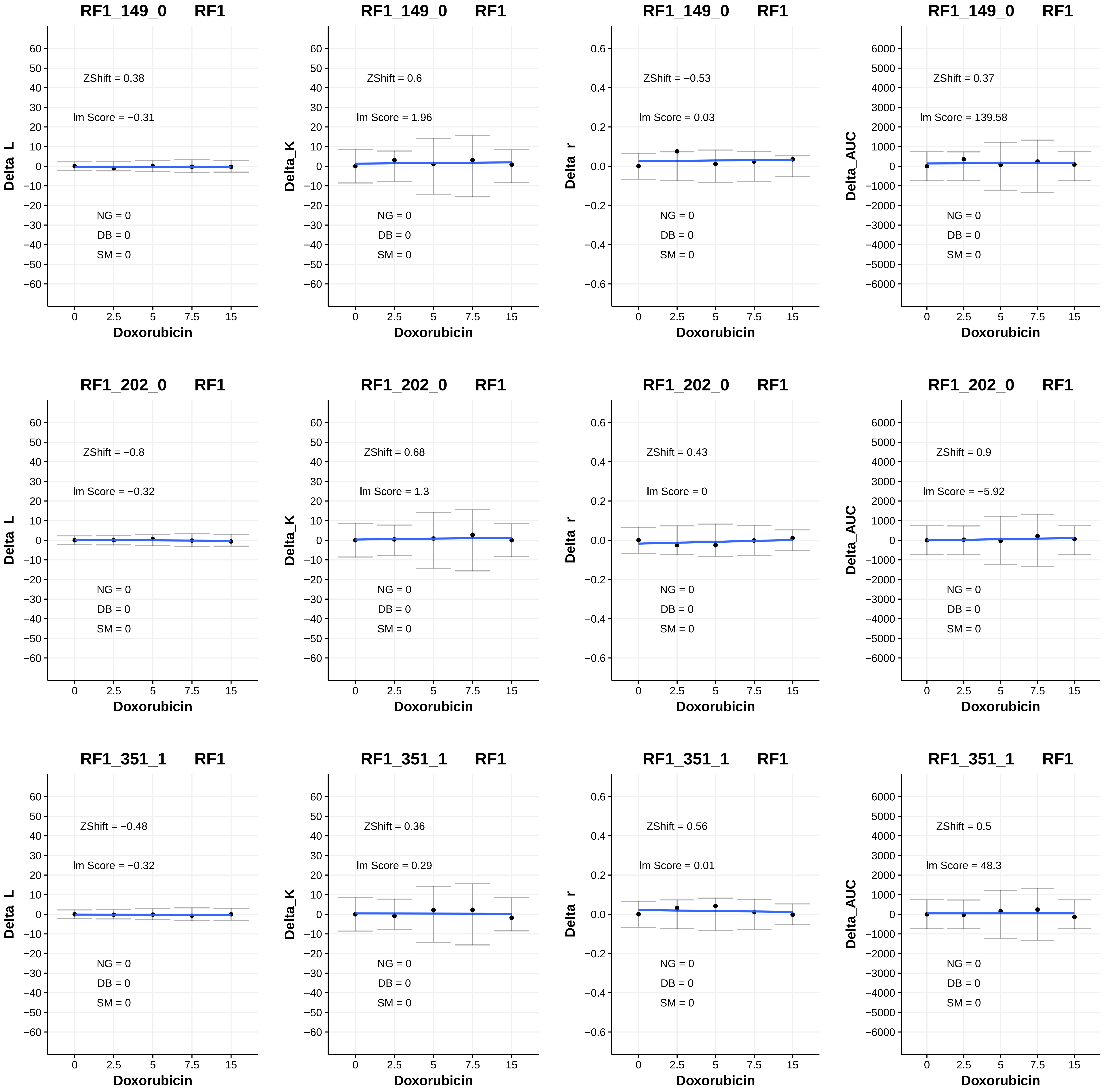
RF1_149_0 RF1
ZShift = 0.38
lm Score = −0.31
NG = 0
DB = 0
SM = 0
Delta_L
Doxorubicin
RF1_149_0 RF1
ZShift = 0.6
lm Score = 1.96
NG = 0
DB = 0
SM = 0
Delta_K
Doxorubicin
RF1_149_0 RF1
ZShift = −0.53
lm Score = 0.03
NG = 0
DB = 0
SM = 0
Delta_r
Doxorubicin
RF1_149_0 RF1
ZShift = 0.37
lm Score = 139.58
NG = 0
DB = 0
SM = 0
Delta_AUC
Doxorubicin
RF1_202_0 RF1
ZShift = −0.8
lm Score = −0.32
NG = 0
DB = 0
SM = 0
Delta_L
Doxorubicin
RF1_202_0 RF1
ZShift = 0.68
lm Score = 1.3
NG = 0
DB = 0
SM = 0
Delta_K
Doxorubicin
RF1_202_0 RF1
ZShift = 0.43
lm Score = 0
NG = 0
DB = 0
SM = 0
Delta_r
Doxorubicin
RF1_202_0 RF1
ZShift = 0.9
lm Score = −5.92
NG = 0
DB = 0
SM = 0
Delta_AUC
Doxorubicin
RF1_351_1 RF1
ZShift = −0.48
lm Score = −0.32
NG = 0
DB = 0
SM = 0
Delta_L
Doxorubicin
RF1_351_1 RF1
ZShift = 0.36
lm Score = 0.29
NG = 0
DB = 0
SM = 0
Delta_K
Doxorubicin
RF1_351_1 RF1
ZShift = 0.56
lm Score = 0.01
NG = 0
DB = 0
SM = 0
Delta_r
Doxorubicin
RF1_351_1 RF1
ZShift = 0.5
lm Score = 48.3
NG = 0
DB = 0
SM = 0
Delta_AUC
Doxorubicin

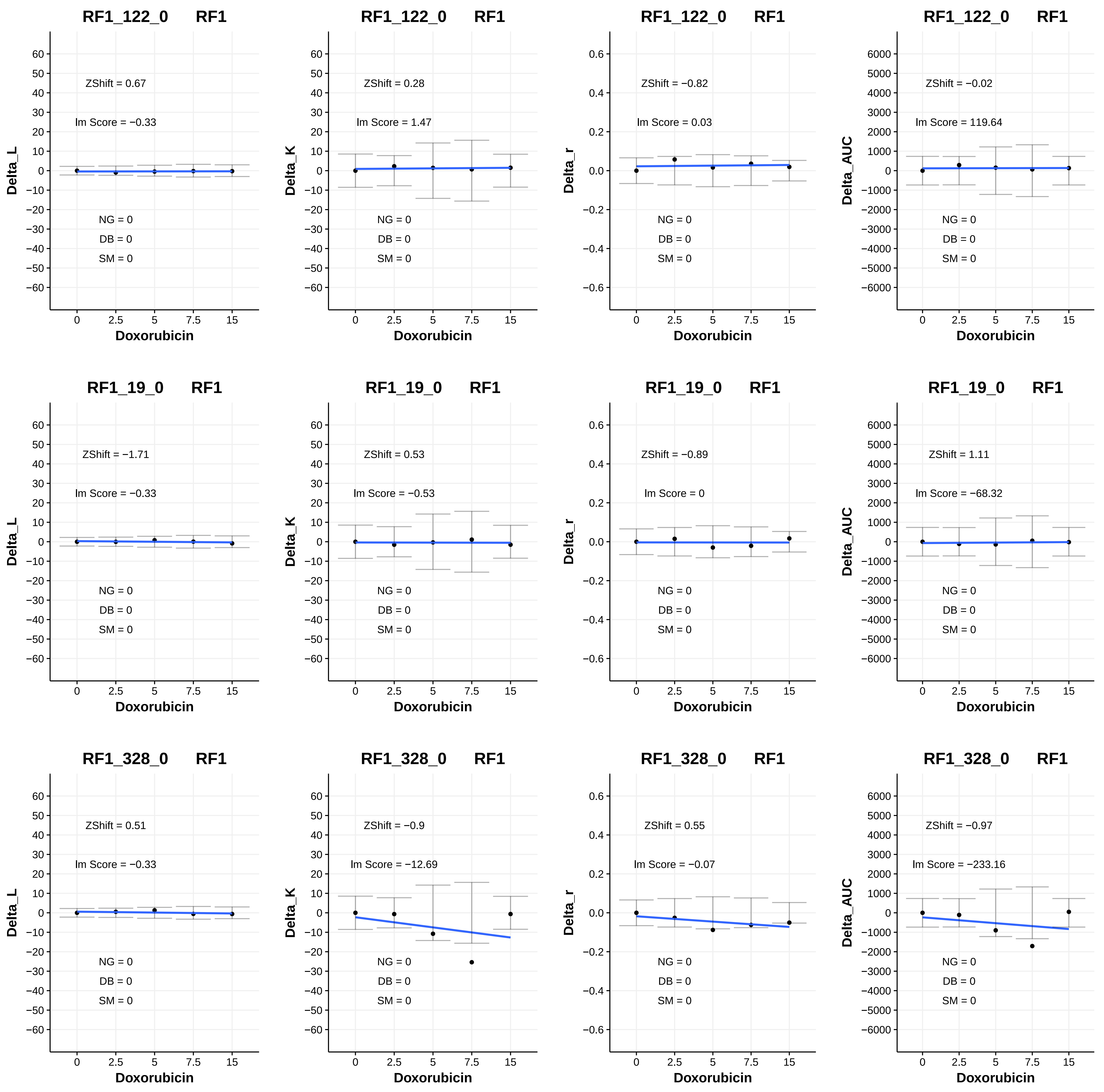
RF1_122_0 RF1
ZShift = 0.67
lm Score = −0.33
NG = 0
DB = 0
SM = 0
Delta_L
Doxorubicin
RF1_122_0 RF1
ZShift = 0.28
lm Score = 1.47
NG = 0
DB = 0
SM = 0
Delta_K
Doxorubicin
RF1_122_0 RF1
ZShift = −0.82
lm Score = 0.03
NG = 0
DB = 0
SM = 0
Delta_r
Doxorubicin
RF1_122_0 RF1
ZShift = −0.02
lm Score = 119.64
NG = 0
DB = 0
SM = 0
Delta_AUC
Doxorubicin
RF1_19_0 RF1
ZShift = −1.71
lm Score = −0.33
NG = 0
DB = 0
SM = 0
Delta_L
Doxorubicin
RF1_19_0 RF1
ZShift = 0.53
lm Score = −0.53
NG = 0
DB = 0
SM = 0
Delta_K
Doxorubicin
RF1_19_0 RF1
ZShift = −0.89
lm Score = 0
NG = 0
DB = 0
SM = 0
Delta_r
Doxorubicin
RF1_19_0 RF1
ZShift = 1.11
lm Score = −68.32
NG = 0
DB = 0
SM = 0
Delta_AUC
Doxorubicin
RF1_328_0 RF1
ZShift = 0.51
lm Score = −0.33
NG = 0
DB = 0
SM = 0
Delta_L
Doxorubicin
RF1_328_0 RF1
ZShift = −0.9
lm Score = −12.69
NG = 0
DB = 0
SM = 0
Delta_K
Doxorubicin
RF1_328_0 RF1
ZShift = 0.55
lm Score = −0.07
NG = 0
DB = 0
SM = 0
Delta_r
Doxorubicin
RF1_328_0 RF1
ZShift = −0.97
lm Score = −233.16
NG = 0
DB = 0
SM = 0
Delta_AUC
Doxorubicin

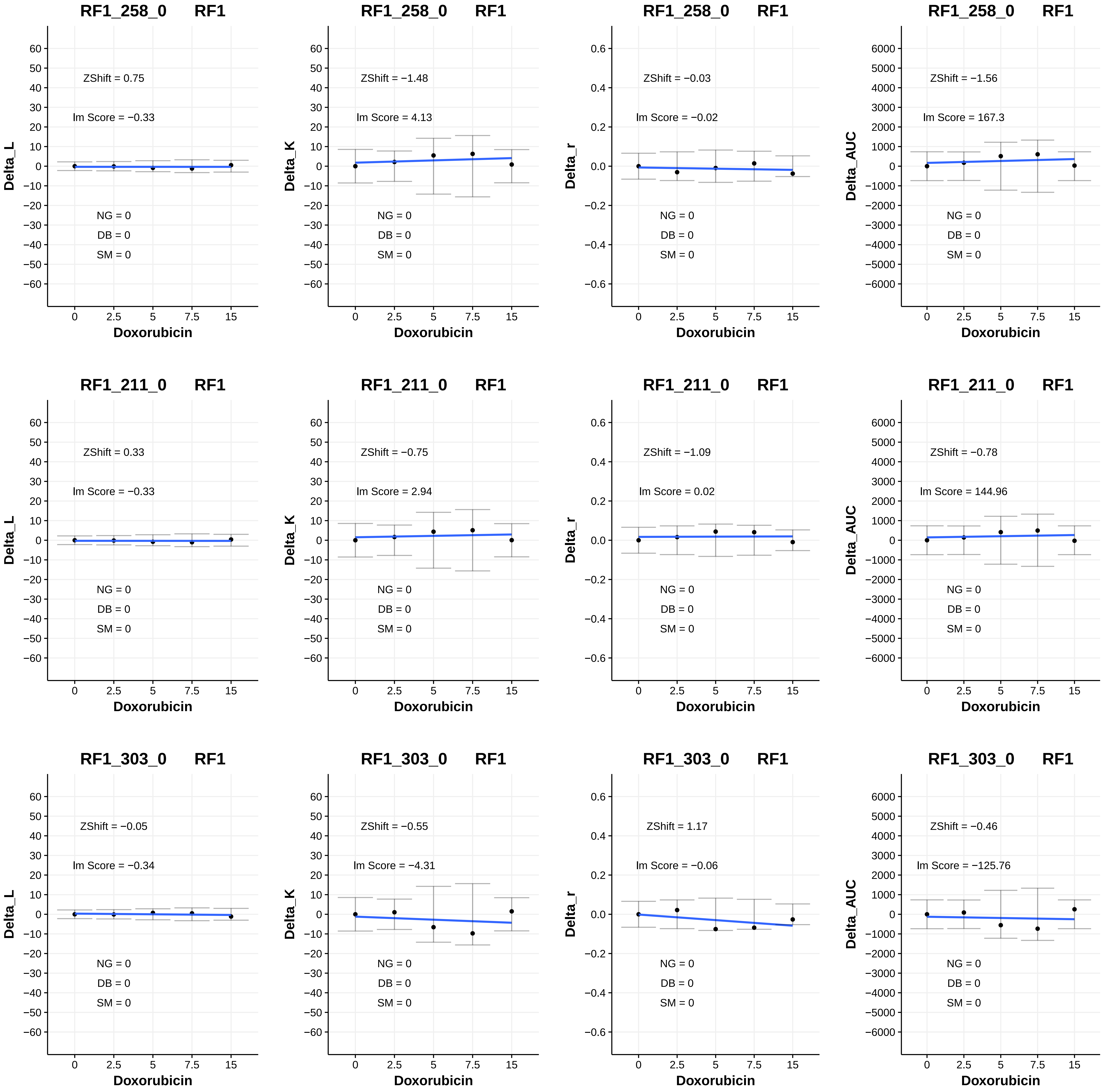

RF1_258_0 RF1
ZShift = 0.75
Im Score = −0.33
NG = 0
DB = 0
SM = 0
Delta_L
Doxorubicin
RF1_258_0 RF1
ZShift = −1.48
Im Score = 4.13
NG = 0
DB = 0
SM = 0
Delta_K
Doxorubicin
RF1_258_0 RF1
ZShift = −0.03
Im Score = −0.02
NG = 0
DB = 0
SM = 0
Delta_r
Doxorubicin
RF1_258_0 RF1
ZShift = −1.56
Im Score = 167.3
NG = 0
DB = 0
SM = 0
Delta_AUC
Doxorubicin
RF1_211_0 RF1
ZShift = 0.33
Im Score = −0.33
NG = 0
DB = 0
SM = 0
Delta_L
Doxorubicin
RF1_211_0 RF1
ZShift = −0.75
Im Score = 2.94
NG = 0
DB = 0
SM = 0
Delta_K
Doxorubicin
RF1_211_0 RF1
ZShift = −1.09
Im Score = 0.02
NG = 0
DB = 0
SM = 0
Delta_r
Doxorubicin
RF1_211_0 RF1
ZShift = −0.78
Im Score = 144.96
NG = 0
DB = 0
SM = 0
Delta_AUC
Doxorubicin
RF1_303_0 RF1
ZShift = −0.05
Im Score = −0.34
NG = 0
DB = 0
SM = 0
Delta_L
Doxorubicin
RF1_303_0 RF1
ZShift = −0.55
Im Score = −4.31
NG = 0
DB = 0
SM = 0
Delta_K
Doxorubicin
RF1_303_0 RF1
ZShift = 1.17
Im Score = −0.06
NG = 0
DB = 0
SM = 0
Delta_r
Doxorubicin
RF1_303_0 RF1
ZShift = −0.46
Im Score = −125.76
NG = 0
DB = 0
SM = 0
Delta_AUC
Doxorubicin

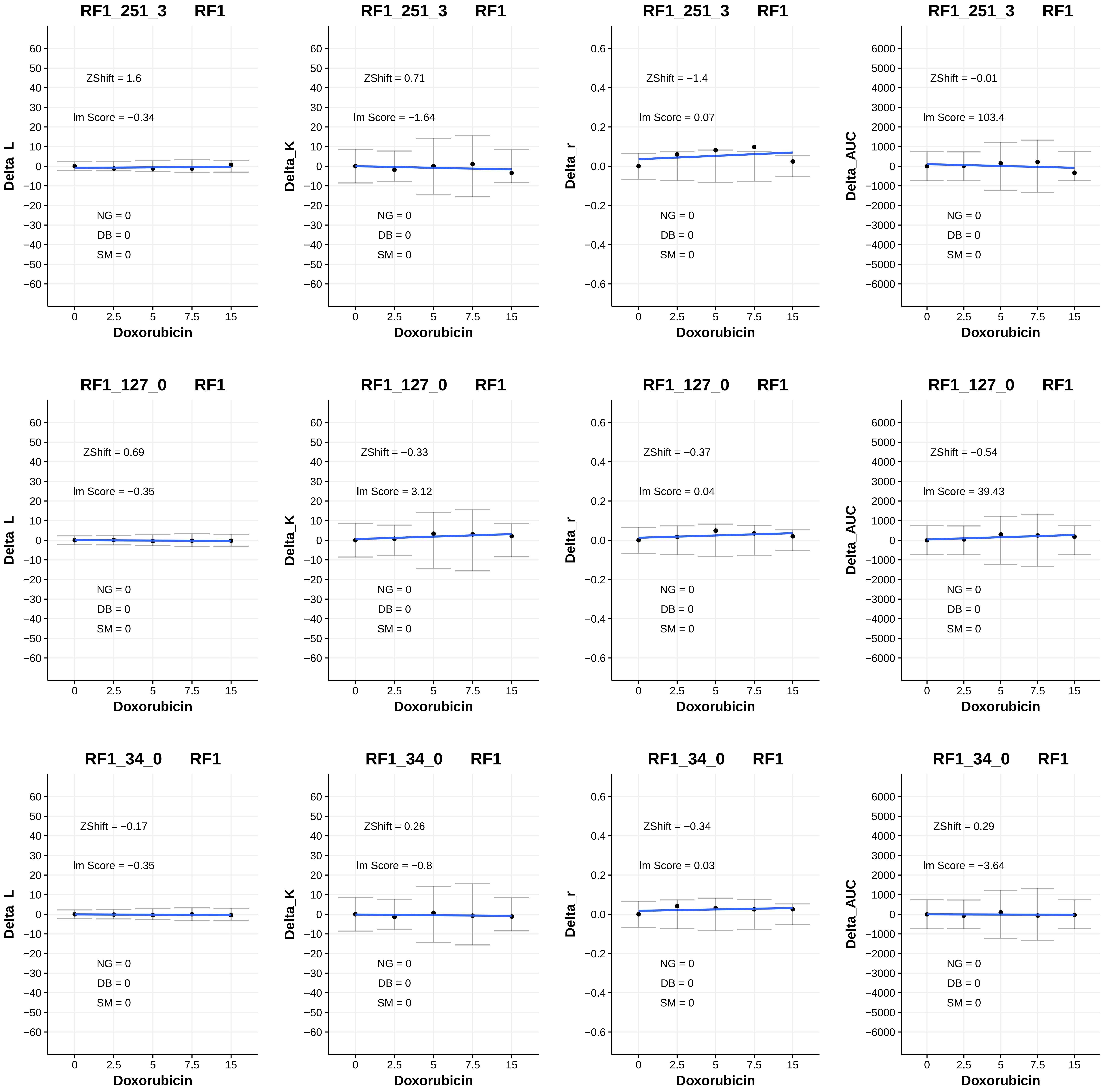
RF1_251_3 RF1
Delta_L
ZShift = 1.6
lm Score = −0.34
NG = 0
DB = 0
SM = 0
Doxorubicin
RF1_251_3 RF1
Delta_K
ZShift = 0.71
lm Score = −1.64
NG = 0
DB = 0
SM = 0
Doxorubicin
RF1_251_3 RF1
Delta_r
ZShift = −1.4
lm Score = 0.07
NG = 0
DB = 0
SM = 0
Doxorubicin
RF1_251_3 RF1
Delta_AUC
ZShift = −0.01
lm Score = 103.4
NG = 0
DB = 0
SM = 0
Doxorubicin
RF1_127_0 RF1
Delta_L
ZShift = 0.69
lm Score = −0.35
NG = 0
DB = 0
SM = 0
Doxorubicin
RF1_127_0 RF1
Delta_K
ZShift = −0.33
lm Score = 3.12
NG = 0
DB = 0
SM = 0
Doxorubicin
RF1_127_0 RF1
Delta_r
ZShift = −0.37
lm Score = 0.04
NG = 0
DB = 0
SM = 0
Doxorubicin
RF1_127_0 RF1
Delta_AUC
ZShift = −0.54
lm Score = 39.43
NG = 0
DB = 0
SM = 0
Doxorubicin
RF1_34_0 RF1
Delta_L
ZShift = −0.17
lm Score = −0.35
NG = 0
DB = 0
SM = 0
Doxorubicin
RF1_34_0 RF1
Delta_K
ZShift = 0.26
lm Score = −0.8
NG = 0
DB = 0
SM = 0
Doxorubicin
RF1_34_0 RF1
Delta_r
ZShift = −0.34
lm Score = 0.03
NG = 0
DB = 0
SM = 0
Doxorubicin
RF1_34_0 RF1
Delta_AUC
ZShift = 0.29
lm Score = −3.64
NG = 0
DB = 0
SM = 0
Doxorubicin

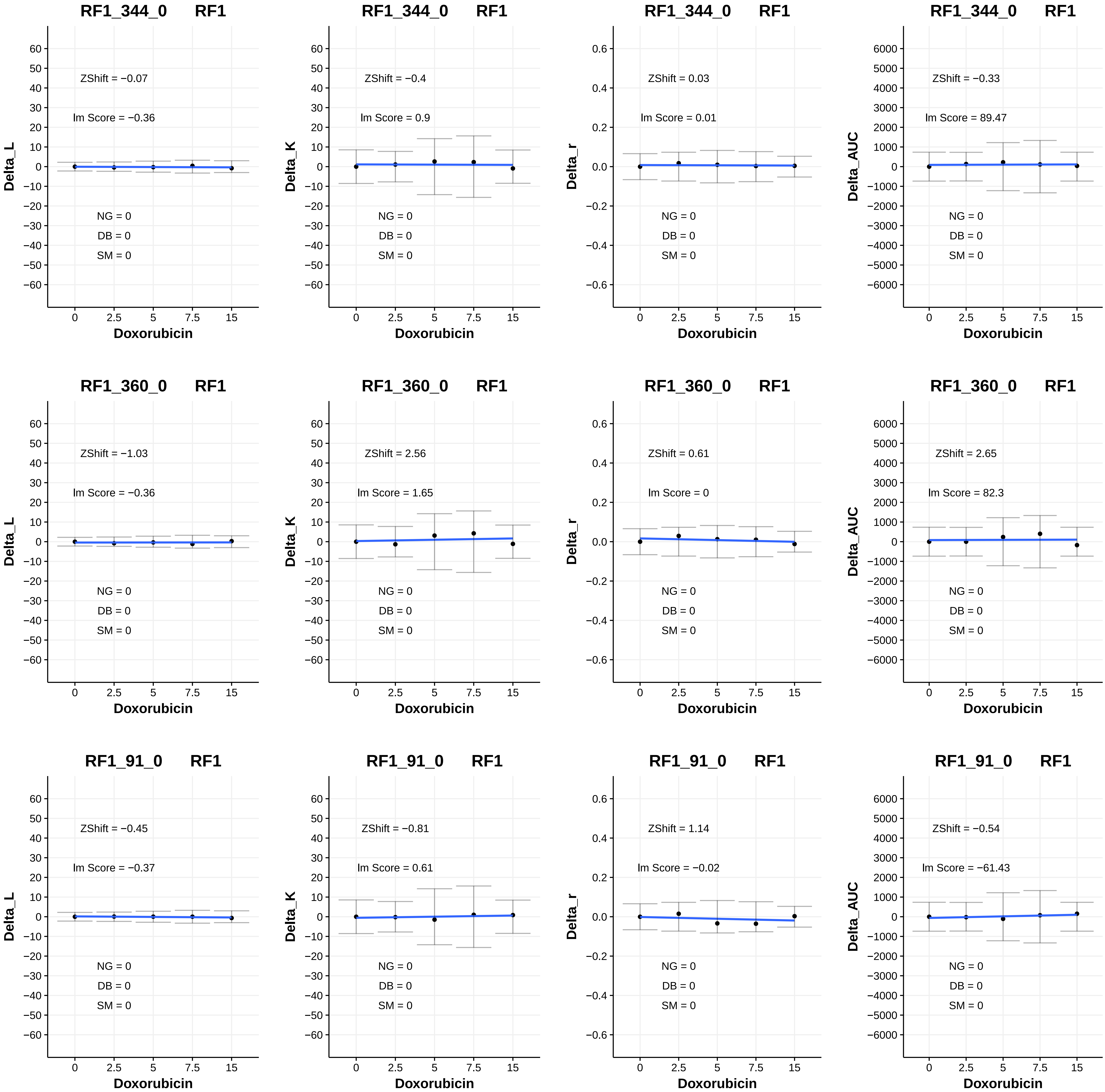
RF1_344_0 RF1
Delta_L
ZShift = −0.07
lm Score = −0.36
NG = 0
DB = 0
SM = 0
Doxorubicin
RF1_344_0 RF1
Delta_K
ZShift = −0.4
lm Score = 0.9
NG = 0
DB = 0
SM = 0
Doxorubicin
RF1_344_0 RF1
Delta_r
ZShift = 0.03
lm Score = 0.01
NG = 0
DB = 0
SM = 0
Doxorubicin
RF1_344_0 RF1
Delta_AUC
ZShift = −0.33
lm Score = 89.47
NG = 0
DB = 0
SM = 0
Doxorubicin
RF1_360_0 RF1
Delta_L
ZShift = −1.03
lm Score = −0.36
NG = 0
DB = 0
SM = 0
Doxorubicin
RF1_360_0 RF1
Delta_K
ZShift = 2.56
lm Score = 1.65
NG = 0
DB = 0
SM = 0
Doxorubicin
RF1_360_0 RF1
Delta_r
ZShift = 0.61
lm Score = 0
NG = 0
DB = 0
SM = 0
Doxorubicin
RF1_360_0 RF1
Delta_AUC
ZShift = 2.65
lm Score = 82.3
NG = 0
DB = 0
SM = 0
Doxorubicin
RF1_91_0 RF1
Delta_L
ZShift = −0.45
lm Score = −0.37
NG = 0
DB = 0
SM = 0
Doxorubicin
RF1_91_0 RF1
Delta_K
ZShift = −0.81
lm Score = 0.61
NG = 0
DB = 0
SM = 0
Doxorubicin
RF1_91_0 RF1
Delta_r
ZShift = 1.14
lm Score = −0.02
NG = 0
DB = 0
SM = 0
Doxorubicin
RF1_91_0 RF1
Delta_AUC
ZShift = −0.54
lm Score = −61.43
NG = 0
DB = 0
SM = 0
Doxorubicin

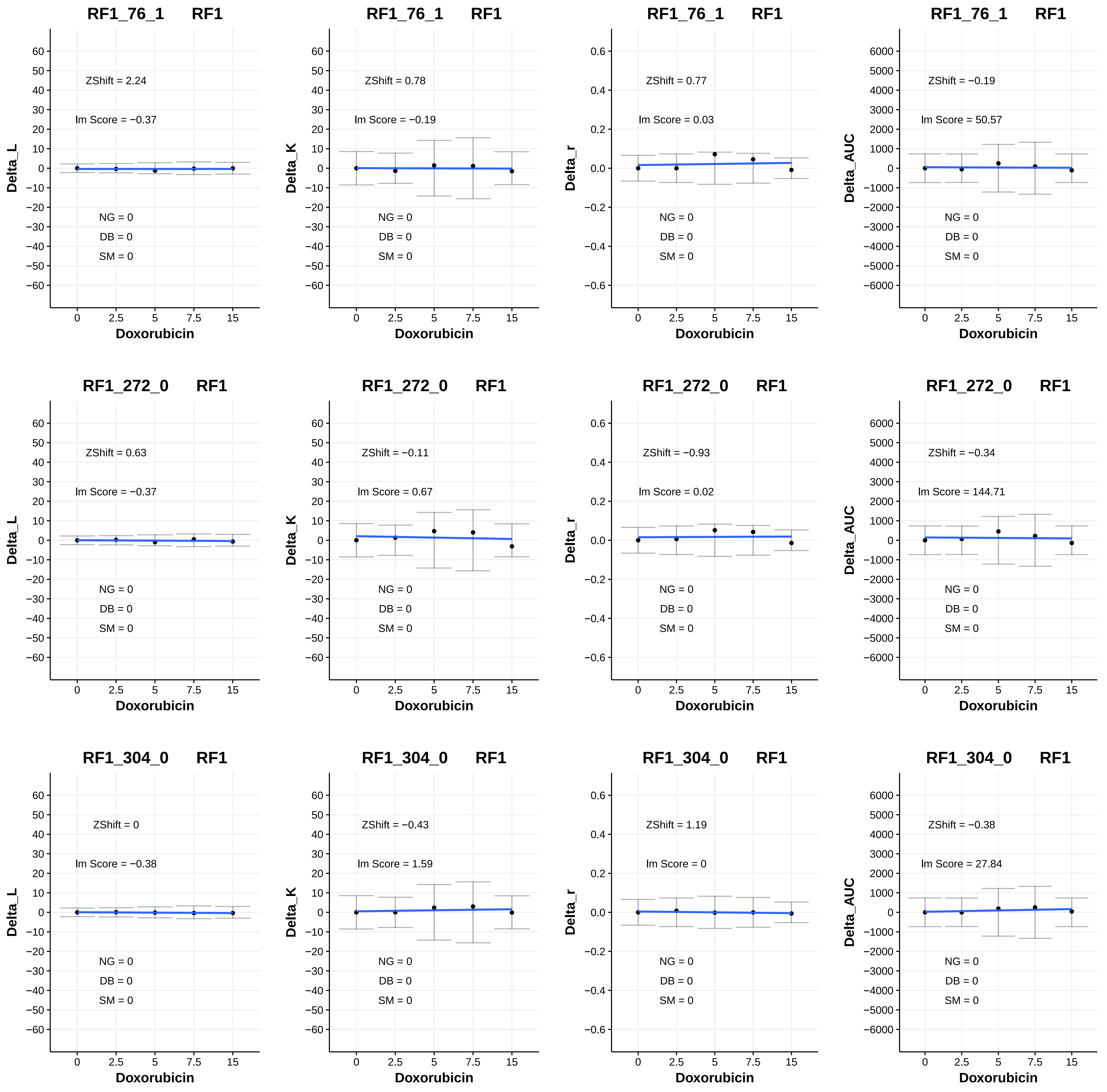
RF1_76_1 RF1
ZShift = 2.24
lm Score = −0.37
NG = 0
DB = 0
SM = 0
Delta_L
Doxorubicin
RF1_76_1 RF1
ZShift = 0.78
lm Score = −0.19
NG = 0
DB = 0
SM = 0
Delta_K
Doxorubicin
RF1_76_1 RF1
ZShift = 0.77
lm Score = 0.03
NG = 0
DB = 0
SM = 0
Delta_r
Doxorubicin
RF1_76_1 RF1
ZShift = −0.19
lm Score = 50.57
NG = 0
DB = 0
SM = 0
Delta_AUC
Doxorubicin
RF1_272_0 RF1
ZShift = 0.63
lm Score = −0.37
NG = 0
DB = 0
SM = 0
Delta_L
Doxorubicin
RF1_272_0 RF1
ZShift = −0.11
lm Score = 0.67
NG = 0
DB = 0
SM = 0
Delta_K
Doxorubicin
RF1_272_0 RF1
ZShift = −0.93
lm Score = 0.02
NG = 0
DB = 0
SM = 0
Delta_r
Doxorubicin
RF1_272_0 RF1
ZShift = −0.34
lm Score = 144.71
NG = 0
DB = 0
SM = 0
Delta_AUC
Doxorubicin
RF1_304_0 RF1
ZShift = 0
lm Score = −0.38
NG = 0
DB = 0
SM = 0
Delta_L
Doxorubicin
RF1_304_0 RF1
ZShift = −0.43
lm Score = 1.59
NG = 0
DB = 0
SM = 0
Delta_K
Doxorubicin
RF1_304_0 RF1
ZShift = 1.19
lm Score = 0
NG = 0
DB = 0
SM = 0
Delta_r
Doxorubicin
RF1_304_0 RF1
ZShift = −0.38
lm Score = 27.84
NG = 0
DB = 0
SM = 0
Delta_AUC
Doxorubicin
0 2.5 5 7.5 15

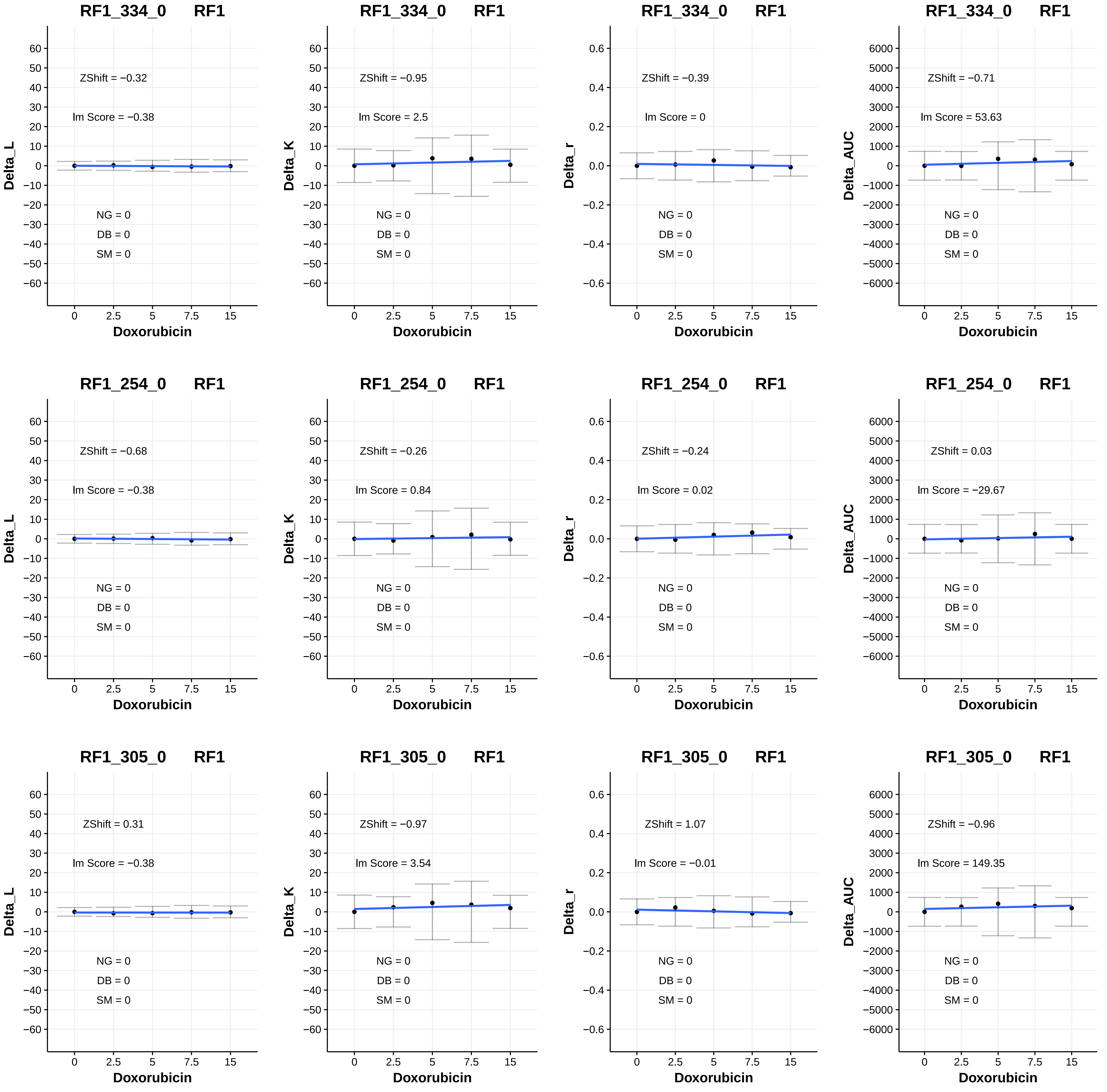
RF1_334_0 RF1
ZShift = −0.32
lm Score = −0.38
NG = 0
DB = 0
SM = 0
Delta_L
Doxorubicin
RF1_334_0 RF1
ZShift = −0.95
lm Score = 2.5
NG = 0
DB = 0
SM = 0
Delta_K
Doxorubicin
RF1_334_0 RF1
ZShift = −0.39
lm Score = 0
NG = 0
DB = 0
SM = 0
Delta_r
Doxorubicin
RF1_334_0 RF1
ZShift = −0.71
lm Score = 53.63
NG = 0
DB = 0
SM = 0
Delta_AUC
Doxorubicin
RF1_254_0 RF1
ZShift = −0.68
lm Score = −0.38
NG = 0
DB = 0
SM = 0
Delta_L
Doxorubicin
RF1_254_0 RF1
ZShift = −0.26
lm Score = 0.84
NG = 0
DB = 0
SM = 0
Delta_K
Doxorubicin
RF1_254_0 RF1
ZShift = −0.24
lm Score = 0.02
NG = 0
DB = 0
SM = 0
Delta_r
Doxorubicin
RF1_254_0 RF1
ZShift = 0.03
lm Score = −29.67
NG = 0
DB = 0
SM = 0
Delta_AUC
Doxorubicin
RF1_305_0 RF1
ZShift = 0.31
lm Score = −0.38
NG = 0
DB = 0
SM = 0
Delta_L
Doxorubicin
RF1_305_0 RF1
ZShift = −0.97
lm Score = 3.54
NG = 0
DB = 0
SM = 0
Delta_K
Doxorubicin
RF1_305_0 RF1
ZShift = 1.07
lm Score = −0.01
NG = 0
DB = 0
SM = 0
Delta_r
Doxorubicin
RF1_305_0 RF1
ZShift = −0.96
lm Score = 149.35
NG = 0
DB = 0
SM = 0
Delta_AUC
Doxorubicin

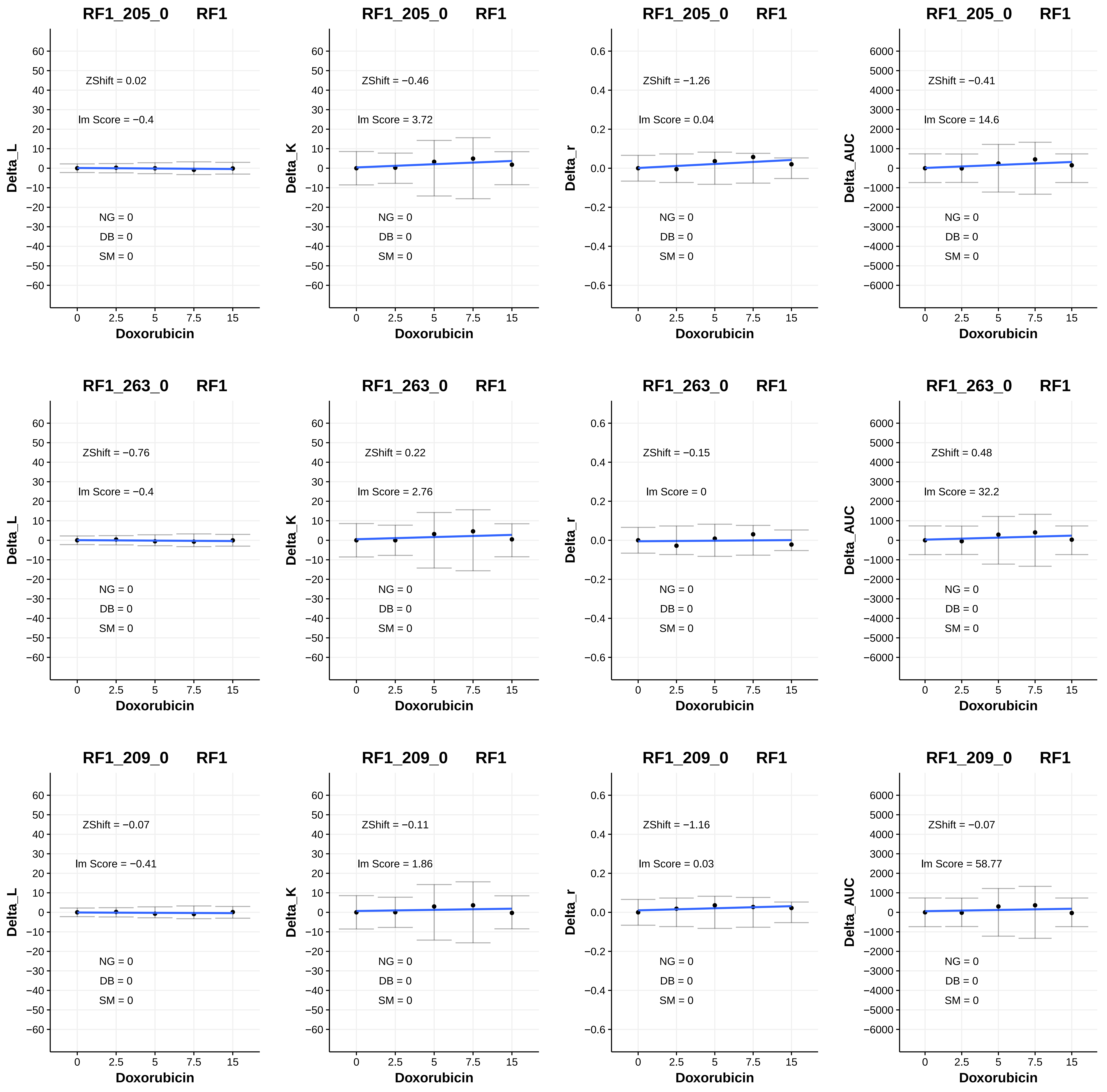
RF1_205_0 RF1
ZShift = 0.02
lm Score = −0.4
NG = 0
DB = 0
SM = 0
Delta_L
Doxorubicin
RF1_205_0 RF1
ZShift = −0.46
lm Score = 3.72
NG = 0
DB = 0
SM = 0
Delta_K
Doxorubicin
RF1_205_0 RF1
ZShift = −1.26
lm Score = 0.04
NG = 0
DB = 0
SM = 0
Delta_r
Doxorubicin
RF1_205_0 RF1
ZShift = −0.41
lm Score = 14.6
NG = 0
DB = 0
SM = 0
Delta_AUC
Doxorubicin
RF1_263_0 RF1
ZShift = −0.76
lm Score = −0.4
NG = 0
DB = 0
SM = 0
Delta_L
Doxorubicin
RF1_263_0 RF1
ZShift = 0.22
lm Score = 2.76
NG = 0
DB = 0
SM = 0
Delta_K
Doxorubicin
RF1_263_0 RF1
ZShift = −0.15
lm Score = 0
NG = 0
DB = 0
SM = 0
Delta_r
Doxorubicin
RF1_263_0 RF1
ZShift = 0.48
lm Score = 32.2
NG = 0
DB = 0
SM = 0
Delta_AUC
Doxorubicin
RF1_209_0 RF1
ZShift = −0.07
lm Score = −0.41
NG = 0
DB = 0
SM = 0
Delta_L
Doxorubicin
RF1_209_0 RF1
ZShift = −0.11
lm Score = 1.86
NG = 0
DB = 0
SM = 0
Delta_K
Doxorubicin
RF1_209_0 RF1
ZShift = −1.16
lm Score = 0.03
NG = 0
DB = 0
SM = 0
Delta_r
Doxorubicin
RF1_209_0 RF1
ZShift = −0.07
lm Score = 58.77
NG = 0
DB = 0
SM = 0
Delta_AUC
Doxorubicin

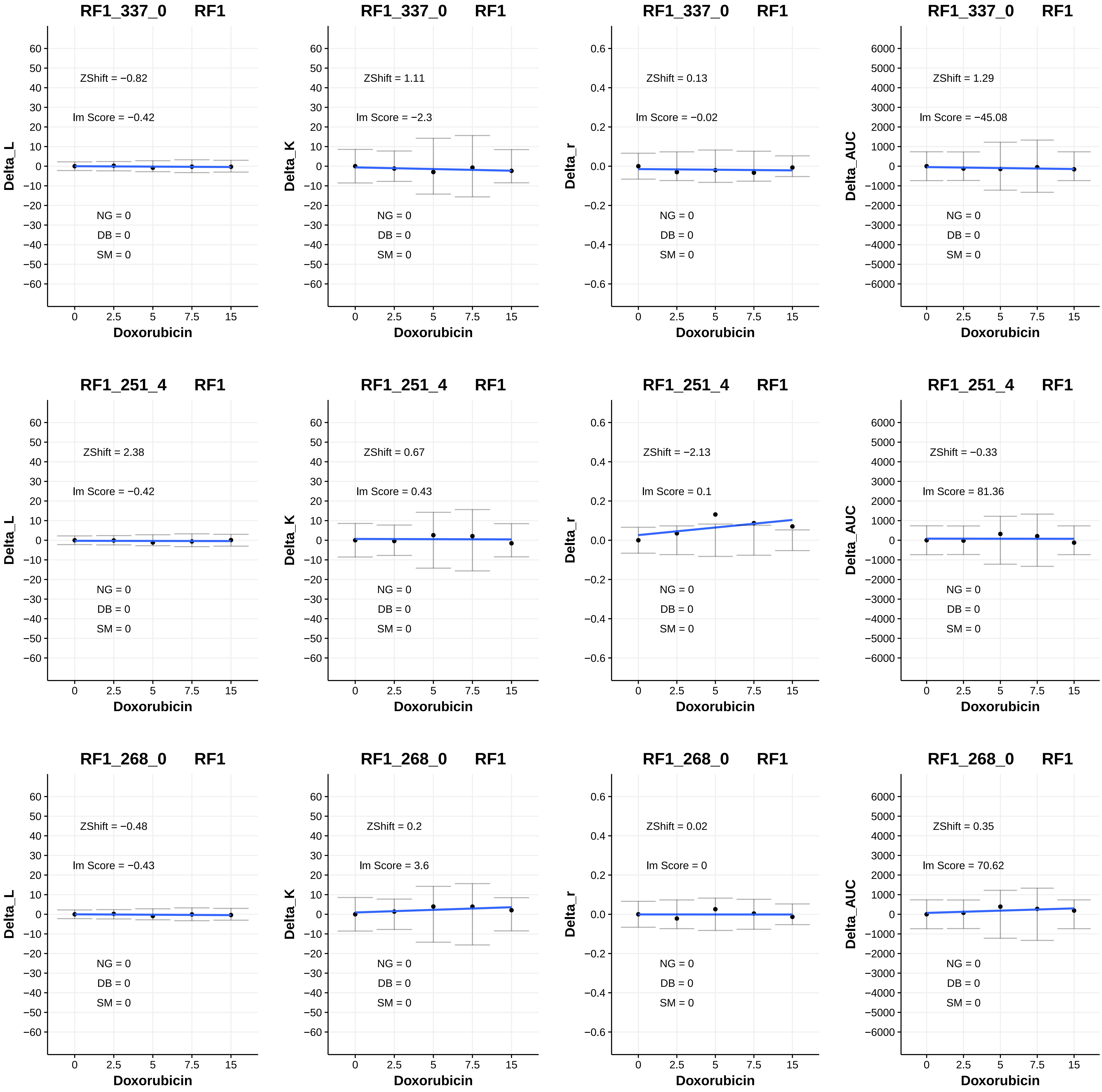
RF1_337_0 RF1
ZShift = −0.82
lm Score = −0.42
NG = 0
DB = 0
SM = 0
Delta_L
Doxorubicin
RF1_337_0 RF1
ZShift = 1.11
lm Score = −2.3
NG = 0
DB = 0
SM = 0
Delta_K
Doxorubicin
RF1_337_0 RF1
ZShift = 0.13
lm Score = −0.02
NG = 0
DB = 0
SM = 0
Delta_r
Doxorubicin
RF1_337_0 RF1
ZShift = 1.29
lm Score = −45.08
NG = 0
DB = 0
SM = 0
Delta_AUC
Doxorubicin
RF1_251_4 RF1
ZShift = 2.38
lm Score = −0.42
NG = 0
DB = 0
SM = 0
Delta_L
Doxorubicin
RF1_251_4 RF1
ZShift = 0.67
lm Score = 0.43
NG = 0
DB = 0
SM = 0
Delta_K
Doxorubicin
RF1_251_4 RF1
ZShift = −2.13
lm Score = 0.1
NG = 0
DB = 0
SM = 0
Delta_r
Doxorubicin
RF1_251_4 RF1
ZShift = −0.33
lm Score = 81.36
NG = 0
DB = 0
SM = 0
Delta_AUC
Doxorubicin
RF1_268_0 RF1
ZShift = −0.48
lm Score = −0.43
NG = 0
DB = 0
SM = 0
Delta_L
Doxorubicin
RF1_268_0 RF1
ZShift = 0.2
lm Score = 3.6
NG = 0
DB = 0
SM = 0
Delta_K
Doxorubicin
RF1_268_0 RF1
ZShift = 0.02
lm Score = 0
NG = 0
DB = 0
SM = 0
Delta_r
Doxorubicin
RF1_268_0 RF1
ZShift = 0.35
lm Score = 70.62
NG = 0
DB = 0
SM = 0
Delta_AUC
Doxorubicin

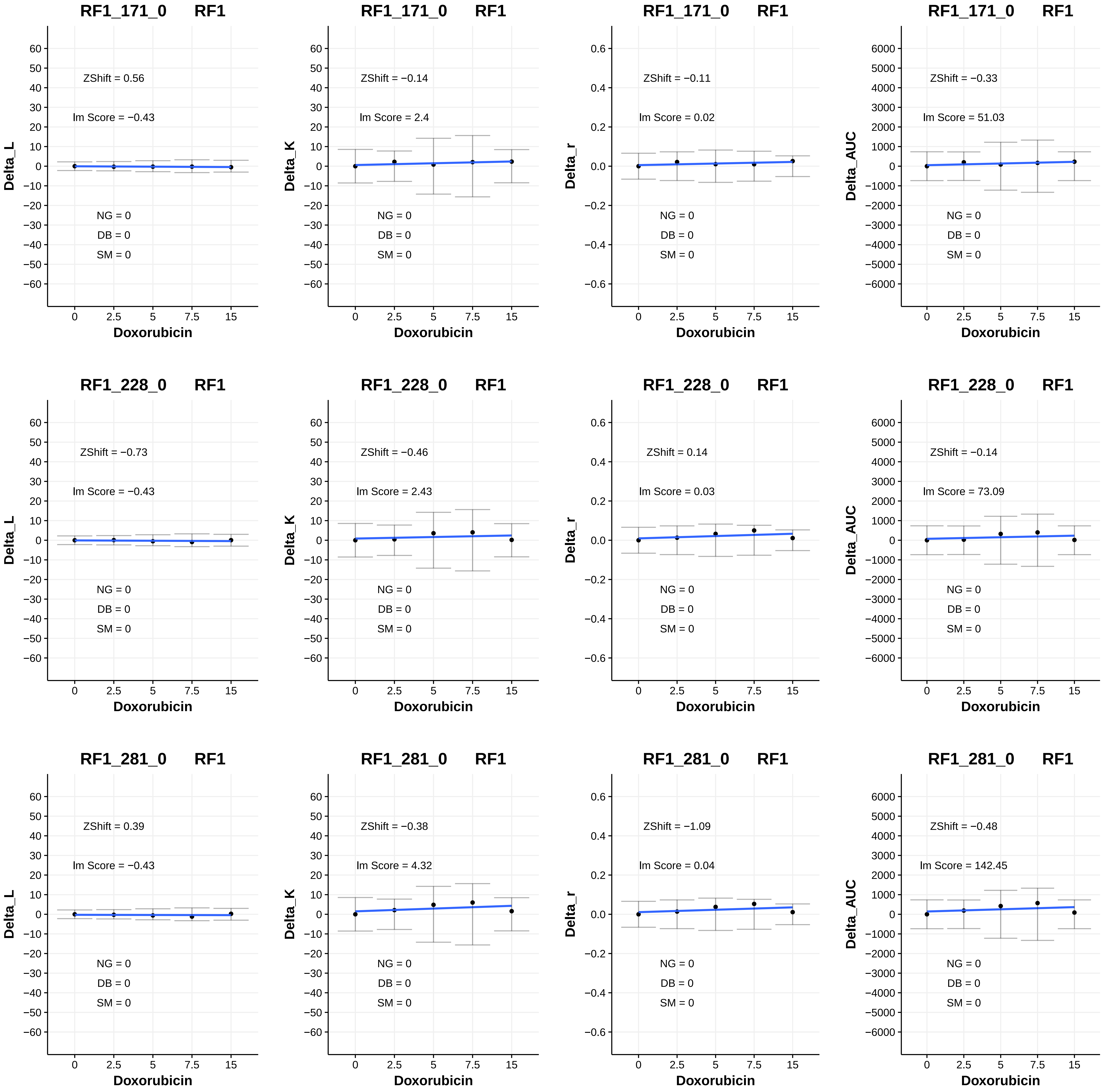
RF1_171_0 RF1
ZShift = 0.56
lm Score = −0.43
NG = 0
DB = 0
SM = 0
Delta_L
Doxorubicin
RF1_171_0 RF1
ZShift = −0.14
lm Score = 2.4
NG = 0
DB = 0
SM = 0
Delta_K
Doxorubicin
RF1_171_0 RF1
ZShift = −0.11
lm Score = 0.02
NG = 0
DB = 0
SM = 0
Delta_r
Doxorubicin
RF1_171_0 RF1
ZShift = −0.33
lm Score = 51.03
NG = 0
DB = 0
SM = 0
Delta_AUC
Doxorubicin
RF1_228_0 RF1
ZShift = −0.73
lm Score = −0.43
NG = 0
DB = 0
SM = 0
Delta_L
Doxorubicin
RF1_228_0 RF1
ZShift = −0.46
lm Score = 2.43
NG = 0
DB = 0
SM = 0
Delta_K
Doxorubicin
RF1_228_0 RF1
ZShift = 0.14
lm Score = 0.03
NG = 0
DB = 0
SM = 0
Delta_r
Doxorubicin
RF1_228_0 RF1
ZShift = −0.14
lm Score = 73.09
NG = 0
DB = 0
SM = 0
Delta_AUC
Doxorubicin
RF1_281_0 RF1
ZShift = 0.39
lm Score = −0.43
NG = 0
DB = 0
SM = 0
Delta_L
Doxorubicin
RF1_281_0 RF1
ZShift = −0.38
lm Score = 4.32
NG = 0
DB = 0
SM = 0
Delta_K
Doxorubicin
RF1_281_0 RF1
ZShift = −1.09
lm Score = 0.04
NG = 0
DB = 0
SM = 0
Delta_r
Doxorubicin
RF1_281_0 RF1
ZShift = −0.48
lm Score = 142.45
NG = 0
DB = 0
SM = 0
Delta_AUC
Doxorubicin

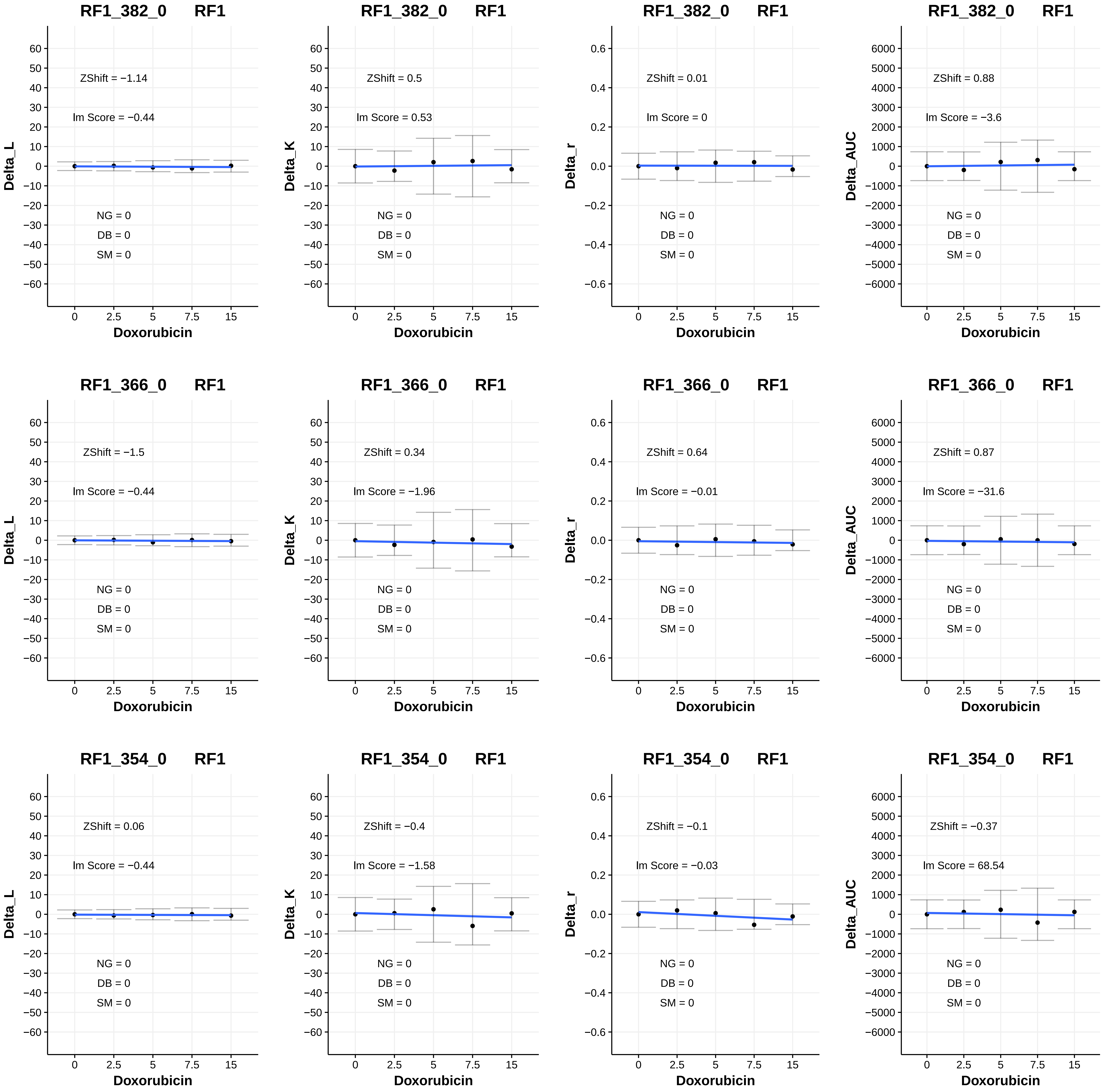
RF1_382_0 RF1
ZShift = −1.14
lm Score = −0.44
NG = 0
DB = 0
SM = 0
Delta_L
Doxorubicin
RF1_382_0 RF1
ZShift = 0.5
lm Score = 0.53
NG = 0
DB = 0
SM = 0
Delta_K
Doxorubicin
RF1_382_0 RF1
ZShift = 0.01
lm Score = 0
NG = 0
DB = 0
SM = 0
Delta_r
Doxorubicin
RF1_382_0 RF1
ZShift = 0.88
lm Score = −3.6
NG = 0
DB = 0
SM = 0
Delta_AUC
Doxorubicin
RF1_366_0 RF1
ZShift = −1.5
lm Score = −0.44
NG = 0
DB = 0
SM = 0
Delta_L
Doxorubicin
RF1_366_0 RF1
ZShift = 0.34
lm Score = −1.96
NG = 0
DB = 0
SM = 0
Delta_K
Doxorubicin
RF1_366_0 RF1
ZShift = 0.64
lm Score = −0.01
NG = 0
DB = 0
SM = 0
Delta_r
Doxorubicin
RF1_366_0 RF1
ZShift = 0.87
lm Score = −31.6
NG = 0
DB = 0
SM = 0
Delta_AUC
Doxorubicin
RF1_354_0 RF1
ZShift = 0.06
lm Score = −0.44
NG = 0
DB = 0
SM = 0
Delta_L
Doxorubicin
RF1_354_0 RF1
ZShift = −0.4
lm Score = −1.58
NG = 0
DB = 0
SM = 0
Delta_K
Doxorubicin
RF1_354_0 RF1
ZShift = −0.1
lm Score = −0.03
NG = 0
DB = 0
SM = 0
Delta_r
Doxorubicin
RF1_354_0 RF1
ZShift = −0.37
lm Score = 68.54
NG = 0
DB = 0
SM = 0
Delta_AUC
Doxorubicin

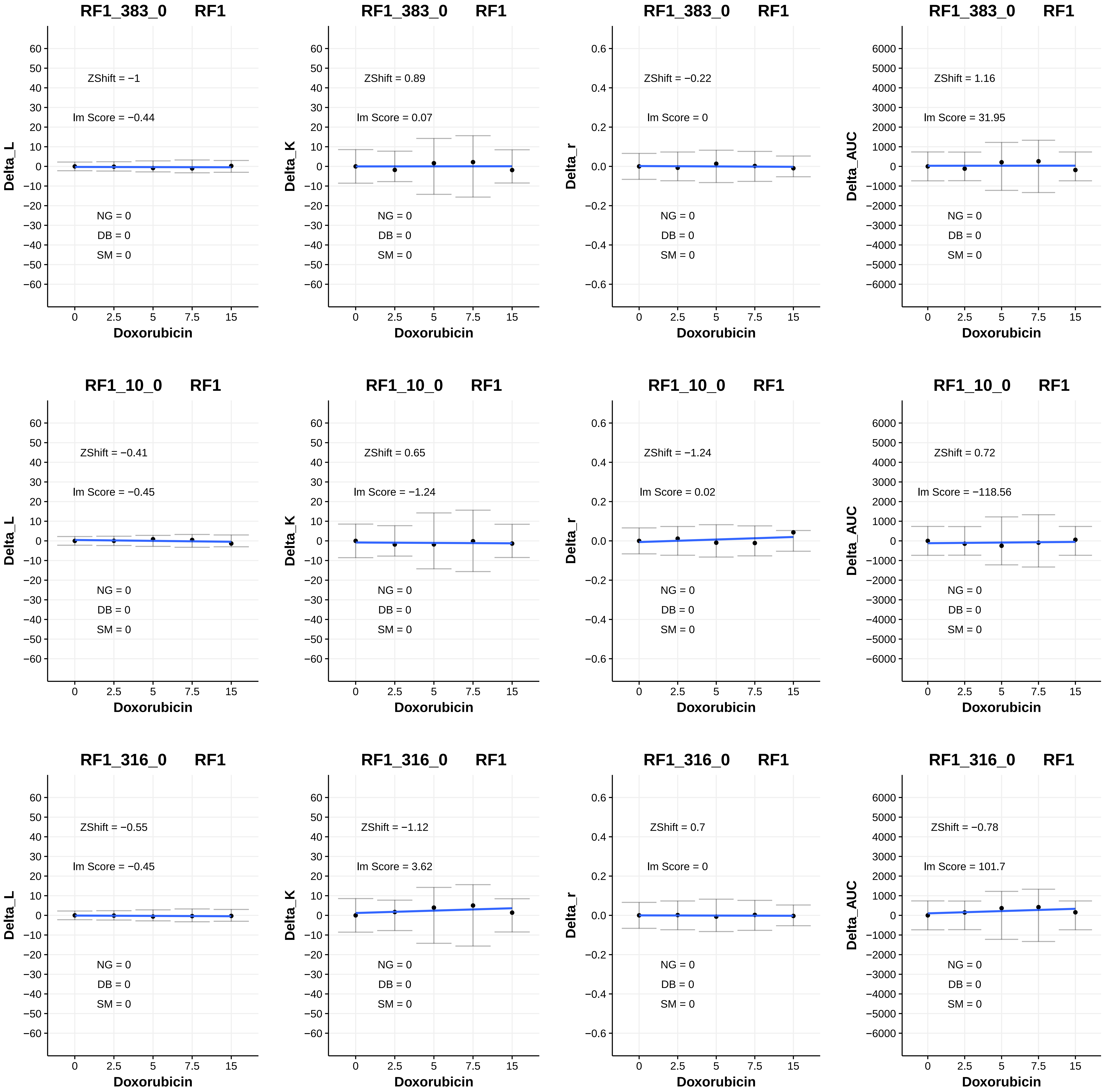

RF1_383_0 RF1
ZShift = −1
lm Score = −0.44
NG = 0
DB = 0
SM = 0
Delta_L
Doxorubicin
RF1_383_0 RF1
ZShift = 0.89
lm Score = 0.07
NG = 0
DB = 0
SM = 0
Delta_K
Doxorubicin
RF1_383_0 RF1
ZShift = −0.22
lm Score = 0
NG = 0
DB = 0
SM = 0
Delta_r
Doxorubicin
RF1_383_0 RF1
ZShift = 1.16
lm Score = 31.95
NG = 0
DB = 0
SM = 0
Delta_AUC
Doxorubicin
RF1_10_0 RF1
ZShift = −0.41
lm Score = −0.45
NG = 0
DB = 0
SM = 0
Delta_L
Doxorubicin
RF1_10_0 RF1
ZShift = 0.65
lm Score = −1.24
NG = 0
DB = 0
SM = 0
Delta_K
Doxorubicin
RF1_10_0 RF1
ZShift = −1.24
lm Score = 0.02
NG = 0
DB = 0
SM = 0
Delta_r
Doxorubicin
RF1_10_0 RF1
ZShift = 0.72
lm Score = −118.56
NG = 0
DB = 0
SM = 0
Delta_AUC
Doxorubicin
RF1_316_0 RF1
ZShift = −0.55
lm Score = −0.45
NG = 0
DB = 0
SM = 0
Delta_L
Doxorubicin
RF1_316_0 RF1
ZShift = −1.12
lm Score = 3.62
NG = 0
DB = 0
SM = 0
Delta_K
Doxorubicin
RF1_316_0 RF1
ZShift = 0.7
lm Score = 0
NG = 0
DB = 0
SM = 0
Delta_r
Doxorubicin
RF1_316_0 RF1
ZShift = −0.78
lm Score = 101.7
NG = 0
DB = 0
SM = 0
Delta_AUC
Doxorubicin

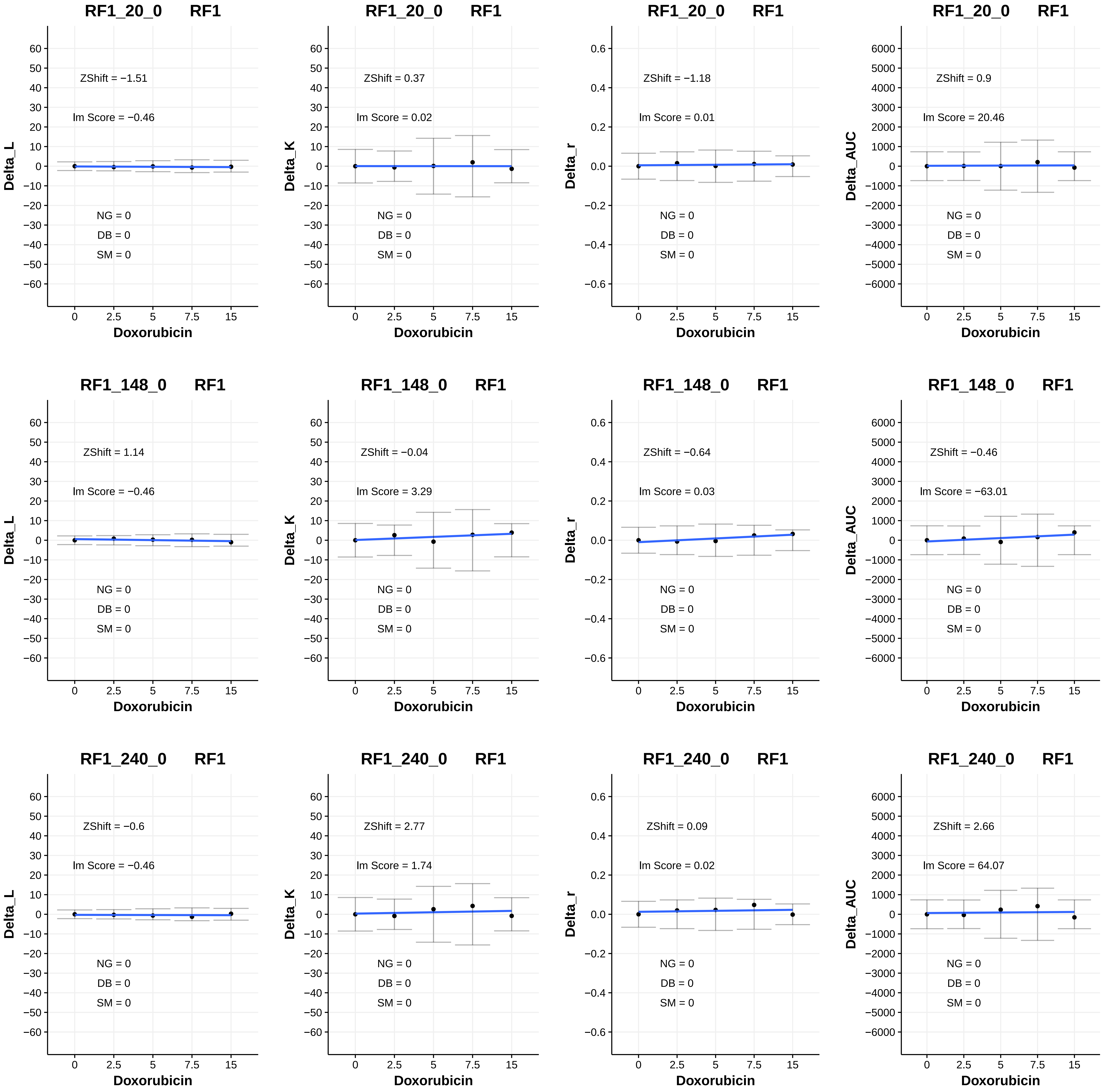
RF1_20_0 RF1
ZShift = −1.51
lm Score = −0.46
NG = 0
DB = 0
SM = 0
Delta_L
Doxorubicin
RF1_20_0 RF1
ZShift = 0.37
lm Score = 0.02
NG = 0
DB = 0
SM = 0
Delta_K
Doxorubicin
RF1_20_0 RF1
ZShift = −1.18
lm Score = 0.01
NG = 0
DB = 0
SM = 0
Delta_r
Doxorubicin
RF1_20_0 RF1
ZShift = 0.9
lm Score = 20.46
NG = 0
DB = 0
SM = 0
Delta_AUC
Doxorubicin
RF1_148_0 RF1
ZShift = 1.14
lm Score = −0.46
NG = 0
DB = 0
SM = 0
Delta_L
Doxorubicin
RF1_148_0 RF1
ZShift = −0.04
lm Score = 3.29
NG = 0
DB = 0
SM = 0
Delta_K
Doxorubicin
RF1_148_0 RF1
ZShift = −0.64
lm Score = 0.03
NG = 0
DB = 0
SM = 0
Delta_r
Doxorubicin
RF1_148_0 RF1
ZShift = −0.46
lm Score = −63.01
NG = 0
DB = 0
SM = 0
Delta_AUC
Doxorubicin
RF1_240_0 RF1
ZShift = −0.6
lm Score = −0.46
NG = 0
DB = 0
SM = 0
Delta_L
Doxorubicin
RF1_240_0 RF1
ZShift = 2.77
lm Score = 1.74
NG = 0
DB = 0
SM = 0
Delta_K
Doxorubicin
RF1_240_0 RF1
ZShift = 0.09
lm Score = 0.02
NG = 0
DB = 0
SM = 0
Delta_r
Doxorubicin
RF1_240_0 RF1
ZShift = 2.66
lm Score = 64.07
NG = 0
DB = 0
SM = 0
Delta_AUC
Doxorubicin

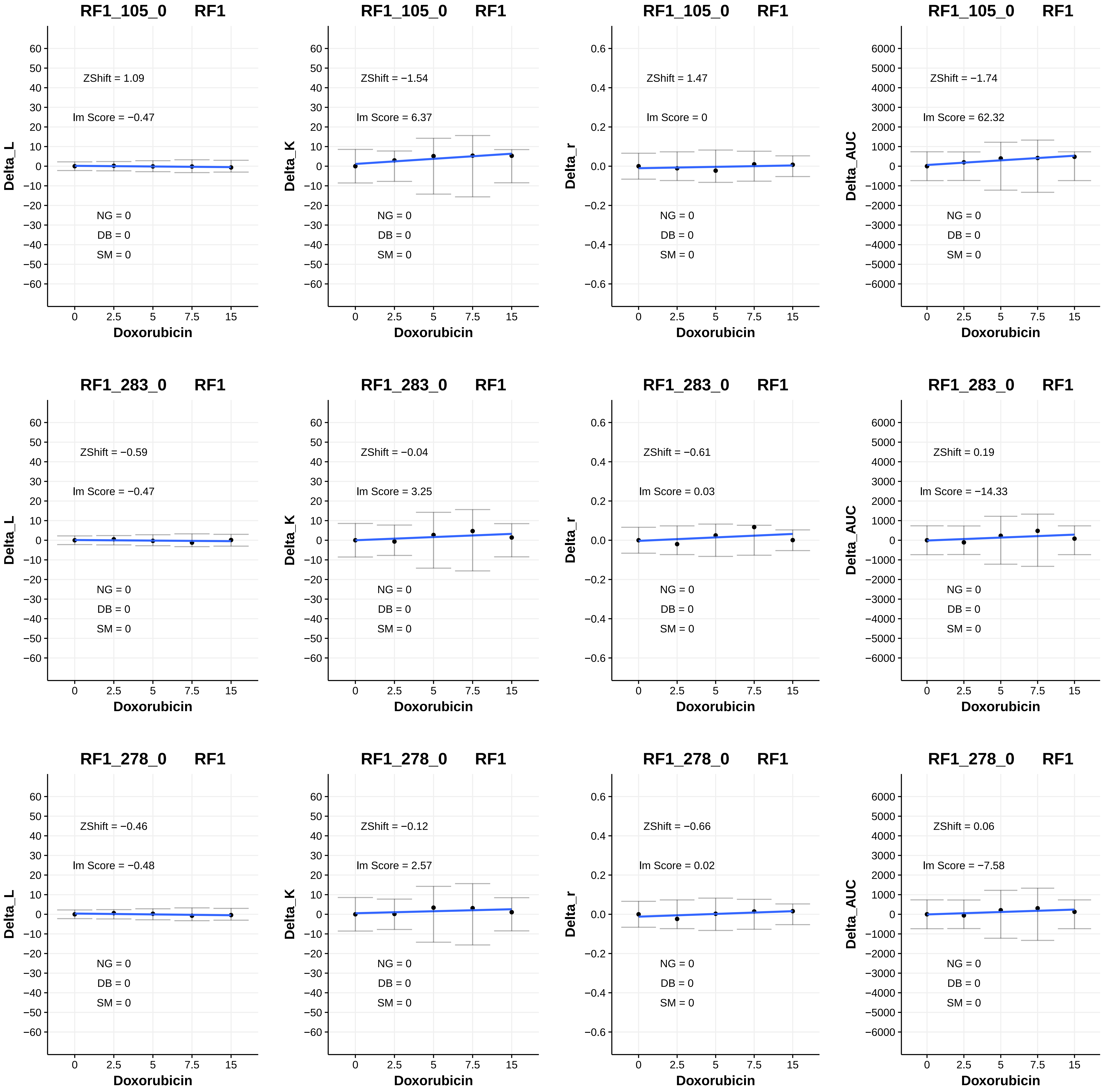

RF1_105_0 RF1
ZShift = 1.09
Im Score = −0.47
NG = 0
DB = 0
SM = 0
Delta_L
Doxorubicin
RF1_105_0 RF1
ZShift = −1.54
Im Score = 6.37
NG = 0
DB = 0
SM = 0
Delta_K
Doxorubicin
RF1_105_0 RF1
ZShift = 1.47
Im Score = 0
NG = 0
DB = 0
SM = 0
Delta_r
Doxorubicin
RF1_105_0 RF1
ZShift = −1.74
Im Score = 62.32
NG = 0
DB = 0
SM = 0
Delta_AUC
Doxorubicin
RF1_283_0 RF1
ZShift = −0.59
Im Score = −0.47
NG = 0
DB = 0
SM = 0
Delta_L
Doxorubicin
RF1_283_0 RF1
ZShift = −0.04
Im Score = 3.25
NG = 0
DB = 0
SM = 0
Delta_K
Doxorubicin
RF1_283_0 RF1
ZShift = −0.61
Im Score = 0.03
NG = 0
DB = 0
SM = 0
Delta_r
Doxorubicin
RF1_283_0 RF1
ZShift = 0.19
Im Score = −14.33
NG = 0
DB = 0
SM = 0
Delta_AUC
Doxorubicin
RF1_278_0 RF1
ZShift = −0.46
Im Score = −0.48
NG = 0
DB = 0
SM = 0
Delta_L
Doxorubicin
RF1_278_0 RF1
ZShift = −0.12
Im Score = 2.57
NG = 0
DB = 0
SM = 0
Delta_K
Doxorubicin
RF1_278_0 RF1
ZShift = −0.66
Im Score = 0.02
NG = 0
DB = 0
SM = 0
Delta_r
Doxorubicin
RF1_278_0 RF1
ZShift = 0.06
Im Score = −7.58
NG = 0
DB = 0
SM = 0
Delta_AUC
Doxorubicin

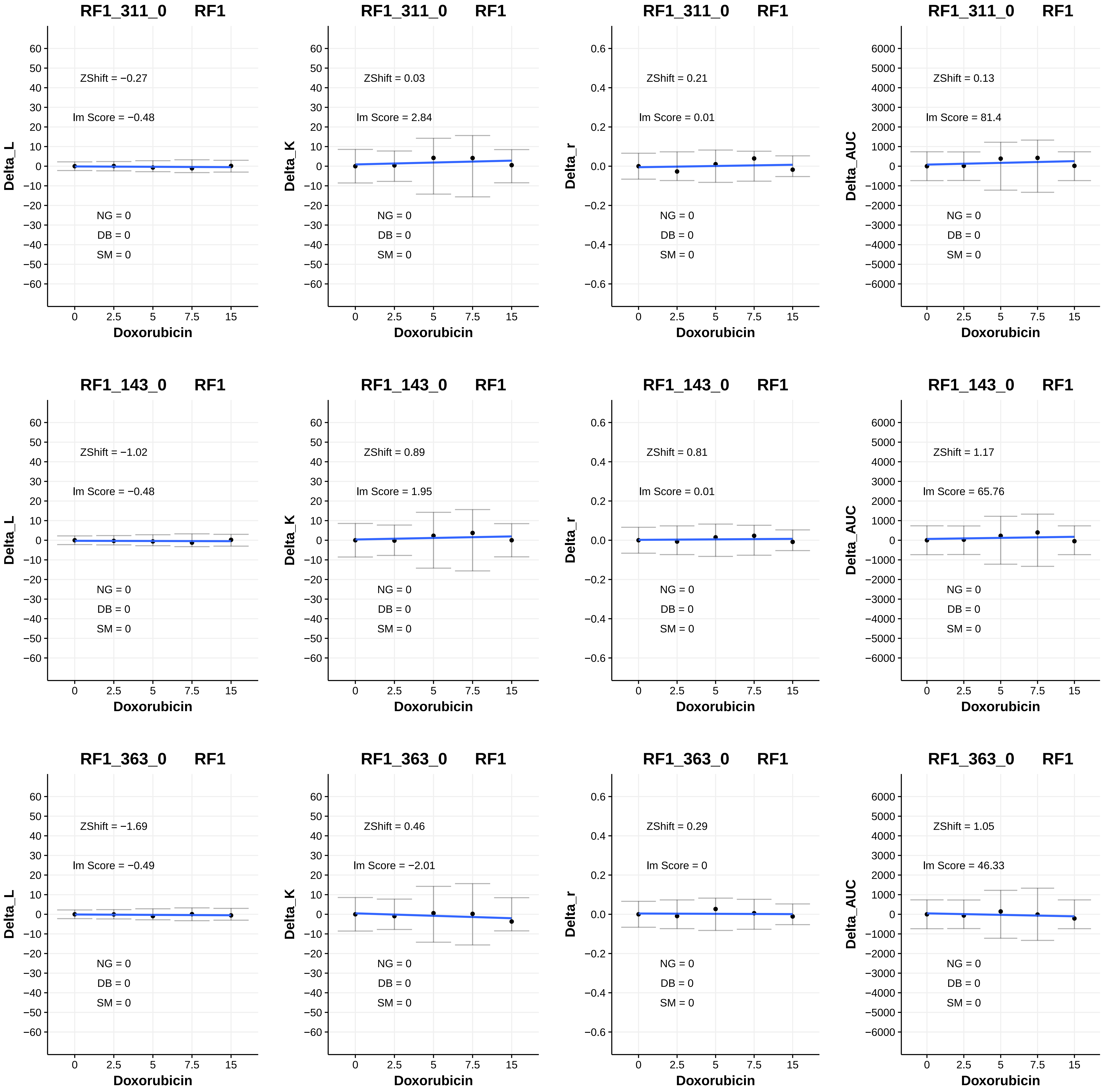
RF1_311_0 RF1
ZShift = −0.27
Im Score = −0.48
NG = 0
DB = 0
SM = 0
Delta_L
Doxorubicin
RF1_311_0 RF1
ZShift = 0.03
Im Score = 2.84
NG = 0
DB = 0
SM = 0
Delta_K
Doxorubicin
RF1_311_0 RF1
ZShift = 0.21
Im Score = 0.01
NG = 0
DB = 0
SM = 0
Delta_r
Doxorubicin
RF1_311_0 RF1
ZShift = 0.13
Im Score = 81.4
NG = 0
DB = 0
SM = 0
Delta_AUC
Doxorubicin
RF1_143_0 RF1
ZShift = −1.02
Im Score = −0.48
NG = 0
DB = 0
SM = 0
Delta_L
Doxorubicin
RF1_143_0 RF1
ZShift = 0.89
Im Score = 1.95
NG = 0
DB = 0
SM = 0
Delta_K
Doxorubicin
RF1_143_0 RF1
ZShift = 0.81
Im Score = 0.01
NG = 0
DB = 0
SM = 0
Delta_r
Doxorubicin
RF1_143_0 RF1
ZShift = 1.17
Im Score = 65.76
NG = 0
DB = 0
SM = 0
Delta_AUC
Doxorubicin
RF1_363_0 RF1
ZShift = −1.69
Im Score = −0.49
NG = 0
DB = 0
SM = 0
Delta_L
Doxorubicin
RF1_363_0 RF1
ZShift = 0.46
Im Score = −2.01
NG = 0
DB = 0
SM = 0
Delta_K
Doxorubicin
RF1_363_0 RF1
ZShift = 0.29
Im Score = 0
NG = 0
DB = 0
SM = 0
Delta_r
Doxorubicin
RF1_363_0 RF1
ZShift = 1.05
Im Score = 46.33
NG = 0
DB = 0
SM = 0
Delta_AUC
Doxorubicin

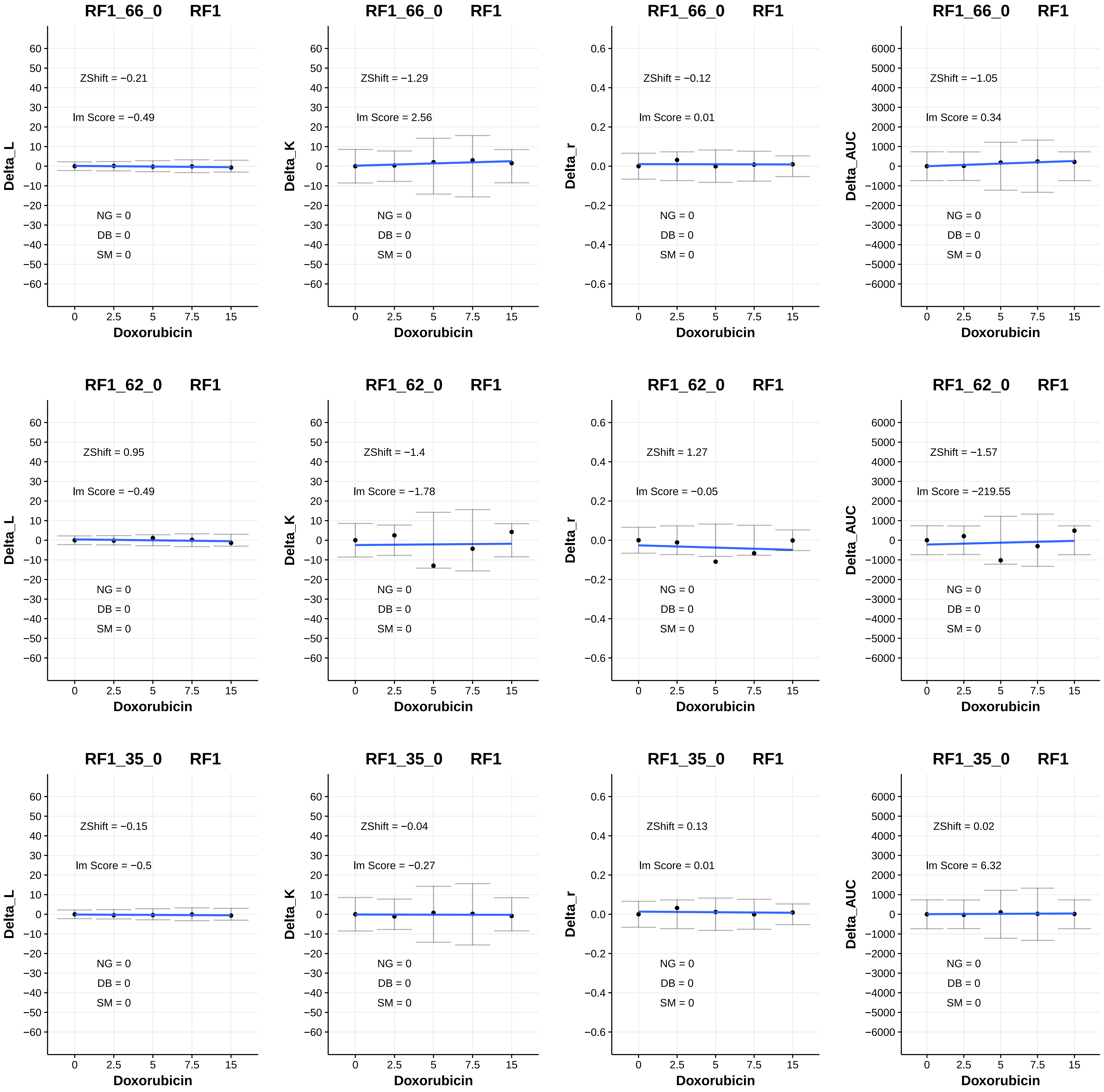
RF1_66_0 RF1
ZShift = −0.21
lm Score = −0.49
NG = 0
DB = 0
SM = 0
Delta_L
Doxorubicin
RF1_66_0 RF1
ZShift = −1.29
lm Score = 2.56
NG = 0
DB = 0
SM = 0
Delta_K
Doxorubicin
RF1_66_0 RF1
ZShift = −0.12
lm Score = 0.01
NG = 0
DB = 0
SM = 0
Delta_r
Doxorubicin
RF1_66_0 RF1
ZShift = −1.05
lm Score = 0.34
NG = 0
DB = 0
SM = 0
Delta_AUC
Doxorubicin
RF1_62_0 RF1
ZShift = 0.95
lm Score = −0.49
NG = 0
DB = 0
SM = 0
Delta_L
Doxorubicin
RF1_62_0 RF1
ZShift = −1.4
lm Score = −1.78
NG = 0
DB = 0
SM = 0
Delta_K
Doxorubicin
RF1_62_0 RF1
ZShift = 1.27
lm Score = −0.05
NG = 0
DB = 0
SM = 0
Delta_r
Doxorubicin
RF1_62_0 RF1
ZShift = −1.57
lm Score = −219.55
NG = 0
DB = 0
SM = 0
Delta_AUC
Doxorubicin
RF1_35_0 RF1
ZShift = −0.15
lm Score = −0.5
NG = 0
DB = 0
SM = 0
Delta_L
Doxorubicin
RF1_35_0 RF1
ZShift = −0.04
lm Score = −0.27
NG = 0
DB = 0
SM = 0
Delta_K
Doxorubicin
RF1_35_0 RF1
ZShift = 0.13
lm Score = 0.01
NG = 0
DB = 0
SM = 0
Delta_r
Doxorubicin
RF1_35_0 RF1
ZShift = 0.02
lm Score = 6.32
NG = 0
DB = 0
SM = 0
Delta_AUC
Doxorubicin

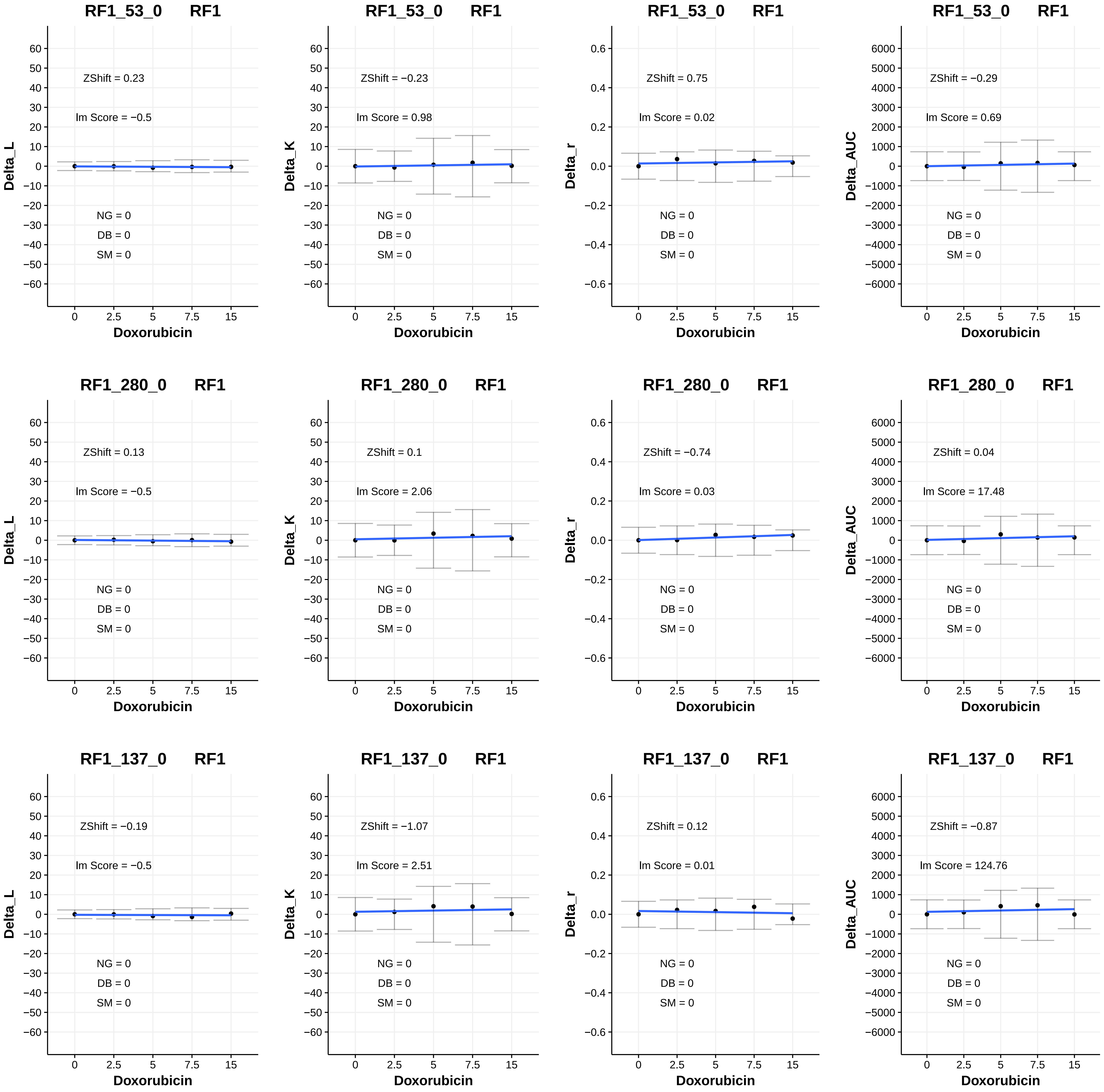
RF1_53_0 RF1
ZShift = 0.23
lm Score = −0.5
NG = 0
DB = 0
SM = 0
Delta_L
Doxorubicin
RF1_53_0 RF1
ZShift = −0.23
lm Score = 0.98
NG = 0
DB = 0
SM = 0
Delta_K
Doxorubicin
RF1_53_0 RF1
ZShift = 0.75
lm Score = 0.02
NG = 0
DB = 0
SM = 0
Delta_r
Doxorubicin
RF1_53_0 RF1
ZShift = −0.29
lm Score = 0.69
NG = 0
DB = 0
SM = 0
Delta_AUC
Doxorubicin
RF1_280_0 RF1
ZShift = 0.13
lm Score = −0.5
NG = 0
DB = 0
SM = 0
Delta_L
Doxorubicin
RF1_280_0 RF1
ZShift = 0.1
lm Score = 2.06
NG = 0
DB = 0
SM = 0
Delta_K
Doxorubicin
RF1_280_0 RF1
ZShift = −0.74
lm Score = 0.03
NG = 0
DB = 0
SM = 0
Delta_r
Doxorubicin
RF1_280_0 RF1
ZShift = 0.04
lm Score = 17.48
NG = 0
DB = 0
SM = 0
Delta_AUC
Doxorubicin
RF1_137_0 RF1
ZShift = −0.19
lm Score = −0.5
NG = 0
DB = 0
SM = 0
Delta_L
Doxorubicin
RF1_137_0 RF1
ZShift = −1.07
lm Score = 2.51
NG = 0
DB = 0
SM = 0
Delta_K
Doxorubicin
RF1_137_0 RF1
ZShift = 0.12
lm Score = 0.01
NG = 0
DB = 0
SM = 0
Delta_r
Doxorubicin
RF1_137_0 RF1
ZShift = −0.87
lm Score = 124.76
NG = 0
DB = 0
SM = 0
Delta_AUC
Doxorubicin

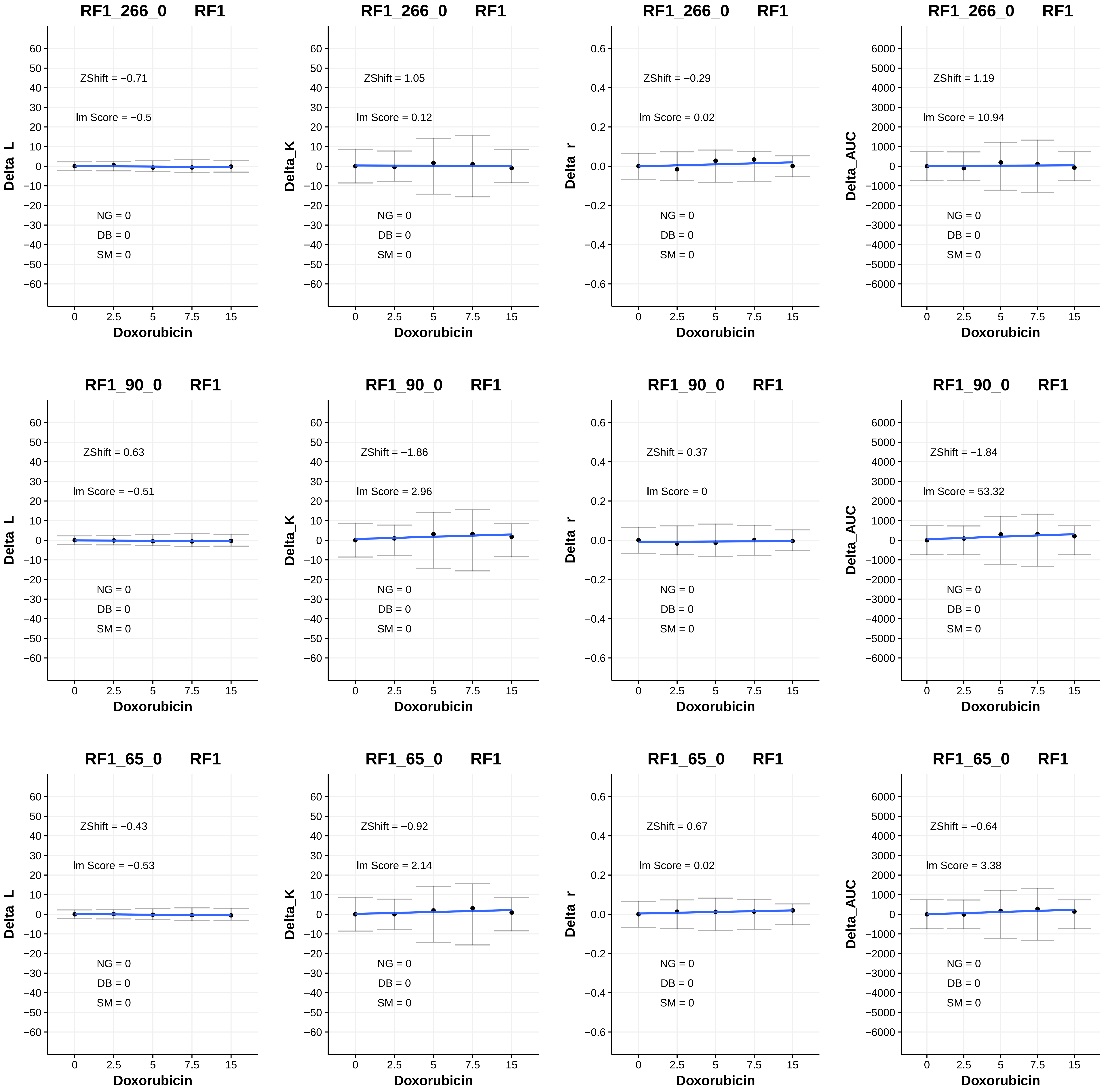

RF1_266_0 RF1
ZShift = −0.71
lm Score = −0.5
NG = 0
DB = 0
SM = 0
Delta_L
Doxorubicin
RF1_266_0 RF1
ZShift = 1.05
lm Score = 0.12
NG = 0
DB = 0
SM = 0
Delta_K
Doxorubicin
RF1_266_0 RF1
ZShift = −0.29
lm Score = 0.02
NG = 0
DB = 0
SM = 0
Delta_r
Doxorubicin
RF1_266_0 RF1
ZShift = 1.19
lm Score = 10.94
NG = 0
DB = 0
SM = 0
Delta_AUC
Doxorubicin
RF1_90_0 RF1
ZShift = 0.63
lm Score = −0.51
NG = 0
DB = 0
SM = 0
Delta_L
Doxorubicin
RF1_90_0 RF1
ZShift = −1.86
lm Score = 2.96
NG = 0
DB = 0
SM = 0
Delta_K
Doxorubicin
RF1_90_0 RF1
ZShift = 0.37
lm Score = 0
NG = 0
DB = 0
SM = 0
Delta_r
Doxorubicin
RF1_90_0 RF1
ZShift = −1.84
lm Score = 53.32
NG = 0
DB = 0
SM = 0
Delta_AUC
Doxorubicin
RF1_65_0 RF1
ZShift = −0.43
lm Score = −0.53
NG = 0
DB = 0
SM = 0
Delta_L
Doxorubicin
RF1_65_0 RF1
ZShift = −0.92
lm Score = 2.14
NG = 0
DB = 0
SM = 0
Delta_K
Doxorubicin
RF1_65_0 RF1
ZShift = 0.67
lm Score = 0.02
NG = 0
DB = 0
SM = 0
Delta_r
Doxorubicin
RF1_65_0 RF1
ZShift = −0.64
lm Score = 3.38
NG = 0
DB = 0
SM = 0
Delta_AUC
Doxorubicin

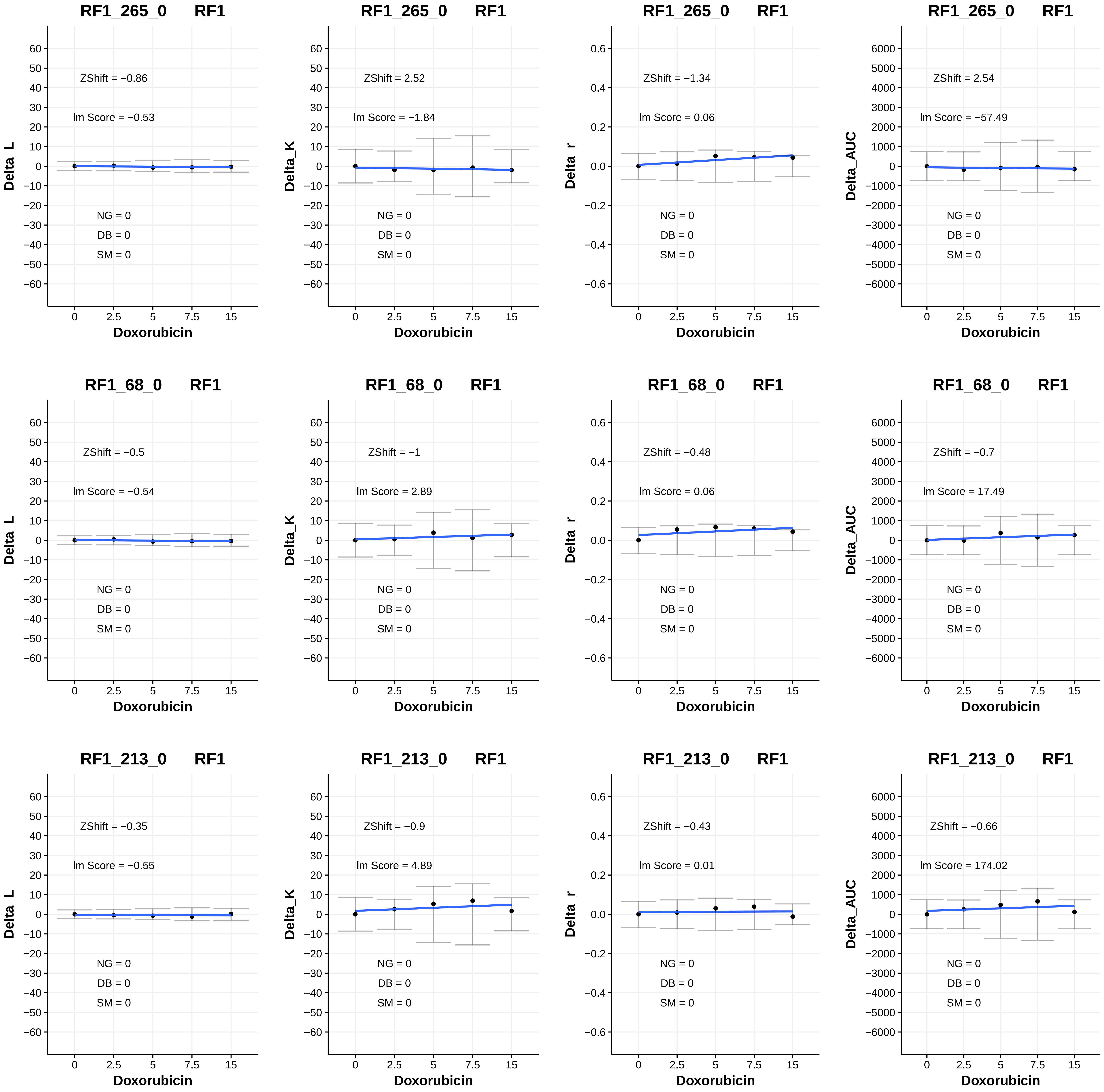
RF1_265_0 RF1
ZShift = −0.86
lm Score = −0.53
NG = 0
DB = 0
SM = 0
Delta_L
Doxorubicin
RF1_265_0 RF1
ZShift = 2.52
lm Score = −1.84
NG = 0
DB = 0
SM = 0
Delta_K
Doxorubicin
RF1_265_0 RF1
ZShift = −1.34
lm Score = 0.06
NG = 0
DB = 0
SM = 0
Delta_r
Doxorubicin
RF1_265_0 RF1
ZShift = 2.54
lm Score = −57.49
NG = 0
DB = 0
SM = 0
Delta_AUC
Doxorubicin
RF1_68_0 RF1
ZShift = −0.5
lm Score = −0.54
NG = 0
DB = 0
SM = 0
Delta_L
Doxorubicin
RF1_68_0 RF1
ZShift = −1
lm Score = 2.89
NG = 0
DB = 0
SM = 0
Delta_K
Doxorubicin
RF1_68_0 RF1
ZShift = −0.48
lm Score = 0.06
NG = 0
DB = 0
SM = 0
Delta_r
Doxorubicin
RF1_68_0 RF1
ZShift = −0.7
lm Score = 17.49
NG = 0
DB = 0
SM = 0
Delta_AUC
Doxorubicin
RF1_213_0 RF1
ZShift = −0.35
lm Score = −0.55
NG = 0
DB = 0
SM = 0
Delta_L
Doxorubicin
RF1_213_0 RF1
ZShift = −0.9
lm Score = 4.89
NG = 0
DB = 0
SM = 0
Delta_K
Doxorubicin
RF1_213_0 RF1
ZShift = −0.43
lm Score = 0.01
NG = 0
DB = 0
SM = 0
Delta_r
Doxorubicin
RF1_213_0 RF1
ZShift = −0.66
lm Score = 174.02
NG = 0
DB = 0
SM = 0
Delta_AUC
Doxorubicin

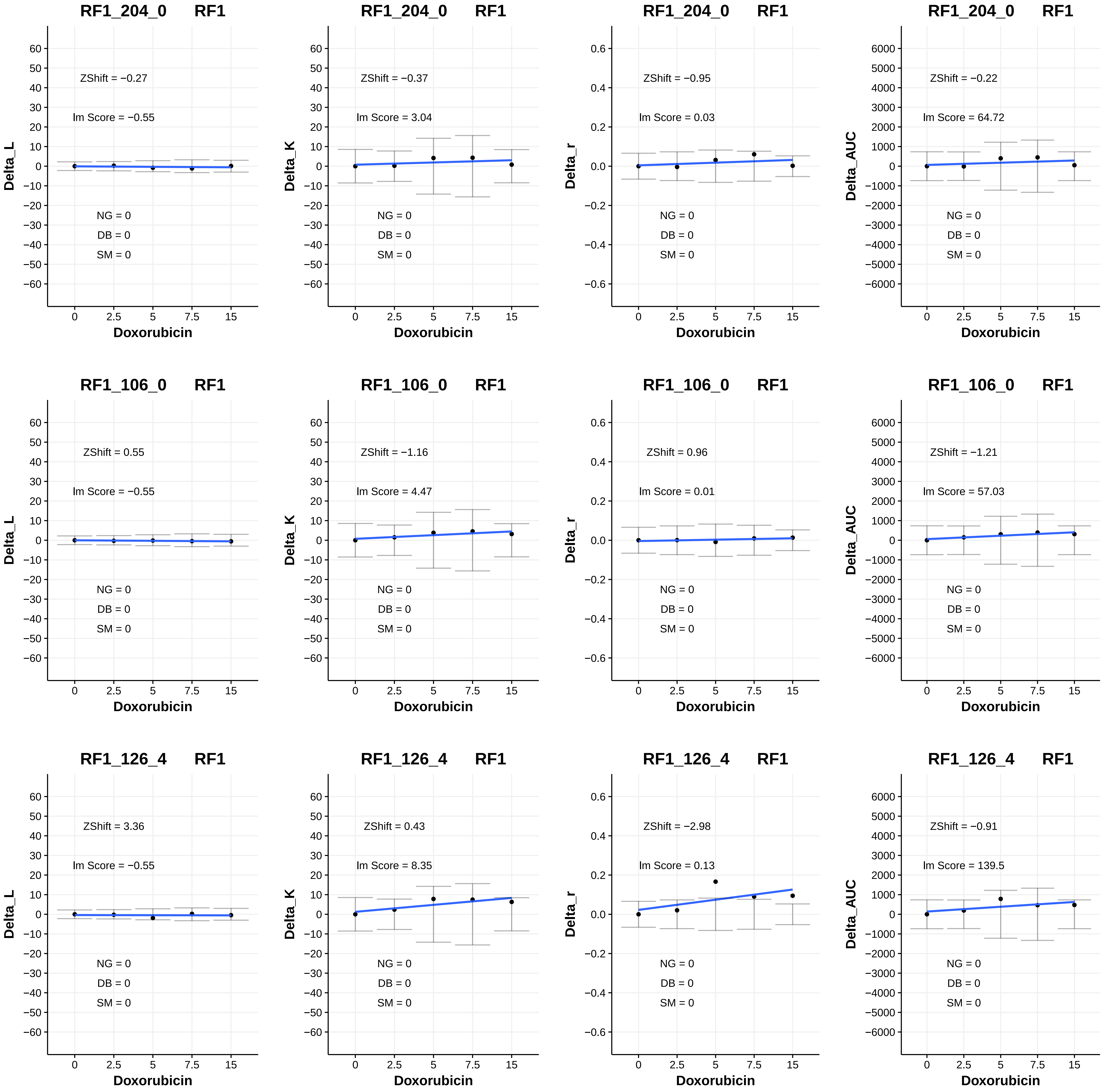

RF1_204_0 RF1
ZShift = −0.27
lm Score = −0.55
NG = 0
DB = 0
SM = 0
Delta_L
Doxorubicin
RF1_204_0 RF1
ZShift = −0.37
lm Score = 3.04
NG = 0
DB = 0
SM = 0
Delta_K
Doxorubicin
RF1_204_0 RF1
ZShift = −0.95
lm Score = 0.03
NG = 0
DB = 0
SM = 0
Delta_r
Doxorubicin
RF1_204_0 RF1
ZShift = −0.22
lm Score = 64.72
NG = 0
DB = 0
SM = 0
Delta_AUC
Doxorubicin
RF1_106_0 RF1
ZShift = 0.55
lm Score = −0.55
NG = 0
DB = 0
SM = 0
Delta_L
Doxorubicin
RF1_106_0 RF1
ZShift = −1.16
lm Score = 4.47
NG = 0
DB = 0
SM = 0
Delta_K
Doxorubicin
RF1_106_0 RF1
ZShift = 0.96
lm Score = 0.01
NG = 0
DB = 0
SM = 0
Delta_r
Doxorubicin
RF1_106_0 RF1
ZShift = −1.21
lm Score = 57.03
NG = 0
DB = 0
SM = 0
Delta_AUC
Doxorubicin
RF1_126_4 RF1
ZShift = 3.36
lm Score = −0.55
NG = 0
DB = 0
SM = 0
Delta_L
Doxorubicin
RF1_126_4 RF1
ZShift = 0.43
lm Score = 8.35
NG = 0
DB = 0
SM = 0
Delta_K
Doxorubicin
RF1_126_4 RF1
ZShift = −2.98
lm Score = 0.13
NG = 0
DB = 0
SM = 0
Delta_r
Doxorubicin
RF1_126_4 RF1
ZShift = −0.91
lm Score = 139.5
NG = 0
DB = 0
SM = 0
Delta_AUC
Doxorubicin

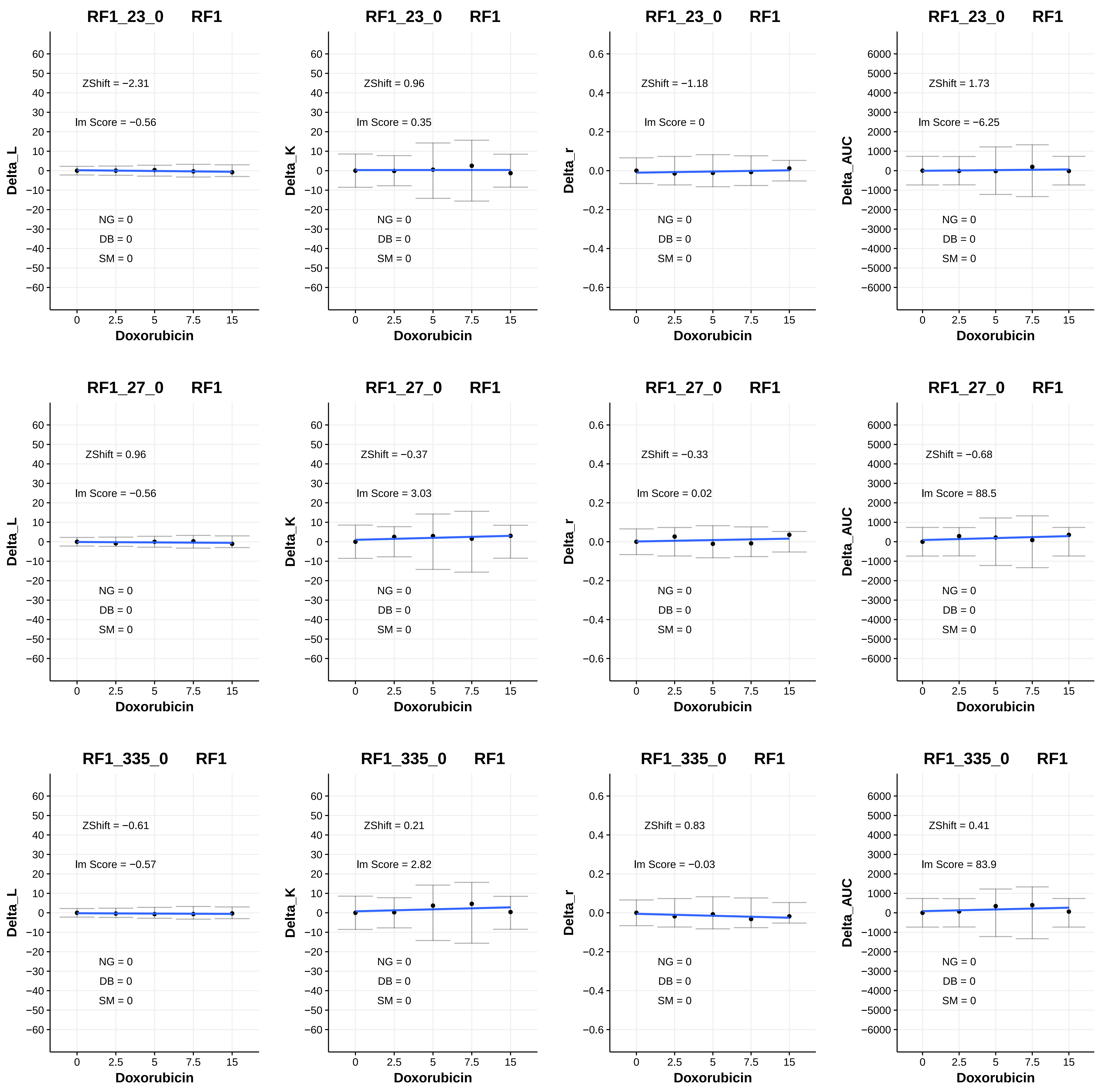
RF1_23_0 RF1
ZShift = −2.31
Im Score = −0.56
NG = 0
DB = 0
SM = 0
Delta_L
Doxorubicin
RF1_23_0 RF1
ZShift = 0.96
Im Score = 0.35
NG = 0
DB = 0
SM = 0
Delta_K
Doxorubicin
RF1_23_0 RF1
ZShift = −1.18
Im Score = 0
NG = 0
DB = 0
SM = 0
Delta_r
Doxorubicin
RF1_23_0 RF1
ZShift = 1.73
Im Score = −6.25
NG = 0
DB = 0
SM = 0
Delta_AUC
Doxorubicin
RF1_27_0 RF1
ZShift = 0.96
Im Score = −0.56
NG = 0
DB = 0
SM = 0
Delta_L
Doxorubicin
RF1_27_0 RF1
ZShift = −0.37
Im Score = 3.03
NG = 0
DB = 0
SM = 0
Delta_K
Doxorubicin
RF1_27_0 RF1
ZShift = −0.33
Im Score = 0.02
NG = 0
DB = 0
SM = 0
Delta_r
Doxorubicin
RF1_27_0 RF1
ZShift = −0.68
Im Score = 88.5
NG = 0
DB = 0
SM = 0
Delta_AUC
Doxorubicin
RF1_335_0 RF1
ZShift = −0.61
Im Score = −0.57
NG = 0
DB = 0
SM = 0
Delta_L
Doxorubicin
RF1_335_0 RF1
ZShift = 0.21
Im Score = 2.82
NG = 0
DB = 0
SM = 0
Delta_K
Doxorubicin
RF1_335_0 RF1
ZShift = 0.83
Im Score = −0.03
NG = 0
DB = 0
SM = 0
Delta_r
Doxorubicin
RF1_335_0 RF1
ZShift = 0.41
Im Score = 83.9
NG = 0
DB = 0
SM = 0
Delta_AUC
Doxorubicin

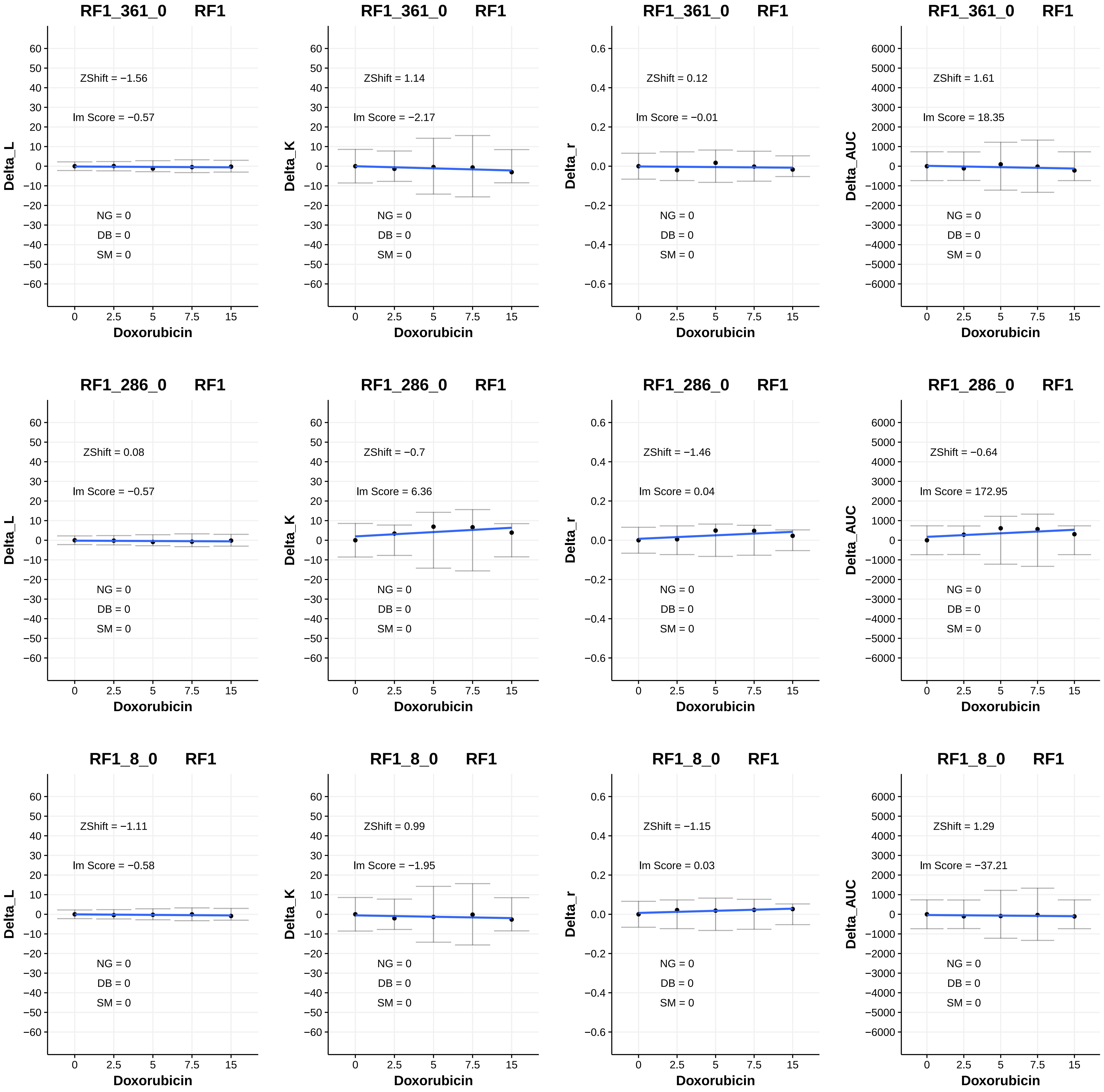
RF1_361_0 RF1
ZShift = −1.56
lm Score = −0.57
NG = 0
DB = 0
SM = 0
Delta_L
Doxorubicin
RF1_361_0 RF1
ZShift = 1.14
lm Score = −2.17
NG = 0
DB = 0
SM = 0
Delta_K
Doxorubicin
RF1_361_0 RF1
ZShift = 0.12
lm Score = −0.01
NG = 0
DB = 0
SM = 0
Delta_r
Doxorubicin
RF1_361_0 RF1
ZShift = 1.61
lm Score = 18.35
NG = 0
DB = 0
SM = 0
Delta_AUC
Doxorubicin
RF1_286_0 RF1
ZShift = 0.08
lm Score = −0.57
NG = 0
DB = 0
SM = 0
Delta_L
Doxorubicin
RF1_286_0 RF1
ZShift = −0.7
lm Score = 6.36
NG = 0
DB = 0
SM = 0
Delta_K
Doxorubicin
RF1_286_0 RF1
ZShift = −1.46
lm Score = 0.04
NG = 0
DB = 0
SM = 0
Delta_r
Doxorubicin
RF1_286_0 RF1
ZShift = −0.64
lm Score = 172.95
NG = 0
DB = 0
SM = 0
Delta_AUC
Doxorubicin
RF1_8_0 RF1
ZShift = −1.11
lm Score = −0.58
NG = 0
DB = 0
SM = 0
Delta_L
Doxorubicin
RF1_8_0 RF1
ZShift = 0.99
lm Score = −1.95
NG = 0
DB = 0
SM = 0
Delta_K
Doxorubicin
RF1_8_0 RF1
ZShift = −1.15
lm Score = 0.03
NG = 0
DB = 0
SM = 0
Delta_r
Doxorubicin
RF1_8_0 RF1
ZShift = 1.29
lm Score = −37.21
NG = 0
DB = 0
SM = 0
Delta_AUC
Doxorubicin

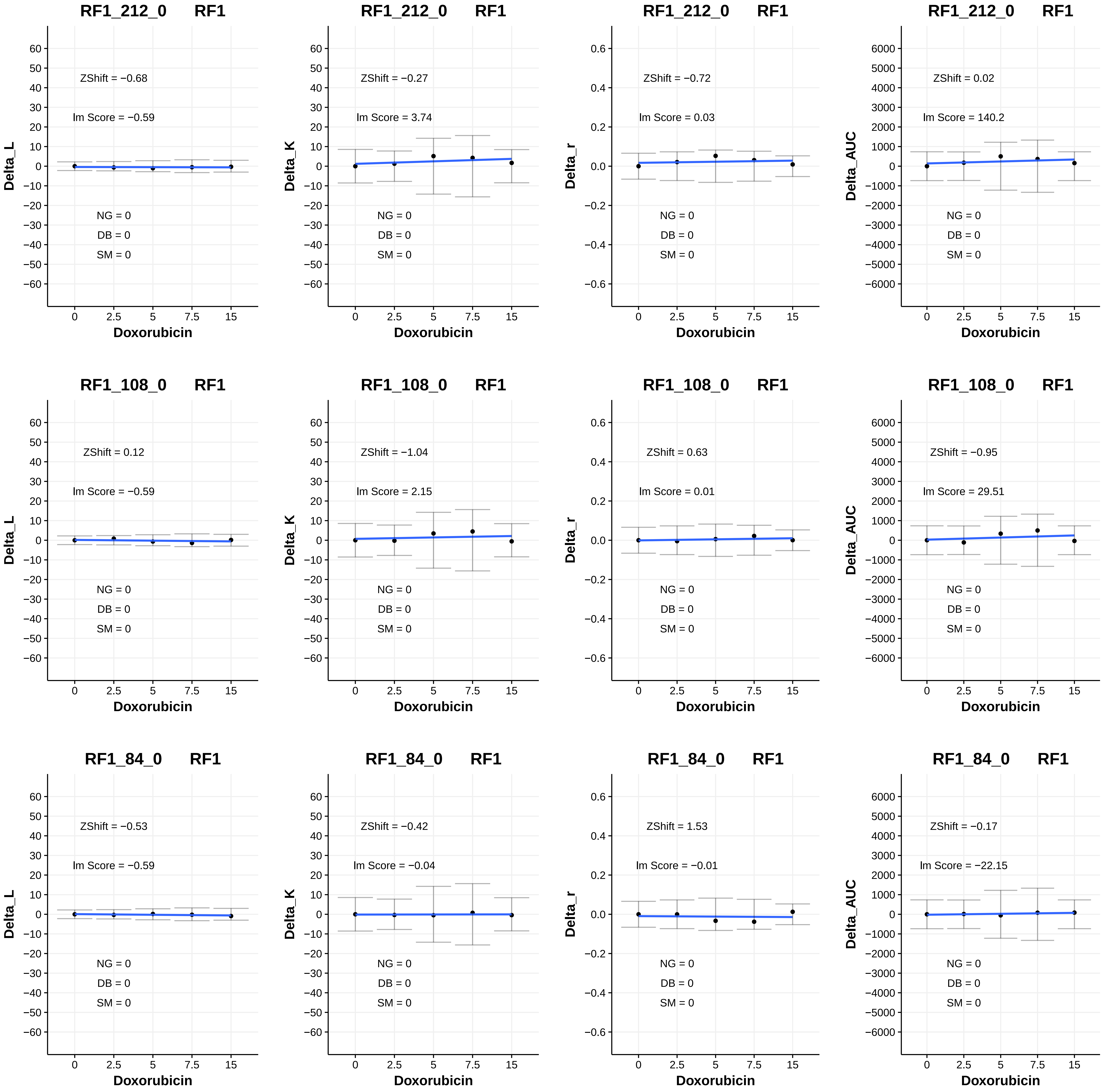

RF1_212_0 RF1
ZShift = −0.68
Im Score = −0.59
NG = 0
DB = 0
SM = 0
Delta_L
Doxorubicin
RF1_212_0 RF1
ZShift = −0.27
Im Score = 3.74
NG = 0
DB = 0
SM = 0
Delta_K
Doxorubicin
RF1_212_0 RF1
ZShift = −0.72
Im Score = 0.03
NG = 0
DB = 0
SM = 0
Delta_r
Doxorubicin
RF1_212_0 RF1
ZShift = 0.02
Im Score = 140.2
NG = 0
DB = 0
SM = 0
Delta_AUC
Doxorubicin
RF1_108_0 RF1
ZShift = 0.12
Im Score = −0.59
NG = 0
DB = 0
SM = 0
Delta_L
Doxorubicin
RF1_108_0 RF1
ZShift = −1.04
Im Score = 2.15
NG = 0
DB = 0
SM = 0
Delta_K
Doxorubicin
RF1_108_0 RF1
ZShift = 0.63
Im Score = 0.01
NG = 0
DB = 0
SM = 0
Delta_r
Doxorubicin
RF1_108_0 RF1
ZShift = −0.95
Im Score = 29.51
NG = 0
DB = 0
SM = 0
Delta_AUC
Doxorubicin
RF1_84_0 RF1
ZShift = −0.53
Im Score = −0.59
NG = 0
DB = 0
SM = 0
Delta_L
Doxorubicin
RF1_84_0 RF1
ZShift = −0.42
Im Score = −0.04
NG = 0
DB = 0
SM = 0
Delta_K
Doxorubicin
RF1_84_0 RF1
ZShift = 1.53
Im Score = −0.01
NG = 0
DB = 0
SM = 0
Delta_r
Doxorubicin
RF1_84_0 RF1
ZShift = −0.17
Im Score = −22.15
NG = 0
DB = 0
SM = 0
Delta_AUC
Doxorubicin

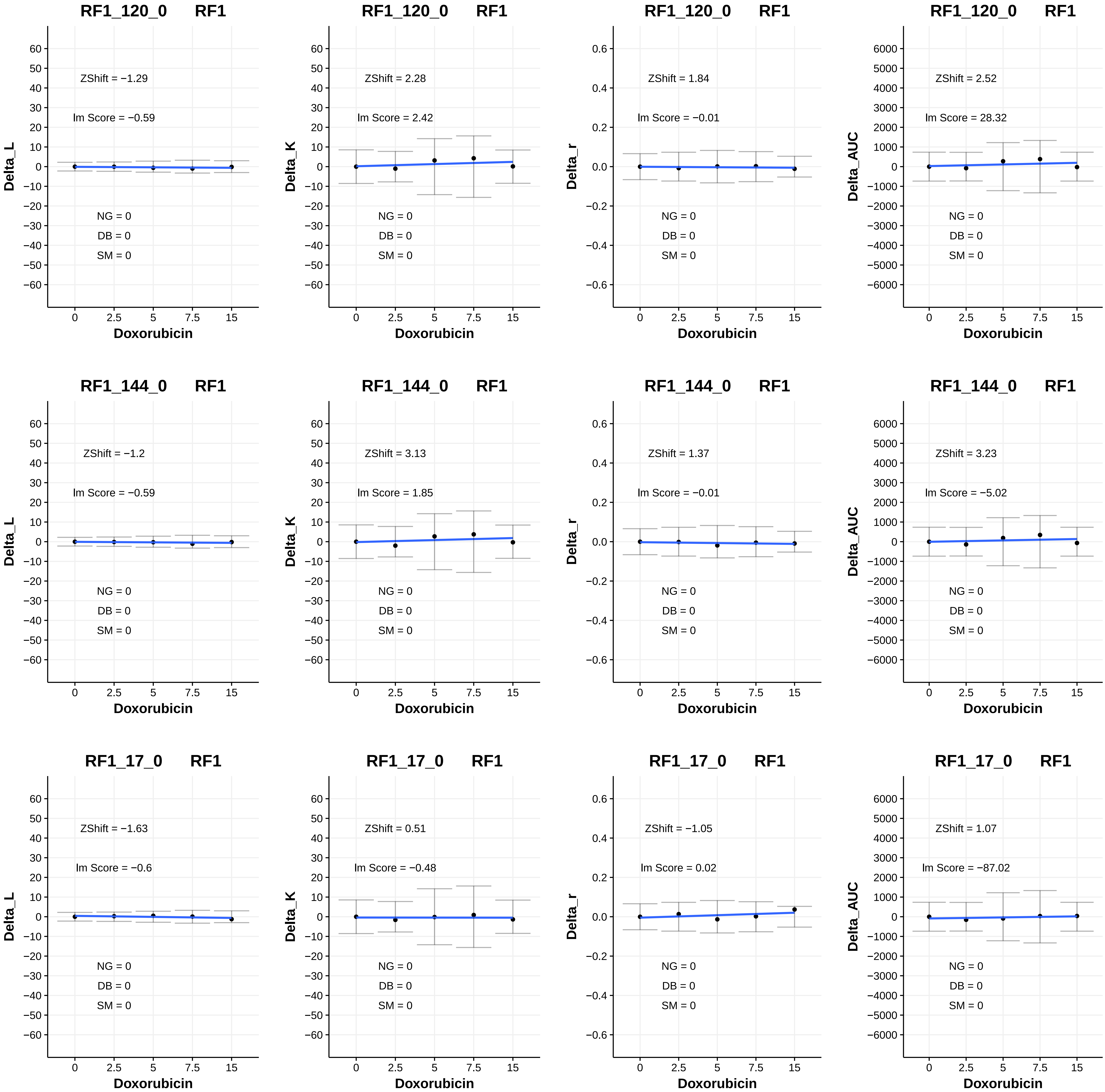

RF1_120_0 RF1
ZShift = −1.29
Im Score = −0.59
NG = 0
DB = 0
SM = 0
Delta_L
Doxorubicin
RF1_120_0 RF1
ZShift = 2.28
Im Score = 2.42
NG = 0
DB = 0
SM = 0
Delta_K
Doxorubicin
RF1_120_0 RF1
ZShift = 1.84
Im Score = −0.01
NG = 0
DB = 0
SM = 0
Delta_r
Doxorubicin
RF1_120_0 RF1
ZShift = 2.52
Im Score = 28.32
NG = 0
DB = 0
SM = 0
Delta_AUC
Doxorubicin
RF1_144_0 RF1
ZShift = −1.2
Im Score = −0.59
NG = 0
DB = 0
SM = 0
Delta_L
Doxorubicin
RF1_144_0 RF1
ZShift = 3.13
Im Score = 1.85
NG = 0
DB = 0
SM = 0
Delta_K
Doxorubicin
RF1_144_0 RF1
ZShift = 1.37
Im Score = −0.01
NG = 0
DB = 0
SM = 0
Delta_r
Doxorubicin
RF1_144_0 RF1
ZShift = 3.23
Im Score = −5.02
NG = 0
DB = 0
SM = 0
Delta_AUC
Doxorubicin
RF1_17_0 RF1
ZShift = −1.63
Im Score = −0.6
NG = 0
DB = 0
SM = 0
Delta_L
Doxorubicin
RF1_17_0 RF1
ZShift = 0.51
Im Score = −0.48
NG = 0
DB = 0
SM = 0
Delta_K
Doxorubicin
RF1_17_0 RF1
ZShift = −1.05
Im Score = 0.02
NG = 0
DB = 0
SM = 0
Delta_r
Doxorubicin
RF1_17_0 RF1
ZShift = 1.07
Im Score = −87.02
NG = 0
DB = 0
SM = 0
Delta_AUC
Doxorubicin

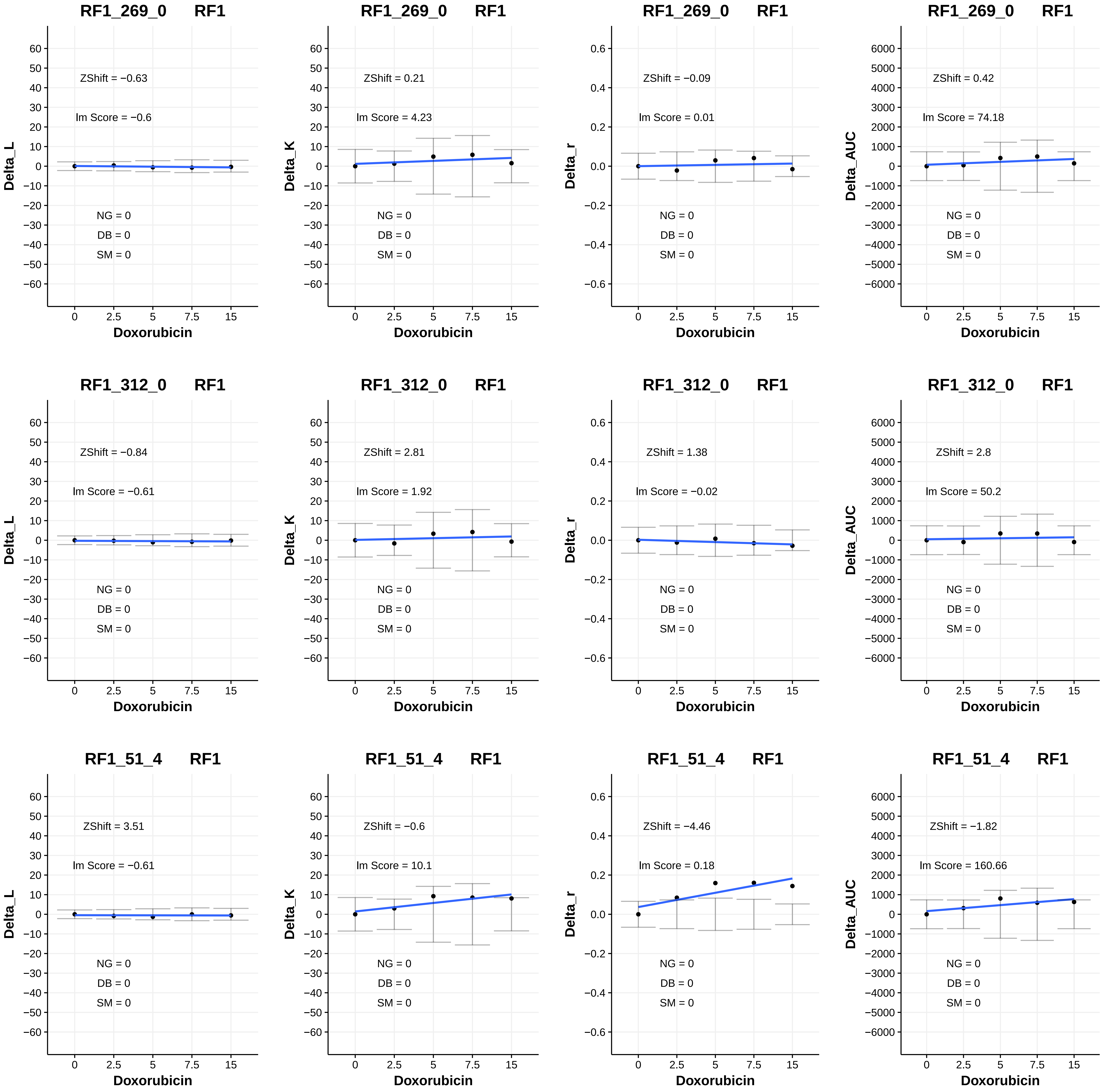
RF1_269_0 RF1
ZShift = −0.63
lm Score = −0.6
NG = 0
DB = 0
SM = 0
Delta_L
Doxorubicin
RF1_269_0 RF1
ZShift = 0.21
lm Score = 4.23
NG = 0
DB = 0
SM = 0
Delta_K
Doxorubicin
RF1_269_0 RF1
ZShift = −0.09
lm Score = 0.01
NG = 0
DB = 0
SM = 0
Delta_r
Doxorubicin
RF1_269_0 RF1
ZShift = 0.42
lm Score = 74.18
NG = 0
DB = 0
SM = 0
Delta_AUC
Doxorubicin
RF1_312_0 RF1
ZShift = −0.84
lm Score = −0.61
NG = 0
DB = 0
SM = 0
Delta_L
Doxorubicin
RF1_312_0 RF1
ZShift = 2.81
lm Score = 1.92
NG = 0
DB = 0
SM = 0
Delta_K
Doxorubicin
RF1_312_0 RF1
ZShift = 1.38
lm Score = −0.02
NG = 0
DB = 0
SM = 0
Delta_r
Doxorubicin
RF1_312_0 RF1
ZShift = 2.8
lm Score = 50.2
NG = 0
DB = 0
SM = 0
Delta_AUC
Doxorubicin
RF1_51_4 RF1
ZShift = 3.51
lm Score = −0.61
NG = 0
DB = 0
SM = 0
Delta_L
Doxorubicin
RF1_51_4 RF1
ZShift = −0.6
lm Score = 10.1
NG = 0
DB = 0
SM = 0
Delta_K
Doxorubicin
RF1_51_4 RF1
ZShift = −4.46
lm Score = 0.18
NG = 0
DB = 0
SM = 0
Delta_r
Doxorubicin
RF1_51_4 RF1
ZShift = −1.82
lm Score = 160.66
NG = 0
DB = 0
SM = 0
Delta_AUC
Doxorubicin

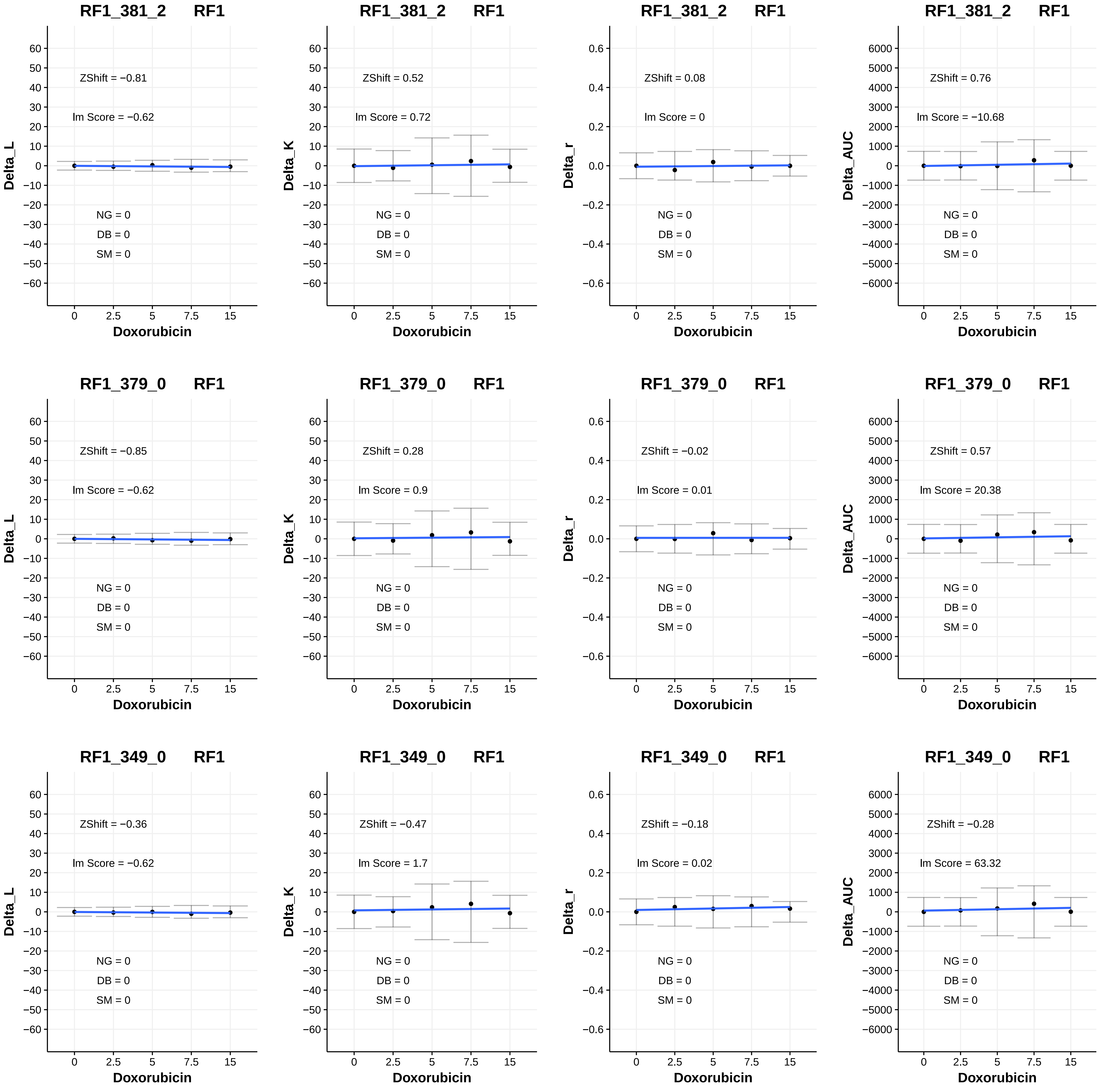
RF1_381_2 RF1
Delta_L
ZShift = −0.81
lm Score = −0.62
NG = 0
DB = 0
SM = 0
Doxorubicin
RF1_381_2 RF1
Delta_K
ZShift = 0.52
lm Score = 0.72
NG = 0
DB = 0
SM = 0
Doxorubicin
RF1_381_2 RF1
Delta_r
ZShift = 0.08
lm Score = 0
NG = 0
DB = 0
SM = 0
Doxorubicin
RF1_381_2 RF1
Delta_AUC
ZShift = 0.76
lm Score = −10.68
NG = 0
DB = 0
SM = 0
Doxorubicin
RF1_379_0 RF1
Delta_L
ZShift = −0.85
lm Score = −0.62
NG = 0
DB = 0
SM = 0
Doxorubicin
RF1_379_0 RF1
Delta_K
ZShift = 0.28
lm Score = 0.9
NG = 0
DB = 0
SM = 0
Doxorubicin
RF1_379_0 RF1
Delta_r
ZShift = −0.02
lm Score = 0.01
NG = 0
DB = 0
SM = 0
Doxorubicin
RF1_379_0 RF1
Delta_AUC
ZShift = 0.57
lm Score = 20.38
NG = 0
DB = 0
SM = 0
Doxorubicin
RF1_349_0 RF1
Delta_L
ZShift = −0.36
lm Score = −0.62
NG = 0
DB = 0
SM = 0
Doxorubicin
RF1_349_0 RF1
Delta_K
ZShift = −0.47
lm Score = 1.7
NG = 0
DB = 0
SM = 0
Doxorubicin
RF1_349_0 RF1
Delta_r
ZShift = −0.18
lm Score = 0.02
NG = 0
DB = 0
SM = 0
Doxorubicin
RF1_349_0 RF1
Delta_AUC
ZShift = −0.28
lm Score = 63.32
NG = 0
DB = 0
SM = 0
Doxorubicin

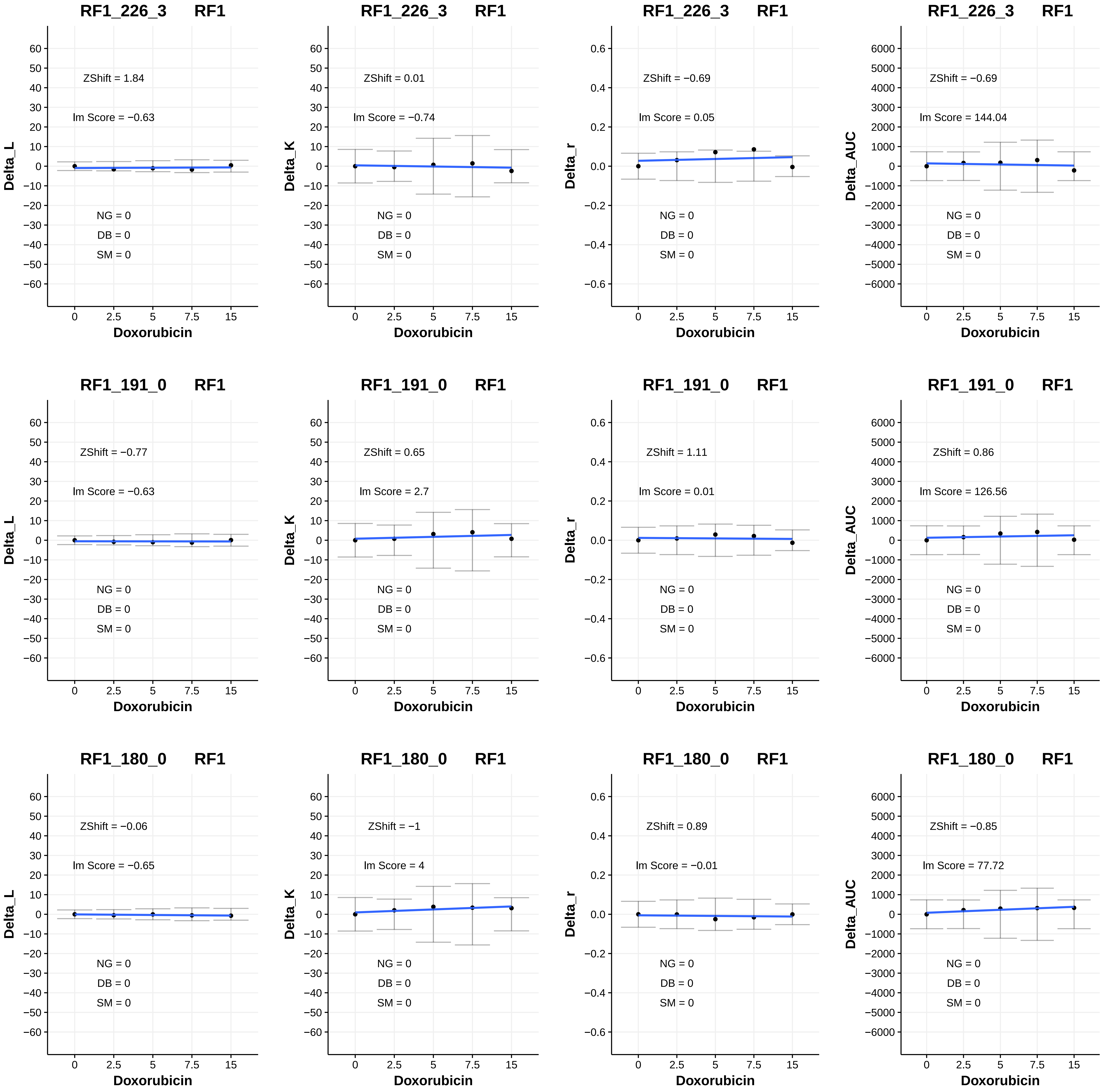
RF1_226_3 RF1
ZShift = 1.84
lm Score = −0.63
NG = 0
DB = 0
SM = 0
Delta_L
Doxorubicin
RF1_226_3 RF1
ZShift = 0.01
lm Score = −0.74
NG = 0
DB = 0
SM = 0
Delta_K
Doxorubicin
RF1_226_3 RF1
ZShift = −0.69
lm Score = 0.05
NG = 0
DB = 0
SM = 0
Delta_r
Doxorubicin
RF1_226_3 RF1
ZShift = −0.69
lm Score = 144.04
NG = 0
DB = 0
SM = 0
Delta_AUC
Doxorubicin
RF1_191_0 RF1
ZShift = −0.77
lm Score = −0.63
NG = 0
DB = 0
SM = 0
Delta_L
Doxorubicin
RF1_191_0 RF1
ZShift = 0.65
lm Score = 2.7
NG = 0
DB = 0
SM = 0
Delta_K
Doxorubicin
RF1_191_0 RF1
ZShift = 1.11
lm Score = 0.01
NG = 0
DB = 0
SM = 0
Delta_r
Doxorubicin
RF1_191_0 RF1
ZShift = 0.86
lm Score = 126.56
NG = 0
DB = 0
SM = 0
Delta_AUC
Doxorubicin
RF1_180_0 RF1
ZShift = −0.06
lm Score = −0.65
NG = 0
DB = 0
SM = 0
Delta_L
Doxorubicin
RF1_180_0 RF1
ZShift = −1
lm Score = 4
NG = 0
DB = 0
SM = 0
Delta_K
Doxorubicin
RF1_180_0 RF1
ZShift = 0.89
lm Score = −0.01
NG = 0
DB = 0
SM = 0
Delta_r
Doxorubicin
RF1_180_0 RF1
ZShift = −0.85
lm Score = 77.72
NG = 0
DB = 0
SM = 0
Delta_AUC
Doxorubicin

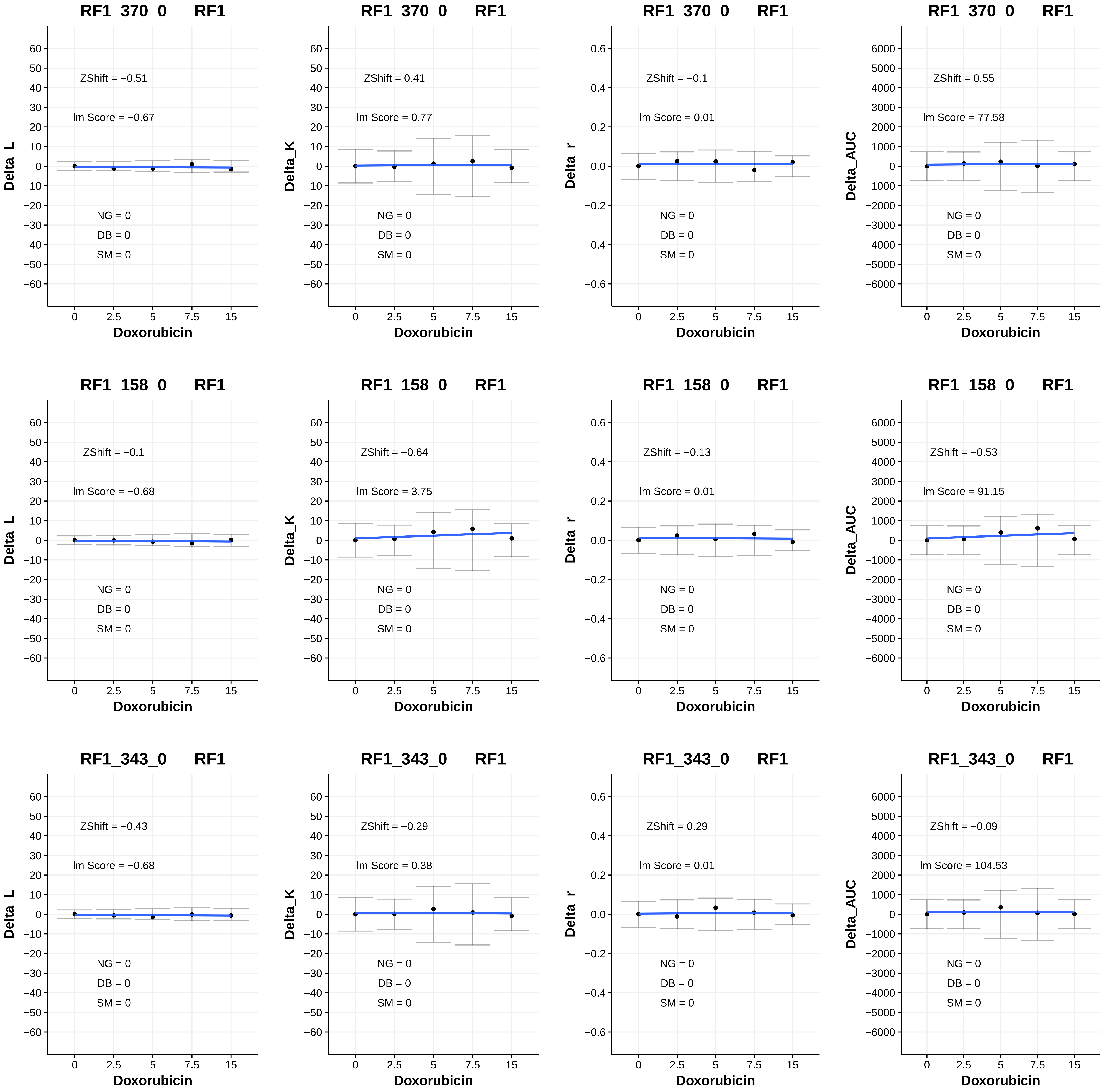

RF1_370_0 RF1
ZShift = −0.51
lm Score = −0.67
NG = 0
DB = 0
SM = 0
Delta_L
Doxorubicin
RF1_370_0 RF1
ZShift = 0.41
lm Score = 0.77
NG = 0
DB = 0
SM = 0
Delta_K
Doxorubicin
RF1_370_0 RF1
ZShift = −0.1
lm Score = 0.01
NG = 0
DB = 0
SM = 0
Delta_r
Doxorubicin
RF1_370_0 RF1
ZShift = 0.55
lm Score = 77.58
NG = 0
DB = 0
SM = 0
Delta_AUC
Doxorubicin
RF1_158_0 RF1
ZShift = −0.1
lm Score = −0.68
NG = 0
DB = 0
SM = 0
Delta_L
Doxorubicin
RF1_158_0 RF1
ZShift = −0.64
lm Score = 3.75
NG = 0
DB = 0
SM = 0
Delta_K
Doxorubicin
RF1_158_0 RF1
ZShift = −0.13
lm Score = 0.01
NG = 0
DB = 0
SM = 0
Delta_r
Doxorubicin
RF1_158_0 RF1
ZShift = −0.53
lm Score = 91.15
NG = 0
DB = 0
SM = 0
Delta_AUC
Doxorubicin
RF1_343_0 RF1
ZShift = −0.43
lm Score = −0.68
NG = 0
DB = 0
SM = 0
Delta_L
Doxorubicin
RF1_343_0 RF1
ZShift = −0.29
lm Score = 0.38
NG = 0
DB = 0
SM = 0
Delta_K
Doxorubicin
RF1_343_0 RF1
ZShift = 0.29
lm Score = 0.01
NG = 0
DB = 0
SM = 0
Delta_r
Doxorubicin
RF1_343_0 RF1
ZShift = −0.09
lm Score = 104.53
NG = 0
DB = 0
SM = 0
Delta_AUC
Doxorubicin

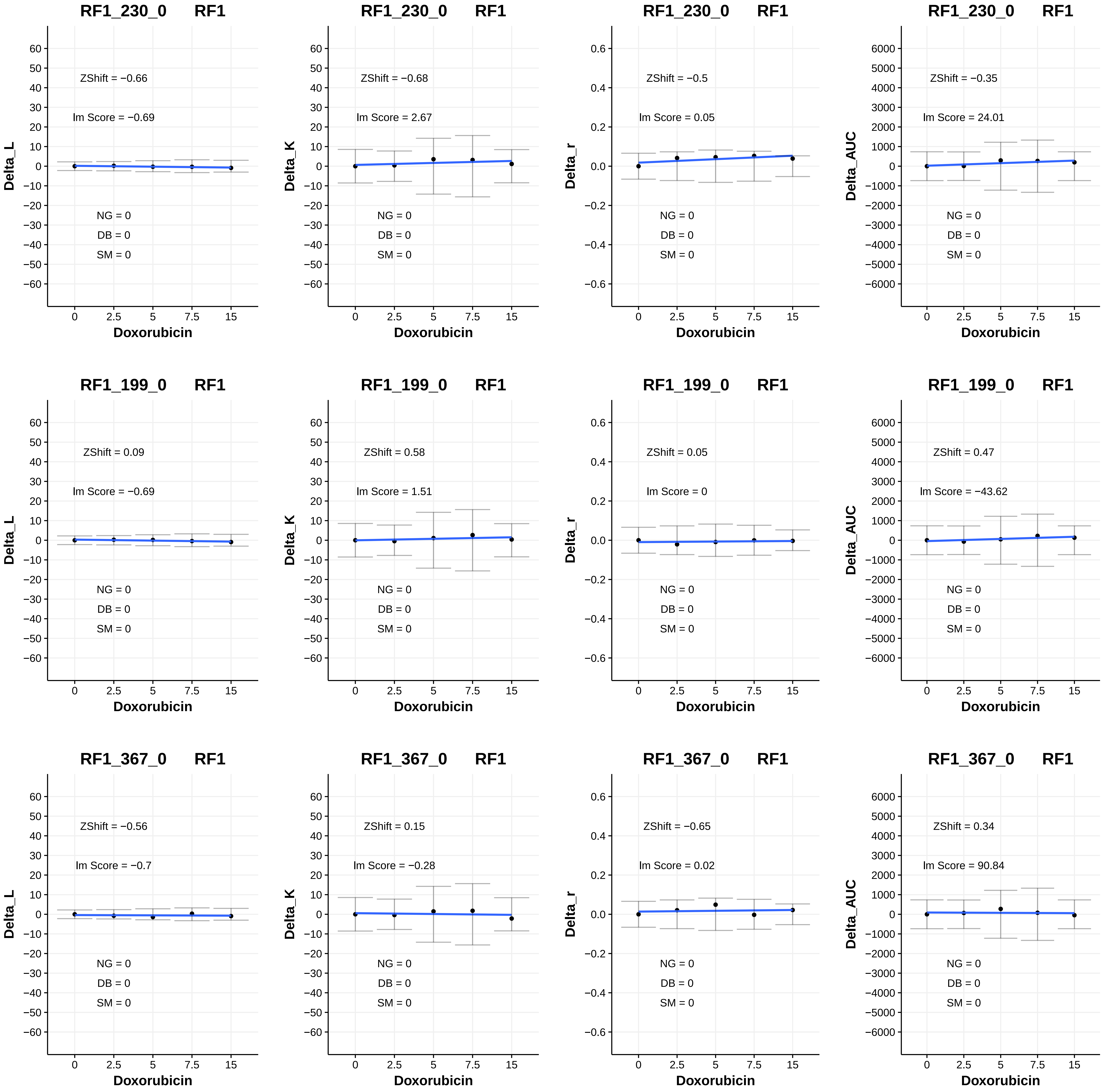
RF1_230_0 RF1
ZShift = −0.66
lm Score = −0.69
NG = 0
DB = 0
SM = 0
Delta_L
Doxorubicin
RF1_230_0 RF1
ZShift = −0.68
lm Score = 2.67
NG = 0
DB = 0
SM = 0
Delta_K
Doxorubicin
RF1_230_0 RF1
ZShift = −0.5
lm Score = 0.05
NG = 0
DB = 0
SM = 0
Delta_r
Doxorubicin
RF1_230_0 RF1
ZShift = −0.35
lm Score = 24.01
NG = 0
DB = 0
SM = 0
Delta_AUC
Doxorubicin
RF1_199_0 RF1
ZShift = 0.09
lm Score = −0.69
NG = 0
DB = 0
SM = 0
Delta_L
Doxorubicin
RF1_199_0 RF1
ZShift = 0.58
lm Score = 1.51
NG = 0
DB = 0
SM = 0
Delta_K
Doxorubicin
RF1_199_0 RF1
ZShift = 0.05
lm Score = 0
NG = 0
DB = 0
SM = 0
Delta_r
Doxorubicin
RF1_199_0 RF1
ZShift = 0.47
lm Score = −43.62
NG = 0
DB = 0
SM = 0
Delta_AUC
Doxorubicin
RF1_367_0 RF1
ZShift = −0.56
lm Score = −0.7
NG = 0
DB = 0
SM = 0
Delta_L
Doxorubicin
RF1_367_0 RF1
ZShift = 0.15
lm Score = −0.28
NG = 0
DB = 0
SM = 0
Delta_K
Doxorubicin
RF1_367_0 RF1
ZShift = −0.65
lm Score = 0.02
NG = 0
DB = 0
SM = 0
Delta_r
Doxorubicin
RF1_367_0 RF1
ZShift = 0.34
lm Score = 90.84
NG = 0
DB = 0
SM = 0
Delta_AUC
Doxorubicin

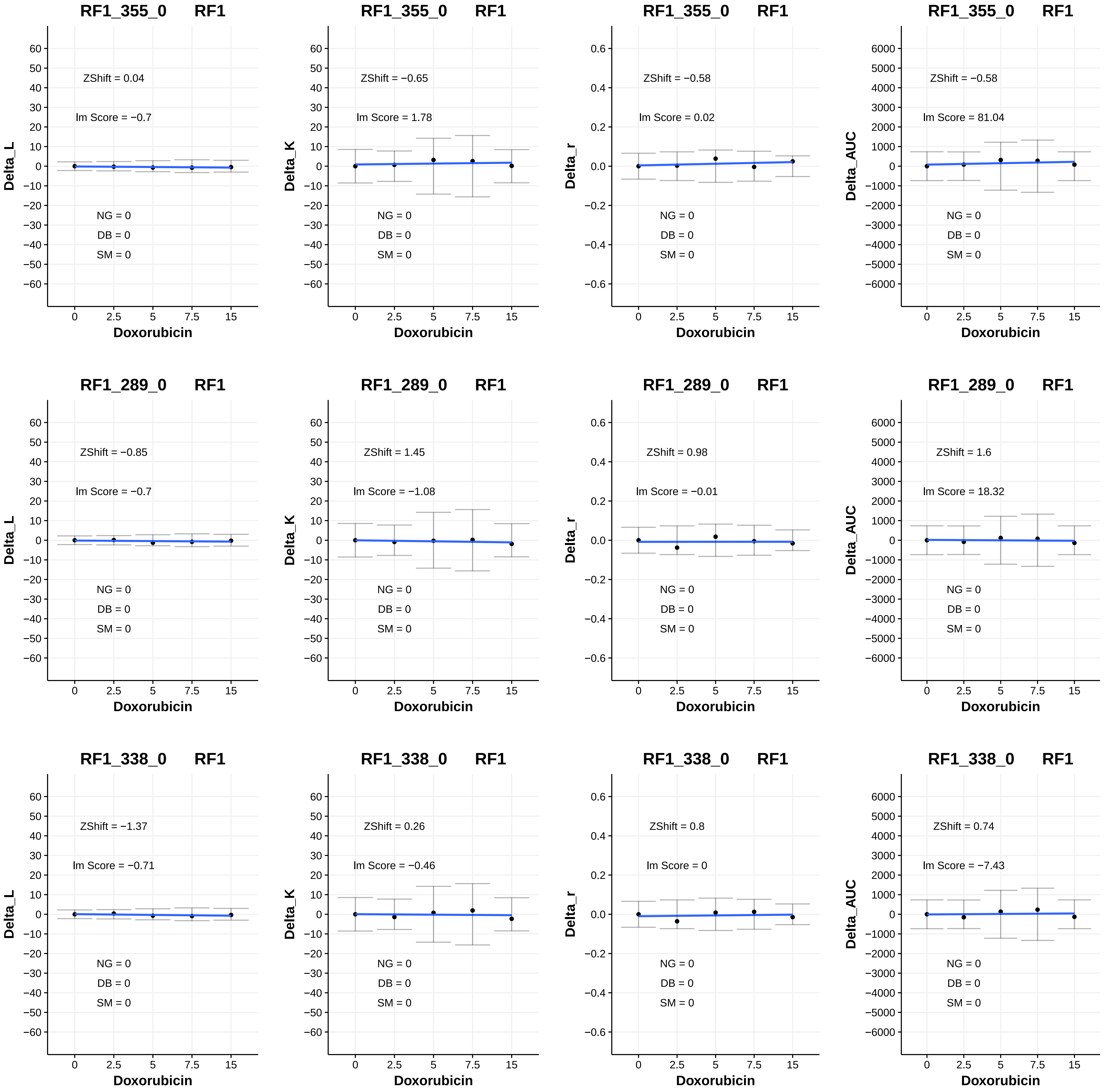
RF1_355_0 RF1
ZShift = 0.04
lm Score = −0.7
NG = 0
DB = 0
SM = 0
Delta_L
Doxorubicin
RF1_355_0 RF1
ZShift = −0.65
lm Score = 1.78
NG = 0
DB = 0
SM = 0
Delta_K
Doxorubicin
RF1_355_0 RF1
ZShift = −0.58
lm Score = 0.02
NG = 0
DB = 0
SM = 0
Delta_r
Doxorubicin
RF1_355_0 RF1
ZShift = −0.58
lm Score = 81.04
NG = 0
DB = 0
SM = 0
Delta_AUC
Doxorubicin
RF1_289_0 RF1
ZShift = −0.85
lm Score = −0.7
NG = 0
DB = 0
SM = 0
Delta_L
Doxorubicin
RF1_289_0 RF1
ZShift = 1.45
lm Score = −1.08
NG = 0
DB = 0
SM = 0
Delta_K
Doxorubicin
RF1_289_0 RF1
ZShift = 0.98
lm Score = −0.01
NG = 0
DB = 0
SM = 0
Delta_r
Doxorubicin
RF1_289_0 RF1
ZShift = 1.6
lm Score = 18.32
NG = 0
DB = 0
SM = 0
Delta_AUC
Doxorubicin
RF1_338_0 RF1
ZShift = −1.37
lm Score = −0.71
NG = 0
DB = 0
SM = 0
Delta_L
Doxorubicin
RF1_338_0 RF1
ZShift = 0.26
lm Score = −0.46
NG = 0
DB = 0
SM = 0
Delta_K
Doxorubicin
RF1_338_0 RF1
ZShift = 0.8
lm Score = 0
NG = 0
DB = 0
SM = 0
Delta_r
Doxorubicin
RF1_338_0 RF1
ZShift = 0.74
lm Score = −7.43
NG = 0
DB = 0
SM = 0
Delta_AUC
Doxorubicin

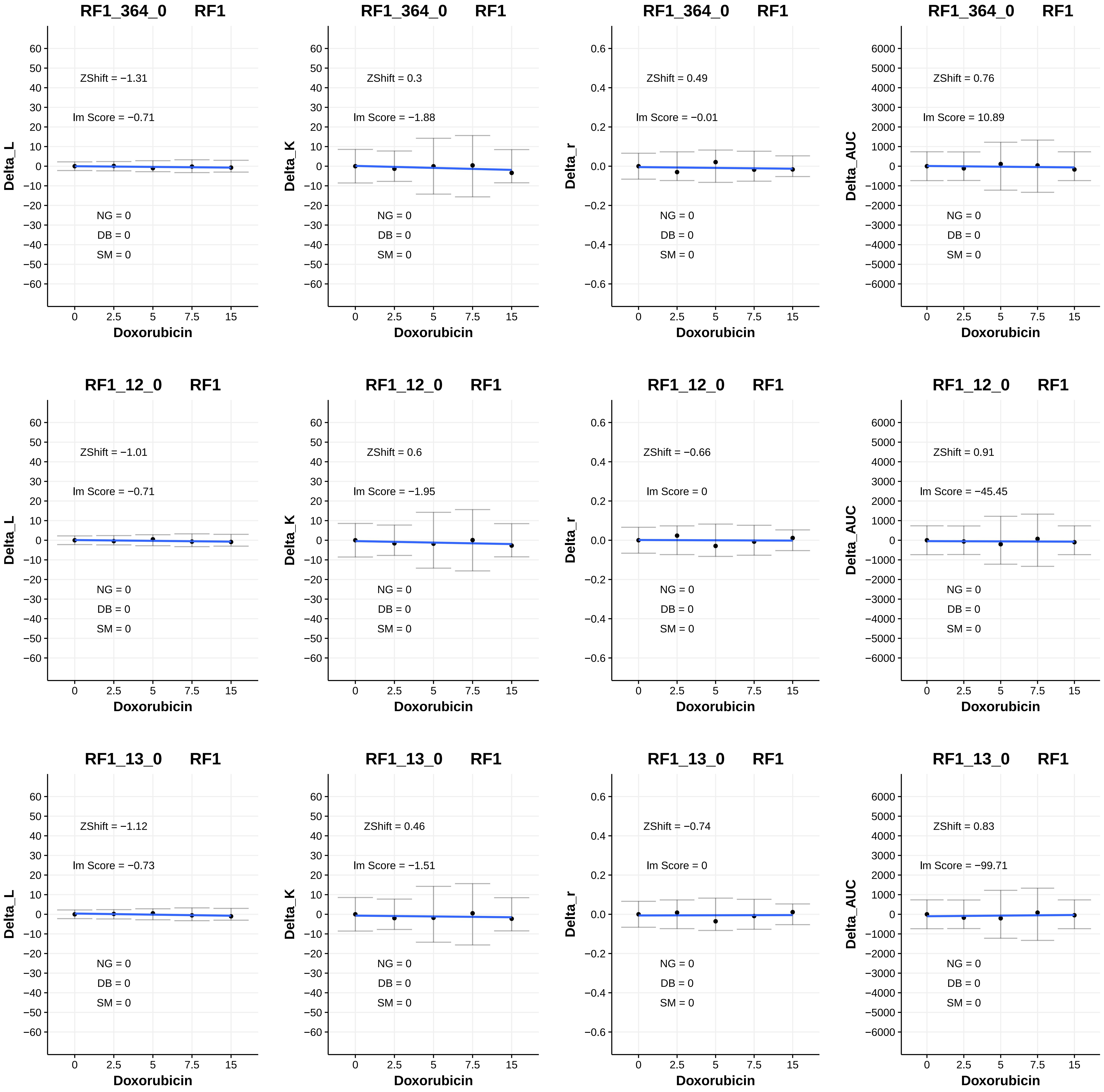
RF1_364_0 RF1
ZShift = −1.31
lm Score = −0.71
NG = 0
DB = 0
SM = 0
Delta_L
Doxorubicin
RF1_364_0 RF1
ZShift = 0.3
lm Score = −1.88
NG = 0
DB = 0
SM = 0
Delta_K
Doxorubicin
RF1_364_0 RF1
ZShift = 0.49
lm Score = −0.01
NG = 0
DB = 0
SM = 0
Delta_r
Doxorubicin
RF1_364_0 RF1
ZShift = 0.76
lm Score = 10.89
NG = 0
DB = 0
SM = 0
Delta_AUC
Doxorubicin
RF1_12_0 RF1
ZShift = −1.01
lm Score = −0.71
NG = 0
DB = 0
SM = 0
Delta_L
Doxorubicin
RF1_12_0 RF1
ZShift = 0.6
lm Score = −1.95
NG = 0
DB = 0
SM = 0
Delta_K
Doxorubicin
RF1_12_0 RF1
ZShift = −0.66
lm Score = 0
NG = 0
DB = 0
SM = 0
Delta_r
Doxorubicin
RF1_12_0 RF1
ZShift = 0.91
lm Score = −45.45
NG = 0
DB = 0
SM = 0
Delta_AUC
Doxorubicin
RF1_13_0 RF1
ZShift = −1.12
lm Score = −0.73
NG = 0
DB = 0
SM = 0
Delta_L
Doxorubicin
RF1_13_0 RF1
ZShift = 0.46
lm Score = −1.51
NG = 0
DB = 0
SM = 0
Delta_K
Doxorubicin
RF1_13_0 RF1
ZShift = −0.74
lm Score = 0
NG = 0
DB = 0
SM = 0
Delta_r
Doxorubicin
RF1_13_0 RF1
ZShift = 0.83
lm Score = −99.71
NG = 0
DB = 0
SM = 0
Delta_AUC
Doxorubicin
0
2.5
5
7.5
15

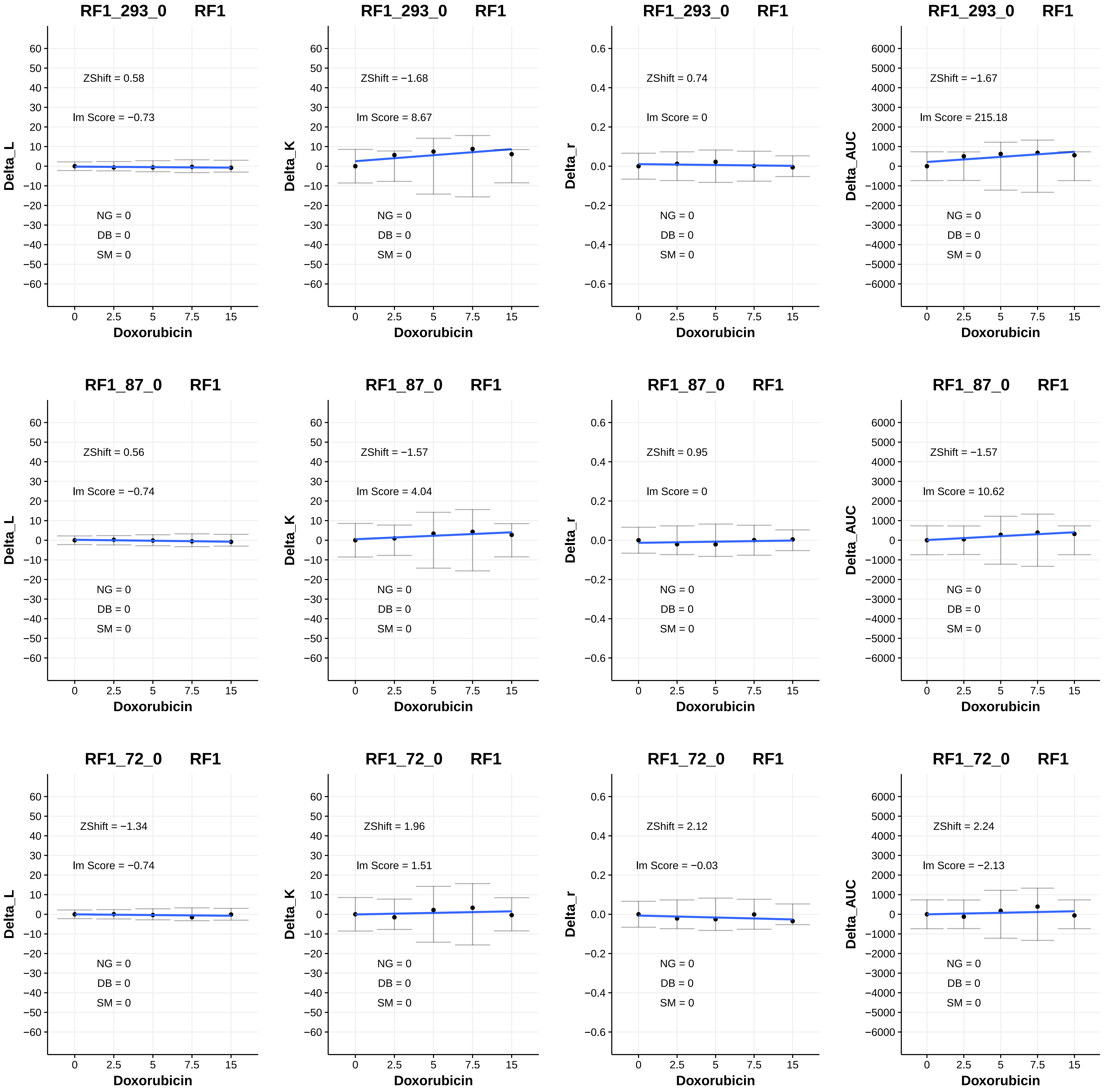

RF1_293_0 RF1
ZShift = 0.58
Im Score = −0.73
NG = 0
DB = 0
SM = 0
Delta_L
Doxorubicin
RF1_293_0 RF1
ZShift = −1.68
Im Score = 8.67
NG = 0
DB = 0
SM = 0
Delta_K
Doxorubicin
RF1_293_0 RF1
ZShift = 0.74
Im Score = 0
NG = 0
DB = 0
SM = 0
Delta_r
Doxorubicin
RF1_293_0 RF1
ZShift = −1.67
Im Score = 215.18
NG = 0
DB = 0
SM = 0
Delta_AUC
Doxorubicin
RF1_87_0 RF1
ZShift = 0.56
Im Score = −0.74
NG = 0
DB = 0
SM = 0
Delta_L
Doxorubicin
RF1_87_0 RF1
ZShift = −1.57
Im Score = 4.04
NG = 0
DB = 0
SM = 0
Delta_K
Doxorubicin
RF1_87_0 RF1
ZShift = 0.95
Im Score = 0
NG = 0
DB = 0
SM = 0
Delta_r
Doxorubicin
RF1_87_0 RF1
ZShift = −1.57
Im Score = 10.62
NG = 0
DB = 0
SM = 0
Delta_AUC
Doxorubicin
RF1_72_0 RF1
ZShift = −1.34
Im Score = −0.74
NG = 0
DB = 0
SM = 0
Delta_L
Doxorubicin
RF1_72_0 RF1
ZShift = 1.96
Im Score = 1.51
NG = 0
DB = 0
SM = 0
Delta_K
Doxorubicin
RF1_72_0 RF1
ZShift = 2.12
Im Score = −0.03
NG = 0
DB = 0
SM = 0
Delta_r
Doxorubicin
RF1_72_0 RF1
ZShift = 2.24
Im Score = −2.13
NG = 0
DB = 0
SM = 0
Delta_AUC
Doxorubicin

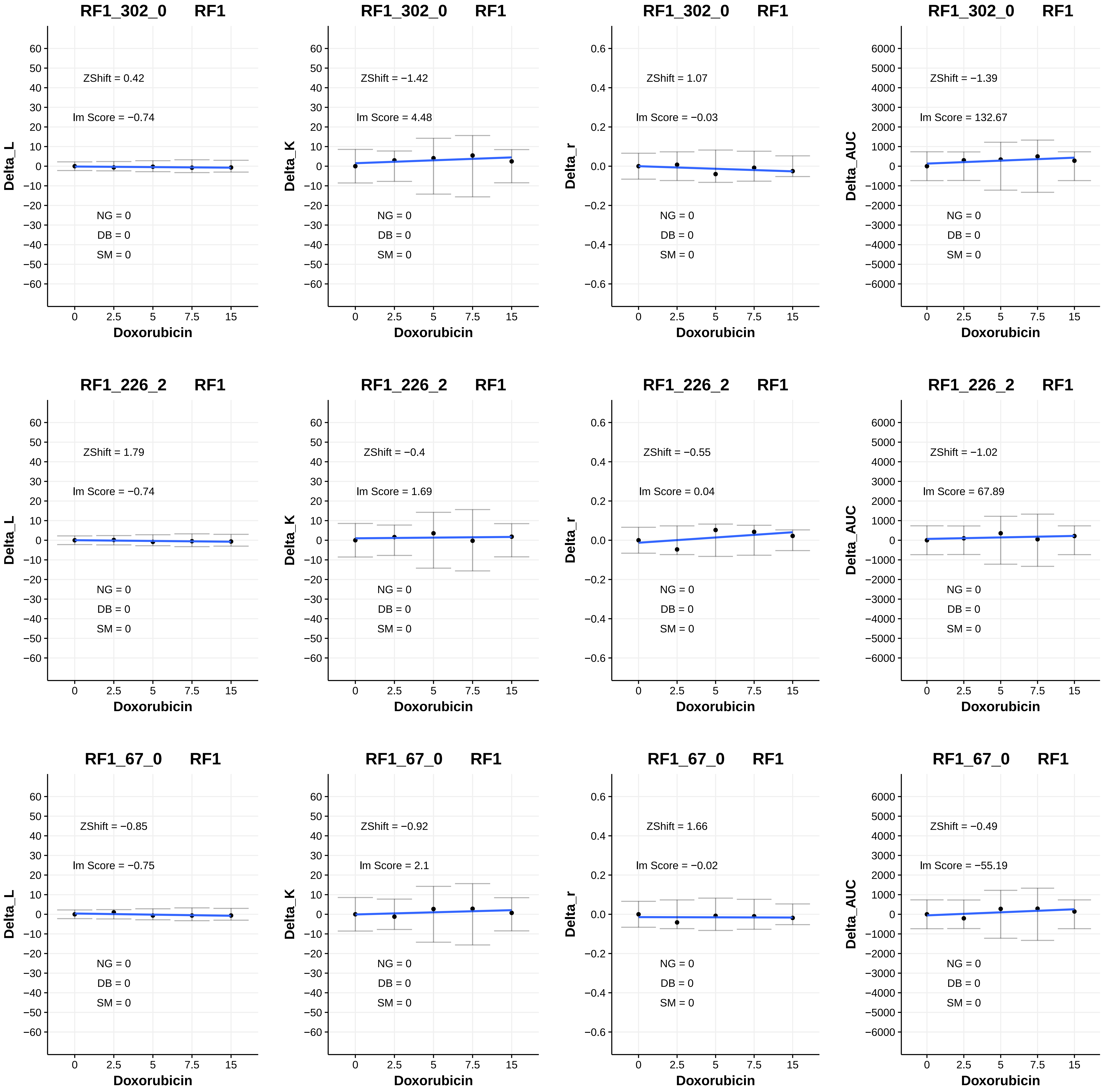
RF1_302_0 RF1
ZShift = 0.42
lm Score = −0.74
NG = 0
DB = 0
SM = 0
Delta_L
Doxorubicin
RF1_302_0 RF1
ZShift = −1.42
lm Score = 4.48
NG = 0
DB = 0
SM = 0
Delta_K
Doxorubicin
RF1_302_0 RF1
ZShift = 1.07
lm Score = −0.03
NG = 0
DB = 0
SM = 0
Delta_r
Doxorubicin
RF1_302_0 RF1
ZShift = −1.39
lm Score = 132.67
NG = 0
DB = 0
SM = 0
Delta_AUC
Doxorubicin
RF1_226_2 RF1
ZShift = 1.79
lm Score = −0.74
NG = 0
DB = 0
SM = 0
Delta_L
Doxorubicin
RF1_226_2 RF1
ZShift = −0.4
lm Score = 1.69
NG = 0
DB = 0
SM = 0
Delta_K
Doxorubicin
RF1_226_2 RF1
ZShift = −0.55
lm Score = 0.04
NG = 0
DB = 0
SM = 0
Delta_r
Doxorubicin
RF1_226_2 RF1
ZShift = −1.02
lm Score = 67.89
NG = 0
DB = 0
SM = 0
Delta_AUC
Doxorubicin
RF1_67_0 RF1
ZShift = −0.85
lm Score = −0.75
NG = 0
DB = 0
SM = 0
Delta_L
Doxorubicin
RF1_67_0 RF1
ZShift = −0.92
lm Score = 2.1
NG = 0
DB = 0
SM = 0
Delta_K
Doxorubicin
RF1_67_0 RF1
ZShift = 1.66
lm Score = −0.02
NG = 0
DB = 0
SM = 0
Delta_r
Doxorubicin
RF1_67_0 RF1
ZShift = −0.49
lm Score = −55.19
NG = 0
DB = 0
SM = 0
Delta_AUC
Doxorubicin

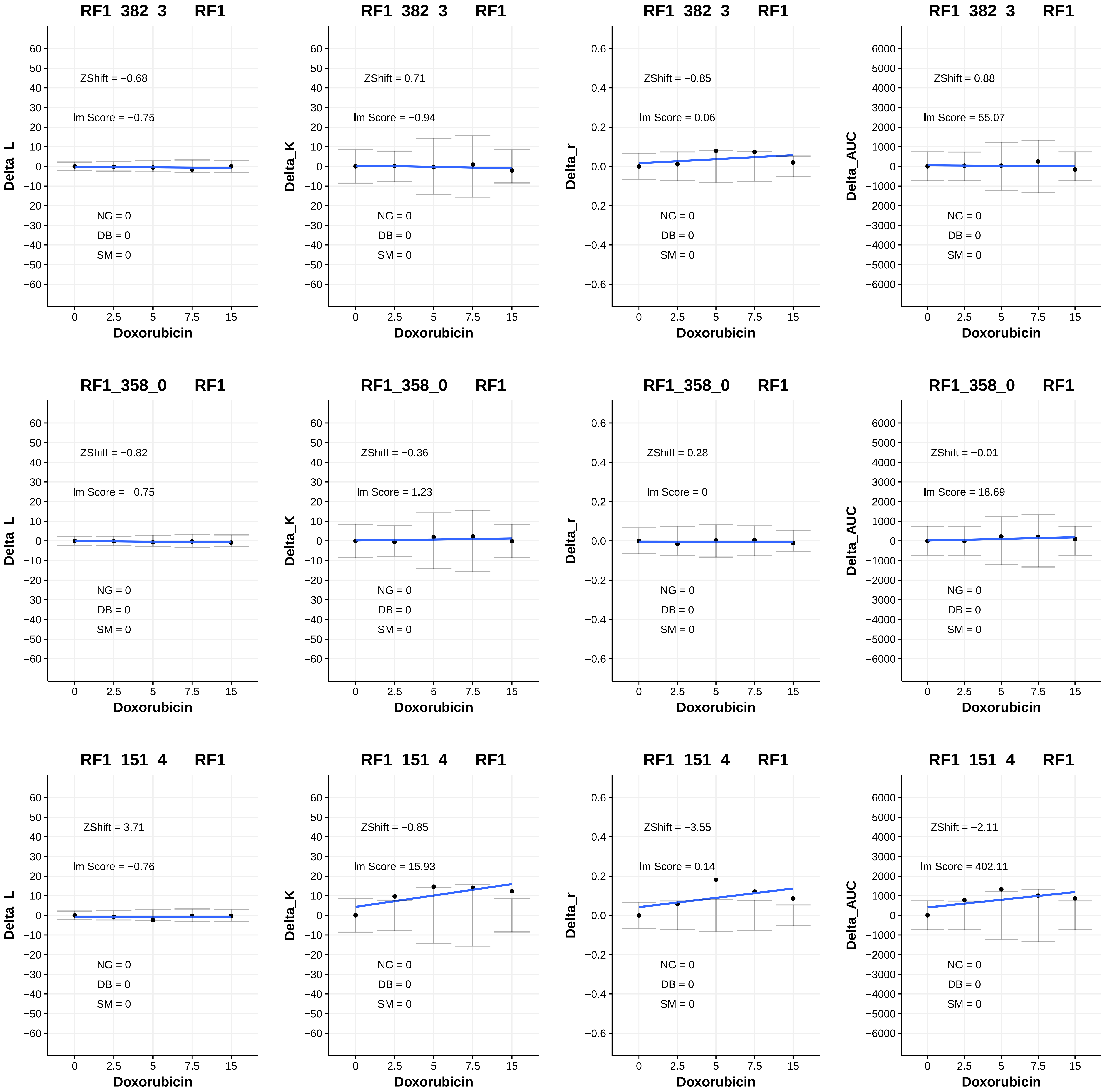
RF1_382_3 RF1
ZShift = −0.68
lm Score = −0.75
NG = 0
DB = 0
SM = 0
Delta_L
Doxorubicin
RF1_382_3 RF1
ZShift = 0.71
lm Score = −0.94
NG = 0
DB = 0
SM = 0
Delta_K
Doxorubicin
RF1_382_3 RF1
ZShift = −0.85
lm Score = 0.06
NG = 0
DB = 0
SM = 0
Delta_r
Doxorubicin
RF1_382_3 RF1
ZShift = 0.88
lm Score = 55.07
NG = 0
DB = 0
SM = 0
Delta_AUC
Doxorubicin
RF1_358_0 RF1
ZShift = −0.82
lm Score = −0.75
NG = 0
DB = 0
SM = 0
Delta_L
Doxorubicin
RF1_358_0 RF1
ZShift = −0.36
lm Score = 1.23
NG = 0
DB = 0
SM = 0
Delta_K
Doxorubicin
RF1_358_0 RF1
ZShift = 0.28
lm Score = 0
NG = 0
DB = 0
SM = 0
Delta_r
Doxorubicin
RF1_358_0 RF1
ZShift = −0.01
lm Score = 18.69
NG = 0
DB = 0
SM = 0
Delta_AUC
Doxorubicin
RF1_151_4 RF1
ZShift = 3.71
lm Score = −0.76
NG = 0
DB = 0
SM = 0
Delta_L
Doxorubicin
RF1_151_4 RF1
ZShift = −0.85
lm Score = 15.93
NG = 0
DB = 0
SM = 0
Delta_K
Doxorubicin
RF1_151_4 RF1
ZShift = −3.55
lm Score = 0.14
NG = 0
DB = 0
SM = 0
Delta_r
Doxorubicin
RF1_151_4 RF1
ZShift = −2.11
lm Score = 402.11
NG = 0
DB = 0
SM = 0
Delta_AUC
Doxorubicin

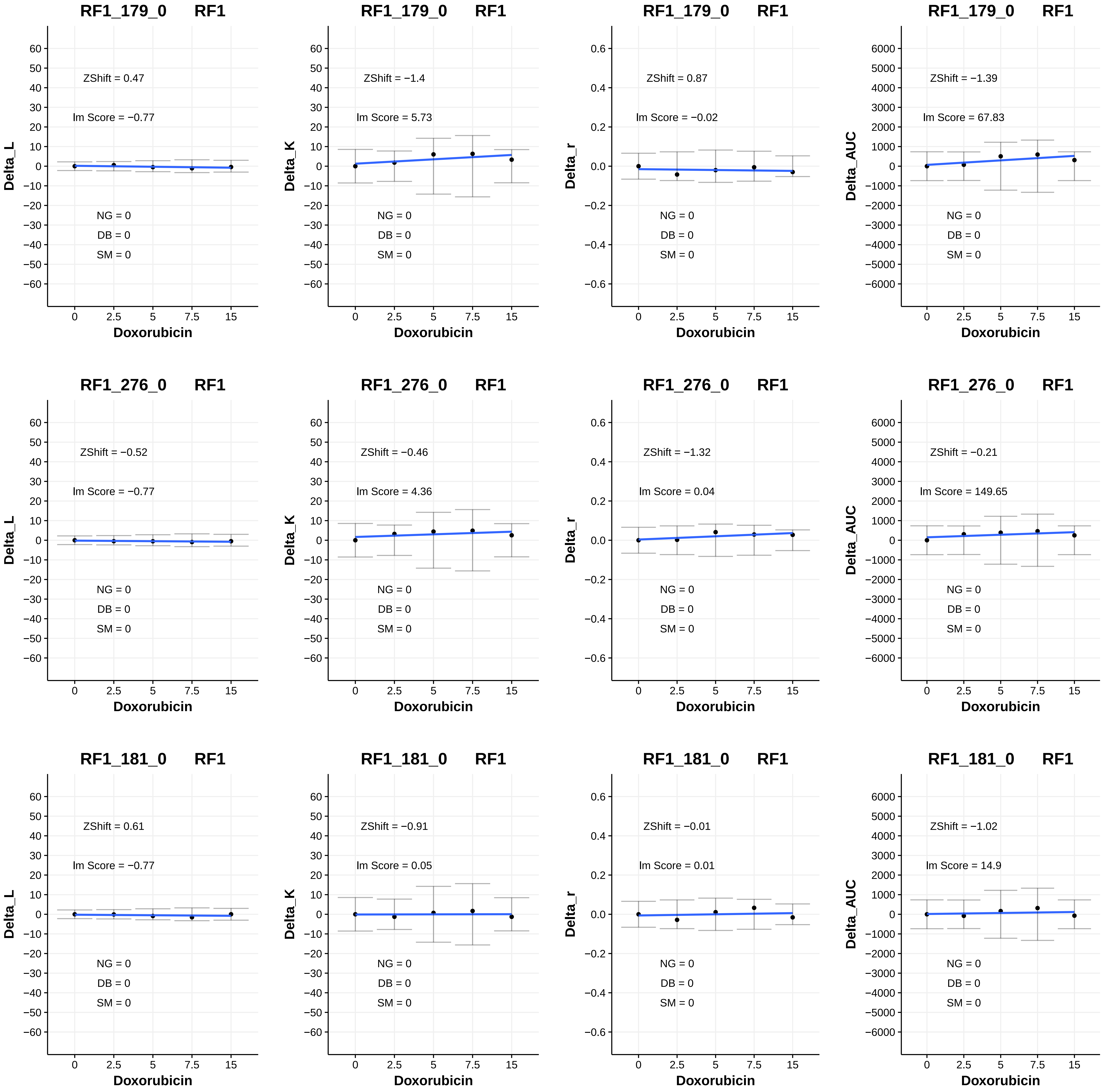
RF1_179_0 RF1
ZShift = 0.47
lm Score = −0.77
NG = 0
DB = 0
SM = 0
Delta_L
Doxorubicin
RF1_179_0 RF1
ZShift = −1.4
lm Score = 5.73
NG = 0
DB = 0
SM = 0
Delta_K
Doxorubicin
RF1_179_0 RF1
ZShift = 0.87
lm Score = −0.02
NG = 0
DB = 0
SM = 0
Delta_r
Doxorubicin
RF1_179_0 RF1
ZShift = −1.39
lm Score = 67.83
NG = 0
DB = 0
SM = 0
Delta_AUC
Doxorubicin
RF1_276_0 RF1
ZShift = −0.52
lm Score = −0.77
NG = 0
DB = 0
SM = 0
Delta_L
Doxorubicin
RF1_276_0 RF1
ZShift = −0.46
lm Score = 4.36
NG = 0
DB = 0
SM = 0
Delta_K
Doxorubicin
RF1_276_0 RF1
ZShift = −1.32
lm Score = 0.04
NG = 0
DB = 0
SM = 0
Delta_r
Doxorubicin
RF1_276_0 RF1
ZShift = −0.21
lm Score = 149.65
NG = 0
DB = 0
SM = 0
Delta_AUC
Doxorubicin
RF1_181_0 RF1
ZShift = 0.61
lm Score = −0.77
NG = 0
DB = 0
SM = 0
Delta_L
Doxorubicin
RF1_181_0 RF1
ZShift = −0.91
lm Score = 0.05
NG = 0
DB = 0
SM = 0
Delta_K
Doxorubicin
RF1_181_0 RF1
ZShift = −0.01
lm Score = 0.01
NG = 0
DB = 0
SM = 0
Delta_r
Doxorubicin
RF1_181_0 RF1
ZShift = −1.02
lm Score = 14.9
NG = 0
DB = 0
SM = 0
Delta_AUC
Doxorubicin

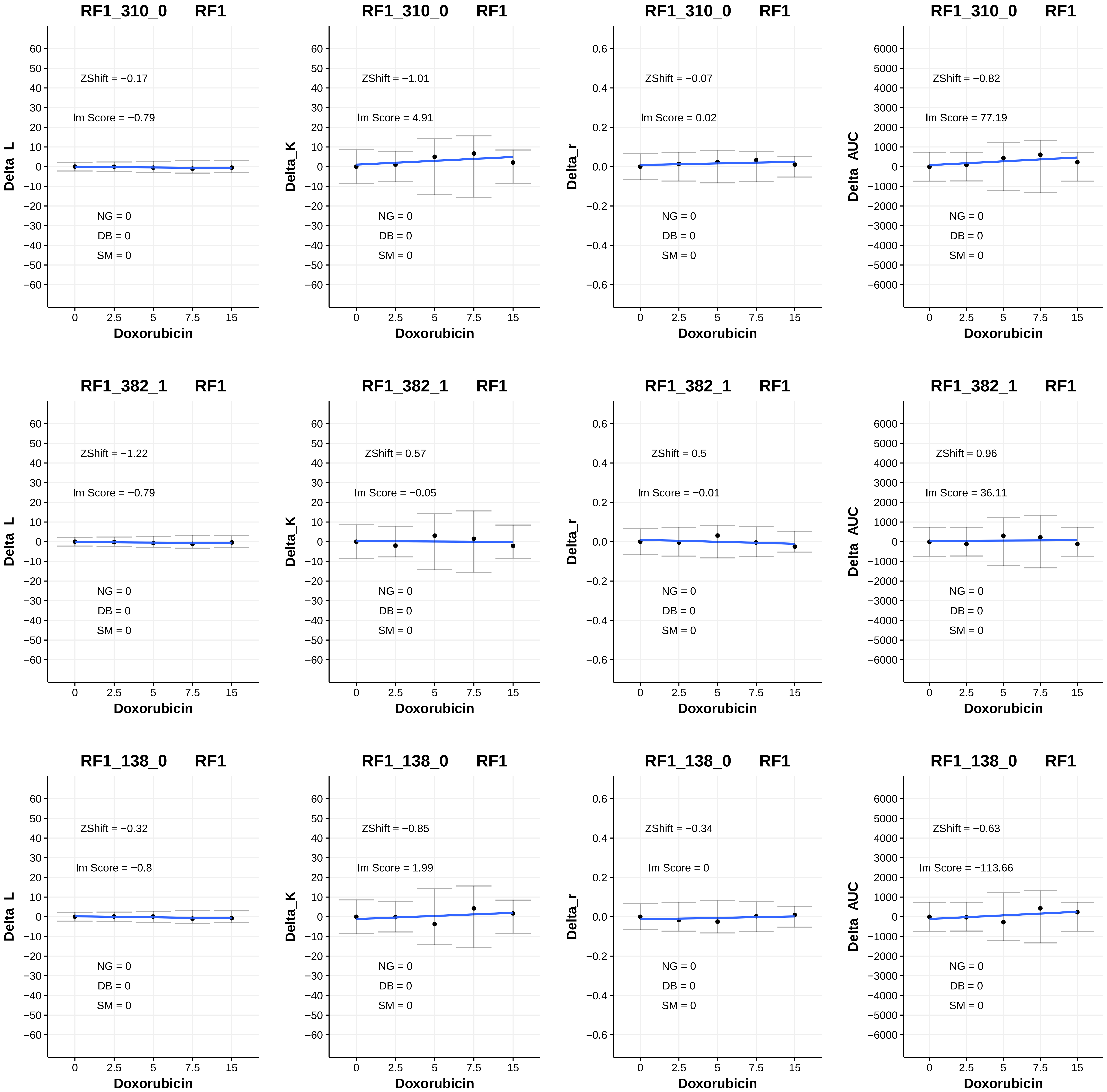
RF1_310_0 RF1
ZShift = −0.17
lm Score = −0.79
NG = 0
DB = 0
SM = 0
Delta_L
Doxorubicin
RF1_310_0 RF1
ZShift = −1.01
lm Score = 4.91
NG = 0
DB = 0
SM = 0
Delta_K
Doxorubicin
RF1_310_0 RF1
ZShift = −0.07
lm Score = 0.02
NG = 0
DB = 0
SM = 0
Delta_r
Doxorubicin
RF1_310_0 RF1
ZShift = −0.82
lm Score = 77.19
NG = 0
DB = 0
SM = 0
Delta_AUC
Doxorubicin
RF1_382_1 RF1
ZShift = −1.22
lm Score = −0.79
NG = 0
DB = 0
SM = 0
Delta_L
Doxorubicin
RF1_382_1 RF1
ZShift = 0.57
lm Score = −0.05
NG = 0
DB = 0
SM = 0
Delta_K
Doxorubicin
RF1_382_1 RF1
ZShift = 0.5
lm Score = −0.01
NG = 0
DB = 0
SM = 0
Delta_r
Doxorubicin
RF1_382_1 RF1
ZShift = 0.96
lm Score = 36.11
NG = 0
DB = 0
SM = 0
Delta_AUC
Doxorubicin
RF1_138_0 RF1
ZShift = −0.32
lm Score = −0.8
NG = 0
DB = 0
SM = 0
Delta_L
Doxorubicin
RF1_138_0 RF1
ZShift = −0.85
lm Score = 1.99
NG = 0
DB = 0
SM = 0
Delta_K
Doxorubicin
RF1_138_0 RF1
ZShift = −0.34
lm Score = 0
NG = 0
DB = 0
SM = 0
Delta_r
Doxorubicin
RF1_138_0 RF1
ZShift = −0.63
lm Score = −113.66
NG = 0
DB = 0
SM = 0
Delta_AUC
Doxorubicin

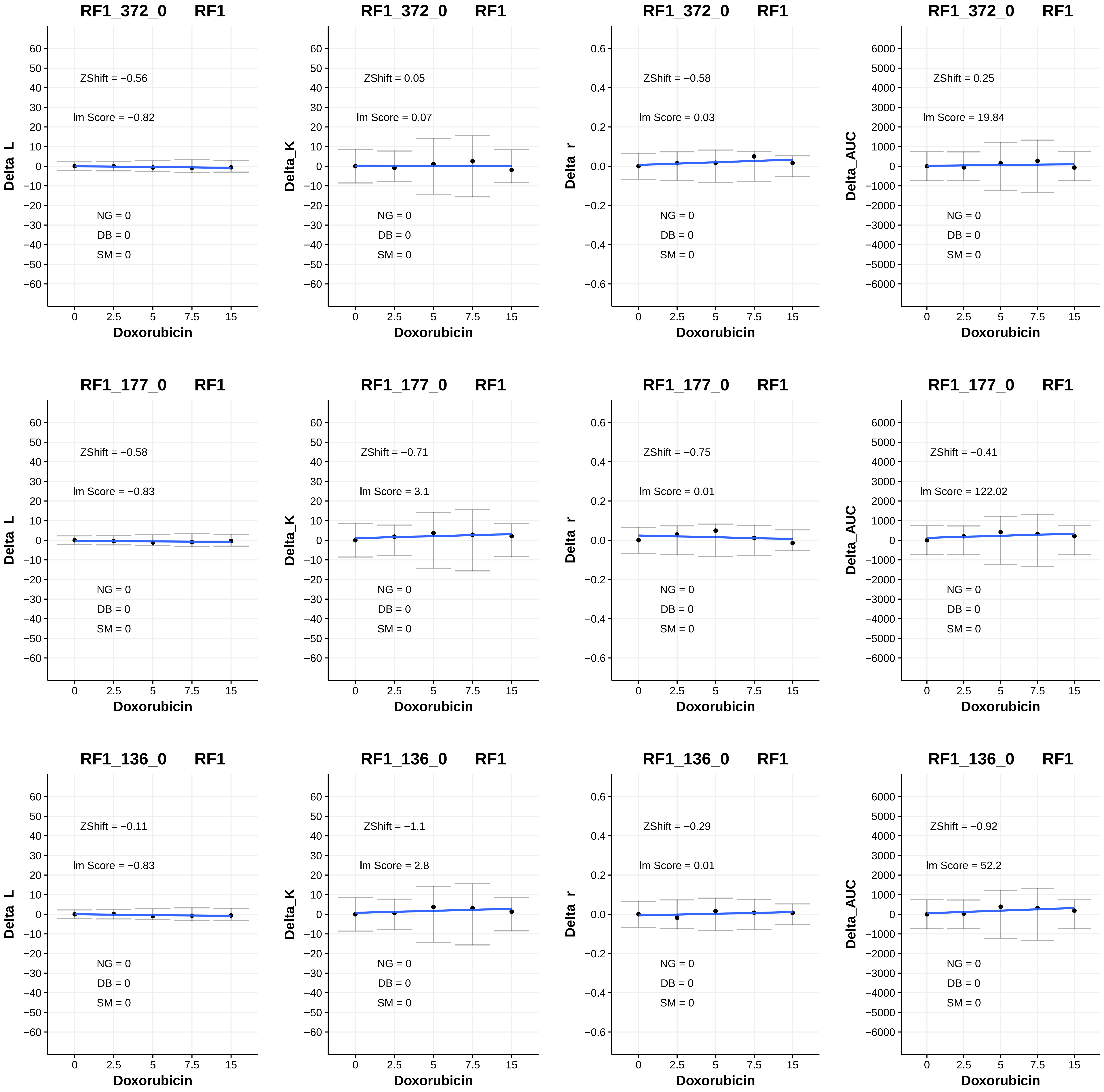

RF1_372_0 RF1
ZShift = −0.56
lm Score = −0.82
NG = 0
DB = 0
SM = 0
Delta_L
Doxorubicin
RF1_372_0 RF1
ZShift = 0.05
lm Score = 0.07
NG = 0
DB = 0
SM = 0
Delta_K
Doxorubicin
RF1_372_0 RF1
ZShift = −0.58
lm Score = 0.03
NG = 0
DB = 0
SM = 0
Delta_r
Doxorubicin
RF1_372_0 RF1
ZShift = 0.25
lm Score = 19.84
NG = 0
DB = 0
SM = 0
Delta_AUC
Doxorubicin
RF1_177_0 RF1
ZShift = −0.58
lm Score = −0.83
NG = 0
DB = 0
SM = 0
Delta_L
Doxorubicin
RF1_177_0 RF1
ZShift = −0.71
lm Score = 3.1
NG = 0
DB = 0
SM = 0
Delta_K
Doxorubicin
RF1_177_0 RF1
ZShift = −0.75
lm Score = 0.01
NG = 0
DB = 0
SM = 0
Delta_r
Doxorubicin
RF1_177_0 RF1
ZShift = −0.41
lm Score = 122.02
NG = 0
DB = 0
SM = 0
Delta_AUC
Doxorubicin
RF1_136_0 RF1
ZShift = −0.11
lm Score = −0.83
NG = 0
DB = 0
SM = 0
Delta_L
Doxorubicin
RF1_136_0 RF1
ZShift = −1.1
lm Score = 2.8
NG = 0
DB = 0
SM = 0
Delta_K
Doxorubicin
RF1_136_0 RF1
ZShift = −0.29
lm Score = 0.01
NG = 0
DB = 0
SM = 0
Delta_r
Doxorubicin
RF1_136_0 RF1
ZShift = −0.92
lm Score = 52.2
NG = 0
DB = 0
SM = 0
Delta_AUC
Doxorubicin

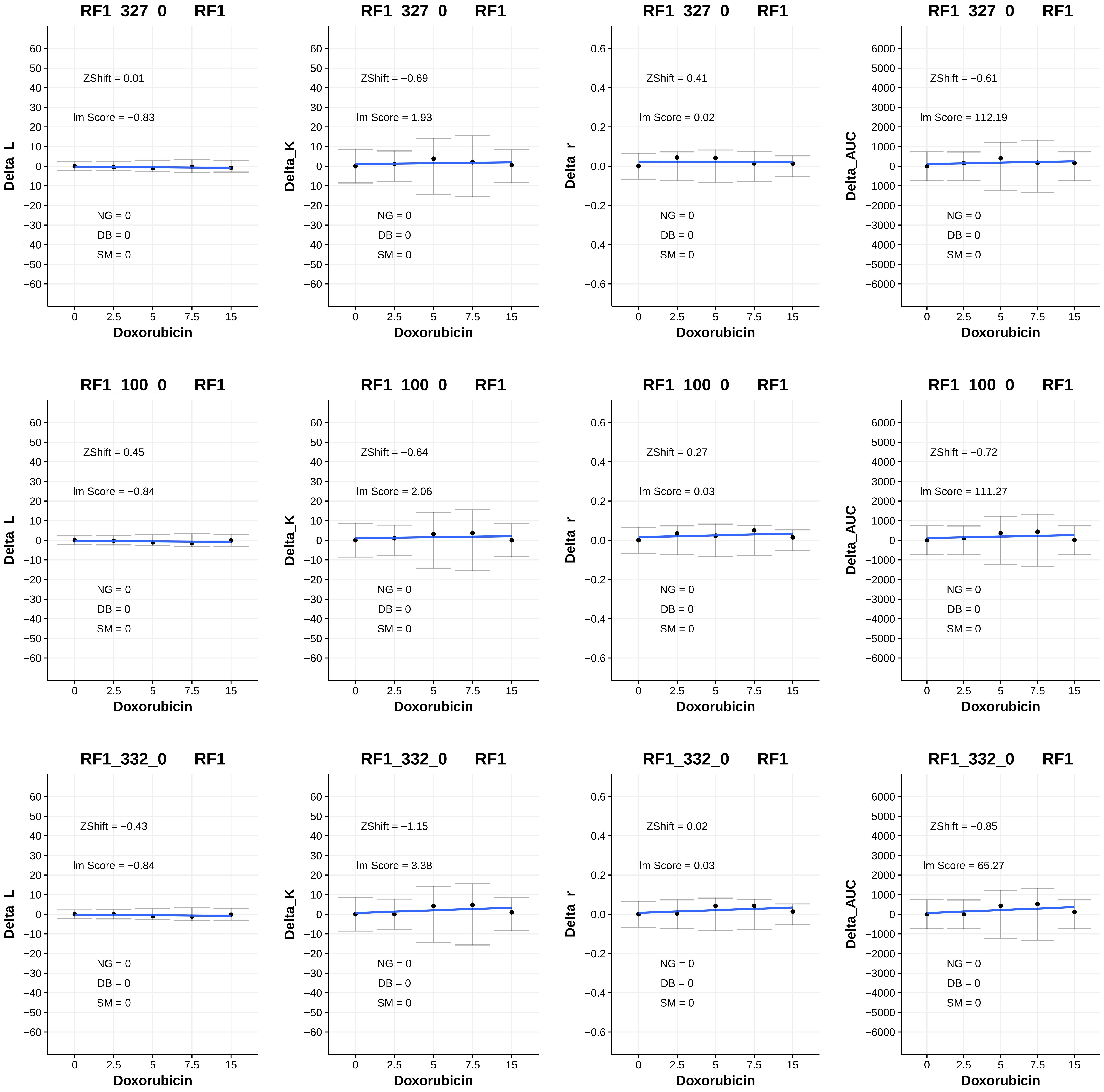
RF1_327_0 RF1
ZShift = 0.01
lm Score = −0.83
NG = 0
DB = 0
SM = 0
Delta_L
Doxorubicin
RF1_327_0 RF1
ZShift = −0.69
lm Score = 1.93
NG = 0
DB = 0
SM = 0
Delta_K
Doxorubicin
RF1_327_0 RF1
ZShift = 0.41
lm Score = 0.02
NG = 0
DB = 0
SM = 0
Delta_r
Doxorubicin
RF1_327_0 RF1
ZShift = −0.61
lm Score = 112.19
NG = 0
DB = 0
SM = 0
Delta_AUC
Doxorubicin
RF1_100_0 RF1
ZShift = 0.45
lm Score = −0.84
NG = 0
DB = 0
SM = 0
Delta_L
Doxorubicin
RF1_100_0 RF1
ZShift = −0.64
lm Score = 2.06
NG = 0
DB = 0
SM = 0
Delta_K
Doxorubicin
RF1_100_0 RF1
ZShift = 0.27
lm Score = 0.03
NG = 0
DB = 0
SM = 0
Delta_r
Doxorubicin
RF1_100_0 RF1
ZShift = −0.72
lm Score = 111.27
NG = 0
DB = 0
SM = 0
Delta_AUC
Doxorubicin
RF1_332_0 RF1
ZShift = −0.43
lm Score = −0.84
NG = 0
DB = 0
SM = 0
Delta_L
Doxorubicin
RF1_332_0 RF1
ZShift = −1.15
lm Score = 3.38
NG = 0
DB = 0
SM = 0
Delta_K
Doxorubicin
RF1_332_0 RF1
ZShift = 0.02
lm Score = 0.03
NG = 0
DB = 0
SM = 0
Delta_r
Doxorubicin
RF1_332_0 RF1
ZShift = −0.85
lm Score = 65.27
NG = 0
DB = 0
SM = 0
Delta_AUC
Doxorubicin

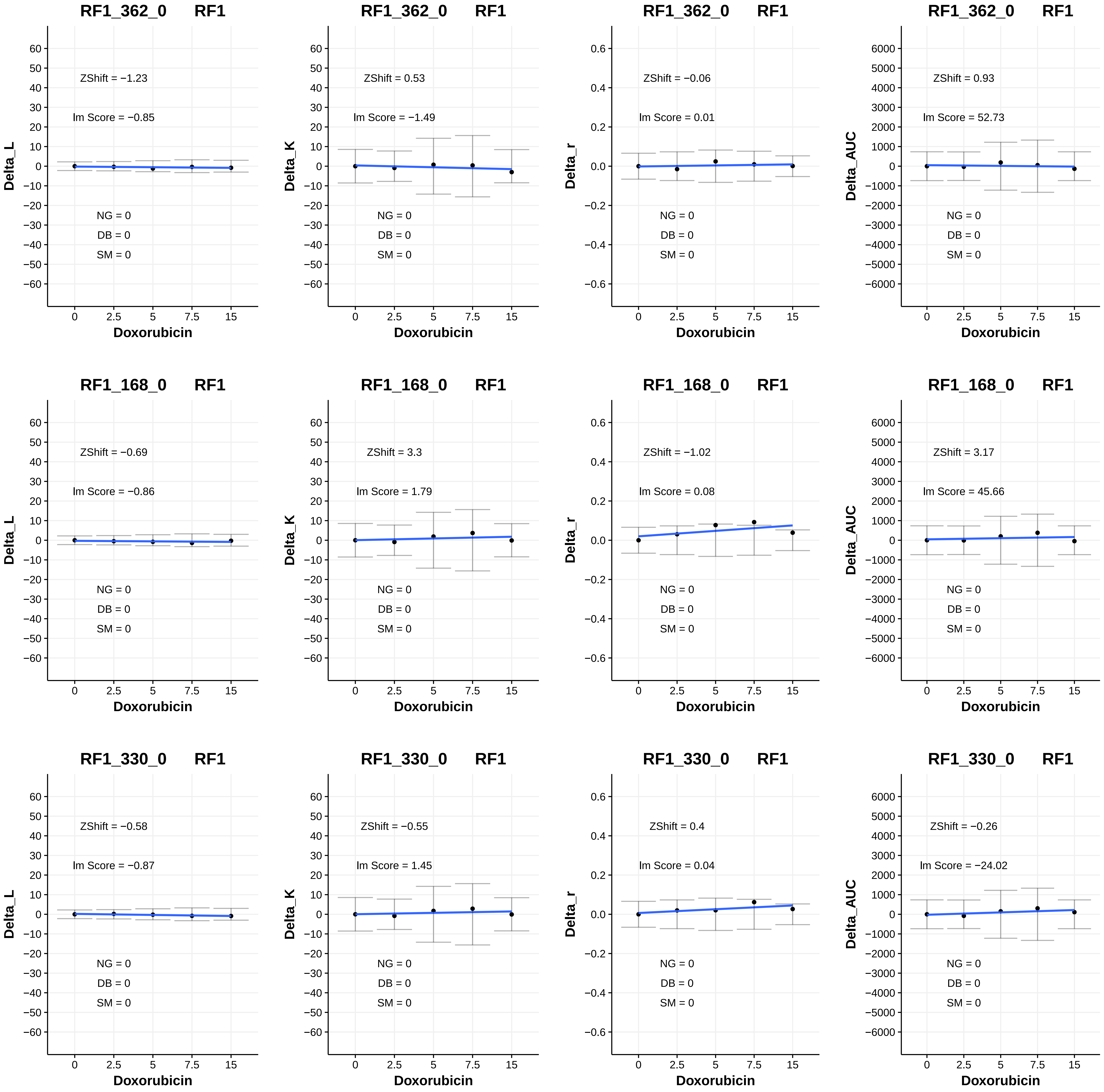
RF1_362_0 RF1
ZShift = −1.23
lm Score = −0.85
NG = 0
DB = 0
SM = 0
Delta_L
Doxorubicin
RF1_362_0 RF1
ZShift = 0.53
lm Score = −1.49
NG = 0
DB = 0
SM = 0
Delta_K
Doxorubicin
RF1_362_0 RF1
ZShift = −0.06
lm Score = 0.01
NG = 0
DB = 0
SM = 0
Delta_r
Doxorubicin
RF1_362_0 RF1
ZShift = 0.93
lm Score = 52.73
NG = 0
DB = 0
SM = 0
Delta_AUC
Doxorubicin
RF1_168_0 RF1
ZShift = −0.69
lm Score = −0.86
NG = 0
DB = 0
SM = 0
Delta_L
Doxorubicin
RF1_168_0 RF1
ZShift = 3.3
lm Score = 1.79
NG = 0
DB = 0
SM = 0
Delta_K
Doxorubicin
RF1_168_0 RF1
ZShift = −1.02
lm Score = 0.08
NG = 0
DB = 0
SM = 0
Delta_r
Doxorubicin
RF1_168_0 RF1
ZShift = 3.17
lm Score = 45.66
NG = 0
DB = 0
SM = 0
Delta_AUC
Doxorubicin
RF1_330_0 RF1
ZShift = −0.58
lm Score = −0.87
NG = 0
DB = 0
SM = 0
Delta_L
Doxorubicin
RF1_330_0 RF1
ZShift = −0.55
lm Score = 1.45
NG = 0
DB = 0
SM = 0
Delta_K
Doxorubicin
RF1_330_0 RF1
ZShift = 0.4
lm Score = 0.04
NG = 0
DB = 0
SM = 0
Delta_r
Doxorubicin
RF1_330_0 RF1
ZShift = −0.26
lm Score = −24.02
NG = 0
DB = 0
SM = 0
Delta_AUC
Doxorubicin

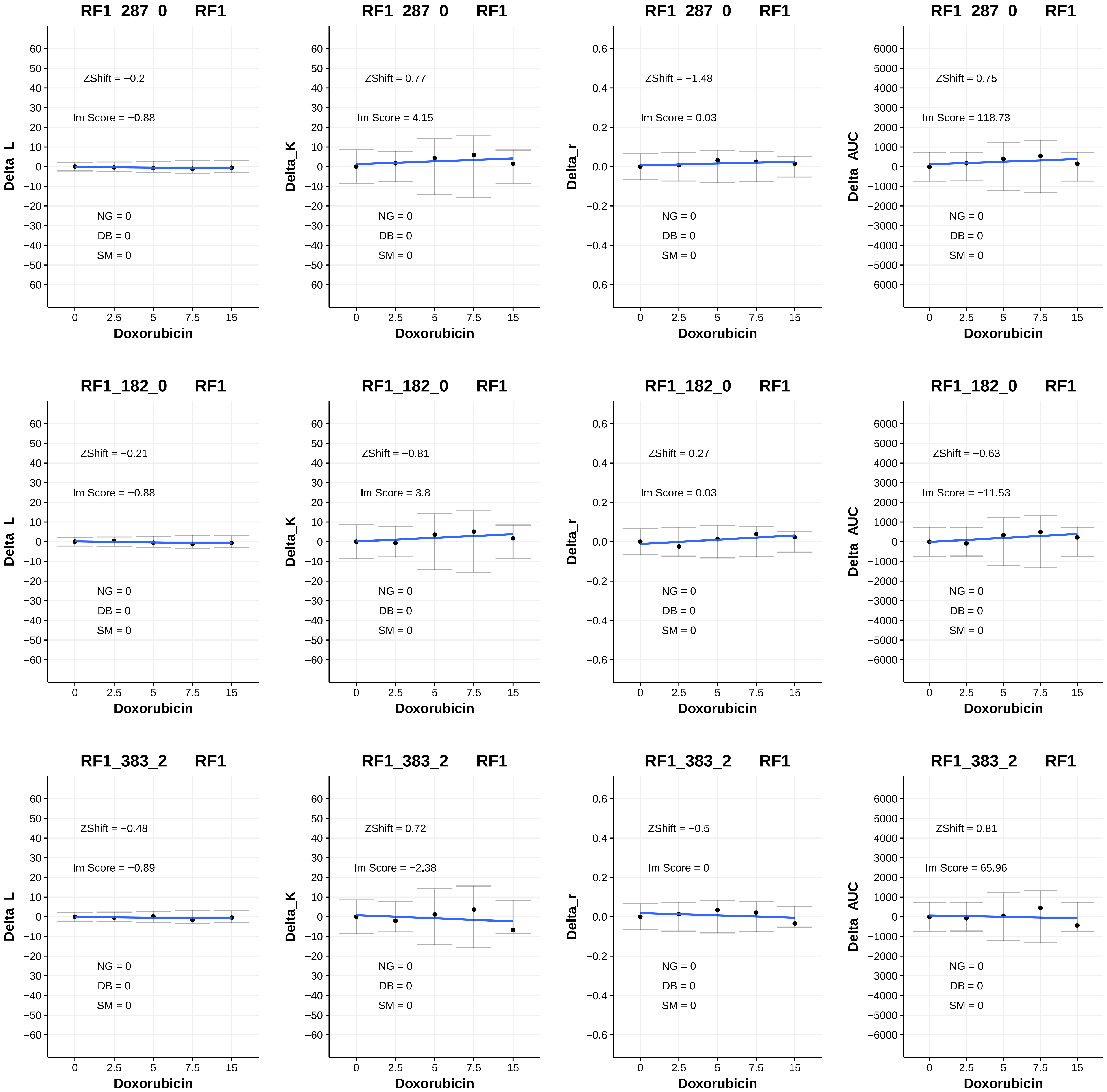
RF1_287_0 RF1
ZShift = −0.2
lm Score = −0.88
NG = 0
DB = 0
SM = 0
Delta_L
Doxorubicin
RF1_287_0 RF1
ZShift = 0.77
lm Score = 4.15
NG = 0
DB = 0
SM = 0
Delta_K
Doxorubicin
RF1_287_0 RF1
ZShift = −1.48
lm Score = 0.03
NG = 0
DB = 0
SM = 0
Delta_r
Doxorubicin
RF1_287_0 RF1
ZShift = 0.75
lm Score = 118.73
NG = 0
DB = 0
SM = 0
Delta_AUC
Doxorubicin
RF1_182_0 RF1
ZShift = −0.21
lm Score = −0.88
NG = 0
DB = 0
SM = 0
Delta_L
Doxorubicin
RF1_182_0 RF1
ZShift = −0.81
lm Score = 3.8
NG = 0
DB = 0
SM = 0
Delta_K
Doxorubicin
RF1_182_0 RF1
ZShift = 0.27
lm Score = 0.03
NG = 0
DB = 0
SM = 0
Delta_r
Doxorubicin
RF1_182_0 RF1
ZShift = −0.63
lm Score = −11.53
NG = 0
DB = 0
SM = 0
Delta_AUC
Doxorubicin
RF1_383_2 RF1
ZShift = −0.48
lm Score = −0.89
NG = 0
DB = 0
SM = 0
Delta_L
Doxorubicin
RF1_383_2 RF1
ZShift = 0.72
lm Score = −2.38
NG = 0
DB = 0
SM = 0
Delta_K
Doxorubicin
RF1_383_2 RF1
ZShift = −0.5
lm Score = 0
NG = 0
DB = 0
SM = 0
Delta_r
Doxorubicin
RF1_383_2 RF1
ZShift = 0.81
lm Score = 65.96
NG = 0
DB = 0
SM = 0
Delta_AUC
Doxorubicin

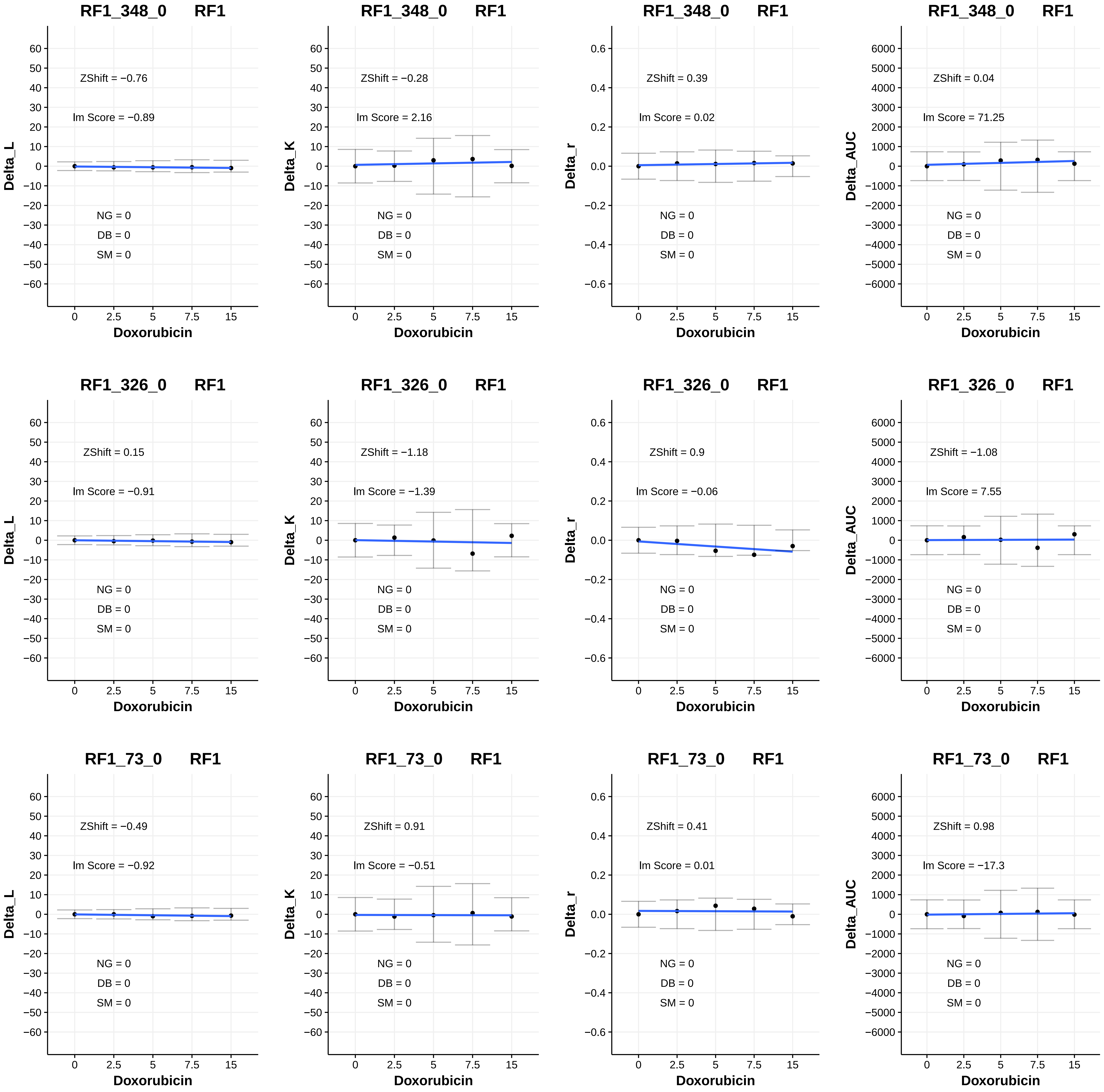

RF1_348_0 RF1
Delta_L
ZShift = −0.76
Im Score = −0.89
NG = 0
DB = 0
SM = 0
Doxorubicin
RF1_348_0 RF1
Delta_K
ZShift = −0.28
Im Score = 2.16
NG = 0
DB = 0
SM = 0
Doxorubicin
RF1_348_0 RF1
Delta_r
ZShift = 0.39
Im Score = 0.02
NG = 0
DB = 0
SM = 0
Doxorubicin
RF1_348_0 RF1
Delta_AUC
ZShift = 0.04
Im Score = 71.25
NG = 0
DB = 0
SM = 0
Doxorubicin
RF1_326_0 RF1
Delta_L
ZShift = 0.15
Im Score = −0.91
NG = 0
DB = 0
SM = 0
Doxorubicin
RF1_326_0 RF1
Delta_K
ZShift = −1.18
Im Score = −1.39
NG = 0
DB = 0
SM = 0
Doxorubicin
RF1_326_0 RF1
Delta_r
ZShift = 0.9
Im Score = −0.06
NG = 0
DB = 0
SM = 0
Doxorubicin
RF1_326_0 RF1
Delta_AUC
ZShift = −1.08
Im Score = 7.55
NG = 0
DB = 0
SM = 0
Doxorubicin
RF1_73_0 RF1
Delta_L
ZShift = −0.49
Im Score = −0.92
NG = 0
DB = 0
SM = 0
Doxorubicin
RF1_73_0 RF1
Delta_K
ZShift = 0.91
Im Score = −0.51
NG = 0
DB = 0
SM = 0
Doxorubicin
RF1_73_0 RF1
Delta_r
ZShift = 0.41
Im Score = 0.01
NG = 0
DB = 0
SM = 0
Doxorubicin
RF1_73_0 RF1
Delta_AUC
ZShift = 0.98
Im Score = −17.3
NG = 0
DB = 0
SM = 0
Doxorubicin

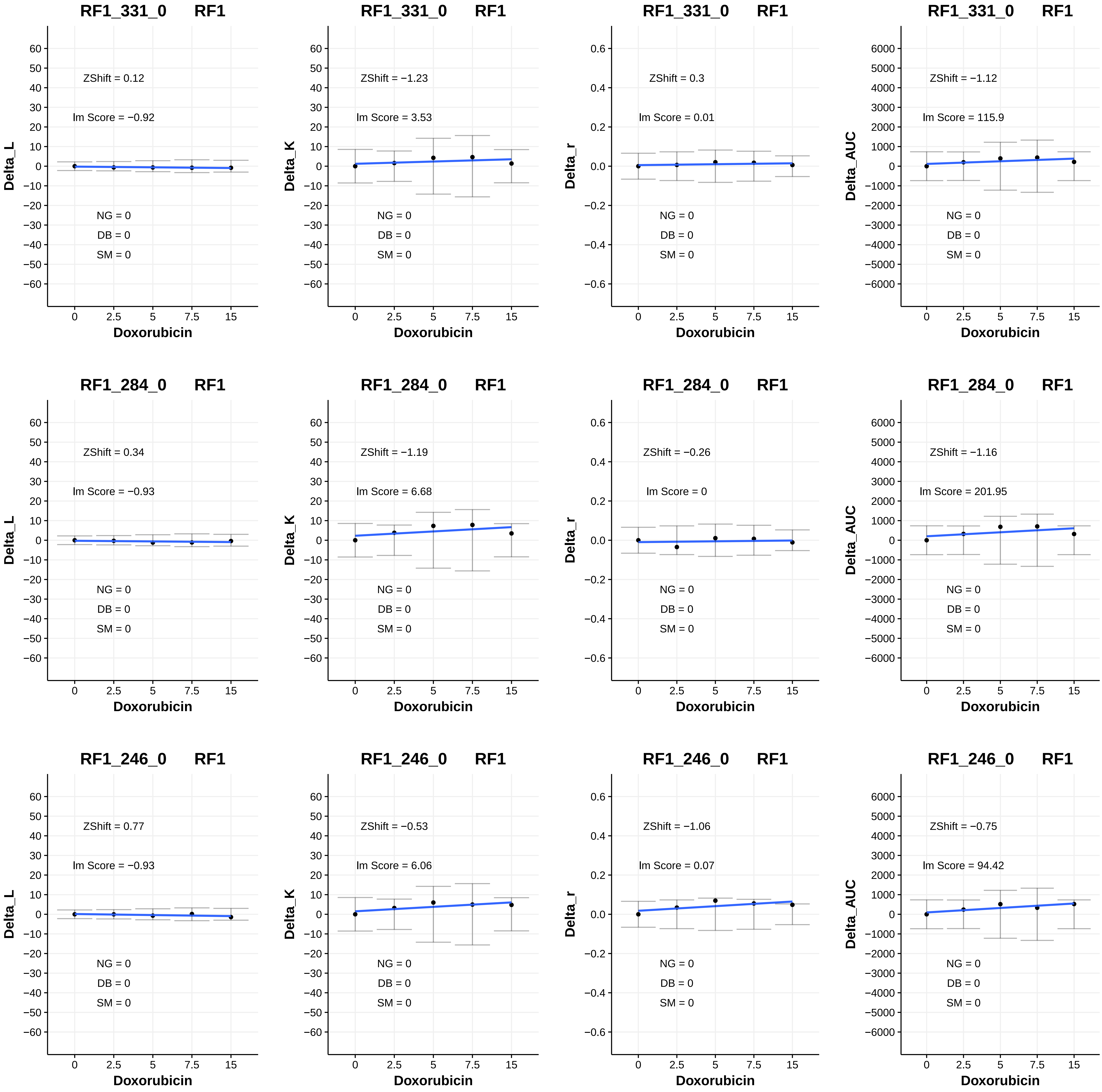
RF1_331_0 RF1
ZShift = 0.12
lm Score = −0.92
NG = 0
DB = 0
SM = 0
Delta_L
Doxorubicin
RF1_331_0 RF1
ZShift = −1.23
lm Score = 3.53
NG = 0
DB = 0
SM = 0
Delta_K
Doxorubicin
RF1_331_0 RF1
ZShift = 0.3
lm Score = 0.01
NG = 0
DB = 0
SM = 0
Delta_r
Doxorubicin
RF1_331_0 RF1
ZShift = −1.12
lm Score = 115.9
NG = 0
DB = 0
SM = 0
Delta_AUC
Doxorubicin
RF1_284_0 RF1
ZShift = 0.34
lm Score = −0.93
NG = 0
DB = 0
SM = 0
Delta_L
Doxorubicin
RF1_284_0 RF1
ZShift = −1.19
lm Score = 6.68
NG = 0
DB = 0
SM = 0
Delta_K
Doxorubicin
RF1_284_0 RF1
ZShift = −0.26
lm Score = 0
NG = 0
DB = 0
SM = 0
Delta_r
Doxorubicin
RF1_284_0 RF1
ZShift = −1.16
lm Score = 201.95
NG = 0
DB = 0
SM = 0
Delta_AUC
Doxorubicin
RF1_246_0 RF1
ZShift = 0.77
lm Score = −0.93
NG = 0
DB = 0
SM = 0
Delta_L
Doxorubicin
RF1_246_0 RF1
ZShift = −0.53
lm Score = 6.06
NG = 0
DB = 0
SM = 0
Delta_K
Doxorubicin
RF1_246_0 RF1
ZShift = −1.06
lm Score = 0.07
NG = 0
DB = 0
SM = 0
Delta_r
Doxorubicin
RF1_246_0 RF1
ZShift = −0.75
lm Score = 94.42
NG = 0
DB = 0
SM = 0
Delta_AUC
Doxorubicin

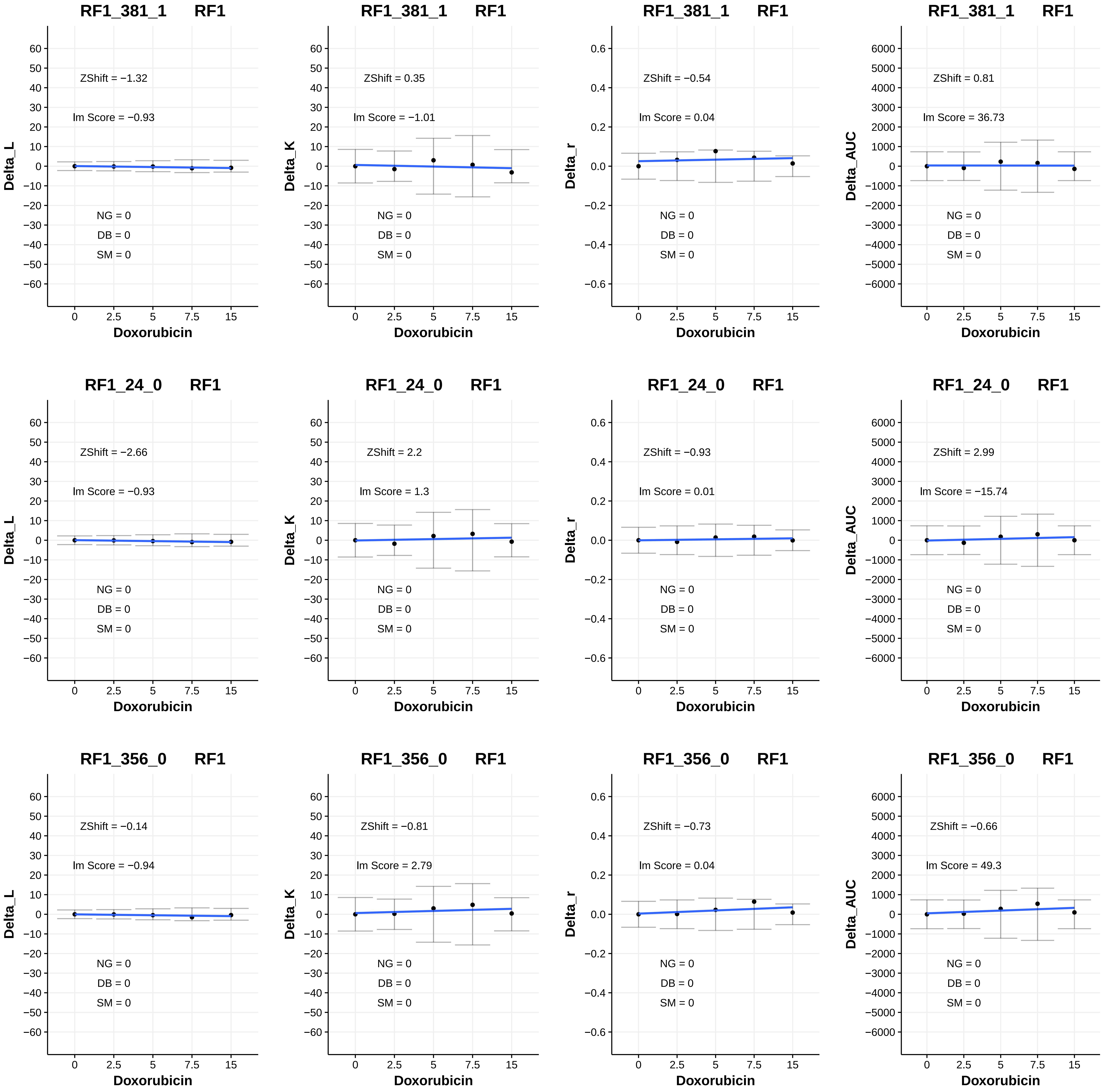
RF1_381_1 RF1
ZShift = −1.32
lm Score = −0.93
NG = 0
DB = 0
SM = 0
Delta_L
Doxorubicin
RF1_381_1 RF1
ZShift = 0.35
lm Score = −1.01
NG = 0
DB = 0
SM = 0
Delta_K
Doxorubicin
RF1_381_1 RF1
ZShift = −0.54
lm Score = 0.04
NG = 0
DB = 0
SM = 0
Delta_r
Doxorubicin
RF1_381_1 RF1
ZShift = 0.81
lm Score = 36.73
NG = 0
DB = 0
SM = 0
Delta_AUC
Doxorubicin
RF1_24_0 RF1
ZShift = −2.66
lm Score = −0.93
NG = 0
DB = 0
SM = 0
Delta_L
Doxorubicin
RF1_24_0 RF1
ZShift = 2.2
lm Score = 1.3
NG = 0
DB = 0
SM = 0
Delta_K
Doxorubicin
RF1_24_0 RF1
ZShift = −0.93
lm Score = 0.01
NG = 0
DB = 0
SM = 0
Delta_r
Doxorubicin
RF1_24_0 RF1
ZShift = 2.99
lm Score = −15.74
NG = 0
DB = 0
SM = 0
Delta_AUC
Doxorubicin
RF1_356_0 RF1
ZShift = −0.14
lm Score = −0.94
NG = 0
DB = 0
SM = 0
Delta_L
Doxorubicin
RF1_356_0 RF1
ZShift = −0.81
lm Score = 2.79
NG = 0
DB = 0
SM = 0
Delta_K
Doxorubicin
RF1_356_0 RF1
ZShift = −0.73
lm Score = 0.04
NG = 0
DB = 0
SM = 0
Delta_r
Doxorubicin
RF1_356_0 RF1
ZShift = −0.66
lm Score = 49.3
NG = 0
DB = 0
SM = 0
Delta_AUC
Doxorubicin

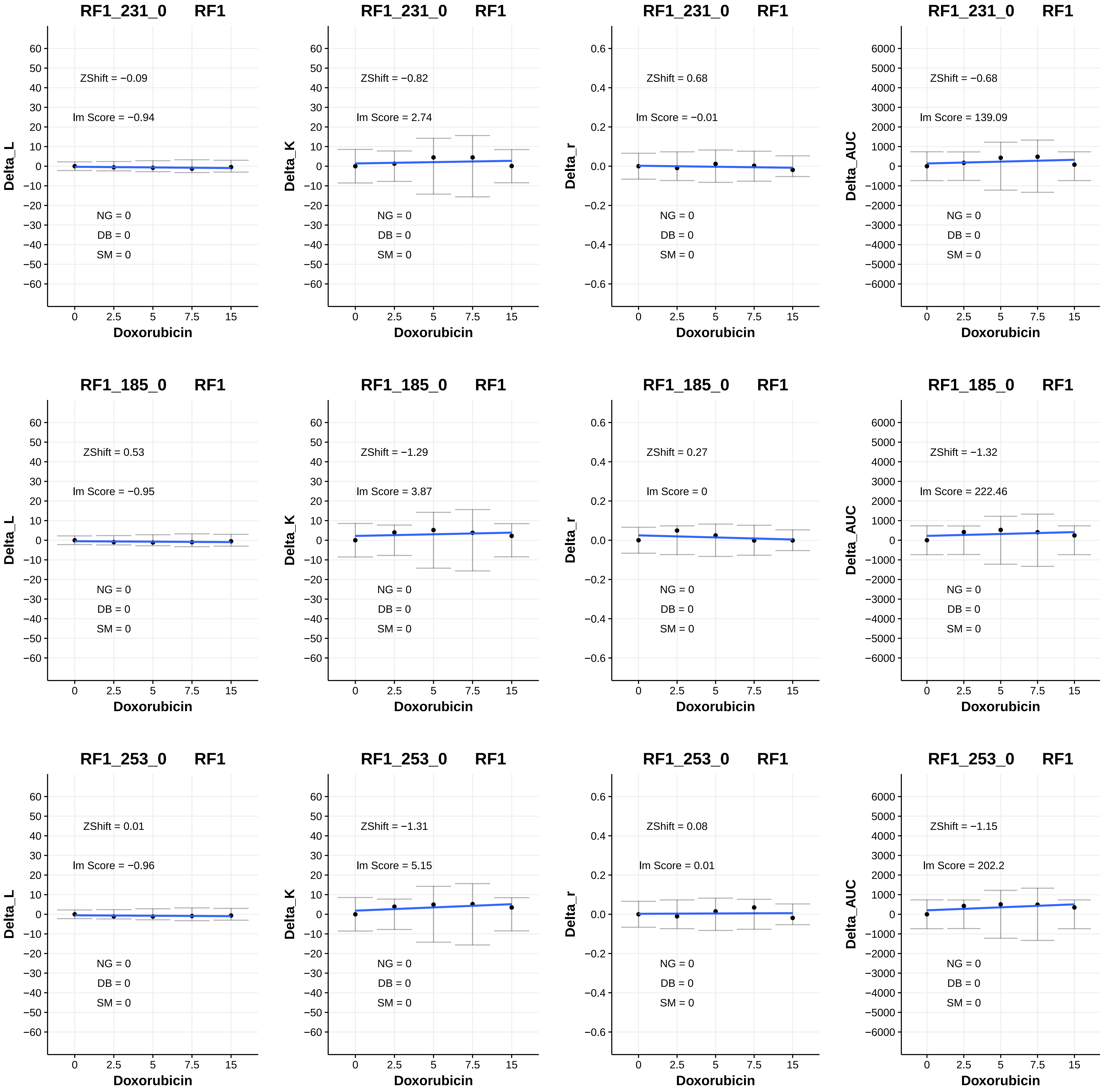
RF1_231_0 RF1
ZShift = −0.09
Im Score = −0.94
NG = 0
DB = 0
SM = 0
Delta_L
Doxorubicin
RF1_231_0 RF1
ZShift = −0.82
Im Score = 2.74
NG = 0
DB = 0
SM = 0
Delta_K
Doxorubicin
RF1_231_0 RF1
ZShift = 0.68
Im Score = −0.01
NG = 0
DB = 0
SM = 0
Delta_r
Doxorubicin
RF1_231_0 RF1
ZShift = −0.68
Im Score = 139.09
NG = 0
DB = 0
SM = 0
Delta_AUC
Doxorubicin
RF1_185_0 RF1
ZShift = 0.53
Im Score = −0.95
NG = 0
DB = 0
SM = 0
Delta_L
Doxorubicin
RF1_185_0 RF1
ZShift = −1.29
Im Score = 3.87
NG = 0
DB = 0
SM = 0
Delta_K
Doxorubicin
RF1_185_0 RF1
ZShift = 0.27
Im Score = 0
NG = 0
DB = 0
SM = 0
Delta_r
Doxorubicin
RF1_185_0 RF1
ZShift = −1.32
Im Score = 222.46
NG = 0
DB = 0
SM = 0
Delta_AUC
Doxorubicin
RF1_253_0 RF1
ZShift = 0.01
Im Score = −0.96
NG = 0
DB = 0
SM = 0
Delta_L
Doxorubicin
RF1_253_0 RF1
ZShift = −1.31
Im Score = 5.15
NG = 0
DB = 0
SM = 0
Delta_K
Doxorubicin
RF1_253_0 RF1
ZShift = 0.08
Im Score = 0.01
NG = 0
DB = 0
SM = 0
Delta_r
Doxorubicin
RF1_253_0 RF1
ZShift = −1.15
Im Score = 202.2
NG = 0
DB = 0
SM = 0
Delta_AUC
Doxorubicin

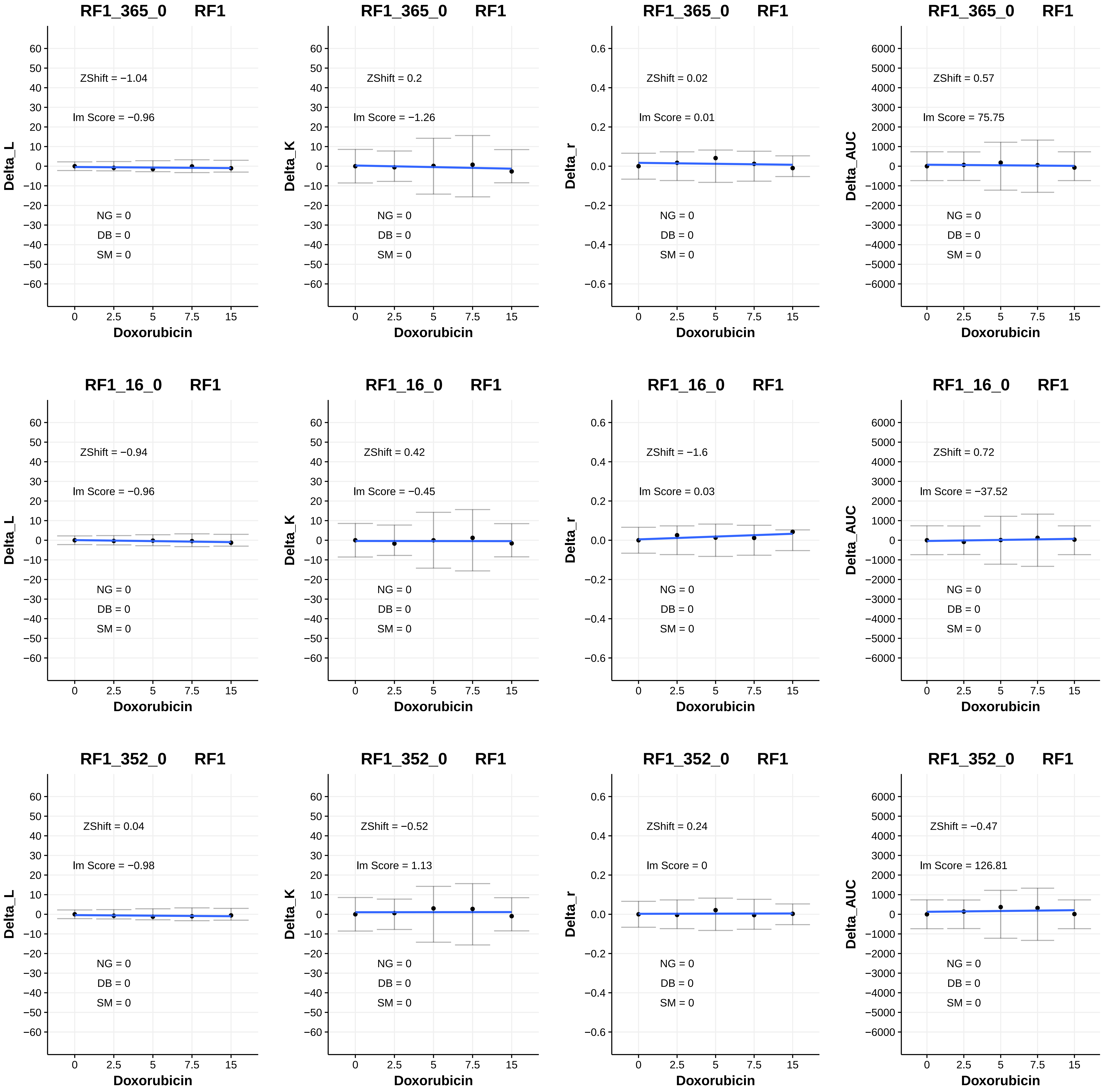
RF1_365_0 RF1
ZShift = −1.04
lm Score = −0.96
NG = 0
DB = 0
SM = 0
Delta_L
Doxorubicin
RF1_365_0 RF1
ZShift = 0.2
lm Score = −1.26
NG = 0
DB = 0
SM = 0
Delta_K
Doxorubicin
RF1_365_0 RF1
ZShift = 0.02
lm Score = 0.01
NG = 0
DB = 0
SM = 0
Delta_r
Doxorubicin
RF1_365_0 RF1
ZShift = 0.57
lm Score = 75.75
NG = 0
DB = 0
SM = 0
Delta_AUC
Doxorubicin
RF1_16_0 RF1
ZShift = −0.94
lm Score = −0.96
NG = 0
DB = 0
SM = 0
Delta_L
Doxorubicin
RF1_16_0 RF1
ZShift = 0.42
lm Score = −0.45
NG = 0
DB = 0
SM = 0
Delta_K
Doxorubicin
RF1_16_0 RF1
ZShift = −1.6
lm Score = 0.03
NG = 0
DB = 0
SM = 0
Delta_r
Doxorubicin
RF1_16_0 RF1
ZShift = 0.72
lm Score = −37.52
NG = 0
DB = 0
SM = 0
Delta_AUC
Doxorubicin
RF1_352_0 RF1
ZShift = 0.04
lm Score = −0.98
NG = 0
DB = 0
SM = 0
Delta_L
Doxorubicin
RF1_352_0 RF1
ZShift = −0.52
lm Score = 1.13
NG = 0
DB = 0
SM = 0
Delta_K
Doxorubicin
RF1_352_0 RF1
ZShift = 0.24
lm Score = 0
NG = 0
DB = 0
SM = 0
Delta_r
Doxorubicin
RF1_352_0 RF1
ZShift = −0.47
lm Score = 126.81
NG = 0
DB = 0
SM = 0
Delta_AUC
Doxorubicin

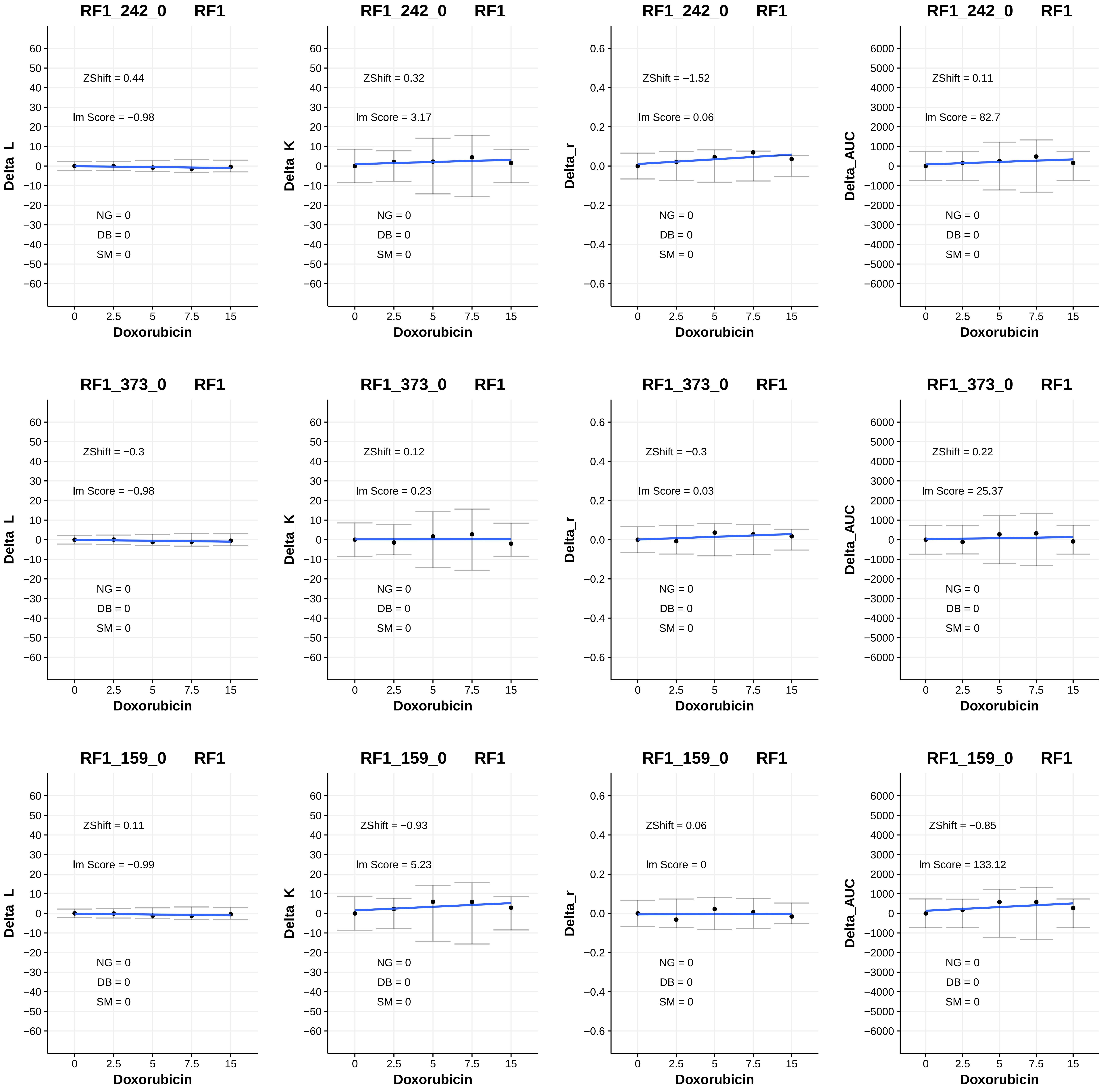
RF1_242_0 RF1
ZShift = 0.44
lm Score = −0.98
NG = 0
DB = 0
SM = 0
Delta_L
Doxorubicin
RF1_242_0 RF1
ZShift = 0.32
lm Score = 3.17
NG = 0
DB = 0
SM = 0
Delta_K
Doxorubicin
RF1_242_0 RF1
ZShift = −1.52
lm Score = 0.06
NG = 0
DB = 0
SM = 0
Delta_r
Doxorubicin
RF1_242_0 RF1
ZShift = 0.11
lm Score = 82.7
NG = 0
DB = 0
SM = 0
Delta_AUC
Doxorubicin
RF1_373_0 RF1
ZShift = −0.3
lm Score = −0.98
NG = 0
DB = 0
SM = 0
Delta_L
Doxorubicin
RF1_373_0 RF1
ZShift = 0.12
lm Score = 0.23
NG = 0
DB = 0
SM = 0
Delta_K
Doxorubicin
RF1_373_0 RF1
ZShift = −0.3
lm Score = 0.03
NG = 0
DB = 0
SM = 0
Delta_r
Doxorubicin
RF1_373_0 RF1
ZShift = 0.22
lm Score = 25.37
NG = 0
DB = 0
SM = 0
Delta_AUC
Doxorubicin
RF1_159_0 RF1
ZShift = 0.11
lm Score = −0.99
NG = 0
DB = 0
SM = 0
Delta_L
Doxorubicin
RF1_159_0 RF1
ZShift = −0.93
lm Score = 5.23
NG = 0
DB = 0
SM = 0
Delta_K
Doxorubicin
RF1_159_0 RF1
ZShift = 0.06
lm Score = 0
NG = 0
DB = 0
SM = 0
Delta_r
Doxorubicin
RF1_159_0 RF1
ZShift = −0.85
lm Score = 133.12
NG = 0
DB = 0
SM = 0
Delta_AUC
Doxorubicin

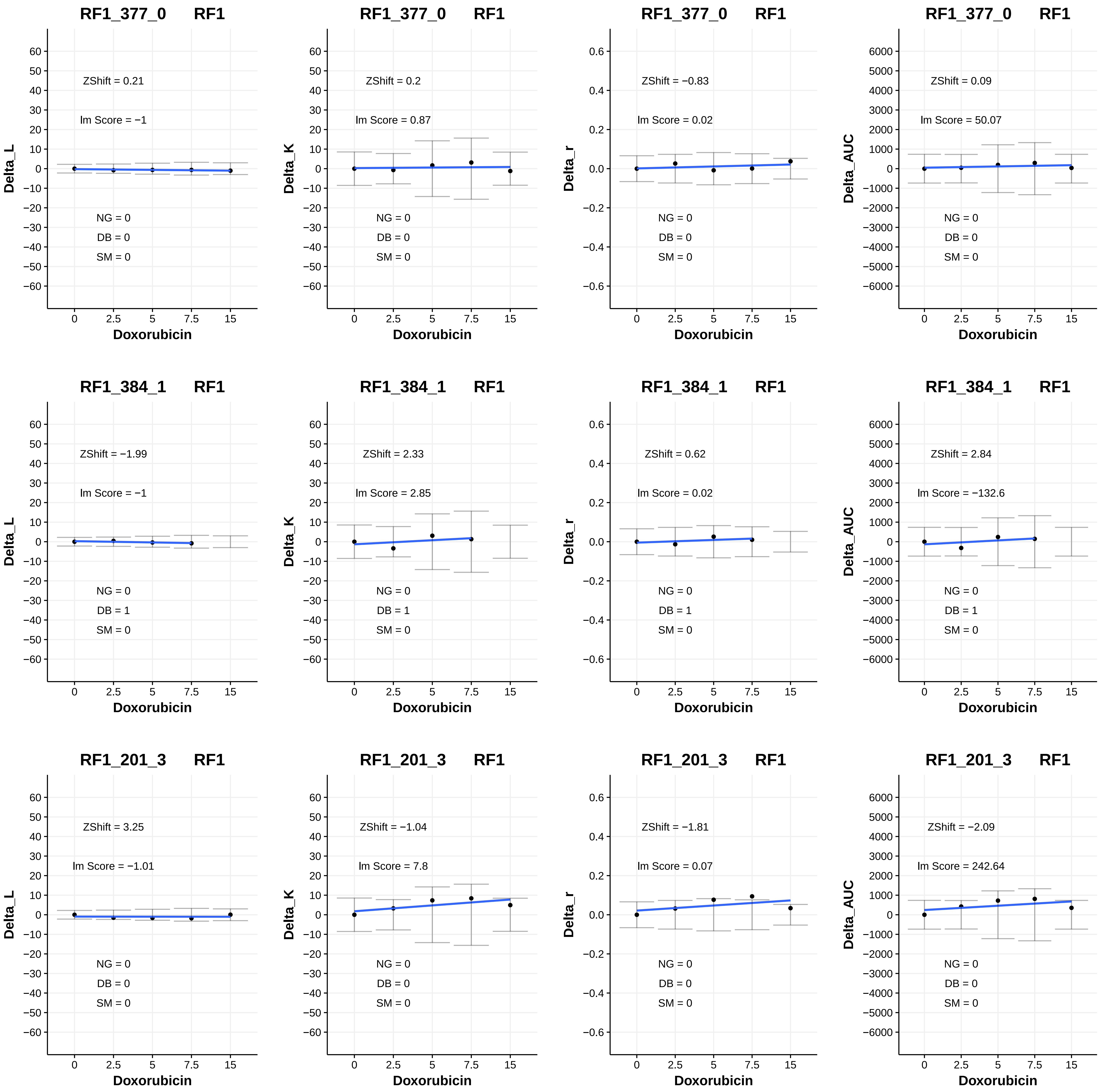

RF1_377_0 RF1
ZShift = 0.21
lm Score = −1
NG = 0
DB = 0
SM = 0
Delta_L
Doxorubicin
RF1_377_0 RF1
ZShift = 0.2
lm Score = 0.87
NG = 0
DB = 0
SM = 0
Delta_K
Doxorubicin
RF1_377_0 RF1
ZShift = −0.83
lm Score = 0.02
NG = 0
DB = 0
SM = 0
Delta_r
Doxorubicin
RF1_377_0 RF1
ZShift = 0.09
lm Score = 50.07
NG = 0
DB = 0
SM = 0
Delta_AUC
Doxorubicin
RF1_384_1 RF1
ZShift = −1.99
lm Score = −1
NG = 0
DB = 1
SM = 0
Delta_L
Doxorubicin
RF1_384_1 RF1
ZShift = 2.33
lm Score = 2.85
NG = 0
DB = 1
SM = 0
Delta_K
Doxorubicin
RF1_384_1 RF1
ZShift = 0.62
lm Score = 0.02
NG = 0
DB = 1
SM = 0
Delta_r
Doxorubicin
RF1_384_1 RF1
ZShift = 2.84
lm Score = −132.6
NG = 0
DB = 1
SM = 0
Delta_AUC
Doxorubicin
RF1_201_3 RF1
ZShift = 3.25
lm Score = −1.01
NG = 0
DB = 0
SM = 0
Delta_L
Doxorubicin
RF1_201_3 RF1
ZShift = −1.04
lm Score = 7.8
NG = 0
DB = 0
SM = 0
Delta_K
Doxorubicin
RF1_201_3 RF1
ZShift = −1.81
lm Score = 0.07
NG = 0
DB = 0
SM = 0
Delta_r
Doxorubicin
RF1_201_3 RF1
ZShift = −2.09
lm Score = 242.64
NG = 0
DB = 0
SM = 0
Delta_AUC
Doxorubicin

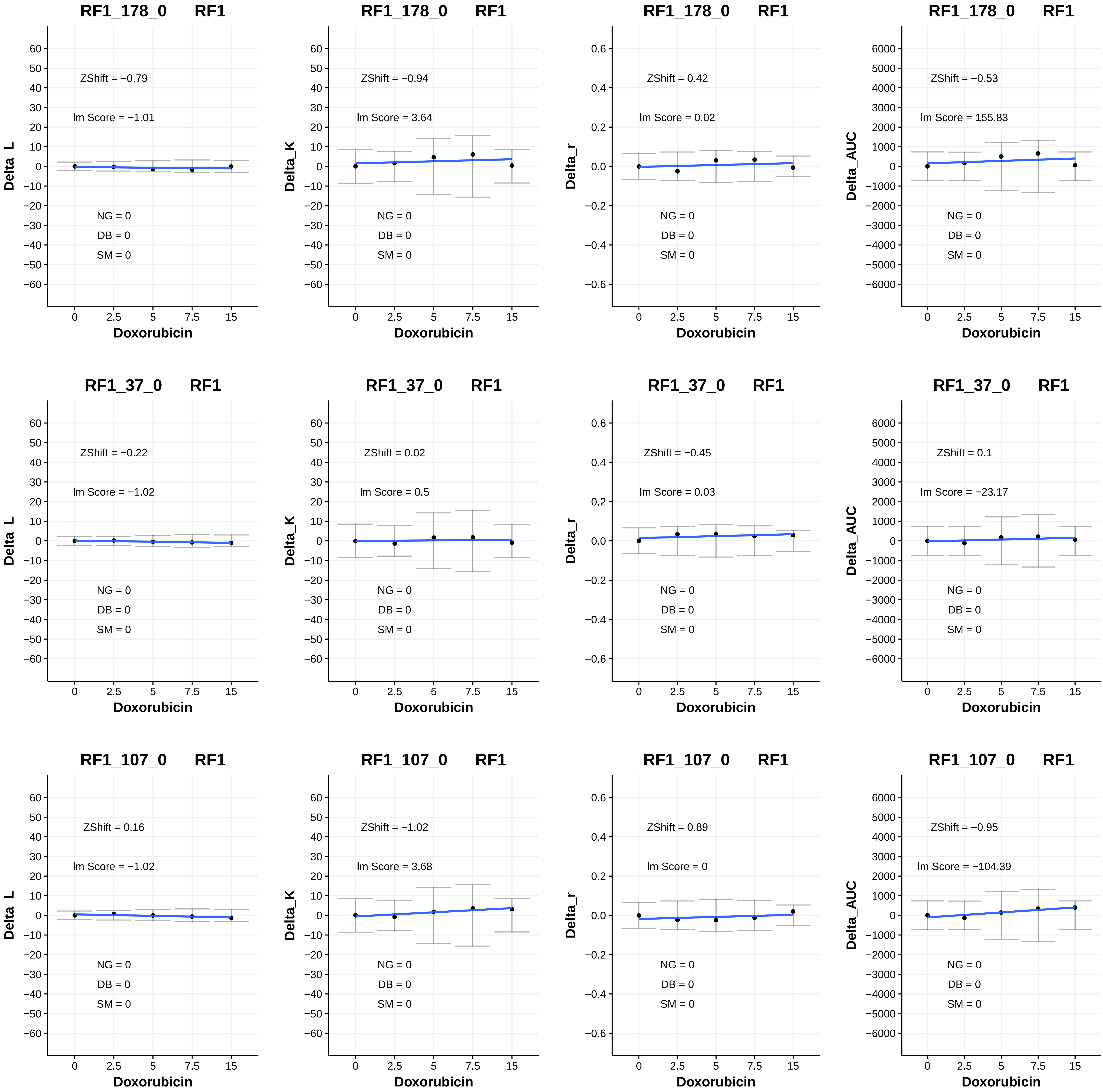
RF1_178_0 RF1
ZShift = −0.79
Im Score = −1.01
NG = 0
DB = 0
SM = 0
Delta_L
Doxorubicin
RF1_178_0 RF1
ZShift = −0.94
Im Score = 3.64
NG = 0
DB = 0
SM = 0
Delta_K
Doxorubicin
RF1_178_0 RF1
ZShift = 0.42
Im Score = 0.02
NG = 0
DB = 0
SM = 0
Delta_r
Doxorubicin
RF1_178_0 RF1
ZShift = −0.53
Im Score = 155.83
NG = 0
DB = 0
SM = 0
Delta_AUC
Doxorubicin
RF1_37_0 RF1
ZShift = −0.22
Im Score = −1.02
NG = 0
DB = 0
SM = 0
Delta_L
Doxorubicin
RF1_37_0 RF1
ZShift = 0.02
Im Score = 0.5
NG = 0
DB = 0
SM = 0
Delta_K
Doxorubicin
RF1_37_0 RF1
ZShift = −0.45
Im Score = 0.03
NG = 0
DB = 0
SM = 0
Delta_r
Doxorubicin
RF1_37_0 RF1
ZShift = 0.1
Im Score = −23.17
NG = 0
DB = 0
SM = 0
Delta_AUC
Doxorubicin
RF1_107_0 RF1
ZShift = 0.16
Im Score = −1.02
NG = 0
DB = 0
SM = 0
Delta_L
Doxorubicin
RF1_107_0 RF1
ZShift = −1.02
Im Score = 3.68
NG = 0
DB = 0
SM = 0
Delta_K
Doxorubicin
RF1_107_0 RF1
ZShift = 0.89
Im Score = 0
NG = 0
DB = 0
SM = 0
Delta_r
Doxorubicin
RF1_107_0 RF1
ZShift = −0.95
Im Score = −104.39
NG = 0
DB = 0
SM = 0
Delta_AUC
Doxorubicin

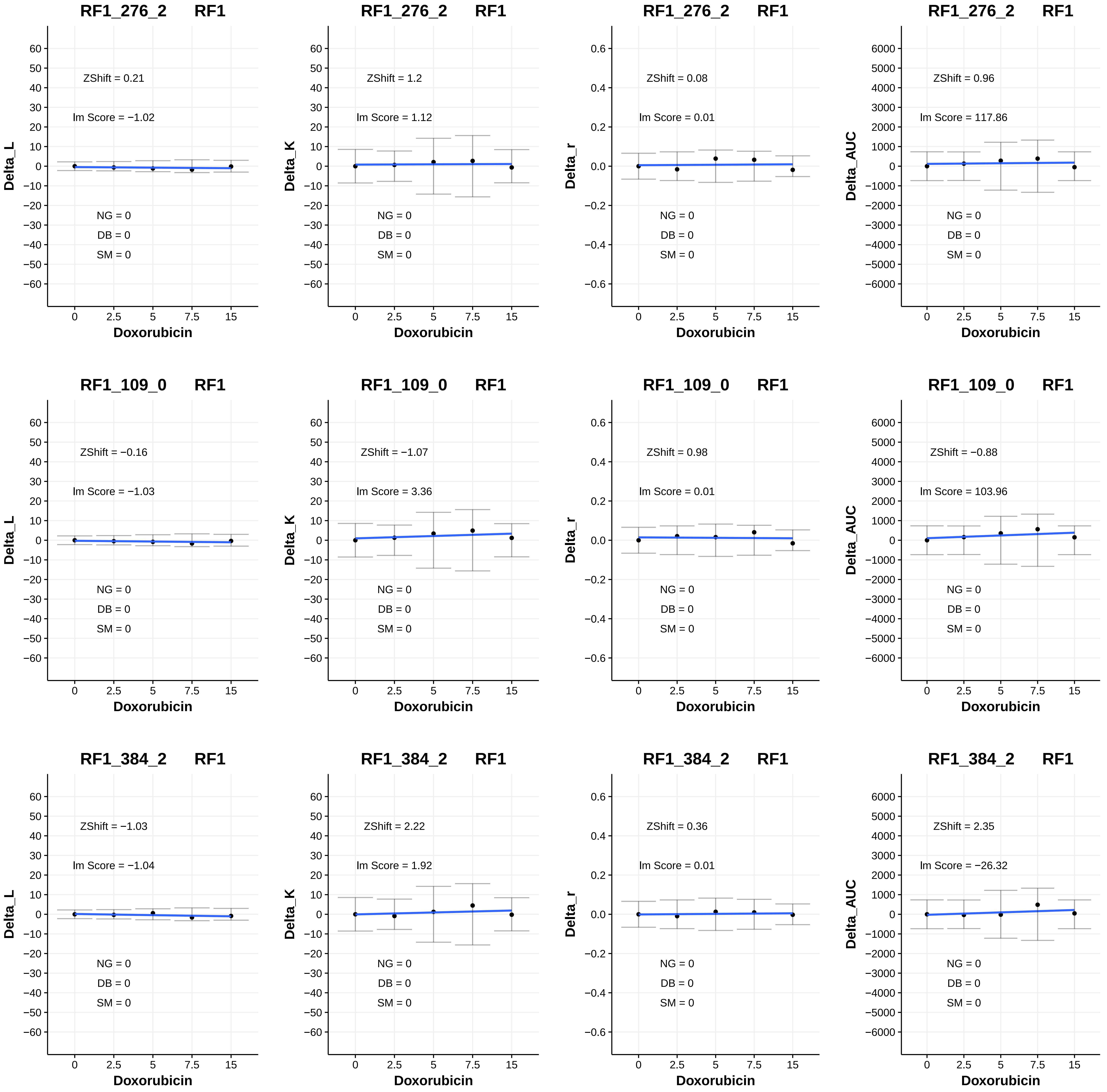

RF1_276_2 RF1
ZShift = 0.21
Im Score = −1.02
NG = 0
DB = 0
SM = 0
Delta_L
Doxorubicin
RF1_276_2 RF1
ZShift = 1.2
Im Score = 1.12
NG = 0
DB = 0
SM = 0
Delta_K
Doxorubicin
RF1_276_2 RF1
ZShift = 0.08
Im Score = 0.01
NG = 0
DB = 0
SM = 0
Delta_r
Doxorubicin
RF1_276_2 RF1
ZShift = 0.96
Im Score = 117.86
NG = 0
DB = 0
SM = 0
Delta_AUC
Doxorubicin
RF1_109_0 RF1
ZShift = −0.16
Im Score = −1.03
NG = 0
DB = 0
SM = 0
Delta_L
Doxorubicin
RF1_109_0 RF1
ZShift = −1.07
Im Score = 3.36
NG = 0
DB = 0
SM = 0
Delta_K
Doxorubicin
RF1_109_0 RF1
ZShift = 0.98
Im Score = 0.01
NG = 0
DB = 0
SM = 0
Delta_r
Doxorubicin
RF1_109_0 RF1
ZShift = −0.88
Im Score = 103.96
NG = 0
DB = 0
SM = 0
Delta_AUC
Doxorubicin
RF1_384_2 RF1
ZShift = −1.03
Im Score = −1.04
NG = 0
DB = 0
SM = 0
Delta_L
Doxorubicin
RF1_384_2 RF1
ZShift = 2.22
Im Score = 1.92
NG = 0
DB = 0
SM = 0
Delta_K
Doxorubicin
RF1_384_2 RF1
ZShift = 0.36
Im Score = 0.01
NG = 0
DB = 0
SM = 0
Delta_r
Doxorubicin
RF1_384_2 RF1
ZShift = 2.35
Im Score = −26.32
NG = 0
DB = 0
SM = 0
Delta_AUC
Doxorubicin

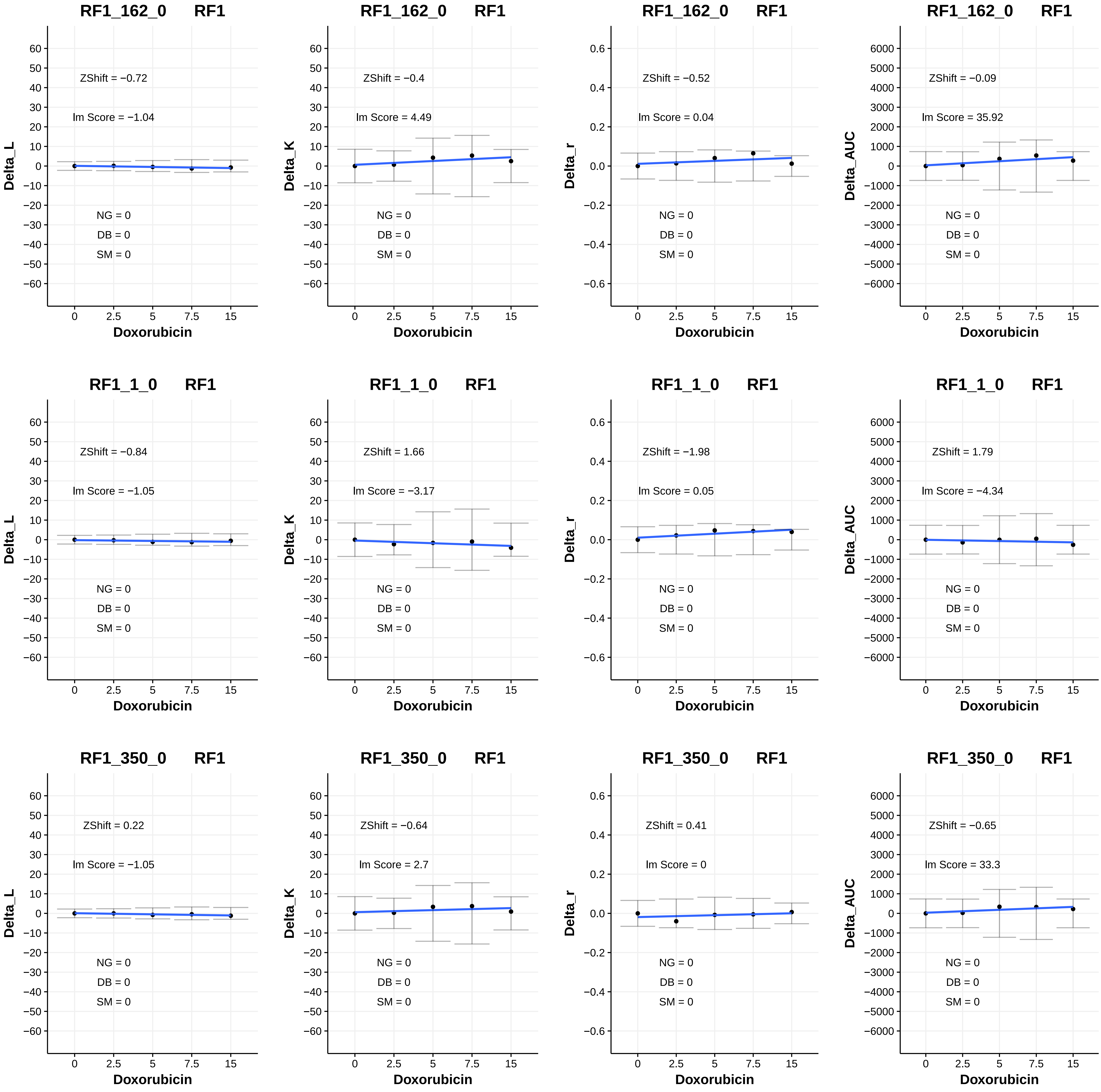
RF1_162_0 RF1
ZShift = −0.72
Im Score = −1.04
NG = 0
DB = 0
SM = 0
Delta_L
Doxorubicin
RF1_162_0 RF1
ZShift = −0.4
Im Score = 4.49
NG = 0
DB = 0
SM = 0
Delta_K
Doxorubicin
RF1_162_0 RF1
ZShift = −0.52
Im Score = 0.04
NG = 0
DB = 0
SM = 0
Delta_r
Doxorubicin
RF1_162_0 RF1
ZShift = −0.09
Im Score = 35.92
NG = 0
DB = 0
SM = 0
Delta_AUC
Doxorubicin
RF1_1_0 RF1
ZShift = −0.84
Im Score = −1.05
NG = 0
DB = 0
SM = 0
Delta_L
Doxorubicin
RF1_1_0 RF1
ZShift = 1.66
Im Score = −3.17
NG = 0
DB = 0
SM = 0
Delta_K
Doxorubicin
RF1_1_0 RF1
ZShift = −1.98
Im Score = 0.05
NG = 0
DB = 0
SM = 0
Delta_r
Doxorubicin
RF1_1_0 RF1
ZShift = 1.79
Im Score = −4.34
NG = 0
DB = 0
SM = 0
Delta_AUC
Doxorubicin
RF1_350_0 RF1
ZShift = 0.22
Im Score = −1.05
NG = 0
DB = 0
SM = 0
Delta_L
Doxorubicin
RF1_350_0 RF1
ZShift = −0.64
Im Score = 2.7
NG = 0
DB = 0
SM = 0
Delta_K
Doxorubicin
RF1_350_0 RF1
ZShift = 0.41
Im Score = 0
NG = 0
DB = 0
SM = 0
Delta_r
Doxorubicin
RF1_350_0 RF1
ZShift = −0.65
Im Score = 33.3
NG = 0
DB = 0
SM = 0
Delta_AUC
Doxorubicin

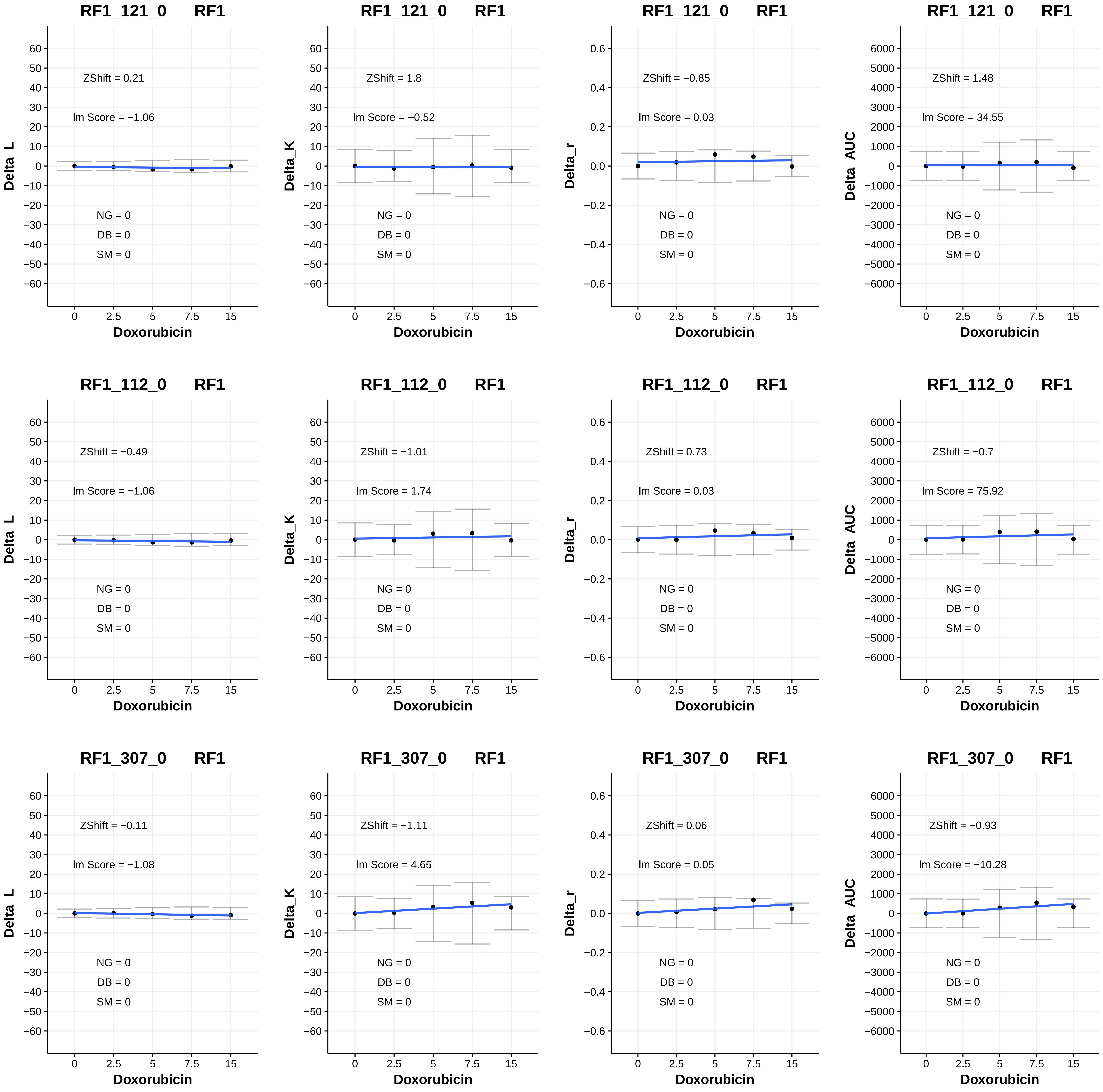
RF1_121_0 RF1
ZShift = 0.21
lm Score = −1.06
NG = 0
DB = 0
SM = 0
Delta_L
Doxorubicin
RF1_121_0 RF1
ZShift = 1.8
lm Score = −0.52
NG = 0
DB = 0
SM = 0
Delta_K
Doxorubicin
RF1_121_0 RF1
ZShift = −0.85
lm Score = 0.03
NG = 0
DB = 0
SM = 0
Delta_r
Doxorubicin
RF1_121_0 RF1
ZShift = 1.48
lm Score = 34.55
NG = 0
DB = 0
SM = 0
Delta_AUC
Doxorubicin
RF1_112_0 RF1
ZShift = −0.49
lm Score = −1.06
NG = 0
DB = 0
SM = 0
Delta_L
Doxorubicin
RF1_112_0 RF1
ZShift = −1.01
lm Score = 1.74
NG = 0
DB = 0
SM = 0
Delta_K
Doxorubicin
RF1_112_0 RF1
ZShift = 0.73
lm Score = 0.03
NG = 0
DB = 0
SM = 0
Delta_r
Doxorubicin
RF1_112_0 RF1
ZShift = −0.7
lm Score = 75.92
NG = 0
DB = 0
SM = 0
Delta_AUC
Doxorubicin
RF1_307_0 RF1
ZShift = −0.11
lm Score = −1.08
NG = 0
DB = 0
SM = 0
Delta_L
Doxorubicin
RF1_307_0 RF1
ZShift = −1.11
lm Score = 4.65
NG = 0
DB = 0
SM = 0
Delta_K
Doxorubicin
RF1_307_0 RF1
ZShift = 0.06
lm Score = 0.05
NG = 0
DB = 0
SM = 0
Delta_r
Doxorubicin
RF1_307_0 RF1
ZShift = −0.93
lm Score = −10.28
NG = 0
DB = 0
SM = 0
Delta_AUC
Doxorubicin

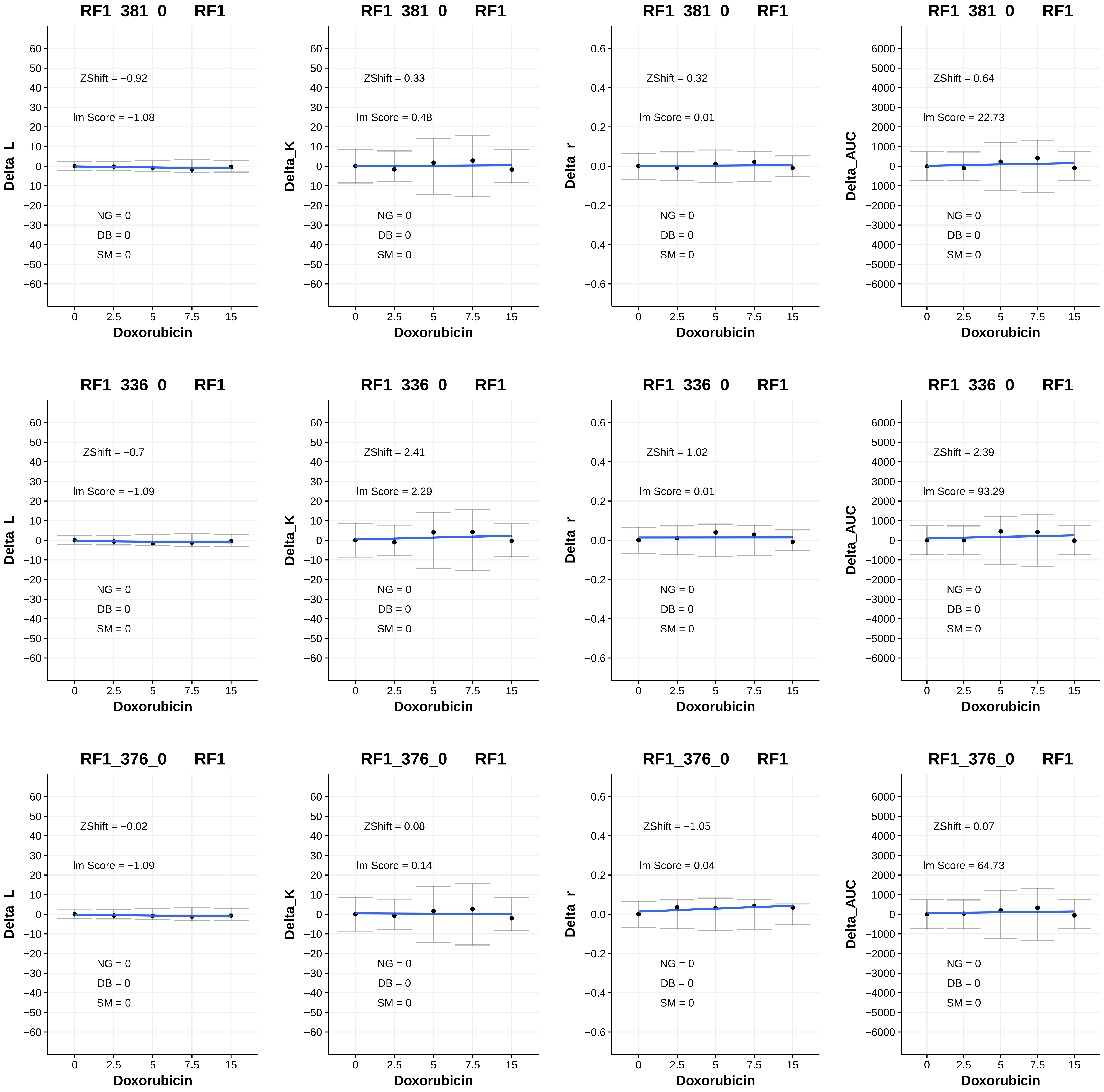
RF1_381_0 RF1
ZShift = −0.92
lm Score = −1.08
NG = 0
DB = 0
SM = 0
Delta_L
Doxorubicin
RF1_381_0 RF1
ZShift = 0.33
lm Score = 0.48
NG = 0
DB = 0
SM = 0
Delta_K
Doxorubicin
RF1_381_0 RF1
ZShift = 0.32
lm Score = 0.01
NG = 0
DB = 0
SM = 0
Delta_r
Doxorubicin
RF1_381_0 RF1
ZShift = 0.64
lm Score = 22.73
NG = 0
DB = 0
SM = 0
Delta_AUC
Doxorubicin
RF1_336_0 RF1
ZShift = −0.7
lm Score = −1.09
NG = 0
DB = 0
SM = 0
Delta_L
Doxorubicin
RF1_336_0 RF1
ZShift = 2.41
lm Score = 2.29
NG = 0
DB = 0
SM = 0
Delta_K
Doxorubicin
RF1_336_0 RF1
ZShift = 1.02
lm Score = 0.01
NG = 0
DB = 0
SM = 0
Delta_r
Doxorubicin
RF1_336_0 RF1
ZShift = 2.39
lm Score = 93.29
NG = 0
DB = 0
SM = 0
Delta_AUC
Doxorubicin
RF1_376_0 RF1
ZShift = −0.02
lm Score = −1.09
NG = 0
DB = 0
SM = 0
Delta_L
Doxorubicin
RF1_376_0 RF1
ZShift = 0.08
lm Score = 0.14
NG = 0
DB = 0
SM = 0
Delta_K
Doxorubicin
RF1_376_0 RF1
ZShift = −1.05
lm Score = 0.04
NG = 0
DB = 0
SM = 0
Delta_r
Doxorubicin
RF1_376_0 RF1
ZShift = 0.07
lm Score = 64.73
NG = 0
DB = 0
SM = 0
Delta_AUC
Doxorubicin

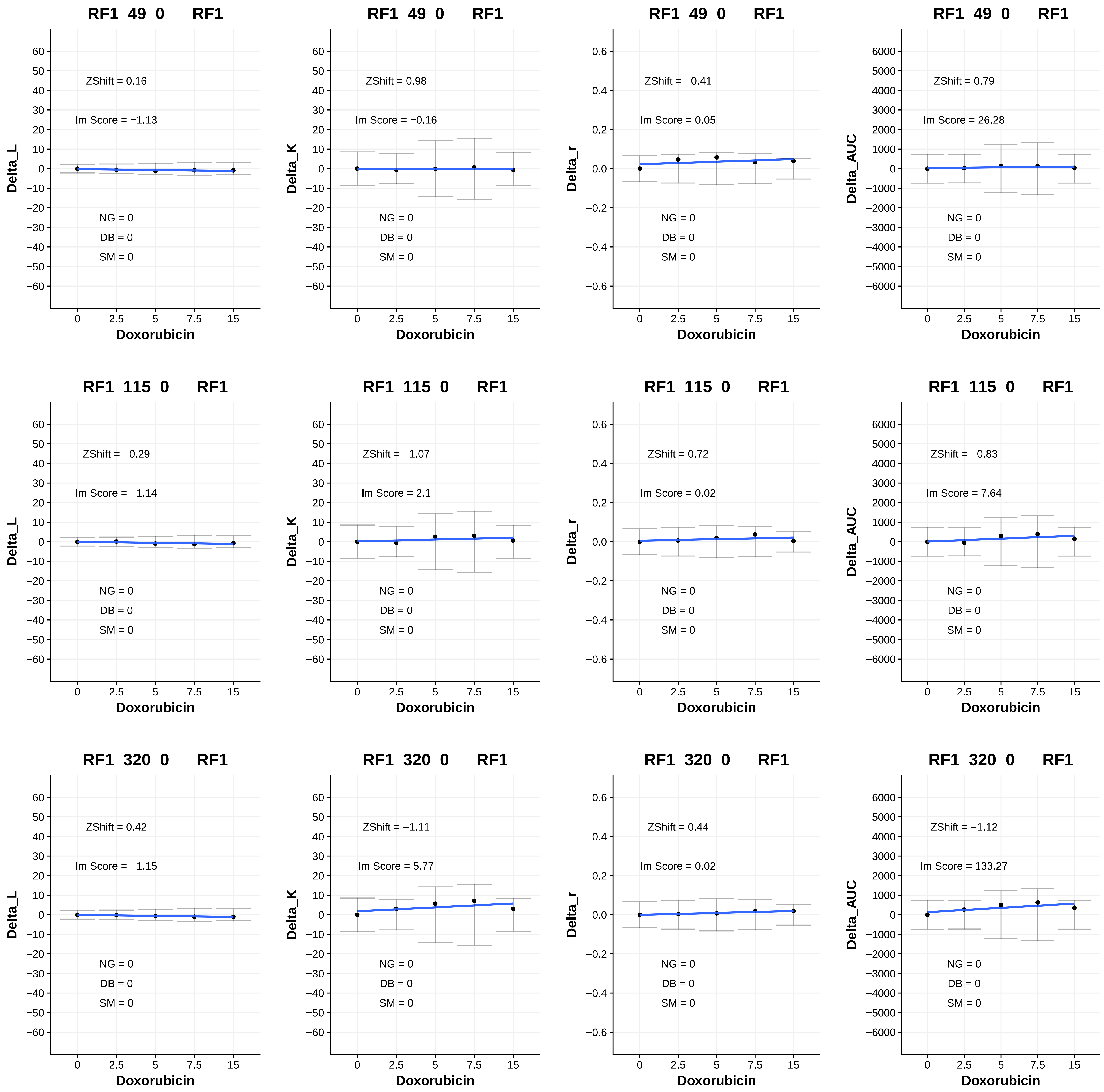
RF1_49_0 RF1
ZShift = 0.16
lm Score = −1.13
NG = 0
DB = 0
SM = 0
Delta_L
Doxorubicin
RF1_49_0 RF1
ZShift = 0.98
lm Score = −0.16
NG = 0
DB = 0
SM = 0
Delta_K
Doxorubicin
RF1_49_0 RF1
ZShift = −0.41
lm Score = 0.05
NG = 0
DB = 0
SM = 0
Delta_r
Doxorubicin
RF1_49_0 RF1
ZShift = 0.79
lm Score = 26.28
NG = 0
DB = 0
SM = 0
Delta_AUC
Doxorubicin
RF1_115_0 RF1
ZShift = −0.29
lm Score = −1.14
NG = 0
DB = 0
SM = 0
Delta_L
Doxorubicin
RF1_115_0 RF1
ZShift = −1.07
lm Score = 2.1
NG = 0
DB = 0
SM = 0
Delta_K
Doxorubicin
RF1_115_0 RF1
ZShift = 0.72
lm Score = 0.02
NG = 0
DB = 0
SM = 0
Delta_r
Doxorubicin
RF1_115_0 RF1
ZShift = −0.83
lm Score = 7.64
NG = 0
DB = 0
SM = 0
Delta_AUC
Doxorubicin
RF1_320_0 RF1
ZShift = 0.42
lm Score = −1.15
NG = 0
DB = 0
SM = 0
Delta_L
Doxorubicin
RF1_320_0 RF1
ZShift = −1.11
lm Score = 5.77
NG = 0
DB = 0
SM = 0
Delta_K
Doxorubicin
RF1_320_0 RF1
ZShift = 0.44
lm Score = 0.02
NG = 0
DB = 0
SM = 0
Delta_r
Doxorubicin
RF1_320_0 RF1
ZShift = −1.12
lm Score = 133.27
NG = 0
DB = 0
SM = 0
Delta_AUC
Doxorubicin

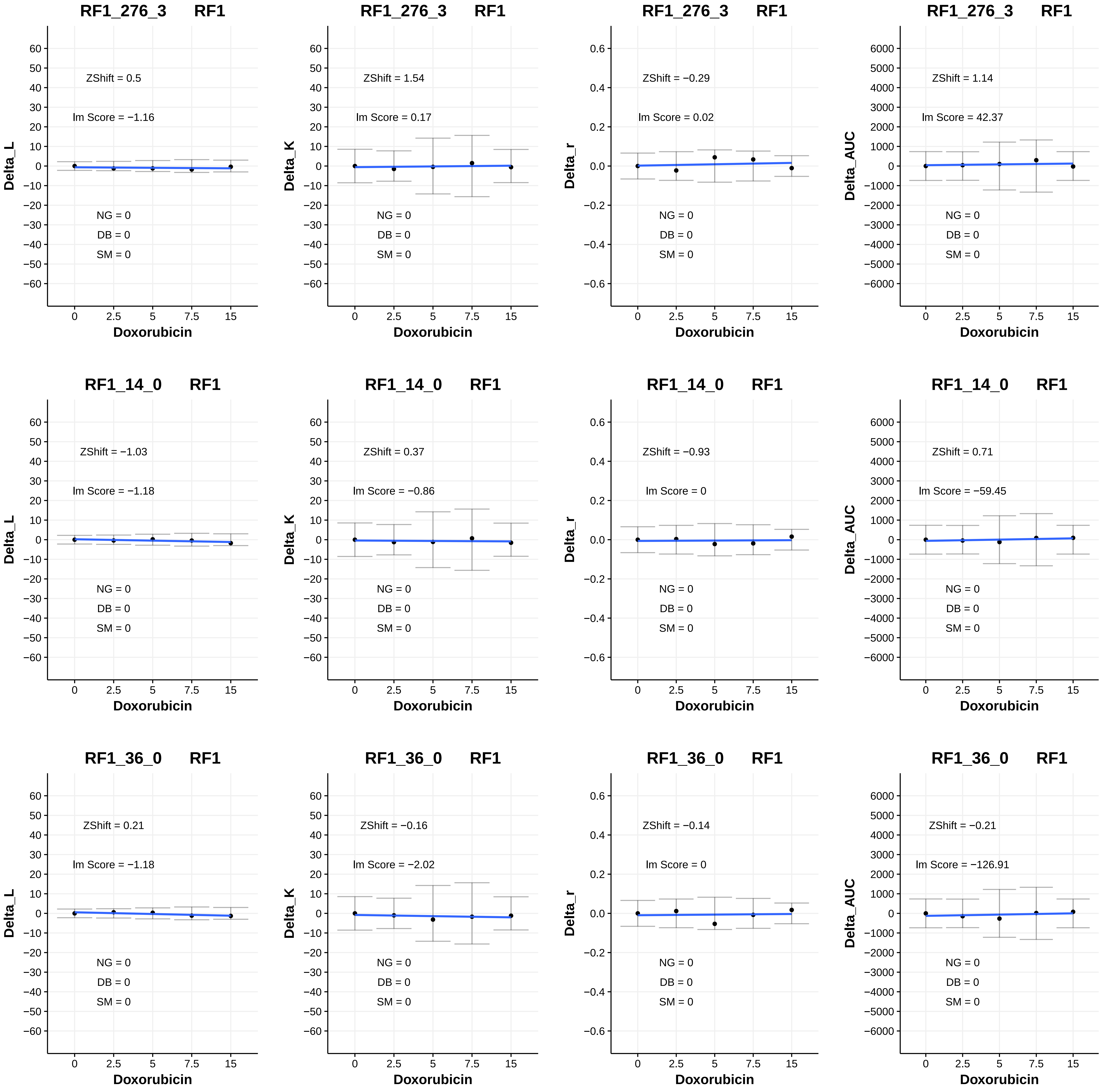

RF1_276_3 RF1
ZShift = 0.5
lm Score = −1.16
NG = 0
DB = 0
SM = 0
Delta_L
Doxorubicin
RF1_276_3 RF1
ZShift = 1.54
lm Score = 0.17
NG = 0
DB = 0
SM = 0
Delta_K
Doxorubicin
RF1_276_3 RF1
ZShift = −0.29
lm Score = 0.02
NG = 0
DB = 0
SM = 0
Delta_r
Doxorubicin
RF1_276_3 RF1
ZShift = 1.14
lm Score = 42.37
NG = 0
DB = 0
SM = 0
Delta_AUC
Doxorubicin
RF1_14_0 RF1
ZShift = −1.03
lm Score = −1.18
NG = 0
DB = 0
SM = 0
Delta_L
Doxorubicin
RF1_14_0 RF1
ZShift = 0.37
lm Score = −0.86
NG = 0
DB = 0
SM = 0
Delta_K
Doxorubicin
RF1_14_0 RF1
ZShift = −0.93
lm Score = 0
NG = 0
DB = 0
SM = 0
Delta_r
Doxorubicin
RF1_14_0 RF1
ZShift = 0.71
lm Score = −59.45
NG = 0
DB = 0
SM = 0
Delta_AUC
Doxorubicin
RF1_36_0 RF1
ZShift = 0.21
lm Score = −1.18
NG = 0
DB = 0
SM = 0
Delta_L
Doxorubicin
RF1_36_0 RF1
ZShift = −0.16
lm Score = −2.02
NG = 0
DB = 0
SM = 0
Delta_K
Doxorubicin
RF1_36_0 RF1
ZShift = −0.14
lm Score = 0
NG = 0
DB = 0
SM = 0
Delta_r
Doxorubicin
RF1_36_0 RF1
ZShift = −0.21
lm Score = −126.91
NG = 0
DB = 0
SM = 0
Delta_AUC
Doxorubicin

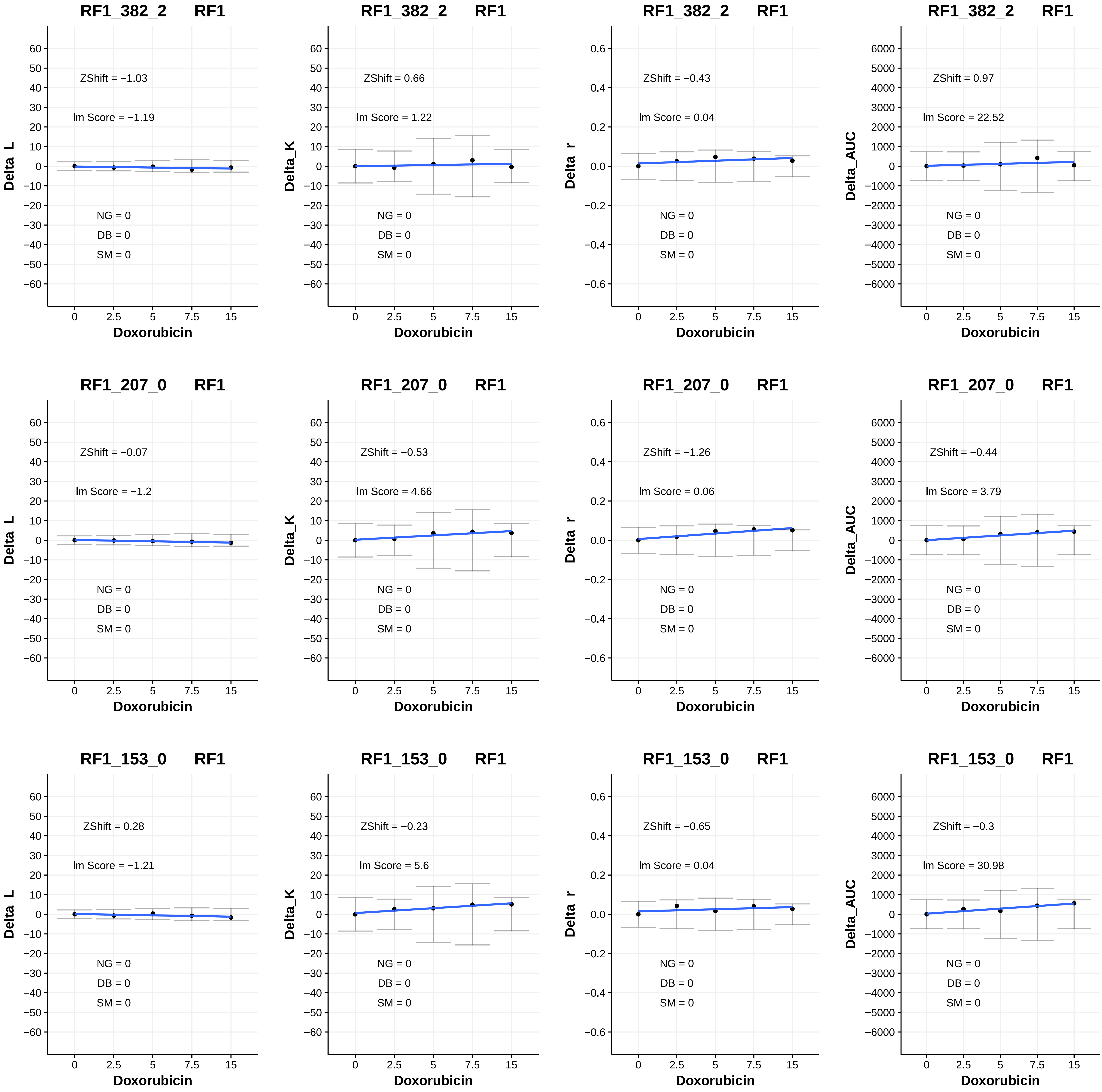
RF1_382_2 RF1
ZShift = −1.03
lm Score = −1.19
NG = 0
DB = 0
SM = 0
Delta_L
Doxorubicin
RF1_382_2 RF1
ZShift = 0.66
lm Score = 1.22
NG = 0
DB = 0
SM = 0
Delta_K
Doxorubicin
RF1_382_2 RF1
ZShift = −0.43
lm Score = 0.04
NG = 0
DB = 0
SM = 0
Delta_r
Doxorubicin
RF1_382_2 RF1
ZShift = 0.97
lm Score = 22.52
NG = 0
DB = 0
SM = 0
Delta_AUC
Doxorubicin
RF1_207_0 RF1
ZShift = −0.07
lm Score = −1.2
NG = 0
DB = 0
SM = 0
Delta_L
Doxorubicin
RF1_207_0 RF1
ZShift = −0.53
lm Score = 4.66
NG = 0
DB = 0
SM = 0
Delta_K
Doxorubicin
RF1_207_0 RF1
ZShift = −1.26
lm Score = 0.06
NG = 0
DB = 0
SM = 0
Delta_r
Doxorubicin
RF1_207_0 RF1
ZShift = −0.44
lm Score = 3.79
NG = 0
DB = 0
SM = 0
Delta_AUC
Doxorubicin
RF1_153_0 RF1
ZShift = 0.28
lm Score = −1.21
NG = 0
DB = 0
SM = 0
Delta_L
Doxorubicin
RF1_153_0 RF1
ZShift = −0.23
lm Score = 5.6
NG = 0
DB = 0
SM = 0
Delta_K
Doxorubicin
RF1_153_0 RF1
ZShift = −0.65
lm Score = 0.04
NG = 0
DB = 0
SM = 0
Delta_r
Doxorubicin
RF1_153_0 RF1
ZShift = −0.3
lm Score = 30.98
NG = 0
DB = 0
SM = 0
Delta_AUC
Doxorubicin

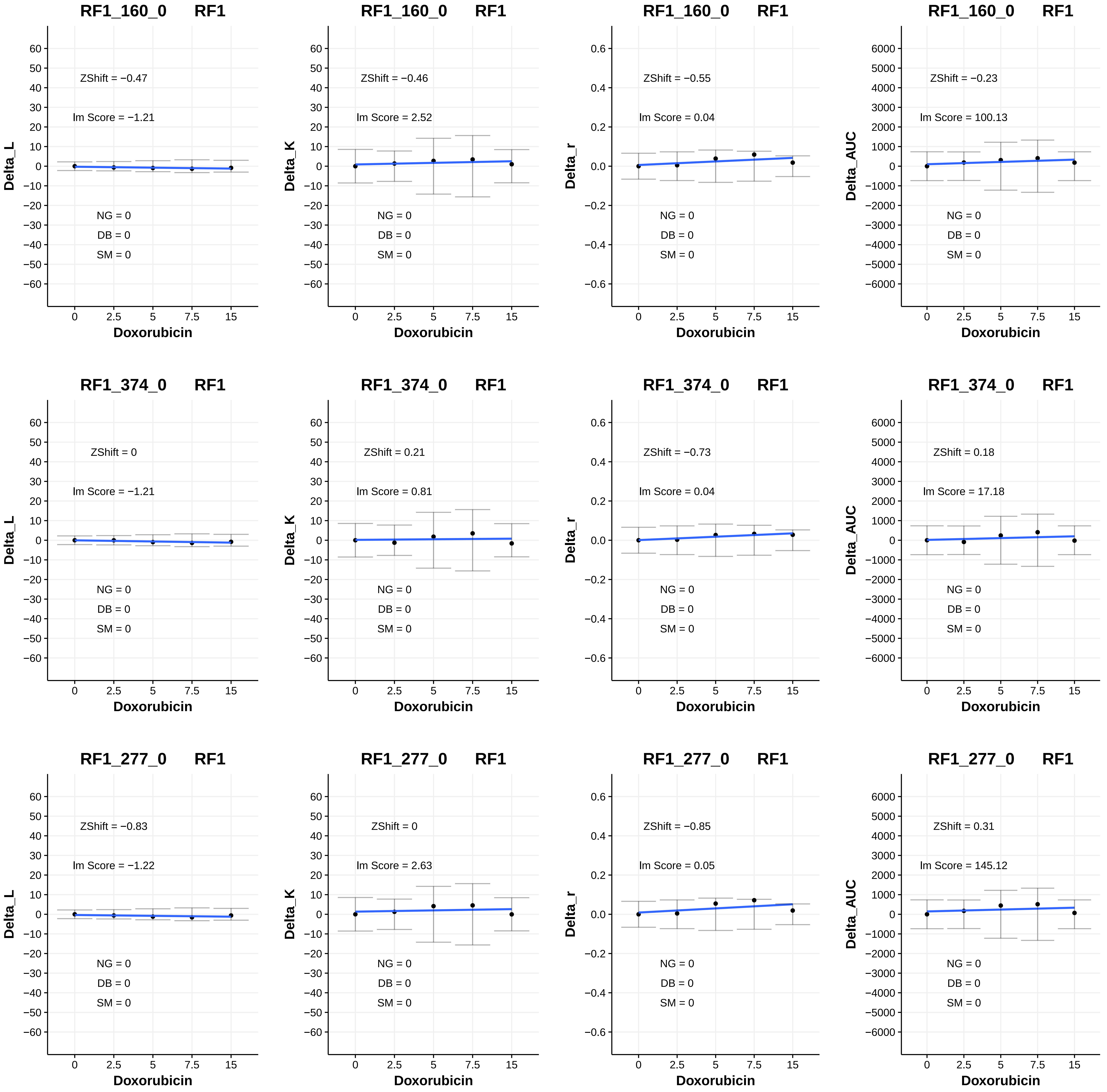

RF1_160_0 RF1
ZShift = −0.47
lm Score = −1.21
NG = 0
DB = 0
SM = 0
Delta_L
Doxorubicin
RF1_160_0 RF1
ZShift = −0.46
lm Score = 2.52
NG = 0
DB = 0
SM = 0
Delta_K
Doxorubicin
RF1_160_0 RF1
ZShift = −0.55
lm Score = 0.04
NG = 0
DB = 0
SM = 0
Delta_r
Doxorubicin
RF1_160_0 RF1
ZShift = −0.23
lm Score = 100.13
NG = 0
DB = 0
SM = 0
Delta_AUC
Doxorubicin
RF1_374_0 RF1
ZShift = 0
lm Score = −1.21
NG = 0
DB = 0
SM = 0
Delta_L
Doxorubicin
RF1_374_0 RF1
ZShift = 0.21
lm Score = 0.81
NG = 0
DB = 0
SM = 0
Delta_K
Doxorubicin
RF1_374_0 RF1
ZShift = −0.73
lm Score = 0.04
NG = 0
DB = 0
SM = 0
Delta_r
Doxorubicin
RF1_374_0 RF1
ZShift = 0.18
lm Score = 17.18
NG = 0
DB = 0
SM = 0
Delta_AUC
Doxorubicin
RF1_277_0 RF1
ZShift = −0.83
lm Score = −1.22
NG = 0
DB = 0
SM = 0
Delta_L
Doxorubicin
RF1_277_0 RF1
ZShift = 0
lm Score = 2.63
NG = 0
DB = 0
SM = 0
Delta_K
Doxorubicin
RF1_277_0 RF1
ZShift = −0.85
lm Score = 0.05
NG = 0
DB = 0
SM = 0
Delta_r
Doxorubicin
RF1_277_0 RF1
ZShift = 0.31
lm Score = 145.12
NG = 0
DB = 0
SM = 0
Delta_AUC
Doxorubicin

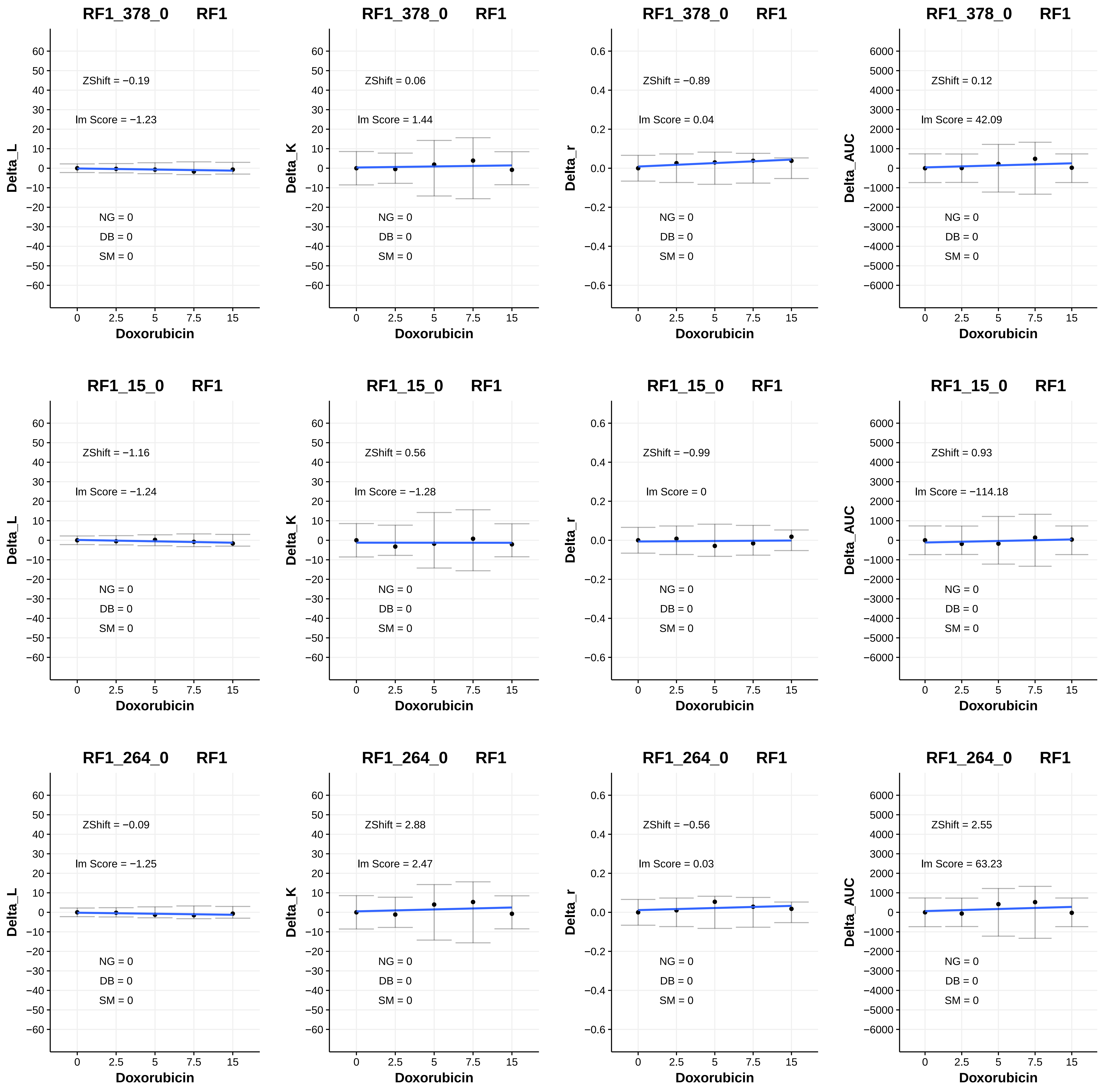

RF1_378_0 RF1
ZShift = −0.19
Im Score = −1.23
NG = 0
DB = 0
SM = 0
Delta_L
Doxorubicin
RF1_378_0 RF1
ZShift = 0.06
Im Score = 1.44
NG = 0
DB = 0
SM = 0
Delta_K
Doxorubicin
RF1_378_0 RF1
ZShift = −0.89
Im Score = 0.04
NG = 0
DB = 0
SM = 0
Delta_r
Doxorubicin
RF1_378_0 RF1
ZShift = 0.12
Im Score = 42.09
NG = 0
DB = 0
SM = 0
Delta_AUC
Doxorubicin
RF1_15_0 RF1
ZShift = −1.16
Im Score = −1.24
NG = 0
DB = 0
SM = 0
Delta_L
Doxorubicin
RF1_15_0 RF1
ZShift = 0.56
Im Score = −1.28
NG = 0
DB = 0
SM = 0
Delta_K
Doxorubicin
RF1_15_0 RF1
ZShift = −0.99
Im Score = 0
NG = 0
DB = 0
SM = 0
Delta_r
Doxorubicin
RF1_15_0 RF1
ZShift = 0.93
Im Score = −114.18
NG = 0
DB = 0
SM = 0
Delta_AUC
Doxorubicin
RF1_264_0 RF1
ZShift = −0.09
Im Score = −1.25
NG = 0
DB = 0
SM = 0
Delta_L
Doxorubicin
RF1_264_0 RF1
ZShift = 2.88
Im Score = 2.47
NG = 0
DB = 0
SM = 0
Delta_K
Doxorubicin
RF1_264_0 RF1
ZShift = −0.56
Im Score = 0.03
NG = 0
DB = 0
SM = 0
Delta_r
Doxorubicin
RF1_264_0 RF1
ZShift = 2.55
Im Score = 63.23
NG = 0
DB = 0
SM = 0
Delta_AUC
Doxorubicin

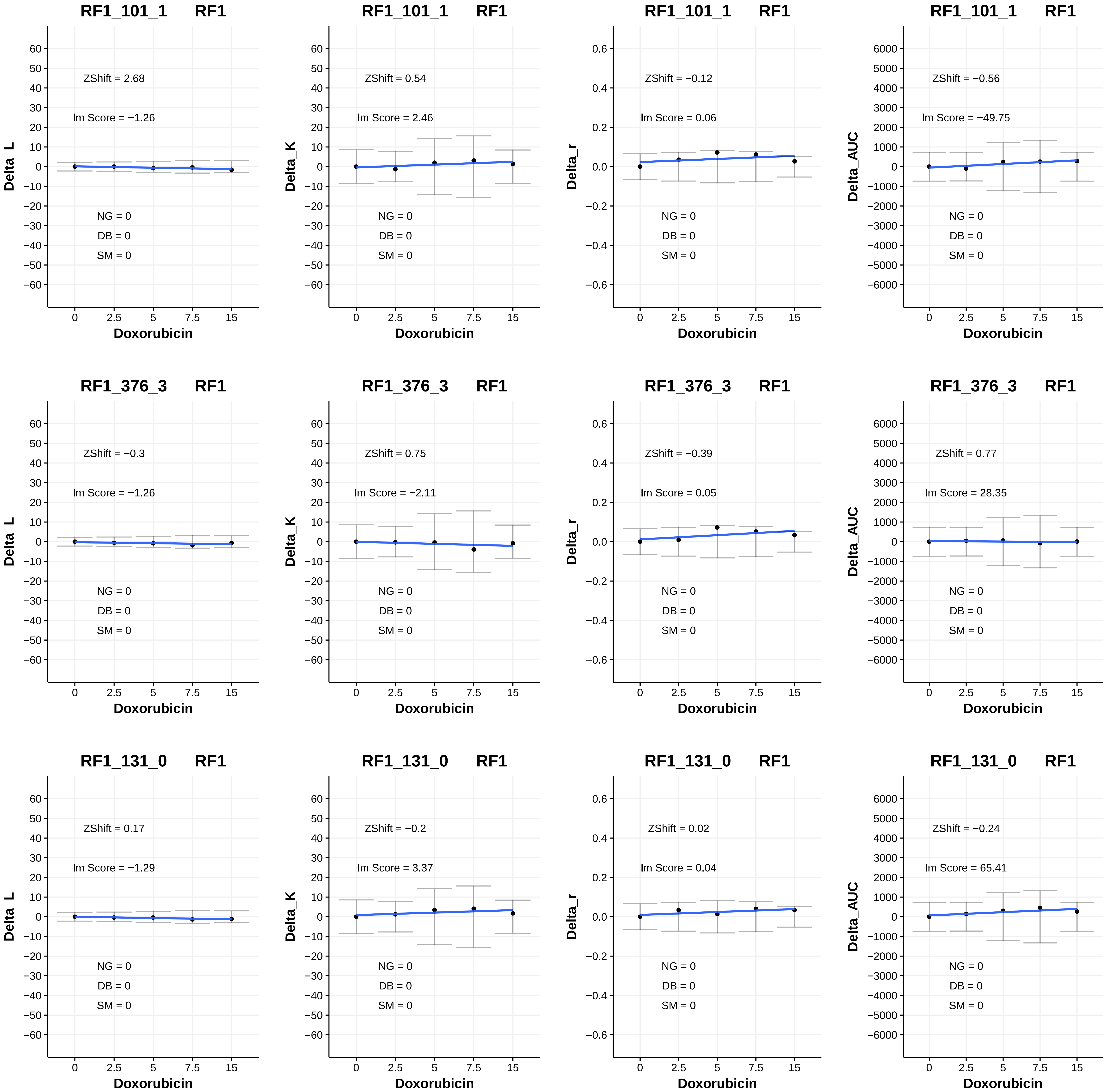
RF1_101_1 RF1
ZShift = 2.68
lm Score = −1.26
NG = 0
DB = 0
SM = 0
Delta_L
Doxorubicin
RF1_101_1 RF1
ZShift = 0.54
lm Score = 2.46
NG = 0
DB = 0
SM = 0
Delta_K
Doxorubicin
RF1_101_1 RF1
ZShift = −0.12
lm Score = 0.06
NG = 0
DB = 0
SM = 0
Delta_r
Doxorubicin
RF1_101_1 RF1
ZShift = −0.56
lm Score = −49.75
NG = 0
DB = 0
SM = 0
Delta_AUC
Doxorubicin
RF1_376_3 RF1
ZShift = −0.3
lm Score = −1.26
NG = 0
DB = 0
SM = 0
Delta_L
Doxorubicin
RF1_376_3 RF1
ZShift = 0.75
lm Score = −2.11
NG = 0
DB = 0
SM = 0
Delta_K
Doxorubicin
RF1_376_3 RF1
ZShift = −0.39
lm Score = 0.05
NG = 0
DB = 0
SM = 0
Delta_r
Doxorubicin
RF1_376_3 RF1
ZShift = 0.77
lm Score = 28.35
NG = 0
DB = 0
SM = 0
Delta_AUC
Doxorubicin
RF1_131_0 RF1
ZShift = 0.17
lm Score = −1.29
NG = 0
DB = 0
SM = 0
Delta_L
Doxorubicin
RF1_131_0 RF1
ZShift = −0.2
lm Score = 3.37
NG = 0
DB = 0
SM = 0
Delta_K
Doxorubicin
RF1_131_0 RF1
ZShift = 0.02
lm Score = 0.04
NG = 0
DB = 0
SM = 0
Delta_r
Doxorubicin
RF1_131_0 RF1
ZShift = −0.24
lm Score = 65.41
NG = 0
DB = 0
SM = 0
Delta_AUC
Doxorubicin

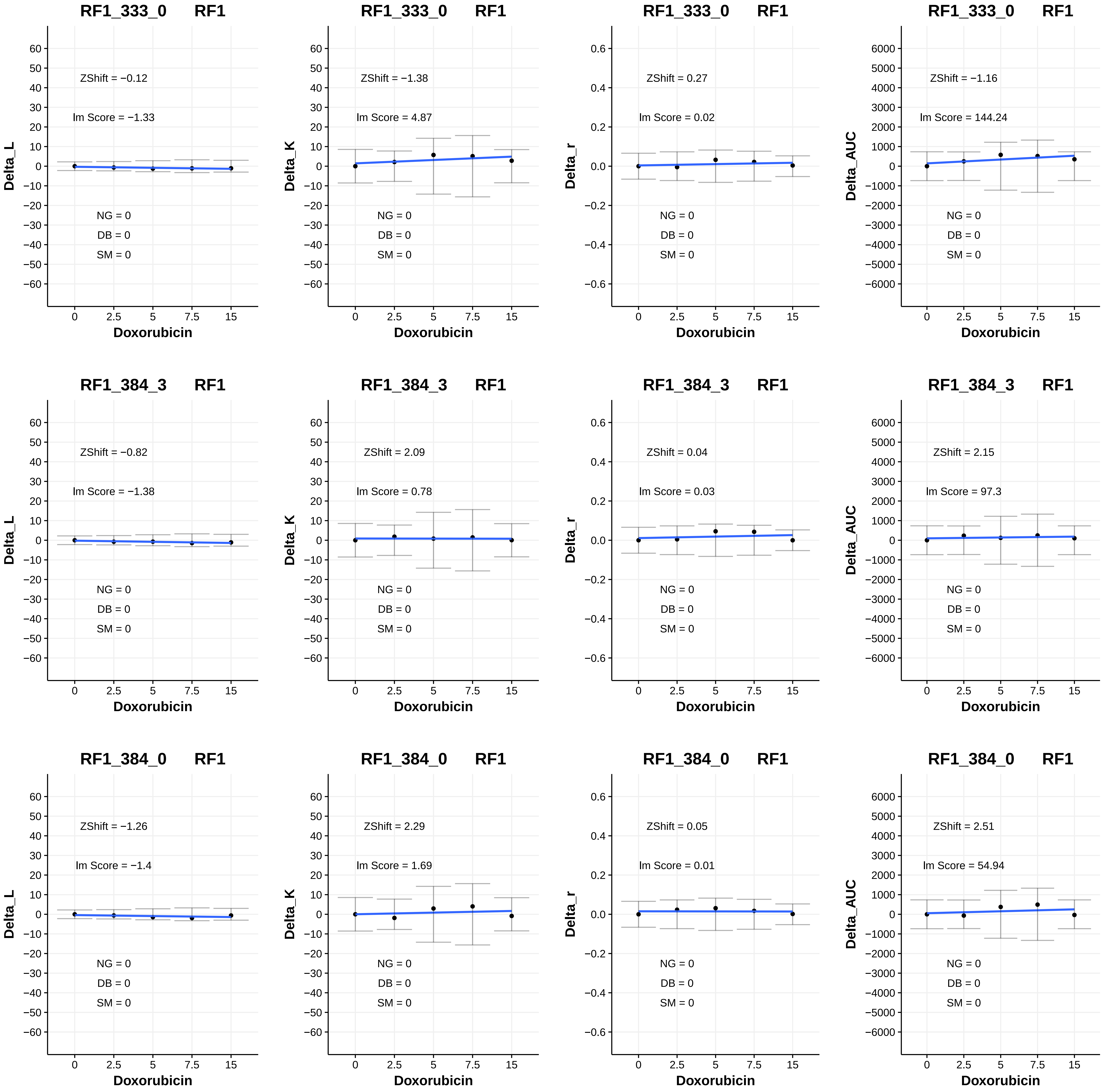
RF1_333_0 RF1
ZShift = −0.12
lm Score = −1.33
NG = 0
DB = 0
SM = 0
Delta_L
Doxorubicin
RF1_333_0 RF1
ZShift = −1.38
lm Score = 4.87
NG = 0
DB = 0
SM = 0
Delta_K
Doxorubicin
RF1_333_0 RF1
ZShift = 0.27
lm Score = 0.02
NG = 0
DB = 0
SM = 0
Delta_r
Doxorubicin
RF1_333_0 RF1
ZShift = −1.16
lm Score = 144.24
NG = 0
DB = 0
SM = 0
Delta_AUC
Doxorubicin
RF1_384_3 RF1
ZShift = −0.82
lm Score = −1.38
NG = 0
DB = 0
SM = 0
Delta_L
Doxorubicin
RF1_384_3 RF1
ZShift = 2.09
lm Score = 0.78
NG = 0
DB = 0
SM = 0
Delta_K
Doxorubicin
RF1_384_3 RF1
ZShift = 0.04
lm Score = 0.03
NG = 0
DB = 0
SM = 0
Delta_r
Doxorubicin
RF1_384_3 RF1
ZShift = 2.15
lm Score = 97.3
NG = 0
DB = 0
SM = 0
Delta_AUC
Doxorubicin
RF1_384_0 RF1
ZShift = −1.26
lm Score = −1.4
NG = 0
DB = 0
SM = 0
Delta_L
Doxorubicin
RF1_384_0 RF1
ZShift = 2.29
lm Score = 1.69
NG = 0
DB = 0
SM = 0
Delta_K
Doxorubicin
RF1_384_0 RF1
ZShift = 0.05
lm Score = 0.01
NG = 0
DB = 0
SM = 0
Delta_r
Doxorubicin
RF1_384_0 RF1
ZShift = 2.51
lm Score = 54.94
NG = 0
DB = 0
SM = 0
Delta_AUC
Doxorubicin

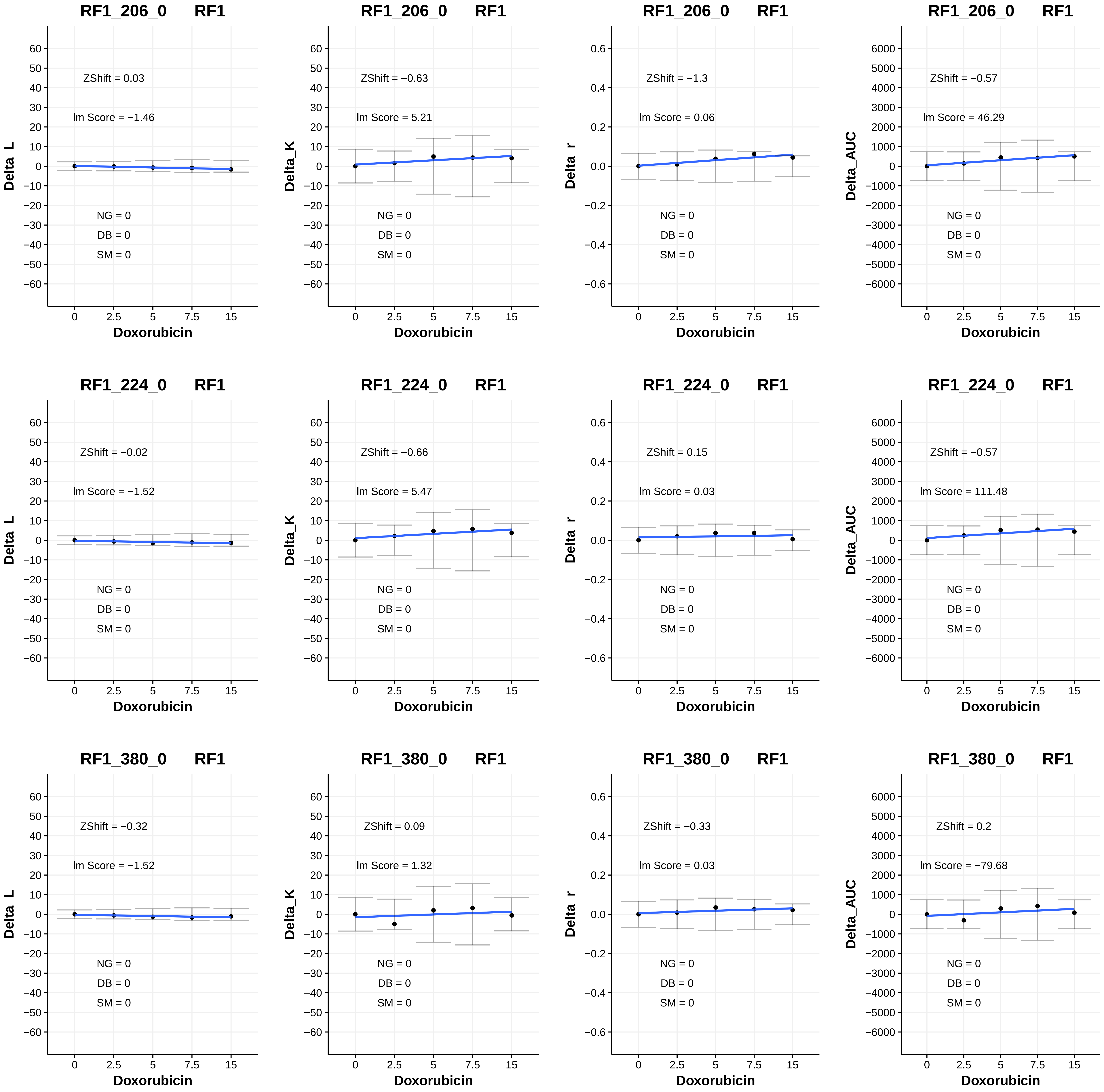

RF1_206_0 RF1
ZShift = 0.03
lm Score = −1.46
Delta_L
NG = 0
DB = 0
SM = 0
Doxorubicin
RF1_206_0 RF1
ZShift = −0.63
lm Score = 5.21
Delta_K
NG = 0
DB = 0
SM = 0
Doxorubicin
RF1_206_0 RF1
ZShift = −1.3
lm Score = 0.06
Delta_r
NG = 0
DB = 0
SM = 0
Doxorubicin
RF1_206_0 RF1
ZShift = −0.57
lm Score = 46.29
Delta_AUC
NG = 0
DB = 0
SM = 0
Doxorubicin
RF1_224_0 RF1
ZShift = −0.02
lm Score = −1.52
Delta_L
NG = 0
DB = 0
SM = 0
Doxorubicin
RF1_224_0 RF1
ZShift = −0.66
lm Score = 5.47
Delta_K
NG = 0
DB = 0
SM = 0
Doxorubicin
RF1_224_0 RF1
ZShift = 0.15
lm Score = 0.03
Delta_r
NG = 0
DB = 0
SM = 0
Doxorubicin
RF1_224_0 RF1
ZShift = −0.57
lm Score = 111.48
Delta_AUC
NG = 0
DB = 0
SM = 0
Doxorubicin
RF1_380_0 RF1
ZShift = −0.32
lm Score = −1.52
Delta_L
NG = 0
DB = 0
SM = 0
Doxorubicin
RF1_380_0 RF1
ZShift = 0.09
lm Score = 1.32
Delta_K
NG = 0
DB = 0
SM = 0
Doxorubicin
RF1_380_0 RF1
ZShift = −0.33
lm Score = 0.03
Delta_r
NG = 0
DB = 0
SM = 0
Doxorubicin
RF1_380_0 RF1
ZShift = 0.2
lm Score = −79.68
Delta_AUC
NG = 0
DB = 0
SM = 0
Doxorubicin

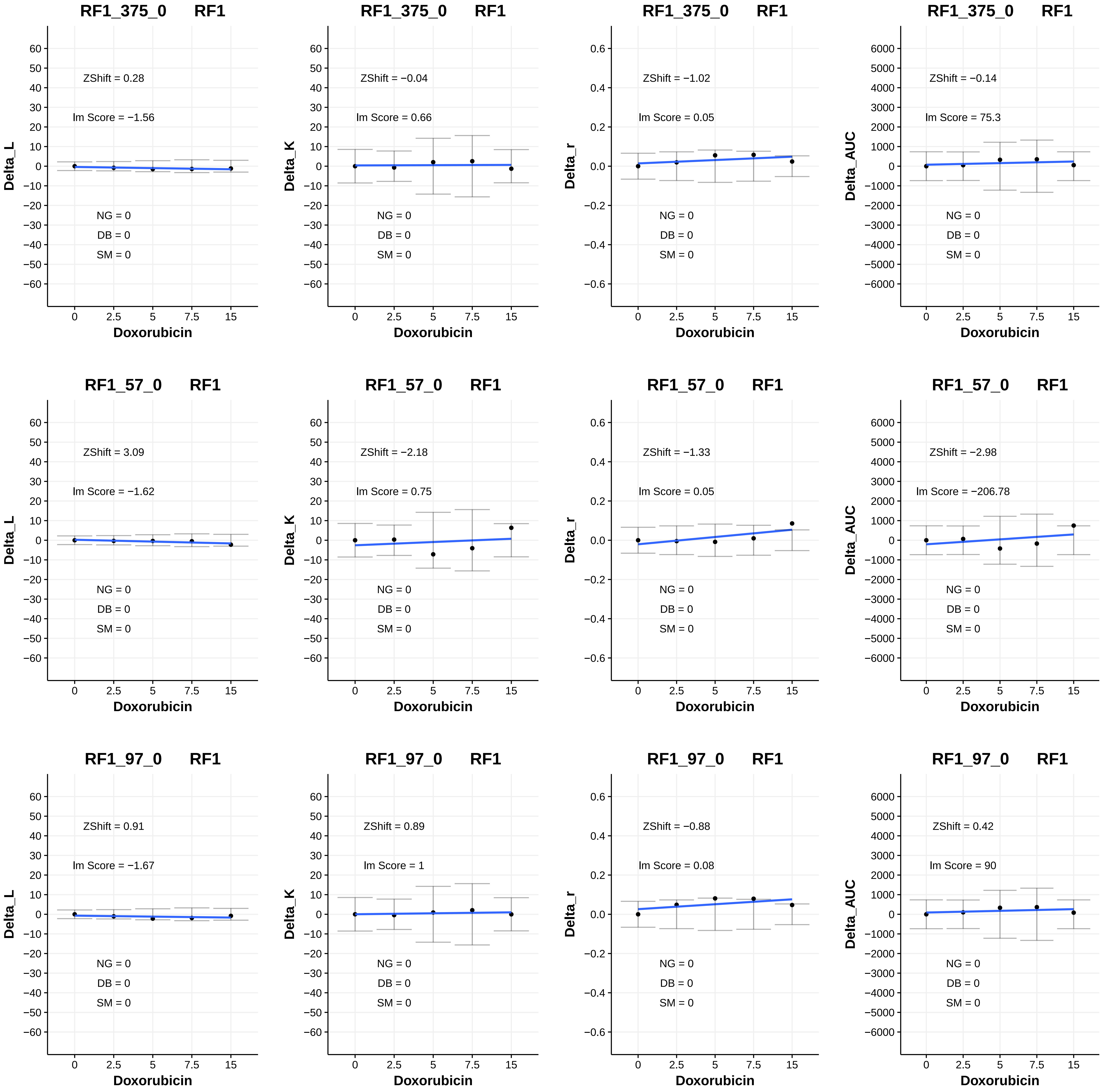

RF1_375_0 RF1
ZShift = 0.28
Im Score = −1.56
NG = 0
DB = 0
SM = 0
Delta_L
RF1_375_0 RF1
ZShift = −0.04
Im Score = 0.66
NG = 0
DB = 0
SM = 0
Delta_K
RF1_375_0 RF1
ZShift = −1.02
Im Score = 0.05
NG = 0
DB = 0
SM = 0
Delta_r
RF1_375_0 RF1
ZShift = −0.14
Im Score = 75.3
NG = 0
DB = 0
SM = 0
Delta_AUC
RF1_57_0 RF1
ZShift = 3.09
Im Score = −1.62
NG = 0
DB = 0
SM = 0
Delta_L
RF1_57_0 RF1
ZShift = −2.18
Im Score = 0.75
NG = 0
DB = 0
SM = 0
Delta_K
RF1_57_0 RF1
ZShift = −1.33
Im Score = 0.05
NG = 0
DB = 0
SM = 0
Delta_r
RF1_57_0 RF1
ZShift = −2.98
Im Score = −206.78
NG = 0
DB = 0
SM = 0
Delta_AUC
RF1_97_0 RF1
ZShift = 0.91
Im Score = −1.67
NG = 0
DB = 0
SM = 0
Delta_L
RF1_97_0 RF1
ZShift = 0.89
Im Score = 1
NG = 0
DB = 0
SM = 0
Delta_K
RF1_97_0 RF1
ZShift = −0.88
Im Score = 0.08
NG = 0
DB = 0
SM = 0
Delta_r
RF1_97_0 RF1
ZShift = 0.42
Im Score = 90
NG = 0
DB = 0
SM = 0
Delta_AUC
Doxorubicin
0
2.5
5
7.5
15

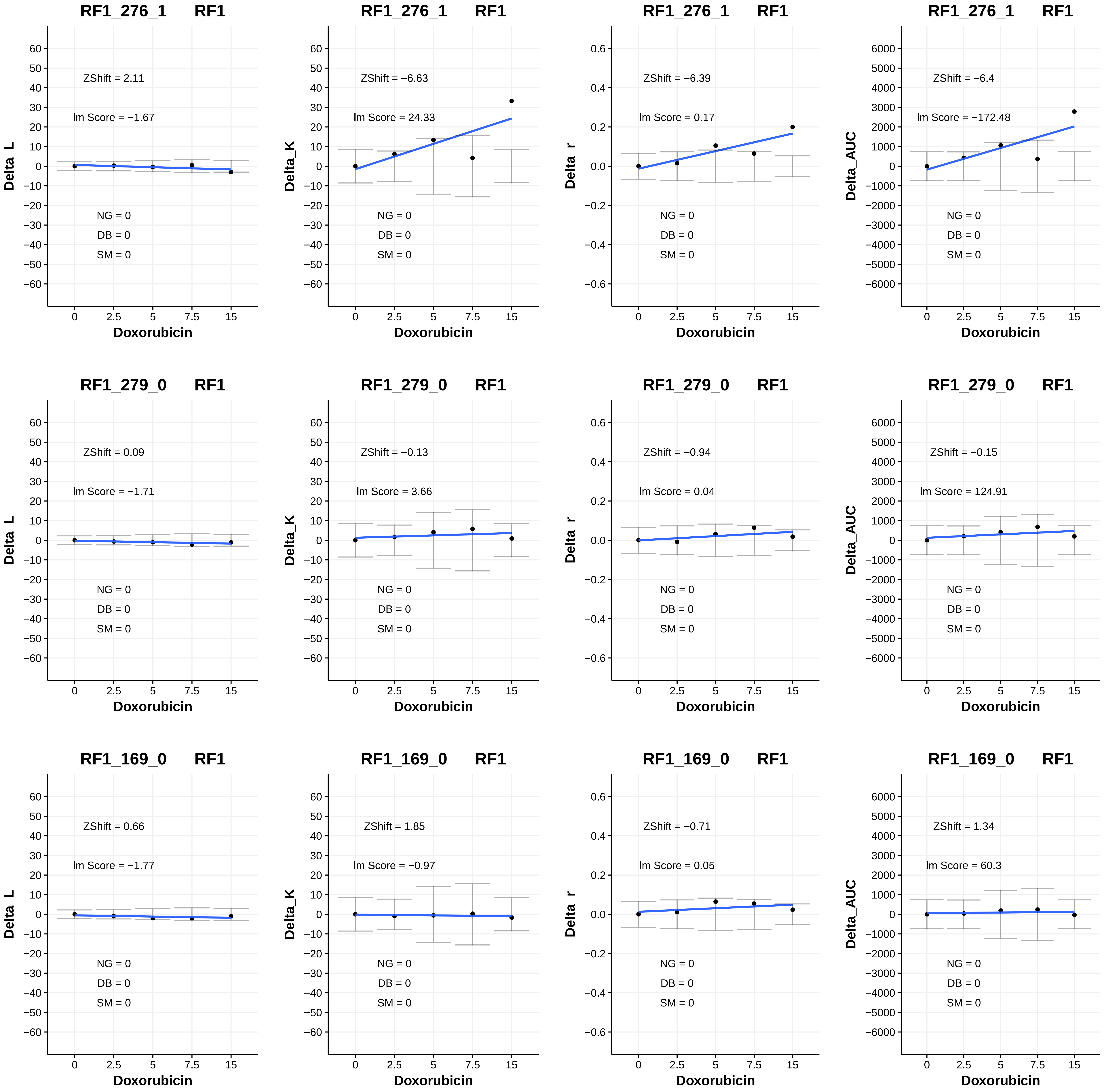

RF1_276_1 RF1
ZShift = 2.11
lm Score = −1.67
NG = 0
DB = 0
SM = 0
Delta_L
Doxorubicin
RF1_276_1 RF1
ZShift = −6.63
lm Score = 24.33
NG = 0
DB = 0
SM = 0
Delta_K
Doxorubicin
RF1_276_1 RF1
ZShift = −6.39
lm Score = 0.17
NG = 0
DB = 0
SM = 0
Delta_r
Doxorubicin
RF1_276_1 RF1
ZShift = −6.4
lm Score = −172.48
NG = 0
DB = 0
SM = 0
Delta_AUC
Doxorubicin
RF1_279_0 RF1
ZShift = 0.09
lm Score = −1.71
NG = 0
DB = 0
SM = 0
Delta_L
Doxorubicin
RF1_279_0 RF1
ZShift = −0.13
lm Score = 3.66
NG = 0
DB = 0
SM = 0
Delta_K
Doxorubicin
RF1_279_0 RF1
ZShift = −0.94
lm Score = 0.04
NG = 0
DB = 0
SM = 0
Delta_r
Doxorubicin
RF1_279_0 RF1
ZShift = −0.15
lm Score = 124.91
NG = 0
DB = 0
SM = 0
Delta_AUC
Doxorubicin
RF1_169_0 RF1
ZShift = 0.66
lm Score = −1.77
NG = 0
DB = 0
SM = 0
Delta_L
Doxorubicin
RF1_169_0 RF1
ZShift = 1.85
lm Score = −0.97
NG = 0
DB = 0
SM = 0
Delta_K
Doxorubicin
RF1_169_0 RF1
ZShift = −0.71
lm Score = 0.05
NG = 0
DB = 0
SM = 0
Delta_r
Doxorubicin
RF1_169_0 RF1
ZShift = 1.34
lm Score = 60.3
NG = 0
DB = 0
SM = 0
Delta_AUC
Doxorubicin

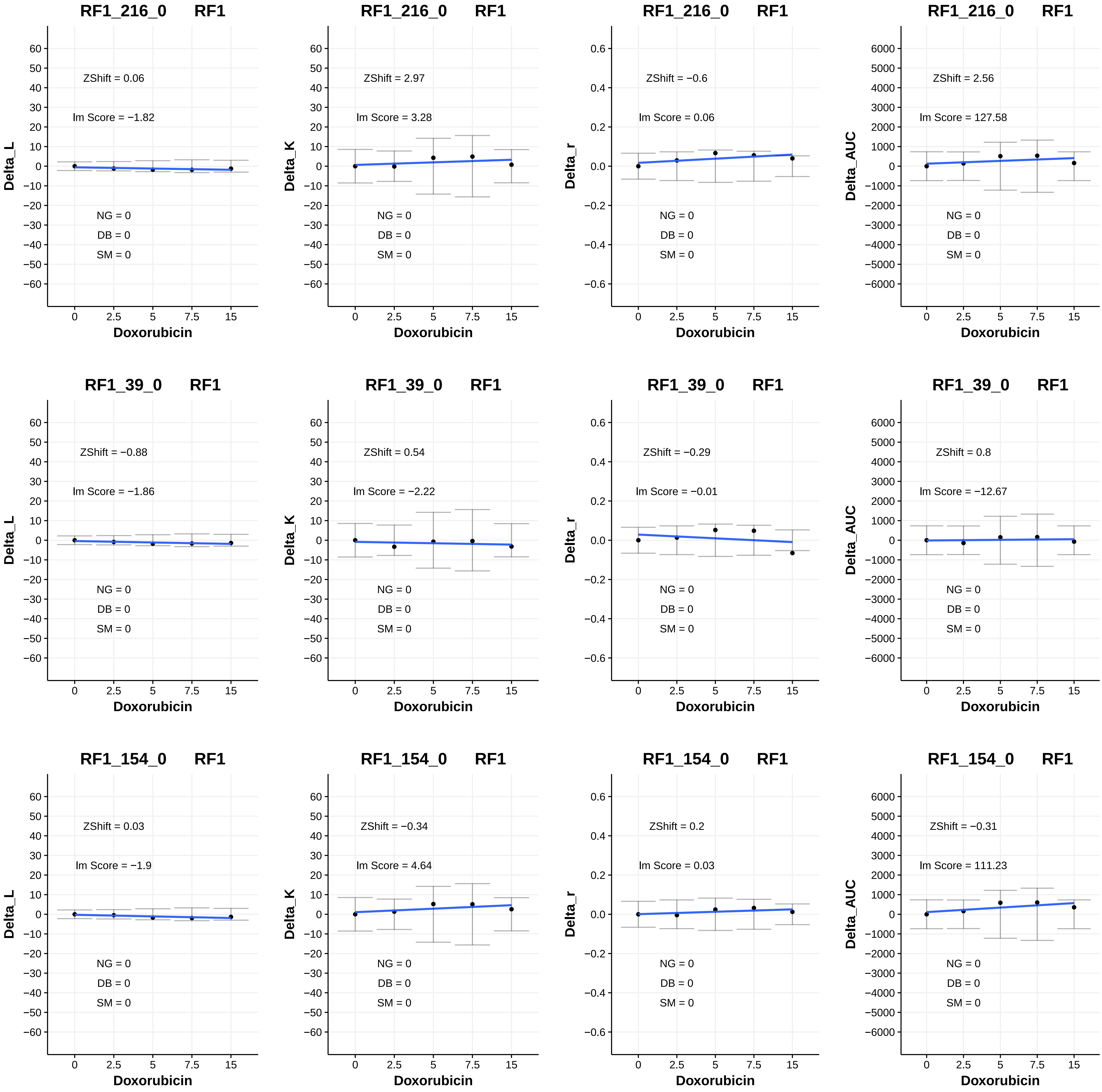

RF1_216_0 RF1
ZShift = 0.06
lm Score = −1.82
NG = 0
DB = 0
SM = 0
Delta_L
Doxorubicin
RF1_216_0 RF1
ZShift = 2.97
lm Score = 3.28
NG = 0
DB = 0
SM = 0
Delta_K
Doxorubicin
RF1_216_0 RF1
ZShift = −0.6
lm Score = 0.06
NG = 0
DB = 0
SM = 0
Delta_r
Doxorubicin
RF1_216_0 RF1
ZShift = 2.56
lm Score = 127.58
NG = 0
DB = 0
SM = 0
Delta_AUC
Doxorubicin
RF1_39_0 RF1
ZShift = −0.88
lm Score = −1.86
NG = 0
DB = 0
SM = 0
Delta_L
Doxorubicin
RF1_39_0 RF1
ZShift = 0.54
lm Score = −2.22
NG = 0
DB = 0
SM = 0
Delta_K
Doxorubicin
RF1_39_0 RF1
ZShift = −0.29
lm Score = −0.01
NG = 0
DB = 0
SM = 0
Delta_r
Doxorubicin
RF1_39_0 RF1
ZShift = 0.8
lm Score = −12.67
NG = 0
DB = 0
SM = 0
Delta_AUC
Doxorubicin
RF1_154_0 RF1
ZShift = 0.03
lm Score = −1.9
NG = 0
DB = 0
SM = 0
Delta_L
Doxorubicin
RF1_154_0 RF1
ZShift = −0.34
lm Score = 4.64
NG = 0
DB = 0
SM = 0
Delta_K
Doxorubicin
RF1_154_0 RF1
ZShift = 0.2
lm Score = 0.03
NG = 0
DB = 0
SM = 0
Delta_r
Doxorubicin
RF1_154_0 RF1
ZShift = −0.31
lm Score = 111.23
NG = 0
DB = 0
SM = 0
Delta_AUC
Doxorubicin

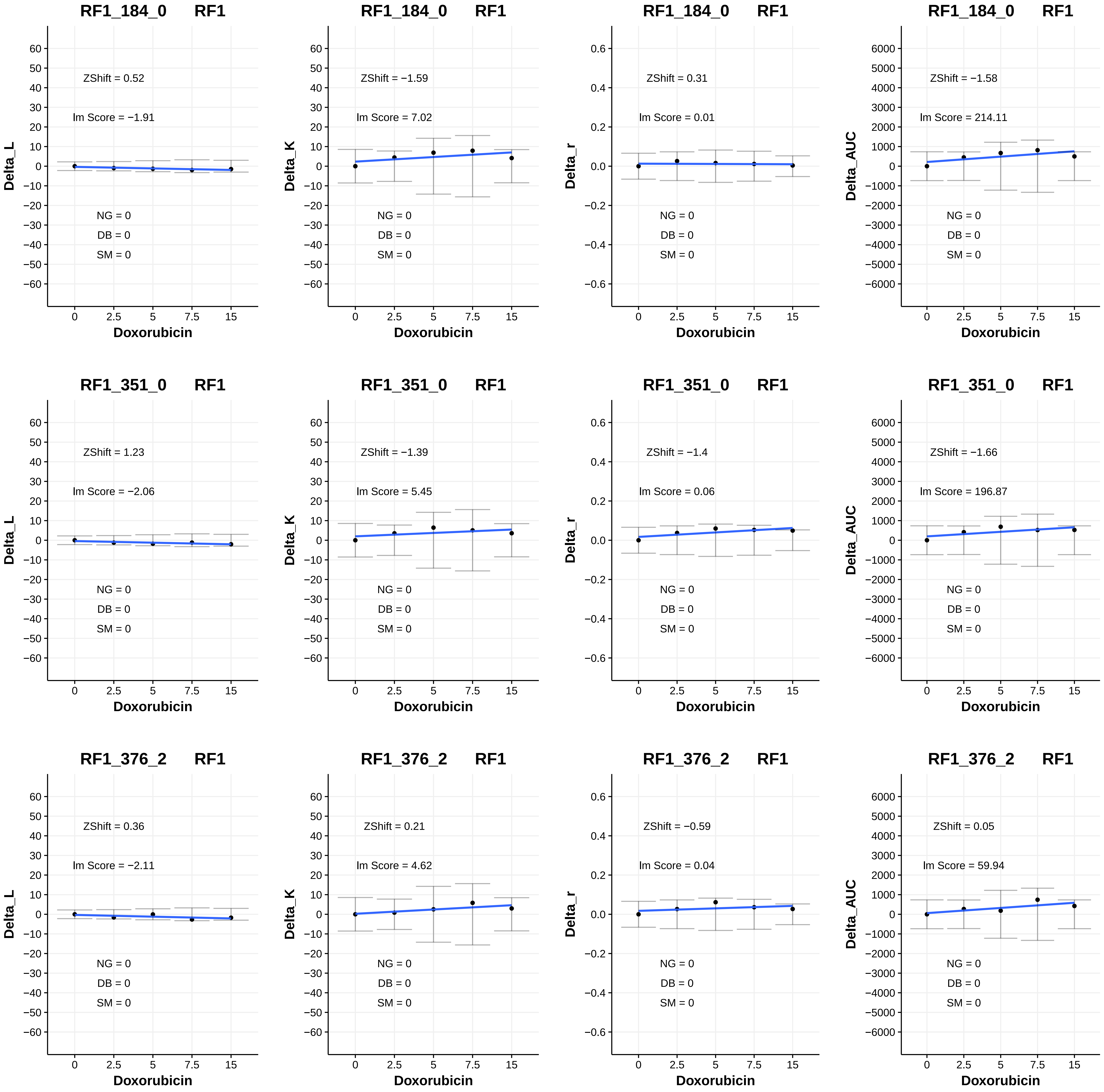
RF1_184_0 RF1
ZShift = 0.52
lm Score = −1.91
NG = 0
DB = 0
SM = 0
Delta_L
Doxorubicin
RF1_184_0 RF1
ZShift = −1.59
lm Score = 7.02
NG = 0
DB = 0
SM = 0
Delta_K
Doxorubicin
RF1_184_0 RF1
ZShift = 0.31
lm Score = 0.01
NG = 0
DB = 0
SM = 0
Delta_r
Doxorubicin
RF1_184_0 RF1
ZShift = −1.58
lm Score = 214.11
NG = 0
DB = 0
SM = 0
Delta_AUC
Doxorubicin
RF1_351_0 RF1
ZShift = 1.23
lm Score = −2.06
NG = 0
DB = 0
SM = 0
Delta_L
Doxorubicin
RF1_351_0 RF1
ZShift = −1.39
lm Score = 5.45
NG = 0
DB = 0
SM = 0
Delta_K
Doxorubicin
RF1_351_0 RF1
ZShift = −1.4
lm Score = 0.06
NG = 0
DB = 0
SM = 0
Delta_r
Doxorubicin
RF1_351_0 RF1
ZShift = −1.66
lm Score = 196.87
NG = 0
DB = 0
SM = 0
Delta_AUC
Doxorubicin
RF1_376_2 RF1
ZShift = 0.36
lm Score = −2.11
NG = 0
DB = 0
SM = 0
Delta_L
Doxorubicin
RF1_376_2 RF1
ZShift = 0.21
lm Score = 4.62
NG = 0
DB = 0
SM = 0
Delta_K
Doxorubicin
RF1_376_2 RF1
ZShift = −0.59
lm Score = 0.04
NG = 0
DB = 0
SM = 0
Delta_r
Doxorubicin
RF1_376_2 RF1
ZShift = 0.05
lm Score = 59.94
NG = 0
DB = 0
SM = 0
Delta_AUC
Doxorubicin

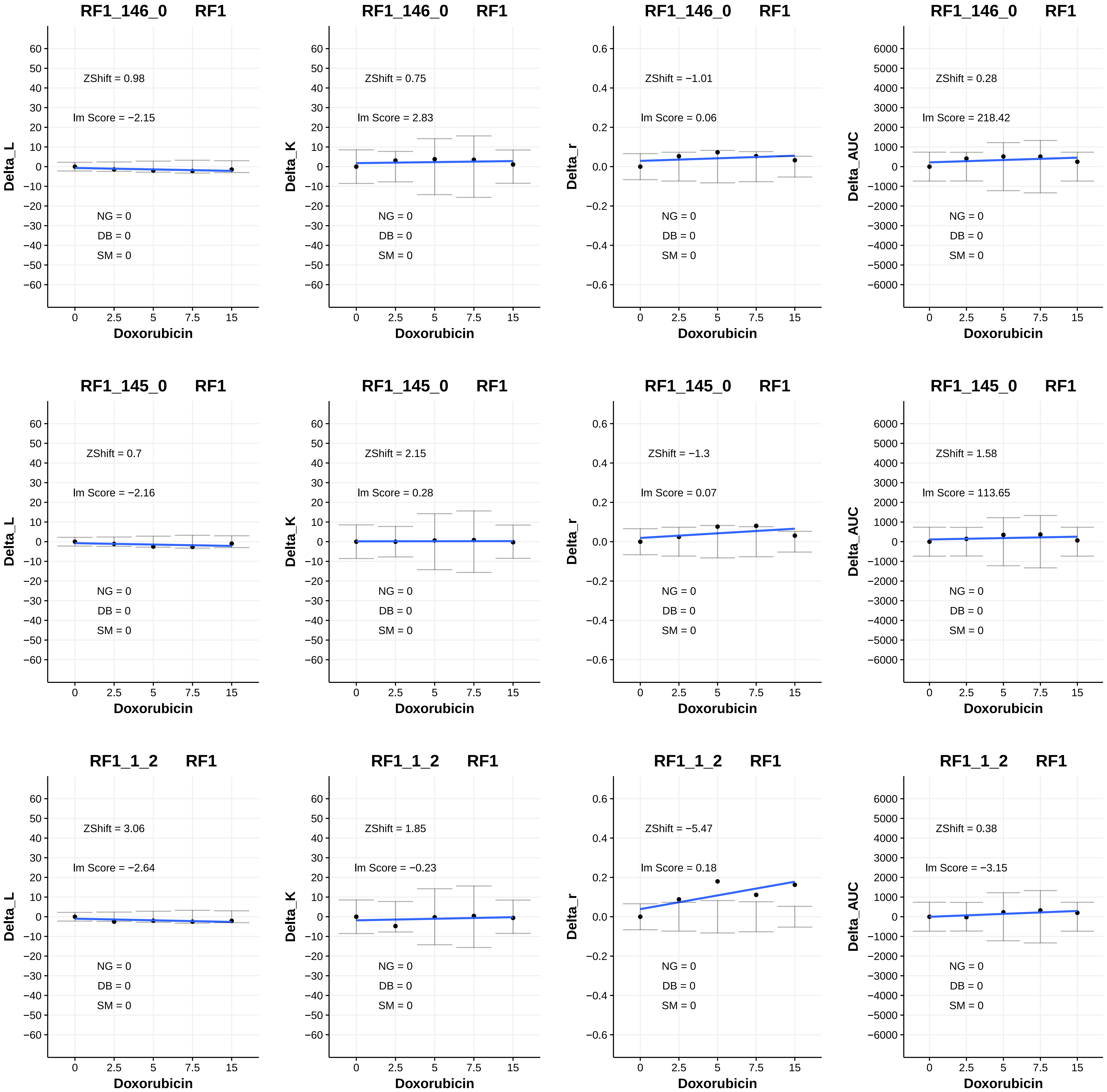
RF1_146_0 RF1
ZShift = 0.98
lm Score = −2.15
NG = 0
DB = 0
SM = 0
Delta_L
Doxorubicin
RF1_146_0 RF1
ZShift = 0.75
lm Score = 2.83
NG = 0
DB = 0
SM = 0
Delta_K
Doxorubicin
RF1_146_0 RF1
ZShift = −1.01
lm Score = 0.06
NG = 0
DB = 0
SM = 0
Delta_r
Doxorubicin
RF1_146_0 RF1
ZShift = 0.28
lm Score = 218.42
NG = 0
DB = 0
SM = 0
Delta_AUC
Doxorubicin
RF1_145_0 RF1
ZShift = 0.7
lm Score = −2.16
NG = 0
DB = 0
SM = 0
Delta_L
Doxorubicin
RF1_145_0 RF1
ZShift = 2.15
lm Score = 0.28
NG = 0
DB = 0
SM = 0
Delta_K
Doxorubicin
RF1_145_0 RF1
ZShift = −1.3
lm Score = 0.07
NG = 0
DB = 0
SM = 0
Delta_r
Doxorubicin
RF1_145_0 RF1
ZShift = 1.58
lm Score = 113.65
NG = 0
DB = 0
SM = 0
Delta_AUC
Doxorubicin
RF1_1_2 RF1
ZShift = 3.06
lm Score = −2.64
NG = 0
DB = 0
SM = 0
Delta_L
Doxorubicin
RF1_1_2 RF1
ZShift = 1.85
lm Score = −0.23
NG = 0
DB = 0
SM = 0
Delta_K
Doxorubicin
RF1_1_2 RF1
ZShift = −5.47
lm Score = 0.18
NG = 0
DB = 0
SM = 0
Delta_r
Doxorubicin
RF1_1_2 RF1
ZShift = 0.38
lm Score = −3.15
NG = 0
DB = 0
SM = 0
Delta_AUC
Doxorubicin

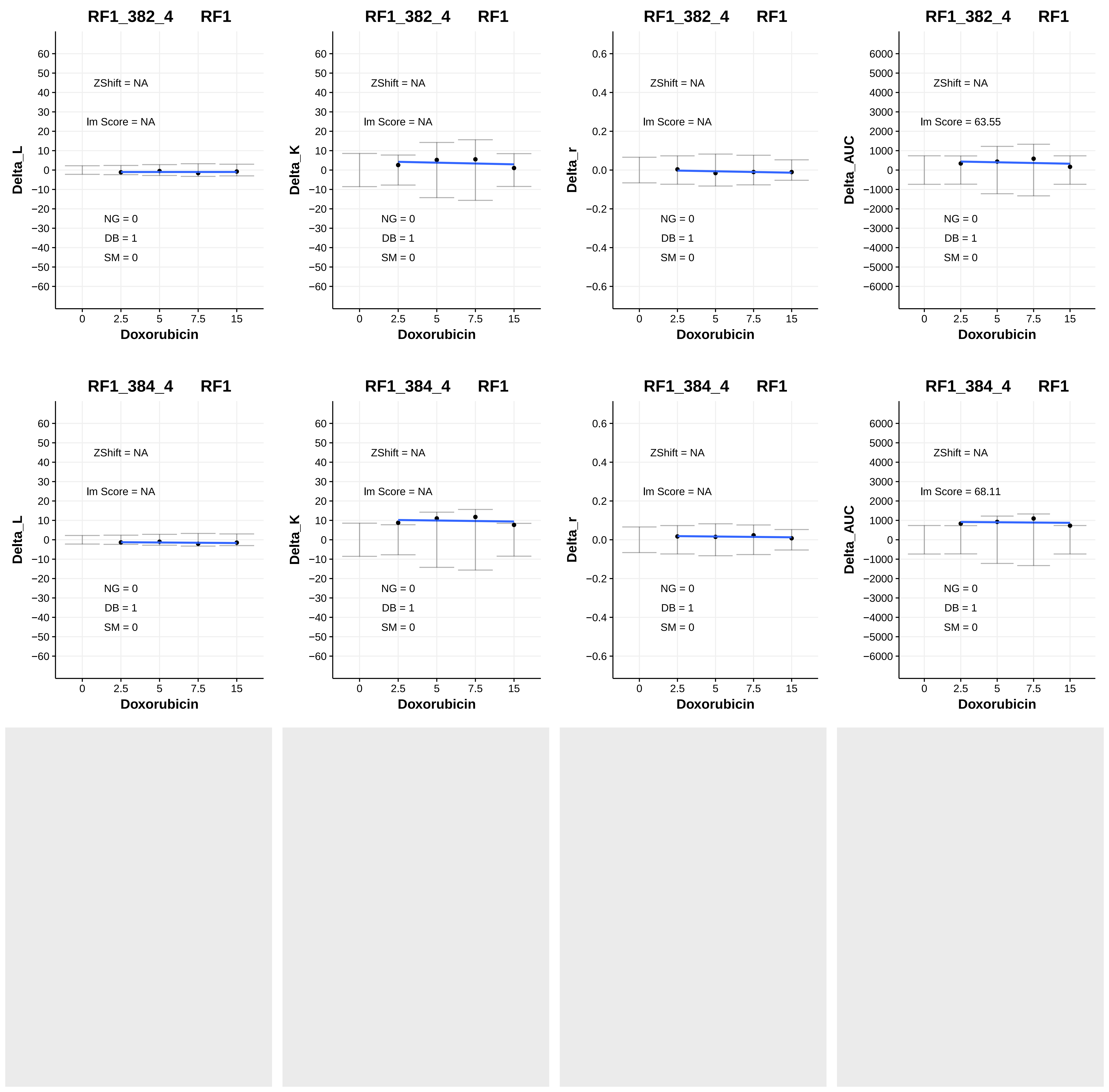

RF1_382_4 RF1
ZShift = NA
lm Score = NA
NG = 0
DB = 1
SM = 0
Delta_L
Doxorubicin
RF1_382_4 RF1
ZShift = NA
lm Score = NA
NG = 0
DB = 1
SM = 0
Delta_K
Doxorubicin
RF1_382_4 RF1
ZShift = NA
lm Score = NA
NG = 0
DB = 1
SM = 0
Delta_r
Doxorubicin
RF1_382_4 RF1
ZShift = NA
lm Score = 63.55
NG = 0
DB = 1
SM = 0
Delta_AUC
Doxorubicin
RF1_384_4 RF1
ZShift = NA
lm Score = NA
NG = 0
DB = 1
SM = 0
Delta_L
Doxorubicin
RF1_384_4 RF1
ZShift = NA
lm Score = NA
NG = 0
DB = 1
SM = 0
Delta_K
Doxorubicin
RF1_384_4 RF1
ZShift = NA
lm Score = NA
NG = 0
DB = 1
SM = 0
Delta_r
Doxorubicin
RF1_384_4 RF1
ZShift = NA
lm Score = 68.11
NG = 0
DB = 1
SM = 0
Delta_AUC
Doxorubicin
