## Additional File 5 for "A yeast phenomic model for the influence of Warburg metabolism on genetic buffering of doxorubicin": B - Heatmaps.pdf

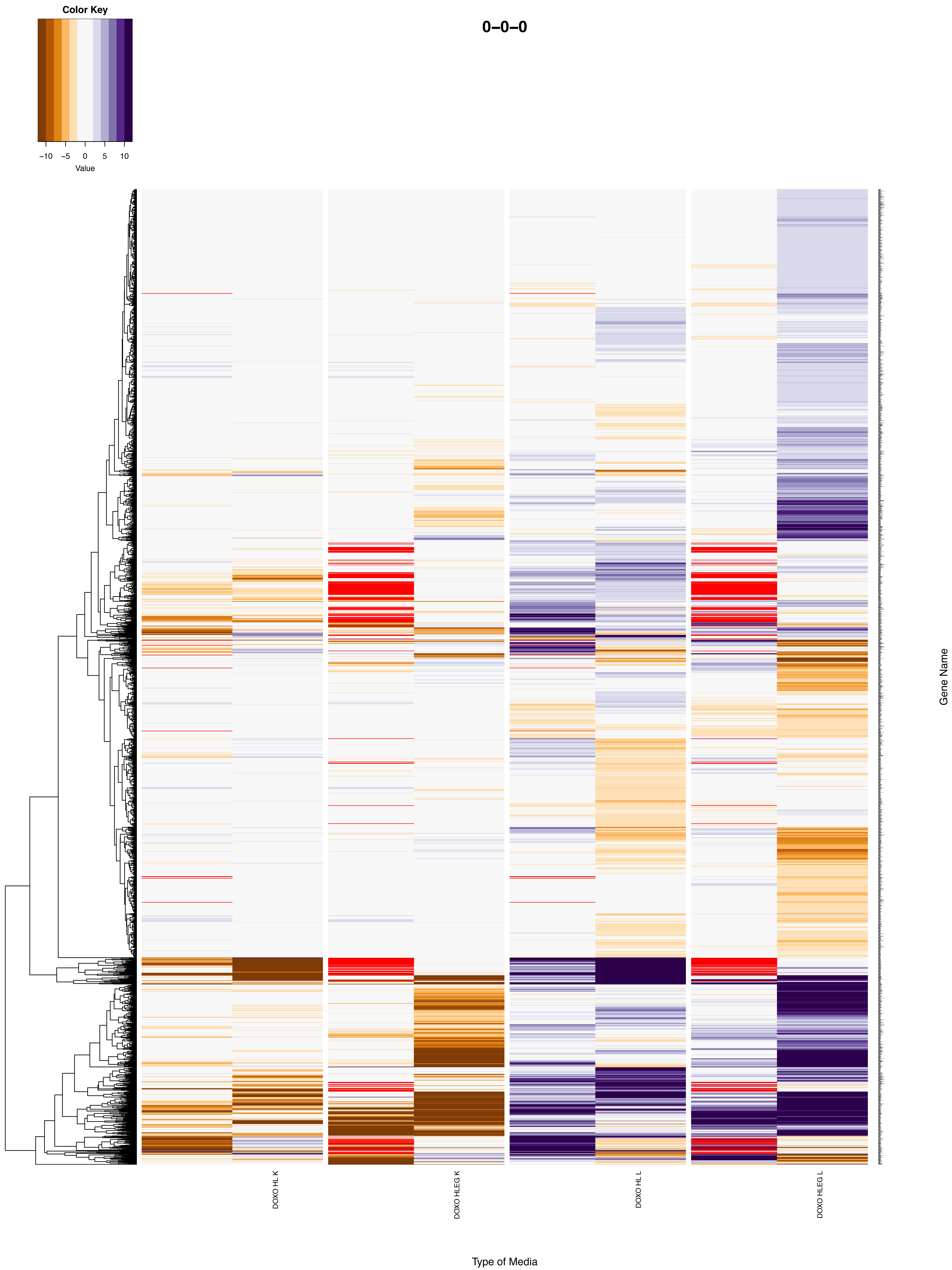
0–0–0
Color Key
−10
−5
0
5
10
Value
Gene Name
DOXO HL K
DOXO HLEG K
DOXO HL L
DOXO HLEG L
Type of Media

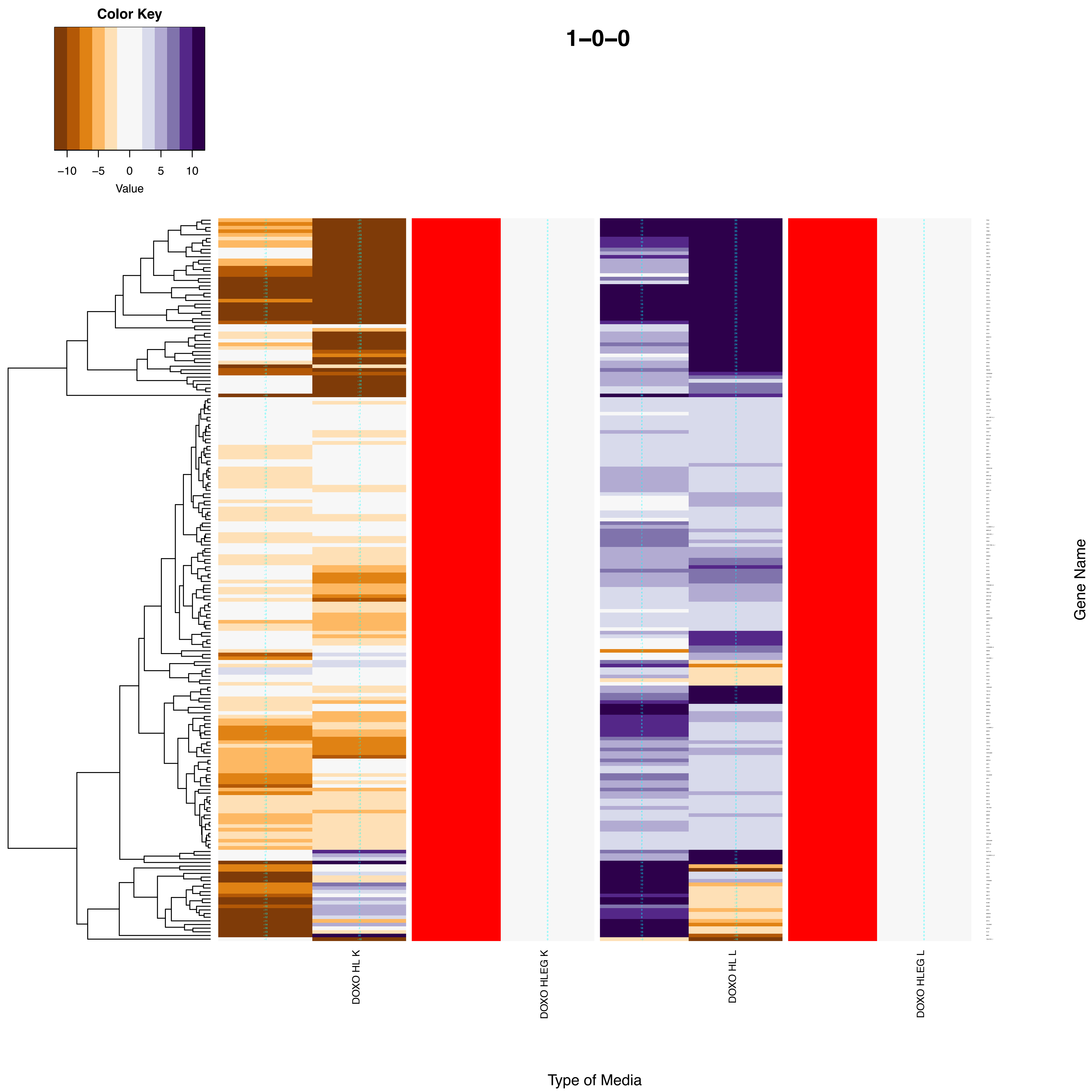

Color Key
−10
−5
0
5
10
Value
1−0−0
DOXO HL K
DOXO HLEG K
DOXO HL L
DOXO HLEG L
Type of Media
Gene Name

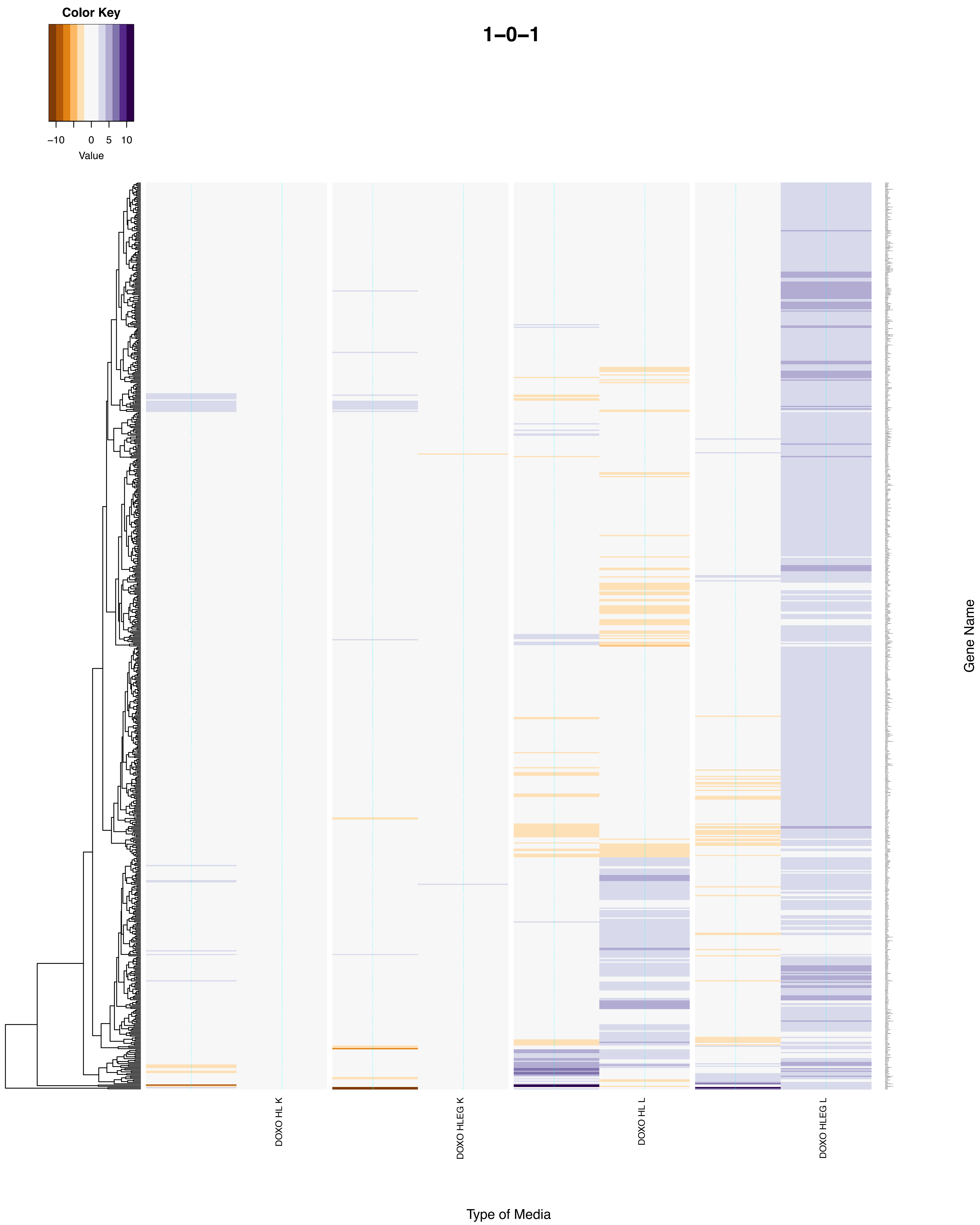
Color Key
−10
0
5
10
Value
1−0−1
DOXO HL K
DOXO HLEG K
DOXO HL L
DOXO HLEG L
Type of Media
Gene Name

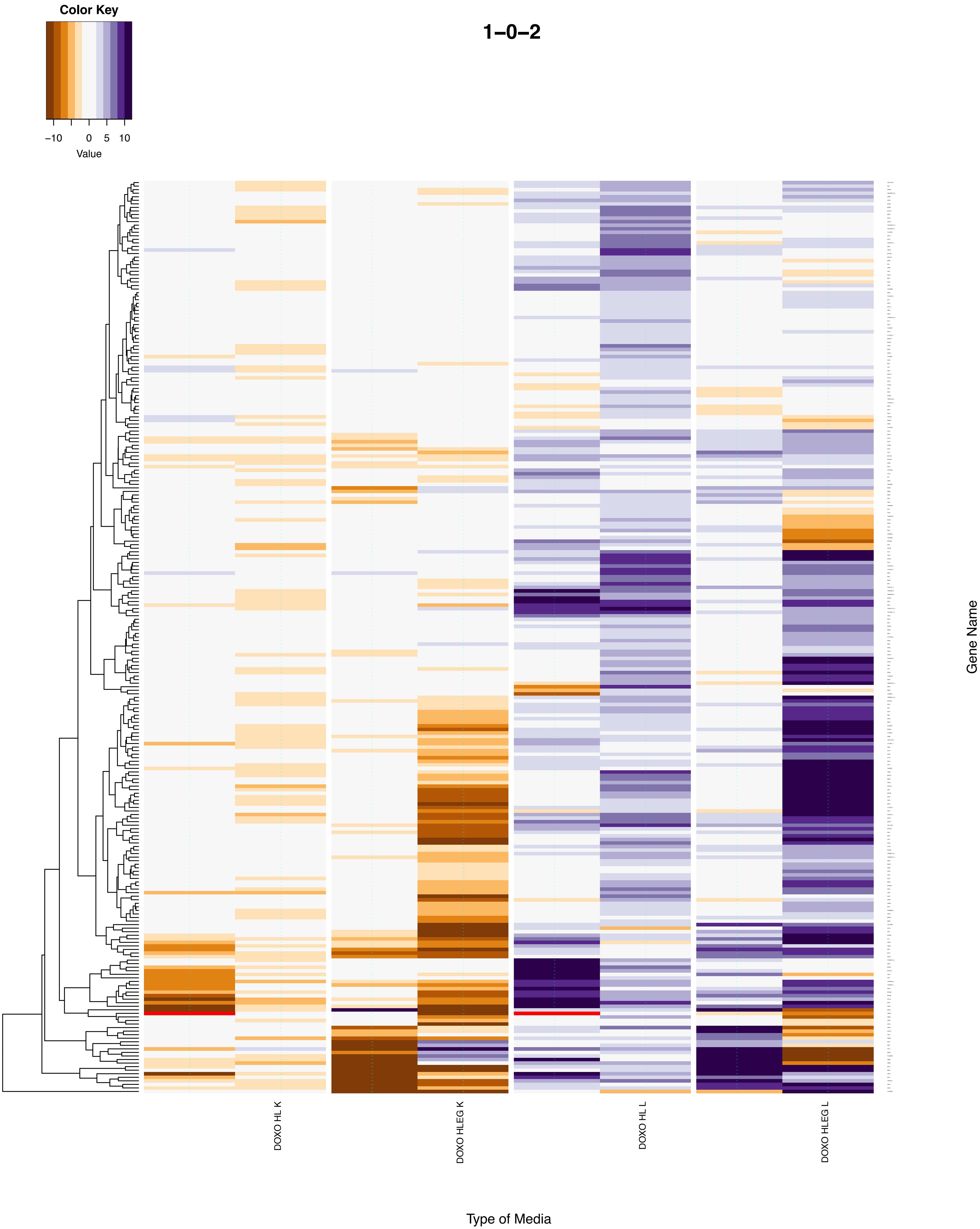

Color Key
−10
0
5
10
Value
1−0−2
DOXO HL K
DOXO HLEG K
DOXO HL L
DOXO HLEG L
Gene Name
Type of Media

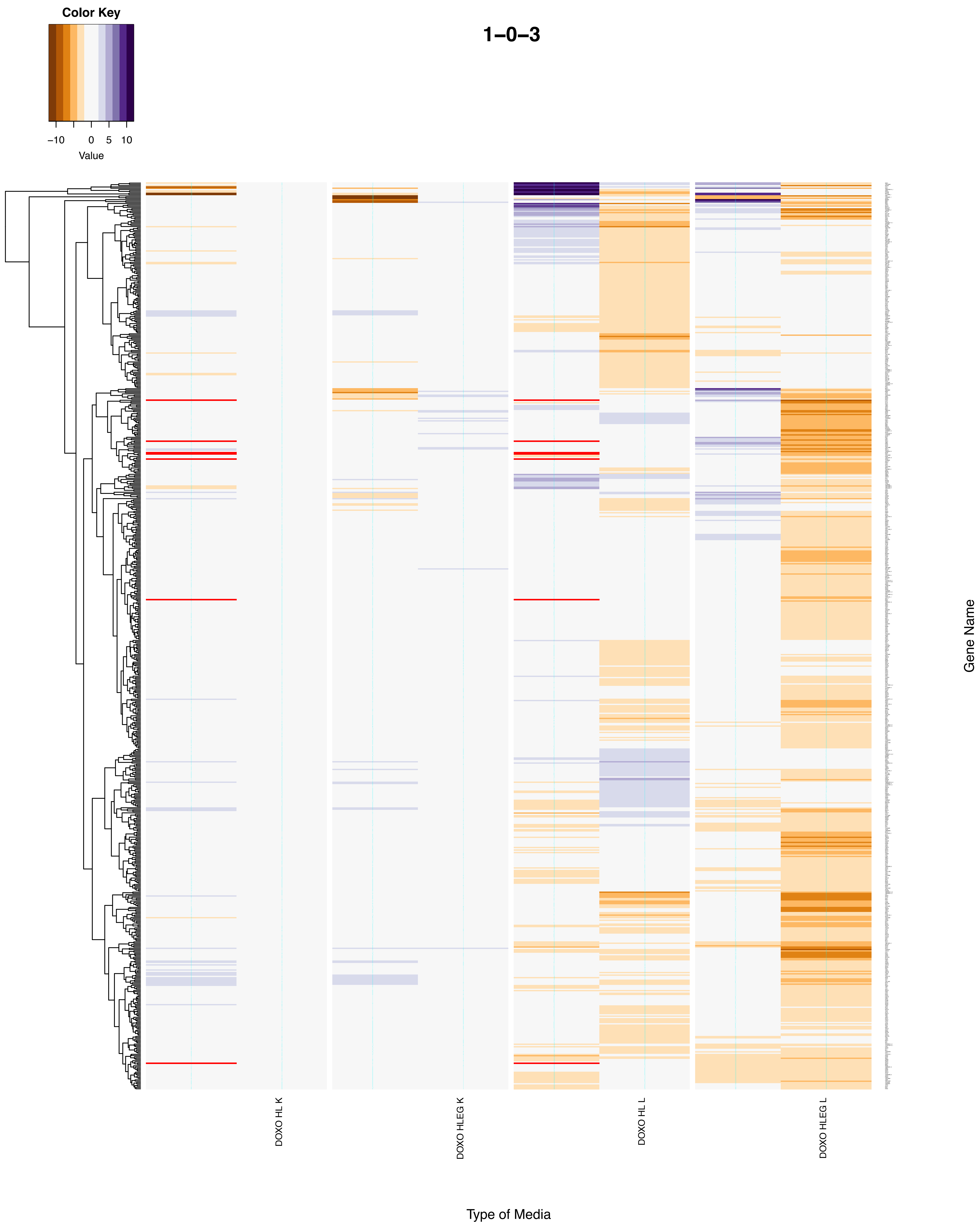

Color Key
−10
0
5
10
Value
1−0−3
DOXO HL K
DOXO HLEG K
DOXO HL L
DOXO HLEG L
Gene Name
Type of Media

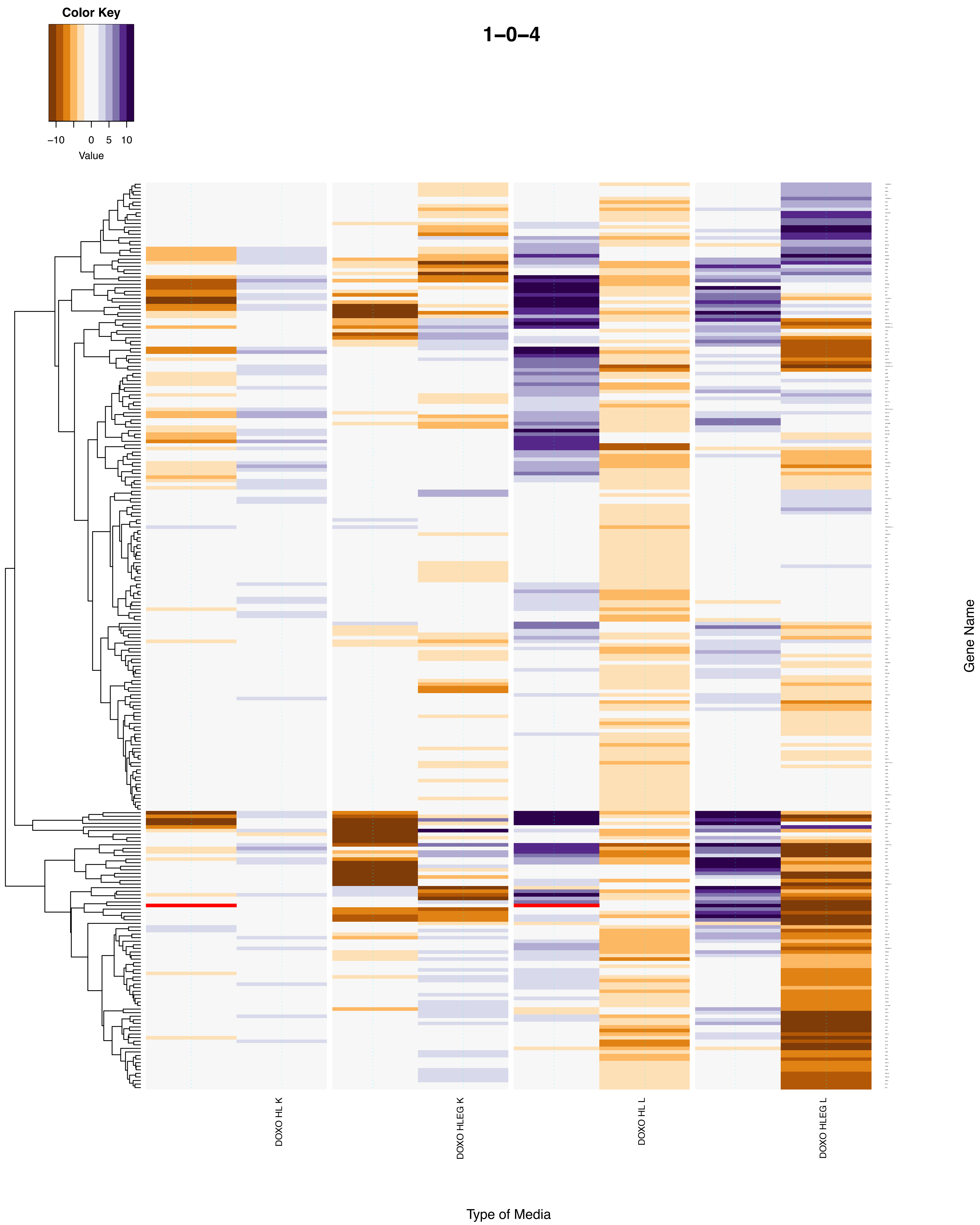
Color Key
−10
0
5
10
Value
1−0−4
DOXO HL K
DOXO HLEG K
DOXO HL L
DOXO HLEG L
Type of Media
Gene Name

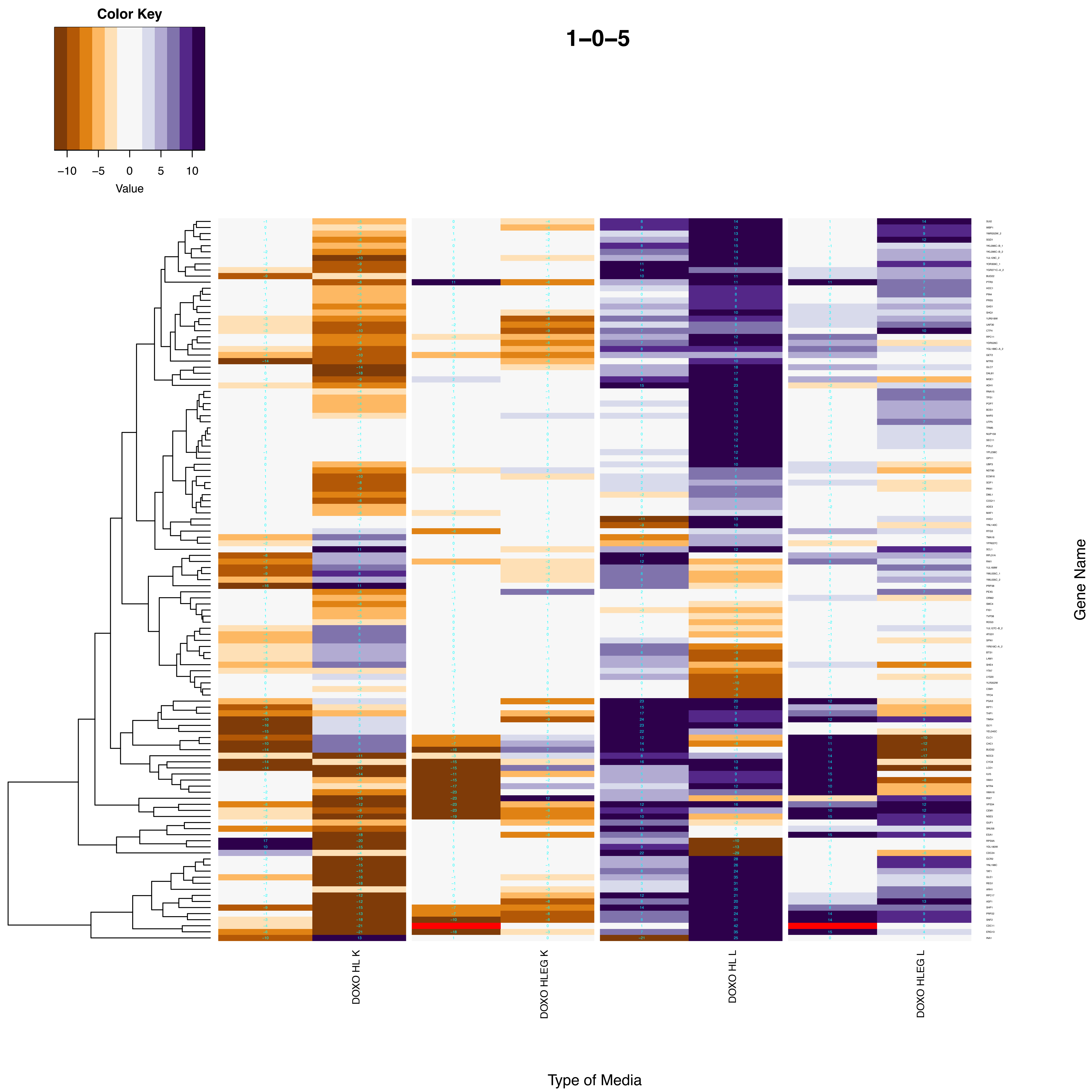

Color Key
−10
−5
0
5
10
Value
1−0−5
DOXO HL K
DOXO HLEG K
DOXO HL L
DOXO HLEG L
Type of Media
Gene Name

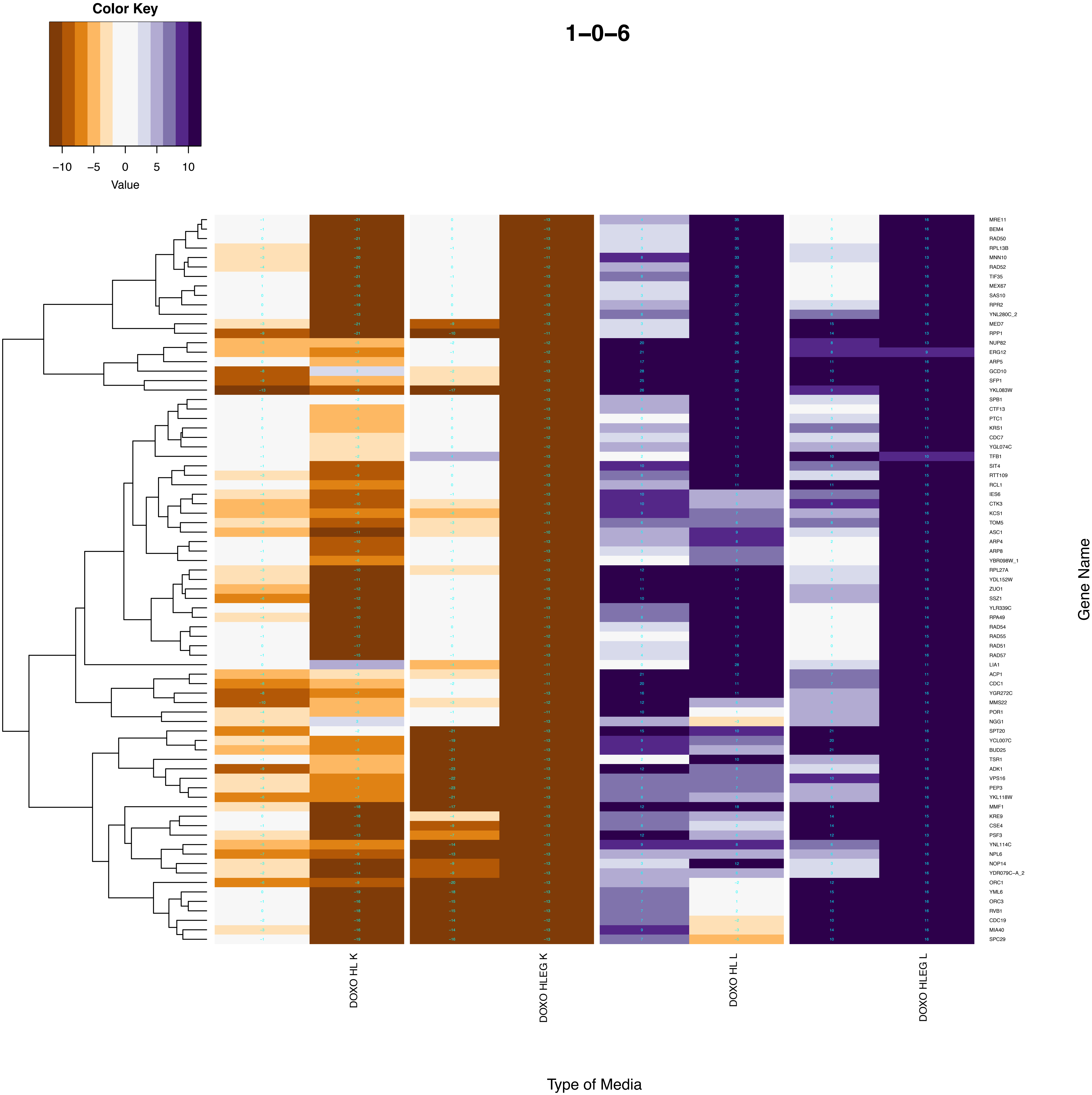
Color Key
−10
−5
0
5
10
Value
1−0−6
MRE11
BEM4
RAD50
RPL13B
MNN10
RAD52
TIF35
MEX67
SAS10
RPR2
YNL280C_2
MED7
RPP1
NUP82
ERG12
ARP5
GCD10
SFP1
YKL083W
SPB1
CTF13
PTC1
KRS1
CDC7
YGL074C
TFB1
SIT4
RTT109
RCL1
IES6
CTK3
KCS1
TOM5
ASC1
ARP4
ARP8
YBR098W_1
RPL27A
YDL152W
ZUO1
SSZ1
YLR339C
RPA49
RAD54
RAD55
RAD51
RAD57
LIA1
ACP1
CDC1
YGR272C
MMS22
POR1
NGG1
SPT20
YCL007C
BUD25
TSR1
ADK1
VPS16
PEP3
YKL118W
MMF1
KRE9
CSE4
PSF3
YNL114C
NPL6
NOP14
YDR079C−A_2
ORC1
YML6
ORC3
RVB1
CDC19
MIA40
SPC29
Gene Name
DOXO HL K
DOXO HLEG K
DOXO HL L
DOXO HLEG L
Type of Media

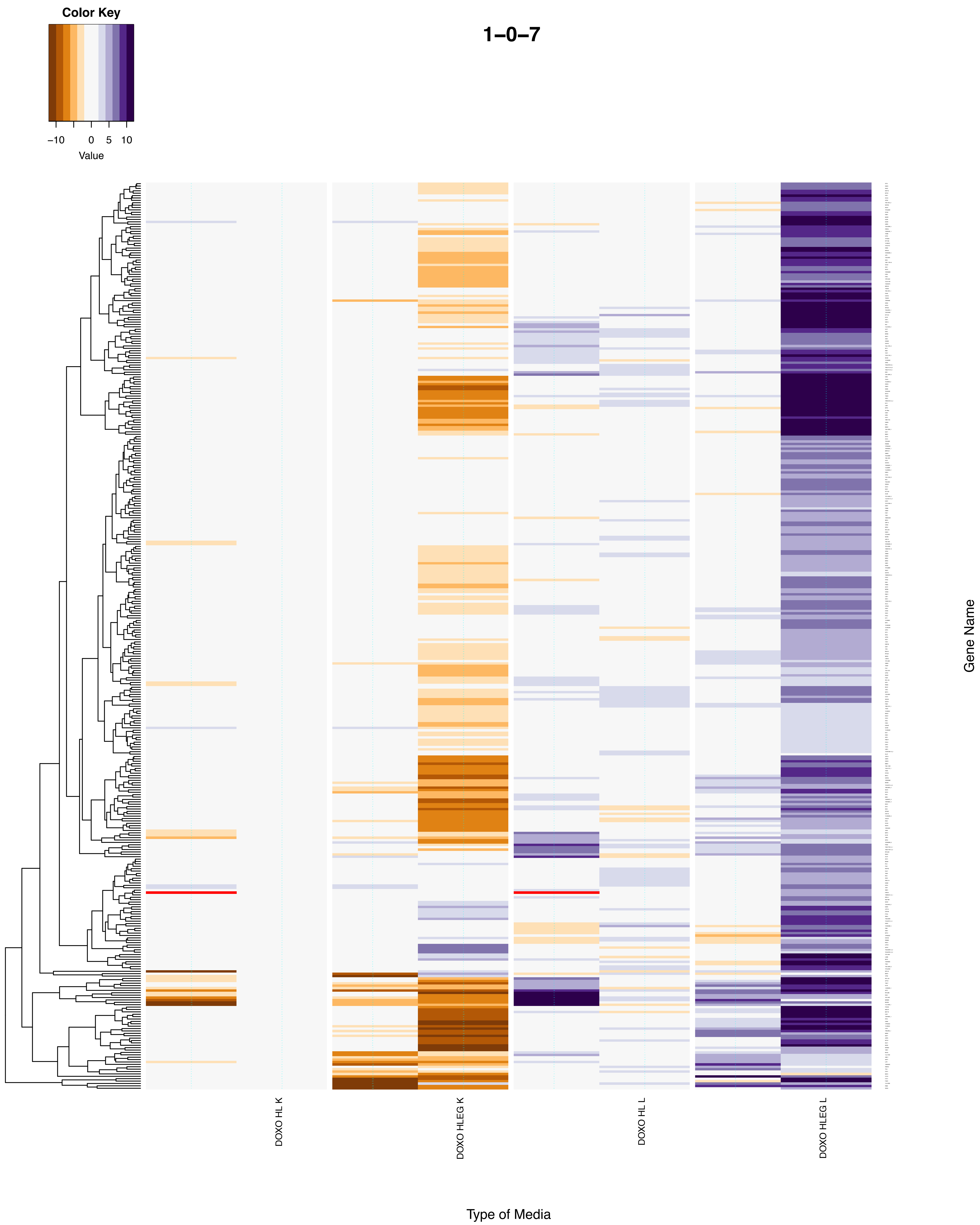

Color Key
-10
0
5
10
Value
1-0-7
DOXO HL K
DOXO HLEG K
DOXO HL L
DOXO HLEG L
Gene Name
Type of Media

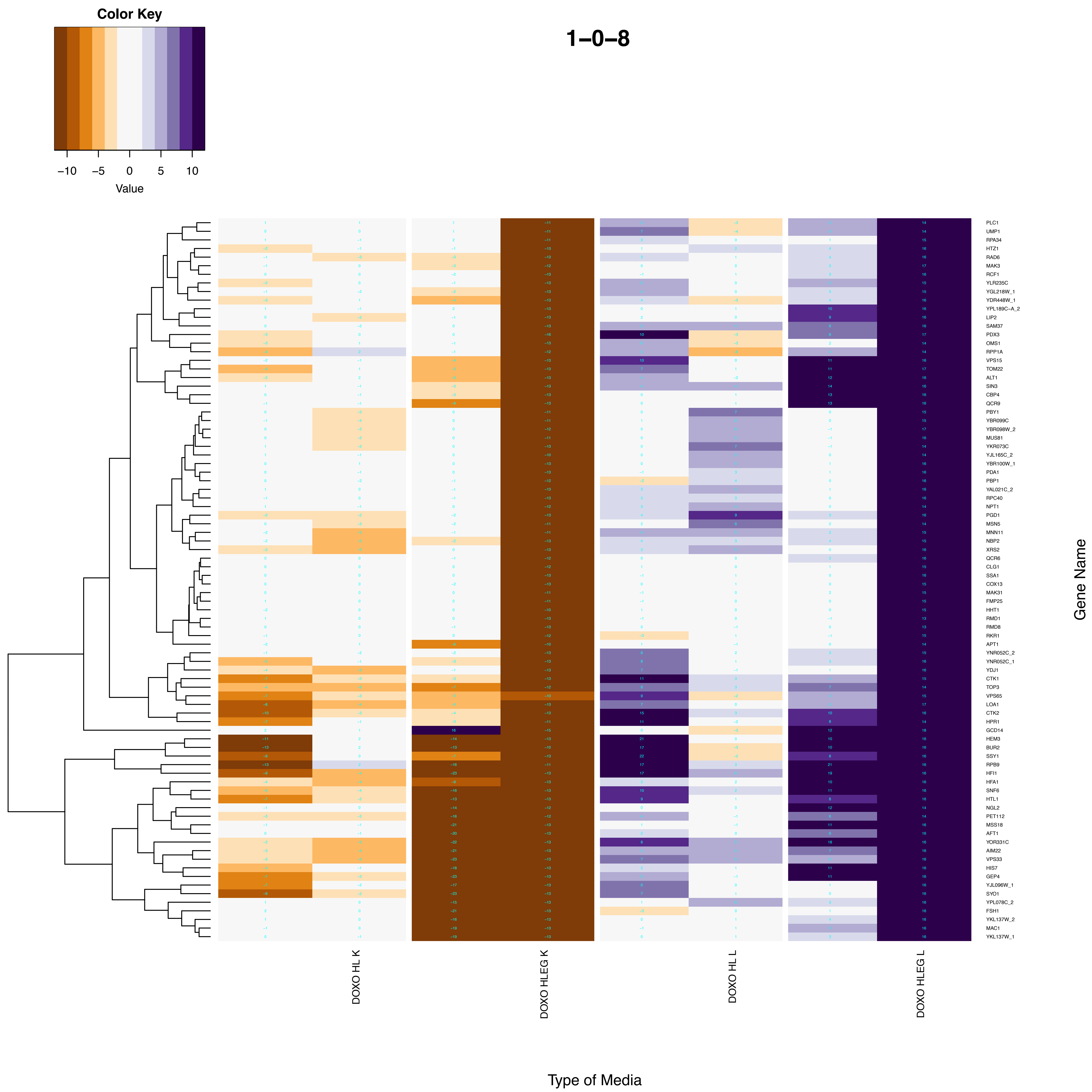

Color Key
−10
−5
0
5
10
Value
1−0−8
PLC1
UMP1
RPA34
HTZ1
RAD6
MAK3
RCF1
YLR235C
YGL218W_1
YDR448W_1
YPL189C−A_2
LIP2
SAM37
PDX3
OMS1
RPP1A
VPS15
TOM22
ALT1
SIN3
CBP4
QCR9
PBY1
YBR099C
YBR098W_2
MUS81
YKR073C
YJL165C_2
YBR100W_1
PDA1
PBP1
YAL021C_2
RPC40
NPT1
PGD1
MSN5
MNN11
NBP2
XRS2
QCR6
CLG1
SSA1
COX13
MAK31
FMP25
HHT1
RMD1
RMD8
RKR1
APT1
YNR052C_2
YNR052C_1
YDJ1
CTK1
TOP3
VPS65
LOA1
CTK2
HPR1
GCD14
HEM3
BUR2
SSY1
RPB9
HFI1
HFA1
SNF6
HTL1
NGL2
PET112
MSS18
AFT1
YOR331C
AIM22
VPS33
HIS7
GEP4
YJL096W_1
SYO1
YPL078C_2
FSH1
YKL137W_2
MAC1
YKL137W_1
Gene Name
DOXO HL K
DOXO HLEG K
DOXO HL L
DOXO HLEG L
Type of Media

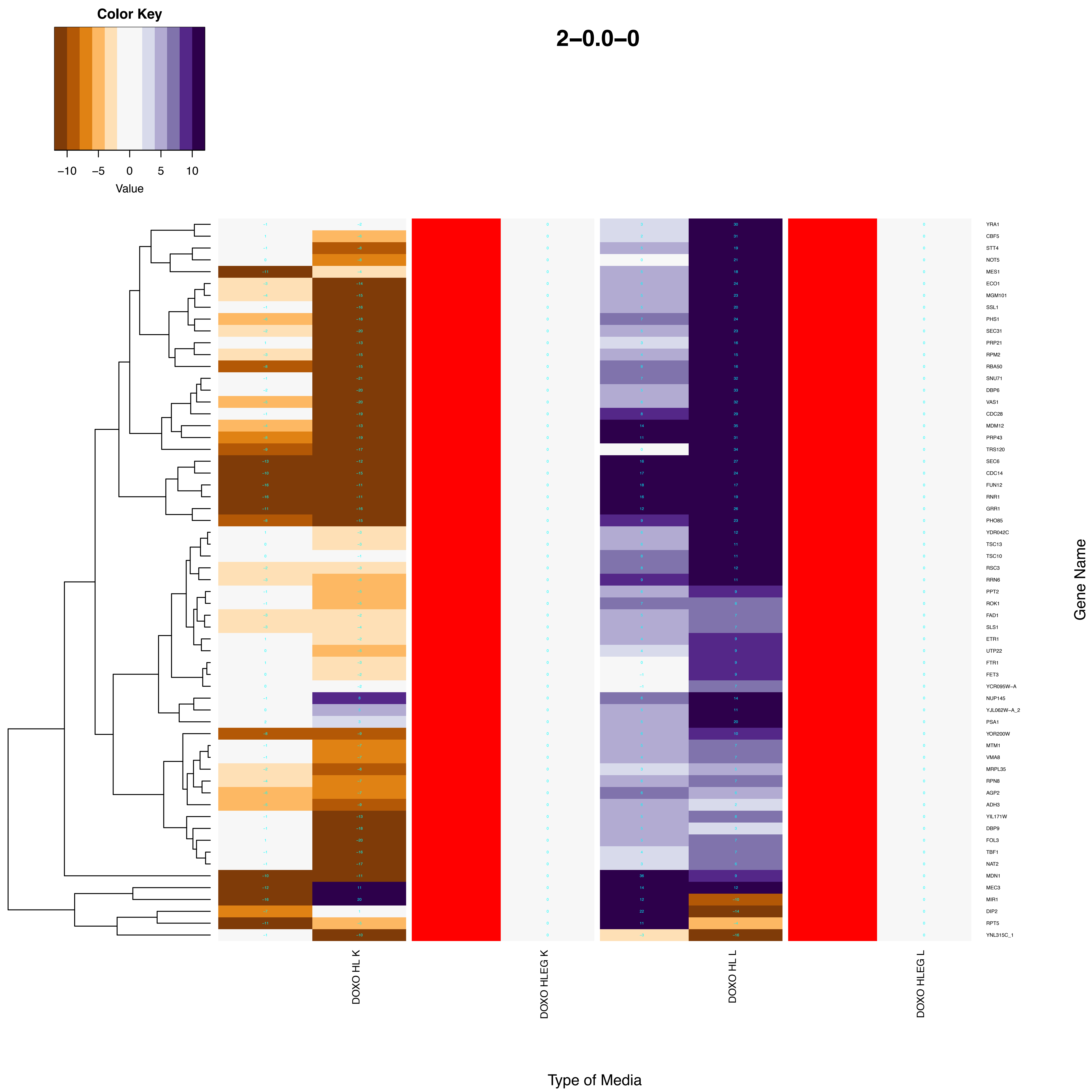
Color Key
−10
−5
0
5
10
Value
2−0.0−0
YRA1
CBF5
STT4
NOT5
MES1
ECO1
MGM101
SSL1
PHS1
SEC31
PRP21
RPM2
RBA50
SNU71
DBP6
VAS1
CDC28
MDM12
PRP43
TRS120
SEC6
CDC14
FUN12
RNR1
GRR1
PHO85
YDR042C
TSC13
TSC10
RSC3
RRN6
PPT2
ROK1
FAD1
SLS1
ETR1
UTP22
FTR1
FET3
YCR095W−A
NUP145
YJL062W−A_2
PSA1
YOR200W
MTM1
VMA8
MRPL35
RPN8
AGP2
ADH3
YIL171W
DBP9
FOL3
TBF1
NAT2
MDN1
MEC3
MIR1
DIP2
RPT5
YNL315C_1
Gene Name
DOXO HL K
DOXO HLEG K
DOXO HL L
DOXO HLEG L
Type of Media

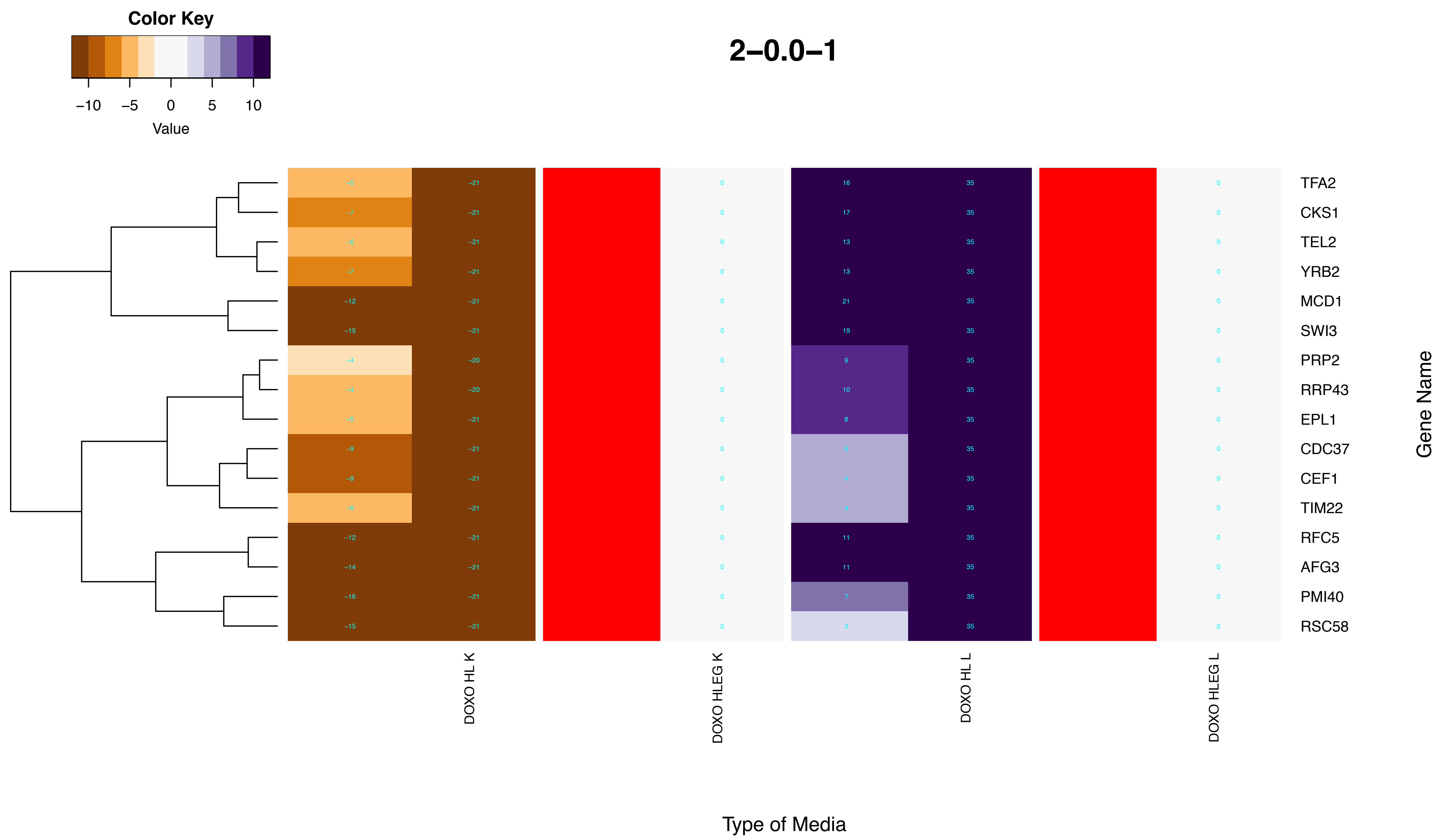
Color Key
−10
−5
0
5
10
Value
2−0.0−1
TFA2
CKS1
TEL2
YRB2
MCD1
SWI3
PRP2
RRP43
EPL1
CDC37
CEF1
TIM22
RFC5
AFG3
PMI40
RSC58
Gene Name
DOXO HL K
DOXO HLEG K
DOXO HL L
DOXO HLEG L
Type of Media

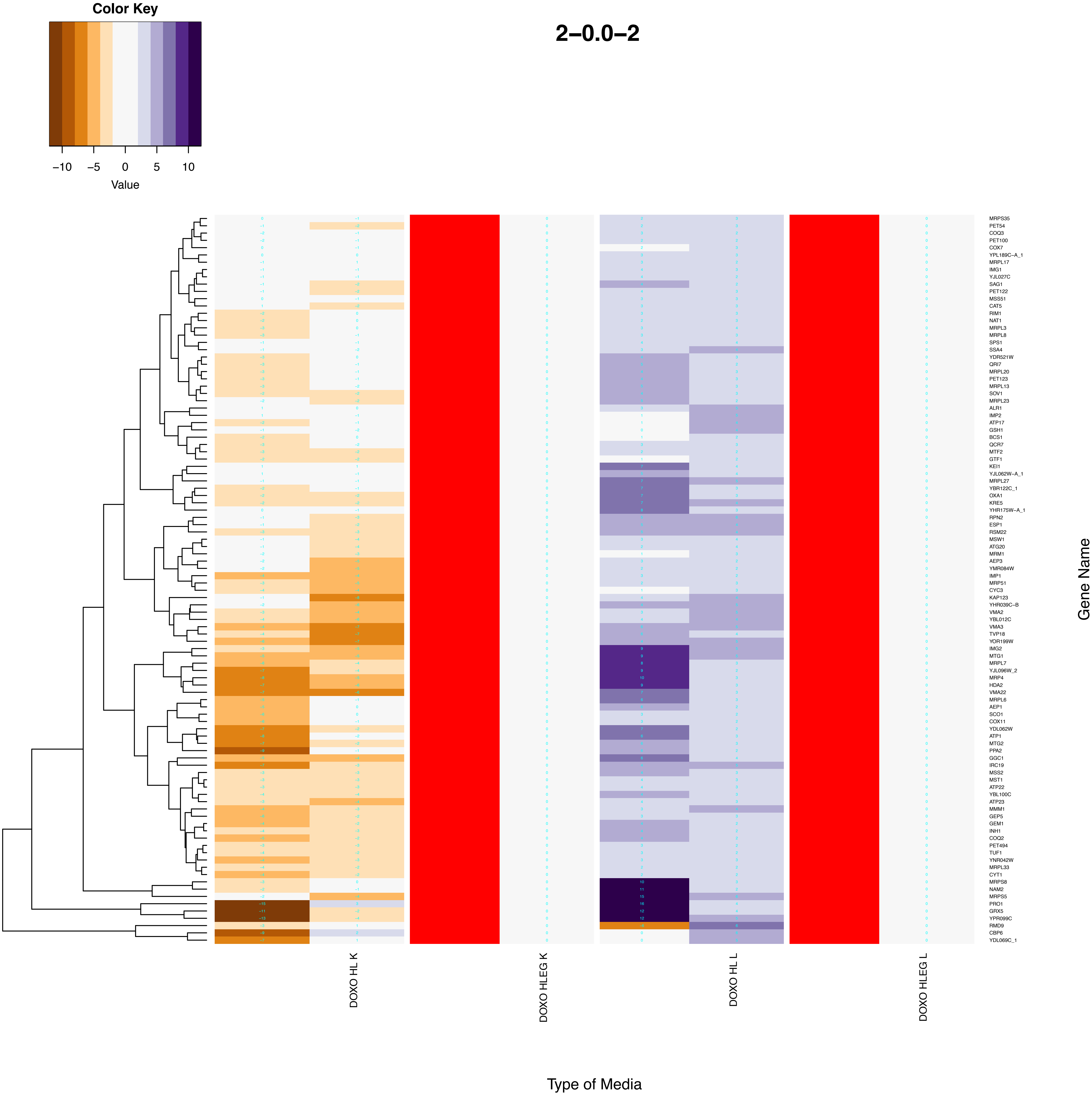

Color Key
−10
−5
0
5
10
Value
2–0.0–2
MRPS35
PET54
COQ3
PET100
COX7
YPL189C–A_1
MRPL17
IMG1
YJL027C
SAG1
PET122
MSS51
CAT5
RIM1
NAT1
MRPL3
MRPL8
SPS1
SSA4
YDR521W
QRI7
MRPL20
PET123
MRPL13
SOV1
MRPL23
ALR1
IMP2
ATP17
GSH1
BCS1
QCR7
MTF2
GTF1
KEI1
YJL062W–A_1
MRPL27
YBR122C_1
OXA1
KRE5
YHR175W–A_1
RPN2
ESP1
RSM22
MSW1
ATG20
MRM1
AEP3
YMR084W
IMP1
MRP51
CYC3
KAP123
YHR039C–B
VMA2
YBL012C
VMA3
TVP18
YOR199W
IMG2
MTG1
MRPL7
YJL096W_2
MRP4
HDA2
VMA22
MRPL6
AEP1
SCO1
COX11
YDL062W
ATP1
MTG2
PPA2
GGC1
IRC19
MSS2
MST1
ATP22
YBL100C
ATP23
MMM1
GEP5
GEM1
INH1
COQ2
PET494
TUF1
YNR042W
MRPL33
CYT1
MRPS8
NAM2
MRPS5
PRO1
GRX5
YPR099C
RMD9
CBP6
YDL069C_1
Gene Name
DOXO HL K
DOXO HLEG K
DOXO HL L
DOXO HLEG L
Type of Media

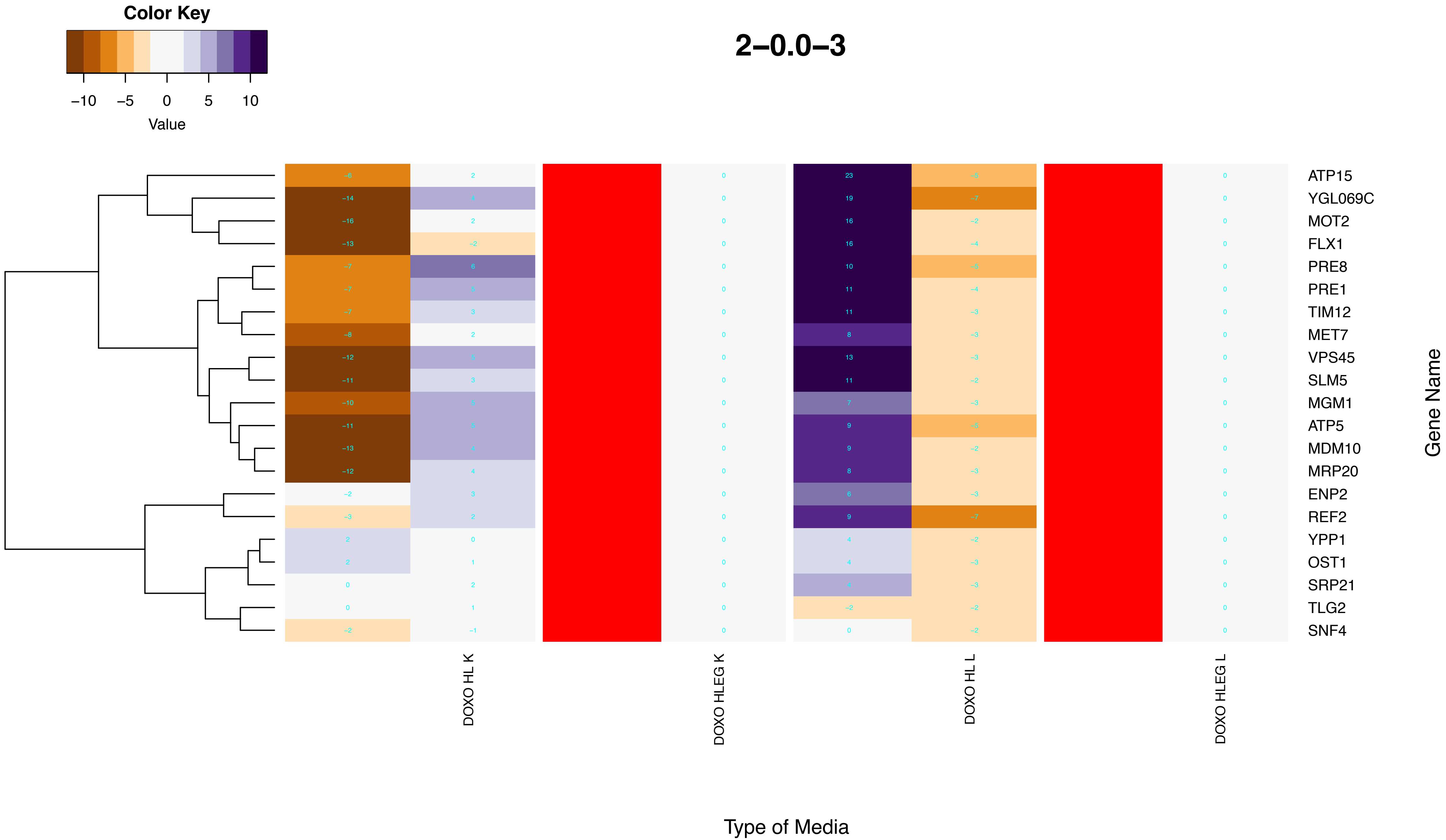
Color Key
−10 −5 0 5 10
Value
2−0.0−3
ATP15
YGL069C
MOT2
FLX1
PRE8
PRE1
TIM12
MET7
VPS45
SLM5
MGM1
ATP5
MDM10
MRP20
ENP2
REF2
YPP1
OST1
SRP21
TLG2
SNF4
Gene Name
DOXO HL K
DOXO HLEG K
DOXO HL L
DOXO HLEG L
Type of Media

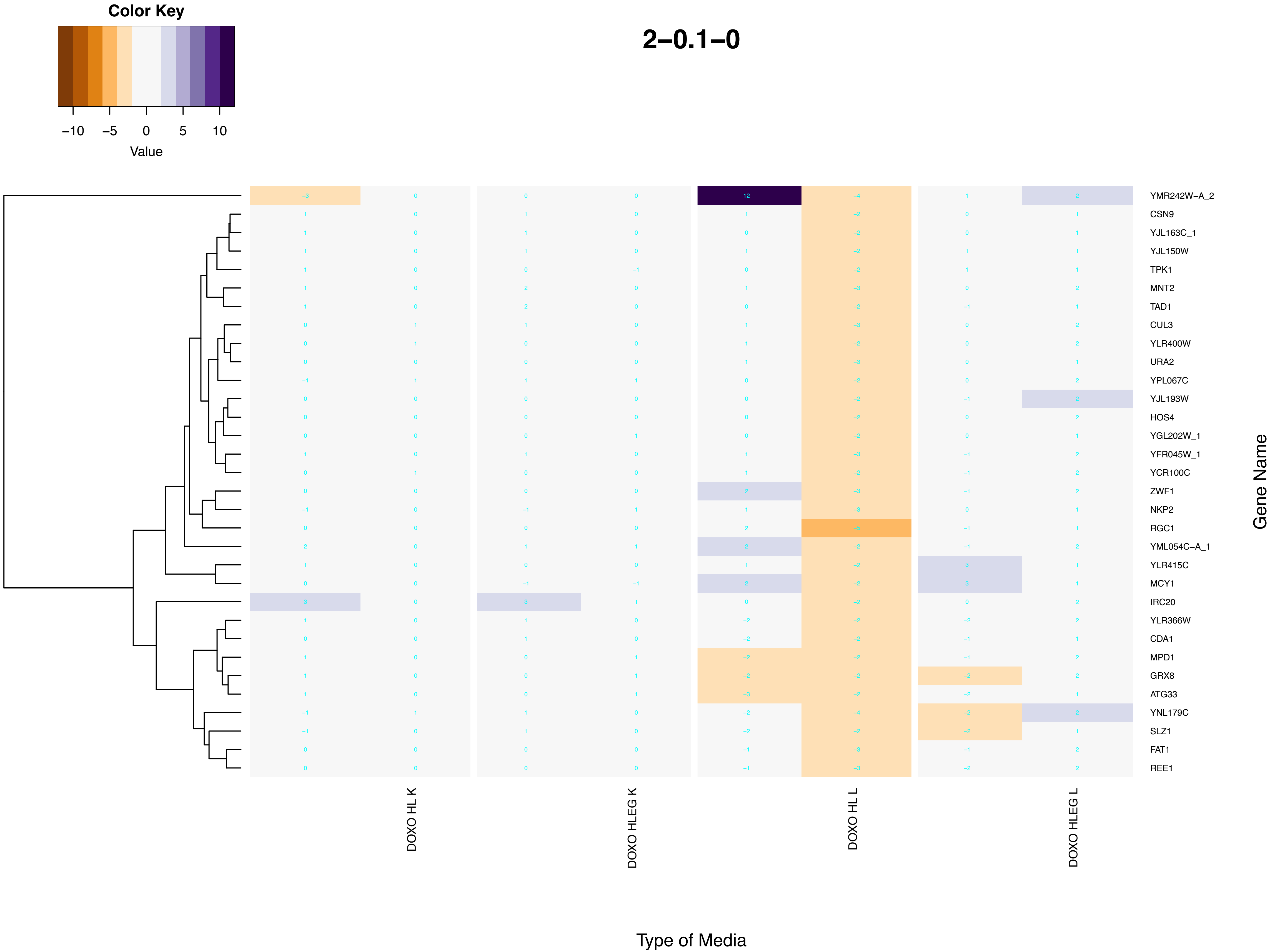

Color Key
−10
−5
0
5
10
Value
2−0.1−0
YMR242W−A_2
CSN9
YJL163C_1
YJL150W
TPK1
MNT2
TAD1
CUL3
YLR400W
URA2
YPL067C
YJL193W
HOS4
YGL202W_1
YFR045W_1
YCR100C
ZWF1
NKP2
RGC1
YML054C−A_1
YLR415C
MCY1
IRC20
YLR366W
CDA1
MPD1
GRX8
ATG33
YNL179C
SLZ1
FAT1
REE1
Gene Name
DOXO HL K
DOXO HLEG K
DOXO HL L
DOXO HLEG L
Type of Media

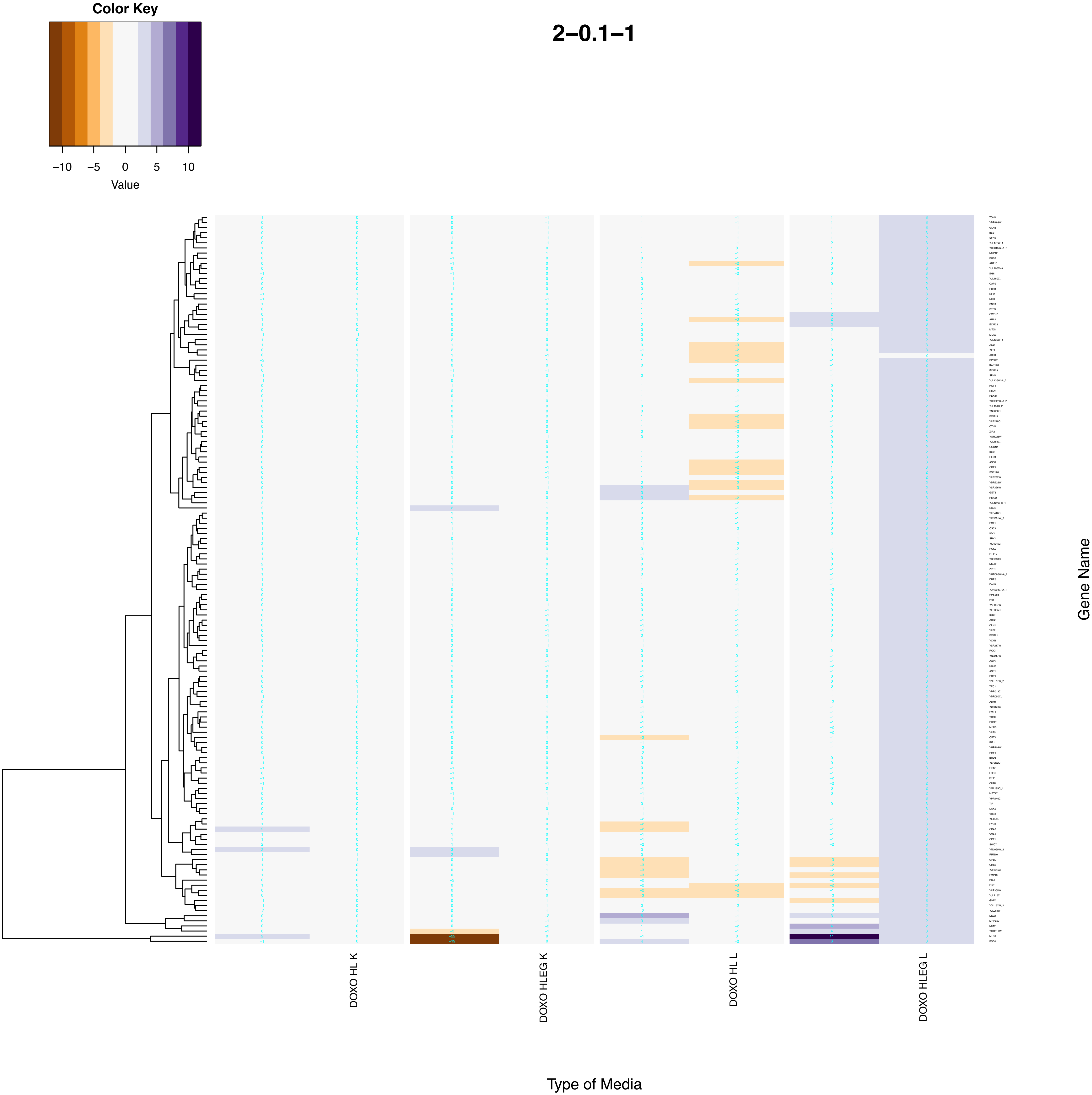
Color Key
−10
−5
0
5
10
Value
2−0.1−1
DOXO HL K
DOXO HLEG K
DOXO HL L
DOXO HLEG L
Type of Media
Gene Name

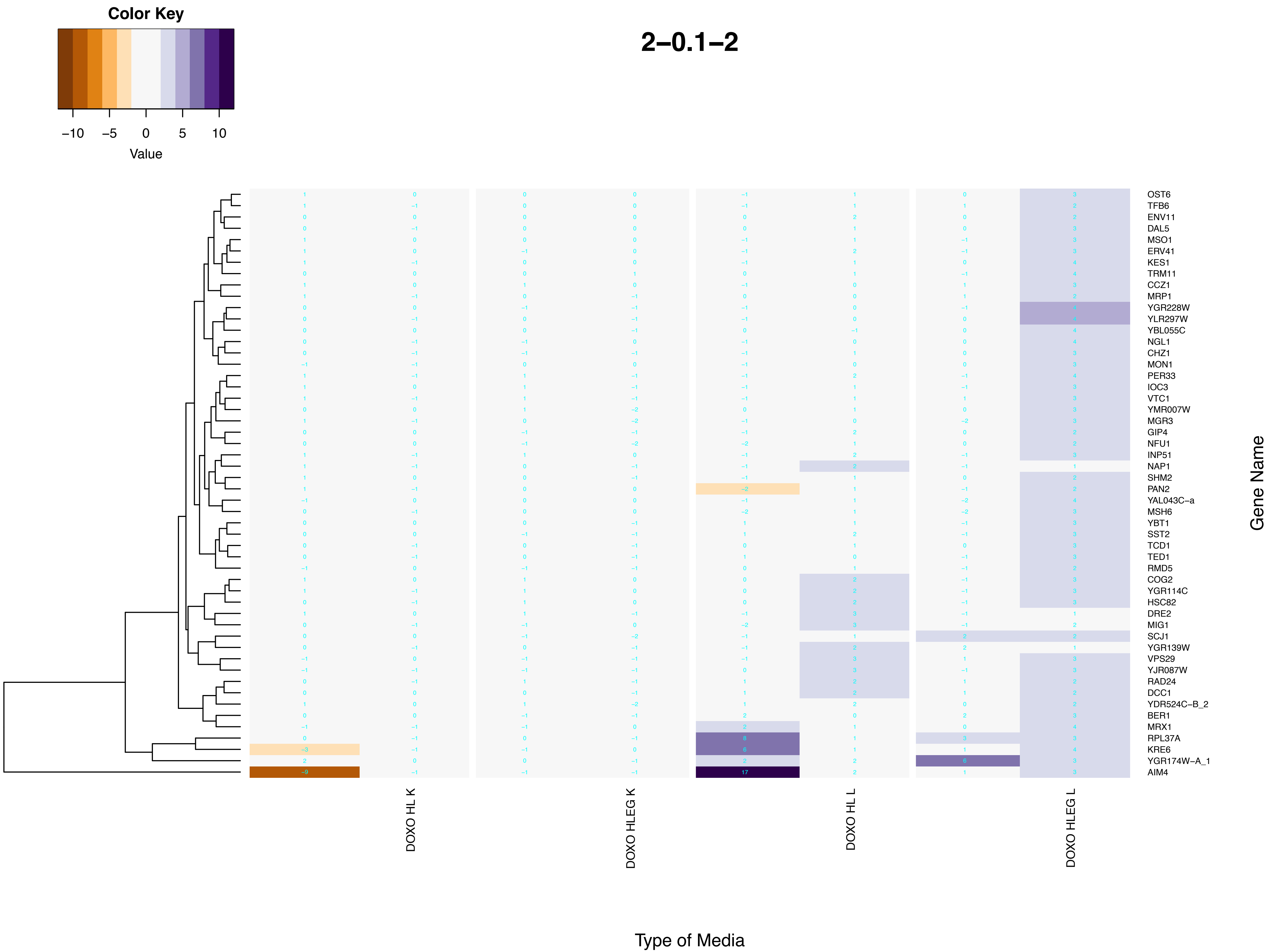
Color Key
−10 −5 0 5 10
Value
2−0.1−2
OST6
TFB6
ENV11
DAL5
MSO1
ERV41
KES1
TRM11
CCZ1
MRP1
YGR228W
YLR297W
YBL055C
NGL1
CHZ1
MON1
PER33
IOC3
VTC1
YMR007W
MGR3
GIP4
NFU1
INP51
NAP1
SHM2
PAN2
YAL043C−a
MSH6
YBT1
SST2
TCD1
TED1
RMD5
COG2
YGR114C
HSC82
DRE2
MIG1
SCJ1
YGR139W
VPS29
YJR087W
RAD24
DCC1
YDR524C−B_2
BER1
MRX1
RPL37A
KRE6
YGR174W−A_1
AIM4
Gene Name
DOXO HL K
DOXO HLEG K
DOXO HL L
DOXO HLEG L
Type of Media

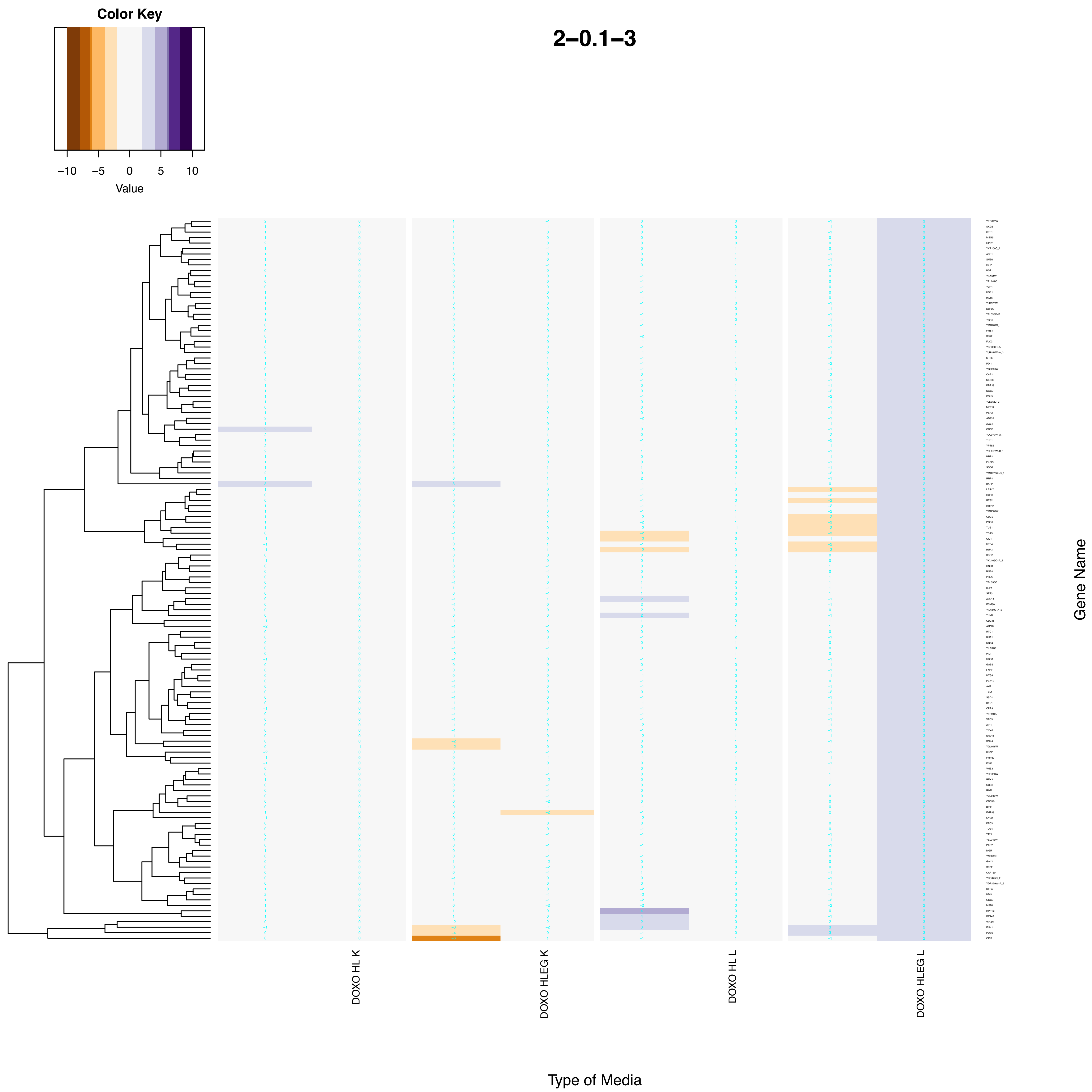
Color Key
−10
−5
0
5
10
Value
2−0.1−3
DOXO HL K
DOXO HLEG K
DOXO HL L
DOXO HLEG L
Type of Media
Gene Name

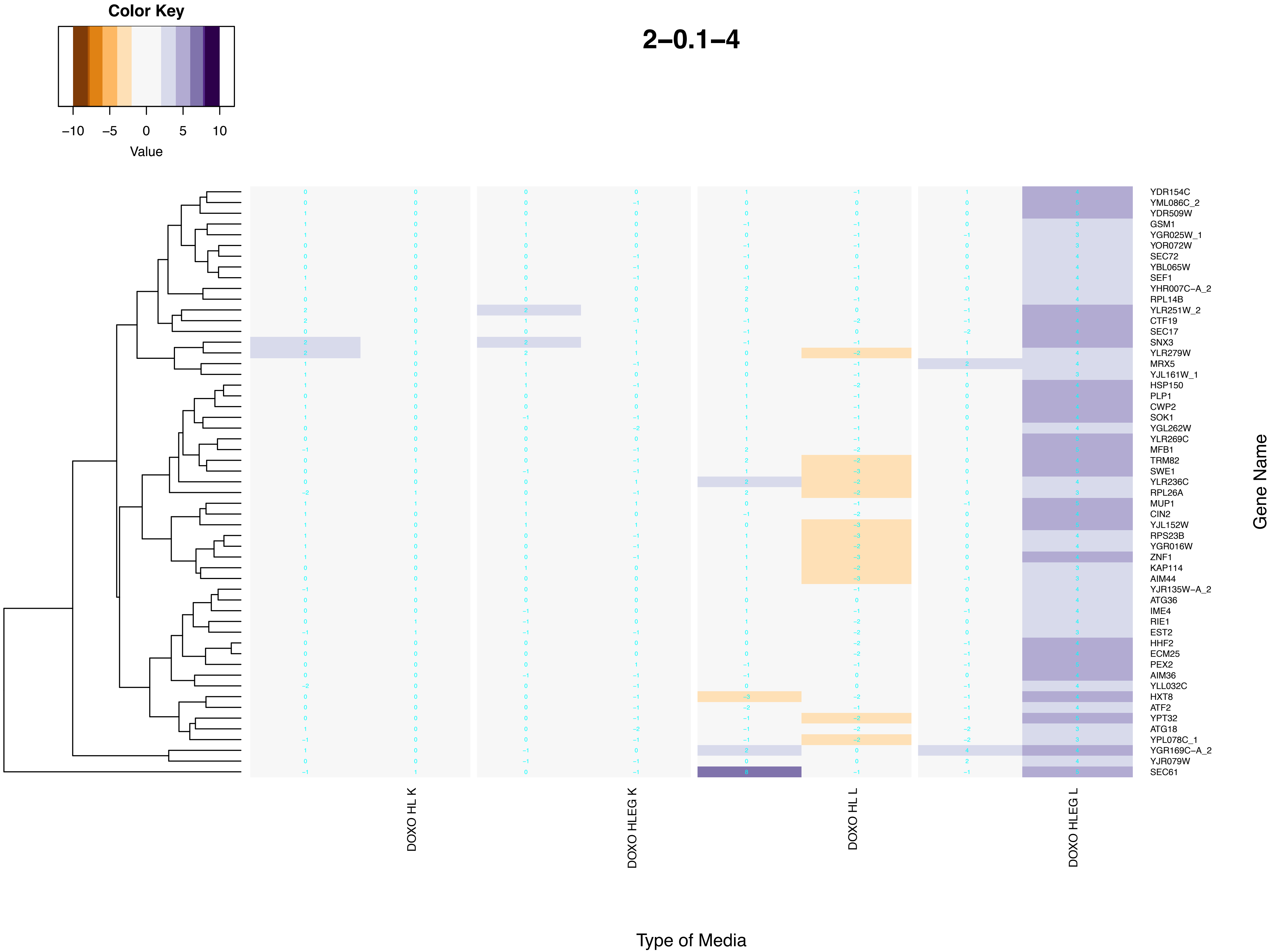

Color Key
−10 −5 0 5 10
Value
2−0.1−4
YDR154C
YML086C_2
YDR509W
GSM1
YGR025W_1
YOR072W
SEC72
YBL065W
SEF1
YHR007C−A_2
RPL14B
YLR251W_2
CTF19
SEC17
SNX3
YLR279W
MRX5
YJL161W_1
HSP150
PLP1
CWP2
SOK1
YGL262W
YLR269C
MFB1
TRM82
SWE1
YLR236C
RPL26A
MUP1
CIN2
YJL152W
RPS23B
YGR016W
ZNF1
KAP114
AIM44
YJR135W−A_2
ATG36
IME4
RIE1
EST2
HHF2
ECM25
PEX2
AIM36
YLL032C
HXT8
ATF2
YPT32
ATG18
YPL078C_1
YGR169C−A_2
YJR079W
SEC61
Gene Name
DOXO HL K
DOXO HLEG K
DOXO HL L
DOXO HLEG L
Type of Media

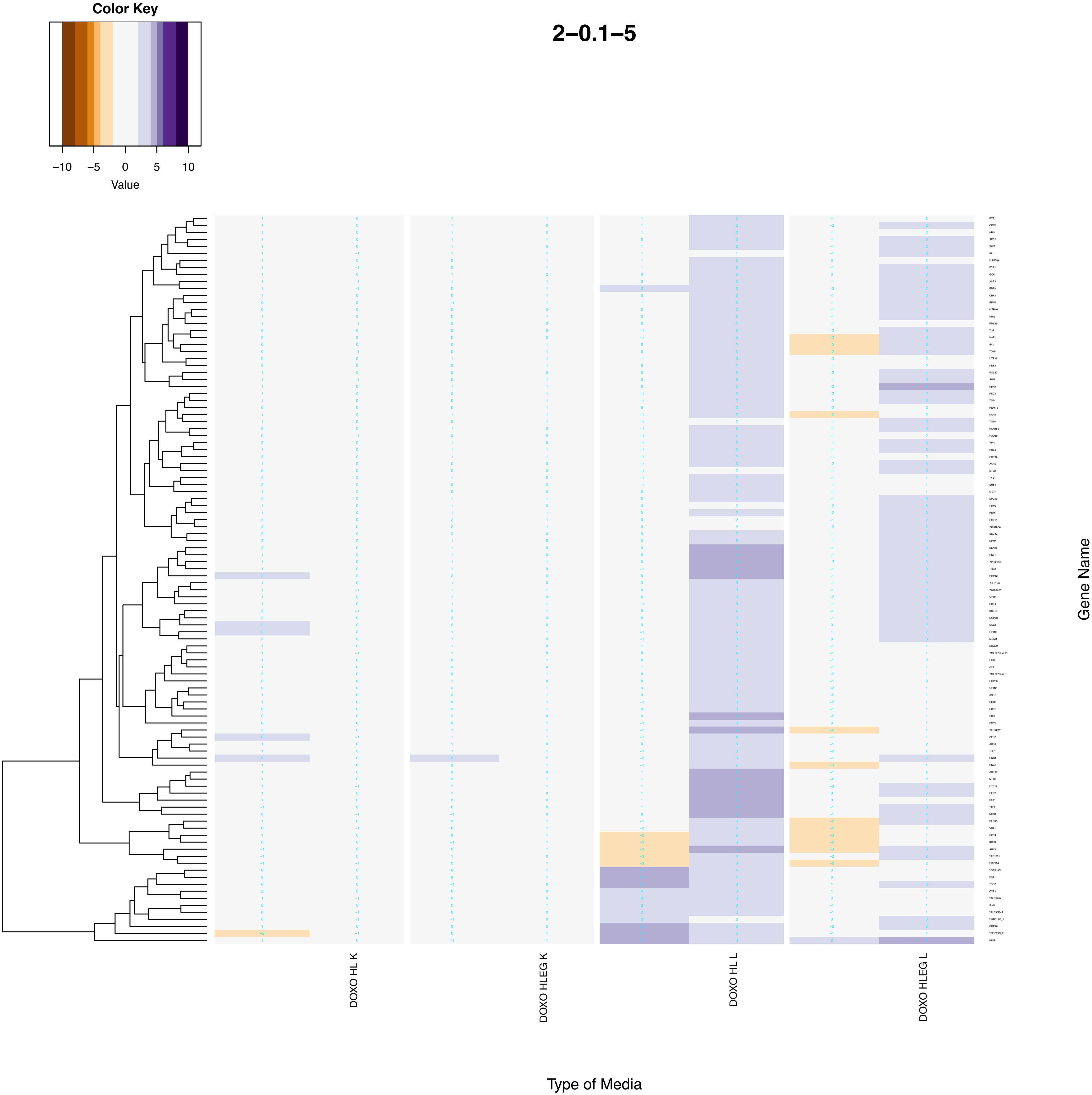
Color Key
−10
−5
0
5
10
Value
2−0.1−5
DOXO HL K
DOXO HLEG K
DOXO HL L
DOXO HLEG L
Gene Name
Type of Media

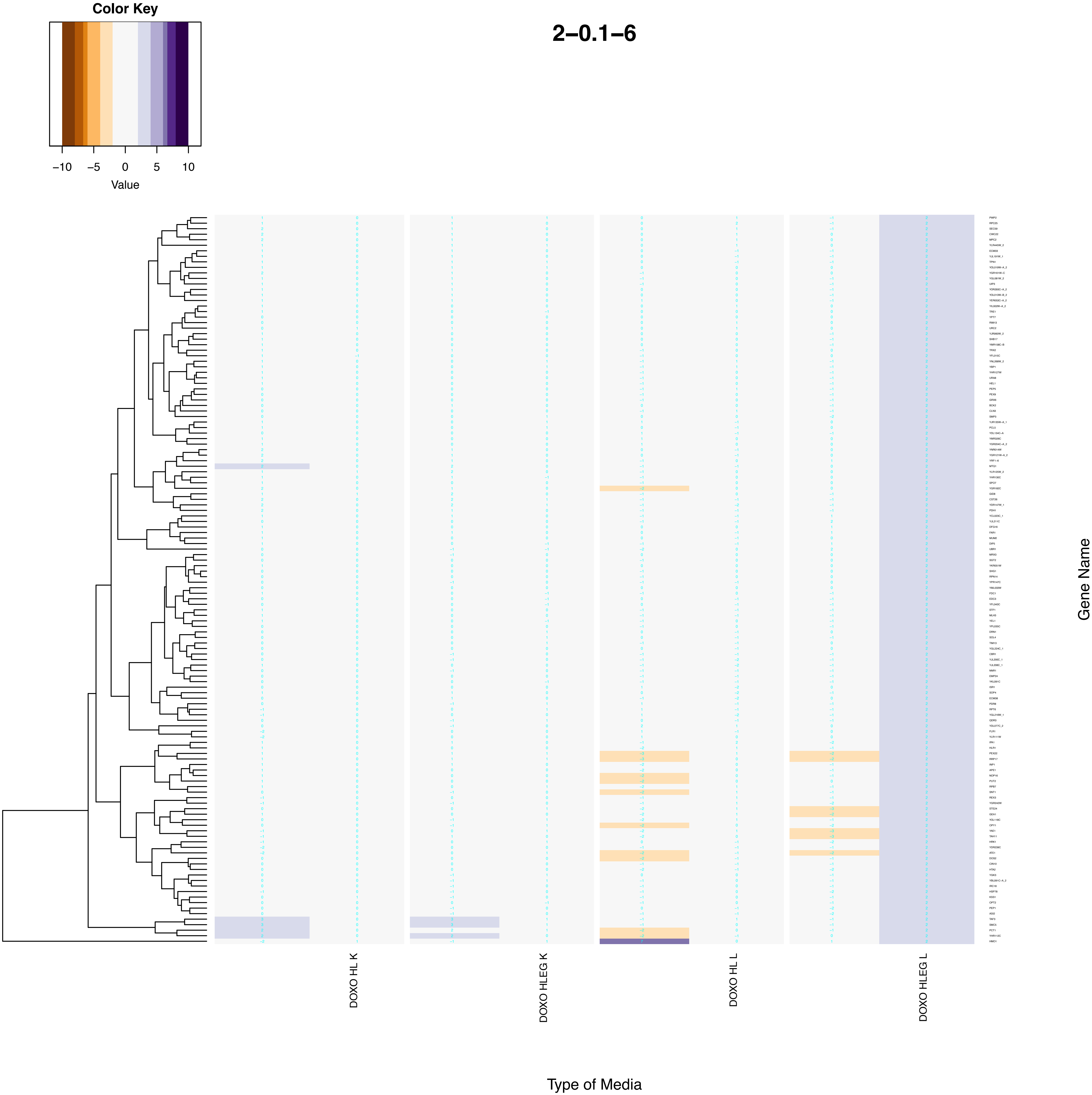
Color Key
−10 −5 0 5 10
Value
2−0.1−6
DOXO HL K
DOXO HLEG K
DOXO HL L
DOXO HLEG L
Gene Name
Type of Media

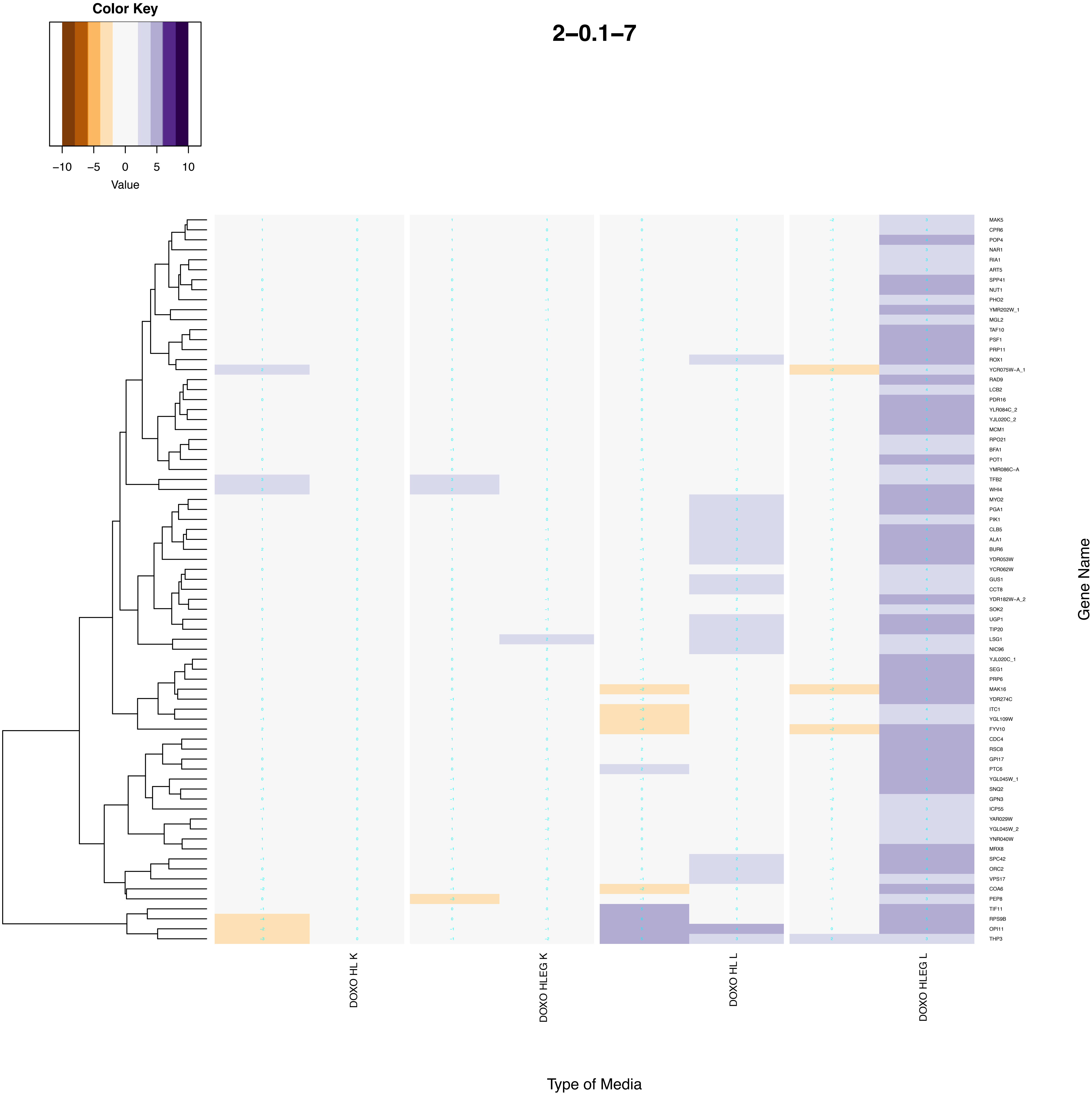
Color Key
−10 −5 0 5 10
Value
2−0.1−7
MAK5
CPR6
POP4
NAR1
RIA1
ART5
SPP41
NUT1
PHO2
YMR202W_1
MGL2
TAF10
PSF1
PRP11
ROX1
YCR075W−A_1
RAD9
LCB2
PDR16
YLR084C_2
YJL020C_2
MCM1
RPO21
BFA1
POT1
YMR086C−A
TFB2
WHI4
MYO2
PGA1
PIK1
CLB5
ALA1
BUR6
YDR053W
YCR062W
GUS1
CCT8
YDR182W−A_2
SOK2
UGP1
TIP20
LSG1
NIC96
YJL020C_1
SEG1
PRP6
MAK16
YDR274C
ITC1
YGL109W
FYV10
CDC4
RSC8
GPI17
PTC6
YGL045W_1
SNQ2
GPN3
ICP55
YAR029W
YGL045W_2
YNR040W
MRX8
SPC42
ORC2
VPS17
COA6
PEP8
TIF11
RPS9B
OPI11
THP3
Gene Name
DOXO HL K
DOXO HLEG K
DOXO HL L
DOXO HLEG L
Type of Media

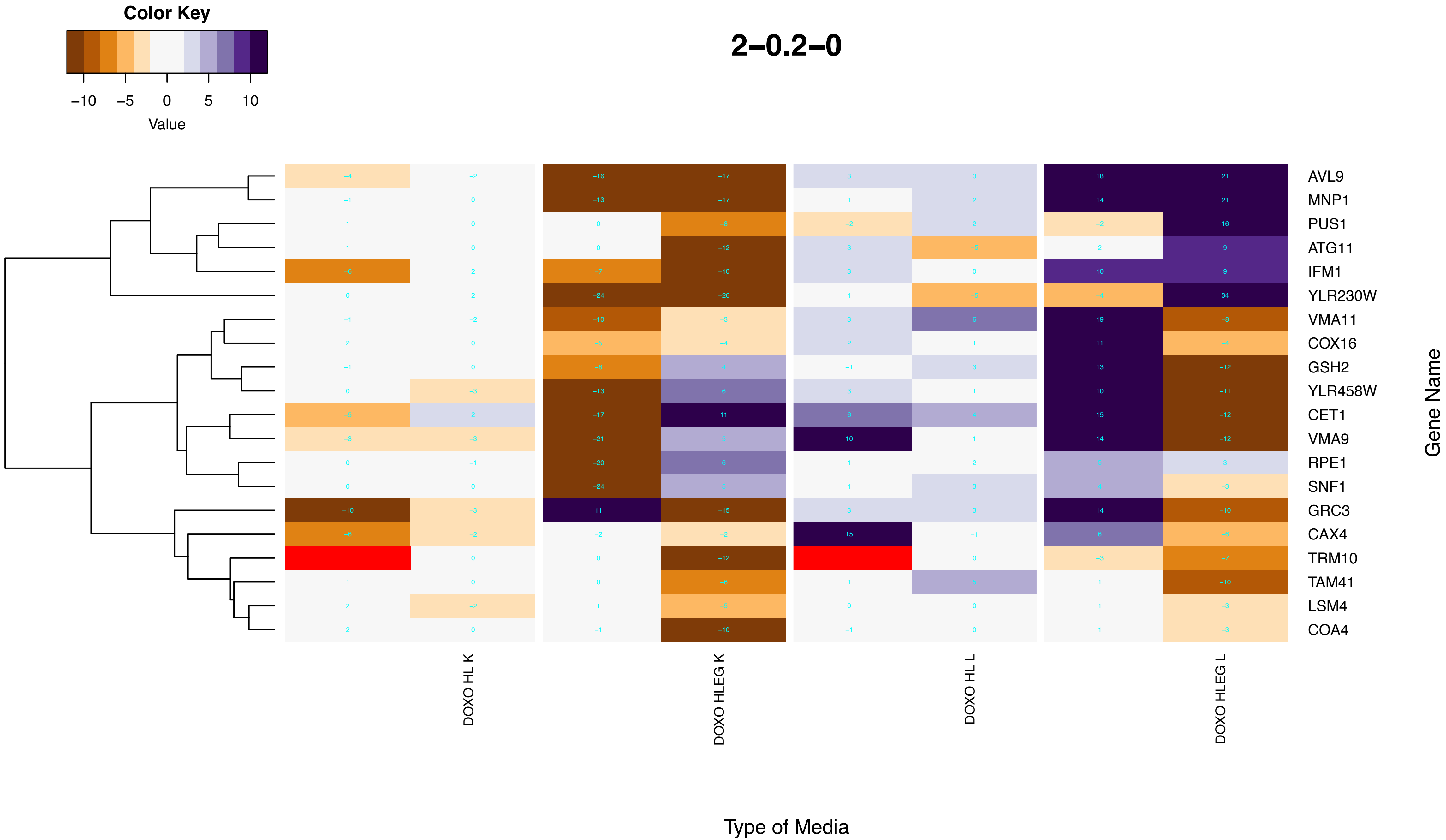
Color Key
−10 −5 0 5 10
Value
2−0.2−0
AVL9
MNP1
PUS1
ATG11
IFM1
YLR230W
VMA11
COX16
GSH2
YLR458W
CET1
VMA9
RPE1
SNF1
GRC3
CAX4
TRM10
TAM41
LSM4
COA4
Gene Name
DOXO HL K
DOXO HLEG K
DOXO HL L
DOXO HLEG L
Type of Media

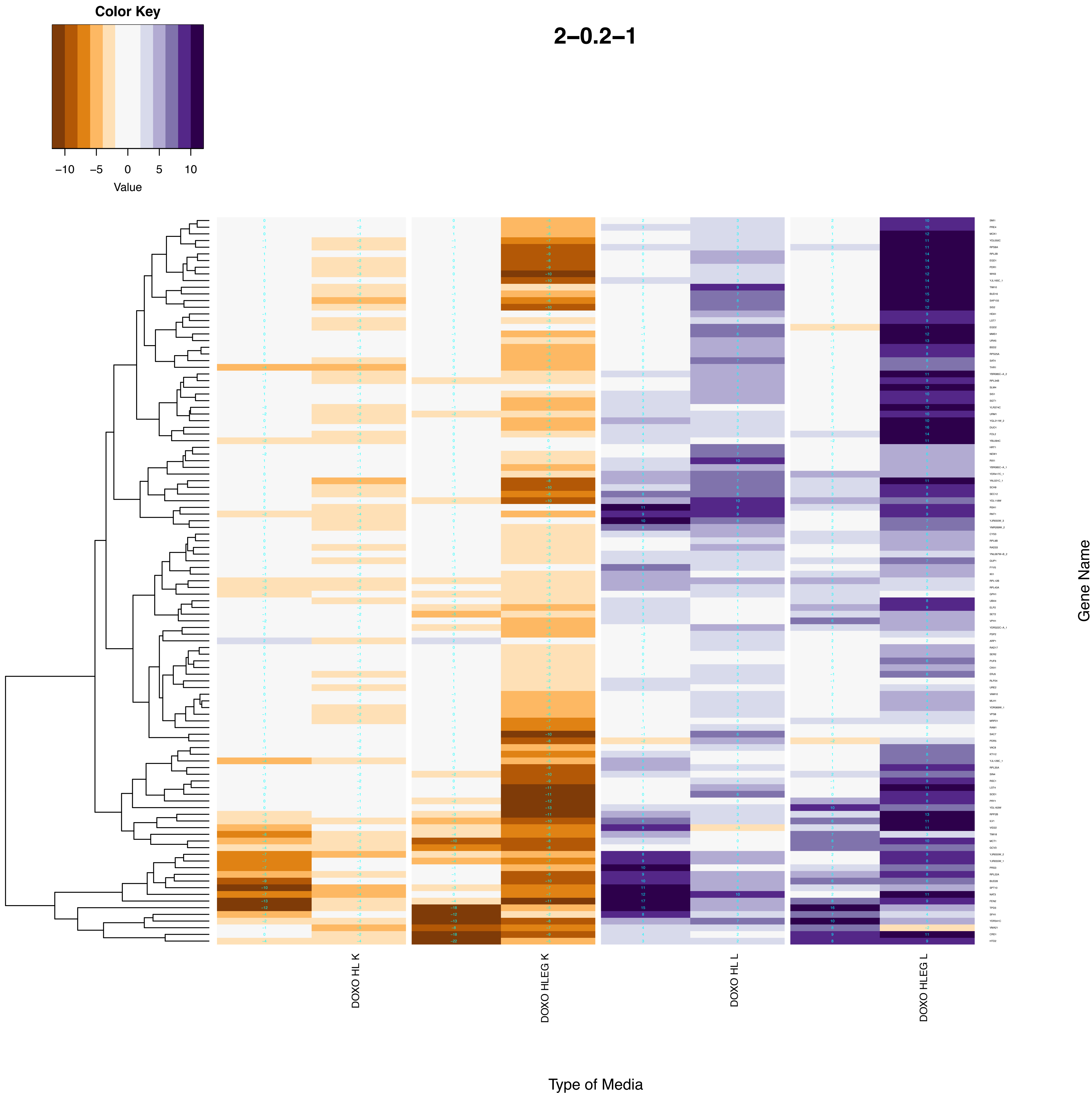
Color Key
−10
−5
0
5
10
Value
2−0.2−1
DOXO HL K
DOXO HLEG K
DOXO HL L
DOXO HLEG L
Gene Name
Type of Media

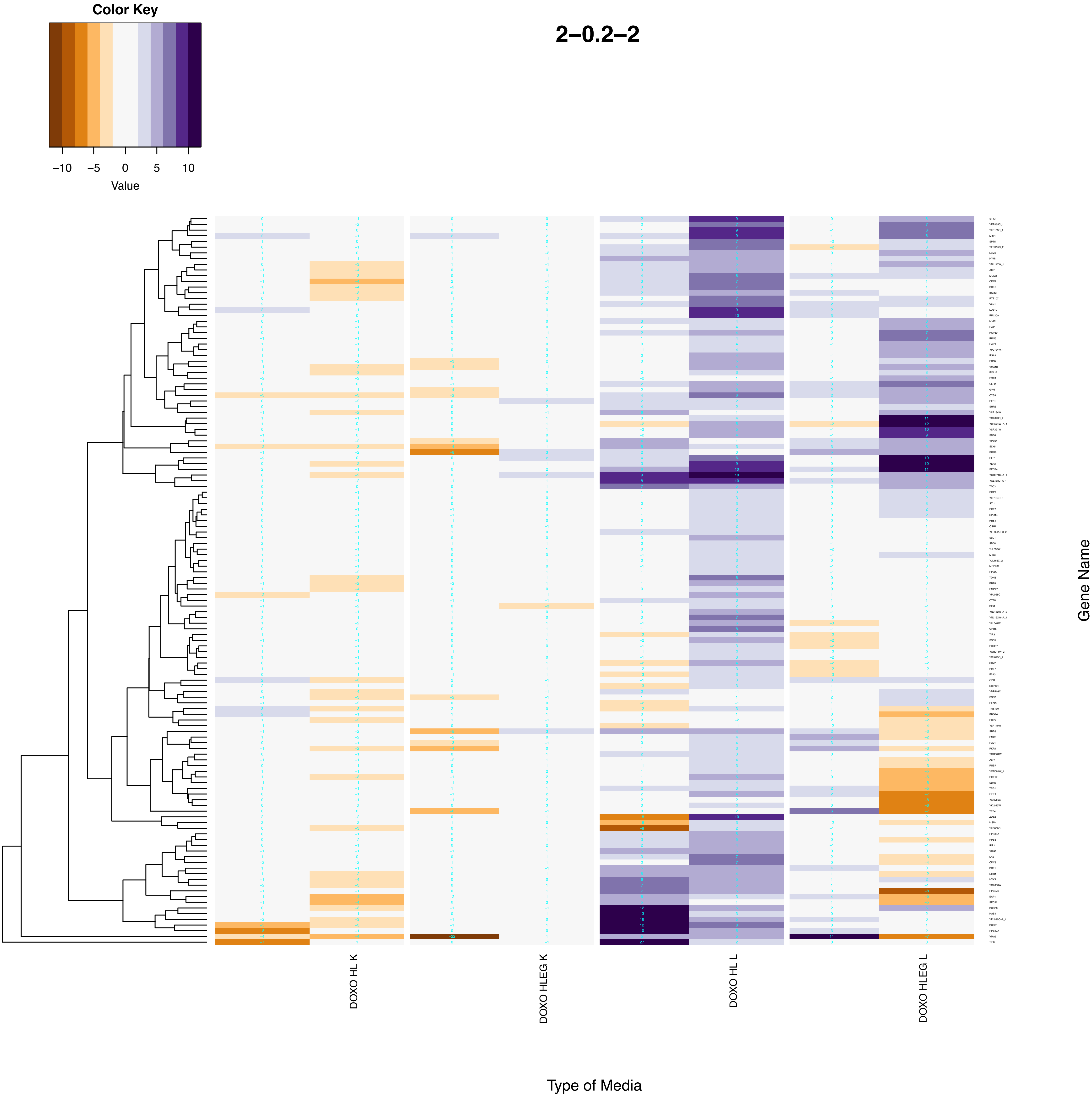

2–0.2–2
Color Key
−10
−5
0
5
10
Value
DOXO HL K
DOXO HLEG K
DOXO HL L
DOXO HLEG L
Type of Media
Gene Name

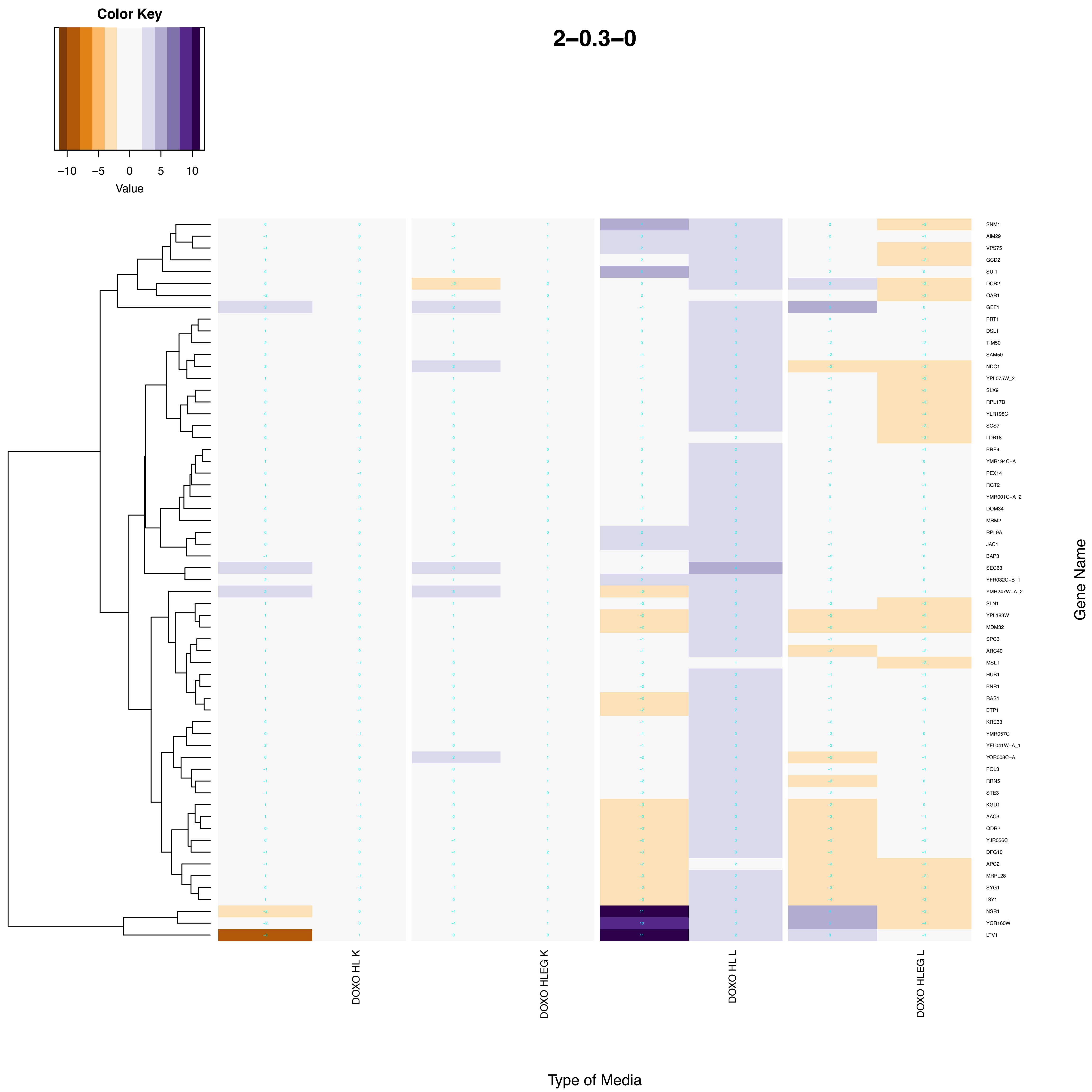

Color Key
−10
−5
0
5
10
Value
2−0.3−0
SNM1
AIM29
VPS75
GCD2
SUI1
DCR2
OAR1
GEF1
PRT1
DSL1
TIM50
SAM50
NDC1
YPL075W_2
SLX9
RPL17B
YLR198C
SCS7
LDB18
BRE4
YMR194C−A
PEX14
RGT2
YMR001C−A_2
DOM34
MRM2
RPL9A
JAC1
BAP3
SEC63
YFR032C−B_1
YMR247W−A_2
SLN1
YPL183W
MDM32
SPC3
ARC40
MSL1
HUB1
BNR1
RAS1
ETP1
KRE33
YMR057C
YFL041W−A_1
YOR008C−A
POL3
RRN5
STE3
KGD1
AAC3
QDR2
YJR056C
DFG10
APC2
MRPL28
SYG1
ISY1
NSR1
YGR160W
LTV1
Gene Name
DOXO HL K
DOXO HLEG K
DOXO HL L
DOXO HLEG L
Type of Media

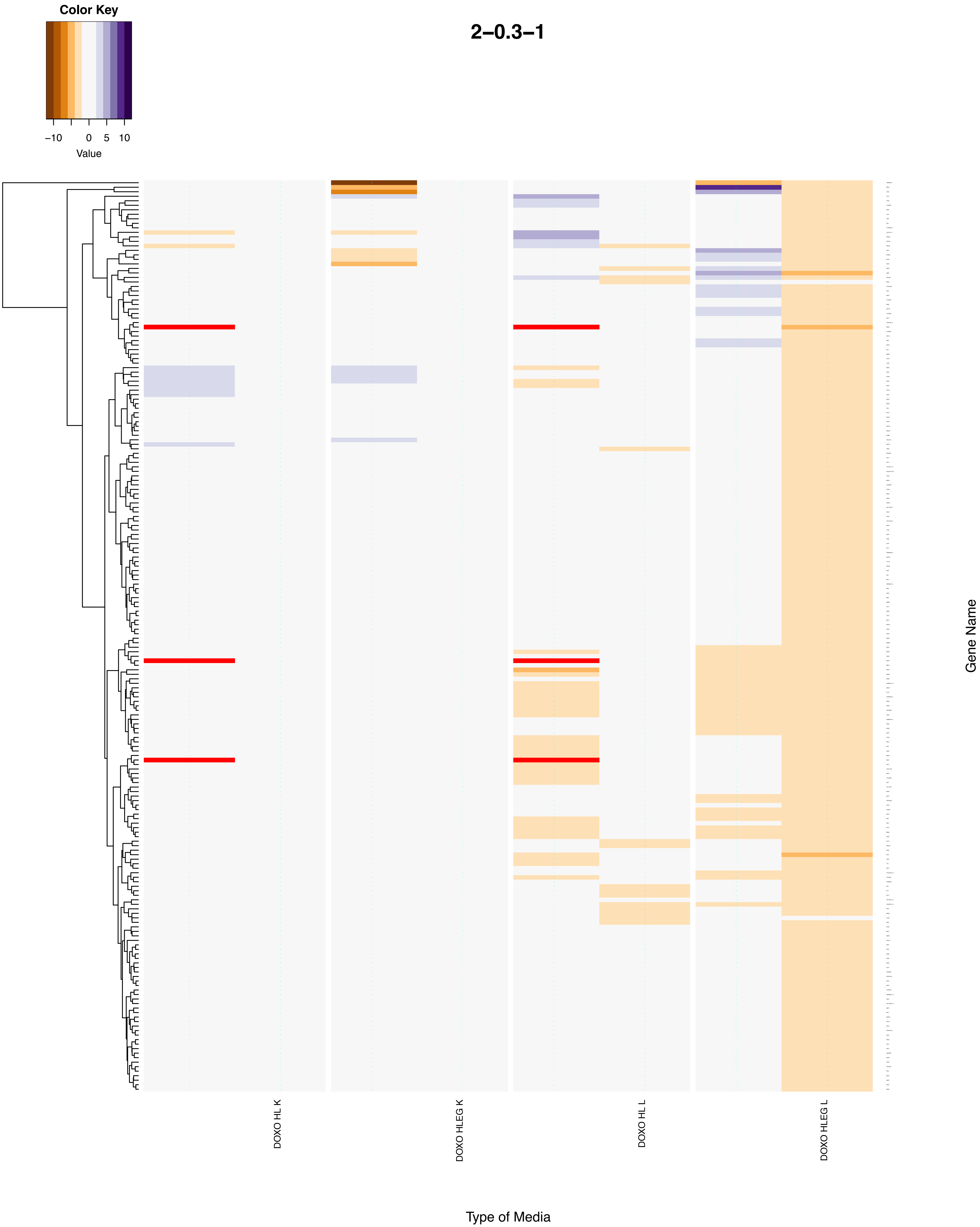

Color Key
−10
0
5
10
Value
2−0.3−1
DOXO HL K
DOXO HLEG K
DOXO HL L
DOXO HLEG L
Gene Name
Type of Media

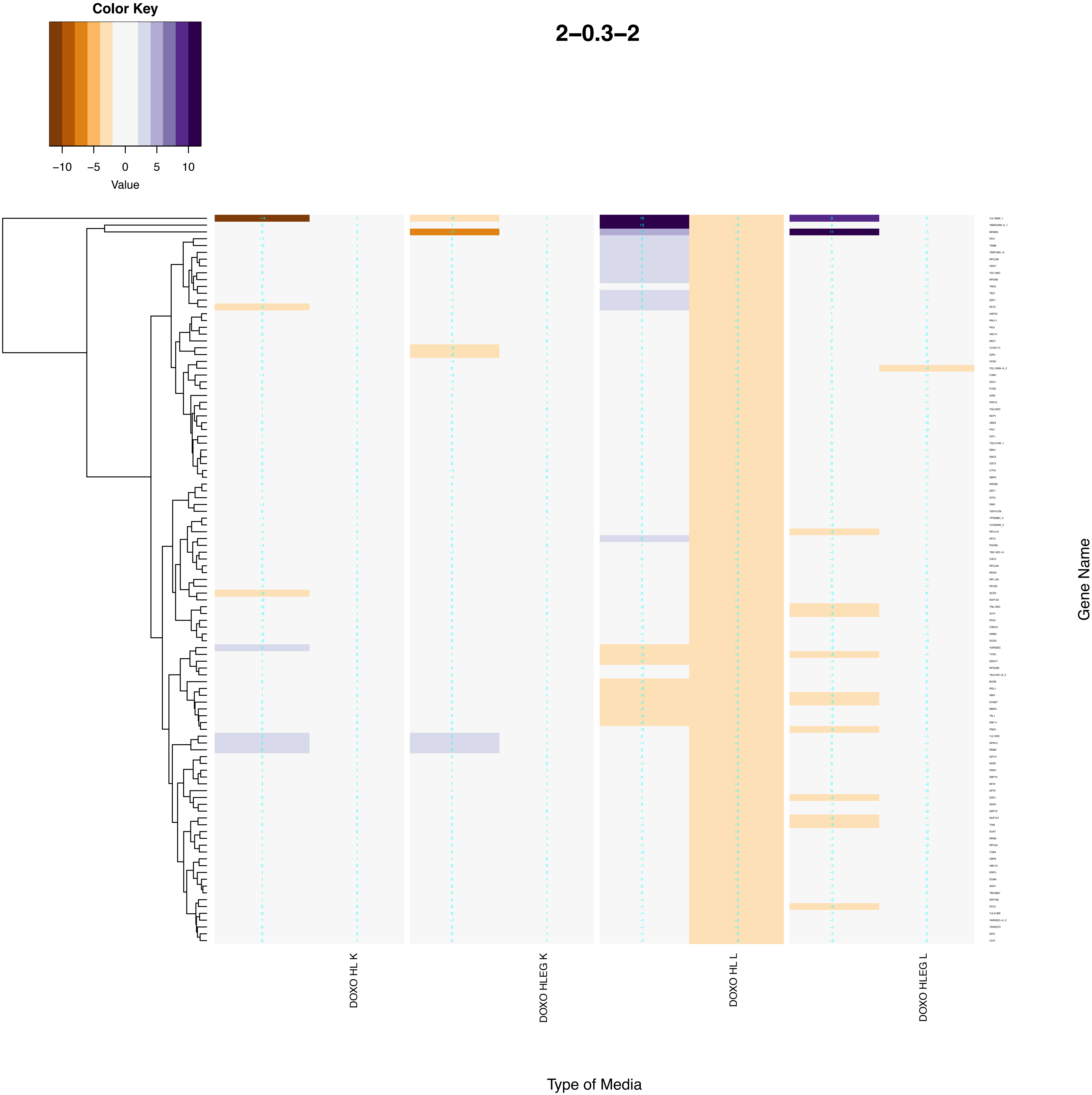
Color Key
−10
−5
0
5
10
Value
2–0.3–2
DOXO HL K
DOXO HLEG K
DOXO HL L
DOXO HLEG L
Type of Media
Gene Name

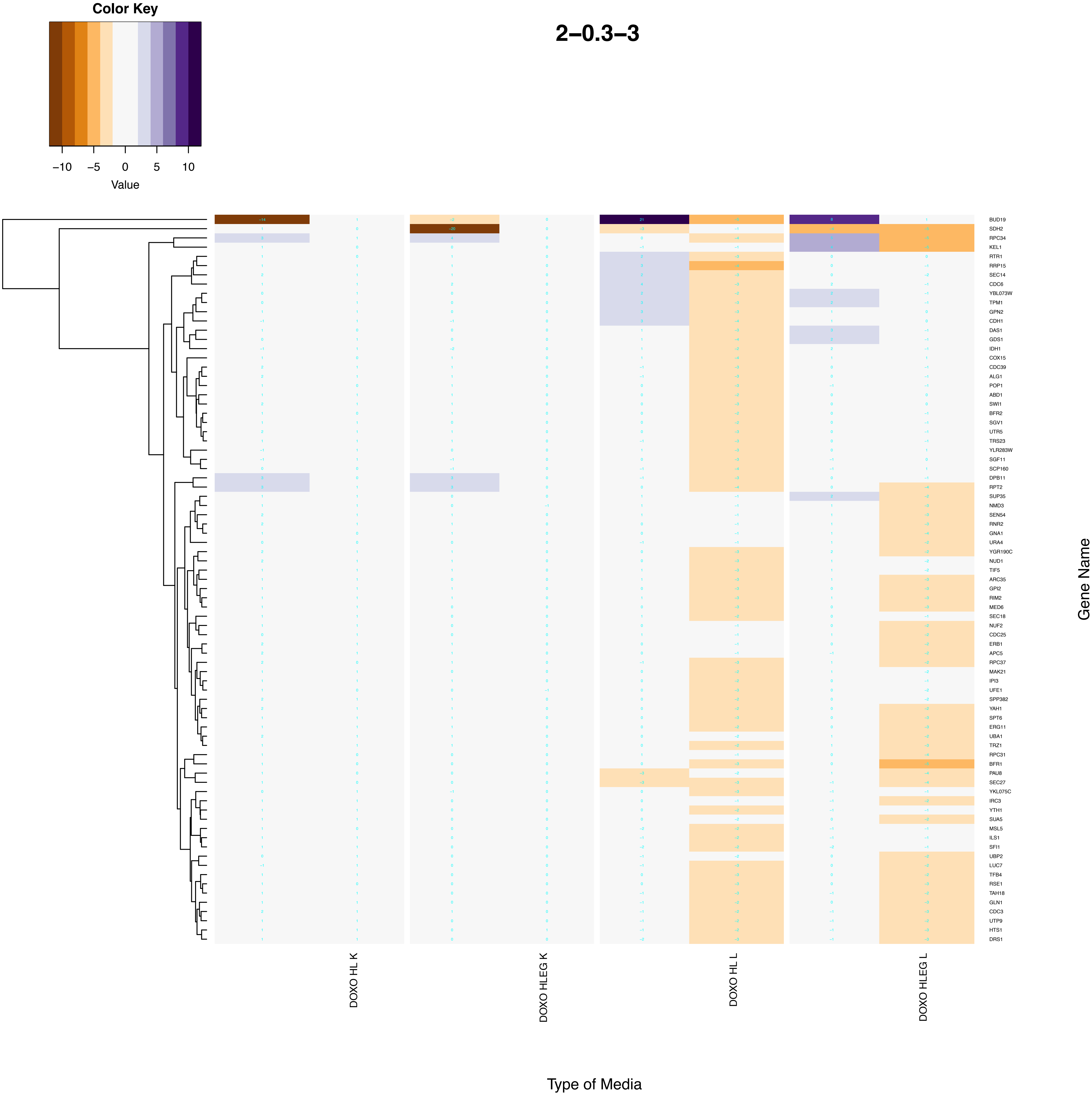

Color Key
−10
−5
0
5
10
Value
2–0.3–3
BUD19
SDH2
RPC34
KEL1
RTR1
RRP15
SEC14
CDC6
YBL073W
TPM1
GPN2
CDH1
DAS1
GDS1
IDH1
COX15
CDC39
ALG1
POP1
ABD1
SWI1
BFR2
SGV1
UTR5
TRS23
YLR283W
SGF11
SCP160
DPB11
RPT2
SUP35
NMD3
SEN54
RNR2
GNA1
URA4
YGR190C
NUD1
TIF5
ARC35
GPI2
RIM2
MED6
SEC18
NUF2
CDC25
ERB1
APC5
RPC37
MAK21
IPI3
UFE1
SPP382
YAH1
SPT6
ERG11
UBA1
TRZ1
RPC31
BFR1
PAU8
SEC27
YKL075C
IRC3
YTH1
SUA5
MSL5
ILS1
SFI1
UBP2
LUC7
TFB4
RSE1
TAH18
GLN1
CDC3
UTP9
HTS1
DRS1
Gene Name
DOXO HL K
DOXO HLEG K
DOXO HL L
DOXO HLEG L
Type of Media

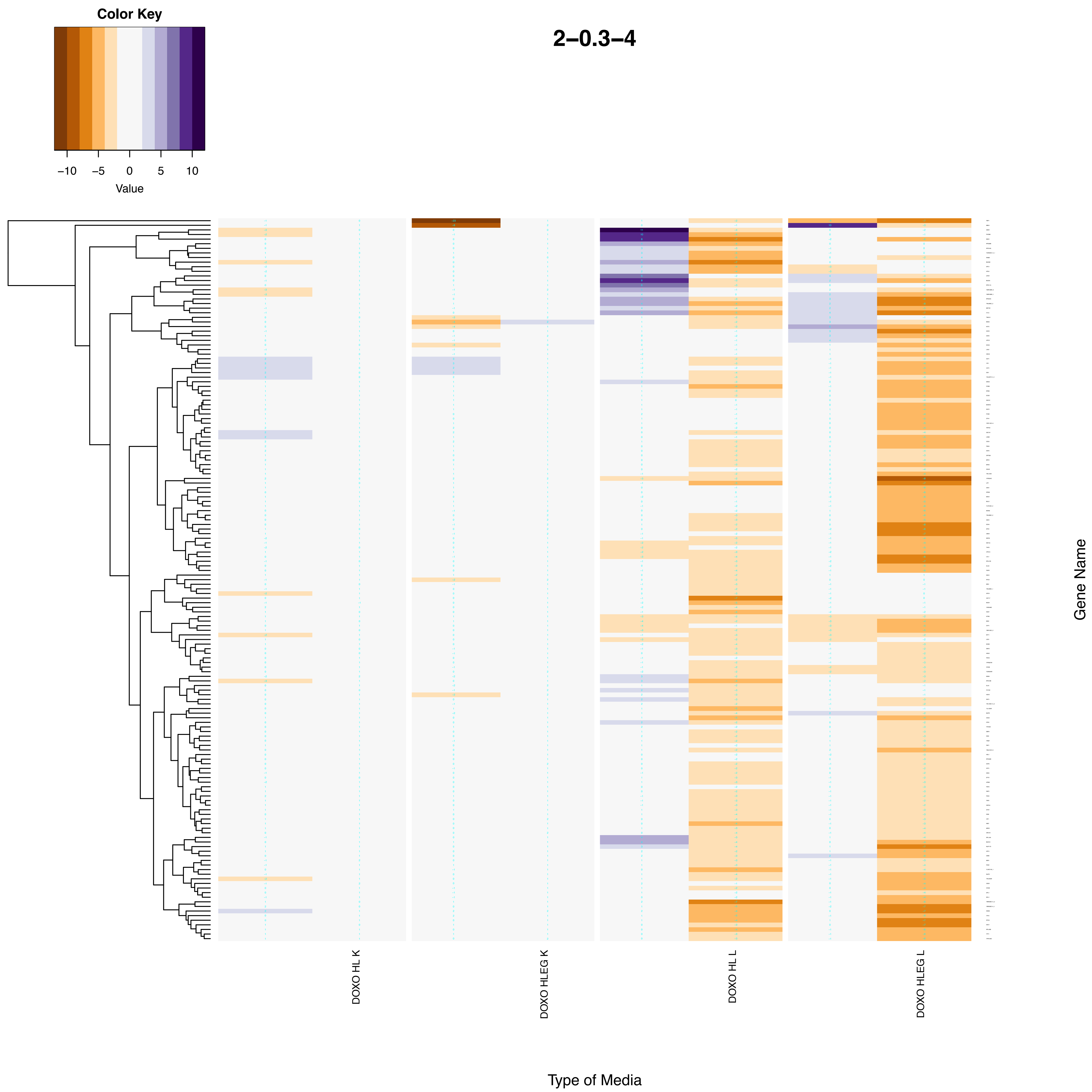
Color Key
−10
−5
0
5
10
Value
2−0.3−4
DOXO HL K
DOXO HLEG K
DOXO HL L
DOXO HLEG L
Gene Name
Type of Media

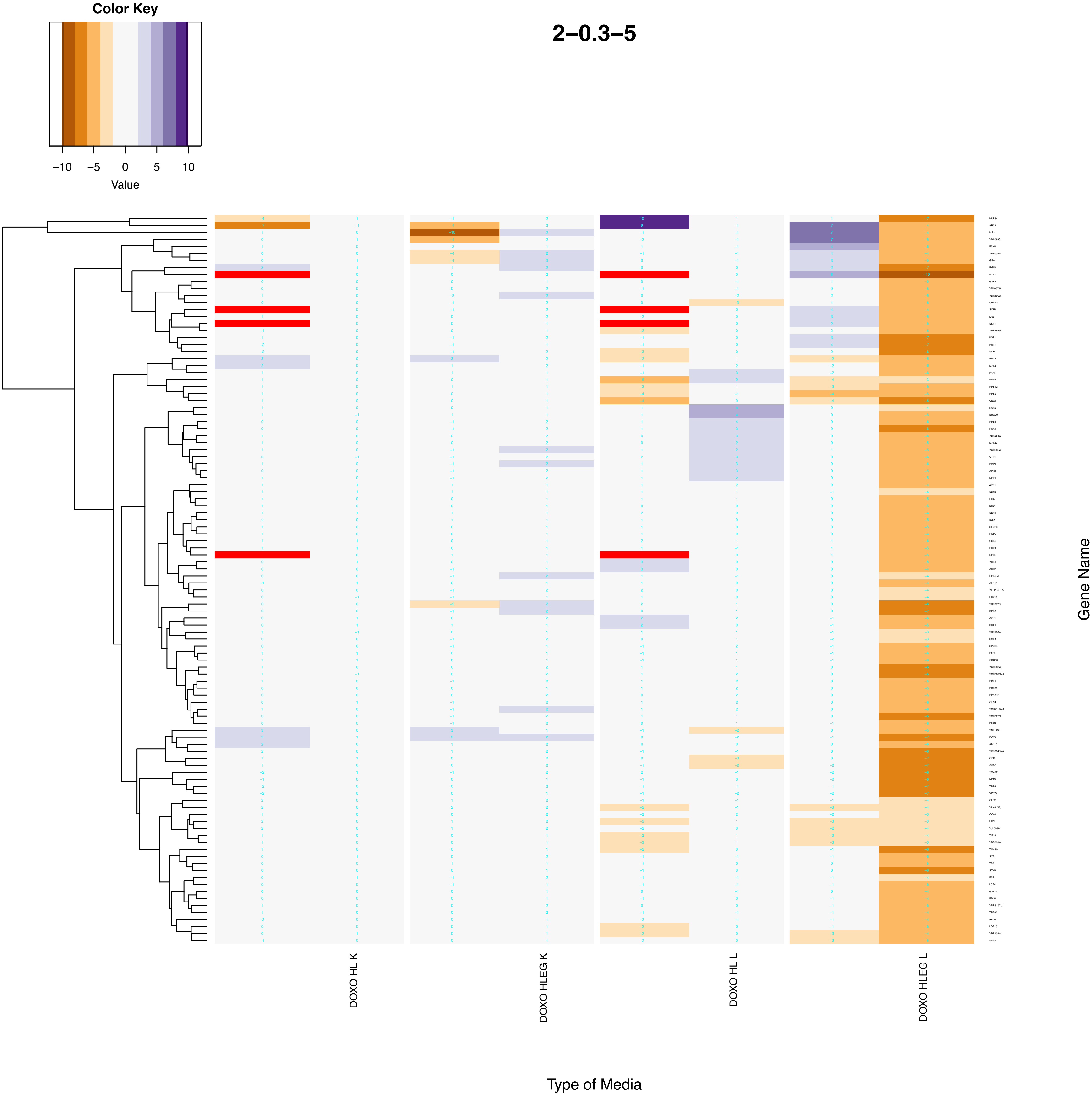
Color Key
−10
−5
0
5
10
Value
2−0.3−5
DOXO HL K
DOXO HLEG K
DOXO HL L
DOXO HLEG L
Type of Media
Gene Name

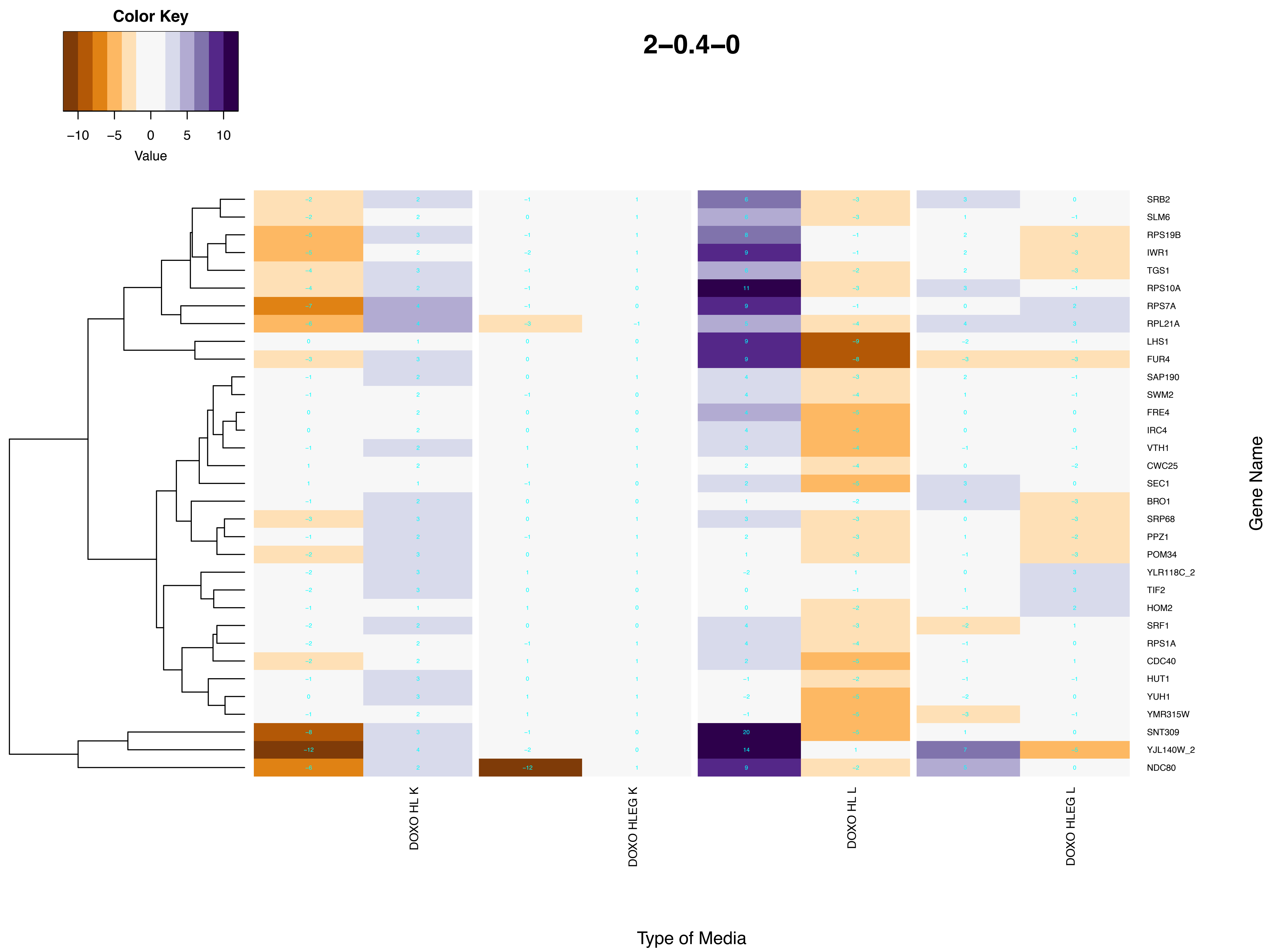
Color Key
−10 −5 0 5 10
Value
2−0.4−0
SRB2
SLM6
RPS19B
IWR1
TGS1
RPS10A
RPS7A
RPL21A
LHS1
FUR4
SAP190
SWM2
FRE4
IRC4
VTH1
CWC25
SEC1
BRO1
SRP68
PPZ1
POM34
YLR118C_2
TIF2
HOM2
SRF1
RPS1A
CDC40
HUT1
YUH1
YMR315W
SNT309
YJL140W_2
NDC80
Gene Name
DOXO HL K
DOXO HLEG K
DOXO HL L
DOXO HLEG L
Type of Media

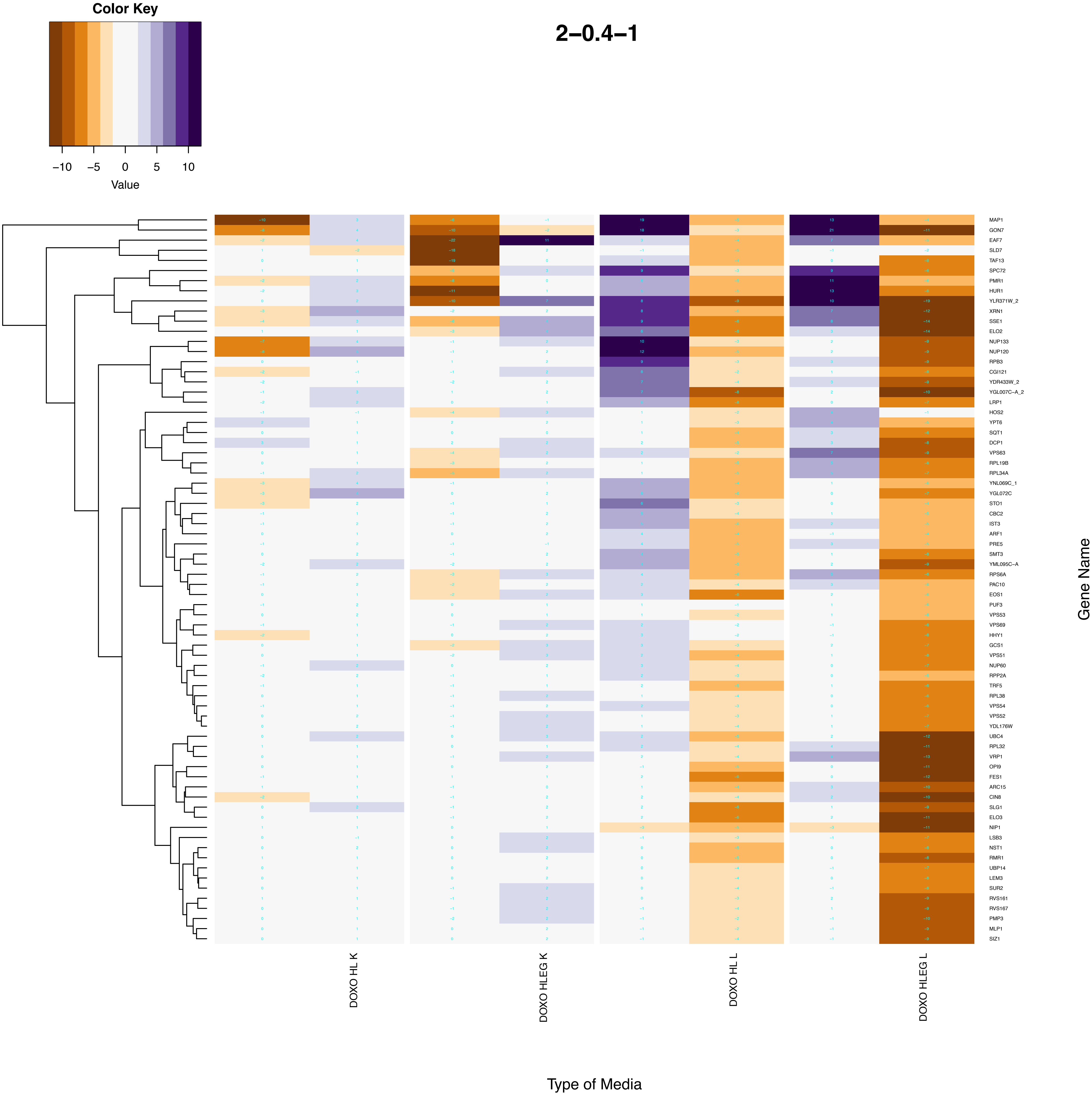
Color Key
−10
−5
0
5
10
Value
2−0.4−1
MAP1
GON7
EAF7
SLD7
TAF13
SPC72
PMR1
HUR1
YLR371W_2
XRN1
SSE1
ELO2
NUP133
NUP120
RPB3
CGI121
YDR433W_2
YGL007C−A_2
LRP1
HOS2
YPT6
SQT1
DCP1
VPS63
RPL19B
RPL34A
YNL069C_1
YGL072C
STO1
CBC2
IST3
ARF1
PRE5
SMT3
YML095C−A
RPS6A
PAC10
EOS1
PUF3
VPS53
VPS69
HHY1
GCS1
VPS51
NUP60
RPP2A
TRF5
RPL38
VPS54
VPS52
YDL176W
UBC4
RPL32
VRP1
OPI9
FES1
ARC15
CIN8
SLG1
ELO3
NIP1
LSB3
NST1
RMR1
UBP14
LEM3
SUR2
RVS161
RVS167
PMP3
MLP1
SIZ1
Gene Name
DOXO HL K
DOXO HLEG K
DOXO HL L
DOXO HLEG L
Type of Media

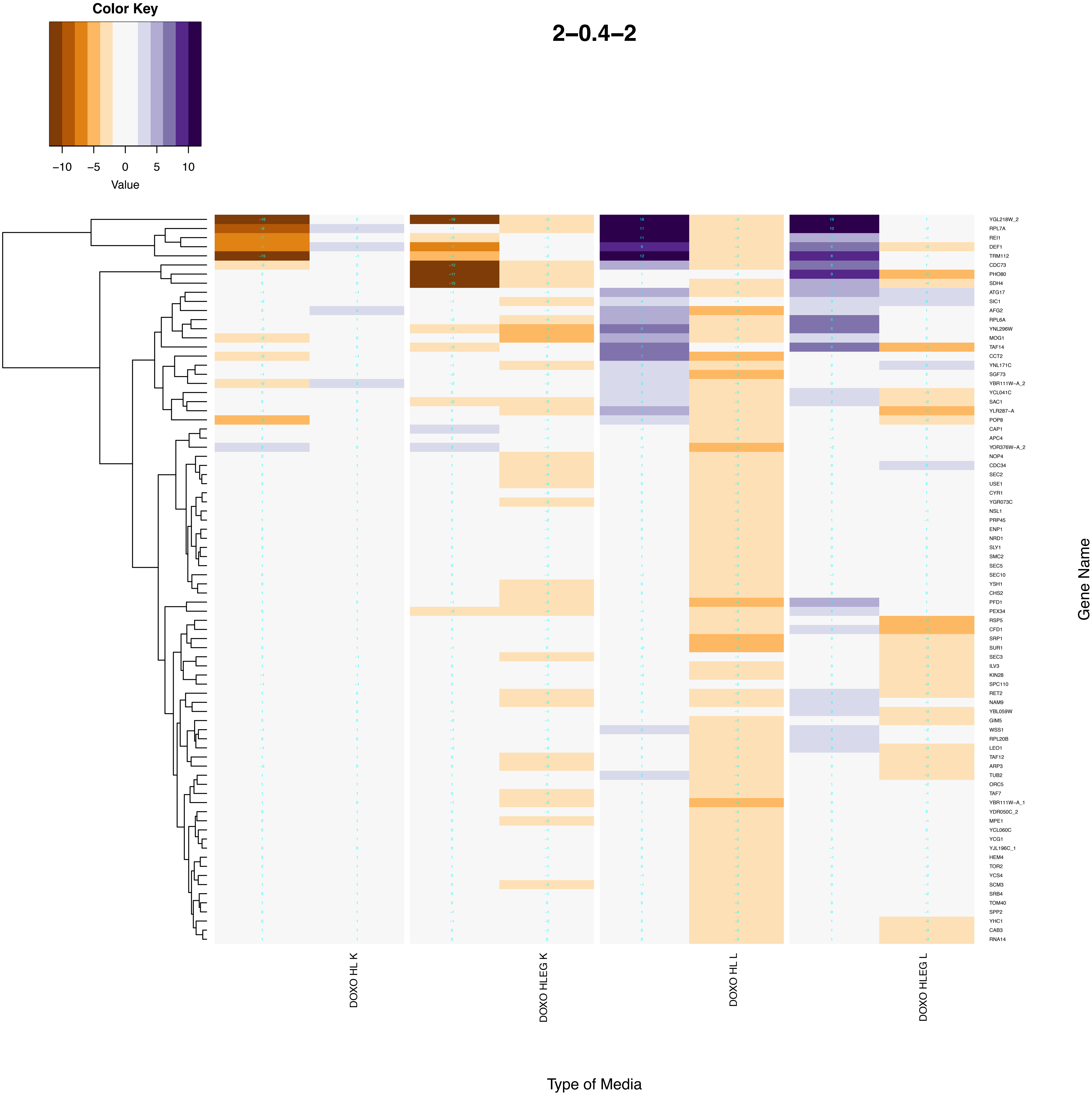

Color Key
−10
−5
0
5
10
Value
2−0.4−2
YGL218W_2
RPL7A
REI1
DEF1
TRM112
CDC73
PHO80
SDH4
ATG17
SIC1
AFG2
RPL6A
YNL296W
MOG1
TAF14
CCT2
YNL171C
SGF73
YBR111W−A_2
YCL041C
SAC1
YLR287−A
POP8
CAP1
APC4
YOR376W−A_2
NOP4
CDC34
SEC2
USE1
CYR1
YGR073C
NSL1
PRP45
ENP1
NRD1
SLY1
SMC2
SEC5
SEC10
YSH1
CHS2
PFD1
PEX34
RSP5
CFD1
SRP1
SUR1
SEC3
ILV3
KIN28
SPC110
RET2
NAM9
YBL059W
GIM5
WSS1
RPL20B
LEO1
TAF12
ARP3
TUB2
ORC5
TAF7
YBR111W−A_1
YDR050C_2
MPE1
YCL060C
YCG1
YJL196C_1
HEM4
TOR2
YCS4
SCM3
SRB4
TOM40
SPP2
YHC1
CAB3
RNA14
Gene Name
DOXO HL K
DOXO HLEG K
DOXO HL L
DOXO HLEG L
Type of Media

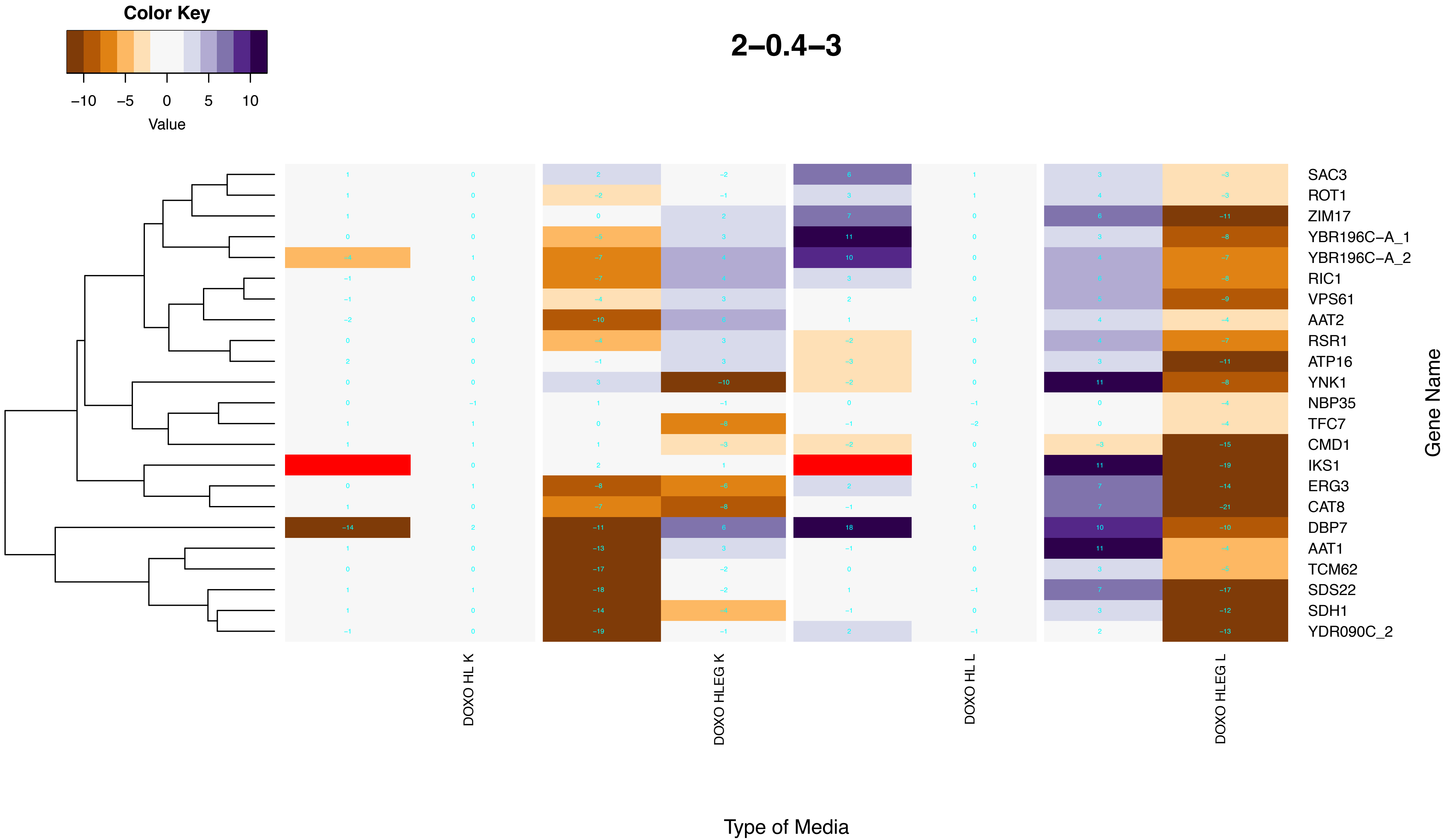
Color Key
−10 −5 0 5 10
Value
2−0.4−3
SAC3
ROT1
ZIM17
YBR196C−A_1
YBR196C−A_2
RIC1
VPS61
AAT2
RSR1
ATP16
YNK1
NBP35
TFC7
CMD1
IKS1
ERG3
CAT8
DBP7
AAT1
TCM62
SDS22
SDH1
YDR090C_2
Gene Name
DOXO HL K
DOXO HLEG K
DOXO HL L
DOXO HLEG L
Type of Media

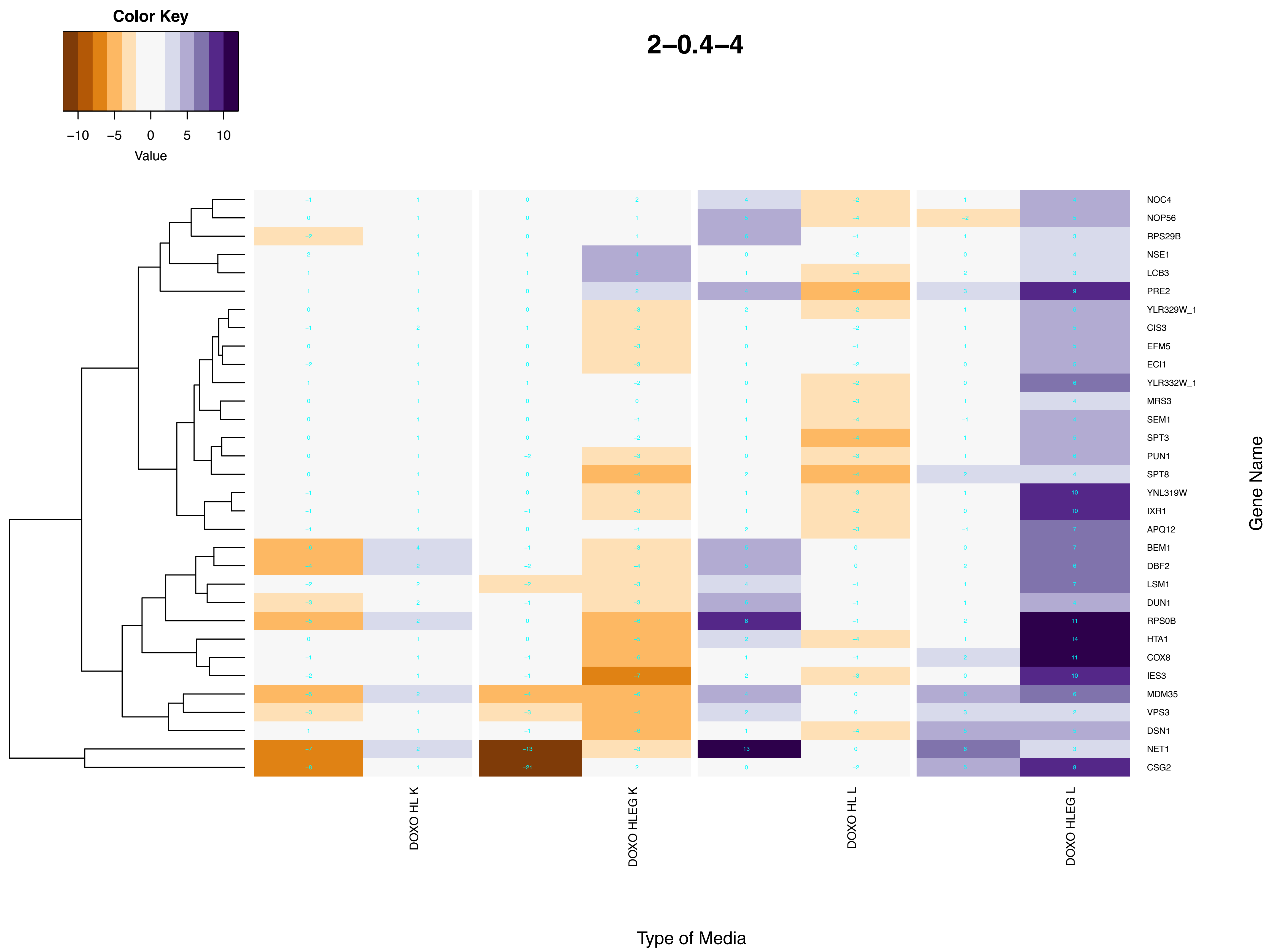
Color Key
−10 −5 0 5 10
Value
2–0.4–4
NOC4
NOP56
RPS29B
NSE1
LCB3
PRE2
YLR329W_1
CIS3
EFM5
ECI1
YLR332W_1
MRS3
SEM1
SPT3
PUN1
SPT8
YNL319W
IXR1
APQ12
BEM1
DBF2
LSM1
DUN1
RPS0B
HTA1
COX8
IES3
MDM35
VPS3
DSN1
NET1
CSG2
Gene Name
DOXO HL K
DOXO HLEG K
DOXO HL L
DOXO HLEG L
Type of Media

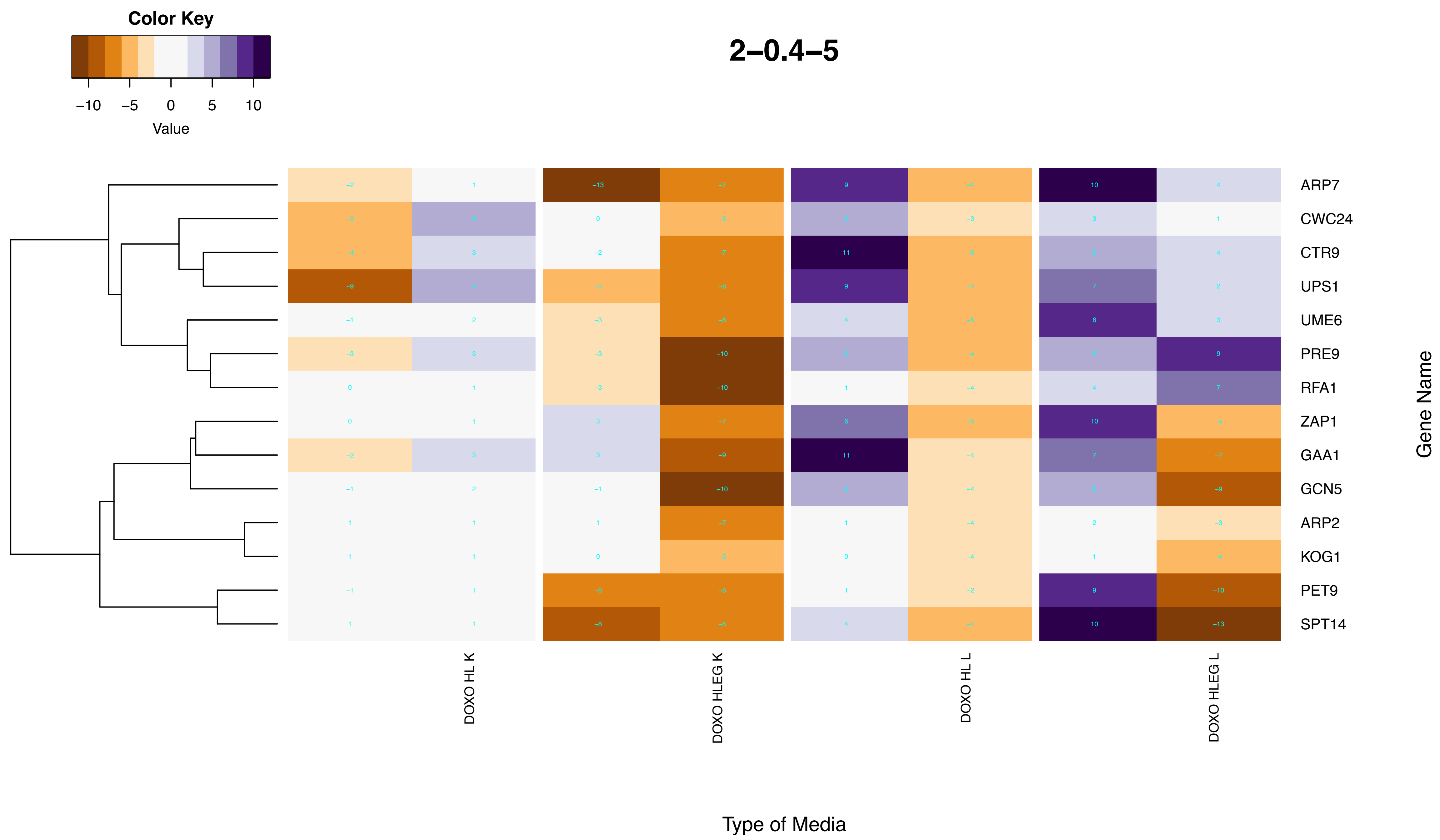
2–0.4–5
Color Key
−10
−5
0
5
10
Value
ARP7
CWC24
CTR9
UPS1
UME6
PRE9
RFA1
ZAP1
GAA1
GCN5
ARP2
KOG1
PET9
SPT14
Gene Name
DOXO HL K
DOXO HLEG K
DOXO HL L
DOXO HLEG L
Type of Media

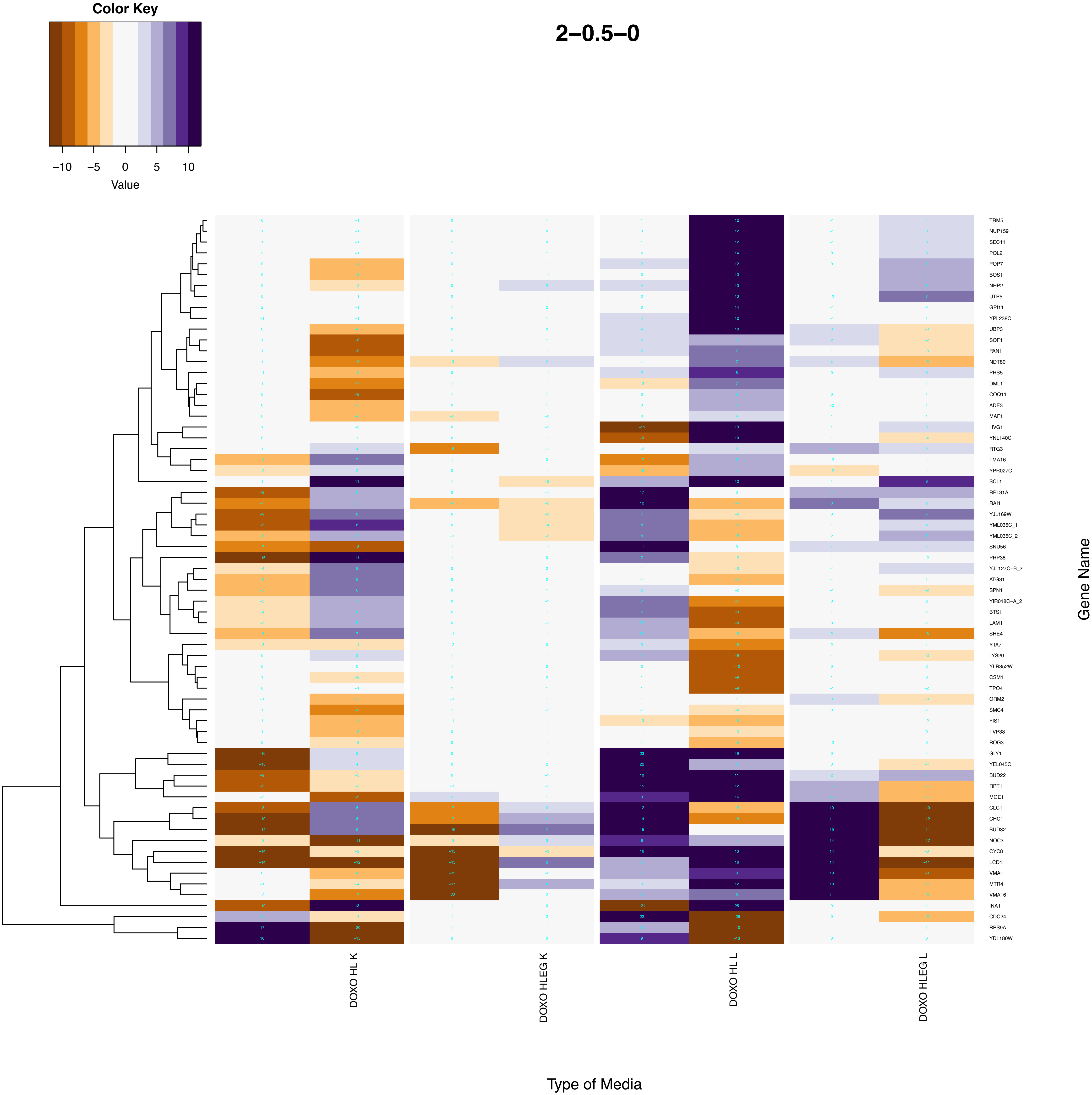

Color Key
−10
−5
0
5
10
Value
2−0.5−0
TRM5
NUP159
SEC11
POL2
POP7
BOS1
NHP2
UTP5
GPI11
YPL238C
UBP3
SOF1
PAN1
NDT80
PRS5
DML1
COQ11
ADE3
MAF1
HVG1
YNL140C
RTG3
TMA16
YPR027C
SCL1
RPL31A
RAI1
YJL169W
YML035C_1
YML035C_2
SNU56
PRP38
YJL127C−B_2
ATG31
SPN1
YIR018C−A_2
BTS1
LAM1
SHE4
YTA7
LYS20
YLR352W
CSM1
TPO4
ORM2
SMC4
FIS1
TVP38
ROG3
GLY1
YEL045C
BUD22
RPT1
MGE1
CLC1
CHC1
BUD32
NOC3
CYC8
LCD1
VMA1
MTR4
VMA16
INA1
CDC24
RPS9A
YDL180W
Gene Name
DOXO HL K
DOXO HLEG K
DOXO HL L
DOXO HLEG L
Type of Media

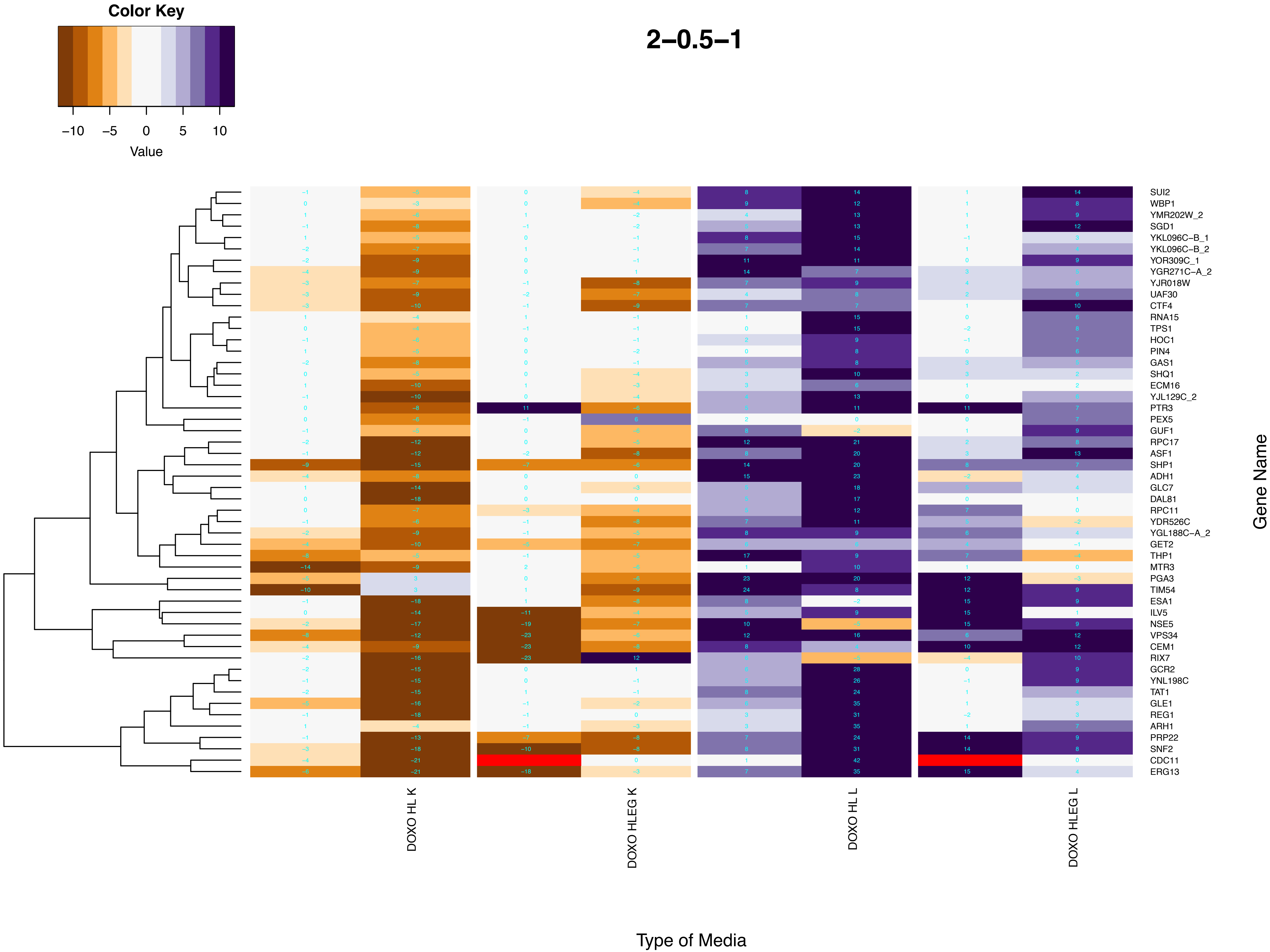

Color Key
−10
−5
0
5
10
Value
2−0.5−1
SUI2
WBP1
YMR202W_2
SGD1
YKL096C−B_1
YKL096C−B_2
YOR309C_1
YGR271C−A_2
YJR018W
UAF30
CTF4
RNA15
TPS1
HOC1
PIN4
GAS1
SHQ1
ECM16
YJL129C_2
PTR3
PEX5
GUF1
RPC17
ASF1
SHP1
ADH1
GLC7
DAL81
RPC11
YDR526C
YGL188C−A_2
GET2
THP1
MTR3
PGA3
TIM54
ESA1
ILV5
NSE5
VPS34
CEM1
RIX7
GCR2
YNL198C
TAT1
GLE1
REG1
ARH1
PRP22
SNF2
CDC11
ERG13
Gene Name
DOXO HL K
DOXO HLEG K
DOXO HL L
DOXO HLEG L
Type of Media

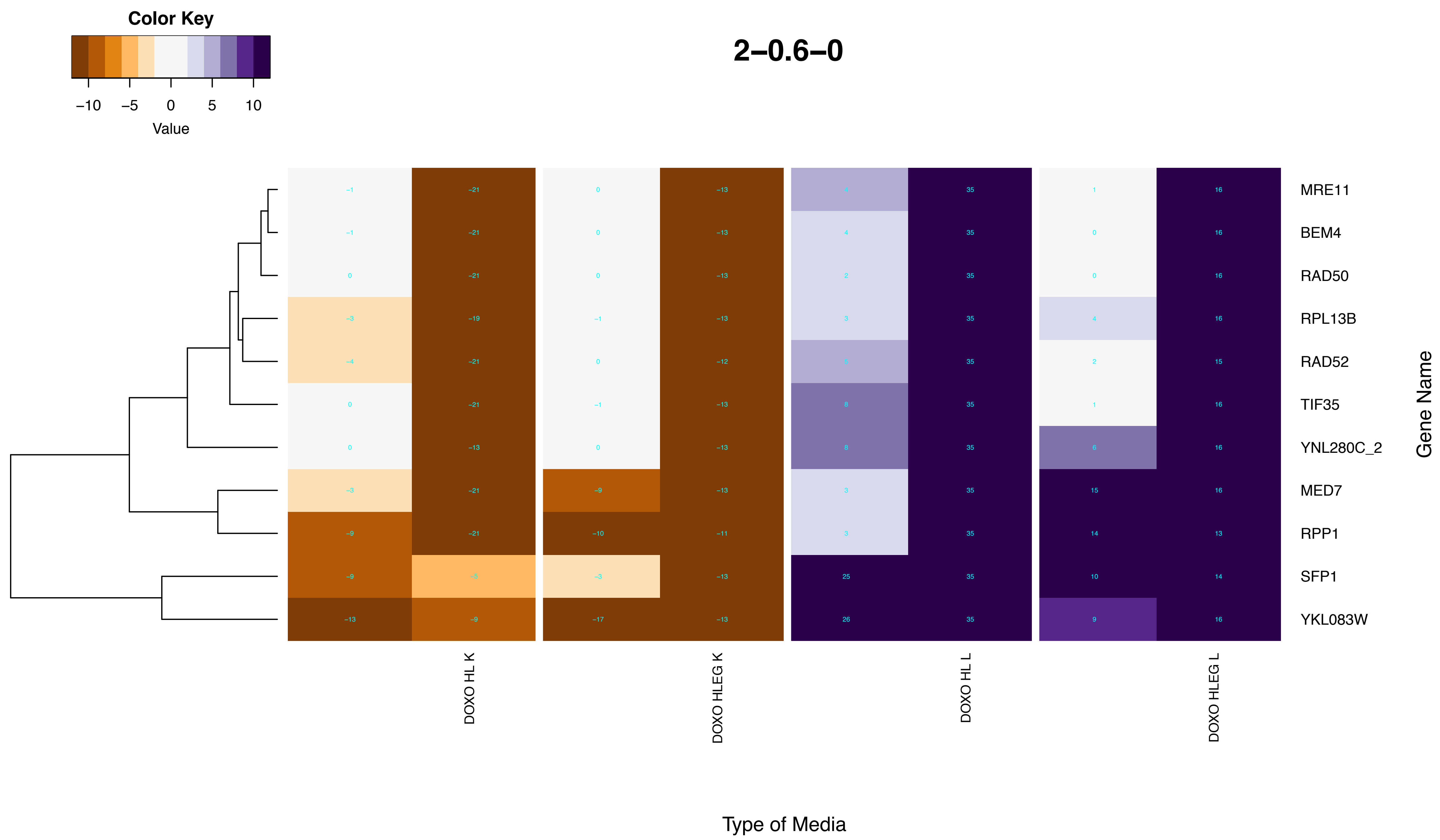
2–0.6–0
Color Key
–10
–5
0
5
10
Value
MRE11
BEM4
RAD50
RPL13B
RAD52
TIF35
YNL280C_2
MED7
RPP1
SFP1
YKL083W
Gene Name
DOXO HL K
DOXO HLEG K
DOXO HL L
DOXO HLEG L
Type of Media

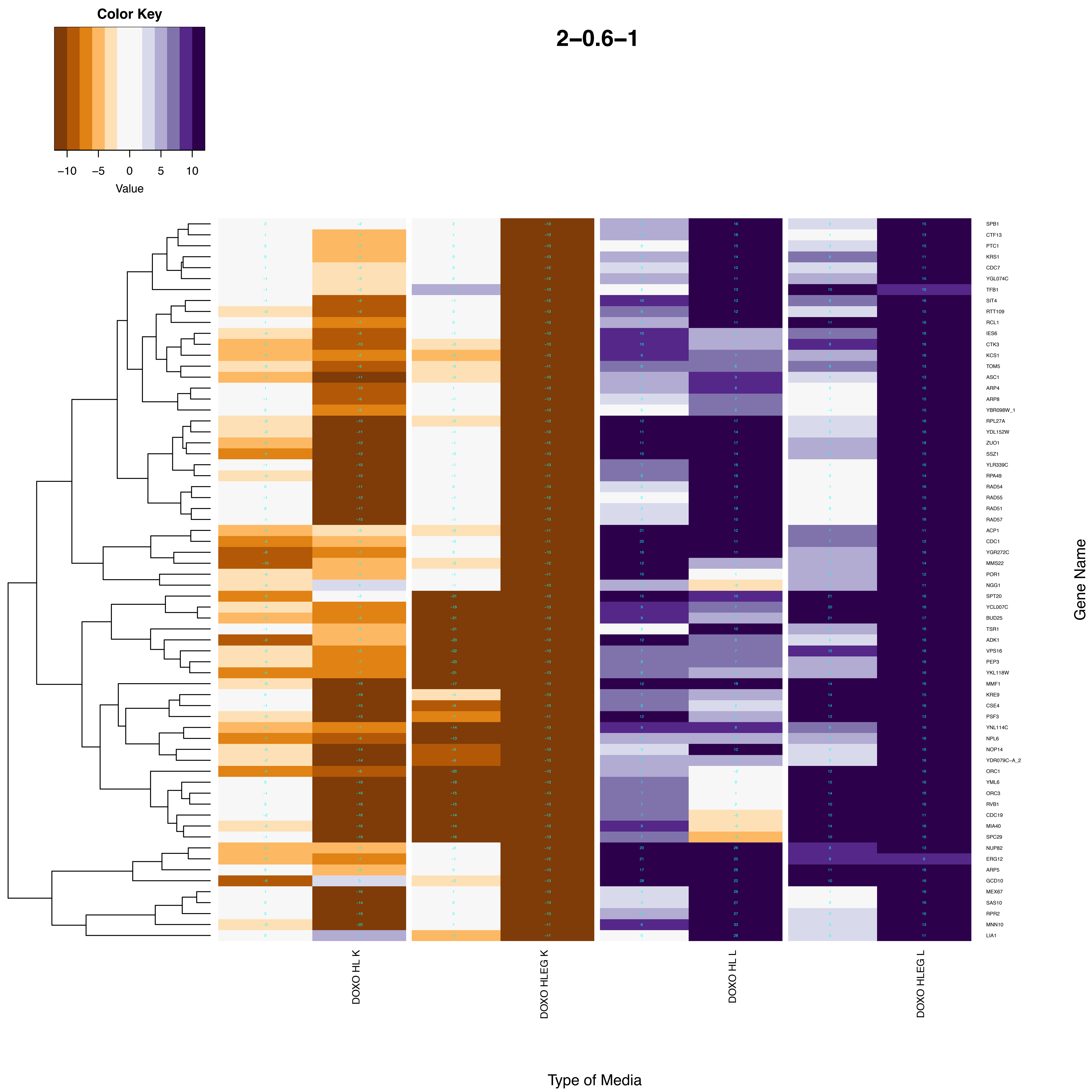

Color Key
−10
−5
0
5
10
Value
2−0.6−1
DOXO HL K
DOXO HLEG K
DOXO HL L
DOXO HLEG L
Type of Media
Gene Name
SPB1
CTF13
PTC1
KRS1
CDC7
YGL074C
TFB1
SIT4
RTT109
RCL1
IES6
CTK3
KCS1
TOM5
ASC1
ARP4
ARP8
YBR098W_1
RPL27A
YDL152W
ZUO1
SSZ1
YLR339C
RPA49
RAD54
RAD55
RAD51
RAD57
ACP1
CDC1
YGR272C
MMS22
POR1
NGG1
SPT20
YCL007C
BUD25
TSR1
ADK1
VPS16
PEP3
YKL118W
MMF1
KRE9
CSE4
PSF3
YNL114C
NPL6
NOP14
YDR079C−A_2
ORC1
YML6
ORC3
RVB1
CDC19
MIA40
SPC29
NUP82
ERG12
ARP5
GCD10
MEX67
SAS10
RPR2
MNN10
LIA1

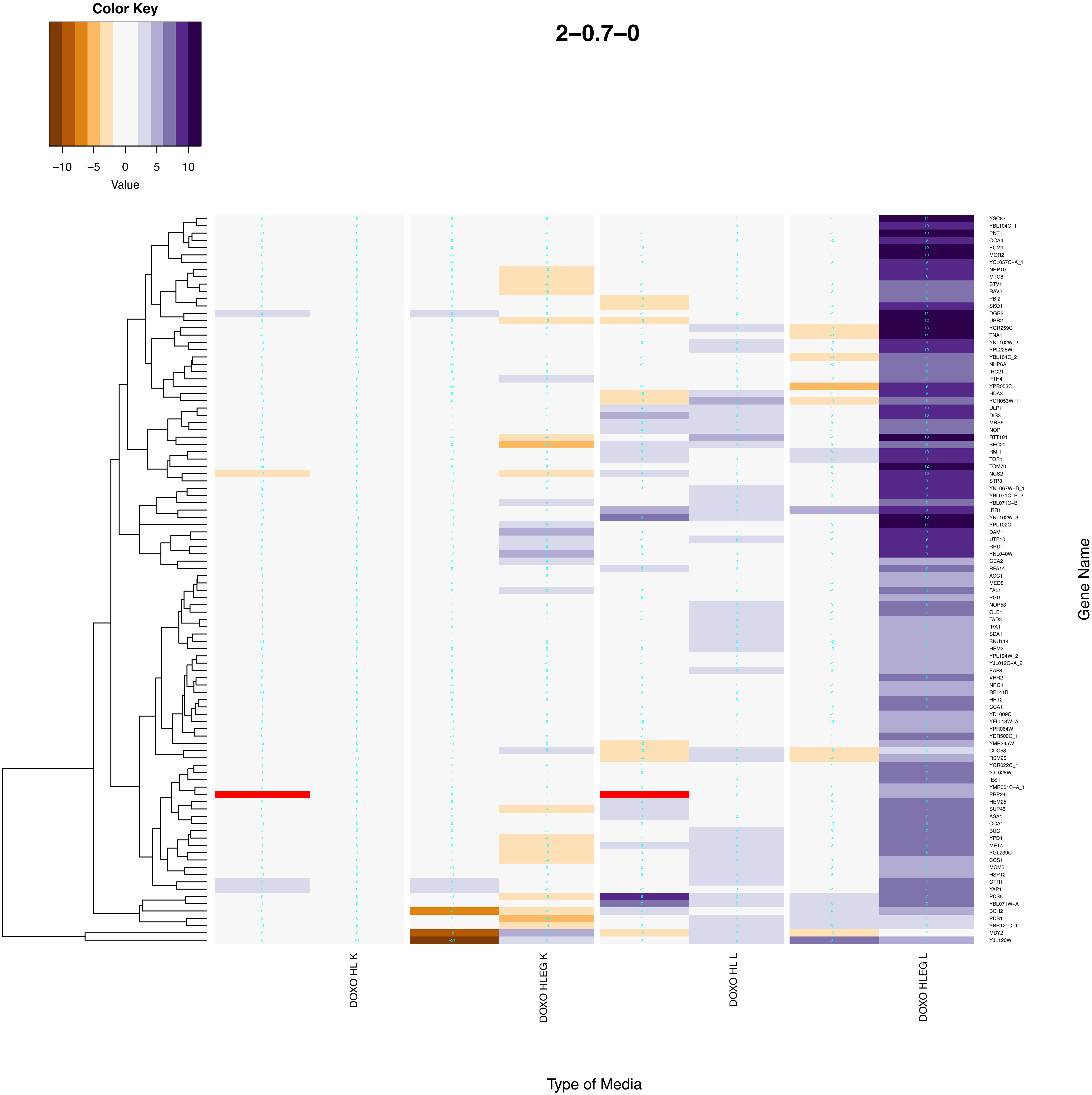

Color Key
−10
−5
0
5
10
Value
2−0.7−0
DOXO HL K
DOXO HLEG K
DOXO HL L
DOXO HLEG L
Type of Media
Gene Name
YSC83
YBL104C_1
PNT1
OCA4
ECM1
MGR2
YCL057C−A_1
NHP10
MTC6
STV1
RAV2
PBI2
SKO1
DGR2
UBR2
YGR259C
TNA1
YNL162W_2
YPL225W
YBL104C_2
NHP6A
IRC21
PTH4
YPR053C
HDA3
YCR053W_1
ULP1
DIS3
MRS6
NOP1
RTT101
SEC20
RMI1
TOP1
TOM70
NCS2
STP3
YNL067W−B_1
YBL071C−B_2
YBL071C−B_1
IRR1
YNL162W_3
YPL102C
DAM1
UTP10
RRD1
YNL040W
GEA2
RPA14
ACC1
MED8
FAL1
PGI1
NOP53
OLE1
TAD3
IRA1
SDA1
SNU114
HEM2
YPL194W_2
YJL012C−A_2
EAF3
VHR2
NRG1
RPL41B
HHT2
CCA1
YDL009C
YFL013W−A
YPR064W
YDR500C_1
YMR245W
CDC53
RSM25
YGR022C_1
YJL028W
IES1
YMR001C−A_1
PRP24
HEM25
SUP45
ASA1
OCA1
BUG1
YPD1
MET4
YGL239C
CCS1
MCM5
HSP12
GTR1
YAP1
PDS5
YBL071W−A_1
BCH2
PDB1
YBR121C_1
MDY2
YJL120W

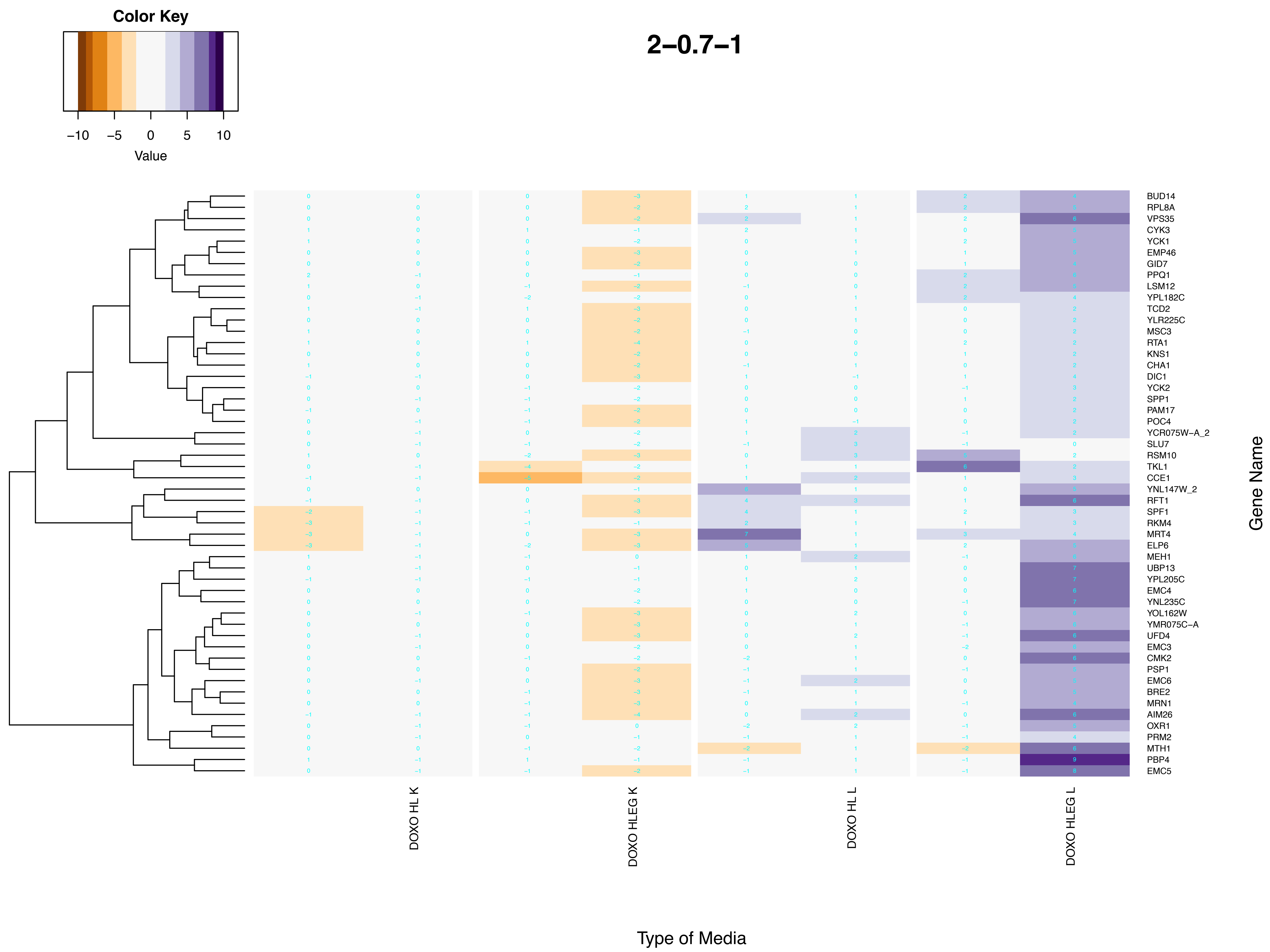
2–0.7–1
Color Key
–10
–5
0
5
10
Value
BUD14
RPL8A
VPS35
CYK3
YCK1
EMP46
GID7
PPQ1
LSM12
YPL182C
TCD2
YLR225C
MSC3
RTA1
KNS1
CHA1
DIC1
YCK2
SPP1
PAM17
POC4
YCR075W–A_2
SLU7
RSM10
TKL1
CCE1
YNL147W_2
RFT1
SPF1
RKM4
MRT4
ELP6
MEH1
UBP13
YPL205C
EMC4
YNL235C
YOL162W
YMR075C–A
UFD4
EMC3
CMK2
PSP1
EMC6
BRE2
MRN1
AIM26
OXR1
PRM2
MTH1
PBP4
EMC5
Gene Name
DOXO HL K
DOXO HLEG K
DOXO HL L
DOXO HLEG L
Type of Media

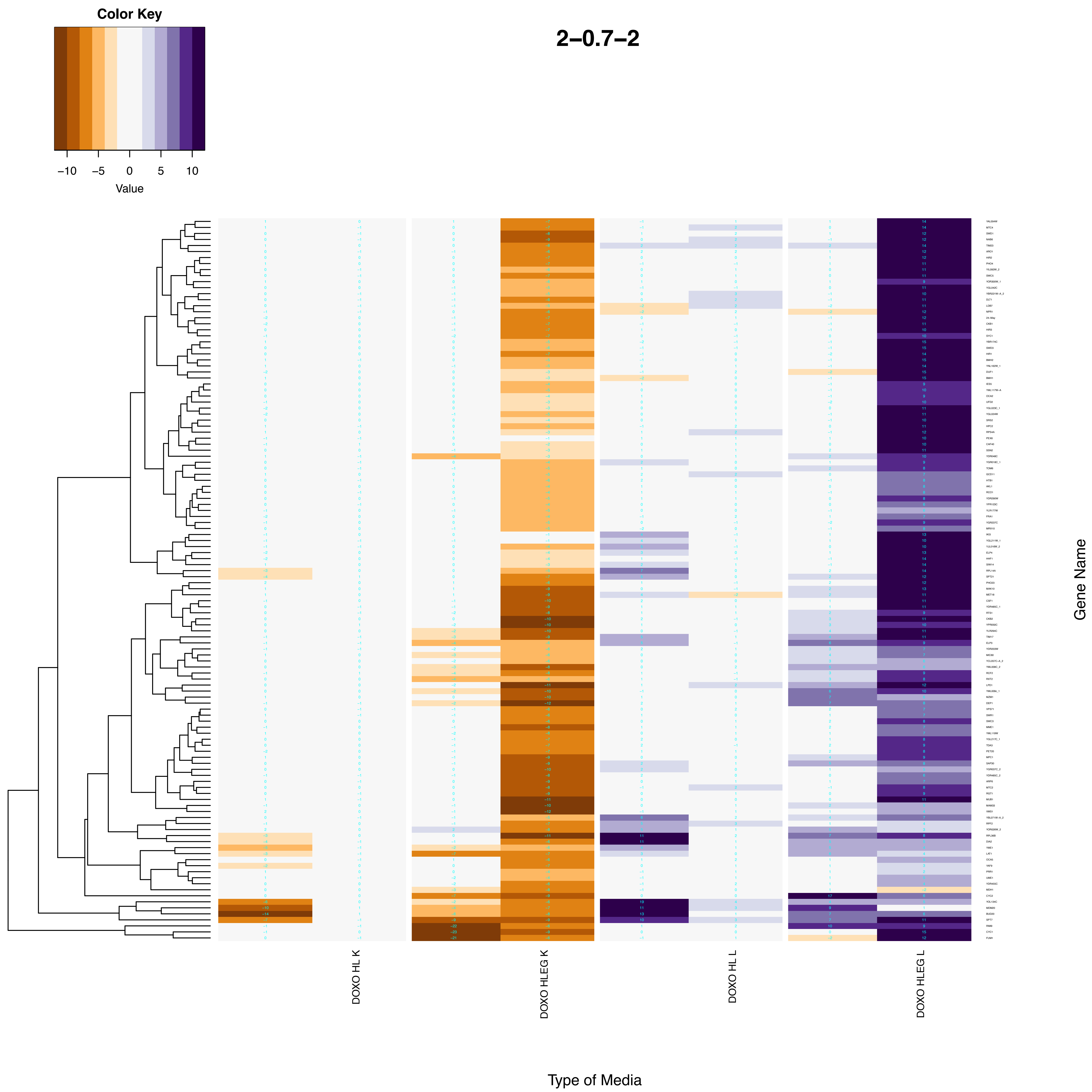

Color Key
−10
−5
0
5
10
Value
2−0.7−2
DOXO HL K
DOXO HLEG K
DOXO HL L
DOXO HLEG L
Gene Name
Type of Media

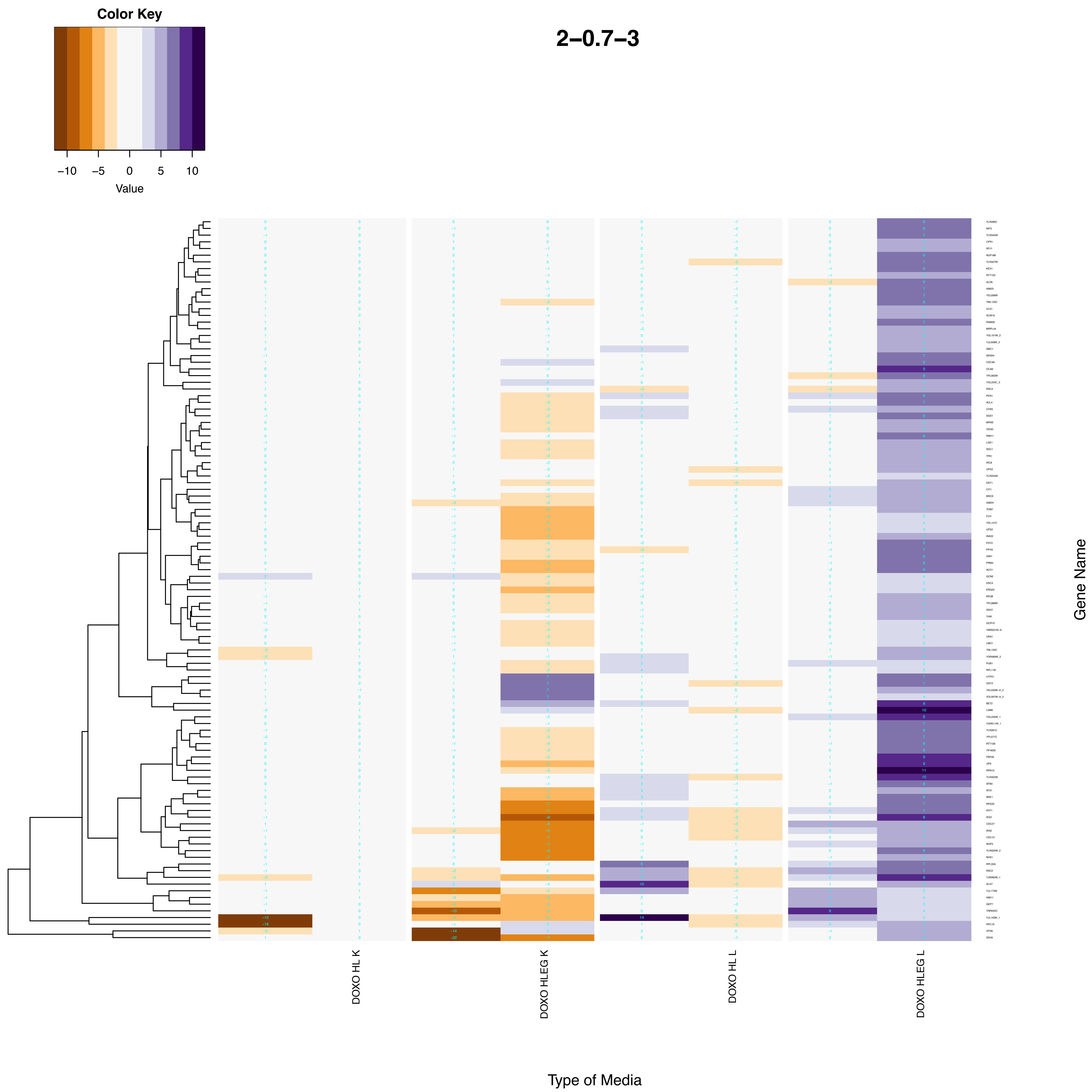
Color Key
−10 −5 0 5 10
Value
2–0.7–3
DOXO HL K
DOXO HLEG K
DOXO HL L
DOXO HLEG L
Gene Name
Type of Media

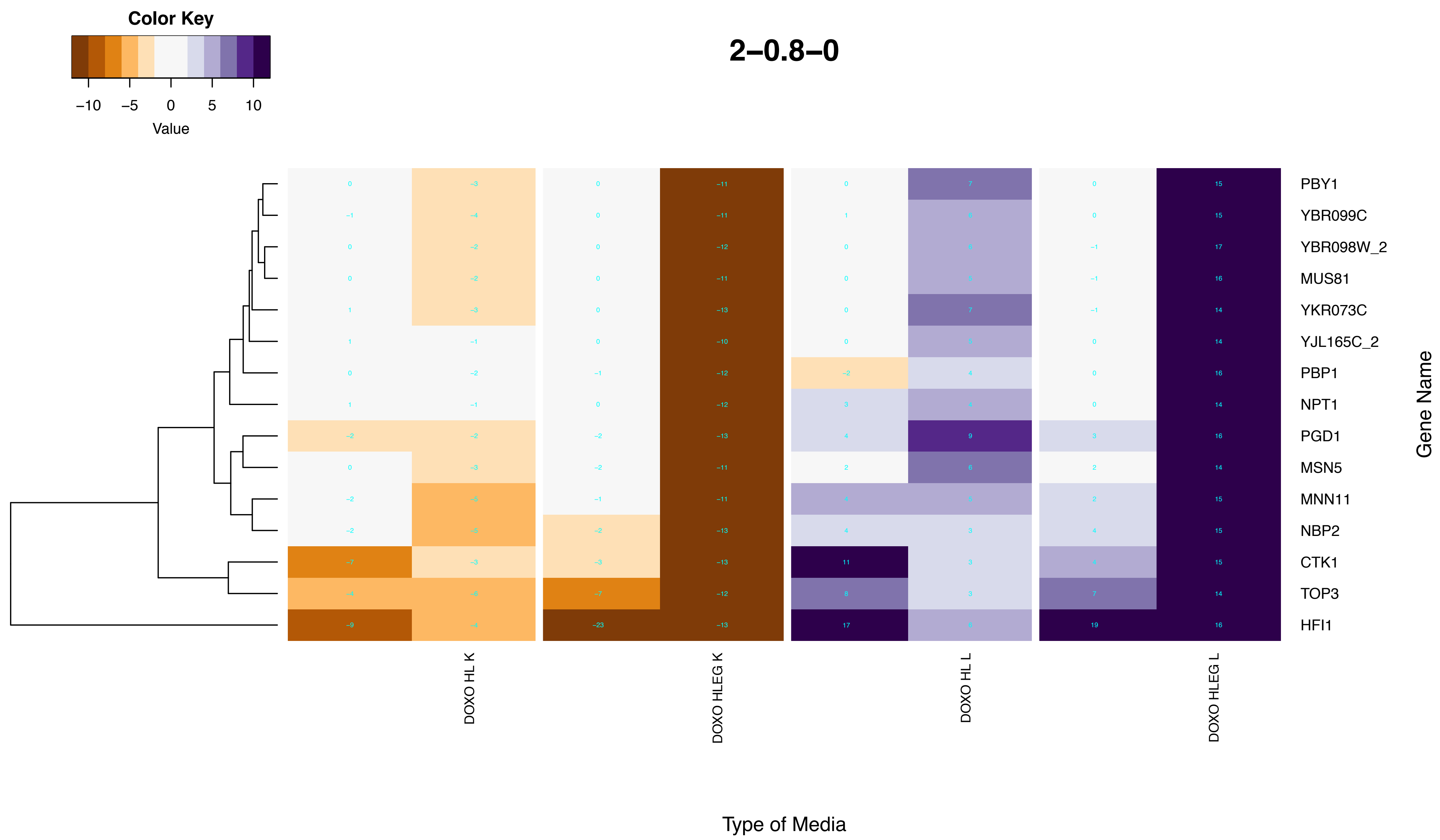
2–0.8–0
Color Key
−10
−5
0
5
10
Value
PBY1
YBR099C
YBR098W_2
MUS81
YKR073C
YJL165C_2
PBP1
NPT1
PGD1
MSN5
MNN11
NBP2
CTK1
TOP3
HFI1
Gene Name
DOXO HL K
DOXO HLEG K
DOXO HL L
DOXO HLEG L
Type of Media

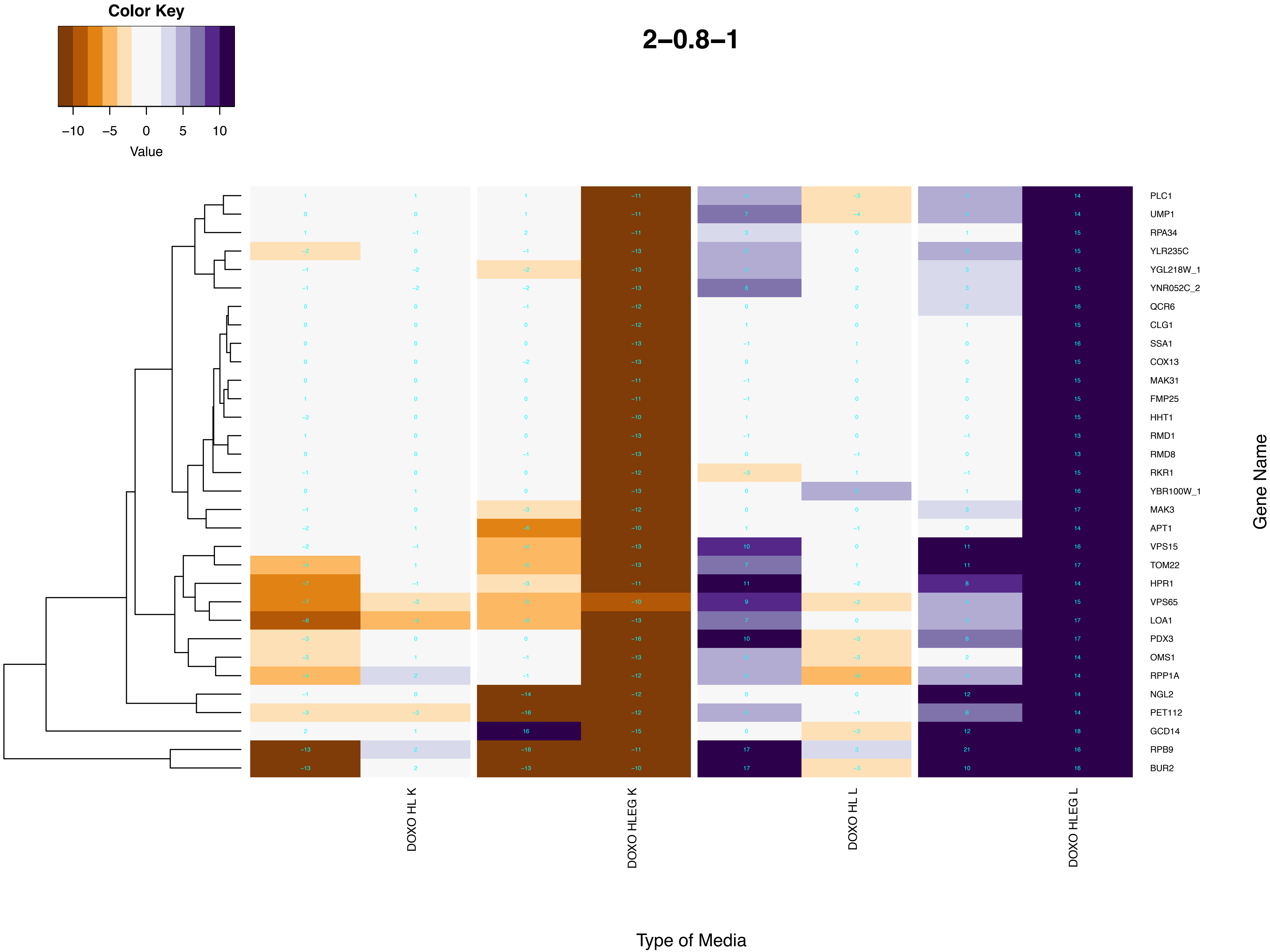
Color Key
−10
−5
0
5
10
Value
2−0.8−1
PLC1
UMP1
RPA34
YLR235C
YGL218W_1
YNR052C_2
QCR6
CLG1
SSA1
COX13
MAK31
FMP25
HHT1
RMD1
RMD8
RKR1
YBR100W_1
MAK3
APT1
VPS15
TOM22
HPR1
VPS65
LOA1
PDX3
OMS1
RPP1A
NGL2
PET112
GCD14
RPB9
BUR2
Gene Name
DOXO HL K
DOXO HLEG K
DOXO HL L
DOXO HLEG L
Type of Media

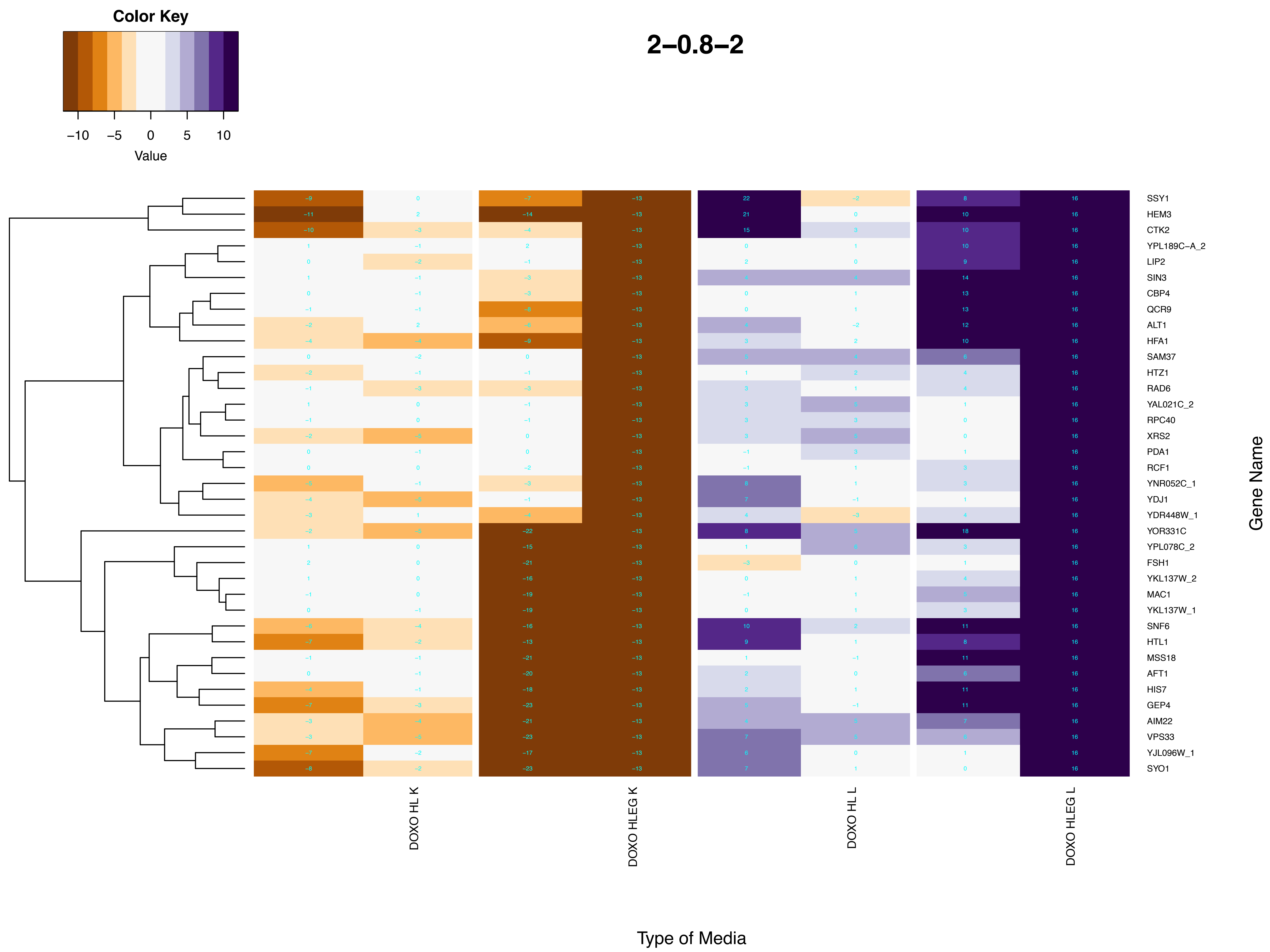

Color Key
−10
−5
0
5
10
Value
2–0.8–2
SSY1
HEM3
CTK2
YPL189C–A_2
LIP2
SIN3
CBP4
QCR9
ALT1
HFA1
SAM37
HTZ1
RAD6
YAL021C_2
RPC40
XRS2
PDA1
RCF1
YNR052C_1
YDJ1
YDR448W_1
YOR331C
YPL078C_2
FSH1
YKL137W_2
MAC1
YKL137W_1
SNF6
HTL1
MSS18
AFT1
HIS7
GEP4
AIM22
VPS33
YJL096W_1
SYO1
Gene Name
DOXO HL K
DOXO HLEG K
DOXO HL L
DOXO HLEG L
Type of Media

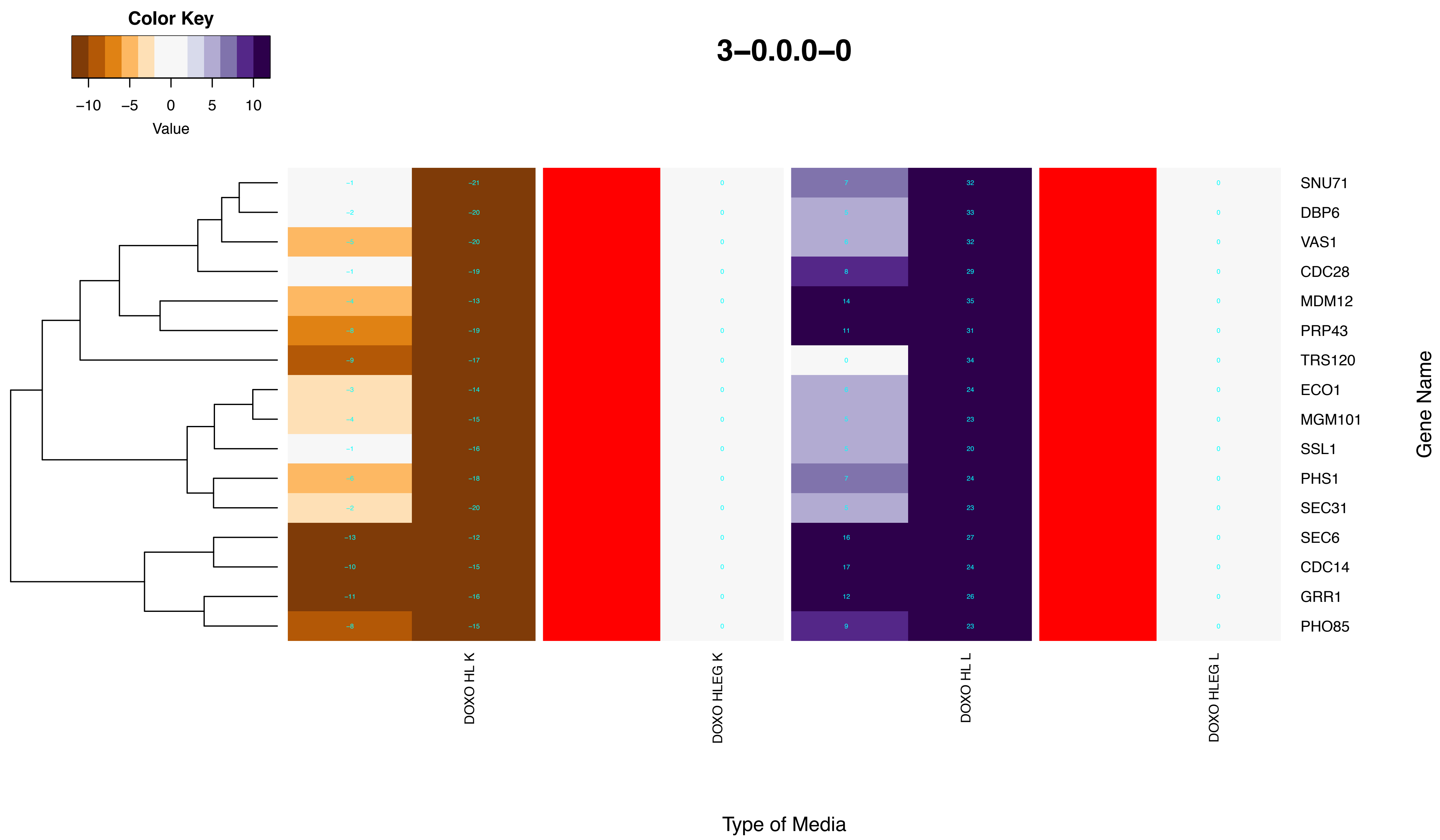

Color Key
−10 −5 0 5 10
Value
3−0.0.0−0
SNU71
DBP6
VAS1
CDC28
MDM12
PRP43
TRS120
ECO1
MGM101
SSL1
PHS1
SEC31
SEC6
CDC14
GRR1
PHO85
Gene Name
DOXO HL K
DOXO HLEG K
DOXO HL L
DOXO HLEG L
Type of Media

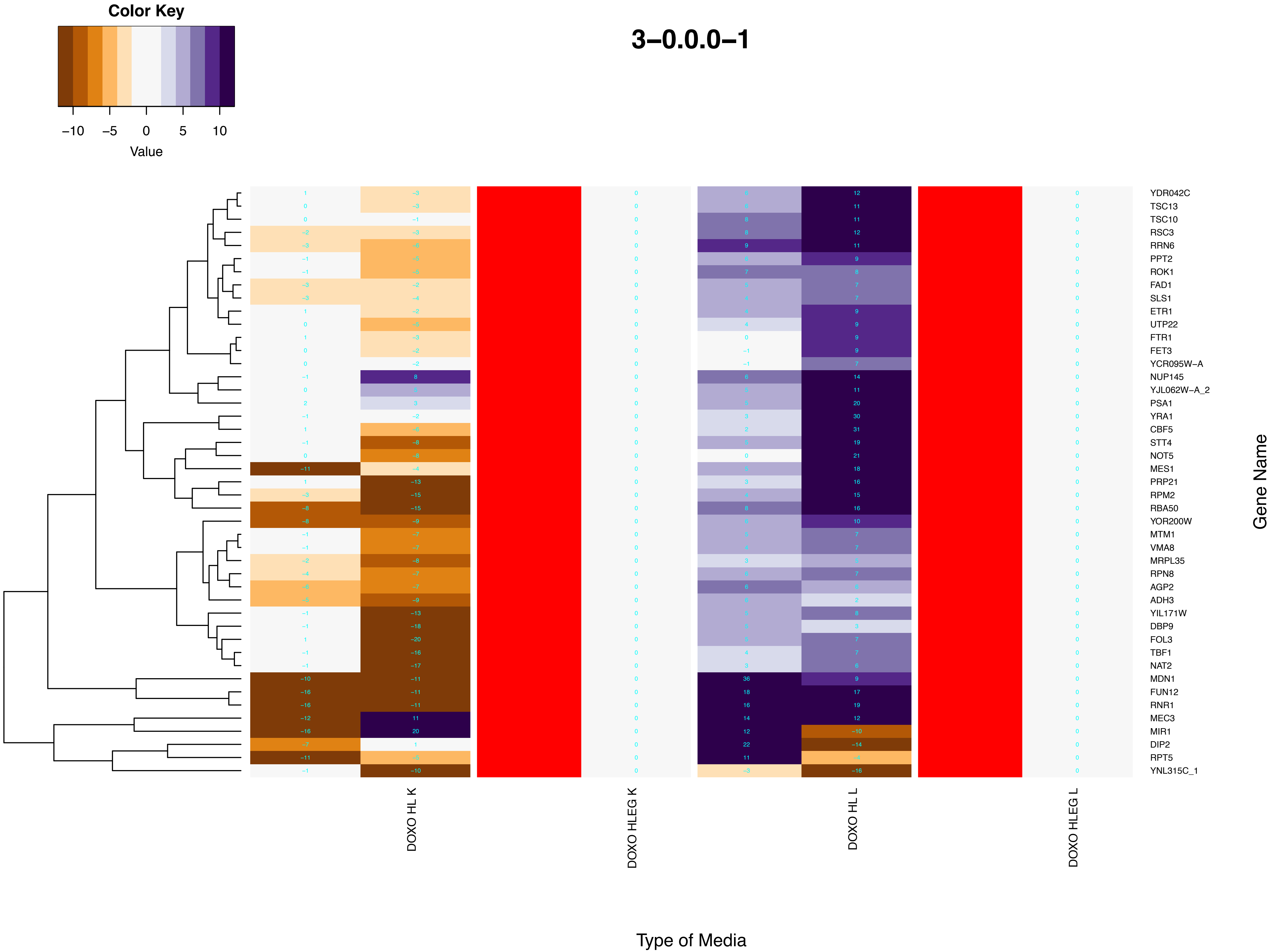

Color Key
−10
−5
0
5
10
Value
3−0.0.0−1
YDR042C
TSC13
TSC10
RSC3
RRN6
PPT2
ROK1
FAD1
SLS1
ETR1
UTP22
FTR1
FET3
YCR095W−A
NUP145
YJL062W−A_2
PSA1
YRA1
CBF5
STT4
NOT5
MES1
PRP21
RPM2
RBA50
YOR200W
MTM1
VMA8
MRPL35
RPN8
AGP2
ADH3
YIL171W
DBP9
FOL3
TBF1
NAT2
MDN1
FUN12
RNR1
MEC3
MIR1
DIP2
RPT5
YNL315C_1
Gene Name
DOXO HL K
DOXO HLEG K
DOXO HL L
DOXO HLEG L
Type of Media

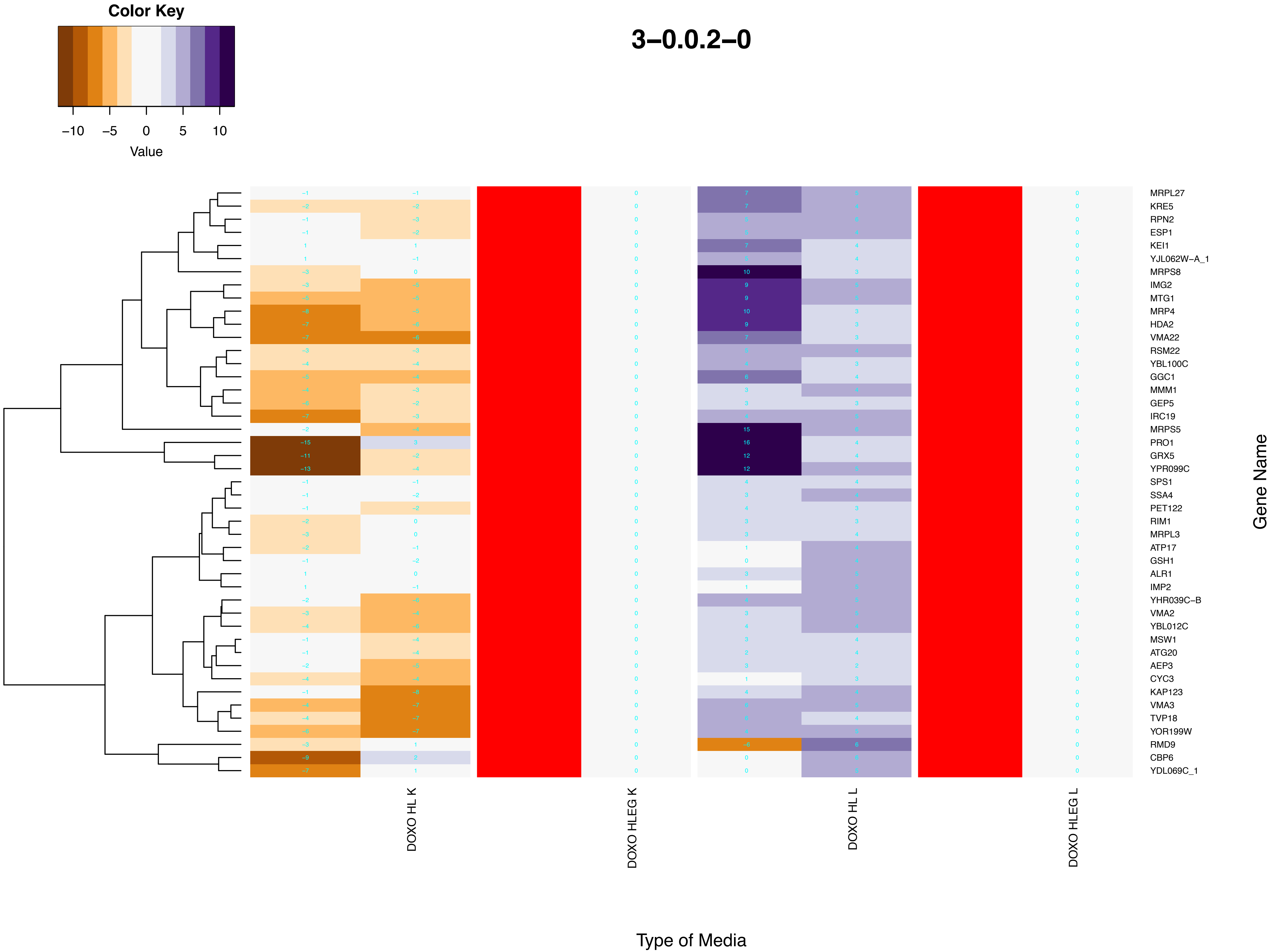

Color Key
−10
−5
0
5
10
Value
3−0.0.2−0
MRPL27
KRE5
RPN2
ESP1
KEI1
YJL062W−A_1
MRPS8
IMG2
MTG1
MRP4
HDA2
VMA22
RSM22
YBL100C
GGC1
MMM1
GEP5
IRC19
MRPS5
PRO1
GRX5
YPR099C
SPS1
SSA4
PET122
RIM1
MRPL3
ATP17
GSH1
ALR1
IMP2
YHR039C−B
VMA2
YBL012C
MSW1
ATG20
AEP3
CYC3
KAP123
VMA3
TVP18
YOR199W
RMD9
CBP6
YDL069C_1
Gene Name
DOXO HL K
DOXO HLEG K
DOXO HL L
DOXO HLEG L
Type of Media

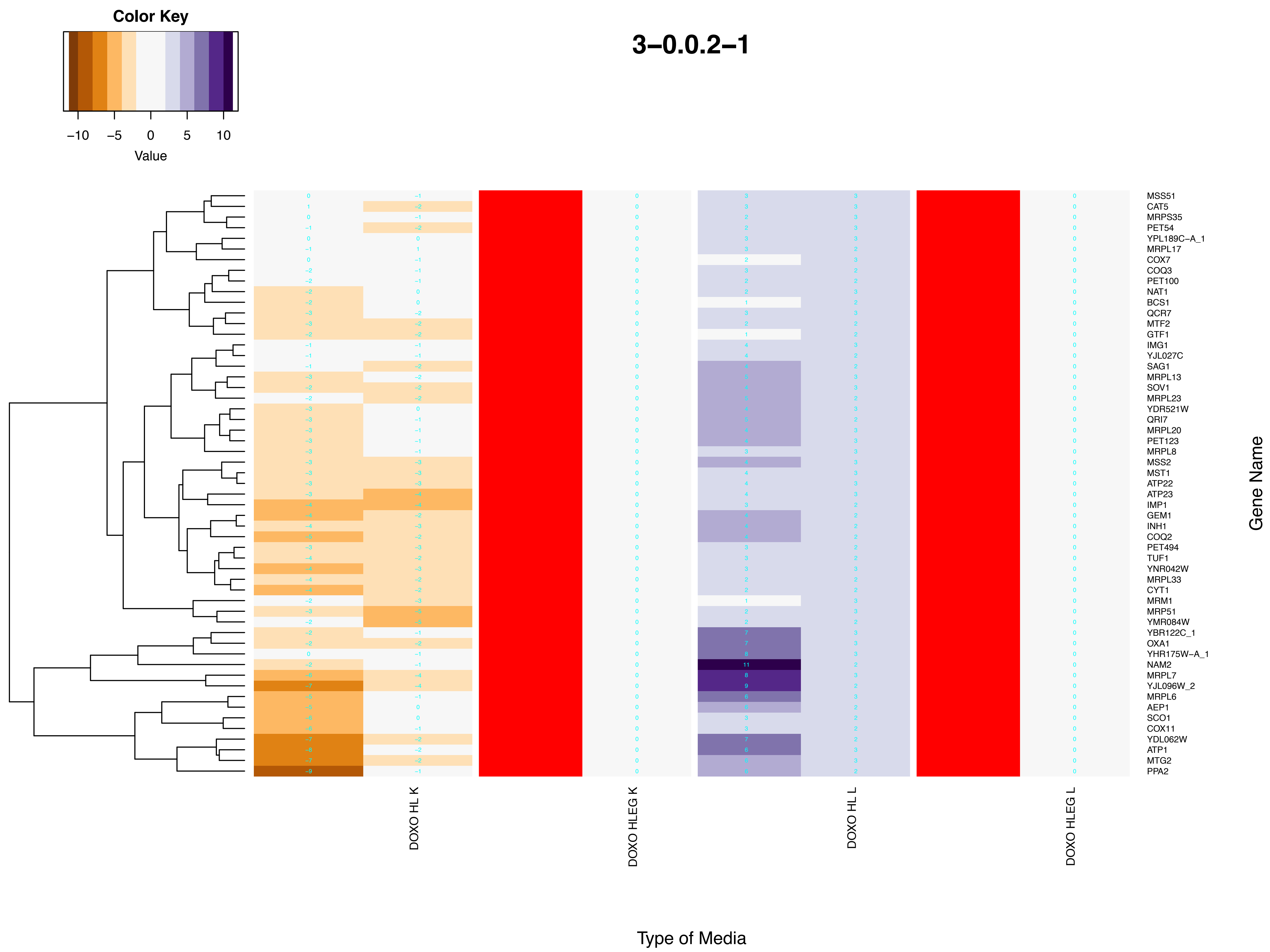

Color Key
−10 −5 0 5 10
Value
3−0.0.2−1
Gene Name
Type of Media
DOXO HL K
DOXO HLEG K
DOXO HL L
DOXO HLEG L
MSS51
CAT5
MRPS35
PET54
YPL189C−A_1
MRPL17
COX7
COQ3
PET100
NAT1
BCS1
QCR7
MTF2
GTF1
IMG1
YJL027C
SAG1
MRPL13
SOV1
MRPL23
YDR521W
QRI7
MRPL20
PET123
MRPL8
MSS2
MST1
ATP22
ATP23
IMP1
GEM1
INH1
COQ2
PET494
TUF1
YNR042W
MRPL33
CYT1
MRM1
MRP51
YMR084W
YBR122C_1
OXA1
YHR175W−A_1
NAM2
MRPL7
YJL096W_2
MRPL6
AEP1
SCO1
COX11
YDL062W
ATP1
MTG2
PPA2

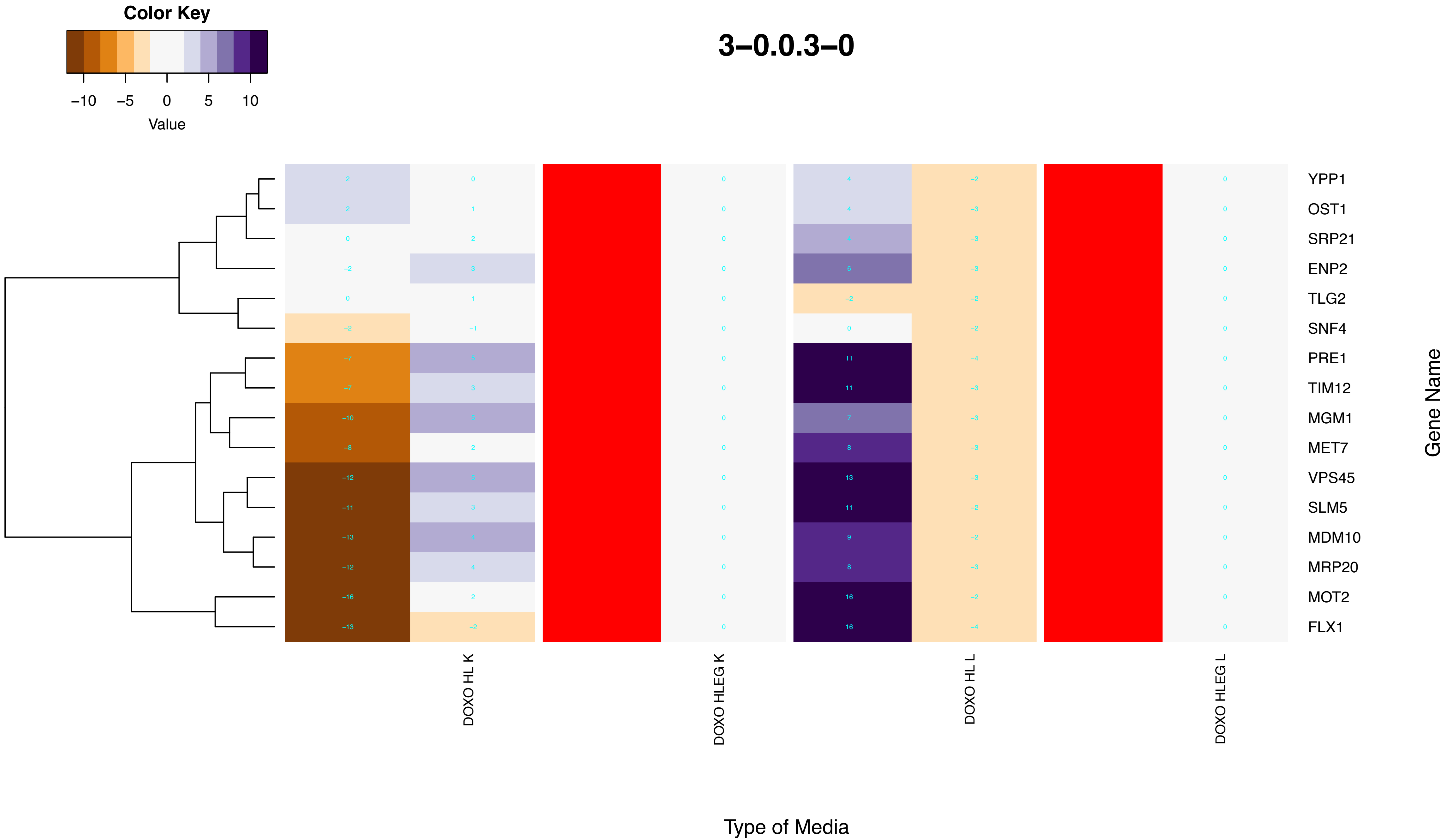
Color Key
−10
−5
0
5
10
Value
3−0.0.3−0
YPP1
OST1
SRP21
ENP2
TLG2
SNF4
PRE1
TIM12
MGM1
MET7
VPS45
SLM5
MDM10
MRP20
MOT2
FLX1
Gene Name
DOXO HL K
DOXO HLEG K
DOXO HL L
DOXO HLEG L
Type of Media

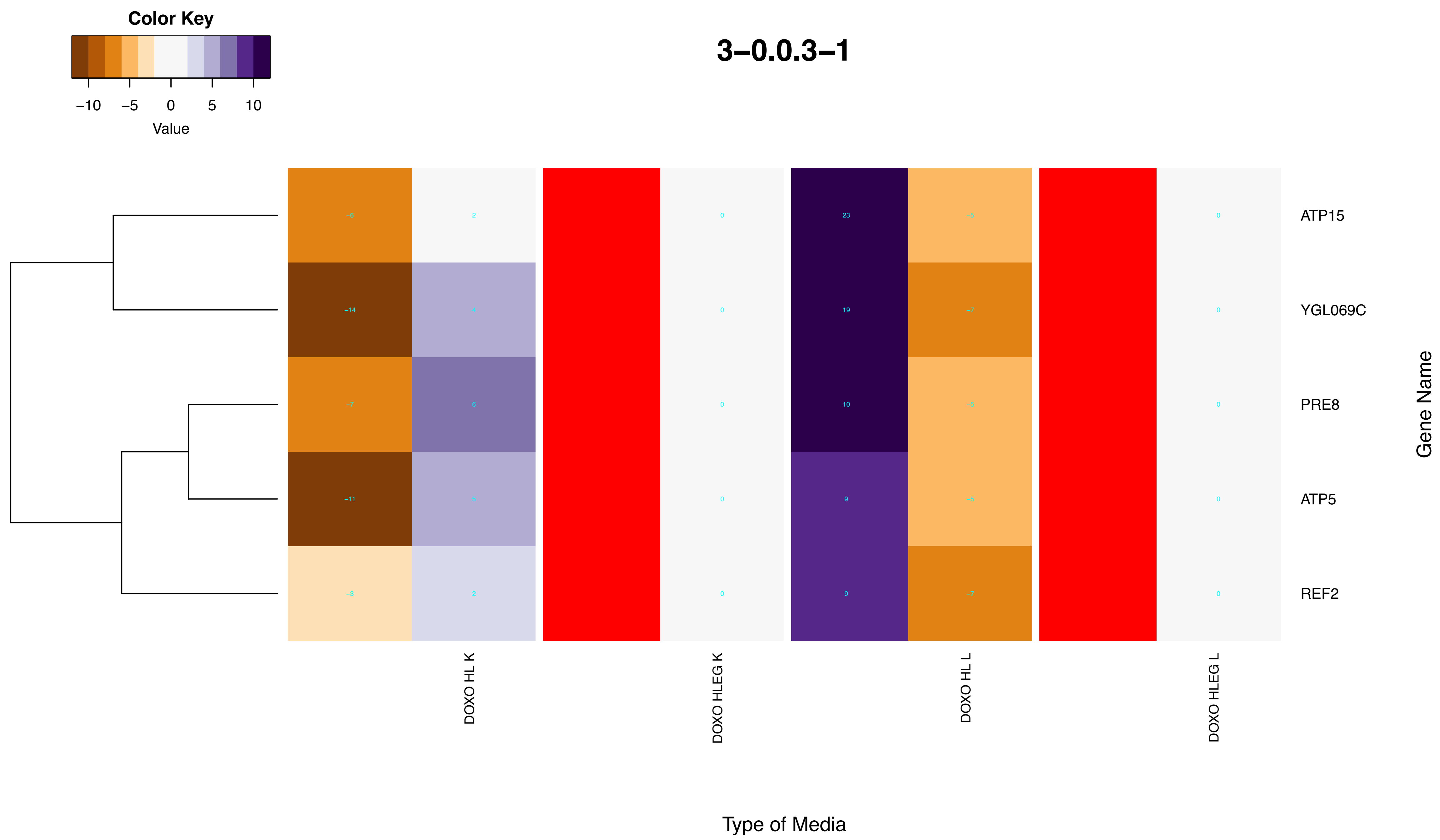
3–0.0.3–1
Color Key
–10
–5
0
5
10
Value
ATP15
YGL069C
PRE8
ATP5
REF2
Gene Name
DOXO HL K
DOXO HLEG K
DOXO HL L
DOXO HLEG L
Type of Media

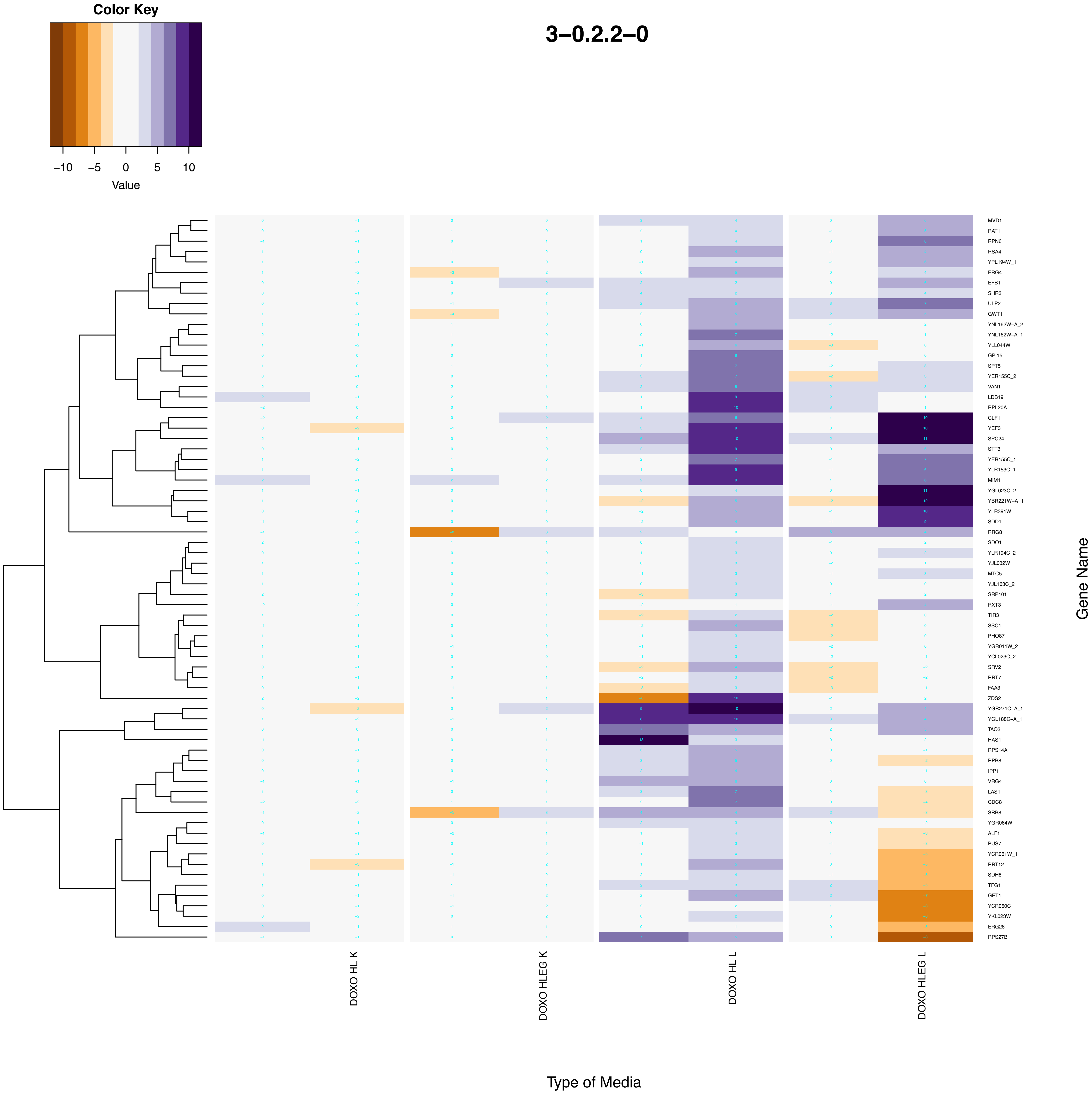
Color Key
−10
−5
0
5
10
Value
3−0.2.2−0
MVD1
RAT1
RPN6
RSA4
YPL194W_1
ERG4
EFB1
SHR3
ULP2
GWT1
YNL162W−A_2
YNL162W−A_1
YLL044W
GPI15
SPT5
YER155C_2
VAN1
LDB19
RPL20A
CLF1
YEF3
SPC24
STT3
YER155C_1
YLR153C_1
MIM1
YGL023C_2
YBR221W−A_1
YLR391W
SDD1
RRG8
SDO1
YLR194C_2
YJL032W
MTC5
YJL163C_2
SRP101
RXT3
TIR3
SSC1
PHO87
YGR011W_2
YCL023C_2
SRV2
RRT7
FAA3
ZDS2
YGR271C−A_1
YGL188C−A_1
TAO3
HAS1
RPS14A
RPB8
IPP1
VRG4
LAS1
CDC8
SRB8
YGR064W
ALF1
PUS7
YCR061W_1
RRT12
SDH8
TFG1
GET1
YCR050C
YKL023W
ERG26
RPS27B
Gene Name
DOXO HL K
DOXO HLEG K
DOXO HL L
DOXO HLEG L
Type of Media

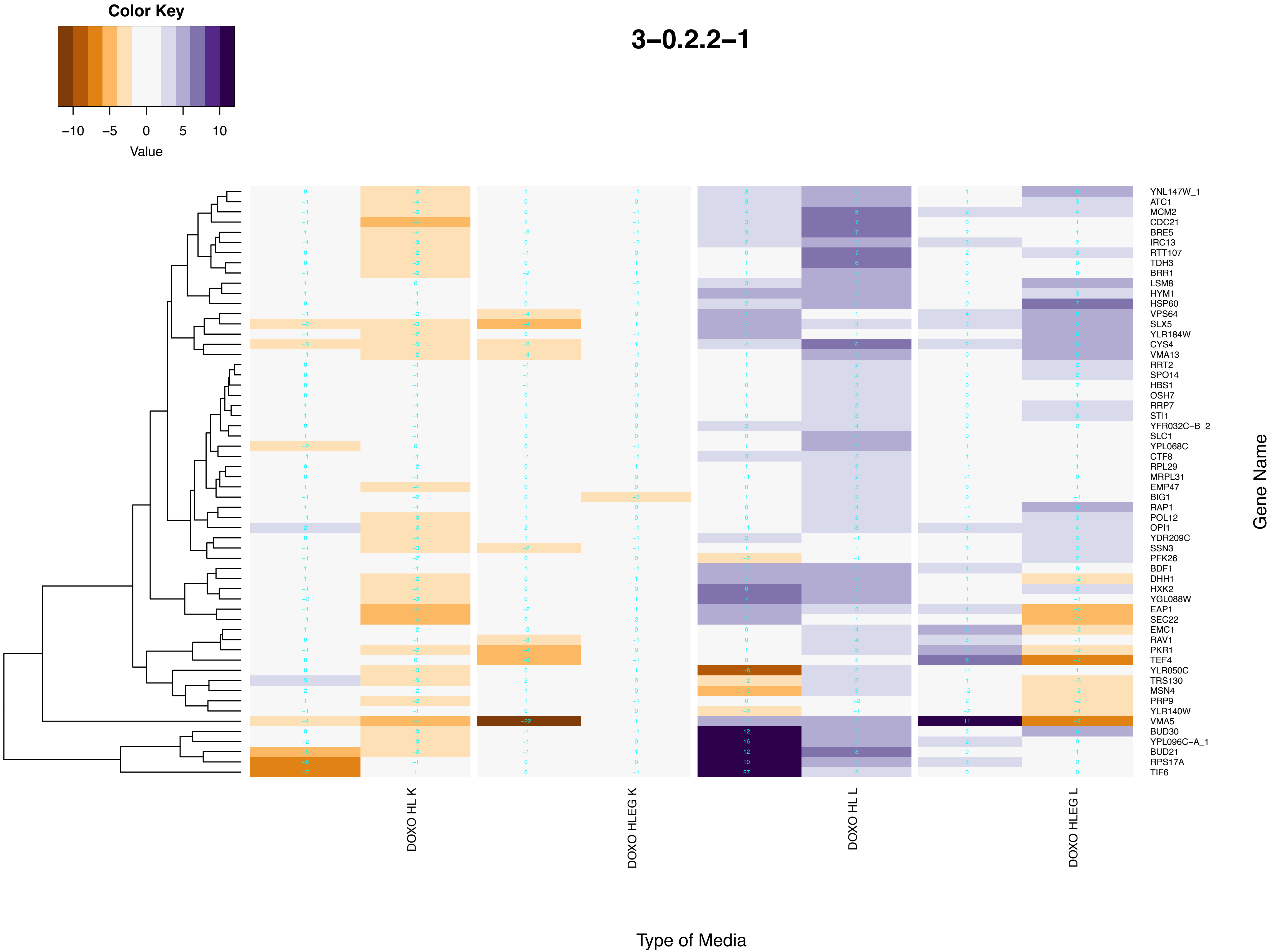
Color Key
−10 −5 0 5 10
Value
3−0.2.2−1
YNL147W_1
ATC1
MCM2
CDC21
BRE5
IRC13
RTT107
TDH3
BRR1
LSM8
HYM1
HSP60
VPS64
SLX5
YLR184W
CYS4
VMA13
RRT2
SPO14
HBS1
OSH7
RRP7
STI1
YFR032C−B_2
SLC1
YPL068C
CTF8
RPL29
MRPL31
EMP47
BIG1
RAP1
POL12
OPI1
YDR209C
SSN3
PFK26
BDF1
DHH1
HXK2
YGL088W
EAP1
SEC22
EMC1
RAV1
PKR1
TEF4
YLR050C
TRS130
MSN4
PRP9
YLR140W
VMA5
BUD30
YPL096C−A_1
BUD21
RPS17A
TIF6
Gene Name
DOXO HL K
DOXO HLEG K
DOXO HL L
DOXO HLEG L
Type of Media

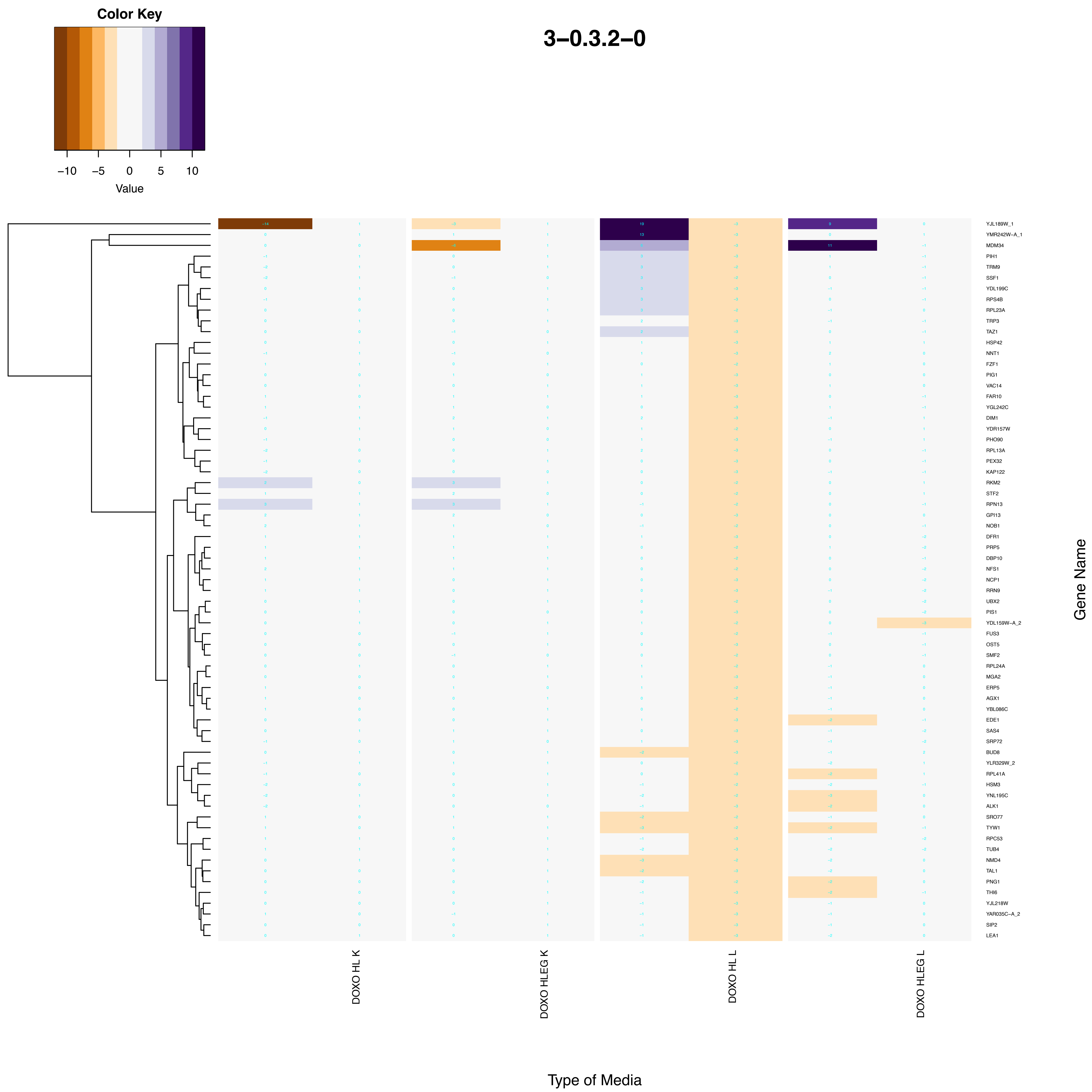

Color Key
−10
−5
0
5
10
Value
3−0.3.2−0
YJL189W_1
YMR242W−A_1
MDM34
PIH1
TRM9
SSF1
YDL199C
RPS4B
RPL23A
TRP3
TAZ1
HSP42
NNT1
FZF1
PIG1
VAC14
FAR10
YGL242C
DIM1
YDR157W
PHO90
RPL13A
PEX32
KAP122
RKM2
STF2
RPN13
GPI13
NOB1
DFR1
PRP5
DBP10
NFS1
NCP1
RRN9
UBX2
PIS1
YDL159W−A_2
FUS3
OST5
SMF2
RPL24A
MGA2
ERP5
AGX1
YBL086C
EDE1
SAS4
SRP72
BUD8
YLR329W_2
RPL41A
HSM3
YNL195C
ALK1
SRO77
TYW1
RPC53
TUB4
NMD4
TAL1
PNG1
THI6
YJL218W
YAR035C−A_2
SIP2
LEA1
Gene Name
DOXO HL K
DOXO HLEG K
DOXO HL L
DOXO HLEG L
Type of Media

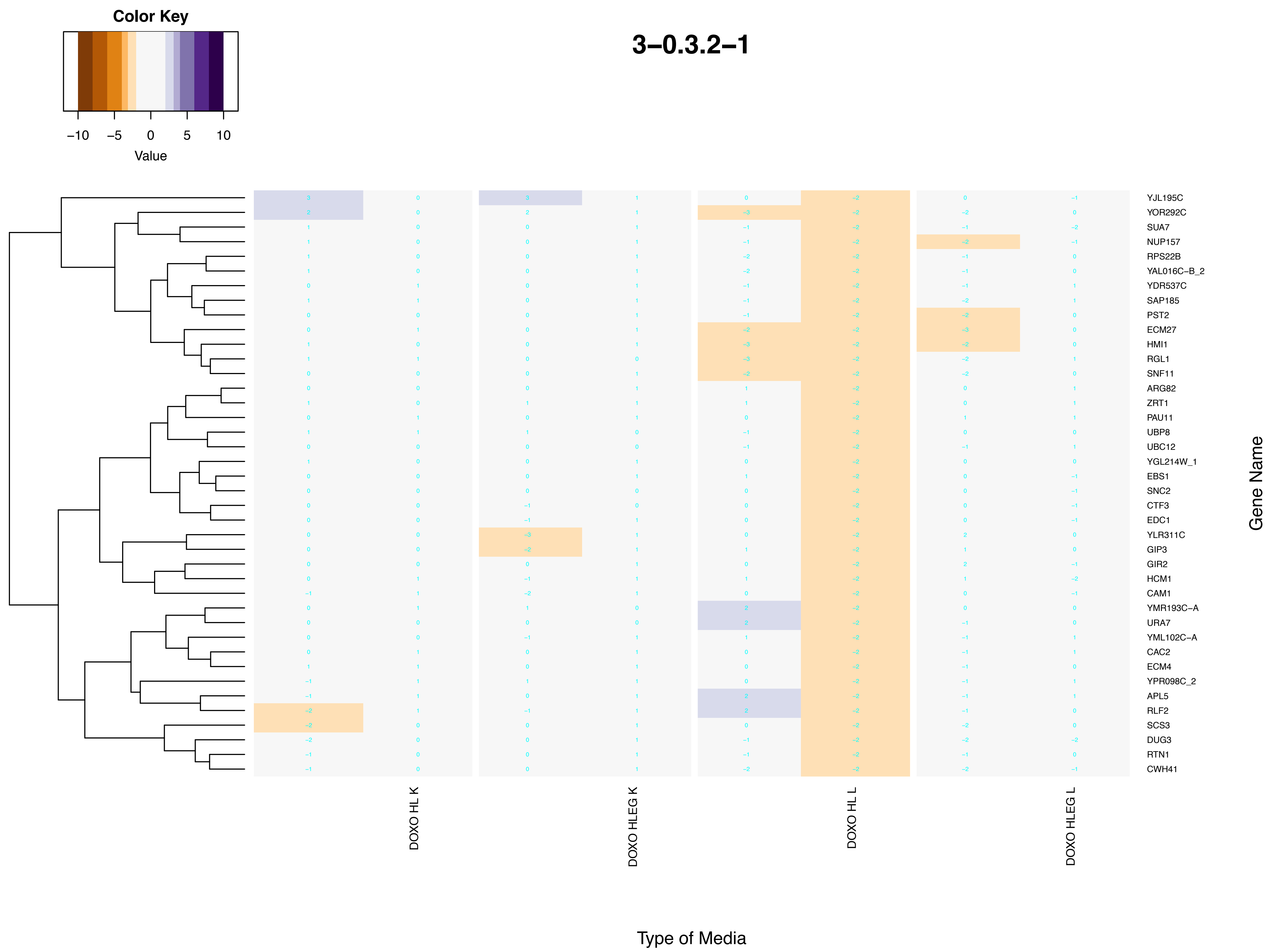
3–0.3.2–1
Color Key
−10 −5 0 5 10
Value
YJL195C
YOR292C
SUA7
NUP157
RPS22B
YAL016C–B_2
YDR537C
SAP185
PST2
ECM27
HMI1
RGL1
SNF11
ARG82
ZRT1
PAU11
UBP8
UBC12
YGL214W_1
EBS1
SNC2
CTF3
EDC1
YLR311C
GIP3
GIR2
HCM1
CAM1
YMR193C–A
URA7
YML102C–A
CAC2
ECM4
YPR098C_2
APL5
RLF2
SCS3
DUG3
RTN1
CWH41
Gene Name
DOXO HL K
DOXO HLEG K
DOXO HL L
DOXO HLEG L
Type of Media

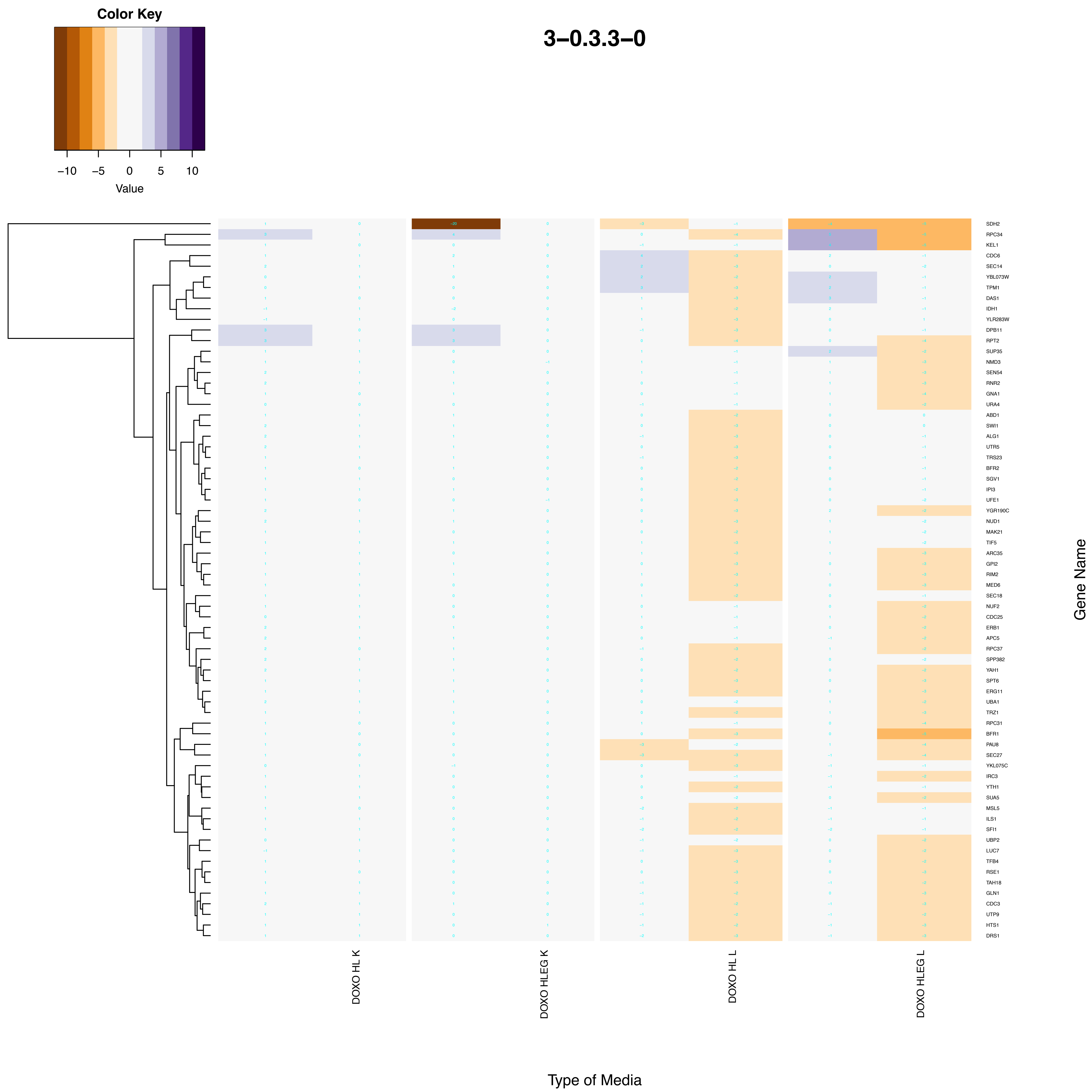

Color Key
−10
−5
0
5
10
Value
3−0.3.3−0
SDH2
RPC34
KEL1
CDC6
SEC14
YBL073W
TPM1
DAS1
IDH1
YLR283W
DPB11
RPT2
SUP35
NMD3
SEN54
RNR2
GNA1
URA4
ABD1
SWI1
ALG1
UTR5
TRS23
BFR2
SGV1
IPI3
UFE1
YGR190C
NUD1
MAK21
TIF5
ARC35
GPI2
RIM2
MED6
SEC18
NUF2
CDC25
ERB1
APC5
RPC37
SPP382
YAH1
SPT6
ERG11
UBA1
TRZ1
RPC31
BFR1
PAU8
SEC27
YKL075C
IRC3
YTH1
SUA5
MSL5
ILS1
SFI1
UBP2
LUC7
TFB4
RSE1
TAH18
GLN1
CDC3
UTP9
HTS1
DRS1
Gene Name
DOXO HL K
DOXO HLEG K
DOXO HL L
DOXO HLEG L
Type of Media

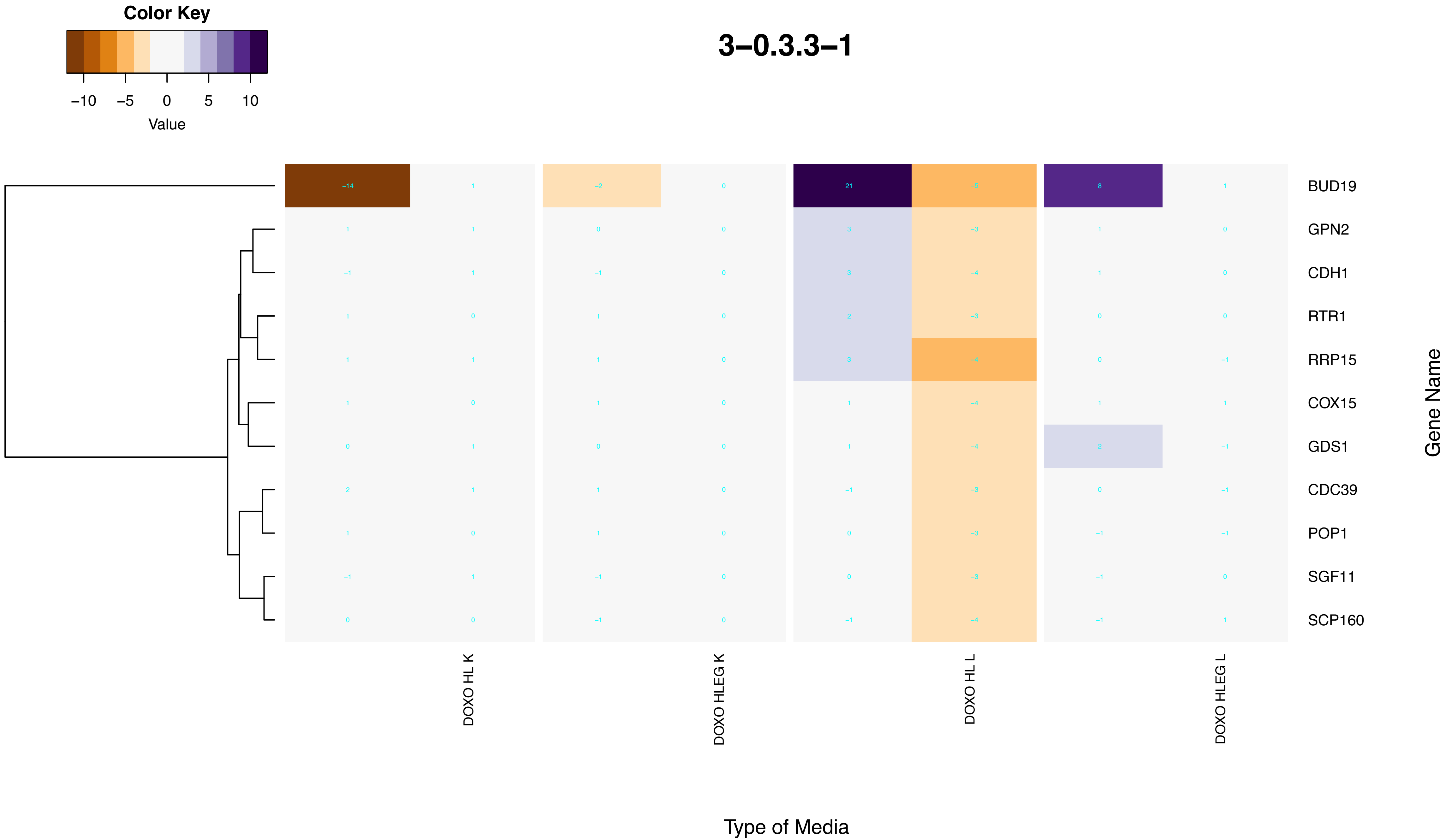
Color Key
−10
−5
0
5
10
Value
3−0.3.3−1
BUD19
GPN2
CDH1
RTR1
RRP15
COX15
GDS1
CDC39
POP1
SGF11
SCP160
Gene Name
DOXO HL K
DOXO HLEG K
DOXO HL L
DOXO HLEG L
Type of Media

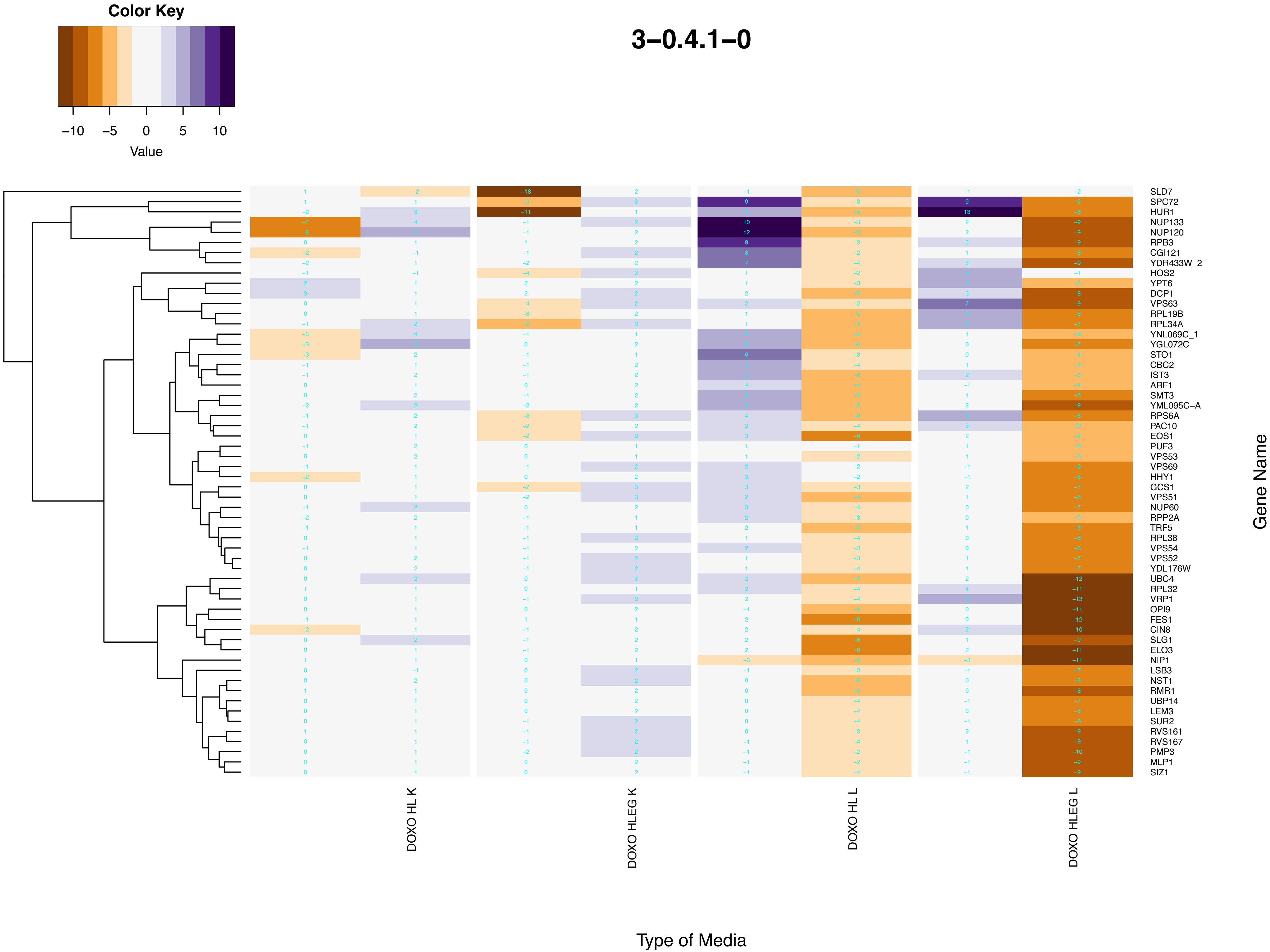
Color Key
−10 −5 0 5 10
Value
3−0.4.1−0
DOXO HL K
DOXO HLEG K
DOXO HL L
DOXO HLEG L
Type of Media
Gene Name
SLD7
SPC72
HUR1
NUP133
NUP120
RPB3
CGI121
YDR433W_2
HOS2
YPT6
DCP1
VPS63
RPL19B
RPL34A
YNL069C_1
YGL072C
STO1
CBC2
IST3
ARF1
SMT3
YML095C−A
RPS6A
PAC10
EOS1
PUF3
VPS53
VPS69
HHY1
GCS1
VPS51
NUP60
RPP2A
TRF5
RPL38
VPS54
VPS52
YDL176W
UBC4
RPL32
VRP1
OPI9
FES1
CIN8
SLG1
ELO3
NIP1
LSB3
NST1
RMR1
UBP14
LEM3
SUR2
RVS161
RVS167
PMP3
MLP1
SIZ1

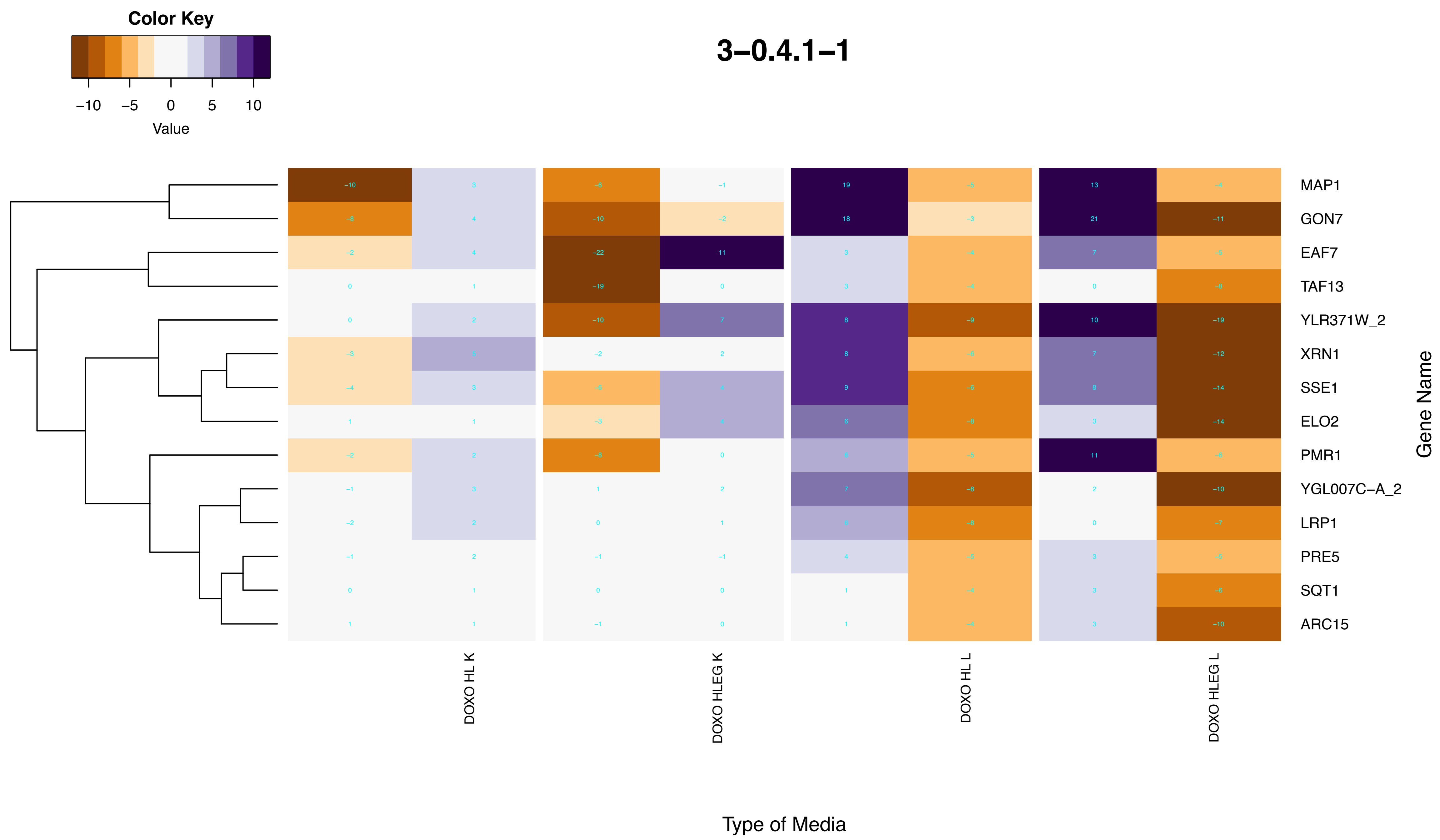
3–0.4.1–1
Color Key
–10
–5
0
5
10
Value
MAP1
GON7
EAF7
TAF13
YLR371W_2
XRN1
SSE1
ELO2
PMR1
YGL007C–A_2
LRP1
PRE5
SQT1
ARC15
Gene Name
DOXO HL K
DOXO HLEG K
DOXO HL L
DOXO HLEG L
Type of Media

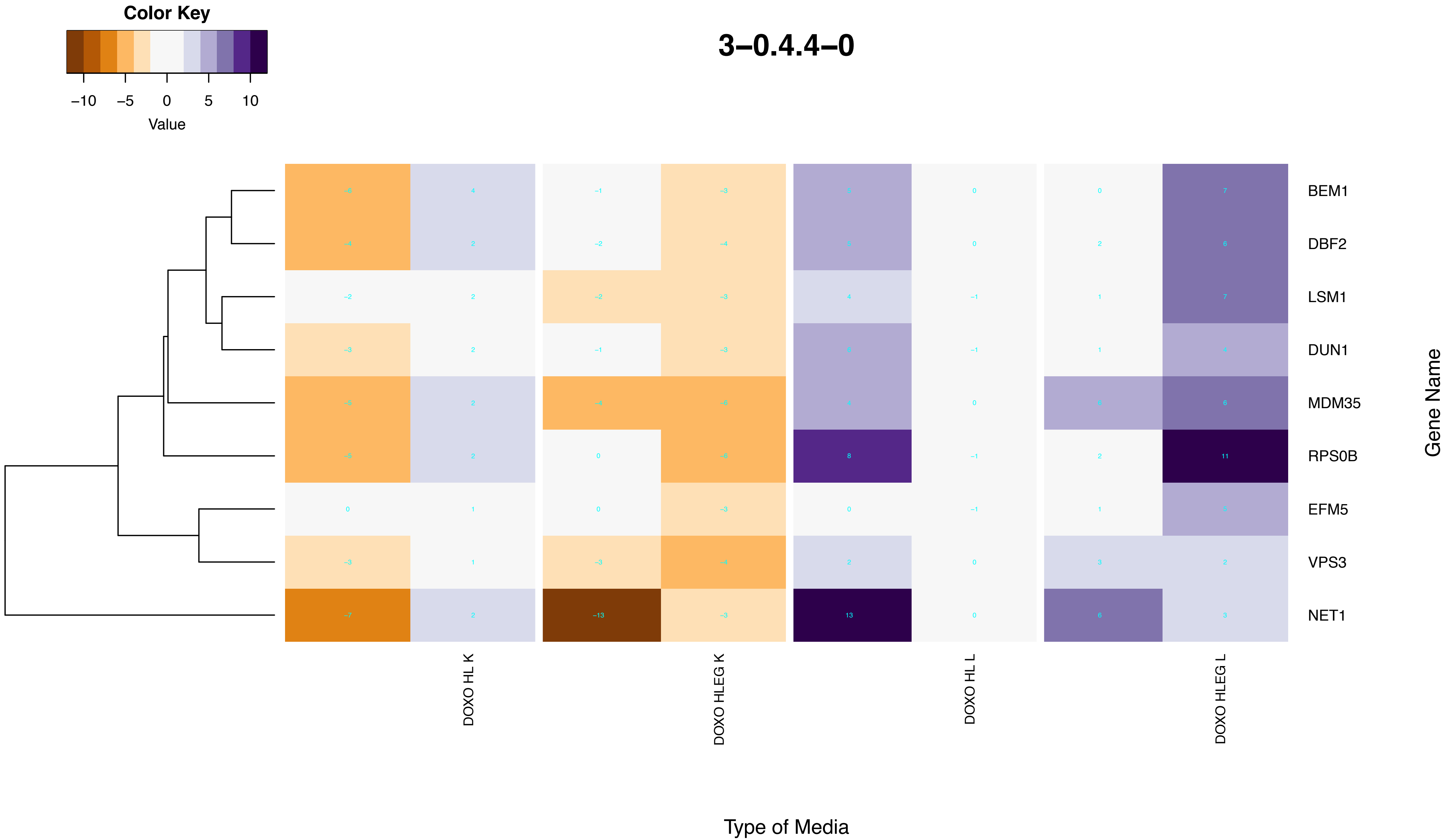
Color Key
–10
–5
0
5
10
Value
3–0.4.4–0
BEM1
DBF2
LSM1
DUN1
MDM35
RPS0B
EFM5
VPS3
NET1
Gene Name
DOXO HL K
DOXO HLEG K
DOXO HL L
DOXO HLEG L
Type of Media

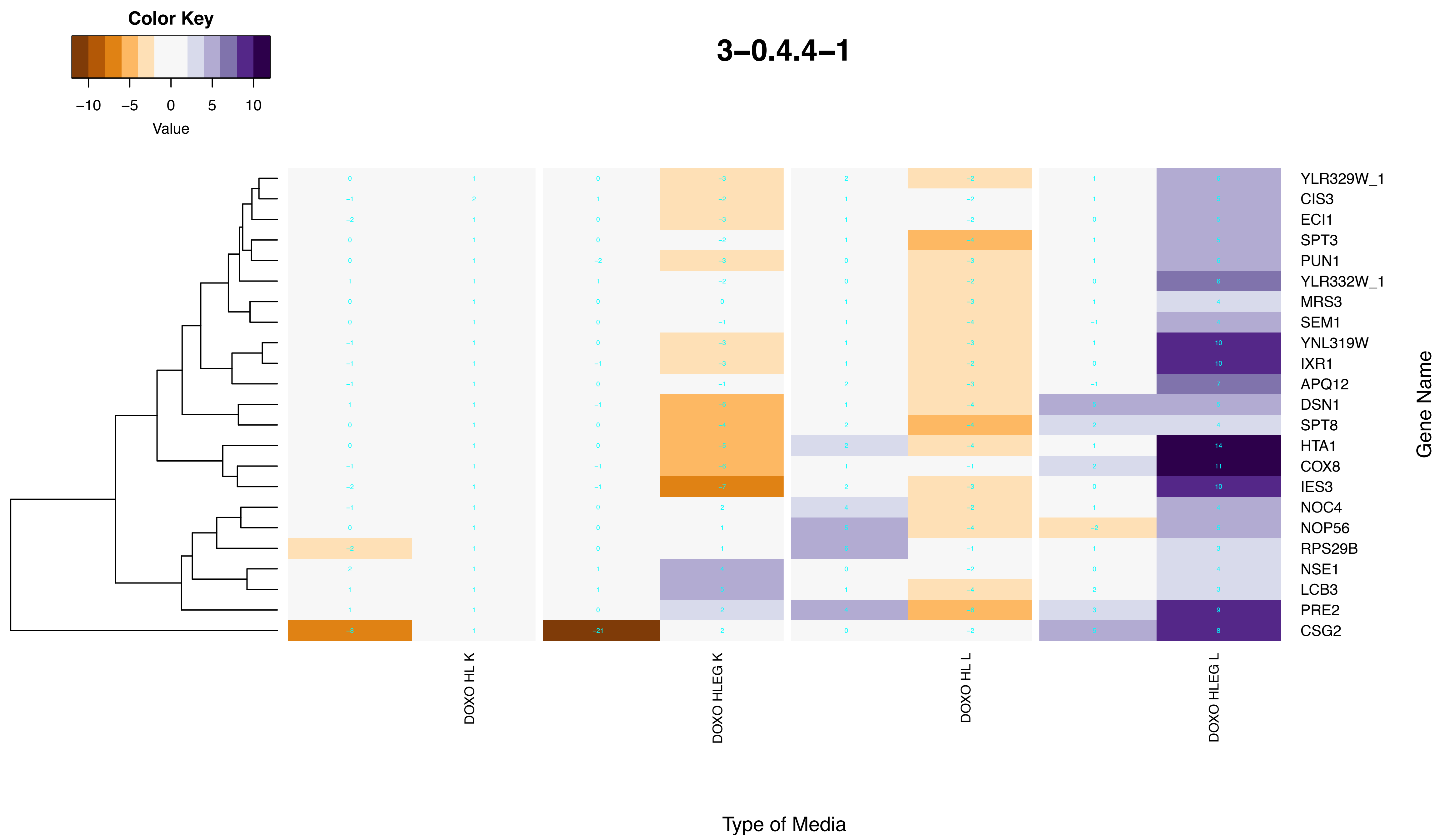
3–0.4.4–1
Color Key
–10 –5 0 5 10
Value
YLR329W_1
CIS3
ECI1
SPT3
PUN1
YLR332W_1
MRS3
SEM1
YNL319W
IXR1
APQ12
DSN1
SPT8
HTA1
COX8
IES3
NOC4
NOP56
RPS29B
NSE1
LCB3
PRE2
CSG2
Gene Name
DOXO HL K
DOXO HLEG K
DOXO HL L
DOXO HLEG L
Type of Media

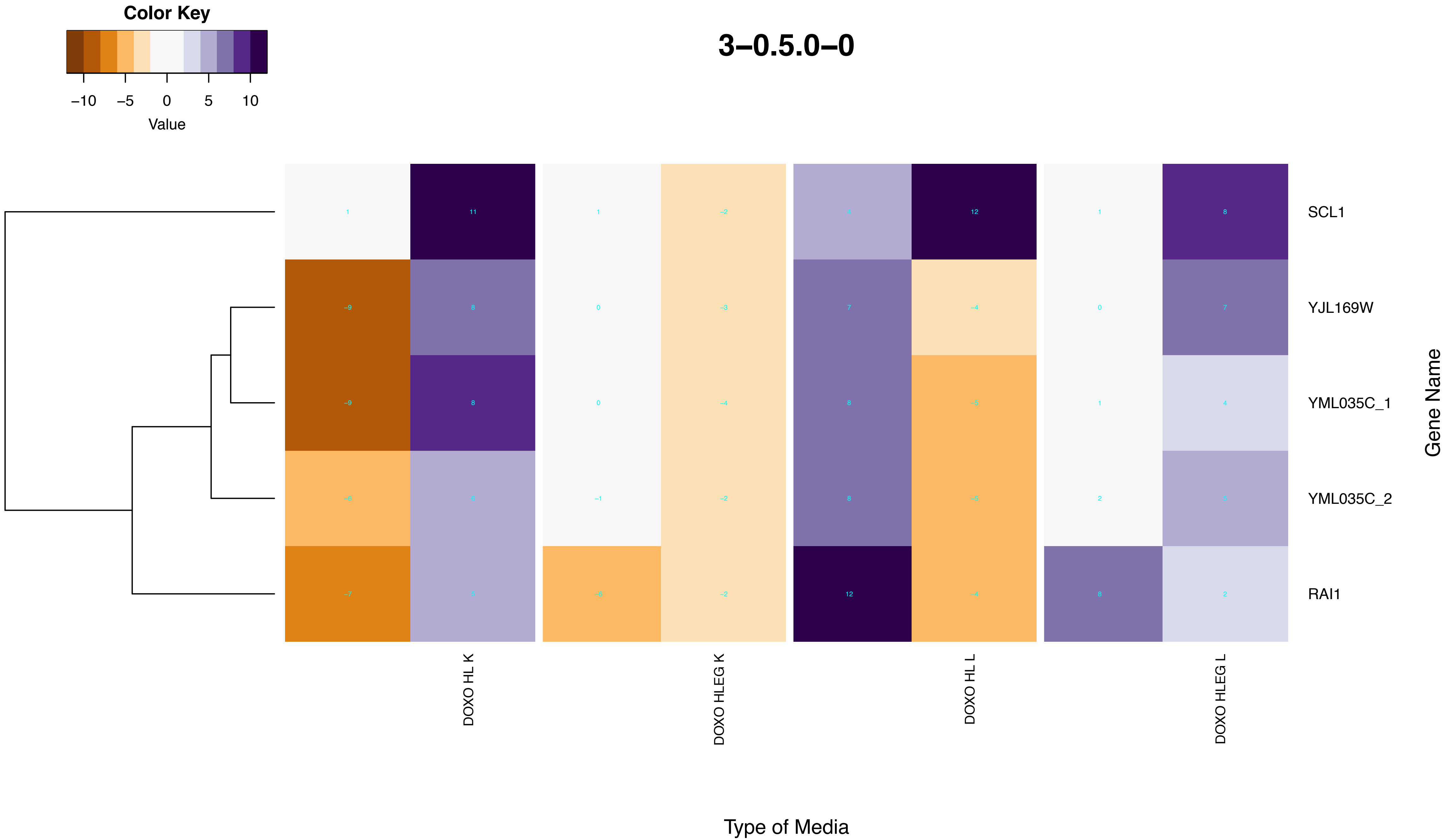

Color Key
−10
−5
0
5
10
Value
3–0.5.0–0
SCL1
YJL169W
YML035C_1
YML035C_2
RAI1
Gene Name
DOXO HL K
DOXO HLEG K
DOXO HL L
DOXO HLEG L
Type of Media

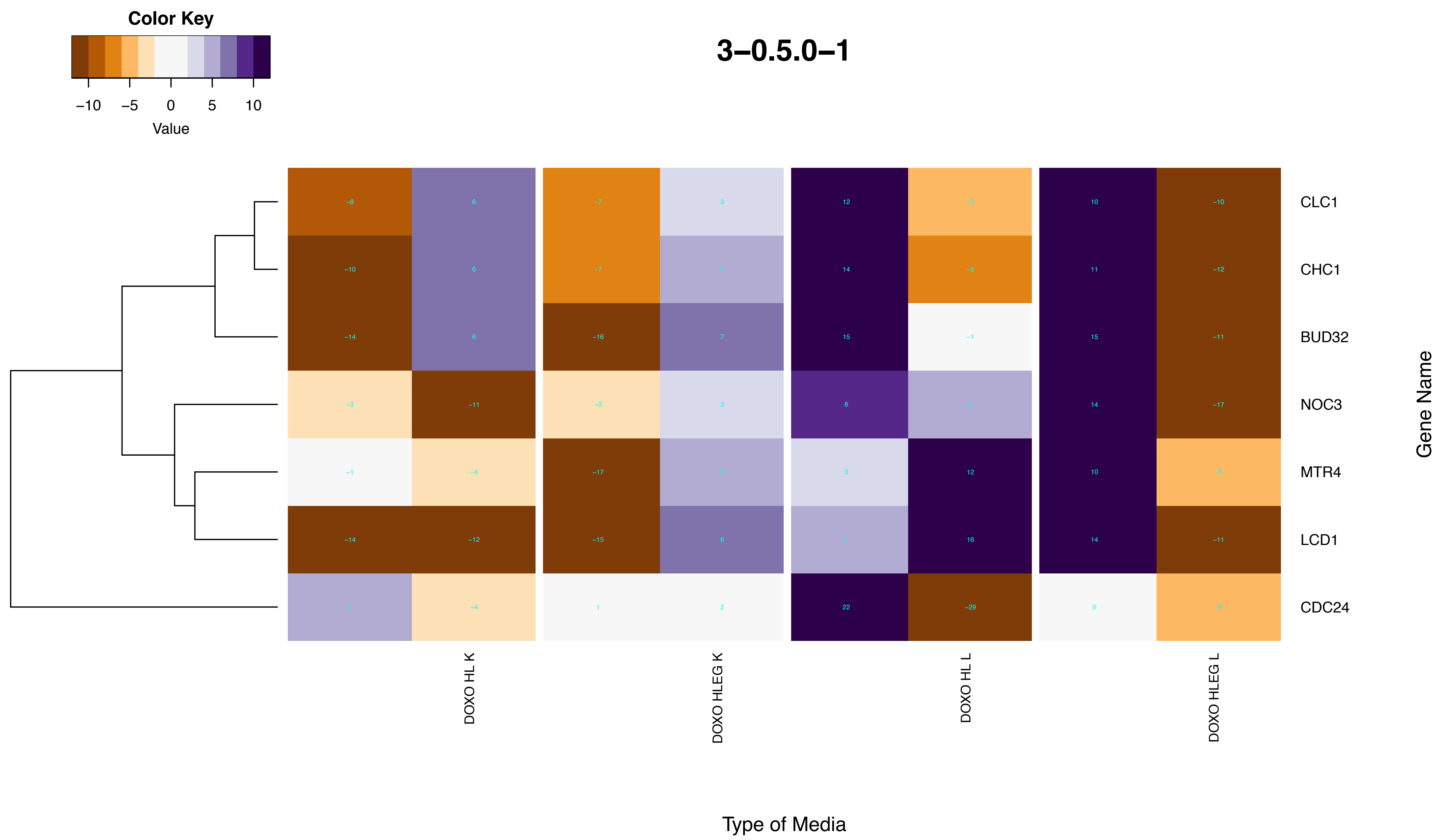
Color Key
−10 −5 0 5 10
Value
3−0.5.0−1
CLC1
CHC1
BUD32
NOC3
MTR4
LCD1
CDC24
Gene Name
DOXO HL K
DOXO HLEG K
DOXO HL L
DOXO HLEG L
Type of Media

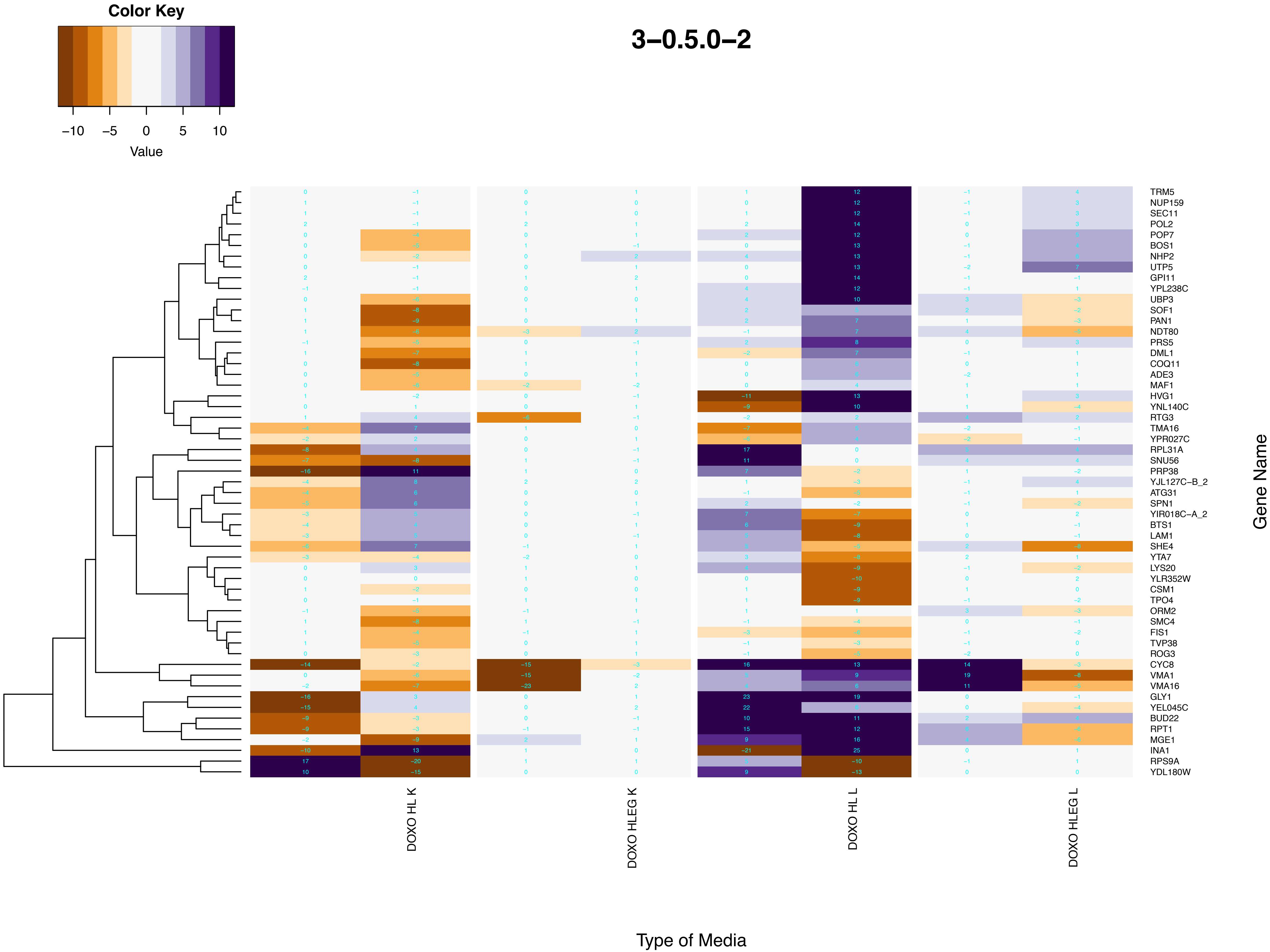
Color Key
−10
−5
0
5
10
Value
3−0.5.0−2
TRM5
NUP159
SEC11
POL2
POP7
BOS1
NHP2
UTP5
GPI11
YPL238C
UBP3
SOF1
PAN1
NDT80
PRS5
DML1
COQ11
ADE3
MAF1
HVG1
YNL140C
RTG3
TMA16
YPR027C
RPL31A
SNU56
PRP38
YJL127C−B_2
ATG31
SPN1
YIR018C−A_2
BTS1
LAM1
SHE4
YTA7
LYS20
YLR352W
CSM1
TPO4
ORM2
SMC4
FIS1
TVP38
ROG3
CYC8
VMA1
VMA16
GLY1
YEL045C
BUD22
RPT1
MGE1
INA1
RPS9A
YDL180W
Gene Name
DOXO HL K
DOXO HLEG K
DOXO HL L
DOXO HLEG L
Type of Media

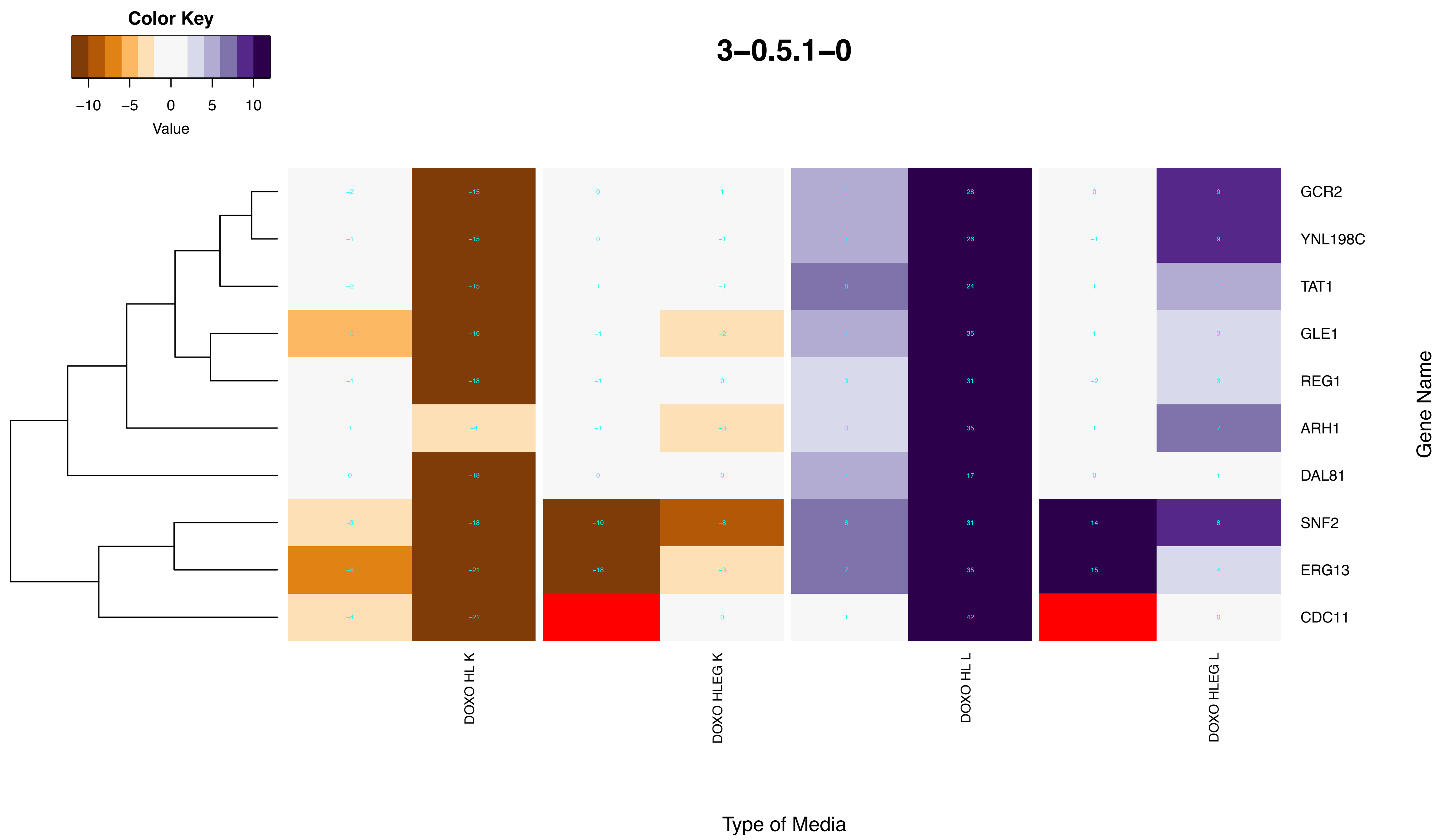
3–0.5.1–0
Color Key
–10 –5 0 5 10
Value
GCR2
YNL198C
TAT1
GLE1
REG1
ARH1
DAL81
SNF2
ERG13
CDC11
Gene Name
DOXO HL K
DOXO HLEG K
DOXO HL L
DOXO HLEG L
Type of Media

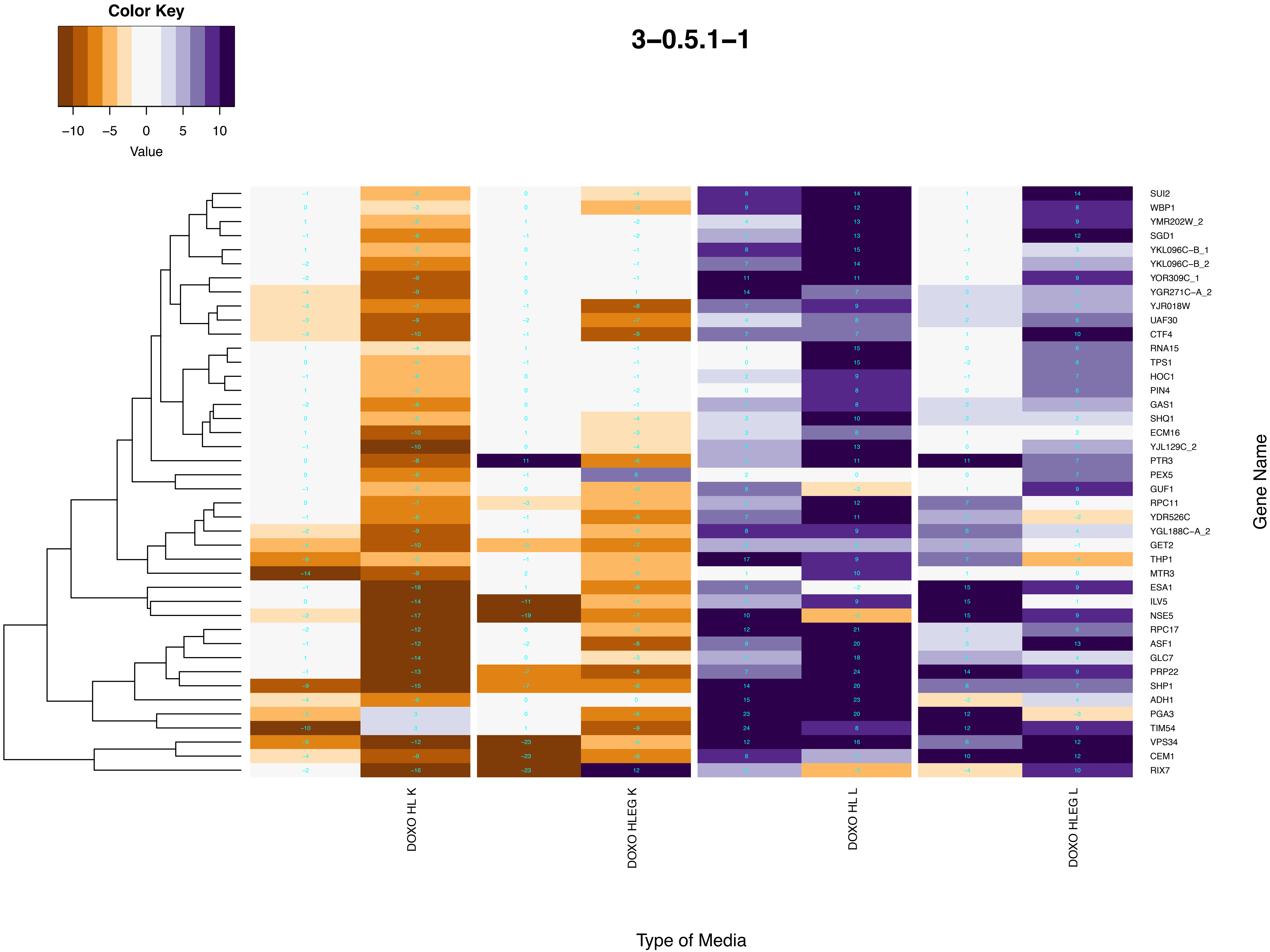

Color Key
−10 −5 0 5 10
Value
3−0.5.1−1
SUI2
WBP1
YMR202W_2
SGD1
YKL096C−B_1
YKL096C−B_2
YOR309C_1
YGR271C−A_2
YJR018W
UAF30
CTF4
RNA15
TPS1
HOC1
PIN4
GAS1
SHQ1
ECM16
YJL129C_2
PTR3
PEX5
GUF1
RPC11
YDR526C
YGL188C−A_2
GET2
THP1
MTR3
ESA1
ILV5
NSE5
RPC17
ASF1
GLC7
PRP22
SHP1
ADH1
PGA3
TIM54
VPS34
CEM1
RIX7
Gene Name
DOXO HL K
DOXO HLEG K
DOXO HL L
DOXO HLEG L
Type of Media

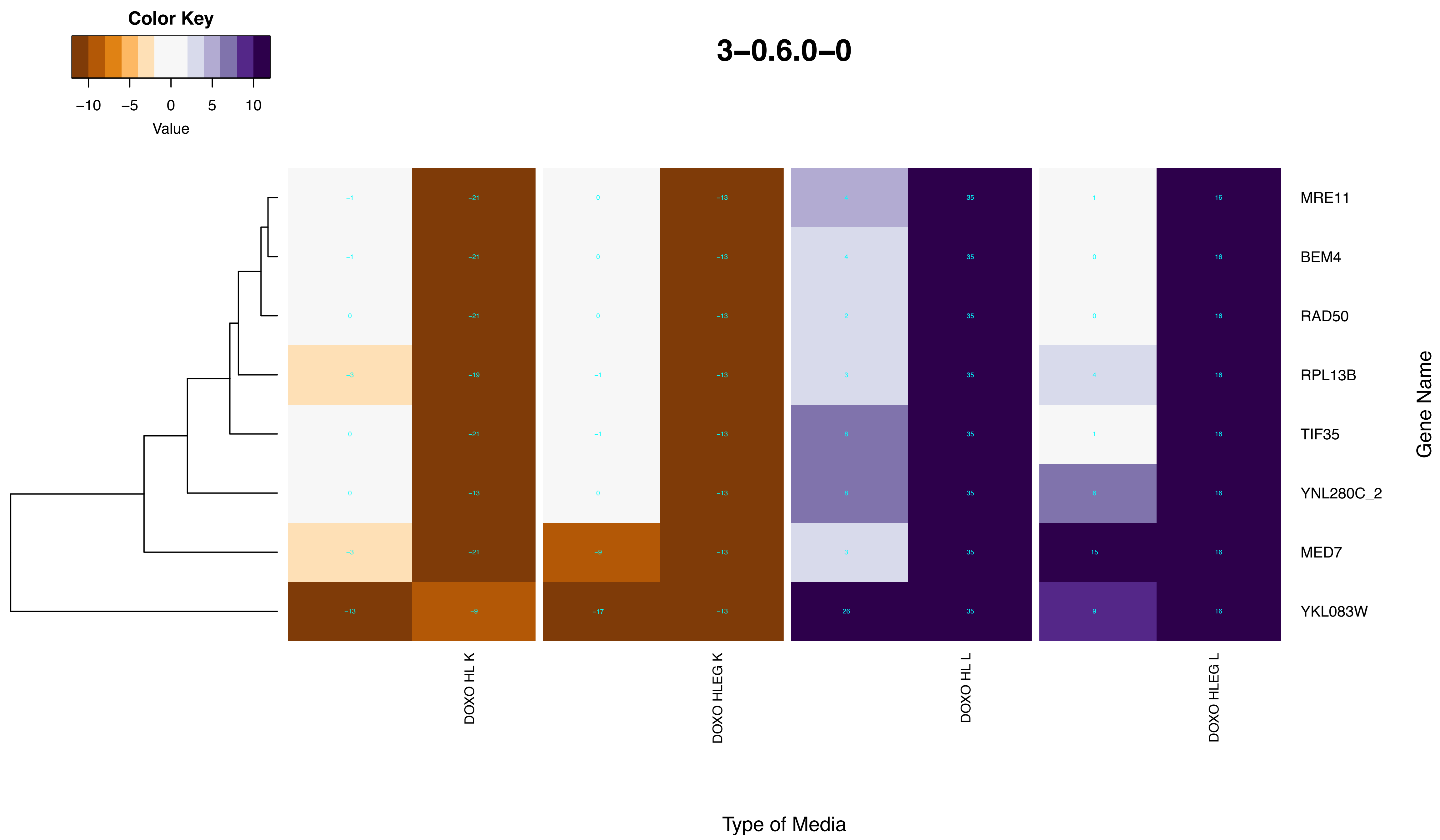
Color Key
−10
−5
0
5
10
Value
3−0.6.0−0
MRE11
BEM4
RAD50
RPL13B
TIF35
YNL280C_2
MED7
YKL083W
Gene Name
DOXO HL K
DOXO HLEG K
DOXO HL L
DOXO HLEG L
Type of Media

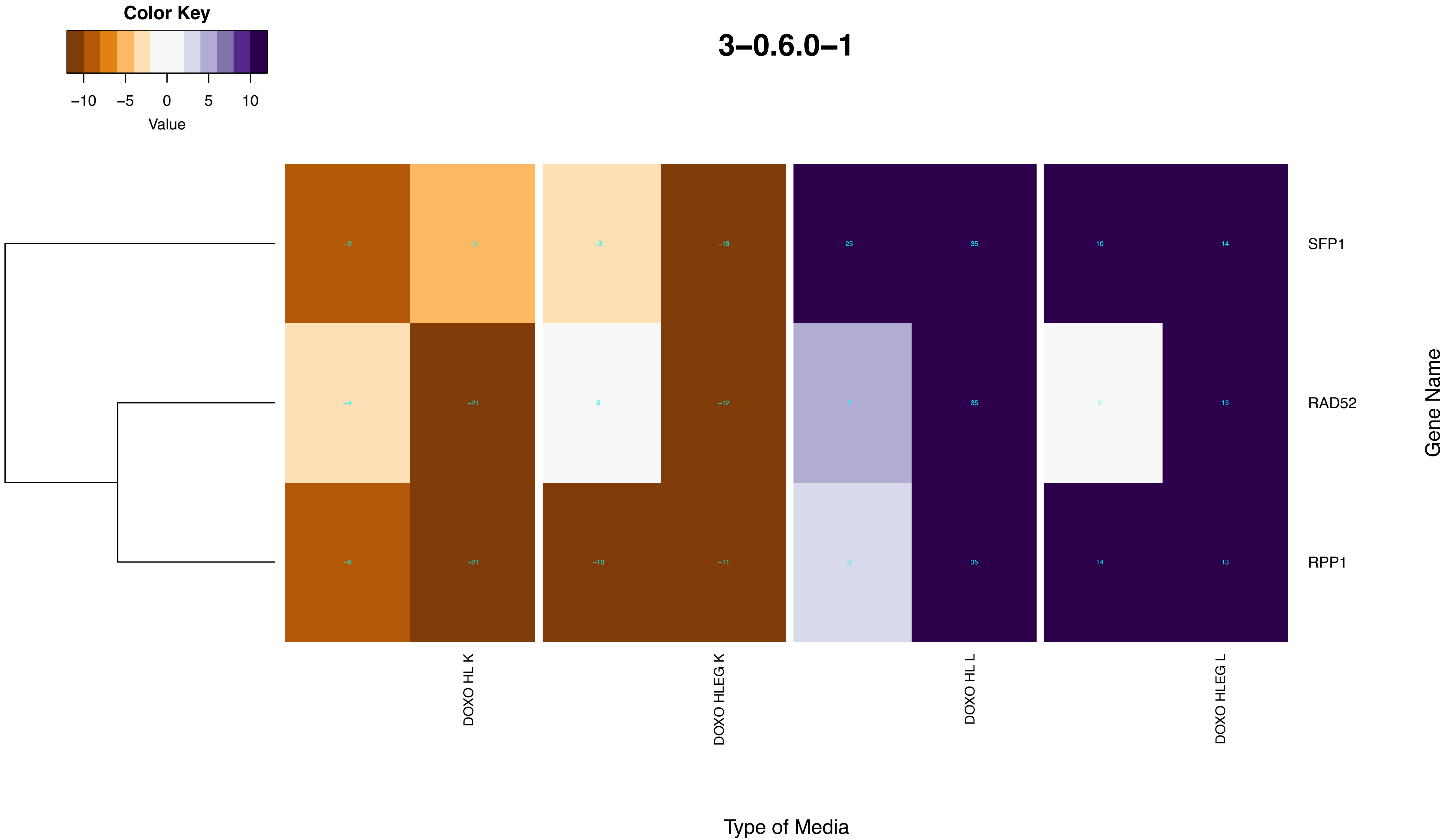
Color Key
−10
−5
0
5
10
Value
3−0.6.0−1
−9
−5
−3
−13
25
35
10
14
SFP1
−4
−21
0
−12
5
35
2
15
RAD52
−9
−21
−10
−11
3
35
14
13
RPP1
Gene Name
DOXO HL K
DOXO HLEG K
DOXO HL L
DOXO HLEG L
Type of Media

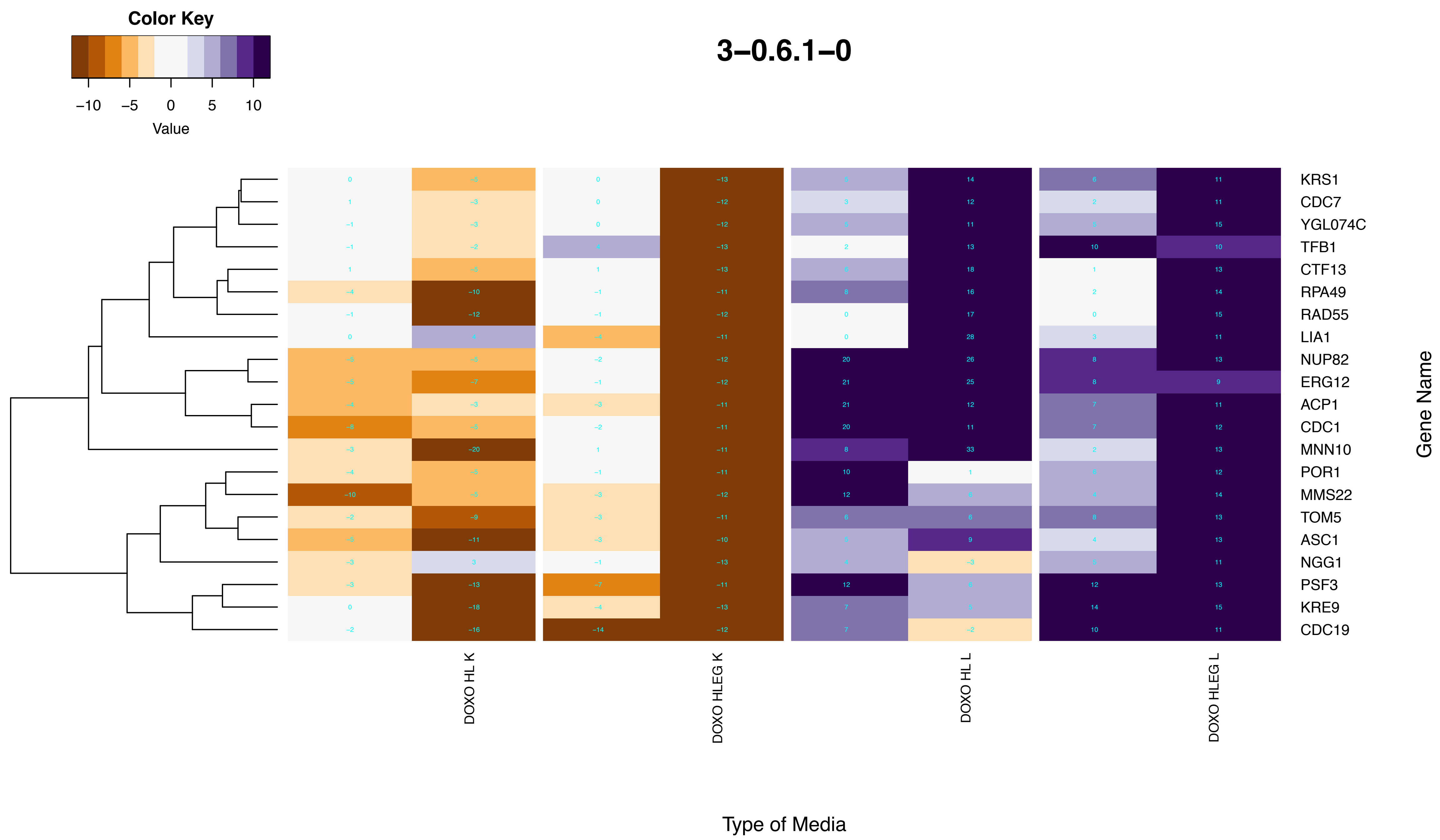
3–0.6.1–0
Color Key
−10 −5 0 5 10
Value
KRS1
CDC7
YGL074C
TFB1
CTF13
RPA49
RAD55
LIA1
NUP82
ERG12
ACP1
CDC1
MNN10
POR1
MMS22
TOM5
ASC1
NGG1
PSF3
KRE9
CDC19
Gene Name
DOXO HL K
DOXO HLEG K
DOXO HL L
DOXO HLEG L
Type of Media

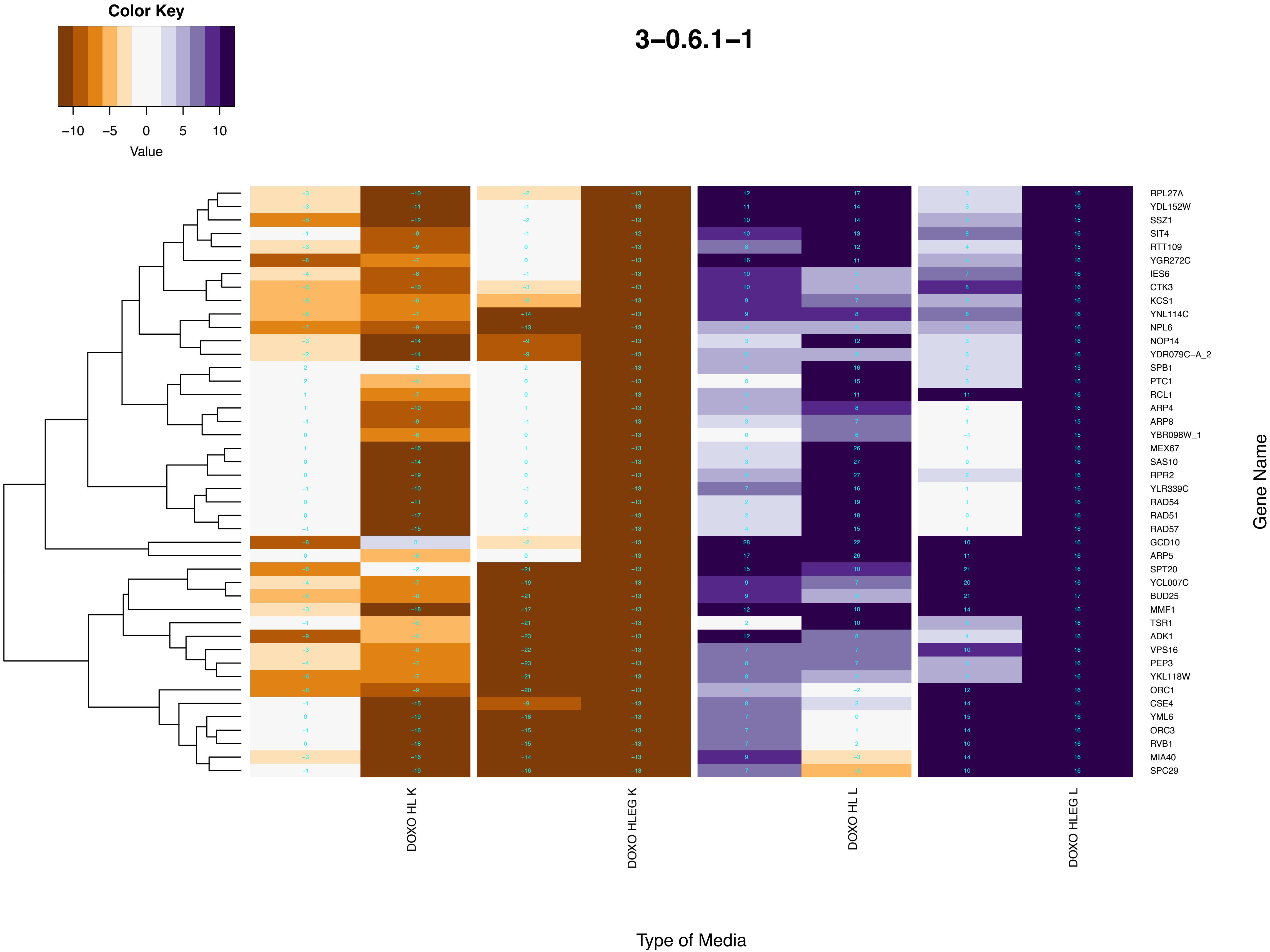
Color Key
−10
−5
0
5
10
Value
3−0.6.1−1
RPL27A
YDL152W
SSZ1
SIT4
RTT109
YGR272C
IES6
CTK3
KCS1
YNL114C
NPL6
NOP14
YDR079C−A_2
SPB1
PTC1
RCL1
ARP4
ARP8
YBR098W_1
MEX67
SAS10
RPR2
YLR339C
RAD54
RAD51
RAD57
GCD10
ARP5
SPT20
YCL007C
BUD25
MMF1
TSR1
ADK1
VPS16
PEP3
YKL118W
ORC1
CSE4
YML6
ORC3
RVB1
MIA40
SPC29
Gene Name
DOXO HL K
DOXO HLEG K
DOXO HL L
DOXO HLEG L
Type of Media

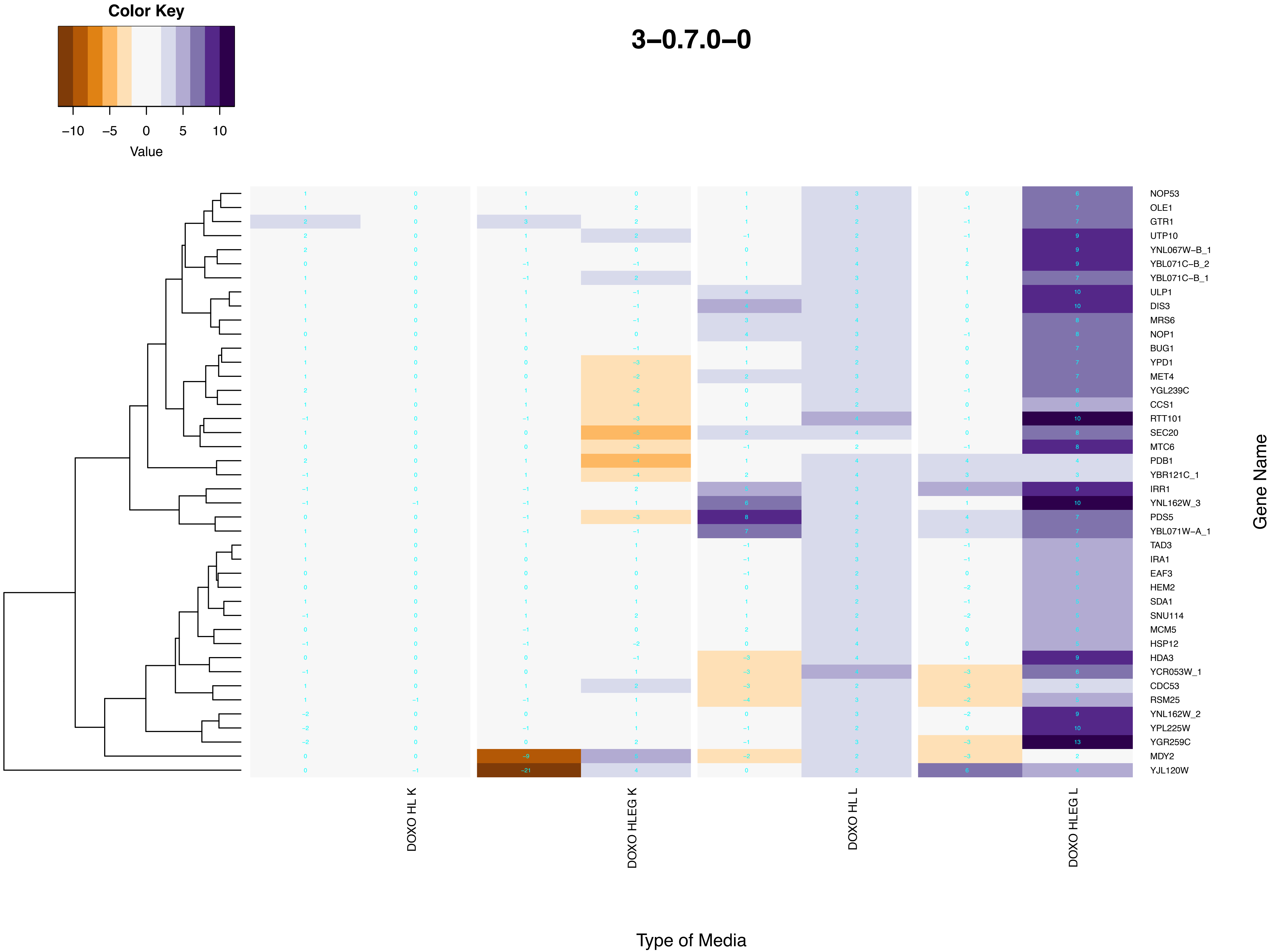

Color Key
−10 −5 0 5 10
Value
3−0.7.0−0
NOP53
OLE1
GTR1
UTP10
YNL067W−B_1
YBL071C−B_2
YBL071C−B_1
ULP1
DIS3
MRS6
NOP1
BUG1
YPD1
MET4
YGL239C
CCS1
RTT101
SEC20
MTC6
PDB1
YBR121C_1
IRR1
YNL162W_3
PDS5
YBL071W−A_1
TAD3
IRA1
EAF3
HEM2
SDA1
SNU114
MCM5
HSP12
HDA3
YCR053W_1
CDC53
RSM25
YNL162W_2
YPL225W
YGR259C
MDY2
YJL120W
Gene Name
DOXO HL K
DOXO HLEG K
DOXO HL L
DOXO HLEG L
Type of Media

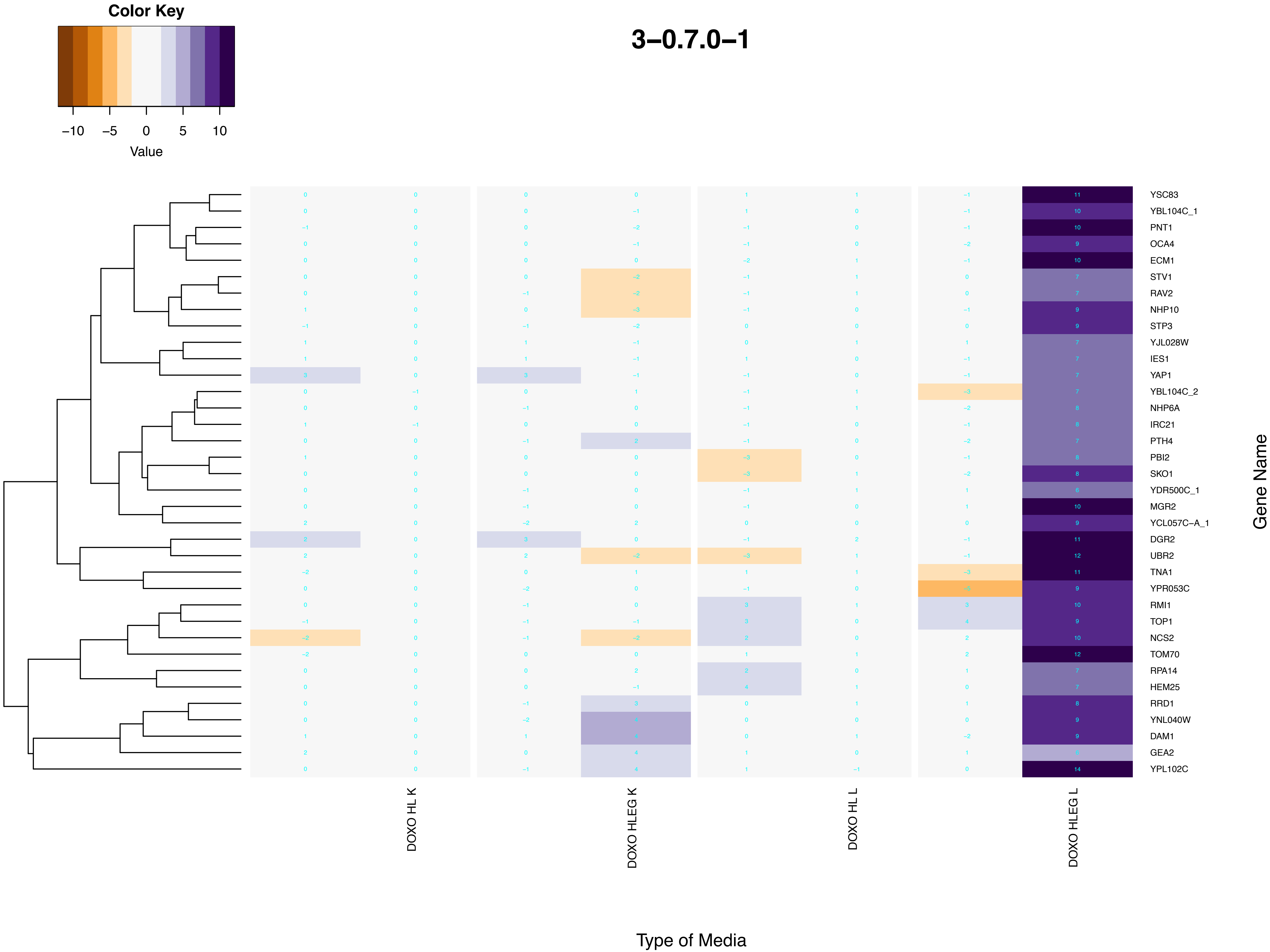
Color Key
−10
−5
0
5
10
Value
3−0.7.0−1
YSC83
YBL104C_1
PNT1
OCA4
ECM1
STV1
RAV2
NHP10
STP3
YJL028W
IES1
YAP1
YBL104C_2
NHP6A
IRC21
PTH4
PBI2
SKO1
YDR500C_1
MGR2
YCL057C−A_1
DGR2
UBR2
TNA1
YPR053C
RMI1
TOP1
NCS2
TOM70
RPA14
HEM25
RRD1
YNL040W
DAM1
GEA2
YPL102C
Gene Name
DOXO HL K
DOXO HLEG K
DOXO HL L
DOXO HLEG L
Type of Media

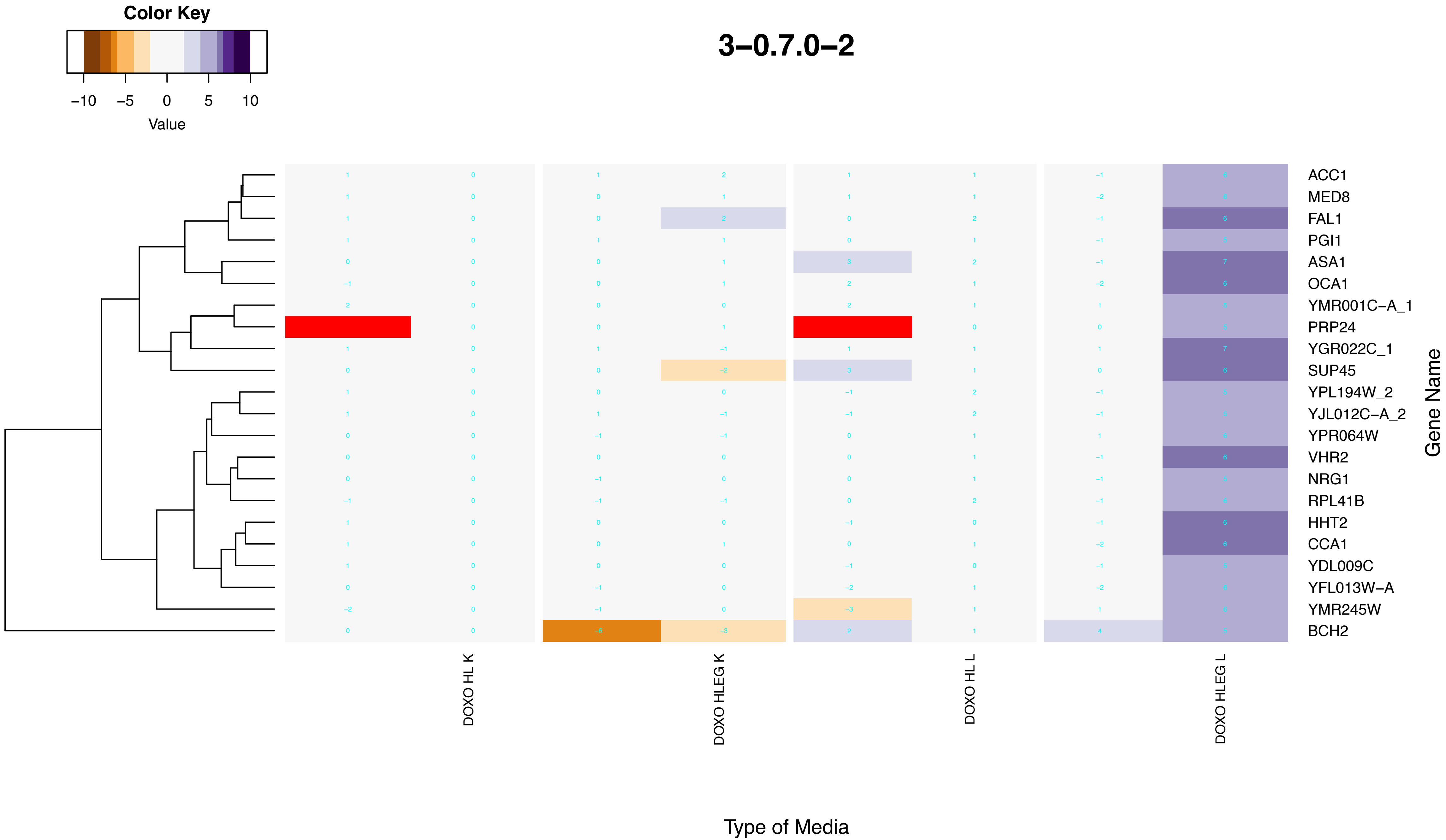

Color Key
−10 −5 0 5 10
Value
3−0.7.0−2
ACC1
MED8
FAL1
PGI1
ASA1
OCA1
YMR001C−A_1
PRP24
YGR022C_1
SUP45
YPL194W_2
YJL012C−A_2
YPR064W
VHR2
NRG1
RPL41B
HHT2
CCA1
YDL009C
YFL013W−A
YMR245W
BCH2
Gene Name
DOXO HL K
DOXO HLEG K
DOXO HL L
DOXO HLEG L
Type of Media

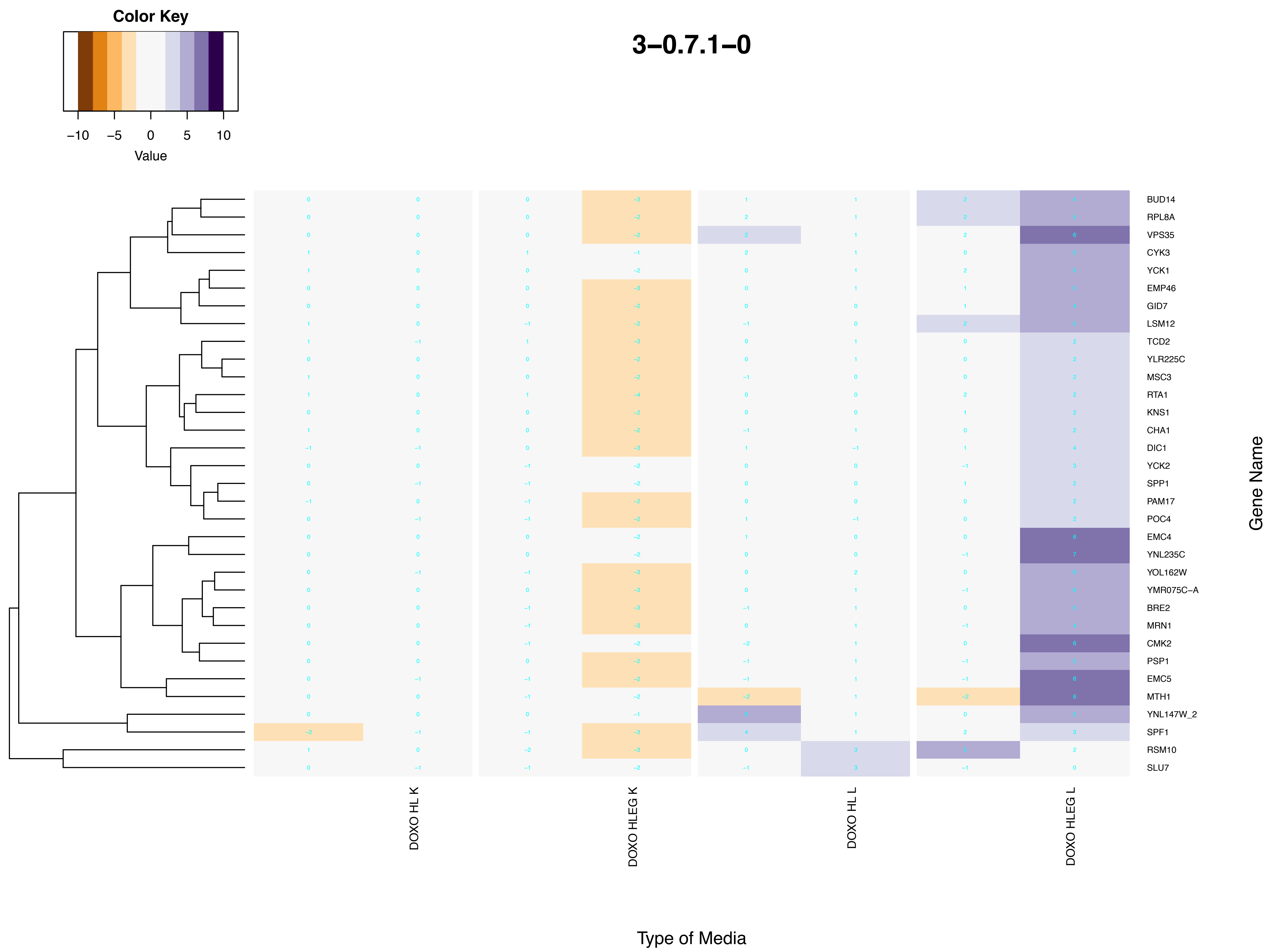

Color Key
−10 −5 0 5 10
Value
3−0.7.1−0
BUD14
RPL8A
VPS35
CYK3
YCK1
EMP46
GID7
LSM12
TCD2
YLR225C
MSC3
RTA1
KNS1
CHA1
DIC1
YCK2
SPP1
PAM17
POC4
EMC4
YNL235C
YOL162W
YMR075C−A
BRE2
MRN1
CMK2
PSP1
EMC5
MTH1
YNL147W_2
SPF1
RSM10
SLU7
Gene Name
DOXO HL K
DOXO HLEG K
DOXO HL L
DOXO HLEG L
Type of Media

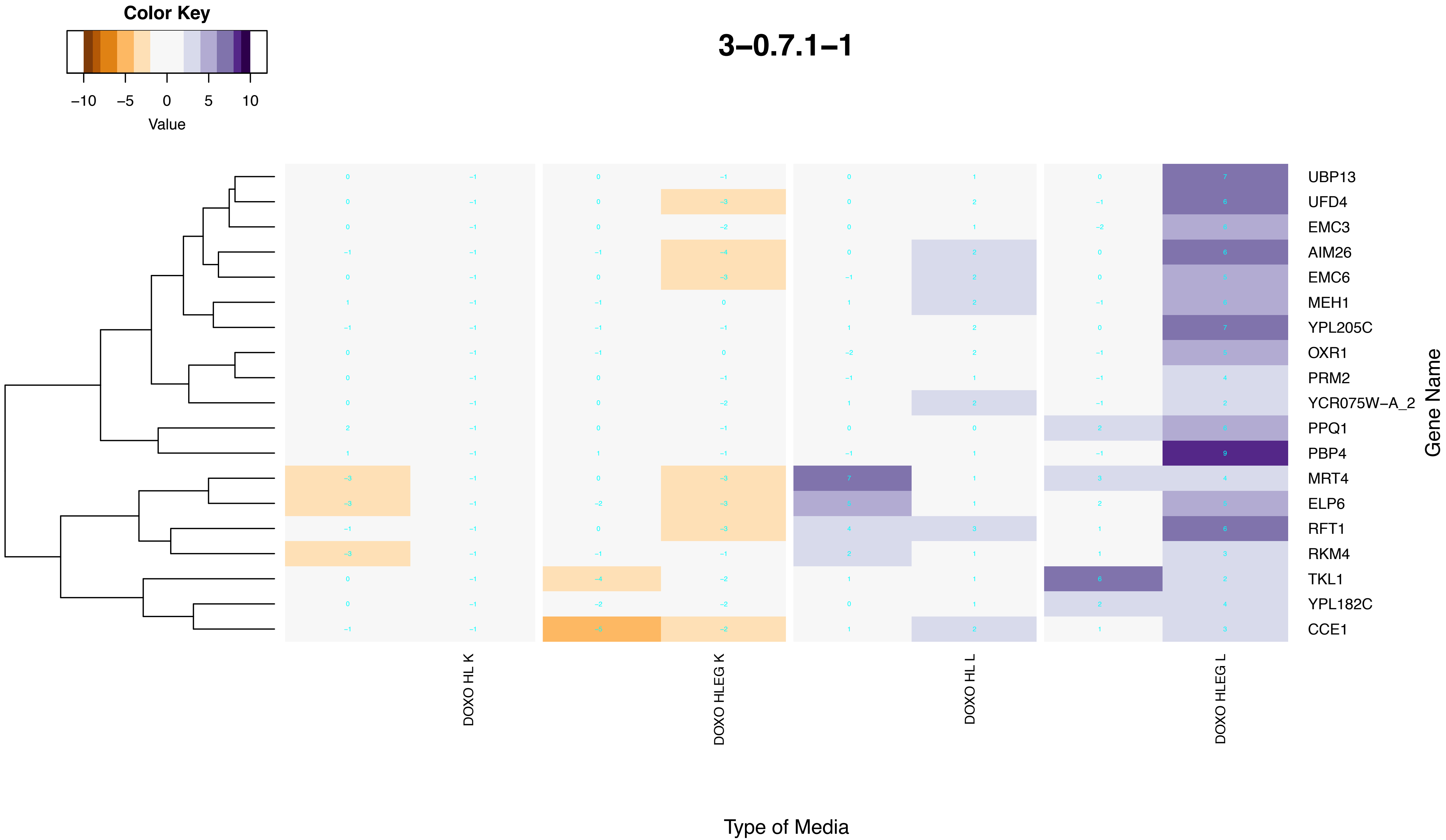

Color Key
−10
−5
0
5
10
Value
3−0.7.1−1
UBP13
UFD4
EMC3
AIM26
EMC6
MEH1
YPL205C
OXR1
PRM2
YCR075W−A_2
PPQ1
PBP4
MRT4
ELP6
RFT1
RKM4
TKL1
YPL182C
CCE1
Gene Name
DOXO HL K
DOXO HLEG K
DOXO HL L
DOXO HLEG L
Type of Media

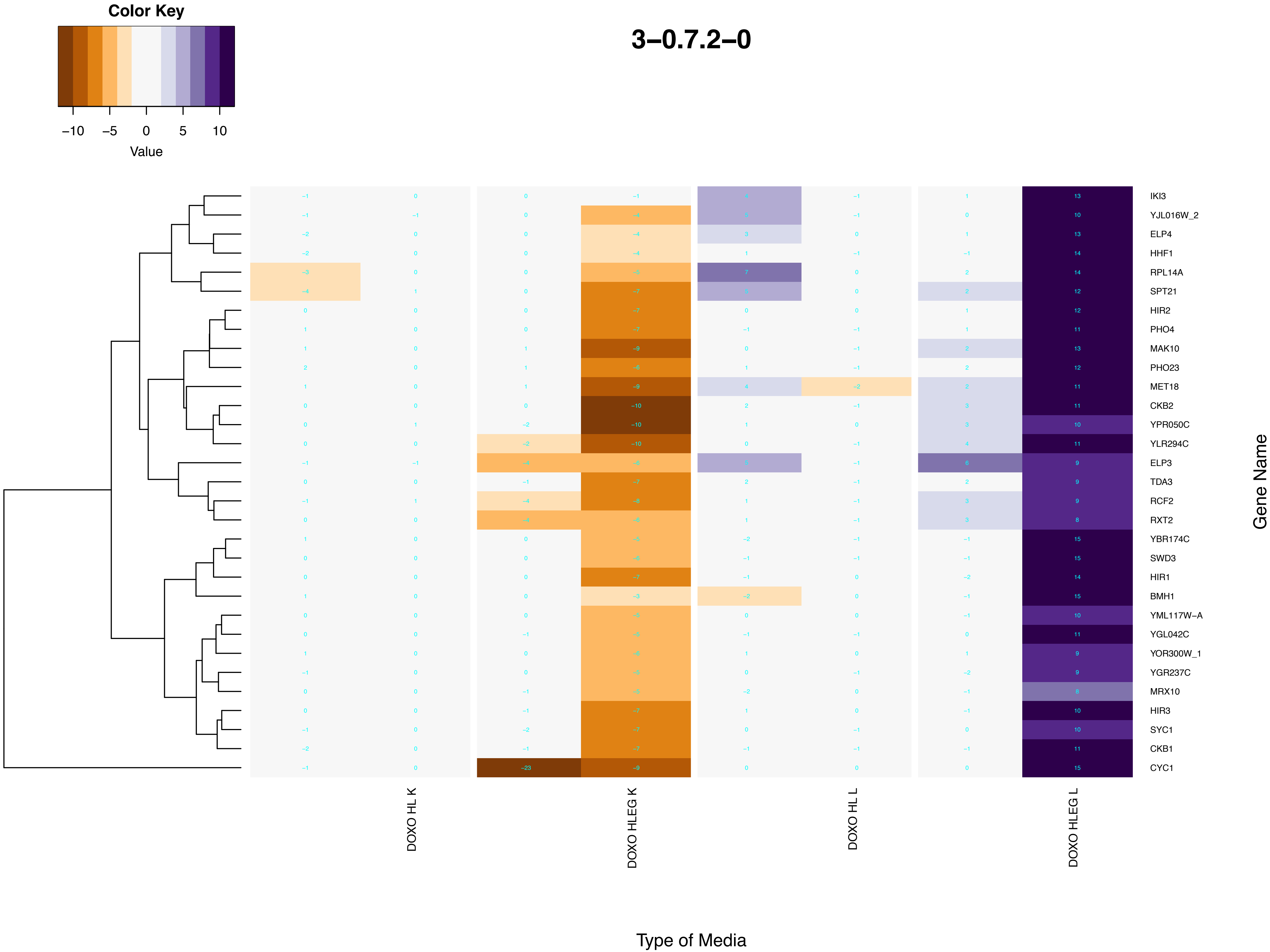

Color Key
−10 −5 0 5 10
Value
3−0.7.2−0
IKI3
YJL016W_2
ELP4
HHF1
RPL14A
SPT21
HIR2
PHO4
MAK10
PHO23
MET18
CKB2
YPR050C
YLR294C
ELP3
TDA3
RCF2
RXT2
YBR174C
SWD3
HIR1
BMH1
YML117W−A
YGL042C
YOR300W_1
YGR237C
MRX10
HIR3
SYC1
CKB1
CYC1
Gene Name
DOXO HL K
DOXO HLEG K
DOXO HL L
DOXO HLEG L
Type of Media

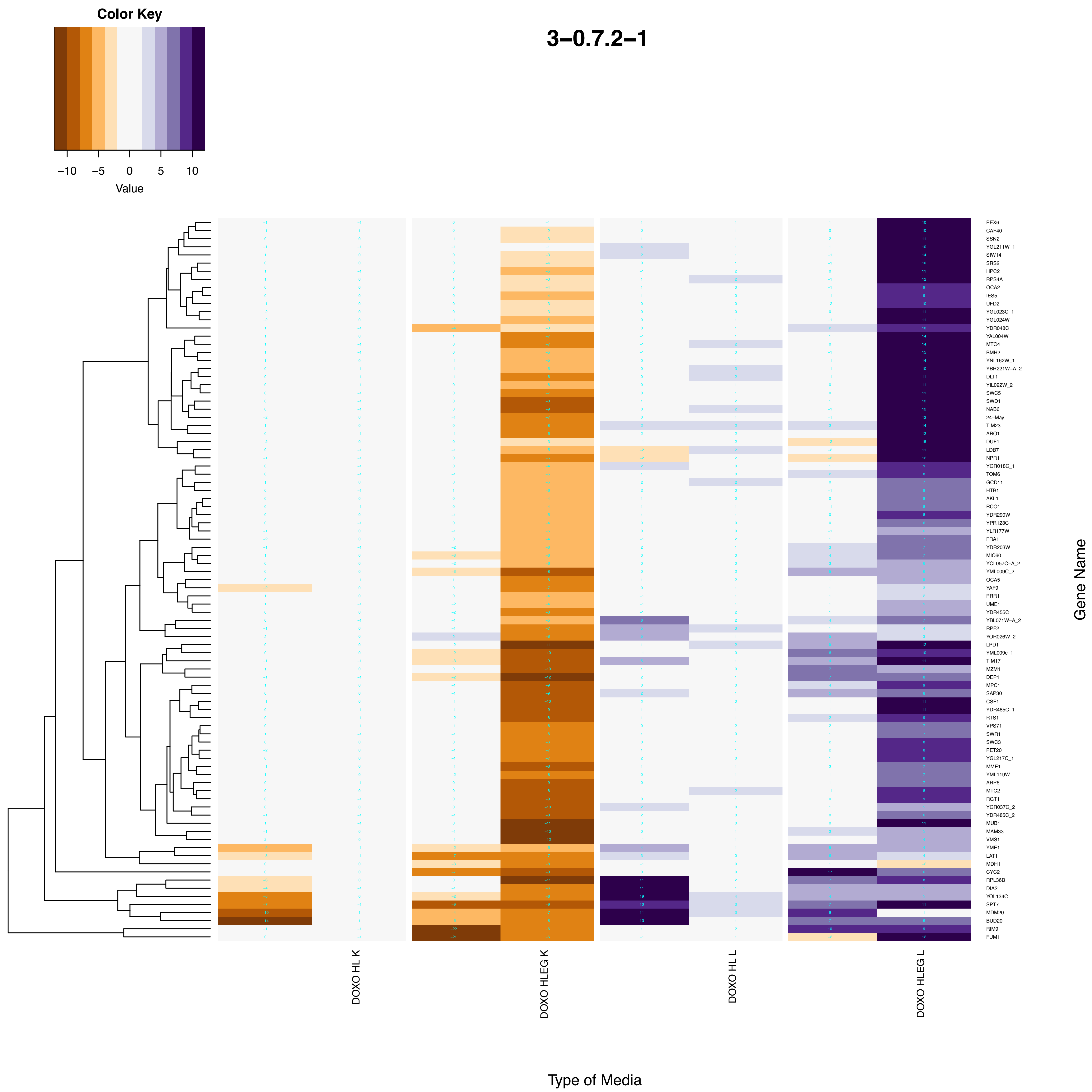

Color Key
−10 −5 0 5 10
Value
3−0.7.2−1
DOXO HL K
DOXO HLEG K
DOXO HL L
DOXO HLEG L
Type of Media
Gene Name
PEX6
CAF40
SSN2
YGL211W_1
SIW14
SRS2
HPC2
RPS4A
OCA2
IES5
UFD2
YGL023C_1
YGL024W
YDR048C
YAL004W
MTC4
BMH2
YNL162W_1
YBR221W−A_2
DLT1
YIL092W_2
SWC5
SWD1
NAB6
24−May
TIM23
ARO1
DUF1
LDB7
NPR1
YGR018C_1
TOM6
GCD11
HTB1
AKL1
RCO1
YDR290W
YPR123C
YLR177W
FRA1
YDR203W
MIC60
YCL057C−A_2
YML009C_2
OCA5
YAF9
PRR1
UME1
YDR455C
YBL071W−A_2
RPF2
YOR026W_2
LPD1
YML009c_1
TIM17
MZM1
DEP1
MPC1
SAP30
CSF1
YDR485C_1
RTS1
VPS71
SWR1
SWC3
PET20
YGL217C_1
MME1
YML119W
ARP6
MTC2
RGT1
YGR037C_2
YDR485C_2
MUB1
MAM33
VMS1
YME1
LAT1
MDH1
CYC2
RPL36B
DIA2
YOL134C
SPT7
MDM20
BUD20
RIM9
FUM1

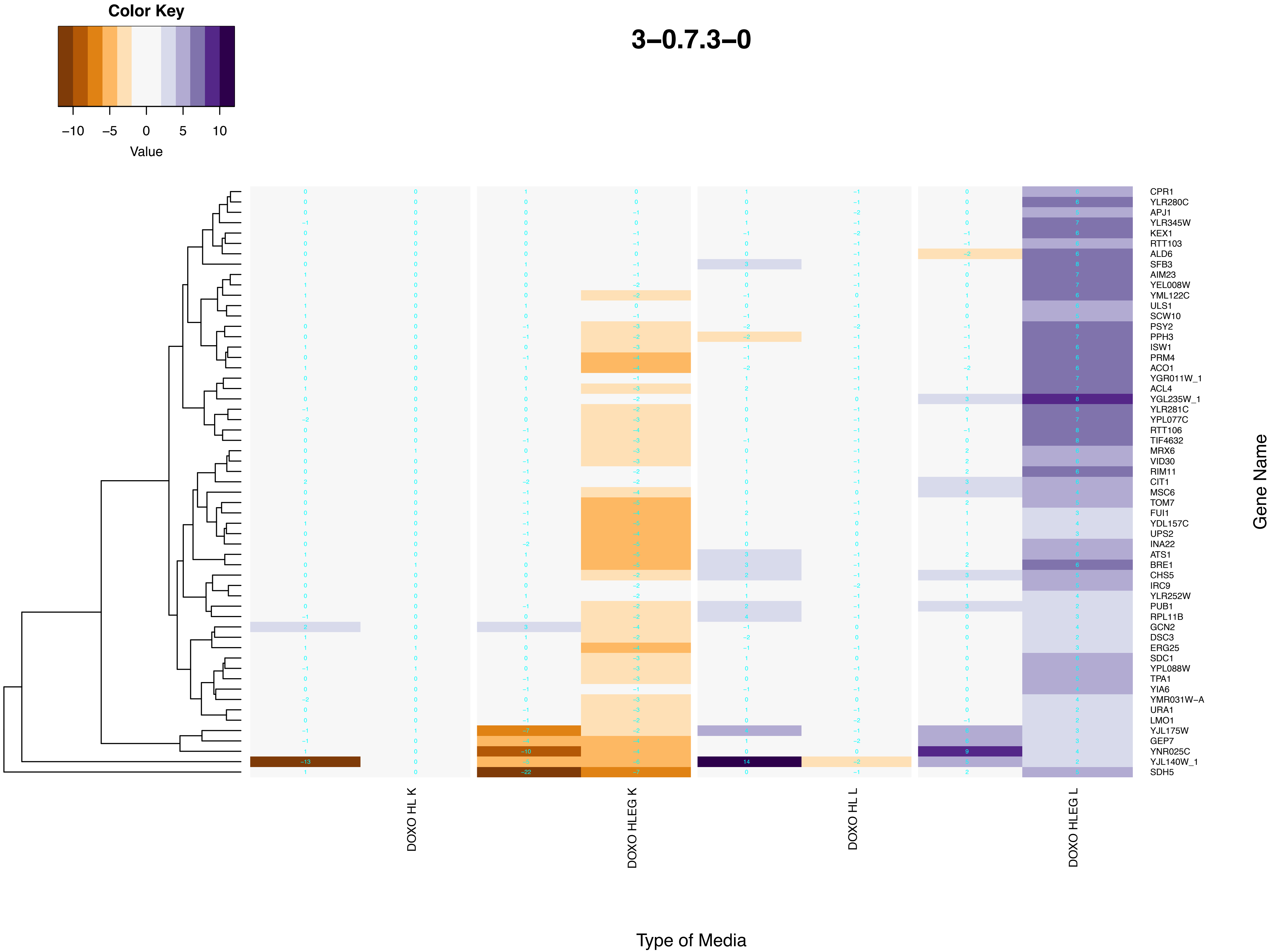

Color Key
−10
−5
0
5
10
Value
3−0.7.3−0
DOXO HL K
DOXO HLEG K
DOXO HL L
DOXO HLEG L
Type of Media
Gene Name
CPR1
YLR280C
APJ1
YLR345W
KEX1
RTT103
ALD6
SFB3
AIM23
YEL008W
YML122C
ULS1
SCW10
PSY2
PPH3
ISW1
PRM4
ACO1
YGR011W_1
ACL4
YGL235W_1
YLR281C
YPL077C
RTT106
TIF4632
MRX6
VID30
RIM11
CIT1
MSC6
TOM7
FUI1
YDL157C
UPS2
INA22
ATS1
BRE1
CHS5
IRC9
YLR252W
PUB1
RPL11B
GCN2
DSC3
ERG25
SDC1
YPL088W
TPA1
YIA6
YMR031W−A
URA1
LMO1
YJL175W
GEP7
YNR025C
YJL140W_1
SDH5

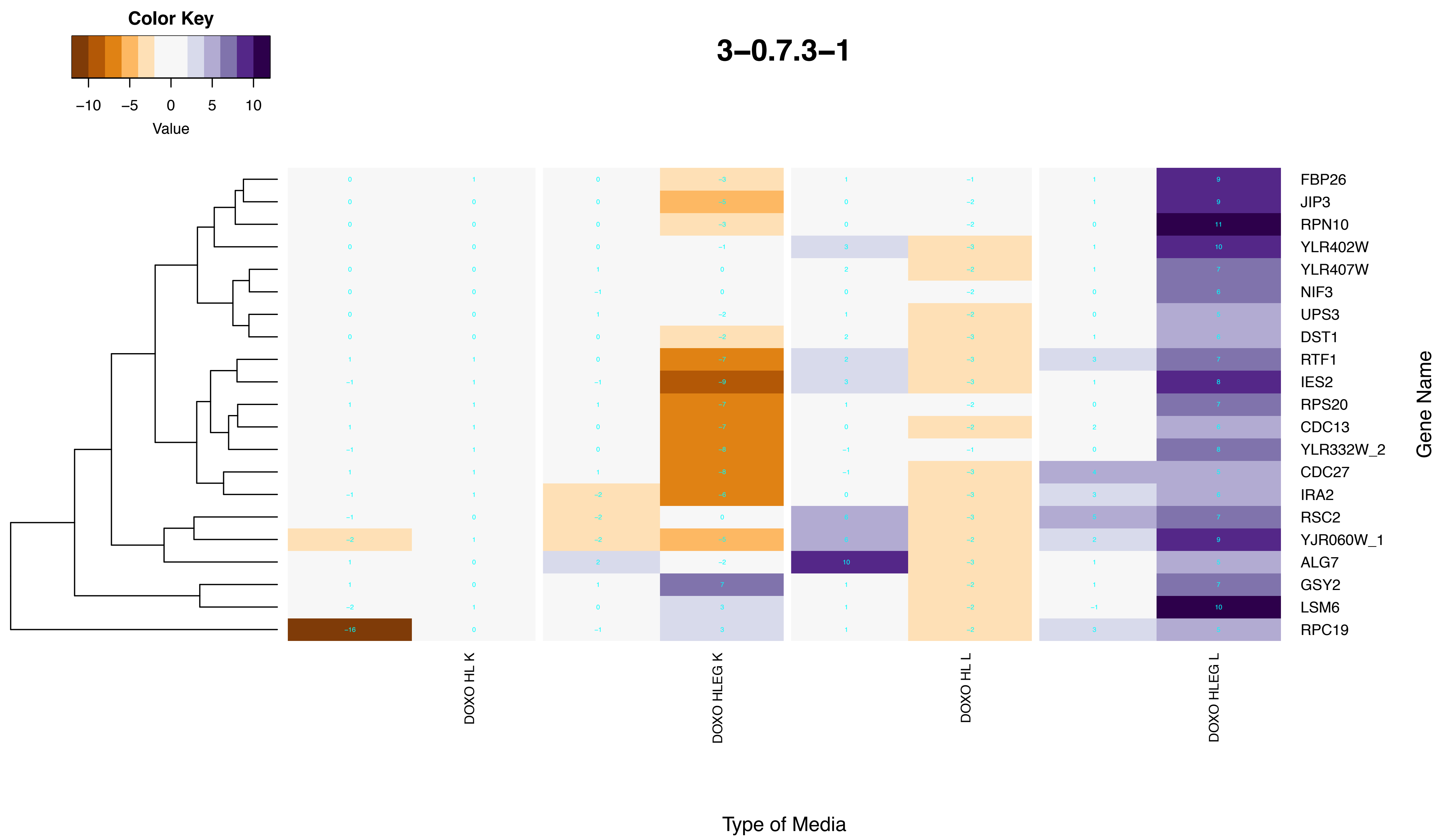
Color Key
−10 −5 0 5 10
Value
3−0.7.3−1
FBP26
JIP3
RPN10
YLR402W
YLR407W
NIF3
UPS3
DST1
RTF1
IES2
RPS20
CDC13
YLR332W_2
CDC27
IRA2
RSC2
YJR060W_1
ALG7
GSY2
LSM6
RPC19
Gene Name
DOXO HL K
DOXO HLEG K
DOXO HL L
DOXO HLEG L
Type of Media

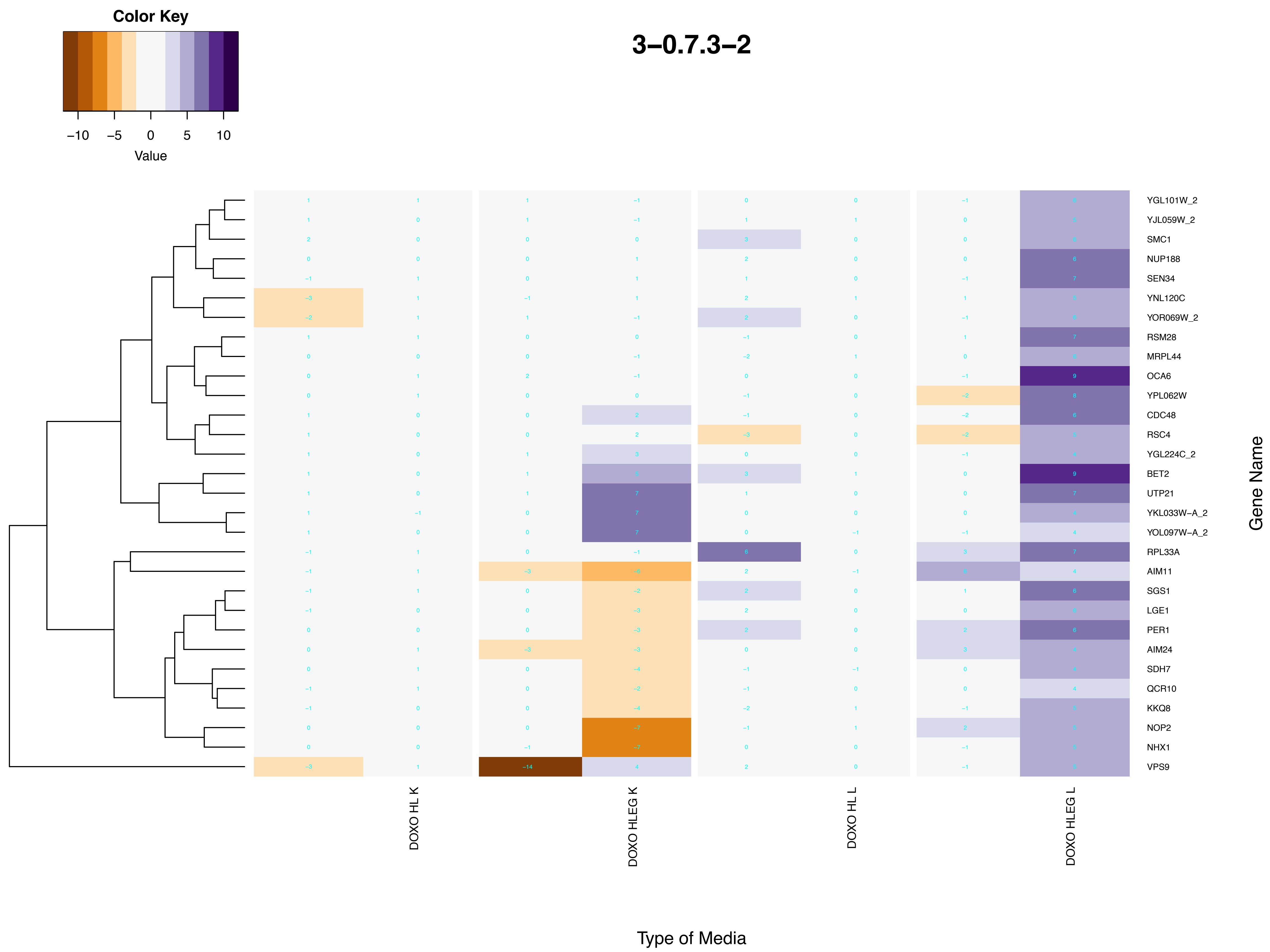
Color Key
−10
−5
0
5
10
Value
3−0.7.3−2
YGL101W_2
YJL059W_2
SMC1
NUP188
SEN34
YNL120C
YOR069W_2
RSM28
MRPL44
OCA6
YPL062W
CDC48
RSC4
YGL224C_2
BET2
UTP21
YKL033W−A_2
YOL097W−A_2
RPL33A
AIM11
SGS1
LGE1
PER1
AIM24
SDH7
QCR10
KKQ8
NOP2
NHX1
VPS9
Gene Name
DOXO HL K
DOXO HLEG K
DOXO HL L
DOXO HLEG L
Type of Media

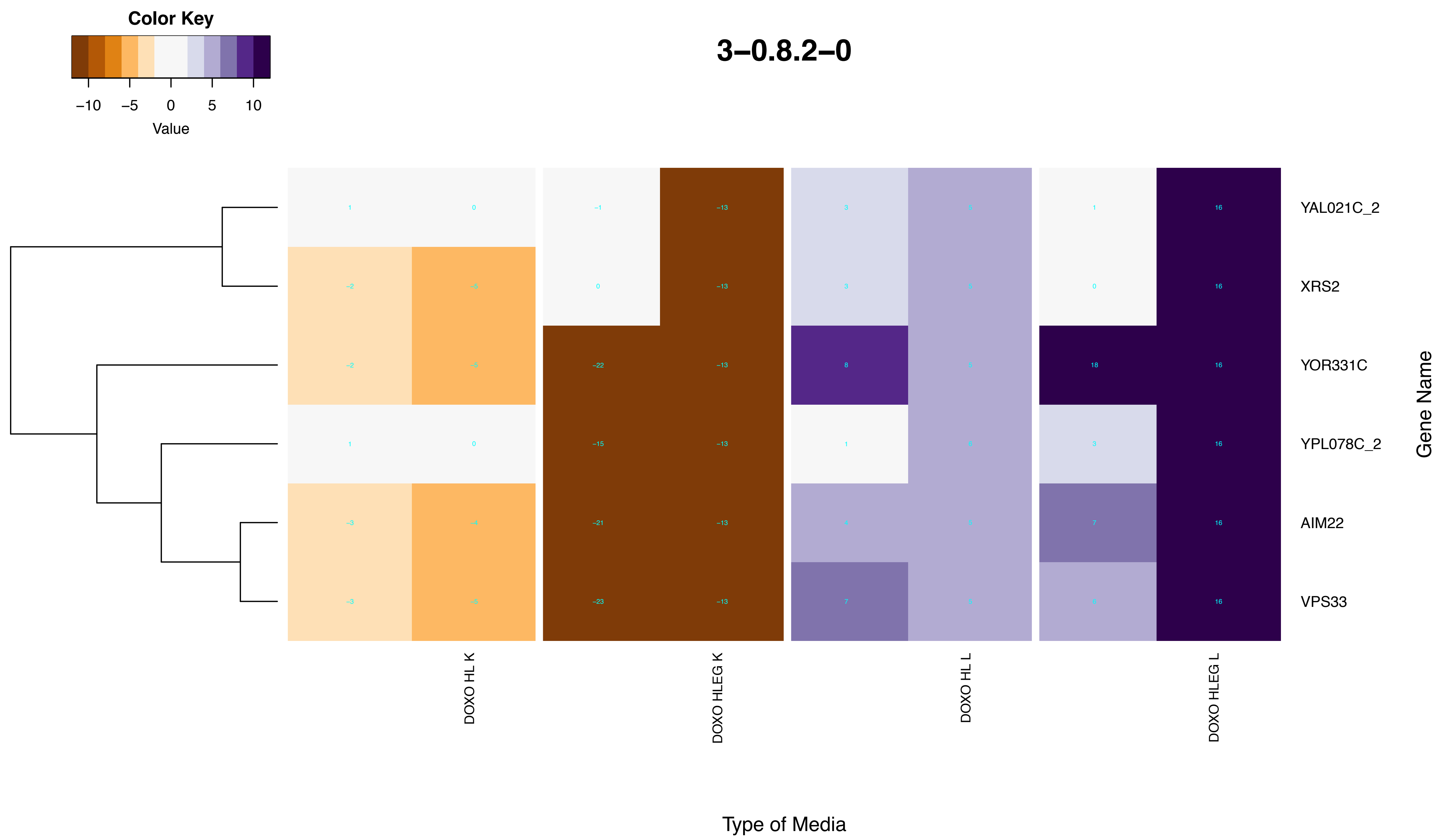

Color Key
−10 −5 0 5 10
Value
3−0.8.2−0
YAL021C_2
XRS2
YOR331C
YPL078C_2
AIM22
VPS33
Gene Name
DOXO HL K
DOXO HLEG K
DOXO HL L
DOXO HLEG L
Type of Media

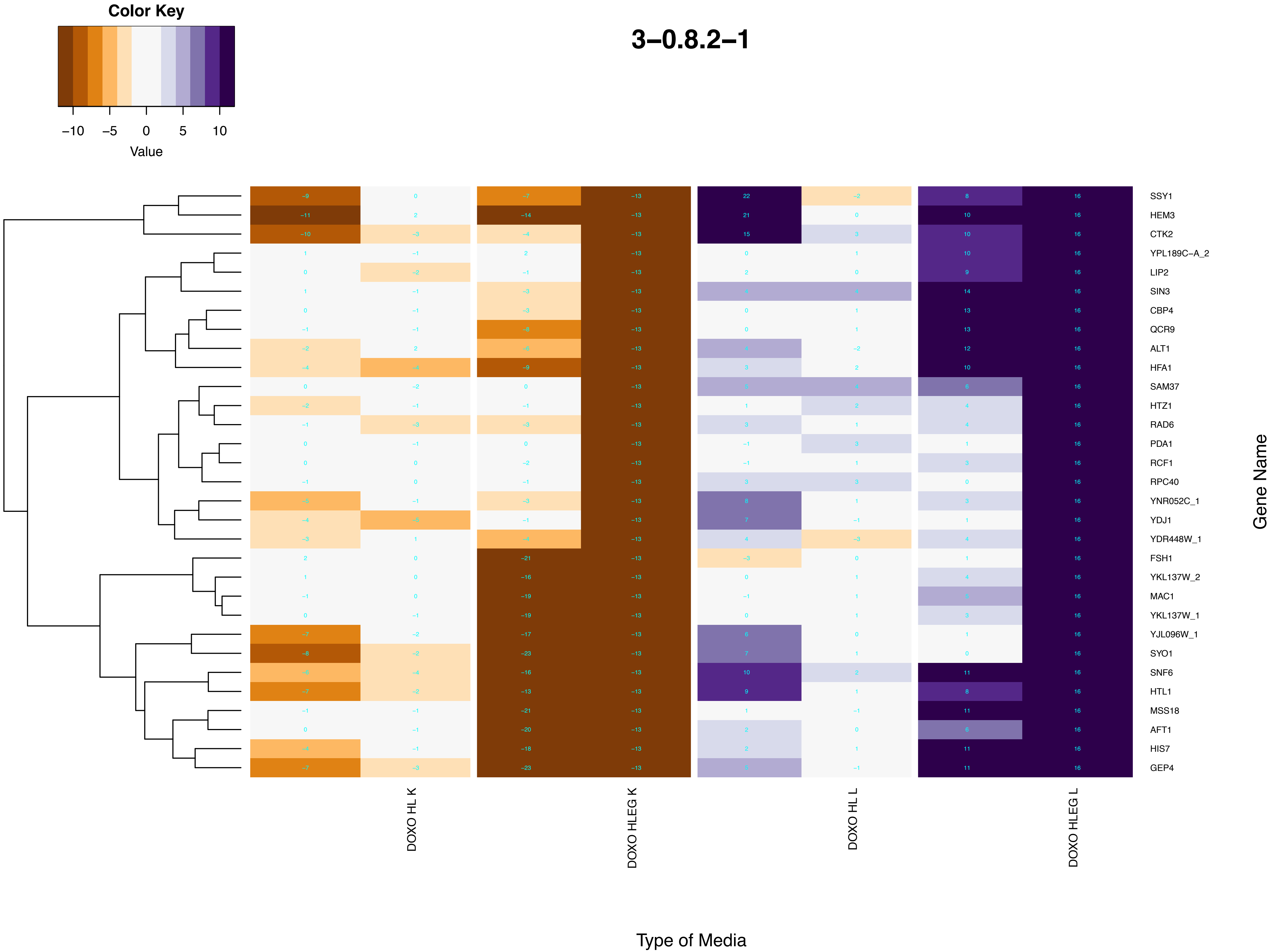

Color Key
−10 −5 0 5 10
Value
3−0.8.2−1
SSY1
HEM3
CTK2
YPL189C−A_2
LIP2
SIN3
CBP4
QCR9
ALT1
HFA1
SAM37
HTZ1
RAD6
PDA1
RCF1
RPC40
YNR052C_1
YDJ1
YDR448W_1
FSH1
YKL137W_2
MAC1
YKL137W_1
YJL096W_1
SYO1
SNF6
HTL1
MSS18
AFT1
HIS7
GEP4
Gene Name
DOXO HL K
DOXO HLEG K
DOXO HL L
DOXO HLEG L
Type of Media

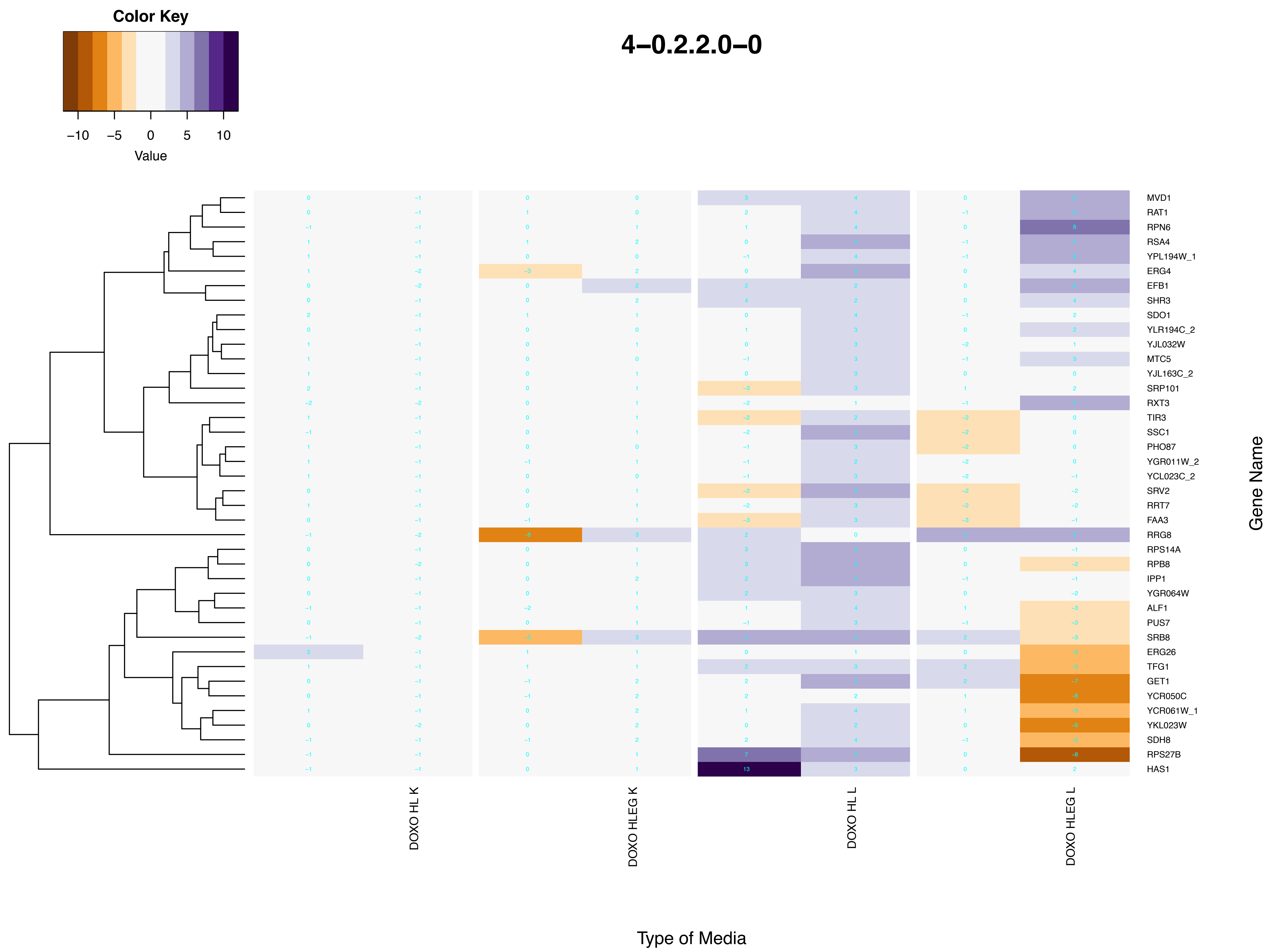
Color Key
−10
−5
0
5
10
Value
4–0.2.2.0–0
MVD1
RAT1
RPN6
RSA4
YPL194W_1
ERG4
EFB1
SHR3
SDO1
YLR194C_2
YJL032W
MTC5
YJL163C_2
SRP101
RXT3
TIR3
SSC1
PHO87
YGR011W_2
YCL023C_2
SRV2
RRT7
FAA3
RRG8
RPS14A
RPB8
IPP1
YGR064W
ALF1
PUS7
SRB8
ERG26
TFG1
GET1
YCR050C
YCR061W_1
YKL023W
SDH8
RPS27B
HAS1
Gene Name
DOXO HL K
DOXO HLEG K
DOXO HL L
DOXO HLEG L
Type of Media

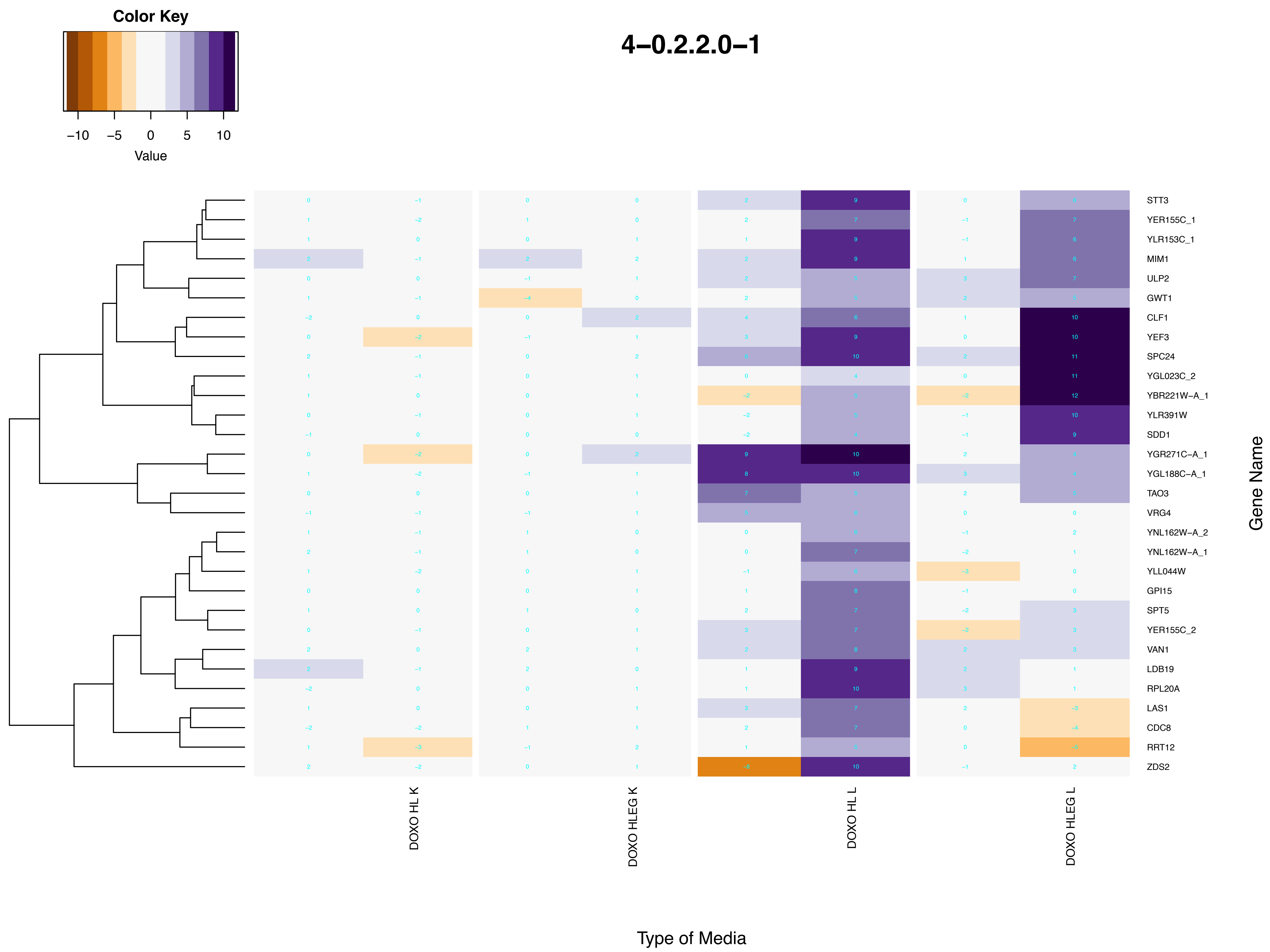

Color Key
−10
−5
0
5
10
Value
4−0.2.2.0−1
STT3
YER155C_1
YLR153C_1
MIM1
ULP2
GWT1
CLF1
YEF3
SPC24
YGL023C_2
YBR221W−A_1
YLR391W
SDD1
YGR271C−A_1
YGL188C−A_1
TAO3
VRG4
YNL162W−A_2
YNL162W−A_1
YLL044W
GPI15
SPT5
YER155C_2
VAN1
LDB19
RPL20A
LAS1
CDC8
RRT12
ZDS2
Gene Name
DOXO HL K
DOXO HLEG K
DOXO HL L
DOXO HLEG L
Type of Media

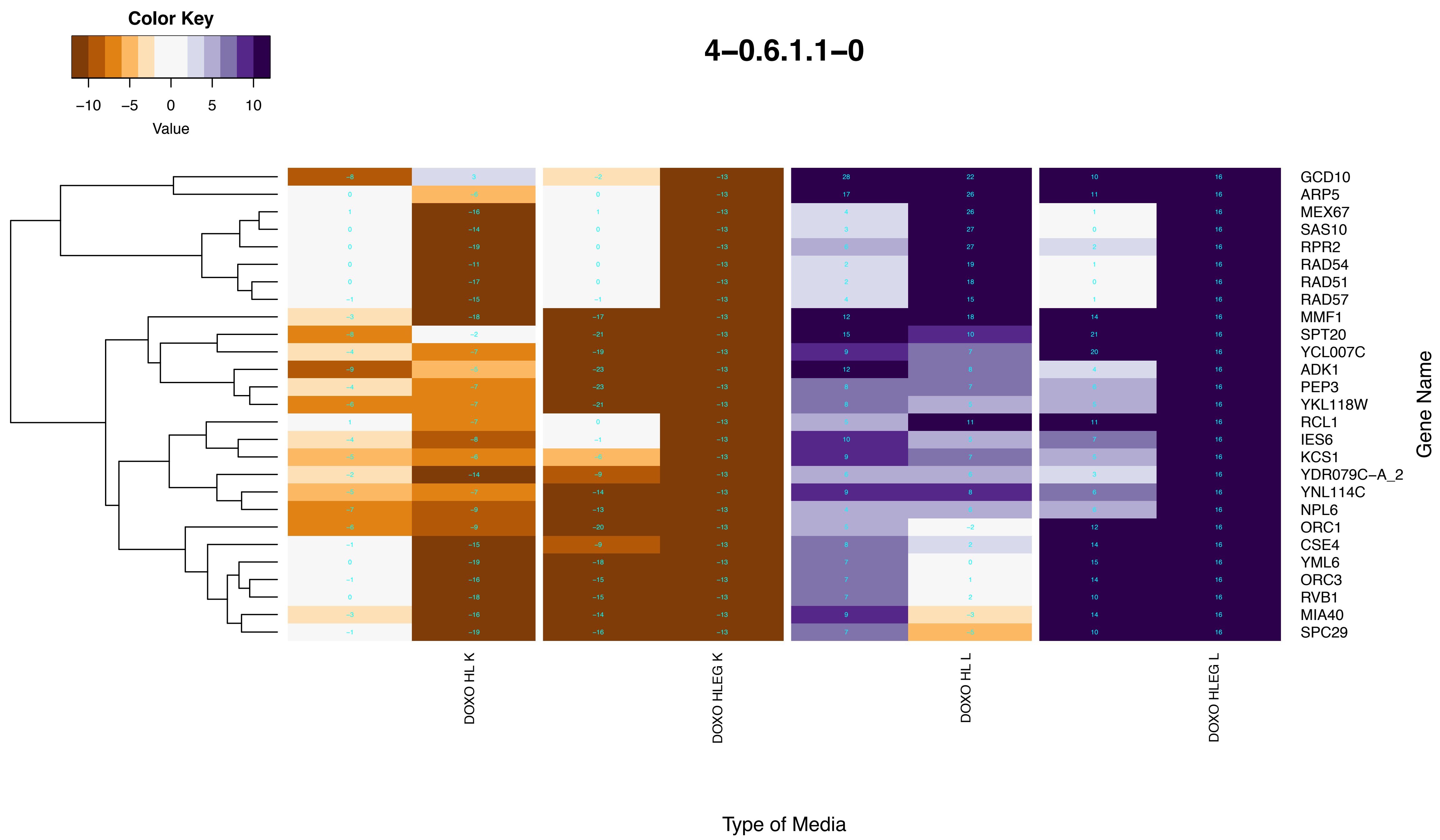
4–0.6.1.1–0
Color Key
−10
−5
0
5
10
Value
GCD10
ARP5
MEX67
SAS10
RPR2
RAD54
RAD51
RAD57
MMF1
SPT20
YCL007C
ADK1
PEP3
YKL118W
RCL1
IES6
KCS1
YDR079C−A_2
YNL114C
NPL6
ORC1
CSE4
YML6
ORC3
RVB1
MIA40
SPC29
Gene Name
DOXO HL K
DOXO HLEG K
DOXO HL L
DOXO HLEG L
Type of Media

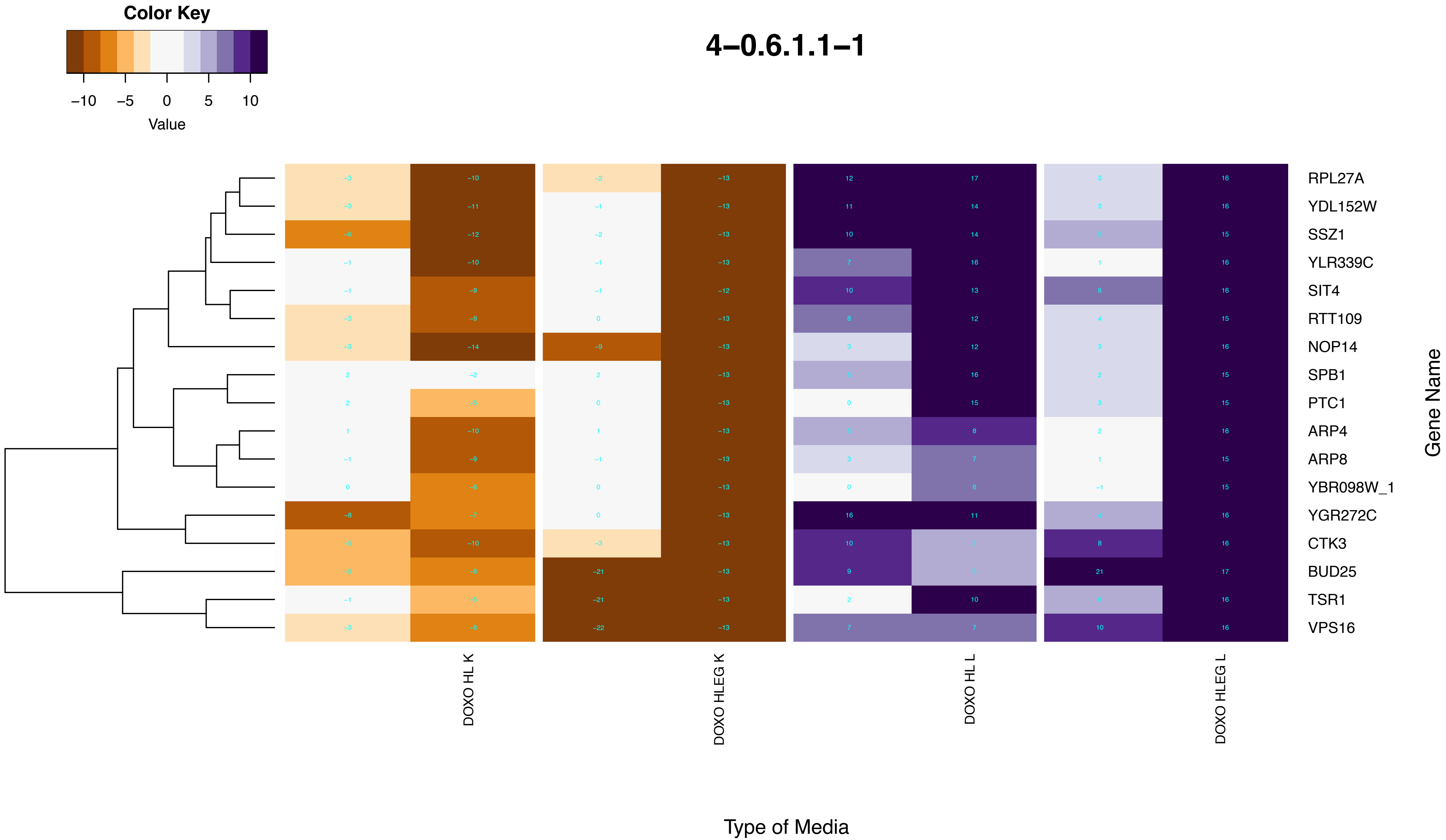

Color Key
−10
−5
0
5
10
Value
4−0.6.1.1−1
RPL27A
YDL152W
SSZ1
YLR339C
SIT4
RTT109
NOP14
SPB1
PTC1
ARP4
ARP8
YBR098W_1
YGR272C
CTK3
BUD25
TSR1
VPS16
Gene Name
DOXO HL K
DOXO HLEG K
DOXO HL L
DOXO HLEG L
Type of Media

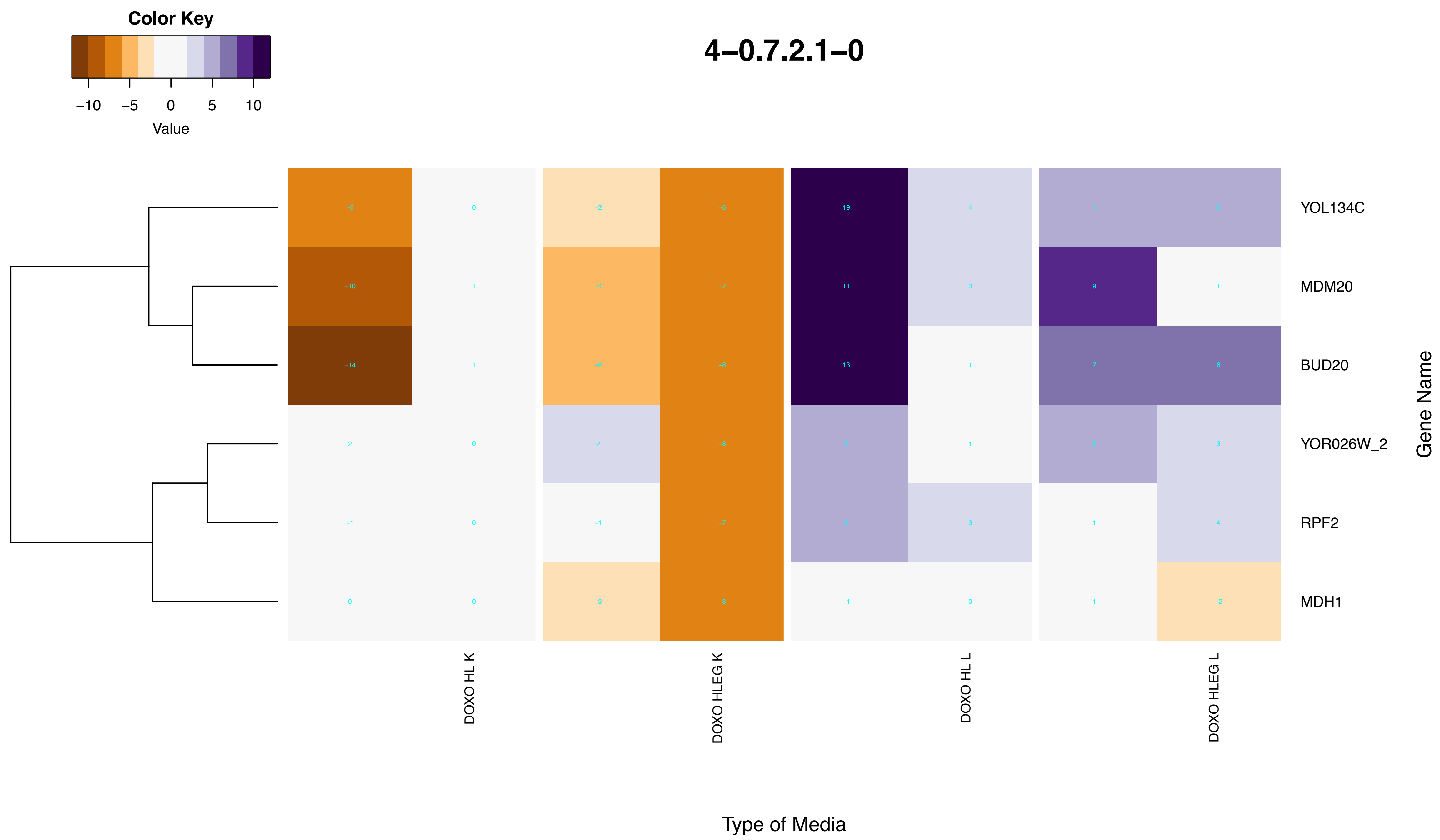
4–0.7.2.1–0
Color Key
−10 −5 0 5 10
Value
YOL134C
MDM20
BUD20
YOR026W_2
RPF2
MDH1
Gene Name
DOXO HL K
DOXO HLEG K
DOXO HL L
DOXO HLEG L
Type of Media

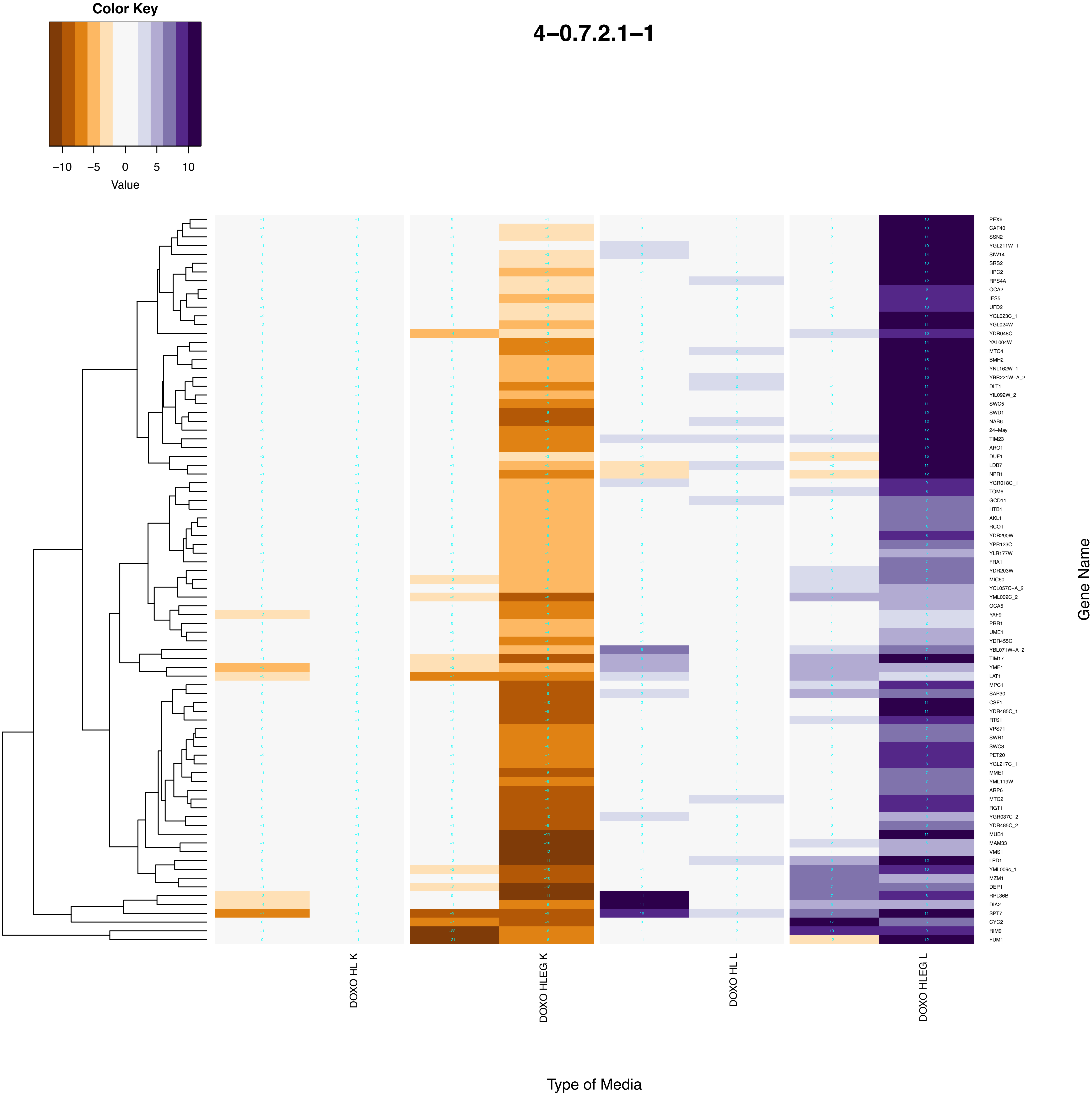
Color Key
Value
−10
−5
0
5
10
4−0.7.2.1−1
DOXO HL K
DOXO HLEG K
DOXO HL L
DOXO HLEG L
Type of Media
Gene Name
PEX6
CAF40
SSN2
YGL211W_1
SIW14
SRS2
HPC2
RPS4A
OCA2
IES5
UFD2
YGL023C_1
YGL024W
YDR048C
YAL004W
MTC4
BMH2
YNL162W_1
YBR221W−A_2
DLT1
YIL092W_2
SWC5
SWD1
NAB6
24−May
TIM23
ARO1
DUF1
LDB7
NPR1
YGR018C_1
TOM6
GCD11
HTB1
AKL1
RCO1
YDR290W
YPR123C
YLR177W
FRA1
YDR203W
MIC60
YCL057C−A_2
YML009C_2
OCA5
YAF9
PRR1
UME1
YDR455C
YBL071W−A_2
TIM17
YME1
LAT1
MPC1
SAP30
CSF1
YDR485C_1
RTS1
VPS71
SWR1
SWC3
PET20
YGL217C_1
MME1
YML119W
ARP6
MTC2
RGT1
YGR037C_2
YDR485C_2
MUB1
MAM33
VMS1
LPD1
YML009c_1
MZM1
DEP1
RPL36B
DIA2
SPT7
CYC2
RIM9
FUM1
