## Supplementary figures and images for "A yeast phenomic model for the influence of Warburg metabolism on genetic buffering of doxorubicin"

### 'de_novo'_cotranslational_protein_folding.pdf

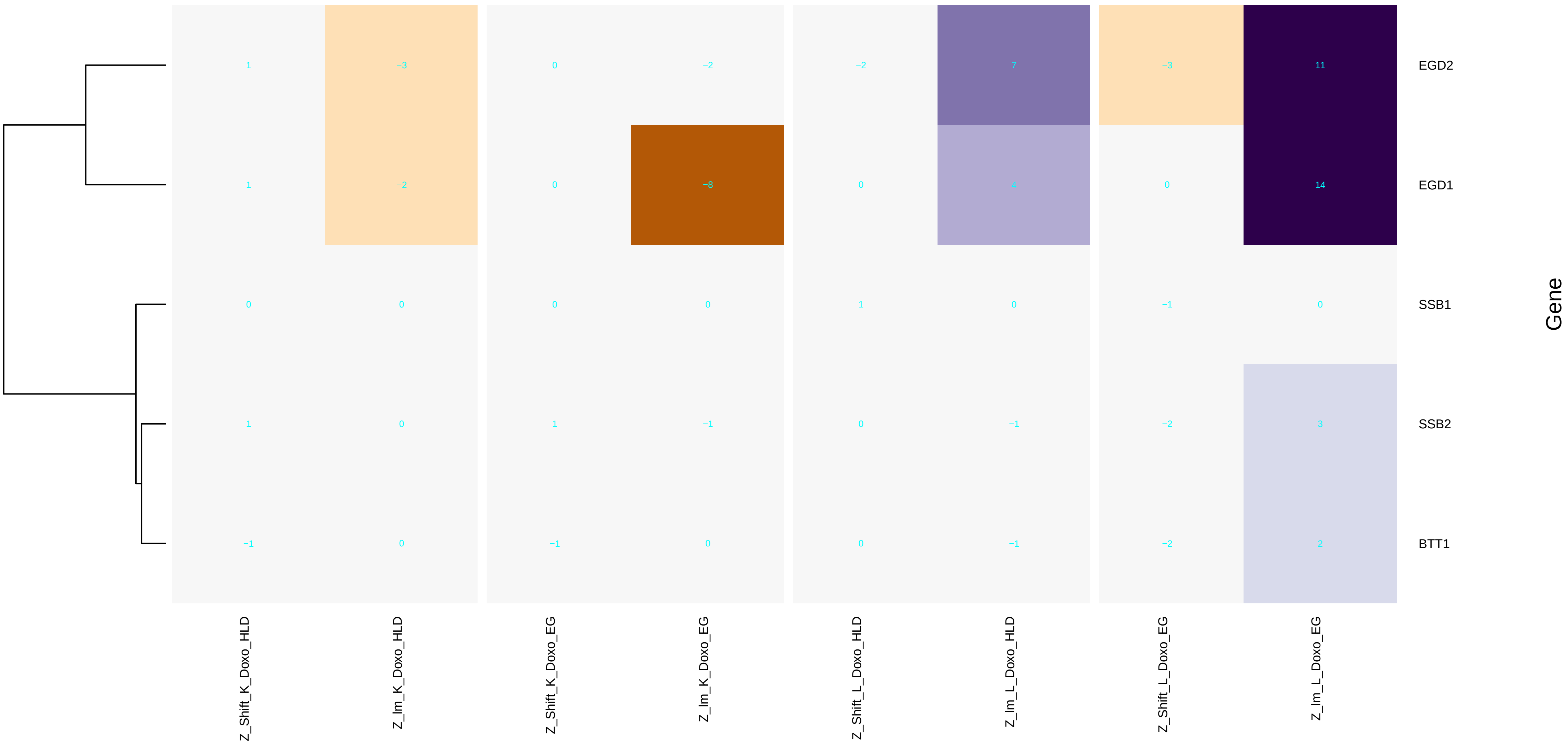

### A - Clust_Scores_by_first_rd_boxplots.pdf

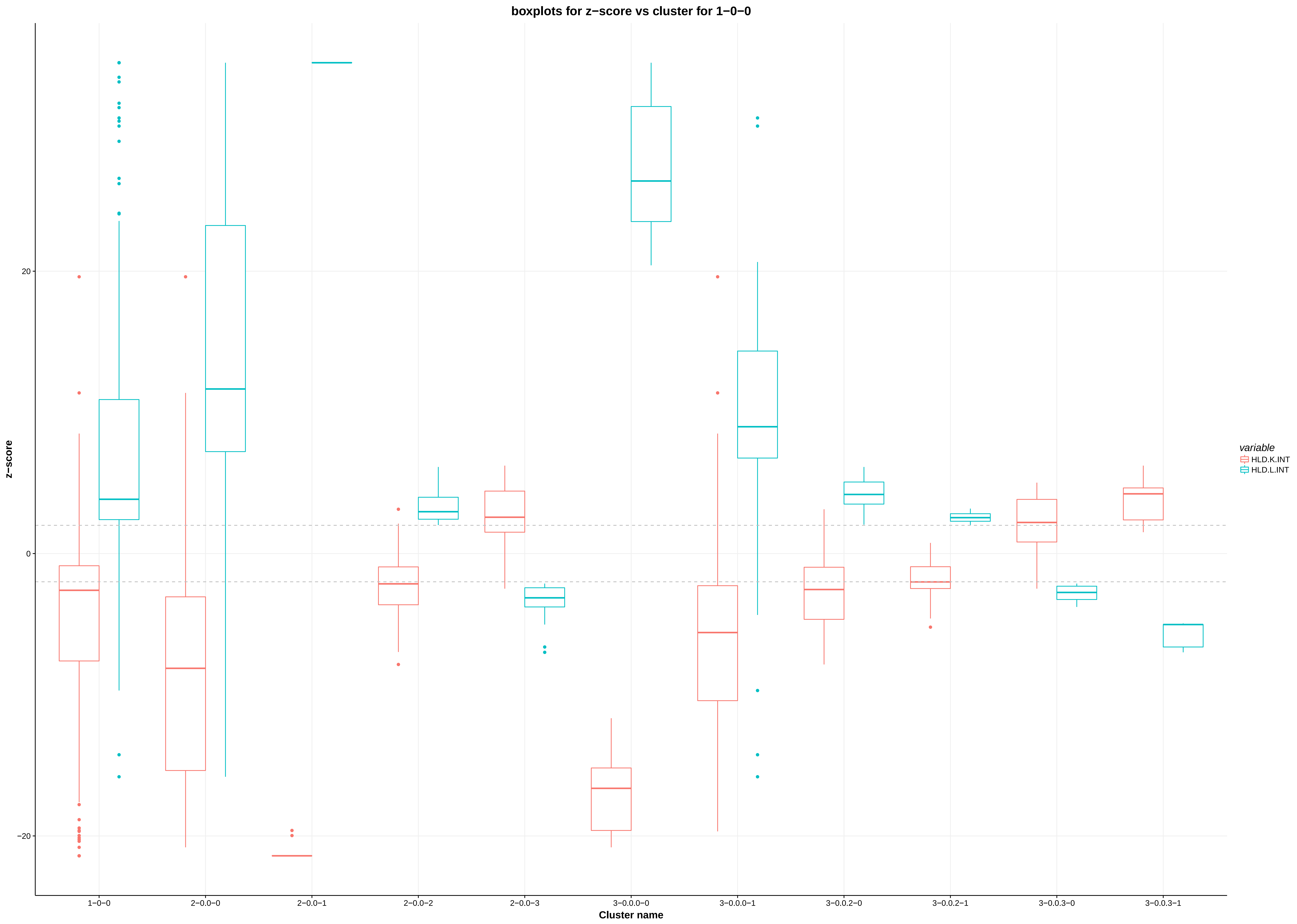

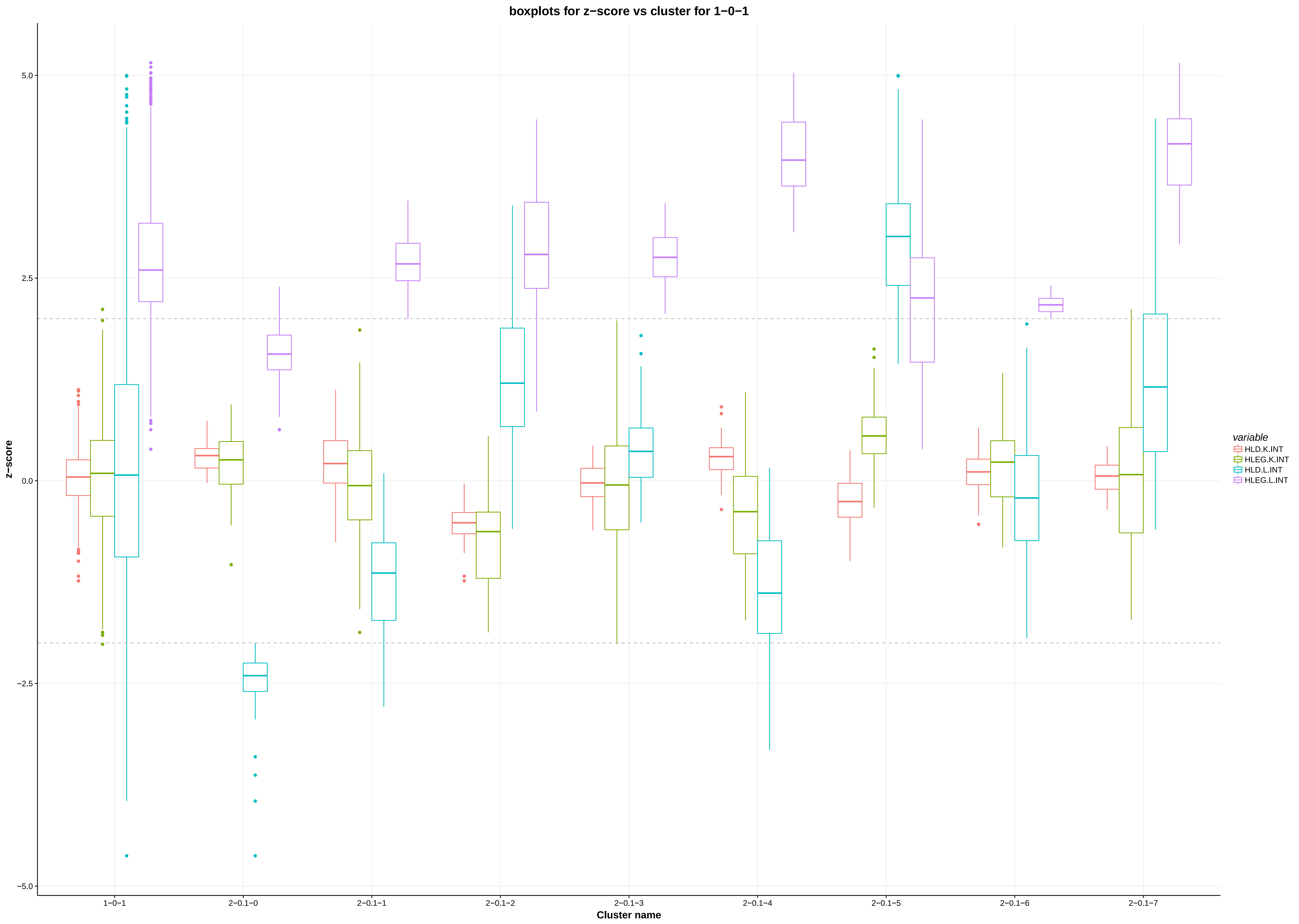

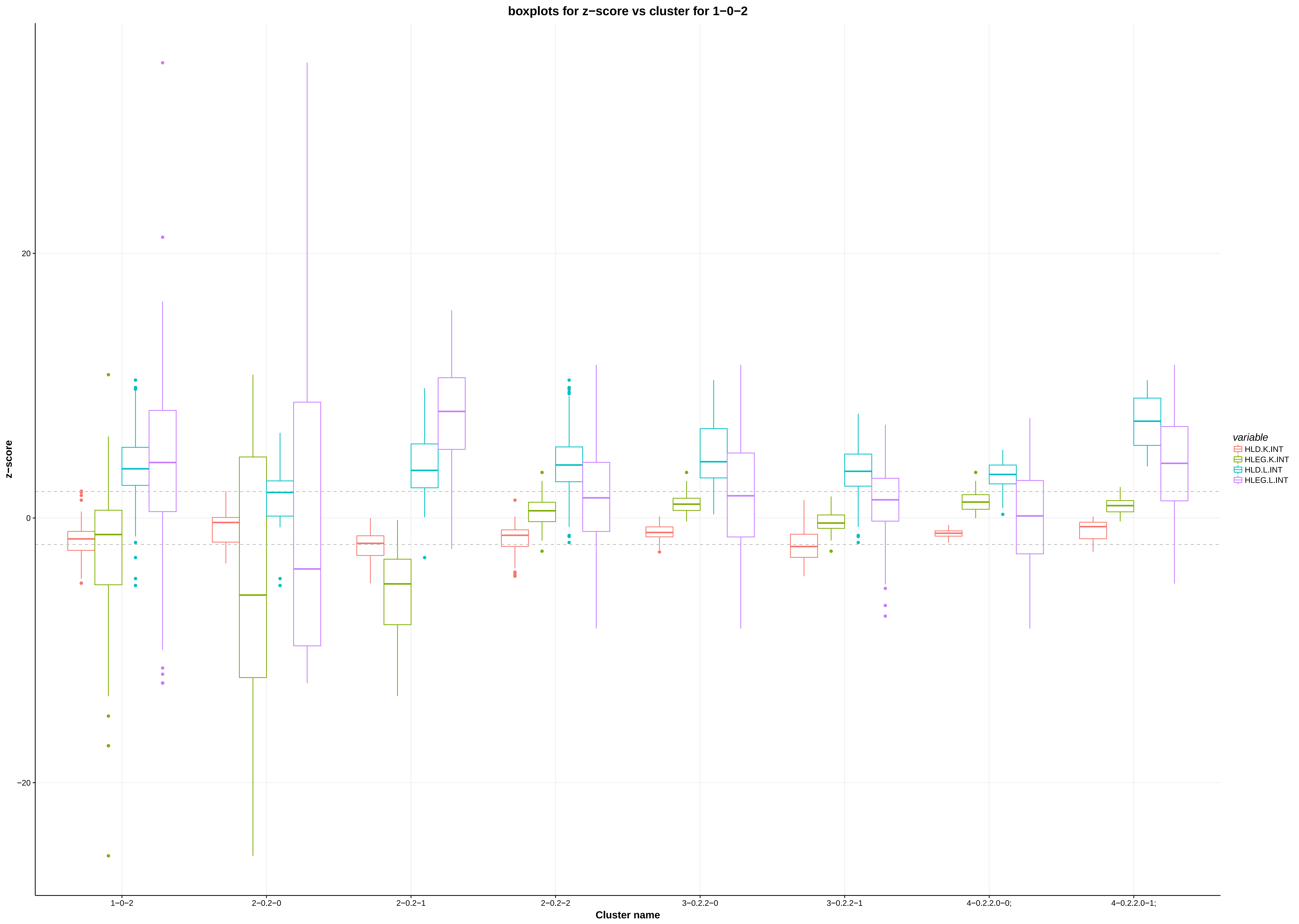

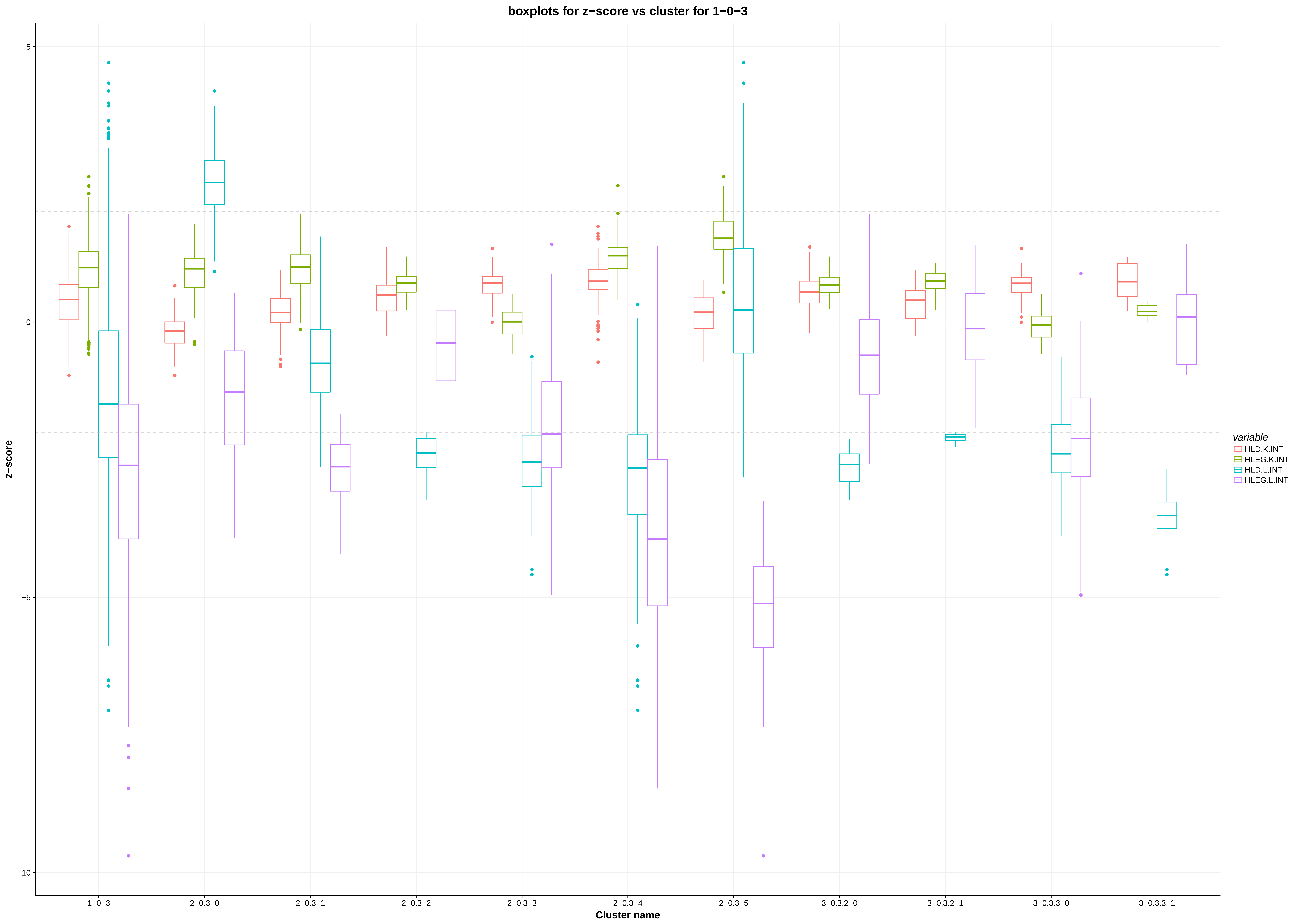

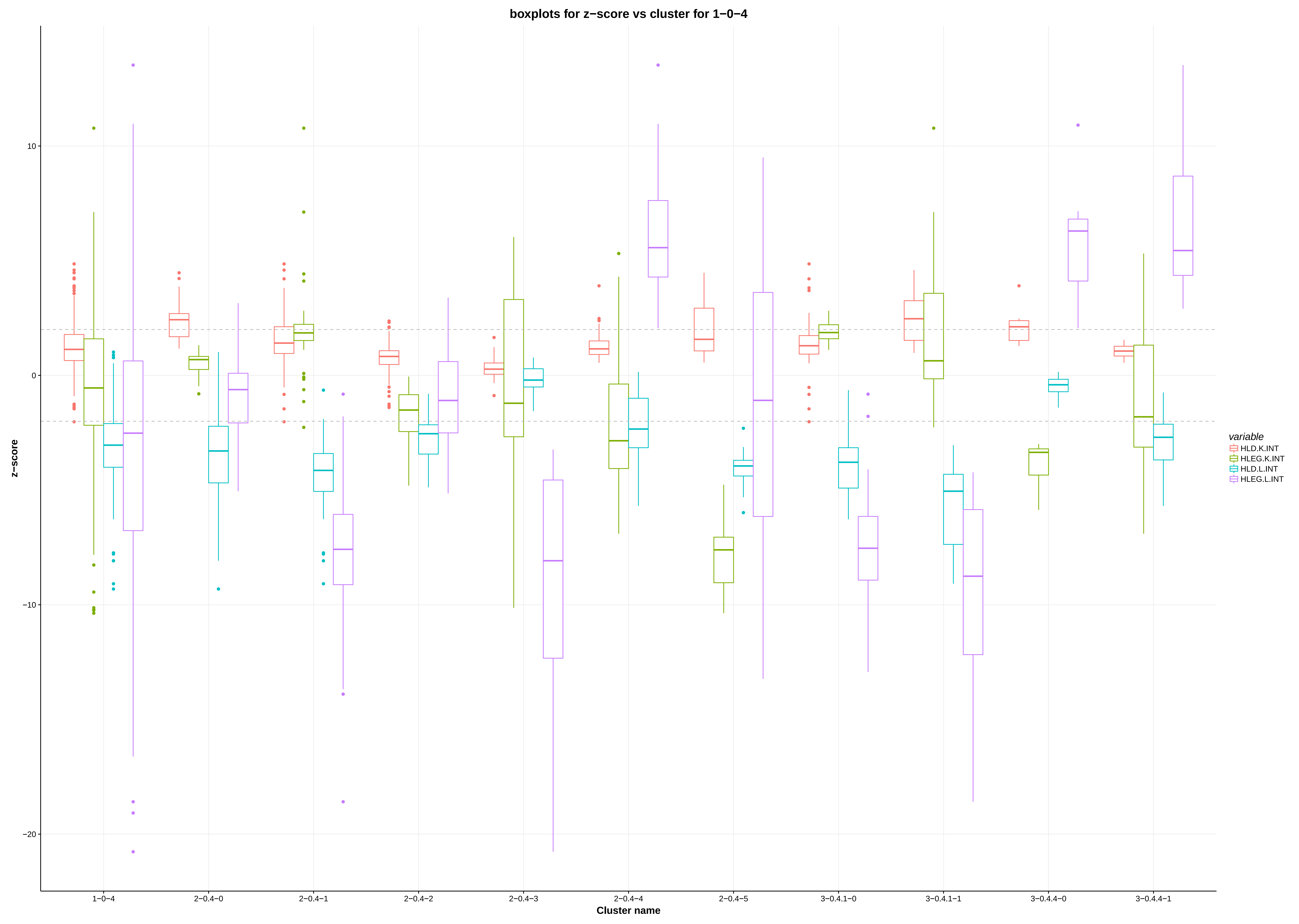

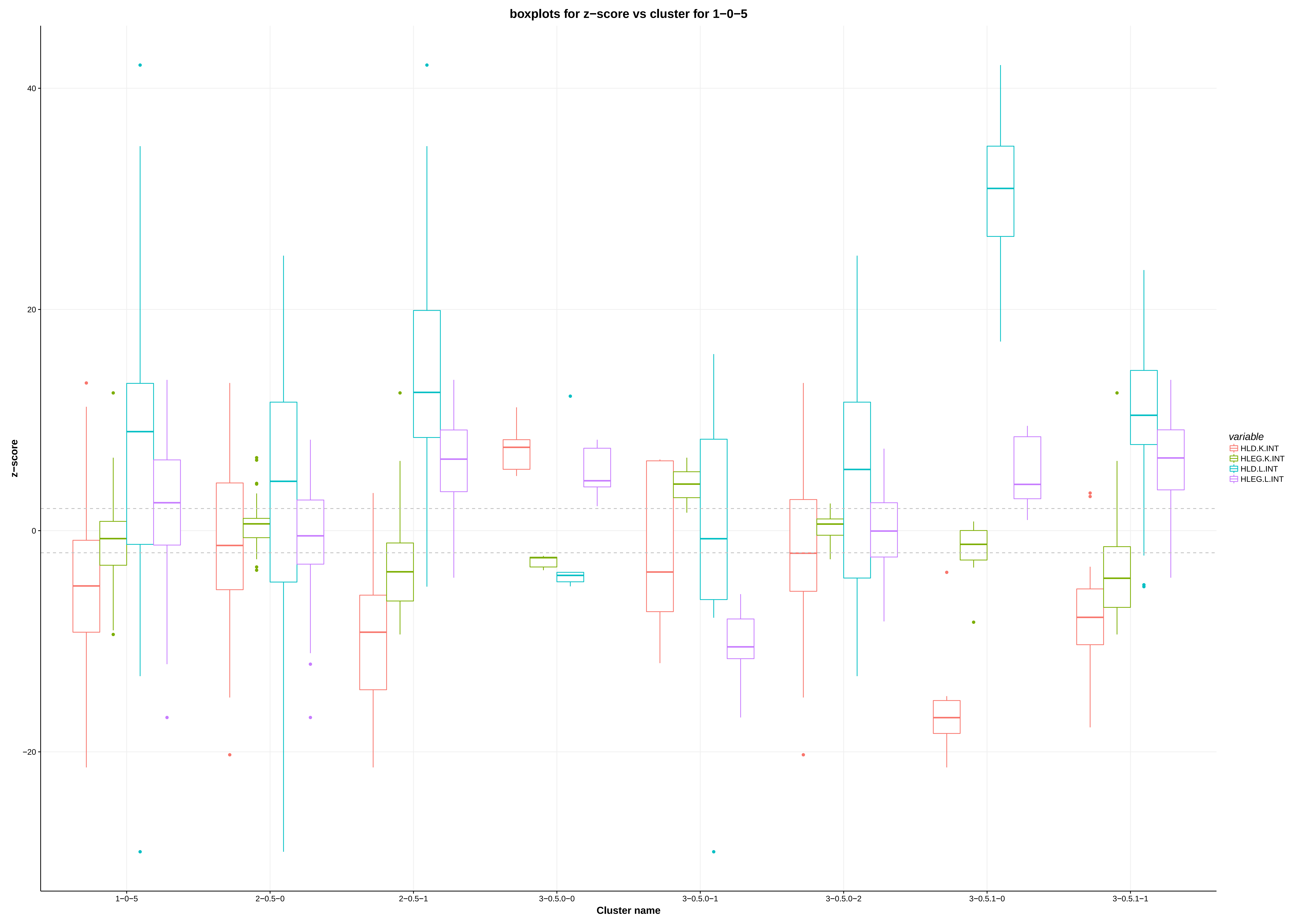

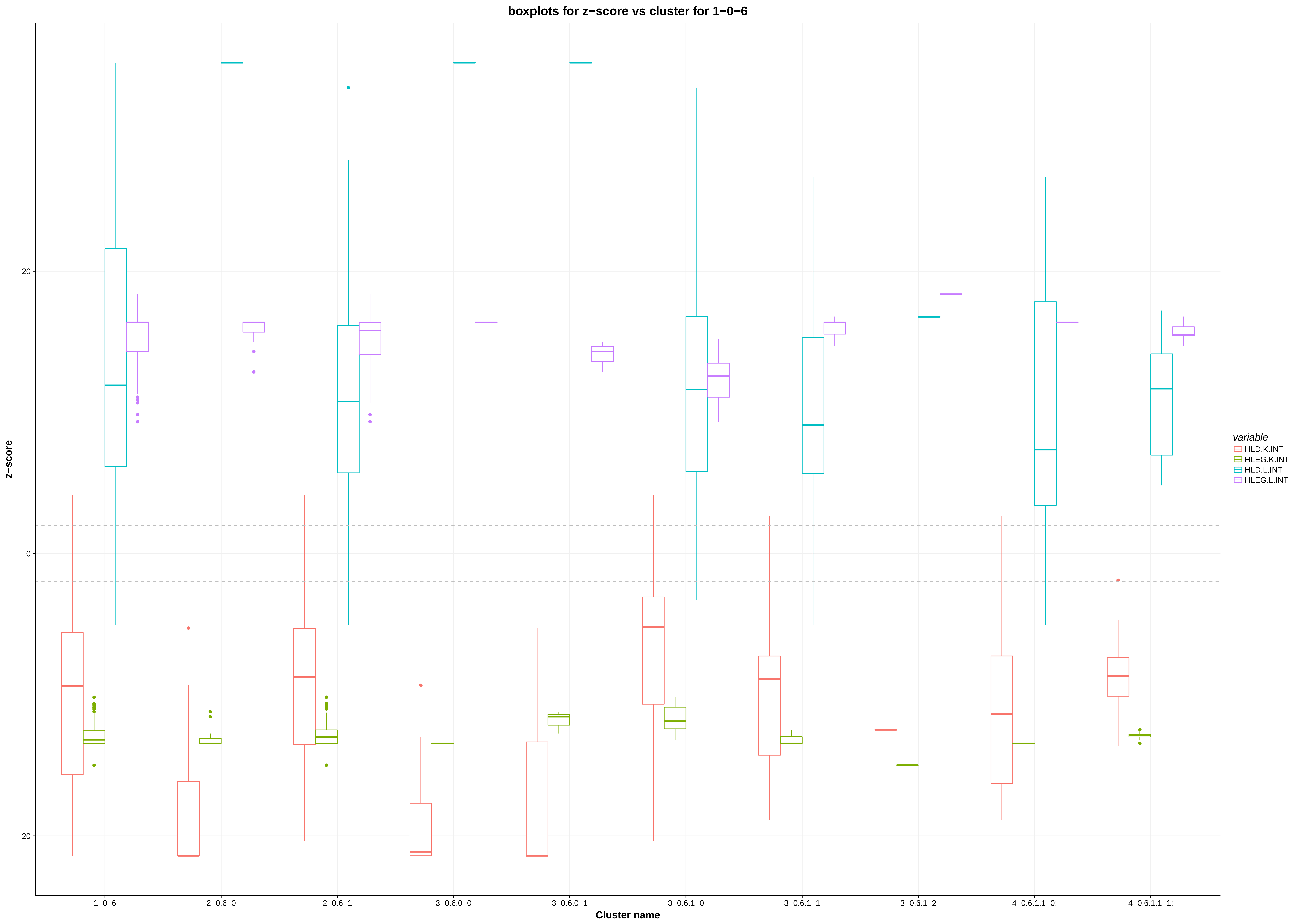

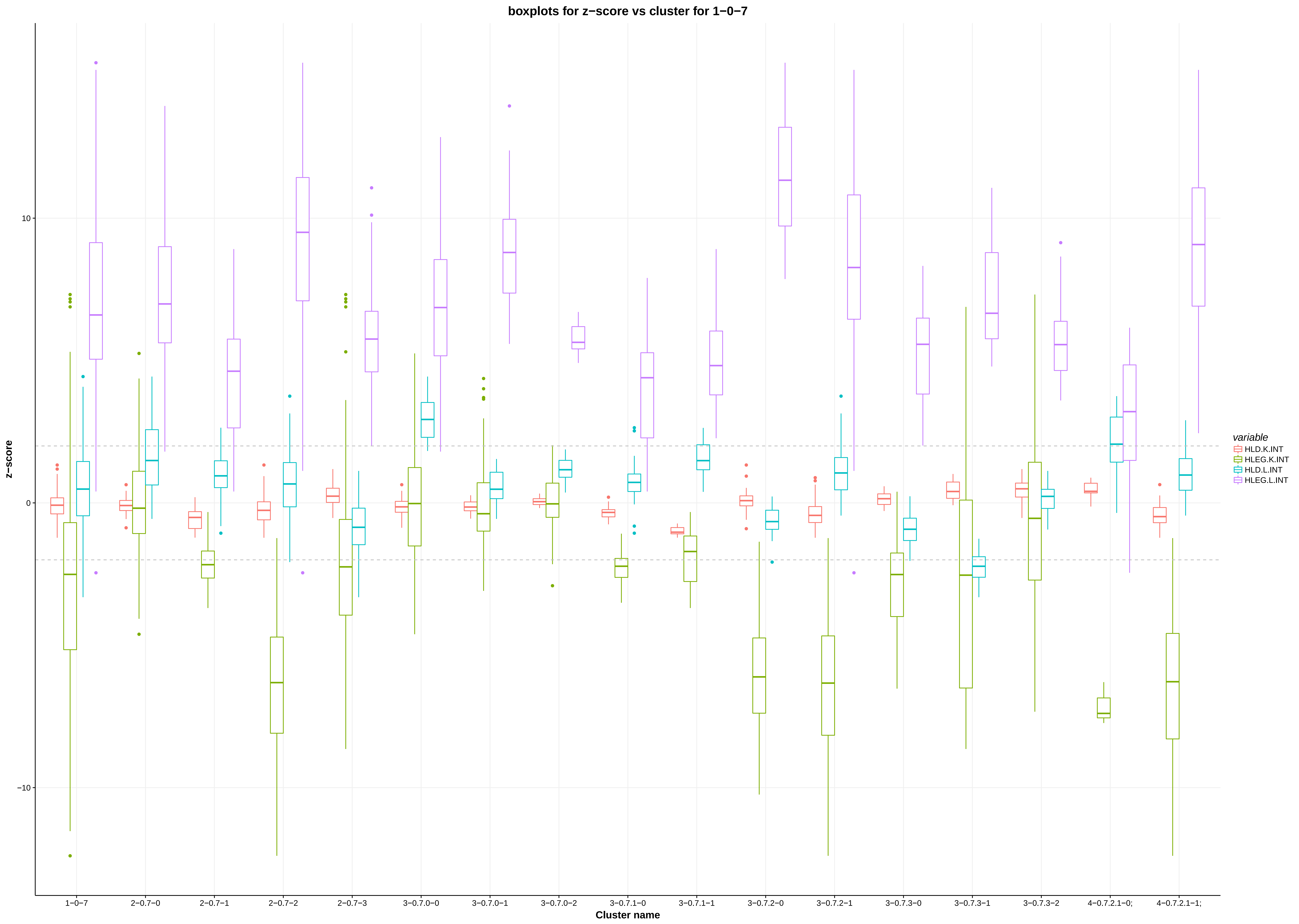

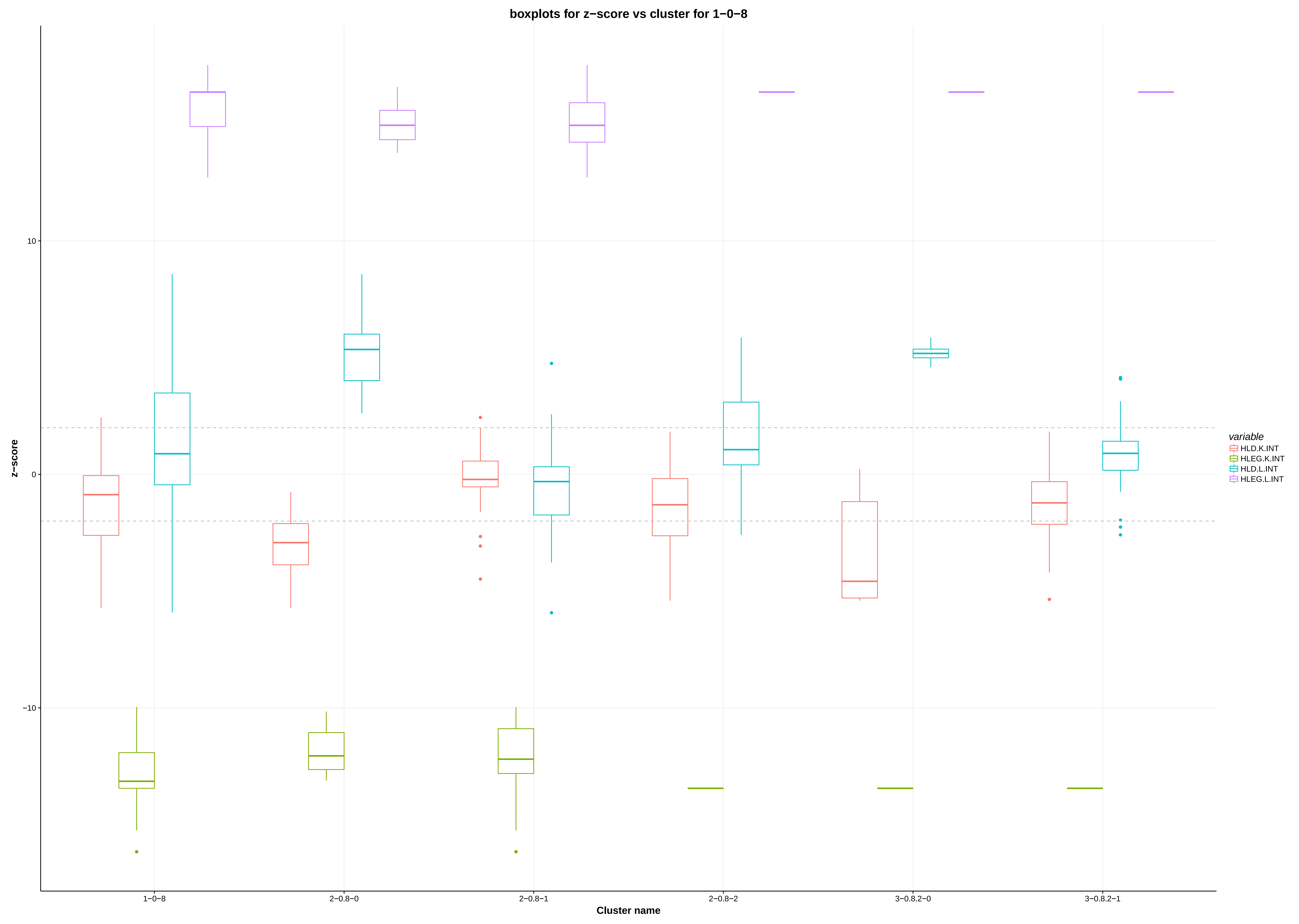

### actin_cortical_patch_localization.pdf

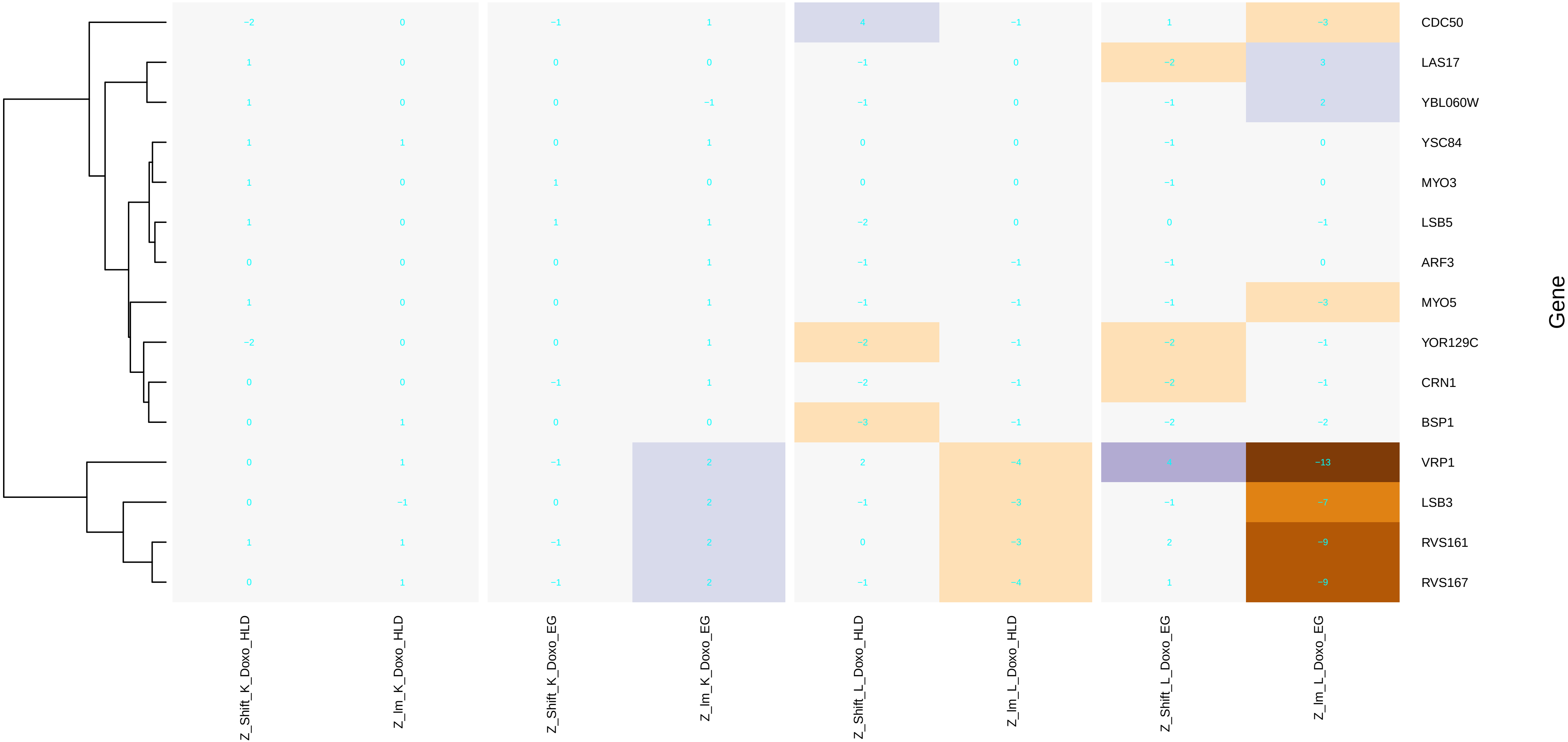

### amino_acid_activation.pdf

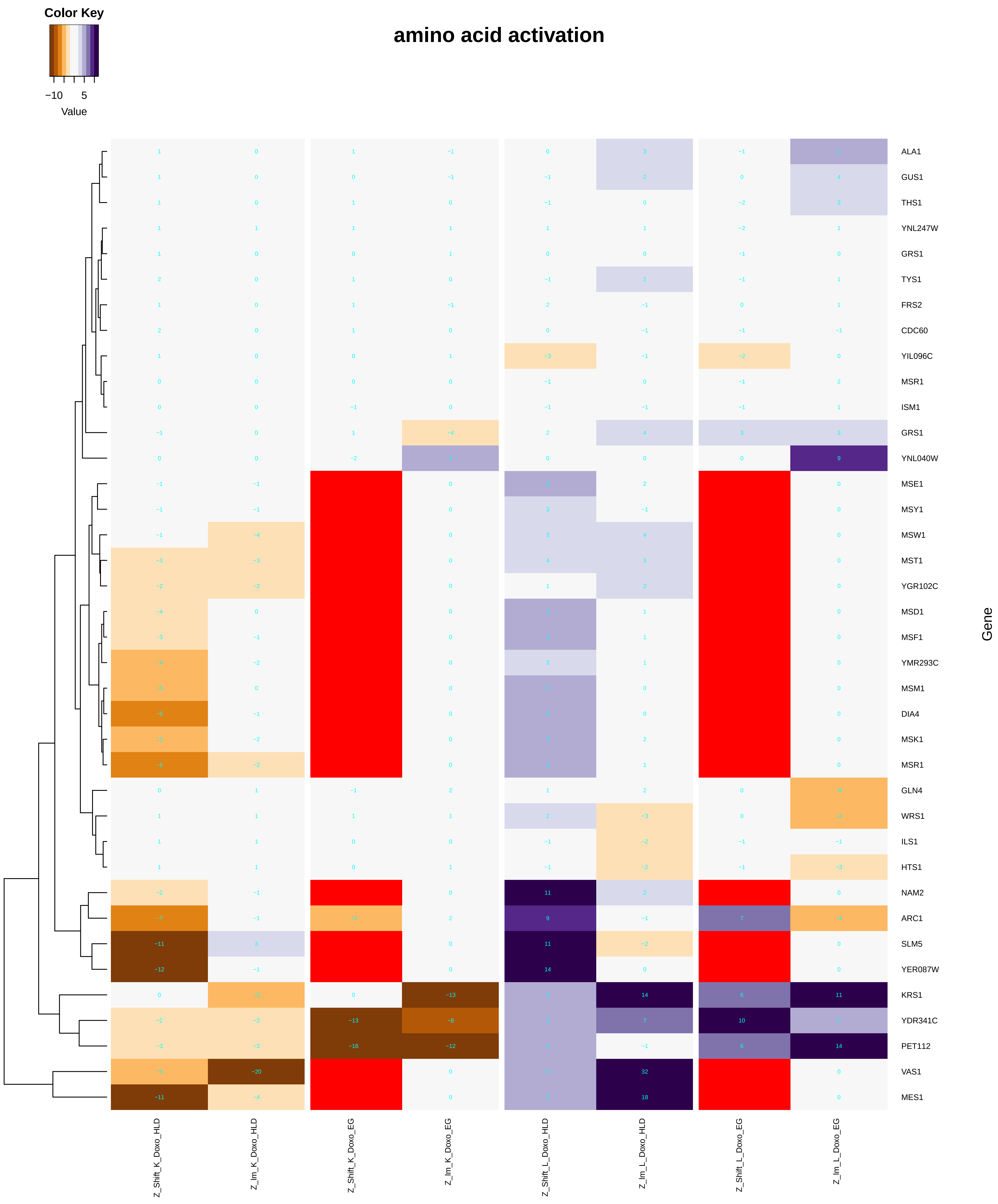

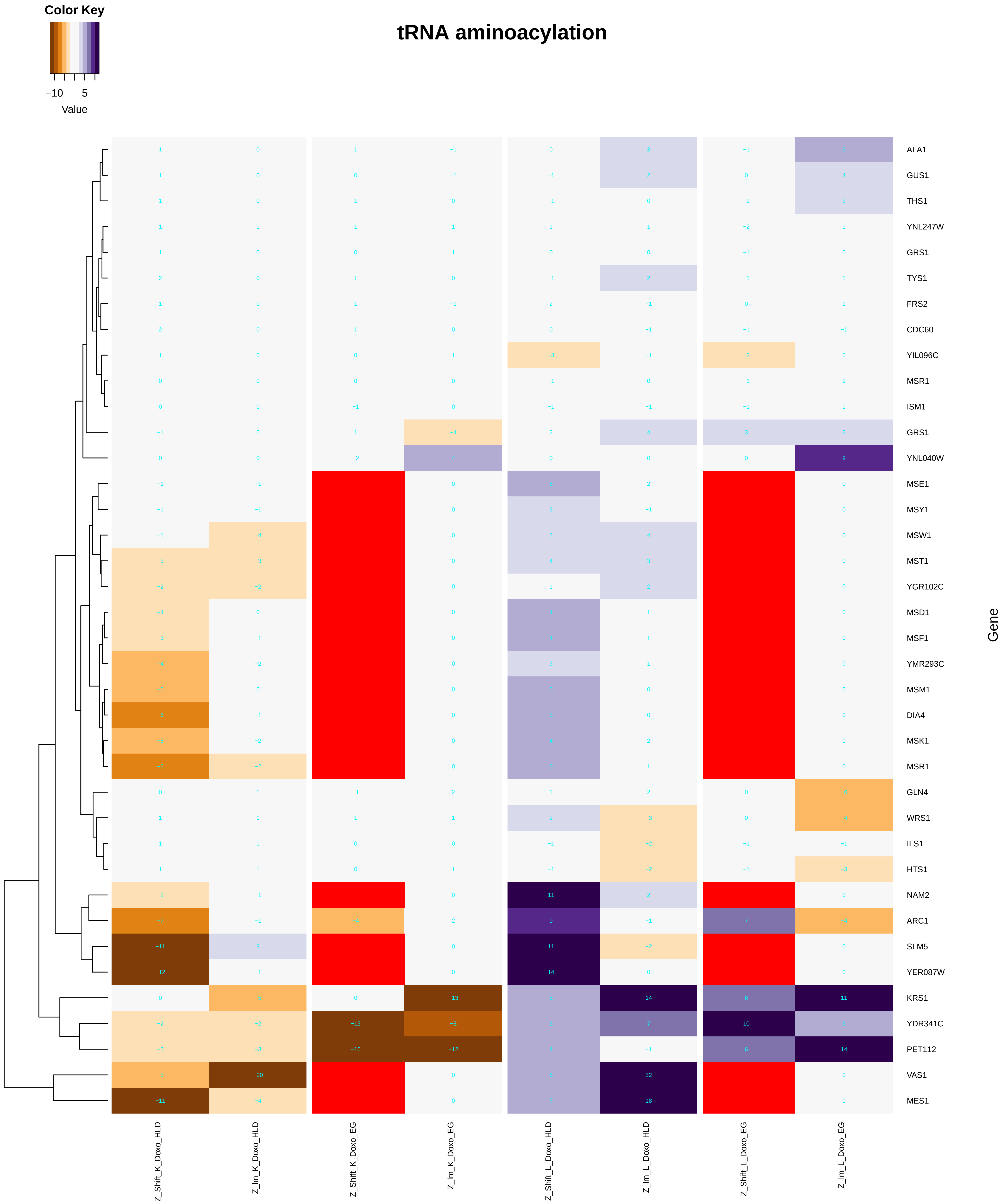

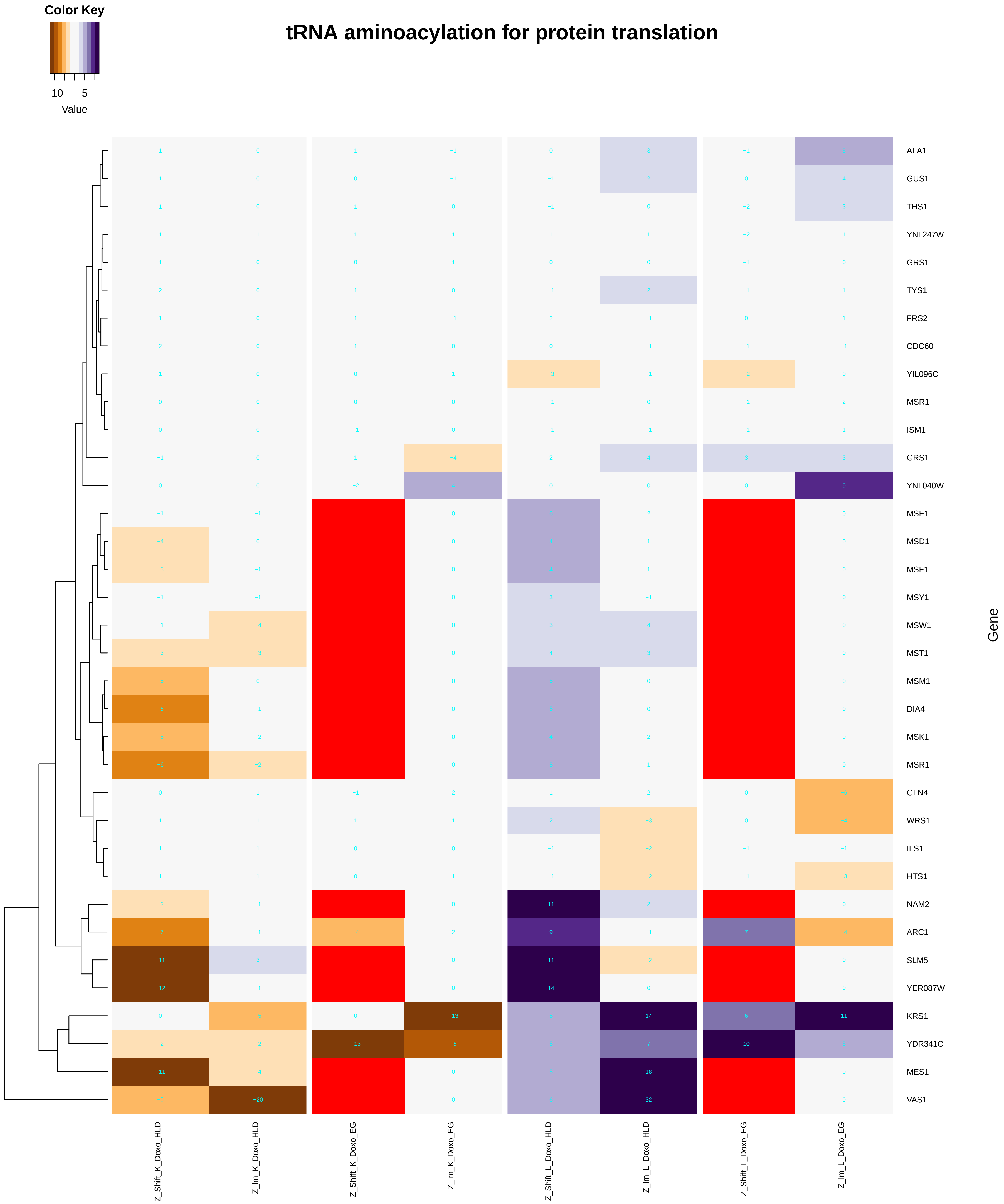

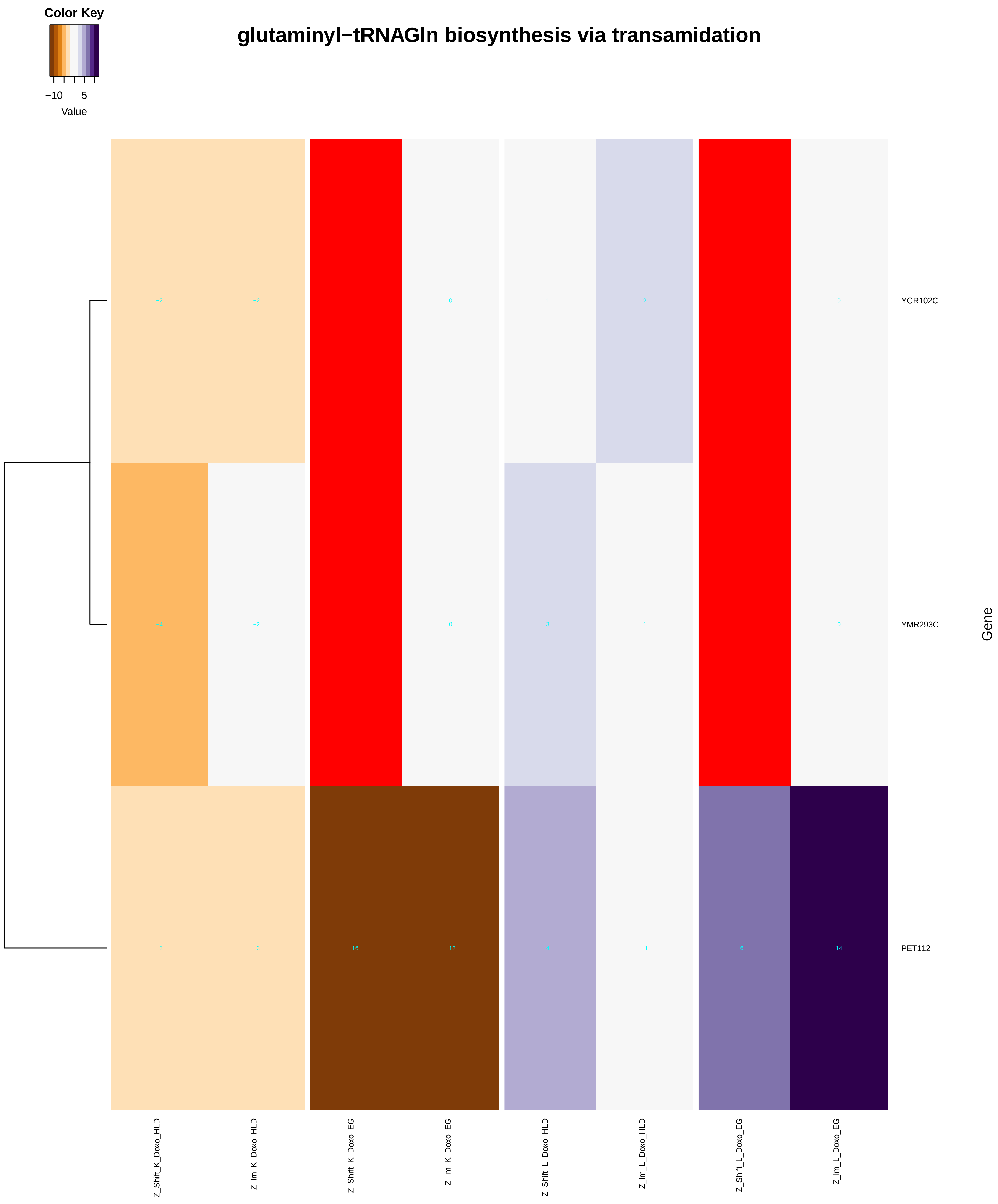

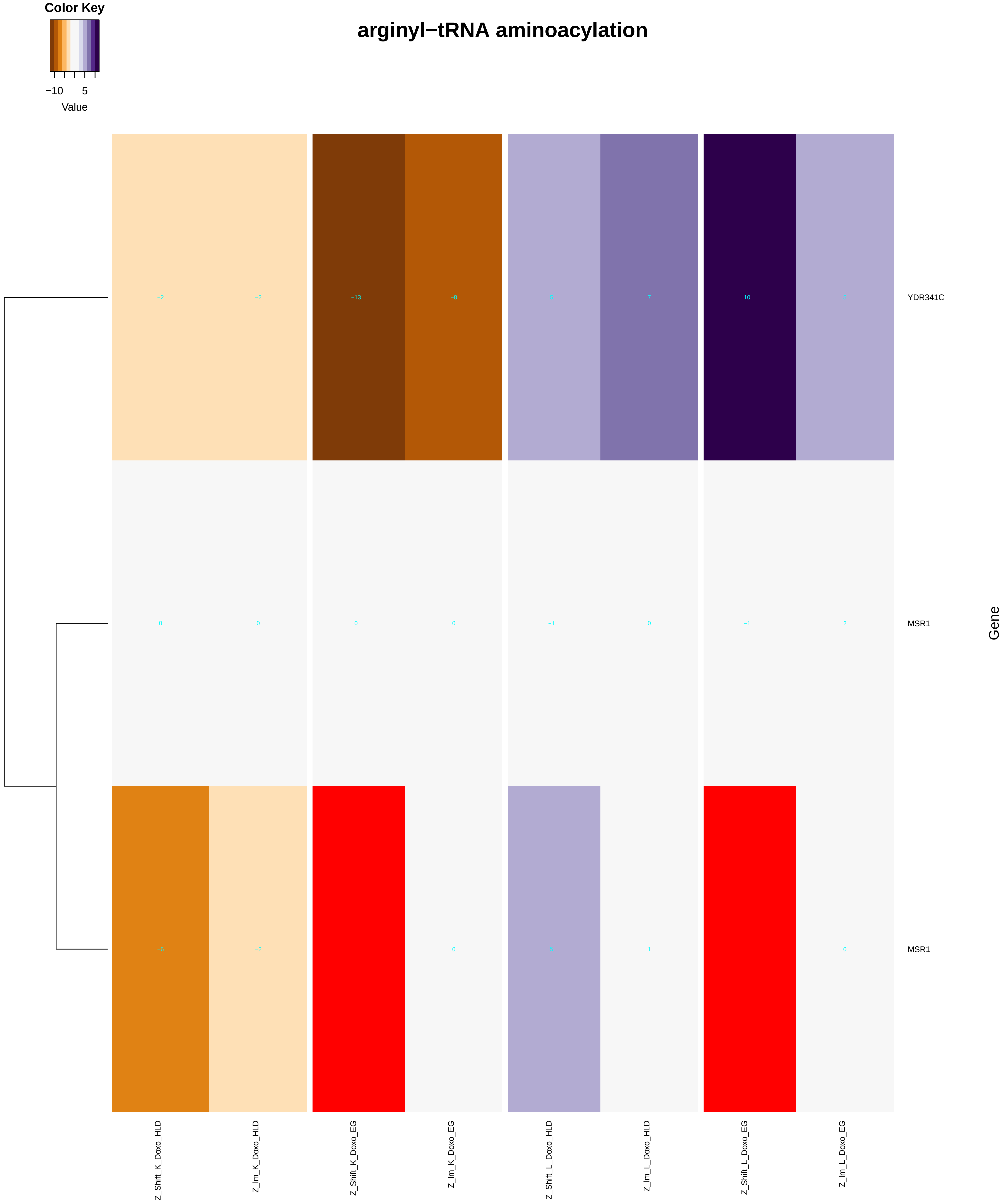

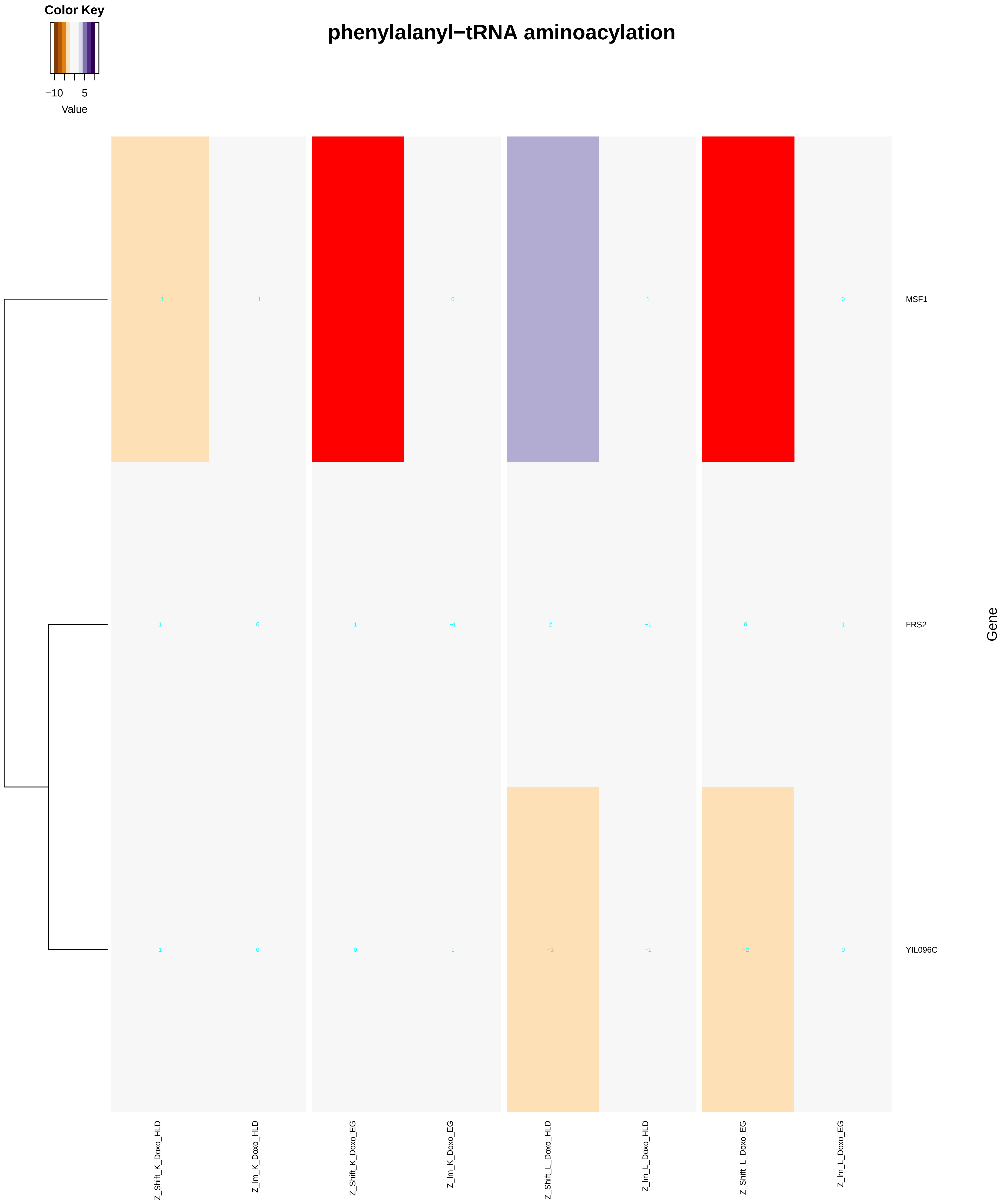

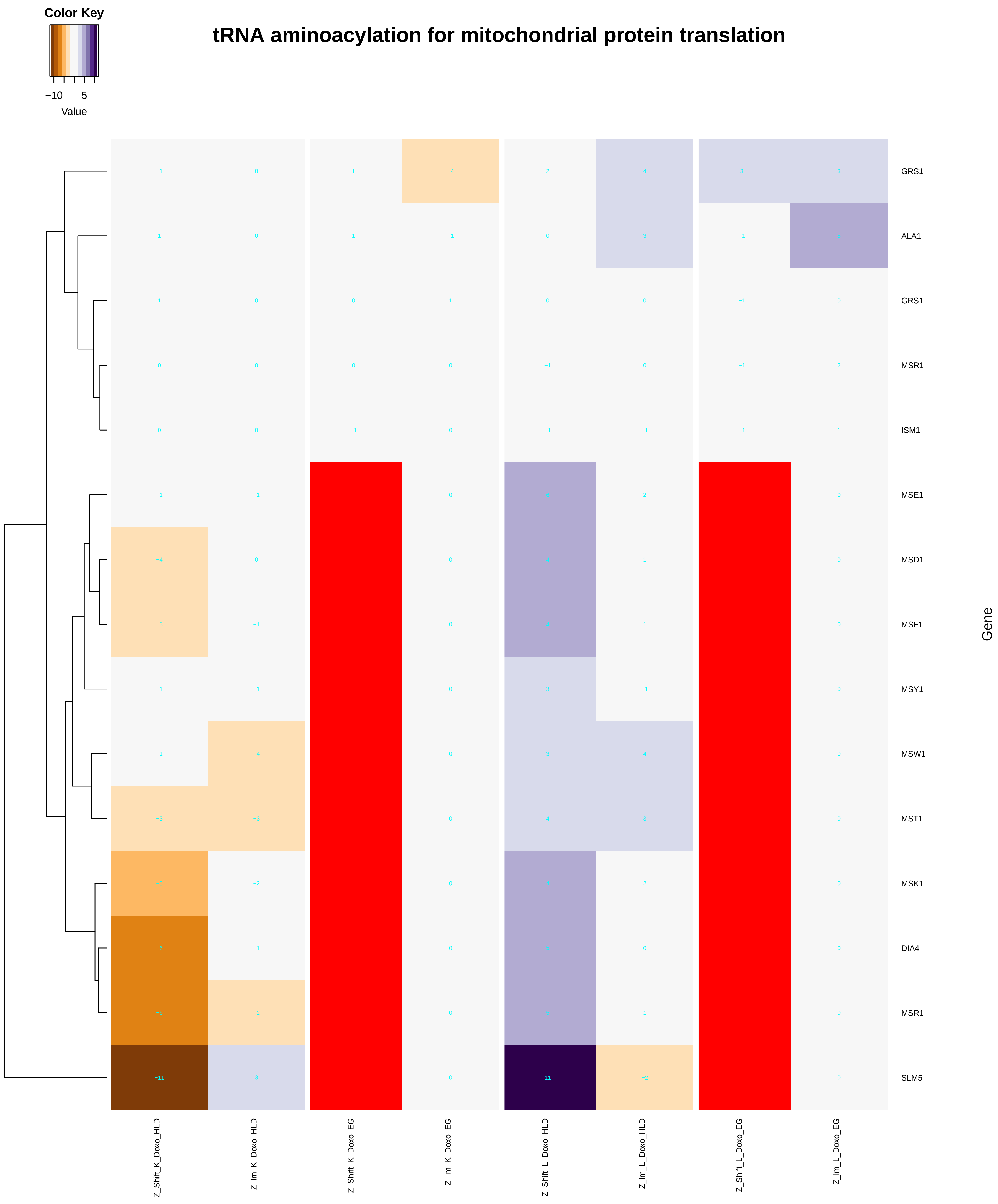

### anatomical_structure_homeostasis.pdf

#### anatomical structure homeostasis

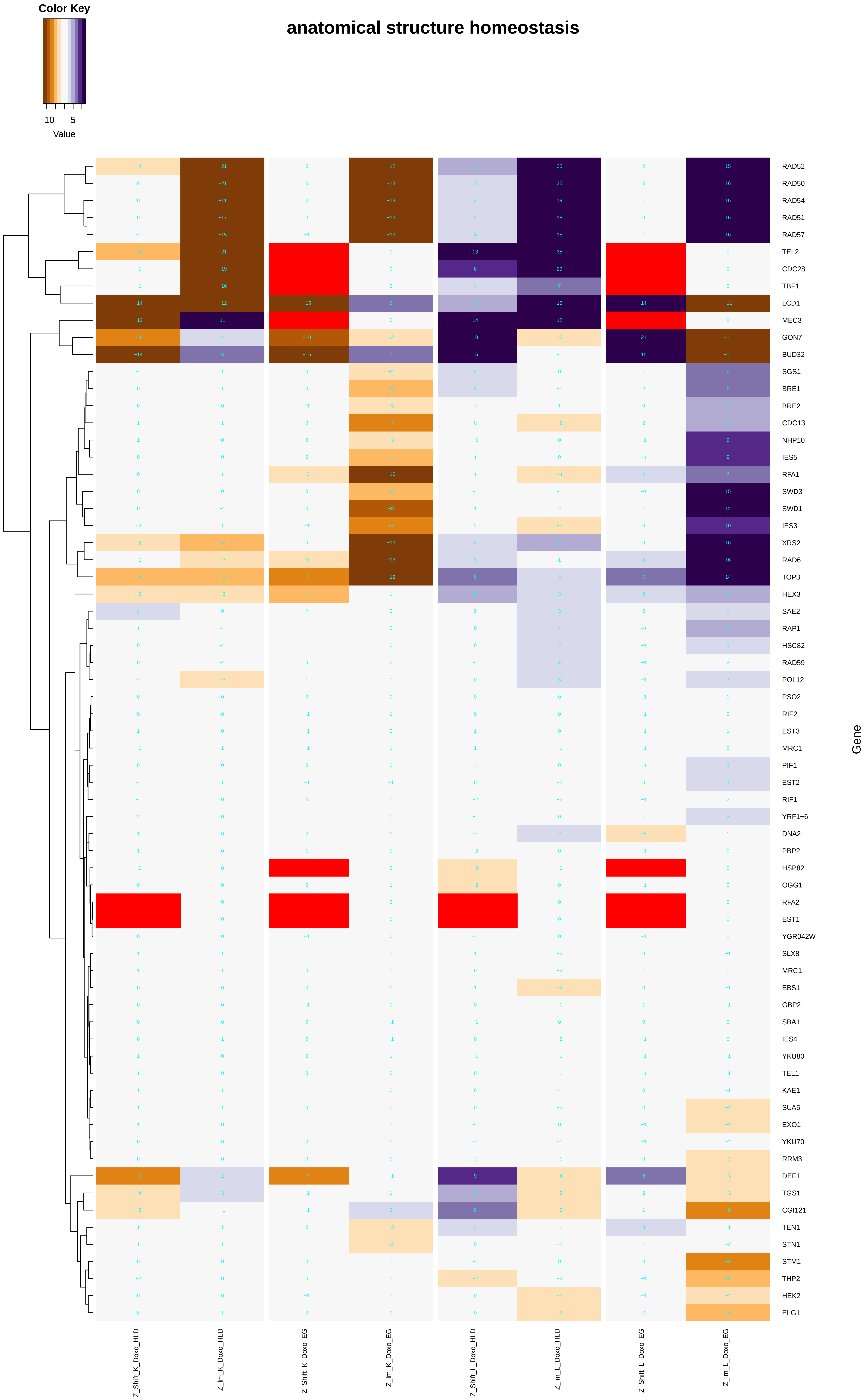

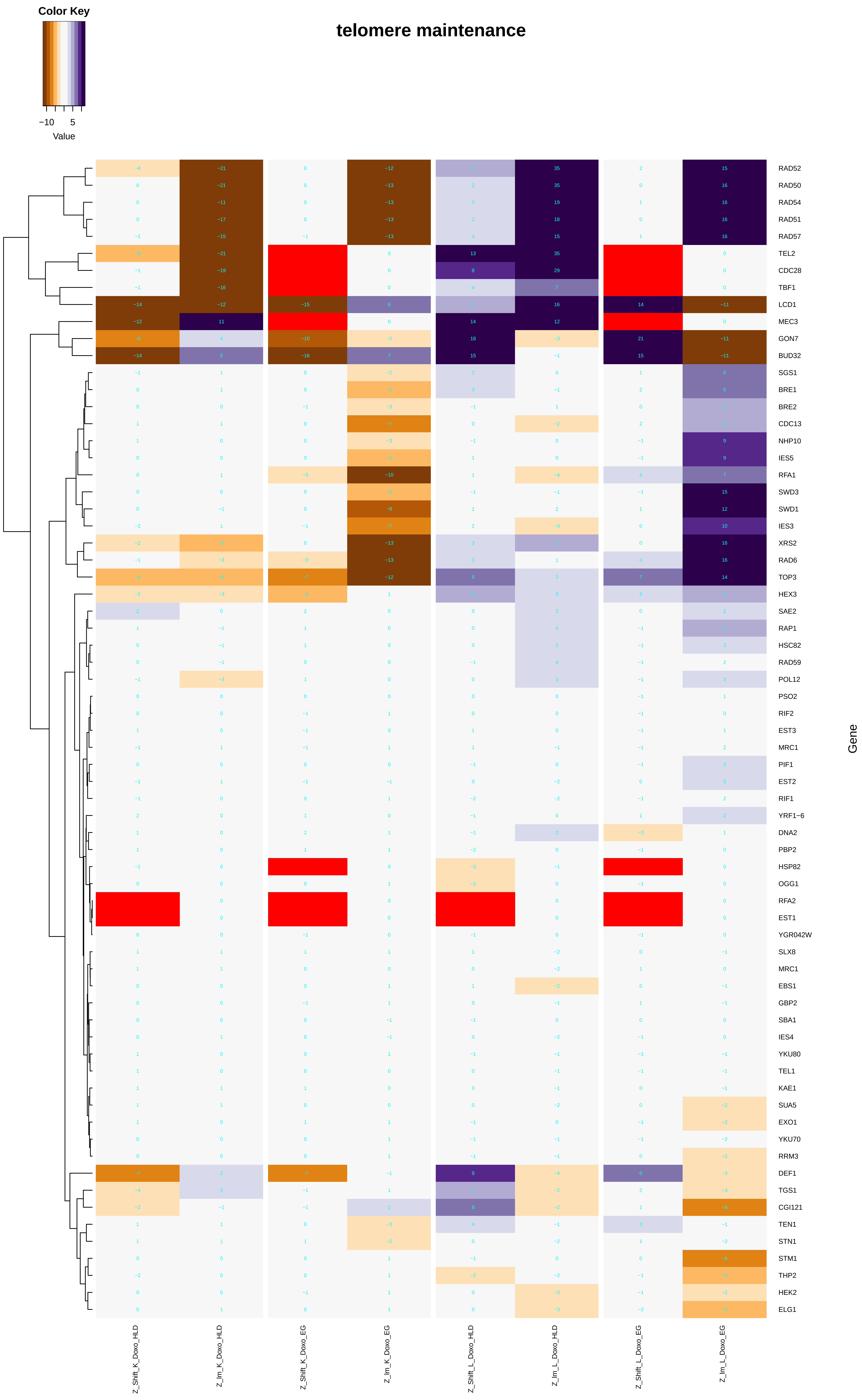

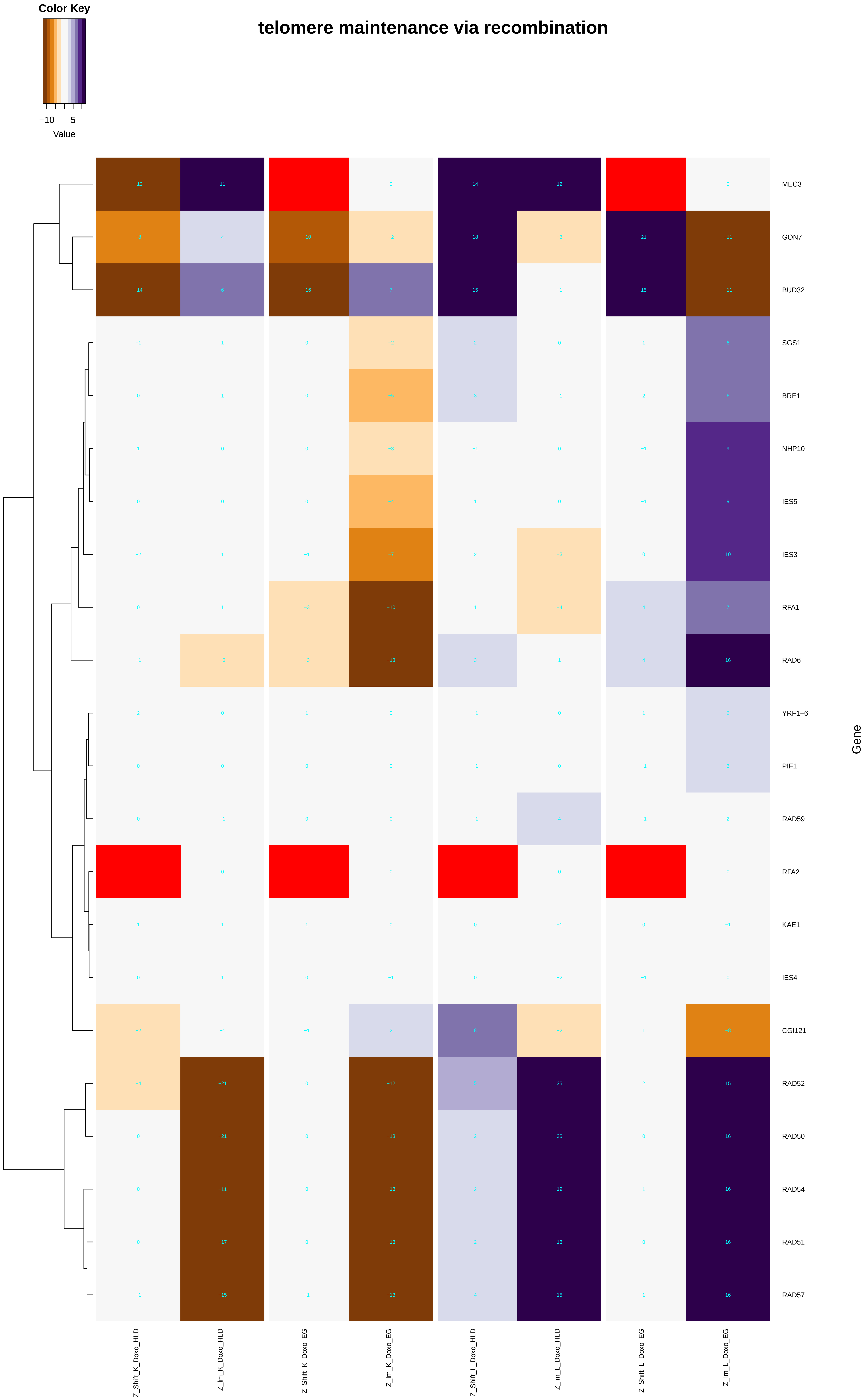

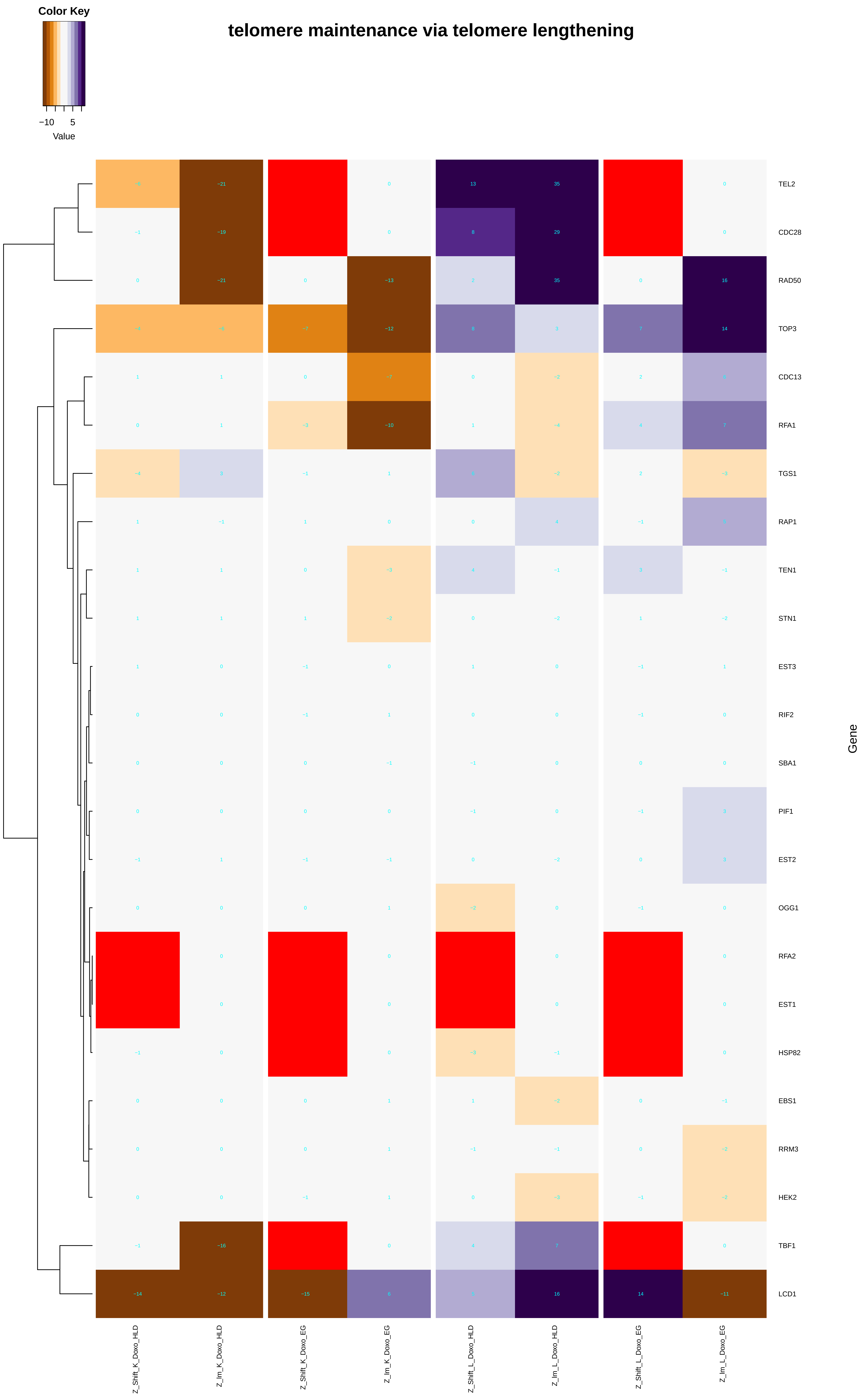

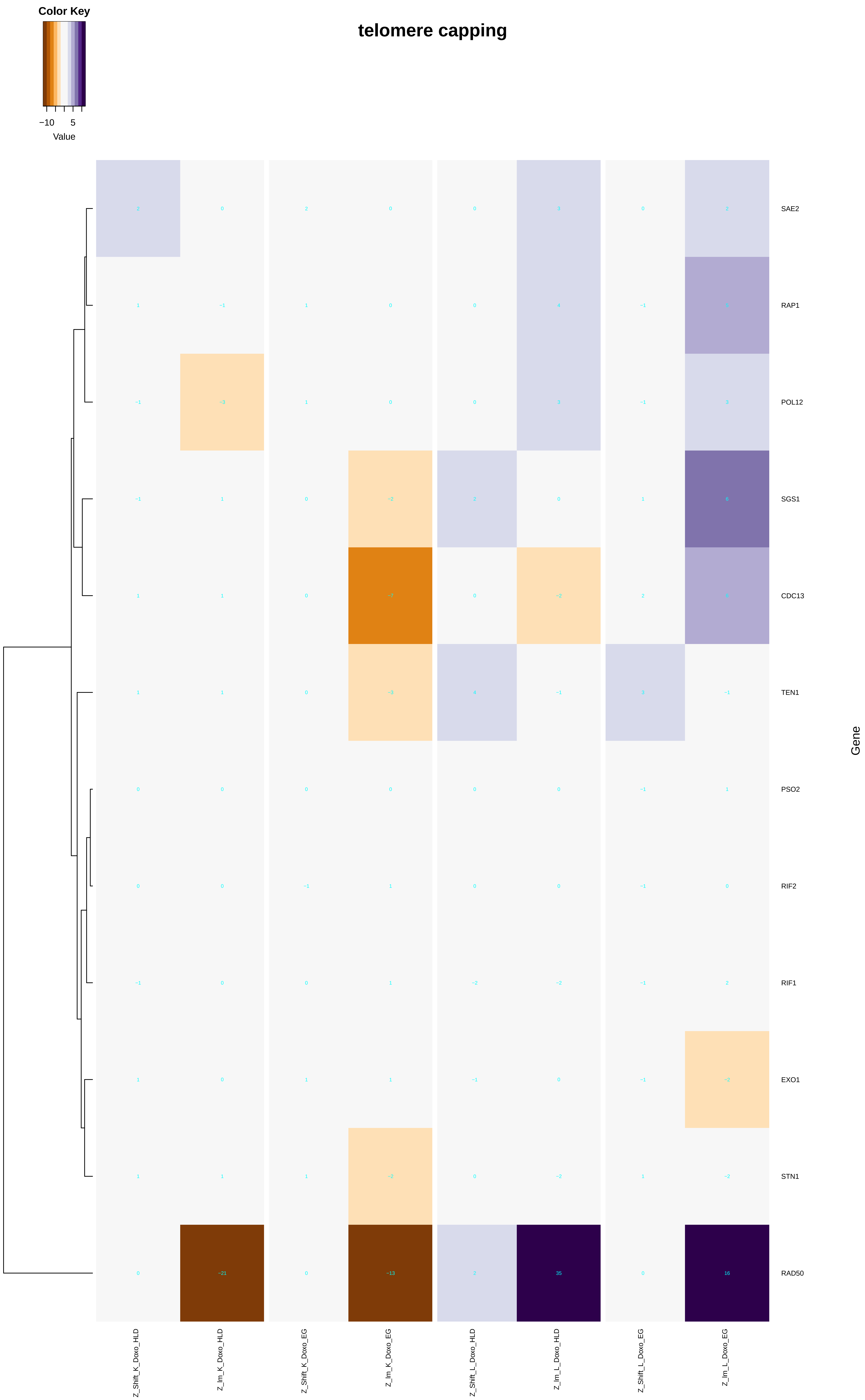

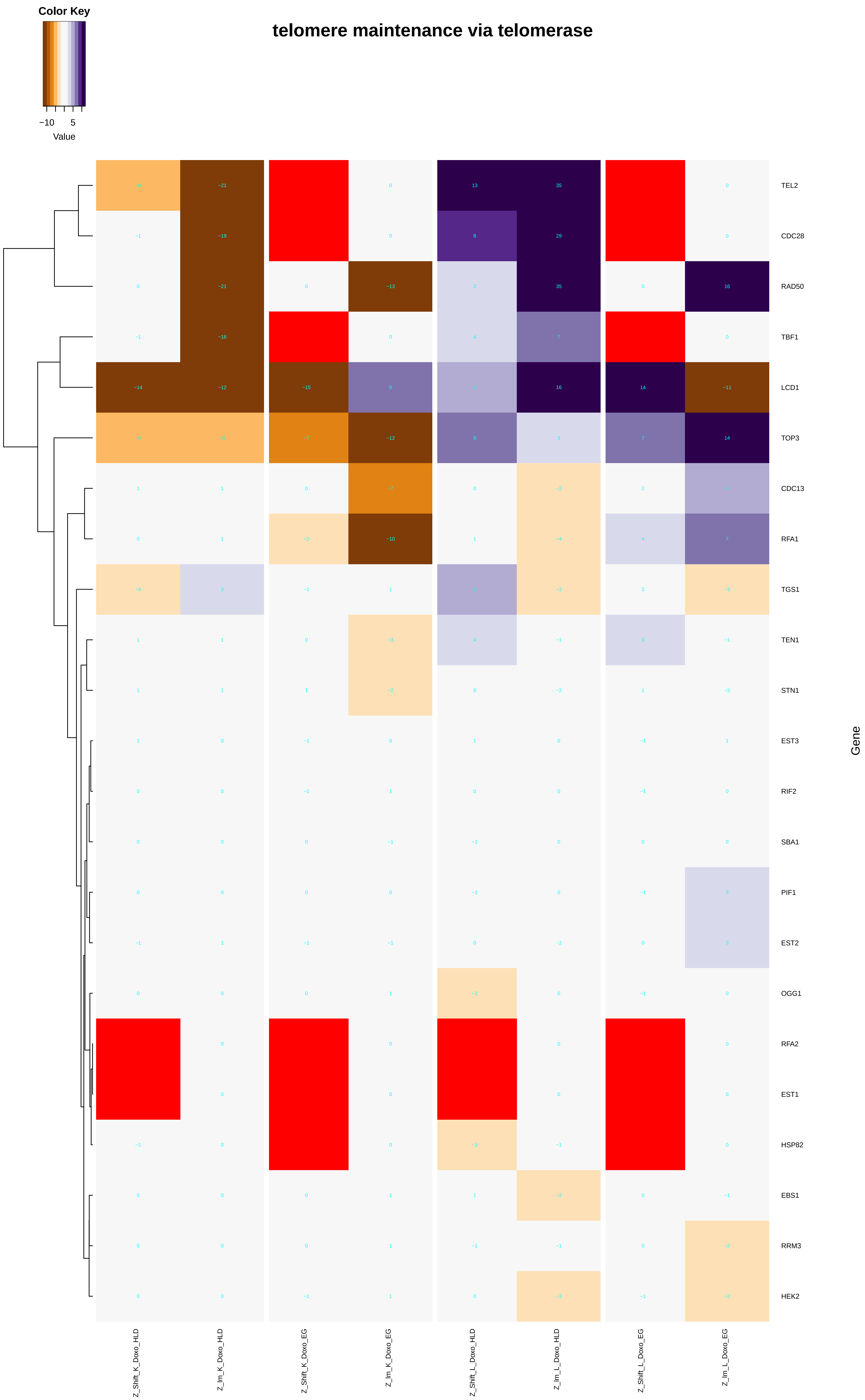

### asparagine_metabolic_process.pdf

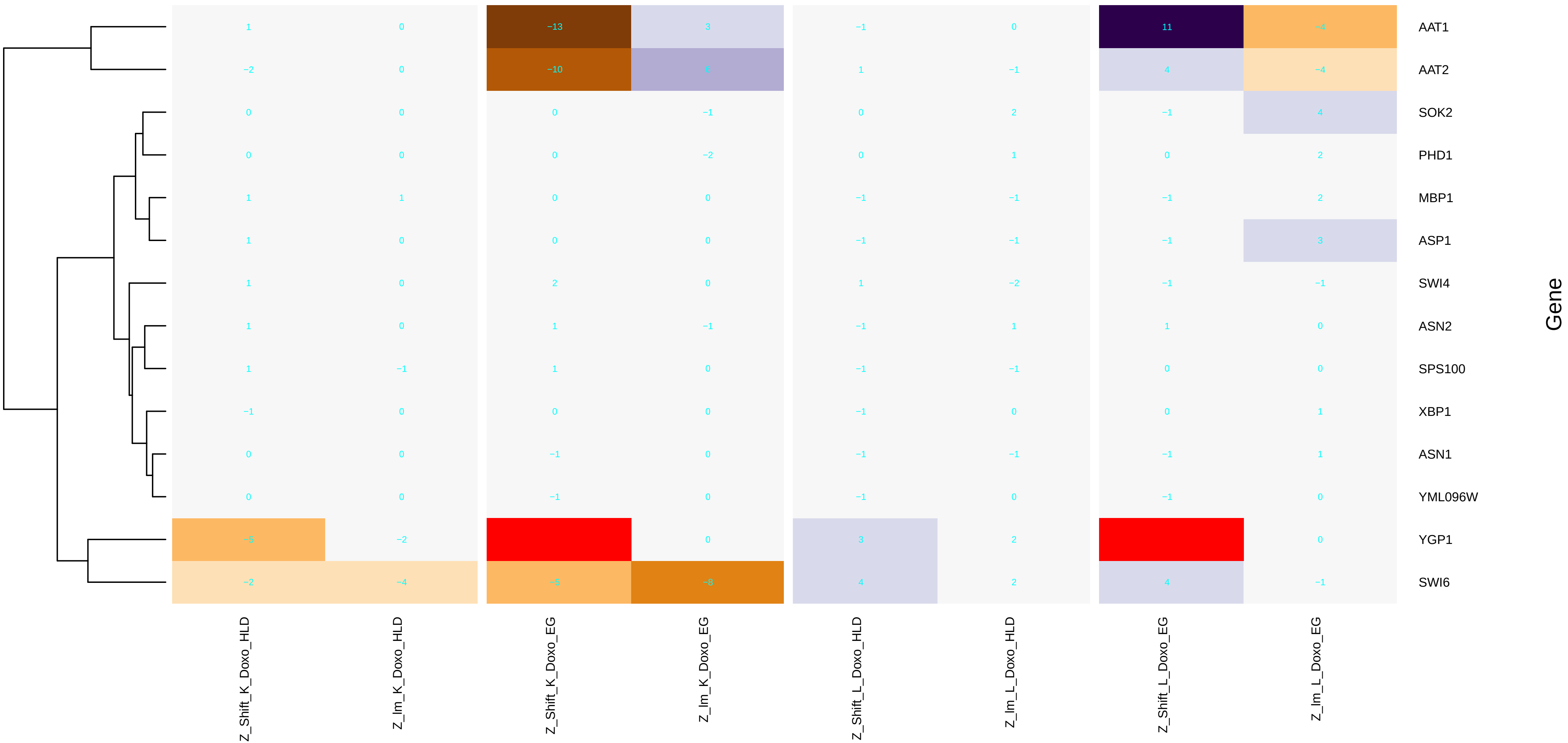

### aspartate_metabolic_process.pdf

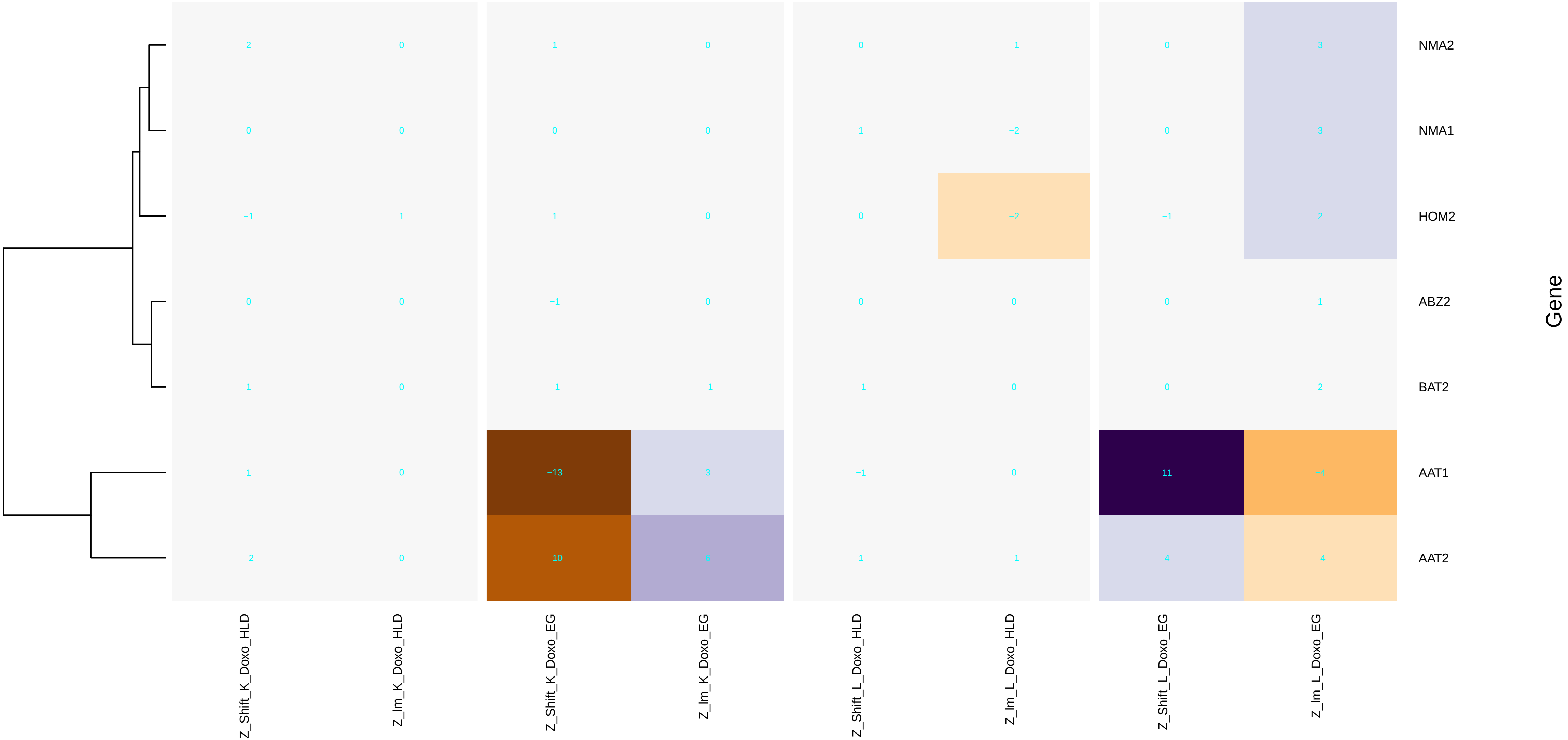

### ATP-dependent_chromatin_remodeling.pdf

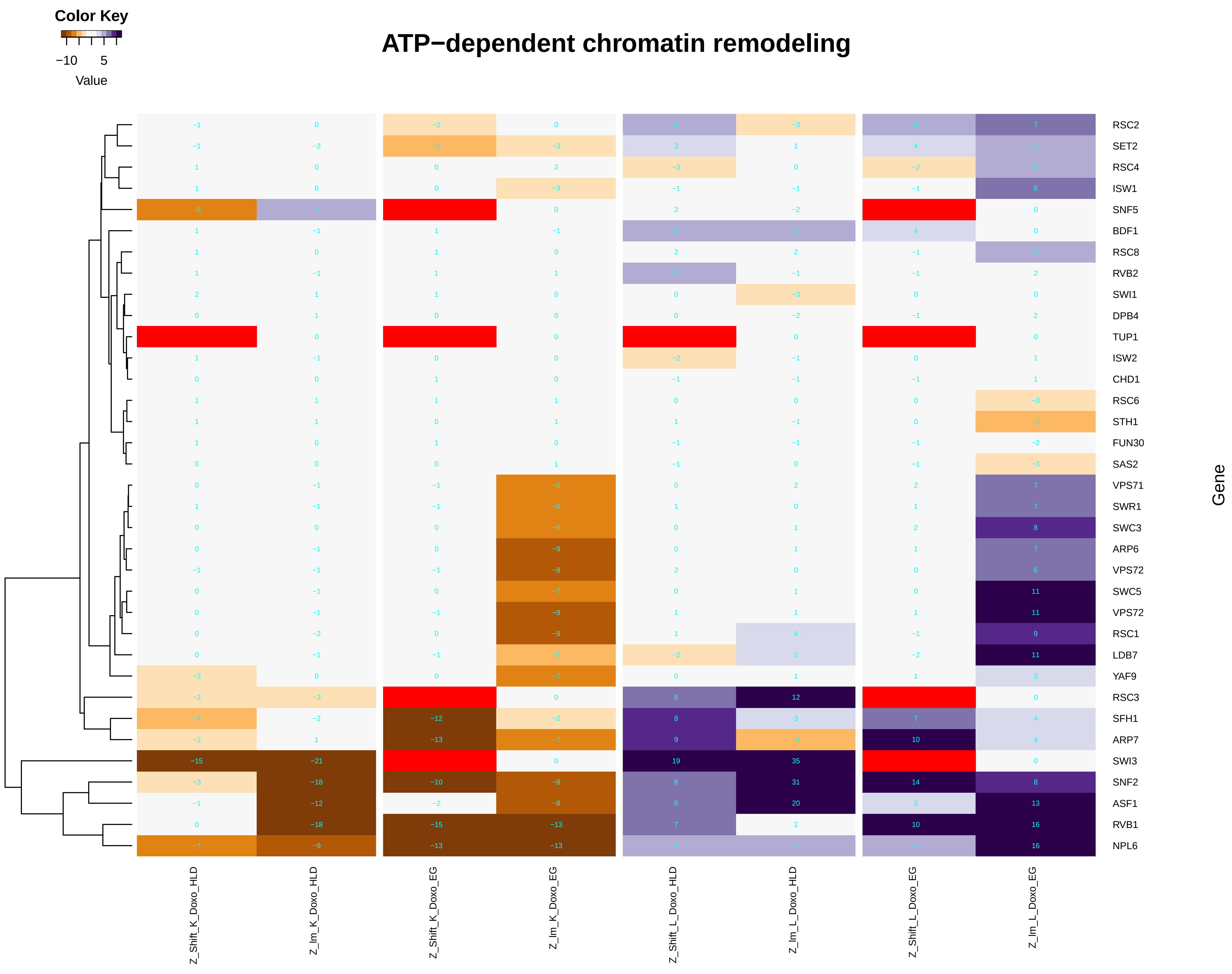

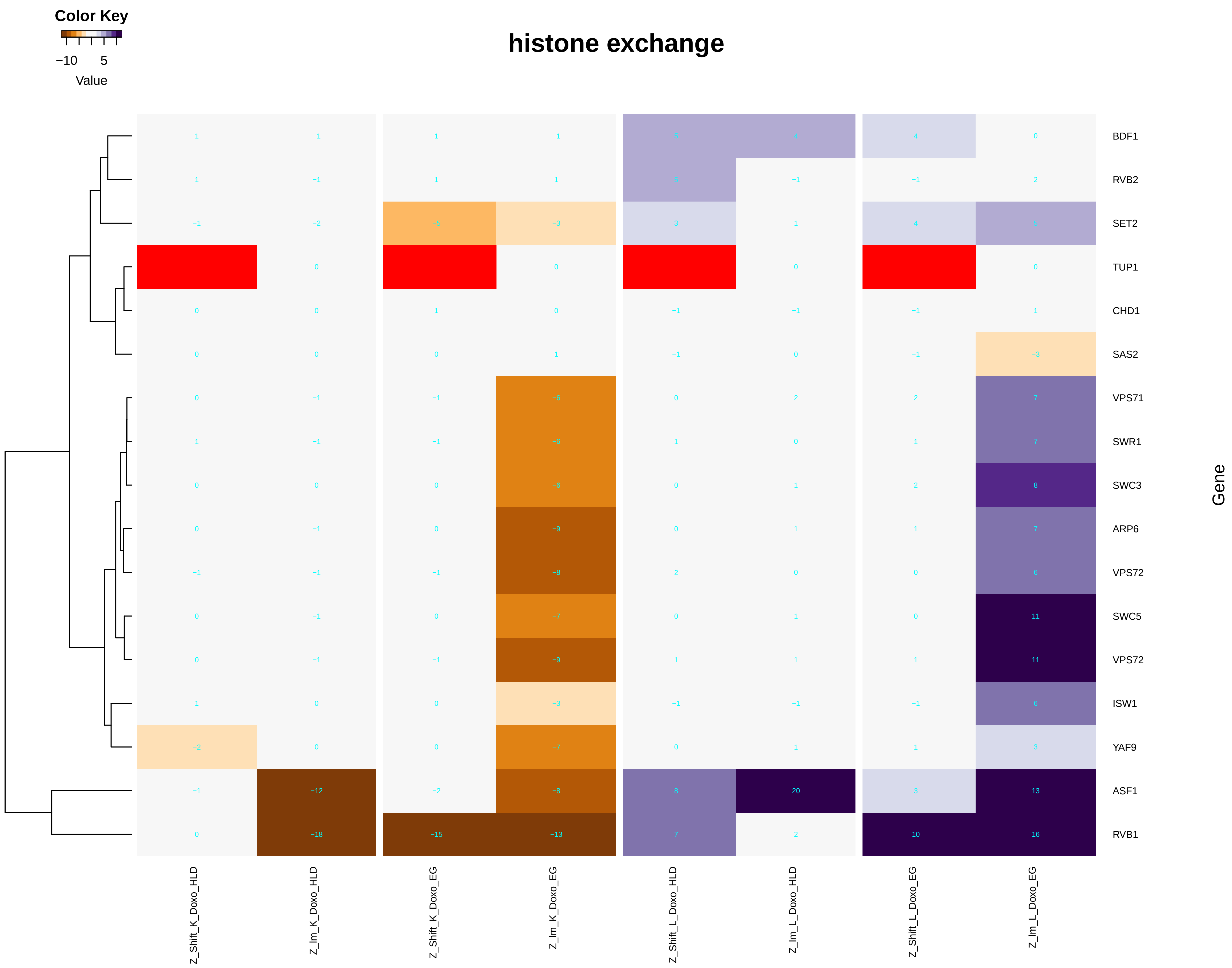

### ATP_biosynthetic_process.pdf

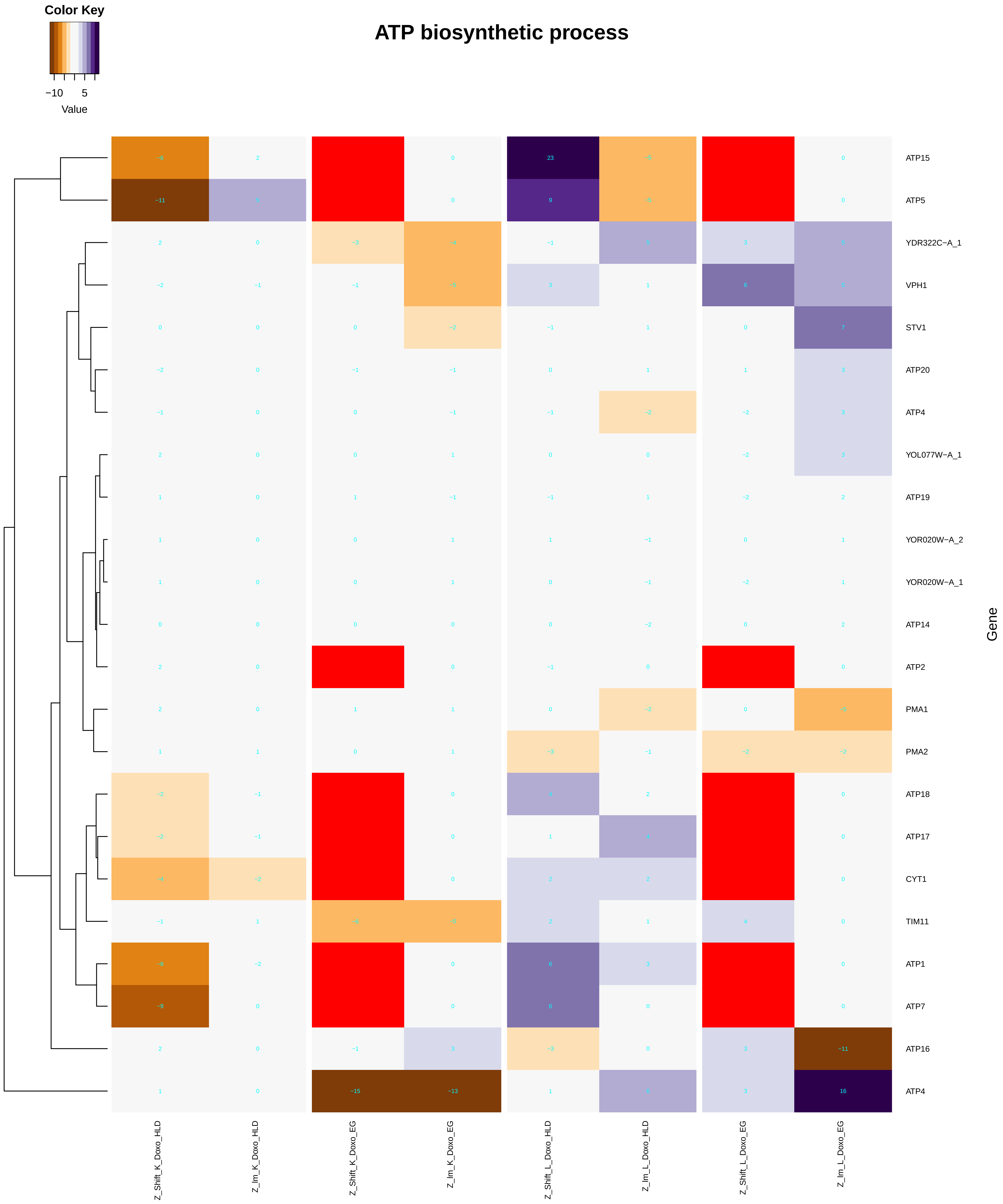

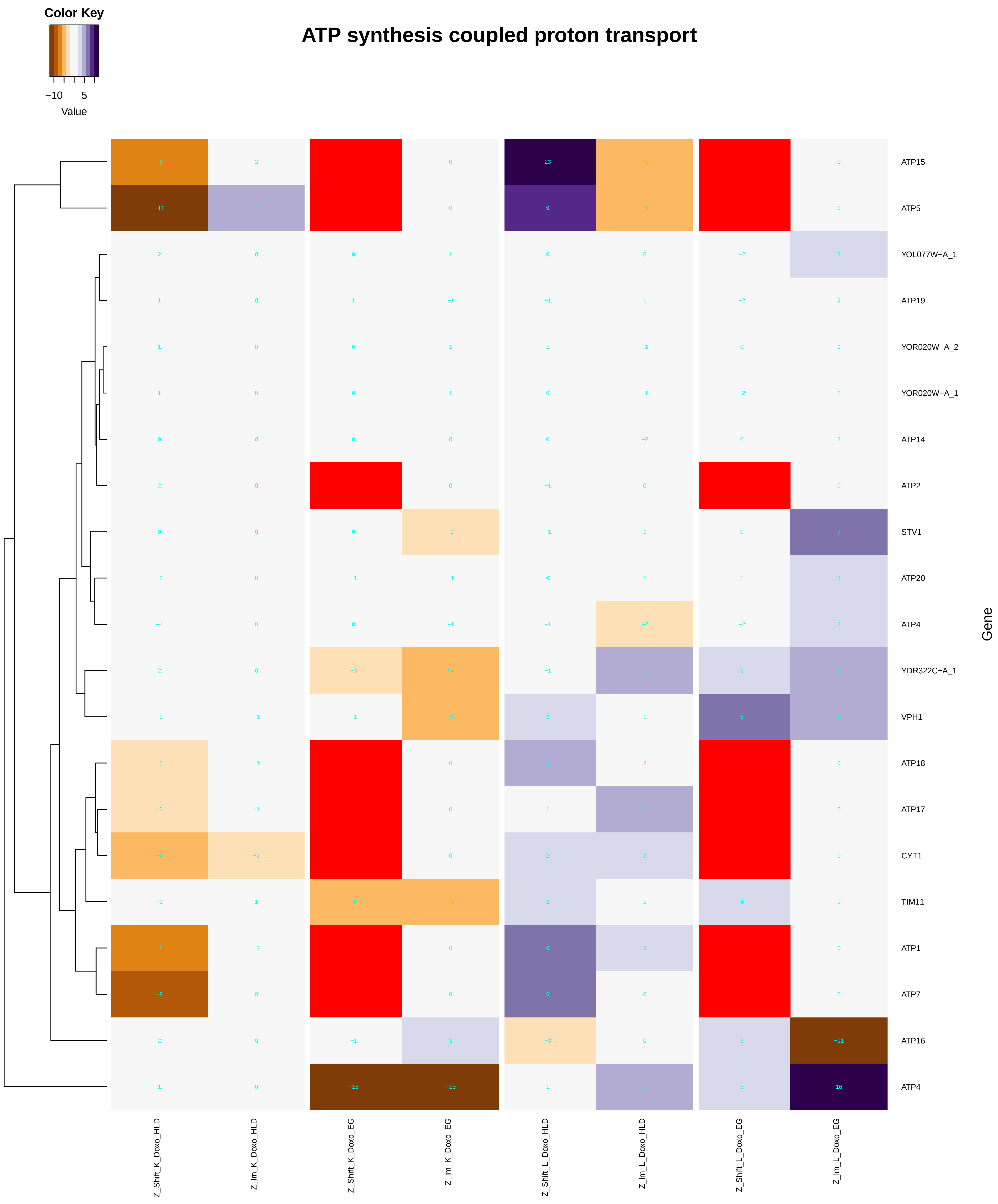

### ATP_hydrolysis_coupled_cation_transmembrane_transport.pdf

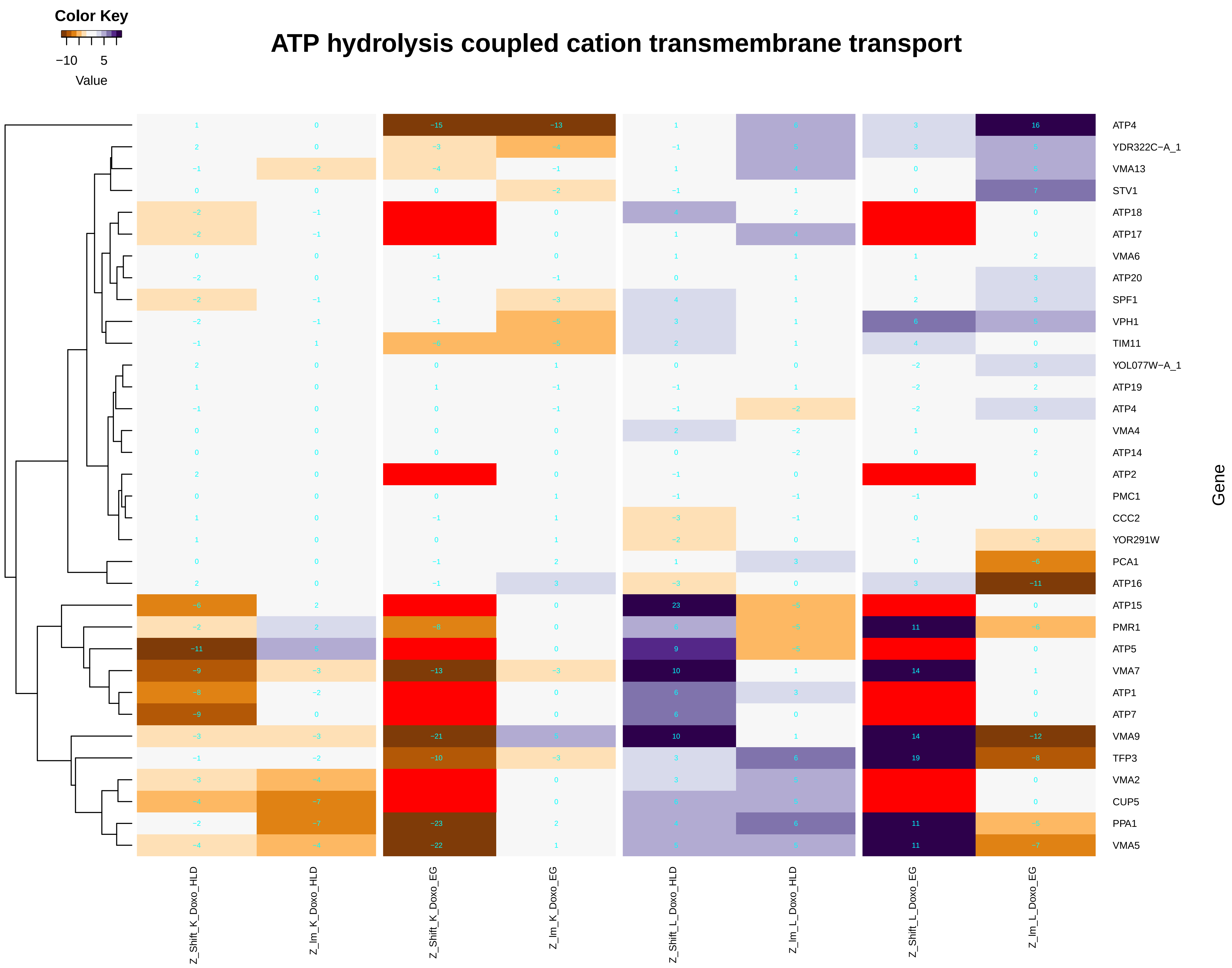

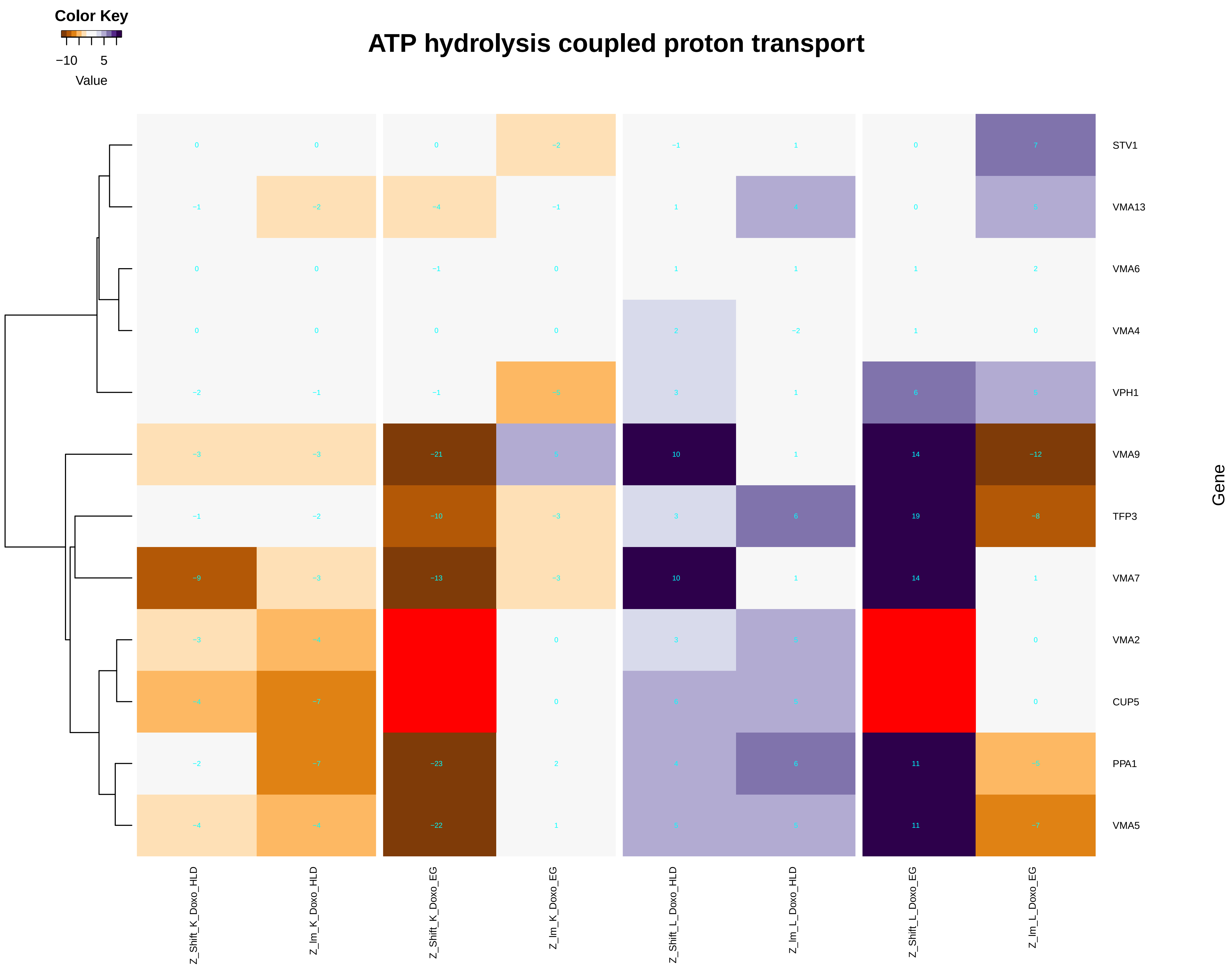

### ATP_hydrolysis_coupled_ion_transmembrane_transport.pdf

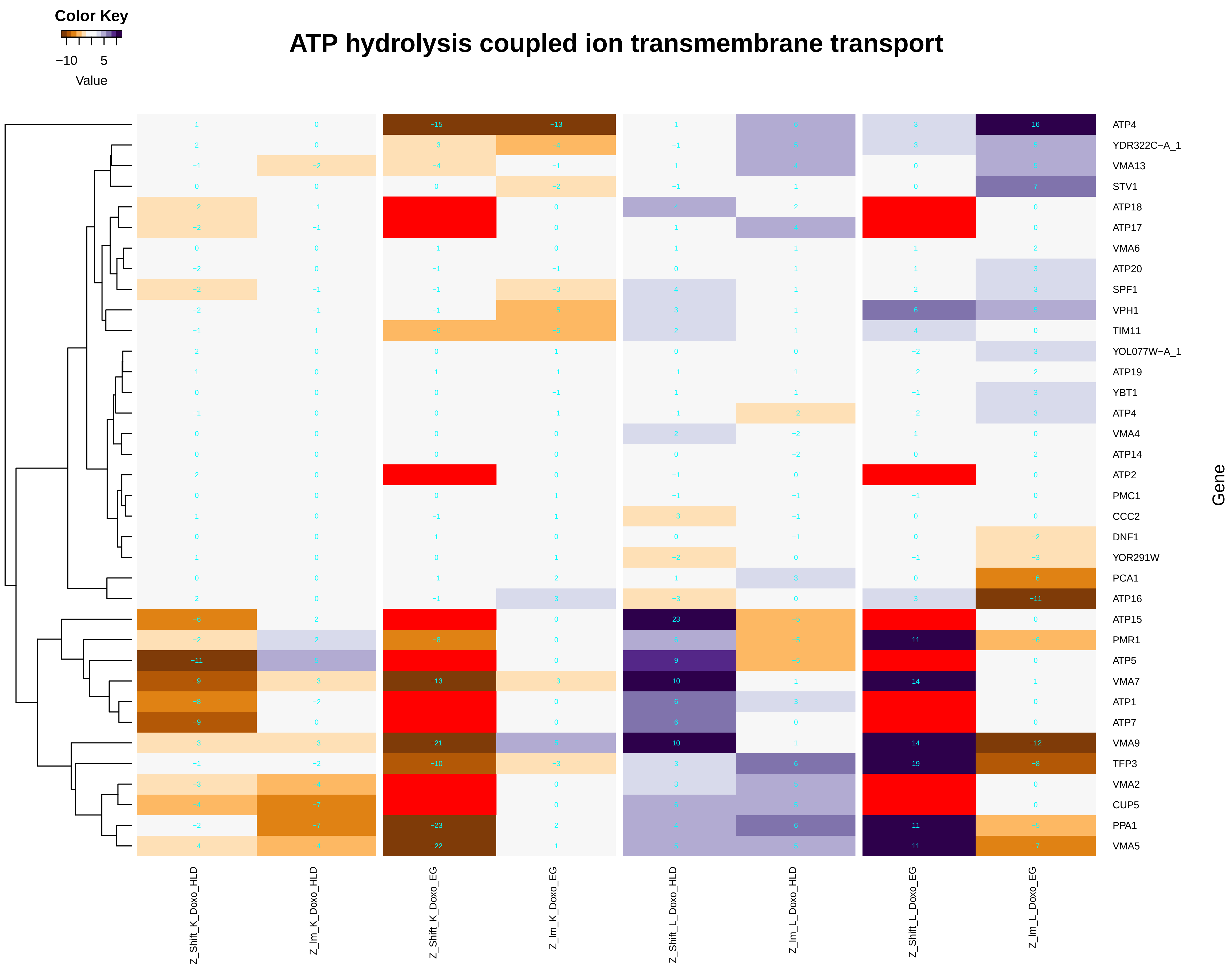

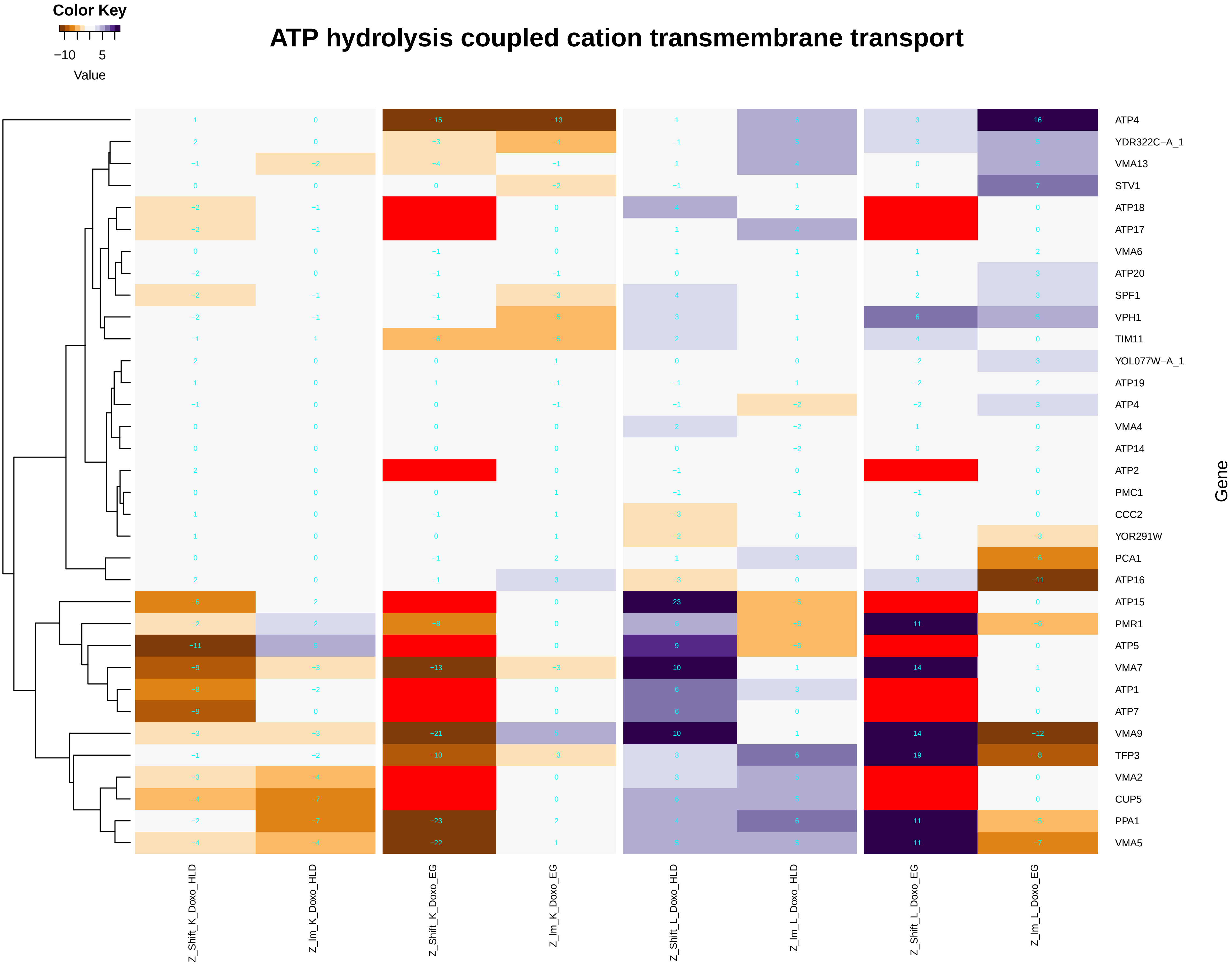

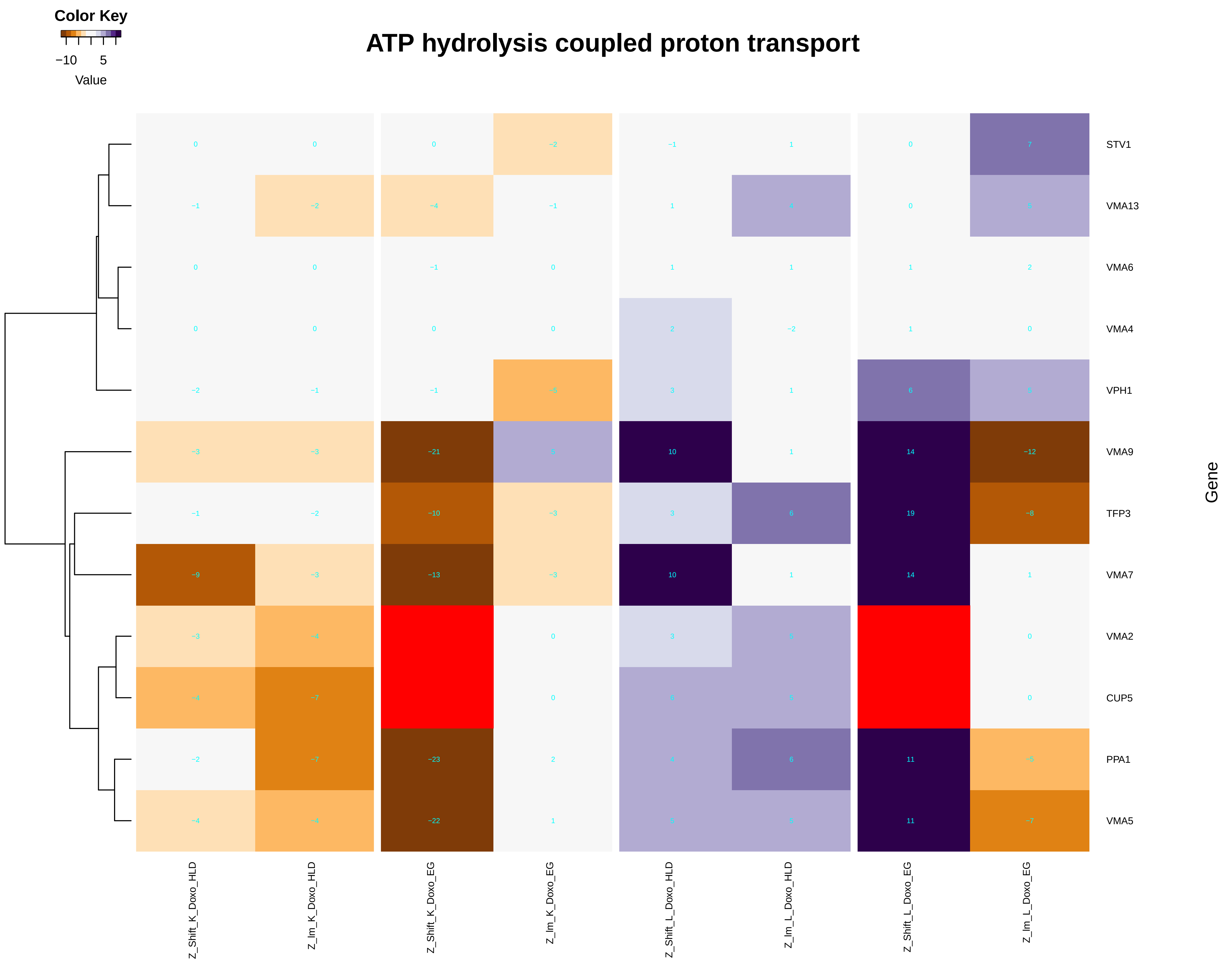

### ATP_hydrolysis_coupled_proton_transport.pdf

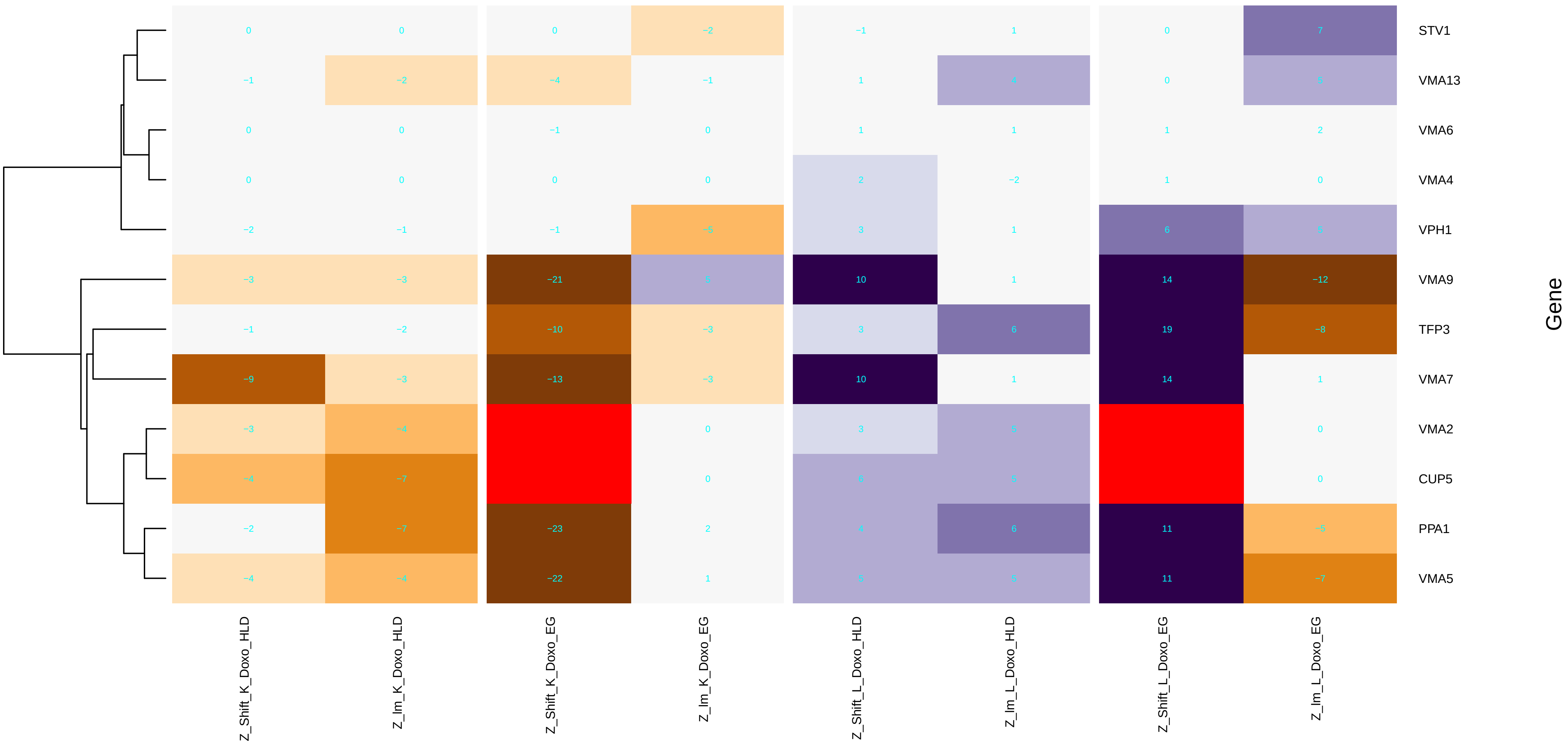

### ATP_hydrolysis_coupled_transmembrane_transport.pdf

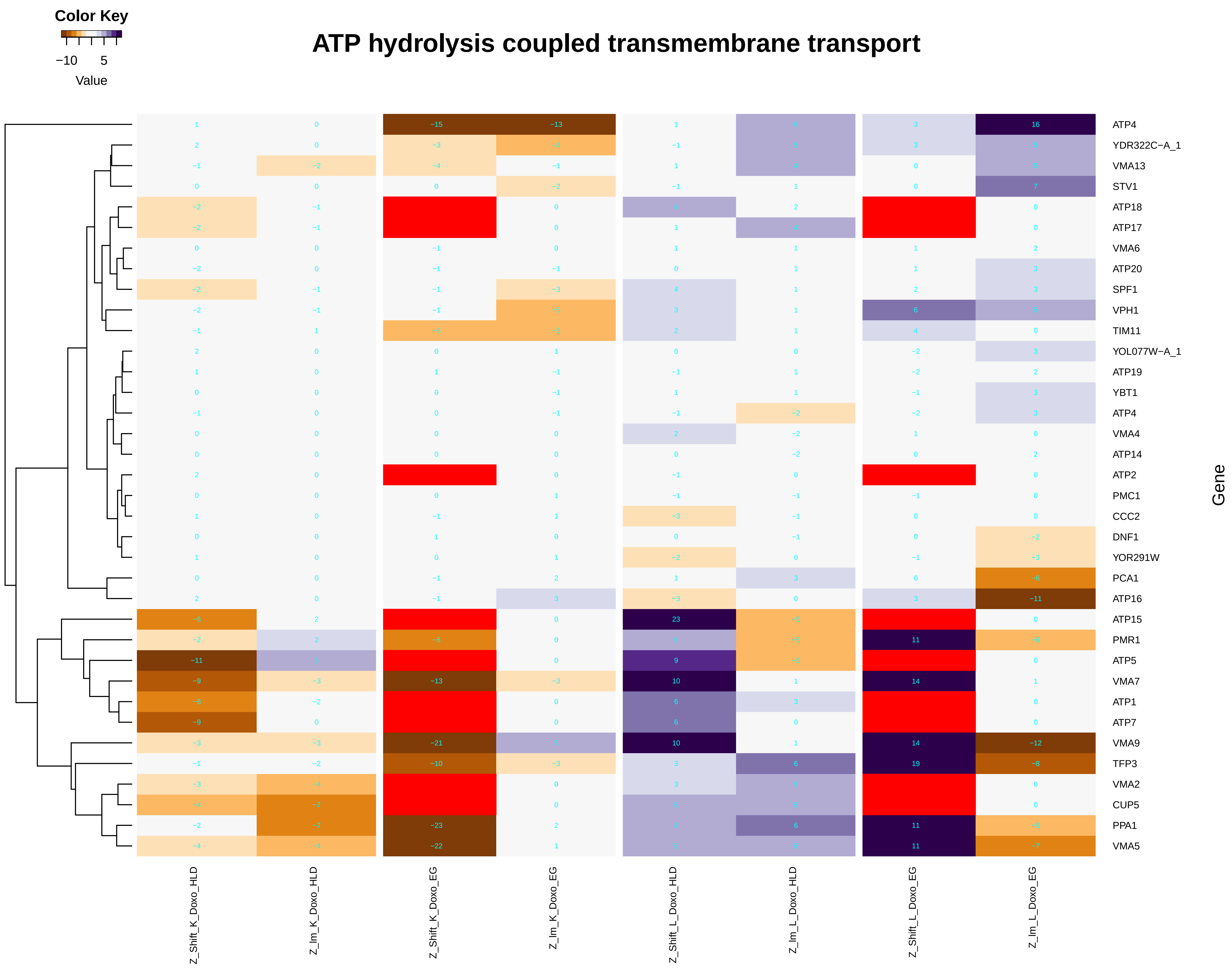

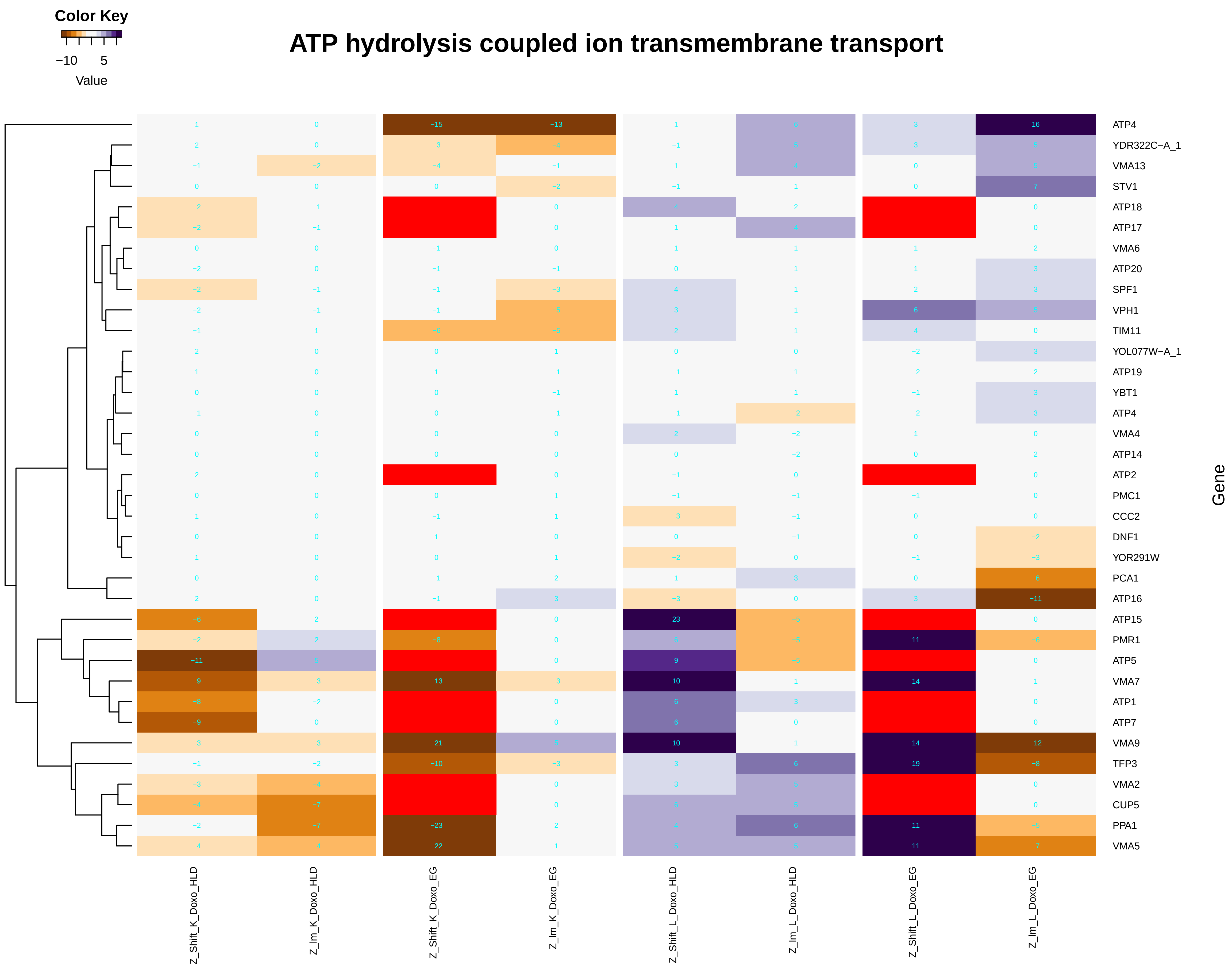

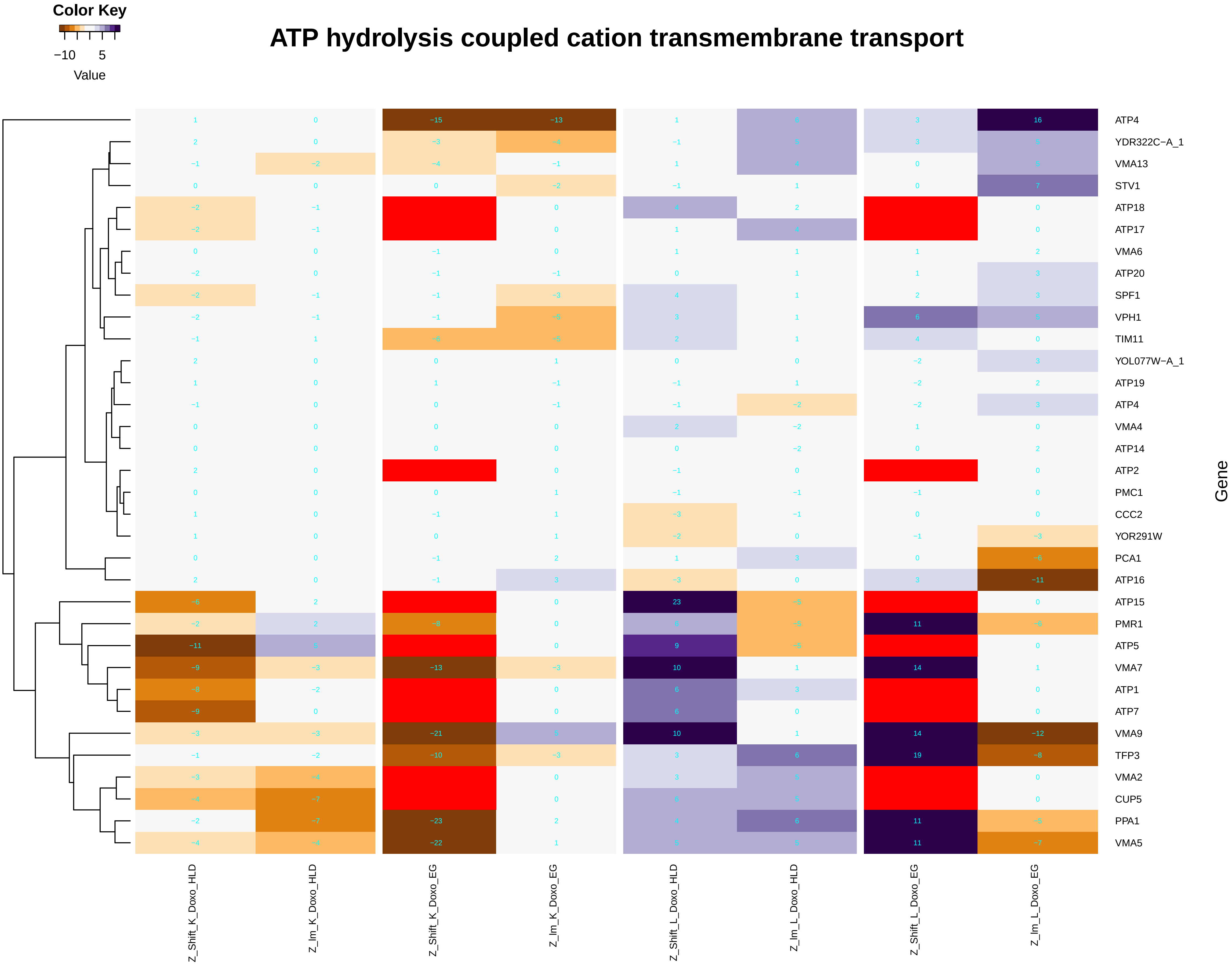

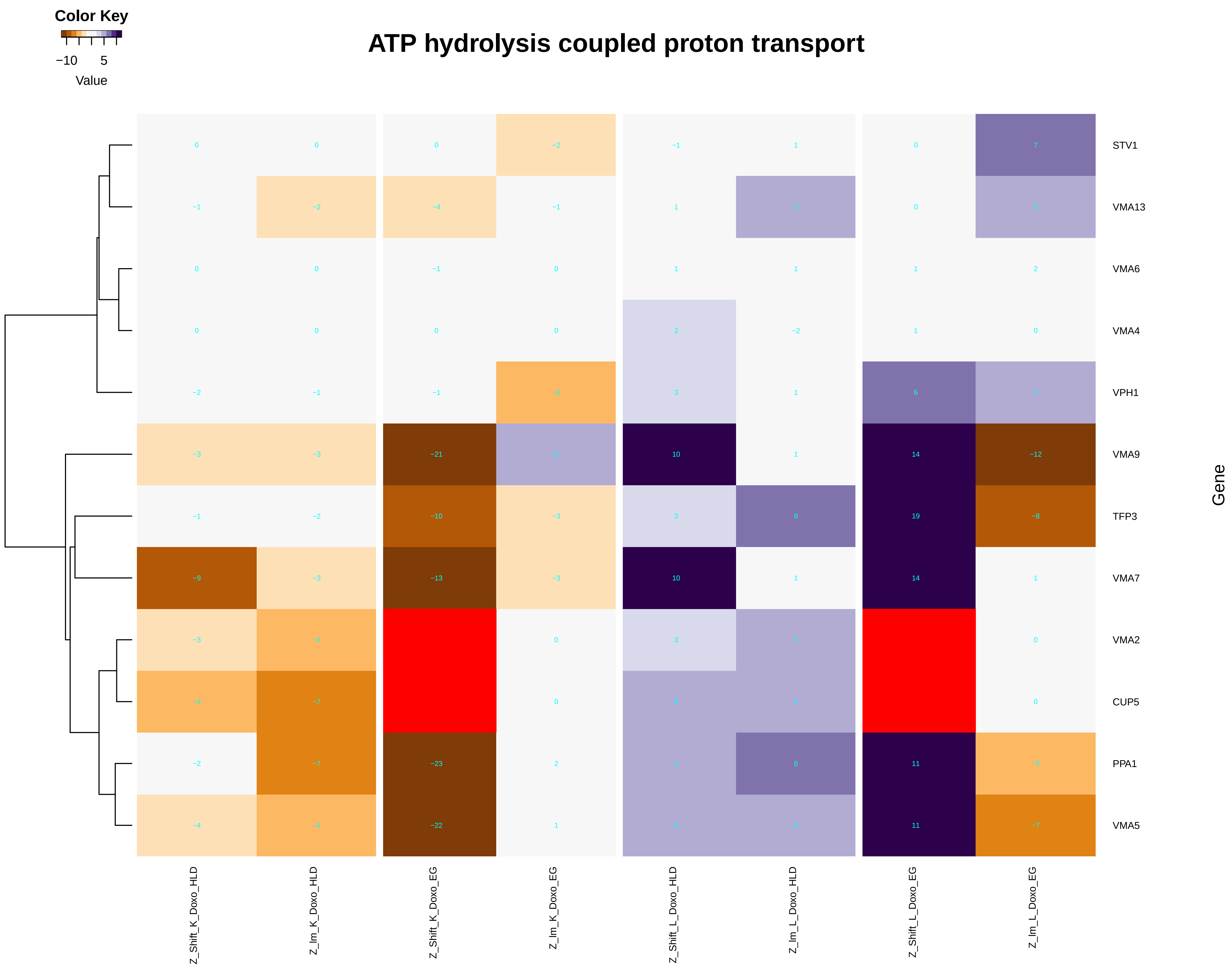

### ATP_metabolic_process.pdf

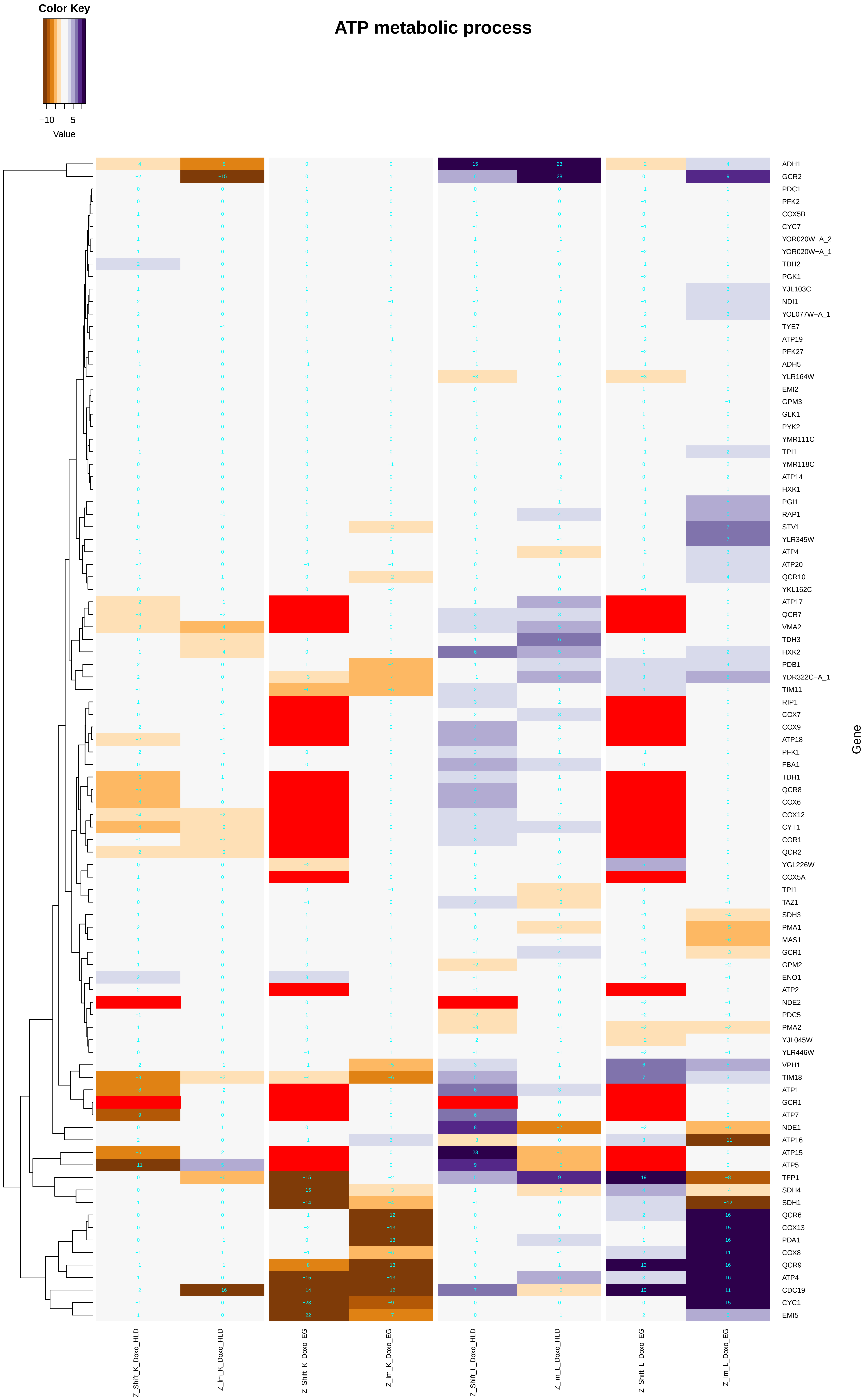

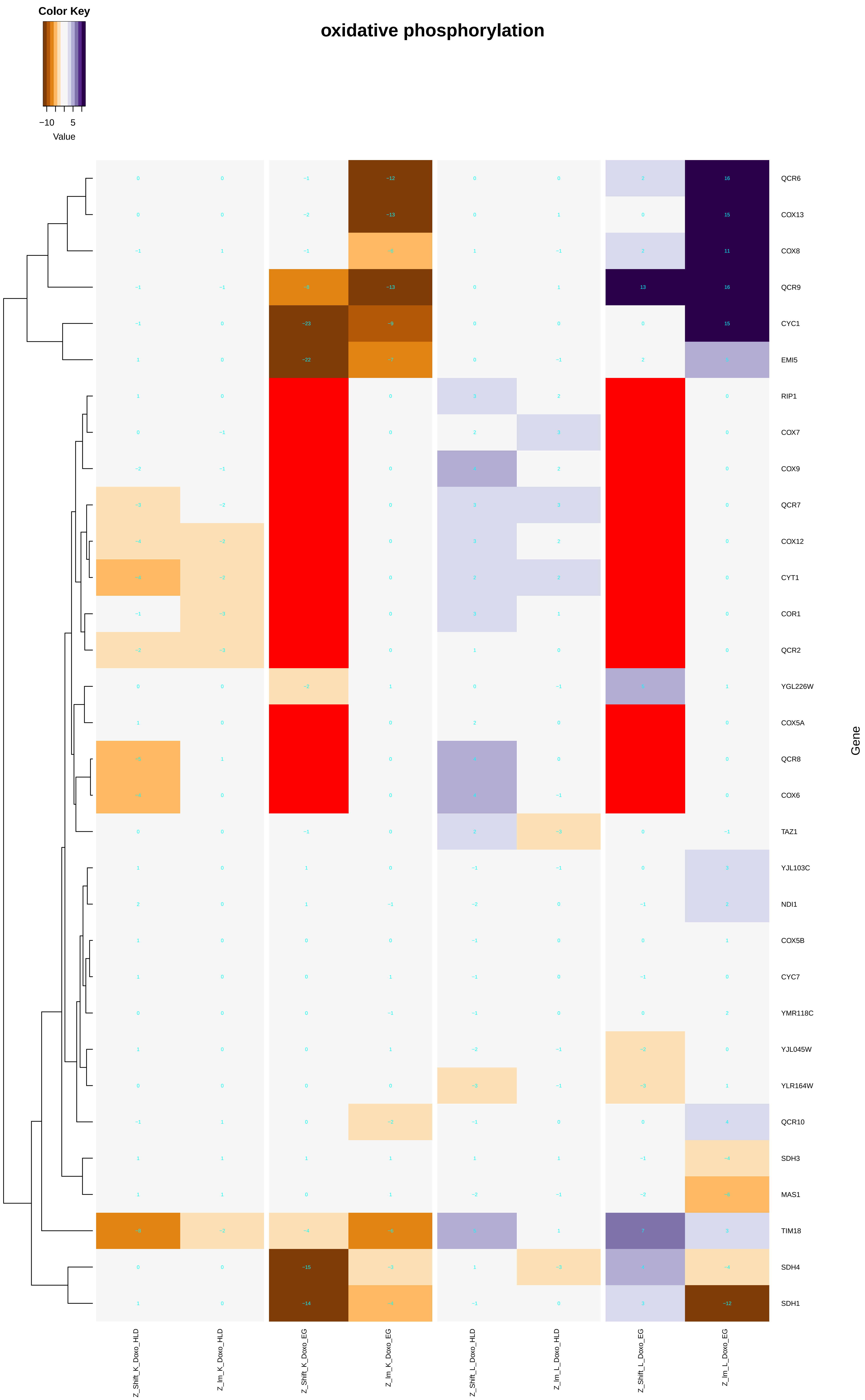

#### ATP biosynthetic process

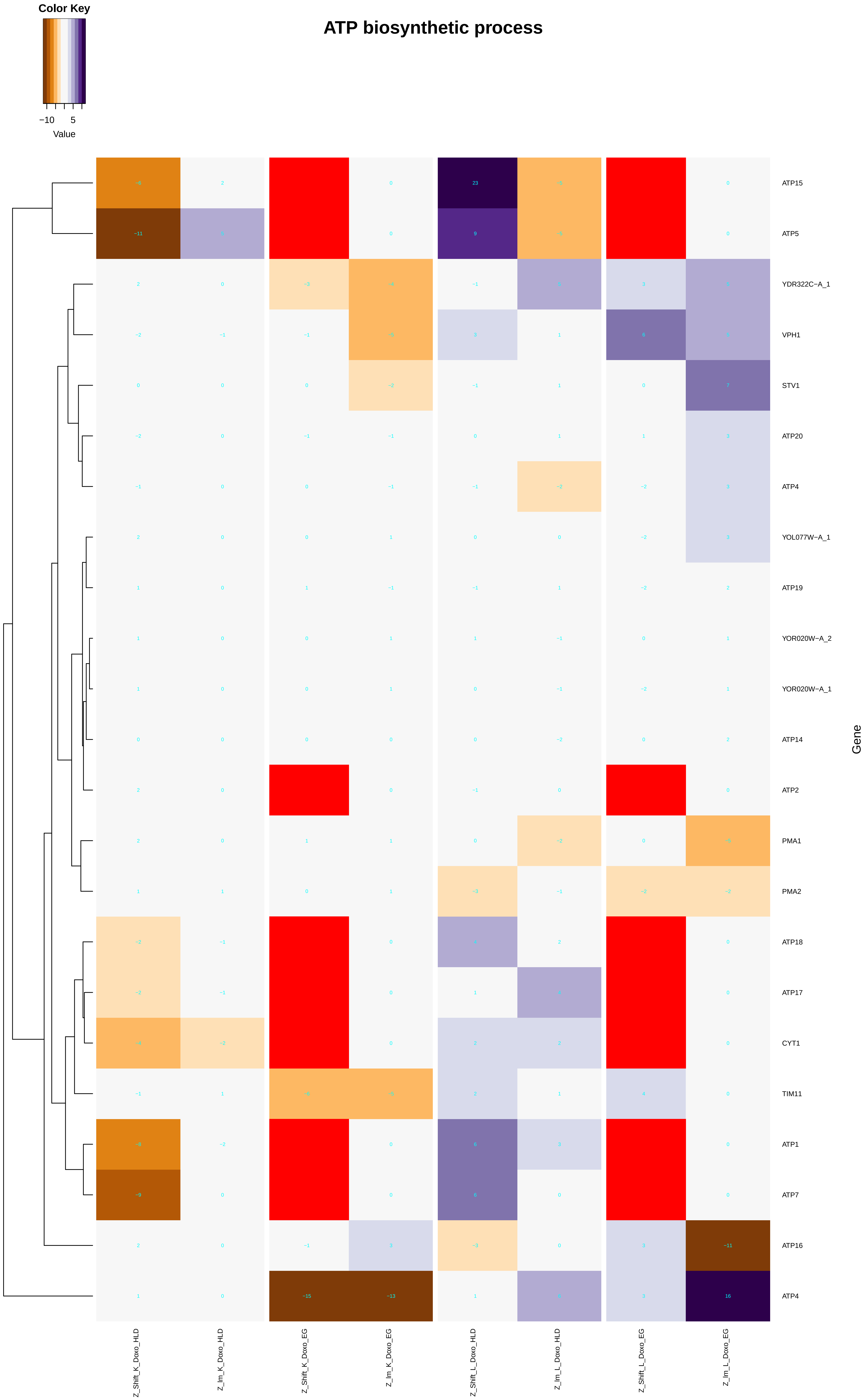

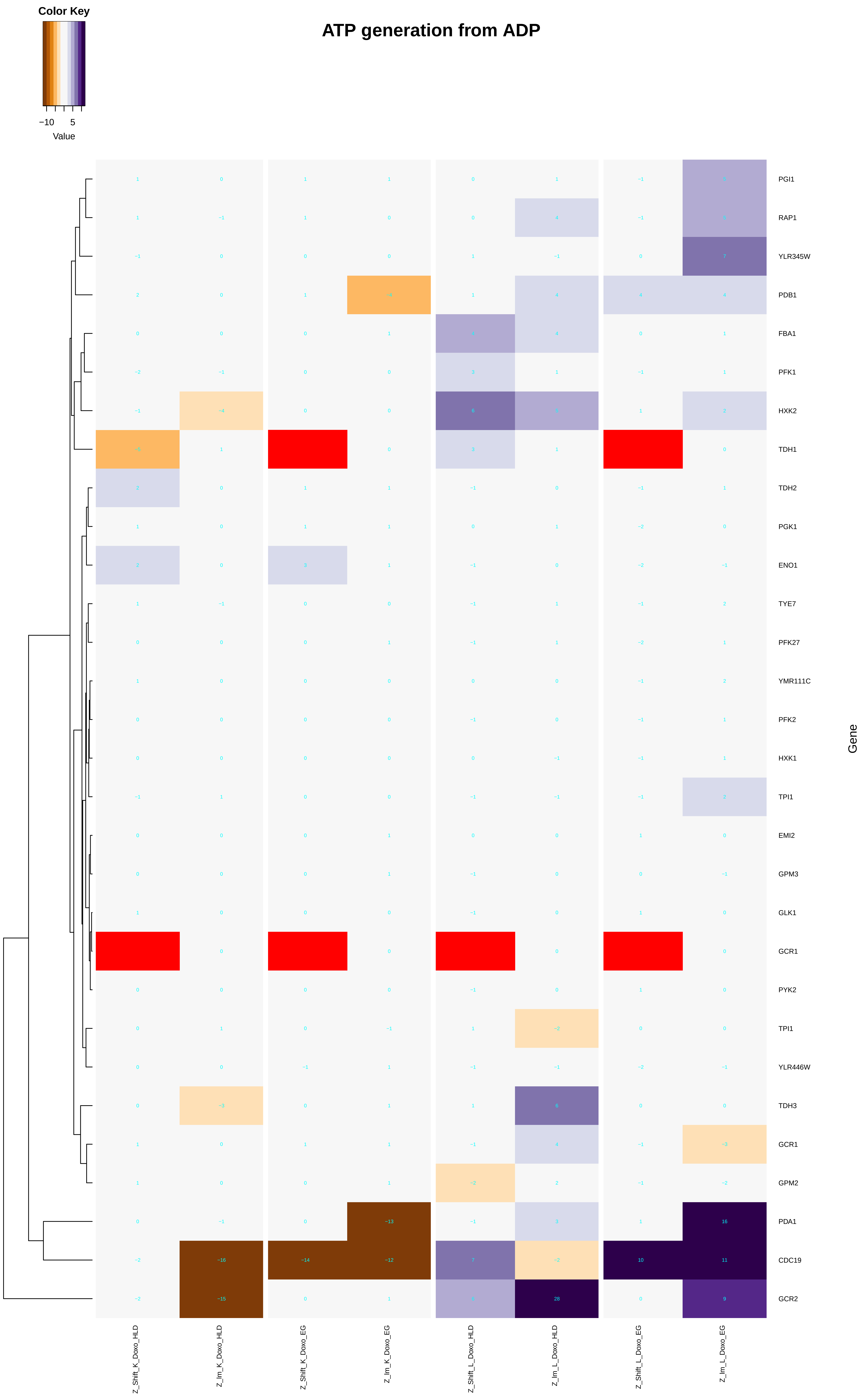

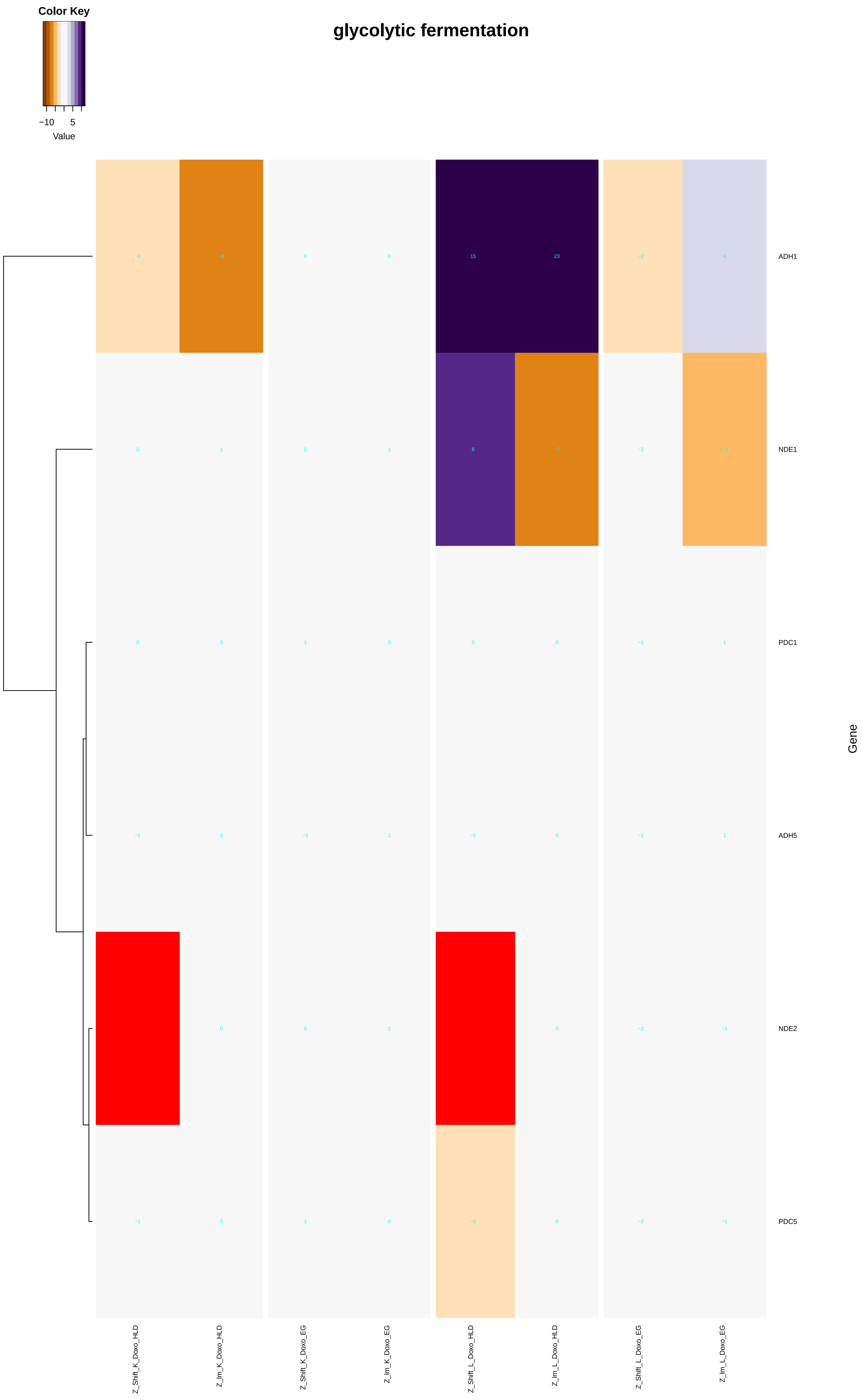

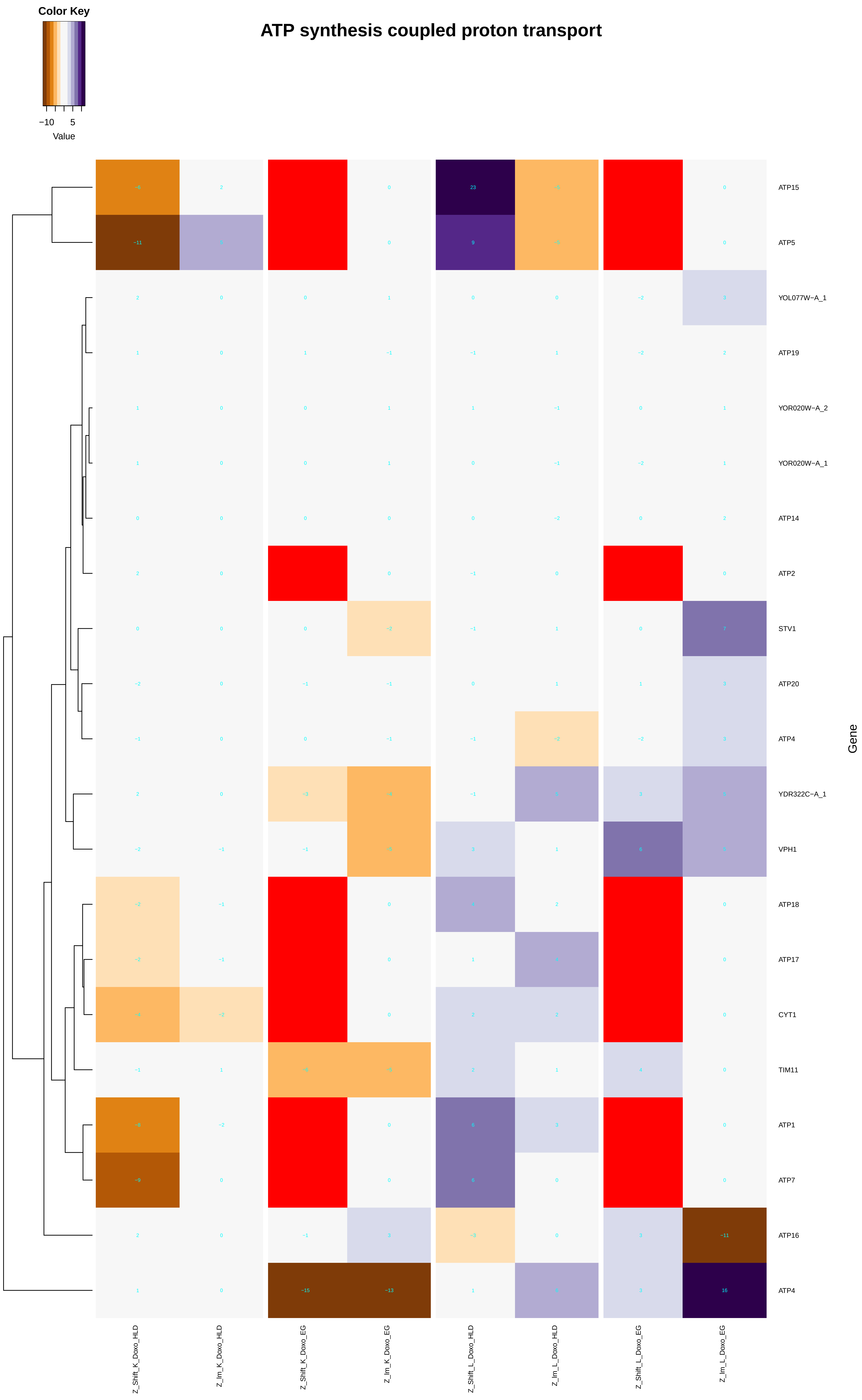

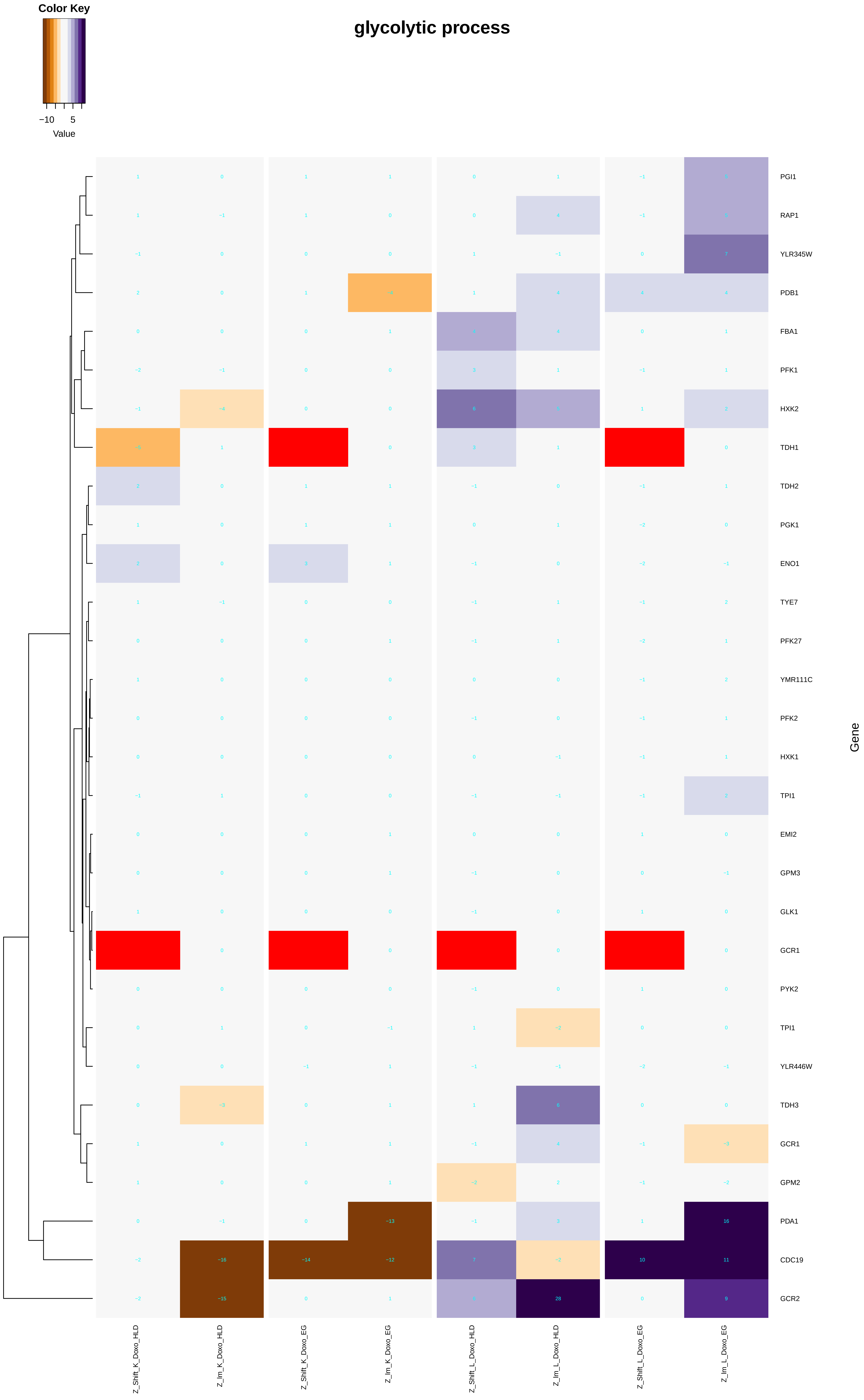

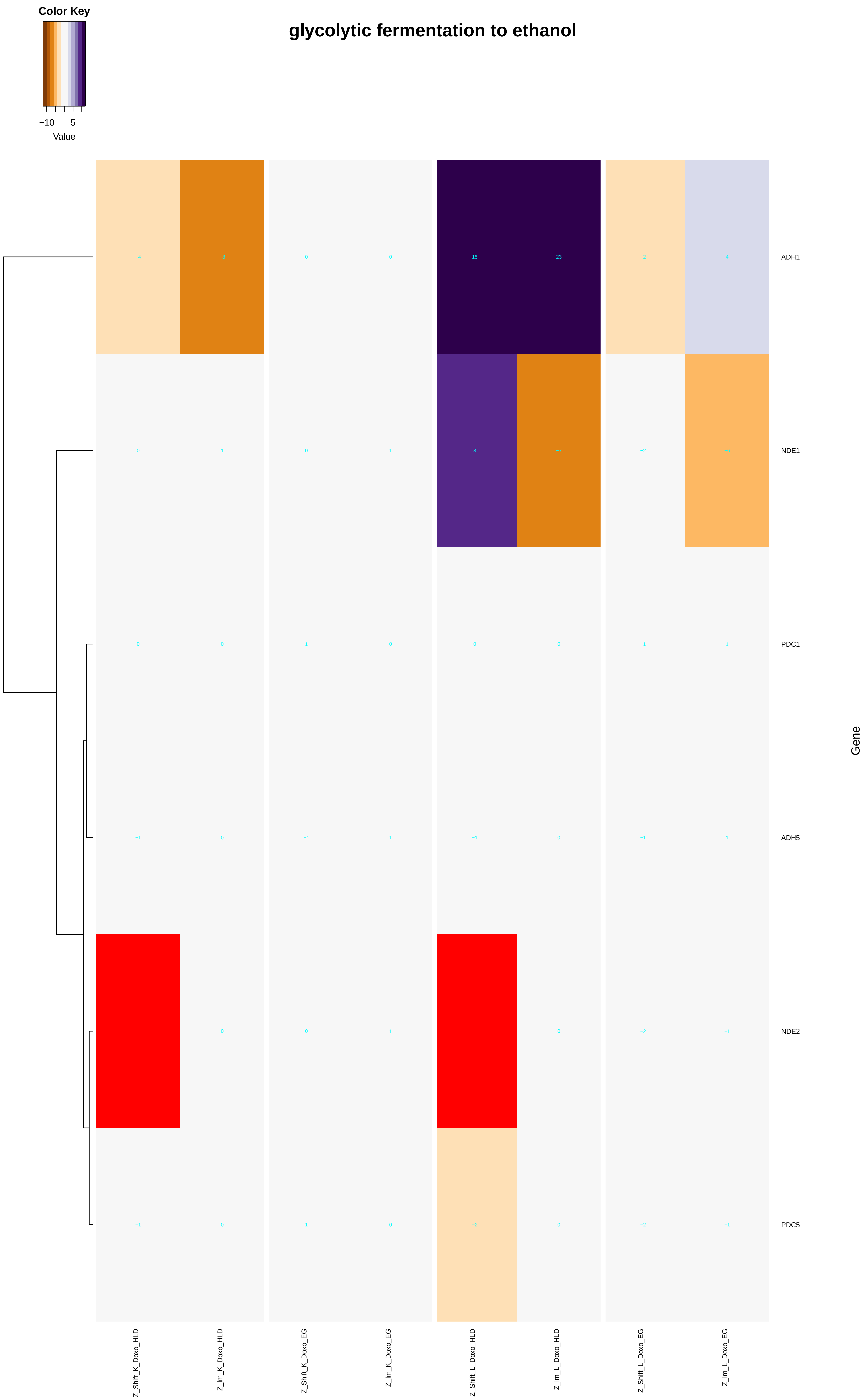

### ATP_synthesis_coupled_proton_transport.pdf

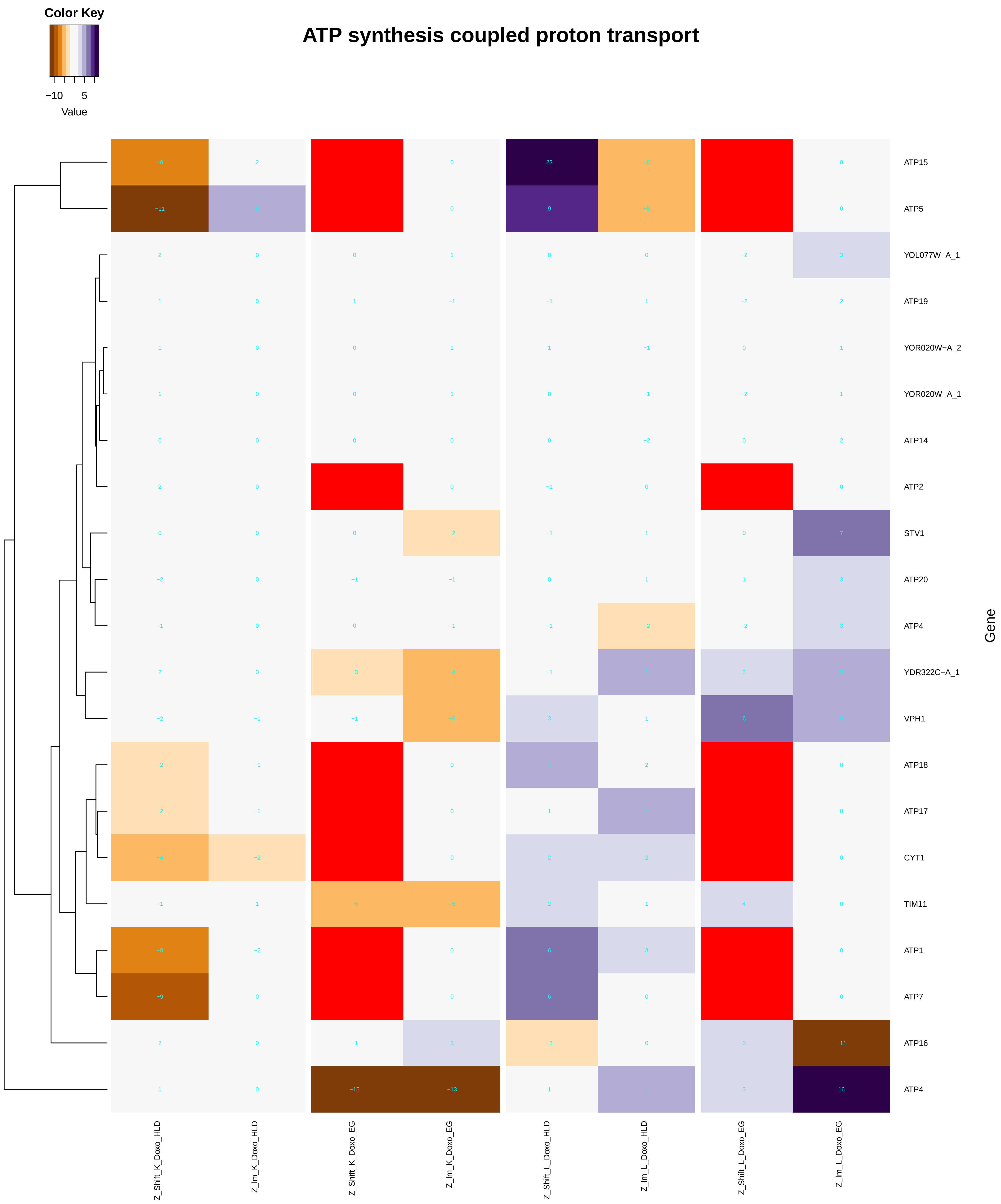

### base-excision_repair.pdf

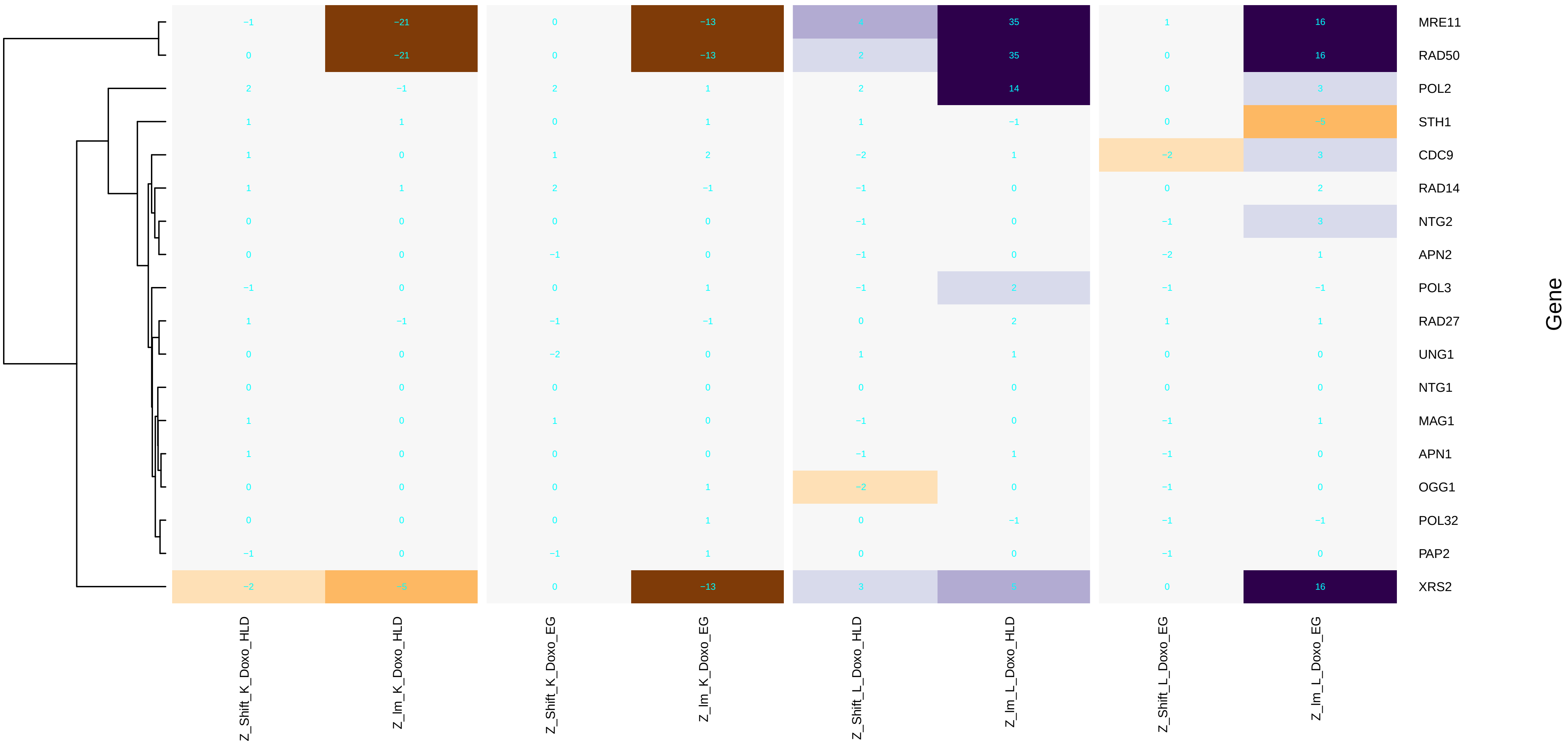

### cell_cycle.pdf

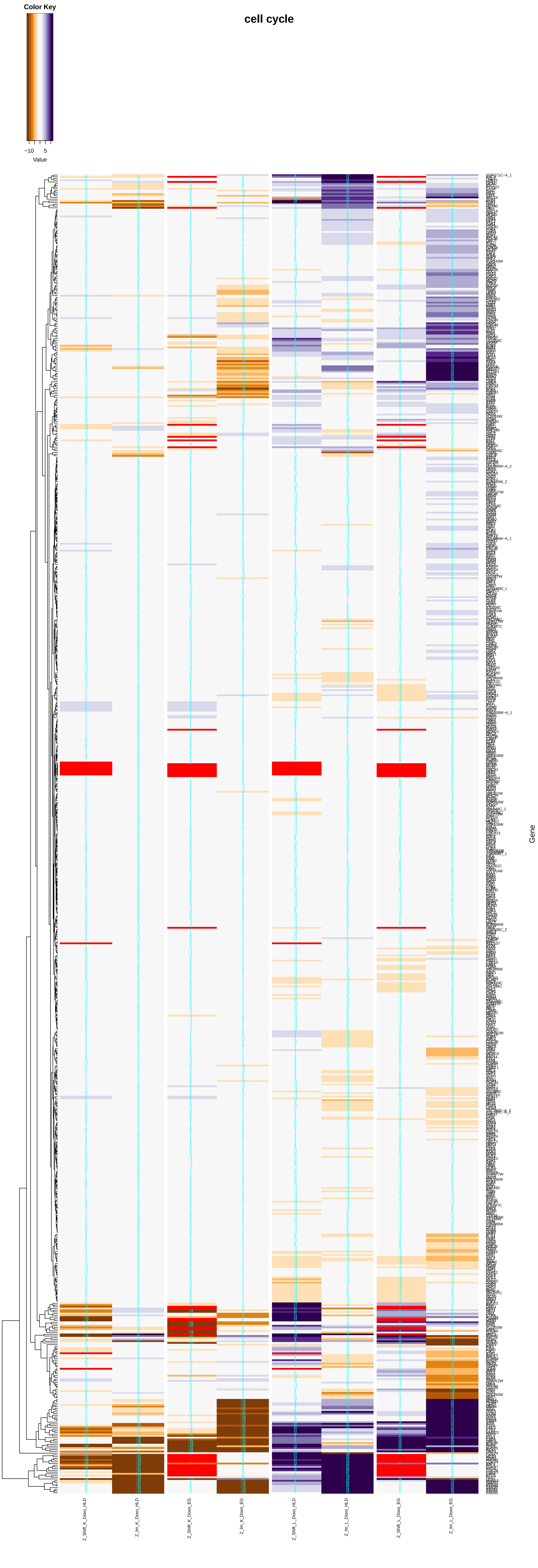

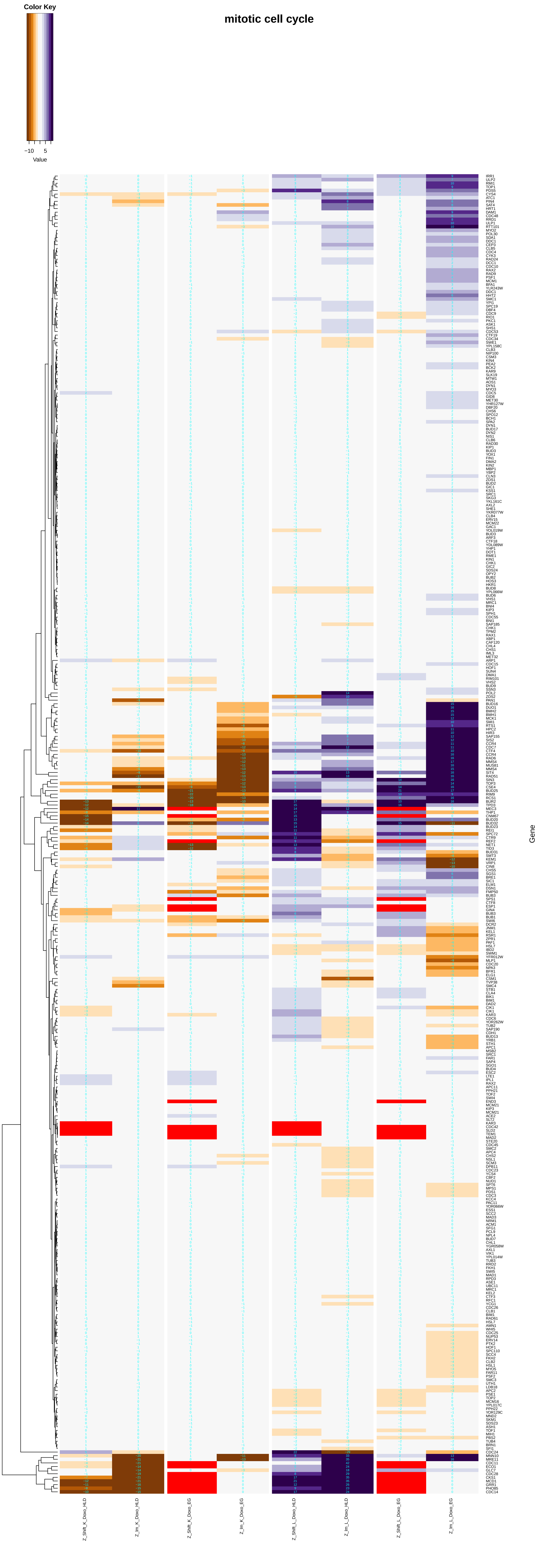

##### meiotic cell cycle

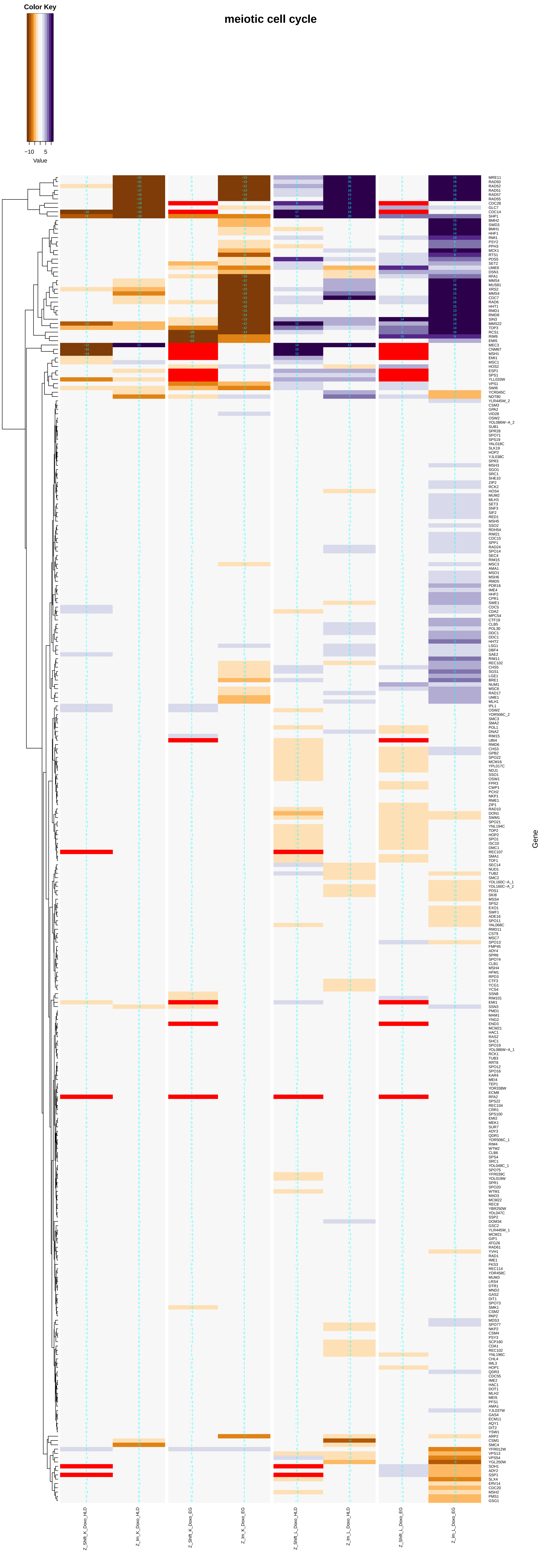

### cell_cycle_arrest.pdf

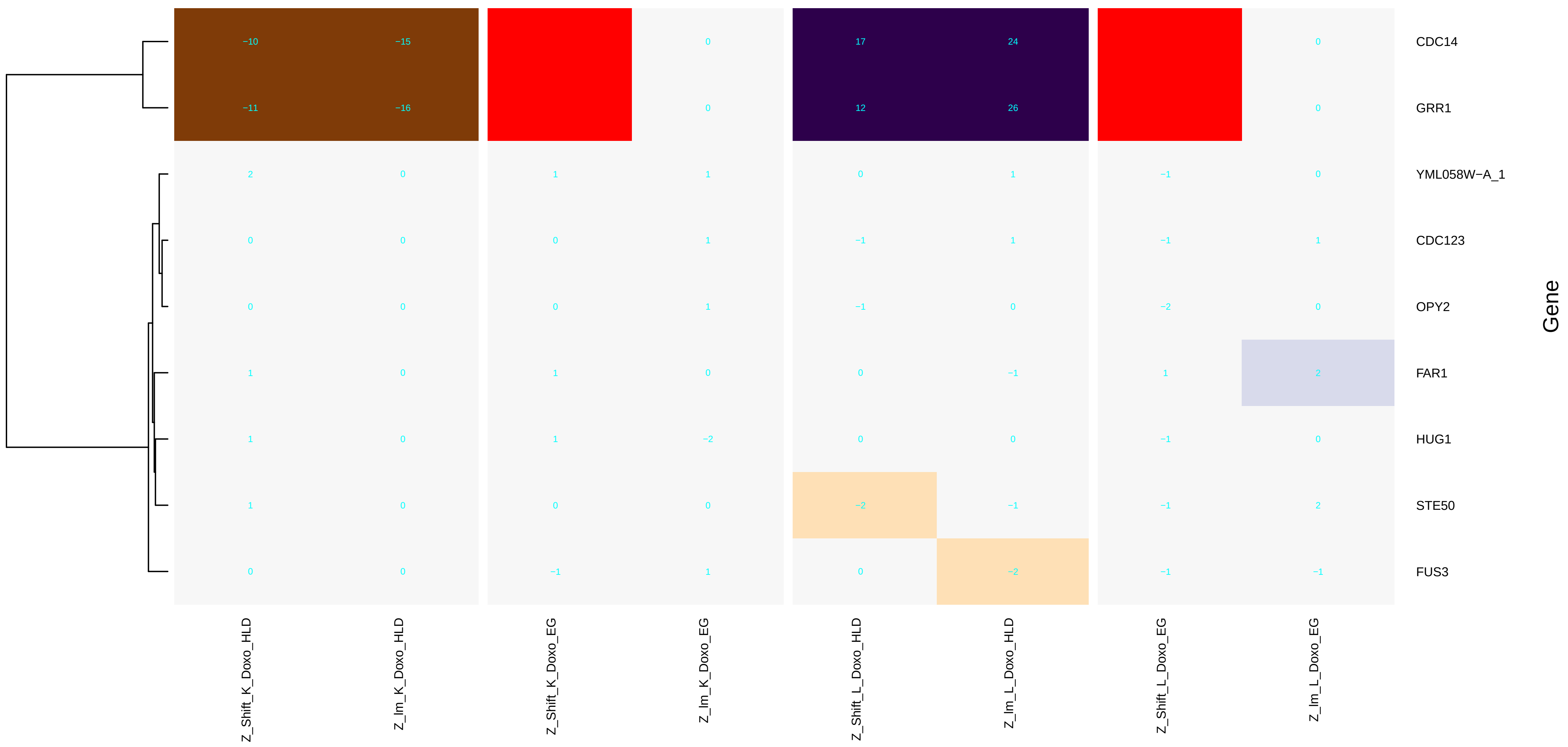

### cell_cycle_DNA_replication.pdf

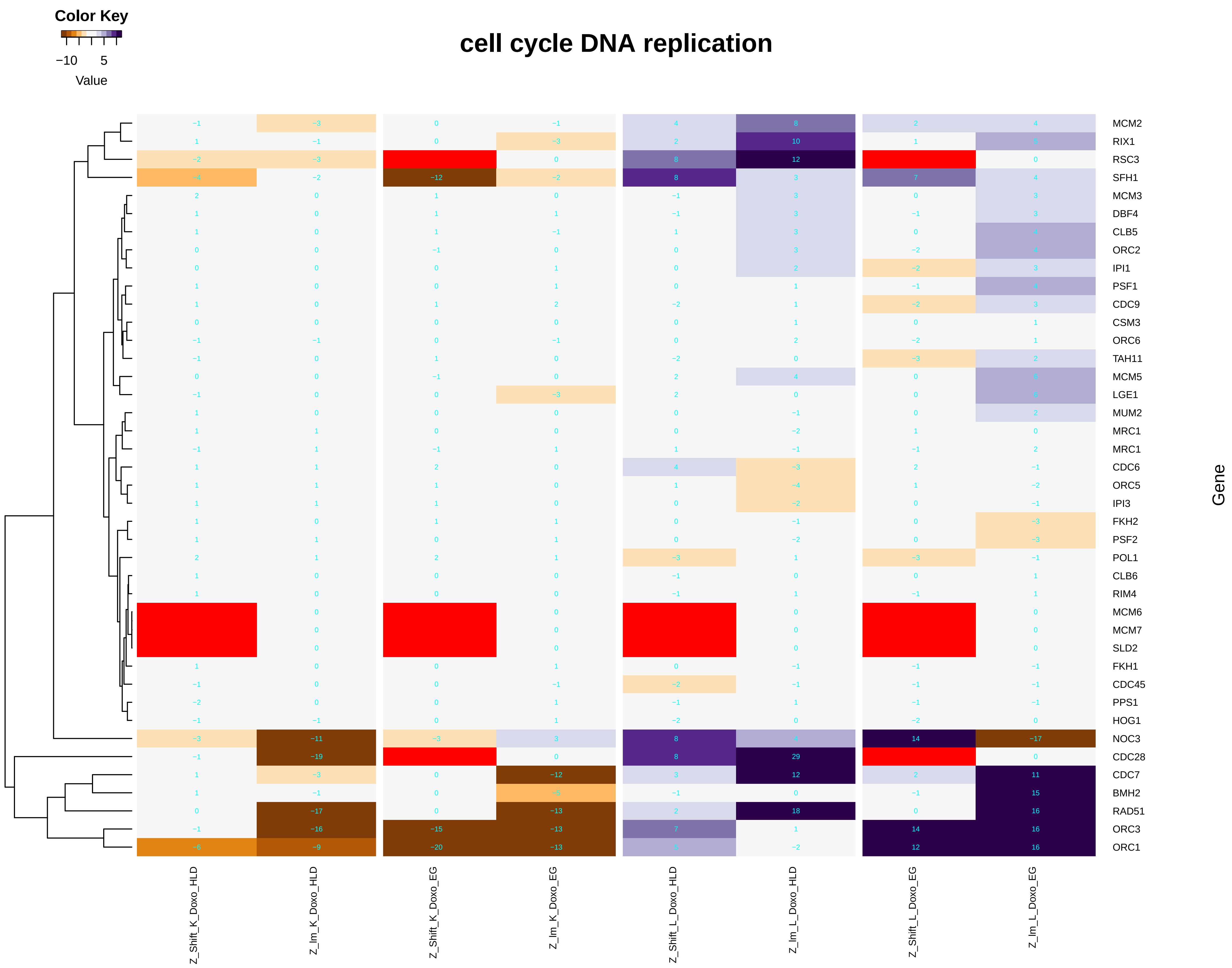

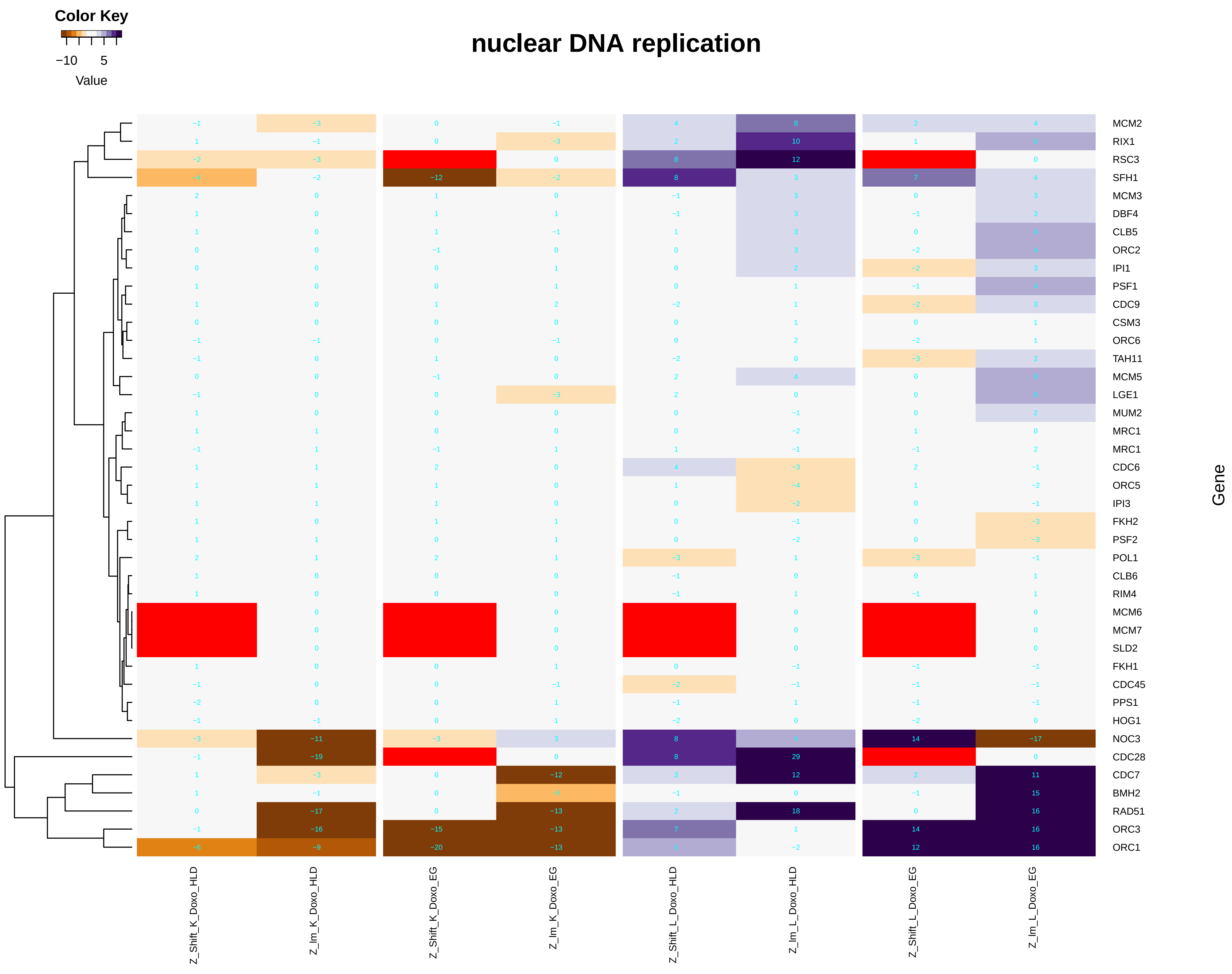

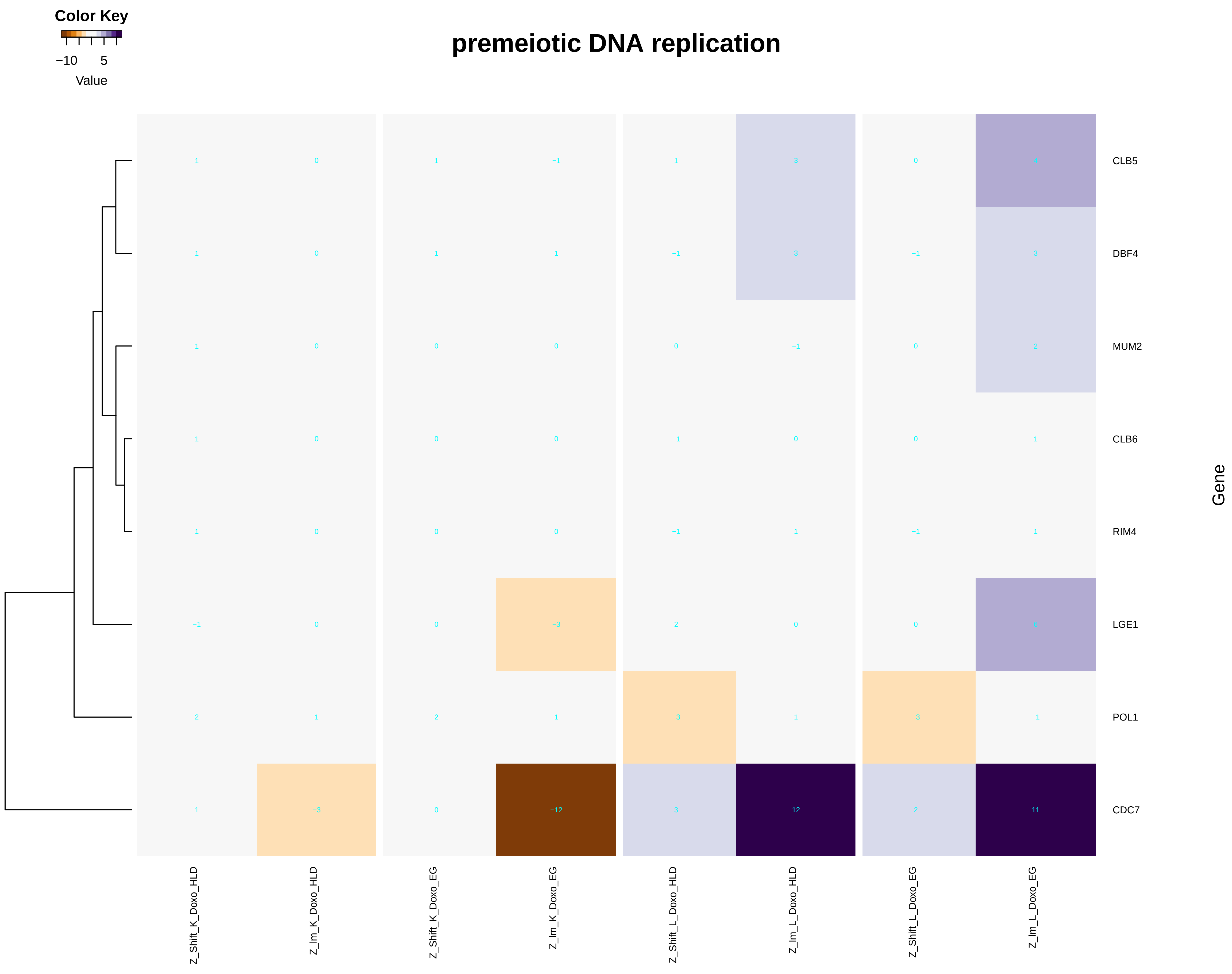

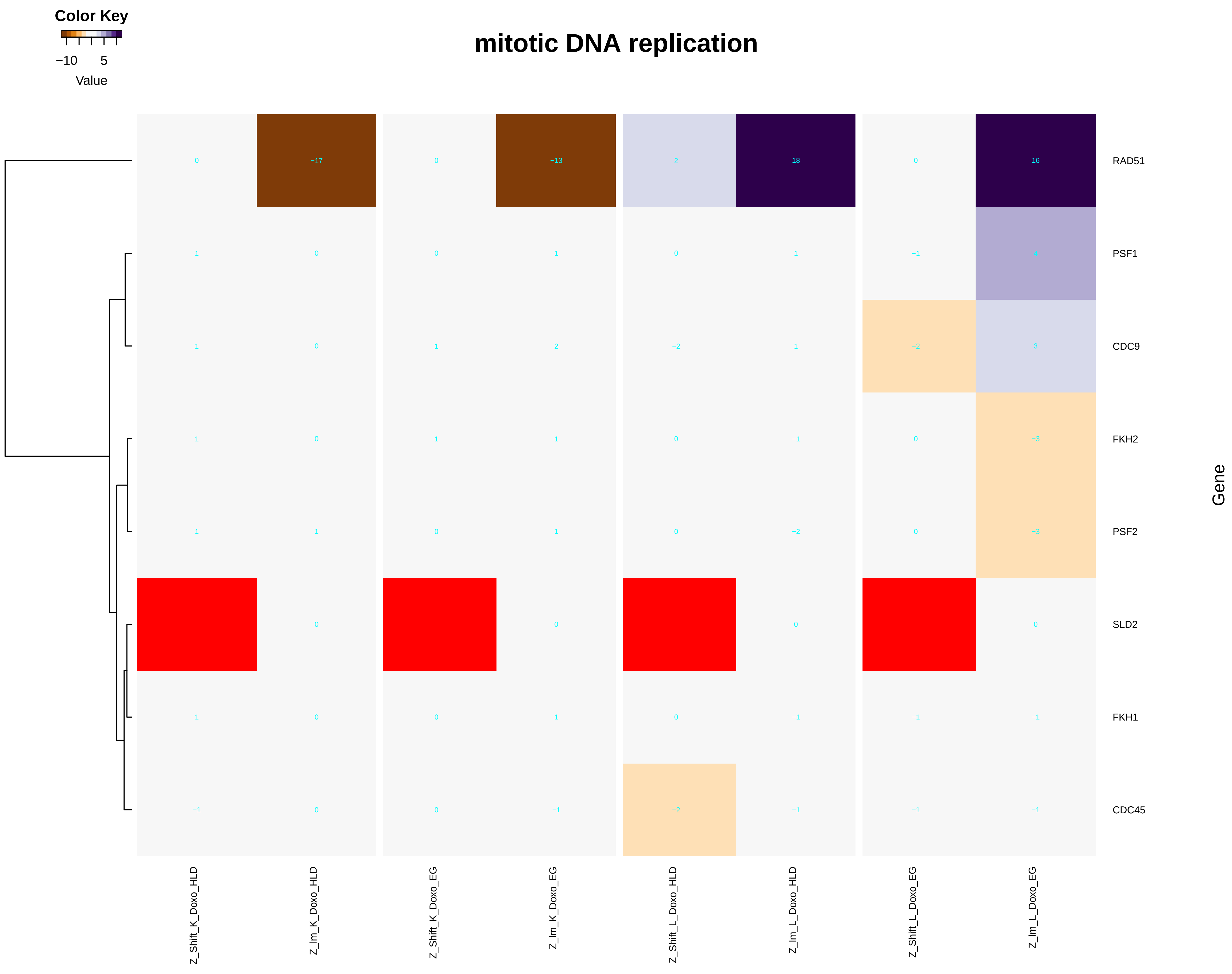

### cell_division.pdf

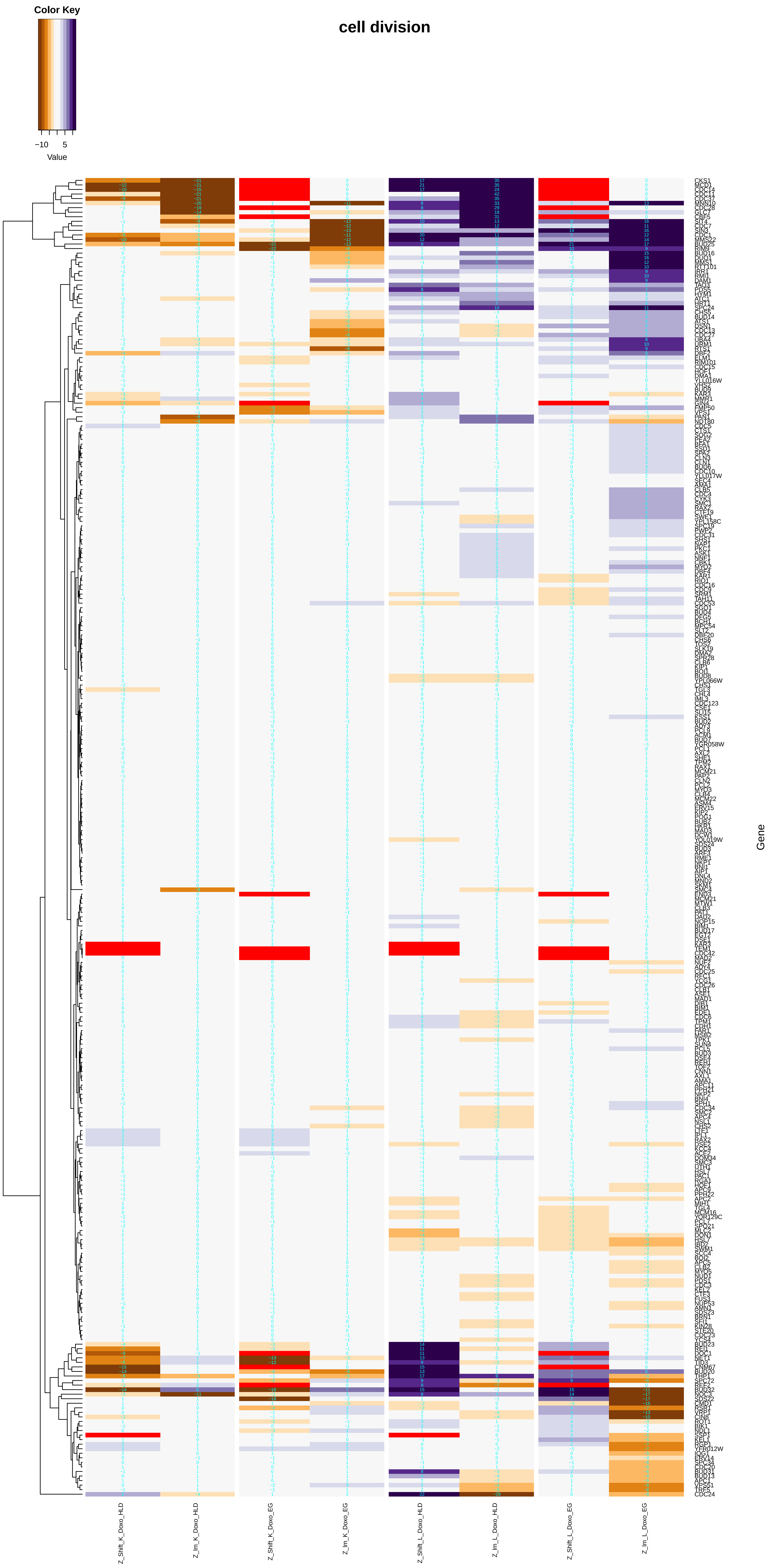

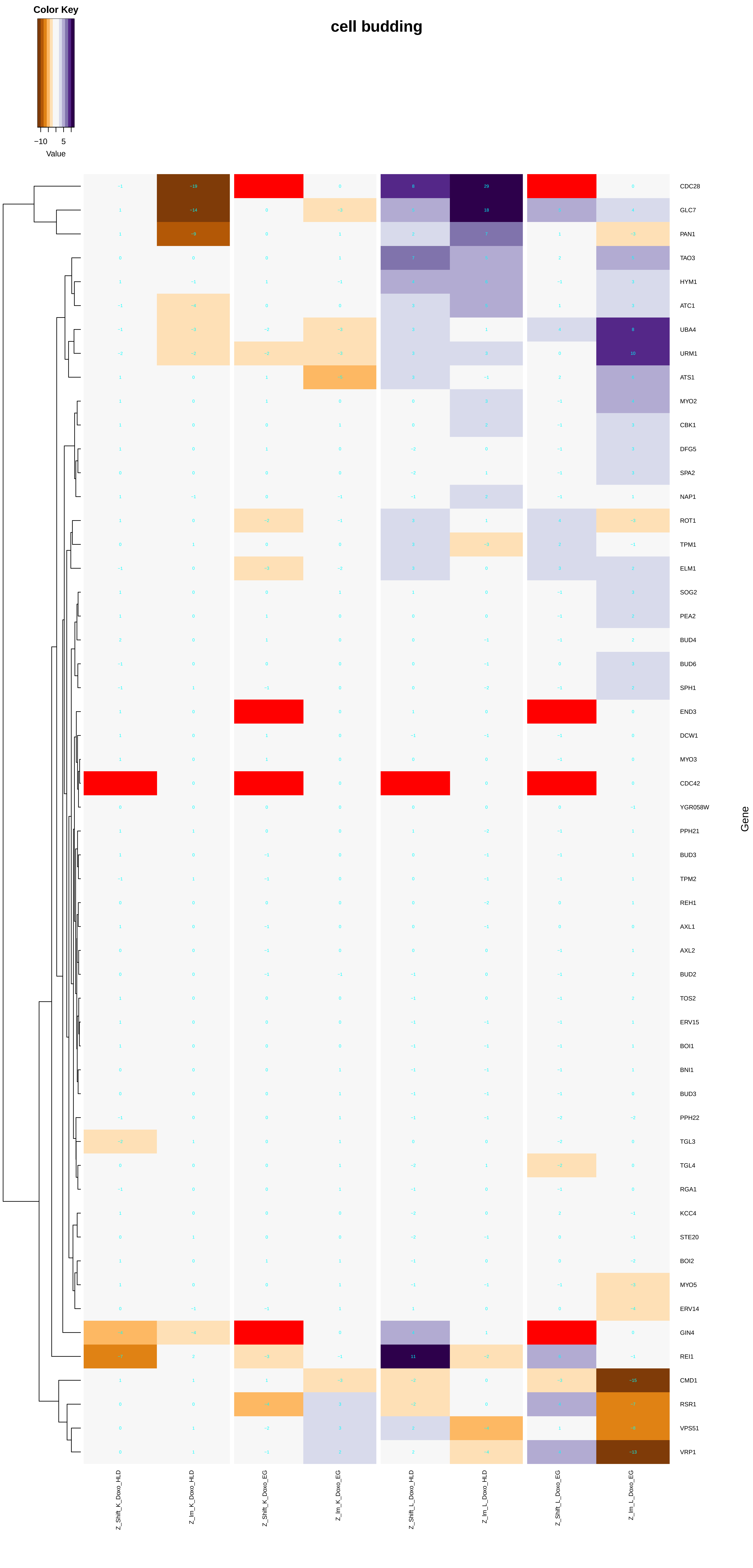

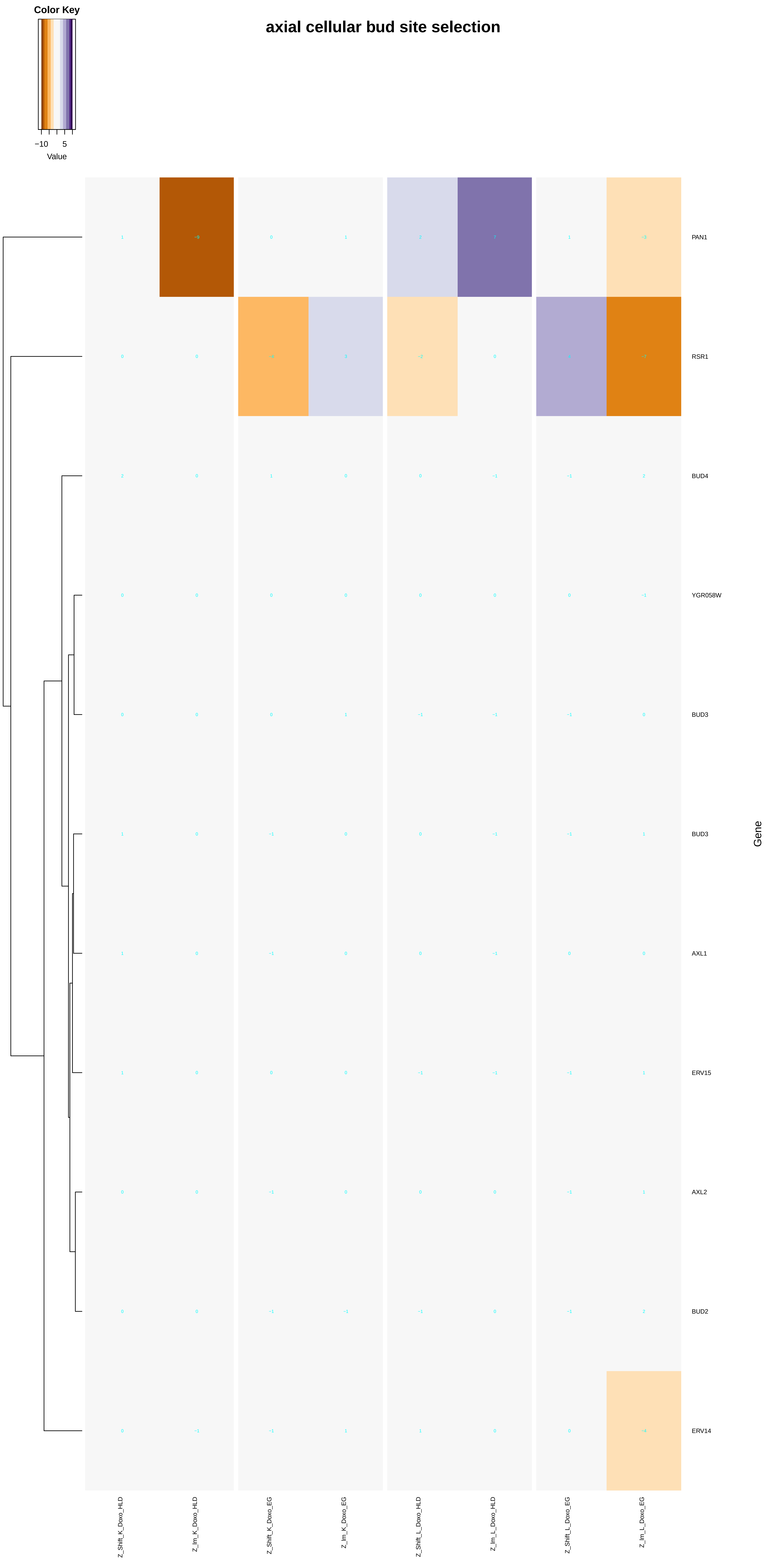

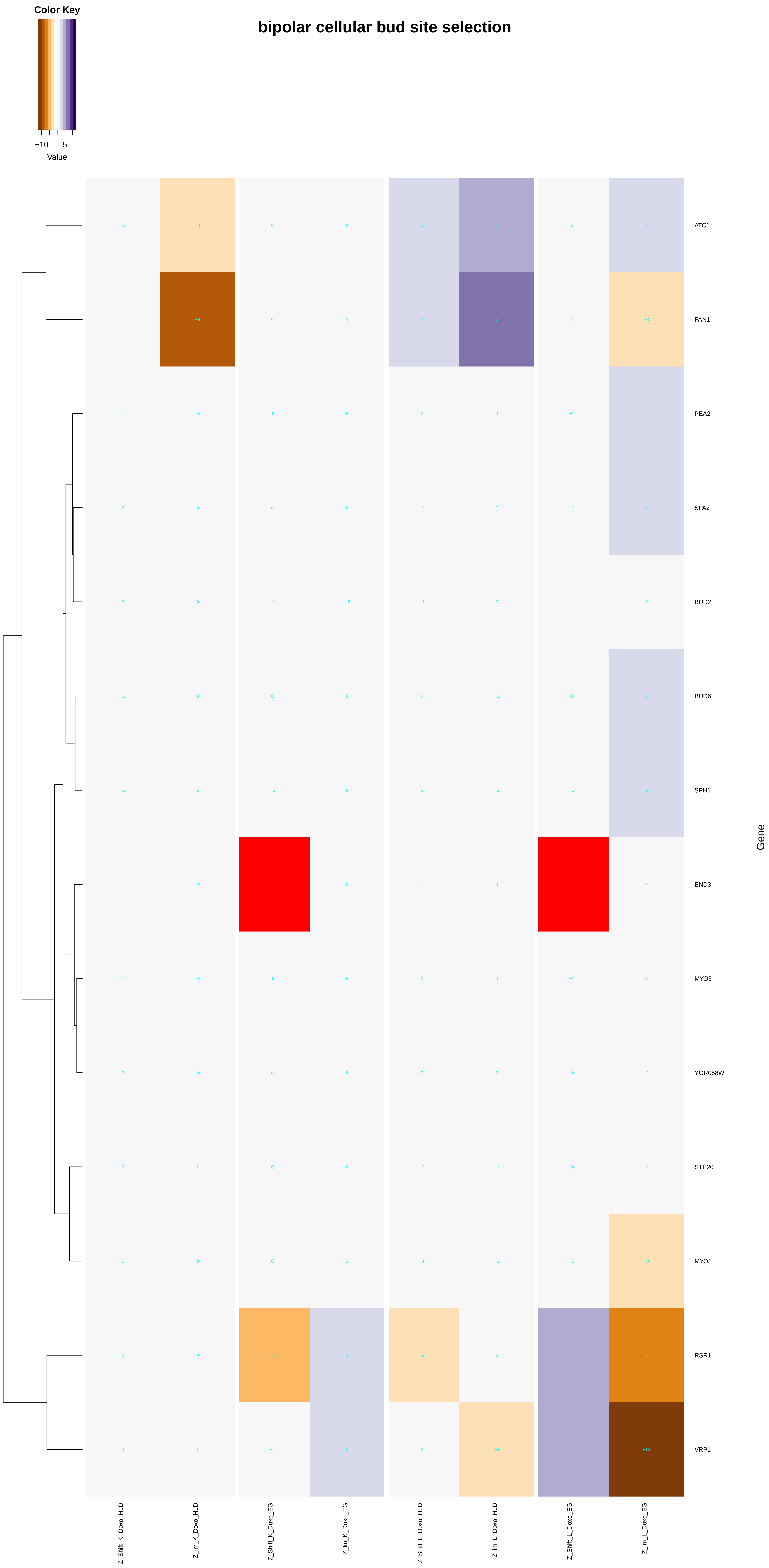

### cell_growth.pdf

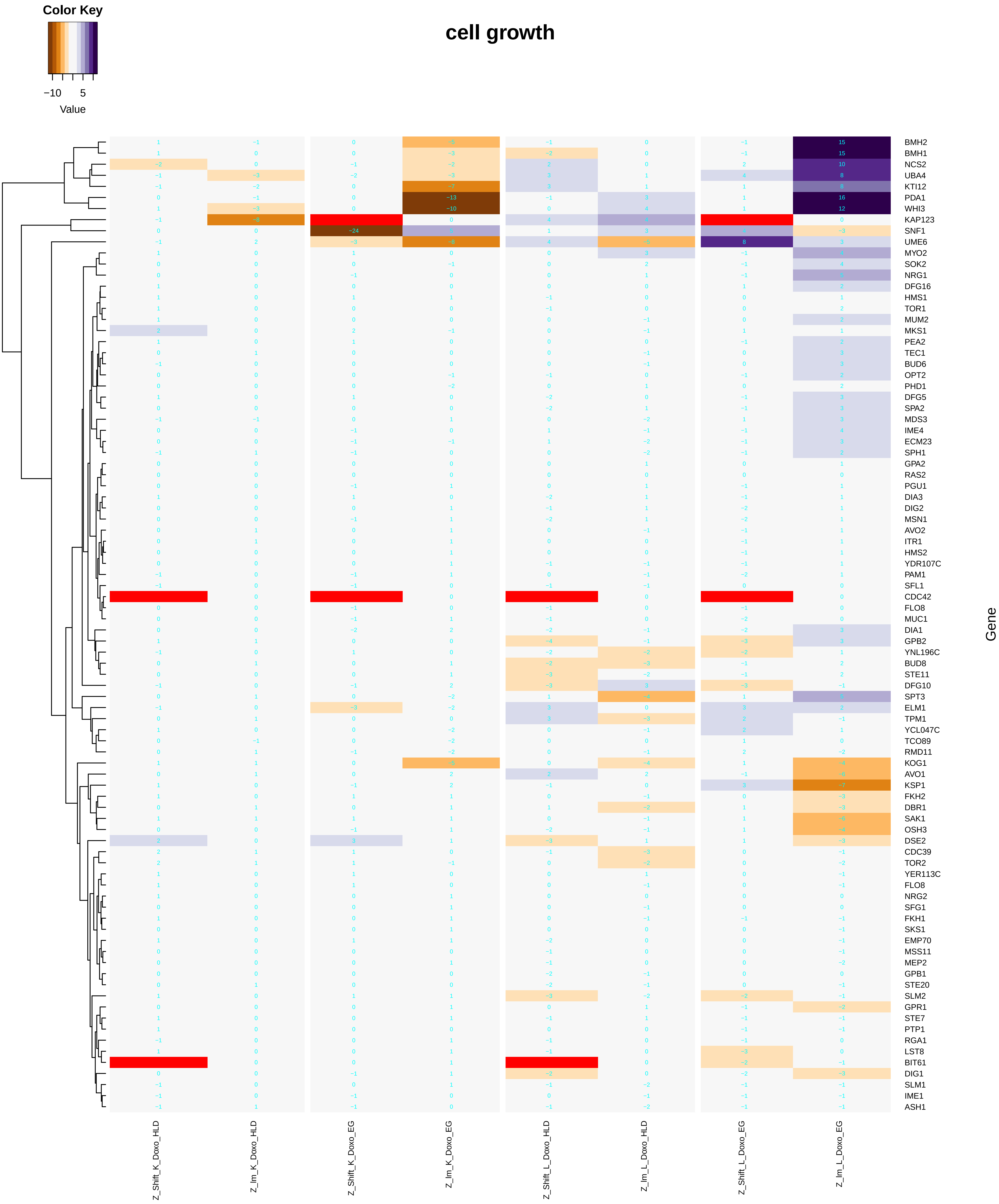

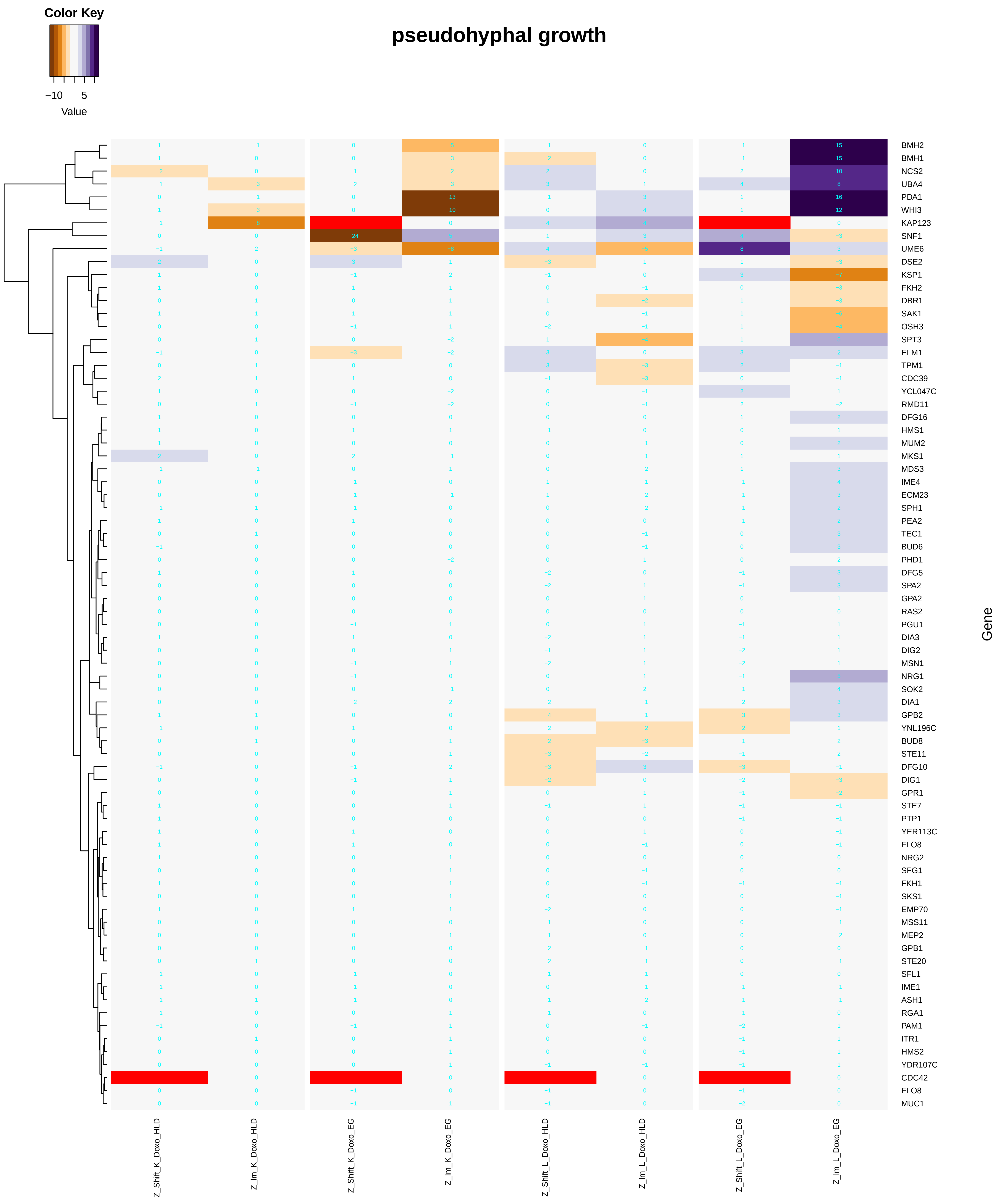

### cellular_monovalent_inorganic_cation_homeostasis.pdf

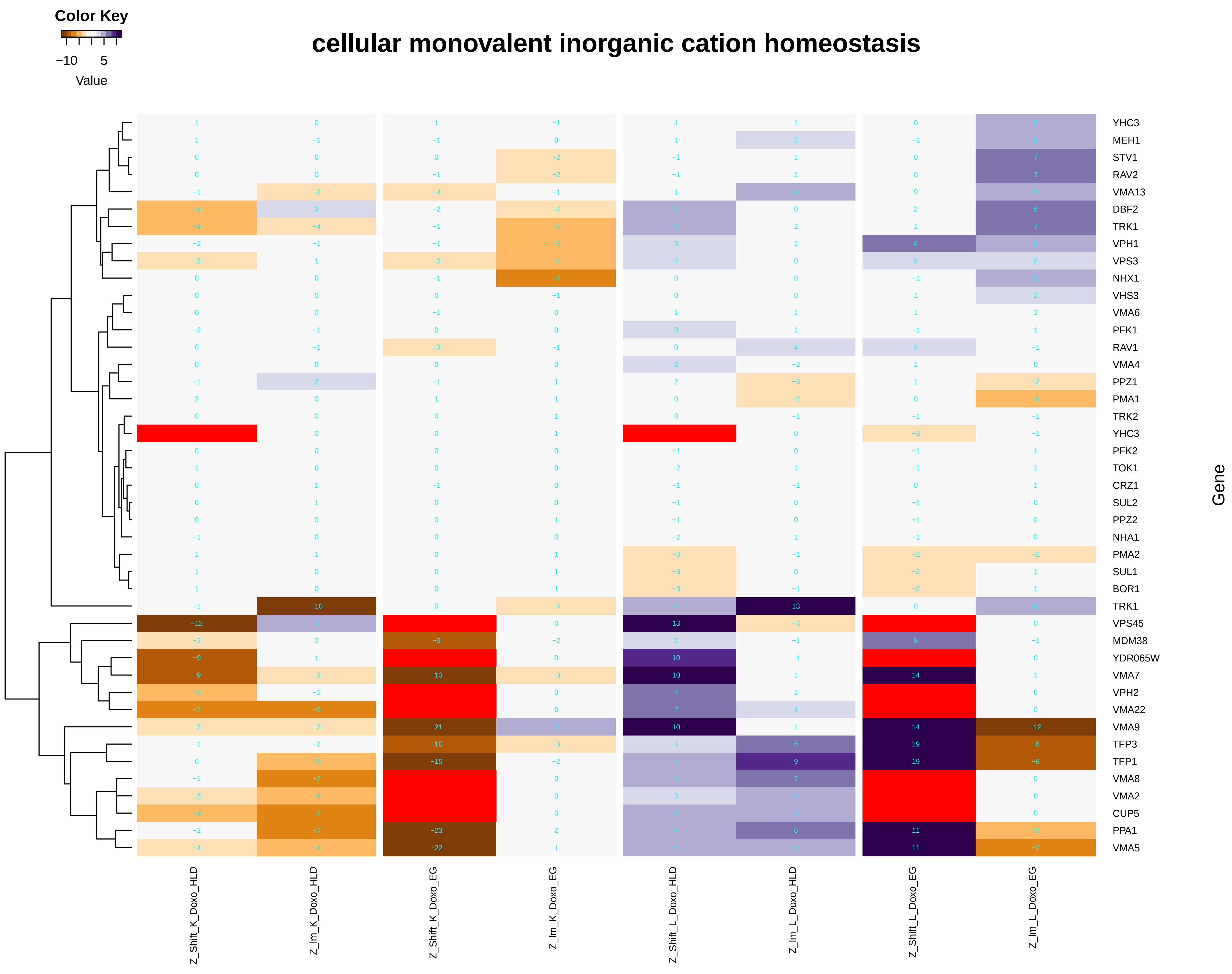

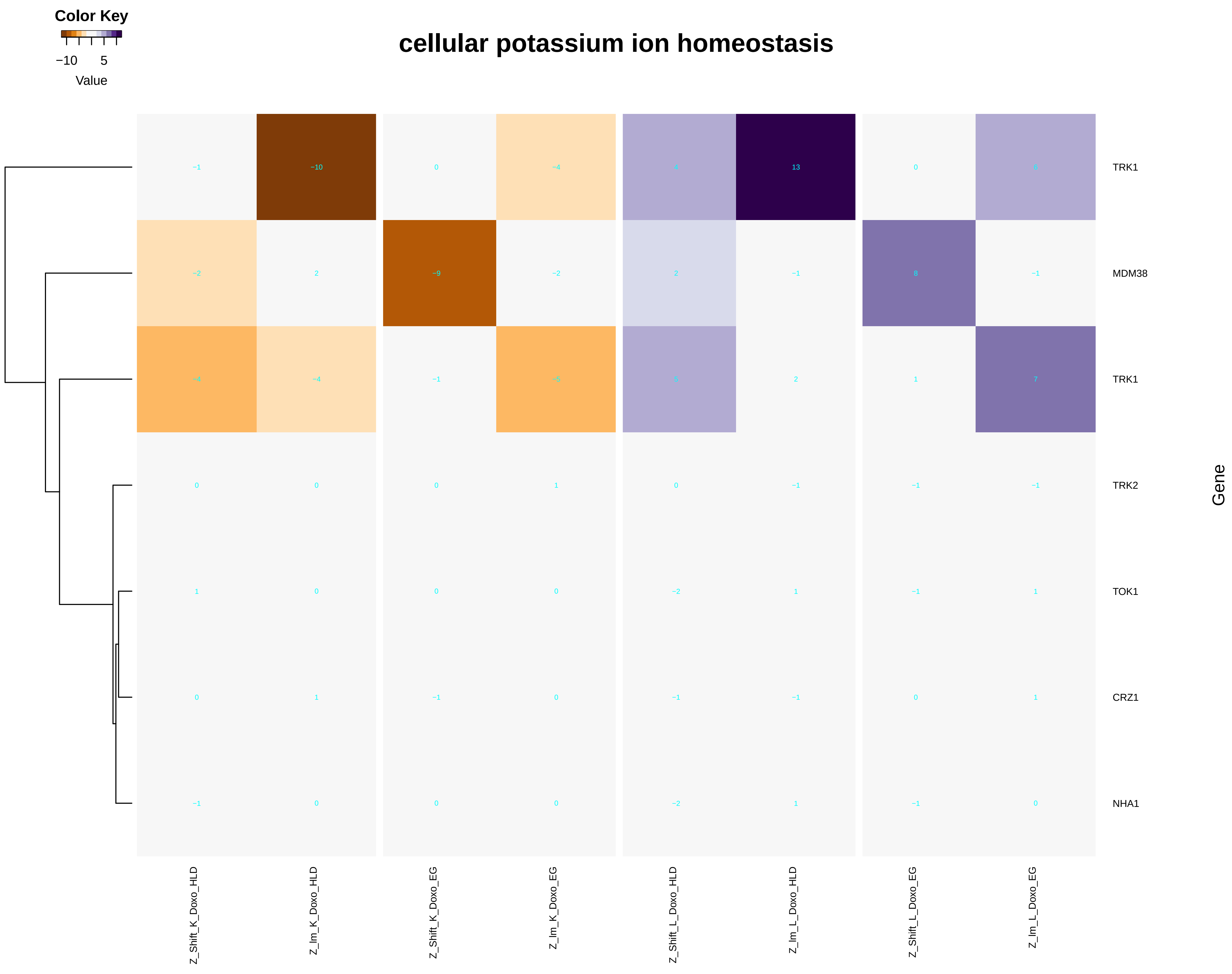

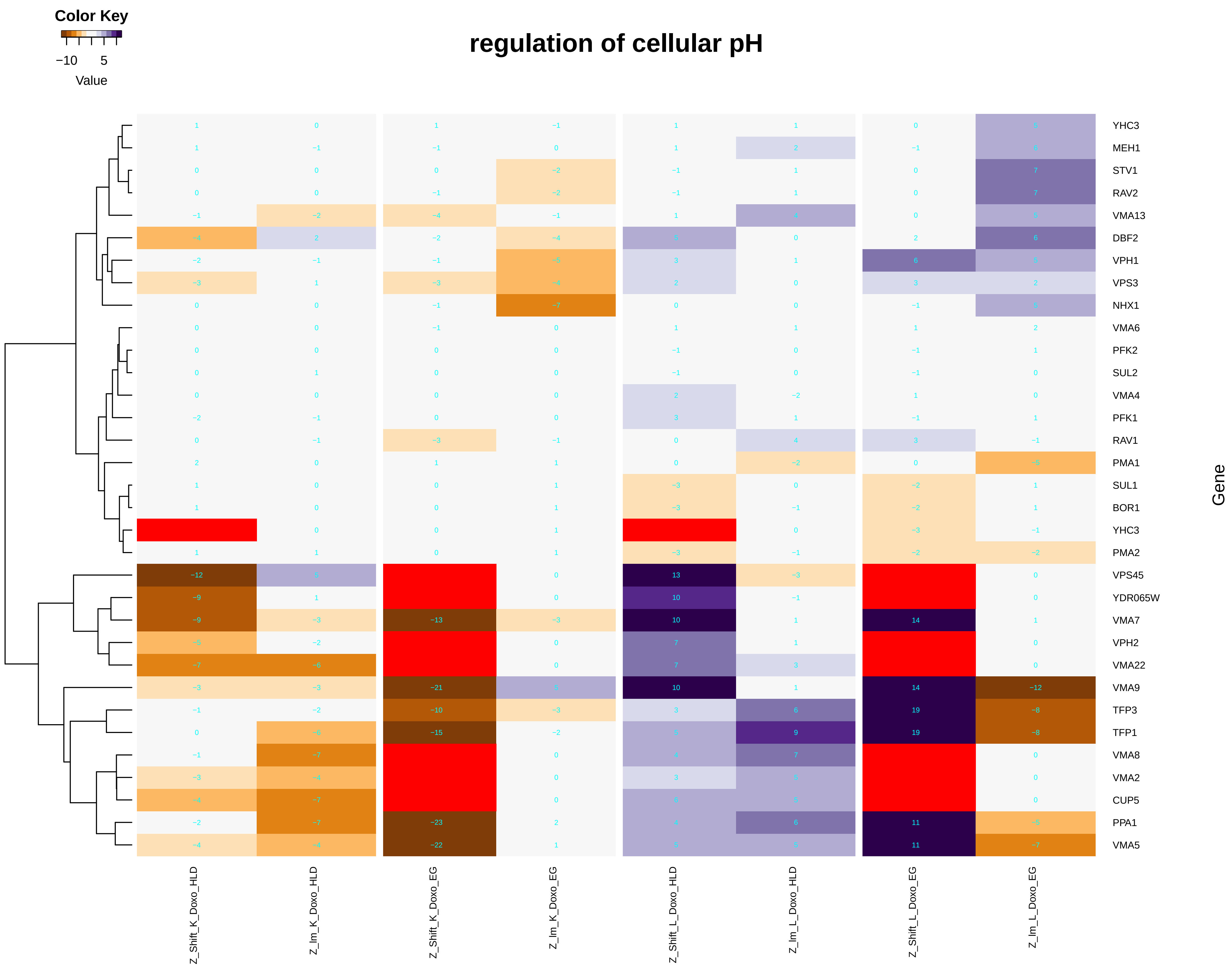

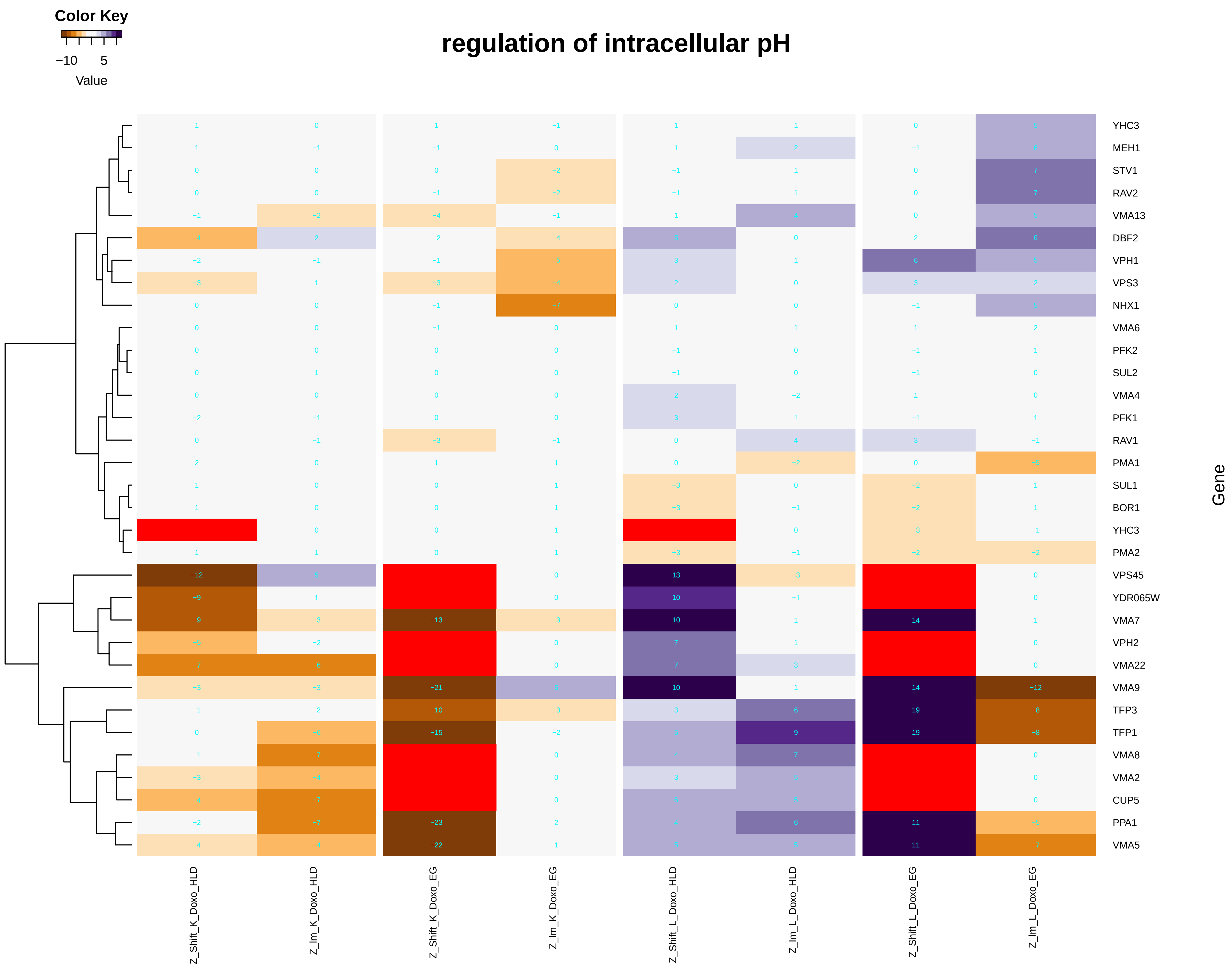

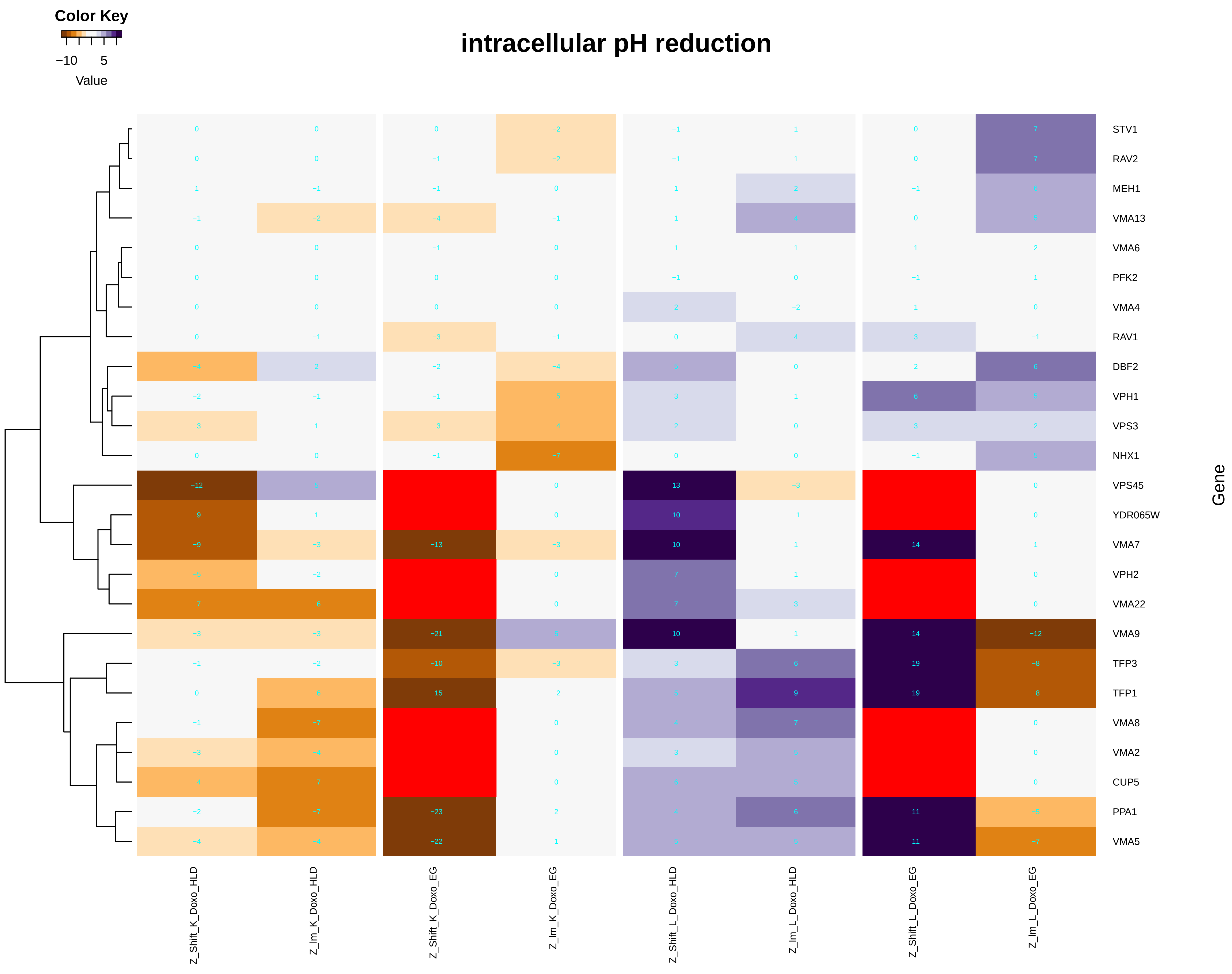

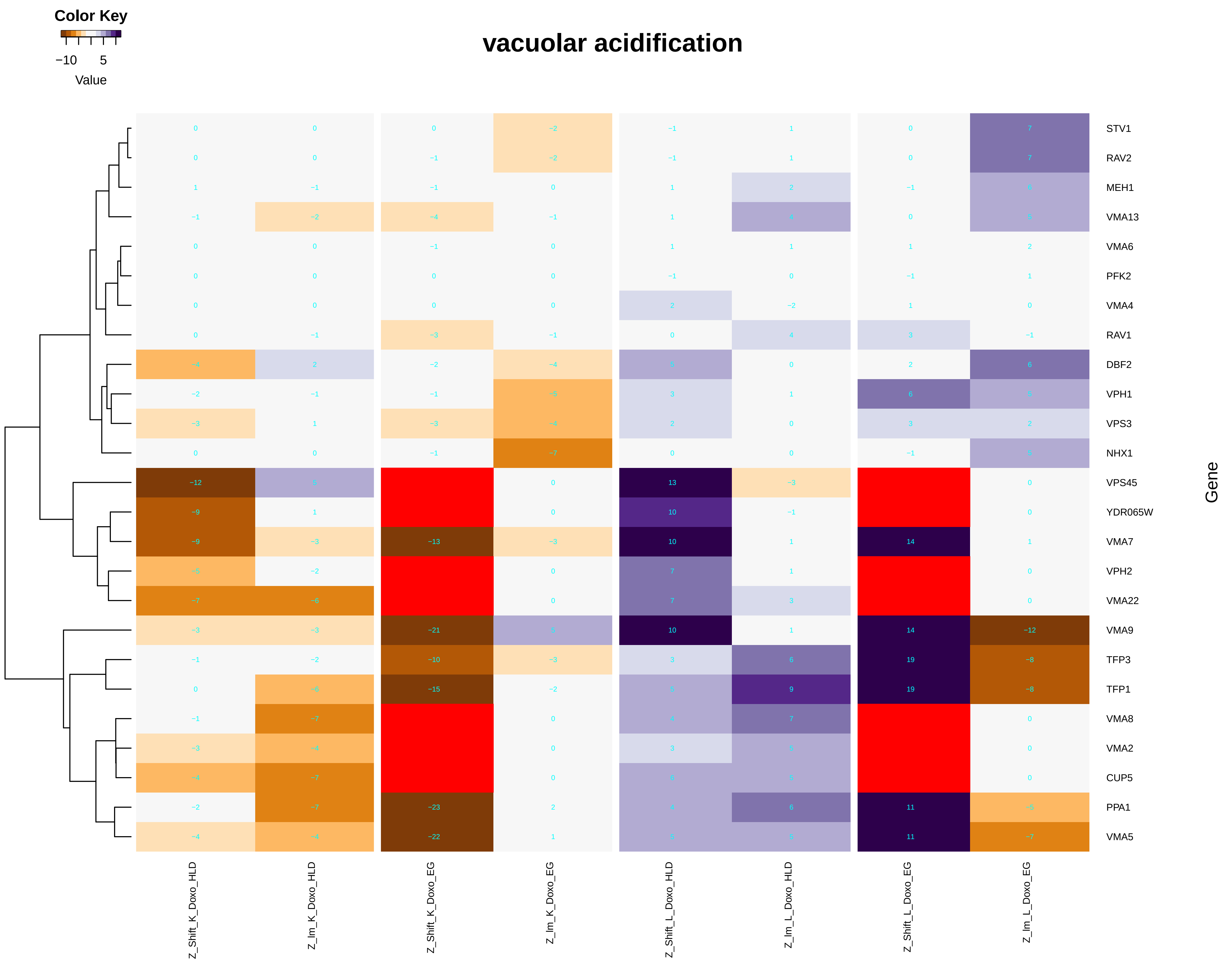

### cellular_protein_complex_localization.pdf

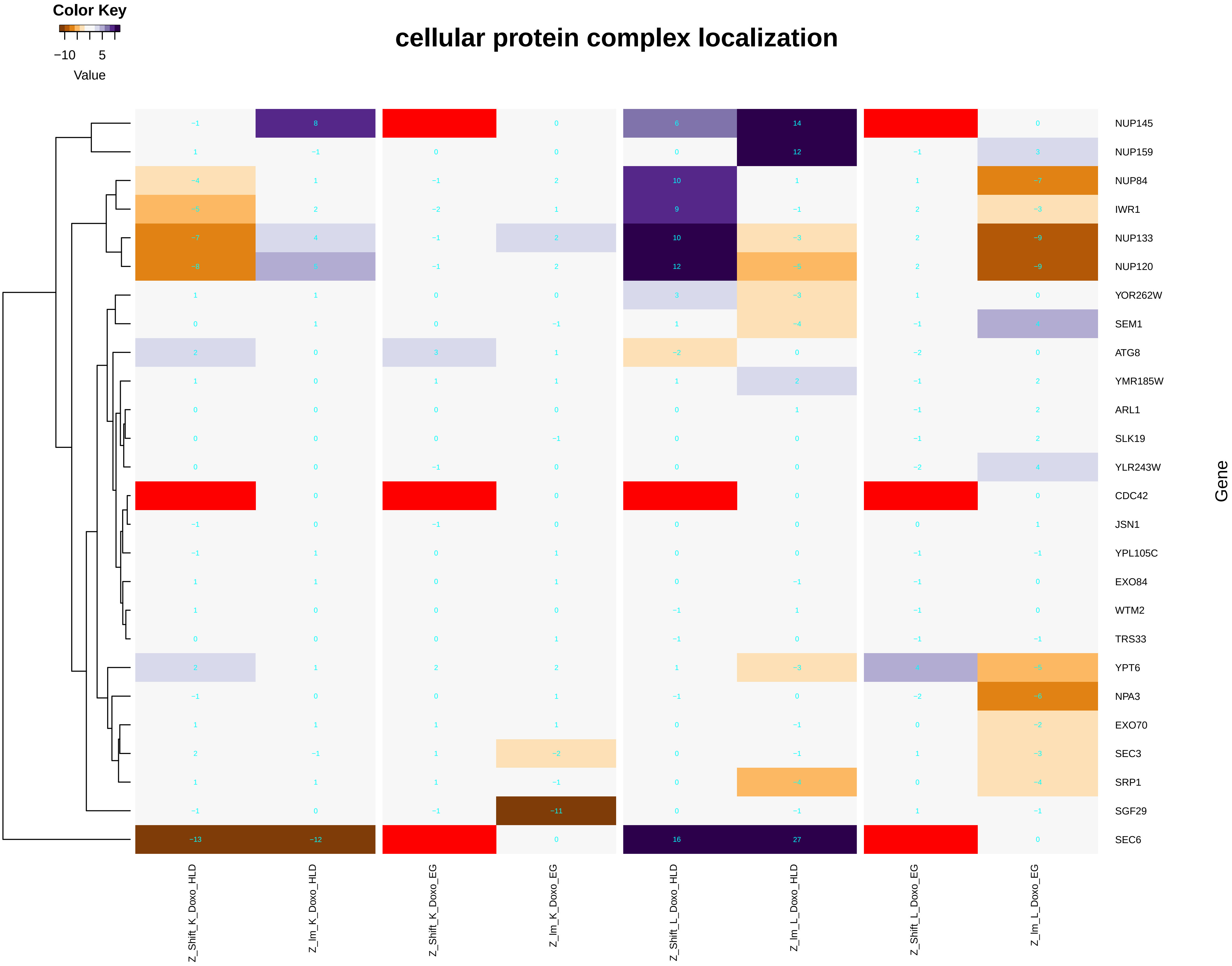

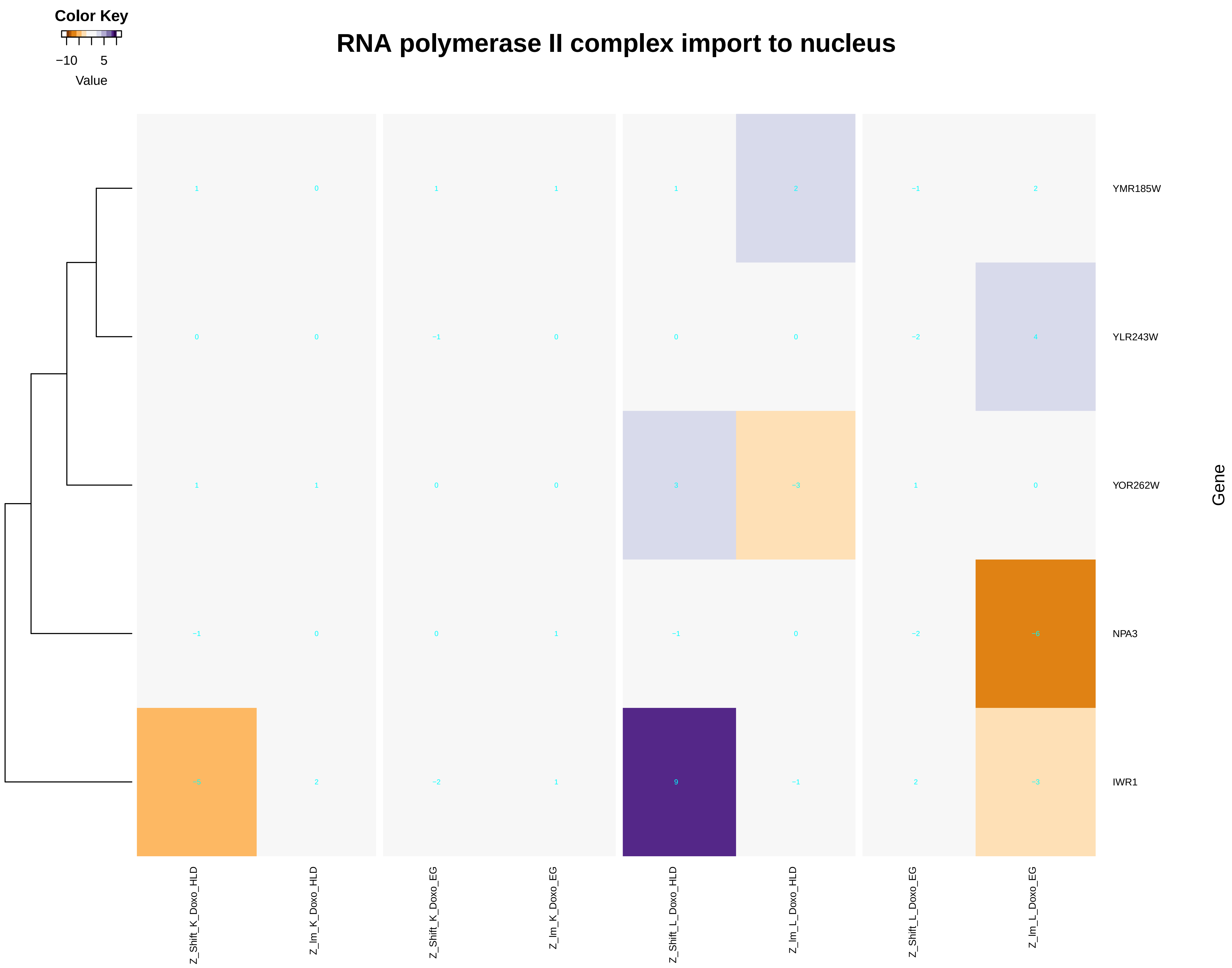

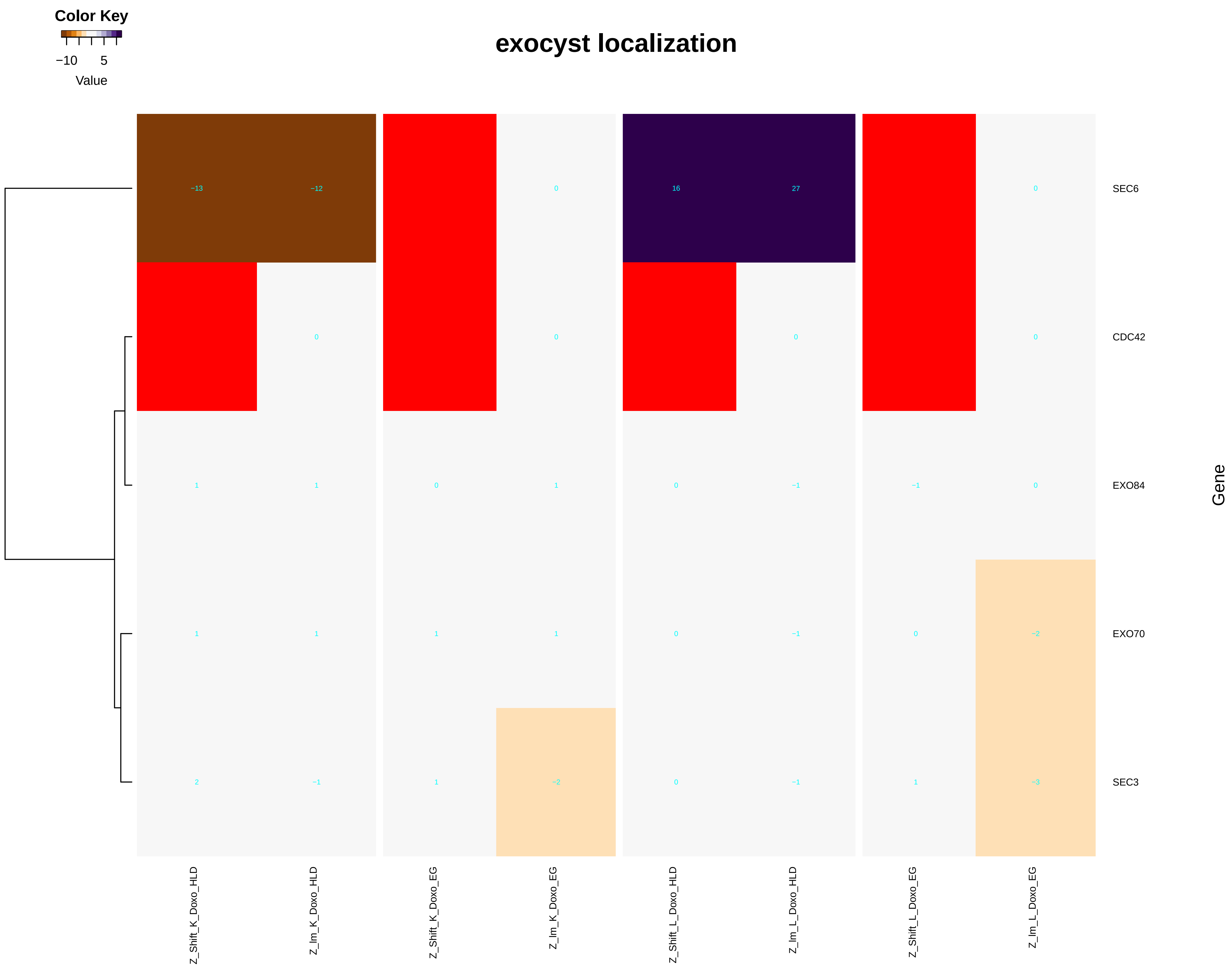

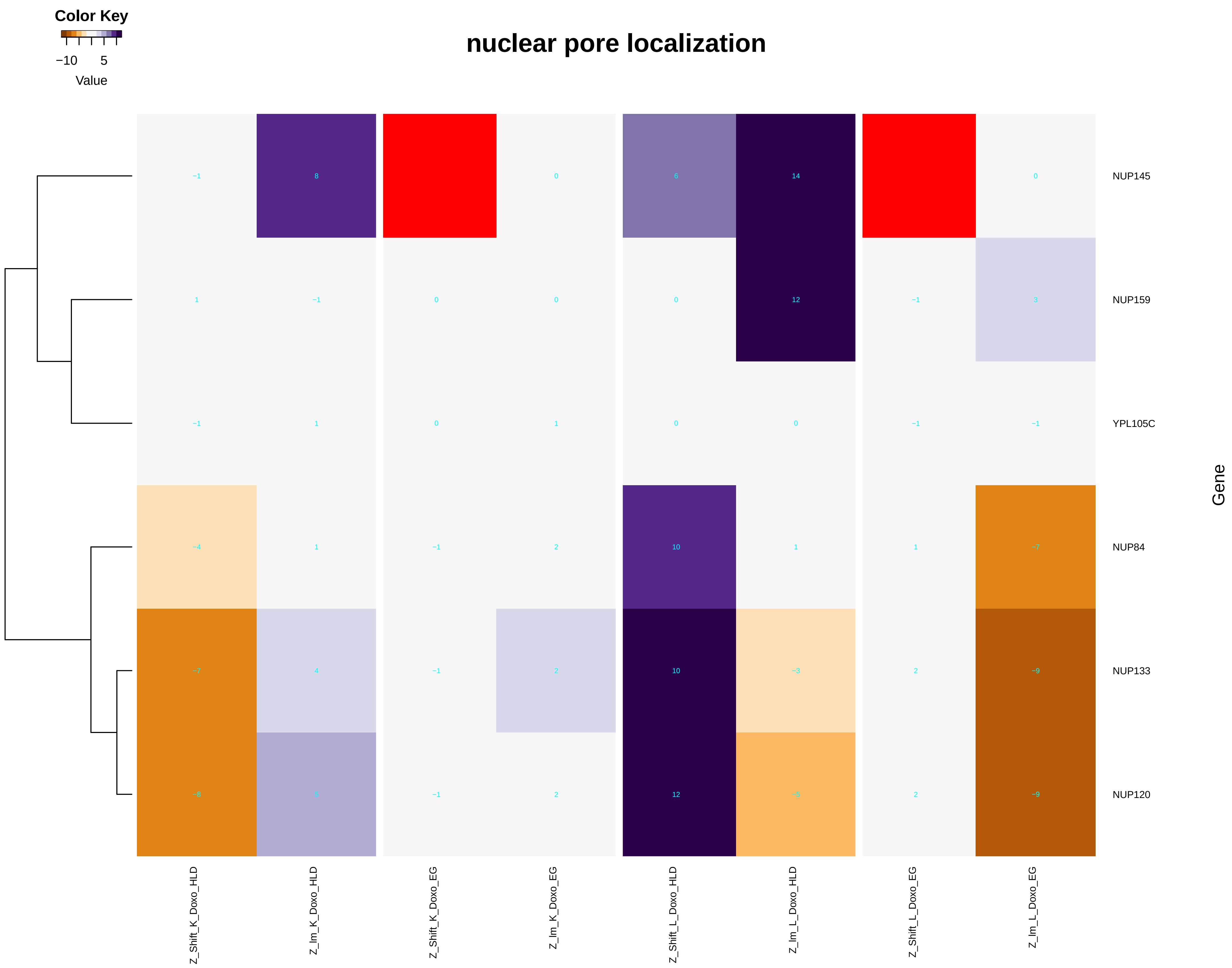

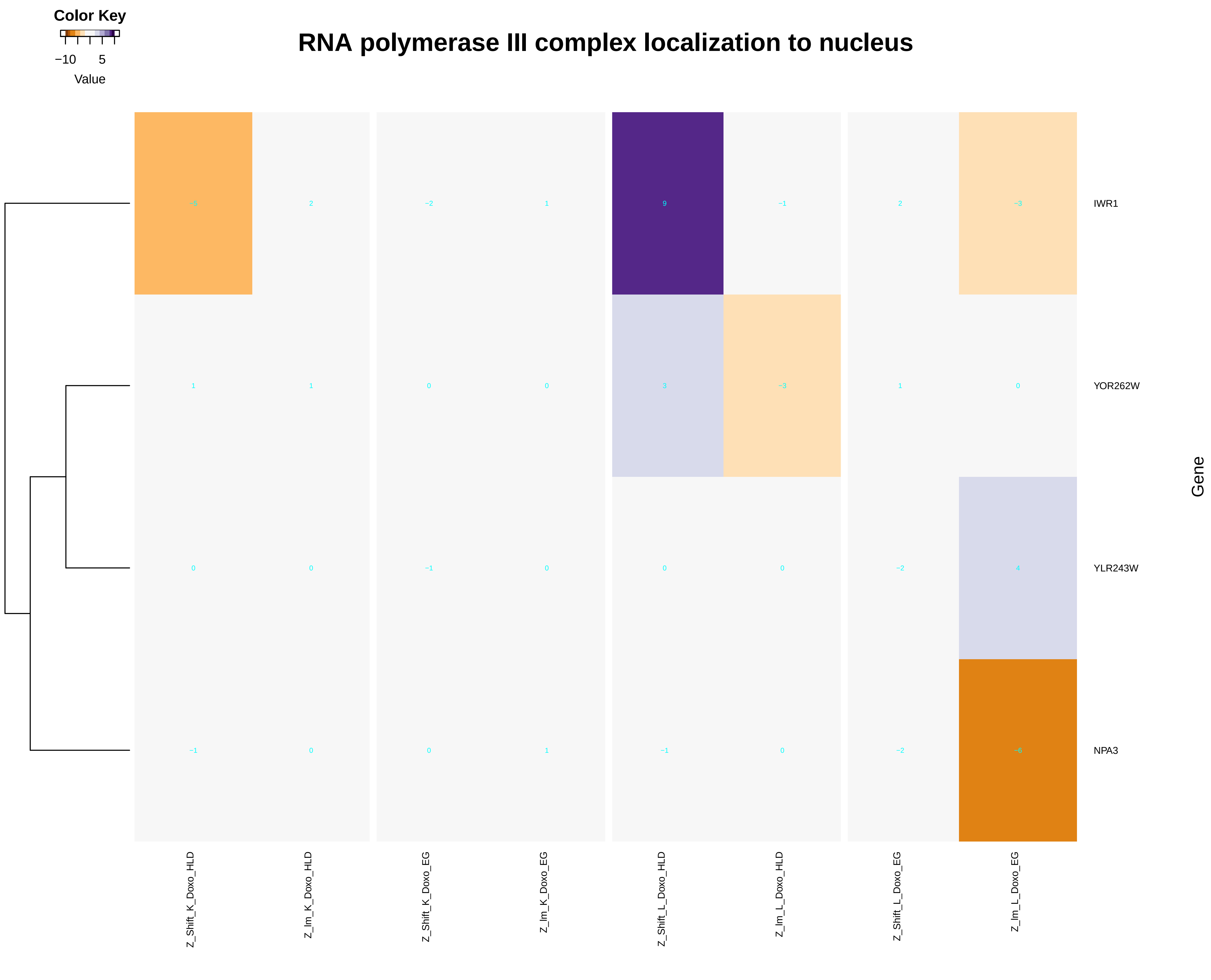

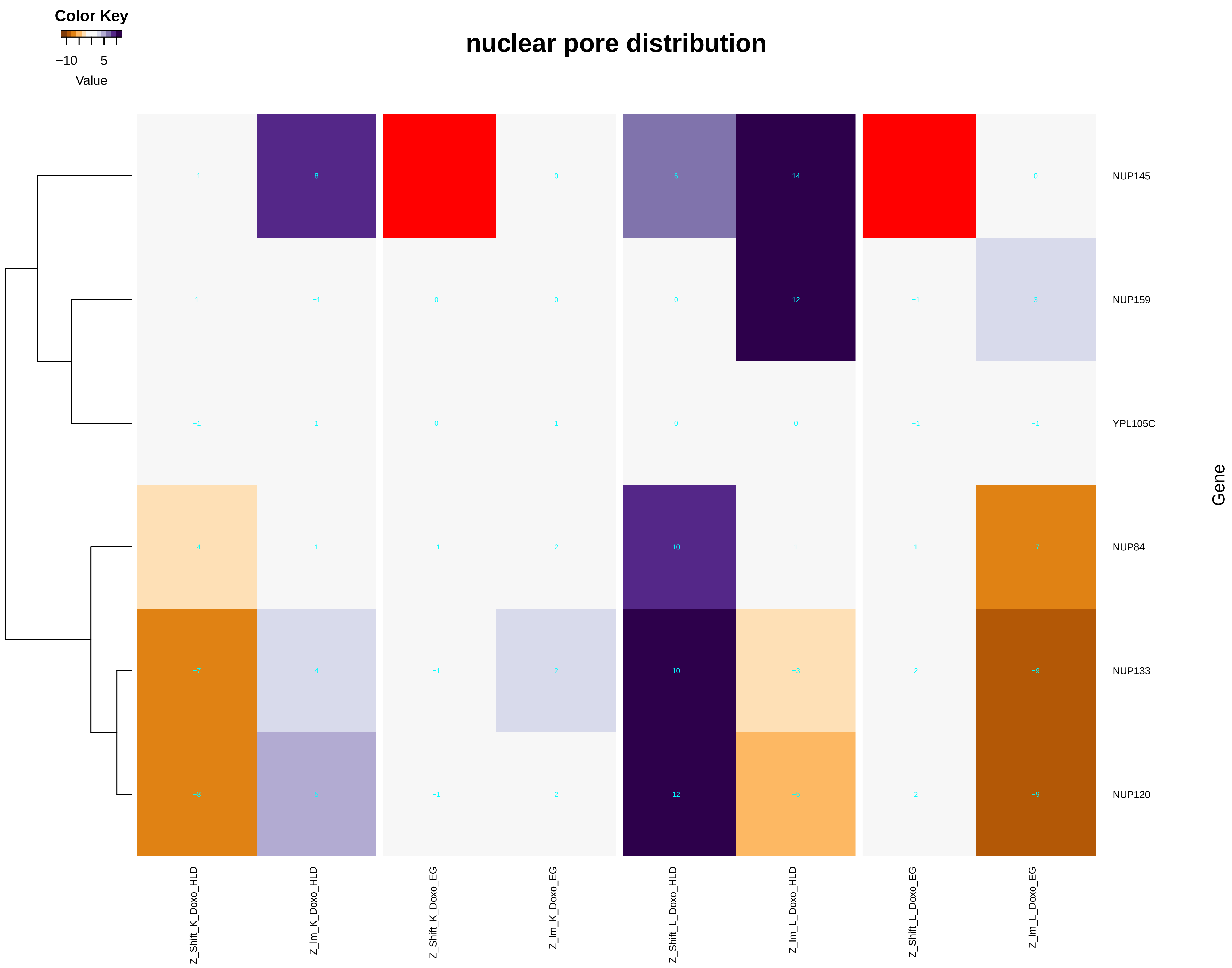

### cellular_response_to_nitrogen_levels.pdf

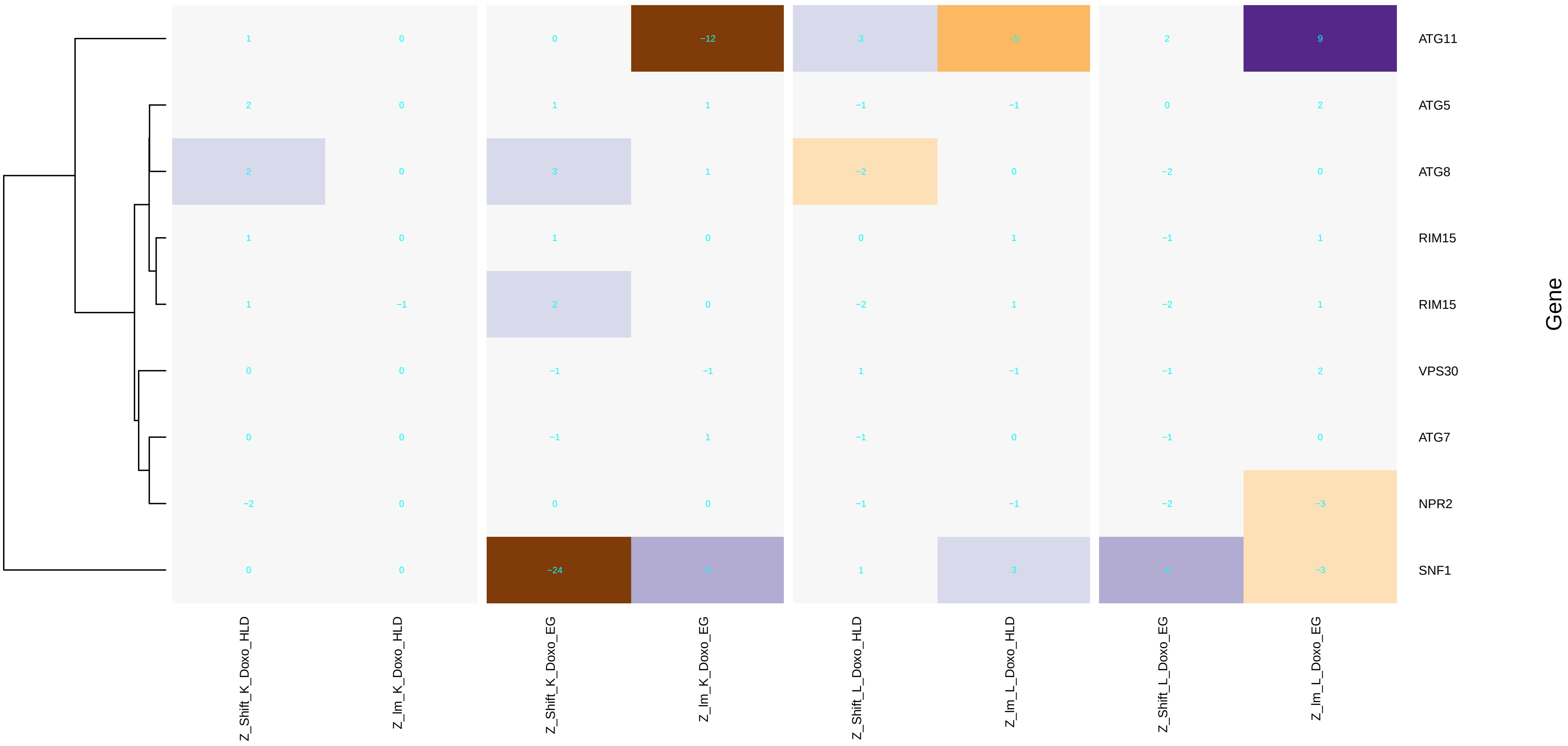

### cellular_response_to_nitrogen_starvation.pdf

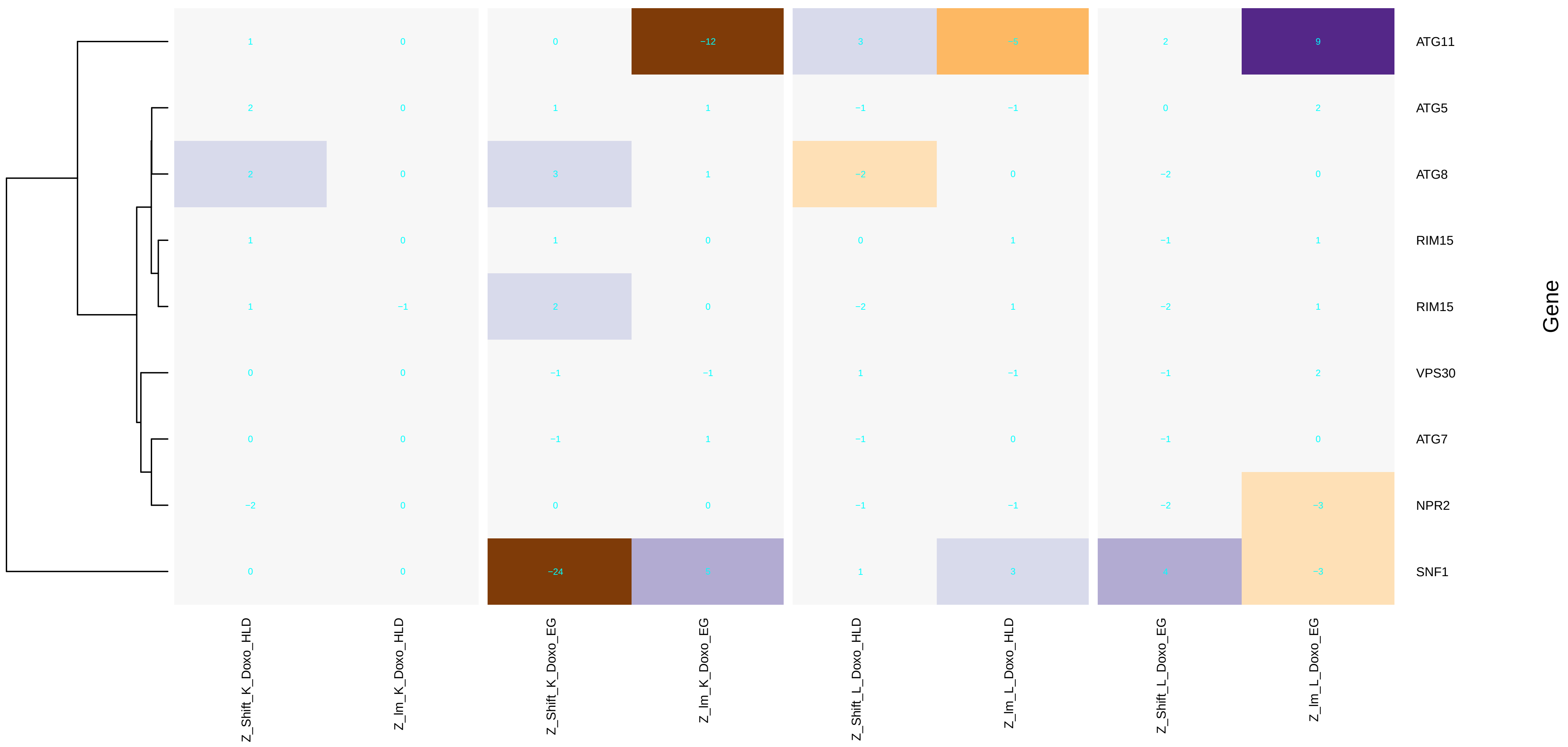

### cellular_sphingolipid_homeostasis.pdf

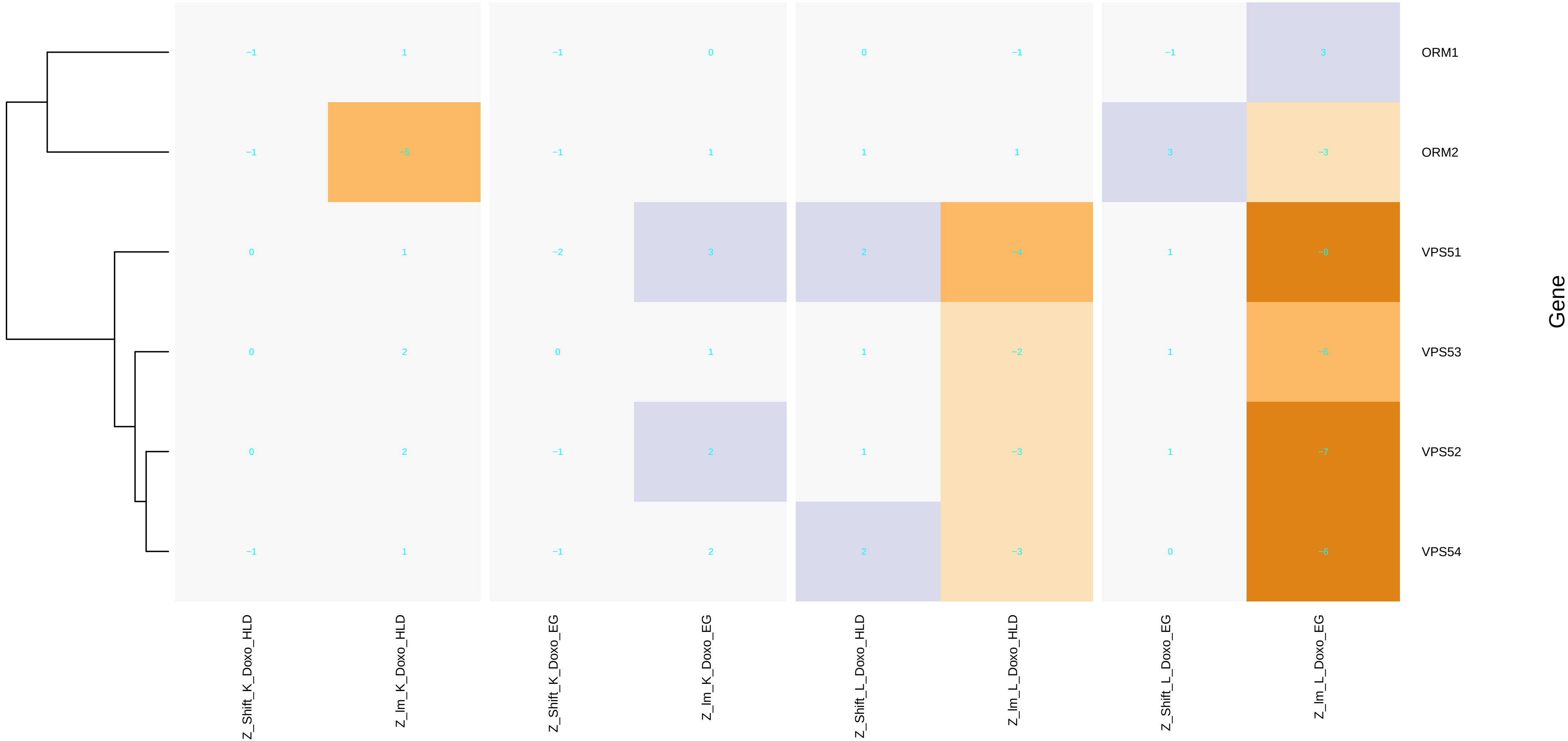

### centromere_complex_assembly.pdf

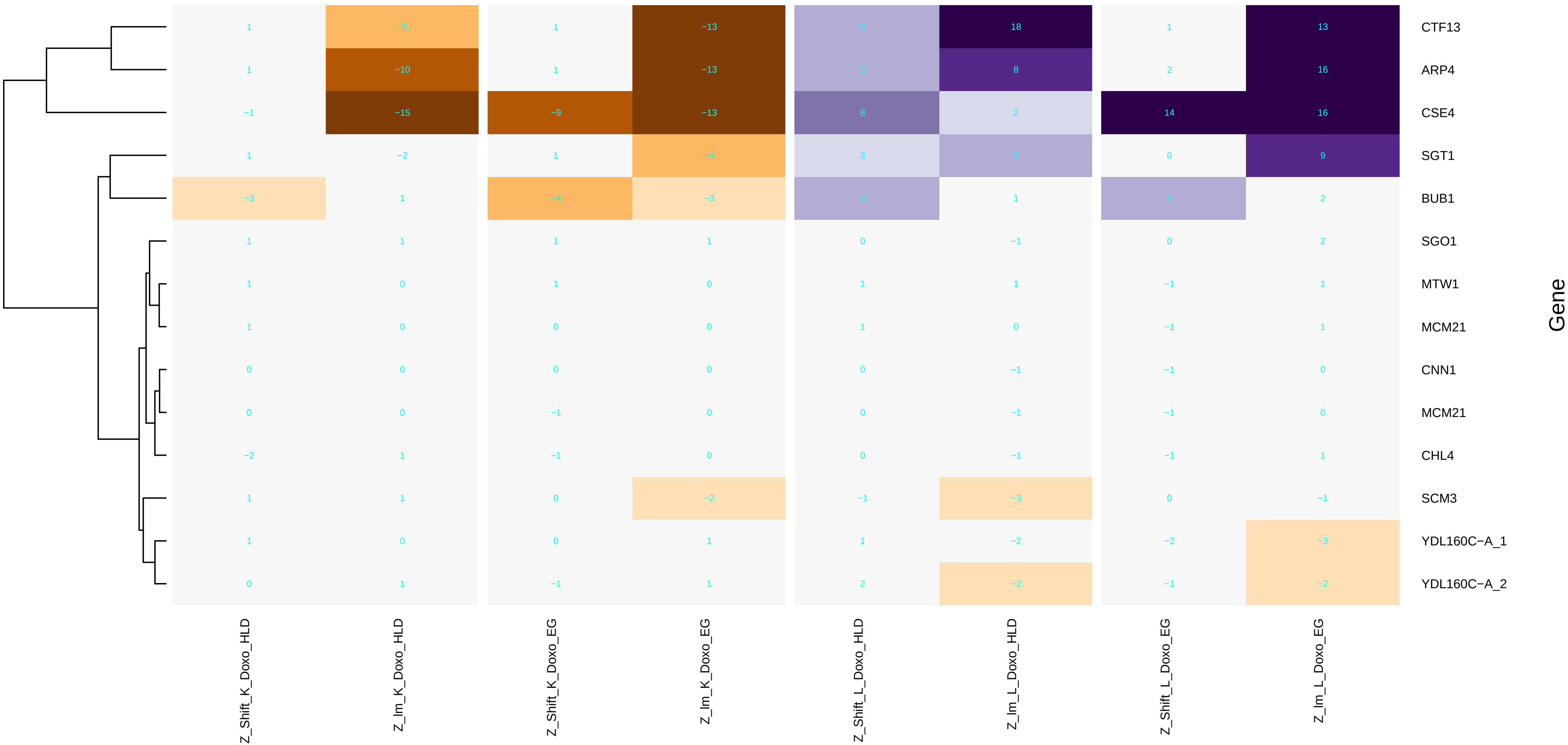

### chromatin_assembly.pdf

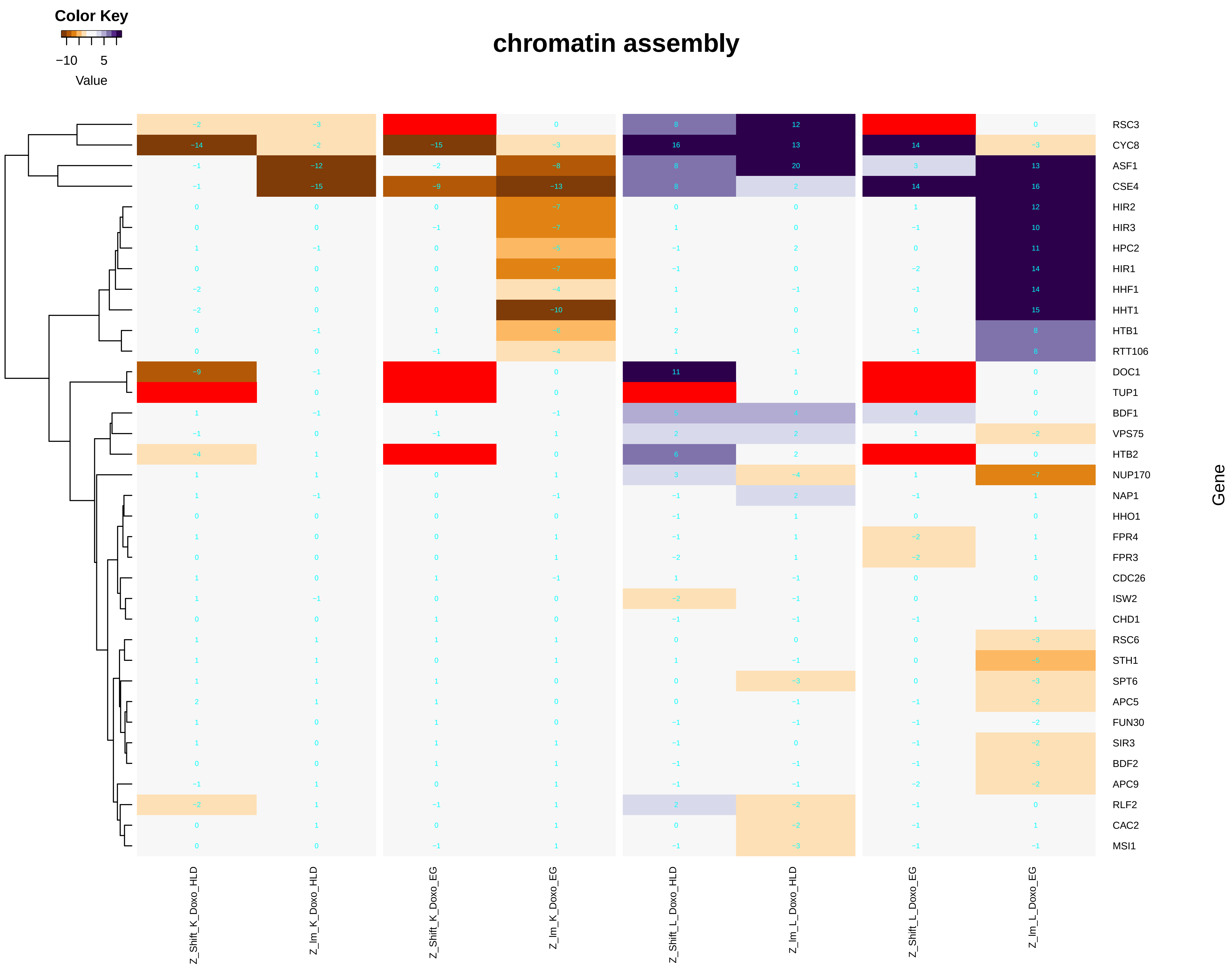

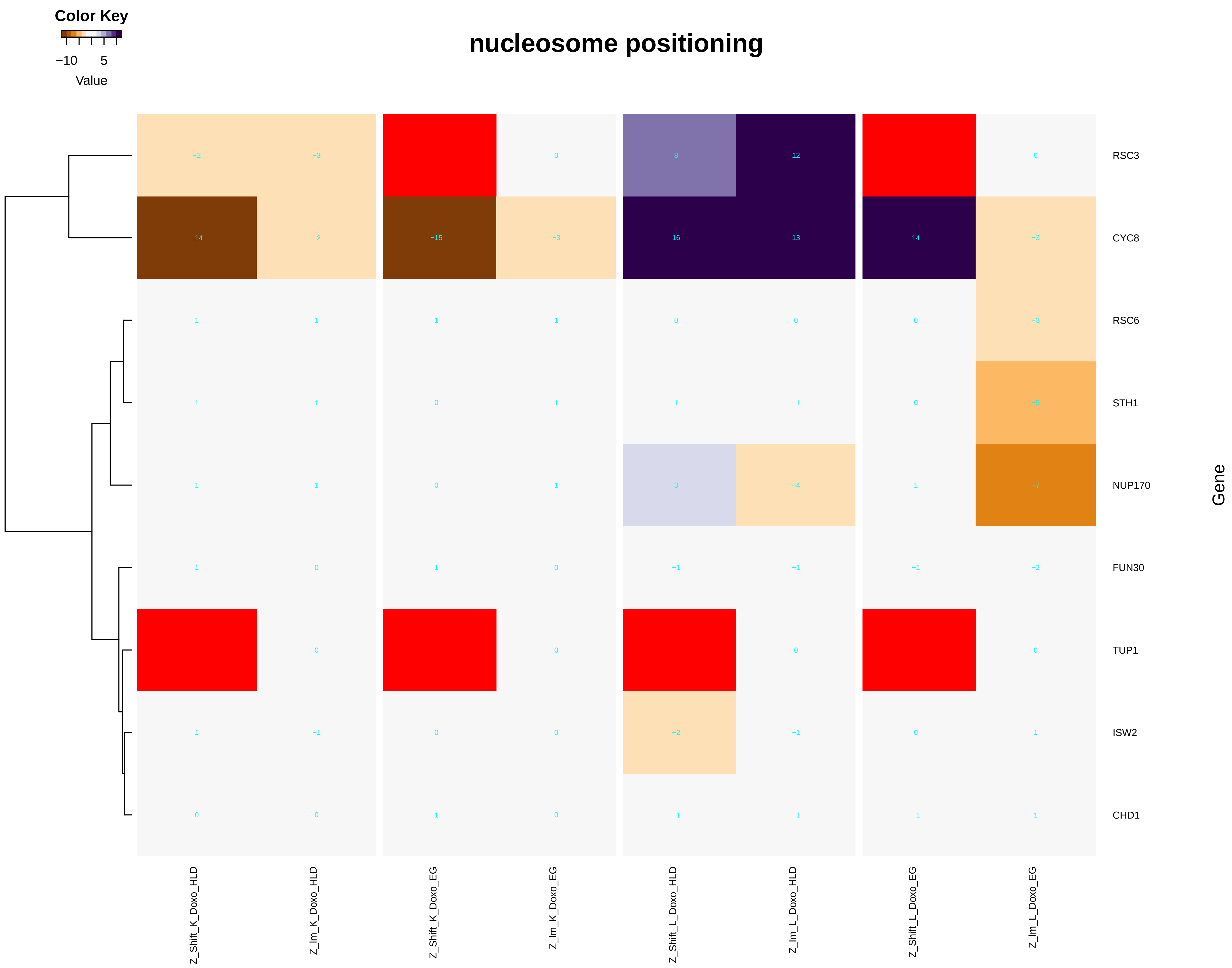

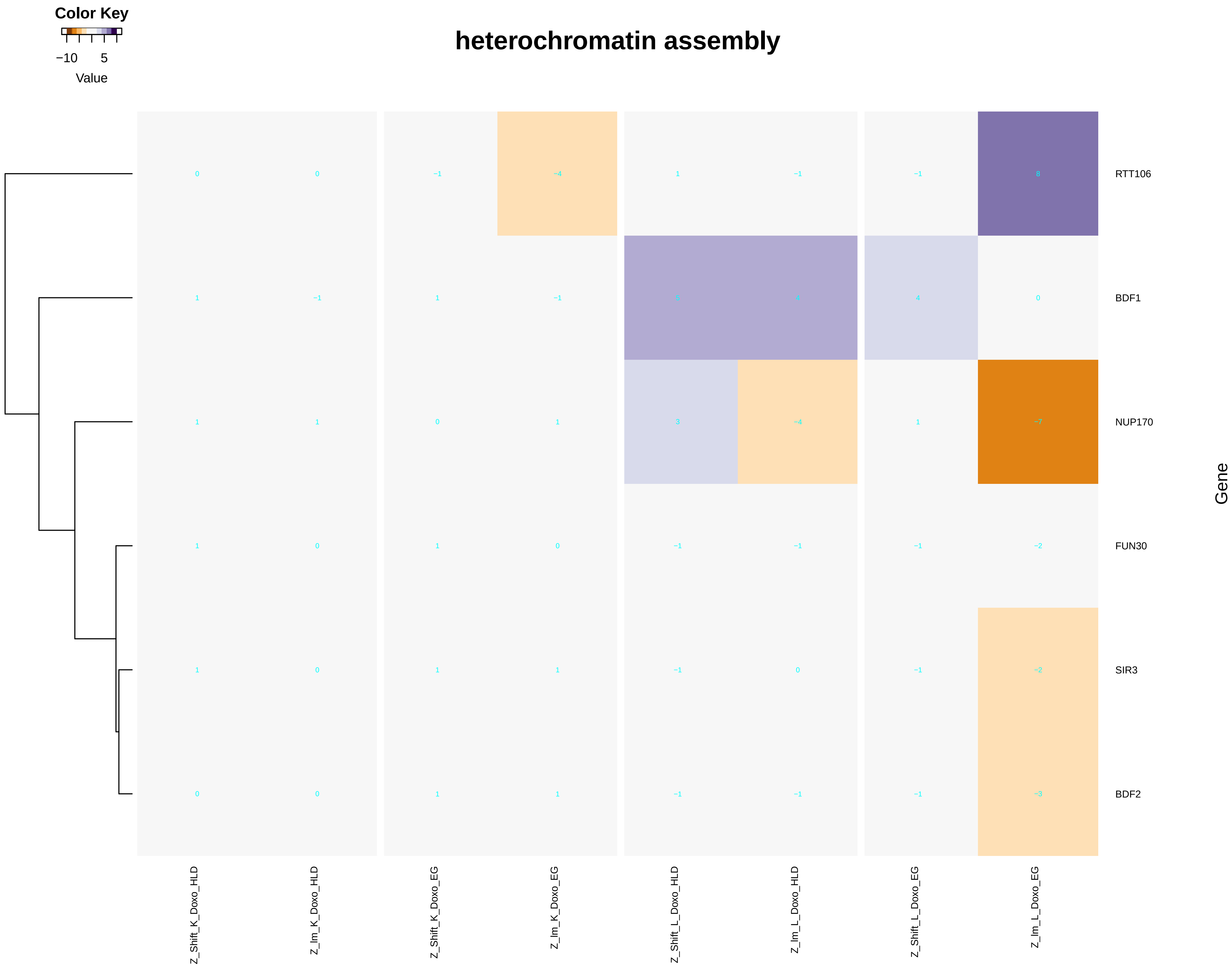

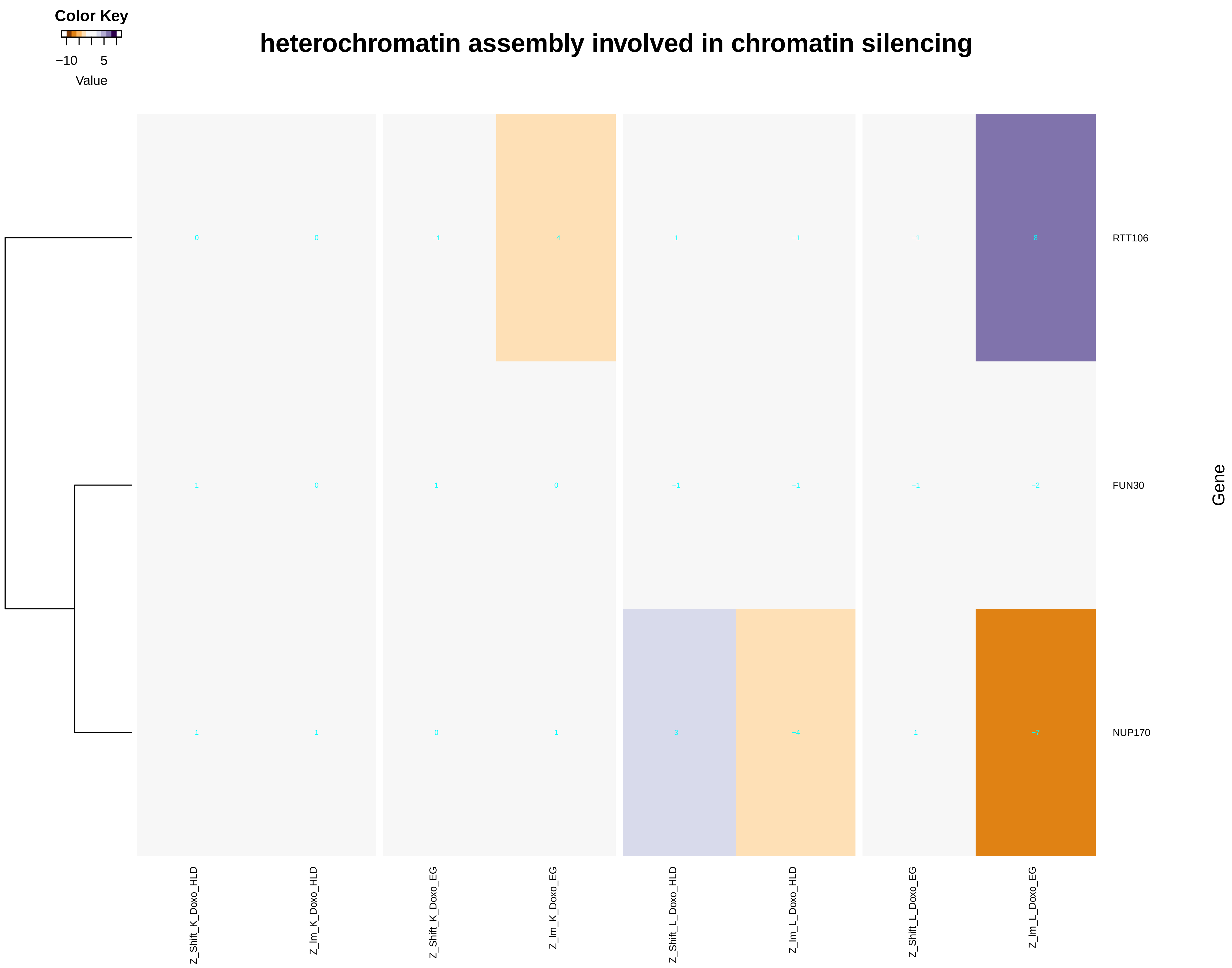

### chromatin_assembly_or_disassembly.pdf

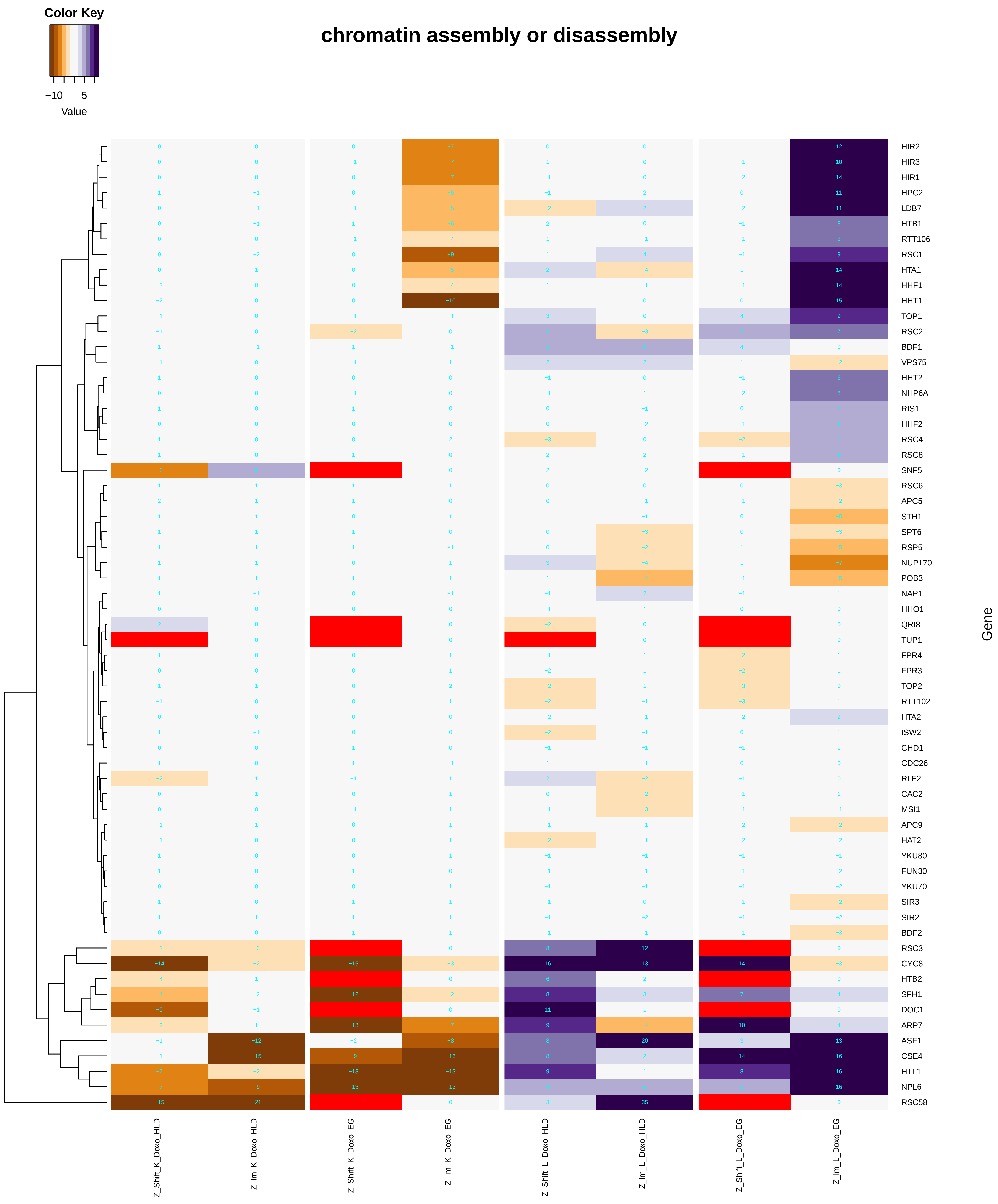

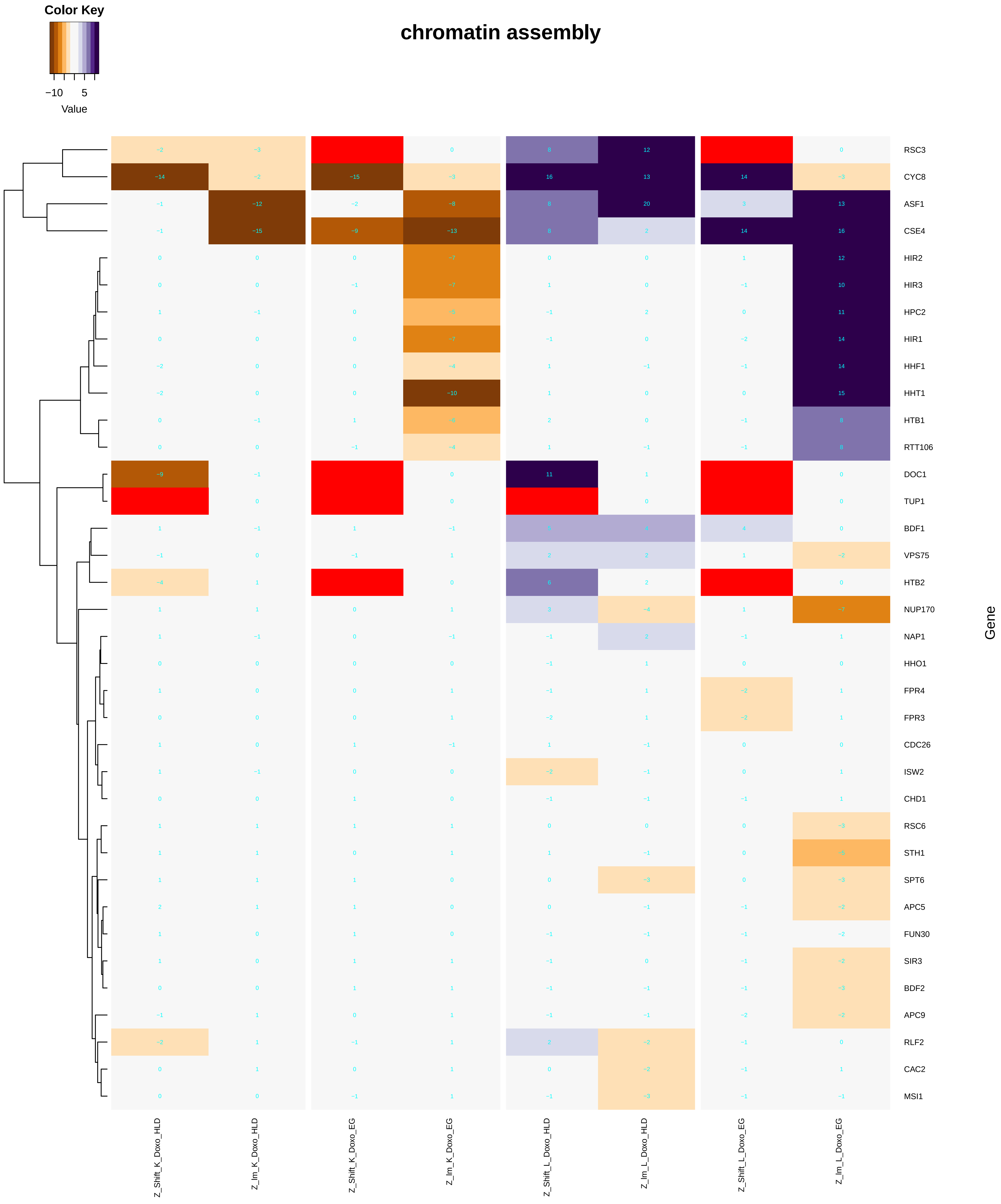

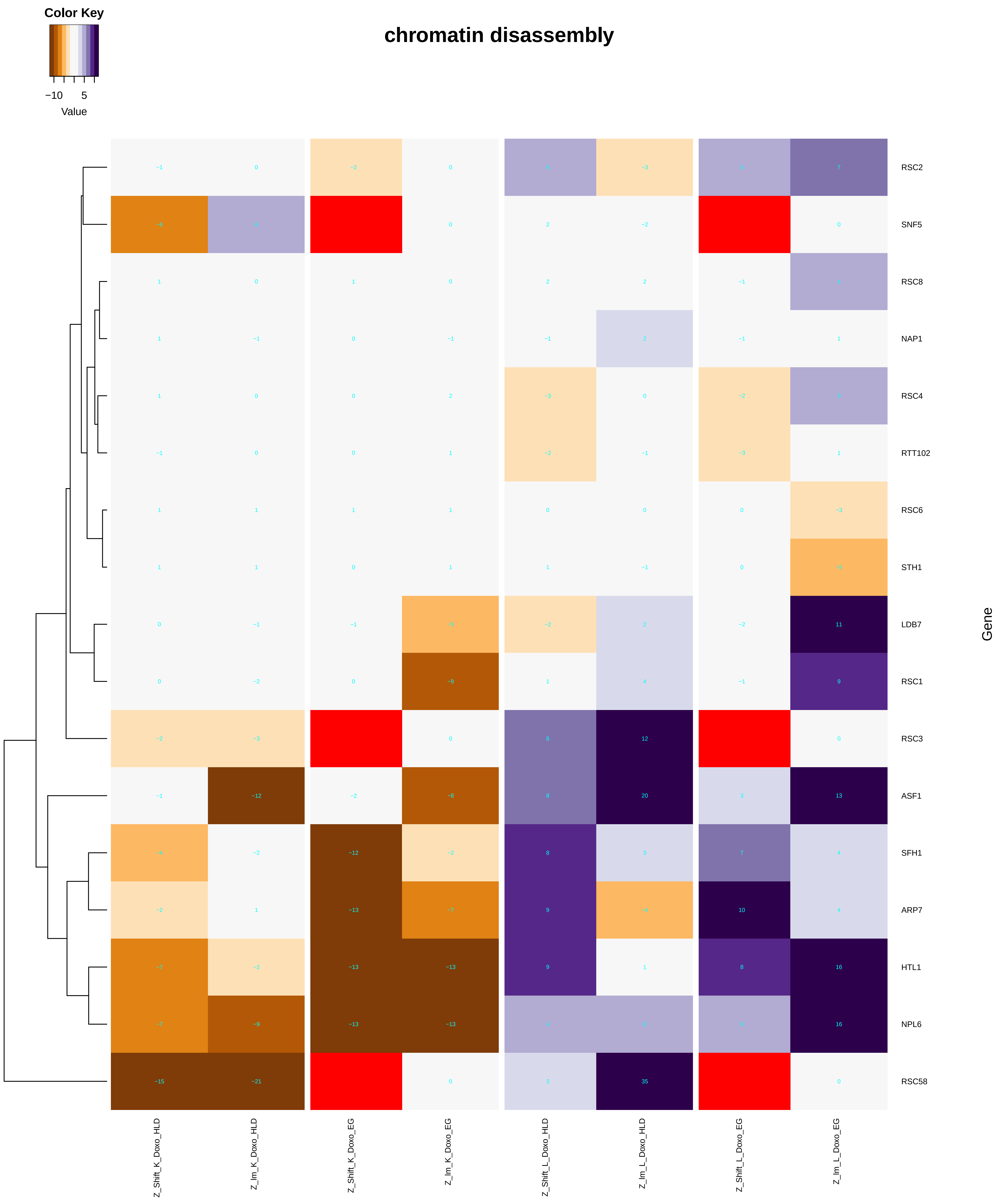

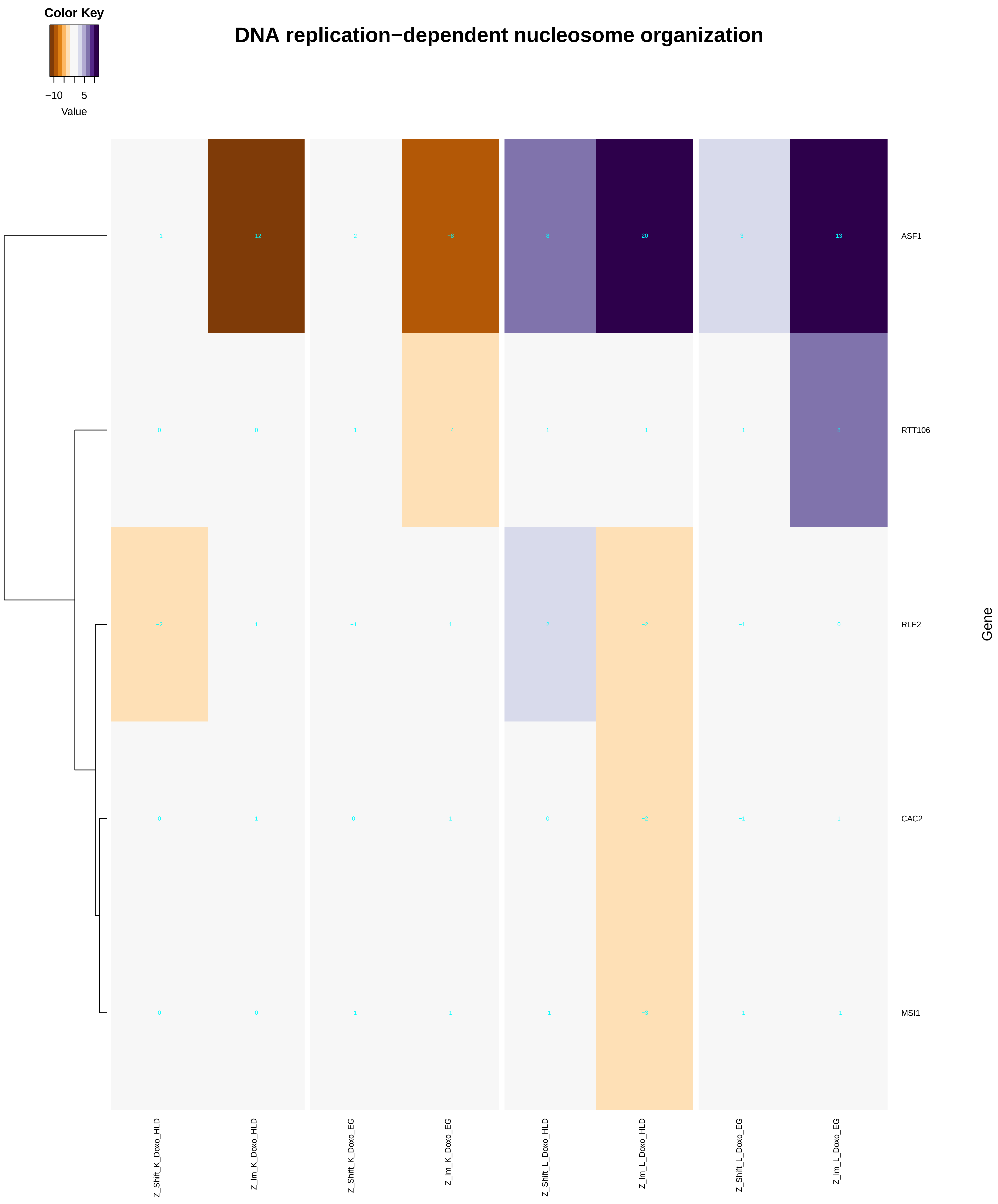

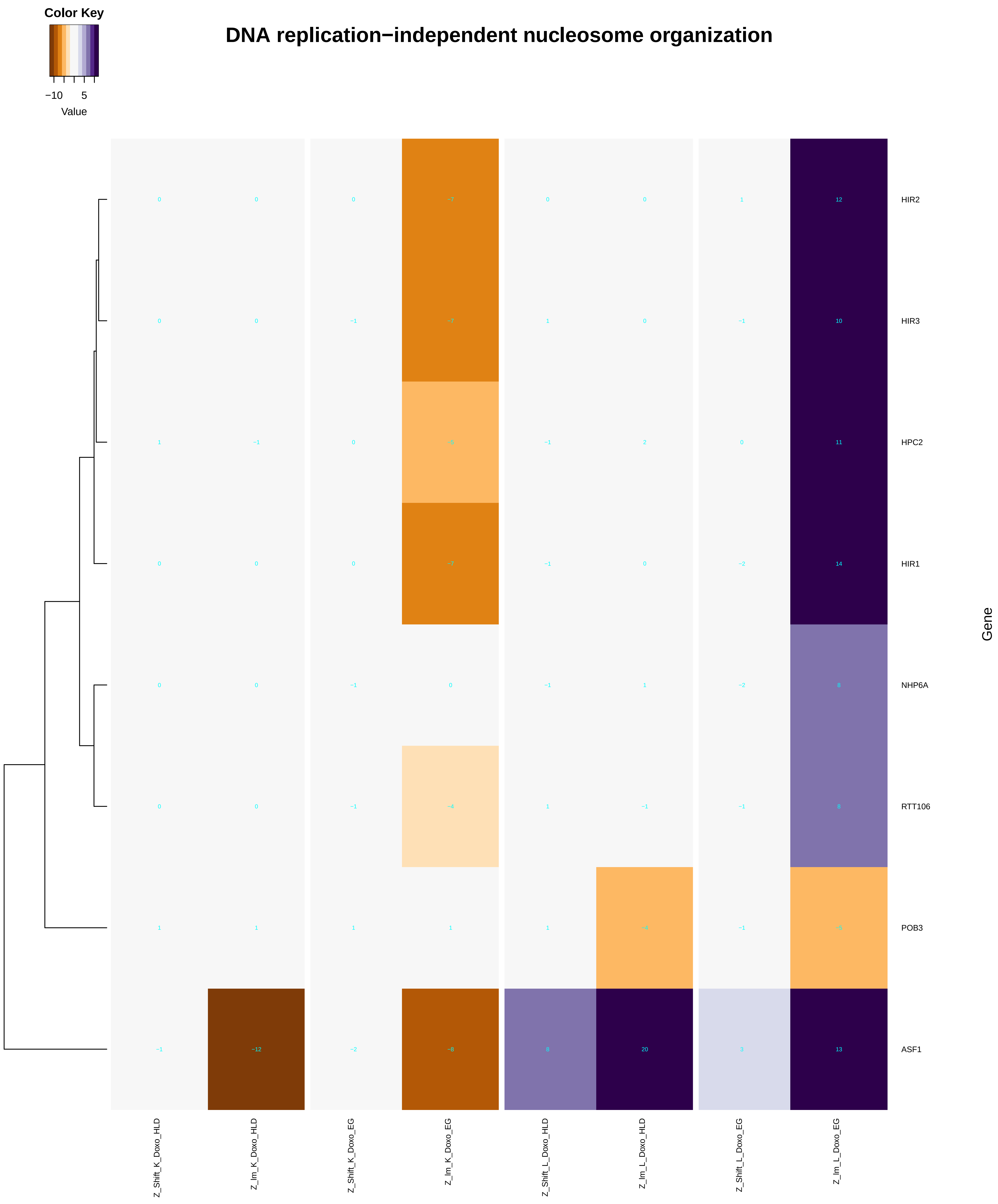

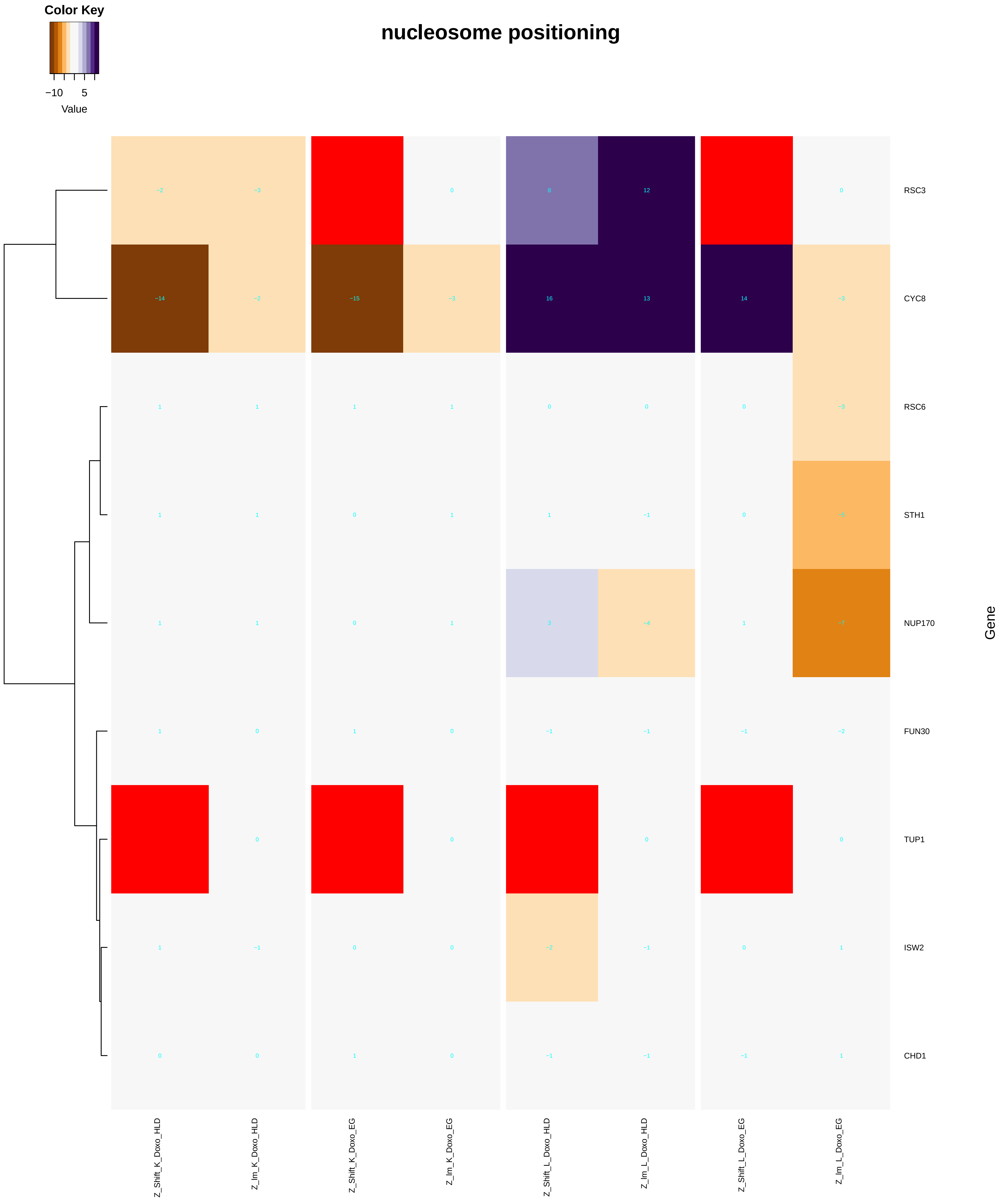

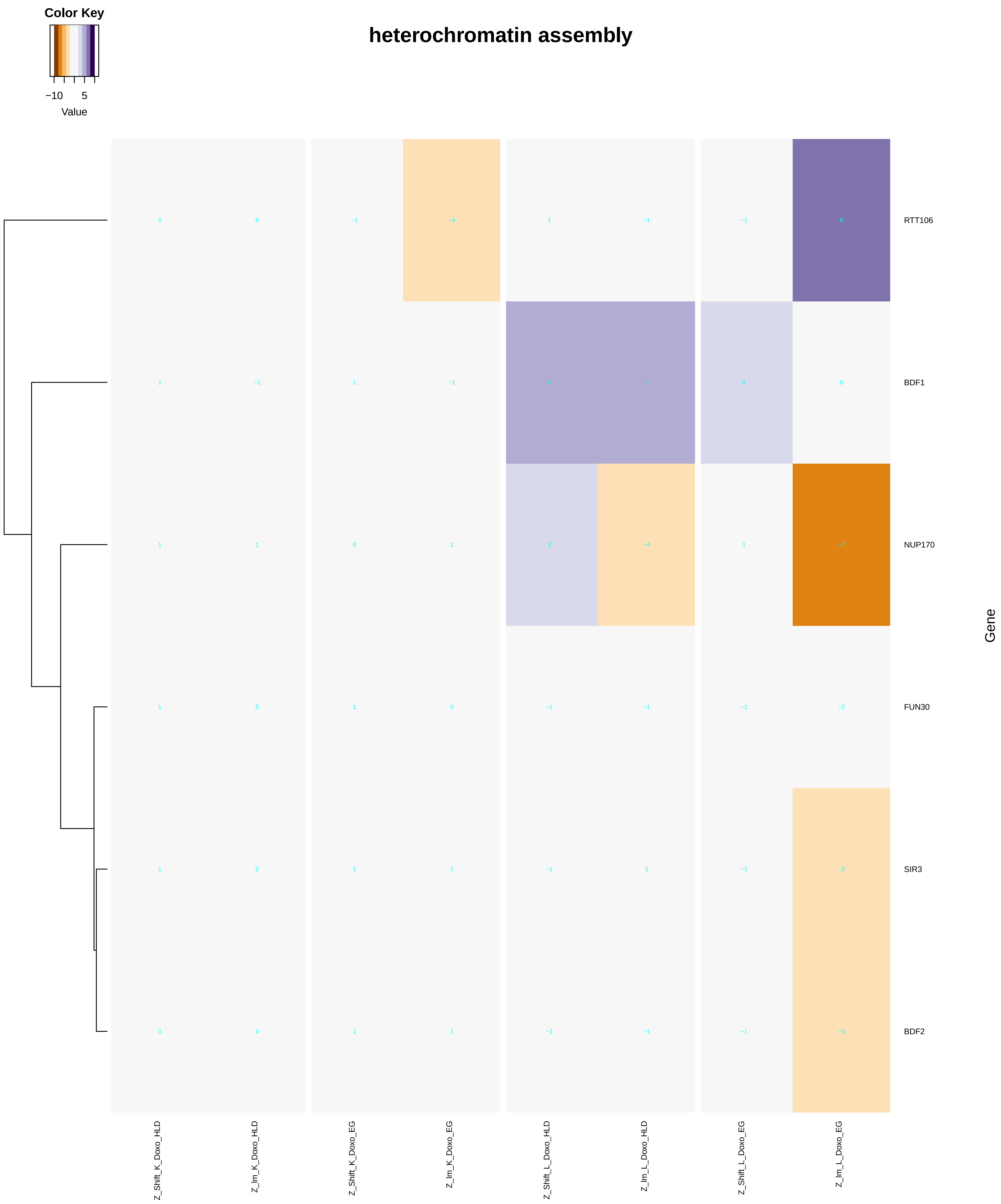

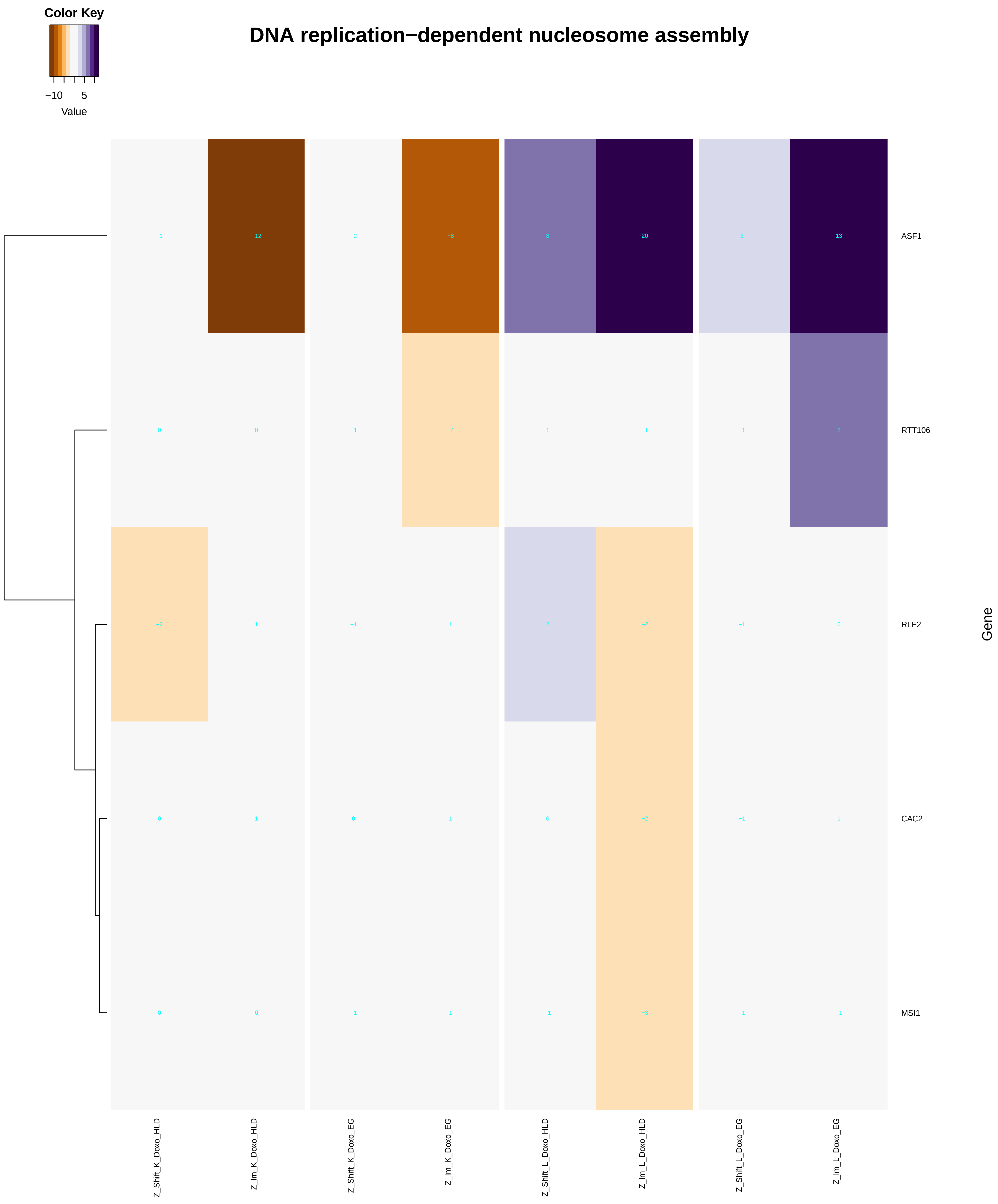

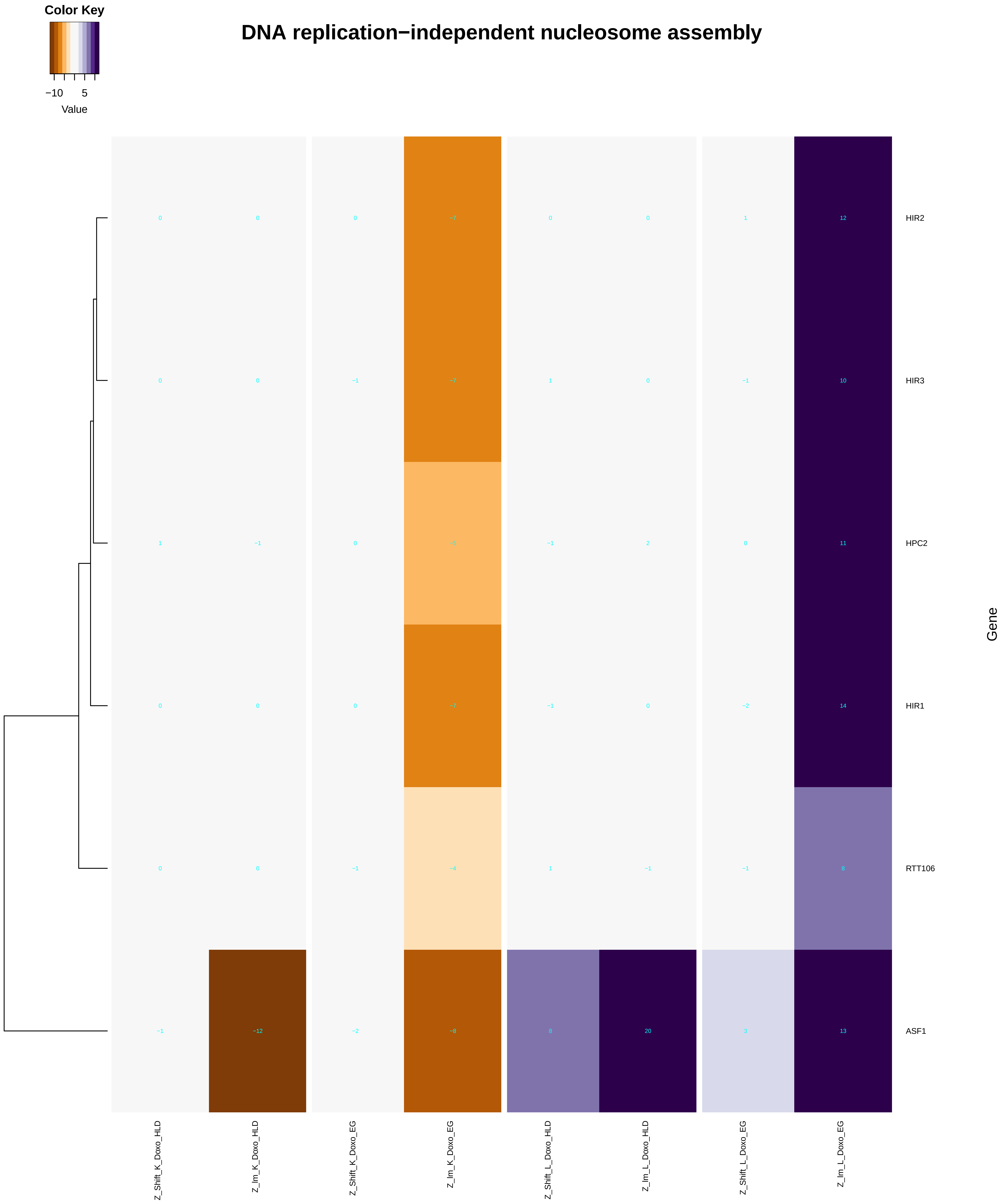

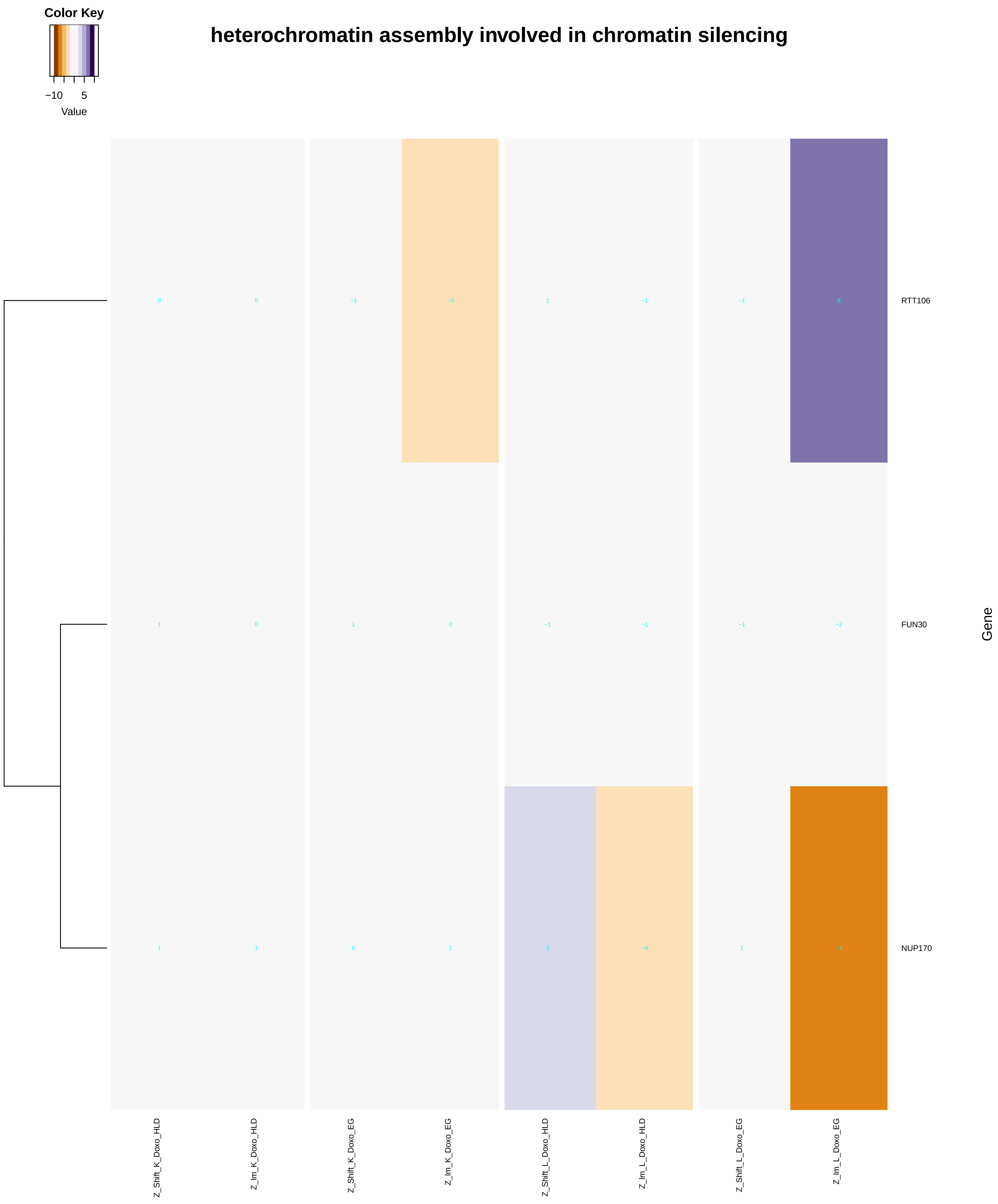

### chromatin_remodeling.pdf

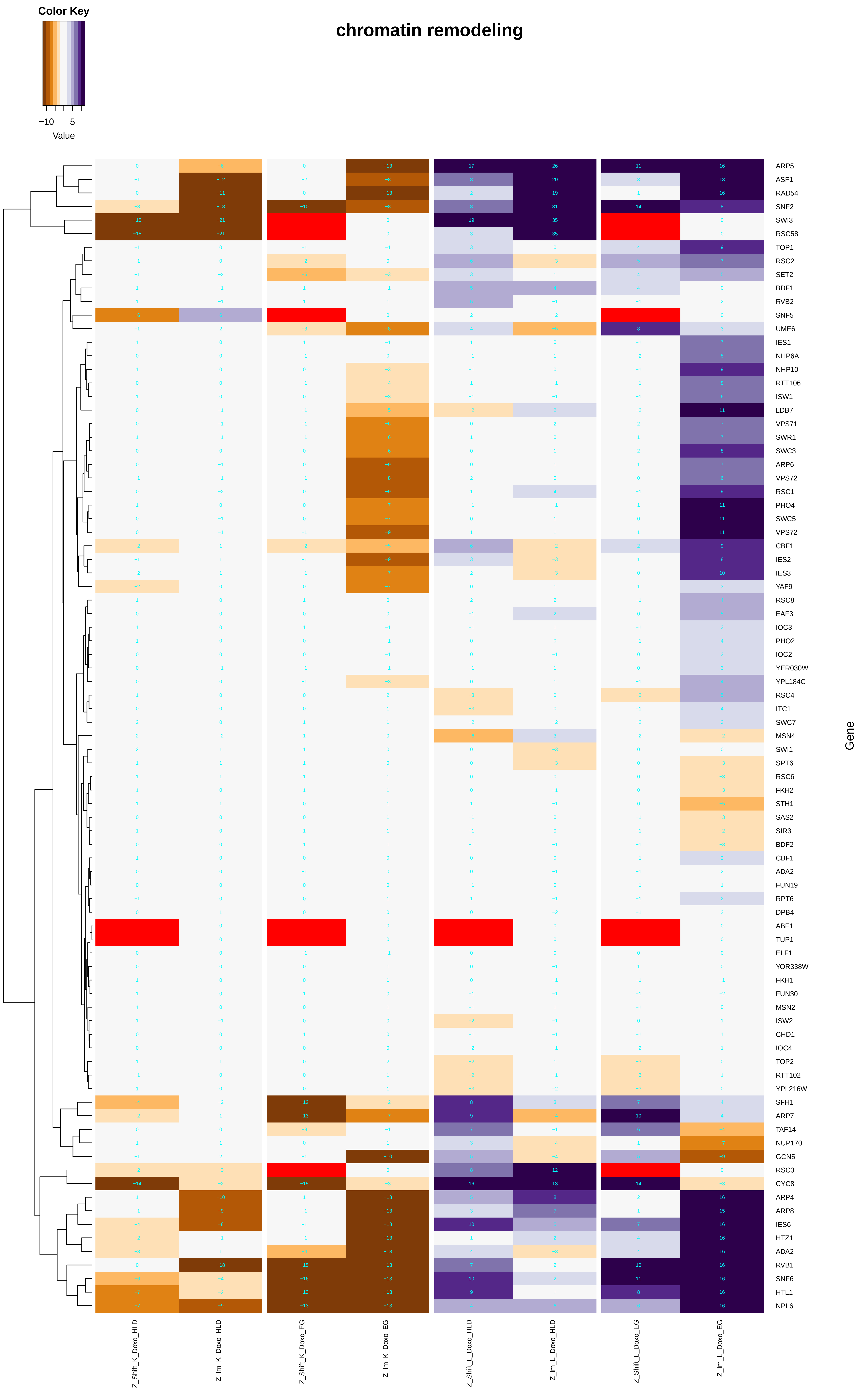

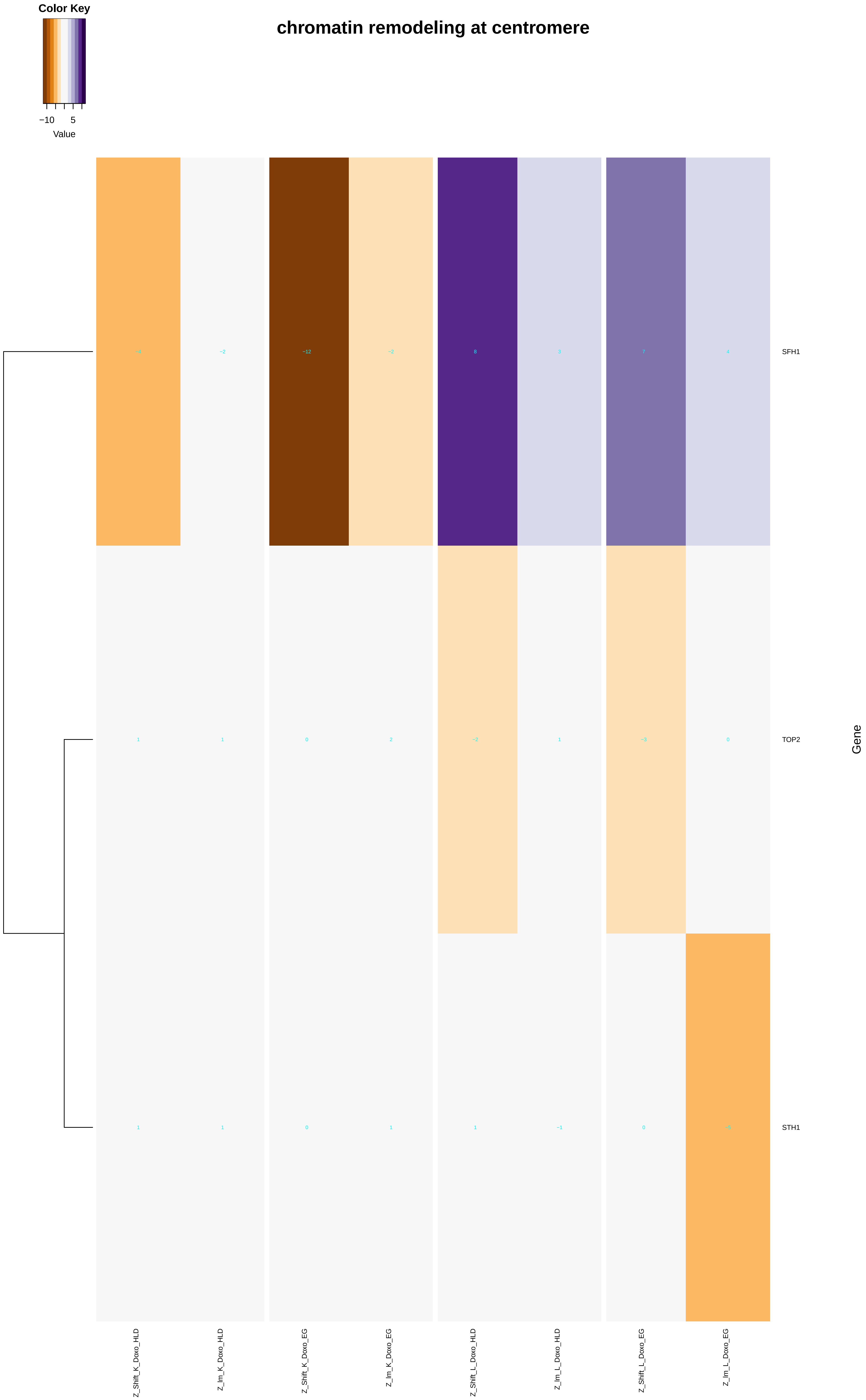

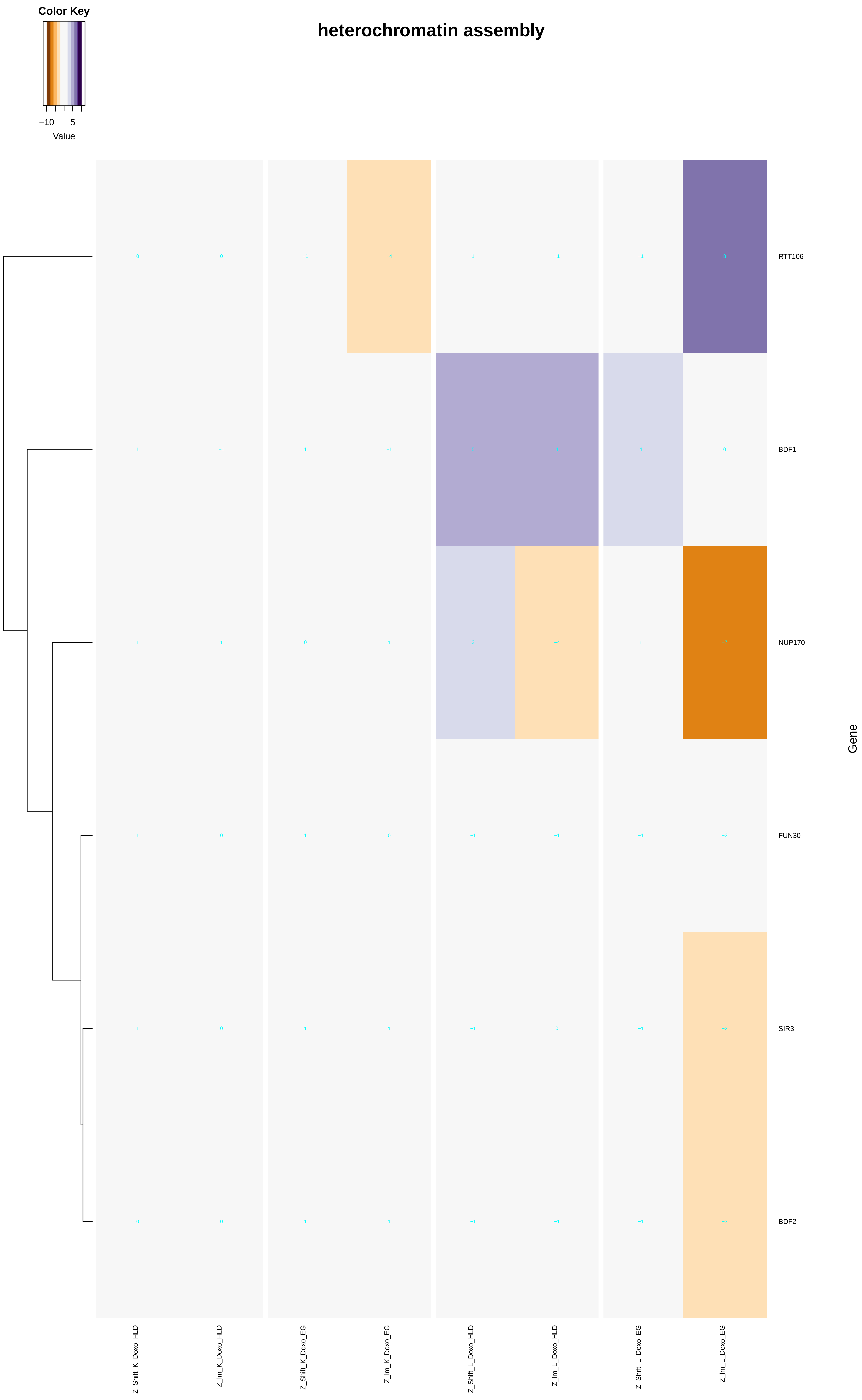

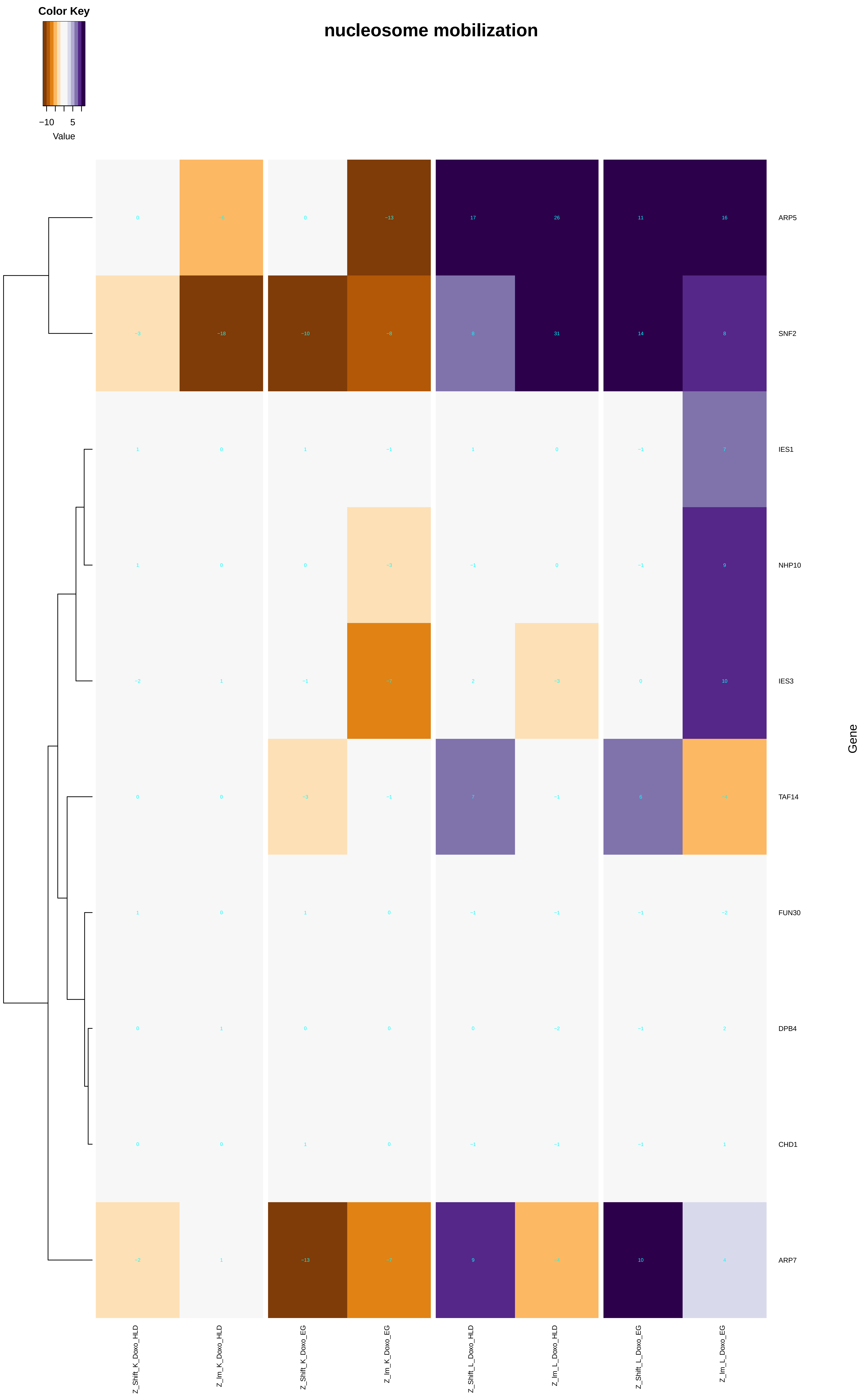

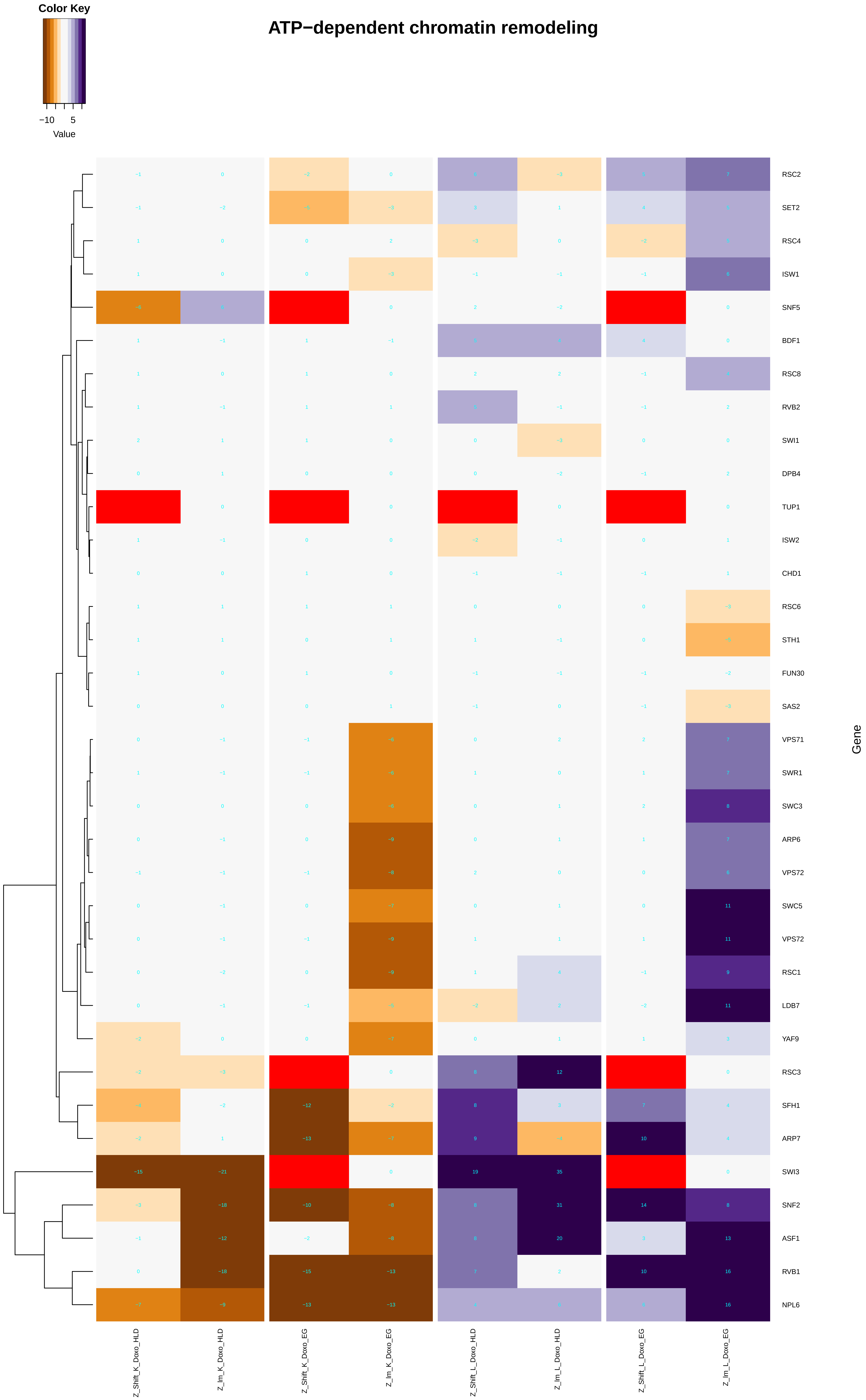

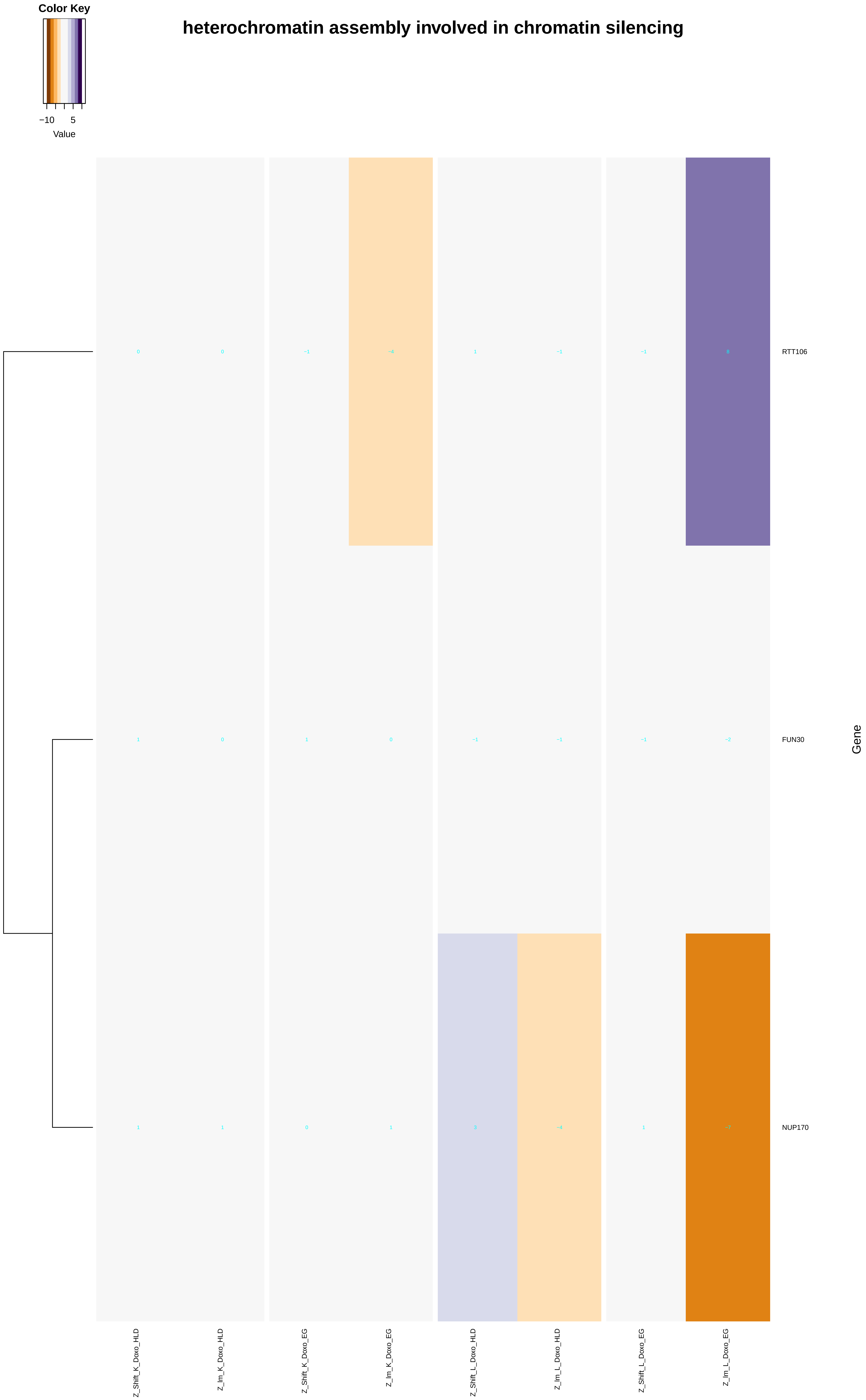

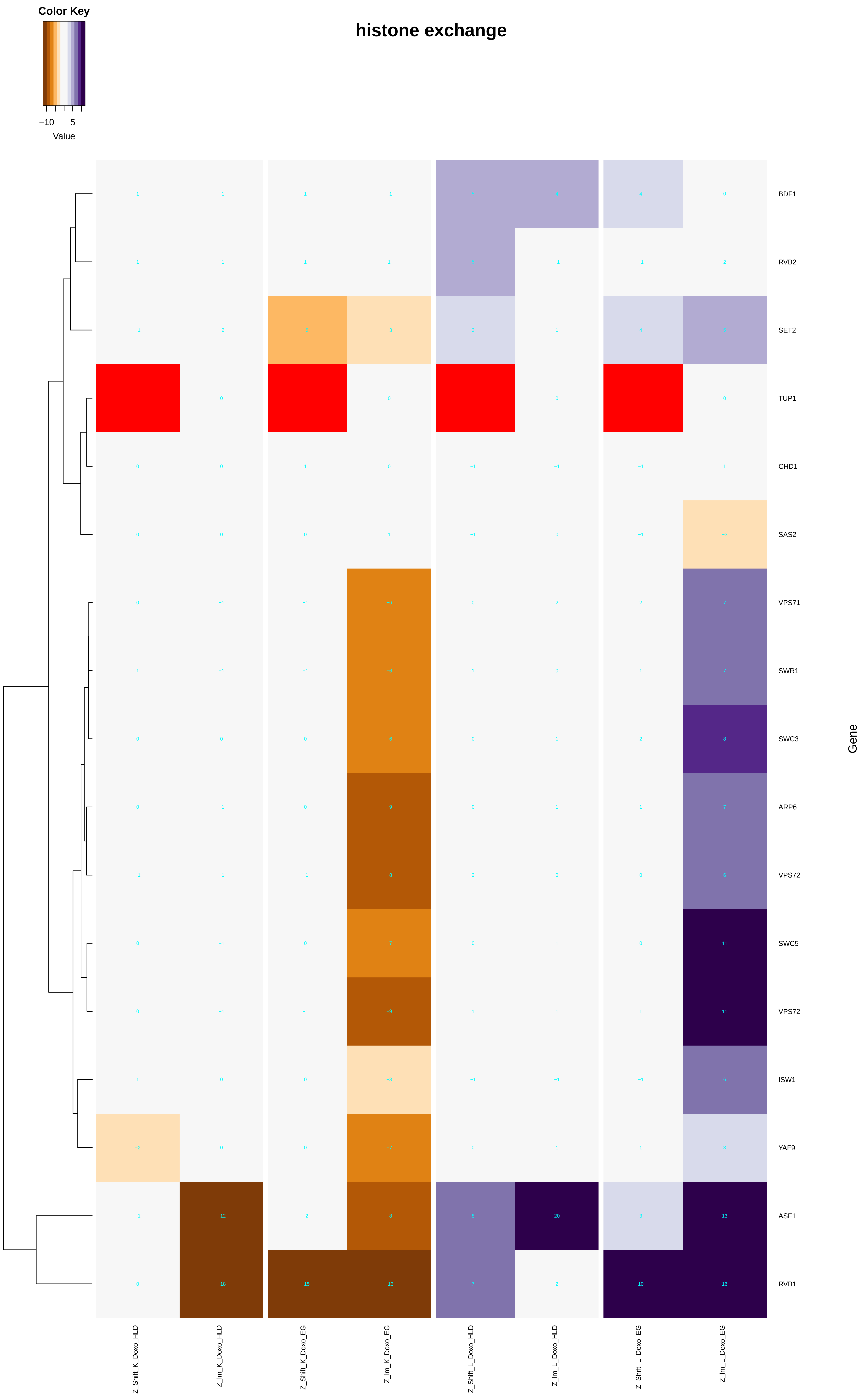

### chromatin_silencing.pdf

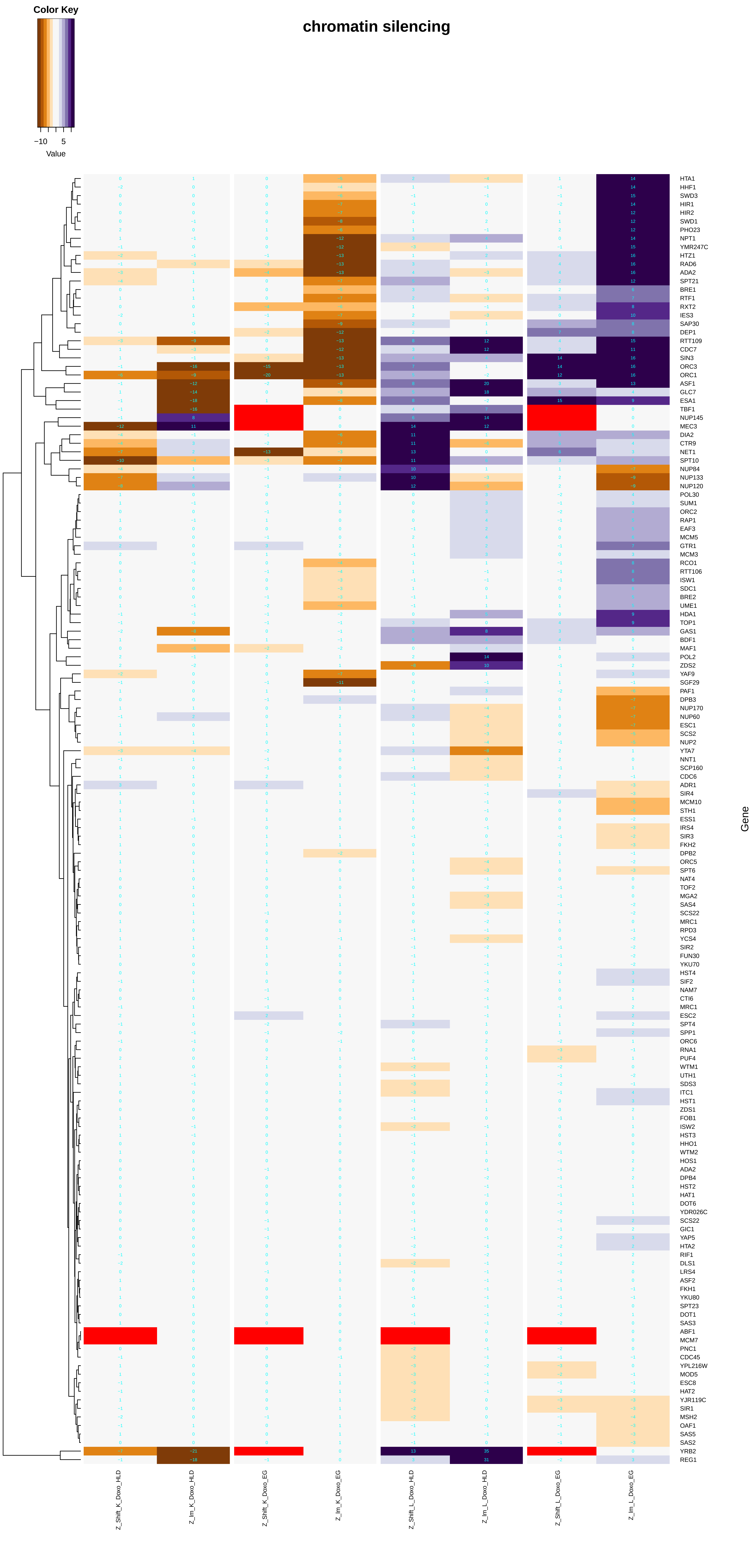

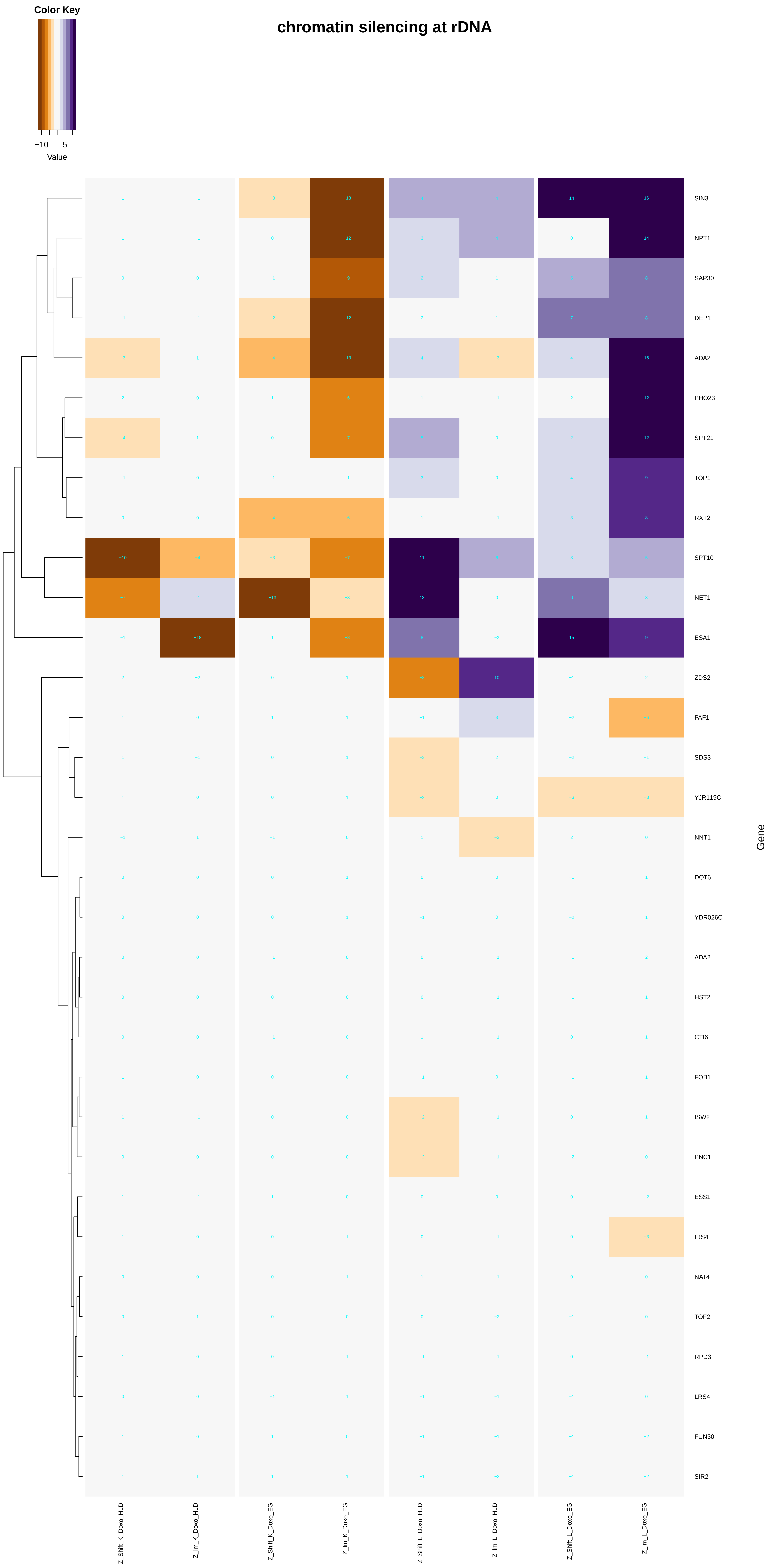

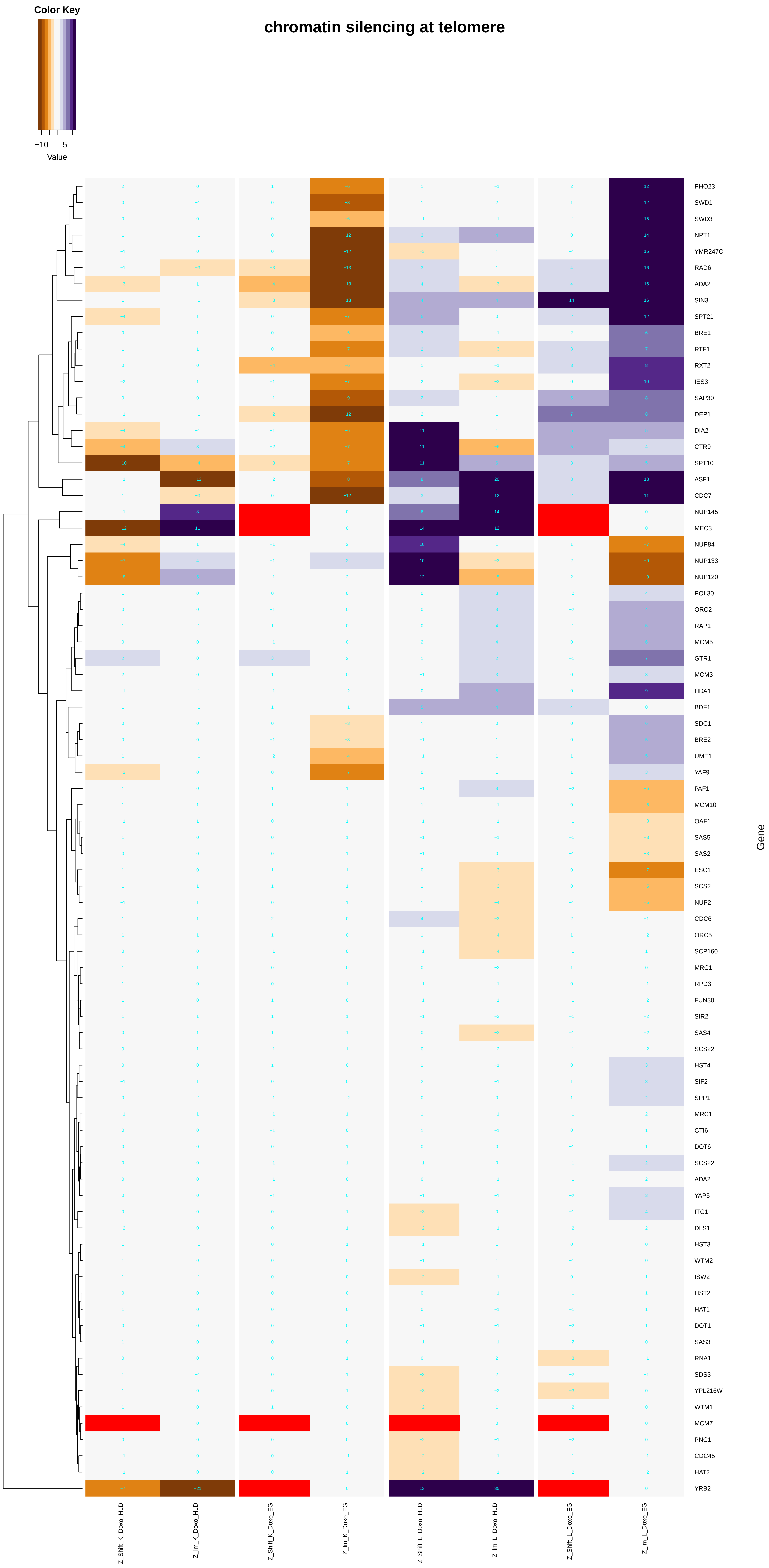

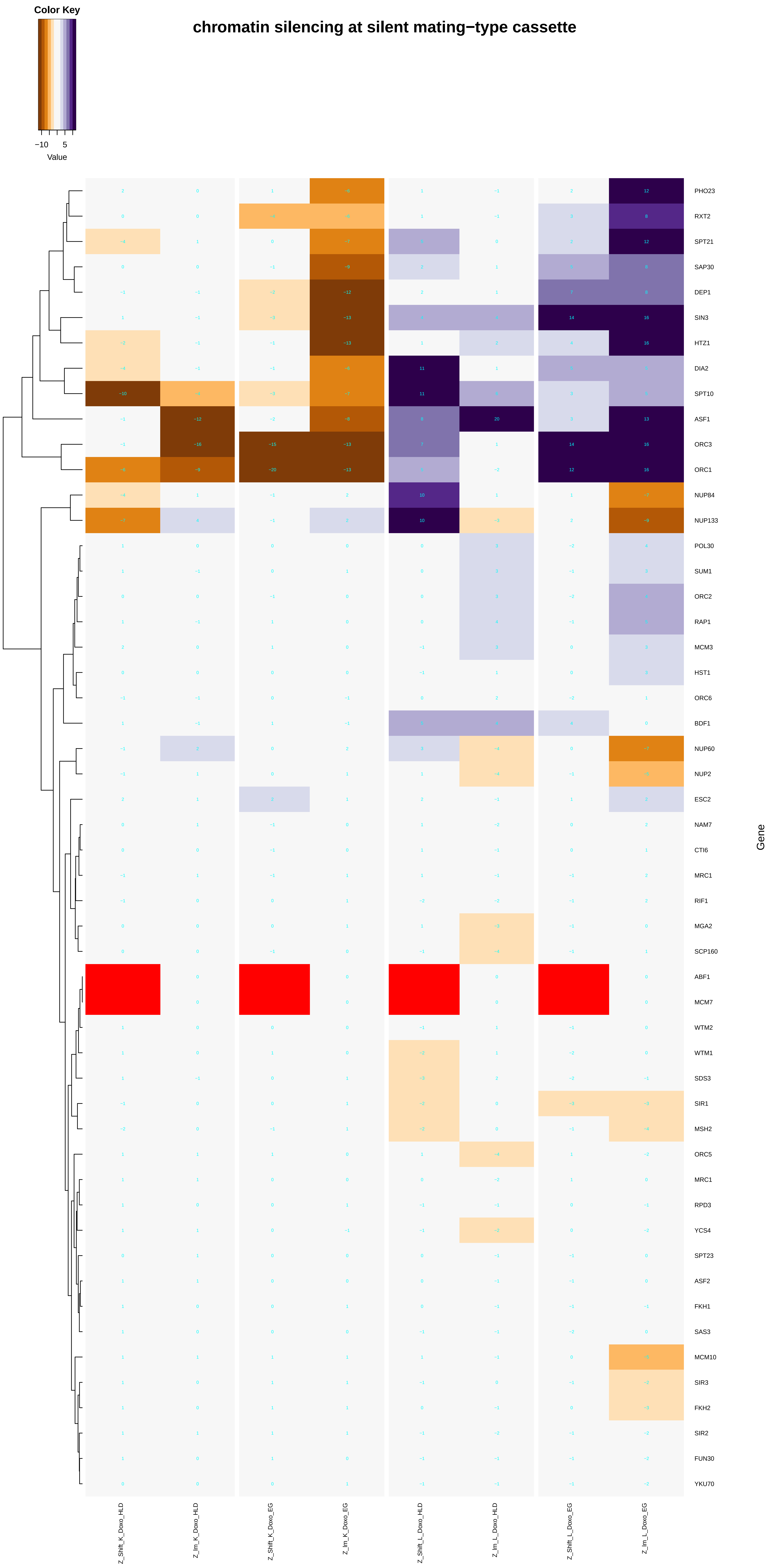

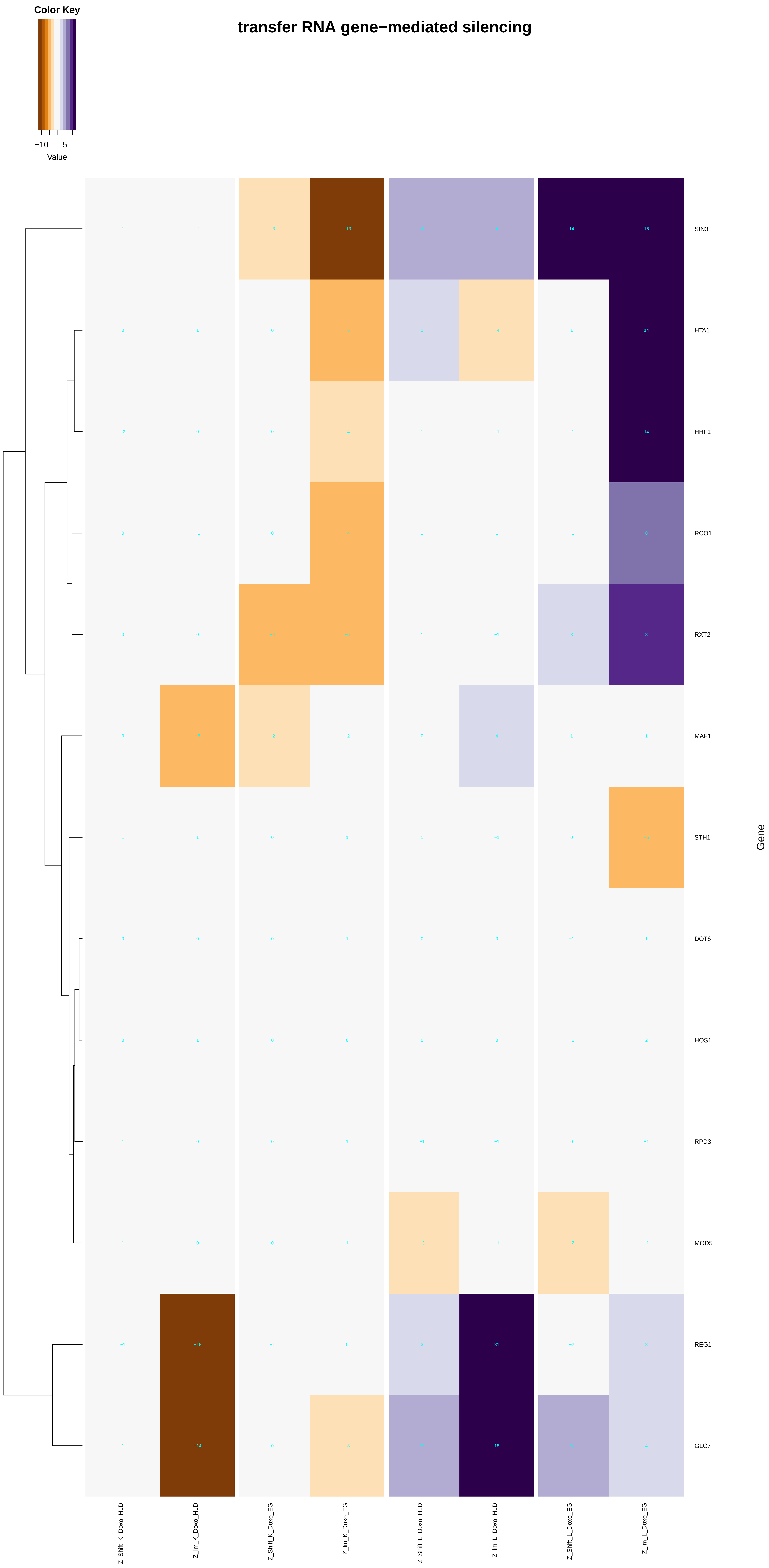

### chromatin_silencing_at_telomere.pdf

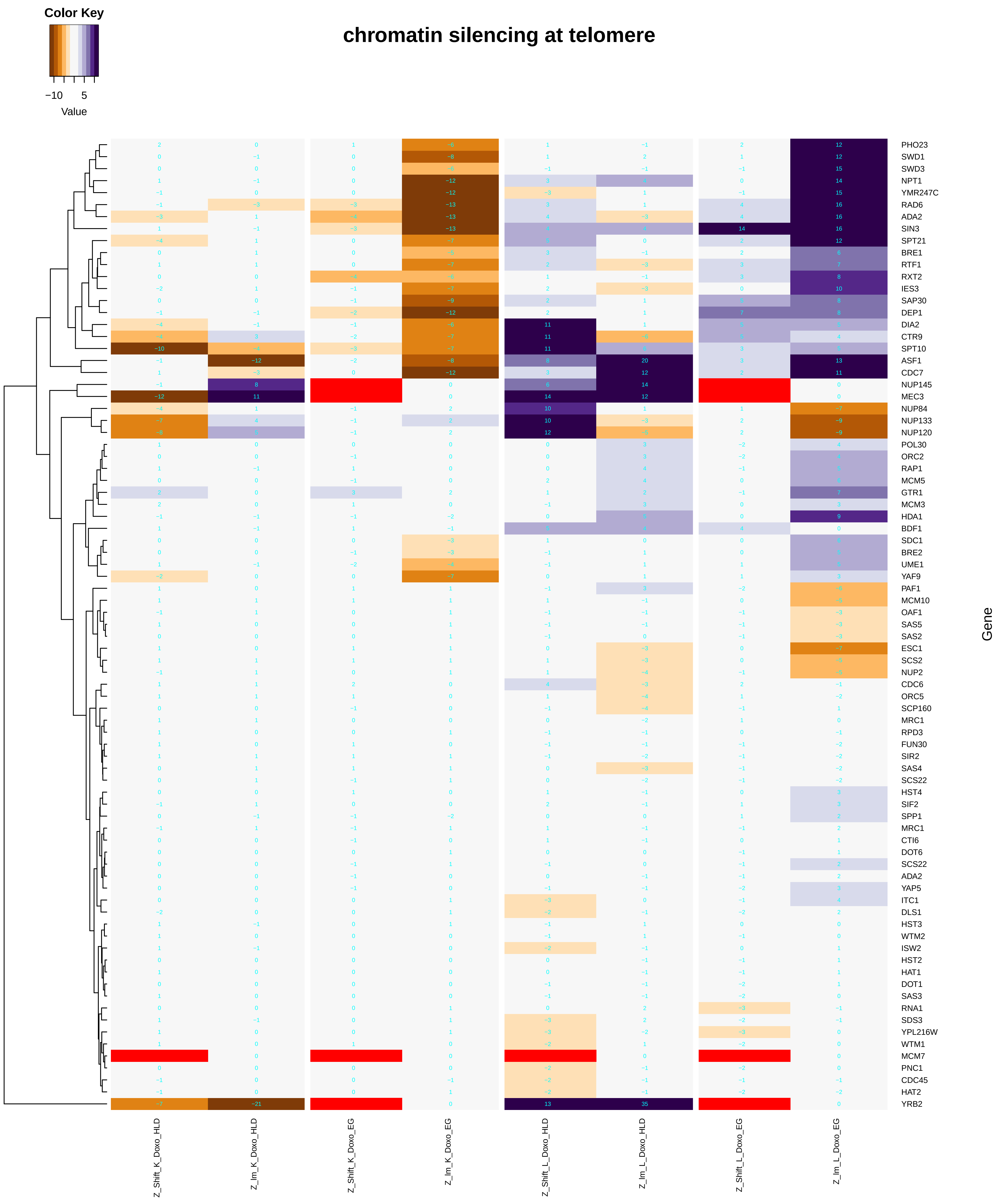

### cleavage_involved_in_rRNA_processing.pdf

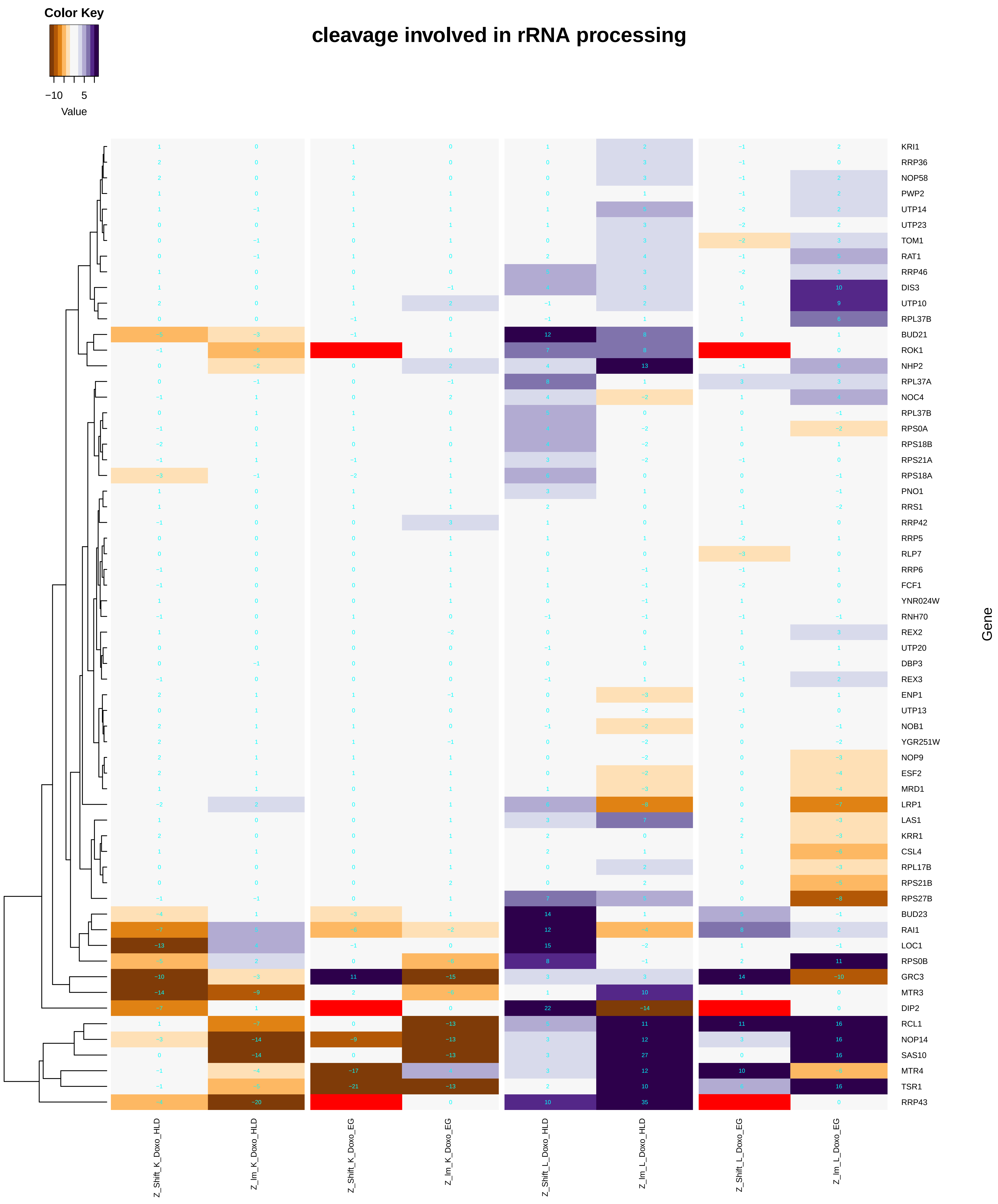

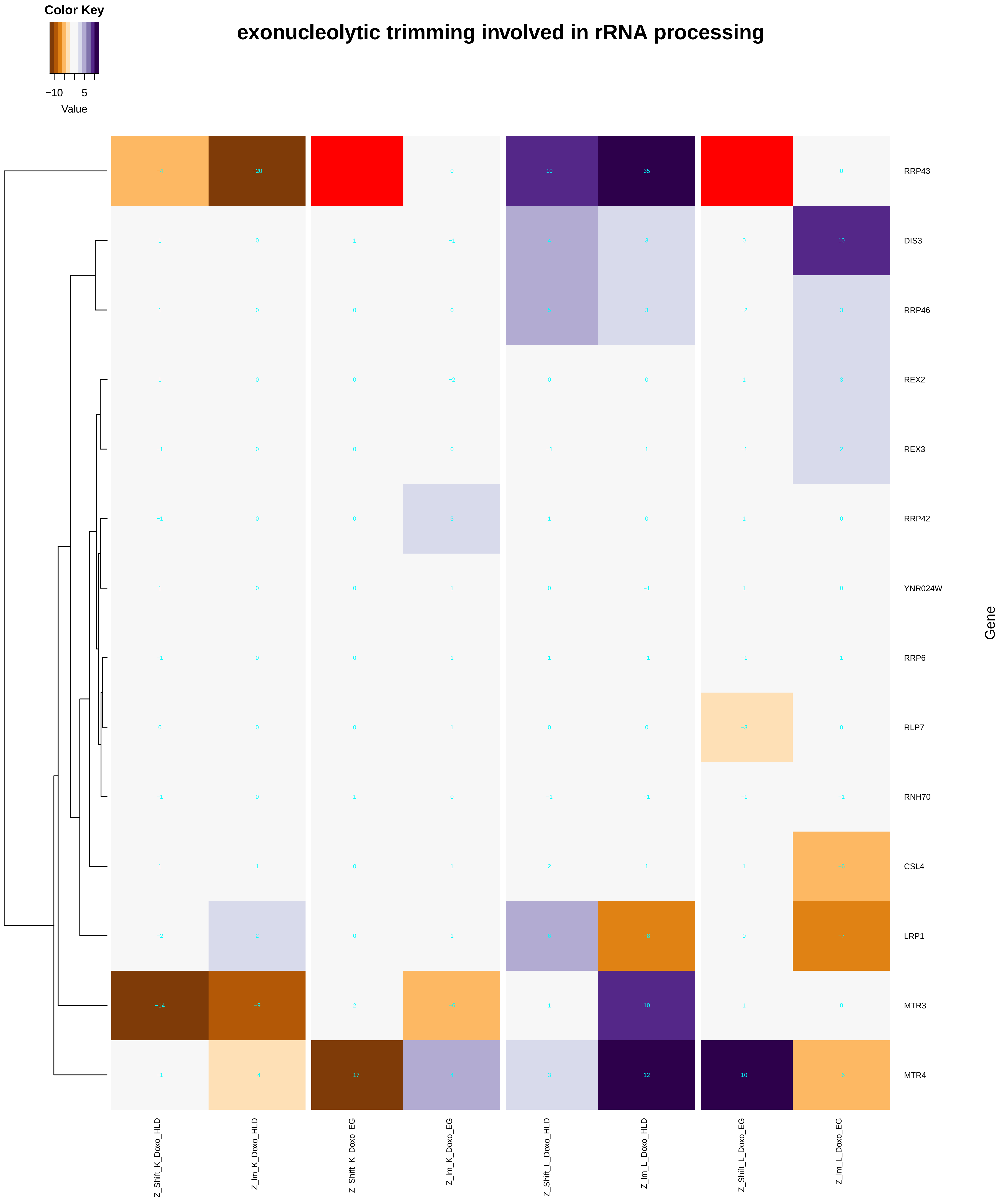

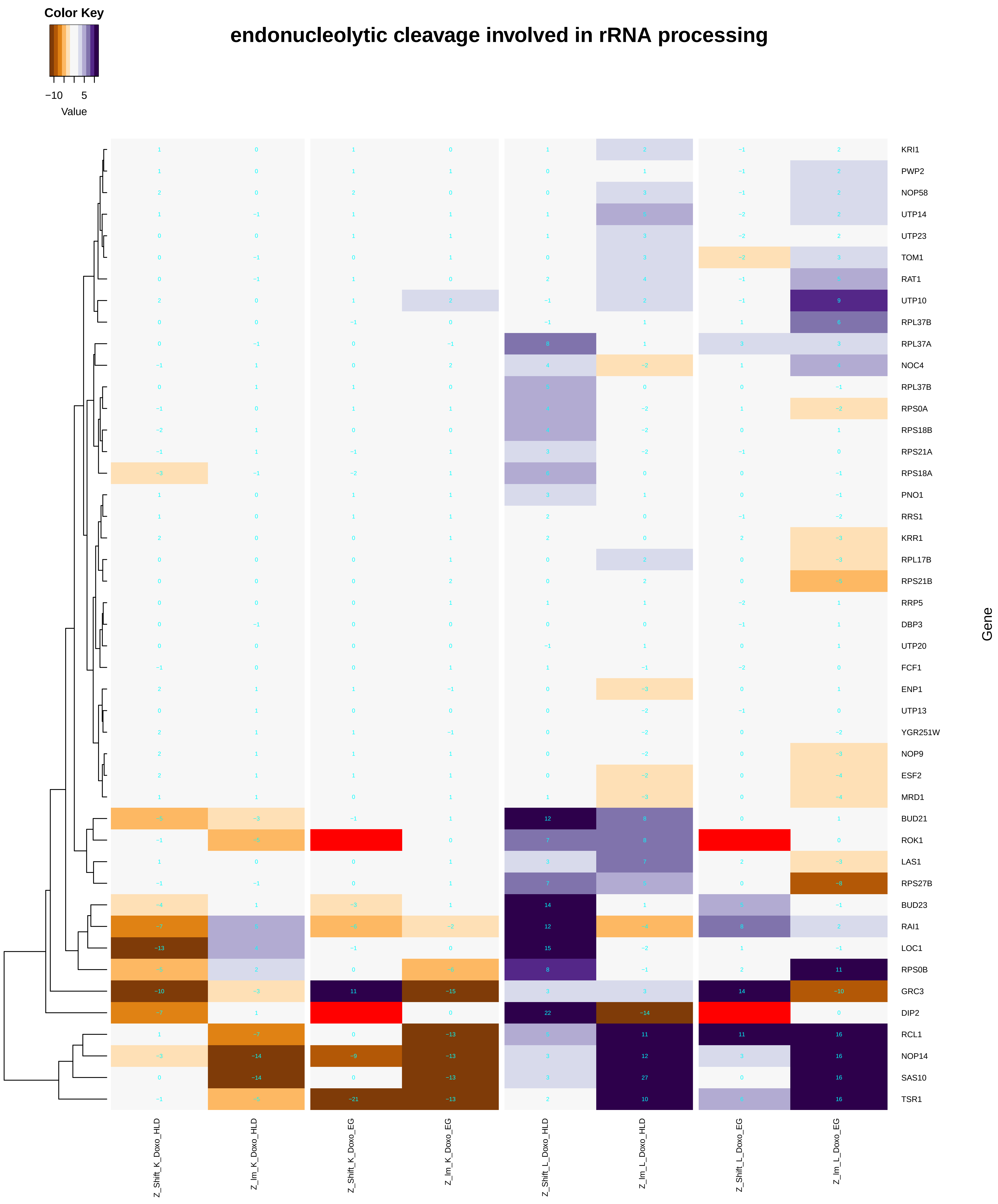

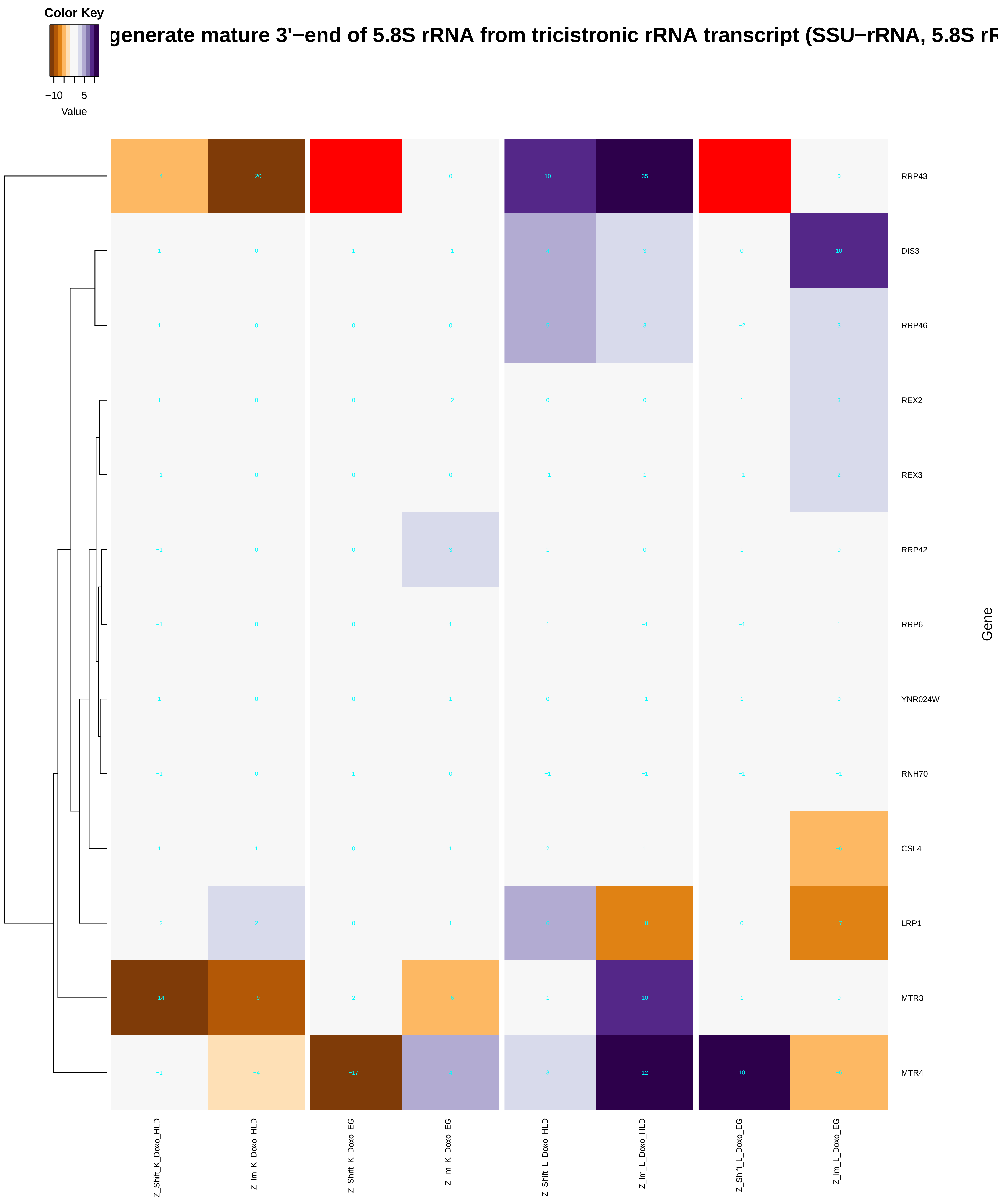

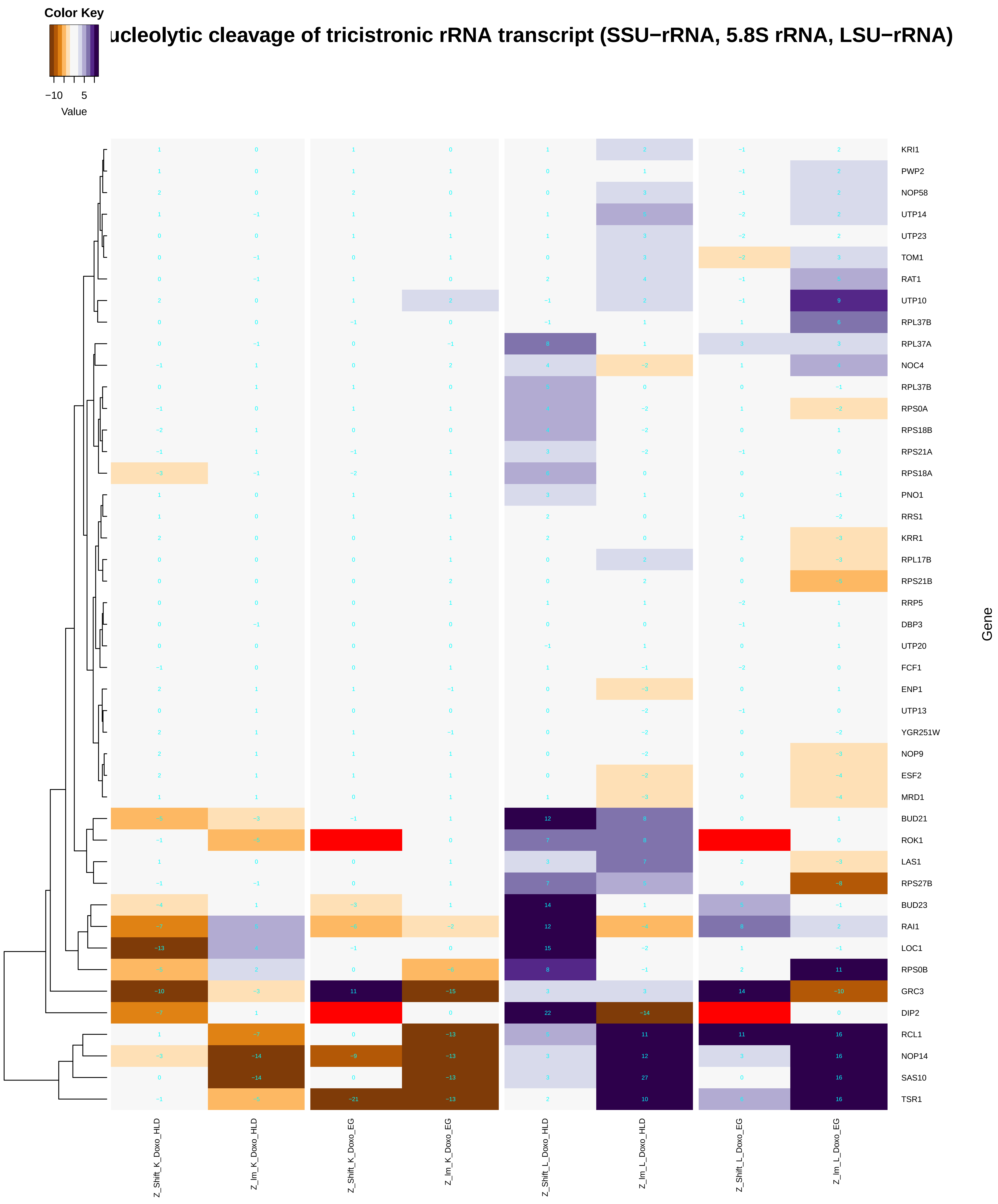

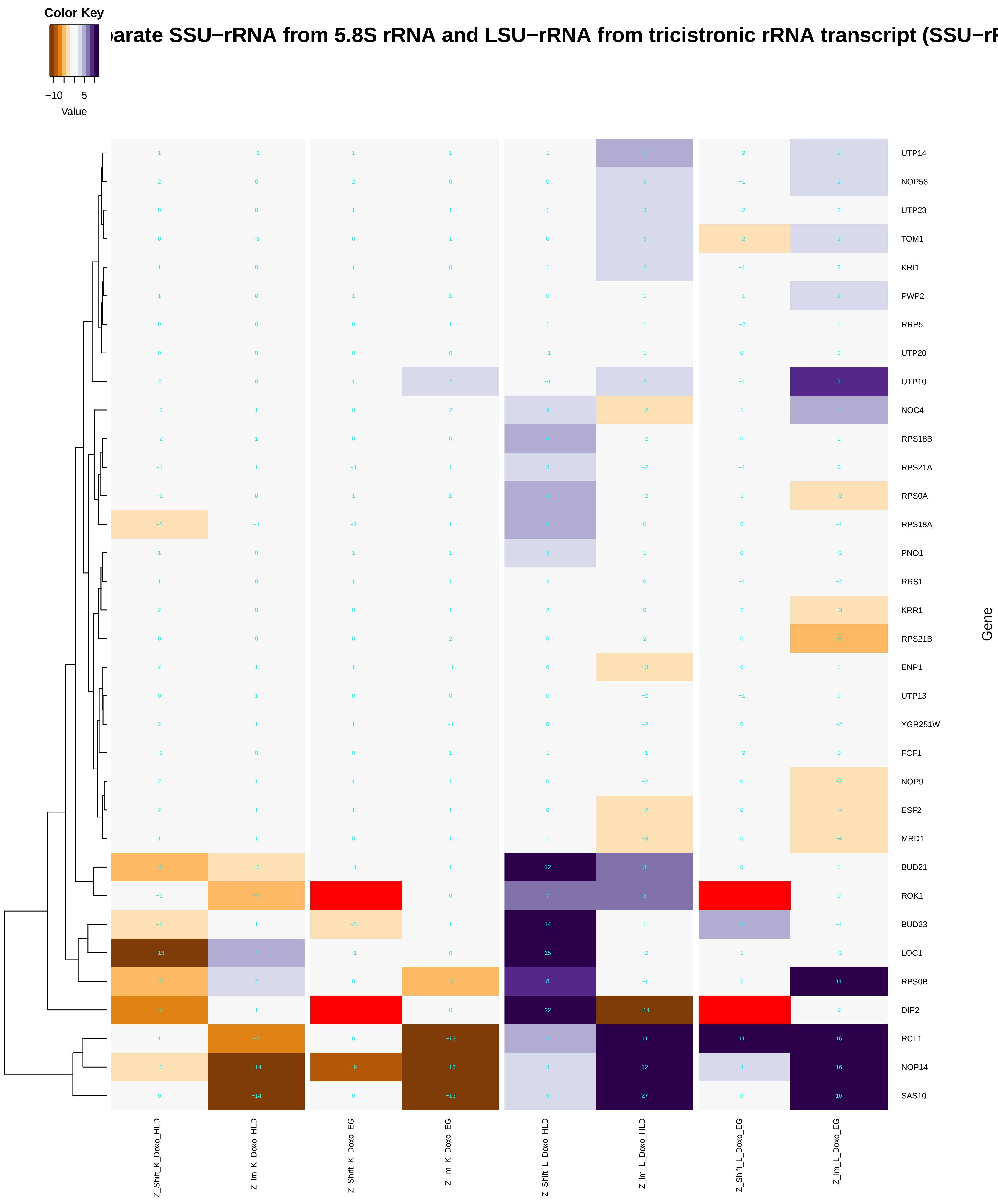

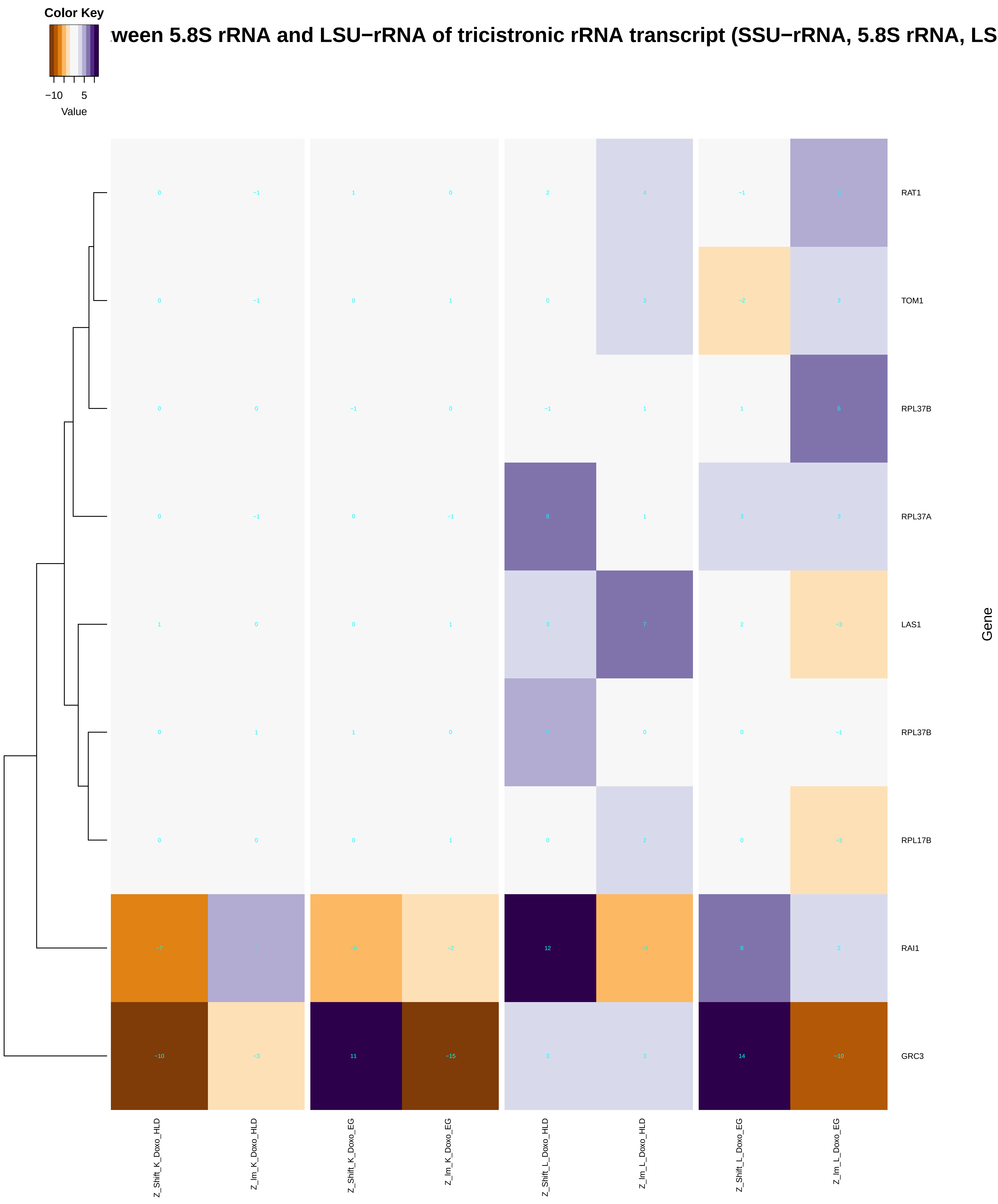

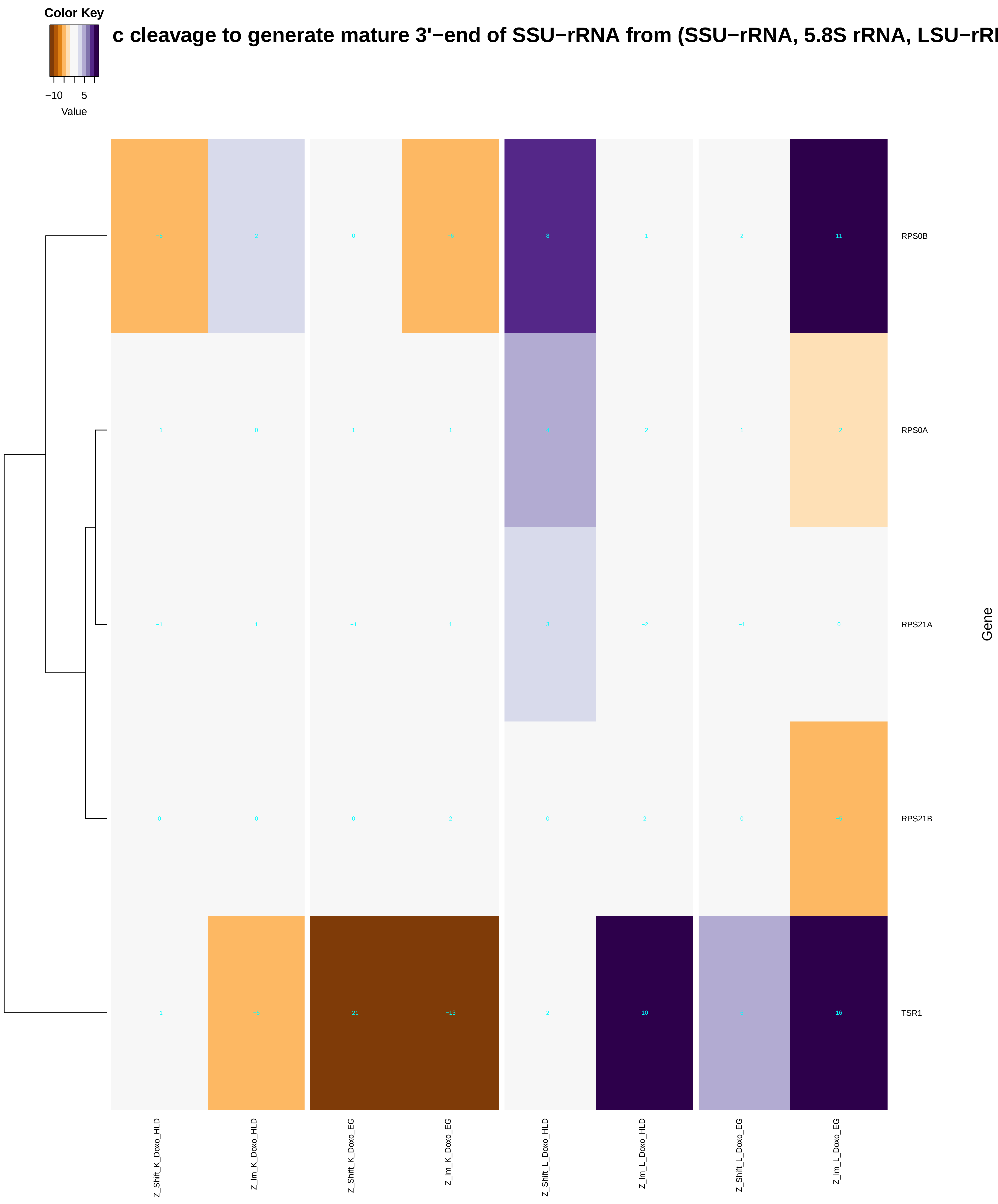

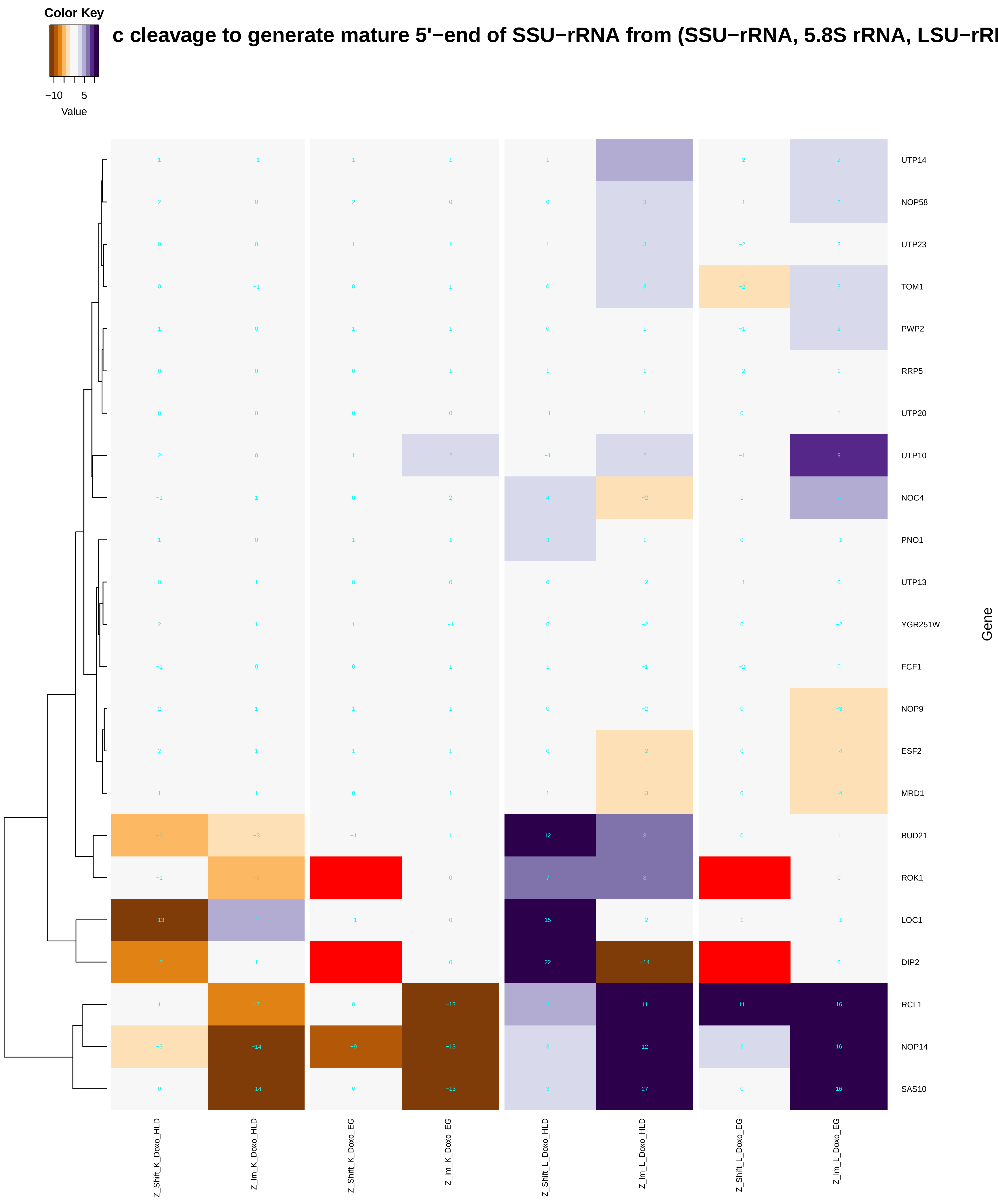

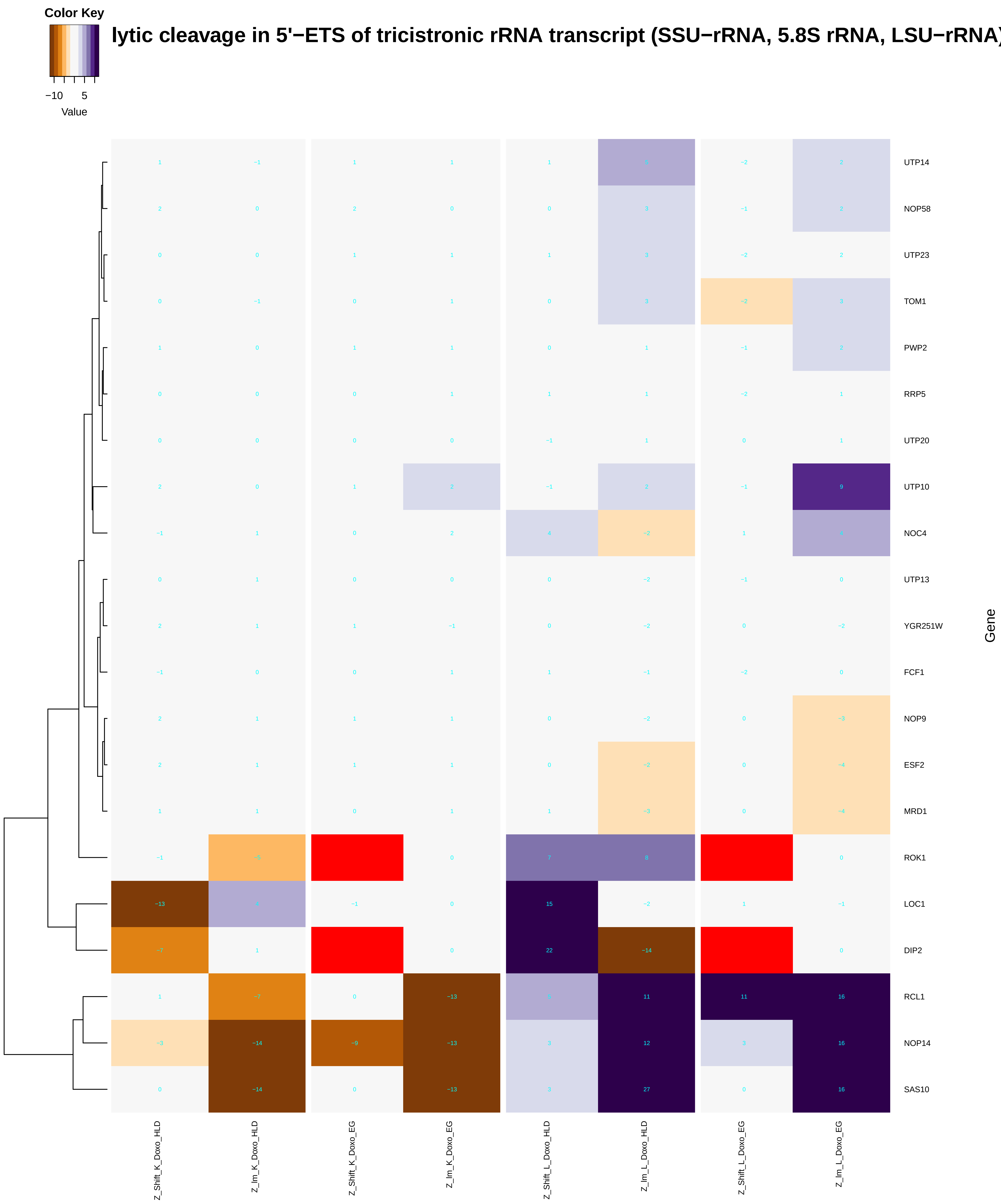

### cytochrome_complex_assembly.pdf

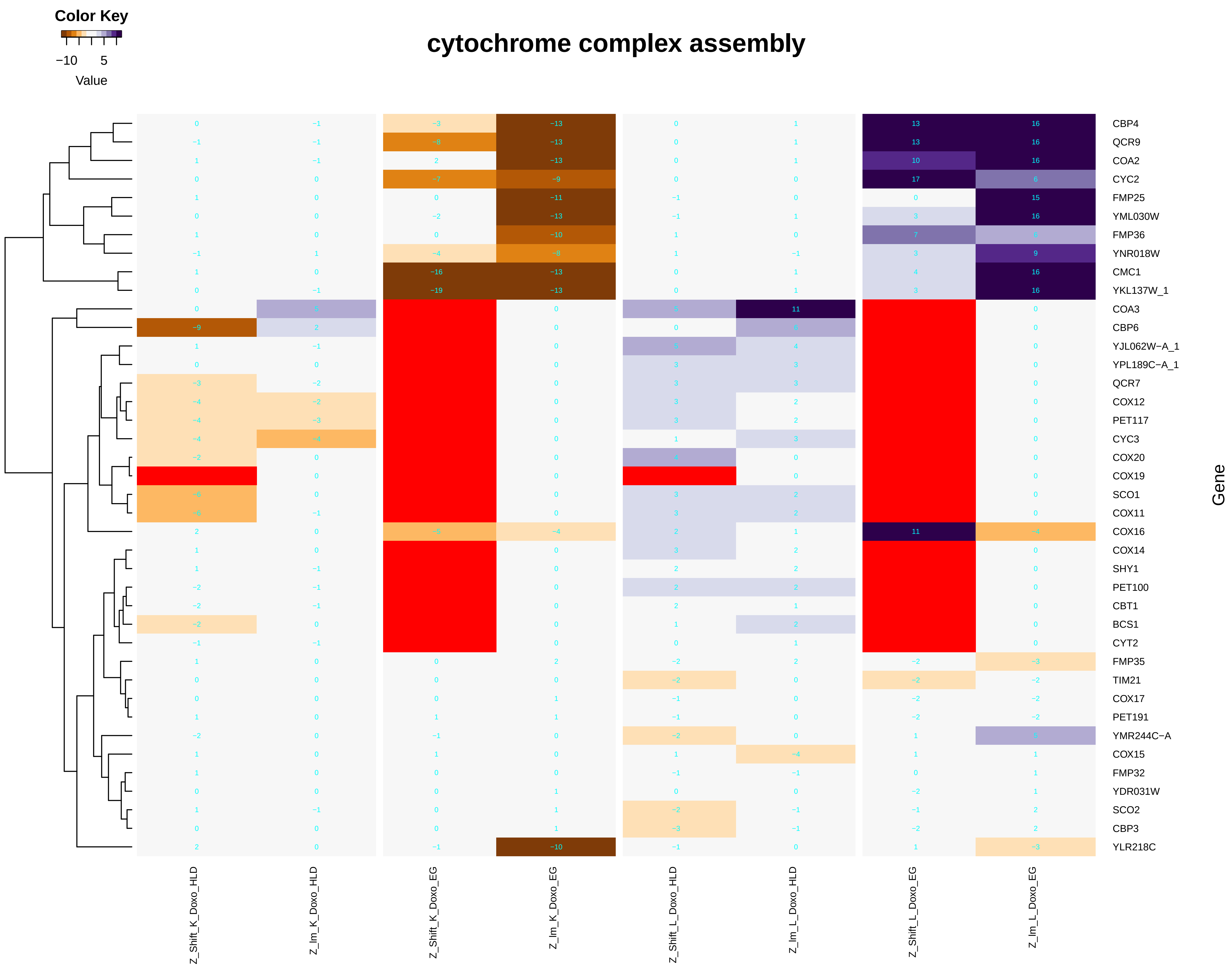

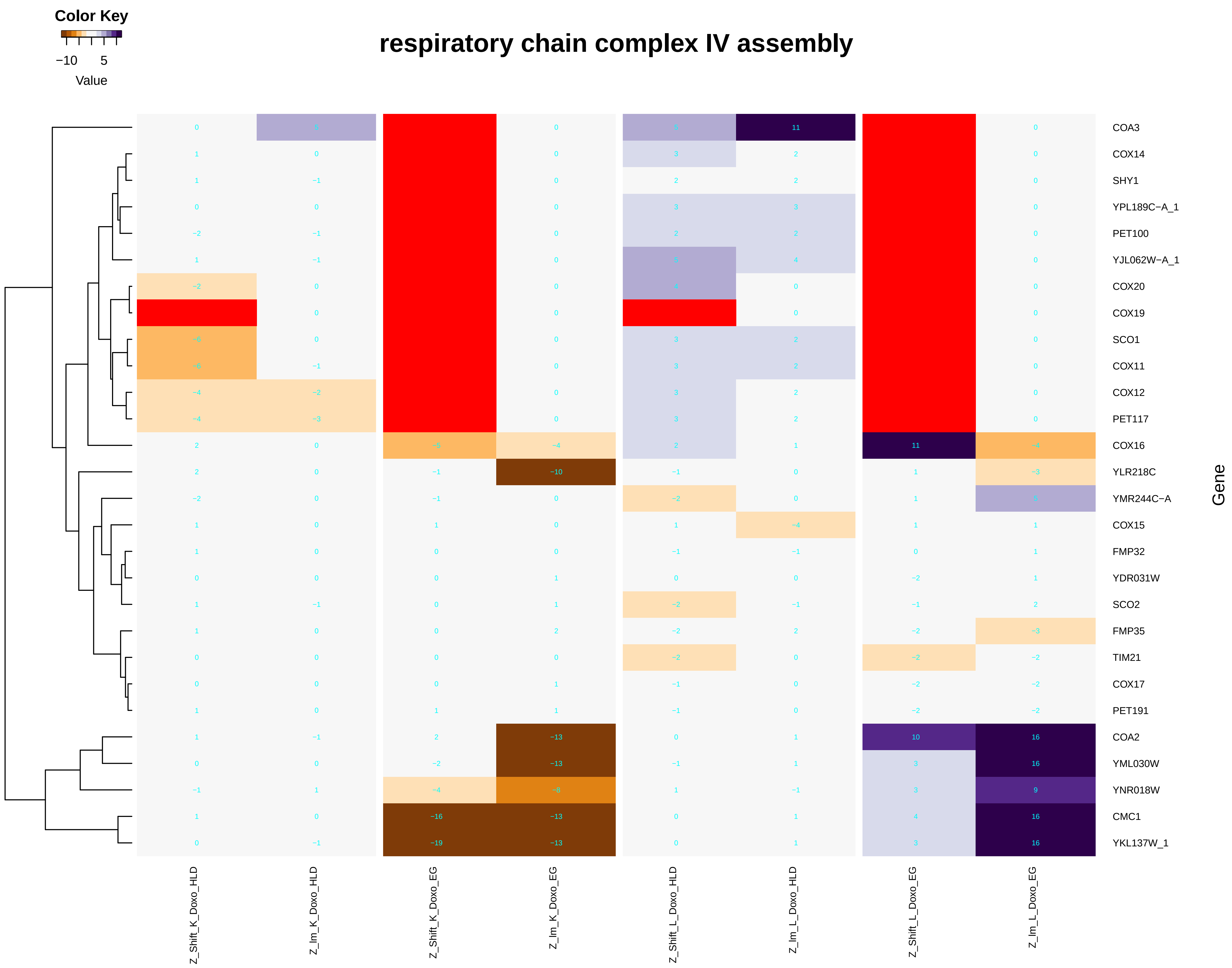

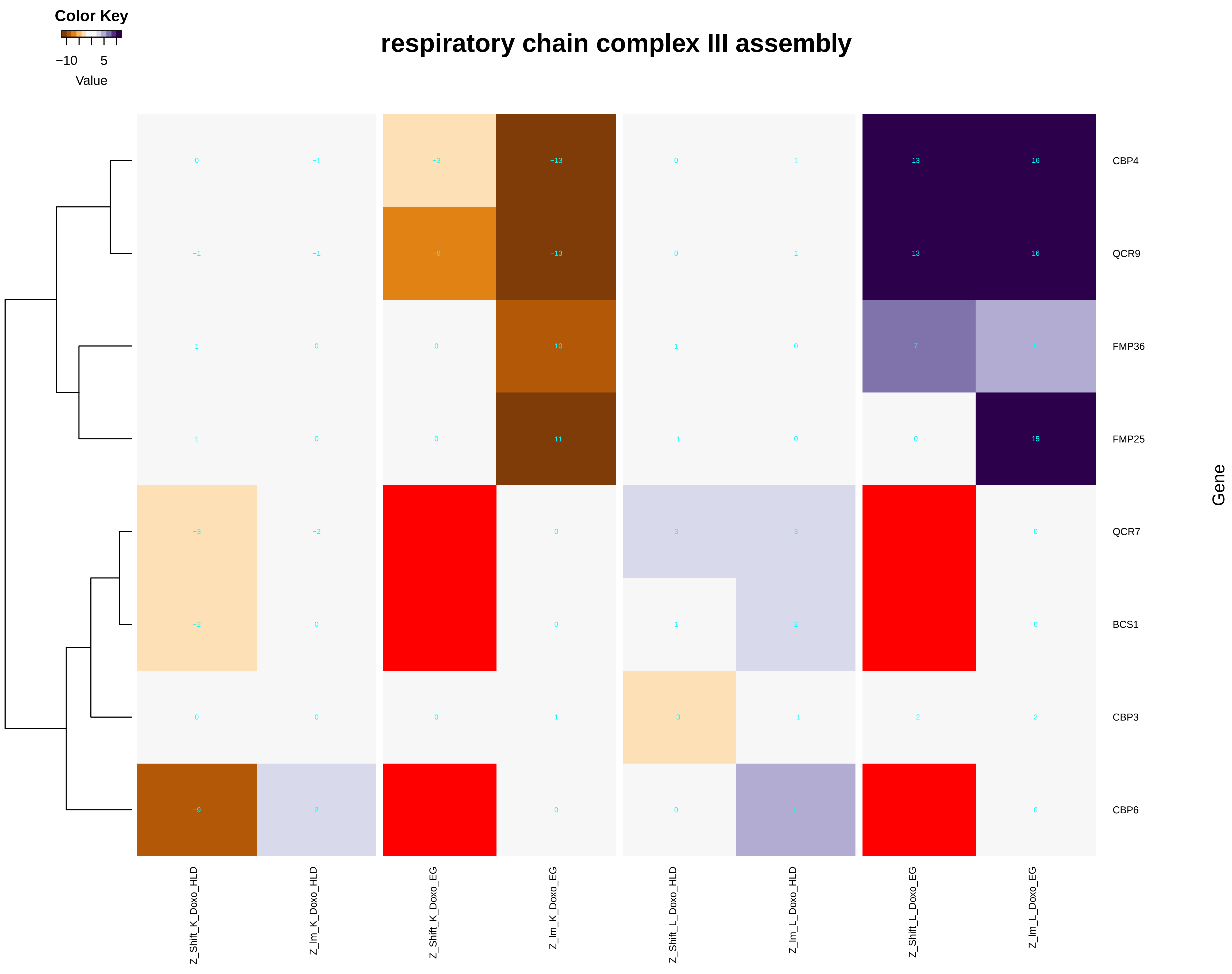

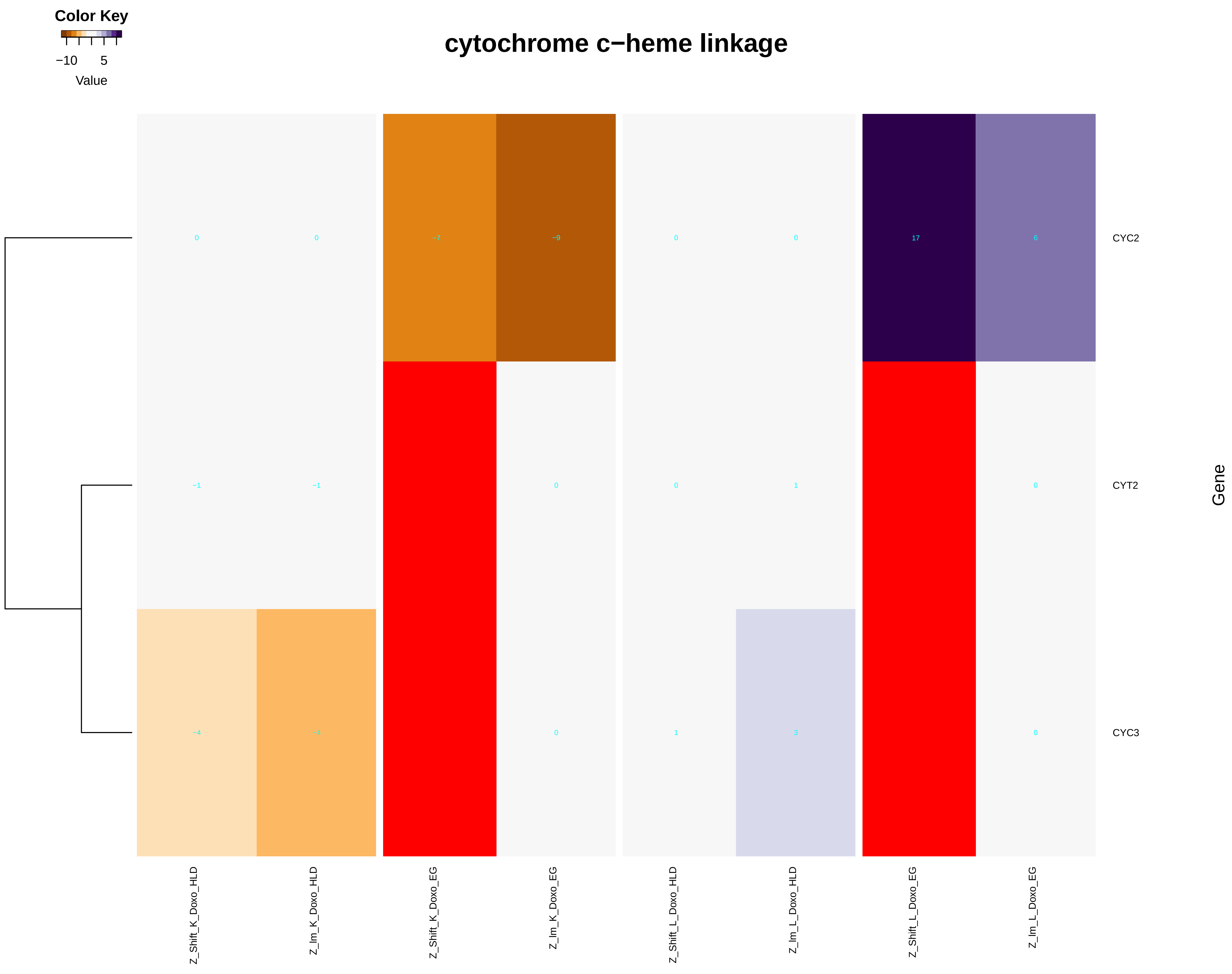

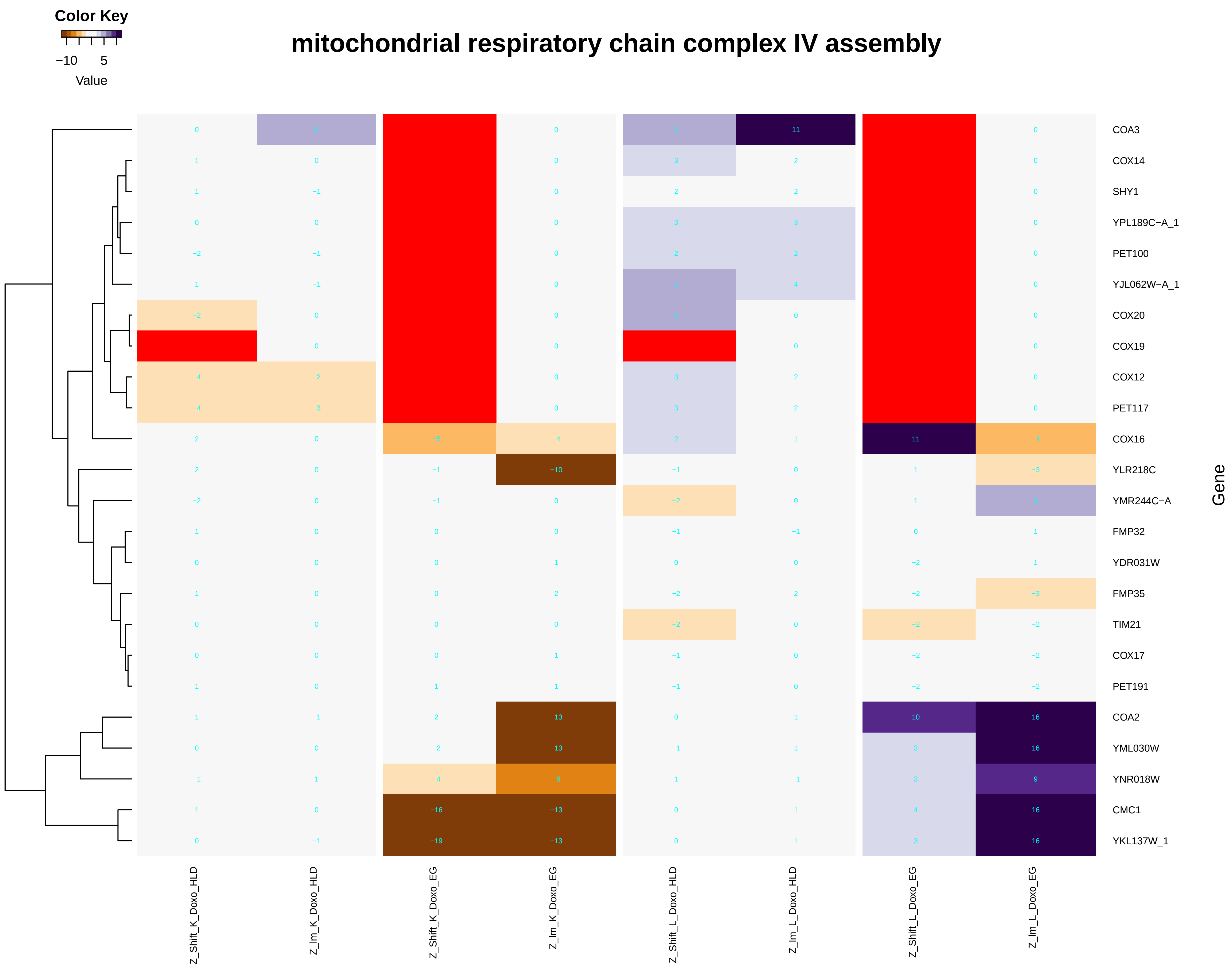

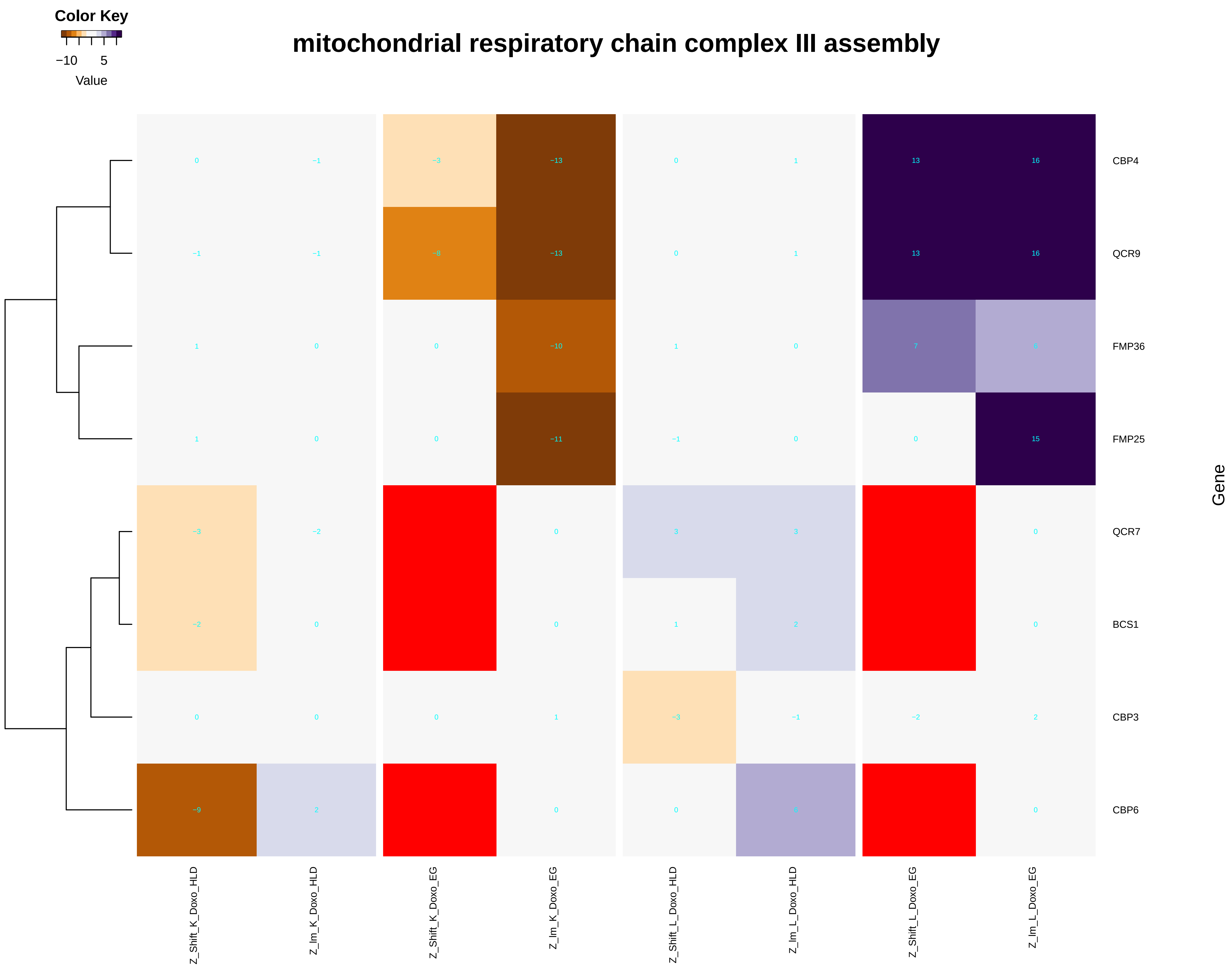

### cytoplasmic_translation.pdf

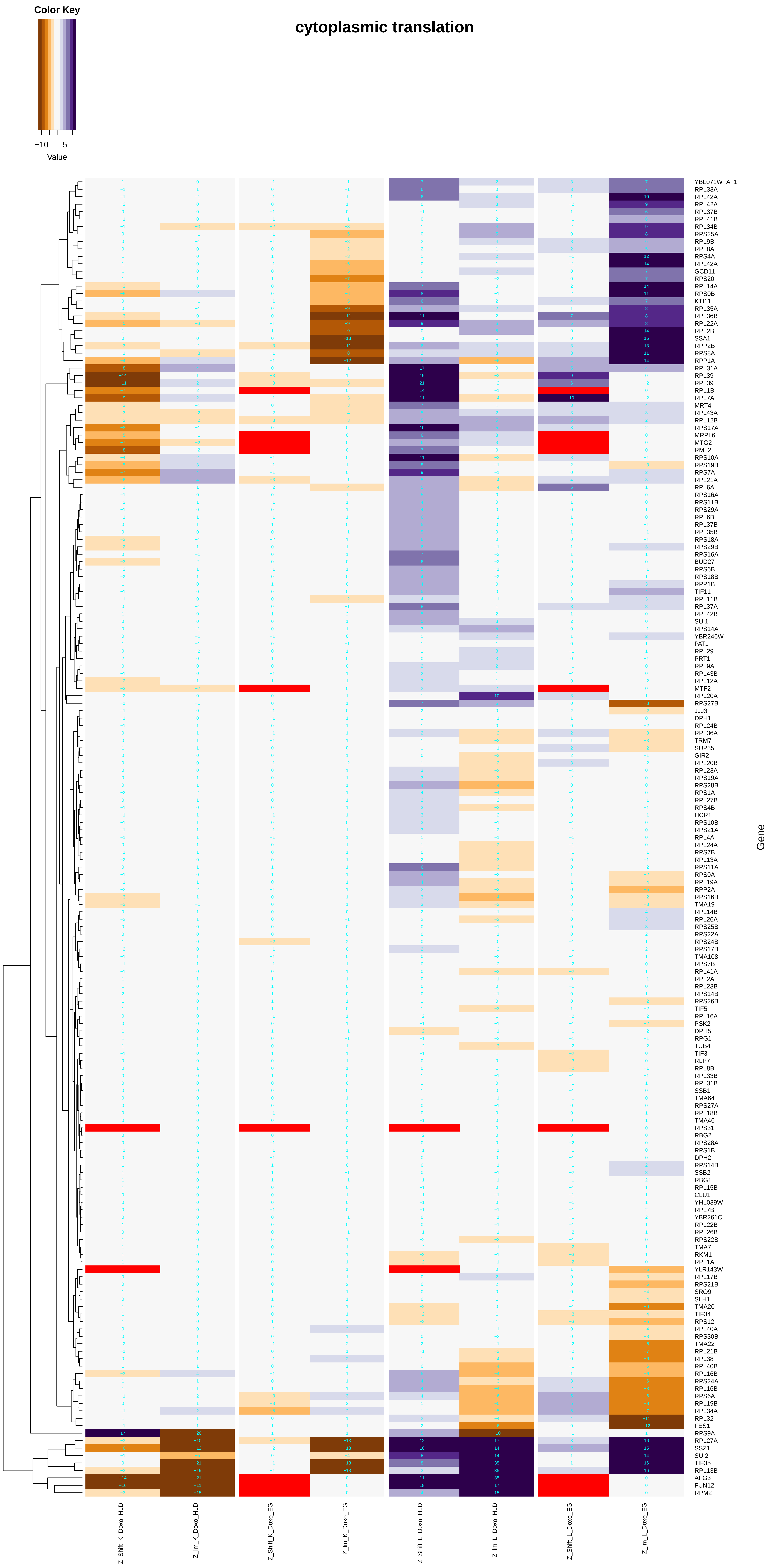

### cytosolic_transport.pdf

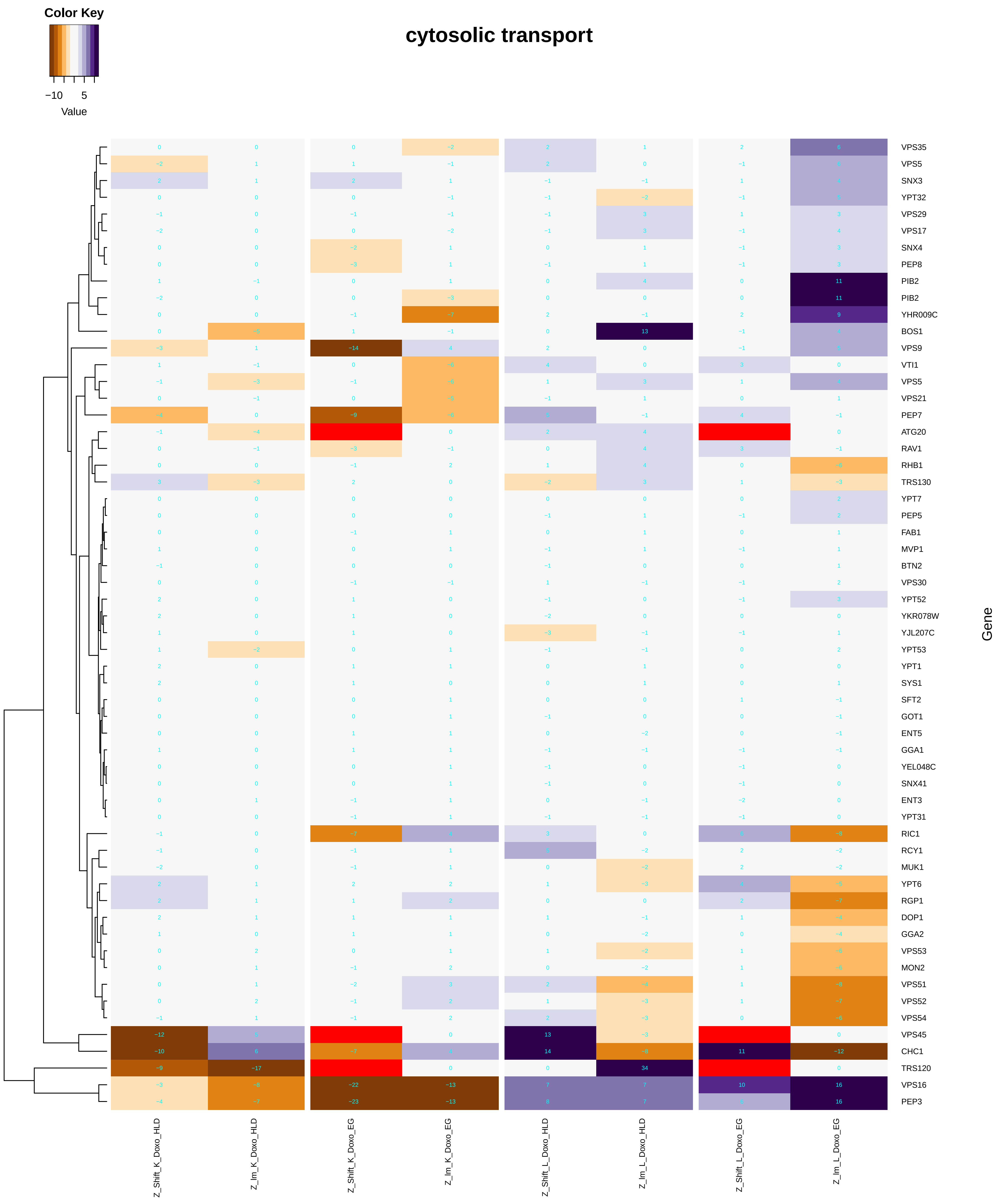

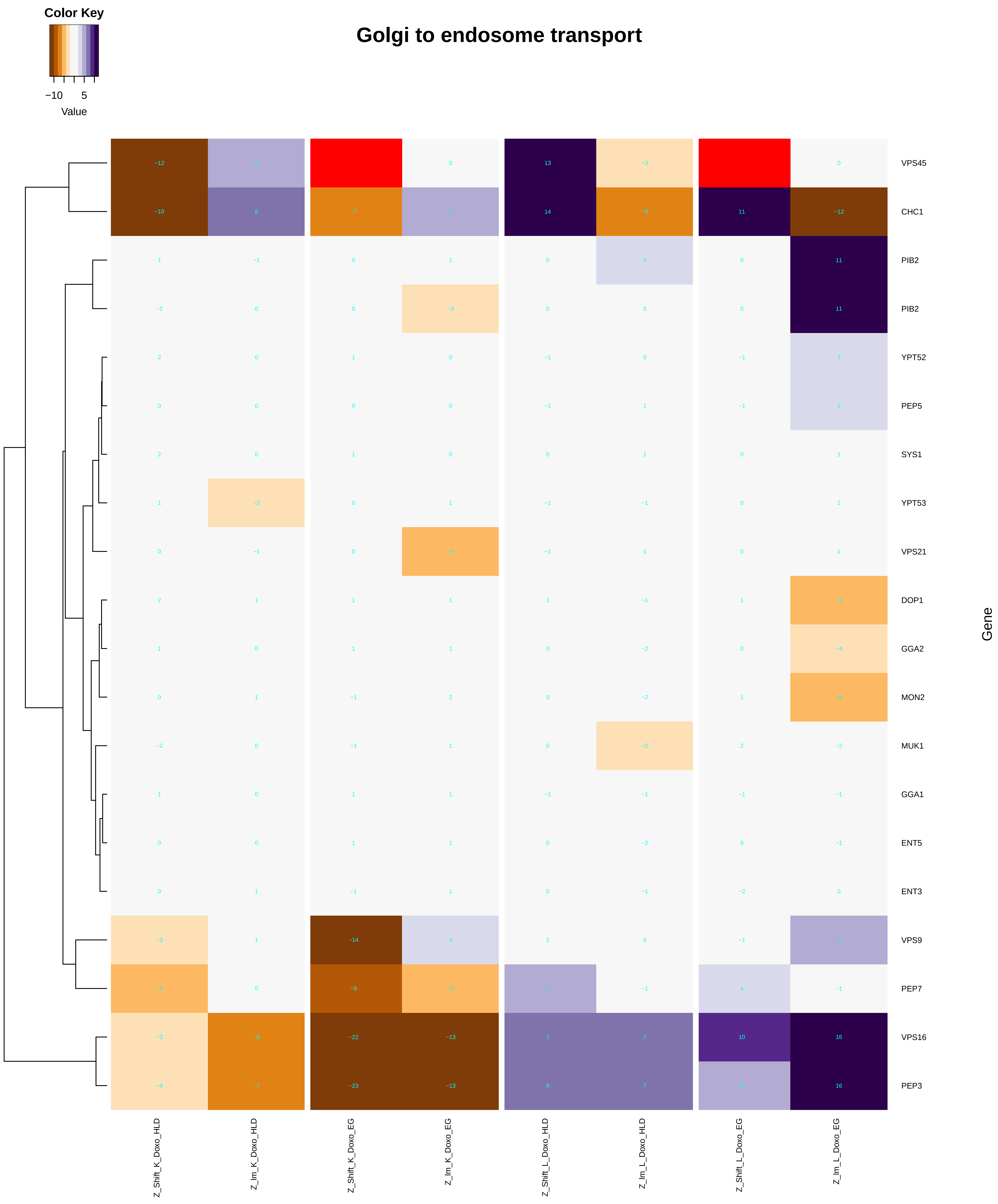

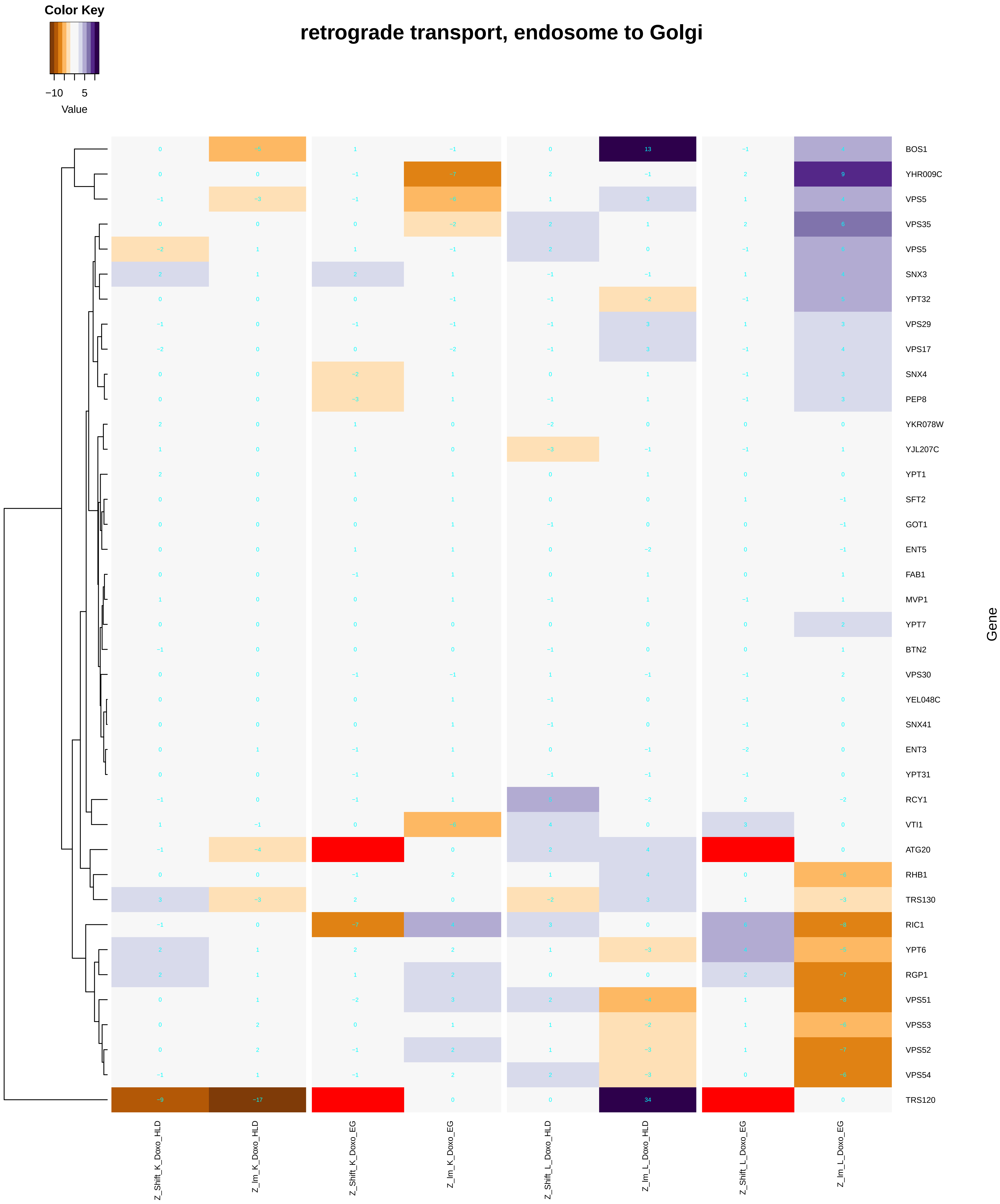

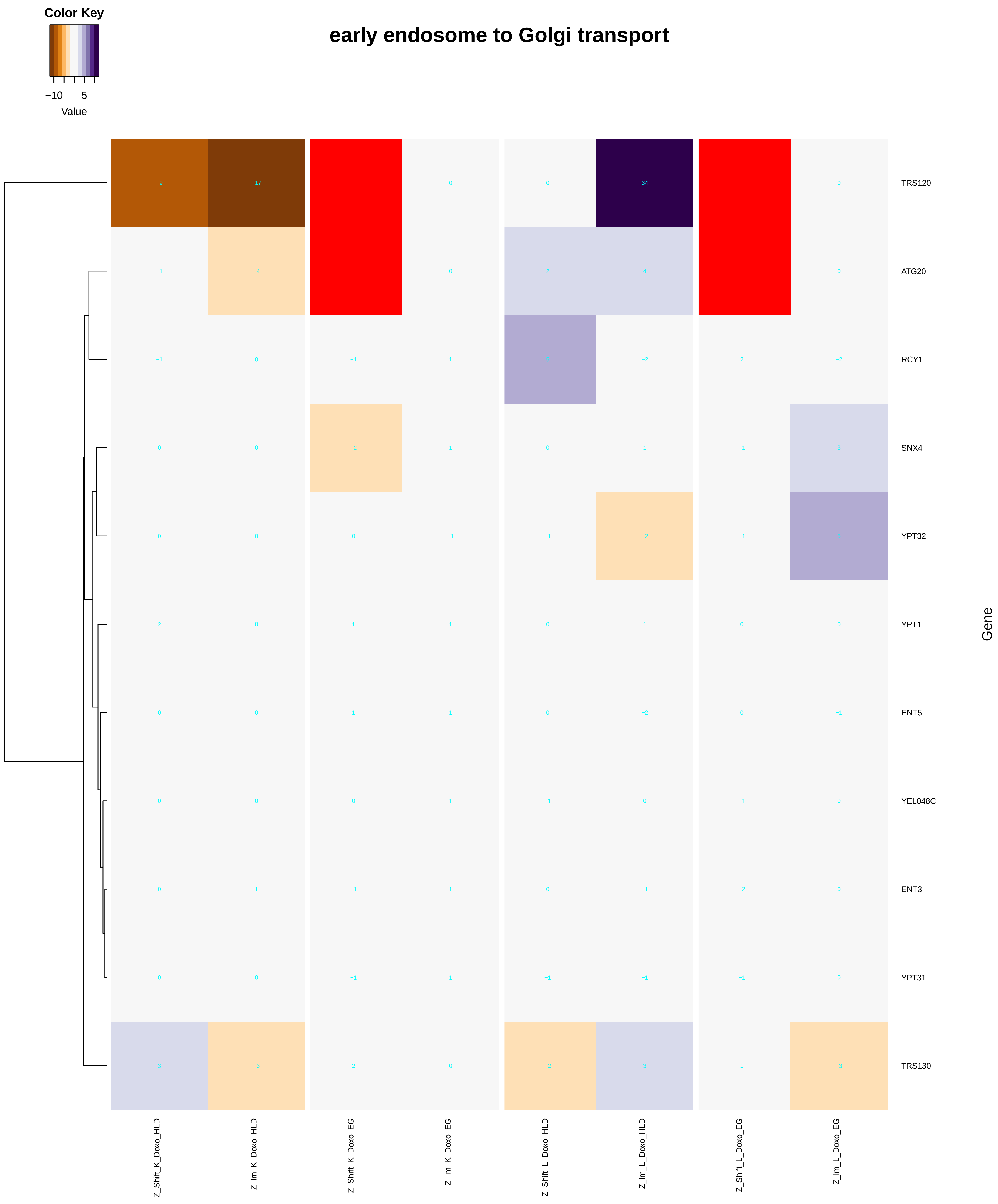

### dicarboxylic_acid_catabolic_process.pdf

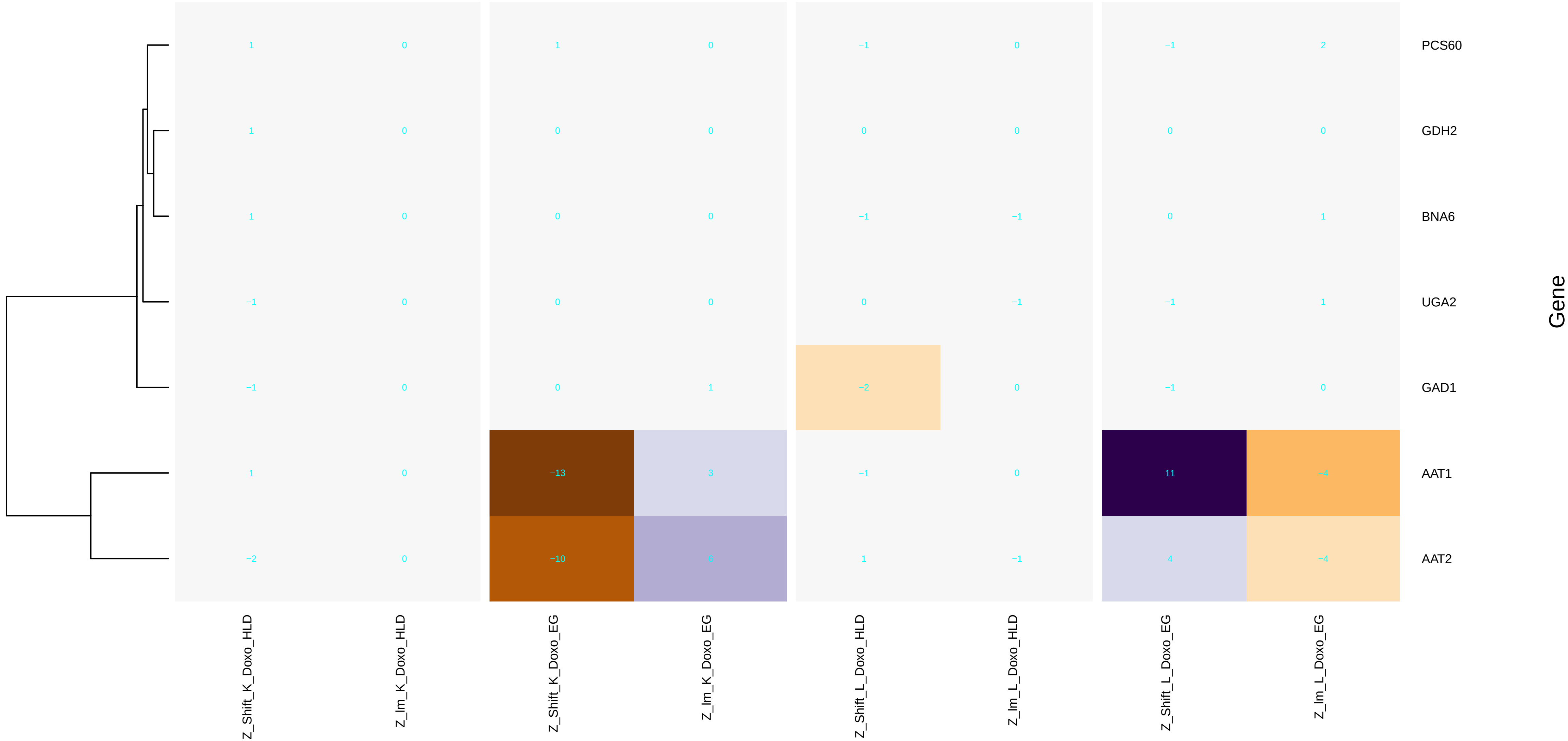

### DNA-dependent_DNA_replication.pdf

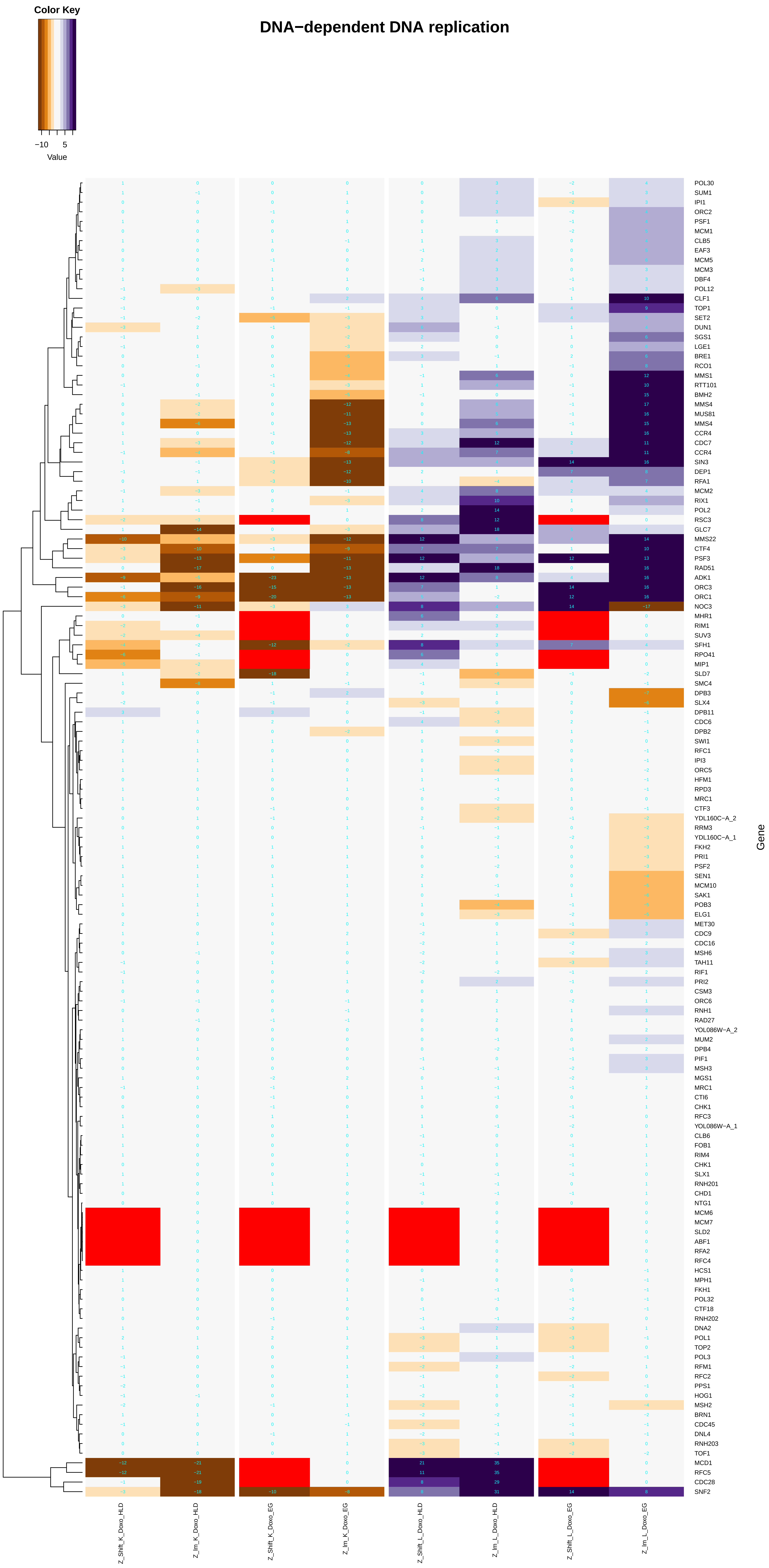

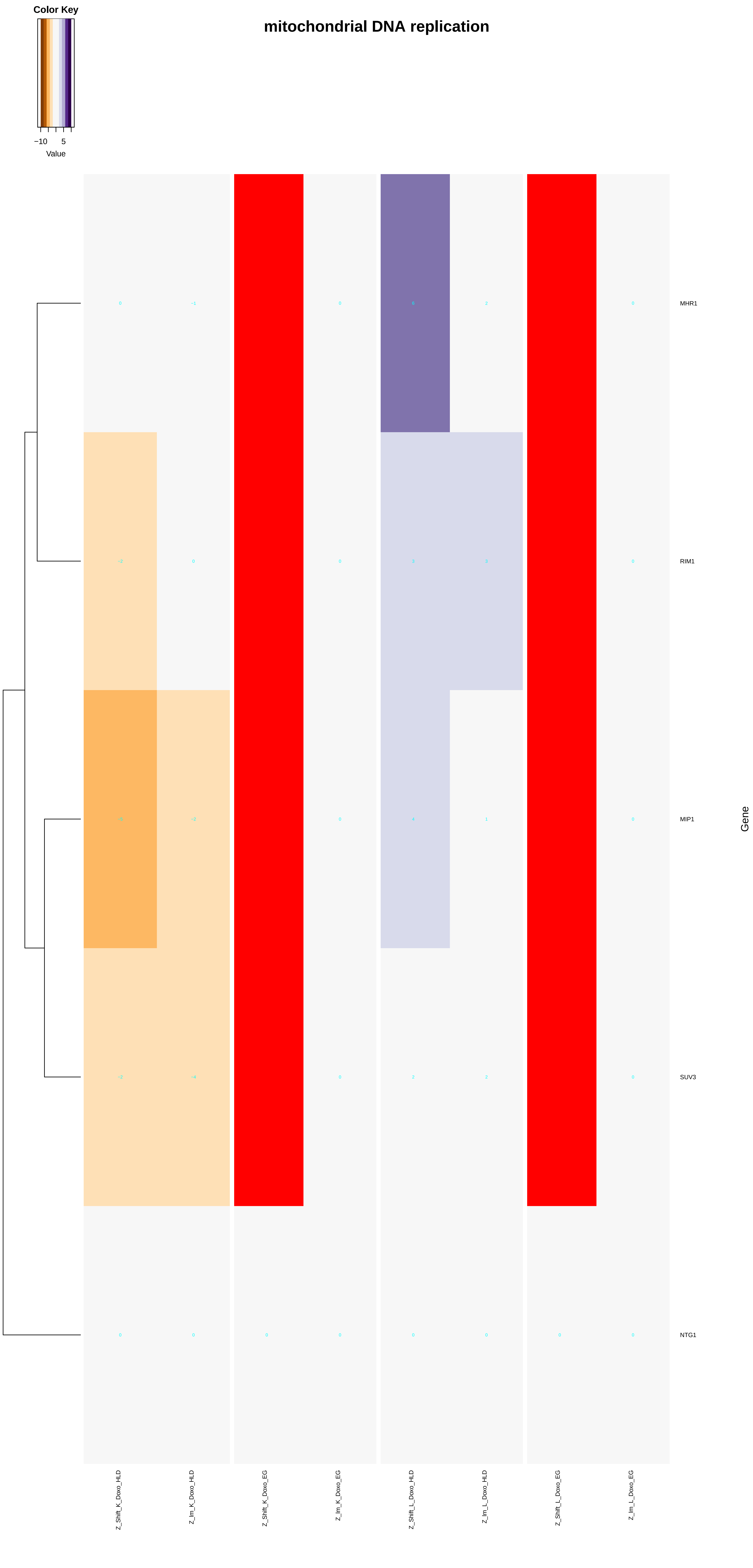

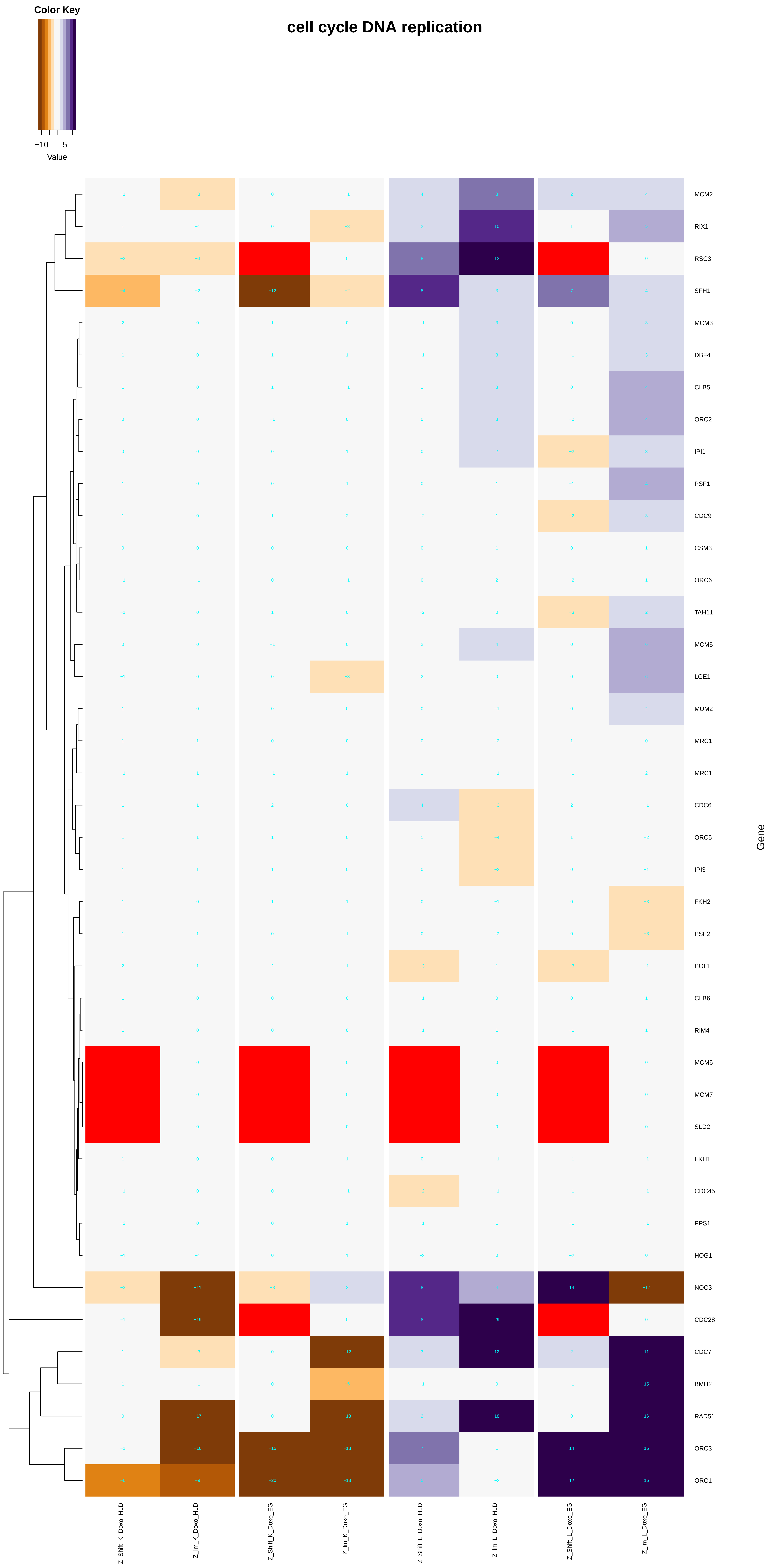

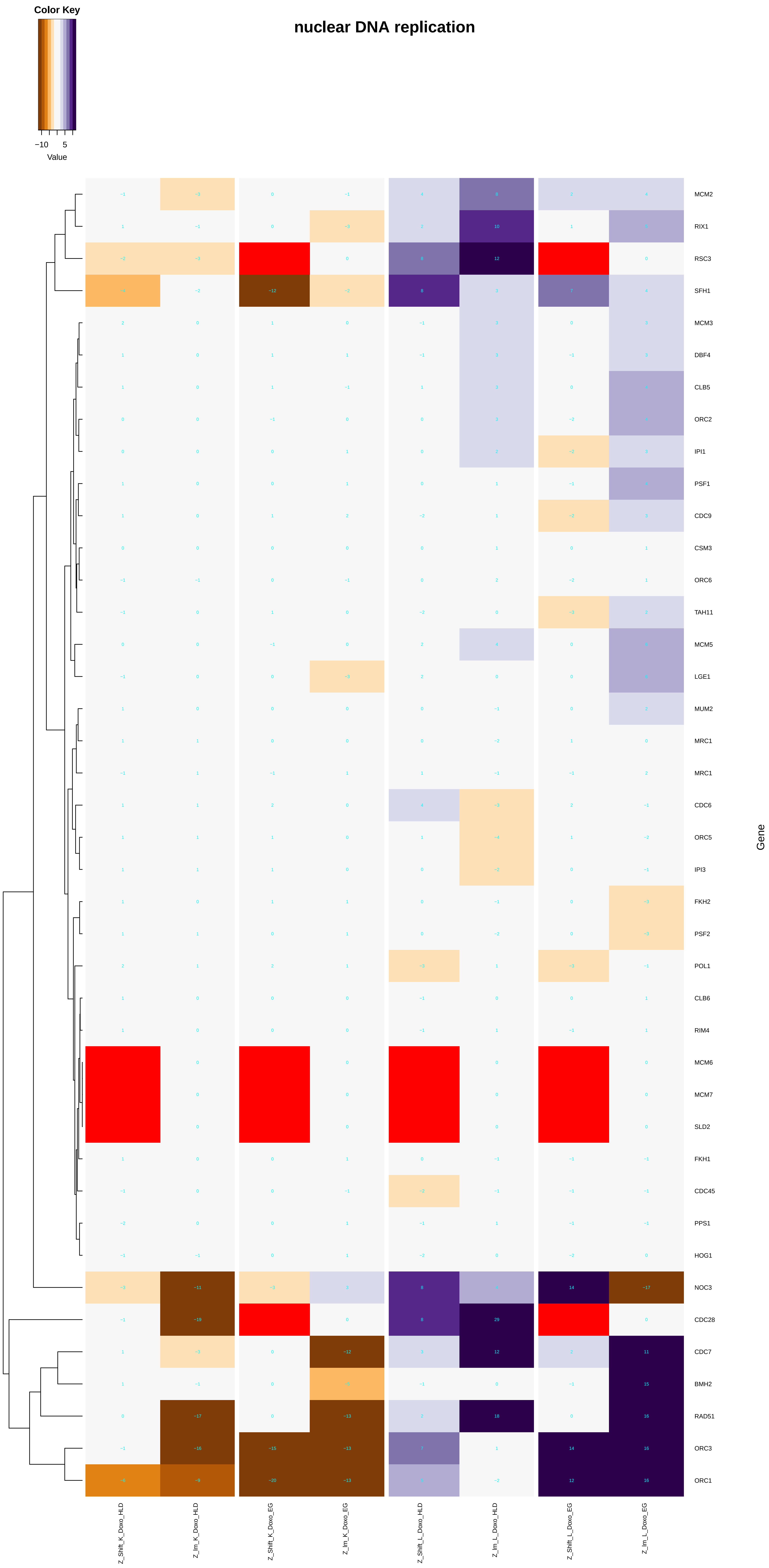

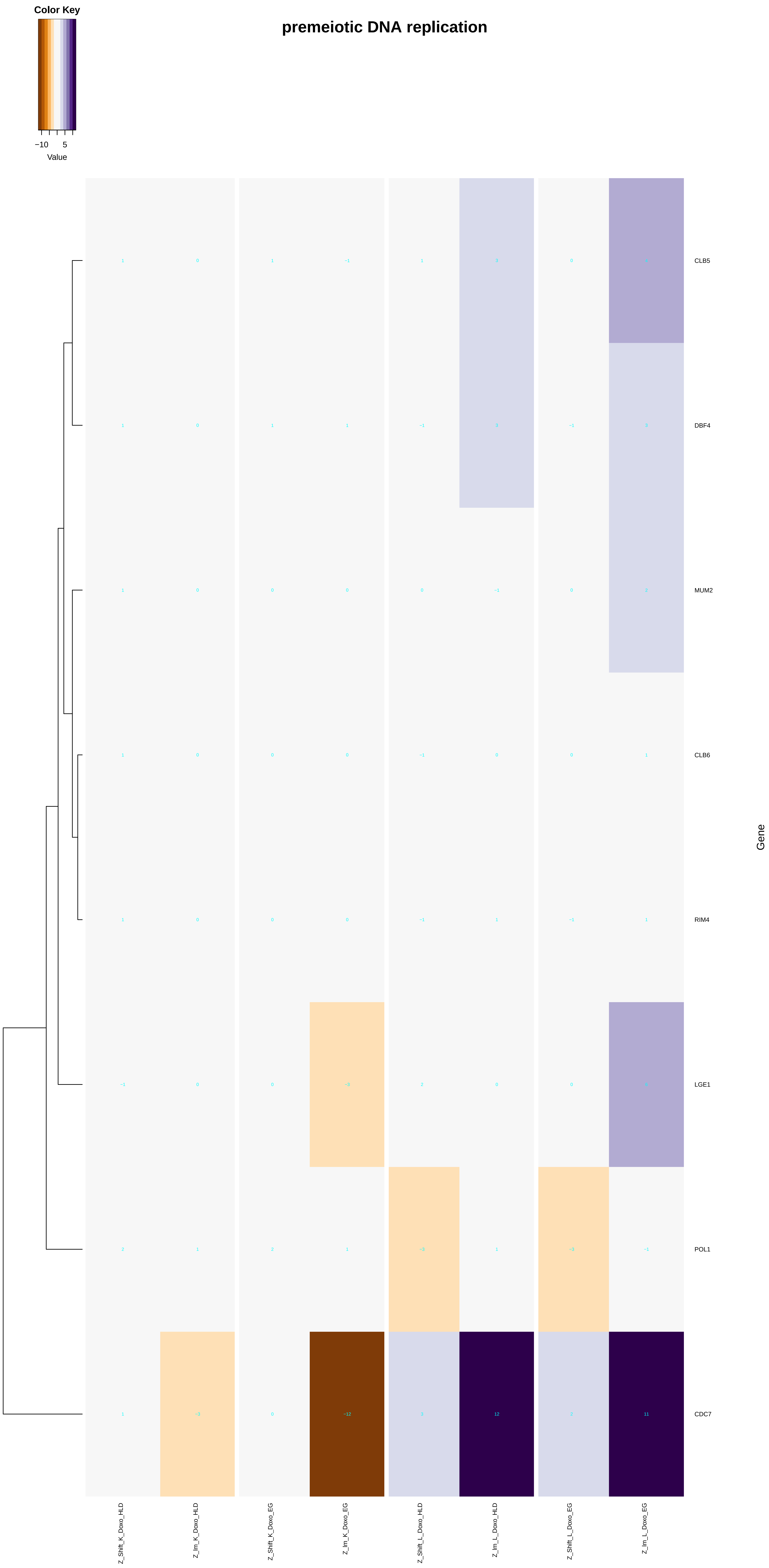

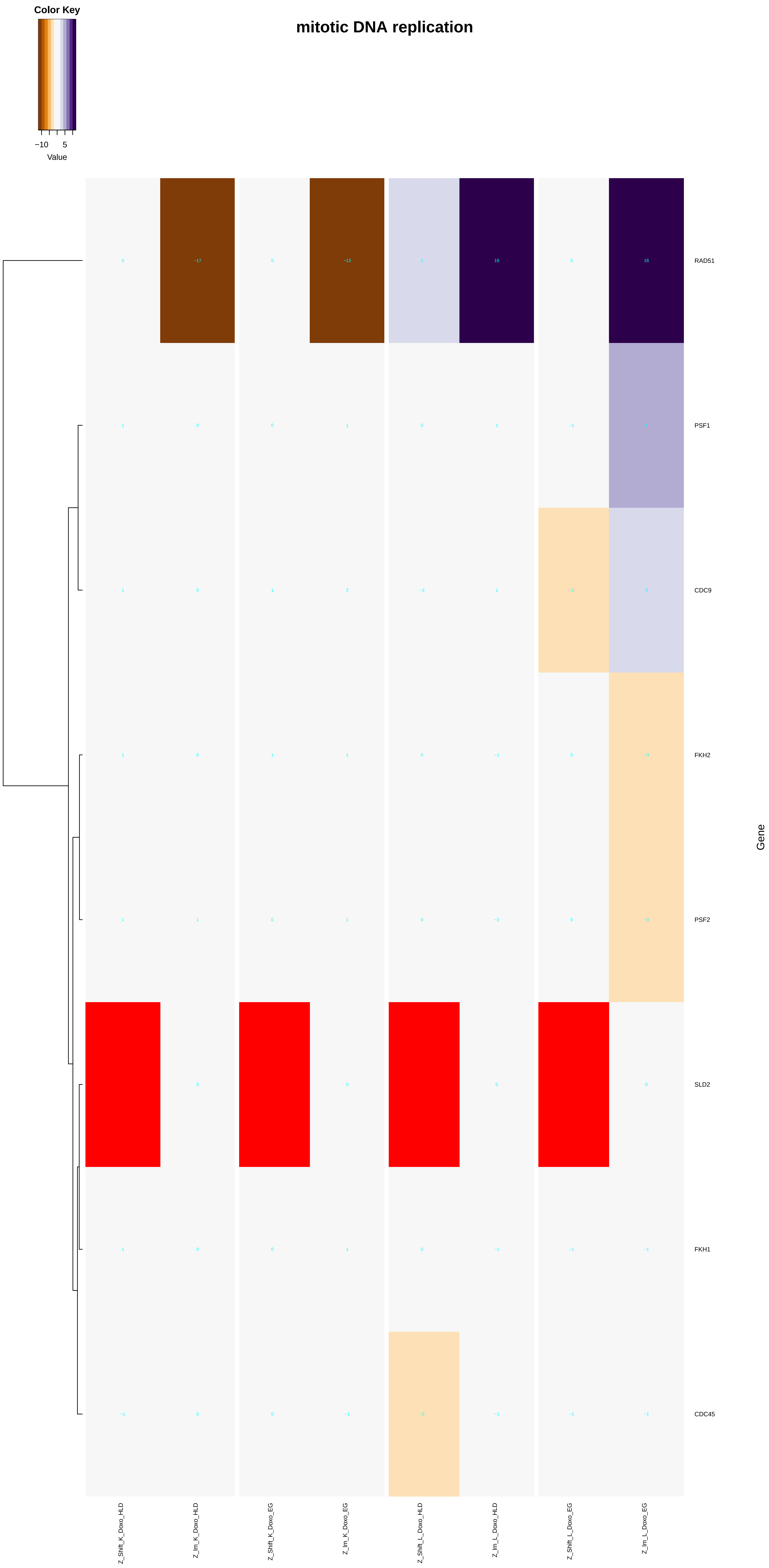

### DNA-templated_transcription,_elongation.pdf

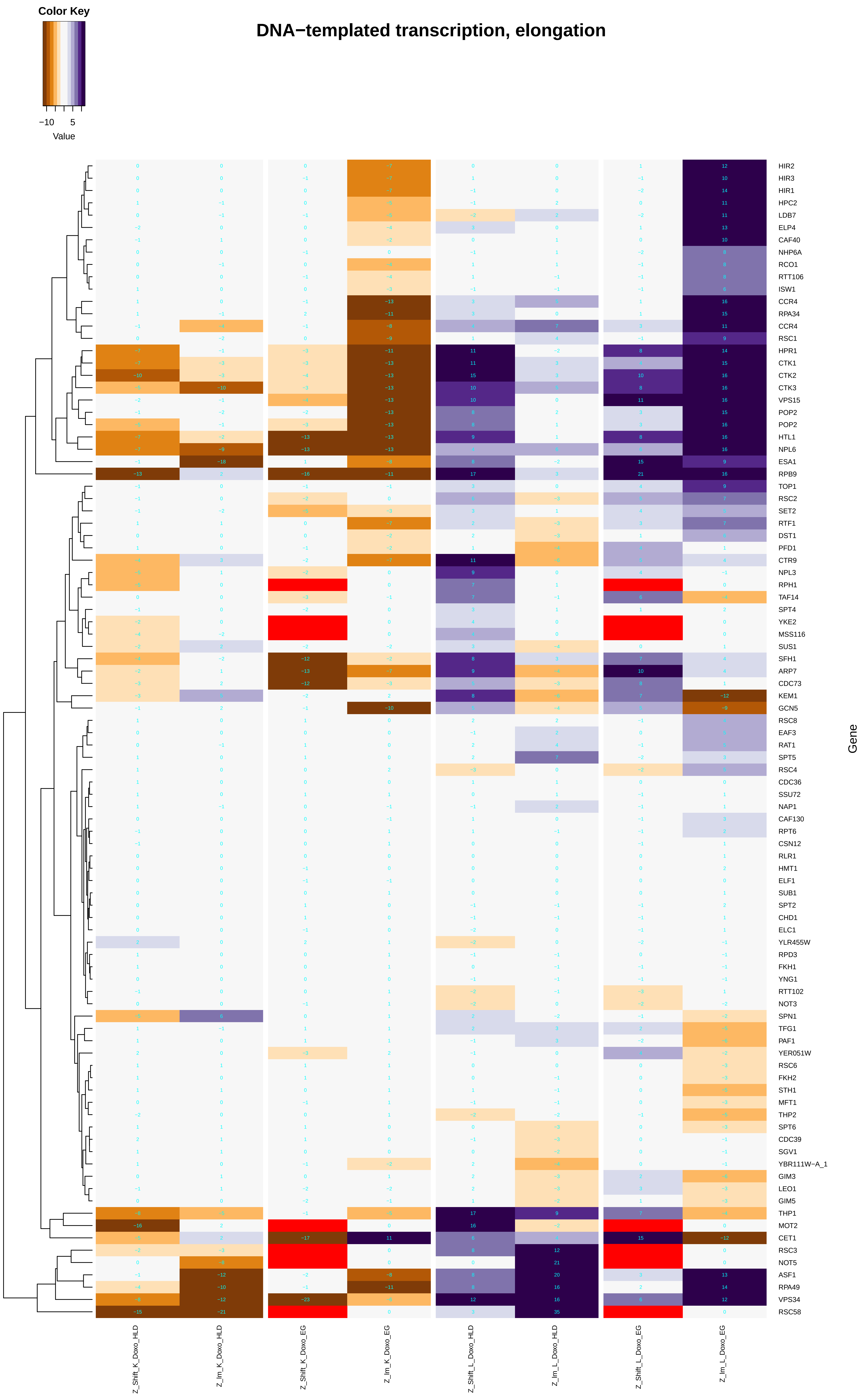

#### transcription elongation from RNA polymerase I promoter

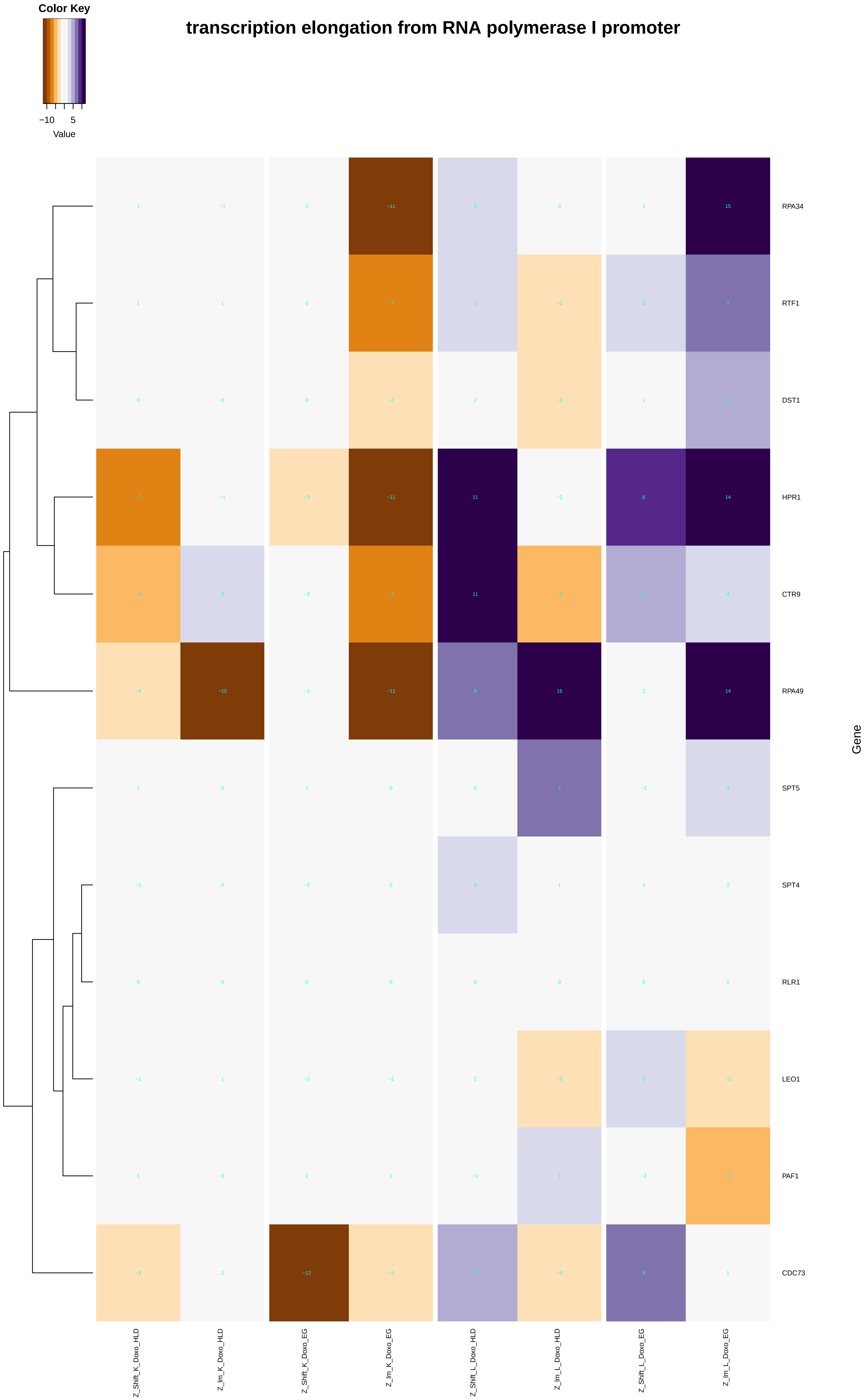

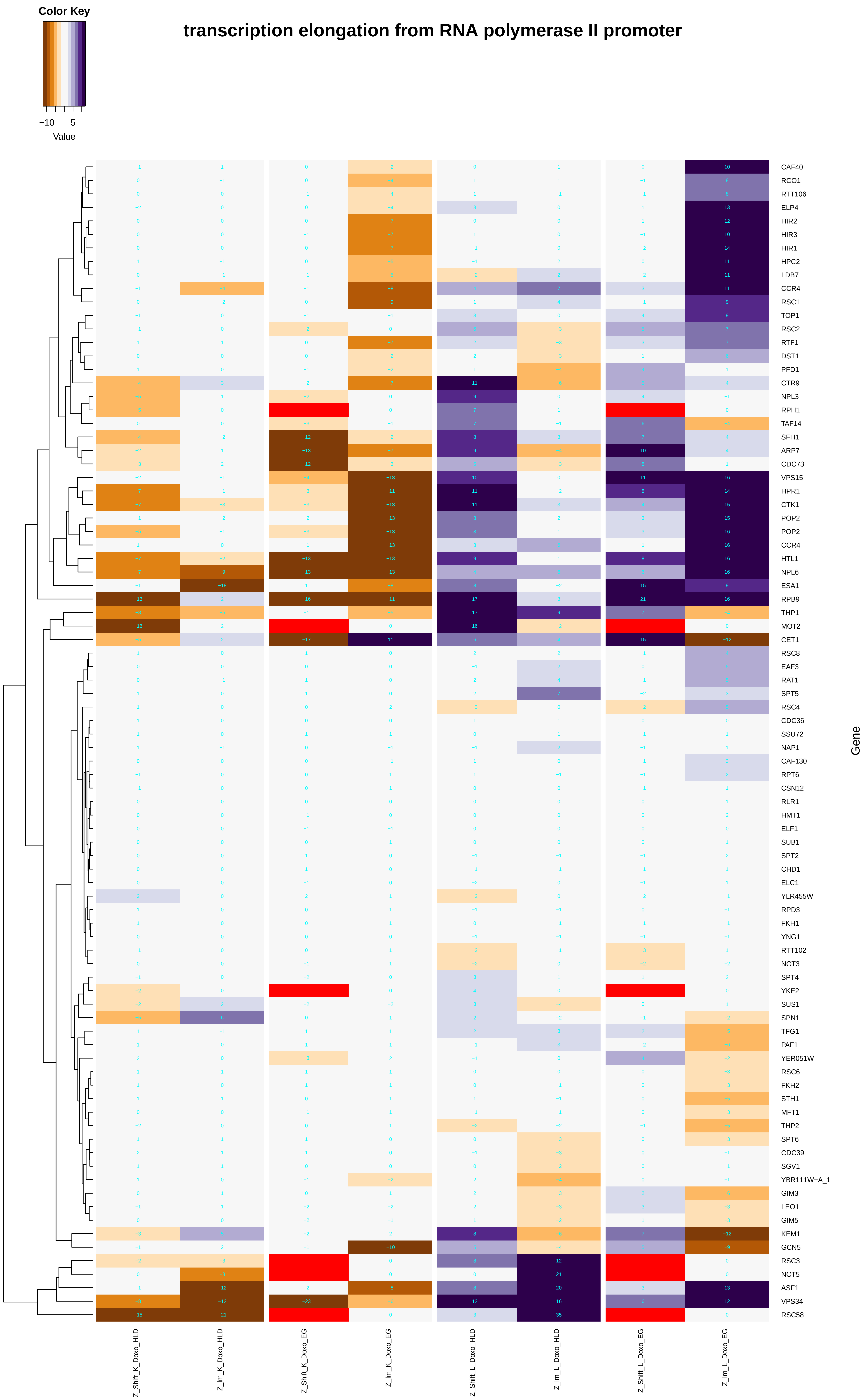

### DNA_double-strand_break_processing.pdf

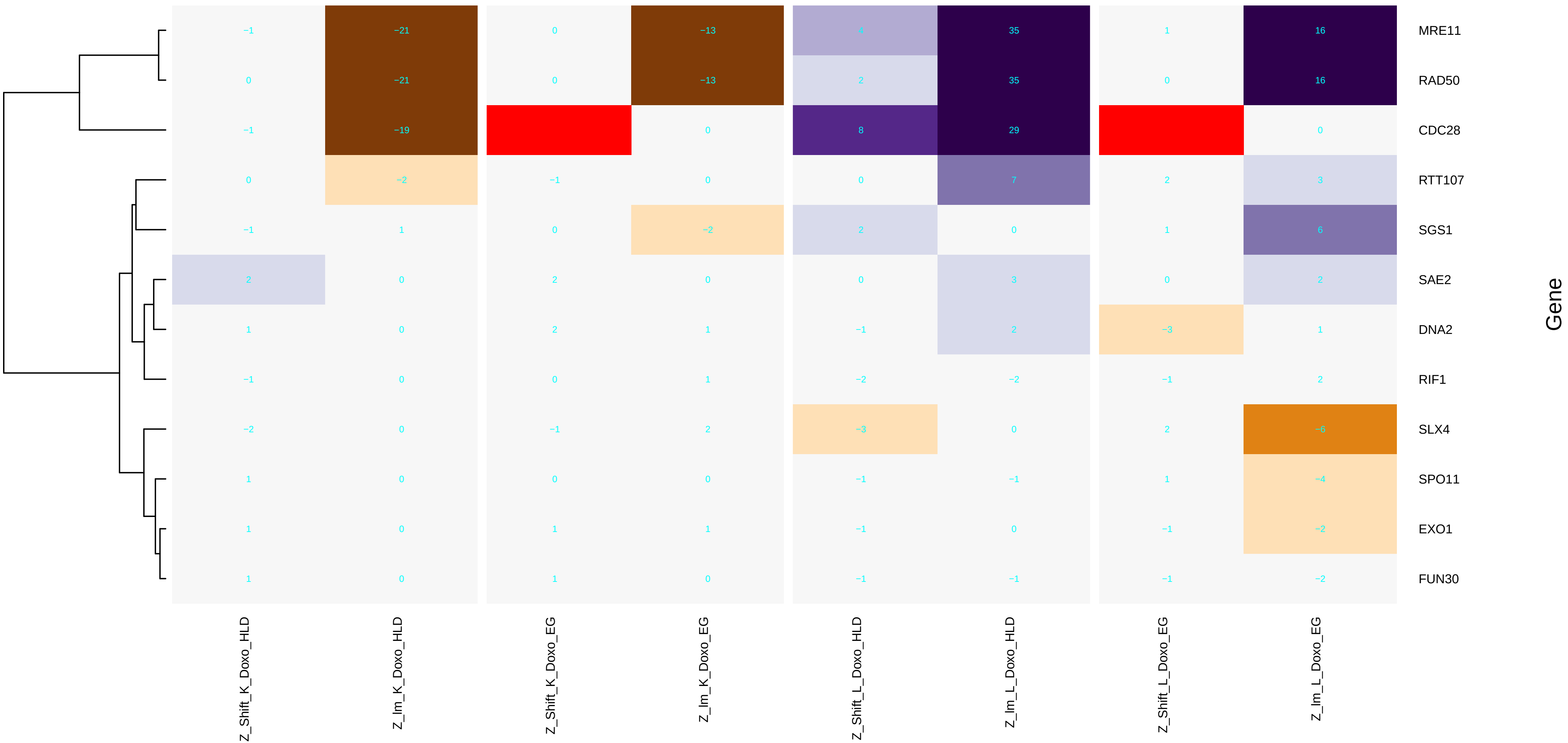

### DNA_duplex_unwinding.pdf

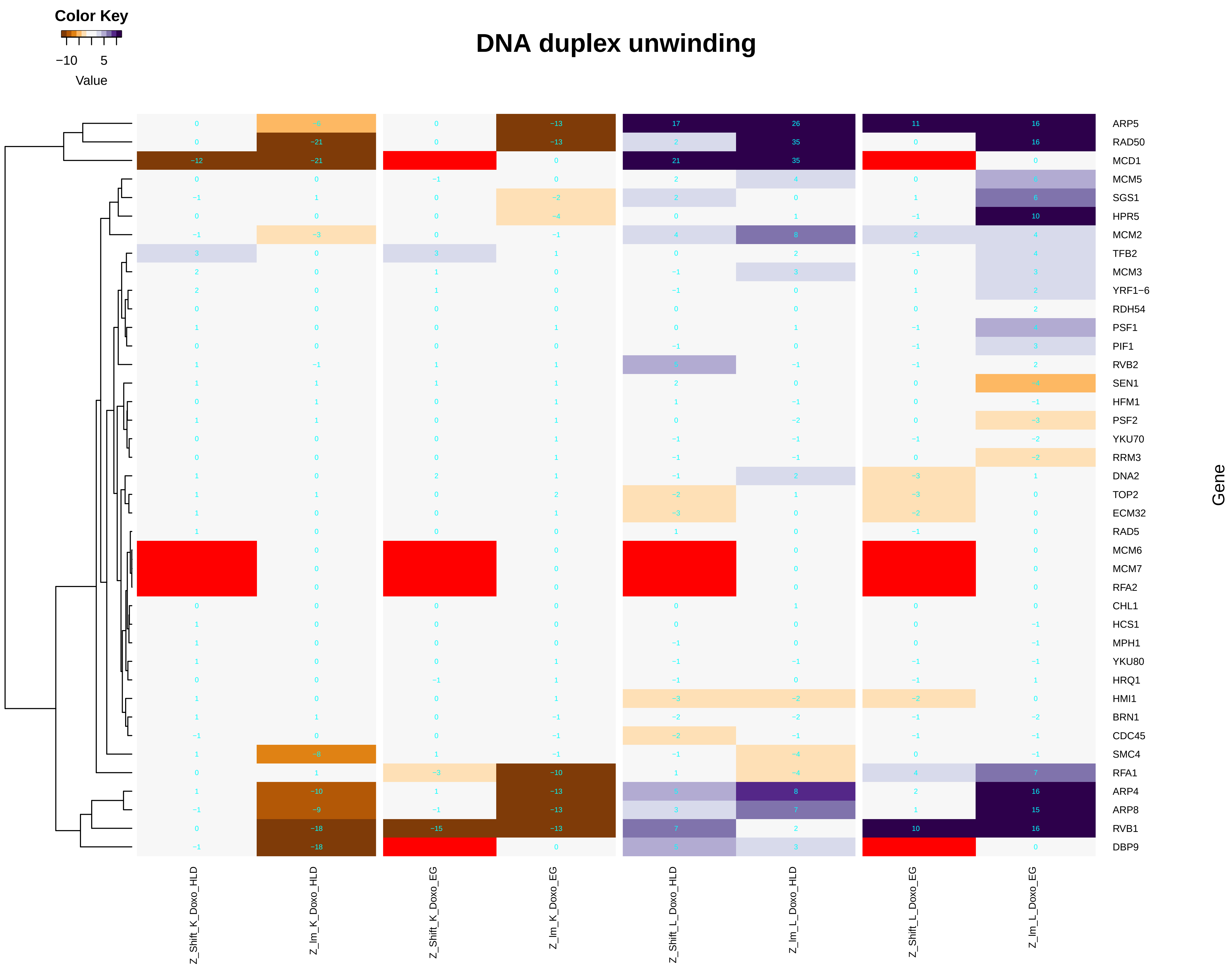

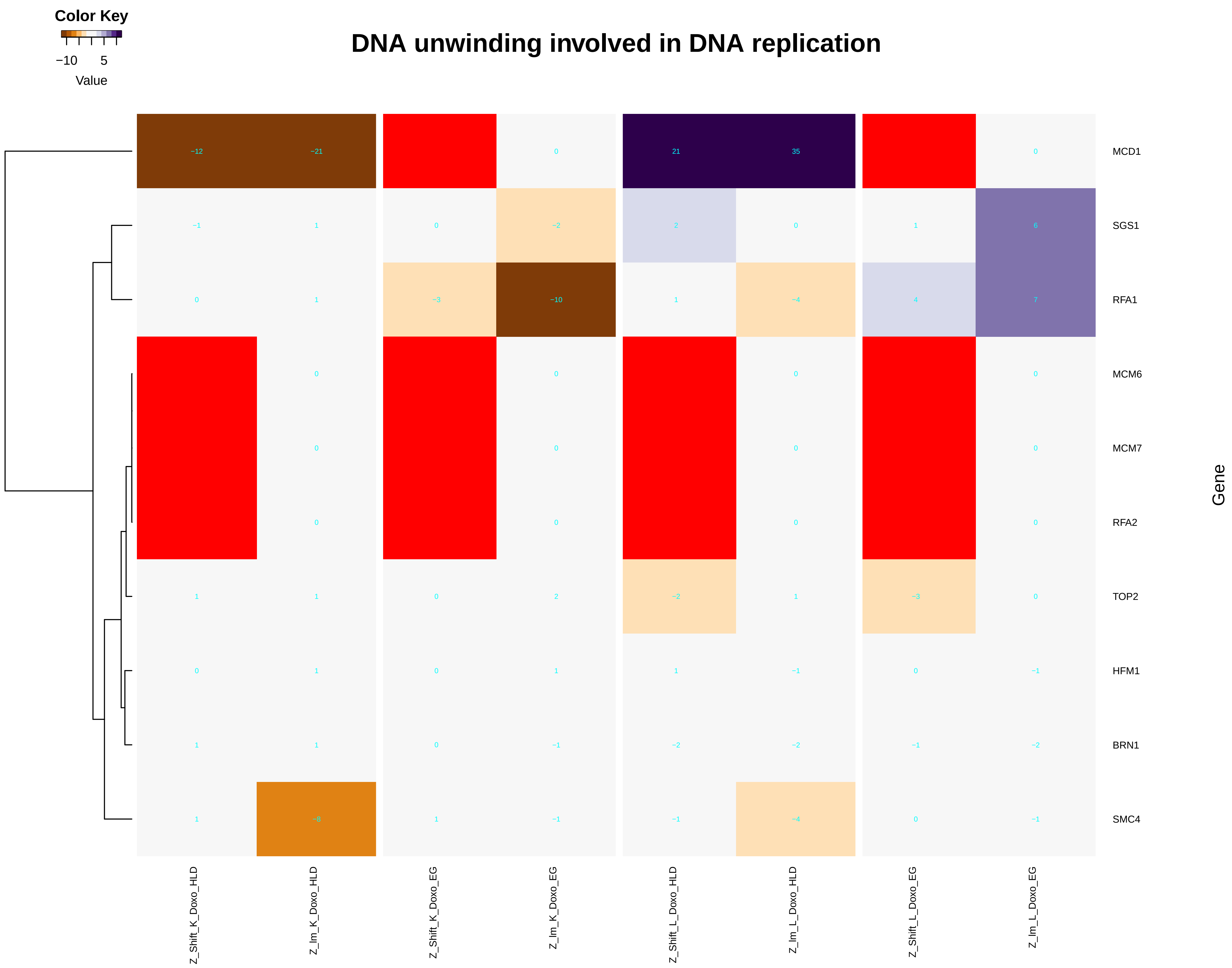

### DNA_geometric_change.pdf

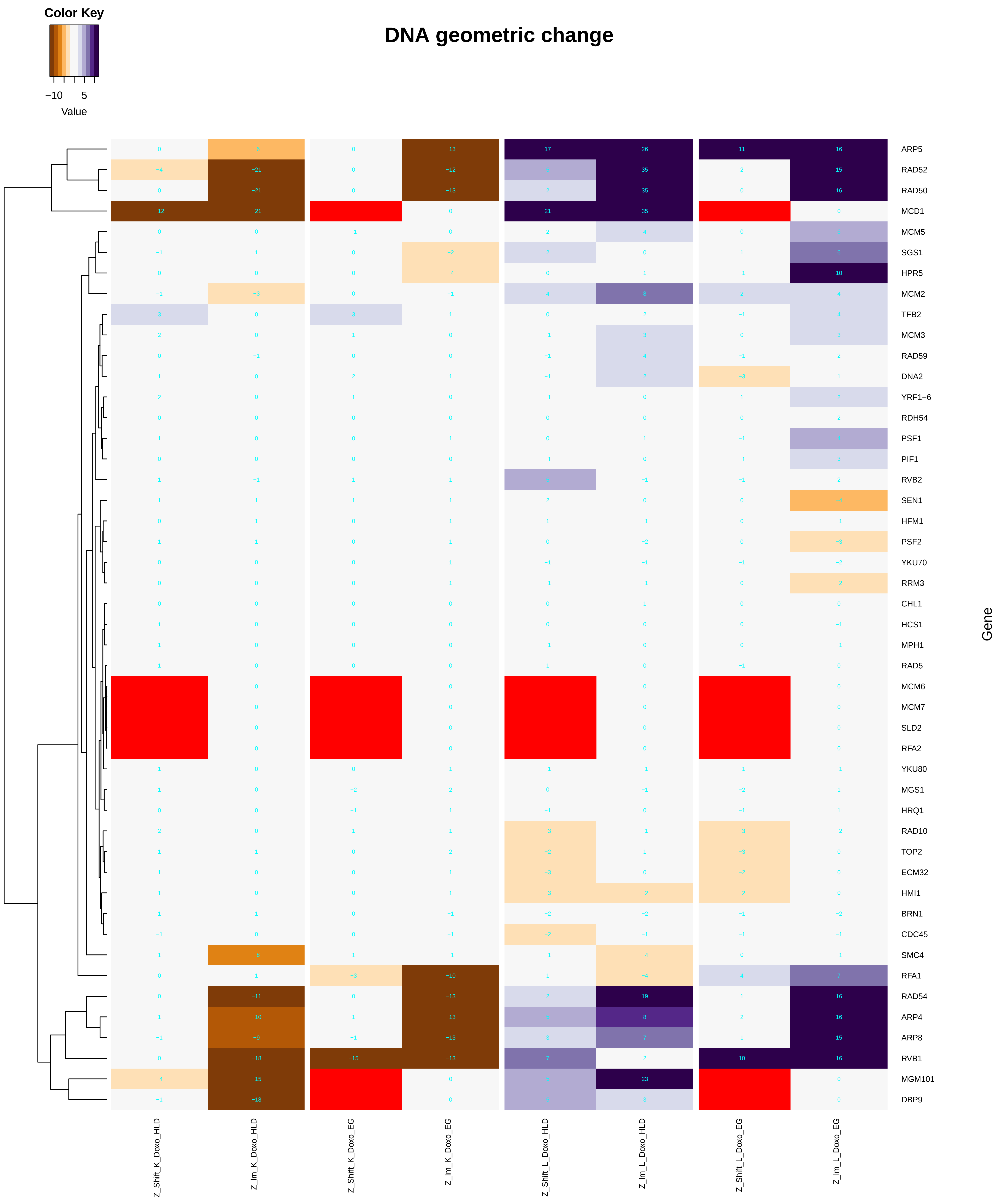

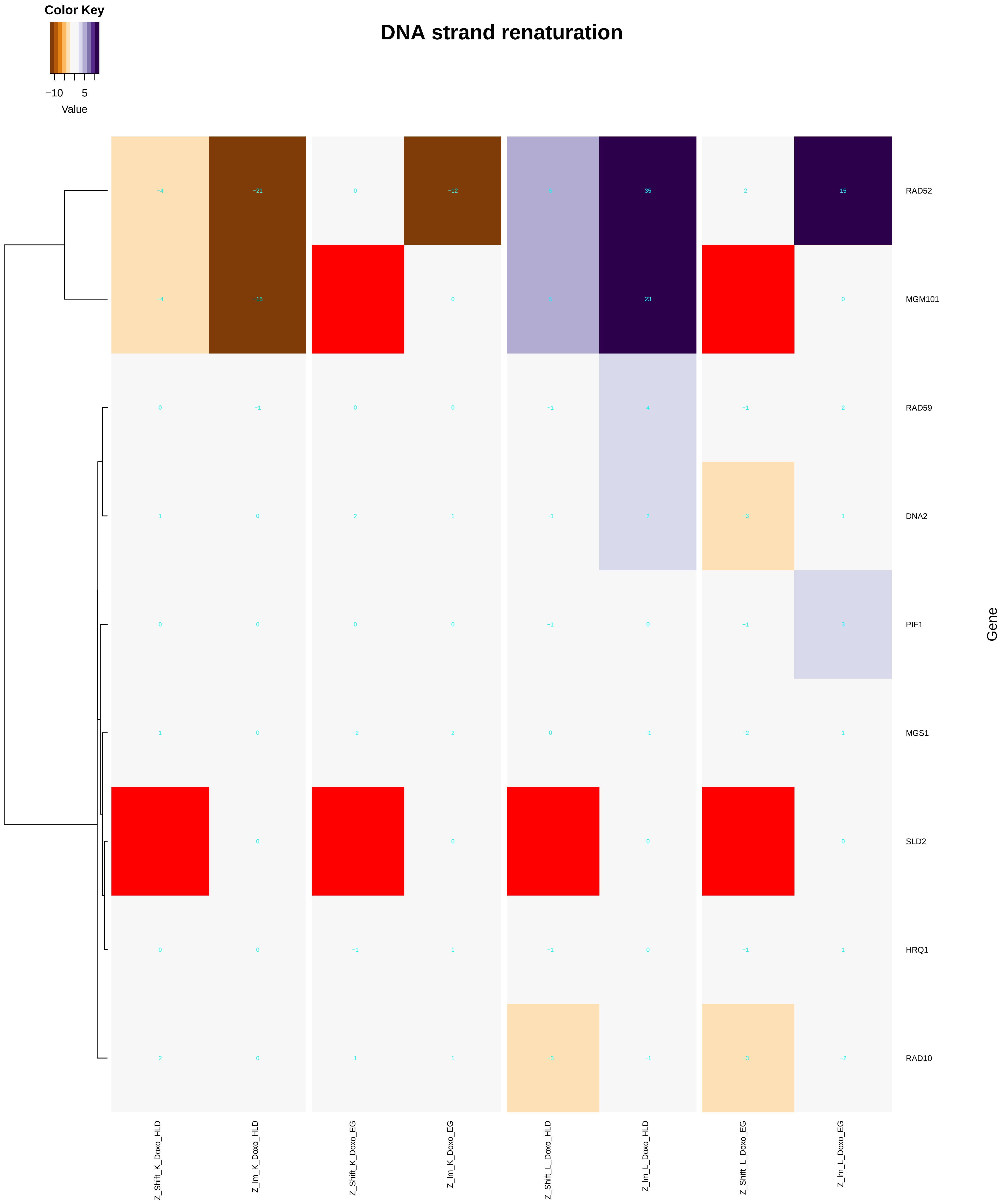

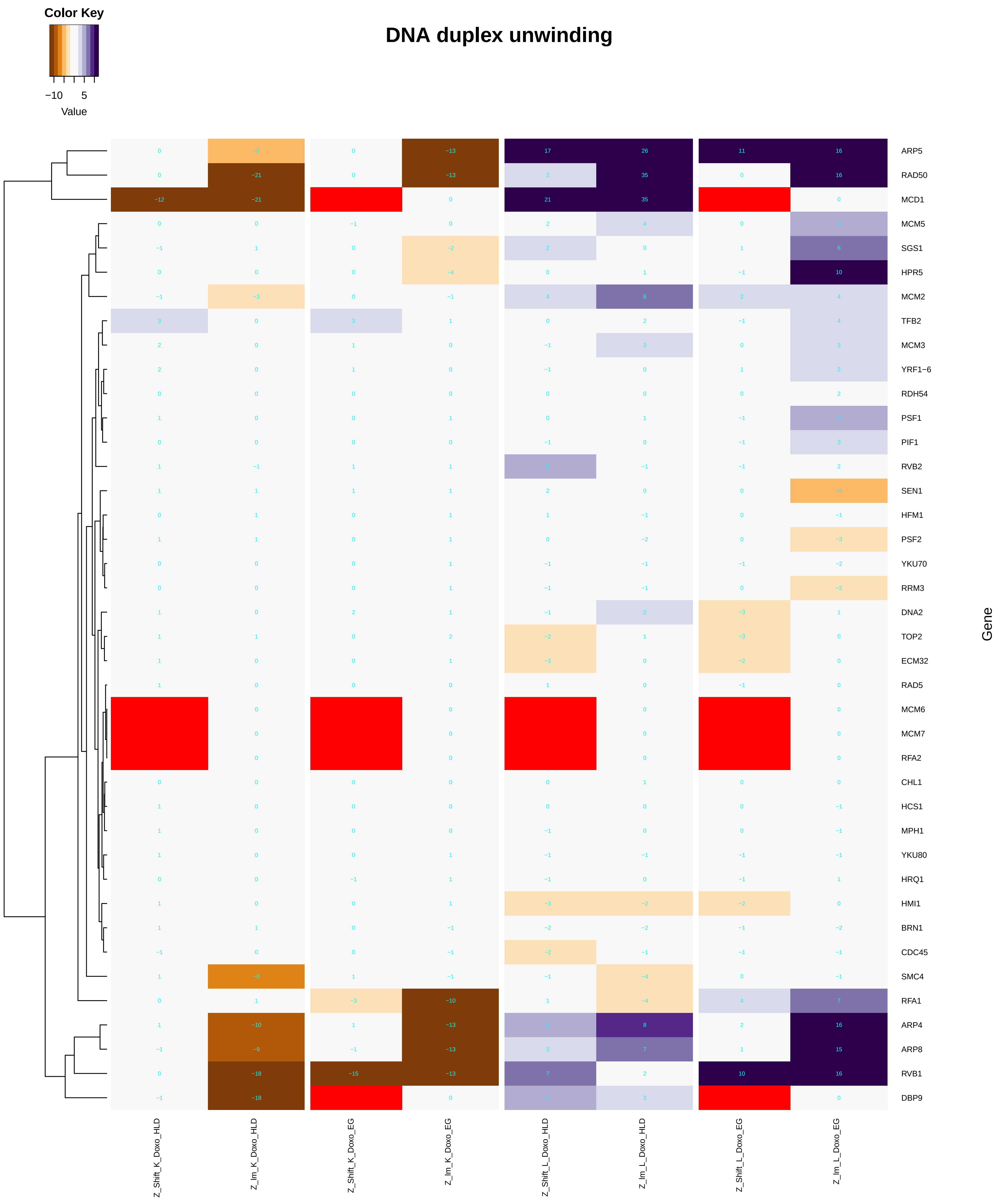

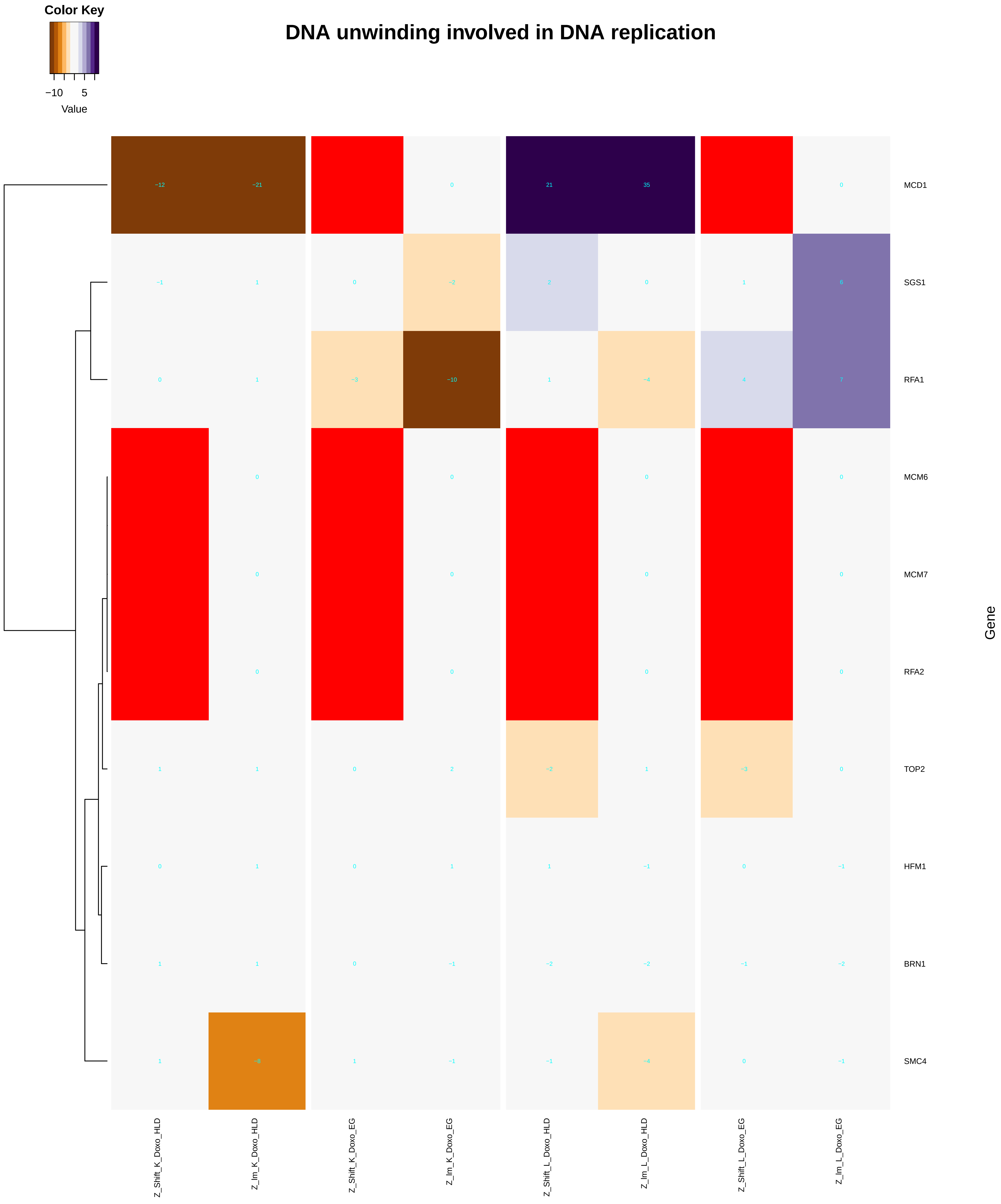

### DNA_packaging.pdf

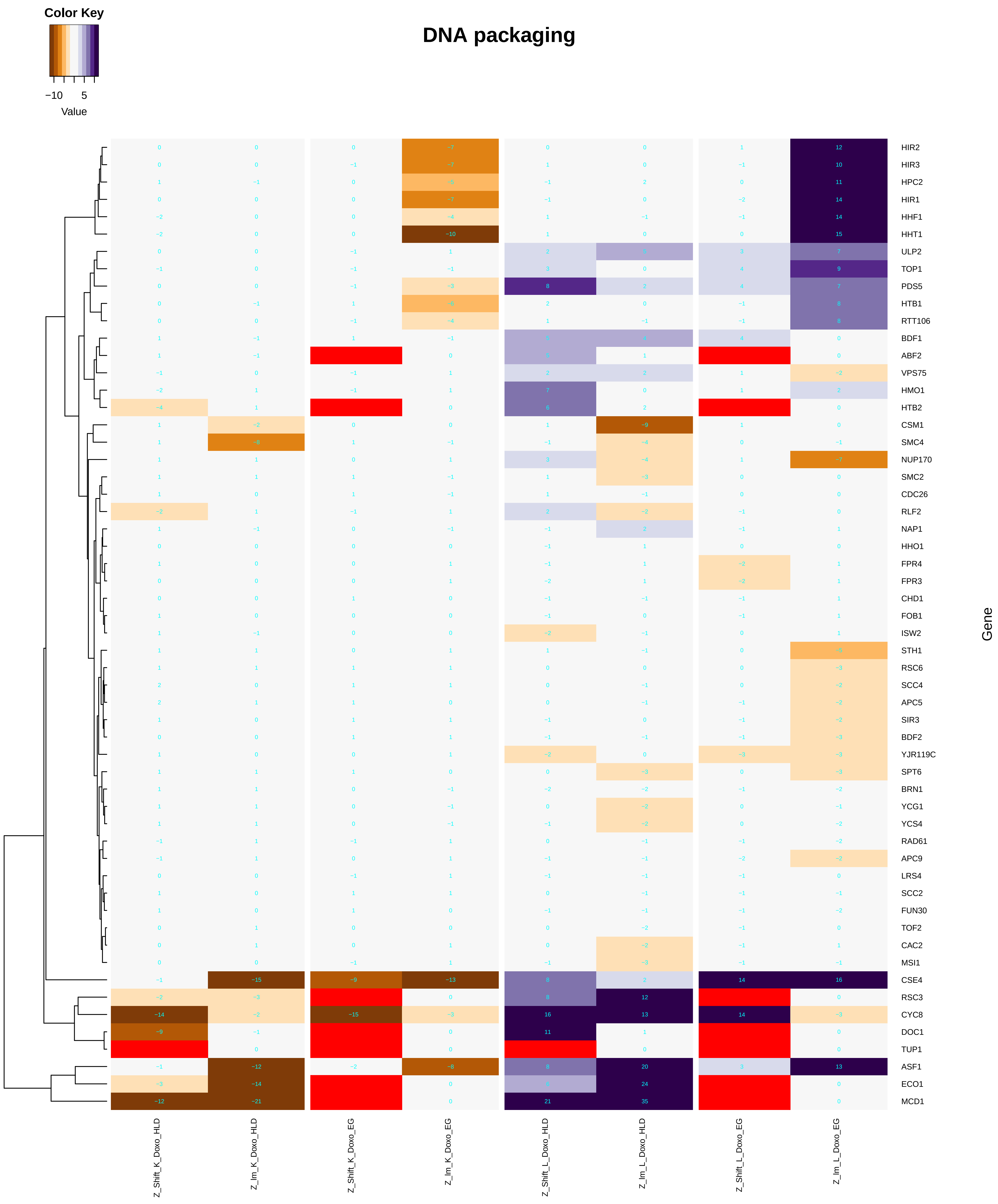

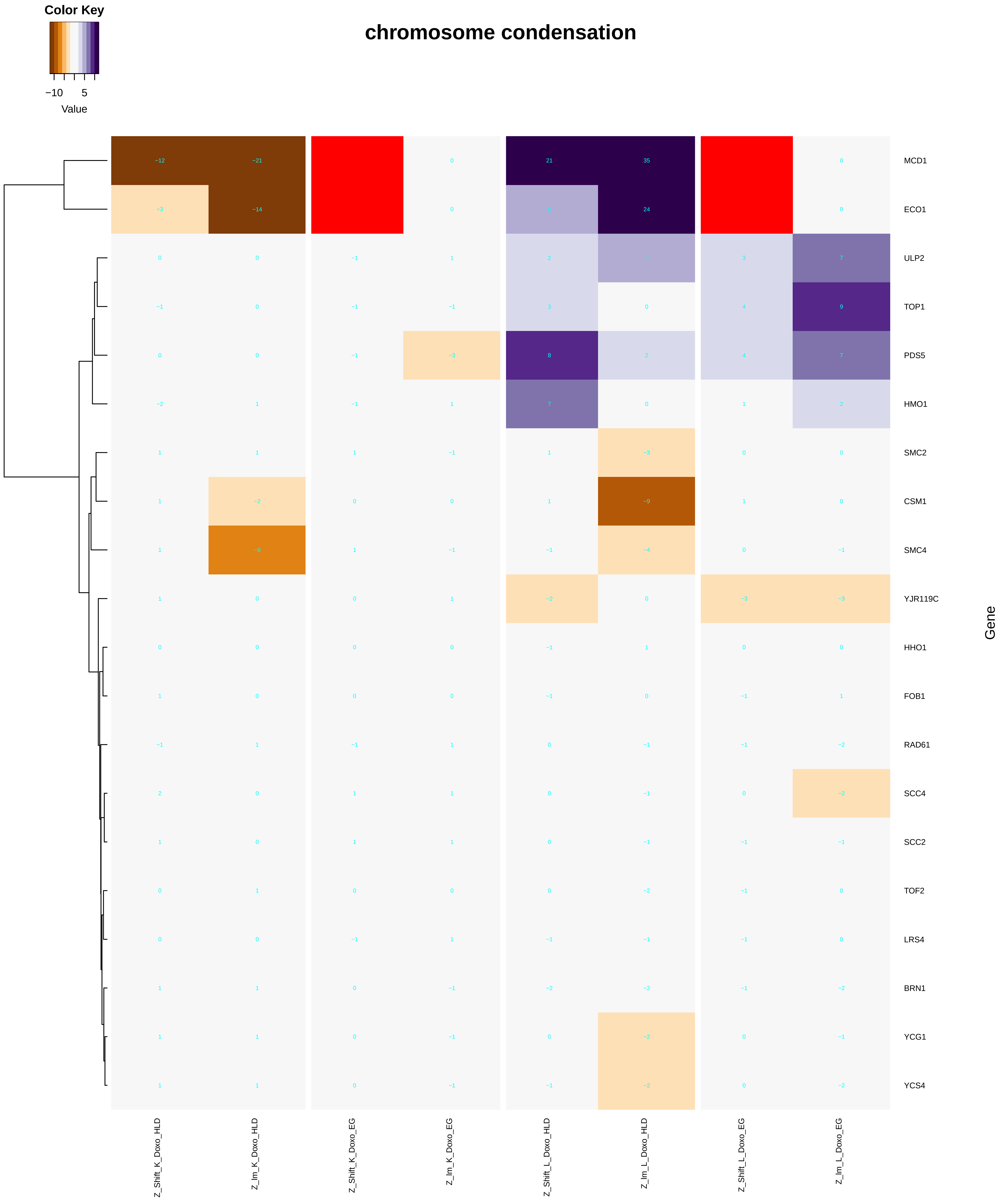

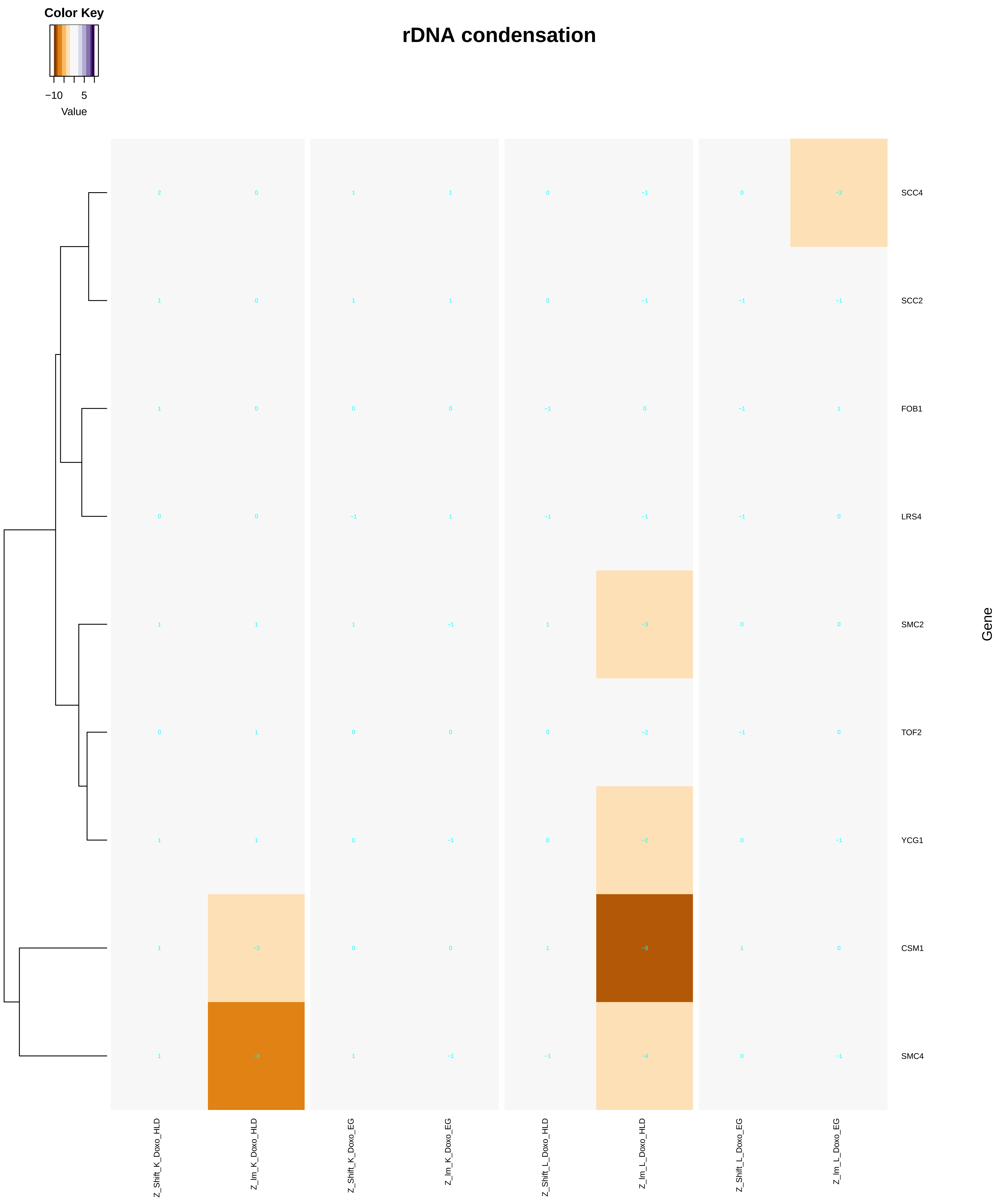

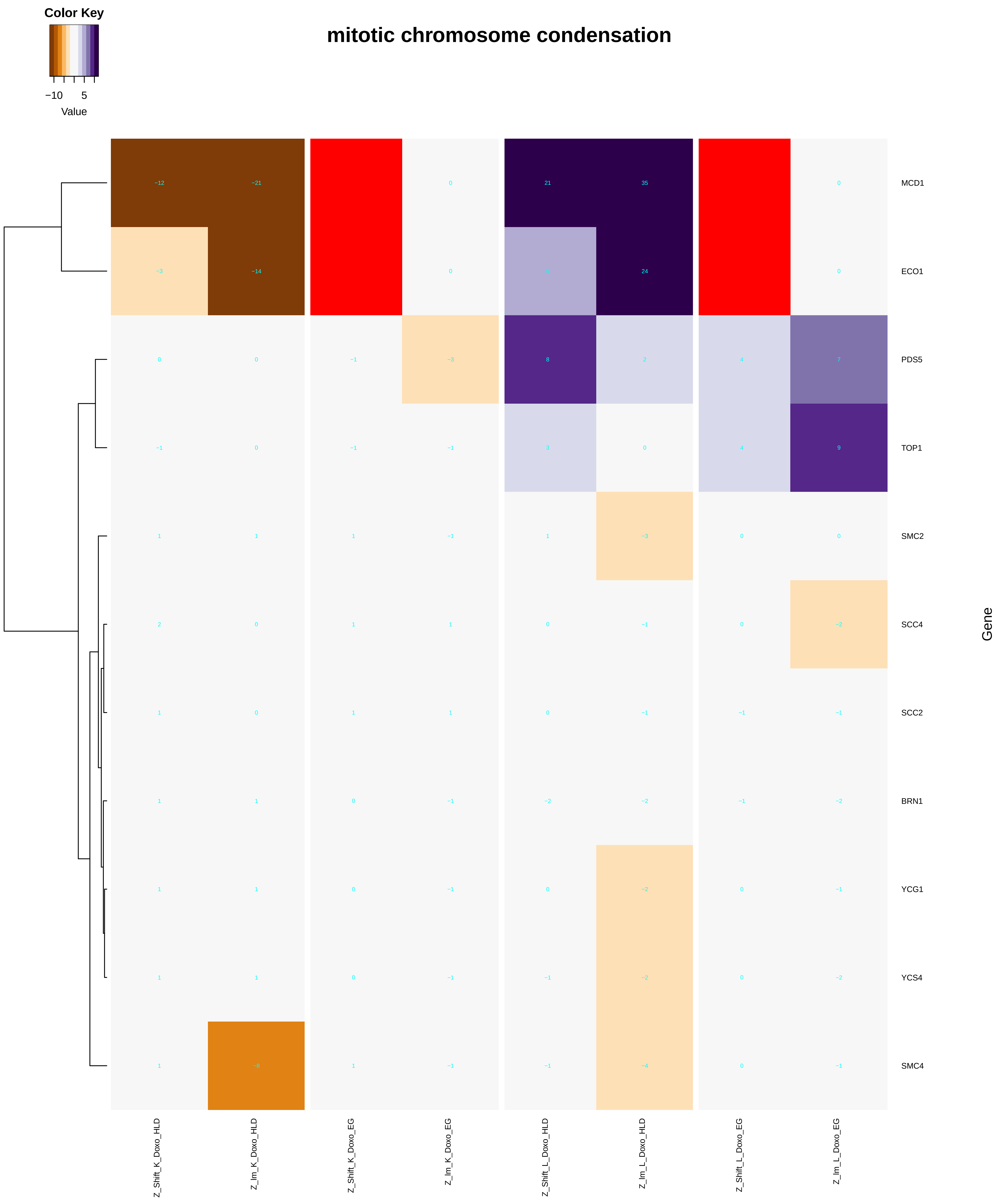

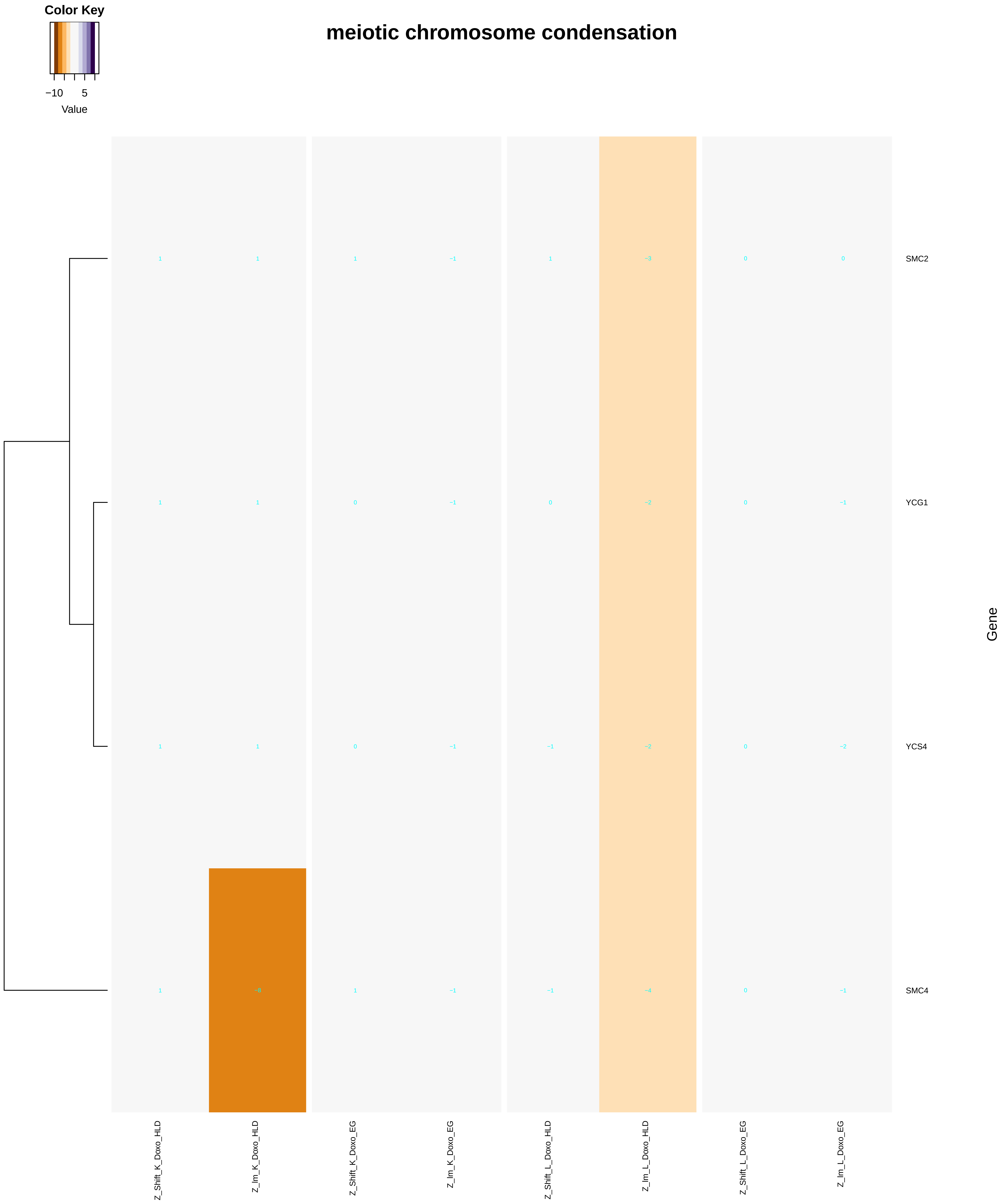

### DNA_recombinase_assembly.pdf

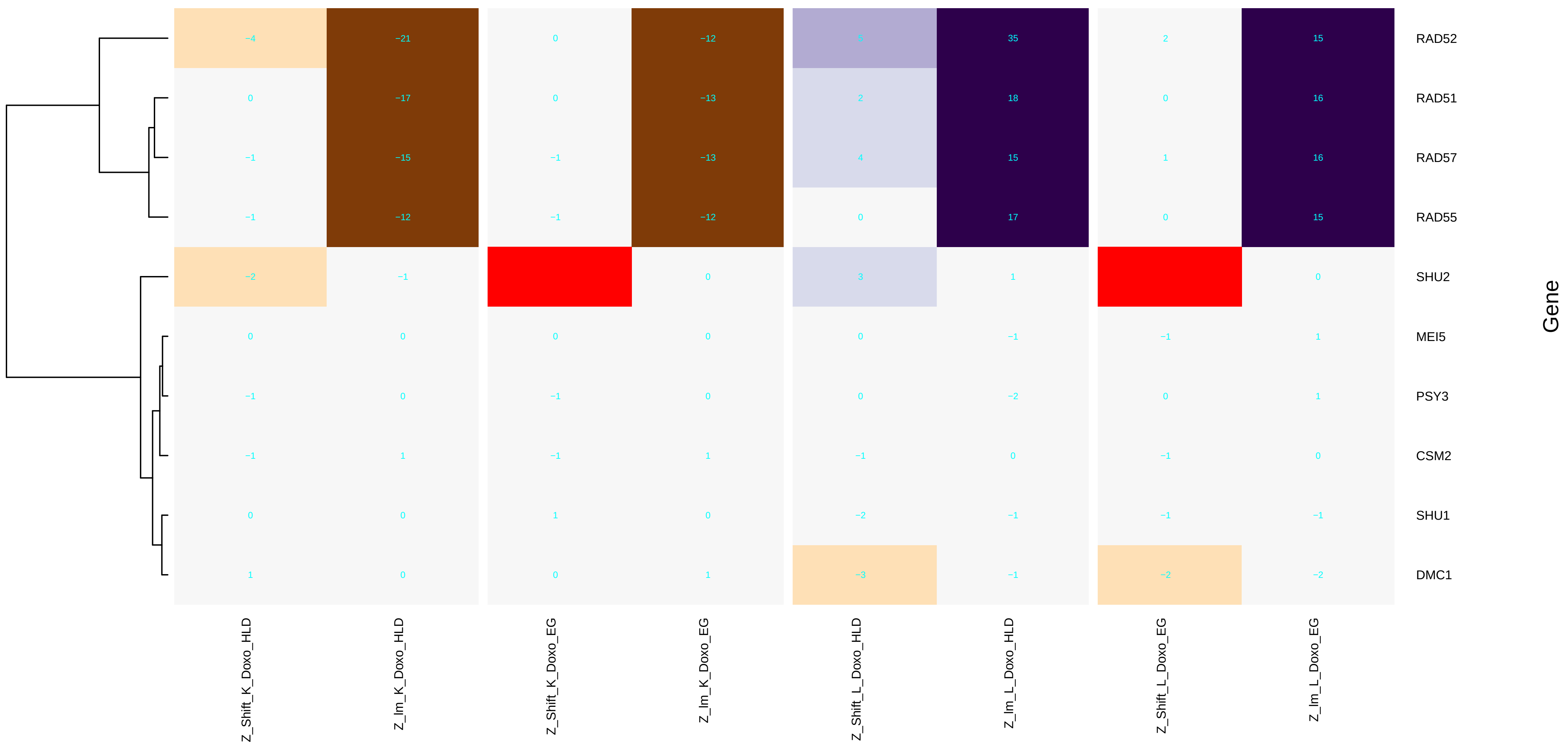

### DNA_repair_complex_assembly.pdf

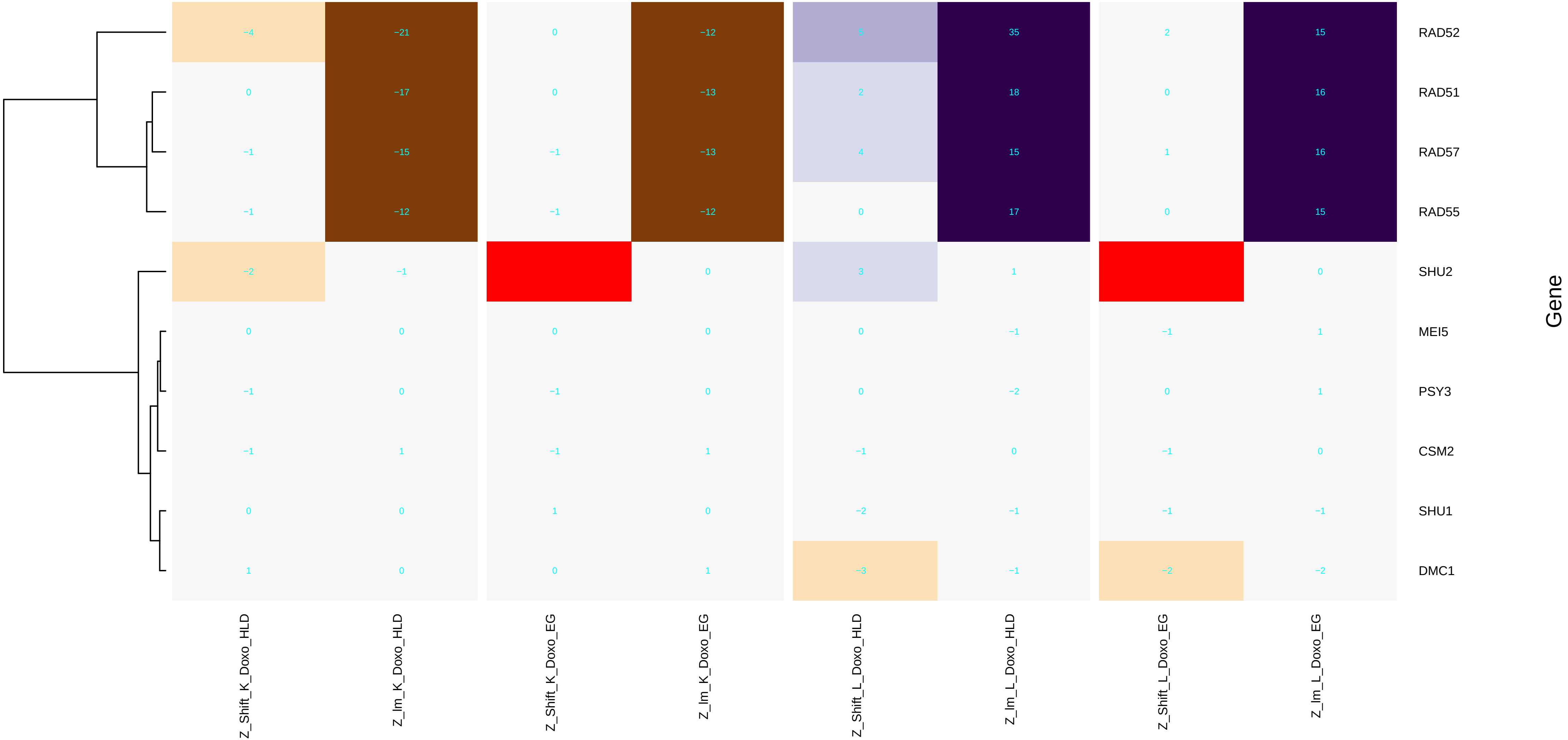

### DNA_replication-dependent_nucleosome_assembly.pdf

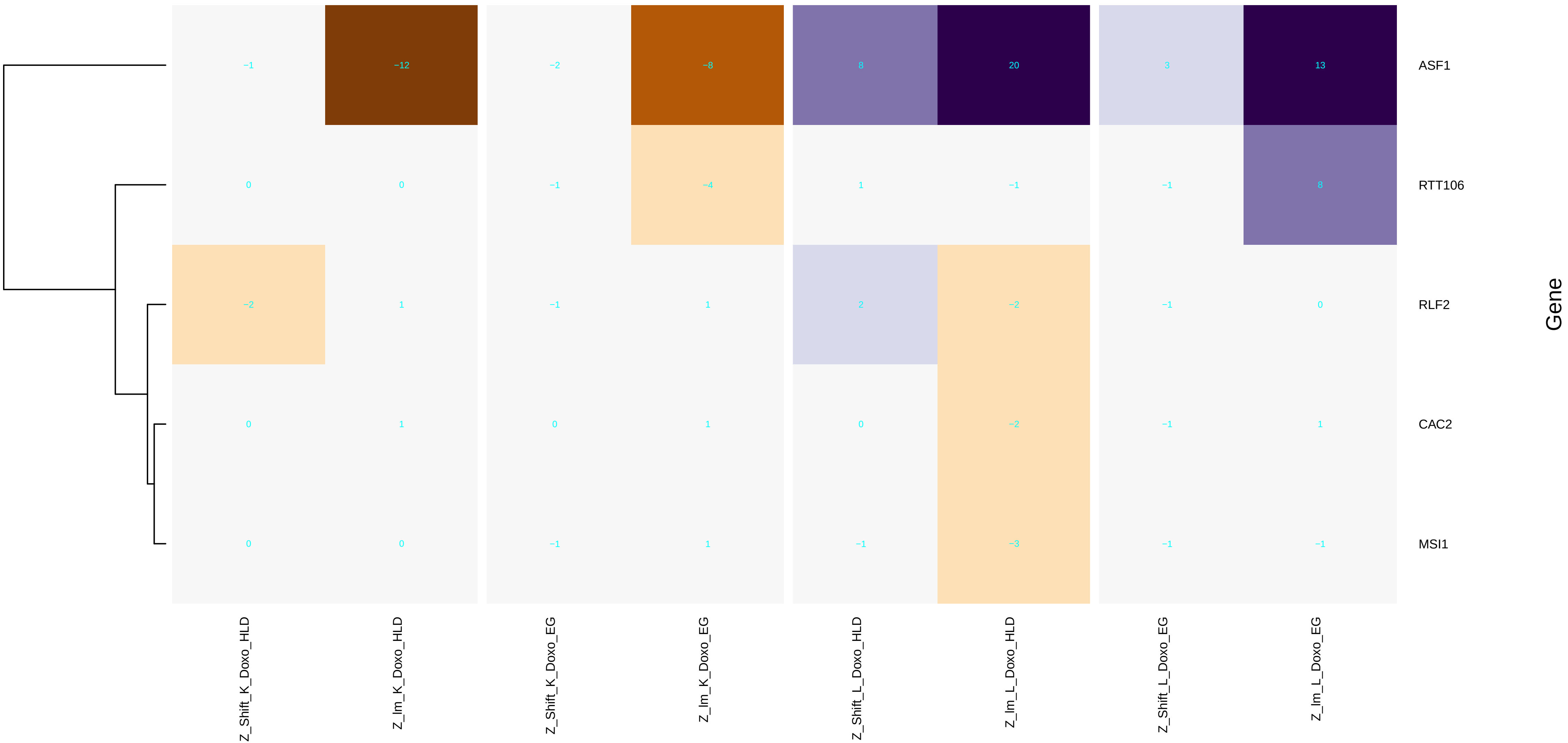

### DNA_replication-dependent_nucleosome_organization.pdf

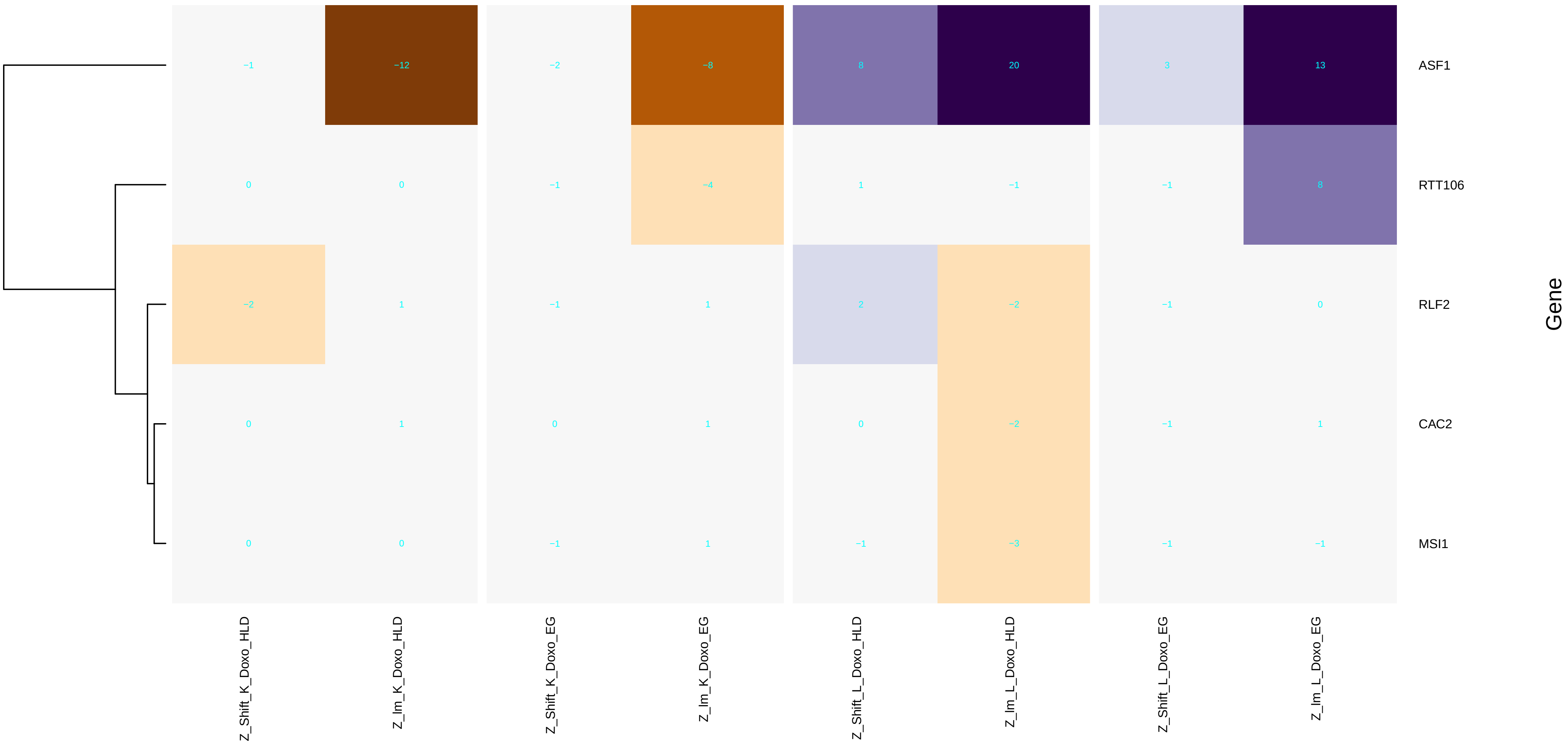

### DNA_replication-independent_nucleosome_assembly.pdf

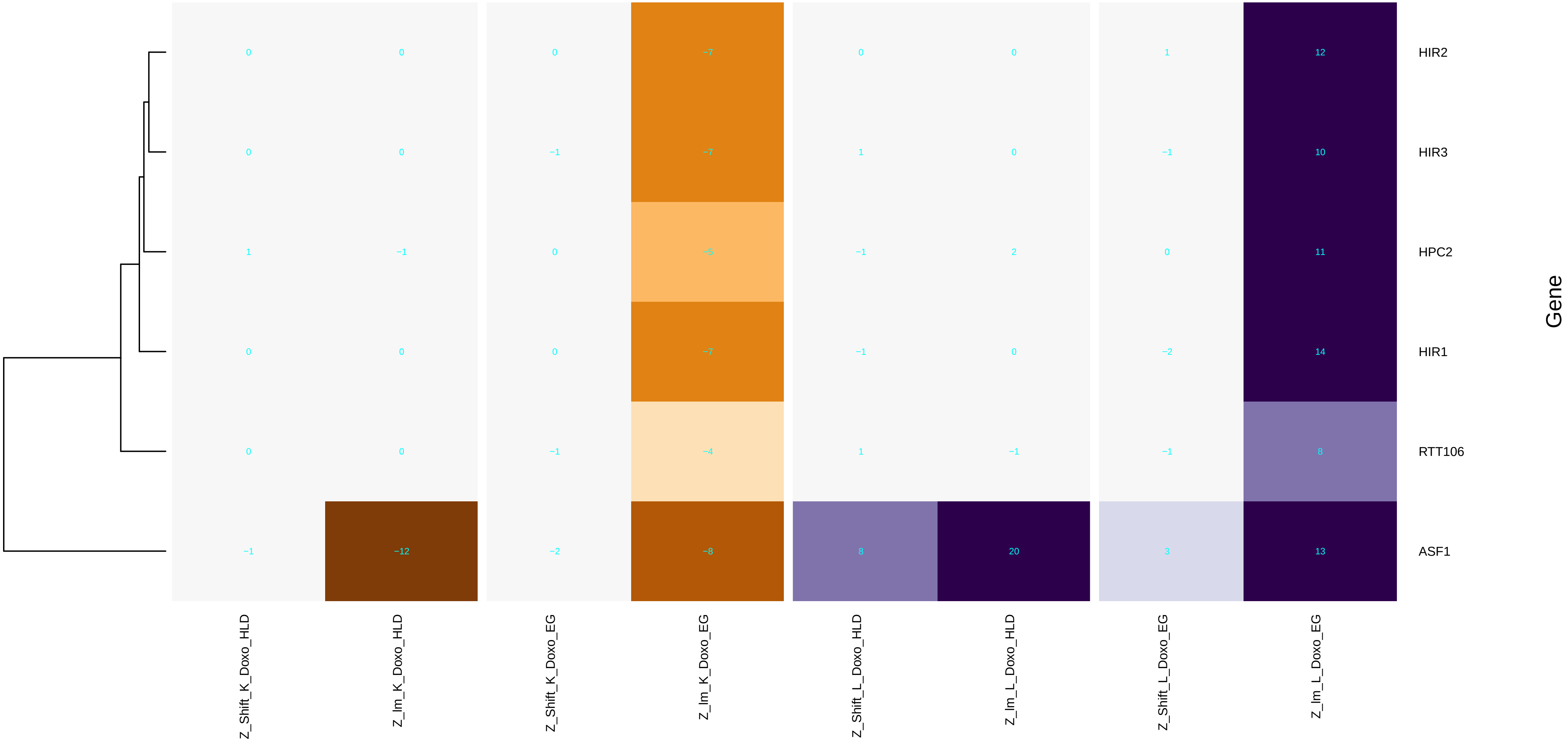

### DNA_replication-independent_nucleosome_organization.pdf

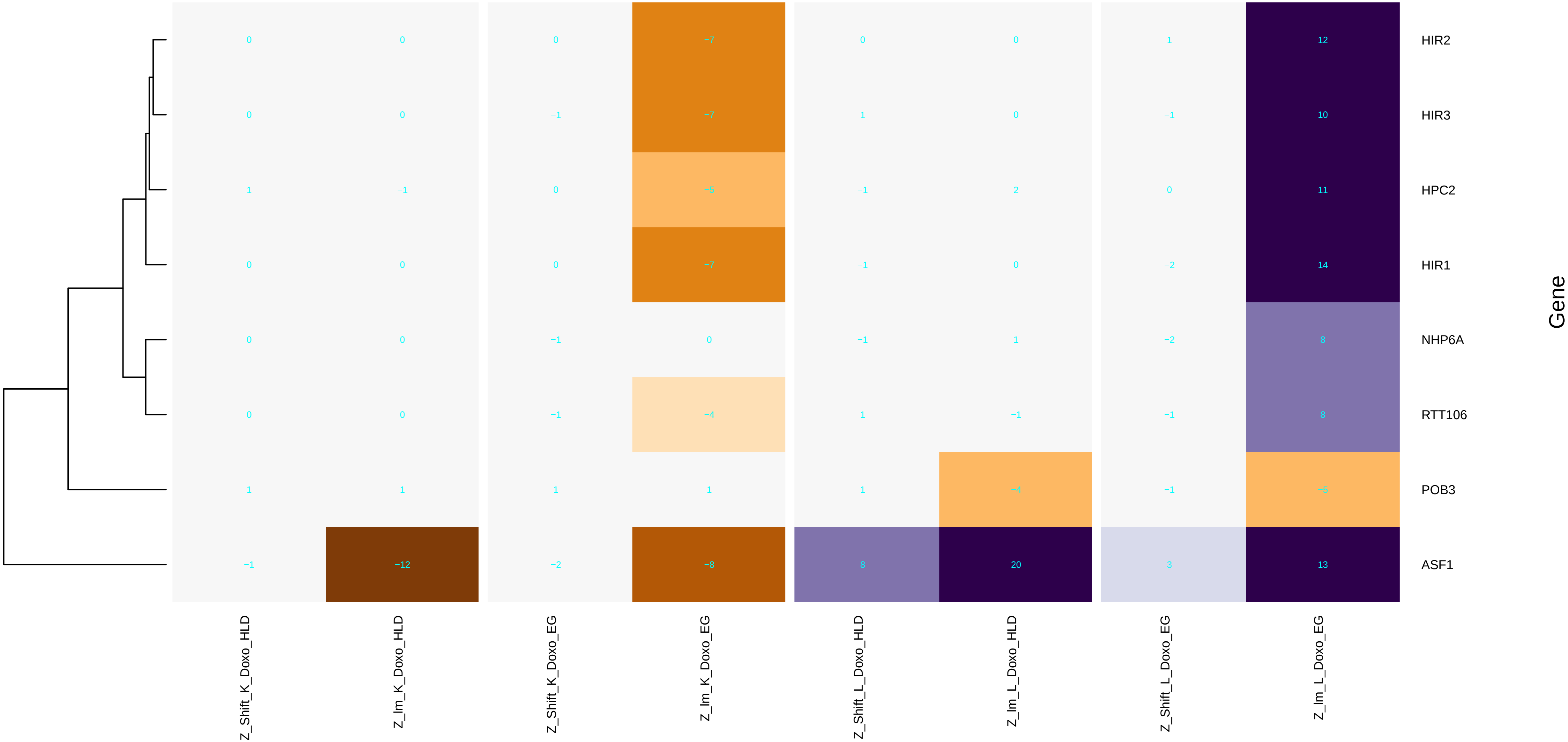

### DNA_replication.pdf

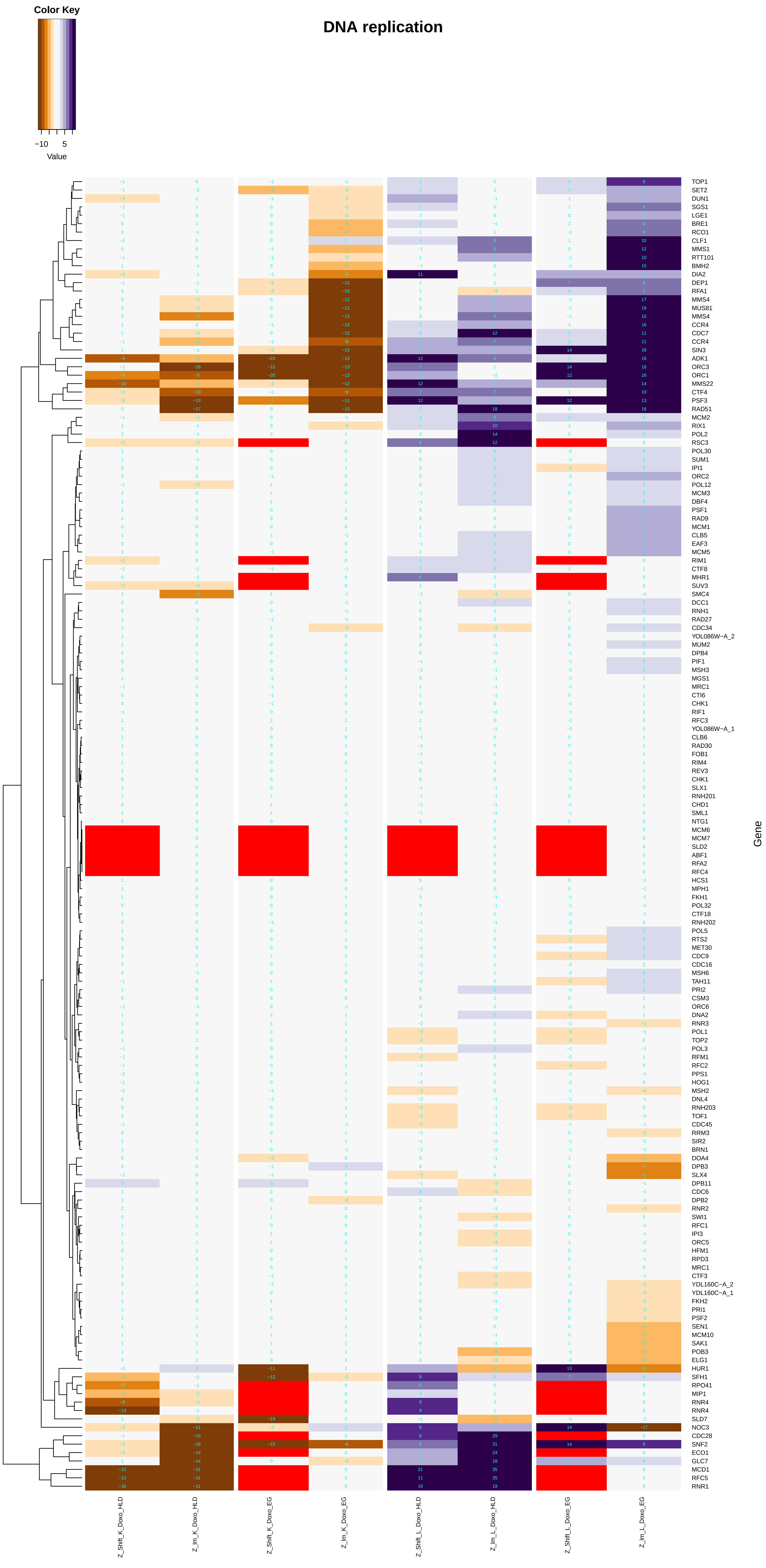

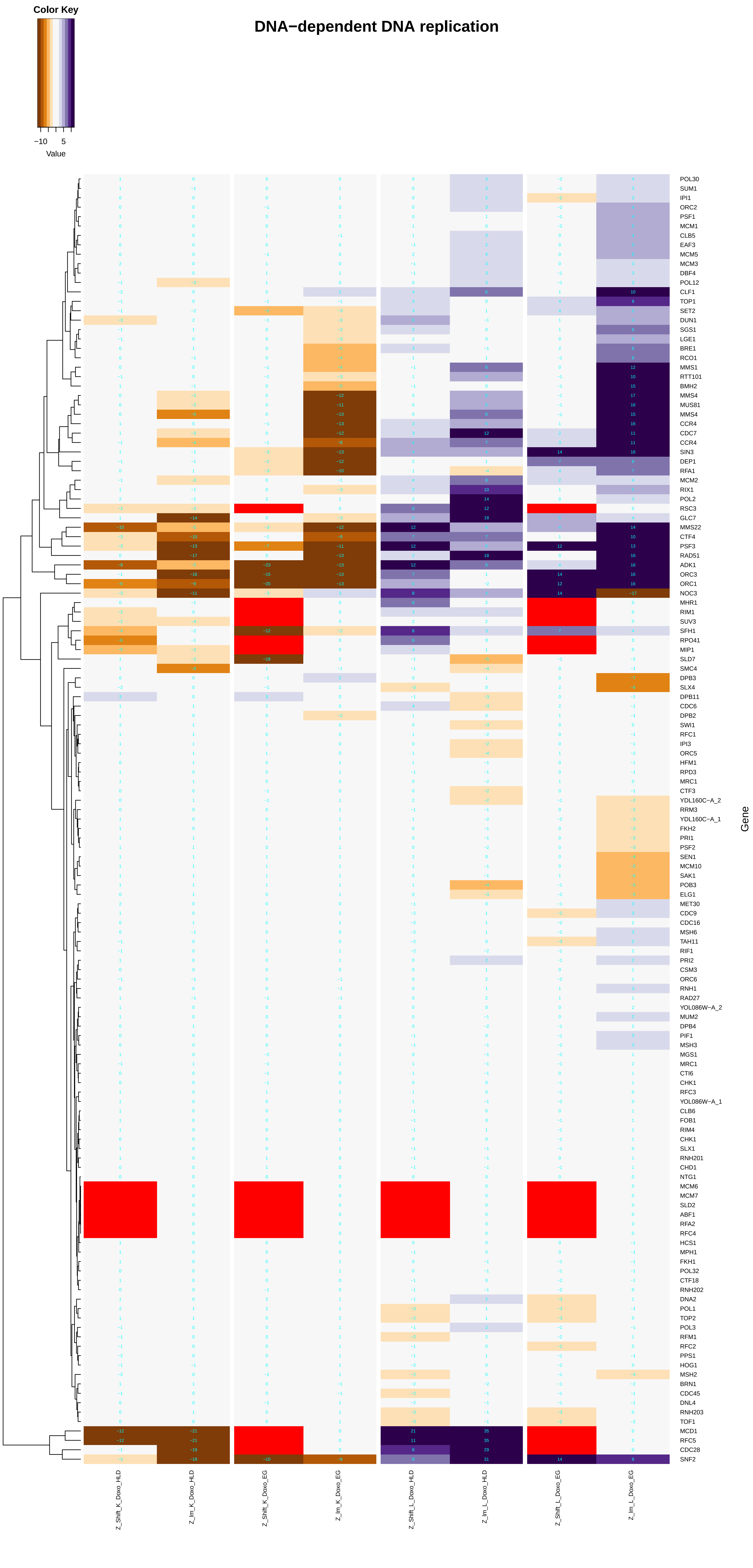

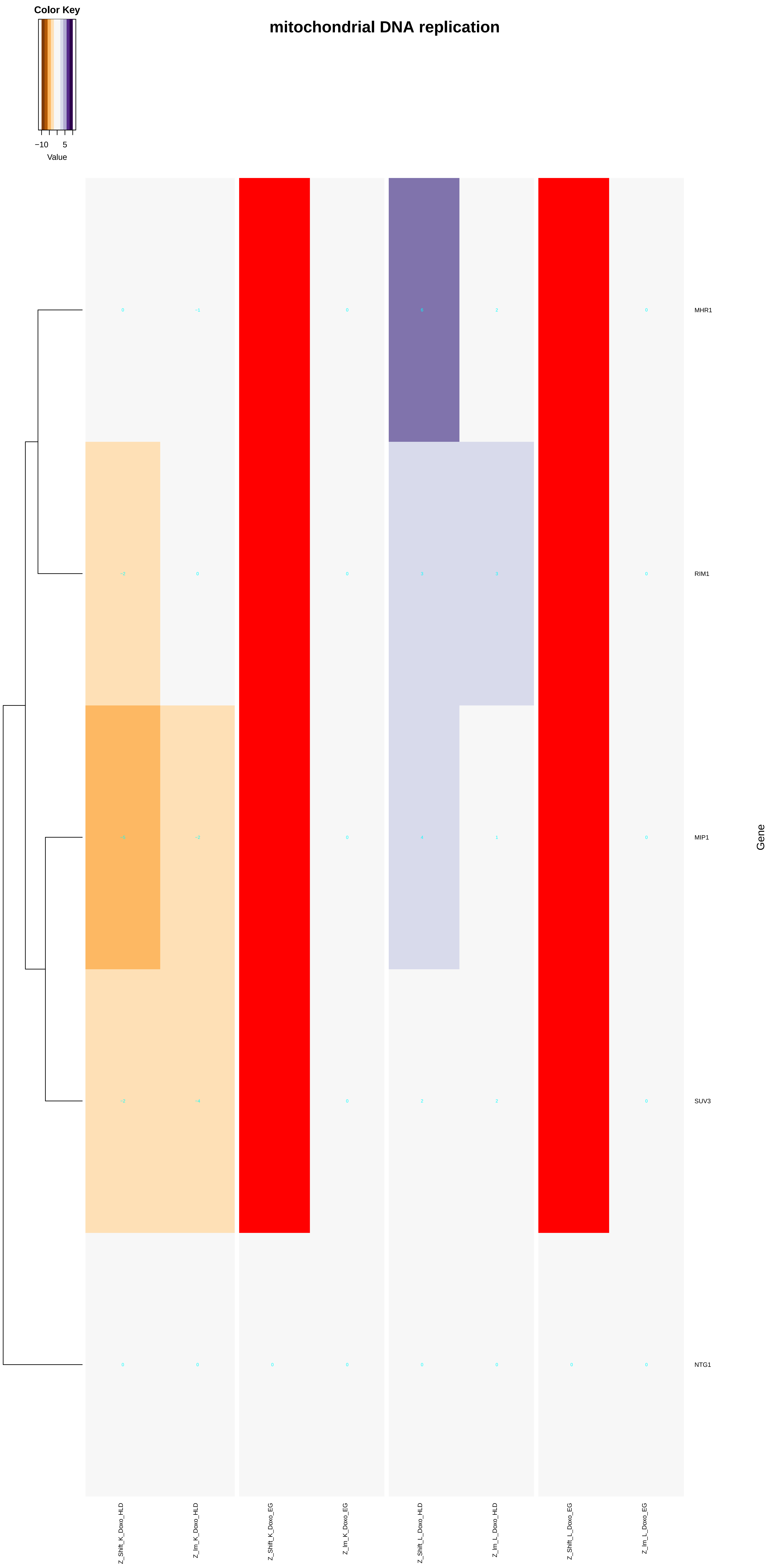

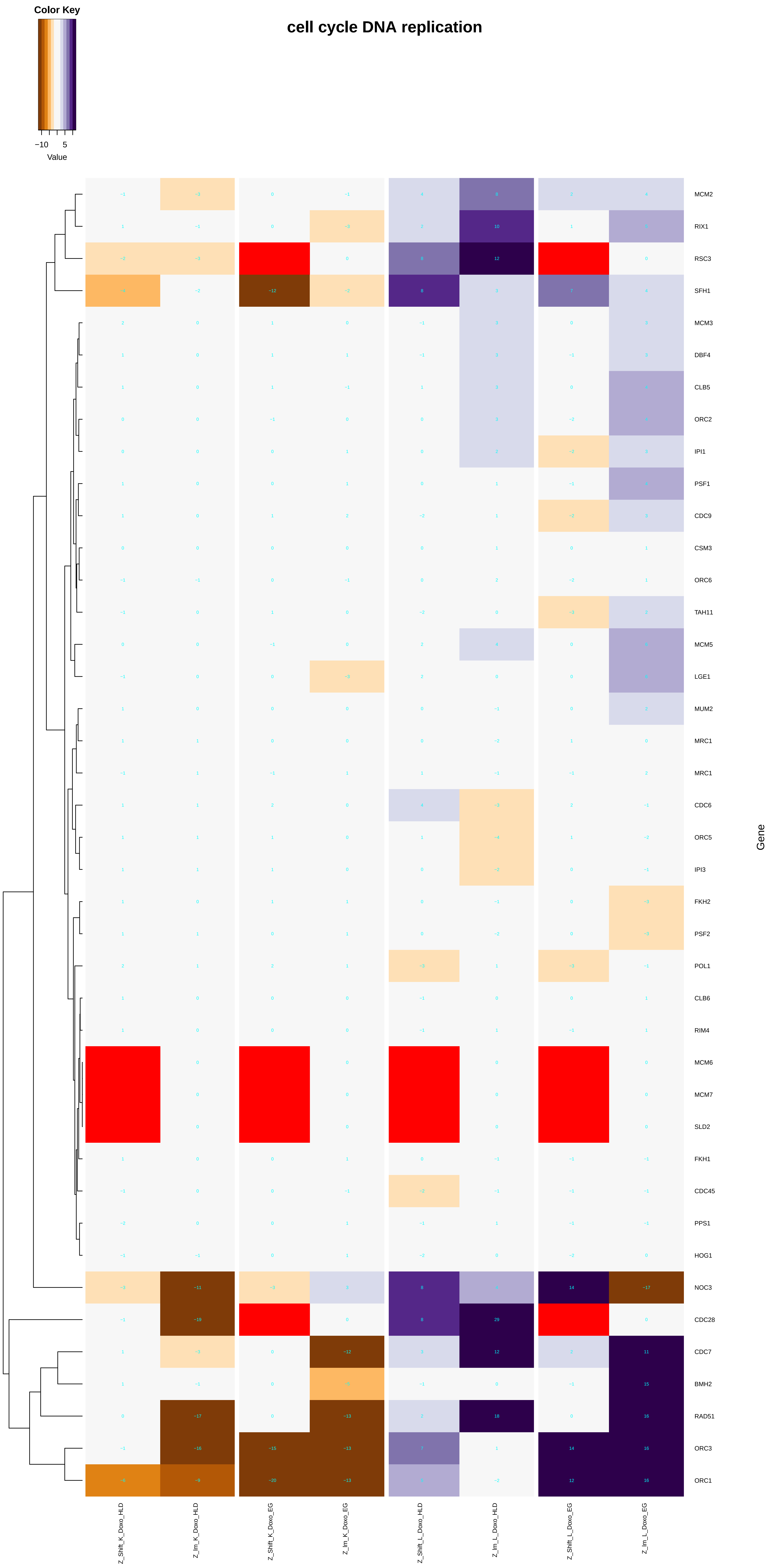

##### nuclear DNA replication

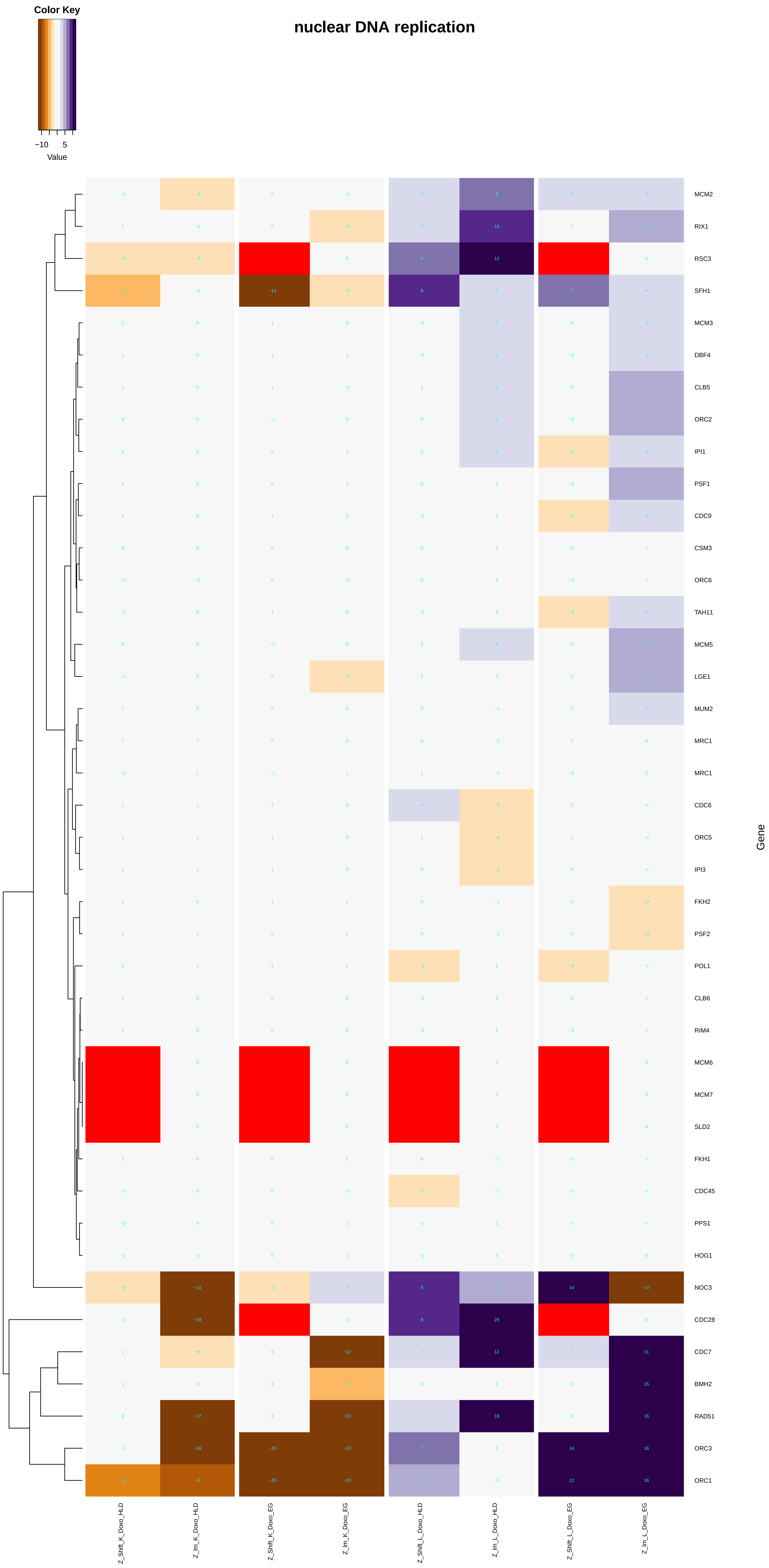

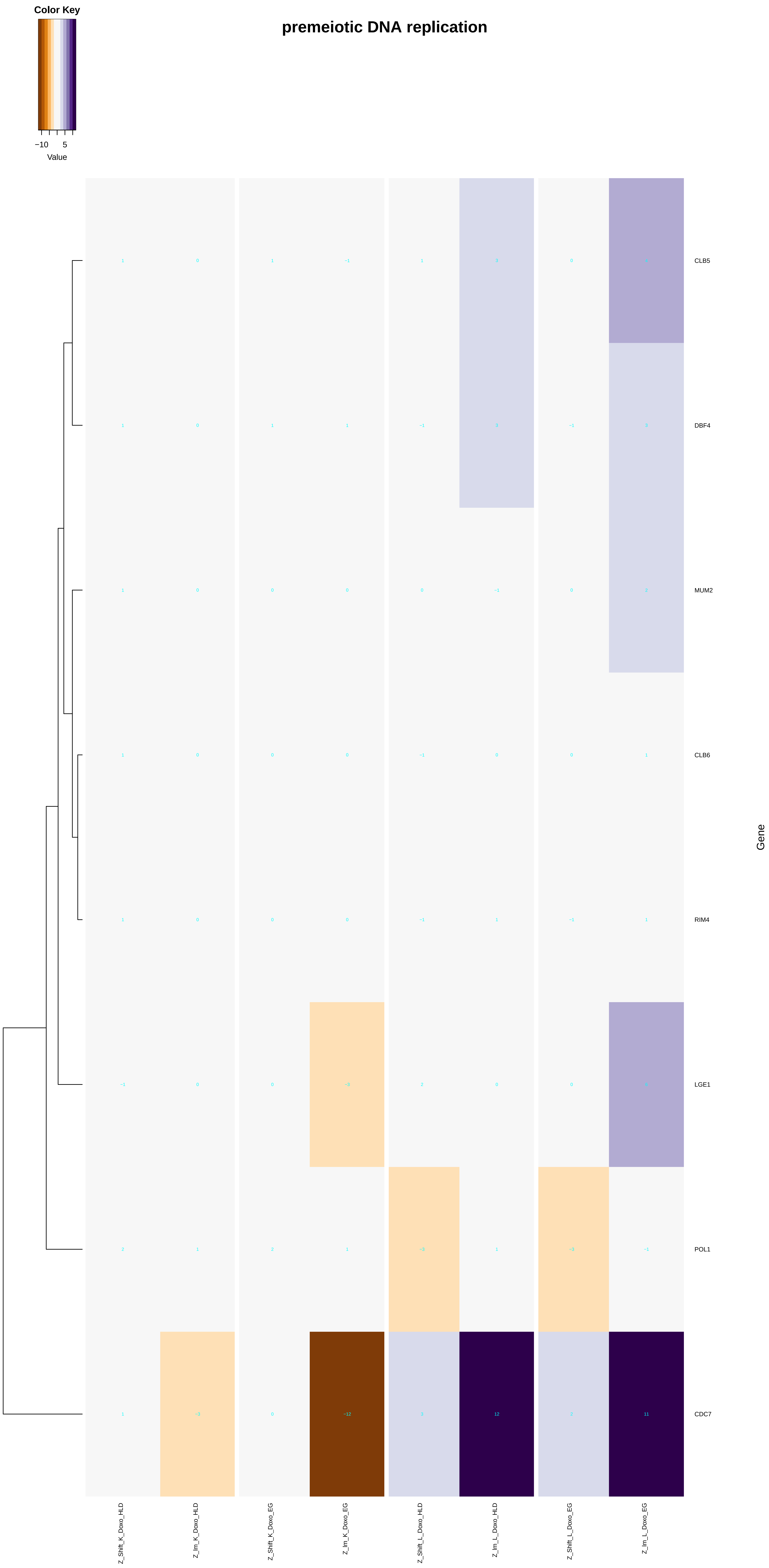

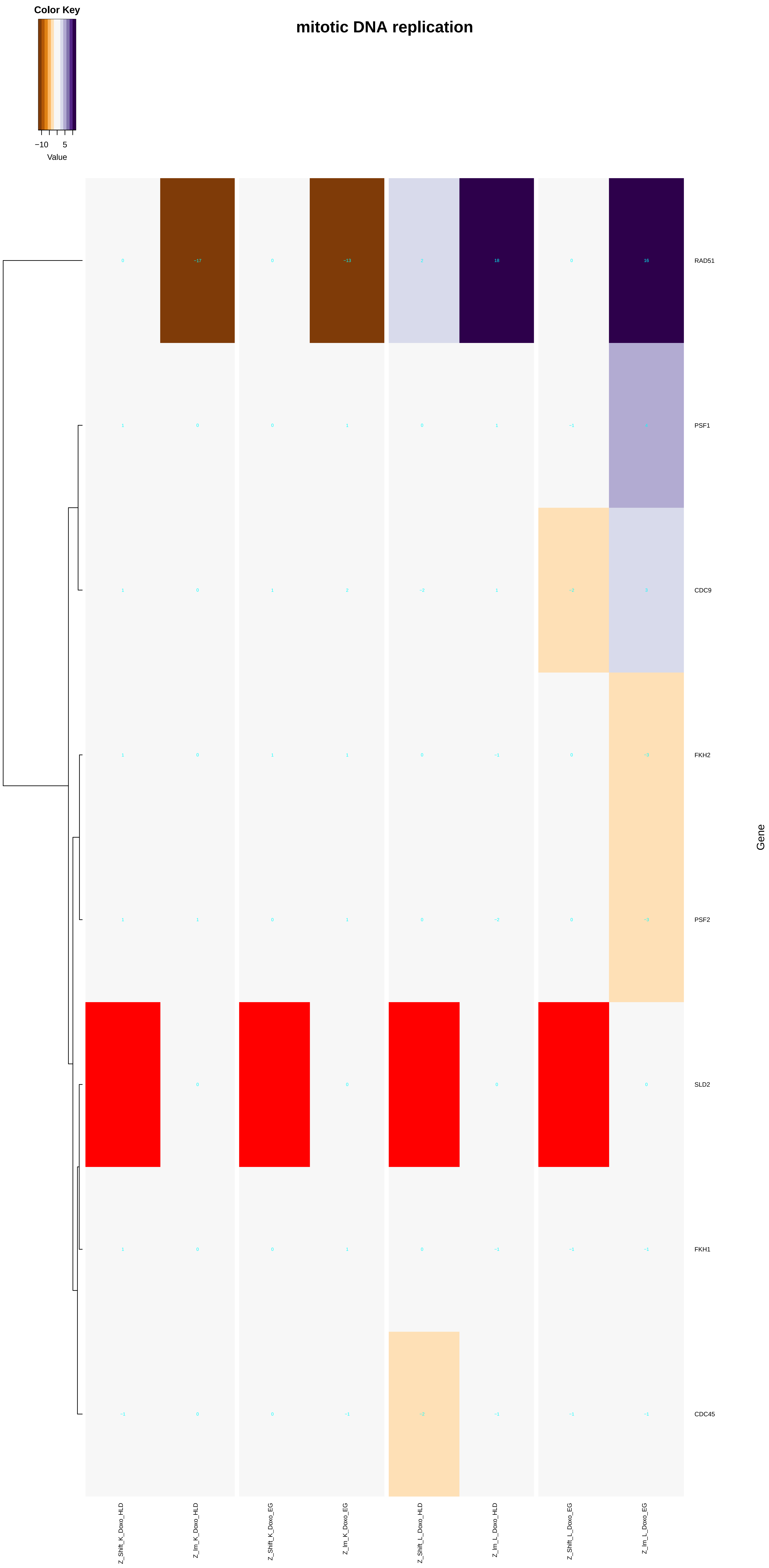

### DNA_replication_initiation.pdf

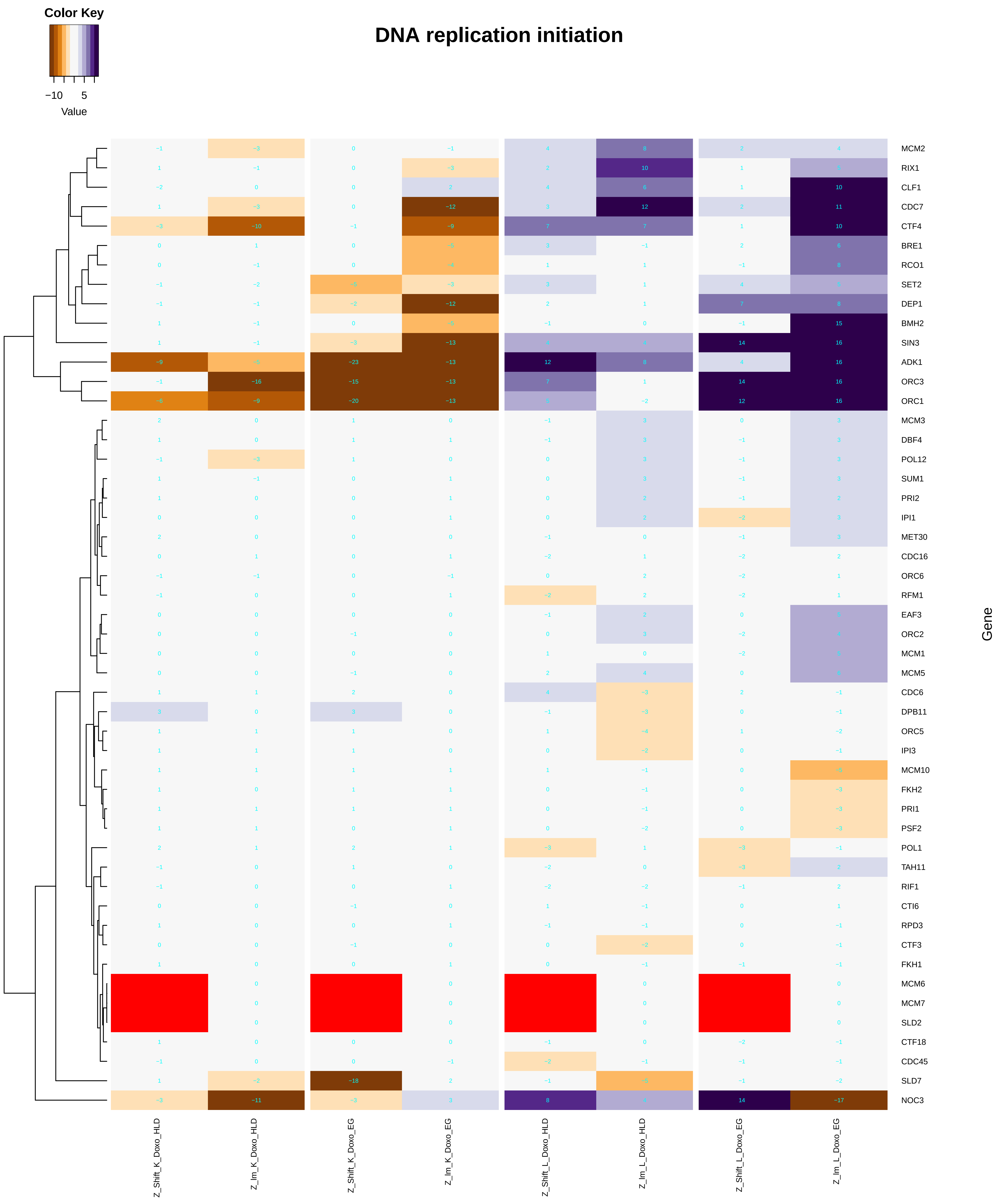

cell cycle DNA replication initiation

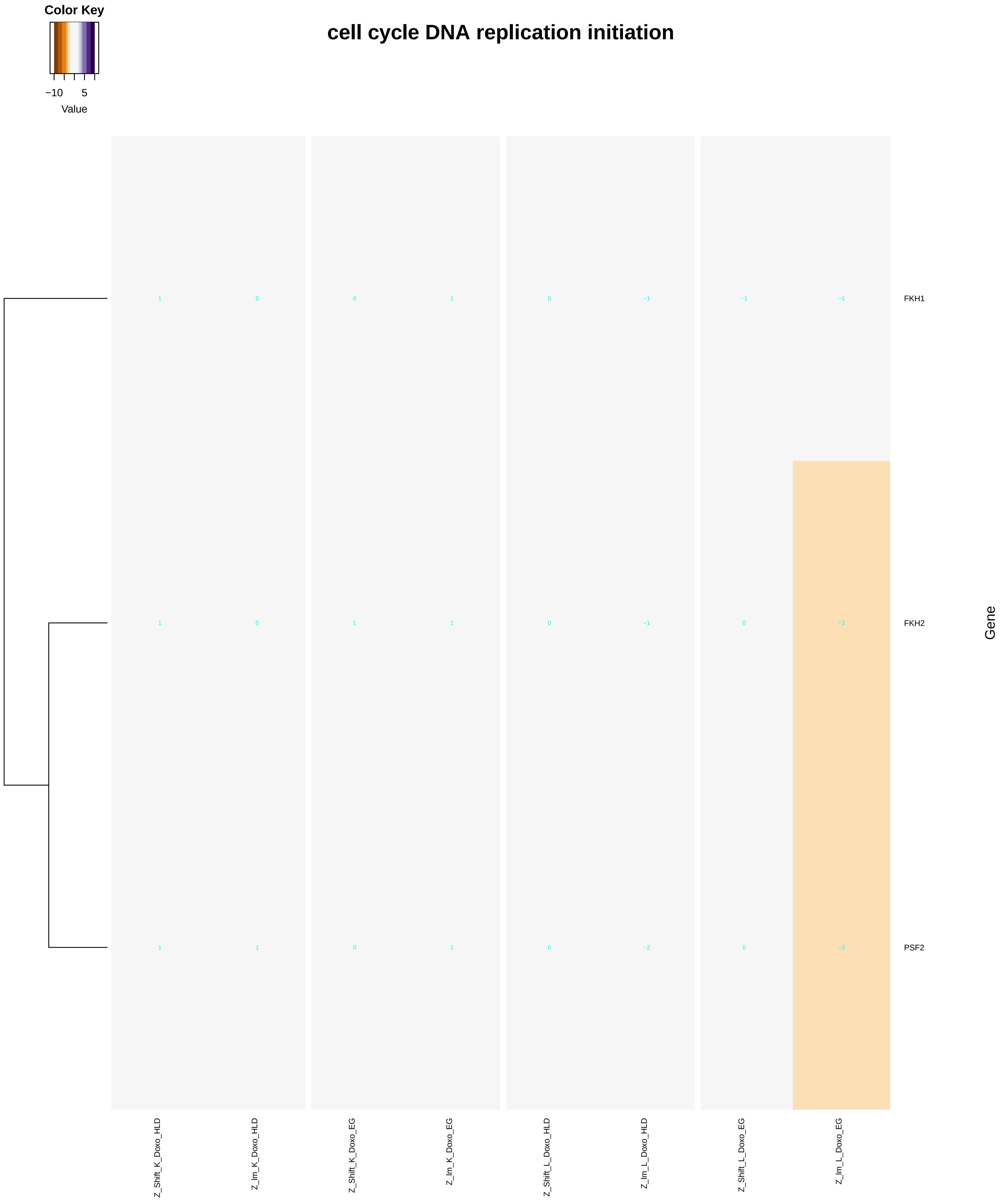

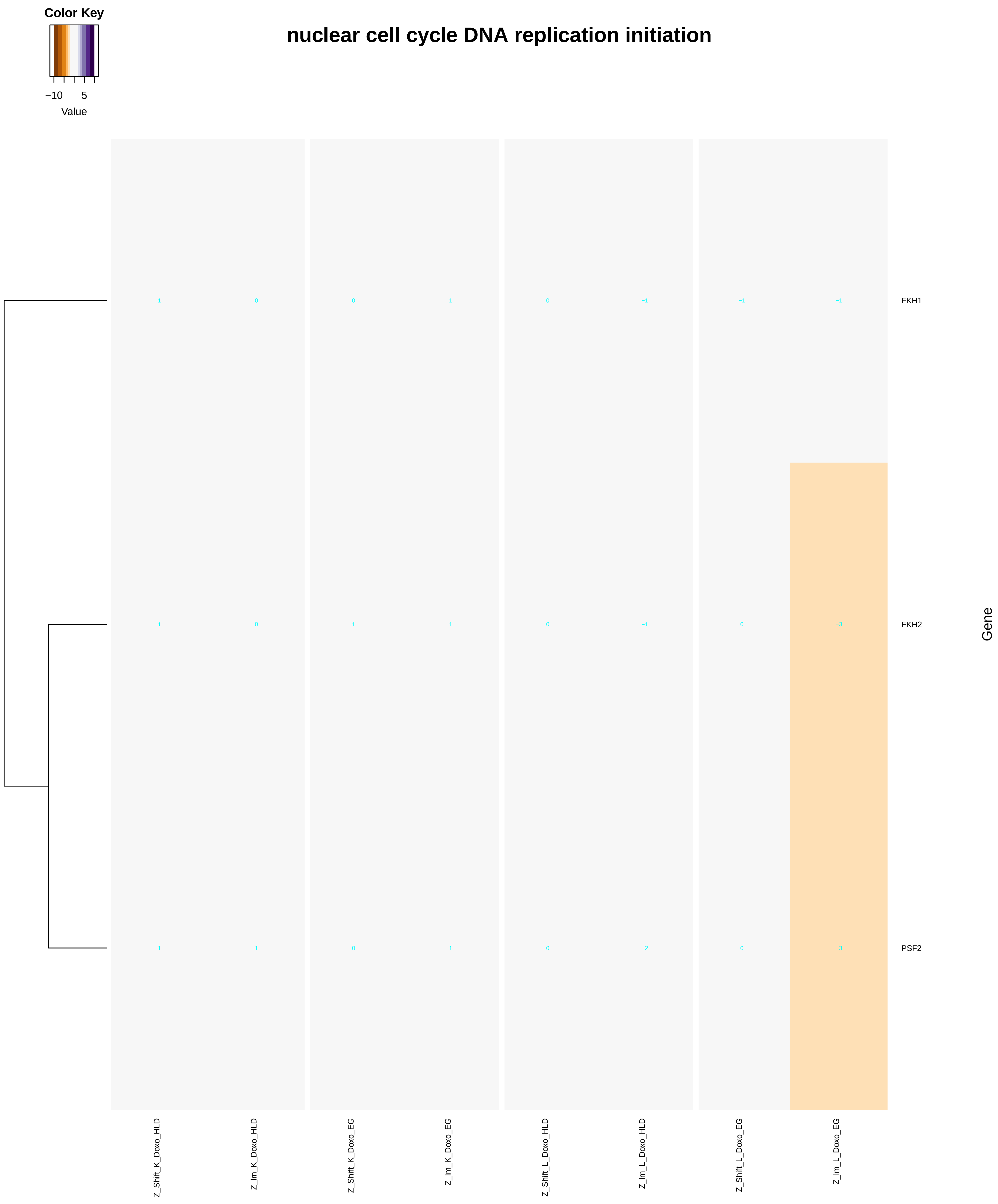

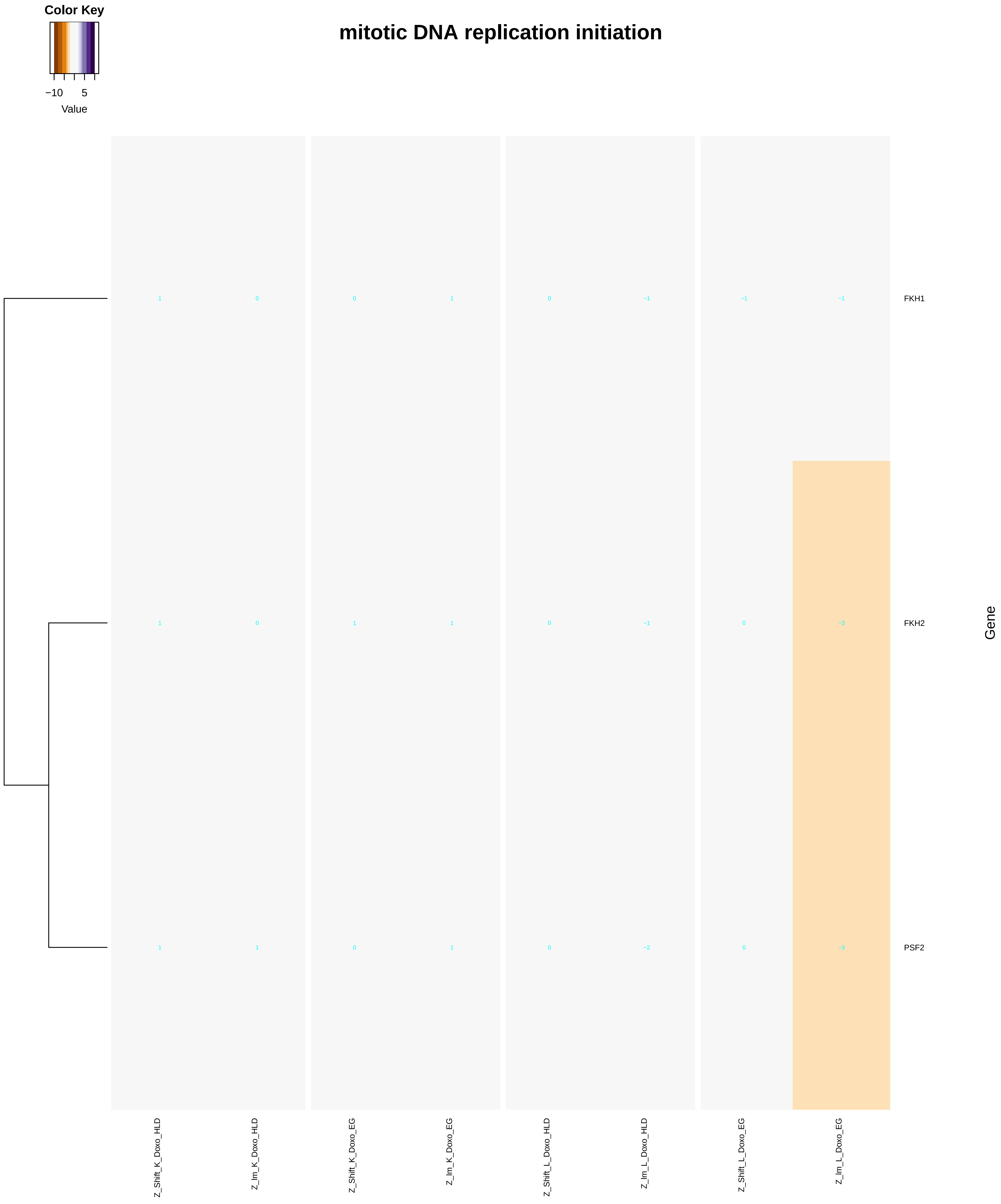

### DNA_topological_change.pdf

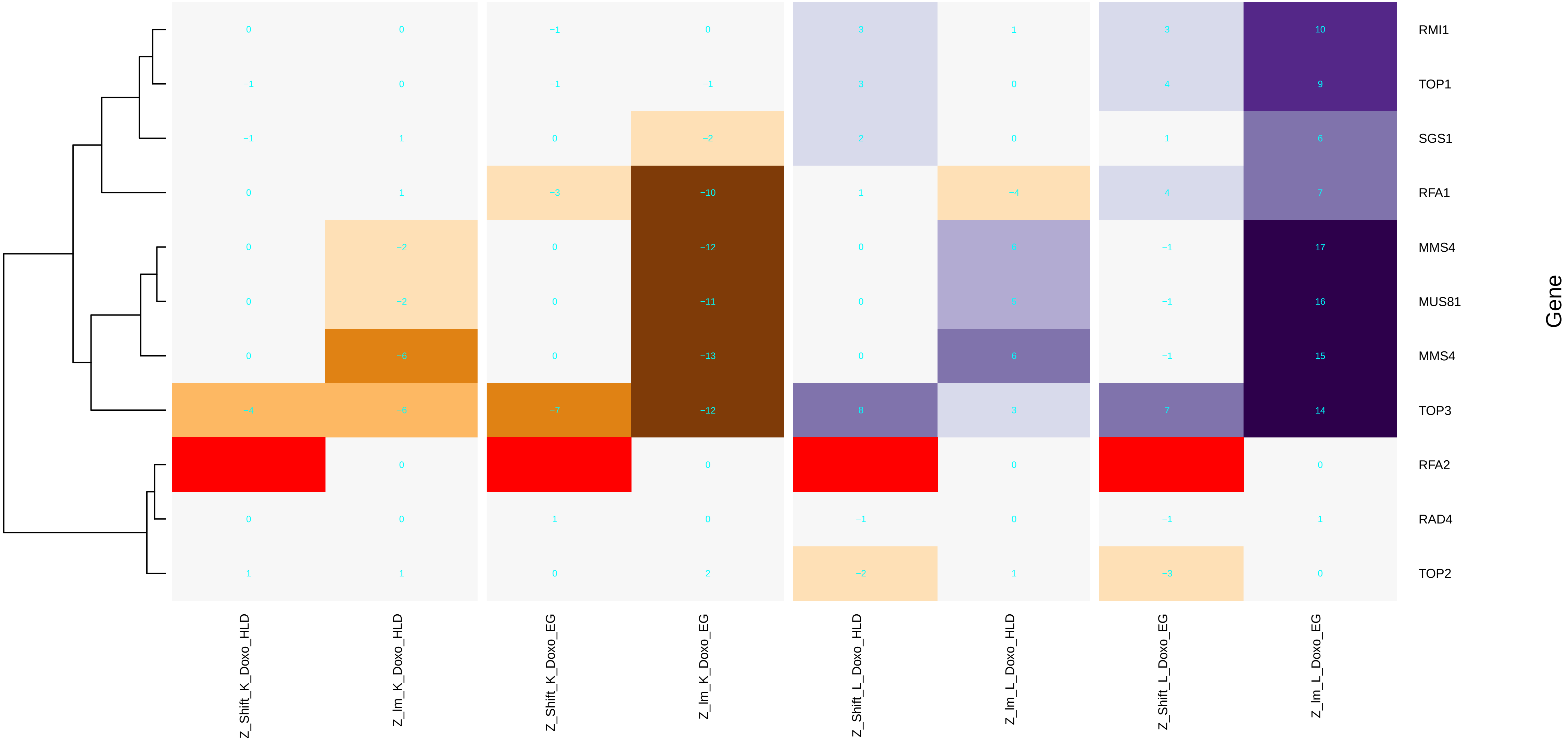

### double-strand_break_repair.pdf

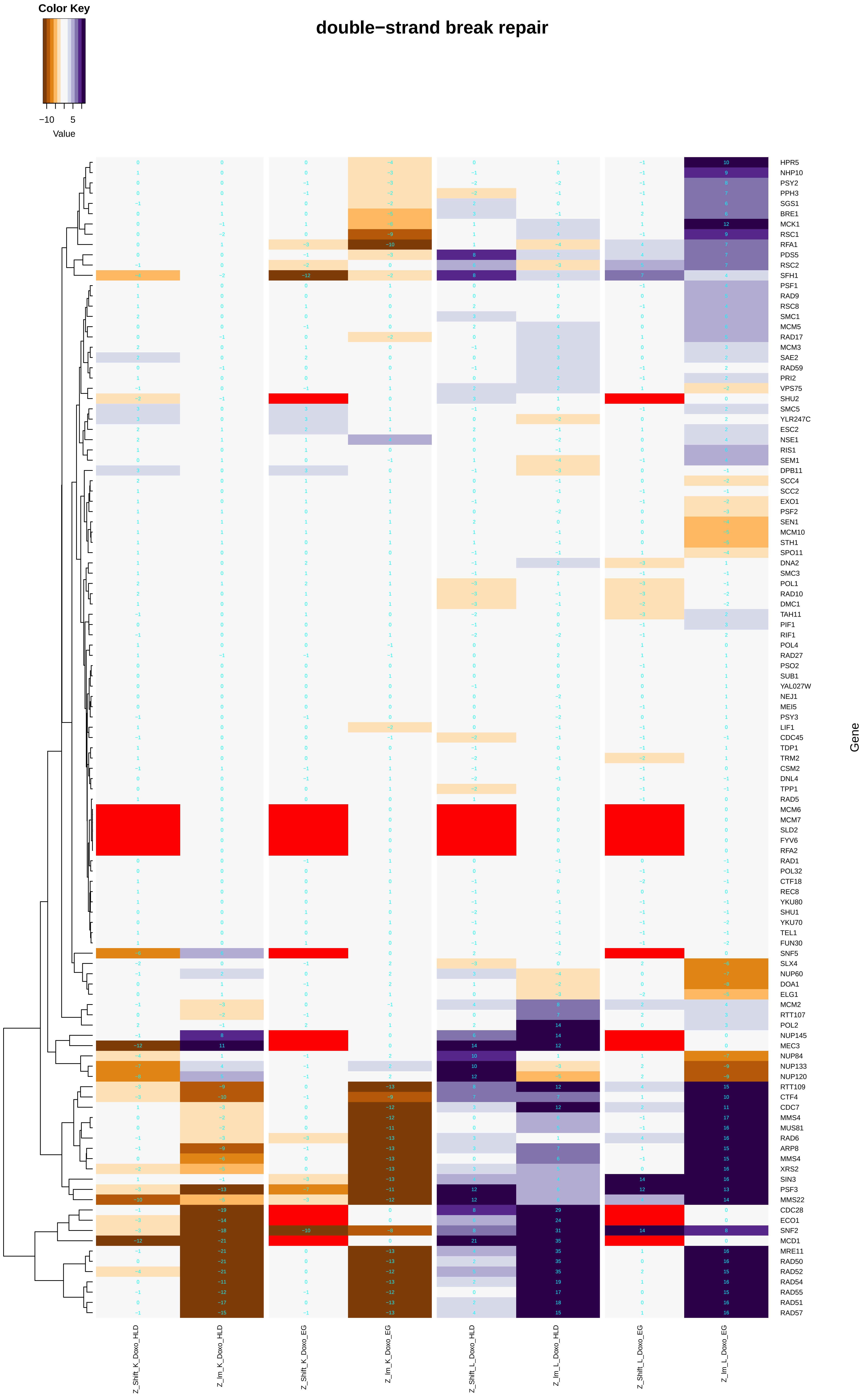

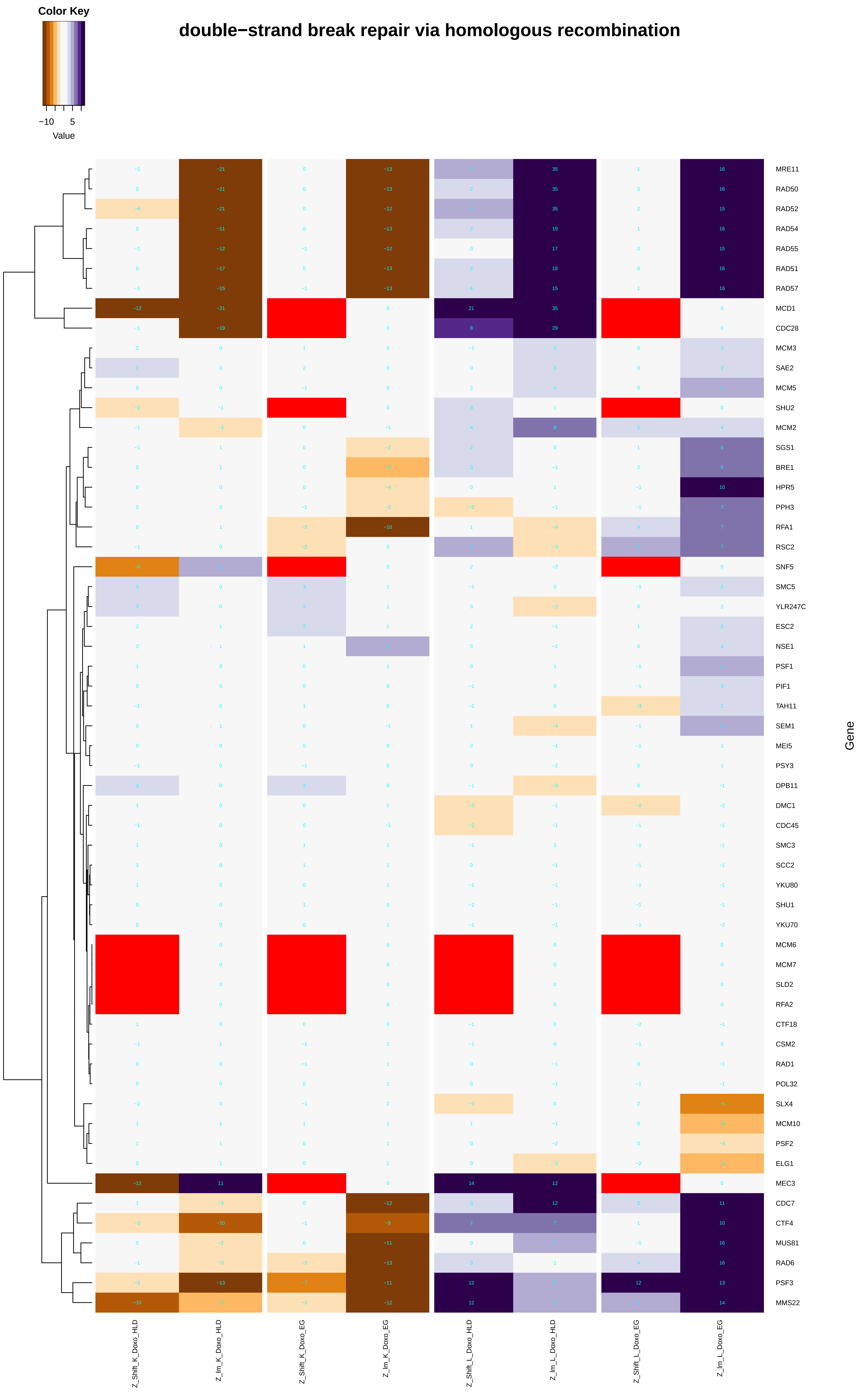

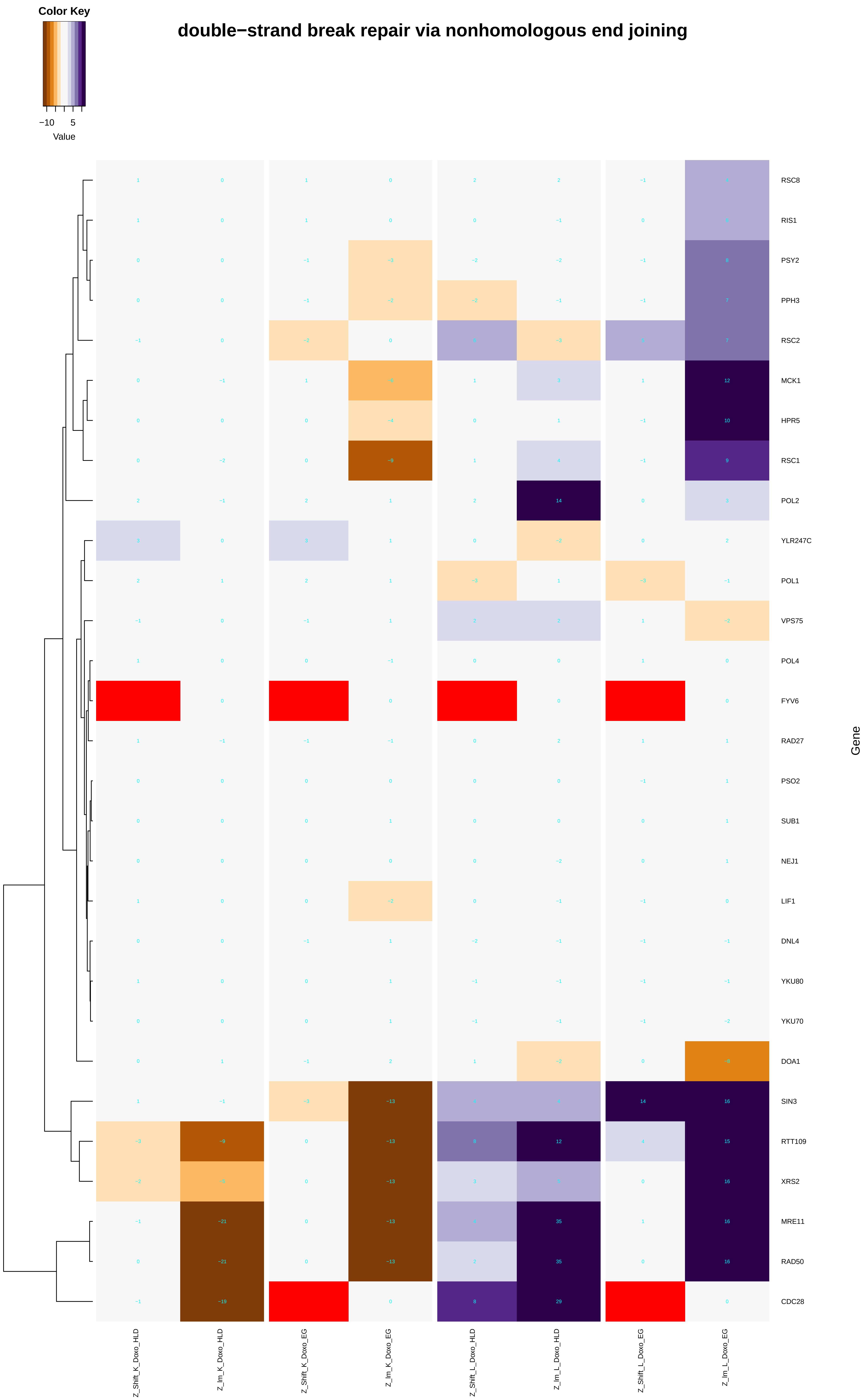

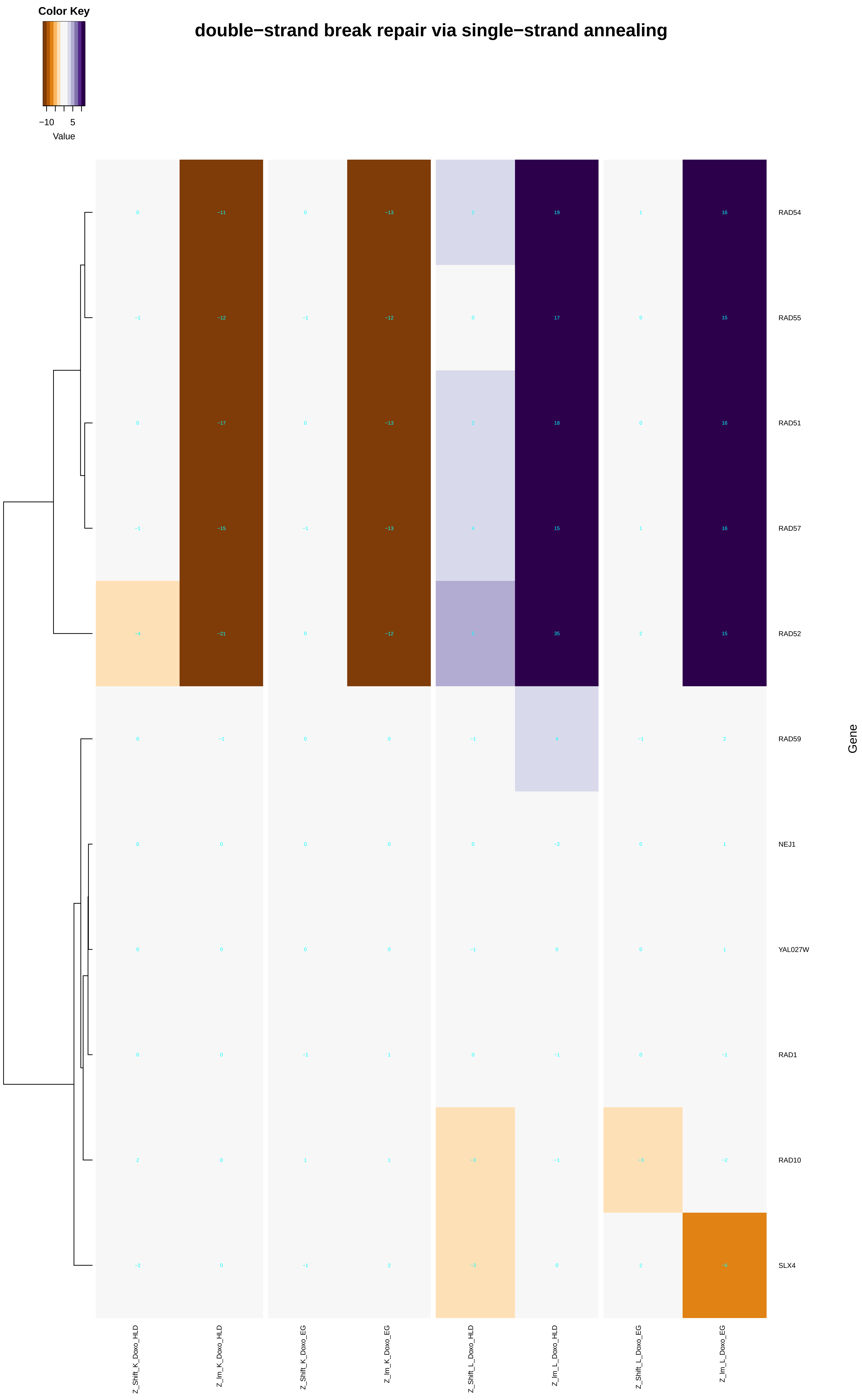

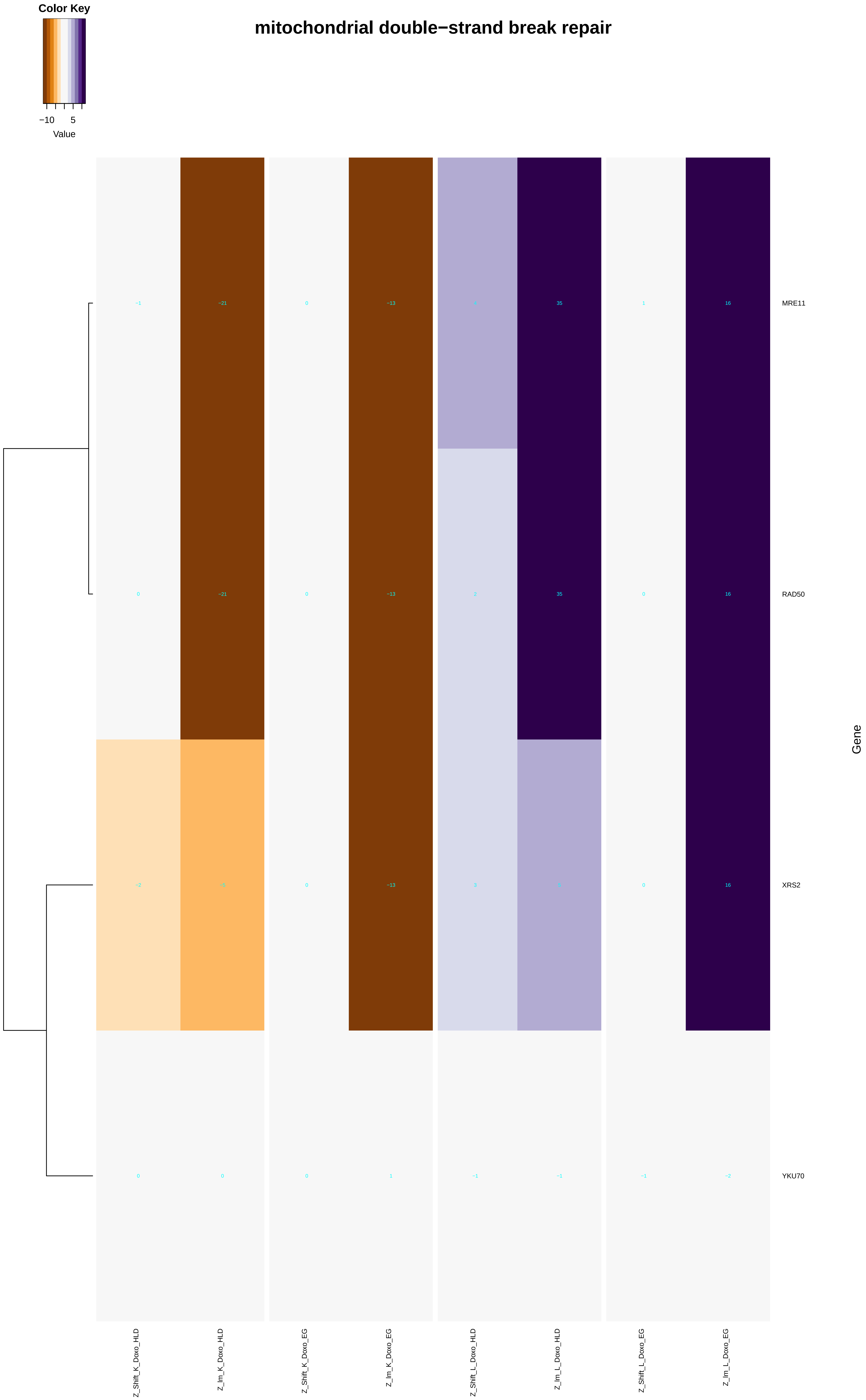

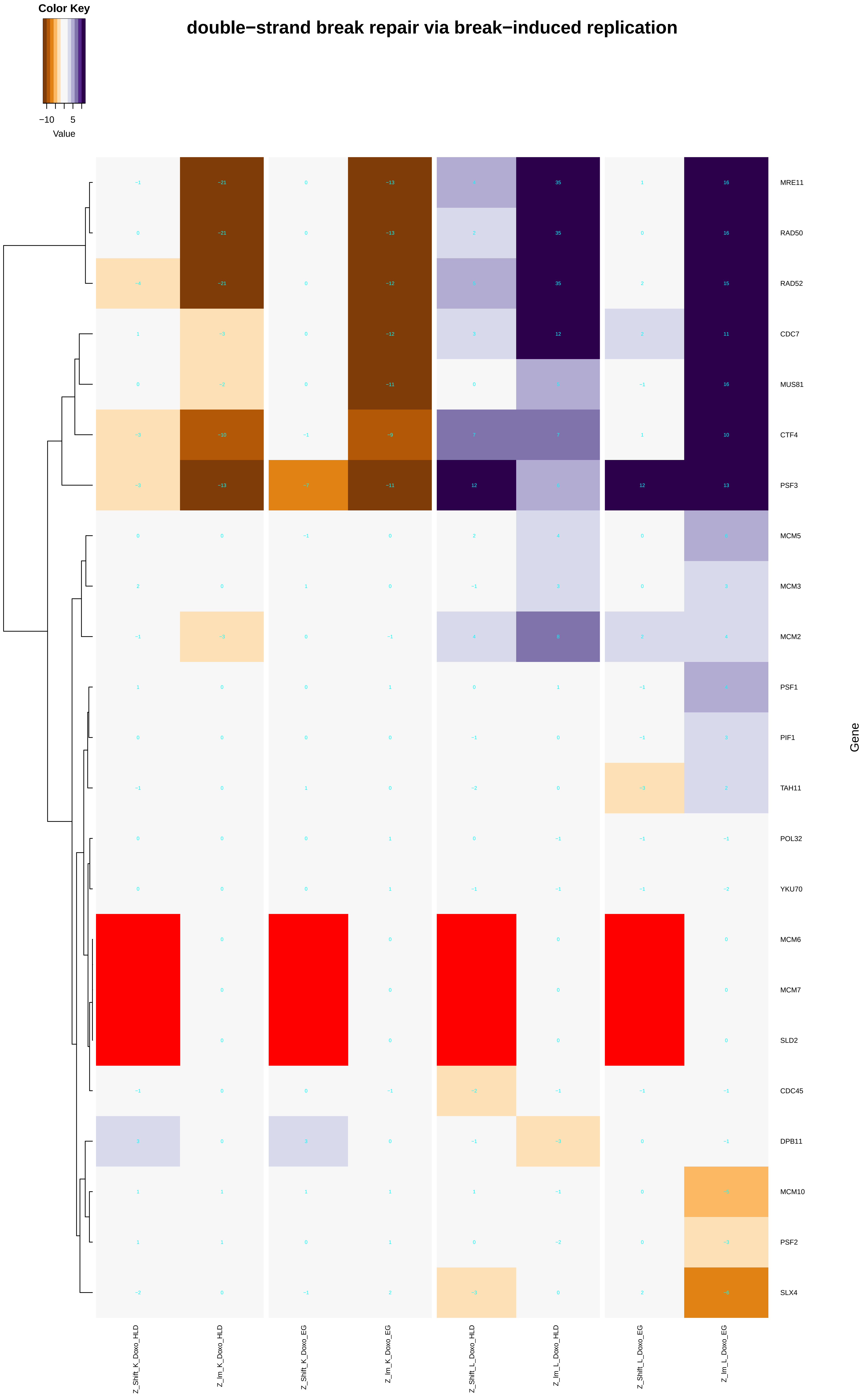

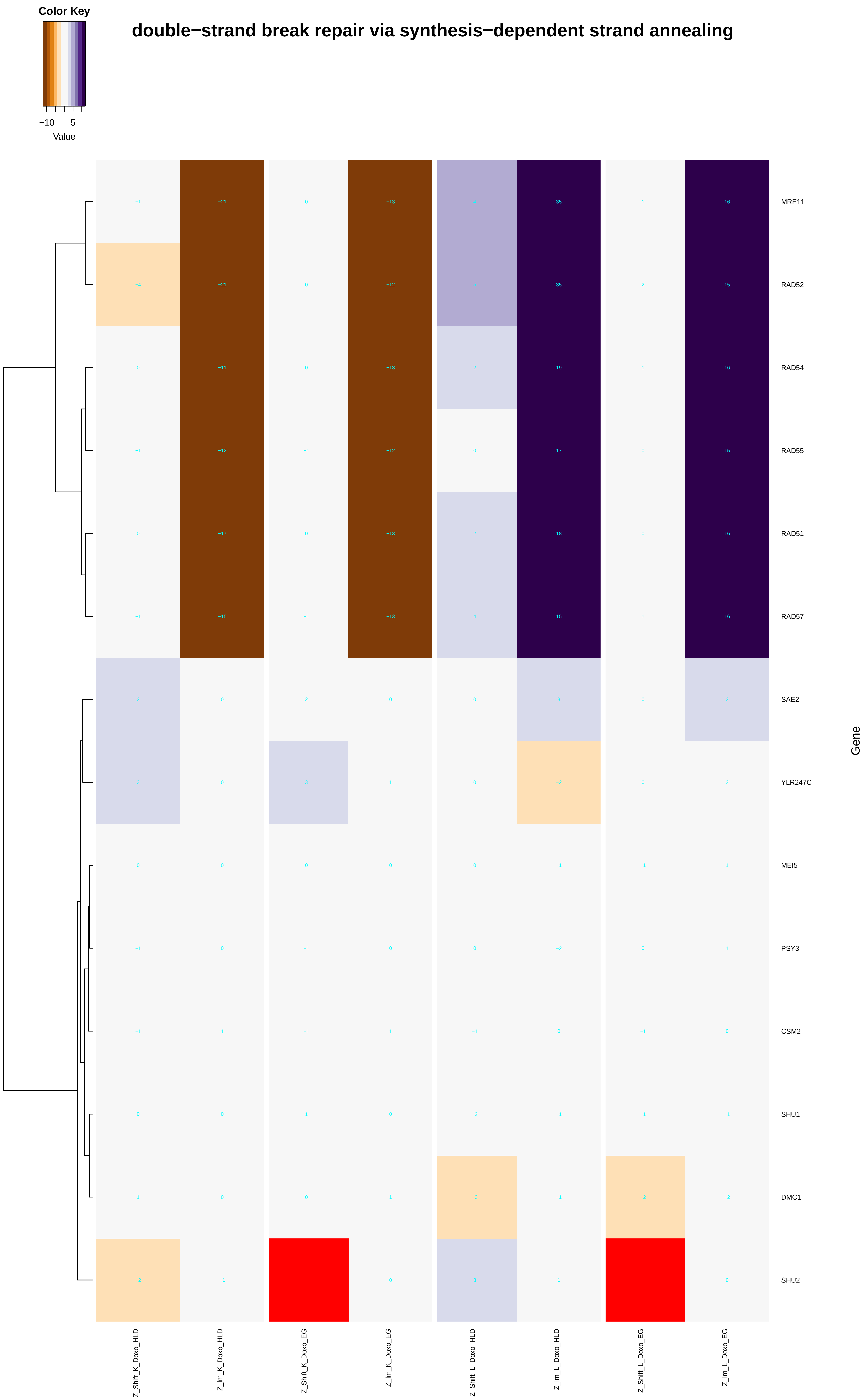

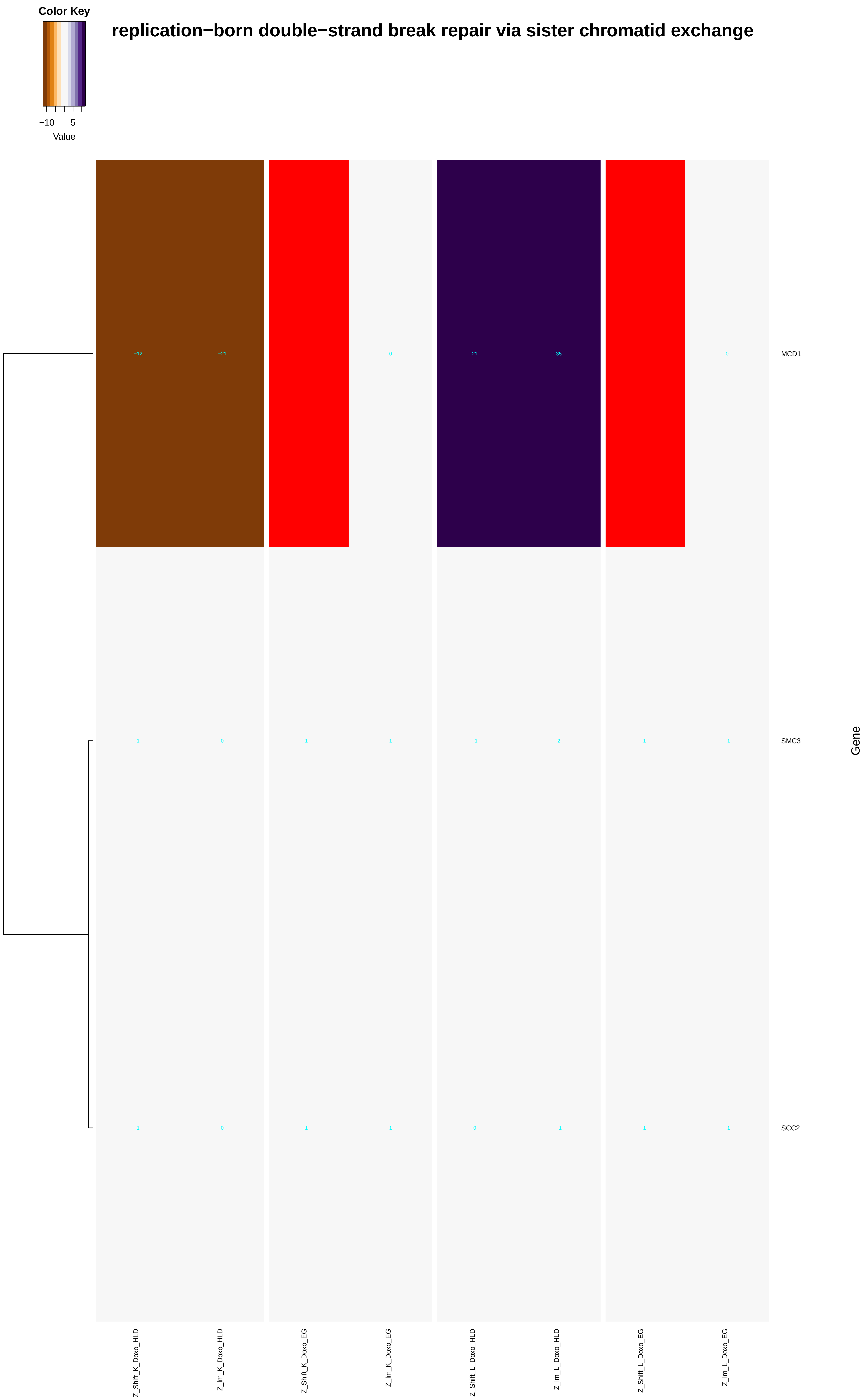

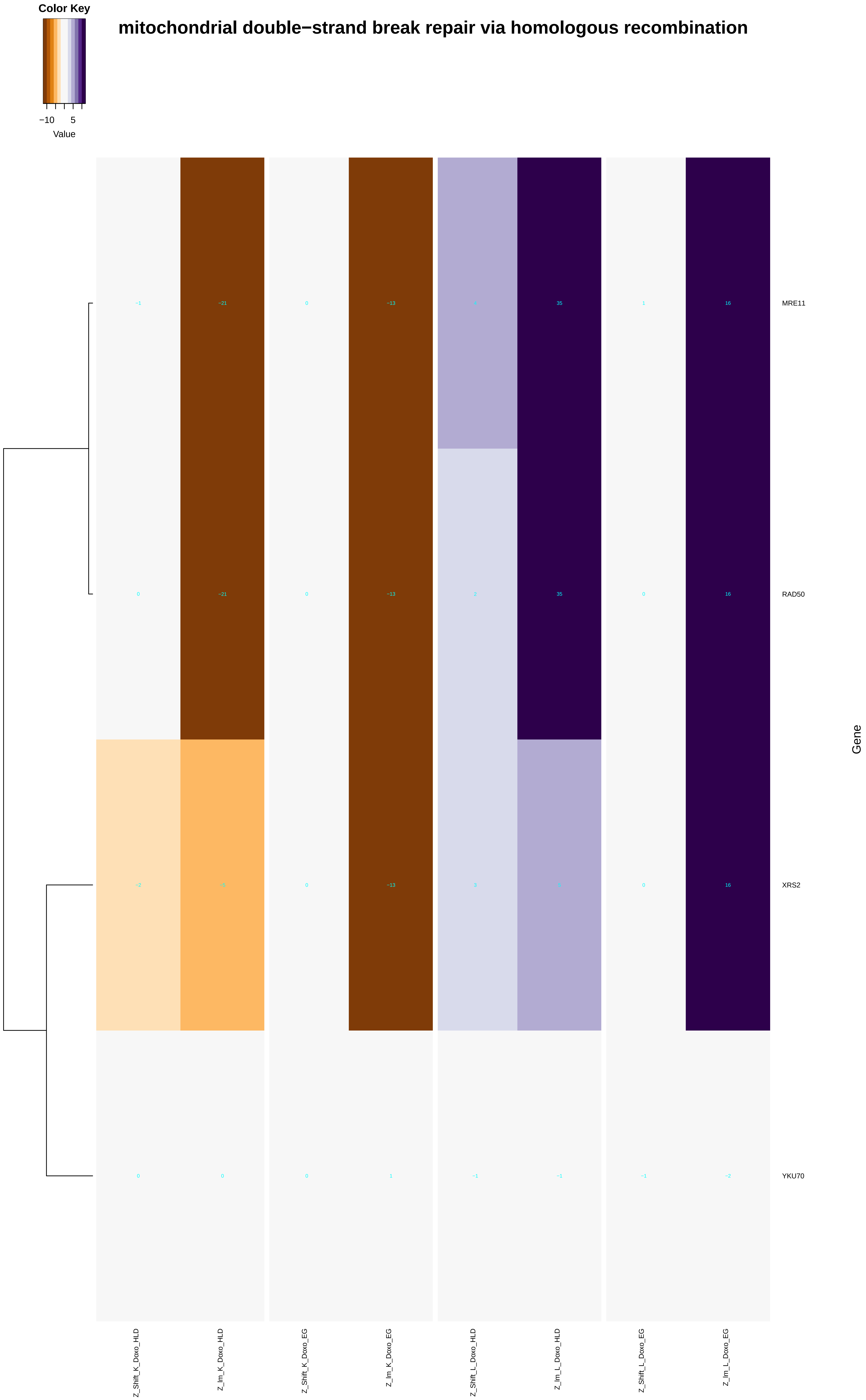

### double-strand_break_repair_via_break-induced_replication.pdf

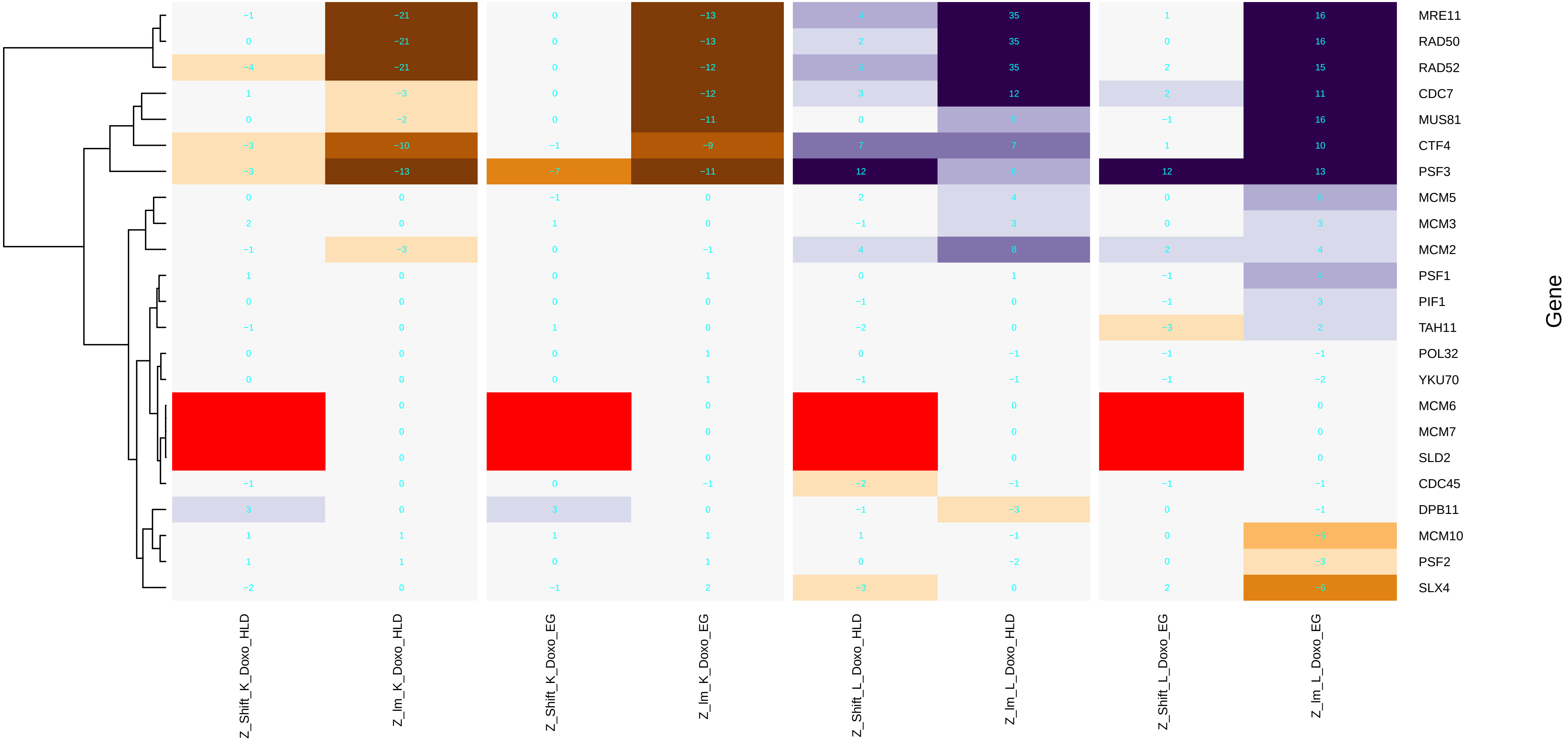

### double-strand_break_repair_via_homologous_recombination.pdf

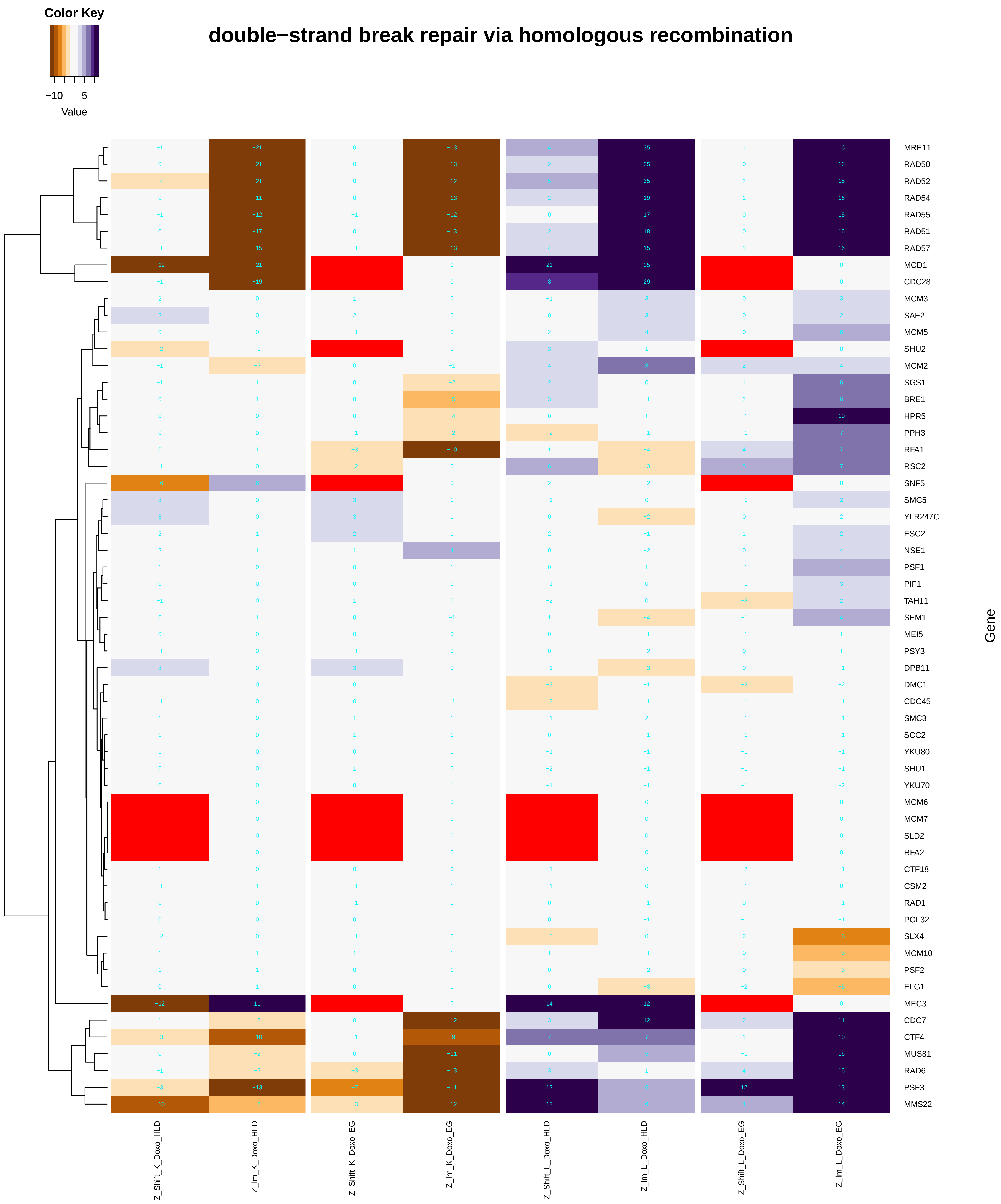

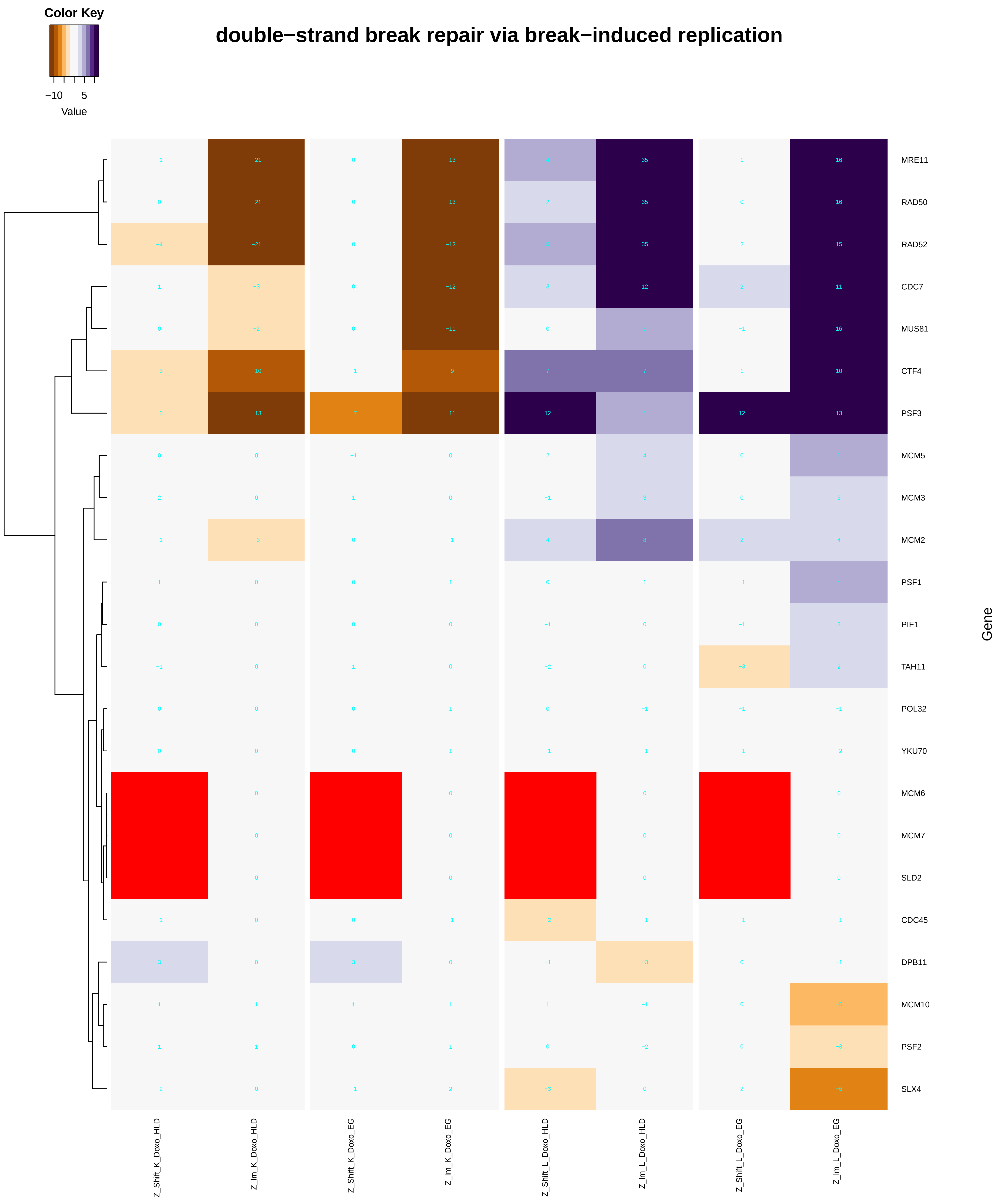

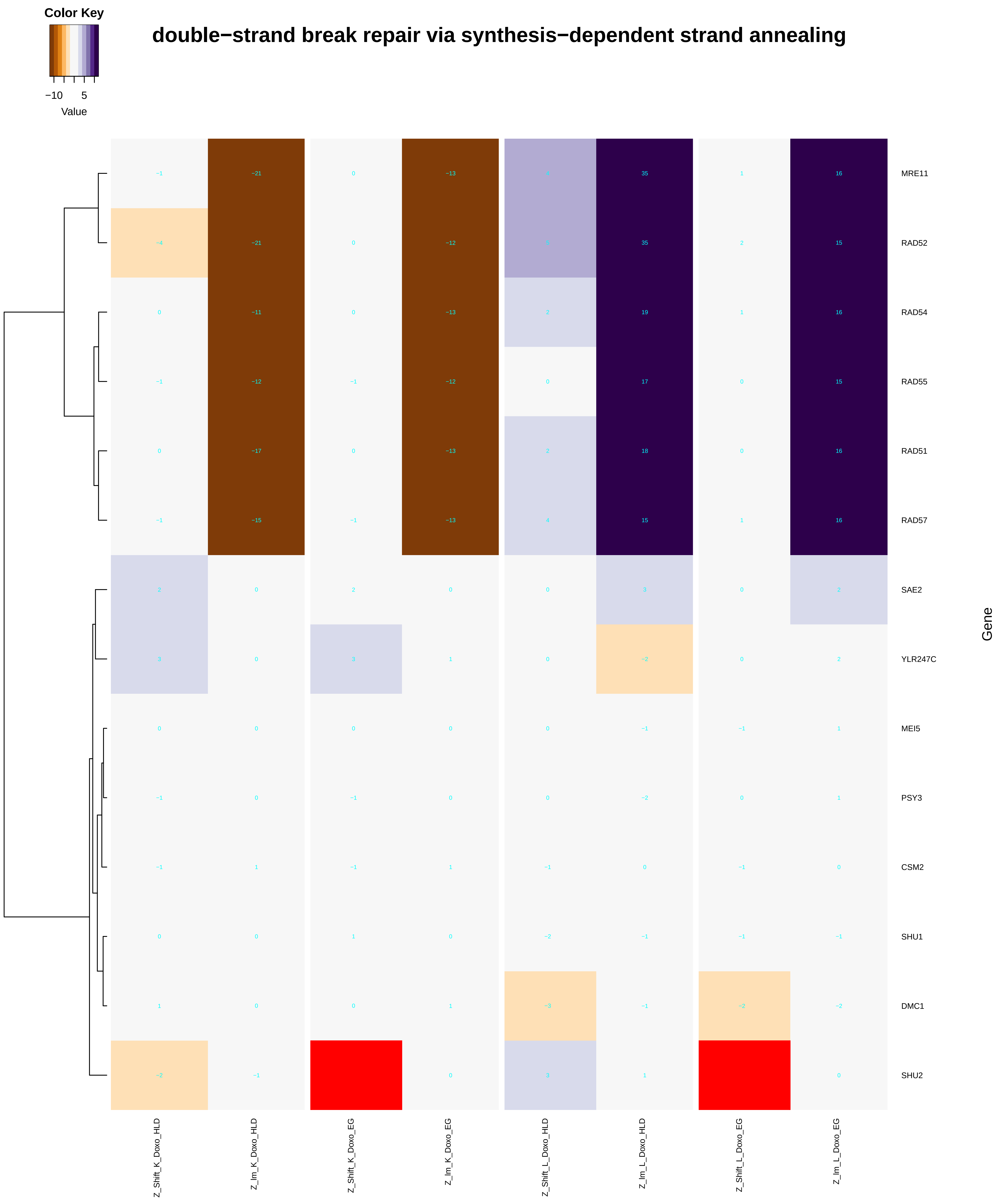

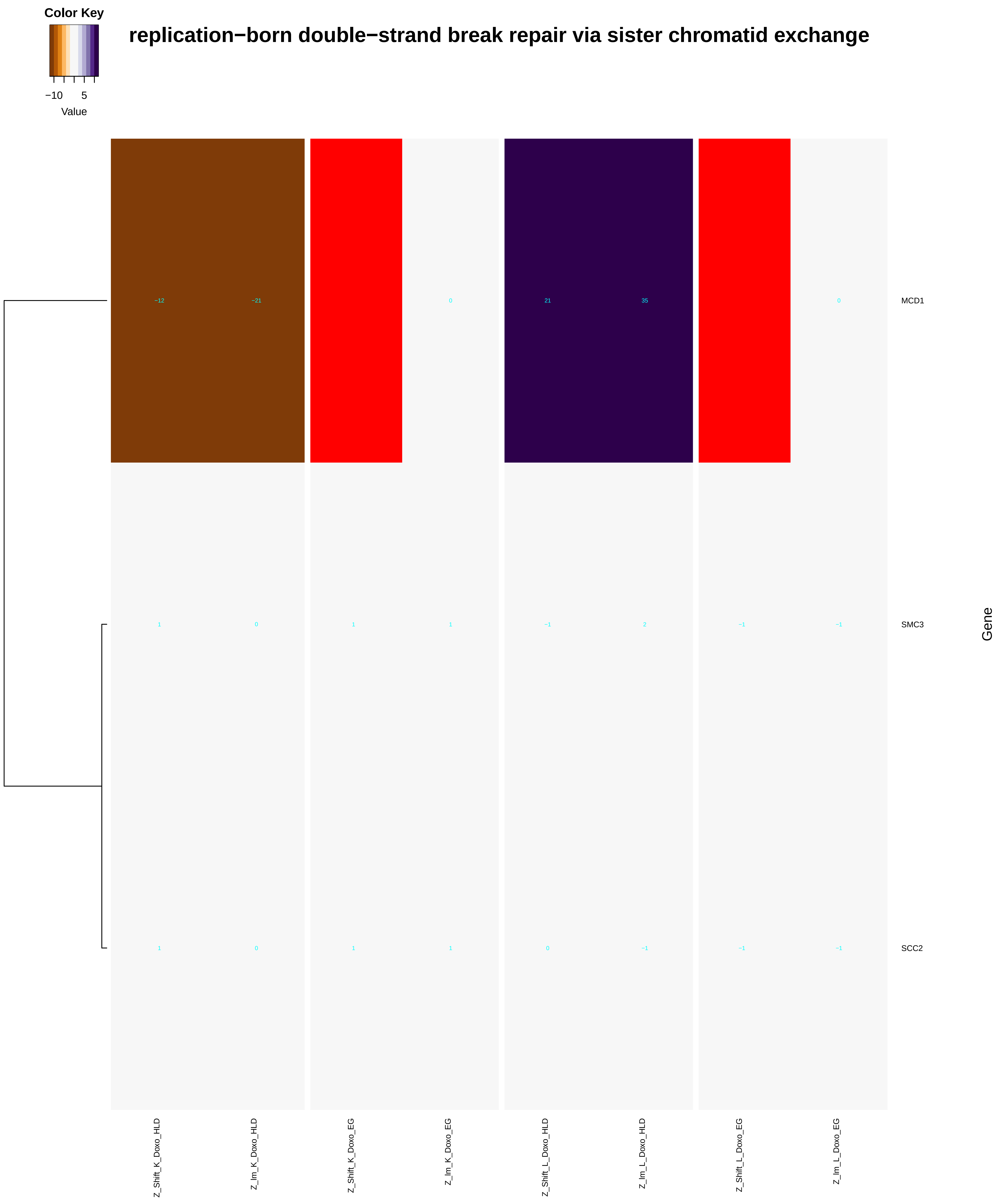

### double-strand_break_repair_via_nonhomologous_end_joining.pdf

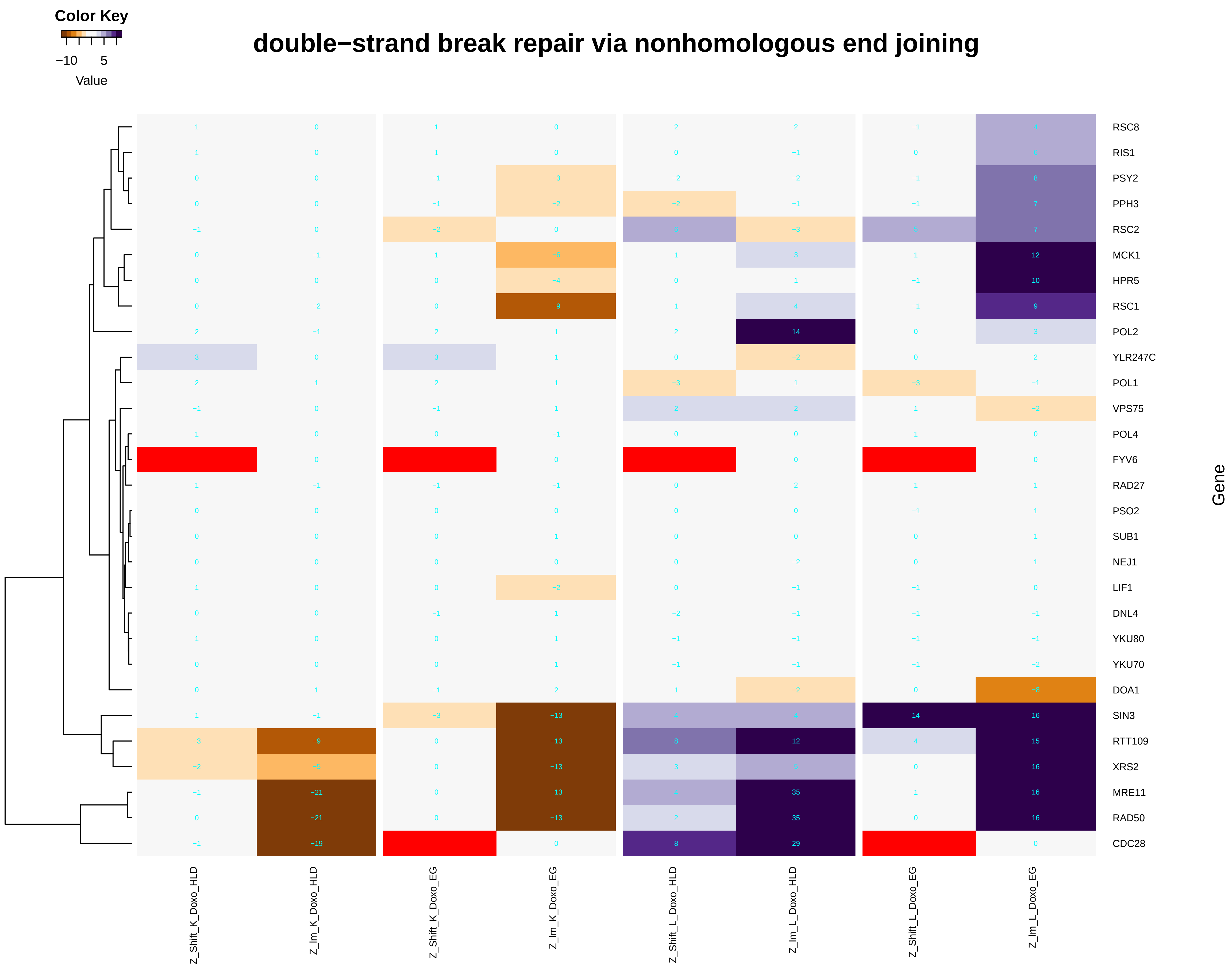

### double-strand_break_repair_via_synthesis-dependent_strand_annealing.pdf

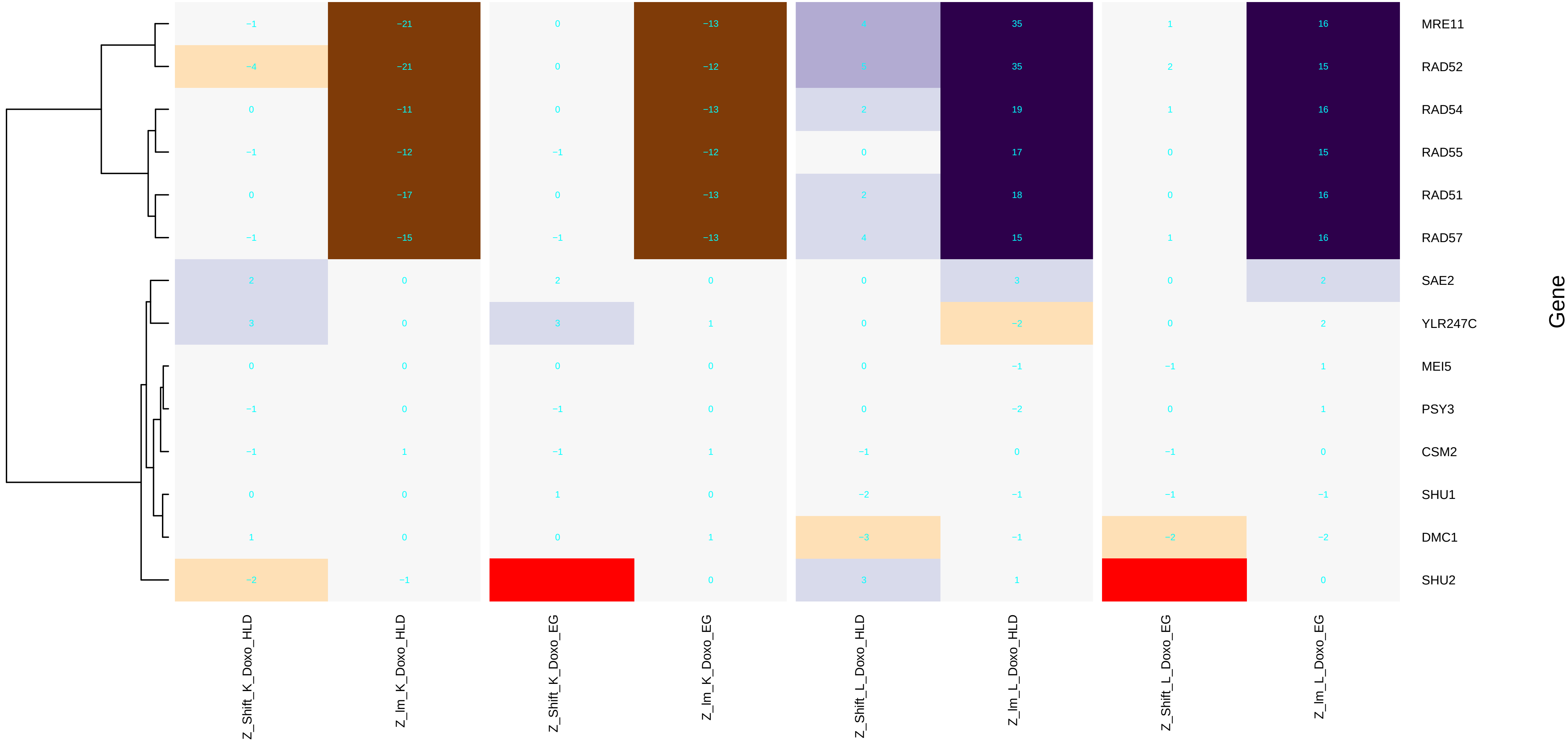

### endonucleolytic_cleavage_in_5'-ETS_of_tricistronic_rRNA_transcript_(SSU-rRNA,_5.8S_rRNA,_LSU-rRNA).pdf

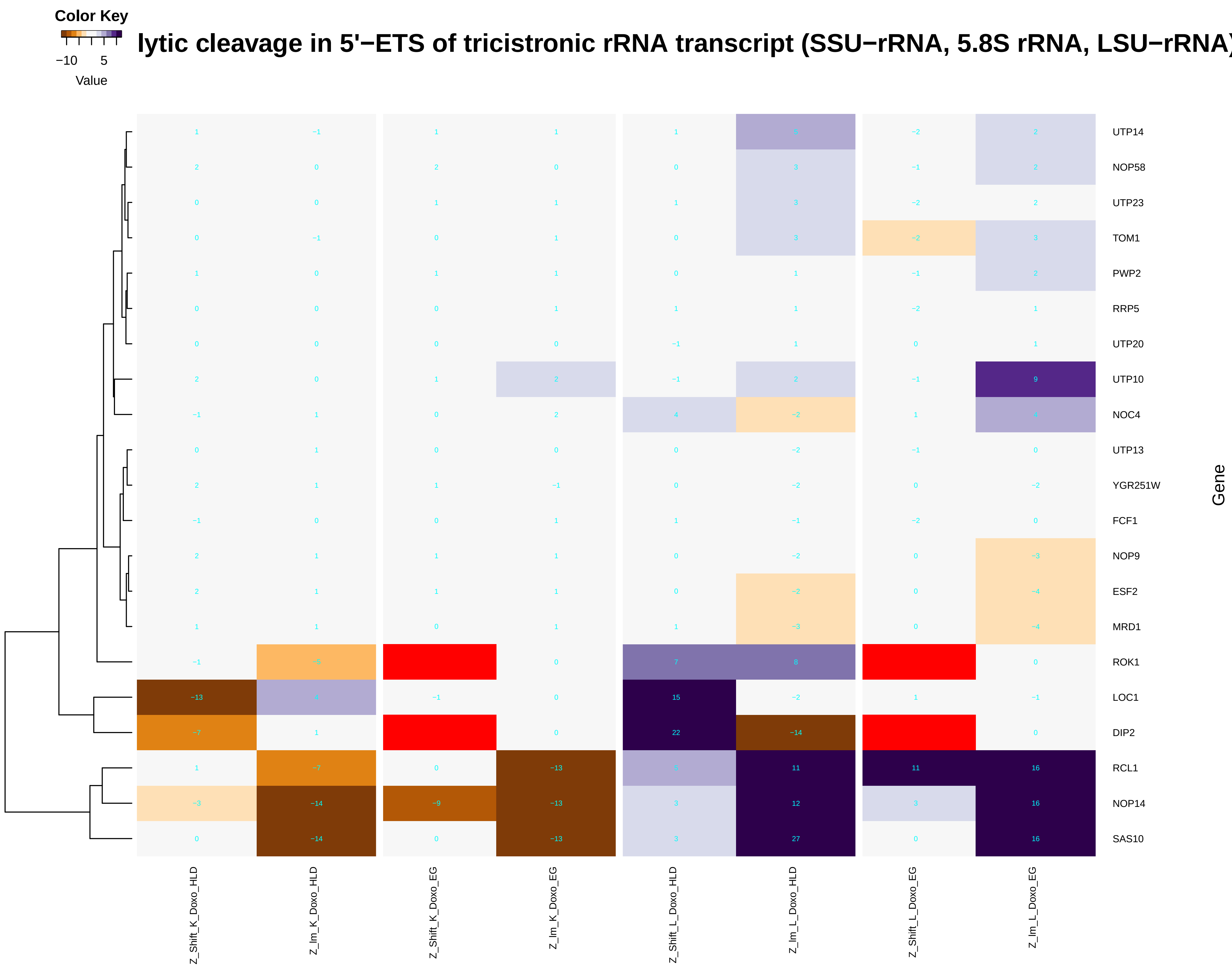

### endonucleolytic_cleavage_involved_in_rRNA_processing.pdf

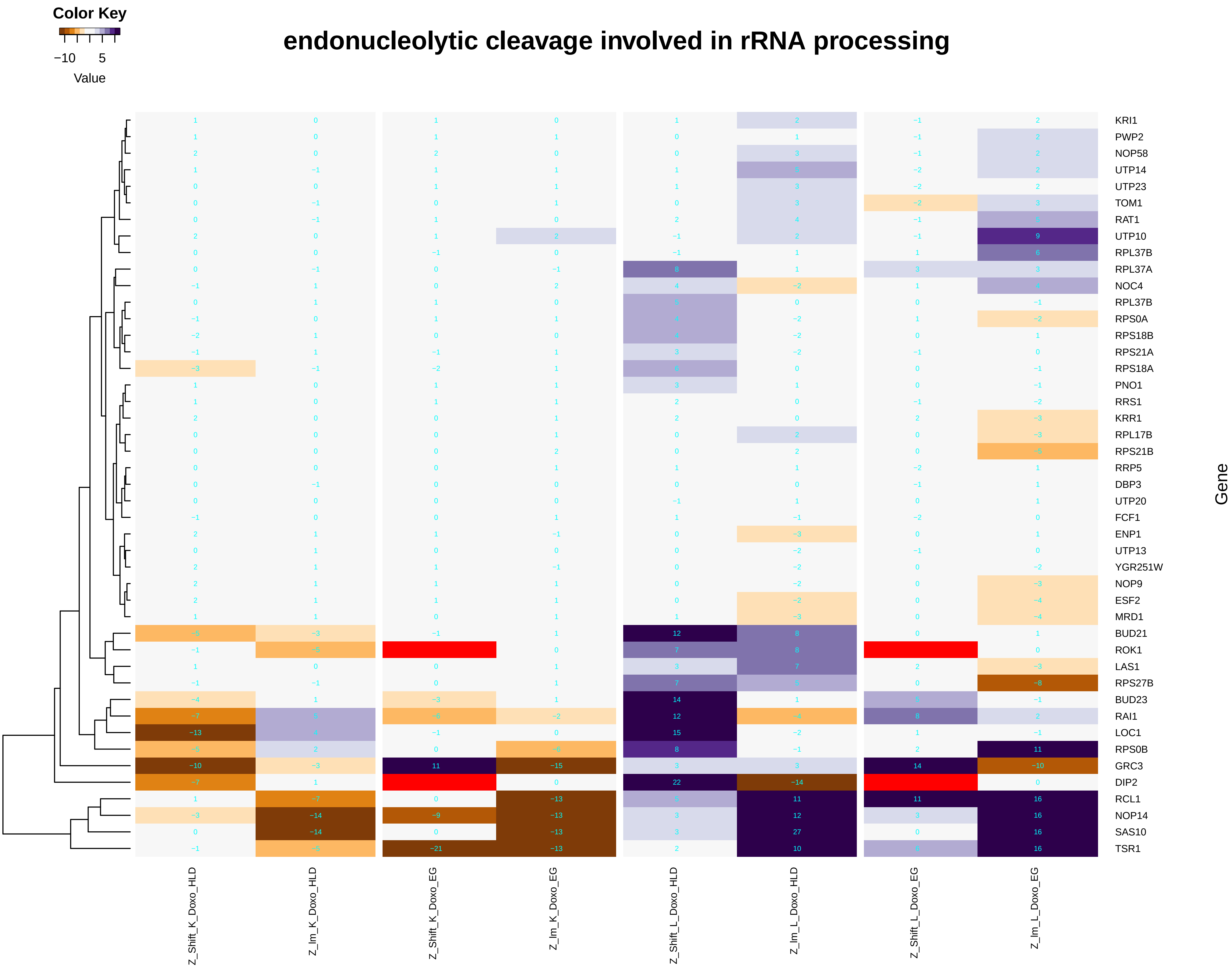

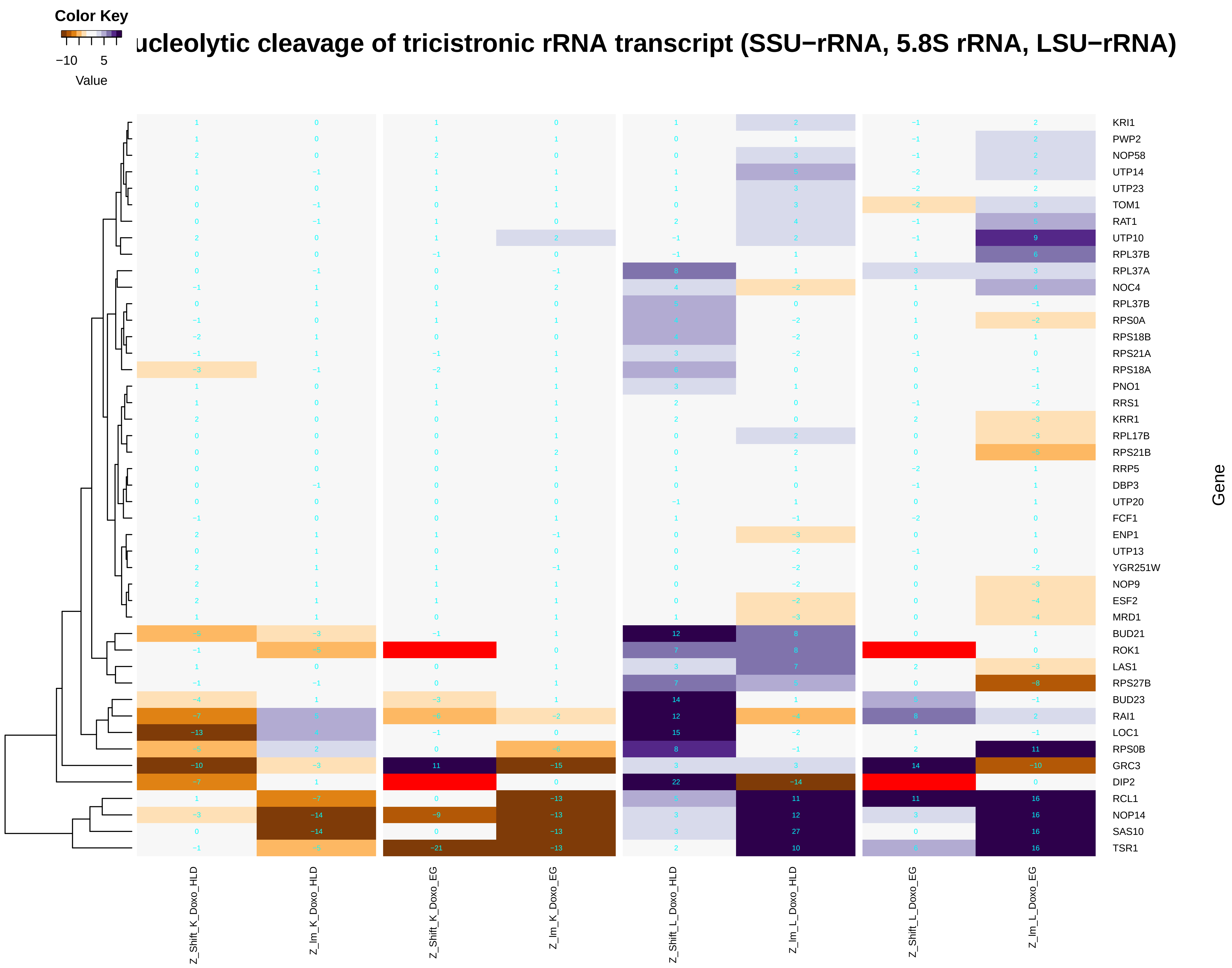

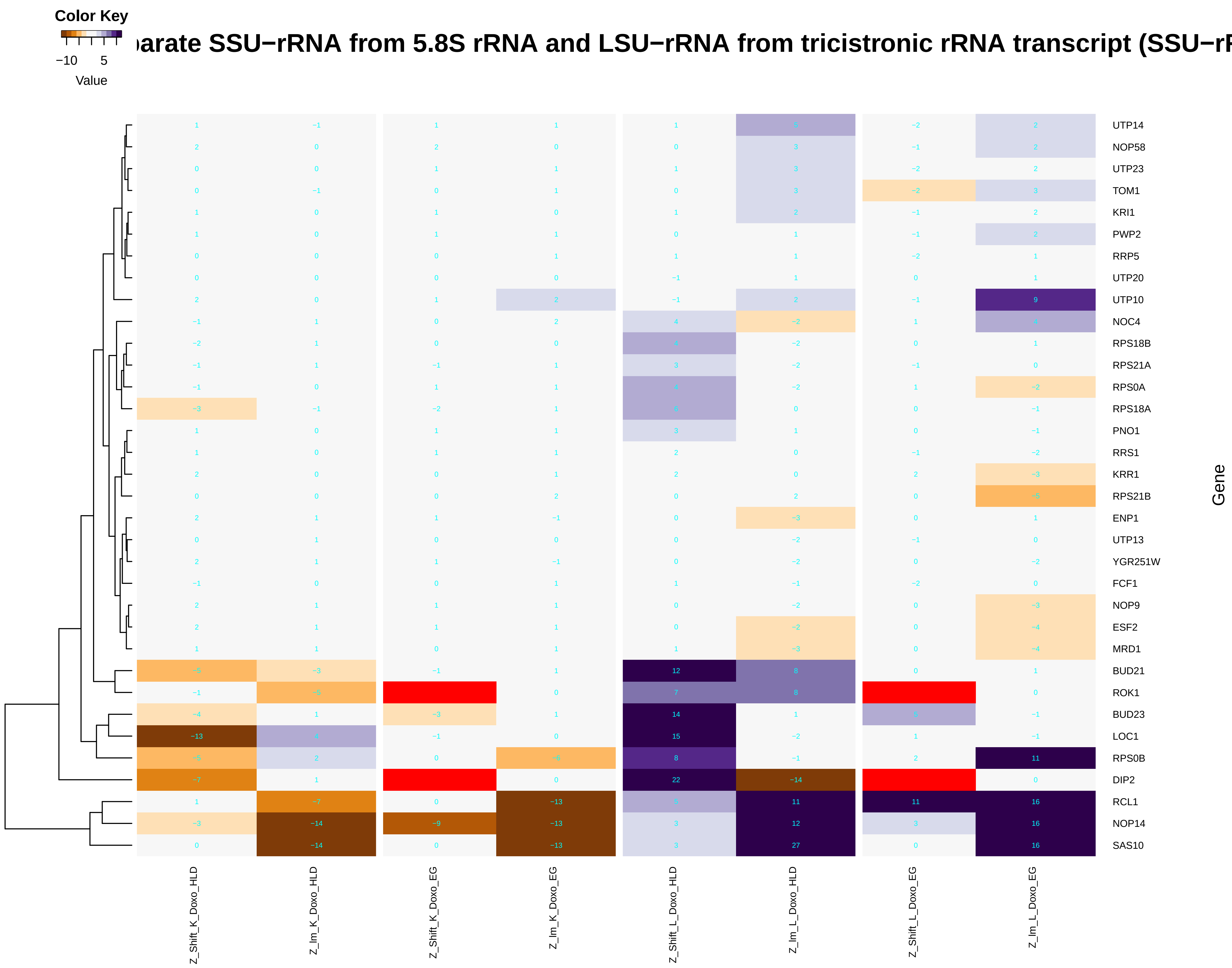

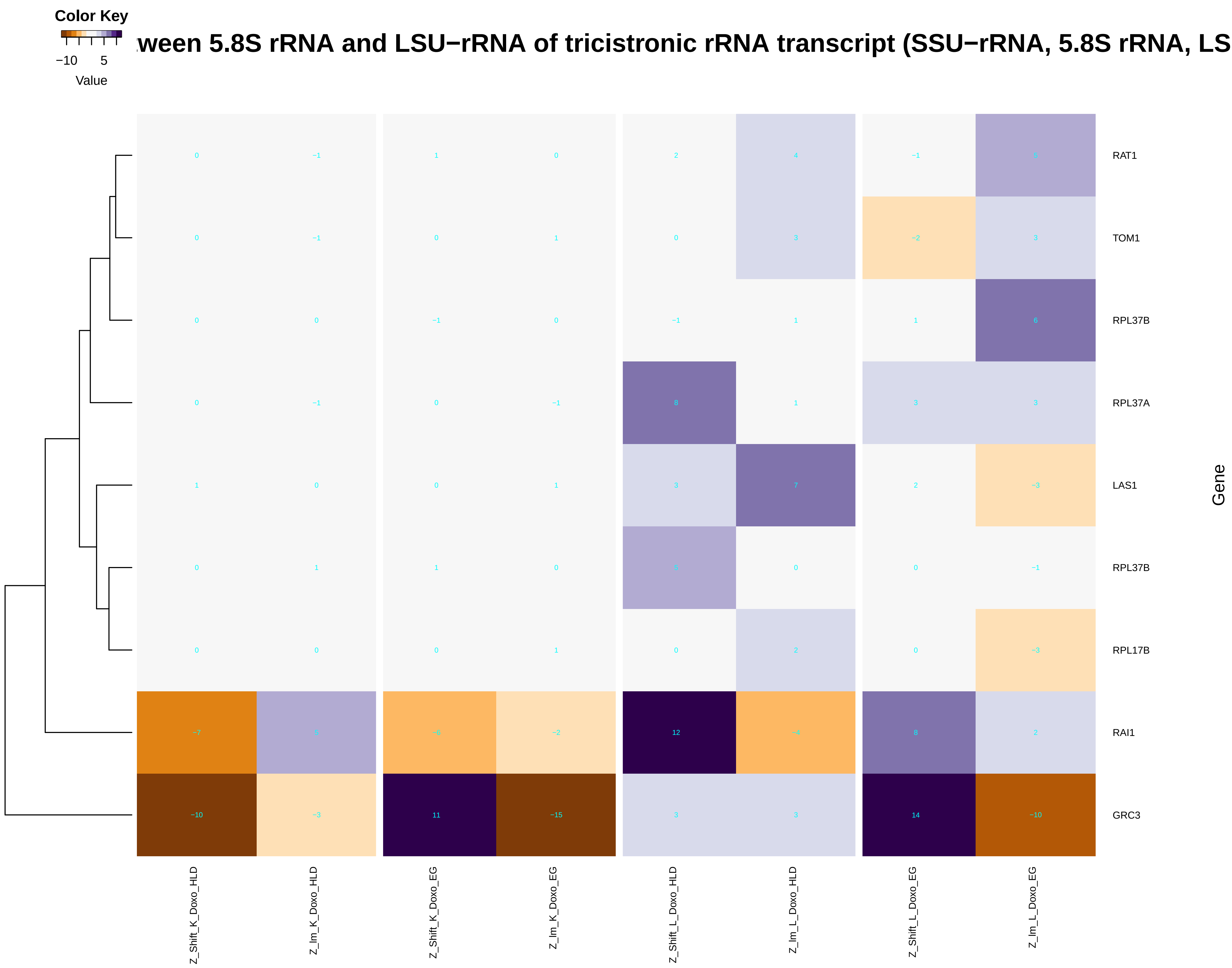

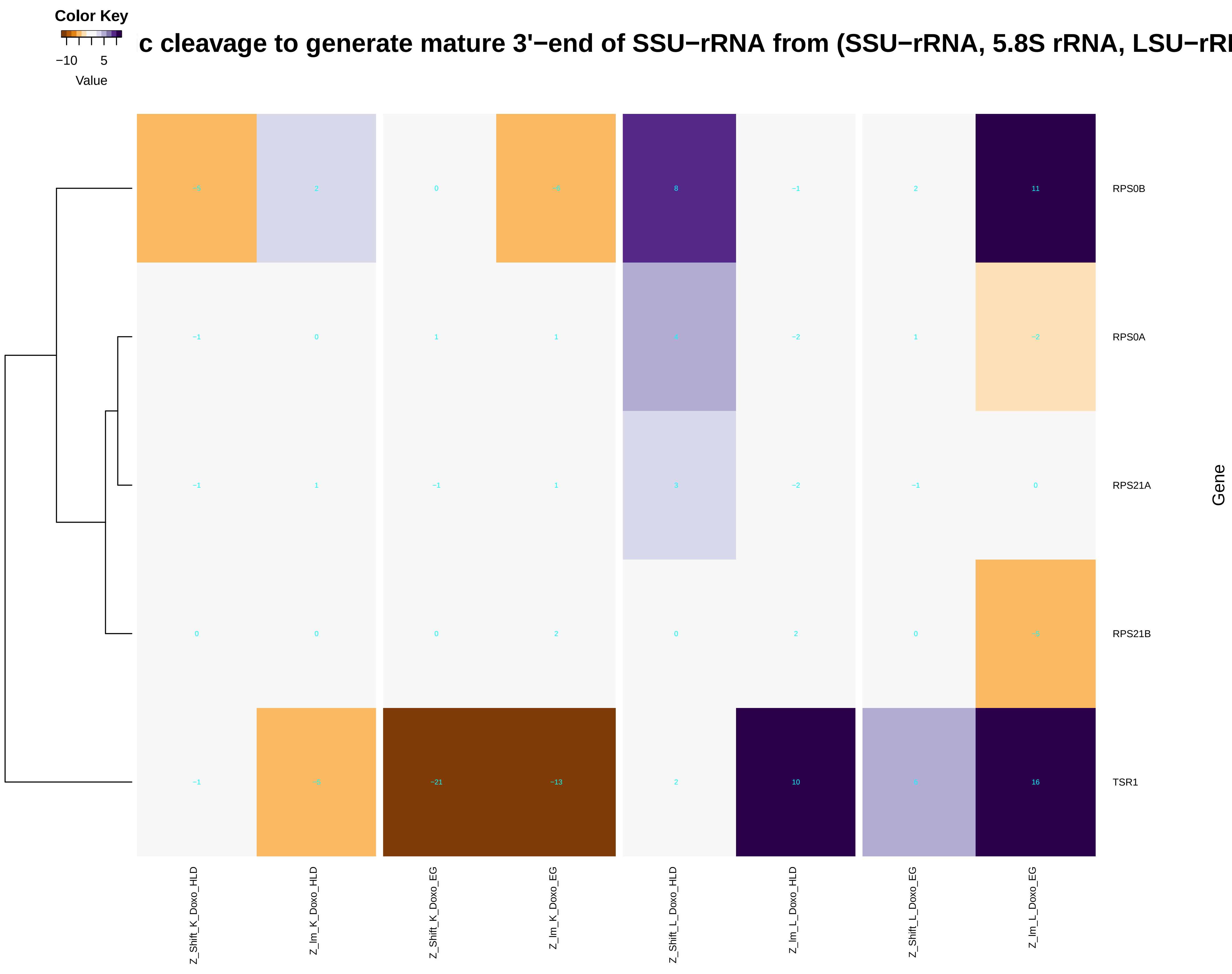

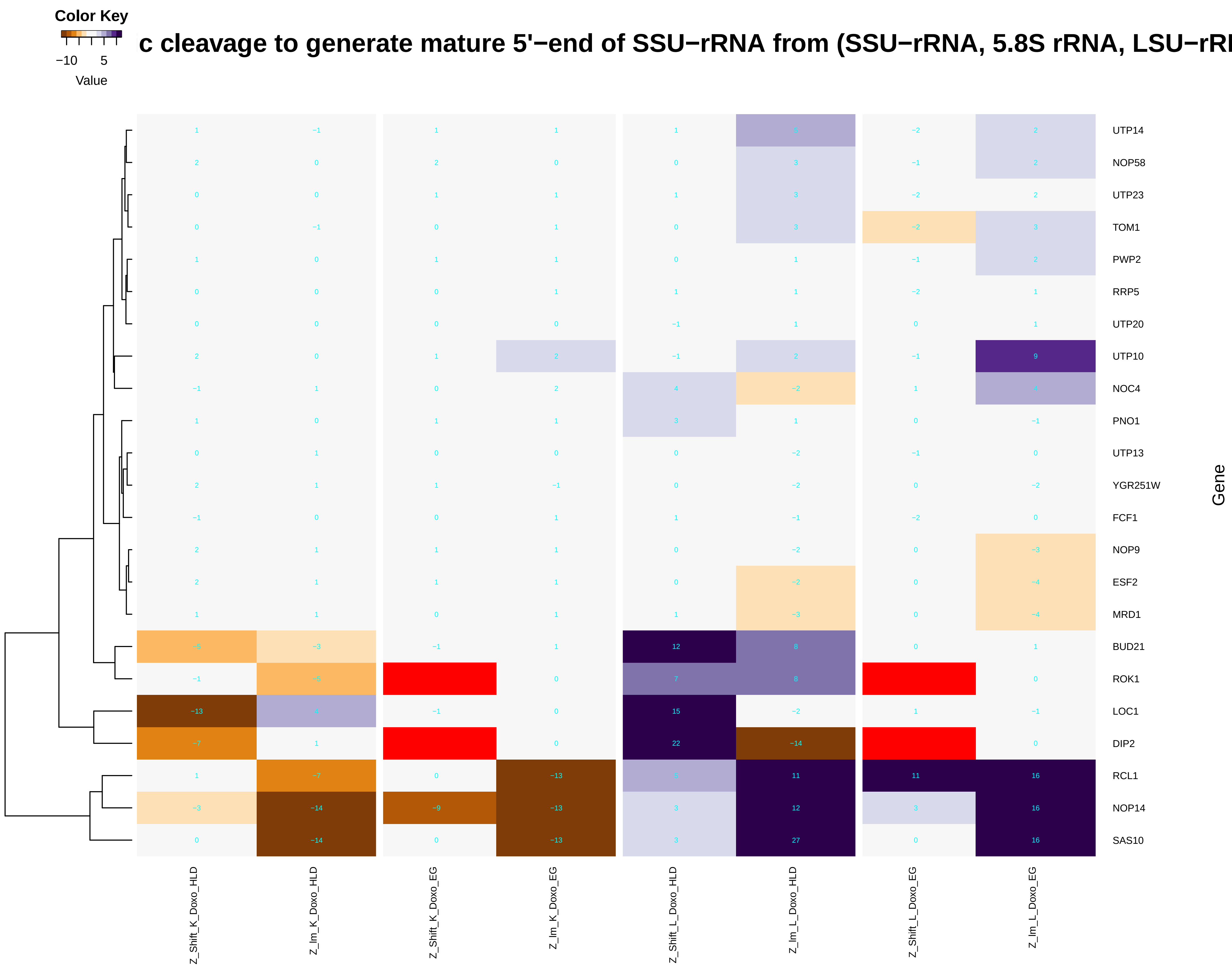

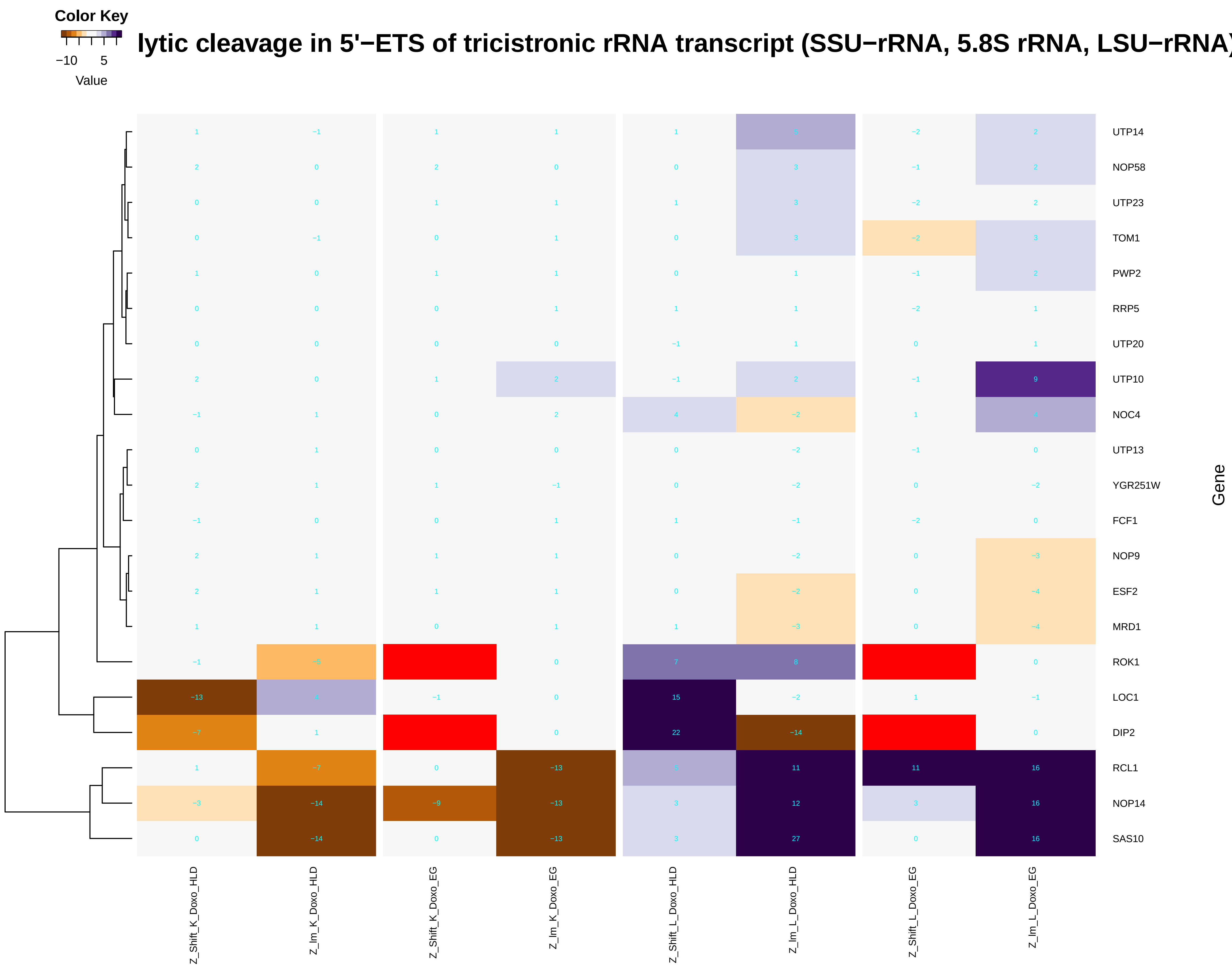

### endonucleolytic_cleavage_of_tricistronic_rRNA_transcript_(SSU-rRNA,_5.8S_rRNA,_LSU-rRNA).pdf

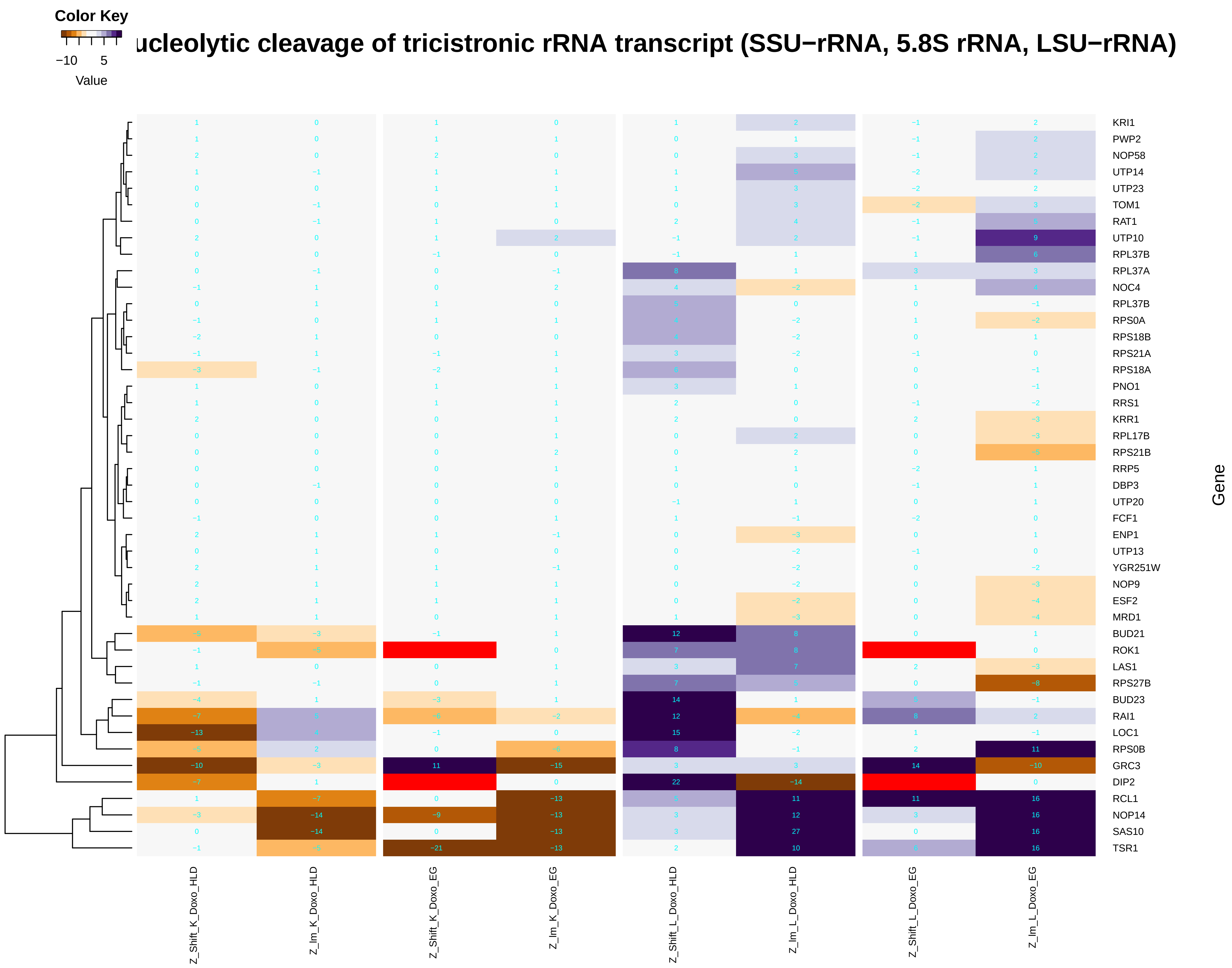

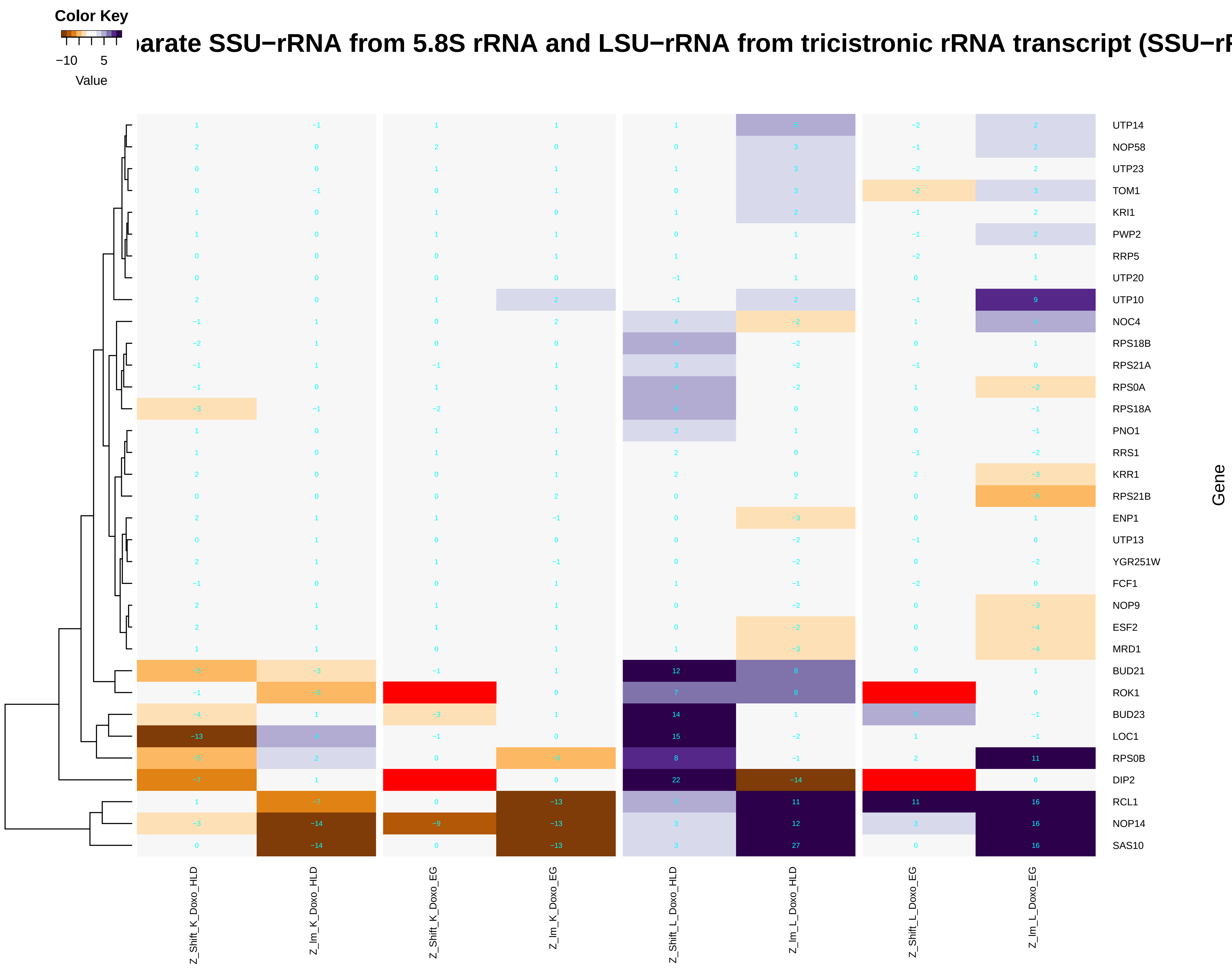

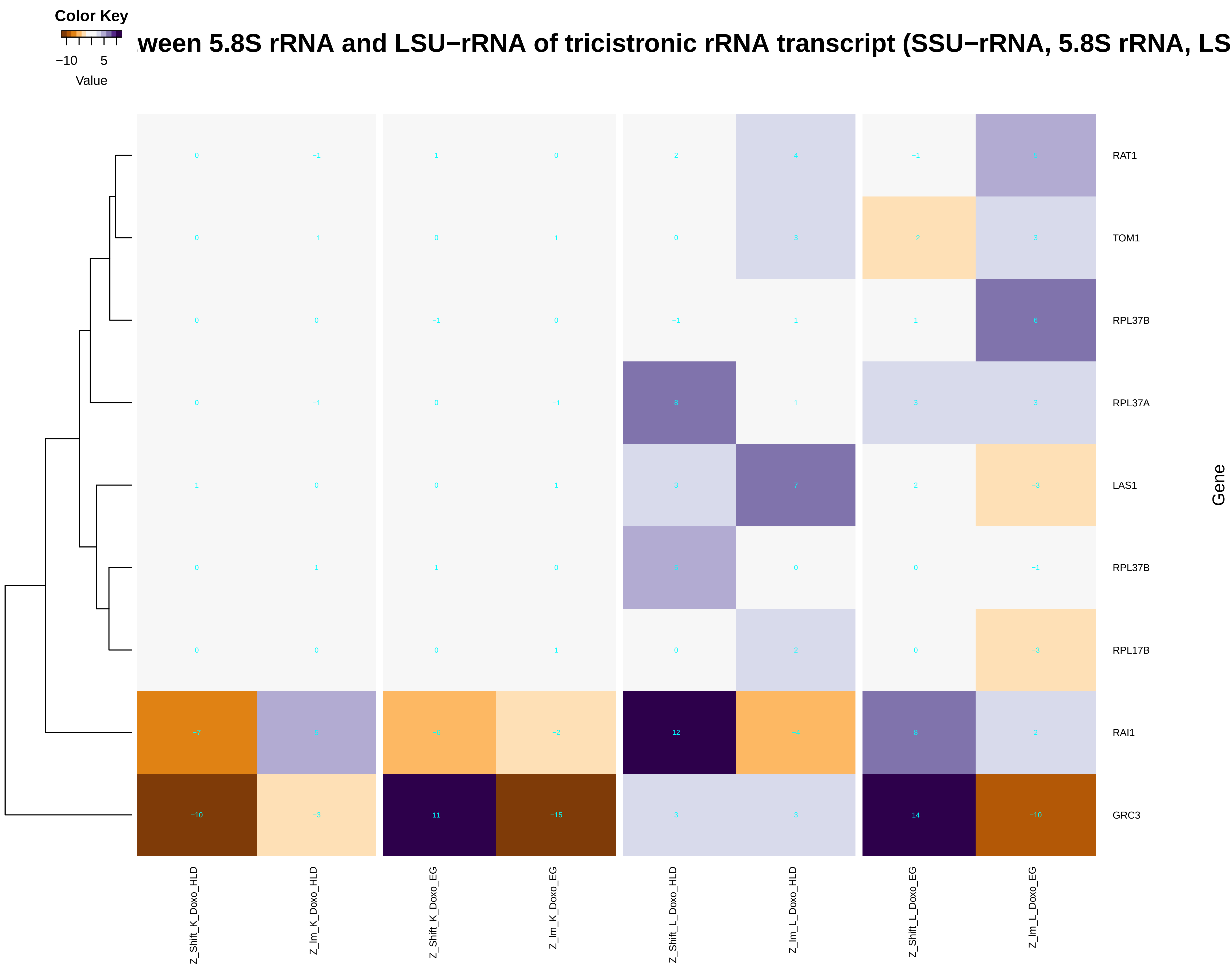

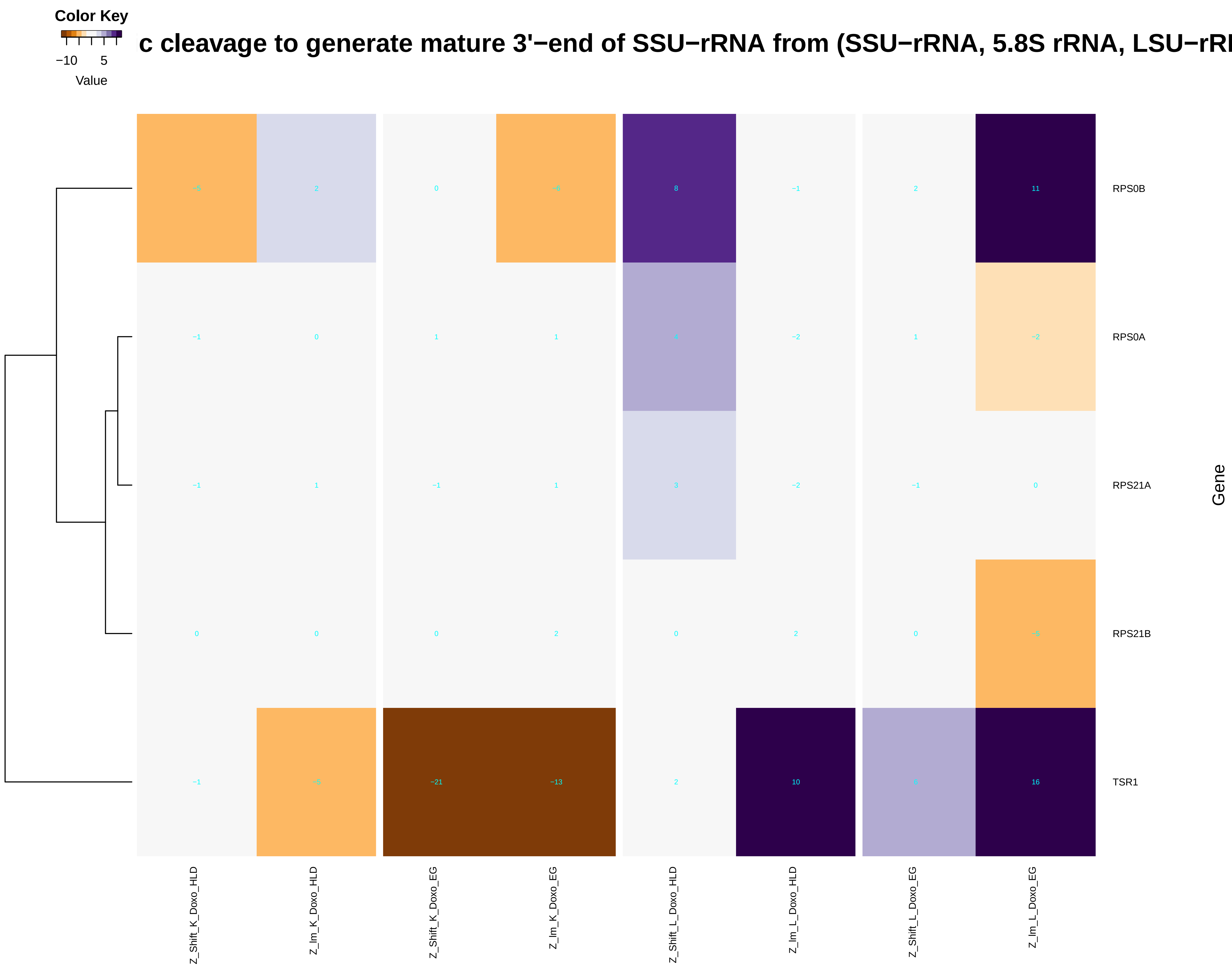

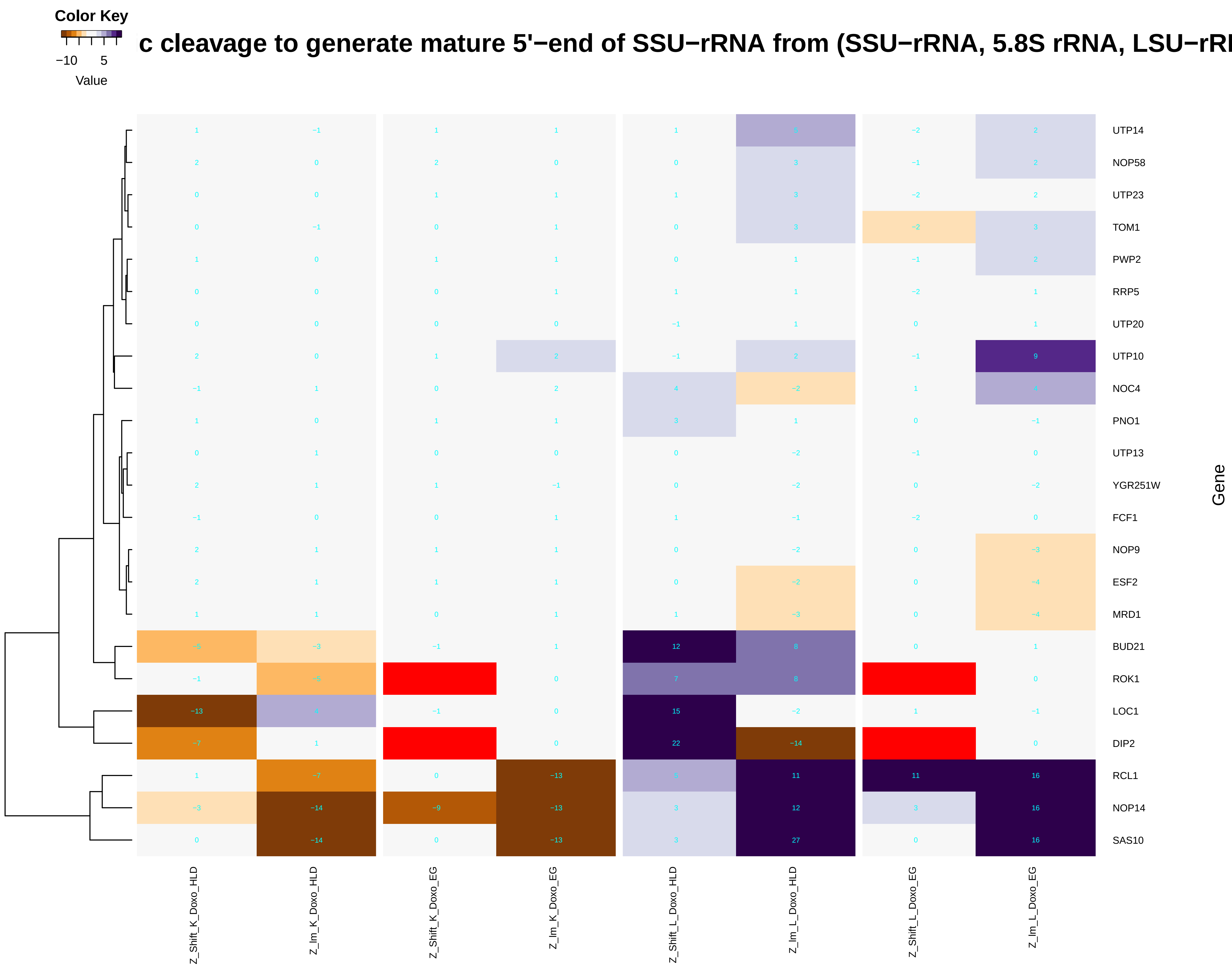

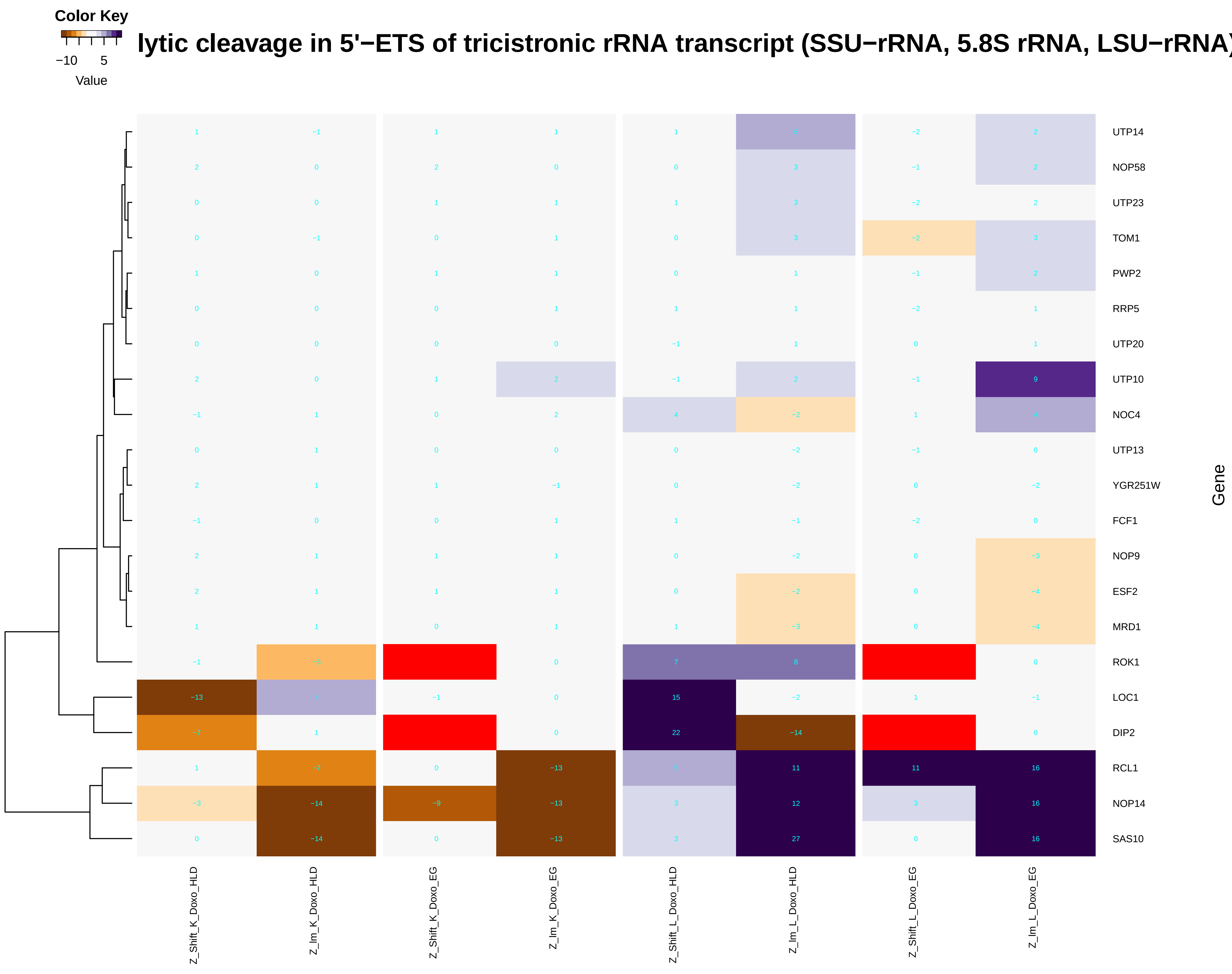

### endosomal_transport.pdf

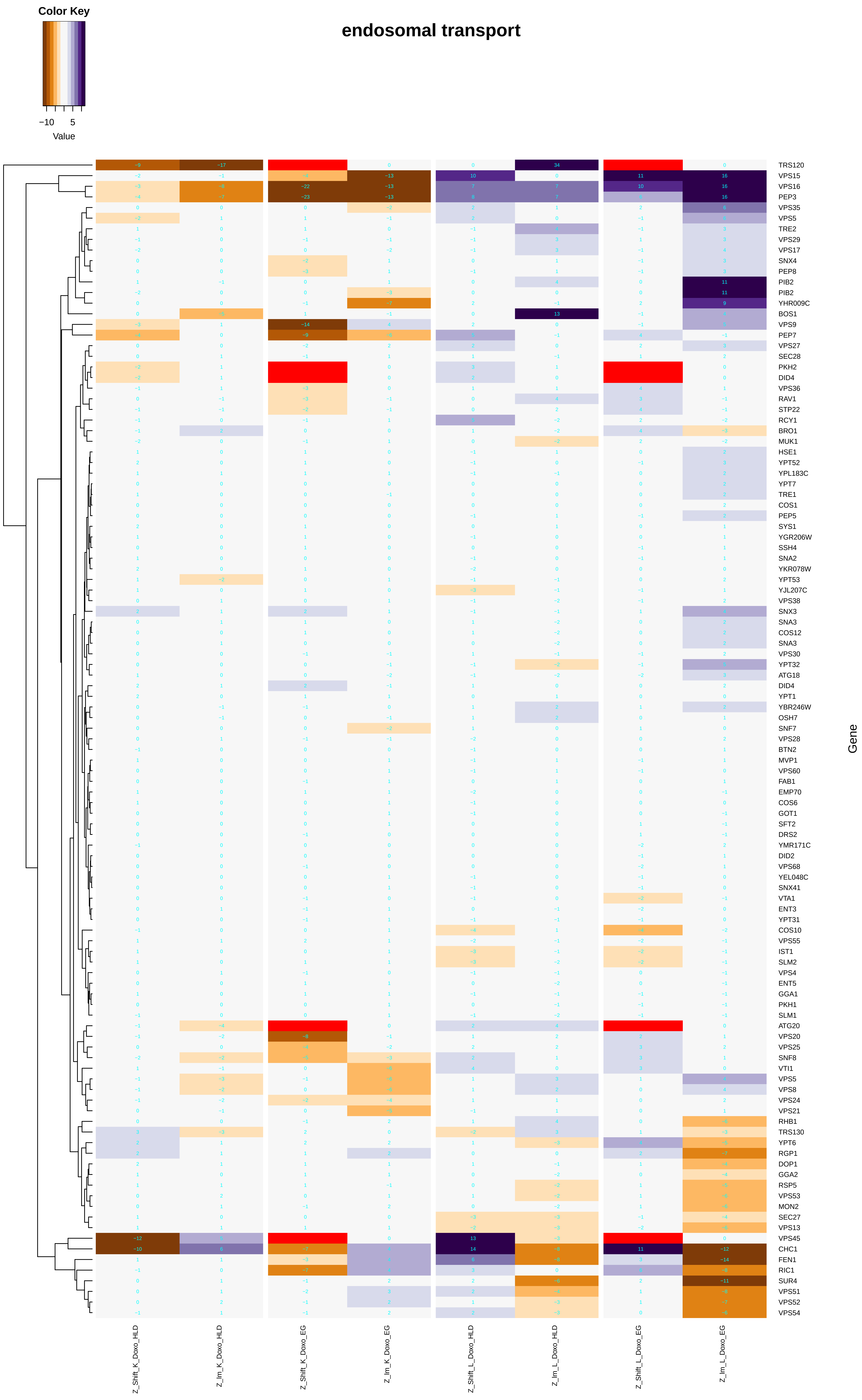

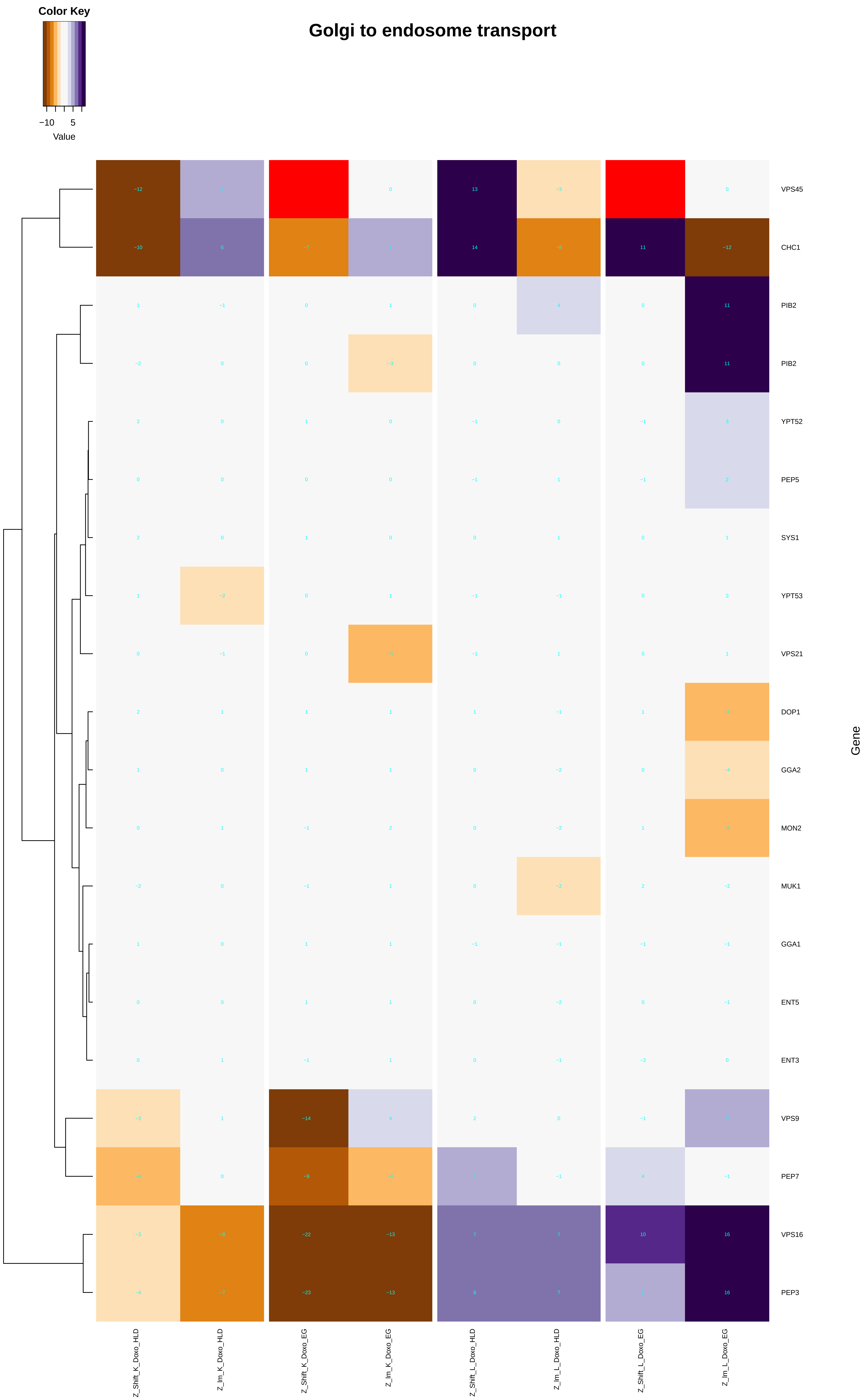

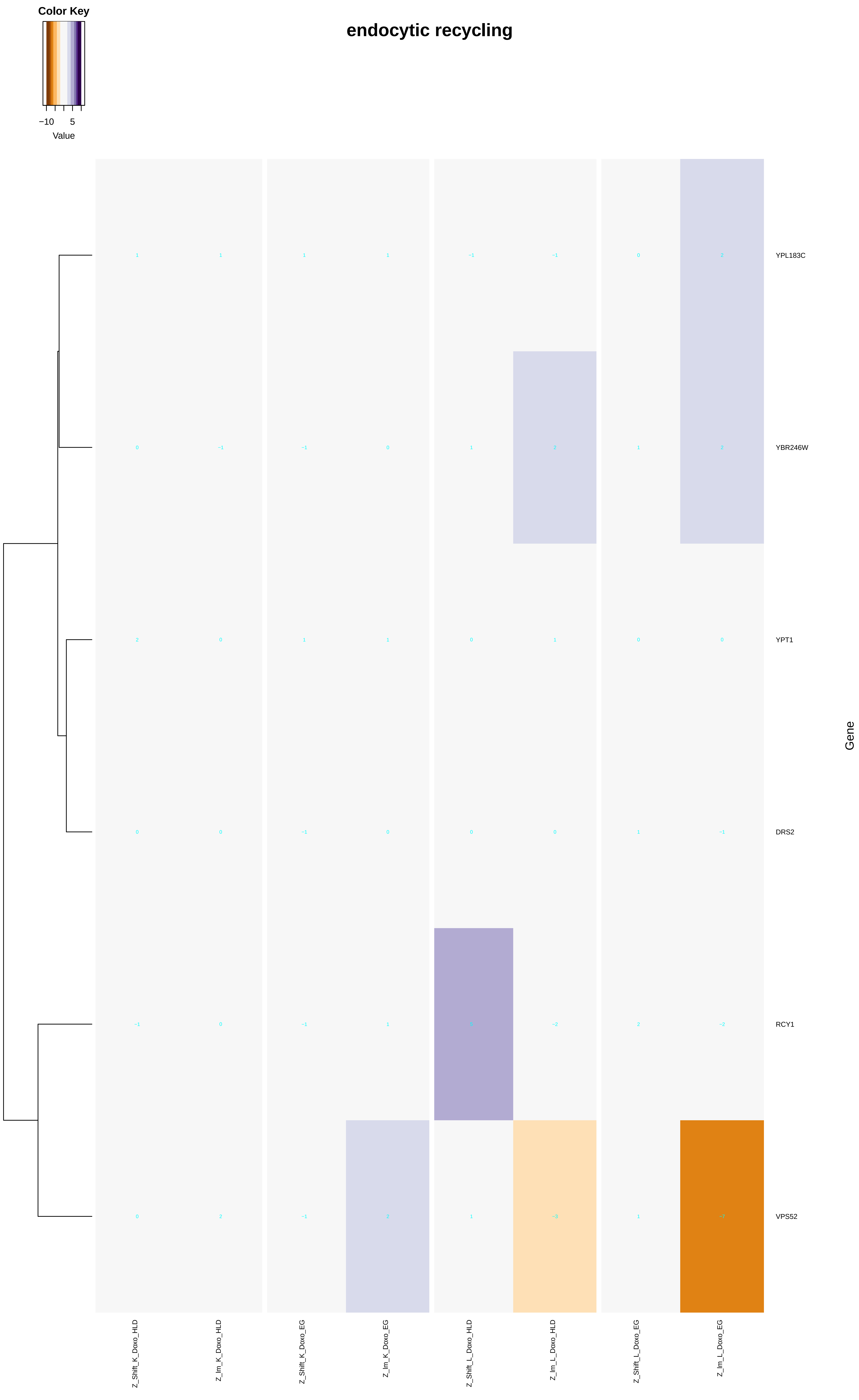

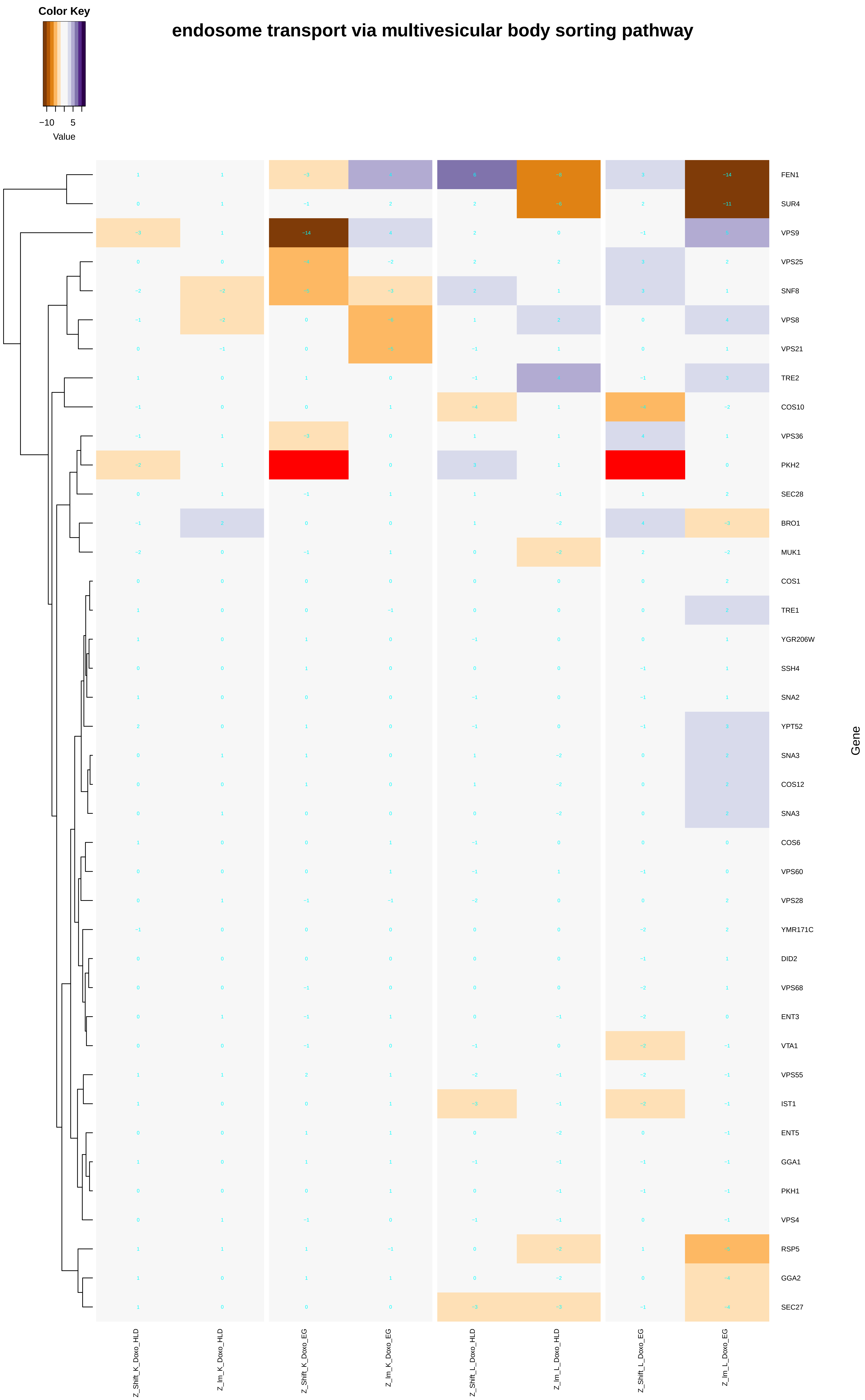

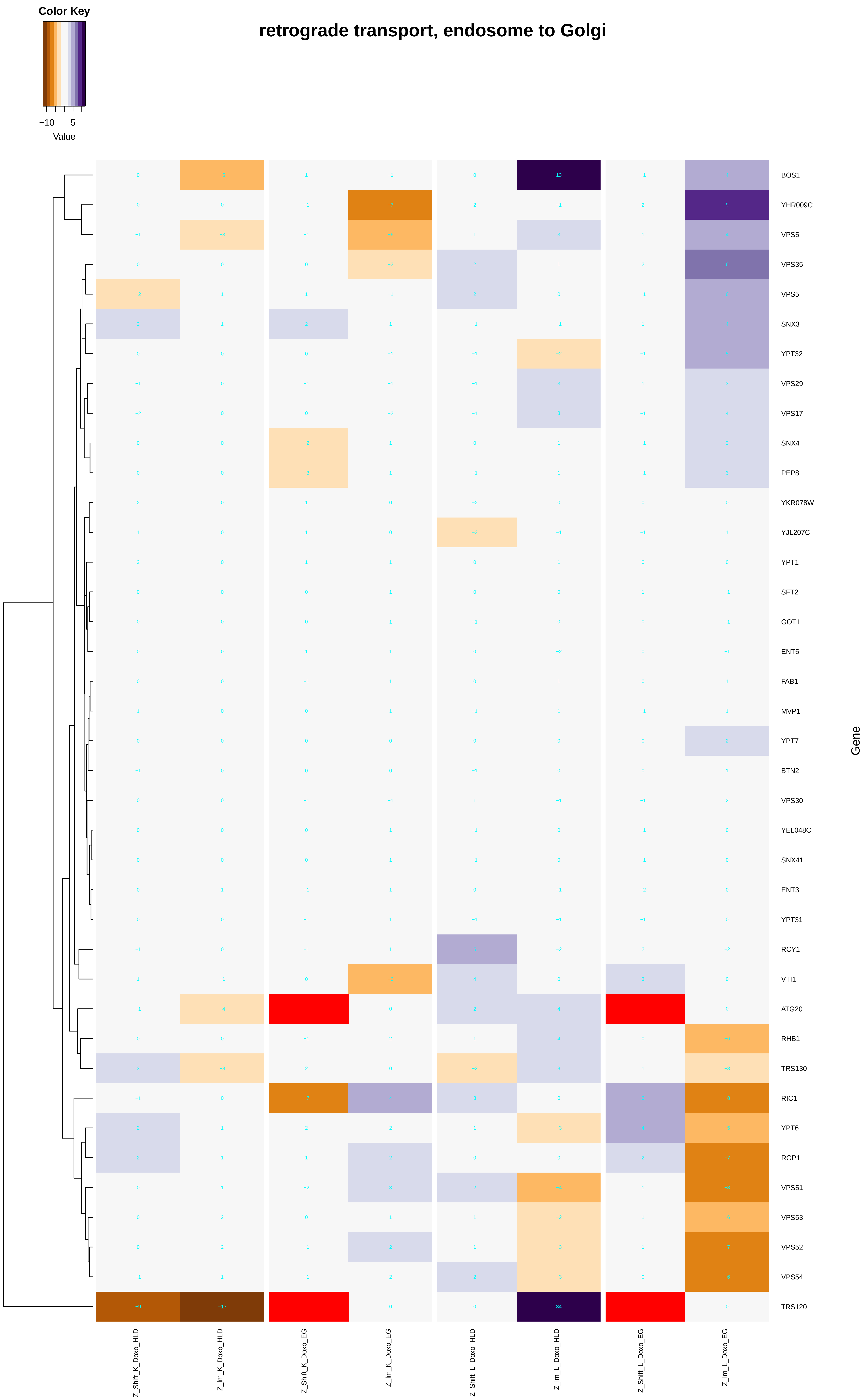

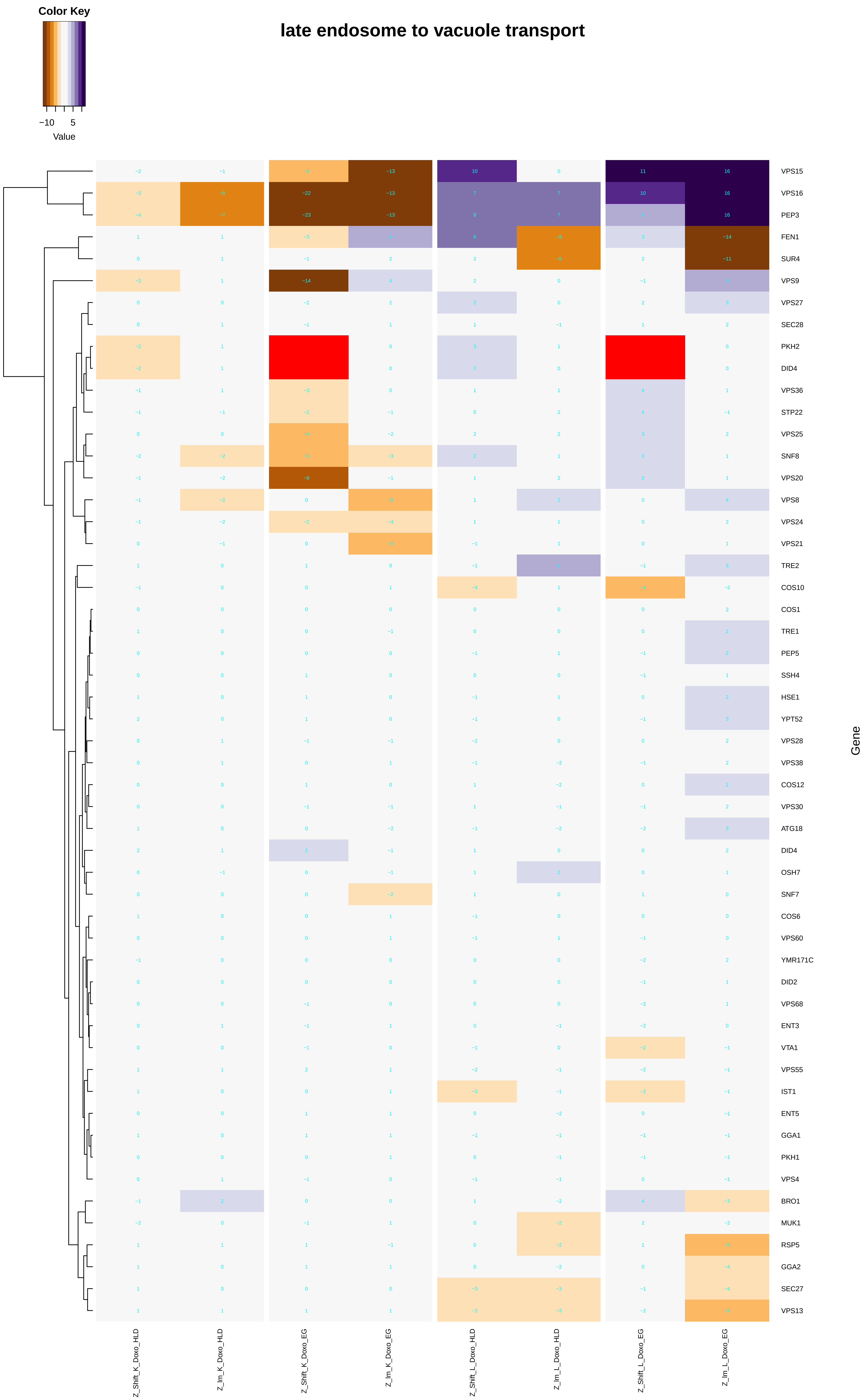

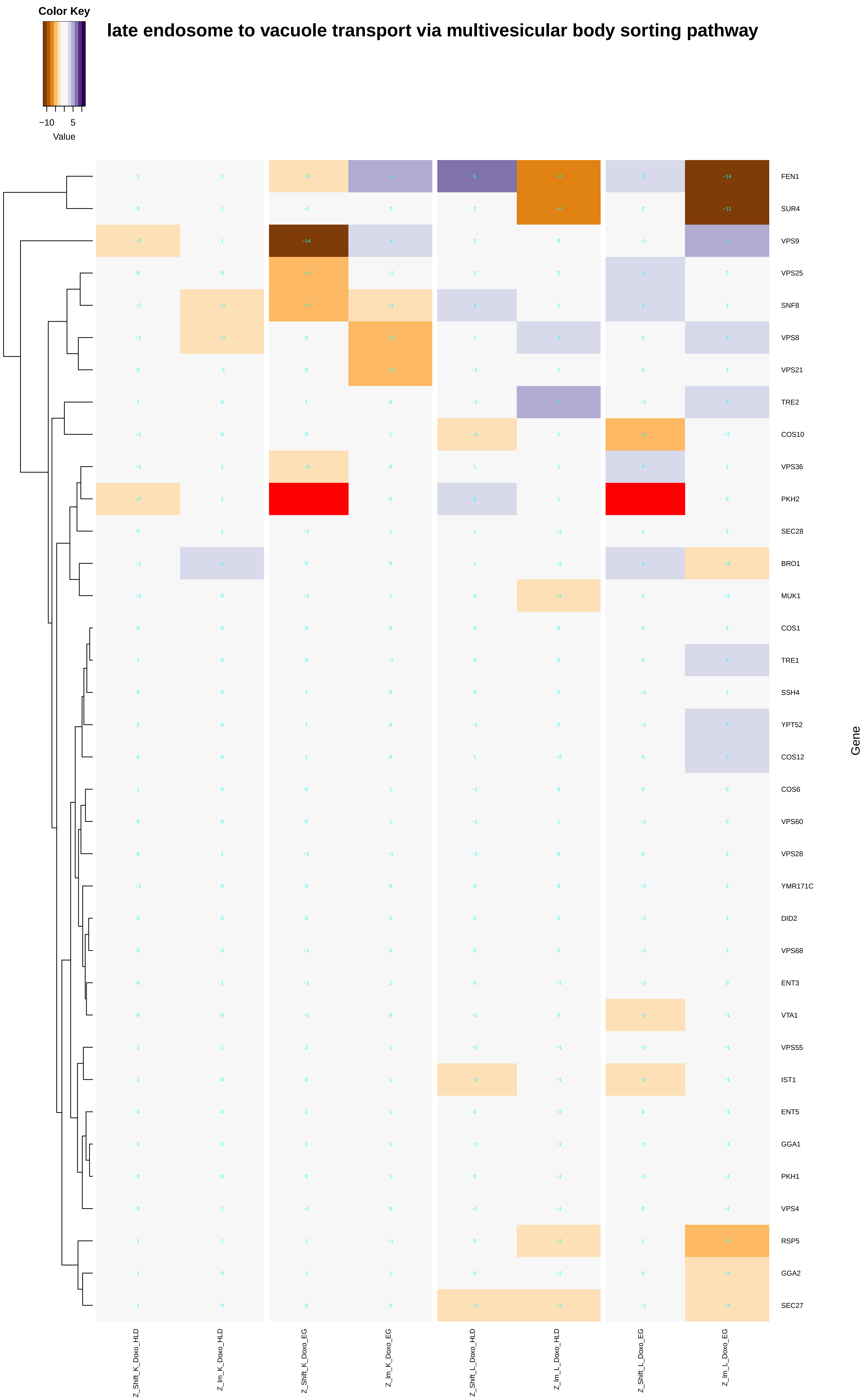

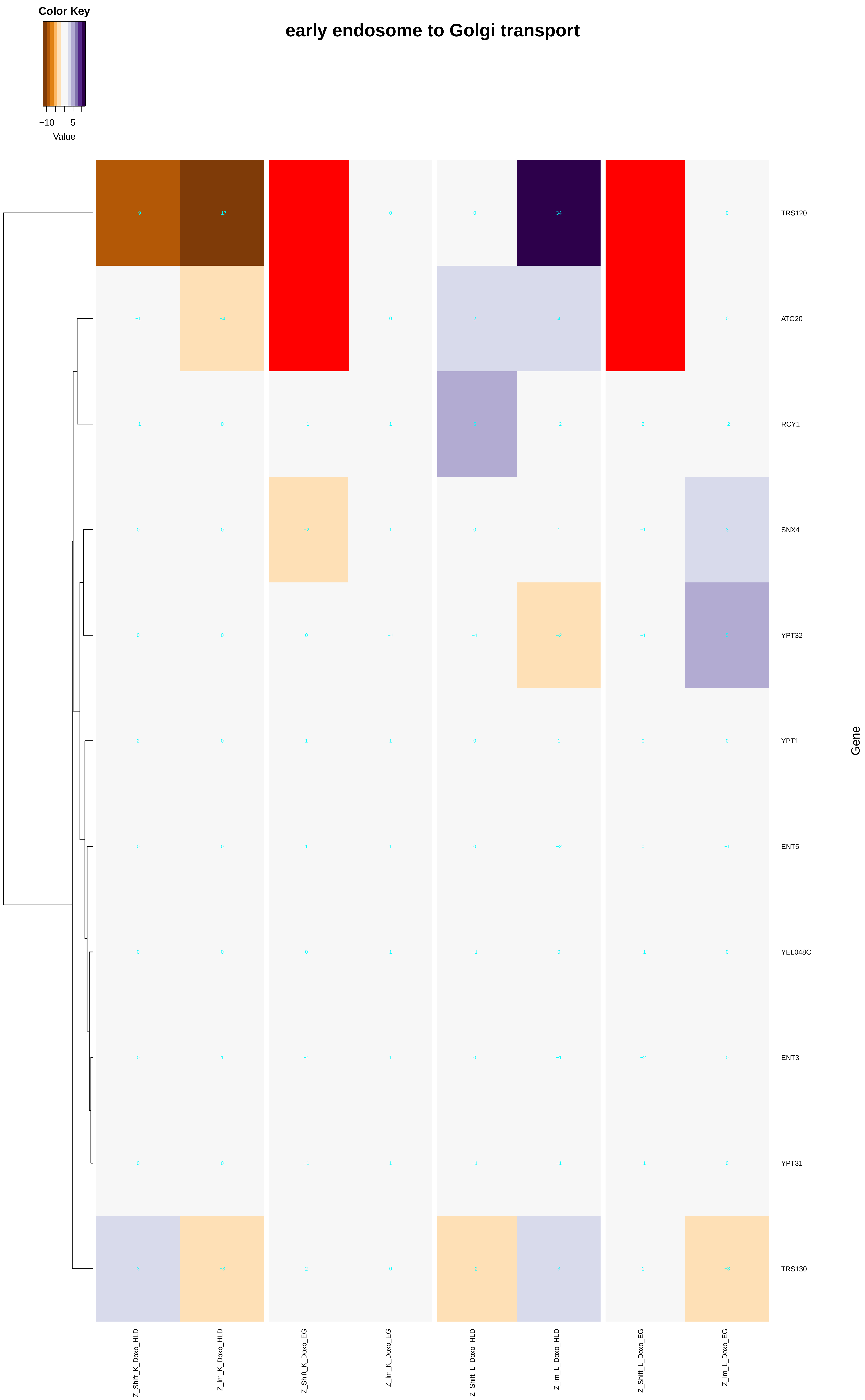

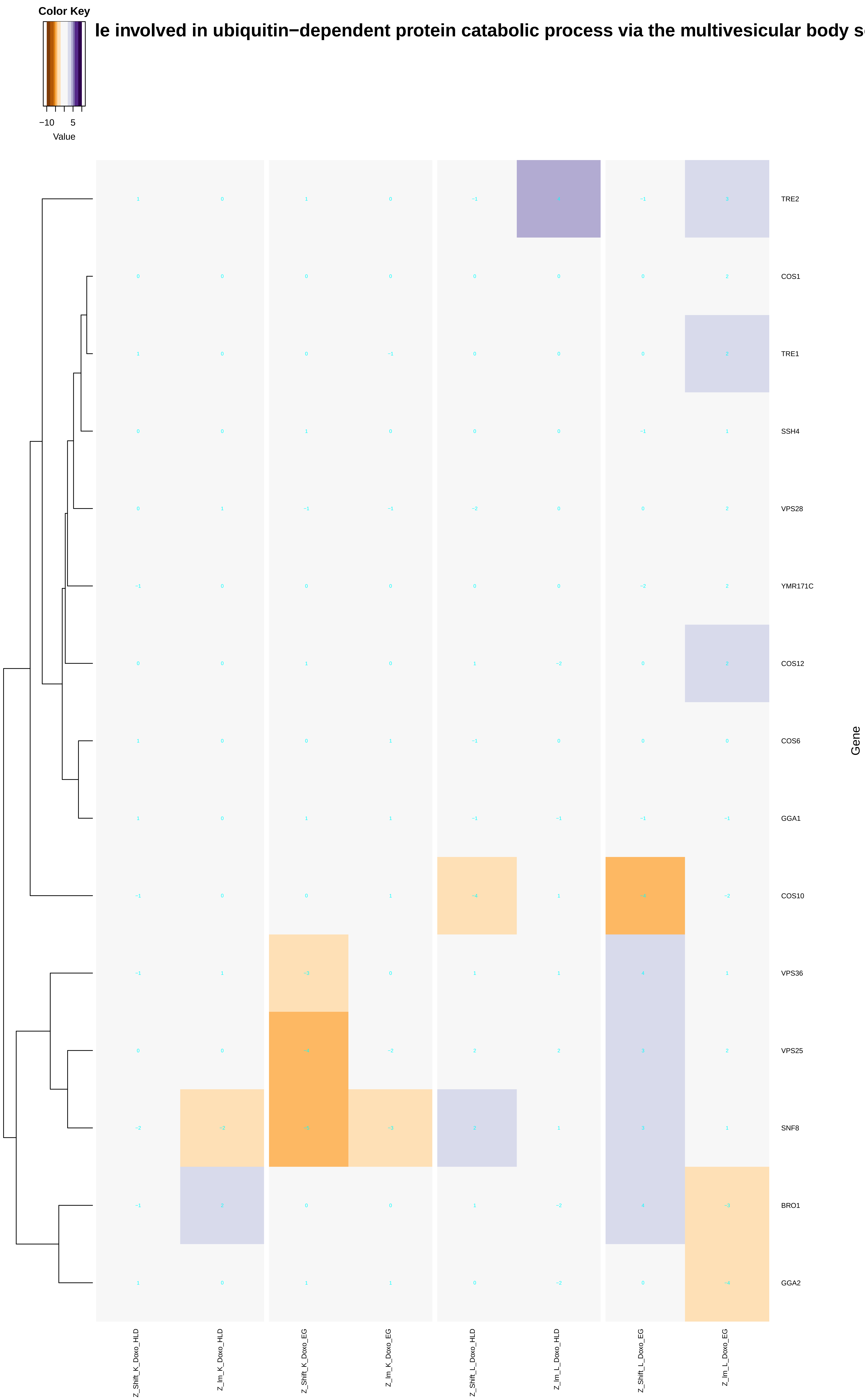

### energy_coupled_proton_transmembrane_transport,_against_electrochemical_gradient.pdf

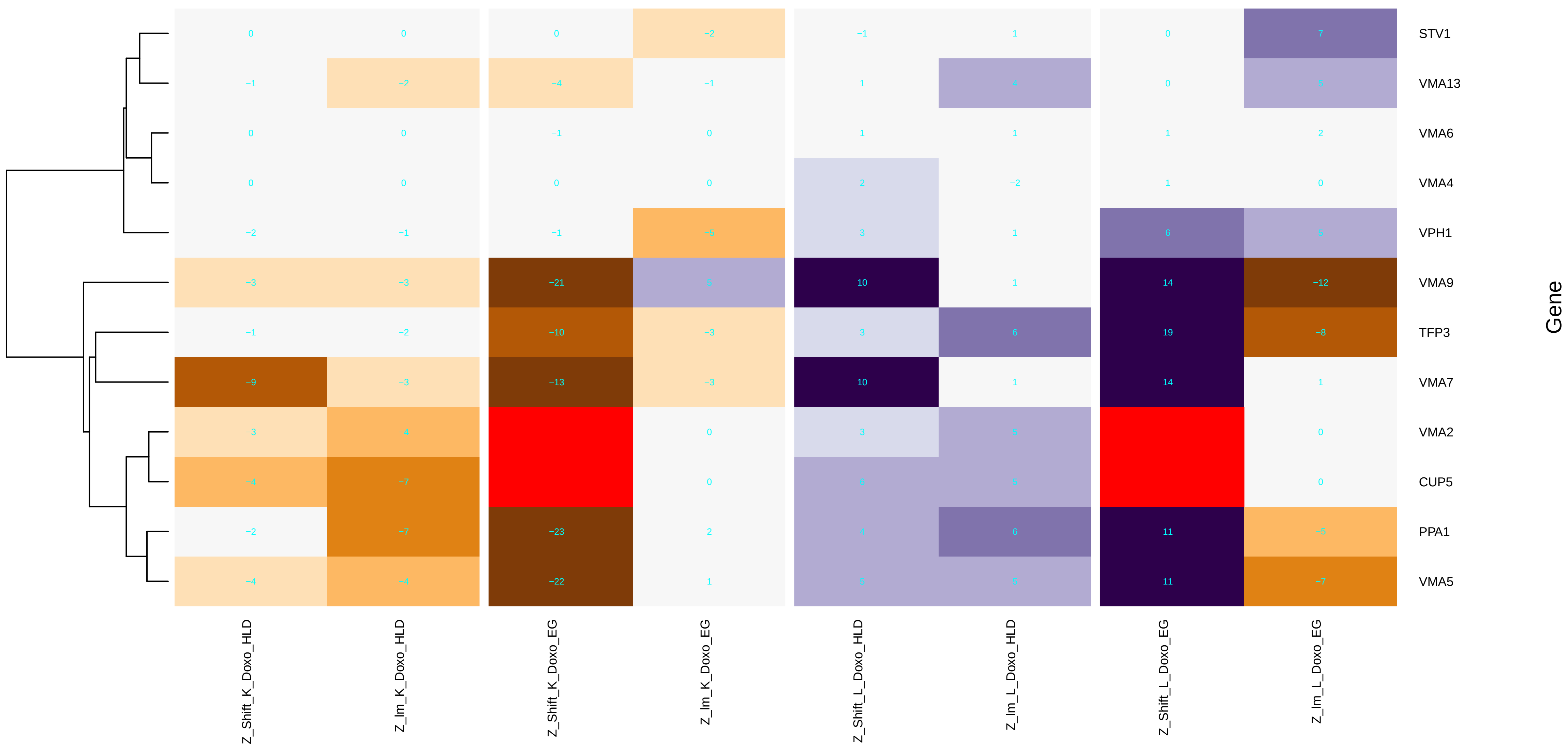

### energy_coupled_proton_transport,_down_electrochemical_gradient.pdf

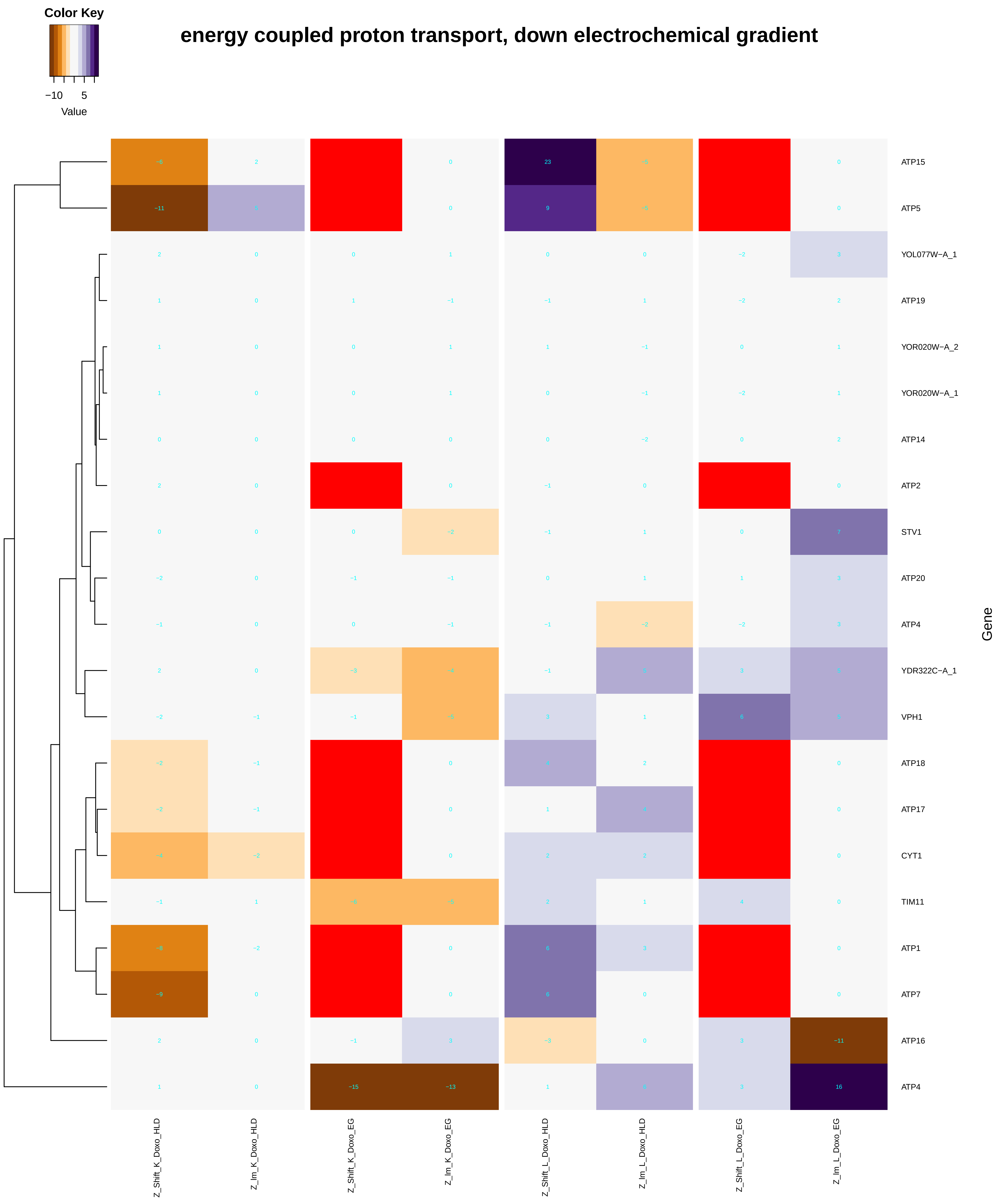

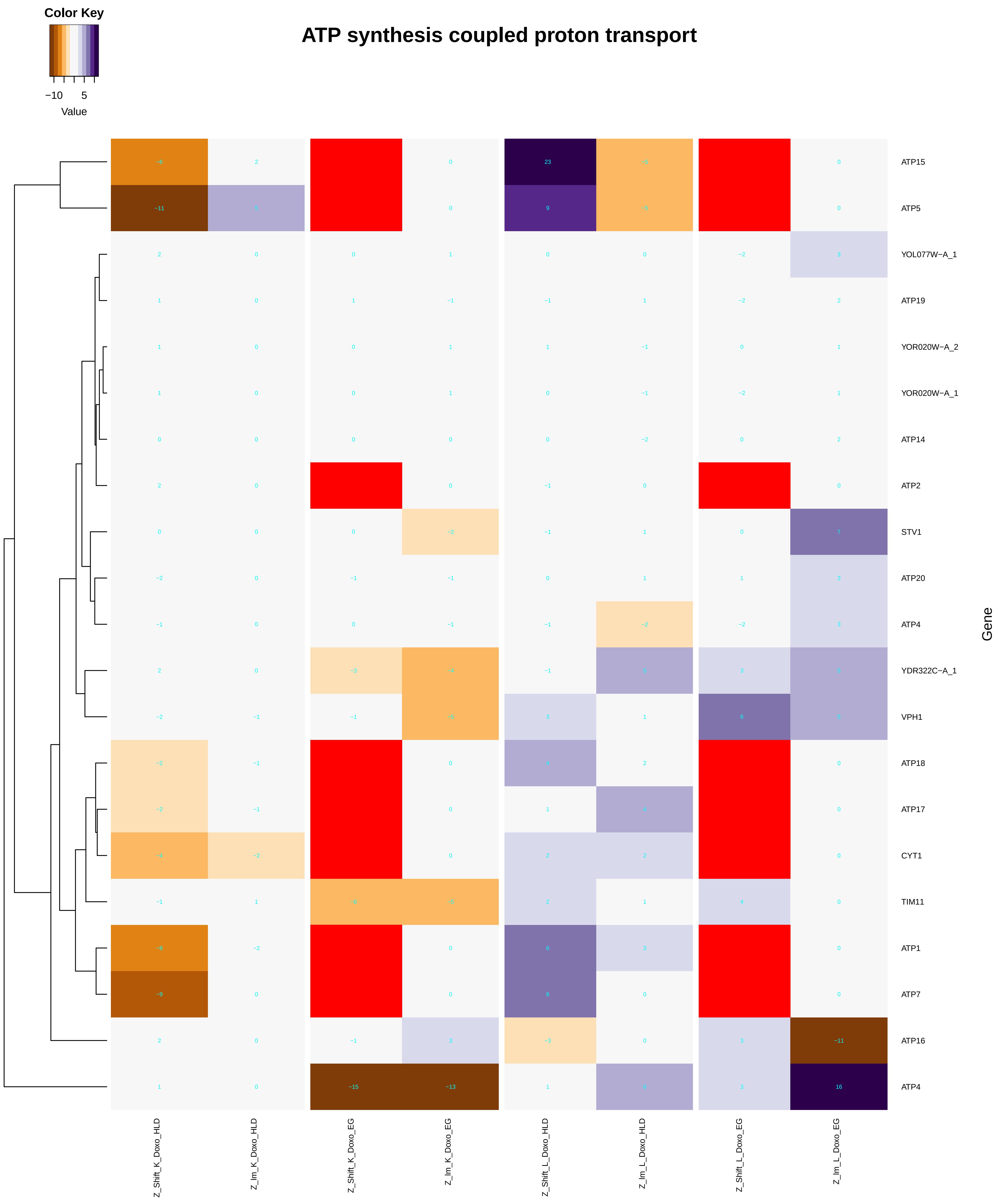

### establishment_of_protein_localization_to_mitochondrial_membrane.pdf

#### establishment of protein localization to mitochondrial membrane

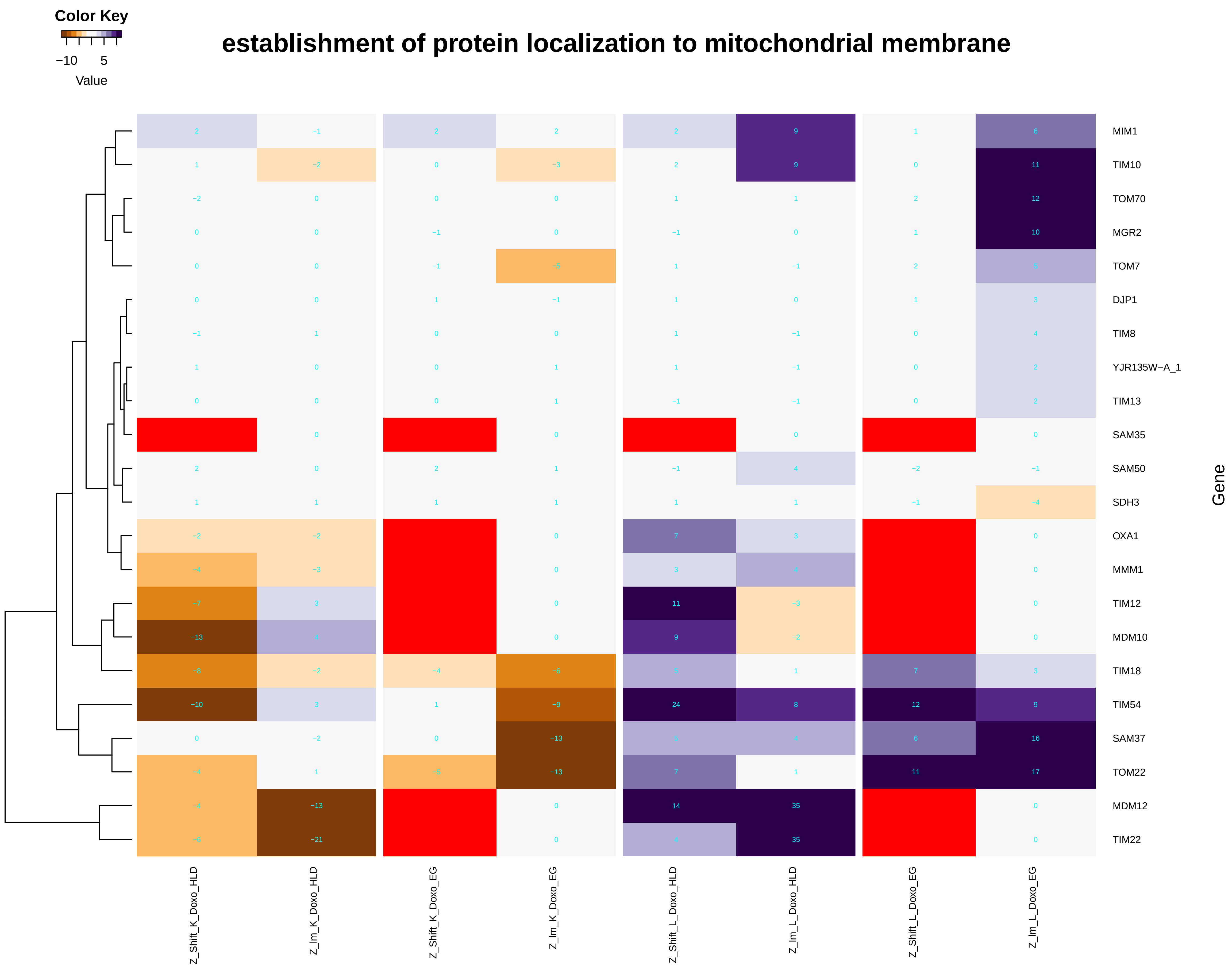

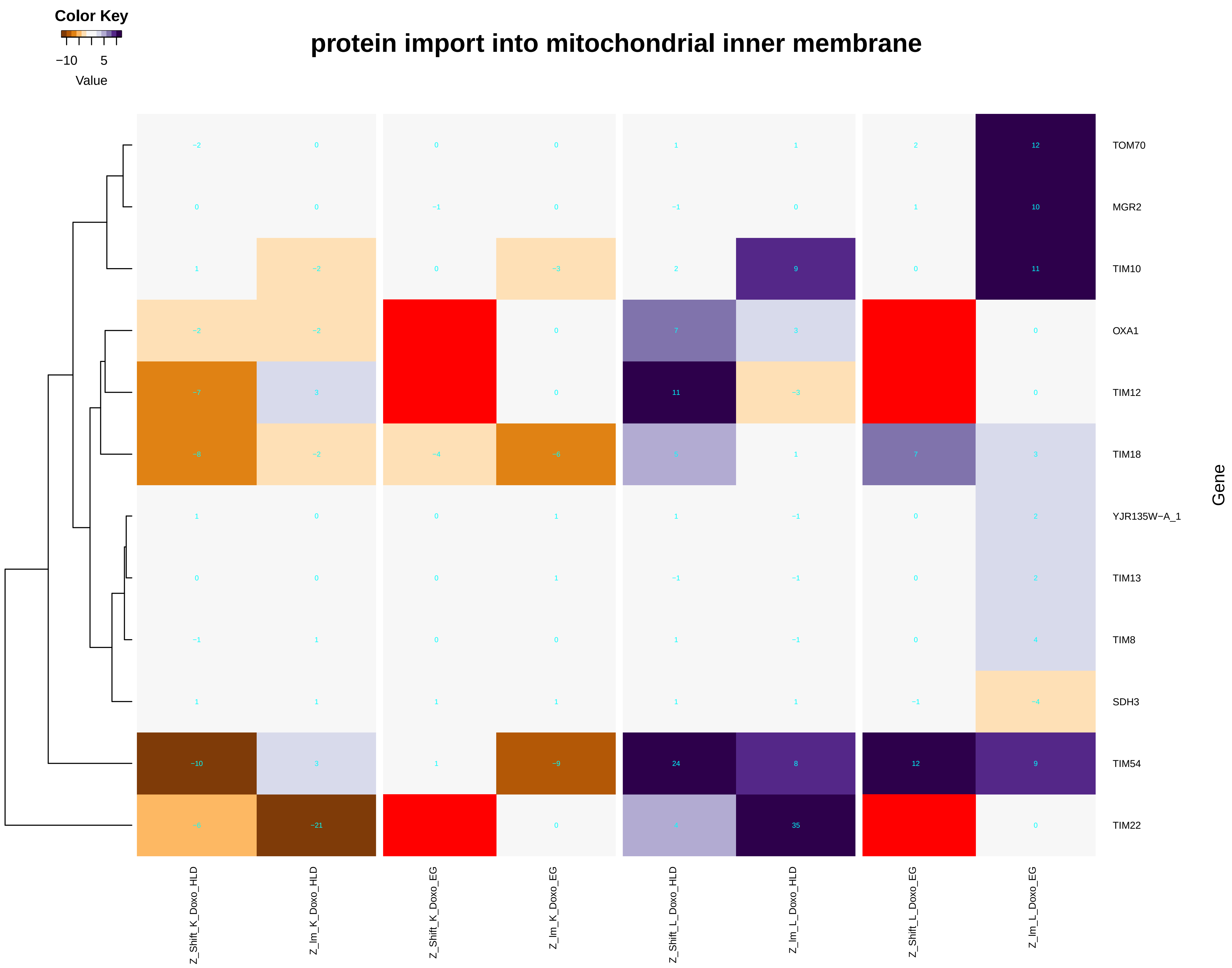

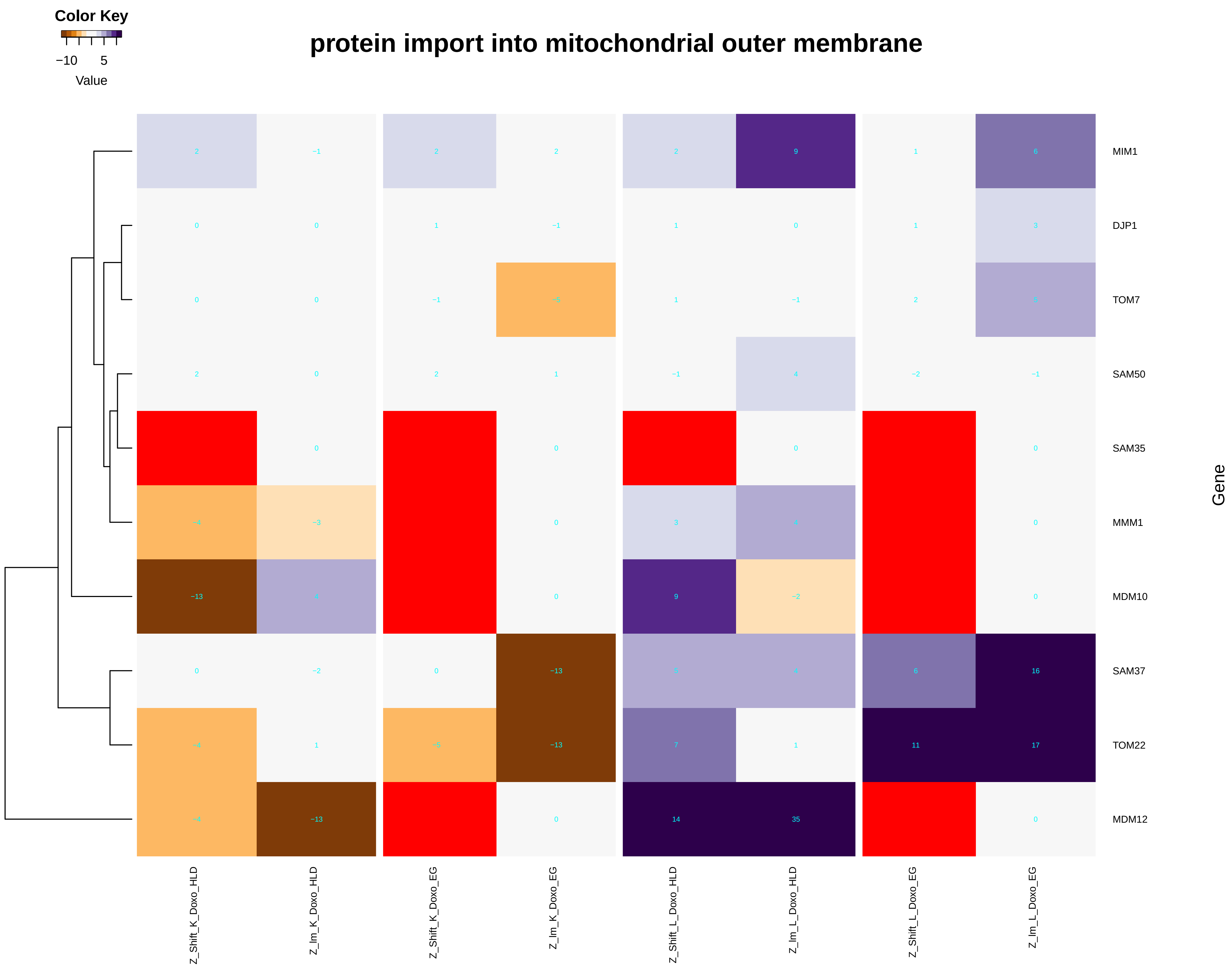

### exocytic_process.pdf

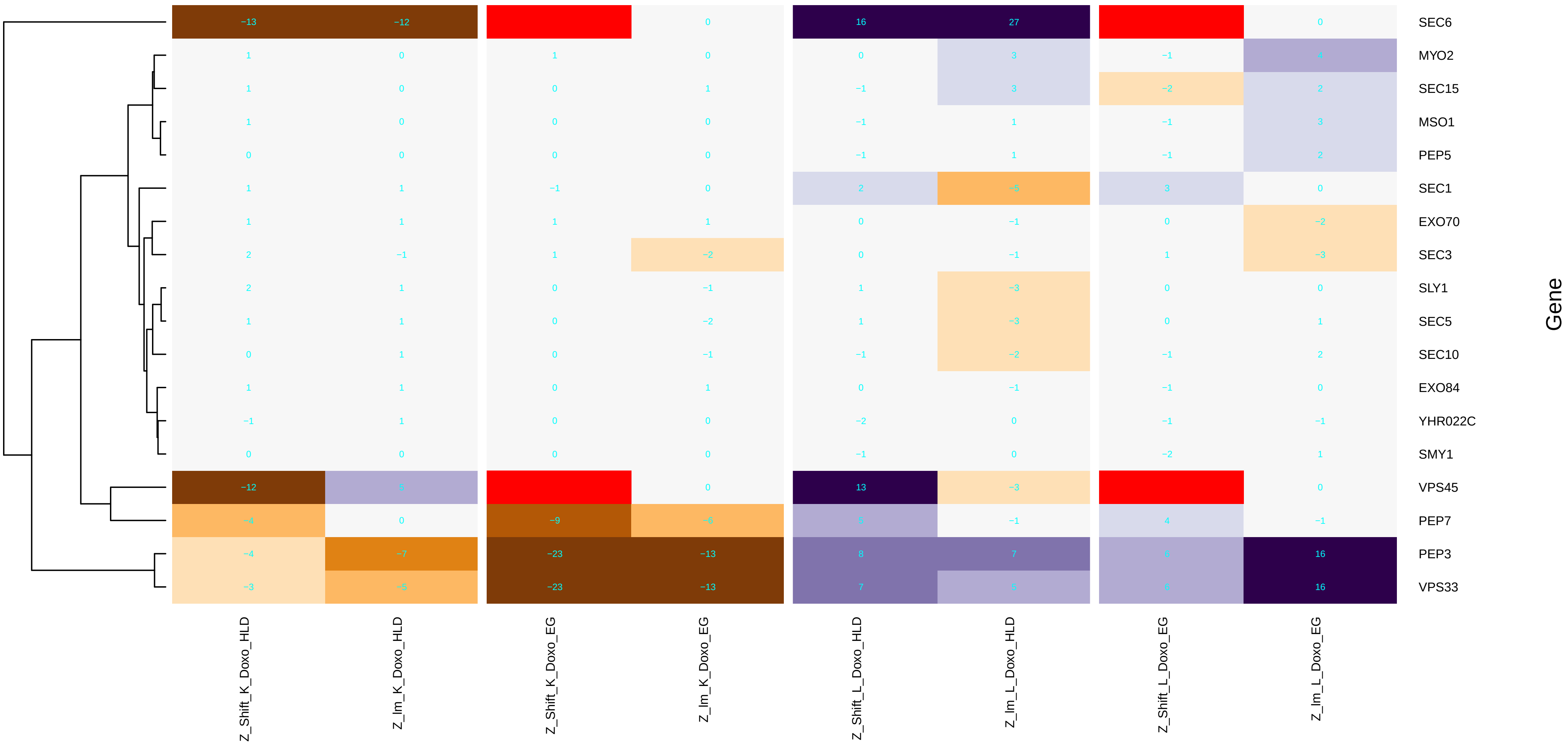

### fatty_acid_biosynthetic_process.pdf

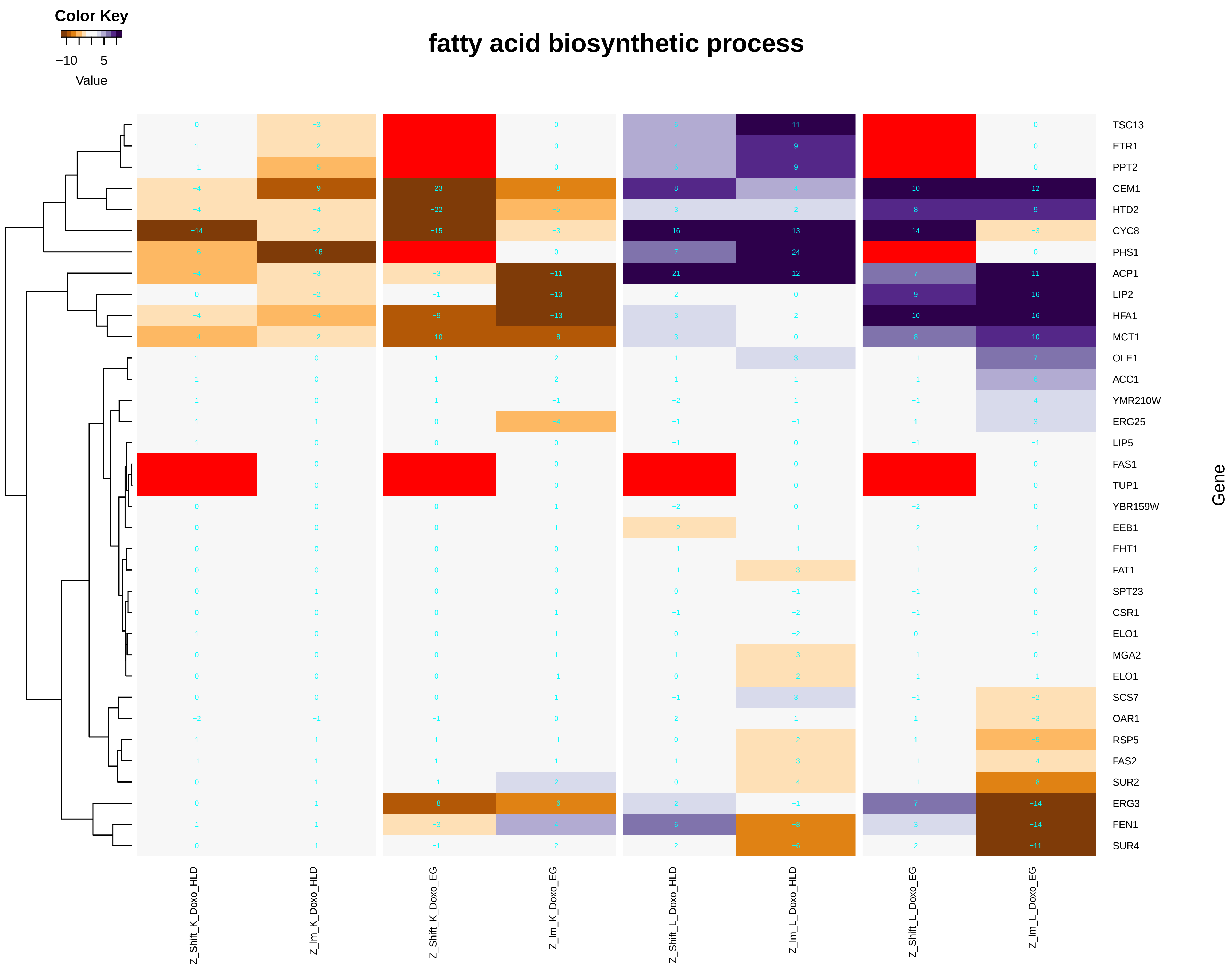

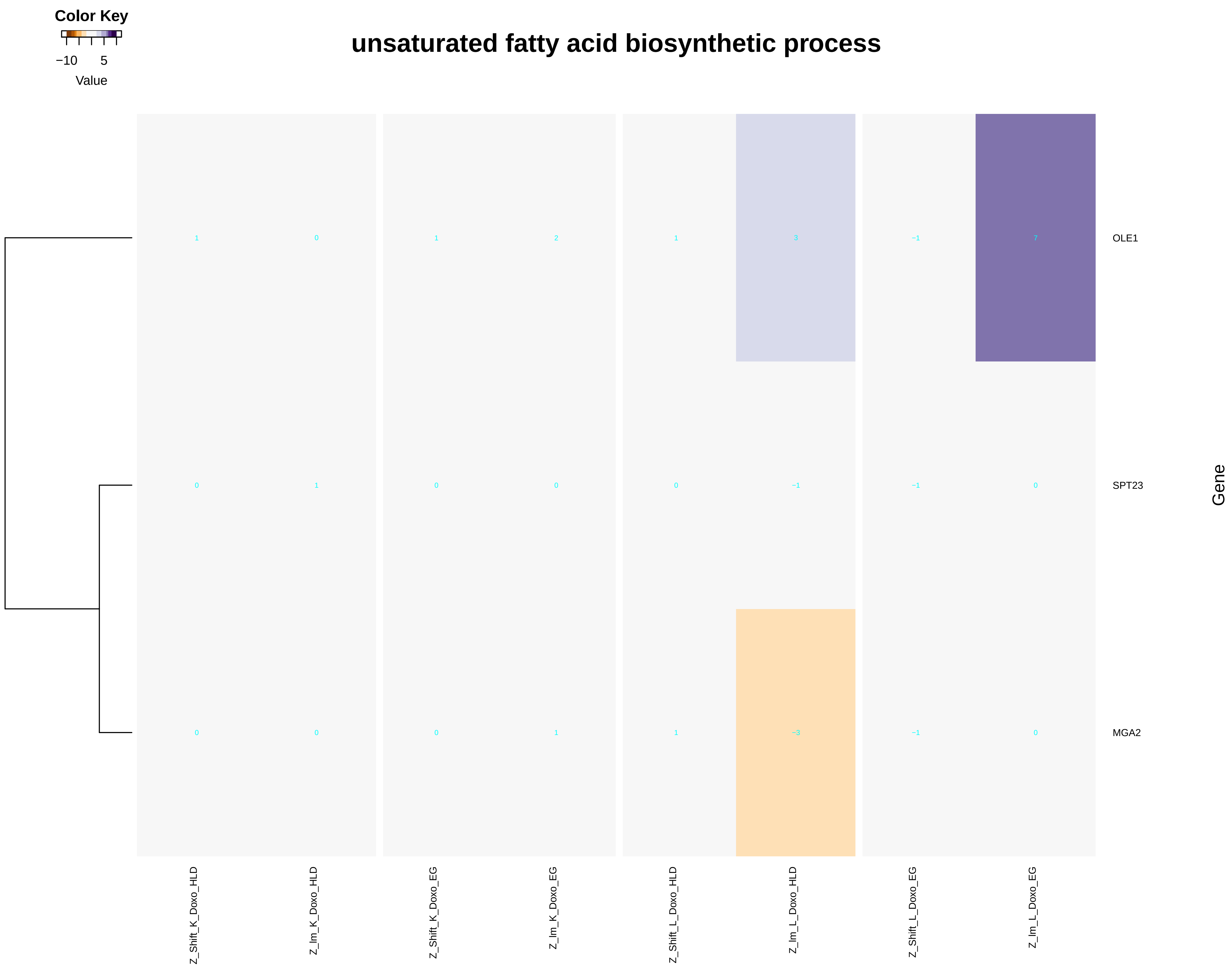

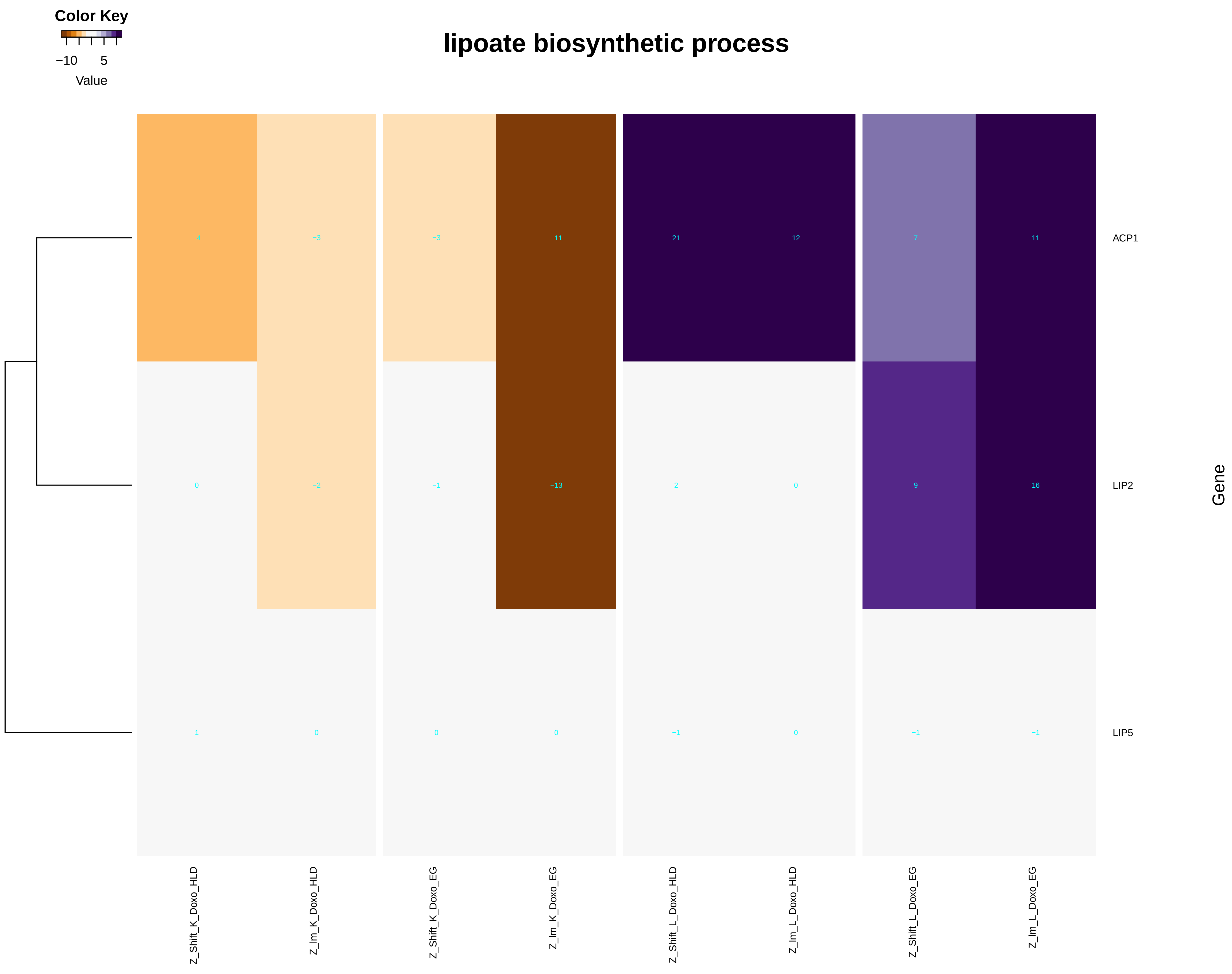

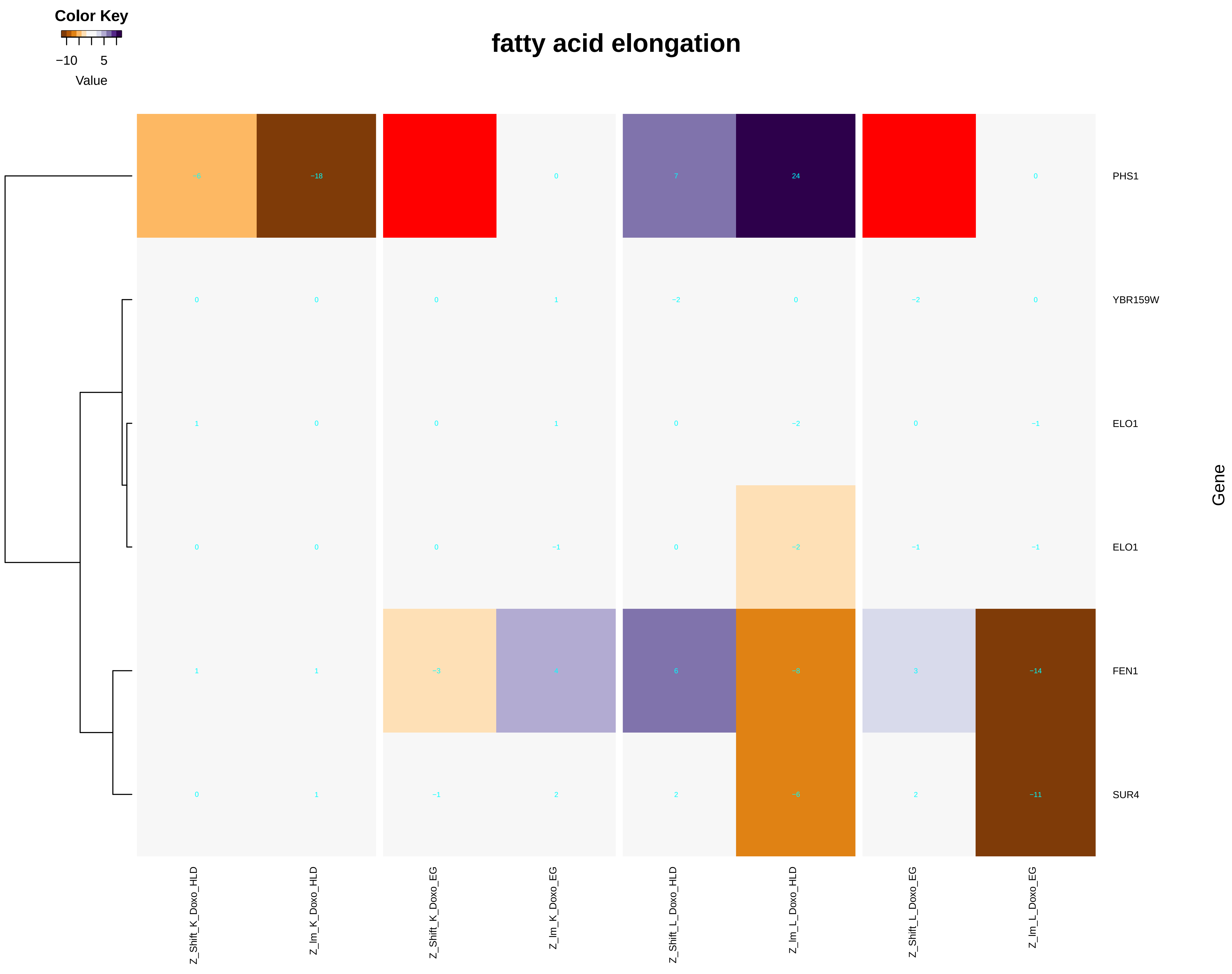

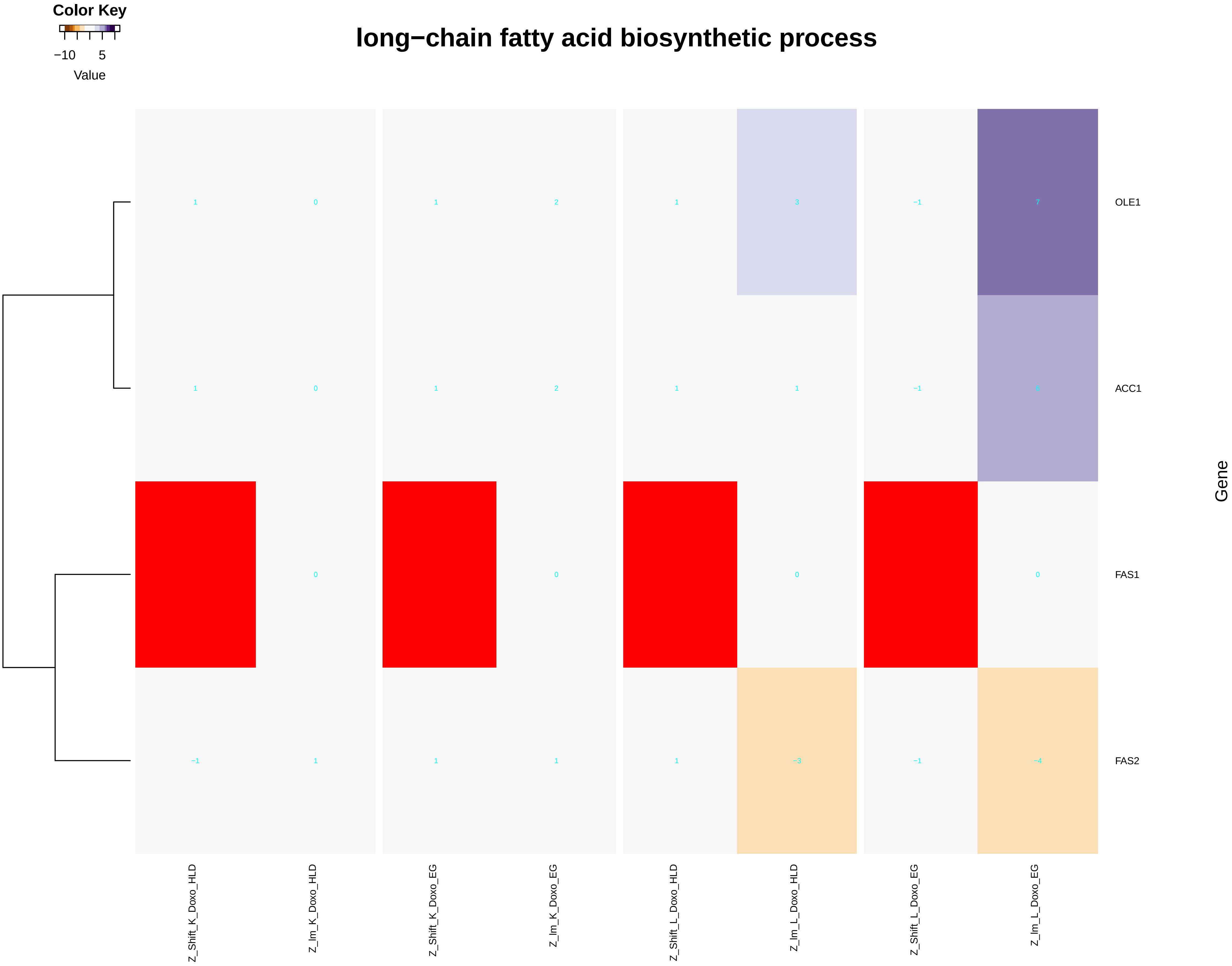

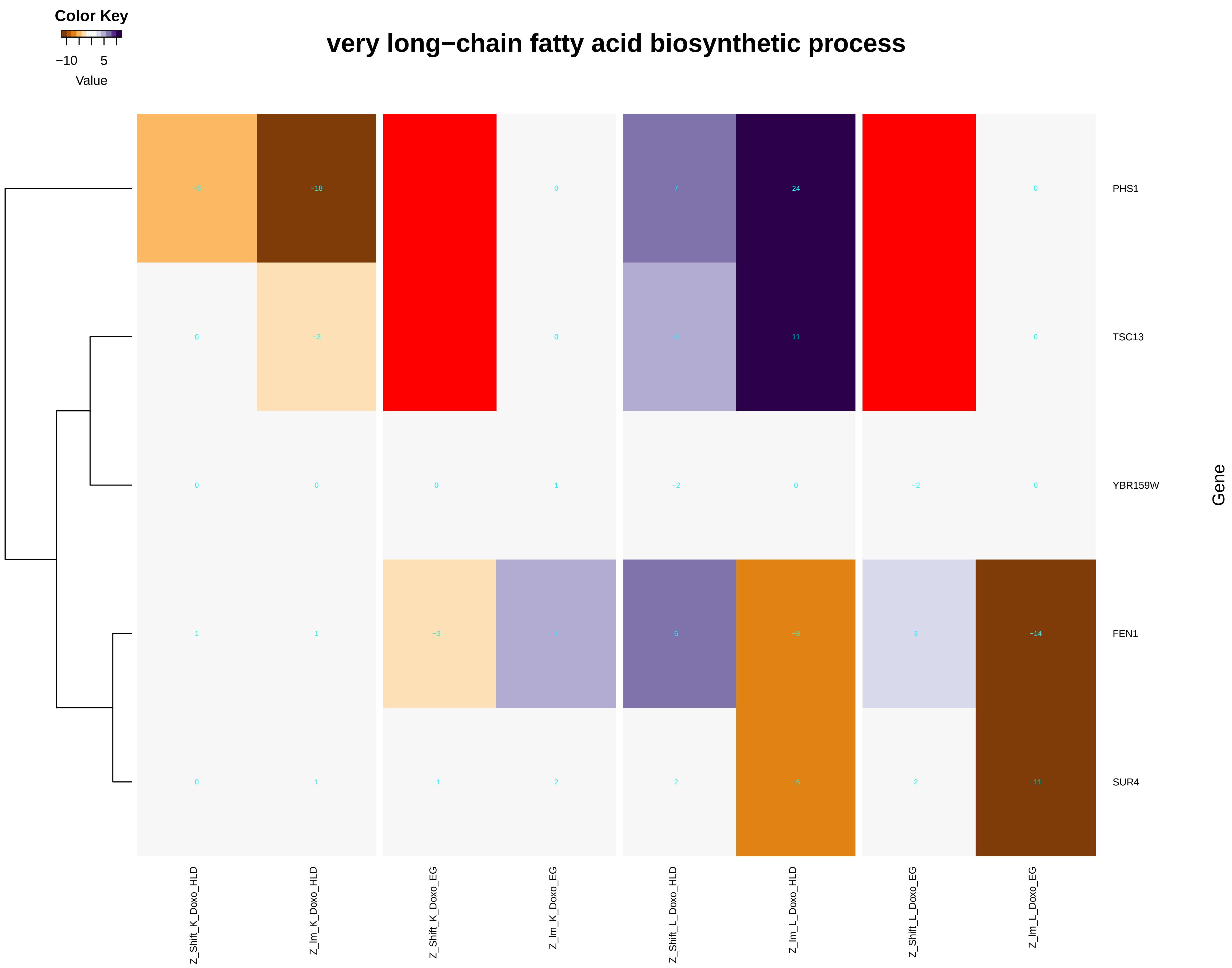

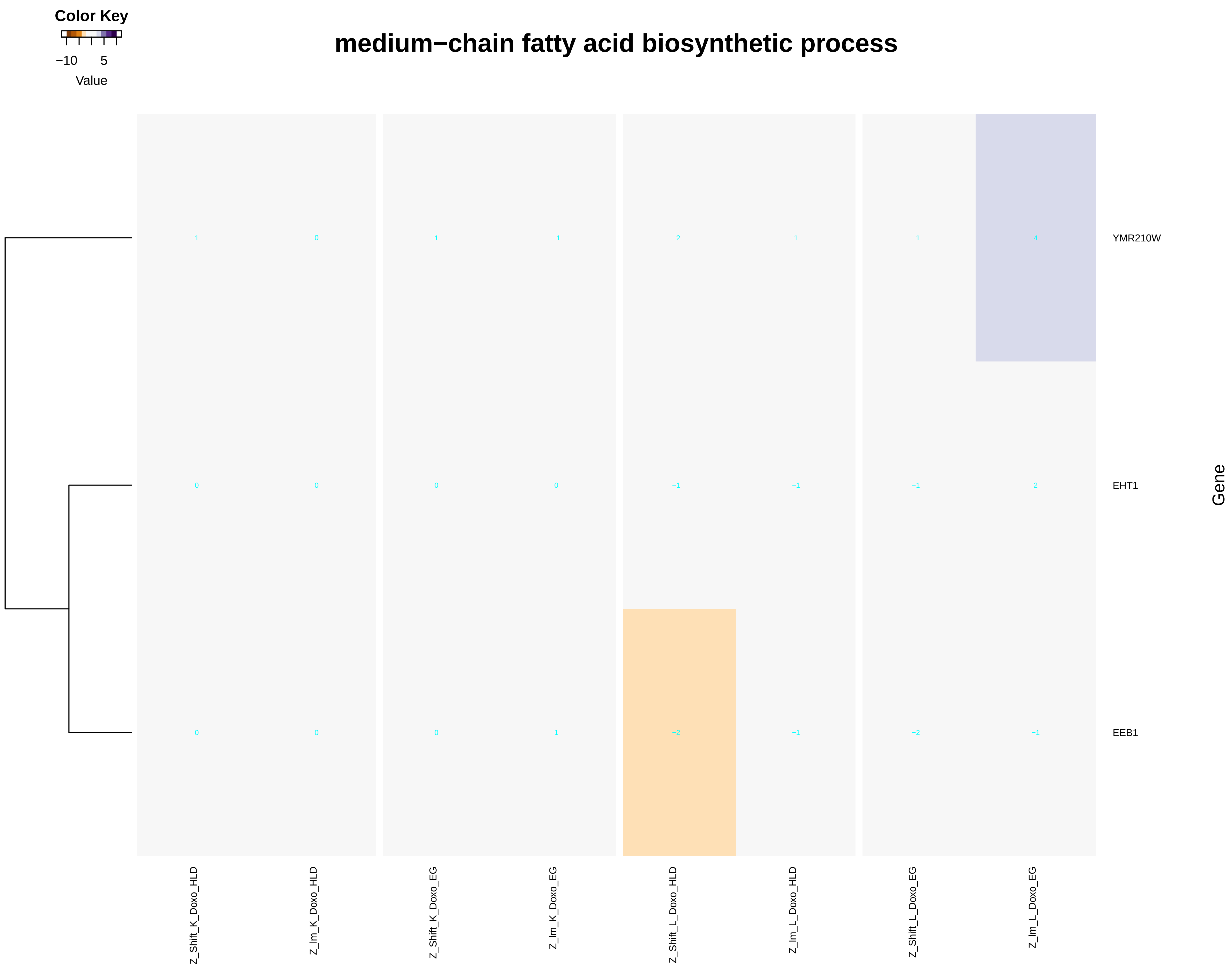

### gene_expression.pdf

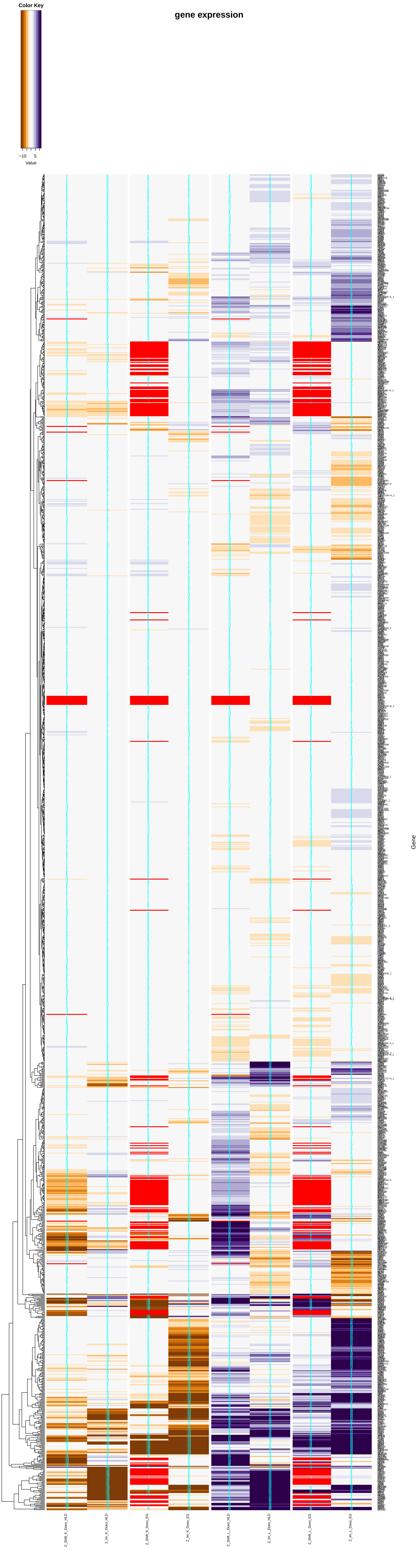

### glycolipid_biosynthetic_process.pdf

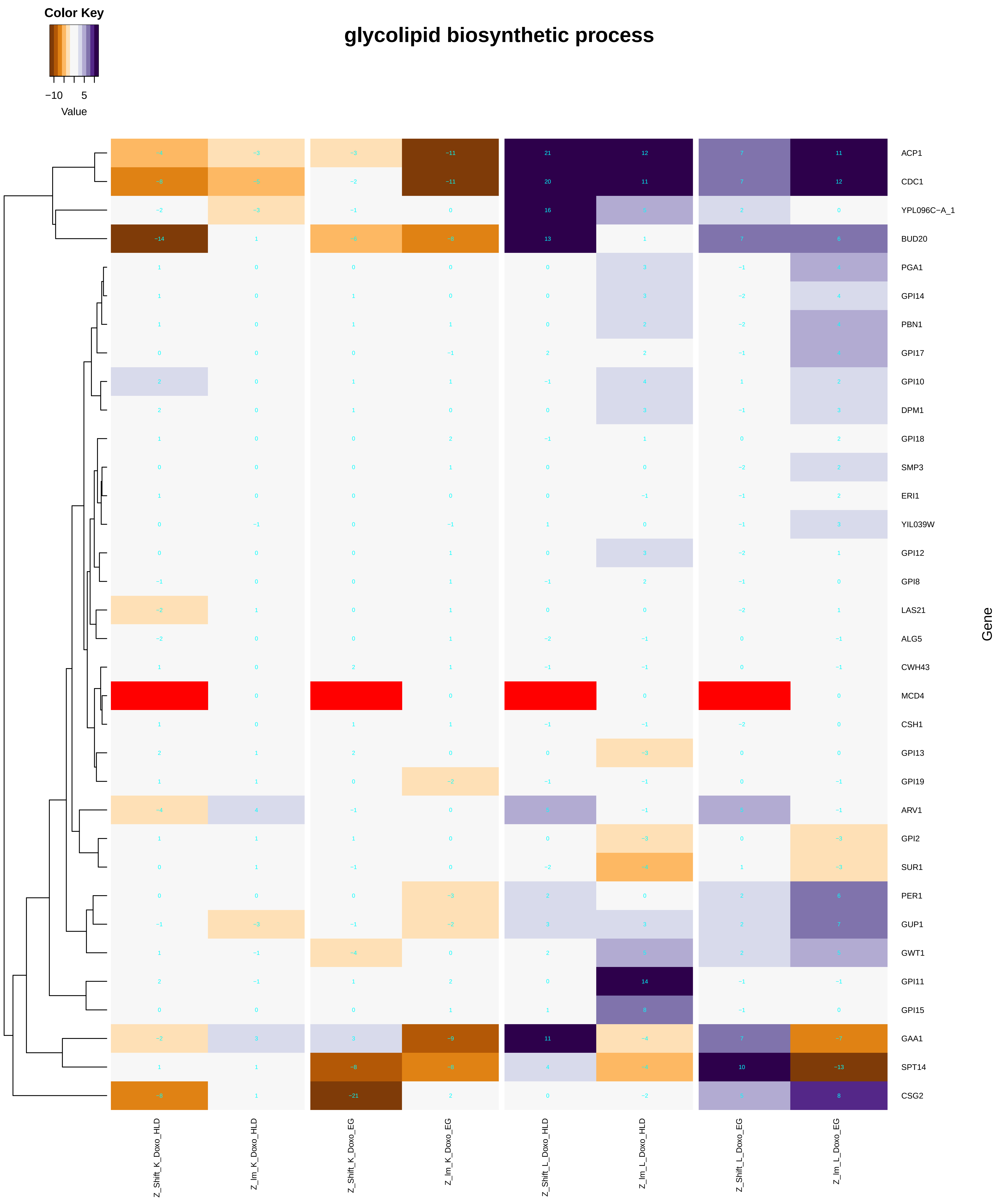

#### GPI anchor biosynthetic process

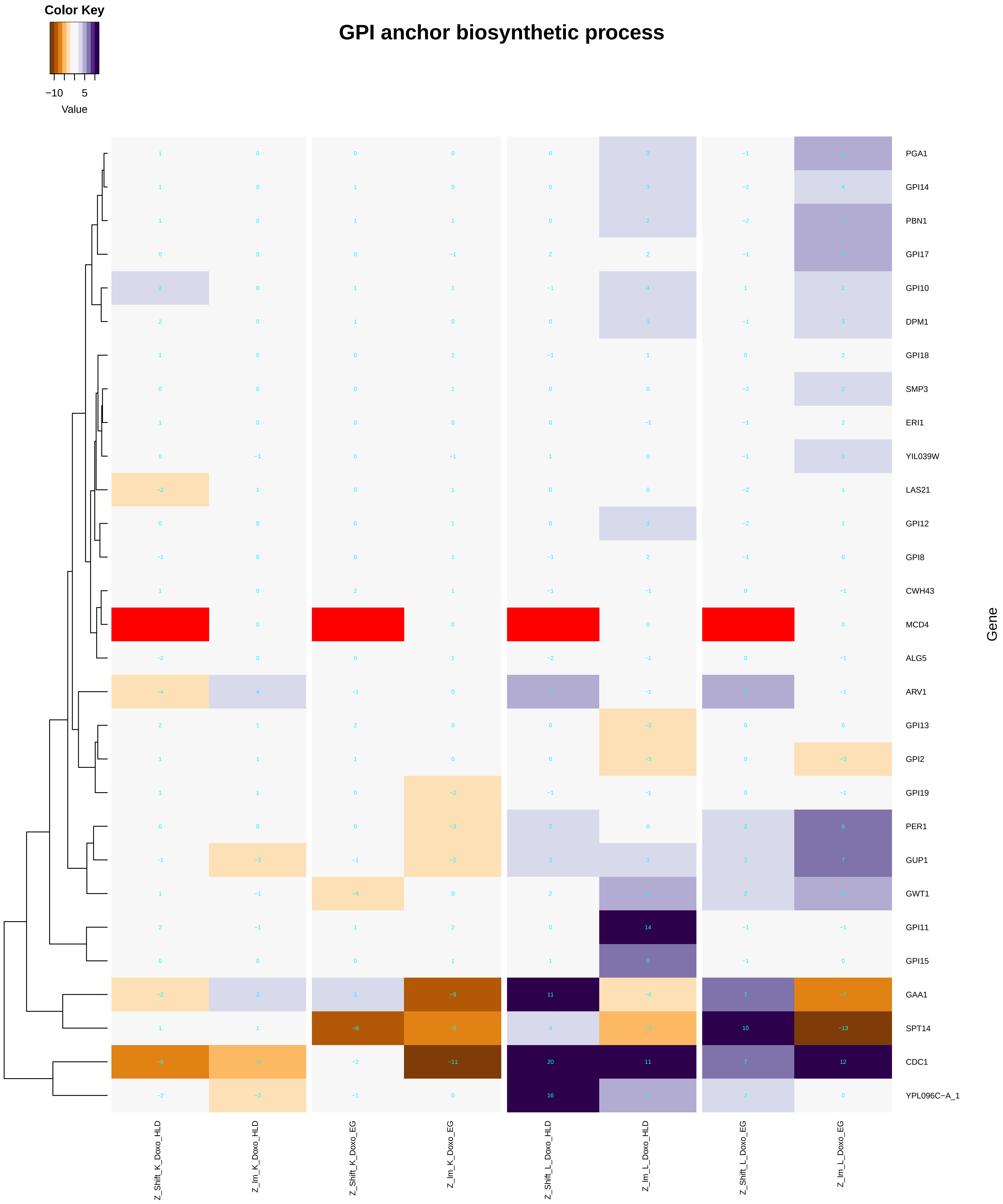

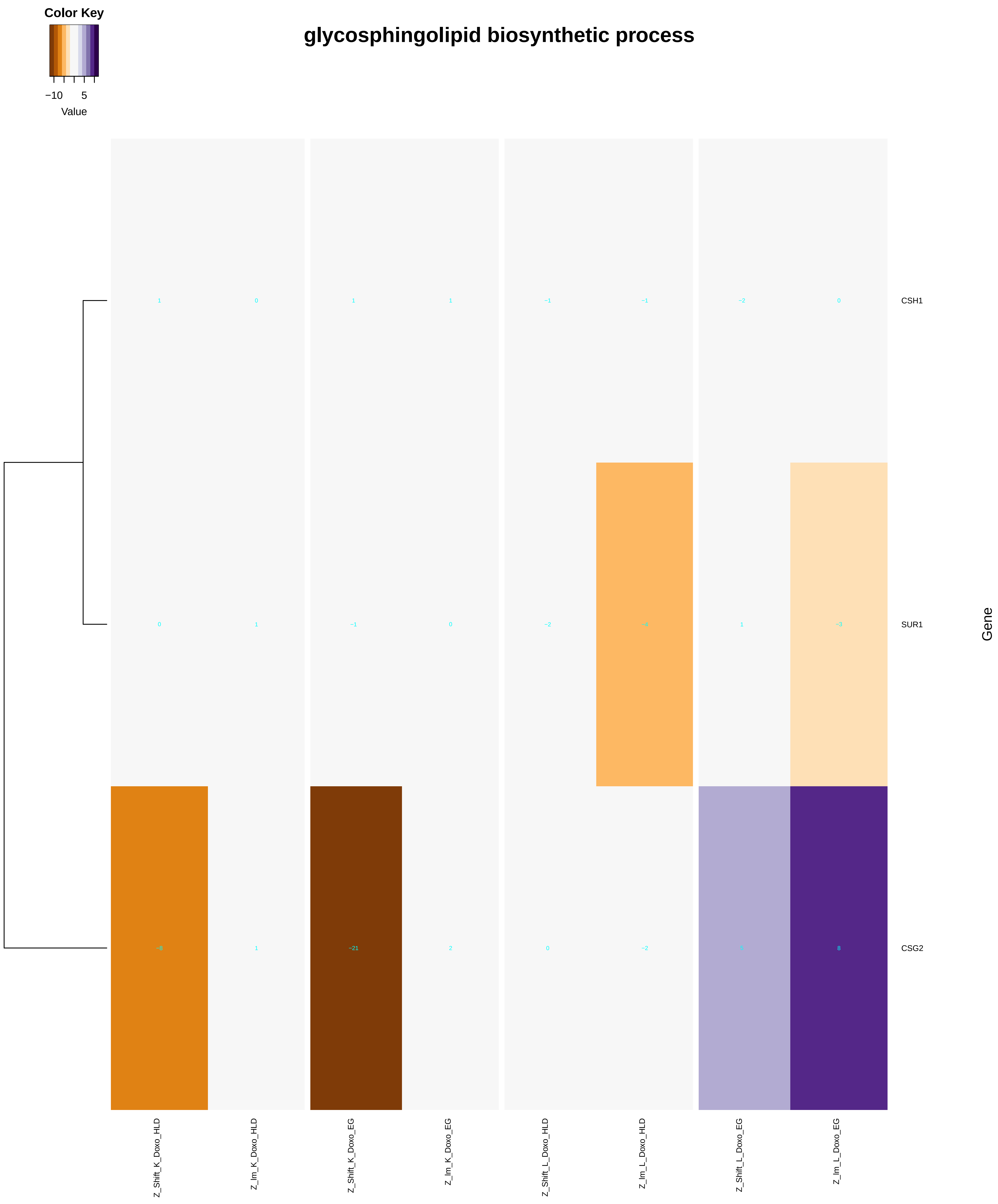

### GPI_anchor_biosynthetic_process.pdf

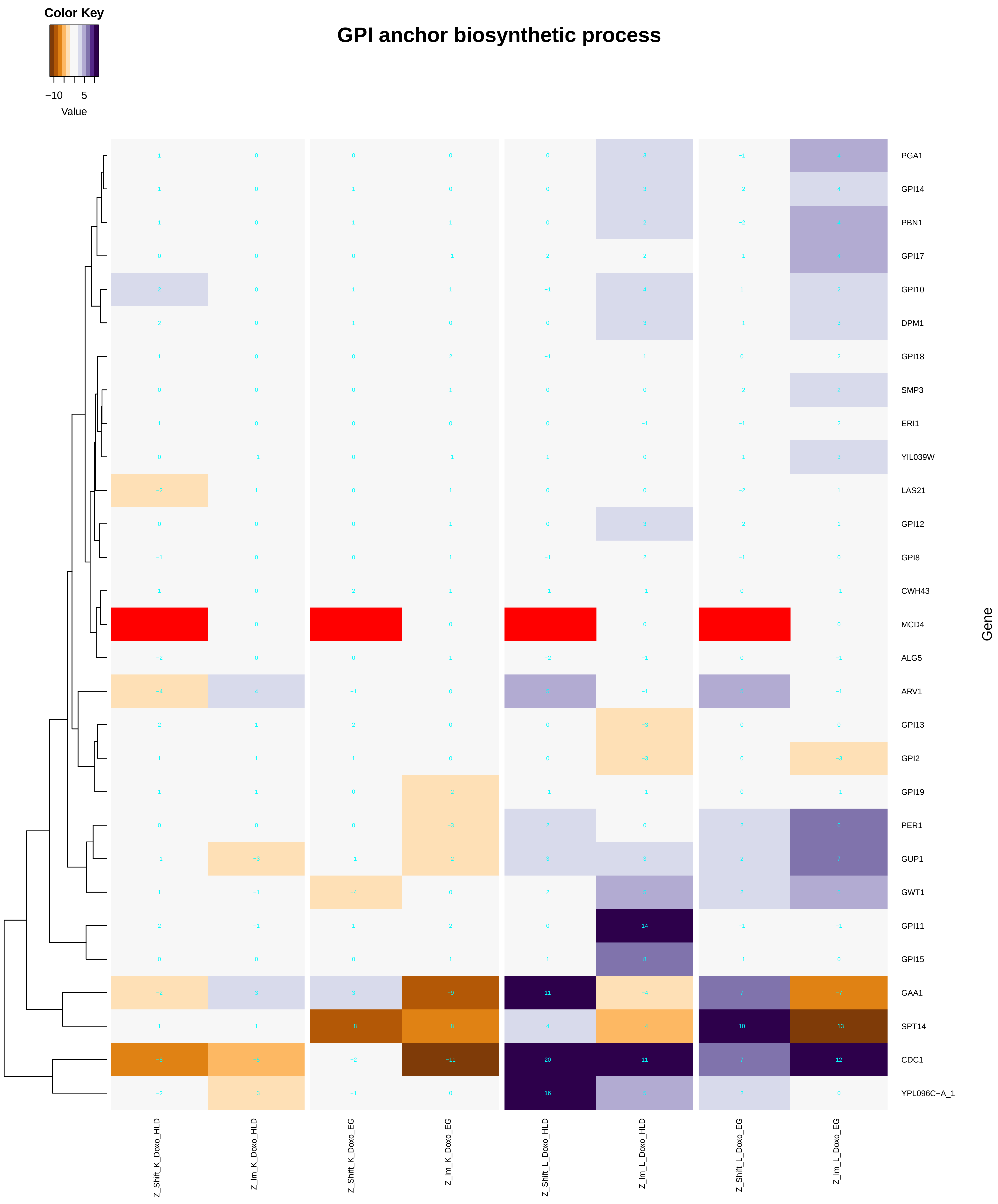

### GPI_anchor_metabolic_process.pdf

#### GPI anchor metabolic process

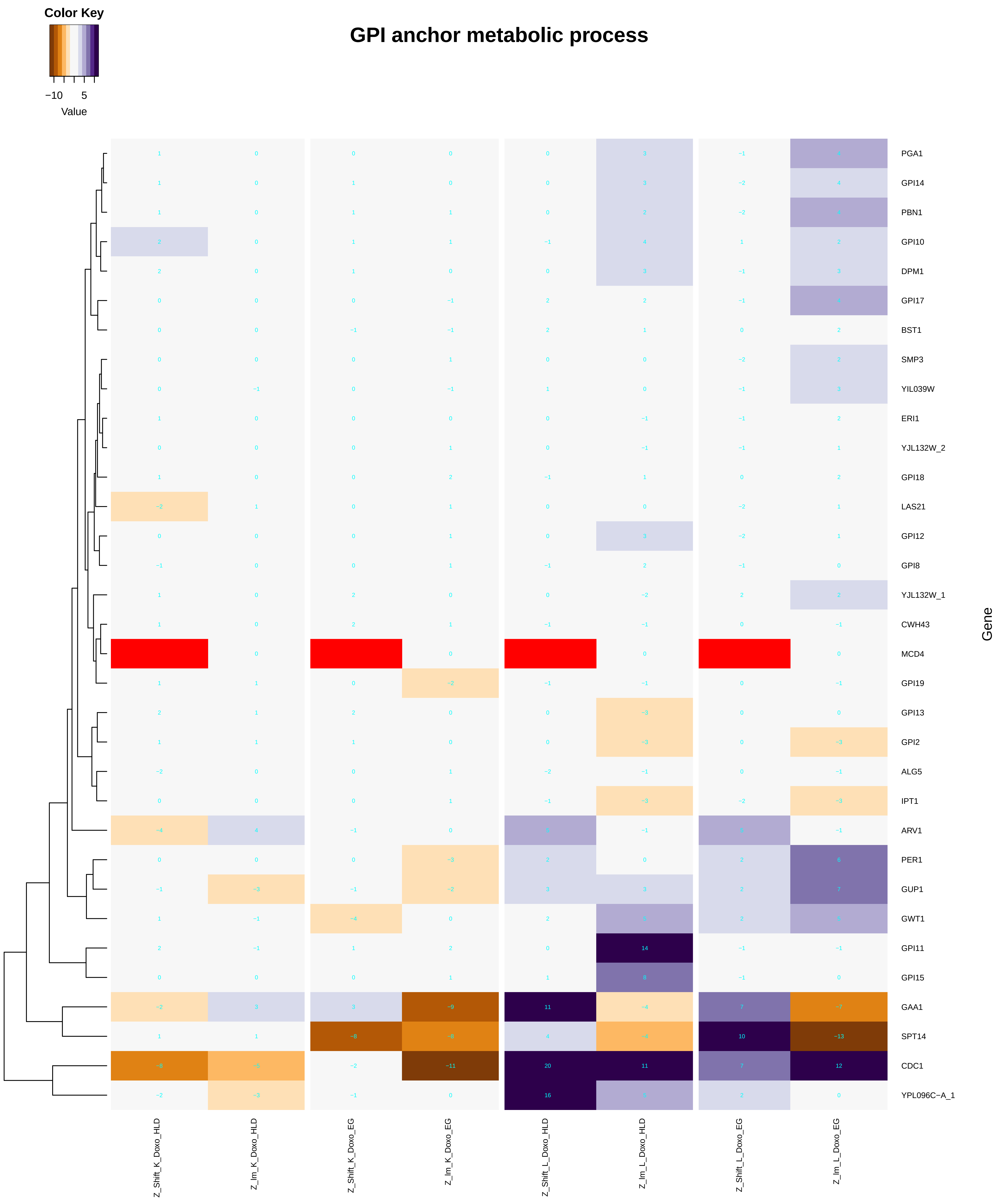

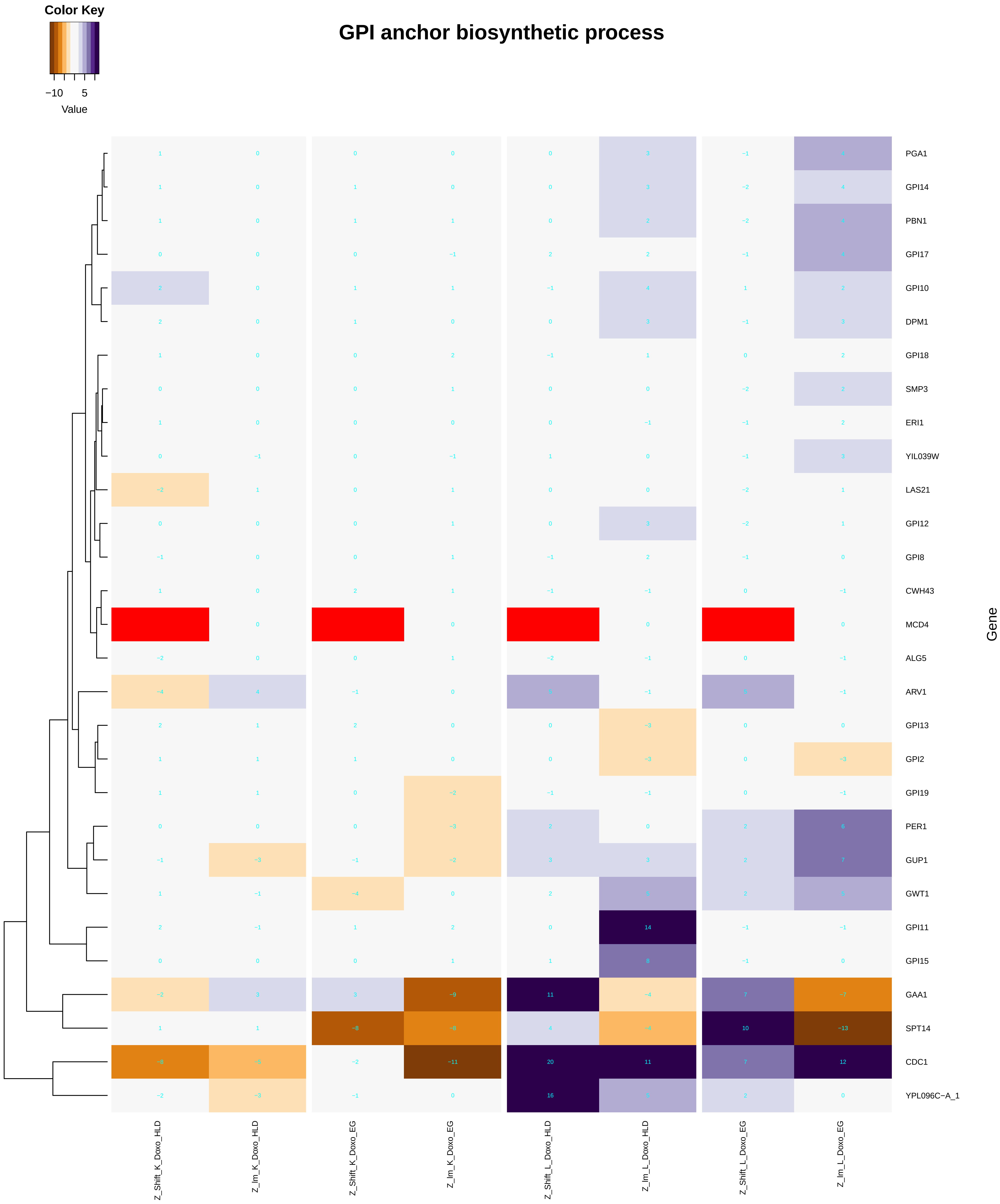

### heteroduplex_formation.pdf

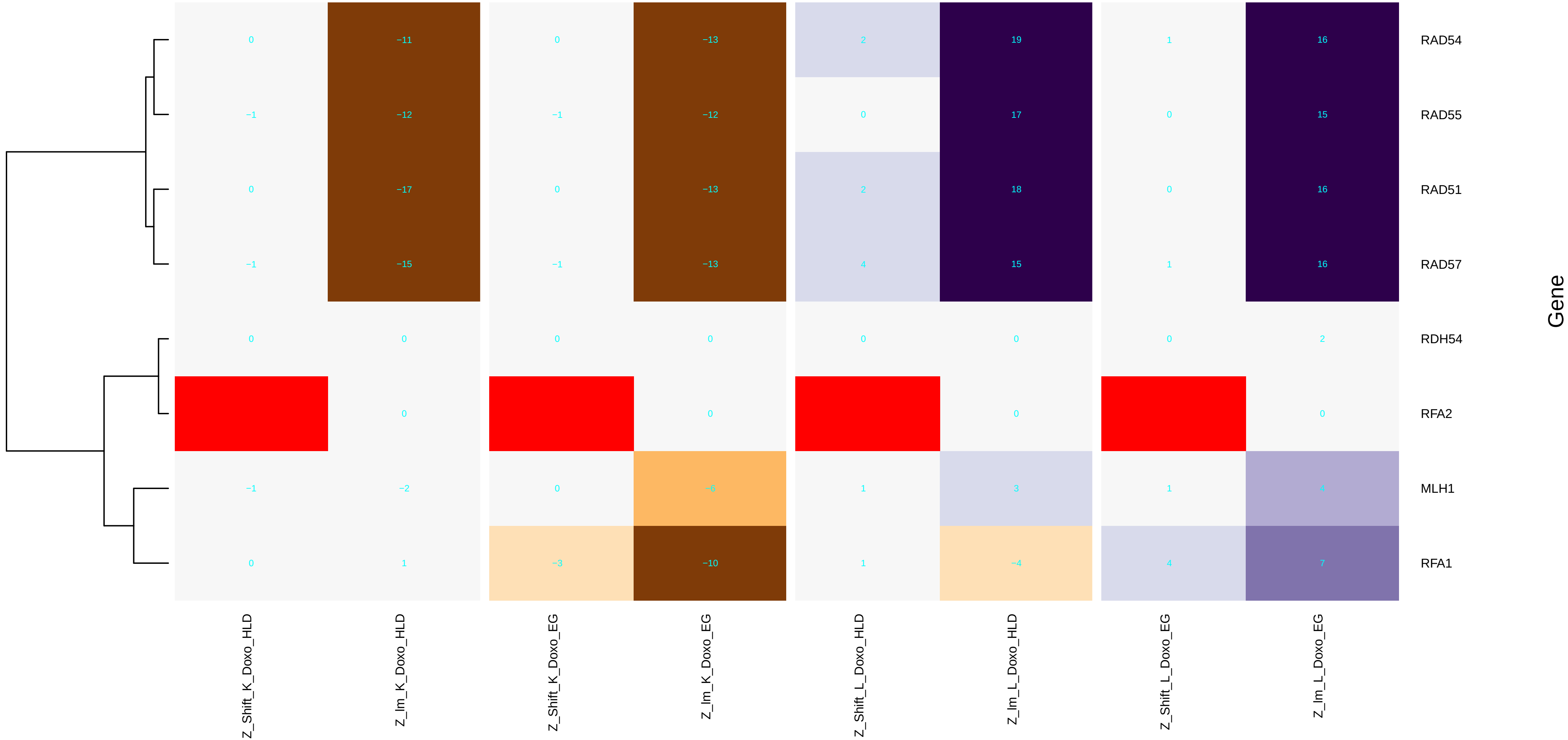

### histone_acetylation.pdf

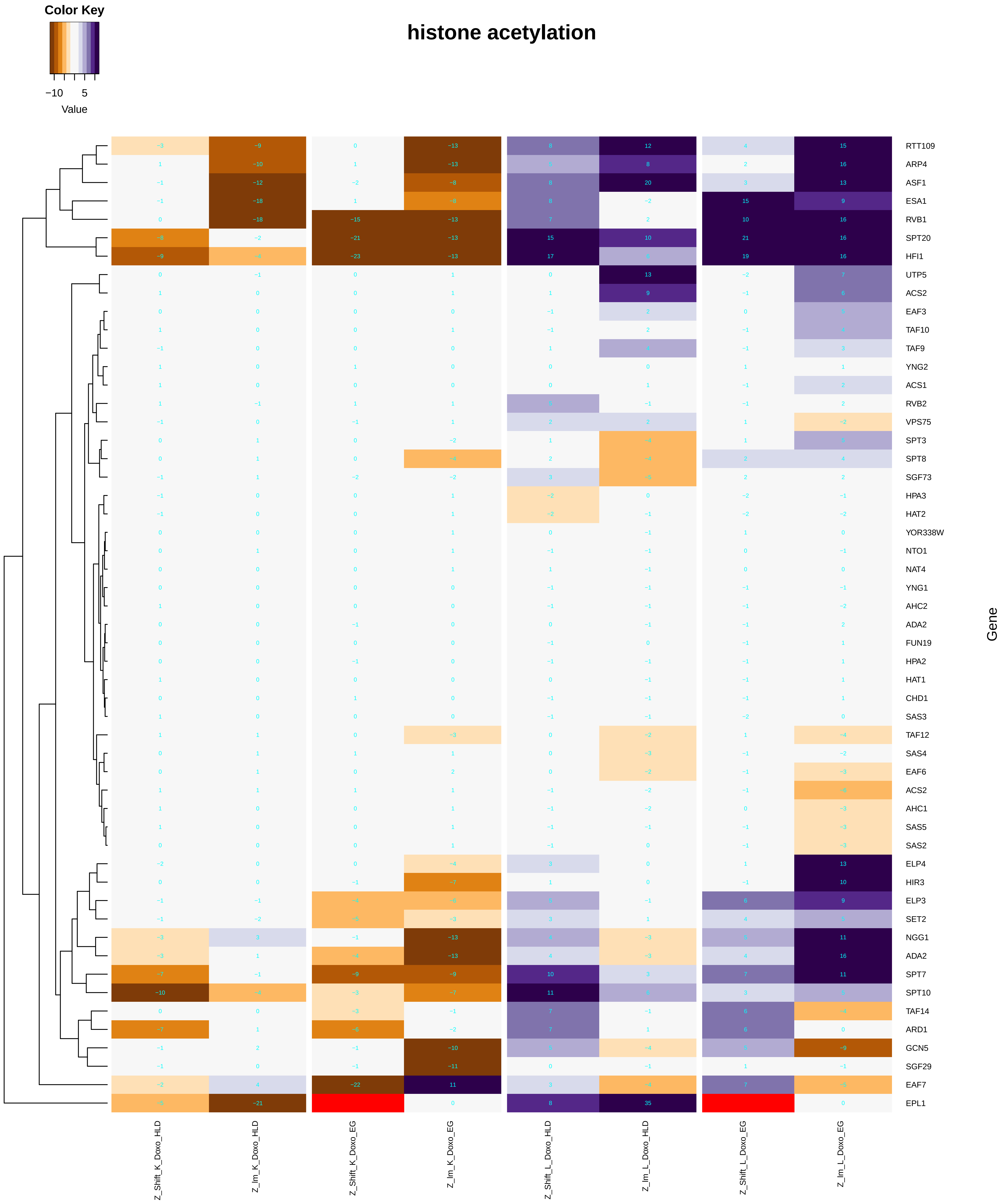

#### histone H3 acetylation

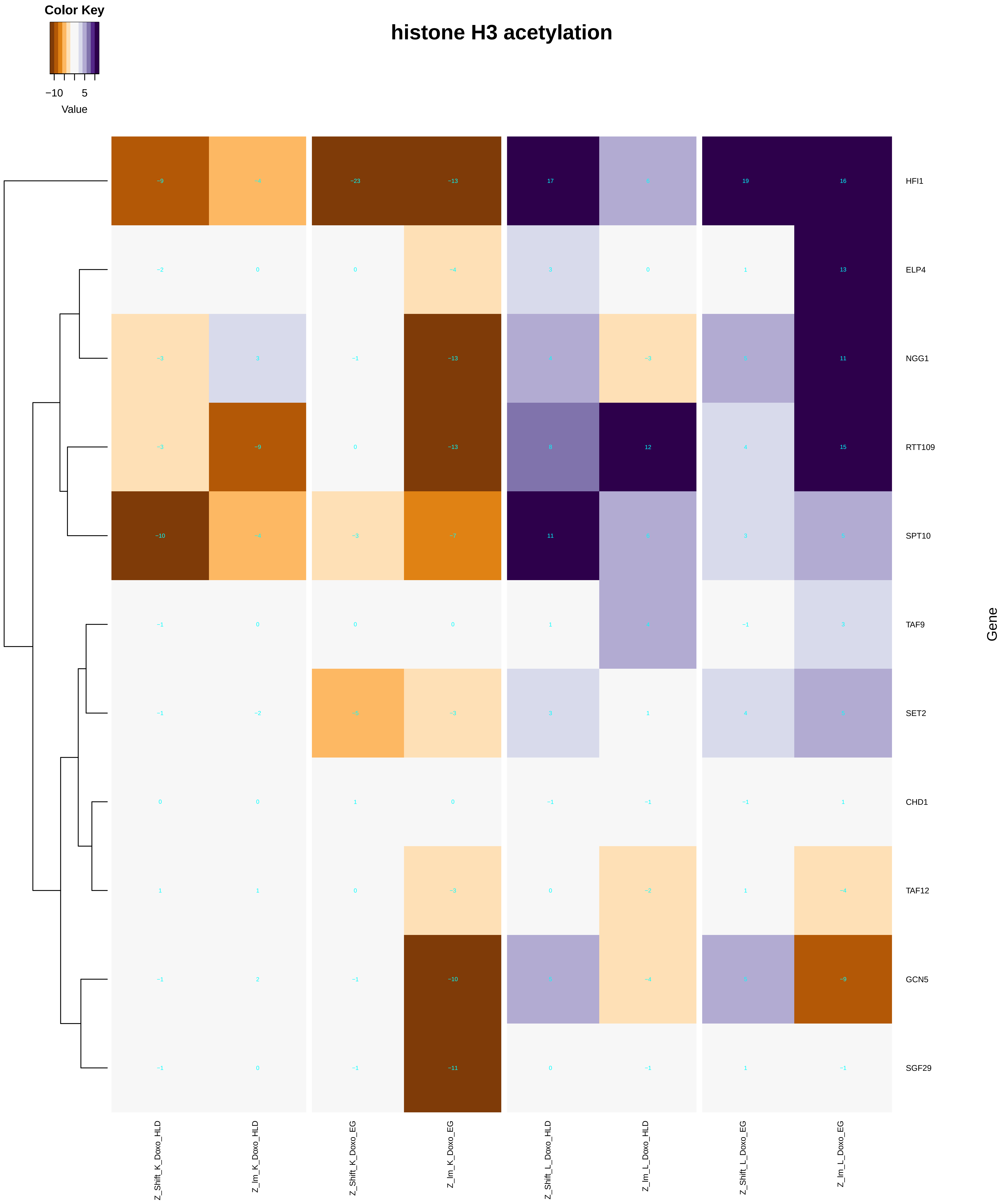

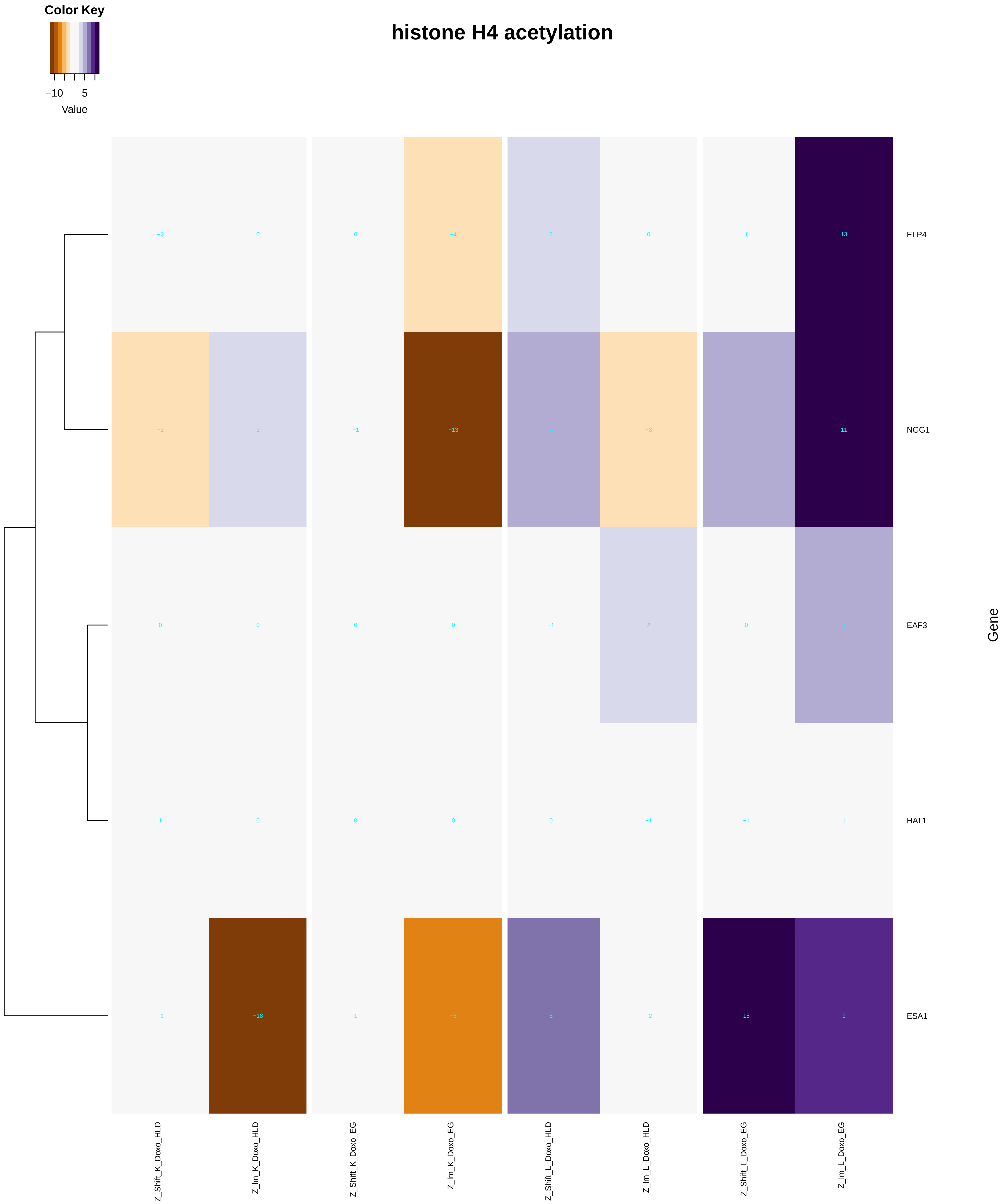

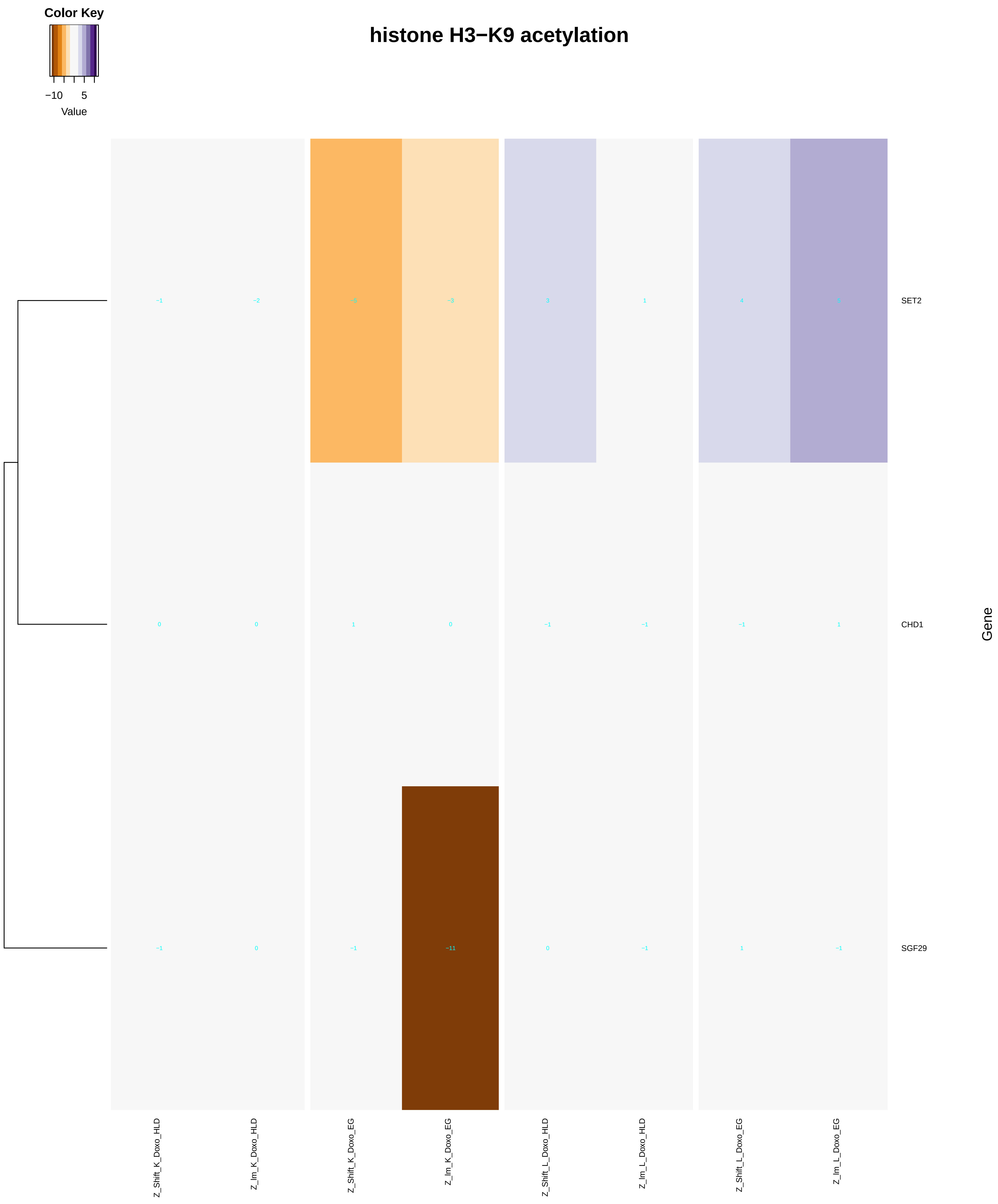

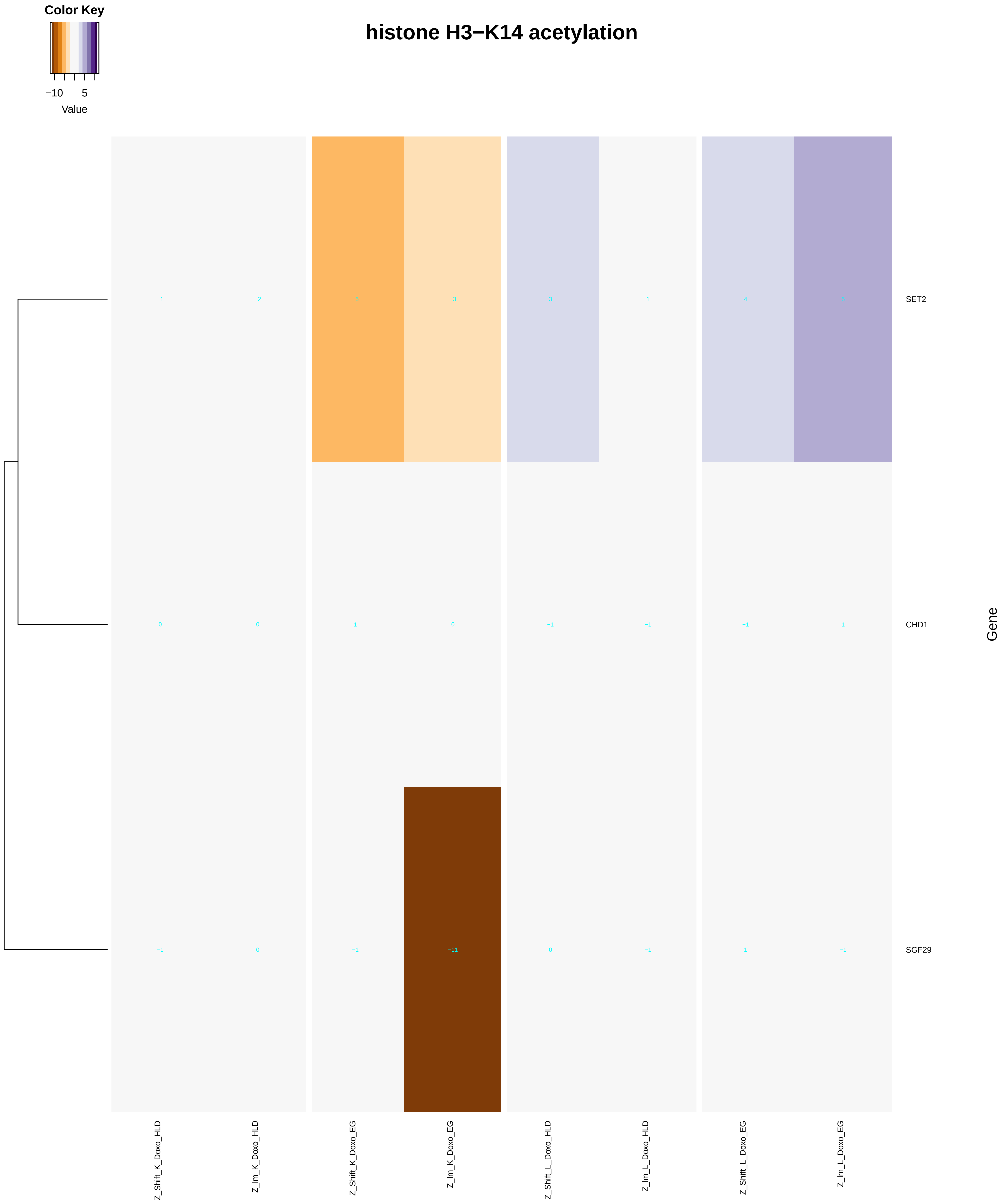

### histone_exchange.pdf

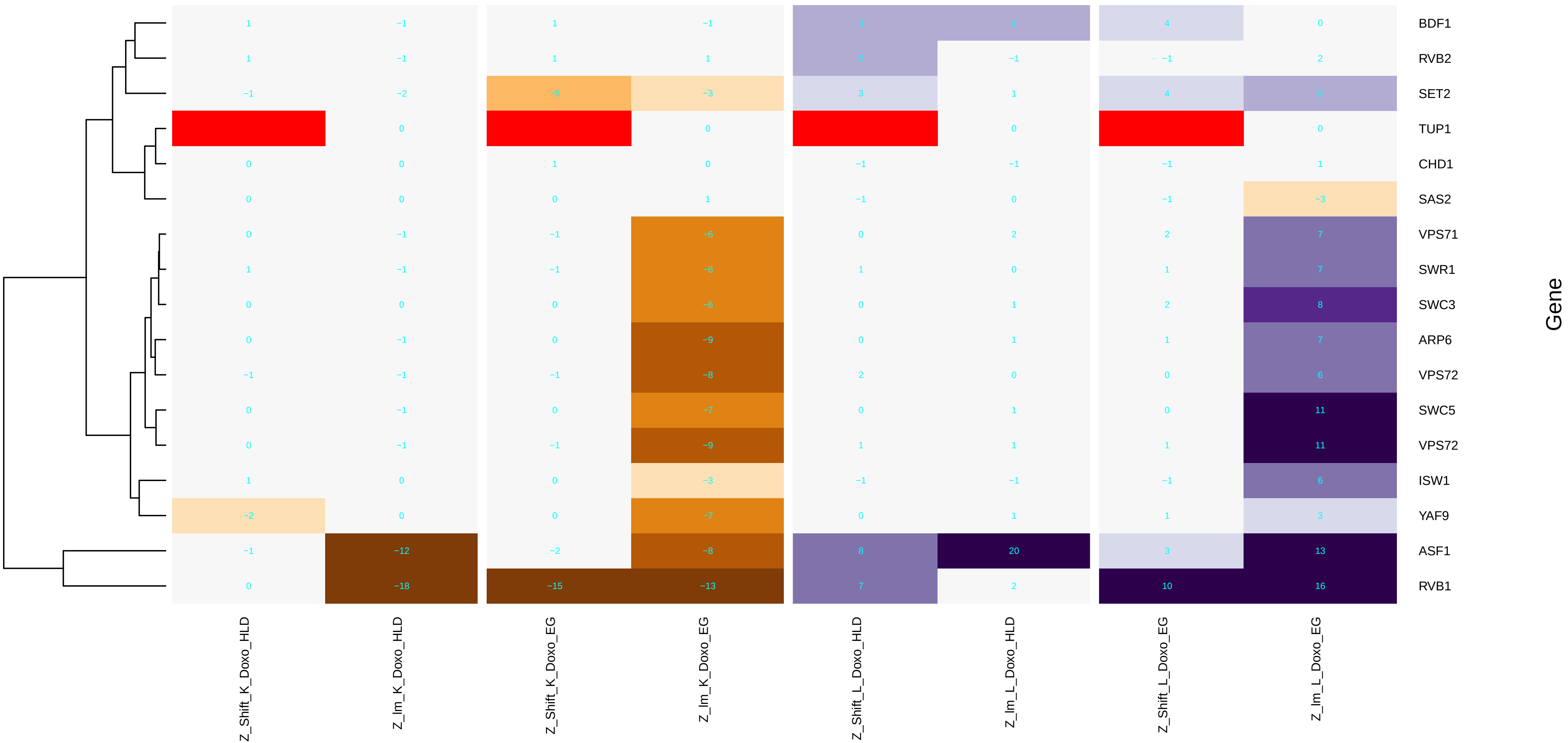

### histone_methylation.pdf

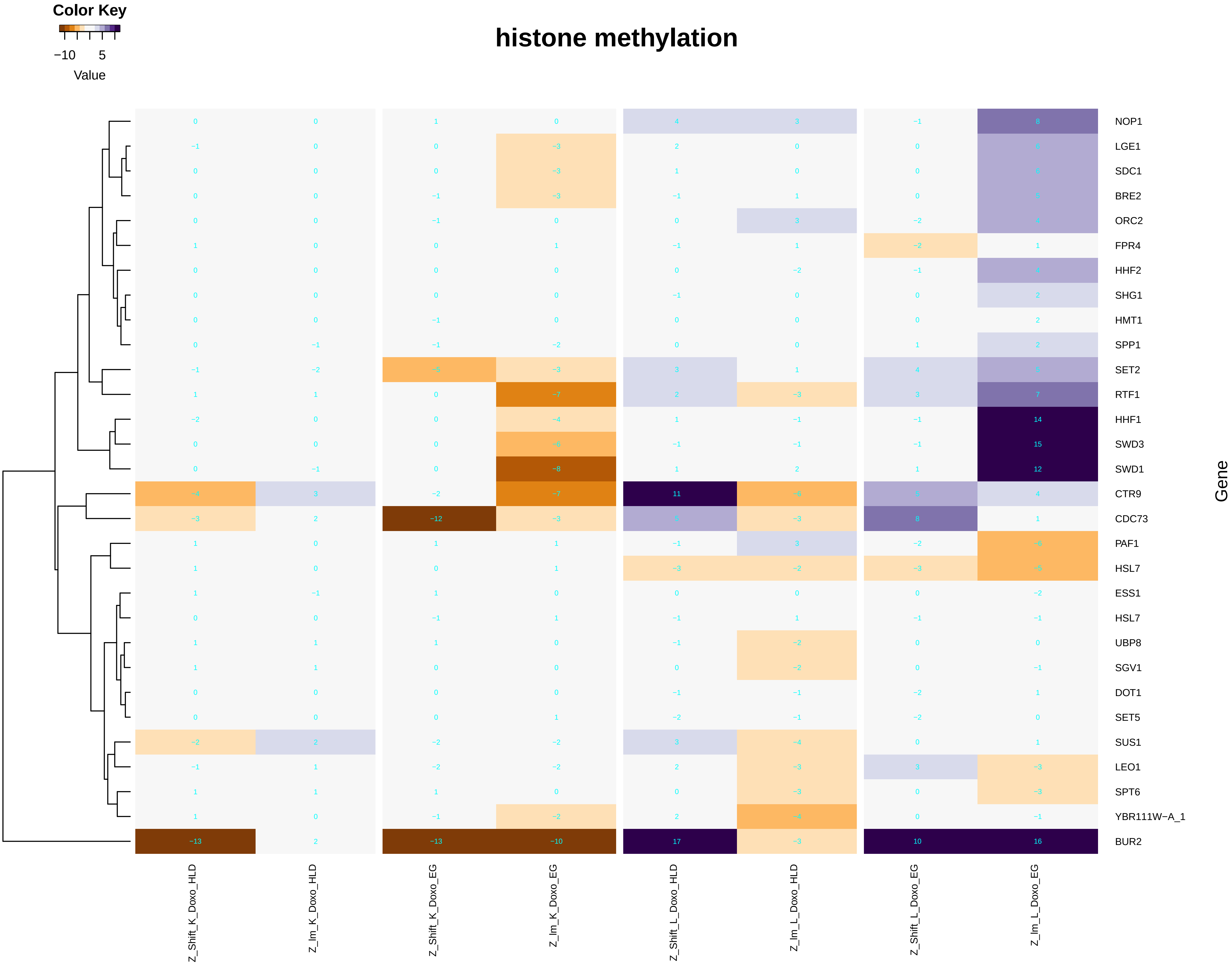

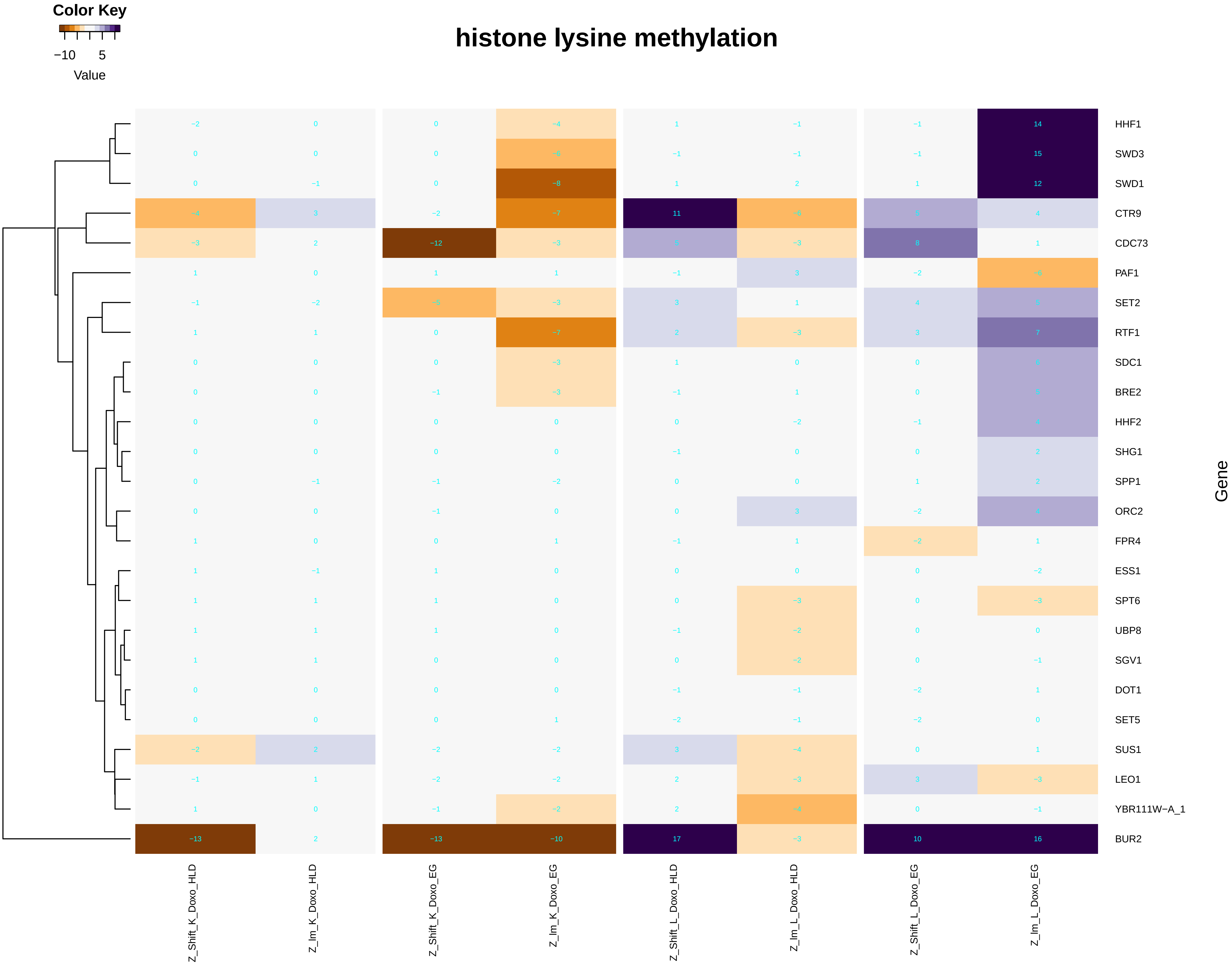

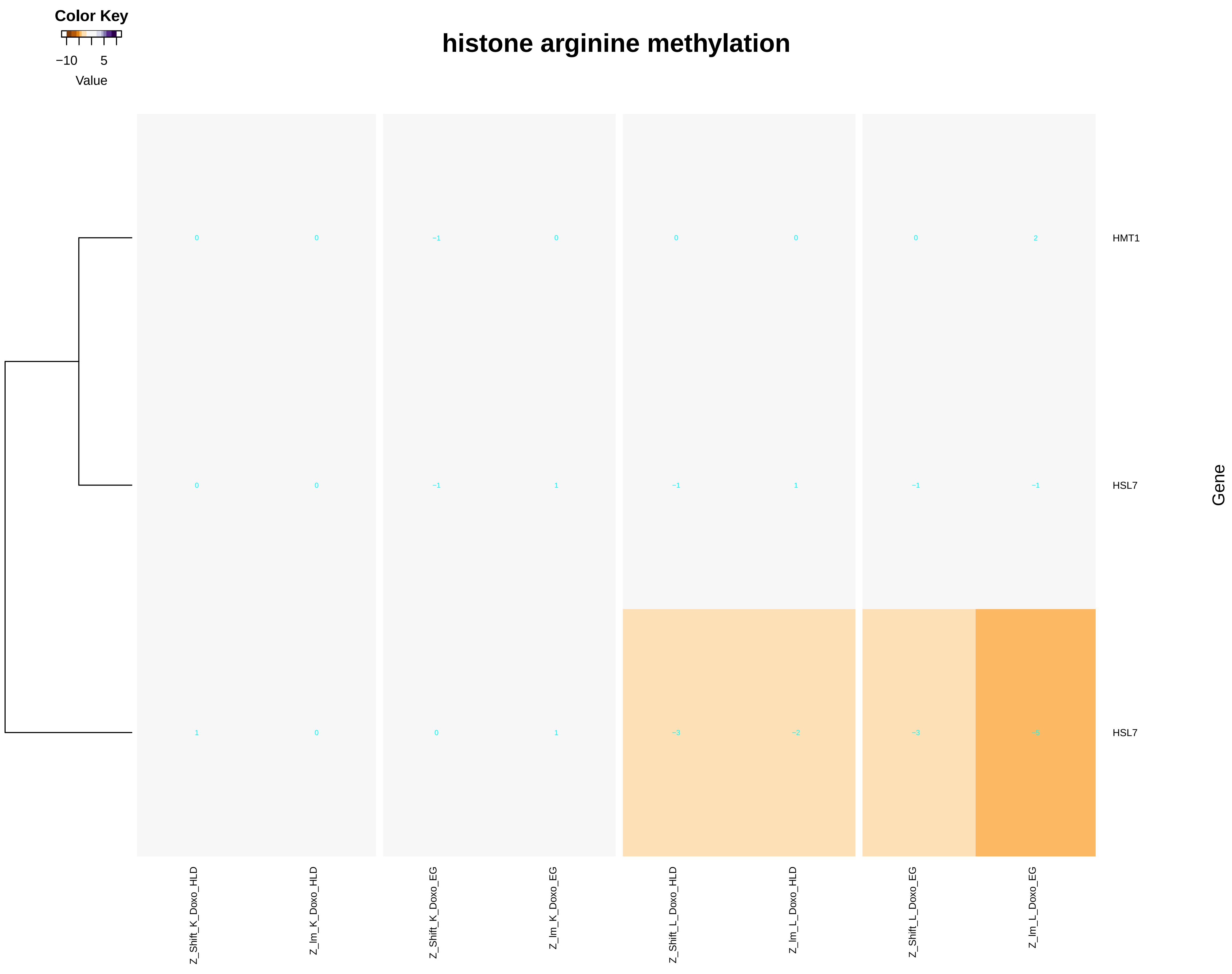

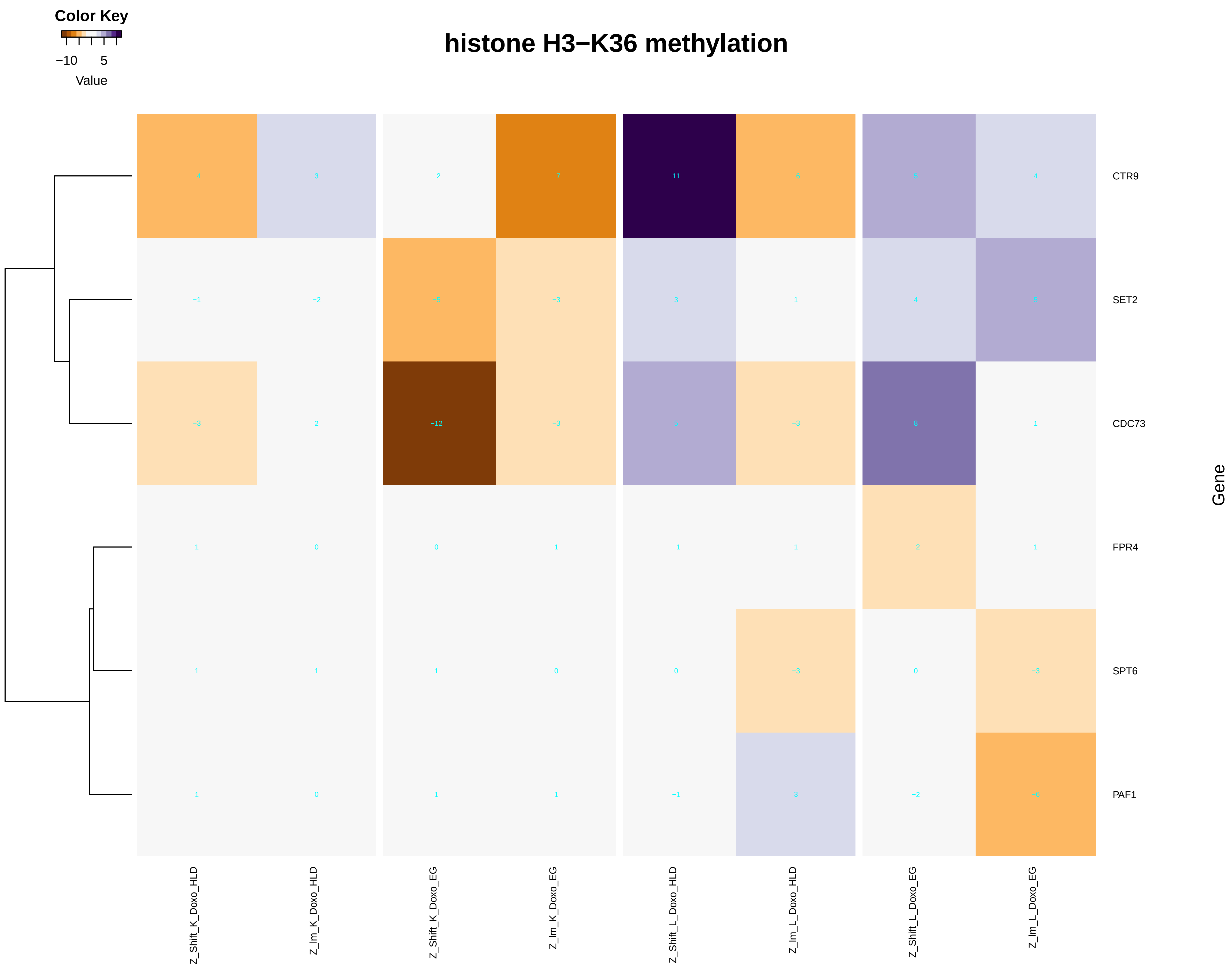

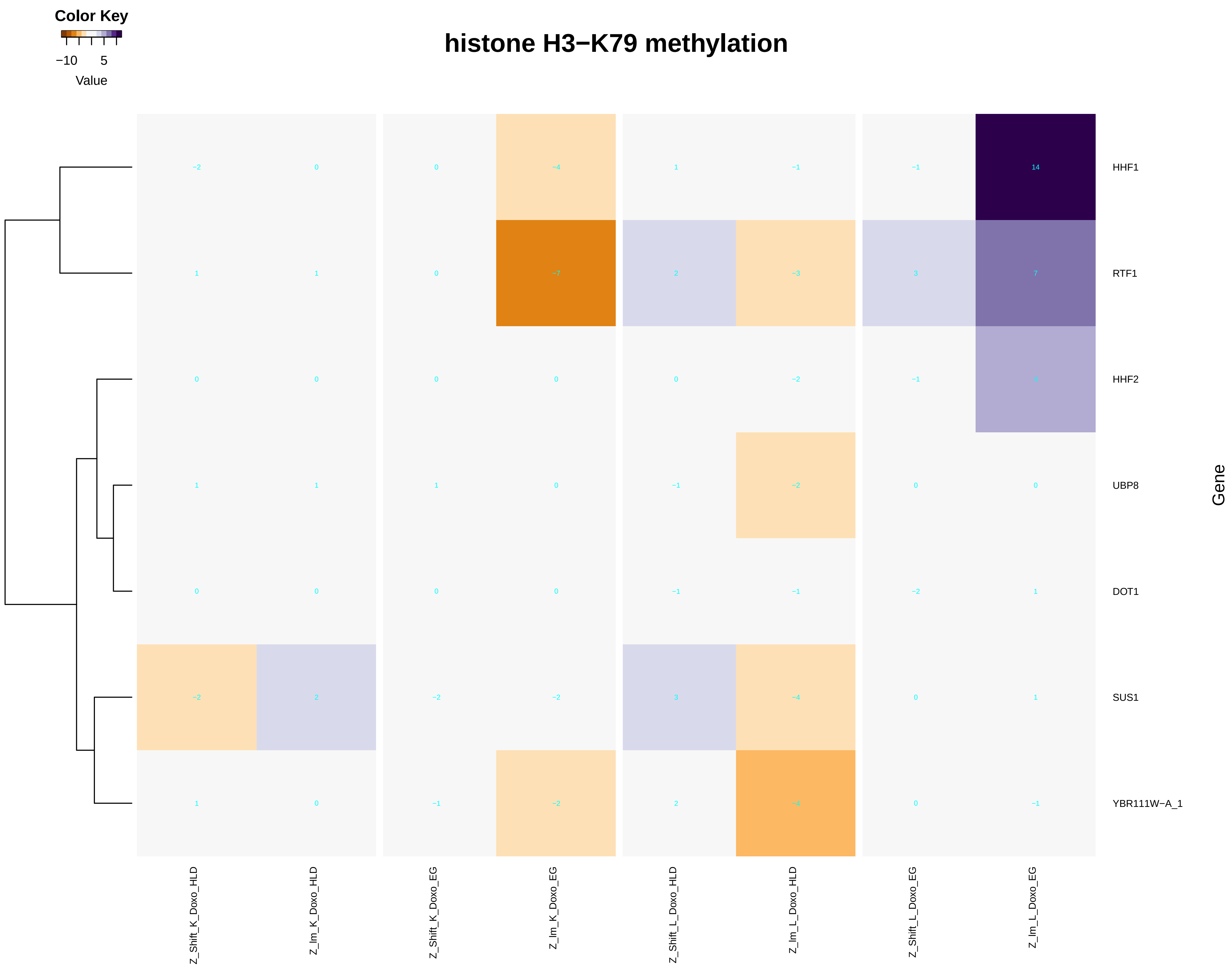

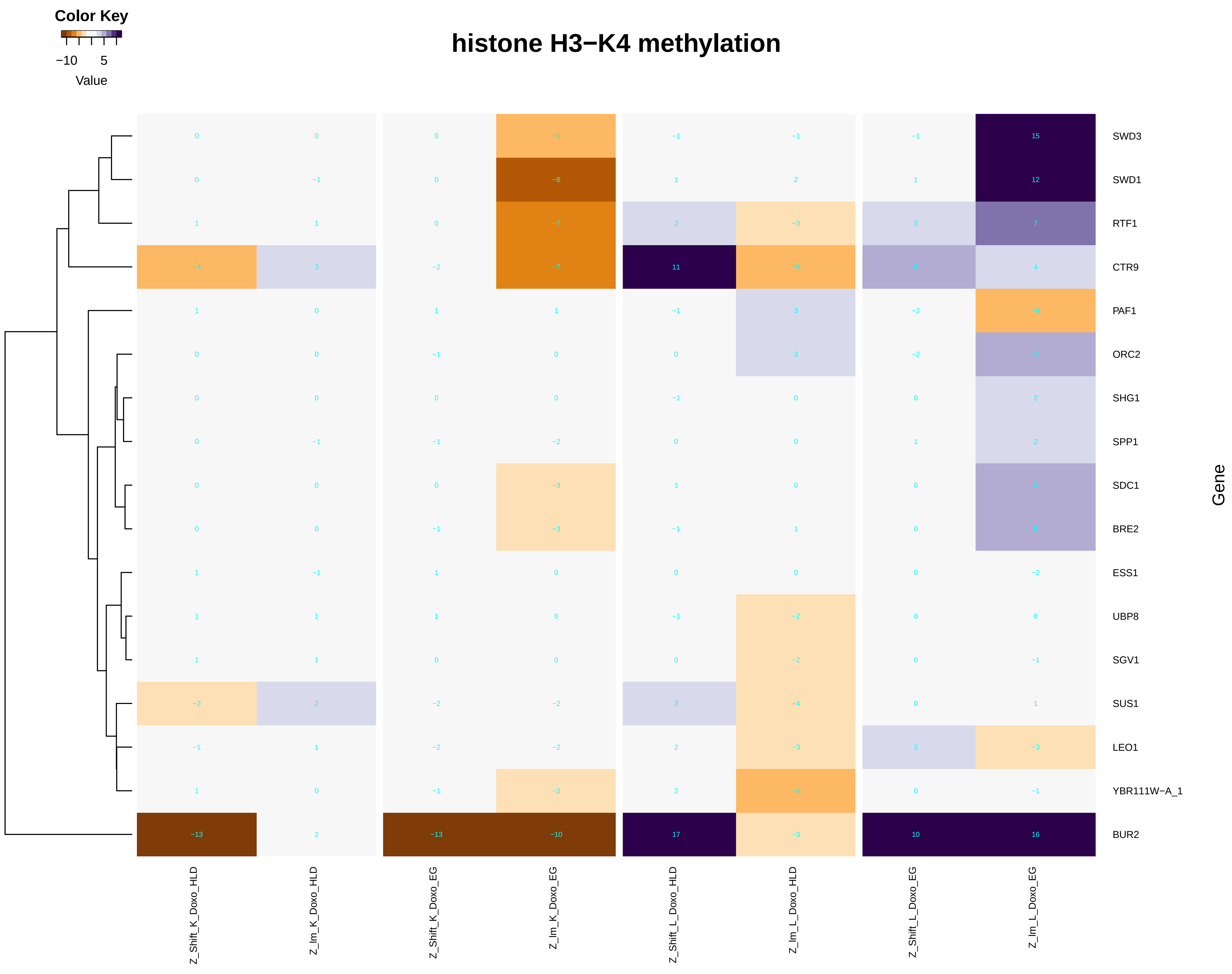

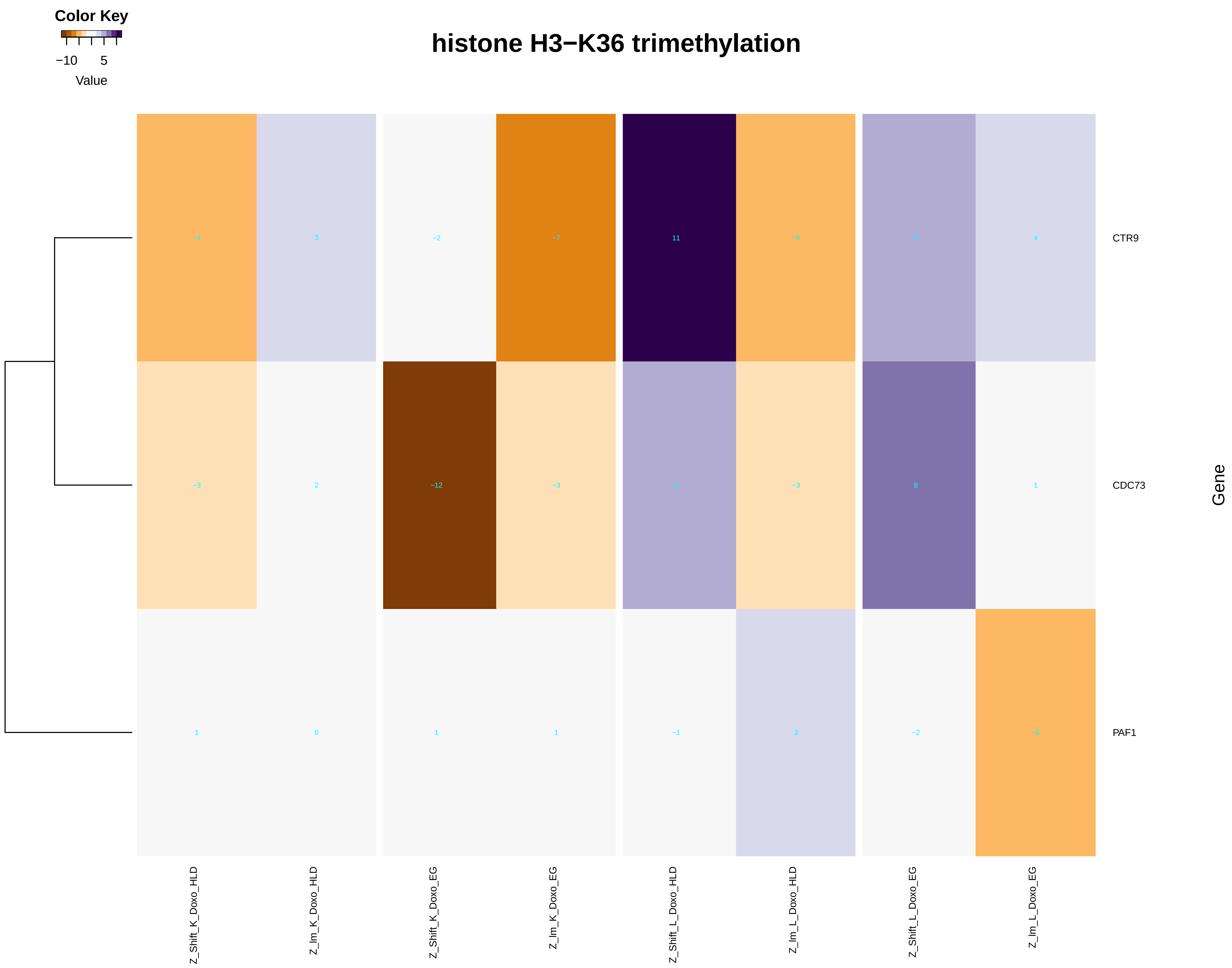

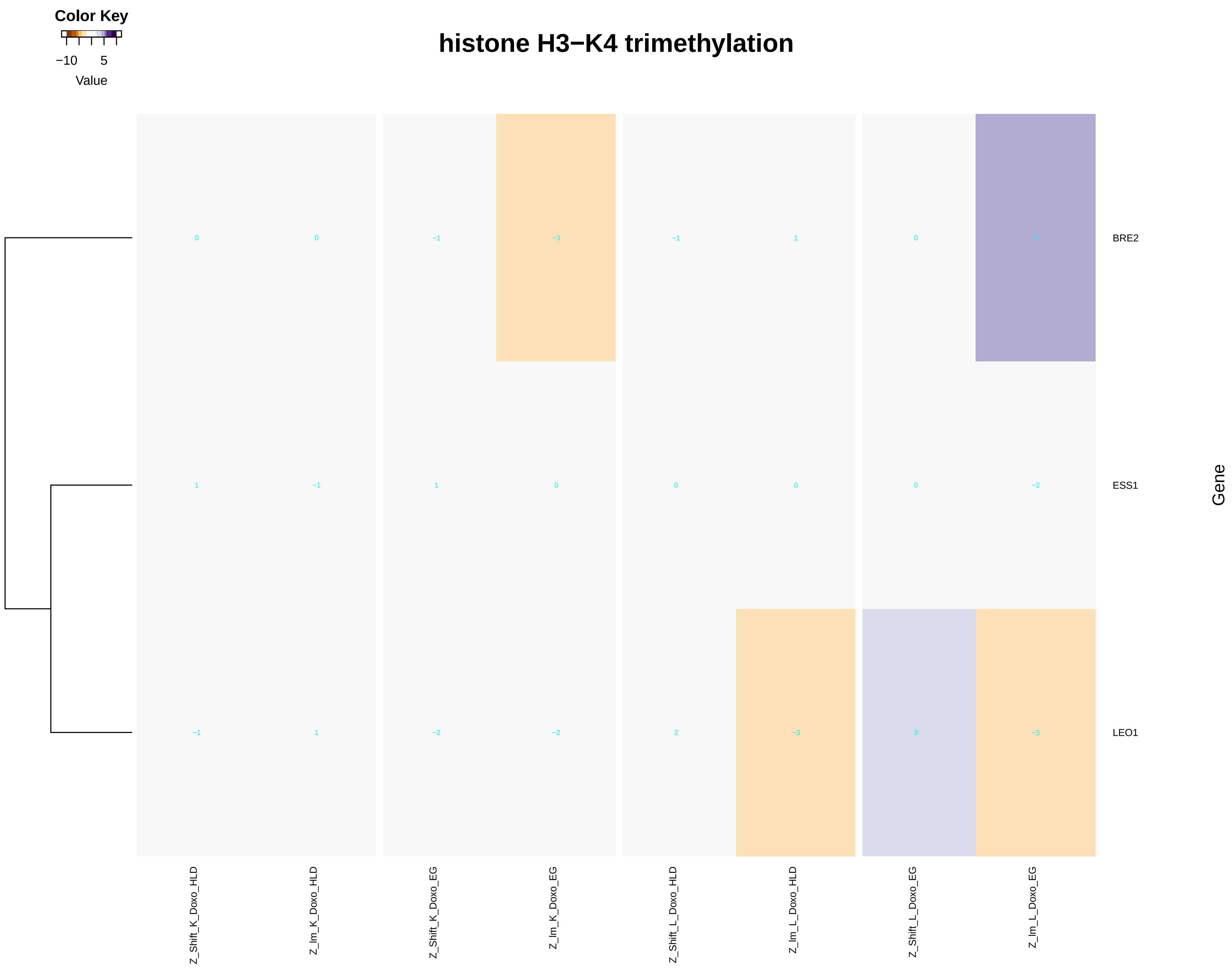

### hydrogen_ion_transmembrane_transport.pdf

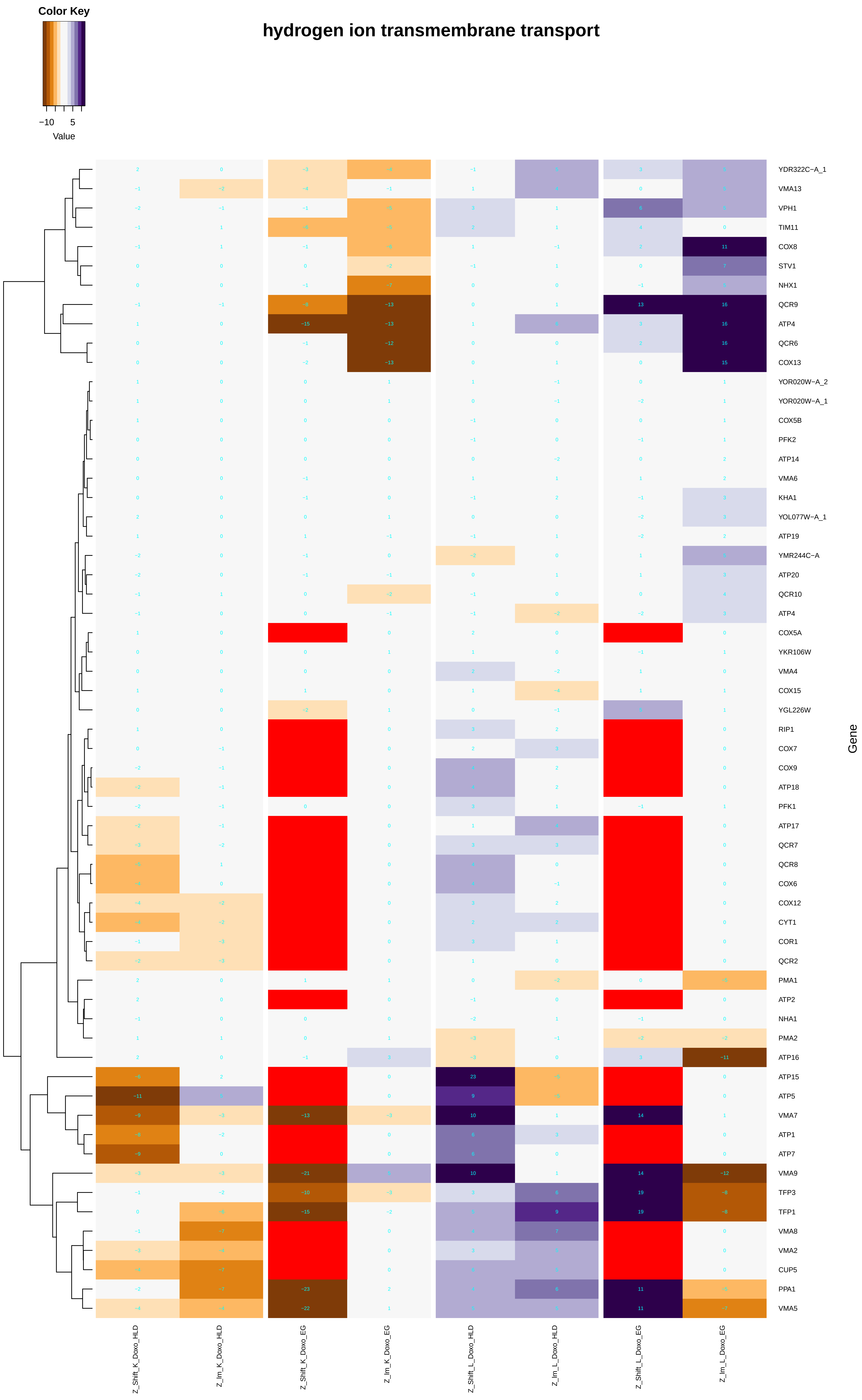

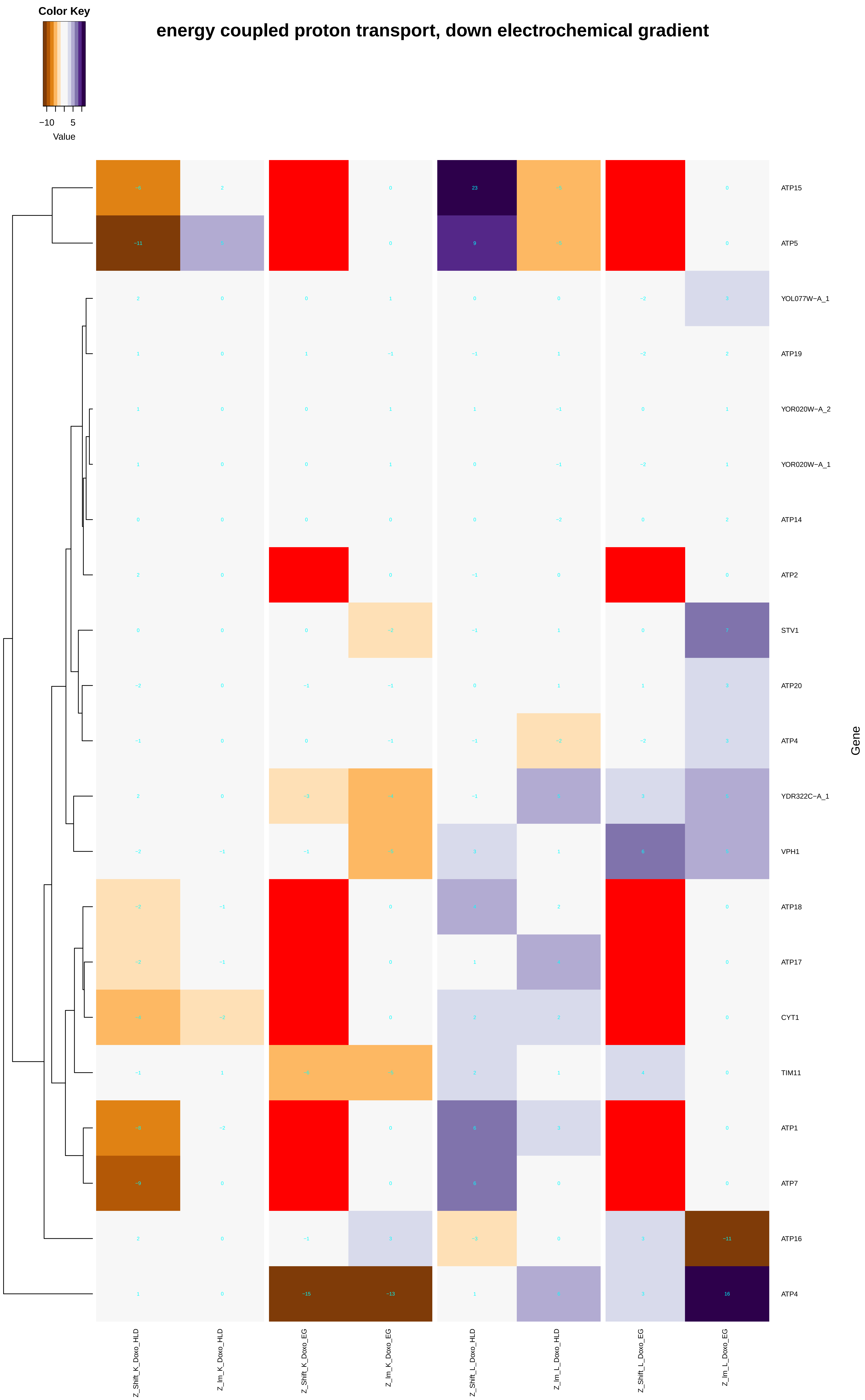

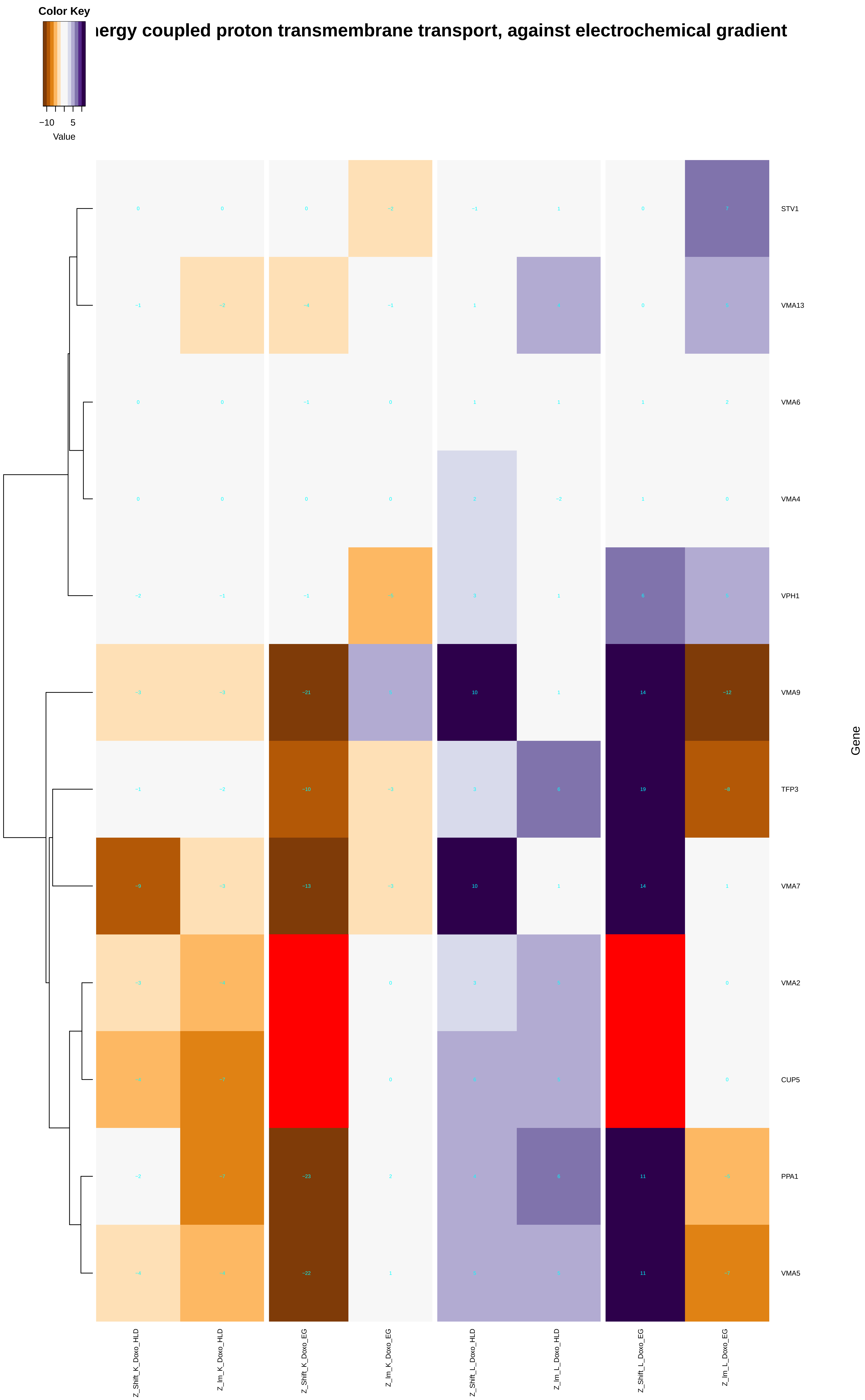

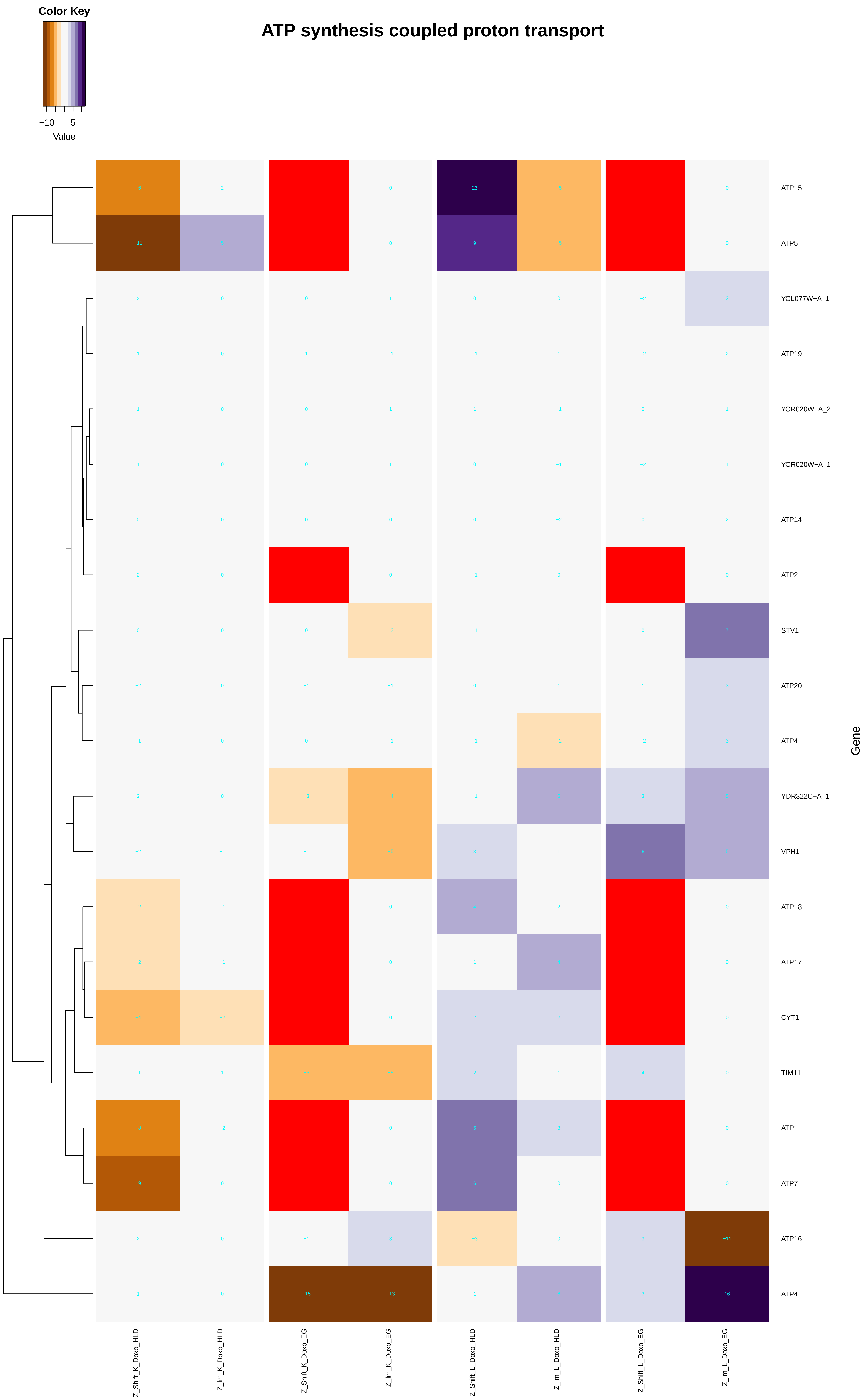

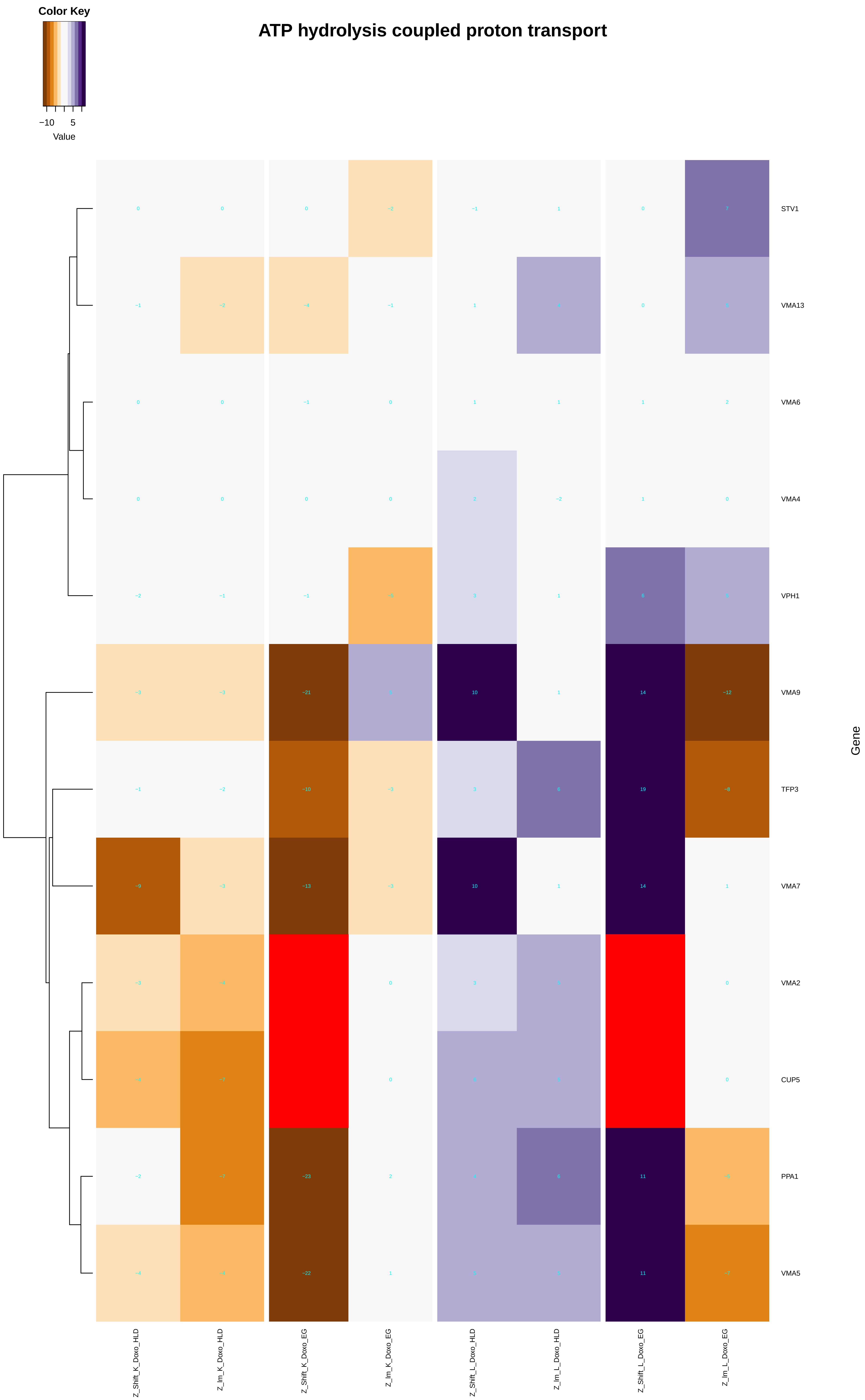

### hydrogen_transport.pdf

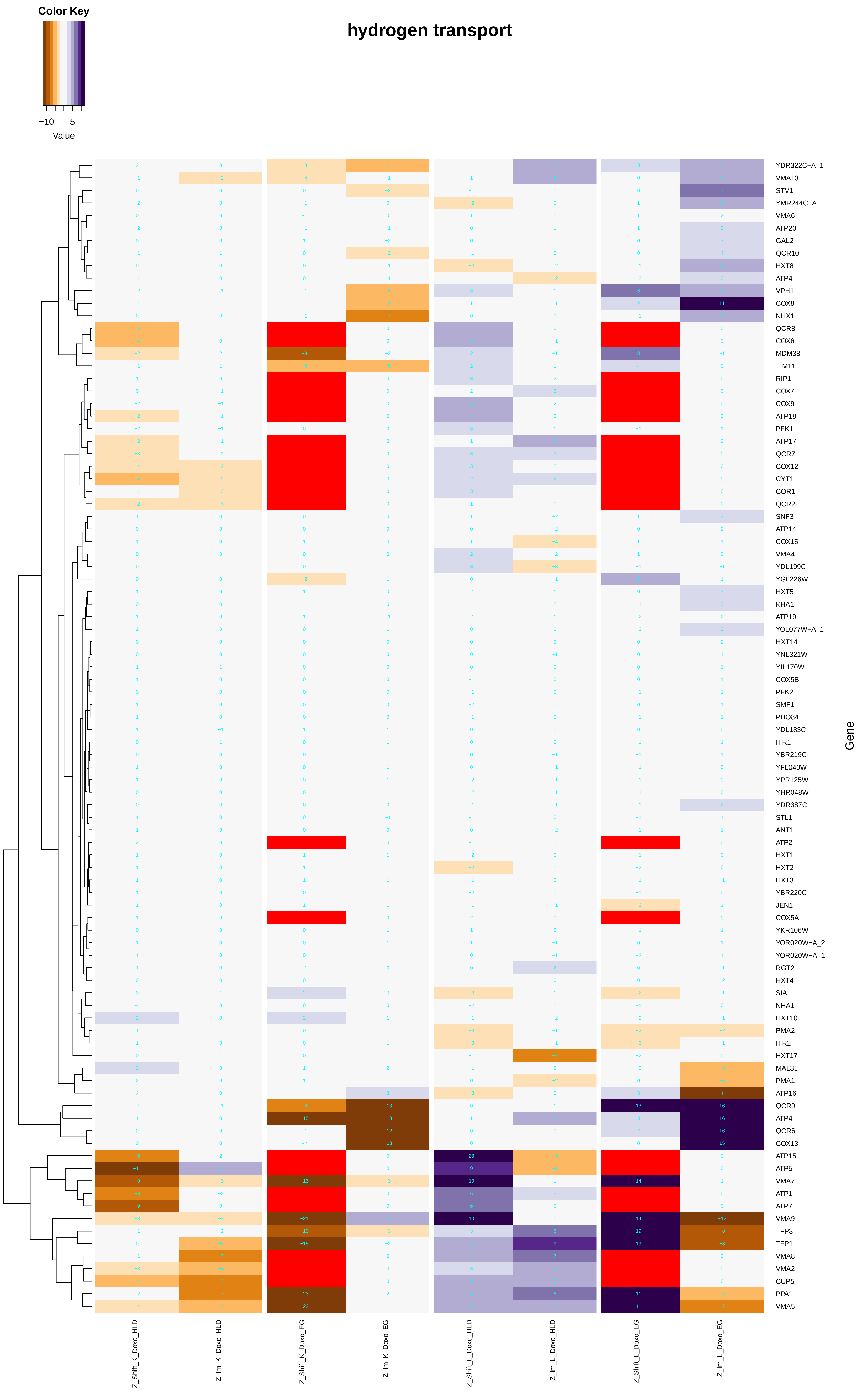

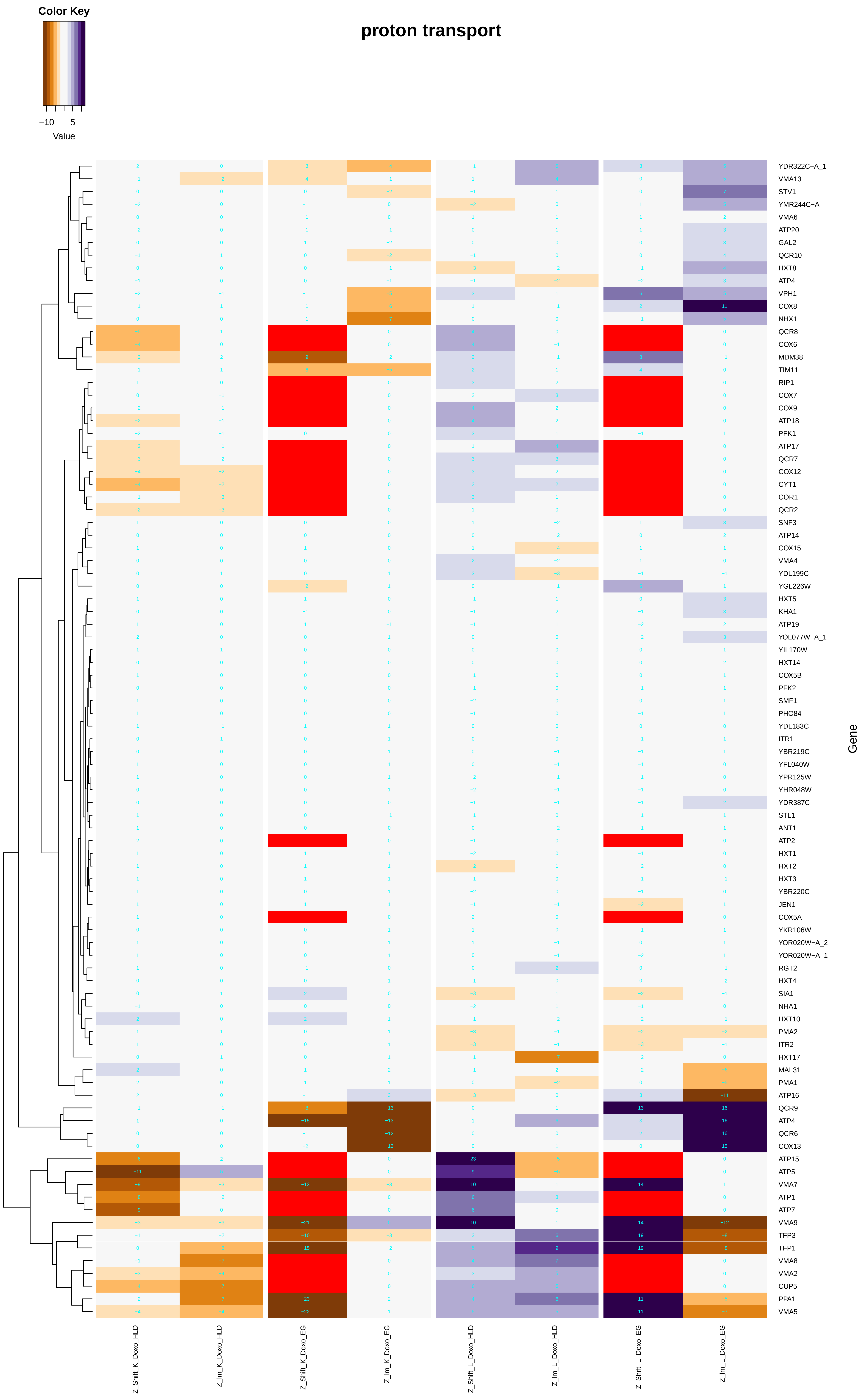

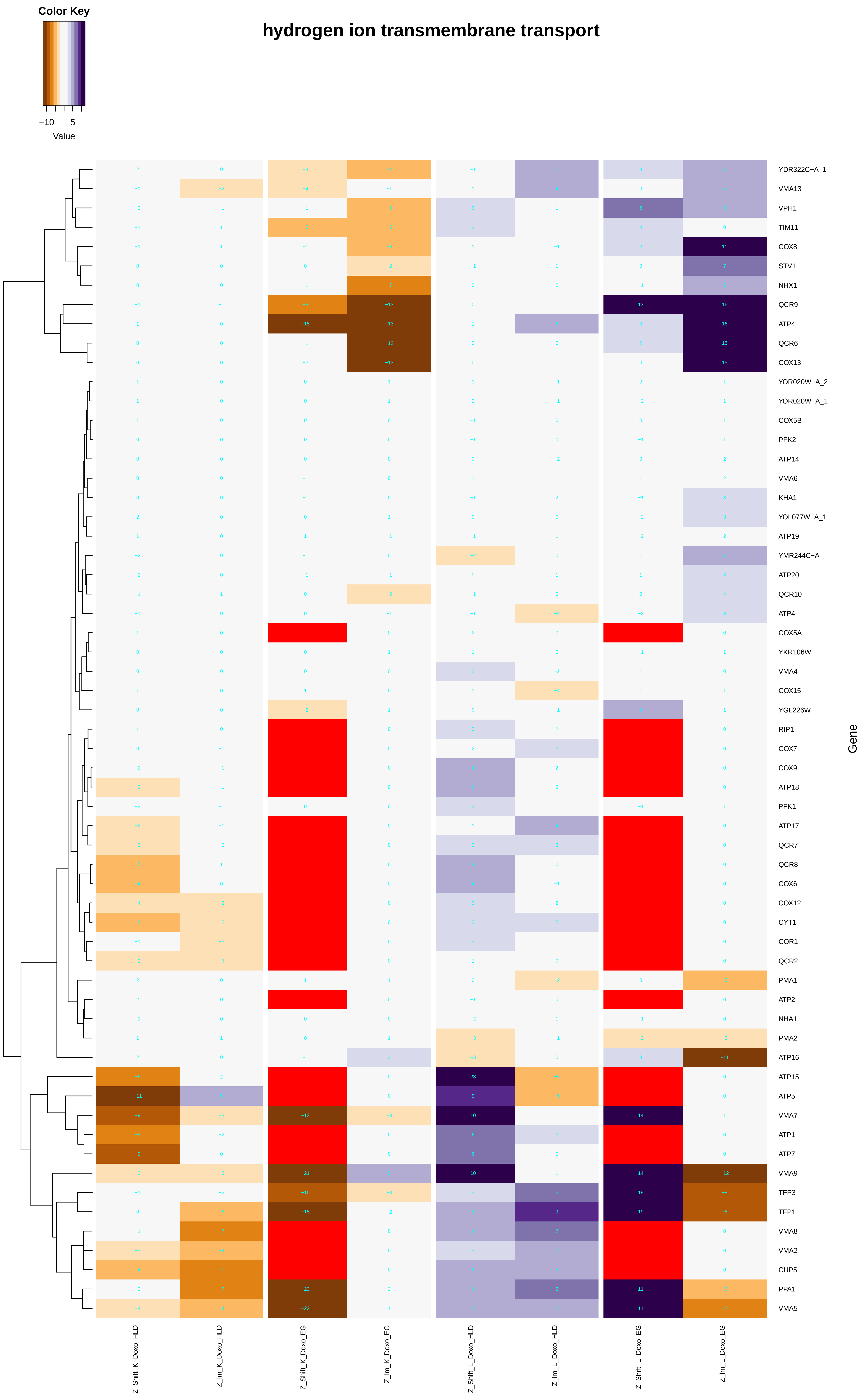

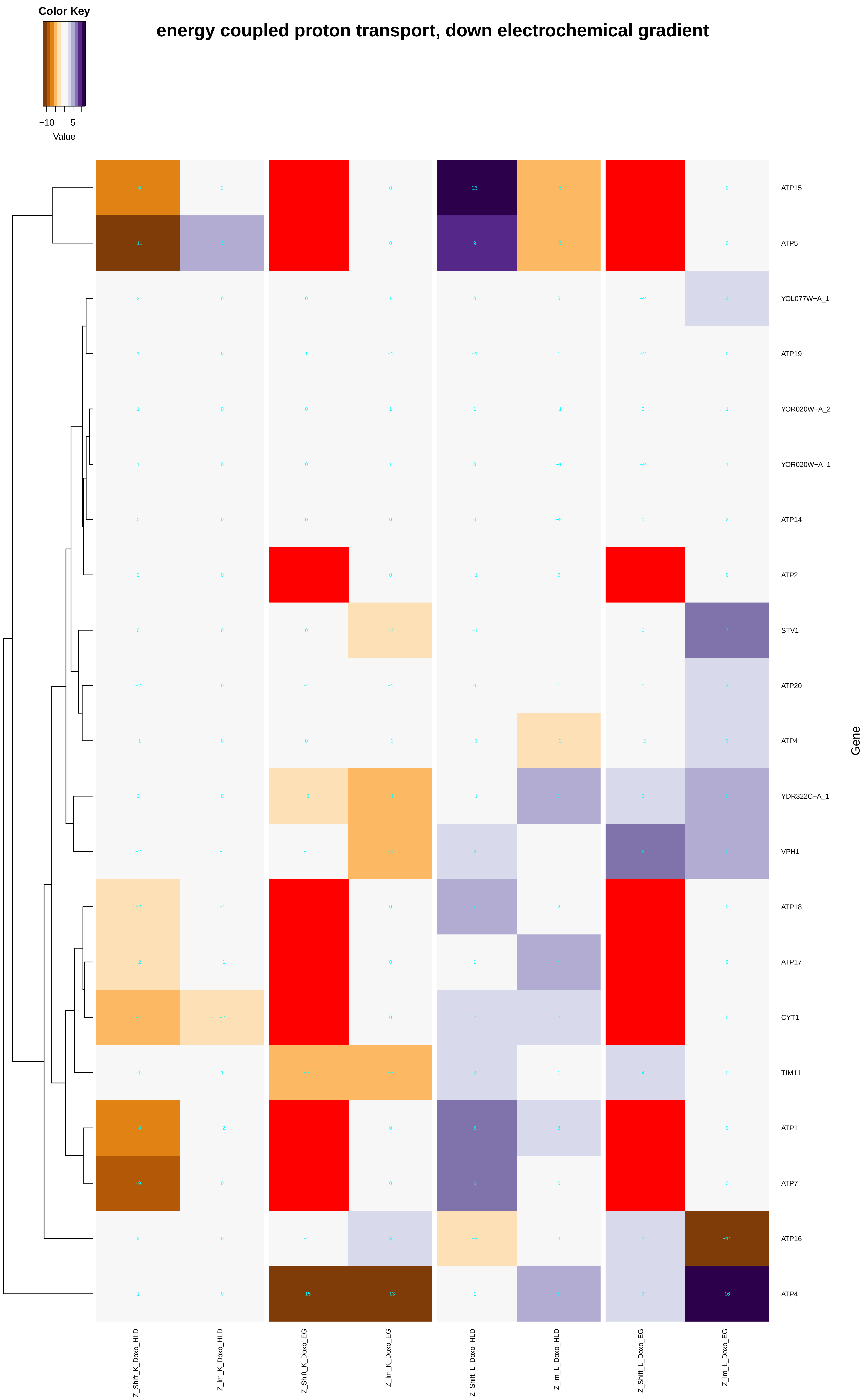

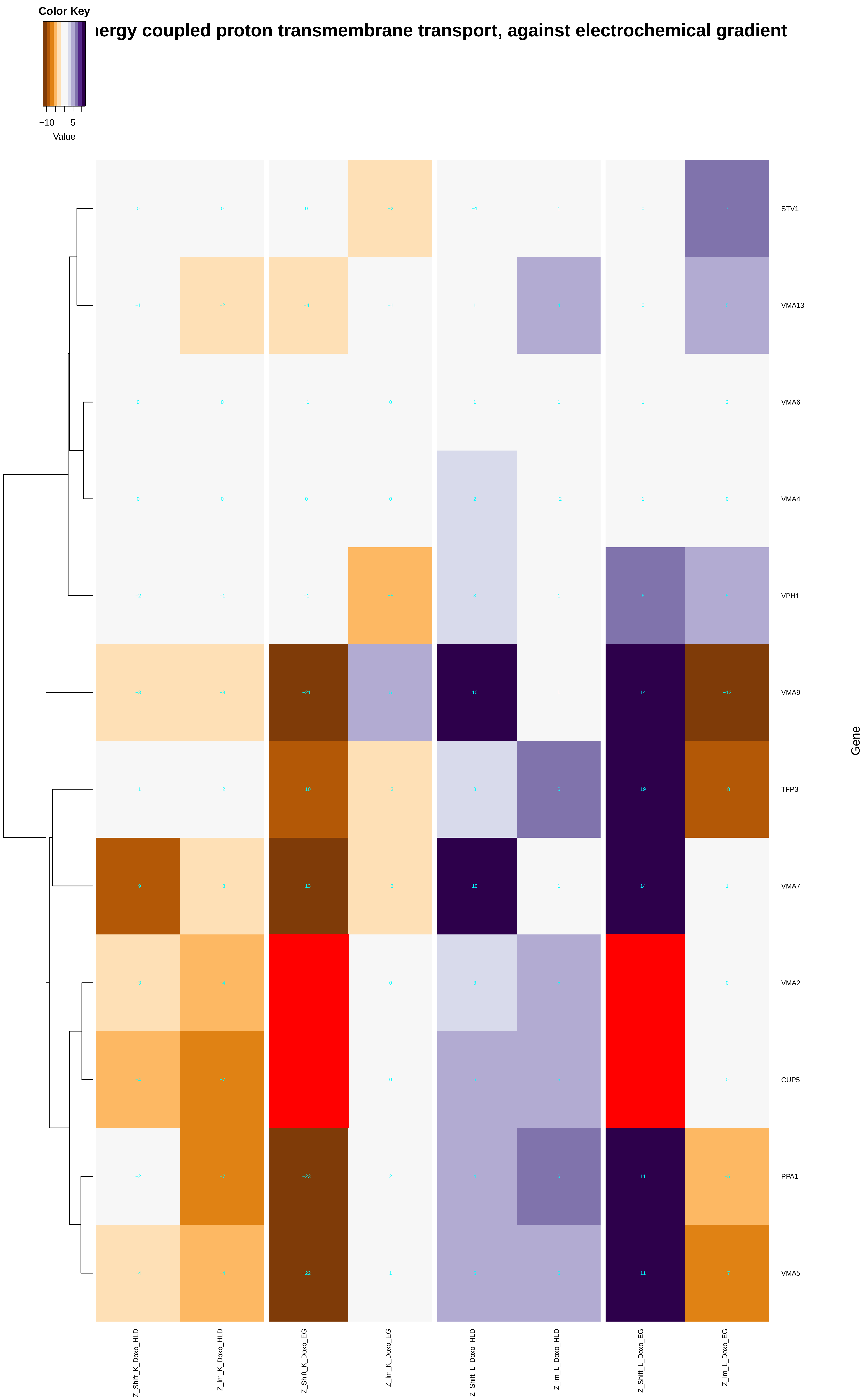

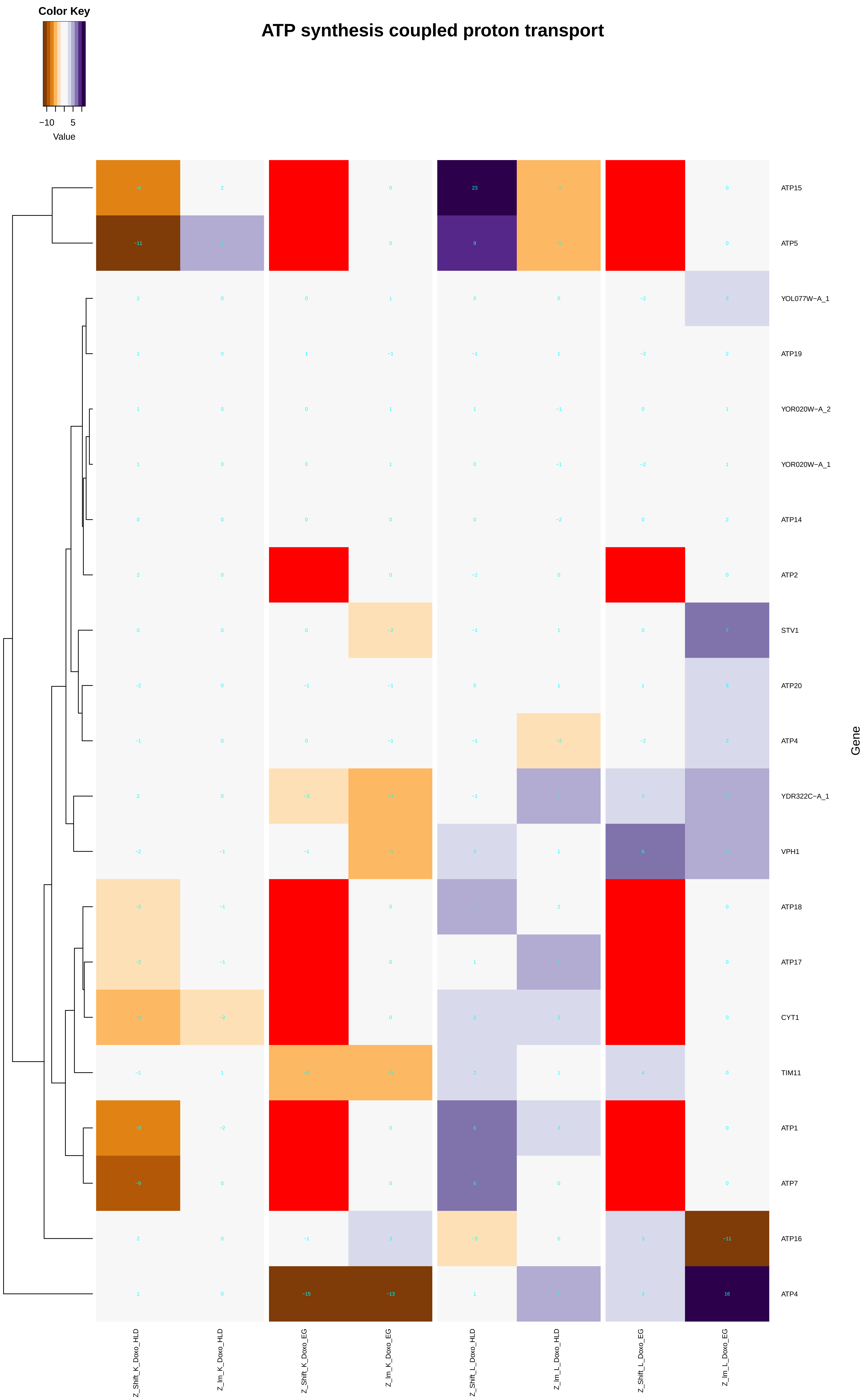

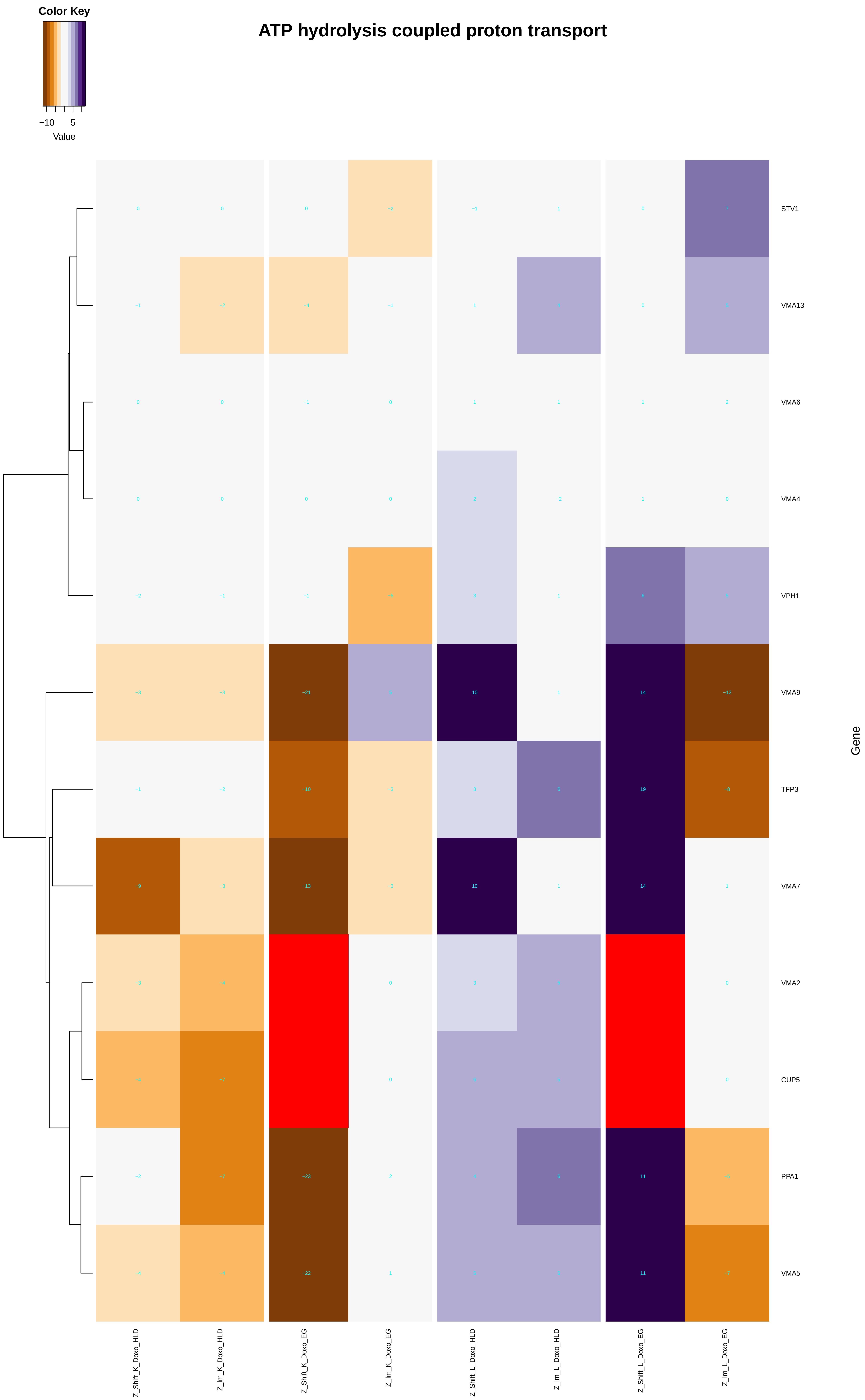

### internal_peptidyl-lysine_acetylation.pdf

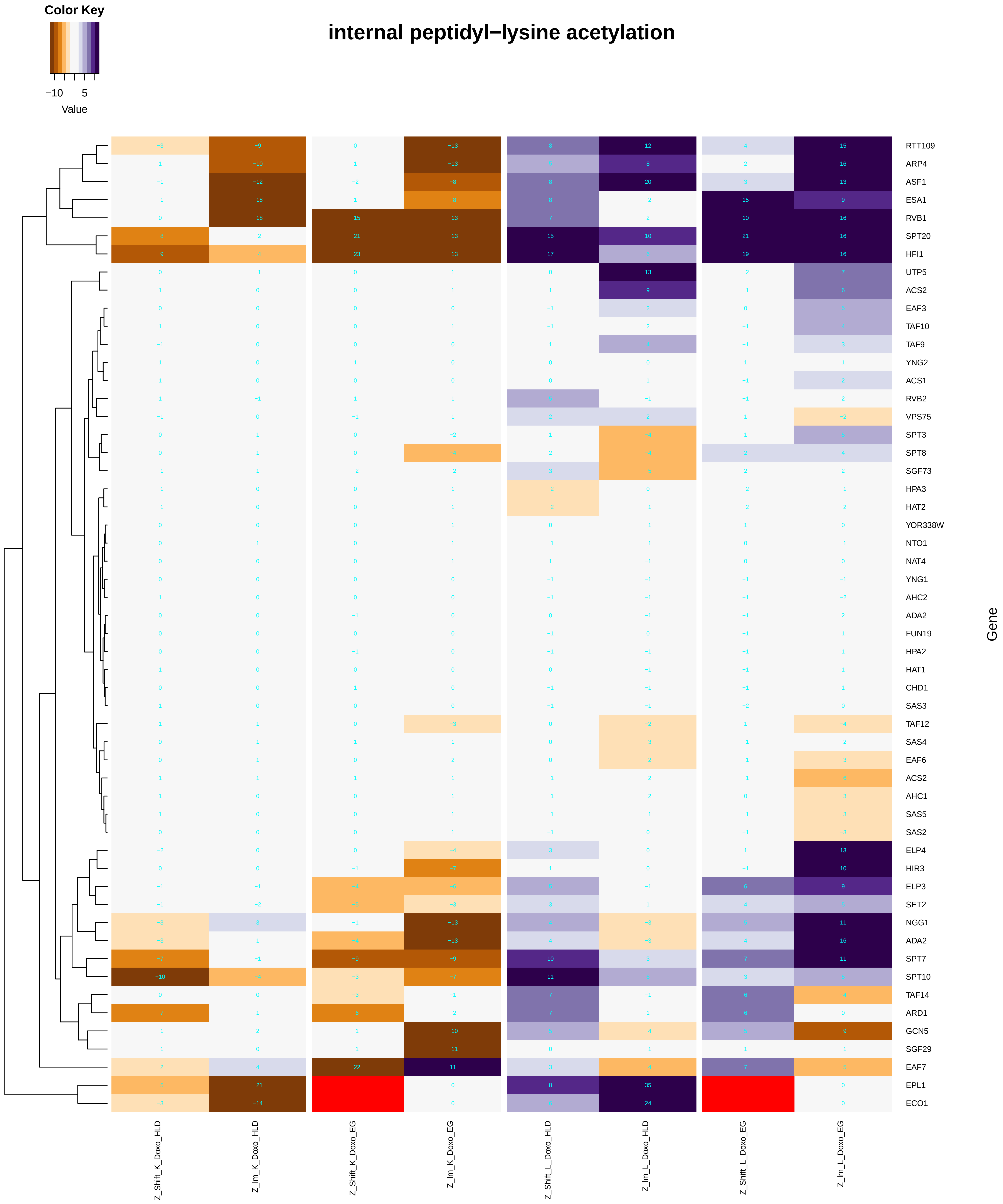

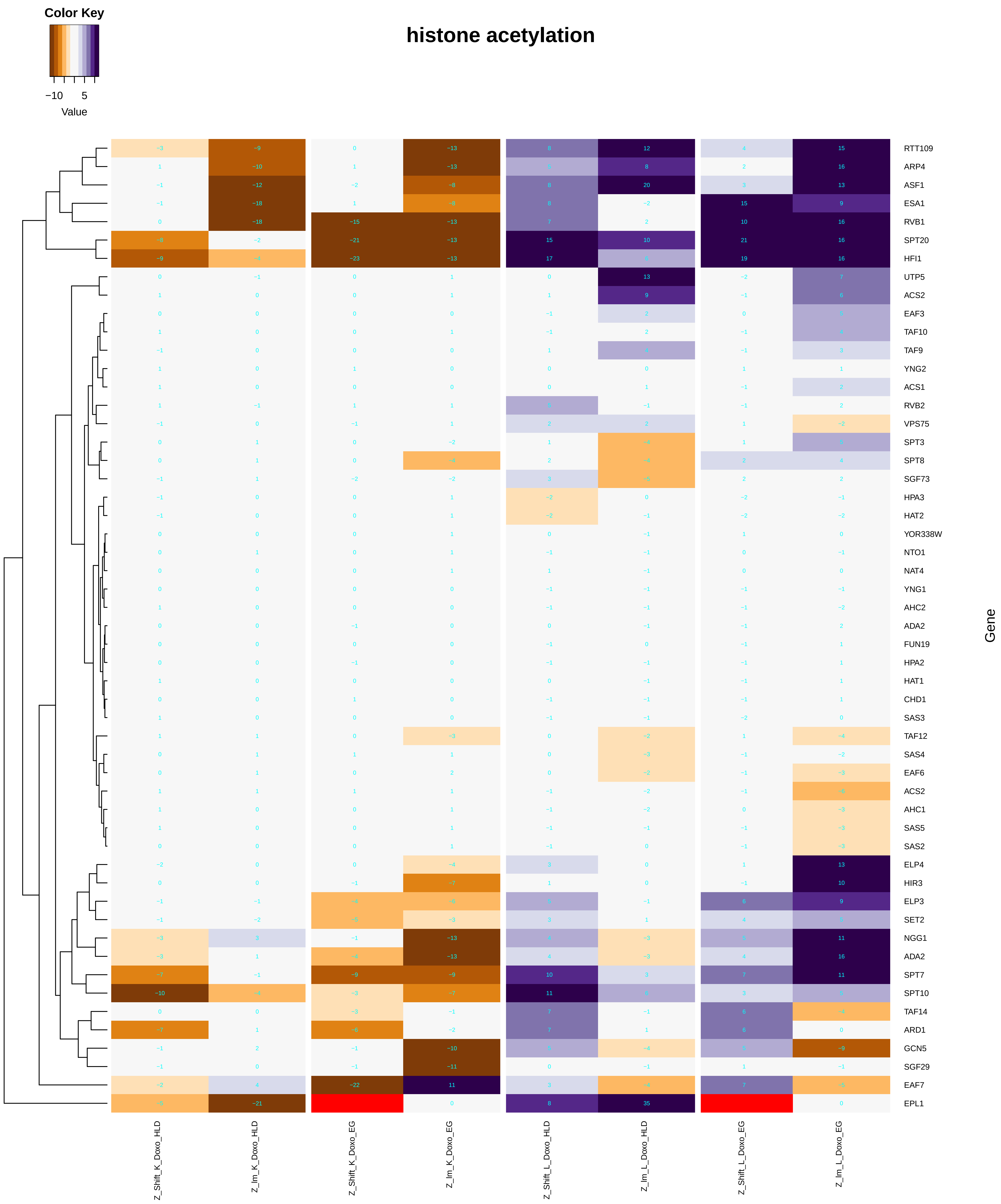

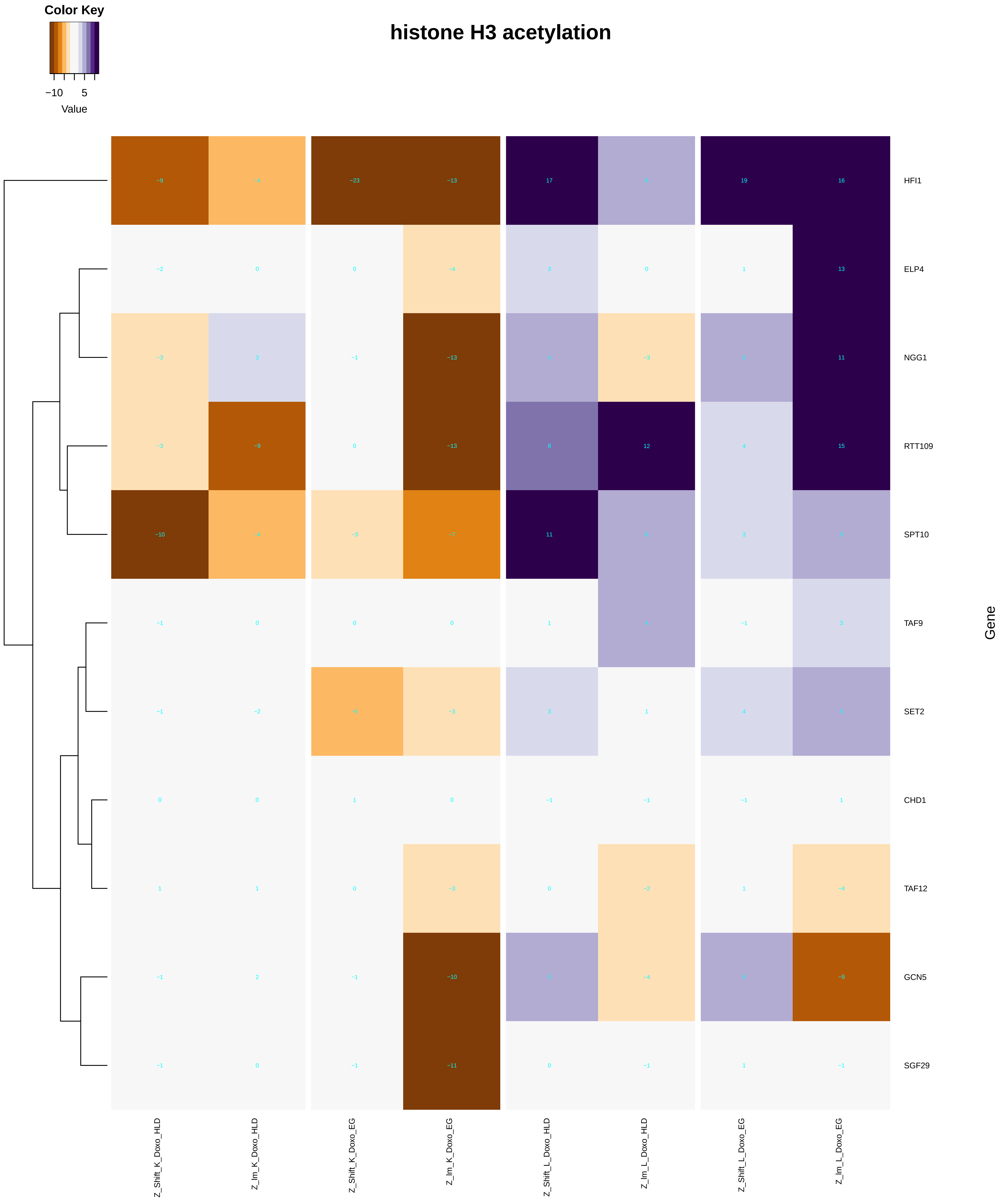

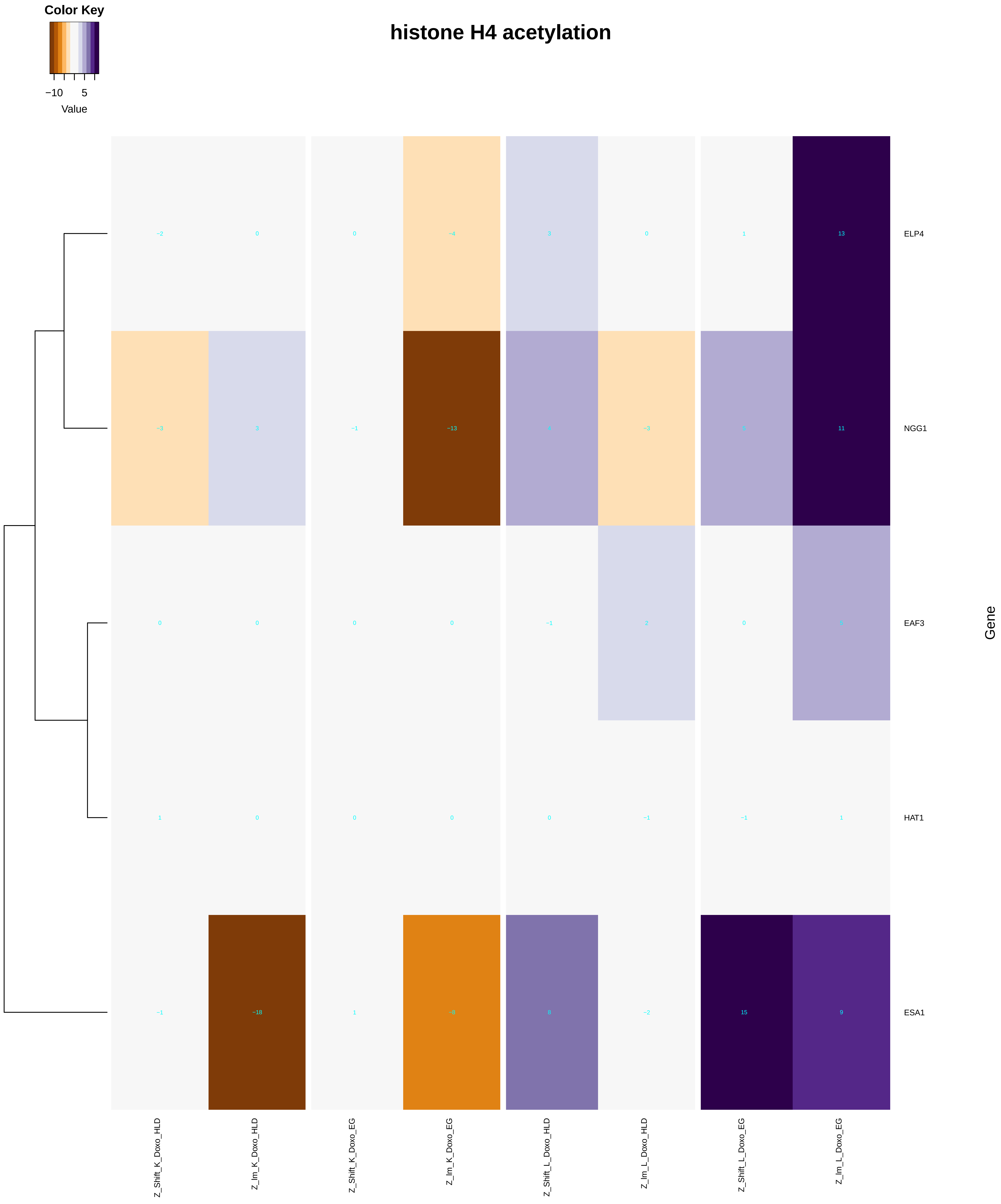

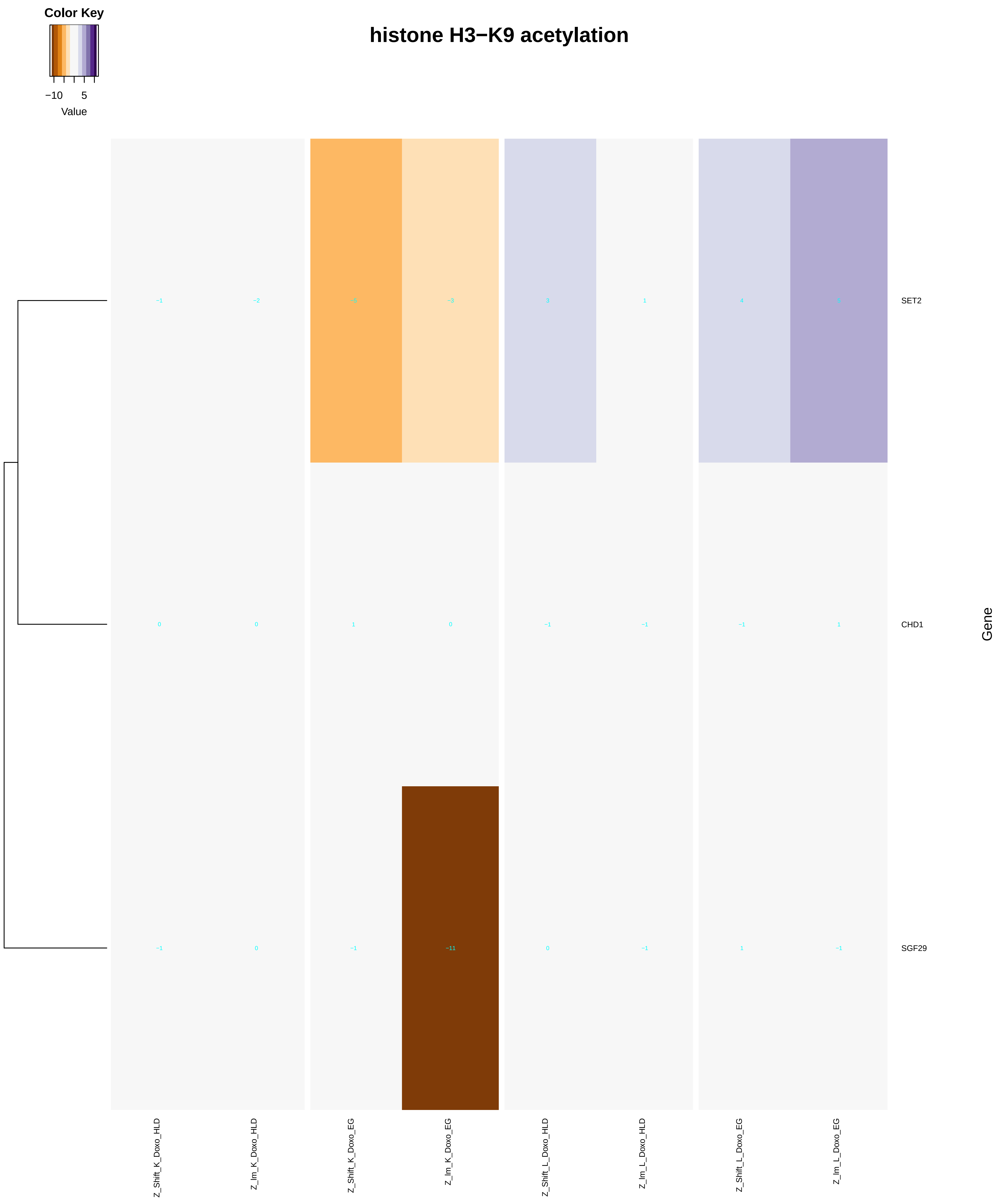

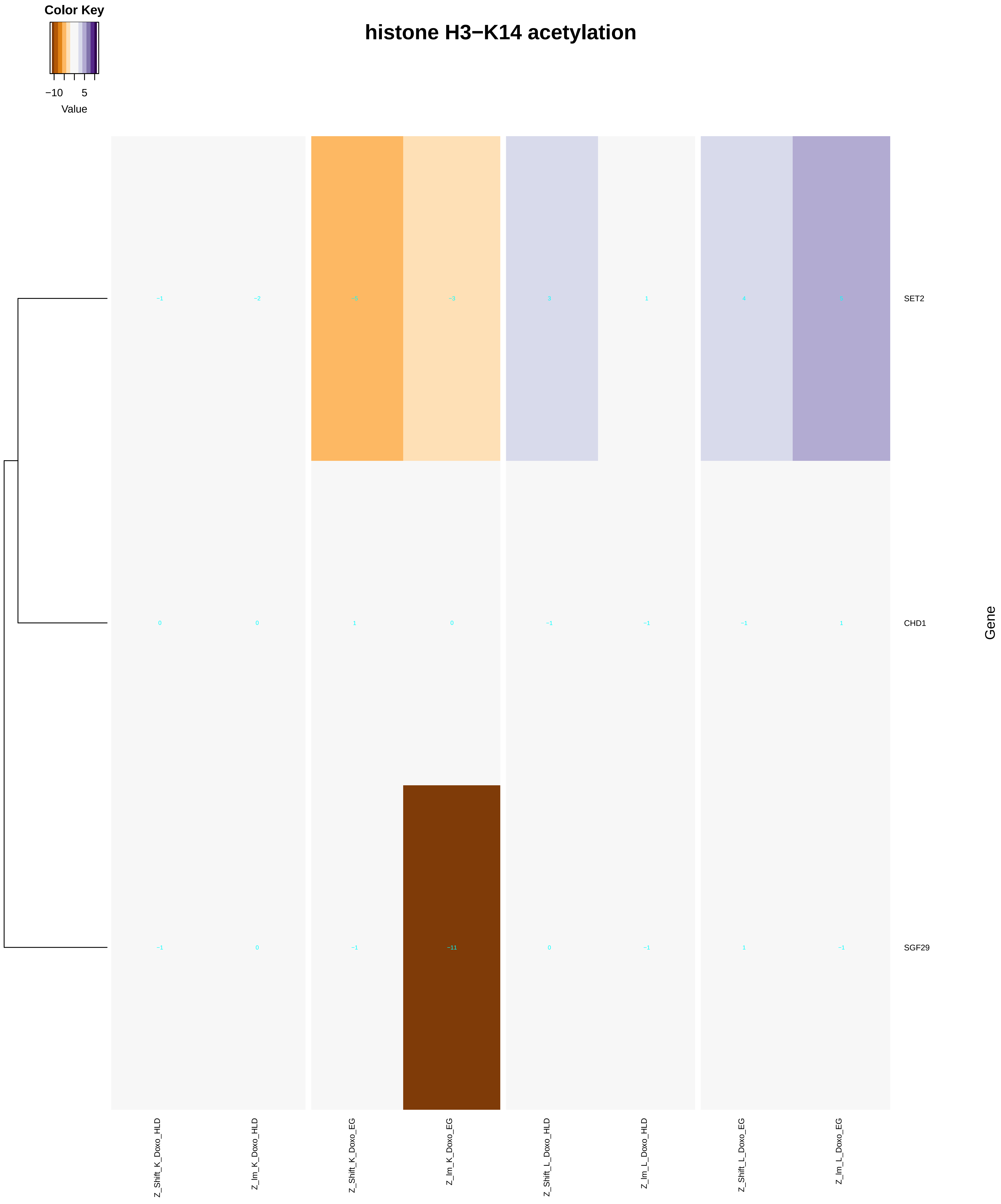

### internal_protein_amino_acid_acetylation.pdf

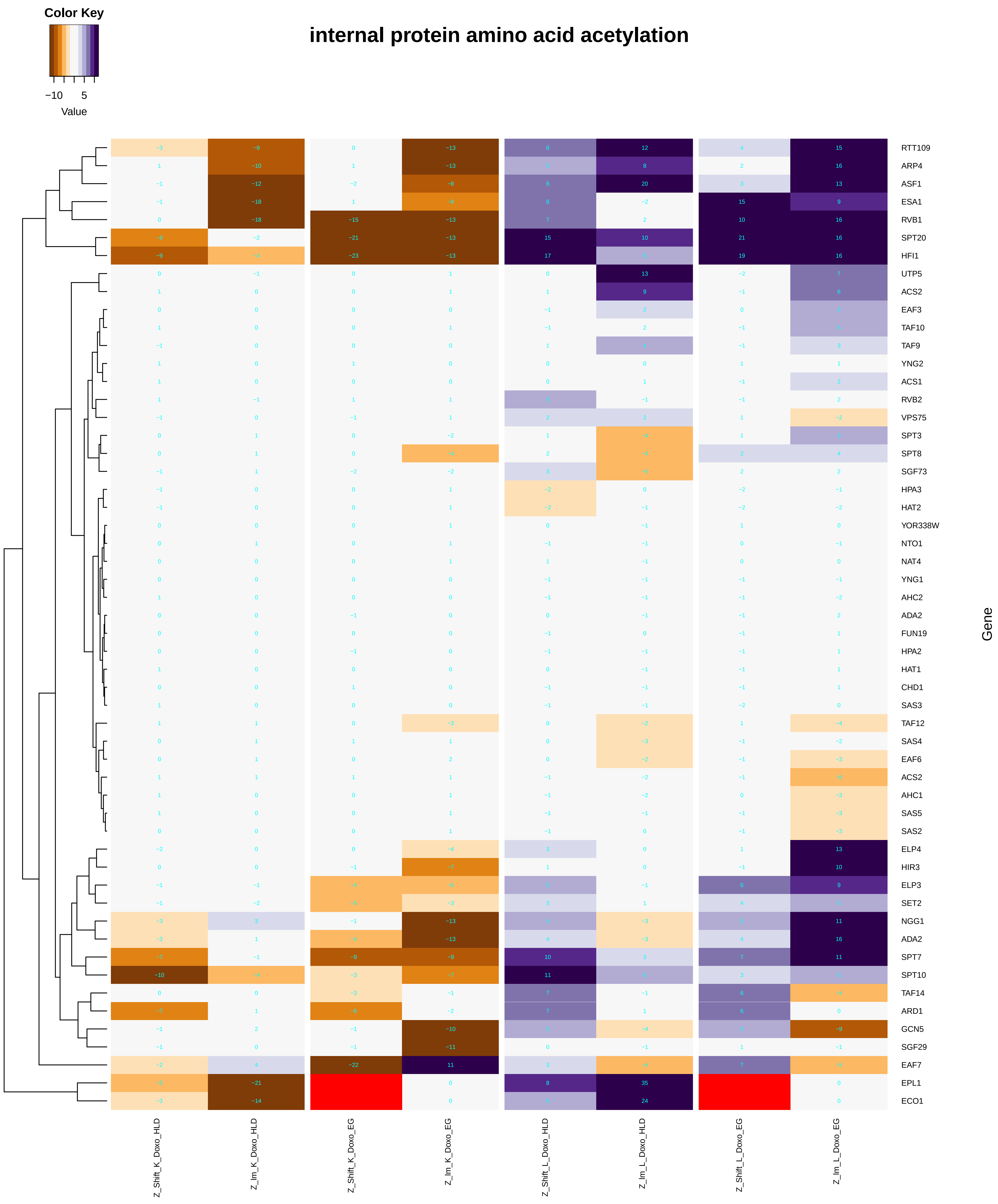

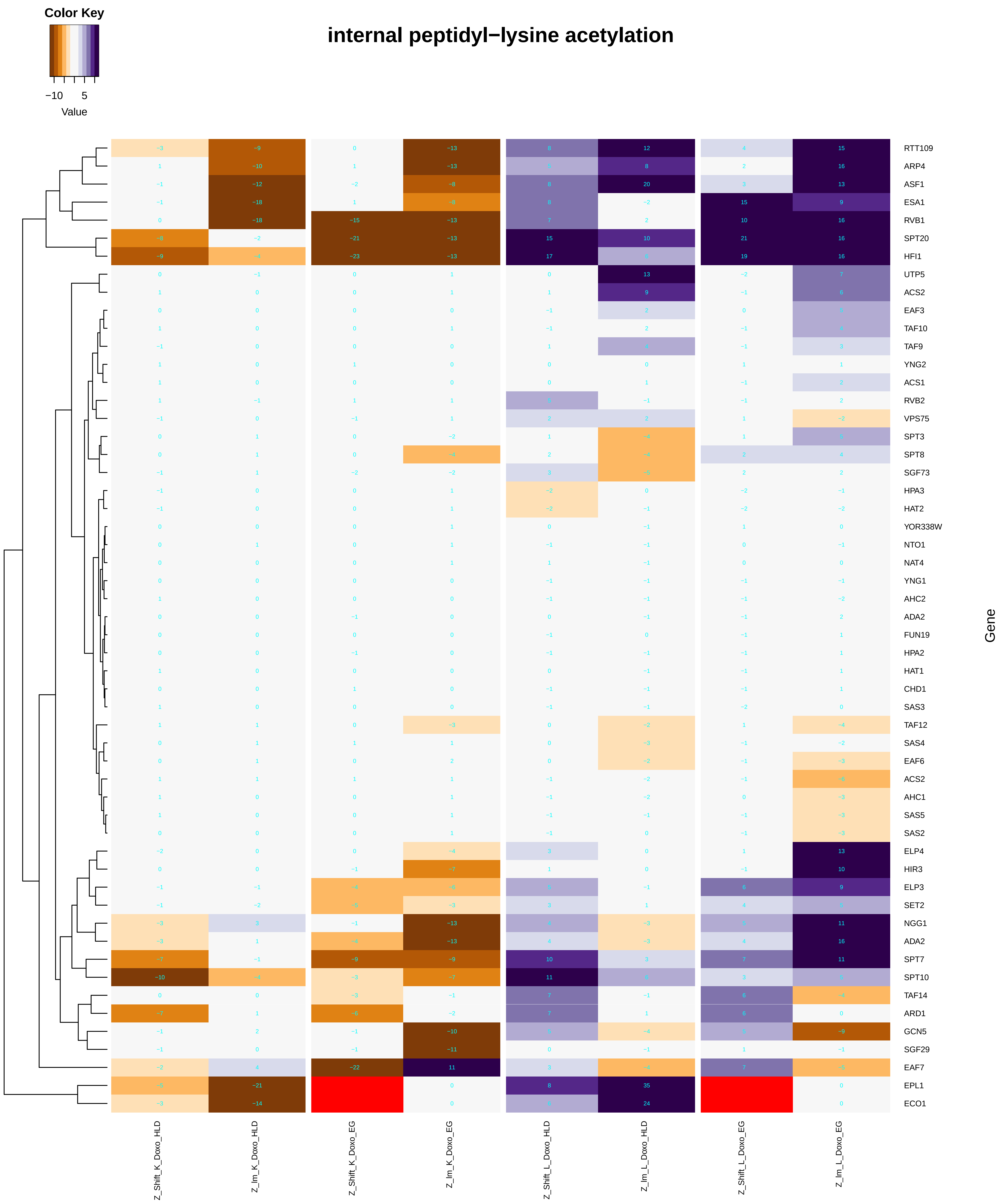

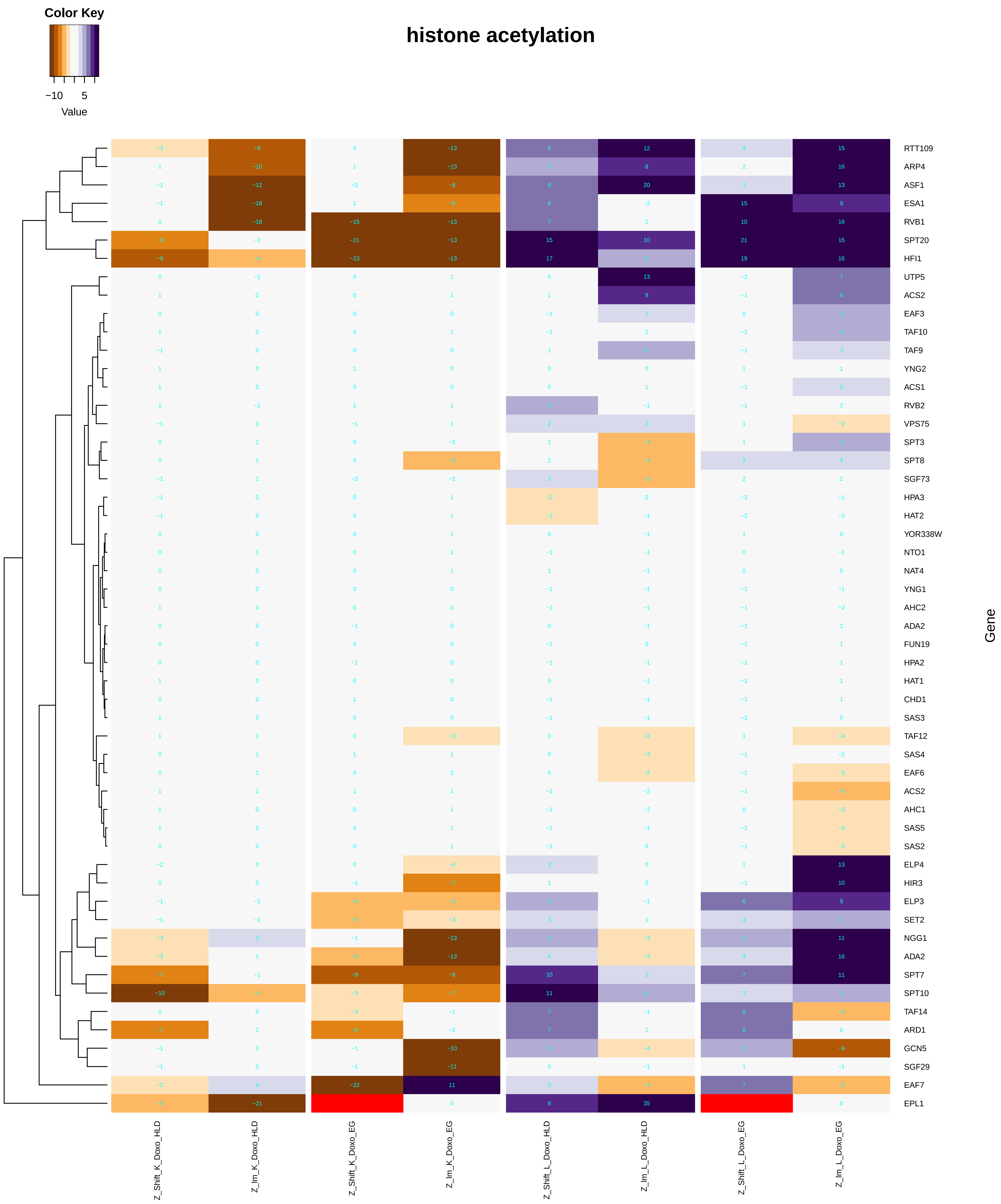

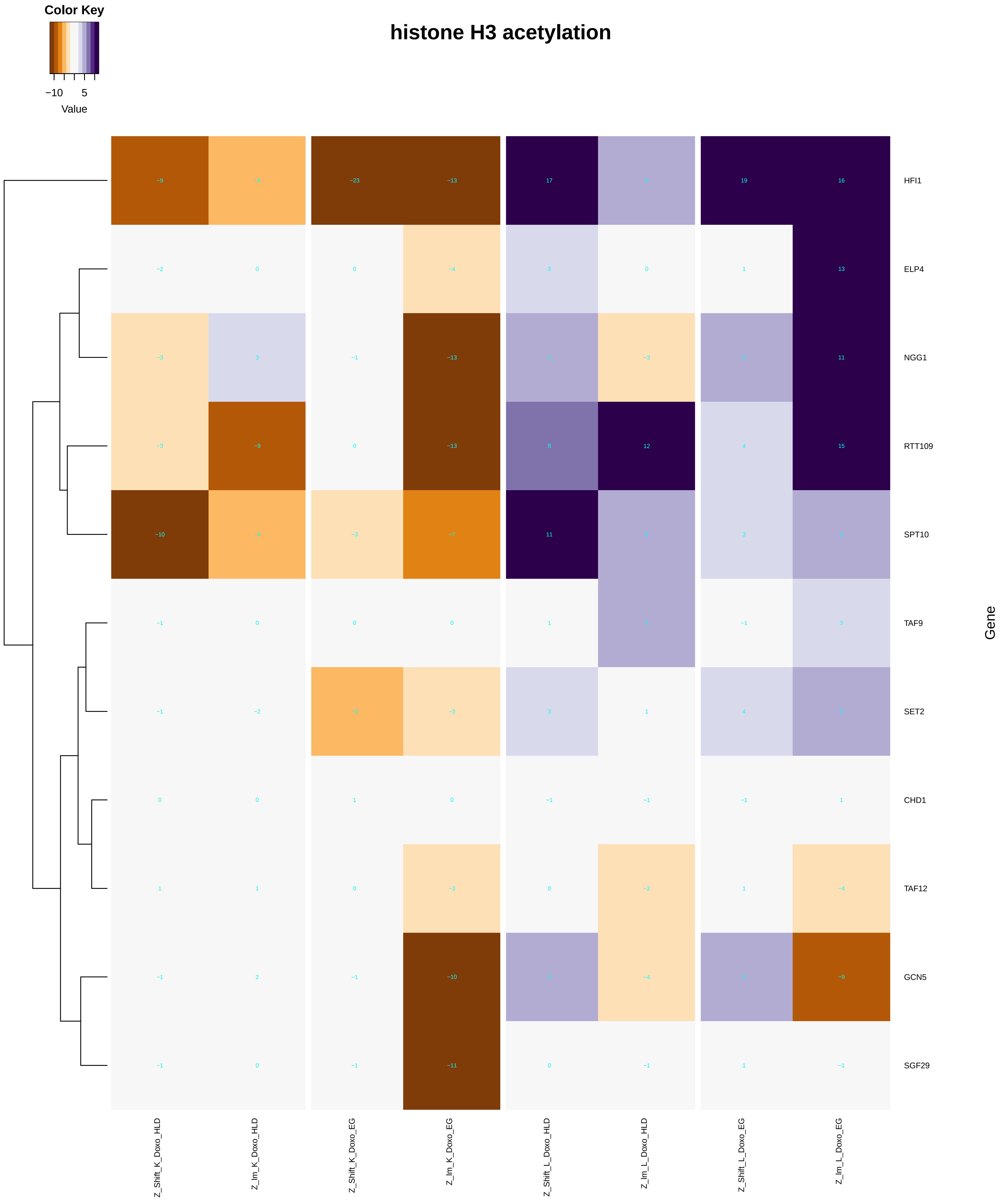

#### histone H4 acetylation

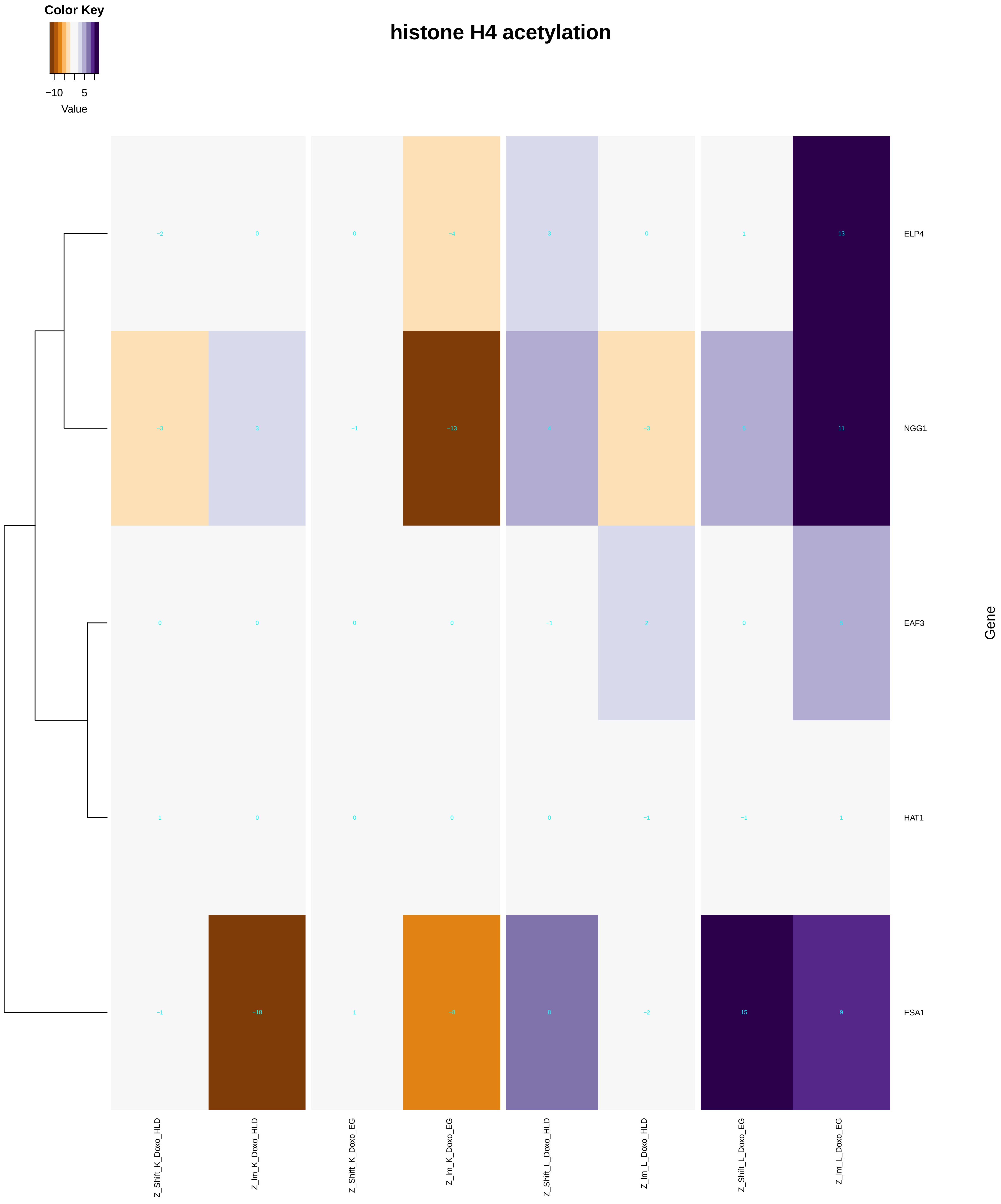

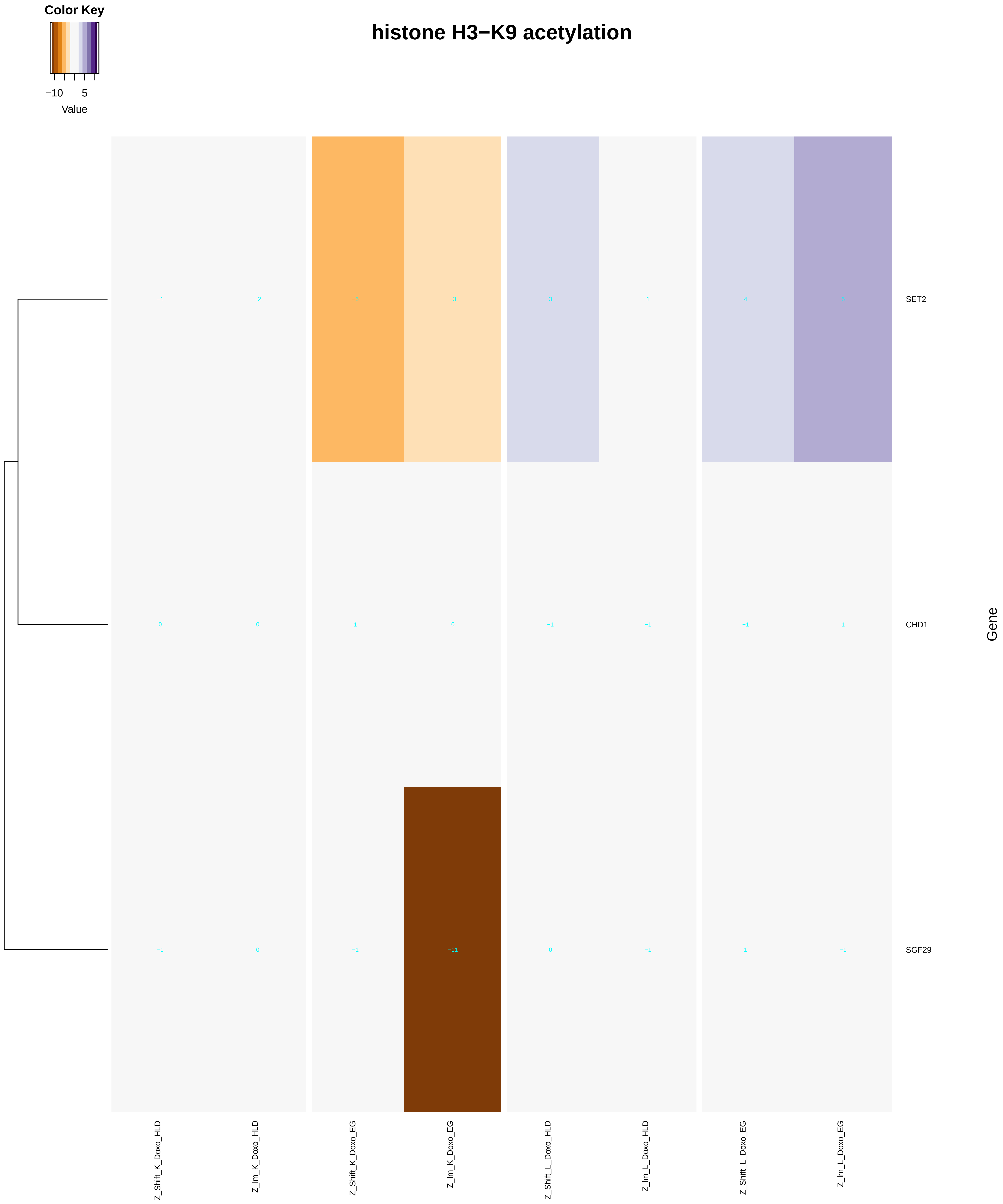

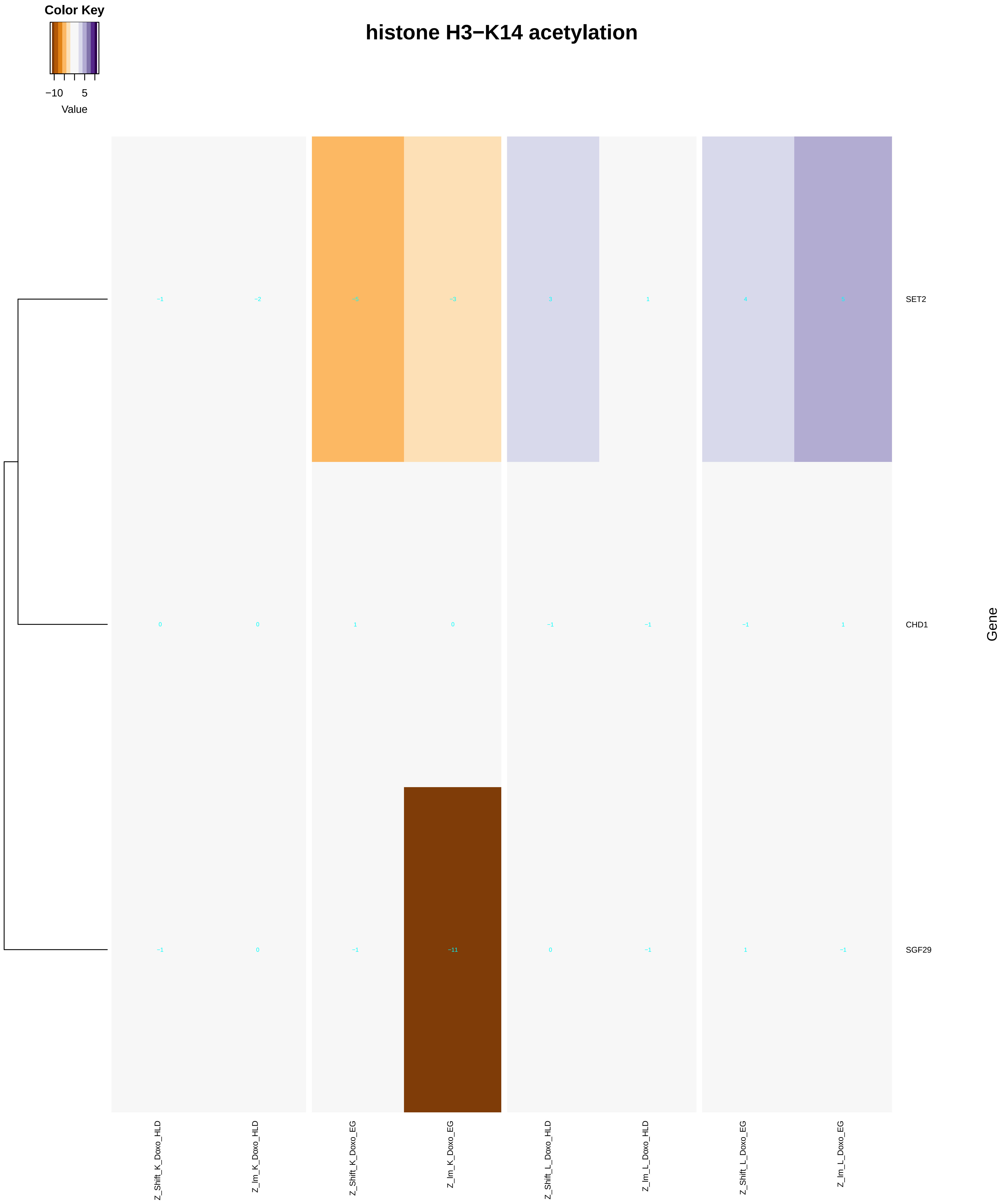

### intracellular_mRNA_localization.pdf

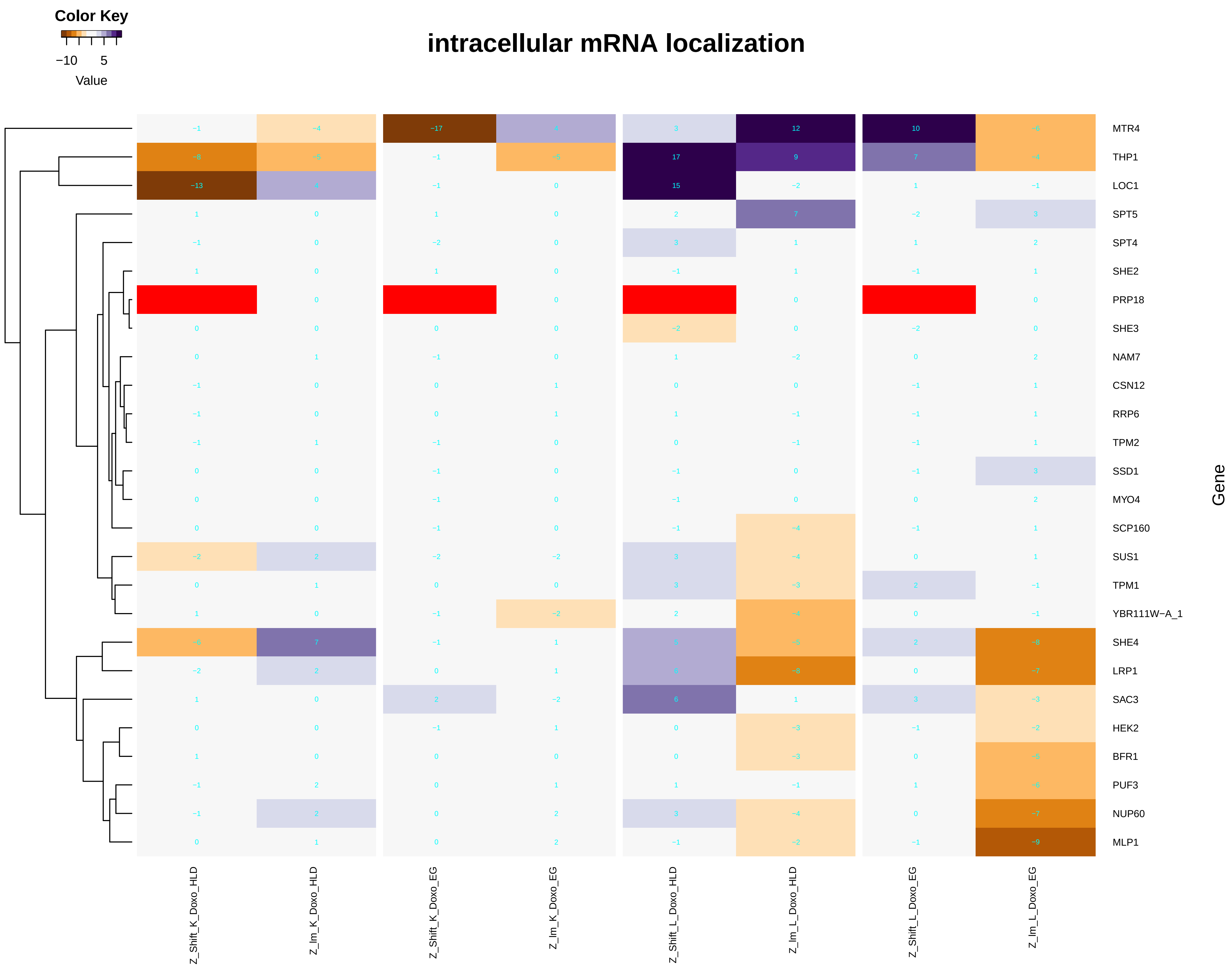

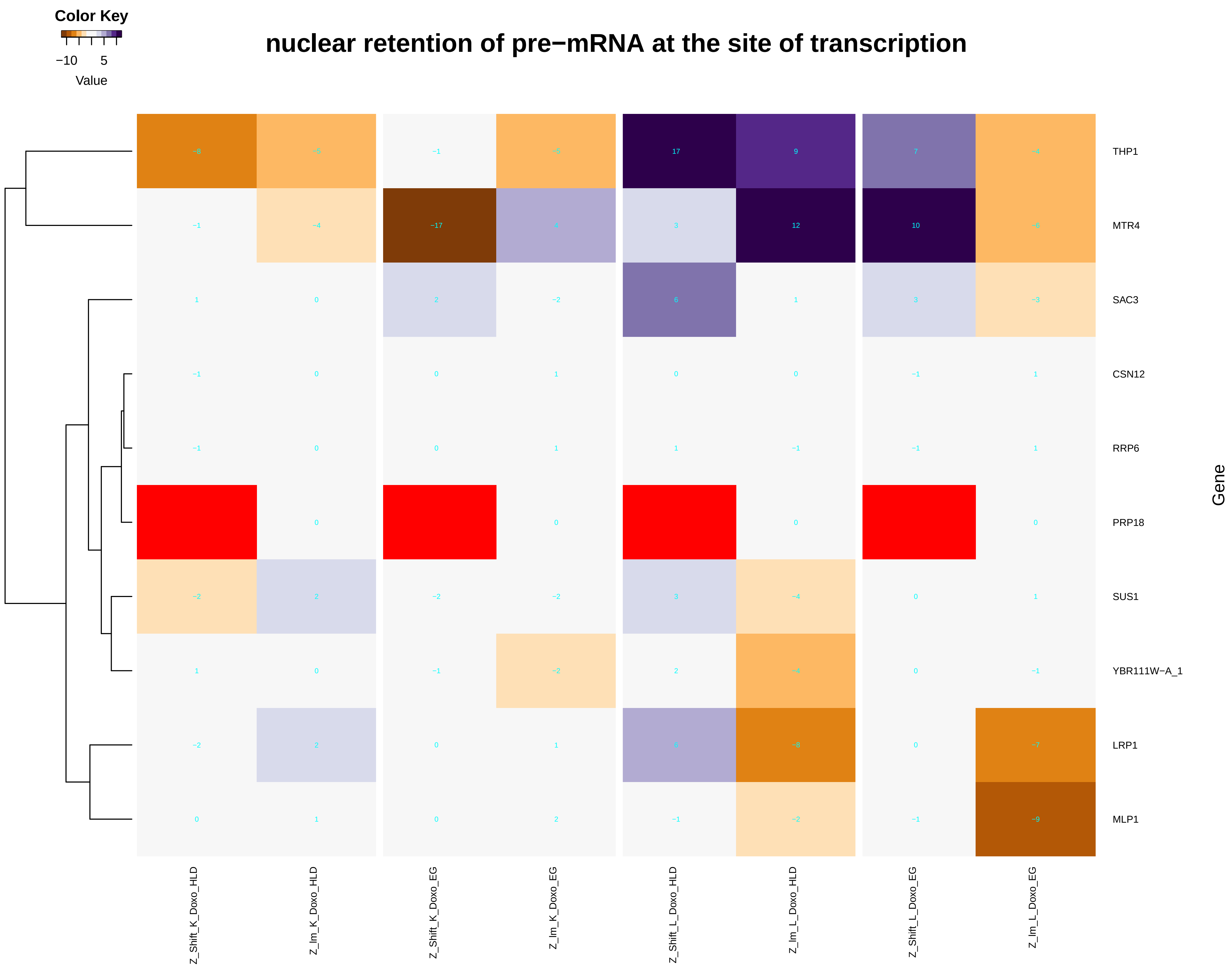

### intracellular_pH_reduction.pdf

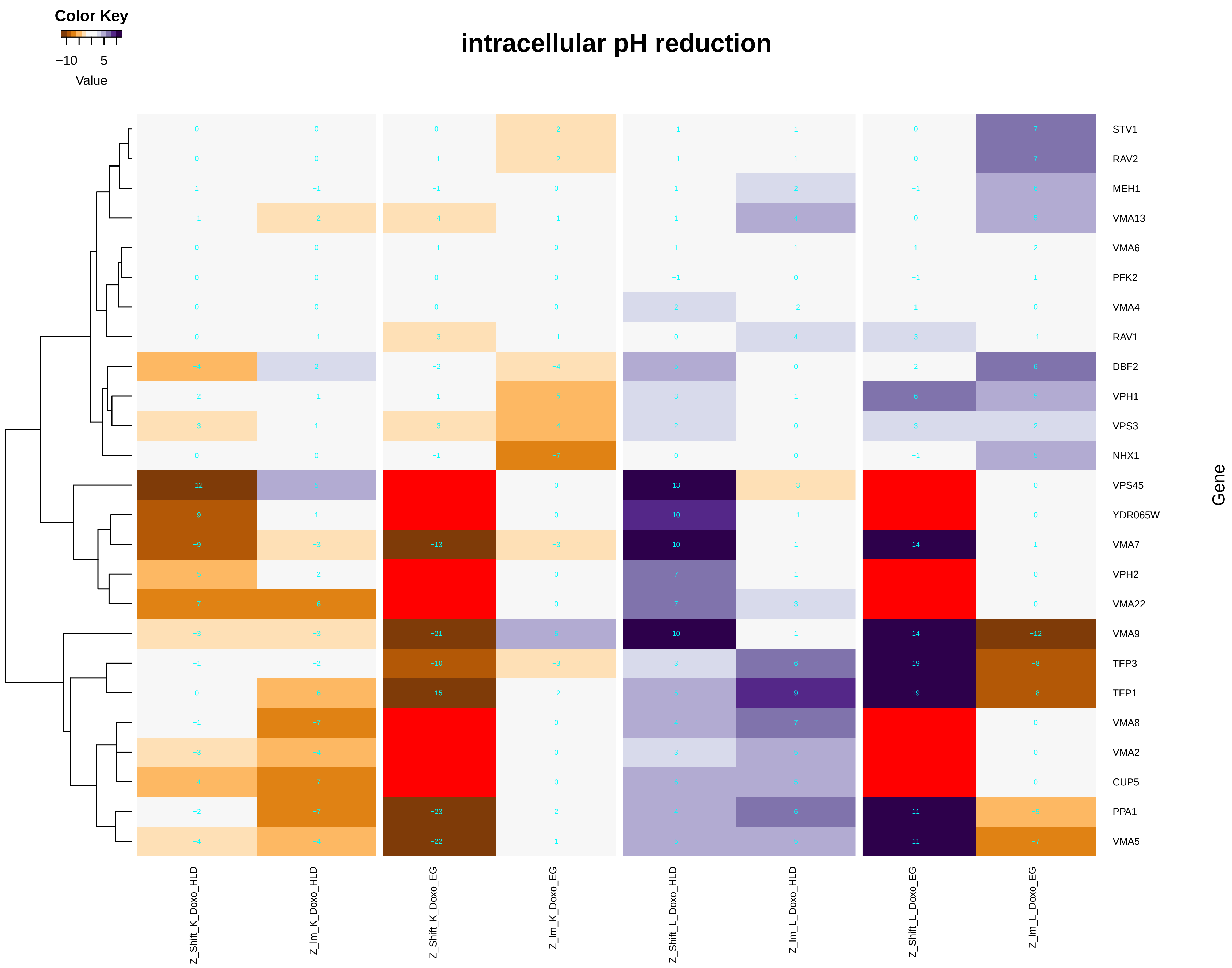

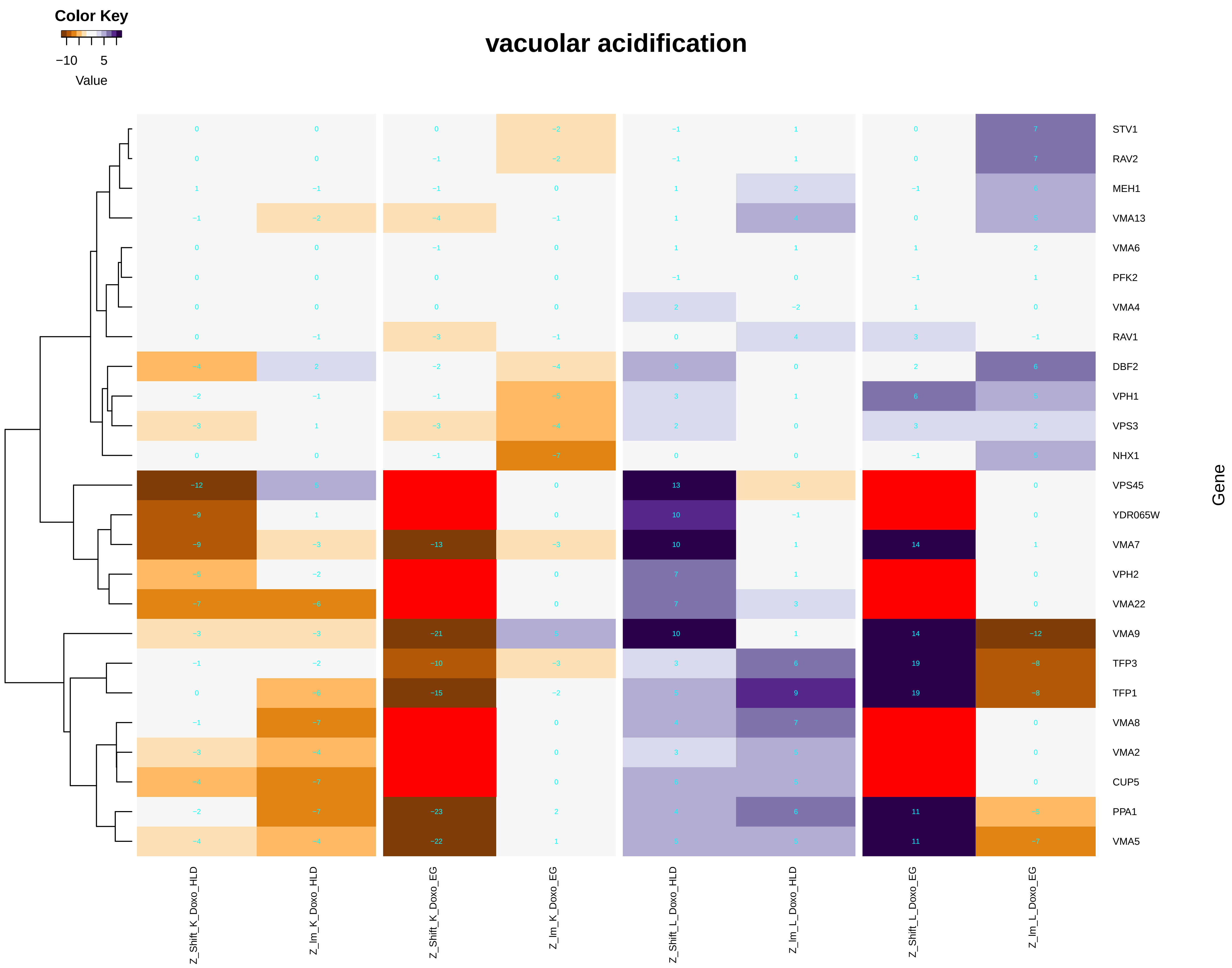

### intracellular_protein_transmembrane_transport.pdf

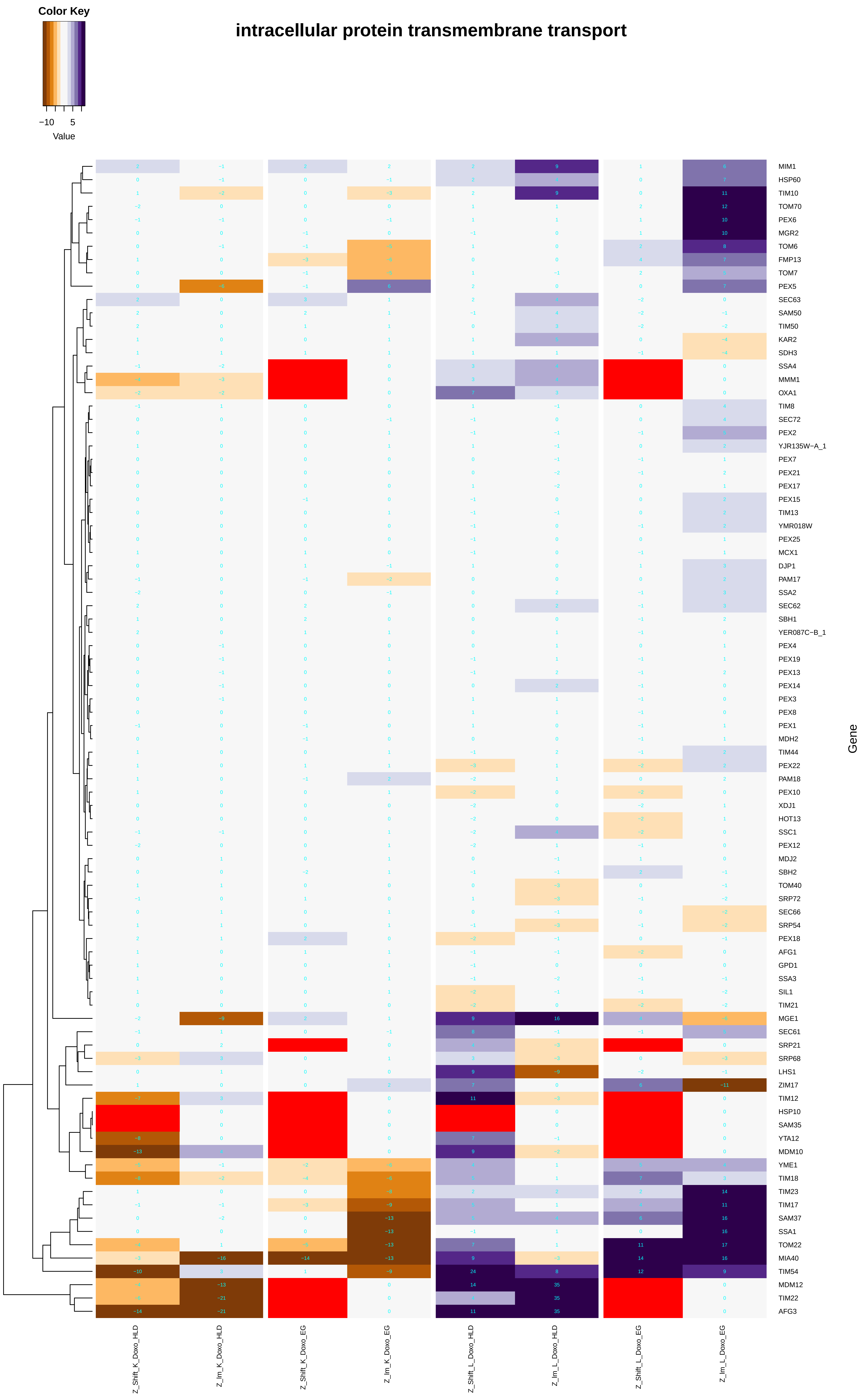

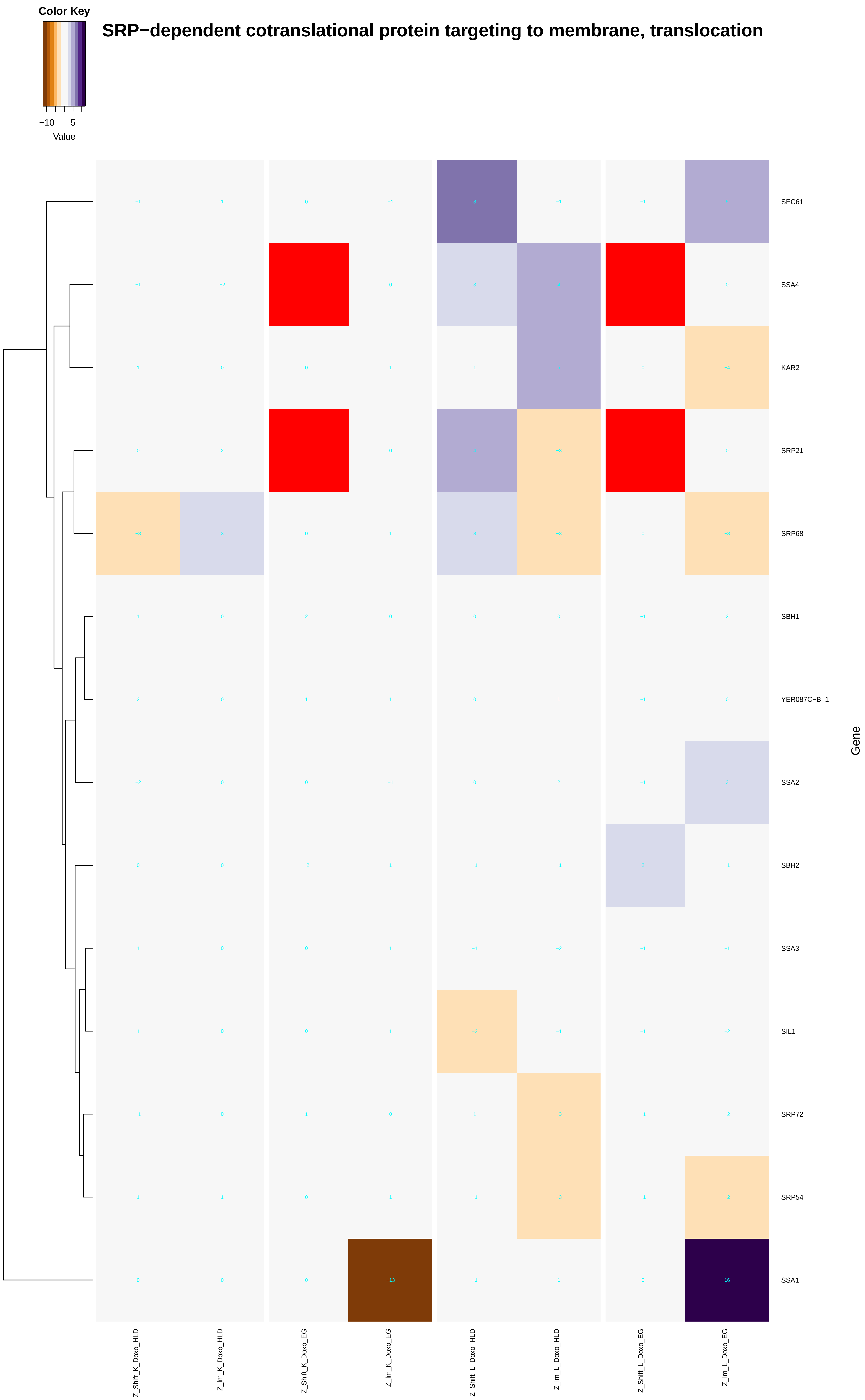

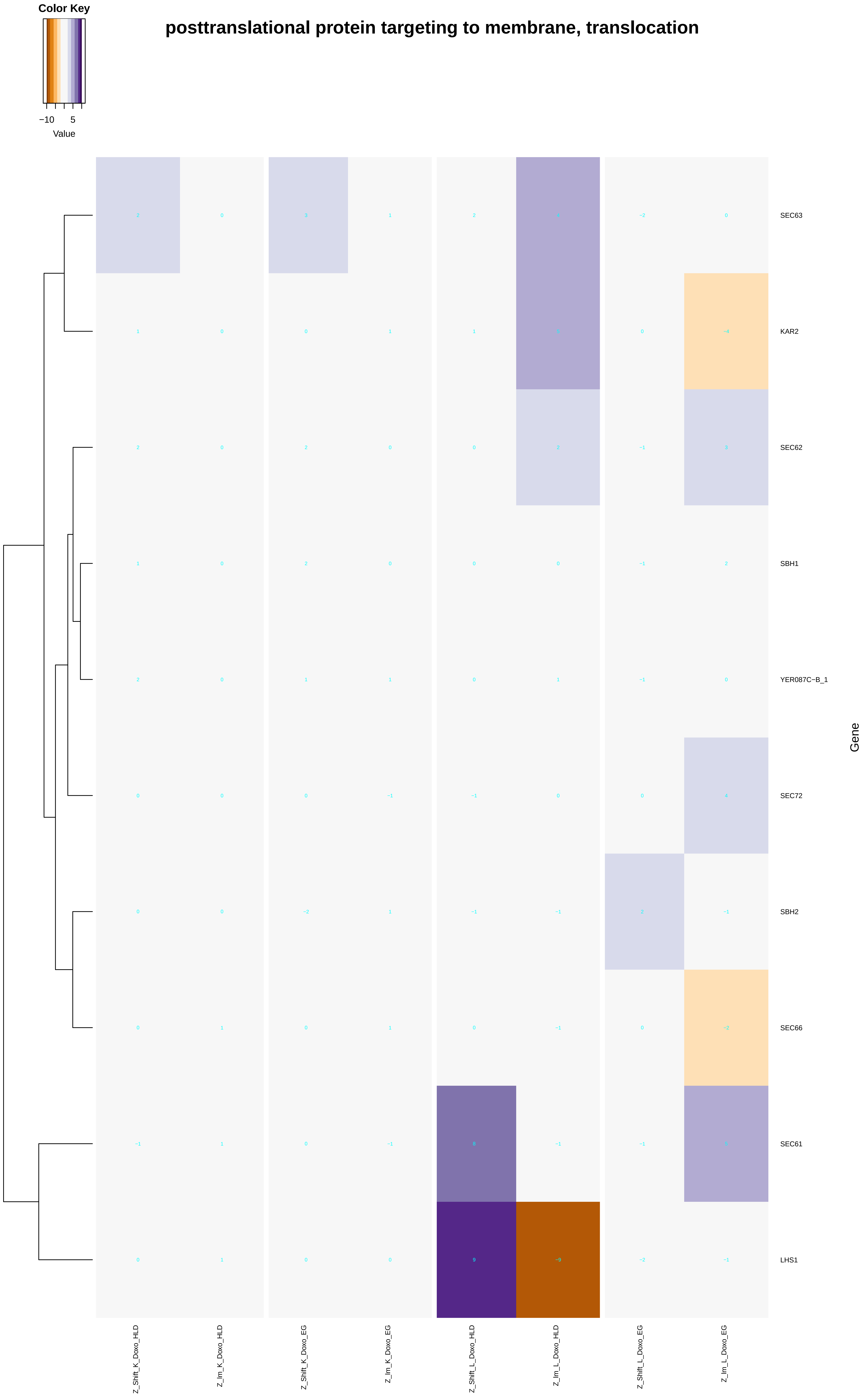

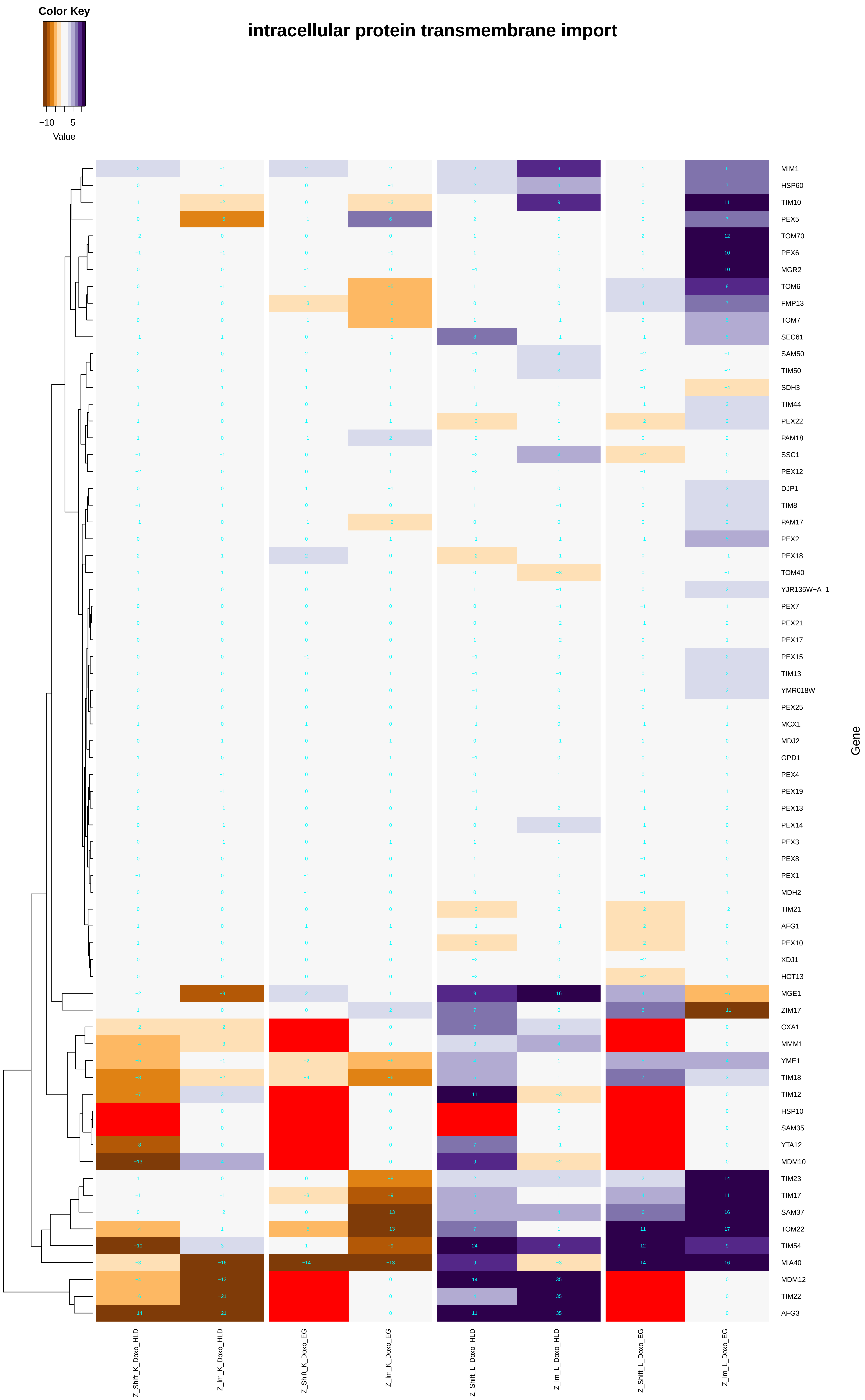

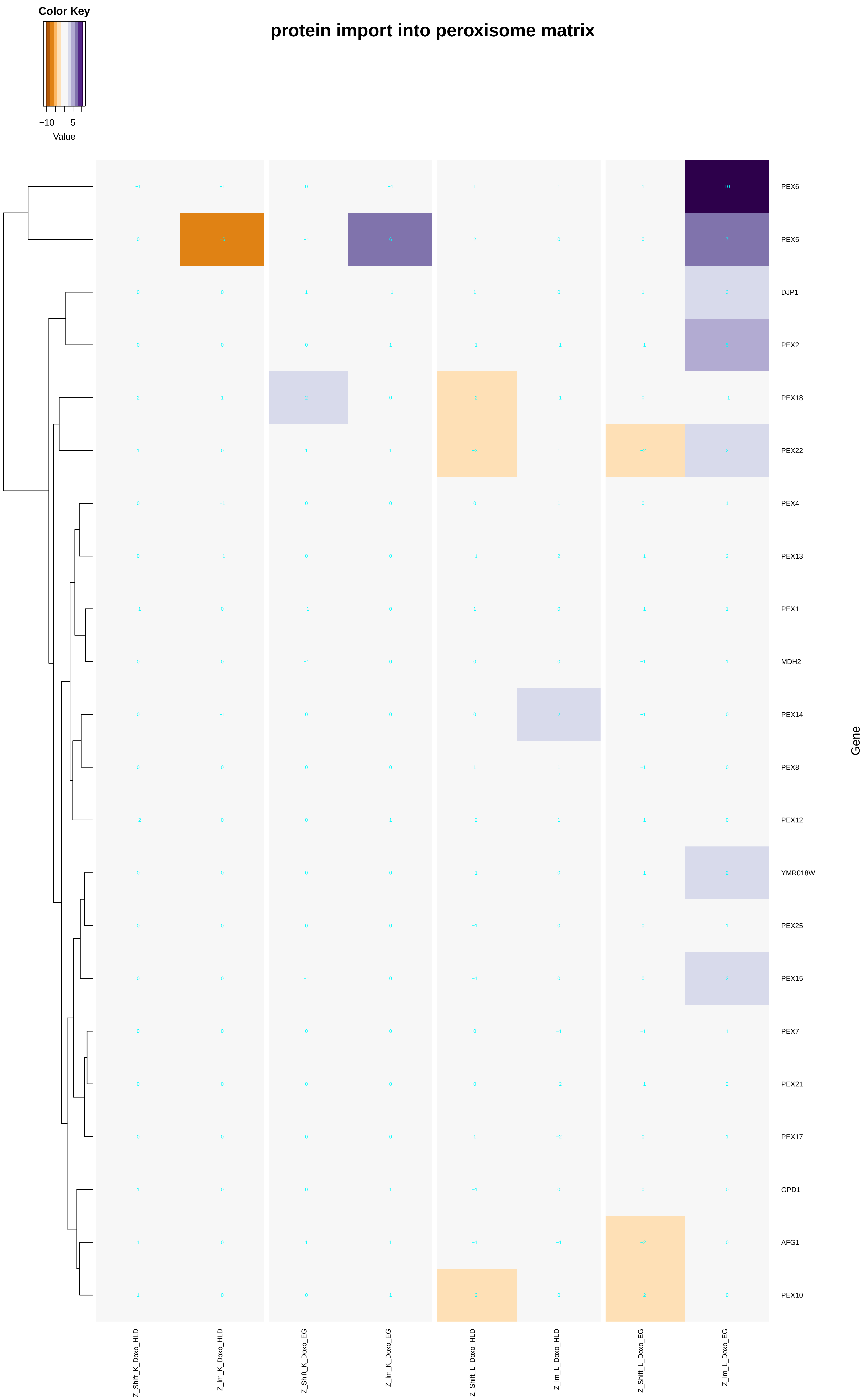

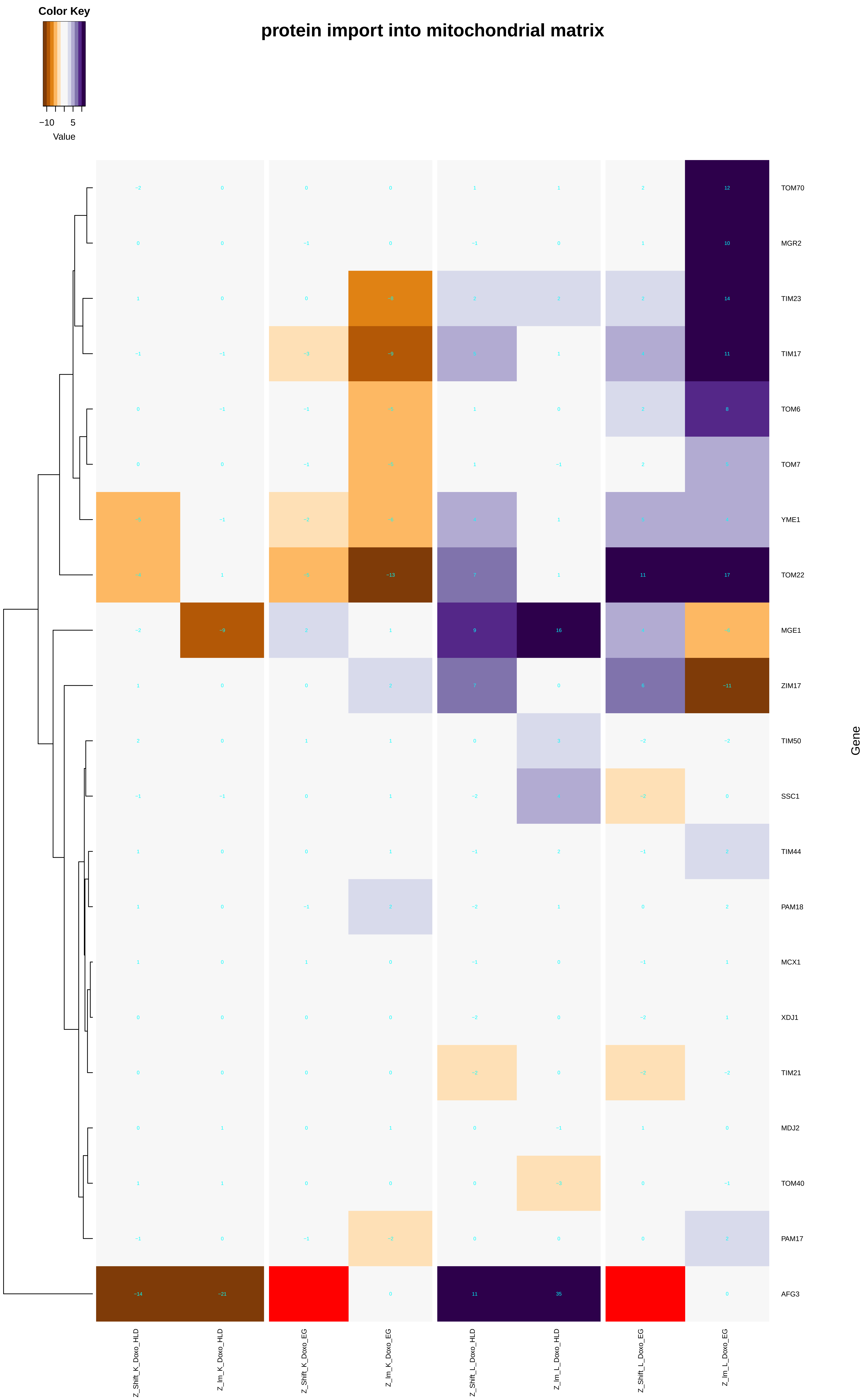

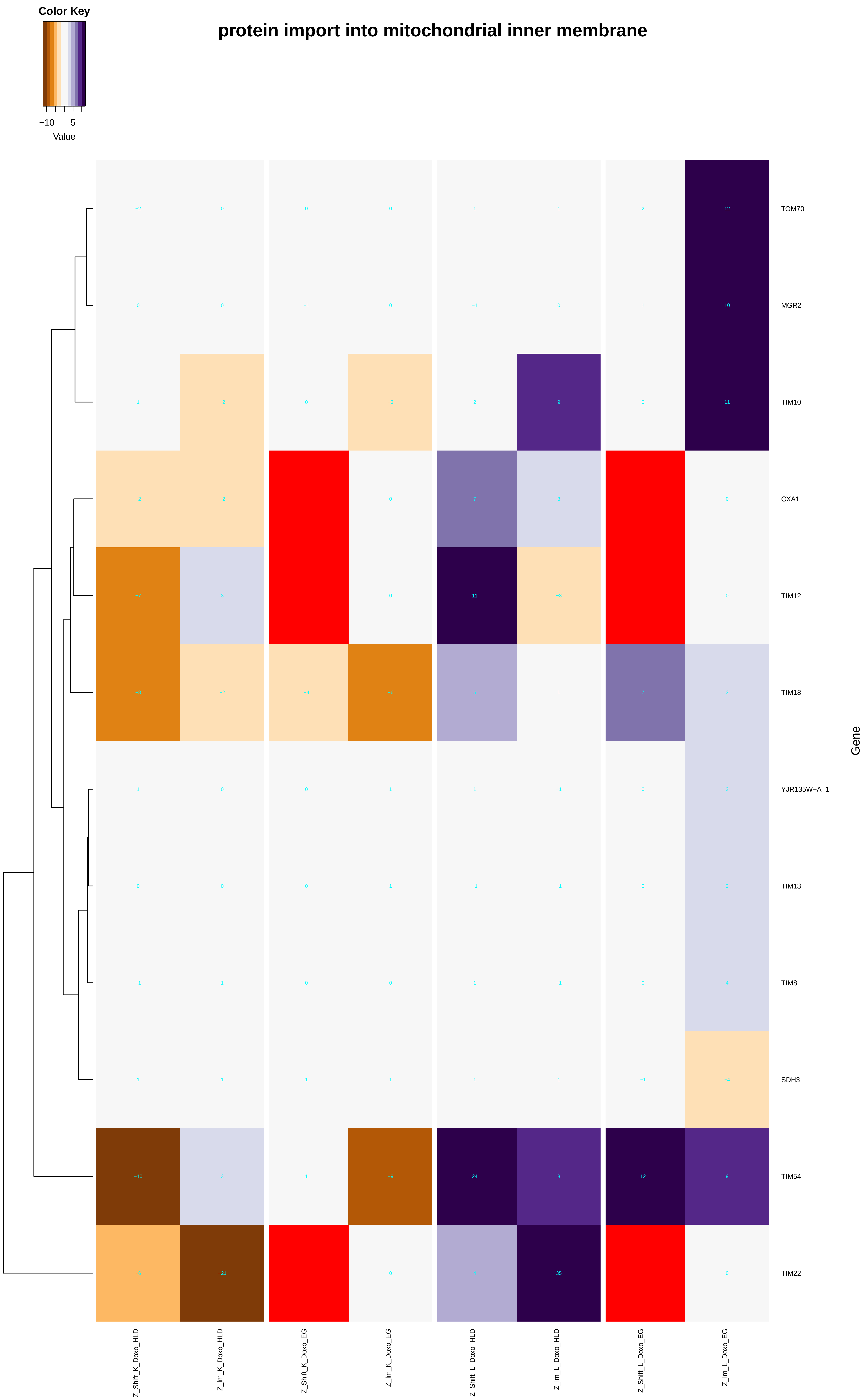

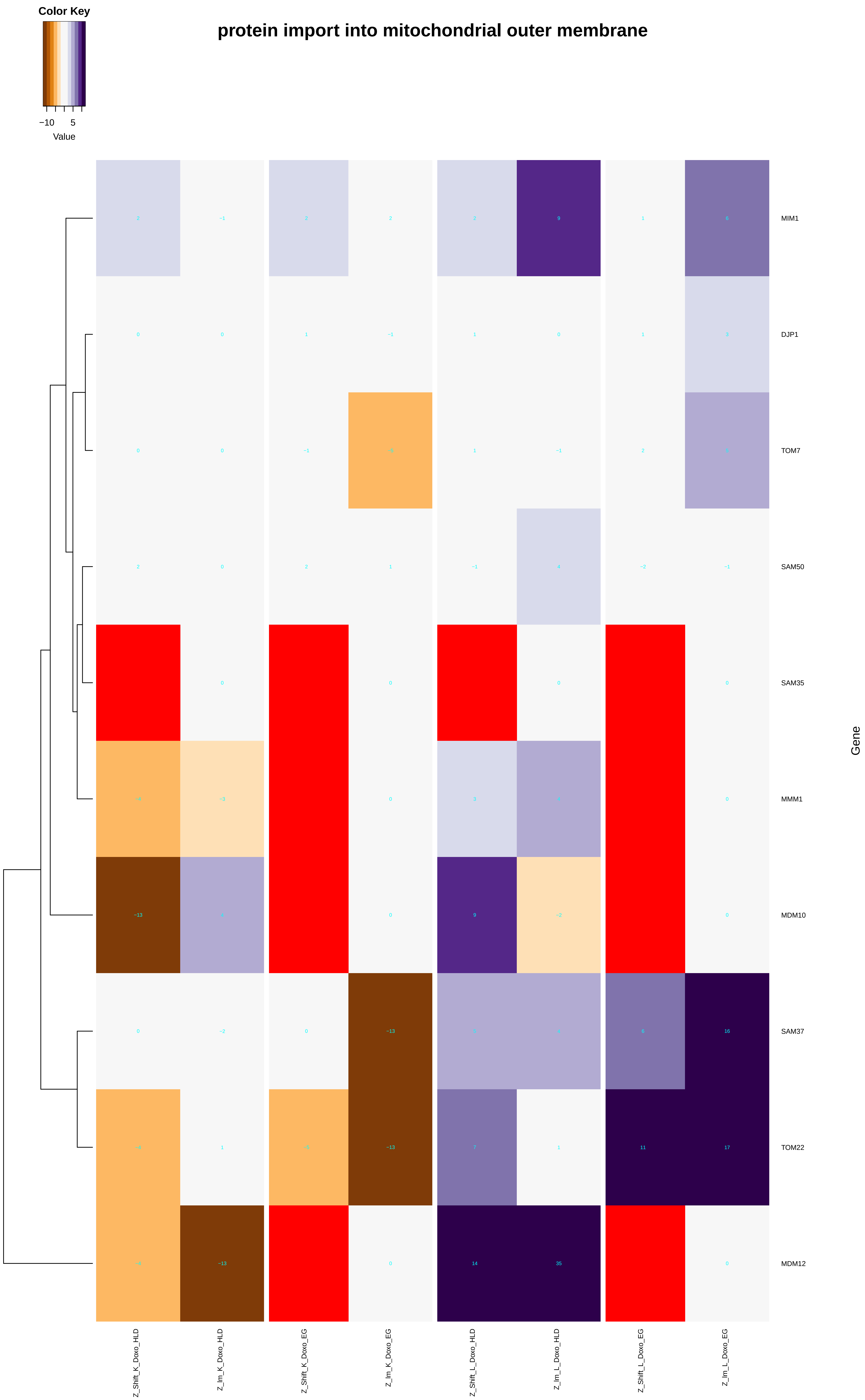

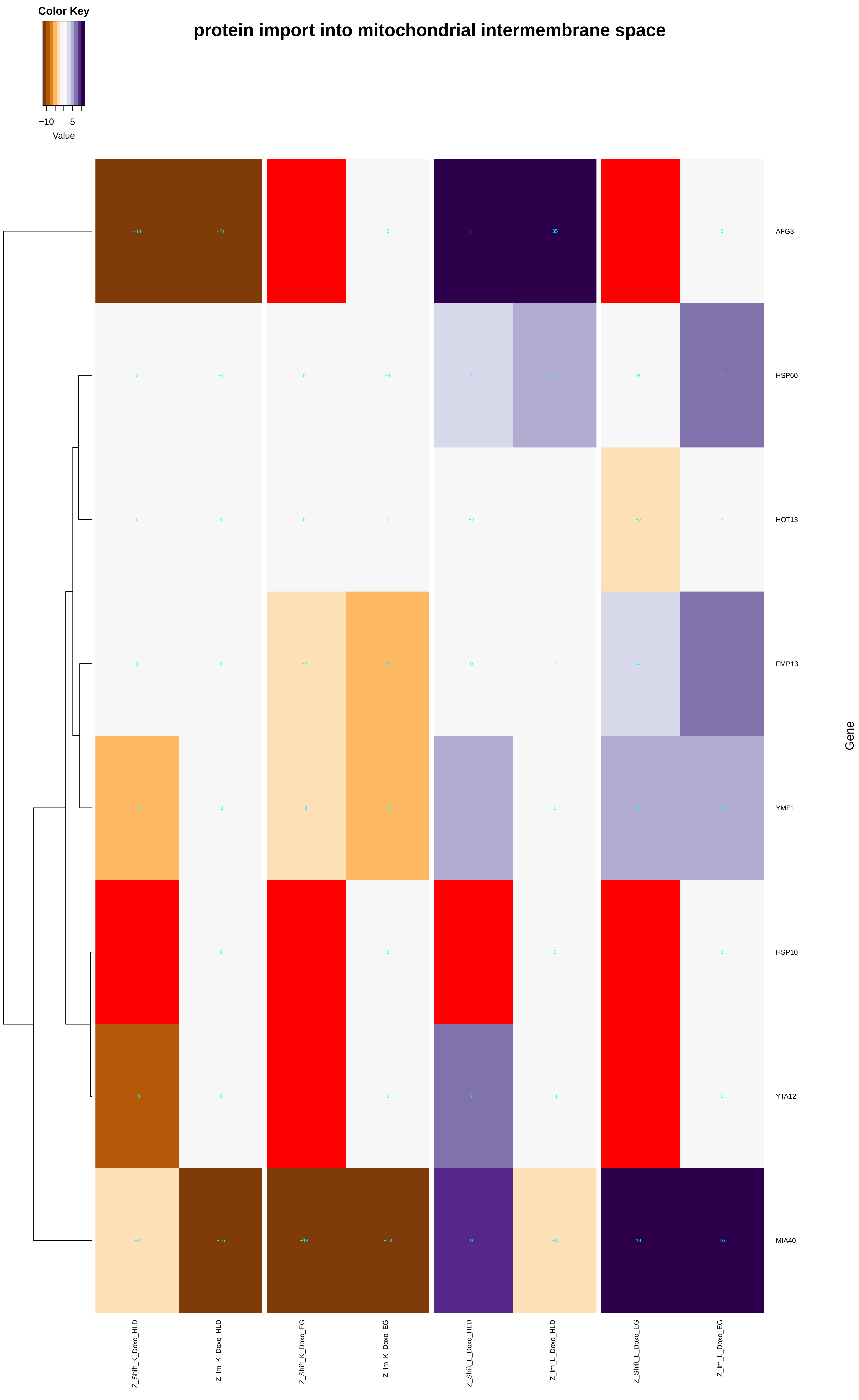

### iron_assimilation.pdf

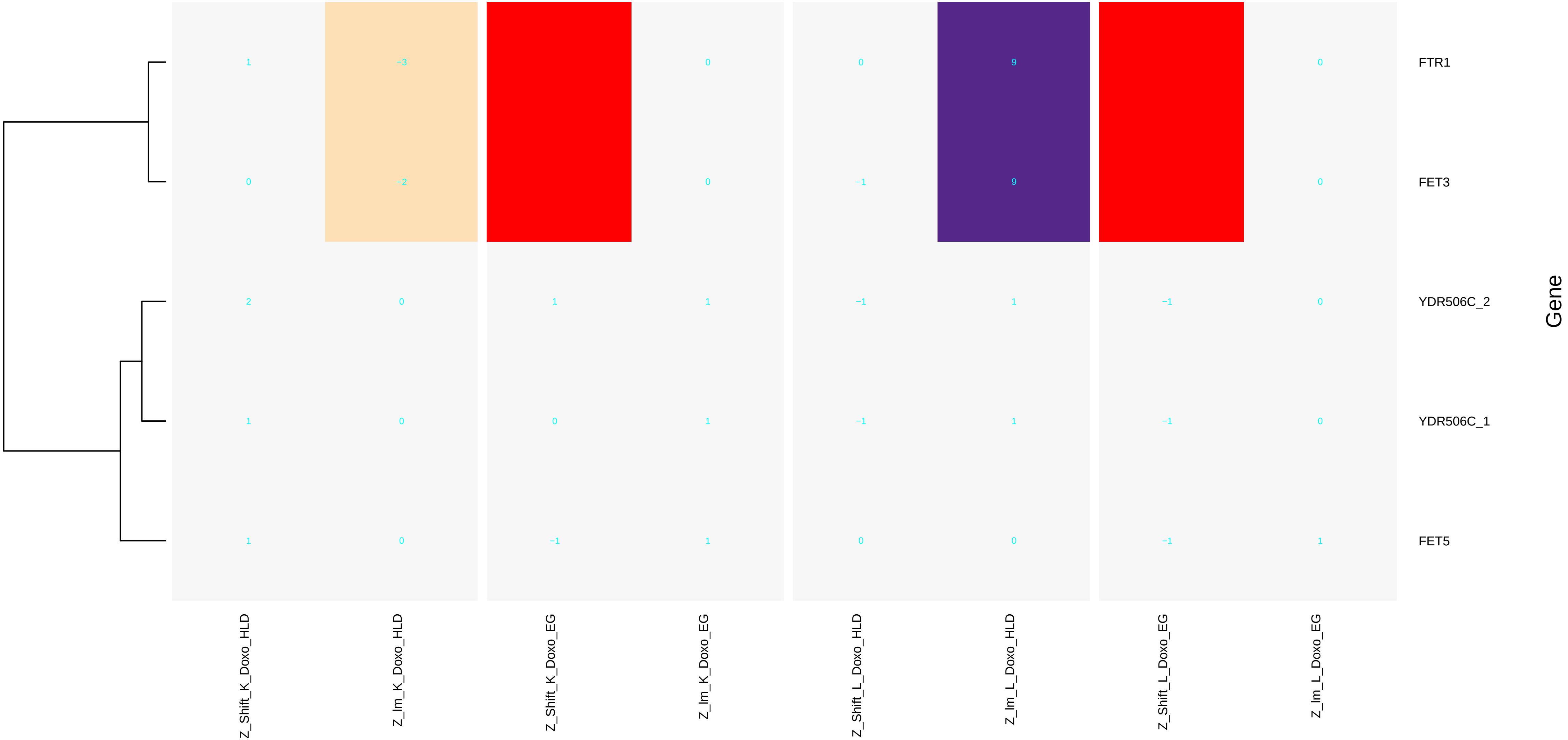

### iron_assimilation_by_reduction_and_transport.pdf

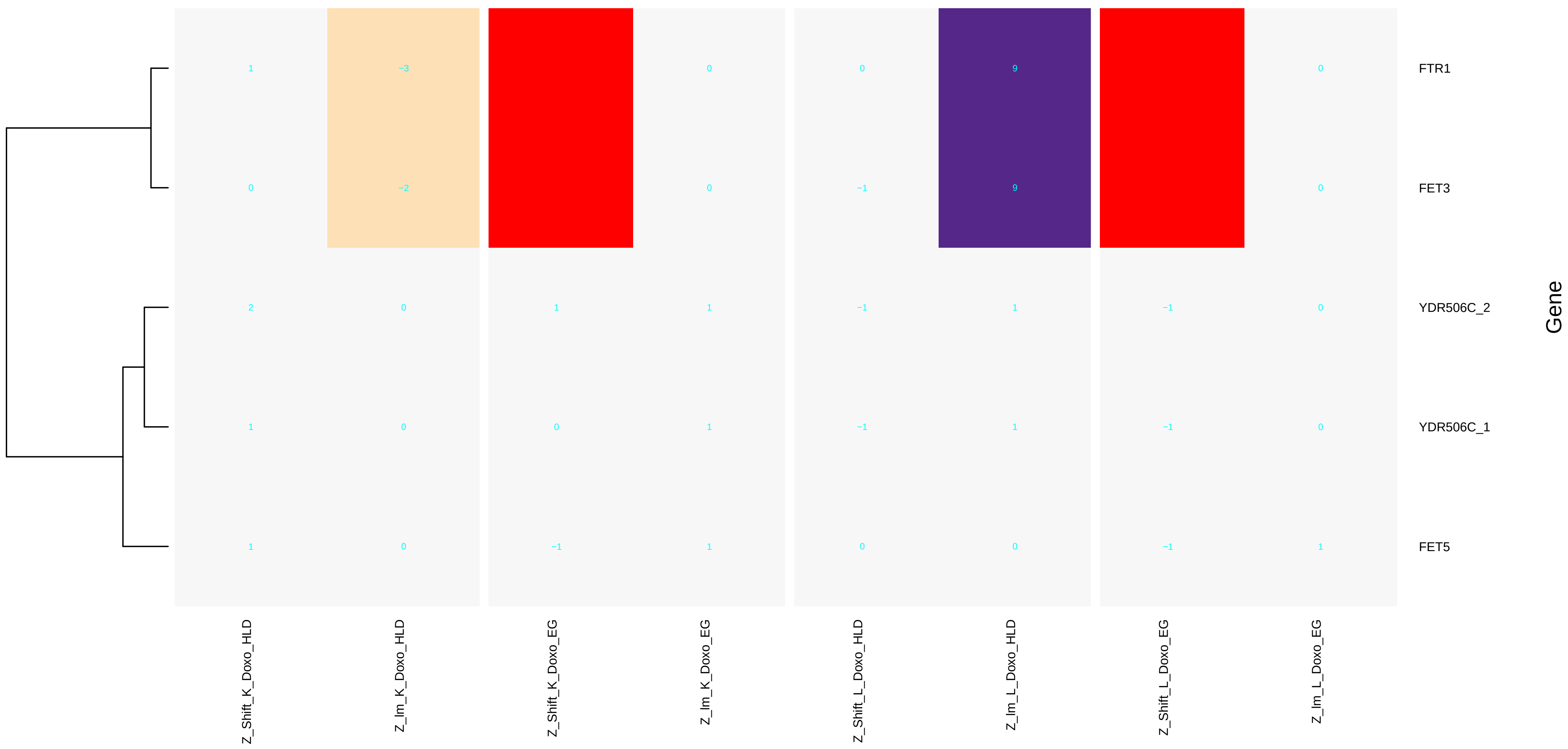

### ketone_biosynthetic_process.pdf

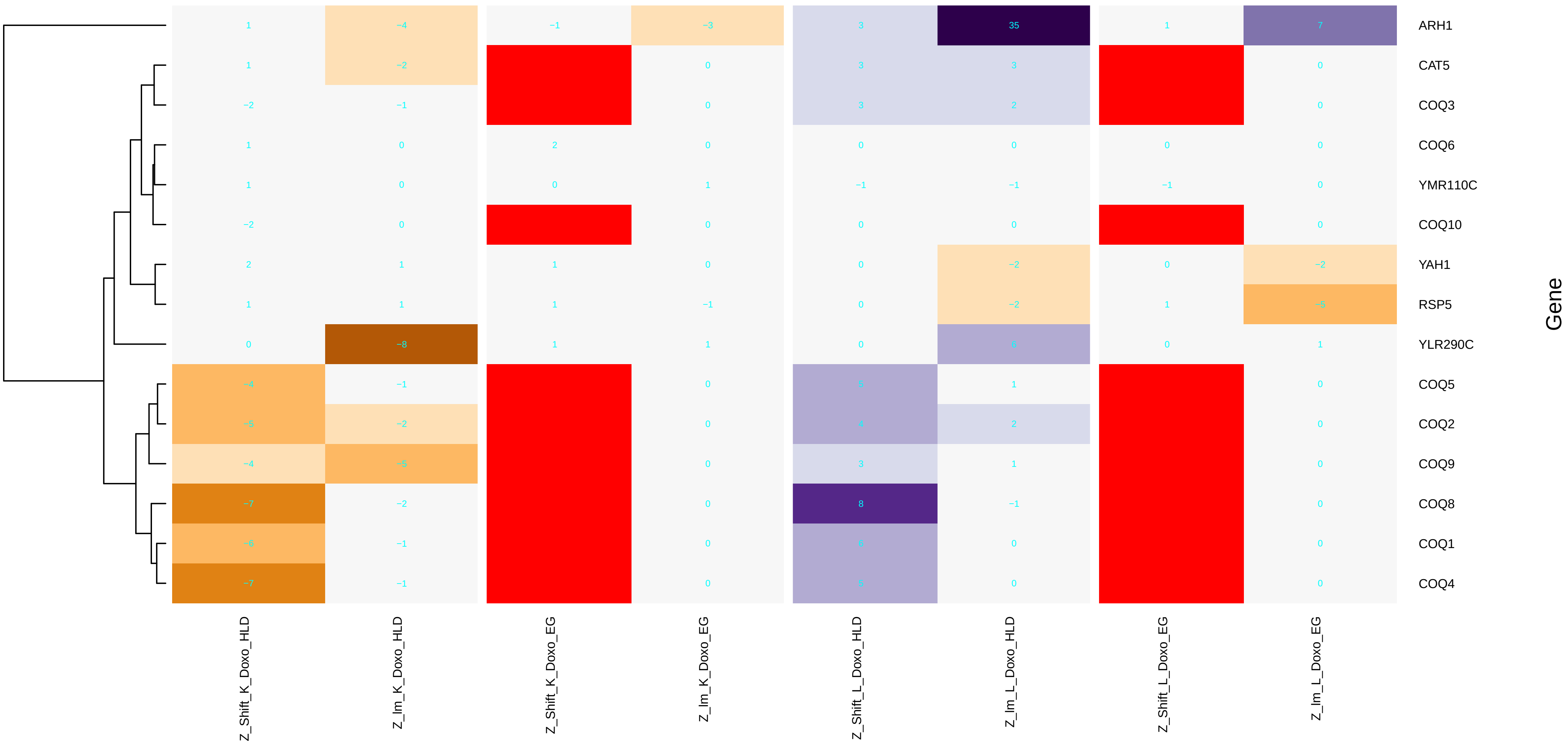

### kinetochore_assembly.pdf

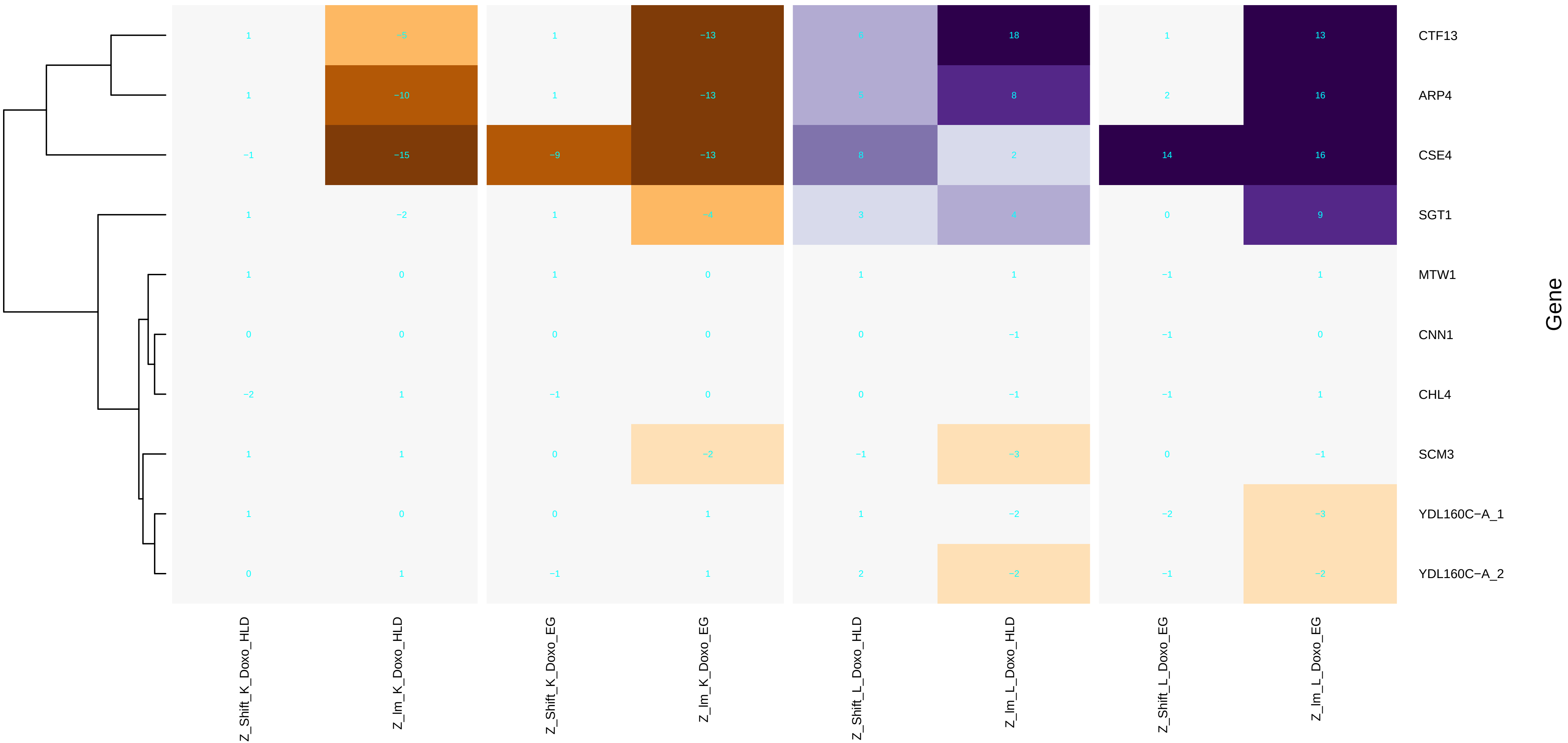

### kinetochore_organization.pdf

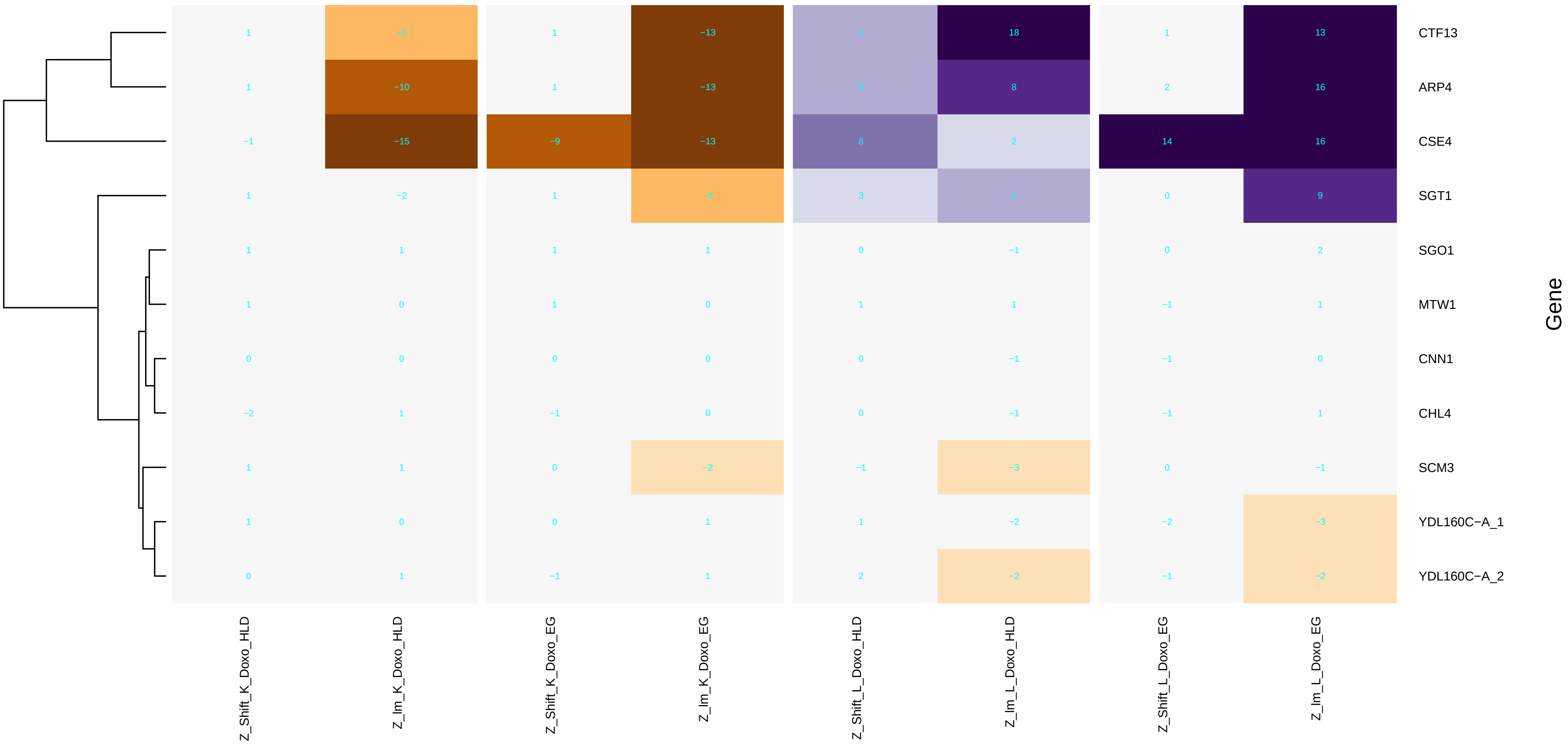

### lipid_homeostasis.pdf

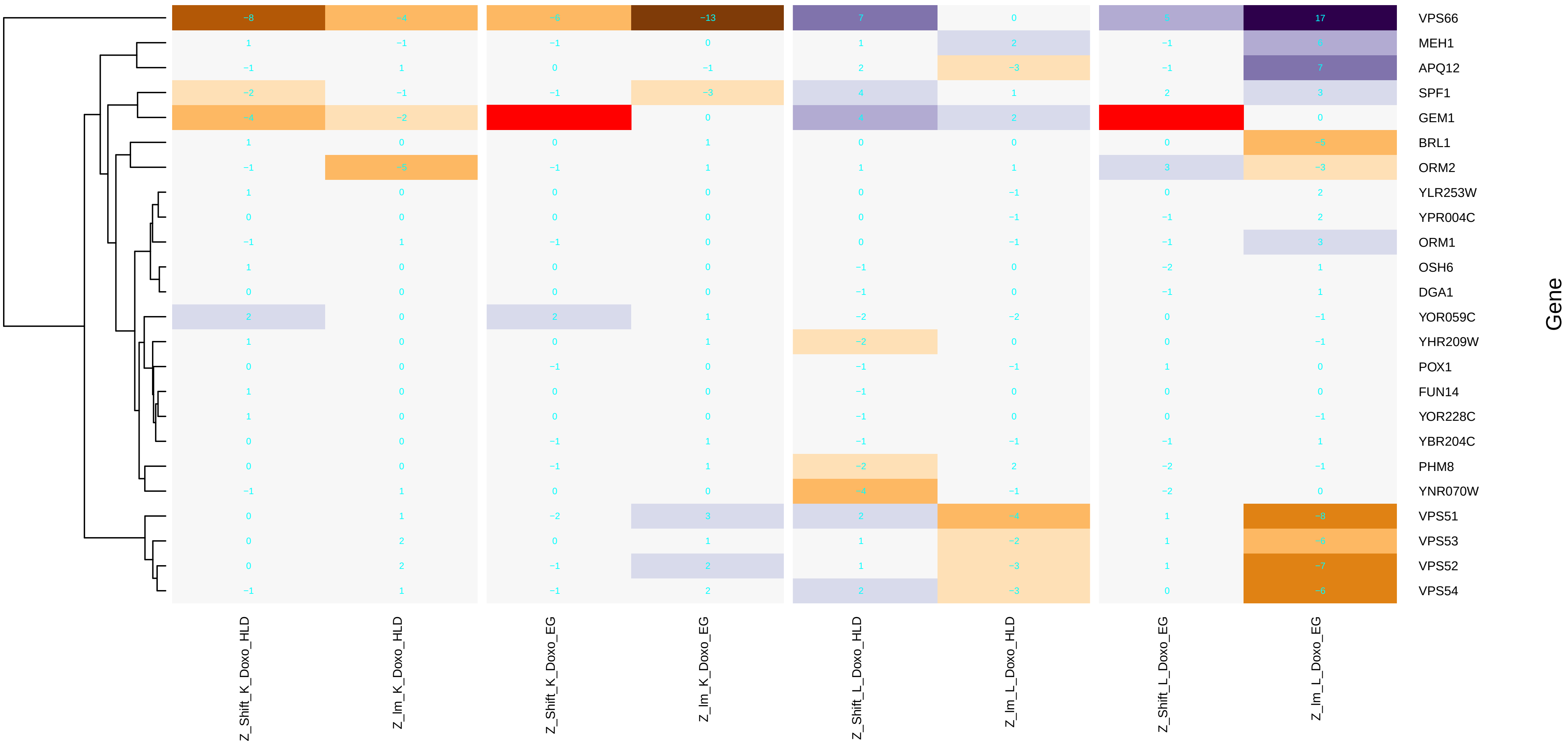

### maintenance_of_DNA_repeat_elements.pdf

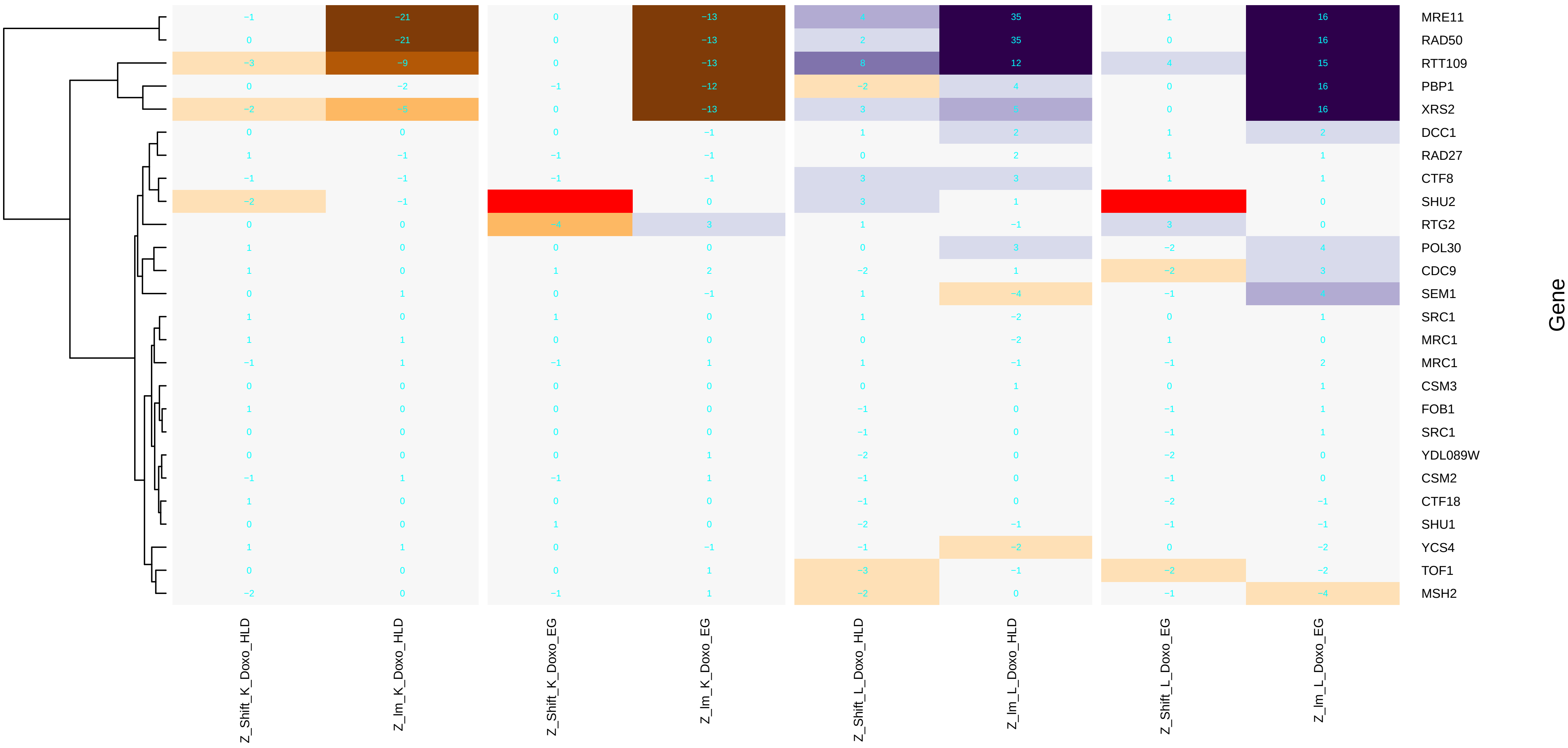

### maintenance_of_DNA_trinucleotide_repeats.pdf

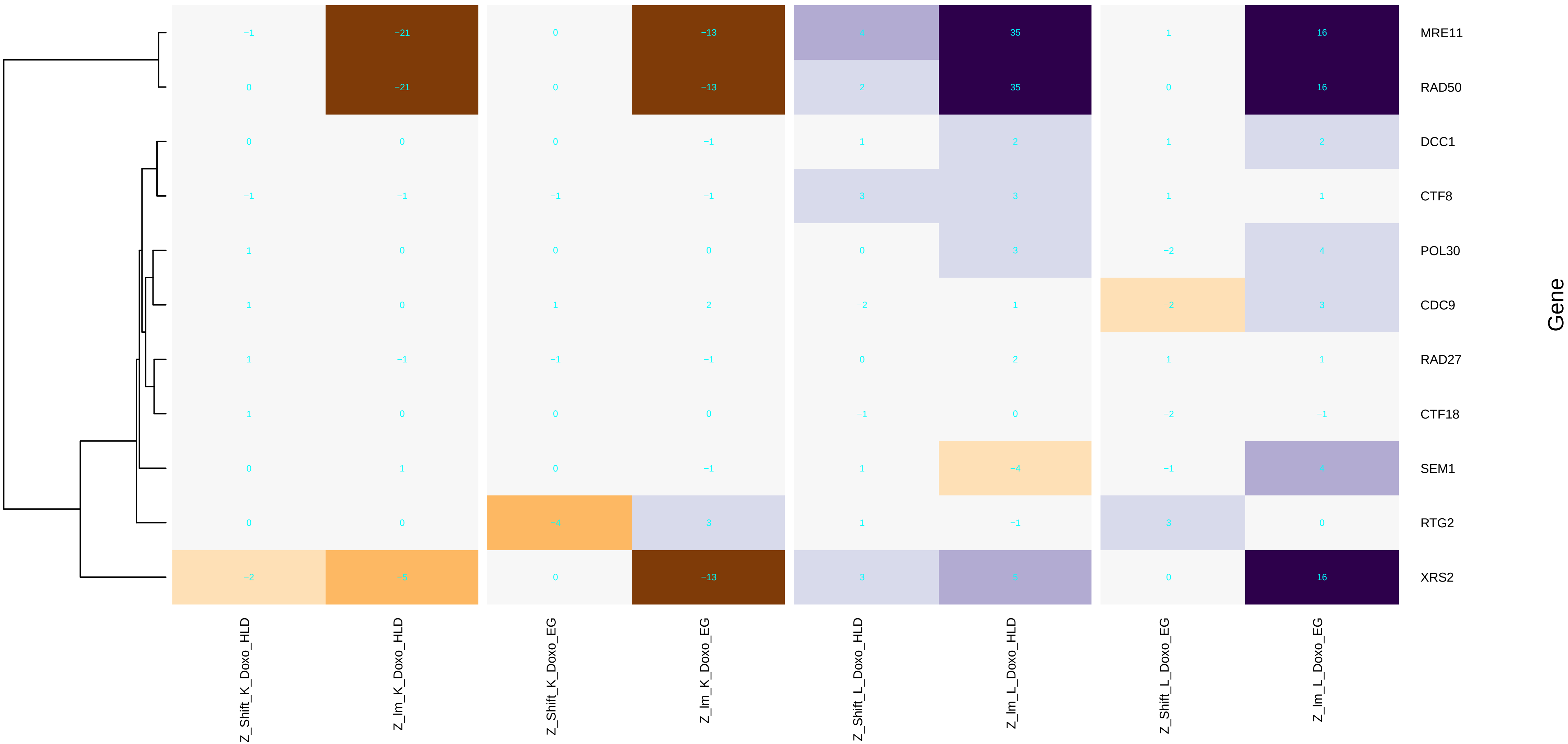

### maintenance_of_organelle_location.pdf

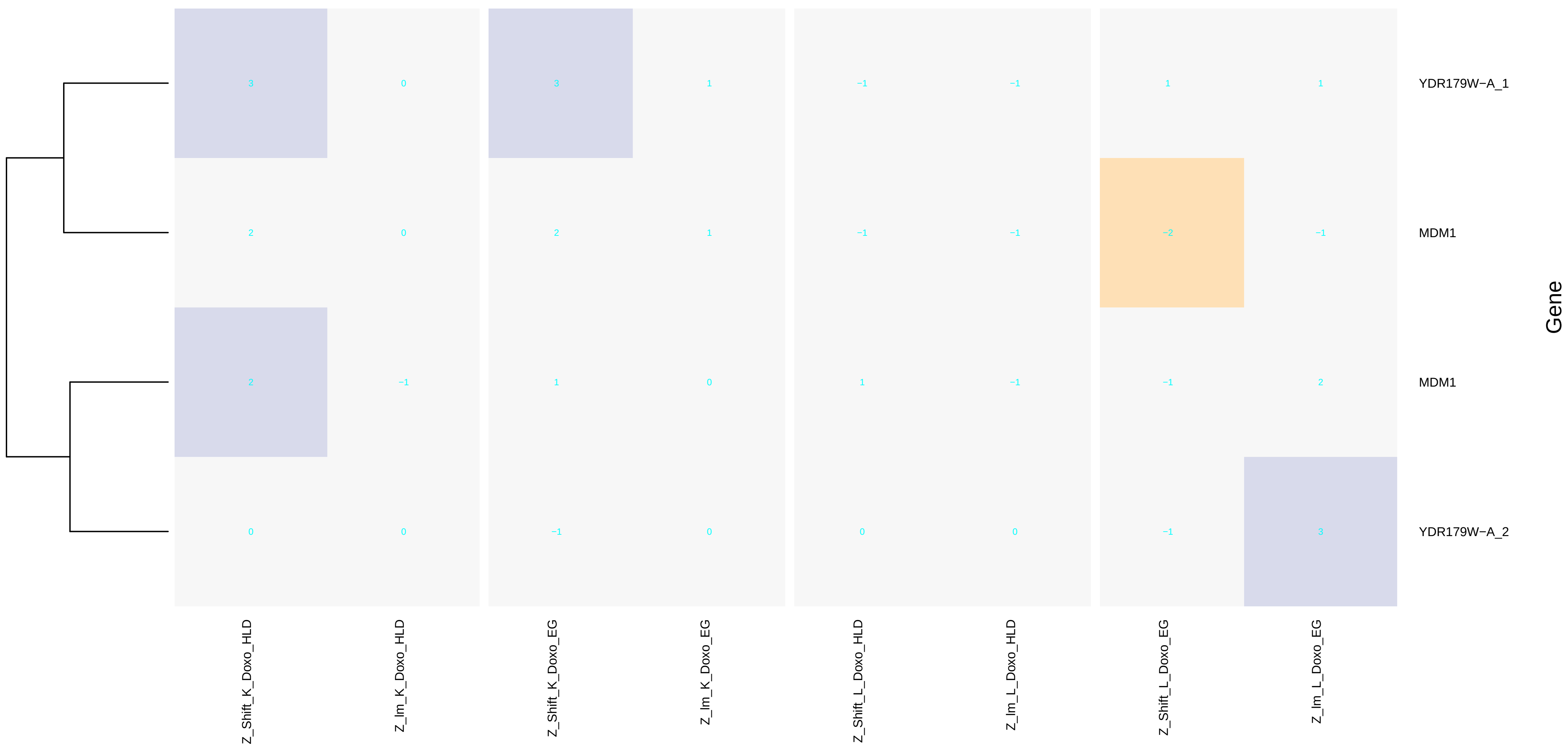

### maturation_of_5.8S_rRNA.pdf

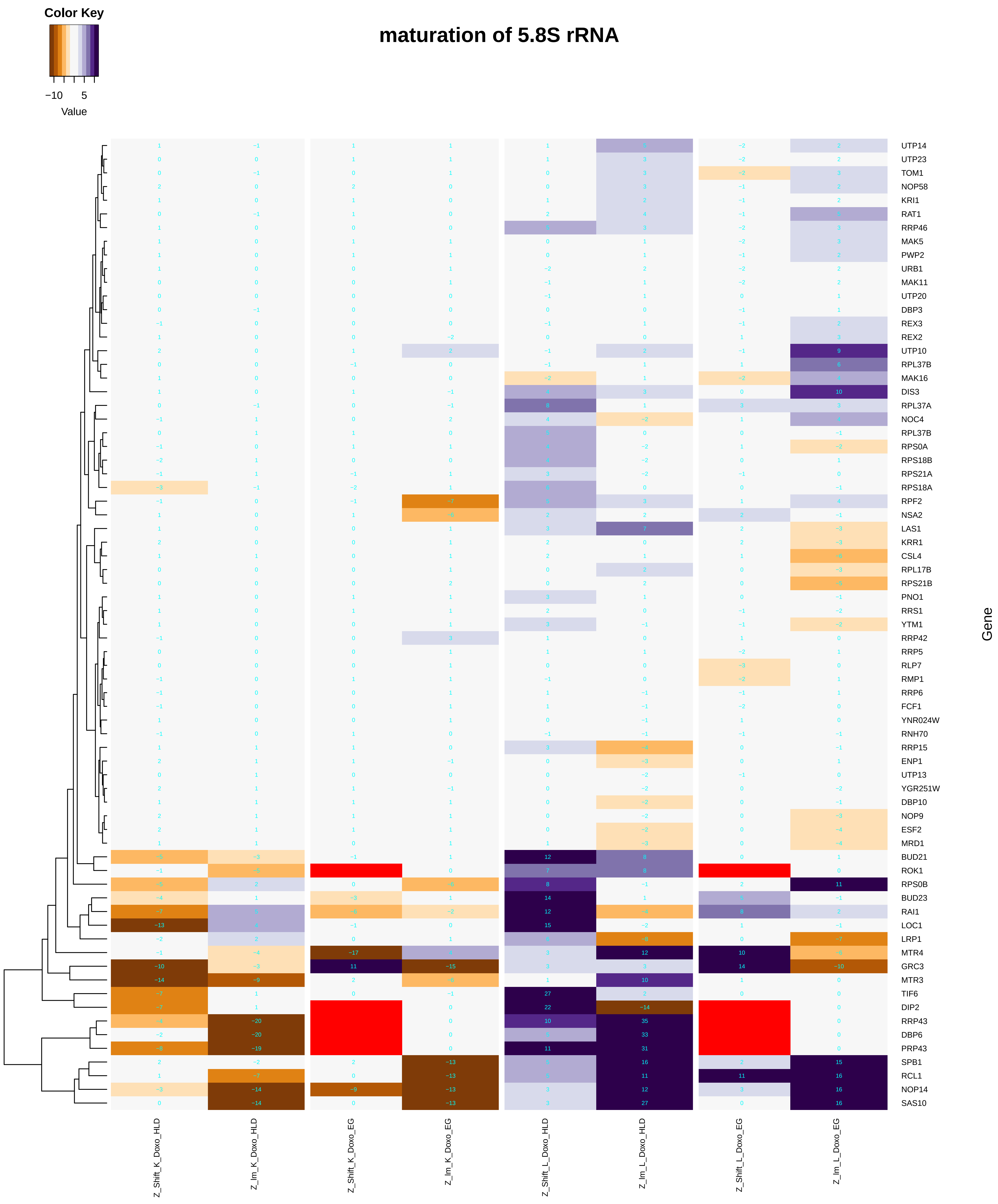

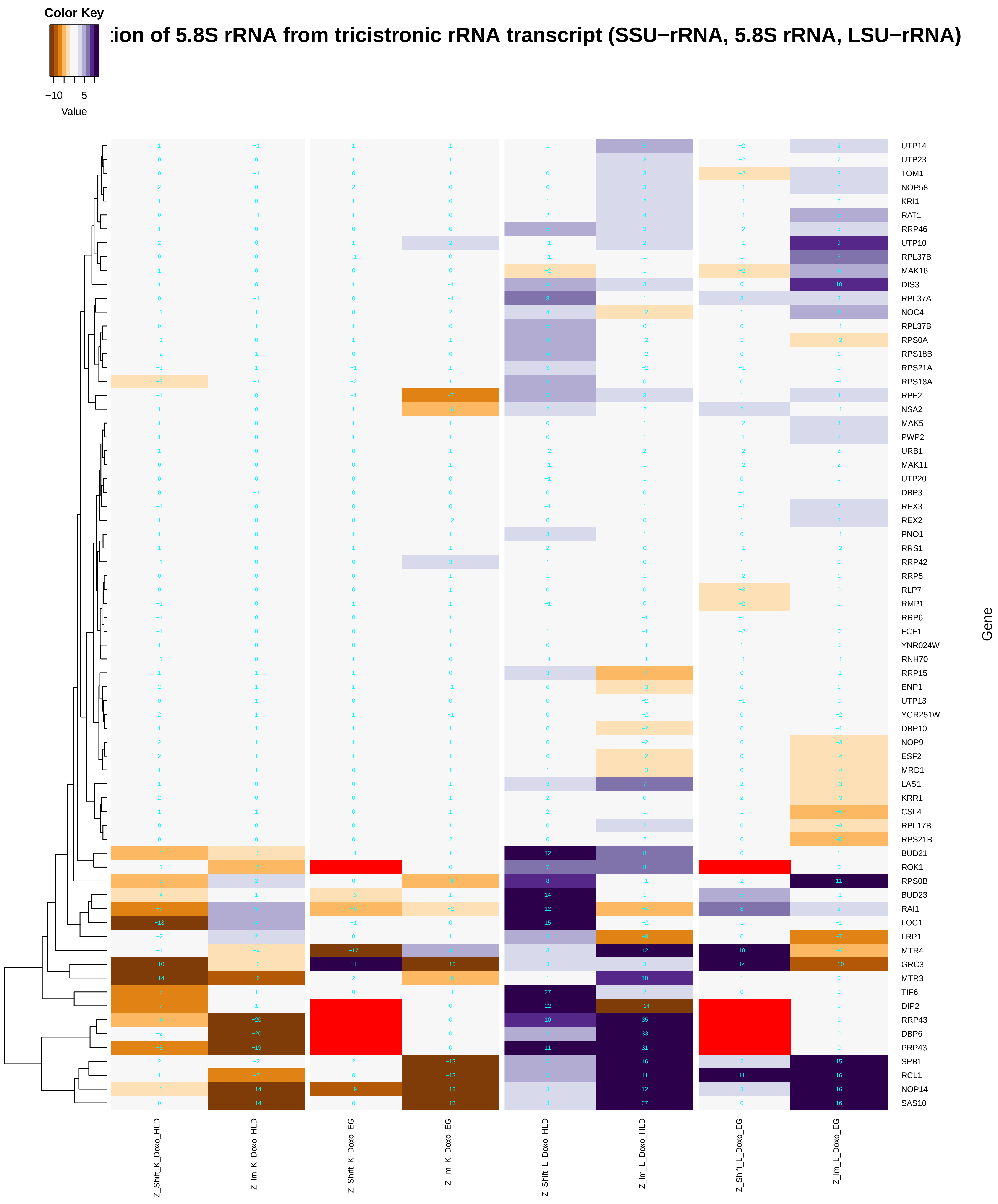

### maturation_of_5.8S_rRNA_from_tricistronic_rRNA_transcript_(SSU-rRNA,_5.8S_rRNA,_LSU-rRNA).pdf

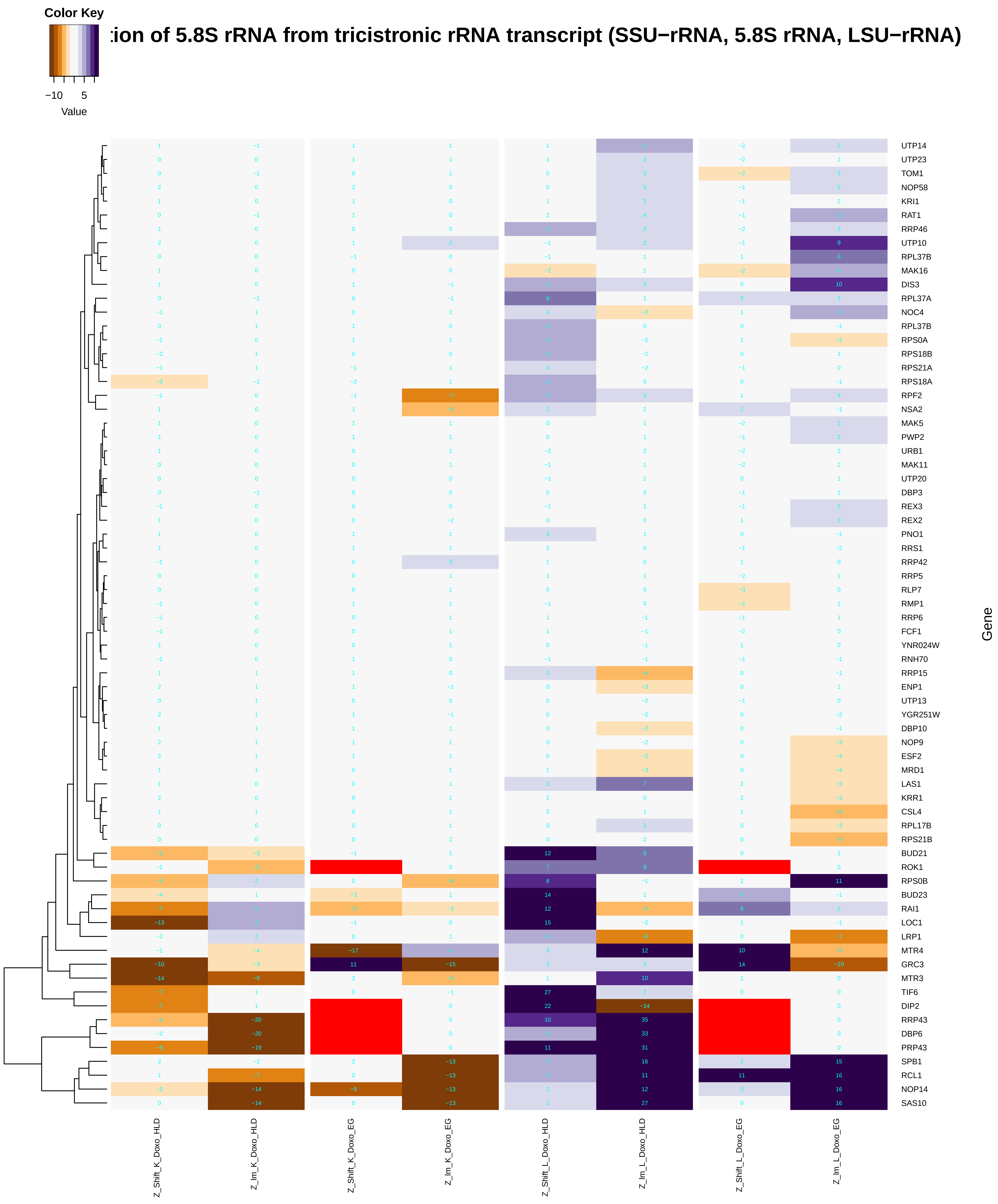

### maturation_of_SSU-rRNA.pdf

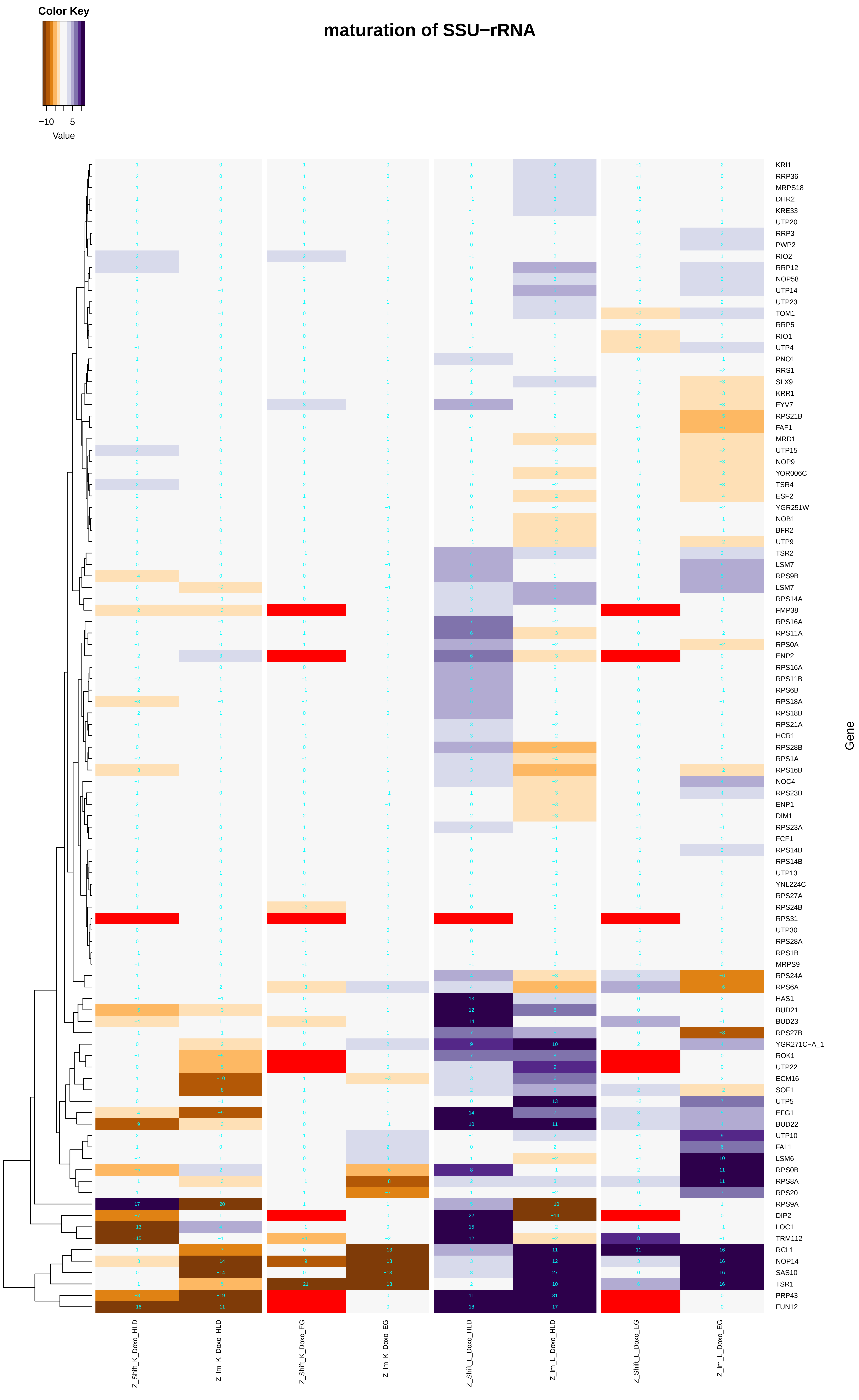

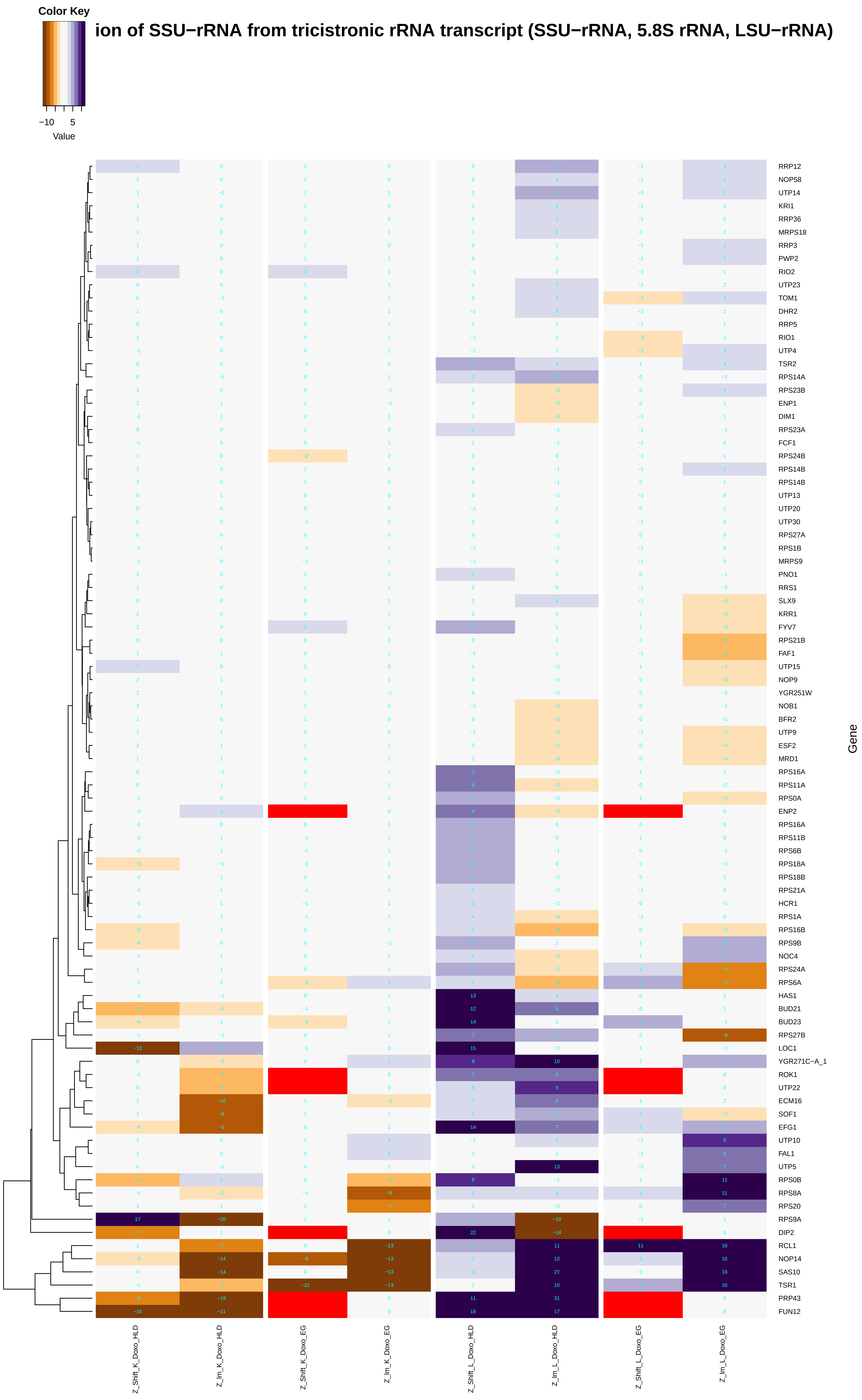

### maturation_of_SSU-rRNA_from_tricistronic_rRNA_transcript_(SSU-rRNA,_5.8S_rRNA,_LSU-rRNA).pdf

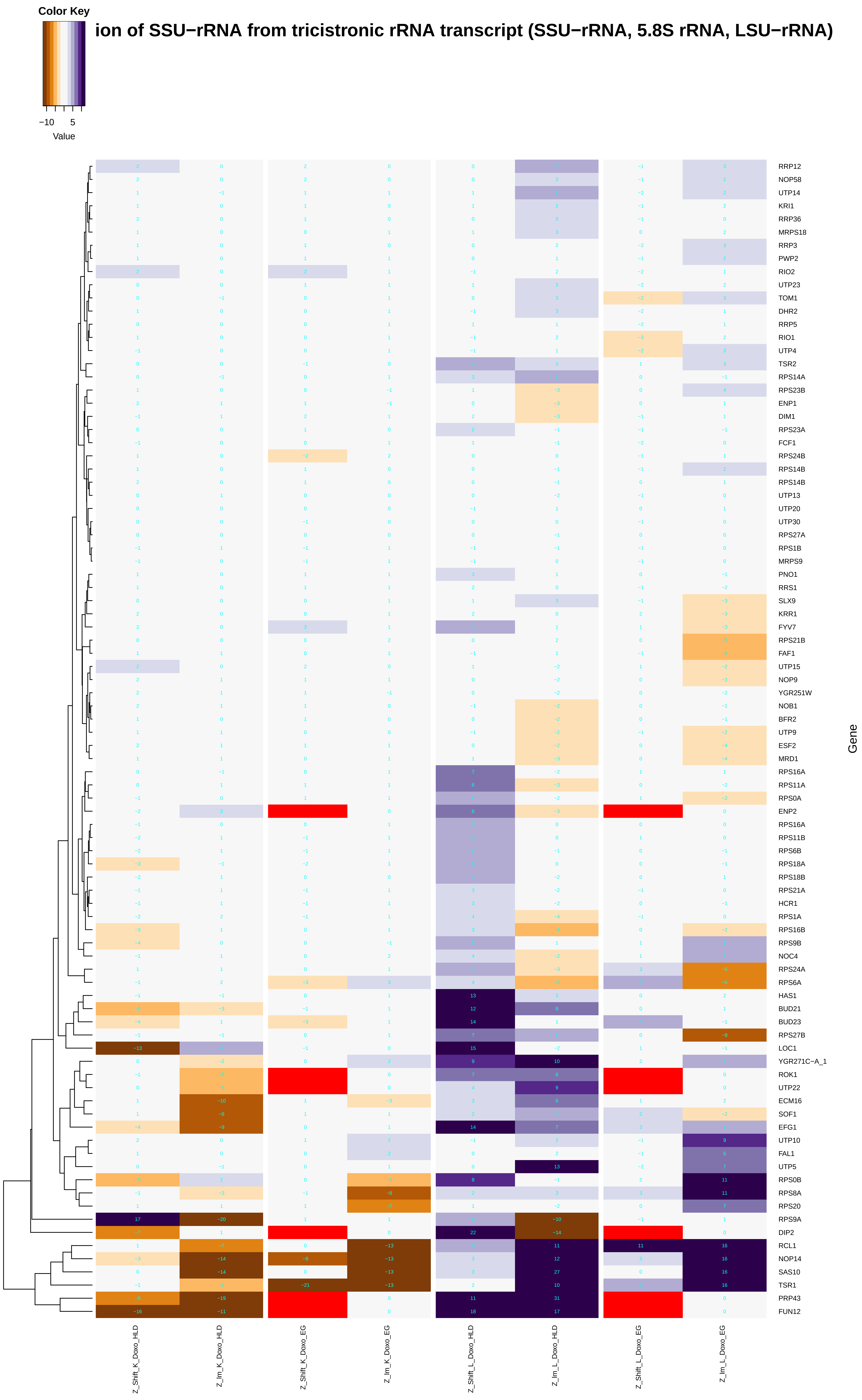

### meiosis_I.pdf

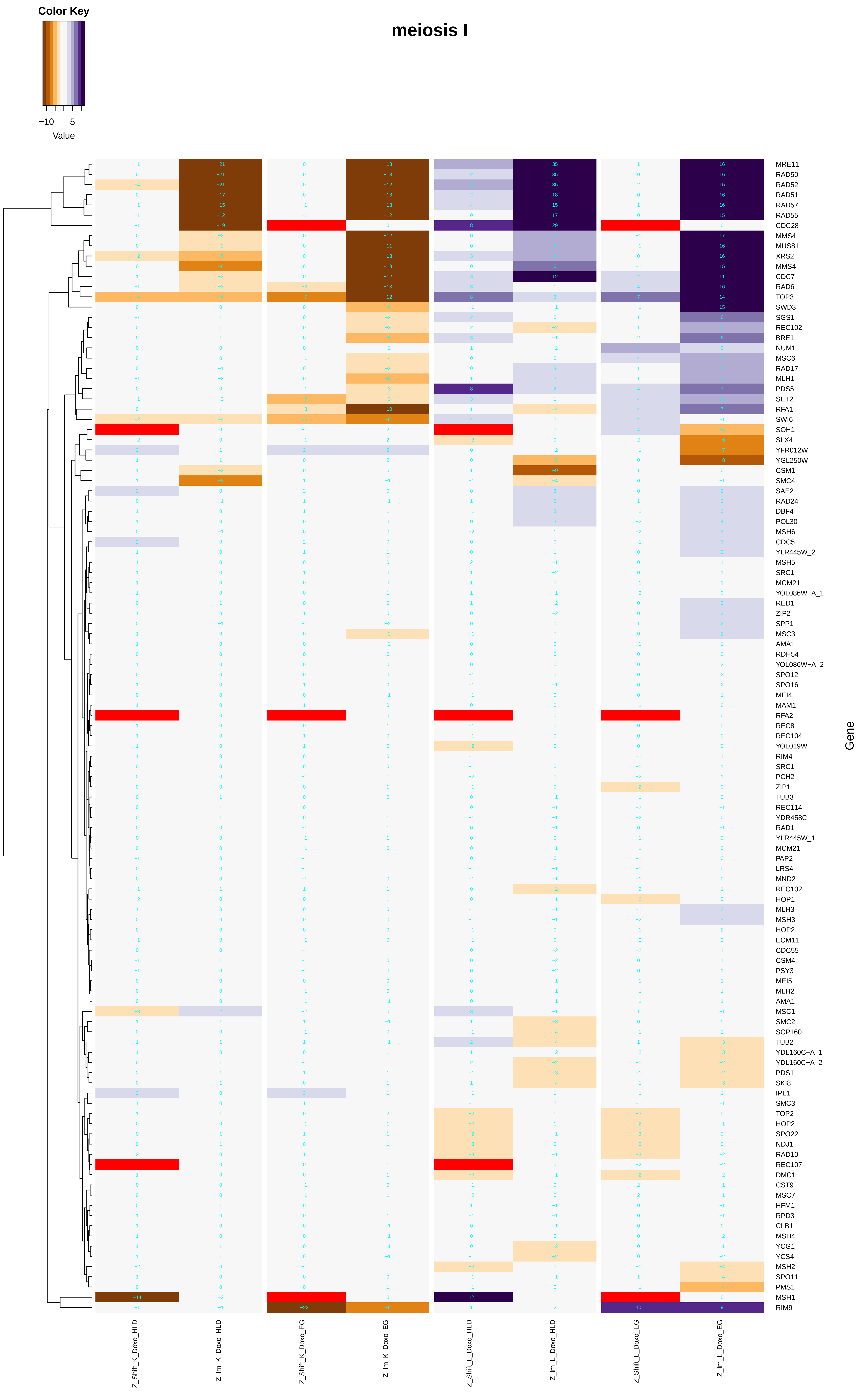

### meiotic_chromosome_condensation.pdf

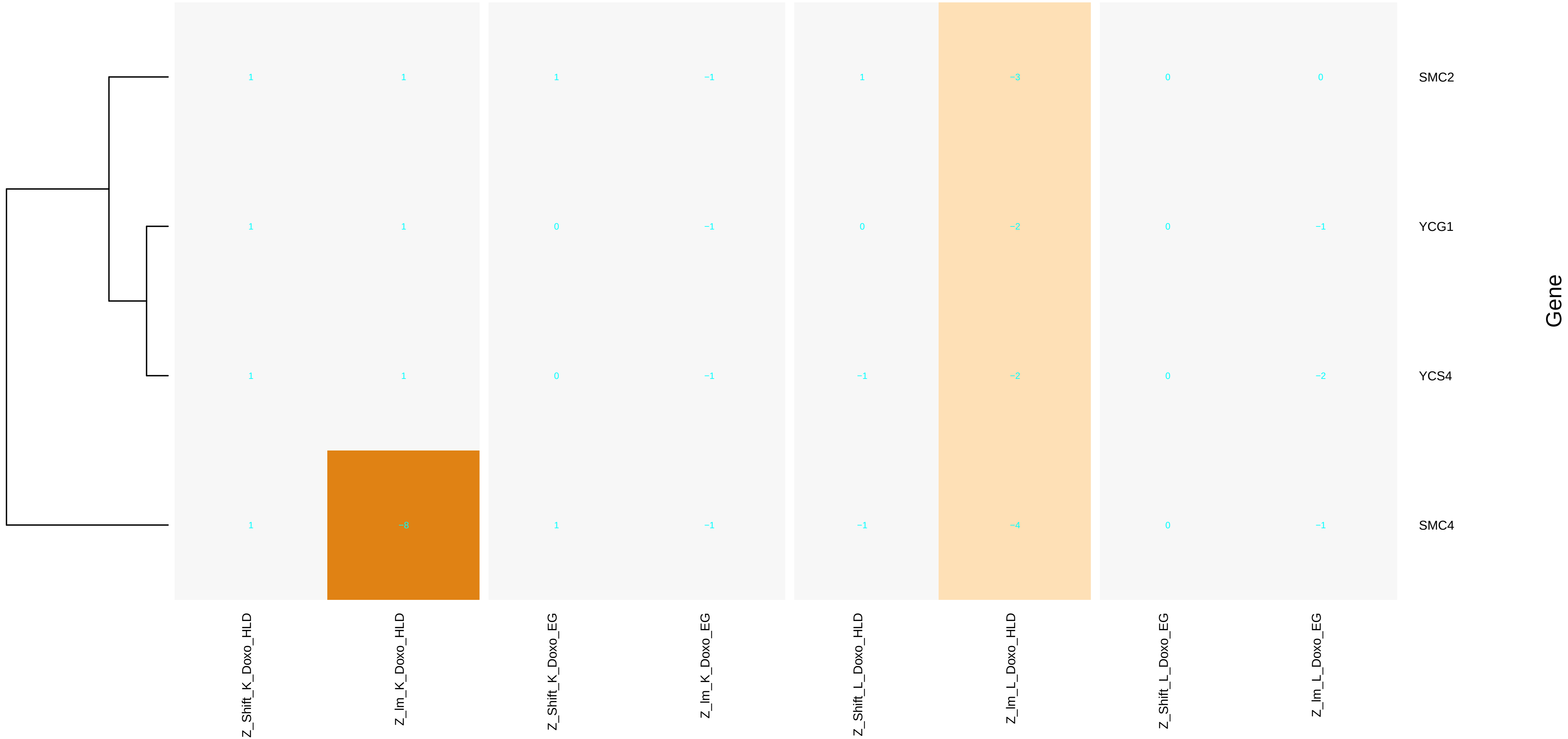

### meiotic_DNA_double-strand_break_formation.pdf

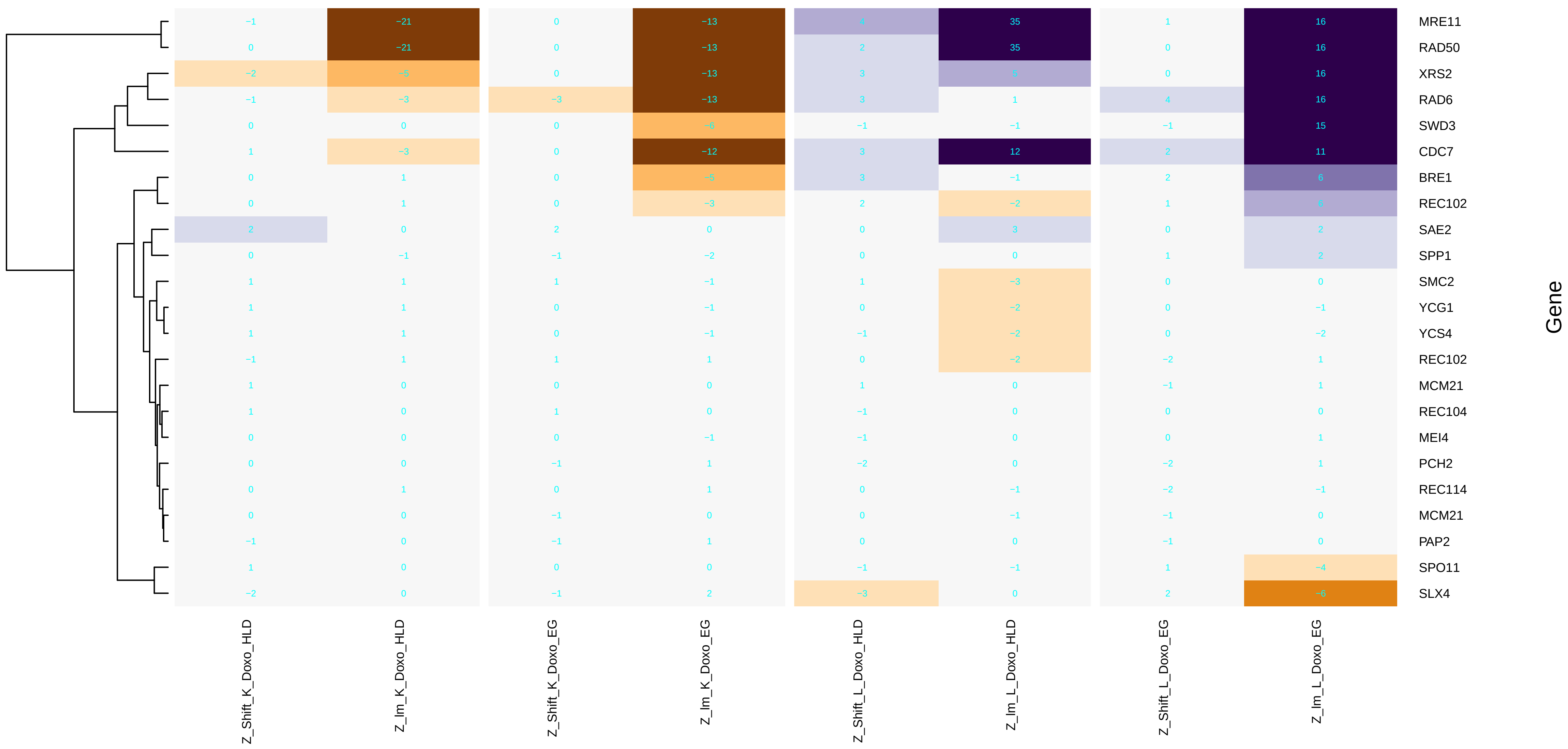

### meiotic_DNA_double-strand_break_processing.pdf

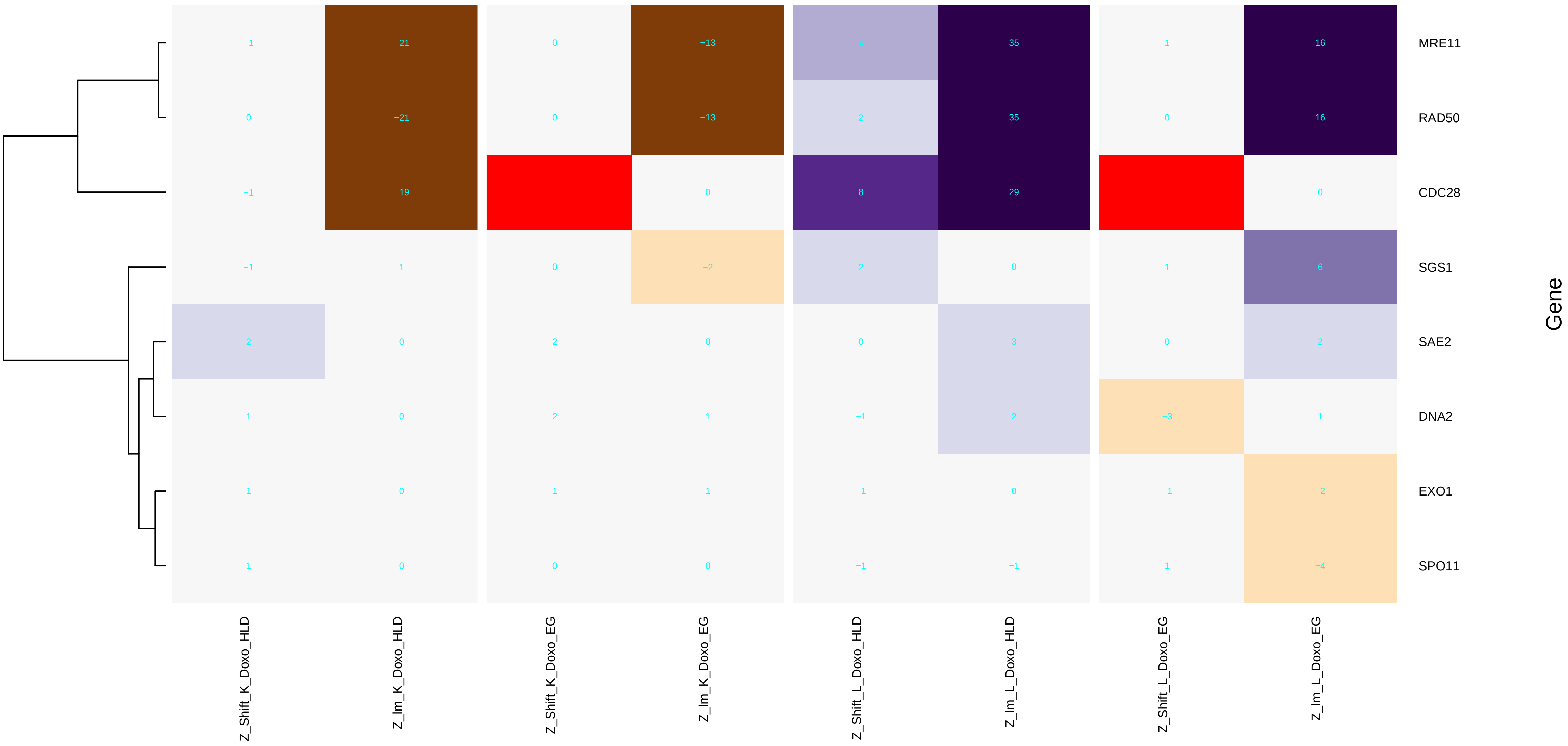

### meiotic_nuclear_division.pdf

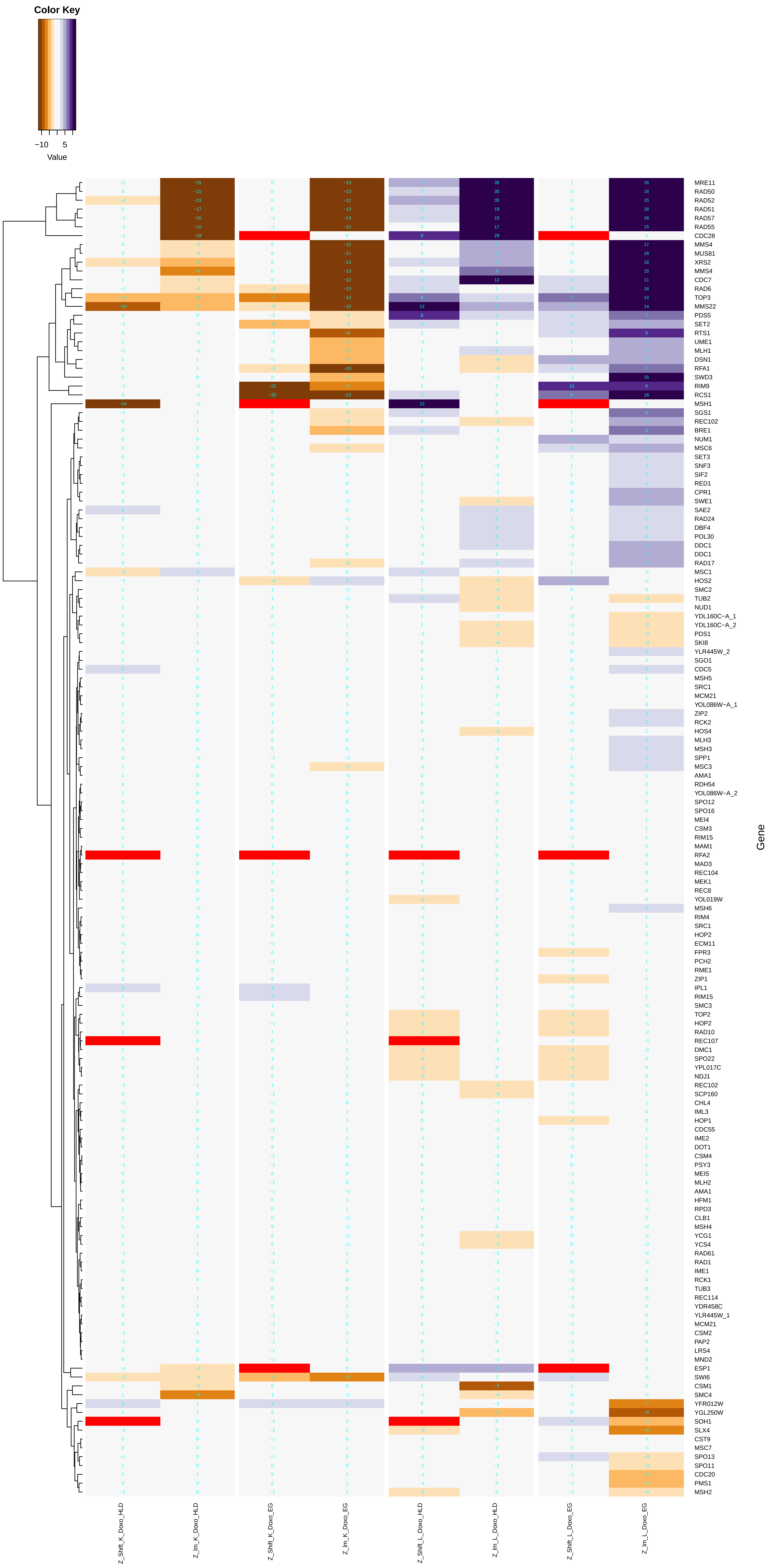

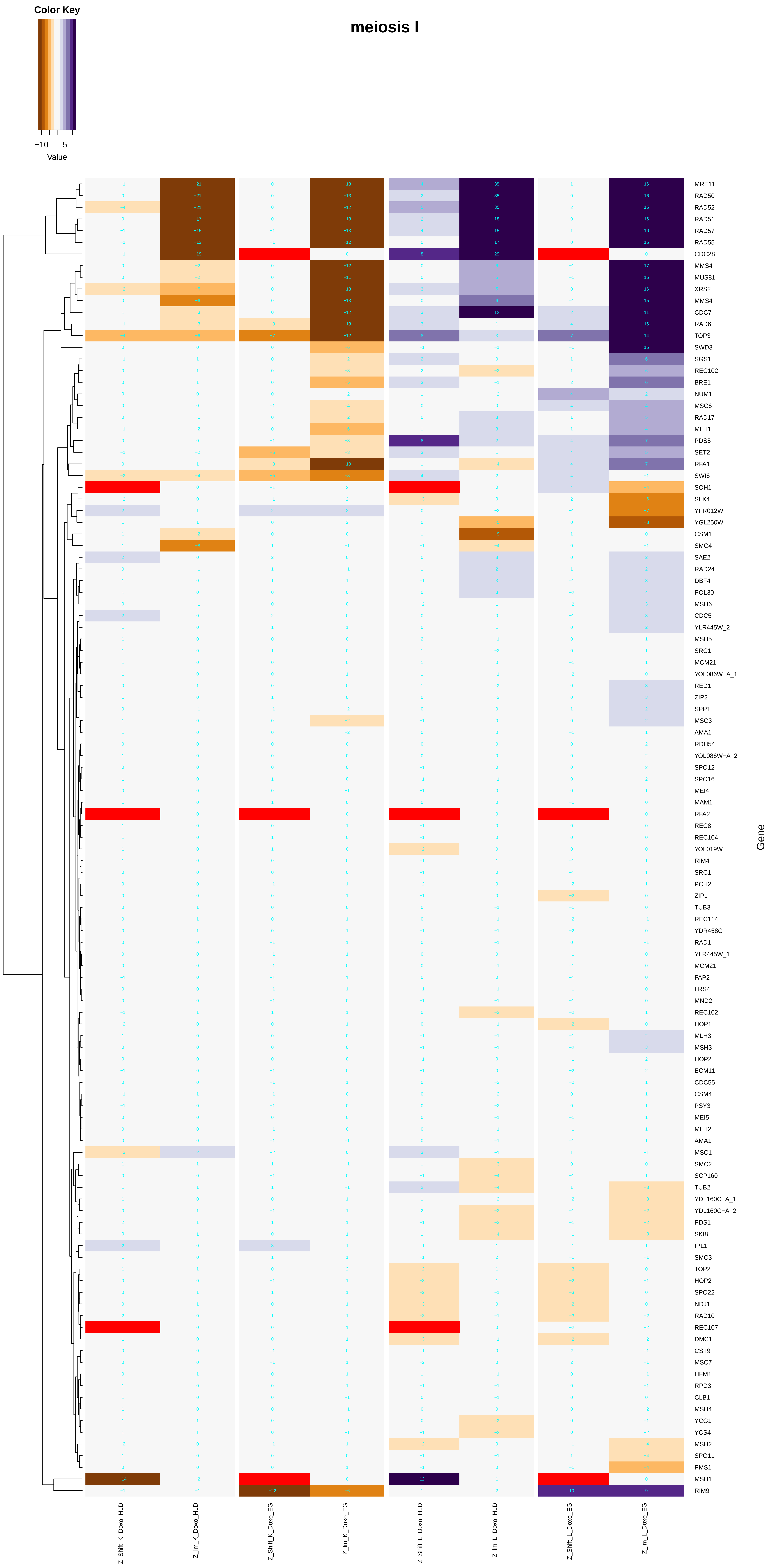

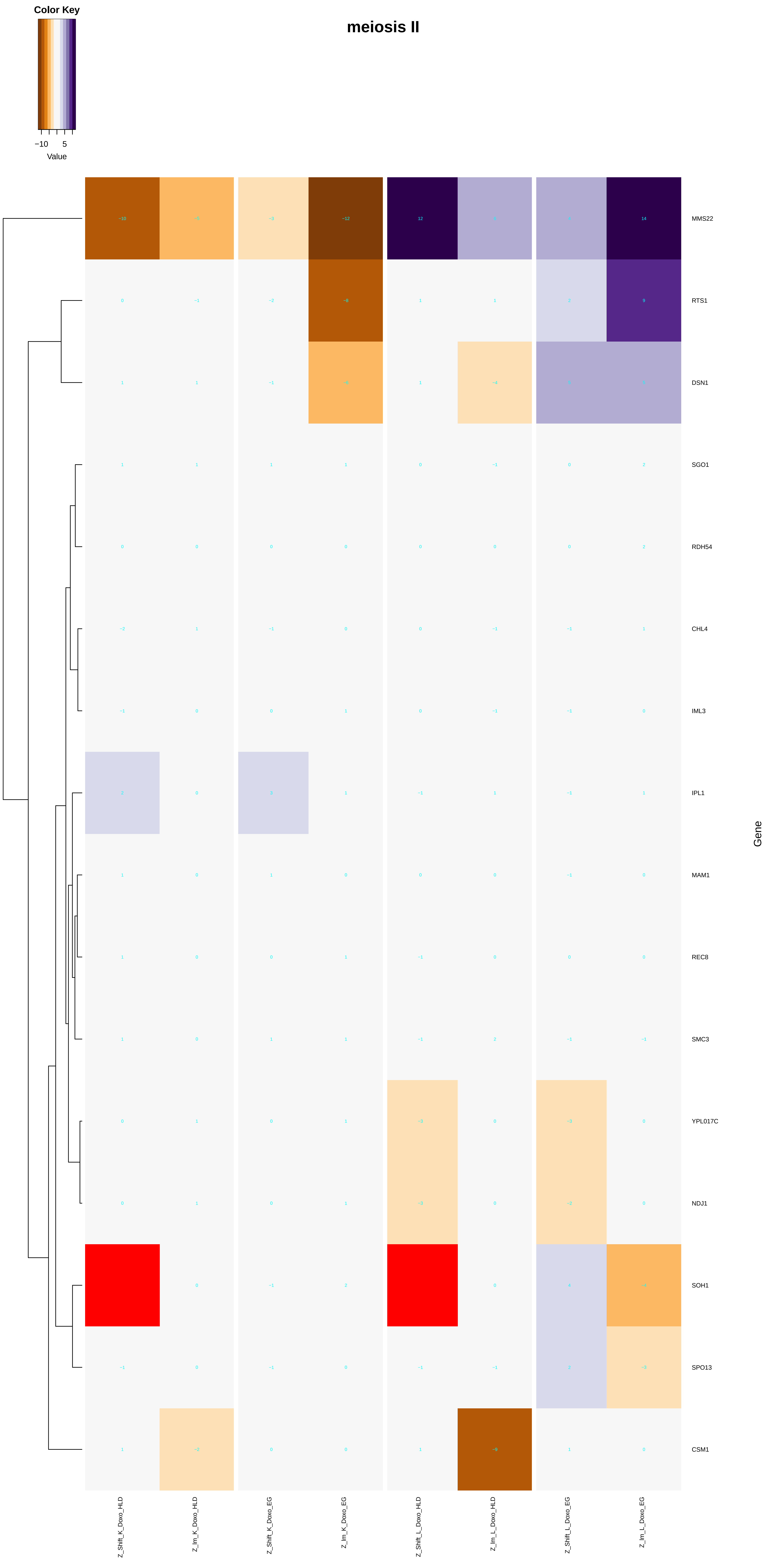

### membrane_lipid_biosynthetic_process.pdf

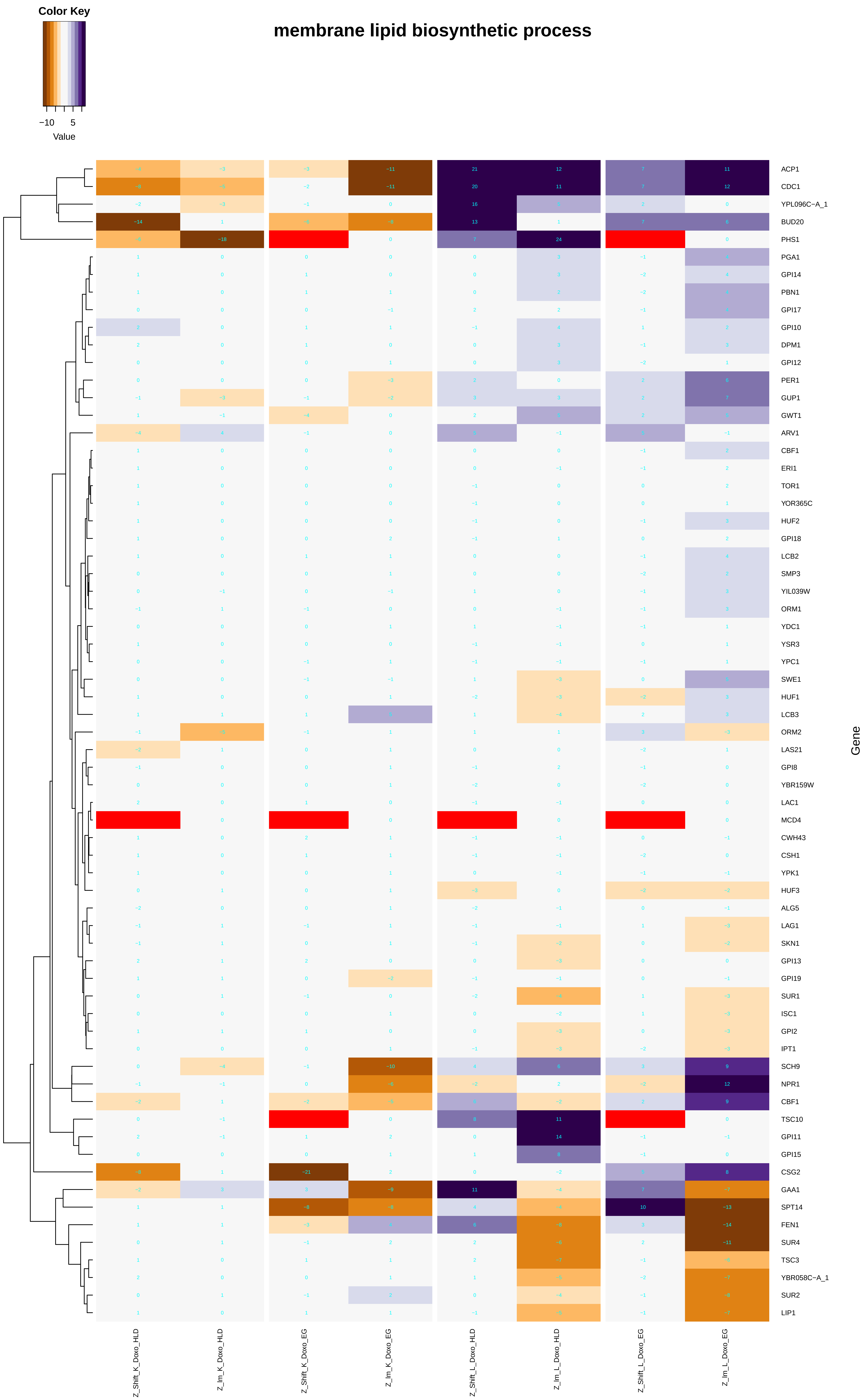

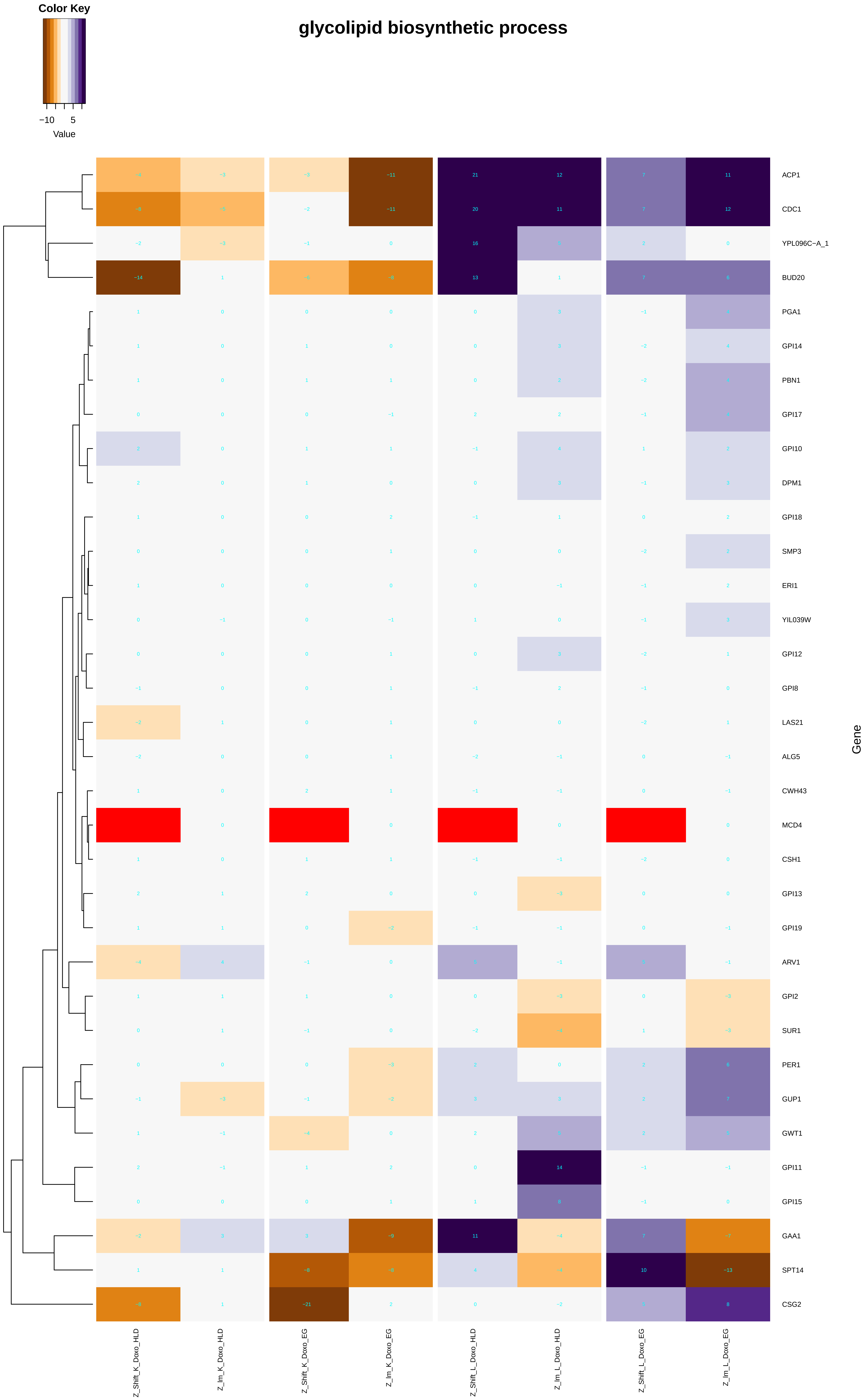

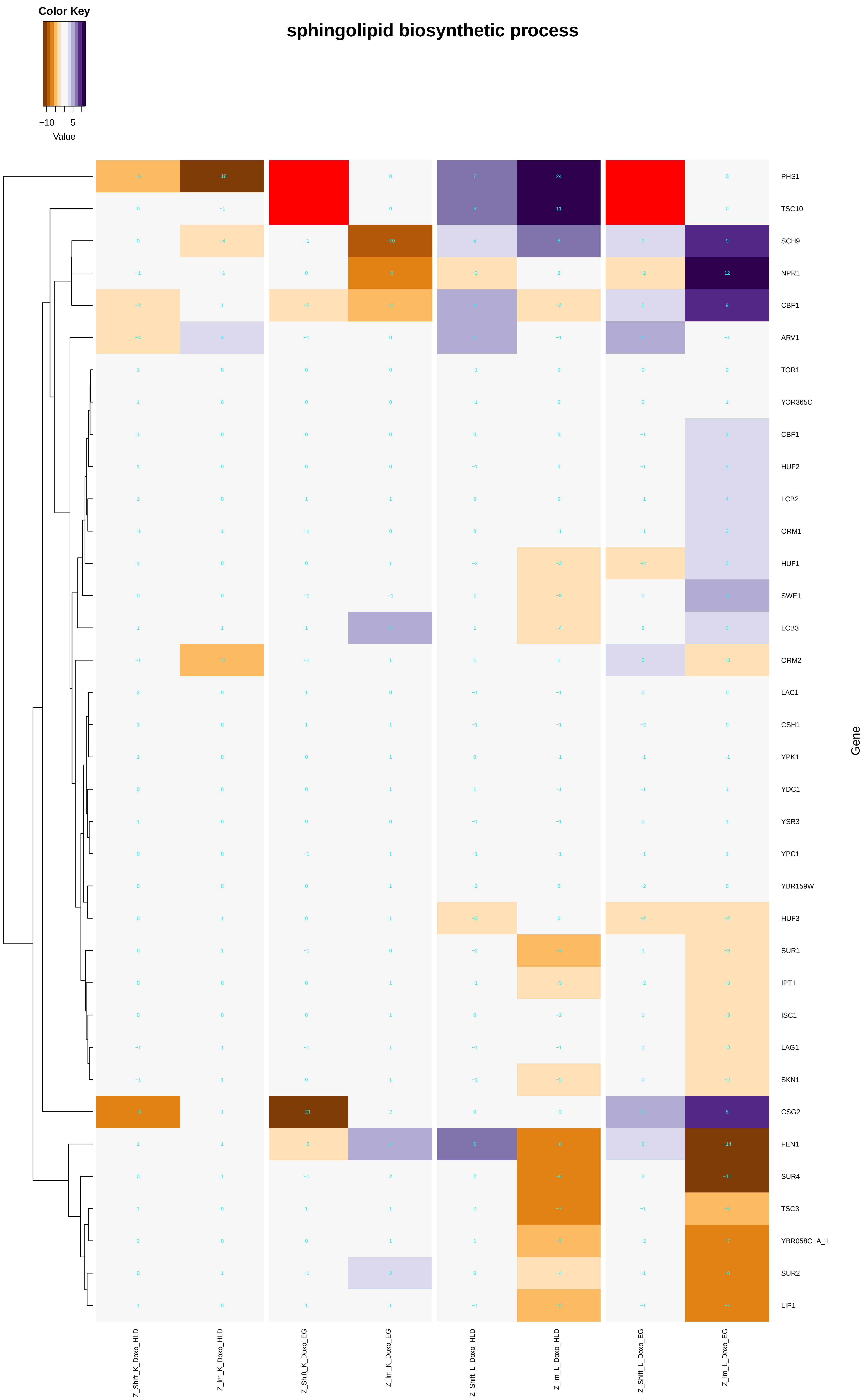

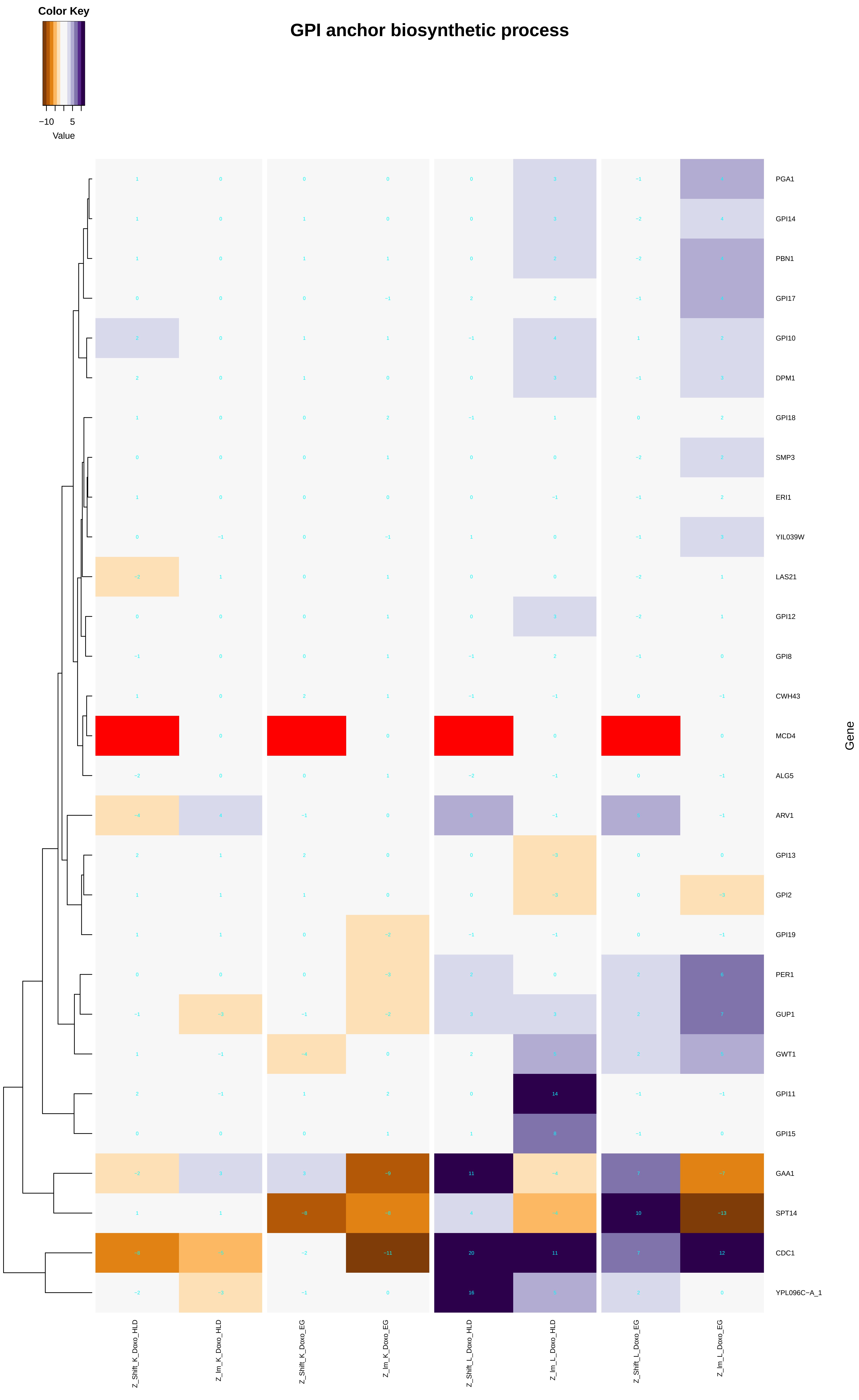

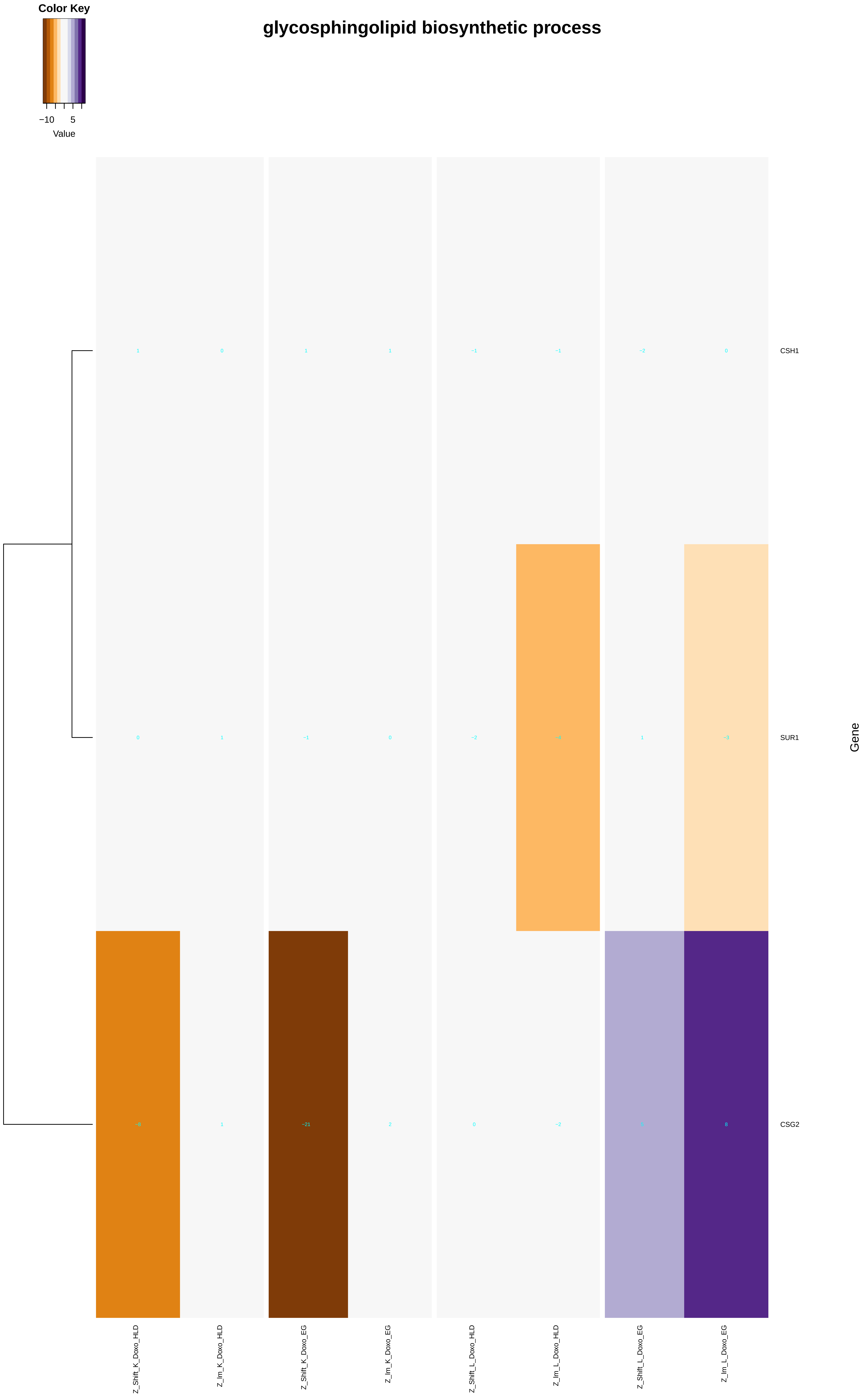

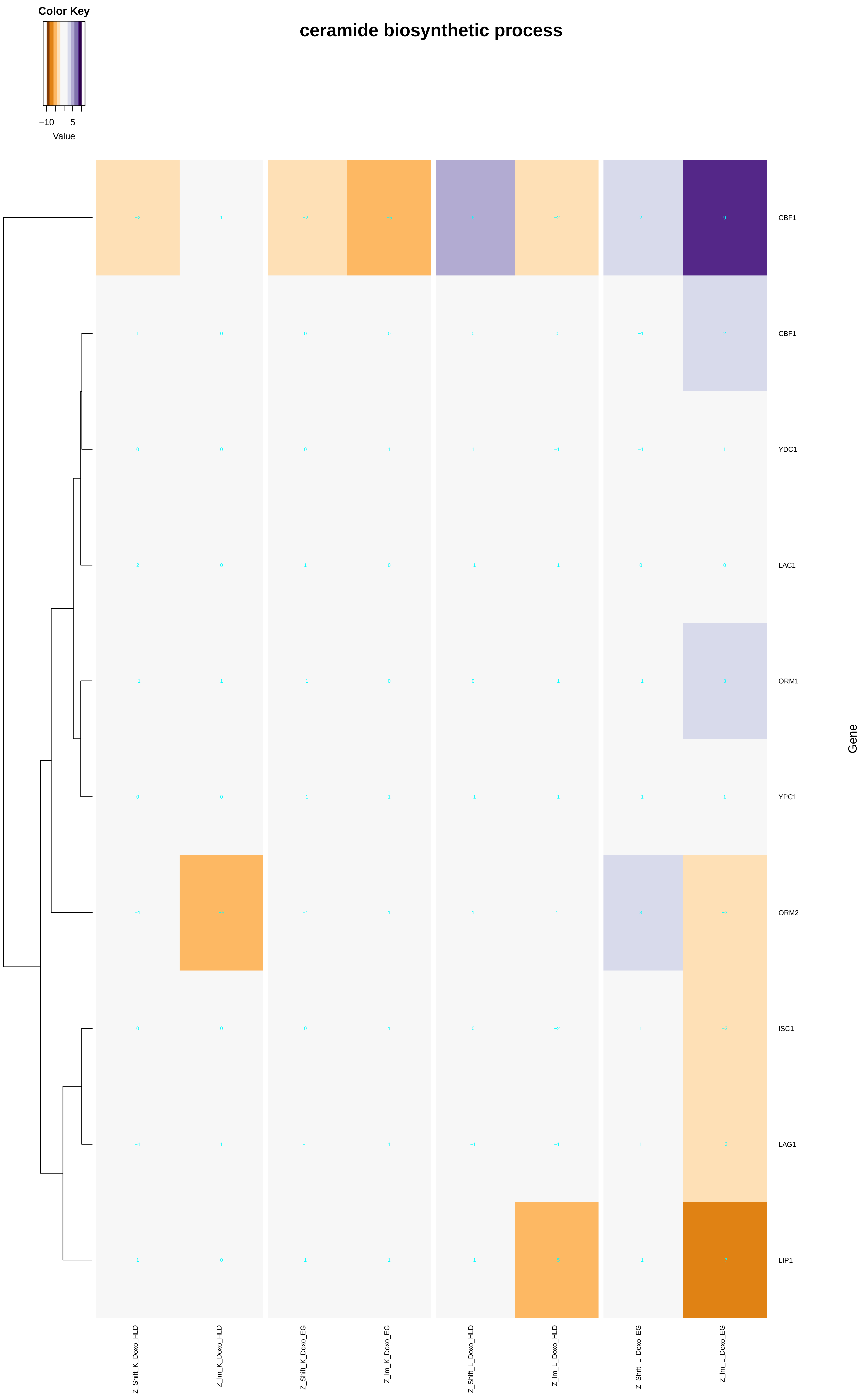
