## Additional File 9 for "A yeast phenomic model for the influence of Warburg metabolism on genetic buffering of doxorubicin": actin_filament-based_process.pdf

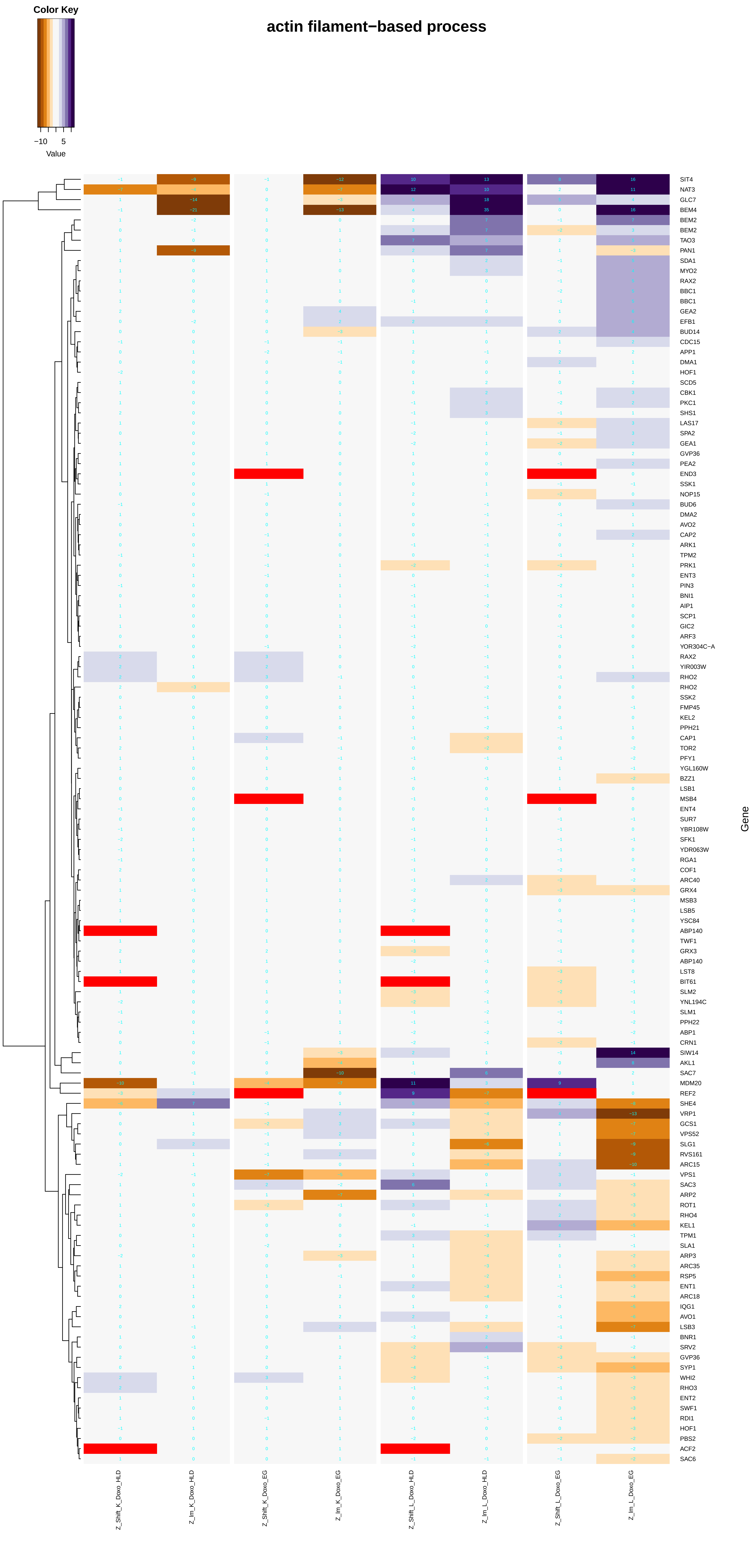

Color Key
−10
5
Value
actin filament−based process
SIT4
NAT3
GLC7
BEM4
BEM2
BEM2
TAO3
PAN1
SDA1
MYO2
RAX2
BBC1
BBC1
GEA2
EFB1
BUD14
CDC15
APP1
DMA1
HOF1
SCD5
CBK1
PKC1
SHS1
LAS17
SPA2
GEA1
GVP36
PEA2
END3
SSK1
NOP15
BUD6
DMA2
AVO2
CAP2
ARK1
TPM2
PRK1
ENT3
PIN3
BNI1
AIP1
SCP1
GIC2
ARF3
YOR304C−A
RAX2
YIR003W
RHO2
RHO2
SSK2
FMP45
KEL2
PPH21
CAP1
TOR2
PFY1
YGL160W
BZZ1
LSB1
MSB4
ENT4
SUR7
YBR108W
SFK1
YDR063W
RGA1
COF1
ARC40
GRX4
MSB3
LSB5
YSC84
ABP140
TWF1
GRX3
ABP140
LST8
BIT61
SLM2
YNL194C
SLM1
PPH22
ABP1
CRN1
SIW14
AKL1
SAC7
MDM20
REF2
SHE4
VRP1
GCS1
VPS52
SLG1
RVS161
ARC15
VPS1
SAC3
ARP2
ROT1
RHO4
KEL1
TPM1
SLA1
ARP3
ARC35
RSP5
ENT1
ARC18
IQG1
AVO1
LSB3
BNR1
SRV2
GVP36
SYP1
WHI2
RHO3
ENT2
SWF1
RDI1
HOF1
PBS2
ACF2
SAC6
Gene
Z_Shift_K_Doxo_HLD
Z_lm_K_Doxo_HLD
Z_Shift_K_Doxo_EG
Z_lm_K_Doxo_EG
Z_Shift_L_Doxo_HLD
Z_lm_L_Doxo_HLD
Z_Shift_L_Doxo_EG
Z_lm_L_Doxo_EG

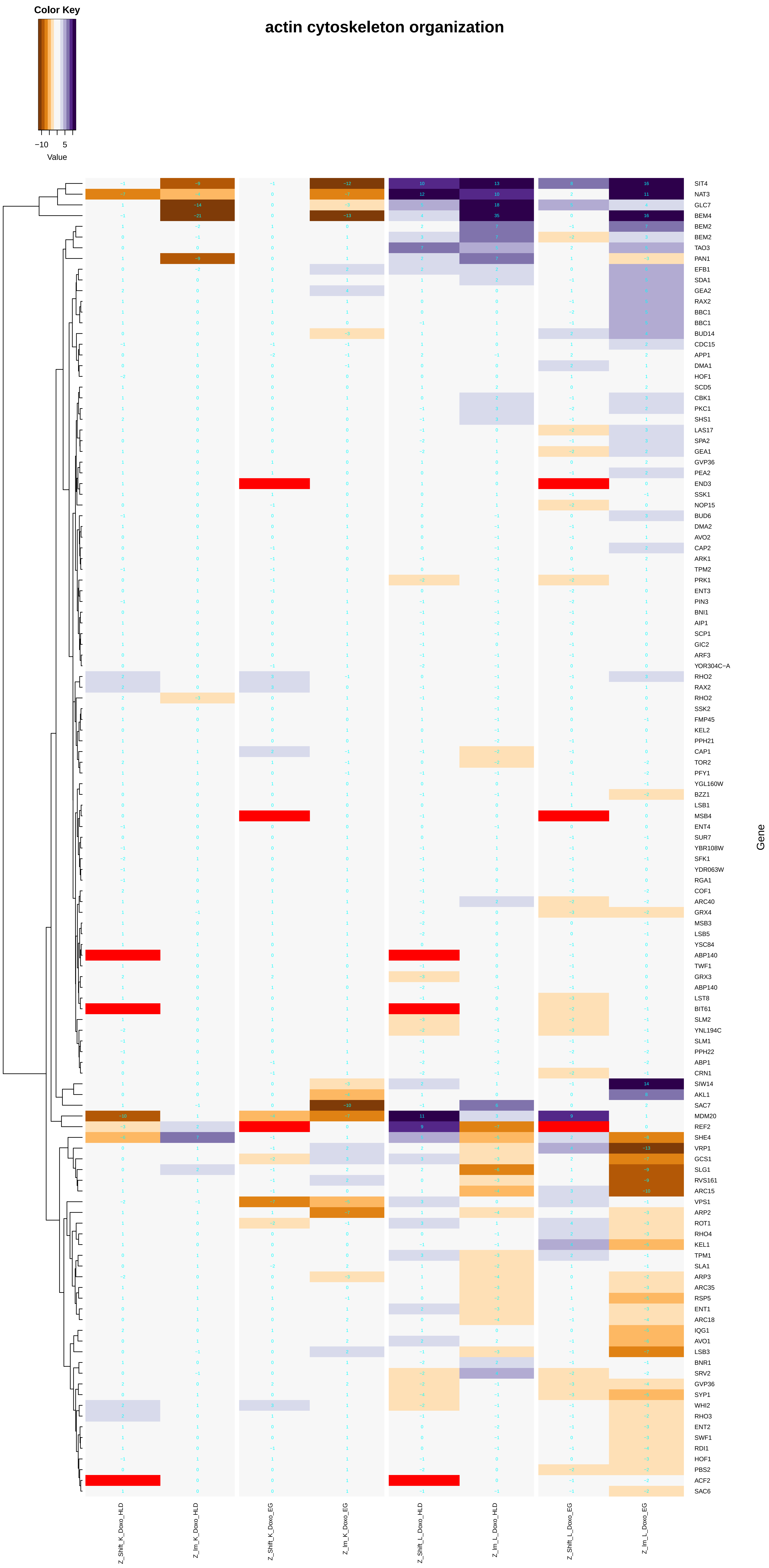

actin cytoskeleton organization
Color Key
−10
5
Value
Gene
SIT4
NAT3
GLC7
BEM4
BEM2
BEM2
TAO3
PAN1
EFB1
SDA1
GEA2
RAX2
BBC1
BBC1
BUD14
CDC15
APP1
DMA1
HOF1
SCD5
CBK1
PKC1
SHS1
LAS17
SPA2
GEA1
GVP36
PEA2
END3
SSK1
NOP15
BUD6
DMA2
AVO2
CAP2
ARK1
TPM2
PRK1
ENT3
PIN3
BNI1
AIP1
SCP1
GIC2
ARF3
YOR304C−A
RHO2
RAX2
RHO2
SSK2
FMP45
KEL2
PPH21
CAP1
TOR2
PFY1
YGL160W
BZZ1
LSB1
MSB4
ENT4
SUR7
YBR108W
SFK1
YDR063W
RGA1
COF1
ARC40
GRX4
MSB3
LSB5
YSC84
ABP140
TWF1
GRX3
ABP140
LST8
BIT61
SLM2
YNL194C
SLM1
PPH22
ABP1
CRN1
SIW14
AKL1
SAC7
MDM20
REF2
SHE4
VRP1
GCS1
SLG1
RVS161
ARC15
VPS1
ARP2
ROT1
RHO4
KEL1
TPM1
SLA1
ARP3
ARC35
RSP5
ENT1
ARC18
IQG1
AVO1
LSB3
BNR1
SRV2
GVP36
SYP1
WHI2
RHO3
ENT2
SWF1
RDI1
HOF1
PBS2
ACF2
SAC6
Z_Shift_K_Doxo_HLD
Z_lm_K_Doxo_HLD
Z_Shift_K_Doxo_EG
Z_lm_K_Doxo_EG
Z_Shift_L_Doxo_HLD
Z_lm_L_Doxo_HLD
Z_Shift_L_Doxo_EG
Z_lm_L_Doxo_EG

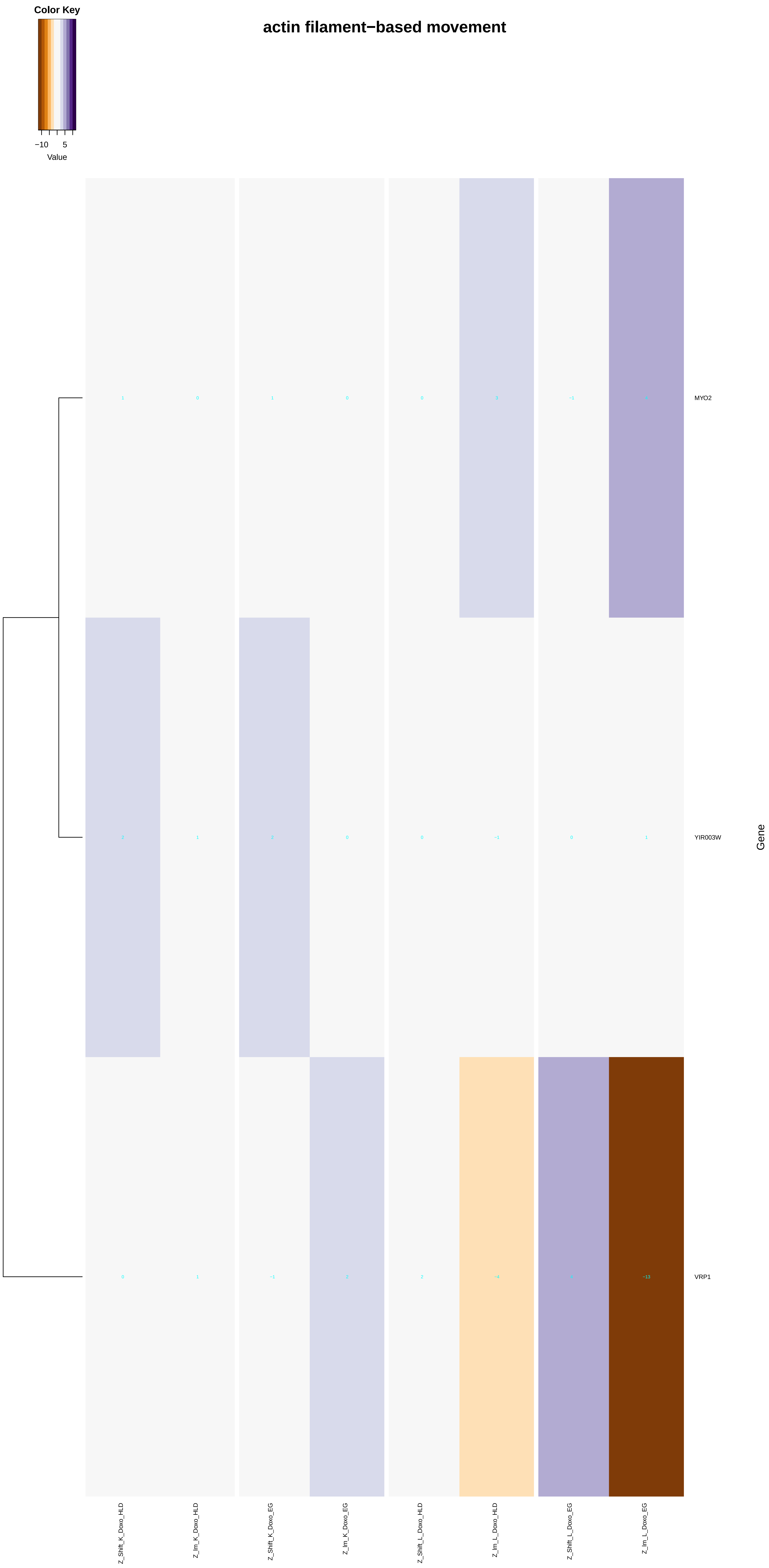

Color Key
−10
5
Value
actin filament−based movement
MYO2
YIR003W
VRP1
Gene
Z_Shift_K_Doxo_HLD
Z_Im_K_Doxo_HLD
Z_Shift_K_Doxo_EG
Z_Im_K_Doxo_EG
Z_Shift_L_Doxo_HLD
Z_Im_L_Doxo_HLD
Z_Shift_L_Doxo_EG
Z_Im_L_Doxo_EG

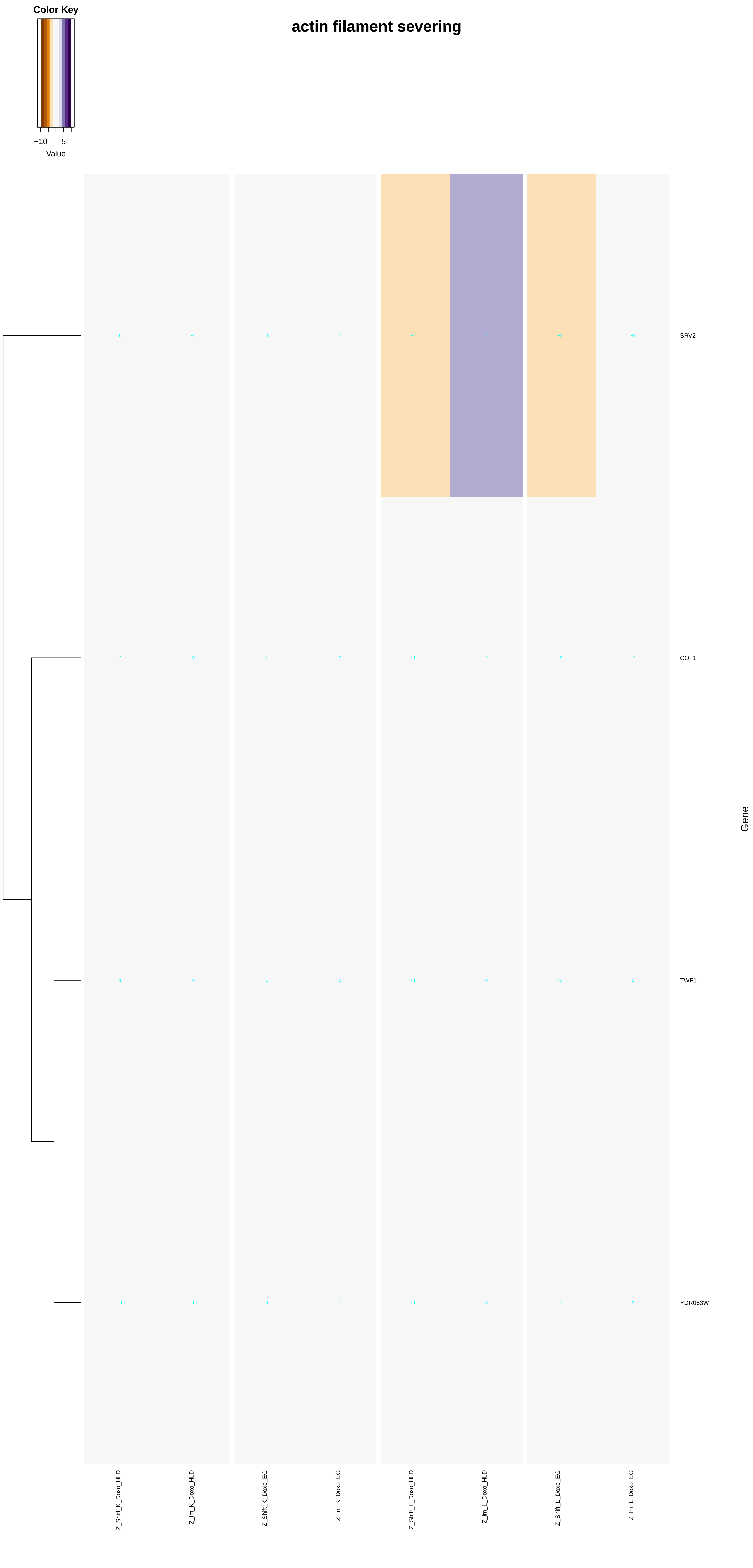

actin filament severing
Color Key
−10
5
Value
SRV2
COF1
TWF1
YDR063W
Gene
Z_Shift_K_Doxo_HLD
Z_lm_K_Doxo_HLD
Z_Shift_K_Doxo_EG
Z_lm_K_Doxo_EG
Z_Shift_L_Doxo_HLD
Z_lm_L_Doxo_HLD
Z_Shift_L_Doxo_EG
Z_lm_L_Doxo_EG

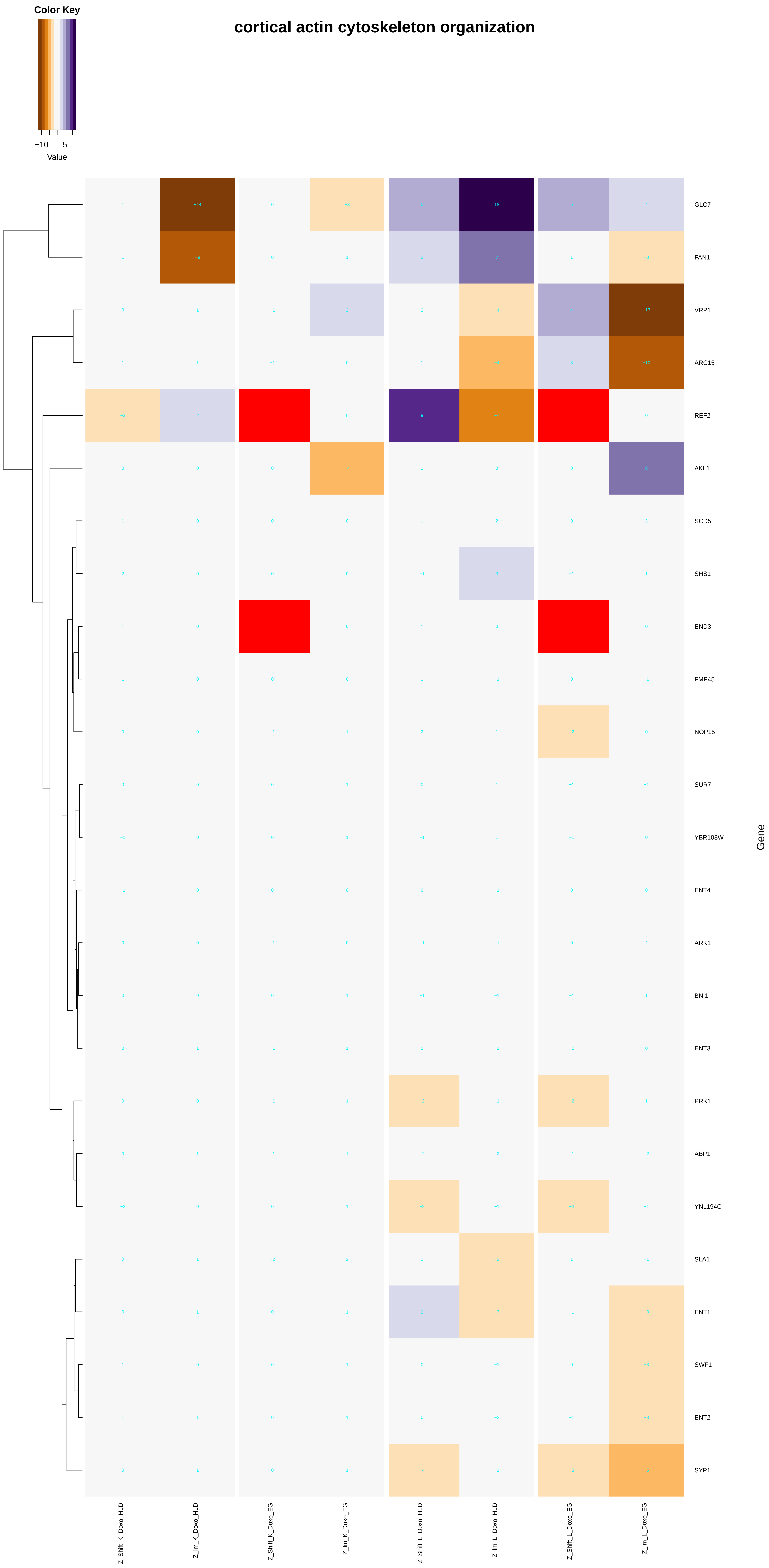

Color Key
−10 5
Value
cortical actin cytoskeleton organization
GLC7
PAN1
VRP1
ARC15
REF2
AKL1
SCD5
SHS1
END3
FMP45
NOP15
SUR7
YBR108W
ENT4
ARK1
BNI1
ENT3
PRK1
ABP1
YNL194C
SLA1
ENT1
SWF1
ENT2
SYP1
Gene
Z_Shift_K_Doxo_HLD
Z_Im_K_Doxo_HLD
Z_Shift_K_Doxo_EG
Z_Im_K_Doxo_EG
Z_Shift_L_Doxo_HLD
Z_Im_L_Doxo_HLD
Z_Shift_L_Doxo_EG
Z_Im_L_Doxo_EG

### establishment or maintenance of actin cytoskeleton polarity

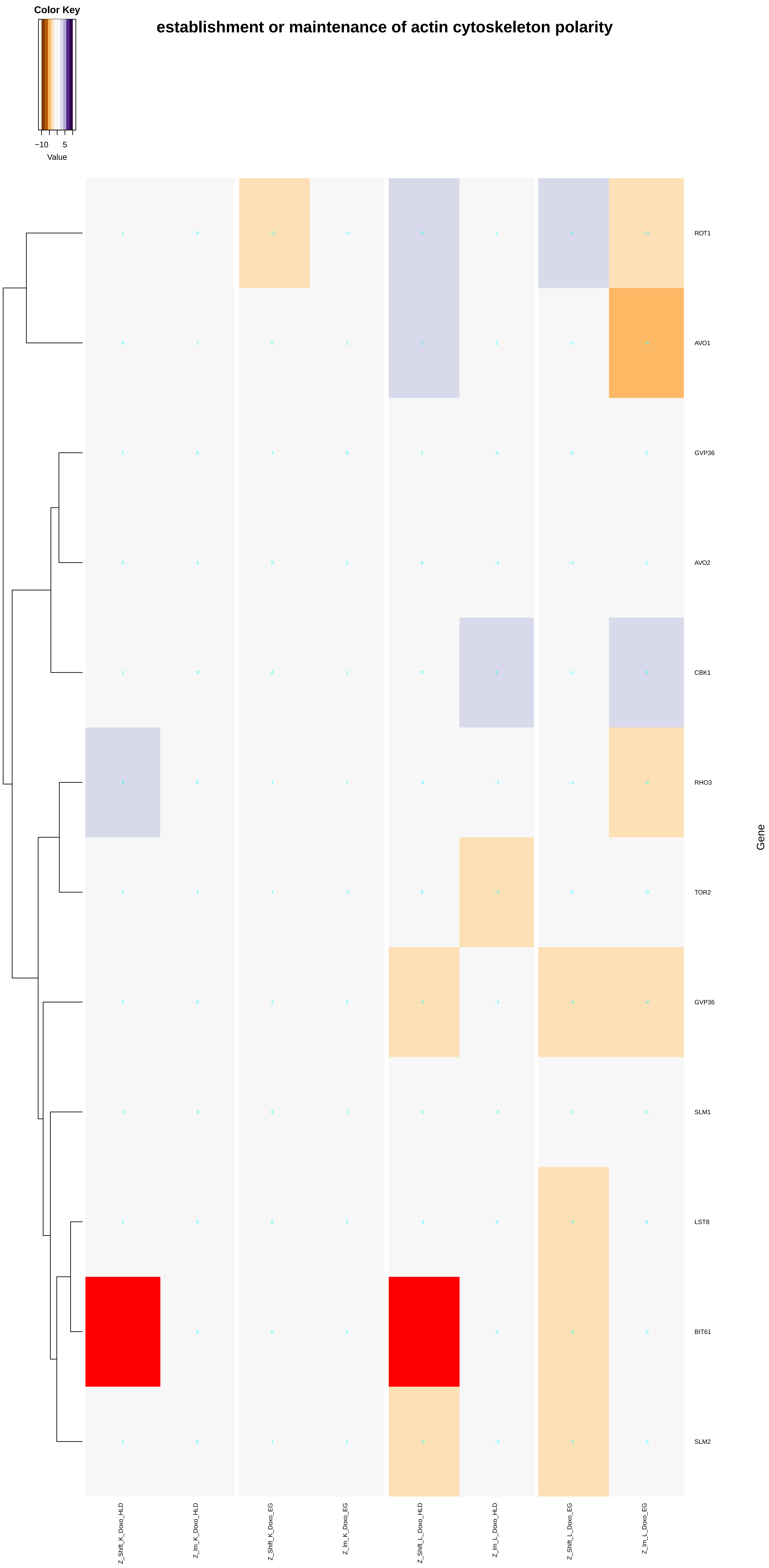

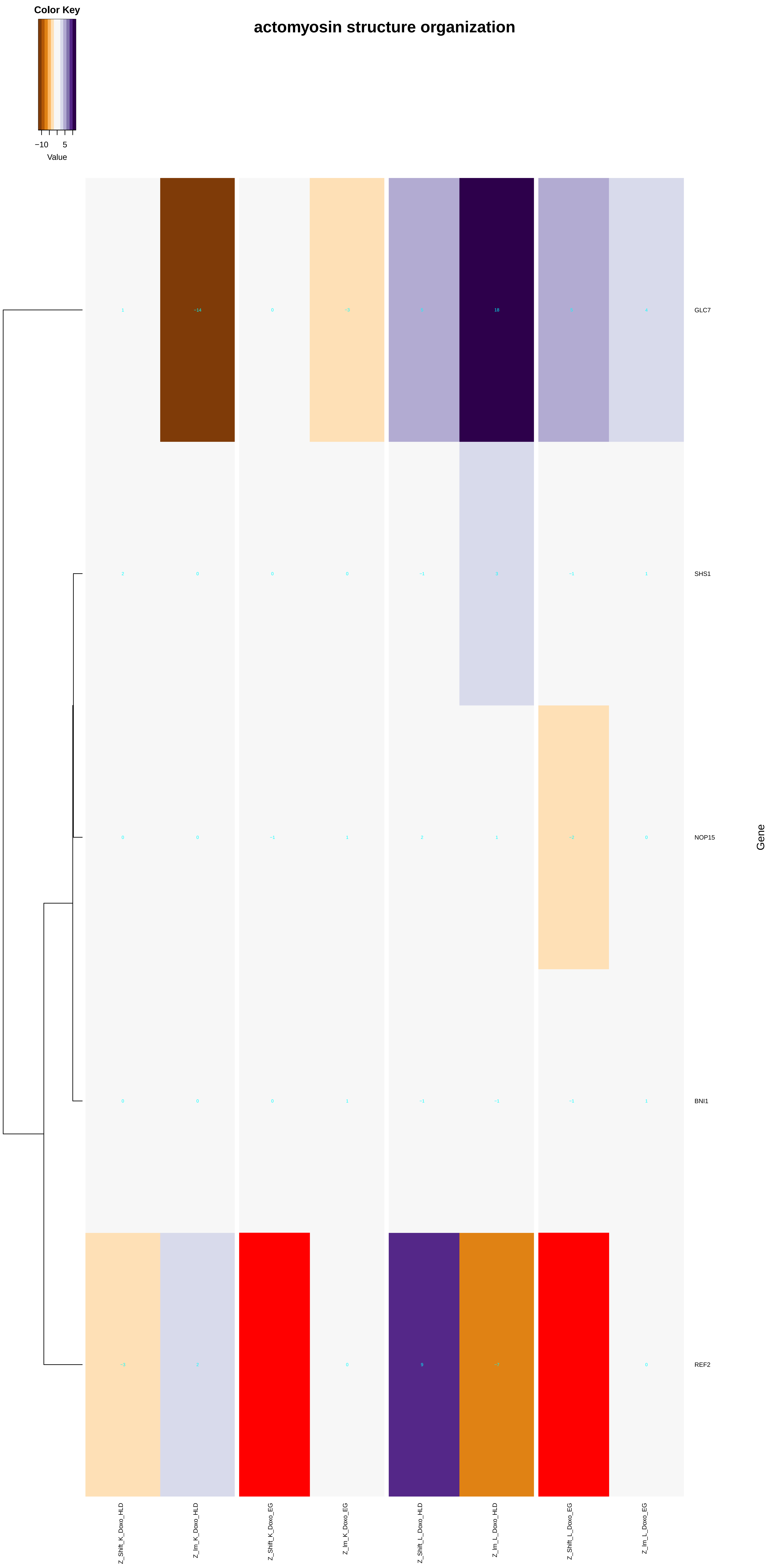
Color Key
−10
5
Value
actomyosin structure organization
GLC7
SHS1
NOP15
BNI1
REF2
Gene
Z_Shift_K_Doxo_HLD
Z_Im_K_Doxo_HLD
Z_Shift_K_Doxo_EG
Z_Im_K_Doxo_EG
Z_Shift_L_Doxo_HLD
Z_Im_L_Doxo_HLD
Z_Shift_L_Doxo_EG
Z_Im_L_Doxo_EG

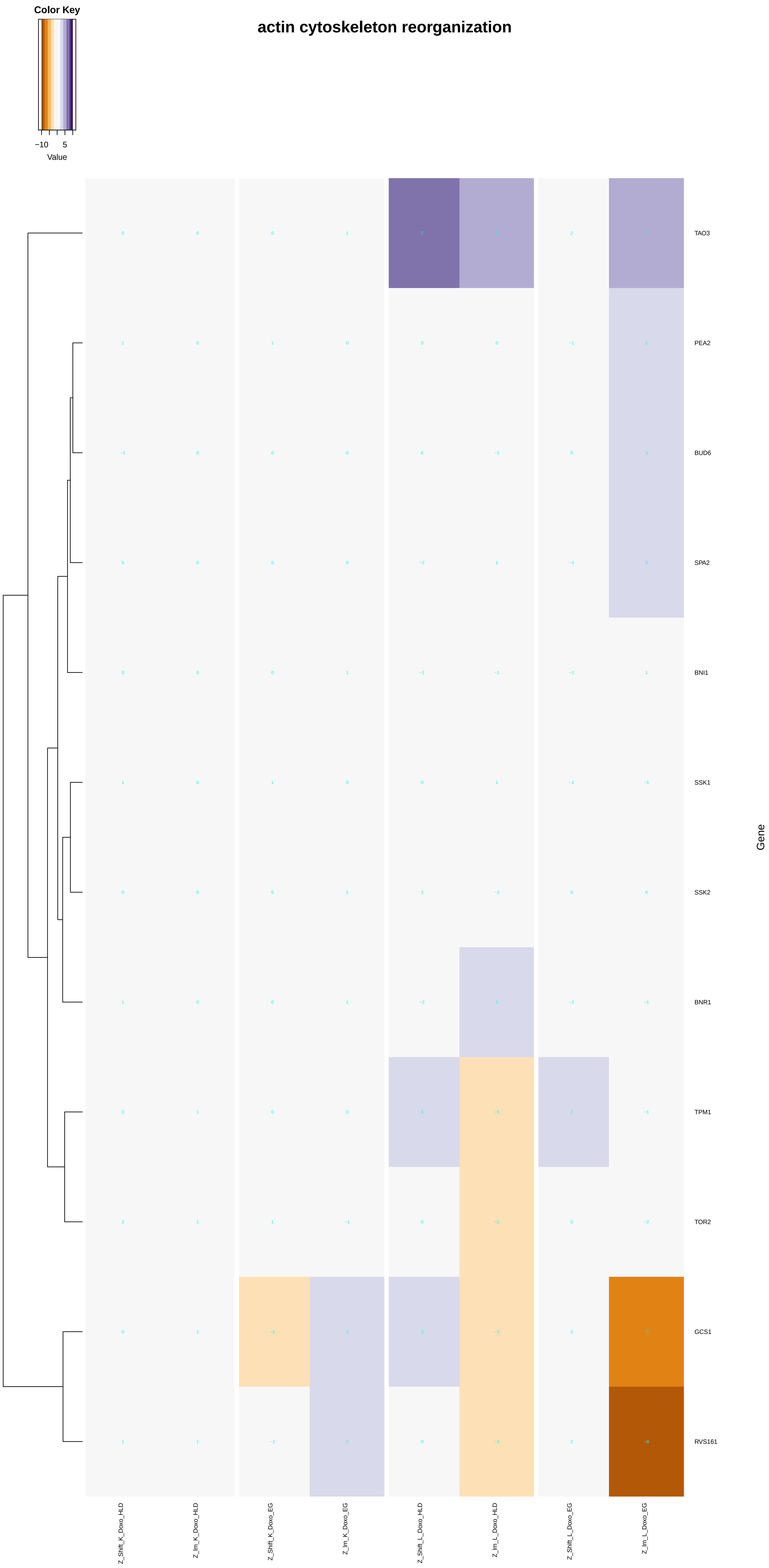

actin cytoskeleton reorganization
Color Key
−10
5
Value
TAO3
PEA2
BUD6
SPA2
BNI1
SSK1
SSK2
BNR1
TPM1
TOR2
GCS1
RVS161
Gene
Z_Shift_K_Doxo_HLD
Z_lm_K_Doxo_HLD
Z_Shift_K_Doxo_EG
Z_lm_K_Doxo_EG
Z_Shift_L_Doxo_HLD
Z_lm_L_Doxo_HLD
Z_Shift_L_Doxo_EG
Z_lm_L_Doxo_EG

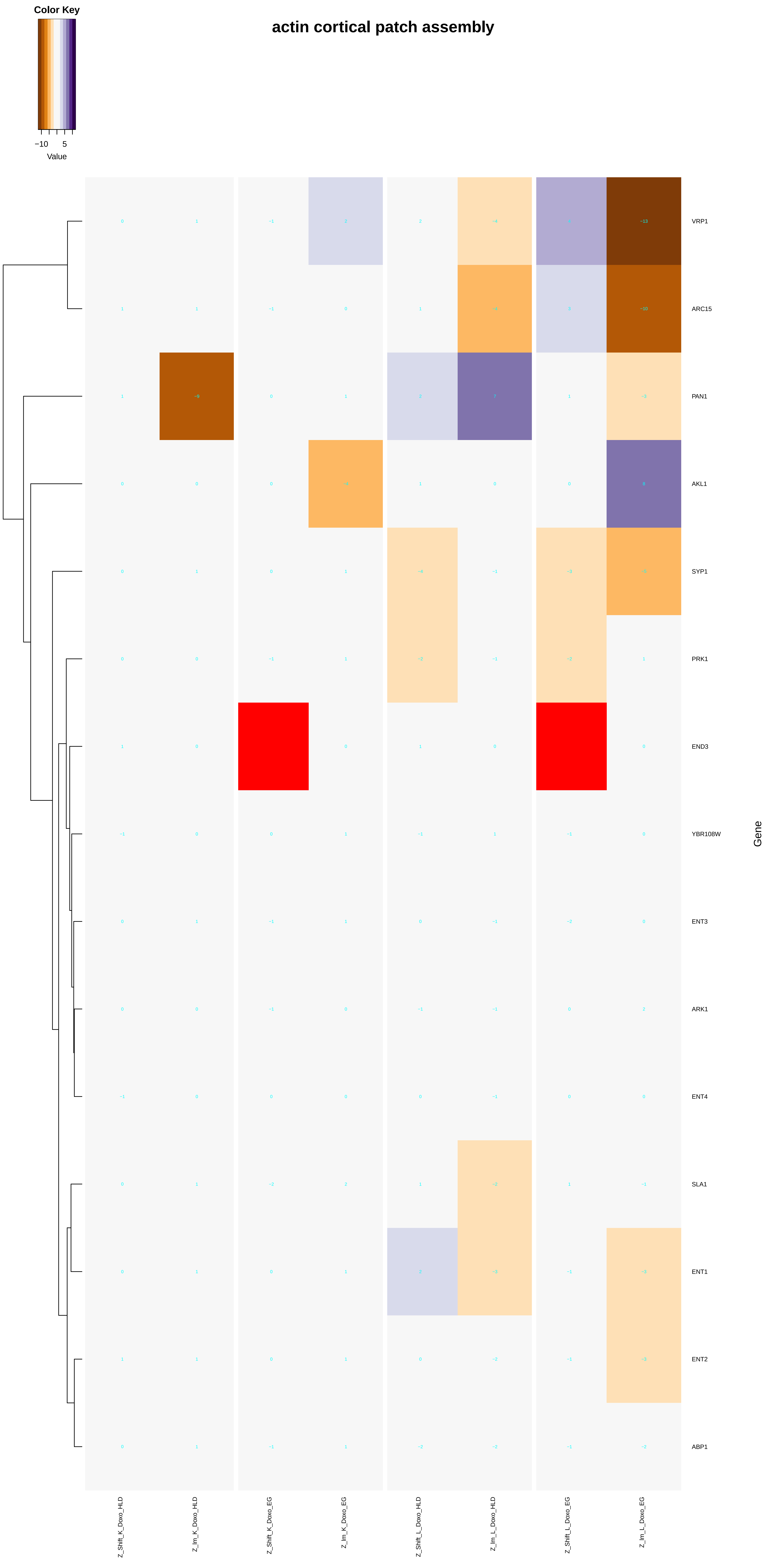
actin cortical patch assembly
Color Key
−10
5
Value
VRP1
ARC15
PAN1
AKL1
SYP1
PRK1
END3
YBR108W
ENT3
ARK1
ENT4
SLA1
ENT1
ENT2
ABP1
Gene
Z_Shift_K_Doxo_HLD
Z_lm_K_Doxo_HLD
Z_Shift_K_Doxo_EG
Z_lm_K_Doxo_EG
Z_Shift_L_Doxo_HLD
Z_lm_L_Doxo_HLD
Z_Shift_L_Doxo_EG
Z_lm_L_Doxo_EG

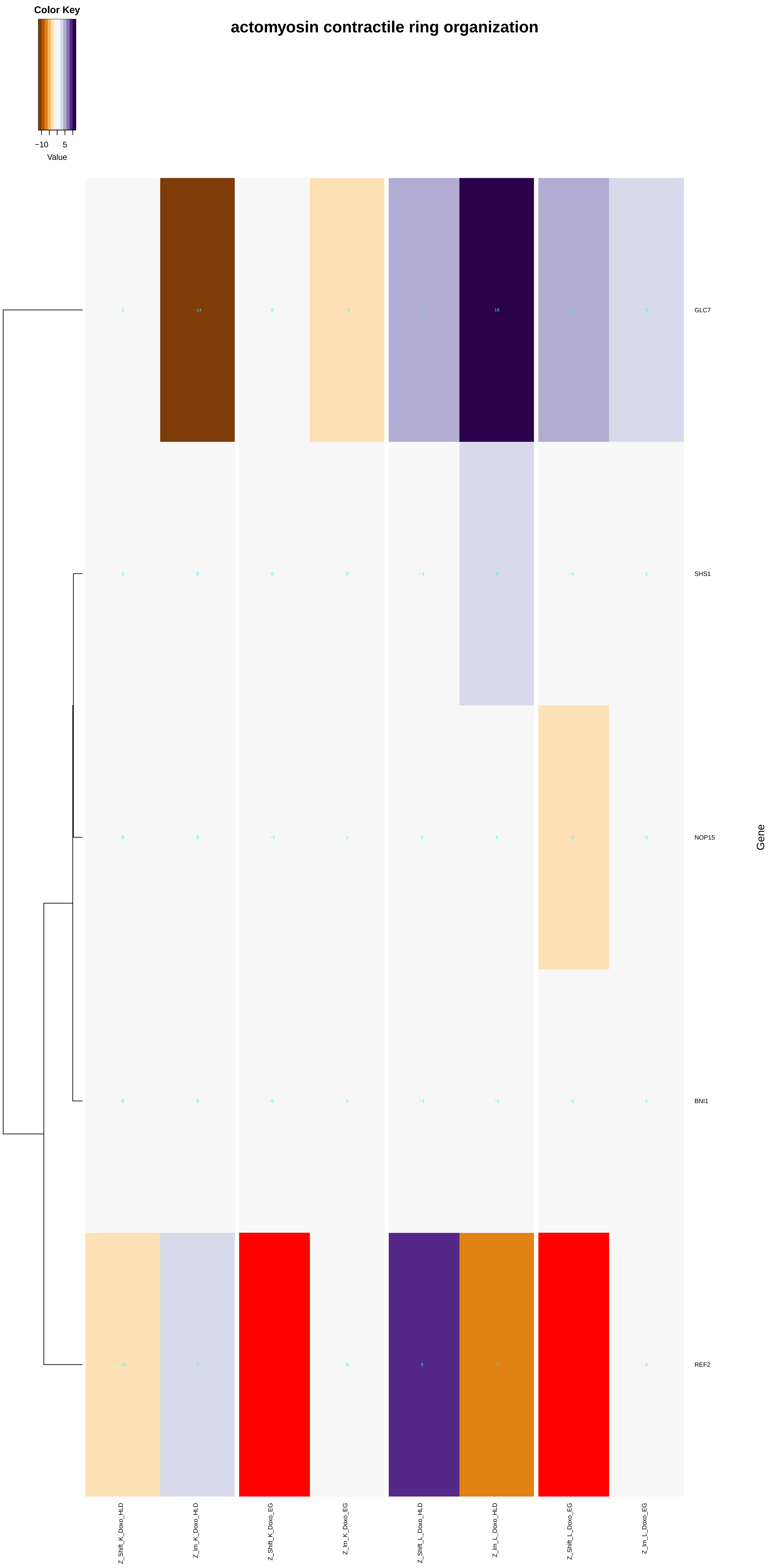
Color Key
−10
5
Value
actomyosin contractile ring organization
GLC7
SHS1
NOP15
BNI1
REF2
Gene
Z_Shift_K_Doxo_HLD
Z_Im_K_Doxo_HLD
Z_Shift_K_Doxo_EG
Z_Im_K_Doxo_EG
Z_Shift_L_Doxo_HLD
Z_Im_L_Doxo_HLD
Z_Shift_L_Doxo_EG
Z_Im_L_Doxo_EG

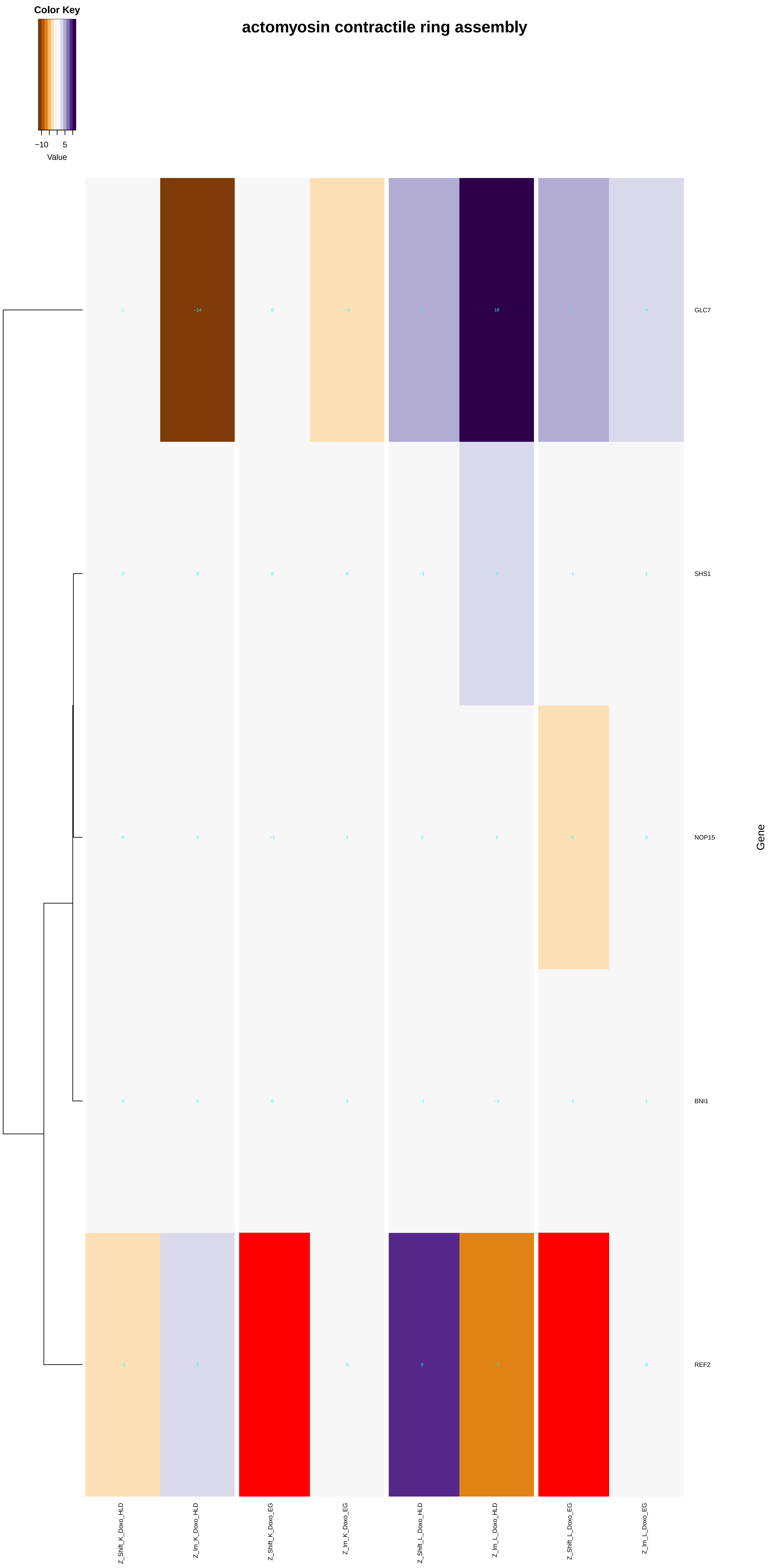

Color Key
−10
5
Value
actomyosin contractile ring assembly
1
−14
0
−3
5
18
5
4
GLC7
2
0
0
0
−1
3
−1
1
SHS1
0
0
−1
1
2
1
−2
0
NOP15
0
0
0
1
−1
−1
−1
1
BNI1
−3
2
0
9
−7
0
REF2
Gene
Z_Shift_K_Doxo_HLD
Z_Im_K_Doxo_HLD
Z_Shift_K_Doxo_EG
Z_Im_K_Doxo_EG
Z_Shift_L_Doxo_HLD
Z_Im_L_Doxo_HLD
Z_Shift_L_Doxo_EG
Z_Im_L_Doxo_EG
