## Additional File 9 for "A yeast phenomic model for the influence of Warburg metabolism on genetic buffering of doxorubicin": cellular_component_biogenesis.pdf

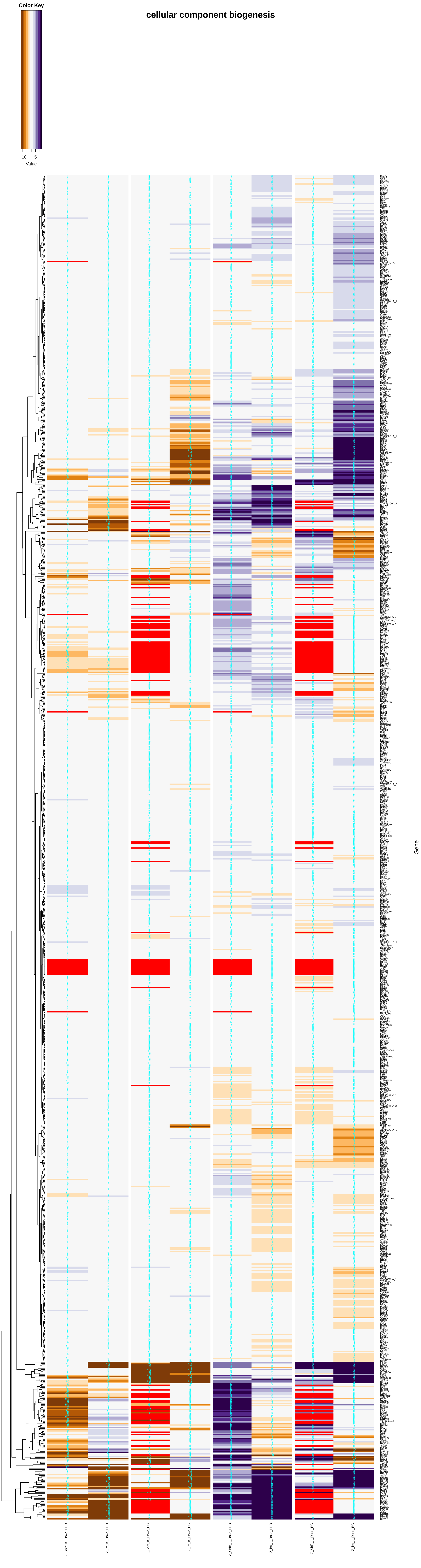

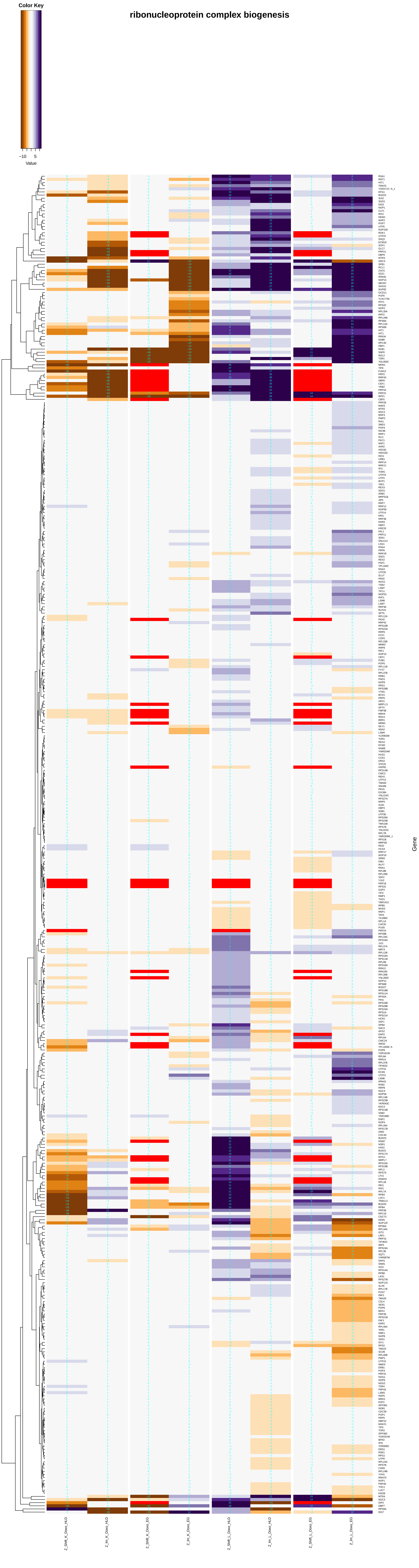
ribonucleoprotein complex biogenesis
Color Key
−10
5
Value
Gene
Z_Shift_K_Doxo_HLD
Z_Im_K_Doxo_HLD
Z_Shift_K_Doxo_EG
Z_Im_K_Doxo_EG
Z_Shift_L_Doxo_HLD
Z_Im_L_Doxo_HLD
Z_Shift_L_Doxo_EG
Z_Im_L_Doxo_EG

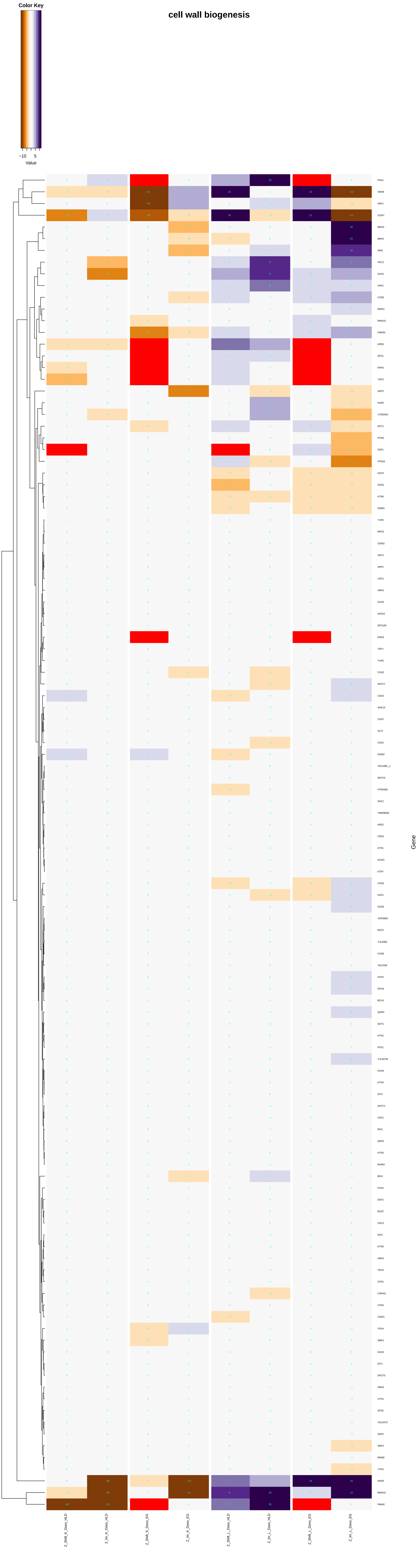

Color Key
−10
5
Value
cell wall biogenesis
Gene

#### membrane biogenesis

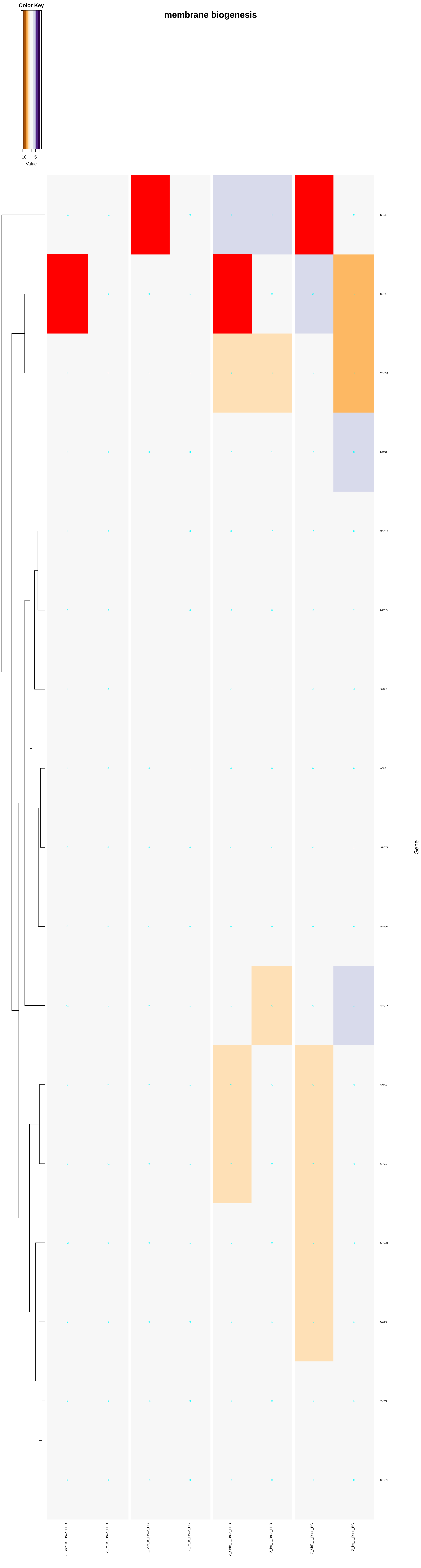

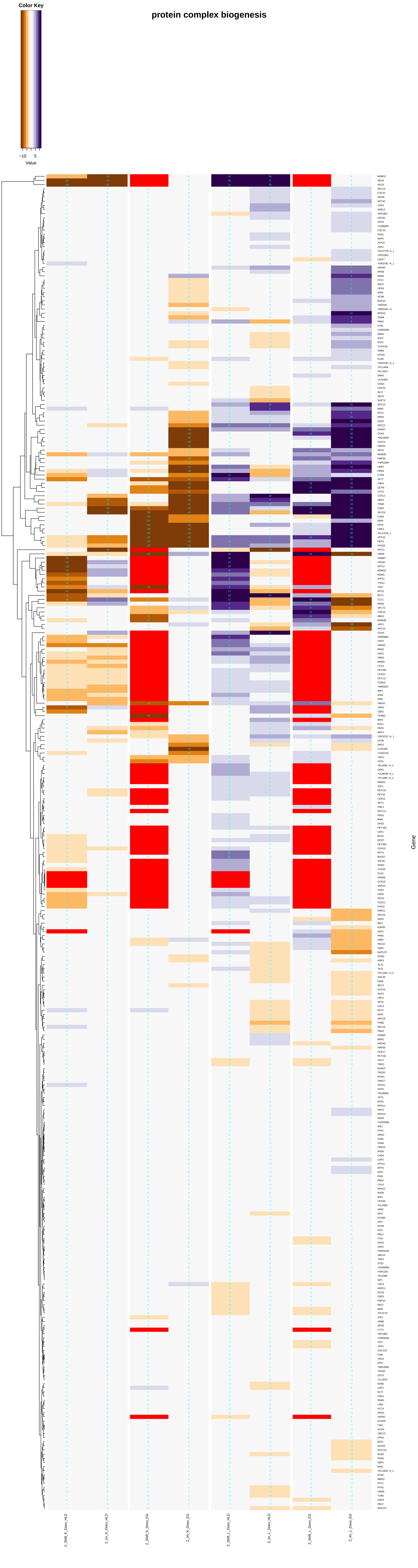

Color Key
-10
5
Value
protein complex biogenesis
Gene
Z_Shift_K_Dose_HLD
Z_Int_K_Dose_HLD
Z_Shift_K_Dose_EG
Z_Int_K_Dose_EG
Z_Shift_L_Dose_HLD
Z_Int_L_Dose_HLD
Z_Shift_L_Dose_EG
Z_Int_L_Dose_EG

### ribosome biogenesis

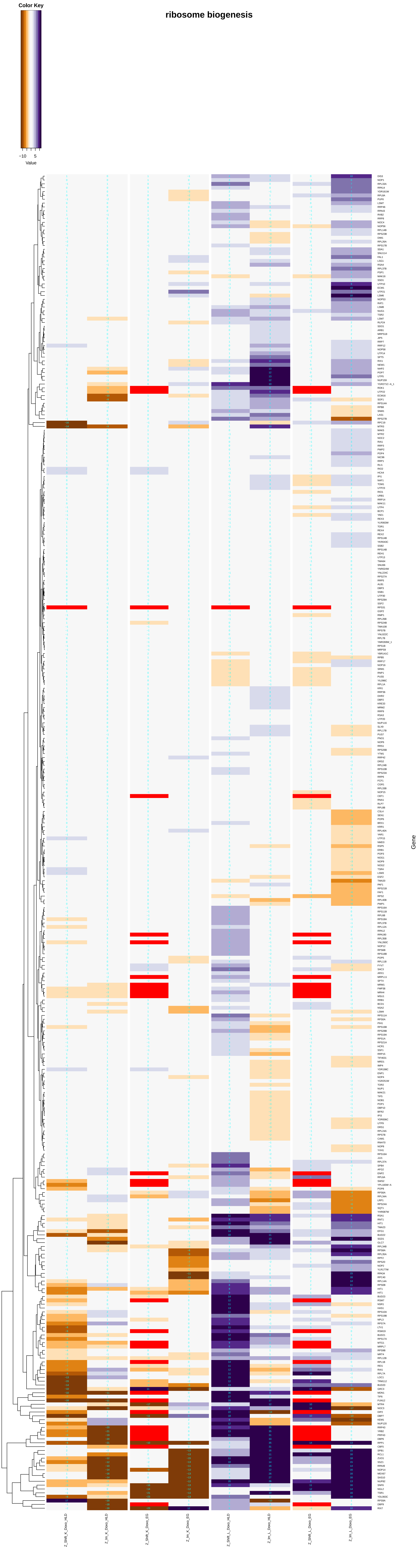

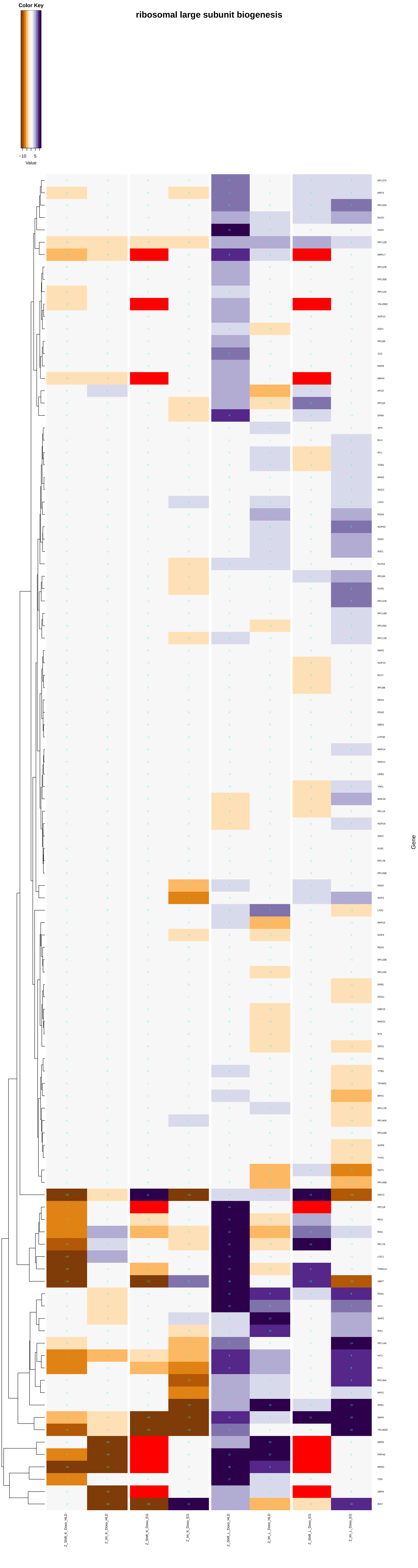
ribosomal large subunit biogenesis
Color Key
−10
5
Value
Gene

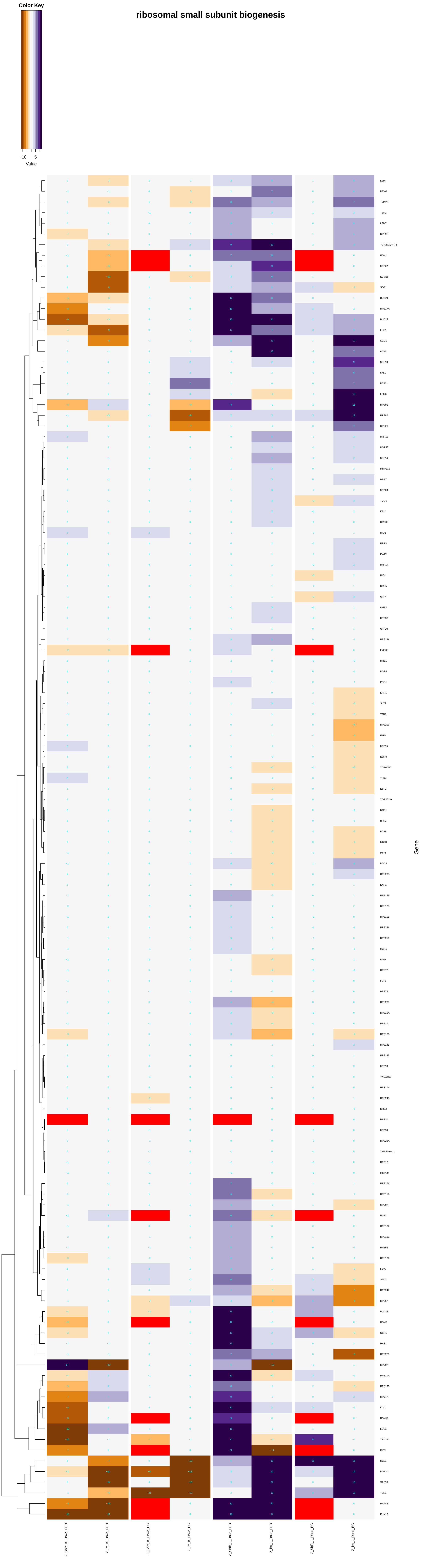

ribosomal small subunit biogenesis
Color Key
−10
5
Value
Gene
Z_Shift_K_Doxo_HLD
Z_Im_K_Doxo_HLD
Z_Shift_K_Doxo_EG
Z_Im_K_Doxo_EG
Z_Shift_L_Doxo_HLD
Z_Im_L_Doxo_HLD
Z_Shift_L_Doxo_EG
Z_Im_L_Doxo_EG

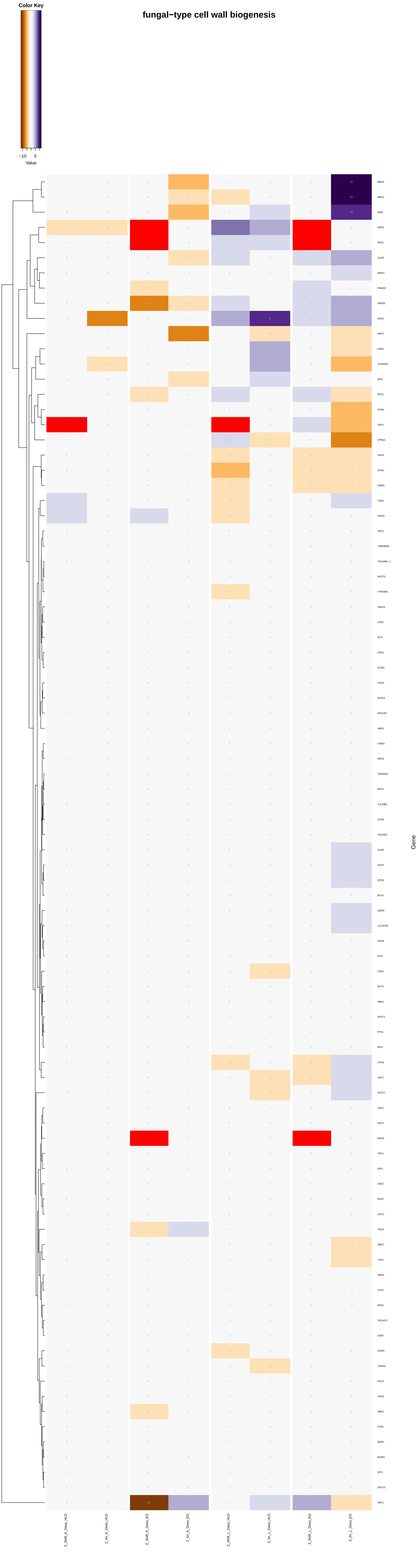

fungal−type cell wall biogenesis
Color Key
−10
5
Value
Gene
Z_Shift_K_Doxo_HLD
Z_Im_K_Doxo_HLD
Z_Shift_K_Doxo_EG
Z_Im_K_Doxo_EG
Z_Shift_L_Doxo_HLD
Z_Im_L_Doxo_HLD
Z_Shift_L_Doxo_EG
Z_Im_L_Doxo_EG

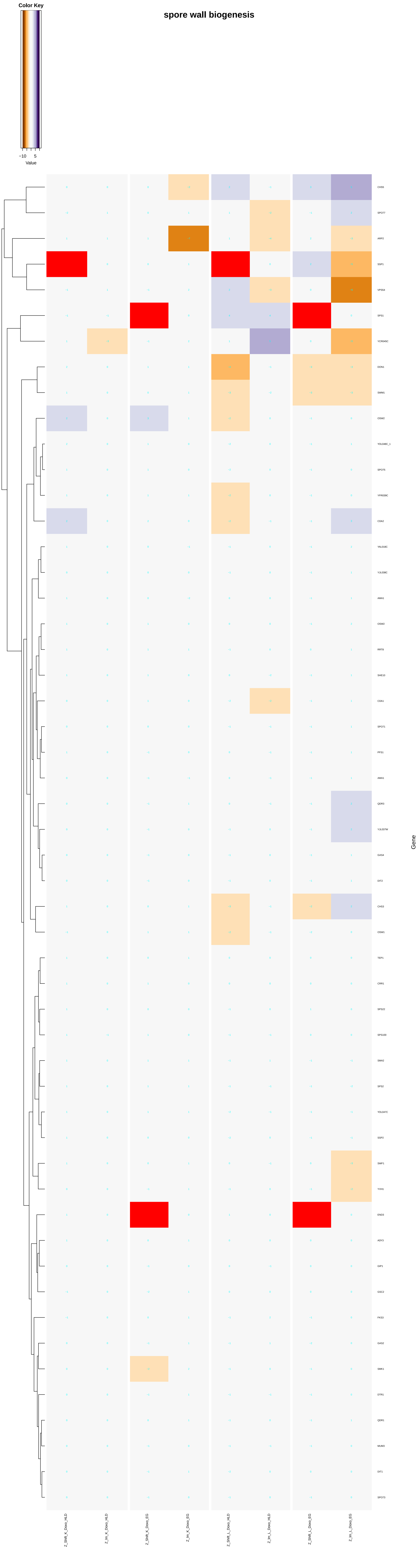

Color Key
−10
5
Value
spore wall biogenesis
CHS5
SPO77
ARP2
SSP1
VPS54
SPS1
YCR045C
DON1
SWM1
OSW2
YOL048C_1
SPO75
YFR039C
CDA2
YAL018C
YJL038C
AMA1
OSW2
RRT8
SHE10
CDA1
SPO71
PFS1
AMA1
QDR3
YJL037W
GAS4
DIT2
CHS3
OSW1
TEP1
CRR1
SPS22
SPS100
SMA2
SPS2
YOL047C
SSP2
SWF1
YVH1
END3
ADY3
GIP1
GSC2
FKS3
GAS2
SMK1
DTR1
QDR1
MUM3
DIT1
SPO73
Gene
Z_Shift_K_Doxo_HLD
Z_Im_K_Doxo_HLD
Z_Shift_K_Doxo_EG
Z_Im_K_Doxo_EG
Z_Shift_L_Doxo_HLD
Z_Im_L_Doxo_HLD
Z_Shift_L_Doxo_EG
Z_Im_L_Doxo_EG

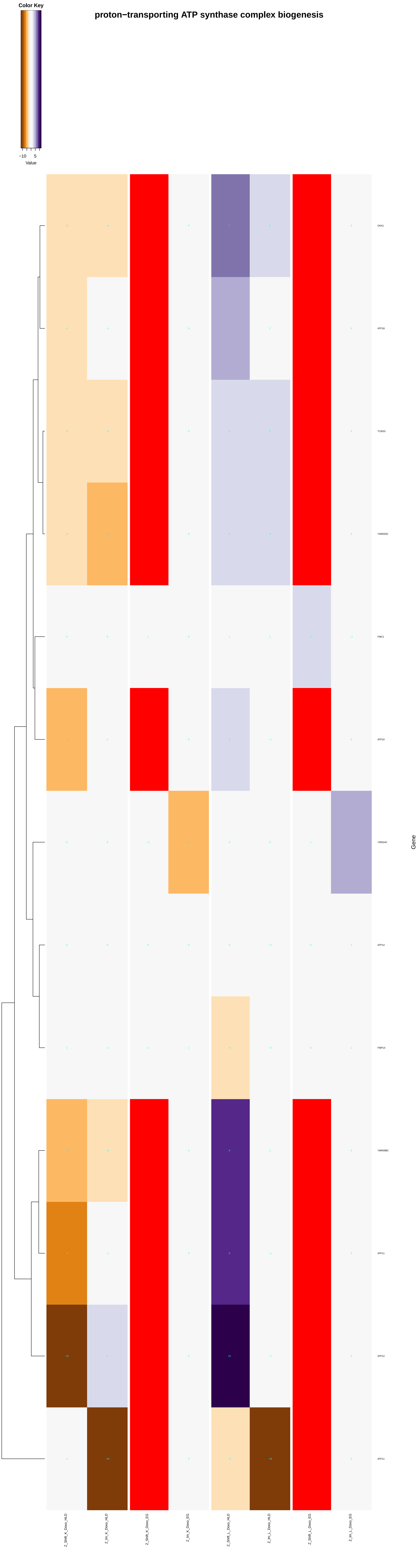
Color Key
−10
5
Value
proton−transporting ATP synthase complex biogenesis
OXA1
ATP18
TCM10
YNR020C
FMC1
ATP10
YIR024C
ATP14
FMP14
YMR098C
ATP11
ATP12
ATP11
Gene
Z_Shift_K_Doxo_HLD
Z_Im_K_Doxo_HLD
Z_Shift_K_Doxo_EG
Z_Im_K_Doxo_EG
Z_Shift_L_Doxo_HLD
Z_Im_L_Doxo_HLD
Z_Shift_L_Doxo_EG
Z_Im_L_Doxo_EG

#### mitochondrial respiratory chain complex II biogenesis

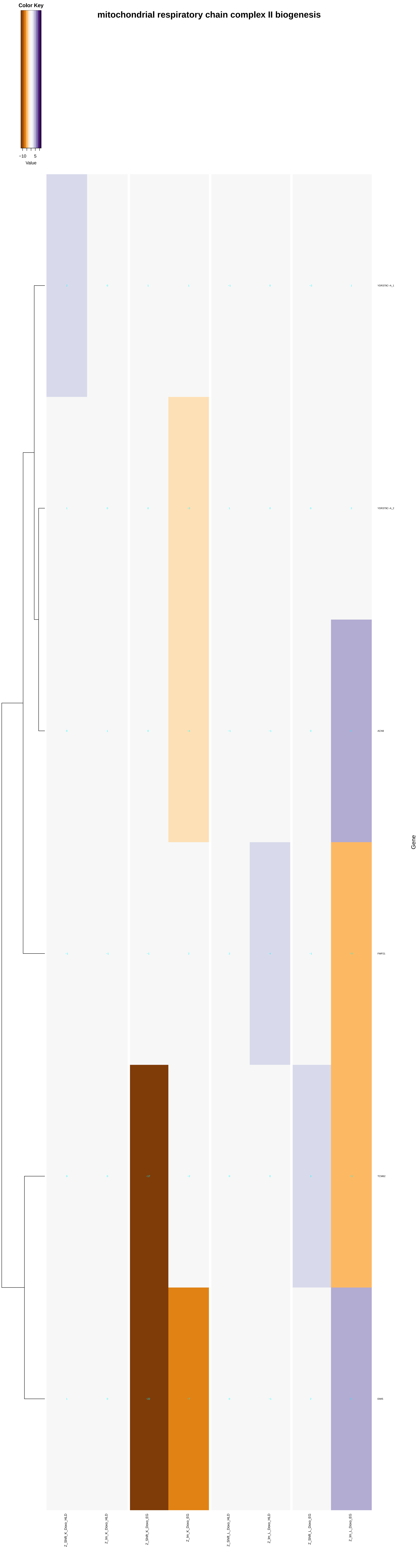

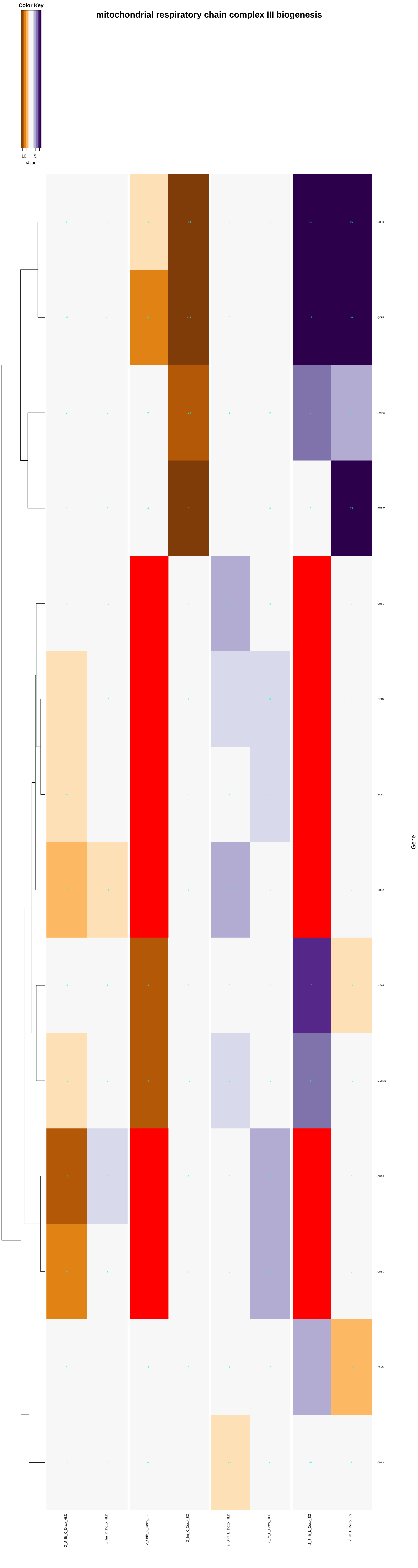
mitochondrial respiratory chain complex III biogenesis
Color Key
−10
5
Value
Gene
CBP4
QCR9
FMP36
FMP25
CBS1
QCR7
BCS1
CBS2
MBA1
MDM38
CBP6
CBS1
PAN5
CBP3
Z_Shift_K_Doxo_HLD
Z_Im_K_Doxo_HLD
Z_Shift_K_Doxo_EG
Z_Im_K_Doxo_EG
Z_Shift_L_Doxo_HLD
Z_Im_L_Doxo_HLD
Z_Shift_L_Doxo_EG
Z_Im_L_Doxo_EG

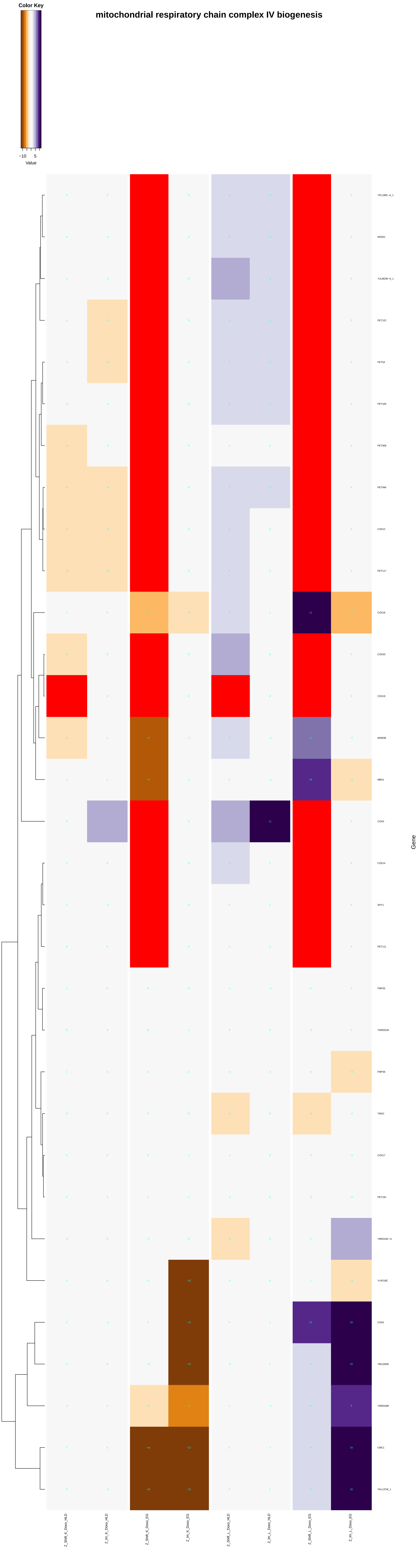

Color Key
−10
5
Value
mitochondrial respiratory chain complex IV biogenesis
YPL189C-A_1
MSS51
YJL062W-A_1
PET122
PET54
PET100
PET309
PET494
COX12
PET117
COX16
COX20
COX19
MDM38
MBA1
COA3
COX14
SHY1
PET111
FMP32
YOR031W
FMP35
TIM21
COX17
PET191
YMR244C-A
YLR218C
COA2
YML030W
YNR018W
CMC1
YKL137W_1
Gene
Z_Shift_K_Doxo_HLD
Z_Im_K_Doxo_HLD
Z_Shift_K_Doxo_EG
Z_Im_K_Doxo_EG
Z_Shift_L_Doxo_HLD
Z_Im_L_Doxo_HLD
Z_Shift_L_Doxo_EG
Z_Im_L_Doxo_EG

### ascospore wall biogenesis

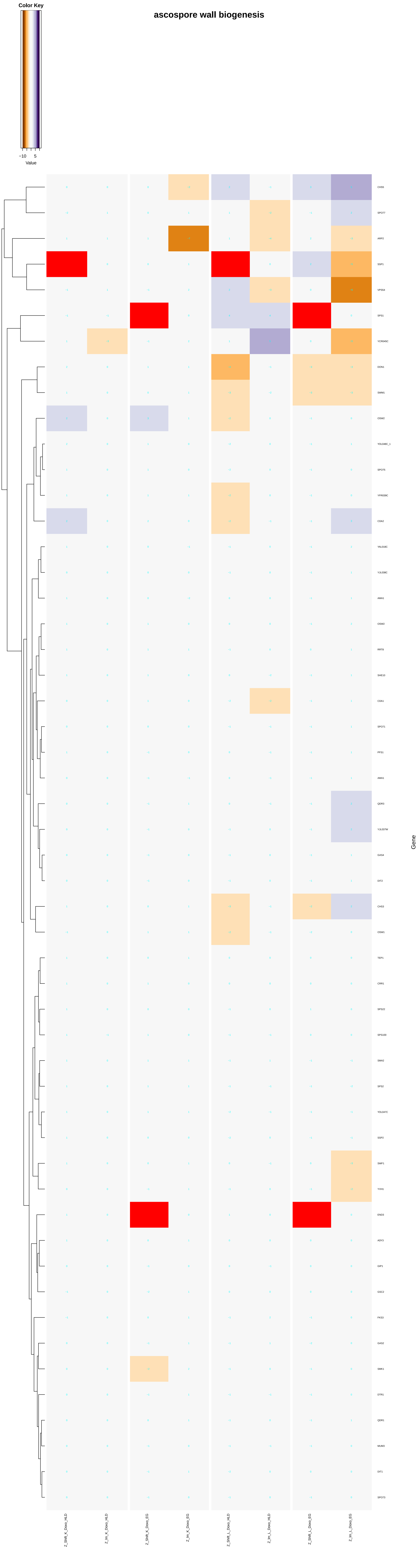

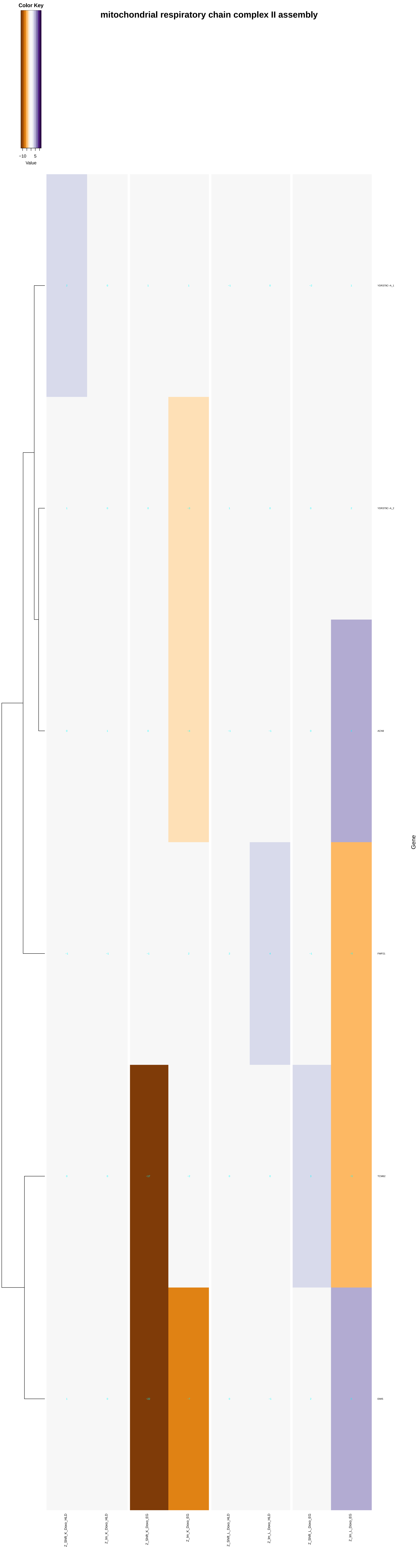
mitochondrial respiratory chain complex II assembly
Color Key
−10
5
Value
YDR379C−A_1
YDR379C−A_2
ACN9
FMP21
TCM62
EMI5
Gene
Z_Shift_K_Doxo_HLD
Z_lm_K_Doxo_HLD
Z_Shift_K_Doxo_EG
Z_lm_K_Doxo_EG
Z_Shift_L_Doxo_HLD
Z_lm_L_Doxo_HLD
Z_Shift_L_Doxo_EG
Z_lm_L_Doxo_EG

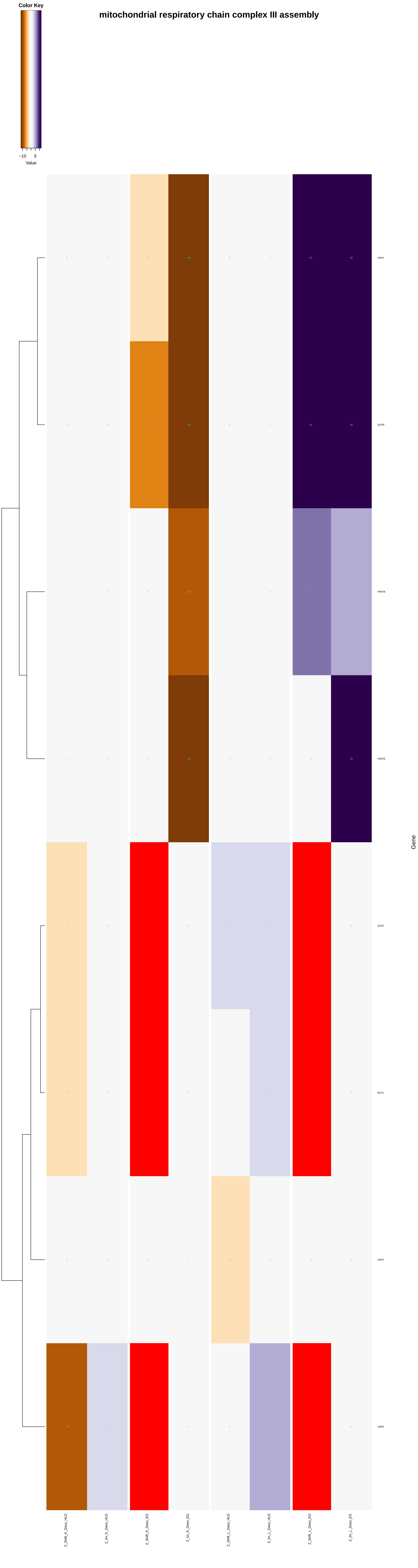
mitochondrial respiratory chain complex III assembly
Color Key
−10
5
Value
CBP4
QCR9
FMP36
FMP25
QCR7
BCS1
CBP3
CBP6
Gene
Z_Shift_K_Doxo_HLD
Z_Im_K_Doxo_HLD
Z_Shift_K_Doxo_EG
Z_Im_K_Doxo_EG
Z_Shift_L_Doxo_HLD
Z_Im_L_Doxo_HLD
Z_Shift_L_Doxo_EG
Z_Im_L_Doxo_EG

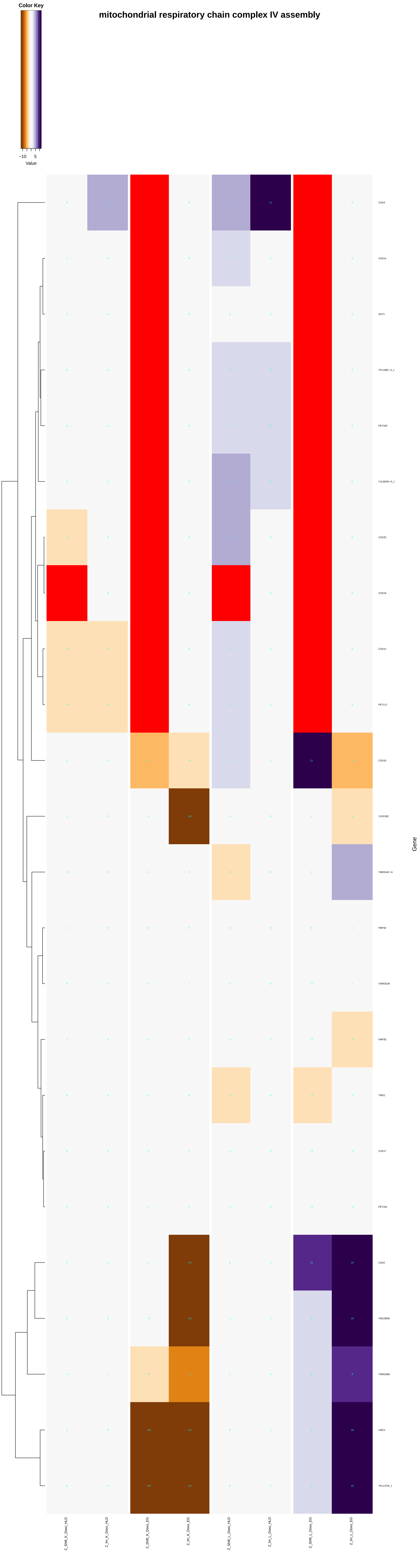

mitochondrial respiratory chain complex IV assembly
Color Key
−10
5
Value
COA3
COX14
SHY1
YPL189C−A_1
PET100
YJL062W−A_1
COX20
COX19
COX12
PET117
COX16
YLR218C
YMR244C−A
FMP32
YDR031W
FMP35
TIM21
COX17
PET191
COA2
YML030W
YNR018W
CMC1
YKL137W_1
Gene
Z_Shift_K_Doxo_HLD
Z_Im_K_Doxo_HLD
Z_Shift_K_Doxo_EG
Z_Im_K_Doxo_EG
Z_Shift_L_Doxo_HLD
Z_Im_L_Doxo_HLD
Z_Shift_L_Doxo_EG
Z_Im_L_Doxo_EG
