## Additional File 9 for "A yeast phenomic model for the influence of Warburg metabolism on genetic buffering of doxorubicin": cellular_response_to_DNA_damage_stimulus.pdf

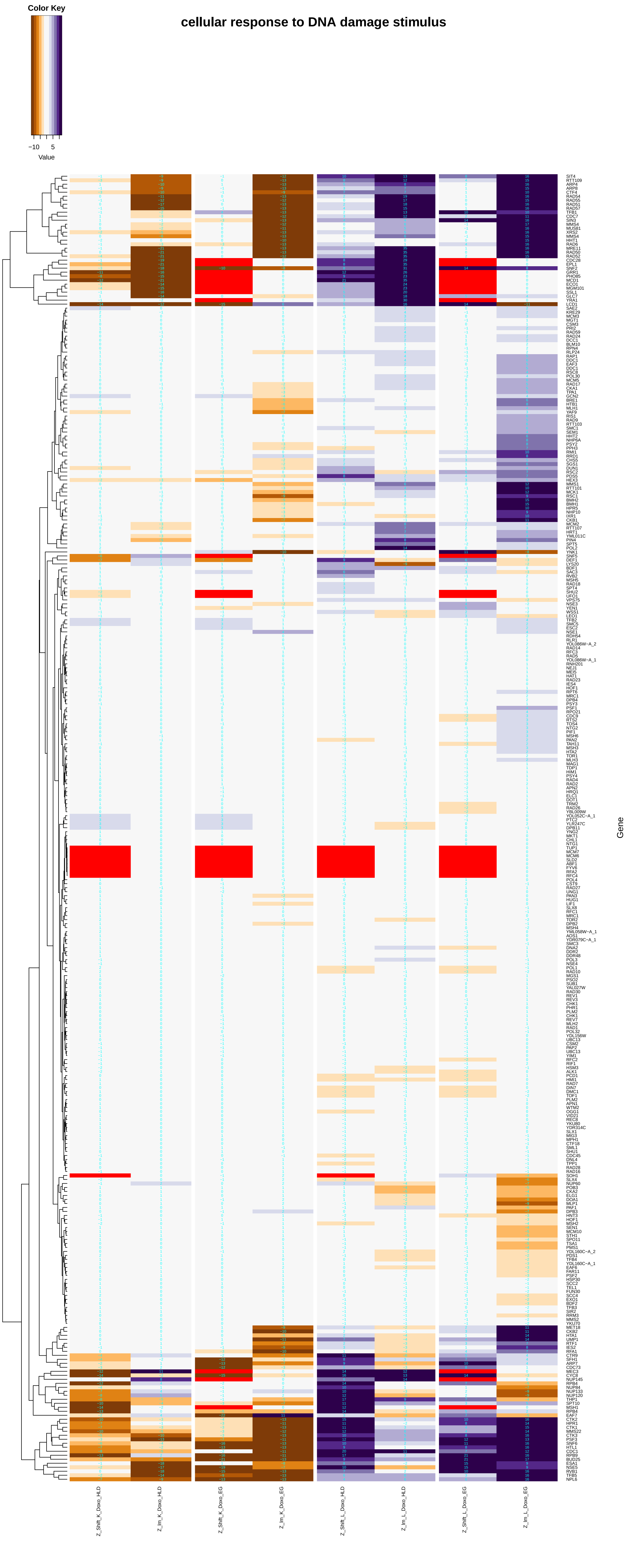
Color Key
−10
5
Value
cellular response to DNA damage stimulus
Gene
Z_Shift_K_Doxo_HLD
Z_lm_K_Doxo_HLD
Z_Shift_K_Doxo_EG
Z_lm_K_Doxo_EG
Z_Shift_L_Doxo_HLD
Z_lm_L_Doxo_HLD
Z_Shift_L_Doxo_EG
Z_lm_L_Doxo_EG

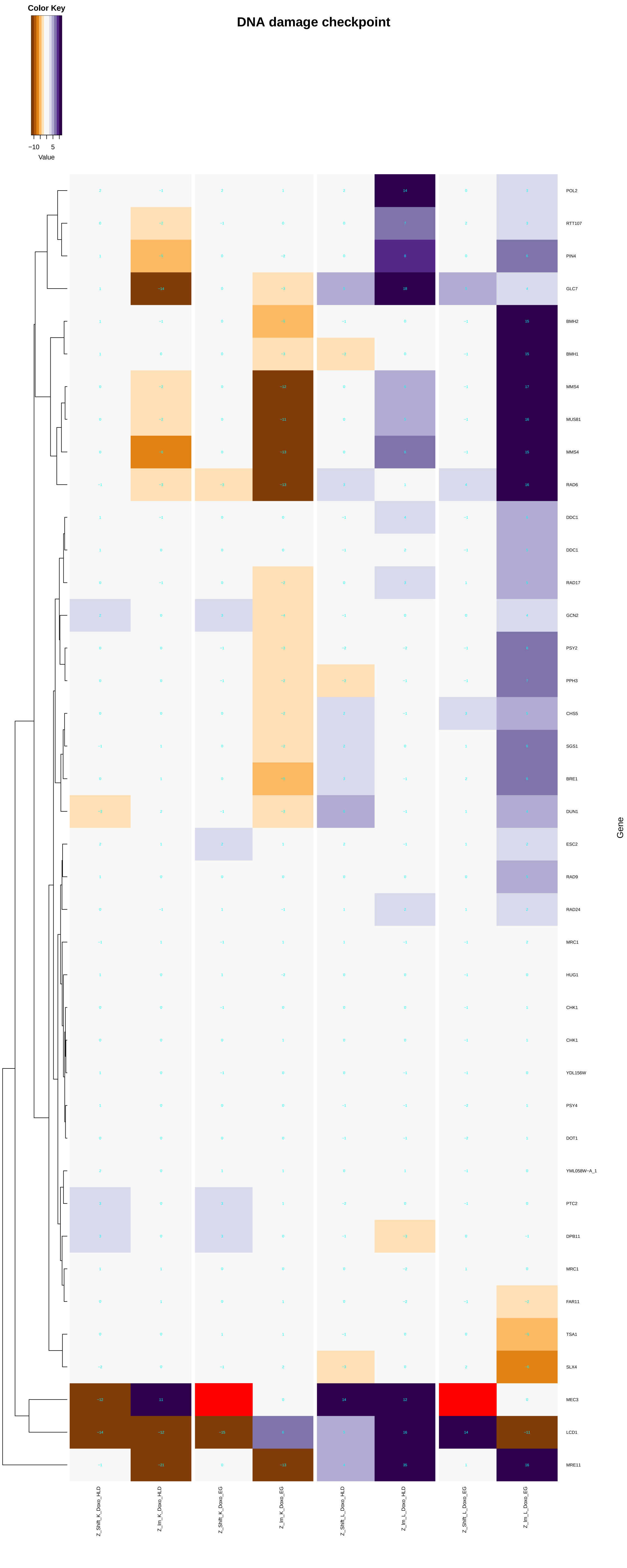

DNA damage checkpoint
Color Key
−10
5
Value
POL2
RTT107
PIN4
GLC7
BMH2
BMH1
MMS4
MUS81
MMS4
RAD6
DDC1
DDC1
RAD17
GCN2
PSY2
PPH3
CHS5
SGS1
BRE1
DUN1
ESC2
RAD9
RAD24
MRC1
HUG1
CHK1
CHK1
YDL156W
PSY4
DOT1
YML058W−A_1
PTC2
DPB11
MRC1
FAR11
TSA1
SLX4
MEC3
LCD1
MRE11
Gene
Z_Shift_K_Doxo_HLD
Z_Im_K_Doxo_HLD
Z_Shift_K_Doxo_EG
Z_Im_K_Doxo_EG
Z_Shift_L_Doxo_HLD
Z_Im_L_Doxo_HLD
Z_Shift_L_Doxo_EG
Z_Im_L_Doxo_EG

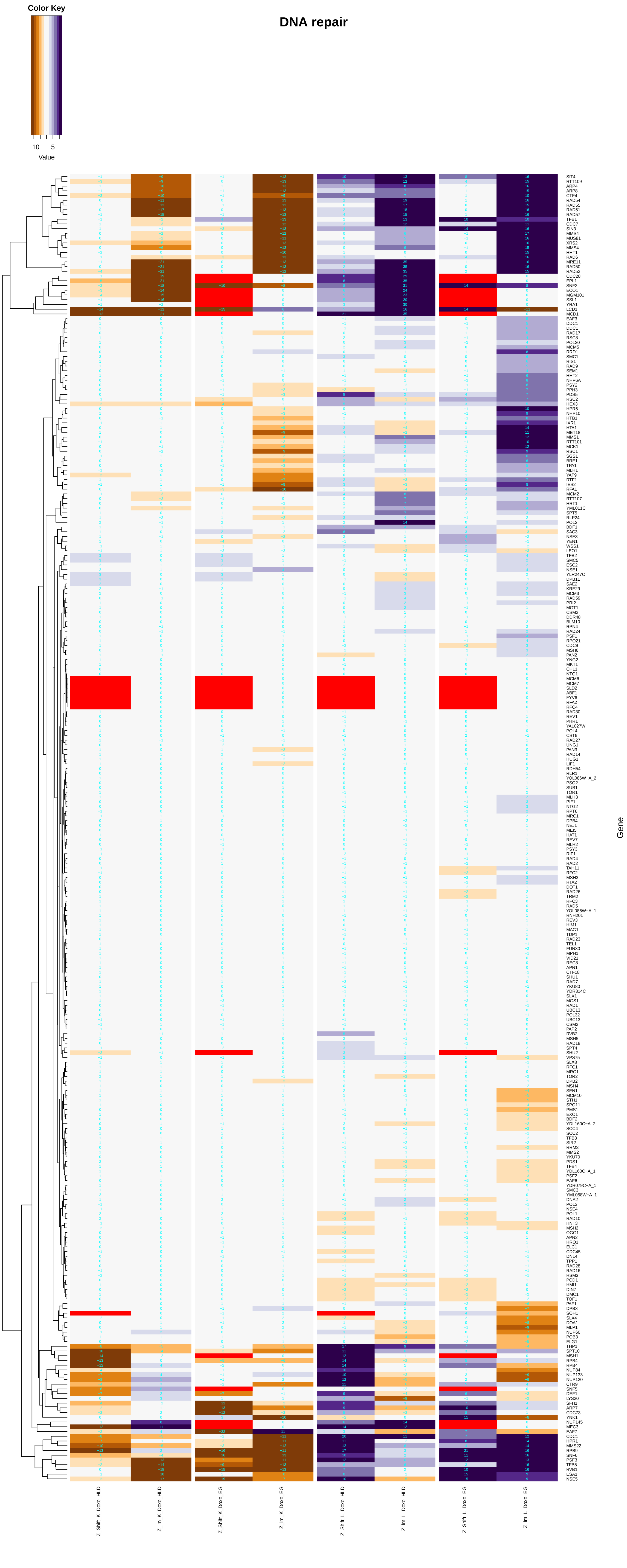
DNA repair
Color Key
−10 5
Value
SIT4
RTT109
ARP4
ARP8
CTF4
RAD54
RAD55
RAD51
RAD57
TFB1
CDC7
SIN3
MMS4
MUS81
XRS2
MMS4
HHT1
RAD6
MRE11
RAD50
RAD52
CDC28
EPL1
SNF2
ECO1
MGM101
SSL1
YRA1
LCD1
MCD1
EAF3
DDC1
DDC1
RAD17
RSC8
POL30
MCM5
RRD1
SMC1
RIS1
RAD9
SEM1
HHT2
NHP6A
PSY2
PPH3
PDS5
RSC2
HEX3
HPR5
NHP10
HTB1
IXR1
HTA1
MET18
MMS1
RTT101
MCK1
RSC1
SGS1
BRE1
TPA1
MLH1
YAF9
RTF1
IES2
RFA1
MCM2
RTT107
HRT1
YML011C
SPT5
RLP24
POL2
BDF1
SAC3
NSE3
YEN1
WSS1
LEO1
TFB2
SMC5
ESC2
NSE1
YLR247C
DPB11
SAE2
KRE29
MCM3
RAD59
PRI2
MGT1
CSM3
DDR48
BLM10
RPN4
RAD24
PSF1
RPO21
CDC9
MSH6
PAN2
YNG2
MKT1
CHL1
NTG1
MCM6
MCM7
SLD2
ABF1
FYV6
RFA2
RFC4
RAD30
REV1
PHR1
YAL027W
POL4
CST9
RAD27
UNG1
PAN3
RAD14
HUG1
LIF1
RDH54
RLR1
YOL086W−A_2
PSO2
SUB1
TOR1
MLH3
PIF1
NTG2
RPT6
MRC1
DPB4
NEJ1
MEI5
HAT1
REV7
MLH2
PSY3
RIF1
RAD4
RAD2
TAH11
RFC2
MSH3
HTA2
DOT1
RAD26
TRM2
RFC3
RAD5
YOL086W−A_1
RNH201
REV3
HIM1
MAG1
TDP1
RAD23
TEL1
FUN30
MPH1
VID21
REC8
APN1
CTF18
SHU1
RAD7
YKU80
YDR314C
SLX1
MGS1
RAD1
UBC13
POL32
UBC13
CSM2
PAP2
RVB2
MSH5
RAD18
SPT4
SHU2
VPS75
SLX8
RFC1
MRC1
TOR2
DPB2
MSH4
SEN1
MCM10
STH1
SPO11
PMS1
EXO1
BDF2
YDL160C−A_2
SCC4
SCC2
TFB3
SIR2
RRM3
MMS2
YKU70
PDS1
TFB4
YDL160C−A_1
PSF2
EAF6
YDR079C−A_1
SMC3
YML058W−A_1
DNA2
POL3
NSE4
POL1
RAD10
HNT3
MSH2
OGG1
APN2
HRQ1
ELC1
CDC45
DNL4
TPP1
RAD28
RAD16
HSM3
PCD1
HMI1
DIN7
DMC1
TOF1
PAF1
DPB3
SOH1
SLX4
DOA1
MLP1
NUP60
POB3
ELG1
THP1
SPT10
MSH1
RPB4
RPB4
NUP84
NUP133
NUP120
CTR9
SNF5
DEF1
LYS20
SFH1
ARP7
CDC73
YNK1
NUP145
MEC3
EAF7
CDC1
HPR1
MMS22
RPB9
SNF6
PSF3
TFB5
RVB1
ESA1
NSE5
Gene
Z_Shift_K_Doxo_HLD
Z_Im_K_Doxo_HLD
Z_Shift_K_Doxo_EG
Z_Im_K_Doxo_EG
Z_Shift_L_Doxo_HLD
Z_Im_L_Doxo_HLD
Z_Shift_L_Doxo_EG
Z_Im_L_Doxo_EG

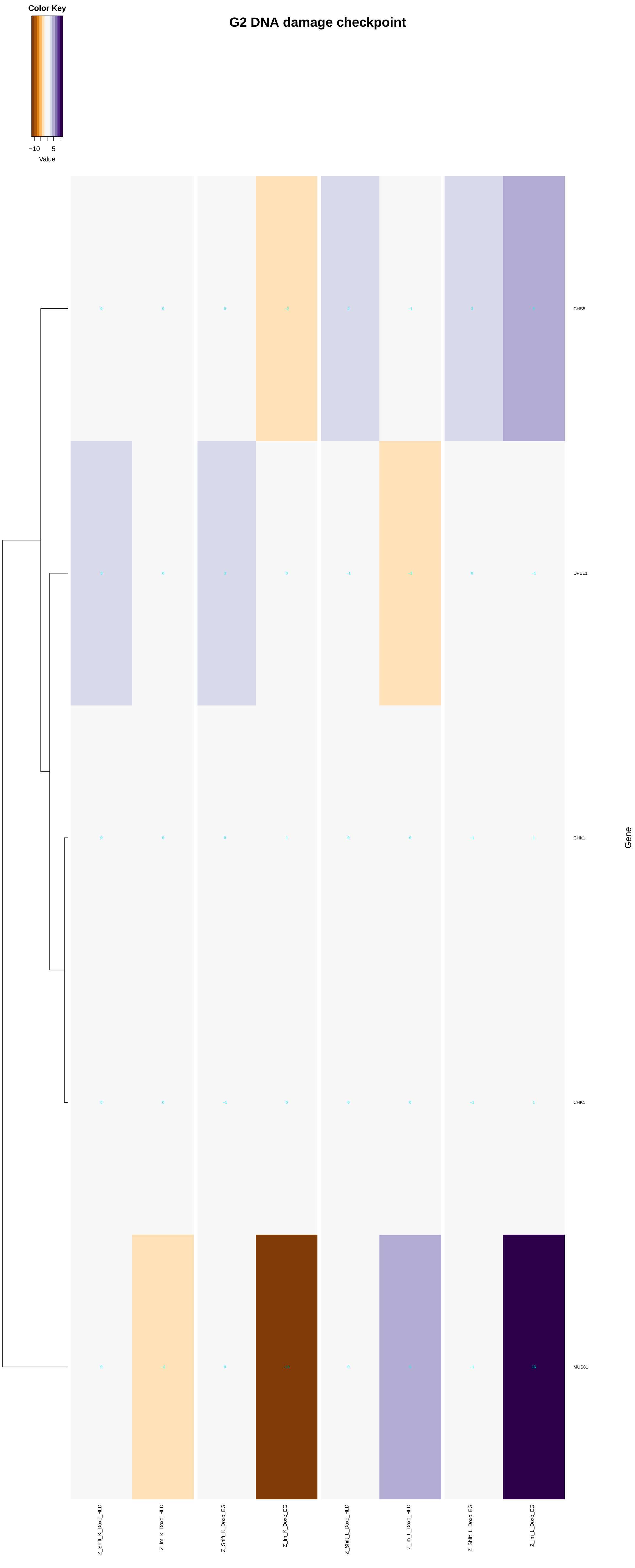
G2 DNA damage checkpoint
Color Key
−10
5
Value
CHS5
DPB11
CHK1
CHK1
MUS81
Gene
Z_Shift_K_Doxo_HLD
Z_Im_K_Doxo_HLD
Z_Shift_K_Doxo_EG
Z_Im_K_Doxo_EG
Z_Shift_L_Doxo_HLD
Z_Im_L_Doxo_HLD
Z_Shift_L_Doxo_EG
Z_Im_L_Doxo_EG

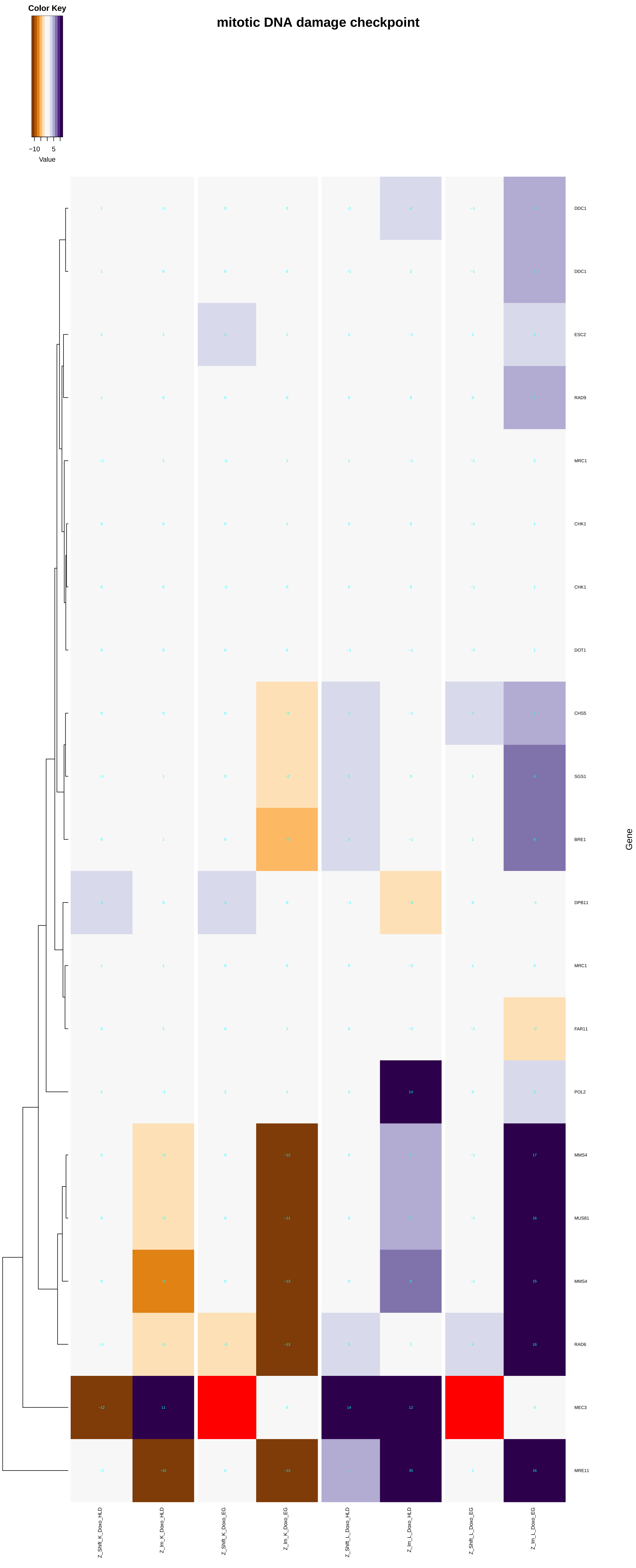

Color Key
−10
5
Value
mitotic DNA damage checkpoint
DDC1
DDC1
ESC2
RAD9
MRC1
CHK1
CHK1
DOT1
CHS5
SGS1
BRE1
DPB11
MRC1
FAR11
POL2
MMS4
MUS81
MMS4
RAD6
MEC3
MRE11
Gene
Z_Shift_K_Doxo_HLD
Z_lm_K_Doxo_HLD
Z_Shift_K_Doxo_EG
Z_lm_K_Doxo_EG
Z_Shift_L_Doxo_HLD
Z_lm_L_Doxo_HLD
Z_Shift_L_Doxo_EG
Z_lm_L_Doxo_EG

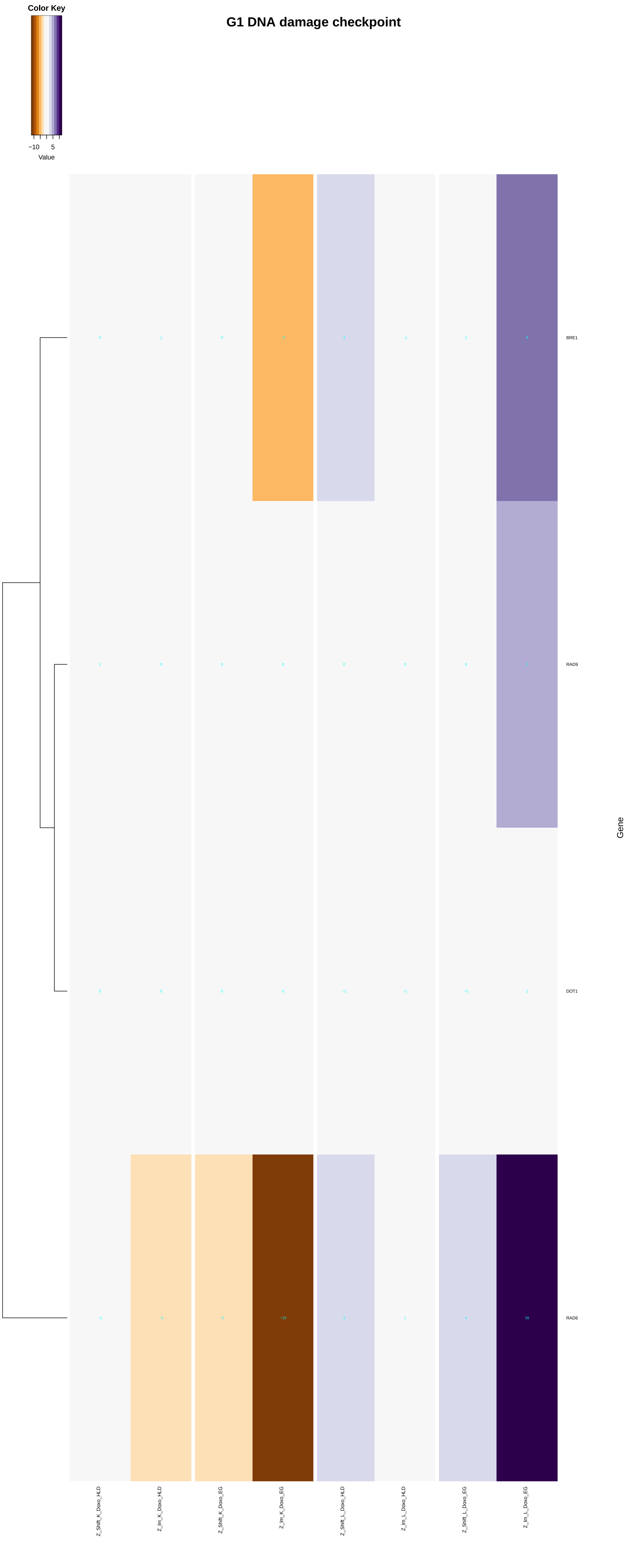

G1 DNA damage checkpoint
Color Key
−10 5
Value
BRE1
RAD9
DOT1
RAD6
Gene
Z_Shift_K_Doxo_HLD
Z_Im_K_Doxo_HLD
Z_Shift_K_Doxo_EG
Z_Im_K_Doxo_EG
Z_Shift_L_Doxo_HLD
Z_Im_L_Doxo_HLD
Z_Shift_L_Doxo_EG
Z_Im_L_Doxo_EG

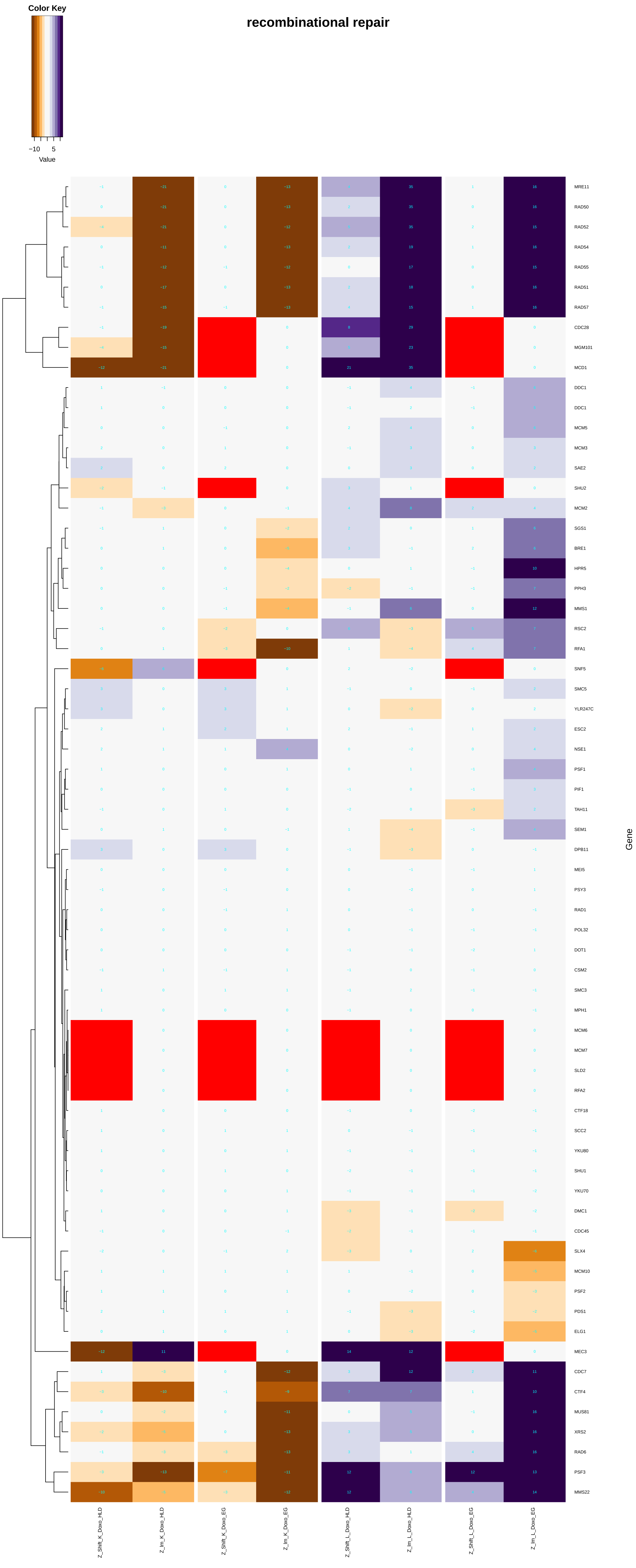

Color Key
−10 5
Value
recombinational repair
Gene
MRE11
RAD50
RAD52
RAD54
RAD55
RAD51
RAD57
CDC28
MGM101
MCD1
DDC1
DDC1
MCM5
MCM3
SAE2
SHU2
MCM2
SGS1
BRE1
HPR5
PPH3
MMS1
RSC2
RFA1
SNF5
SMC5
YLR247C
ESC2
NSE1
PSF1
PIF1
TAH11
SEM1
DPB11
MEI5
PSY3
RAD1
POL32
DOT1
CSM2
SMC3
MPH1
MCM6
MCM7
SLD2
RFA2
CTF18
SCC2
YKU80
SHU1
YKU70
DMC1
CDC45
SLX4
MCM10
PSF2
PDS1
ELG1
MEC3
CDC7
CTF4
MUS81
XRS2
RAD6
PSF3
MMS22
Z_Shift_K_Doxo_HLD
Z_Im_K_Doxo_HLD
Z_Shift_K_Doxo_EG
Z_Im_K_Doxo_EG
Z_Shift_L_Doxo_HLD
Z_Im_L_Doxo_HLD
Z_Shift_L_Doxo_EG
Z_Im_L_Doxo_EG

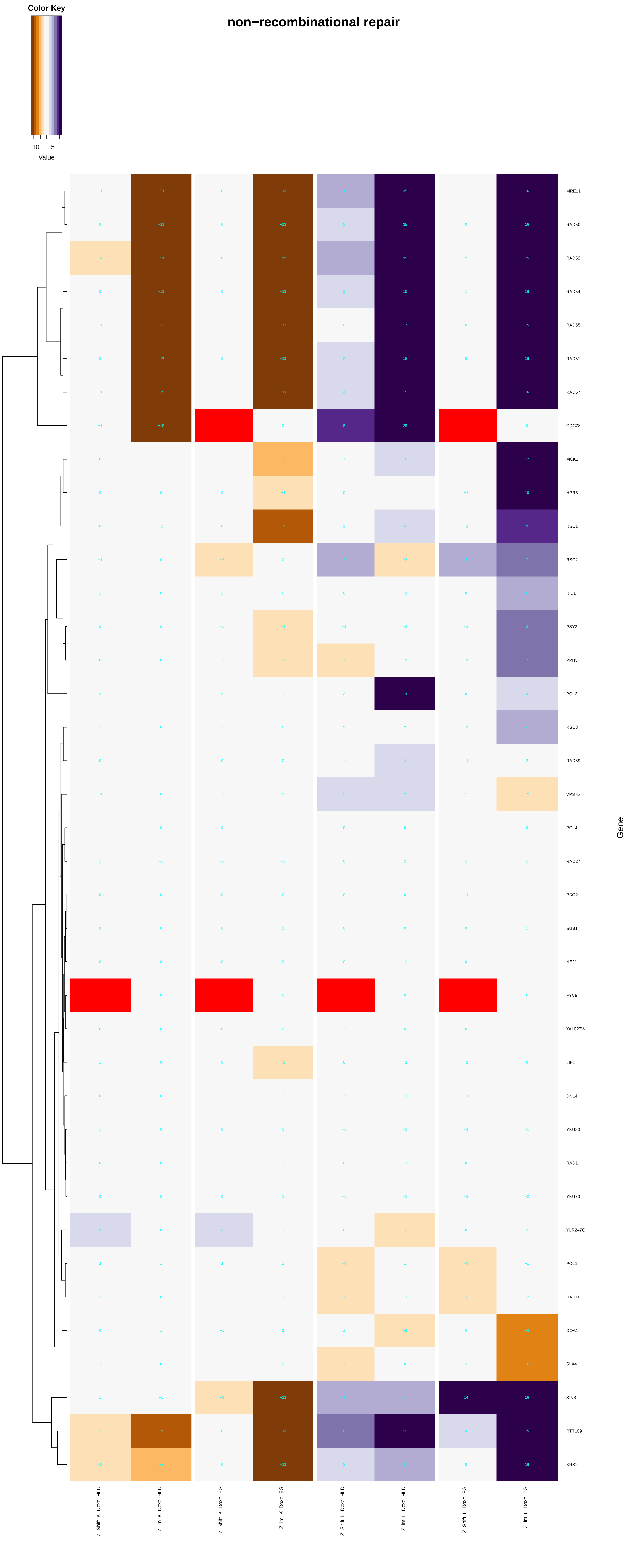
non−recombinational repair
Color Key
−10
5
Value
MRE11
RAD50
RAD52
RAD54
RAD55
RAD51
RAD57
CDC28
MCK1
HPR5
RSC1
RSC2
RIS1
PSY2
PPH3
POL2
RSC8
RAD59
VPS75
POL4
RAD27
PSO2
SUB1
NEJ1
FYV6
YAL027W
LIF1
DNL4
YKU80
RAD1
YKU70
YLR247C
POL1
RAD10
DOA1
SLX4
SIN3
RTT109
XRS2
Gene
Z_Shift_K_Doxo_HLD
Z_Im_K_Doxo_HLD
Z_Shift_K_Doxo_EG
Z_Im_K_Doxo_EG
Z_Shift_L_Doxo_HLD
Z_Im_L_Doxo_HLD
Z_Shift_L_Doxo_EG
Z_Im_L_Doxo_EG

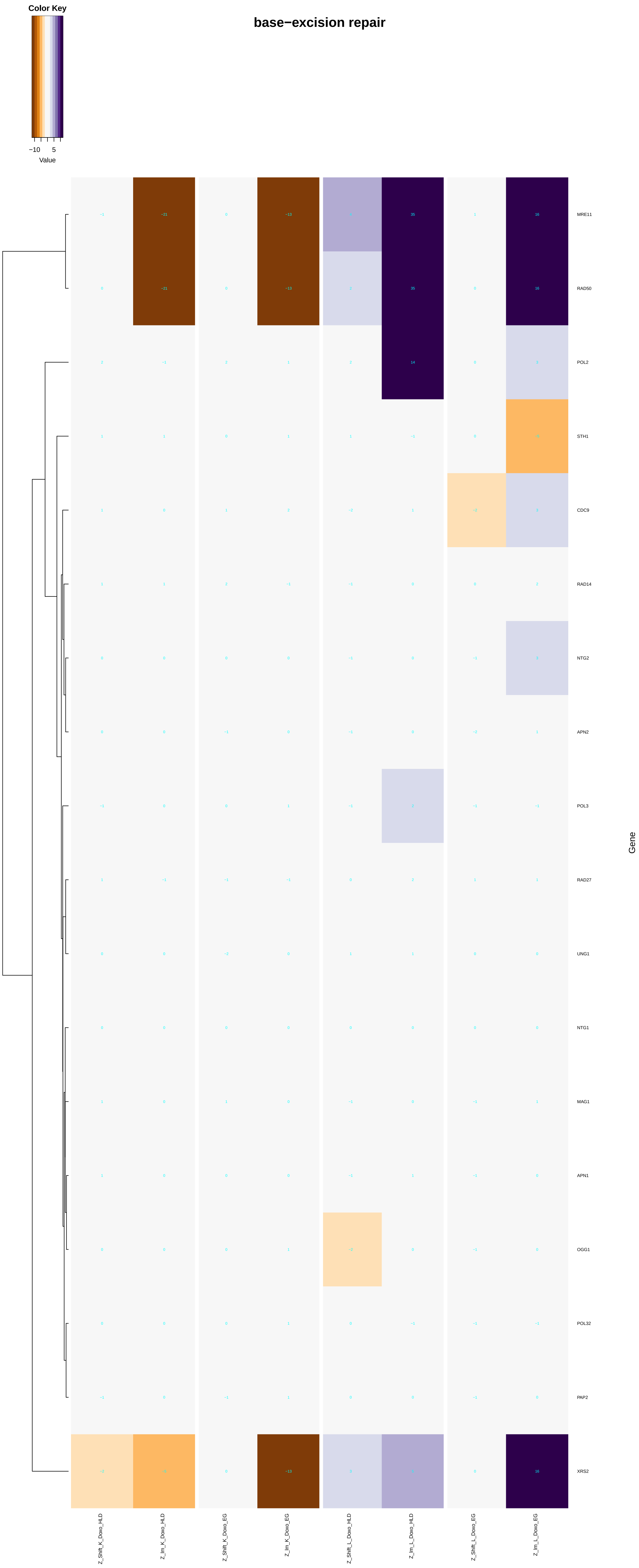
Color Key
−10
5
Value
base−excision repair
MRE11
RAD50
POL2
STH1
CDC9
RAD14
NTG2
APN2
POL3
RAD27
UNG1
NTG1
MAG1
APN1
OGG1
POL32
PAP2
XRS2
Gene
Z_Shift_K_Doxo_HLD
Z_Im_K_Doxo_HLD
Z_Shift_K_Doxo_EG
Z_Im_K_Doxo_EG
Z_Shift_L_Doxo_HLD
Z_Im_L_Doxo_HLD
Z_Shift_L_Doxo_EG
Z_Im_L_Doxo_EG

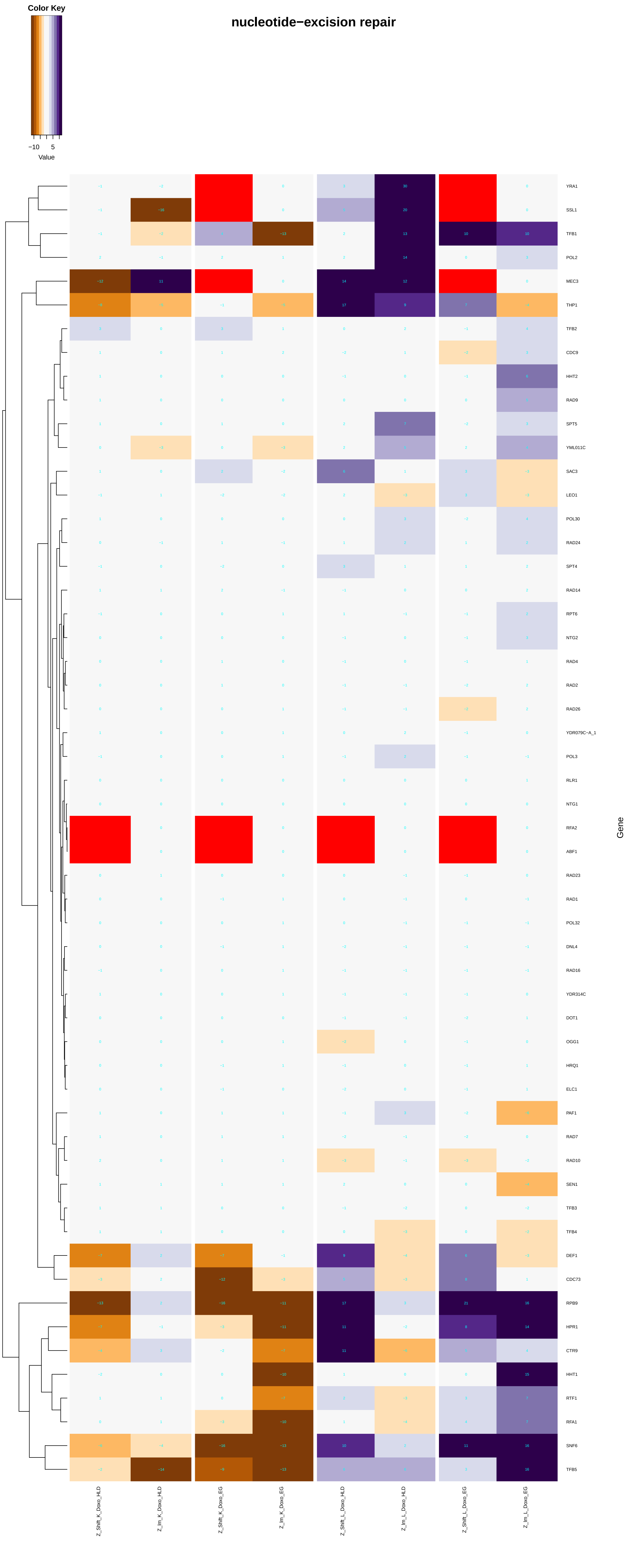

nucleotide−excision repair
Color Key
−10
5
Value
YRA1
SSL1
TFB1
POL2
MEC3
THP1
TFB2
CDC9
HHT2
RAD9
SPT5
YML011C
SAC3
LEO1
POL30
RAD24
SPT4
RAD14
RPT6
NTG2
RAD4
RAD2
RAD26
YDR079C−A_1
POL3
RLR1
NTG1
RFA2
ABF1
RAD23
RAD1
POL32
DNL4
RAD16
YDR314C
DOT1
OGG1
HRQ1
ELC1
PAF1
RAD7
RAD10
SEN1
TFB3
TFB4
DEF1
CDC73
RPB9
HPR1
CTR9
HHT1
RTF1
RFA1
SNF6
TFB5
Gene
Z_Shift_K_Doxo_HLD
Z_Im_K_Doxo_HLD
Z_Shift_K_Doxo_EG
Z_Im_K_Doxo_EG
Z_Shift_L_Doxo_HLD
Z_Im_L_Doxo_HLD
Z_Shift_L_Doxo_EG
Z_Im_L_Doxo_EG

mismatch repair

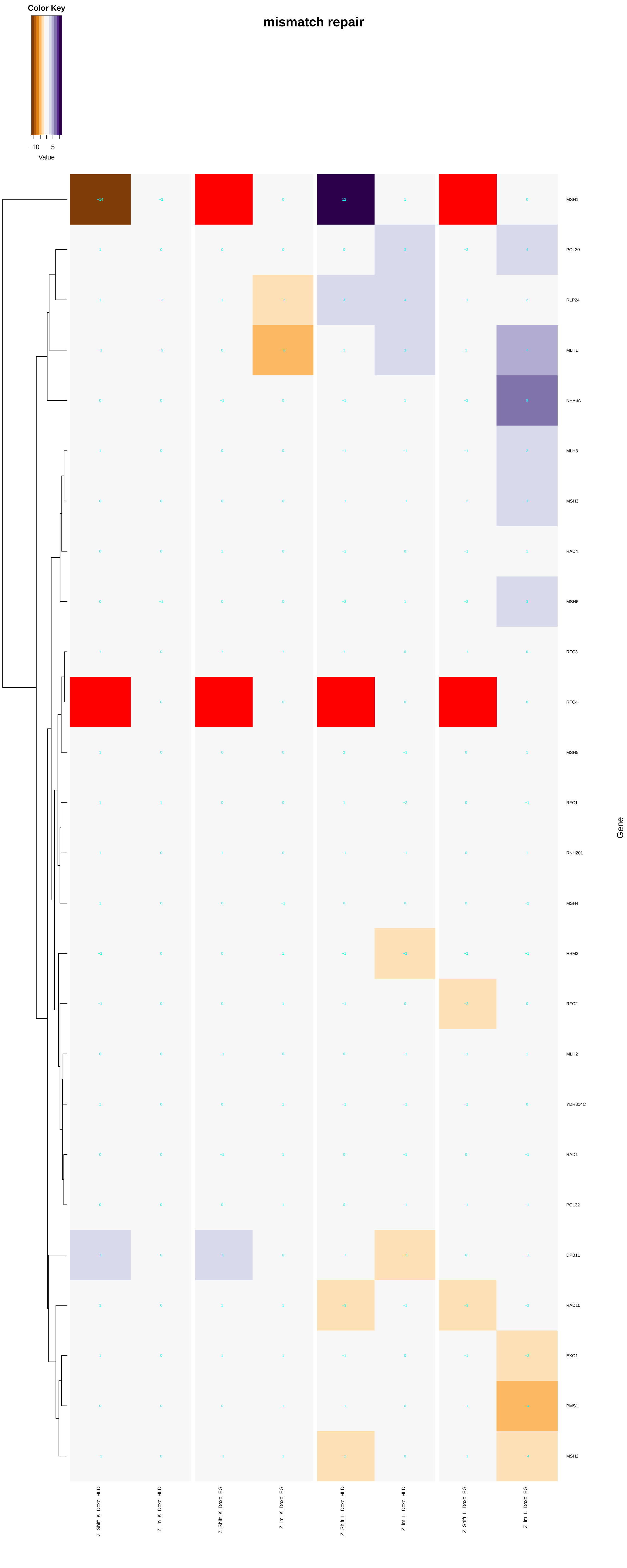

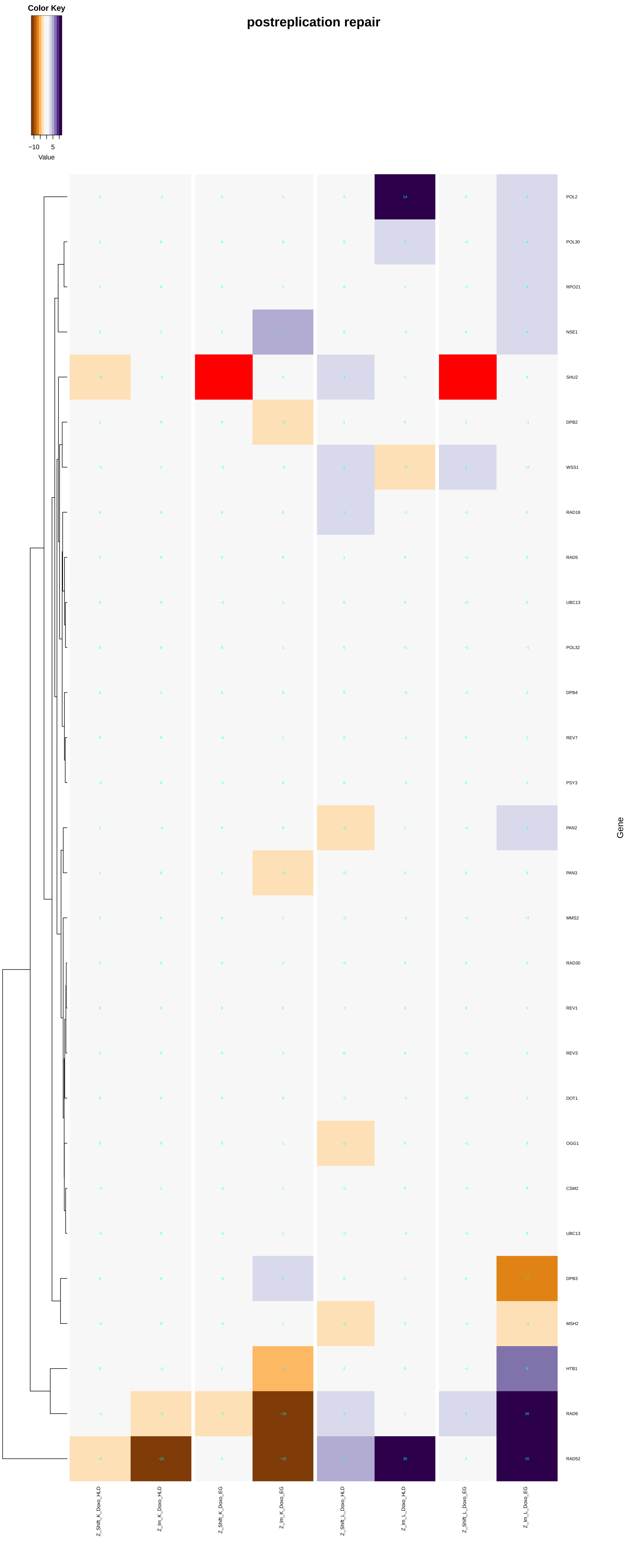
Color Key
−10
5
Value
postreplication repair
POL2
POL30
RPO21
NSE1
SHU2
DPB2
WSS1
RAD18
RAD5
UBC13
POL32
DPB4
REV7
PSY3
PAN2
PAN3
MMS2
RAD30
REV1
REV3
DOT1
OGG1
CSM2
UBC13
DPB3
MSH2
HTB1
RAD6
RAD52
Gene
Z_Shift_K_Doxo_HLD
Z_Im_K_Doxo_HLD
Z_Shift_K_Doxo_EG
Z_Im_K_Doxo_EG
Z_Shift_L_Doxo_HLD
Z_Im_L_Doxo_HLD
Z_Shift_L_Doxo_EG
Z_Im_L_Doxo_EG

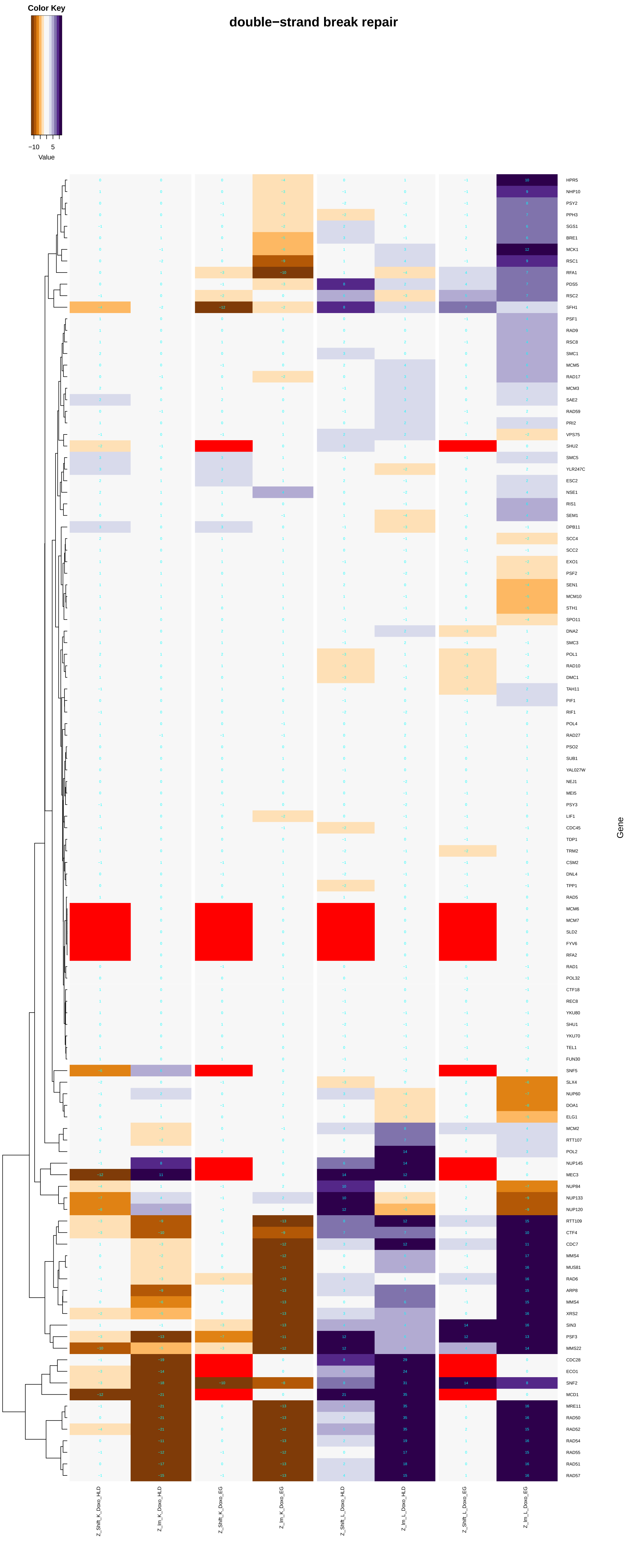
Color Key
−10 5
Value
double−strand break repair
Gene
HPR5
NHP10
PSY2
PPH3
SGS1
BRE1
MCK1
RSC1
RFA1
PDS5
RSC2
SFH1
PSF1
RAD9
RSC8
SMC1
MCM5
RAD17
MCM3
SAE2
RAD59
PRI2
VPS75
SHU2
SMC5
YLR247C
ESC2
NSE1
RIS1
SEM1
DPB11
SCC4
SCC2
EXO1
PSF2
SEN1
MCM10
STH1
SPO11
DNA2
SMC3
POL1
RAD10
DMC1
TAH11
PIF1
RIF1
POL4
RAD27
PSO2
SUB1
YAL027W
NEJ1
MEI5
PSY3
LIF1
CDC45
TDP1
TRM2
CSM2
DNL4
TPP1
RAD5
MCM6
MCM7
SLD2
FYV6
RFA2
RAD1
POL32
CTF18
REC8
YKU80
SHU1
YKU70
TEL1
FUN30
SNF5
SLX4
NUP60
DOA1
ELG1
MCM2
RTT107
POL2
NUP145
MEC3
NUP84
NUP133
NUP120
RTT109
CTF4
CDC7
MMS4
MUS81
RAD6
ARP8
MMS4
XRS2
SIN3
PSF3
MMS22
CDC28
ECO1
SNF2
MCD1
MRE11
RAD50
RAD52
RAD54
RAD55
RAD51
RAD57
Z_Shift_K_Doxo_HLD
Z_Im_K_Doxo_HLD
Z_Shift_K_Doxo_EG
Z_Im_K_Doxo_EG
Z_Shift_L_Doxo_HLD
Z_Im_L_Doxo_HLD
Z_Shift_L_Doxo_EG
Z_Im_L_Doxo_EG

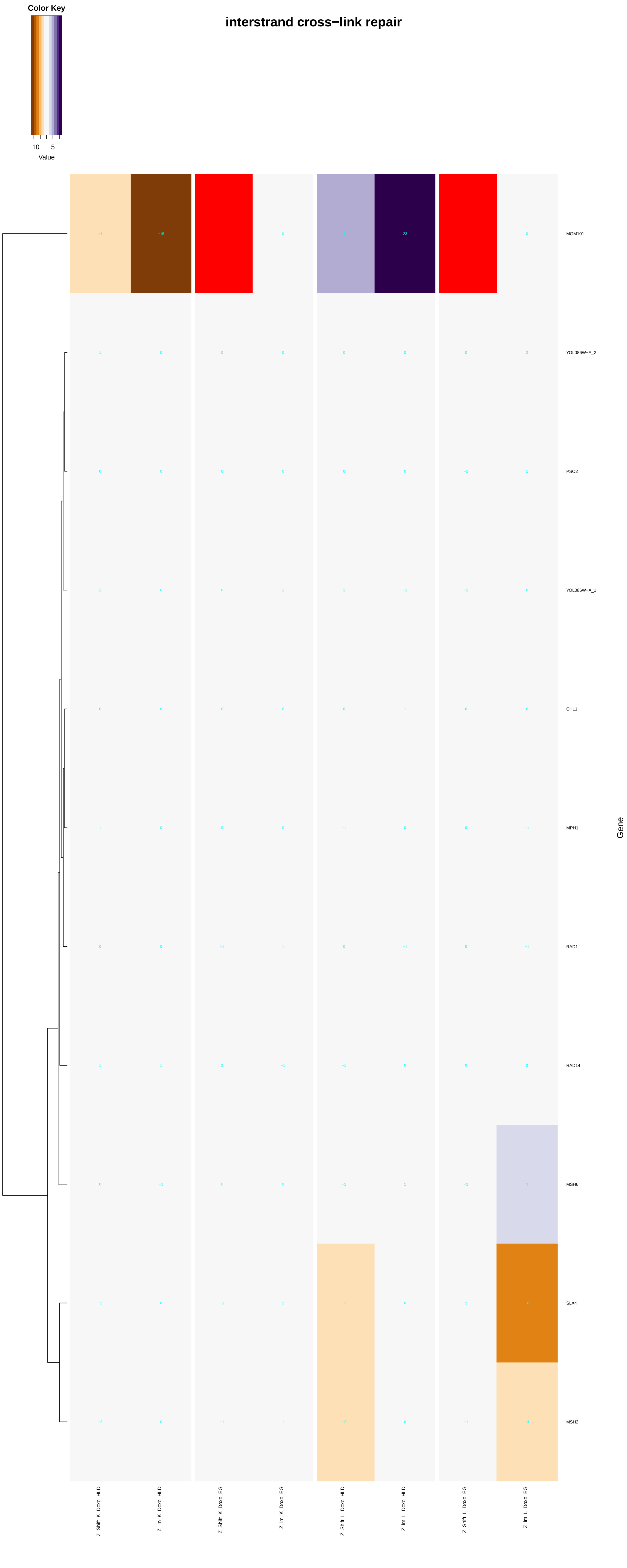

interstrand cross−link repair
Color Key
−10 5
Value
MGM101
YOL086W−A_2
PSO2
YOL086W−A_1
CHL1
MPH1
RAD1
RAD14
MSH6
SLX4
MSH2
Gene
Z_Shift_K_Doxo_HLD
Z_Im_K_Doxo_HLD
Z_Shift_K_Doxo_EG
Z_Im_K_Doxo_EG
Z_Shift_L_Doxo_HLD
Z_Im_L_Doxo_HLD
Z_Shift_L_Doxo_EG
Z_Im_L_Doxo_EG

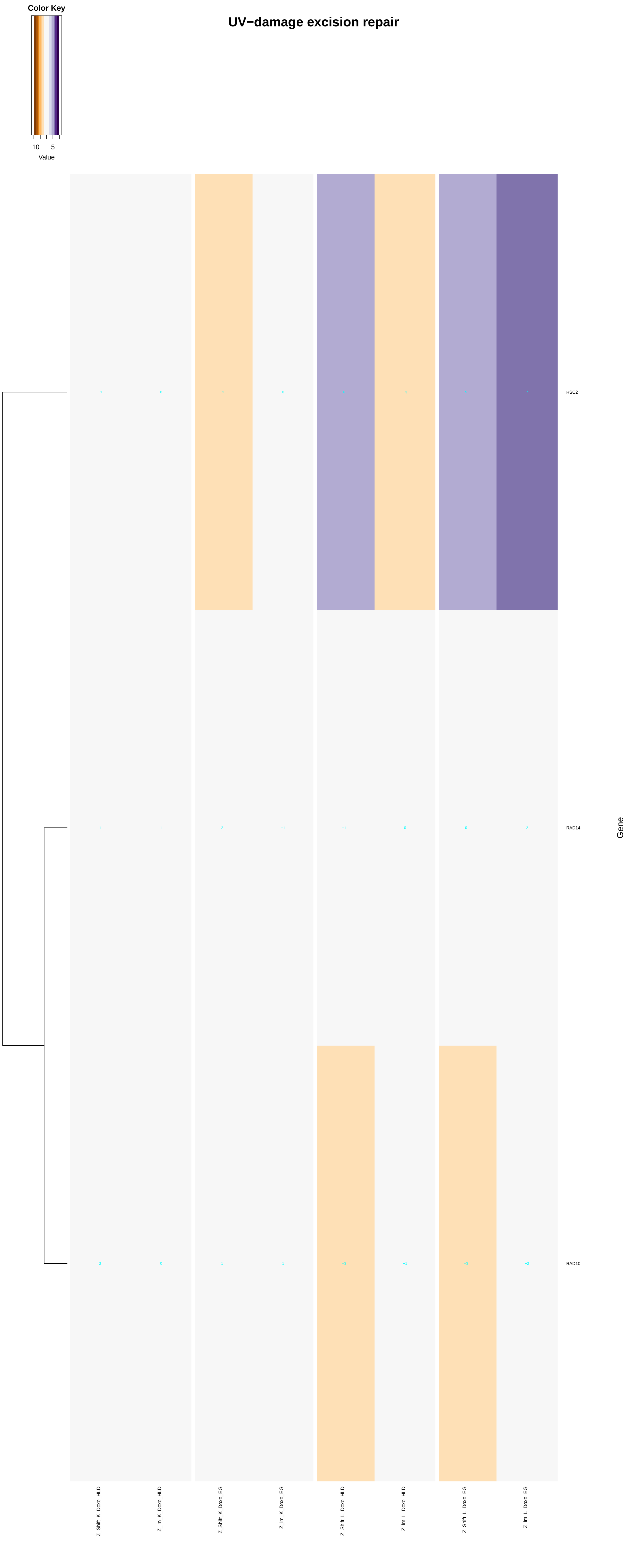
UV−damage excision repair
Color Key
−10
5
Value
RSC2
RAD14
RAD10
Gene
Z_Shift_K_Doxo_HLD
Z_Im_K_Doxo_HLD
Z_Shift_K_Doxo_EG
Z_Im_K_Doxo_EG
Z_Shift_L_Doxo_HLD
Z_Im_L_Doxo_HLD
Z_Shift_L_Doxo_EG
Z_Im_L_Doxo_EG

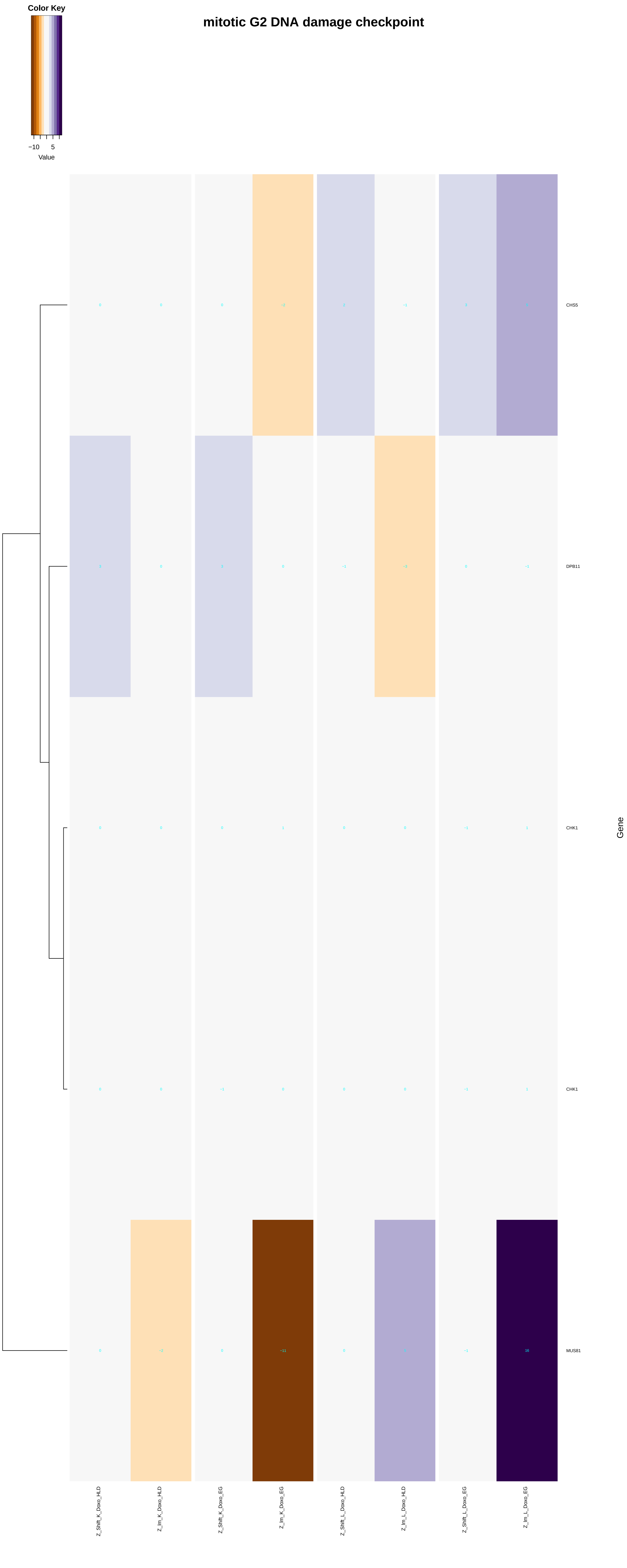

mitotic G2 DNA damage checkpoint
Color Key
−10
5
Value
CHS5
DPB11
CHK1
CHK1
MUS81
Gene
Z_Shift_K_Doxo_HLD
Z_Im_K_Doxo_HLD
Z_Shift_K_Doxo_EG
Z_Im_K_Doxo_EG
Z_Shift_L_Doxo_HLD
Z_Im_L_Doxo_HLD
Z_Shift_L_Doxo_EG
Z_Im_L_Doxo_EG

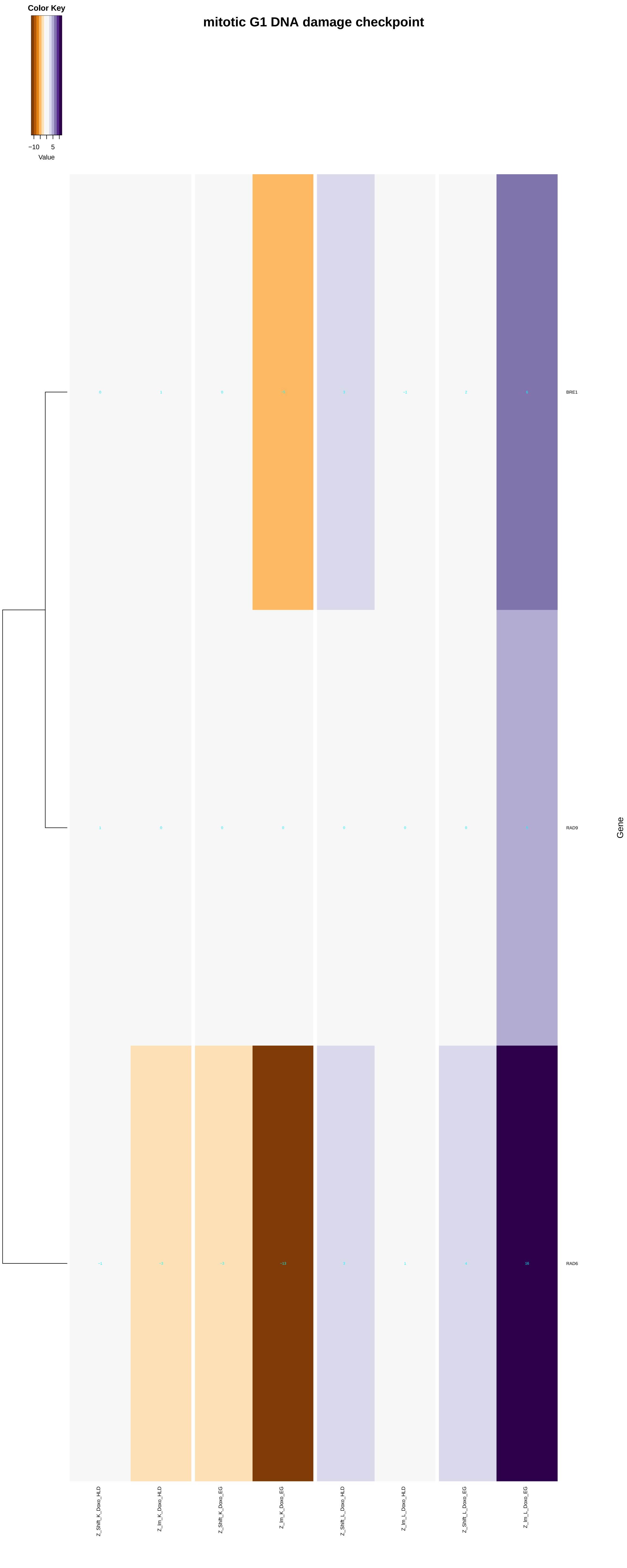

Color Key
−10
5
Value
mitotic G1 DNA damage checkpoint
0
1
0
−5
3
−1
2
6
BRE1
1
0
0
0
0
0
0
5
RAD9
−1
−3
−3
−13
3
1
4
16
RAD6
Gene
Z_Shift_K_Doxo_HLD
Z_Im_K_Doxo_HLD
Z_Shift_K_Doxo_EG
Z_Im_K_Doxo_EG
Z_Shift_L_Doxo_HLD
Z_Im_L_Doxo_HLD
Z_Shift_L_Doxo_EG
Z_Im_L_Doxo_EG

#### intra−S DNA damage checkpoint

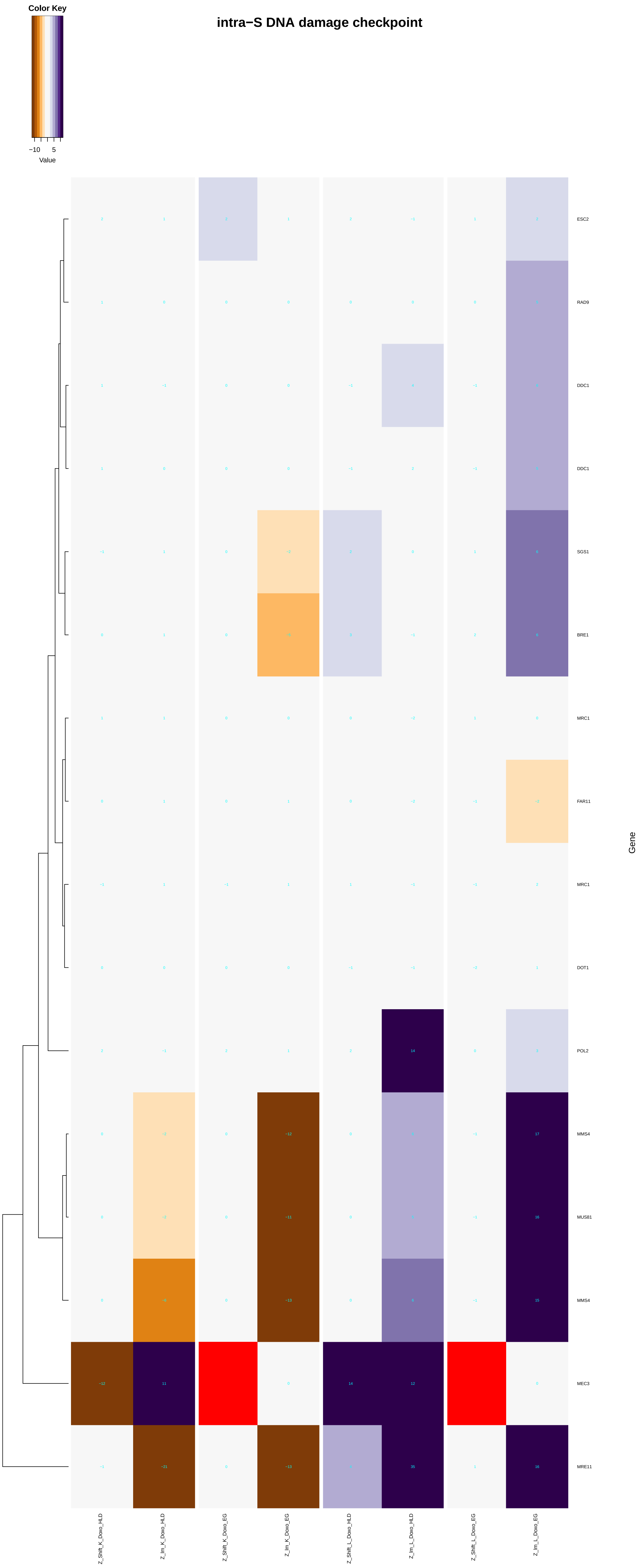

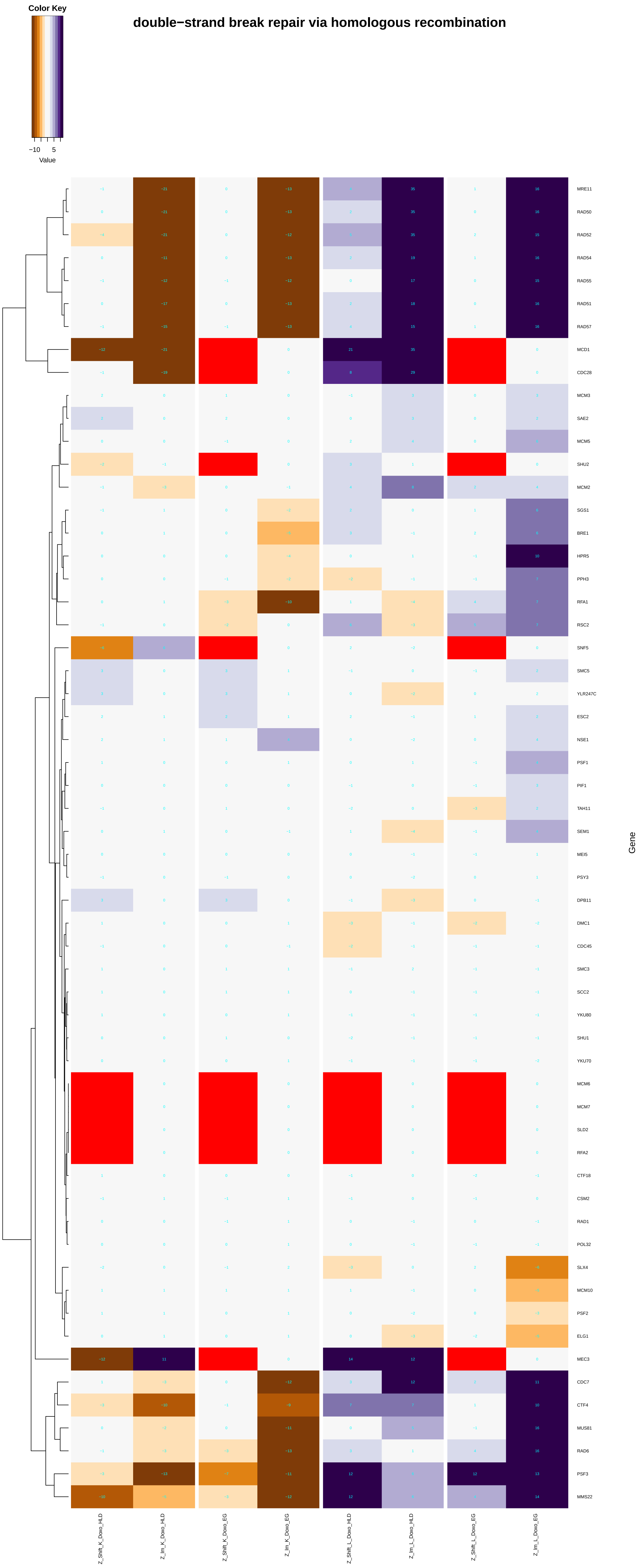
Color Key
−10
5
Value
double−strand break repair via homologous recombination
MRE11
RAD50
RAD52
RAD54
RAD55
RAD51
RAD57
MCD1
CDC28
MCM3
SAE2
MCM5
SHU2
MCM2
SGS1
BRE1
HPR5
PPH3
RFA1
RSC2
SNF5
SMC5
YLR247C
ESC2
NSE1
PSF1
PIF1
TAH11
SEM1
MEI5
PSY3
DPB11
DMC1
CDC45
SMC3
SCC2
YKU80
SHU1
YKU70
MCM6
MCM7
SLD2
RFA2
CTF18
CSM2
RAD1
POL32
SLX4
MCM10
PSF2
ELG1
MEC3
CDC7
CTF4
MUS81
RAD6
PSF3
MMS22
Gene
Z_Shift_K_Doxo_HLD
Z_Im_K_Doxo_HLD
Z_Shift_K_Doxo_EG
Z_Im_K_Doxo_EG
Z_Shift_L_Doxo_HLD
Z_Im_L_Doxo_HLD
Z_Shift_L_Doxo_EG
Z_Im_L_Doxo_EG

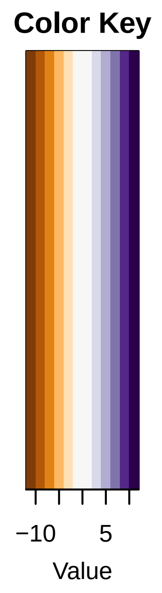


### mitochondrial double−strand break repair via homologous recombination

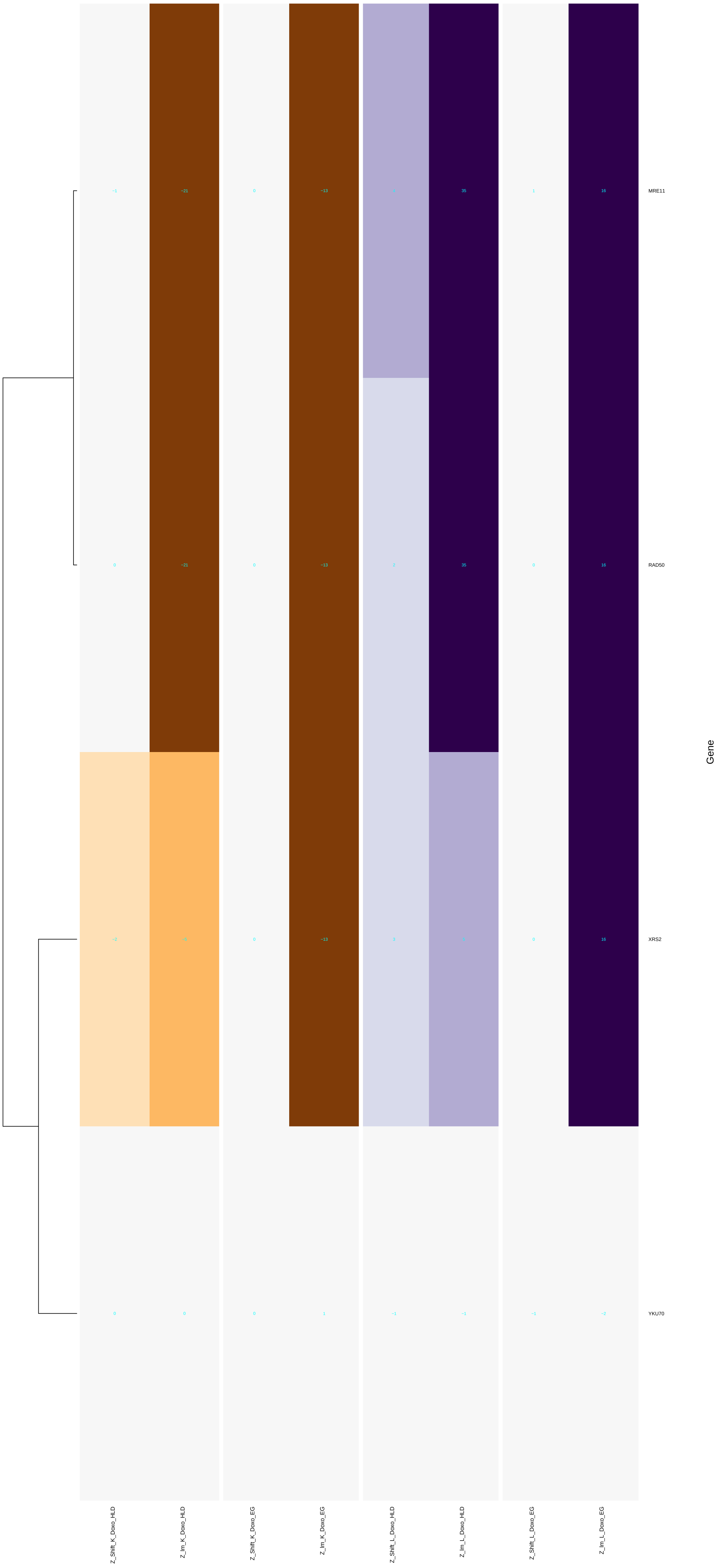

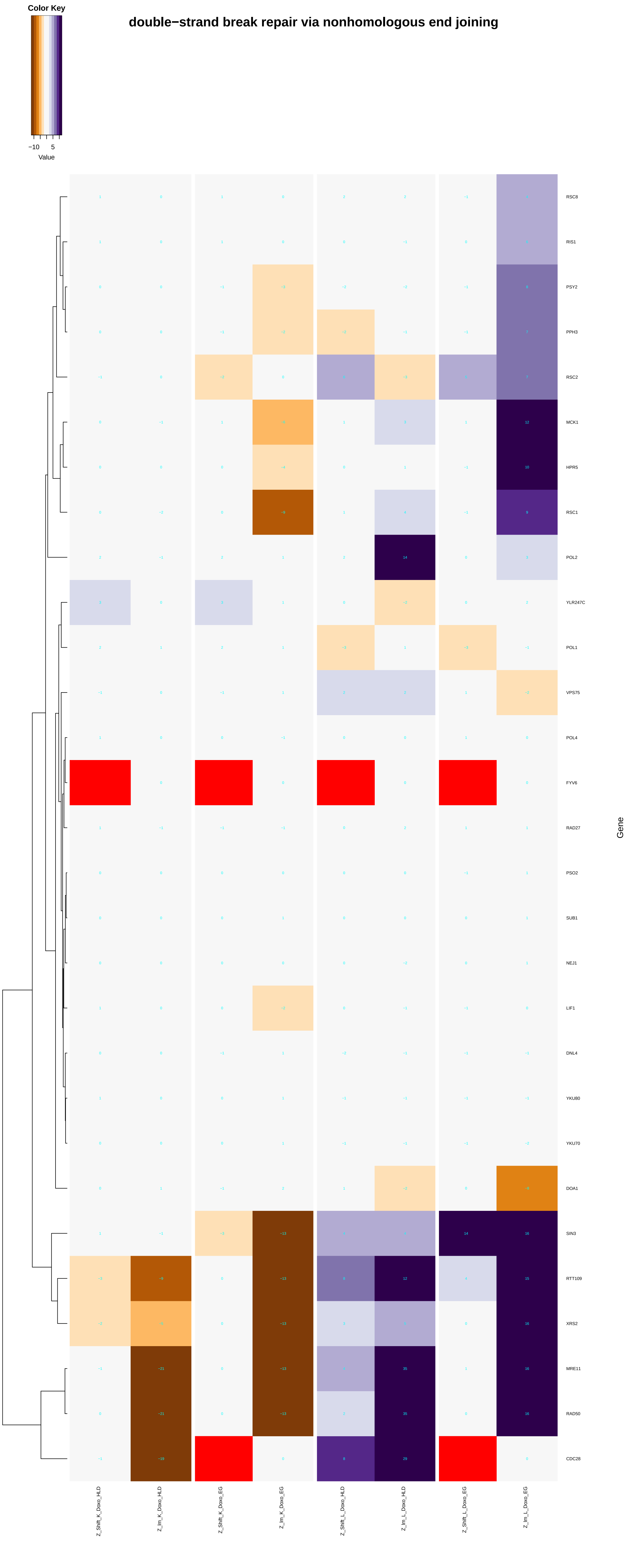
double−strand break repair via nonhomologous end joining
Color Key
−10
5
Value
RSC8
RIS1
PSY2
PPH3
RSC2
MCK1
HPR5
RSC1
POL2
YLR247C
POL1
VPS75
POL4
FYV6
RAD27
PSO2
SUB1
NEJ1
LIF1
DNL4
YKU80
YKU70
DOA1
SIN3
RTT109
XRS2
MRE11
RAD50
CDC28
Gene
Z_Shift_K_Doxo_HLD
Z_Im_K_Doxo_HLD
Z_Shift_K_Doxo_EG
Z_Im_K_Doxo_EG
Z_Shift_L_Doxo_HLD
Z_Im_L_Doxo_HLD
Z_Shift_L_Doxo_EG
Z_Im_L_Doxo_EG

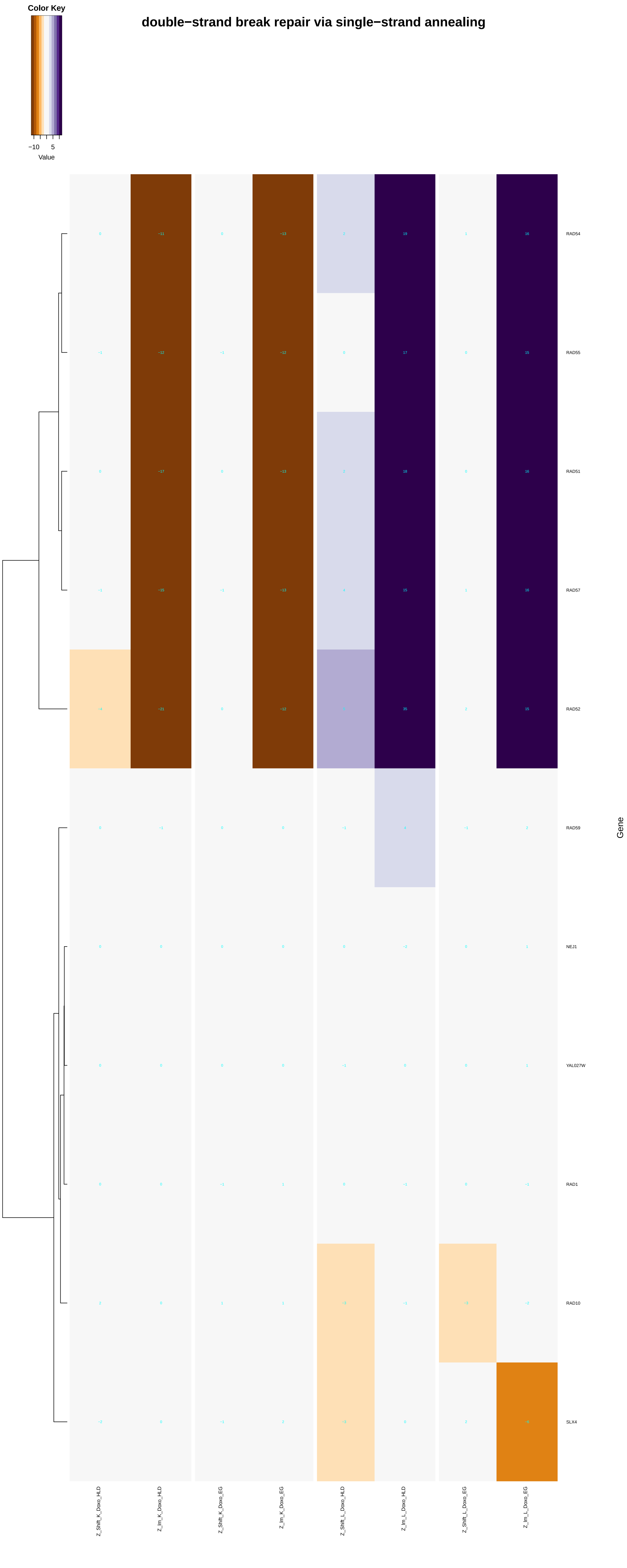
Color Key
−10
5
Value
double−strand break repair via single−strand annealing
RAD54
RAD55
RAD51
RAD57
RAD52
RAD59
NEJ1
YAL027W
RAD1
RAD10
SLX4
Gene
Z_Shift_K_Doxo_HLD
Z_Im_K_Doxo_HLD
Z_Shift_K_Doxo_EG
Z_Im_K_Doxo_EG
Z_Shift_L_Doxo_HLD
Z_Im_L_Doxo_HLD
Z_Shift_L_Doxo_EG
Z_Im_L_Doxo_EG

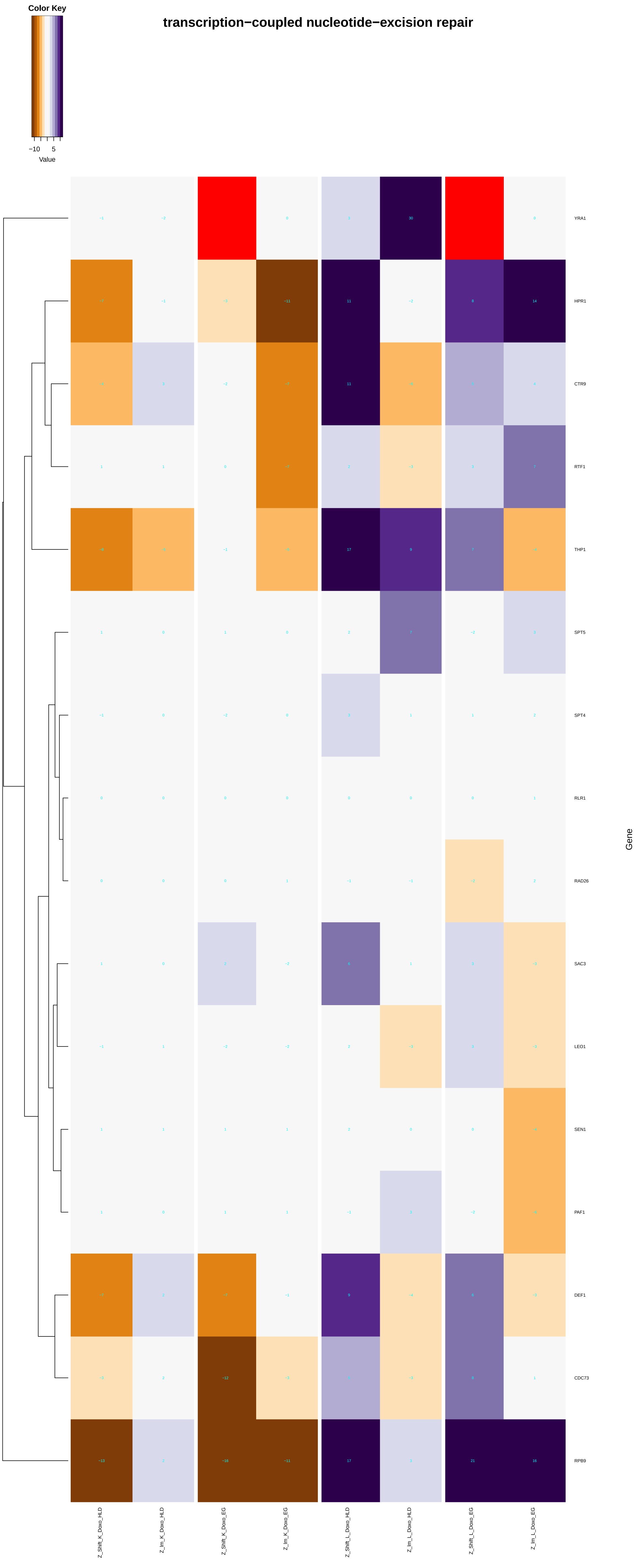
transcription−coupled nucleotide−excision repair
Color Key
−10
5
Value
YRA1
HPR1
CTR9
RTF1
THP1
SPT5
SPT4
RLR1
RAD26
SAC3
LEO1
SEN1
PAF1
DEF1
CDC73
RPB9
Gene
Z_Shift_K_Doxo_HLD
Z_lm_K_Doxo_HLD
Z_Shift_K_Doxo_EG
Z_lm_K_Doxo_EG
Z_Shift_L_Doxo_HLD
Z_lm_L_Doxo_HLD
Z_Shift_L_Doxo_EG
Z_lm_L_Doxo_EG

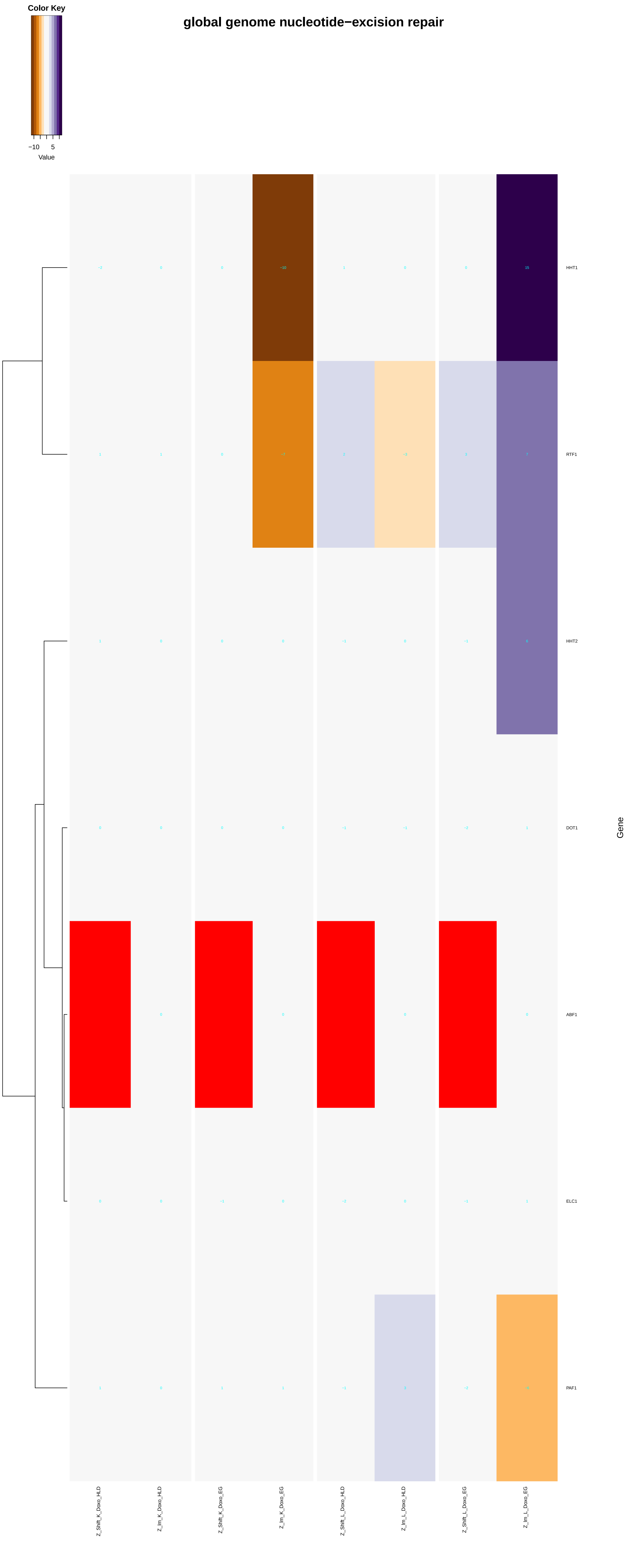
global genome nucleotide−excision repair
Color Key
−10
5
Value
HHT1
RTF1
HHT2
DOT1
ABF1
ELC1
PAF1
Gene
Z_Shift_K_Doxo_HLD
Z_Im_K_Doxo_HLD
Z_Shift_K_Doxo_EG
Z_Im_K_Doxo_EG
Z_Shift_L_Doxo_HLD
Z_Im_L_Doxo_HLD
Z_Shift_L_Doxo_EG
Z_Im_L_Doxo_EG

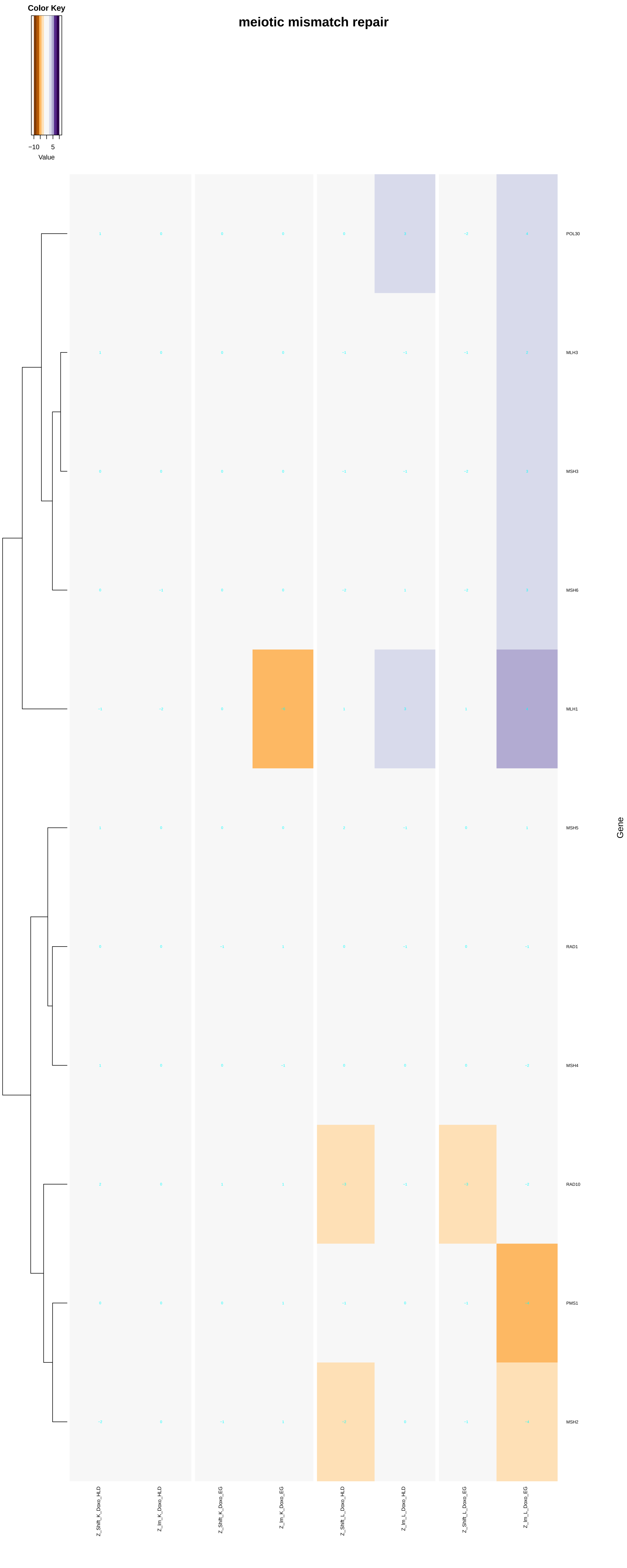

meiotic mismatch repair
Color Key
−10
5
Value
POL30
MLH3
MSH3
MSH6
MLH1
MSH5
RAD1
MSH4
RAD10
PMS1
MSH2
Gene
Z_Shift_K_Doxo_HLD
Z_Im_K_Doxo_HLD
Z_Shift_K_Doxo_EG
Z_Im_K_Doxo_EG
Z_Shift_L_Doxo_HLD
Z_Im_L_Doxo_HLD
Z_Shift_L_Doxo_EG
Z_Im_L_Doxo_EG

### translesion synthesis

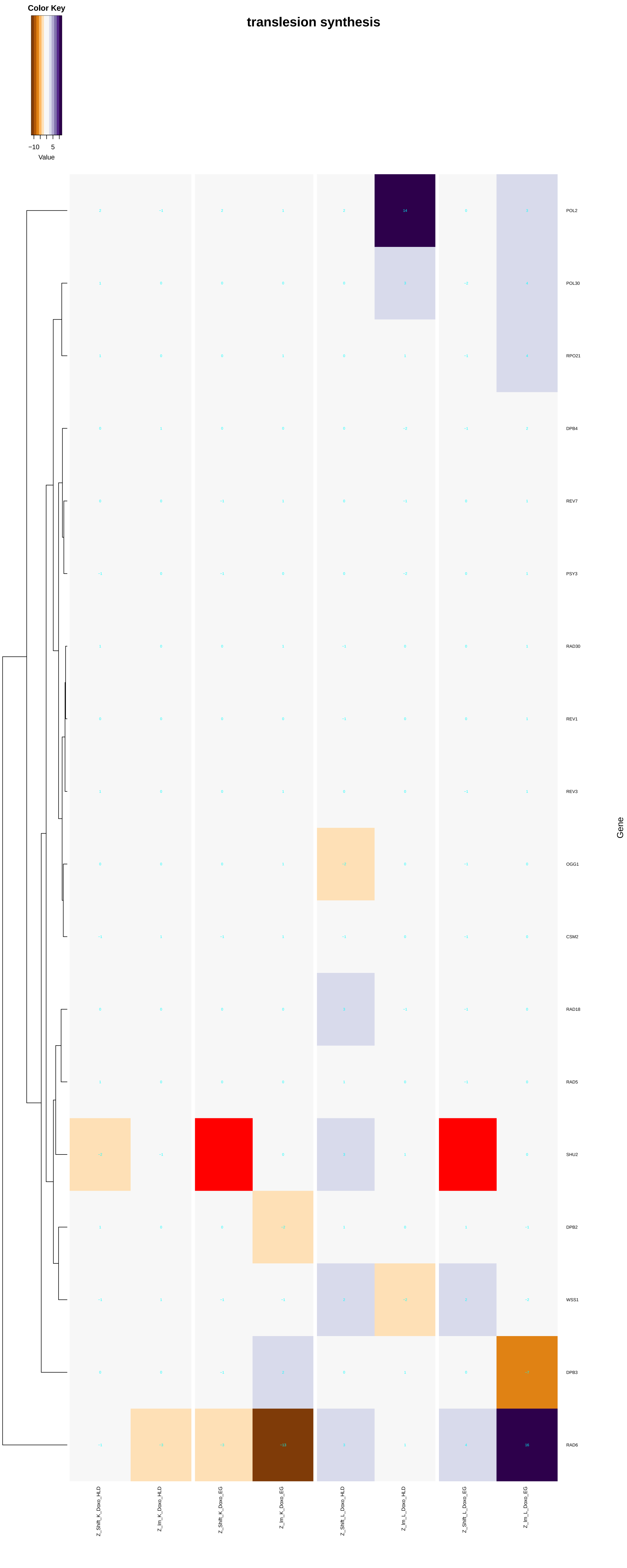

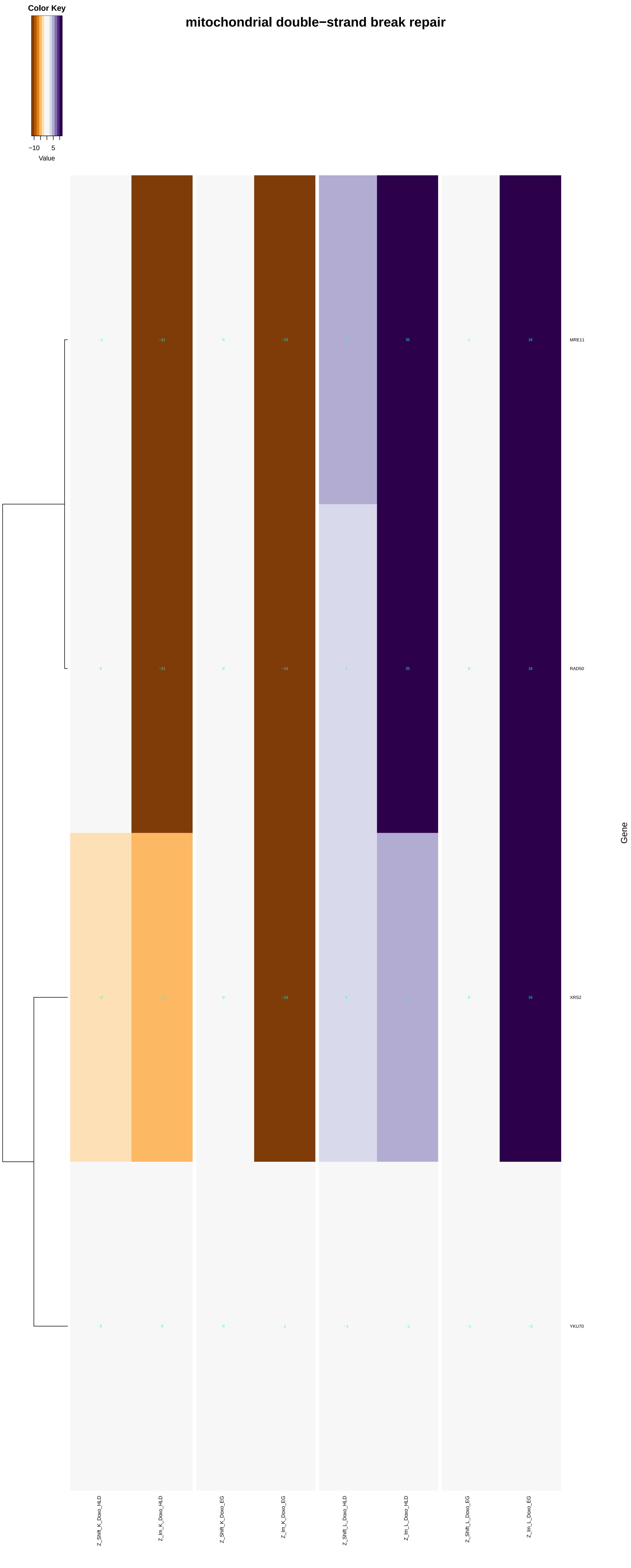

Color Key
−10 5
Value
mitochondrial double−strand break repair
−1 −21 0 −13 4 35 1 16 MRE11
0 −21 0 −13 3 35 0 16 RAD50
−2 −5 0 −13 3 5 0 16 XRS2
0 0 0 1 −1 −1 −1 −2 YKU70
Gene
Z_Shift_K_Doxo_HLD
Z_Im_K_Doxo_HLD
Z_Shift_K_Doxo_EG
Z_Im_K_Doxo_EG
Z_Shift_L_Doxo_HLD
Z_Im_L_Doxo_HLD
Z_Shift_L_Doxo_EG
Z_Im_L_Doxo_EG

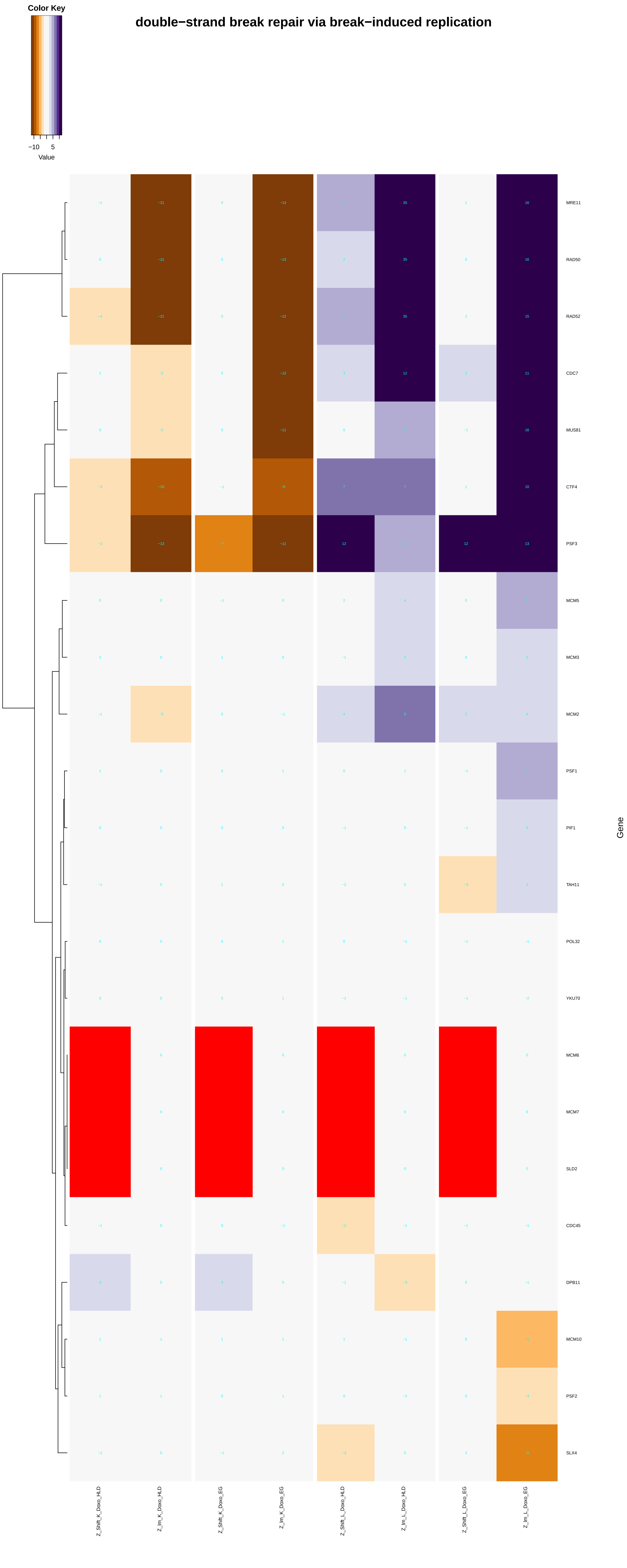
double−strand break repair via break−induced replication
Color Key
−10
5
Value
MRE11
RAD50
RAD52
CDC7
MUS81
CTF4
PSF3
MCM5
MCM3
MCM2
PSF1
PIF1
TAH11
POL32
YKU70
MCM6
MCM7
SLD2
CDC45
DPB11
MCM10
PSF2
SLX4
Gene
Z_Shift_K_Doxo_HLD
Z_Im_K_Doxo_HLD
Z_Shift_K_Doxo_EG
Z_Im_K_Doxo_EG
Z_Shift_L_Doxo_HLD
Z_Im_L_Doxo_HLD
Z_Shift_L_Doxo_EG
Z_Im_L_Doxo_EG

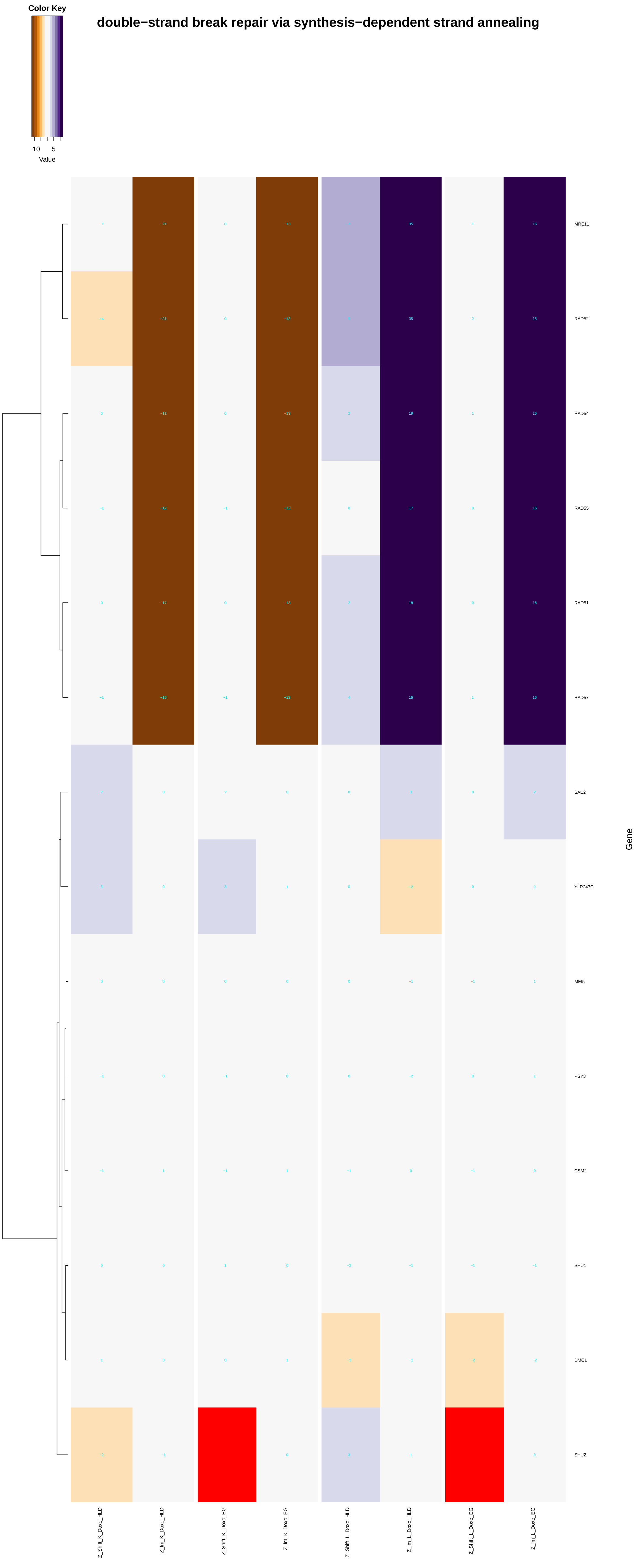
Color Key
−10 5
Value
double−strand break repair via synthesis−dependent strand annealing
MRE11
RAD52
RAD54
RAD55
RAD51
RAD57
SAE2
YLR247C
MEI5
PSY3
CSM2
SHU1
DMC1
SHU2
Gene
Z_Shift_K_Doxo_HLD
Z_Im_K_Doxo_HLD
Z_Shift_K_Doxo_EG
Z_Im_K_Doxo_EG
Z_Shift_L_Doxo_HLD
Z_Im_L_Doxo_HLD
Z_Shift_L_Doxo_EG
Z_Im_L_Doxo_EG

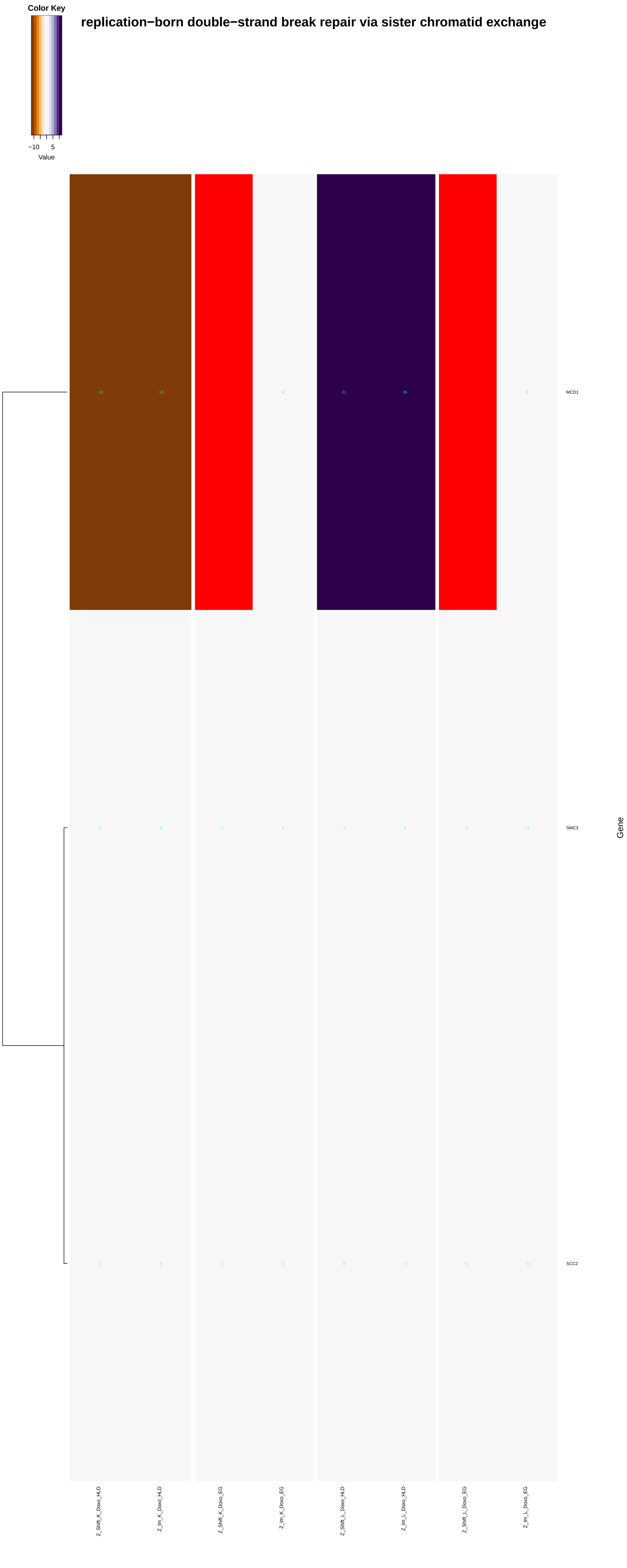

Color Key
−10
5
Value
replication−born double−strand break repair via sister chromatid exchange
−12
−21
0
21
35
0
MCD1
1
0
1
1
−1
2
−1
−1
SMC3
1
0
1
1
0
−1
−1
−1
SCC2
Gene
Z_Shift_K_Doxo_HLD
Z_lm_K_Doxo_HLD
Z_Shift_K_Doxo_EG
Z_lm_K_Doxo_EG
Z_Shift_L_Doxo_HLD
Z_lm_L_Doxo_HLD
Z_Shift_L_Doxo_EG
Z_lm_L_Doxo_EG

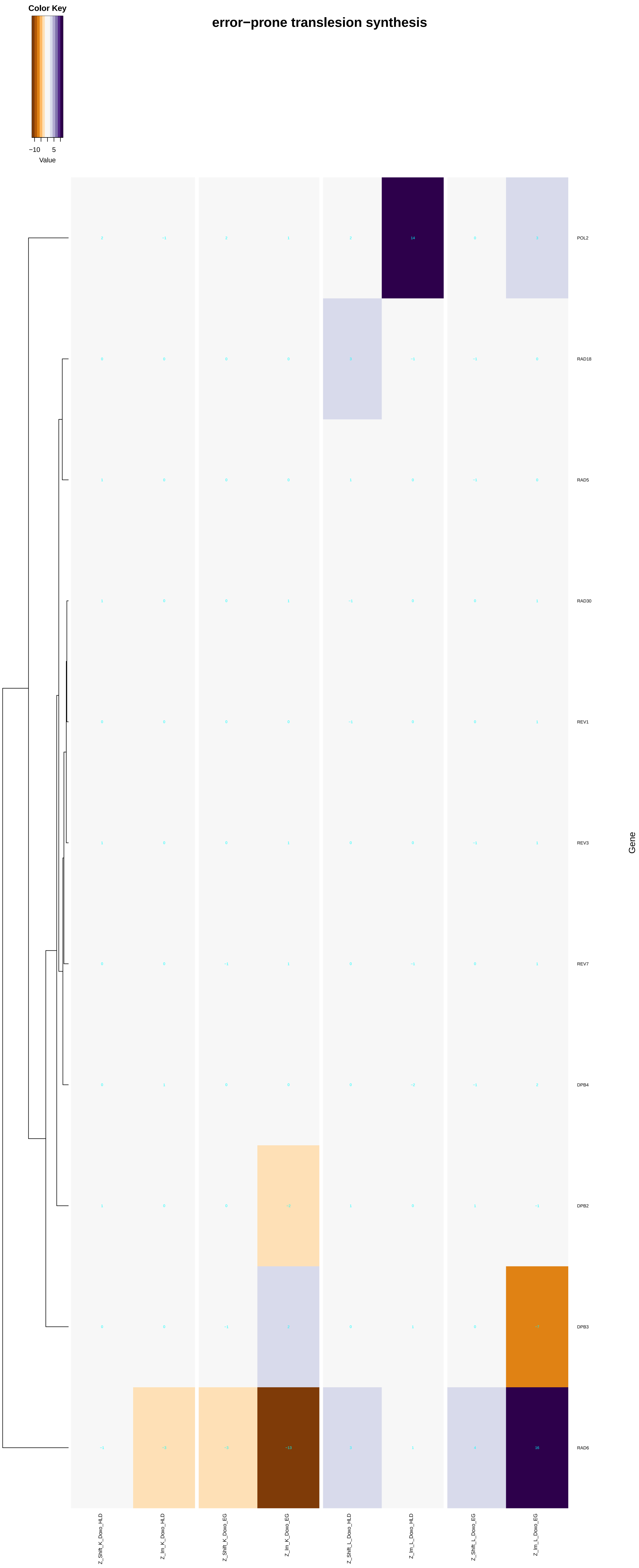
error−prone translesion synthesis
Color Key
−10
5
Value
POL2
RAD18
RAD5
RAD30
REV1
REV3
REV7
DPB4
DPB2
DPB3
RAD6
Gene
Z_Shift_K_Doxo_HLD
Z_Im_K_Doxo_HLD
Z_Shift_K_Doxo_EG
Z_Im_K_Doxo_EG
Z_Shift_L_Doxo_HLD
Z_Im_L_Doxo_HLD
Z_Shift_L_Doxo_EG
Z_Im_L_Doxo_EG

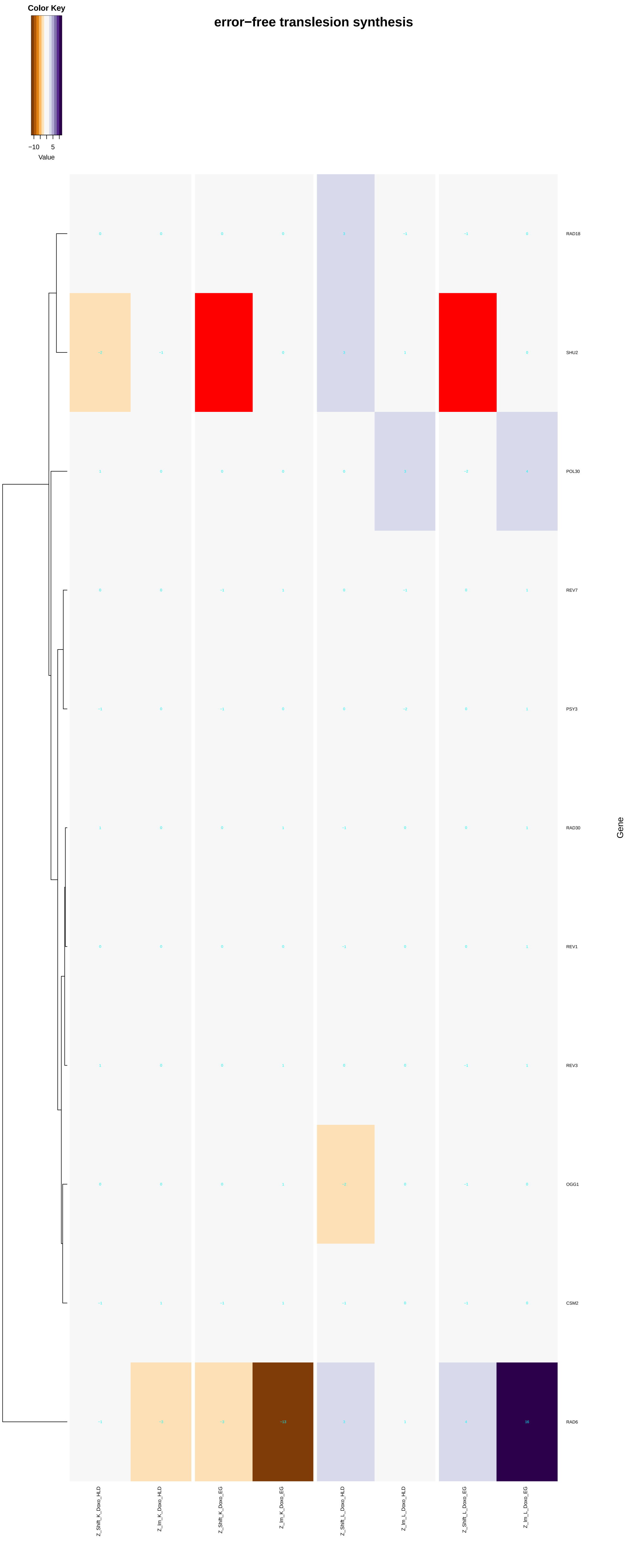

Color Key
−10
5
Value
error−free translesion synthesis
RAD18
SHU2
POL30
REV7
PSY3
RAD30
REV1
REV3
OGG1
CSM2
RAD6
Gene
Z_Shift_K_Doxo_HLD
Z_Im_K_Doxo_HLD
Z_Shift_K_Doxo_EG
Z_Im_K_Doxo_EG
Z_Shift_L_Doxo_HLD
Z_Im_L_Doxo_HLD
Z_Shift_L_Doxo_EG
Z_Im_L_Doxo_EG
