## Additional File 9 for "A yeast phenomic model for the influence of Warburg metabolism on genetic buffering of doxorubicin": cellular_response_to_extracellular_stimulus.pdf

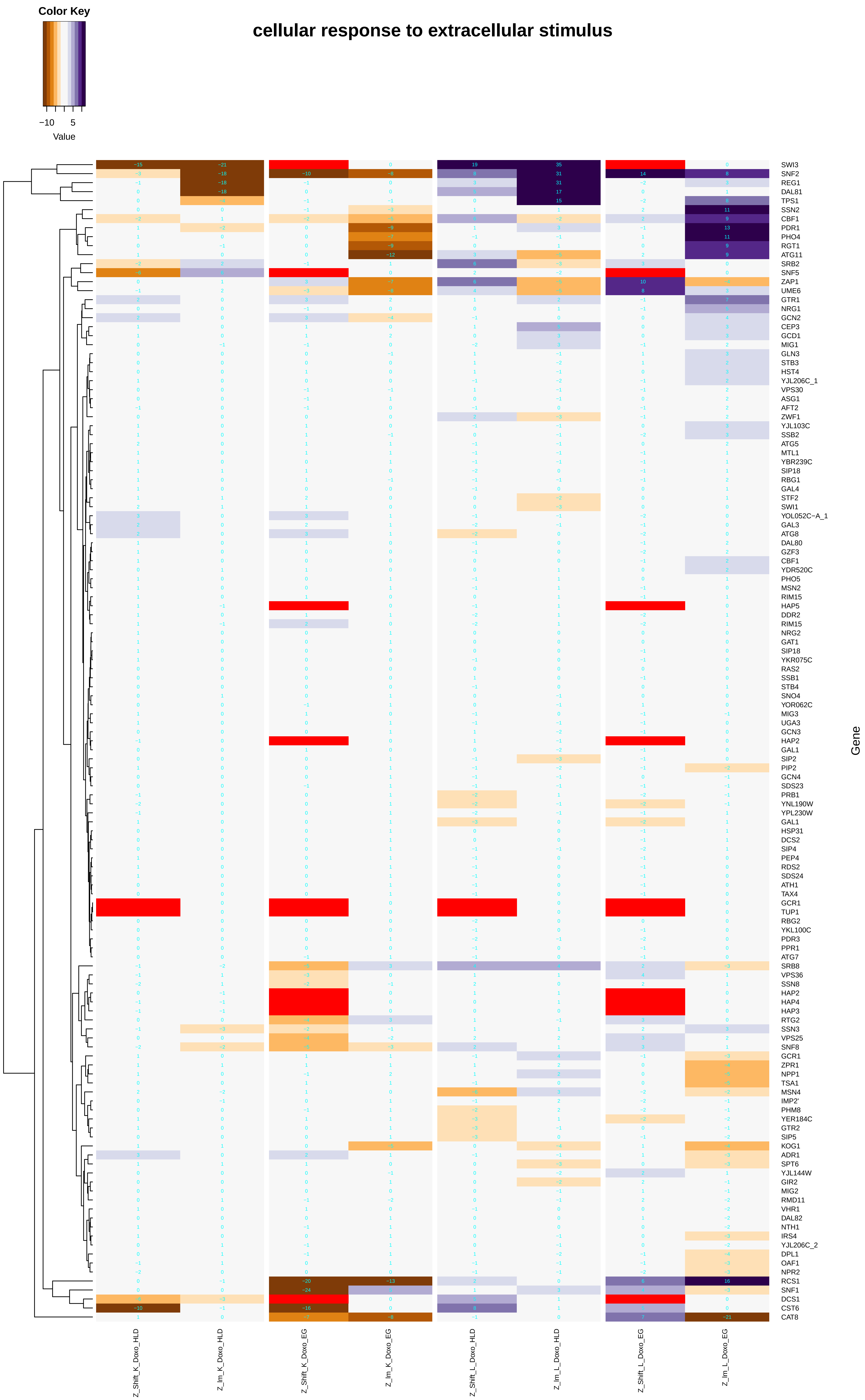

Color Key
−10
5
Value
cellular response to extracellular stimulus
SWI3
SNF2
REG1
DAL81
TPS1
SSN2
CBF1
PDR1
PHO4
RGT1
ATG11
SRB2
SNF5
ZAP1
UME6
GTR1
NRG1
GCN2
CEP3
GCD1
MIG1
GLN3
STB3
HST4
YJL206C_1
VPS30
ASG1
AFT2
ZWF1
YJL103C
SSB2
ATG5
MTL1
YBR239C
SIP18
RBG1
GAL4
STF2
SWI1
YOL052C−A_1
GAL3
ATG8
DAL80
GZF3
CBF1
YDR520C
PHO5
MSN2
RIM15
HAP5
DDR2
RIM15
NRG2
GAT1
SIP18
YKR075C
RAS2
SSB1
STB4
SNO4
YOR062C
MIG3
UGA3
GCN3
HAP2
GAL1
SIP2
PIP2
GCN4
SDS23
PRB1
YNL190W
YPL230W
GAL1
HSP31
DCS2
SIP4
PEP4
RDS2
SDS24
ATH1
TAX4
GCR1
TUP1
RBG2
YKL100C
PDR3
PPR1
ATG7
SRB8
VPS36
SSN8
HAP2
HAP4
HAP3
RTG2
SSN3
VPS25
SNF8
GCR1
ZPR1
NPP1
TSA1
MSN4
IMP2'
PHM8
YER184C
GTR2
SIP5
KOG1
ADR1
SPT6
YJL144W
GIR2
MIG2
RMD11
VHR1
DAL82
NTH1
IRS4
YJL206C_2
DPL1
OAF1
NPR2
RCS1
SNF1
DCS1
CST6
CAT8
Gene
Z_Shift_K_Doxo_HLD
Z_Im_K_Doxo_HLD
Z_Shift_K_Doxo_EG
Z_Im_K_Doxo_EG
Z_Shift_L_Doxo_HLD
Z_Im_L_Doxo_HLD
Z_Shift_L_Doxo_EG
Z_Im_L_Doxo_EG

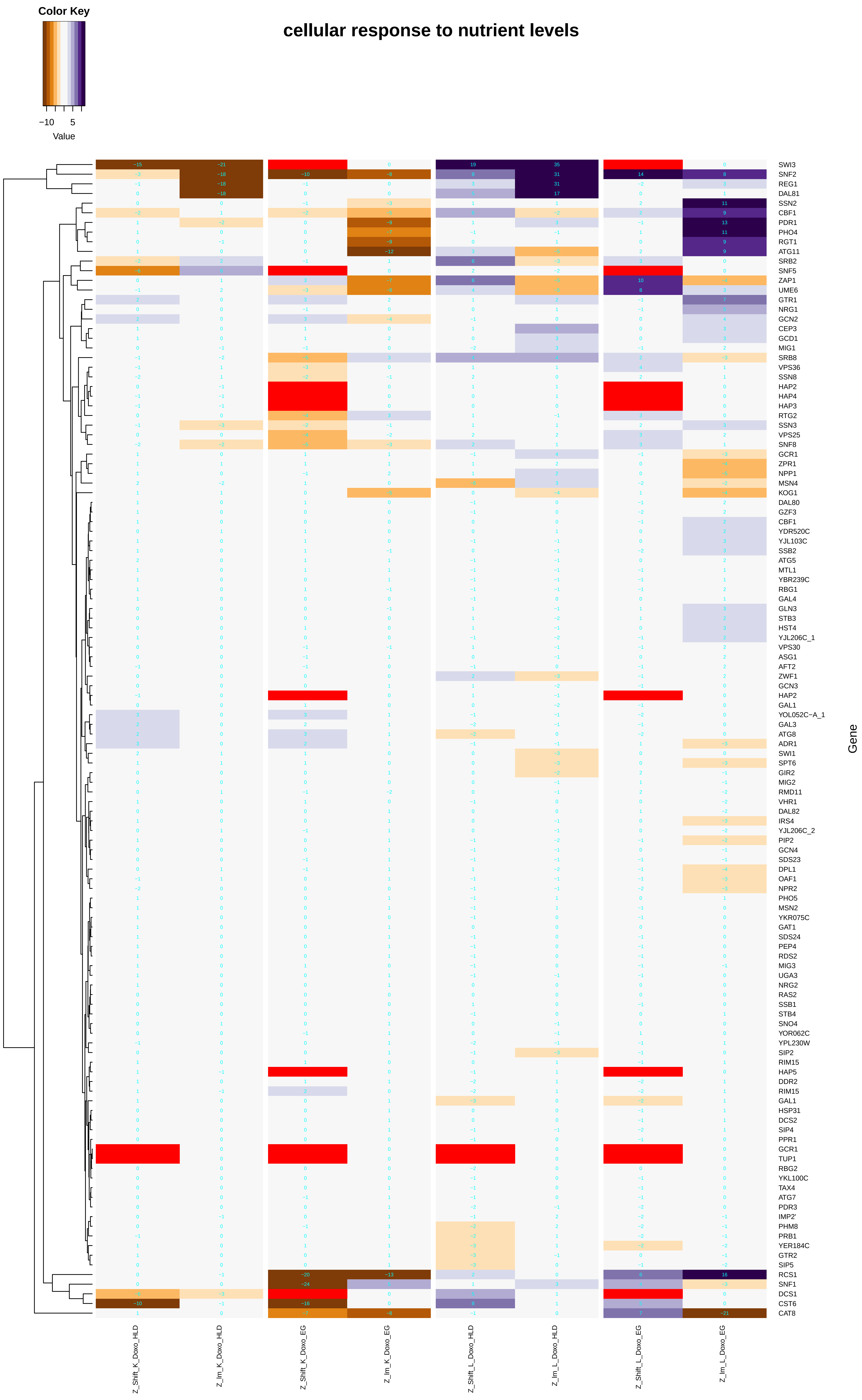
cellular response to nutrient levels
Color Key
−10
5
Value
Gene
Z_Shift_K_Doxo_HLD
Z_Im_K_Doxo_HLD
Z_Shift_K_Doxo_EG
Z_Im_K_Doxo_EG
Z_Shift_L_Doxo_HLD
Z_Im_L_Doxo_HLD
Z_Shift_L_Doxo_EG
Z_Im_L_Doxo_EG
SWI3
SNF2
REG1
DAL81
SSN2
CBF1
PDR1
PHO4
RGT1
ATG11
SRB2
SNF5
ZAP1
UME6
GTR1
NRG1
GCN2
CEP3
GCD1
MIG1
SRB8
VPS36
SSN8
HAP2
HAP4
HAP3
RTG2
SSN3
VPS25
SNF8
GCR1
ZPR1
NPP1
MSN4
KOG1
DAL80
GZF3
CBF1
YDR520C
YJL103C
SSB2
ATG5
MTL1
YBR239C
RBG1
GAL4
GLN3
STB3
HST4
YJL206C_1
VPS30
ASG1
AFT2
ZWF1
GCN3
HAP2
GAL1
YOL052C−A_1
GAL3
ATG8
ADR1
SWI1
SPT6
GIR2
MIG2
RMD11
VHR1
DAL82
IRS4
YJL206C_2
PIP2
GCN4
SDS23
DPL1
OAF1
NPR2
PHO5
MSN2
YKR075C
GAT1
SDS24
PEP4
RDS2
MIG3
UGA3
NRG2
RAS2
SSB1
STB4
SNO4
YOR062C
YPL230W
SIP2
RIM15
HAP5
DDR2
RIM15
GAL1
HSP31
DCS2
SIP4
PPR1
GCR1
TUP1
RBG2
YKL100C
TAX4
ATG7
PDR3
IMP2'
PHM8
PRB1
YER184C
GTR2
SIP5
RCS1
SNF1
DCS1
CST6
CAT8

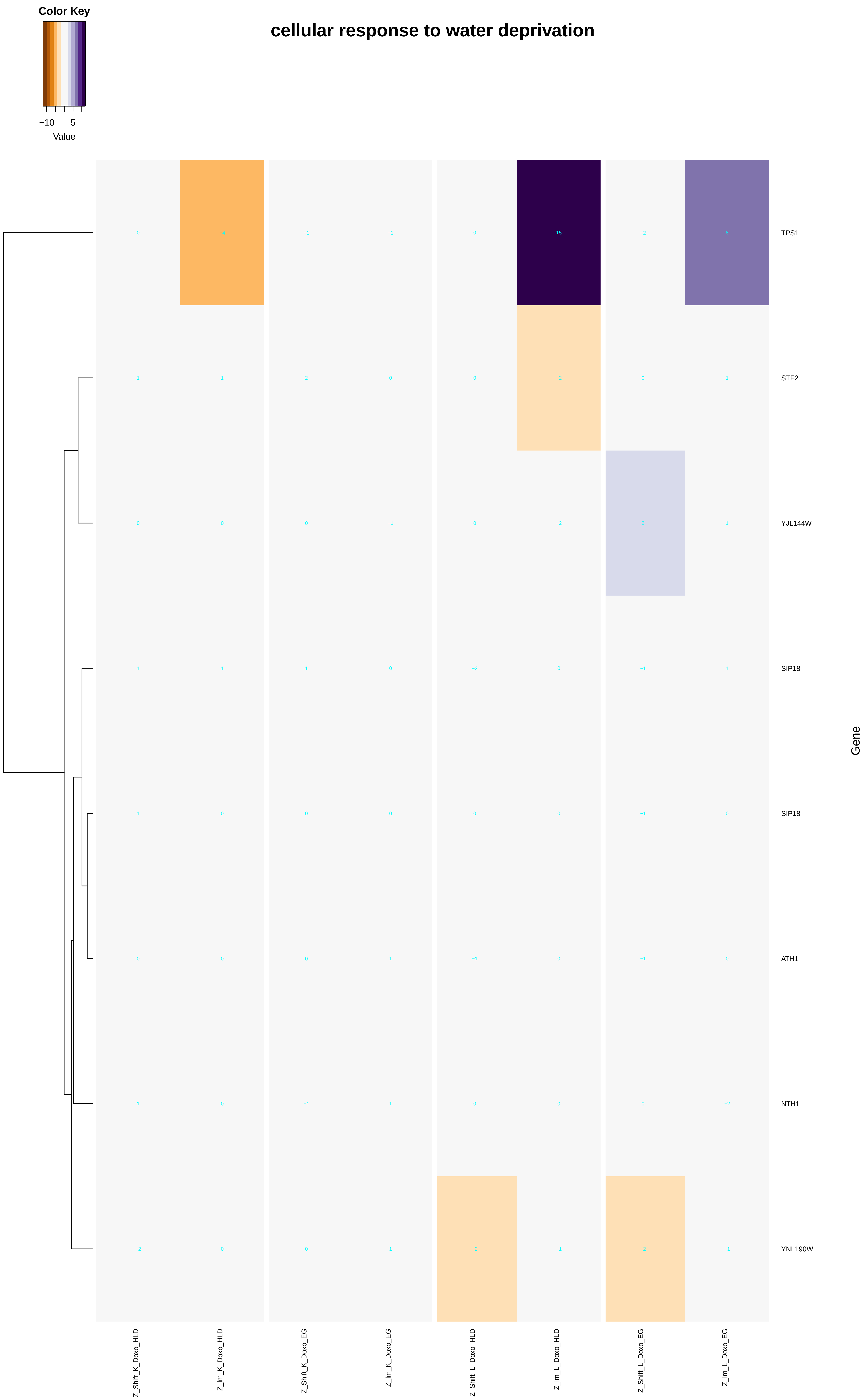

Color Key
−10
5
Value
cellular response to water deprivation
TPS1
STF2
YJL144W
SIP18
SIP18
ATH1
NTH1
YNL190W
Gene
Z_Shift_K_Doxo_HLD
Z_Im_K_Doxo_HLD
Z_Shift_K_Doxo_EG
Z_Im_K_Doxo_EG
Z_Shift_L_Doxo_HLD
Z_Im_L_Doxo_HLD
Z_Shift_L_Doxo_EG
Z_Im_L_Doxo_EG

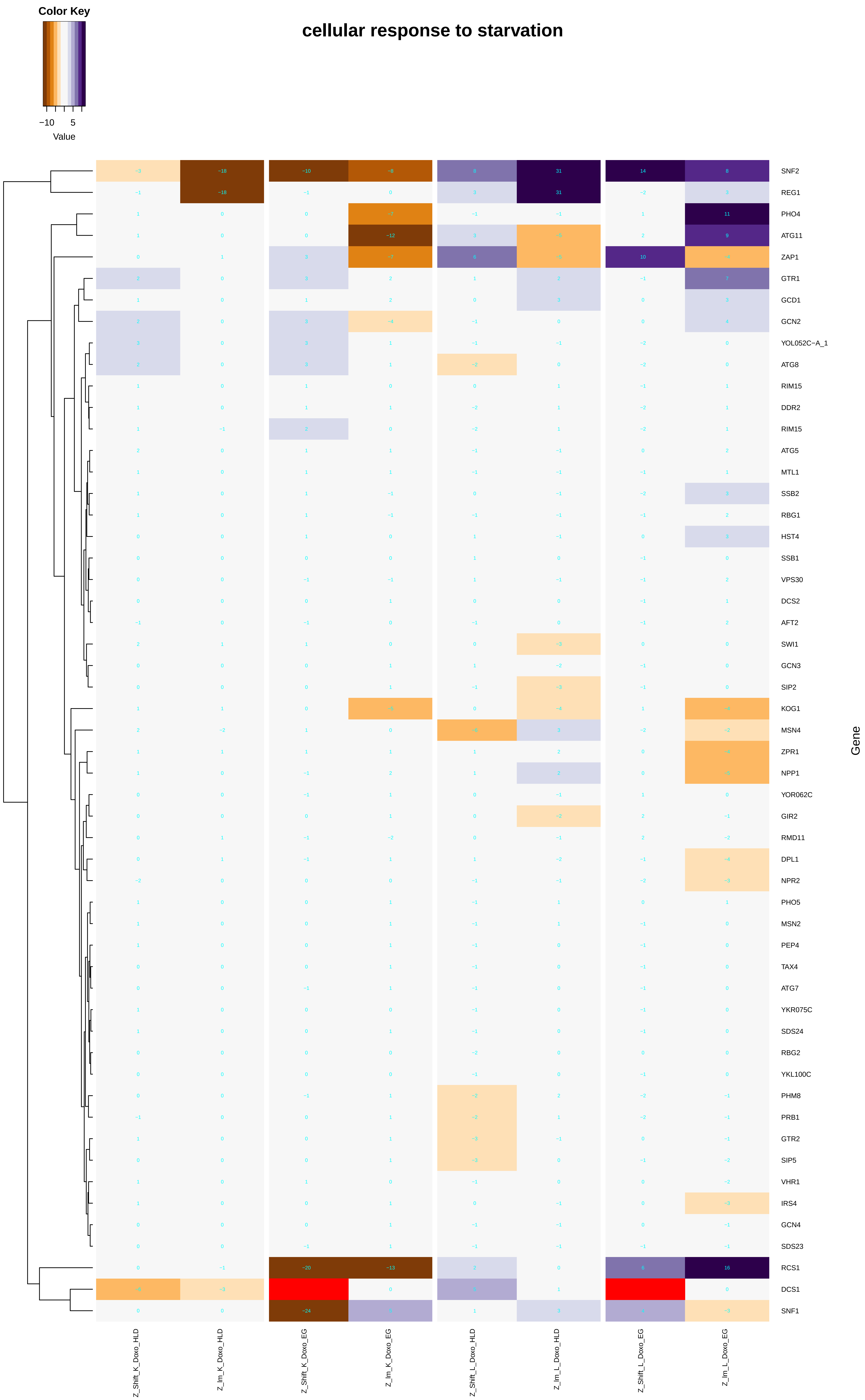
Color Key
−10
5
Value
cellular response to starvation
SNF2
REG1
PHO4
ATG11
ZAP1
GTR1
GCD1
GCN2
YOL052C−A_1
ATG8
RIM15
DDR2
RIM15
ATG5
MTL1
SSB2
RBG1
HST4
SSB1
VPS30
DCS2
AFT2
SWI1
GCN3
SIP2
KOG1
MSN4
ZPR1
NPP1
YOR062C
GIR2
RMD11
DPL1
NPR2
PHO5
MSN2
PEP4
TAX4
ATG7
YKR075C
SDS24
RBG2
YKL100C
PHM8
PRB1
GTR2
SIP5
VHR1
IRS4
GCN4
SDS23
RCS1
DCS1
SNF1
Gene
Z_Shift_K_Doxo_HLD
Z_Im_K_Doxo_HLD
Z_Shift_K_Doxo_EG
Z_Im_K_Doxo_EG
Z_Shift_L_Doxo_HLD
Z_Im_L_Doxo_HLD
Z_Shift_L_Doxo_EG
Z_Im_L_Doxo_EG

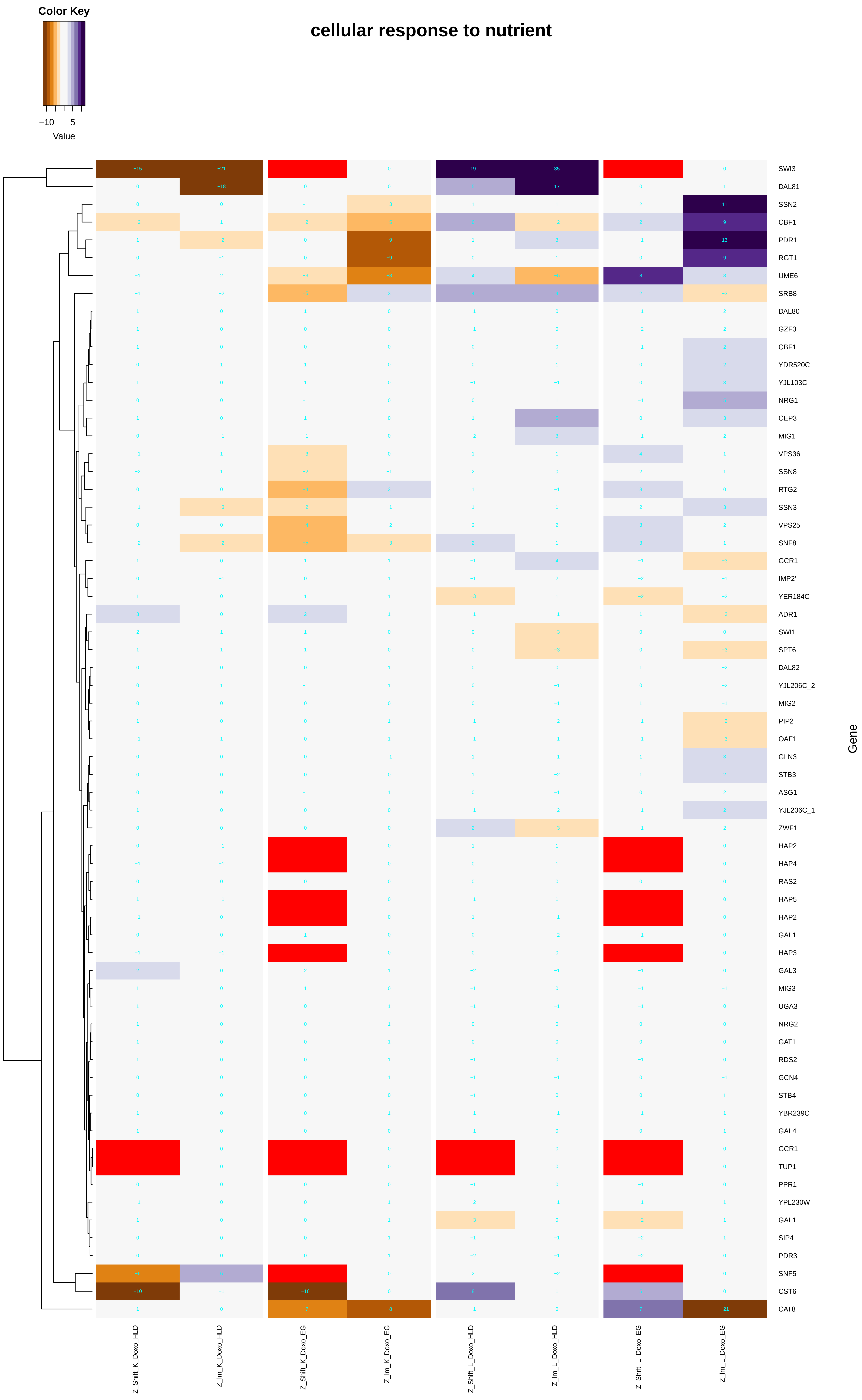
cellular response to nutrient
Color Key
−10
5
Value
SWI3
DAL81
SSN2
CBF1
PDR1
RGT1
UME6
SRB8
DAL80
GZF3
CBF1
YDR520C
YJL103C
NRG1
CEP3
MIG1
VPS36
SSN8
RTG2
SSN3
VPS25
SNF8
GCR1
IMP2'
YER184C
ADR1
SWI1
SPT6
DAL82
YJL206C_2
MIG2
PIP2
OAF1
GLN3
STB3
ASG1
YJL206C_1
ZWF1
HAP2
HAP4
RAS2
HAP5
HAP2
GAL1
HAP3
GAL3
MIG3
UGA3
NRG2
GAT1
RDS2
GCN4
STB4
YBR239C
GAL4
GCR1
TUP1
PPR1
YPL230W
GAL1
SIP4
PDR3
SNF5
CST6
CAT8
Gene
Z_Shift_K_Doxo_HLD
Z_Im_K_Doxo_HLD
Z_Shift_K_Doxo_EG
Z_Im_K_Doxo_EG
Z_Shift_L_Doxo_HLD
Z_Im_L_Doxo_HLD
Z_Shift_L_Doxo_EG
Z_Im_L_Doxo_EG

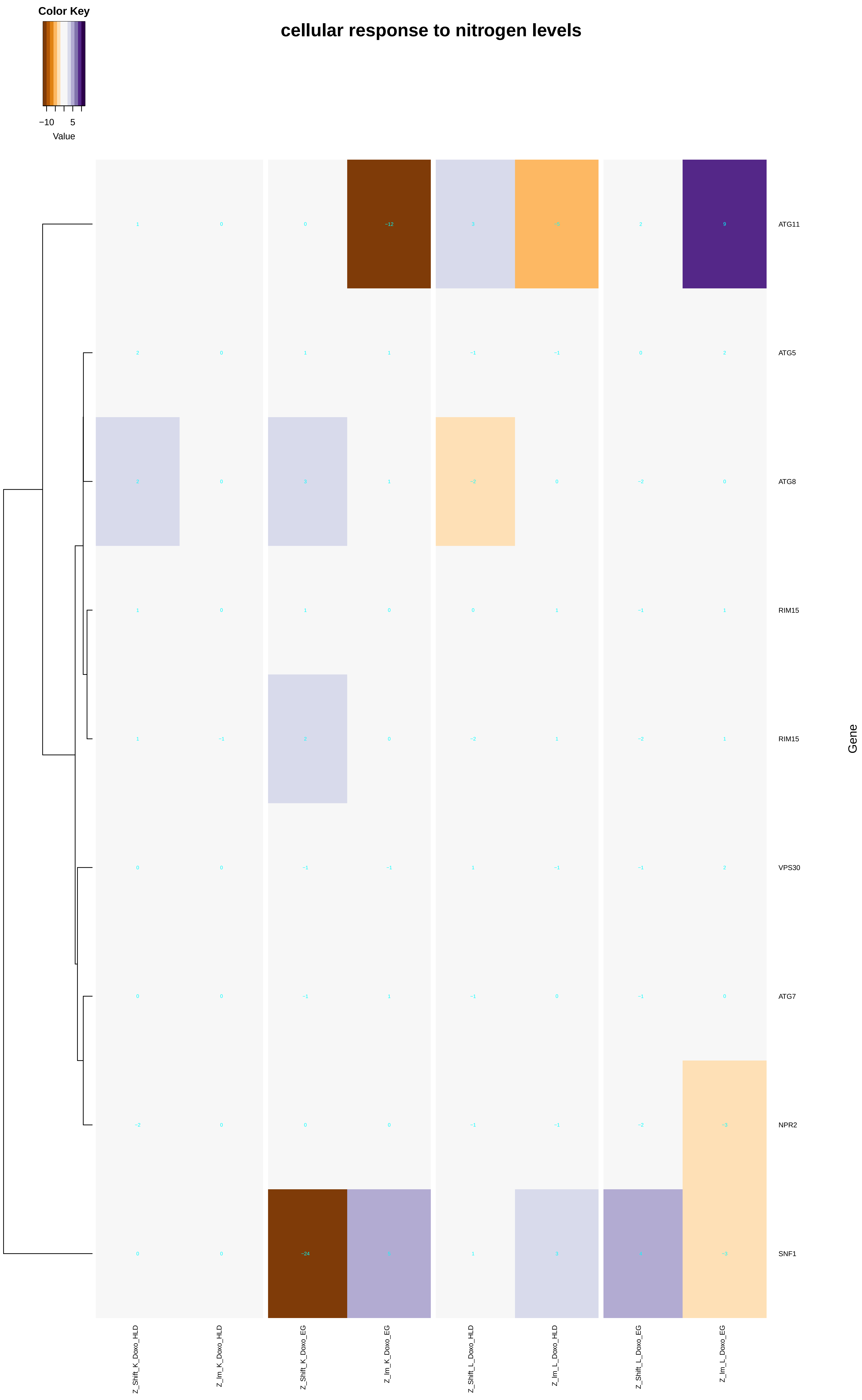

Color Key
−10
5
Value
cellular response to nitrogen levels
ATG11
ATG5
ATG8
RIM15
RIM15
VPS30
ATG7
NPR2
SNF1
Gene
Z_Shift_K_Doxo_HLD
Z_Im_K_Doxo_HLD
Z_Shift_K_Doxo_EG
Z_Im_K_Doxo_EG
Z_Shift_L_Doxo_HLD
Z_Im_L_Doxo_HLD
Z_Shift_L_Doxo_EG
Z_Im_L_Doxo_EG

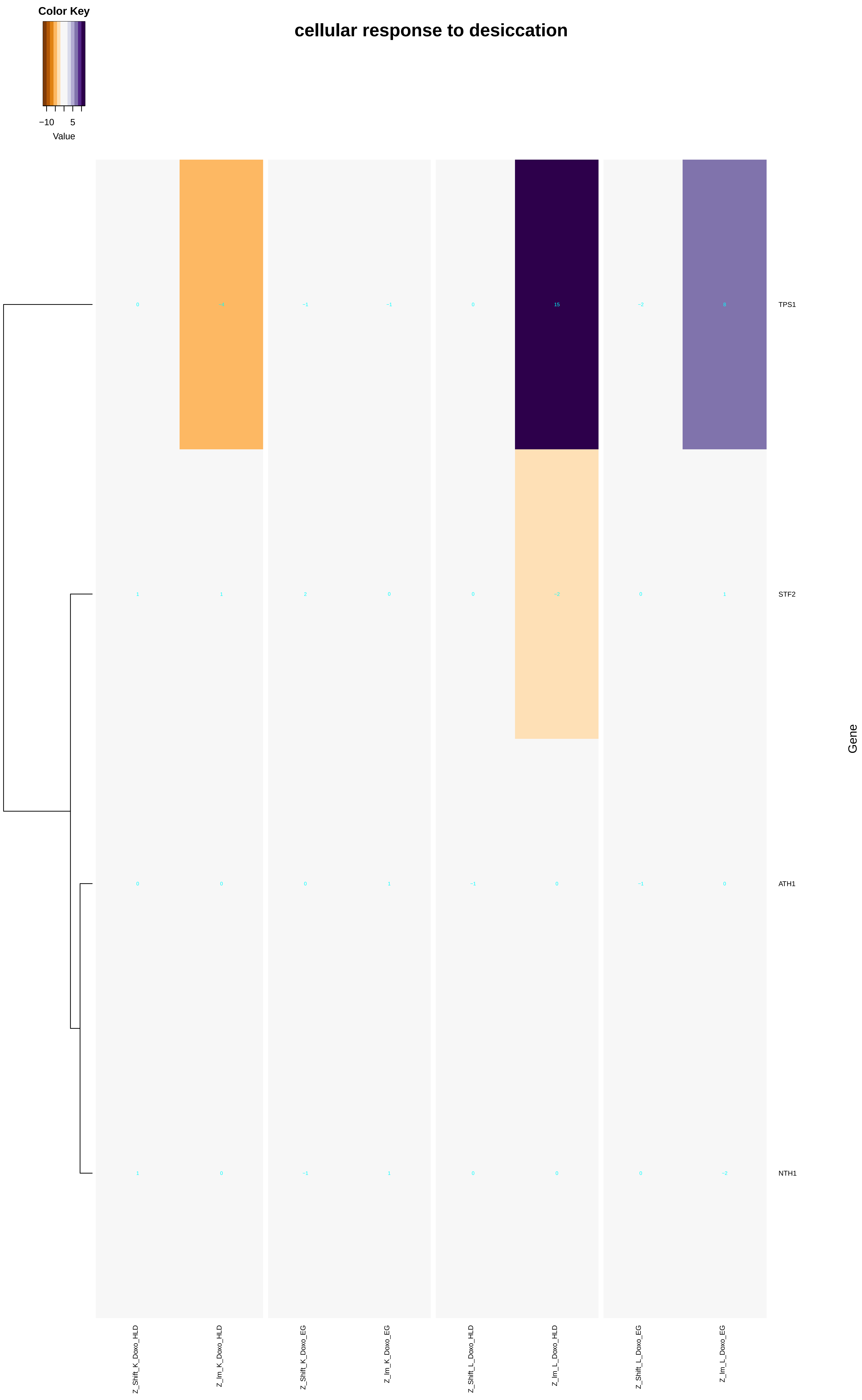
Color Key
−10
5
Value
cellular response to desiccation
TPS1
STF2
ATH1
NTH1
Gene
Z_Shift_K_Doxo_HLD
Z_Im_K_Doxo_HLD
Z_Shift_K_Doxo_EG
Z_Im_K_Doxo_EG
Z_Shift_L_Doxo_HLD
Z_Im_L_Doxo_HLD
Z_Shift_L_Doxo_EG
Z_Im_L_Doxo_EG

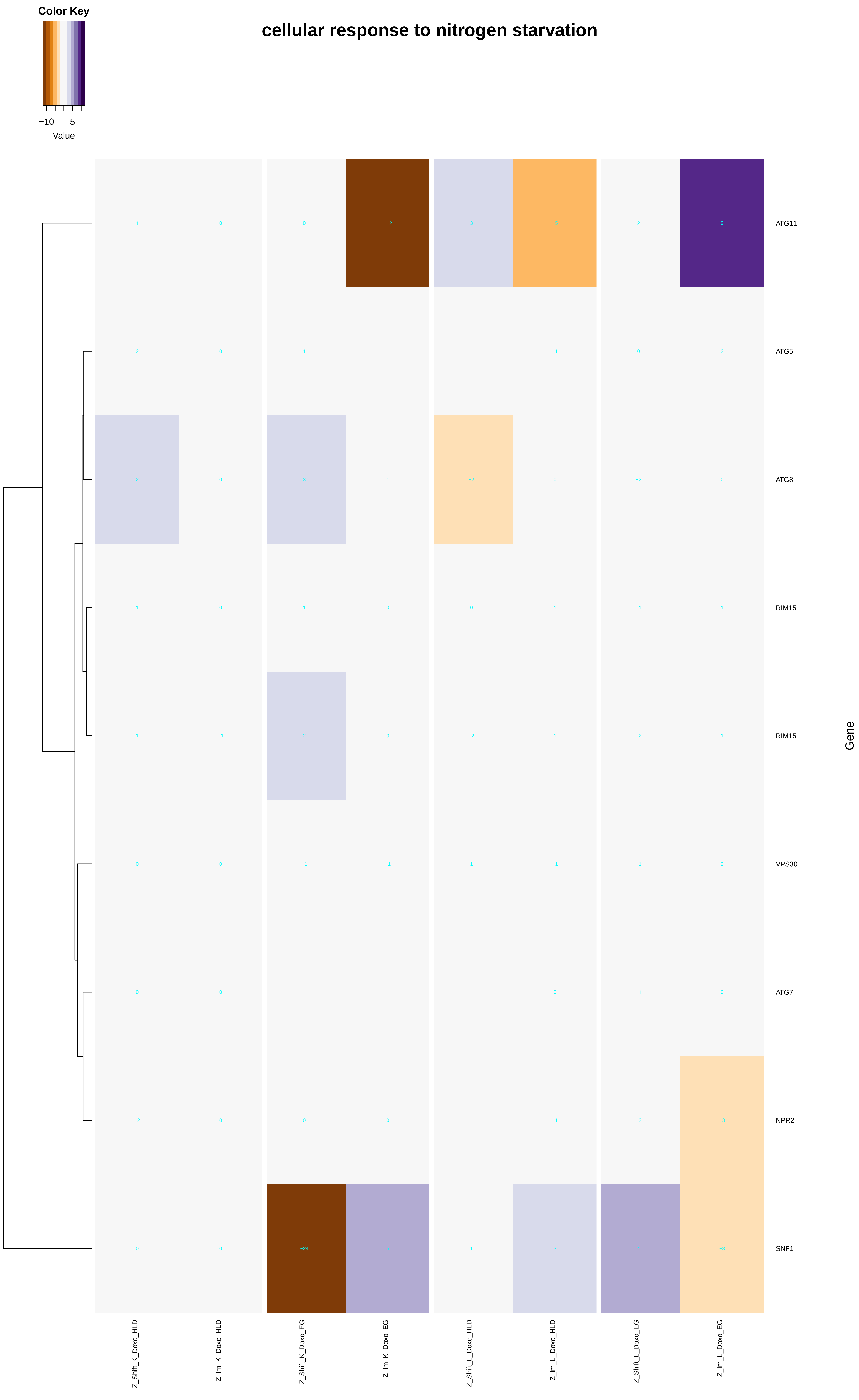
cellular response to nitrogen starvation
Color Key
−10
5
Value
ATG11
ATG5
ATG8
RIM15
RIM15
VPS30
ATG7
NPR2
SNF1
Gene
Z_Shift_K_Doxo_HLD
Z_Im_K_Doxo_HLD
Z_Shift_K_Doxo_EG
Z_Im_K_Doxo_EG
Z_Shift_L_Doxo_HLD
Z_Im_L_Doxo_HLD
Z_Shift_L_Doxo_EG
Z_Im_L_Doxo_EG

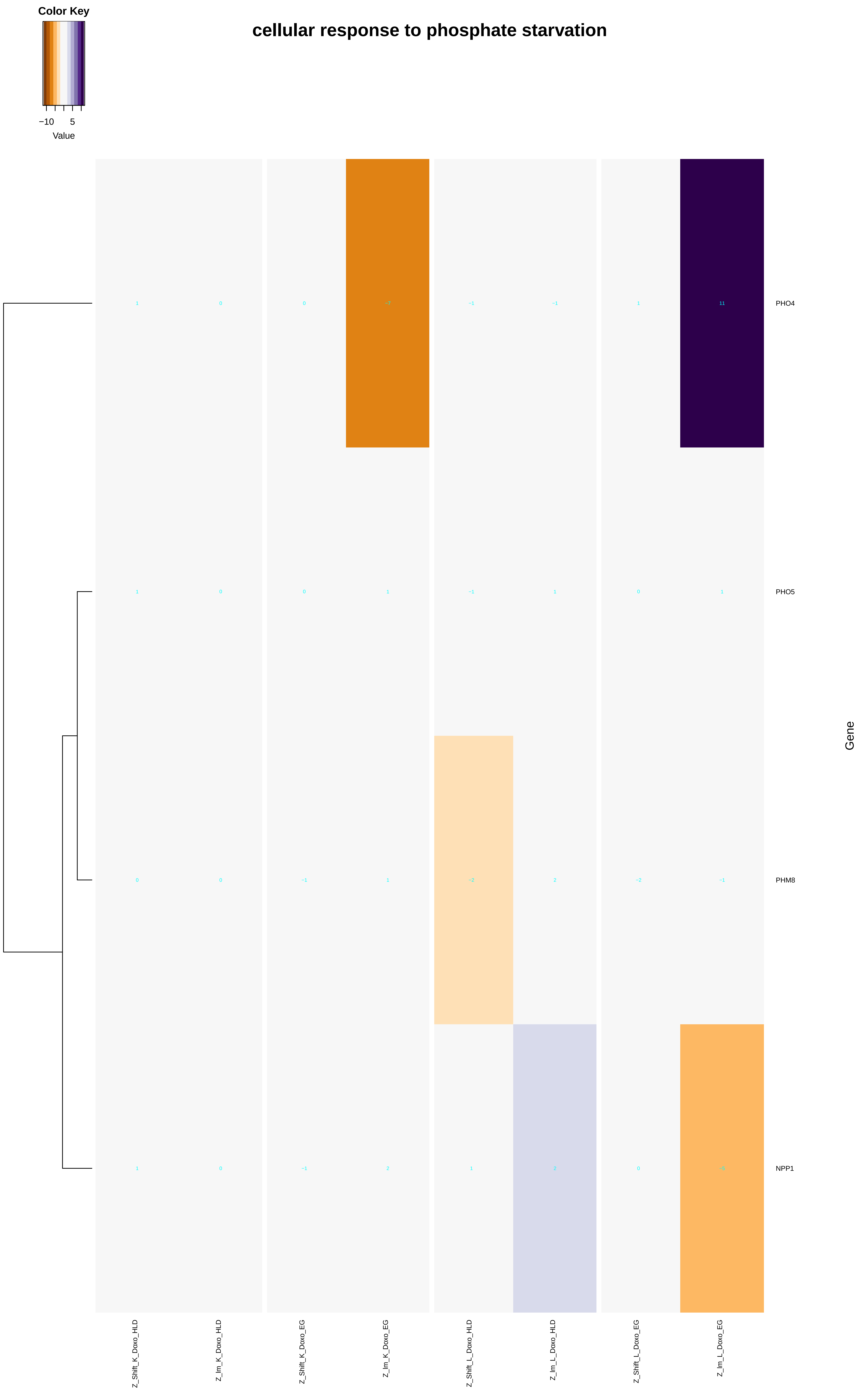
cellular response to phosphate starvation
Color Key
−10
5
Value
PHO4
PHO5
PHM8
NPP1
Gene
Z_Shift_K_Doxo_HLD
Z_Im_K_Doxo_HLD
Z_Shift_K_Doxo_EG
Z_Im_K_Doxo_EG
Z_Shift_L_Doxo_HLD
Z_Im_L_Doxo_HLD
Z_Shift_L_Doxo_EG
Z_Im_L_Doxo_EG

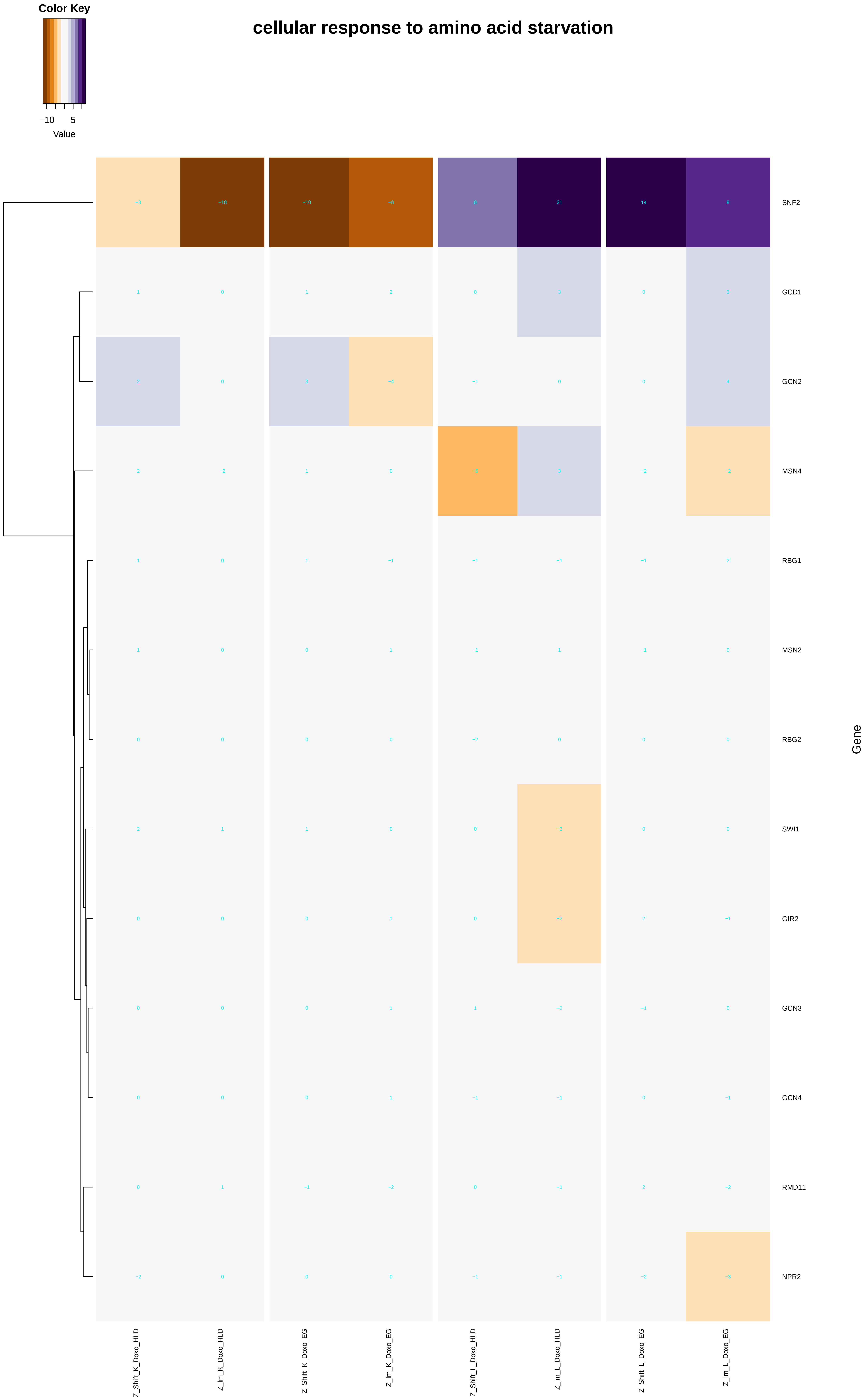

Color Key
−10
5
Value
cellular response to amino acid starvation
SNF2
GCD1
GCN2
MSN4
RBG1
MSN2
RBG2
SWI1
GIR2
GCN3
GCN4
RMD11
NPR2
Gene
Z_Shift_K_Doxo_HLD
Z_Im_K_Doxo_HLD
Z_Shift_K_Doxo_EG
Z_Im_K_Doxo_EG
Z_Shift_L_Doxo_HLD
Z_Im_L_Doxo_HLD
Z_Shift_L_Doxo_EG
Z_Im_L_Doxo_EG

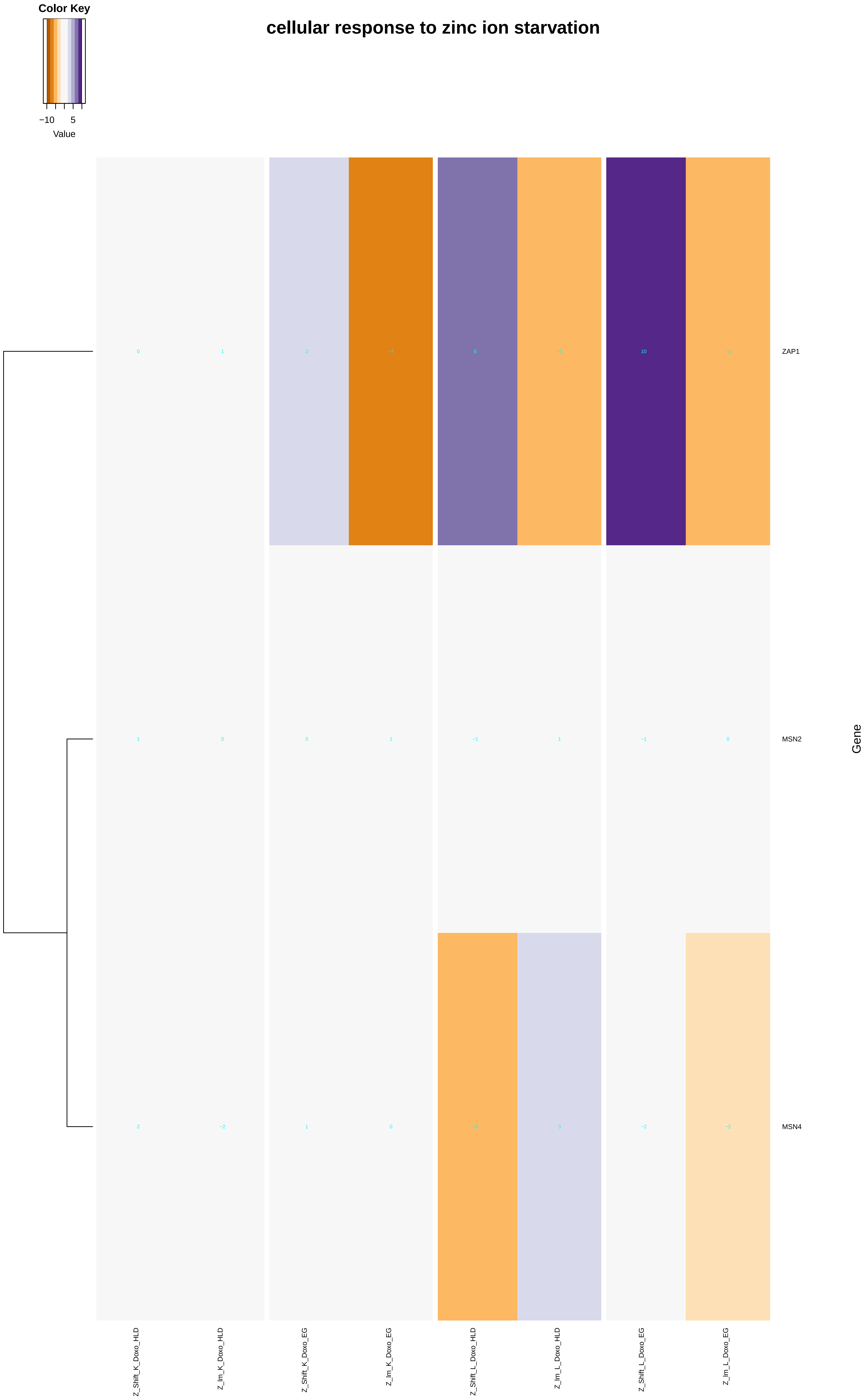

Color Key
−10
5
Value
cellular response to zinc ion starvation
0
1
3
−7
6
−5
10
−4
ZAP1
1
0
0
1
−1
1
−1
0
MSN2
2
−2
1
0
−6
3
−2
−2
MSN4
Gene
Z_Shift_K_Doxo_HLD
Z_Im_K_Doxo_HLD
Z_Shift_K_Doxo_EG
Z_Im_K_Doxo_EG
Z_Shift_L_Doxo_HLD
Z_Im_L_Doxo_HLD
Z_Shift_L_Doxo_EG
Z_Im_L_Doxo_EG

### cellular response to glucose starvation

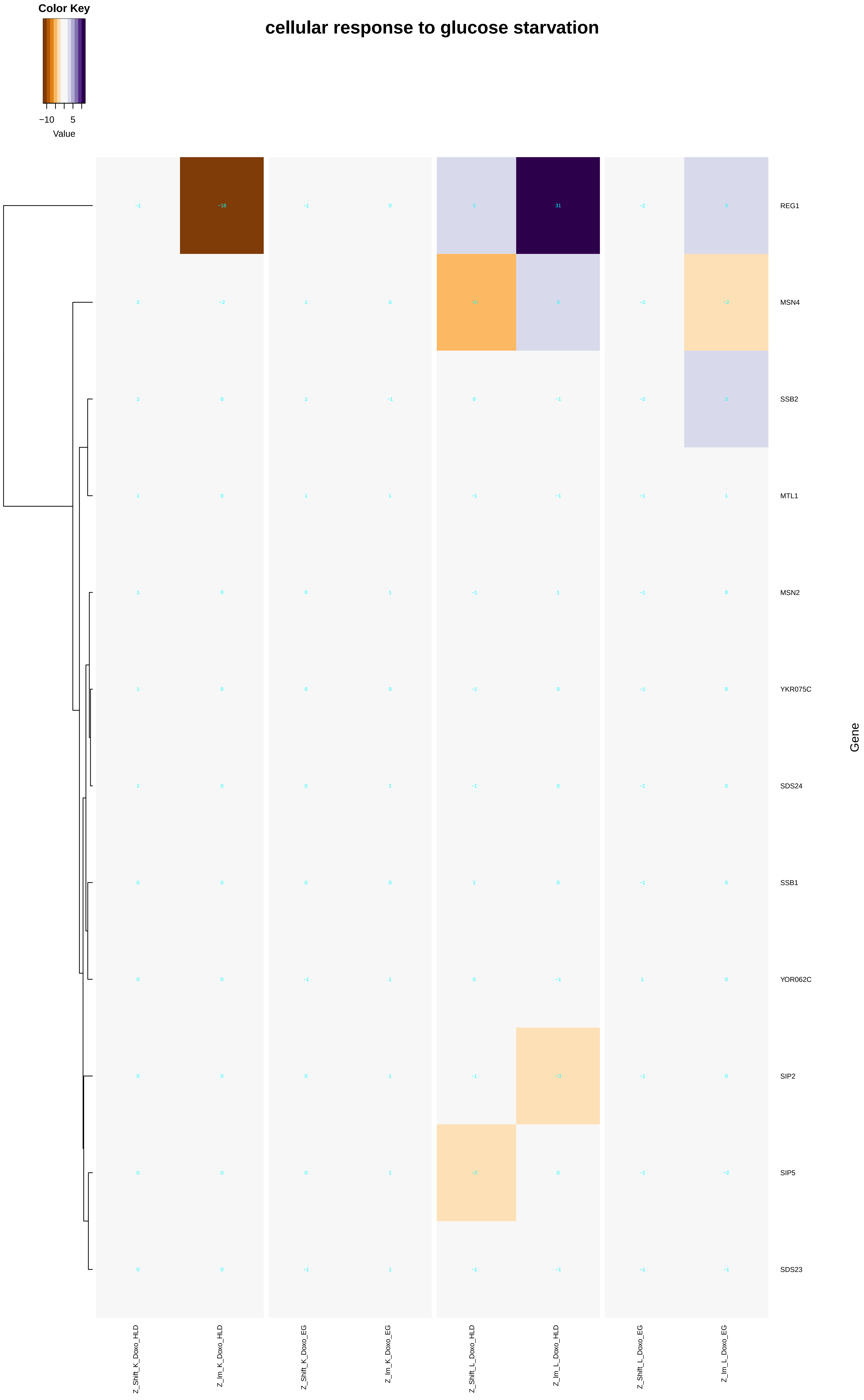

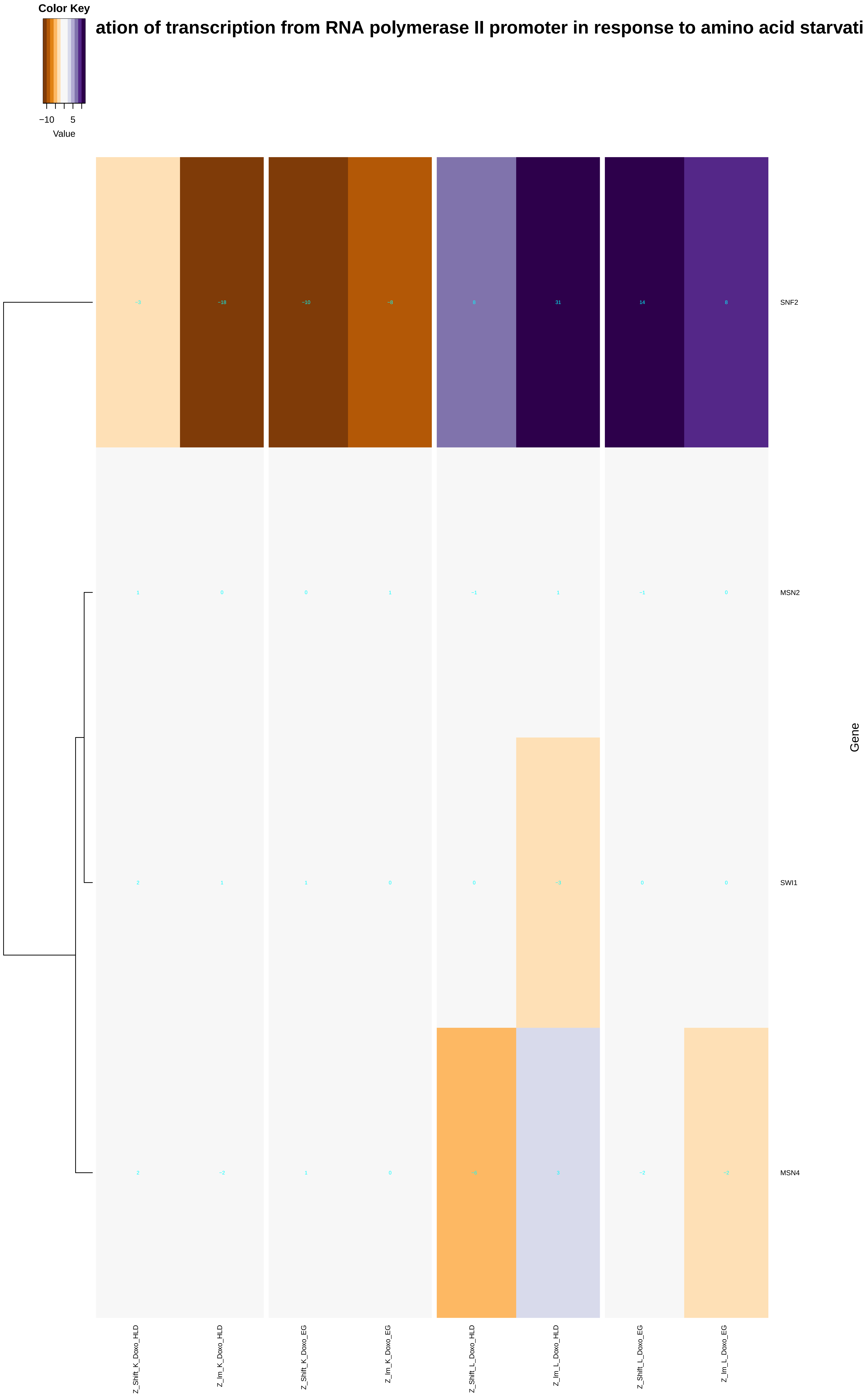
Color Key
−10 5
Value
ation of transcription from RNA polymerase II promoter in response to amino acid starvati
SNF2
MSN2
SWI1
MSN4
Gene
Z_Shift_K_Doxo_HLD
Z_Im_K_Doxo_HLD
Z_Shift_K_Doxo_EG
Z_Im_K_Doxo_EG
Z_Shift_L_Doxo_HLD
Z_Im_L_Doxo_HLD
Z_Shift_L_Doxo_EG
Z_Im_L_Doxo_EG

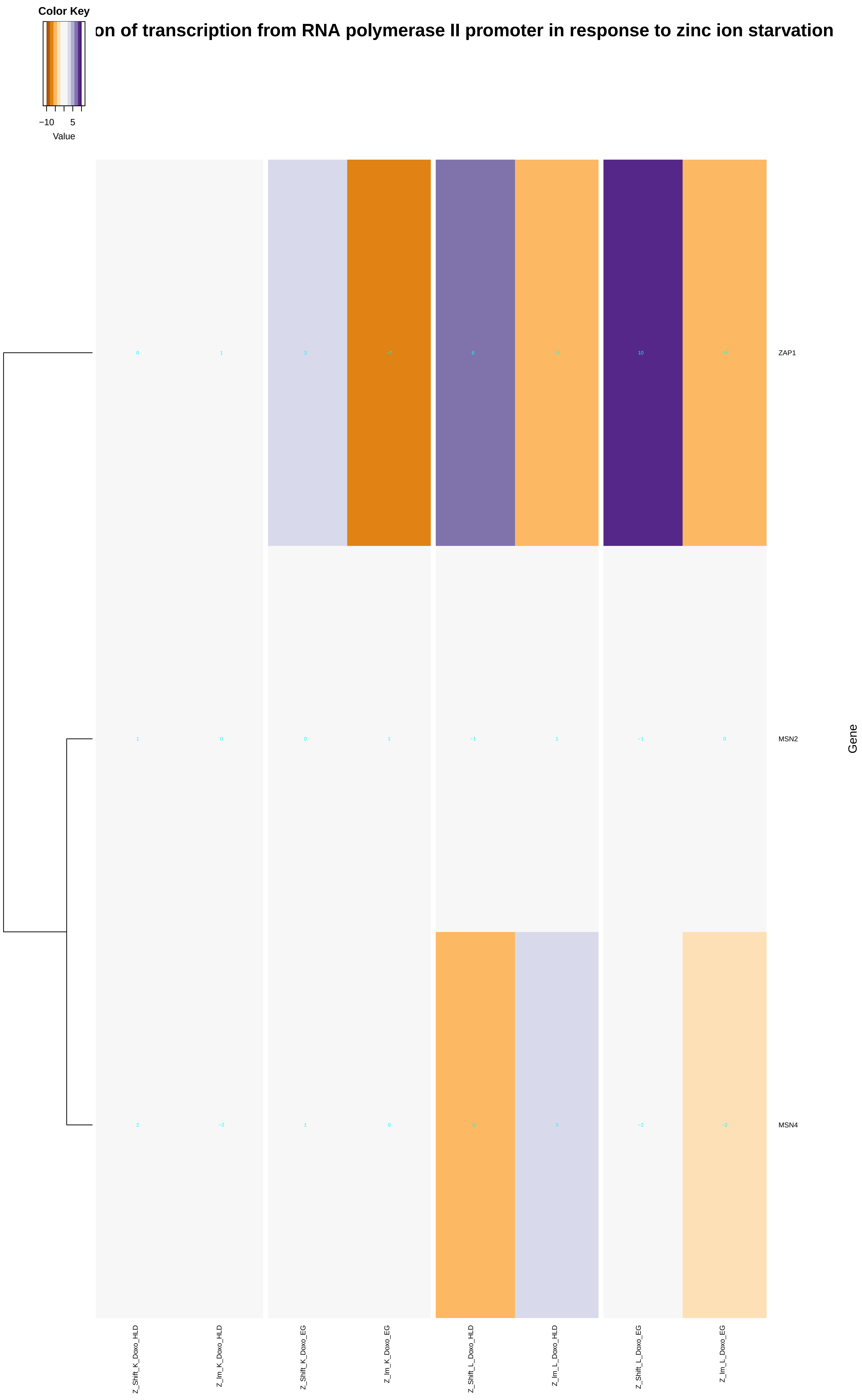
Color Key
−10
5
Value
on of transcription from RNA polymerase II promoter in response to zinc ion starvation
ZAP1
MSN2
MSN4
Gene
Z_Shift_K_Doxo_HLD
Z_Im_K_Doxo_HLD
Z_Shift_K_Doxo_EG
Z_Im_K_Doxo_EG
Z_Shift_L_Doxo_HLD
Z_Im_L_Doxo_HLD
Z_Shift_L_Doxo_EG
Z_Im_L_Doxo_EG

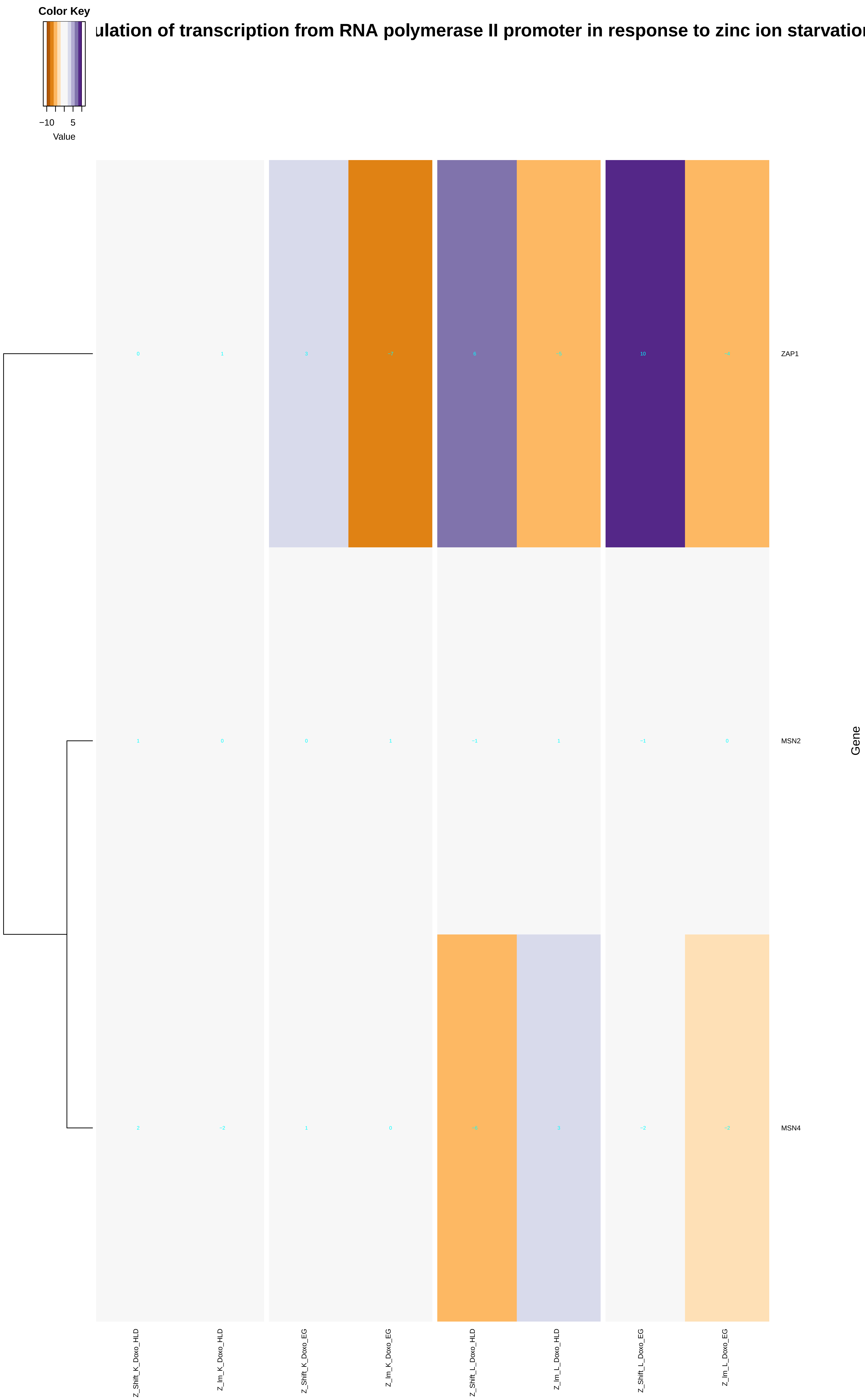

Color Key
−10
5
Value
ulation of transcription from RNA polymerase II promoter in response to zinc ion starvatio
ZAP1
MSN2
MSN4
Gene
Z_Shift_K_Doxo_HLD
Z_lm_K_Doxo_HLD
Z_Shift_K_Doxo_EG
Z_lm_K_Doxo_EG
Z_Shift_L_Doxo_HLD
Z_lm_L_Doxo_HLD
Z_Shift_L_Doxo_EG
Z_lm_L_Doxo_EG
