## Additional File 9 for "A yeast phenomic model for the influence of Warburg metabolism on genetic buffering of doxorubicin": cellular_response_to_nutrient_levels.pdf

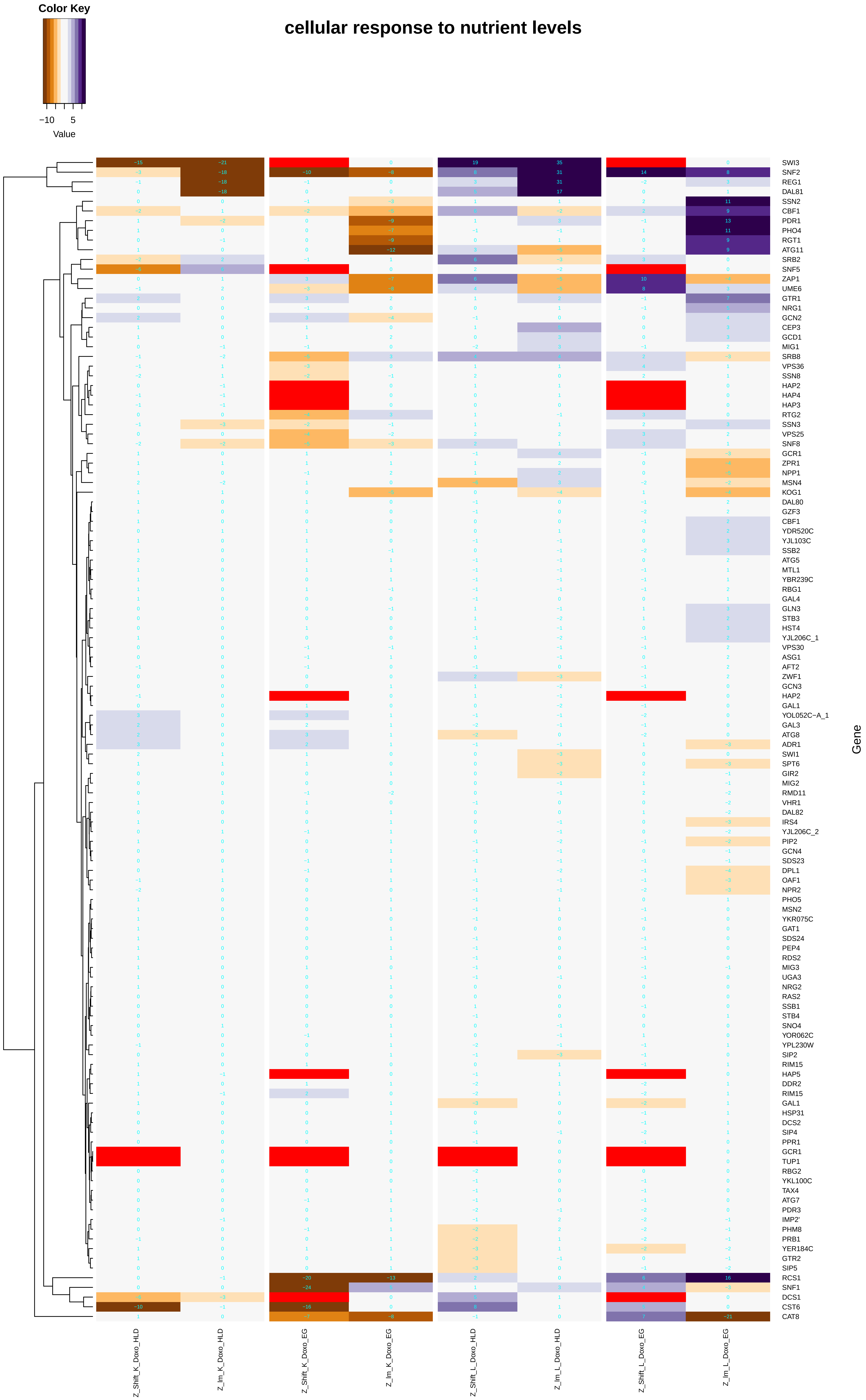

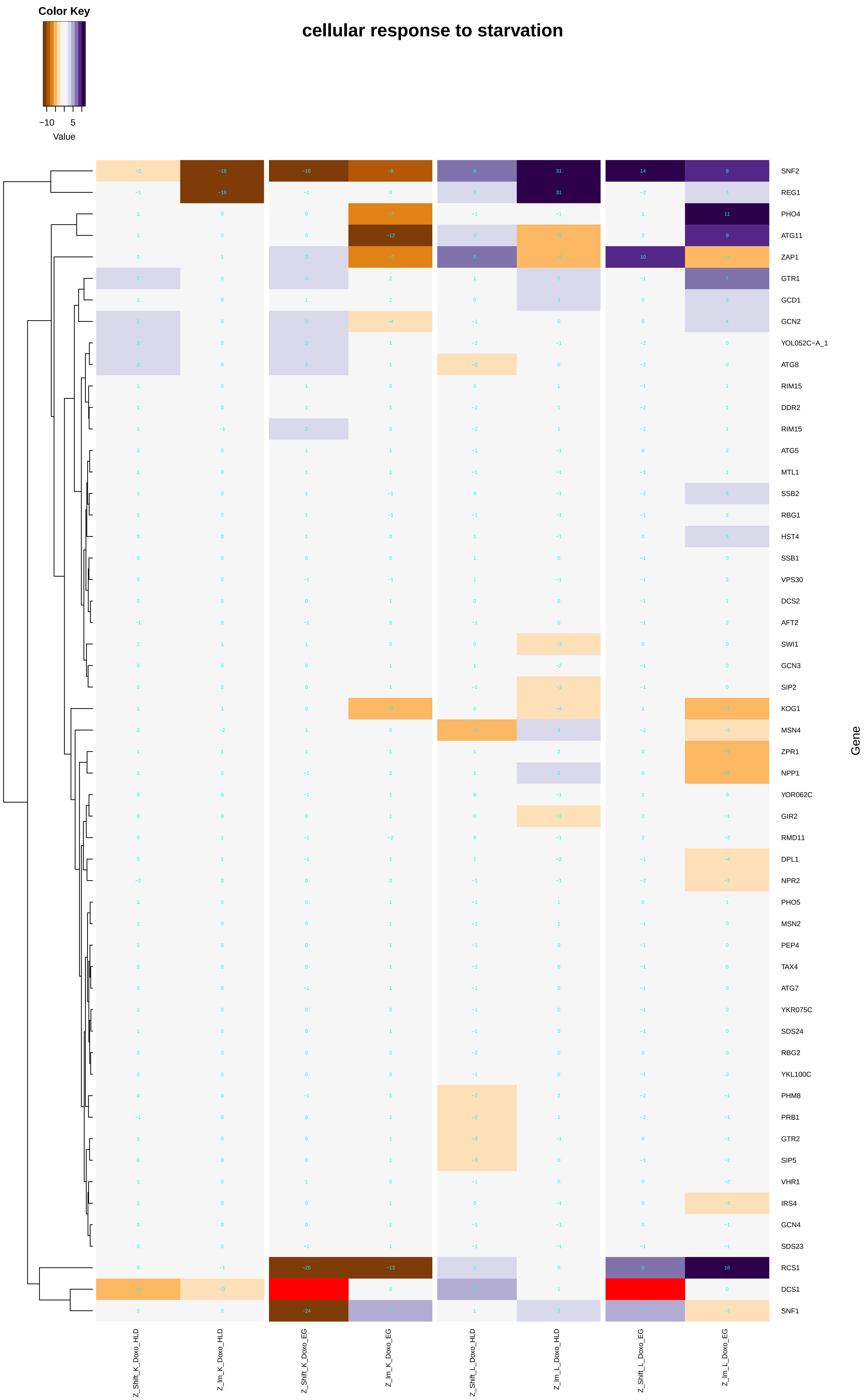

Color Key
−10
5
Value
cellular response to starvation
SNF2
REG1
PHO4
ATG11
ZAP1
GTR1
GCD1
GCN2
YOL052C−A_1
ATG8
RIM15
DDR2
RIM15
ATG5
MTL1
SSB2
RBG1
HST4
SSB1
VPS30
DCS2
AFT2
SWI1
GCN3
SIP2
KOG1
MSN4
ZPR1
NPP1
YOR062C
GIR2
RMD11
DPL1
NPR2
PHO5
MSN2
PEP4
TAX4
ATG7
YKR075C
SDS24
RBG2
YKL100C
PHM8
PRB1
GTR2
SIP5
VHR1
IRS4
GCN4
SDS23
RCS1
DCS1
SNF1
Gene
Z_Shift_K_Doxo_HLD
Z_Im_K_Doxo_HLD
Z_Shift_K_Doxo_EG
Z_Im_K_Doxo_EG
Z_Shift_L_Doxo_HLD
Z_Im_L_Doxo_HLD
Z_Shift_L_Doxo_EG
Z_Im_L_Doxo_EG

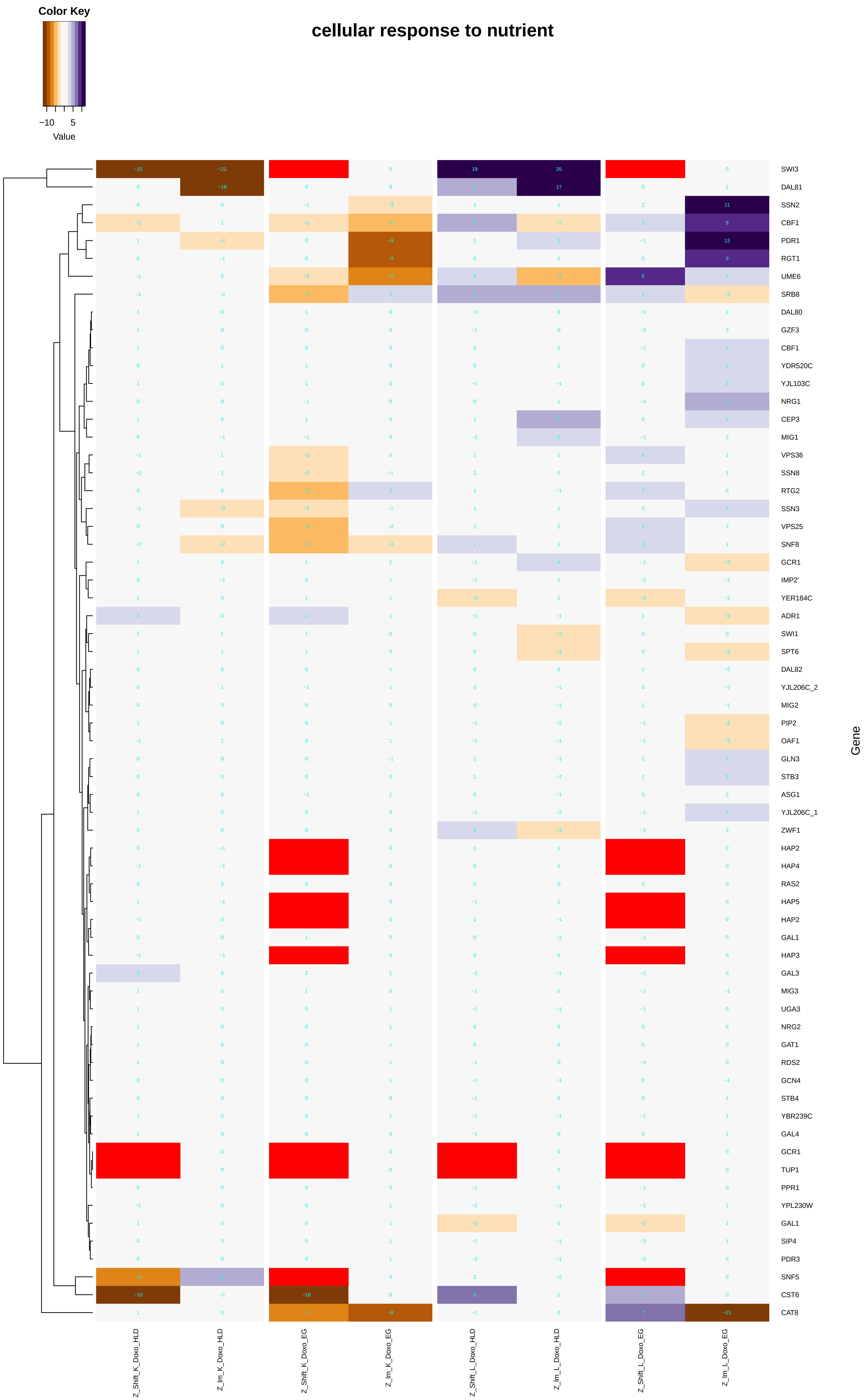

cellular response to nutrient
Color Key
−10
5
Value
Gene
SWI3
DAL81
SSN2
CBF1
PDR1
RGT1
UME6
SRB8
DAL80
GZF3
CBF1
YDR520C
YJL103C
NRG1
CEP3
MIG1
VPS36
SSN8
RTG2
SSN3
VPS25
SNF8
GCR1
IMP2'
YER184C
ADR1
SWI1
SPT6
DAL82
YJL206C_2
MIG2
PIP2
OAF1
GLN3
STB3
ASG1
YJL206C_1
ZWF1
HAP2
HAP4
RAS2
HAP5
HAP2
GAL1
HAP3
GAL3
MIG3
UGA3
NRG2
GAT1
RDS2
GCN4
STB4
YBR239C
GAL4
GCR1
TUP1
PPR1
YPL230W
GAL1
SIP4
PDR3
SNF5
CST6
CAT8
Z_Shift_K_Doxo_HLD
Z_Im_K_Doxo_HLD
Z_Shift_K_Doxo_EG
Z_Im_K_Doxo_EG
Z_Shift_L_Doxo_HLD
Z_Im_L_Doxo_HLD
Z_Shift_L_Doxo_EG
Z_Im_L_Doxo_EG

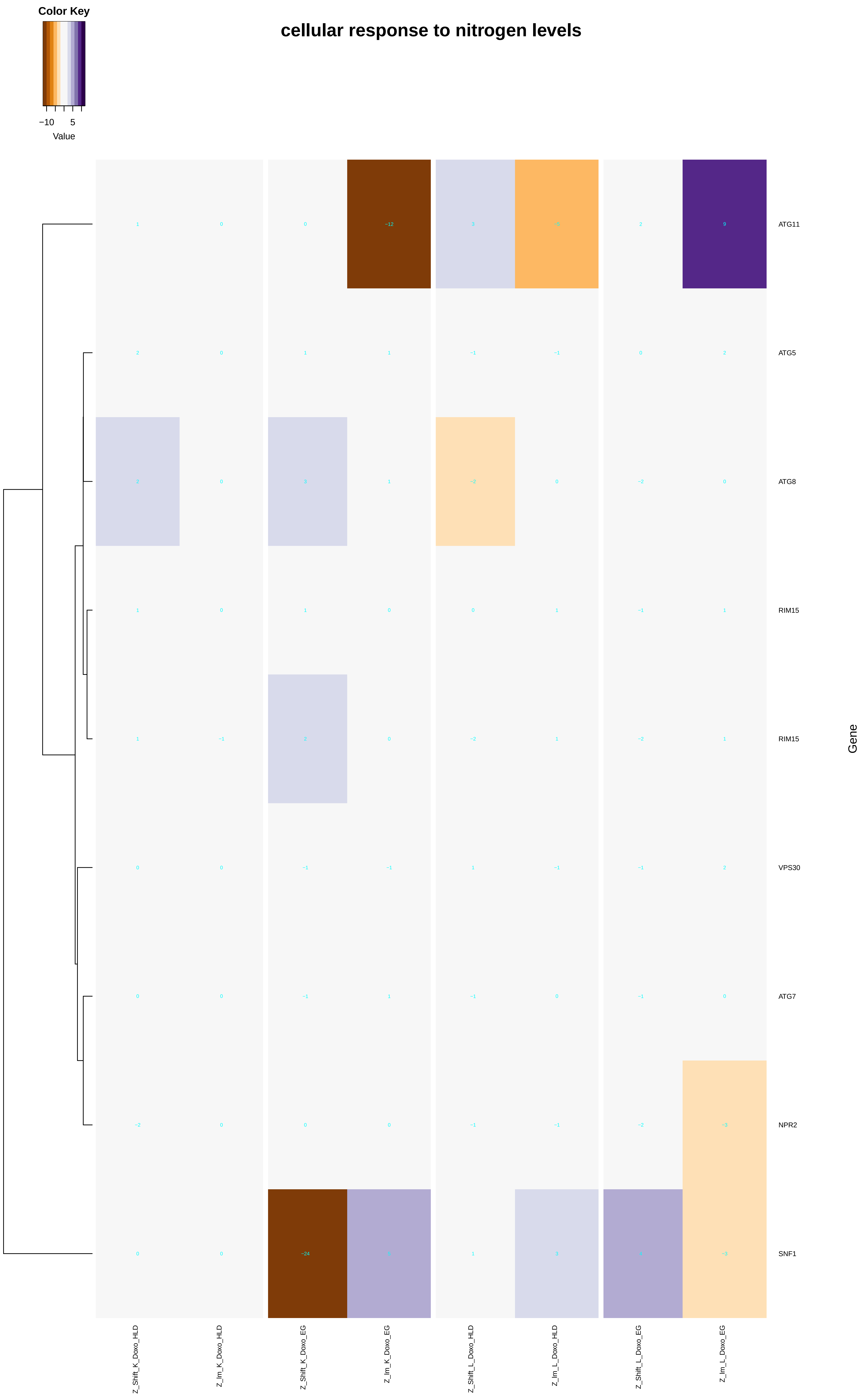
cellular response to nitrogen levels
Color Key
−10
5
Value
ATG11
ATG5
ATG8
RIM15
RIM15
VPS30
ATG7
NPR2
SNF1
Gene
Z_Shift_K_Doxo_HLD
Z_Im_K_Doxo_HLD
Z_Shift_K_Doxo_EG
Z_Im_K_Doxo_EG
Z_Shift_L_Doxo_HLD
Z_Im_L_Doxo_HLD
Z_Shift_L_Doxo_EG
Z_Im_L_Doxo_EG

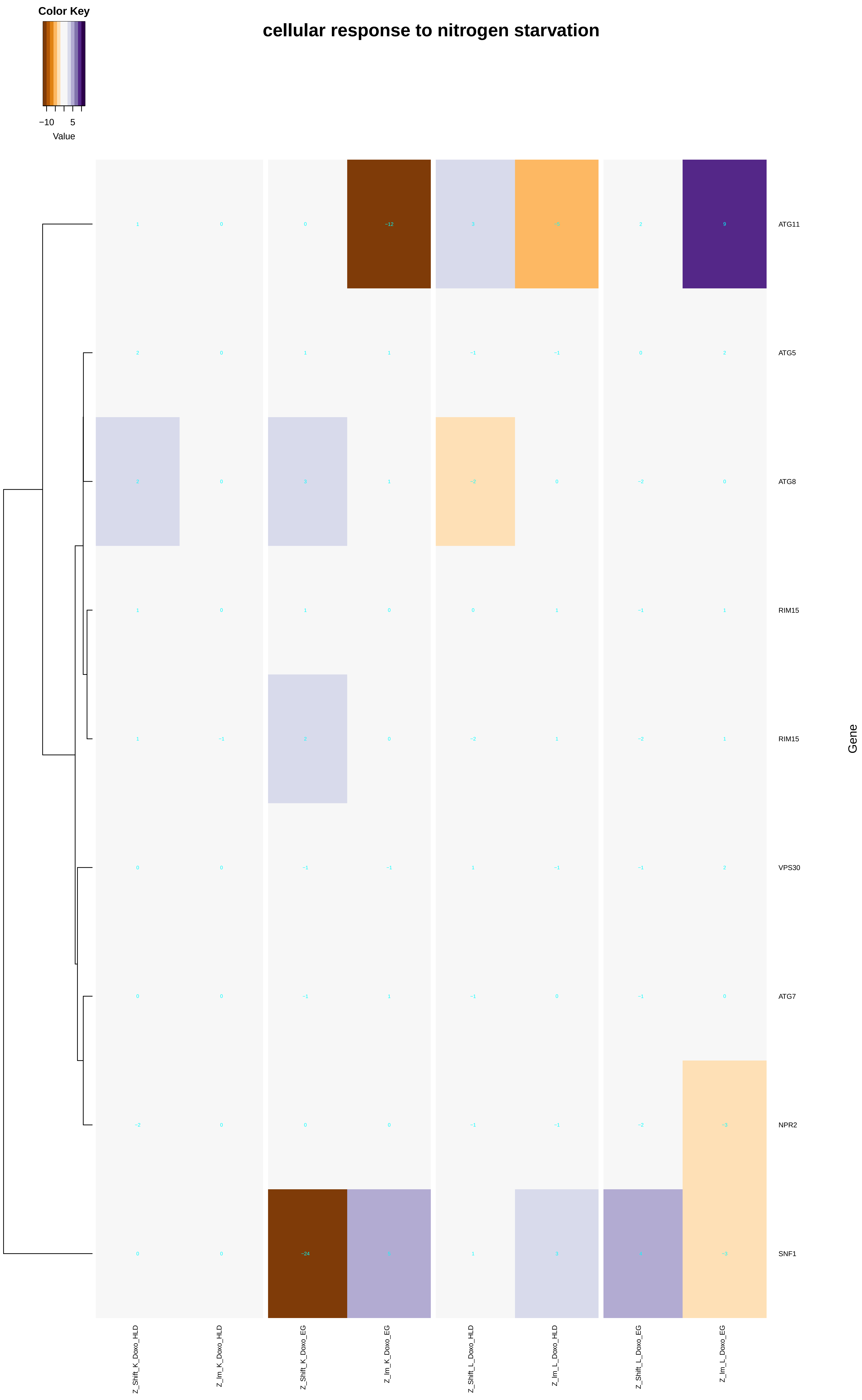
Color Key
−10
5
Value
cellular response to nitrogen starvation
ATG11
ATG5
ATG8
RIM15
RIM15
VPS30
ATG7
NPR2
SNF1
Gene
Z_Shift_K_Doxo_HLD
Z_lm_K_Doxo_HLD
Z_Shift_K_Doxo_EG
Z_lm_K_Doxo_EG
Z_Shift_L_Doxo_HLD
Z_lm_L_Doxo_HLD
Z_Shift_L_Doxo_EG
Z_lm_L_Doxo_EG

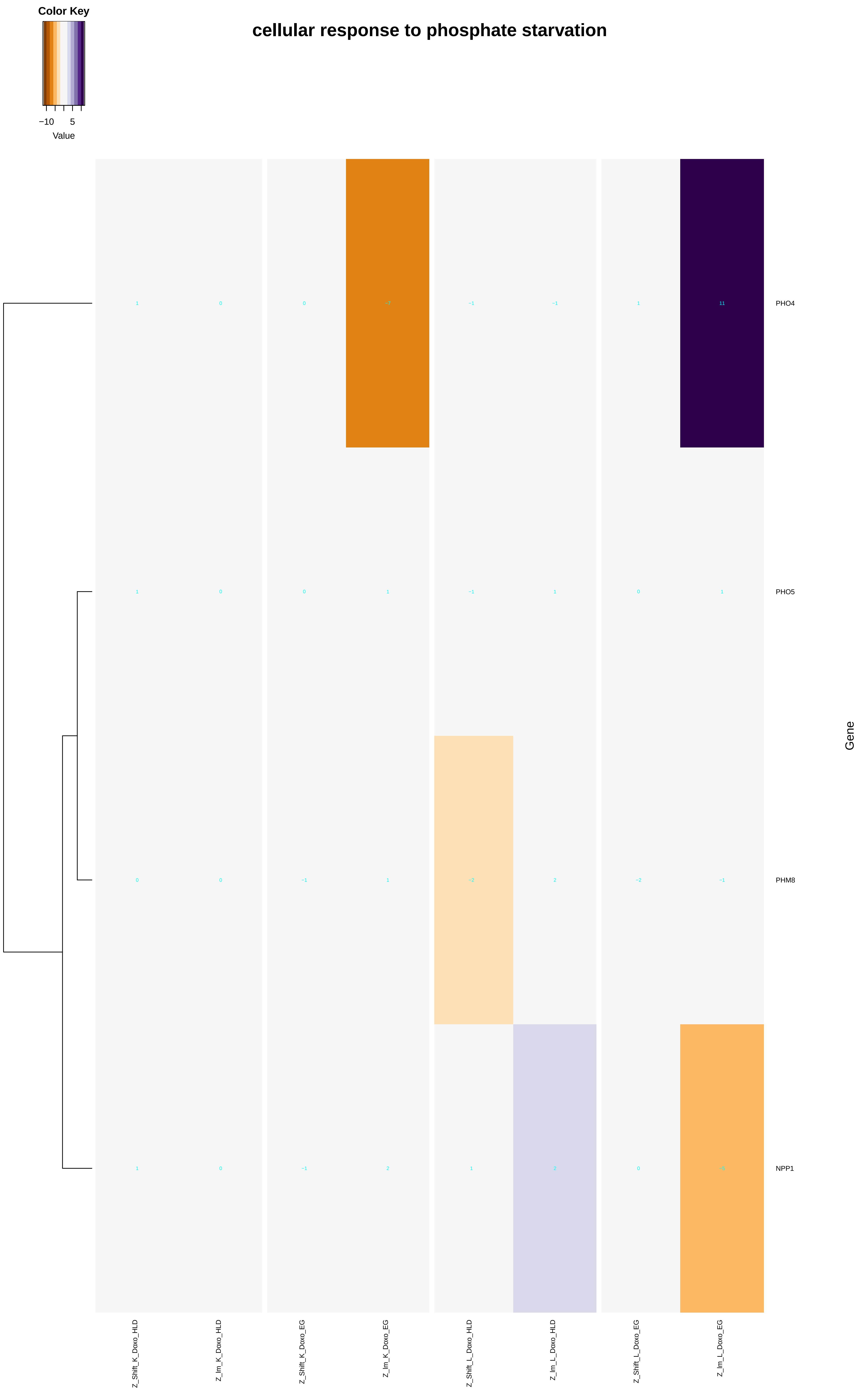
Color Key
−10
5
Value
cellular response to phosphate starvation
PHO4
PHO5
PHM8
NPP1
Gene
Z_Shift_K_Doxo_HLD
Z_Im_K_Doxo_HLD
Z_Shift_K_Doxo_EG
Z_Im_K_Doxo_EG
Z_Shift_L_Doxo_HLD
Z_Im_L_Doxo_HLD
Z_Shift_L_Doxo_EG
Z_Im_L_Doxo_EG

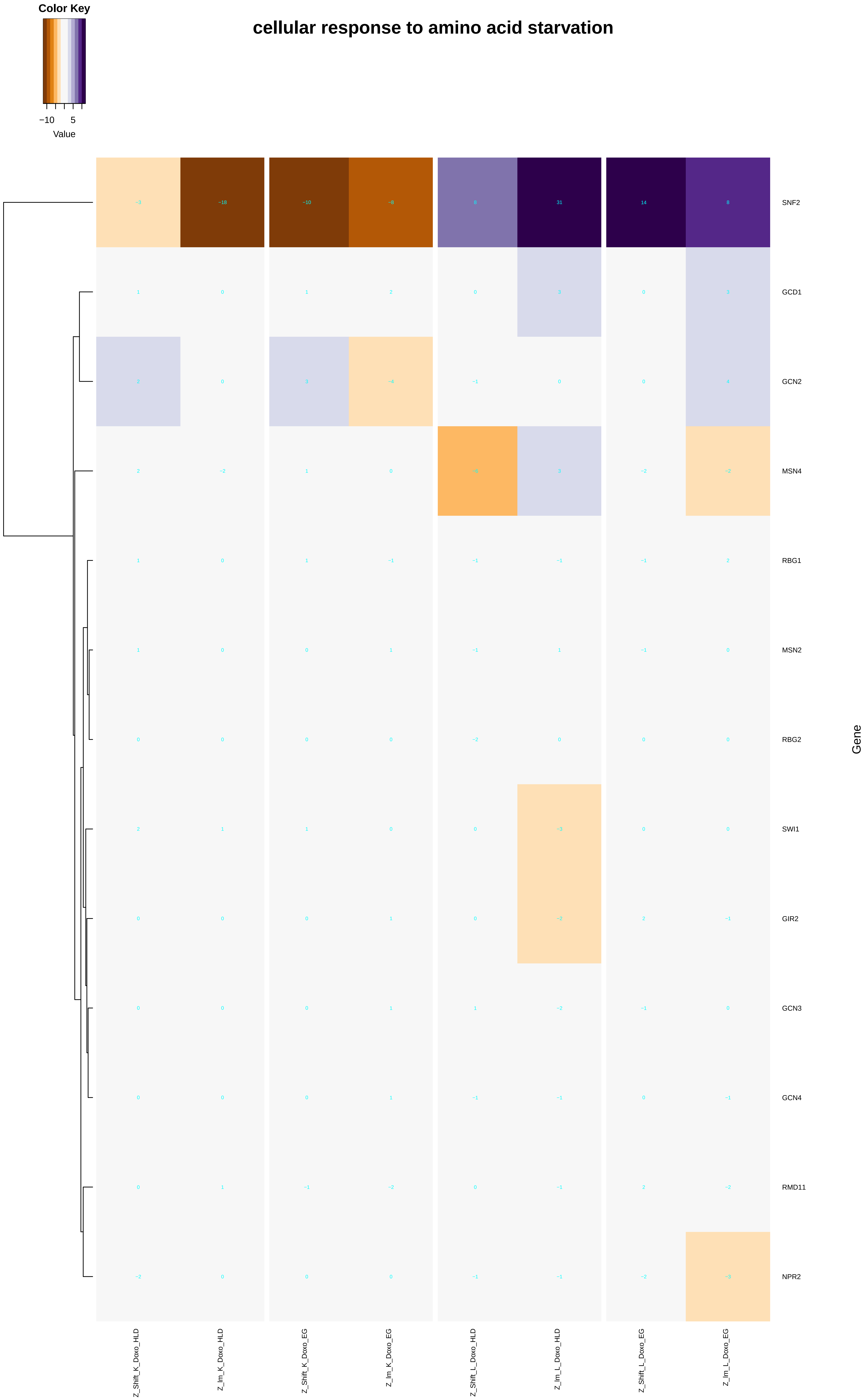
Color Key
−10
5
Value
cellular response to amino acid starvation
SNF2
GCD1
GCN2
MSN4
RBG1
MSN2
RBG2
SWI1
GIR2
GCN3
GCN4
RMD11
NPR2
Gene
Z_Shift_K_Doxo_HLD
Z_Im_K_Doxo_HLD
Z_Shift_K_Doxo_EG
Z_Im_K_Doxo_EG
Z_Shift_L_Doxo_HLD
Z_Im_L_Doxo_HLD
Z_Shift_L_Doxo_EG
Z_Im_L_Doxo_EG

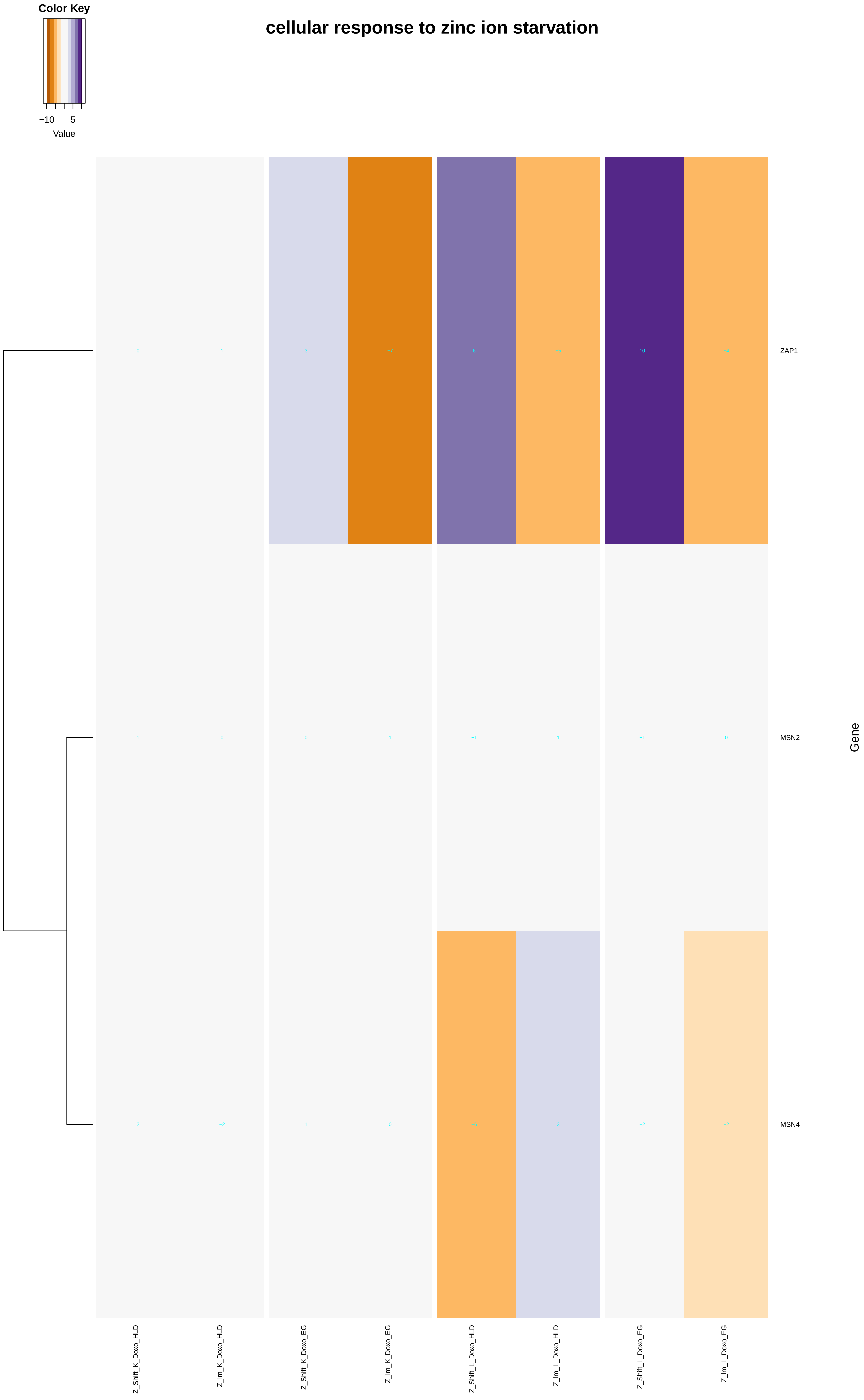
Color Key
−10
5
Value
cellular response to zinc ion starvation
0
1
3
−7
6
−5
10
−4
ZAP1
1
0
0
1
−1
1
−1
0
MSN2
2
−2
1
0
−6
3
−2
−2
MSN4
Gene
Z_Shift_K_Doxo_HLD
Z_lm_K_Doxo_HLD
Z_Shift_K_Doxo_EG
Z_lm_K_Doxo_EG
Z_Shift_L_Doxo_HLD
Z_lm_L_Doxo_HLD
Z_Shift_L_Doxo_EG
Z_lm_L_Doxo_EG

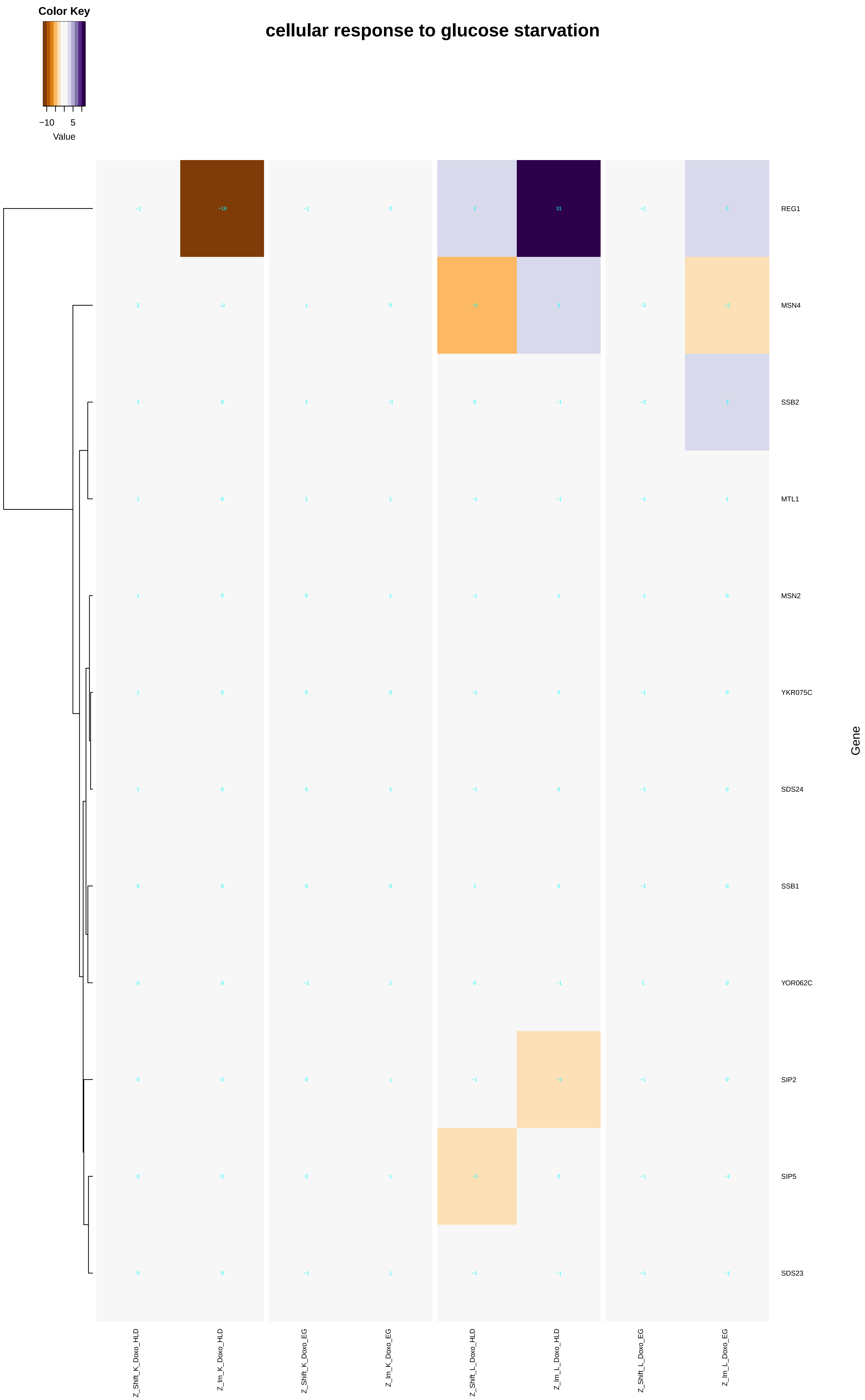

Color Key
−10
5
Value
cellular response to glucose starvation
REG1
MSN4
SSB2
MTL1
MSN2
YKR075C
SDS24
SSB1
YOR062C
SIP2
SIP5
SDS23
Gene
Z_Shift_K_Doxo_HLD
Z_Im_K_Doxo_HLD
Z_Shift_K_Doxo_EG
Z_Im_K_Doxo_EG
Z_Shift_L_Doxo_HLD
Z_Im_L_Doxo_HLD
Z_Shift_L_Doxo_EG
Z_Im_L_Doxo_EG

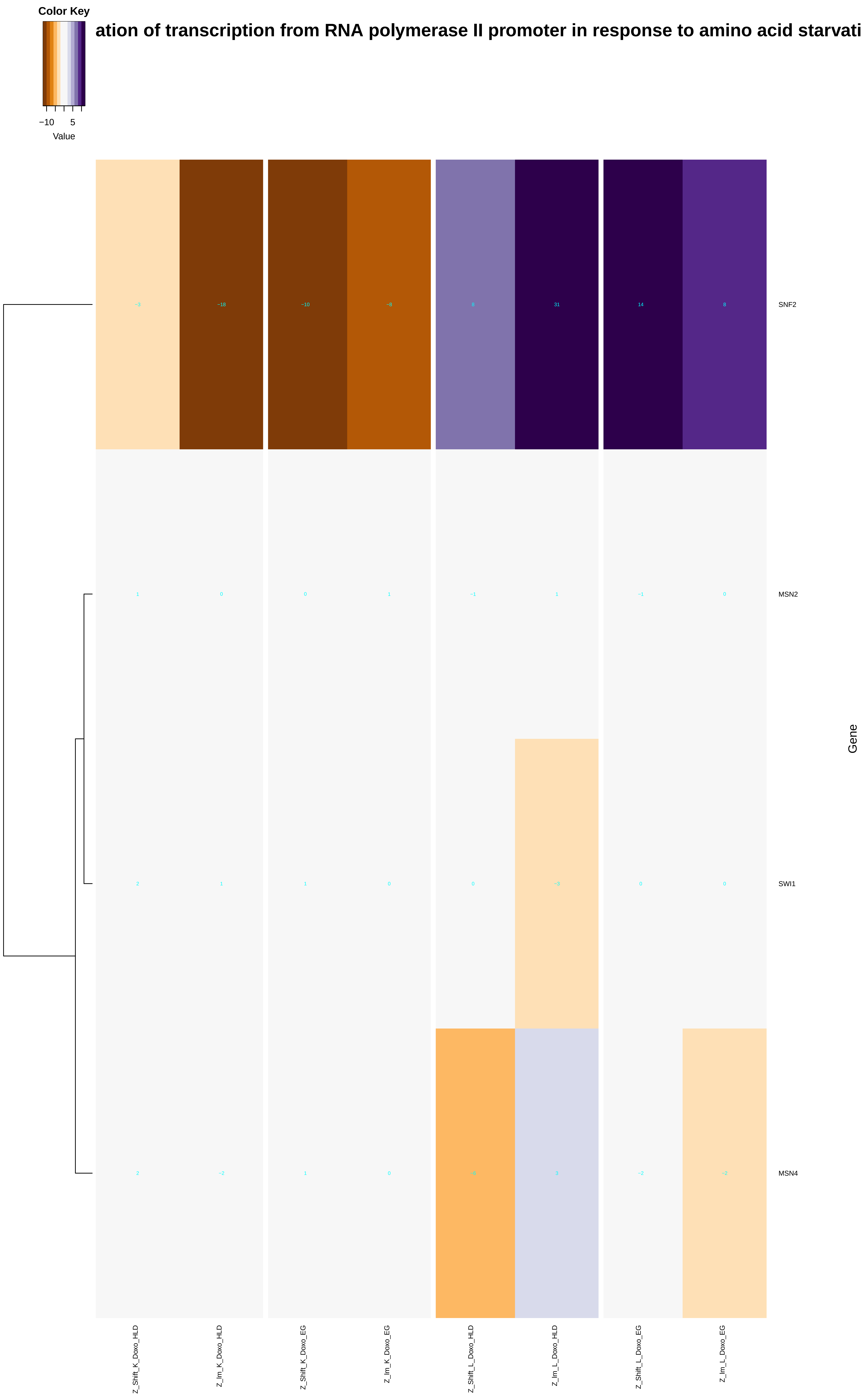

Color Key
−10
5
Value
ation of transcription from RNA polymerase II promoter in response to amino acid starvati
SNF2
MSN2
SWI1
MSN4
Gene
Z_Shift_K_Doxo_HLD
Z_lm_K_Doxo_HLD
Z_Shift_K_Doxo_EG
Z_lm_K_Doxo_EG
Z_Shift_L_Doxo_HLD
Z_lm_L_Doxo_HLD
Z_Shift_L_Doxo_EG
Z_lm_L_Doxo_EG

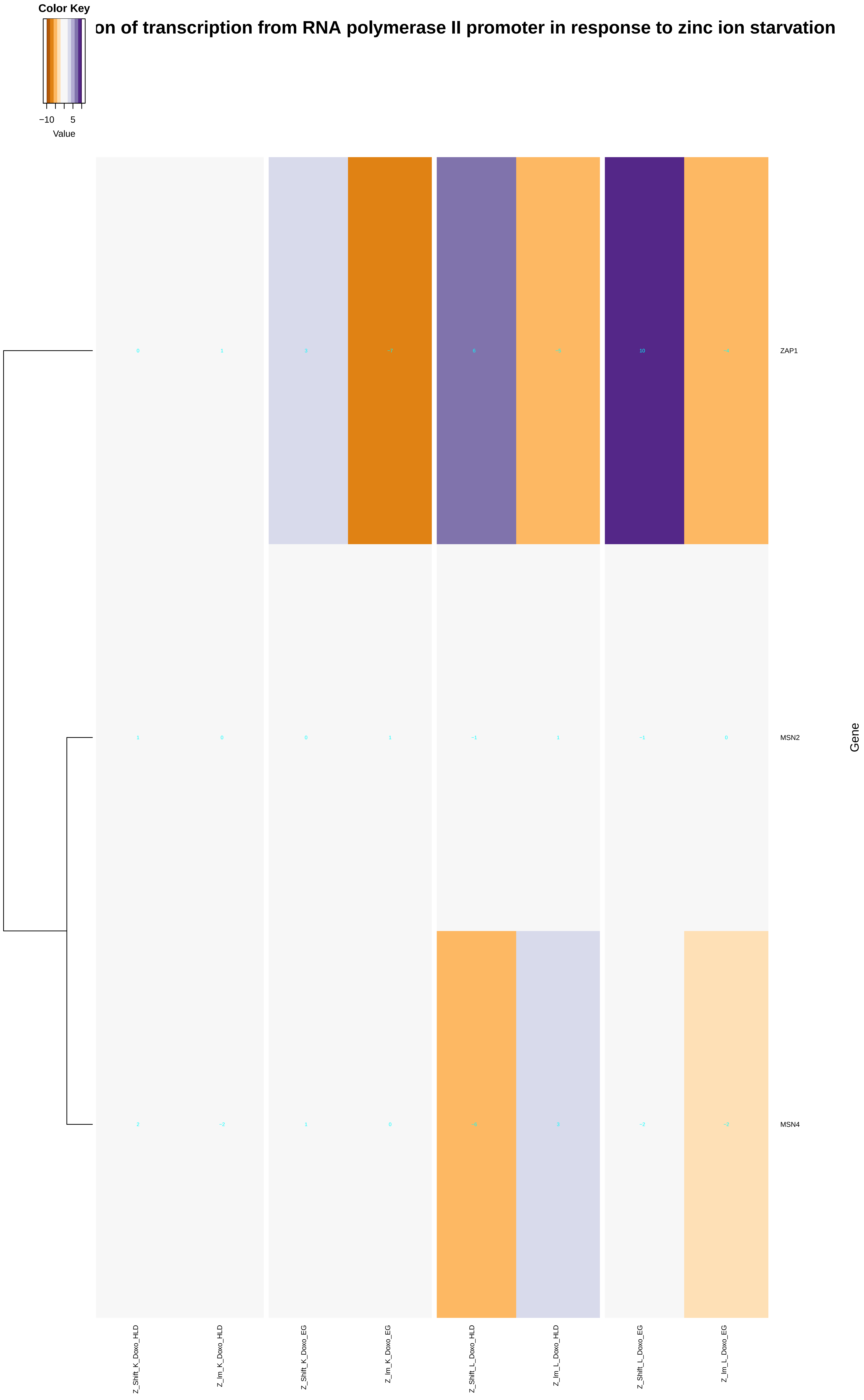

Color Key
−10
5
Value
on of transcription from RNA polymerase II promoter in response to zinc ion starvation
ZAP1
MSN2
MSN4
Gene
Z_Shift_K_Doxo_HLD
Z_Im_K_Doxo_HLD
Z_Shift_K_Doxo_EG
Z_Im_K_Doxo_EG
Z_Shift_L_Doxo_HLD
Z_Im_L_Doxo_HLD
Z_Shift_L_Doxo_EG
Z_Im_L_Doxo_EG

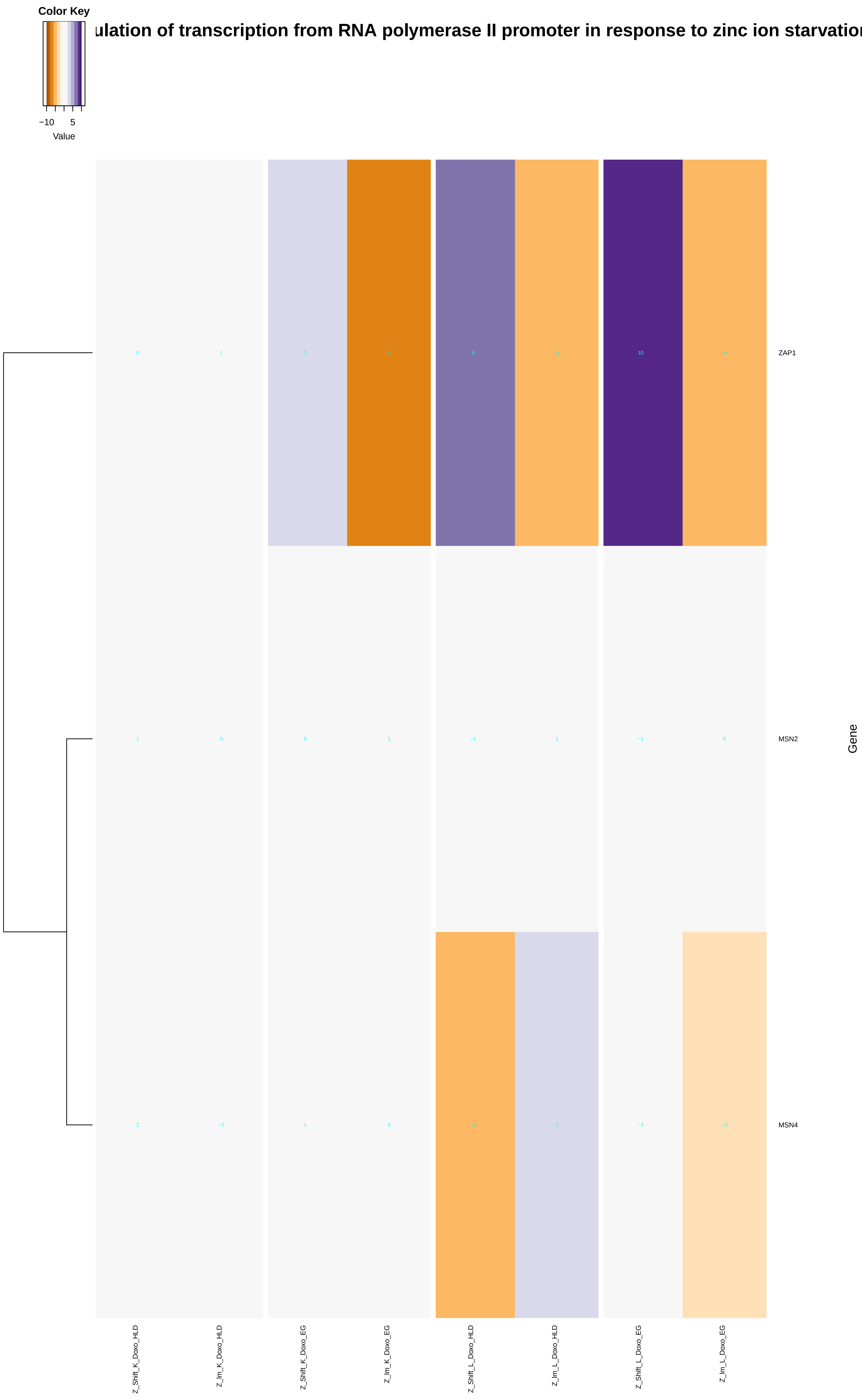
ulation of transcription from RNA polymerase II promoter in response to zinc ion starvatio
Color Key
−10
5
Value
ZAP1
MSN2
MSN4
Gene
Z_Shift_K_Doxo_HLD
Z_Im_K_Doxo_HLD
Z_Shift_K_Doxo_EG
Z_Im_K_Doxo_EG
Z_Shift_L_Doxo_HLD
Z_Im_L_Doxo_HLD
Z_Shift_L_Doxo_EG
Z_Im_L_Doxo_EG
