## Additional File 9 for "A yeast phenomic model for the influence of Warburg metabolism on genetic buffering of doxorubicin": chromatin_organization.pdf

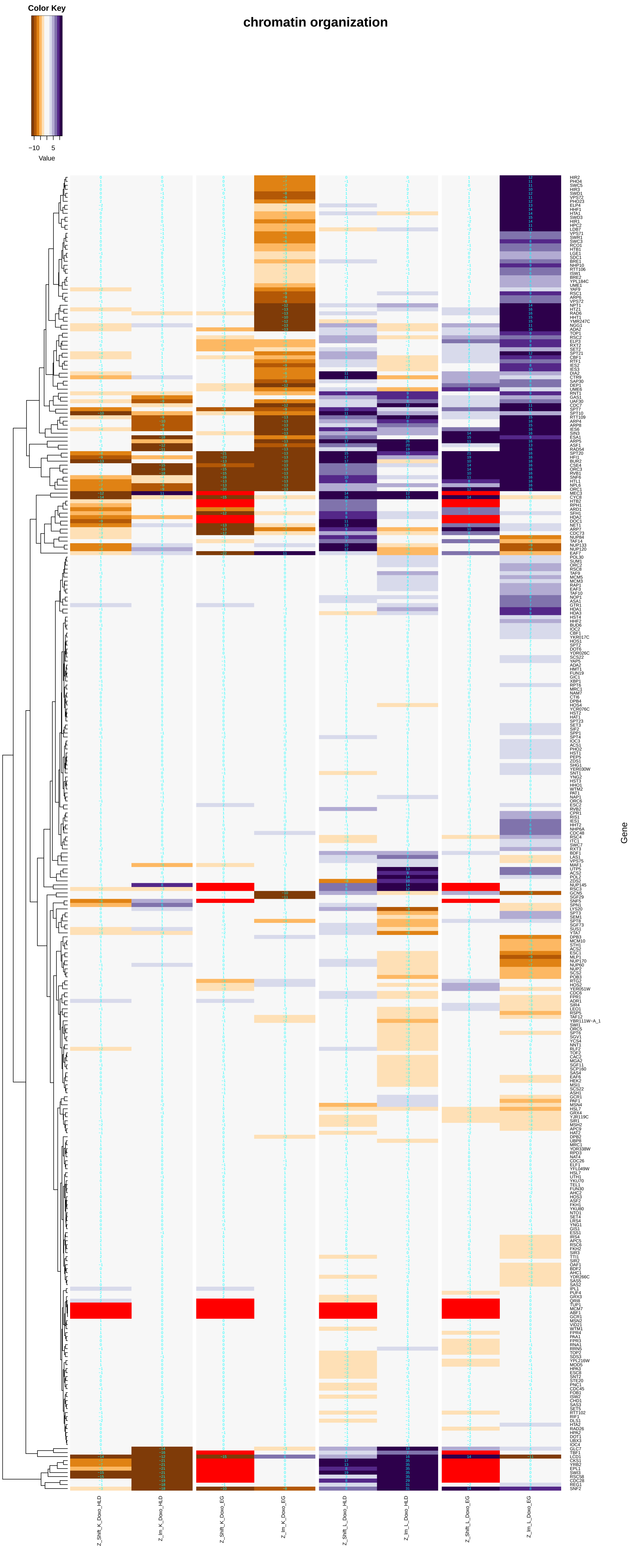

chromatin organization
Color Key
−10
5
Value
Z_Shift_K_Doxo_HLD
Z_Im_K_Doxo_HLD
Z_Shift_K_Doxo_EG
Z_Im_K_Doxo_EG
Z_Shift_L_Doxo_HLD
Z_Im_L_Doxo_HLD
Z_Shift_L_Doxo_EG
Z_Im_L_Doxo_EG
Gene

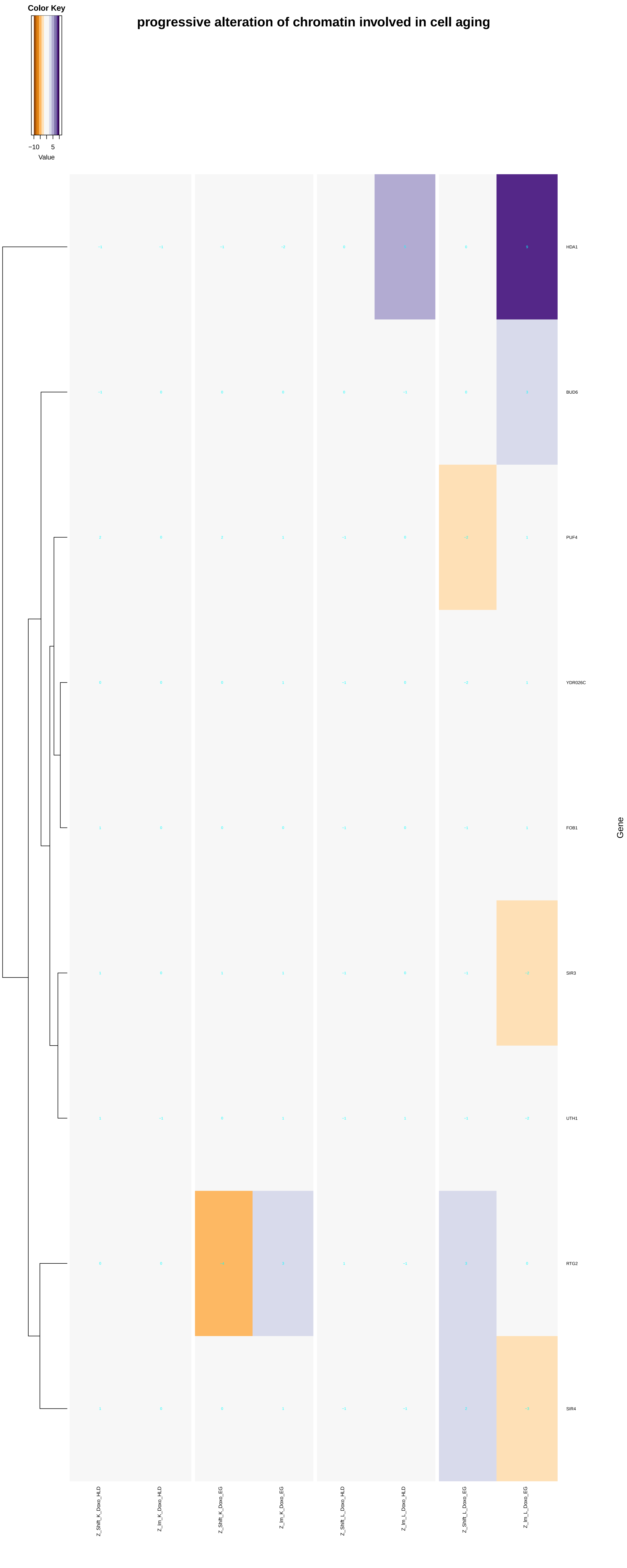

Color Key
−10
5
Value
progressive alteration of chromatin involved in cell aging
HDA1
BUD6
PUF4
YDR026C
FOB1
SIR3
UTH1
RTG2
SIR4
Gene
Z_Shift_K_Doxo_HLD
Z_lm_K_Doxo_HLD
Z_Shift_K_Doxo_EG
Z_lm_K_Doxo_EG
Z_Shift_L_Doxo_HLD
Z_lm_L_Doxo_HLD
Z_Shift_L_Doxo_EG
Z_lm_L_Doxo_EG

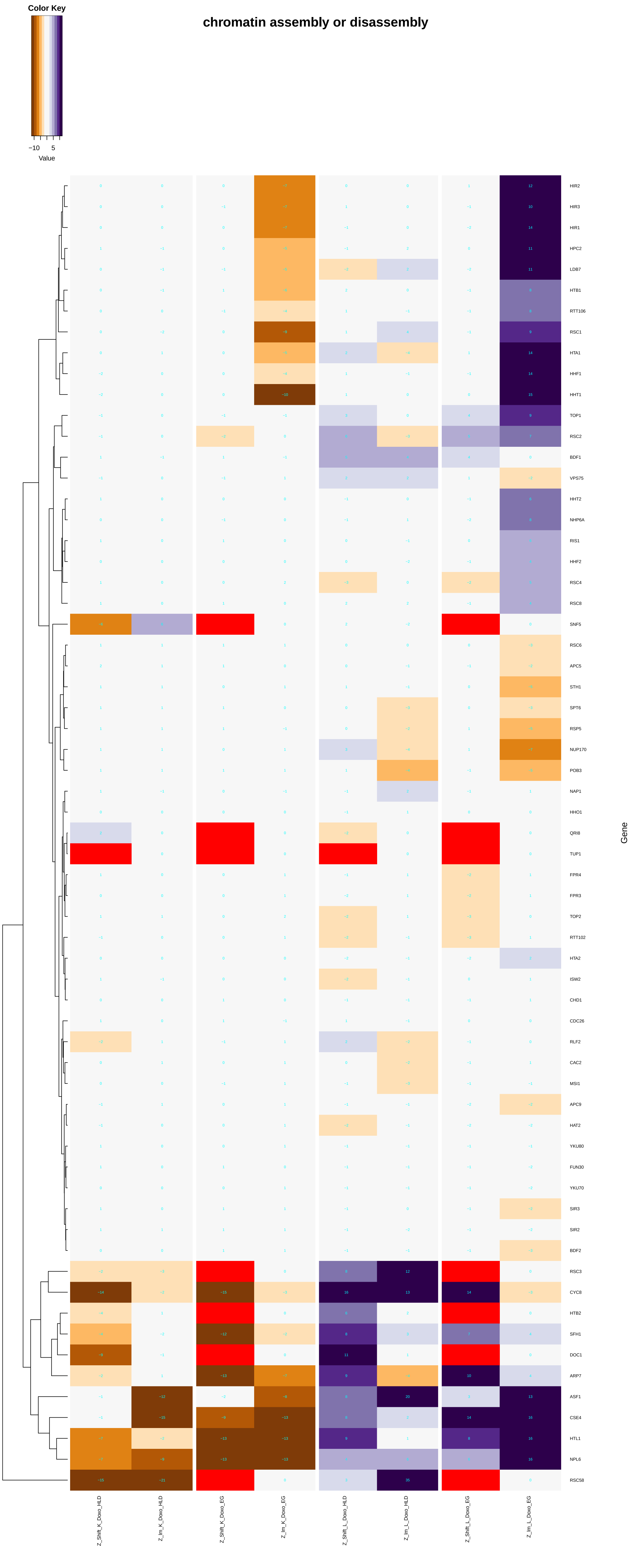
chromatin assembly or disassembly
Color Key
−10
5
Value
HIR2
HIR3
HIR1
HPC2
LDB7
HTB1
RTT106
RSC1
HTA1
HHF1
HHT1
TOP1
RSC2
BDF1
VPS75
HHT2
NHP6A
RIS1
HHF2
RSC4
RSC8
SNF5
RSC6
APC5
STH1
SPT6
RSP5
NUP170
POB3
NAP1
HHO1
QRI8
TUP1
FPR4
FPR3
TOP2
RTT102
HTA2
ISW2
CHD1
CDC26
RLF2
CAC2
MSI1
APC9
HAT2
YKU80
FUN30
YKU70
SIR3
SIR2
BDF2
RSC3
CYC8
HTB2
SFH1
DOC1
ARP7
ASF1
CSE4
HTL1
NPL6
RSC58
Gene
Z_Shift_K_Doxo_HLD
Z_Im_K_Doxo_HLD
Z_Shift_K_Doxo_EG
Z_Im_K_Doxo_EG
Z_Shift_L_Doxo_HLD
Z_Im_L_Doxo_HLD
Z_Shift_L_Doxo_EG
Z_Im_L_Doxo_EG

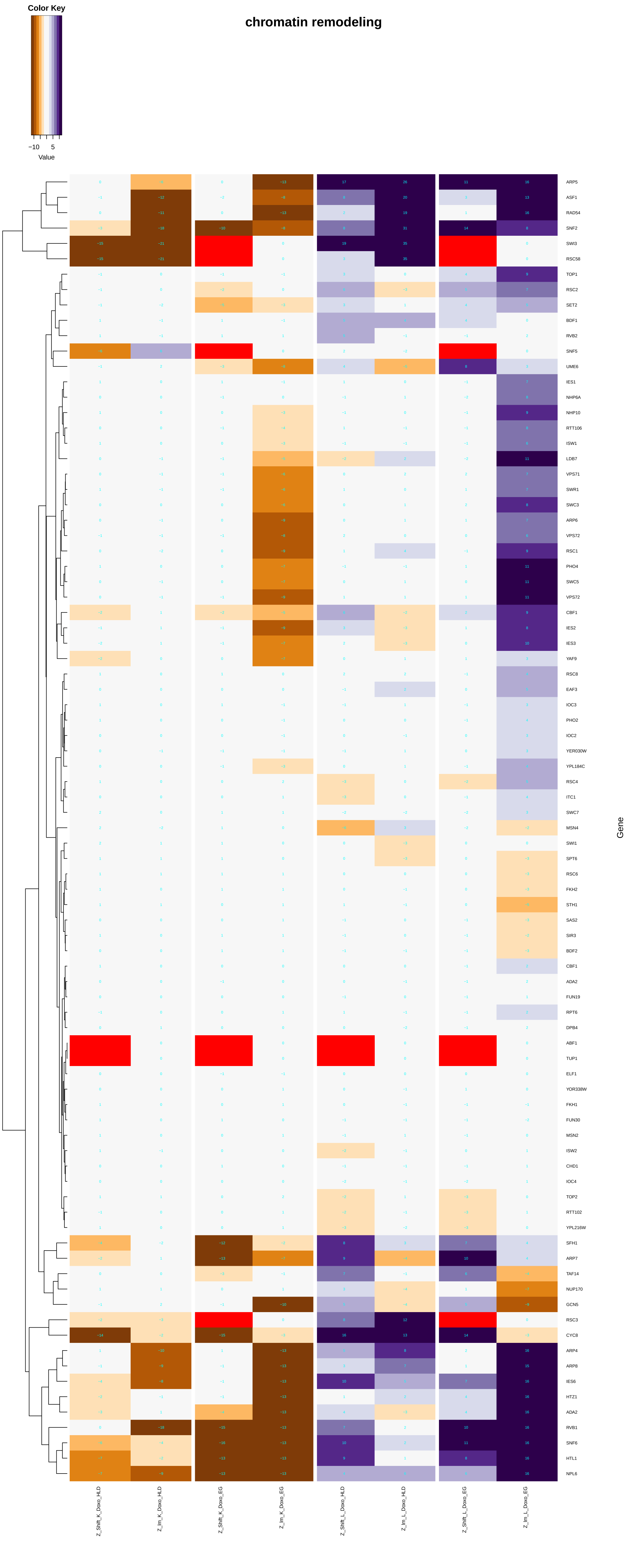
Color Key
−10
5
Value
chromatin remodeling
ARP5
ASF1
RAD54
SNF2
SWI3
RSC58
TOP1
RSC2
SET2
BDF1
RVB2
SNF5
UME6
IES1
NHP6A
NHP10
RTT106
ISW1
LDB7
VPS71
SWR1
SWC3
ARP6
VPS72
RSC1
PHO4
SWC5
VPS72
CBF1
IES2
IES3
YAF9
RSC8
EAF3
IOC3
PHO2
IOC2
YER030W
YPL184C
RSC4
ITC1
SWC7
MSN4
SWI1
SPT6
RSC6
FKH2
STH1
SAS2
SIR3
BDF2
CBF1
ADA2
FUN19
RPT6
DPB4
ABF1
TUP1
ELF1
YOR338W
FKH1
FUN30
MSN2
ISW2
CHD1
IOC4
TOP2
RTT102
YPL216W
SFH1
ARP7
TAF14
NUP170
GCN5
RSC3
CYC8
ARP4
ARP8
IES6
HTZ1
ADA2
RVB1
SNF6
HTL1
NPL6
Gene
Z_Shift_K_Doxo_HLD
Z_lm_K_Doxo_HLD
Z_Shift_K_Doxo_EG
Z_lm_K_Doxo_EG
Z_Shift_L_Doxo_HLD
Z_lm_L_Doxo_HLD
Z_Shift_L_Doxo_EG
Z_lm_L_Doxo_EG

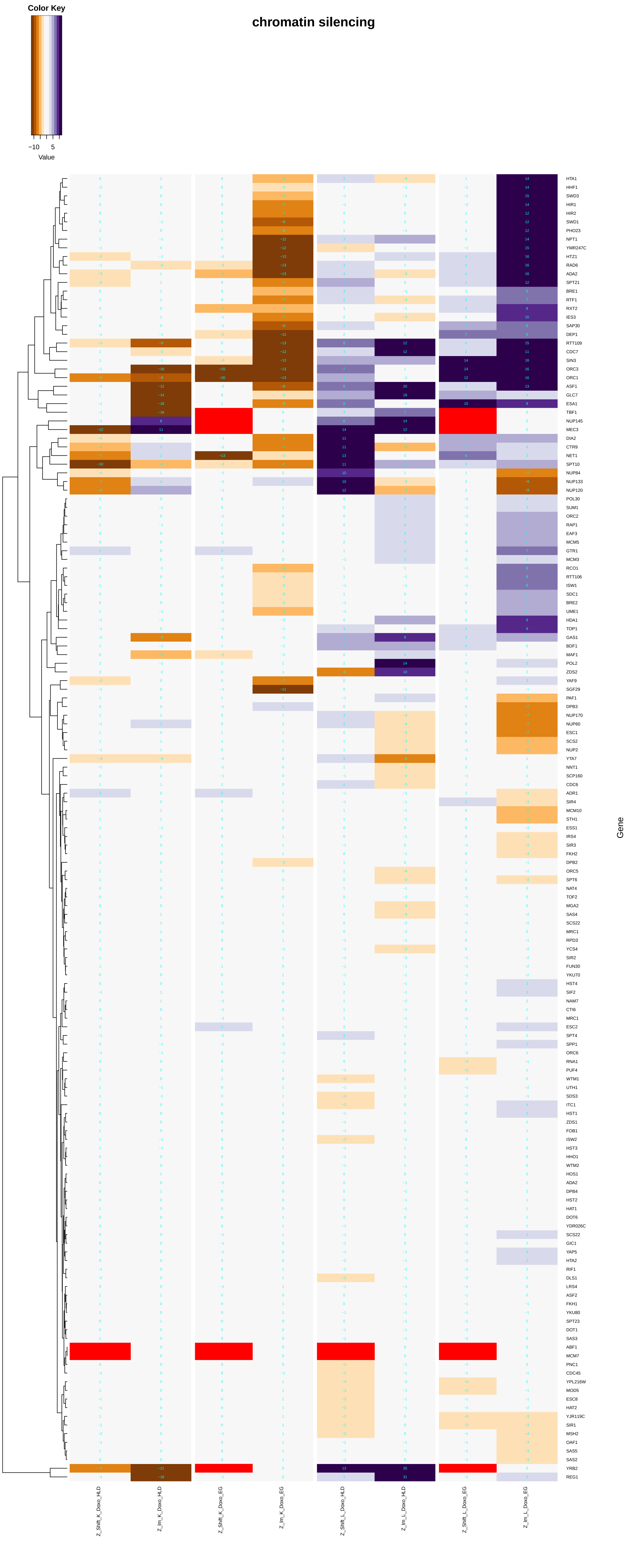

Color Key
−10
5
Value
chromatin silencing
HTA1
HHF1
SWD3
HIR1
HIR2
SWD1
PHO23
NPT1
YMR247C
HTZ1
RAD6
ADA2
SPT21
BRE1
RTF1
RXT2
IES3
SAP30
DEP1
RTT109
CDC7
SIN3
ORC3
ORC1
ASF1
GLC7
ESA1
TBF1
NUP145
MEC3
DIA2
CTR9
NET1
SPT10
NUP84
NUP133
NUP120
POL30
SUM1
ORC2
RAP1
EAF3
MCM5
GTR1
MCM3
RCO1
RTT106
ISW1
SDC1
BRE2
UME1
HDA1
TOP1
GAS1
BDF1
MAF1
POL2
ZDS2
YAF9
SGF29
PAF1
DPB3
NUP170
NUP60
ESC1
SCS2
NUP2
YTA7
NNT1
SCP160
CDC6
ADR1
SIR4
MCM10
STH1
ESS1
IRS4
SIR3
FKH2
DPB2
ORC5
SPT6
NAT4
TOF2
MGA2
SAS4
SCS22
MRC1
RPD3
YCS4
SIR2
FUN30
YKU70
HST4
SIF2
NAM7
CTI6
MRC1
ESC2
SPT4
SPP1
ORC6
RNA1
PUF4
WTM1
UTH1
SDS3
ITC1
HST1
ZDS1
FOB1
ISW2
HST3
HHO1
WTM2
HOS1
ADA2
DPB4
HST2
HAT1
DOT6
YDR026C
SCS22
GIC1
YAP5
HTA2
RIF1
DLS1
LRS4
ASF2
FKH1
YKU80
SPT23
DOT1
SAS3
ABF1
MCM7
PNC1
CDC45
YPL216W
MOD5
ESC8
HAT2
YJR119C
SIR1
MSH2
OAF1
SAS5
SAS2
YRB2
REG1
Gene
Z_Shift_K_Doxo_HLD
Z_Im_K_Doxo_HLD
Z_Shift_K_Doxo_EG
Z_Im_K_Doxo_EG
Z_Shift_L_Doxo_HLD
Z_Im_L_Doxo_HLD
Z_Shift_L_Doxo_EG
Z_Im_L_Doxo_EG

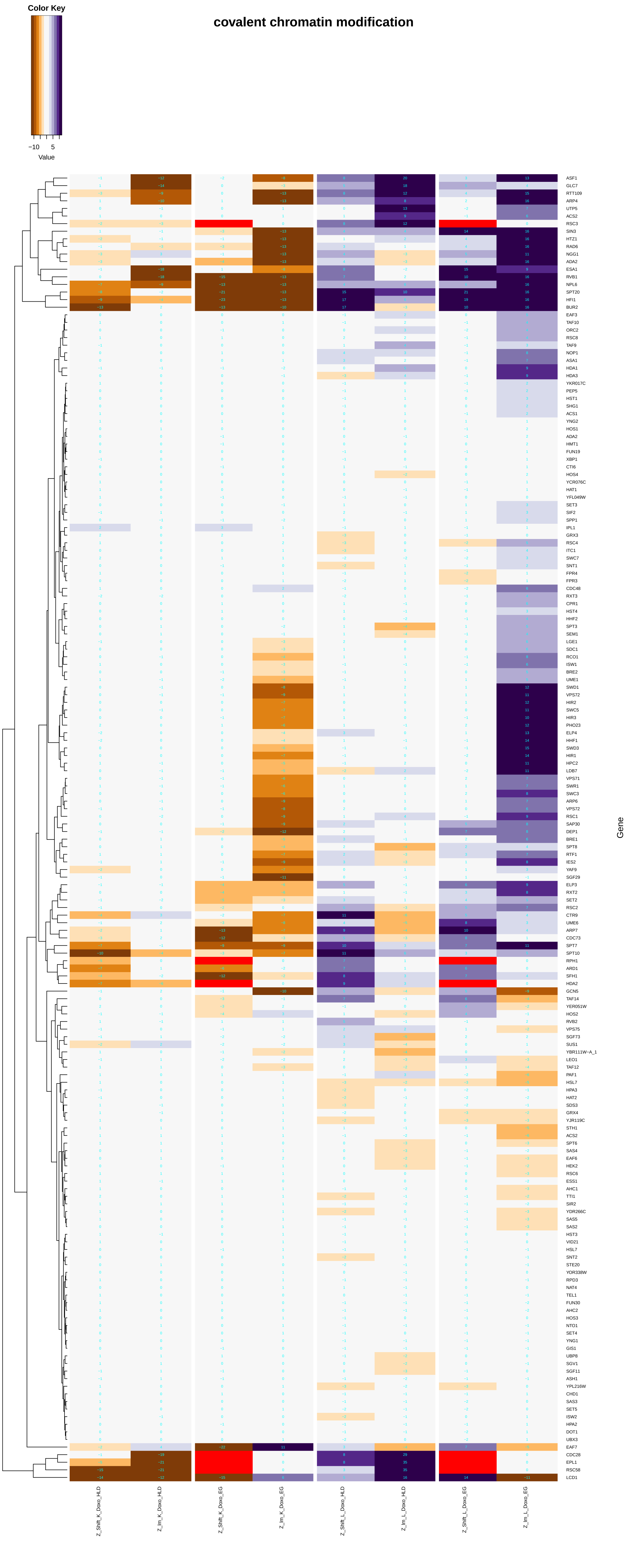
covalent chromatin modification
Color Key
−10
5
Value
Gene
Z_Shift_K_Doxo_HLD
Z_Im_K_Doxo_HLD
Z_Shift_K_Doxo_EG
Z_Im_K_Doxo_EG
Z_Shift_L_Doxo_HLD
Z_Im_L_Doxo_HLD
Z_Shift_L_Doxo_EG
Z_Im_L_Doxo_EG

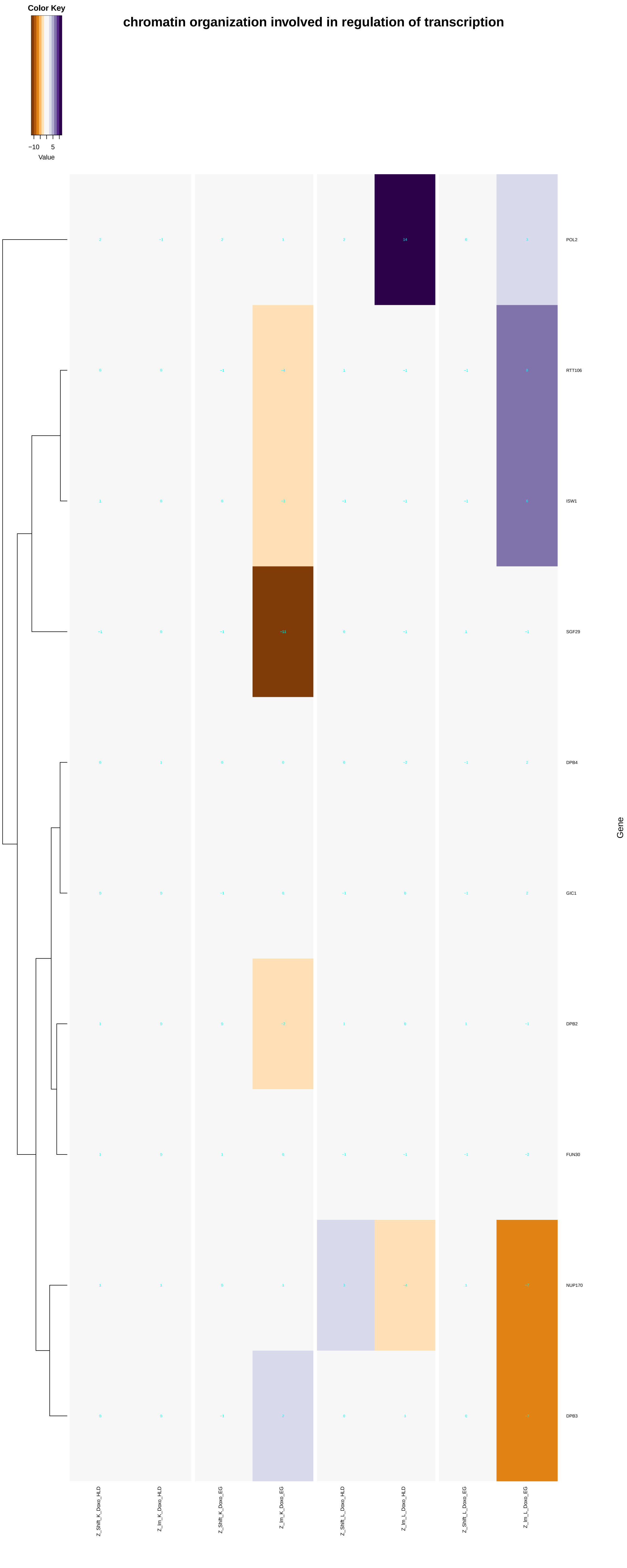
chromatin organization involved in regulation of transcription
Color Key
−10
5
Value
POL2
RTT106
ISW1
SGF29
DPB4
GIC1
DPB2
FUN30
NUP170
DPB3
Gene
Z_Shift_K_Doxo_HLD
Z_Im_K_Doxo_HLD
Z_Shift_K_Doxo_EG
Z_Im_K_Doxo_EG
Z_Shift_L_Doxo_HLD
Z_Im_L_Doxo_HLD
Z_Shift_L_Doxo_EG
Z_Im_L_Doxo_EG

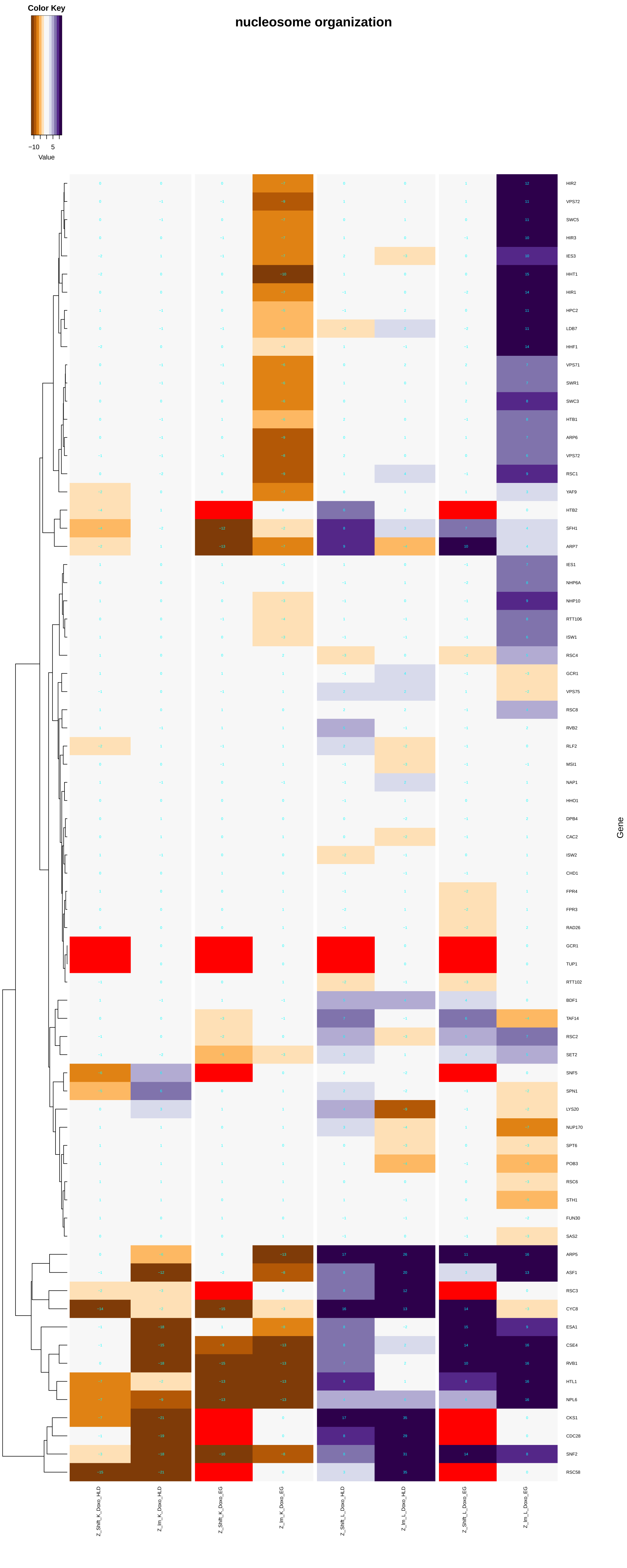

nucleosome organization
Color Key
−10
5
Value
HIR2
VPS72
SWC5
HIR3
IES3
HHT1
HIR1
HPC2
LDB7
HHF1
VPS71
SWR1
SWC3
HTB1
ARP6
VPS72
RSC1
YAF9
HTB2
SFH1
ARP7
IES1
NHP6A
NHP10
RTT106
ISW1
RSC4
GCR1
VPS75
RSC8
RVB2
RLF2
MSI1
NAP1
HHO1
DPB4
CAC2
ISW2
CHD1
FPR4
FPR3
RAD26
GCR1
TUP1
RTT102
BDF1
TAF14
RSC2
SET2
SNF5
SPN1
LYS20
NUP170
SPT6
POB3
RSC6
STH1
FUN30
SAS2
ARP5
ASF1
RSC3
CYC8
ESA1
CSE4
RVB1
HTL1
NPL6
CKS1
CDC28
SNF2
RSC58
Gene
Z_Shift_K_Doxo_HLD
Z_Im_K_Doxo_HLD
Z_Shift_K_Doxo_EG
Z_Im_K_Doxo_EG
Z_Shift_L_Doxo_HLD
Z_Im_L_Doxo_HLD
Z_Shift_L_Doxo_EG
Z_Im_L_Doxo_EG

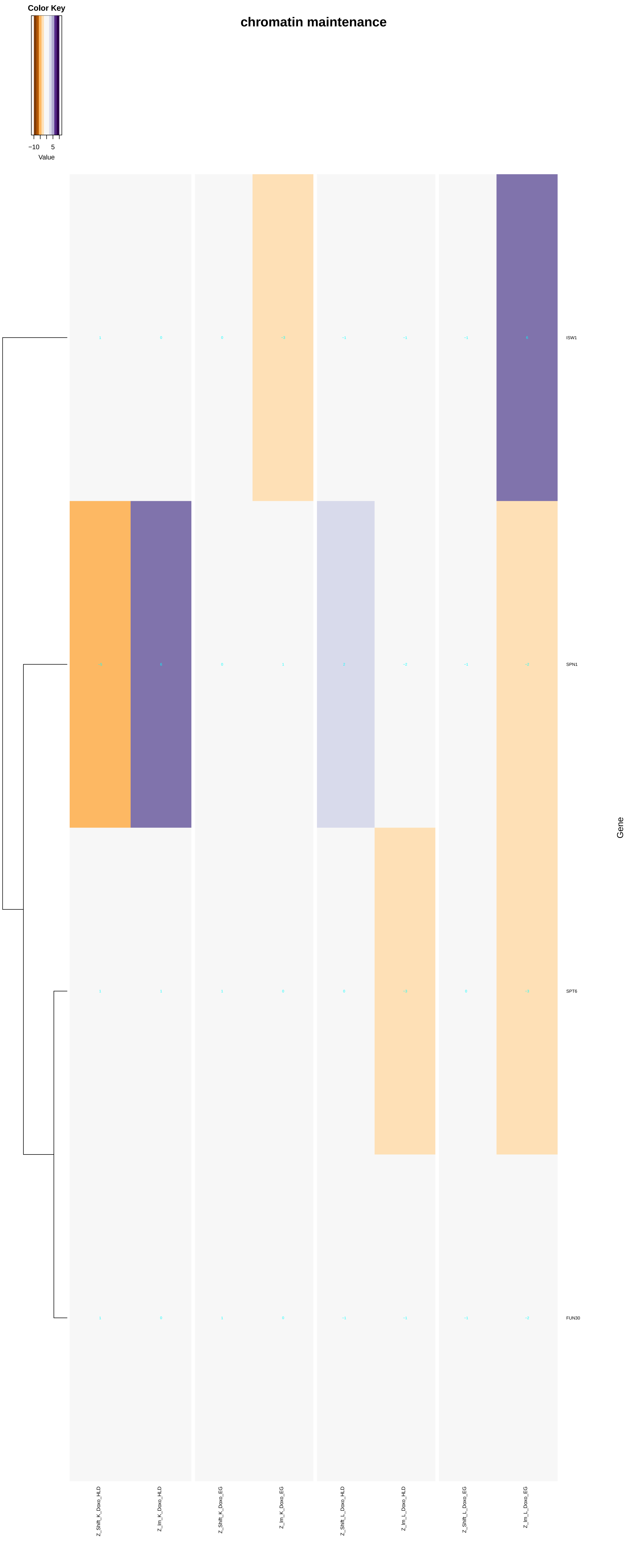

chromatin maintenance
Color Key
−10
5
Value
ISW1
SPN1
SPT6
FUN30
Gene
Z_Shift_K_Doxo_HLD
Z_lm_K_Doxo_HLD
Z_Shift_K_Doxo_EG
Z_lm_K_Doxo_EG
Z_Shift_L_Doxo_HLD
Z_lm_L_Doxo_HLD
Z_Shift_L_Doxo_EG
Z_lm_L_Doxo_EG

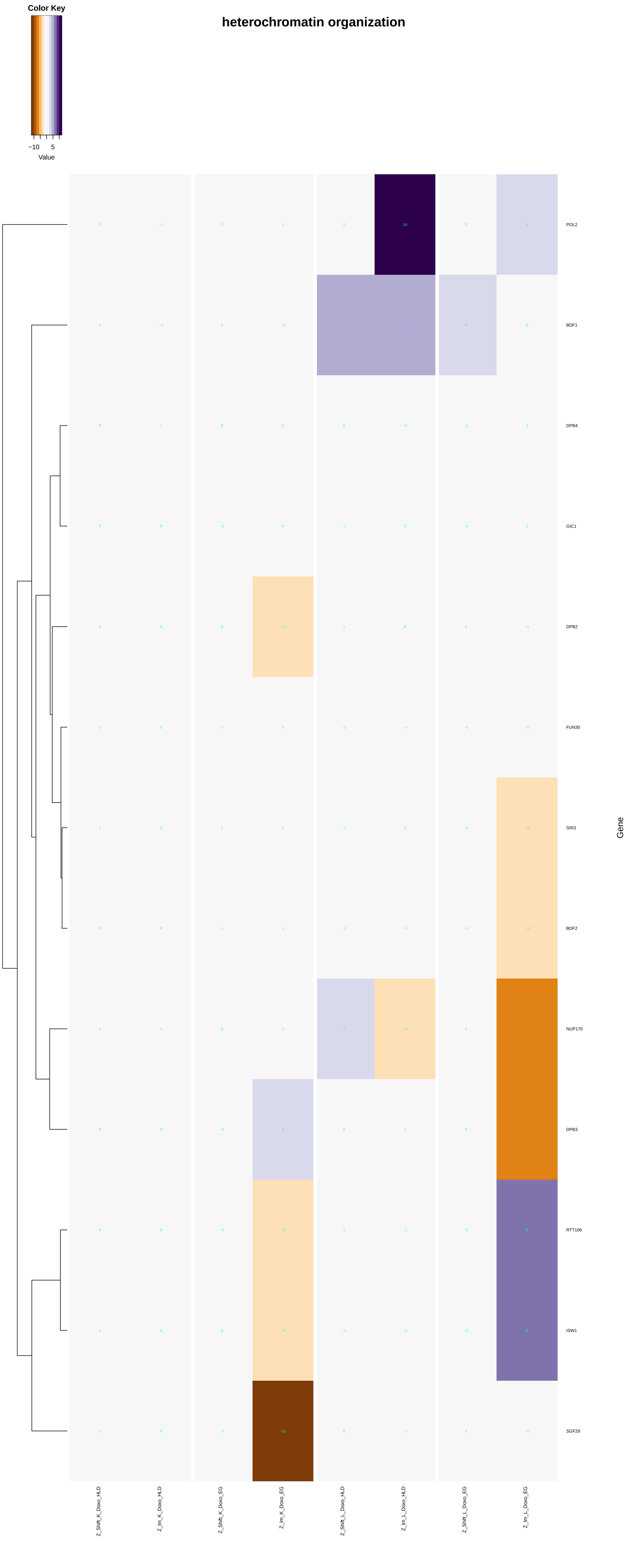
heterochromatin organization
Color Key
−10
5
Value
POL2
BDF1
DPB4
GIC1
DPB2
FUN30
SIR3
BDF2
NUP170
DPB3
RTT106
ISW1
SGF29
Gene
Z_Shift_K_Doxo_HLD
Z_lm_K_Doxo_HLD
Z_Shift_K_Doxo_EG
Z_lm_K_Doxo_EG
Z_Shift_L_Doxo_HLD
Z_lm_L_Doxo_HLD
Z_Shift_L_Doxo_EG
Z_lm_L_Doxo_EG

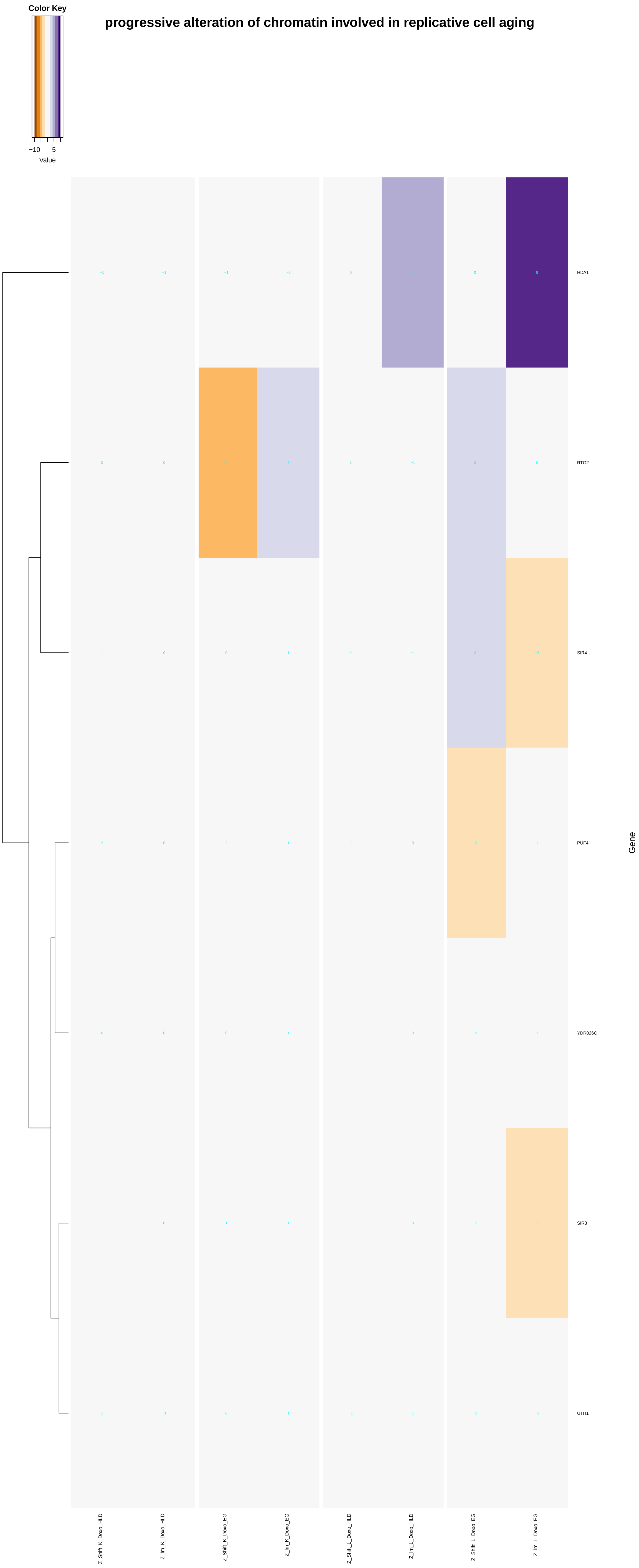

Color Key
−10
5
Value
progressive alteration of chromatin involved in replicative cell aging
HDA1
RTG2
SIR4
PUF4
YDR026C
SIR3
UTH1
Gene
Z_Shift_K_Doxo_HLD
Z_Im_K_Doxo_HLD
Z_Shift_K_Doxo_EG
Z_Im_K_Doxo_EG
Z_Shift_L_Doxo_HLD
Z_Im_L_Doxo_HLD
Z_Shift_L_Doxo_EG
Z_Im_L_Doxo_EG

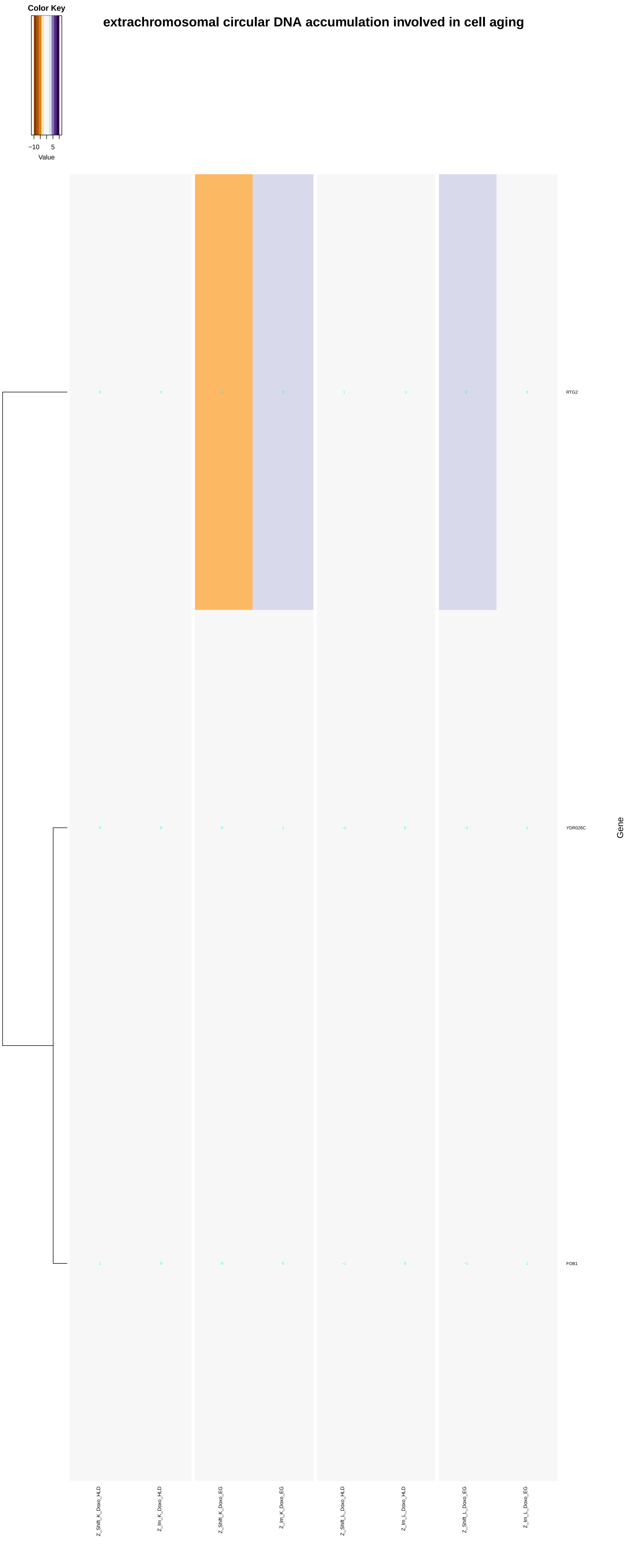
extrachromosomal circular DNA accumulation involved in cell aging
Color Key
−10
5
Value
RTG2
YDR026C
FOB1
Gene
Z_Shift_K_Doxo_HLD
Z_Im_K_Doxo_HLD
Z_Shift_K_Doxo_EG
Z_Im_K_Doxo_EG
Z_Shift_L_Doxo_HLD
Z_Im_L_Doxo_HLD
Z_Shift_L_Doxo_EG
Z_Im_L_Doxo_EG

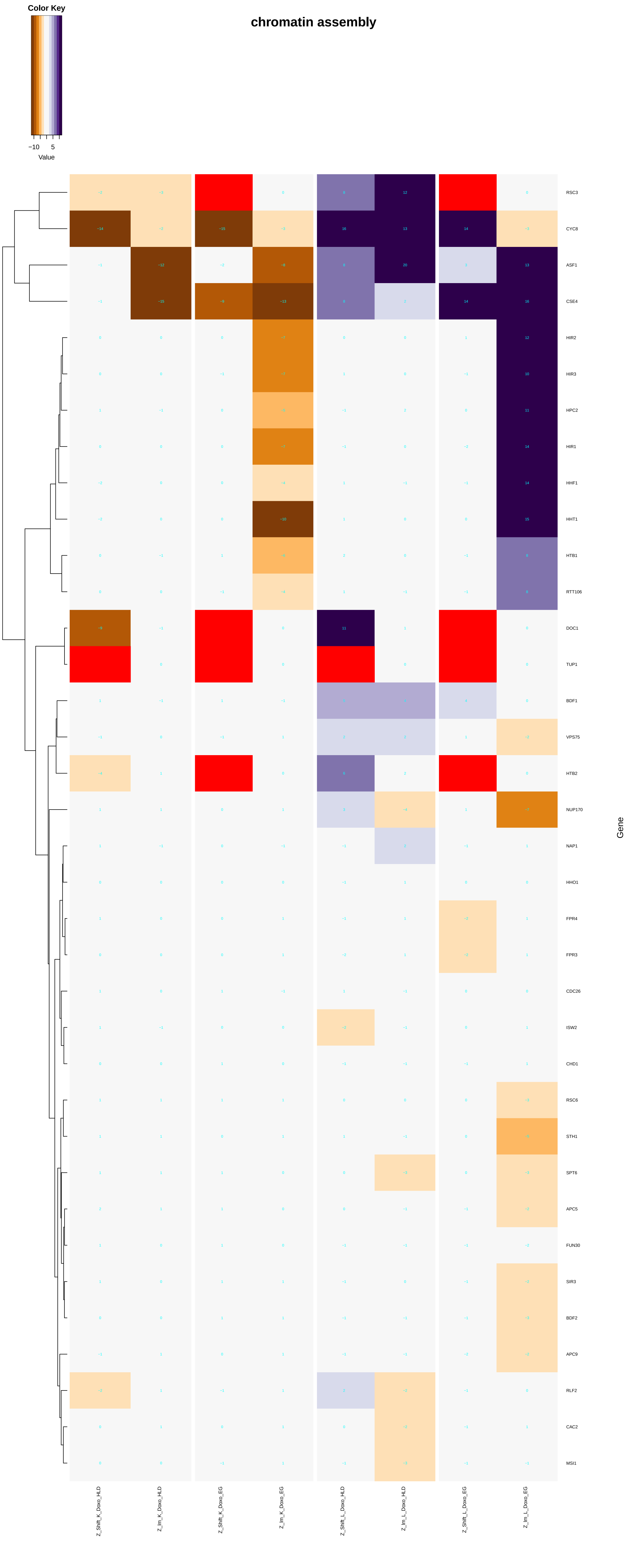
chromatin assembly
Color Key
−10
5
Value
RSC3
CYC8
ASF1
CSE4
HIR2
HIR3
HPC2
HIR1
HHF1
HHT1
HTB1
RTT106
DOC1
TUP1
BDF1
VPS75
HTB2
NUP170
NAP1
HHO1
FPR4
FPR3
CDC26
ISW2
CHD1
RSC6
STH1
SPT6
APC5
FUN30
SIR3
BDF2
APC9
RLF2
CAC2
MSI1
Gene
Z_Shift_K_Doxo_HLD
Z_lm_K_Doxo_HLD
Z_Shift_K_Doxo_EG
Z_lm_K_Doxo_EG
Z_Shift_L_Doxo_HLD
Z_lm_L_Doxo_HLD
Z_Shift_L_Doxo_EG
Z_lm_L_Doxo_EG

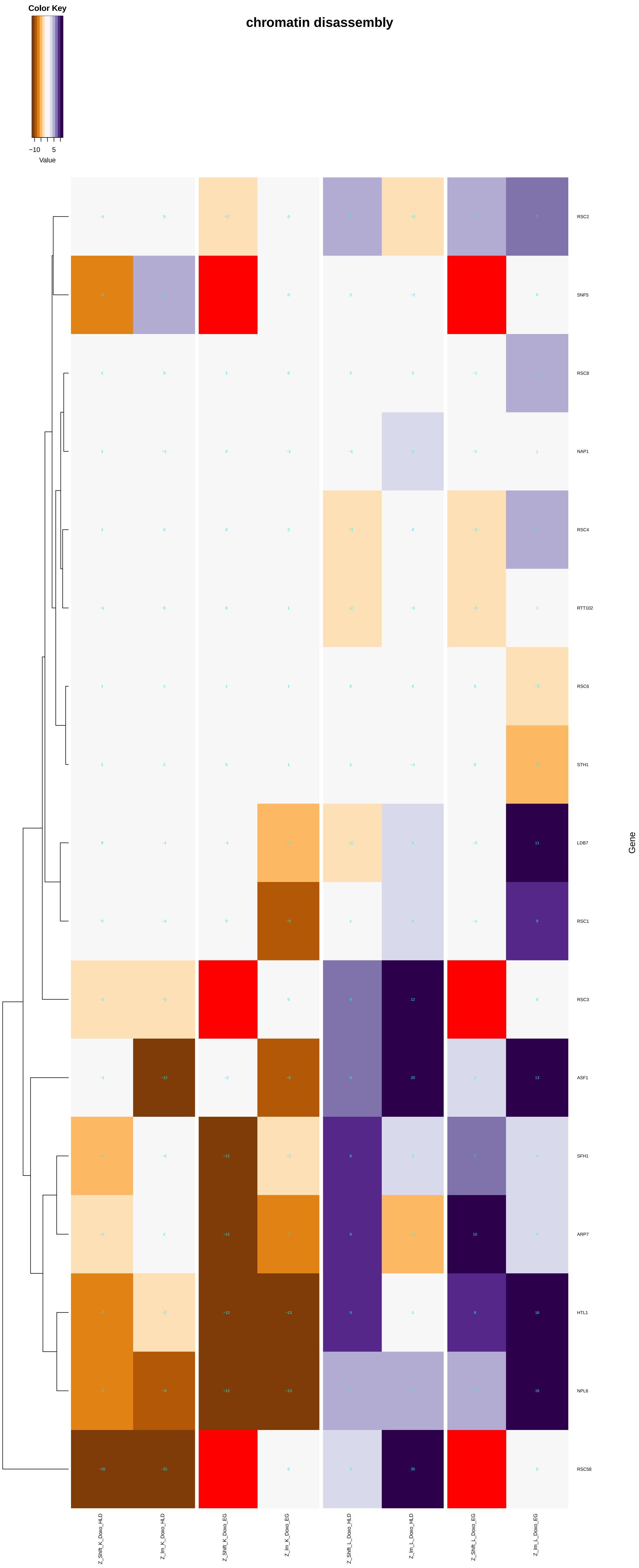
chromatin disassembly
Color Key
−10
5
Value
RSC2
SNF5
RSC8
NAP1
RSC4
RTT102
RSC6
STH1
LDB7
RSC1
RSC3
ASF1
SFH1
ARP7
HTL1
NPL6
RSC58
Gene
Z_Shift_K_Doxo_HLD
Z_Im_K_Doxo_HLD
Z_Shift_K_Doxo_EG
Z_Im_K_Doxo_EG
Z_Shift_L_Doxo_HLD
Z_Im_L_Doxo_HLD
Z_Shift_L_Doxo_EG
Z_Im_L_Doxo_EG

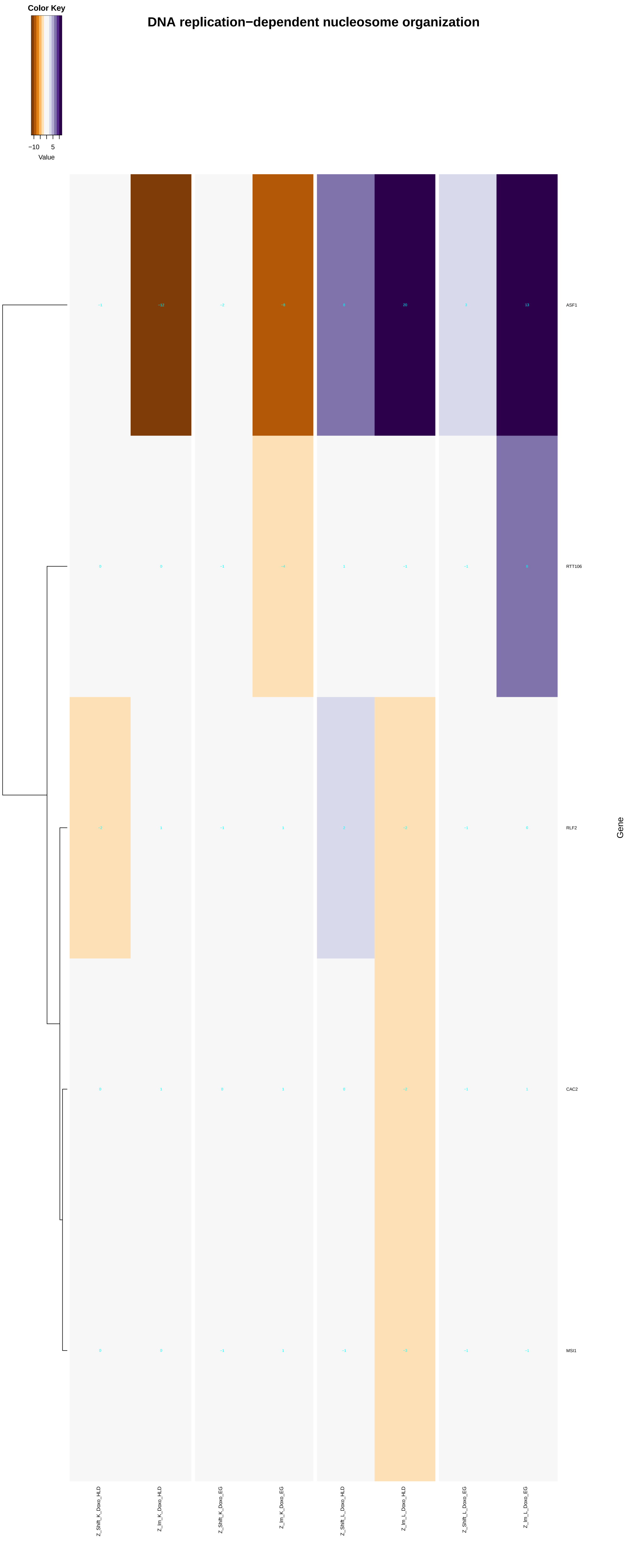

Color Key
−10
5
Value
DNA replication−dependent nucleosome organization
−1
−12
−2
−8
8
20
3
13
ASF1
0
0
−1
−4
1
−1
−1
8
RTT106
−2
1
−1
1
2
−2
−1
0
RLF2
0
1
0
1
0
−2
−1
1
CAC2
0
0
−1
1
−1
−3
−1
−1
MSI1
Gene
Z_Shift_K_Doxo_HLD
Z_Im_K_Doxo_HLD
Z_Shift_K_Doxo_EG
Z_Im_K_Doxo_EG
Z_Shift_L_Doxo_HLD
Z_Im_L_Doxo_HLD
Z_Shift_L_Doxo_EG
Z_Im_L_Doxo_EG

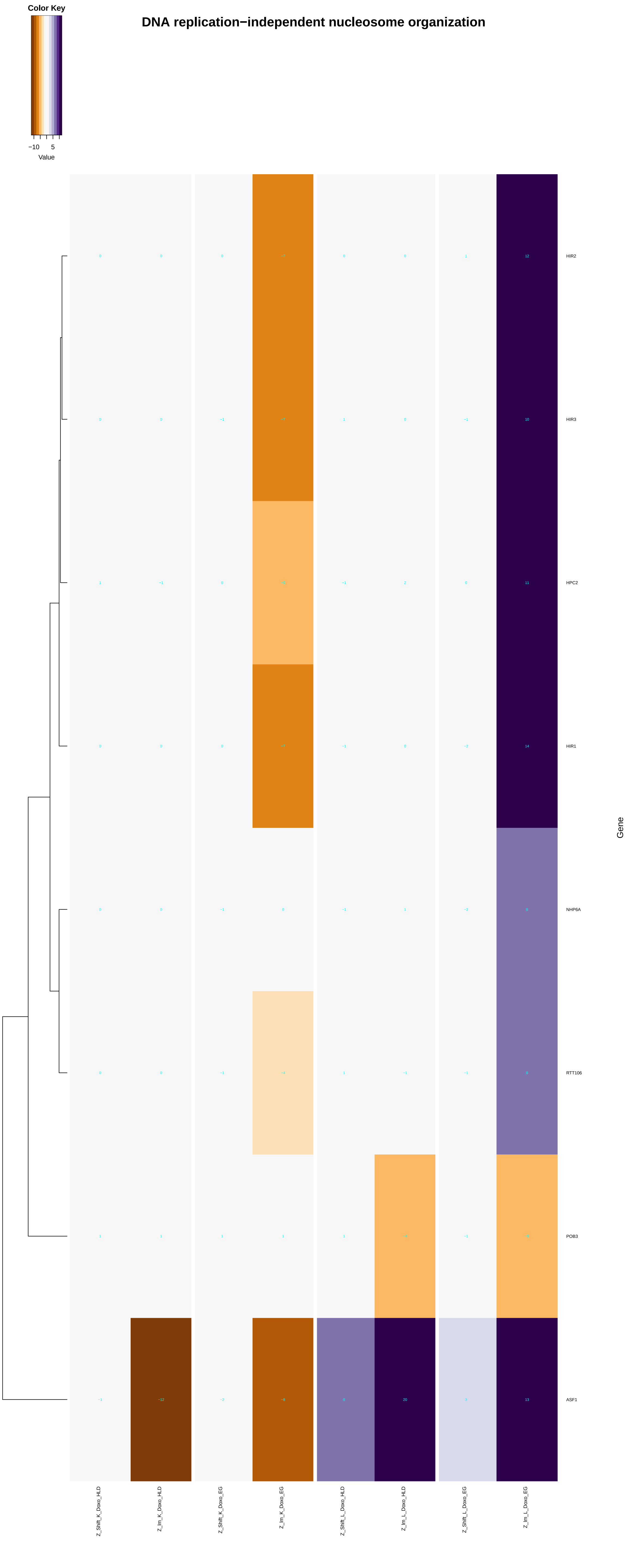
DNA replication−independent nucleosome organization
Color Key
−10
5
Value
HIR2
HIR3
HPC2
HIR1
NHP6A
RTT106
POB3
ASF1
Gene
Z_Shift_K_Doxo_HLD
Z_lm_K_Doxo_HLD
Z_Shift_K_Doxo_EG
Z_lm_K_Doxo_EG
Z_Shift_L_Doxo_HLD
Z_lm_L_Doxo_HLD
Z_Shift_L_Doxo_EG
Z_lm_L_Doxo_EG

### chromatin remodeling at centromere

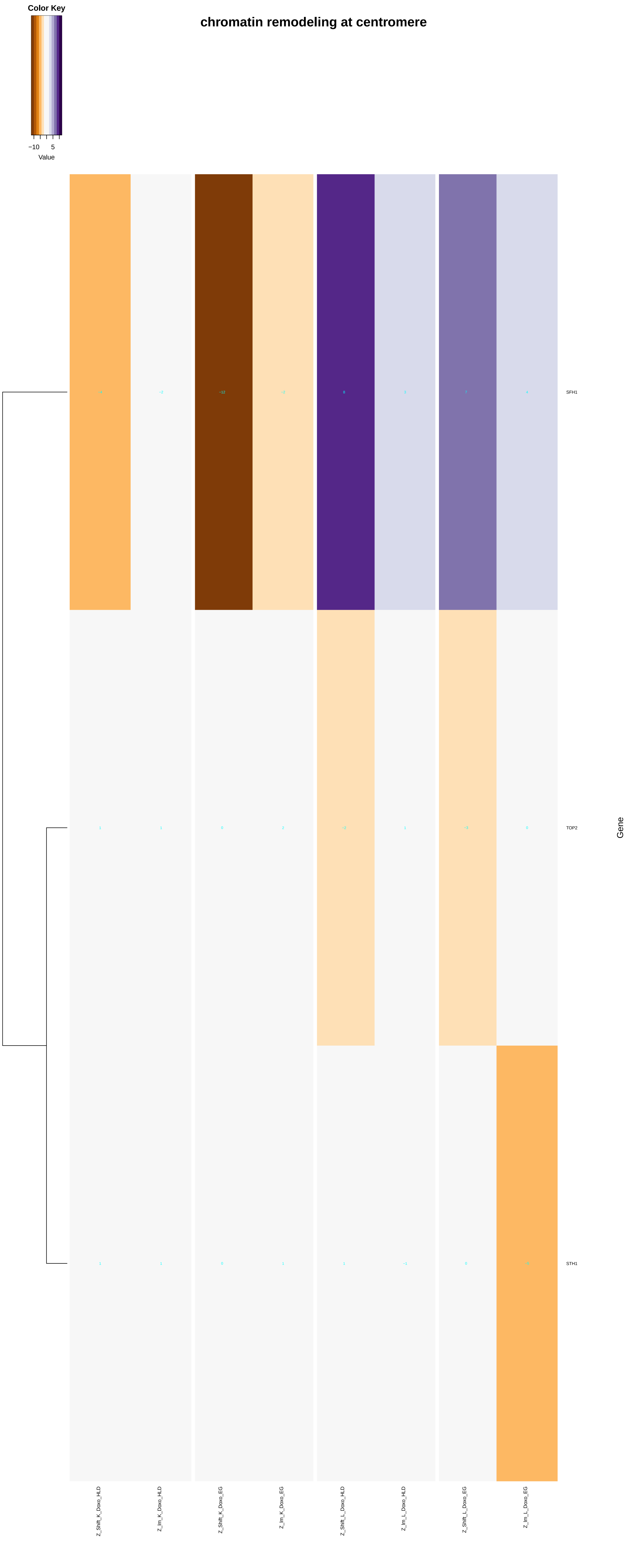

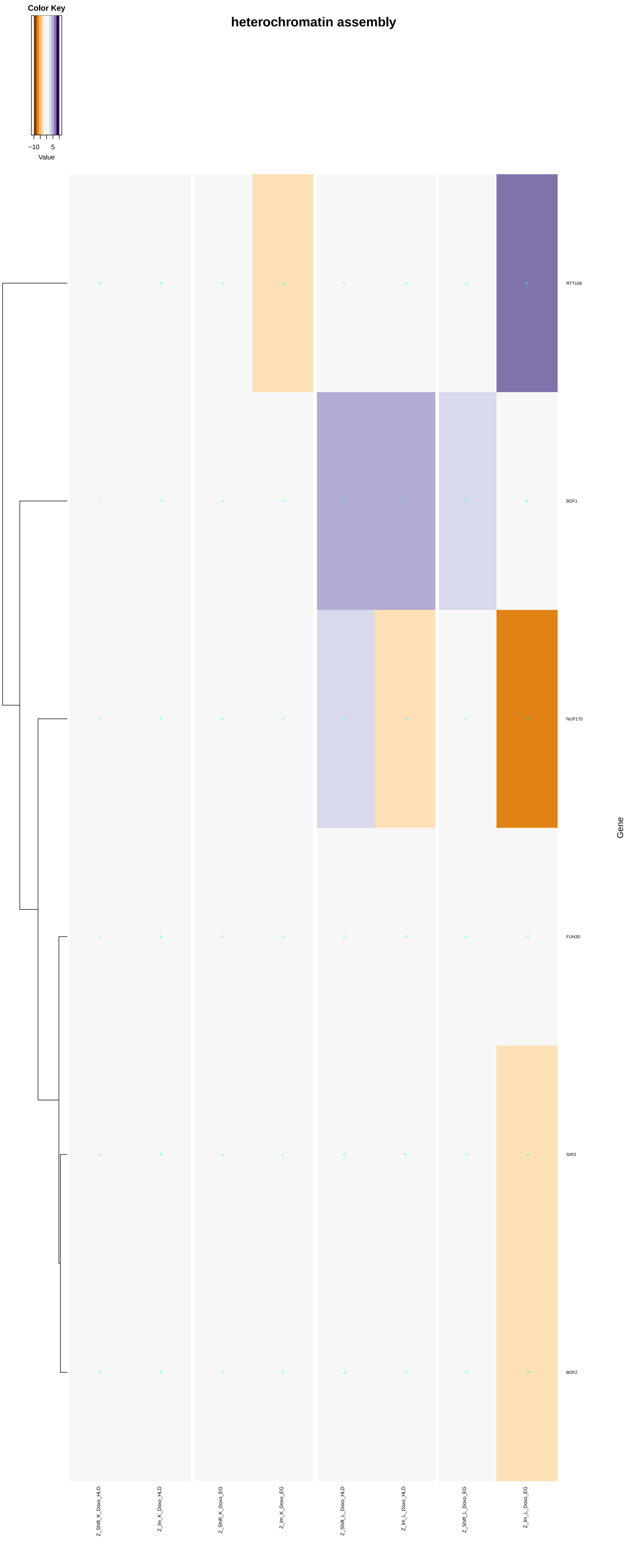

heterochromatin assembly
Color Key
−10
5
Value
RTT106
BDF1
NUP170
FUN30
SIR3
BDF2
Gene
Z_Shift_K_Doxo_HLD
Z_Im_K_Doxo_HLD
Z_Shift_K_Doxo_EG
Z_Im_K_Doxo_EG
Z_Shift_L_Doxo_HLD
Z_Im_L_Doxo_HLD
Z_Shift_L_Doxo_EG
Z_Im_L_Doxo_EG

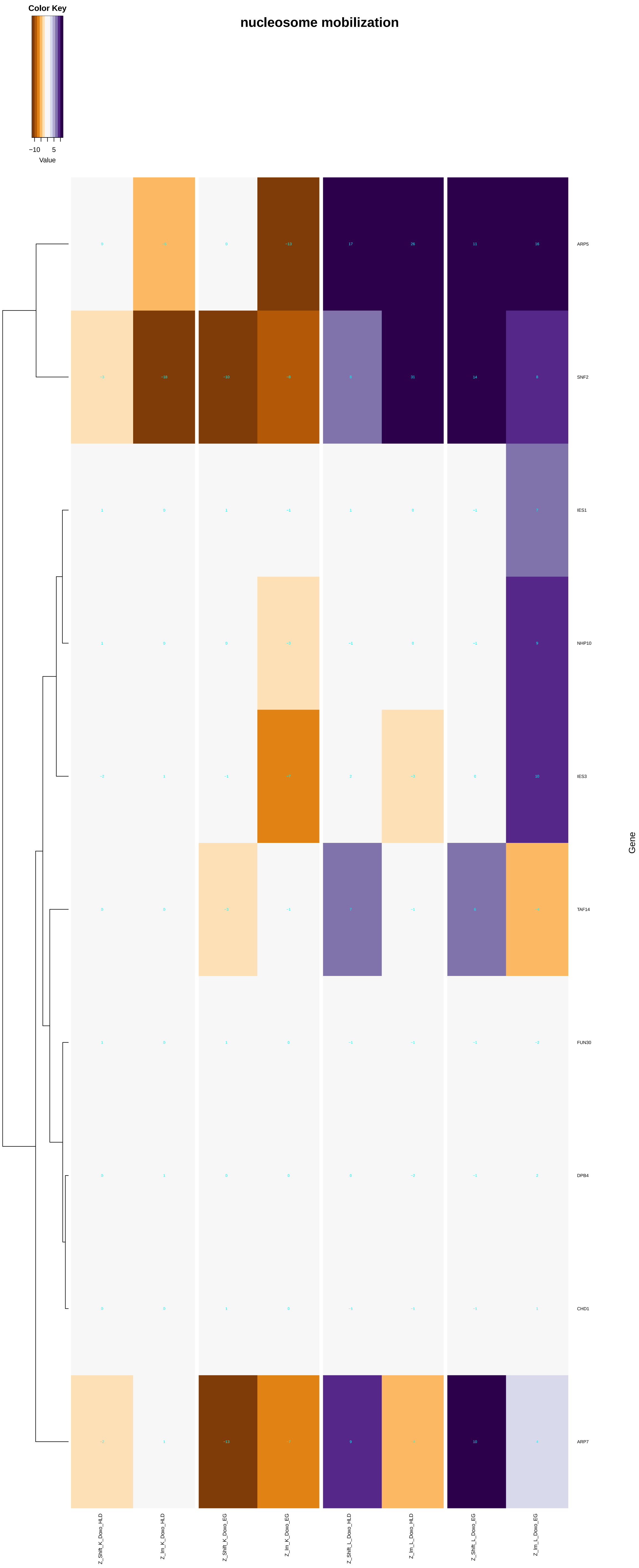

Color Key
−10 5
Value
nucleosome mobilization
ARP5
SNF2
IES1
NHP10
IES3
TAF14
FUN30
DPB4
CHD1
ARP7
Gene
Z_Shift_K_Doxo_HLD
Z_lm_K_Doxo_HLD
Z_Shift_K_Doxo_EG
Z_lm_K_Doxo_EG
Z_Shift_L_Doxo_HLD
Z_lm_L_Doxo_HLD
Z_Shift_L_Doxo_EG
Z_lm_L_Doxo_EG

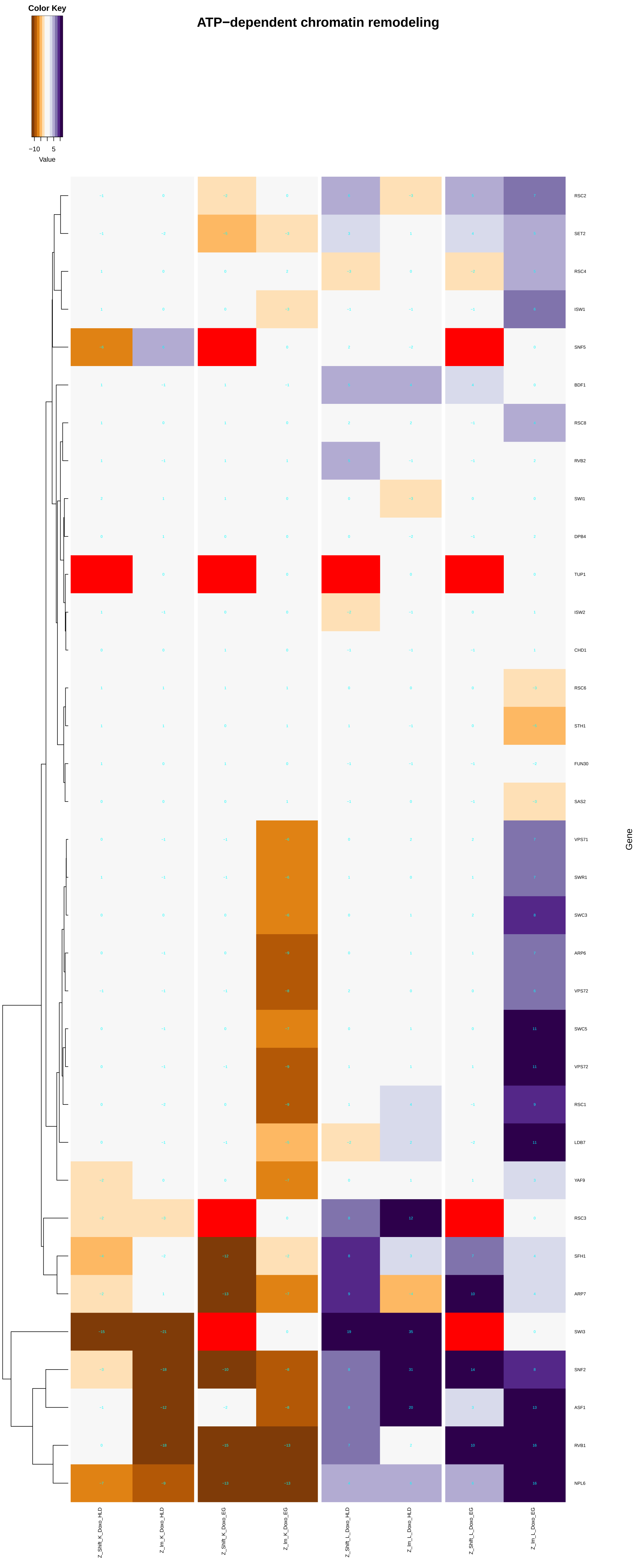

ATP−dependent chromatin remodeling
Color Key
−10
5
Value
RSC2
SET2
RSC4
ISW1
SNF5
BDF1
RSC8
RVB2
SWI1
DPB4
TUP1
ISW2
CHD1
RSC6
STH1
FUN30
SAS2
VPS71
SWR1
SWC3
ARP6
VPS72
SWC5
VPS72
RSC1
LDB7
YAF9
RSC3
SFH1
ARP7
SWI3
SNF2
ASF1
RVB1
NPL6
Gene
Z_Shift_K_Doxo_HLD
Z_Im_K_Doxo_HLD
Z_Shift_K_Doxo_EG
Z_Im_K_Doxo_EG
Z_Shift_L_Doxo_HLD
Z_Im_L_Doxo_HLD
Z_Shift_L_Doxo_EG
Z_Im_L_Doxo_EG

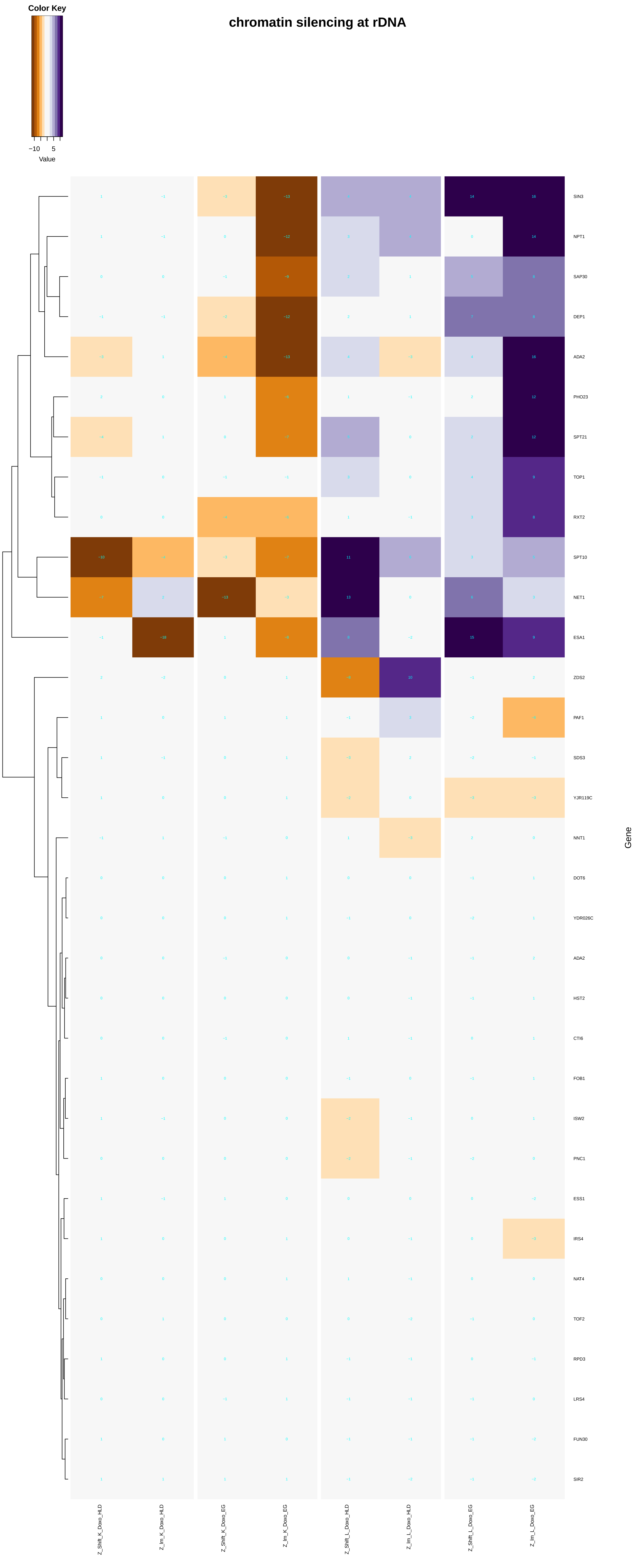
chromatin silencing at rDNA
Color Key
−10
5
Value
SIN3
NPT1
SAP30
DEP1
ADA2
PHO23
SPT21
TOP1
RXT2
SPT10
NET1
ESA1
ZDS2
PAF1
SDS3
YJR119C
NNT1
DOT6
YDR026C
ADA2
HST2
CTI6
FOB1
ISW2
PNC1
ESS1
IRS4
NAT4
TOF2
RPD3
LRS4
FUN30
SIR2
Gene
Z_Shift_K_Doxo_HLD
Z_Im_K_Doxo_HLD
Z_Shift_K_Doxo_EG
Z_Im_K_Doxo_EG
Z_Shift_L_Doxo_HLD
Z_Im_L_Doxo_HLD
Z_Shift_L_Doxo_EG
Z_Im_L_Doxo_EG

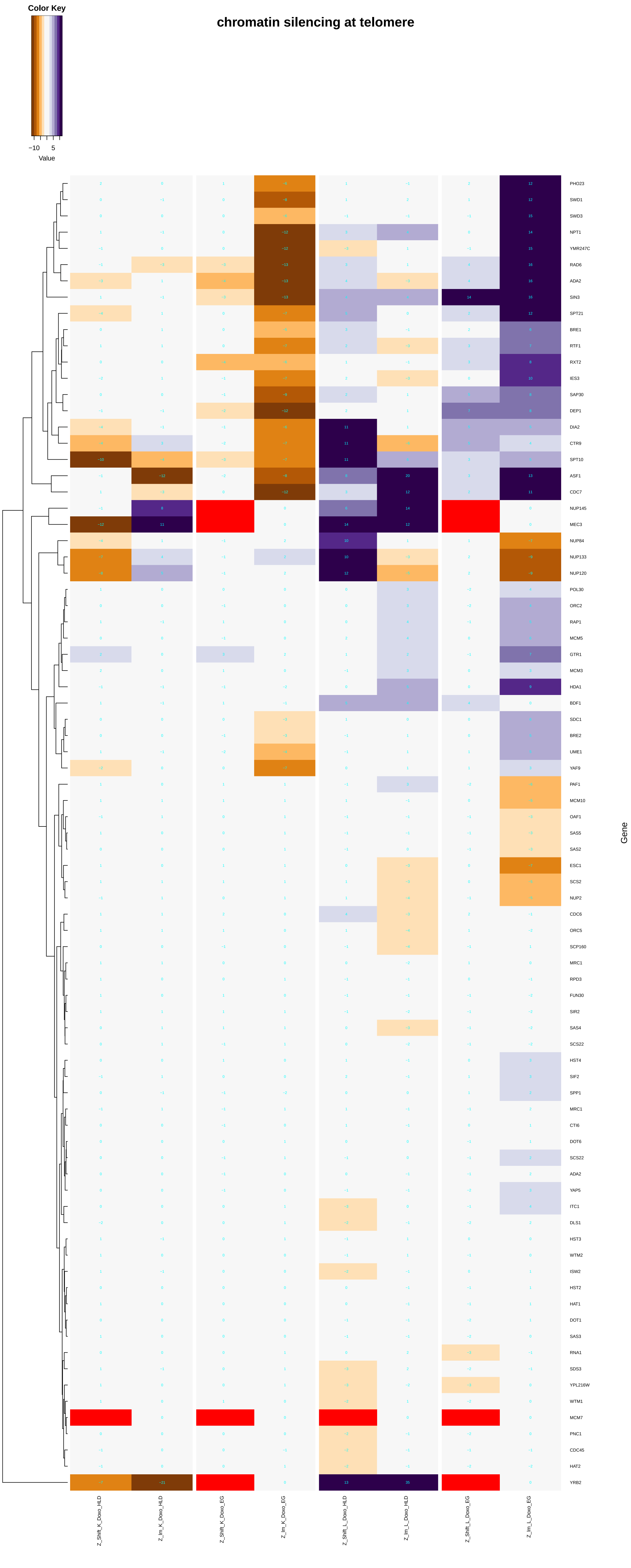
chromatin silencing at telomere
Color Key
−10
5
Value
PHO23
SWD1
SWD3
NPT1
YMR247C
RAD6
ADA2
SIN3
SPT21
BRE1
RTF1
RXT2
IES3
SAP30
DEP1
DIA2
CTR9
SPT10
ASF1
CDC7
NUP145
MEC3
NUP84
NUP133
NUP120
POL30
ORC2
RAP1
MCM5
GTR1
MCM3
HDA1
BDF1
SDC1
BRE2
UME1
YAF9
PAF1
MCM10
OAF1
SAS5
SAS2
ESC1
SCS2
NUP2
CDC6
ORC5
SCP160
MRC1
RPD3
FUN30
SIR2
SAS4
SCS22
HST4
SIF2
SPP1
MRC1
CTI6
DOT6
SCS22
ADA2
YAP5
ITC1
DLS1
HST3
WTM2
ISW2
HST2
HAT1
DOT1
SAS3
RNA1
SDS3
YPL216W
WTM1
MCM7
PNC1
CDC45
HAT2
YRB2
Gene
Z_Shift_K_Doxo_HLD
Z_Im_K_Doxo_HLD
Z_Shift_K_Doxo_EG
Z_Im_K_Doxo_EG
Z_Shift_L_Doxo_HLD
Z_Im_L_Doxo_HLD
Z_Shift_L_Doxo_EG
Z_Im_L_Doxo_EG

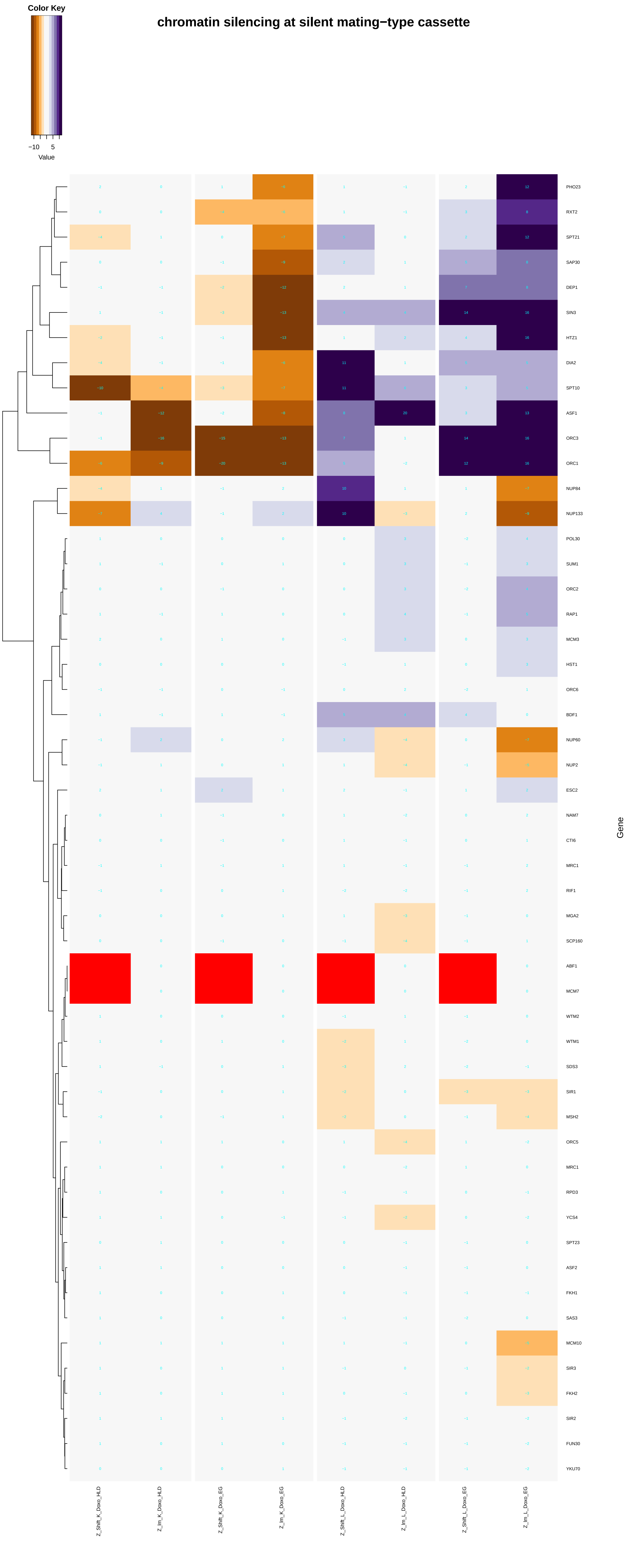
Color Key
−10
5
Value
chromatin silencing at silent mating−type cassette
PHO23
RXT2
SPT21
SAP30
DEP1
SIN3
HTZ1
DIA2
SPT10
ASF1
ORC3
ORC1
NUP84
NUP133
POL30
SUM1
ORC2
RAP1
MCM3
HST1
ORC6
BDF1
NUP60
NUP2
ESC2
NAM7
CTI6
MRC1
RIF1
MGA2
SCP160
ABF1
MCM7
WTM2
WTM1
SDS3
SIR1
MSH2
ORC5
MRC1
RPD3
YCS4
SPT23
ASF2
FKH1
SAS3
MCM10
SIR3
FKH2
SIR2
FUN30
YKU70
Gene
Z_Shift_K_Doxo_HLD
Z_Im_K_Doxo_HLD
Z_Shift_K_Doxo_EG
Z_Im_K_Doxo_EG
Z_Shift_L_Doxo_HLD
Z_Im_L_Doxo_HLD
Z_Shift_L_Doxo_EG
Z_Im_L_Doxo_EG

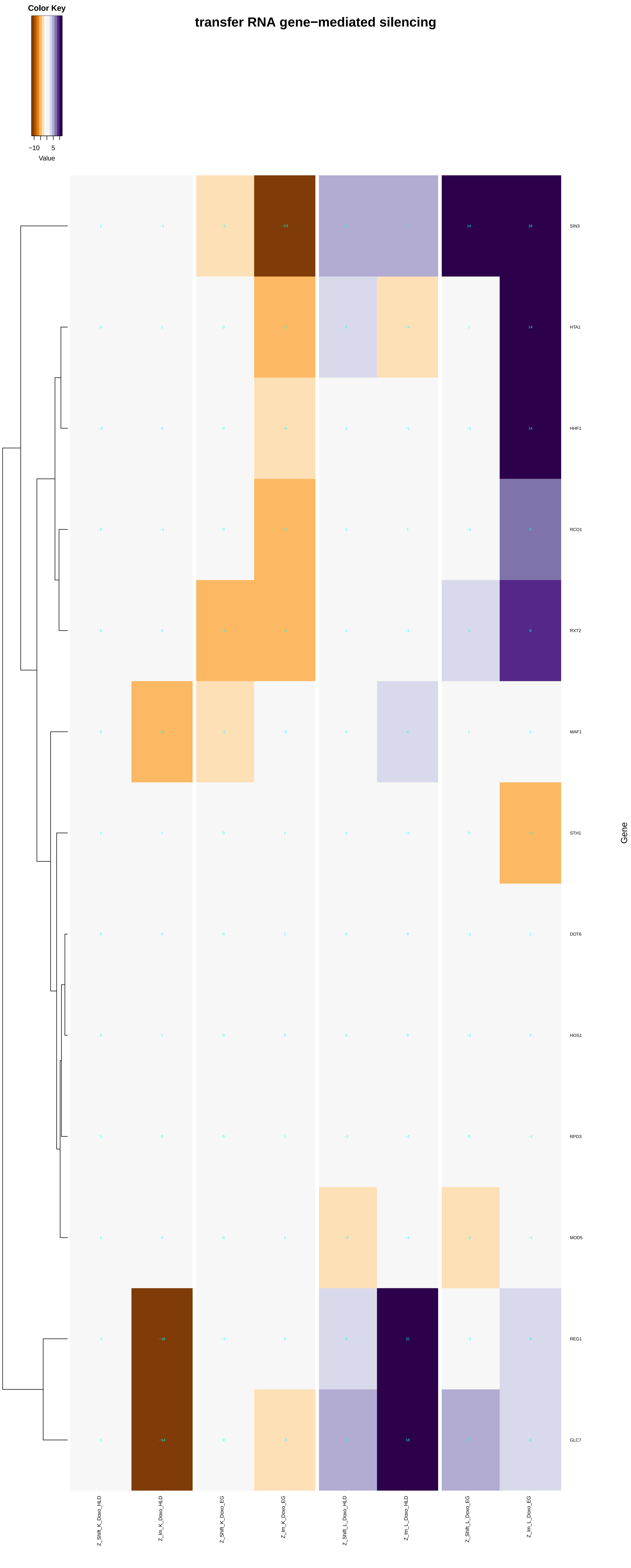
Color Key
−10 5
Value
transfer RNA gene−mediated silencing
SIN3
HTA1
HHF1
RCO1
RXT2
MAF1
STH1
DOT6
HOS1
RPD3
MOD5
REG1
GLC7
Gene
Z_Shift_K_Doxo_HLD
Z_Im_K_Doxo_HLD
Z_Shift_K_Doxo_EG
Z_Im_K_Doxo_EG
Z_Shift_L_Doxo_HLD
Z_Im_L_Doxo_HLD
Z_Shift_L_Doxo_EG
Z_Im_L_Doxo_EG

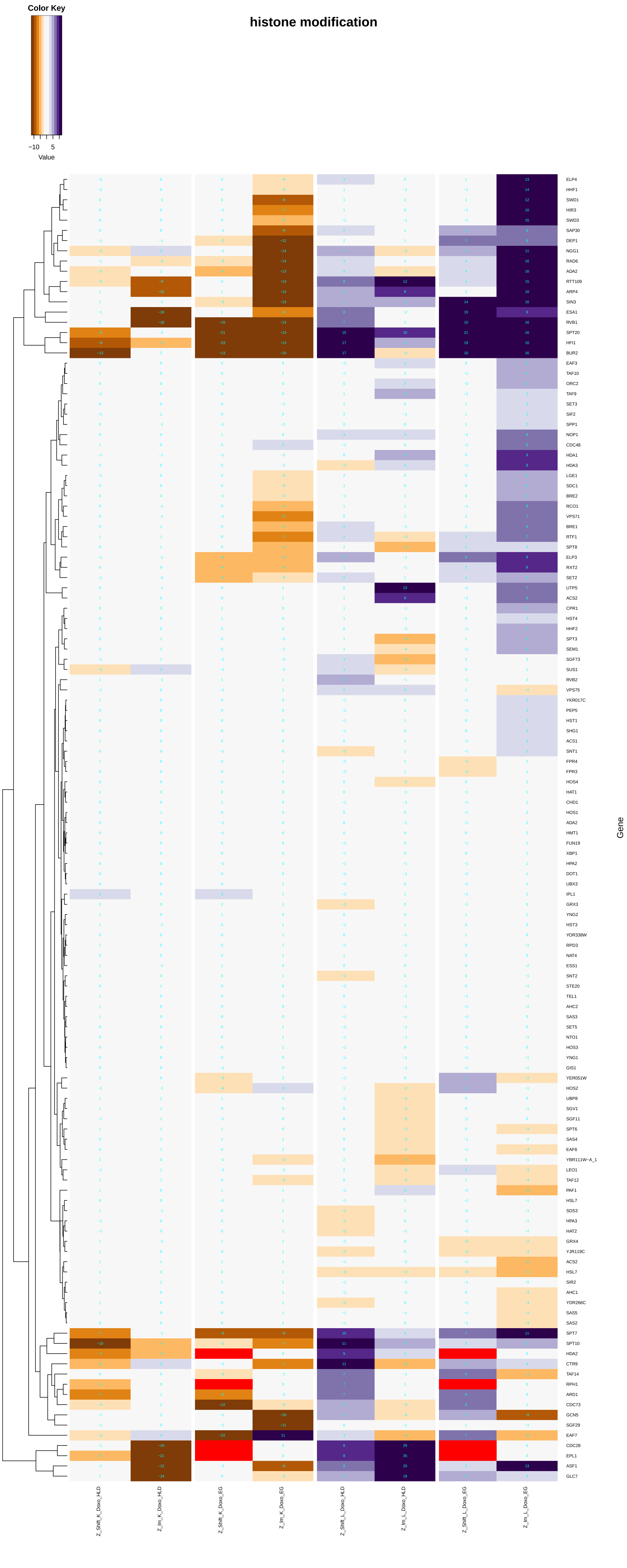
histone modification
Color Key
−10
5
Value
ELP4
HHF1
SWD1
HIR3
SWD3
SAP30
DEP1
NGG1
RAD6
ADA2
RTT109
ARP4
SIN3
ESA1
RVB1
SPT20
HFI1
BUR2
EAF3
TAF10
ORC2
TAF9
SET3
SIF2
SPP1
NOP1
CDC48
HDA1
HDA3
LGE1
SDC1
BRE2
RCO1
VPS71
BRE1
RTF1
SPT8
ELP3
RXT2
SET2
UTP5
ACS2
CPR1
HST4
HHF2
SPT3
SEM1
SGF73
SUS1
RVB2
VPS75
YKR017C
PEP5
HST1
SHG1
ACS1
SNT1
FPR4
FPR3
HOS4
HAT1
CHD1
HOS1
ADA2
HMT1
FUN19
XBP1
HPA2
DOT1
UBX3
IPL1
GRX3
YNG2
HST3
YOR338W
RPD3
NAT4
ESS1
SNT2
STE20
TEL1
AHC2
SAS3
SET5
NTO1
HOS3
YNG1
GIS1
YER051W
HOS2
UBP8
SGV1
SGF11
SPT6
SAS4
EAF6
YBR111W−A_1
LEO1
TAF12
PAF1
HSL7
SDS3
HPA3
HAT2
GRX4
YJR119C
ACS2
HSL7
SIR2
AHC1
YDR266C
SAS5
SAS2
SPT7
SPT10
HDA2
CTR9
TAF14
RPH1
ARD1
CDC73
GCN5
SGF29
EAF7
CDC28
EPL1
ASF1
GLC7
Gene
Z_Shift_K_Doxo_HLD
Z_Im_K_Doxo_HLD
Z_Shift_K_Doxo_EG
Z_Im_K_Doxo_EG
Z_Shift_L_Doxo_HLD
Z_Im_L_Doxo_HLD
Z_Shift_L_Doxo_EG
Z_Im_L_Doxo_EG

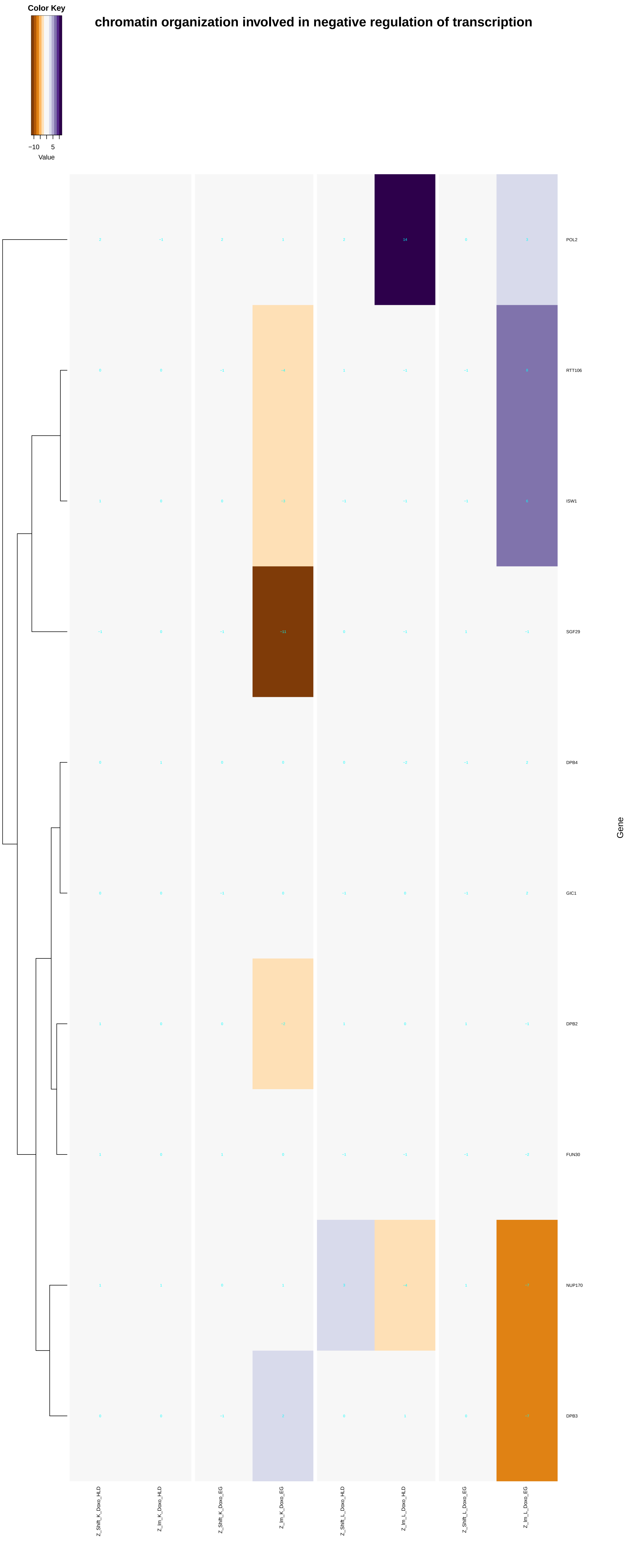

Color Key
−10 5
Value
chromatin organization involved in negative regulation of transcription
POL2
RTT106
ISW1
SGF29
DPB4
GIC1
DPB2
FUN30
NUP170
DPB3
Gene
Z_Shift_K_Doxo_HLD
Z_Im_K_Doxo_HLD
Z_Shift_K_Doxo_EG
Z_Im_K_Doxo_EG
Z_Shift_L_Doxo_HLD
Z_Im_L_Doxo_HLD
Z_Shift_L_Doxo_EG
Z_Im_L_Doxo_EG

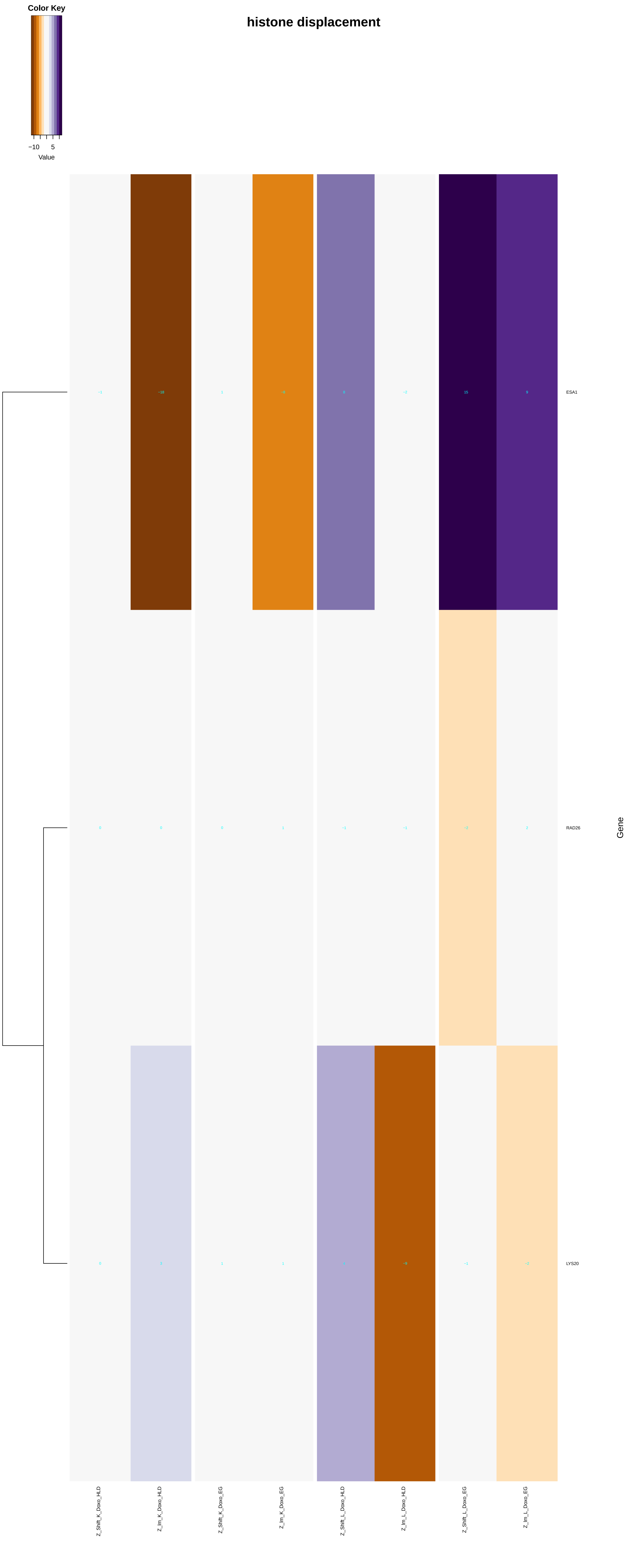
histone displacement
Color Key
−10
5
Value
−1
−18
1
−8
8
−2
15
9
ESA1
0
0
0
1
−1
−1
−2
2
RAD26
0
3
1
1
4
−9
−1
−2
LYS20
Gene
Z_Shift_K_Doxo_HLD
Z_Im_K_Doxo_HLD
Z_Shift_K_Doxo_EG
Z_Im_K_Doxo_EG
Z_Shift_L_Doxo_HLD
Z_Im_L_Doxo_HLD
Z_Shift_L_Doxo_EG
Z_Im_L_Doxo_EG

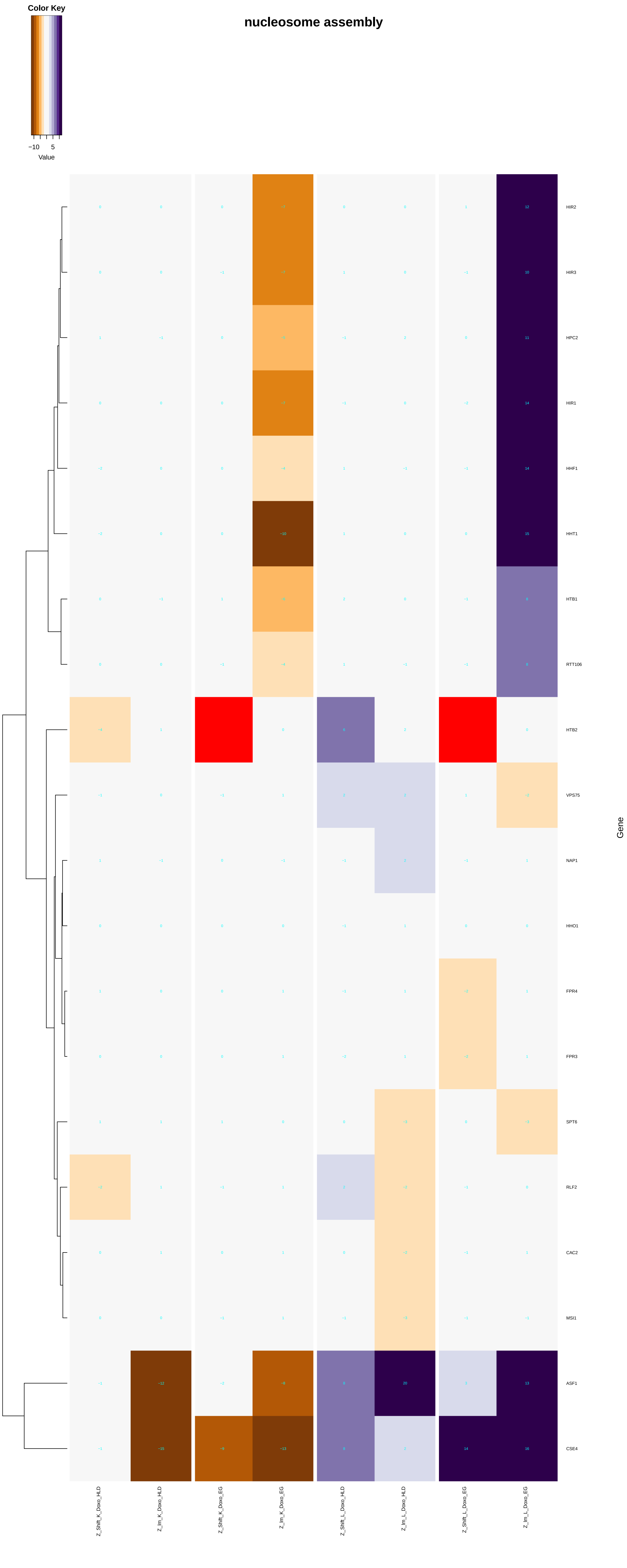

Color Key
−10
5
Value
nucleosome assembly
HIR2
HIR3
HPC2
HIR1
HHF1
HHT1
HTB1
RTT106
HTB2
VPS75
NAP1
HHO1
FPR4
FPR3
SPT6
RLF2
CAC2
MSI1
ASF1
CSE4
Gene
Z_Shift_K_Doxo_HLD
Z_lm_K_Doxo_HLD
Z_Shift_K_Doxo_EG
Z_lm_K_Doxo_EG
Z_Shift_L_Doxo_HLD
Z_lm_L_Doxo_HLD
Z_Shift_L_Doxo_EG
Z_lm_L_Doxo_EG

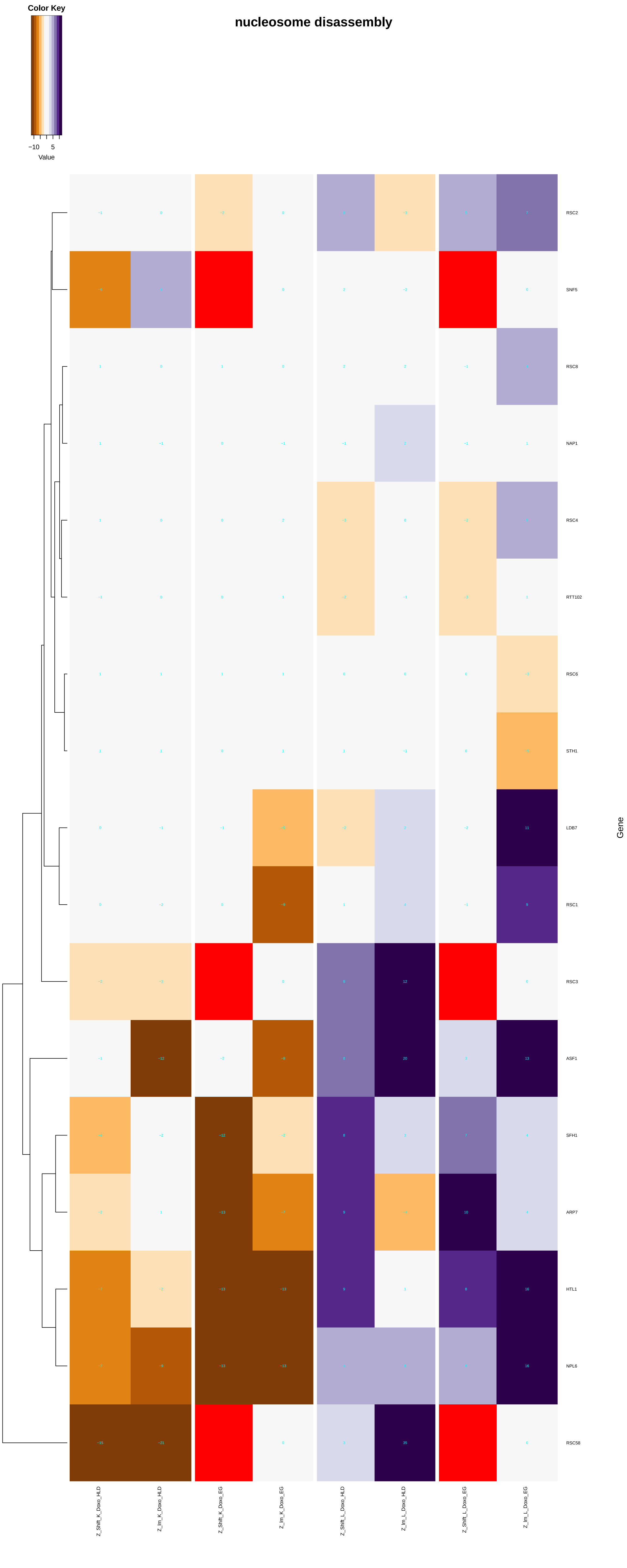

nucleosome disassembly
Color Key
−10
5
Value
RSC2
SNF5
RSC8
NAP1
RSC4
RTT102
RSC6
STH1
LDB7
RSC1
RSC3
ASF1
SFH1
ARP7
HTL1
NPL6
RSC58
Gene
Z_Shift_K_Doxo_HLD
Z_lm_K_Doxo_HLD
Z_Shift_K_Doxo_EG
Z_lm_K_Doxo_EG
Z_Shift_L_Doxo_HLD
Z_lm_L_Doxo_HLD
Z_Shift_L_Doxo_EG
Z_lm_L_Doxo_EG

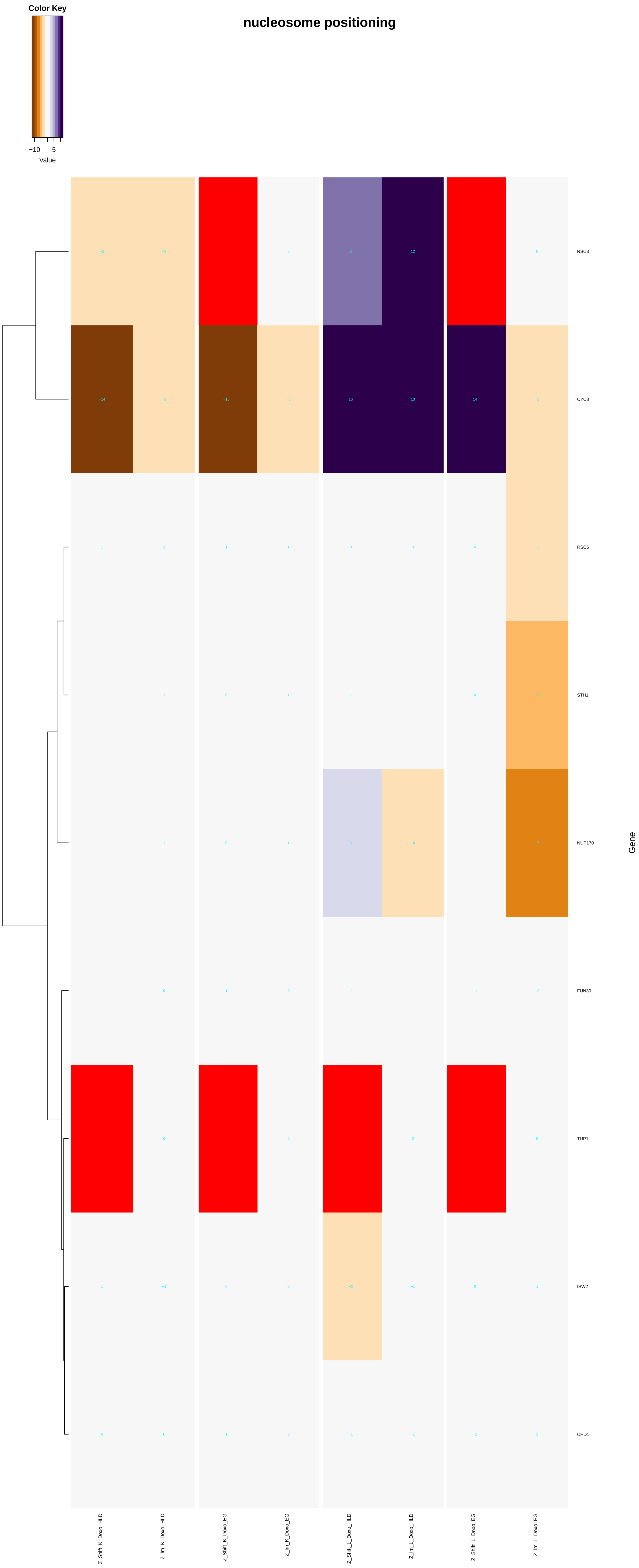
nucleosome positioning
Color Key
−10
5
Value
RSC3
CYC8
RSC6
STH1
NUP170
FUN30
TUP1
ISW2
CHD1
Gene
Z_Shift_K_Doxo_HLD
Z_Im_K_Doxo_HLD
Z_Shift_K_Doxo_EG
Z_Im_K_Doxo_EG
Z_Shift_L_Doxo_HLD
Z_Im_L_Doxo_HLD
Z_Shift_L_Doxo_EG
Z_Im_L_Doxo_EG

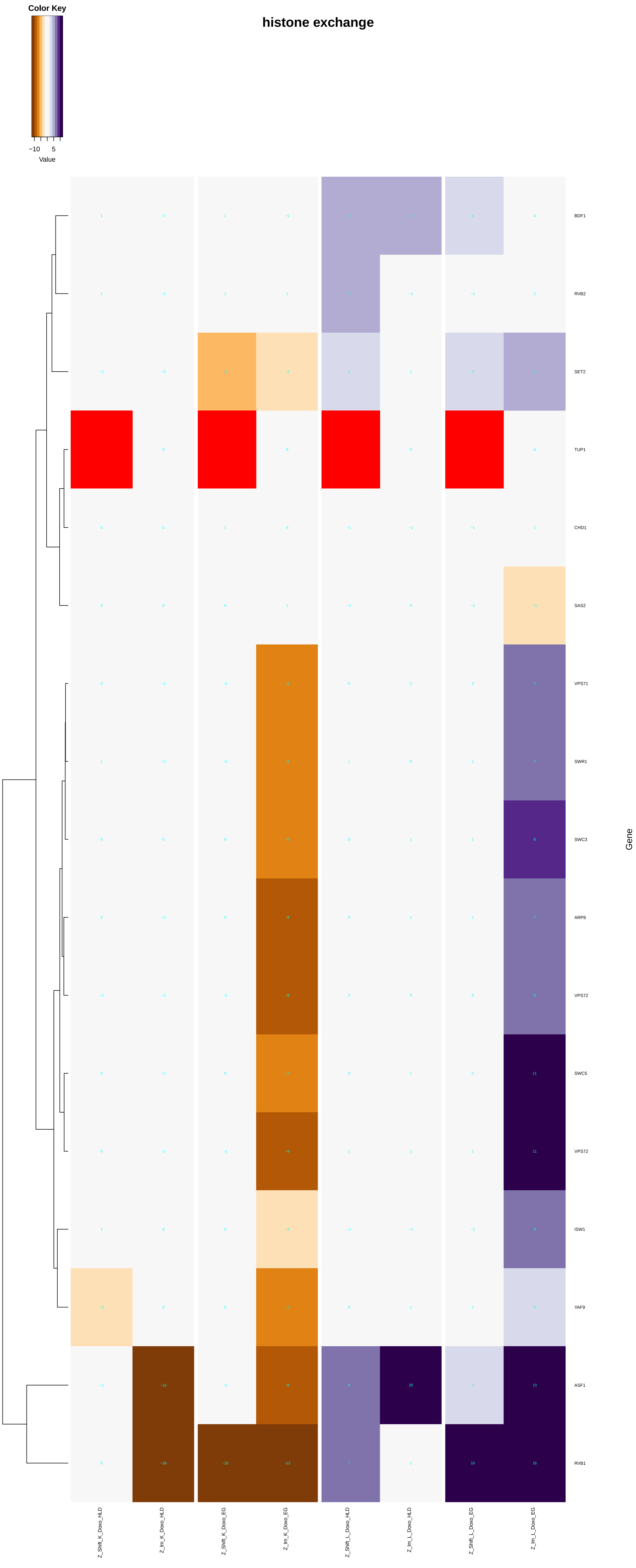

Color Key
−10
5
Value
histone exchange
BDF1
RVB2
SET2
TUP1
CHD1
SAS2
VPS71
SWR1
SWC3
ARP6
VPS72
SWC5
VPS72
ISW1
YAF9
ASF1
RVB1
Gene
Z_Shift_K_Doxo_HLD
Z_Im_K_Doxo_HLD
Z_Shift_K_Doxo_EG
Z_Im_K_Doxo_EG
Z_Shift_L_Doxo_HLD
Z_Im_L_Doxo_HLD
Z_Shift_L_Doxo_EG
Z_Im_L_Doxo_EG

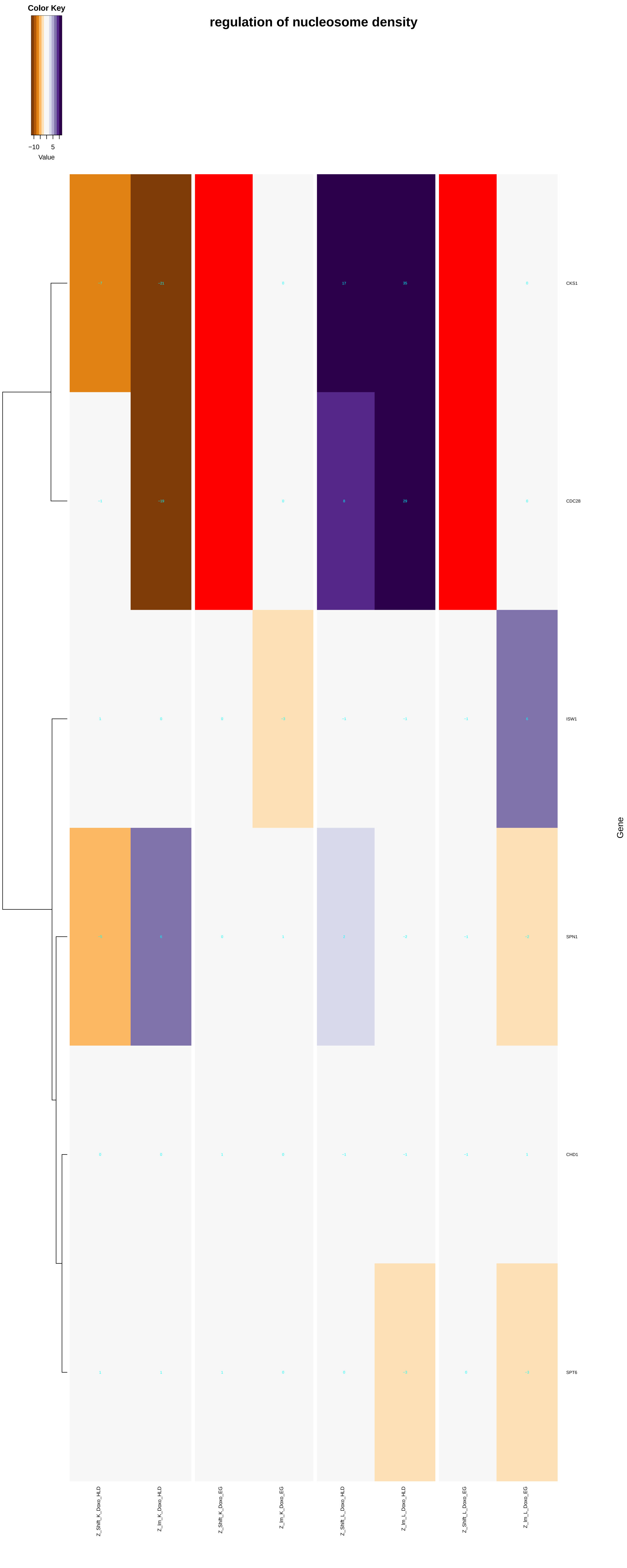
regulation of nucleosome density
Color Key
−10
5
Value
CKS1
CDC28
ISW1
SPN1
CHD1
SPT6
Gene
Z_Shift_K_Doxo_HLD
Z_Im_K_Doxo_HLD
Z_Shift_K_Doxo_EG
Z_Im_K_Doxo_EG
Z_Shift_L_Doxo_HLD
Z_Im_L_Doxo_HLD
Z_Shift_L_Doxo_EG
Z_Im_L_Doxo_EG

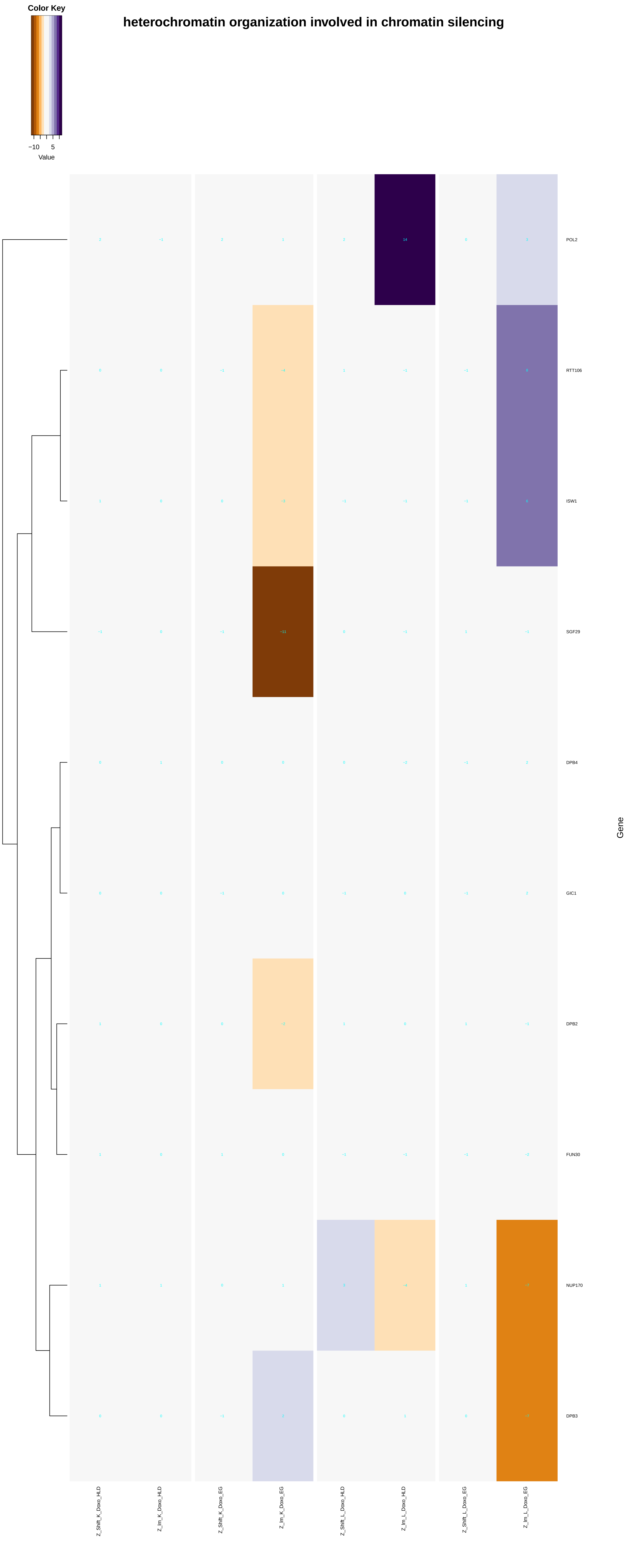

heterochromatin organization involved in chromatin silencing
Color Key
−10
5
Value
POL2
RTT106
ISW1
SGF29
DPB4
GIC1
DPB2
FUN30
NUP170
DPB3
Gene
Z_Shift_K_Doxo_HLD
Z_lm_K_Doxo_HLD
Z_Shift_K_Doxo_EG
Z_lm_K_Doxo_EG
Z_Shift_L_Doxo_HLD
Z_lm_L_Doxo_HLD
Z_Shift_L_Doxo_EG
Z_lm_L_Doxo_EG

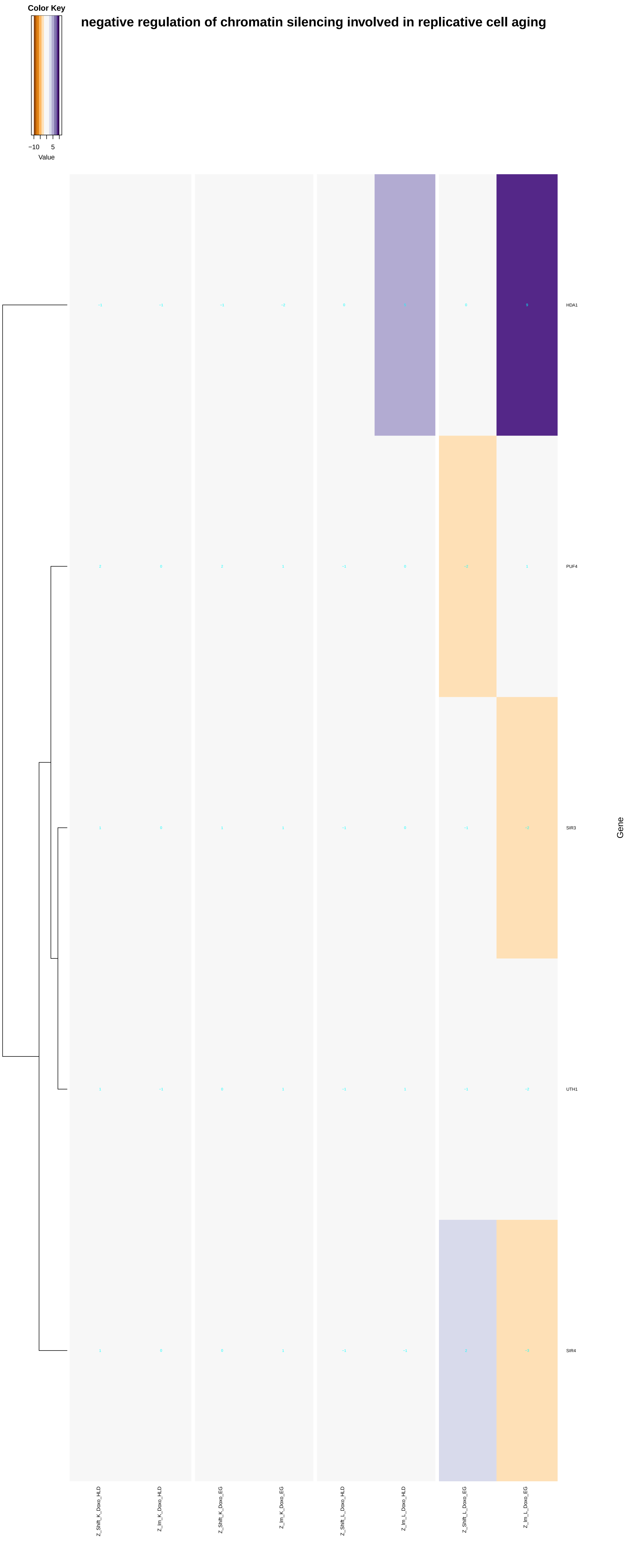
Color Key
−10
5
Value
negative regulation of chromatin silencing involved in replicative cell aging
HDA1
PUF4
SIR3
UTH1
SIR4
Gene
Z_Shift_K_Doxo_HLD
Z_lm_K_Doxo_HLD
Z_Shift_K_Doxo_EG
Z_lm_K_Doxo_EG
Z_Shift_L_Doxo_HLD
Z_lm_L_Doxo_HLD
Z_Shift_L_Doxo_EG
Z_lm_L_Doxo_EG

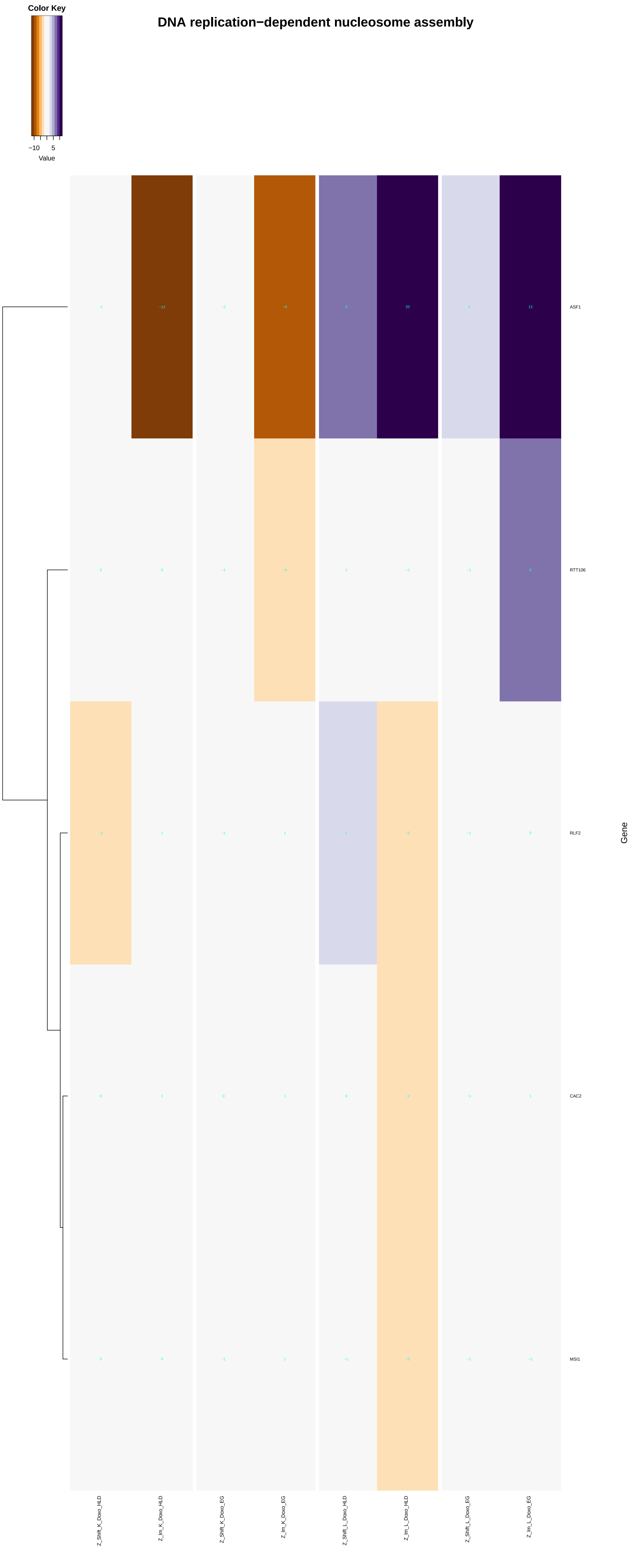

DNA replication−dependent nucleosome assembly
Color Key
−10
5
Value
−1
−12
−2
−8
8
20
3
13
ASF1
0
0
−1
−4
1
−1
−1
8
RTT106
−2
1
−1
1
2
−2
−1
0
RLF2
0
1
0
1
0
−2
−1
1
CAC2
0
0
−1
1
−1
−3
−1
−1
MSI1
Gene
Z_Shift_K_Doxo_HLD
Z_Im_K_Doxo_HLD
Z_Shift_K_Doxo_EG
Z_Im_K_Doxo_EG
Z_Shift_L_Doxo_HLD
Z_Im_L_Doxo_HLD
Z_Shift_L_Doxo_EG
Z_Im_L_Doxo_EG

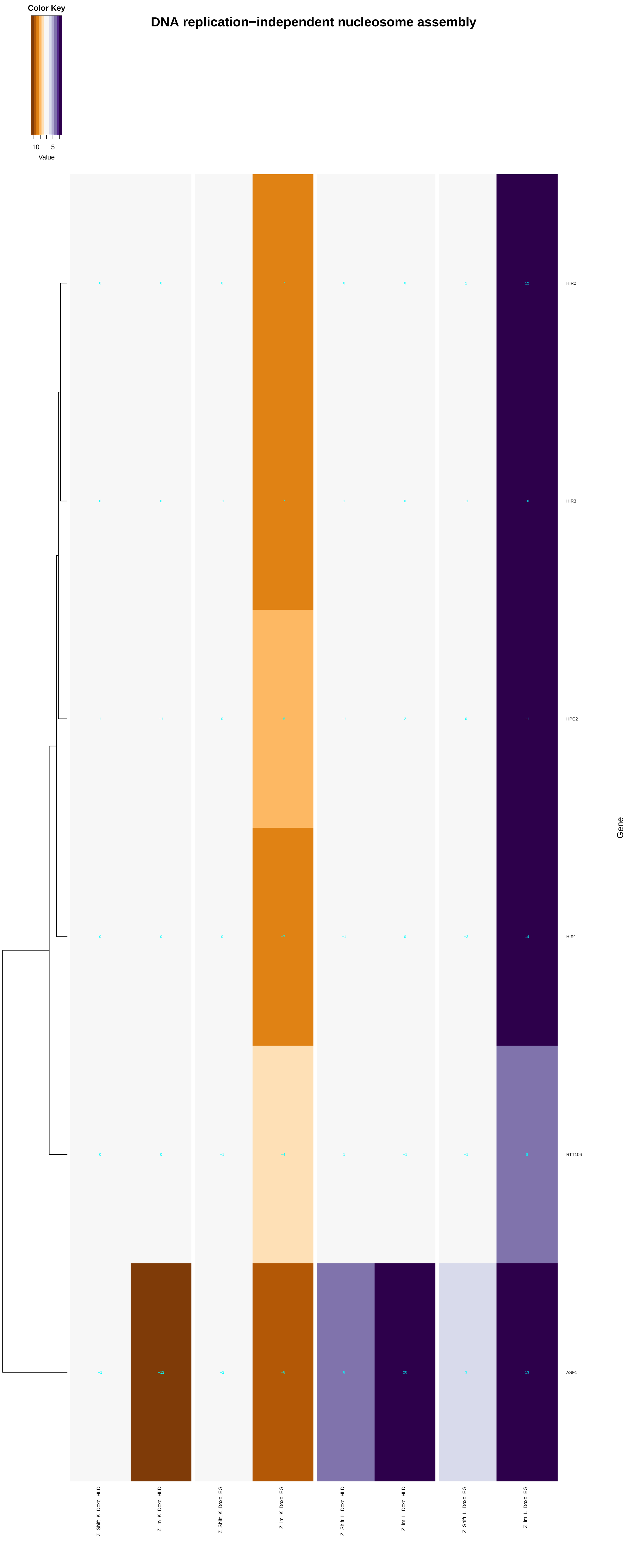

DNA replication−independent nucleosome assembly
Color Key
−10
5
Value
HIR2
HIR3
HPC2
HIR1
RTT106
ASF1
Gene
Z_Shift_K_Doxo_HLD
Z_Im_K_Doxo_HLD
Z_Shift_K_Doxo_EG
Z_Im_K_Doxo_EG
Z_Shift_L_Doxo_HLD
Z_Im_L_Doxo_HLD
Z_Shift_L_Doxo_EG
Z_Im_L_Doxo_EG

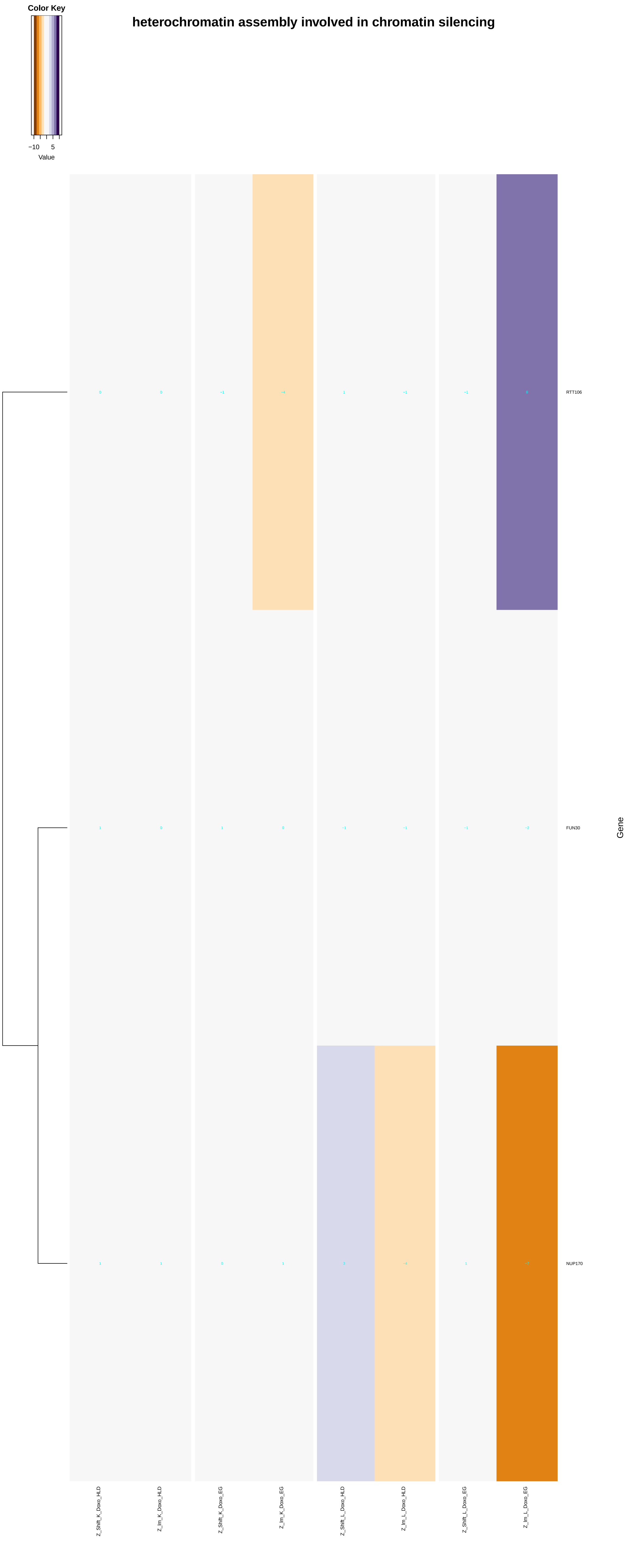
heterochromatin assembly involved in chromatin silencing
Color Key
−10
5
Value
RTT106
FUN30
NUP170
Gene
Z_Shift_K_Doxo_HLD
Z_Im_K_Doxo_HLD
Z_Shift_K_Doxo_EG
Z_Im_K_Doxo_EG
Z_Shift_L_Doxo_HLD
Z_Im_L_Doxo_HLD
Z_Shift_L_Doxo_EG
Z_Im_L_Doxo_EG

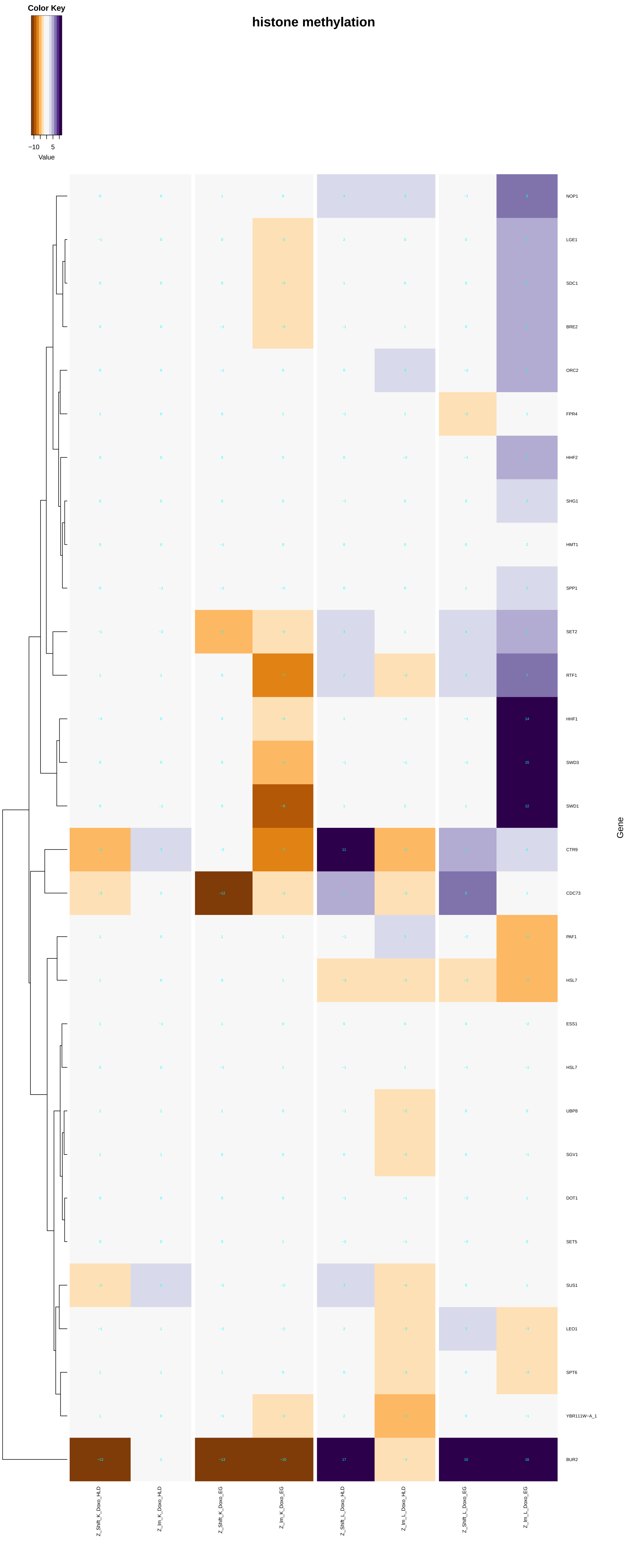

Color Key
−10
5
Value
histone methylation
NOP1
LGE1
SDC1
BRE2
ORC2
FPR4
HHF2
SHG1
HMT1
SPP1
SET2
RTF1
HHF1
SWD3
SWD1
CTR9
CDC73
PAF1
HSL7
ESS1
HSL7
UBP8
SGV1
DOT1
SET5
SUS1
LEO1
SPT6
YBR111W−A_1
BUR2
Gene
Z_Shift_K_Doxo_HLD
Z_lm_K_Doxo_HLD
Z_Shift_K_Doxo_EG
Z_lm_K_Doxo_EG
Z_Shift_L_Doxo_HLD
Z_lm_L_Doxo_HLD
Z_Shift_L_Doxo_EG
Z_lm_L_Doxo_EG

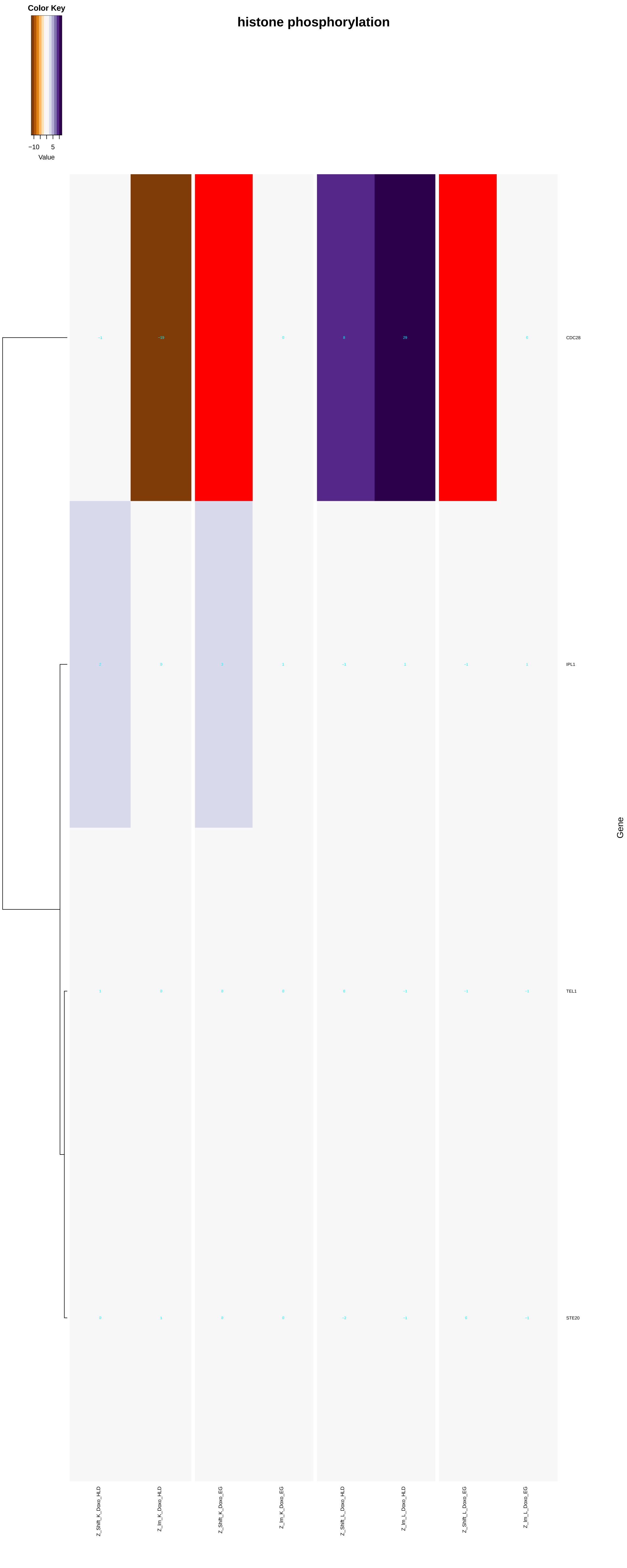
histone phosphorylation
Color Key
−10
5
Value
CDC28
IPL1
TEL1
STE20
Gene
Z_Shift_K_Doxo_HLD
Z_Im_K_Doxo_HLD
Z_Shift_K_Doxo_EG
Z_Im_K_Doxo_EG
Z_Shift_L_Doxo_HLD
Z_Im_L_Doxo_HLD
Z_Shift_L_Doxo_EG
Z_Im_L_Doxo_EG

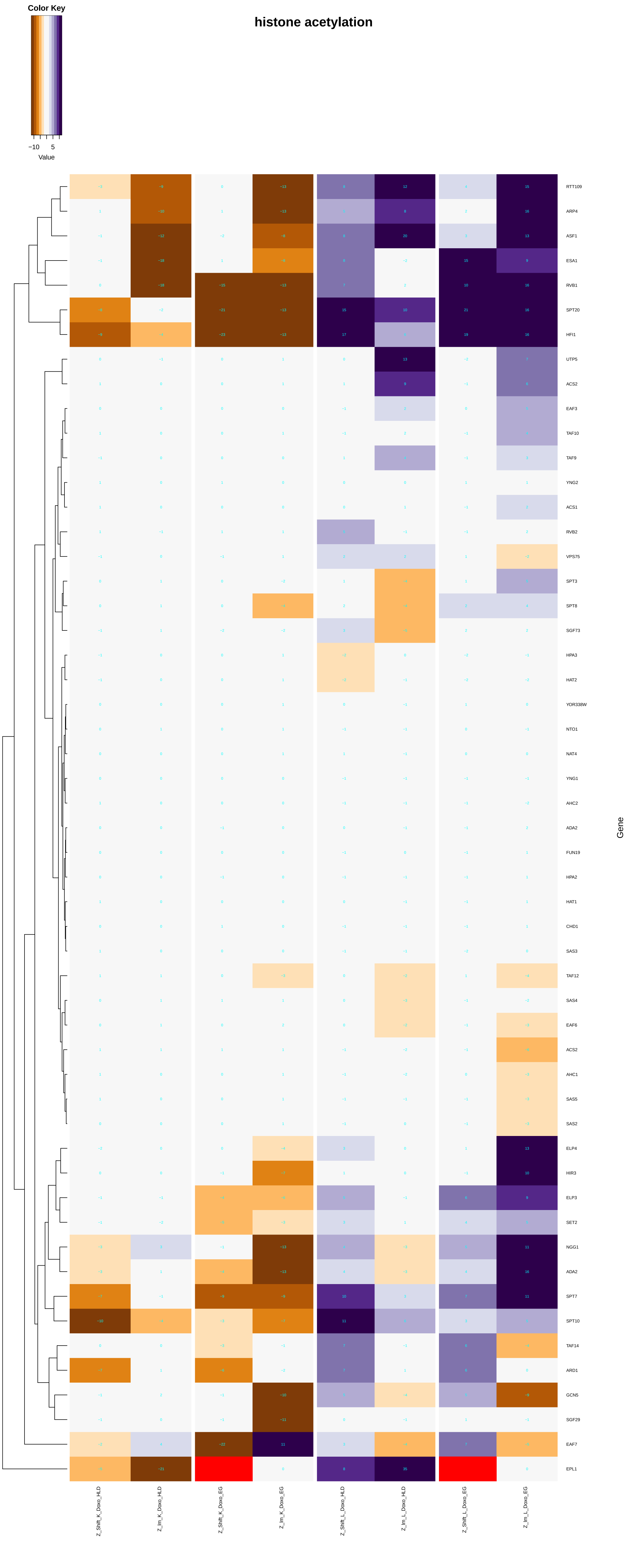

Color Key
−10
5
Value
histone acetylation
RTT109
ARP4
ASF1
ESA1
RVB1
SPT20
HFI1
UTP5
ACS2
EAF3
TAF10
TAF9
YNG2
ACS1
RVB2
VPS75
SPT3
SPT8
SGF73
HPA3
HAT2
YOR338W
NTO1
NAT4
YNG1
AHC2
ADA2
FUN19
HPA2
HAT1
CHD1
SAS3
TAF12
SAS4
EAF6
ACS2
AHC1
SAS5
SAS2
ELP4
HIR3
ELP3
SET2
NGG1
ADA2
SPT7
SPT10
TAF14
ARD1
GCN5
SGF29
EAF7
EPL1
Gene
Z_Shift_K_Doxo_HLD
Z_Im_K_Doxo_HLD
Z_Shift_K_Doxo_EG
Z_Im_K_Doxo_EG
Z_Shift_L_Doxo_HLD
Z_Im_L_Doxo_HLD
Z_Shift_L_Doxo_EG
Z_Im_L_Doxo_EG

### histone ubiquitination

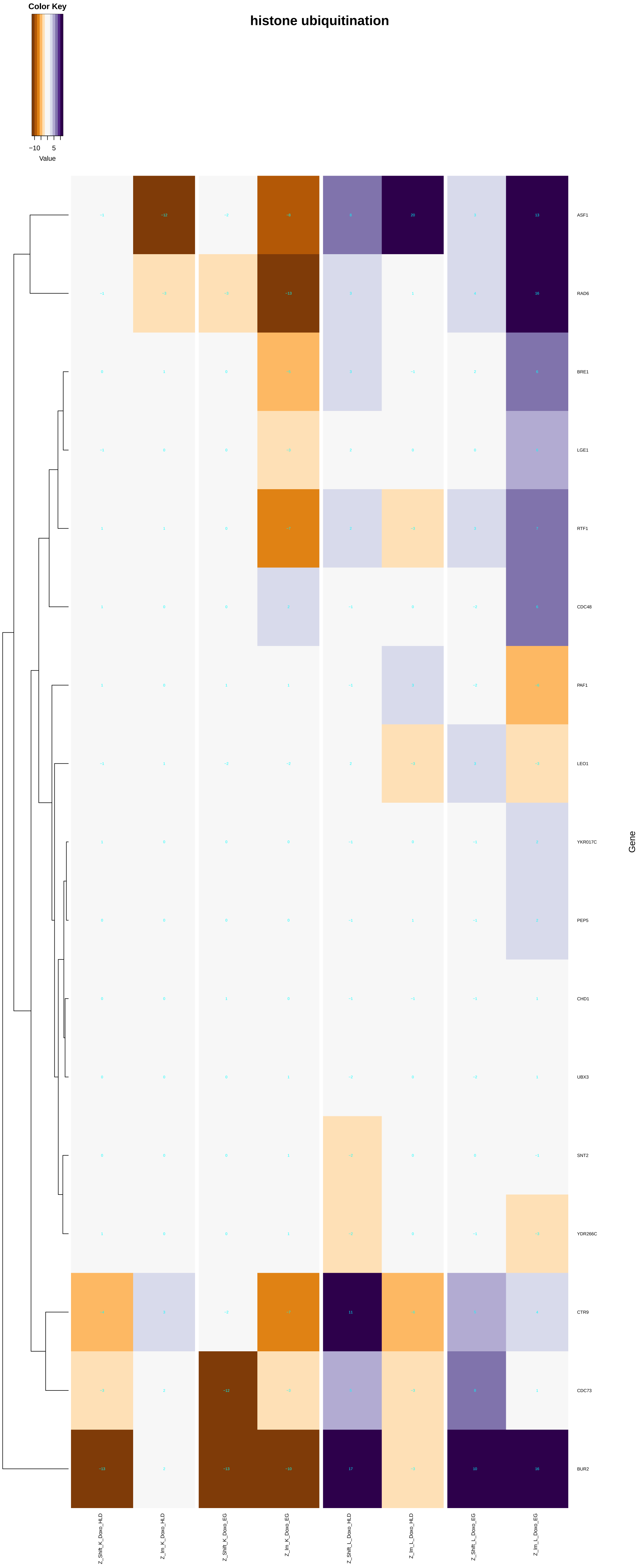

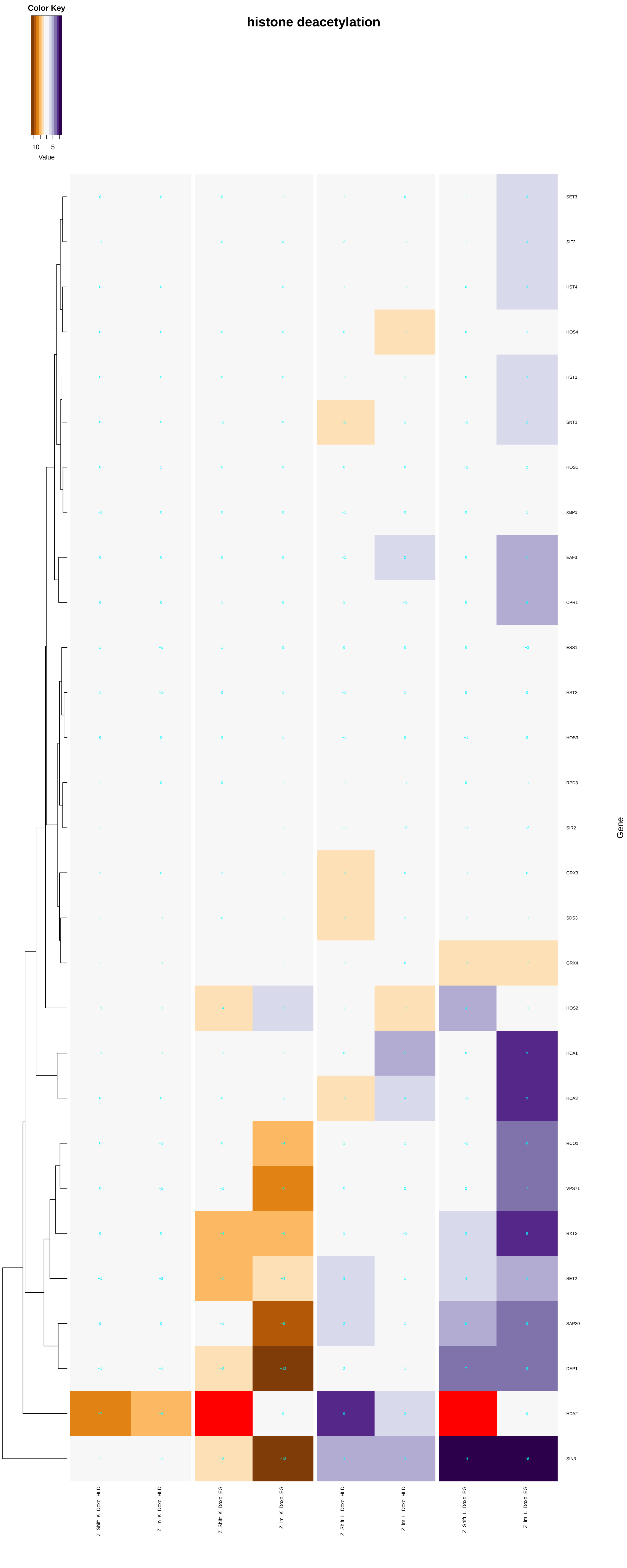

histone deacetylation
Color Key
−10
5
Value
SET3
SIF2
HST4
HOS4
HST1
SNT1
HOS1
XBP1
EAF3
CPR1
ESS1
HST3
HOS3
RPD3
SIR2
GRX3
SDS3
GRX4
HOS2
HDA1
HDA3
RCO1
VPS71
RXT2
SET2
SAP30
DEP1
HDA2
SIN3
Gene
Z_Shift_K_Doxo_HLD
Z_Im_K_Doxo_HLD
Z_Shift_K_Doxo_EG
Z_Im_K_Doxo_EG
Z_Shift_L_Doxo_HLD
Z_Im_L_Doxo_HLD
Z_Shift_L_Doxo_EG
Z_Im_L_Doxo_EG

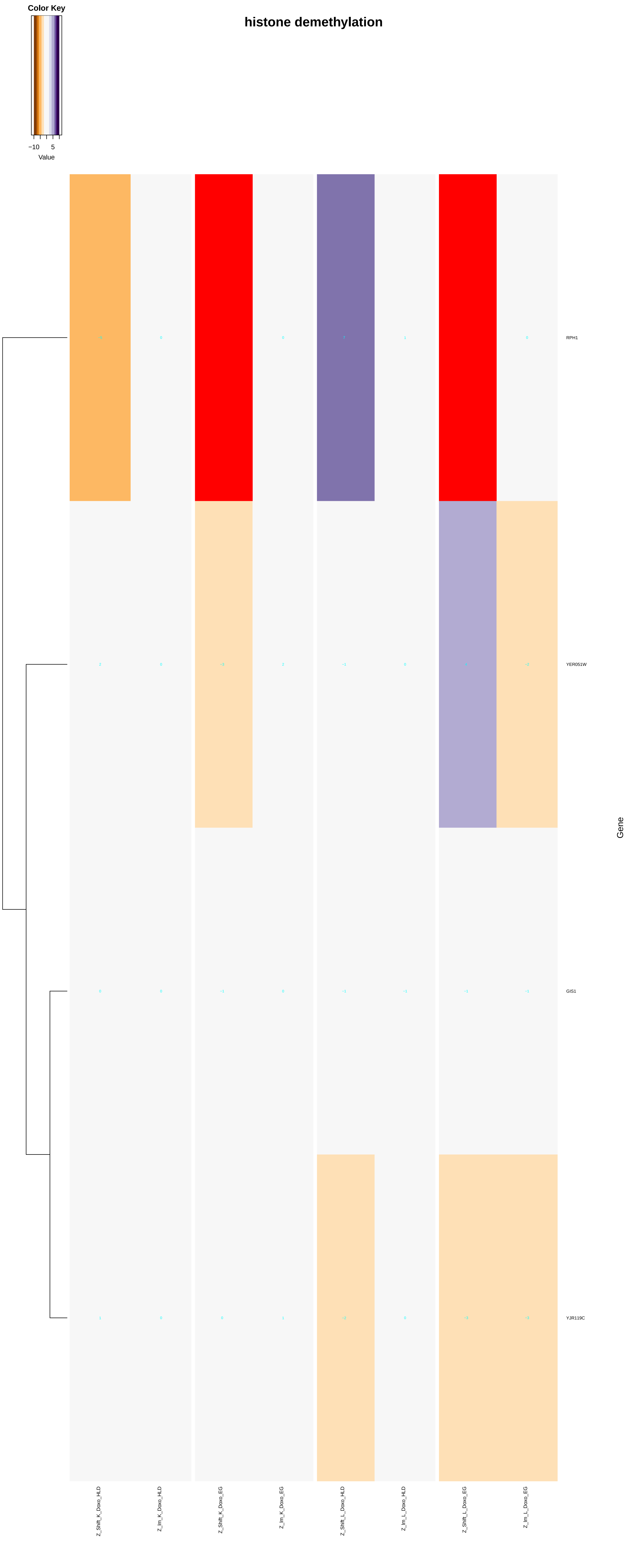
Color Key
−10
5
Value
histone demethylation
RPH1
YER051W
GIS1
YJR119C
Gene
Z_Shift_K_Doxo_HLD
Z_Im_K_Doxo_HLD
Z_Shift_K_Doxo_EG
Z_Im_K_Doxo_EG
Z_Shift_L_Doxo_HLD
Z_Im_L_Doxo_HLD
Z_Shift_L_Doxo_EG
Z_Im_L_Doxo_EG

### histone deubiquitination

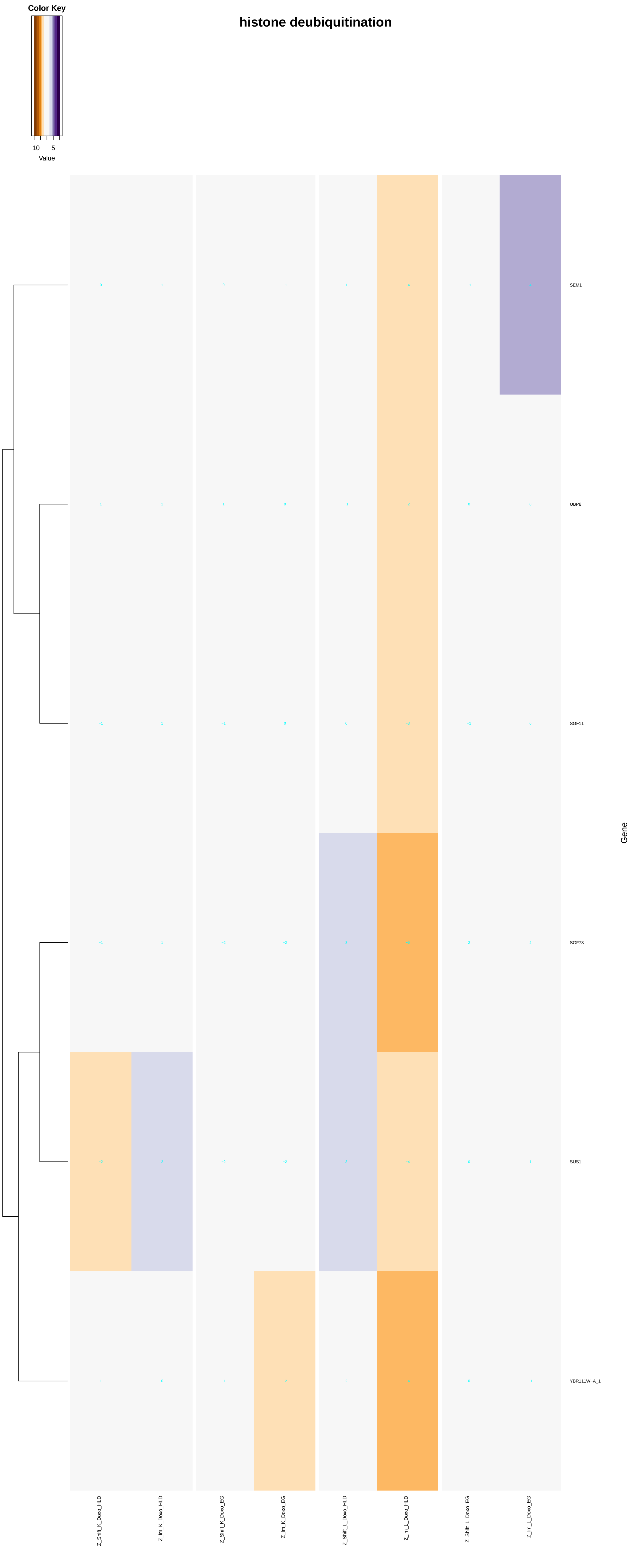

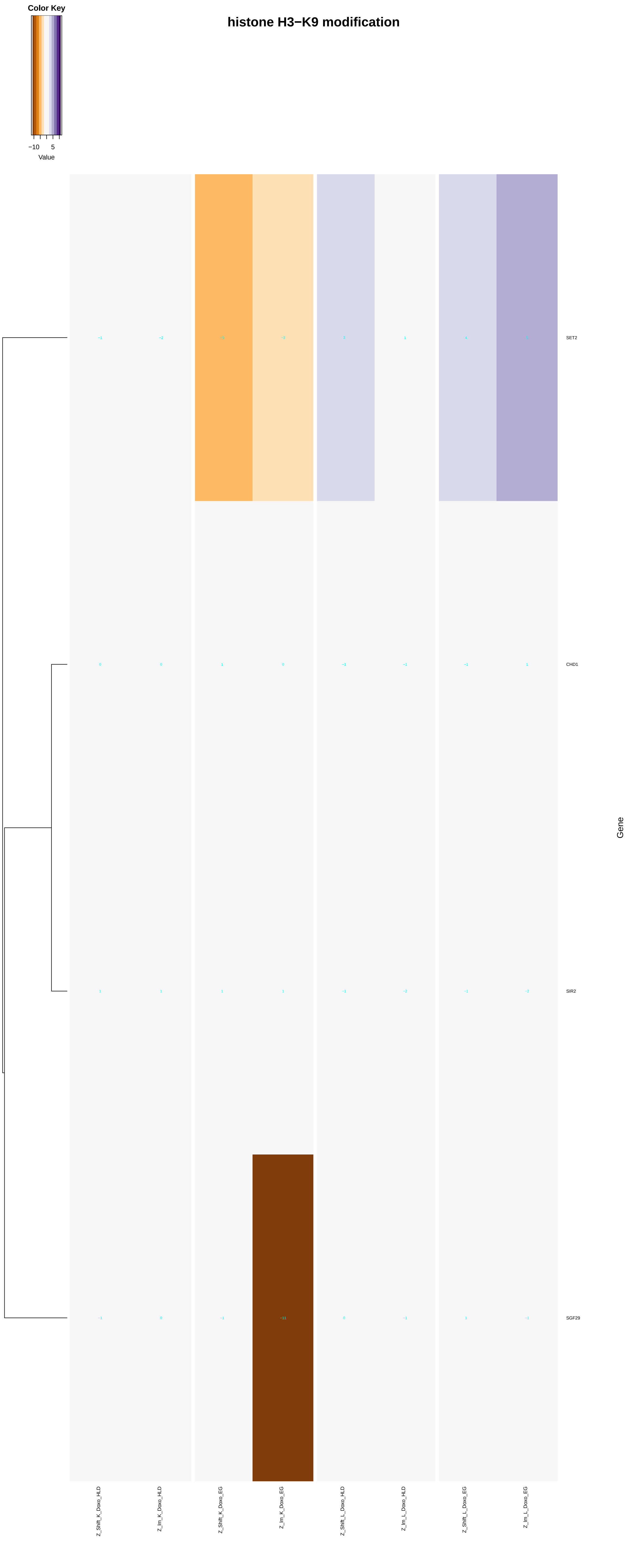
histone H3−K9 modification
Color Key
−10
5
Value
SET2
CHD1
SIR2
SGF29
Gene
Z_Shift_K_Doxo_HLD
Z_Im_K_Doxo_HLD
Z_Shift_K_Doxo_EG
Z_Im_K_Doxo_EG
Z_Shift_L_Doxo_HLD
Z_Im_L_Doxo_HLD
Z_Shift_L_Doxo_EG
Z_Im_L_Doxo_EG

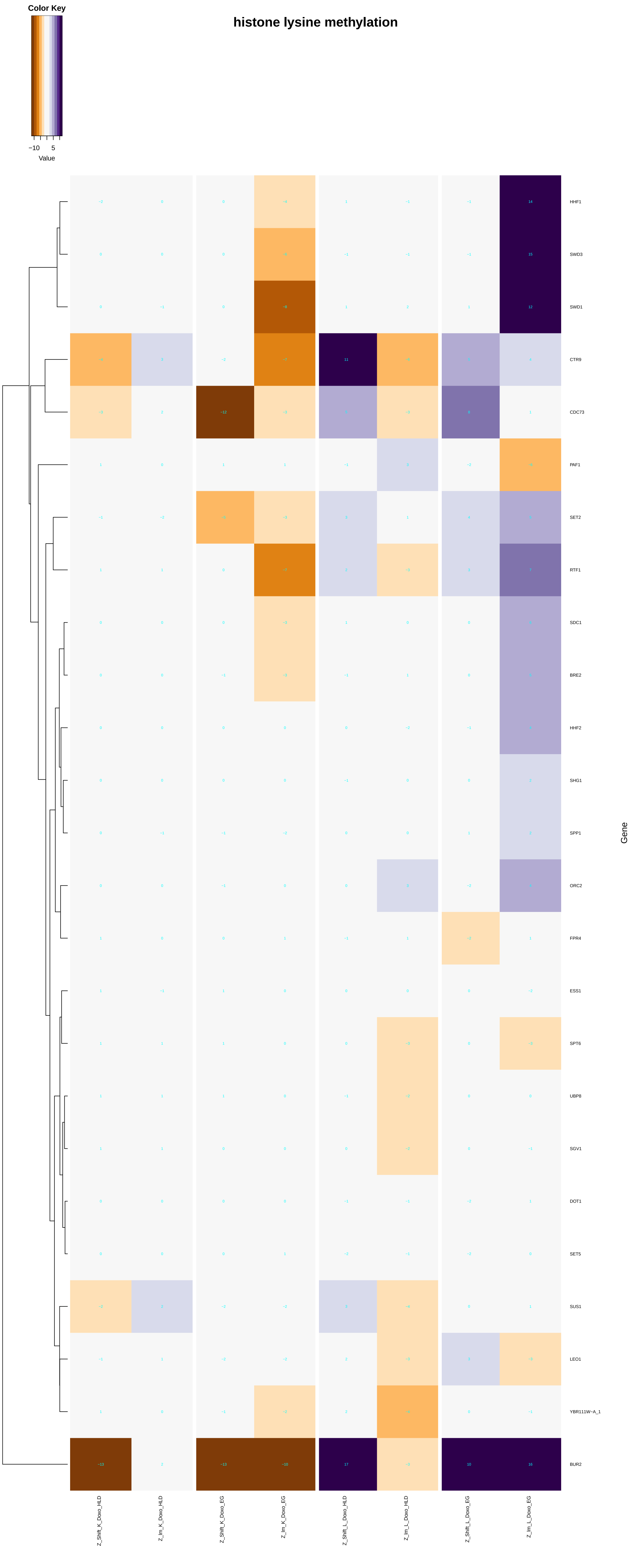
Color Key
−10 5
Value
histone lysine methylation
HHF1
SWD3
SWD1
CTR9
CDC73
PAF1
SET2
RTF1
SDC1
BRE2
HHF2
SHG1
SPP1
ORC2
FPR4
ESS1
SPT6
UBP8
SGV1
DOT1
SET5
SUS1
LEO1
YBR111W−A_1
BUR2
Gene
Z_Shift_K_Doxo_HLD
Z_lm_K_Doxo_HLD
Z_Shift_K_Doxo_EG
Z_lm_K_Doxo_EG
Z_Shift_L_Doxo_HLD
Z_lm_L_Doxo_HLD
Z_Shift_L_Doxo_EG
Z_lm_L_Doxo_EG

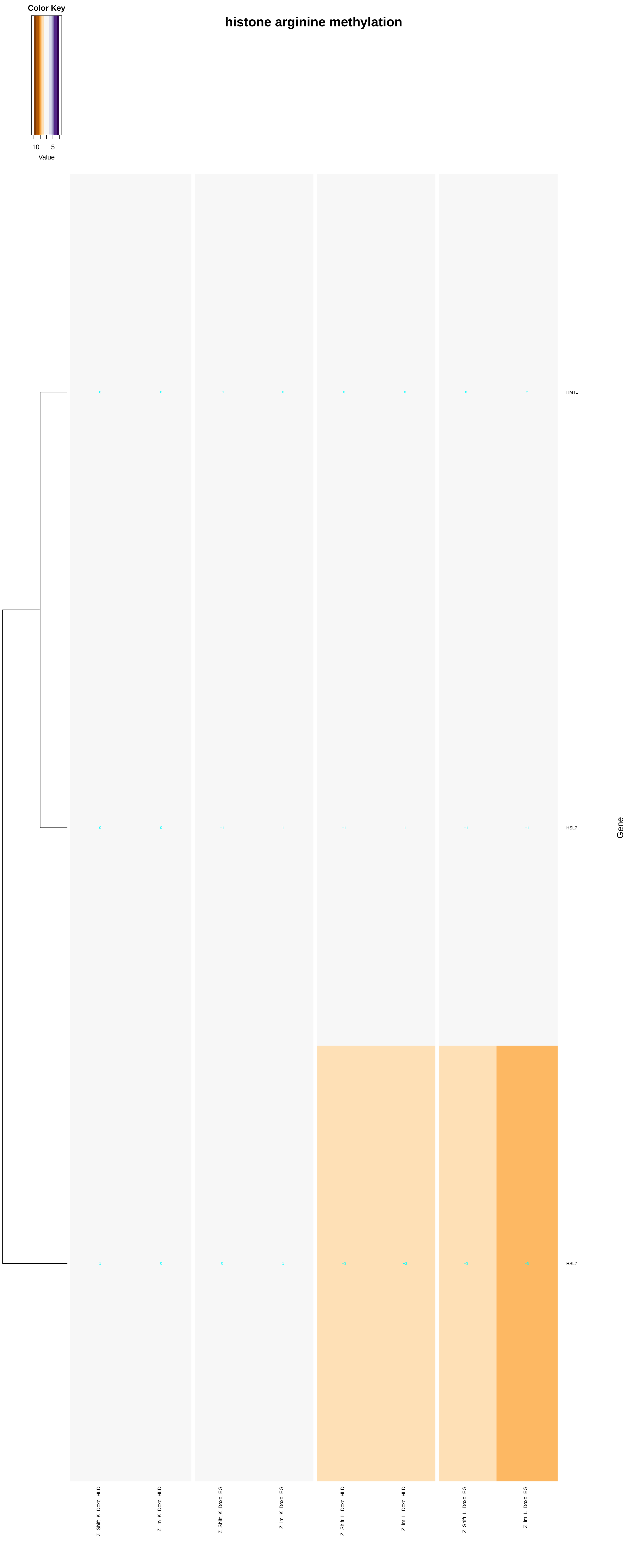

Color Key
−10
5
Value
histone arginine methylation
HMT1
HSL7
HSL7
Gene
Z_Shift_K_Doxo_HLD
Z_lm_K_Doxo_HLD
Z_Shift_K_Doxo_EG
Z_lm_K_Doxo_EG
Z_Shift_L_Doxo_HLD
Z_lm_L_Doxo_HLD
Z_Shift_L_Doxo_EG
Z_lm_L_Doxo_EG

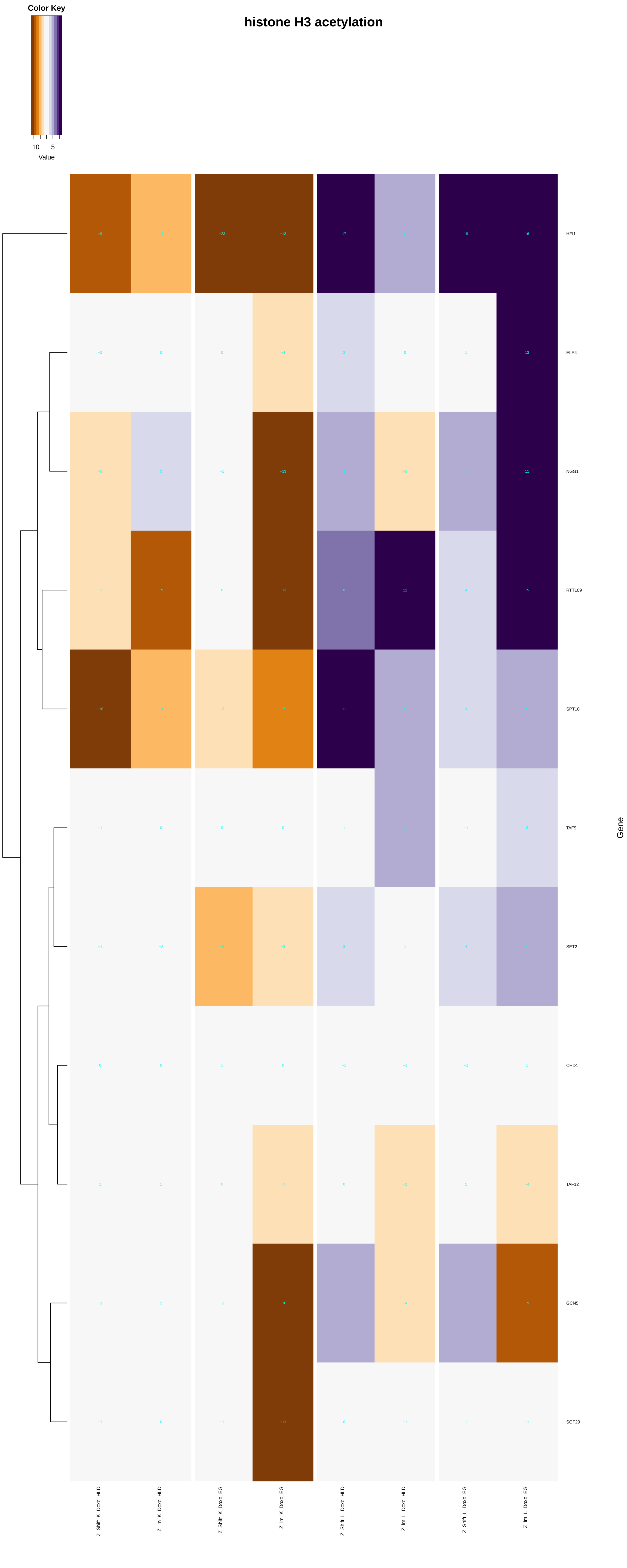

histone H3 acetylation
Color Key
−10
5
Value
HFI1
ELP4
NGG1
RTT109
SPT10
TAF9
SET2
CHD1
TAF12
GCN5
SGF29
Gene
Z_Shift_K_Doxo_HLD
Z_Im_K_Doxo_HLD
Z_Shift_K_Doxo_EG
Z_Im_K_Doxo_EG
Z_Shift_L_Doxo_HLD
Z_Im_L_Doxo_HLD
Z_Shift_L_Doxo_EG
Z_Im_L_Doxo_EG

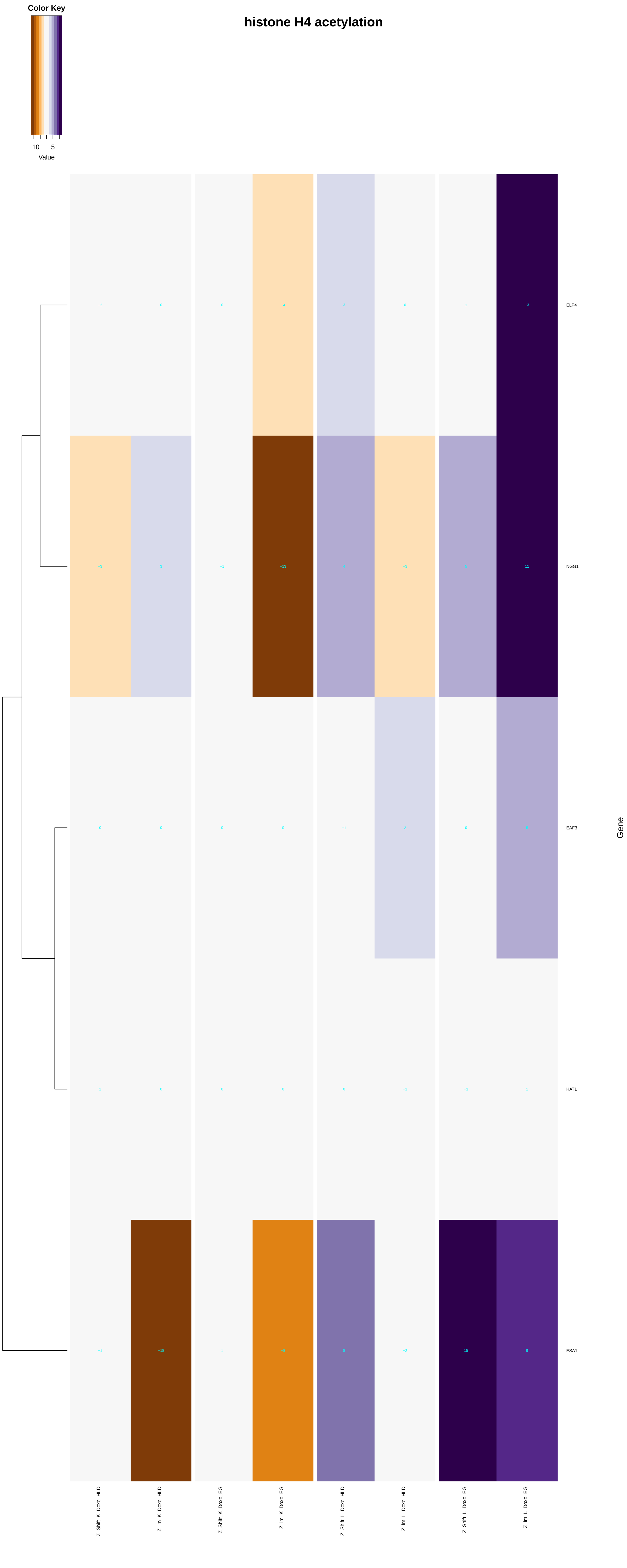
histone H4 acetylation
Color Key
−10
5
Value
ELP4
NGG1
EAF3
HAT1
ESA1
Gene
Z_Shift_K_Doxo_HLD
Z_Im_K_Doxo_HLD
Z_Shift_K_Doxo_EG
Z_Im_K_Doxo_EG
Z_Shift_L_Doxo_HLD
Z_Im_L_Doxo_HLD
Z_Shift_L_Doxo_EG
Z_Im_L_Doxo_EG

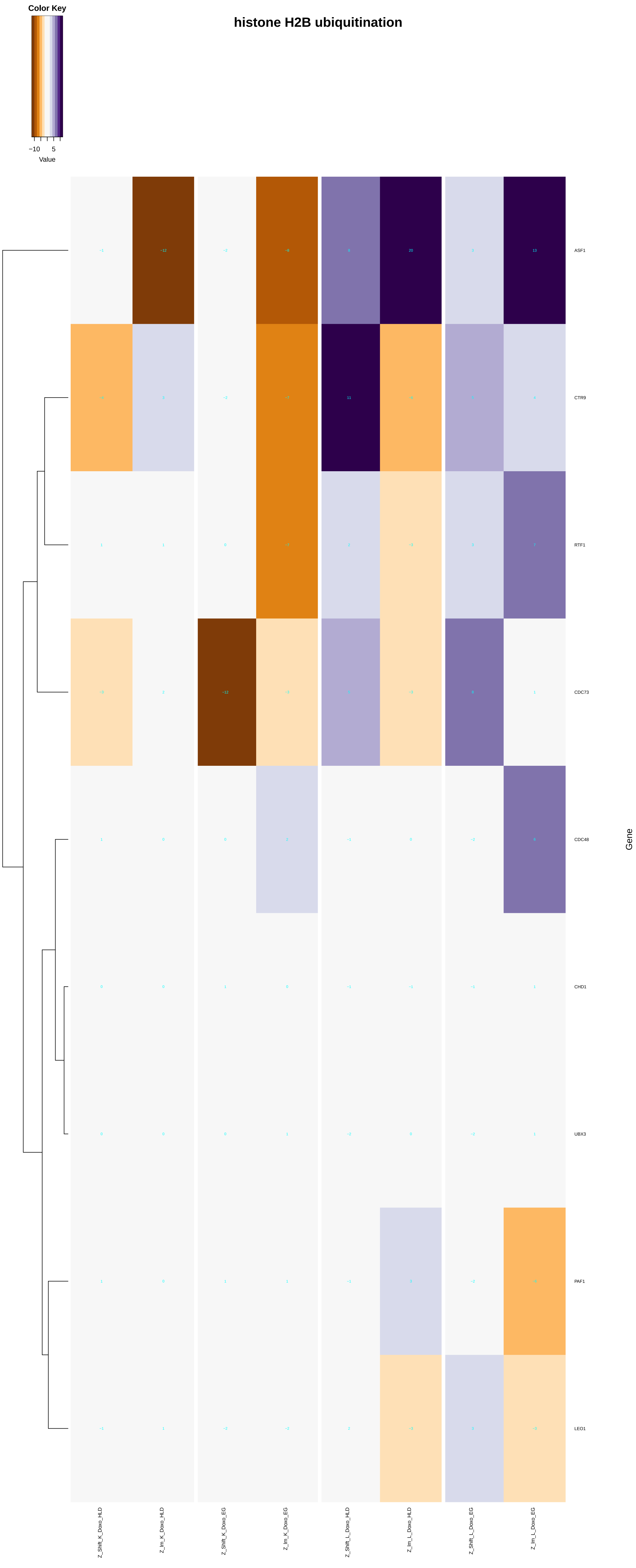

histone H2B ubiquitination
Color Key
−10
5
Value
ASF1
CTR9
RTF1
CDC73
CDC48
CHD1
UBX3
PAF1
LEO1
Gene
Z_Shift_K_Doxo_HLD
Z_Im_K_Doxo_HLD
Z_Shift_K_Doxo_EG
Z_Im_K_Doxo_EG
Z_Shift_L_Doxo_HLD
Z_Im_L_Doxo_HLD
Z_Shift_L_Doxo_EG
Z_Im_L_Doxo_EG

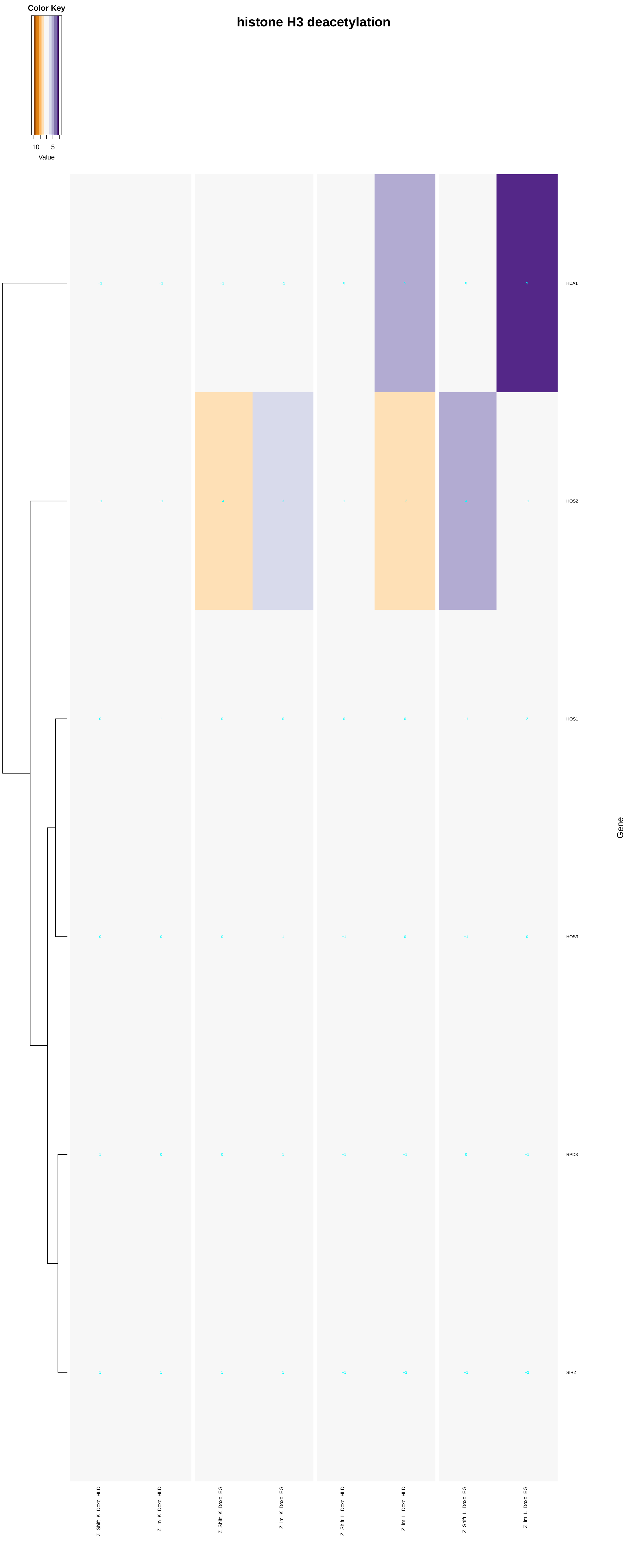

Color Key
−10
5
Value
histone H3 deacetylation
HDA1
HOS2
HOS1
HOS3
RPD3
SIR2
Gene
Z_Shift_K_Doxo_HLD
Z_Im_K_Doxo_HLD
Z_Shift_K_Doxo_EG
Z_Im_K_Doxo_EG
Z_Shift_L_Doxo_HLD
Z_Im_L_Doxo_HLD
Z_Shift_L_Doxo_EG
Z_Im_L_Doxo_EG

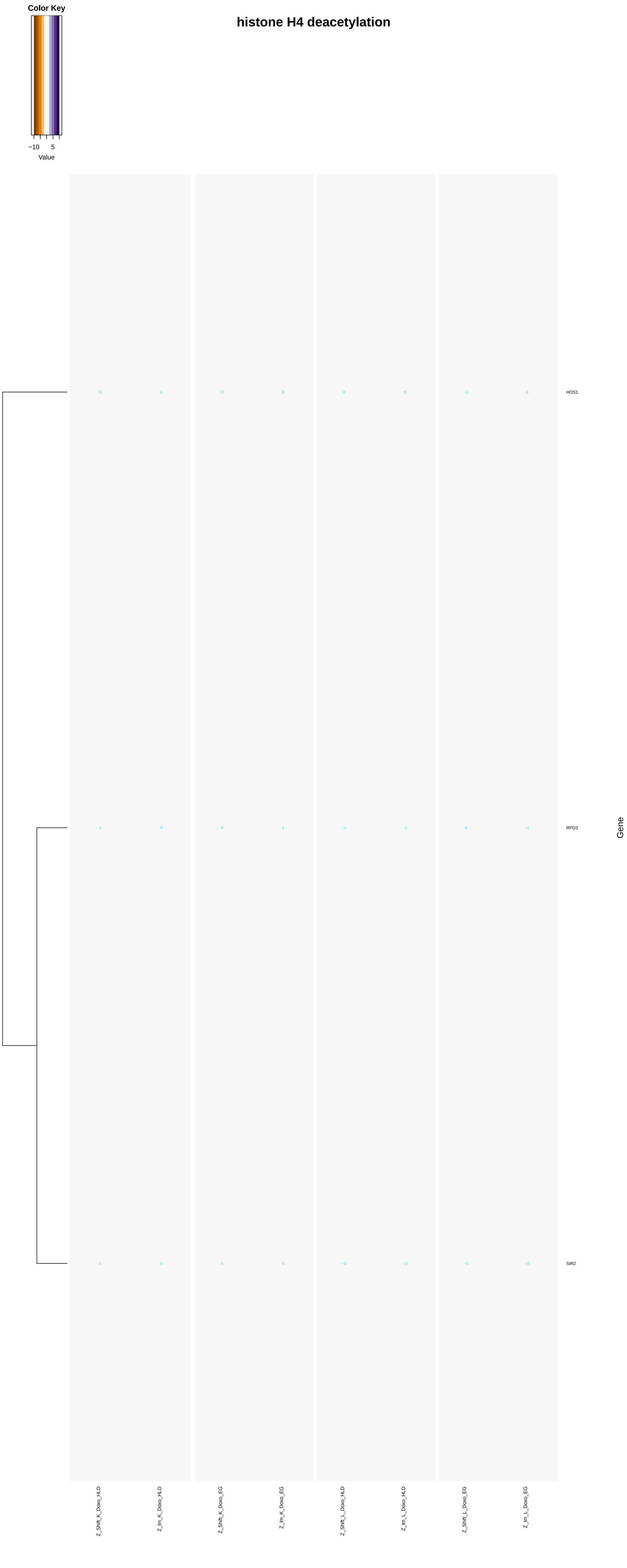
Color Key
−10
5
Value
histone H4 deacetylation
0
1
0
0
0
0
−1
2
HOS1
1
0
0
1
−1
−1
0
−1
RPD3
1
1
1
1
−1
−2
−1
−2
SIR2
Gene
Z_Shift_K_Doxo_HLD
Z_Im_K_Doxo_HLD
Z_Shift_K_Doxo_EG
Z_Im_K_Doxo_EG
Z_Shift_L_Doxo_HLD
Z_Im_L_Doxo_HLD
Z_Shift_L_Doxo_EG
Z_Im_L_Doxo_EG

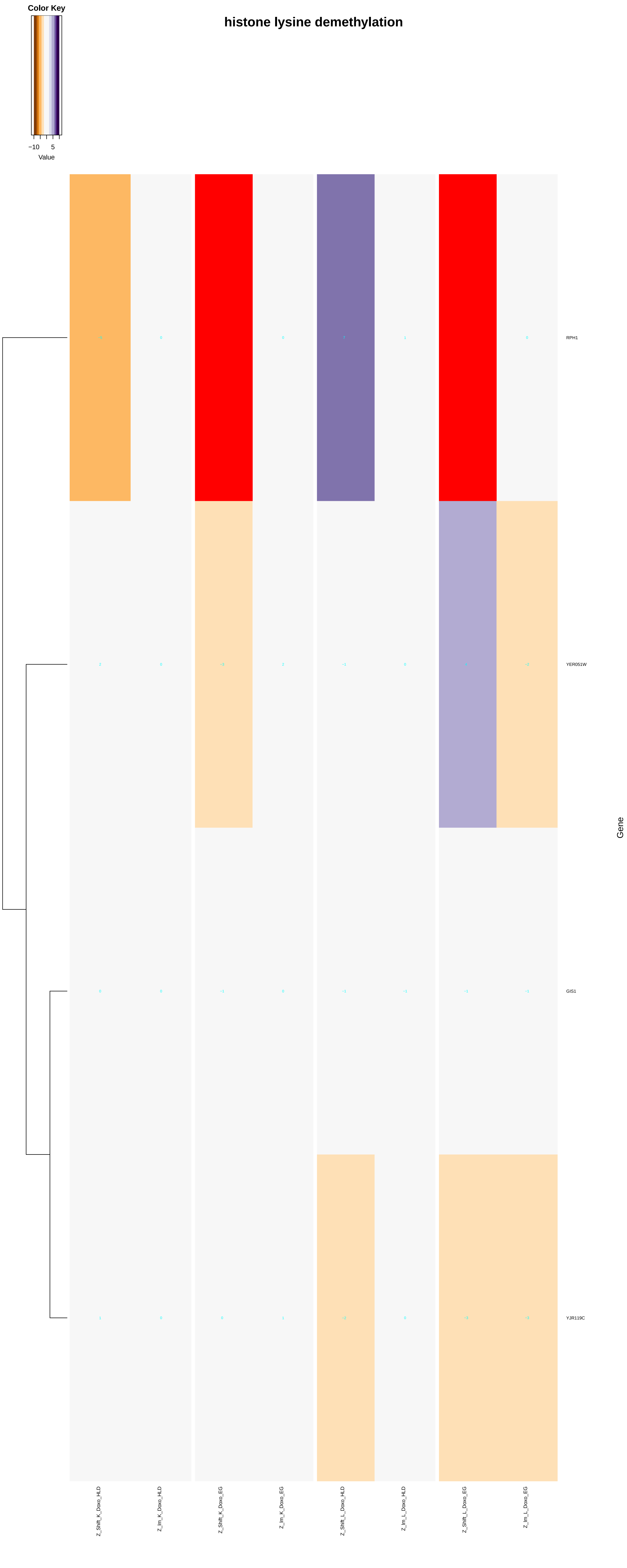

histone lysine demethylation
Color Key
−10
5
Value
RPH1
YER051W
GIS1
YJR119C
Gene
Z_Shift_K_Doxo_HLD
Z_Im_K_Doxo_HLD
Z_Shift_K_Doxo_EG
Z_Im_K_Doxo_EG
Z_Shift_L_Doxo_HLD
Z_Im_L_Doxo_HLD
Z_Shift_L_Doxo_EG
Z_Im_L_Doxo_EG

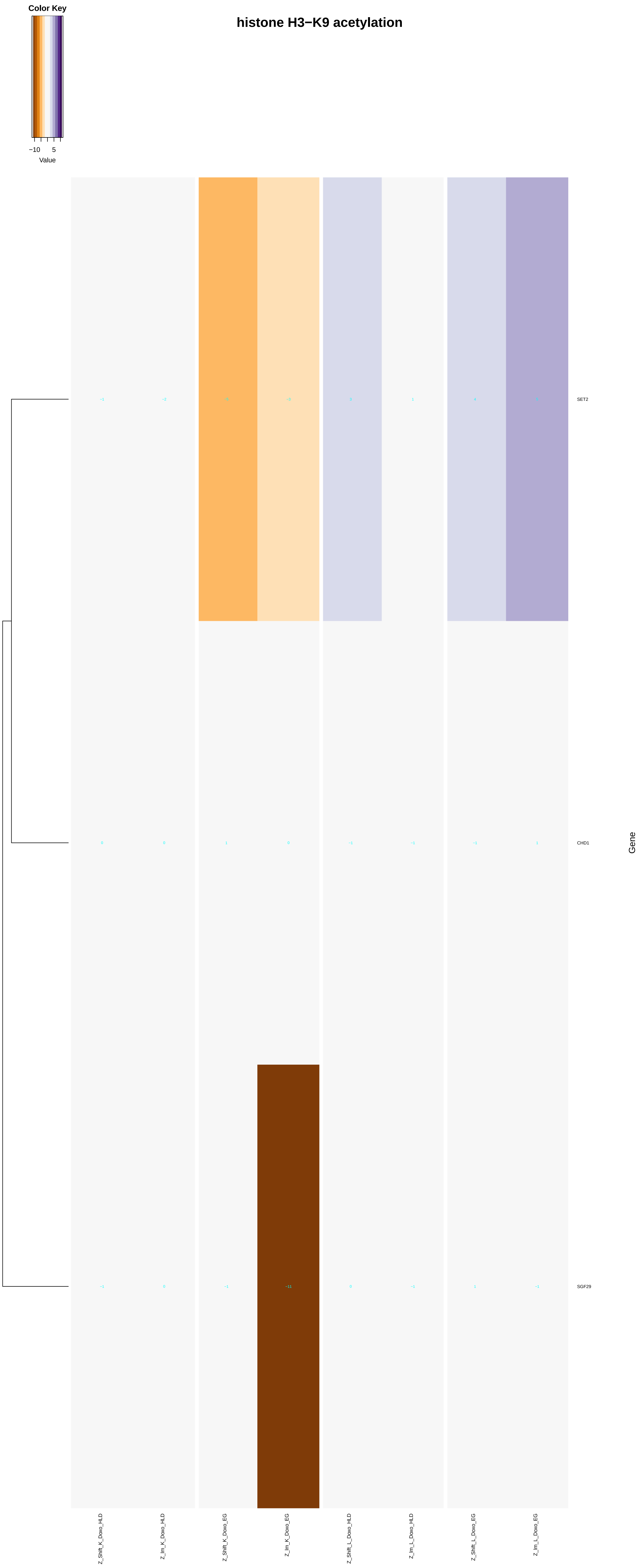
histone H3−K9 acetylation
Color Key
−10
5
Value
SET2
CHD1
SGF29
Gene
Z_Shift_K_Doxo_HLD
Z_Im_K_Doxo_HLD
Z_Shift_K_Doxo_EG
Z_Im_K_Doxo_EG
Z_Shift_L_Doxo_HLD
Z_Im_L_Doxo_HLD
Z_Shift_L_Doxo_EG
Z_Im_L_Doxo_EG

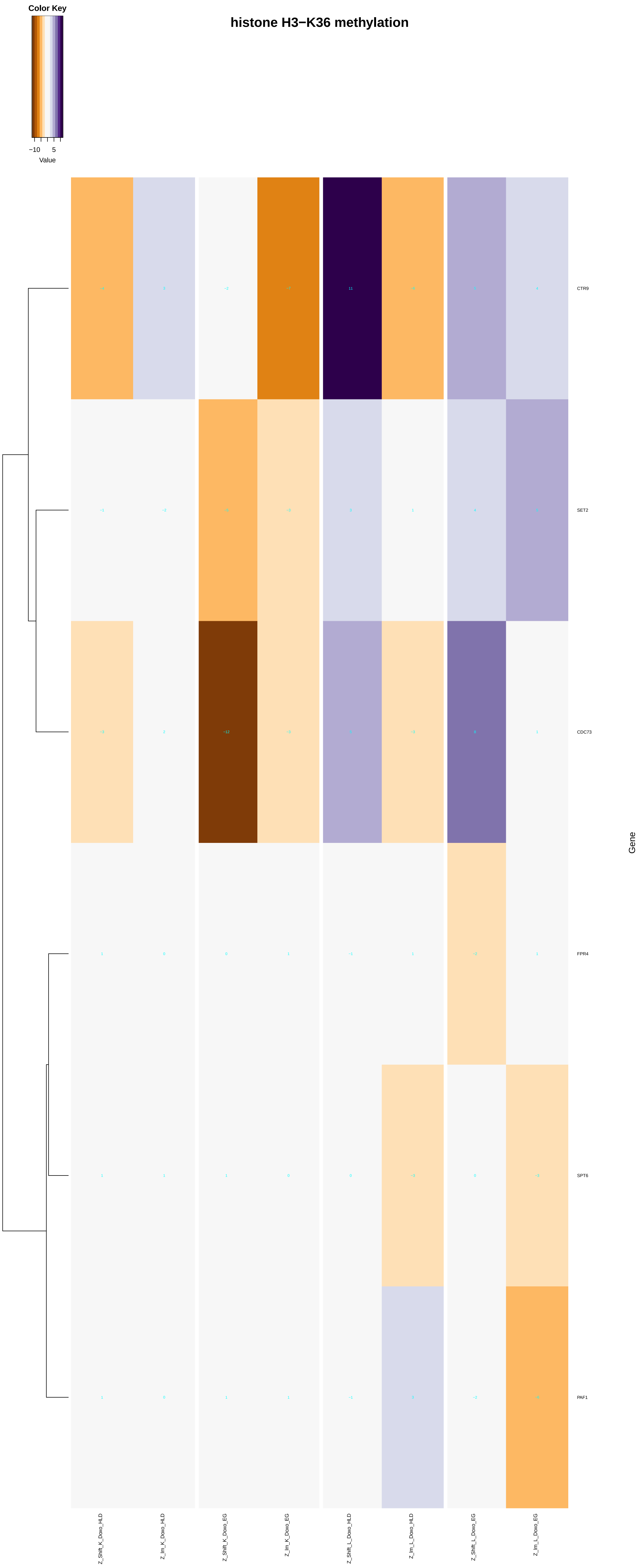
Color Key
−10
5
Value
histone H3−K36 methylation
CTR9
SET2
CDC73
FPR4
SPT6
PAF1
Gene
Z_Shift_K_Doxo_HLD
Z_Im_K_Doxo_HLD
Z_Shift_K_Doxo_EG
Z_Im_K_Doxo_EG
Z_Shift_L_Doxo_HLD
Z_Im_L_Doxo_HLD
Z_Shift_L_Doxo_EG
Z_Im_L_Doxo_EG

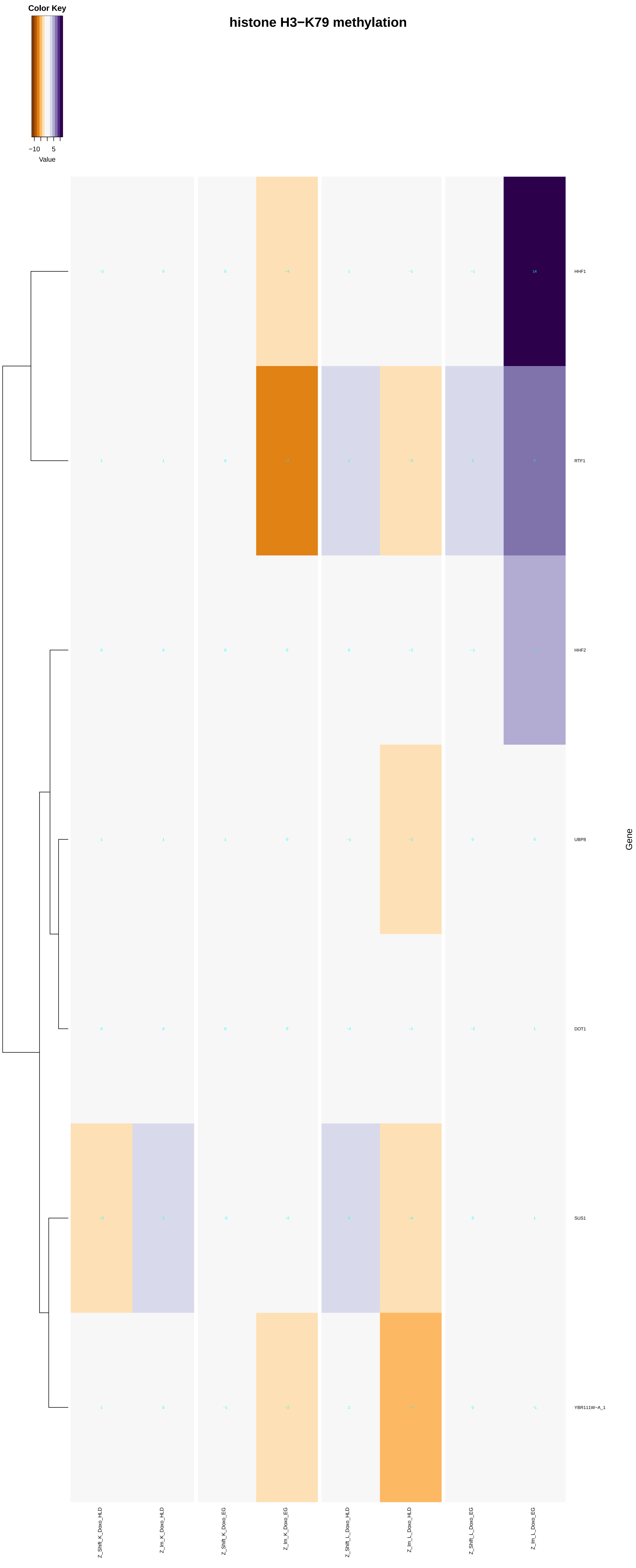
histone H3−K79 methylation
Color Key
−10
5
Value
HHF1
RTF1
HHF2
UBP8
DOT1
SUS1
YBR111W−A_1
Gene
Z_Shift_K_Doxo_HLD
Z_Im_K_Doxo_HLD
Z_Shift_K_Doxo_EG
Z_Im_K_Doxo_EG
Z_Shift_L_Doxo_HLD
Z_Im_L_Doxo_HLD
Z_Shift_L_Doxo_EG
Z_Im_L_Doxo_EG

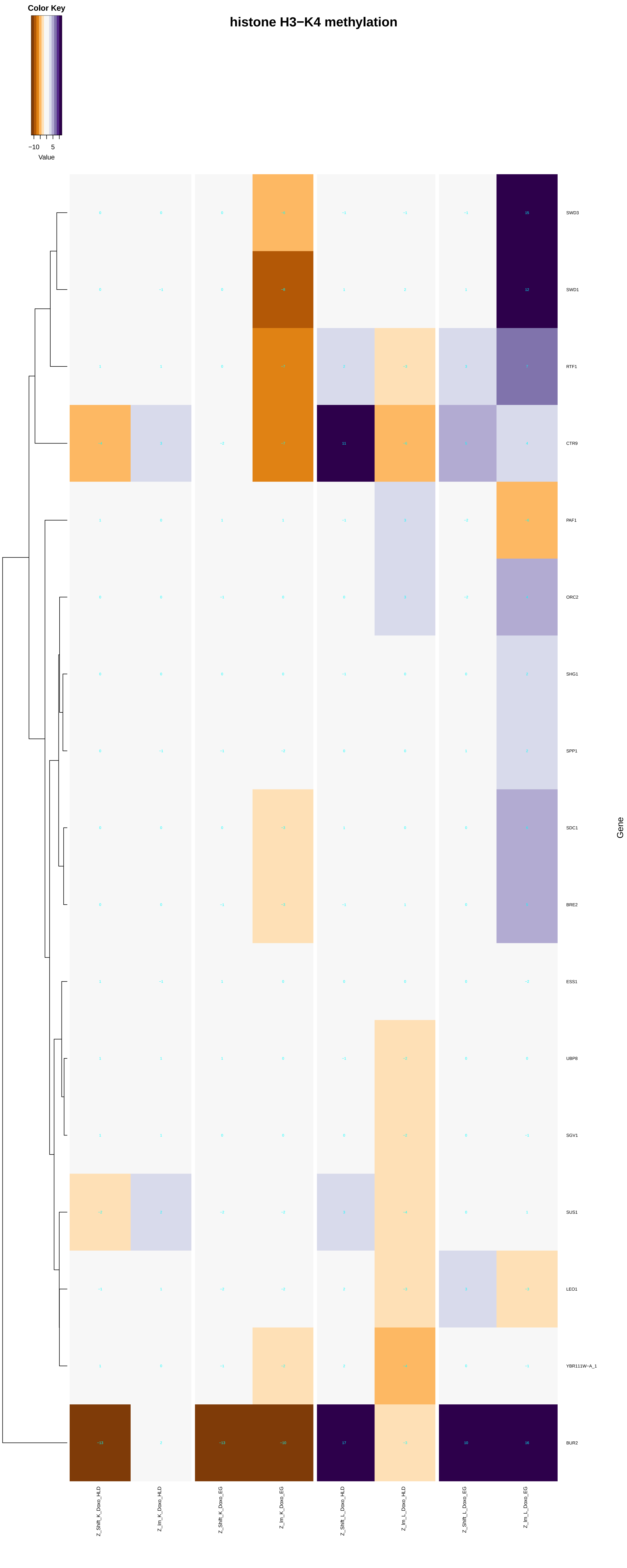

Color Key
−10 5
Value
histone H3−K4 methylation
SWD3
SWD1
RTF1
CTR9
PAF1
ORC2
SHG1
SPP1
SDC1
BRE2
ESS1
UBP8
SGV1
SUS1
LEO1
YBR111W−A_1
BUR2
Gene
Z_Shift_K_Doxo_HLD
Z_Im_K_Doxo_HLD
Z_Shift_K_Doxo_EG
Z_Im_K_Doxo_EG
Z_Shift_L_Doxo_HLD
Z_Im_L_Doxo_HLD
Z_Shift_L_Doxo_EG
Z_Im_L_Doxo_EG

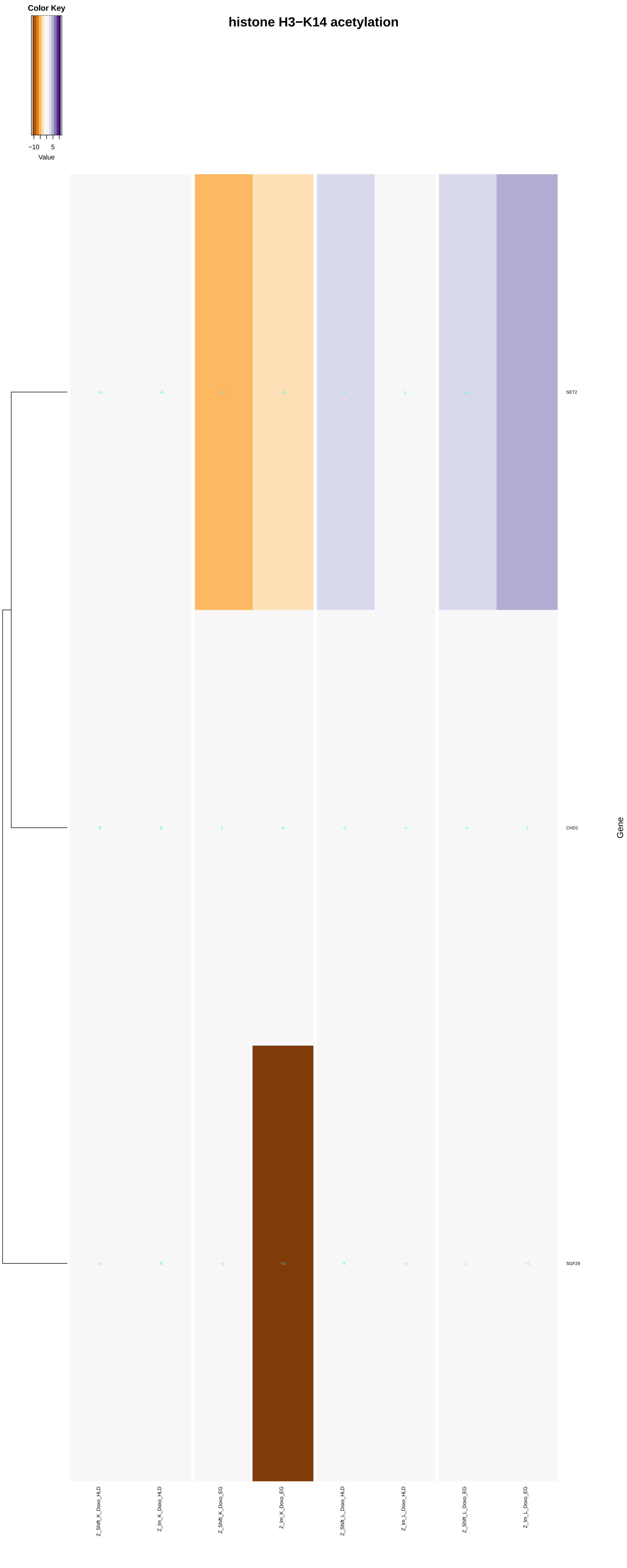
Color Key
−10
5
Value
histone H3−K14 acetylation
SET2
CHD1
SGF29
Gene
Z_Shift_K_Doxo_HLD
Z_Im_K_Doxo_HLD
Z_Shift_K_Doxo_EG
Z_Im_K_Doxo_EG
Z_Shift_L_Doxo_HLD
Z_Im_L_Doxo_HLD
Z_Shift_L_Doxo_EG
Z_Im_L_Doxo_EG

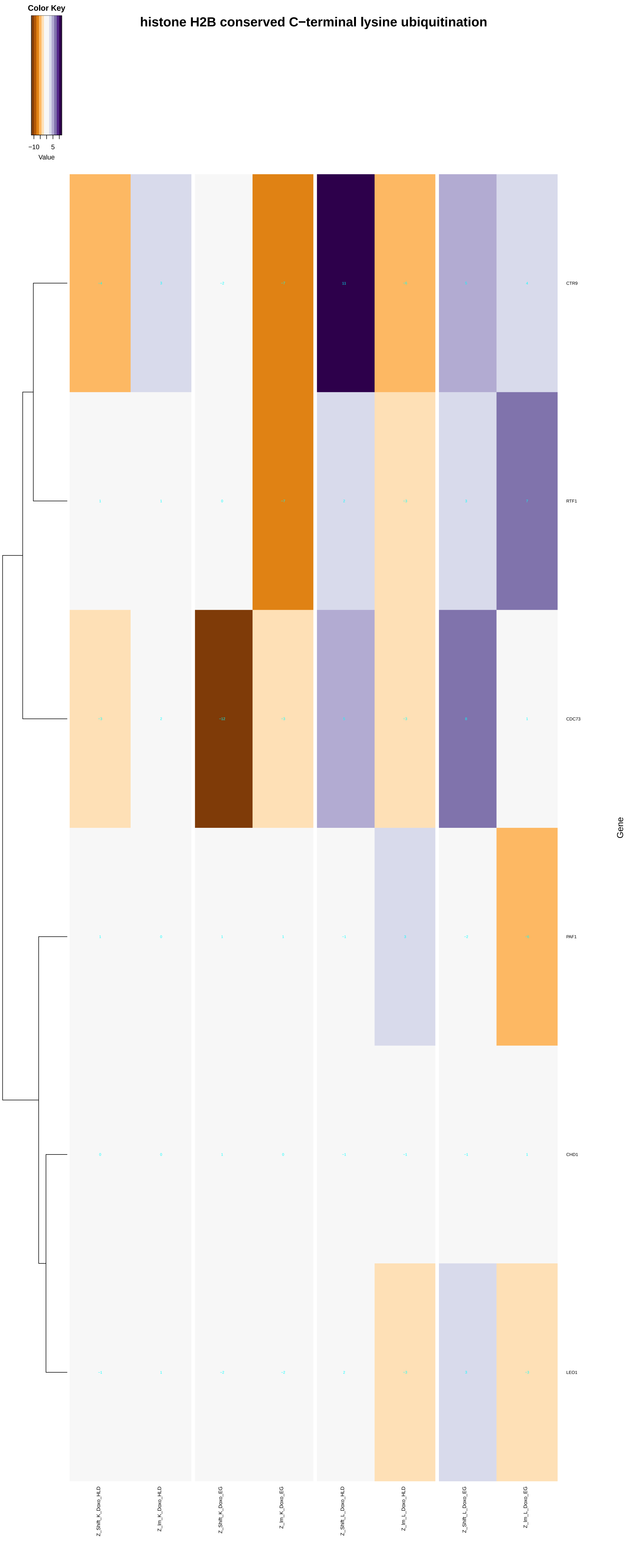

Color Key
−10
5
Value
histone H2B conserved C−terminal lysine ubiquitination
CTR9
RTF1
CDC73
PAF1
CHD1
LEO1
Gene
Z_Shift_K_Doxo_HLD
Z_Im_K_Doxo_HLD
Z_Shift_K_Doxo_EG
Z_Im_K_Doxo_EG
Z_Shift_L_Doxo_HLD
Z_Im_L_Doxo_HLD
Z_Shift_L_Doxo_EG
Z_Im_L_Doxo_EG

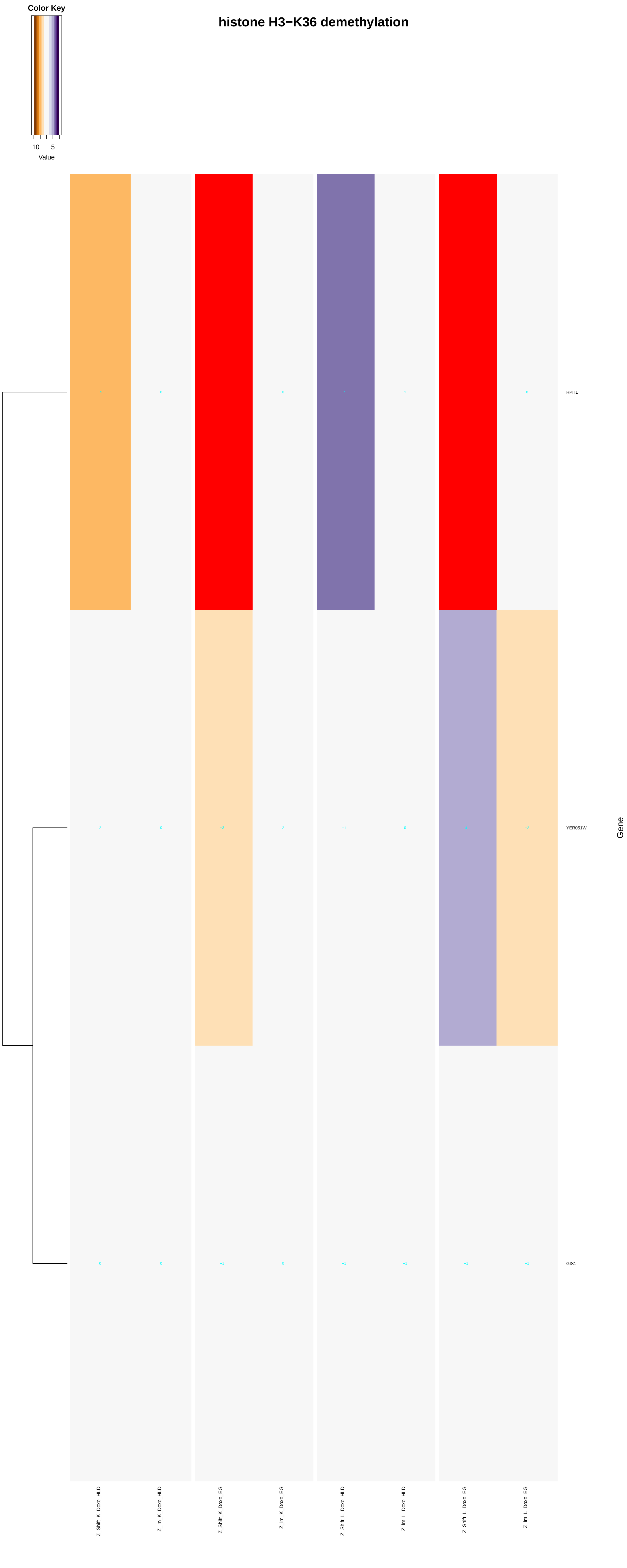
histone H3−K36 demethylation
Color Key
−10
5
Value
RPH1
YER051W
GIS1
Gene
Z_Shift_K_Doxo_HLD
Z_Im_K_Doxo_HLD
Z_Shift_K_Doxo_EG
Z_Im_K_Doxo_EG
Z_Shift_L_Doxo_HLD
Z_Im_L_Doxo_HLD
Z_Shift_L_Doxo_EG
Z_Im_L_Doxo_EG

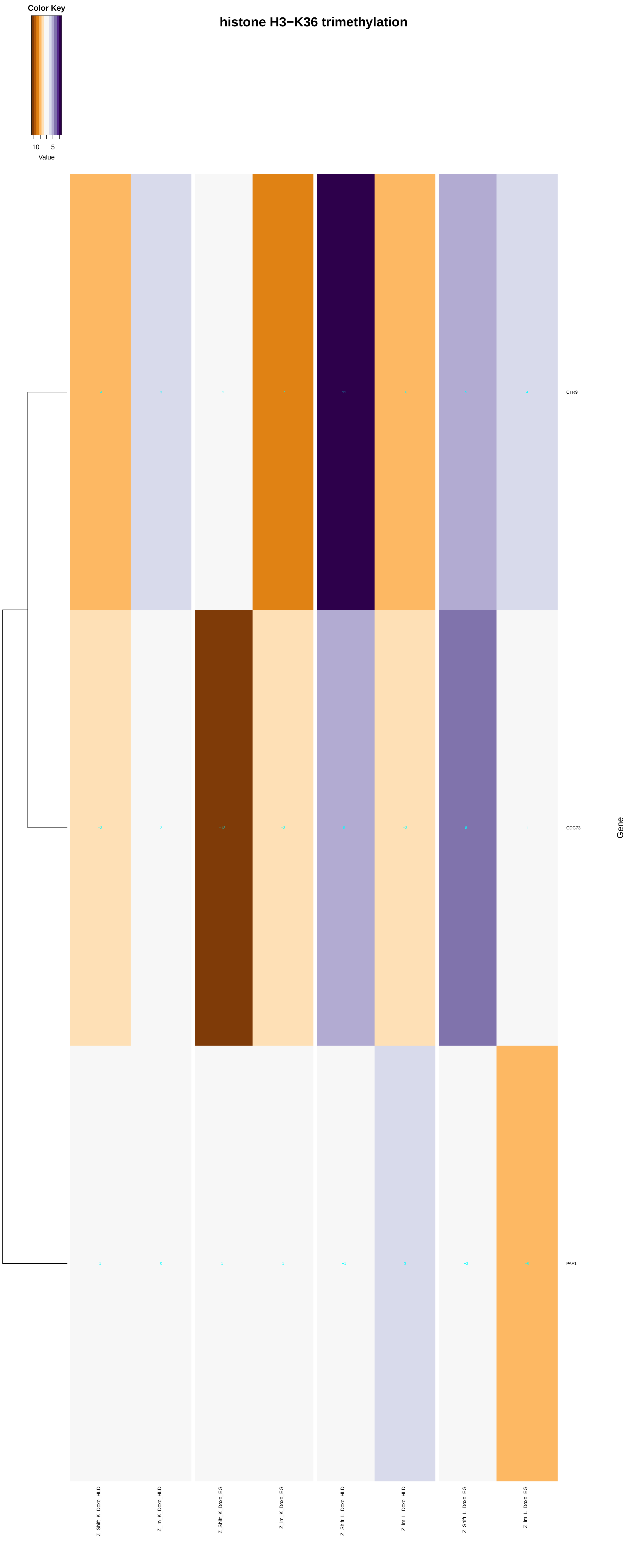

histone H3−K36 trimethylation
Color Key
−10
5
Value
CTR9
CDC73
PAF1
Gene
Z_Shift_K_Doxo_HLD
Z_Im_K_Doxo_HLD
Z_Shift_K_Doxo_EG
Z_Im_K_Doxo_EG
Z_Shift_L_Doxo_HLD
Z_Im_L_Doxo_HLD
Z_Shift_L_Doxo_EG
Z_Im_L_Doxo_EG

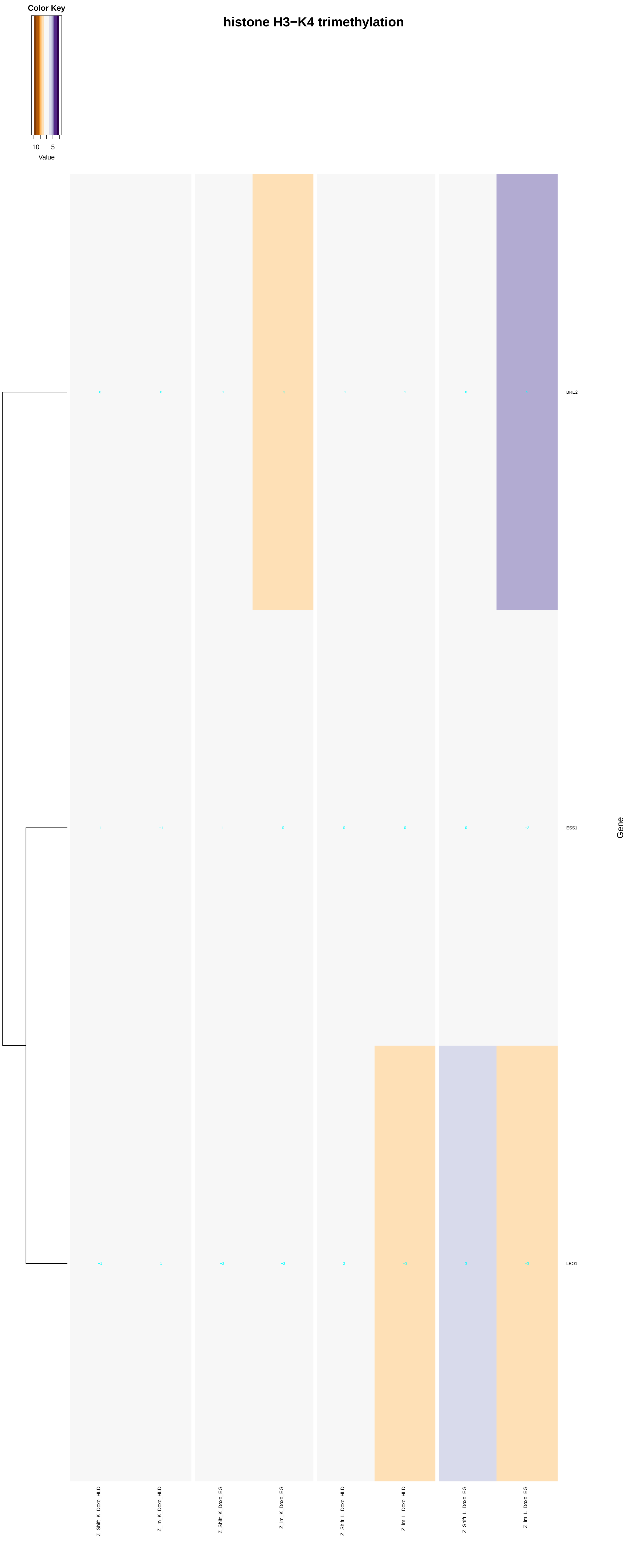

Color Key
−10
5
Value
histone H3−K4 trimethylation
BRE2
ESS1
LEO1
Gene
Z_Shift_K_Doxo_HLD
Z_Im_K_Doxo_HLD
Z_Shift_K_Doxo_EG
Z_Im_K_Doxo_EG
Z_Shift_L_Doxo_HLD
Z_Im_L_Doxo_HLD
Z_Shift_L_Doxo_EG
Z_Im_L_Doxo_EG
