## Additional File 9 for "A yeast phenomic model for the influence of Warburg metabolism on genetic buffering of doxorubicin": chromosome_localization.pdf

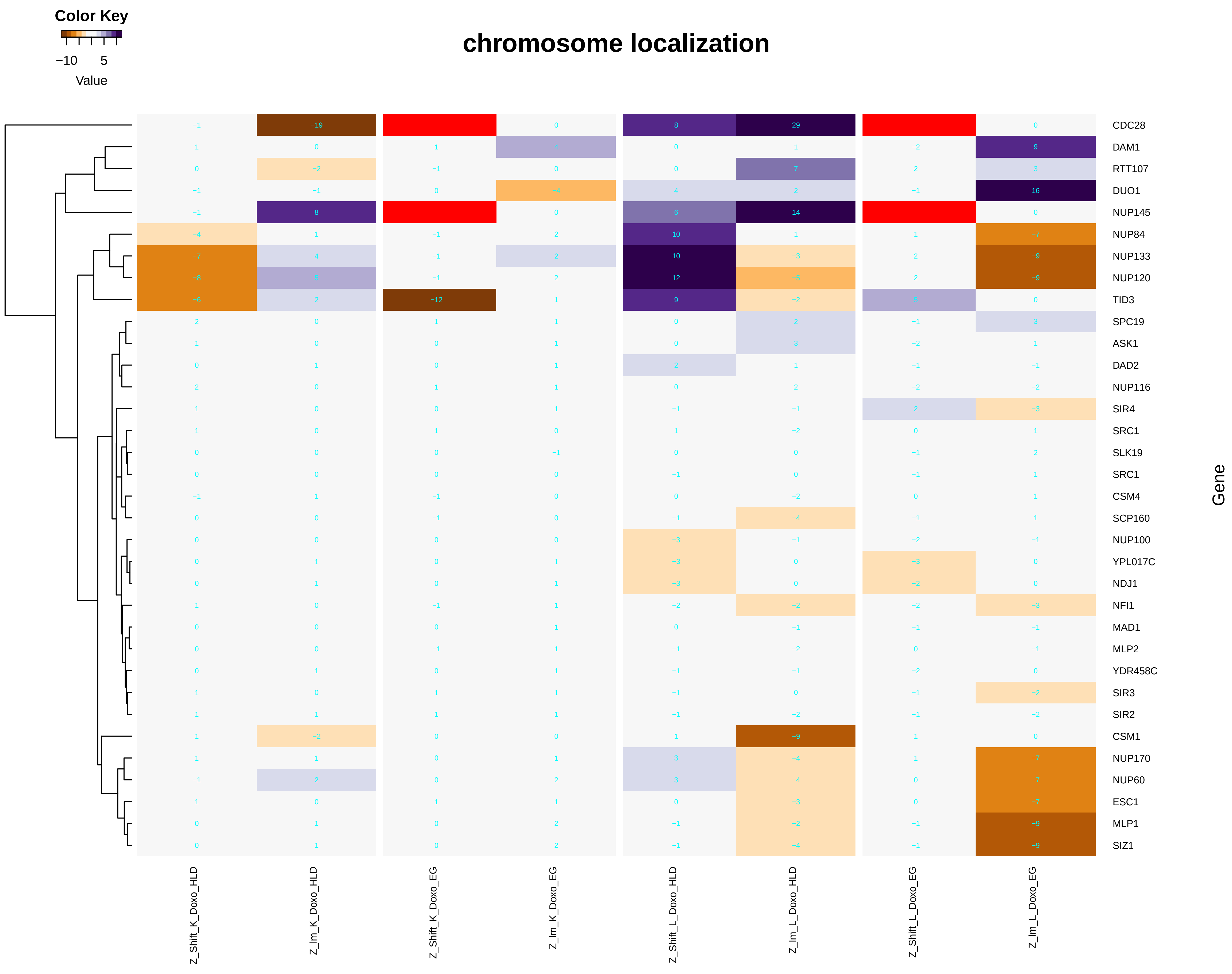

chromosome localization
Color Key
−10
5
Value
Gene
CDC28
DAM1
RTT107
DUO1
NUP145
NUP84
NUP133
NUP120
TID3
SPC19
ASK1
DAD2
NUP116
SIR4
SRC1
SLK19
SRC1
CSM4
SCP160
NUP100
YPL017C
NDJ1
NFI1
MAD1
MLP2
YDR458C
SIR3
SIR2
CSM1
NUP170
NUP60
ESC1
MLP1
SIZ1
Z_Shift_K_Doxo_HLD
Z_Im_K_Doxo_HLD
Z_Shift_K_Doxo_EG
Z_Im_K_Doxo_EG
Z_Shift_L_Doxo_HLD
Z_Im_L_Doxo_HLD
Z_Shift_L_Doxo_EG
Z_Im_L_Doxo_EG

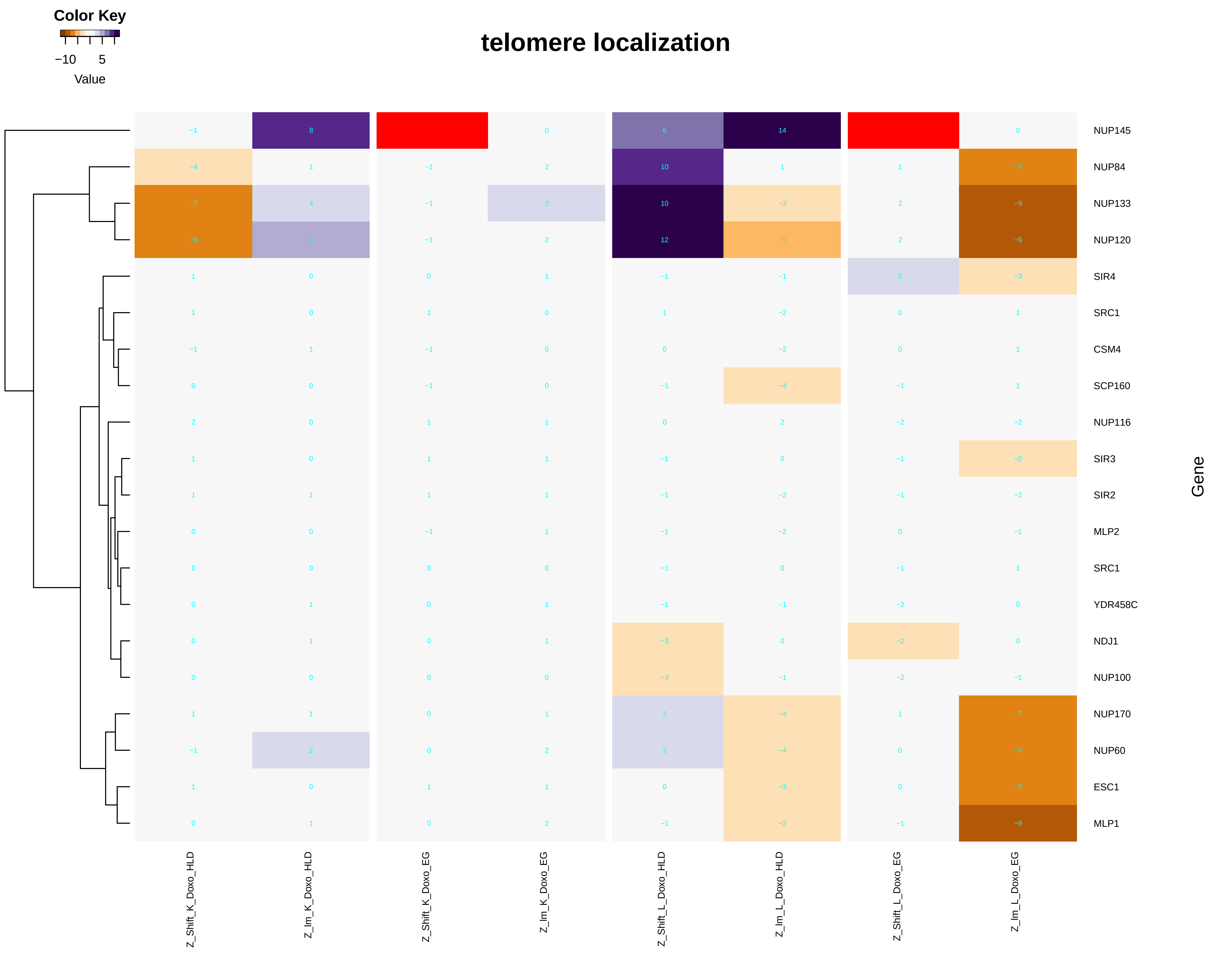

telomere localization
Color Key
−10
5
Value
NUP145
NUP84
NUP133
NUP120
SIR4
SRC1
CSM4
SCP160
NUP116
SIR3
SIR2
MLP2
SRC1
YDR458C
NDJ1
NUP100
NUP170
NUP60
ESC1
MLP1
Gene
Z_Shift_K_Doxo_HLD
Z_Im_K_Doxo_HLD
Z_Shift_K_Doxo_EG
Z_Im_K_Doxo_EG
Z_Shift_L_Doxo_HLD
Z_Im_L_Doxo_HLD
Z_Shift_L_Doxo_EG
Z_Im_L_Doxo_EG

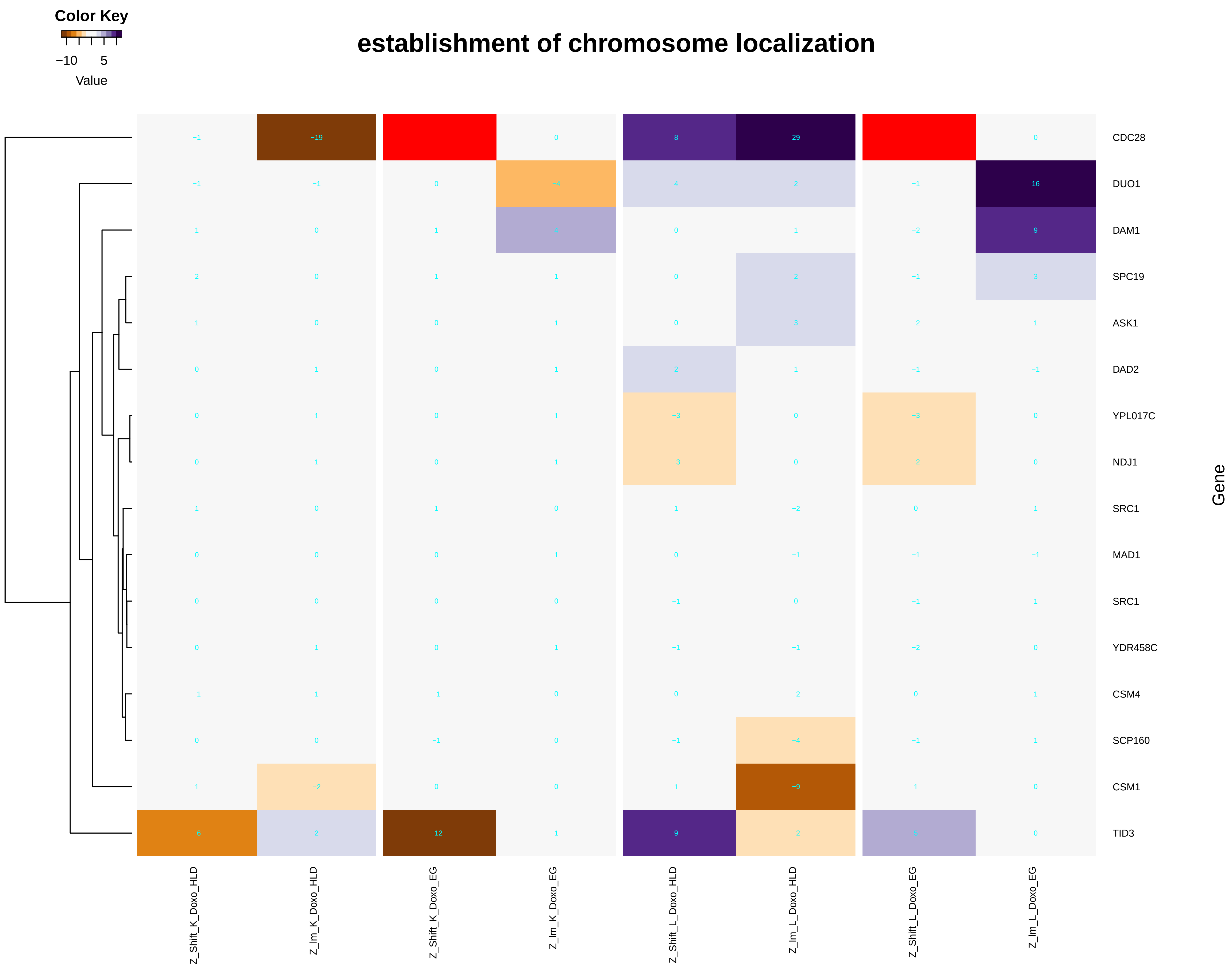
Color Key
−10 5
Value
establishment of chromosome localization
CDC28
DUO1
DAM1
SPC19
ASK1
DAD2
YPL017C
NDJ1
SRC1
MAD1
SRC1
YDR458C
CSM4
SCP160
CSM1
TID3
Gene
Z_Shift_K_Doxo_HLD
Z_Im_K_Doxo_HLD
Z_Shift_K_Doxo_EG
Z_Im_K_Doxo_EG
Z_Shift_L_Doxo_HLD
Z_Im_L_Doxo_HLD
Z_Shift_L_Doxo_EG
Z_Im_L_Doxo_EG

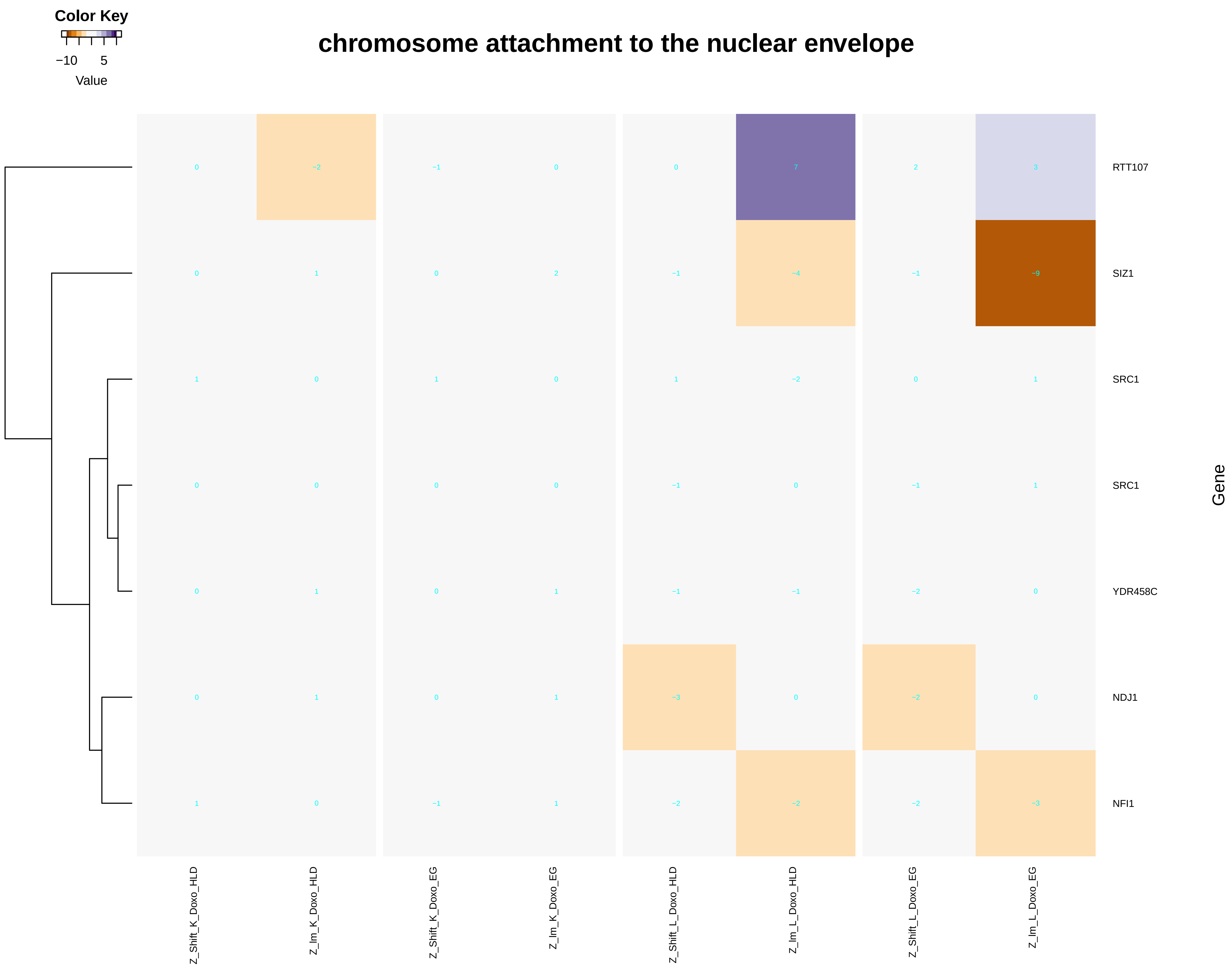

Color Key
−10
5
Value
chromosome attachment to the nuclear envelope
RTT107
SIZ1
SRC1
SRC1
YDR458C
NDJ1
NFI1
Gene
Z_Shift_K_Doxo_HLD
Z_Im_K_Doxo_HLD
Z_Shift_K_Doxo_EG
Z_Im_K_Doxo_EG
Z_Shift_L_Doxo_HLD
Z_Im_L_Doxo_HLD
Z_Shift_L_Doxo_EG
Z_Im_L_Doxo_EG

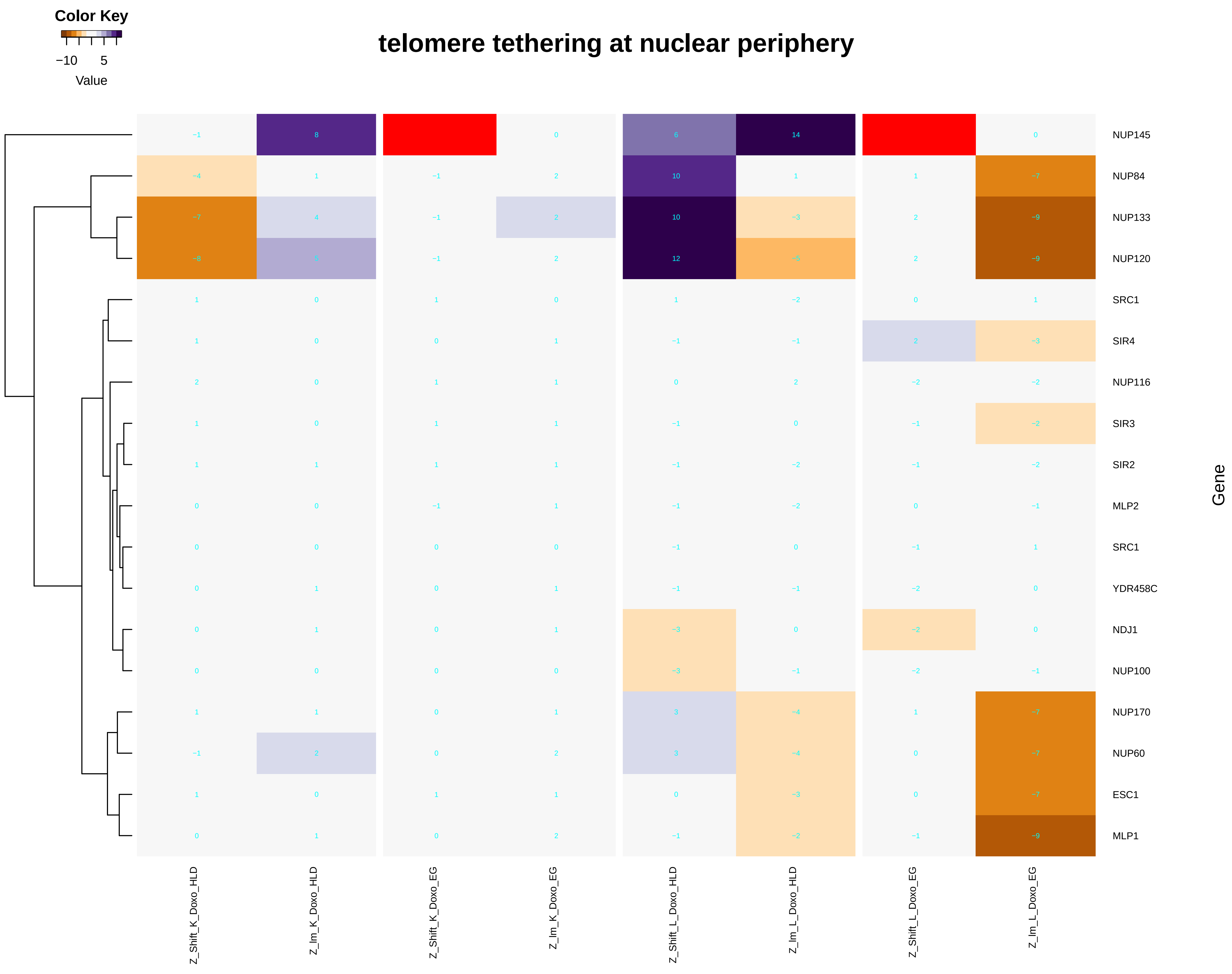
Color Key
−10 5
Value
telomere tethering at nuclear periphery
NUP145 −1 8 0 6 14 0
NUP84 −4 1 −1 2 10 1 1 −7
NUP133 −7 4 −1 2 10 −3 2 −9
NUP120 −8 5 −1 2 12 −5 2 −9
SRC1 1 0 1 0 1 −2 0 1
SIR4 1 0 0 1 −1 −1 2 −3
NUP116 2 0 1 1 0 2 −2 −2
SIR3 1 0 1 1 −1 0 −1 −2
SIR2 1 1 1 1 −1 −2 −1 −2
MLP2 0 0 −1 1 −1 −2 0 −1
SRC1 0 0 0 0 −1 0 −1 1
YDR458C 0 1 0 1 −1 −1 −2 0
NDJ1 0 1 0 1 −3 0 −2 0
NUP100 0 0 0 0 −3 −1 −2 −1
NUP170 1 1 0 1 3 −4 1 −7
NUP60 −1 2 0 2 3 −4 0 −7
ESC1 1 0 1 1 0 −3 0 −7
MLP1 0 1 0 2 −1 −2 −1 −9
Gene
Z_Shift_K_Doxo_HLD
Z_Im_K_Doxo_HLD
Z_Shift_K_Doxo_EG
Z_Im_K_Doxo_EG
Z_Shift_L_Doxo_HLD
Z_Im_L_Doxo_HLD
Z_Shift_L_Doxo_EG
Z_Im_L_Doxo_EG

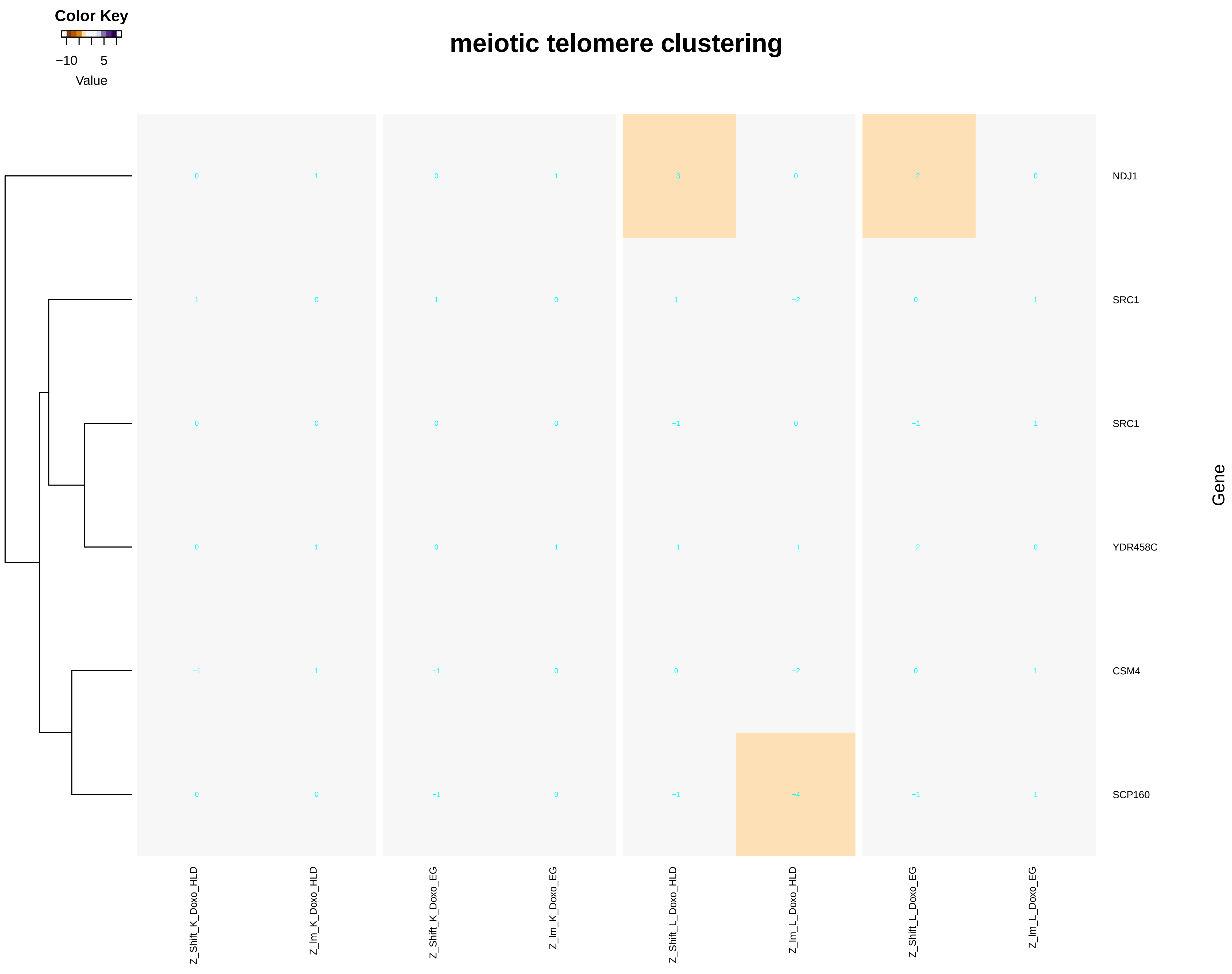

Color Key
−10
5
Value
meiotic telomere clustering
NDJ1
SRC1
SRC1
YDR458C
CSM4
SCP160
Gene
Z_Shift_K_Doxo_HLD
Z_Im_K_Doxo_HLD
Z_Shift_K_Doxo_EG
Z_Im_K_Doxo_EG
Z_Shift_L_Doxo_HLD
Z_Im_L_Doxo_HLD
Z_Shift_L_Doxo_EG
Z_Im_L_Doxo_EG
0 1 0 1 −3 0 −2 0
1 0 1 0 1 −2 0 1
0 0 0 0 −1 0 −1 1
0 1 0 1 −1 −1 −2 0
−1 1 −1 0 0 −2 0 1
0 0 −1 0 −1 −4 −1 1

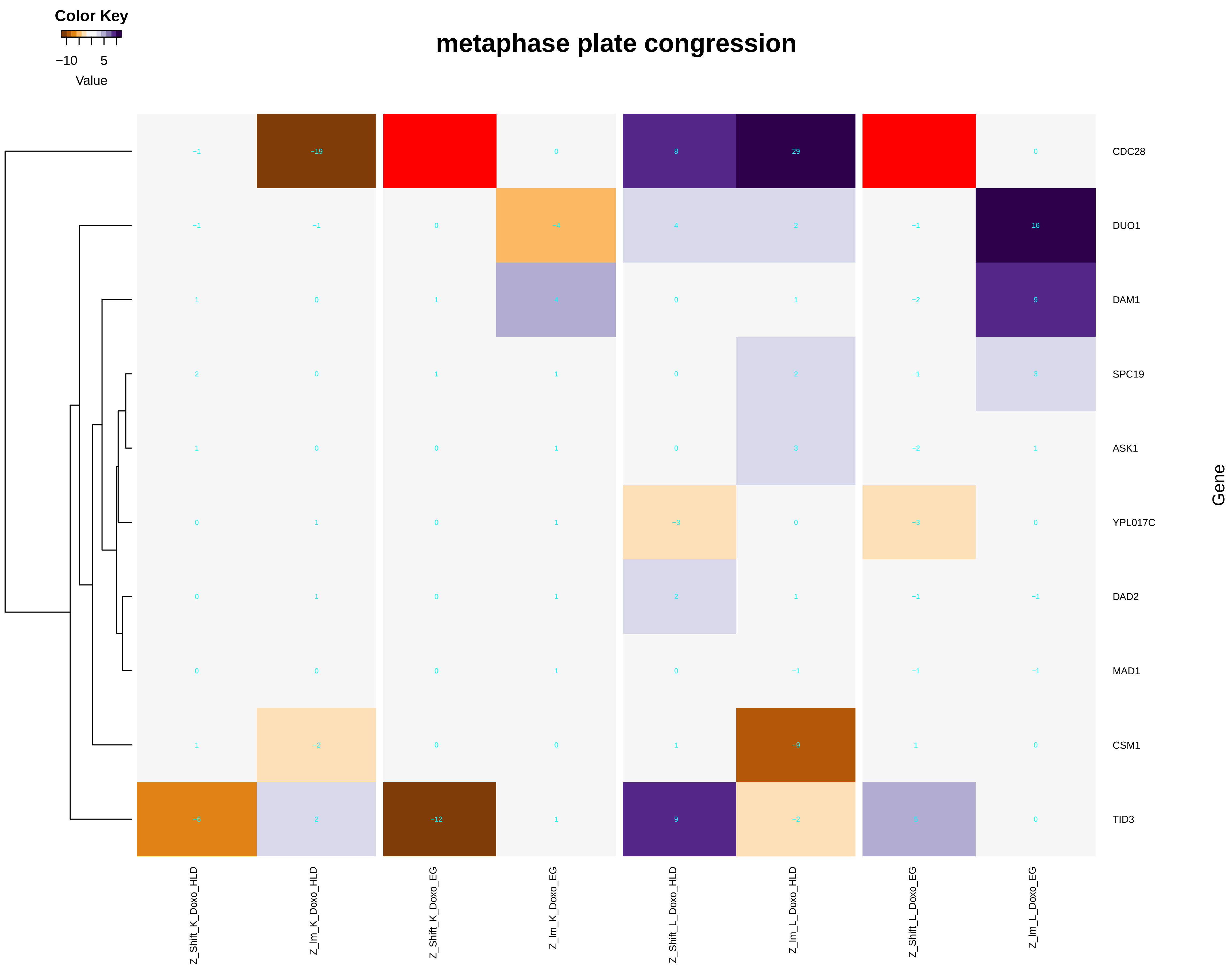
Color Key
−10
5
Value
metaphase plate congression
CDC28
DUO1
DAM1
SPC19
ASK1
YPL017C
DAD2
MAD1
CSM1
TID3
Gene
Z_Shift_K_Doxo_HLD
Z_Im_K_Doxo_HLD
Z_Shift_K_Doxo_EG
Z_Im_K_Doxo_EG
Z_Shift_L_Doxo_HLD
Z_Im_L_Doxo_HLD
Z_Shift_L_Doxo_EG
Z_Im_L_Doxo_EG

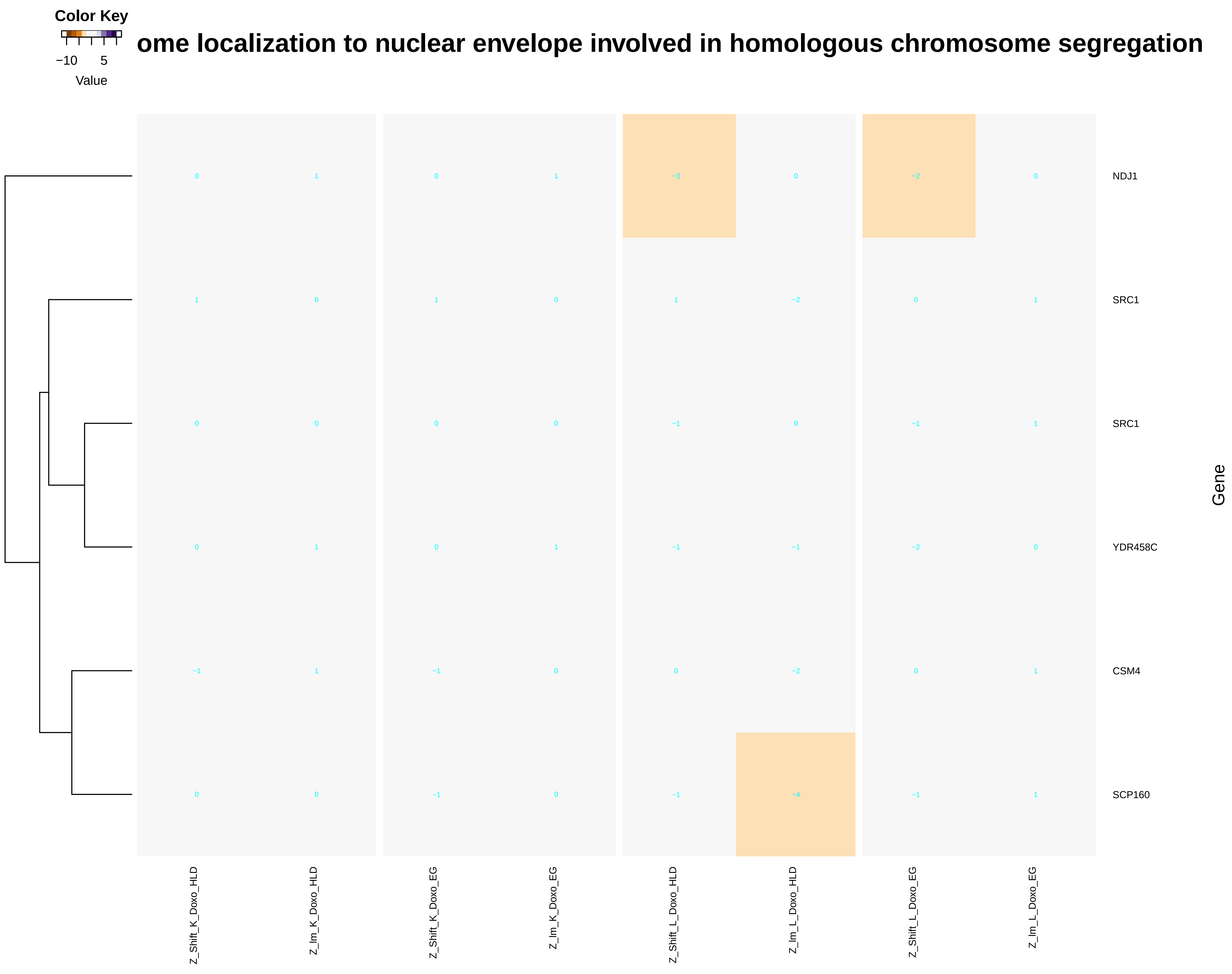
Color Key
−10
5
Value
ome localization to nuclear envelope involved in homologous chromosome segregation
NDJ1
SRC1
SRC1
YDR458C
CSM4
SCP160
Gene
Z_Shift_K_Doxo_HLD
Z_Im_K_Doxo_HLD
Z_Shift_K_Doxo_EG
Z_Im_K_Doxo_EG
Z_Shift_L_Doxo_HLD
Z_Im_L_Doxo_HLD
Z_Shift_L_Doxo_EG
Z_Im_L_Doxo_EG

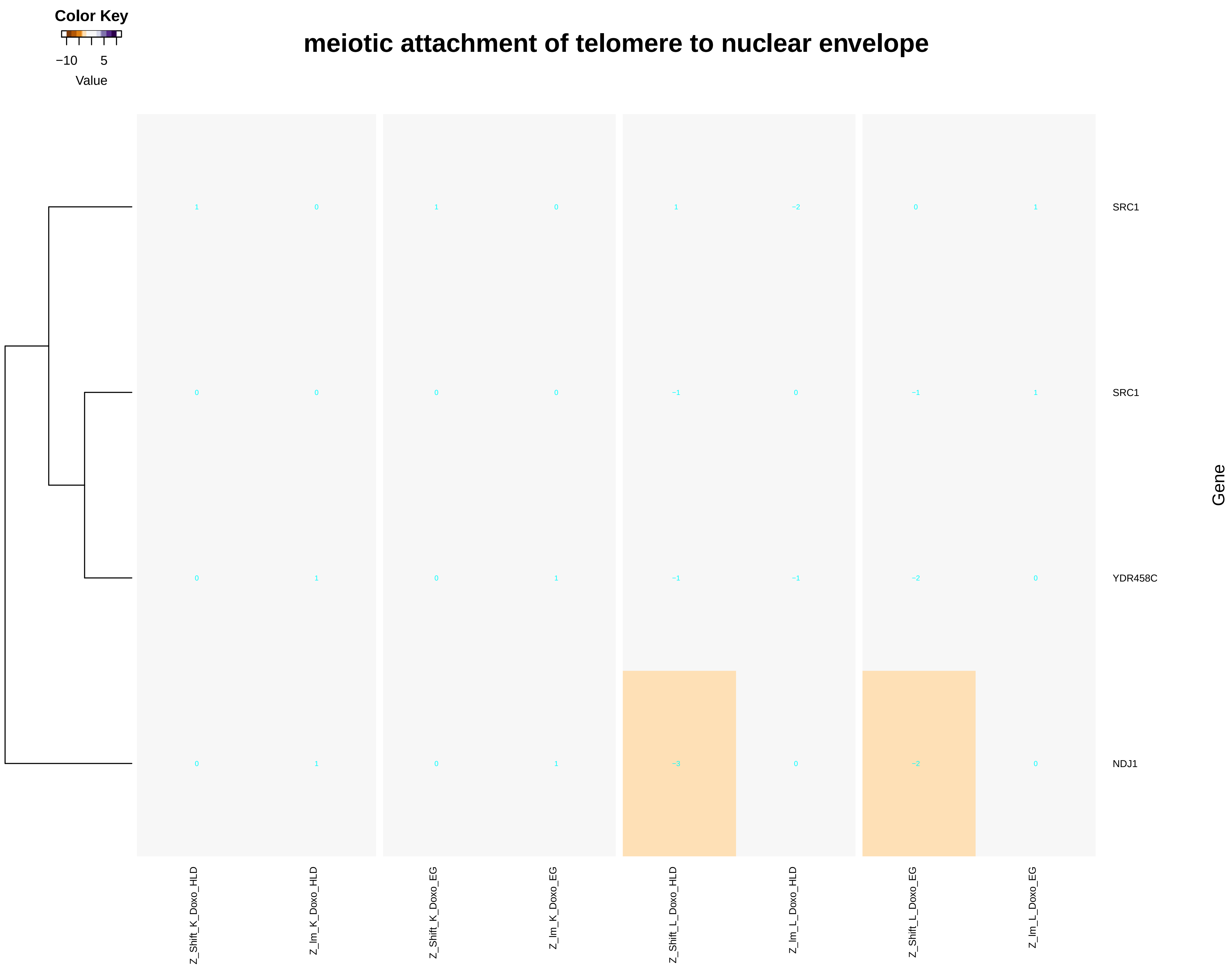

Color Key
−10
5
Value
meiotic attachment of telomere to nuclear envelope
SRC1
SRC1
YDR458C
NDJ1
Gene
Z_Shift_K_Doxo_HLD
Z_Im_K_Doxo_HLD
Z_Shift_K_Doxo_EG
Z_Im_K_Doxo_EG
Z_Shift_L_Doxo_HLD
Z_Im_L_Doxo_HLD
Z_Shift_L_Doxo_EG
Z_Im_L_Doxo_EG

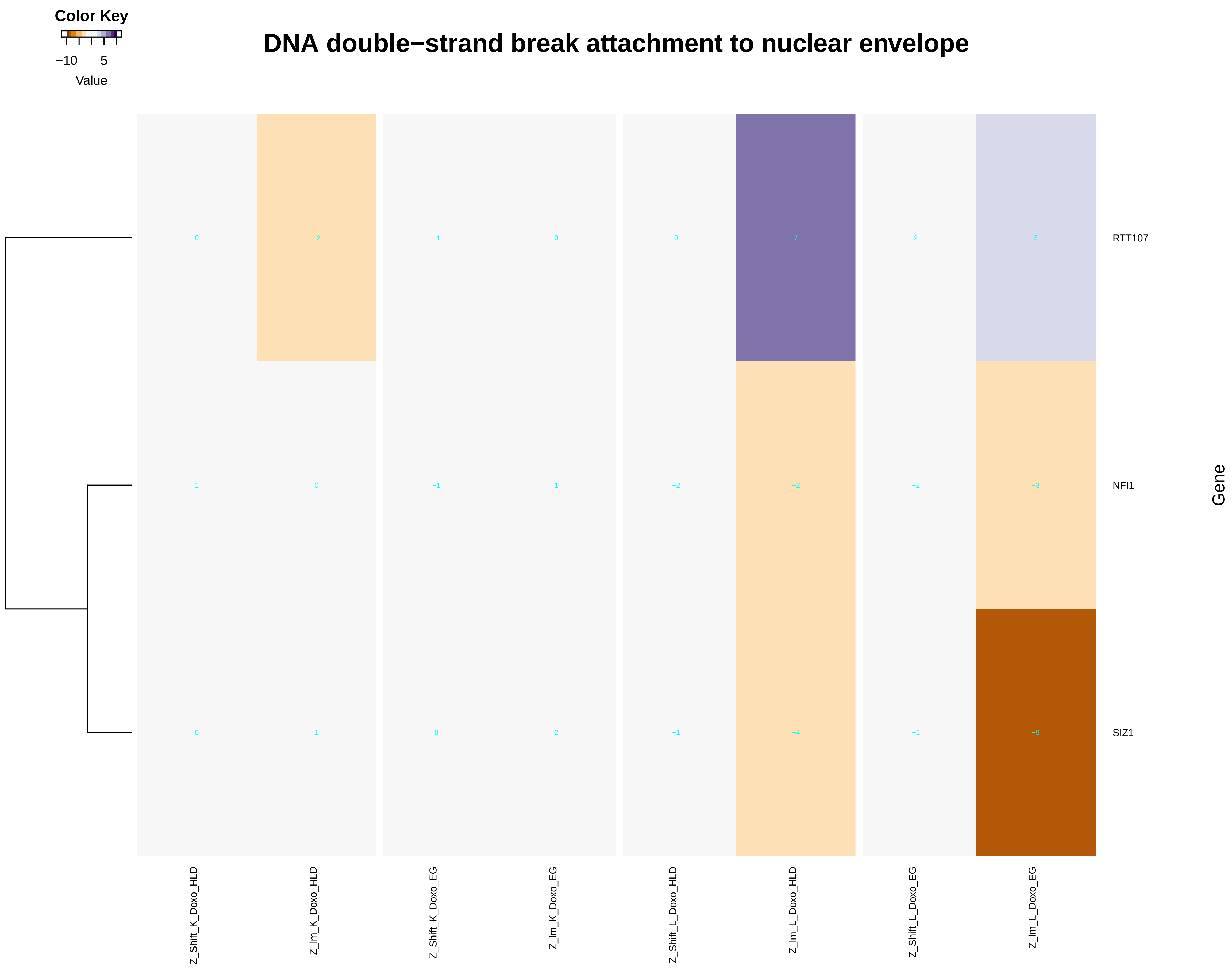
Color Key
−10
5
Value
DNA double−strand break attachment to nuclear envelope
0
−2
−1
0
0
7
2
3
RTT107
1
0
−1
1
−2
−2
−2
−3
NFI1
0
1
0
2
−1
−4
−1
−9
SIZ1
Gene
Z_Shift_K_Doxo_HLD
Z_Im_K_Doxo_HLD
Z_Shift_K_Doxo_EG
Z_Im_K_Doxo_EG
Z_Shift_L_Doxo_HLD
Z_Im_L_Doxo_HLD
Z_Shift_L_Doxo_EG
Z_Im_L_Doxo_EG

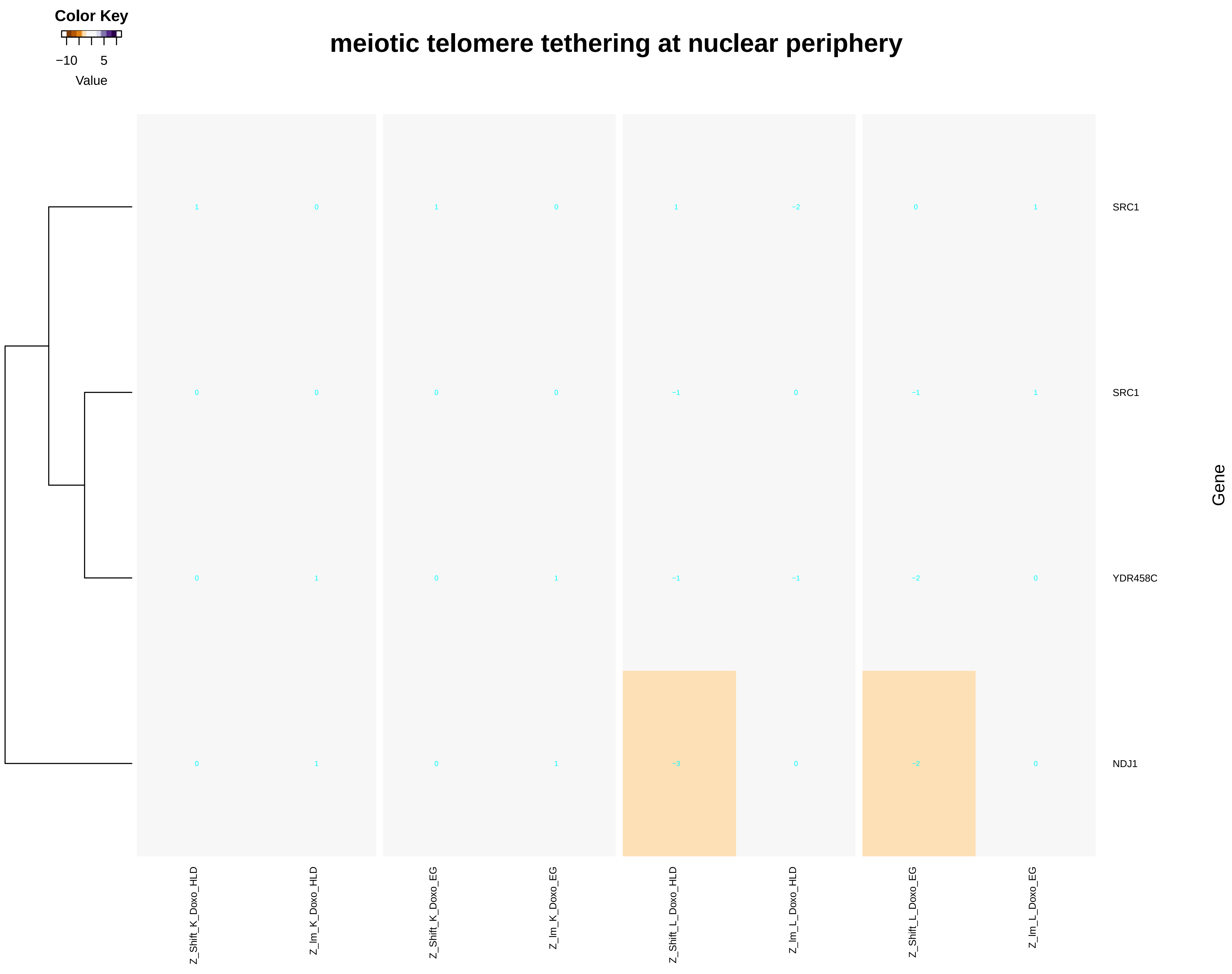
Color Key
−10
5
Value
meiotic telomere tethering at nuclear periphery
SRC1
SRC1
YDR458C
NDJ1
Gene
Z_Shift_K_Doxo_HLD
Z_lm_K_Doxo_HLD
Z_Shift_K_Doxo_EG
Z_lm_K_Doxo_EG
Z_Shift_L_Doxo_HLD
Z_lm_L_Doxo_HLD
Z_Shift_L_Doxo_EG
Z_lm_L_Doxo_EG

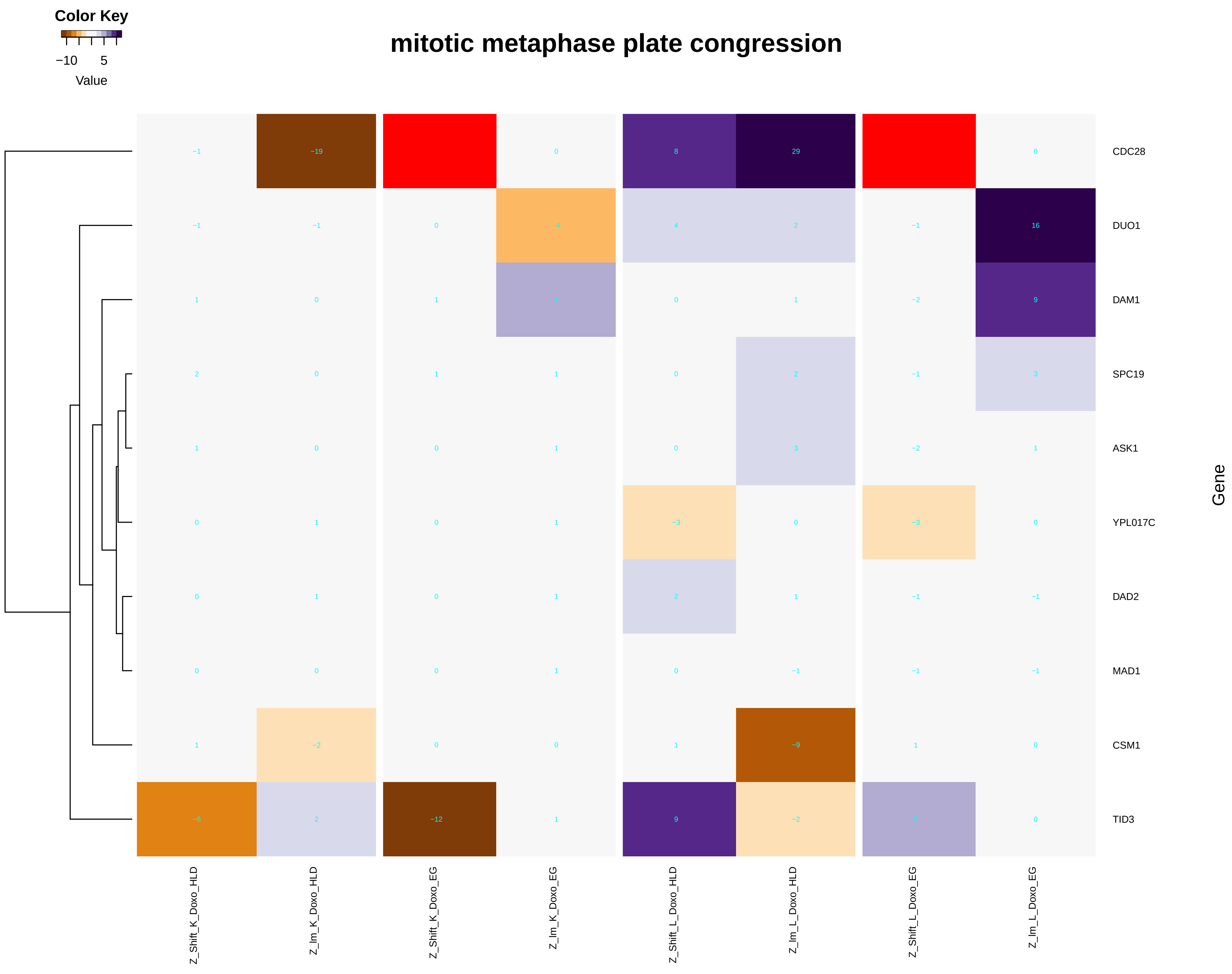
Color Key
−10
5
Value
mitotic metaphase plate congression
CDC28
DUO1
DAM1
SPC19
ASK1
YPL017C
DAD2
MAD1
CSM1
TID3
Gene
Z_Shift_K_Doxo_HLD
Z_Im_K_Doxo_HLD
Z_Shift_K_Doxo_EG
Z_Im_K_Doxo_EG
Z_Shift_L_Doxo_HLD
Z_Im_L_Doxo_HLD
Z_Shift_L_Doxo_EG
Z_Im_L_Doxo_EG
