## Additional File 9 for "A yeast phenomic model for the influence of Warburg metabolism on genetic buffering of doxorubicin": DNA_conformation_change.pdf

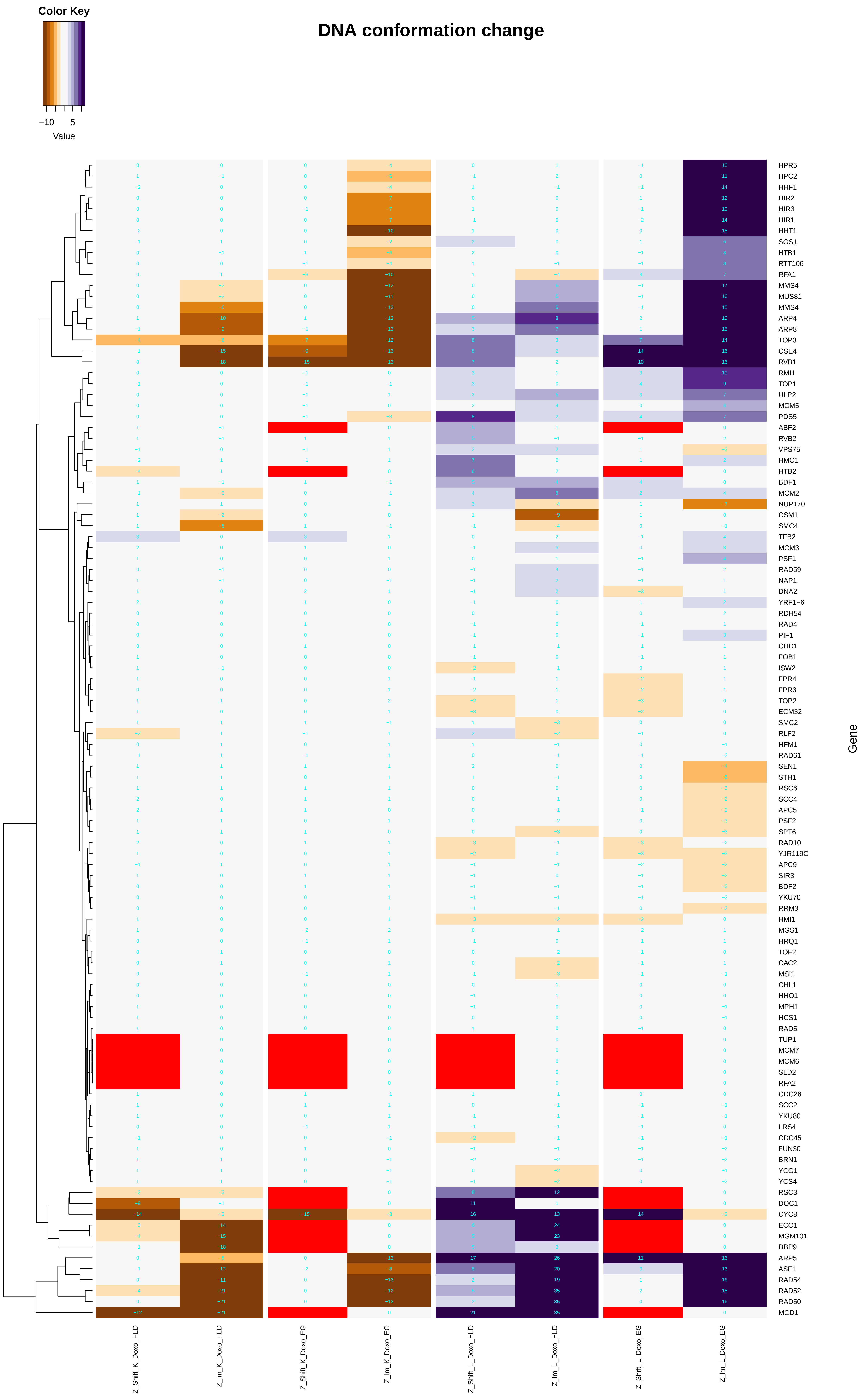

Color Key
−10
5
Value
DNA conformation change
HPR5
HPC2
HHF1
HIR2
HIR3
HIR1
HHT1
SGS1
HTB1
RTT106
RFA1
MMS4
MUS81
MMS4
ARP4
ARP8
TOP3
CSE4
RVB1
RMI1
TOP1
ULP2
MCM5
PDS5
ABF2
RVB2
VPS75
HMO1
HTB2
BDF1
MCM2
NUP170
CSM1
SMC4
TFB2
MCM3
PSF1
RAD59
NAP1
DNA2
YRF1−6
RDH54
RAD4
PIF1
CHD1
FOB1
ISW2
FPR4
FPR3
TOP2
ECM32
SMC2
RLF2
HFM1
RAD61
SEN1
STH1
RSC6
SCC4
APC5
PSF2
SPT6
RAD10
YJR119C
APC9
SIR3
BDF2
YKU70
RRM3
HMI1
MGS1
HRQ1
TOF2
CAC2
MSI1
CHL1
HHO1
MPH1
HCS1
RAD5
TUP1
MCM7
MCM6
SLD2
RFA2
CDC26
SCC2
YKU80
LRS4
CDC45
FUN30
BRN1
YCG1
YCS4
RSC3
DOC1
CYC8
ECO1
MGM101
DBP9
ARP5
ASF1
RAD54
RAD52
RAD50
MCD1
Gene
Z_Shift_K_Doxo_HLD
Z_Im_K_Doxo_HLD
Z_Shift_K_Doxo_EG
Z_Im_K_Doxo_EG
Z_Shift_L_Doxo_HLD
Z_Im_L_Doxo_HLD
Z_Shift_L_Doxo_EG
Z_Im_L_Doxo_EG

### DNA topological change

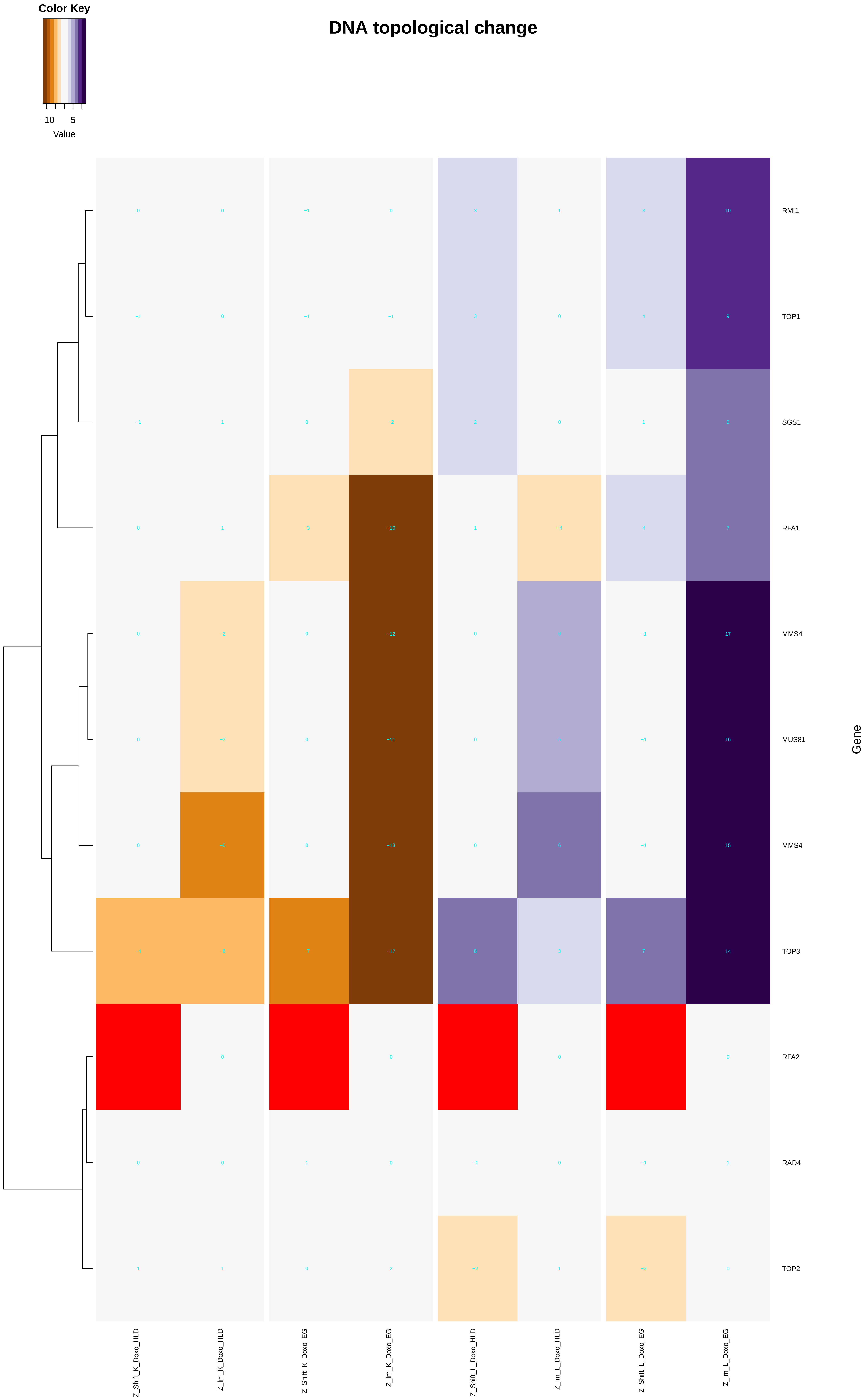

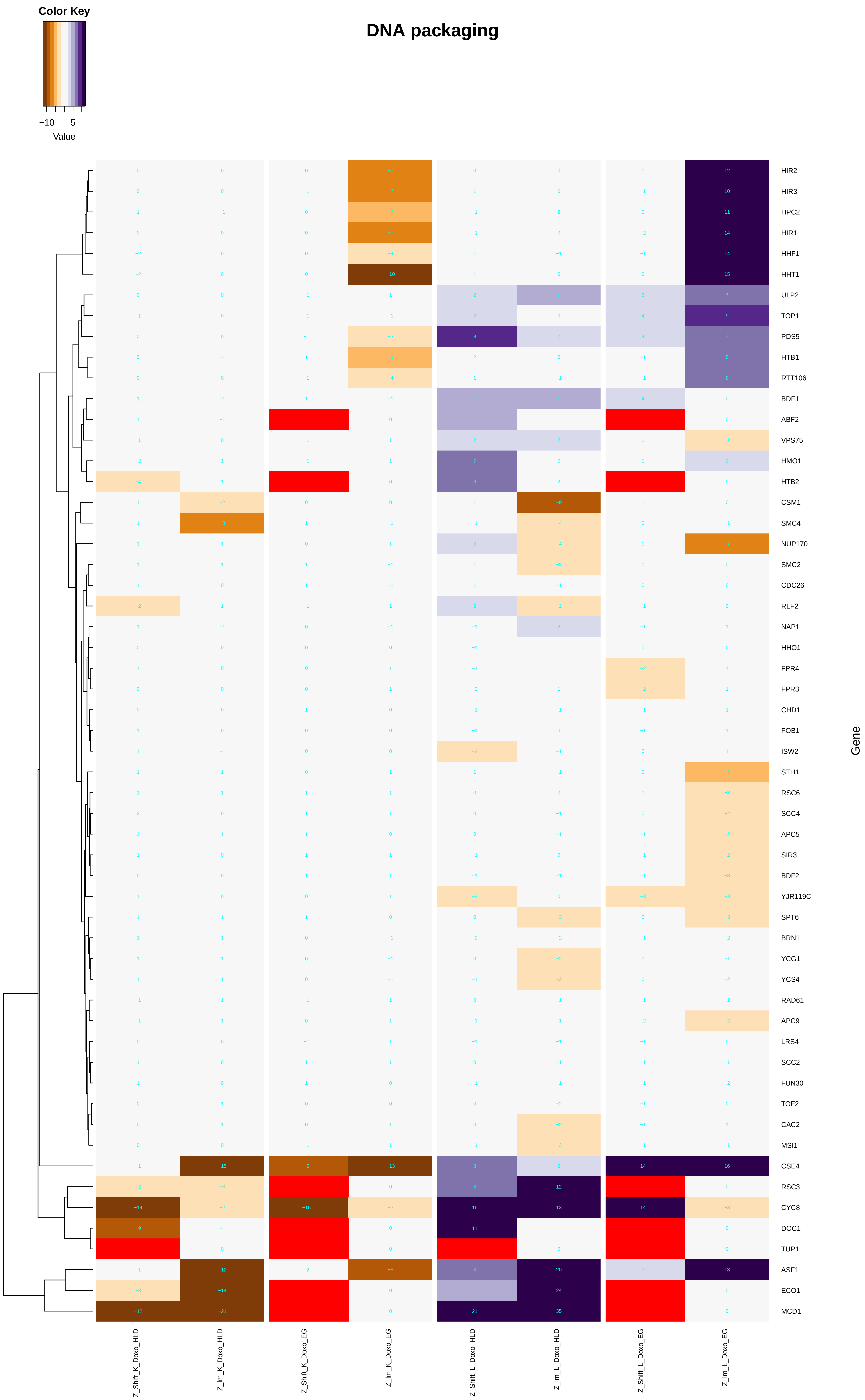

Color Key
−10 5
Value
DNA packaging
HIR2
HIR3
HPC2
HIR1
HHF1
HHT1
ULP2
TOP1
PDS5
HTB1
RTT106
BDF1
ABF2
VPS75
HMO1
HTB2
CSM1
SMC4
NUP170
SMC2
CDC26
RLF2
NAP1
HHO1
FPR4
FPR3
CHD1
FOB1
ISW2
STH1
RSC6
SCC4
APC5
SIR3
BDF2
YJR119C
SPT6
BRN1
YCG1
YCS4
RAD61
APC9
LRS4
SCC2
FUN30
TOF2
CAC2
MSI1
CSE4
RSC3
CYC8
DOC1
TUP1
ASF1
ECO1
MCD1
Gene
Z_Shift_K_Doxo_HLD
Z_Im_K_Doxo_HLD
Z_Shift_K_Doxo_EG
Z_Im_K_Doxo_EG
Z_Shift_L_Doxo_HLD
Z_Im_L_Doxo_HLD
Z_Shift_L_Doxo_EG
Z_Im_L_Doxo_EG

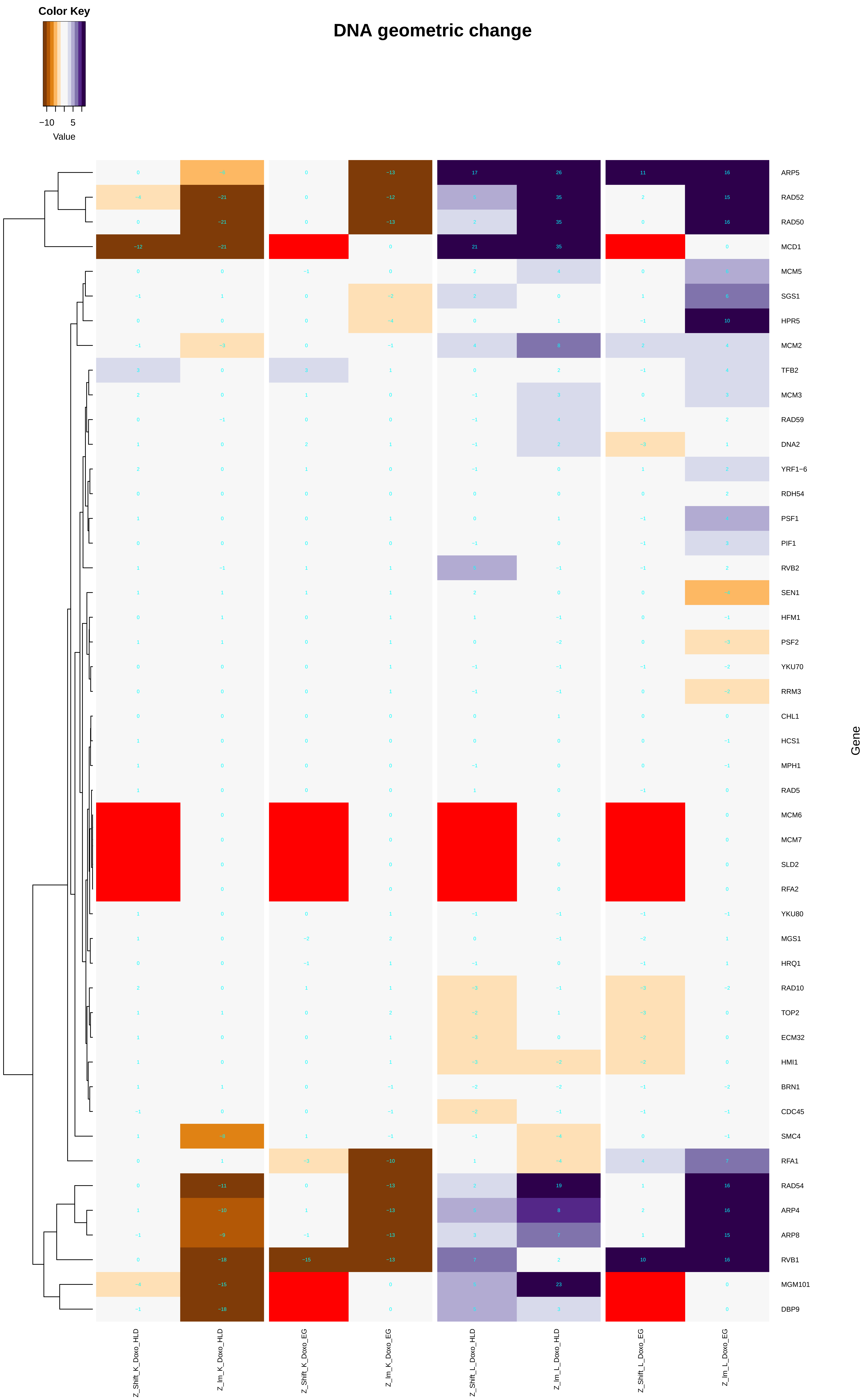

Color Key
−10 5
Value
DNA geometric change
ARP5
RAD52
RAD50
MCD1
MCM5
SGS1
HPR5
MCM2
TFB2
MCM3
RAD59
DNA2
YRF1−6
RDH54
PSF1
PIF1
RVB2
SEN1
HFM1
PSF2
YKU70
RRM3
CHL1
HCS1
MPH1
RAD5
MCM6
MCM7
SLD2
RFA2
YKU80
MGS1
HRQ1
RAD10
TOP2
ECM32
HMI1
BRN1
CDC45
SMC4
RFA1
RAD54
ARP4
ARP8
RVB1
MGM101
DBP9
Gene
Z_Shift_K_Doxo_HLD
Z_Im_K_Doxo_HLD
Z_Shift_K_Doxo_EG
Z_Im_K_Doxo_EG
Z_Shift_L_Doxo_HLD
Z_Im_L_Doxo_HLD
Z_Shift_L_Doxo_EG
Z_Im_L_Doxo_EG

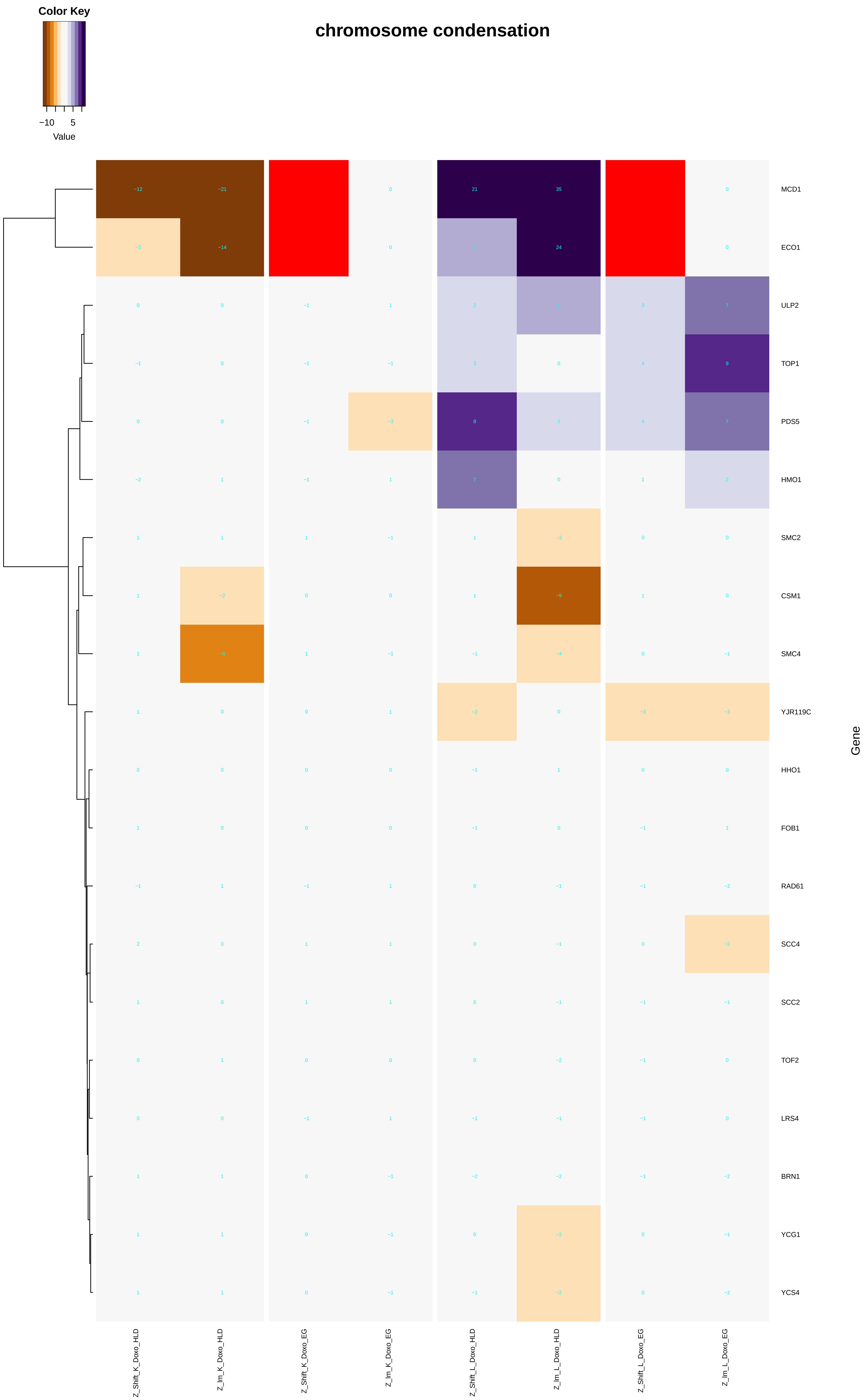

Color Key
−10
5
Value
chromosome condensation
MCD1
ECO1
ULP2
TOP1
PDS5
HMO1
SMC2
CSM1
SMC4
YJR119C
HHO1
FOB1
RAD61
SCC4
SCC2
TOF2
LRS4
BRN1
YCG1
YCS4
Gene
Z_Shift_K_Doxo_HLD
Z_lm_K_Doxo_HLD
Z_Shift_K_Doxo_EG
Z_lm_K_Doxo_EG
Z_Shift_L_Doxo_HLD
Z_lm_L_Doxo_HLD
Z_Shift_L_Doxo_EG
Z_lm_L_Doxo_EG

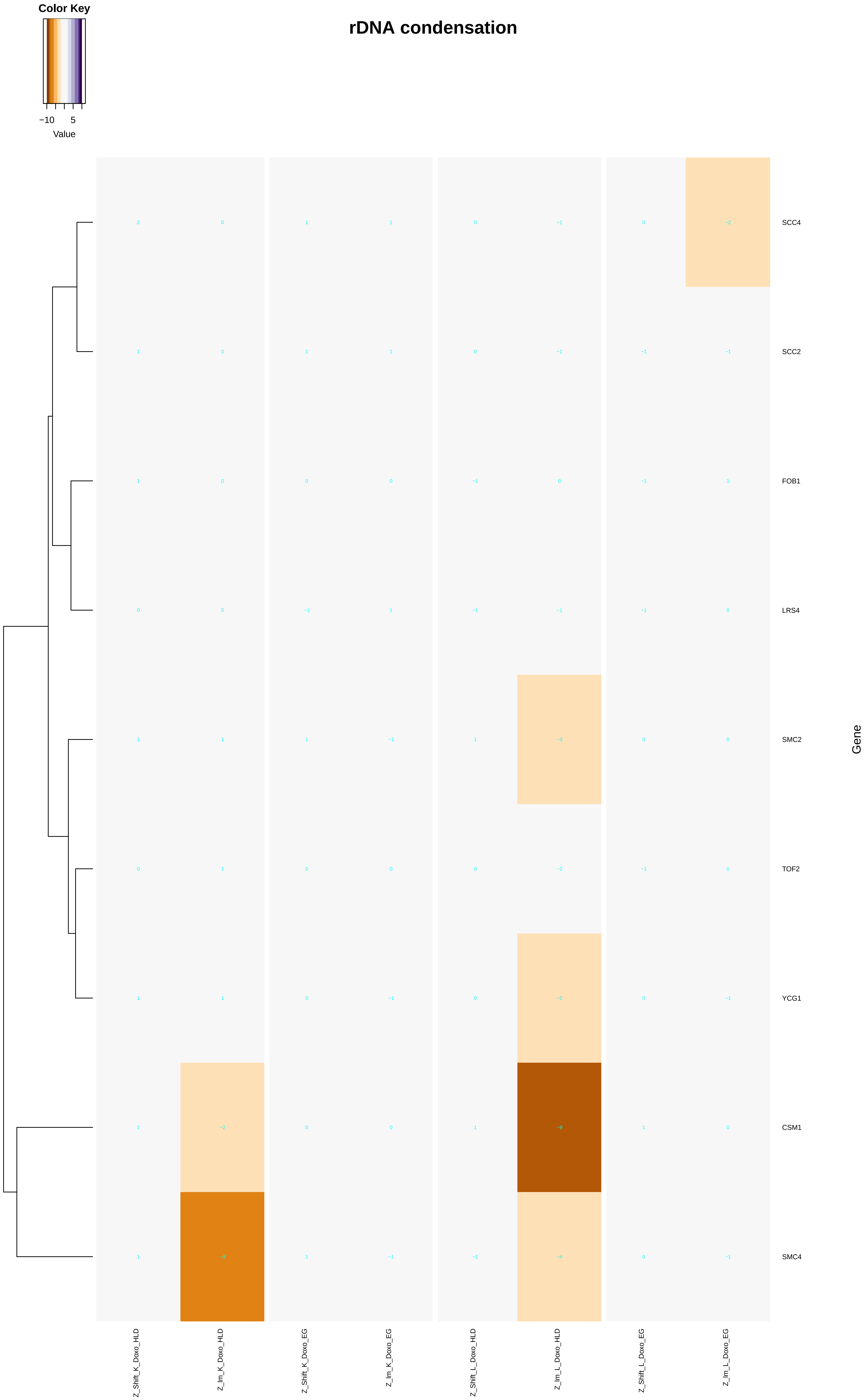
Color Key
−10
5
Value
rDNA condensation
SCC4
SCC2
FOB1
LRS4
SMC2
TOF2
YCG1
CSM1
SMC4
Gene
Z_Shift_K_Doxo_HLD
Z_Im_K_Doxo_HLD
Z_Shift_K_Doxo_EG
Z_Im_K_Doxo_EG
Z_Shift_L_Doxo_HLD
Z_Im_L_Doxo_HLD
Z_Shift_L_Doxo_EG
Z_Im_L_Doxo_EG

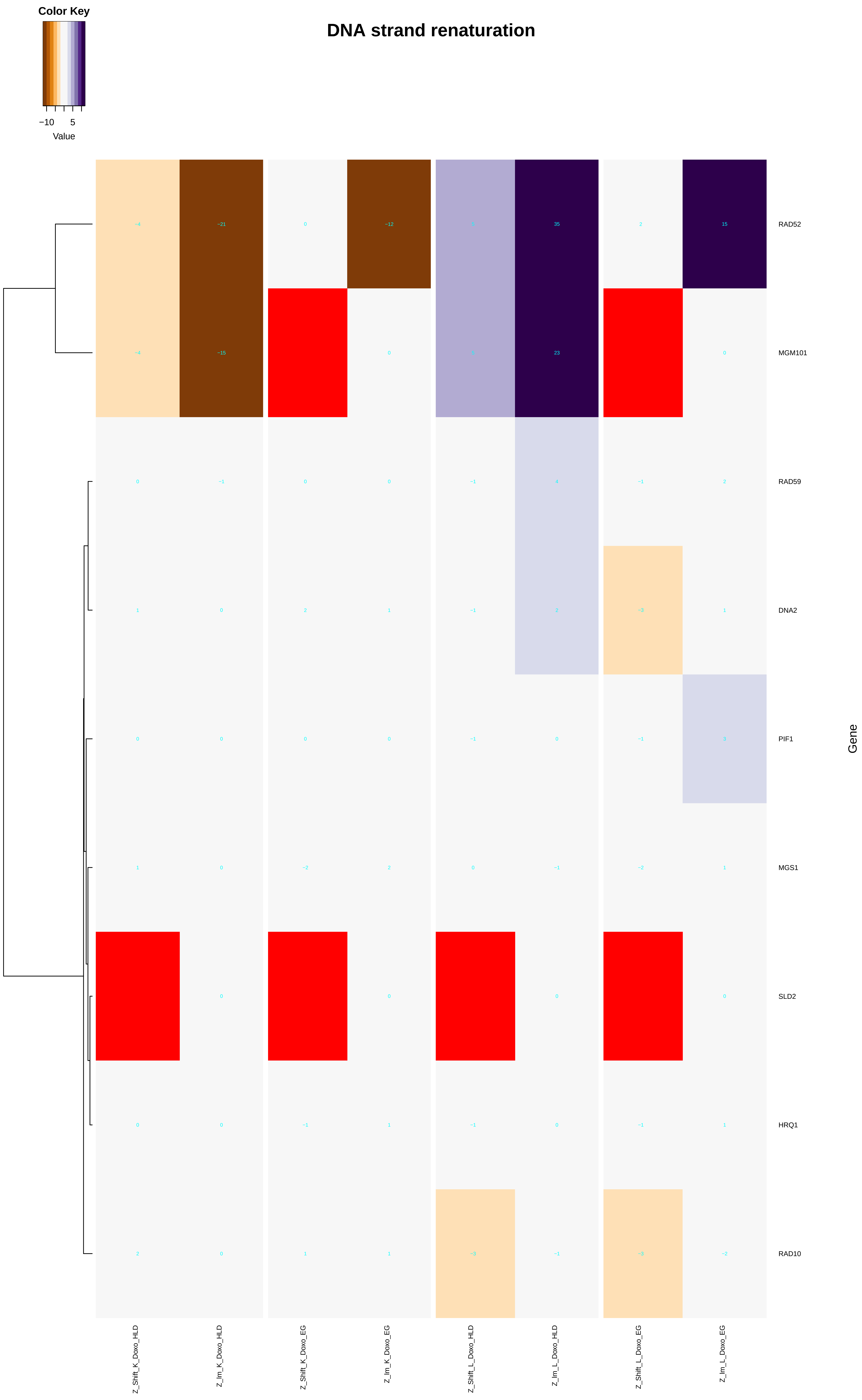

DNA strand renaturation
Color Key
−10
5
Value
RAD52
MGM101
RAD59
DNA2
PIF1
MGS1
SLD2
HRQ1
RAD10
Gene
Z_Shift_K_Doxo_HLD
Z_Im_K_Doxo_HLD
Z_Shift_K_Doxo_EG
Z_Im_K_Doxo_EG
Z_Shift_L_Doxo_HLD
Z_Im_L_Doxo_HLD
Z_Shift_L_Doxo_EG
Z_Im_L_Doxo_EG

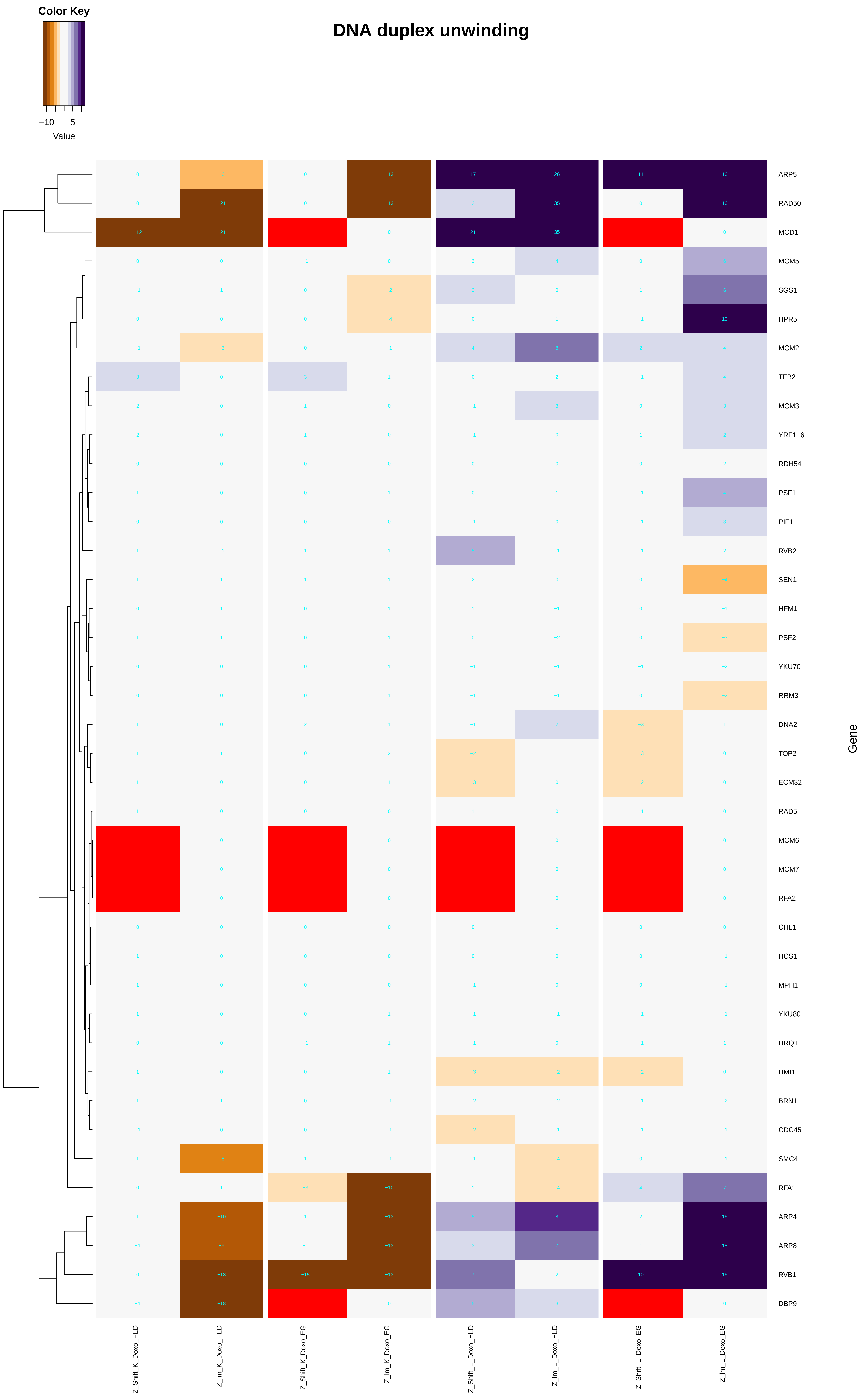
DNA duplex unwinding
Color Key
−10 5
Value
ARP5
RAD50
MCD1
MCM5
SGS1
HPR5
MCM2
TFB2
MCM3
YRF1−6
RDH54
PSF1
PIF1
RVB2
SEN1
HFM1
PSF2
YKU70
RRM3
DNA2
TOP2
ECM32
RAD5
MCM6
MCM7
RFA2
CHL1
HCS1
MPH1
YKU80
HRQ1
HMI1
BRN1
CDC45
SMC4
RFA1
ARP4
ARP8
RVB1
DBP9
Gene
Z_Shift_K_Doxo_HLD
Z_lm_K_Doxo_HLD
Z_Shift_K_Doxo_EG
Z_lm_K_Doxo_EG
Z_Shift_L_Doxo_HLD
Z_lm_L_Doxo_HLD
Z_Shift_L_Doxo_EG
Z_lm_L_Doxo_EG

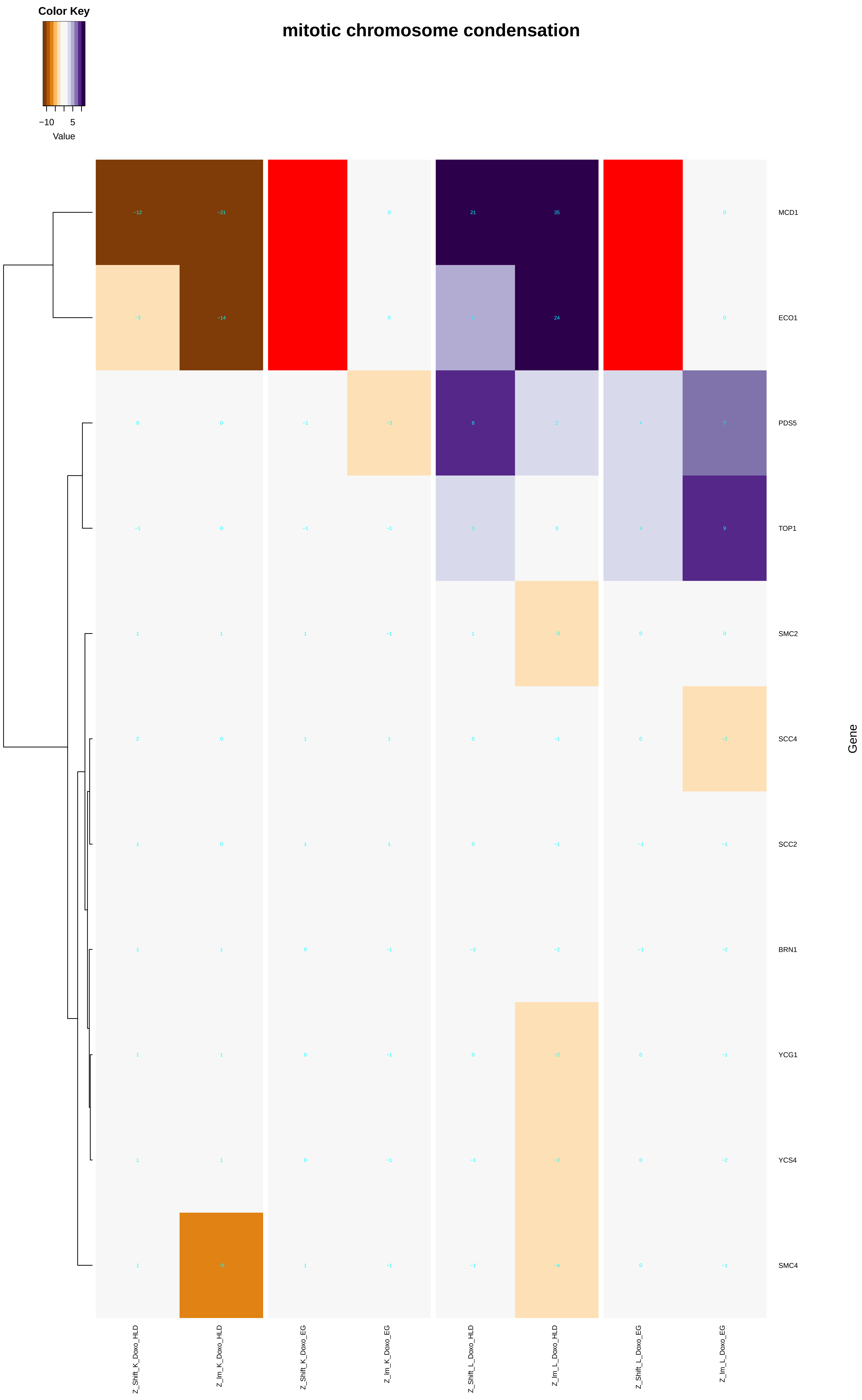
Color Key
−10
5
Value
mitotic chromosome condensation
MCD1
ECO1
PDS5
TOP1
SMC2
SCC4
SCC2
BRN1
YCG1
YCS4
SMC4
Gene
Z_Shift_K_Doxo_HLD
Z_Im_K_Doxo_HLD
Z_Shift_K_Doxo_EG
Z_Im_K_Doxo_EG
Z_Shift_L_Doxo_HLD
Z_Im_L_Doxo_HLD
Z_Shift_L_Doxo_EG
Z_Im_L_Doxo_EG

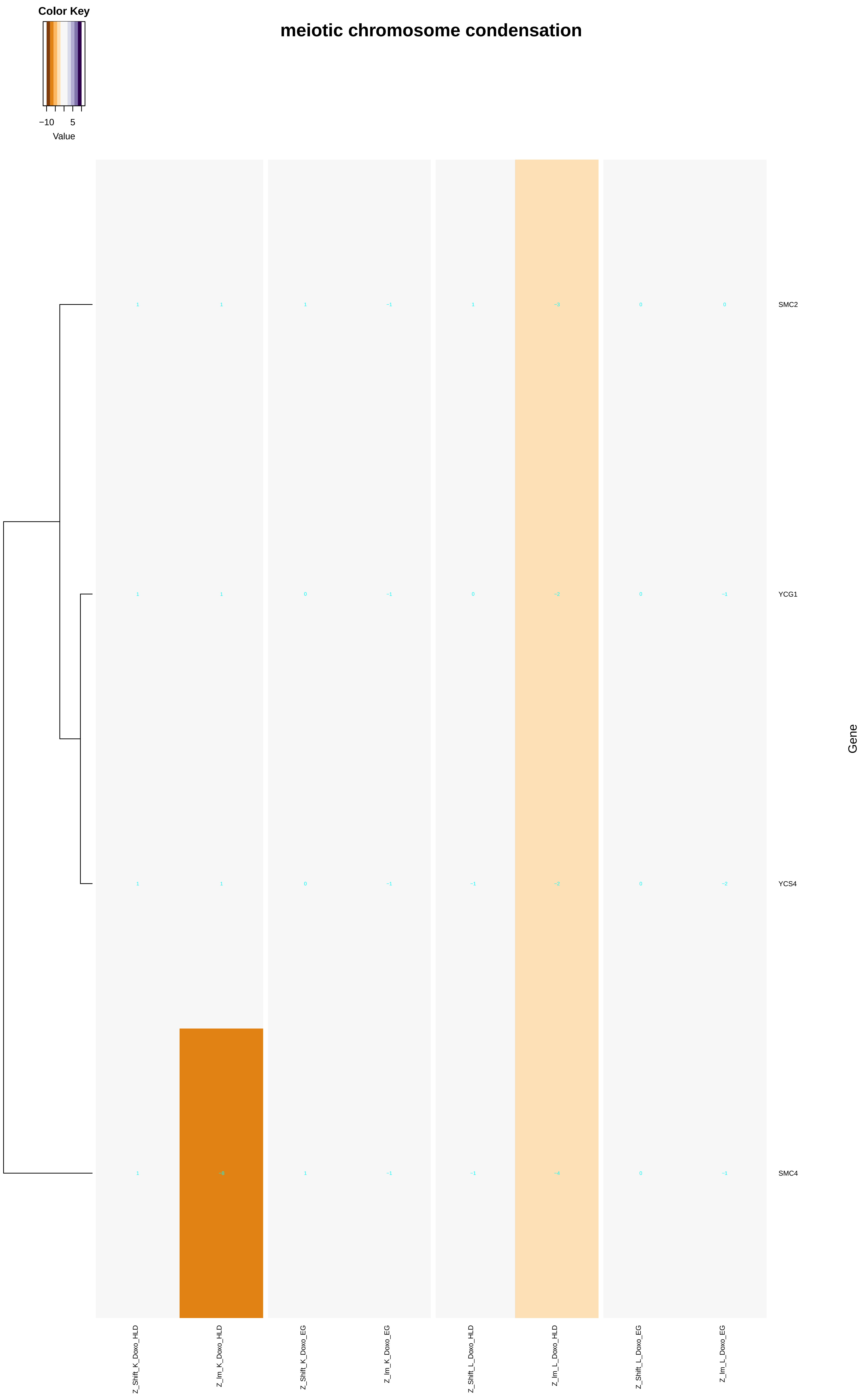
Color Key
−10
5
Value
meiotic chromosome condensation
1
1
1
−1
1
−3
0
0
SMC2
1
1
0
−1
0
−2
0
−1
YCG1
1
1
0
−1
−1
−2
0
−2
YCS4
1
−8
1
−1
−1
−4
0
−1
SMC4
Gene
Z_Shift_K_Doxo_HLD
Z_lm_K_Doxo_HLD
Z_Shift_K_Doxo_EG
Z_lm_K_Doxo_EG
Z_Shift_L_Doxo_HLD
Z_lm_L_Doxo_HLD
Z_Shift_L_Doxo_EG
Z_lm_L_Doxo_EG

### DNA unwinding involved in DNA replication

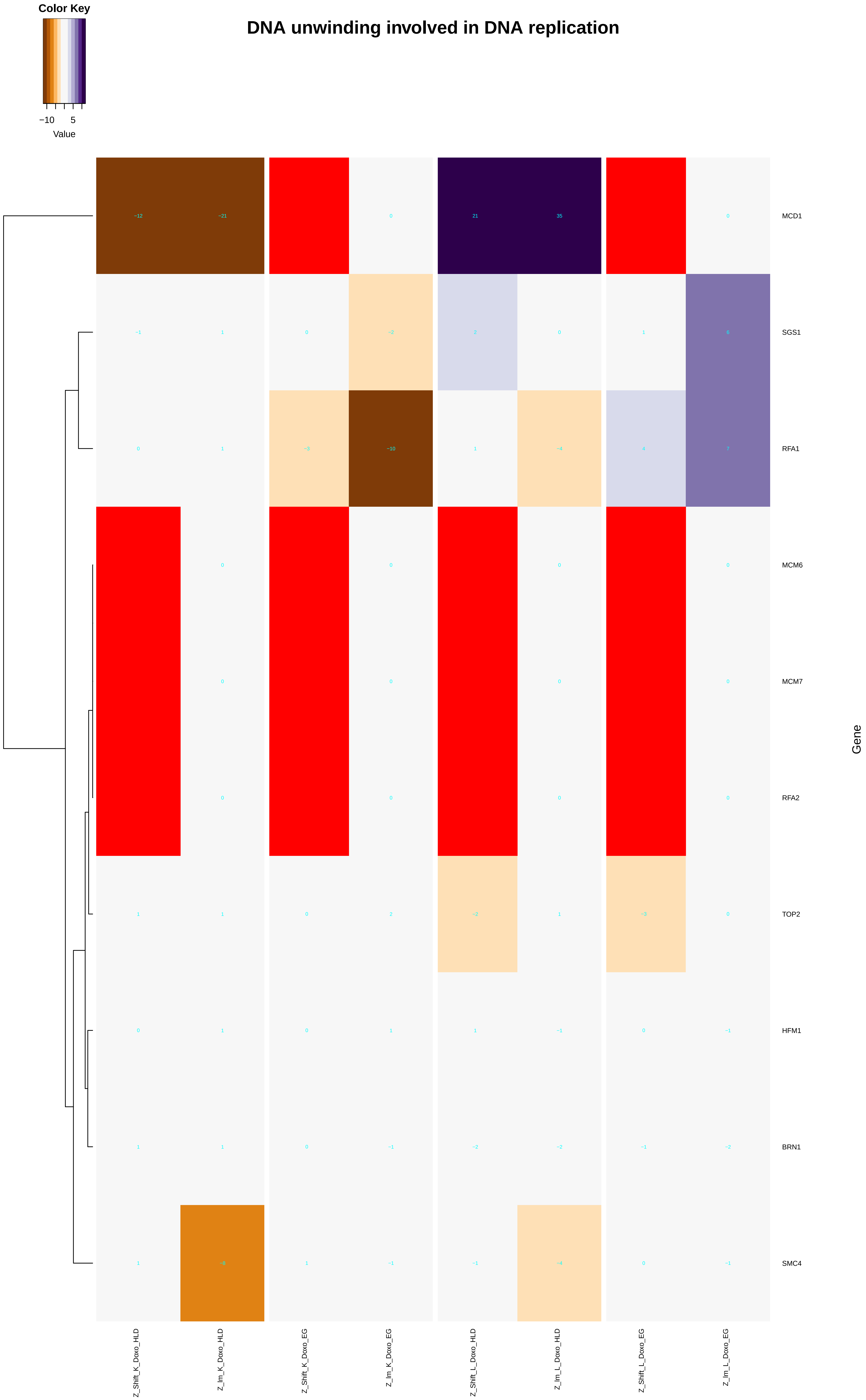
