## Additional File 9 for "A yeast phenomic model for the influence of Warburg metabolism on genetic buffering of doxorubicin": DNA_recombination.pdf

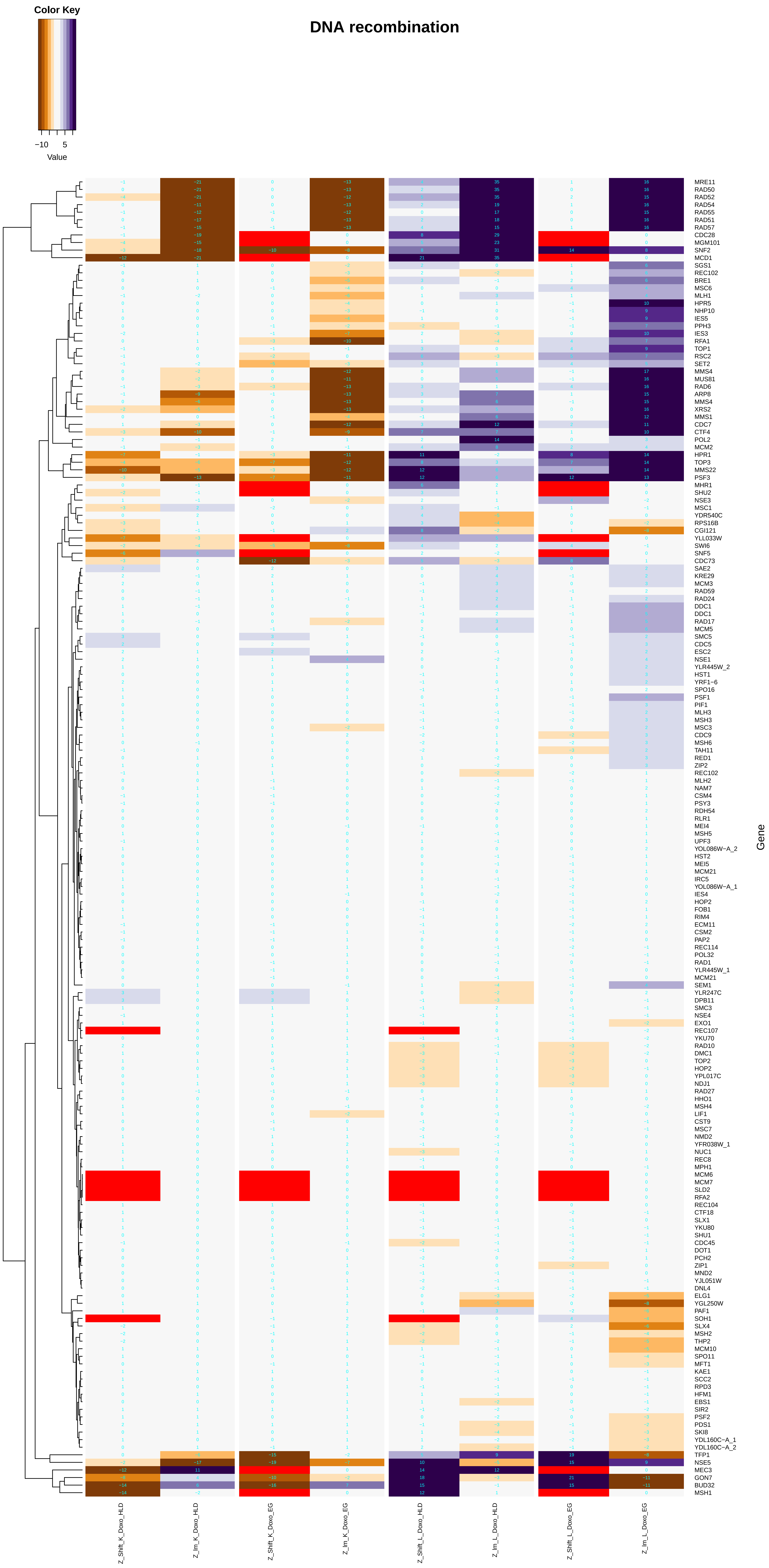

DNA recombination
Color Key
−10
5
Value
Gene
MRE11
RAD50
RAD52
RAD54
RAD55
RAD51
RAD57
CDC28
MGM101
SNF2
MCD1
SGS1
REC102
BRE1
MSC6
MLH1
HPR5
NHP10
IES5
PPH3
IES3
RFA1
TOP1
RSC2
SET2
MMS4
MUS81
RAD6
ARP8
MMS4
XRS2
MMS1
CDC7
CTF4
POL2
MCM2
HPR1
TOP3
MMS22
PSF3
MHR1
SHU2
NSE3
MSC1
YDR540C
RPS16B
CGI121
YLL033W
SWI6
SNF5
CDC73
SAE2
KRE29
MCM3
RAD59
RAD24
DDC1
DDC1
RAD17
MCM5
SMC5
CDC5
ESC2
NSE1
YLR445W_2
HST1
YRF1−6
SPO16
PSF1
PIF1
MLH3
MSH3
MSC3
CDC9
MSH6
TAH11
RED1
ZIP2
REC102
MLH2
NAM7
CSM4
PSY3
RDH54
RLR1
MEI4
MSH5
UPF3
YOL086W−A_2
HST2
MEI5
MCM21
IRC5
YOL086W−A_1
IES4
HOP2
FOB1
RIM4
ECM11
CSM2
PAP2
REC114
POL32
RAD1
YLR445W_1
MCM21
SEM1
YLR247C
DPB11
SMC3
NSE4
EXO1
REC107
YKU70
RAD10
DMC1
TOP2
HOP2
YPL017C
NDJ1
RAD27
HHO1
MSH4
LIF1
CST9
MSC7
NMD2
YFR038W_1
NUC1
REC8
MPH1
MCM6
MCM7
SLD2
RFA2
REC104
CTF18
SLX1
YKU80
SHU1
CDC45
DOT1
PCH2
ZIP1
MND2
YJL051W
DNL4
ELG1
YGL250W
PAF1
SOH1
SLX4
MSH2
THP2
MCM10
SPO11
MFT1
KAE1
SCC2
RPD3
HFM1
EBS1
SIR2
PSF2
PDS1
SKI8
YDL160C−A_1
YDL160C−A_2
TFP1
NSE5
MEC3
GON7
BUD32
MSH1
Z_Shift_K_Doxo_HLD
Z_Im_K_Doxo_HLD
Z_Shift_K_Doxo_EG
Z_Im_K_Doxo_EG
Z_Shift_L_Doxo_HLD
Z_Im_L_Doxo_HLD
Z_Shift_L_Doxo_EG
Z_Im_L_Doxo_EG

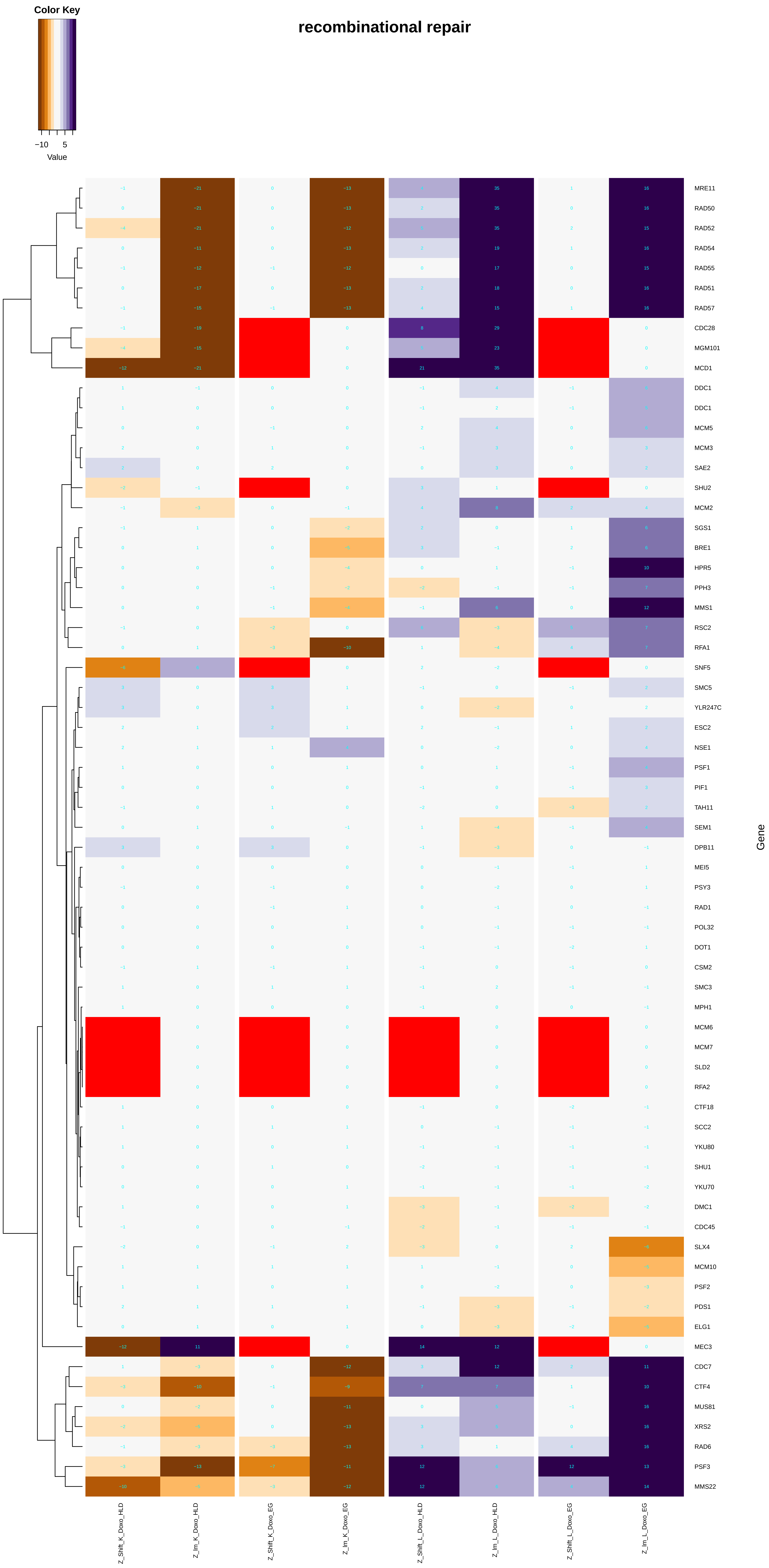
recombinational repair
Color Key
−10
5
Value
MRE11
RAD50
RAD52
RAD54
RAD55
RAD51
RAD57
CDC28
MGM101
MCD1
DDC1
DDC1
MCM5
MCM3
SAE2
SHU2
MCM2
SGS1
BRE1
HPR5
PPH3
MMS1
RSC2
RFA1
SNF5
SMC5
YLR247C
ESC2
NSE1
PSF1
PIF1
TAH11
SEM1
DPB11
MEI5
PSY3
RAD1
POL32
DOT1
CSM2
SMC3
MPH1
MCM6
MCM7
SLD2
RFA2
CTF18
SCC2
YKU80
SHU1
YKU70
DMC1
CDC45
SLX4
MCM10
PSF2
PDS1
ELG1
MEC3
CDC7
CTF4
MUS81
XRS2
RAD6
PSF3
MMS22
Gene
Z_Shift_K_Doxo_HLD
Z_lm_K_Doxo_HLD
Z_Shift_K_Doxo_EG
Z_lm_K_Doxo_EG
Z_Shift_L_Doxo_HLD
Z_lm_L_Doxo_HLD
Z_Shift_L_Doxo_EG
Z_lm_L_Doxo_EG

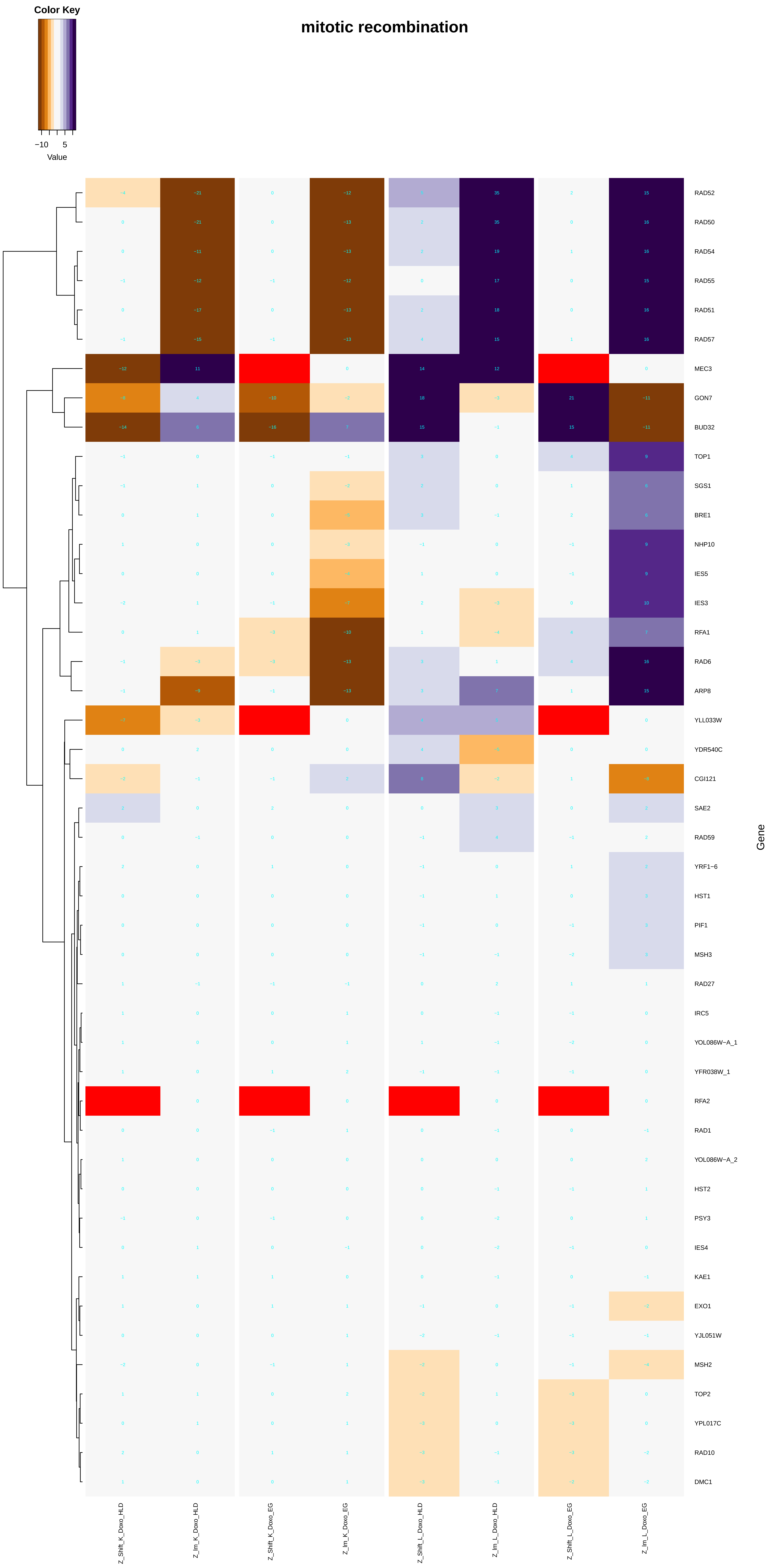
Color Key
−10
5
Value
mitotic recombination
RAD52
RAD50
RAD54
RAD55
RAD51
RAD57
MEC3
GON7
BUD32
TOP1
SGS1
BRE1
NHP10
IES5
IES3
RFA1
RAD6
ARP8
YLL033W
YDR540C
CGI121
SAE2
RAD59
YRF1−6
HST1
PIF1
MSH3
RAD27
IRC5
YOL086W−A_1
YFR038W_1
RFA2
RAD1
YOL086W−A_2
HST2
PSY3
IES4
KAE1
EXO1
YJL051W
MSH2
TOP2
YPL017C
RAD10
DMC1
Gene
Z_Shift_K_Doxo_HLD
Z_Im_K_Doxo_HLD
Z_Shift_K_Doxo_EG
Z_Im_K_Doxo_EG
Z_Shift_L_Doxo_HLD
Z_Im_L_Doxo_HLD
Z_Shift_L_Doxo_EG
Z_Im_L_Doxo_EG

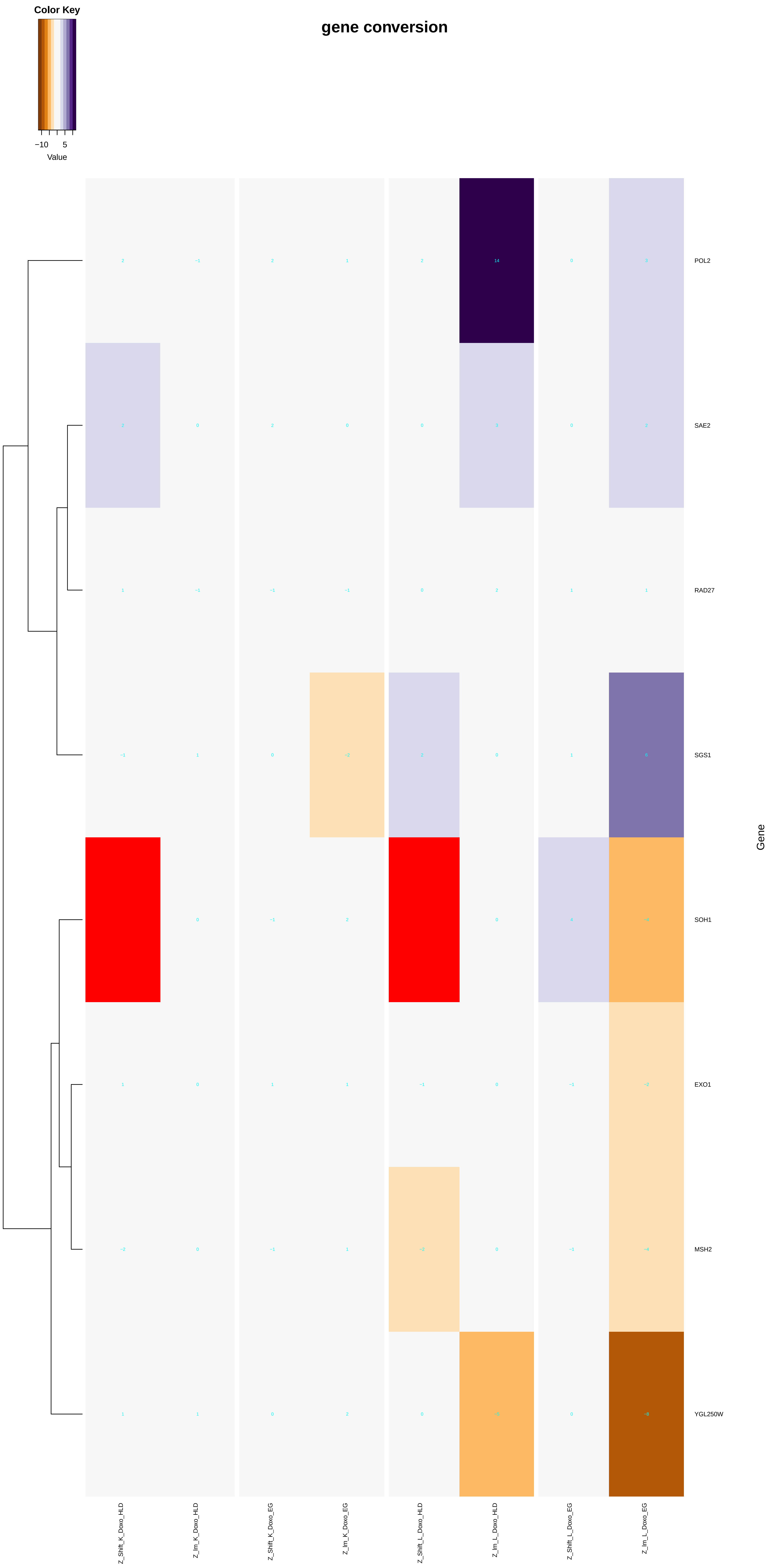
gene conversion
Color Key
−10
5
Value
POL2
SAE2
RAD27
SGS1
SOH1
EXO1
MSH2
YGL250W
Gene
Z_Shift_K_Doxo_HLD
Z_Im_K_Doxo_HLD
Z_Shift_K_Doxo_EG
Z_Im_K_Doxo_EG
Z_Shift_L_Doxo_HLD
Z_Im_L_Doxo_HLD
Z_Shift_L_Doxo_EG
Z_Im_L_Doxo_EG

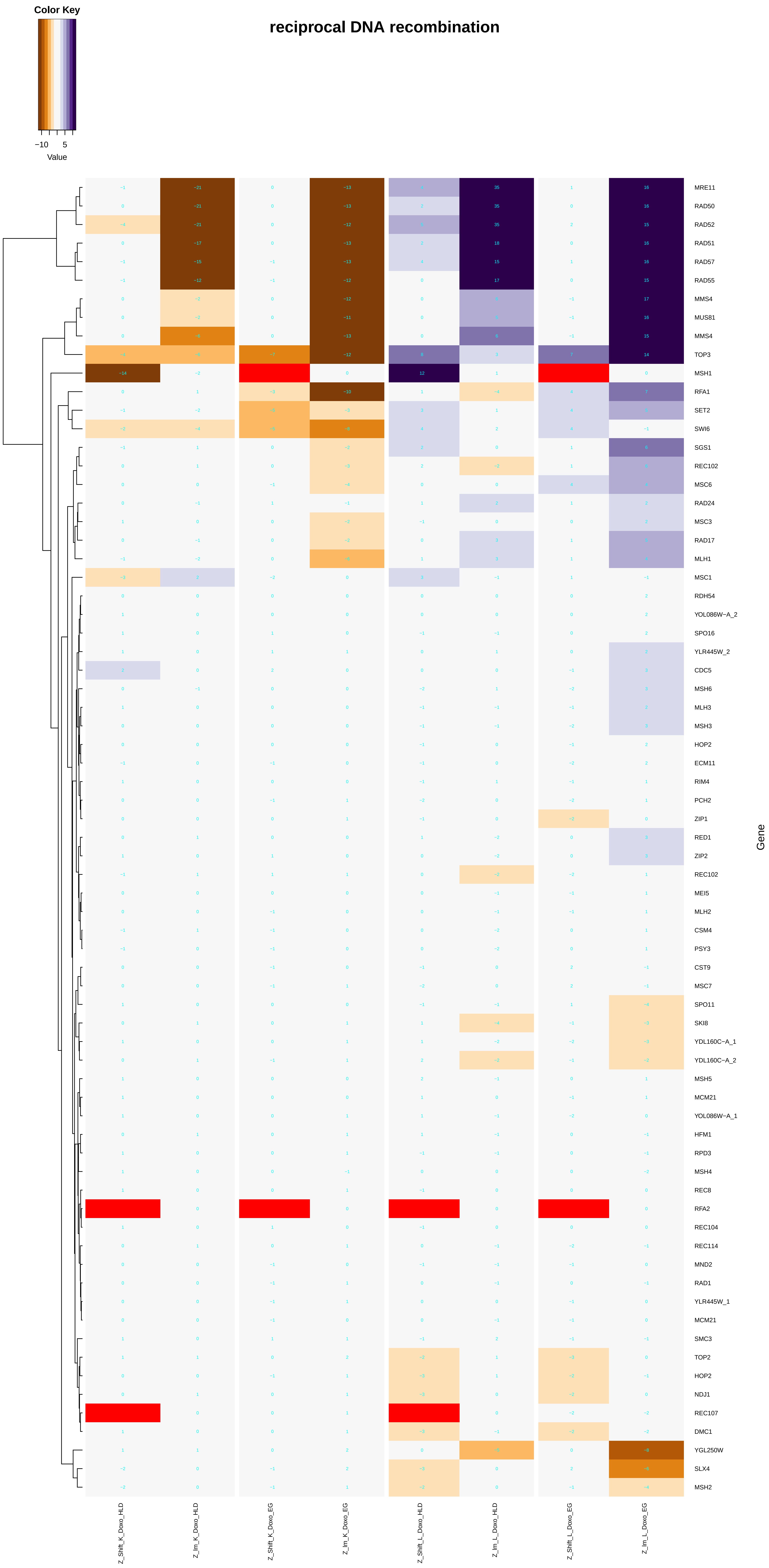

Color Key
−10
5
Value
reciprocal DNA recombination
Gene
MRE11
RAD50
RAD52
RAD51
RAD57
RAD55
MMS4
MUS81
MMS4
TOP3
MSH1
RFA1
SET2
SWI6
SGS1
REC102
MSC6
RAD24
MSC3
RAD17
MLH1
MSC1
RDH54
YOL086W−A_2
SPO16
YLR445W_2
CDC5
MSH6
MLH3
MSH3
HOP2
ECM11
RIM4
PCH2
ZIP1
RED1
ZIP2
REC102
MEI5
MLH2
CSM4
PSY3
CST9
MSC7
SPO11
SKI8
YDL160C−A_1
YDL160C−A_2
MSH5
MCM21
YOL086W−A_1
HFM1
RPD3
MSH4
REC8
RFA2
REC104
REC114
MND2
RAD1
YLR445W_1
MCM21
SMC3
TOP2
HOP2
NDJ1
REC107
DMC1
YGL250W
SLX4
MSH2
Z_Shift_K_Doxo_HLD
Z_Im_K_Doxo_HLD
Z_Shift_K_Doxo_EG
Z_Im_K_Doxo_EG
Z_Shift_L_Doxo_HLD
Z_Im_L_Doxo_HLD
Z_Shift_L_Doxo_EG
Z_Im_L_Doxo_EG

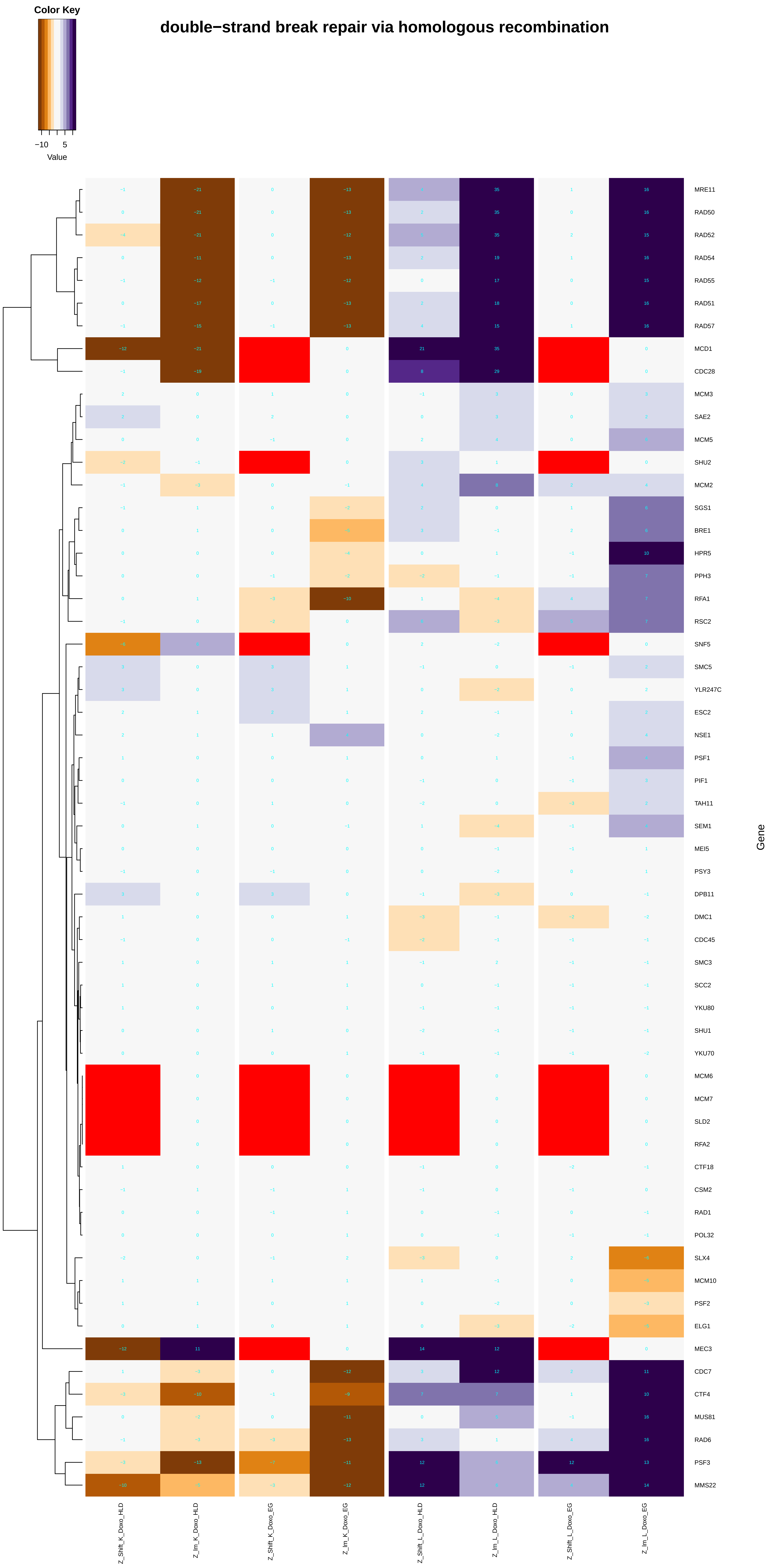
double−strand break repair via homologous recombination
Color Key
−10
5
Value
MRE11
RAD50
RAD52
RAD54
RAD55
RAD51
RAD57
MCD1
CDC28
MCM3
SAE2
MCM5
SHU2
MCM2
SGS1
BRE1
HPR5
PPH3
RFA1
RSC2
SNF5
SMC5
YLR247C
ESC2
NSE1
PSF1
PIF1
TAH11
SEM1
MEI5
PSY3
DPB11
DMC1
CDC45
SMC3
SCC2
YKU80
SHU1
YKU70
MCM6
MCM7
SLD2
RFA2
CTF18
CSM2
RAD1
POL32
SLX4
MCM10
PSF2
ELG1
MEC3
CDC7
CTF4
MUS81
RAD6
PSF3
MMS22
Gene
Z_Shift_K_Doxo_HLD
Z_lm_K_Doxo_HLD
Z_Shift_K_Doxo_EG
Z_lm_K_Doxo_EG
Z_Shift_L_Doxo_HLD
Z_lm_L_Doxo_HLD
Z_Shift_L_Doxo_EG
Z_lm_L_Doxo_EG

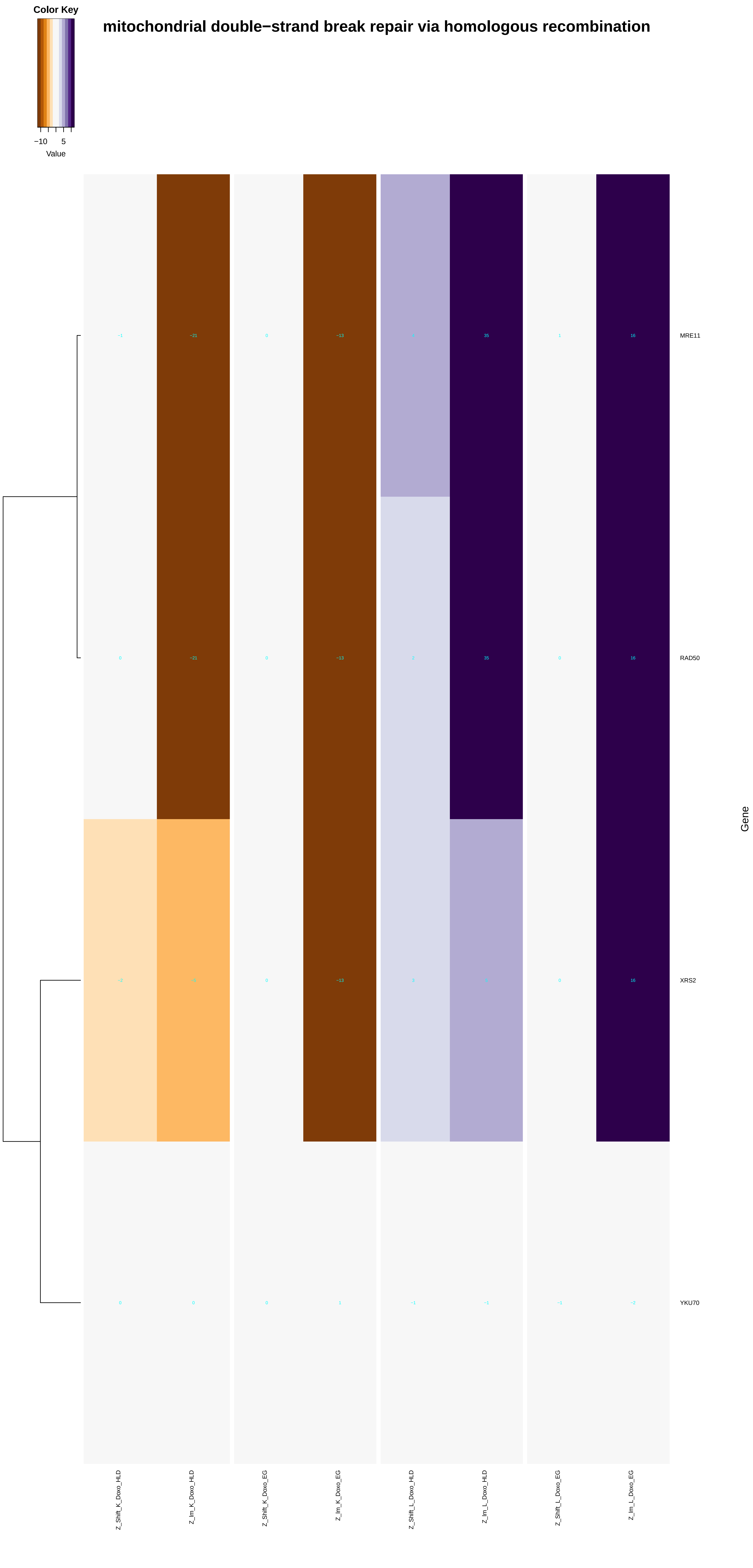

mitochondrial double−strand break repair via homologous recombination
Color Key
−10
5
Value
MRE11
RAD50
XRS2
YKU70
Gene
Z_Shift_K_Doxo_HLD
Z_lm_K_Doxo_HLD
Z_Shift_K_Doxo_EG
Z_lm_K_Doxo_EG
Z_Shift_L_Doxo_HLD
Z_lm_L_Doxo_HLD
Z_Shift_L_Doxo_EG
Z_lm_L_Doxo_EG

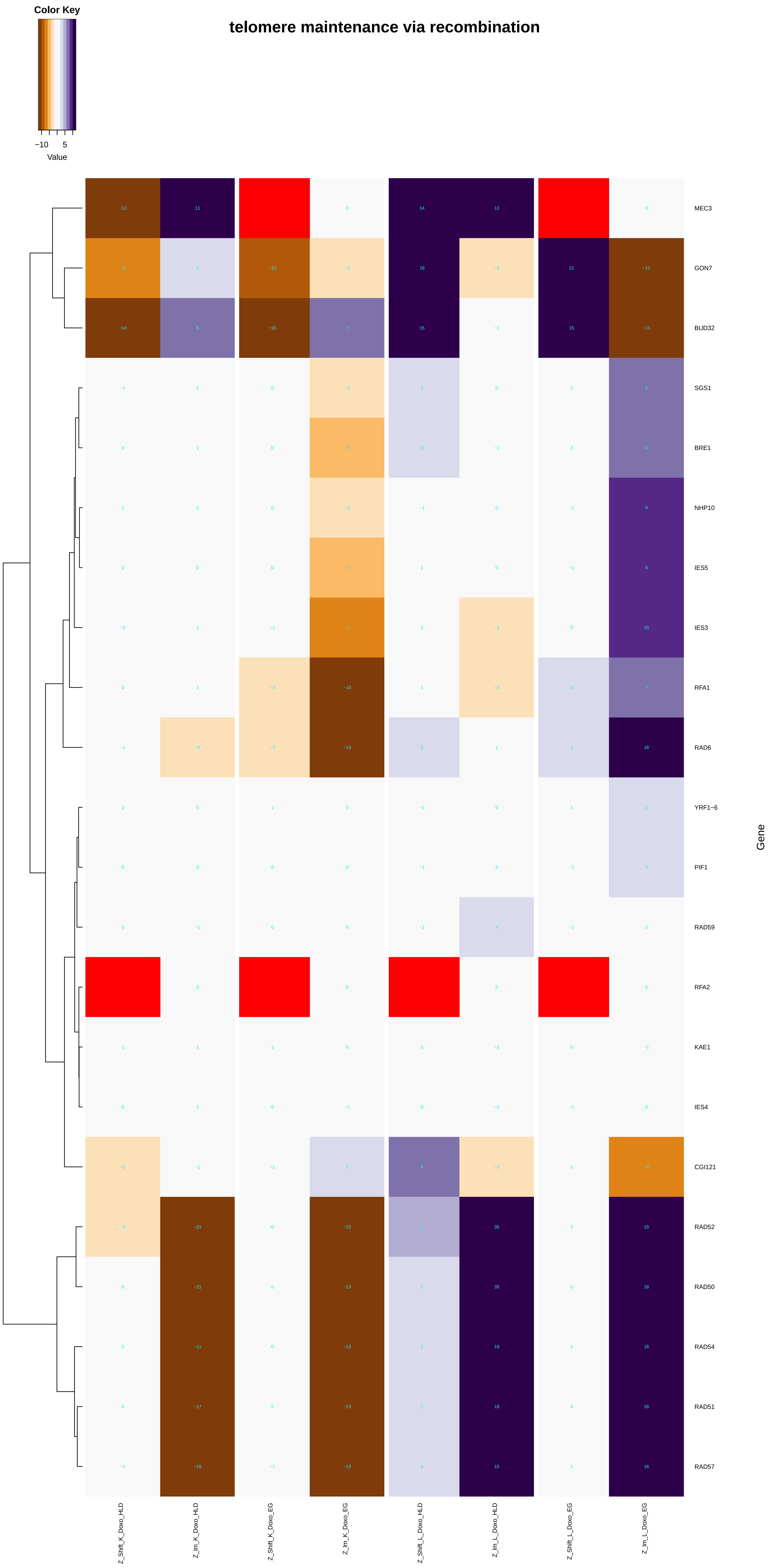

Color Key
−10
5
Value
telomere maintenance via recombination
MEC3
GON7
BUD32
SGS1
BRE1
NHP10
IES5
IES3
RFA1
RAD6
YRF1−6
PIF1
RAD59
RFA2
KAE1
IES4
CGI121
RAD52
RAD50
RAD54
RAD51
RAD57
Gene
Z_Shift_K_Doxo_HLD
Z_lm_K_Doxo_HLD
Z_Shift_K_Doxo_EG
Z_lm_K_Doxo_EG
Z_Shift_L_Doxo_HLD
Z_lm_L_Doxo_HLD
Z_Shift_L_Doxo_EG
Z_lm_L_Doxo_EG

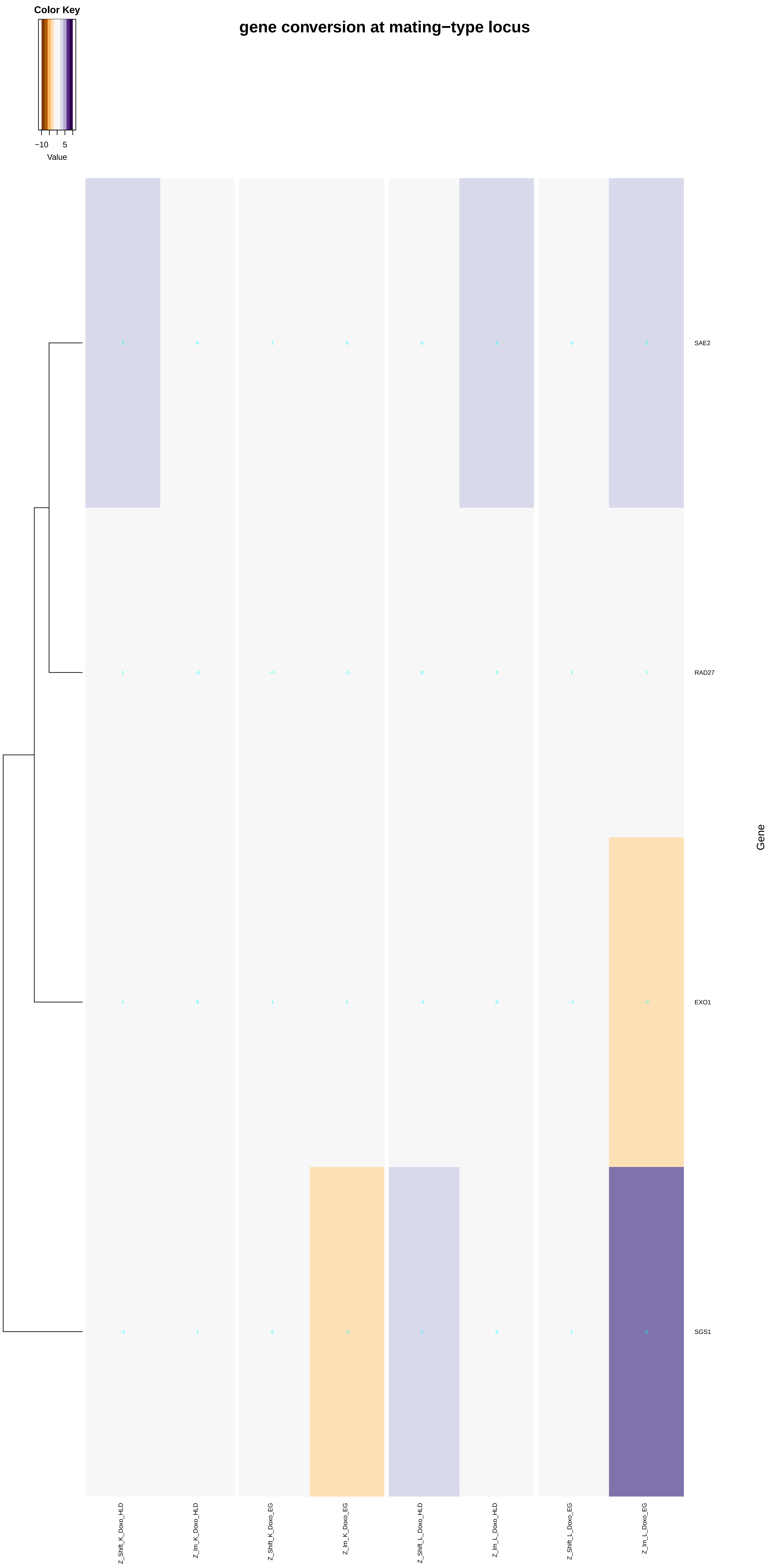
gene conversion at mating−type locus
Color Key
−10
5
Value
SAE2
RAD27
EXO1
SGS1
Gene
Z_Shift_K_Doxo_HLD
Z_Im_K_Doxo_HLD
Z_Shift_K_Doxo_EG
Z_Im_K_Doxo_EG
Z_Shift_L_Doxo_HLD
Z_Im_L_Doxo_HLD
Z_Shift_L_Doxo_EG
Z_Im_L_Doxo_EG

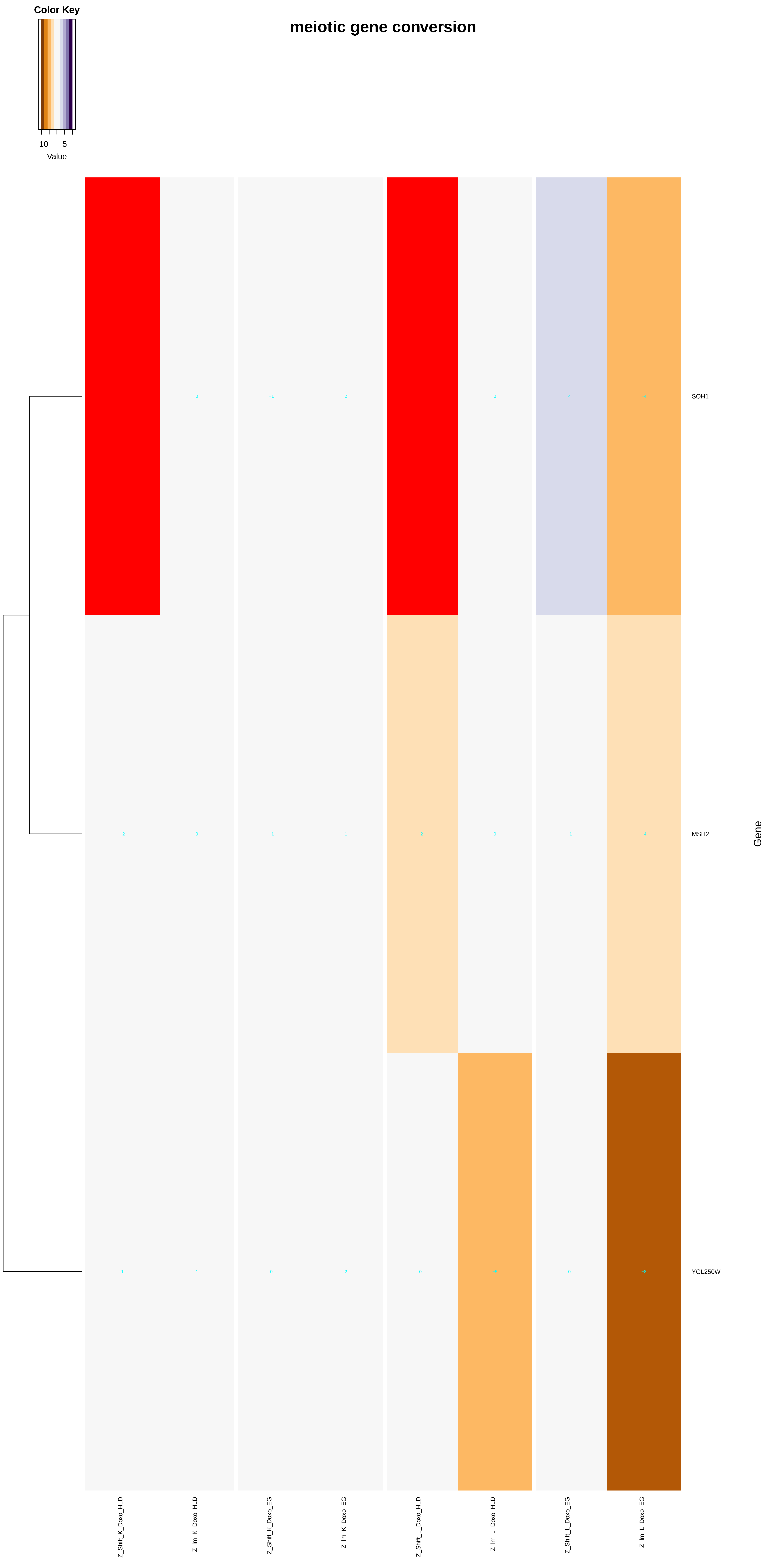
meiotic gene conversion
Color Key
−10
5
Value
SOH1
MSH2
YGL250W
Gene
Z_Shift_K_Doxo_HLD
Z_Im_K_Doxo_HLD
Z_Shift_K_Doxo_EG
Z_Im_K_Doxo_EG
Z_Shift_L_Doxo_HLD
Z_Im_L_Doxo_HLD
Z_Shift_L_Doxo_EG
Z_Im_L_Doxo_EG

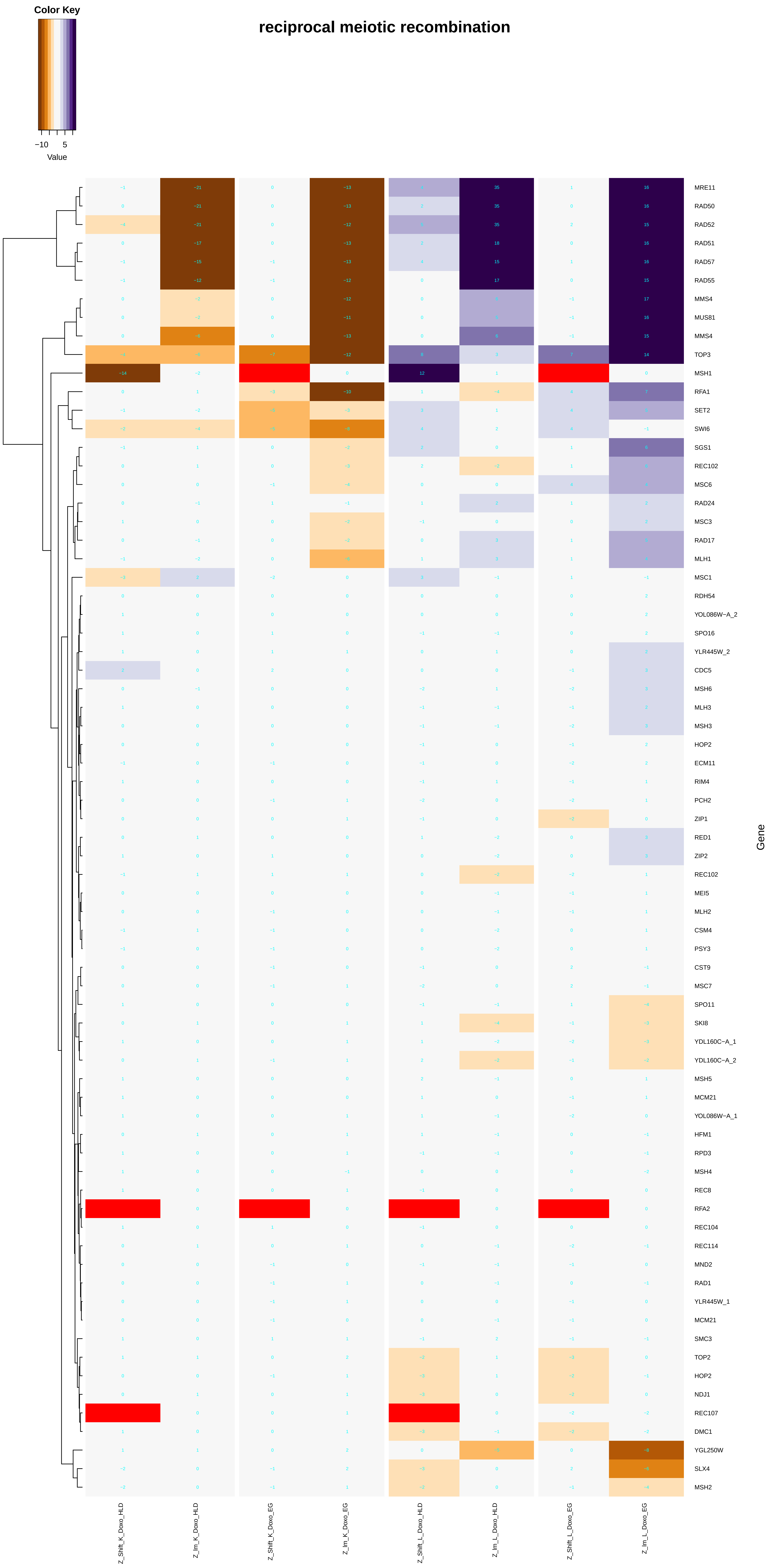
Color Key
−10
5
Value
reciprocal meiotic recombination
Gene
MRE11
RAD50
RAD52
RAD51
RAD57
RAD55
MMS4
MUS81
MMS4
TOP3
MSH1
RFA1
SET2
SWI6
SGS1
REC102
MSC6
RAD24
MSC3
RAD17
MLH1
MSC1
RDH54
YOL086W−A_2
SPO16
YLR445W_2
CDC5
MSH6
MLH3
MSH3
HOP2
ECM11
RIM4
PCH2
ZIP1
RED1
ZIP2
REC102
MEI5
MLH2
CSM4
PSY3
CST9
MSC7
SPO11
SKI8
YDL160C−A_1
YDL160C−A_2
MSH5
MCM21
YOL086W−A_1
HFM1
RPD3
MSH4
REC8
RFA2
REC104
REC114
MND2
RAD1
YLR445W_1
MCM21
SMC3
TOP2
HOP2
NDJ1
REC107
DMC1
YGL250W
SLX4
MSH2
Z_Shift_K_Doxo_HLD
Z_Im_K_Doxo_HLD
Z_Shift_K_Doxo_EG
Z_Im_K_Doxo_EG
Z_Shift_L_Doxo_HLD
Z_Im_L_Doxo_HLD
Z_Shift_L_Doxo_EG
Z_Im_L_Doxo_EG

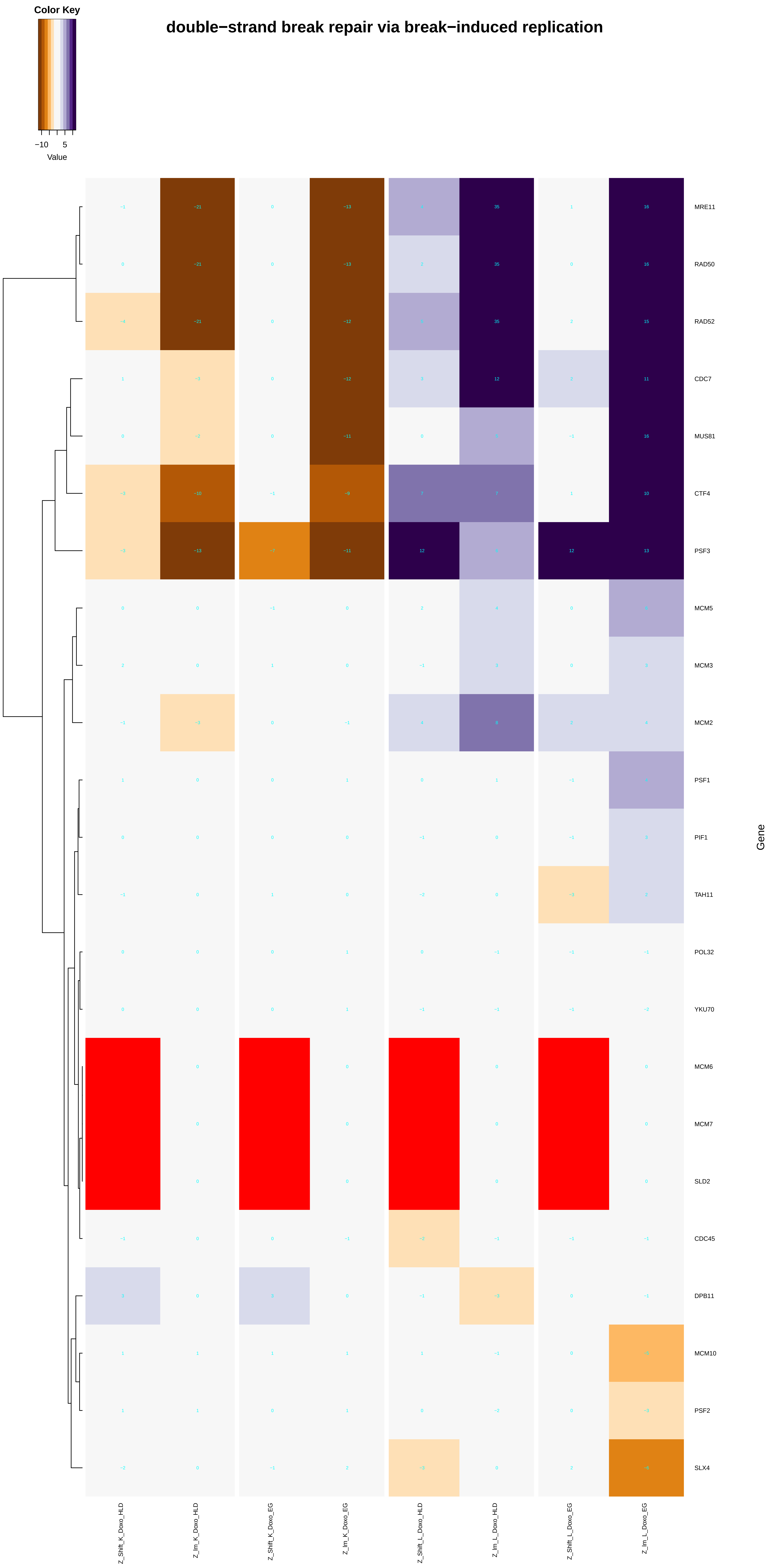
double−strand break repair via break−induced replication
Color Key
−10
5
Value
MRE11
RAD50
RAD52
CDC7
MUS81
CTF4
PSF3
MCM5
MCM3
MCM2
PSF1
PIF1
TAH11
POL32
YKU70
MCM6
MCM7
SLD2
CDC45
DPB11
MCM10
PSF2
SLX4
Gene
Z_Shift_K_Doxo_HLD
Z_Im_K_Doxo_HLD
Z_Shift_K_Doxo_EG
Z_Im_K_Doxo_EG
Z_Shift_L_Doxo_HLD
Z_Im_L_Doxo_HLD
Z_Shift_L_Doxo_EG
Z_Im_L_Doxo_EG

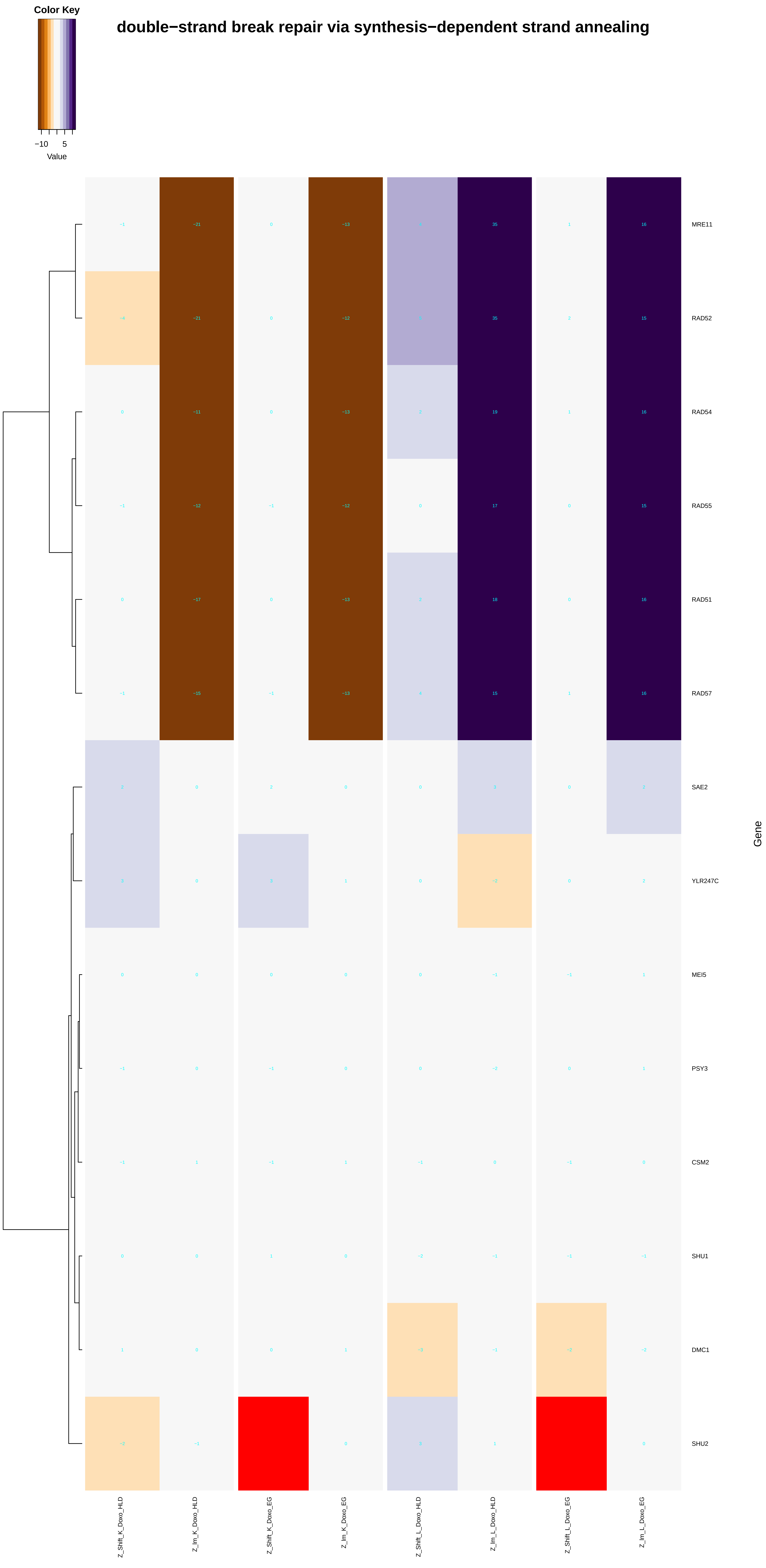

Color Key
−10
5
Value
double−strand break repair via synthesis−dependent strand annealing
MRE11
RAD52
RAD54
RAD55
RAD51
RAD57
SAE2
YLR247C
MEI5
PSY3
CSM2
SHU1
DMC1
SHU2
Gene
Z_Shift_K_Doxo_HLD
Z_Im_K_Doxo_HLD
Z_Shift_K_Doxo_EG
Z_Im_K_Doxo_EG
Z_Shift_L_Doxo_HLD
Z_Im_L_Doxo_HLD
Z_Shift_L_Doxo_EG
Z_Im_L_Doxo_EG

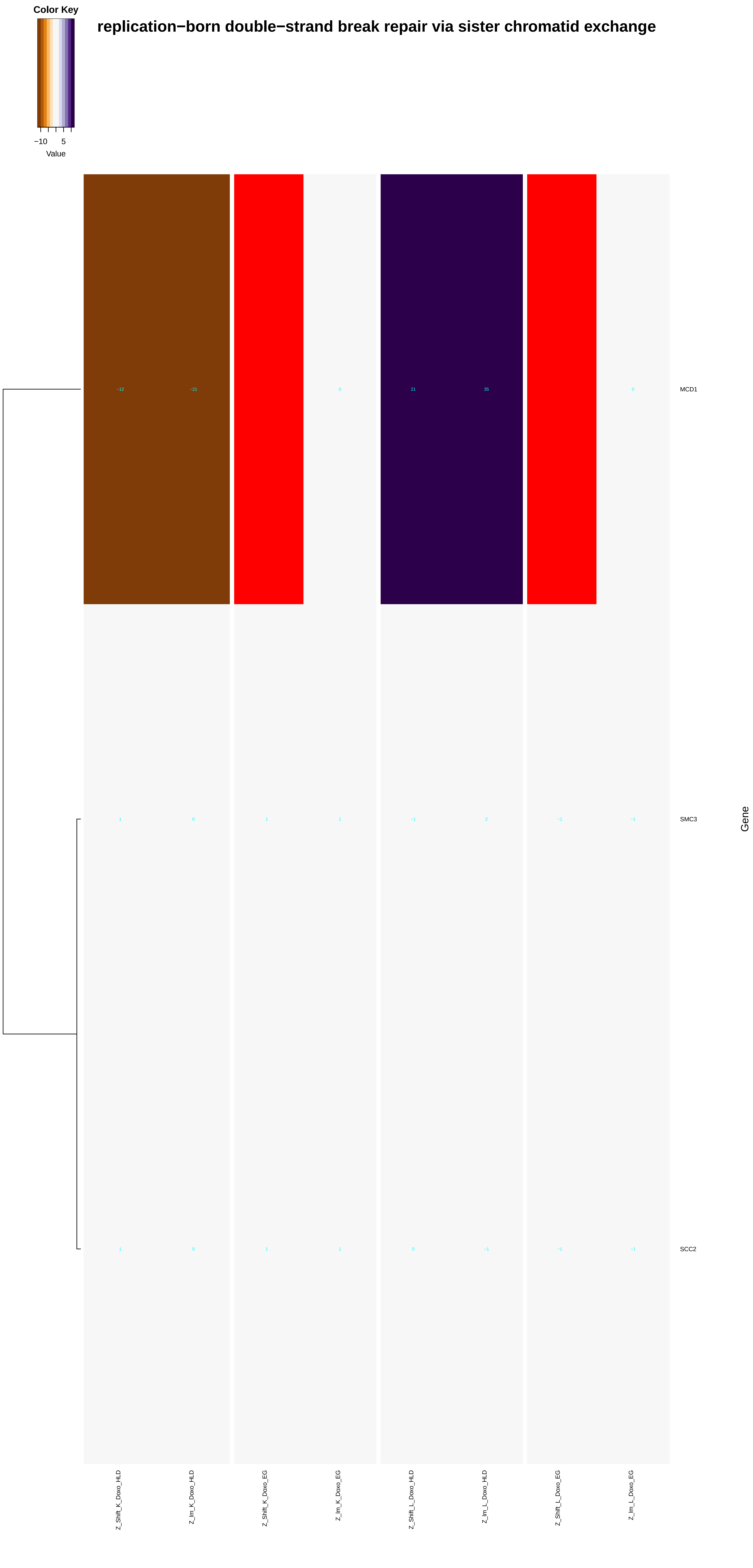
Color Key
−10
5
Value
replication−born double−strand break repair via sister chromatid exchange
MCD1
SMC3
SCC2
Gene
Z_Shift_K_Doxo_HLD
Z_Im_K_Doxo_HLD
Z_Shift_K_Doxo_EG
Z_Im_K_Doxo_EG
Z_Shift_L_Doxo_HLD
Z_Im_L_Doxo_HLD
Z_Shift_L_Doxo_EG
Z_Im_L_Doxo_EG
