## Additional File 9 for "A yeast phenomic model for the influence of Warburg metabolism on genetic buffering of doxorubicin": establishment_of_protein_localization_to_membrane.pdf

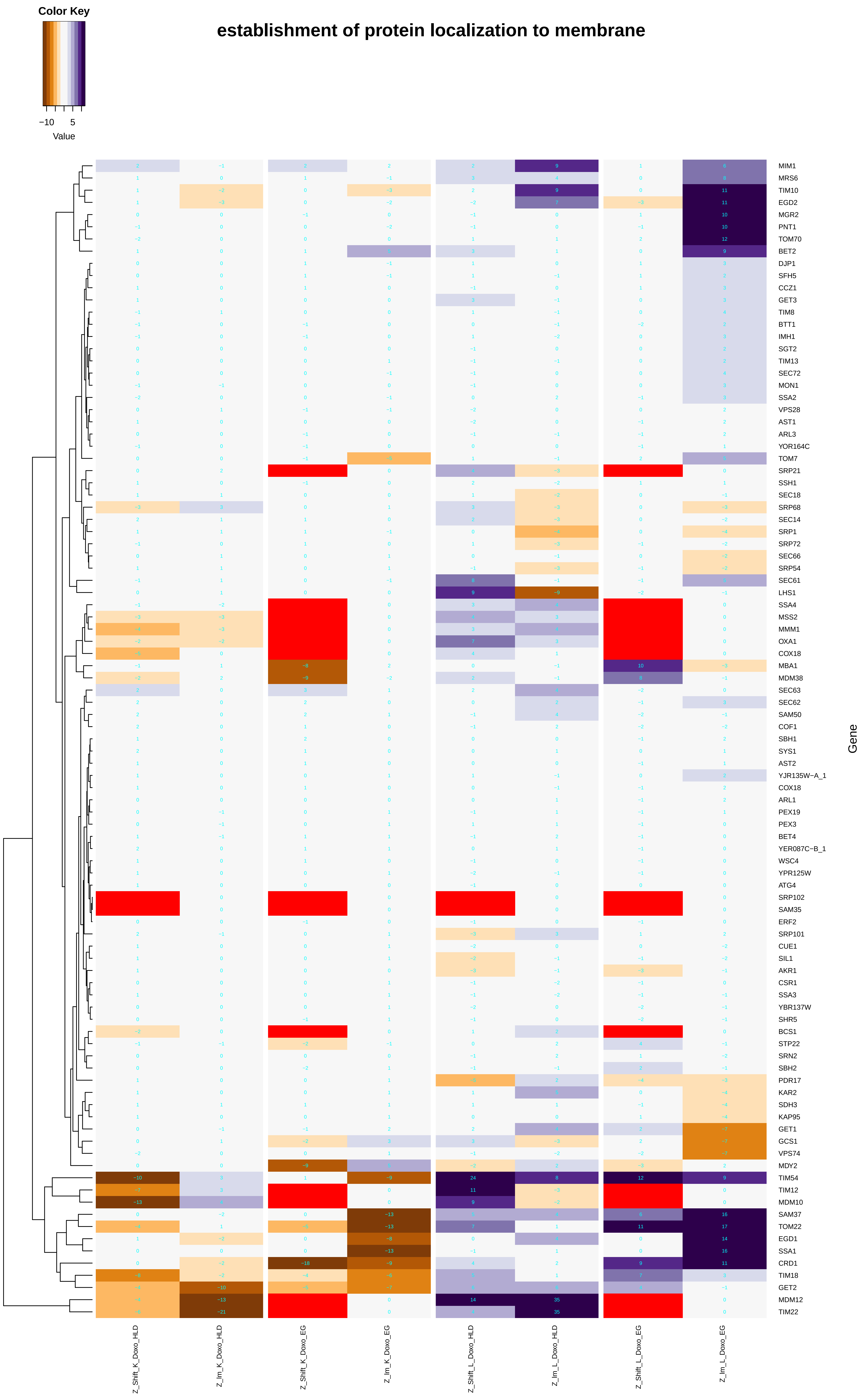

establishment of protein localization to membrane
Color Key
−10
5
Value
Gene
MIM1
MRS6
TIM10
EGD2
MGR2
PNT1
TOM70
BET2
DJP1
SFH5
CCZ1
GET3
TIM8
BTT1
IMH1
SGT2
TIM13
SEC72
MON1
SSA2
VPS28
AST1
ARL3
YOR164C
TOM7
SRP21
SSH1
SEC18
SRP68
SEC14
SRP1
SRP72
SEC66
SRP54
SEC61
LHS1
SSA4
MSS2
MMM1
OXA1
COX18
MBA1
MDM38
SEC63
SEC62
SAM50
COF1
SBH1
SYS1
AST2
YJR135W−A_1
COX18
ARL1
PEX19
PEX3
BET4
YER087C−B_1
WSC4
YPR125W
ATG4
SRP102
SAM35
ERF2
SRP101
CUE1
SIL1
AKR1
CSR1
SSA3
YBR137W
SHR5
BCS1
STP22
SRN2
SBH2
PDR17
KAR2
SDH3
KAP95
GET1
GCS1
VPS74
MDY2
TIM54
TIM12
MDM10
SAM37
TOM22
EGD1
SSA1
CRD1
TIM18
GET2
MDM12
TIM22
Z_Shift_K_Doxo_HLD
Z_Im_K_Doxo_HLD
Z_Shift_K_Doxo_EG
Z_Im_K_Doxo_EG
Z_Shift_L_Doxo_HLD
Z_Im_L_Doxo_HLD
Z_Shift_L_Doxo_EG
Z_Im_L_Doxo_EG

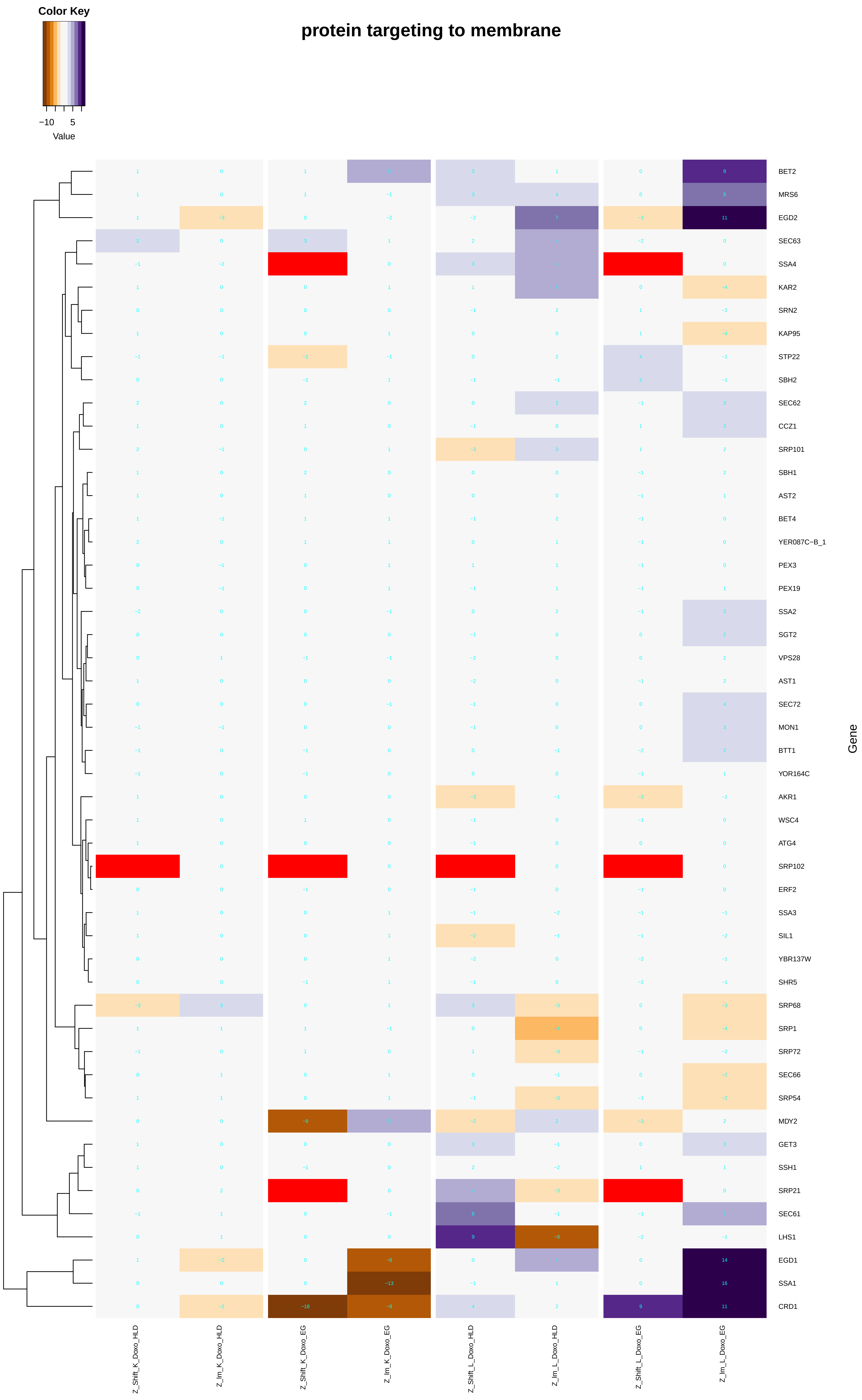

protein targeting to membrane
Color Key
−10
5
Value
BET2
MRS6
EGD2
SEC63
SSA4
KAR2
SRN2
KAP95
STP22
SBH2
SEC62
CCZ1
SRP101
SBH1
AST2
BET4
YER087C−B_1
PEX3
PEX19
SSA2
SGT2
VPS28
AST1
SEC72
MON1
BTT1
YOR164C
AKR1
WSC4
ATG4
SRP102
ERF2
SSA3
SIL1
YBR137W
SHR5
SRP68
SRP1
SRP72
SEC66
SRP54
MDY2
GET3
SSH1
SRP21
SEC61
LHS1
EGD1
SSA1
CRD1
Gene
Z_Shift_K_Doxo_HLD
Z_Im_K_Doxo_HLD
Z_Shift_K_Doxo_EG
Z_Im_K_Doxo_EG
Z_Shift_L_Doxo_HLD
Z_Im_L_Doxo_HLD
Z_Shift_L_Doxo_EG
Z_Im_L_Doxo_EG

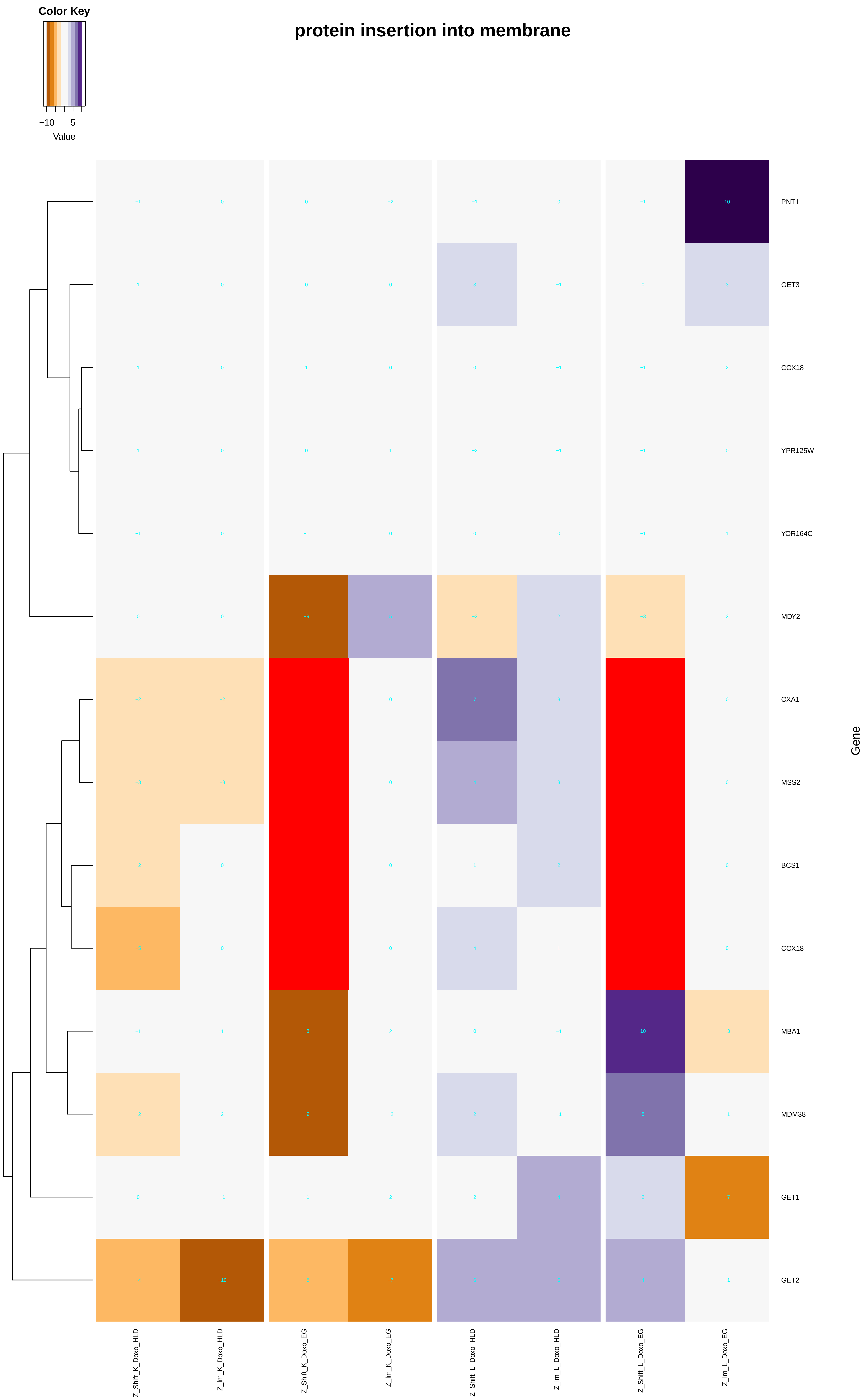
Color Key
−10
5
Value
protein insertion into membrane
PNT1
GET3
COX18
YPR125W
YOR164C
MDY2
OXA1
MSS2
BCS1
COX18
MBA1
MDM38
GET1
GET2
Gene
Z_Shift_K_Doxo_HLD
Z_Im_K_Doxo_HLD
Z_Shift_K_Doxo_EG
Z_Im_K_Doxo_EG
Z_Shift_L_Doxo_HLD
Z_Im_L_Doxo_HLD
Z_Shift_L_Doxo_EG
Z_Im_L_Doxo_EG

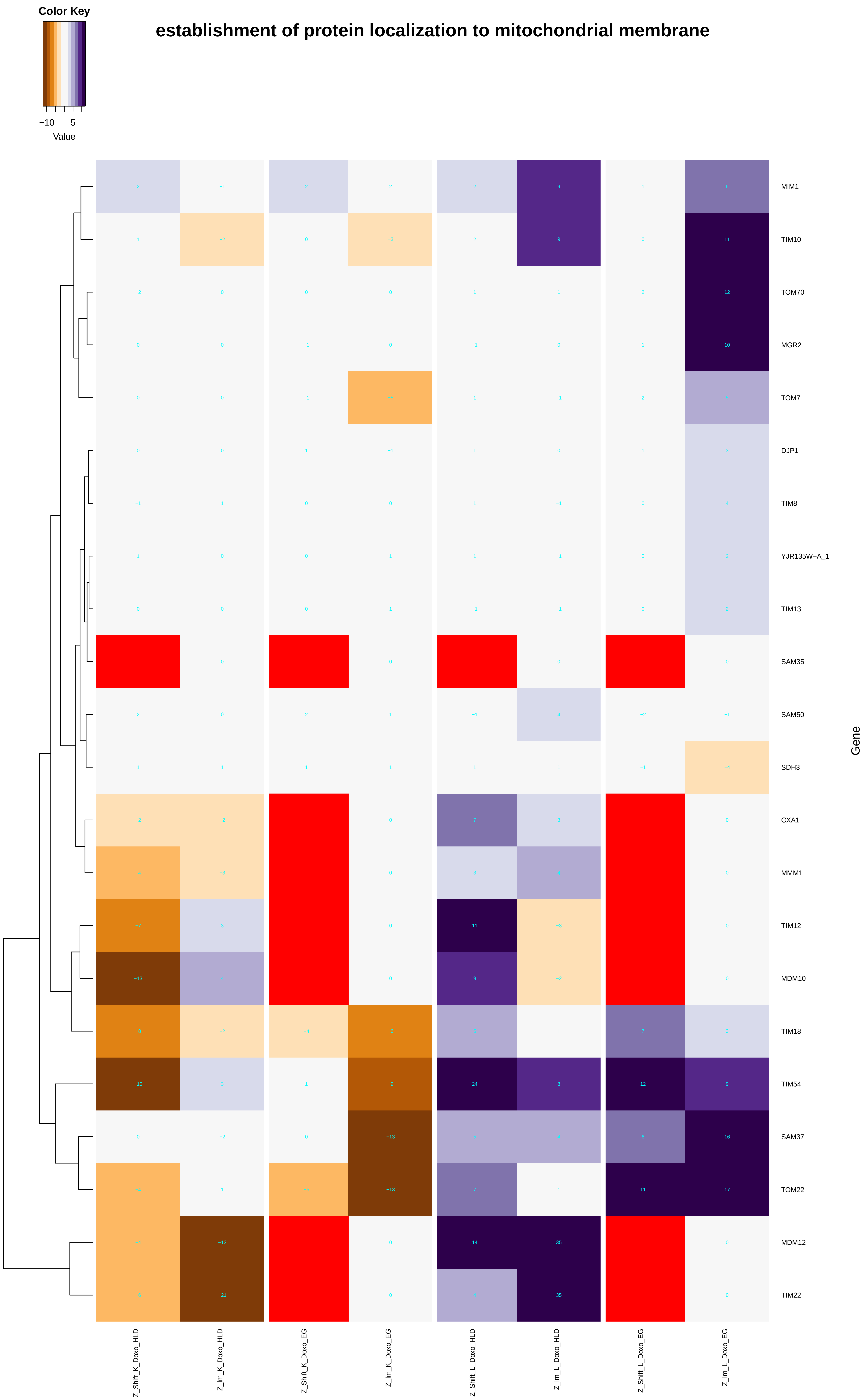
Color Key
−10
5
Value
establishment of protein localization to mitochondrial membrane
MIM1
TIM10
TOM70
MGR2
TOM7
DJP1
TIM8
YJR135W−A_1
TIM13
SAM35
SAM50
SDH3
OXA1
MMM1
TIM12
MDM10
TIM18
TIM54
SAM37
TOM22
MDM12
TIM22
Gene
Z_Shift_K_Doxo_HLD
Z_Im_K_Doxo_HLD
Z_Shift_K_Doxo_EG
Z_Im_K_Doxo_EG
Z_Shift_L_Doxo_HLD
Z_Im_L_Doxo_HLD
Z_Shift_L_Doxo_EG
Z_Im_L_Doxo_EG

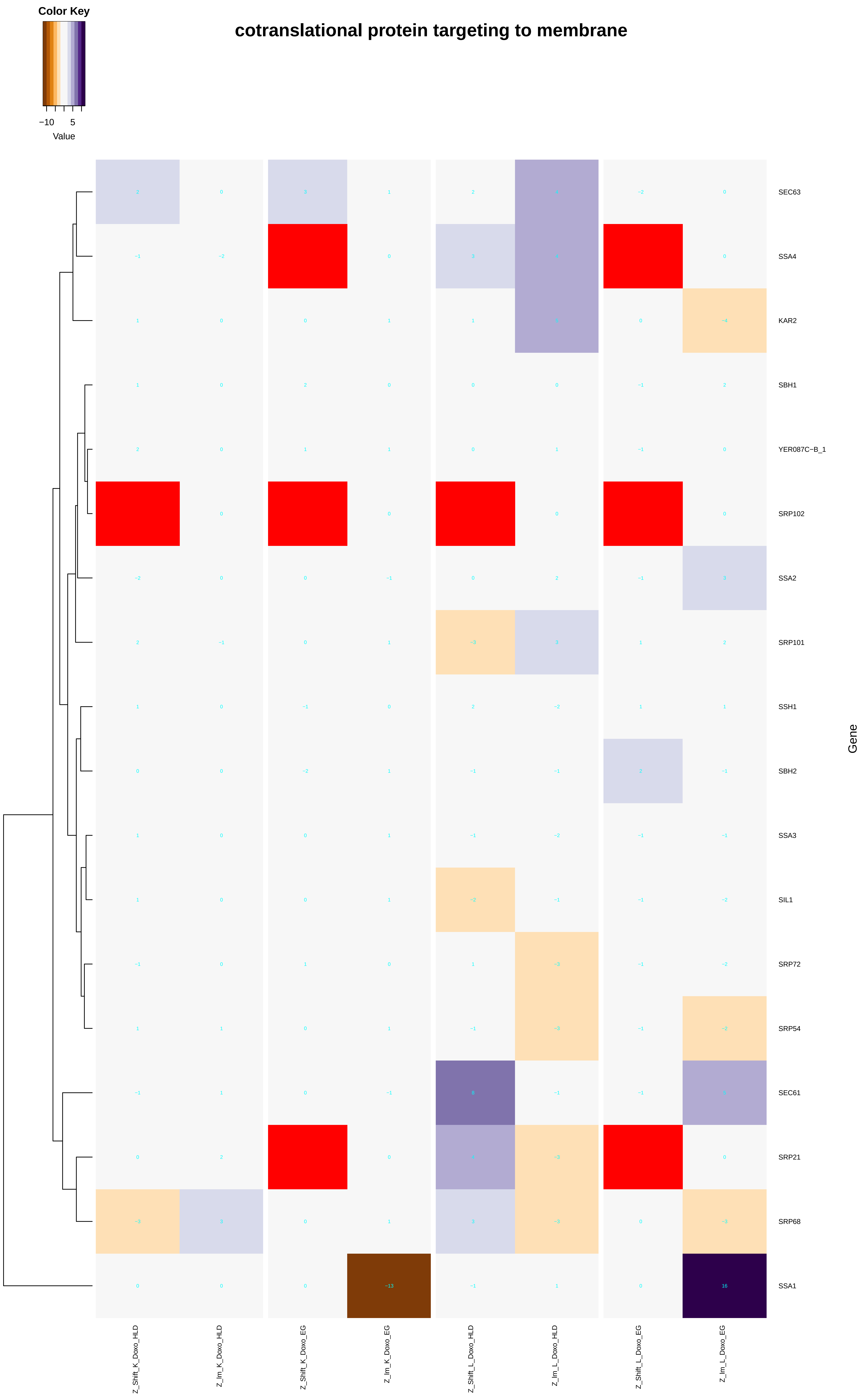

Color Key
−10
5
Value
cotranslational protein targeting to membrane
SEC63
SSA4
KAR2
SBH1
YER087C−B_1
SRP102
SSA2
SRP101
SSH1
SBH2
SSA3
SIL1
SRP72
SRP54
SEC61
SRP21
SRP68
SSA1
Gene
Z_Shift_K_Doxo_HLD
Z_Im_K_Doxo_HLD
Z_Shift_K_Doxo_EG
Z_Im_K_Doxo_EG
Z_Shift_L_Doxo_HLD
Z_Im_L_Doxo_HLD
Z_Shift_L_Doxo_EG
Z_Im_L_Doxo_EG

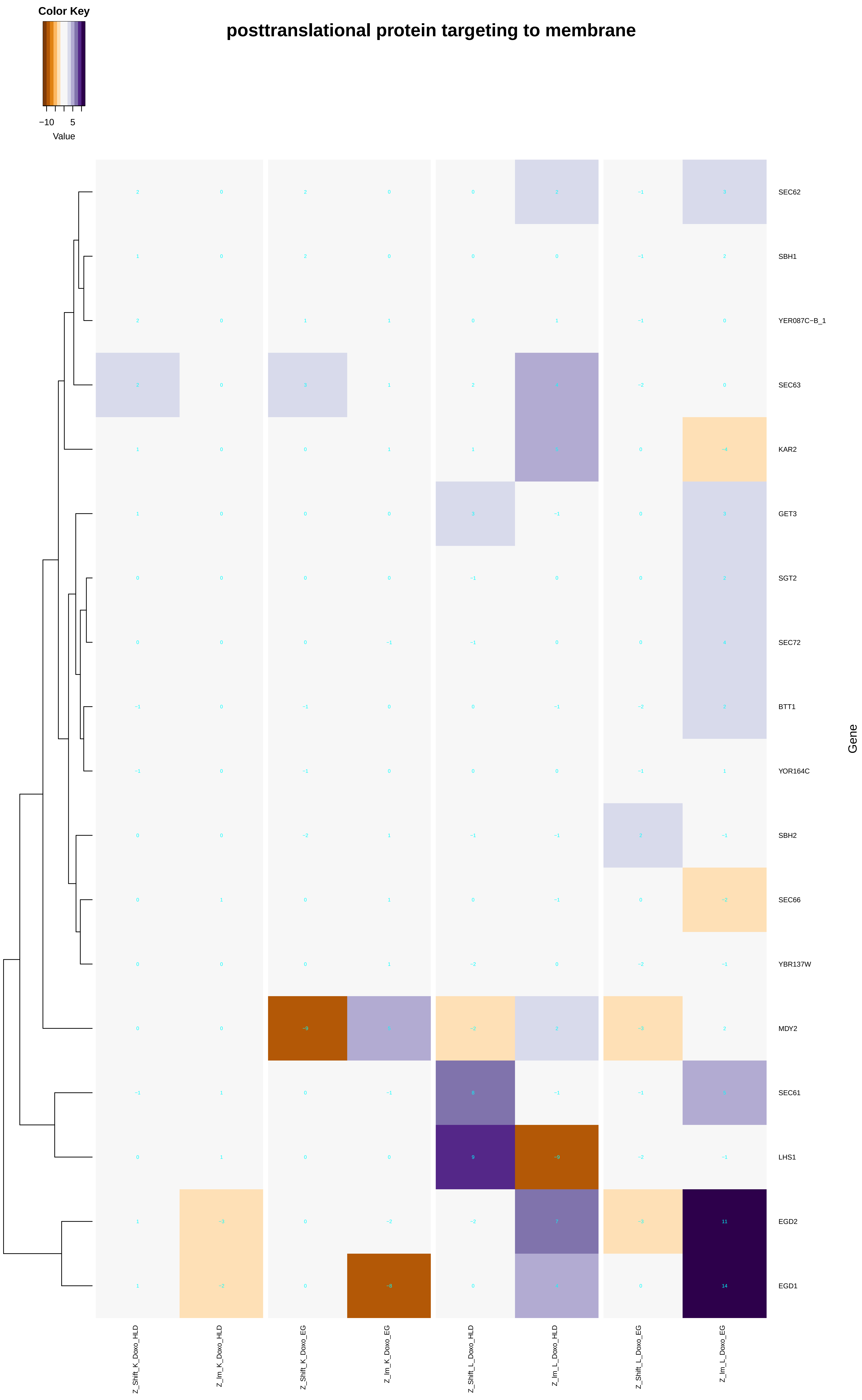
posttranslational protein targeting to membrane
Color Key
−10
5
Value
Gene
SEC62
SBH1
YER087C−B_1
SEC63
KAR2
GET3
SGT2
SEC72
BTT1
YOR164C
SBH2
SEC66
YBR137W
MDY2
SEC61
LHS1
EGD2
EGD1
Z_Shift_K_Doxo_HLD
Z_Im_K_Doxo_HLD
Z_Shift_K_Doxo_EG
Z_Im_K_Doxo_EG
Z_Shift_L_Doxo_HLD
Z_Im_L_Doxo_HLD
Z_Shift_L_Doxo_EG
Z_Im_L_Doxo_EG

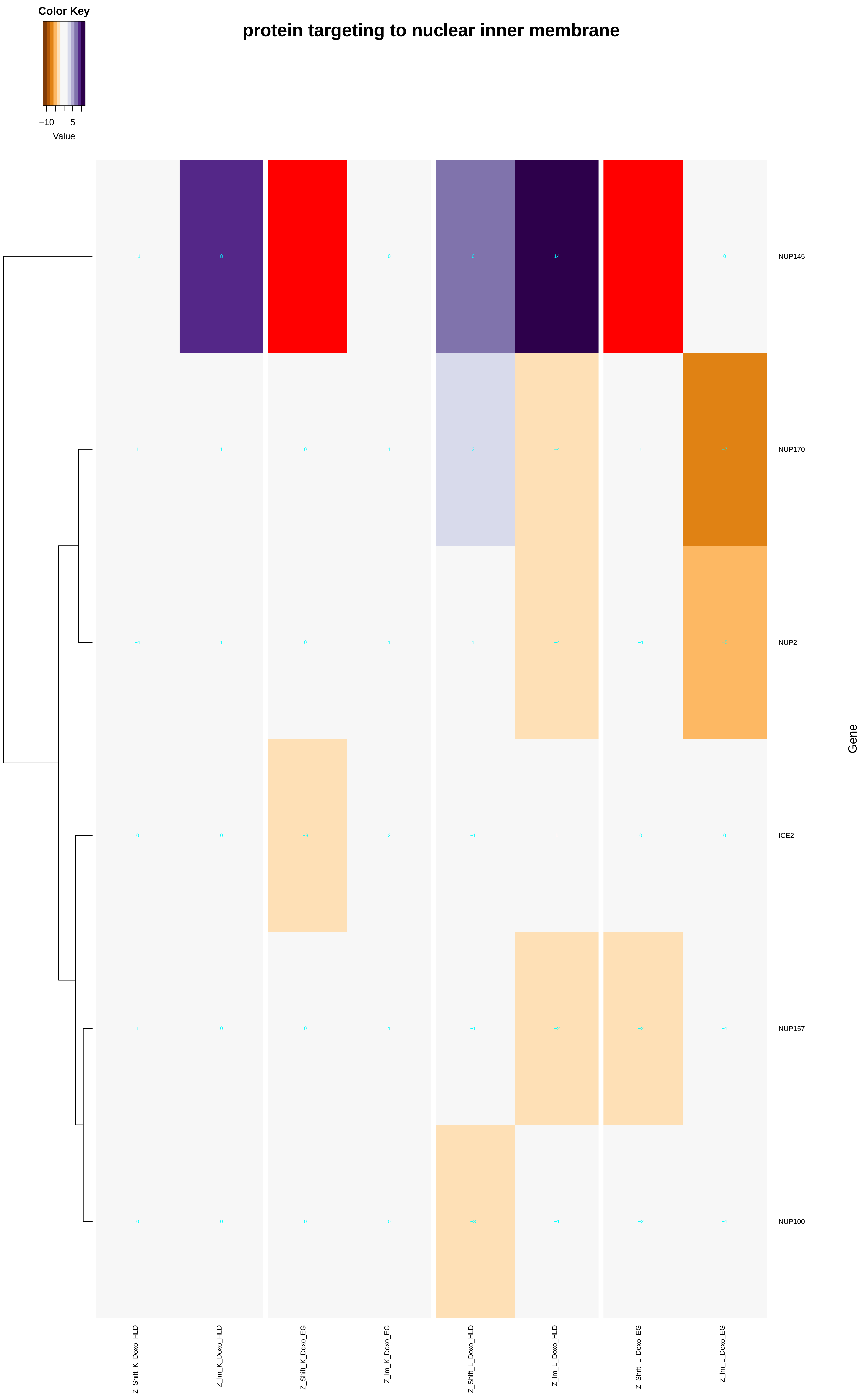
Color Key
−10
5
Value
protein targeting to nuclear inner membrane
NUP145
NUP170
NUP2
ICE2
NUP157
NUP100
Gene
Z_Shift_K_Doxo_HLD
Z_Im_K_Doxo_HLD
Z_Shift_K_Doxo_EG
Z_Im_K_Doxo_EG
Z_Shift_L_Doxo_HLD
Z_Im_L_Doxo_HLD
Z_Shift_L_Doxo_EG
Z_Im_L_Doxo_EG

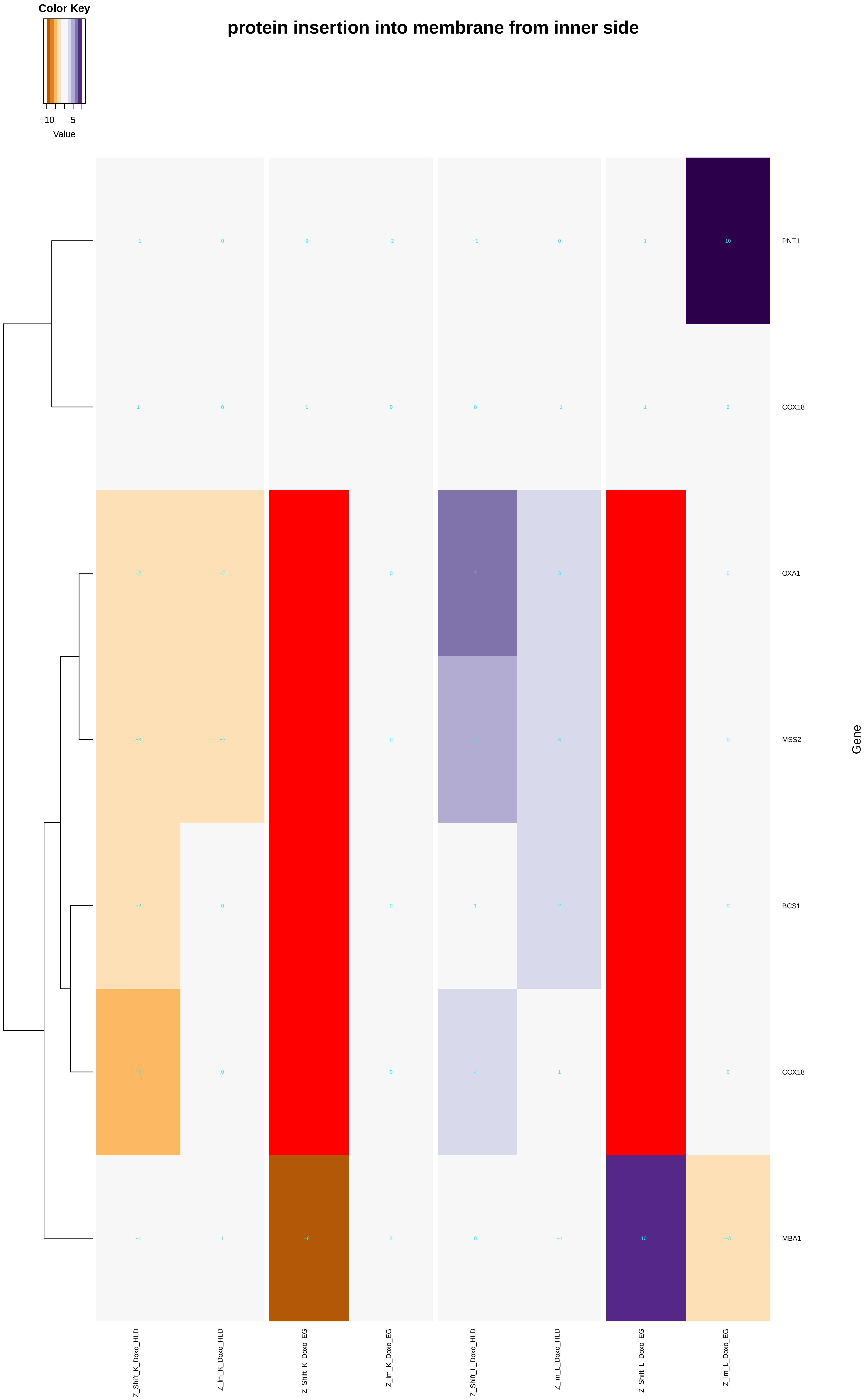
protein insertion into membrane from inner side
Color Key
−10
5
Value
PNT1
COX18
OXA1
MSS2
BCS1
COX18
MBA1
Gene
Z_Shift_K_Doxo_HLD
Z_Im_K_Doxo_HLD
Z_Shift_K_Doxo_EG
Z_Im_K_Doxo_EG
Z_Shift_L_Doxo_HLD
Z_Im_L_Doxo_HLD
Z_Shift_L_Doxo_EG
Z_Im_L_Doxo_EG

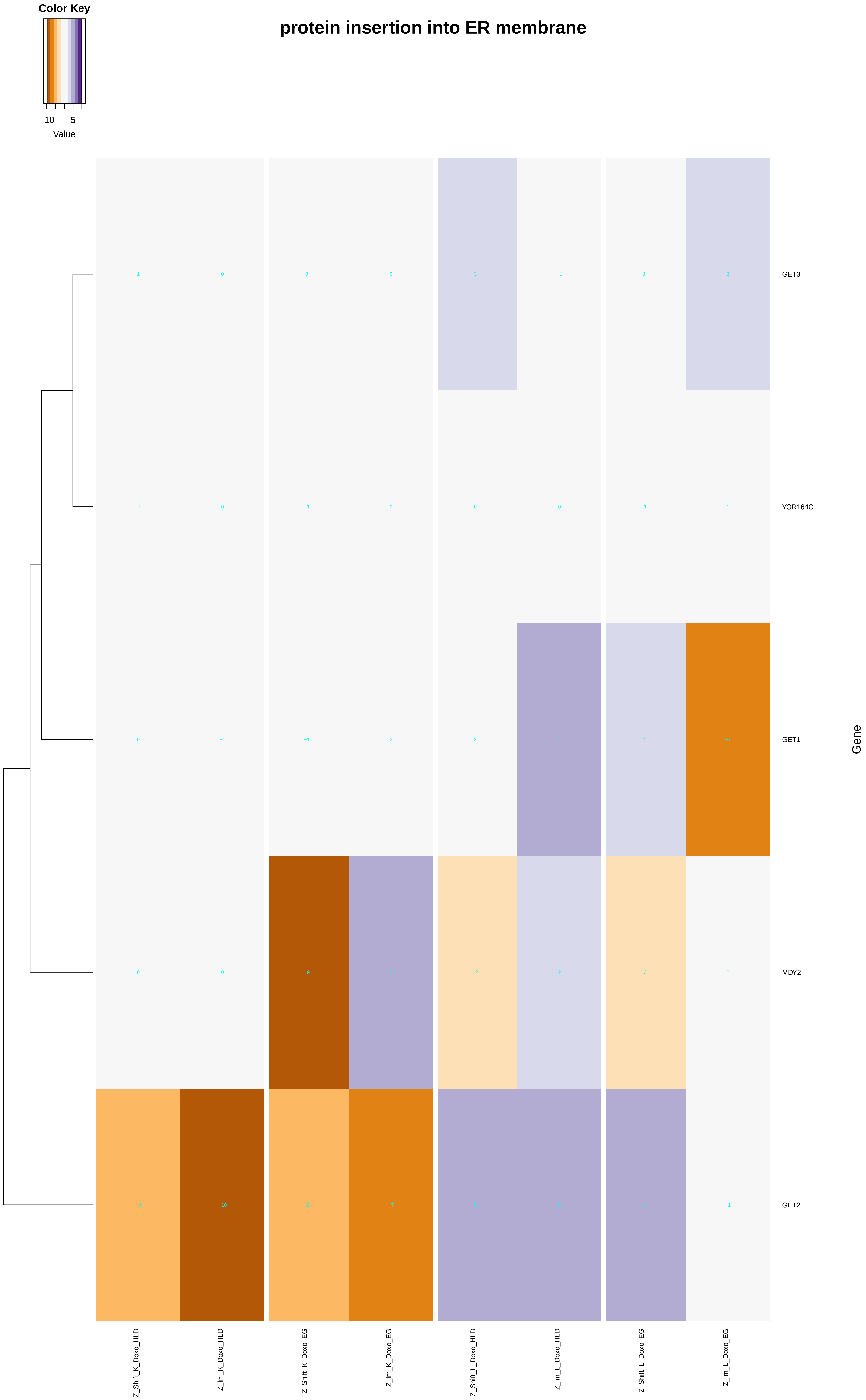
Color Key
−10
5
Value
protein insertion into ER membrane
GET3
YOR164C
GET1
MDY2
GET2
Gene
Z_Shift_K_Doxo_HLD
Z_Im_K_Doxo_HLD
Z_Shift_K_Doxo_EG
Z_Im_K_Doxo_EG
Z_Shift_L_Doxo_HLD
Z_Im_L_Doxo_HLD
Z_Shift_L_Doxo_EG
Z_Im_L_Doxo_EG

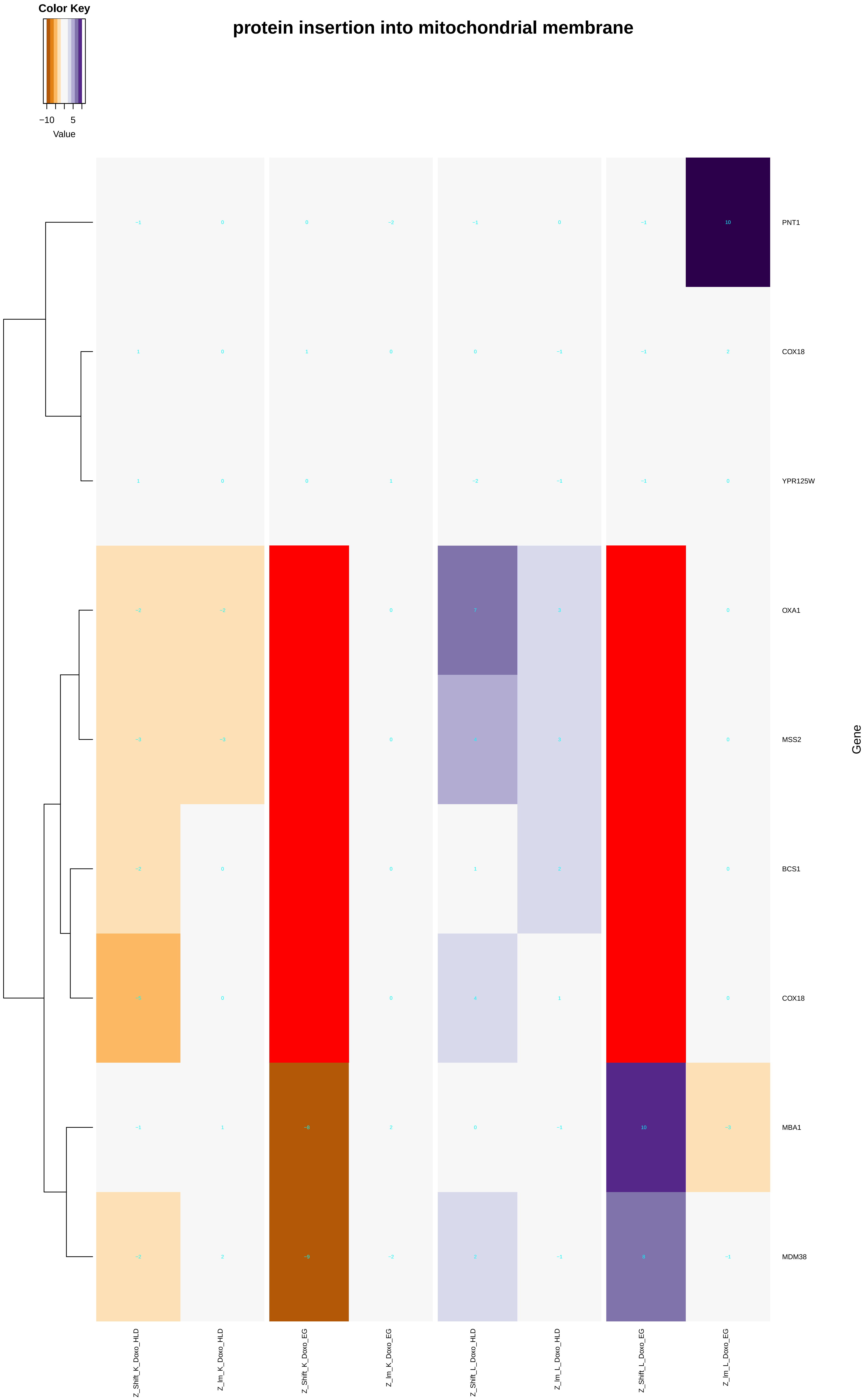
protein insertion into mitochondrial membrane
Color Key
−10
5
Value
PNT1
COX18
YPR125W
OXA1
MSS2
BCS1
COX18
MBA1
MDM38
Gene
Z_Shift_K_Doxo_HLD
Z_Im_K_Doxo_HLD
Z_Shift_K_Doxo_EG
Z_Im_K_Doxo_EG
Z_Shift_L_Doxo_HLD
Z_Im_L_Doxo_HLD
Z_Shift_L_Doxo_EG
Z_Im_L_Doxo_EG

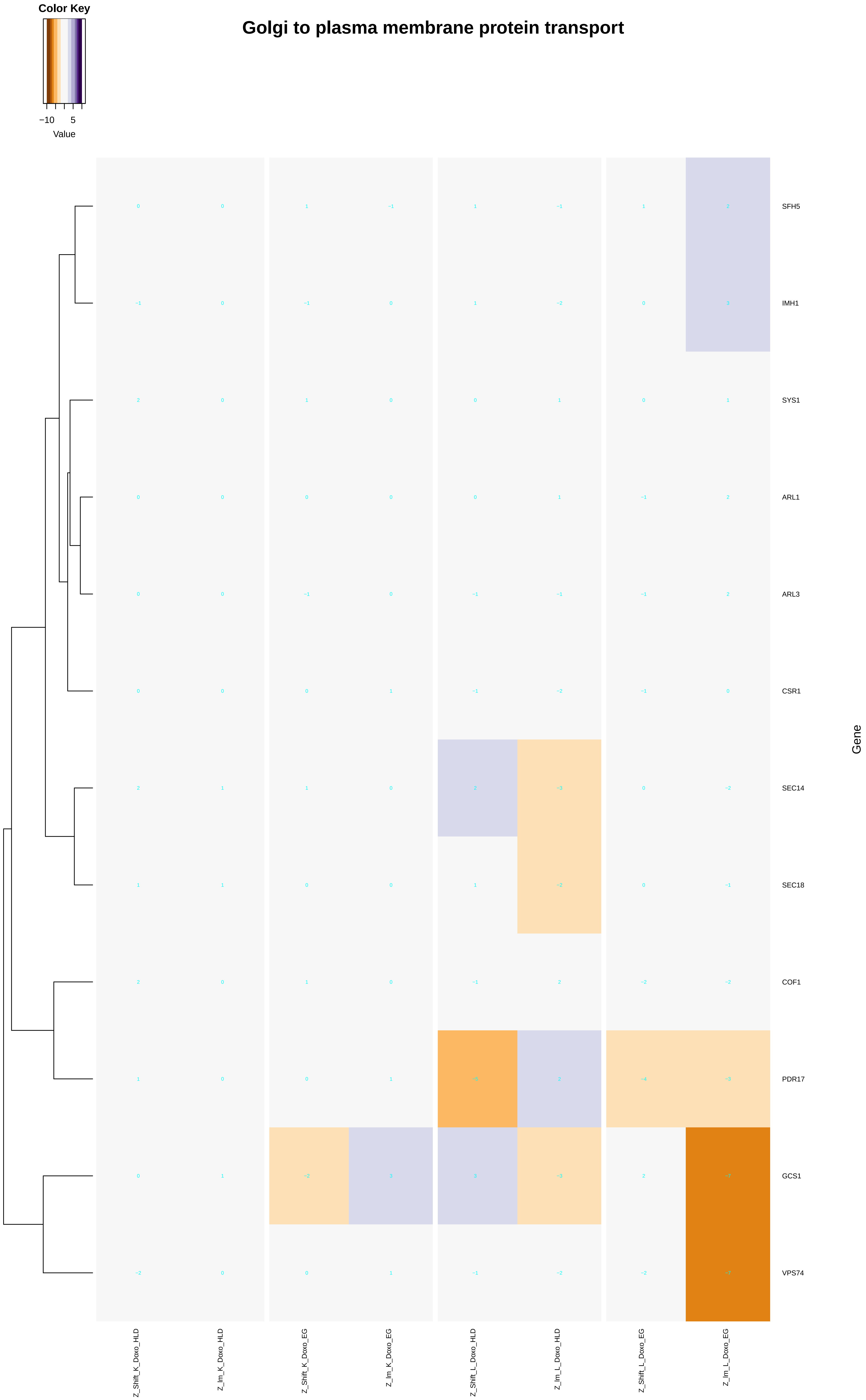
Color Key
−10
5
Value
Golgi to plasma membrane protein transport
SFH5
IMH1
SYS1
ARL1
ARL3
CSR1
SEC14
SEC18
COF1
PDR17
GCS1
VPS74
Gene
Z_Shift_K_Doxo_HLD
Z_Im_K_Doxo_HLD
Z_Shift_K_Doxo_EG
Z_Im_K_Doxo_EG
Z_Shift_L_Doxo_HLD
Z_Im_L_Doxo_HLD
Z_Shift_L_Doxo_EG
Z_Im_L_Doxo_EG

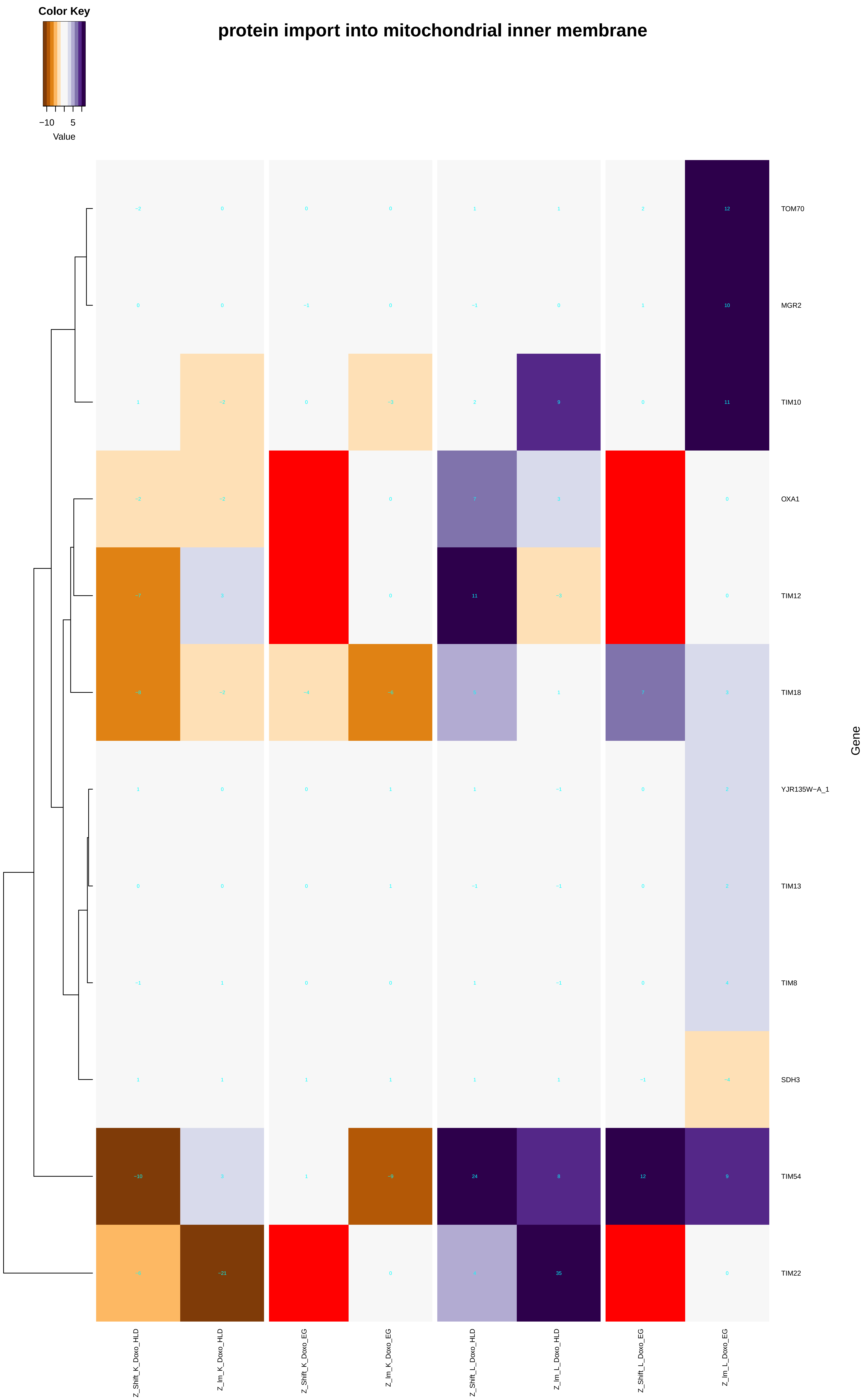

Color Key
−10
5
Value
protein import into mitochondrial inner membrane
TOM70
MGR2
TIM10
OXA1
TIM12
TIM18
YJR135W−A_1
TIM13
TIM8
SDH3
TIM54
TIM22
Gene
Z_Shift_K_Doxo_HLD
Z_Im_K_Doxo_HLD
Z_Shift_K_Doxo_EG
Z_Im_K_Doxo_EG
Z_Shift_L_Doxo_HLD
Z_Im_L_Doxo_HLD
Z_Shift_L_Doxo_EG
Z_Im_L_Doxo_EG

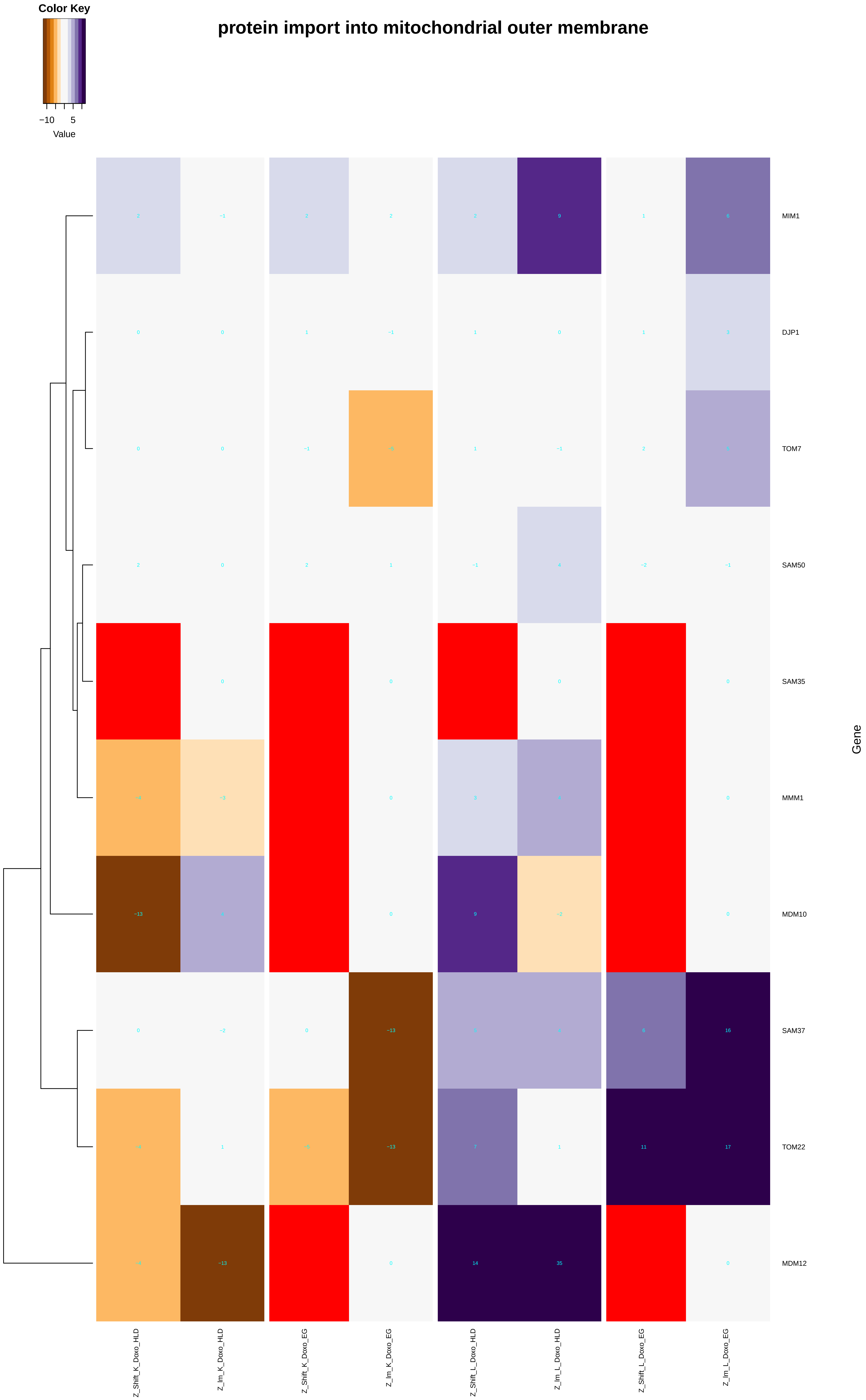
protein import into mitochondrial outer membrane
Color Key
−10
5
Value
MIM1
DJP1
TOM7
SAM50
SAM35
MMM1
MDM10
SAM37
TOM22
MDM12
Gene
Z_Shift_K_Doxo_HLD
Z_Im_K_Doxo_HLD
Z_Shift_K_Doxo_EG
Z_Im_K_Doxo_EG
Z_Shift_L_Doxo_HLD
Z_Im_L_Doxo_HLD
Z_Shift_L_Doxo_EG
Z_Im_L_Doxo_EG

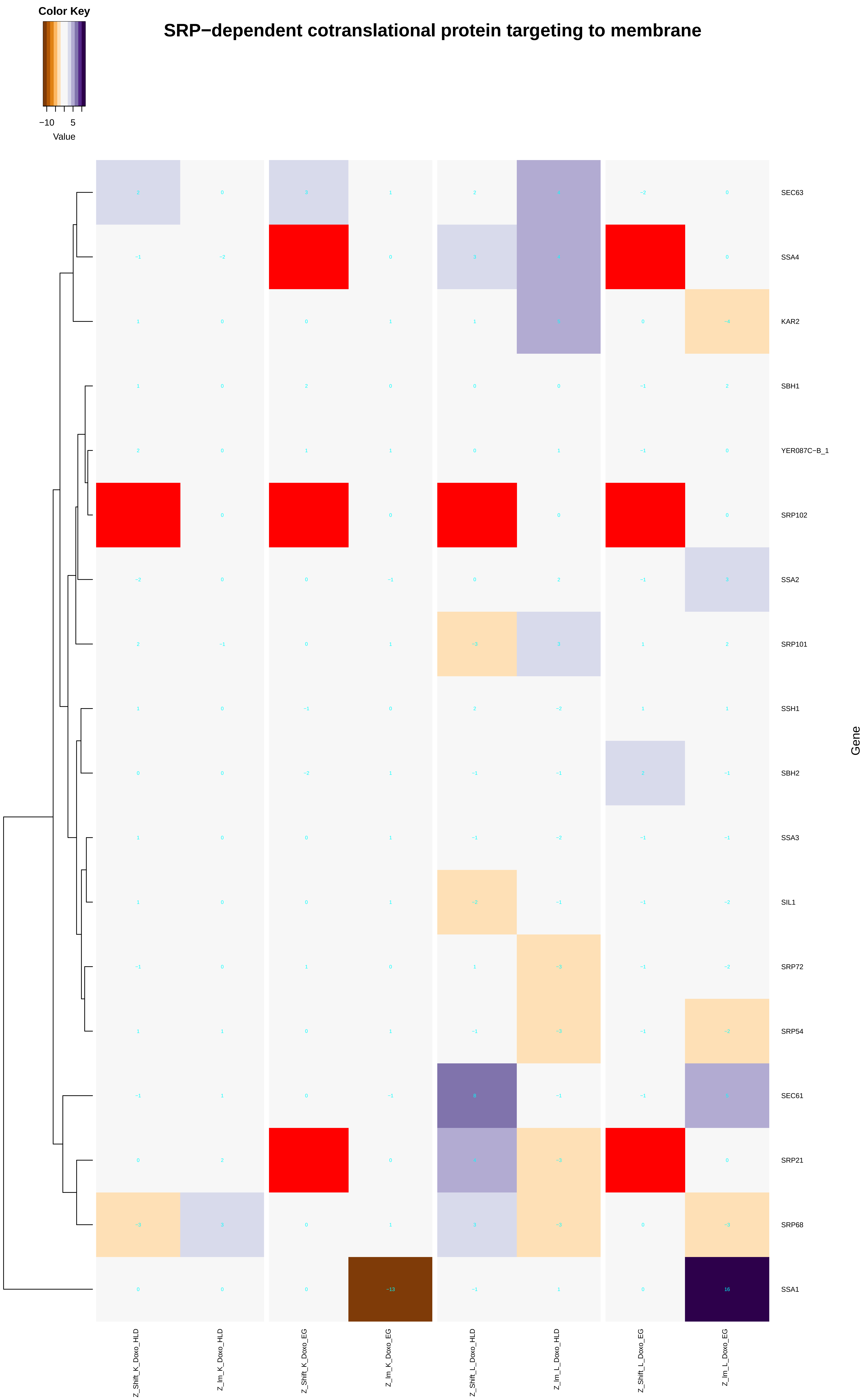

Color Key
−10
5
Value
SRP−dependent cotranslational protein targeting to membrane
SEC63
SSA4
KAR2
SBH1
YER087C−B_1
SRP102
SSA2
SRP101
SSH1
SBH2
SSA3
SIL1
SRP72
SRP54
SEC61
SRP21
SRP68
SSA1
Gene
Z_Shift_K_Doxo_HLD
Z_Im_K_Doxo_HLD
Z_Shift_K_Doxo_EG
Z_Im_K_Doxo_EG
Z_Shift_L_Doxo_HLD
Z_Im_L_Doxo_HLD
Z_Shift_L_Doxo_EG
Z_Im_L_Doxo_EG

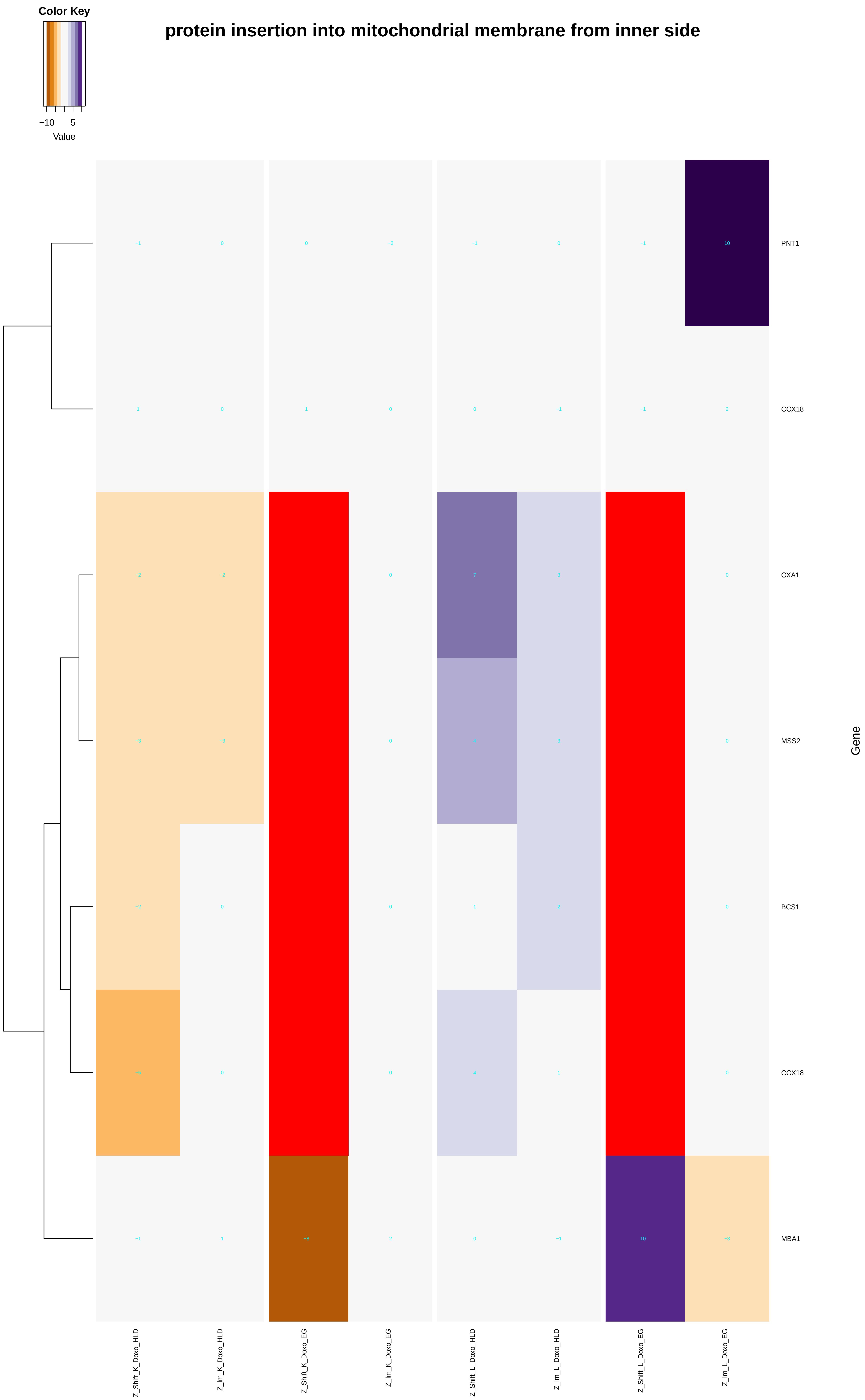

Color Key
−10
5
Value
protein insertion into mitochondrial membrane from inner side
PNT1
COX18
OXA1
MSS2
BCS1
COX18
MBA1
Gene
Z_Shift_K_Doxo_HLD
Z_Im_K_Doxo_HLD
Z_Shift_K_Doxo_EG
Z_Im_K_Doxo_EG
Z_Shift_L_Doxo_HLD
Z_Im_L_Doxo_HLD
Z_Shift_L_Doxo_EG
Z_Im_L_Doxo_EG
