## Additional File 9 for "A yeast phenomic model for the influence of Warburg metabolism on genetic buffering of doxorubicin": establishment_of_protein_localization_to_mitochondrion.pdf

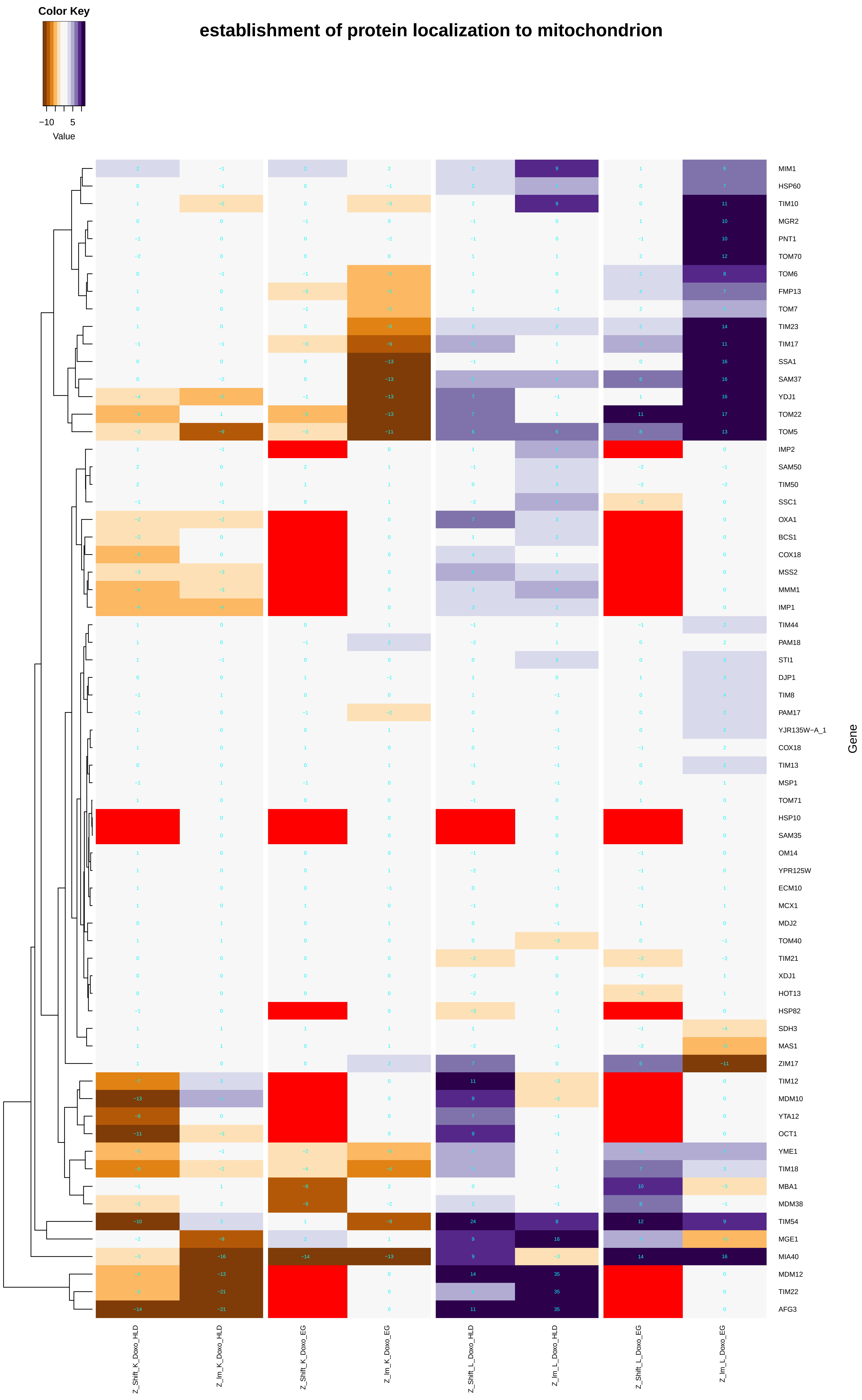
establishment of protein localization to mitochondrion
Color Key
−10
5
Value
Gene
MIM1
HSP60
TIM10
MGR2
PNT1
TOM70
TOM6
FMP13
TOM7
TIM23
TIM17
SSA1
SAM37
YDJ1
TOM22
TOM5
IMP2
SAM50
TIM50
SSC1
OXA1
BCS1
COX18
MSS2
MMM1
IMP1
TIM44
PAM18
STI1
DJP1
TIM8
PAM17
YJR135W−A_1
COX18
TIM13
MSP1
TOM71
HSP10
SAM35
OM14
YPR125W
ECM10
MCX1
MDJ2
TOM40
TIM21
XDJ1
HOT13
HSP82
SDH3
MAS1
ZIM17
TIM12
MDM10
YTA12
OCT1
YME1
TIM18
MBA1
MDM38
TIM54
MGE1
MIA40
MDM12
TIM22
AFG3
Z_Shift_K_Doxo_HLD
Z_lm_K_Doxo_HLD
Z_Shift_K_Doxo_EG
Z_lm_K_Doxo_EG
Z_Shift_L_Doxo_HLD
Z_lm_L_Doxo_HLD
Z_Shift_L_Doxo_EG
Z_lm_L_Doxo_EG

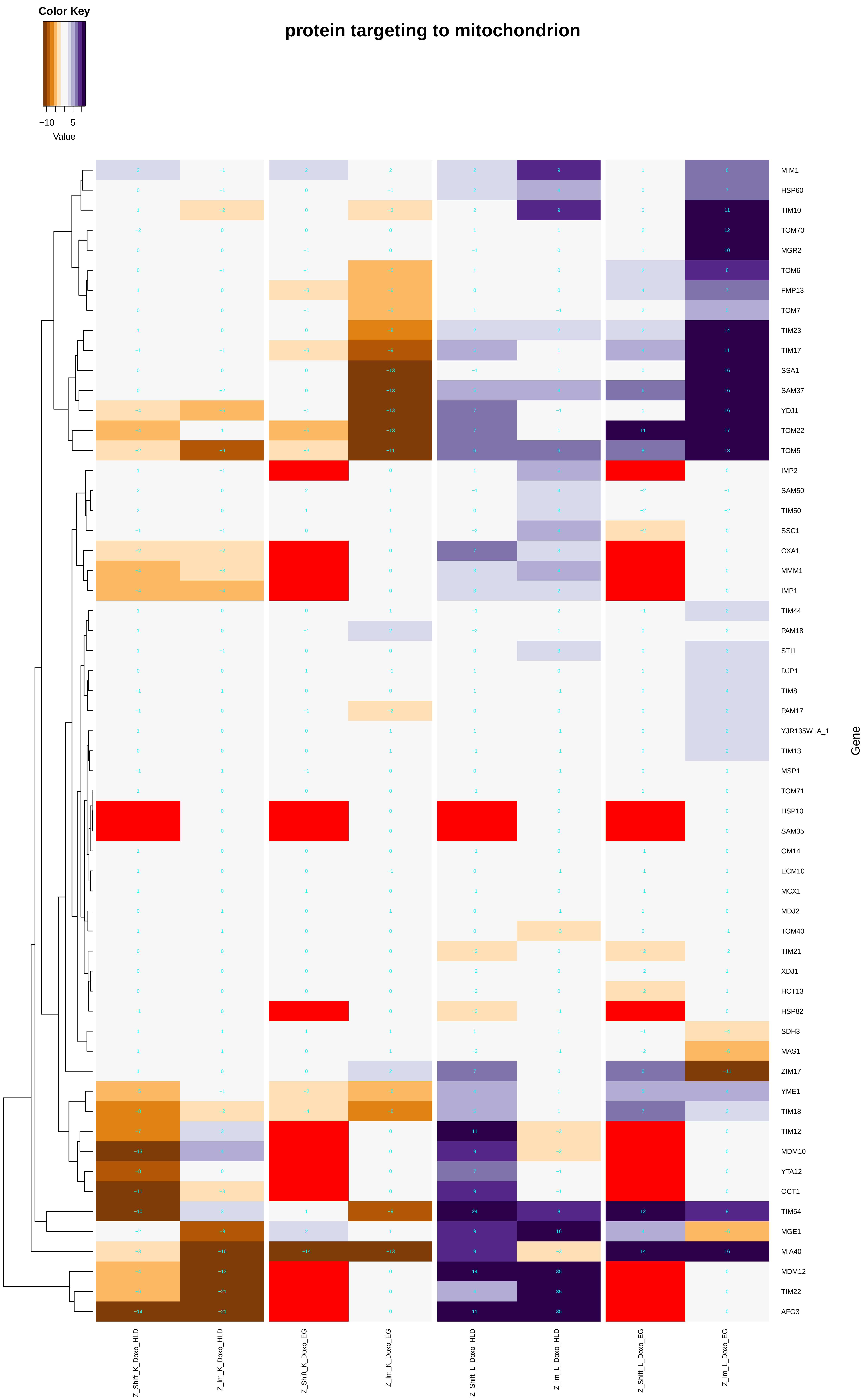
protein targeting to mitochondrion
Color Key
−10
5
Value
MIM1
HSP60
TIM10
TOM70
MGR2
TOM6
FMP13
TOM7
TIM23
TIM17
SSA1
SAM37
YDJ1
TOM22
TOM5
IMP2
SAM50
TIM50
SSC1
OXA1
MMM1
IMP1
TIM44
PAM18
STI1
DJP1
TIM8
PAM17
YJR135W−A_1
TIM13
MSP1
TOM71
HSP10
SAM35
OM14
ECM10
MCX1
MDJ2
TOM40
TIM21
XDJ1
HOT13
HSP82
SDH3
MAS1
ZIM17
YME1
TIM18
TIM12
MDM10
YTA12
OCT1
TIM54
MGE1
MIA40
MDM12
TIM22
AFG3
Gene
Z_Shift_K_Doxo_HLD
Z_lm_K_Doxo_HLD
Z_Shift_K_Doxo_EG
Z_lm_K_Doxo_EG
Z_Shift_L_Doxo_HLD
Z_lm_L_Doxo_HLD
Z_Shift_L_Doxo_EG
Z_lm_L_Doxo_EG

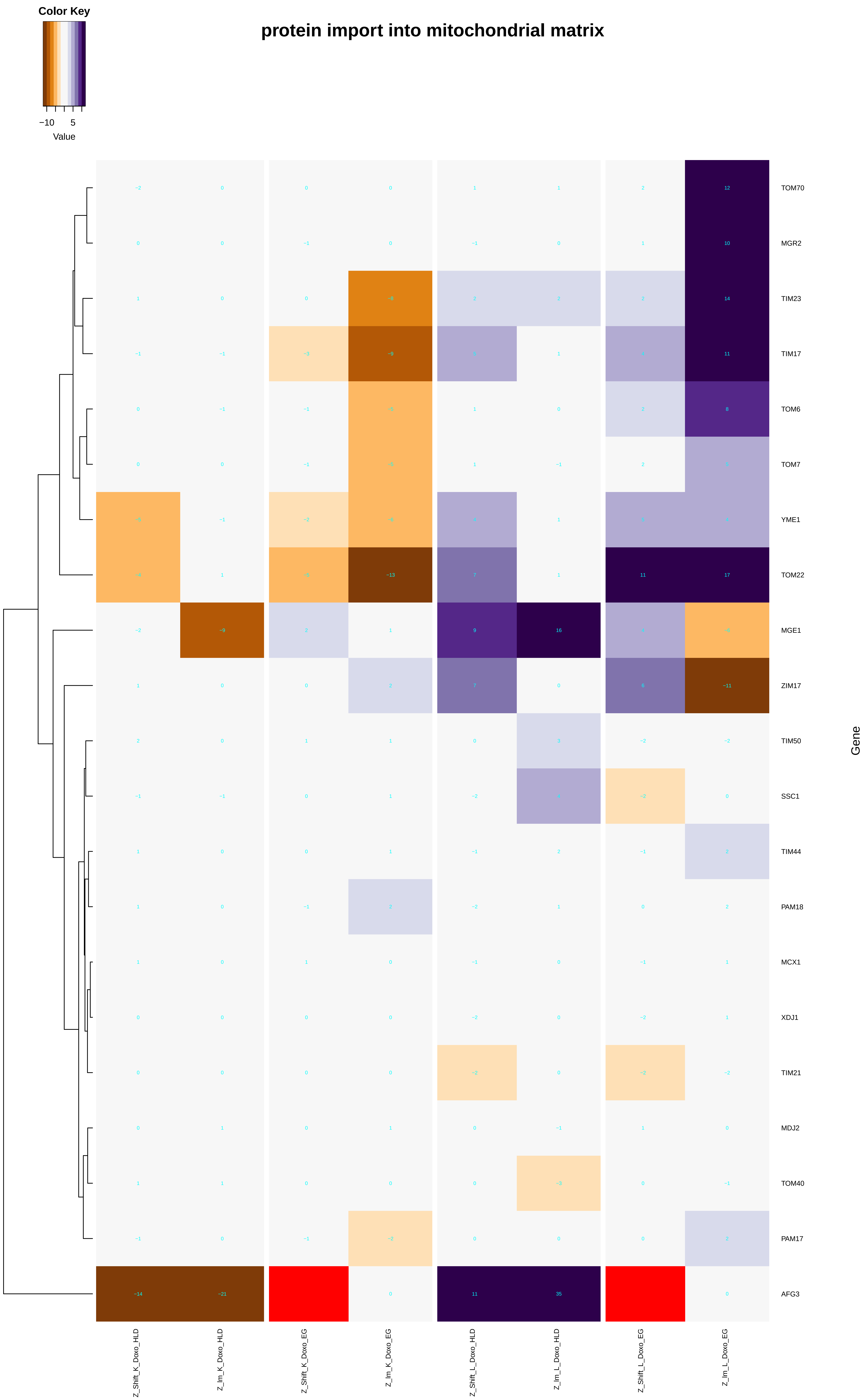
Color Key
−10 5
Value
protein import into mitochondrial matrix
TOM70
MGR2
TIM23
TIM17
TOM6
TOM7
YME1
TOM22
MGE1
ZIM17
TIM50
SSC1
TIM44
PAM18
MCX1
XDJ1
TIM21
MDJ2
TOM40
PAM17
AFG3
Gene
Z_Shift_K_Doxo_HLD
Z_Im_K_Doxo_HLD
Z_Shift_K_Doxo_EG
Z_Im_K_Doxo_EG
Z_Shift_L_Doxo_HLD
Z_Im_L_Doxo_HLD
Z_Shift_L_Doxo_EG
Z_Im_L_Doxo_EG

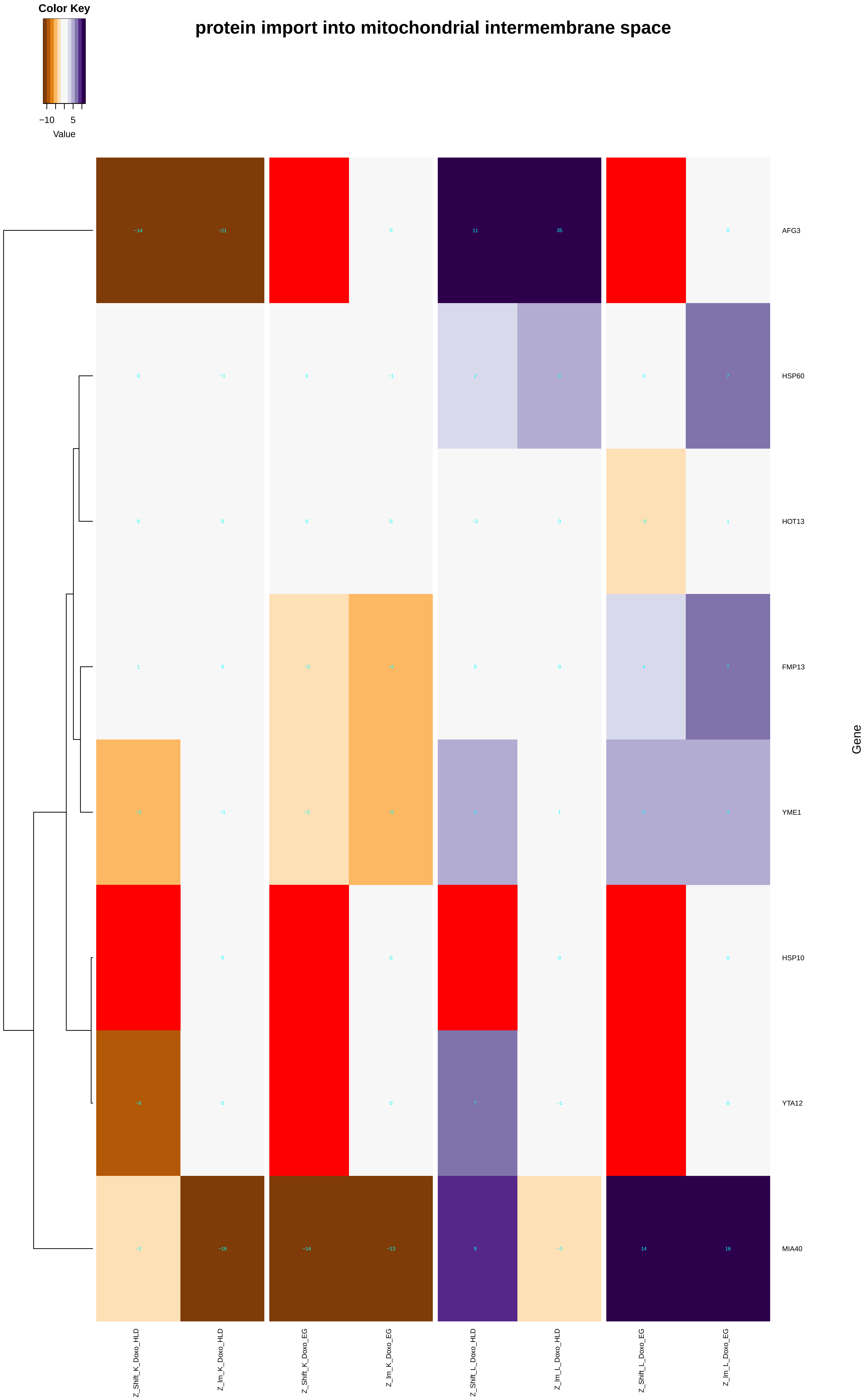
Color Key
−10
5
Value
protein import into mitochondrial intermembrane space
AFG3
HSP60
HOT13
FMP13
YME1
HSP10
YTA12
MIA40
Gene
Z_Shift_K_Doxo_HLD
Z_Im_K_Doxo_HLD
Z_Shift_K_Doxo_EG
Z_Im_K_Doxo_EG
Z_Shift_L_Doxo_HLD
Z_Im_L_Doxo_HLD
Z_Shift_L_Doxo_EG
Z_Im_L_Doxo_EG

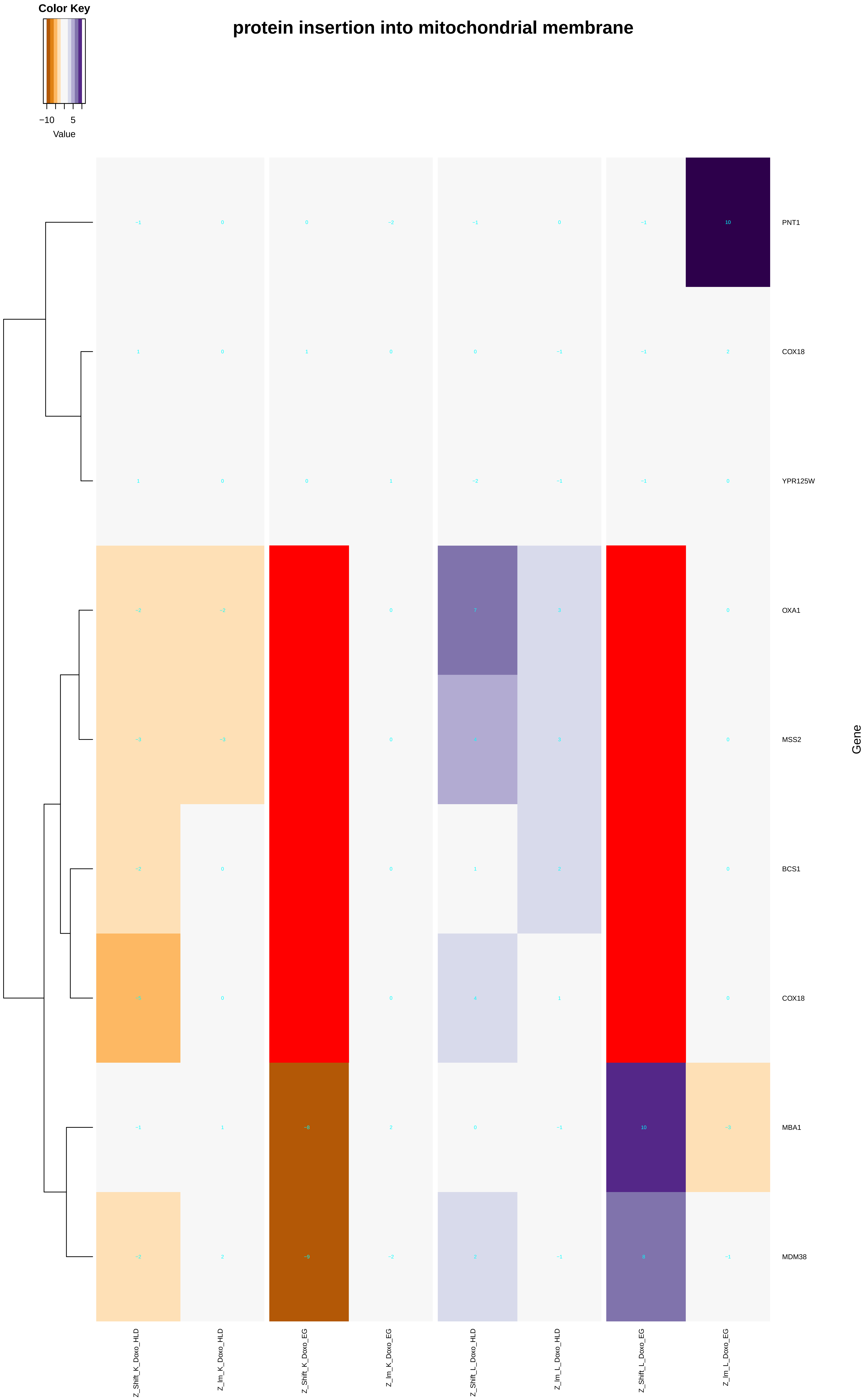
Color Key
−10
5
Value
protein insertion into mitochondrial membrane
PNT1
COX18
YPR125W
OXA1
MSS2
BCS1
COX18
MBA1
MDM38
Gene
Z_Shift_K_Doxo_HLD
Z_Im_K_Doxo_HLD
Z_Shift_K_Doxo_EG
Z_Im_K_Doxo_EG
Z_Shift_L_Doxo_HLD
Z_Im_L_Doxo_HLD
Z_Shift_L_Doxo_EG
Z_Im_L_Doxo_EG

### establishment of protein localization to mitochondrial membrane

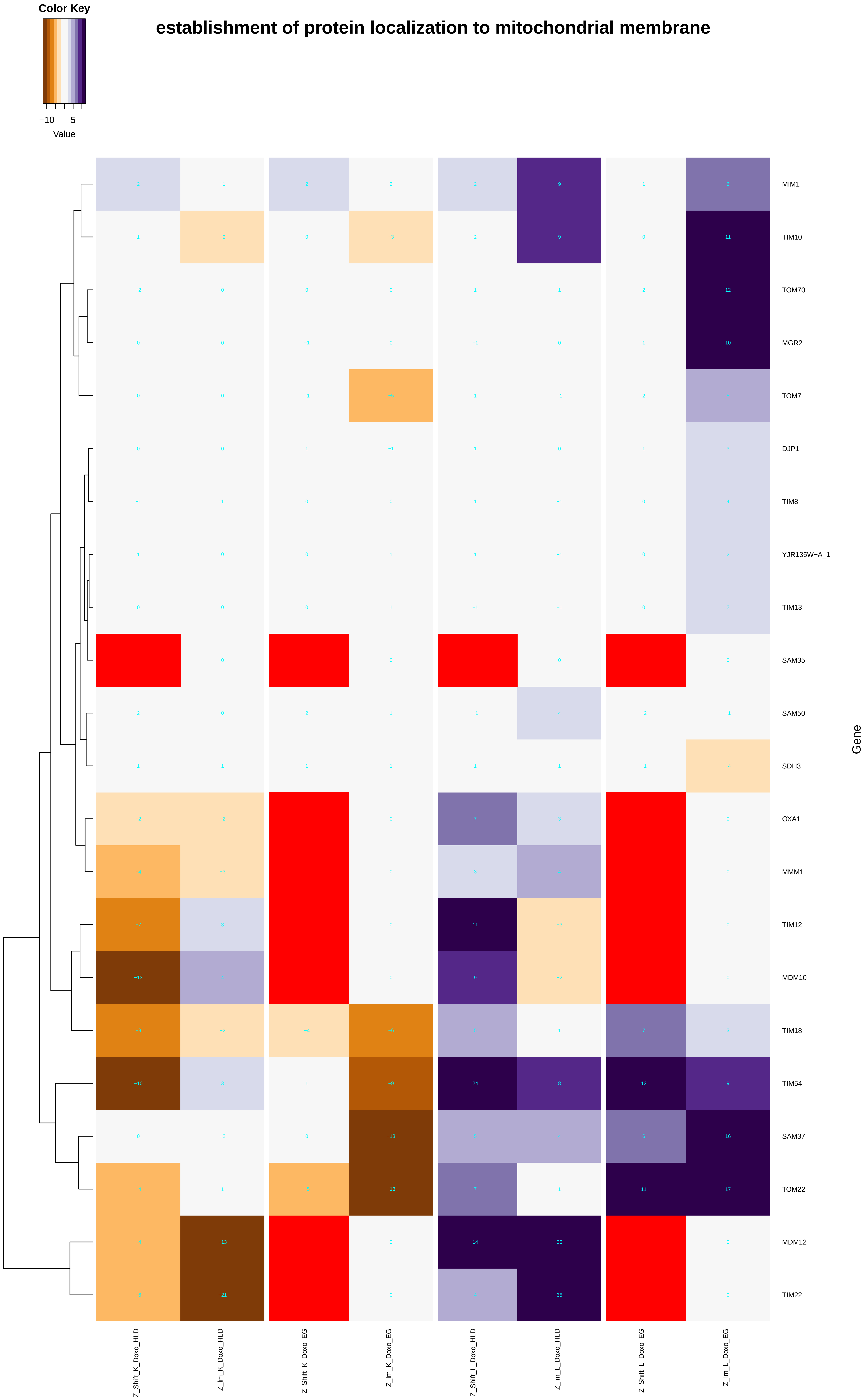

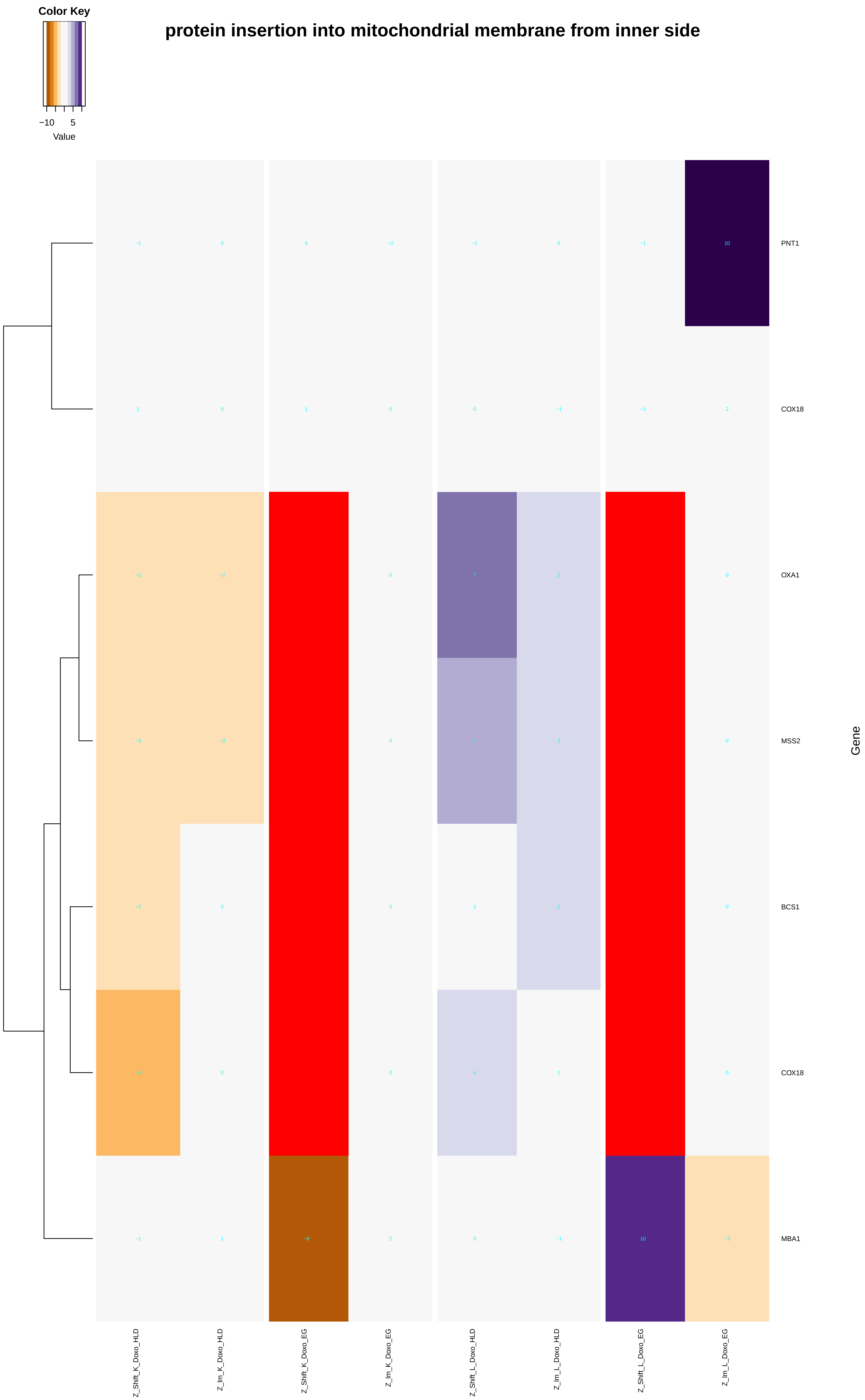
Color Key
−10
5
Value
protein insertion into mitochondrial membrane from inner side
PNT1
COX18
OXA1
MSS2
BCS1
COX18
MBA1
Gene
Z_Shift_K_Doxo_HLD
Z_Im_K_Doxo_HLD
Z_Shift_K_Doxo_EG
Z_Im_K_Doxo_EG
Z_Shift_L_Doxo_HLD
Z_Im_L_Doxo_HLD
Z_Shift_L_Doxo_EG
Z_Im_L_Doxo_EG
−1 0 0 −2 −1 0 −1 10
1 0 1 0 0 −1 −1 2
−2 −2 0 7 3 0
−3 −3 0 3 0
−2 0 0 1 2 0
0 0 4 1 0
−1 1 −8 2 0 −1 10 −3

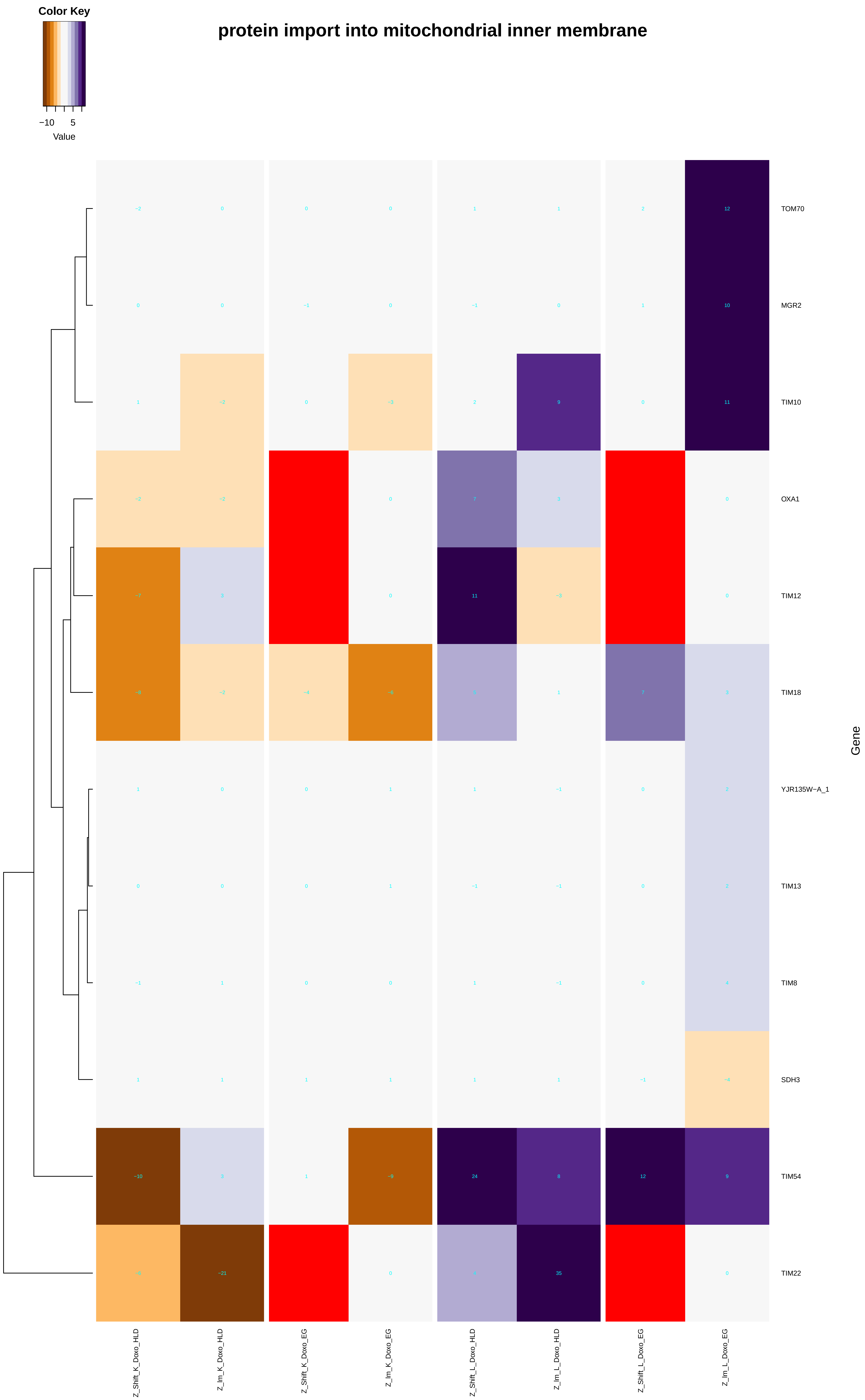

Color Key
−10
5
Value
protein import into mitochondrial inner membrane
TOM70
MGR2
TIM10
OXA1
TIM12
TIM18
YJR135W−A_1
TIM13
TIM8
SDH3
TIM54
TIM22
Gene
Z_Shift_K_Doxo_HLD
Z_Im_K_Doxo_HLD
Z_Shift_K_Doxo_EG
Z_Im_K_Doxo_EG
Z_Shift_L_Doxo_HLD
Z_Im_L_Doxo_HLD
Z_Shift_L_Doxo_EG
Z_Im_L_Doxo_EG

### protein import into mitochondrial outer membrane

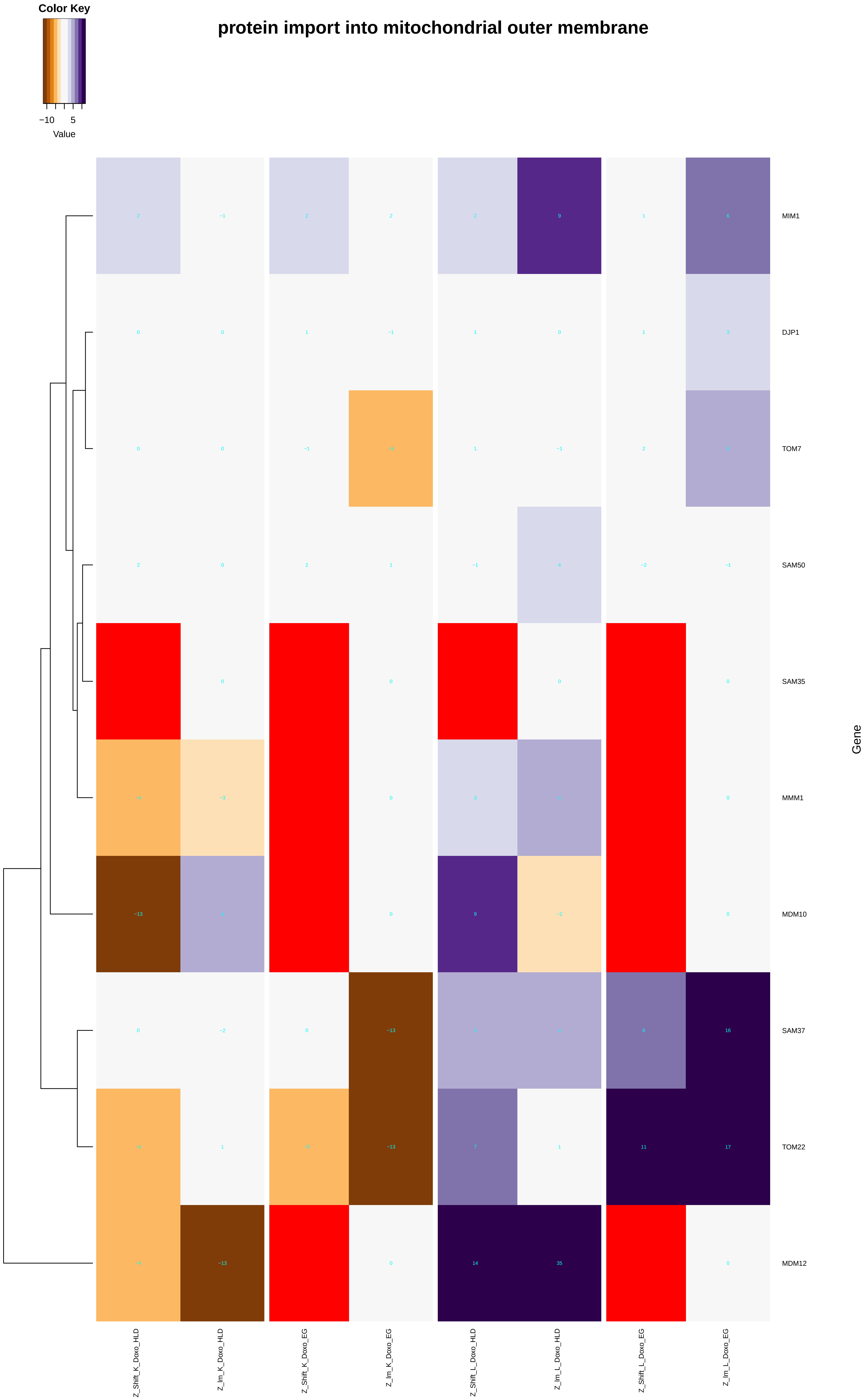
