## Additional File 9 for "A yeast phenomic model for the influence of Warburg metabolism on genetic buffering of doxorubicin": establishment_of_protein_localization_to_organelle.pdf

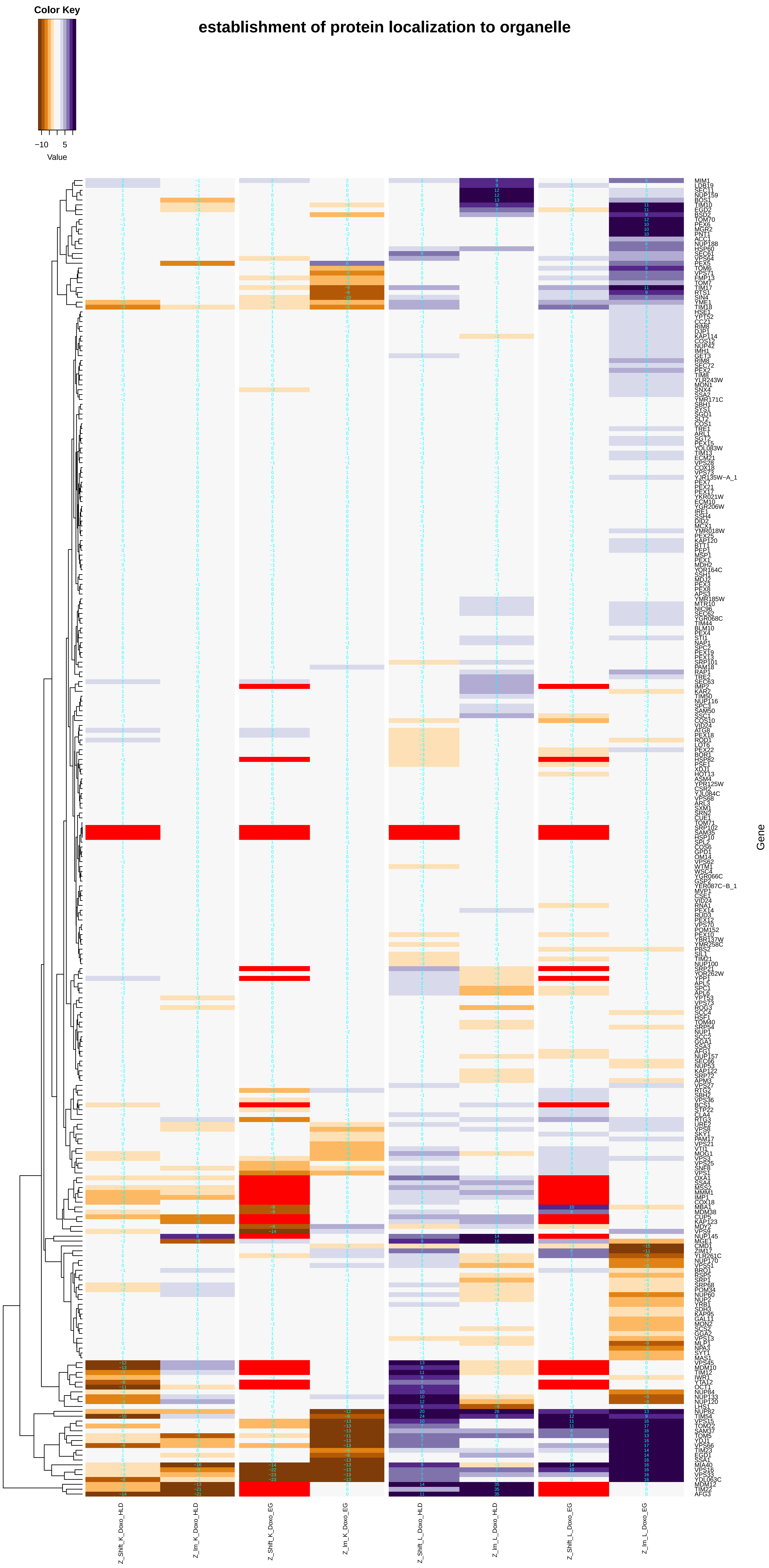

establishment of protein localization to organelle
Color Key
−10
5
Value
Gene
MIM1
LDB19
SEC11
NUP159
BOS1
TIM10
EGD2
BSD2
TOM70
PEX6
MGR2
PNT1
ACC1
NUP188
HSP60
SEC61
VPS64
PEX5
TOM6
VPS71
FMP13
TOM7
TIM17
RTS1
SIN4
YME1
TIM18
HSE1
YPT52
CCZ1
RIM8
DJP1
KAP114
COS12
NUP42
IMH1
GET3
RIM8
SEC72
PEX2
TIM8
YLR243W
MON1
SNX4
SSA2
YMR171C
SBH1
SYS1
SGO1
SLT2
COS1
TRE1
ARL1
SGT2
PEX15
YOL083W
TIM13
ECM21
VPS28
COX18
VPS73
YJR135W−A_1
PEX7
PEX21
PEX17
YKR021W
ECM10
YGR206W
IRE1
SSH4
DID2
MCX1
YMR018W
PEX25
KAP120
BTT1
PEP1
MSP1
PEX1
MDH2
YOR164C
SSH1
MDJ2
PEX3
PEX8
APS3
YMR185W
MTR10
NIC96
SEC62
YGR068C
TIM44
BLM10
PEX4
STI1
NAP1
SPC2
PEX19
PEX13
SRP101
PAM18
RAP1
TRE2
SEC63
IMP2
KAR2
TIM50
NUP116
SPC3
SAM50
SSC1
COS10
VID24
ATG8
PEX18
ROD1
LOT6
PEX22
BOR1
HSP82
PSE1
XDJ1
HOT13
ASM4
YPR125W
CSR2
YJL084C
VPS68
ARL3
SXM1
SRN2
CUE1
TOM71
SRP102
SAM35
HSP10
SPL2
COS6
GPD1
OM14
VPS62
WTM1
WSC4
YGR066C
GSP2
YER087C−B_1
MVP1
CSE1
VID24
RNA1
PEX14
RUD3
PEX12
VPS70
POM152
PEX10
YBR137W
YMR258C
PBS2
SIL1
TIM21
NUP100
SRP21
YOR262W
YPP1
APL5
SPC1
APL6
YPT53
VPS73
ROG3
SCC4
HSF1
TOM40
SRP54
NUP1
SCC2
GGA1
SSA3
AFG1
NUP157
SEC66
NUP53
KAP122
SRP72
APM3
VPS27
RTG2
SBH2
VPS36
BCS1
STP22
CLA4
RTG3
URE2
VPS8
SKY1
PAM17
VPS21
VTI1
MOG1
VPS3
VPS25
SNF8
VPS1
OXA1
SSA4
MSS2
MMM1
IMP1
COX18
MBA1
MDM38
CUP5
KAP123
MDY2
VPS9
NUP145
MGE1
CMD1
ZIM17
YLR261C
NUP170
VPS51
BRO1
RSP5
SRP1
SRP68
POM34
NUP60
NUP2
YRB1
SDH3
KAP95
GAL11
MON2
SCS2
GGA2
VPS13
MLP1
NPA3
SYT1
MAS1
VPS45
MDM10
TIM12
IWR1
YTA12
OCT1
NUP84
NUP133
NUP120
LHS1
NUP82
TIM54
VPS15
TOM22
SAM37
TOM5
YDJ1
VPS66
TIM23
EGD1
SSA1
MIA40
VPS16
VPS33
YDL063C
MDM12
TIM22
AFG3
Z_Shift_K_Doxo_HLD
Z_lm_K_Doxo_HLD
Z_Shift_K_Doxo_EG
Z_lm_K_Doxo_EG
Z_Shift_L_Doxo_HLD
Z_lm_L_Doxo_HLD
Z_Shift_L_Doxo_EG
Z_lm_L_Doxo_EG

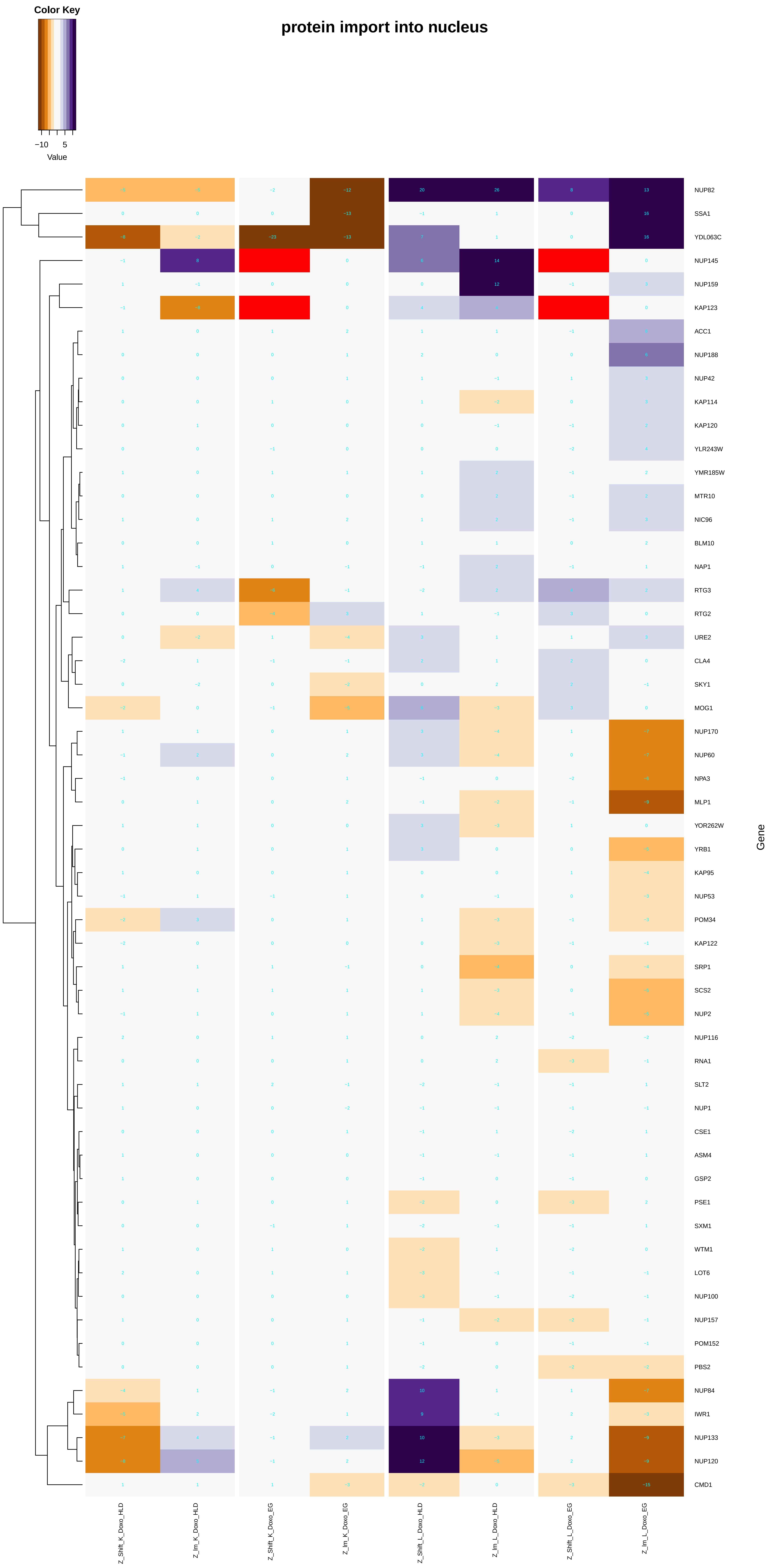

Color Key
−10
5
Value
protein import into nucleus
NUP82
SSA1
YDL063C
NUP145
NUP159
KAP123
ACC1
NUP188
NUP42
KAP114
KAP120
YLR243W
YMR185W
MTR10
NIC96
BLM10
NAP1
RTG3
RTG2
URE2
CLA4
SKY1
MOG1
NUP170
NUP60
NPA3
MLP1
YOR262W
YRB1
KAP95
NUP53
POM34
KAP122
SRP1
SCS2
NUP2
NUP116
RNA1
SLT2
NUP1
CSE1
ASM4
GSP2
PSE1
SXM1
WTM1
LOT6
NUP100
NUP157
POM152
PBS2
NUP84
IWR1
NUP133
NUP120
CMD1
Gene
Z_Shift_K_Doxo_HLD
Z_lm_K_Doxo_HLD
Z_Shift_K_Doxo_EG
Z_lm_K_Doxo_EG
Z_Shift_L_Doxo_HLD
Z_lm_L_Doxo_HLD
Z_Shift_L_Doxo_EG
Z_lm_L_Doxo_EG

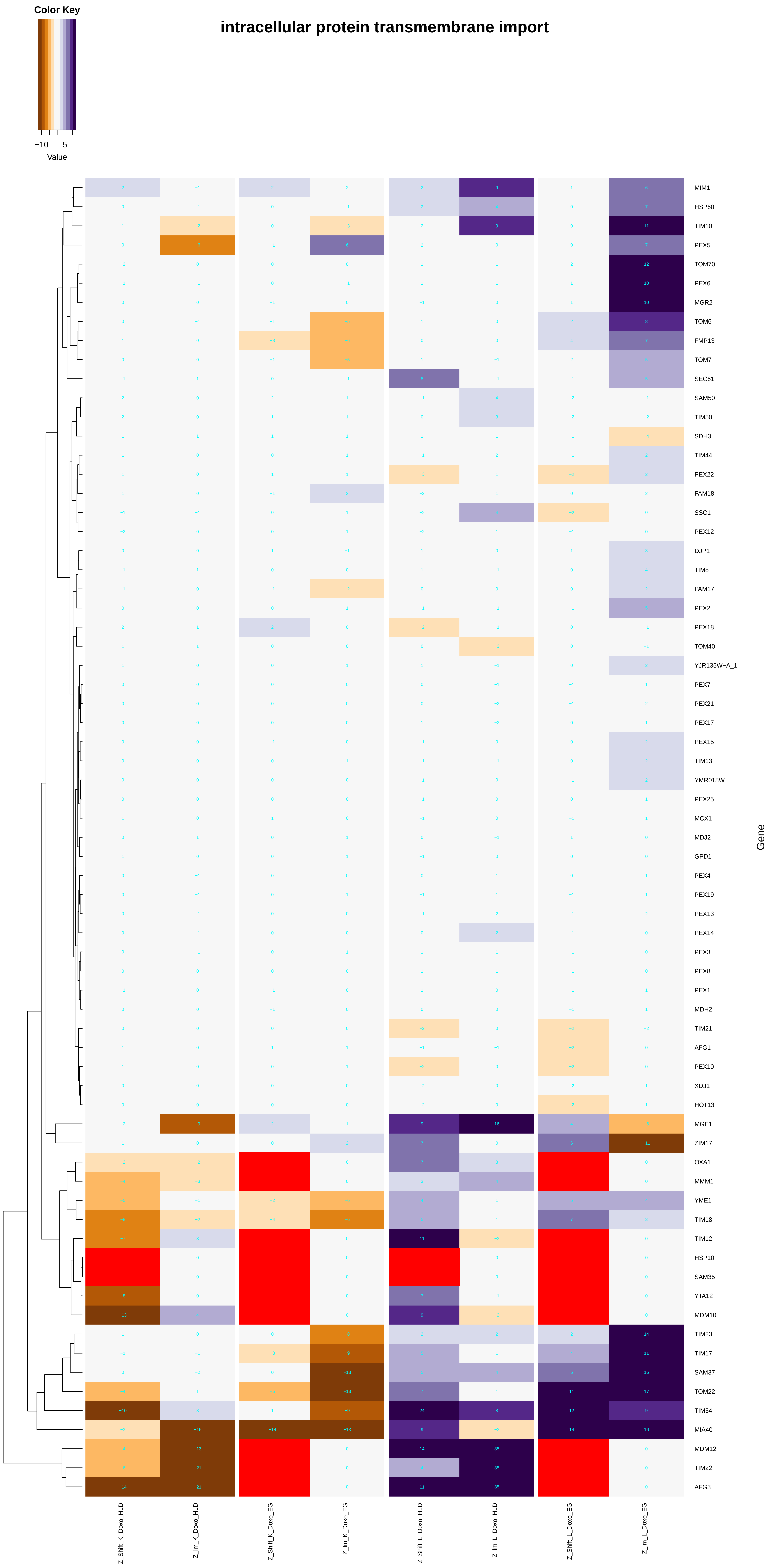

Color Key
−10
5
Value
intracellular protein transmembrane import
MIM1
HSP60
TIM10
PEX5
TOM70
PEX6
MGR2
TOM6
FMP13
TOM7
SEC61
SAM50
TIM50
SDH3
TIM44
PEX22
PAM18
SSC1
PEX12
DJP1
TIM8
PAM17
PEX2
PEX18
TOM40
YJR135W−A_1
PEX7
PEX21
PEX17
PEX15
TIM13
YMR018W
PEX25
MCX1
MDJ2
GPD1
PEX4
PEX19
PEX13
PEX14
PEX3
PEX8
PEX1
MDH2
TIM21
AFG1
PEX10
XDJ1
HOT13
MGE1
ZIM17
OXA1
MMM1
YME1
TIM18
TIM12
HSP10
SAM35
YTA12
MDM10
TIM23
TIM17
SAM37
TOM22
TIM54
MIA40
MDM12
TIM22
AFG3
Gene
Z_Shift_K_Doxo_HLD
Z_lm_K_Doxo_HLD
Z_Shift_K_Doxo_EG
Z_lm_K_Doxo_EG
Z_Shift_L_Doxo_HLD
Z_lm_L_Doxo_HLD
Z_Shift_L_Doxo_EG
Z_lm_L_Doxo_EG

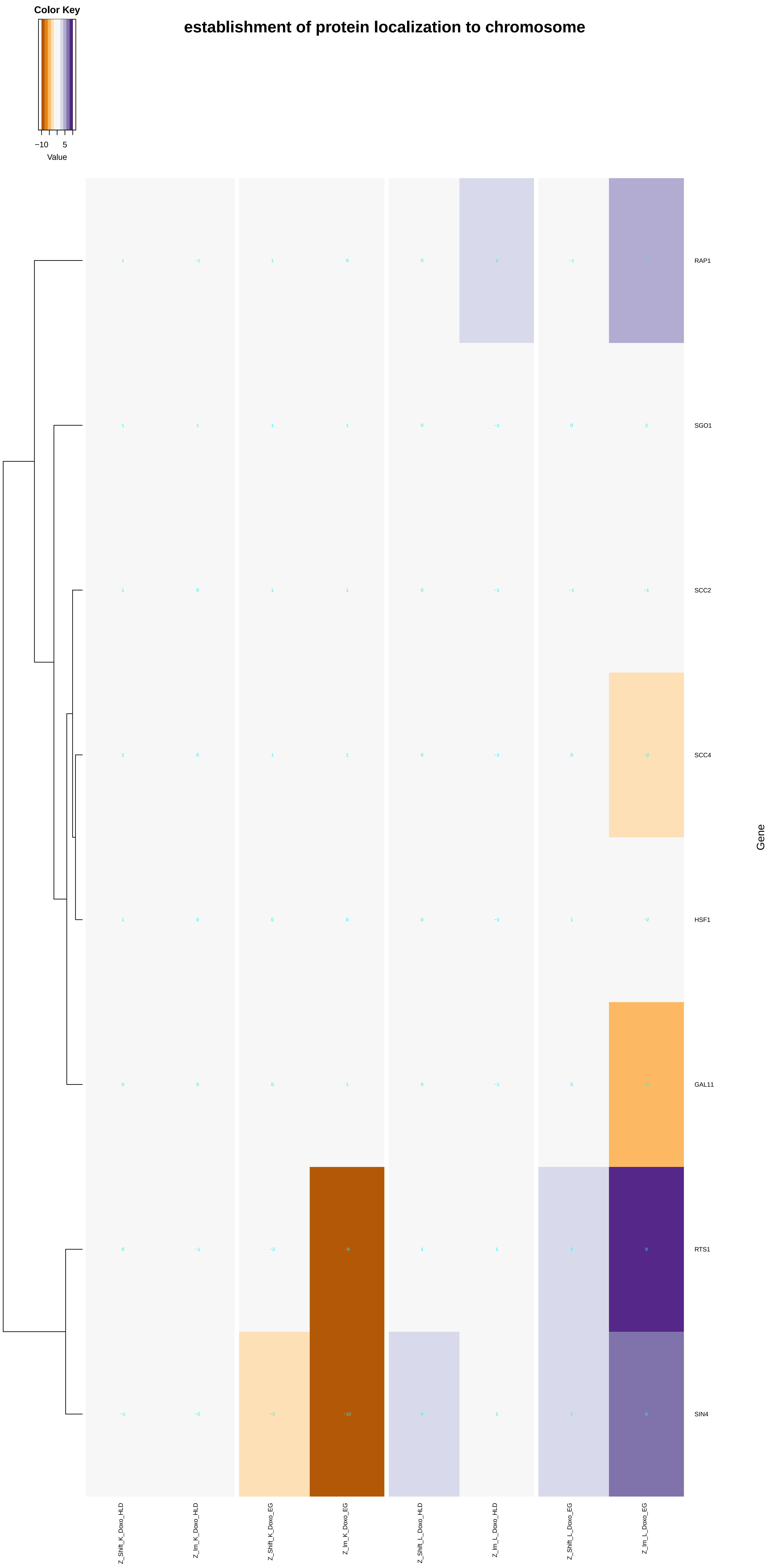
Color Key
−10
5
Value
establishment of protein localization to chromosome
RAP1
SGO1
SCC2
SCC4
HSF1
GAL11
RTS1
SIN4
Gene
Z_Shift_K_Doxo_HLD
Z_lm_K_Doxo_HLD
Z_Shift_K_Doxo_EG
Z_lm_K_Doxo_EG
Z_Shift_L_Doxo_HLD
Z_lm_L_Doxo_HLD
Z_Shift_L_Doxo_EG
Z_lm_L_Doxo_EG

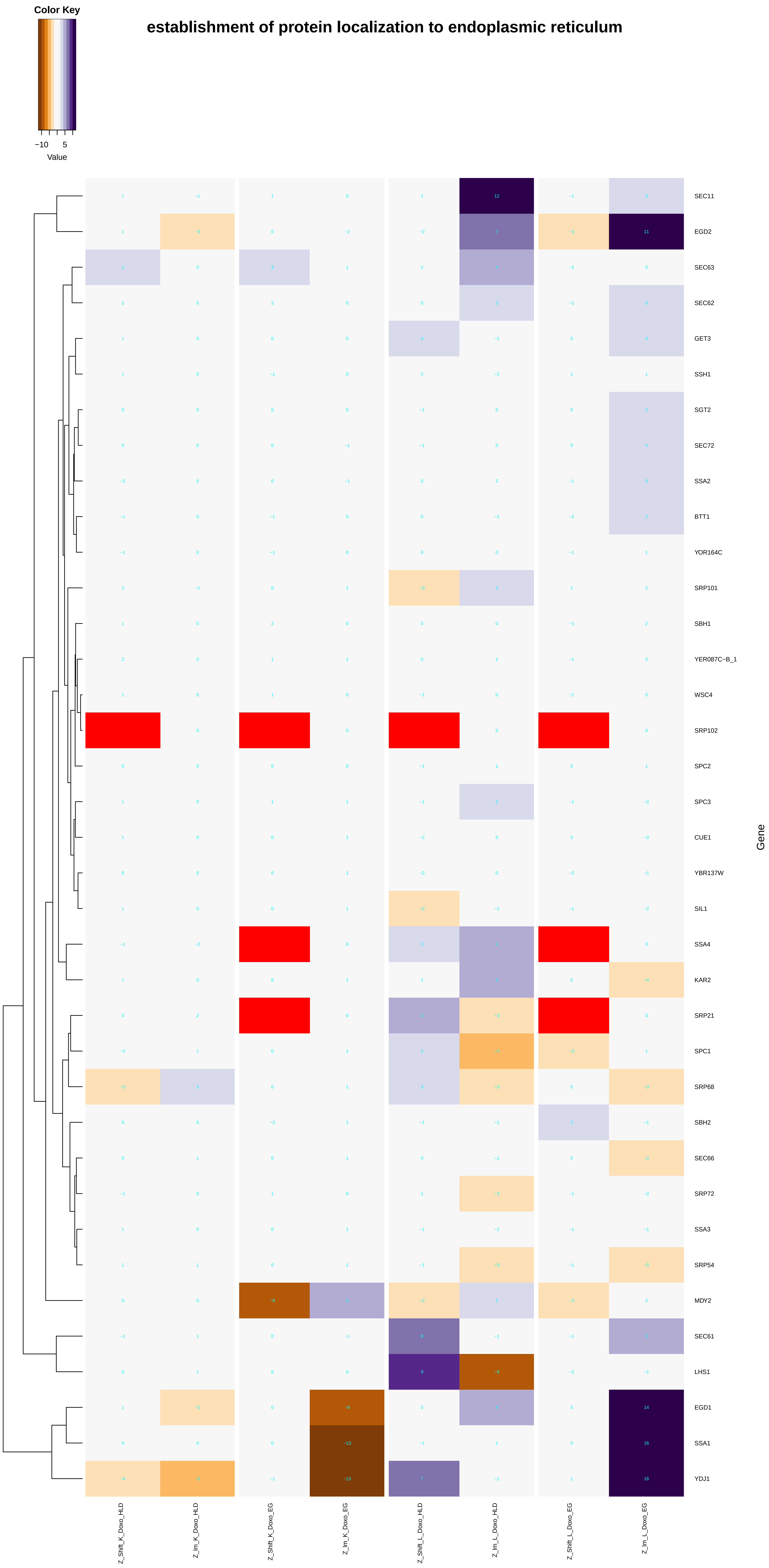
Color Key
−10
5
Value
establishment of protein localization to endoplasmic reticulum
SEC11
EGD2
SEC63
SEC62
GET3
SSH1
SGT2
SEC72
SSA2
BTT1
YOR164C
SRP101
SBH1
YER087C−B_1
WSC4
SRP102
SPC2
SPC3
CUE1
YBR137W
SIL1
SSA4
KAR2
SRP21
SPC1
SRP68
SBH2
SEC66
SRP72
SSA3
SRP54
MDY2
SEC61
LHS1
EGD1
SSA1
YDJ1
Gene
Z_Shift_K_Doxo_HLD
Z_Im_K_Doxo_HLD
Z_Shift_K_Doxo_EG
Z_Im_K_Doxo_EG
Z_Shift_L_Doxo_HLD
Z_Im_L_Doxo_HLD
Z_Shift_L_Doxo_EG
Z_Im_L_Doxo_EG

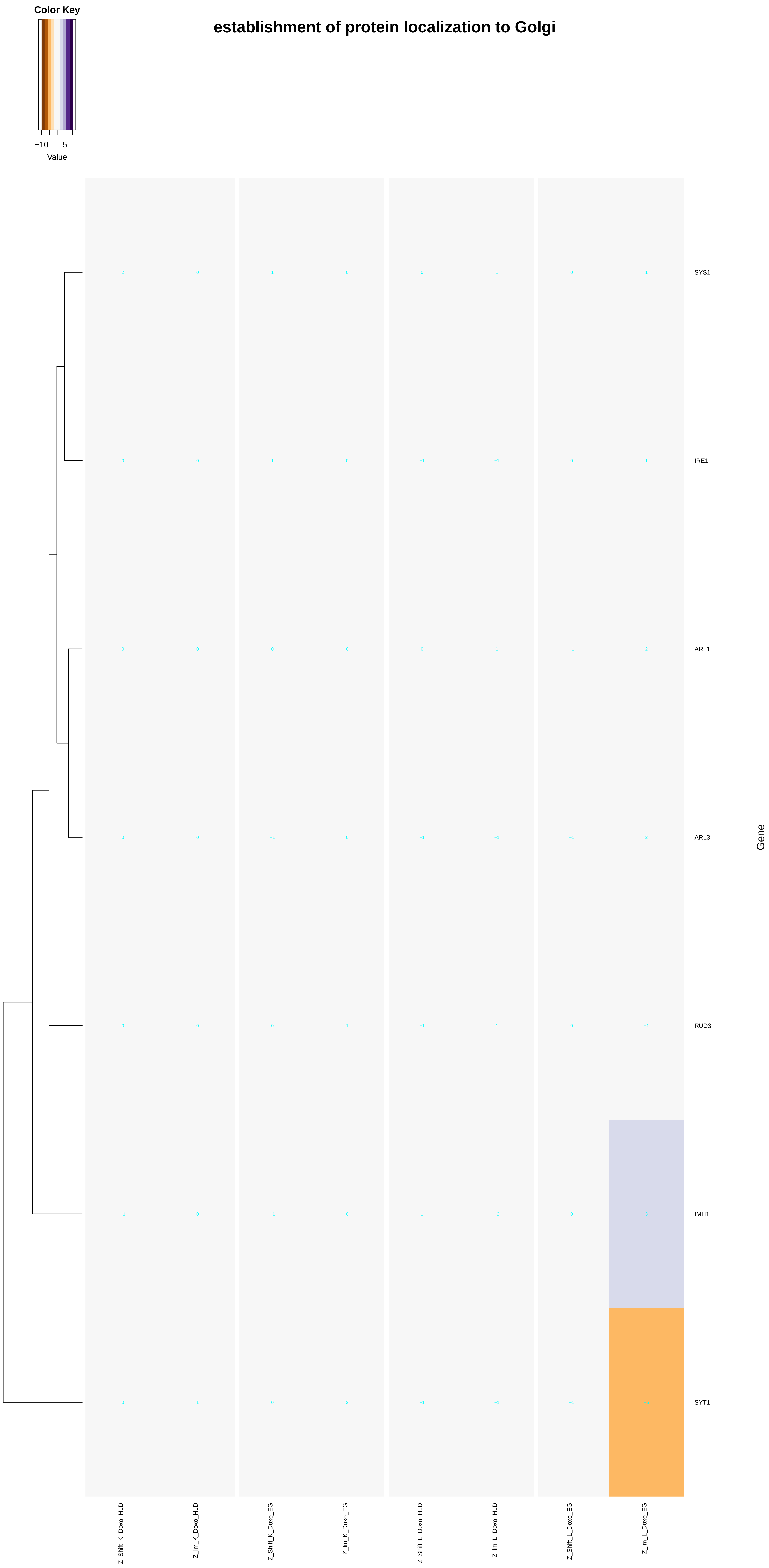
establishment of protein localization to Golgi
Color Key
−10
5
Value
SYS1
IRE1
ARL1
ARL3
RUD3
IMH1
SYT1
Gene
Z_Shift_K_Doxo_HLD
Z_Im_K_Doxo_HLD
Z_Shift_K_Doxo_EG
Z_Im_K_Doxo_EG
Z_Shift_L_Doxo_HLD
Z_Im_L_Doxo_HLD
Z_Shift_L_Doxo_EG
Z_Im_L_Doxo_EG

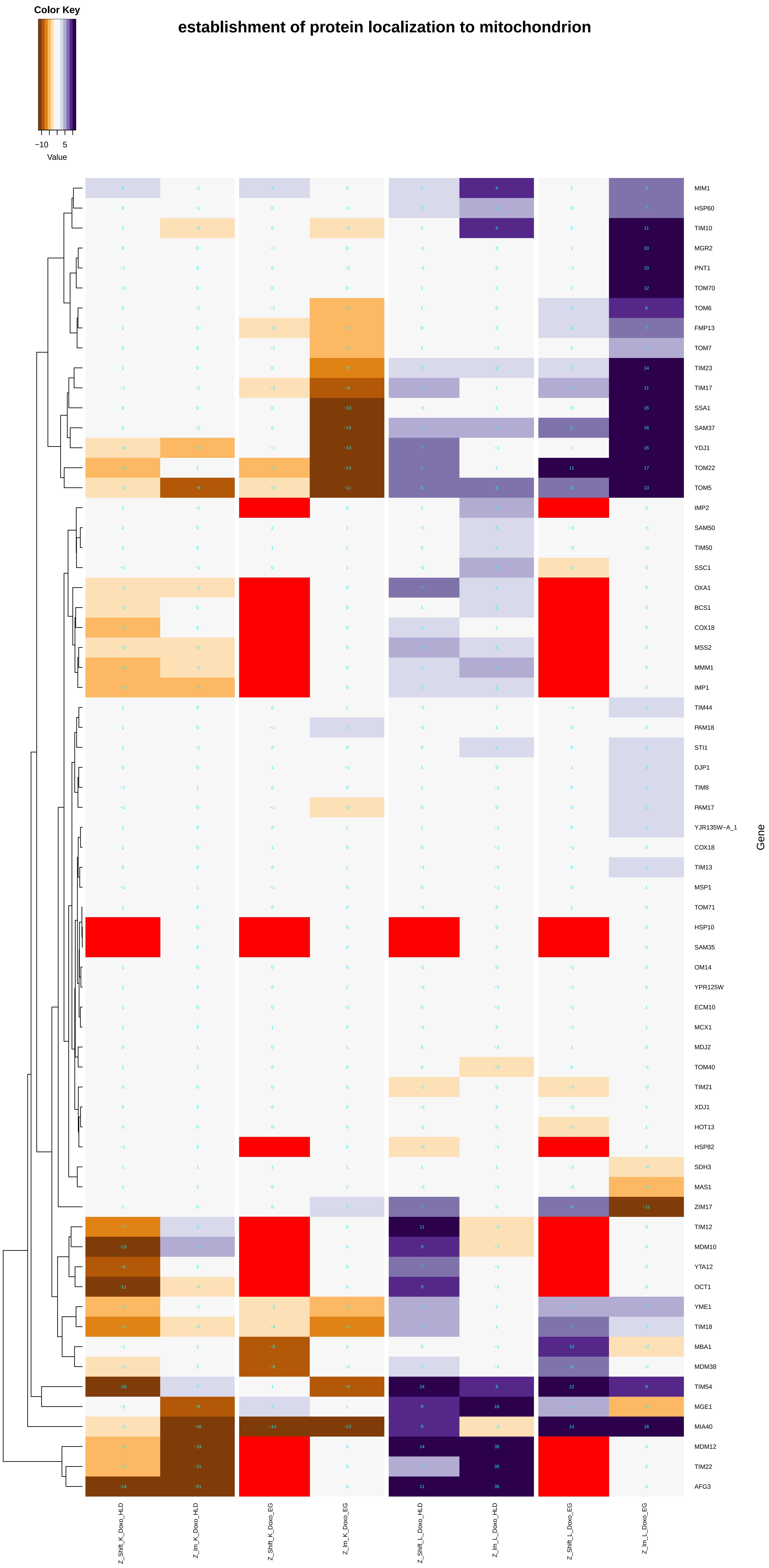
establishment of protein localization to mitochondrion
Color Key
−10 5
Value
MIM1
HSP60
TIM10
MGR2
PNT1
TOM70
TOM6
FMP13
TOM7
TIM23
TIM17
SSA1
SAM37
YDJ1
TOM22
TOM5
IMP2
SAM50
TIM50
SSC1
OXA1
BCS1
COX18
MSS2
MMM1
IMP1
TIM44
PAM18
STI1
DJP1
TIM8
PAM17
YJR135W−A_1
COX18
TIM13
MSP1
TOM71
HSP10
SAM35
OM14
YPR125W
ECM10
MCX1
MDJ2
TOM40
TIM21
XDJ1
HOT13
HSP82
SDH3
MAS1
ZIM17
TIM12
MDM10
YTA12
OCT1
YME1
TIM18
MBA1
MDM38
TIM54
MGE1
MIA40
MDM12
TIM22
AFG3
Gene
Z_Shift_K_Doxo_HLD
Z_lm_K_Doxo_HLD
Z_Shift_K_Doxo_EG
Z_lm_K_Doxo_EG
Z_Shift_L_Doxo_HLD
Z_lm_L_Doxo_HLD
Z_Shift_L_Doxo_EG
Z_lm_L_Doxo_EG

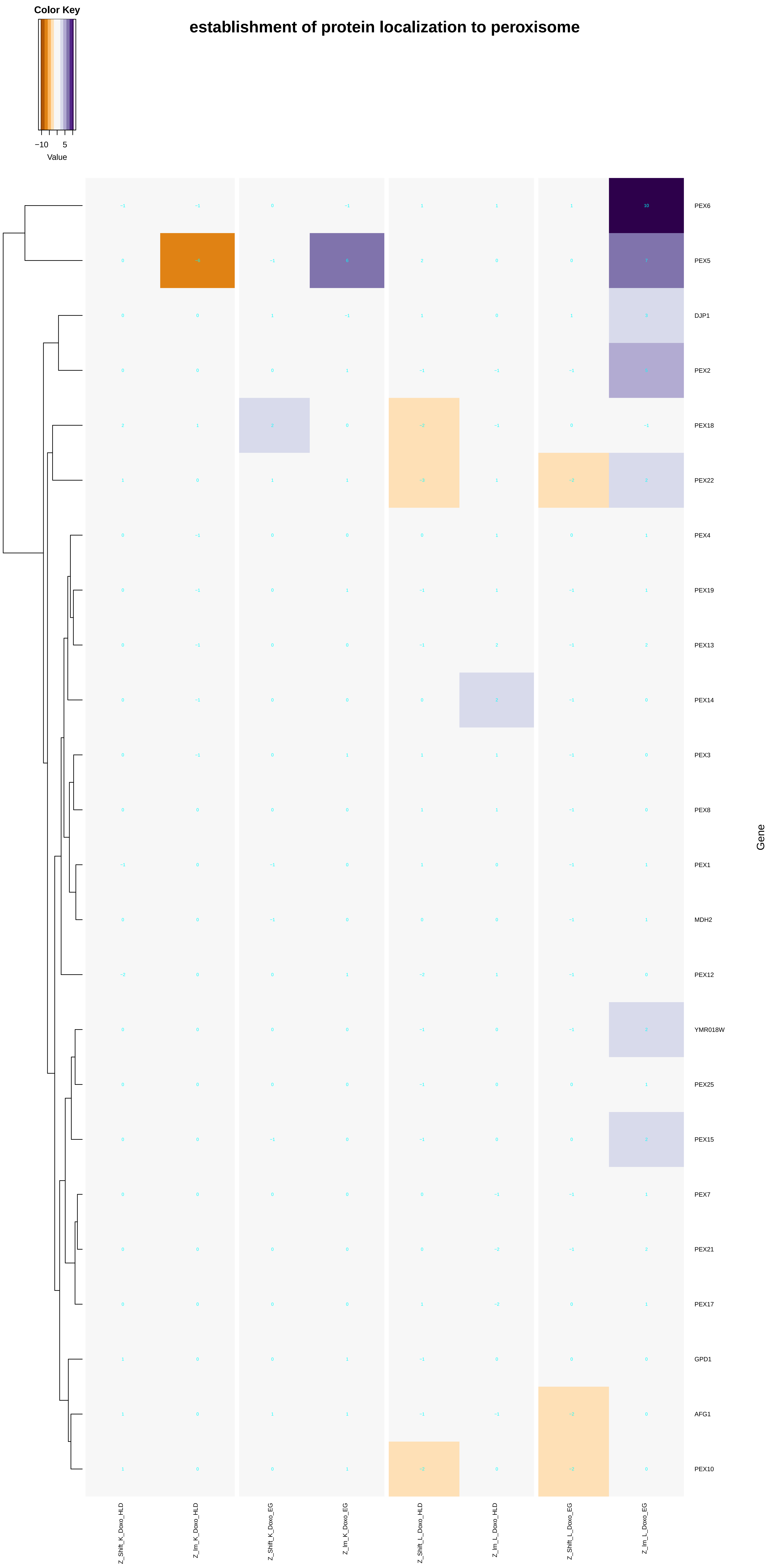
Color Key
−10
5
Value
establishment of protein localization to peroxisome
PEX6
PEX5
DJP1
PEX2
PEX18
PEX22
PEX4
PEX19
PEX13
PEX14
PEX3
PEX8
PEX1
MDH2
PEX12
YMR018W
PEX25
PEX15
PEX7
PEX21
PEX17
GPD1
AFG1
PEX10
Gene
Z_Shift_K_Doxo_HLD
Z_lm_K_Doxo_HLD
Z_Shift_K_Doxo_EG
Z_lm_K_Doxo_EG
Z_Shift_L_Doxo_HLD
Z_lm_L_Doxo_HLD
Z_Shift_L_Doxo_EG
Z_lm_L_Doxo_EG

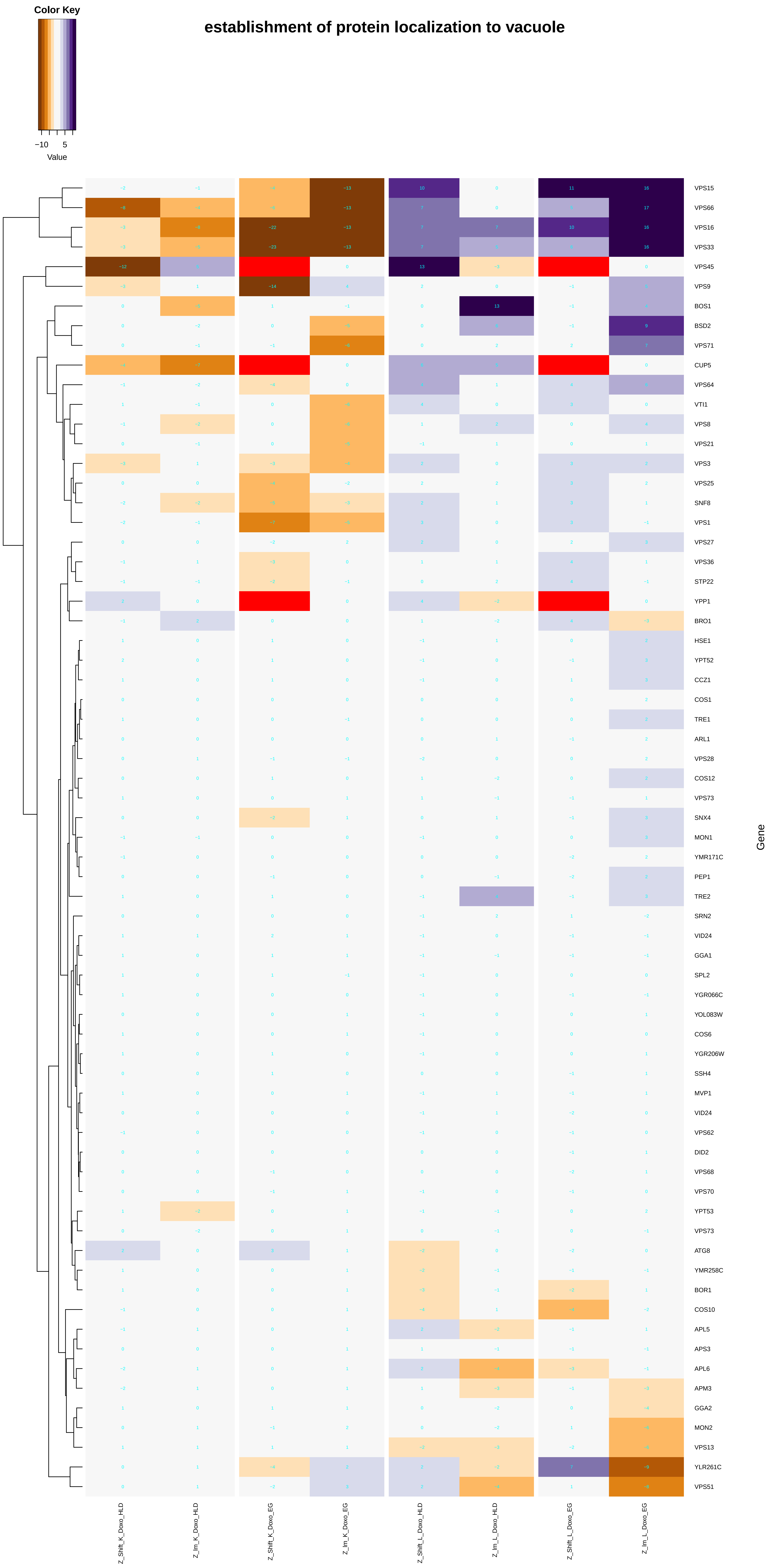
establishment of protein localization to vacuole
Color Key
−10
5
Value
VPS15
VPS66
VPS16
VPS33
VPS45
VPS9
BOS1
BSD2
VPS71
CUP5
VPS64
VTI1
VPS8
VPS21
VPS3
VPS25
SNF8
VPS1
VPS27
VPS36
STP22
YPP1
BRO1
HSE1
YPT52
CCZ1
COS1
TRE1
ARL1
VPS28
COS12
VPS73
SNX4
MON1
YMR171C
PEP1
TRE2
SRN2
VID24
GGA1
SPL2
YGR066C
YOL083W
COS6
YGR206W
SSH4
MVP1
VID24
VPS62
DID2
VPS68
VPS70
YPT53
VPS73
ATG8
YMR258C
BOR1
COS10
APL5
APS3
APL6
APM3
GGA2
MON2
VPS13
YLR261C
VPS51
Gene
Z_Shift_K_Doxo_HLD
Z_lm_K_Doxo_HLD
Z_Shift_K_Doxo_EG
Z_lm_K_Doxo_EG
Z_Shift_L_Doxo_HLD
Z_lm_L_Doxo_HLD
Z_Shift_L_Doxo_EG
Z_lm_L_Doxo_EG

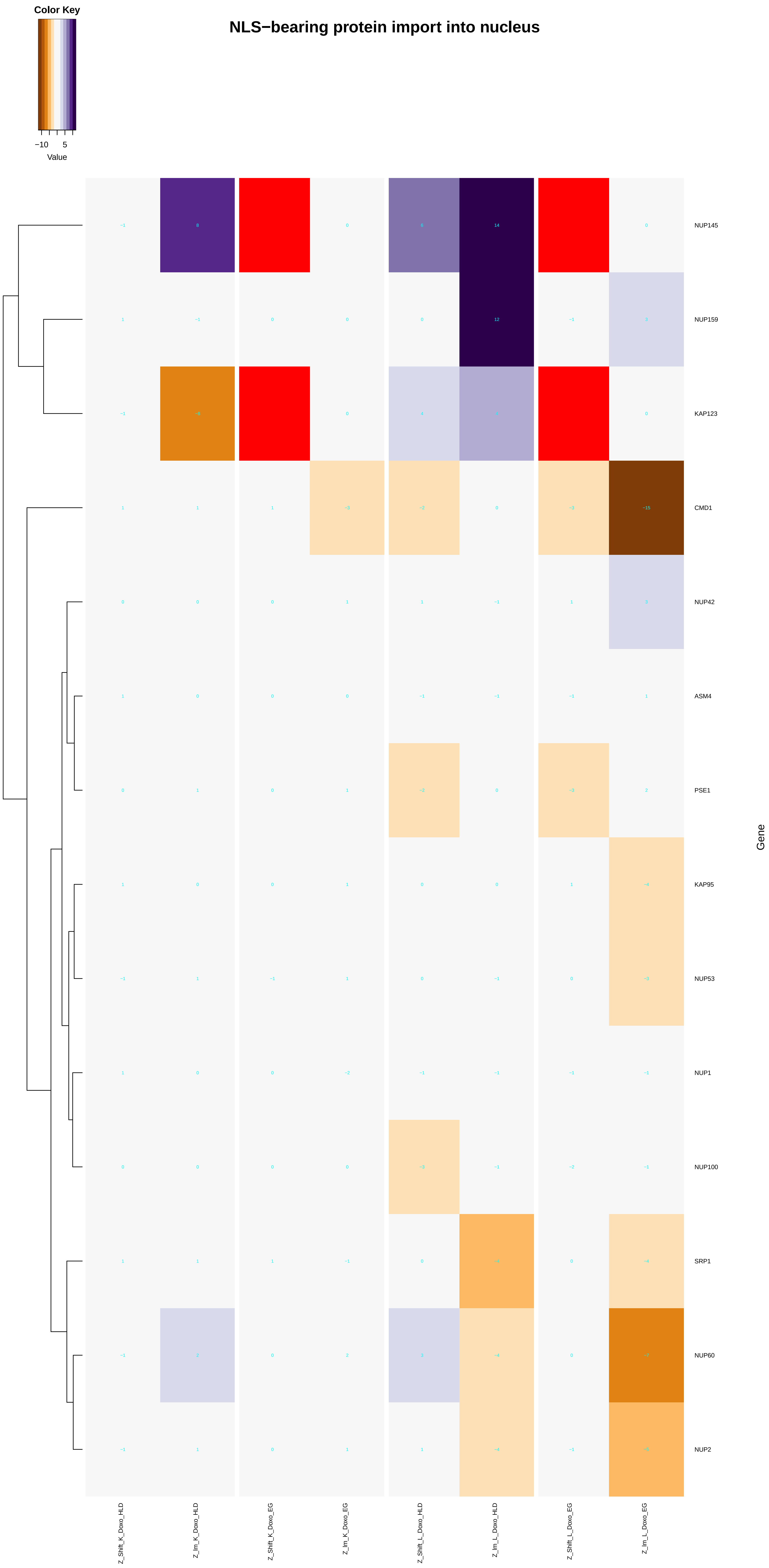
NLS−bearing protein import into nucleus
Color Key
−10
5
Value
NUP145
NUP159
KAP123
CMD1
NUP42
ASM4
PSE1
KAP95
NUP53
NUP1
NUP100
SRP1
NUP60
NUP2
Gene
Z_Shift_K_Doxo_HLD
Z_Im_K_Doxo_HLD
Z_Shift_K_Doxo_EG
Z_Im_K_Doxo_EG
Z_Shift_L_Doxo_HLD
Z_Im_L_Doxo_HLD
Z_Shift_L_Doxo_EG
Z_Im_L_Doxo_EG

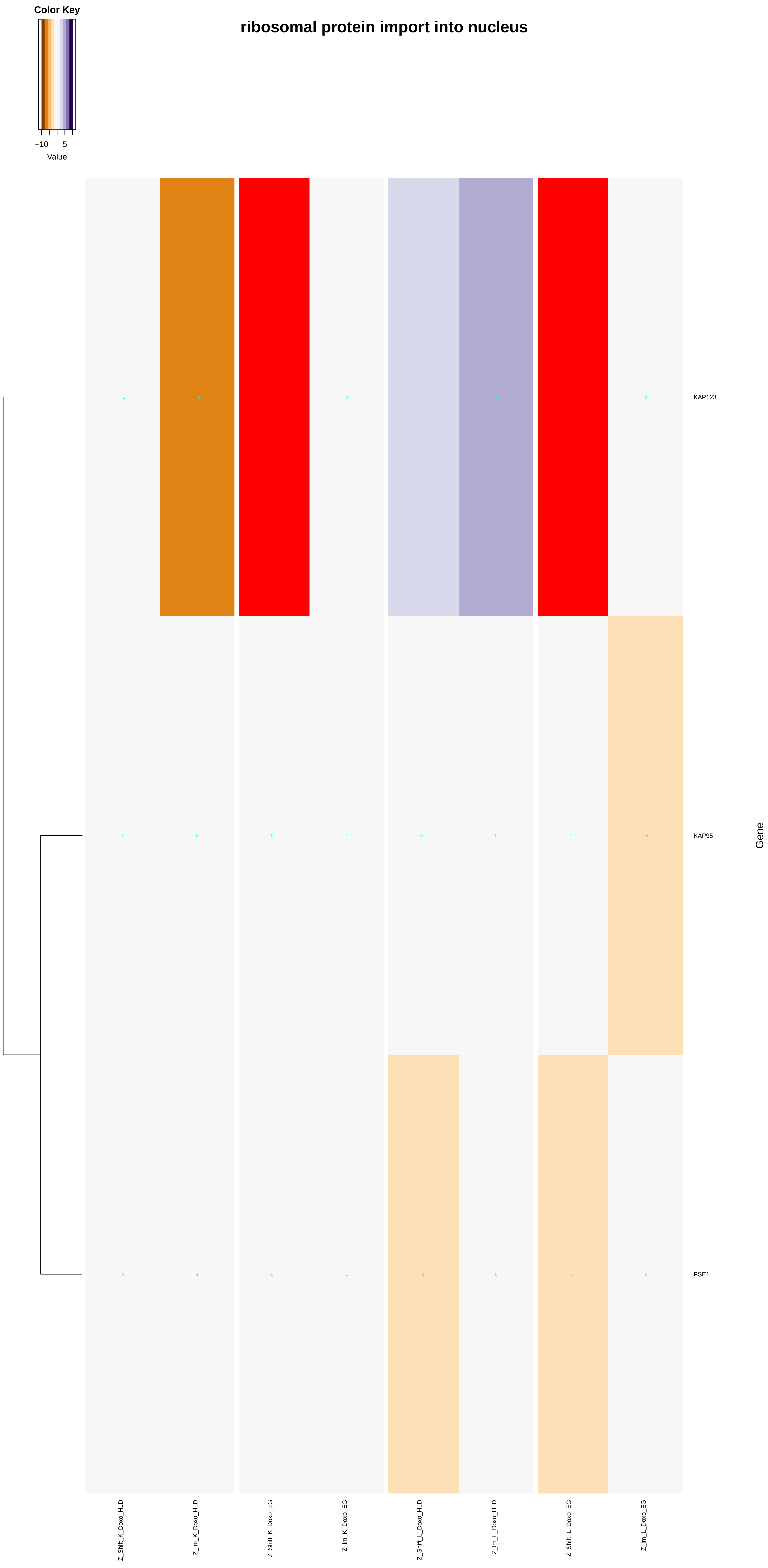
ribosomal protein import into nucleus
Color Key
−10
5
Value
KAP123
KAP95
PSE1
Gene
Z_Shift_K_Doxo_HLD
Z_Im_K_Doxo_HLD
Z_Shift_K_Doxo_EG
Z_Im_K_Doxo_EG
Z_Shift_L_Doxo_HLD
Z_Im_L_Doxo_HLD
Z_Shift_L_Doxo_EG
Z_Im_L_Doxo_EG

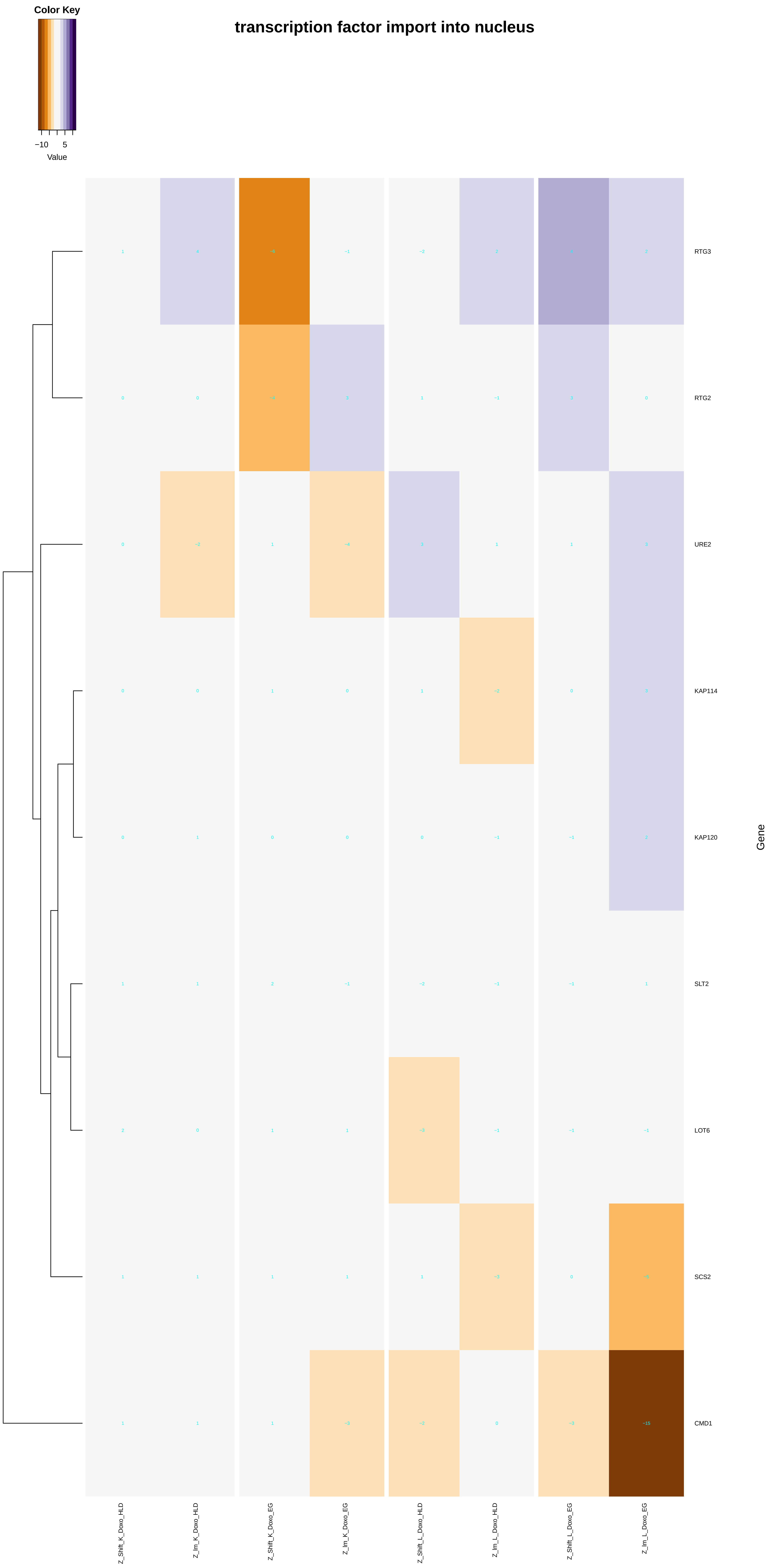
transcription factor import into nucleus
Color Key
−10
5
Value
RTG3
RTG2
URE2
KAP114
KAP120
SLT2
LOT6
SCS2
CMD1
Gene
Z_Shift_K_Doxo_HLD
Z_Im_K_Doxo_HLD
Z_Shift_K_Doxo_EG
Z_Im_K_Doxo_EG
Z_Shift_L_Doxo_HLD
Z_Im_L_Doxo_HLD
Z_Shift_L_Doxo_EG
Z_Im_L_Doxo_EG

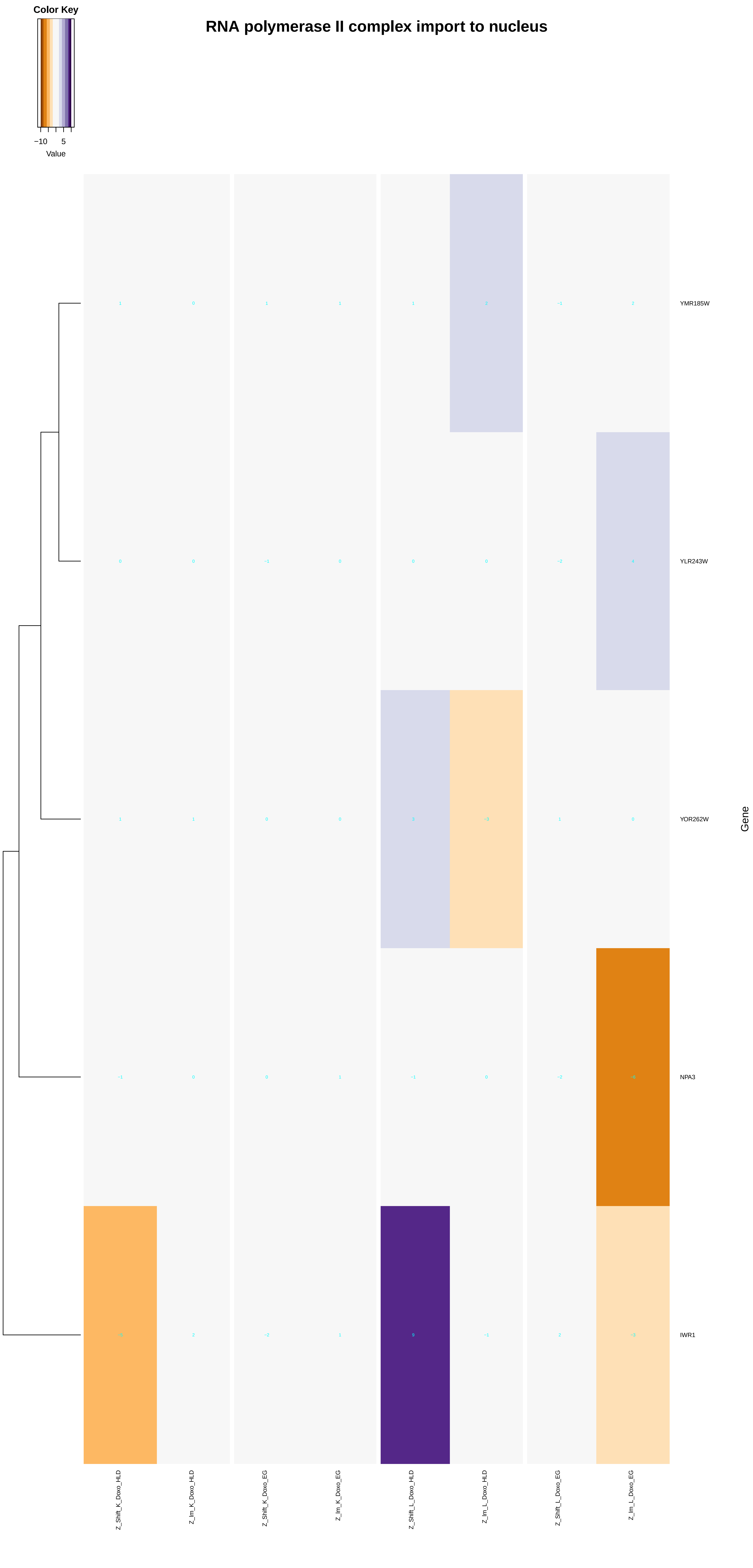
RNA polymerase II complex import to nucleus
Color Key
−10
5
Value
YMR185W
YLR243W
YOR262W
NPA3
IWR1
Gene
Z_Shift_K_Doxo_HLD
Z_lm_K_Doxo_HLD
Z_Shift_K_Doxo_EG
Z_lm_K_Doxo_EG
Z_Shift_L_Doxo_HLD
Z_lm_L_Doxo_HLD
Z_Shift_L_Doxo_EG
Z_lm_L_Doxo_EG

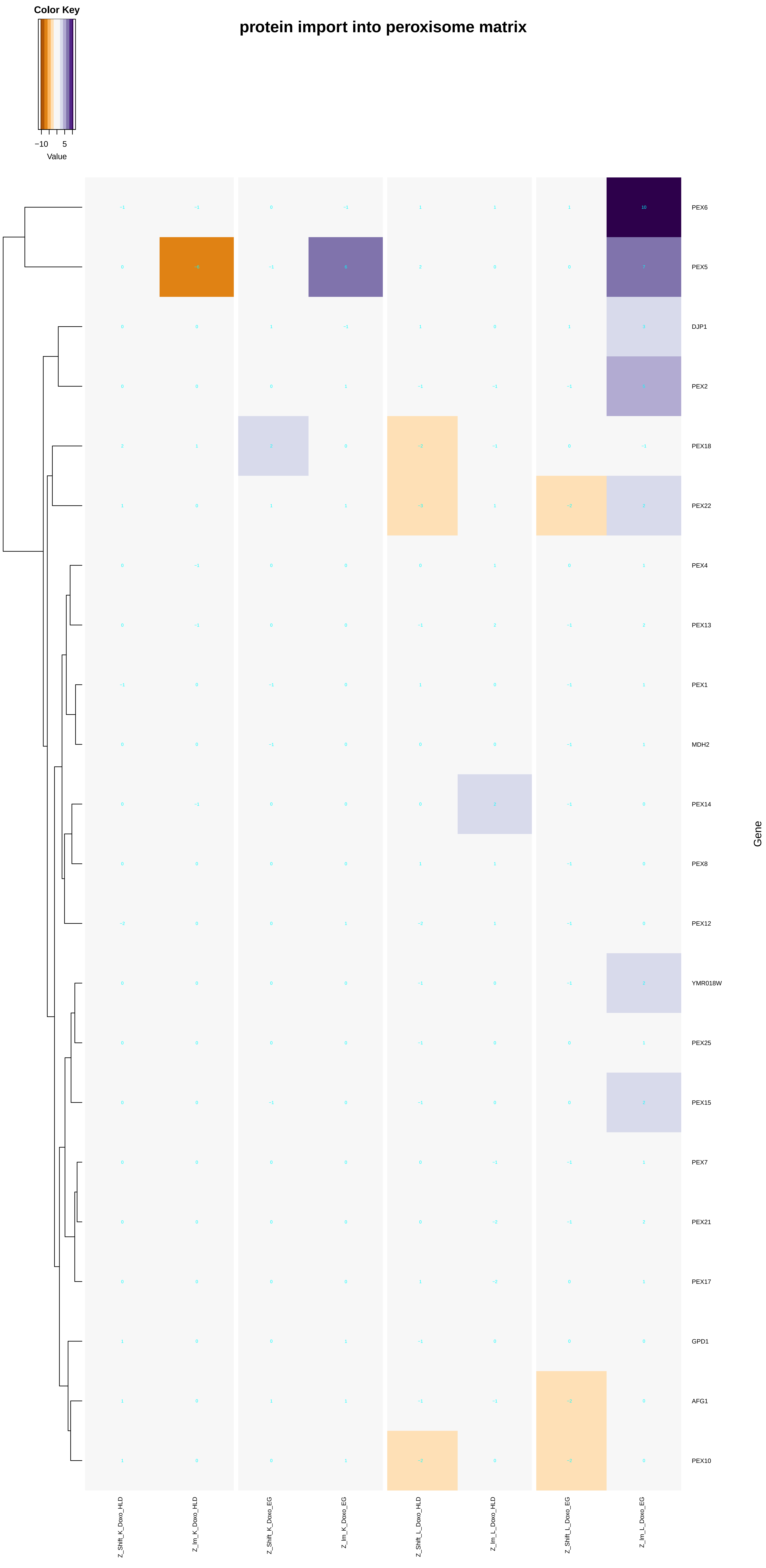

Color Key
−10
5
Value
protein import into peroxisome matrix
PEX6
PEX5
DJP1
PEX2
PEX18
PEX22
PEX4
PEX13
PEX1
MDH2
PEX14
PEX8
PEX12
YMR018W
PEX25
PEX15
PEX7
PEX21
PEX17
GPD1
AFG1
PEX10
Gene
Z_Shift_K_Doxo_HLD
Z_Im_K_Doxo_HLD
Z_Shift_K_Doxo_EG
Z_Im_K_Doxo_EG
Z_Shift_L_Doxo_HLD
Z_Im_L_Doxo_HLD
Z_Shift_L_Doxo_EG
Z_Im_L_Doxo_EG

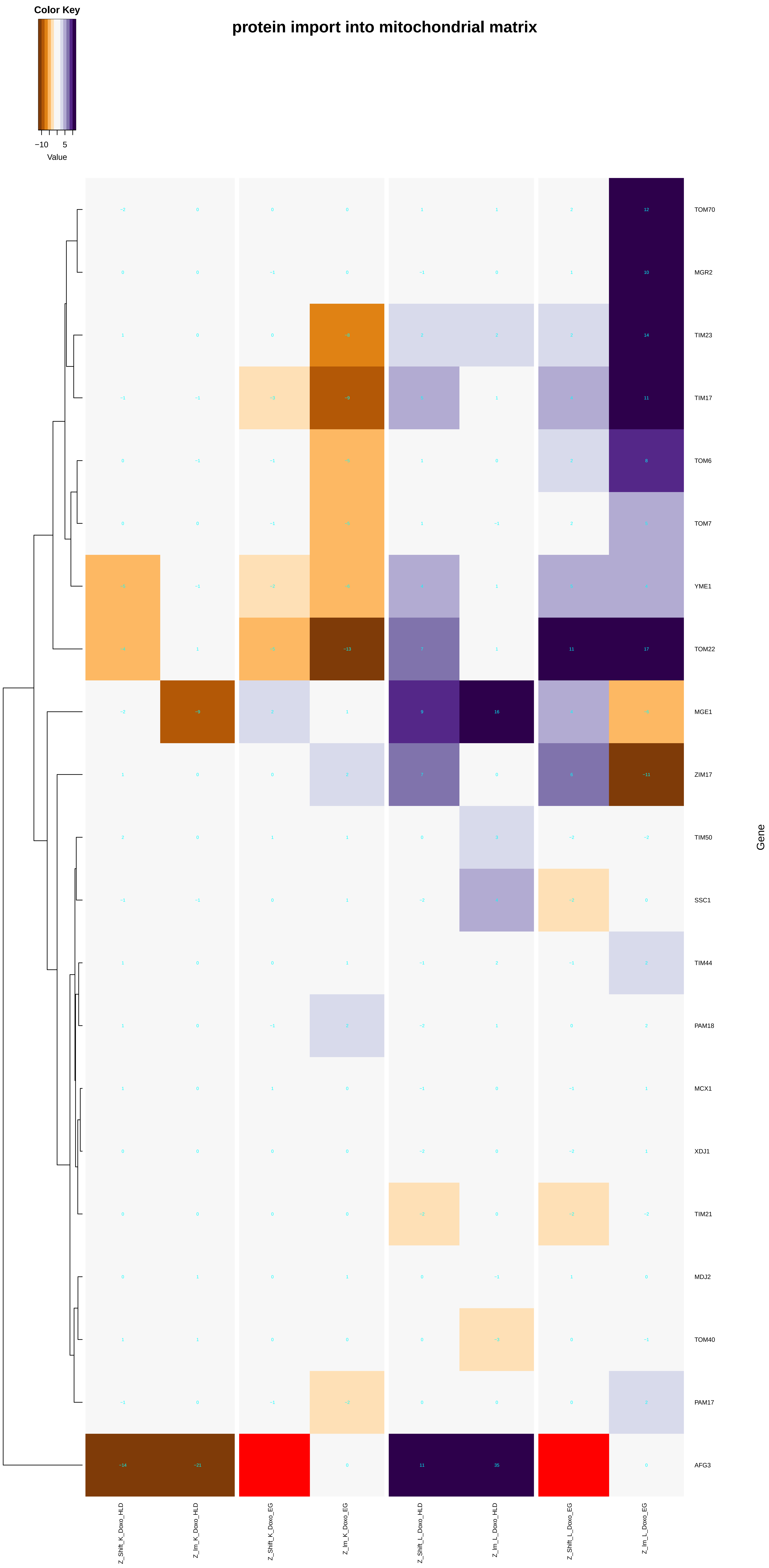
protein import into mitochondrial matrix
Color Key
−10
5
Value
TOM70
MGR2
TIM23
TIM17
TOM6
TOM7
YME1
TOM22
MGE1
ZIM17
TIM50
SSC1
TIM44
PAM18
MCX1
XDJ1
TIM21
MDJ2
TOM40
PAM17
AFG3
Gene
Z_Shift_K_Doxo_HLD
Z_Im_K_Doxo_HLD
Z_Shift_K_Doxo_EG
Z_Im_K_Doxo_EG
Z_Shift_L_Doxo_HLD
Z_Im_L_Doxo_HLD
Z_Shift_L_Doxo_EG
Z_Im_L_Doxo_EG

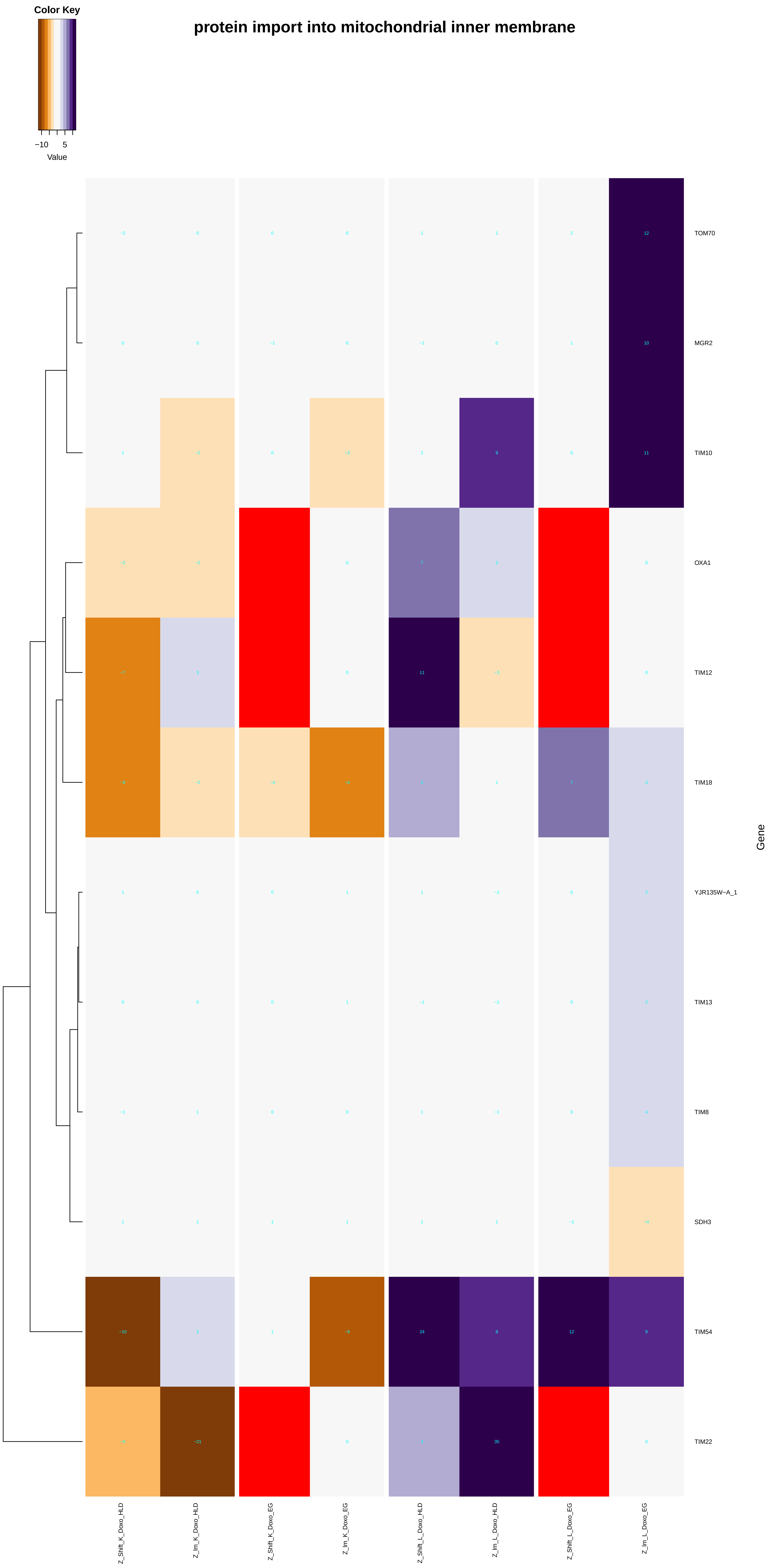
protein import into mitochondrial inner membrane
Color Key
−10
5
Value
Gene
TOM70
MGR2
TIM10
OXA1
TIM12
TIM18
YJR135W−A_1
TIM13
TIM8
SDH3
TIM54
TIM22
Z_Shift_K_Doxo_HLD
Z_Im_K_Doxo_HLD
Z_Shift_K_Doxo_EG
Z_Im_K_Doxo_EG
Z_Shift_L_Doxo_HLD
Z_Im_L_Doxo_HLD
Z_Shift_L_Doxo_EG
Z_Im_L_Doxo_EG

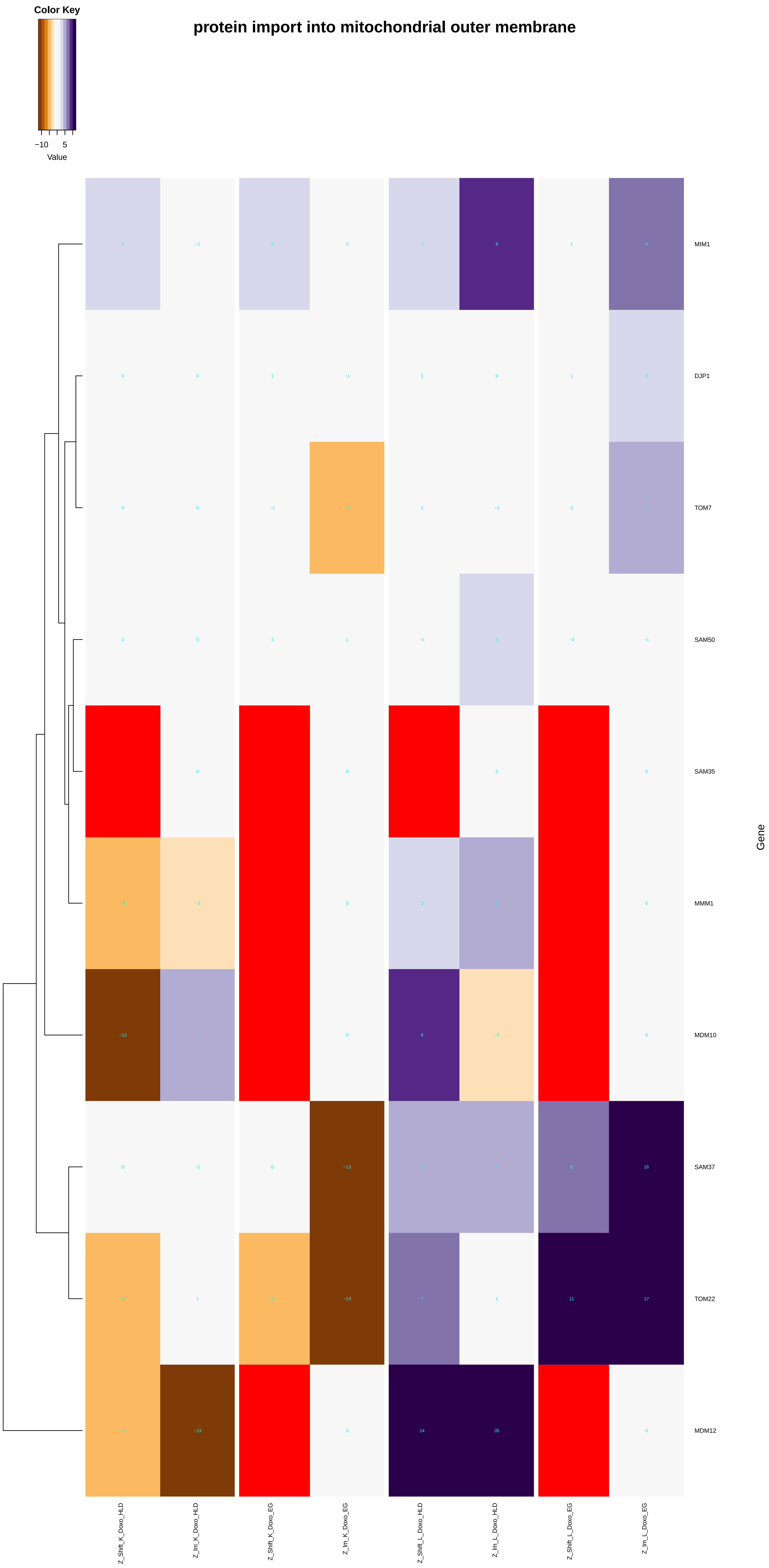

Color Key
−10
5
Value
protein import into mitochondrial outer membrane
MIM1
DJP1
TOM7
SAM50
SAM35
MMM1
MDM10
SAM37
TOM22
MDM12
Gene
Z_Shift_K_Doxo_HLD
Z_Im_K_Doxo_HLD
Z_Shift_K_Doxo_EG
Z_Im_K_Doxo_EG
Z_Shift_L_Doxo_HLD
Z_Im_L_Doxo_HLD
Z_Shift_L_Doxo_EG
Z_Im_L_Doxo_EG

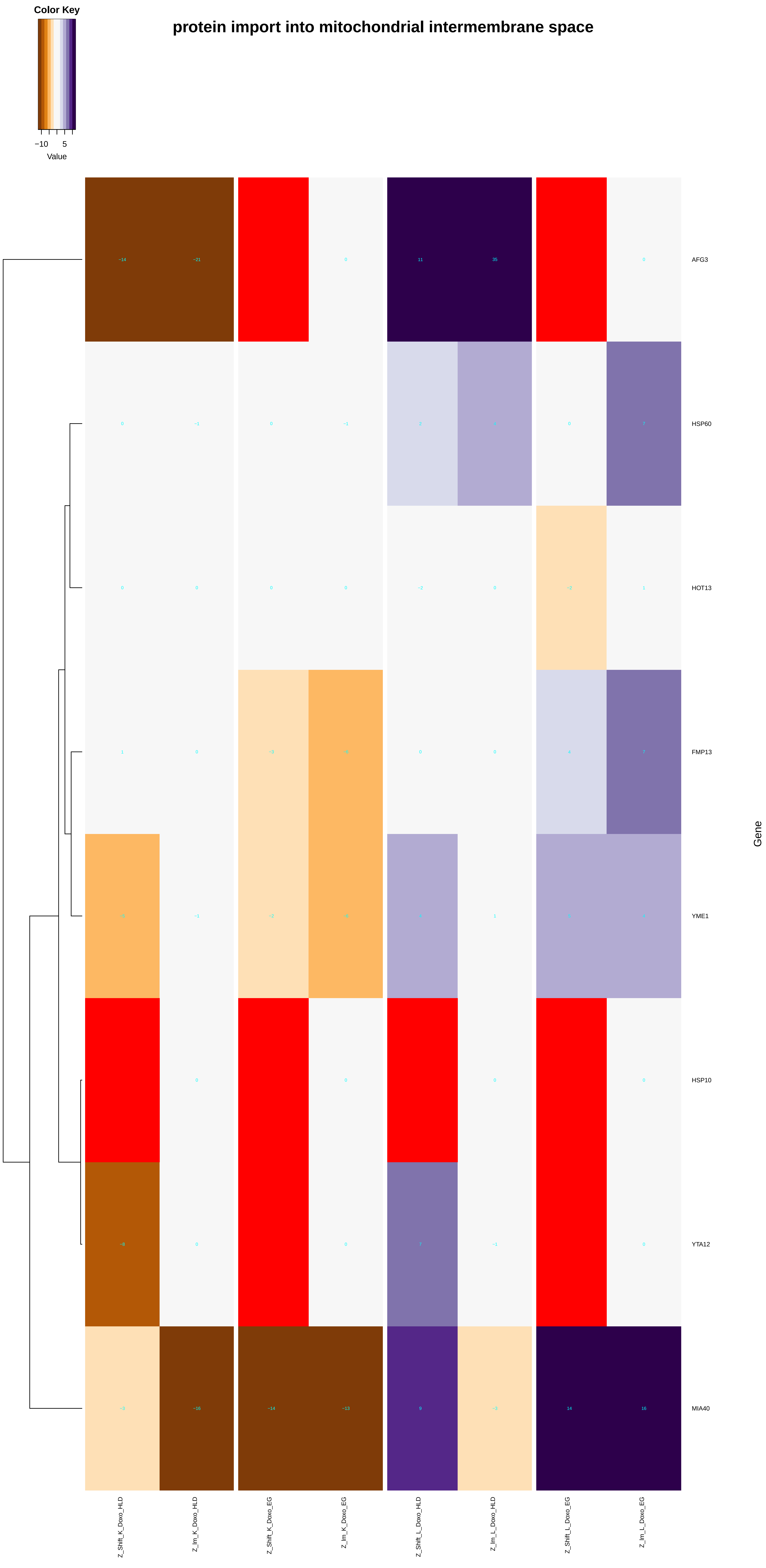

Color Key
−10
5
Value
protein import into mitochondrial intermembrane space
AFG3
HSP60
HOT13
FMP13
YME1
HSP10
YTA12
MIA40
Gene
Z_Shift_K_Doxo_HLD
Z_Im_K_Doxo_HLD
Z_Shift_K_Doxo_EG
Z_Im_K_Doxo_EG
Z_Shift_L_Doxo_HLD
Z_Im_L_Doxo_HLD
Z_Shift_L_Doxo_EG
Z_Im_L_Doxo_EG

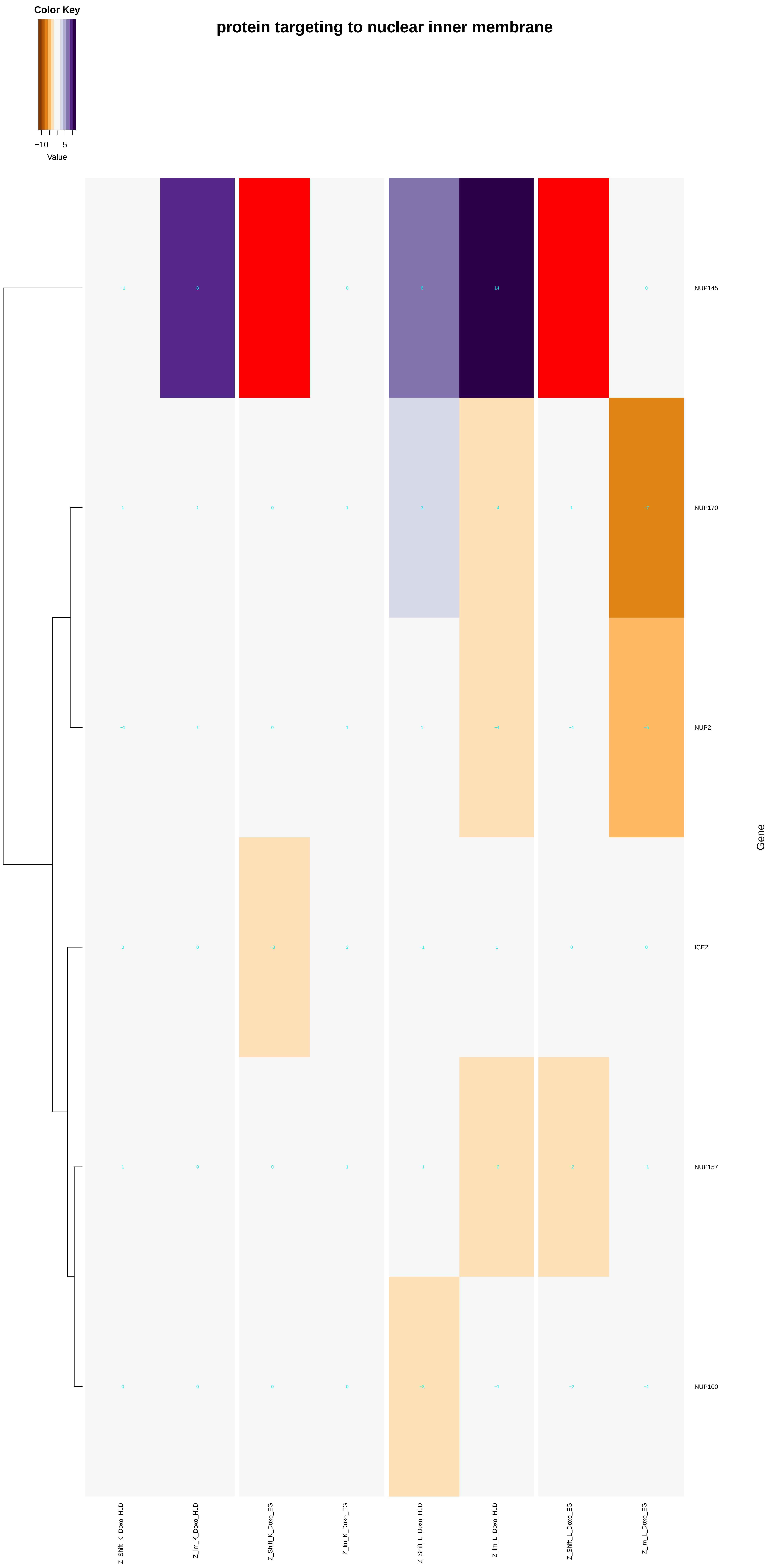
Color Key
−10
5
Value
protein targeting to nuclear inner membrane
NUP145
NUP170
NUP2
ICE2
NUP157
NUP100
Gene
Z_Shift_K_Doxo_HLD
Z_lm_K_Doxo_HLD
Z_Shift_K_Doxo_EG
Z_lm_K_Doxo_EG
Z_Shift_L_Doxo_HLD
Z_lm_L_Doxo_HLD
Z_Shift_L_Doxo_EG
Z_lm_L_Doxo_EG

### establishment of protein localization to chromatin

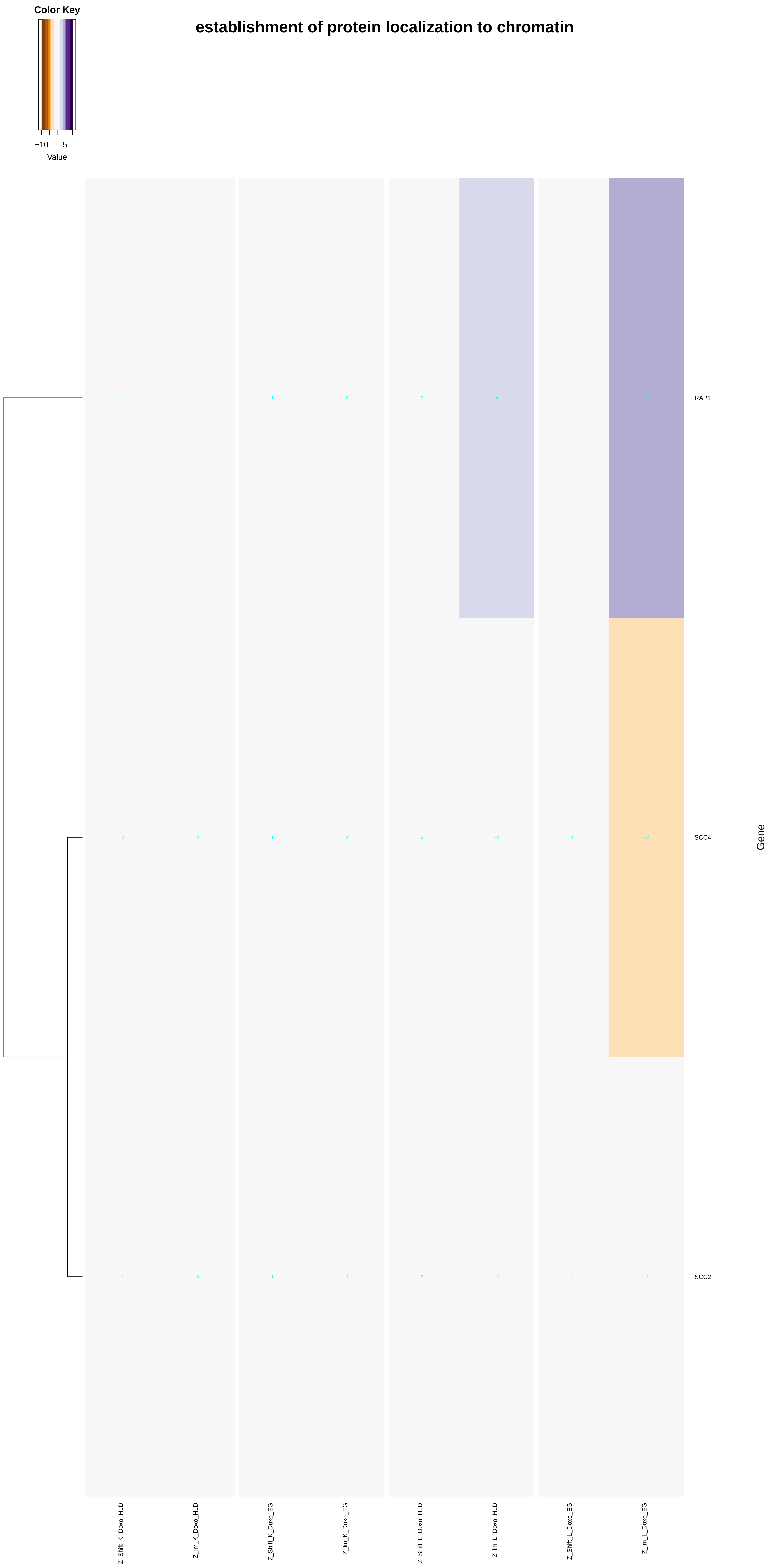

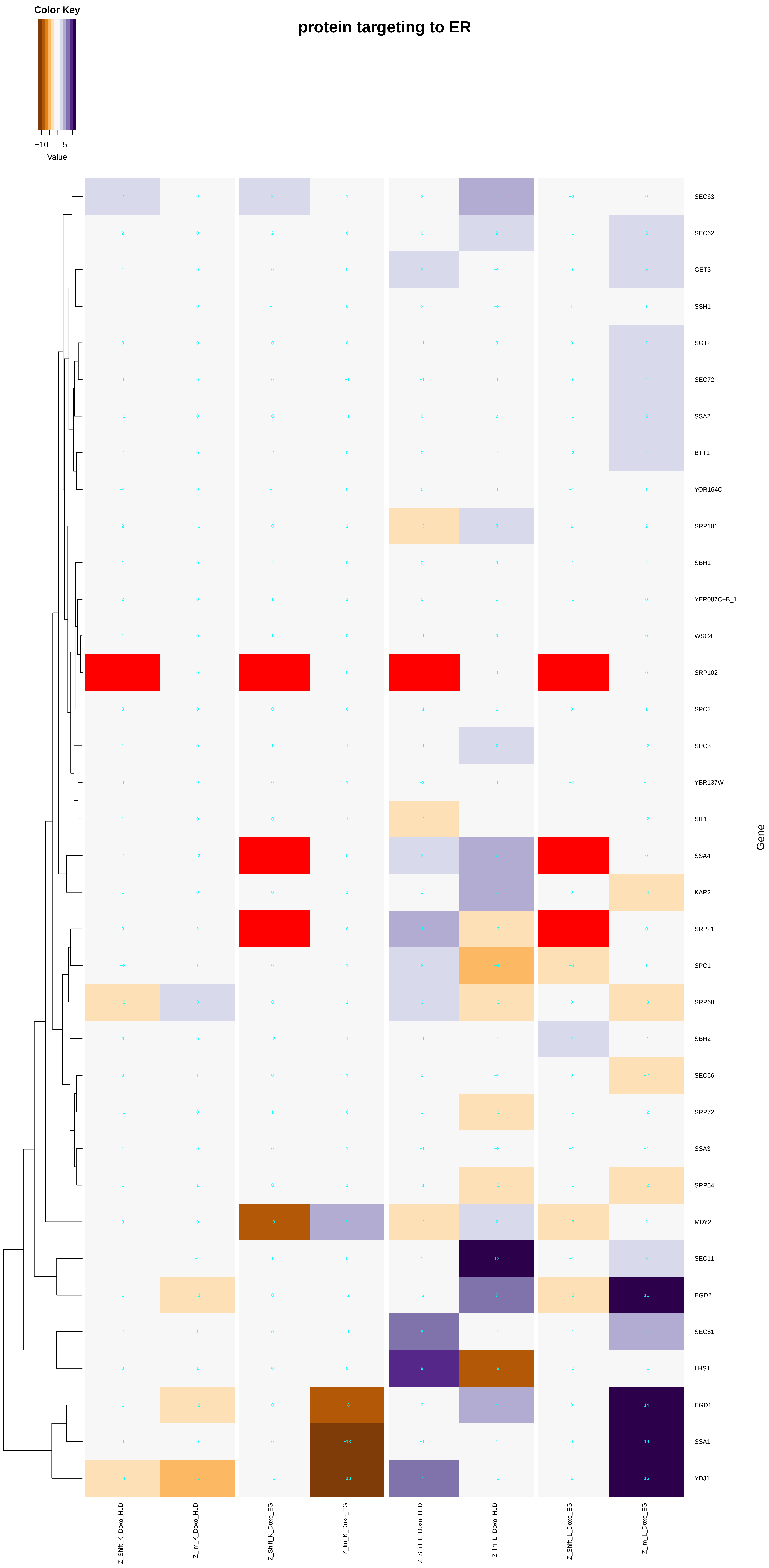

Color Key
−10
5
Value
protein targeting to ER
SEC63
SEC62
GET3
SSH1
SGT2
SEC72
SSA2
BTT1
YOR164C
SRP101
SBH1
YER087C−B_1
WSC4
SRP102
SPC2
SPC3
YBR137W
SIL1
SSA4
KAR2
SRP21
SPC1
SRP68
SBH2
SEC66
SRP72
SSA3
SRP54
MDY2
SEC11
EGD2
SEC61
LHS1
EGD1
SSA1
YDJ1
Gene
Z_Shift_K_Doxo_HLD
Z_lm_K_Doxo_HLD
Z_Shift_K_Doxo_EG
Z_lm_K_Doxo_EG
Z_Shift_L_Doxo_HLD
Z_lm_L_Doxo_HLD
Z_Shift_L_Doxo_EG
Z_lm_L_Doxo_EG

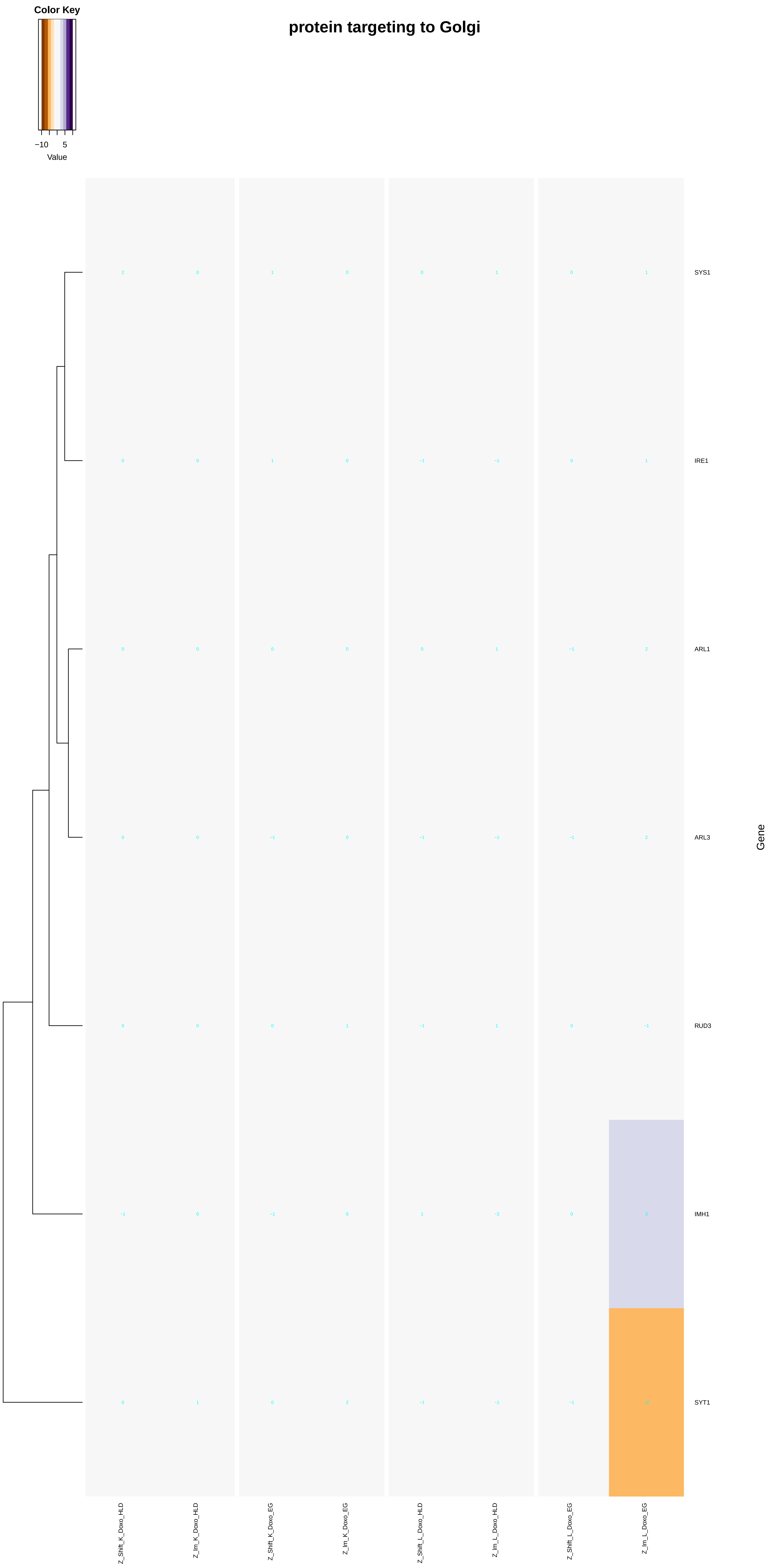
protein targeting to Golgi
Color Key
−10
5
Value
SYS1
IRE1
ARL1
ARL3
RUD3
IMH1
SYT1
Gene
Z_Shift_K_Doxo_HLD
Z_lm_K_Doxo_HLD
Z_Shift_K_Doxo_EG
Z_lm_K_Doxo_EG
Z_Shift_L_Doxo_HLD
Z_lm_L_Doxo_HLD
Z_Shift_L_Doxo_EG
Z_lm_L_Doxo_EG

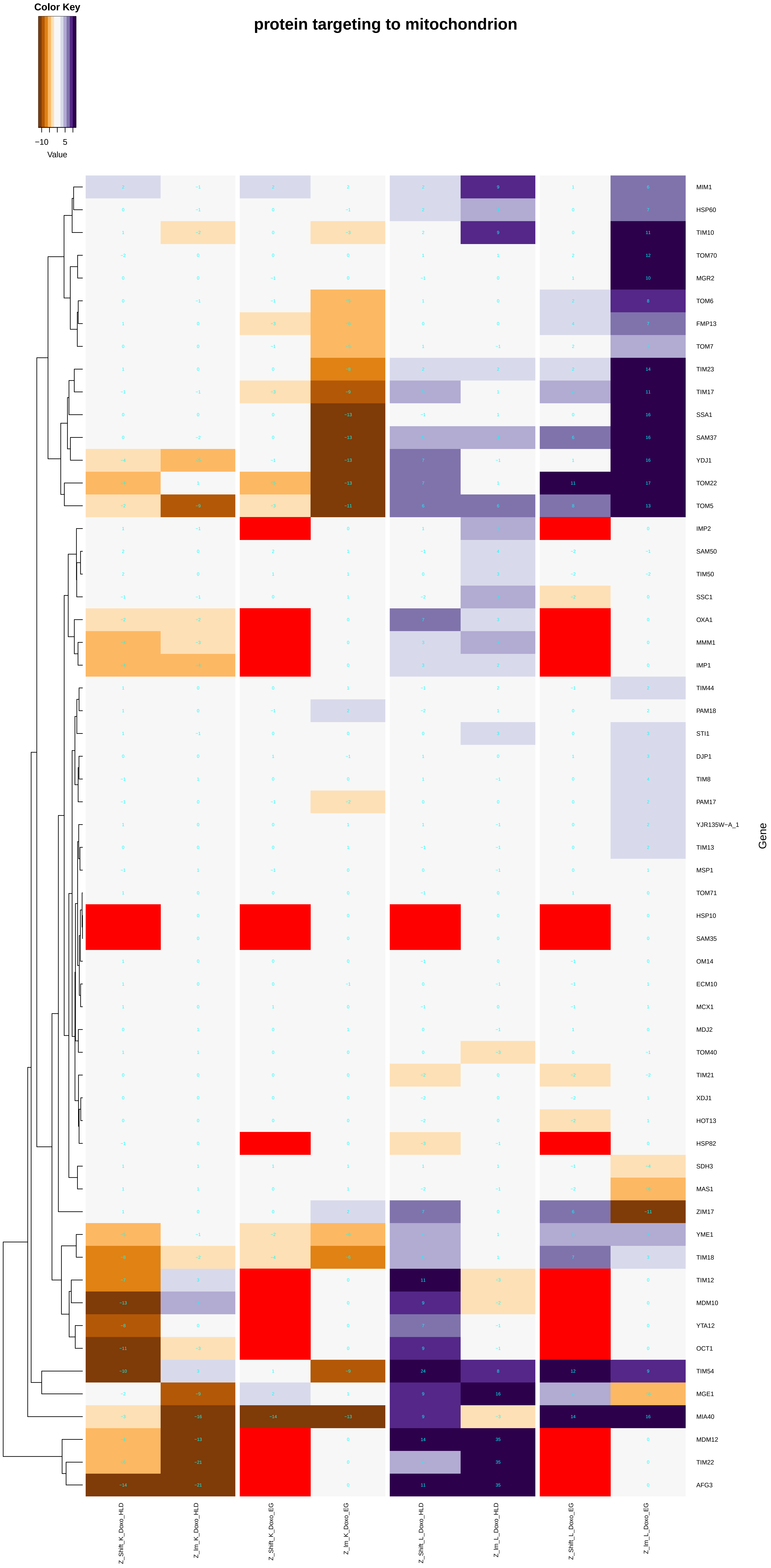
protein targeting to mitochondrion
Color Key
−10
5
Value
Gene
MIM1
HSP60
TIM10
TOM70
MGR2
TOM6
FMP13
TOM7
TIM23
TIM17
SSA1
SAM37
YDJ1
TOM22
TOM5
IMP2
SAM50
TIM50
SSC1
OXA1
MMM1
IMP1
TIM44
PAM18
STI1
DJP1
TIM8
PAM17
YJR135W−A_1
TIM13
MSP1
TOM71
HSP10
SAM35
OM14
ECM10
MCX1
MDJ2
TOM40
TIM21
XDJ1
HOT13
HSP82
SDH3
MAS1
ZIM17
YME1
TIM18
TIM12
MDM10
YTA12
OCT1
TIM54
MGE1
MIA40
MDM12
TIM22
AFG3
Z_Shift_K_Doxo_HLD
Z_lm_K_Doxo_HLD
Z_Shift_K_Doxo_EG
Z_lm_K_Doxo_EG
Z_Shift_L_Doxo_HLD
Z_lm_L_Doxo_HLD
Z_Shift_L_Doxo_EG
Z_lm_L_Doxo_EG

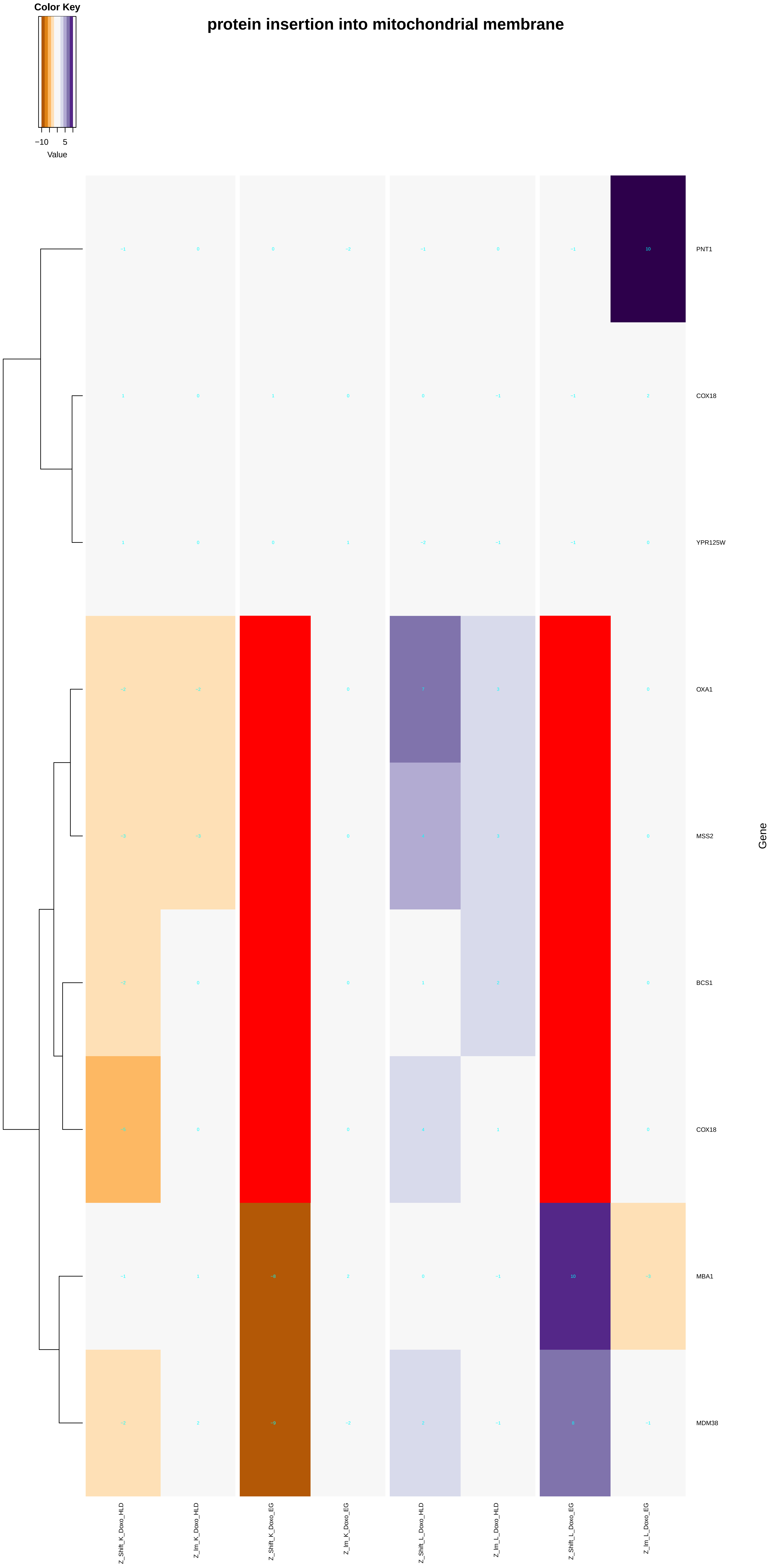
protein insertion into mitochondrial membrane
Color Key
−10
5
Value
PNT1
COX18
YPR125W
OXA1
MSS2
BCS1
COX18
MBA1
MDM38
Gene
Z_Shift_K_Doxo_HLD
Z_lm_K_Doxo_HLD
Z_Shift_K_Doxo_EG
Z_lm_K_Doxo_EG
Z_Shift_L_Doxo_HLD
Z_lm_L_Doxo_HLD
Z_Shift_L_Doxo_EG
Z_lm_L_Doxo_EG

### establishment of protein localization to mitochondrial membrane

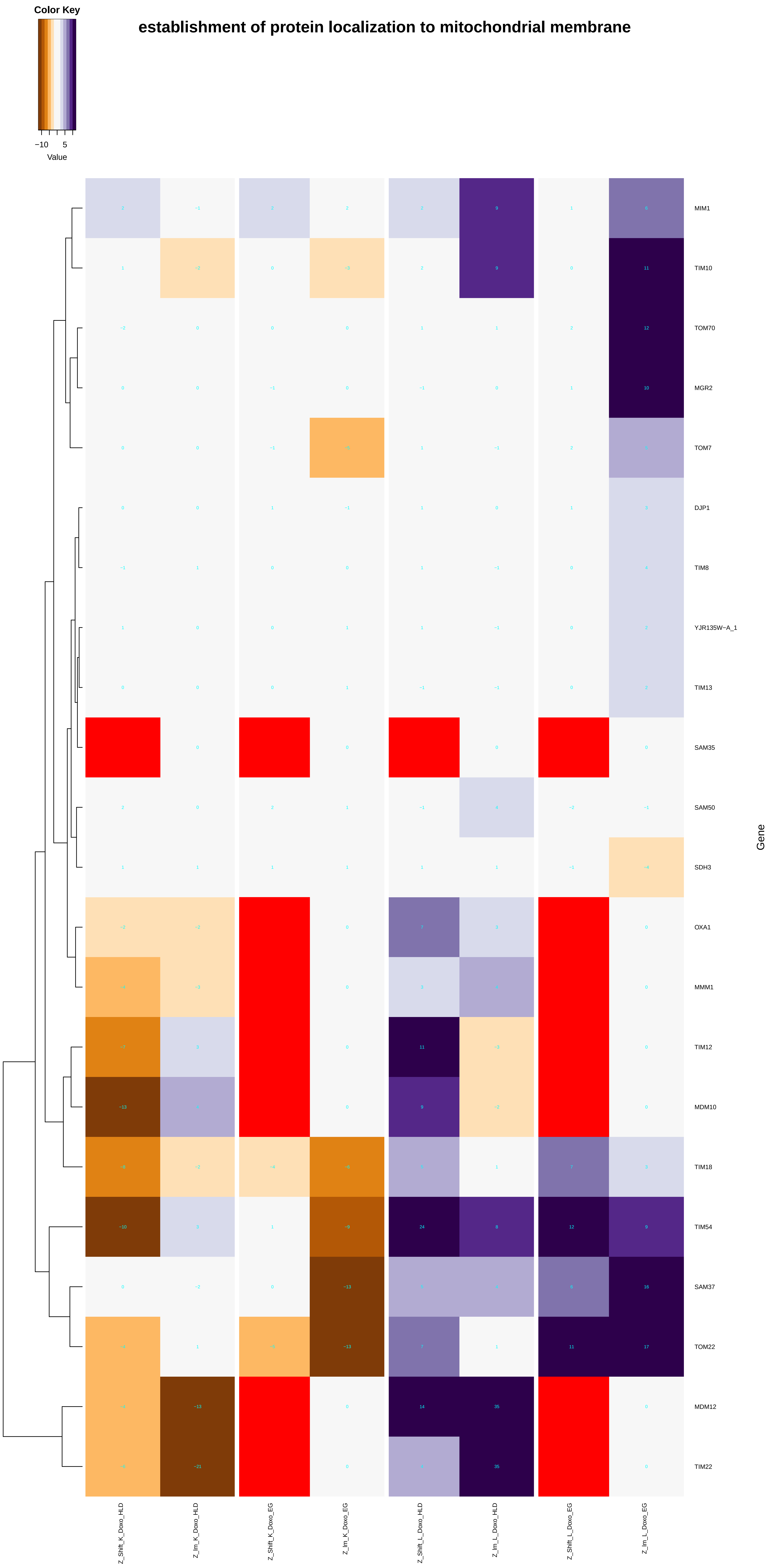

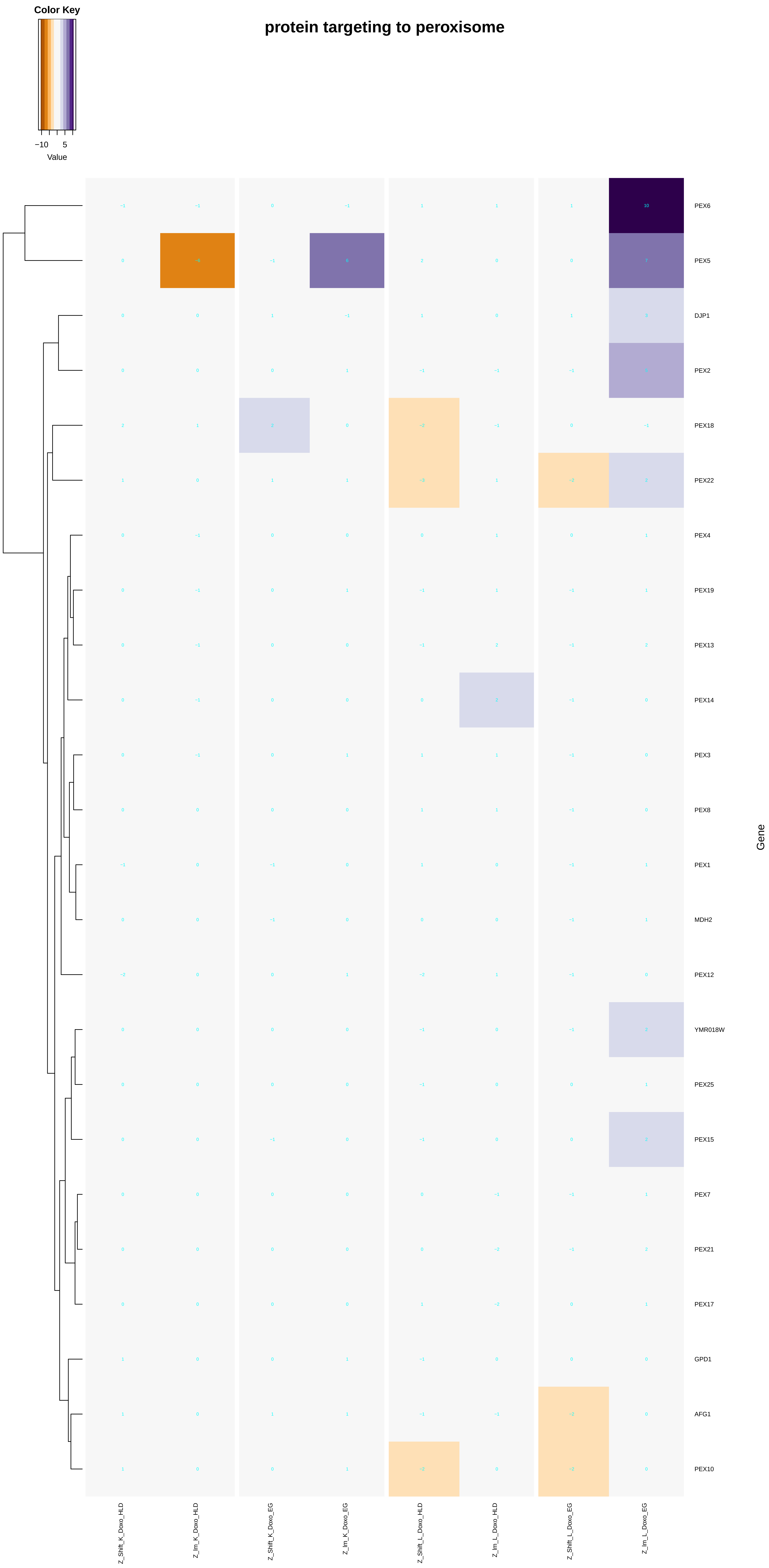
protein targeting to peroxisome
Color Key
−10
5
Value
PEX6
PEX5
DJP1
PEX2
PEX18
PEX22
PEX4
PEX19
PEX13
PEX14
PEX3
PEX8
PEX1
MDH2
PEX12
YMR018W
PEX25
PEX15
PEX7
PEX21
PEX17
GPD1
AFG1
PEX10
Gene
Z_Shift_K_Doxo_HLD
Z_Im_K_Doxo_HLD
Z_Shift_K_Doxo_EG
Z_Im_K_Doxo_EG
Z_Shift_L_Doxo_HLD
Z_Im_L_Doxo_HLD
Z_Shift_L_Doxo_EG
Z_Im_L_Doxo_EG

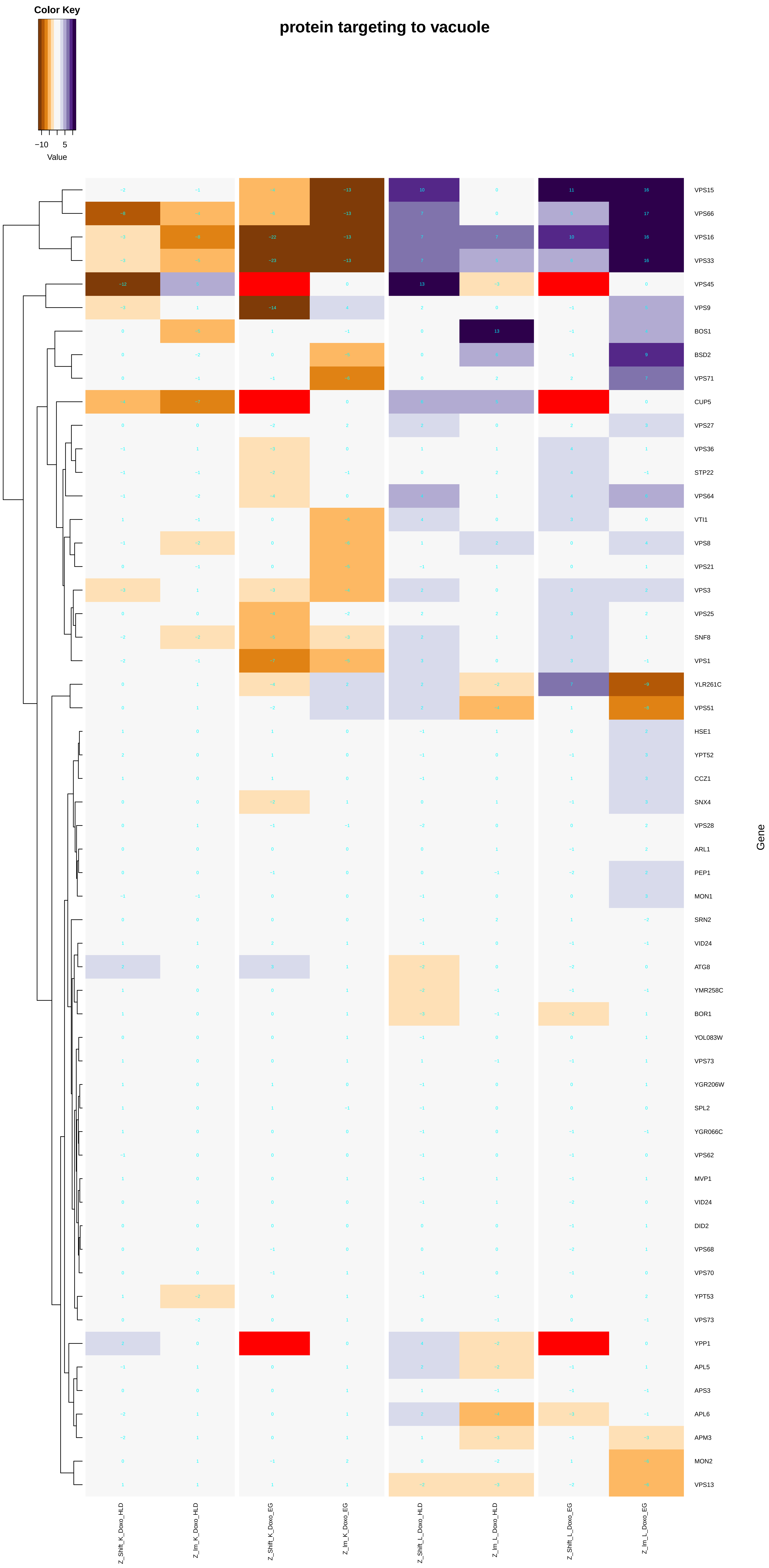

Color Key
−10
5
Value
protein targeting to vacuole
VPS15
VPS66
VPS16
VPS33
VPS45
VPS9
BOS1
BSD2
VPS71
CUP5
VPS27
VPS36
STP22
VPS64
VTI1
VPS8
VPS21
VPS3
VPS25
SNF8
VPS1
YLR261C
VPS51
HSE1
YPT52
CCZ1
SNX4
VPS28
ARL1
PEP1
MON1
SRN2
VID24
ATG8
YMR258C
BOR1
YOL083W
VPS73
YGR206W
SPL2
YGR066C
VPS62
MVP1
VID24
DID2
VPS68
VPS70
YPT53
VPS73
YPP1
APL5
APS3
APL6
APM3
MON2
VPS13
Gene
Z_Shift_K_Doxo_HLD
Z_lm_K_Doxo_HLD
Z_Shift_K_Doxo_EG
Z_lm_K_Doxo_EG
Z_Shift_L_Doxo_HLD
Z_lm_L_Doxo_HLD
Z_Shift_L_Doxo_EG
Z_lm_L_Doxo_EG

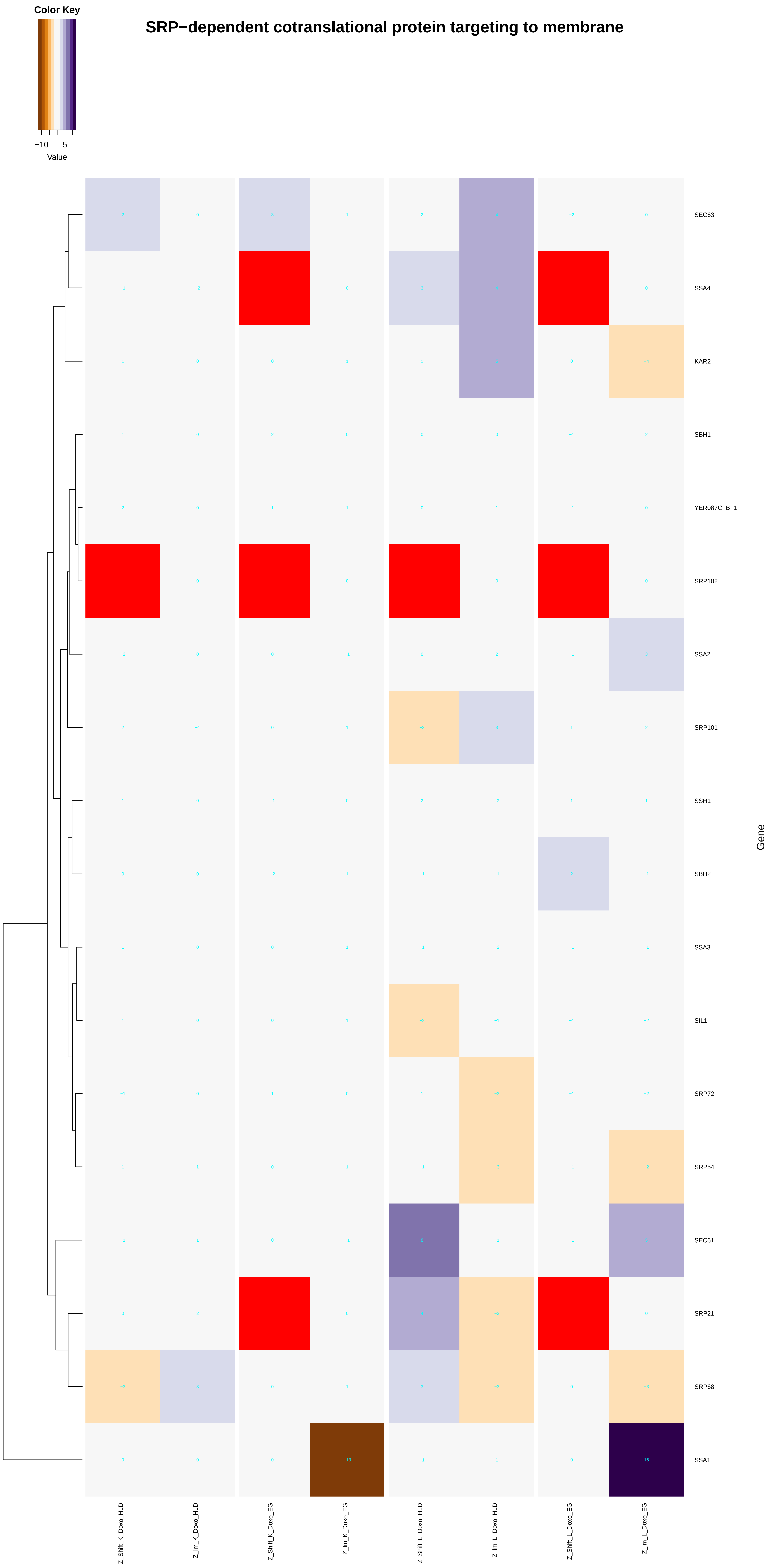
Color Key
−10
5
Value
SRP−dependent cotranslational protein targeting to membrane
SEC63
SSA4
KAR2
SBH1
YER087C−B_1
SRP102
SSA2
SRP101
SSH1
SBH2
SSA3
SIL1
SRP72
SRP54
SEC61
SRP21
SRP68
SSA1
Gene
Z_Shift_K_Doxo_HLD
Z_lm_K_Doxo_HLD
Z_Shift_K_Doxo_EG
Z_lm_K_Doxo_EG
Z_Shift_L_Doxo_HLD
Z_lm_L_Doxo_HLD
Z_Shift_L_Doxo_EG
Z_lm_L_Doxo_EG
−13
16

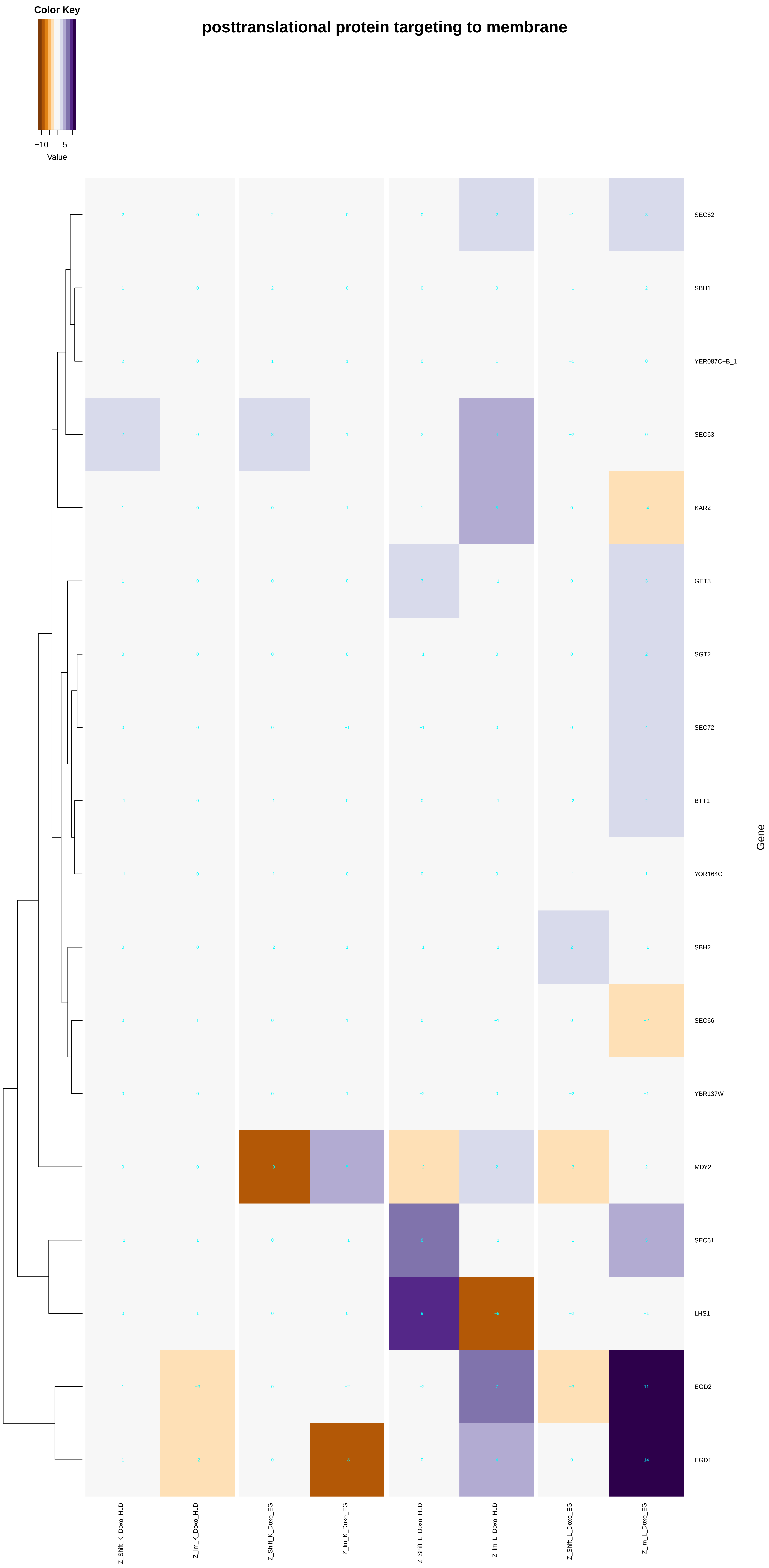
Color Key
−10 5
Value
posttranslational protein targeting to membrane
SEC62
SBH1
YER087C−B_1
SEC63
KAR2
GET3
SGT2
SEC72
BTT1
YOR164C
SBH2
SEC66
YBR137W
MDY2
SEC61
LHS1
EGD2
EGD1
Gene
Z_Shift_K_Doxo_HLD
Z_Im_K_Doxo_HLD
Z_Shift_K_Doxo_EG
Z_Im_K_Doxo_EG
Z_Shift_L_Doxo_HLD
Z_Im_L_Doxo_HLD
Z_Shift_L_Doxo_EG
Z_Im_L_Doxo_EG

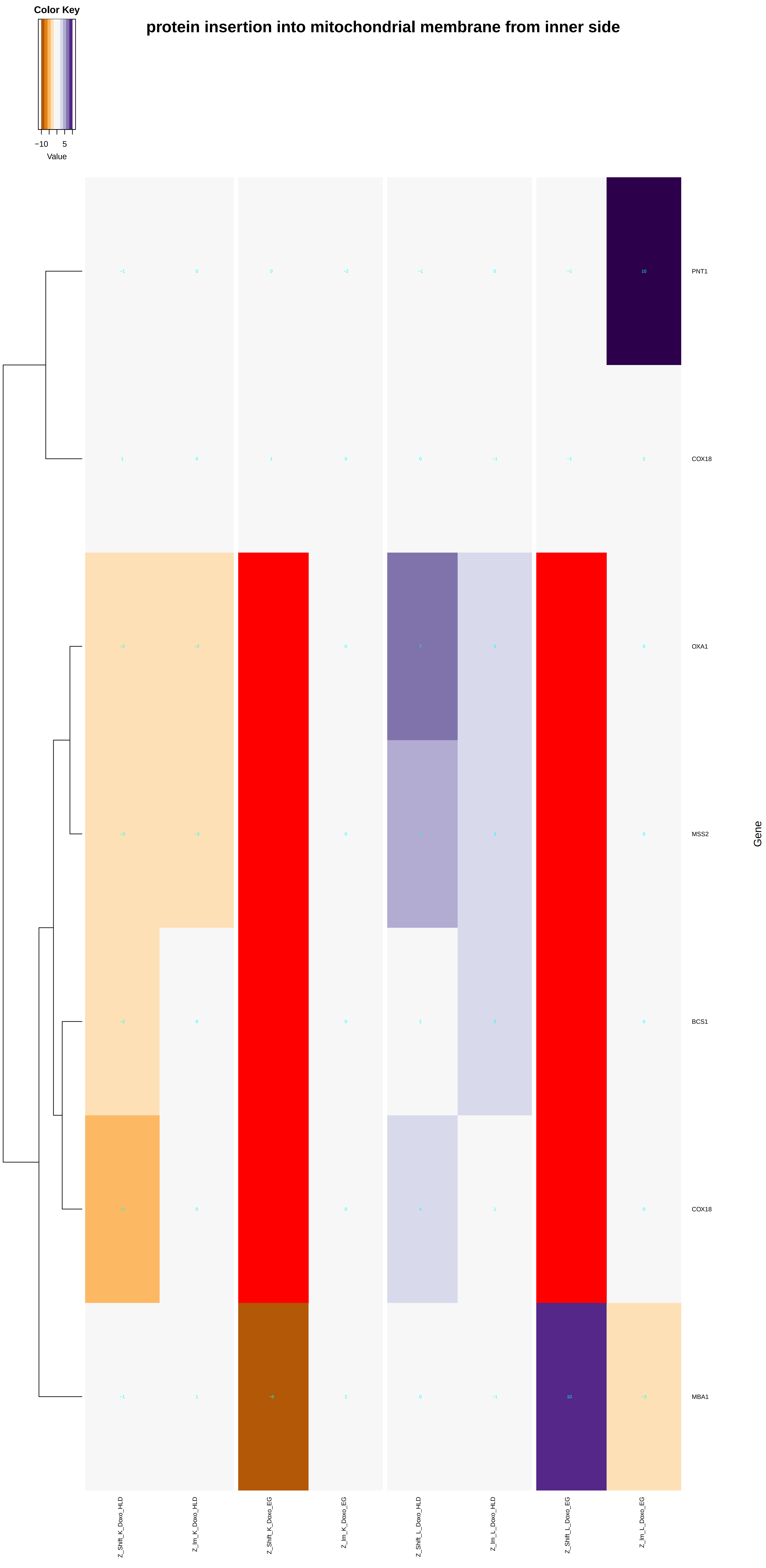

protein insertion into mitochondrial membrane from inner side
Color Key
−10
5
Value
PNT1
COX18
OXA1
MSS2
BCS1
COX18
MBA1
Gene
Z_Shift_K_Doxo_HLD
Z_lm_K_Doxo_HLD
Z_Shift_K_Doxo_EG
Z_lm_K_Doxo_EG
Z_Shift_L_Doxo_HLD
Z_lm_L_Doxo_HLD
Z_Shift_L_Doxo_EG
Z_lm_L_Doxo_EG

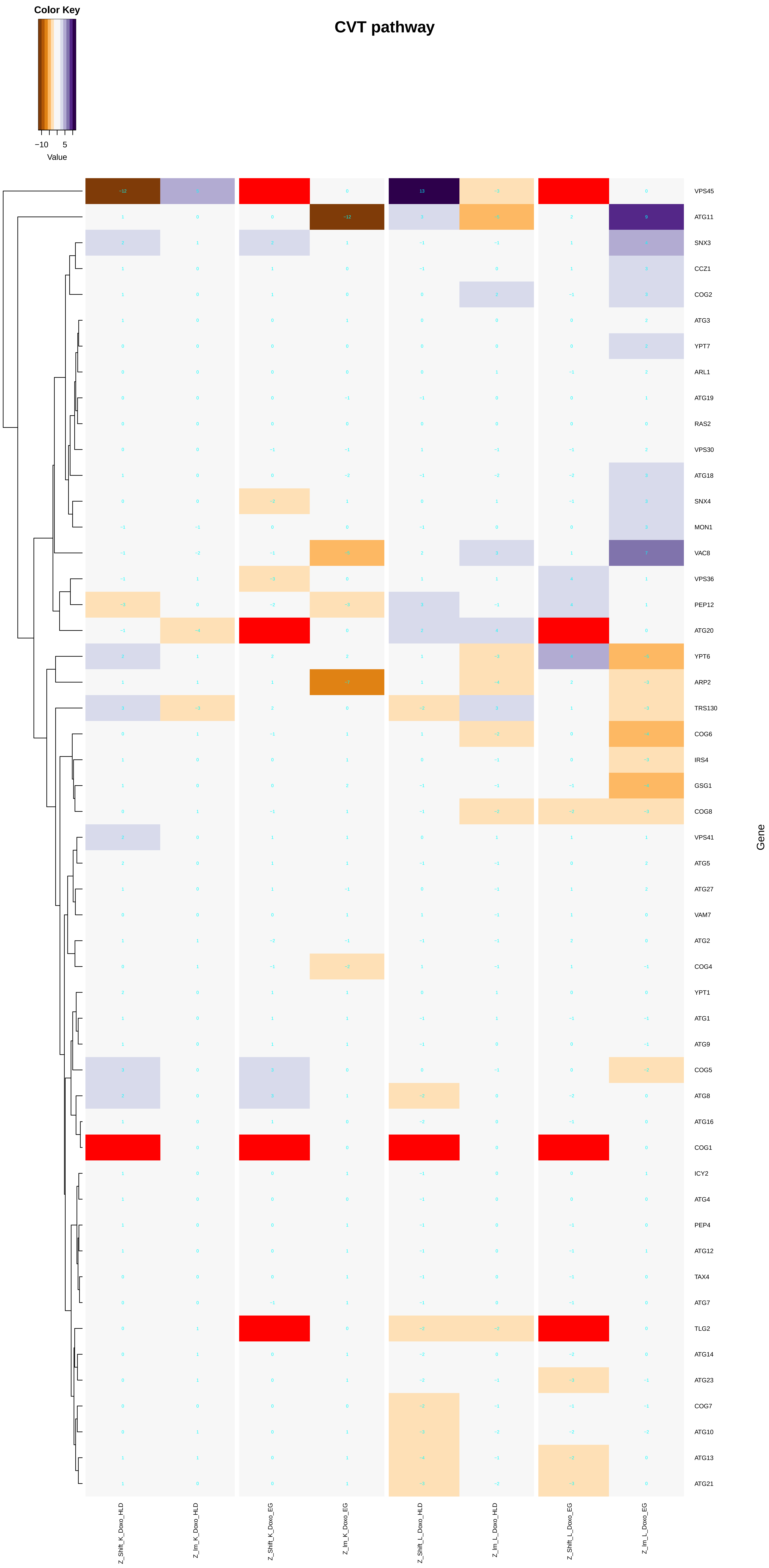
CVT pathway
Color Key
−10
5
Value
VPS45
ATG11
SNX3
CCZ1
COG2
ATG3
YPT7
ARL1
ATG19
RAS2
VPS30
ATG18
SNX4
MON1
VAC8
VPS36
PEP12
ATG20
YPT6
ARP2
TRS130
COG6
IRS4
GSG1
COG8
VPS41
ATG5
ATG27
VAM7
ATG2
COG4
YPT1
ATG1
ATG9
COG5
ATG8
ATG16
COG1
ICY2
ATG4
PEP4
ATG12
TAX4
ATG7
TLG2
ATG14
ATG23
COG7
ATG10
ATG13
ATG21
Gene
Z_Shift_K_Doxo_HLD
Z_Im_K_Doxo_HLD
Z_Shift_K_Doxo_EG
Z_Im_K_Doxo_EG
Z_Shift_L_Doxo_HLD
Z_Im_L_Doxo_HLD
Z_Shift_L_Doxo_EG
Z_Im_L_Doxo_EG

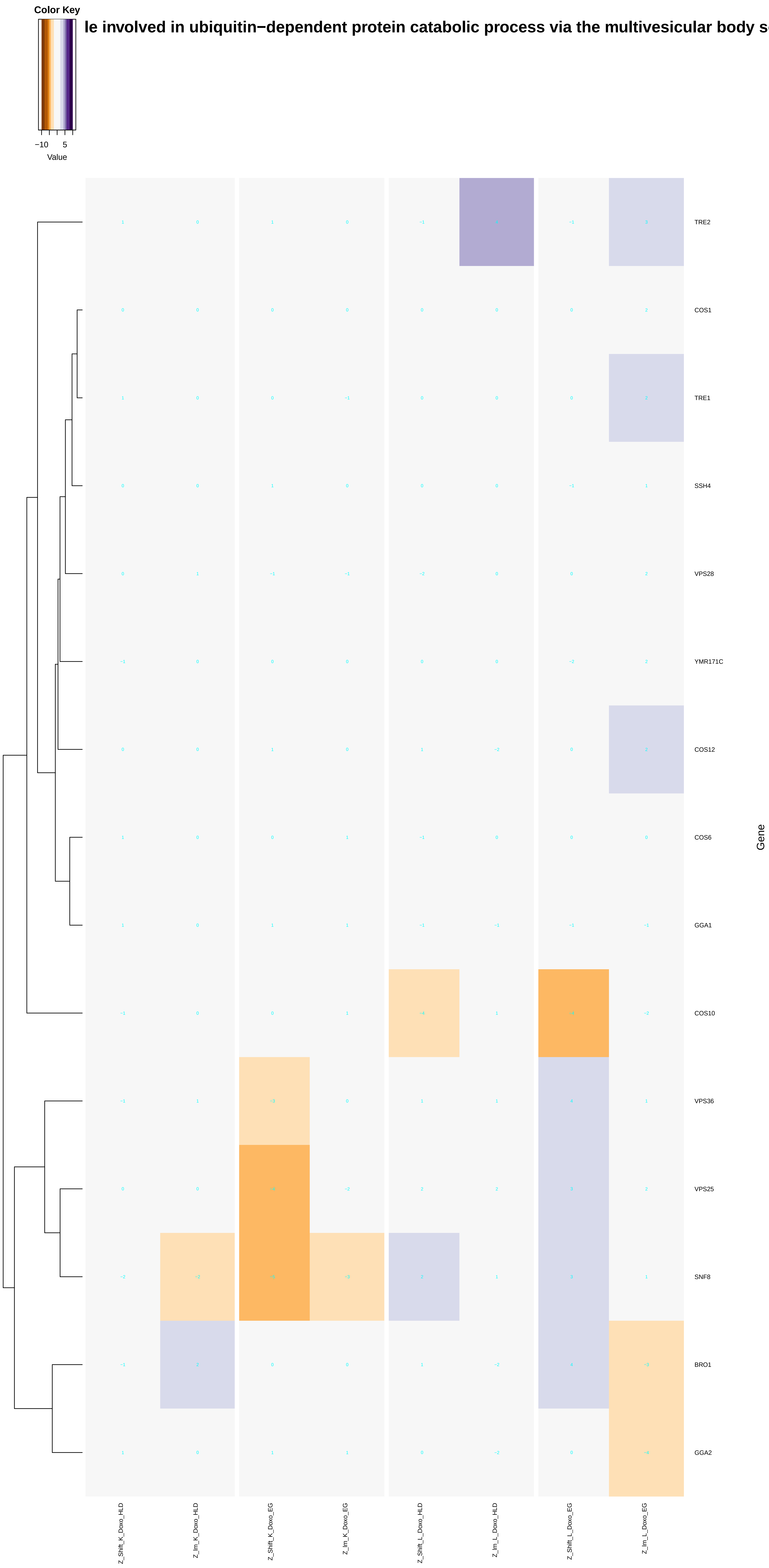
Color Key
−10
5
Value
le involved in ubiquitin−dependent protein catabolic process via the multivesicular body s
TRE2
COS1
TRE1
SSH4
VPS28
YMR171C
COS12
COS6
GGA1
COS10
VPS36
VPS25
SNF8
BRO1
GGA2
Gene
Z_Shift_K_Doxo_HLD
Z_Im_K_Doxo_HLD
Z_Shift_K_Doxo_EG
Z_Im_K_Doxo_EG
Z_Shift_L_Doxo_HLD
Z_Im_L_Doxo_HLD
Z_Shift_L_Doxo_EG
Z_Im_L_Doxo_EG
