## Additional File 9 for "A yeast phenomic model for the influence of Warburg metabolism on genetic buffering of doxorubicin": gene_silencing.pdf

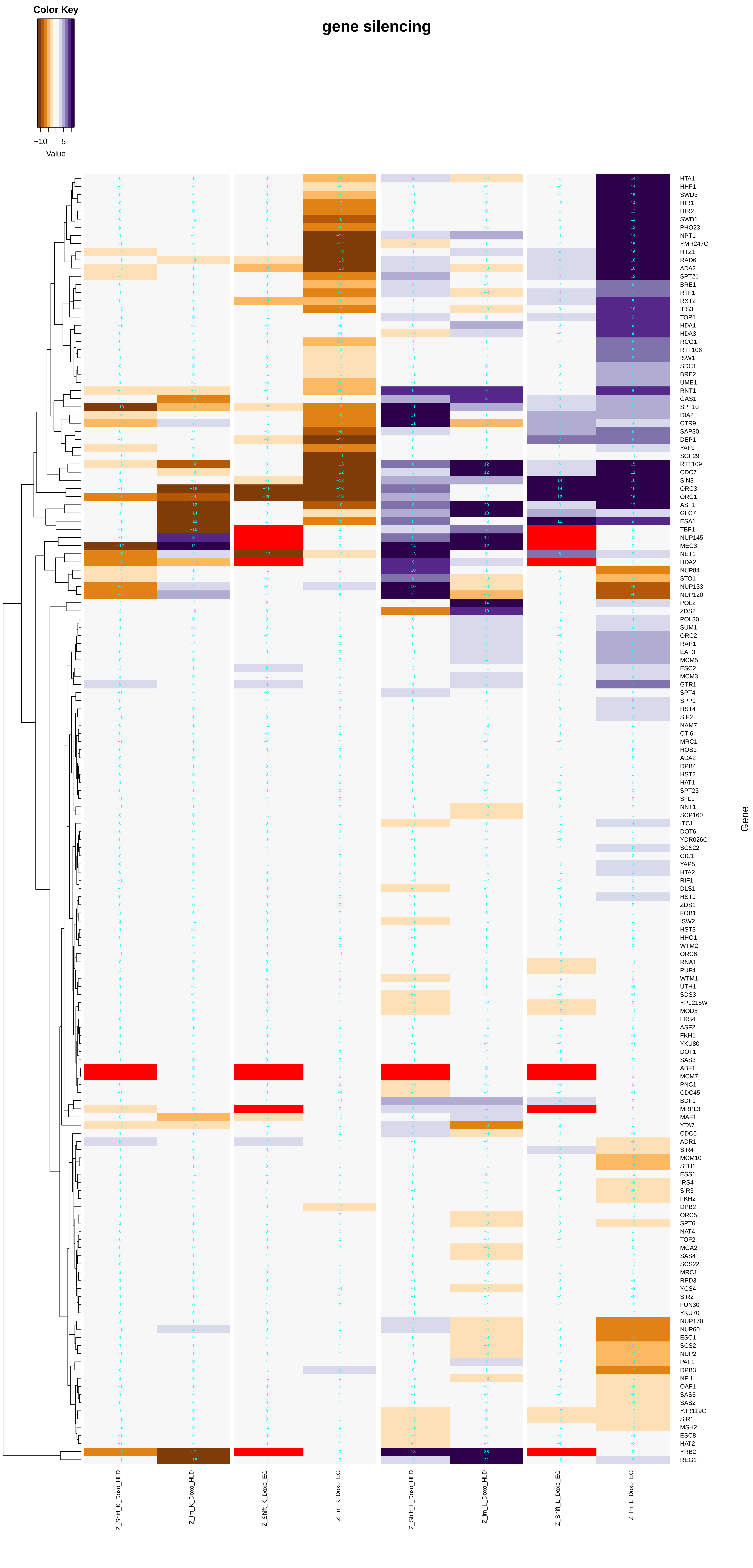

Color Key
−10
5
Value
gene silencing
HTA1
HHF1
SWD3
HIR1
HIR2
SWD1
PHO23
NPT1
YMR247C
HTZ1
RAD6
ADA2
SPT21
BRE1
RTF1
RXT2
IES3
TOP1
HDA1
HDA3
RCO1
RTT106
ISW1
SDC1
BRE2
UME1
RNT1
GAS1
SPT10
DIA2
CTR9
SAP30
DEP1
YAF9
SGF29
RTT109
CDC7
SIN3
ORC3
ORC1
ASF1
GLC7
ESA1
TBF1
NUP145
MEC3
NET1
HDA2
NUP84
STO1
NUP133
NUP120
POL2
ZDS2
POL30
SUM1
ORC2
RAP1
EAF3
MCM5
ESC2
MCM3
GTR1
SPT4
SPP1
HST4
SIF2
NAM7
CTI6
MRC1
HOS1
ADA2
DPB4
HST2
HAT1
SPT23
SFL1
NNT1
SCP160
ITC1
DOT6
YDR026C
SCS22
GIC1
YAP5
HTA2
RIF1
DLS1
HST1
ZDS1
FOB1
ISW2
HST3
HHO1
WTM2
ORC6
RNA1
PUF4
WTM1
UTH1
SDS3
YPL216W
MOD5
LRS4
ASF2
FKH1
YKU80
DOT1
SAS3
ABF1
MCM7
PNC1
CDC45
BDF1
MRPL3
MAF1
YTA7
CDC6
ADR1
SIR4
MCM10
STH1
ESS1
IRS4
SIR3
FKH2
DPB2
ORC5
SPT6
NAT4
TOF2
MGA2
SAS4
SCS22
MRC1
RPD3
YCS4
SIR2
FUN30
YKU70
NUP170
NUP60
ESC1
SCS2
NUP2
PAF1
DPB3
NFI1
OAF1
SAS5
SAS2
YJR119C
SIR1
MSH2
ESC8
HAT2
YRB2
REG1
Gene
Z_Shift_K_Doxo_HLD
Z_Im_K_Doxo_HLD
Z_Shift_K_Doxo_EG
Z_Im_K_Doxo_EG
Z_Shift_L_Doxo_HLD
Z_Im_L_Doxo_HLD
Z_Shift_L_Doxo_EG
Z_Im_L_Doxo_EG

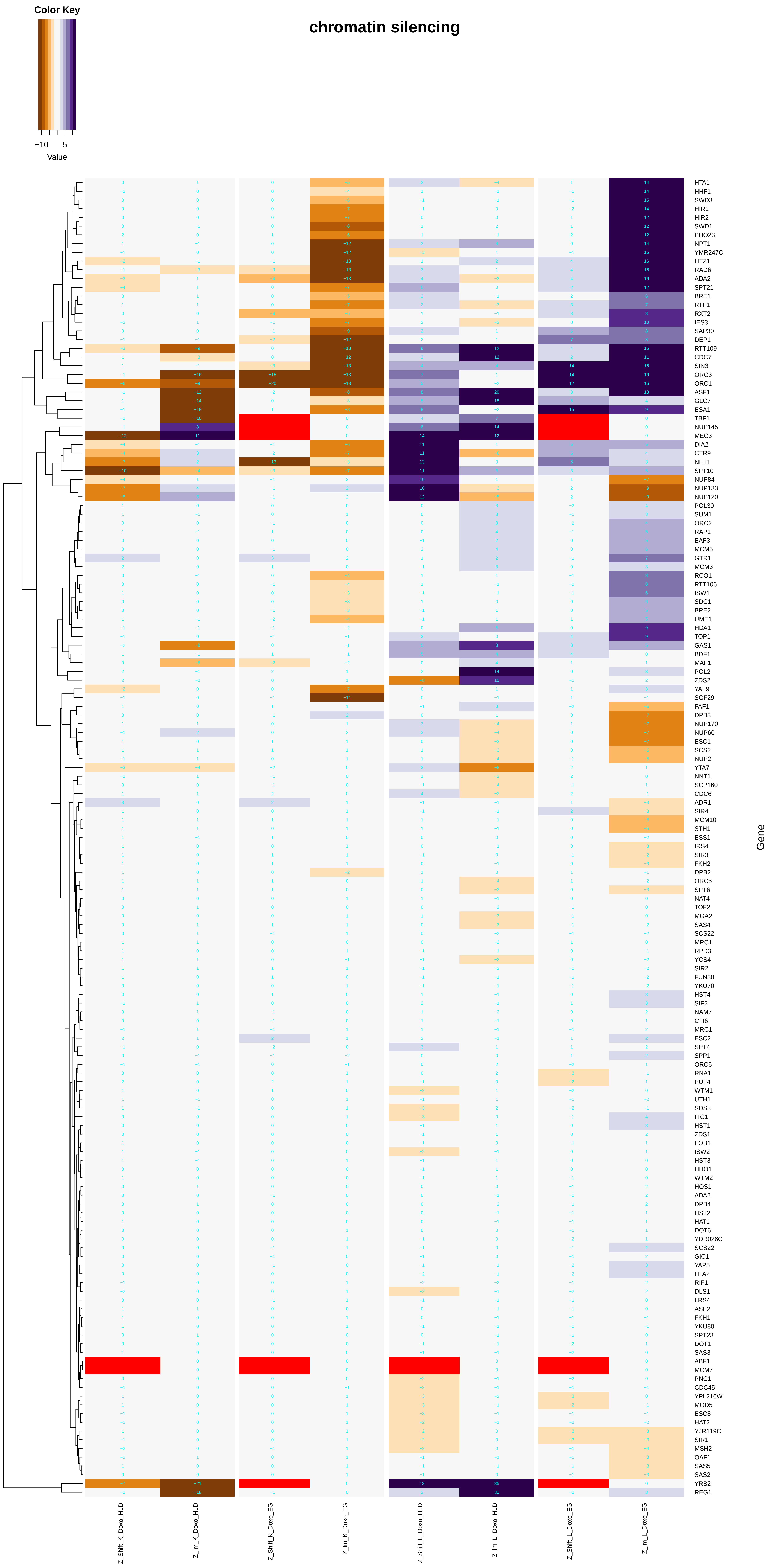
Color Key
−10 5
Value
chromatin silencing
Gene
Z_Shift_K_Doxo_HLD
Z_Im_K_Doxo_HLD
Z_Shift_K_Doxo_EG
Z_Im_K_Doxo_EG
Z_Shift_L_Doxo_HLD
Z_Im_L_Doxo_HLD
Z_Shift_L_Doxo_EG
Z_Im_L_Doxo_EG

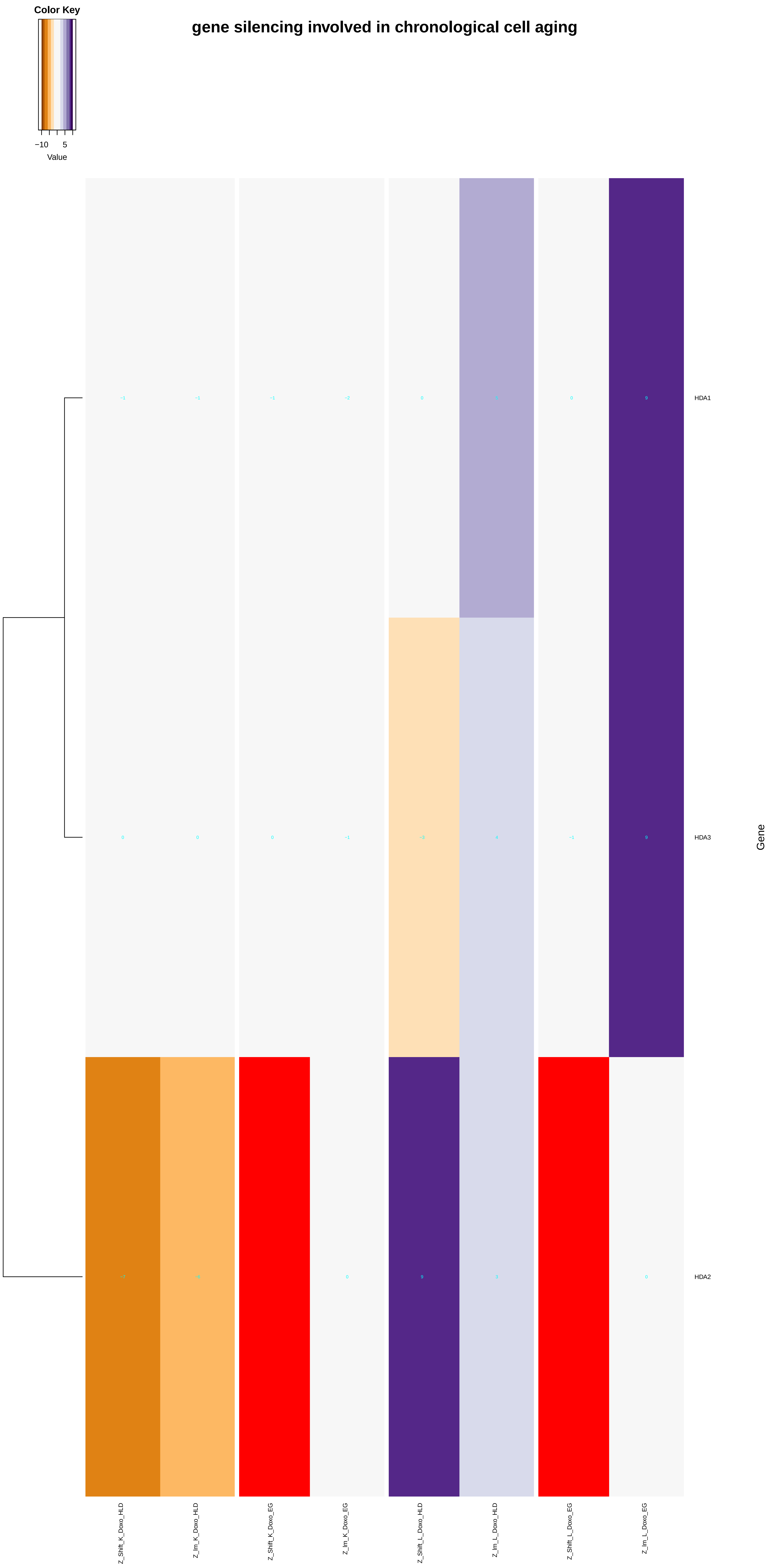
Color Key
−10
5
Value
gene silencing involved in chronological cell aging
HDA1
HDA3
HDA2
Gene
Z_Shift_K_Doxo_HLD
Z_lm_K_Doxo_HLD
Z_Shift_K_Doxo_EG
Z_lm_K_Doxo_EG
Z_Shift_L_Doxo_HLD
Z_lm_L_Doxo_HLD
Z_Shift_L_Doxo_EG
Z_lm_L_Doxo_EG

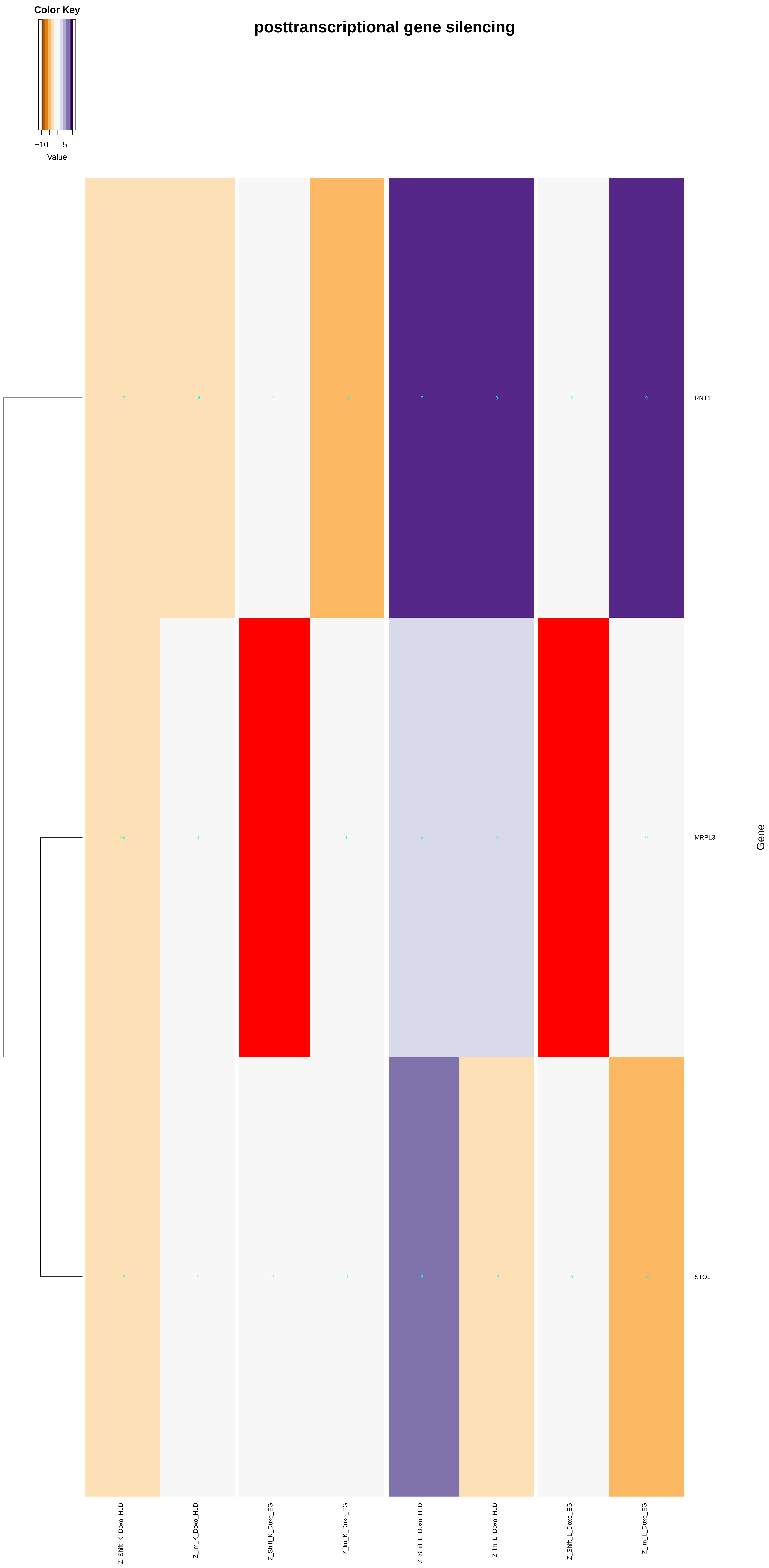
Color Key
−10
5
Value
posttranscriptional gene silencing
RNT1
MRPL3
STO1
Gene
Z_Shift_K_Doxo_HLD
Z_lm_K_Doxo_HLD
Z_Shift_K_Doxo_EG
Z_lm_K_Doxo_EG
Z_Shift_L_Doxo_HLD
Z_lm_L_Doxo_HLD
Z_Shift_L_Doxo_EG
Z_lm_L_Doxo_EG

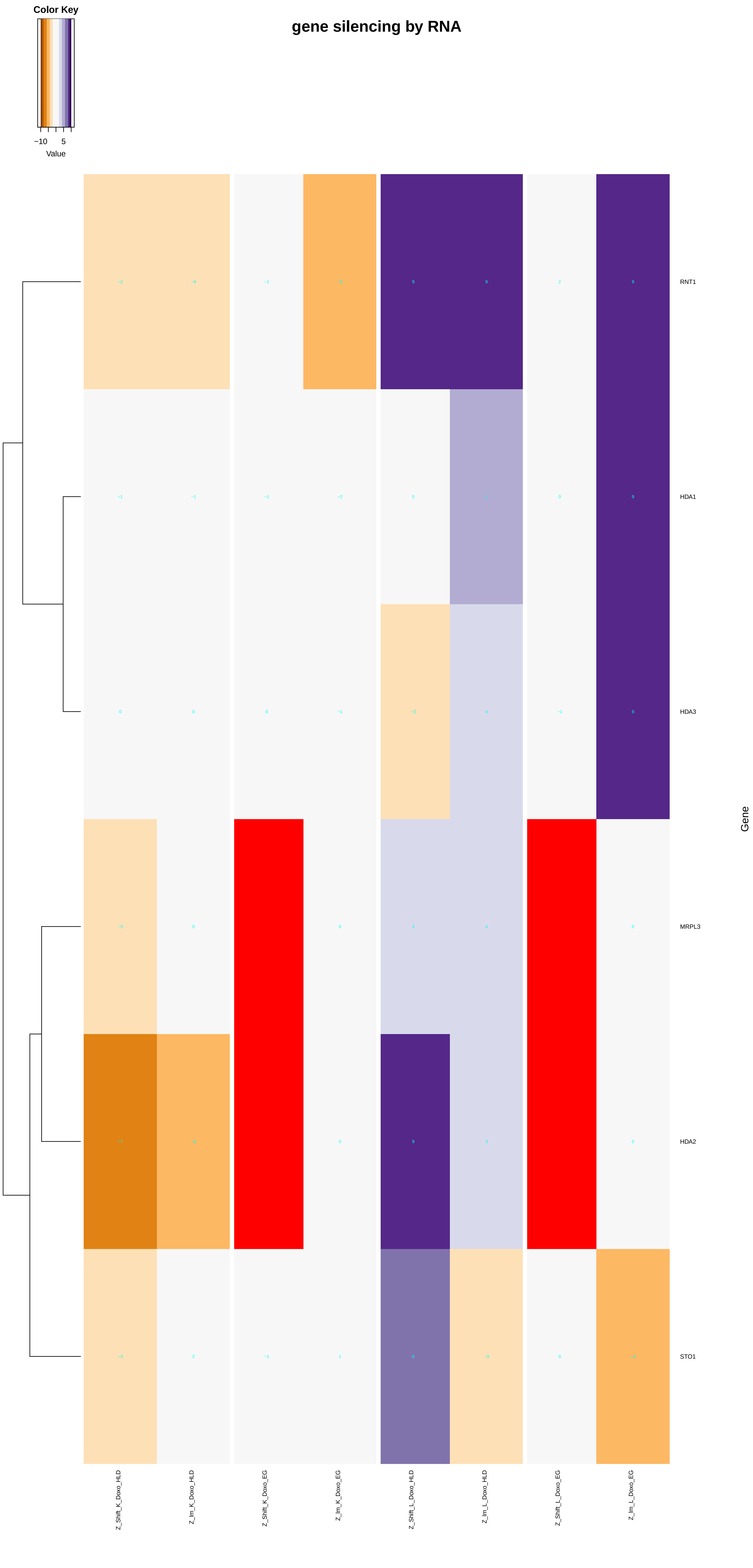
Color Key
−10
5
Value
gene silencing by RNA
RNT1
HDA1
HDA3
MRPL3
HDA2
STO1
Gene
Z_Shift_K_Doxo_HLD
Z_Im_K_Doxo_HLD
Z_Shift_K_Doxo_EG
Z_Im_K_Doxo_EG
Z_Shift_L_Doxo_HLD
Z_Im_L_Doxo_HLD
Z_Shift_L_Doxo_EG
Z_Im_L_Doxo_EG

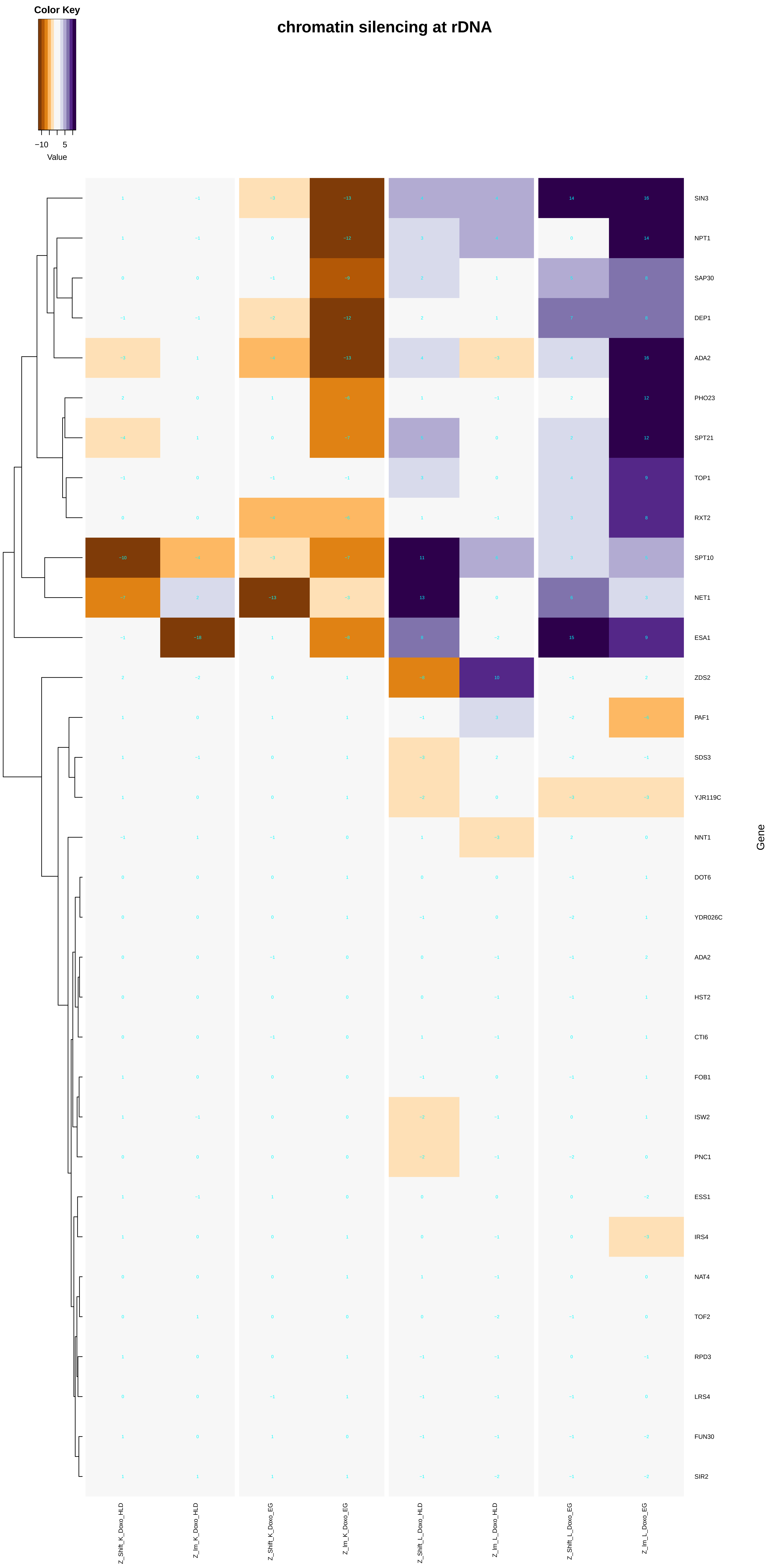

Color Key
−10
5
Value
chromatin silencing at rDNA
SIN3
NPT1
SAP30
DEP1
ADA2
PHO23
SPT21
TOP1
RXT2
SPT10
NET1
ESA1
ZDS2
PAF1
SDS3
YJR119C
NNT1
DOT6
YDR026C
ADA2
HST2
CTI6
FOB1
ISW2
PNC1
ESS1
IRS4
NAT4
TOF2
RPD3
LRS4
FUN30
SIR2
Gene
Z_Shift_K_Doxo_HLD
Z_Im_K_Doxo_HLD
Z_Shift_K_Doxo_EG
Z_Im_K_Doxo_EG
Z_Shift_L_Doxo_HLD
Z_Im_L_Doxo_HLD
Z_Shift_L_Doxo_EG
Z_Im_L_Doxo_EG

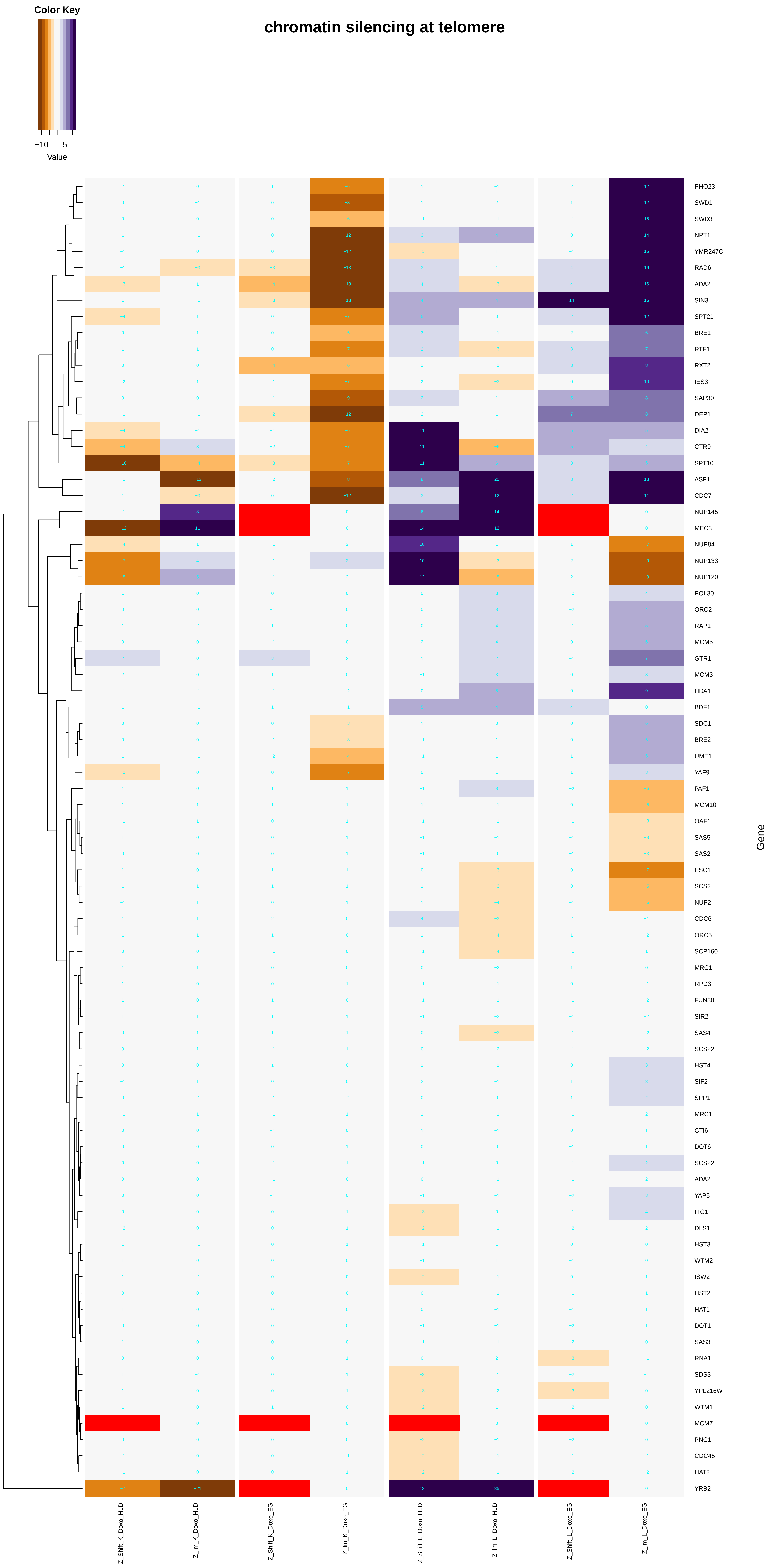
chromatin silencing at telomere
Color Key
−10
5
Value
PHO23
SWD1
SWD3
NPT1
YMR247C
RAD6
ADA2
SIN3
SPT21
BRE1
RTF1
RXT2
IES3
SAP30
DEP1
DIA2
CTR9
SPT10
ASF1
CDC7
NUP145
MEC3
NUP84
NUP133
NUP120
POL30
ORC2
RAP1
MCM5
GTR1
MCM3
HDA1
BDF1
SDC1
BRE2
UME1
YAF9
PAF1
MCM10
OAF1
SAS5
SAS2
ESC1
SCS2
NUP2
CDC6
ORC5
SCP160
MRC1
RPD3
FUN30
SIR2
SAS4
SCS22
HST4
SIF2
SPP1
MRC1
CTI6
DOT6
SCS22
ADA2
YAP5
ITC1
DLS1
HST3
WTM2
ISW2
HST2
HAT1
DOT1
SAS3
RNA1
SDS3
YPL216W
WTM1
MCM7
PNC1
CDC45
HAT2
YRB2
Gene
Z_Shift_K_Doxo_HLD
Z_Im_K_Doxo_HLD
Z_Shift_K_Doxo_EG
Z_Im_K_Doxo_EG
Z_Shift_L_Doxo_HLD
Z_Im_L_Doxo_HLD
Z_Shift_L_Doxo_EG
Z_Im_L_Doxo_EG

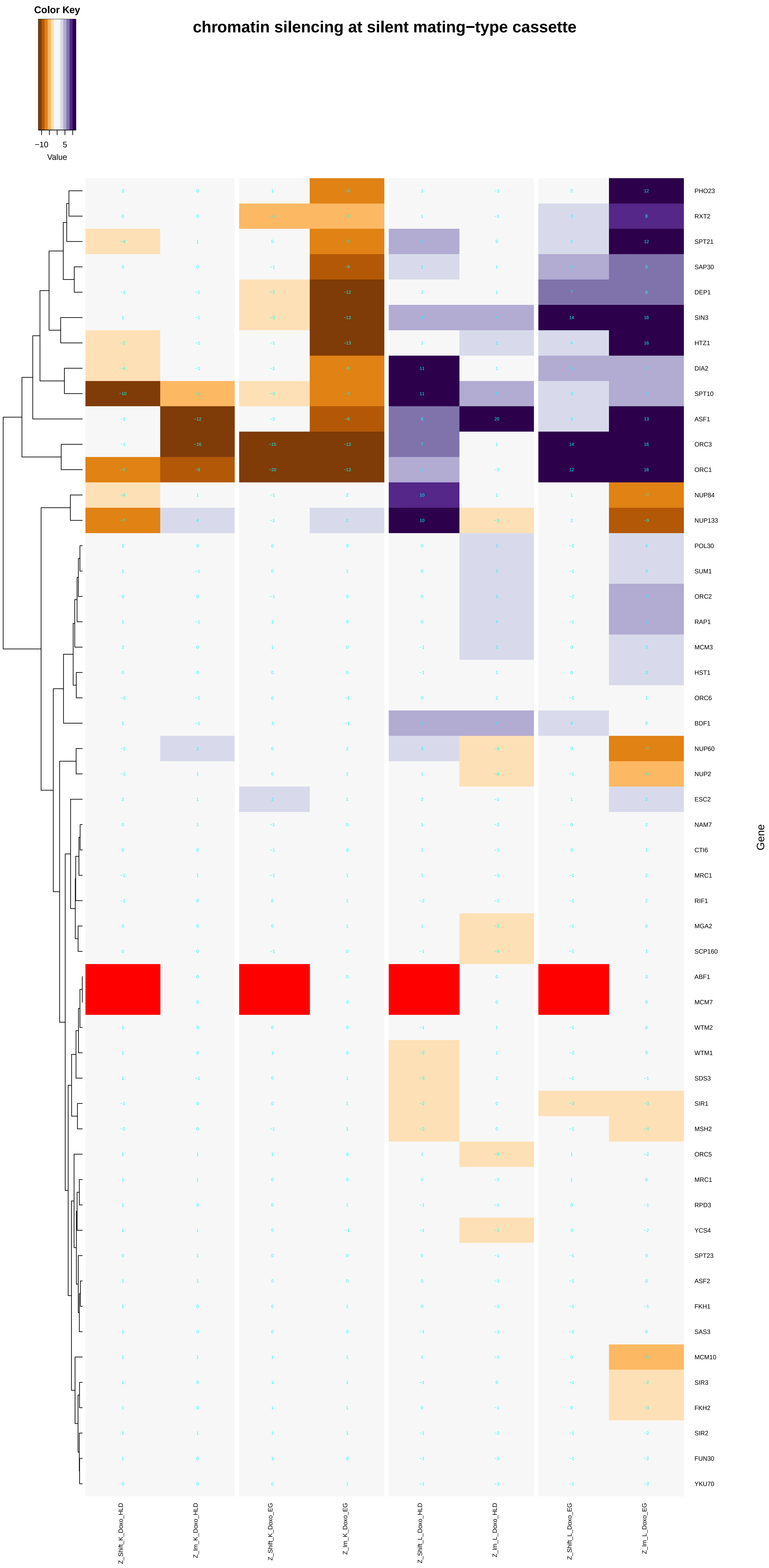

Color Key
−10
5
Value
chromatin silencing at silent mating−type cassette
PHO23
RXT2
SPT21
SAP30
DEP1
SIN3
HTZ1
DIA2
SPT10
ASF1
ORC3
ORC1
NUP84
NUP133
POL30
SUM1
ORC2
RAP1
MCM3
HST1
ORC6
BDF1
NUP60
NUP2
ESC2
NAM7
CTI6
MRC1
RIF1
MGA2
SCP160
ABF1
MCM7
WTM2
WTM1
SDS3
SIR1
MSH2
ORC5
MRC1
RPD3
YCS4
SPT23
ASF2
FKH1
SAS3
MCM10
SIR3
FKH2
SIR2
FUN30
YKU70
Gene
Z_Shift_K_Doxo_HLD
Z_Im_K_Doxo_HLD
Z_Shift_K_Doxo_EG
Z_Im_K_Doxo_EG
Z_Shift_L_Doxo_HLD
Z_Im_L_Doxo_HLD
Z_Shift_L_Doxo_EG
Z_Im_L_Doxo_EG

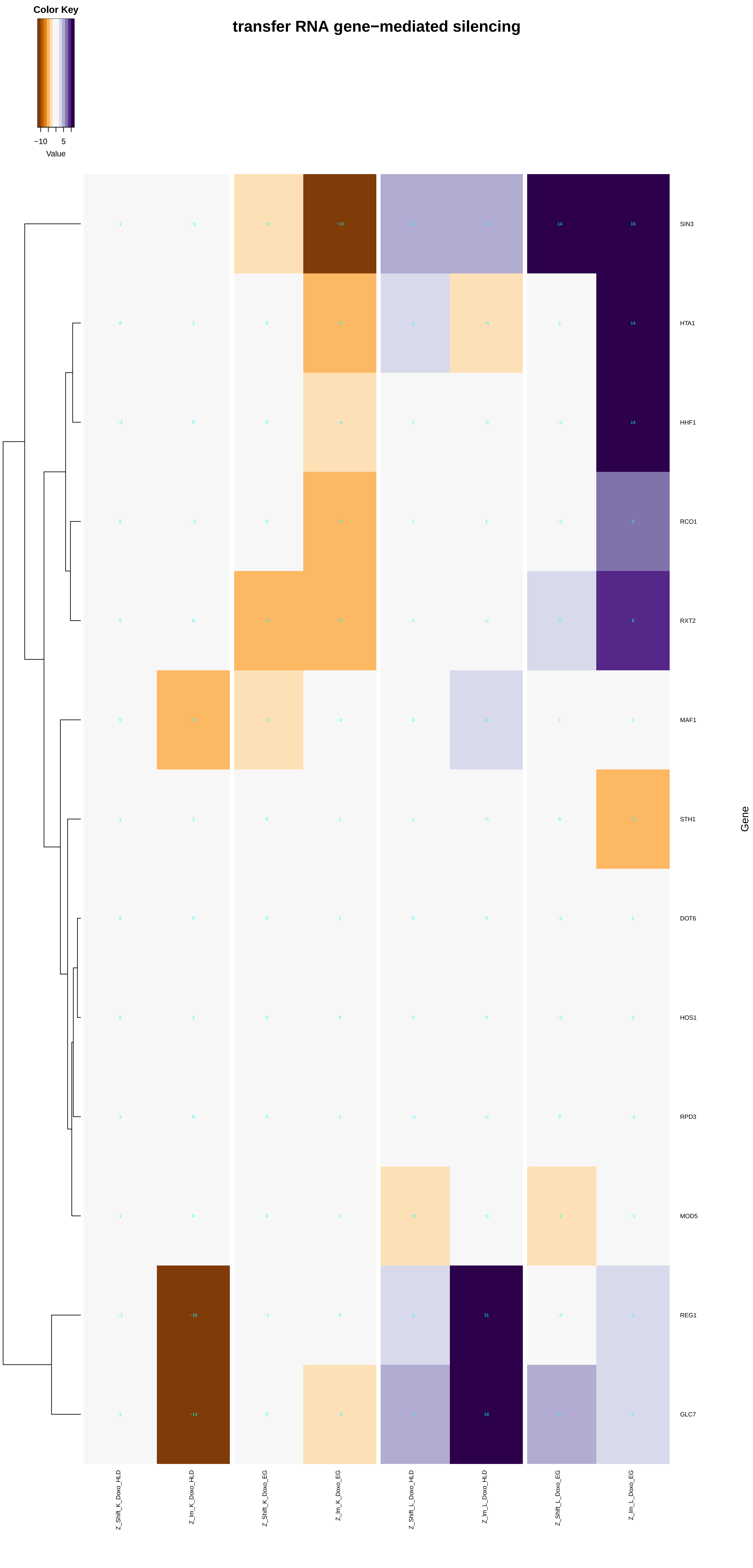
transfer RNA gene−mediated silencing
Color Key
−10
5
Value
SIN3
HTA1
HHF1
RCO1
RXT2
MAF1
STH1
DOT6
HOS1
RPD3
MOD5
REG1
GLC7
Gene
Z_Shift_K_Doxo_HLD
Z_lm_K_Doxo_HLD
Z_Shift_K_Doxo_EG
Z_lm_K_Doxo_EG
Z_Shift_L_Doxo_HLD
Z_lm_L_Doxo_HLD
Z_Shift_L_Doxo_EG
Z_lm_L_Doxo_EG

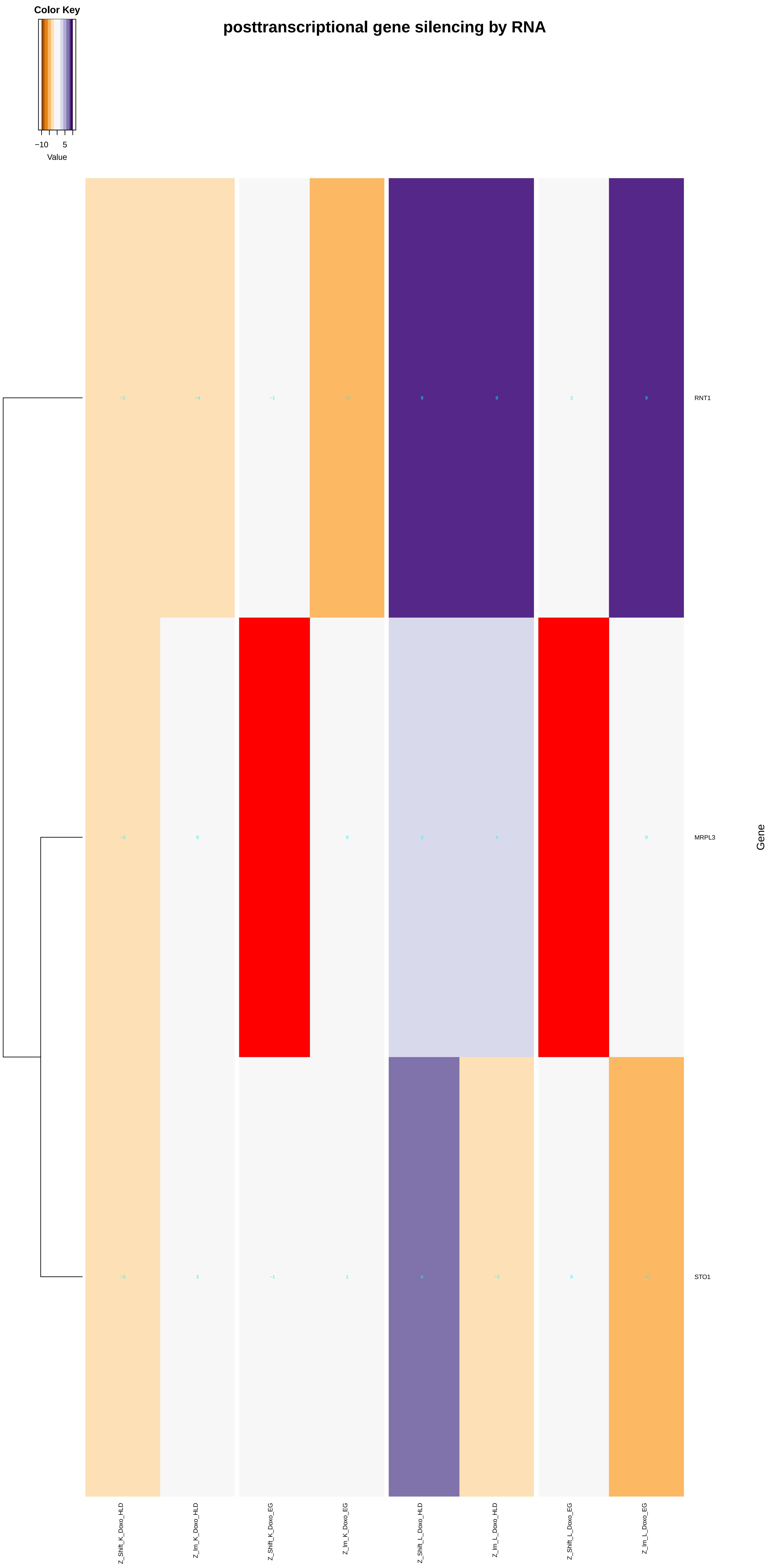

posttranscriptional gene silencing by RNA
Color Key
−10
5
Value
RNT1
MRPL3
STO1
Gene
Z_Shift_K_Doxo_HLD
Z_Im_K_Doxo_HLD
Z_Shift_K_Doxo_EG
Z_Im_K_Doxo_EG
Z_Shift_L_Doxo_HLD
Z_Im_L_Doxo_HLD
Z_Shift_L_Doxo_EG
Z_Im_L_Doxo_EG

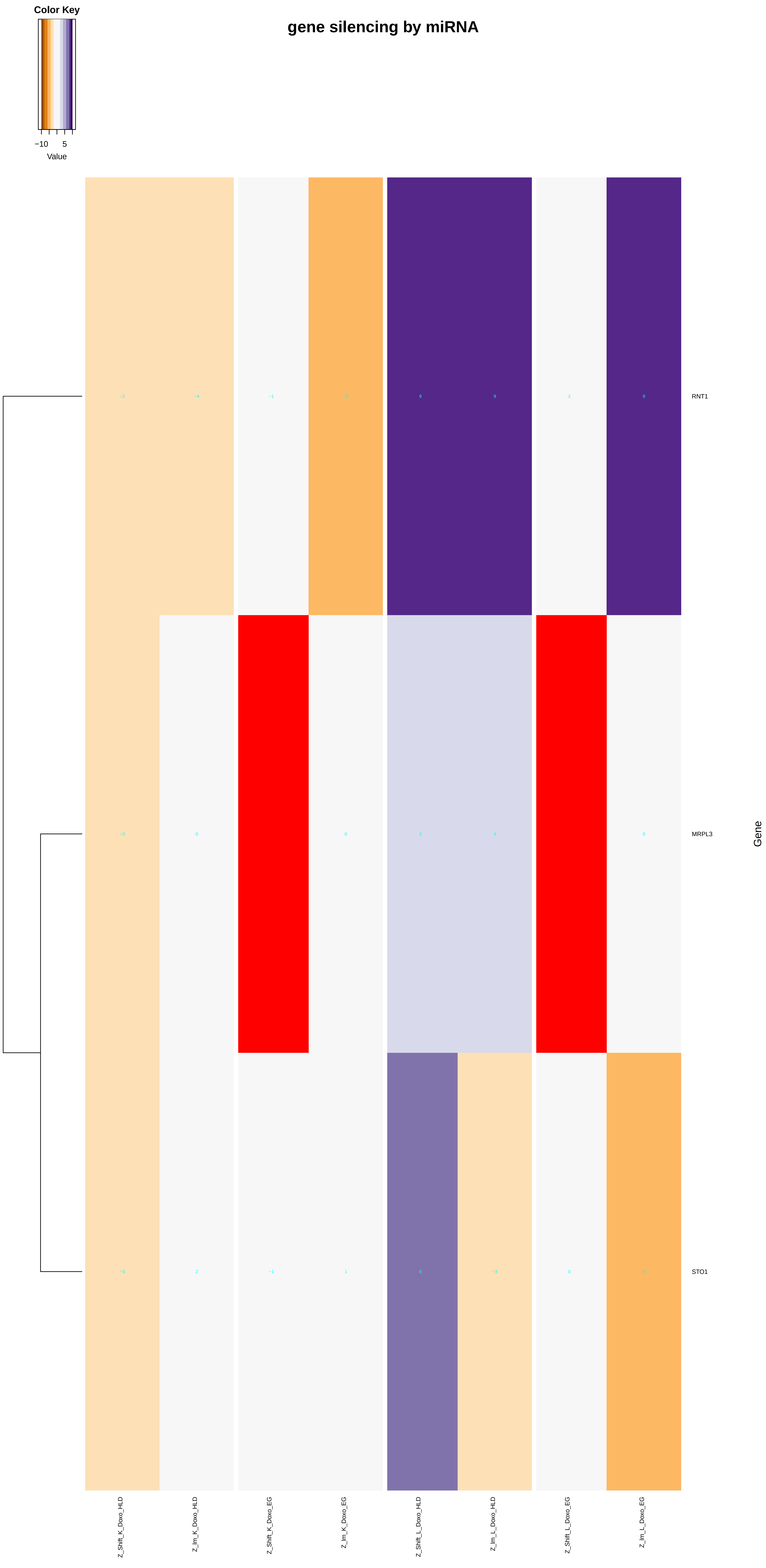

gene silencing by miRNA
Color Key
−10
5
Value
RNT1
MRPL3
STO1
Gene
Z_Shift_K_Doxo_HLD
Z_Im_K_Doxo_HLD
Z_Shift_K_Doxo_EG
Z_Im_K_Doxo_EG
Z_Shift_L_Doxo_HLD
Z_Im_L_Doxo_HLD
Z_Shift_L_Doxo_EG
Z_Im_L_Doxo_EG
