## Additional File 9 for "A yeast phenomic model for the influence of Warburg metabolism on genetic buffering of doxorubicin": Golgi_vesicle_transport.pdf

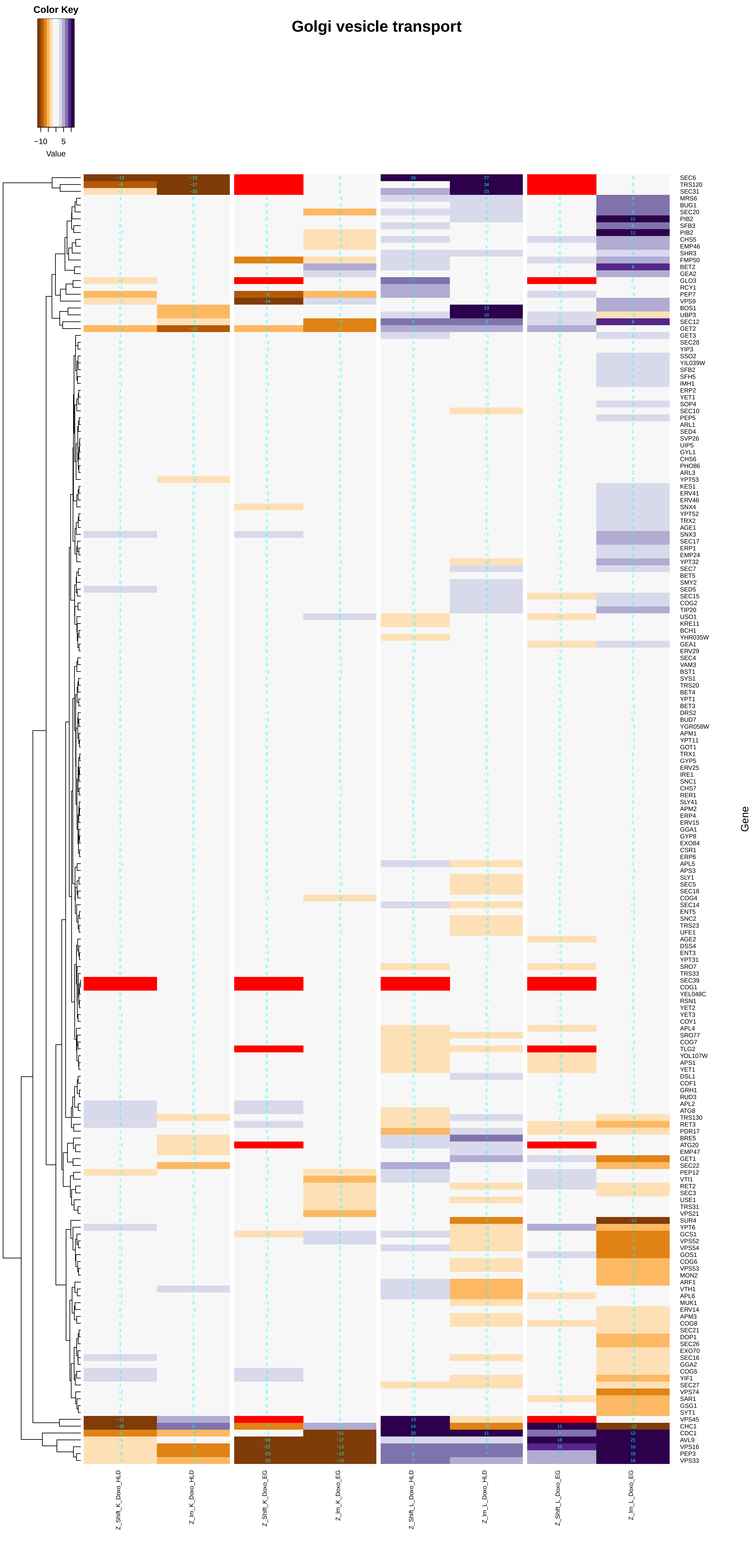
Golgi vesicle transport
Color Key
−10
5
Value
Gene
SEC6
TRS120
SEC31
MRS6
BUG1
SEC20
PIB2
SFB3
PIB2
CHS5
EMP46
SHR3
FMP50
BET2
GEA2
GLO3
RCY1
PEP7
VPS9
BOS1
UBP3
SEC12
GET2
GET3
SEC28
YIP3
SSO2
YIL039W
SFB2
SFH5
IMH1
ERP2
YET1
SOP4
SEC10
PEP5
ARL1
SED4
SVP26
UIP5
GYL1
CHS6
PHO86
ARL3
YPT53
KES1
ERV41
ERV46
SNX4
YPT52
TRX2
AGE1
SNX3
SEC17
ERP1
EMP24
YPT32
SEC7
BET5
SMY2
SED5
SEC15
COG2
TIP20
USO1
KRE11
BCH1
YHR035W
GEA1
ERV29
SEC4
VAM3
BST1
SYS1
TRS20
BET4
YPT1
BET3
DRS2
BUD7
YGR058W
APM1
YPT11
GOT1
TRX1
GYP5
ERV25
IRE1
SNC1
CHS7
RER1
SLY41
APM2
ERP4
ERV15
GGA1
GYP8
EXO84
CSR1
ERP6
APL5
APS3
SLY1
SEC5
SEC18
COG4
SEC14
ENT5
SNC2
TRS23
UFE1
AGE2
DSS4
ENT3
YPT31
SRO7
TRS33
SEC39
COG1
YEL048C
RSN1
YET2
YET3
COY1
APL4
SRO77
COG7
TLG2
YOL107W
APS1
YET1
DSL1
COF1
GRH1
RUD3
APL2
ATG8
TRS130
RET3
PDR17
BRE5
ATG20
EMP47
GET1
SEC22
PEP12
VTI1
RET2
SEC3
USE1
TRS31
VPS21
SUR4
YPT6
GCS1
VPS52
VPS54
GOS1
COG6
VPS53
MON2
ARF1
VTH1
APL6
MUK1
ERV14
APM3
COG8
SEC21
DOP1
SEC26
EXO70
SEC16
GGA2
COG5
YIF1
SEC27
VPS74
SAR1
GSG1
SYT1
VPS45
CHC1
CDC1
AVL9
VPS16
PEP3
VPS33
Z_Shift_K_Doxo_HLD
Z_lm_K_Doxo_HLD
Z_Shift_K_Doxo_EG
Z_lm_K_Doxo_EG
Z_Shift_L_Doxo_HLD
Z_lm_L_Doxo_HLD
Z_Shift_L_Doxo_EG
Z_lm_L_Doxo_EG

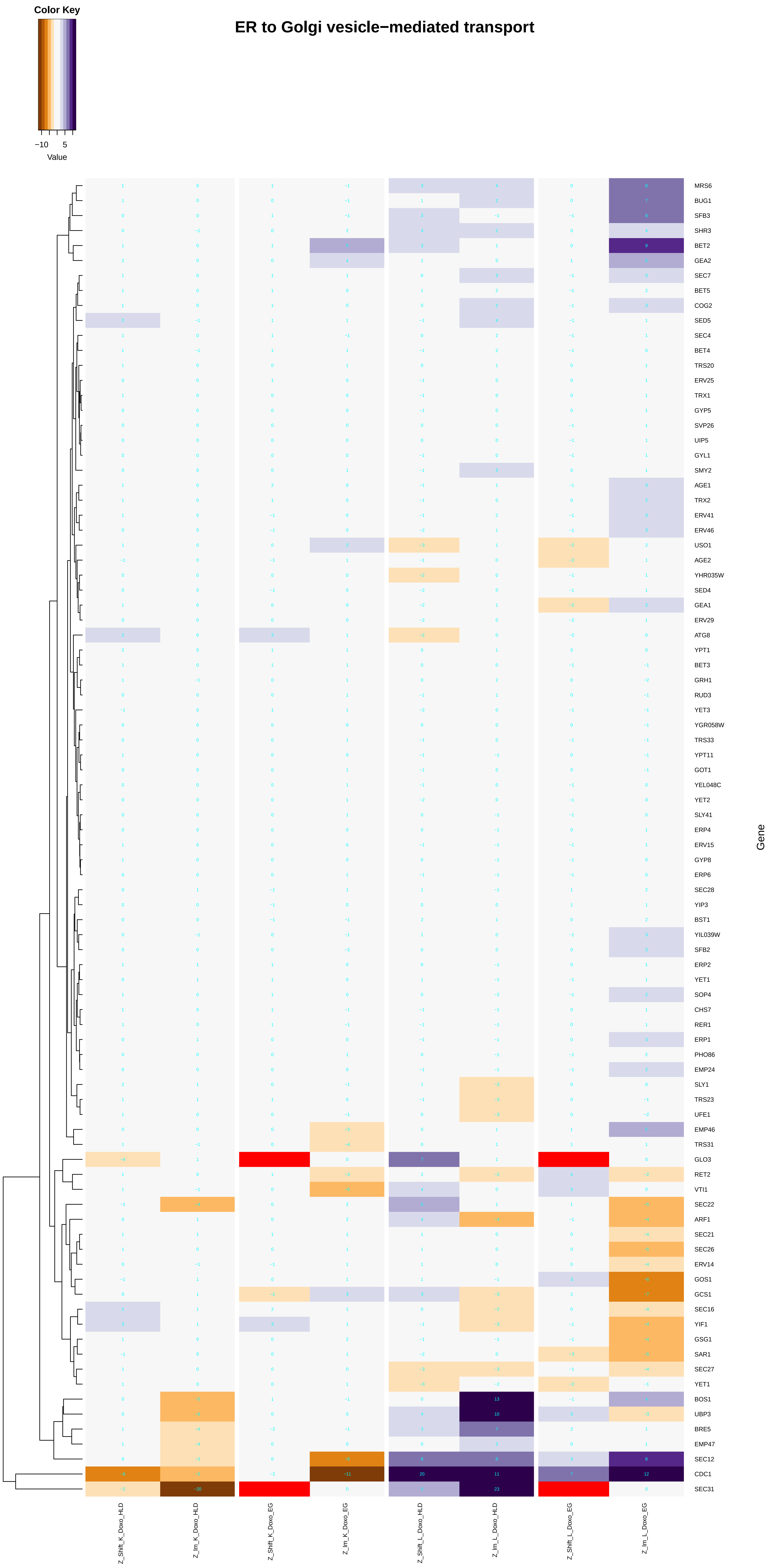

Color Key
−10
5
Value
ER to Golgi vesicle−mediated transport
MRS6
BUG1
SFB3
SHR3
BET2
GEA2
SEC7
BET5
COG2
SED5
SEC4
BET4
TRS20
ERV25
TRX1
GYP5
SVP26
UIP5
GYL1
SMY2
AGE1
TRX2
ERV41
ERV46
USO1
AGE2
YHR035W
SED4
GEA1
ERV29
ATG8
YPT1
BET3
GRH1
RUD3
YET3
YGR058W
TRS33
YPT11
GOT1
YEL048C
YET2
SLY41
ERP4
ERV15
GYP8
ERP6
SEC28
YIP3
BST1
YIL039W
SFB2
ERP2
YET1
SOP4
CHS7
RER1
ERP1
PHO86
EMP24
SLY1
TRS23
UFE1
EMP46
TRS31
GLO3
RET2
VTI1
SEC22
ARF1
SEC21
SEC26
ERV14
GOS1
GCS1
SEC16
YIF1
GSG1
SAR1
SEC27
YET1
BOS1
UBP3
BRE5
EMP47
SEC12
CDC1
SEC31
Gene
Z_Shift_K_Doxo_HLD
Z_lm_K_Doxo_HLD
Z_Shift_K_Doxo_EG
Z_lm_K_Doxo_EG
Z_Shift_L_Doxo_HLD
Z_lm_L_Doxo_HLD
Z_Shift_L_Doxo_EG
Z_lm_L_Doxo_EG

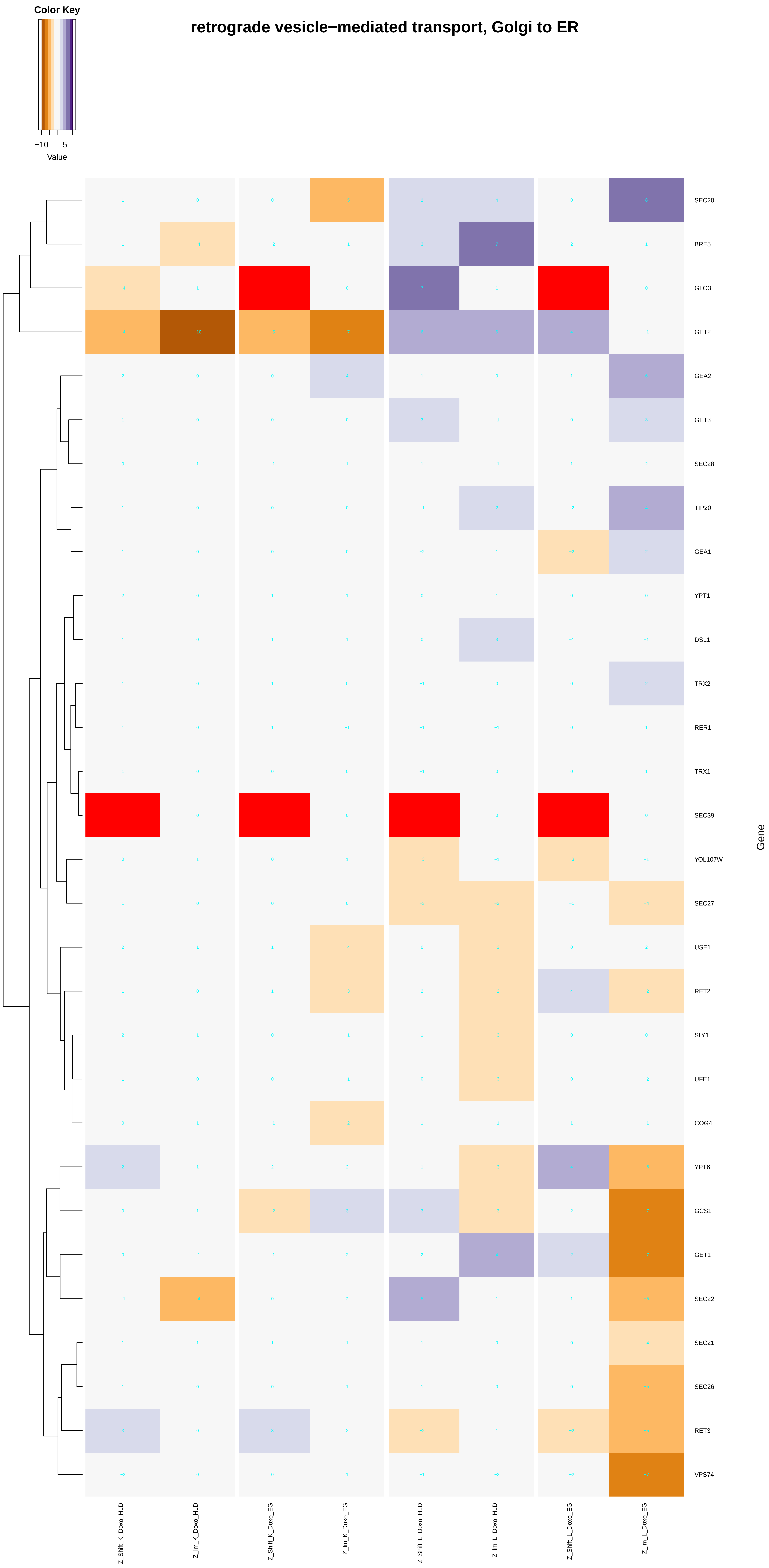
Color Key
−10
5
Value
retrograde vesicle−mediated transport, Golgi to ER
SEC20
BRE5
GLO3
GET2
GEA2
GET3
SEC28
TIP20
GEA1
YPT1
DSL1
TRX2
RER1
TRX1
SEC39
YOL107W
SEC27
USE1
RET2
SLY1
UFE1
COG4
YPT6
GCS1
GET1
SEC22
SEC21
SEC26
RET3
VPS74
Gene
Z_Shift_K_Doxo_HLD
Z_Im_K_Doxo_HLD
Z_Shift_K_Doxo_EG
Z_Im_K_Doxo_EG
Z_Shift_L_Doxo_HLD
Z_Im_L_Doxo_HLD
Z_Shift_L_Doxo_EG
Z_Im_L_Doxo_EG

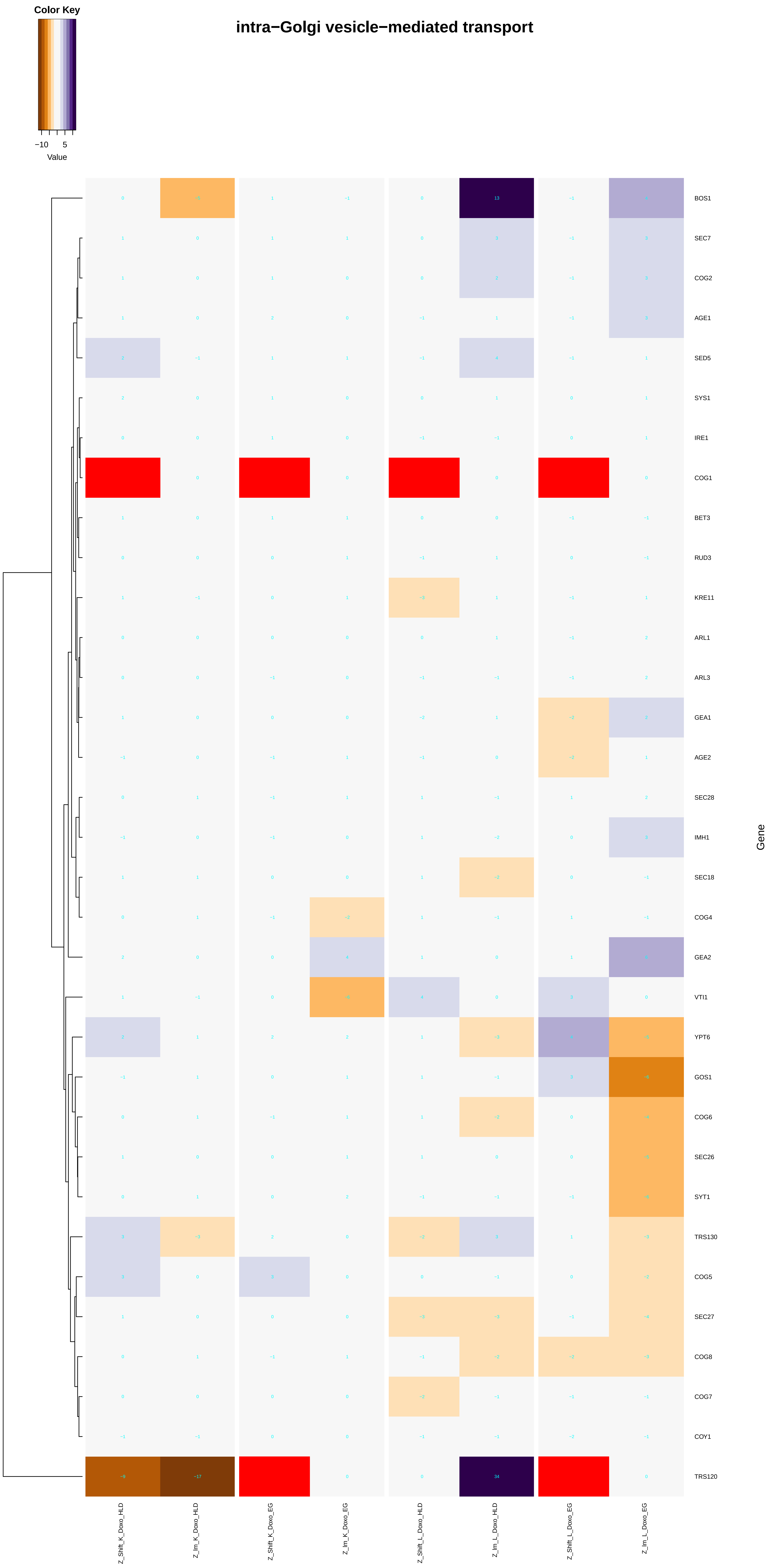
Color Key
−10
5
Value
intra−Golgi vesicle−mediated transport
BOS1
SEC7
COG2
AGE1
SED5
SYS1
IRE1
COG1
BET3
RUD3
KRE11
ARL1
ARL3
GEA1
AGE2
SEC28
IMH1
SEC18
COG4
GEA2
VTI1
YPT6
GOS1
COG6
SEC26
SYT1
TRS130
COG5
SEC27
COG8
COG7
COY1
TRS120
Gene
Z_Shift_K_Doxo_HLD
Z_lm_K_Doxo_HLD
Z_Shift_K_Doxo_EG
Z_lm_K_Doxo_EG
Z_Shift_L_Doxo_HLD
Z_lm_L_Doxo_HLD
Z_Shift_L_Doxo_EG
Z_lm_L_Doxo_EG

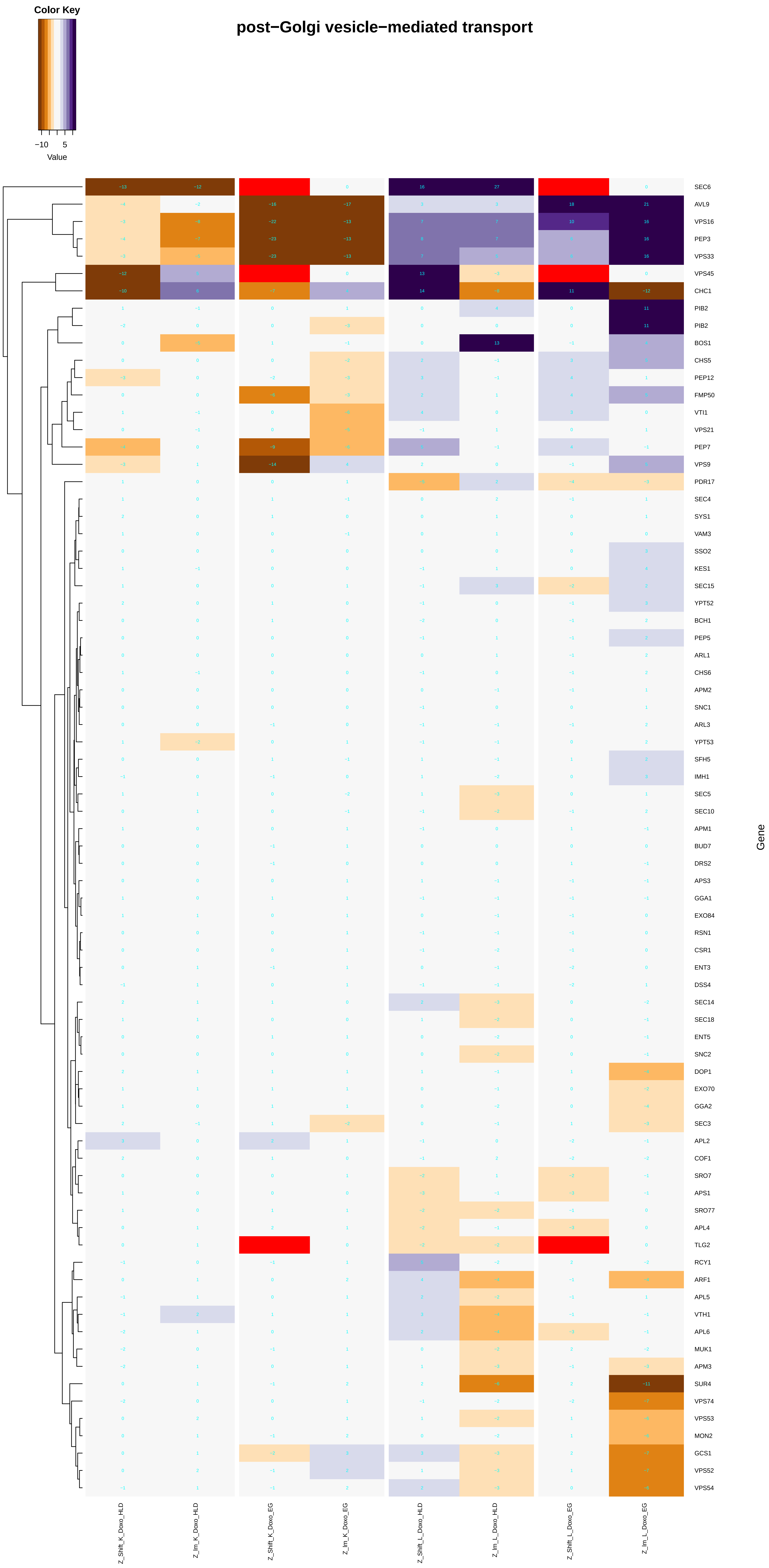
Color Key
−10
5
Value
post−Golgi vesicle−mediated transport
SEC6
AVL9
VPS16
PEP3
VPS33
VPS45
CHC1
PIB2
PIB2
BOS1
CHS5
PEP12
FMP50
VTI1
VPS21
PEP7
VPS9
PDR17
SEC4
SYS1
VAM3
SSO2
KES1
SEC15
YPT52
BCH1
PEP5
ARL1
CHS6
APM2
SNC1
ARL3
YPT53
SFH5
IMH1
SEC5
SEC10
APM1
BUD7
DRS2
APS3
GGA1
EXO84
RSN1
CSR1
ENT3
DSS4
SEC14
SEC18
ENT5
SNC2
DOP1
EXO70
GGA2
SEC3
APL2
COF1
SRO7
APS1
SRO77
APL4
TLG2
RCY1
ARF1
APL5
VTH1
APL6
MUK1
APM3
SUR4
VPS74
VPS53
MON2
GCS1
VPS52
VPS54
Gene
Z_Shift_K_Doxo_HLD
Z_Im_K_Doxo_HLD
Z_Shift_K_Doxo_EG
Z_Im_K_Doxo_EG
Z_Shift_L_Doxo_HLD
Z_Im_L_Doxo_HLD
Z_Shift_L_Doxo_EG
Z_Im_L_Doxo_EG

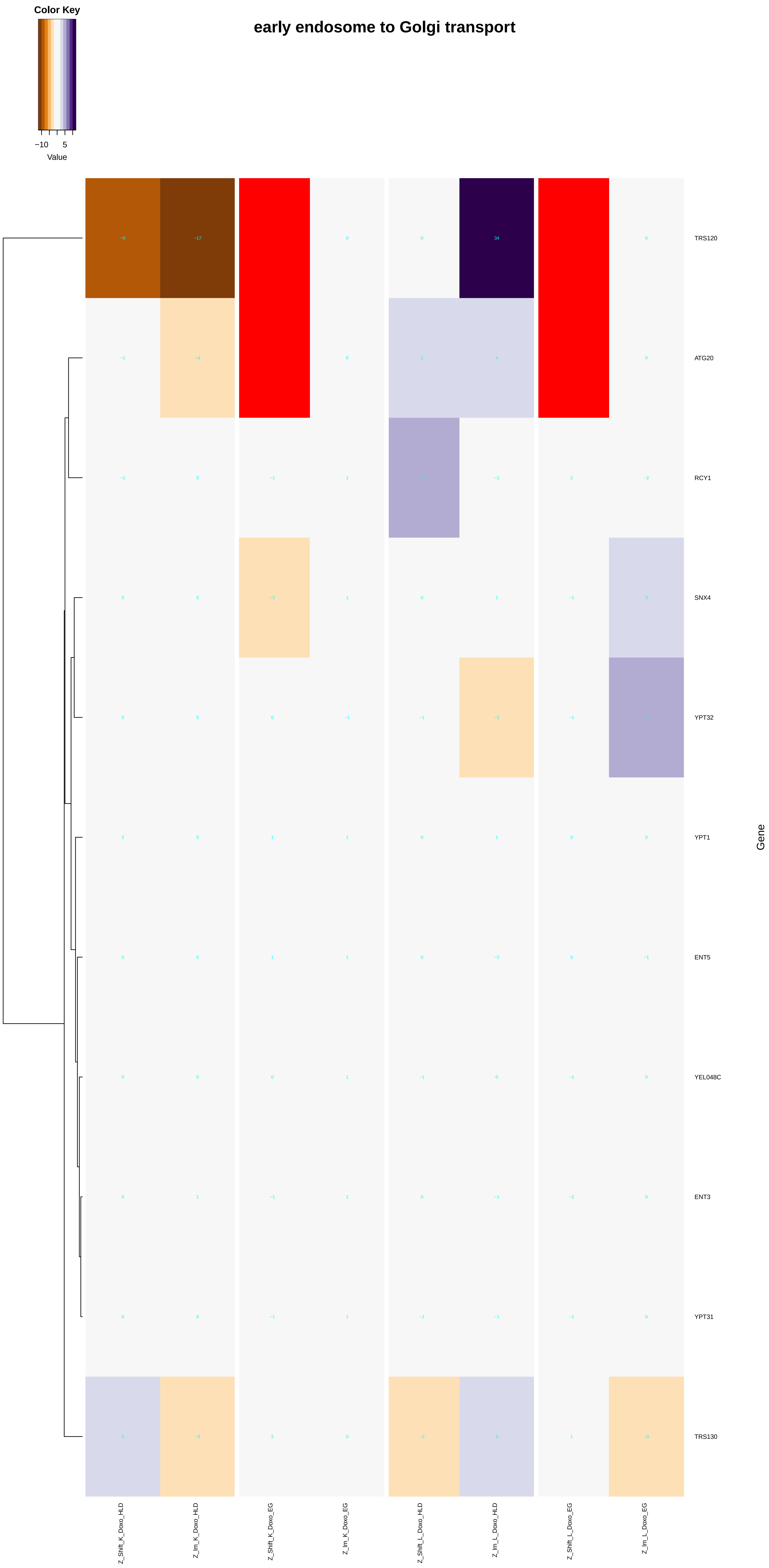
Color Key
−10
5
Value
early endosome to Golgi transport
TRS120
ATG20
RCY1
SNX4
YPT32
YPT1
ENT5
YEL048C
ENT3
YPT31
TRS130
Gene
Z_Shift_K_Doxo_HLD
Z_Im_K_Doxo_HLD
Z_Shift_K_Doxo_EG
Z_Im_K_Doxo_EG
Z_Shift_L_Doxo_HLD
Z_Im_L_Doxo_HLD
Z_Shift_L_Doxo_EG
Z_Im_L_Doxo_EG

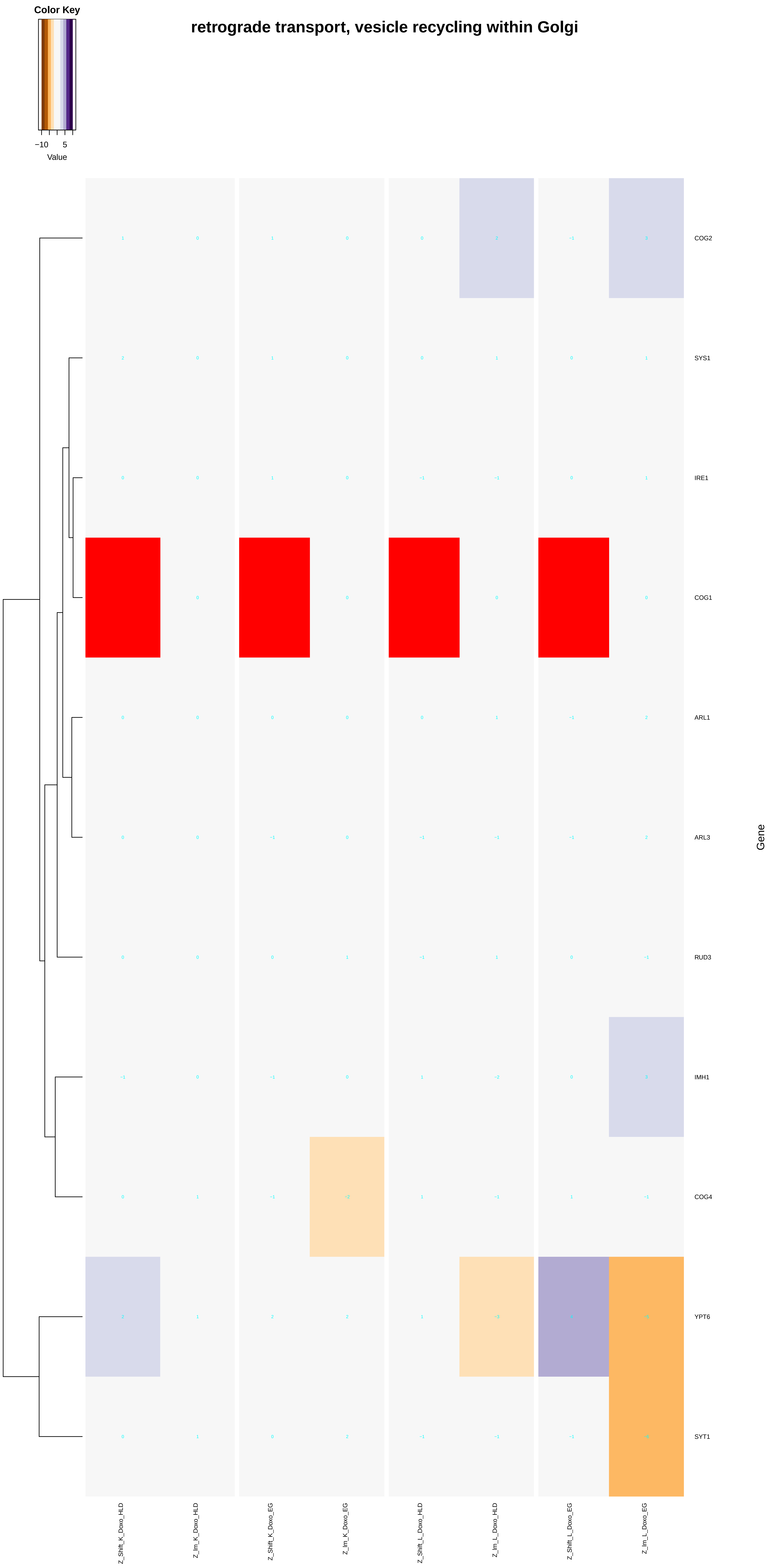
Color Key
−10
5
Value
retrograde transport, vesicle recycling within Golgi
COG2
SYS1
IRE1
COG1
ARL1
ARL3
RUD3
IMH1
COG4
YPT6
SYT1
Gene
Z_Shift_K_Doxo_HLD
Z_Im_K_Doxo_HLD
Z_Shift_K_Doxo_EG
Z_Im_K_Doxo_EG
Z_Shift_L_Doxo_HLD
Z_Im_L_Doxo_HLD
Z_Shift_L_Doxo_EG
Z_Im_L_Doxo_EG

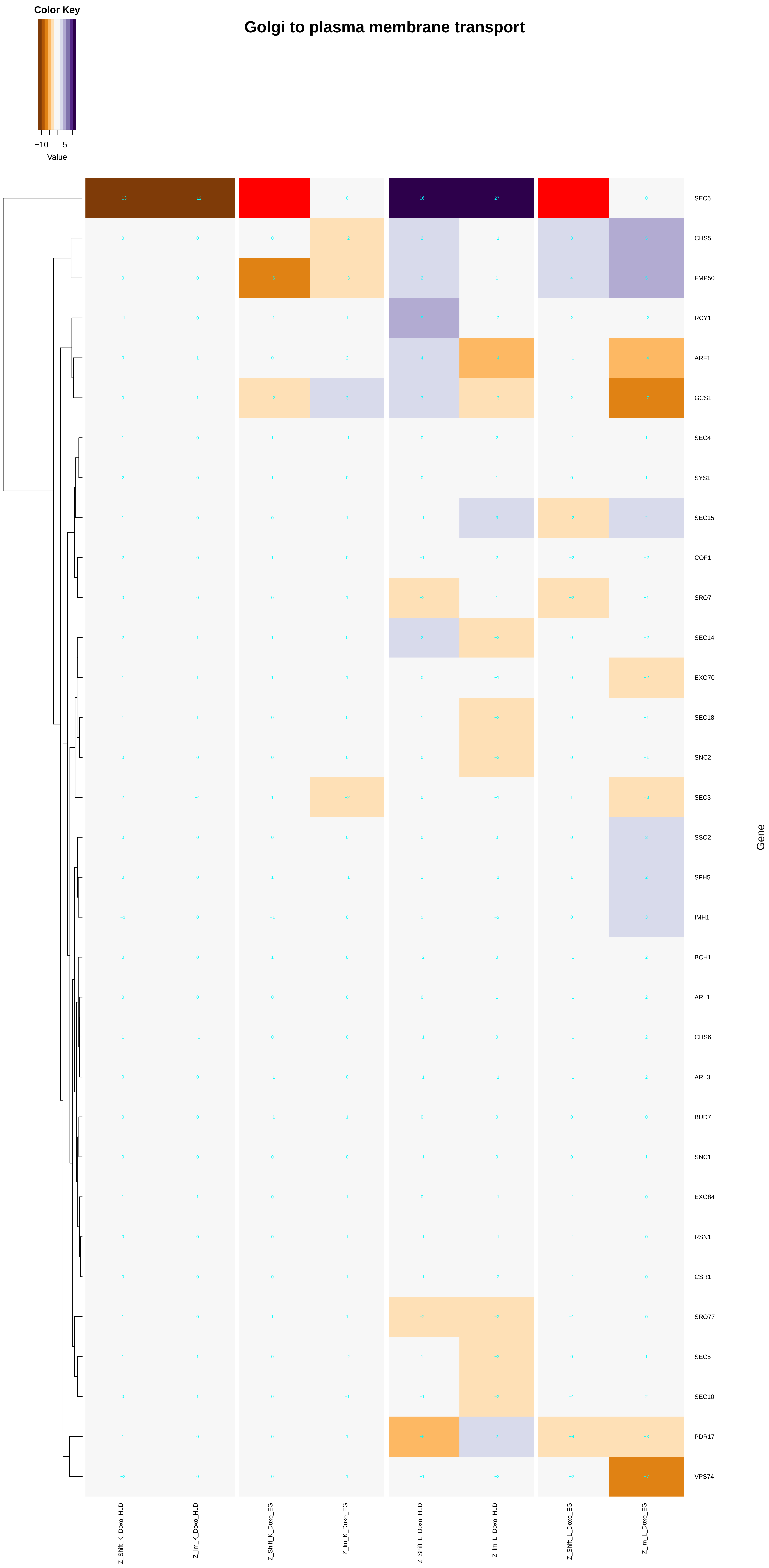
Color Key
−10
5
Value
Golgi to plasma membrane transport
SEC6
CHS5
FMP50
RCY1
ARF1
GCS1
SEC4
SYS1
SEC15
COF1
SRO7
SEC14
EXO70
SEC18
SNC2
SEC3
SSO2
SFH5
IMH1
BCH1
ARL1
CHS6
ARL3
BUD7
SNC1
EXO84
RSN1
CSR1
SRO77
SEC5
SEC10
PDR17
VPS74
Gene
Z_Shift_K_Doxo_HLD
Z_Im_K_Doxo_HLD
Z_Shift_K_Doxo_EG
Z_Im_K_Doxo_EG
Z_Shift_L_Doxo_HLD
Z_Im_L_Doxo_HLD
Z_Shift_L_Doxo_EG
Z_Im_L_Doxo_EG

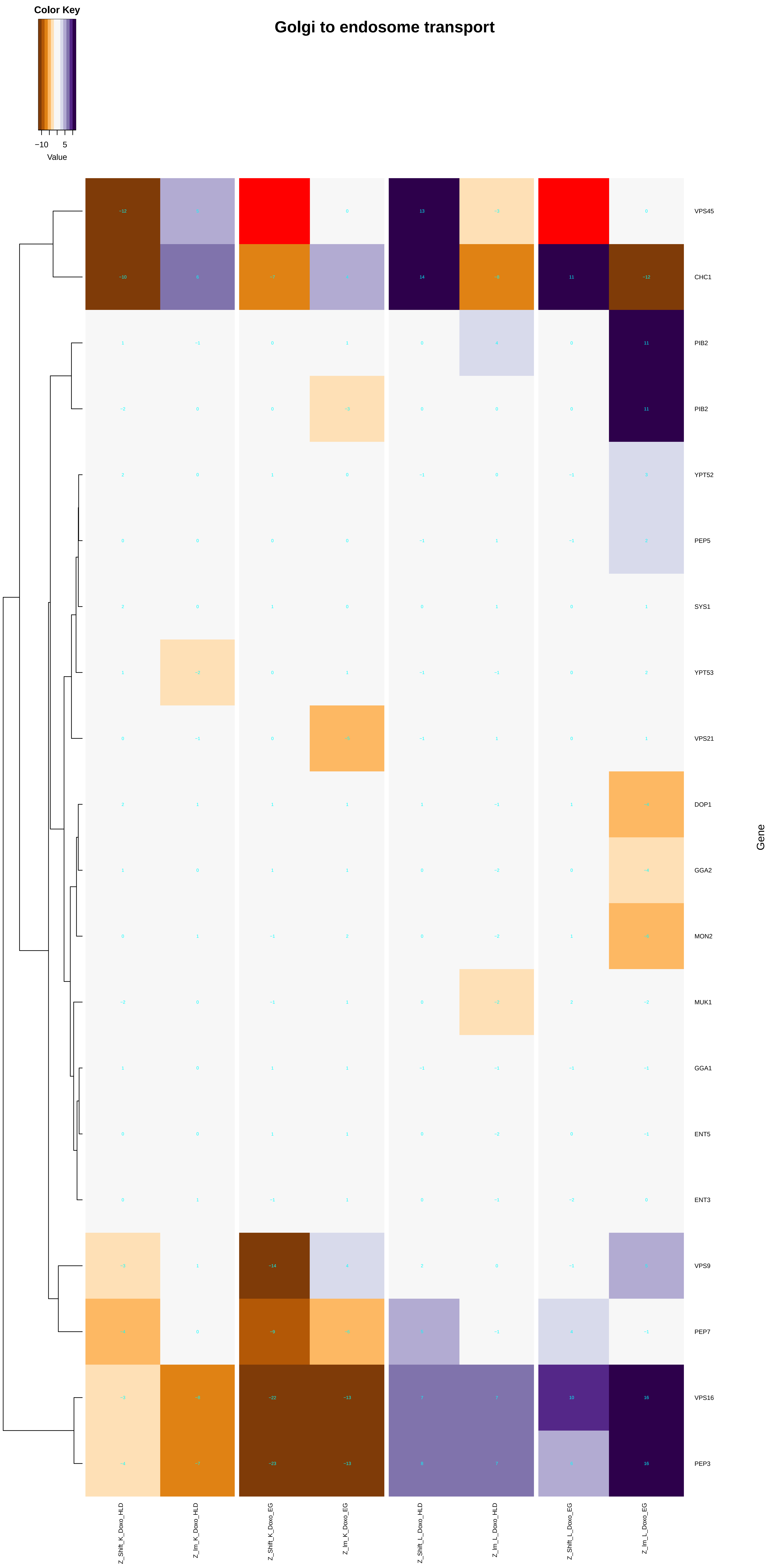

Color Key
−10 5
Value
Golgi to endosome transport
VPS45
CHC1
PIB2
PIB2
YPT52
PEP5
SYS1
YPT53
VPS21
DOP1
GGA2
MON2
MUK1
GGA1
ENT5
ENT3
VPS9
PEP7
VPS16
PEP3
Gene
Z_Shift_K_Doxo_HLD
Z_Im_K_Doxo_HLD
Z_Shift_K_Doxo_EG
Z_Im_K_Doxo_EG
Z_Shift_L_Doxo_HLD
Z_Im_L_Doxo_HLD
Z_Shift_L_Doxo_EG
Z_Im_L_Doxo_EG

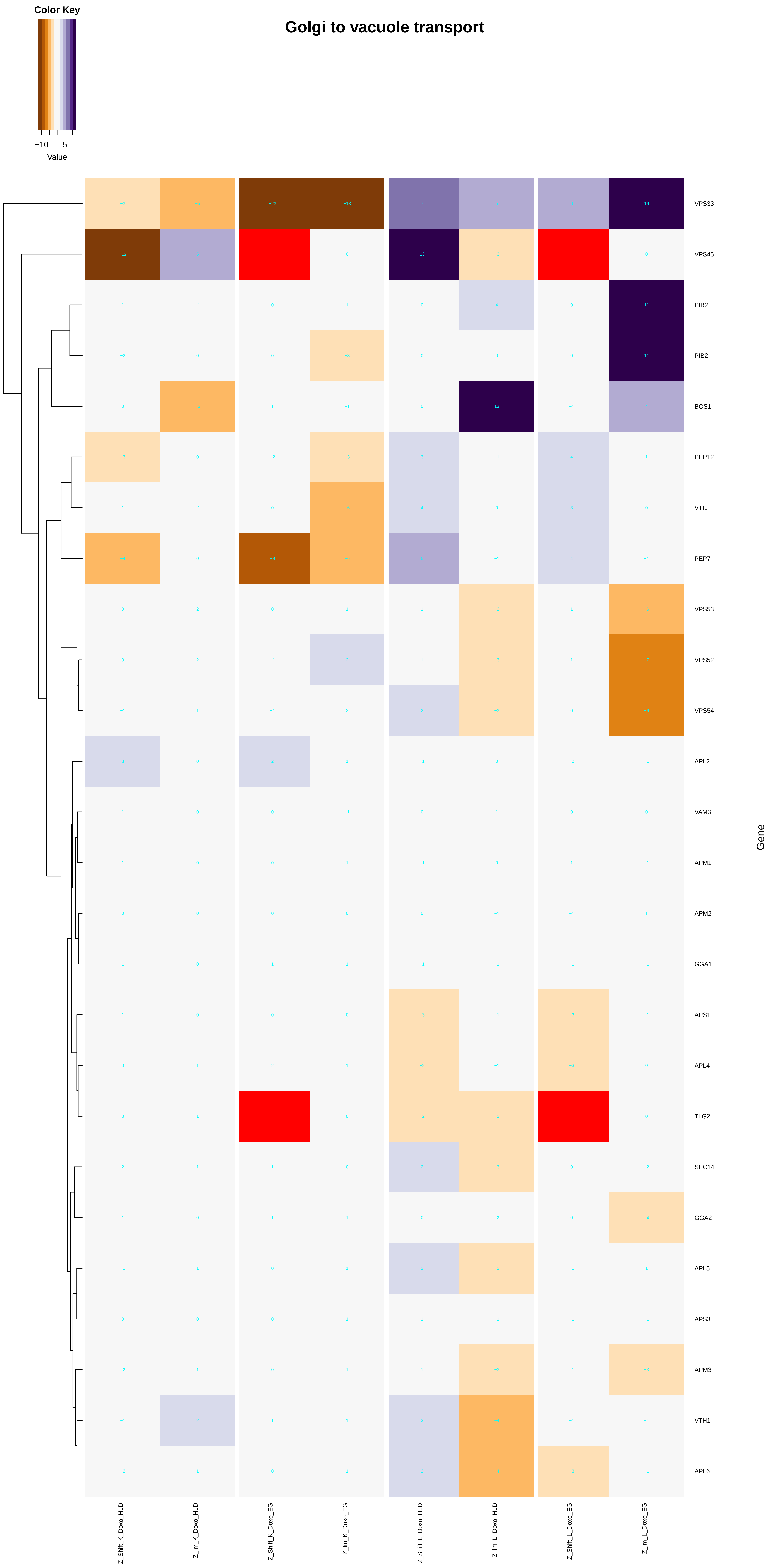

Color Key
−10
5
Value
Golgi to vacuole transport
VPS33
VPS45
PIB2
PIB2
BOS1
PEP12
VTI1
PEP7
VPS53
VPS52
VPS54
APL2
VAM3
APM1
APM2
GGA1
APS1
APL4
TLG2
SEC14
GGA2
APL5
APS3
APM3
VTH1
APL6
Gene
Z_Shift_K_Doxo_HLD
Z_lm_K_Doxo_HLD
Z_Shift_K_Doxo_EG
Z_lm_K_Doxo_EG
Z_Shift_L_Doxo_HLD
Z_lm_L_Doxo_HLD
Z_Shift_L_Doxo_EG
Z_lm_L_Doxo_EG

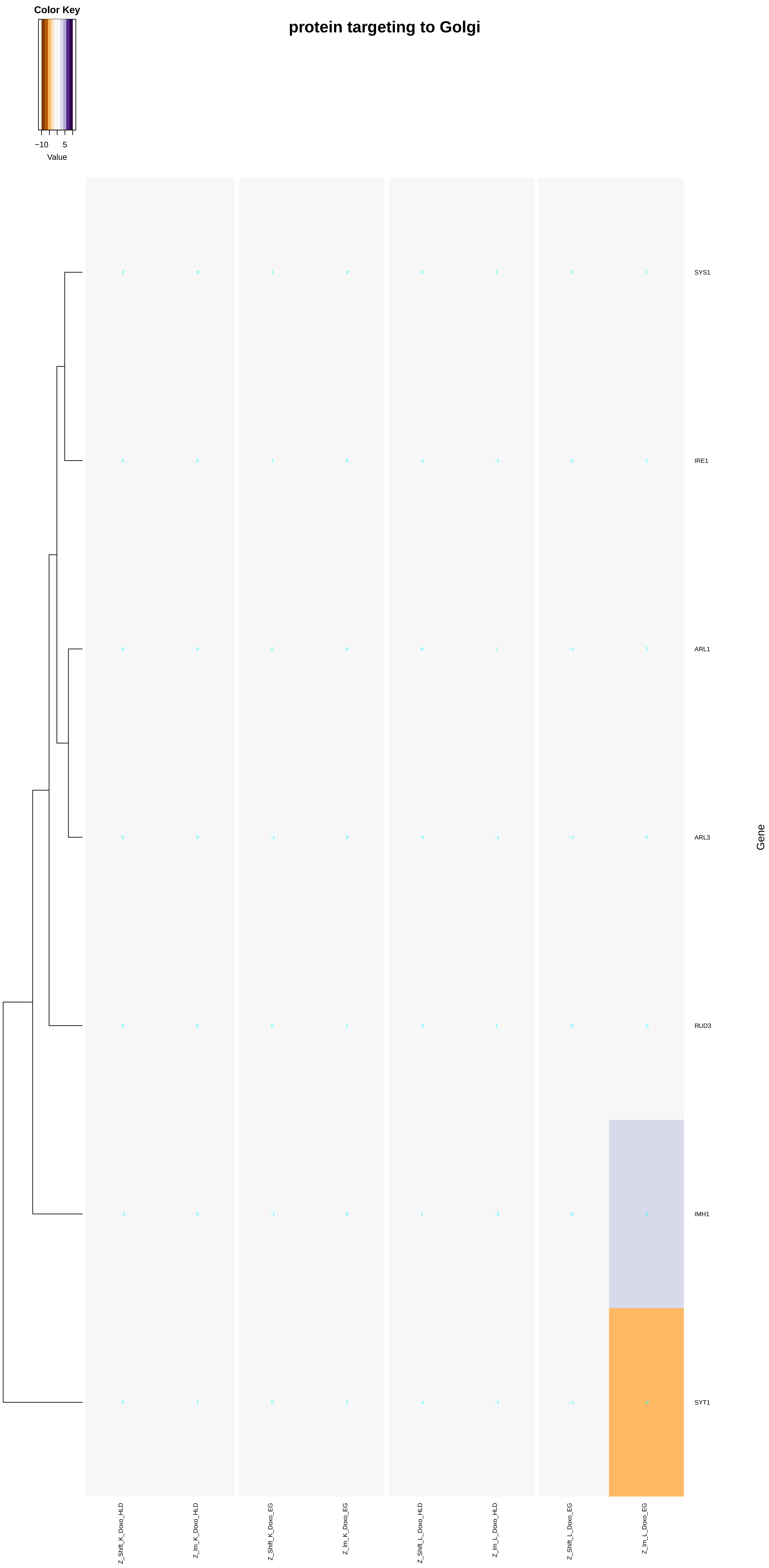

Color Key
−10
5
Value
protein targeting to Golgi
SYS1
IRE1
ARL1
ARL3
RUD3
IMH1
SYT1
Gene
Z_Shift_K_Doxo_HLD
Z_Im_K_Doxo_HLD
Z_Shift_K_Doxo_EG
Z_Im_K_Doxo_EG
Z_Shift_L_Doxo_HLD
Z_Im_L_Doxo_HLD
Z_Shift_L_Doxo_EG
Z_Im_L_Doxo_EG

### Golgi to plasma membrane protein transport

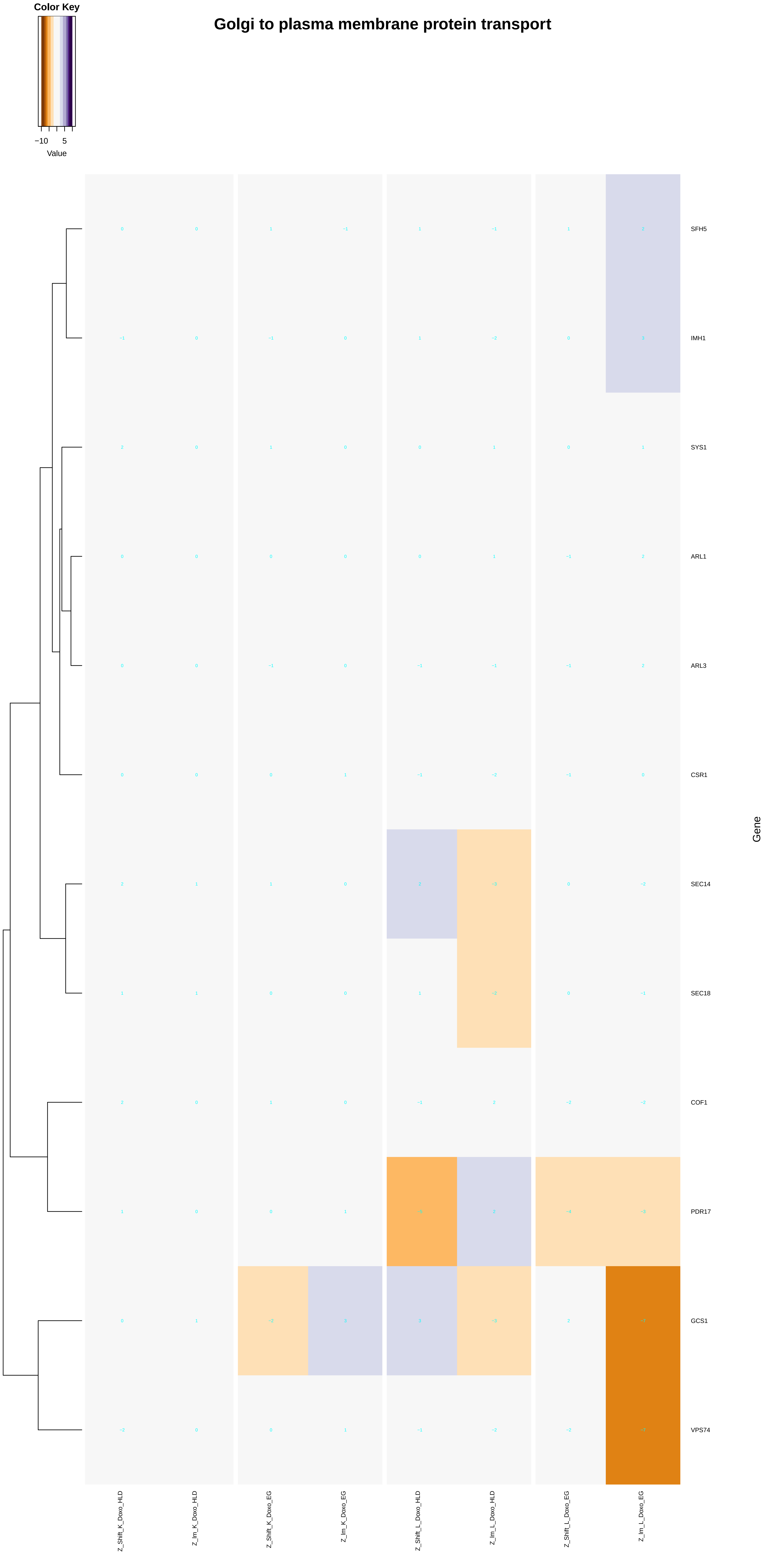
