## Additional File 9 for "A yeast phenomic model for the influence of Warburg metabolism on genetic buffering of doxorubicin": inorganic_ion_transmembrane_transport.pdf

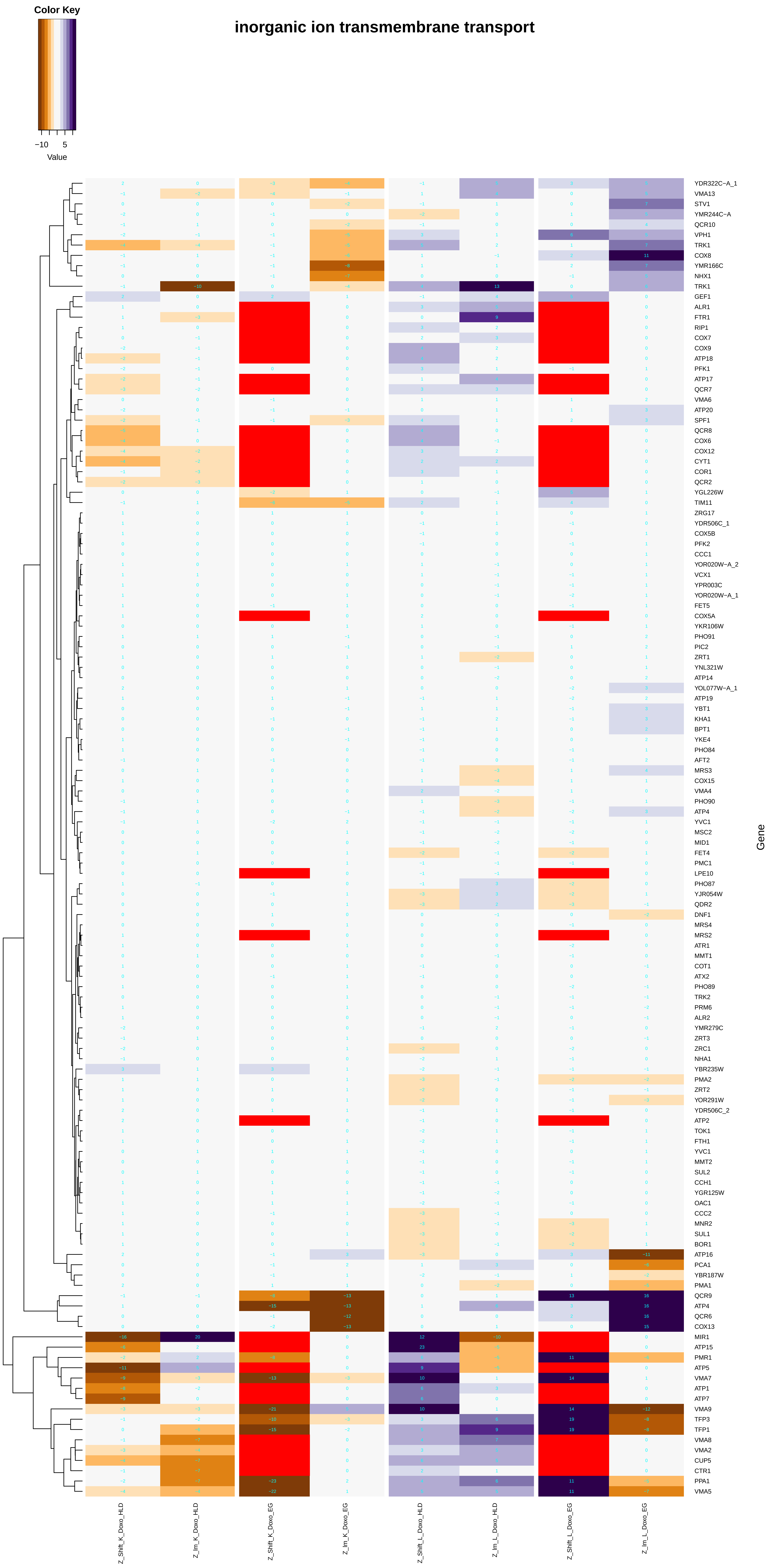
Color Key
−10
5
Value
inorganic ion transmembrane transport
YDR322C−A_1
VMA13
STV1
YMR244C−A
QCR10
VPH1
TRK1
COX8
YMR166C
NHX1
TRK1
GEF1
ALR1
FTR1
RIP1
COX7
COX9
ATP18
PFK1
ATP17
QCR7
VMA6
ATP20
SPF1
QCR8
COX6
COX12
CYT1
COR1
QCR2
YGL226W
TIM11
ZRG17
YDR506C_1
COX5B
PFK2
CCC1
YOR020W−A_2
VCX1
YPR003C
YOR020W−A_1
FET5
COX5A
YKR106W
PHO91
PIC2
ZRT1
YNL321W
ATP14
YOL077W−A_1
ATP19
YBT1
KHA1
BPT1
YKE4
PHO84
AFT2
MRS3
COX15
VMA4
PHO90
ATP4
YVC1
MSC2
MID1
FET4
PMC1
LPE10
PHO87
YJR054W
QDR2
DNF1
MRS4
MRS2
ATR1
MMT1
COT1
ATX2
PHO89
TRK2
PRM6
ALR2
YMR279C
ZRT3
ZRC1
NHA1
YBR235W
PMA2
ZRT2
YOR291W
YDR506C_2
ATP2
TOK1
FTH1
YVC1
MMT2
SUL2
CCH1
YGR125W
OAC1
CCC2
MNR2
SUL1
BOR1
ATP16
PCA1
YBR187W
PMA1
QCR9
ATP4
QCR6
COX13
MIR1
ATP15
PMR1
ATP5
VMA7
ATP1
ATP7
VMA9
TFP3
TFP1
VMA8
VMA2
CUP5
CTR1
PPA1
VMA5
Gene
Z_Shift_K_Doxo_HLD
Z_lm_K_Doxo_HLD
Z_Shift_K_Doxo_EG
Z_lm_K_Doxo_EG
Z_Shift_L_Doxo_HLD
Z_lm_L_Doxo_HLD
Z_Shift_L_Doxo_EG
Z_lm_L_Doxo_EG

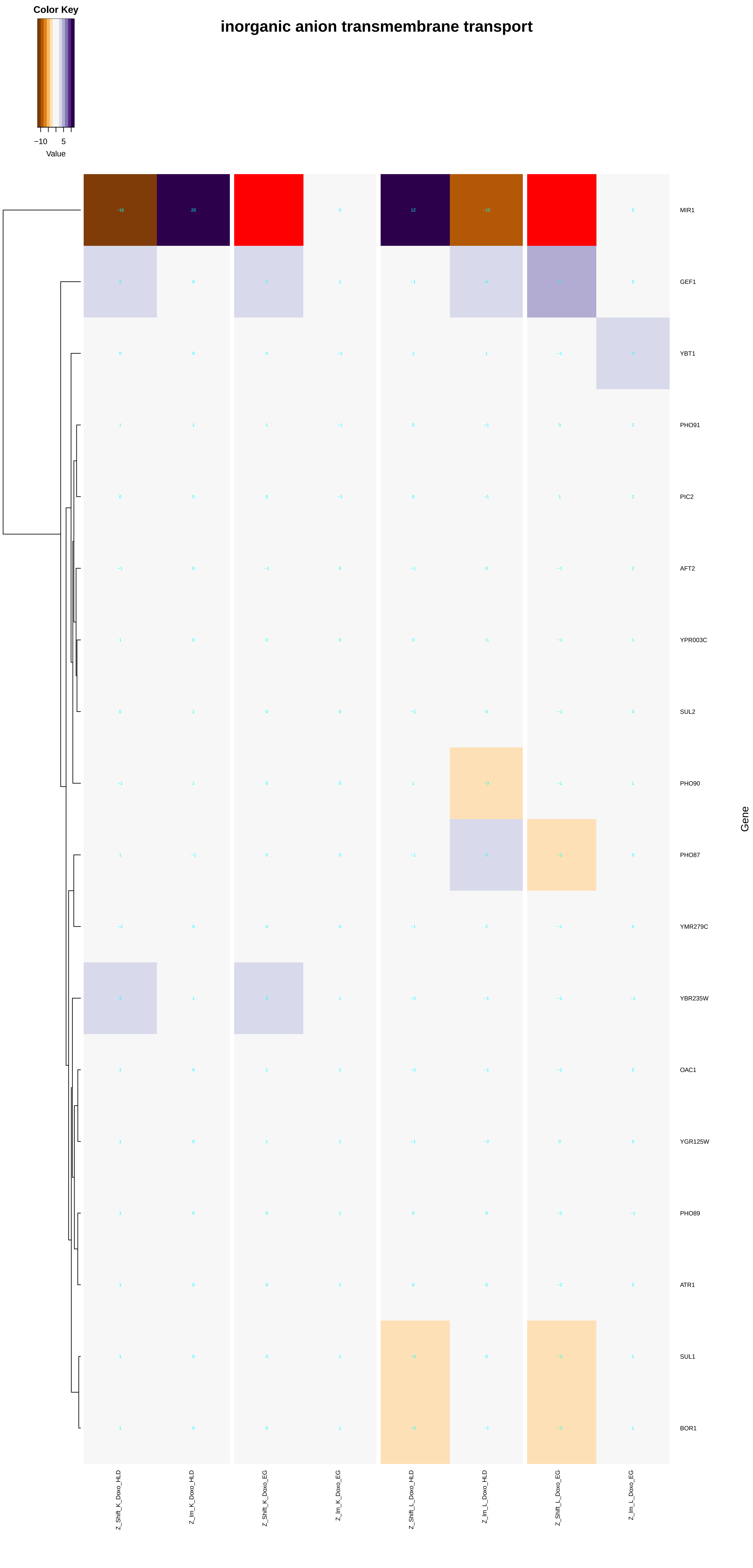
Color Key
−10
5
Value
inorganic anion transmembrane transport
MIR1
GEF1
YBT1
PHO91
PIC2
AFT2
YPR003C
SUL2
PHO90
PHO87
YMR279C
YBR235W
OAC1
YGR125W
PHO89
ATR1
SUL1
BOR1
Gene
Z_Shift_K_Doxo_HLD
Z_Im_K_Doxo_HLD
Z_Shift_K_Doxo_EG
Z_Im_K_Doxo_EG
Z_Shift_L_Doxo_HLD
Z_Im_L_Doxo_HLD
Z_Shift_L_Doxo_EG
Z_Im_L_Doxo_EG

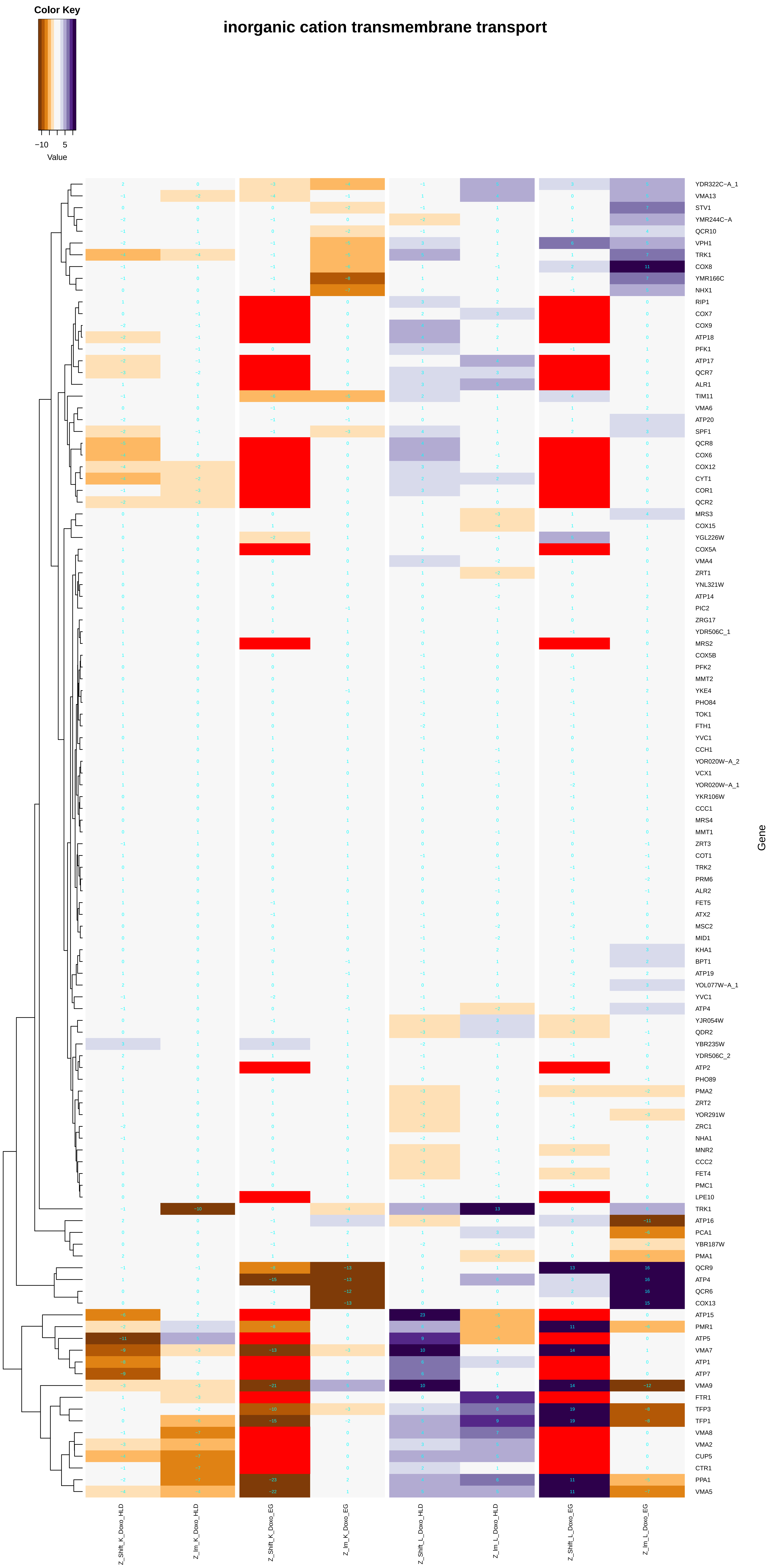

inorganic cation transmembrane transport
Color Key
−10
5
Value
YDR322C−A_1
VMA13
STV1
YMR244C−A
QCR10
VPH1
TRK1
COX8
YMR166C
NHX1
RIP1
COX7
COX9
ATP18
PFK1
ATP17
QCR7
ALR1
TIM11
VMA6
ATP20
SPF1
QCR8
COX6
COX12
CYT1
COR1
QCR2
MRS3
COX15
YGL226W
COX5A
VMA4
ZRT1
YNL321W
ATP14
PIC2
ZRG17
YDR506C_1
MRS2
COX5B
PFK2
MMT2
YKE4
PHO84
TOK1
FTH1
YVC1
CCH1
YOR020W−A_2
VCX1
YOR020W−A_1
YKR106W
CCC1
MRS4
MMT1
ZRT3
COT1
TRK2
PRM6
ALR2
FET5
ATX2
MSC2
MID1
KHA1
BPT1
ATP19
YOL077W−A_1
YVC1
ATP4
YJR054W
QDR2
YBR235W
YDR506C_2
ATP2
PHO89
PMA2
ZRT2
YOR291W
ZRC1
NHA1
MNR2
CCC2
FET4
PMC1
LPE10
TRK1
ATP16
PCA1
YBR187W
PMA1
QCR9
ATP4
QCR6
COX13
ATP15
PMR1
ATP5
VMA7
ATP1
ATP7
VMA9
FTR1
TFP3
TFP1
VMA8
VMA2
CUP5
CTR1
PPA1
VMA5
Gene
Z_Shift_K_Doxo_HLD
Z_lm_K_Doxo_HLD
Z_Shift_K_Doxo_EG
Z_lm_K_Doxo_EG
Z_Shift_L_Doxo_HLD
Z_lm_L_Doxo_HLD
Z_Shift_L_Doxo_EG
Z_lm_L_Doxo_EG

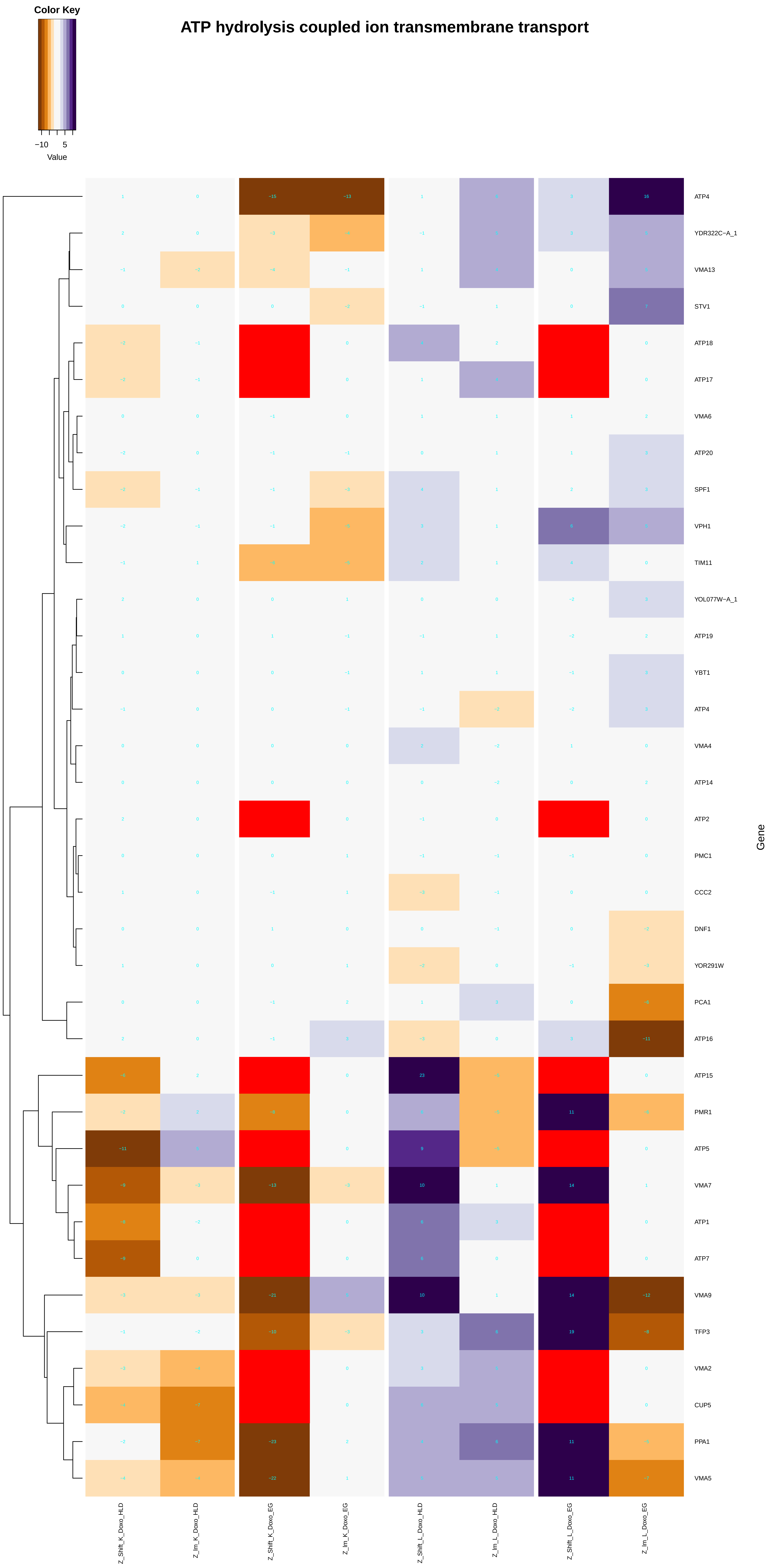
ATP hydrolysis coupled ion transmembrane transport
Color Key
−10
5
Value
ATP4
YDR322C−A_1
VMA13
STV1
ATP18
ATP17
VMA6
ATP20
SPF1
VPH1
TIM11
YOL077W−A_1
ATP19
YBT1
ATP4
VMA4
ATP14
ATP2
PMC1
CCC2
DNF1
YOR291W
PCA1
ATP16
ATP15
PMR1
ATP5
VMA7
ATP1
ATP7
VMA9
TFP3
VMA2
CUP5
PPA1
VMA5
Gene
Z_Shift_K_Doxo_HLD
Z_Im_K_Doxo_HLD
Z_Shift_K_Doxo_EG
Z_Im_K_Doxo_EG
Z_Shift_L_Doxo_HLD
Z_Im_L_Doxo_HLD
Z_Shift_L_Doxo_EG
Z_Im_L_Doxo_EG

### phosphate ion transmembrane transport

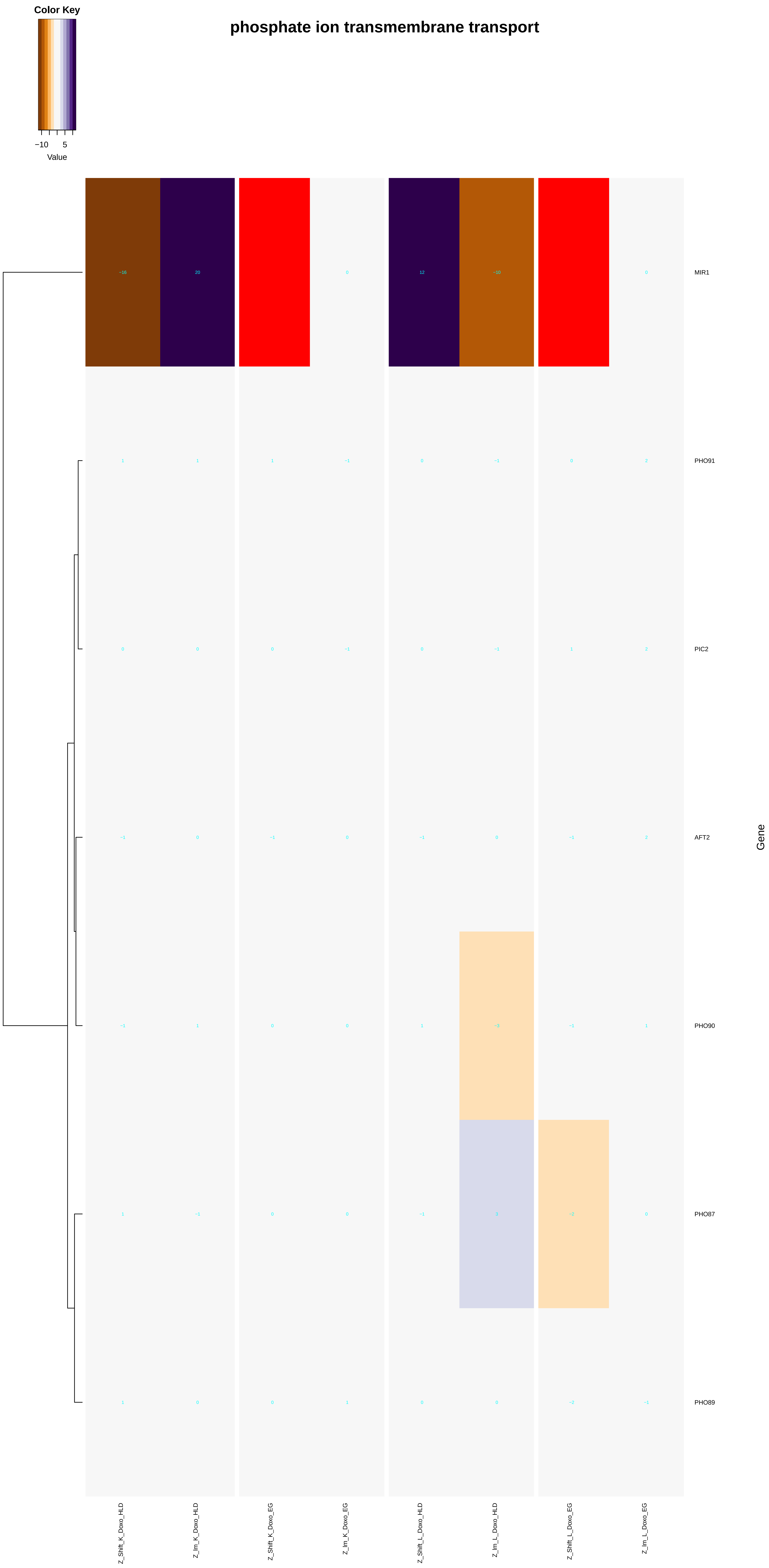

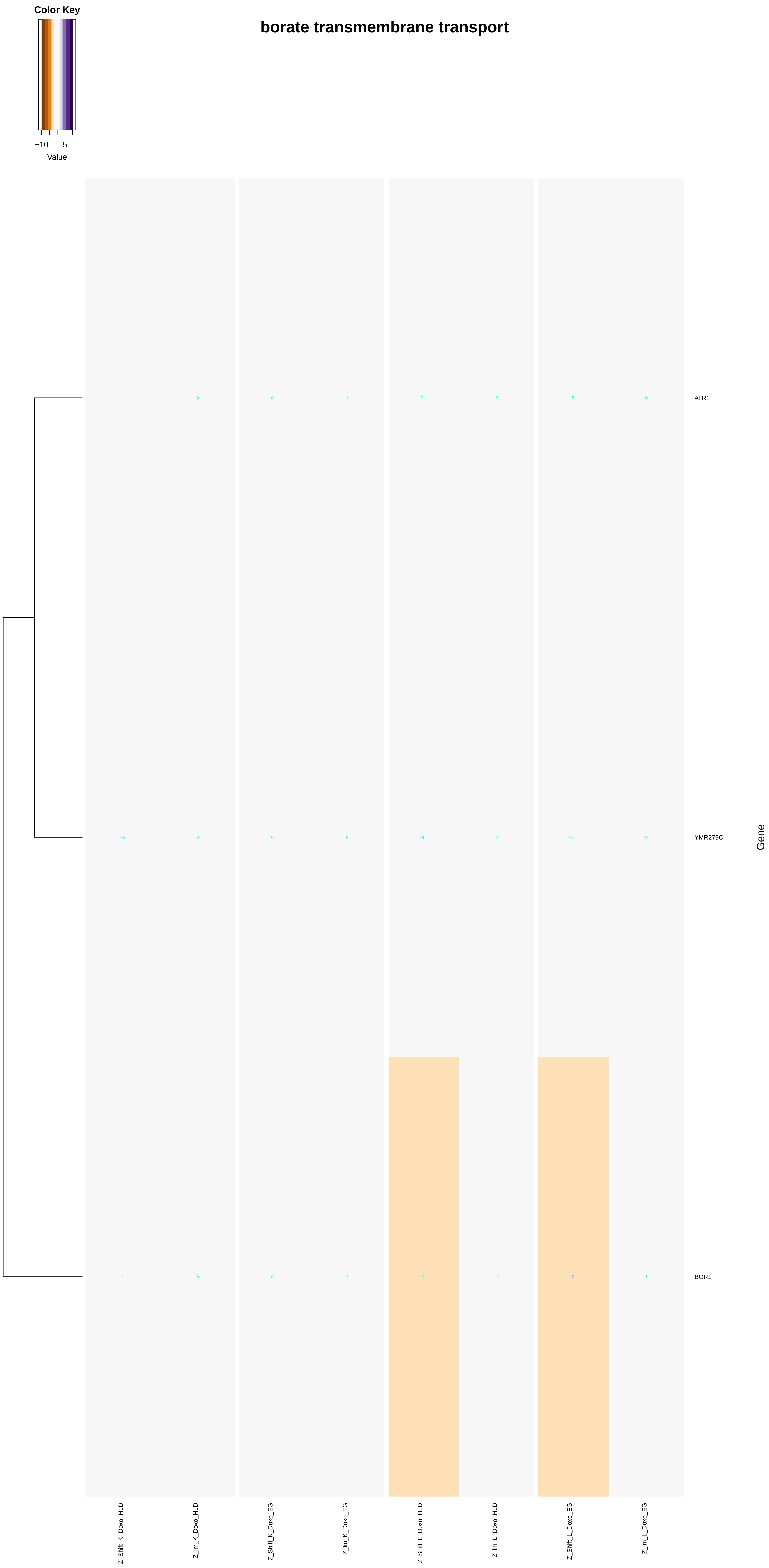
Color Key
−10
5
Value
borate transmembrane transport
1 0 0 1 0 0 −2 0
ATR1
−2 0 0 0 −1 2 −1 0
YMR279C
1 0 0 1 −3 −1 −2 1
BOR1
Gene
Z_Shift_K_Doxo_HLD
Z_lm_K_Doxo_HLD
Z_Shift_K_Doxo_EG
Z_lm_K_Doxo_EG
Z_Shift_L_Doxo_HLD
Z_lm_L_Doxo_HLD
Z_Shift_L_Doxo_EG
Z_lm_L_Doxo_EG

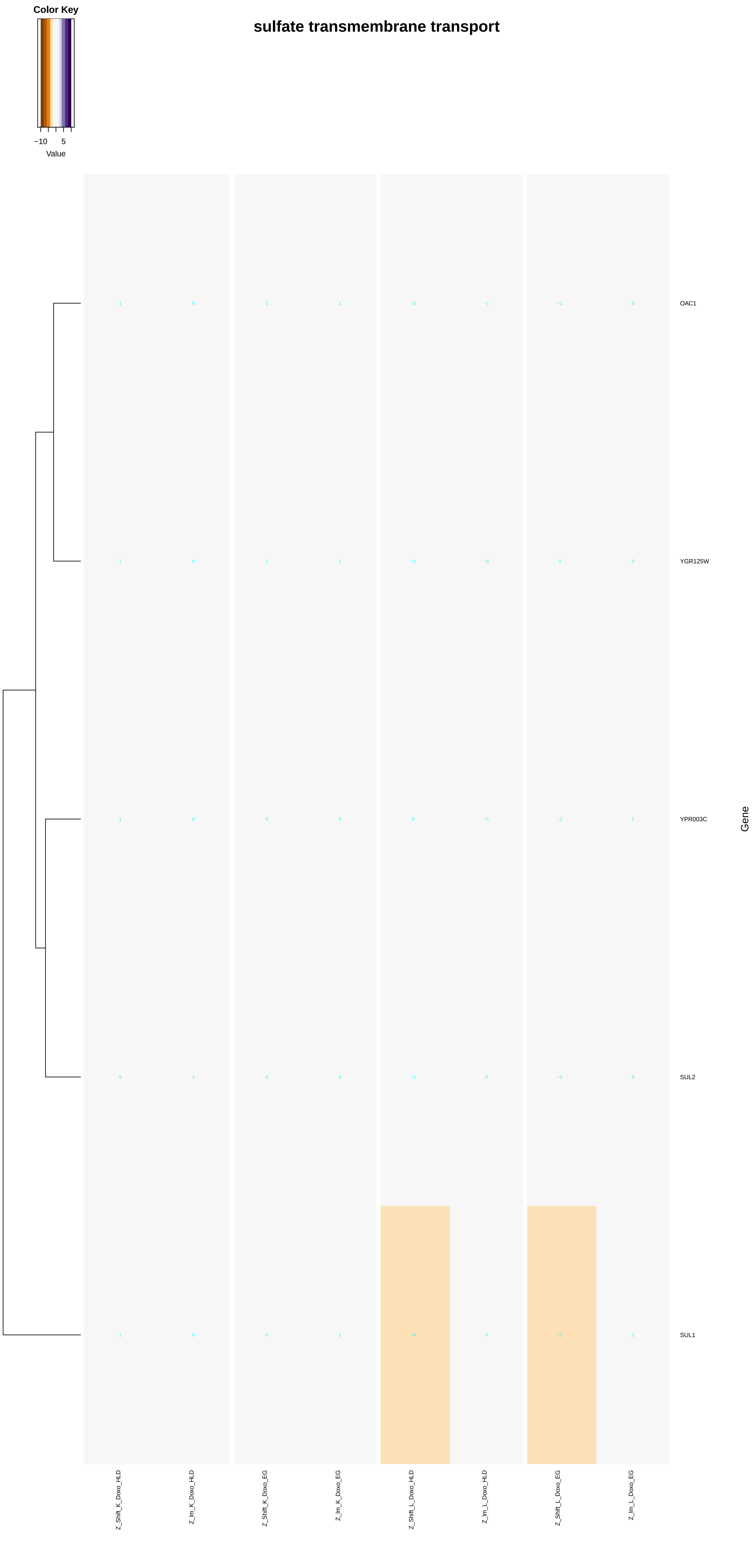
Color Key
−10
5
Value
sulfate transmembrane transport
OAC1
YGR125W
YPR003C
SUL2
SUL1
Gene
Z_Shift_K_Doxo_HLD
Z_lm_K_Doxo_HLD
Z_Shift_K_Doxo_EG
Z_lm_K_Doxo_EG
Z_Shift_L_Doxo_HLD
Z_lm_L_Doxo_HLD
Z_Shift_L_Doxo_EG
Z_lm_L_Doxo_EG

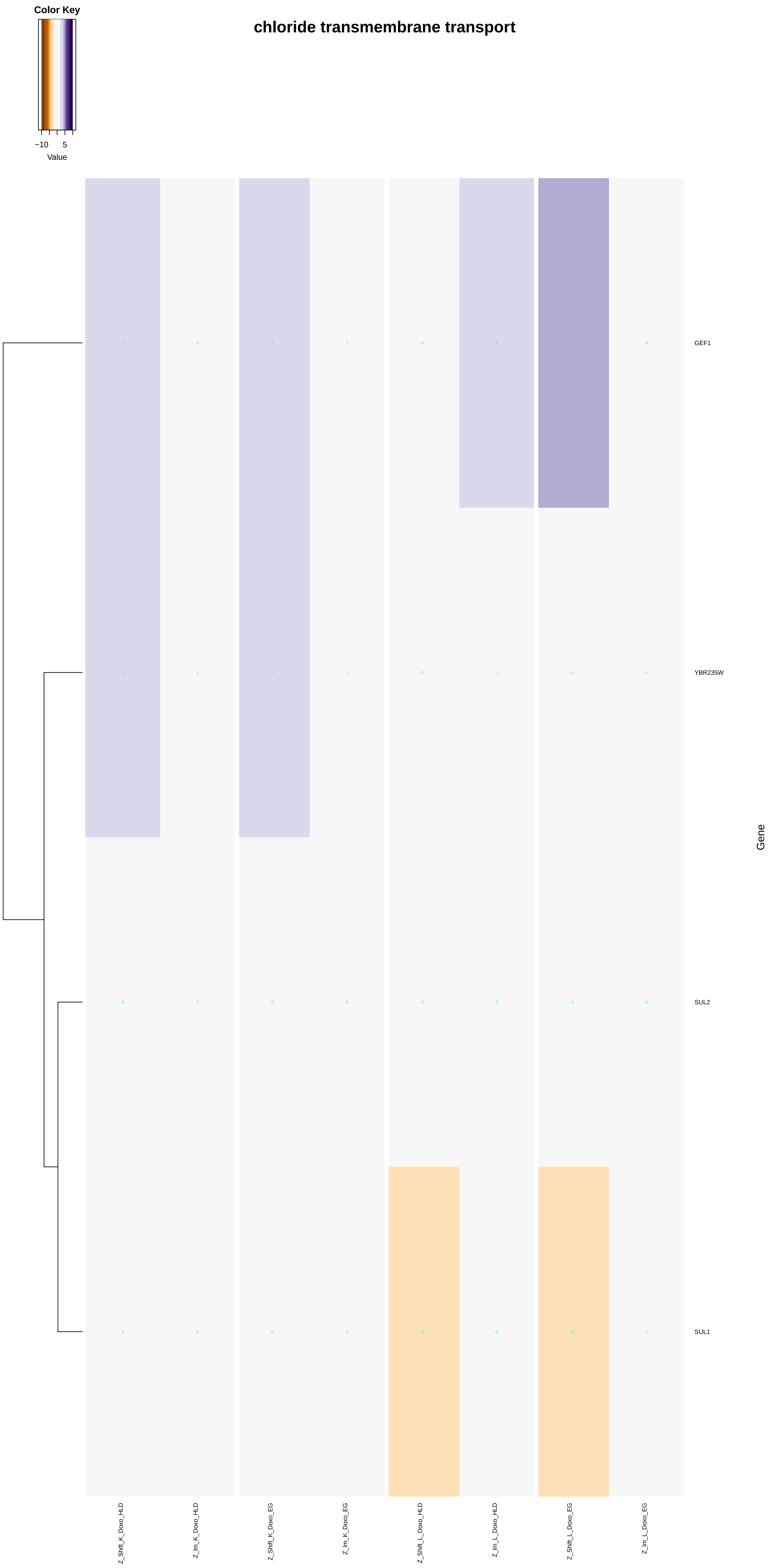
chloride transmembrane transport
Color Key
−10
5
Value
GEF1
YBR235W
SUL2
SUL1
Gene
Z_Shift_K_Doxo_HLD
Z_lm_K_Doxo_HLD
Z_Shift_K_Doxo_EG
Z_lm_K_Doxo_EG
Z_Shift_L_Doxo_HLD
Z_lm_L_Doxo_HLD
Z_Shift_L_Doxo_EG
Z_lm_L_Doxo_EG

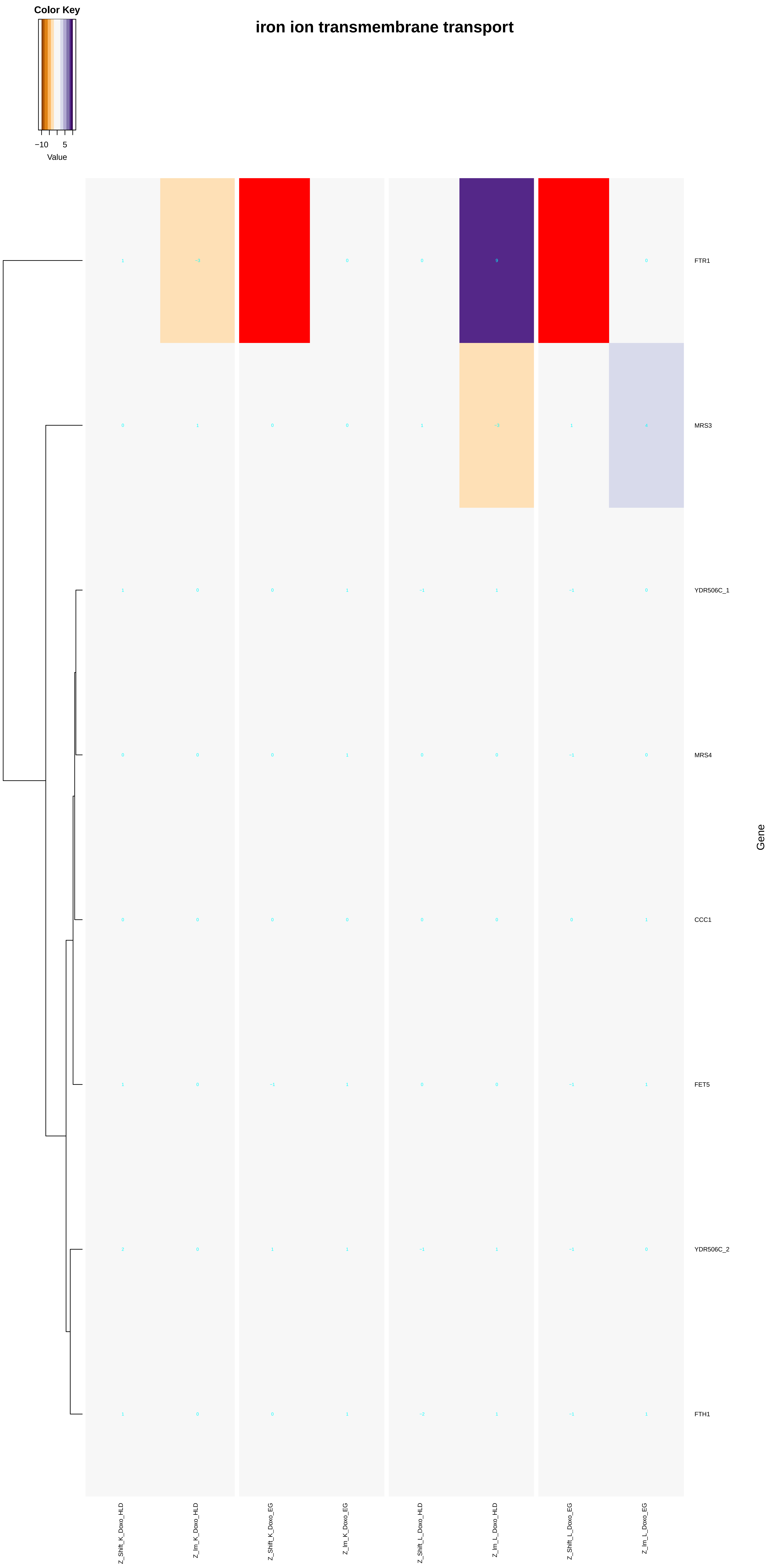
Color Key
−10
5
Value
iron ion transmembrane transport
FTR1
MRS3
YDR506C_1
MRS4
CCC1
FET5
YDR506C_2
FTH1
Gene
Z_Shift_K_Doxo_HLD
Z_lm_K_Doxo_HLD
Z_Shift_K_Doxo_EG
Z_lm_K_Doxo_EG
Z_Shift_L_Doxo_HLD
Z_lm_L_Doxo_HLD
Z_Shift_L_Doxo_EG
Z_lm_L_Doxo_EG

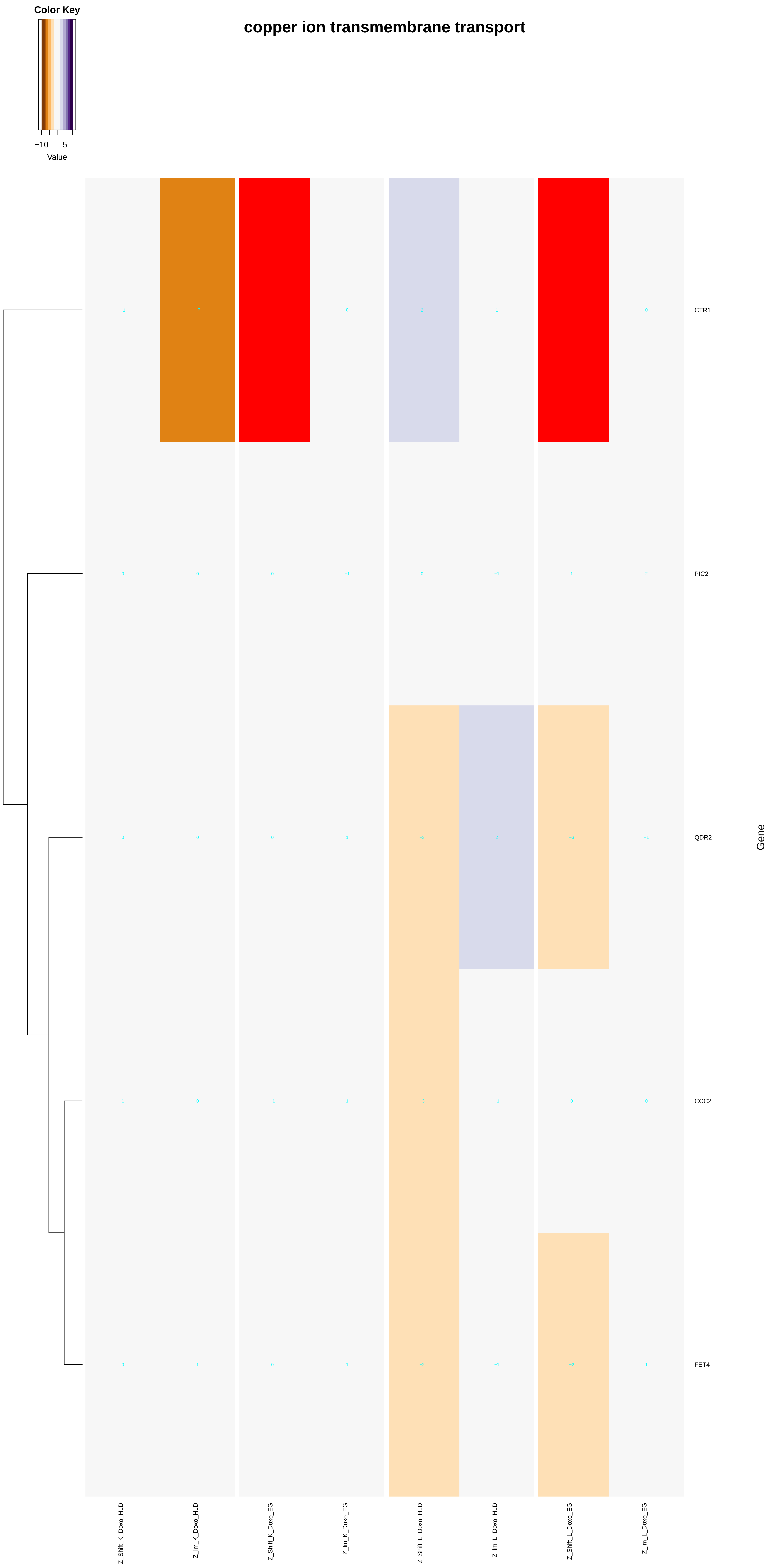

Color Key
−10
5
Value
copper ion transmembrane transport
CTR1
PIC2
QDR2
CCC2
FET4
Gene
Z_Shift_K_Doxo_HLD
Z_Im_K_Doxo_HLD
Z_Shift_K_Doxo_EG
Z_Im_K_Doxo_EG
Z_Shift_L_Doxo_HLD
Z_Im_L_Doxo_HLD
Z_Shift_L_Doxo_EG
Z_Im_L_Doxo_EG

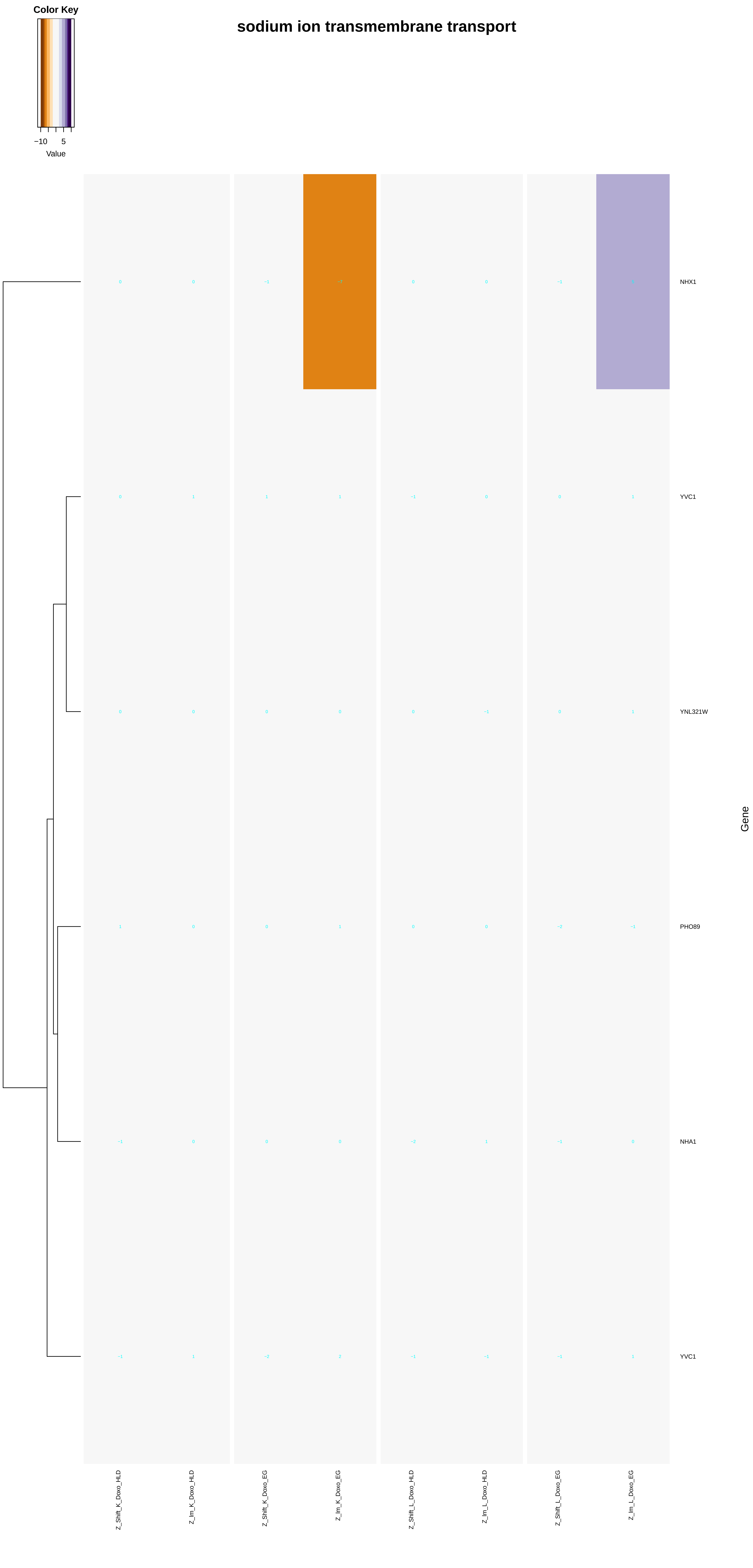

Color Key
−10
5
Value
sodium ion transmembrane transport
NHX1
YVC1
YNL321W
PHO89
NHA1
YVC1
Gene
Z_Shift_K_Doxo_HLD
Z_Im_K_Doxo_HLD
Z_Shift_K_Doxo_EG
Z_Im_K_Doxo_EG
Z_Shift_L_Doxo_HLD
Z_Im_L_Doxo_HLD
Z_Shift_L_Doxo_EG
Z_Im_L_Doxo_EG

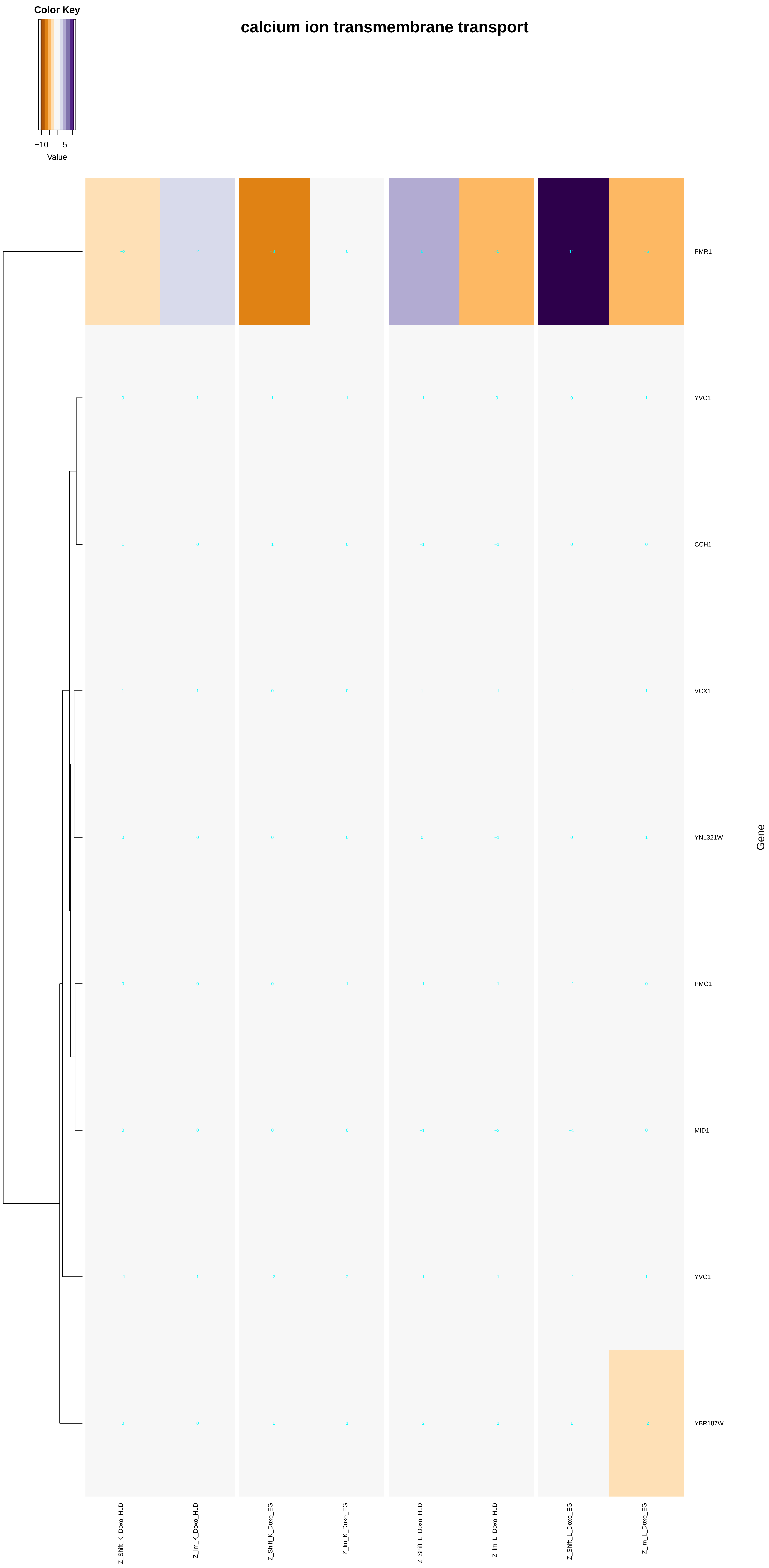
Color Key
−10
5
Value
calcium ion transmembrane transport
PMR1
YVC1
CCH1
VCX1
YNL321W
PMC1
MID1
YVC1
YBR187W
Gene
Z_Shift_K_Doxo_HLD
Z_Im_K_Doxo_HLD
Z_Shift_K_Doxo_EG
Z_Im_K_Doxo_EG
Z_Shift_L_Doxo_HLD
Z_Im_L_Doxo_HLD
Z_Shift_L_Doxo_EG
Z_Im_L_Doxo_EG

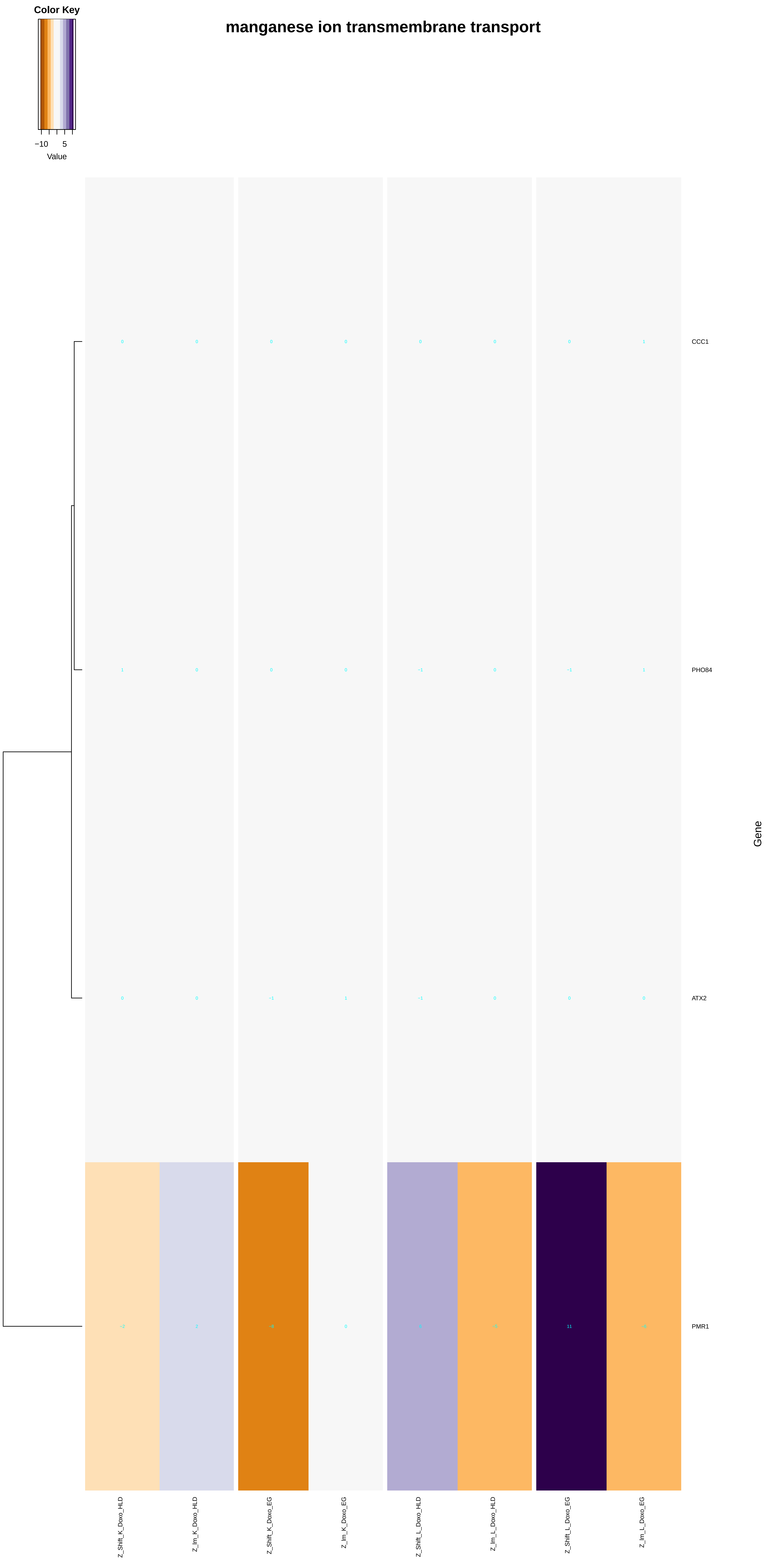
manganese ion transmembrane transport
Color Key
−10
5
Value
CCC1
PHO84
ATX2
PMR1
Gene
Z_Shift_K_Doxo_HLD
Z_Im_K_Doxo_HLD
Z_Shift_K_Doxo_EG
Z_Im_K_Doxo_EG
Z_Shift_L_Doxo_HLD
Z_Im_L_Doxo_HLD
Z_Shift_L_Doxo_EG
Z_Im_L_Doxo_EG

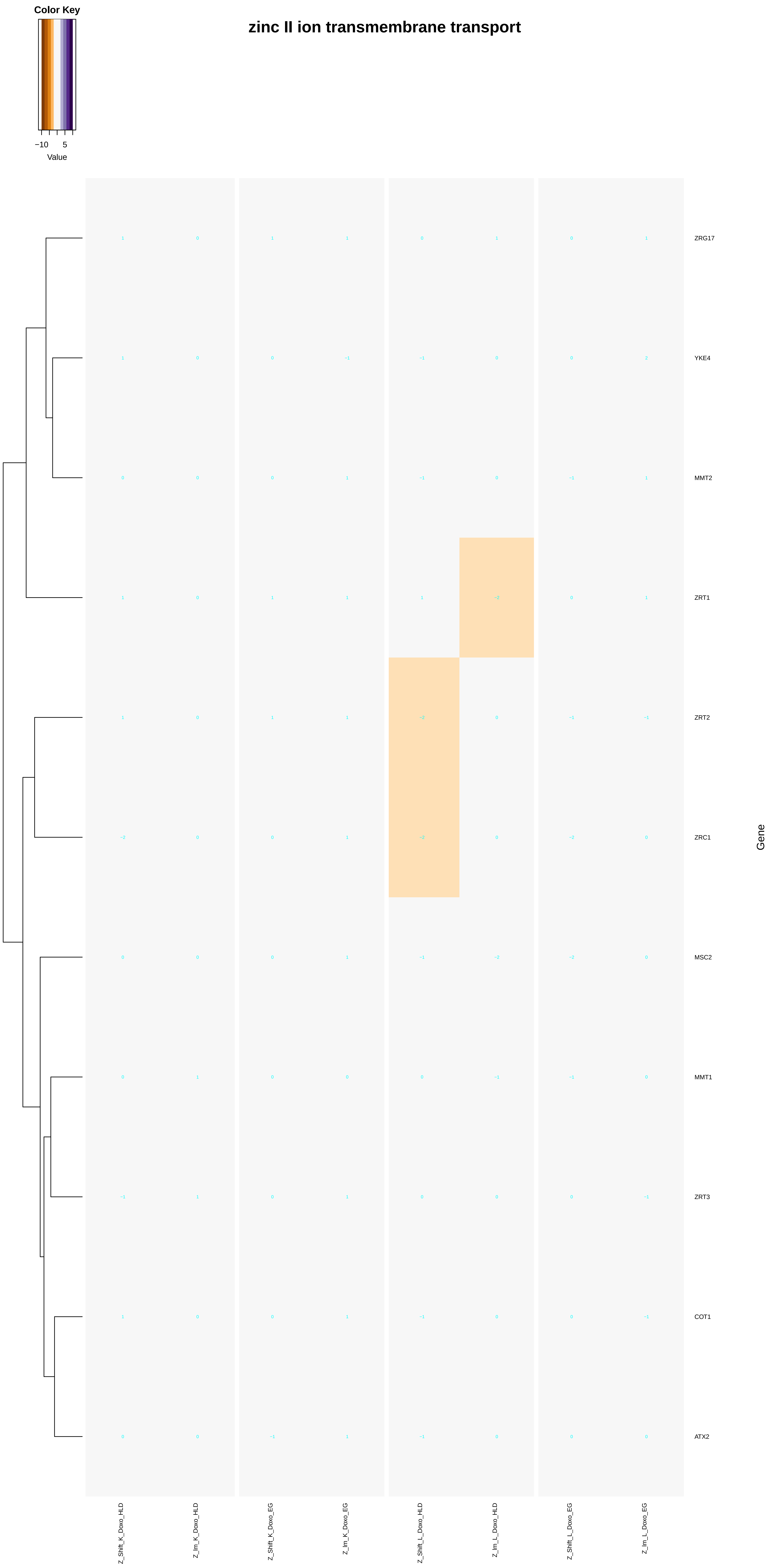

Color Key
−10
5
Value
zinc II ion transmembrane transport
ZRG17
YKE4
MMT2
ZRT1
ZRT2
ZRC1
MSC2
MMT1
ZRT3
COT1
ATX2
Gene
Z_Shift_K_Doxo_HLD
Z_Im_K_Doxo_HLD
Z_Shift_K_Doxo_EG
Z_Im_K_Doxo_EG
Z_Shift_L_Doxo_HLD
Z_Im_L_Doxo_HLD
Z_Shift_L_Doxo_EG
Z_Im_L_Doxo_EG

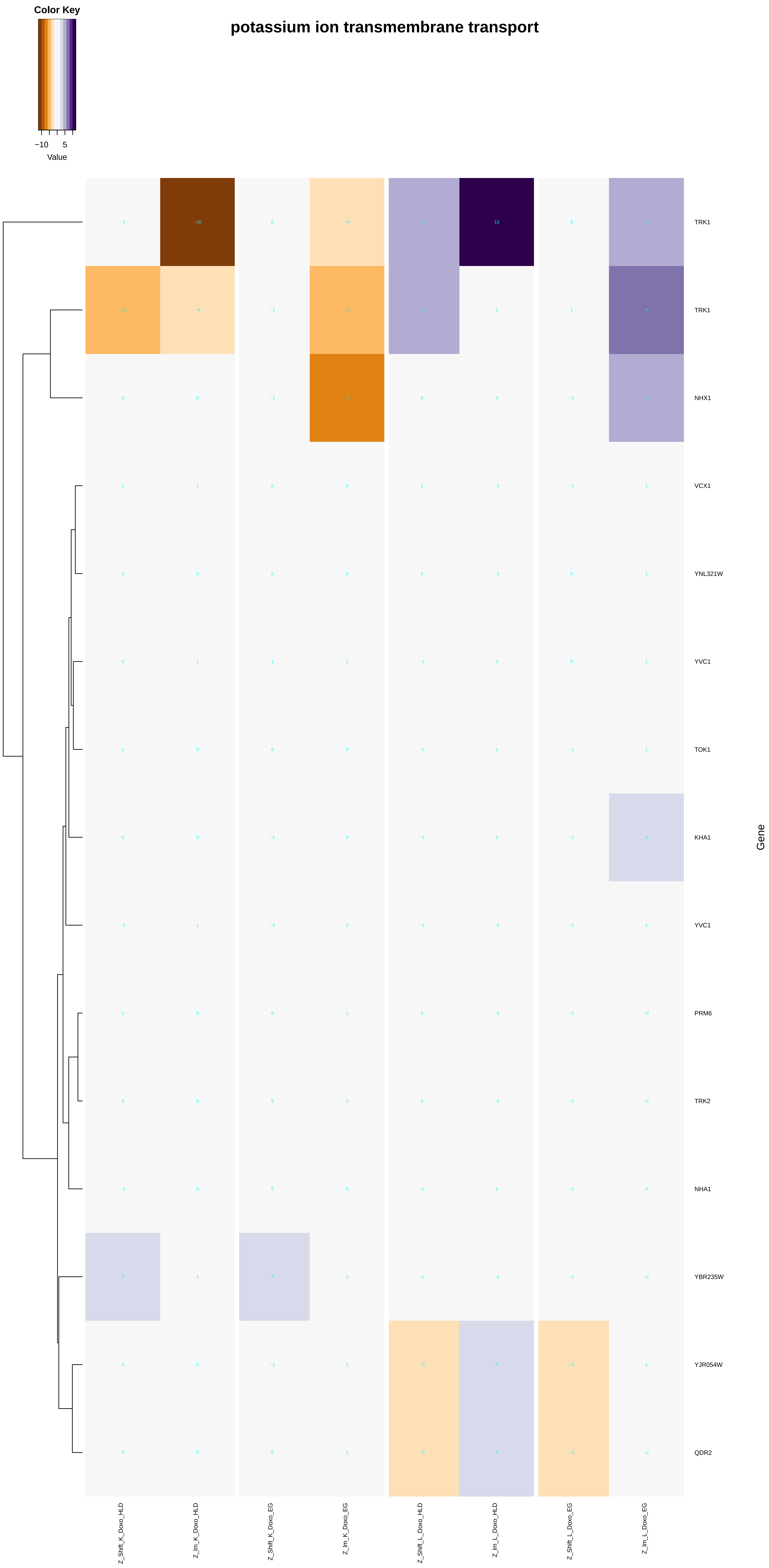

Color Key
−10
5
Value
potassium ion transmembrane transport
TRK1
TRK1
NHX1
VCX1
YNL321W
YVC1
TOK1
KHA1
YVC1
PRM6
TRK2
NHA1
YBR235W
YJR054W
QDR2
Gene
Z_Shift_K_Doxo_HLD
Z_Im_K_Doxo_HLD
Z_Shift_K_Doxo_EG
Z_Im_K_Doxo_EG
Z_Shift_L_Doxo_HLD
Z_Im_L_Doxo_HLD
Z_Shift_L_Doxo_EG
Z_Im_L_Doxo_EG

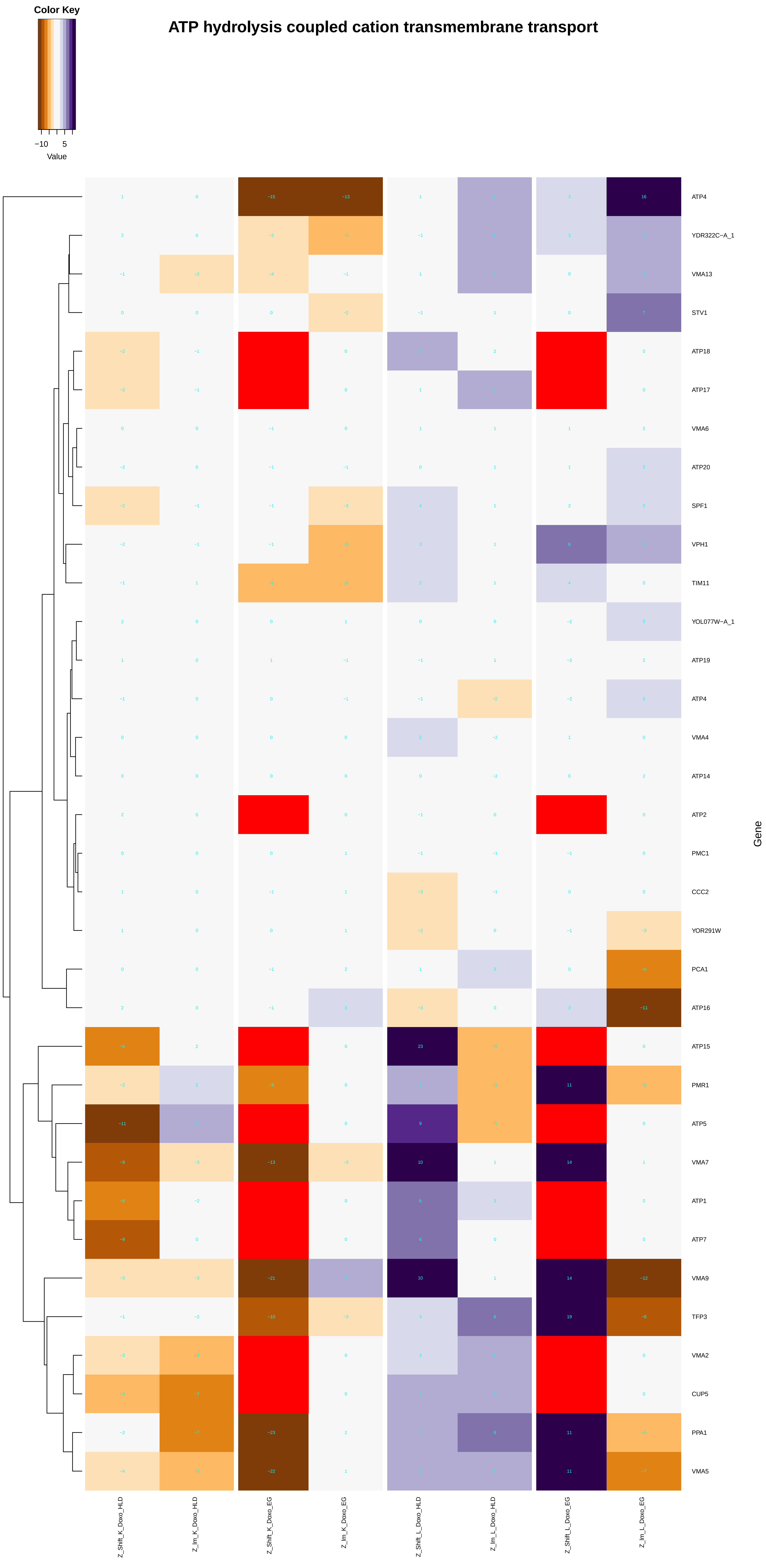
Color Key
−10
5
Value
ATP hydrolysis coupled cation transmembrane transport
ATP4
YDR322C−A_1
VMA13
STV1
ATP18
ATP17
VMA6
ATP20
SPF1
VPH1
TIM11
YOL077W−A_1
ATP19
ATP4
VMA4
ATP14
ATP2
PMC1
CCC2
YOR291W
PCA1
ATP16
ATP15
PMR1
ATP5
VMA7
ATP1
ATP7
VMA9
TFP3
VMA2
CUP5
PPA1
VMA5
Gene
Z_Shift_K_Doxo_HLD
Z_lm_K_Doxo_HLD
Z_Shift_K_Doxo_EG
Z_lm_K_Doxo_EG
Z_Shift_L_Doxo_HLD
Z_lm_L_Doxo_HLD
Z_Shift_L_Doxo_EG
Z_lm_L_Doxo_EG

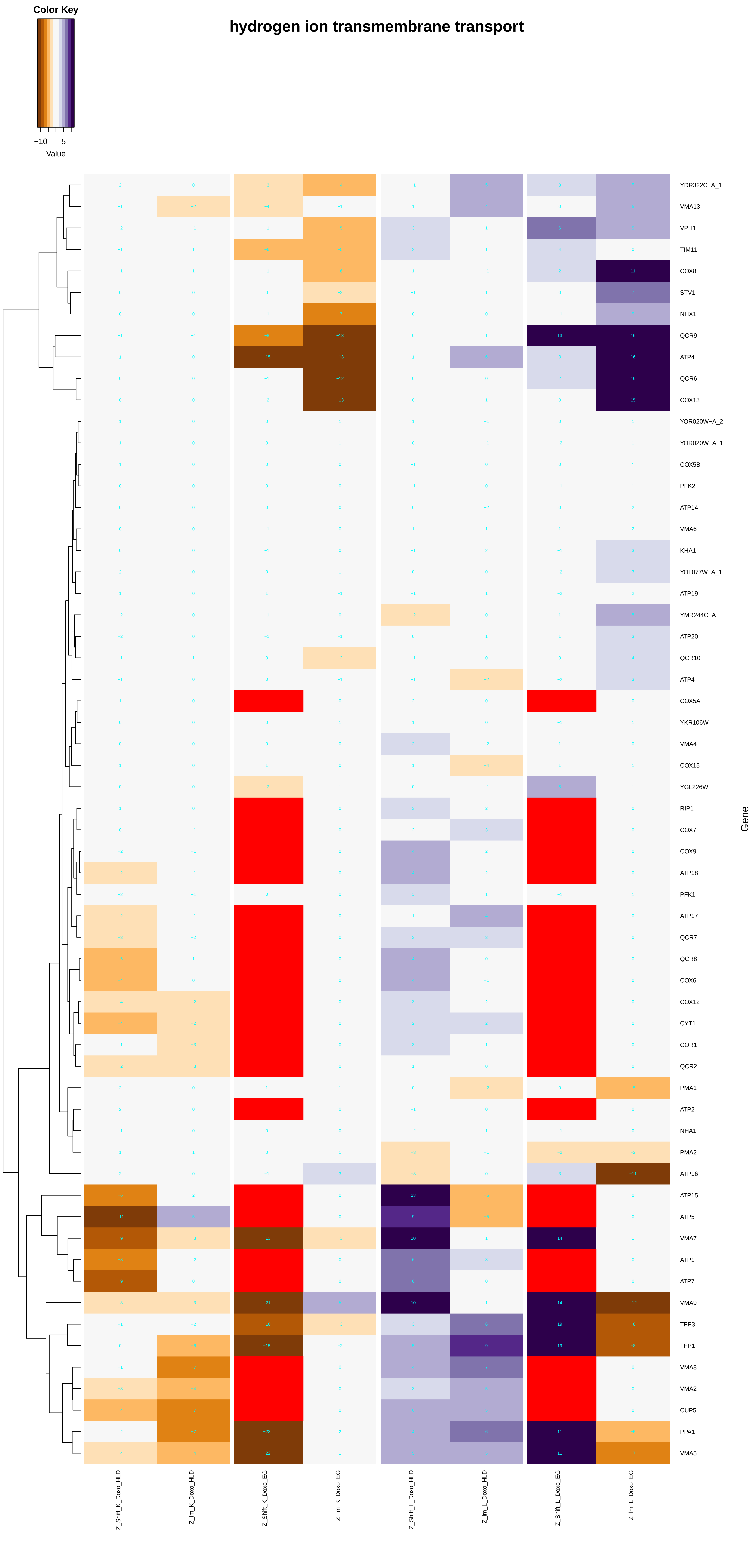

Color Key
−10
5
Value
hydrogen ion transmembrane transport
YDR322C−A_1
VMA13
VPH1
TIM11
COX8
STV1
NHX1
QCR9
ATP4
QCR6
COX13
YOR020W−A_2
YOR020W−A_1
COX5B
PFK2
ATP14
VMA6
KHA1
YOL077W−A_1
ATP19
YMR244C−A
ATP20
QCR10
ATP4
COX5A
YKR106W
VMA4
COX15
YGL226W
RIP1
COX7
COX9
ATP18
PFK1
ATP17
QCR7
QCR8
COX6
COX12
CYT1
COR1
QCR2
PMA1
ATP2
NHA1
PMA2
ATP16
ATP15
ATP5
VMA7
ATP1
ATP7
VMA9
TFP3
TFP1
VMA8
VMA2
CUP5
PPA1
VMA5
Gene
Z_Shift_K_Doxo_HLD
Z_lm_K_Doxo_HLD
Z_Shift_K_Doxo_EG
Z_lm_K_Doxo_EG
Z_Shift_L_Doxo_HLD
Z_lm_L_Doxo_HLD
Z_Shift_L_Doxo_EG
Z_lm_L_Doxo_EG

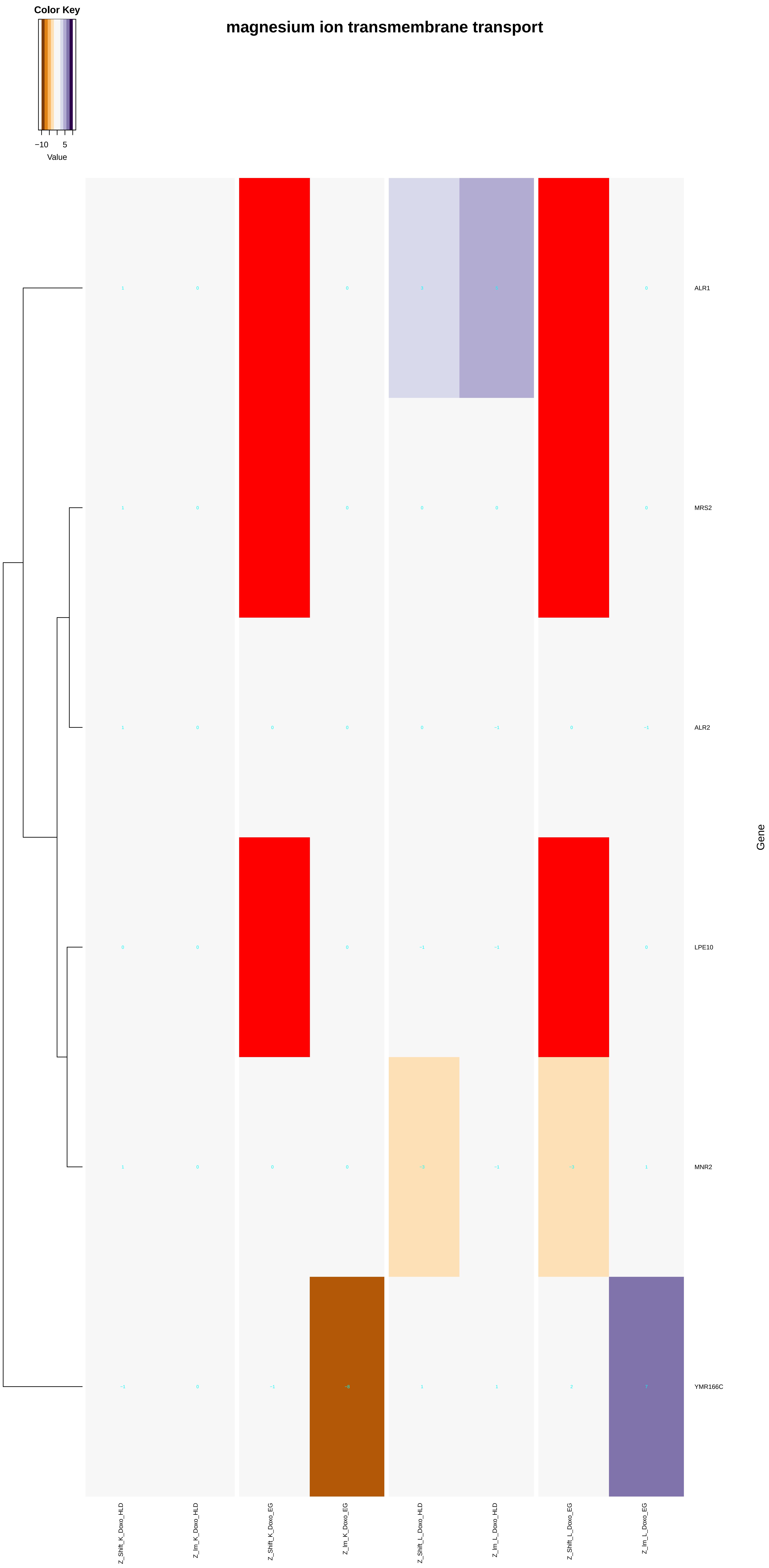
magnesium ion transmembrane transport
Color Key
−10
5
Value
ALR1
MRS2
ALR2
LPE10
MNR2
YMR166C
Gene
Z_Shift_K_Doxo_HLD
Z_lm_K_Doxo_HLD
Z_Shift_K_Doxo_EG
Z_lm_K_Doxo_EG
Z_Shift_L_Doxo_HLD
Z_lm_L_Doxo_HLD
Z_Shift_L_Doxo_EG
Z_lm_L_Doxo_EG

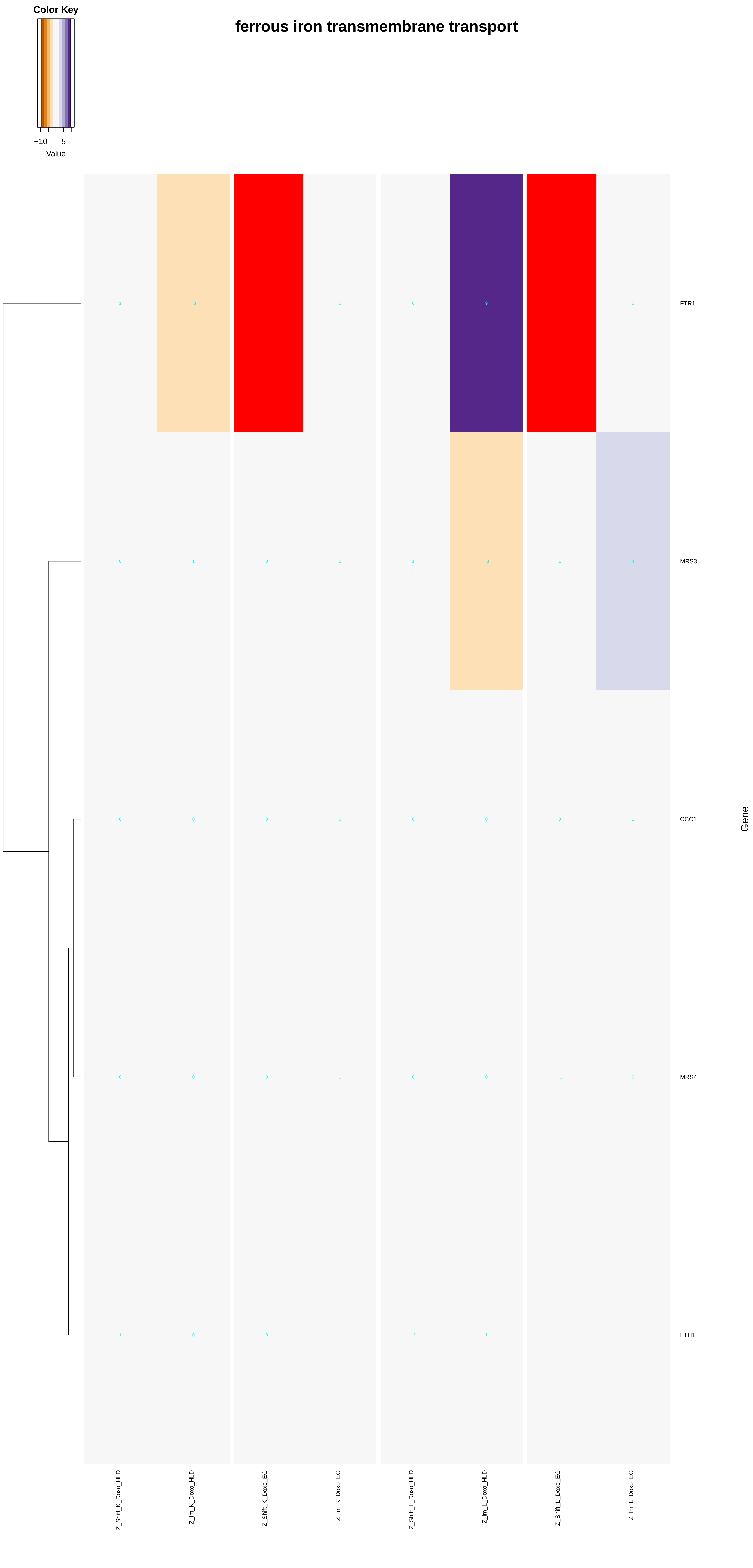
ferrous iron transmembrane transport
Color Key
−10
5
Value
FTR1
MRS3
CCC1
MRS4
FTH1
Gene
Z_Shift_K_Doxo_HLD
Z_Im_K_Doxo_HLD
Z_Shift_K_Doxo_EG
Z_Im_K_Doxo_EG
Z_Shift_L_Doxo_HLD
Z_Im_L_Doxo_HLD
Z_Shift_L_Doxo_EG
Z_Im_L_Doxo_EG

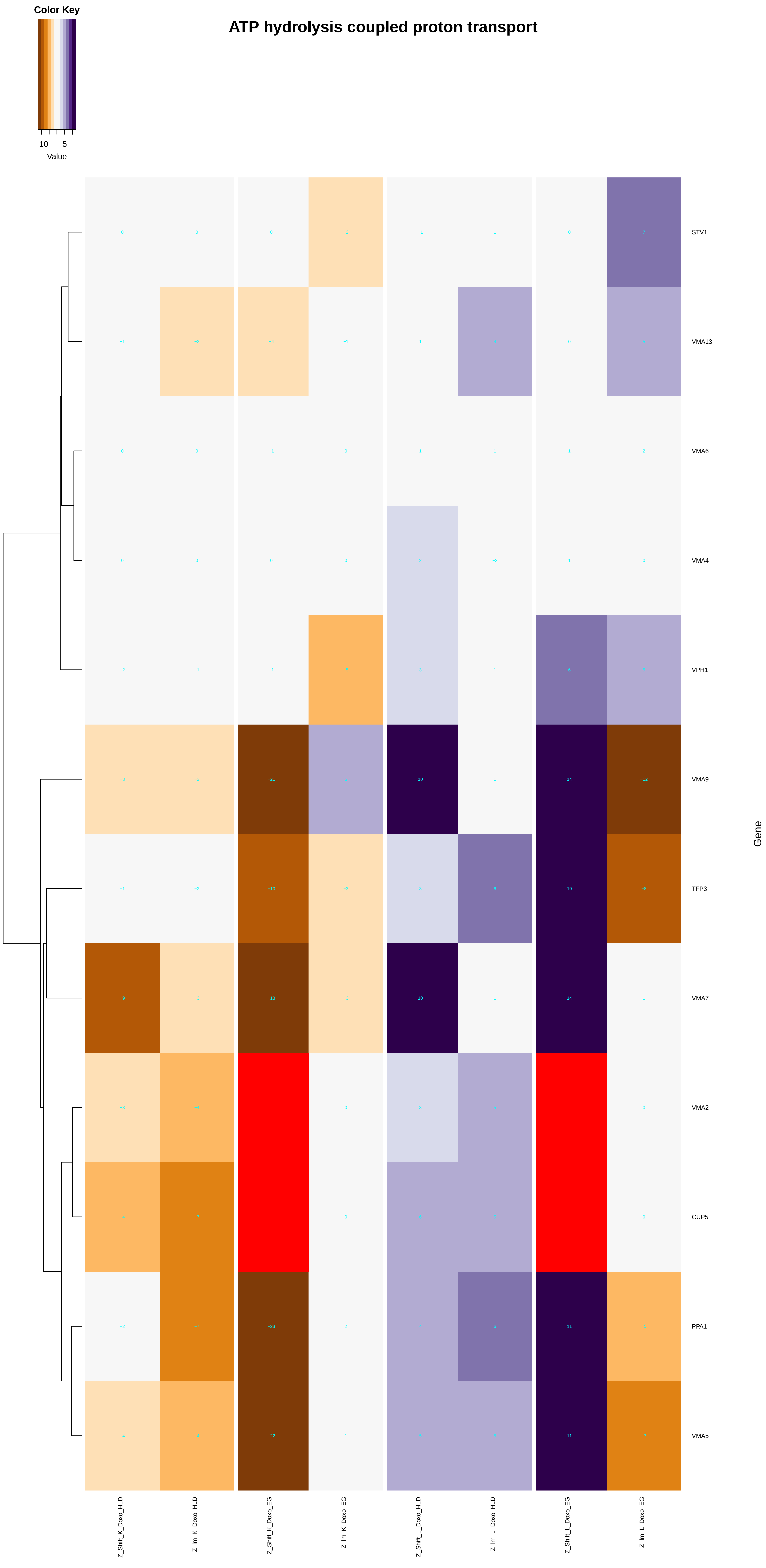
Color Key
−10
5
Value
ATP hydrolysis coupled proton transport
STV1
VMA13
VMA6
VMA4
VPH1
VMA9
TFP3
VMA7
VMA2
CUP5
PPA1
VMA5
Gene
Z_Shift_K_Doxo_HLD
Z_Im_K_Doxo_HLD
Z_Shift_K_Doxo_EG
Z_Im_K_Doxo_EG
Z_Shift_L_Doxo_HLD
Z_Im_L_Doxo_HLD
Z_Shift_L_Doxo_EG
Z_Im_L_Doxo_EG

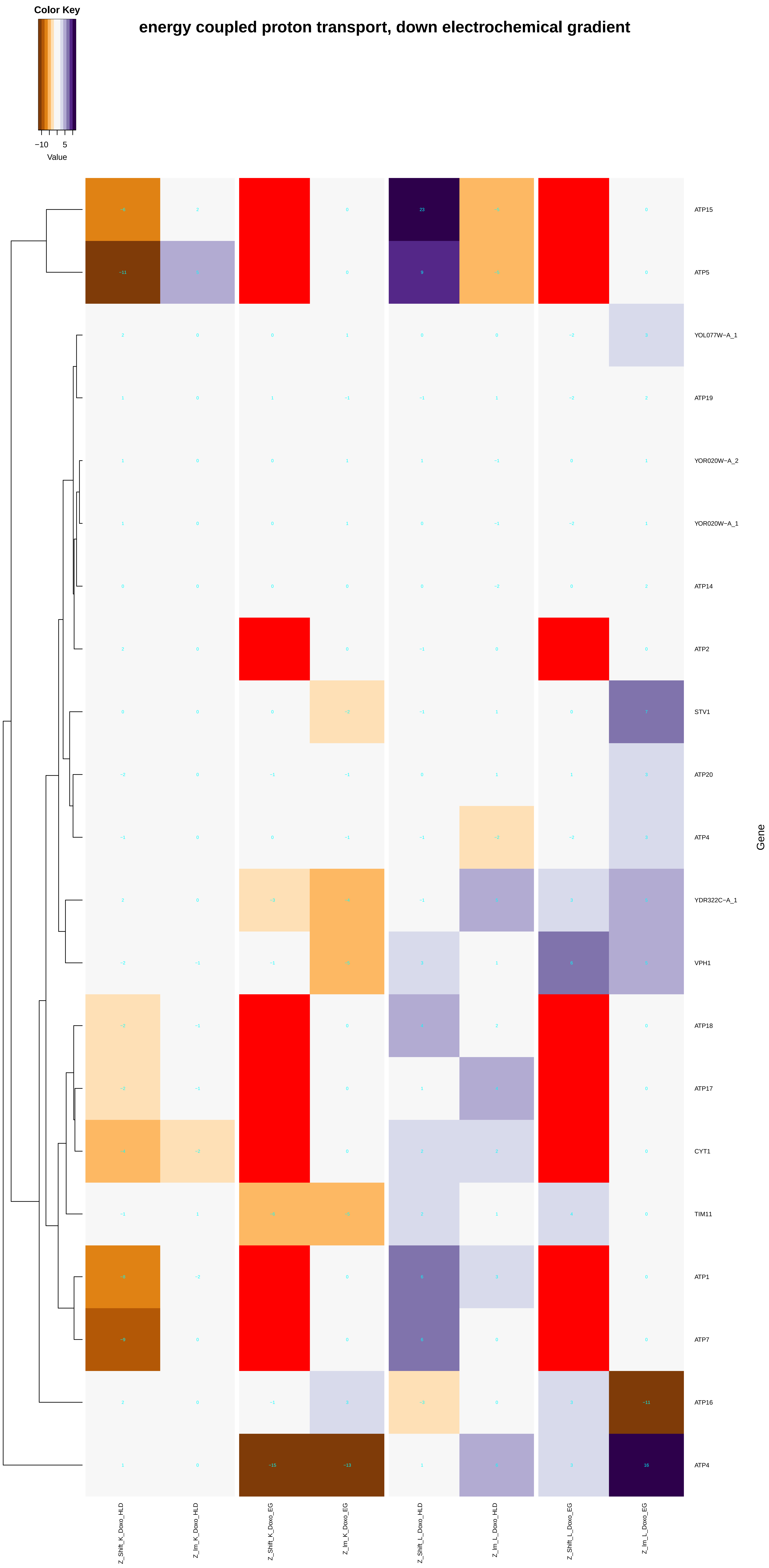
Color Key
−10
5
Value
energy coupled proton transport, down electrochemical gradient
ATP15
ATP5
YOL077W−A_1
ATP19
YOR020W−A_2
YOR020W−A_1
ATP14
ATP2
STV1
ATP20
ATP4
YDR322C−A_1
VPH1
ATP18
ATP17
CYT1
TIM11
ATP1
ATP7
ATP16
ATP4
Gene
Z_Shift_K_Doxo_HLD
Z_Im_K_Doxo_HLD
Z_Shift_K_Doxo_EG
Z_Im_K_Doxo_EG
Z_Shift_L_Doxo_HLD
Z_Im_L_Doxo_HLD
Z_Shift_L_Doxo_EG
Z_Im_L_Doxo_EG

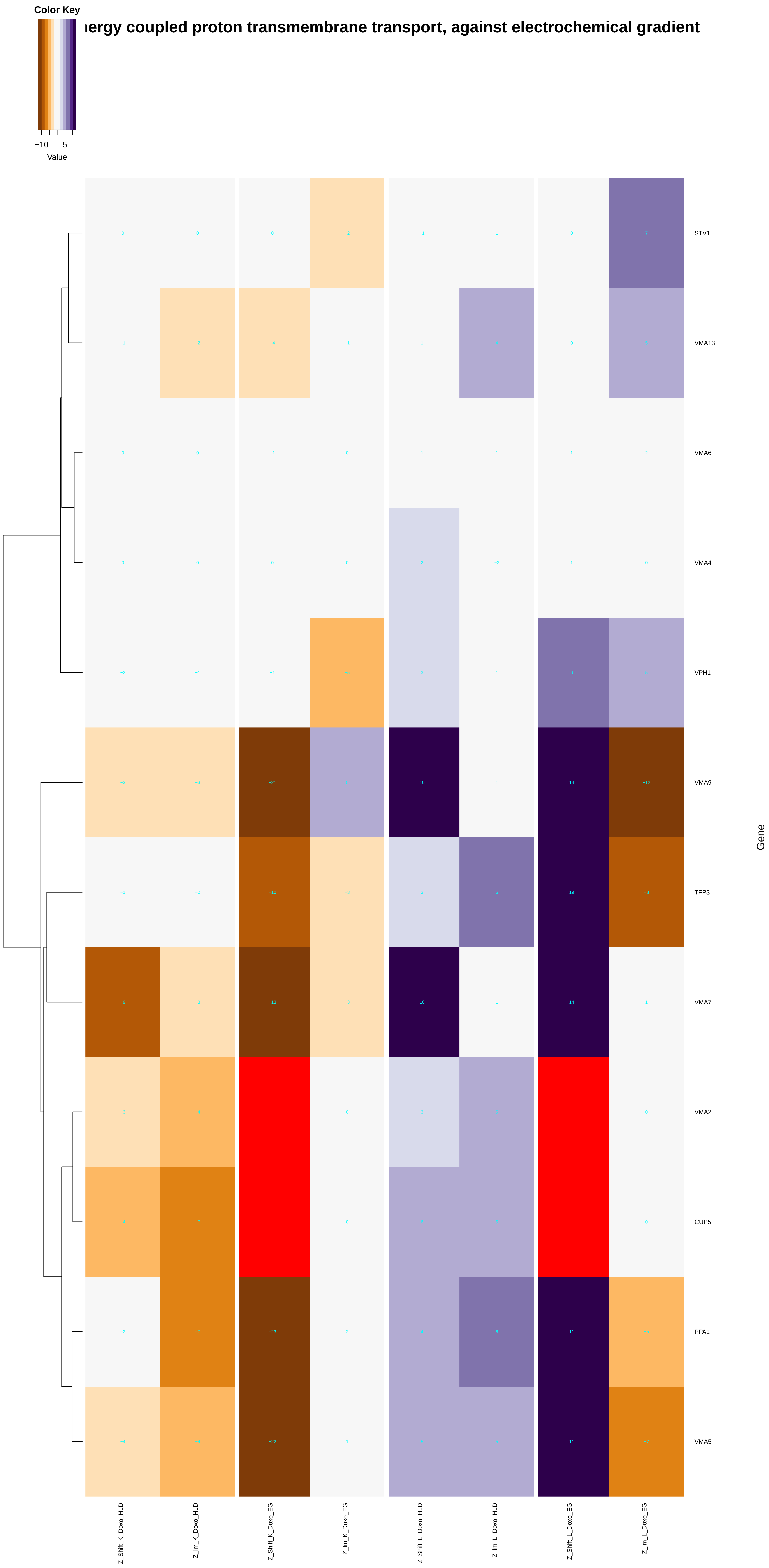

Color Key
−10
5
Value
ıergy coupled proton transmembrane transport, against electrochemical gradient
STV1
VMA13
VMA6
VMA4
VPH1
VMA9
TFP3
VMA7
VMA2
CUP5
PPA1
VMA5
Gene
Z_Shift_K_Doxo_HLD
Z_lm_K_Doxo_HLD
Z_Shift_K_Doxo_EG
Z_lm_K_Doxo_EG
Z_Shift_L_Doxo_HLD
Z_lm_L_Doxo_HLD
Z_Shift_L_Doxo_EG
Z_lm_L_Doxo_EG

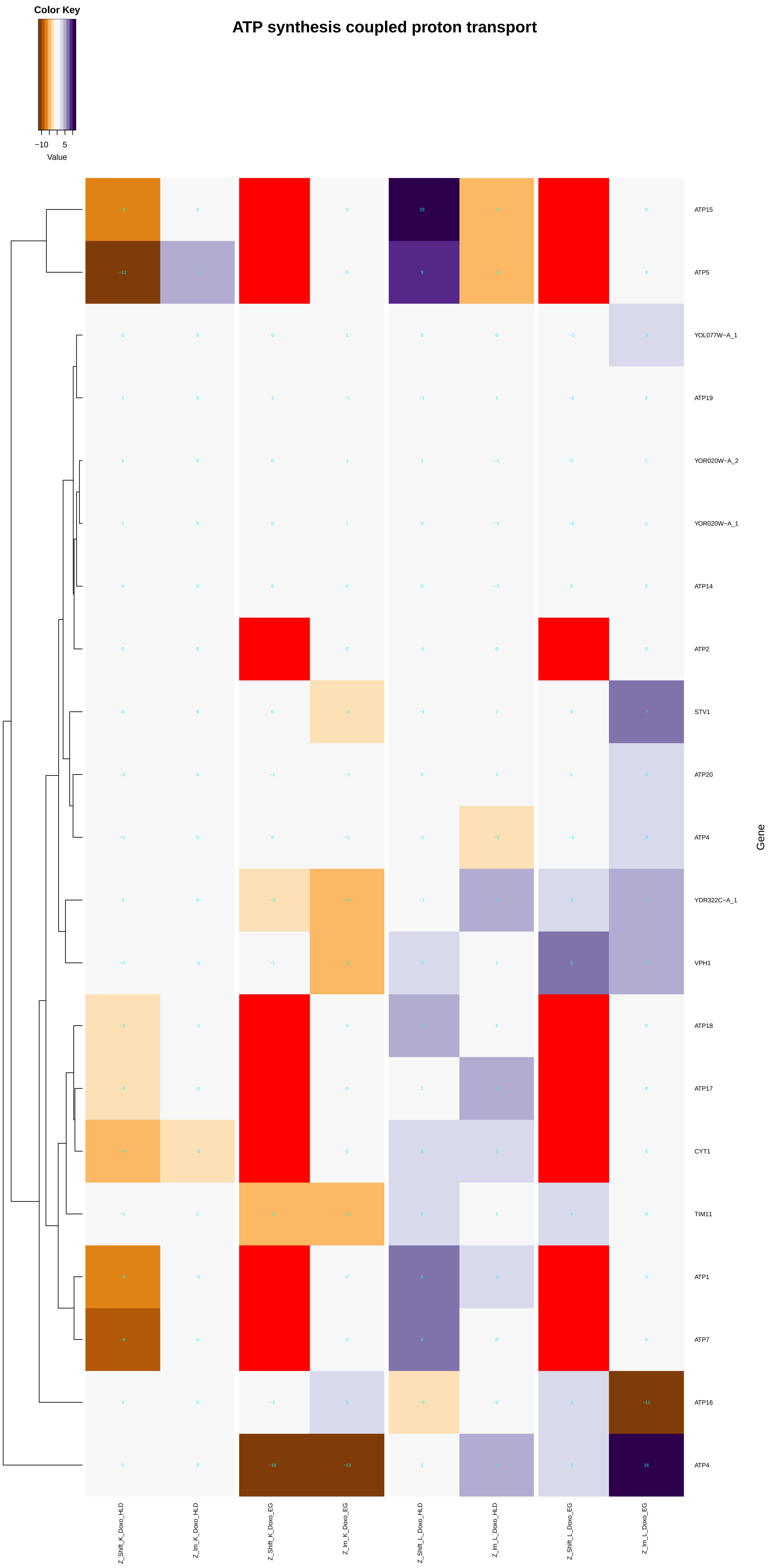

Color Key
−10
5
Value
ATP synthesis coupled proton transport
ATP15
ATP5
YOL077W−A_1
ATP19
YOR020W−A_2
YOR020W−A_1
ATP14
ATP2
STV1
ATP20
ATP4
YDR322C−A_1
VPH1
ATP18
ATP17
CYT1
TIM11
ATP1
ATP7
ATP16
ATP4
Gene
Z_Shift_K_Doxo_HLD
Z_Im_K_Doxo_HLD
Z_Shift_K_Doxo_EG
Z_Im_K_Doxo_EG
Z_Shift_L_Doxo_HLD
Z_Im_L_Doxo_HLD
Z_Shift_L_Doxo_EG
Z_Im_L_Doxo_EG
