## Additional File 9 for "A yeast phenomic model for the influence of Warburg metabolism on genetic buffering of doxorubicin": macromolecular_complex_disassembly.pdf

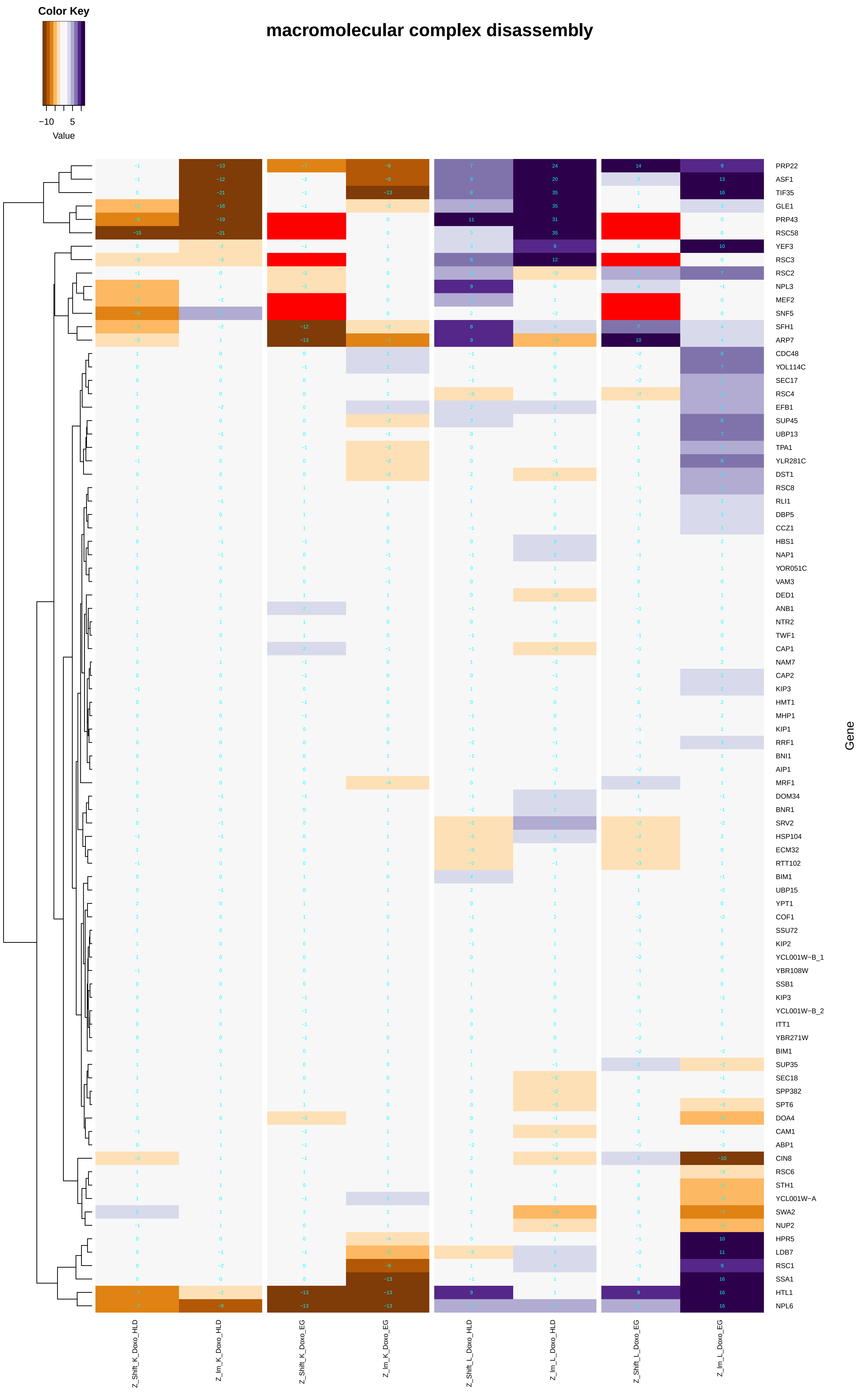
Color Key
−10 5
Value
macromolecular complex disassembly
Gene
Z_Shift_K_Doxo_HLD
Z_Im_K_Doxo_HLD
Z_Shift_K_Doxo_EG
Z_Im_K_Doxo_EG
Z_Shift_L_Doxo_HLD
Z_Im_L_Doxo_HLD
Z_Shift_L_Doxo_EG
Z_Im_L_Doxo_EG
PRP22
ASF1
TIF35
GLE1
PRP43
RSC58
YEF3
RSC3
RSC2
NPL3
MEF2
SNF5
SFH1
ARP7
CDC48
YOL114C
SEC17
RSC4
EFB1
SUP45
UBP13
TPA1
YLR281C
DST1
RSC8
RLI1
DBP5
CCZ1
HBS1
NAP1
YOR051C
VAM3
DED1
ANB1
NTR2
TWF1
CAP1
NAM7
CAP2
KIP3
HMT1
MHP1
KIP1
RRF1
BNI1
AIP1
MRF1
DOM34
BNR1
SRV2
HSP104
ECM32
RTT102
BIM1
UBP15
YPT1
COF1
SSU72
KIP2
YCL001W−B_1
YBR108W
SSB1
KIP3
YCL001W−B_2
ITT1
YBR271W
BIM1
SUP35
SEC18
SPP382
SPT6
DOA4
CAM1
ABP1
CIN8
RSC6
STH1
YCL001W−A
SWA2
NUP2
HPR5
LDB7
RSC1
SSA1
HTL1
NPL6

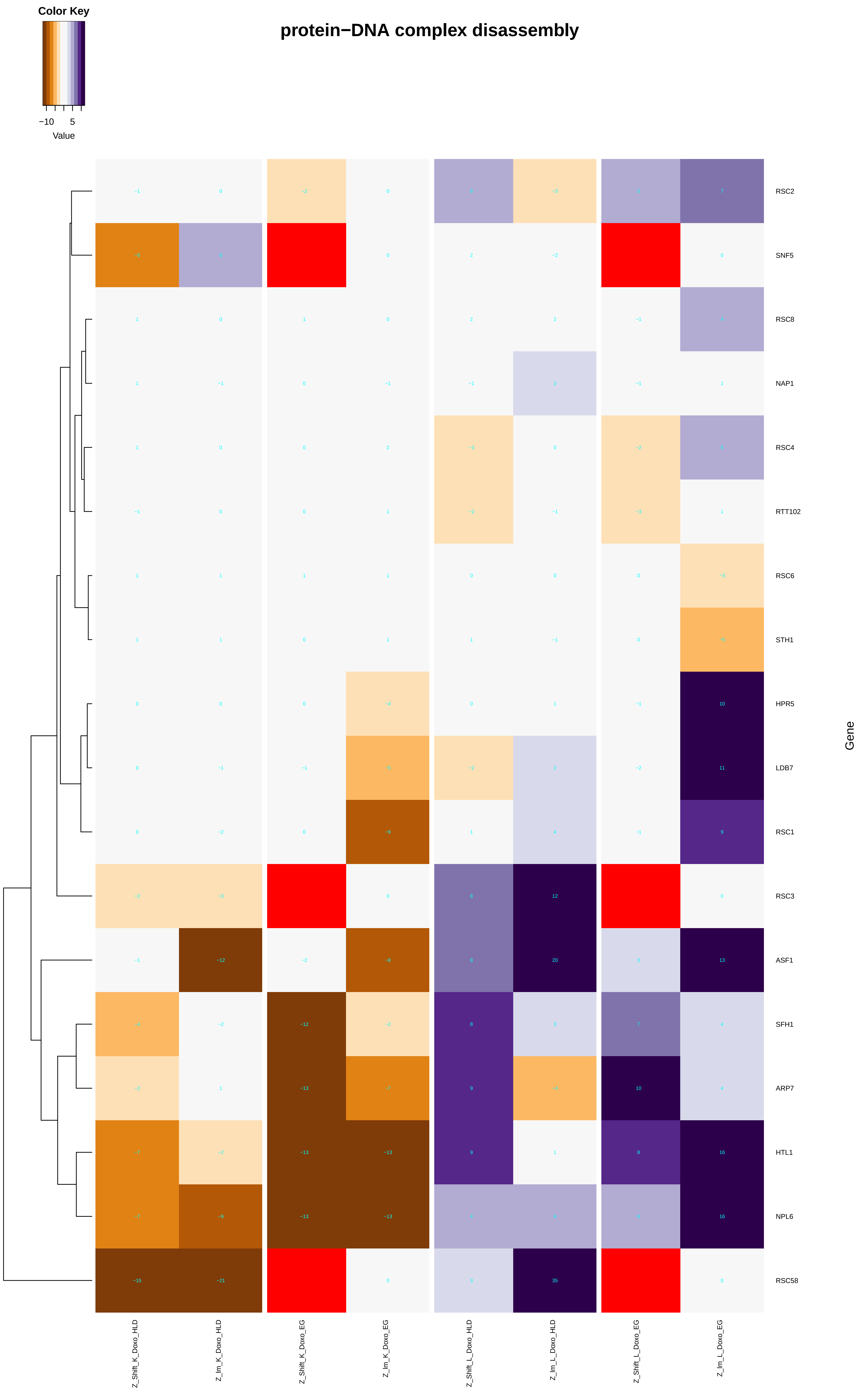
Color Key
−10
5
Value
protein−DNA complex disassembly
RSC2
SNF5
RSC8
NAP1
RSC4
RTT102
RSC6
STH1
HPR5
LDB7
RSC1
RSC3
ASF1
SFH1
ARP7
HTL1
NPL6
RSC58
Gene
Z_Shift_K_Doxo_HLD
Z_Im_K_Doxo_HLD
Z_Shift_K_Doxo_EG
Z_Im_K_Doxo_EG
Z_Shift_L_Doxo_HLD
Z_Im_L_Doxo_HLD
Z_Shift_L_Doxo_EG
Z_Im_L_Doxo_EG

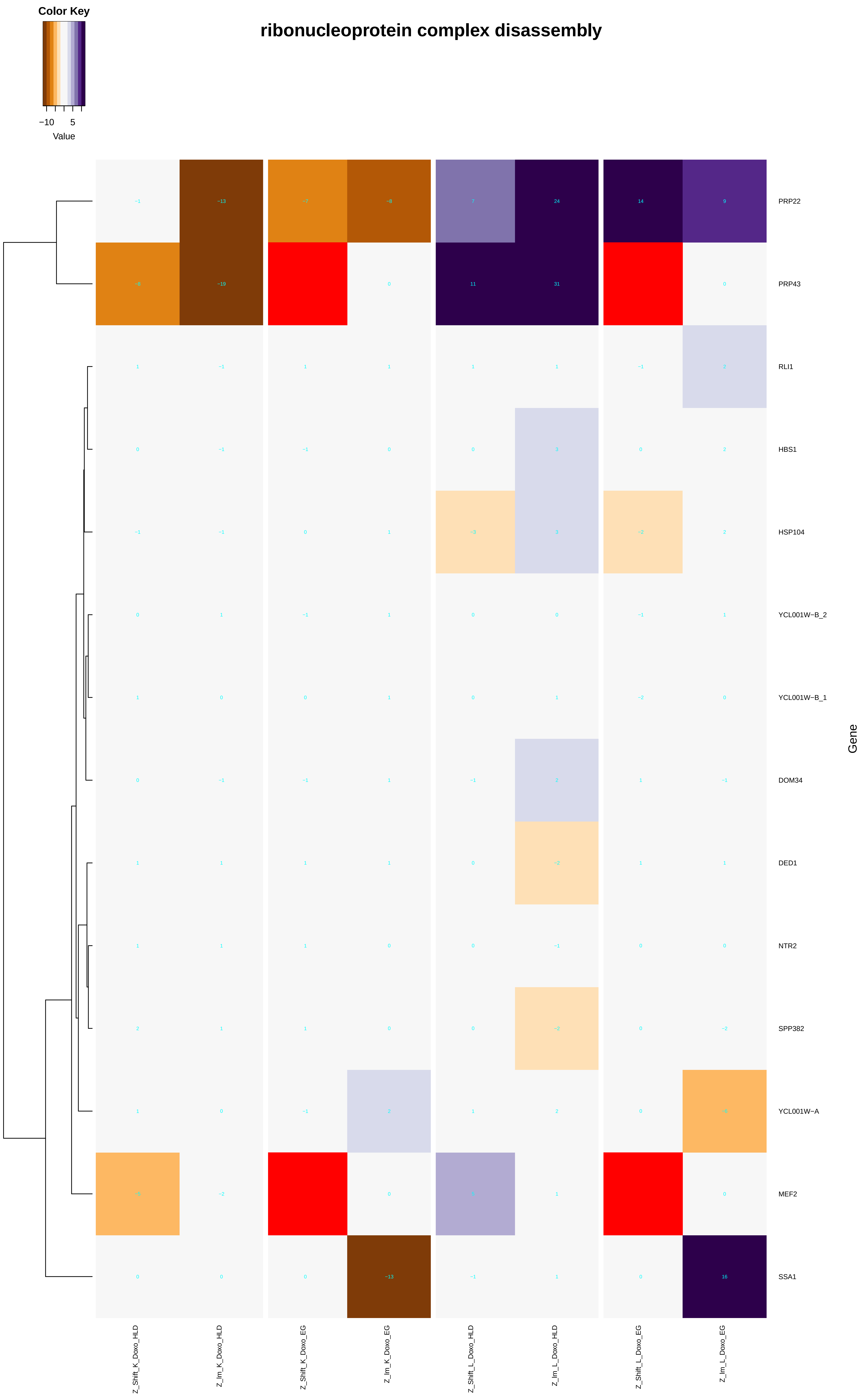

ribonucleoprotein complex disassembly
Color Key
−10
5
Value
PRP22
PRP43
RLI1
HBS1
HSP104
YCL001W−B_2
YCL001W−B_1
DOM34
DED1
NTR2
SPP382
YCL001W−A
MEF2
SSA1
Gene
Z_Shift_K_Doxo_HLD
Z_Im_K_Doxo_HLD
Z_Shift_K_Doxo_EG
Z_Im_K_Doxo_EG
Z_Shift_L_Doxo_HLD
Z_Im_L_Doxo_HLD
Z_Shift_L_Doxo_EG
Z_Im_L_Doxo_EG

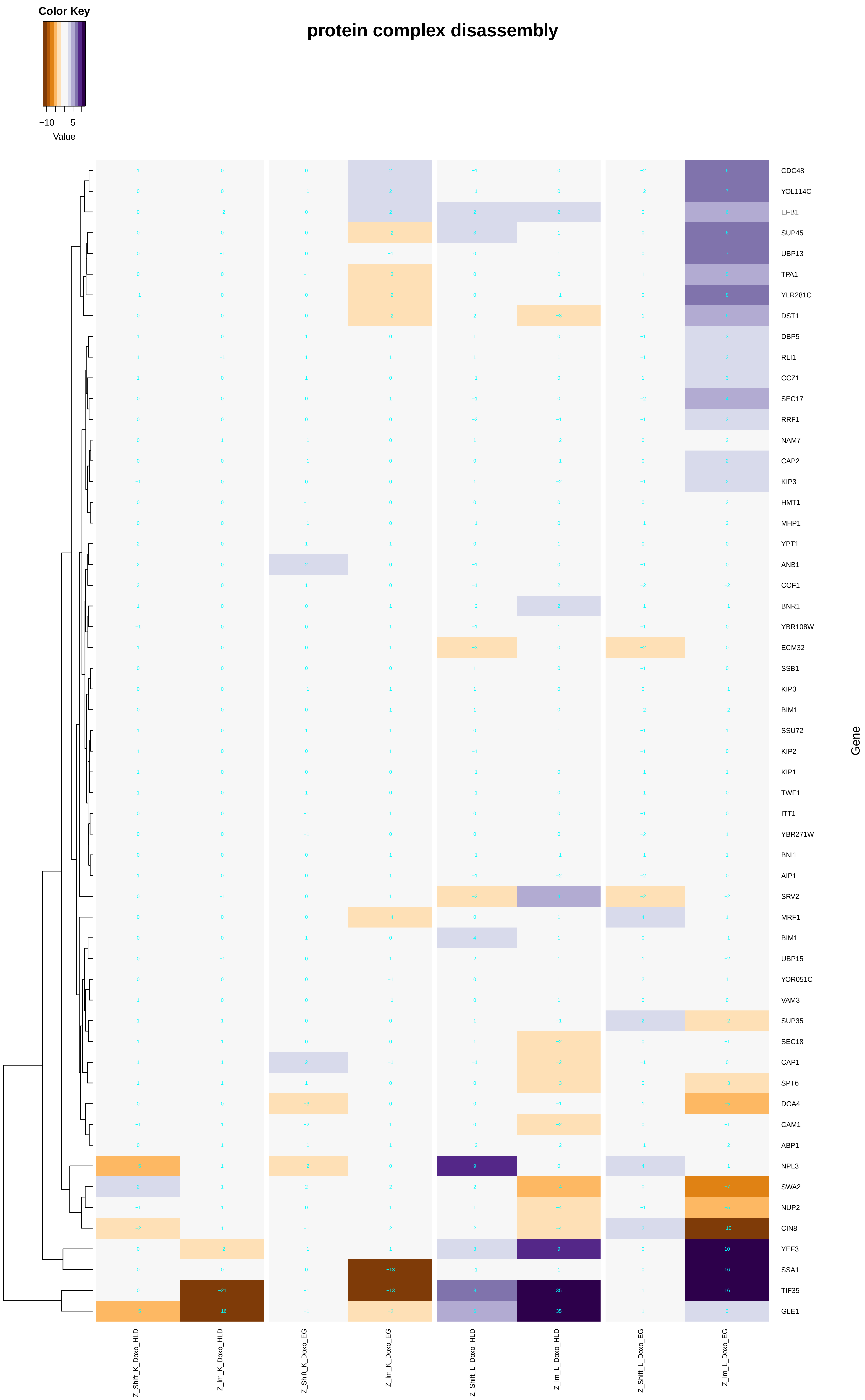
protein complex disassembly
Color Key
−10
5
Value
Gene
CDC48
YOL114C
EFB1
SUP45
UBP13
TPA1
YLR281C
DST1
DBP5
RLI1
CCZ1
SEC17
RRF1
NAM7
CAP2
KIP3
HMT1
MHP1
YPT1
ANB1
COF1
BNR1
YBR108W
ECM32
SSB1
KIP3
BIM1
SSU72
KIP2
KIP1
TWF1
ITT1
YBR271W
BNI1
AIP1
SRV2
MRF1
BIM1
UBP15
YOR051C
VAM3
SUP35
SEC18
CAP1
SPT6
DOA4
CAM1
ABP1
NPL3
SWA2
NUP2
CIN8
YEF3
SSA1
TIF35
GLE1
Z_Shift_K_Doxo_HLD
Z_Im_K_Doxo_HLD
Z_Shift_K_Doxo_EG
Z_Im_K_Doxo_EG
Z_Shift_L_Doxo_HLD
Z_Im_L_Doxo_HLD
Z_Shift_L_Doxo_EG
Z_Im_L_Doxo_EG

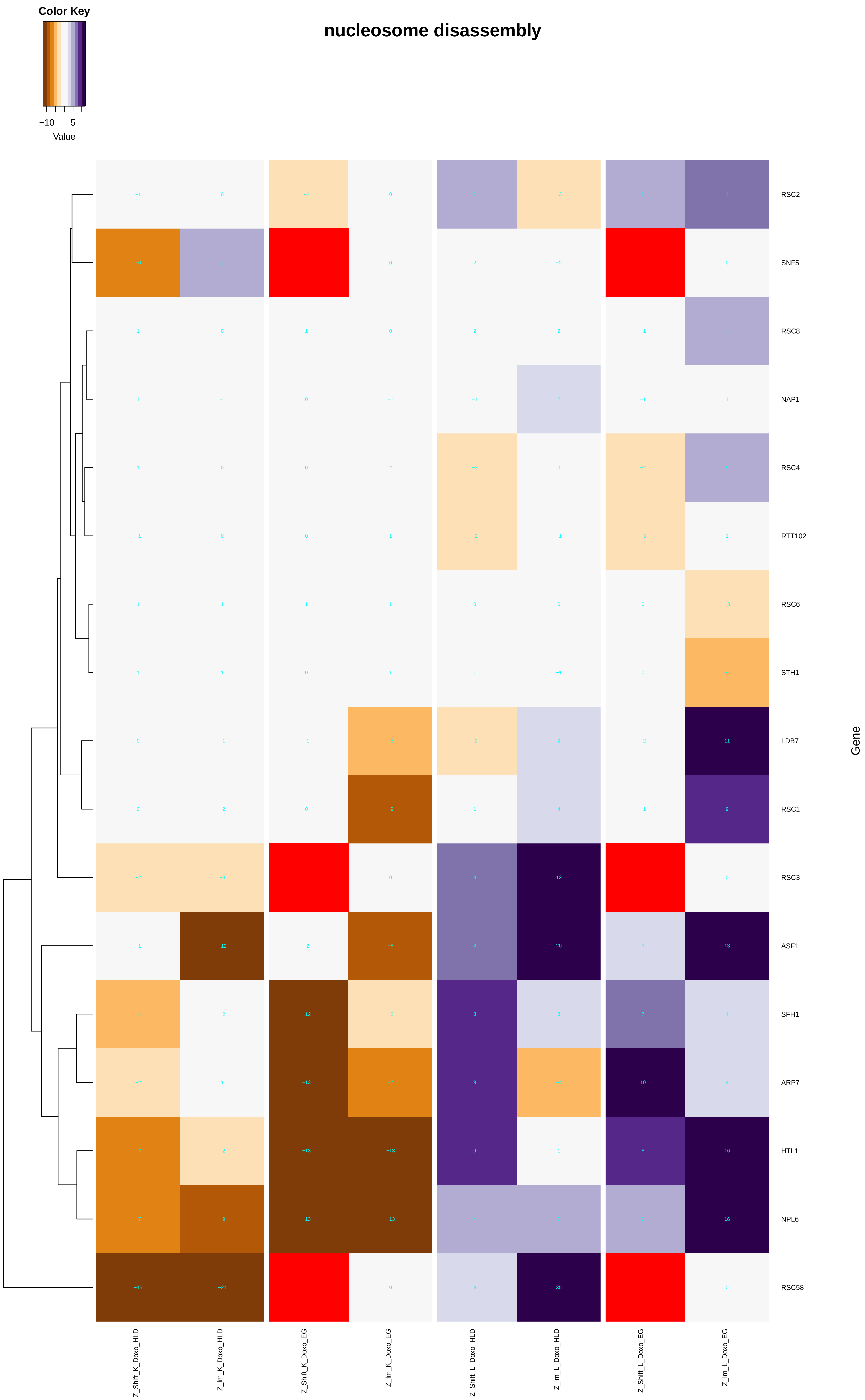
nucleosome disassembly
Color Key
−10
5
Value
RSC2
SNF5
RSC8
NAP1
RSC4
RTT102
RSC6
STH1
LDB7
RSC1
RSC3
ASF1
SFH1
ARP7
HTL1
NPL6
RSC58
Gene
Z_Shift_K_Doxo_HLD
Z_Im_K_Doxo_HLD
Z_Shift_K_Doxo_EG
Z_Im_K_Doxo_EG
Z_Shift_L_Doxo_HLD
Z_Im_L_Doxo_HLD
Z_Shift_L_Doxo_EG
Z_Im_L_Doxo_EG

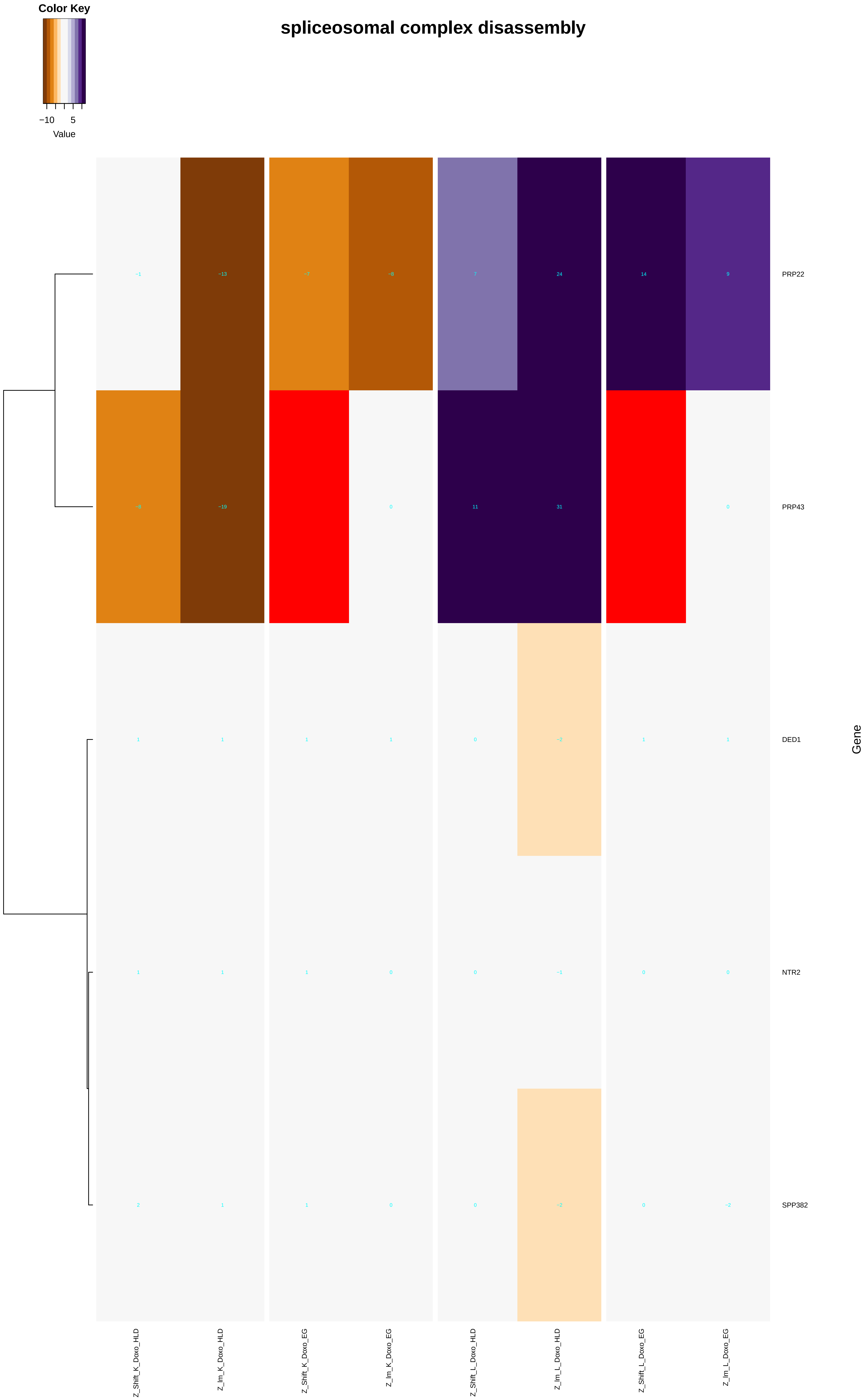
Color Key
−10
5
Value
spliceosomal complex disassembly
PRP22
PRP43
DED1
NTR2
SPP382
Gene
Z_Shift_K_Doxo_HLD
Z_lm_K_Doxo_HLD
Z_Shift_K_Doxo_EG
Z_lm_K_Doxo_EG
Z_Shift_L_Doxo_HLD
Z_lm_L_Doxo_HLD
Z_Shift_L_Doxo_EG
Z_lm_L_Doxo_EG

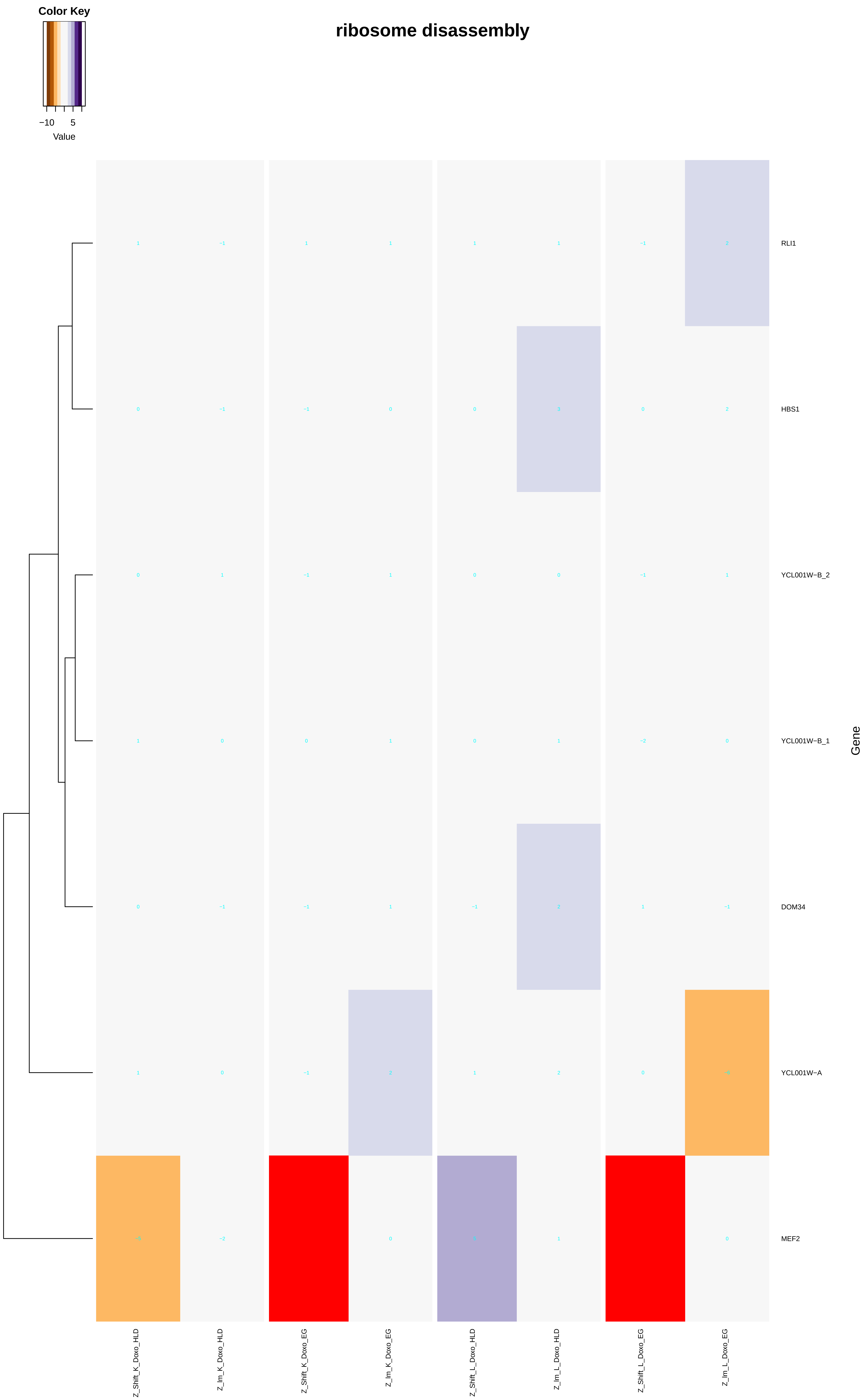
Color Key
−10
5
Value
ribosome disassembly
RLI1
HBS1
YCL001W−B_2
YCL001W−B_1
DOM34
YCL001W−A
MEF2
Gene
Z_Shift_K_Doxo_HLD
Z_Im_K_Doxo_HLD
Z_Shift_K_Doxo_EG
Z_Im_K_Doxo_EG
Z_Shift_L_Doxo_HLD
Z_Im_L_Doxo_HLD
Z_Shift_L_Doxo_EG
Z_Im_L_Doxo_EG

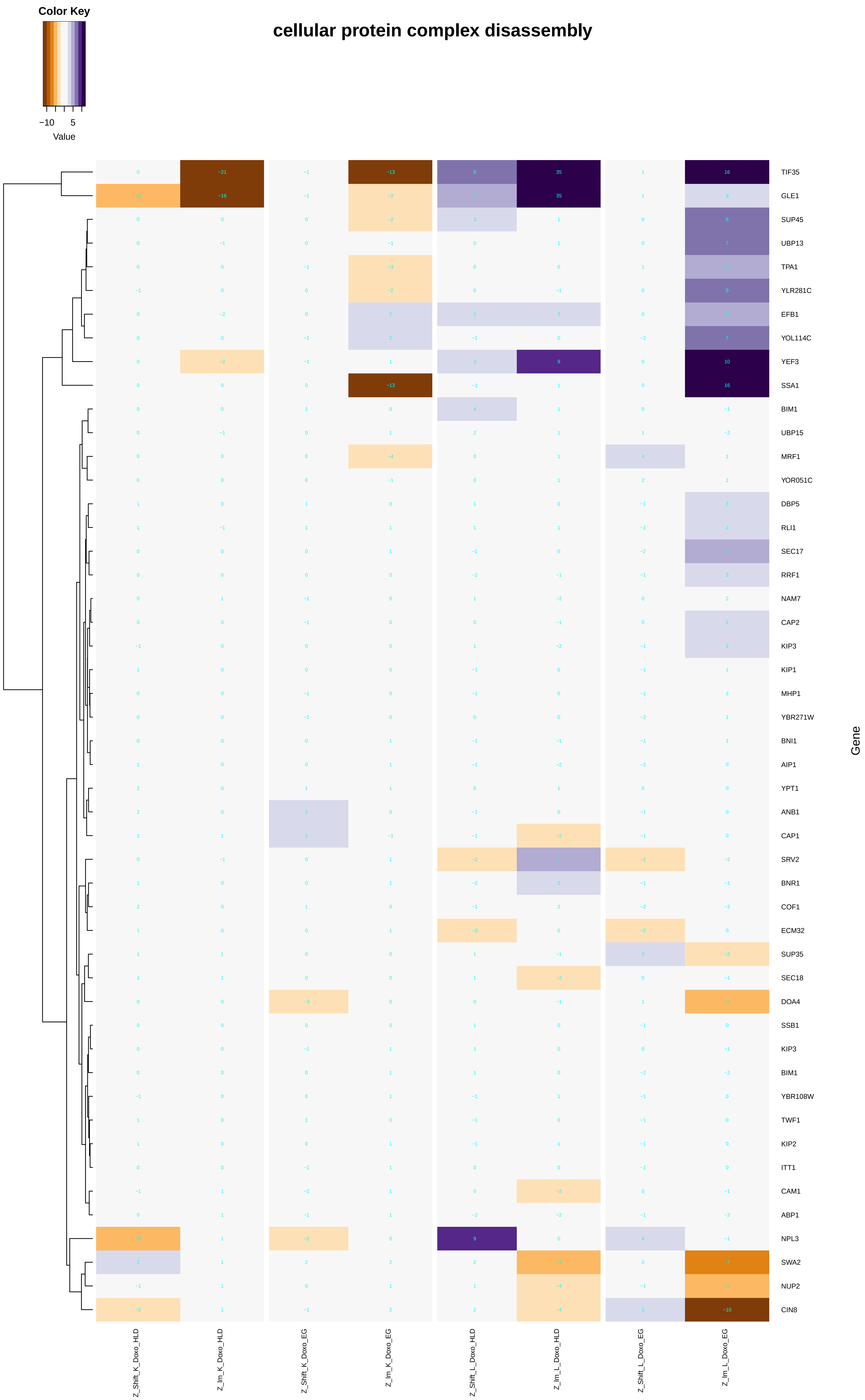

Color Key
−10
5
Value
cellular protein complex disassembly
TIF35
GLE1
SUP45
UBP13
TPA1
YLR281C
EFB1
YOL114C
YEF3
SSA1
BIM1
UBP15
MRF1
YOR051C
DBP5
RLI1
SEC17
RRF1
NAM7
CAP2
KIP3
KIP1
MHP1
YBR271W
BNI1
AIP1
YPT1
ANB1
CAP1
SRV2
BNR1
COF1
ECM32
SUP35
SEC18
DOA4
SSB1
KIP3
BIM1
YBR108W
TWF1
KIP2
ITT1
CAM1
ABP1
NPL3
SWA2
NUP2
CIN8
Gene
Z_Shift_K_Doxo_HLD
Z_lm_K_Doxo_HLD
Z_Shift_K_Doxo_EG
Z_lm_K_Doxo_EG
Z_Shift_L_Doxo_HLD
Z_lm_L_Doxo_HLD
Z_Shift_L_Doxo_EG
Z_lm_L_Doxo_EG

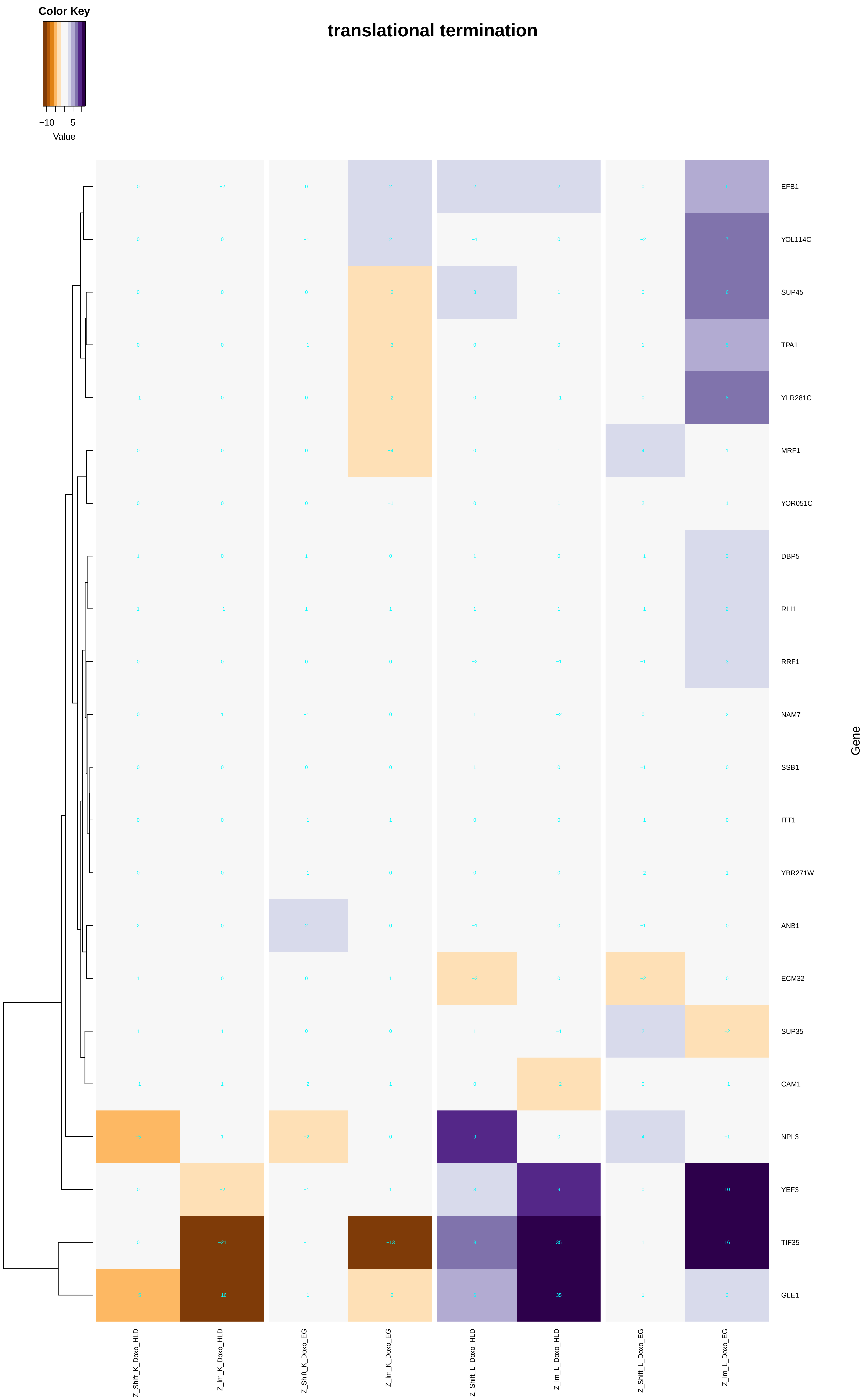
translational termination
Color Key
−10
5
Value
EFB1
YOL114C
SUP45
TPA1
YLR281C
MRF1
YOR051C
DBP5
RLI1
RRF1
NAM7
SSB1
ITT1
YBR271W
ANB1
ECM32
SUP35
CAM1
NPL3
YEF3
TIF35
GLE1
Gene
Z_Shift_K_Doxo_HLD
Z_Im_K_Doxo_HLD
Z_Shift_K_Doxo_EG
Z_Im_K_Doxo_EG
Z_Shift_L_Doxo_HLD
Z_Im_L_Doxo_HLD
Z_Shift_L_Doxo_EG
Z_Im_L_Doxo_EG

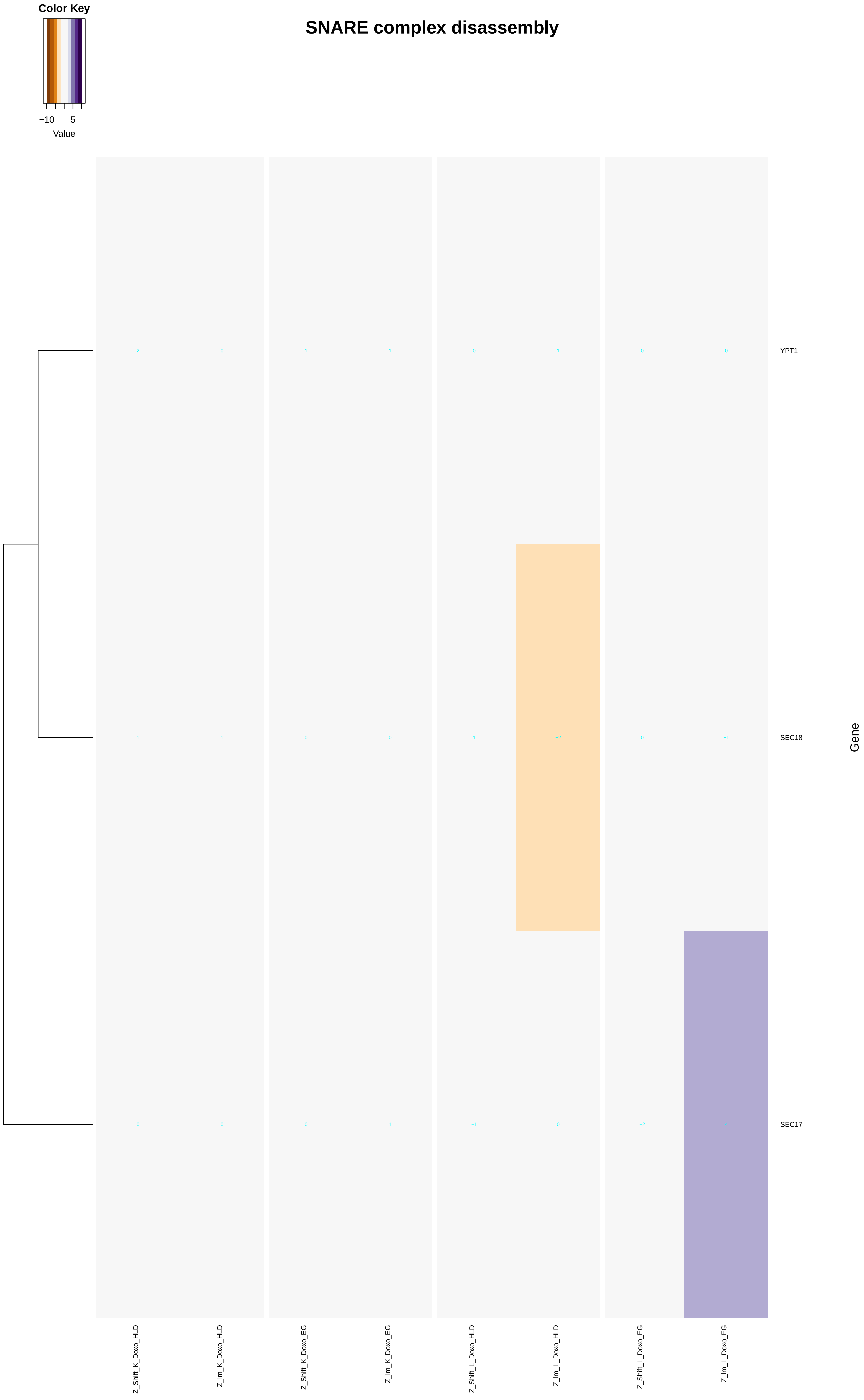
SNARE complex disassembly
Color Key
−10
5
Value
YPT1
SEC18
SEC17
Gene
Z_Shift_K_Doxo_HLD
Z_Im_K_Doxo_HLD
Z_Shift_K_Doxo_EG
Z_Im_K_Doxo_EG
Z_Shift_L_Doxo_HLD
Z_Im_L_Doxo_HLD
Z_Shift_L_Doxo_EG
Z_Im_L_Doxo_EG

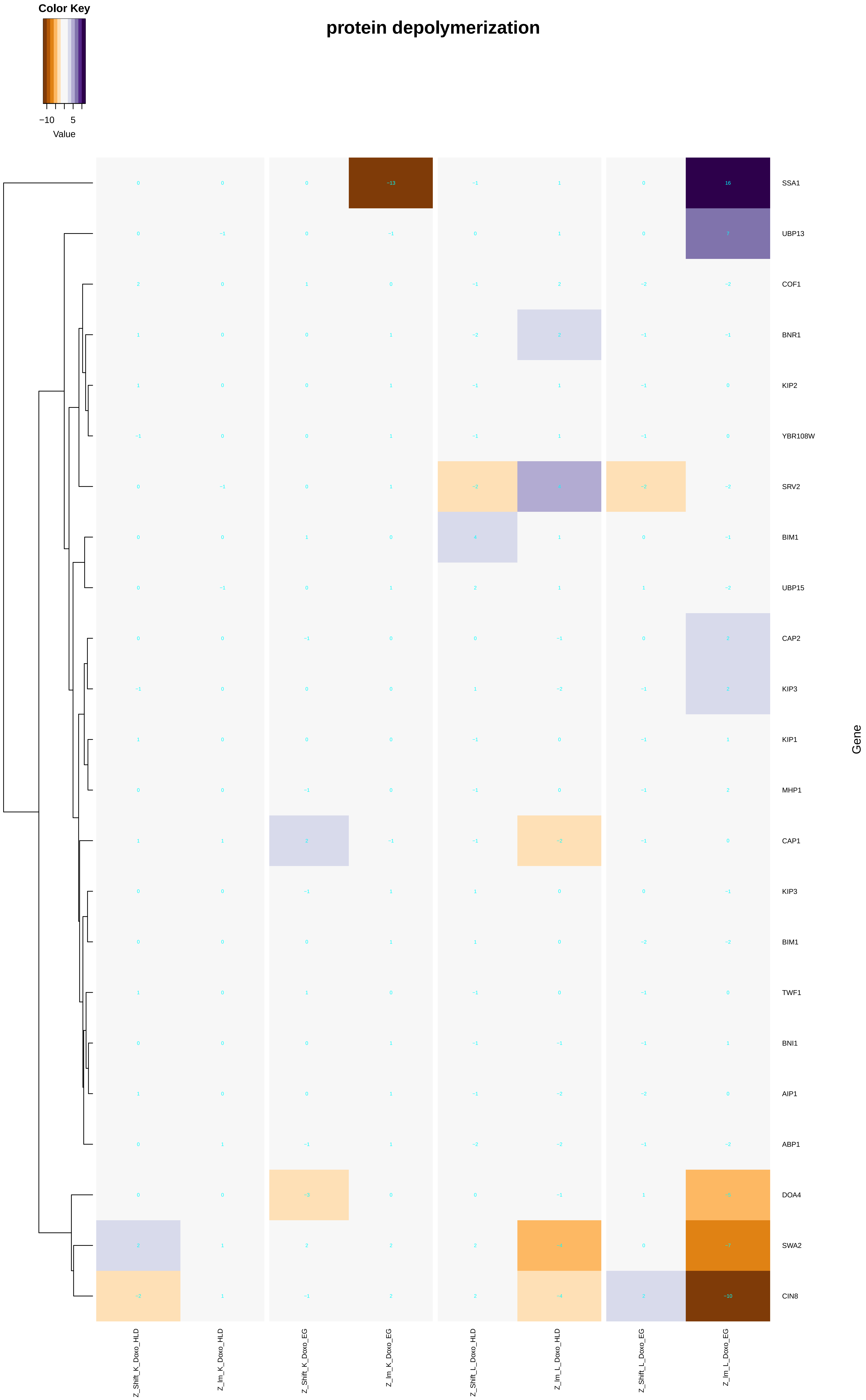

Color Key
−10
5
Value
protein depolymerization
SSA1
UBP13
COF1
BNR1
KIP2
YBR108W
SRV2
BIM1
UBP15
CAP2
KIP3
KIP1
MHP1
CAP1
KIP3
BIM1
TWF1
BNI1
AIP1
ABP1
DOA4
SWA2
CIN8
Gene
Z_Shift_K_Doxo_HLD
Z_Im_K_Doxo_HLD
Z_Shift_K_Doxo_EG
Z_Im_K_Doxo_EG
Z_Shift_L_Doxo_HLD
Z_Im_L_Doxo_HLD
Z_Shift_L_Doxo_EG
Z_Im_L_Doxo_EG

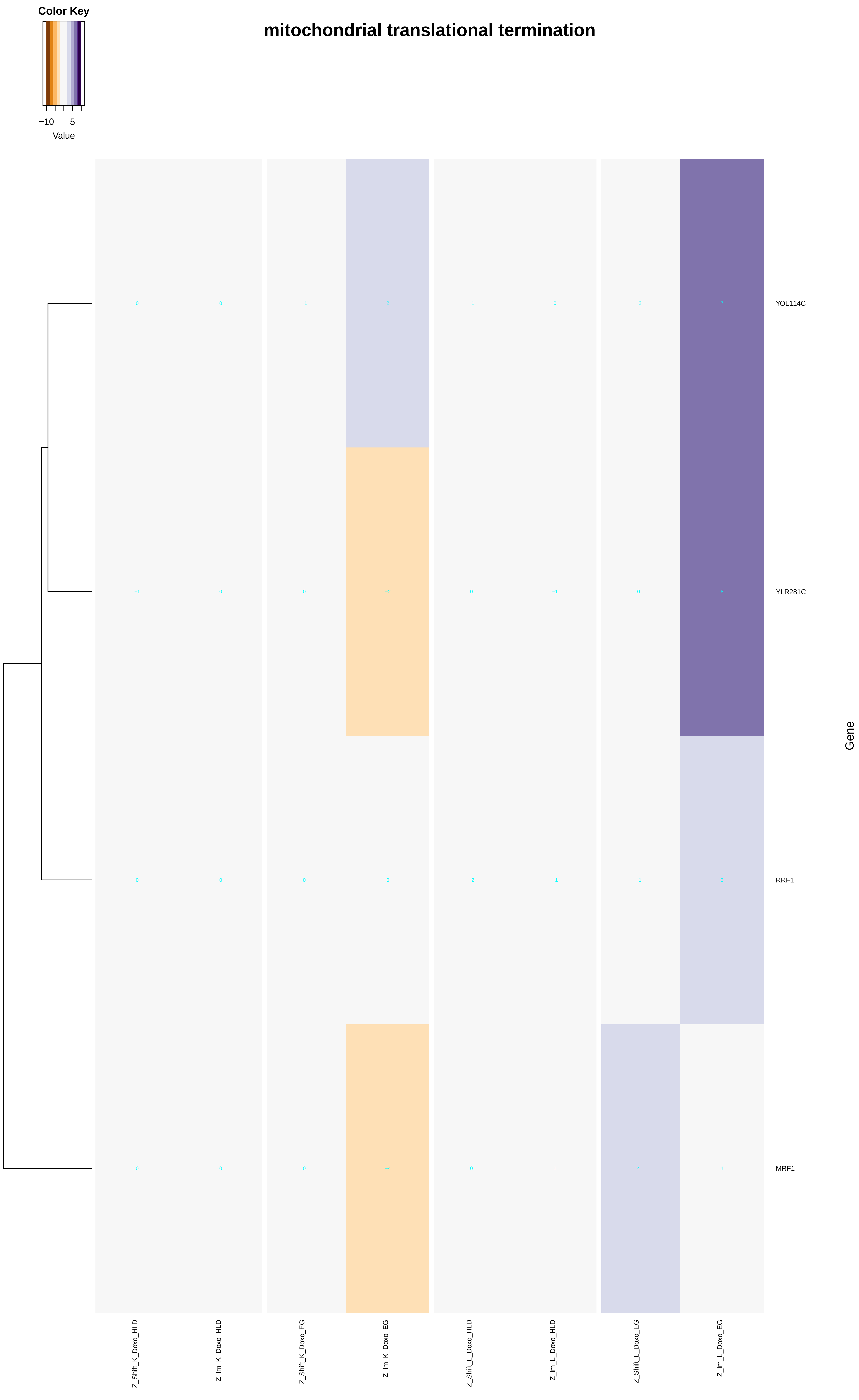

mitochondrial translational termination
Color Key
−10
5
Value
YOL114C
YLR281C
RRF1
MRF1
Gene
0 0 −1 2 −1 0 −2 7
−1 0 0 −2 0 −1 0 8
0 0 0 0 −2 −1 −1 3
0 0 0 −4 0 1 4 1
Z_Shift_K_Doxo_HLD
Z_Im_K_Doxo_HLD
Z_Shift_K_Doxo_EG
Z_Im_K_Doxo_EG
Z_Shift_L_Doxo_HLD
Z_Im_L_Doxo_HLD
Z_Shift_L_Doxo_EG
Z_Im_L_Doxo_EG

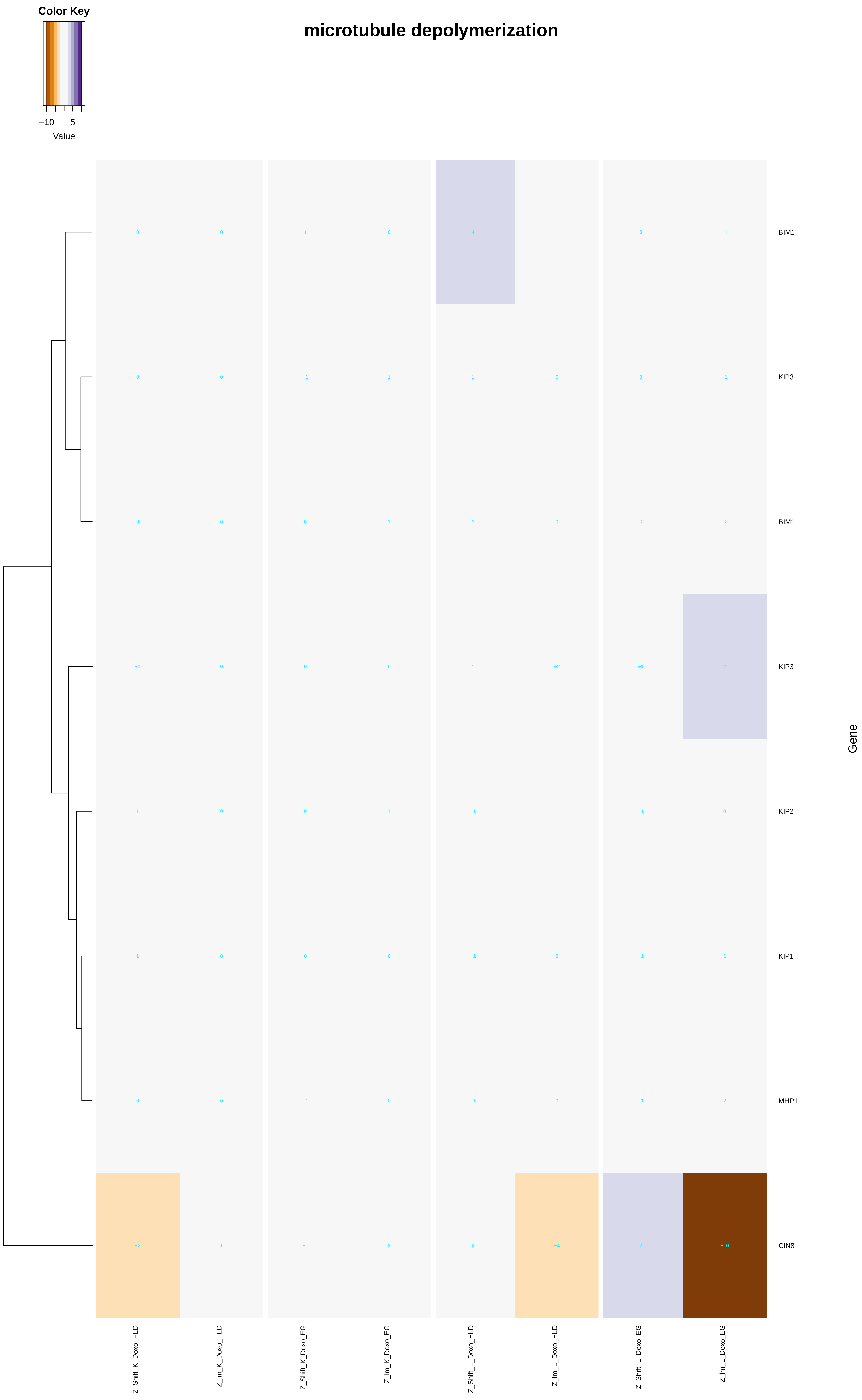

Color Key
−10
5
Value
microtubule depolymerization
BIM1
KIP3
BIM1
KIP3
KIP2
KIP1
MHP1
CIN8
Gene
Z_Shift_K_Doxo_HLD
Z_Im_K_Doxo_HLD
Z_Shift_K_Doxo_EG
Z_Im_K_Doxo_EG
Z_Shift_L_Doxo_HLD
Z_Im_L_Doxo_HLD
Z_Shift_L_Doxo_EG
Z_Im_L_Doxo_EG
0 0 1 0 4 1 0 −1
0 0 −1 1 1 0 0 −1
0 0 0 1 1 0 −2 −2
−1 0 0 0 1 −2 −1 2
1 0 0 1 −1 1 −1 0
1 0 0 0 −1 0 −1 1
0 0 −1 0 −1 0 −1 2
−2 1 −1 2 2 −4 2 −10

### free ubiquitin chain depolymerization

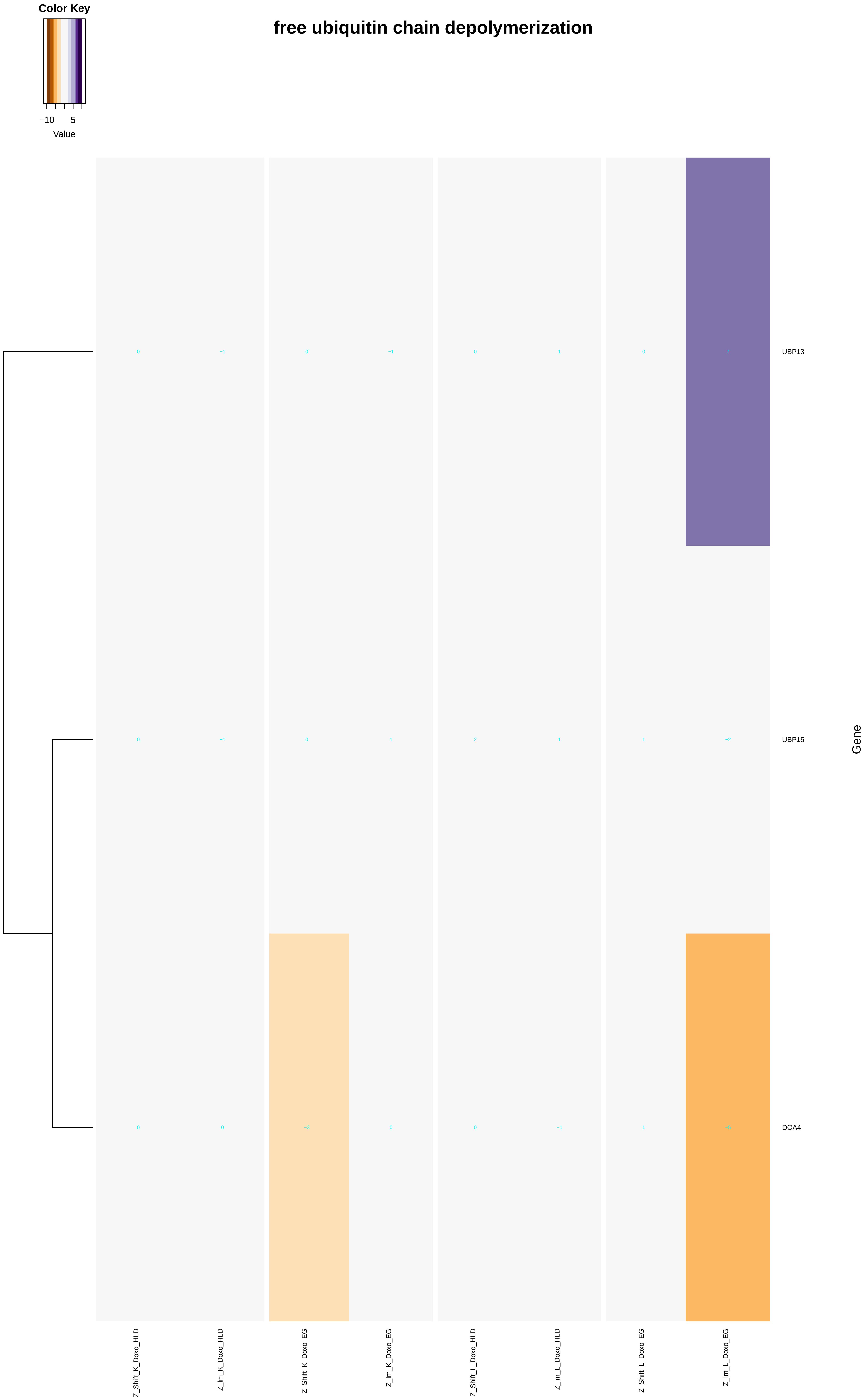

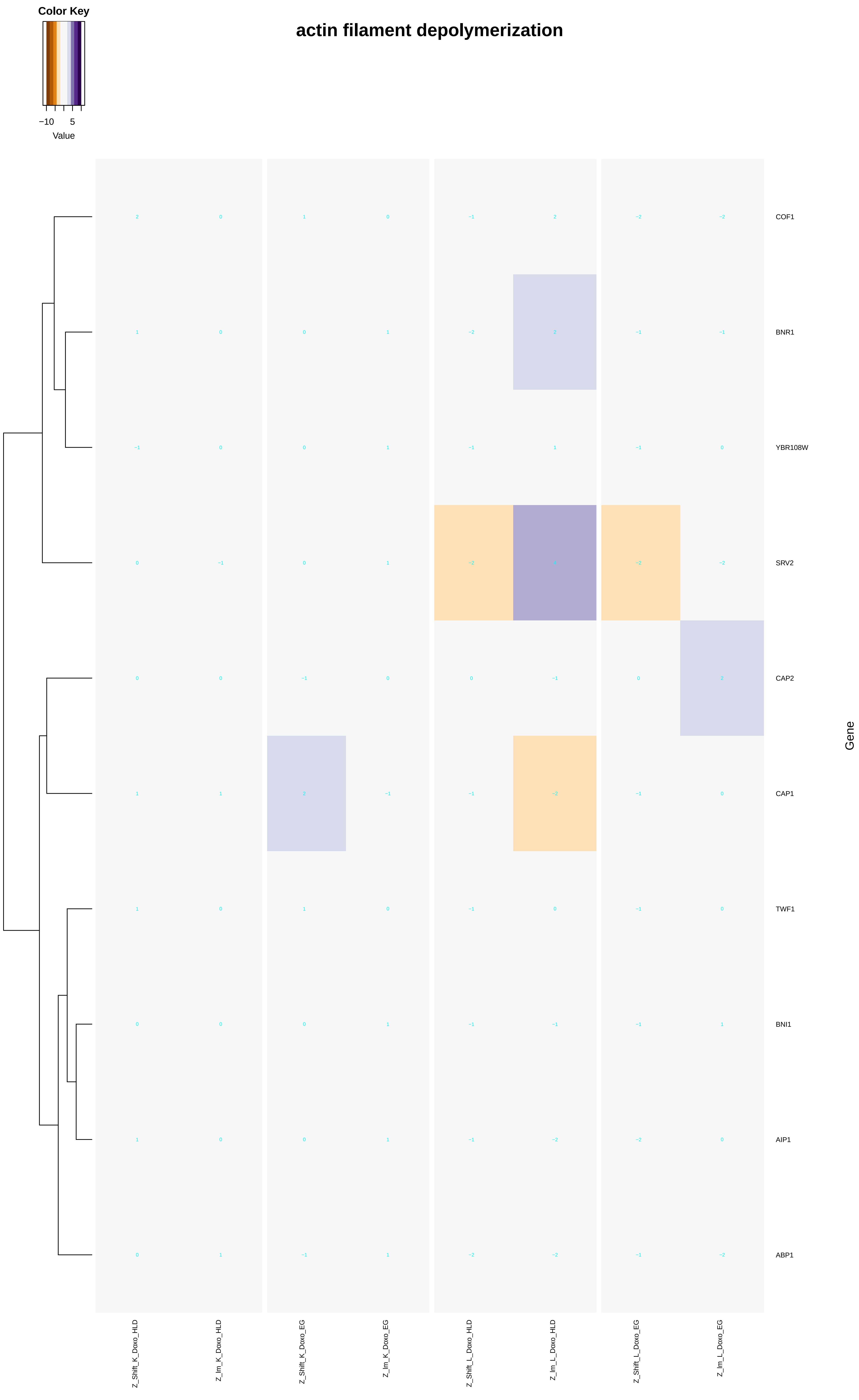
Color Key
−10
5
Value
actin filament depolymerization
COF1
BNR1
YBR108W
SRV2
CAP2
CAP1
TWF1
BNI1
AIP1
ABP1
Gene
Z_Shift_K_Doxo_HLD
Z_Im_K_Doxo_HLD
Z_Shift_K_Doxo_EG
Z_Im_K_Doxo_EG
Z_Shift_L_Doxo_HLD
Z_Im_L_Doxo_HLD
Z_Shift_L_Doxo_EG
Z_Im_L_Doxo_EG
