## Additional File 9 for "A yeast phenomic model for the influence of Warburg metabolism on genetic buffering of doxorubicin": mitochondrion_organization.pdf

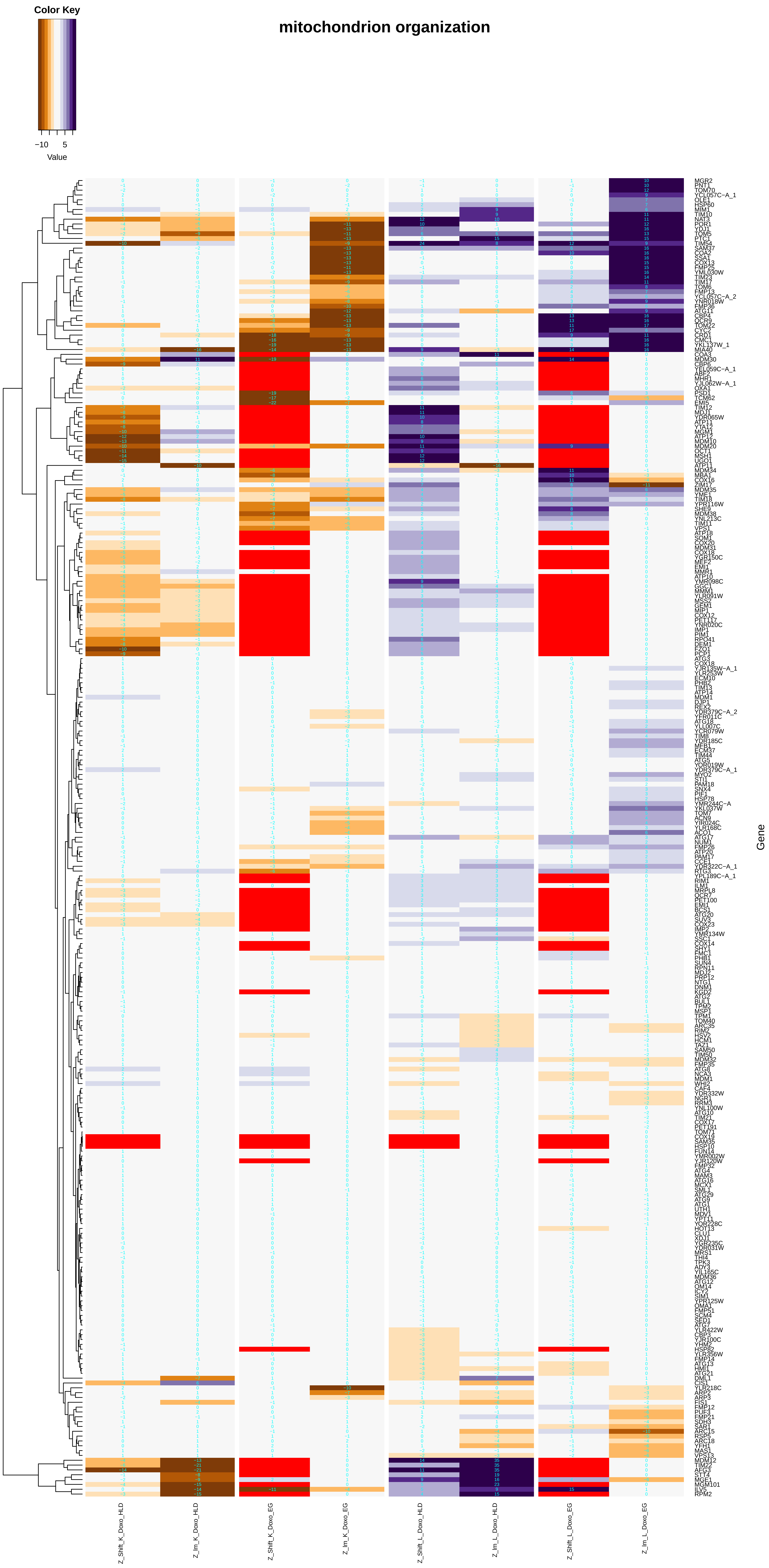

mitochondrion organization
Color Key
−10
5
Value
Gene
Z_Shift_K_Doxo_HLD
Z_Im_K_Doxo_HLD
Z_Shift_K_Doxo_EG
Z_Im_K_Doxo_EG
Z_Shift_L_Doxo_HLD
Z_Im_L_Doxo_HLD
Z_Shift_L_Doxo_EG
Z_Im_L_Doxo_EG

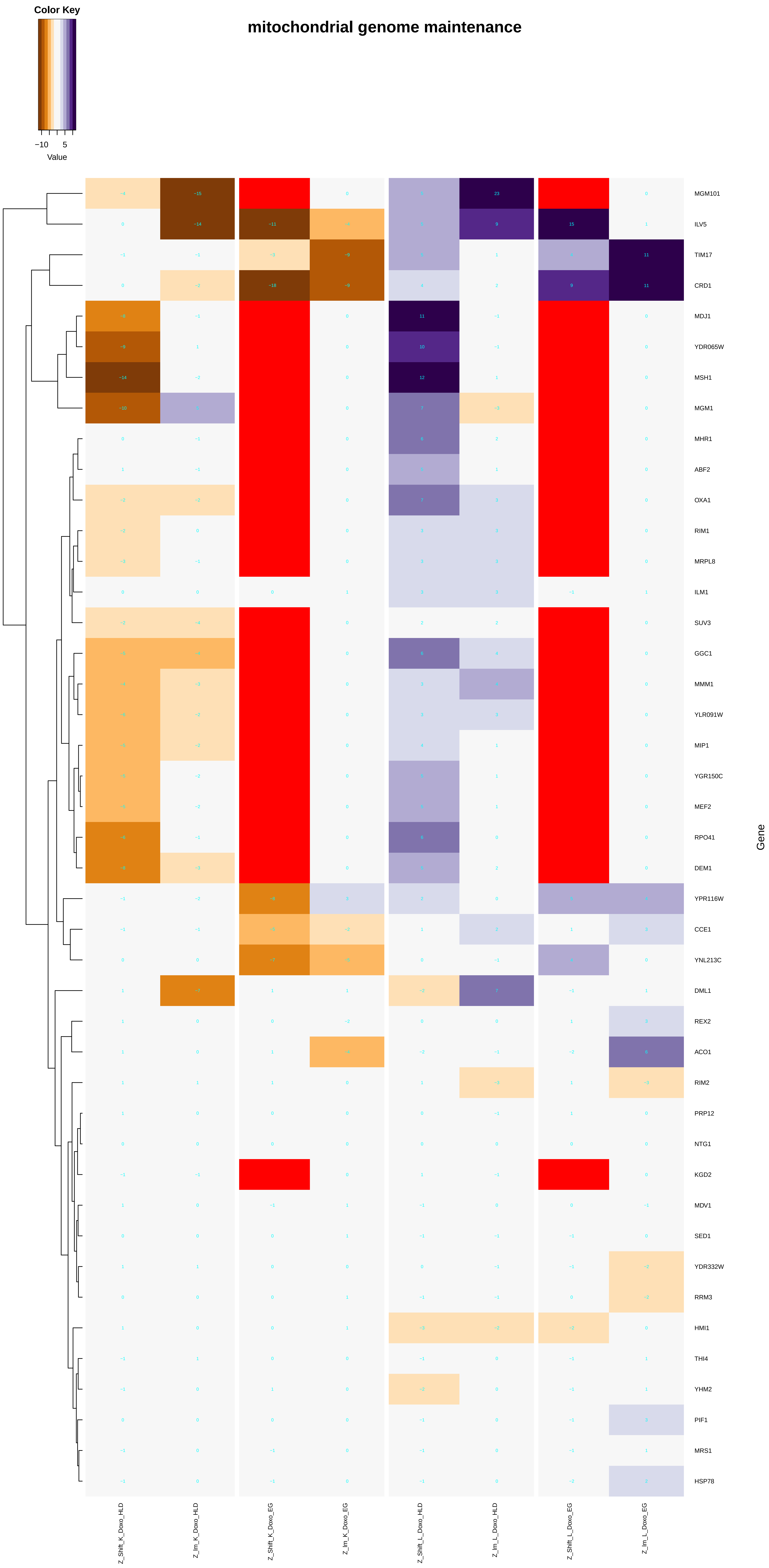
Color Key
−10
5
Value
mitochondrial genome maintenance
MGM101
ILV5
TIM17
CRD1
MDJ1
YDR065W
MSH1
MGM1
MHR1
ABF2
OXA1
RIM1
MRPL8
ILM1
SUV3
GGC1
MMM1
YLR091W
MIP1
YGR150C
MEF2
RPO41
DEM1
YPR116W
CCE1
YNL213C
DML1
REX2
ACO1
RIM2
PRP12
NTG1
KGD2
MDV1
SED1
YDR332W
RRM3
HMI1
THI4
YHM2
PIF1
MRS1
HSP78
Gene
Z_Shift_K_Doxo_HLD
Z_lm_K_Doxo_HLD
Z_Shift_K_Doxo_EG
Z_lm_K_Doxo_EG
Z_Shift_L_Doxo_HLD
Z_lm_L_Doxo_HLD
Z_Shift_L_Doxo_EG
Z_lm_L_Doxo_EG

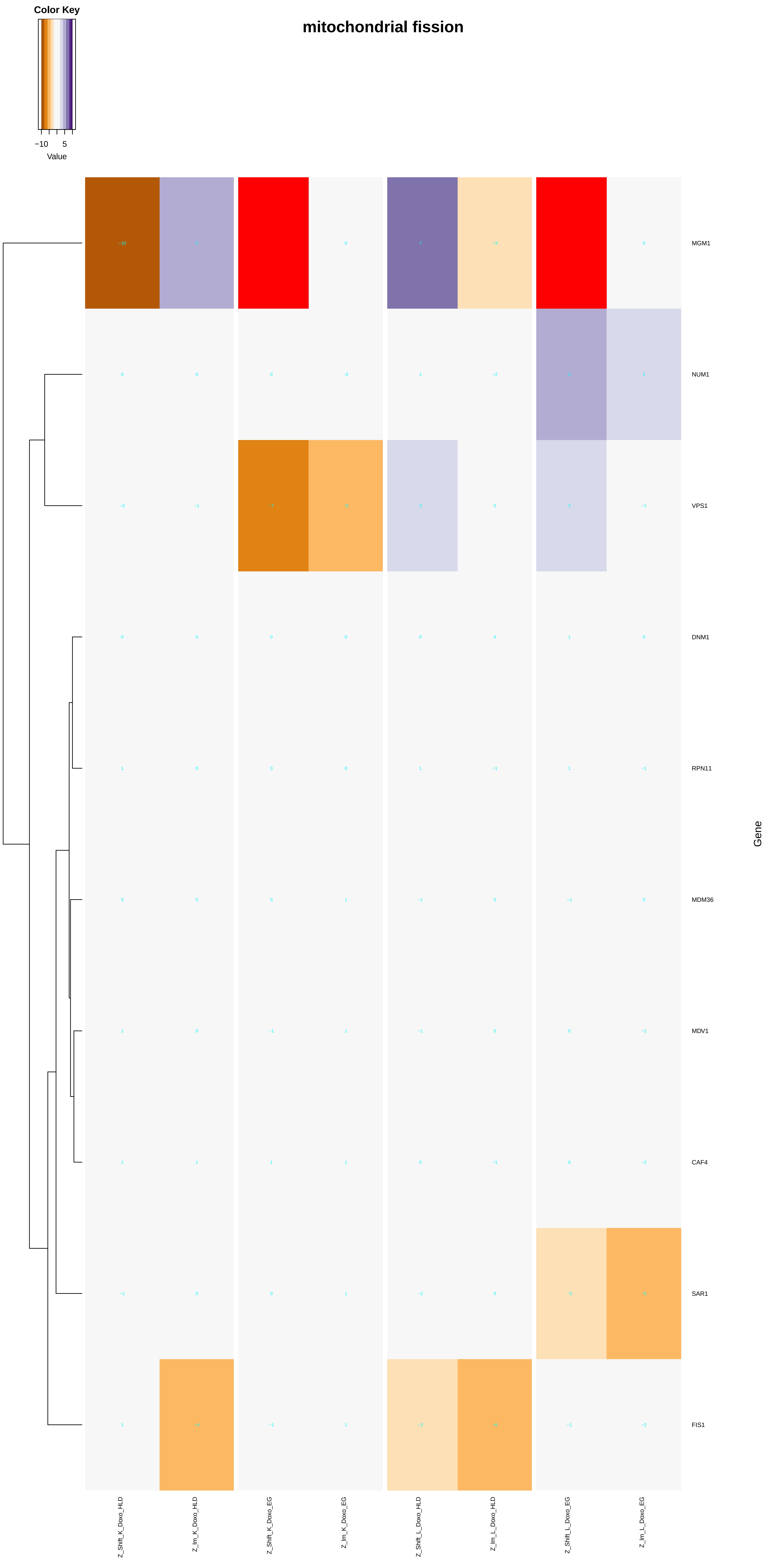
Color Key
−10
5
Value
mitochondrial fission
MGM1
NUM1
VPS1
DNM1
RPN11
MDM36
MDV1
CAF4
SAR1
FIS1
Gene
Z_Shift_K_Doxo_HLD
Z_Im_K_Doxo_HLD
Z_Shift_K_Doxo_EG
Z_Im_K_Doxo_EG
Z_Shift_L_Doxo_HLD
Z_Im_L_Doxo_HLD
Z_Shift_L_Doxo_EG
Z_Im_L_Doxo_EG

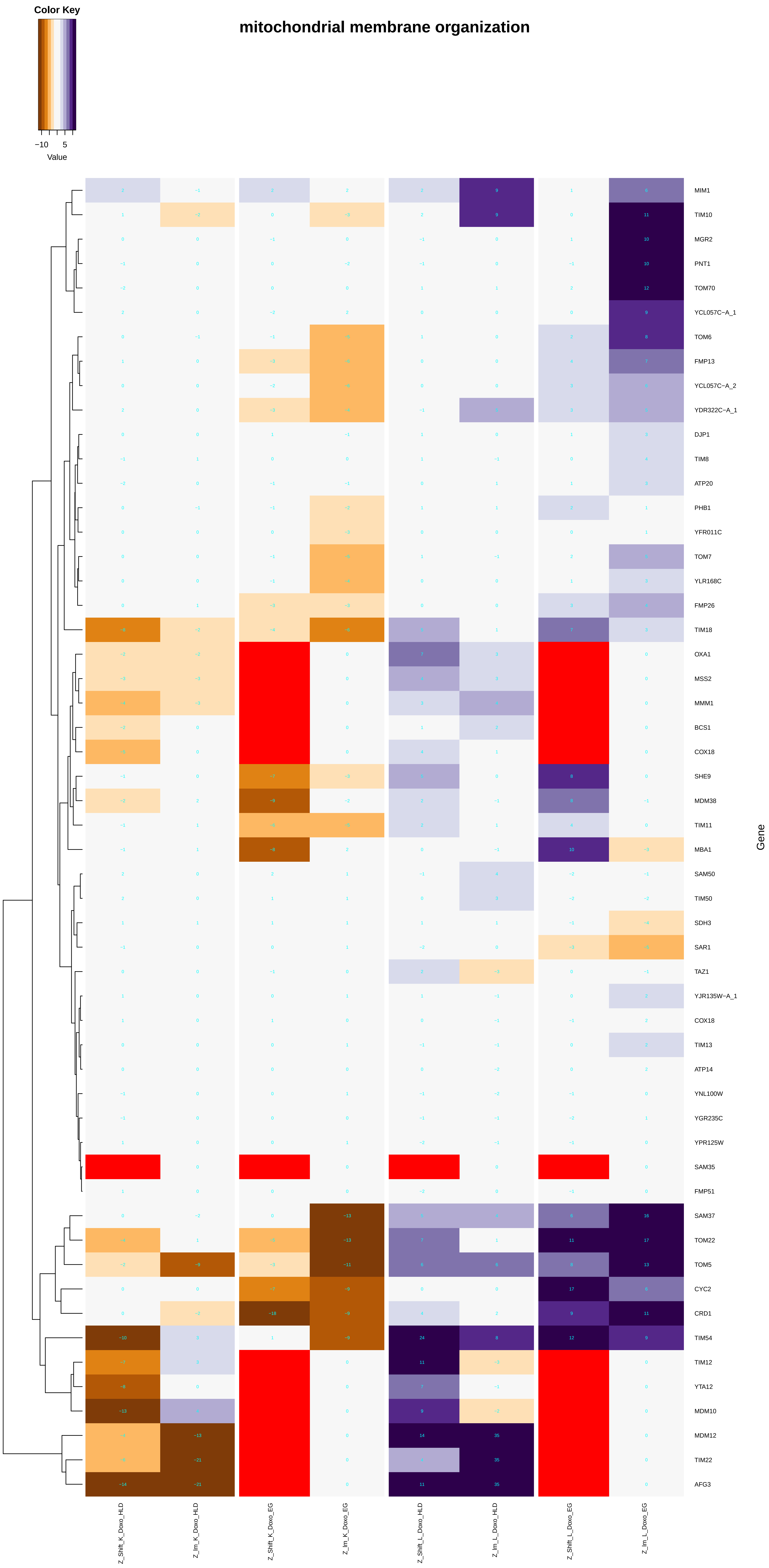

mitochondrial membrane organization
Color Key
−10
5
Value
MIM1
TIM10
MGR2
PNT1
TOM70
YCL057C−A_1
TOM6
FMP13
YCL057C−A_2
YDR322C−A_1
DJP1
TIM8
ATP20
PHB1
YFR011C
TOM7
YLR168C
FMP26
TIM18
OXA1
MSS2
MMM1
BCS1
COX18
SHE9
MDM38
TIM11
MBA1
SAM50
TIM50
SDH3
SAR1
TAZ1
YJR135W−A_1
COX18
TIM13
ATP14
YNL100W
YGR235C
YPR125W
SAM35
FMP51
SAM37
TOM22
TOM5
CYC2
CRD1
TIM54
TIM12
YTA12
MDM10
MDM12
TIM22
AFG3
Gene
Z_Shift_K_Doxo_HLD
Z_Im_K_Doxo_HLD
Z_Shift_K_Doxo_EG
Z_Im_K_Doxo_EG
Z_Shift_L_Doxo_HLD
Z_Im_L_Doxo_HLD
Z_Shift_L_Doxo_EG
Z_Im_L_Doxo_EG

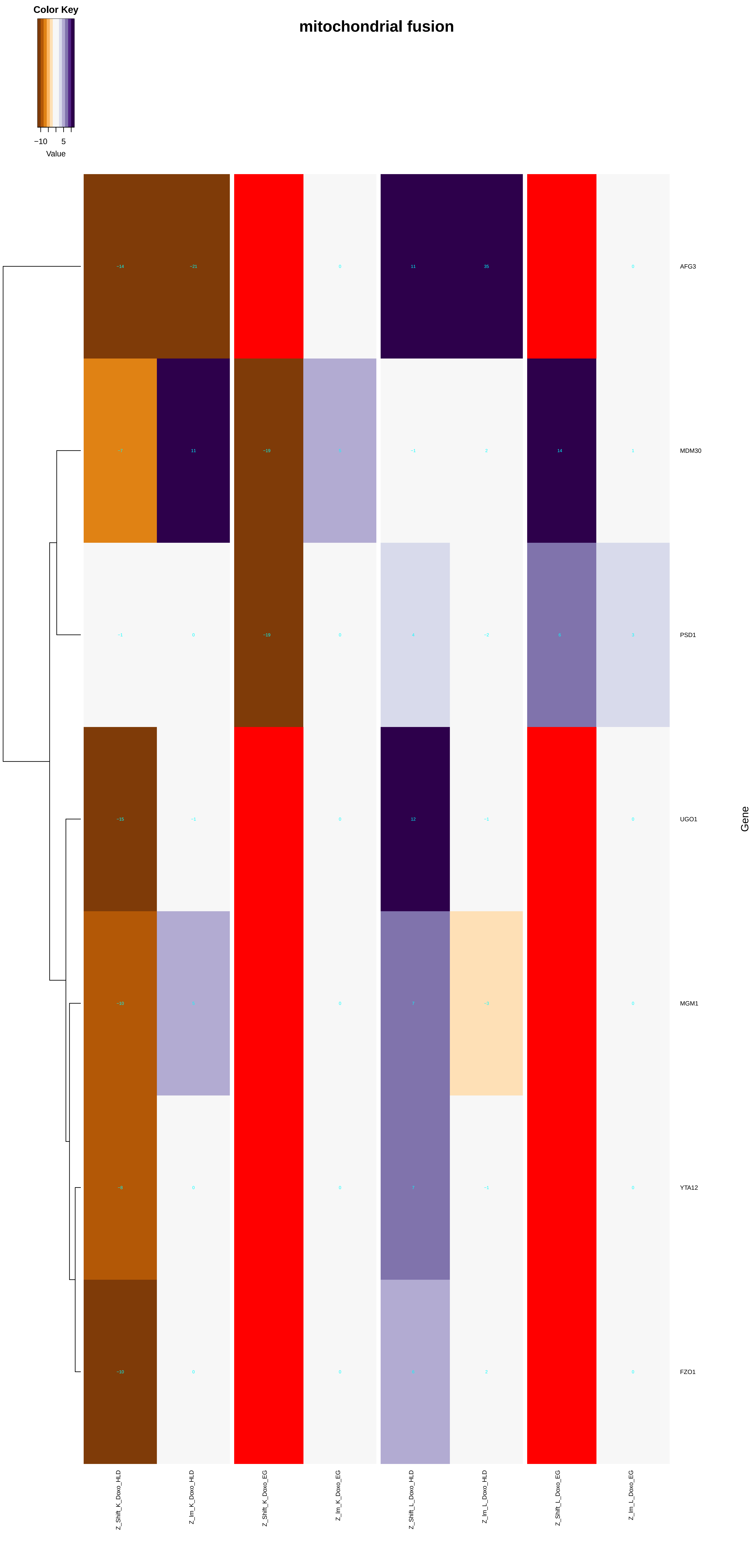

Color Key
−10
5
Value
mitochondrial fusion
AFG3
MDM30
PSD1
UGO1
MGM1
YTA12
FZO1
Gene
Z_Shift_K_Doxo_HLD
Z_Im_K_Doxo_HLD
Z_Shift_K_Doxo_EG
Z_Im_K_Doxo_EG
Z_Shift_L_Doxo_HLD
Z_Im_L_Doxo_HLD
Z_Shift_L_Doxo_EG
Z_Im_L_Doxo_EG

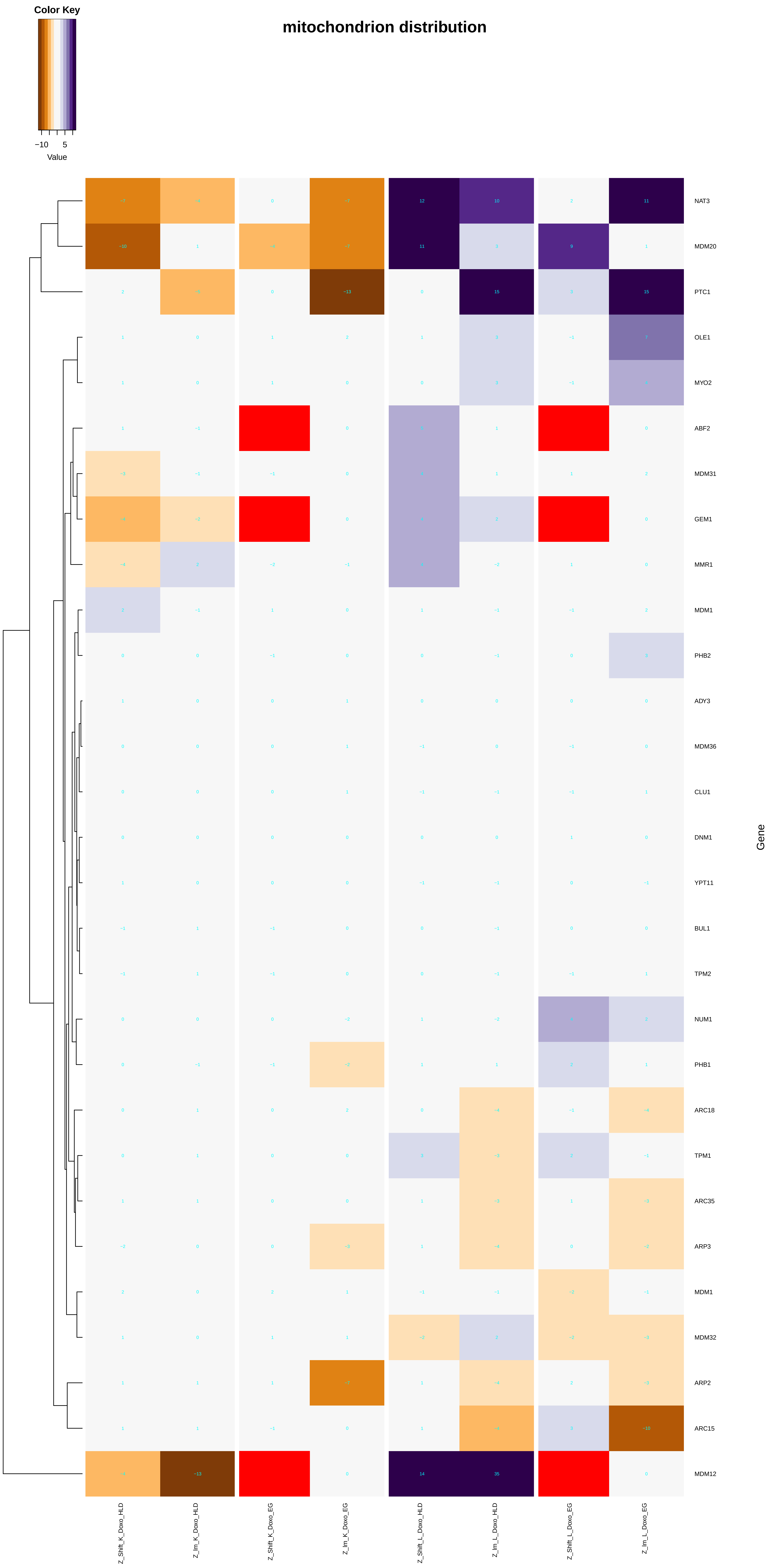

Color Key
−10
5
Value
mitochondrion distribution
NAT3
MDM20
PTC1
OLE1
MYO2
ABF2
MDM31
GEM1
MMR1
MDM1
PHB2
ADY3
MDM36
CLU1
DNM1
YPT11
BUL1
TPM2
NUM1
PHB1
ARC18
TPM1
ARC35
ARP3
MDM1
MDM32
ARP2
ARC15
MDM12
Gene
Z_Shift_K_Doxo_HLD
Z_Im_K_Doxo_HLD
Z_Shift_K_Doxo_EG
Z_Im_K_Doxo_EG
Z_Shift_L_Doxo_HLD
Z_Im_L_Doxo_HLD
Z_Shift_L_Doxo_EG
Z_Im_L_Doxo_EG

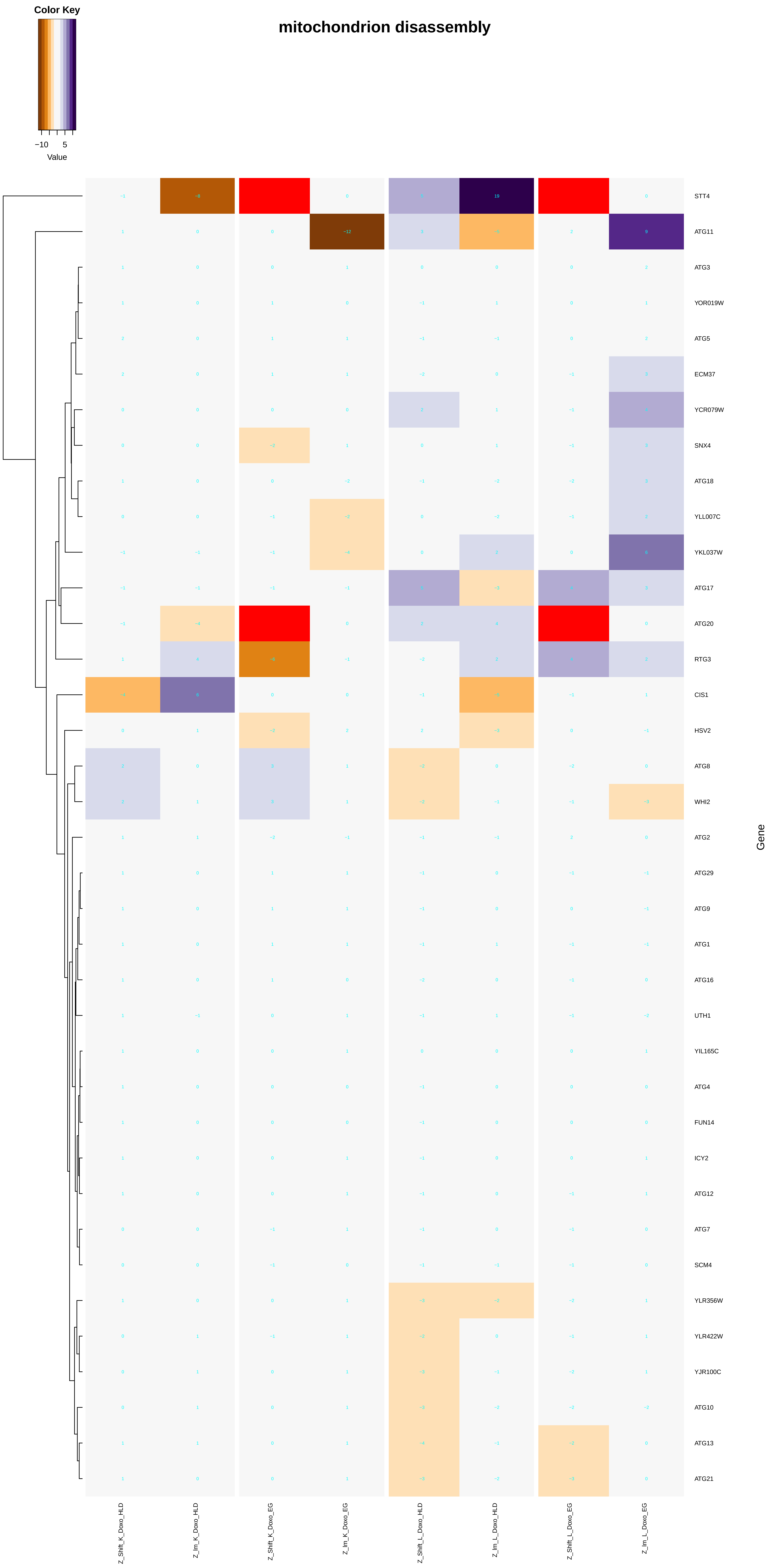

mitochondrion disassembly
Color Key
−10
5
Value
STT4
ATG11
ATG3
YOR019W
ATG5
ECM37
YCR079W
SNX4
ATG18
YLL007C
YKL037W
ATG17
ATG20
RTG3
CIS1
HSV2
ATG8
WHI2
ATG2
ATG29
ATG9
ATG1
ATG16
UTH1
YIL165C
ATG4
FUN14
ICY2
ATG12
ATG7
SCM4
YLR356W
YLR422W
YJR100C
ATG10
ATG13
ATG21
Gene
Z_Shift_K_Doxo_HLD
Z_lm_K_Doxo_HLD
Z_Shift_K_Doxo_EG
Z_lm_K_Doxo_EG
Z_Shift_L_Doxo_HLD
Z_lm_L_Doxo_HLD
Z_Shift_L_Doxo_EG
Z_lm_L_Doxo_EG

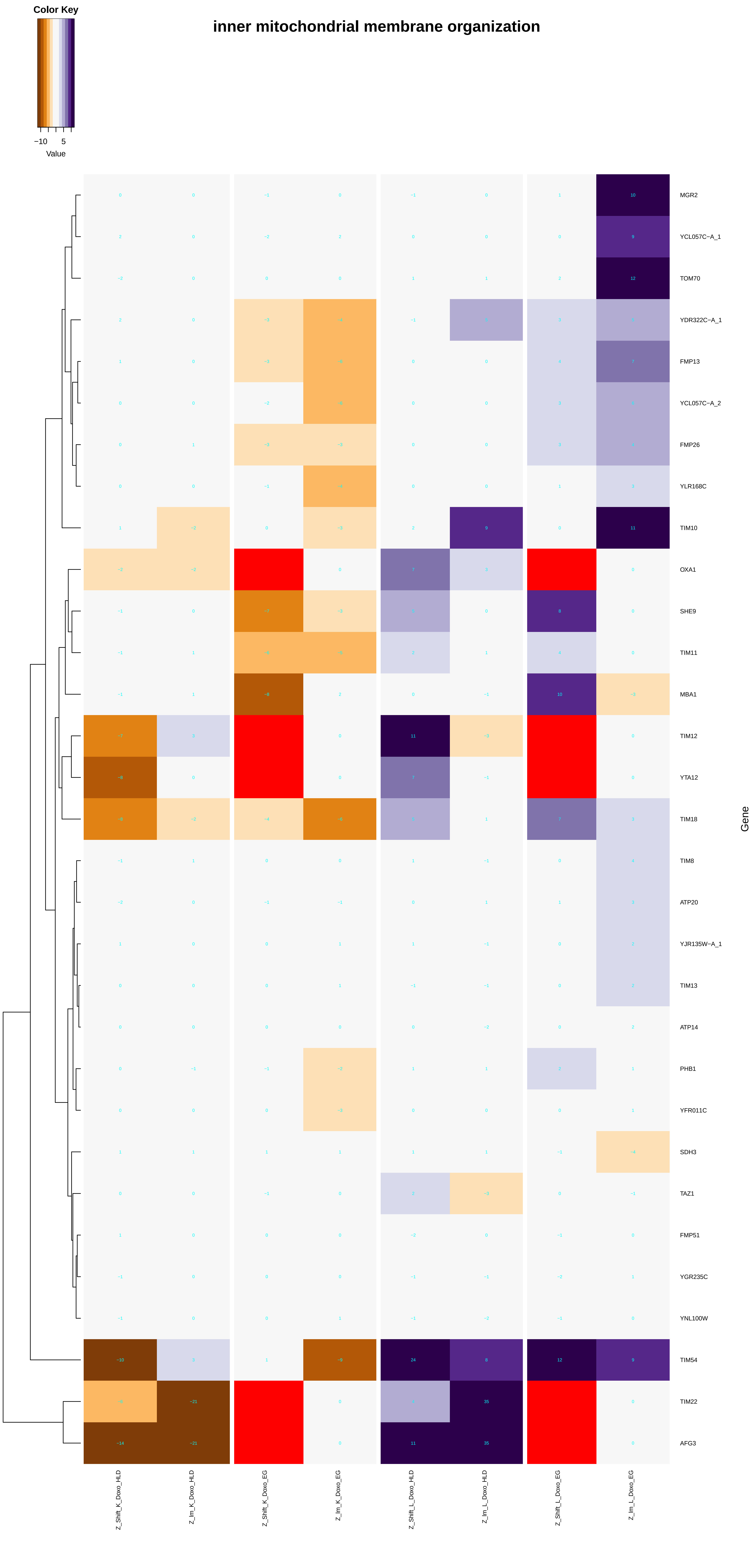

Color Key
−10
5
Value
inner mitochondrial membrane organization
MGR2
YCL057C−A_1
TOM70
YDR322C−A_1
FMP13
YCL057C−A_2
FMP26
YLR168C
TIM10
OXA1
SHE9
TIM11
MBA1
TIM12
YTA12
TIM18
TIM8
ATP20
YJR135W−A_1
TIM13
ATP14
PHB1
YFR011C
SDH3
TAZ1
FMP51
YGR235C
YNL100W
TIM54
TIM22
AFG3
Gene
Z_Shift_K_Doxo_HLD
Z_Im_K_Doxo_HLD
Z_Shift_K_Doxo_EG
Z_Im_K_Doxo_EG
Z_Shift_L_Doxo_HLD
Z_Im_L_Doxo_HLD
Z_Shift_L_Doxo_EG
Z_Im_L_Doxo_EG

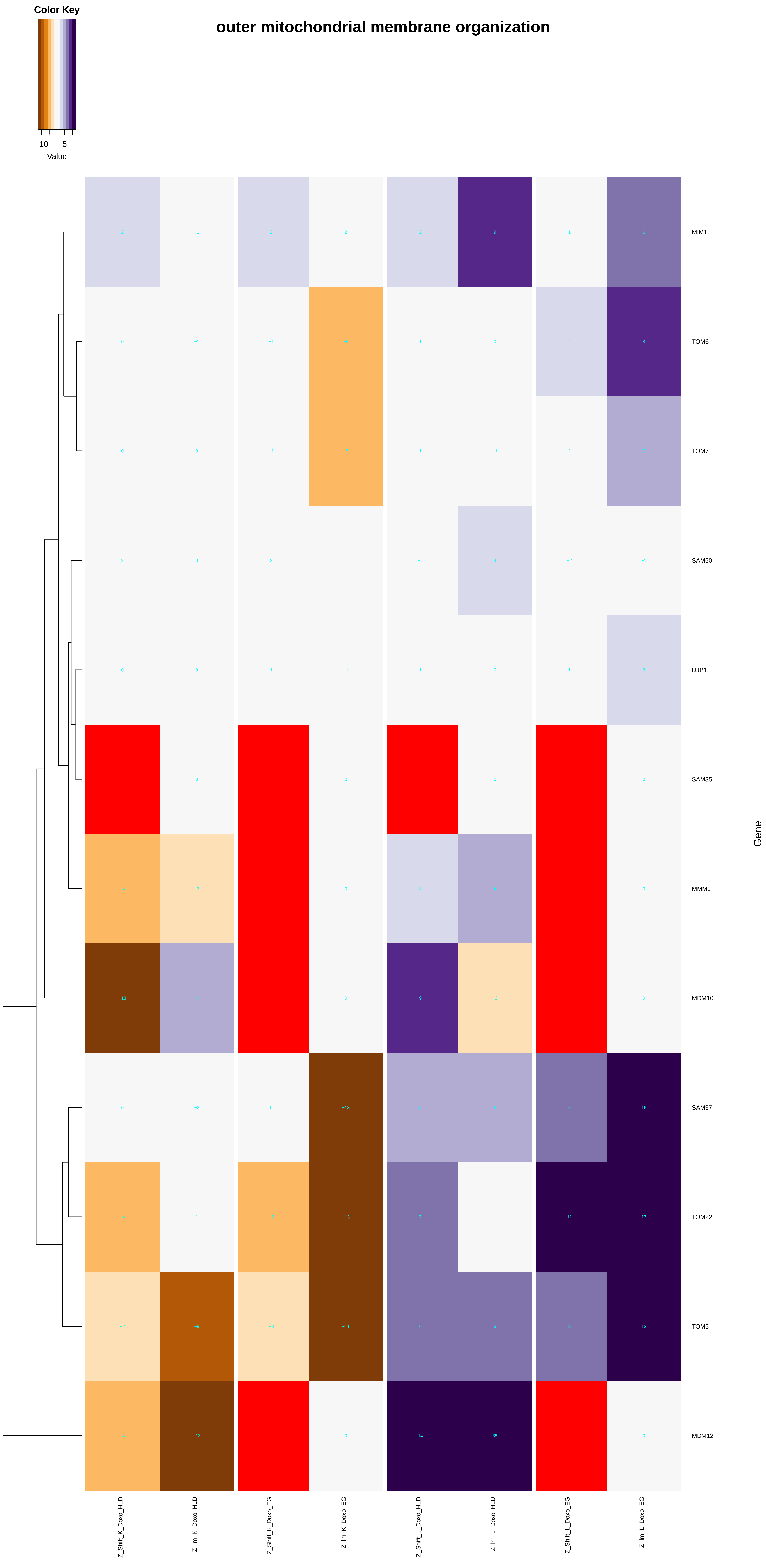
Color Key
−10
5
Value
outer mitochondrial membrane organization
MIM1
TOM6
TOM7
SAM50
DJP1
SAM35
MMM1
MDM10
SAM37
TOM22
TOM5
MDM12
Gene
Z_Shift_K_Doxo_HLD
Z_Im_K_Doxo_HLD
Z_Shift_K_Doxo_EG
Z_Im_K_Doxo_EG
Z_Shift_L_Doxo_HLD
Z_Im_L_Doxo_HLD
Z_Shift_L_Doxo_EG
Z_Im_L_Doxo_EG

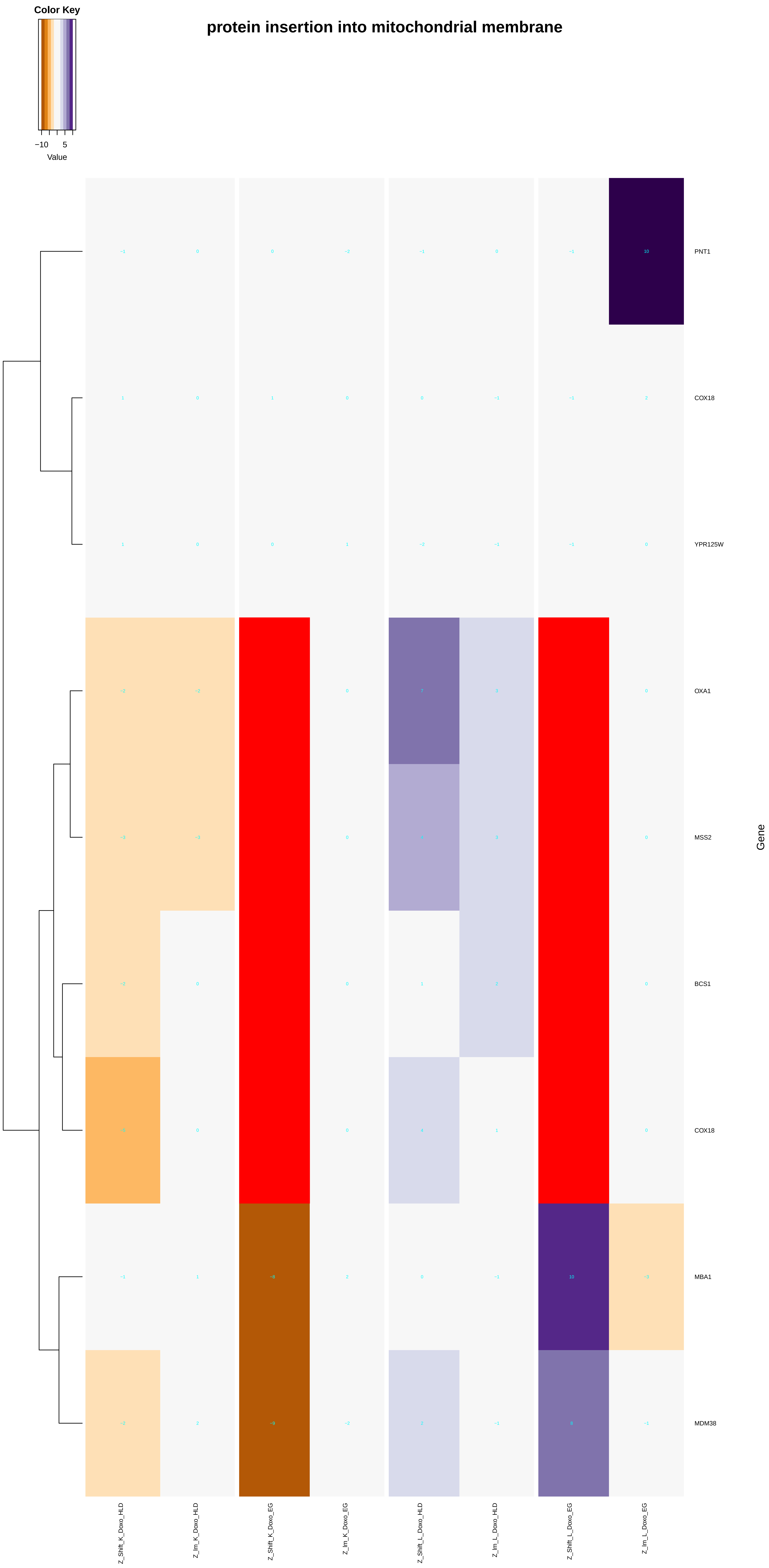
protein insertion into mitochondrial membrane
Color Key
−10
5
Value
PNT1
COX18
YPR125W
OXA1
MSS2
BCS1
COX18
MBA1
MDM38
Gene
Z_Shift_K_Doxo_HLD
Z_lm_K_Doxo_HLD
Z_Shift_K_Doxo_EG
Z_lm_K_Doxo_EG
Z_Shift_L_Doxo_HLD
Z_lm_L_Doxo_HLD
Z_Shift_L_Doxo_EG
Z_lm_L_Doxo_EG

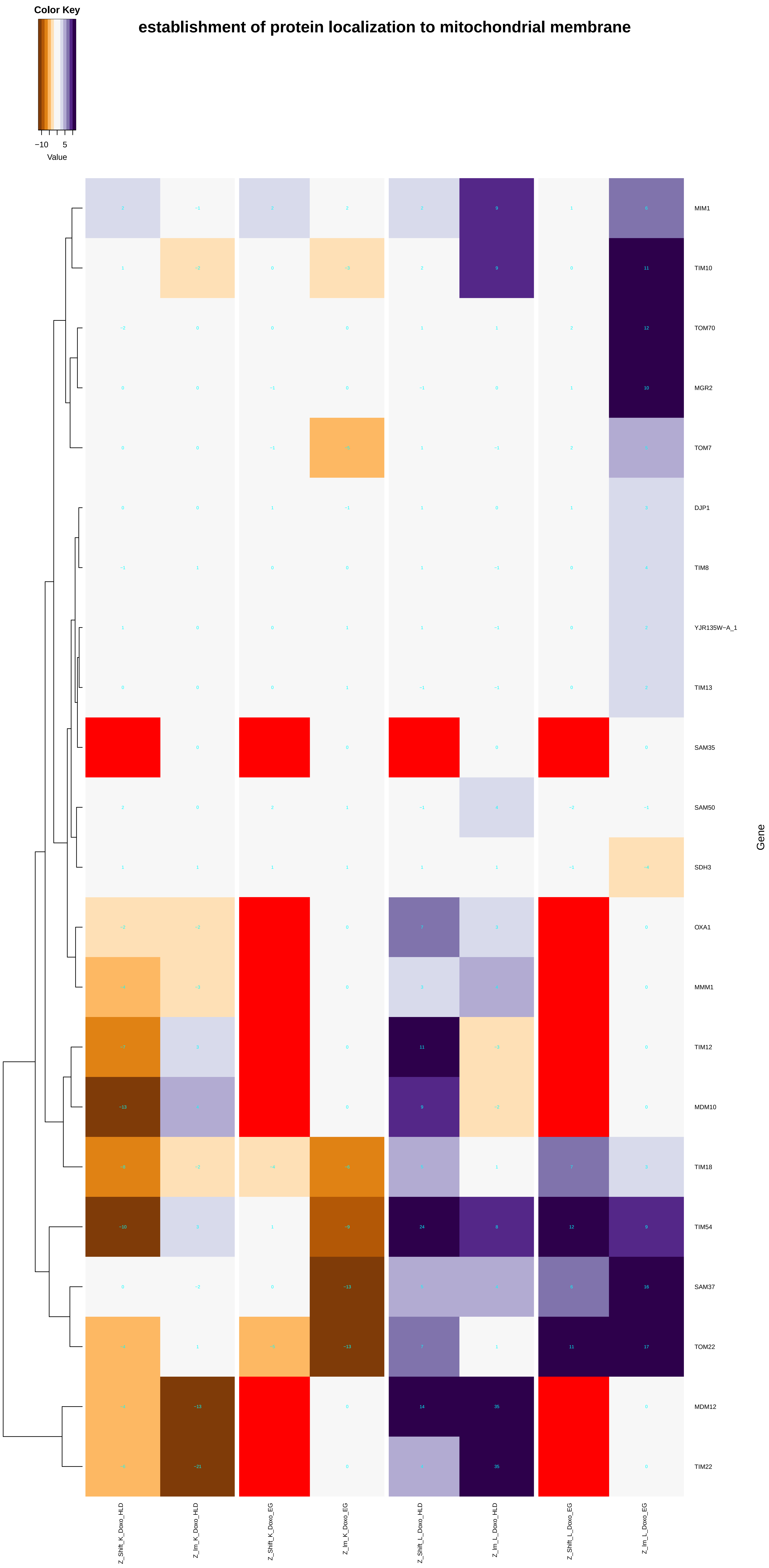

Color Key
−10
5
Value
establishment of protein localization to mitochondrial membrane
MIM1
TIM10
TOM70
MGR2
TOM7
DJP1
TIM8
YJR135W−A_1
TIM13
SAM35
SAM50
SDH3
OXA1
MMM1
TIM12
MDM10
TIM18
TIM54
SAM37
TOM22
MDM12
TIM22
Gene
Z_Shift_K_Doxo_HLD
Z_Im_K_Doxo_HLD
Z_Shift_K_Doxo_EG
Z_Im_K_Doxo_EG
Z_Shift_L_Doxo_HLD
Z_Im_L_Doxo_HLD
Z_Shift_L_Doxo_EG
Z_Im_L_Doxo_EG

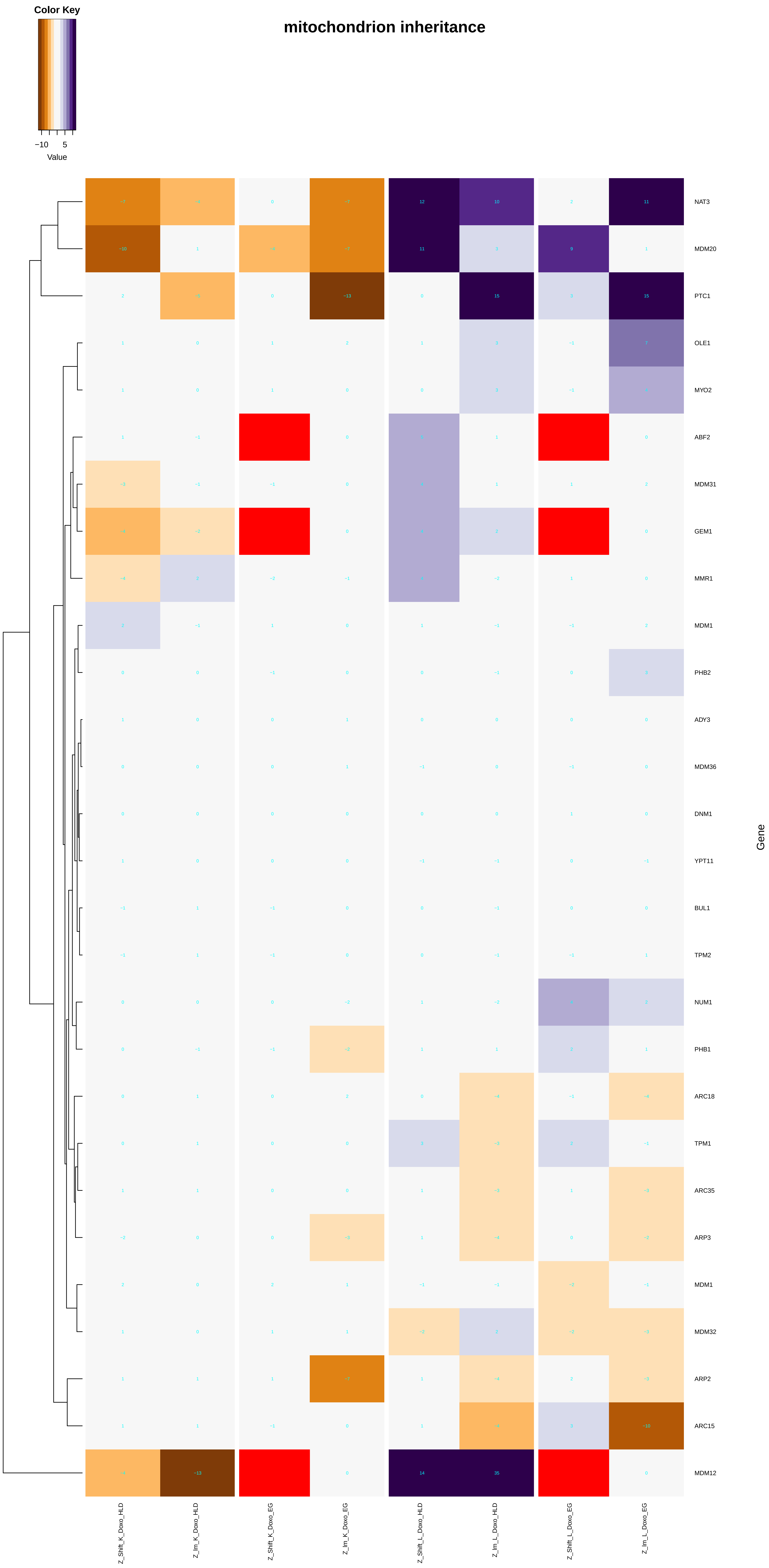

mitochondrion inheritance
Color Key
−10
5
Value
Gene
NAT3
MDM20
PTC1
OLE1
MYO2
ABF2
MDM31
GEM1
MMR1
MDM1
PHB2
ADY3
MDM36
DNM1
YPT11
BUL1
TPM2
NUM1
PHB1
ARC18
TPM1
ARC35
ARP3
MDM1
MDM32
ARP2
ARC15
MDM12
Z_Shift_K_Doxo_HLD
Z_Im_K_Doxo_HLD
Z_Shift_K_Doxo_EG
Z_Im_K_Doxo_EG
Z_Shift_L_Doxo_HLD
Z_Im_L_Doxo_HLD
Z_Shift_L_Doxo_EG
Z_Im_L_Doxo_EG

mitophagy

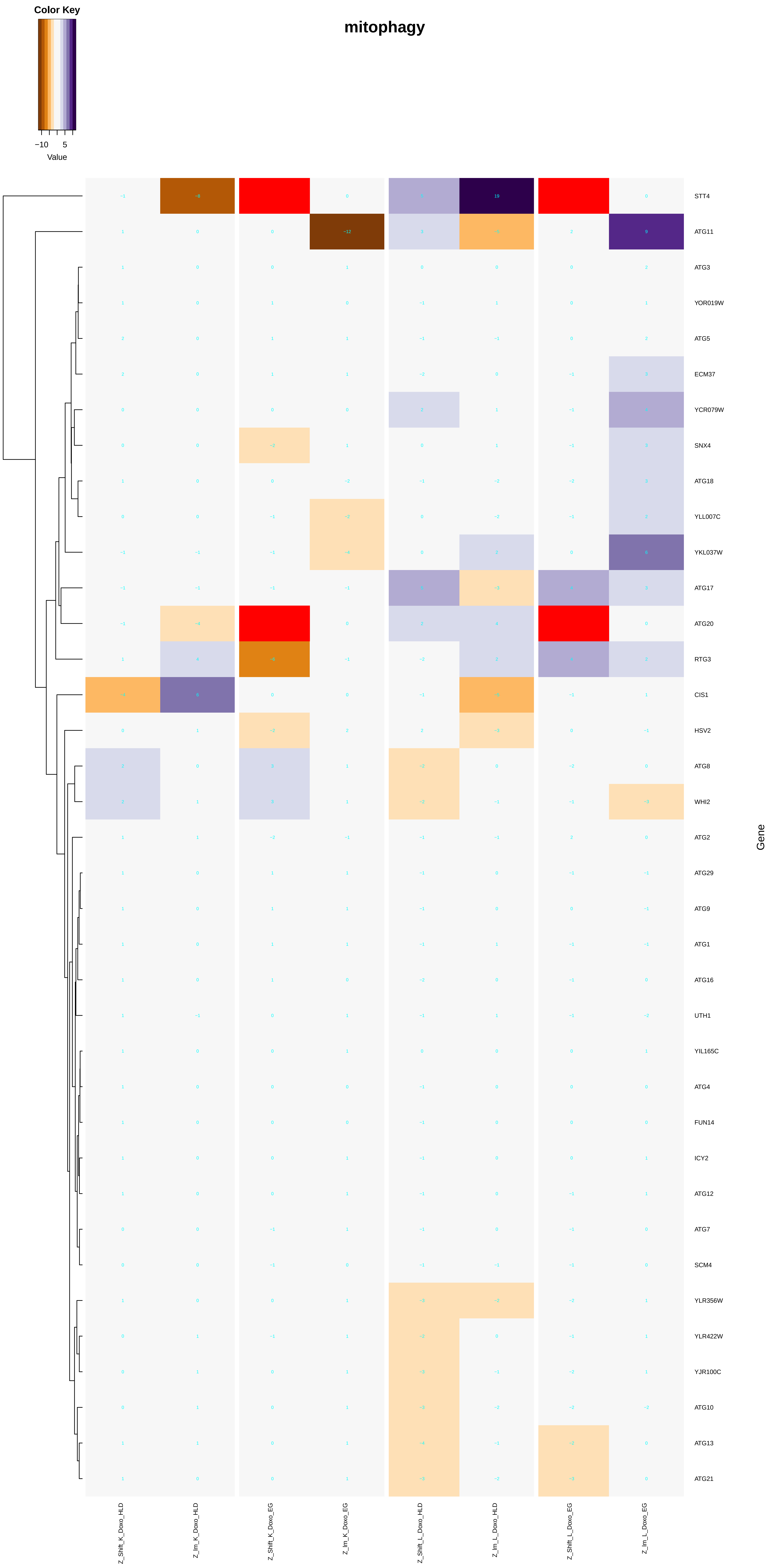

Color Key
−10
5
Value
STT4
ATG11
ATG3
YOR019W
ATG5
ECM37
YCR079W
SNX4
ATG18
YLL007C
YKL037W
ATG17
ATG20
RTG3
CIS1
HSV2
ATG8
WHI2
ATG2
ATG29
ATG9
ATG1
ATG16
UTH1
YIL165C
ATG4
FUN14
ICY2
ATG12
ATG7
SCM4
YLR356W
YLR422W
YJR100C
ATG10
ATG13
ATG21
Gene
Z_Shift_K_Doxo_HLD
Z_Im_K_Doxo_HLD
Z_Shift_K_Doxo_EG
Z_Im_K_Doxo_EG
Z_Shift_L_Doxo_HLD
Z_Im_L_Doxo_HLD
Z_Shift_L_Doxo_EG
Z_Im_L_Doxo_EG

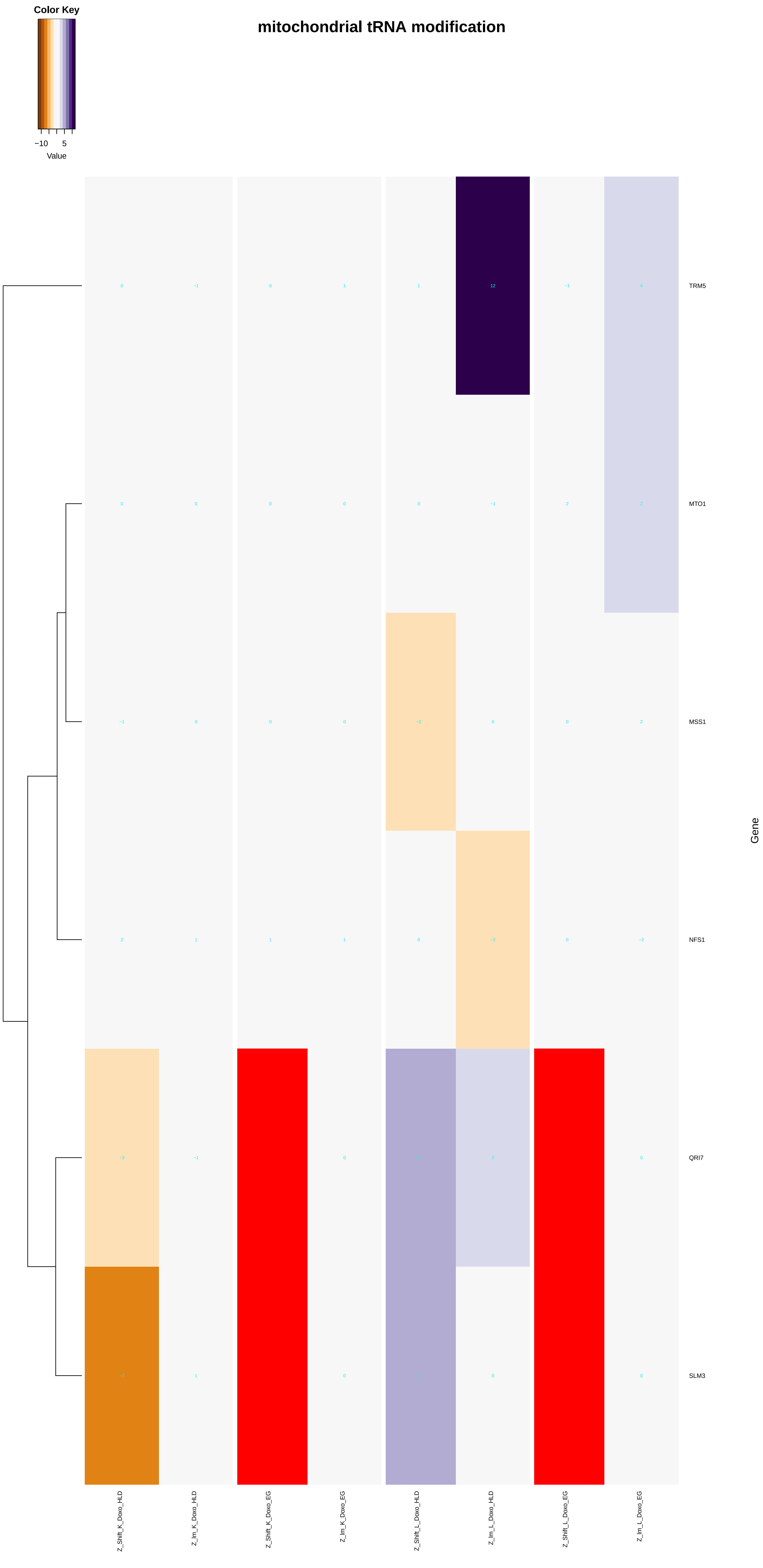

mitochondrial tRNA modification
Color Key
−10
5
Value
TRM5
MTO1
MSS1
NFS1
QRI7
SLM3
Gene
Z_Shift_K_Doxo_HLD
Z_Im_K_Doxo_HLD
Z_Shift_K_Doxo_EG
Z_Im_K_Doxo_EG
Z_Shift_L_Doxo_HLD
Z_Im_L_Doxo_HLD
Z_Shift_L_Doxo_EG
Z_Im_L_Doxo_EG

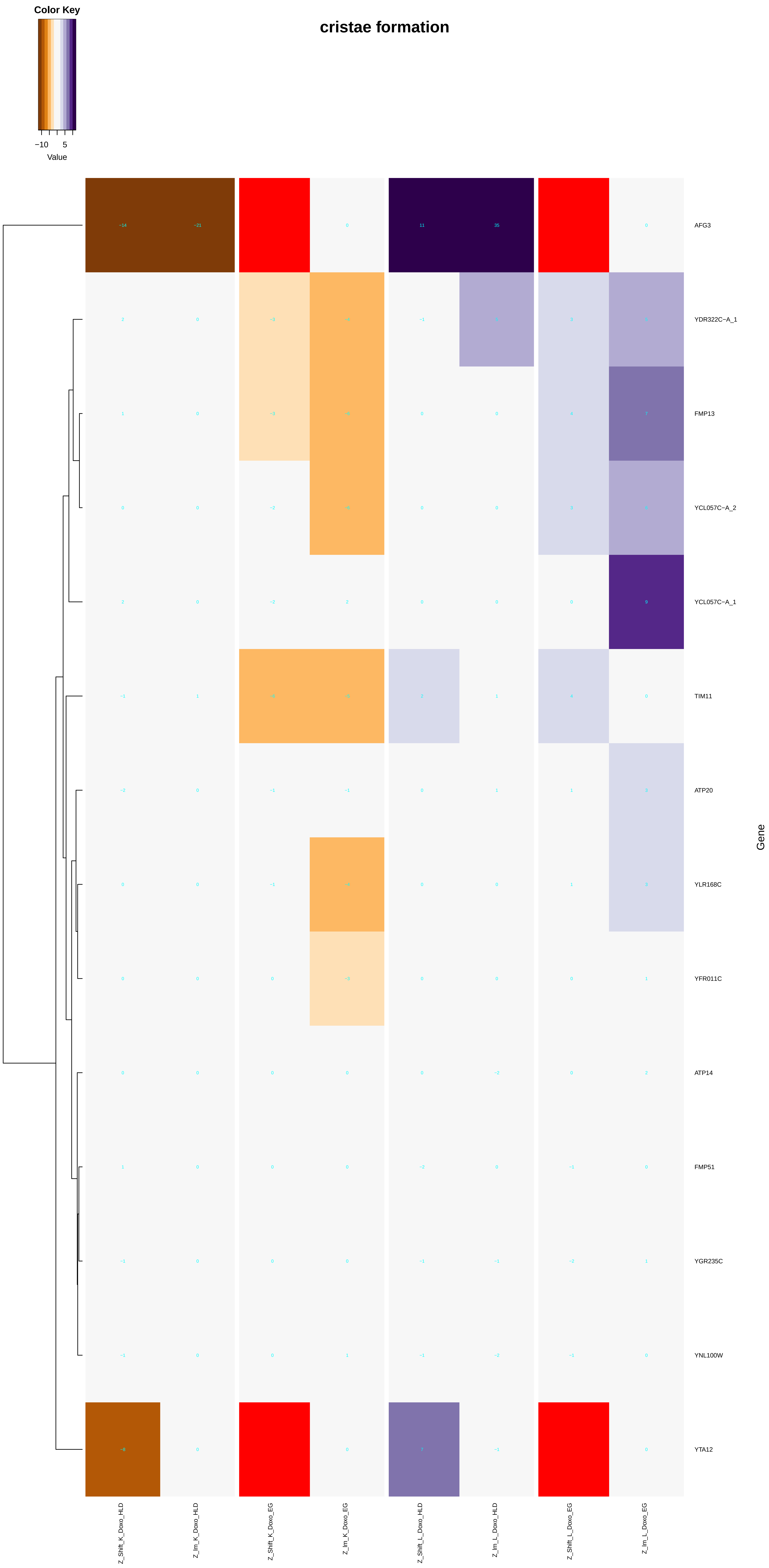

cristae formation
Color Key
−10
5
Value
AFG3
YDR322C−A_1
FMP13
YCL057C−A_2
YCL057C−A_1
TIM11
ATP20
YLR168C
YFR011C
ATP14
FMP51
YGR235C
YNL100W
YTA12
Gene
Z_Shift_K_Doxo_HLD
Z_Im_K_Doxo_HLD
Z_Shift_K_Doxo_EG
Z_Im_K_Doxo_EG
Z_Shift_L_Doxo_HLD
Z_Im_L_Doxo_HLD
Z_Shift_L_Doxo_EG
Z_Im_L_Doxo_EG

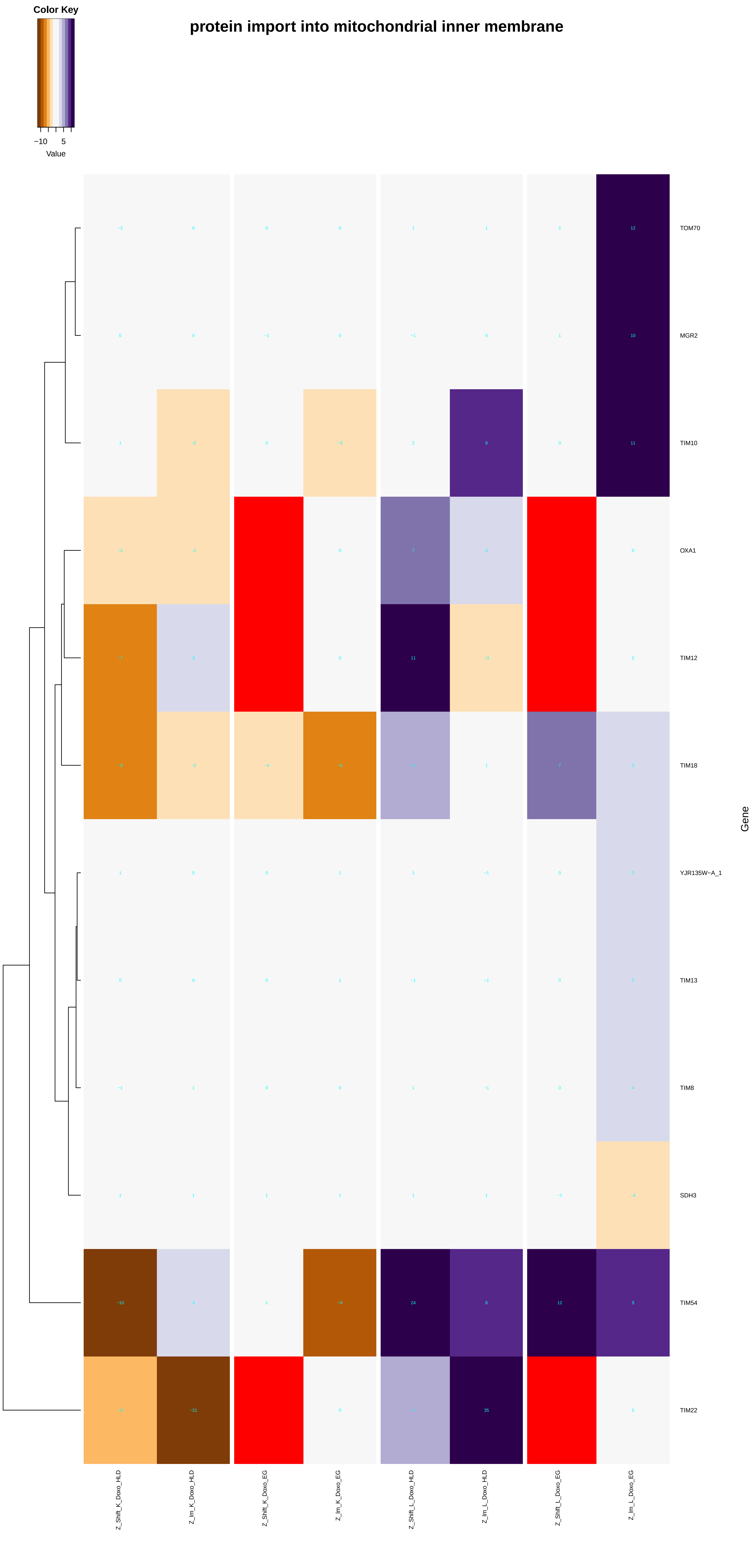
Color Key
−10
5
Value
protein import into mitochondrial inner membrane
TOM70
MGR2
TIM10
OXA1
TIM12
TIM18
YJR135W−A_1
TIM13
TIM8
SDH3
TIM54
TIM22
Gene
Z_Shift_K_Doxo_HLD
Z_Im_K_Doxo_HLD
Z_Shift_K_Doxo_EG
Z_Im_K_Doxo_EG
Z_Shift_L_Doxo_HLD
Z_Im_L_Doxo_HLD
Z_Shift_L_Doxo_EG
Z_Im_L_Doxo_EG

### protein import into mitochondrial outer membrane

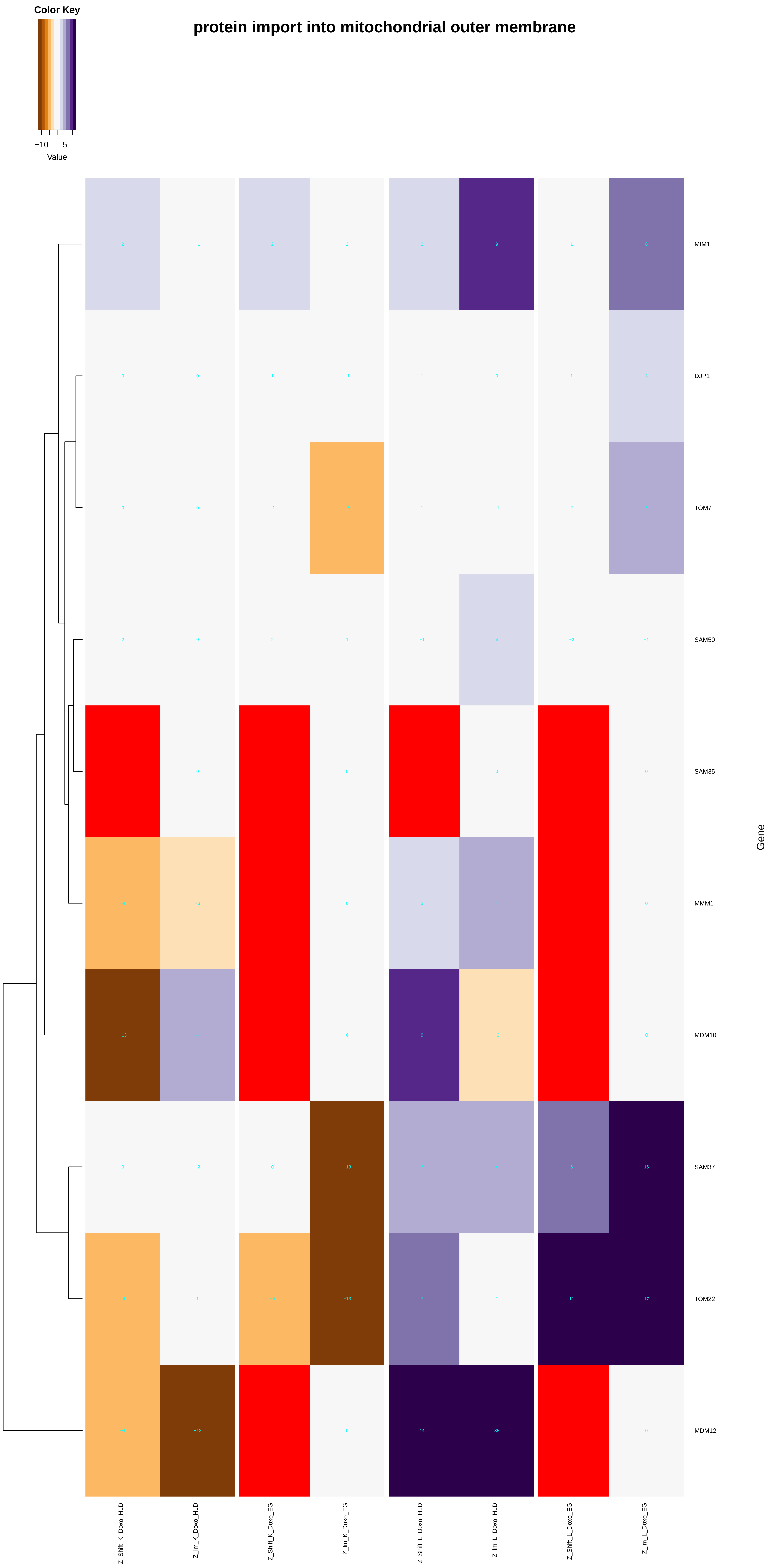

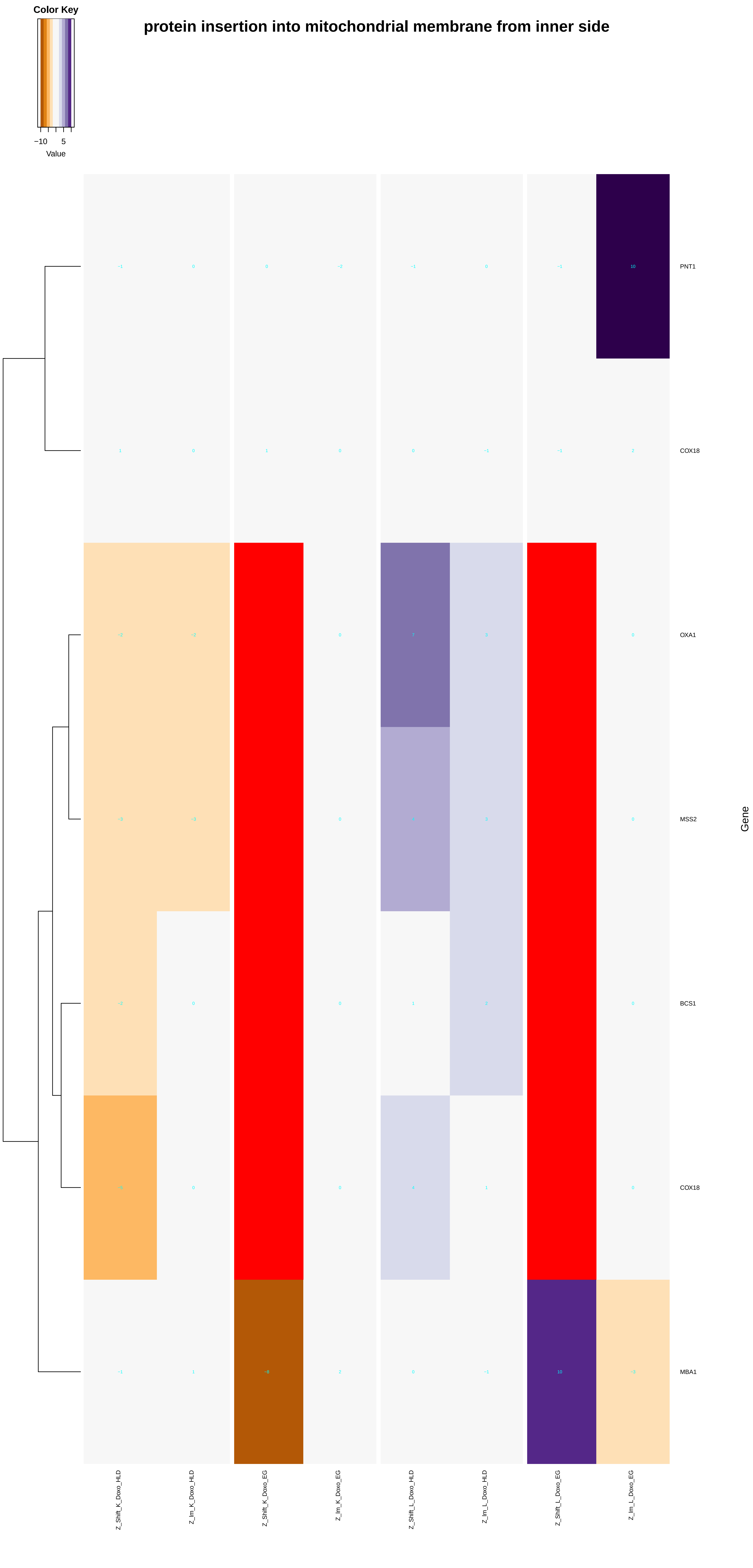

Color Key
−10
5
Value
protein insertion into mitochondrial membrane from inner side
PNT1
COX18
OXA1
MSS2
BCS1
COX18
MBA1
Gene
Z_Shift_K_Doxo_HLD
Z_Im_K_Doxo_HLD
Z_Shift_K_Doxo_EG
Z_Im_K_Doxo_EG
Z_Shift_L_Doxo_HLD
Z_Im_L_Doxo_HLD
Z_Shift_L_Doxo_EG
Z_Im_L_Doxo_EG

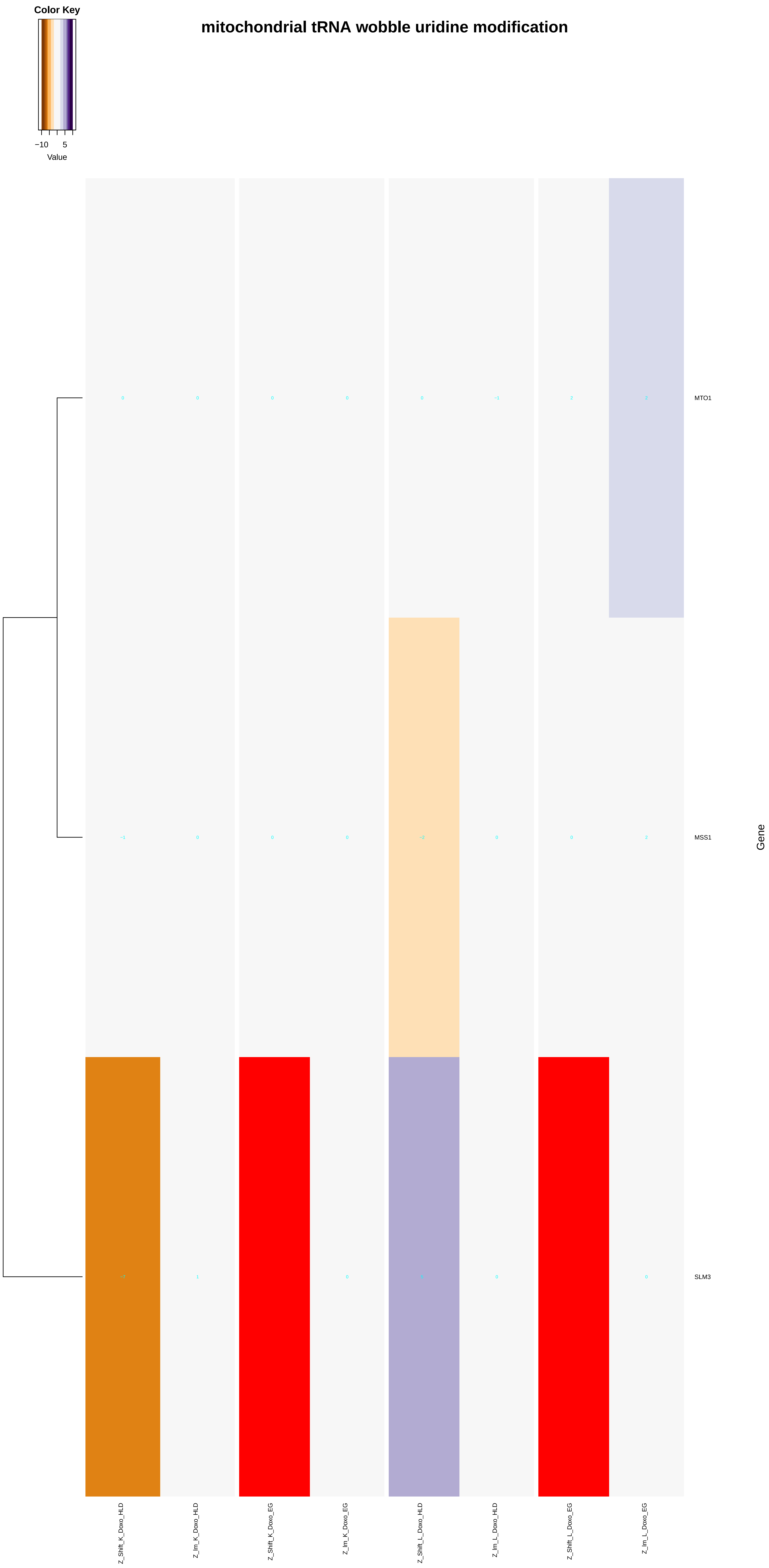

mitochondrial tRNA wobble uridine modification
Color Key
−10
5
Value
MTO1
MSS1
SLM3
Gene
Z_Shift_K_Doxo_HLD
Z_Im_K_Doxo_HLD
Z_Shift_K_Doxo_EG
Z_Im_K_Doxo_EG
Z_Shift_L_Doxo_HLD
Z_Im_L_Doxo_HLD
Z_Shift_L_Doxo_EG
Z_Im_L_Doxo_EG
