## Additional File 9 for "A yeast phenomic model for the influence of Warburg metabolism on genetic buffering of doxorubicin": modification-dependent_macromolecule_catabolic_process.pdf

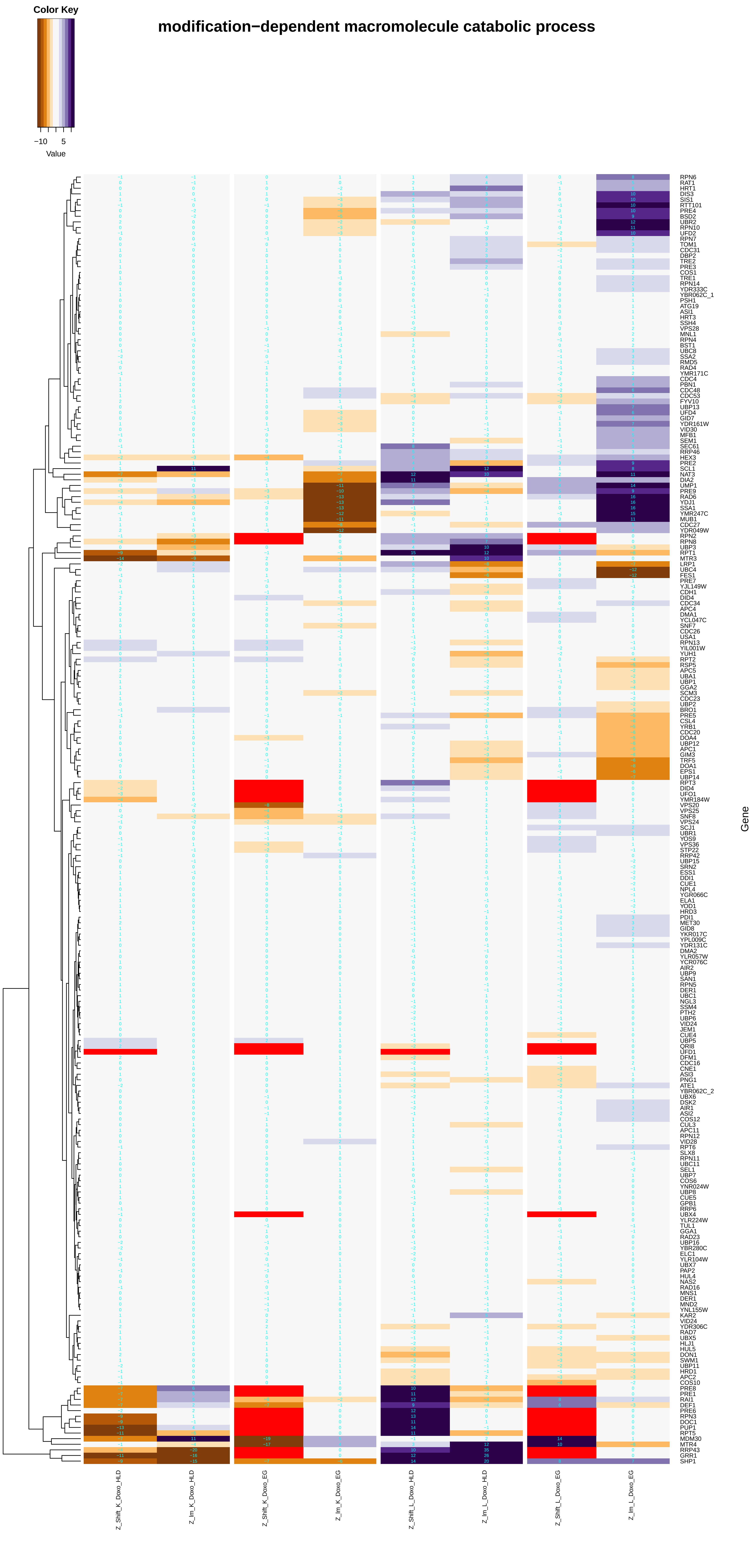
modification−dependent macromolecule catabolic process
Color Key
−10
5
Value
RPN6
RAT1
HRT1
DIS3
SIS1
RTT101
PRE4
BSD2
UBR2
RPN10
UFD2
RPN7
TOM1
CDC31
DBP2
TRE2
PRE3
COS1
TRE1
RPN14
YDR333C
YBR062C_1
PSH1
ATG19
ASI1
HRT3
SSH4
VPS28
MNL1
RPN4
BST1
UBC8
SSA2
RMD5
RAD4
YMR171C
CDC4
PBN1
CDC48
CDC53
FYV10
UBP13
UFD4
GID7
YDR161W
VID30
MFB1
SEM1
SEC61
RRP46
HEX3
PRE2
SCL1
NAT3
DIA2
UMP1
PRE9
RAD6
YDJ1
SSA1
YMR247C
MUB1
CDC27
YDR049W
RPN2
RPN8
UBP3
RPT1
MTR3
LRP1
UBC4
FES1
PRE7
YJL149W
CDH1
DID4
CDC34
APC4
DMA1
YCL047C
SNF7
CDC26
USA1
RPN13
YIL001W
YUH1
RPT2
RSP5
APC5
UBA1
UBP1
GGA2
SCM3
CDC23
UBP2
BRO1
PRE5
CSL4
YRB1
CDC20
DOA4
UBP12
APC1
GIM3
TRF5
DOA1
EPS1
UBP14
RPT3
DID4
UFO1
YMR184W
VPS20
VPS25
SNF8
VPS24
SCJ1
UBR1
YOS9
VPS36
STP22
RRP42
UBP15
SRN2
ESS1
DDI1
CUE1
NPL4
YGR066C
ELA1
YOD1
HRD3
PDI1
MET30
GID8
YKR017C
YPL009C
YDR131C
DMA2
YLR057W
YCR076C
AIR2
UBP9
SAN1
RPN5
DER1
UBC1
NGL3
SSM4
PTH2
UBP6
VID24
JEM1
CUE4
UBP5
QRI8
UFD1
DFM1
CDC16
CNE1
ASI3
PNG1
ATE1
YBR062C_2
UBX6
DSK2
AIR1
ASI2
COS12
CUL3
APC11
RPN12
VID28
RPT6
SLX8
RPN11
UBC11
SEL1
UBP7
COS6
YNR024W
UBP8
CUE5
GPB1
RRP6
UBX4
YLR224W
TUL1
GGA1
RAD23
UBP16
YBR280C
ELC1
YLR104W
UBX7
PAP2
HUL4
NAS2
RAD16
MNS1
DER1
MND2
YNL155W
KAR2
VID24
YDR306C
RAD7
UBX5
HLJ1
HUL5
DON1
SWM1
UBP11
HRD1
APC2
COS10
PRE8
PRE1
RAI1
DEF1
PRE6
RPN3
DOC1
PUP1
RPT5
MDM30
MTR4
RRP43
GRR1
SHP1
Gene
Z_Shift_K_Doxo_HLD
Z_lm_K_Doxo_HLD
Z_Shift_K_Doxo_EG
Z_lm_K_Doxo_EG
Z_Shift_L_Doxo_HLD
Z_lm_L_Doxo_HLD
Z_Shift_L_Doxo_EG
Z_lm_L_Doxo_EG

### modification−dependent protein catabolic process

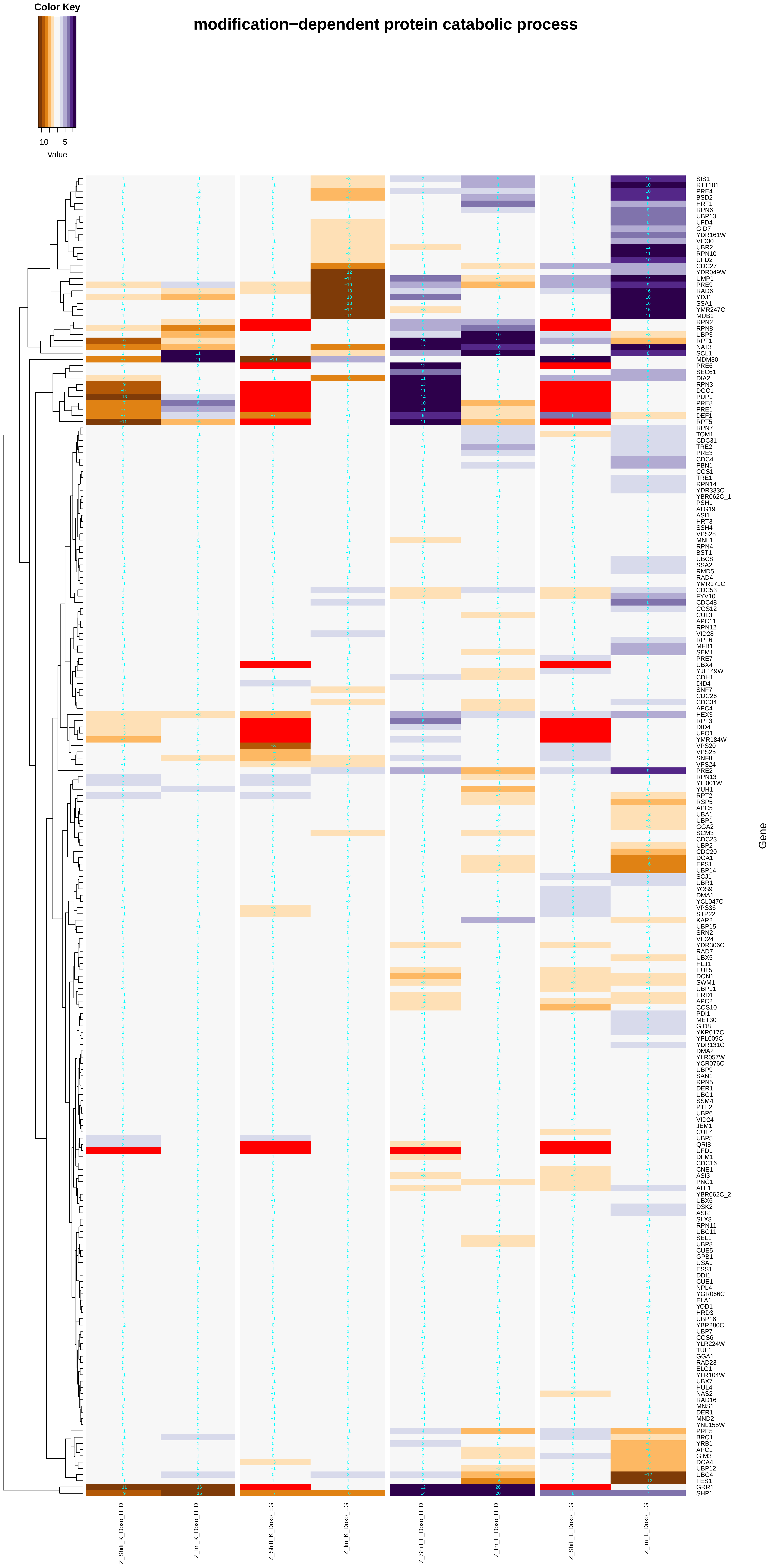

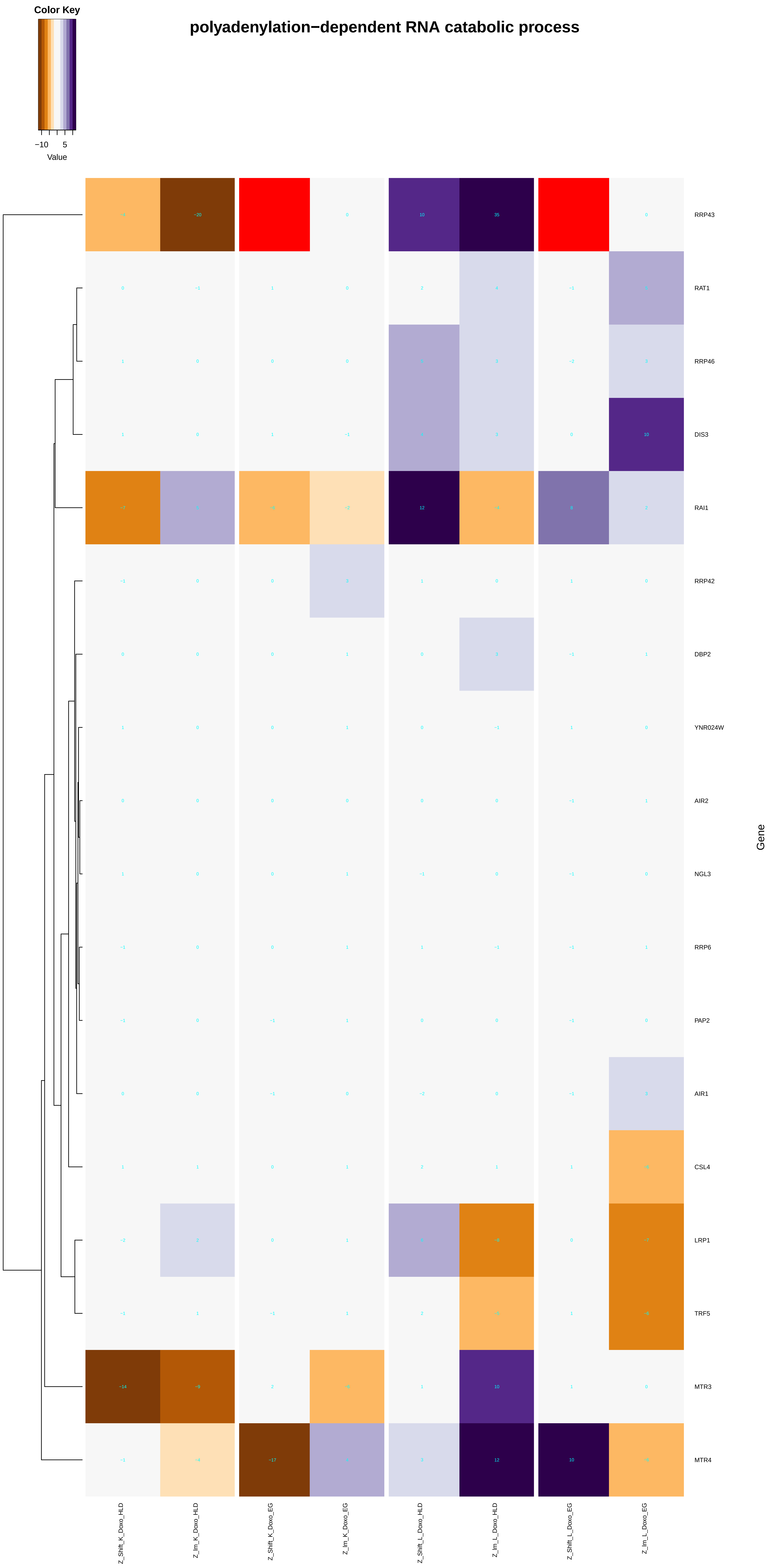
Color Key
−10
5
Value
polyadenylation−dependent RNA catabolic process
RRP43
RAT1
RRP46
DIS3
RAI1
RRP42
DBP2
YNR024W
AIR2
NGL3
RRP6
PAP2
AIR1
CSL4
LRP1
TRF5
MTR3
MTR4
Gene
Z_Shift_K_Doxo_HLD
Z_Im_K_Doxo_HLD
Z_Shift_K_Doxo_EG
Z_Im_K_Doxo_EG
Z_Shift_L_Doxo_HLD
Z_Im_L_Doxo_HLD
Z_Shift_L_Doxo_EG
Z_Im_L_Doxo_EG

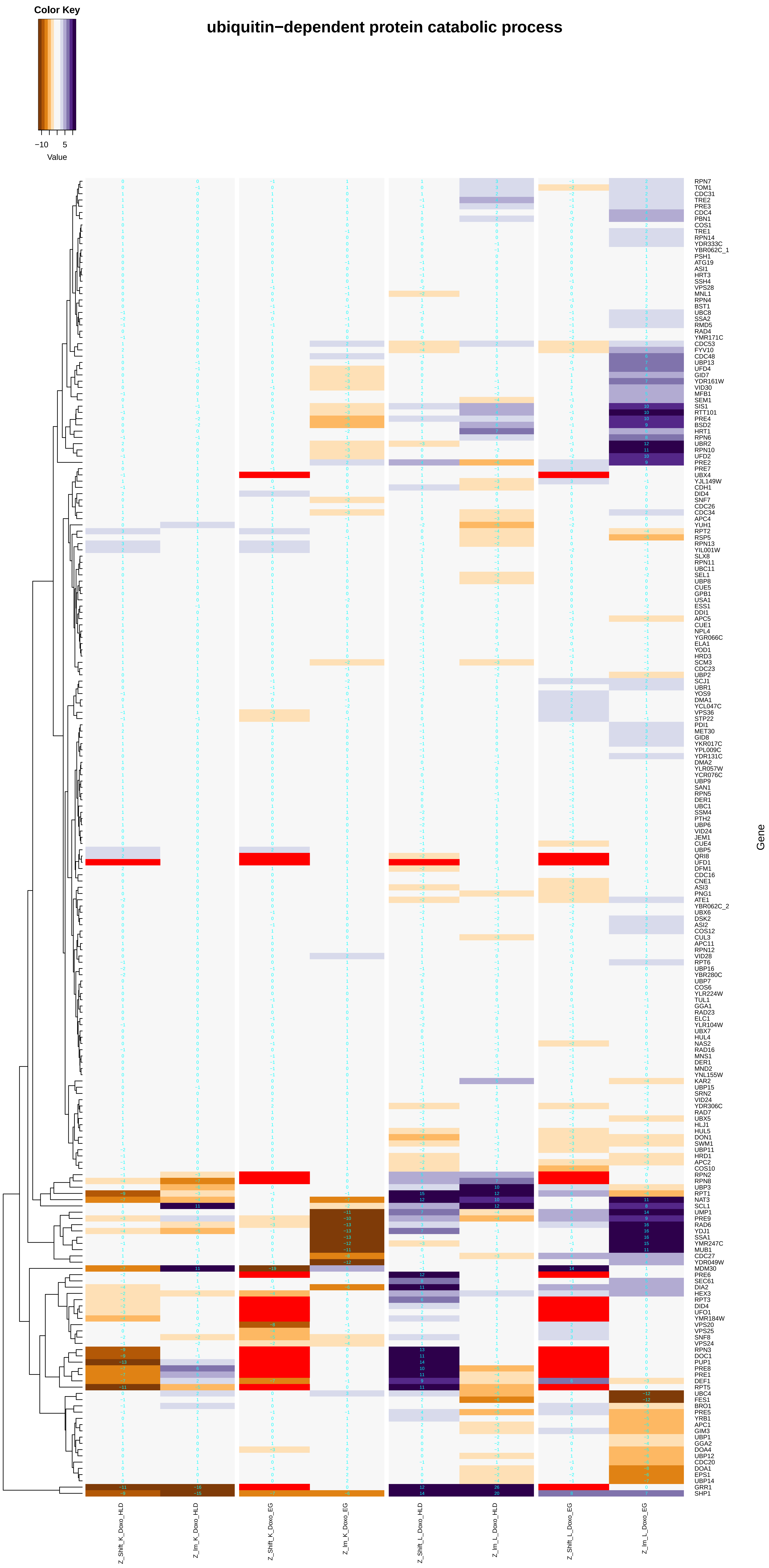
ubiquitin−dependent protein catabolic process
Color Key
−10
5
Value
Z_Shift_K_Doxo_HLD
Z_Im_K_Doxo_HLD
Z_Shift_K_Doxo_EG
Z_Im_K_Doxo_EG
Z_Shift_L_Doxo_HLD
Z_Im_L_Doxo_HLD
Z_Shift_L_Doxo_EG
Z_Im_L_Doxo_EG
Gene
RPN7
TOM1
CDC31
TRE2
PRE3
CDC4
PBN1
COS1
TRE1
RPN14
YDR333C
YBR062C_1
PSH1
ATG19
ASI1
HRT3
SSH4
VPS28
MNL1
RPN4
BST1
UBC8
SSA2
RMD5
RAD4
YMR171C
CDC53
FYV10
CDC48
UBP13
UFD4
GID7
YDR161W
VID30
MFB1
SEM1
SIS1
RTT101
PRE4
BSD2
HRT1
RPN6
UBR2
RPN10
UFD2
PRE2
PRE7
UBX4
YJL149W
CDH1
DID4
SNF7
CDC26
CDC34
APC4
YUH1
RPT2
RSP5
RPN13
YIL001W
SLX8
RPN11
UBC11
SEL1
UBP8
CUE5
GPB1
USA1
ESS1
DDI1
APC5
CUE1
NPL4
YGR066C
ELA1
YOD1
HRD3
SCM3
CDC23
UBP2
SCJ1
UBR1
YOS9
DMA1
YCL047C
VPS36
STP22
PDI1
MET30
GID8
YKR017C
YPL009C
YDR131C
DMA2
YLR057W
YCR076C
UBP9
SAN1
RPN5
DER1
UBC1
SSM4
PTH2
UBP6
VID24
JEM1
CUE4
UBP5
QRI8
UFD1
DFM1
CDC16
CNE1
ASI3
PNG1
ATE1
YBR062C_2
UBX6
DSK2
ASI2
COS12
CUL3
APC11
RPN12
VID28
RPT6
UBP16
YBR280C
UBP7
COS6
YLR224W
TUL1
GGA1
RAD23
ELC1
YLR104W
UBX7
HUL4
NAS2
RAD16
MNS1
DER1
MND2
YNL155W
KAR2
UBP15
SRN2
VID24
YDR306C
RAD7
UBX5
HLJ1
HUL5
DON1
SWM1
UBP11
HRD1
APC2
COS10
RPN2
RPN8
UBP3
RPT1
NAT3
SCL1
UMP1
PRE9
RAD6
YDJ1
SSA1
YMR247C
MUB1
CDC27
YDR049W
MDM30
PRE6
SEC61
DIA2
HEX3
RPT3
DID4
UFO1
YMR184W
VPS20
VPS25
SNF8
VPS24
RPN3
DOC1
PUP1
PRE8
PRE1
DEF1
RPT5
UBC4
FES1
BRO1
PRE5
YRB1
APC1
GIM3
UBP1
GGA2
DOA4
UBP12
CDC20
DOA1
EPS1
UBP14
GRR1
SHP1

### polyadenylation−dependent ncRNA catabolic process

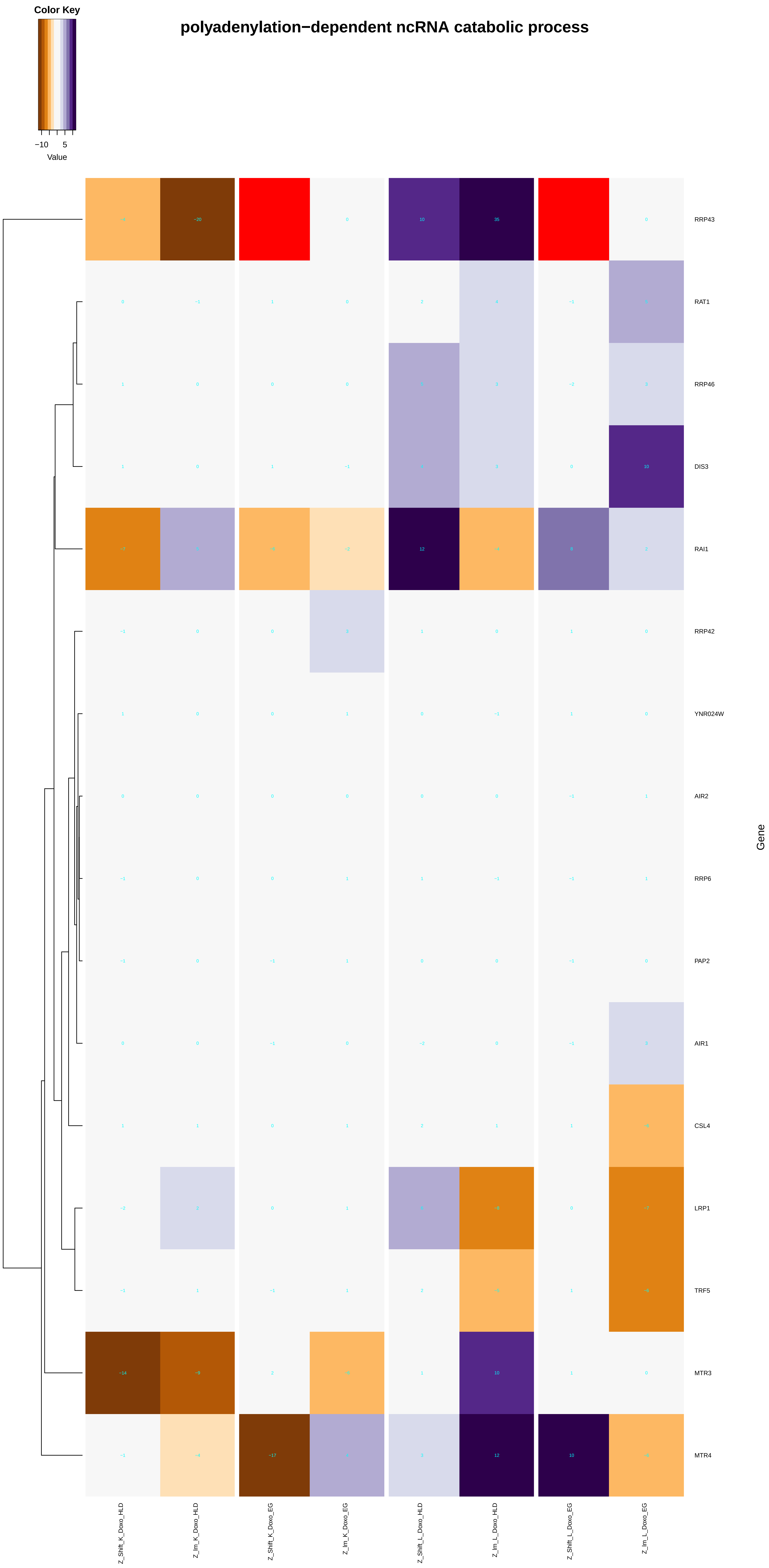

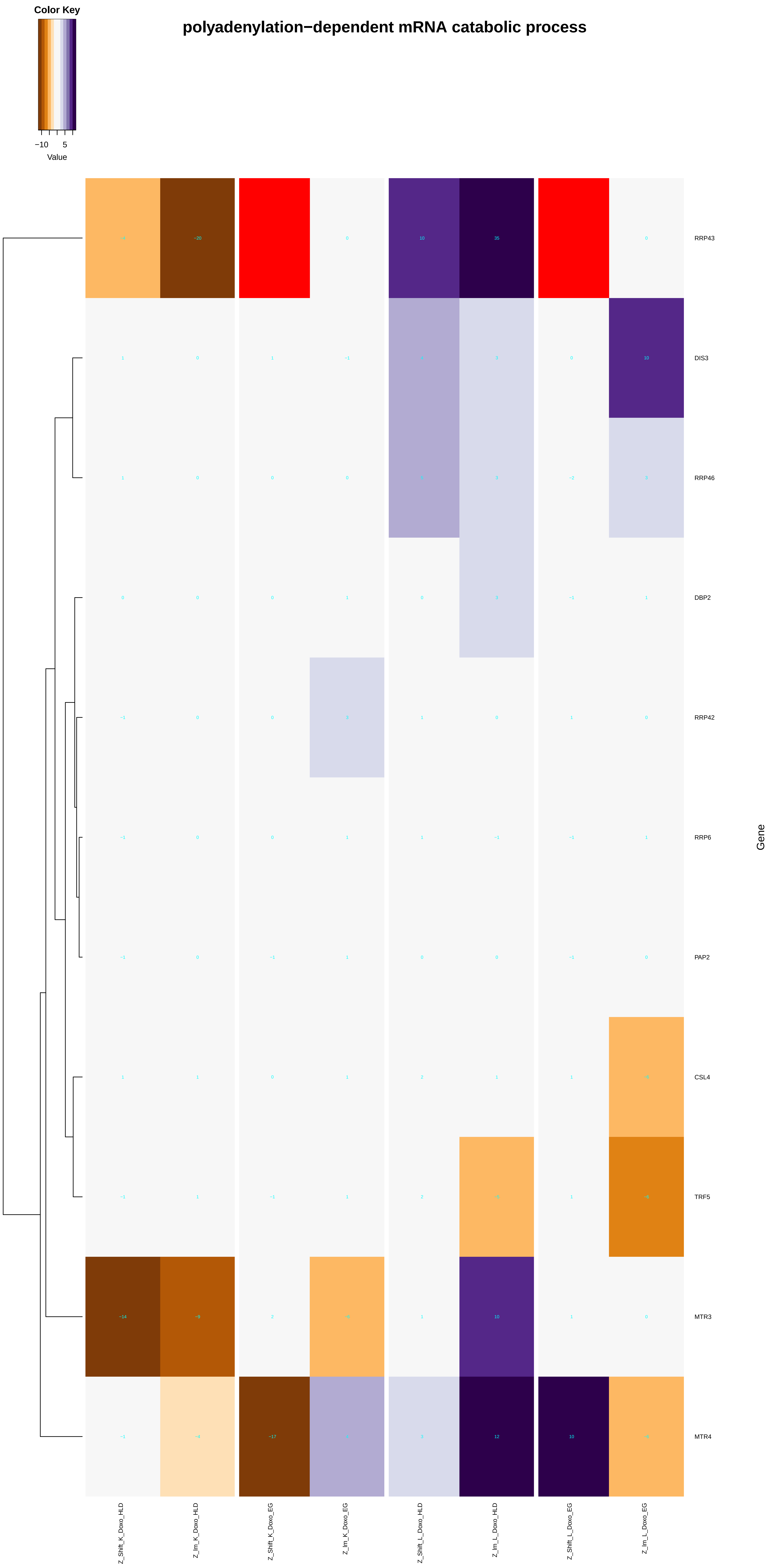

Color Key
−10
5
Value
polyadenylation−dependent mRNA catabolic process
RRP43
DIS3
RRP46
DBP2
RRP42
RRP6
PAP2
CSL4
TRF5
MTR3
MTR4
Gene
Z_Shift_K_Doxo_HLD
Z_Im_K_Doxo_HLD
Z_Shift_K_Doxo_EG
Z_Im_K_Doxo_EG
Z_Shift_L_Doxo_HLD
Z_Im_L_Doxo_HLD
Z_Shift_L_Doxo_EG
Z_Im_L_Doxo_EG

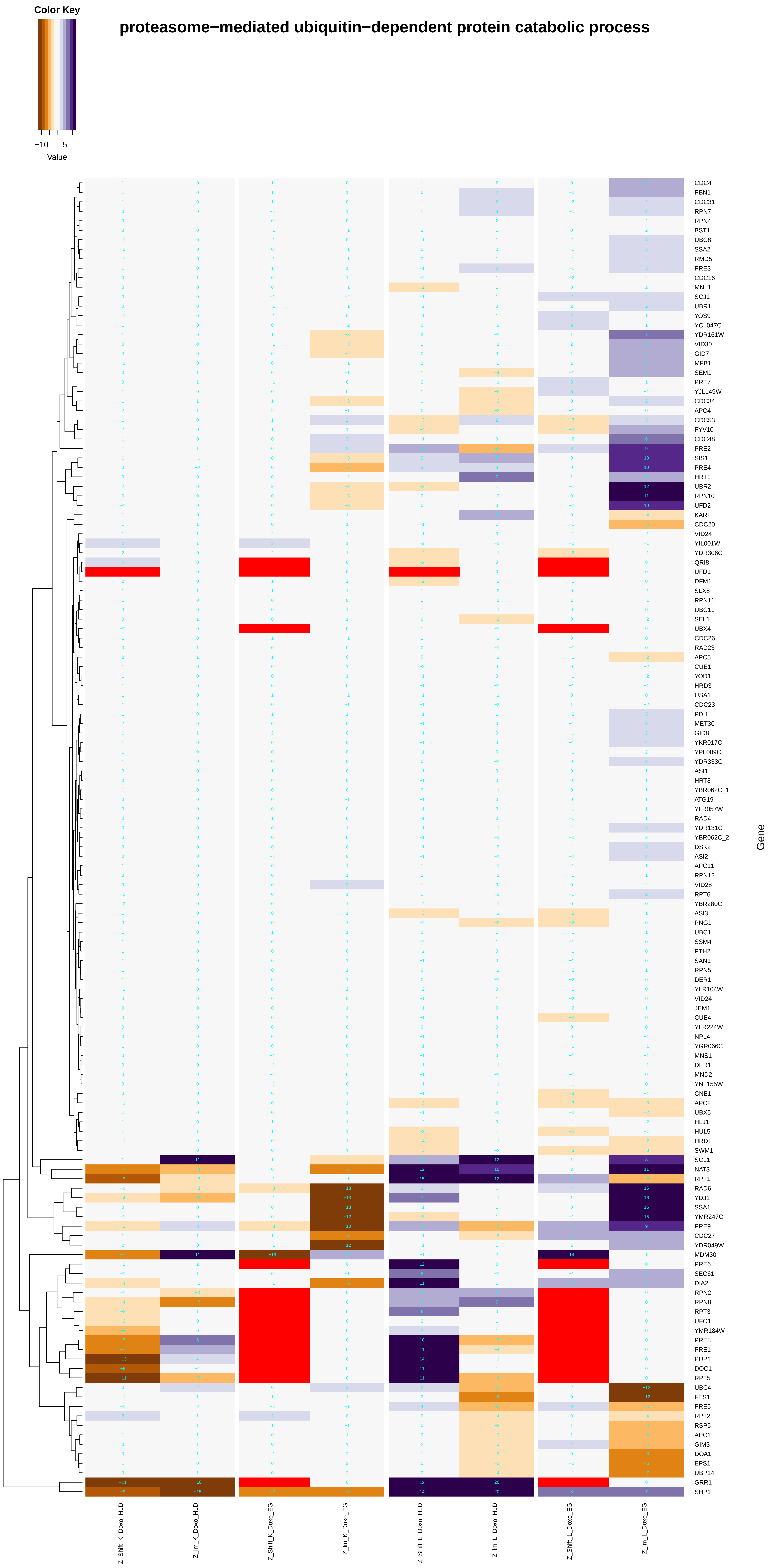

Color Key
−10
5
Value
proteasome−mediated ubiquitin−dependent protein catabolic process
CDC4
PBN1
CDC31
RPN7
RPN4
BST1
UBC8
SSA2
RMD5
PRE3
CDC16
MNL1
SCJ1
UBR1
YOS9
YCL047C
YDR161W
VID30
GID7
MFB1
SEM1
PRE7
YJL149W
CDC34
APC4
CDC53
FYV10
CDC48
PRE2
SIS1
PRE4
HRT1
UBR2
RPN10
UFD2
KAR2
CDC20
VID24
YIL001W
YDR306C
QRI8
UFD1
DFM1
SLX8
RPN11
UBC11
SEL1
UBX4
CDC26
RAD23
APC5
CUE1
YOD1
HRD3
USA1
CDC23
PDI1
MET30
GID8
YKR017C
YPL009C
YDR333C
ASI1
HRT3
YBR062C_1
ATG19
YLR057W
RAD4
YDR131C
YBR062C_2
DSK2
ASI2
APC11
RPN12
VID28
RPT6
YBR280C
ASI3
PNG1
UBC1
SSM4
PTH2
SAN1
RPN5
DER1
YLR104W
VID24
JEM1
CUE4
YLR224W
NPL4
YGR066C
MNS1
DER1
MND2
YNL155W
CNE1
APC2
UBX5
HLJ1
HUL5
HRD1
SWM1
SCL1
NAT3
RPT1
RAD6
YDJ1
SSA1
YMR247C
PRE9
CDC27
YDR049W
MDM30
PRE6
SEC61
DIA2
RPN2
RPN8
RPT3
UFO1
YMR184W
PRE8
PRE1
PUP1
DOC1
RPT5
UBC4
FES1
PRE5
RPT2
RSP5
APC1
GIM3
DOA1
EPS1
UBP14
GRR1
SHP1
Gene
Z_Shift_K_Doxo_HLD
Z_lm_K_Doxo_HLD
Z_Shift_K_Doxo_EG
Z_lm_K_Doxo_EG
Z_Shift_L_Doxo_HLD
Z_lm_L_Doxo_HLD
Z_Shift_L_Doxo_EG
Z_lm_L_Doxo_EG

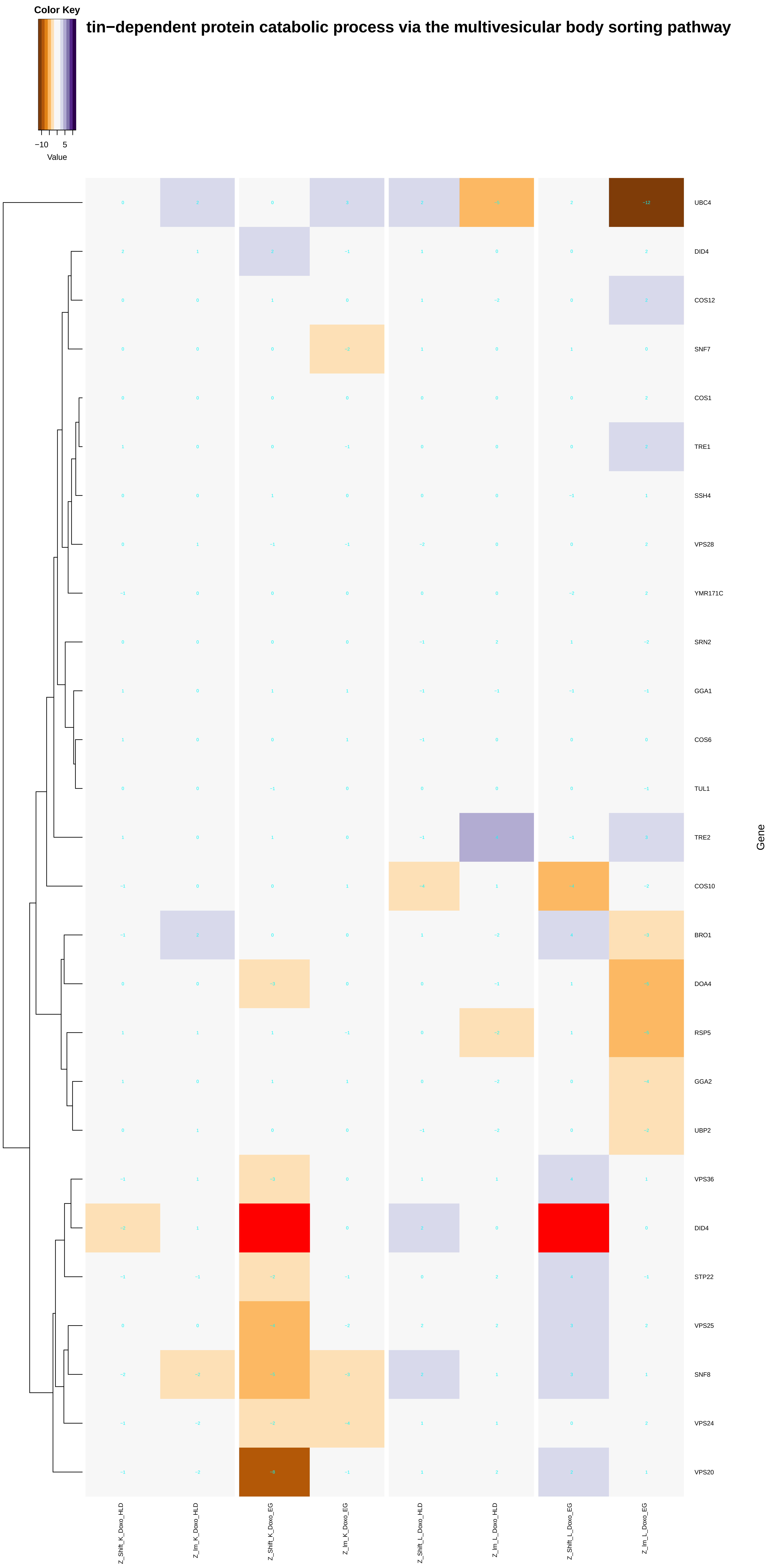
Color Key
−10
5
Value
tin−dependent protein catabolic process via the multivesicular body sorting pathway
UBC4
DID4
COS12
SNF7
COS1
TRE1
SSH4
VPS28
YMR171C
SRN2
GGA1
COS6
TUL1
TRE2
COS10
BRO1
DOA4
RSP5
GGA2
UBP2
VPS36
DID4
STP22
VPS25
SNF8
VPS24
VPS20
Gene
Z_Shift_K_Doxo_HLD
Z_Im_K_Doxo_HLD
Z_Shift_K_Doxo_EG
Z_Im_K_Doxo_EG
Z_Shift_L_Doxo_HLD
Z_Im_L_Doxo_HLD
Z_Shift_L_Doxo_EG
Z_Im_L_Doxo_EG

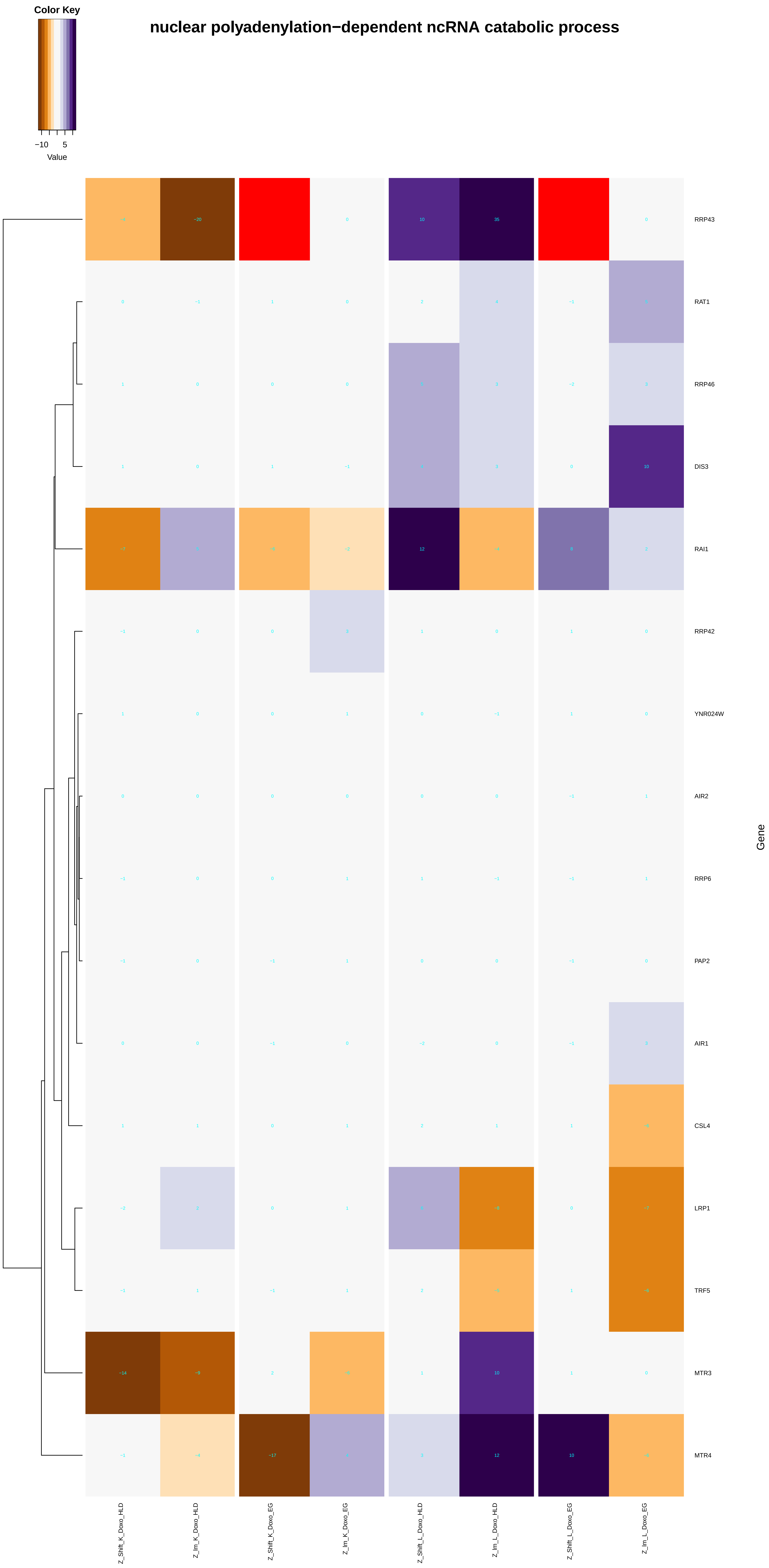
nuclear polyadenylation−dependent ncRNA catabolic process
Color Key
−10
5
Value
RRP43
RAT1
RRP46
DIS3
RAI1
RRP42
YNR024W
AIR2
RRP6
PAP2
AIR1
CSL4
LRP1
TRF5
MTR3
MTR4
Gene
Z_Shift_K_Doxo_HLD
Z_Im_K_Doxo_HLD
Z_Shift_K_Doxo_EG
Z_Im_K_Doxo_EG
Z_Shift_L_Doxo_HLD
Z_Im_L_Doxo_HLD
Z_Shift_L_Doxo_EG
Z_Im_L_Doxo_EG

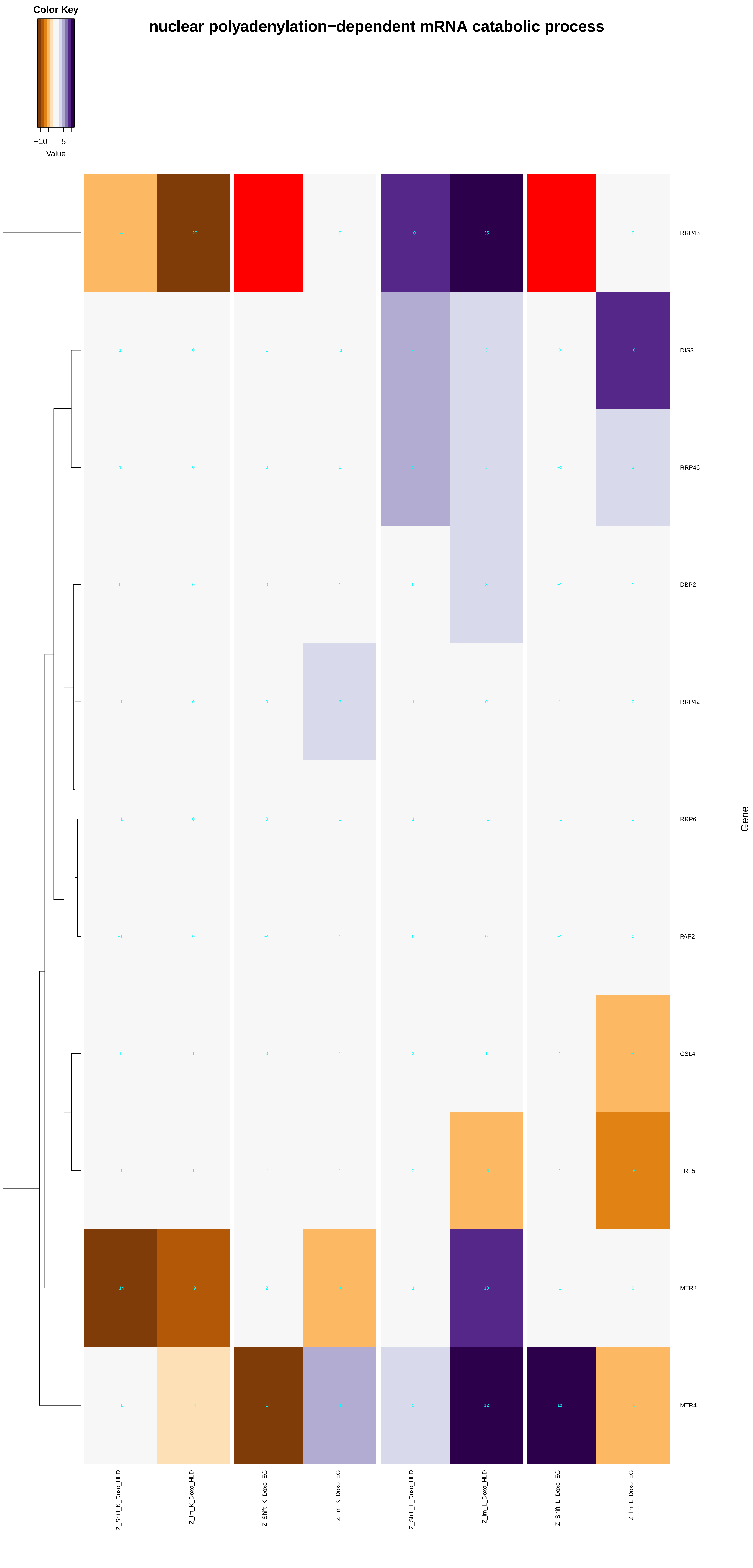
nuclear polyadenylation−dependent mRNA catabolic process
Color Key
−10
5
Value
RRP43
DIS3
RRP46
DBP2
RRP42
RRP6
PAP2
CSL4
TRF5
MTR3
MTR4
Gene
Z_Shift_K_Doxo_HLD
Z_Im_K_Doxo_HLD
Z_Shift_K_Doxo_EG
Z_Im_K_Doxo_EG
Z_Shift_L_Doxo_HLD
Z_Im_L_Doxo_HLD
Z_Shift_L_Doxo_EG
Z_Im_L_Doxo_EG

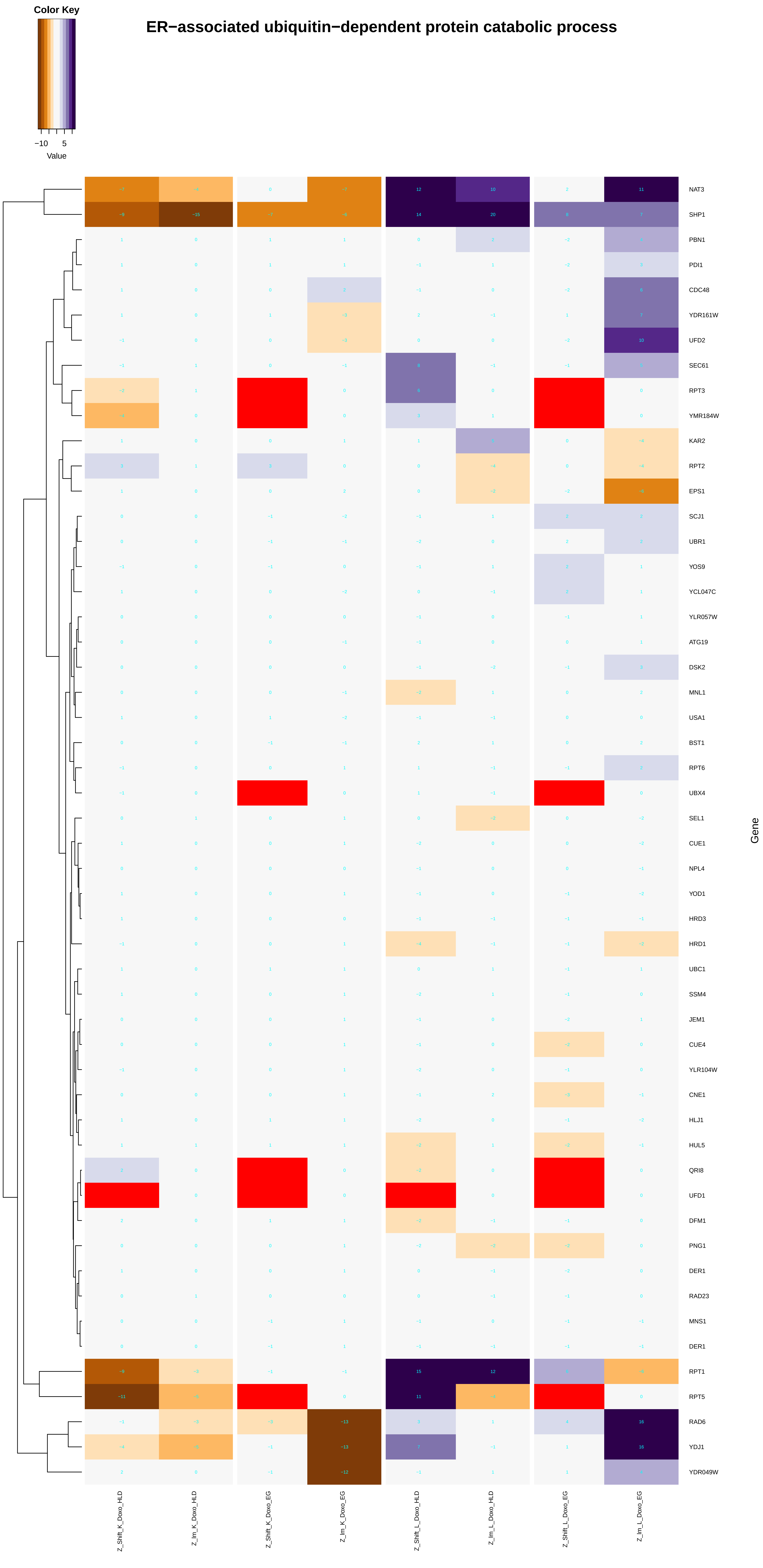
Color Key
−10
5
Value
ER−associated ubiquitin−dependent protein catabolic process
NAT3
SHP1
PBN1
PDI1
CDC48
YDR161W
UFD2
SEC61
RPT3
YMR184W
KAR2
RPT2
EPS1
SCJ1
UBR1
YOS9
YCL047C
YLR057W
ATG19
DSK2
MNL1
USA1
BST1
RPT6
UBX4
SEL1
CUE1
NPL4
YOD1
HRD3
HRD1
UBC1
SSM4
JEM1
CUE4
YLR104W
CNE1
HLJ1
HUL5
QRI8
UFD1
DFM1
PNG1
DER1
RAD23
MNS1
DER1
RPT1
RPT5
RAD6
YDJ1
YDR049W
Gene
Z_Shift_K_Doxo_HLD
Z_Im_K_Doxo_HLD
Z_Shift_K_Doxo_EG
Z_Im_K_Doxo_EG
Z_Shift_L_Doxo_HLD
Z_Im_L_Doxo_HLD
Z_Shift_L_Doxo_EG
Z_Im_L_Doxo_EG

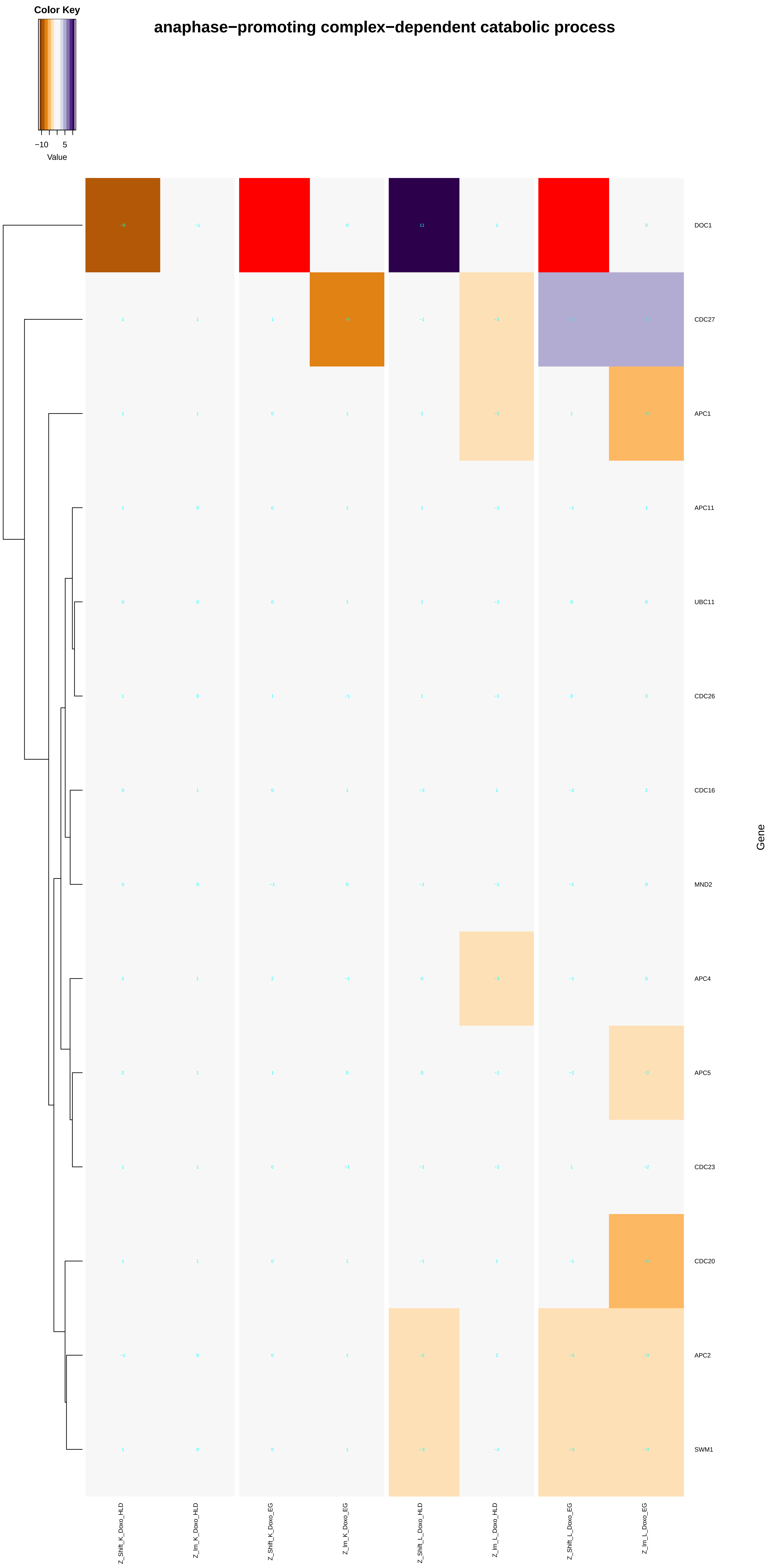
Color Key
−10
5
Value
anaphase−promoting complex−dependent catabolic process
DOC1
CDC27
APC1
APC11
UBC11
CDC26
CDC16
MND2
APC4
APC5
CDC23
CDC20
APC2
SWM1
Gene
Z_Shift_K_Doxo_HLD
Z_Im_K_Doxo_HLD
Z_Shift_K_Doxo_EG
Z_Im_K_Doxo_EG
Z_Shift_L_Doxo_HLD
Z_Im_L_Doxo_HLD
Z_Shift_L_Doxo_EG
Z_Im_L_Doxo_EG

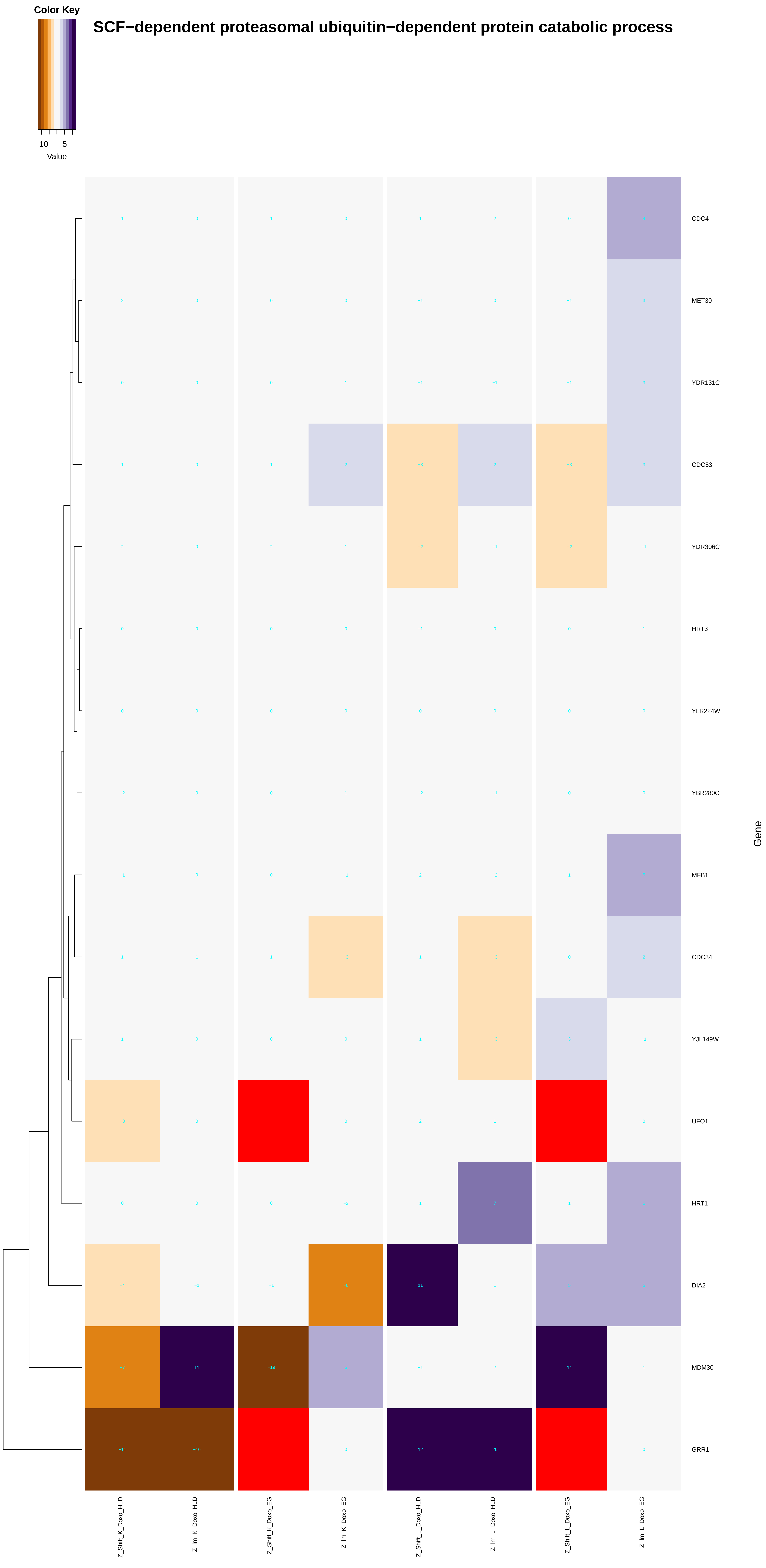
Color Key
−10 5
Value
SCF−dependent proteasomal ubiquitin−dependent protein catabolic process
CDC4
MET30
YDR131C
CDC53
YDR306C
HRT3
YLR224W
YBR280C
MFB1
CDC34
YJL149W
UFO1
HRT1
DIA2
MDM30
GRR1
Gene
Z_Shift_K_Doxo_HLD
Z_Im_K_Doxo_HLD
Z_Shift_K_Doxo_EG
Z_Im_K_Doxo_EG
Z_Shift_L_Doxo_HLD
Z_Im_L_Doxo_HLD
Z_Shift_L_Doxo_EG
Z_Im_L_Doxo_EG

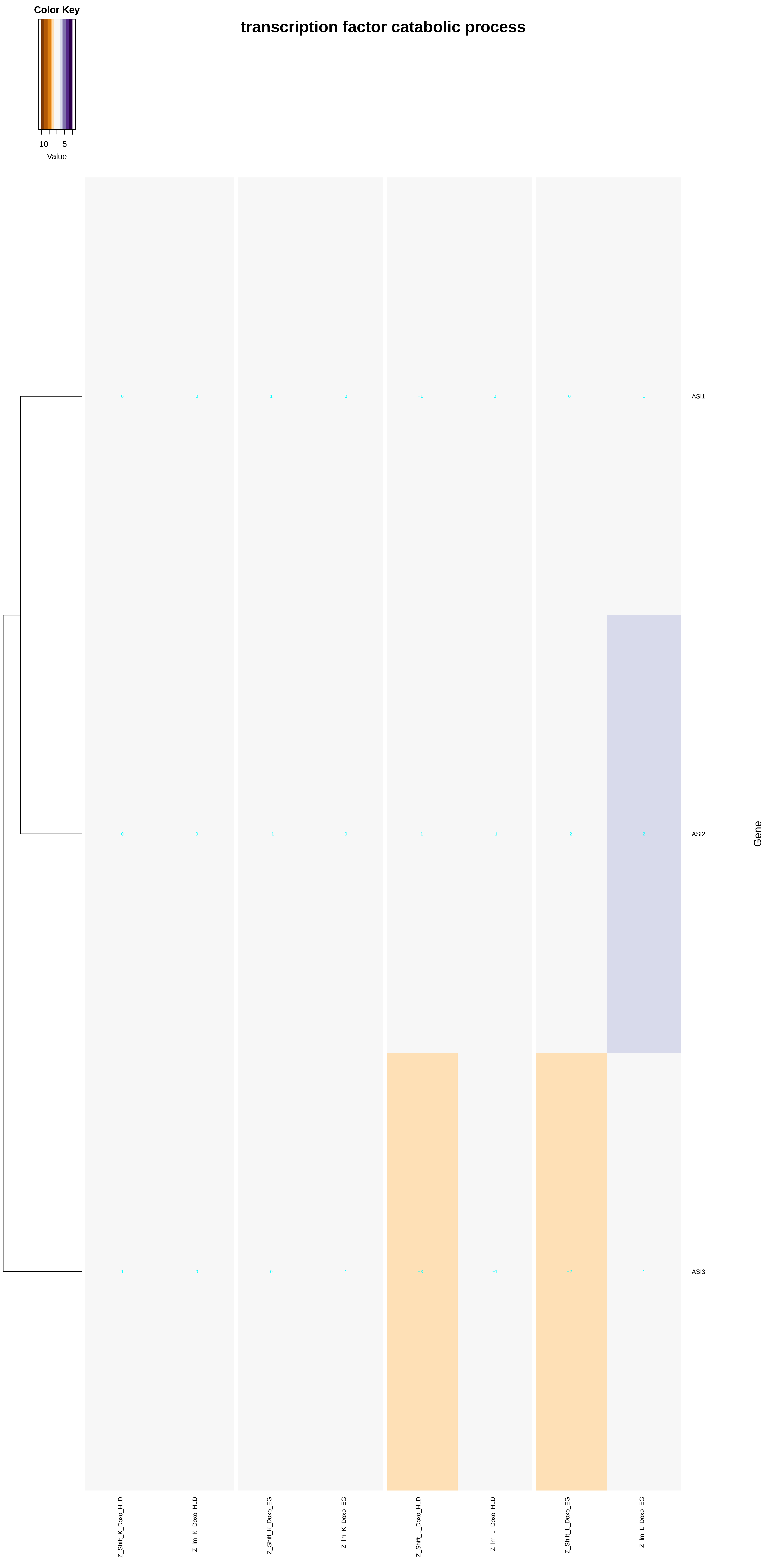

Color Key
−10
5
Value
transcription factor catabolic process
ASI1
ASI2
ASI3
Gene
Z_Shift_K_Doxo_HLD
Z_Im_K_Doxo_HLD
Z_Shift_K_Doxo_EG
Z_Im_K_Doxo_EG
Z_Shift_L_Doxo_HLD
Z_Im_L_Doxo_HLD
Z_Shift_L_Doxo_EG
Z_Im_L_Doxo_EG

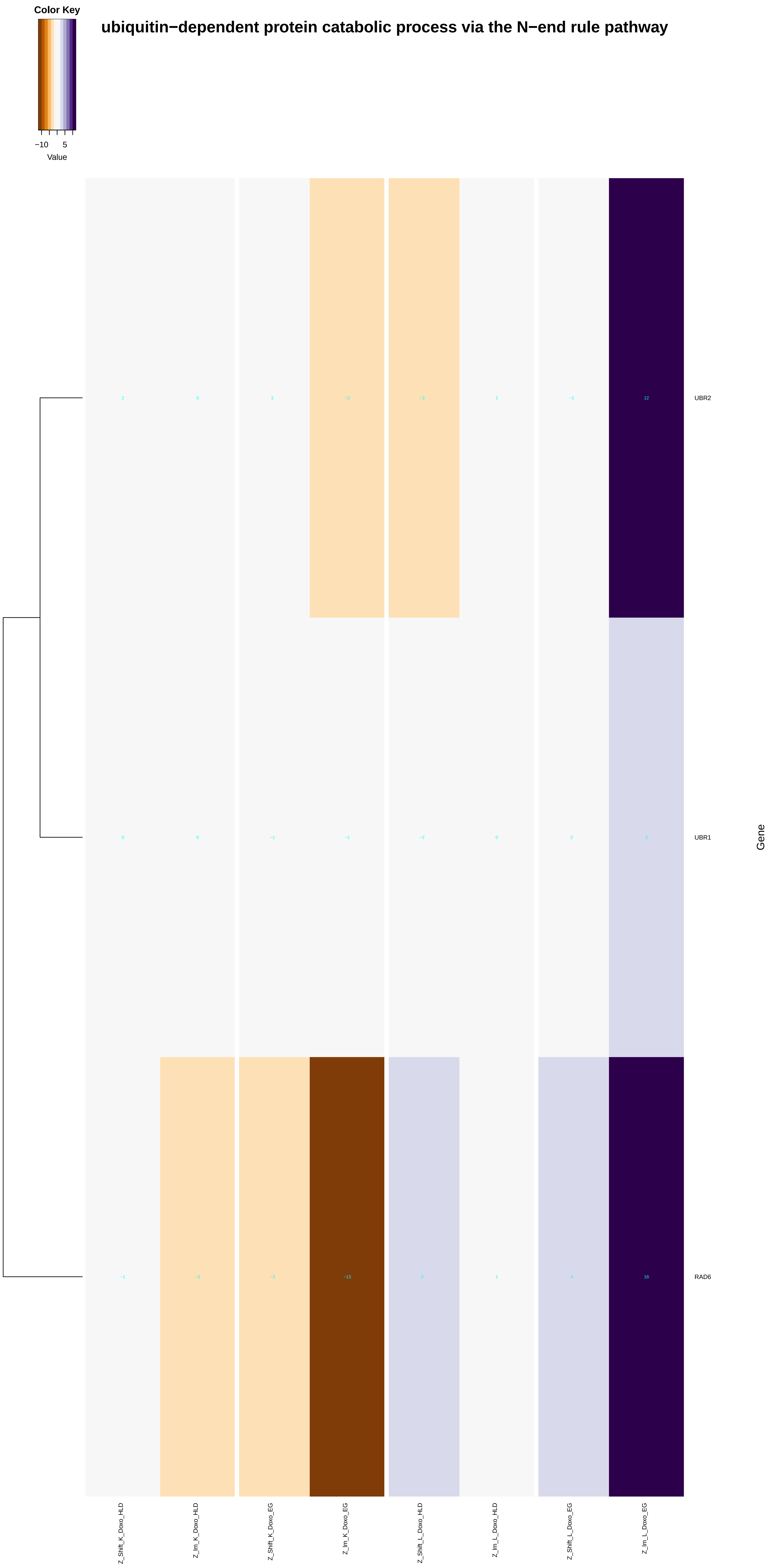
ubiquitin−dependent protein catabolic process via the N−end rule pathway
Color Key
−10
5
Value
UBR2
UBR1
RAD6
Gene
Z_Shift_K_Doxo_HLD
Z_Im_K_Doxo_HLD
Z_Shift_K_Doxo_EG
Z_Im_K_Doxo_EG
Z_Shift_L_Doxo_HLD
Z_Im_L_Doxo_HLD
Z_Shift_L_Doxo_EG
Z_Im_L_Doxo_EG

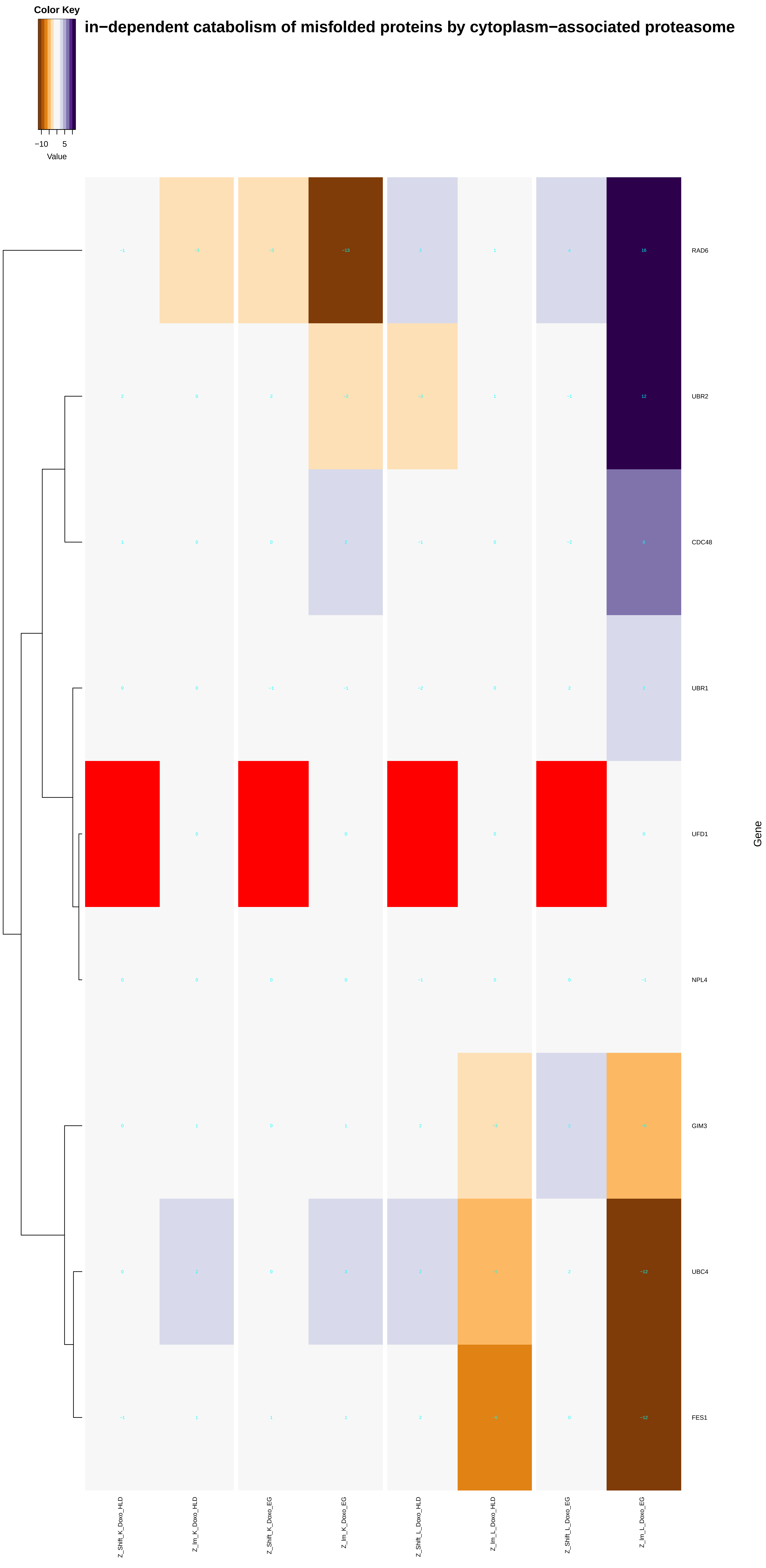

in−dependent catabolism of misfolded proteins by cytoplasm−associated proteasome
Color Key
−10
5
Value
RAD6
UBR2
CDC48
UBR1
UFD1
NPL4
GIM3
UBC4
FES1
Gene
Z_Shift_K_Doxo_HLD
Z_Im_K_Doxo_HLD
Z_Shift_K_Doxo_EG
Z_Im_K_Doxo_EG
Z_Shift_L_Doxo_HLD
Z_Im_L_Doxo_HLD
Z_Shift_L_Doxo_EG
Z_Im_L_Doxo_EG

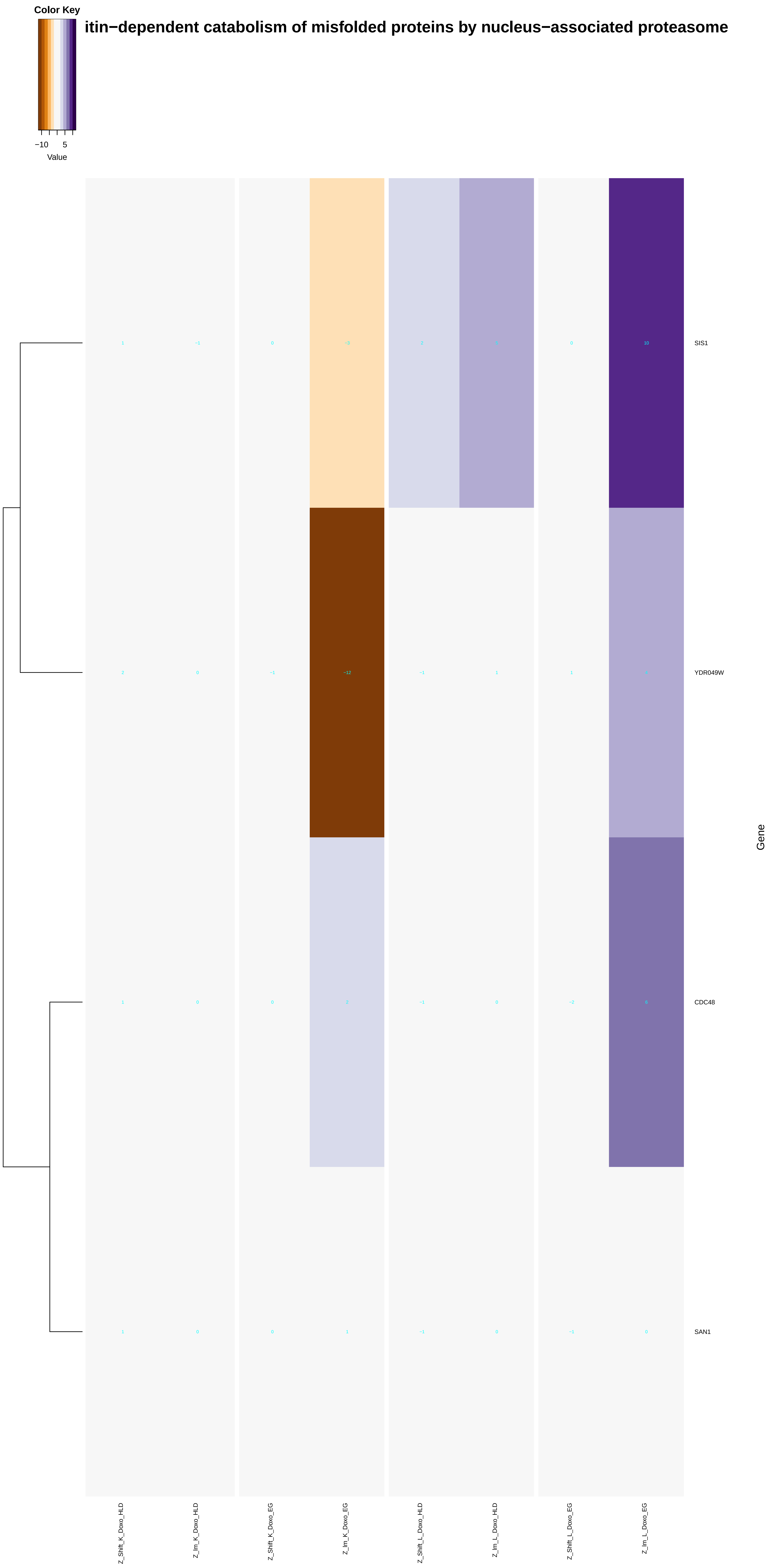
Color Key
−10 5
Value
itin−dependent catabolism of misfolded proteins by nucleus−associated proteasome
SIS1
YDR049W
CDC48
SAN1
Gene
Z_Shift_K_Doxo_HLD
Z_Im_K_Doxo_HLD
Z_Shift_K_Doxo_EG
Z_Im_K_Doxo_EG
Z_Shift_L_Doxo_HLD
Z_Im_L_Doxo_HLD
Z_Shift_L_Doxo_EG
Z_Im_L_Doxo_EG

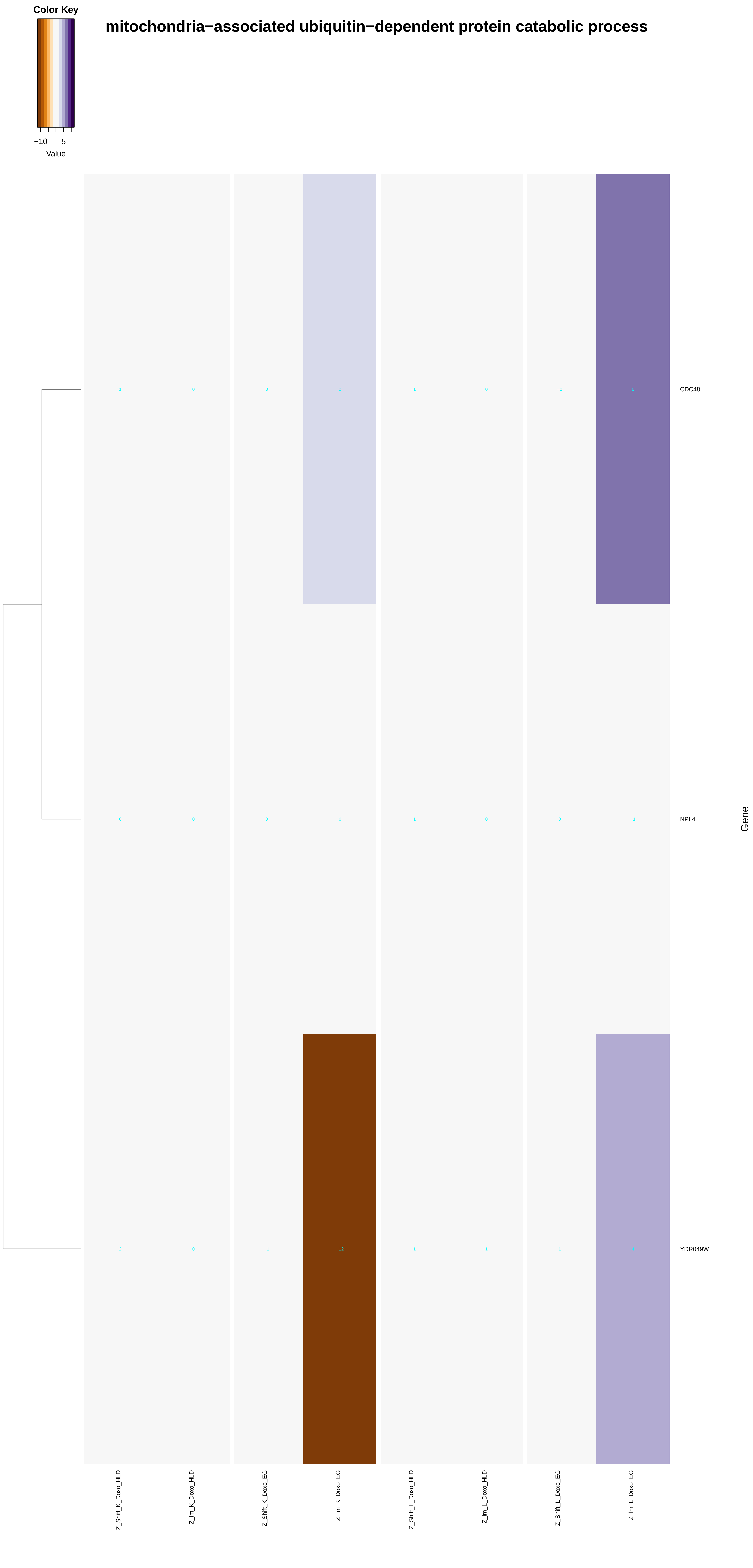
Color Key
−10
5
Value
mitochondria−associated ubiquitin−dependent protein catabolic process
CDC48
NPL4
YDR049W
Gene
Z_Shift_K_Doxo_HLD
Z_lm_K_Doxo_HLD
Z_Shift_K_Doxo_EG
Z_lm_K_Doxo_EG
Z_Shift_L_Doxo_HLD
Z_lm_L_Doxo_HLD
Z_Shift_L_Doxo_EG
Z_lm_L_Doxo_EG

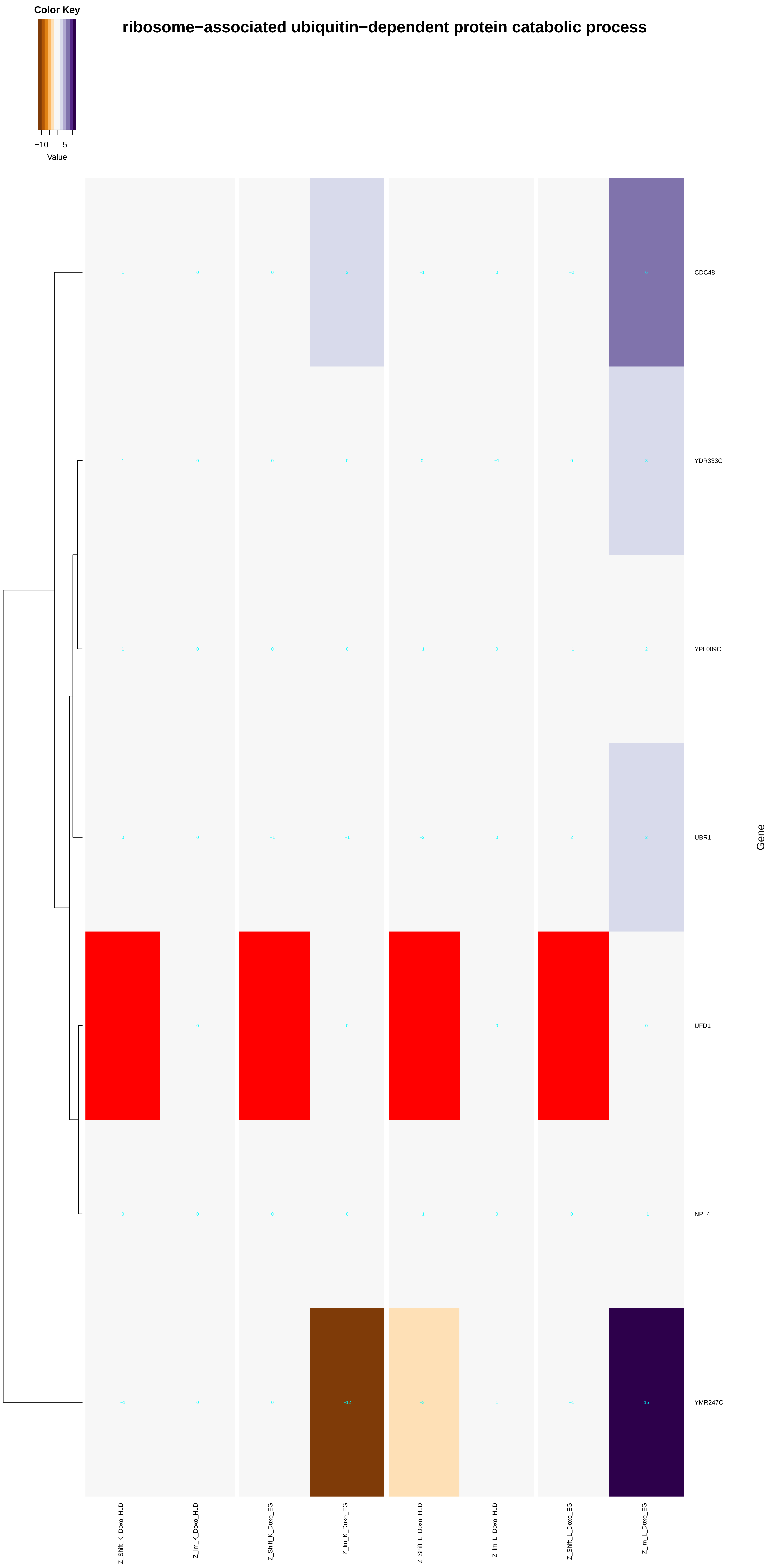
ribosome−associated ubiquitin−dependent protein catabolic process
Color Key
−10
5
Value
CDC48
YDR333C
YPL009C
UBR1
UFD1
NPL4
YMR247C
Gene
Z_Shift_K_Doxo_HLD
Z_Im_K_Doxo_HLD
Z_Shift_K_Doxo_EG
Z_Im_K_Doxo_EG
Z_Shift_L_Doxo_HLD
Z_Im_L_Doxo_HLD
Z_Shift_L_Doxo_EG
Z_Im_L_Doxo_EG

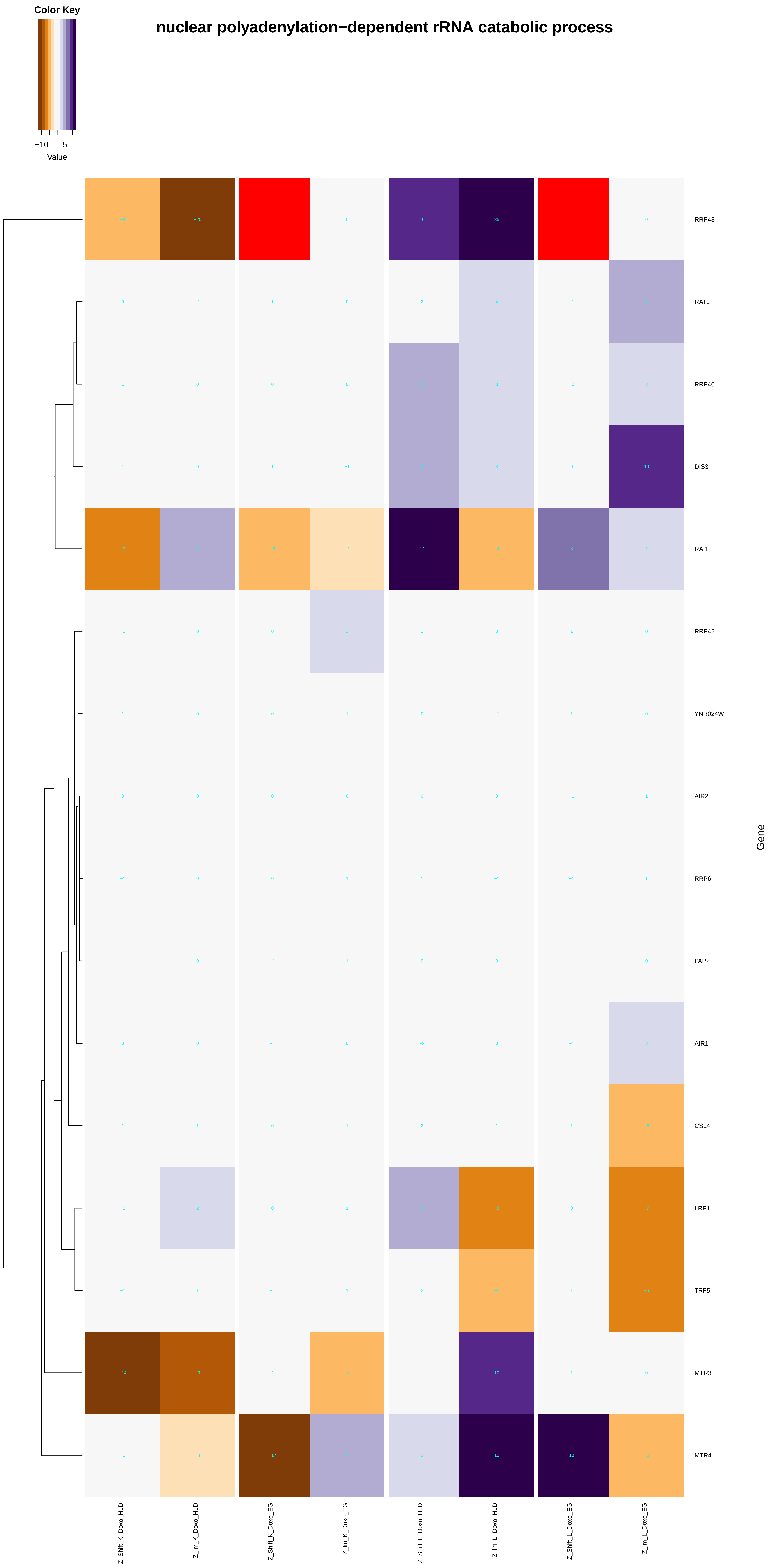

Color Key
−10
5
Value
nuclear polyadenylation−dependent rRNA catabolic process
RRP43
RAT1
RRP46
DIS3
RAI1
RRP42
YNR024W
AIR2
RRP6
PAP2
AIR1
CSL4
LRP1
TRF5
MTR3
MTR4
Gene
Z_Shift_K_Doxo_HLD
Z_Im_K_Doxo_HLD
Z_Shift_K_Doxo_EG
Z_Im_K_Doxo_EG
Z_Shift_L_Doxo_HLD
Z_Im_L_Doxo_HLD
Z_Shift_L_Doxo_EG
Z_Im_L_Doxo_EG

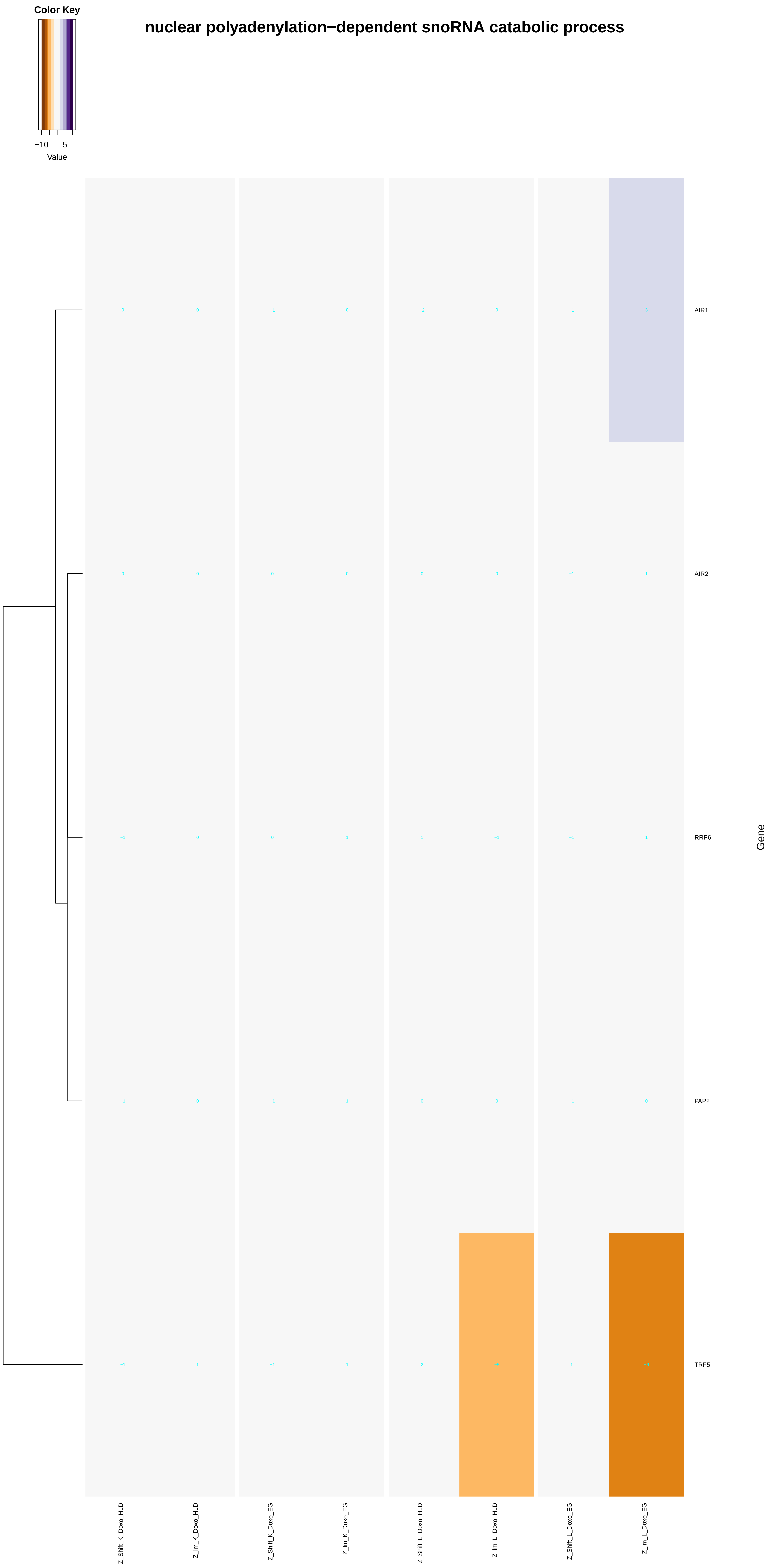
Color Key
−10
5
Value
nuclear polyadenylation−dependent snoRNA catabolic process
AIR1
AIR2
RRP6
PAP2
TRF5
Gene
Z_Shift_K_Doxo_HLD
Z_lm_K_Doxo_HLD
Z_Shift_K_Doxo_EG
Z_lm_K_Doxo_EG
Z_Shift_L_Doxo_HLD
Z_lm_L_Doxo_HLD
Z_Shift_L_Doxo_EG
Z_lm_L_Doxo_EG

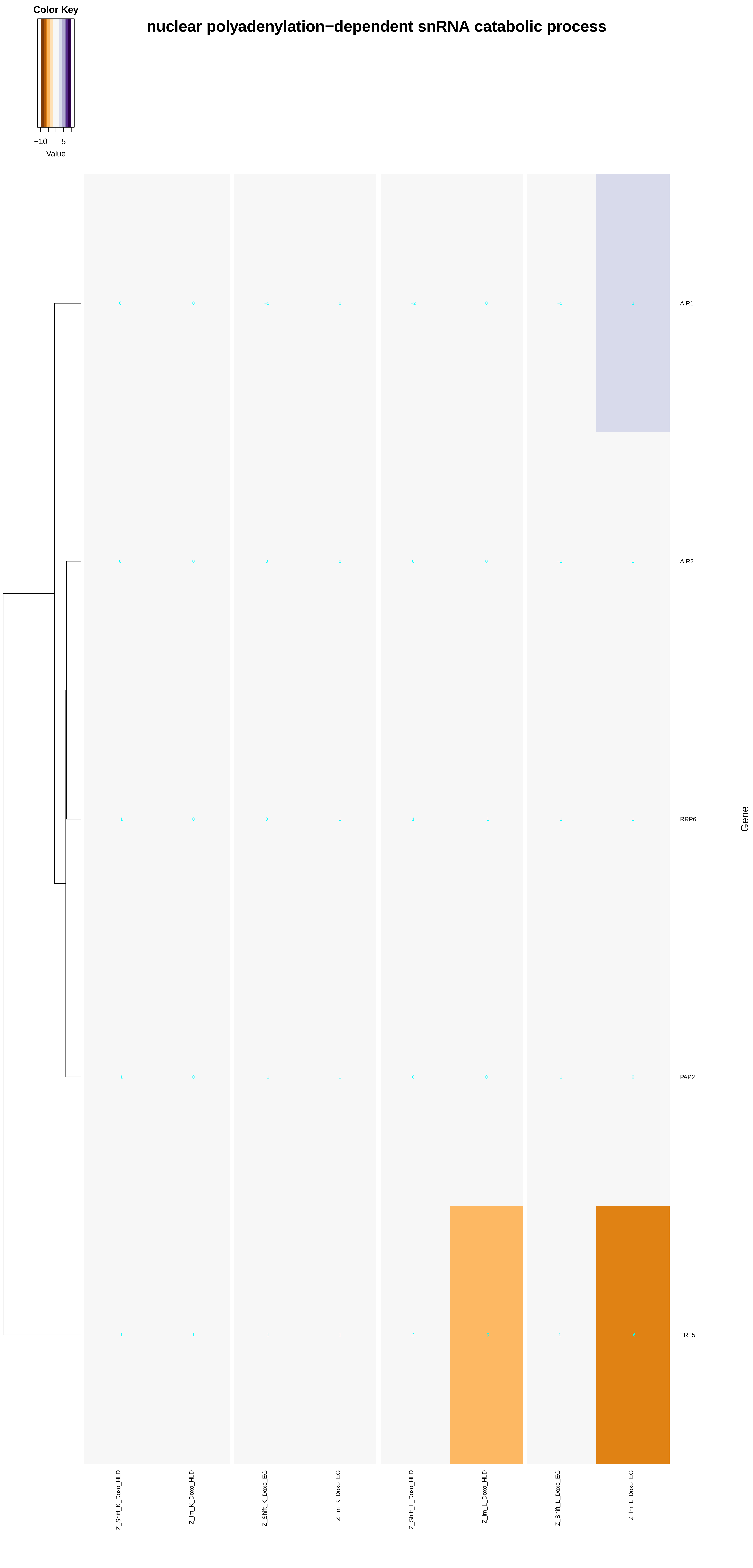
Color Key
−10
5
Value
nuclear polyadenylation−dependent snRNA catabolic process
AIR1
AIR2
RRP6
PAP2
TRF5
Gene
Z_Shift_K_Doxo_HLD
Z_Im_K_Doxo_HLD
Z_Shift_K_Doxo_EG
Z_Im_K_Doxo_EG
Z_Shift_L_Doxo_HLD
Z_Im_L_Doxo_HLD
Z_Shift_L_Doxo_EG
Z_Im_L_Doxo_EG

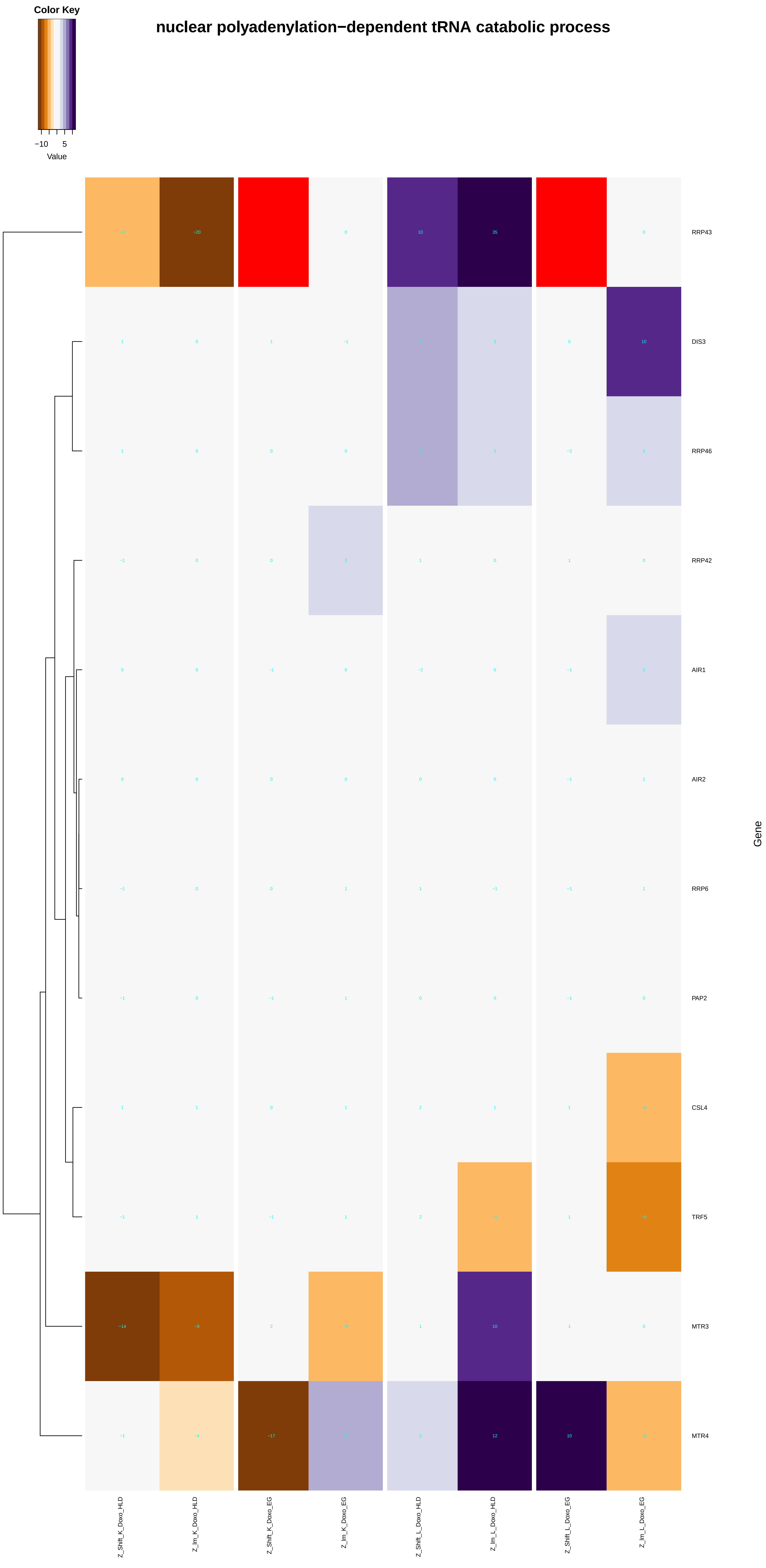
nuclear polyadenylation−dependent tRNA catabolic process
Color Key
−10
5
Value
RRP43
DIS3
RRP46
RRP42
AIR1
AIR2
RRP6
PAP2
CSL4
TRF5
MTR3
MTR4
Gene
Z_Shift_K_Doxo_HLD
Z_lm_K_Doxo_HLD
Z_Shift_K_Doxo_EG
Z_lm_K_Doxo_EG
Z_Shift_L_Doxo_HLD
Z_lm_L_Doxo_HLD
Z_Shift_L_Doxo_EG
Z_lm_L_Doxo_EG

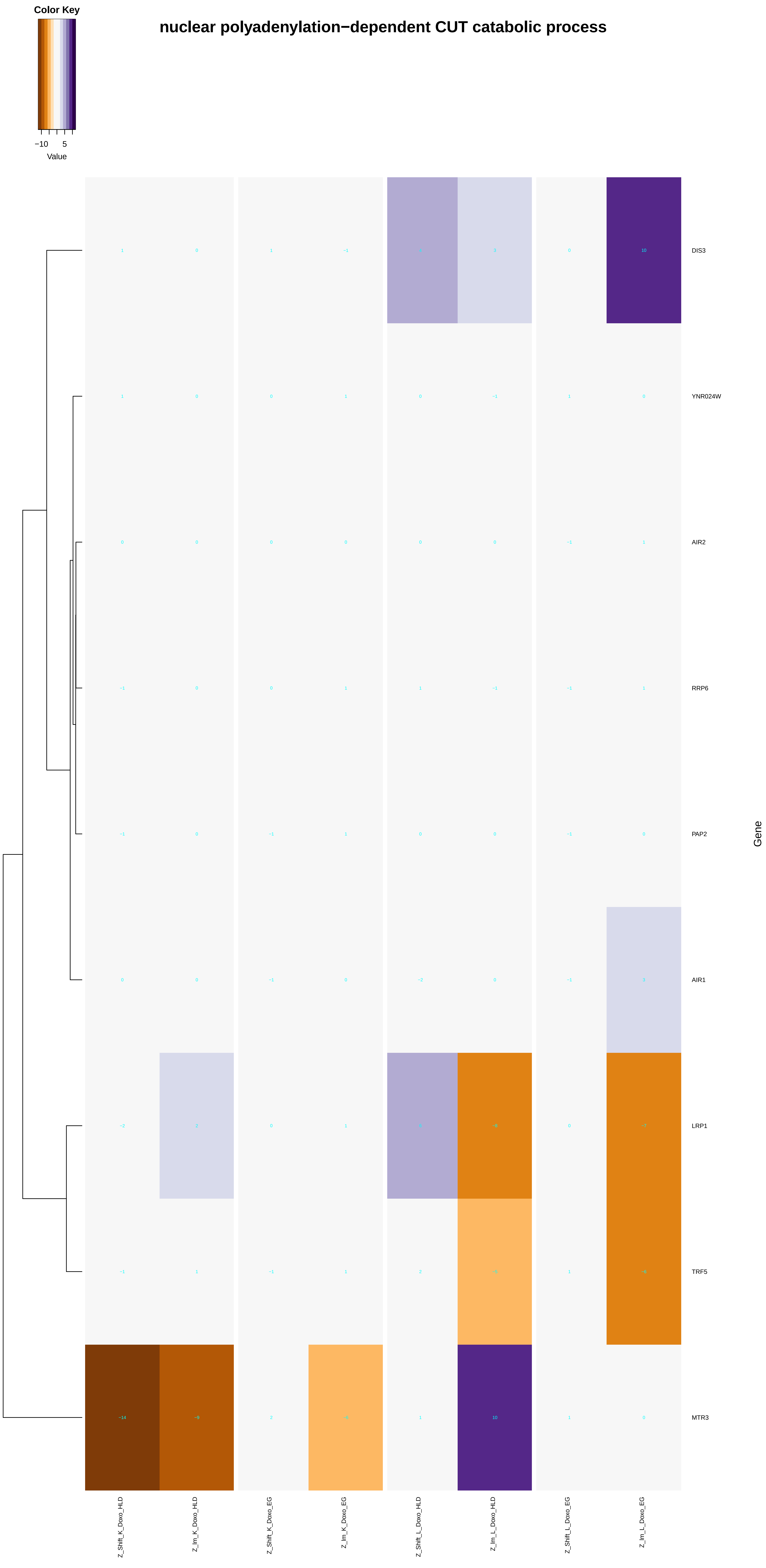

Color Key
−10
5
Value
nuclear polyadenylation−dependent CUT catabolic process
DIS3
YNR024W
AIR2
RRP6
PAP2
AIR1
LRP1
TRF5
MTR3
Gene
Z_Shift_K_Doxo_HLD
Z_Im_K_Doxo_HLD
Z_Shift_K_Doxo_EG
Z_Im_K_Doxo_EG
Z_Shift_L_Doxo_HLD
Z_Im_L_Doxo_HLD
Z_Shift_L_Doxo_EG
Z_Im_L_Doxo_EG

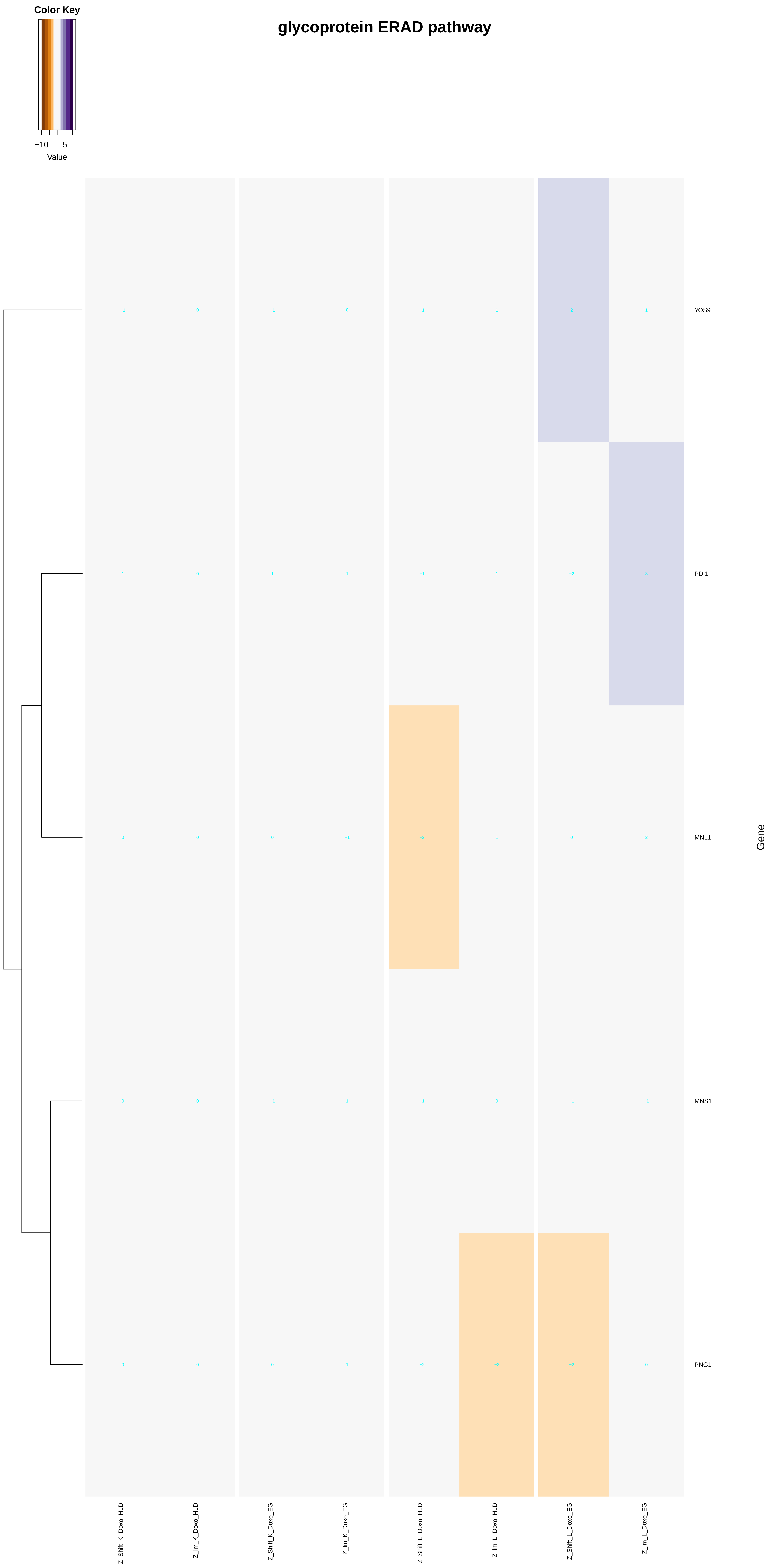
Color Key
−10
5
Value
glycoprotein ERAD pathway
YOS9
PDI1
MNL1
MNS1
PNG1
Gene
Z_Shift_K_Doxo_HLD
Z_Im_K_Doxo_HLD
Z_Shift_K_Doxo_EG
Z_Im_K_Doxo_EG
Z_Shift_L_Doxo_HLD
Z_Im_L_Doxo_HLD
Z_Shift_L_Doxo_EG
Z_Im_L_Doxo_EG
