## Additional File 9 for "A yeast phenomic model for the influence of Warburg metabolism on genetic buffering of doxorubicin": monovalent_inorganic_cation_transport.pdf

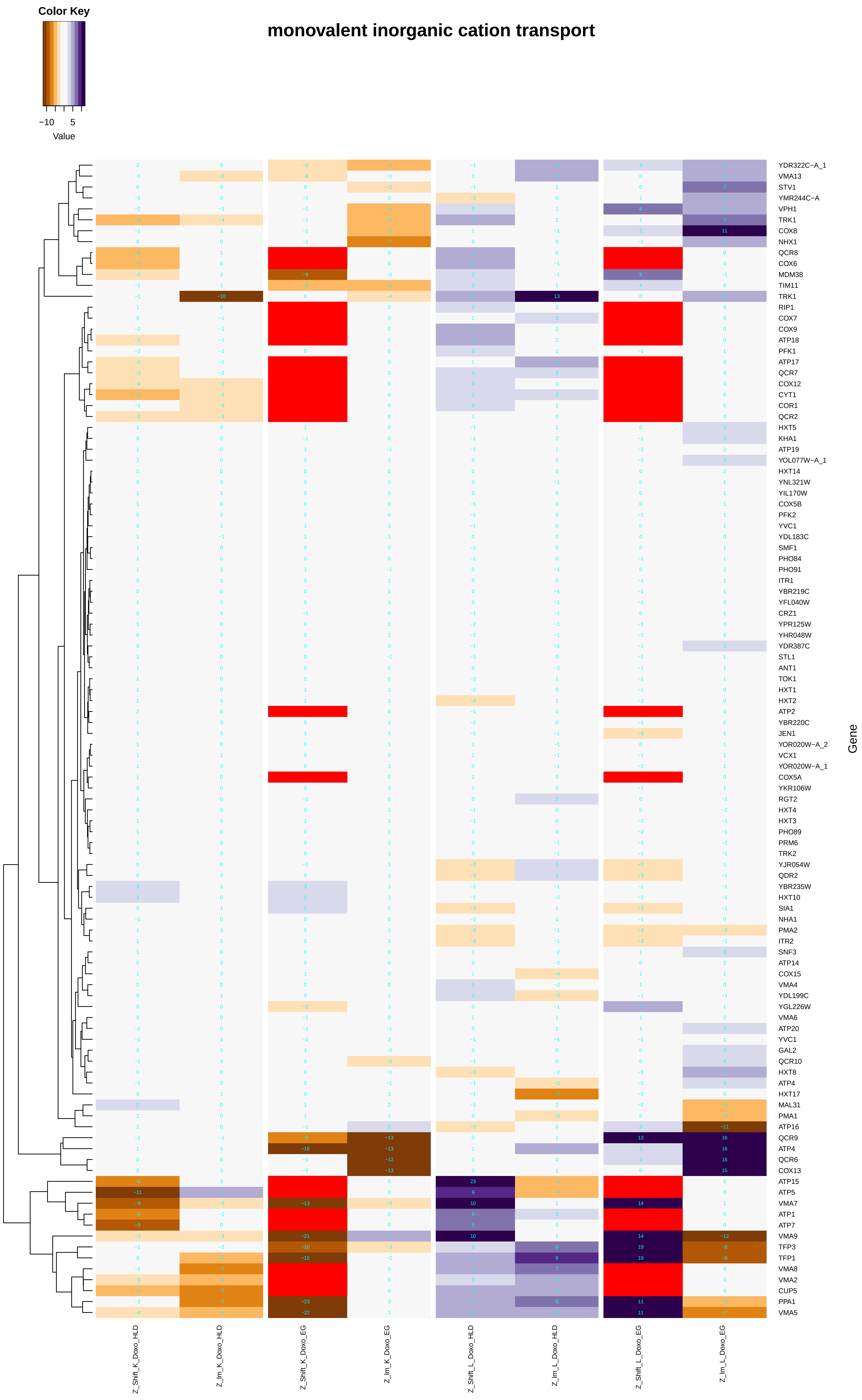

monovalent inorganic cation transport
Color Key
−10
5
Value
YDR322C−A_1
VMA13
STV1
YMR244C−A
VPH1
TRK1
COX8
NHX1
QCR8
COX6
MDM38
TIM11
TRK1
RIP1
COX7
COX9
ATP18
PFK1
ATP17
QCR7
COX12
CYT1
COR1
QCR2
HXT5
KHA1
ATP19
YOL077W−A_1
HXT14
YNL321W
YIL170W
COX5B
PFK2
YVC1
YDL183C
SMF1
PHO84
PHO91
ITR1
YBR219C
YFL040W
CRZ1
YPR125W
YHR048W
YDR387C
STL1
ANT1
TOK1
HXT1
HXT2
ATP2
YBR220C
JEN1
YOR020W−A_2
VCX1
YOR020W−A_1
COX5A
YKR106W
RGT2
HXT4
HXT3
PHO89
PRM6
TRK2
YJR054W
QDR2
YBR235W
HXT10
SIA1
NHA1
PMA2
ITR2
SNF3
ATP14
COX15
VMA4
YDL199C
YGL226W
VMA6
ATP20
YVC1
GAL2
QCR10
HXT8
ATP4
HXT17
MAL31
PMA1
ATP16
QCR9
ATP4
QCR6
COX13
ATP15
ATP5
VMA7
ATP1
ATP7
VMA9
TFP3
TFP1
VMA8
VMA2
CUP5
PPA1
VMA5
Gene
Z_Shift_K_Doxo_HLD
Z_lm_K_Doxo_HLD
Z_Shift_K_Doxo_EG
Z_lm_K_Doxo_EG
Z_Shift_L_Doxo_HLD
Z_lm_L_Doxo_HLD
Z_Shift_L_Doxo_EG
Z_lm_L_Doxo_EG

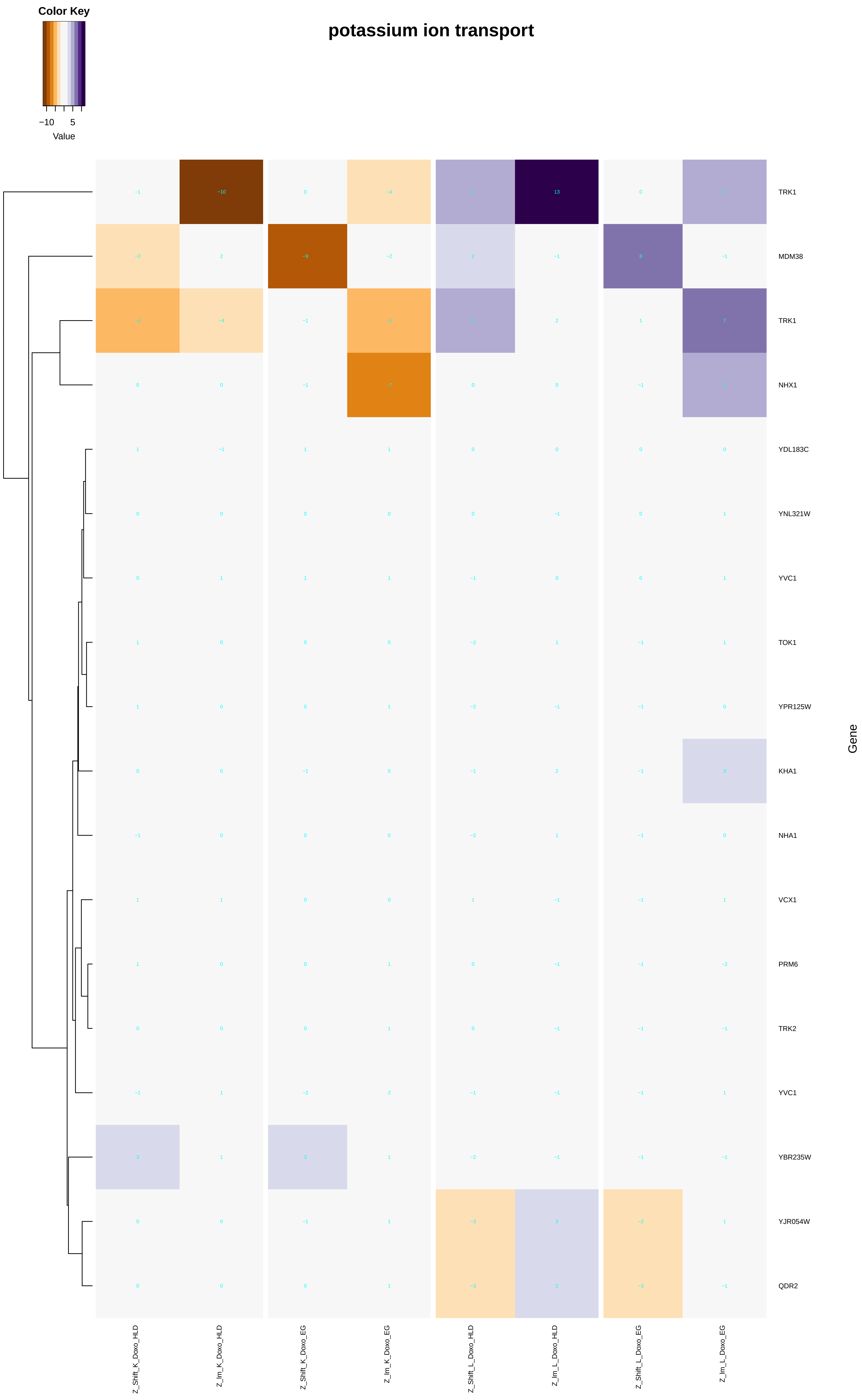
potassium ion transport
Color Key
−10
5
Value
Gene
TRK1
MDM38
TRK1
NHX1
YDL183C
YNL321W
YVC1
TOK1
YPR125W
KHA1
NHA1
VCX1
PRM6
TRK2
YVC1
YBR235W
YJR054W
QDR2
Z_Shift_K_Doxo_HLD
Z_Im_K_Doxo_HLD
Z_Shift_K_Doxo_EG
Z_Im_K_Doxo_EG
Z_Shift_L_Doxo_HLD
Z_Im_L_Doxo_HLD
Z_Shift_L_Doxo_EG
Z_Im_L_Doxo_EG

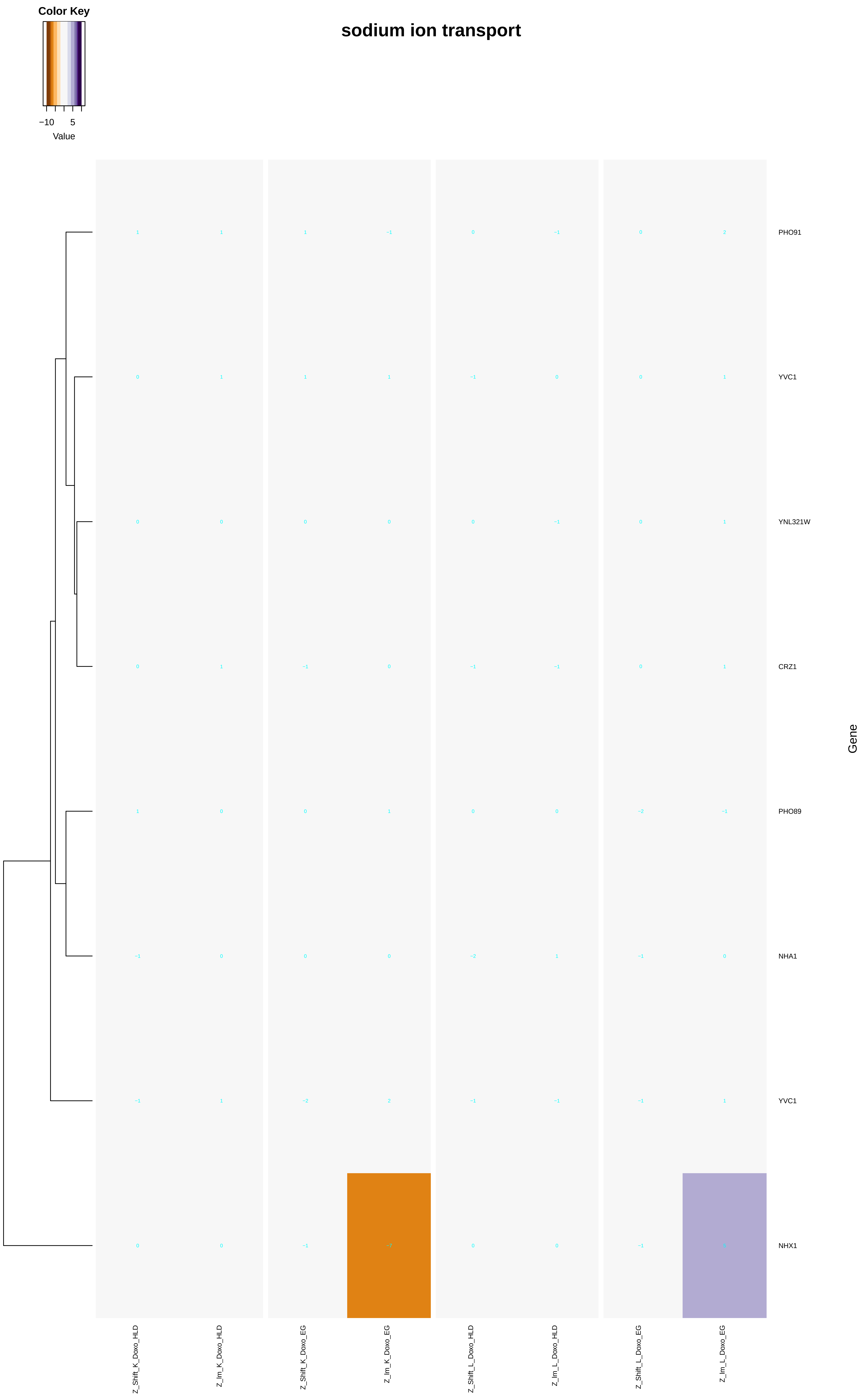
Color Key
−10
5
Value
sodium ion transport
PHO91
YVC1
YNL321W
CRZ1
PHO89
NHA1
YVC1
NHX1
Gene
Z_Shift_K_Doxo_HLD
Z_Im_K_Doxo_HLD
Z_Shift_K_Doxo_EG
Z_Im_K_Doxo_EG
Z_Shift_L_Doxo_HLD
Z_Im_L_Doxo_HLD
Z_Shift_L_Doxo_EG
Z_Im_L_Doxo_EG

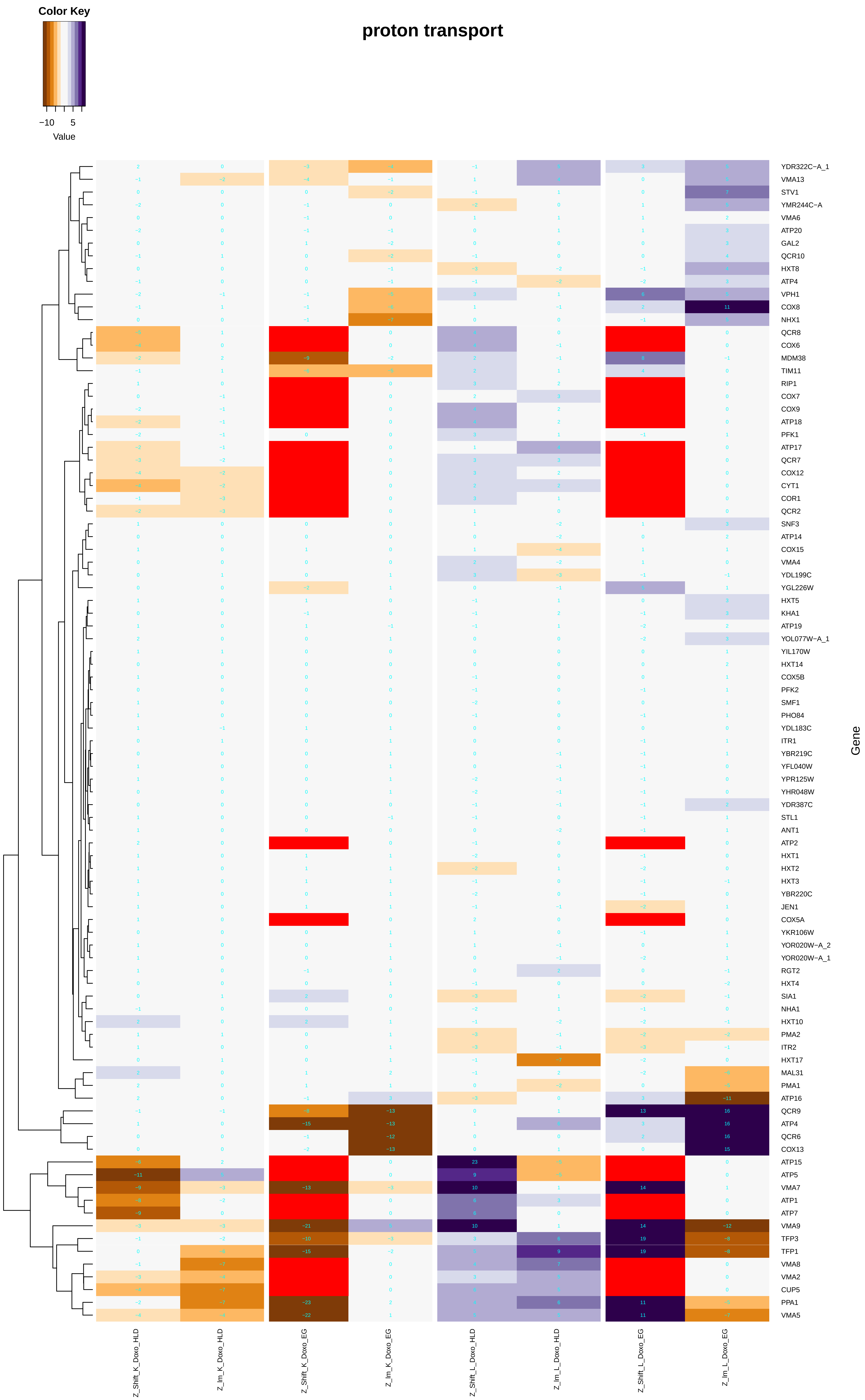

proton transport
Color Key
−10
5
Value
Gene
YDR322C−A_1
VMA13
STV1
YMR244C−A
VMA6
ATP20
GAL2
QCR10
HXT8
ATP4
VPH1
COX8
NHX1
QCR8
COX6
MDM38
TIM11
RIP1
COX7
COX9
ATP18
PFK1
ATP17
QCR7
COX12
CYT1
COR1
QCR2
SNF3
ATP14
COX15
VMA4
YDL199C
YGL226W
HXT5
KHA1
ATP19
YOL077W−A_1
YIL170W
HXT14
COX5B
PFK2
SMF1
PHO84
YDL183C
ITR1
YBR219C
YFL040W
YPR125W
YHR048W
YDR387C
STL1
ANT1
ATP2
HXT1
HXT2
HXT3
YBR220C
JEN1
COX5A
YKR106W
YOR020W−A_2
YOR020W−A_1
RGT2
HXT4
SIA1
NHA1
HXT10
PMA2
ITR2
HXT17
MAL31
PMA1
ATP16
QCR9
ATP4
QCR6
COX13
ATP15
ATP5
VMA7
ATP1
ATP7
VMA9
TFP3
TFP1
VMA8
VMA2
CUP5
PPA1
VMA5
Z_Shift_K_Doxo_HLD
Z_lm_K_Doxo_HLD
Z_Shift_K_Doxo_EG
Z_lm_K_Doxo_EG
Z_Shift_L_Doxo_HLD
Z_lm_L_Doxo_HLD
Z_Shift_L_Doxo_EG
Z_lm_L_Doxo_EG

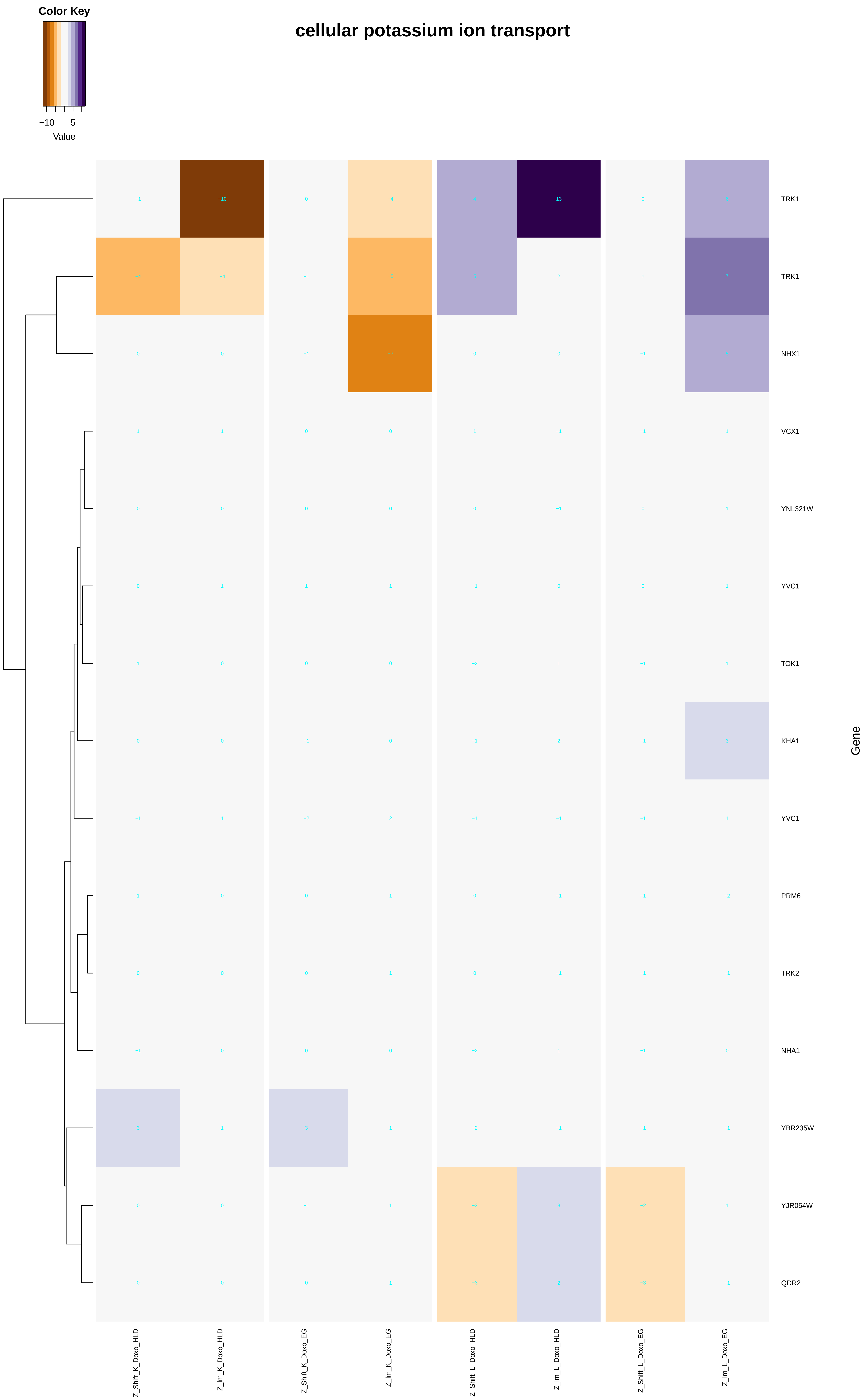
cellular potassium ion transport
Color Key
−10
5
Value
TRK1
TRK1
NHX1
VCX1
YNL321W
YVC1
TOK1
KHA1
YVC1
PRM6
TRK2
NHA1
YBR235W
YJR054W
QDR2
Gene
Z_Shift_K_Doxo_HLD
Z_Im_K_Doxo_HLD
Z_Shift_K_Doxo_EG
Z_Im_K_Doxo_EG
Z_Shift_L_Doxo_HLD
Z_Im_L_Doxo_HLD
Z_Shift_L_Doxo_EG
Z_Im_L_Doxo_EG

### sodium ion transmembrane transport

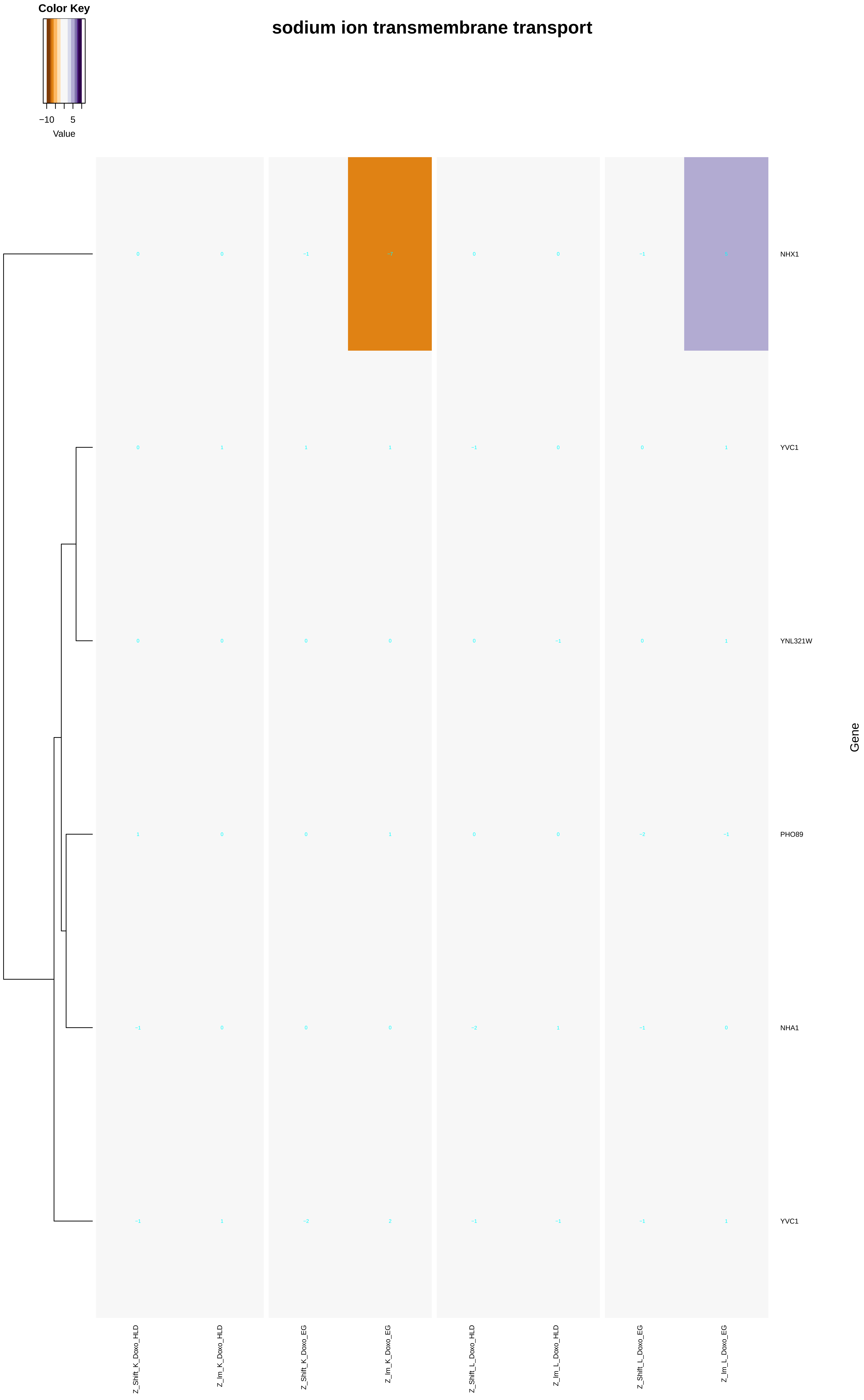

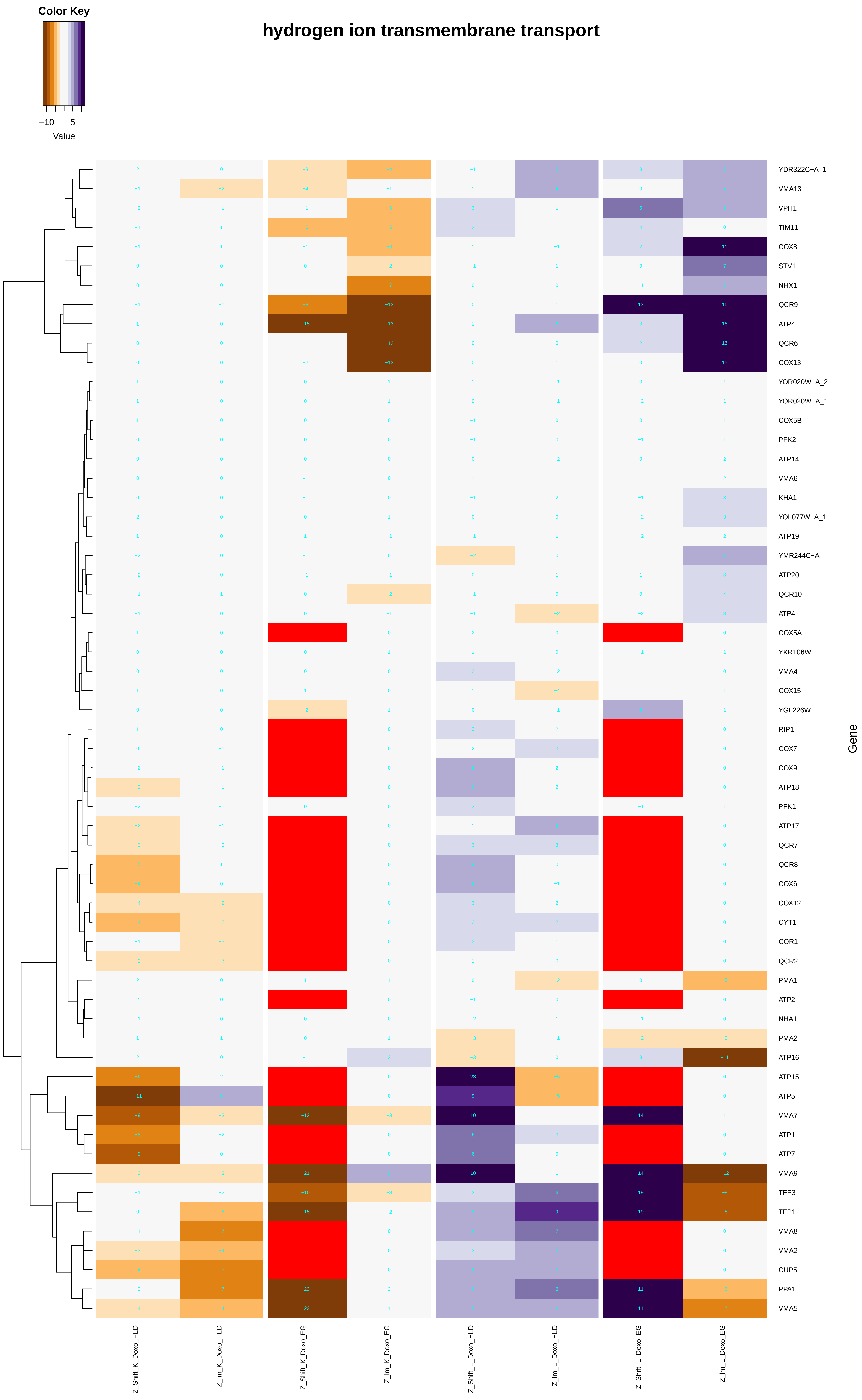
Color Key
−10
5
Value
hydrogen ion transmembrane transport
YDR322C−A_1
VMA13
VPH1
TIM11
COX8
STV1
NHX1
QCR9
ATP4
QCR6
COX13
YOR020W−A_2
YOR020W−A_1
COX5B
PFK2
ATP14
VMA6
KHA1
YOL077W−A_1
ATP19
YMR244C−A
ATP20
QCR10
ATP4
COX5A
YKR106W
VMA4
COX15
YGL226W
RIP1
COX7
COX9
ATP18
PFK1
ATP17
QCR7
QCR8
COX6
COX12
CYT1
COR1
QCR2
PMA1
ATP2
NHA1
PMA2
ATP16
ATP15
ATP5
VMA7
ATP1
ATP7
VMA9
TFP3
TFP1
VMA8
VMA2
CUP5
PPA1
VMA5
Gene
Z_Shift_K_Doxo_HLD
Z_Im_K_Doxo_HLD
Z_Shift_K_Doxo_EG
Z_Im_K_Doxo_EG
Z_Shift_L_Doxo_HLD
Z_Im_L_Doxo_HLD
Z_Shift_L_Doxo_EG
Z_Im_L_Doxo_EG

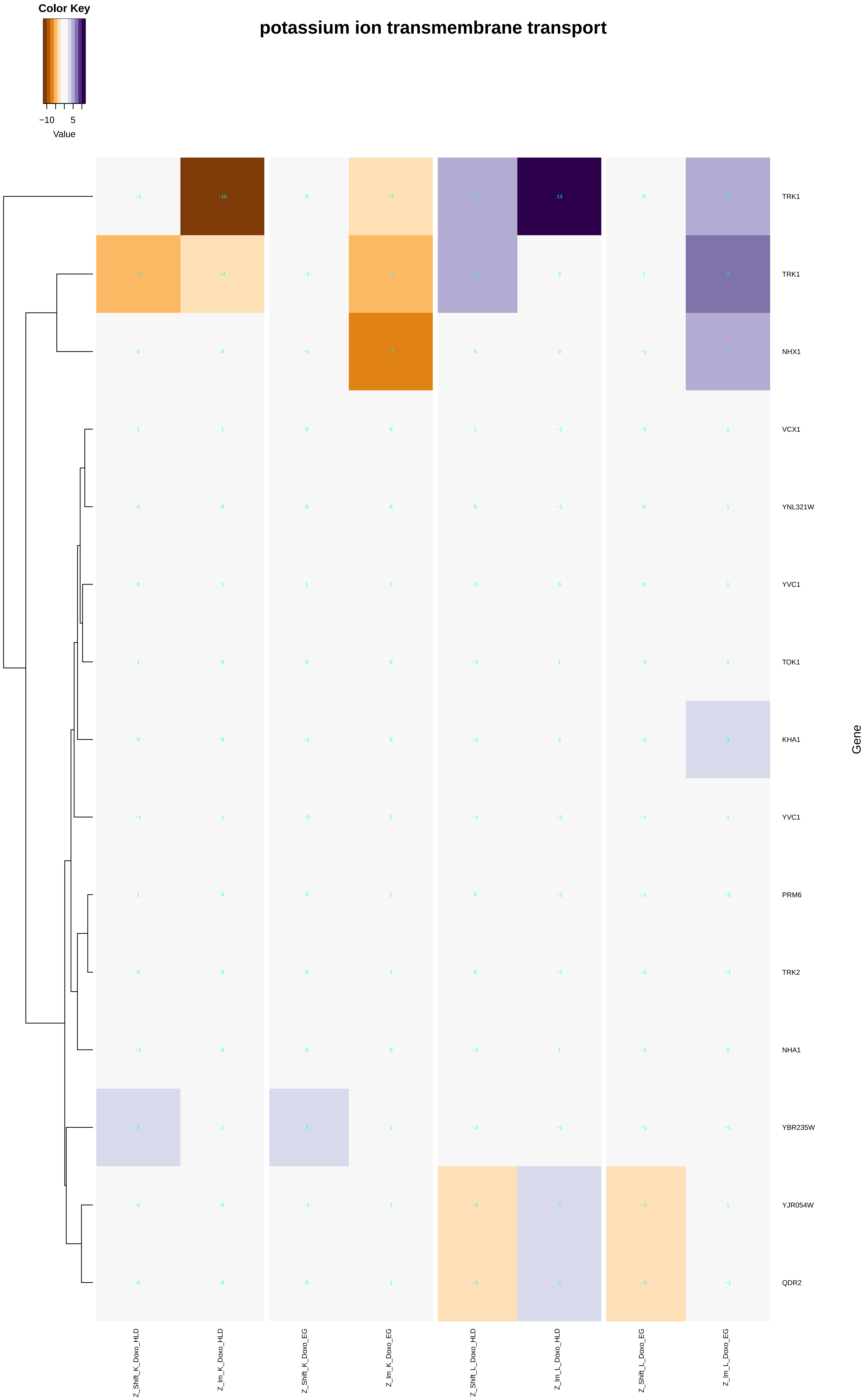

Color Key
−10
5
Value
potassium ion transmembrane transport
TRK1
TRK1
NHX1
VCX1
YNL321W
YVC1
TOK1
KHA1
YVC1
PRM6
TRK2
NHA1
YBR235W
YJR054W
QDR2
Gene
Z_Shift_K_Doxo_HLD
Z_Im_K_Doxo_HLD
Z_Shift_K_Doxo_EG
Z_Im_K_Doxo_EG
Z_Shift_L_Doxo_HLD
Z_Im_L_Doxo_HLD
Z_Shift_L_Doxo_EG
Z_Im_L_Doxo_EG

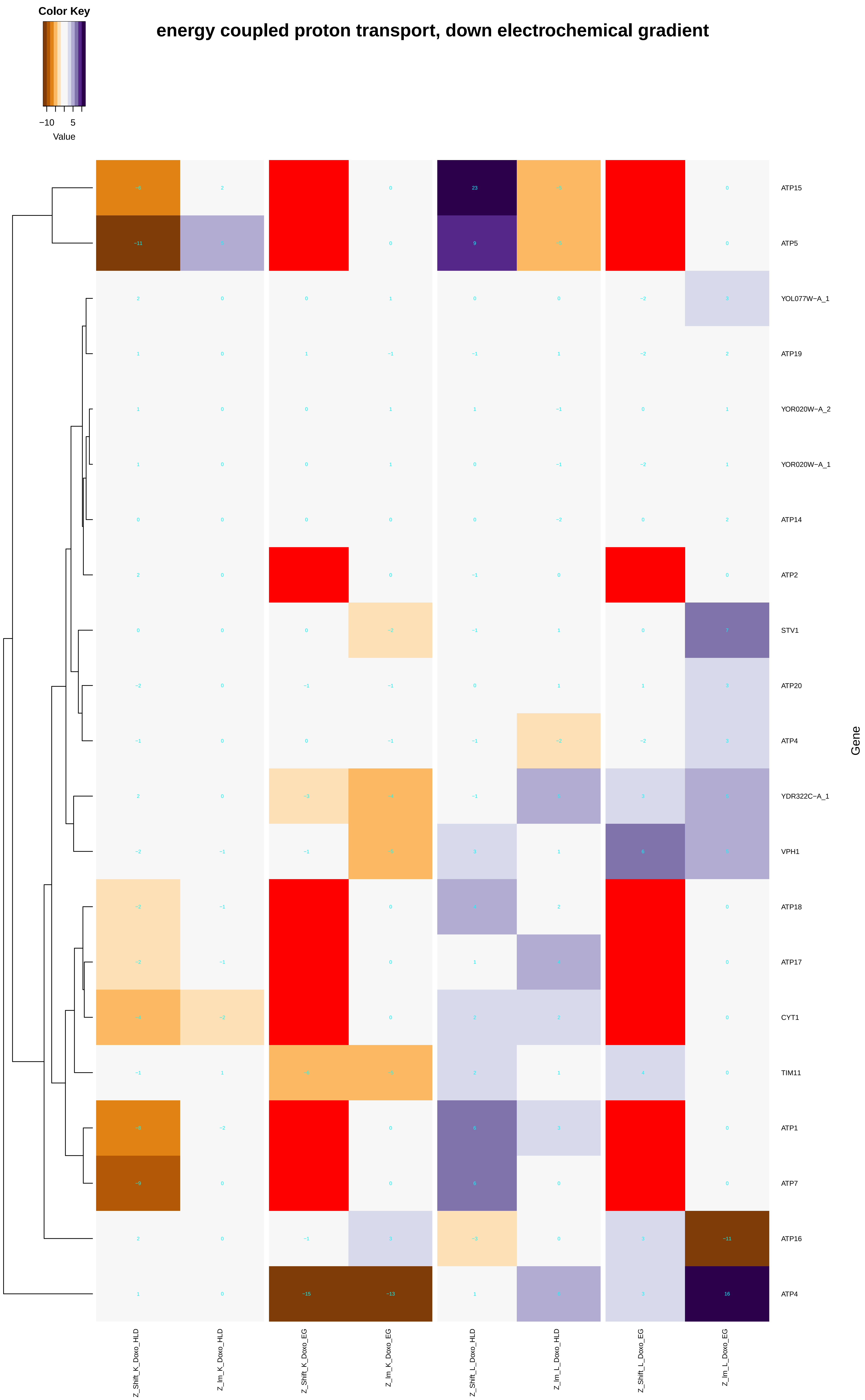
Color Key
−10
5
Value
energy coupled proton transport, down electrochemical gradient
ATP15
ATP5
YOL077W−A_1
ATP19
YOR020W−A_2
YOR020W−A_1
ATP14
ATP2
STV1
ATP20
ATP4
YDR322C−A_1
VPH1
ATP18
ATP17
CYT1
TIM11
ATP1
ATP7
ATP16
ATP4
Gene
Z_Shift_K_Doxo_HLD
Z_Im_K_Doxo_HLD
Z_Shift_K_Doxo_EG
Z_Im_K_Doxo_EG
Z_Shift_L_Doxo_HLD
Z_Im_L_Doxo_HLD
Z_Shift_L_Doxo_EG
Z_Im_L_Doxo_EG

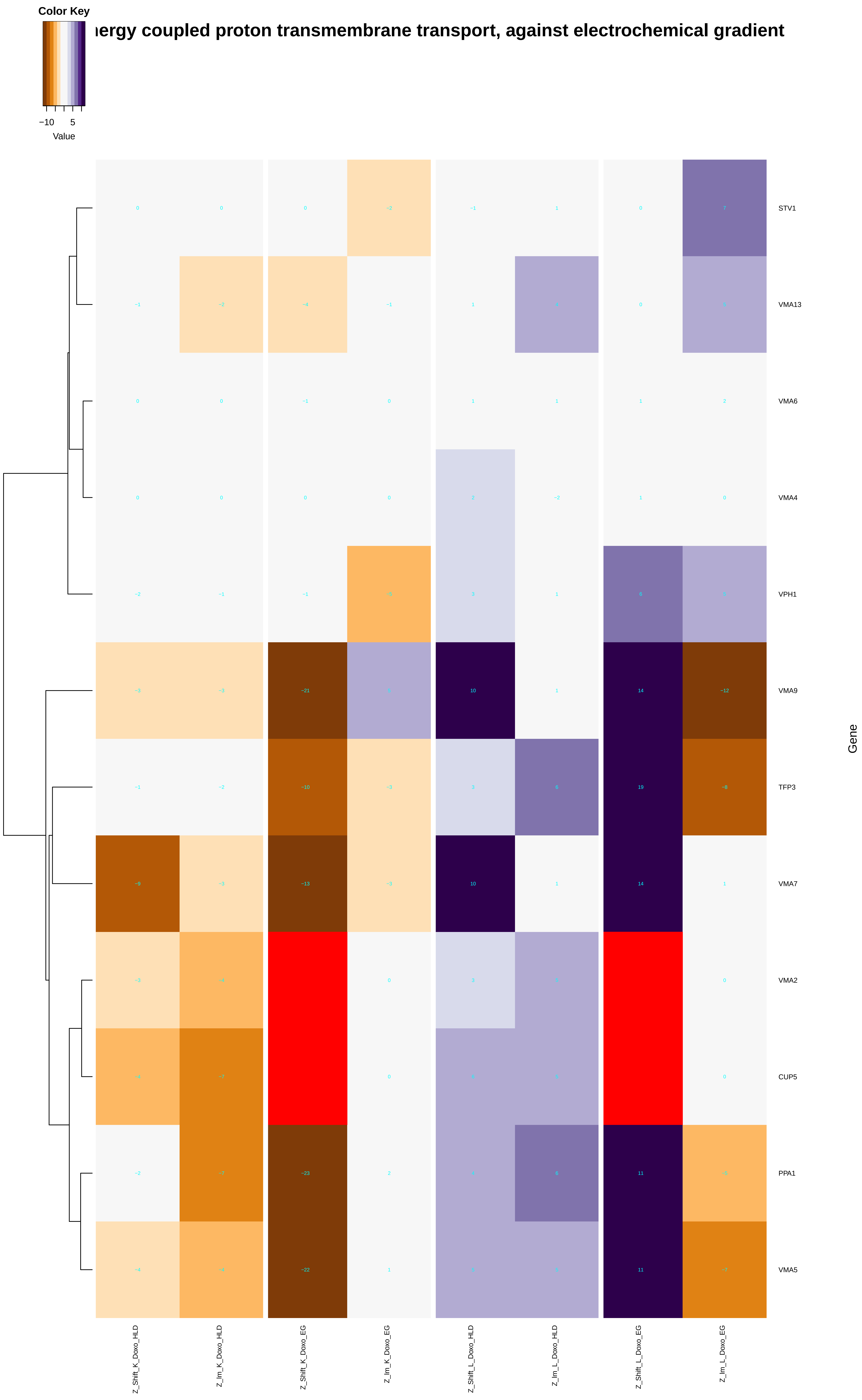
Color Key
−10
5
Value
ergy coupled proton transmembrane transport, against electrochemical gradient
STV1
VMA13
VMA6
VMA4
VPH1
VMA9
TFP3
VMA7
VMA2
CUP5
PPA1
VMA5
Gene
Z_Shift_K_Doxo_HLD
Z_Im_K_Doxo_HLD
Z_Shift_K_Doxo_EG
Z_Im_K_Doxo_EG
Z_Shift_L_Doxo_HLD
Z_Im_L_Doxo_HLD
Z_Shift_L_Doxo_EG
Z_Im_L_Doxo_EG

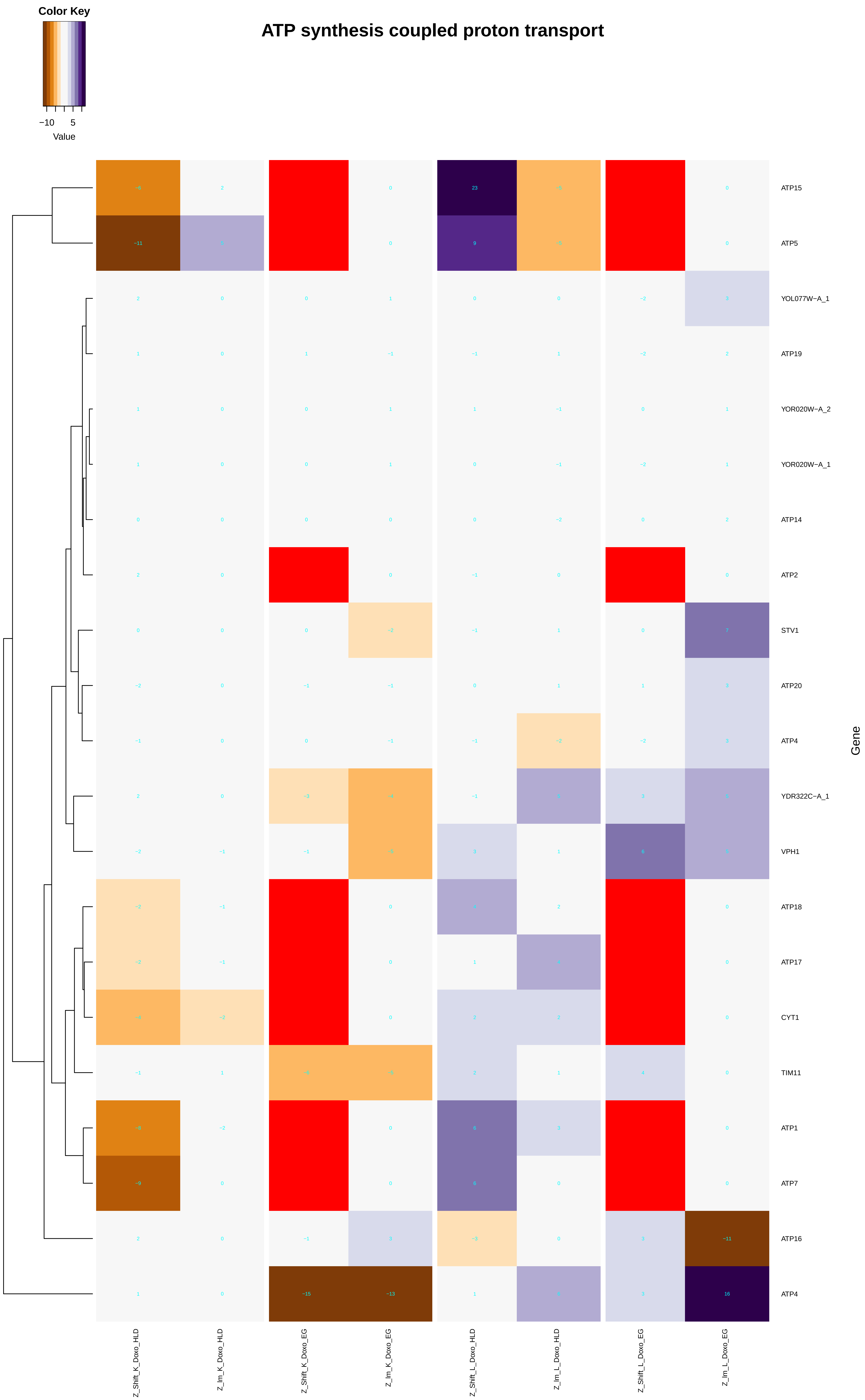

Color Key
−10
5
Value
ATP synthesis coupled proton transport
ATP15
ATP5
YOL077W−A_1
ATP19
YOR020W−A_2
YOR020W−A_1
ATP14
ATP2
STV1
ATP20
ATP4
YDR322C−A_1
VPH1
ATP18
ATP17
CYT1
TIM11
ATP1
ATP7
ATP16
ATP4
Gene
Z_Shift_K_Doxo_HLD
Z_Im_K_Doxo_HLD
Z_Shift_K_Doxo_EG
Z_Im_K_Doxo_EG
Z_Shift_L_Doxo_HLD
Z_Im_L_Doxo_HLD
Z_Shift_L_Doxo_EG
Z_Im_L_Doxo_EG

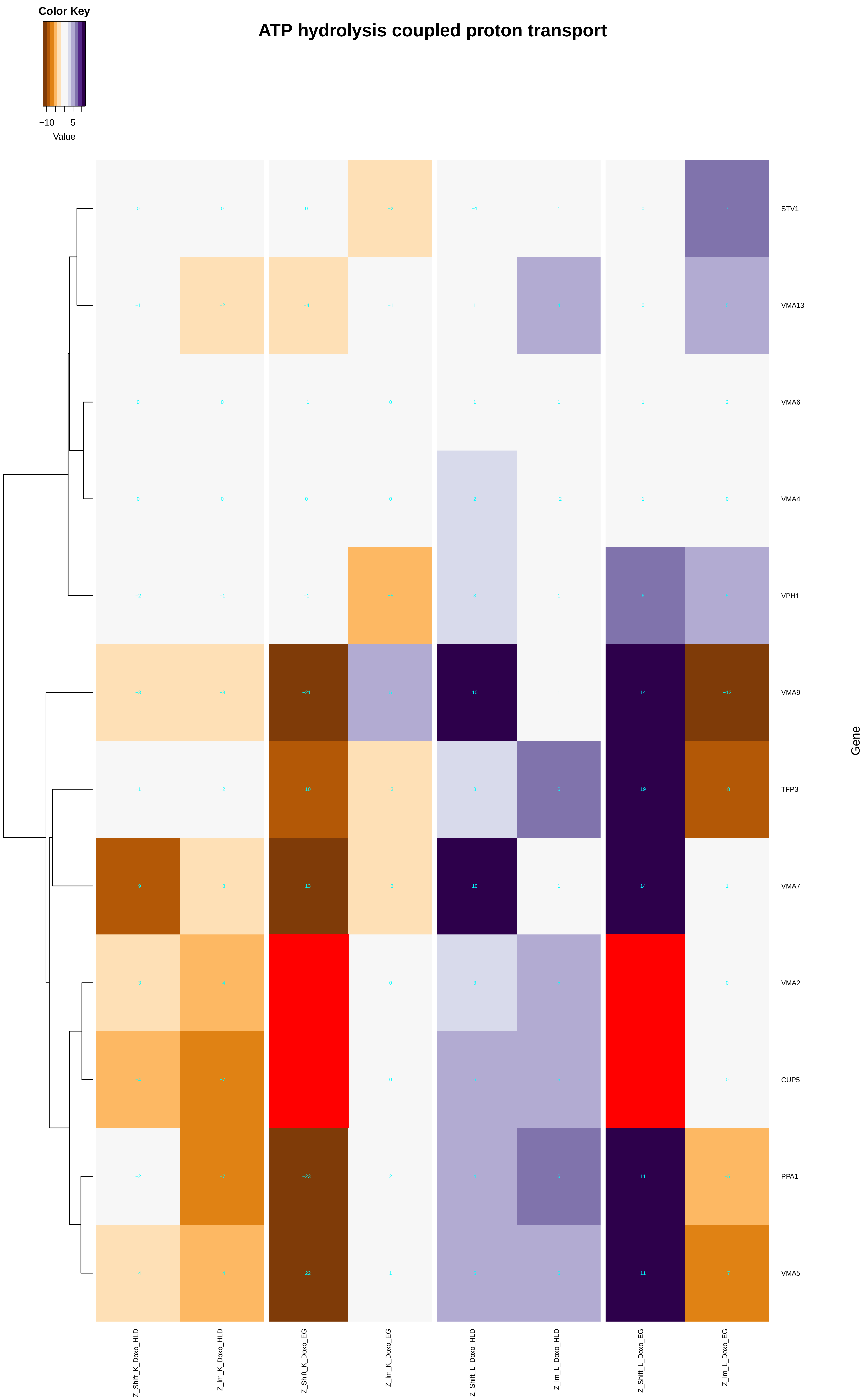
ATP hydrolysis coupled proton transport
Color Key
−10
5
Value
STV1
VMA13
VMA6
VMA4
VPH1
VMA9
TFP3
VMA7
VMA2
CUP5
PPA1
VMA5
Gene
Z_Shift_K_Doxo_HLD
Z_Im_K_Doxo_HLD
Z_Shift_K_Doxo_EG
Z_Im_K_Doxo_EG
Z_Shift_L_Doxo_HLD
Z_Im_L_Doxo_HLD
Z_Shift_L_Doxo_EG
Z_Im_L_Doxo_EG
