## Additional File 9 for "A yeast phenomic model for the influence of Warburg metabolism on genetic buffering of doxorubicin": mRNA_metabolic_process.pdf

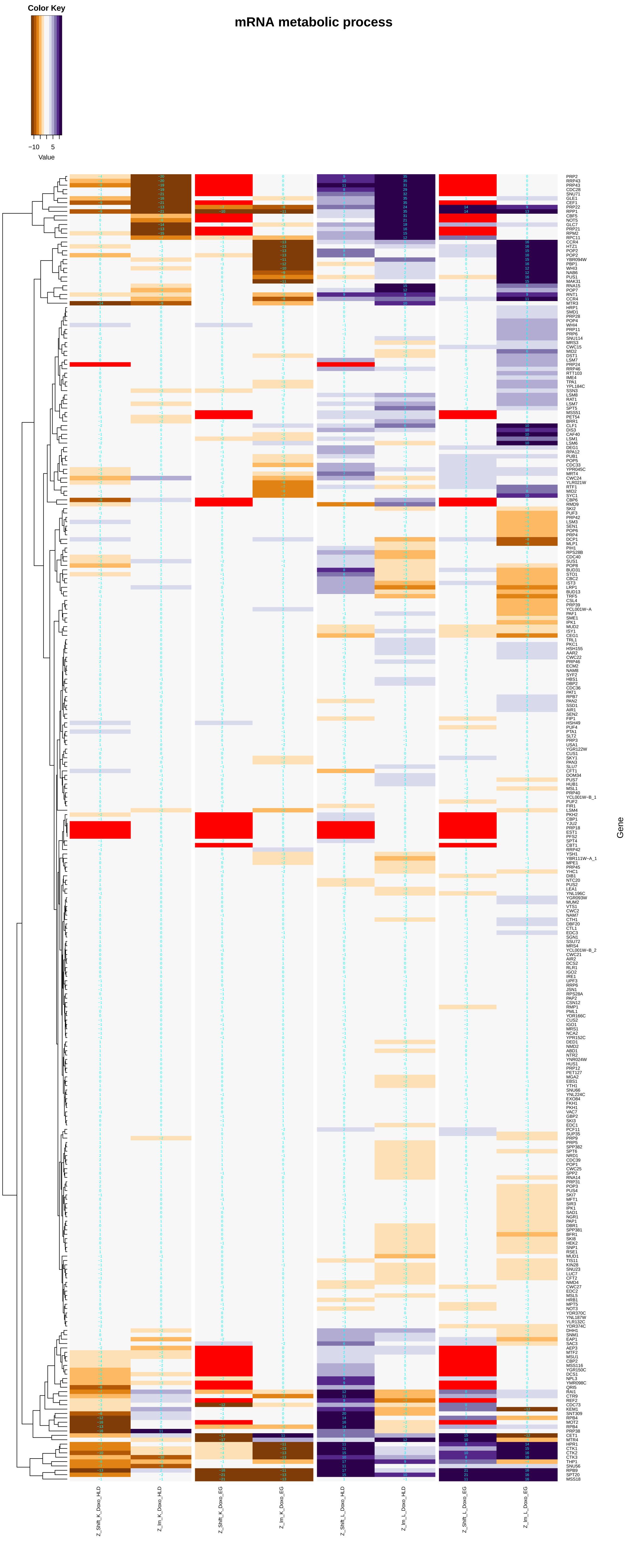
mRNA metabolic process
Color Key
−10 5
Value
Gene
Z_Shift_K_Doxo_HLD
Z_Im_K_Doxo_HLD
Z_Shift_K_Doxo_EG
Z_Im_K_Doxo_EG
Z_Shift_L_Doxo_HLD
Z_Im_L_Doxo_HLD
Z_Shift_L_Doxo_EG
Z_Im_L_Doxo_EG

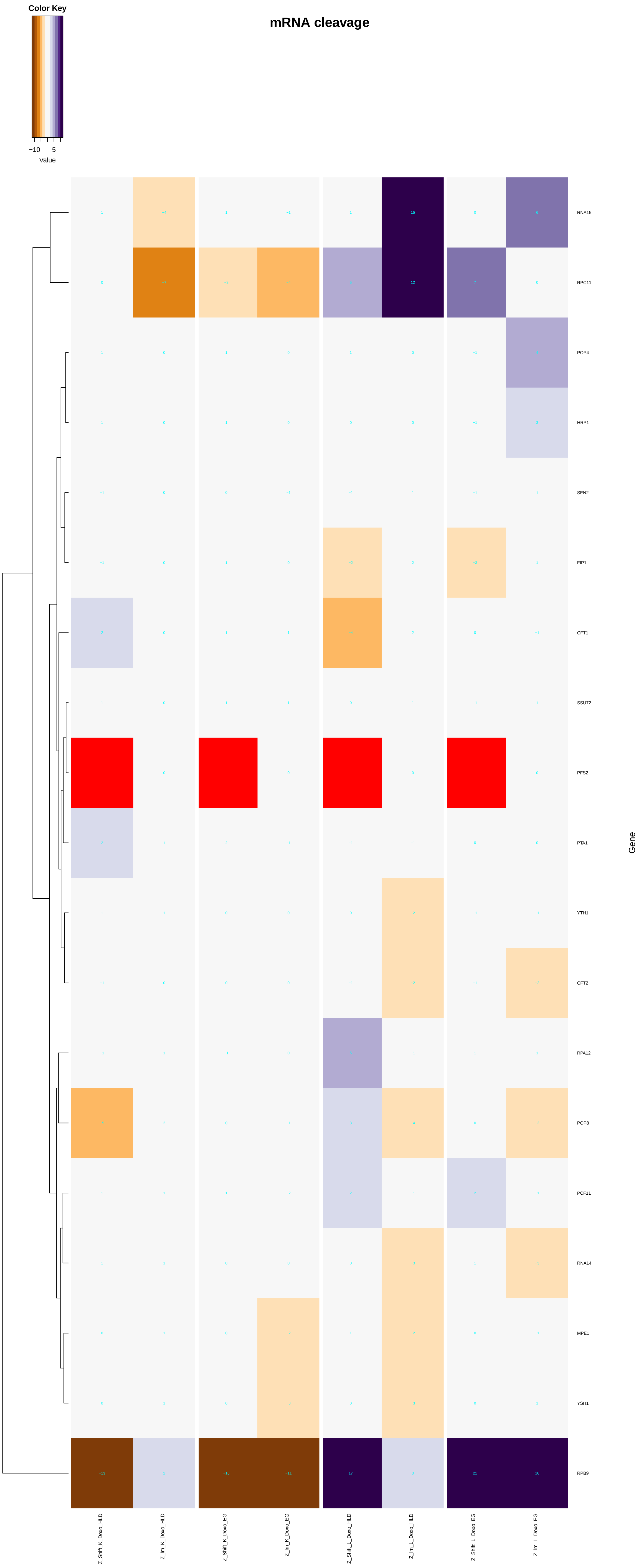

Color Key
−10
5
Value
mRNA cleavage
RNA15
RPC11
POP4
HRP1
SEN2
FIP1
CFT1
SSU72
PFS2
PTA1
YTH1
CFT2
RPA12
POP8
PCF11
RNA14
MPE1
YSH1
RPB9
Gene
Z_Shift_K_Doxo_HLD
Z_Im_K_Doxo_HLD
Z_Shift_K_Doxo_EG
Z_Im_K_Doxo_EG
Z_Shift_L_Doxo_HLD
Z_Im_L_Doxo_HLD
Z_Shift_L_Doxo_EG
Z_Im_L_Doxo_EG

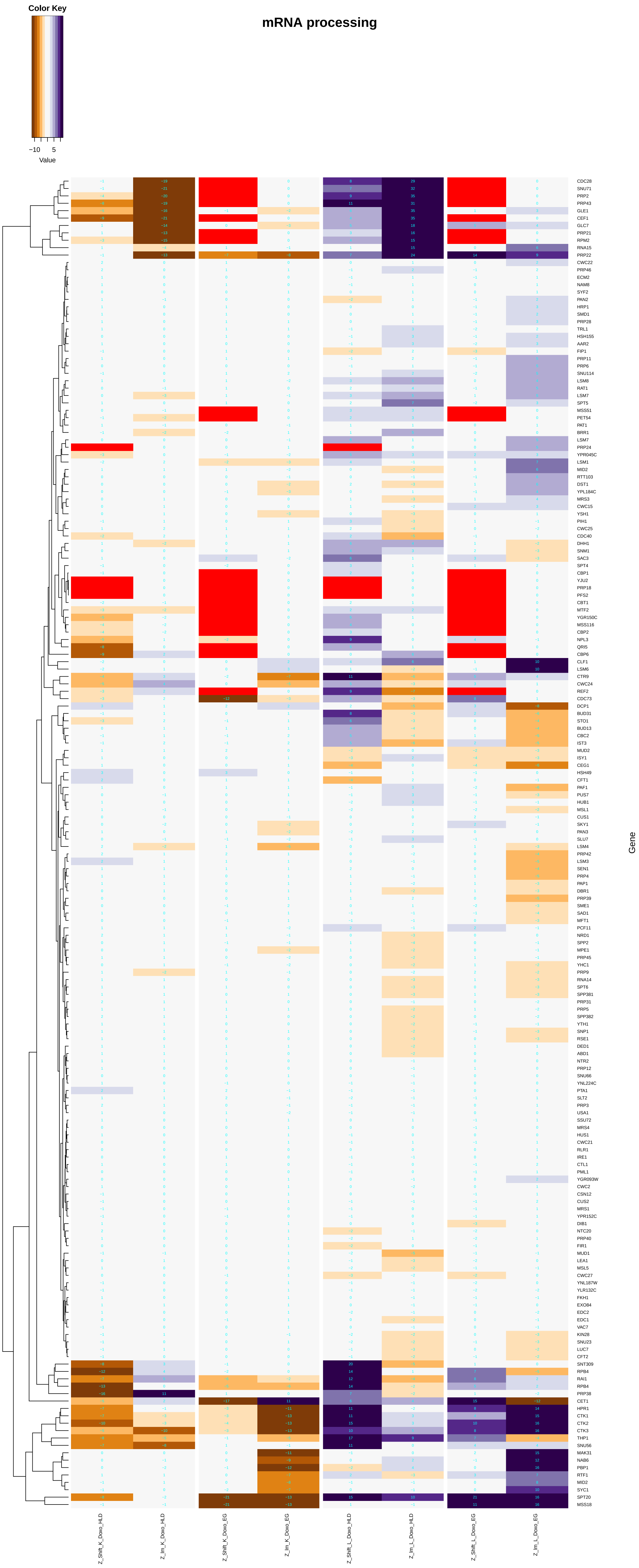

mRNA processing
Color Key
−10 5
Value
Gene
CDC28
SNU71
PRP2
PRP43
GLE1
CEF1
GLC7
PRP21
RPM2
RNA15
PRP22
CWC22
PRP46
ECM2
NAM8
SYF2
PAN2
HRP1
SMD1
PRP28
TRL1
HSH155
AAR2
FIP1
PRP11
PRP6
SNU114
LSM8
RAT1
LSM7
SPT5
MSS51
PET54
PAT1
BRR1
LSM7
PRP24
YPR045C
LSM1
MID2
RTT103
DST1
YPL184C
MRS3
CWC15
YSH1
PIH1
CWC25
CDC40
DHH1
SNM1
SAC3
SPT4
CBP1
YJU2
PRP18
PFS2
CBT1
MTF2
YGR150C
MSS116
CBP2
NPL3
QRI5
CBP6
CLF1
LSM6
CTR9
CWC24
REF2
CDC73
DCP1
BUD31
STO1
BUD13
CBC2
IST3
MUD2
ISY1
CEG1
HSH49
CFT1
PAF1
PUS7
HUB1
MSL1
CUS1
SKY1
PAN3
SLU7
LSM4
PRP42
LSM3
SEN1
PRP4
PAP1
DBR1
PRP39
SME1
SAD1
MFT1
PCF11
NRD1
SPP2
MPE1
PRP45
YHC1
PRP9
RNA14
SPT6
SPP381
PRP31
PRP5
SPP382
YTH1
SNP1
RSE1
DED1
ABD1
NTR2
PRP12
SNU66
YNL224C
PTA1
SLT2
PRP3
USA1
SSU72
MRS4
HUS1
CWC21
RLR1
IRE1
CTL1
PML1
YGR093W
CWC2
CSN12
CUS2
MRS1
YPR152C
DIB1
NTC20
PRP40
FIR1
MUD1
LEA1
MSL5
CWC27
YNL187W
YLR132C
FKH1
EXO84
EDC2
EDC1
VAC7
KIN28
SNU23
LUC7
CFT2
SNT309
RPB4
RAI1
RPB4
PRP38
CET1
HPR1
CTK1
CTK2
CTK3
THP1
SNU56
MAK31
NAB6
PBP1
RTF1
MID2
SYC1
SPT20
MSS18
Z_Shift_K_Doxo_HLD
Z_Im_K_Doxo_HLD
Z_Shift_K_Doxo_EG
Z_Im_K_Doxo_EG
Z_Shift_L_Doxo_HLD
Z_Im_L_Doxo_HLD
Z_Shift_L_Doxo_EG
Z_Im_L_Doxo_EG

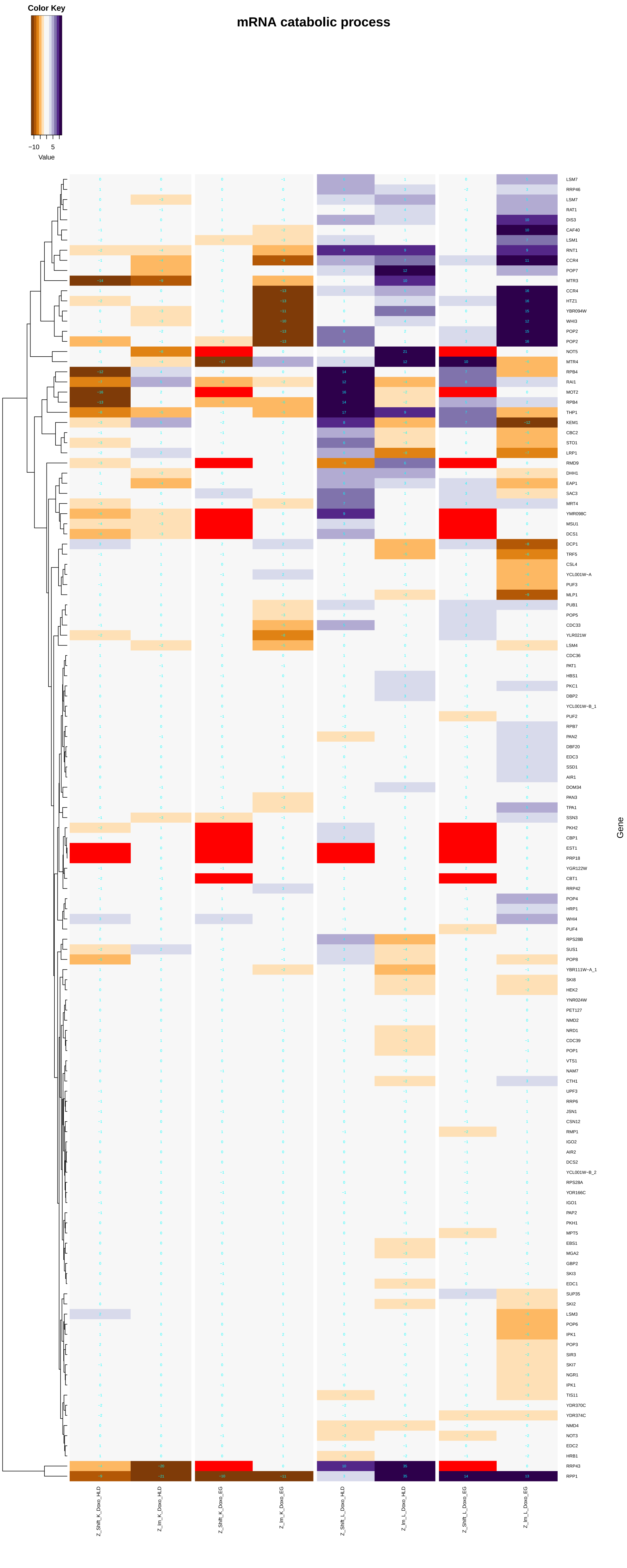
mRNA catabolic process
Color Key
−10
5
Value
LSM7
RRP46
LSM7
RAT1
DIS3
CAF40
LSM1
RNT1
CCR4
POP7
MTR3
CCR4
HTZ1
YBR094W
WHI3
POP2
POP2
NOT5
MTR4
RPB4
RAI1
MOT2
RPB4
THP1
KEM1
CBC2
STO1
LRP1
RMD9
DHH1
EAP1
SAC3
MRT4
YMR098C
MSU1
DCS1
DCP1
TRF5
CSL4
YCL001W−A
PUF3
MLP1
PUB1
POP5
CDC33
YLR021W
LSM4
CDC36
PAT1
HBS1
PKC1
DBP2
YCL001W−B_1
PUF2
RPB7
PAN2
DBF20
EDC3
SSD1
AIR1
DOM34
PAN3
TPA1
SSN3
PKH2
CBP1
EST1
PRP18
YGR122W
CBT1
RRP42
POP4
HRP1
WHI4
PUF4
RPS28B
SUS1
POP8
YBR111W−A_1
SKI8
HEK2
YNR024W
PET127
NMD2
NRD1
CDC39
POP1
VTS1
NAM7
CTH1
UPF3
RRP6
JSN1
CSN12
RMP1
IGO2
AIR2
DCS2
YCL001W−B_2
RPS28A
YOR166C
IGO1
PAP2
PKH1
MPT5
EBS1
MGA2
GBP2
SKI3
EDC1
SUP35
SKI2
LSM3
POP6
IPK1
POP3
SIR3
SKI7
NGR1
IPK1
TIS11
YDR370C
YDR374C
NMD4
NOT3
EDC2
HRB1
RRP43
RPP1
Gene
Z_Shift_K_Doxo_HLD
Z_lm_K_Doxo_HLD
Z_Shift_K_Doxo_EG
Z_lm_K_Doxo_EG
Z_Shift_L_Doxo_HLD
Z_lm_L_Doxo_HLD
Z_Shift_L_Doxo_EG
Z_lm_L_Doxo_EG

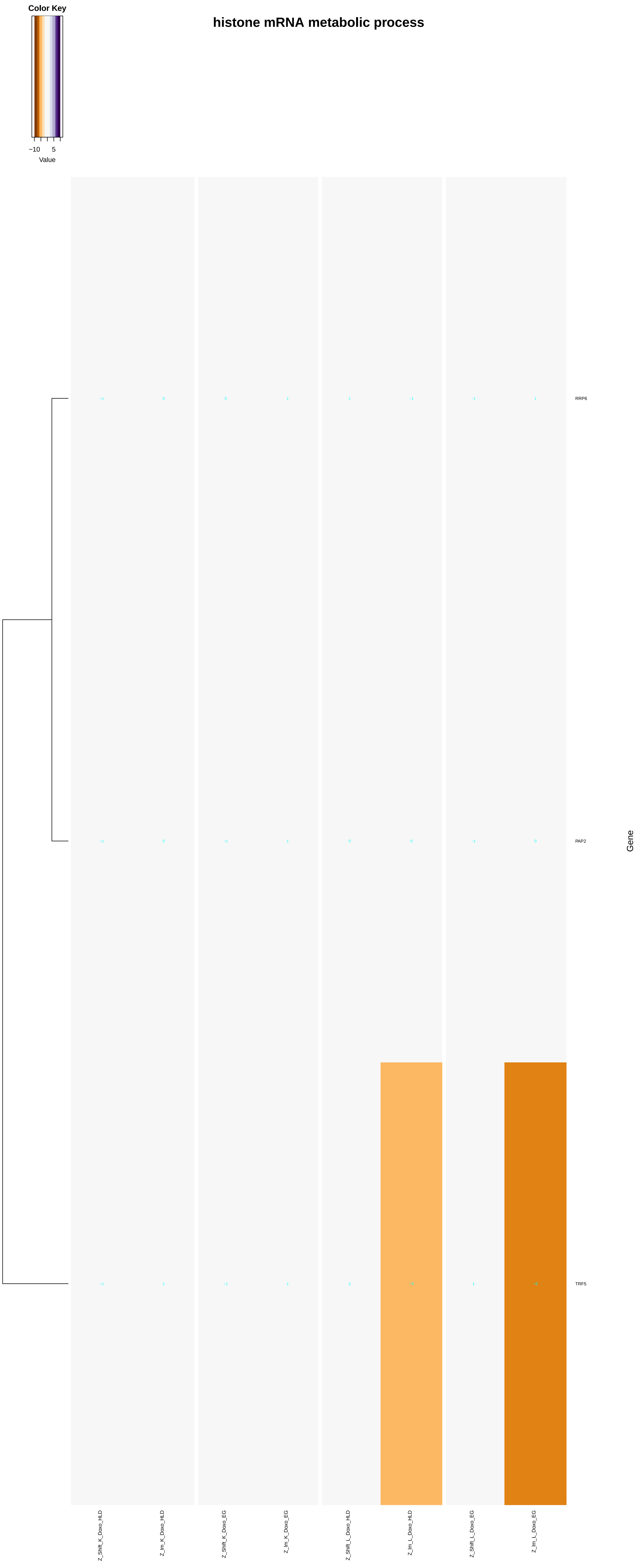

histone mRNA metabolic process
Color Key
−10
5
Value
RRP6
PAP2
TRF5
Gene
Z_Shift_K_Doxo_HLD
Z_Im_K_Doxo_HLD
Z_Shift_K_Doxo_EG
Z_Im_K_Doxo_EG
Z_Shift_L_Doxo_HLD
Z_Im_L_Doxo_HLD
Z_Shift_L_Doxo_EG
Z_Im_L_Doxo_EG

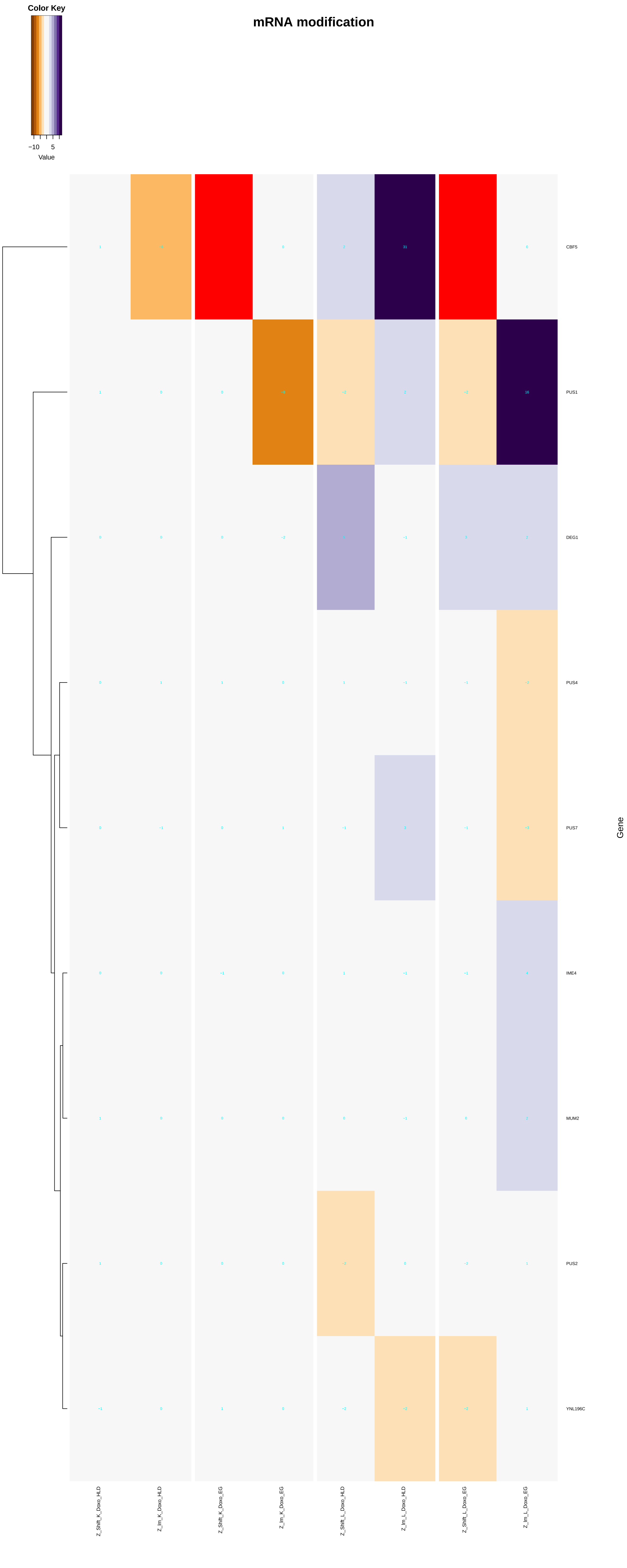
Color Key
−10
5
Value
mRNA modification
CBF5
PUS1
DEG1
PUS4
PUS7
IME4
MUM2
PUS2
YNL196C
Gene
Z_Shift_K_Doxo_HLD
Z_Im_K_Doxo_HLD
Z_Shift_K_Doxo_EG
Z_Im_K_Doxo_EG
Z_Shift_L_Doxo_HLD
Z_Im_L_Doxo_HLD
Z_Shift_L_Doxo_EG
Z_Im_L_Doxo_EG

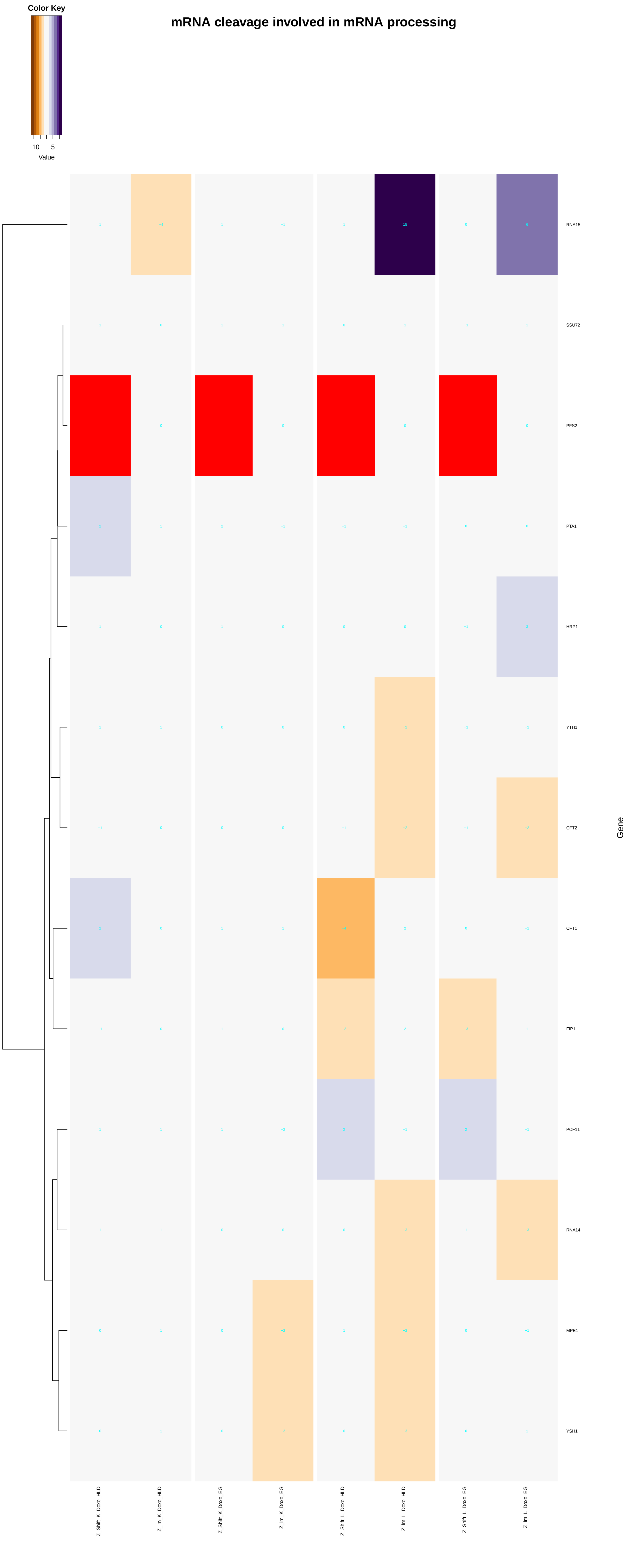

mRNA cleavage involved in mRNA processing
Color Key
−10
5
Value
RNA15
SSU72
PFS2
PTA1
HRP1
YTH1
CFT2
CFT1
FIP1
PCF11
RNA14
MPE1
YSH1
Gene
Z_Shift_K_Doxo_HLD
Z_Im_K_Doxo_HLD
Z_Shift_K_Doxo_EG
Z_Im_K_Doxo_EG
Z_Shift_L_Doxo_HLD
Z_Im_L_Doxo_HLD
Z_Shift_L_Doxo_EG
Z_Im_L_Doxo_EG

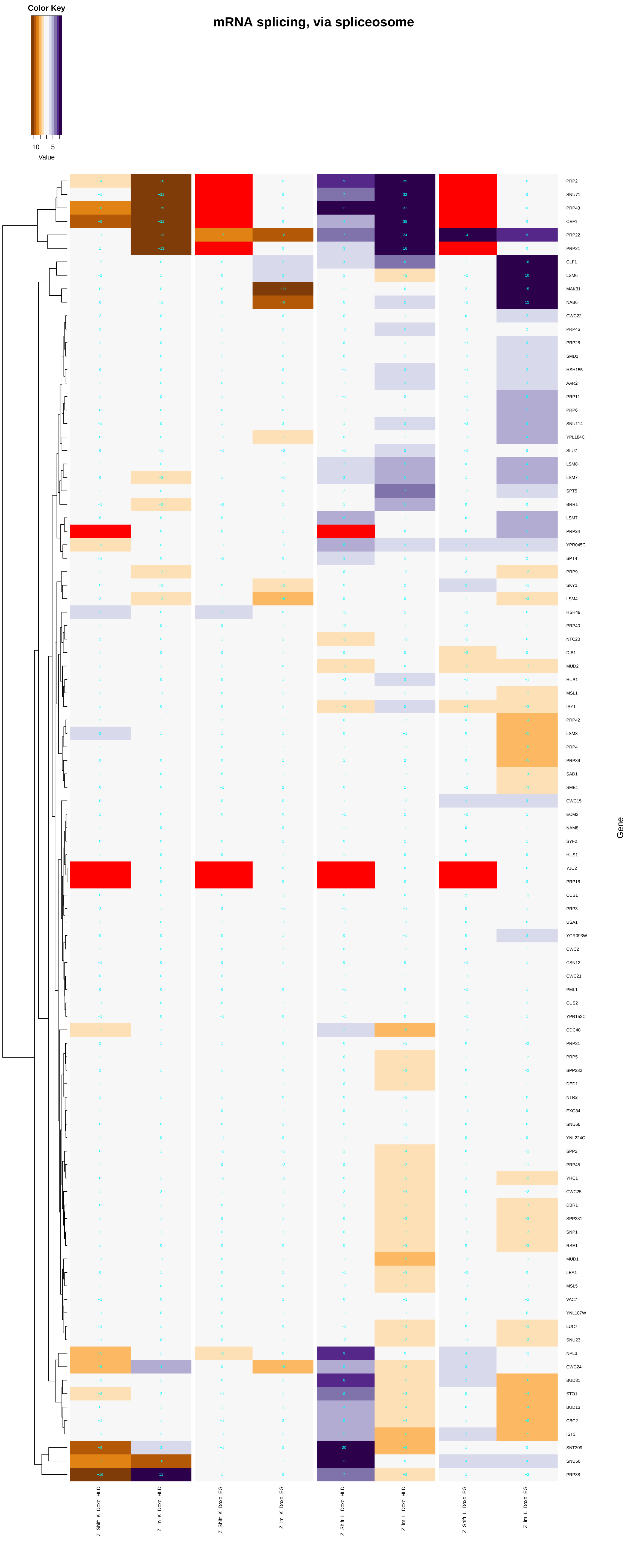
Color Key
−10
5
Value
mRNA splicing, via spliceosome
PRP2
SNU71
PRP43
CEF1
PRP22
PRP21
CLF1
LSM6
MAK31
NAB6
CWC22
PRP46
PRP28
SMD1
HSH155
AAR2
PRP11
PRP6
SNU114
YPL184C
SLU7
LSM8
LSM7
SPT5
BRR1
LSM7
PRP24
YPR045C
SPT4
PRP9
SKY1
LSM4
HSH49
PRP40
NTC20
DIB1
MUD2
HUB1
MSL1
ISY1
PRP42
LSM3
PRP4
PRP39
SAD1
SME1
CWC15
ECM2
NAM8
SYF2
HUS1
YJU2
PRP18
CUS1
PRP3
USA1
YGR093W
CWC2
CSN12
CWC21
PML1
CUS2
YPR152C
CDC40
PRP31
PRP5
SPP382
DED1
NTR2
EXO84
SNU66
YNL224C
SPP2
PRP45
YHC1
CWC25
DBR1
SPP381
SNP1
RSE1
MUD1
LEA1
MSL5
VAC7
YNL187W
LUC7
SNU23
NPL3
CWC24
BUD31
STO1
BUD13
CBC2
IST3
SNT309
SNU56
PRP38
Gene
Z_Shift_K_Doxo_HLD
Z_Im_K_Doxo_HLD
Z_Shift_K_Doxo_EG
Z_Im_K_Doxo_EG
Z_Shift_L_Doxo_HLD
Z_Im_L_Doxo_HLD
Z_Shift_L_Doxo_EG
Z_Im_L_Doxo_EG

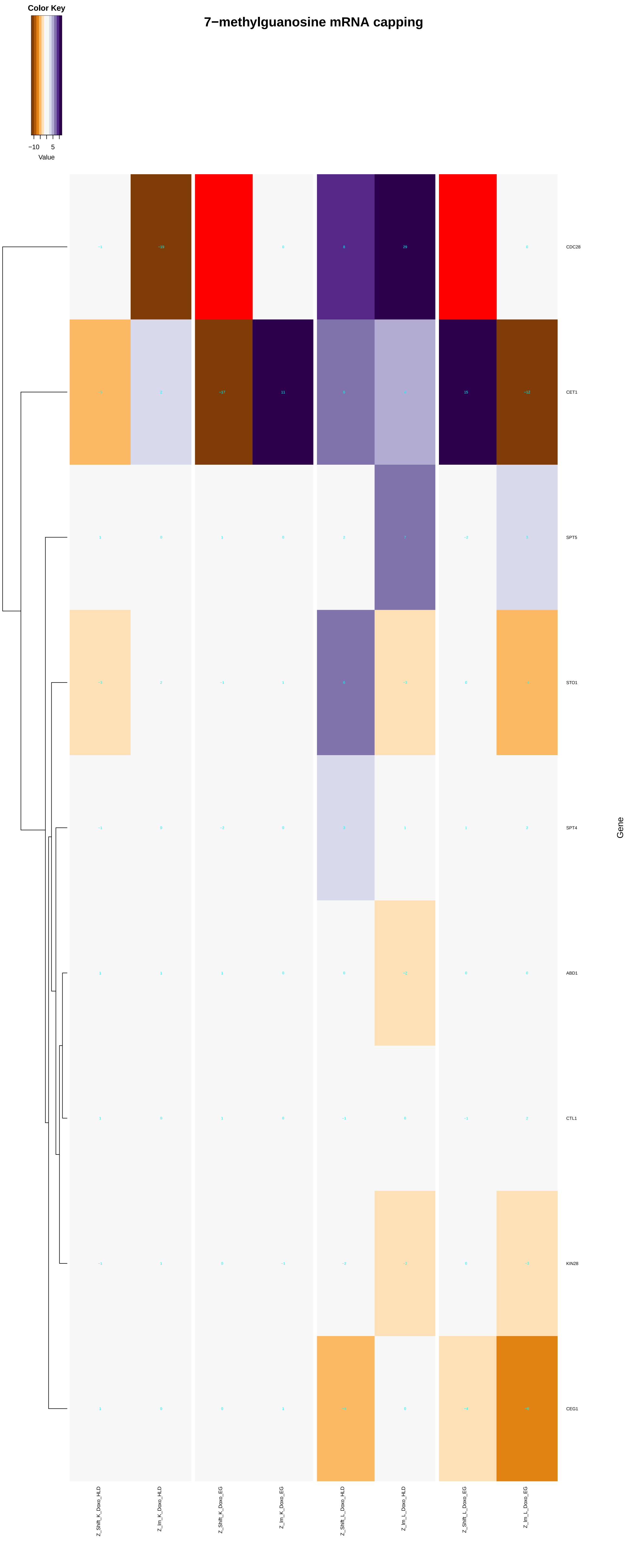

Color Key
−10
5
Value
7−methylguanosine mRNA capping
CDC28
CET1
SPT5
STO1
SPT4
ABD1
CTL1
KIN28
CEG1
Gene
Z_Shift_K_Doxo_HLD
Z_Im_K_Doxo_HLD
Z_Shift_K_Doxo_EG
Z_Im_K_Doxo_EG
Z_Shift_L_Doxo_HLD
Z_Im_L_Doxo_HLD
Z_Shift_L_Doxo_EG
Z_Im_L_Doxo_EG

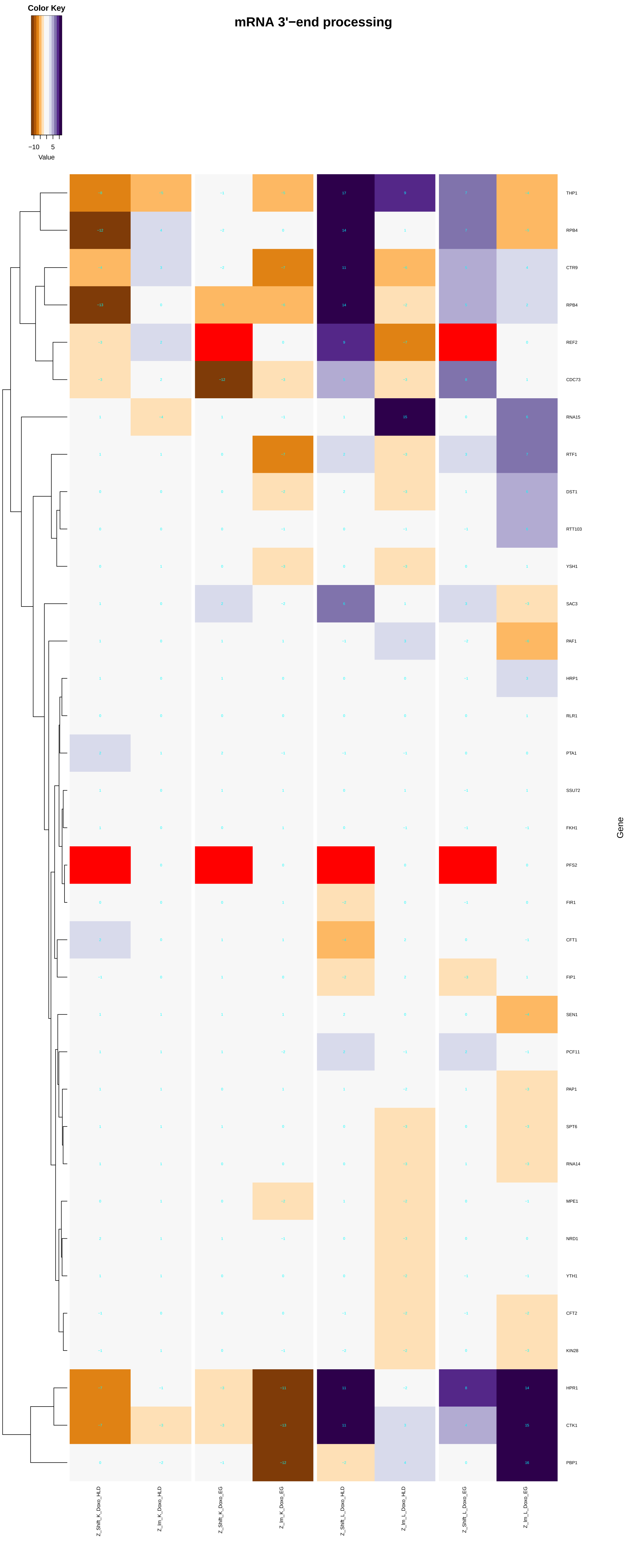
mRNA 3'−end processing
Color Key
−10
5
Value
THP1
RPB4
CTR9
RPB4
REF2
CDC73
RNA15
RTF1
DST1
RTT103
YSH1
SAC3
PAF1
HRP1
RLR1
PTA1
SSU72
FKH1
PFS2
FIR1
CFT1
FIP1
SEN1
PCF11
PAP1
SPT6
RNA14
MPE1
NRD1
YTH1
CFT2
KIN28
HPR1
CTK1
PBP1
Gene
Z_Shift_K_Doxo_HLD
Z_Im_K_Doxo_HLD
Z_Shift_K_Doxo_EG
Z_Im_K_Doxo_EG
Z_Shift_L_Doxo_HLD
Z_Im_L_Doxo_HLD
Z_Shift_L_Doxo_EG
Z_Im_L_Doxo_EG

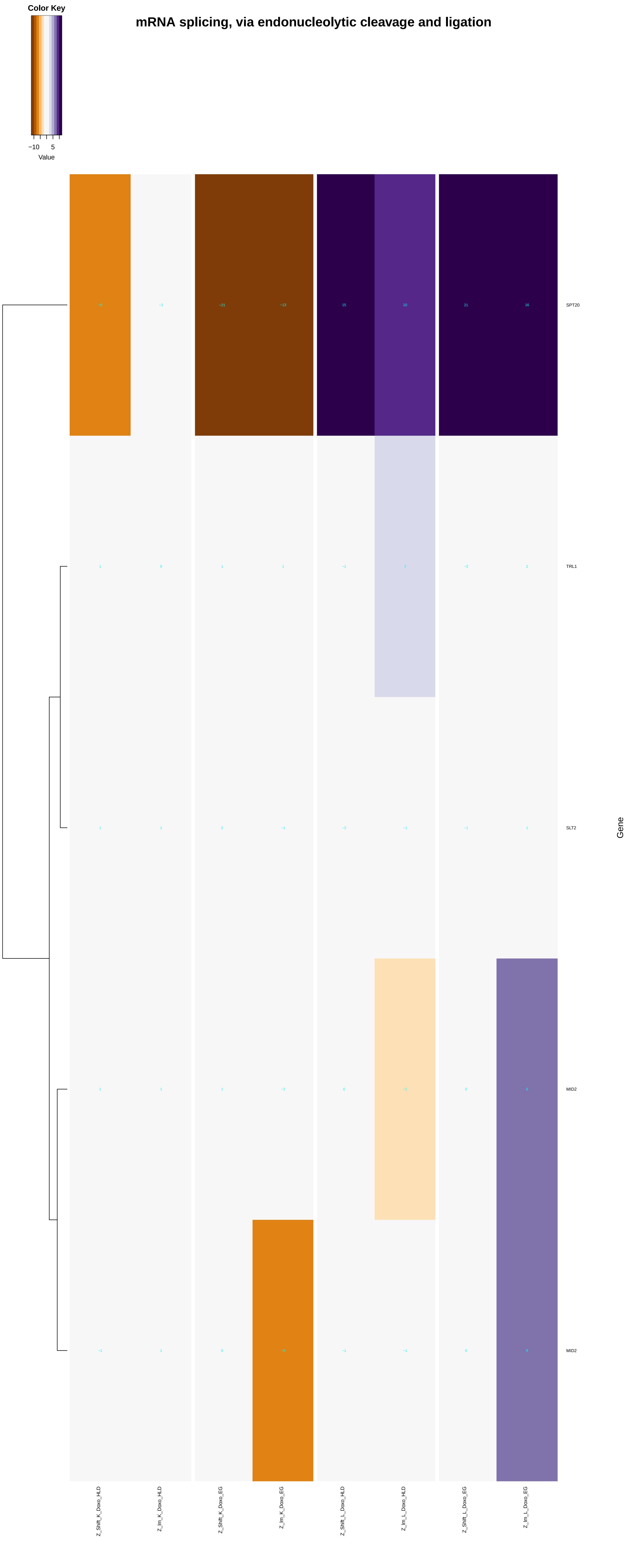
mRNA splicing, via endonucleolytic cleavage and ligation
Color Key
−10
5
Value
SPT20
TRL1
SLT2
MID2
MID2
Gene
Z_Shift_K_Doxo_HLD
Z_Im_K_Doxo_HLD
Z_Shift_K_Doxo_EG
Z_Im_K_Doxo_EG
Z_Shift_L_Doxo_HLD
Z_Im_L_Doxo_HLD
Z_Shift_L_Doxo_EG
Z_Im_L_Doxo_EG

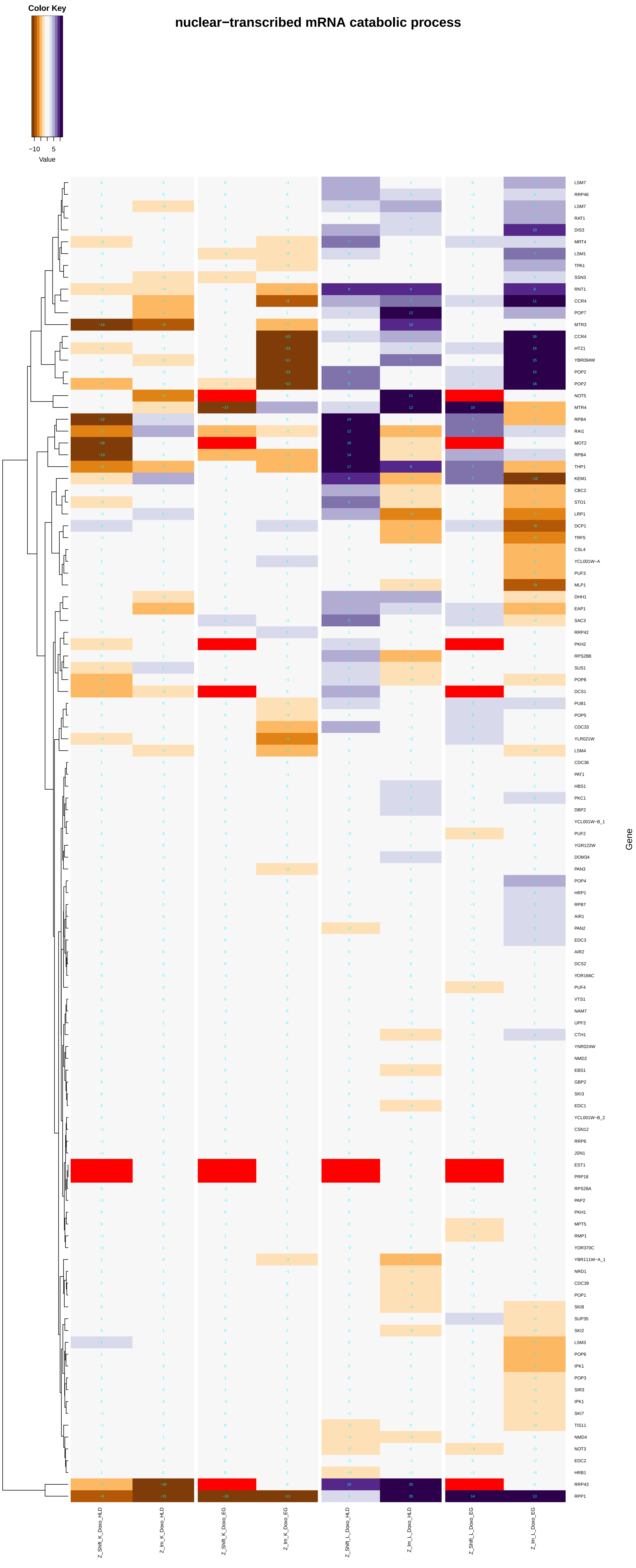
nuclear−transcribed mRNA catabolic process
Color Key
−10
5
Value
Gene
LSM7
RRP46
LSM7
RAT1
DIS3
MRT4
LSM1
TPA1
SSN3
RNT1
CCR4
POP7
MTR3
CCR4
HTZ1
YBR094W
POP2
POP2
NOT5
MTR4
RPB4
RAI1
MOT2
RPB4
THP1
KEM1
CBC2
STO1
LRP1
DCP1
TRF5
CSL4
YCL001W−A
PUF3
MLP1
DHH1
EAP1
SAC3
RRP42
PKH2
RPS28B
SUS1
POP8
DCS1
PUB1
POP5
CDC33
YLR021W
LSM4
CDC36
PAT1
HBS1
PKC1
DBP2
YCL001W−B_1
PUF2
YGR122W
DOM34
PAN3
POP4
HRP1
RPB7
AIR1
PAN2
EDC3
AIR2
DCS2
YOR166C
PUF4
VTS1
NAM7
UPF3
CTH1
YNR024W
NMD2
EBS1
GBP2
SKI3
EDC1
YCL001W−B_2
CSN12
RRP6
JSN1
EST1
PRP18
RPS28A
PAP2
PKH1
MPT5
RMP1
YDR370C
YBR111W−A_1
NRD1
CDC39
POP1
SKI8
SUP35
SKI2
LSM3
POP6
IPK1
POP3
SIR3
IPK1
SKI7
TIS11
NMD4
NOT3
EDC2
HRB1
RRP43
RPP1
Z_Shift_K_Doxo_HLD
Z_Im_K_Doxo_HLD
Z_Shift_K_Doxo_EG
Z_Im_K_Doxo_EG
Z_Shift_L_Doxo_HLD
Z_Im_L_Doxo_HLD
Z_Shift_L_Doxo_EG
Z_Im_L_Doxo_EG

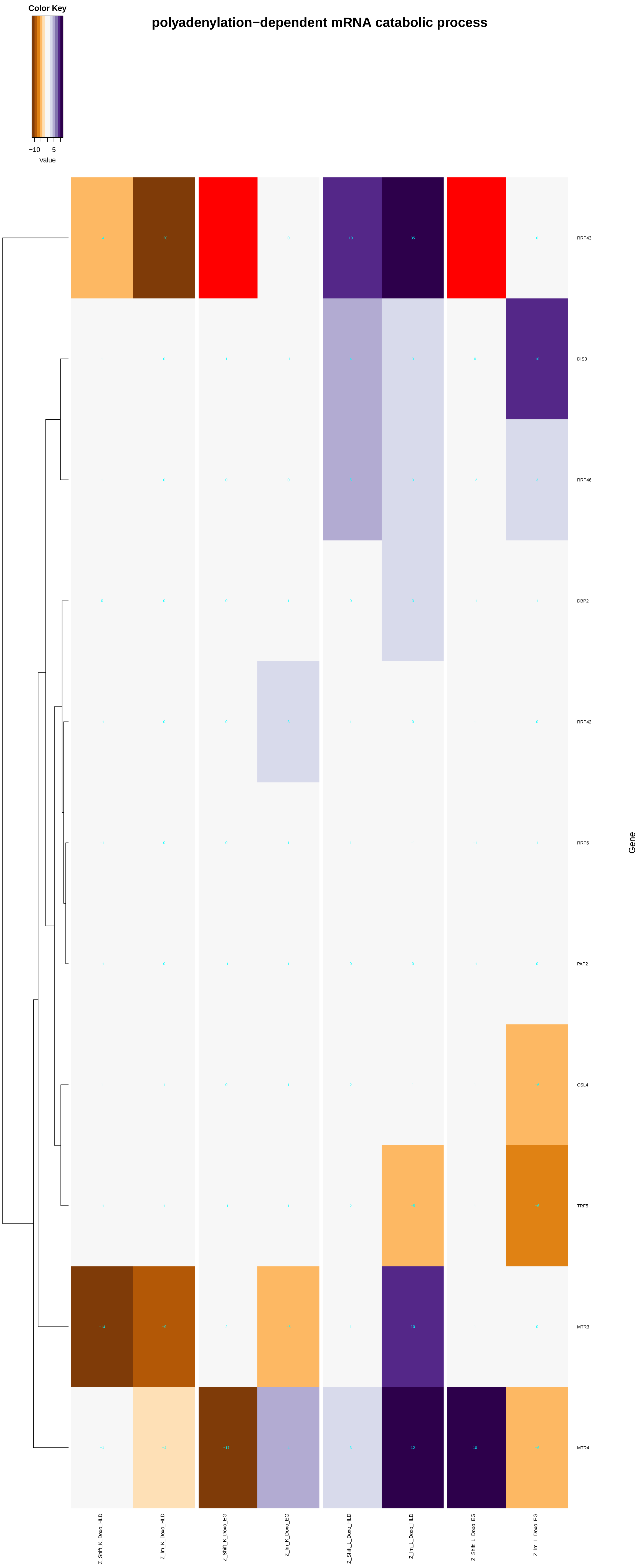
Color Key
−10
5
Value
polyadenylation−dependent mRNA catabolic process
RRP43
DIS3
RRP46
DBP2
RRP42
RRP6
PAP2
CSL4
TRF5
MTR3
MTR4
Gene
Z_Shift_K_Doxo_HLD
Z_Im_K_Doxo_HLD
Z_Shift_K_Doxo_EG
Z_Im_K_Doxo_EG
Z_Shift_L_Doxo_HLD
Z_Im_L_Doxo_HLD
Z_Shift_L_Doxo_EG
Z_Im_L_Doxo_EG

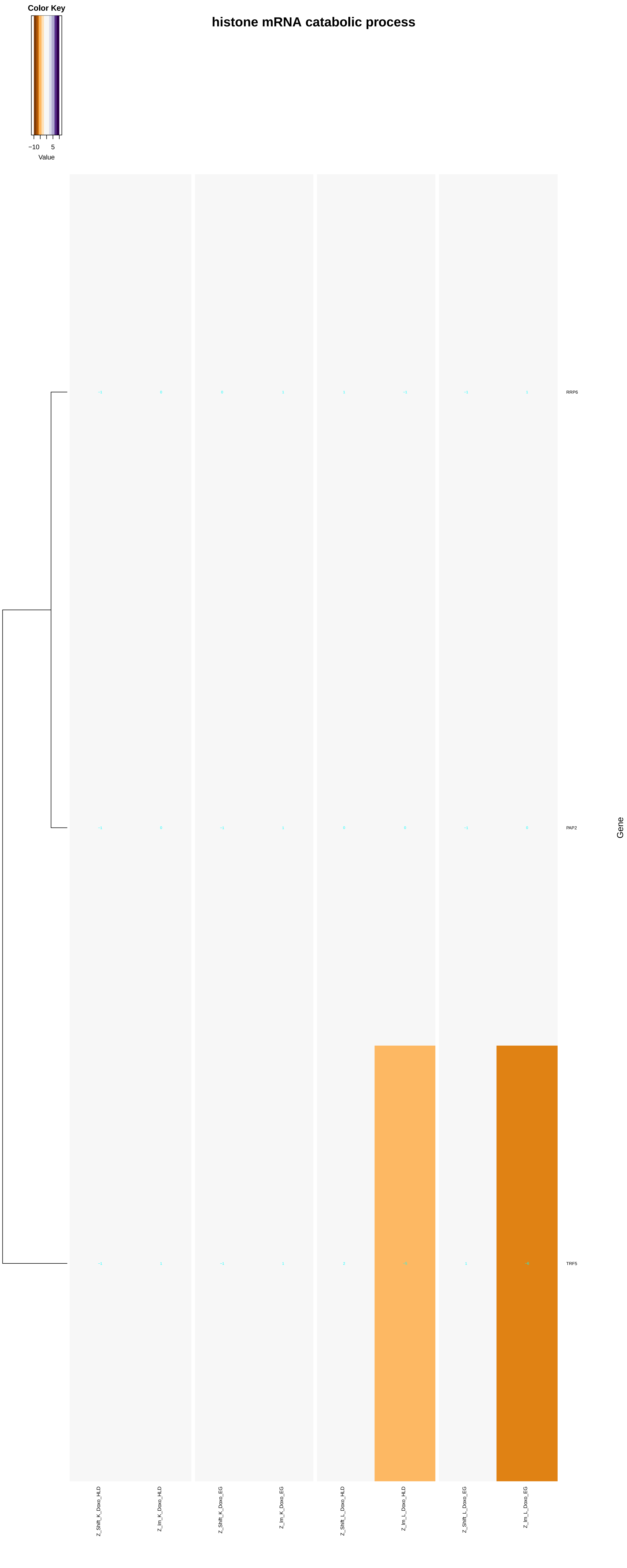
histone mRNA catabolic process
Color Key
−10
5
Value
RRP6
PAP2
TRF5
Gene
Z_Shift_K_Doxo_HLD
Z_Im_K_Doxo_HLD
Z_Shift_K_Doxo_EG
Z_Im_K_Doxo_EG
Z_Shift_L_Doxo_HLD
Z_Im_L_Doxo_HLD
Z_Shift_L_Doxo_EG
Z_Im_L_Doxo_EG

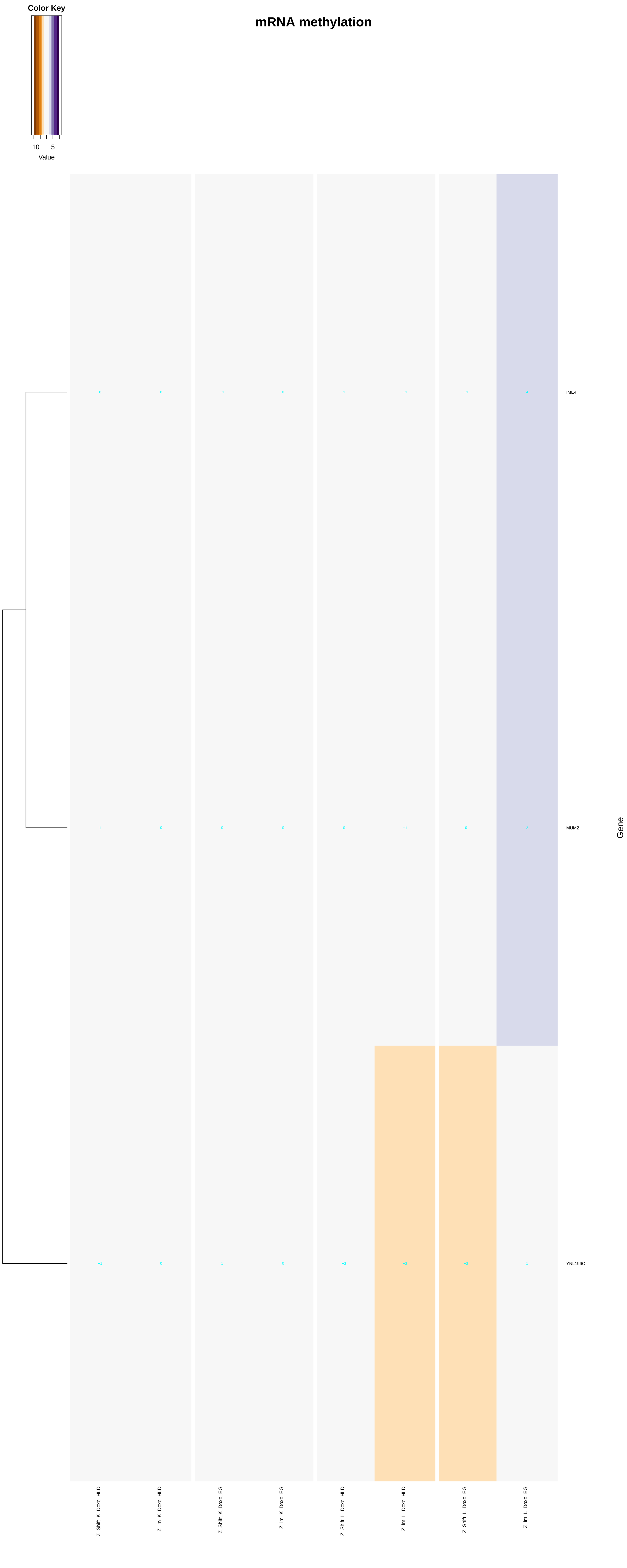

Color Key
−10
5
Value
mRNA methylation
IME4
MUM2
YNL196C
Gene
Z_Shift_K_Doxo_HLD
Z_Im_K_Doxo_HLD
Z_Shift_K_Doxo_EG
Z_Im_K_Doxo_EG
Z_Shift_L_Doxo_HLD
Z_Im_L_Doxo_HLD
Z_Shift_L_Doxo_EG
Z_Im_L_Doxo_EG

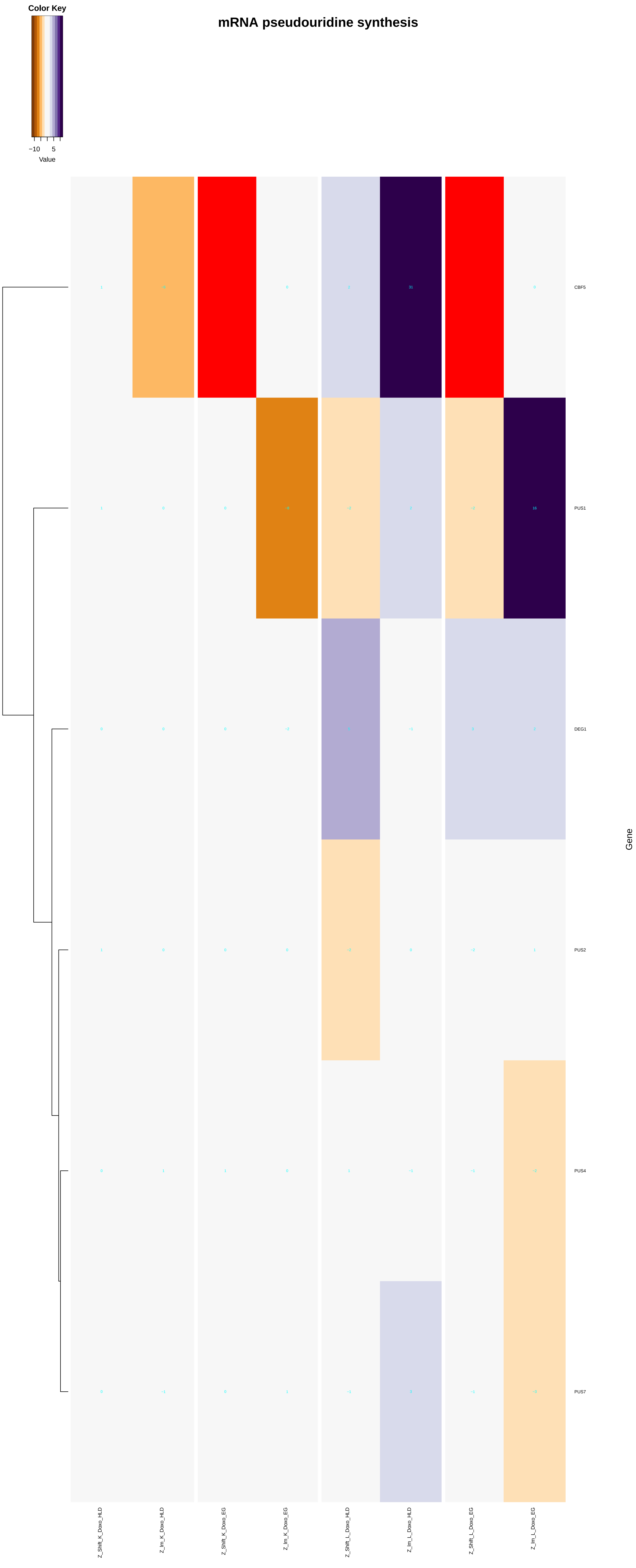
mRNA pseudouridine synthesis
Color Key
−10
5
Value
CBF5
PUS1
DEG1
PUS2
PUS4
PUS7
Gene
Z_Shift_K_Doxo_HLD
Z_lm_K_Doxo_HLD
Z_Shift_K_Doxo_EG
Z_lm_K_Doxo_EG
Z_Shift_L_Doxo_HLD
Z_lm_L_Doxo_HLD
Z_Shift_L_Doxo_EG
Z_lm_L_Doxo_EG

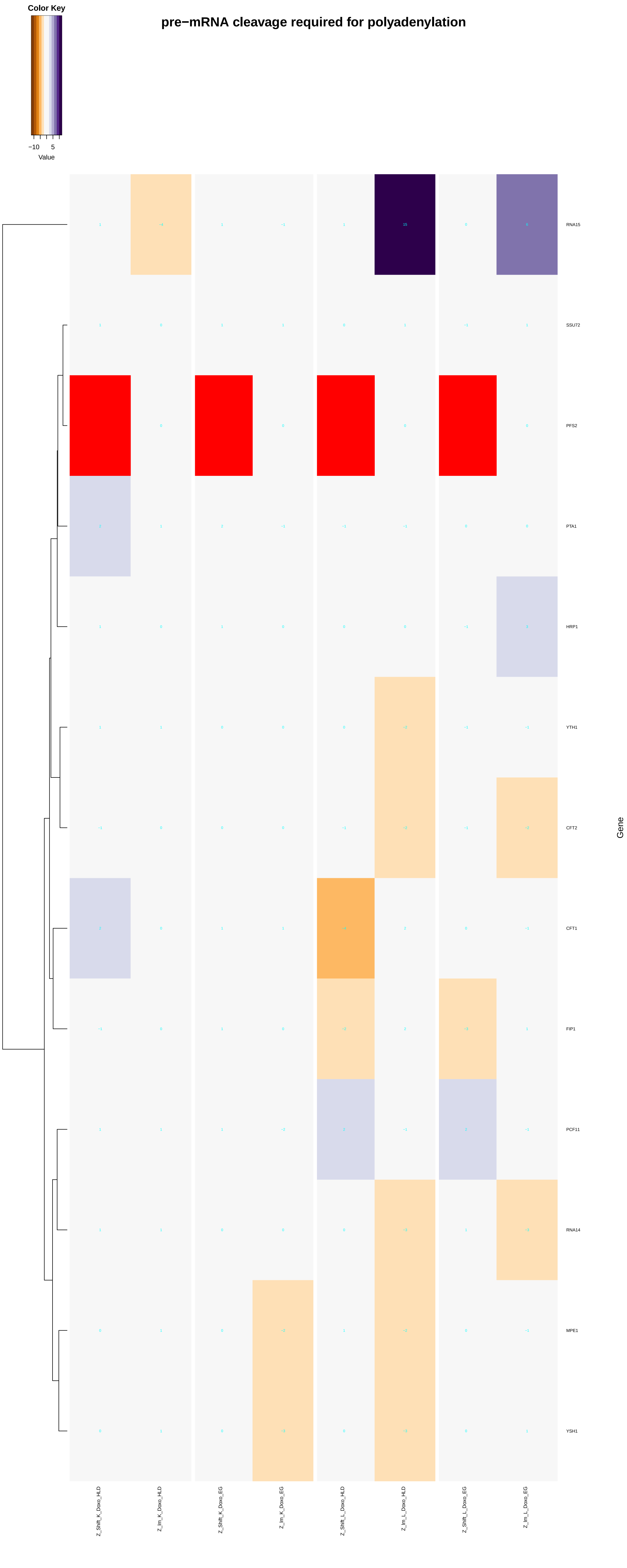
pre−mRNA cleavage required for polyadenylation
Color Key
−10
5
Value
RNA15
SSU72
PFS2
PTA1
HRP1
YTH1
CFT2
CFT1
FIP1
PCF11
RNA14
MPE1
YSH1
Gene
Z_Shift_K_Doxo_HLD
Z_Im_K_Doxo_HLD
Z_Shift_K_Doxo_EG
Z_Im_K_Doxo_EG
Z_Shift_L_Doxo_HLD
Z_Im_L_Doxo_HLD
Z_Shift_L_Doxo_EG
Z_Im_L_Doxo_EG

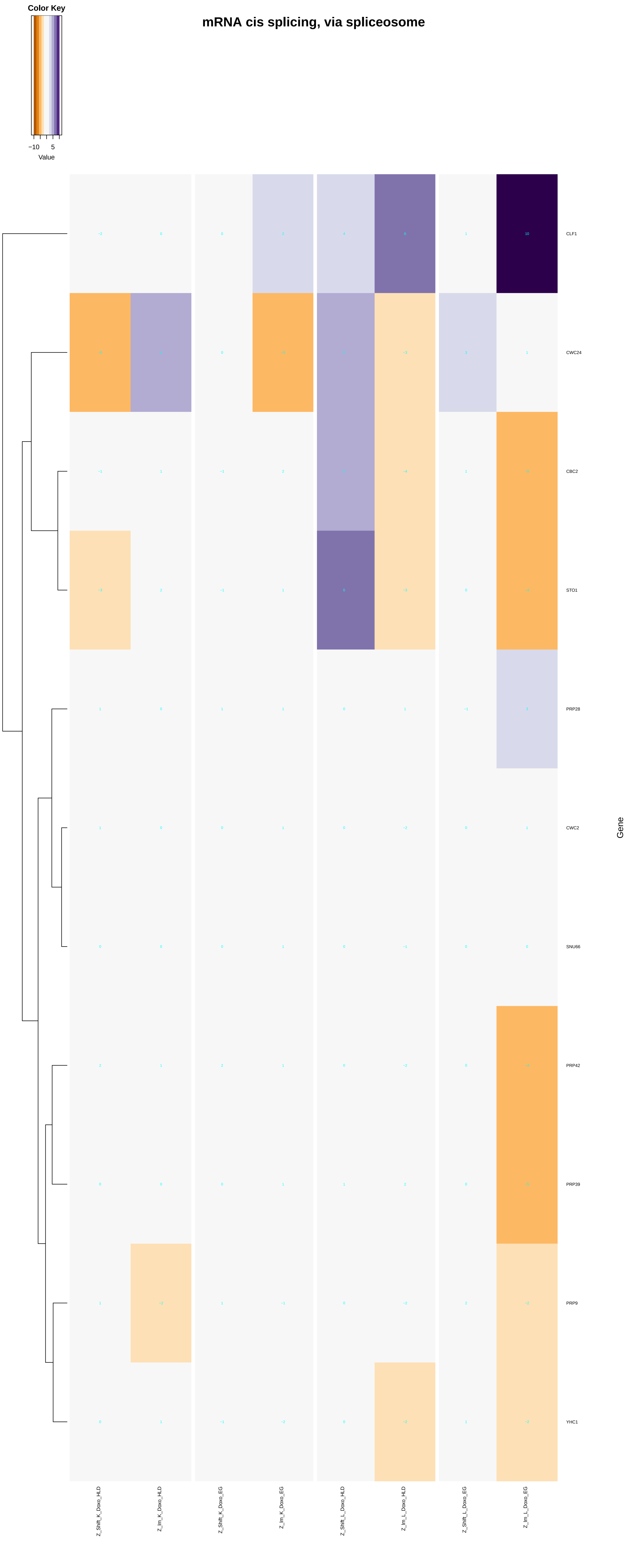
mRNA cis splicing, via spliceosome
Color Key
−10
5
Value
CLF1
CWC24
CBC2
STO1
PRP28
CWC2
SNU66
PRP42
PRP39
PRP9
YHC1
Gene
Z_Shift_K_Doxo_HLD
Z_Im_K_Doxo_HLD
Z_Shift_K_Doxo_EG
Z_Im_K_Doxo_EG
Z_Shift_L_Doxo_HLD
Z_Im_L_Doxo_HLD
Z_Shift_L_Doxo_EG
Z_Im_L_Doxo_EG

#### mRNA polyadenylation

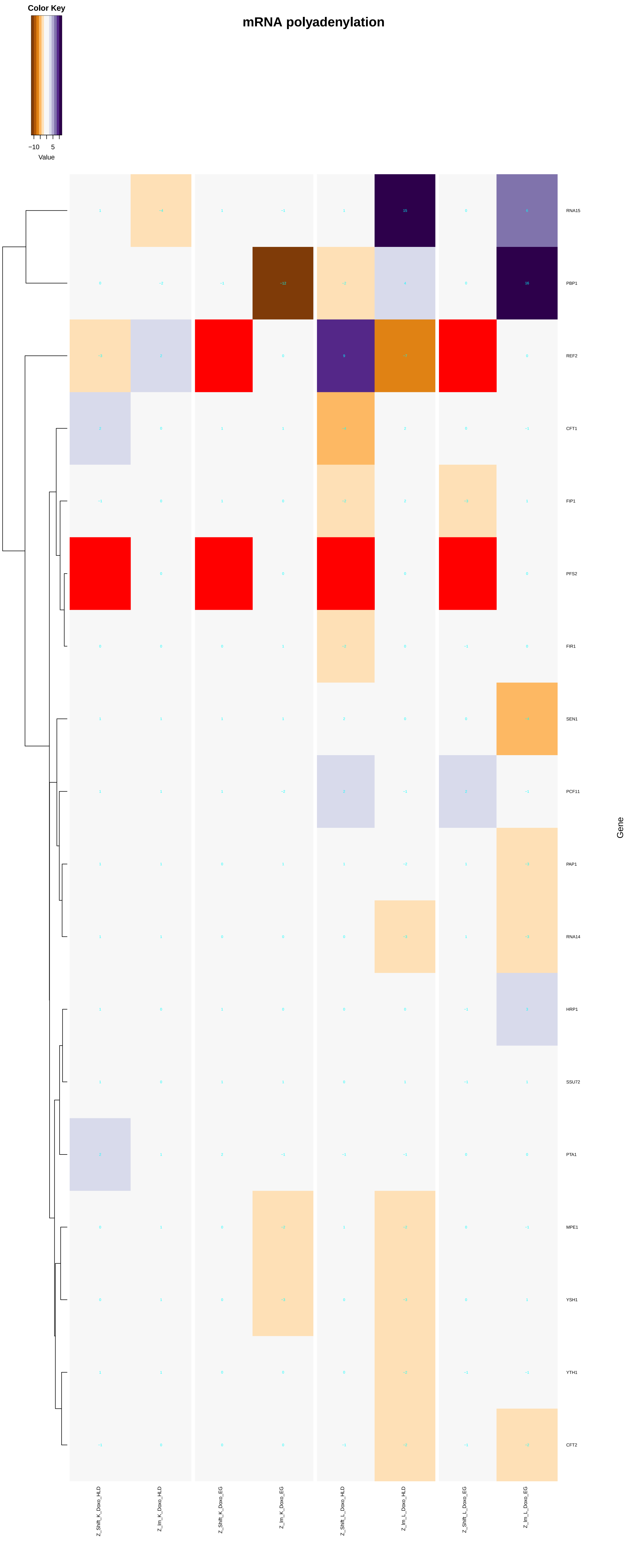

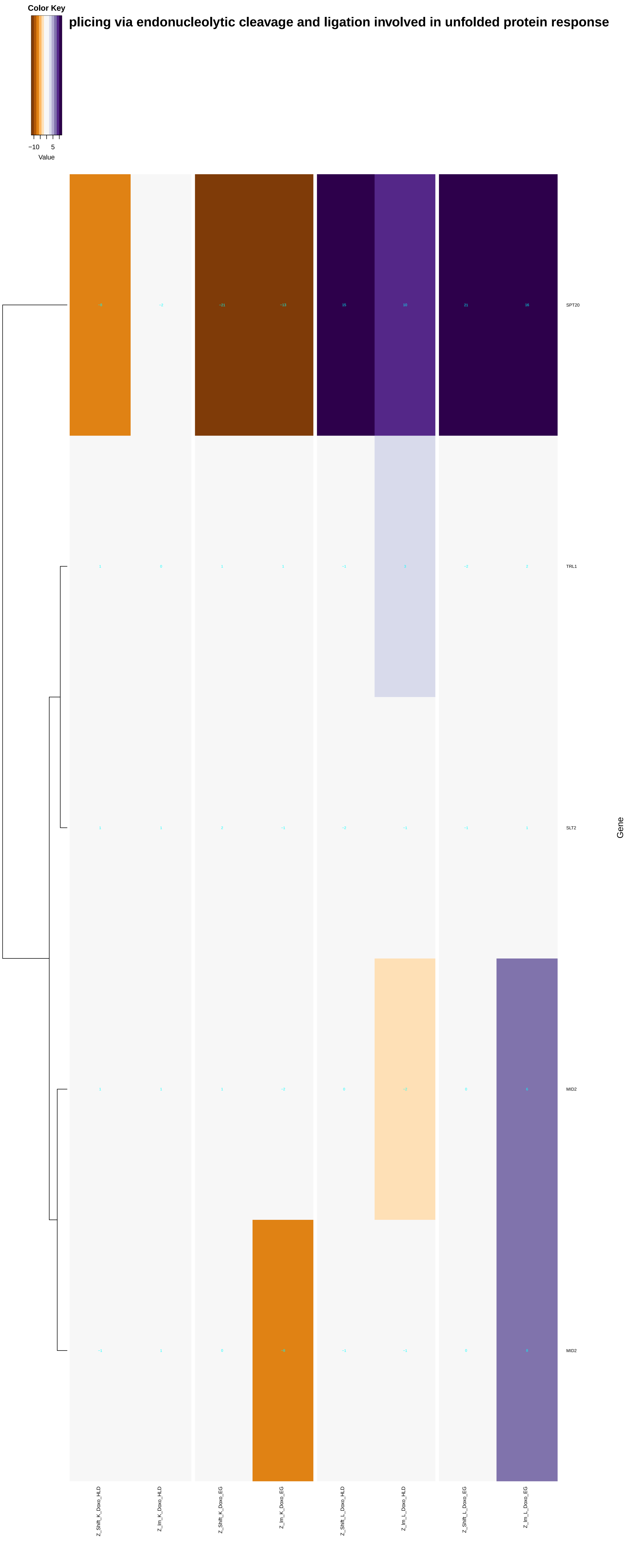
Color Key
−10
5
Value
plicing via endonucleolytic cleavage and ligation involved in unfolded protein response
SPT20
TRL1
SLT2
MID2
MID2
Gene
Z_Shift_K_Doxo_HLD
Z_Im_K_Doxo_HLD
Z_Shift_K_Doxo_EG
Z_Im_K_Doxo_EG
Z_Shift_L_Doxo_HLD
Z_Im_L_Doxo_HLD
Z_Shift_L_Doxo_EG
Z_Im_L_Doxo_EG

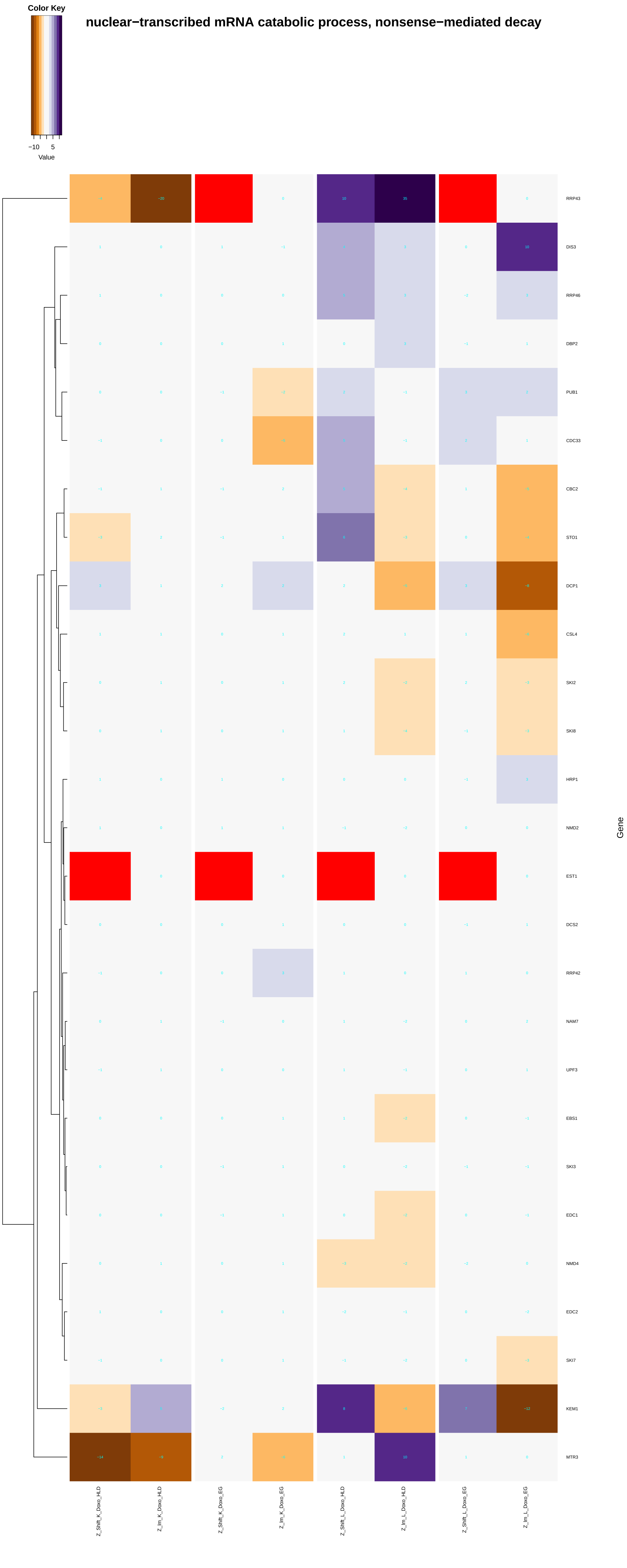
nuclear−transcribed mRNA catabolic process, nonsense−mediated decay
Color Key
−10
5
Value
RRP43
DIS3
RRP46
DBP2
PUB1
CDC33
CBC2
STO1
DCP1
CSL4
SKI2
SKI8
HRP1
NMD2
EST1
DCS2
RRP42
NAM7
UPF3
EBS1
SKI3
EDC1
NMD4
EDC2
SKI7
KEM1
MTR3
Gene
Z_Shift_K_Doxo_HLD
Z_Im_K_Doxo_HLD
Z_Shift_K_Doxo_EG
Z_Im_K_Doxo_EG
Z_Shift_L_Doxo_HLD
Z_Im_L_Doxo_HLD
Z_Shift_L_Doxo_EG
Z_Im_L_Doxo_EG

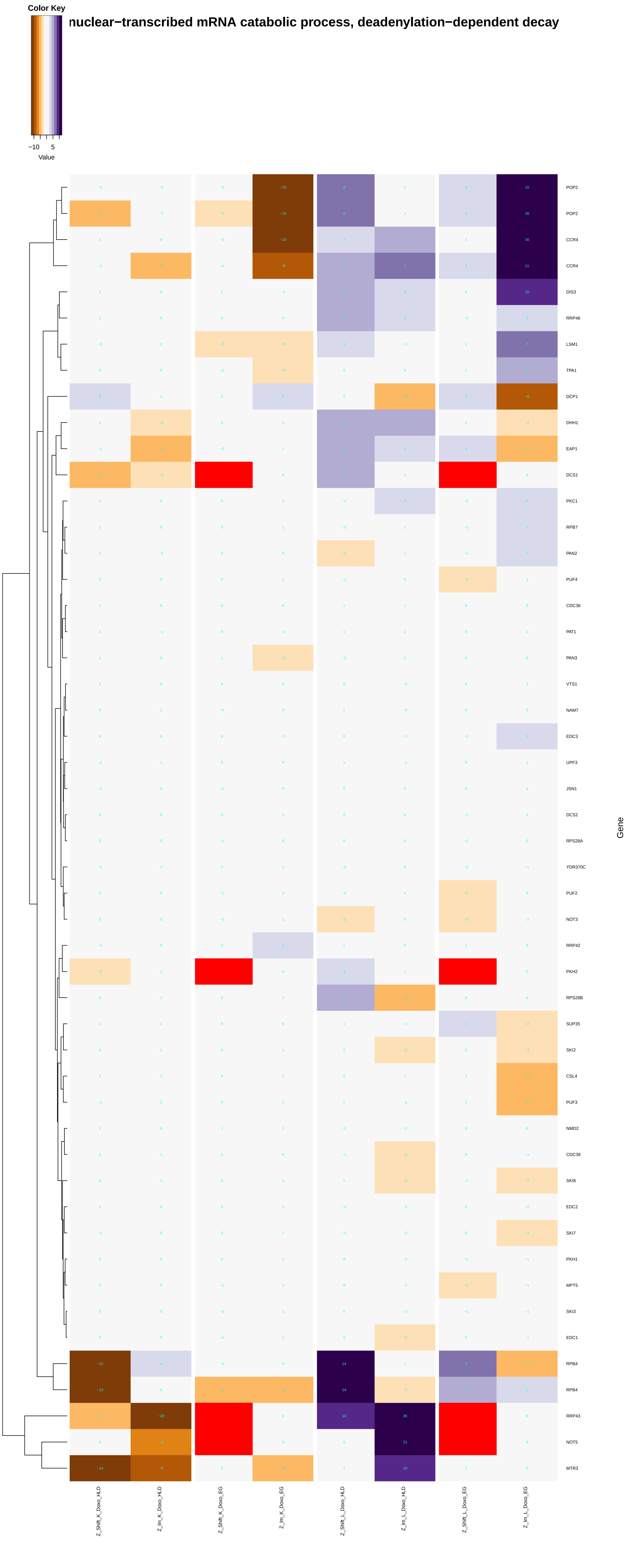

nuclear−transcribed mRNA catabolic process, deadenylation−dependent decay
Color Key
−10 5
Value
POP2
POP2
CCR4
CCR4
DIS3
RRP46
LSM1
TPA1
DCP1
DHH1
EAP1
DCS1
PKC1
RPB7
PAN2
PUF4
CDC36
PAT1
PAN3
VTS1
NAM7
EDC3
UPF3
JSN1
DCS2
RPS28A
YDR370C
PUF2
NOT3
RRP42
PKH2
RPS28B
SUP35
SKI2
CSL4
PUF3
NMD2
CDC39
SKI8
EDC2
SKI7
PKH1
MPT5
SKI3
EDC1
RPB4
RPB4
RRP43
NOT5
MTR3
Gene
Z_Shift_K_Doxo_HLD
Z_Im_K_Doxo_HLD
Z_Shift_K_Doxo_EG
Z_Im_K_Doxo_EG
Z_Shift_L_Doxo_HLD
Z_Im_L_Doxo_HLD
Z_Shift_L_Doxo_EG
Z_Im_L_Doxo_EG

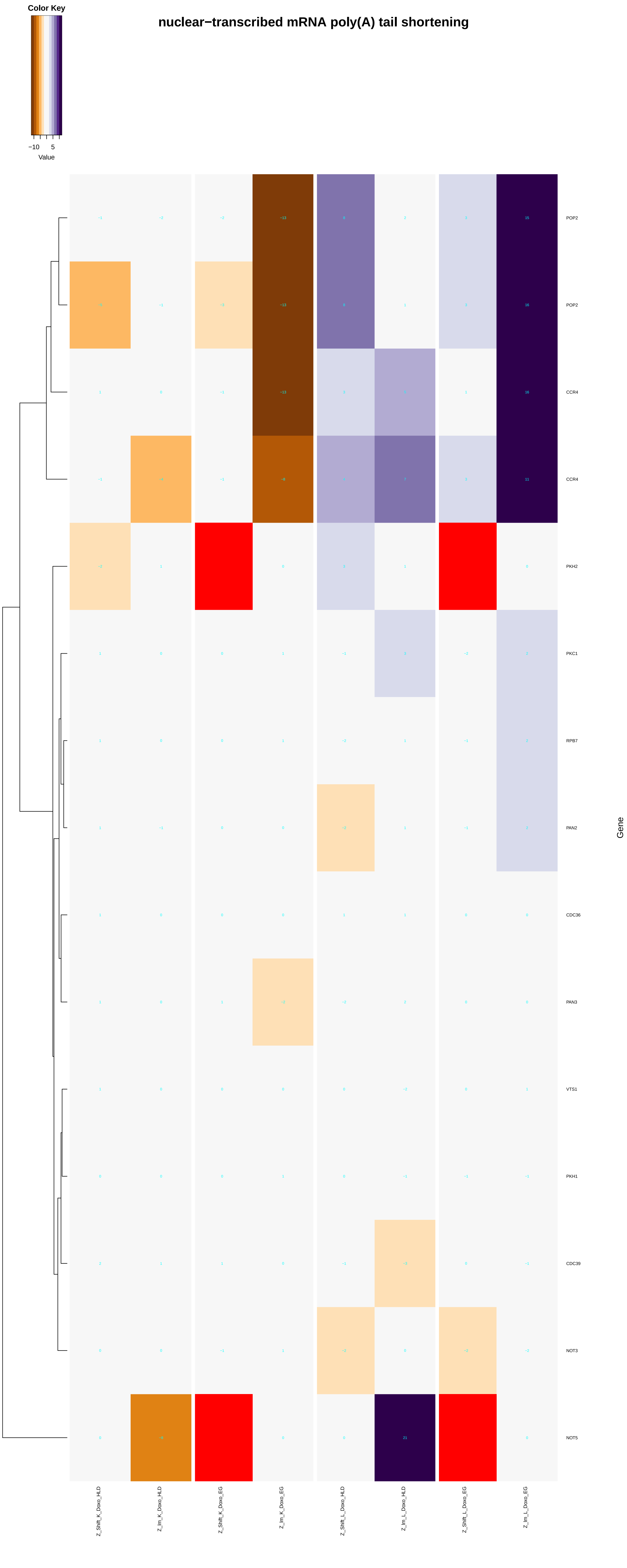

Color Key
−10
5
Value
nuclear−transcribed mRNA poly(A) tail shortening
POP2
POP2
CCR4
CCR4
PKH2
PKC1
RPB7
PAN2
CDC36
PAN3
VTS1
PKH1
CDC39
NOT3
NOT5
Gene
Z_Shift_K_Doxo_HLD
Z_Im_K_Doxo_HLD
Z_Shift_K_Doxo_EG
Z_Im_K_Doxo_EG
Z_Shift_L_Doxo_HLD
Z_Im_L_Doxo_HLD
Z_Shift_L_Doxo_EG
Z_Im_L_Doxo_EG

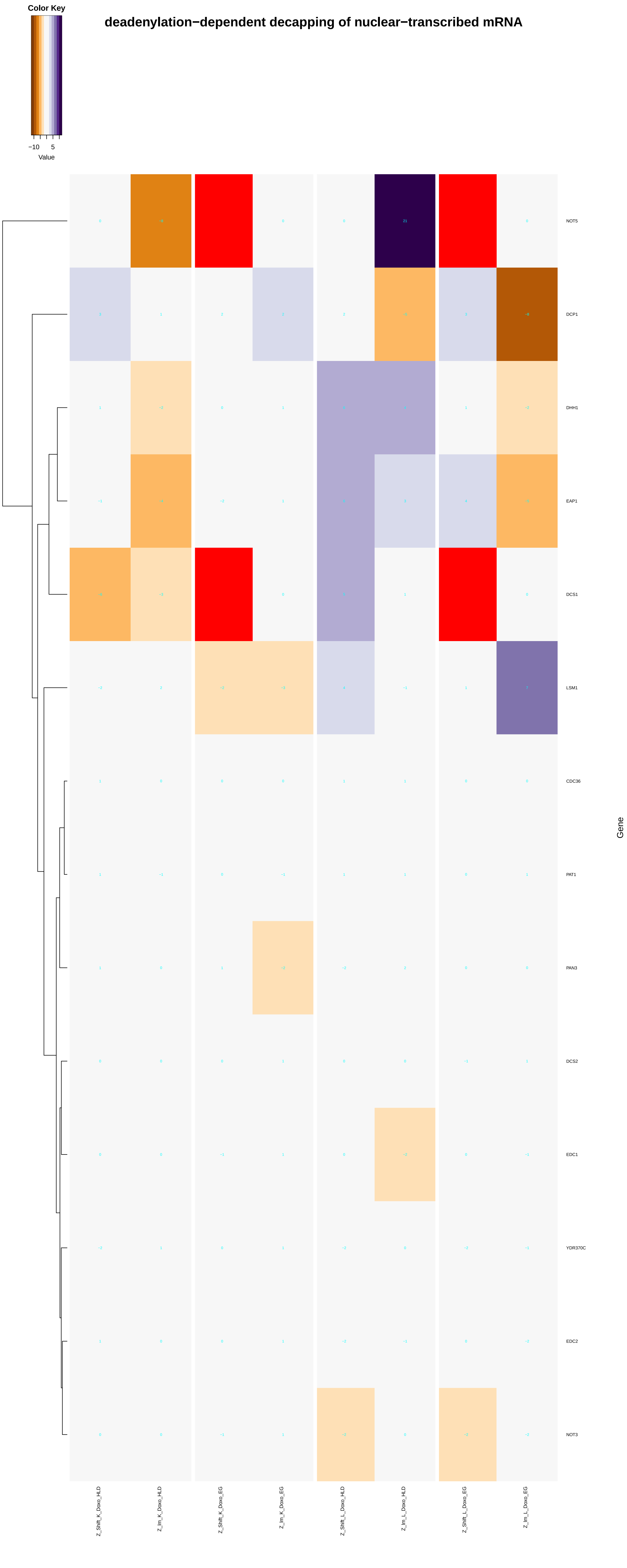

deadenylation−dependent decapping of nuclear−transcribed mRNA
Color Key
−10
5
Value
NOT5
DCP1
DHH1
EAP1
DCS1
LSM1
CDC36
PAT1
PAN3
DCS2
EDC1
YDR370C
EDC2
NOT3
Gene
Z_Shift_K_Doxo_HLD
Z_Im_K_Doxo_HLD
Z_Shift_K_Doxo_EG
Z_Im_K_Doxo_EG
Z_Shift_L_Doxo_HLD
Z_Im_L_Doxo_HLD
Z_Shift_L_Doxo_EG
Z_Im_L_Doxo_EG

### nuclear−transcribed mRNA catabolic process, exonucleolytic

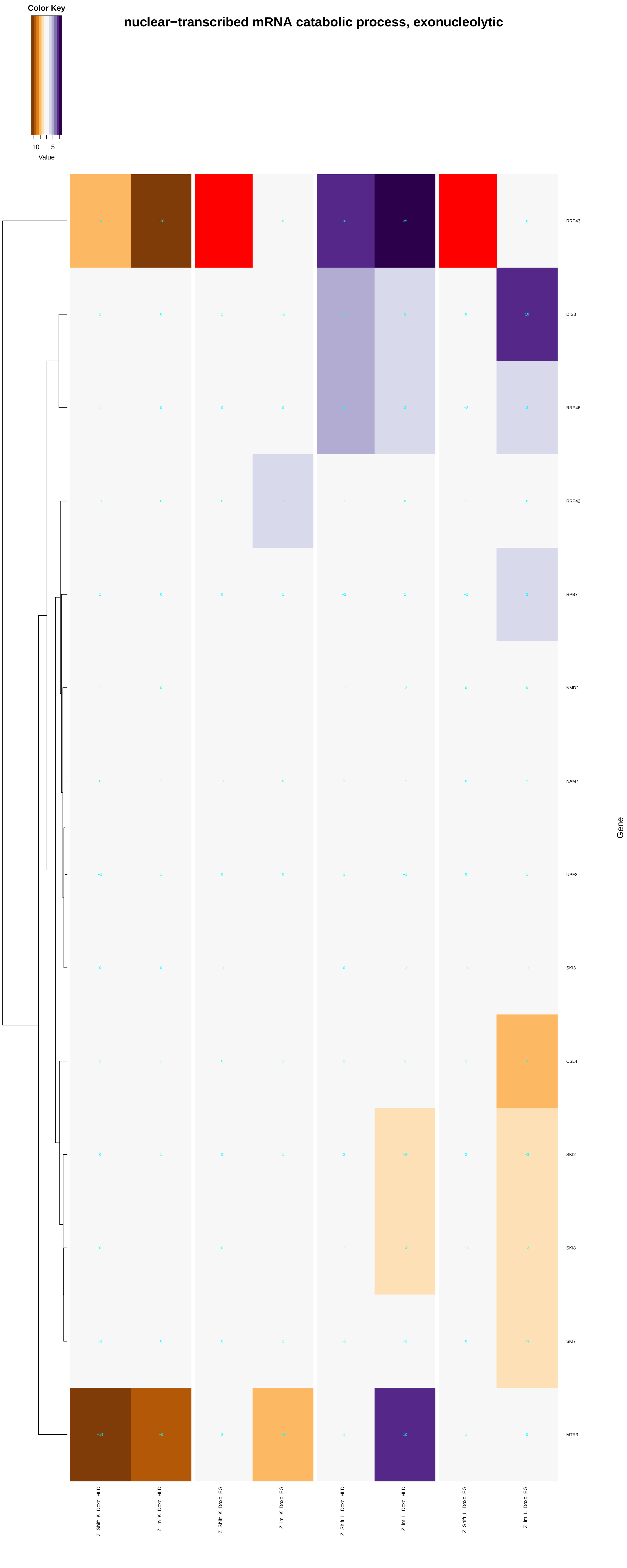

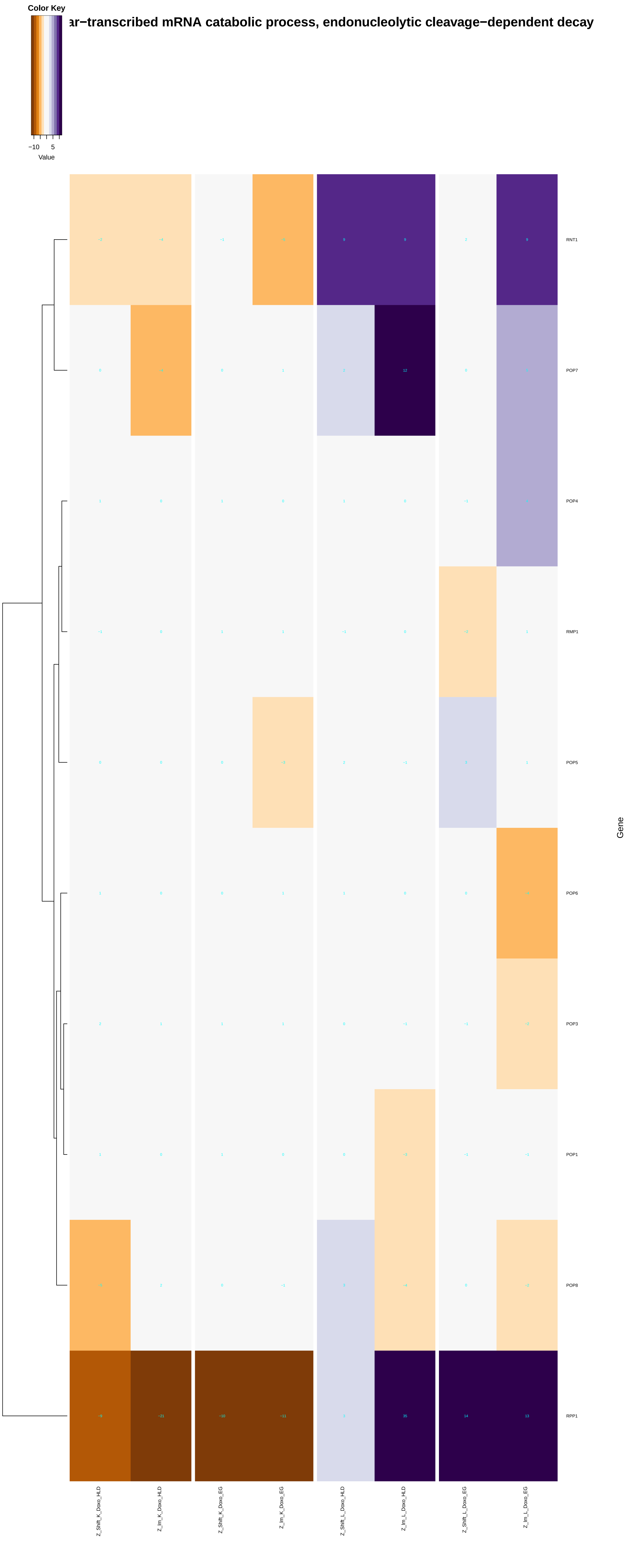

Color Key
−10
5
Value
ar−transcribed mRNA catabolic process, endonucleolytic cleavage−dependent decay
RNT1
POP7
POP4
RMP1
POP5
POP6
POP3
POP1
POP8
RPP1
Gene
Z_Shift_K_Doxo_HLD
Z_Im_K_Doxo_HLD
Z_Shift_K_Doxo_EG
Z_Im_K_Doxo_EG
Z_Shift_L_Doxo_HLD
Z_Im_L_Doxo_HLD
Z_Shift_L_Doxo_EG
Z_Im_L_Doxo_EG

#### uclear−transcribed mRNA catabolic process, deadenylation−independent decay

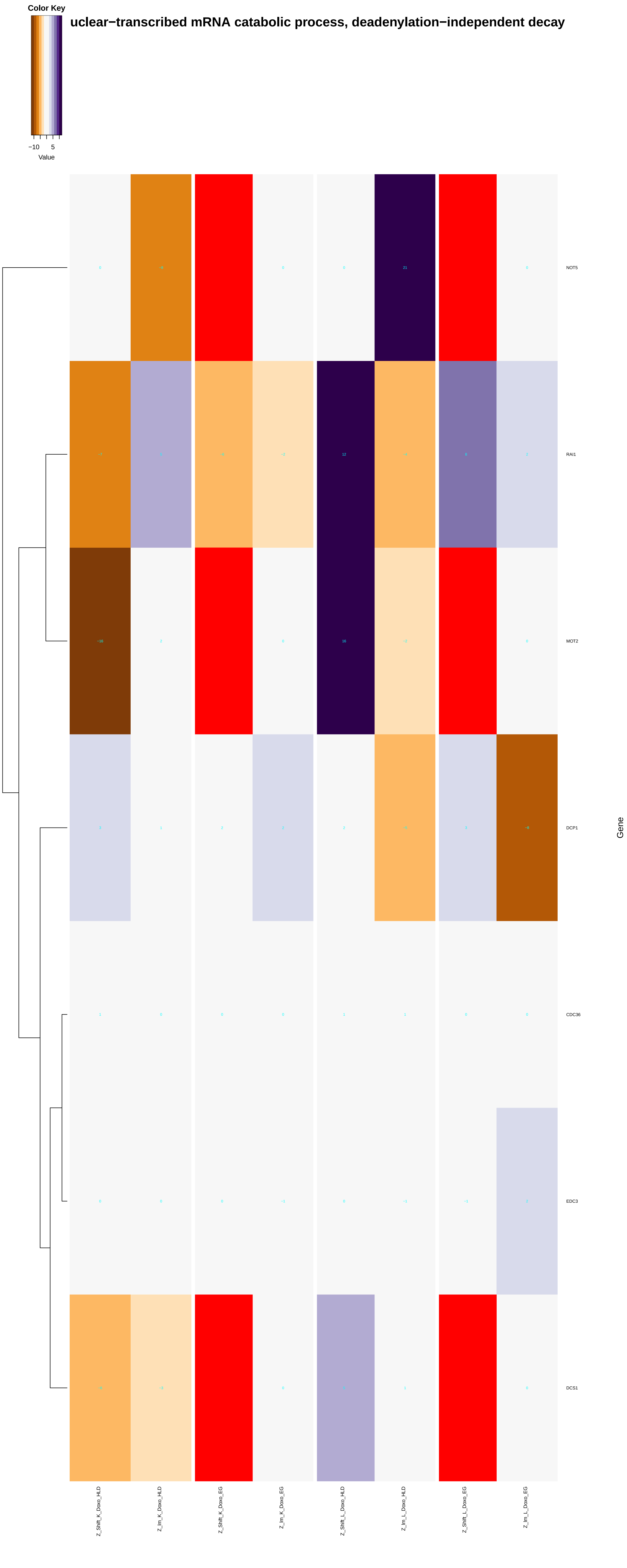

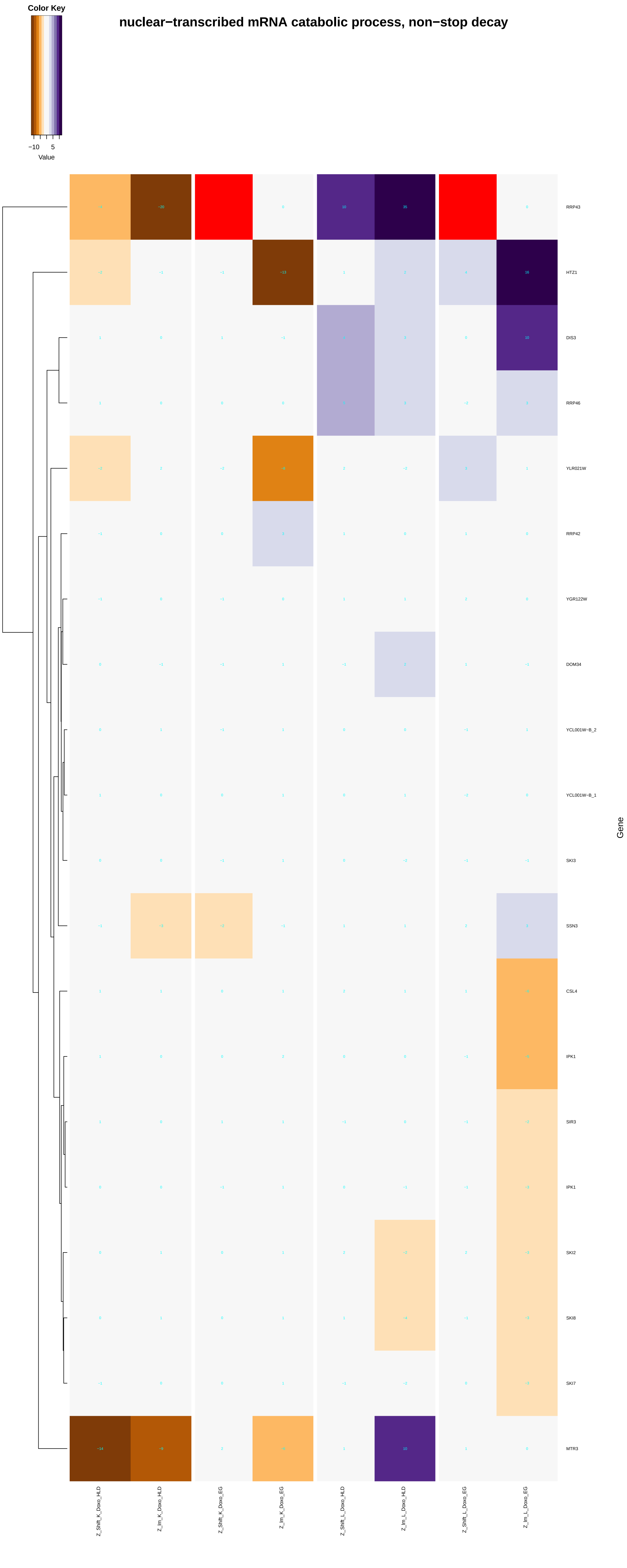

Color Key
−10
5
Value
nuclear−transcribed mRNA catabolic process, non−stop decay
RRP43
HTZ1
DIS3
RRP46
YLR021W
RRP42
YGR122W
DOM34
YCL001W−B_2
YCL001W−B_1
SKI3
SSN3
CSL4
IPK1
SIR3
IPK1
SKI2
SKI8
SKI7
MTR3
Gene
Z_Shift_K_Doxo_HLD
Z_Im_K_Doxo_HLD
Z_Shift_K_Doxo_EG
Z_Im_K_Doxo_EG
Z_Shift_L_Doxo_HLD
Z_Im_L_Doxo_HLD
Z_Shift_L_Doxo_EG
Z_Im_L_Doxo_EG

### nuclear−transcribed mRNA catabolic process, no−go decay

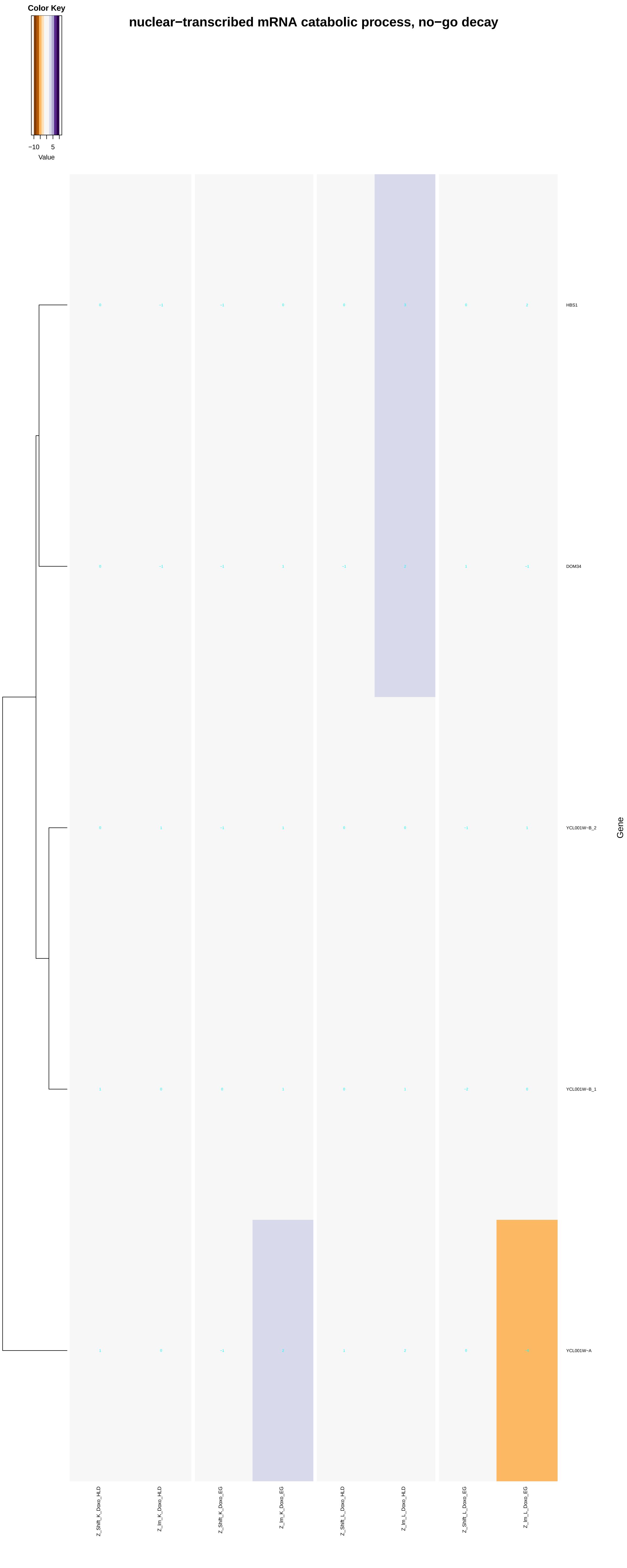

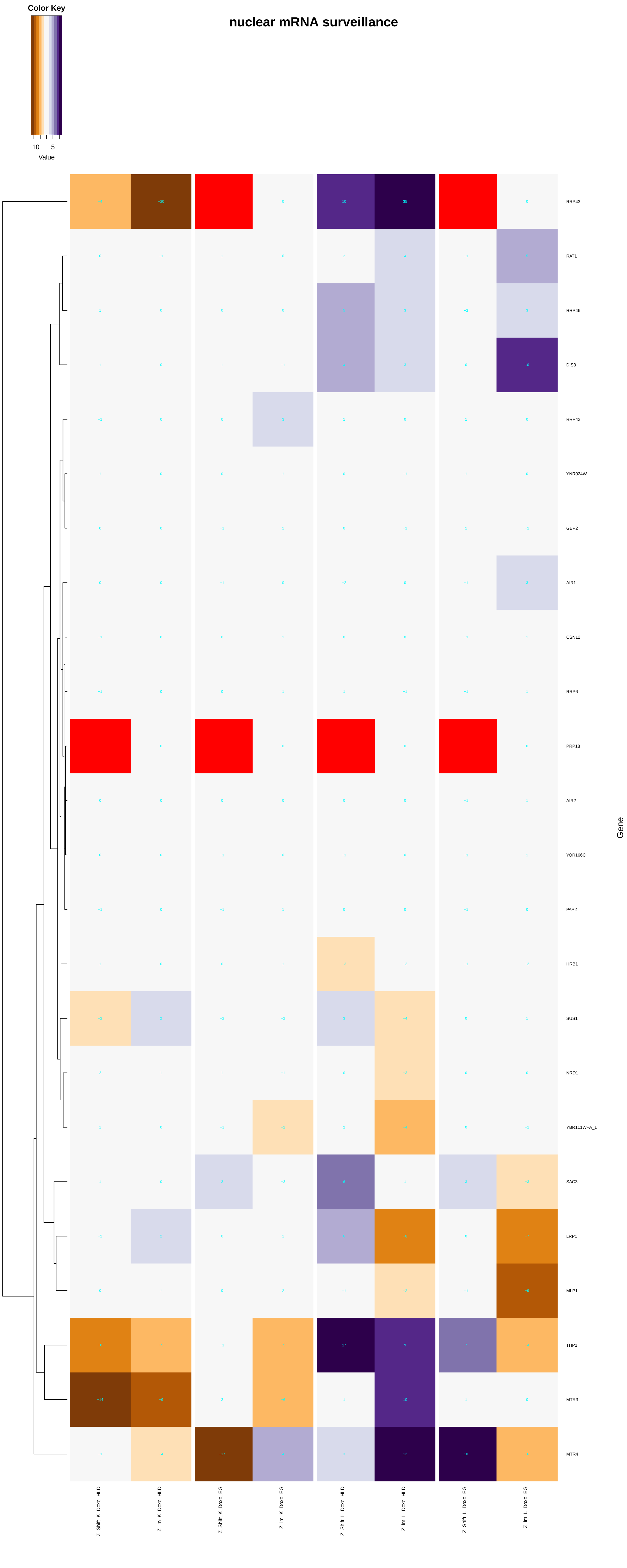
Color Key
Value
nuclear mRNA surveillance
RRP43
RAT1
RRP46
DIS3
RRP42
YNR024W
GBP2
AIR1
CSN12
RRP6
PRP18
AIR2
YOR166C
PAP2
HRB1
SUS1
NRD1
YBR111W-A_1
SAC3
LRP1
MLP1
THP1
MTR3
MTR4
Gene
Z_Shift_K_Doxo_HLD
Z_Im_K_Doxo_HLD
Z_Shift_K_Doxo_EG
Z_Im_K_Doxo_EG
Z_Shift_L_Doxo_HLD
Z_Im_L_Doxo_HLD
Z_Shift_L_Doxo_EG
Z_Im_L_Doxo_EG

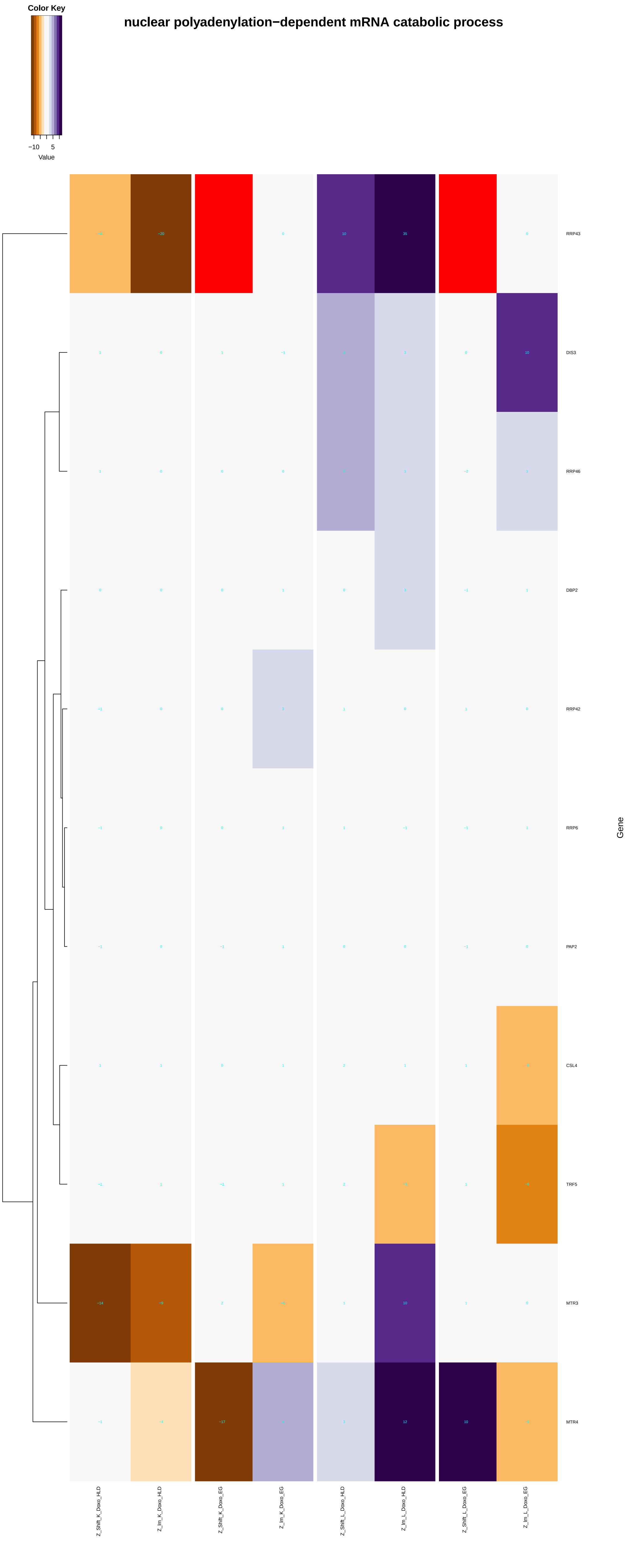

Color Key
−10
5
Value
nuclear polyadenylation−dependent mRNA catabolic process
RRP43
DIS3
RRP46
DBP2
RRP42
RRP6
PAP2
CSL4
TRF5
MTR3
MTR4
Gene
Z_Shift_K_Doxo_HLD
Z_Im_K_Doxo_HLD
Z_Shift_K_Doxo_EG
Z_Im_K_Doxo_EG
Z_Shift_L_Doxo_HLD
Z_Im_L_Doxo_HLD
Z_Shift_L_Doxo_EG
Z_Im_L_Doxo_EG

−transcribed mRNA catabolic process, 3'−5' exonucleolytic nonsense−mediated decay

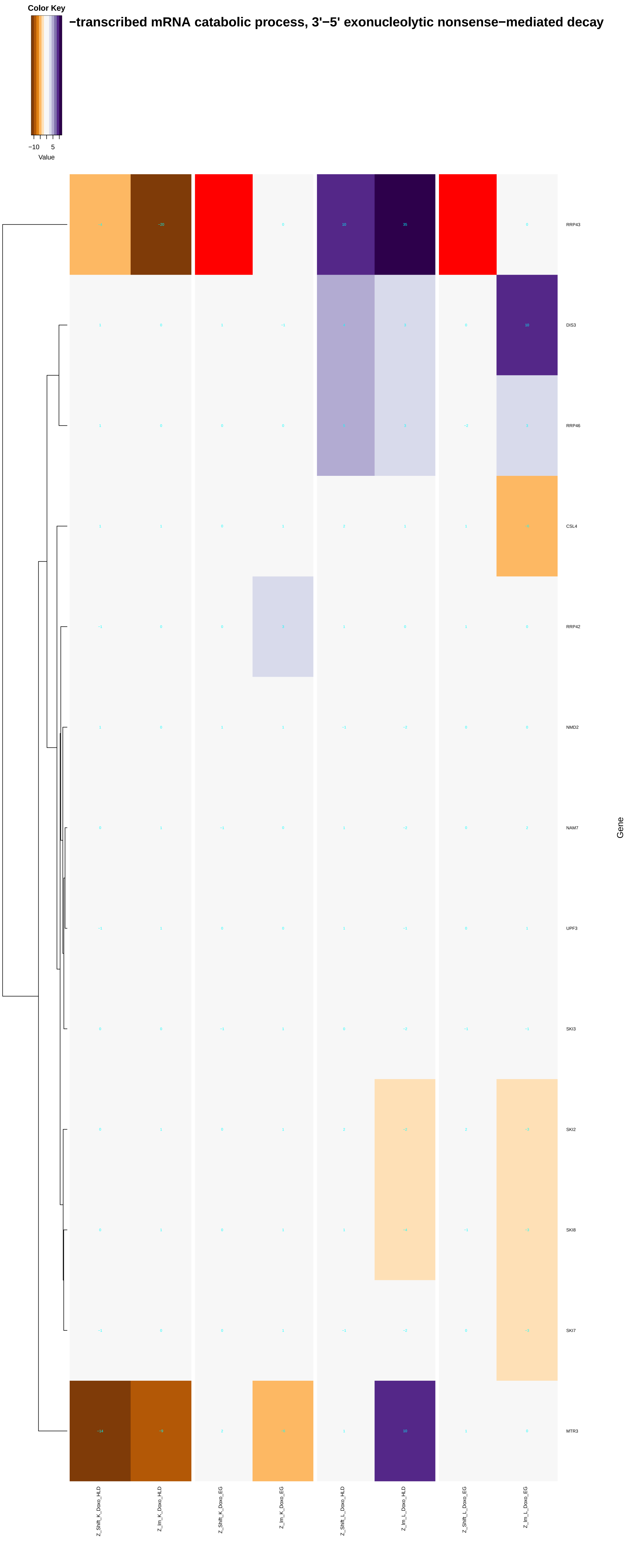

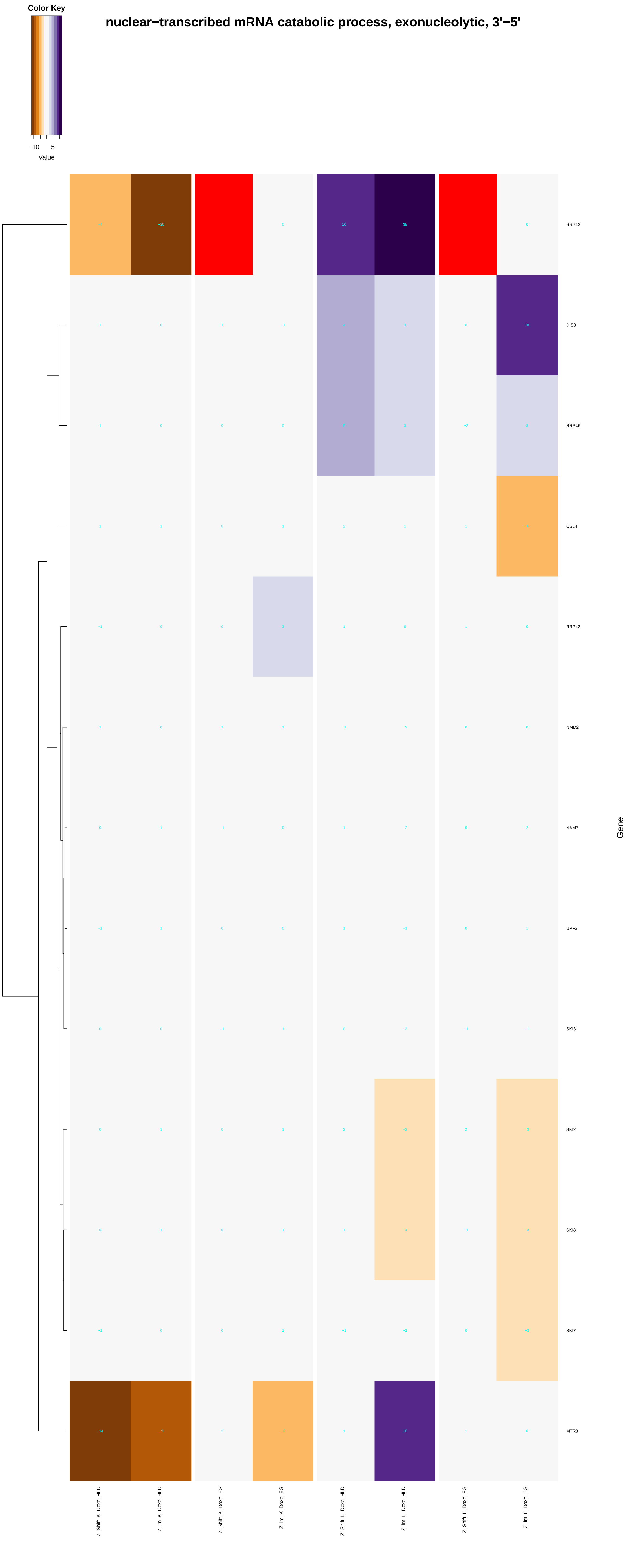

Color Key
−10
5
Value
nuclear−transcribed mRNA catabolic process, exonucleolytic, 3'−5'
RRP43
DIS3
RRP46
CSL4
RRP42
NMD2
NAM7
UPF3
SKI3
SKI2
SKI8
SKI7
MTR3
Gene
Z_Shift_K_Doxo_HLD
Z_Im_K_Doxo_HLD
Z_Shift_K_Doxo_EG
Z_Im_K_Doxo_EG
Z_Shift_L_Doxo_HLD
Z_Im_L_Doxo_HLD
Z_Shift_L_Doxo_EG
Z_Im_L_Doxo_EG

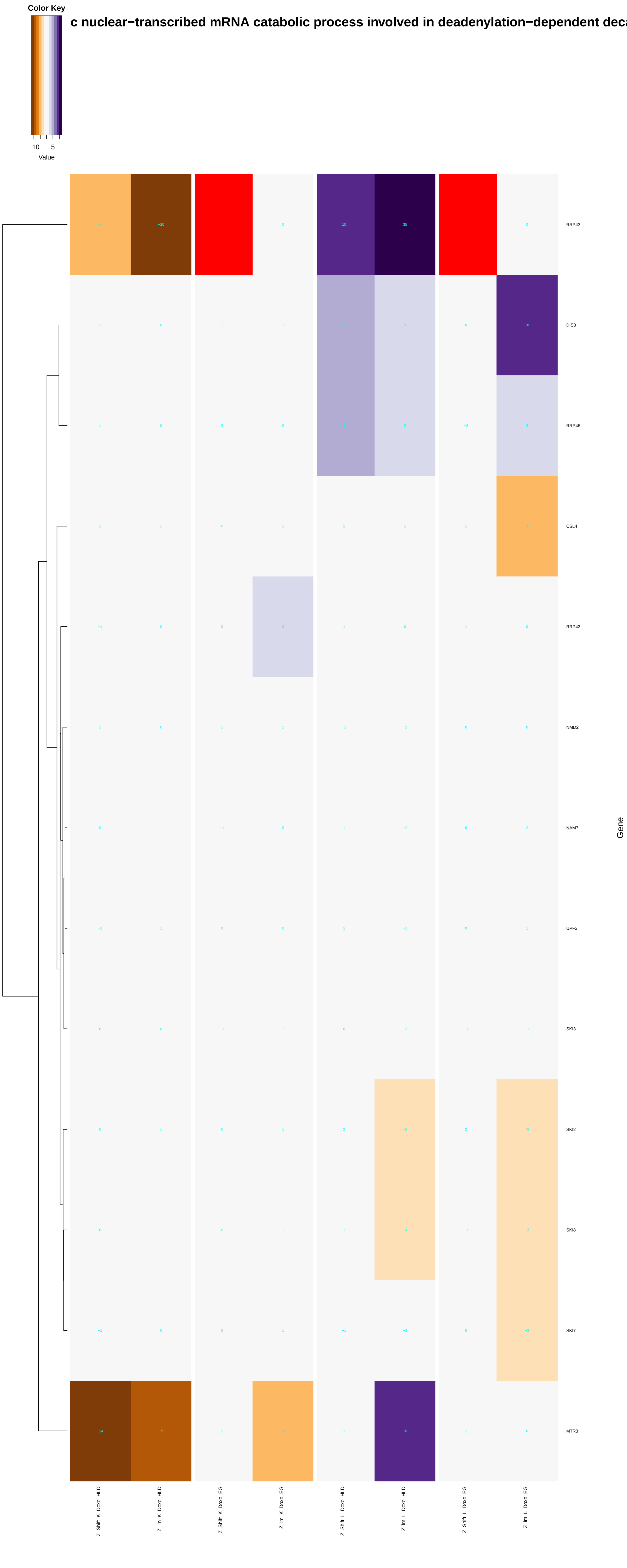
Color Key
−10 5
Value
c nuclear−transcribed mRNA catabolic process involved in deadenylation−dependent deca
RRP43
DIS3
RRP46
CSL4
RRP42
NMD2
NAM7
UPF3
SKI3
SKI2
SKI8
SKI7
MTR3
Gene
Z_Shift_K_Doxo_HLD
Z_lm_K_Doxo_HLD
Z_Shift_K_Doxo_EG
Z_lm_K_Doxo_EG
Z_Shift_L_Doxo_HLD
Z_lm_L_Doxo_HLD
Z_Shift_L_Doxo_EG
Z_lm_L_Doxo_EG

#### nuclear mRNA surveillance of spliceosomal pre−mRNA splicing

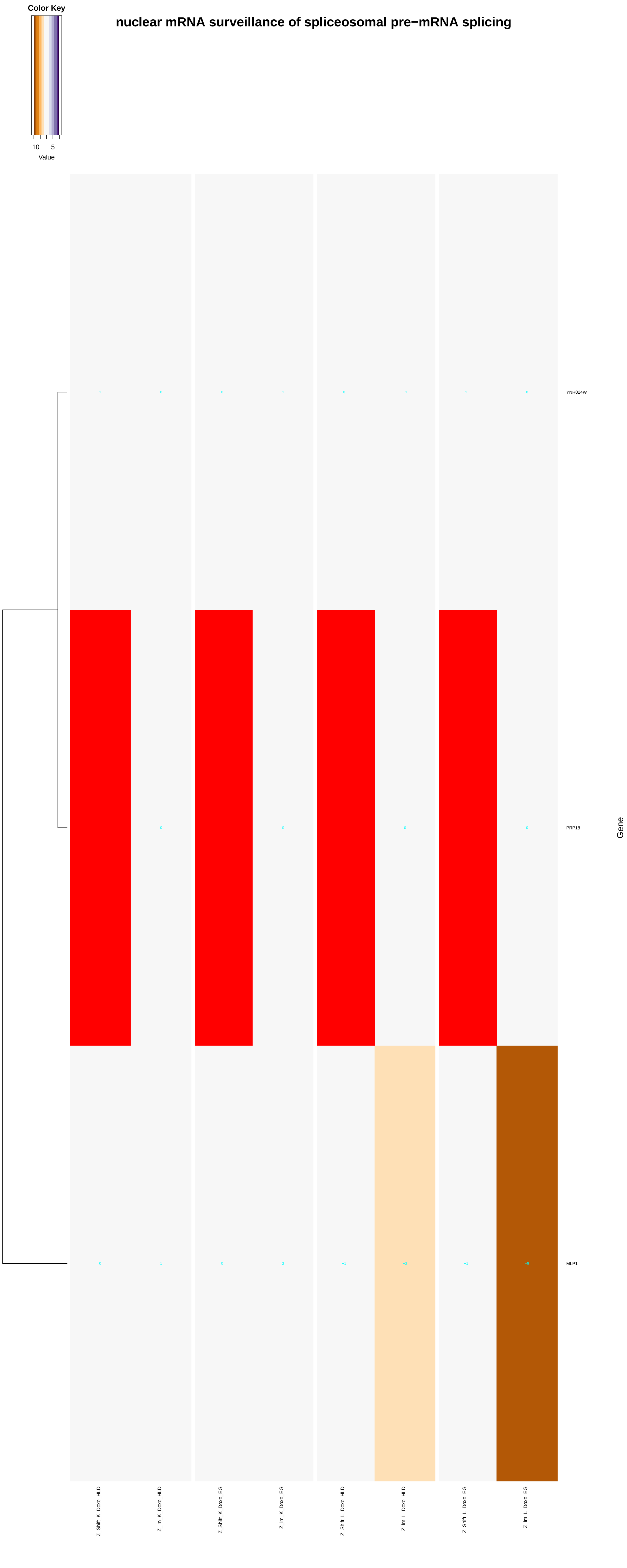

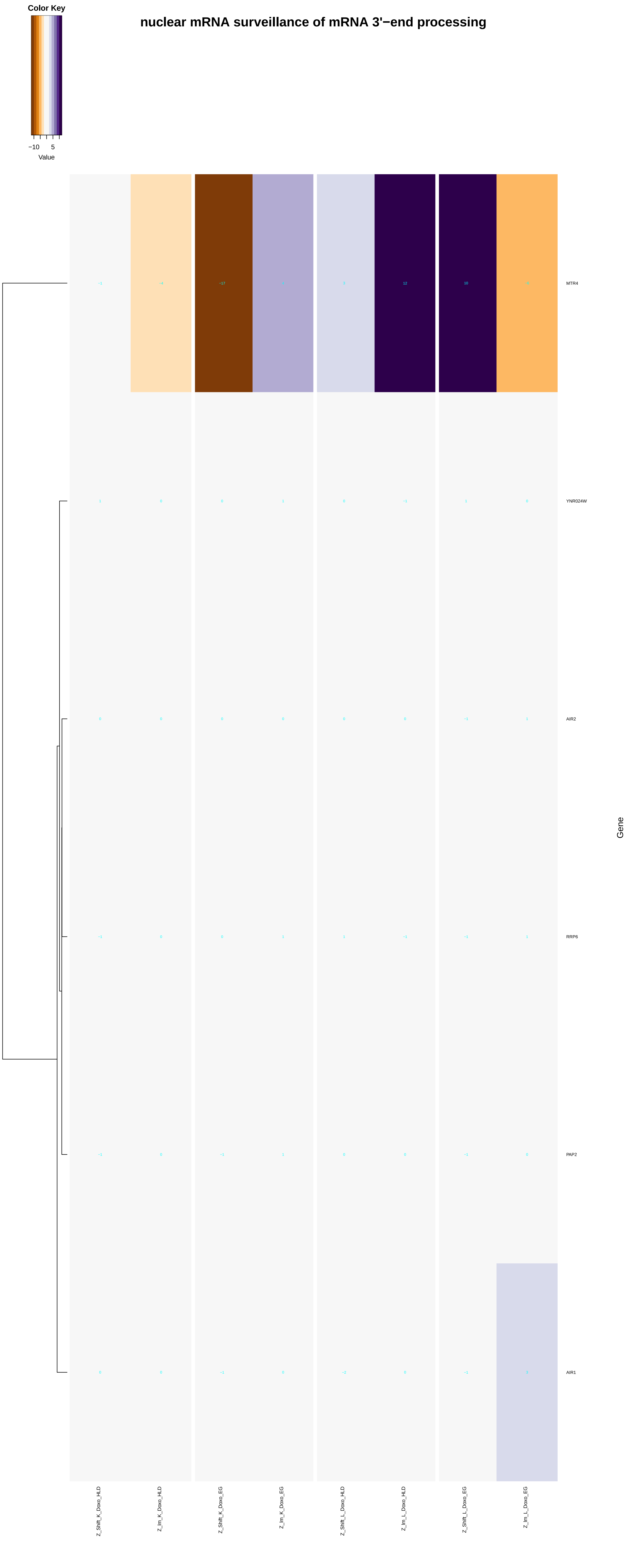
nuclear mRNA surveillance of mRNA 3'−end processing
Color Key
−10
5
Value
MTR4
YNR024W
AIR2
RRP6
PAP2
AIR1
Gene
Z_Shift_K_Doxo_HLD
Z_Im_K_Doxo_HLD
Z_Shift_K_Doxo_EG
Z_Im_K_Doxo_EG
Z_Shift_L_Doxo_HLD
Z_Im_L_Doxo_HLD
Z_Shift_L_Doxo_EG
Z_Im_L_Doxo_EG
