## Additional File 9 for "A yeast phenomic model for the influence of Warburg metabolism on genetic buffering of doxorubicin": ncRNA_3'-end_processing.pdf

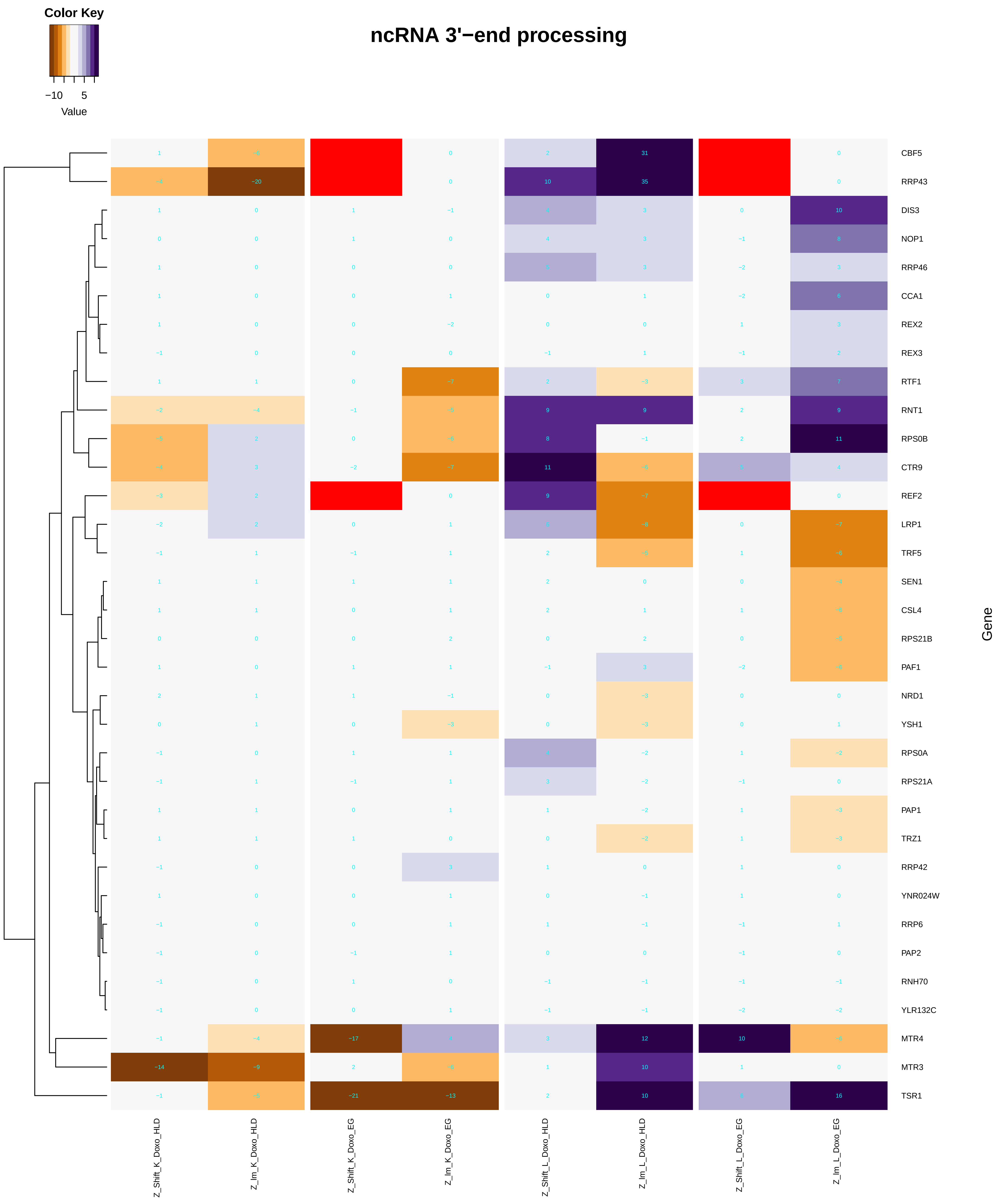
ncRNA 3'−end processing
Color Key
−10
5
Value
Gene
CBF5
RRP43
DIS3
NOP1
RRP46
CCA1
REX2
REX3
RTF1
RNT1
RPS0B
CTR9
REF2
LRP1
TRF5
SEN1
CSL4
RPS21B
PAF1
NRD1
YSH1
RPS0A
RPS21A
PAP1
TRZ1
RRP42
YNR024W
RRP6
PAP2
RNH70
YLR132C
MTR4
MTR3
TSR1
Z_Shift_K_Doxo_HLD
Z_lm_K_Doxo_HLD
Z_Shift_K_Doxo_EG
Z_lm_K_Doxo_EG
Z_Shift_L_Doxo_HLD
Z_lm_L_Doxo_HLD
Z_Shift_L_Doxo_EG
Z_lm_L_Doxo_EG

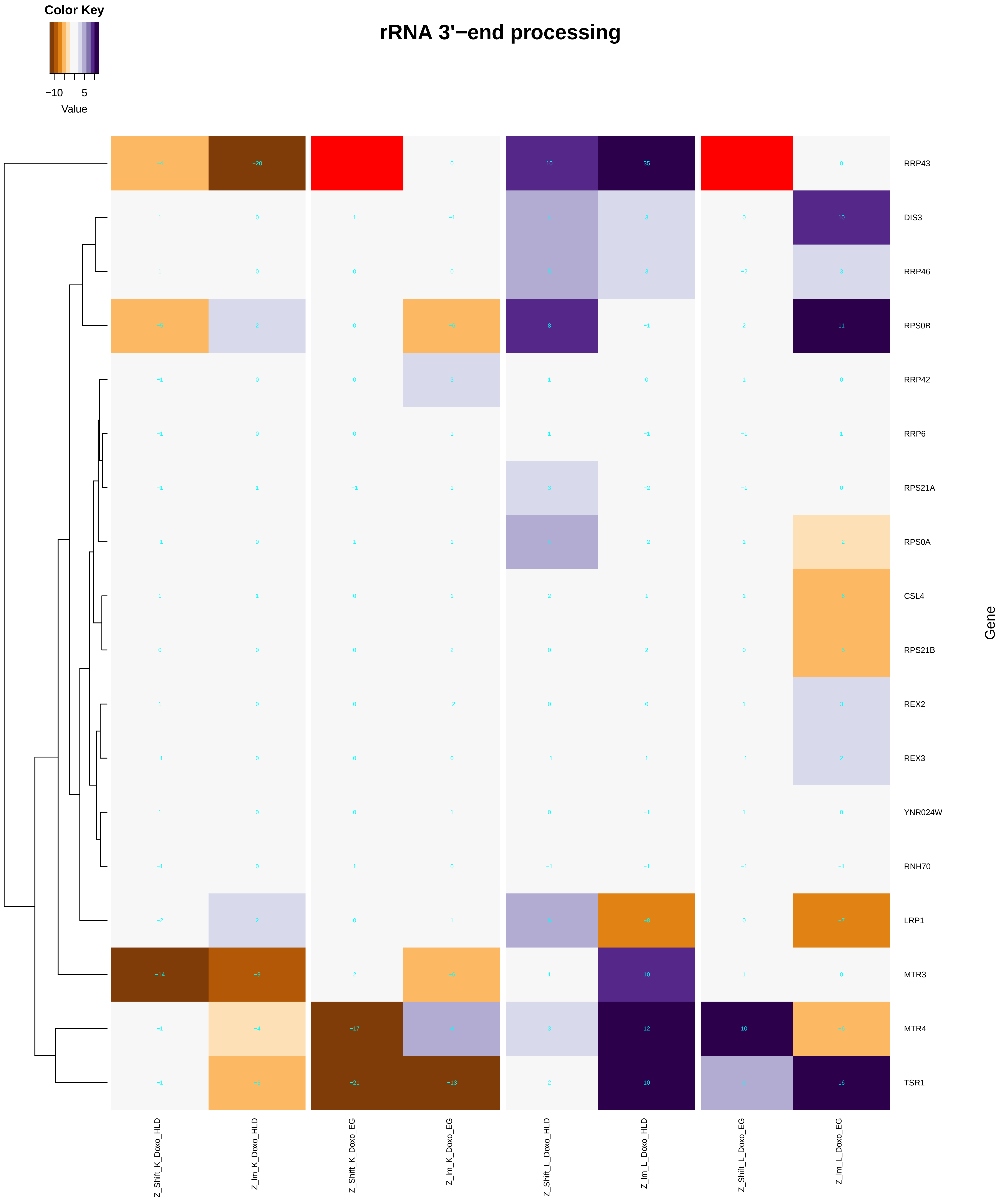
Color Key
−10 5
Value
rRNA 3'−end processing
RRP43
DIS3
RRP46
RPS0B
RRP42
RRP6
RPS21A
RPS0A
CSL4
RPS21B
REX2
REX3
YNR024W
RNH70
LRP1
MTR3
MTR4
TSR1
Gene
Z_Shift_K_Doxo_HLD
Z_lm_K_Doxo_HLD
Z_Shift_K_Doxo_EG
Z_lm_K_Doxo_EG
Z_Shift_L_Doxo_HLD
Z_lm_L_Doxo_HLD
Z_Shift_L_Doxo_EG
Z_lm_L_Doxo_EG

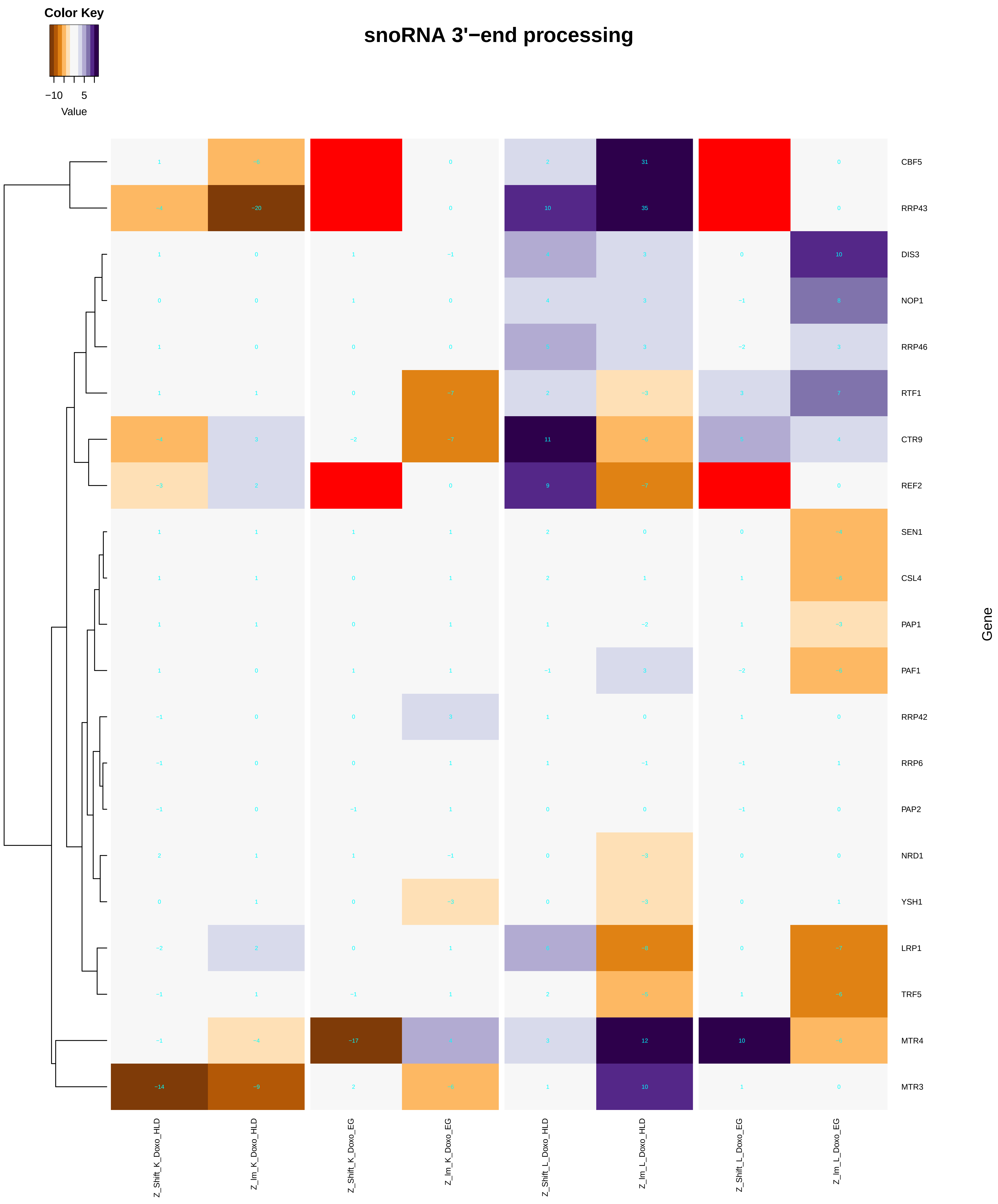
Color Key
−10 5
Value
snoRNA 3'−end processing
CBF5
RRP43
DIS3
NOP1
RRP46
RTF1
CTR9
REF2
SEN1
CSL4
PAP1
PAF1
RRP42
RRP6
PAP2
NRD1
YSH1
LRP1
TRF5
MTR4
MTR3
Gene
Z_Shift_K_Doxo_HLD
Z_Im_K_Doxo_HLD
Z_Shift_K_Doxo_EG
Z_Im_K_Doxo_EG
Z_Shift_L_Doxo_HLD
Z_Im_L_Doxo_HLD
Z_Shift_L_Doxo_EG
Z_Im_L_Doxo_EG

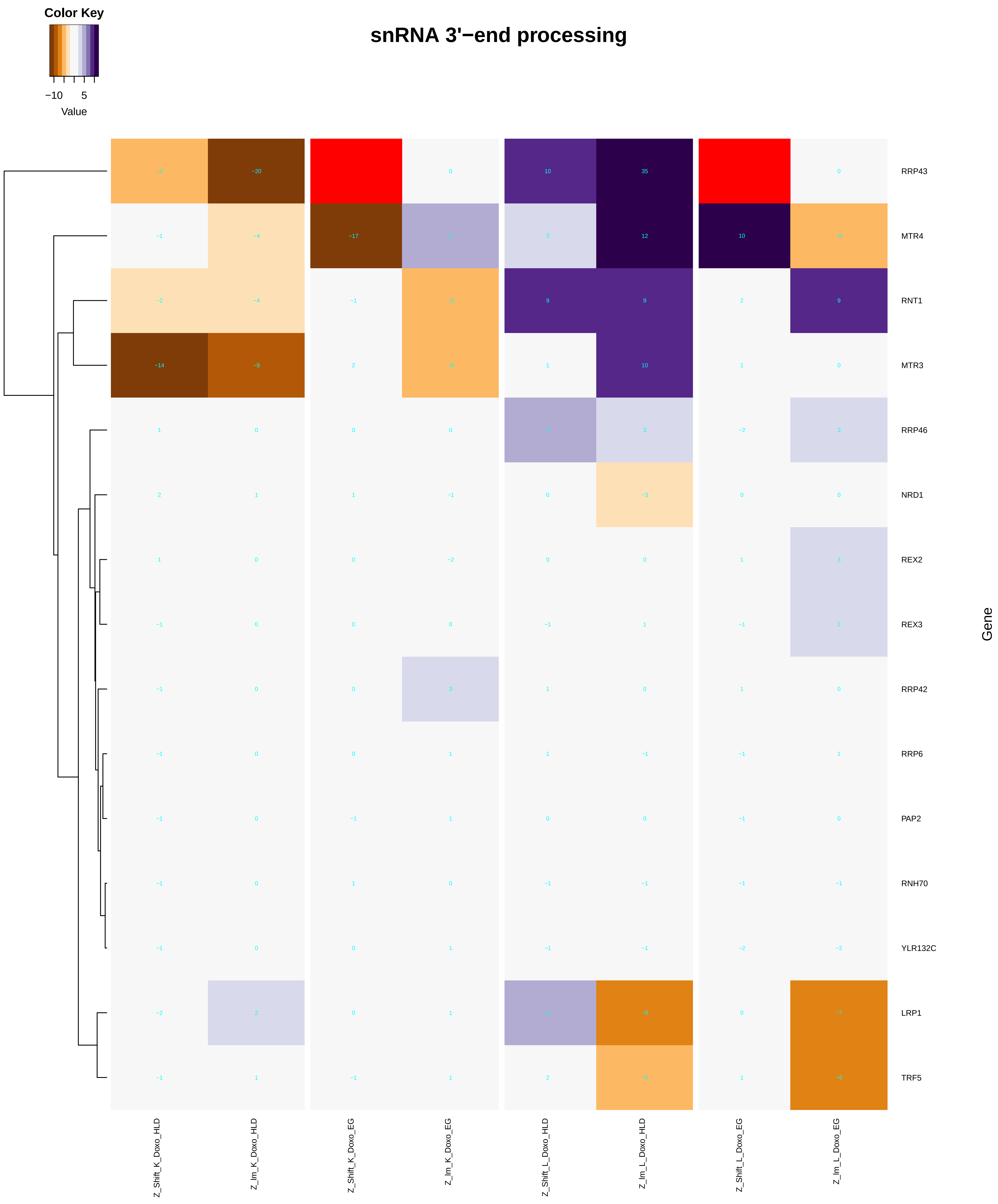
Color Key
−10
5
Value
snRNA 3'−end processing
RRP43
MTR4
RNT1
MTR3
RRP46
NRD1
REX2
REX3
RRP42
RRP6
PAP2
RNH70
YLR132C
LRP1
TRF5
Gene
Z_Shift_K_Doxo_HLD
Z_Im_K_Doxo_HLD
Z_Shift_K_Doxo_EG
Z_Im_K_Doxo_EG
Z_Shift_L_Doxo_HLD
Z_Im_L_Doxo_HLD
Z_Shift_L_Doxo_EG
Z_Im_L_Doxo_EG

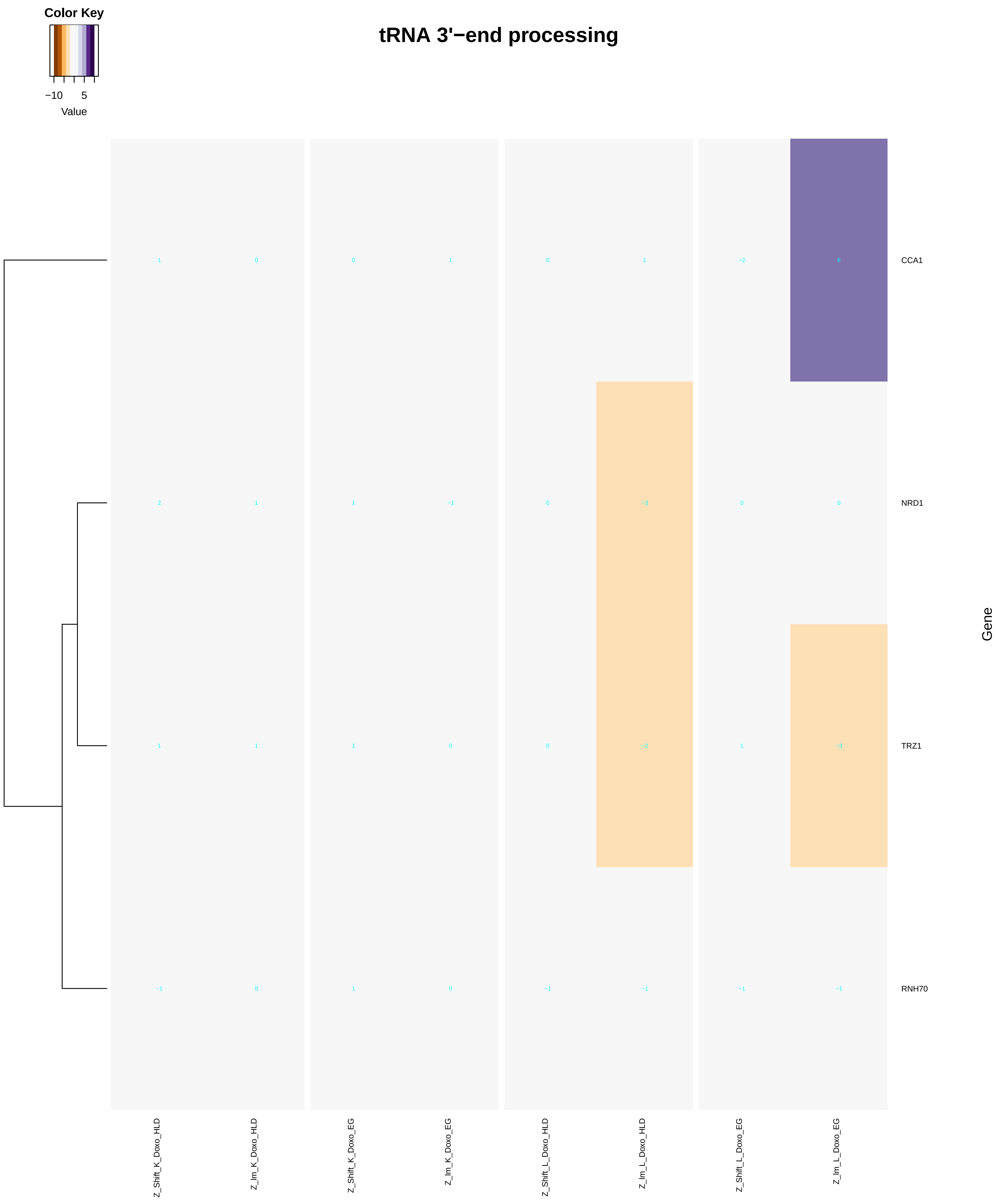
Color Key
−10
5
Value
tRNA 3'−end processing
1
0
0
1
0
1
−2
6
CCA1
2
1
1
−1
0
−3
0
0
NRD1
1
1
1
0
0
−2
1
−3
TRZ1
−1
0
1
0
−1
−1
−1
−1
RNH70
Gene
Z_Shift_K_Doxo_HLD
Z_lm_K_Doxo_HLD
Z_Shift_K_Doxo_EG
Z_lm_K_Doxo_EG
Z_Shift_L_Doxo_HLD
Z_lm_L_Doxo_HLD
Z_Shift_L_Doxo_EG
Z_lm_L_Doxo_EG

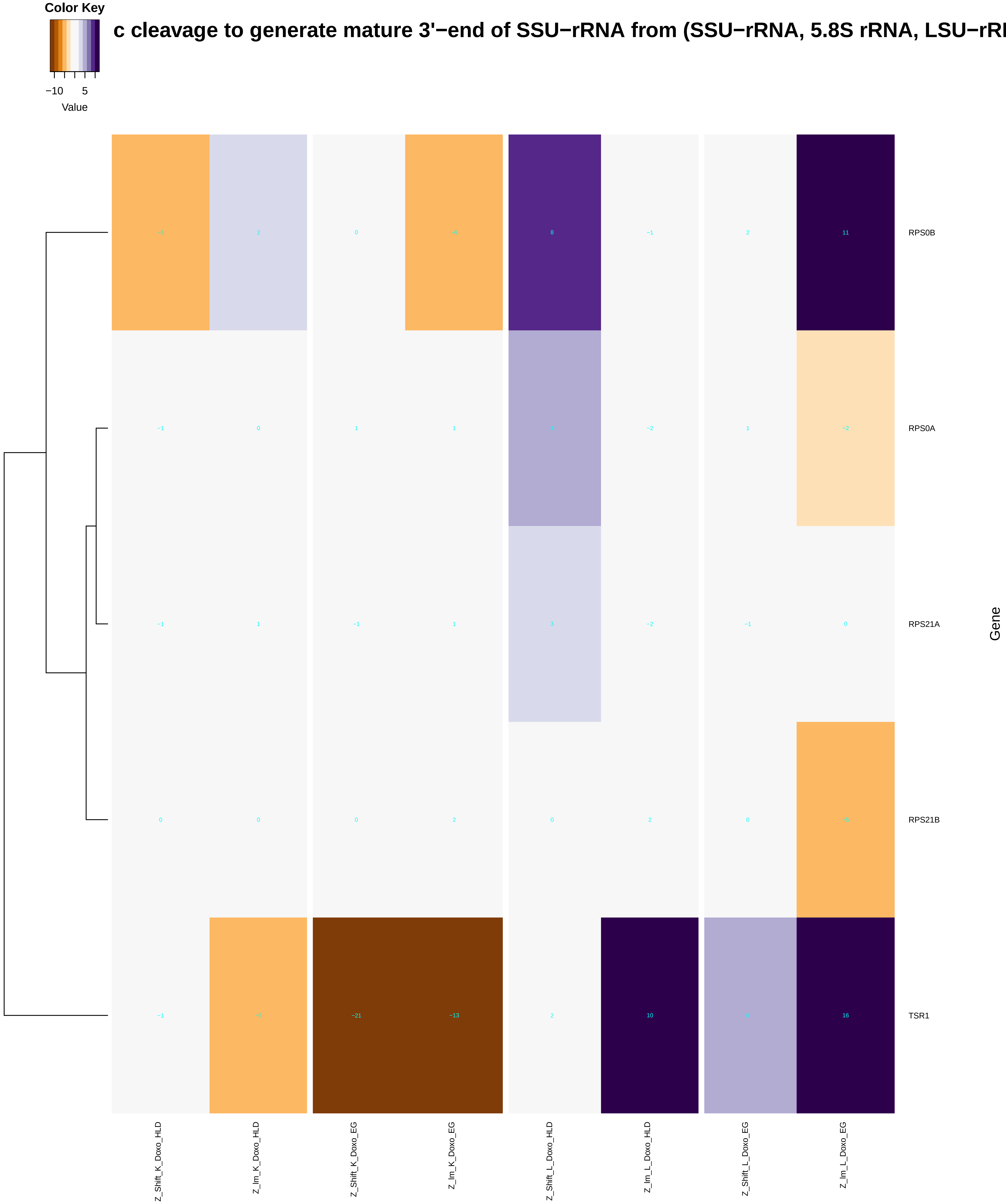

c cleavage to generate mature 3'−end of SSU−rRNA from (SSU−rRNA, 5.8S rRNA, LSU−rR
Color Key
−10
5
Value
−5
2
0
−6
8
−1
2
11
RPS0B
−1
0
1
1
−2
1
−2
RPS0A
−1
1
−1
1
3
−2
−1
0
RPS21A
0
0
0
2
0
2
0
−5
RPS21B
−1
−5
−21
−13
2
10
16
TSR1
Gene
Z_Shift_K_Doxo_HLD
Z_Im_K_Doxo_HLD
Z_Shift_K_Doxo_EG
Z_Im_K_Doxo_EG
Z_Shift_L_Doxo_HLD
Z_Im_L_Doxo_HLD
Z_Shift_L_Doxo_EG
Z_Im_L_Doxo_EG

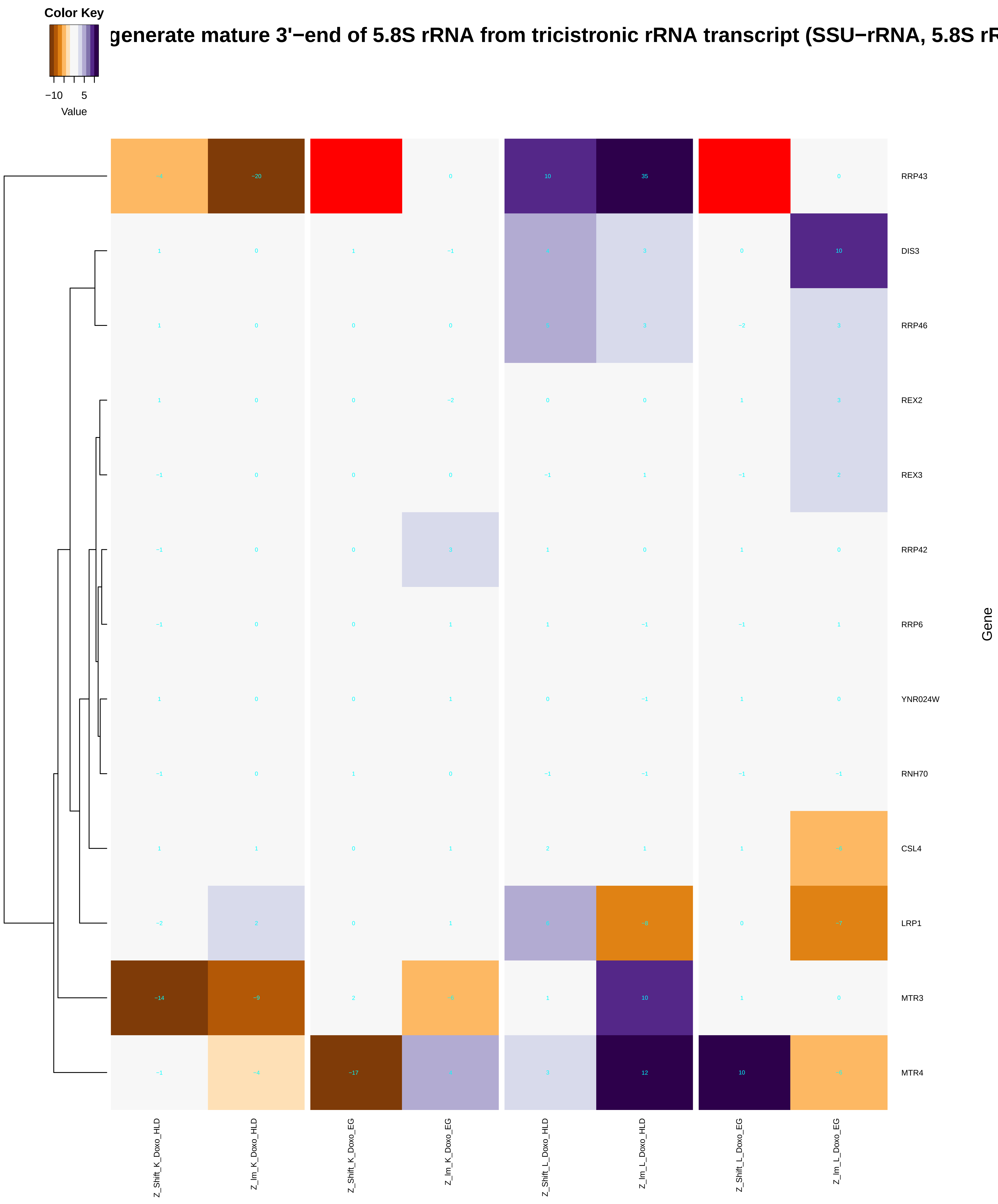
Color Key
−10
5
Value
generate mature 3'−end of 5.8S rRNA from tricistronic rRNA transcript (SSU−rRNA, 5.8S rR
RRP43
DIS3
RRP46
REX2
REX3
RRP42
RRP6
YNR024W
RNH70
CSL4
LRP1
MTR3
MTR4
Gene
Z_Shift_K_Doxo_HLD
Z_lm_K_Doxo_HLD
Z_Shift_K_Doxo_EG
Z_lm_K_Doxo_EG
Z_Shift_L_Doxo_HLD
Z_lm_L_Doxo_HLD
Z_Shift_L_Doxo_EG
Z_lm_L_Doxo_EG

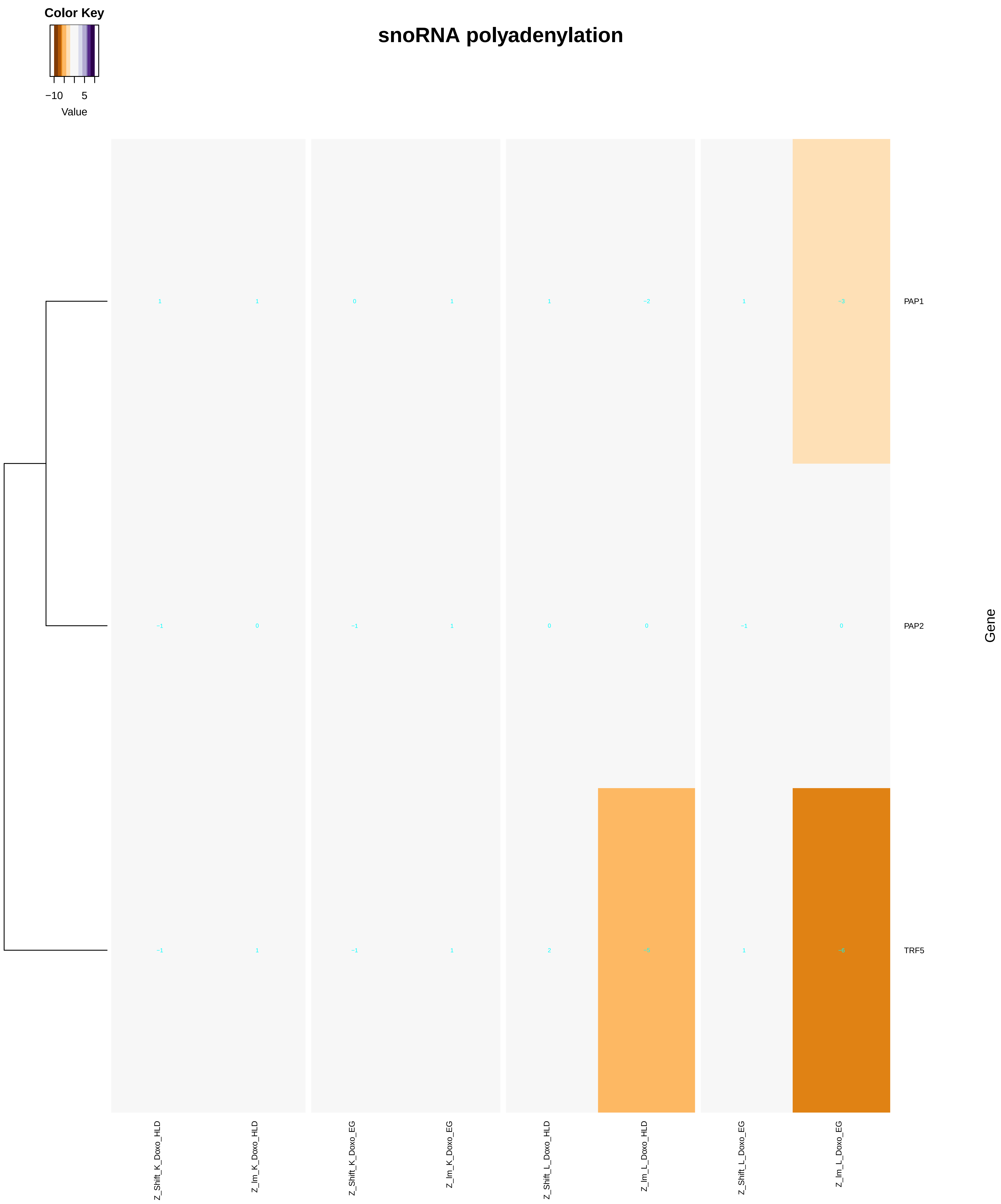
snoRNA polyadenylation
Color Key
−10
5
Value
1
1
0
1
1
−2
1
−3
PAP1
−1
0
−1
1
0
0
−1
0
PAP2
−1
1
−1
1
2
−5
1
−6
TRF5
Gene
Z_Shift_K_Doxo_HLD
Z_Im_K_Doxo_HLD
Z_Shift_K_Doxo_EG
Z_Im_K_Doxo_EG
Z_Shift_L_Doxo_HLD
Z_Im_L_Doxo_HLD
Z_Shift_L_Doxo_EG
Z_Im_L_Doxo_EG

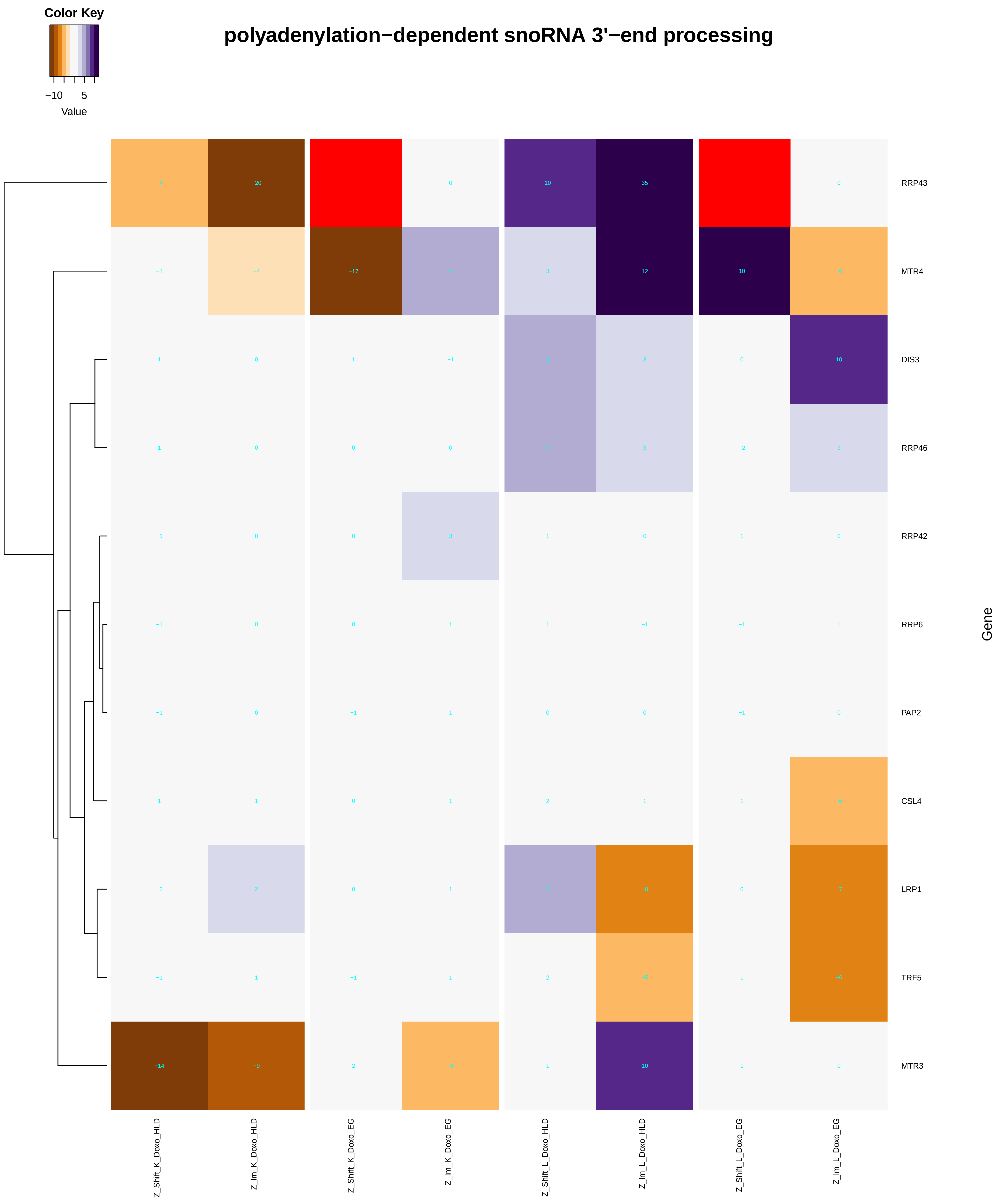
Color Key
−10
5
Value
polyadenylation−dependent snoRNA 3'−end processing
RRP43
MTR4
DIS3
RRP46
RRP42
RRP6
PAP2
CSL4
LRP1
TRF5
MTR3
Gene
Z_Shift_K_Doxo_HLD
Z_lm_K_Doxo_HLD
Z_Shift_K_Doxo_EG
Z_lm_K_Doxo_EG
Z_Shift_L_Doxo_HLD
Z_lm_L_Doxo_HLD
Z_Shift_L_Doxo_EG
Z_lm_L_Doxo_EG

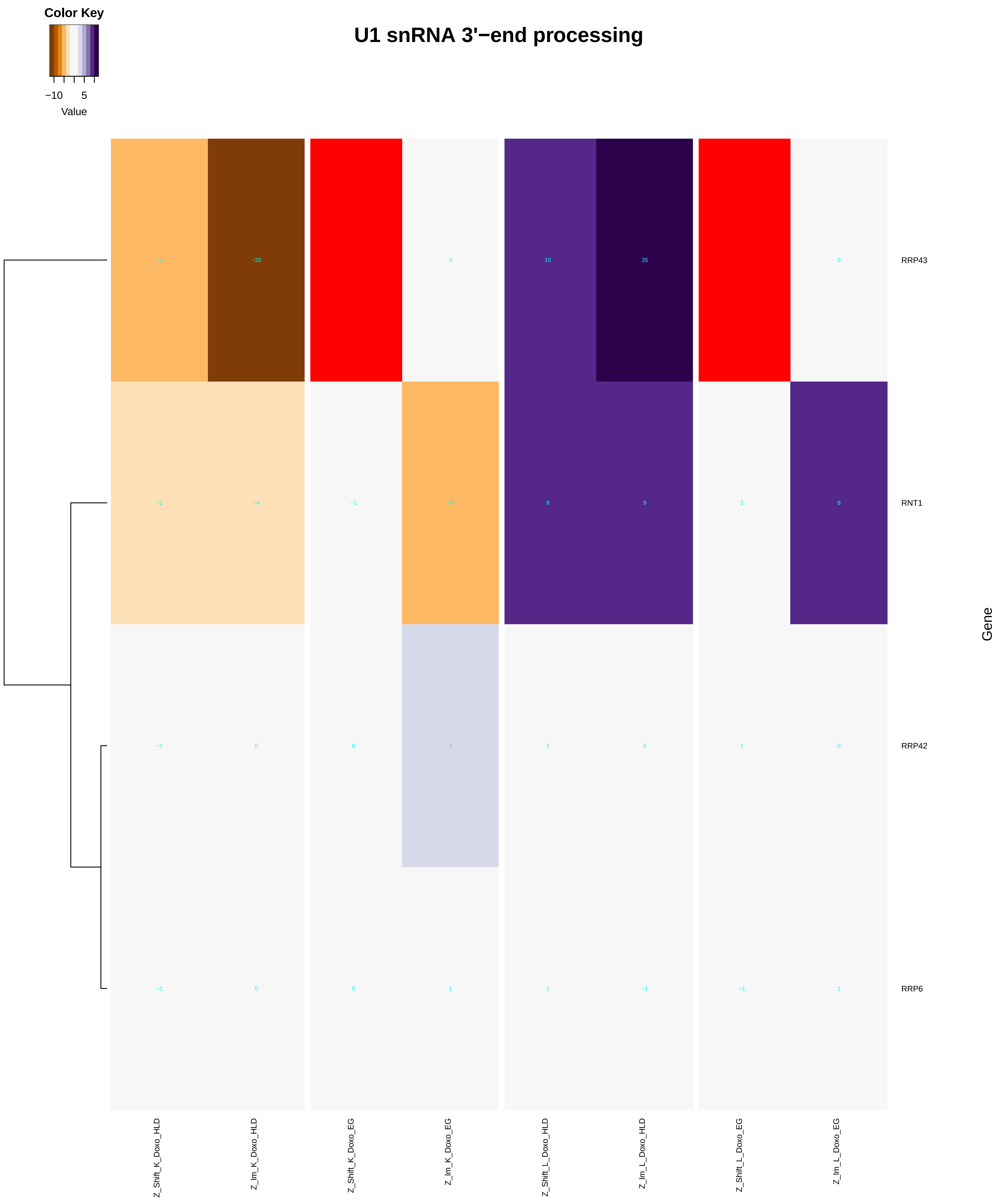

Color Key
−10
5
Value
U1 snRNA 3'−end processing
RRP43
RNT1
RRP42
RRP6
Gene
Z_Shift_K_Doxo_HLD
Z_Im_K_Doxo_HLD
Z_Shift_K_Doxo_EG
Z_Im_K_Doxo_EG
Z_Shift_L_Doxo_HLD
Z_Im_L_Doxo_HLD
Z_Shift_L_Doxo_EG
Z_Im_L_Doxo_EG

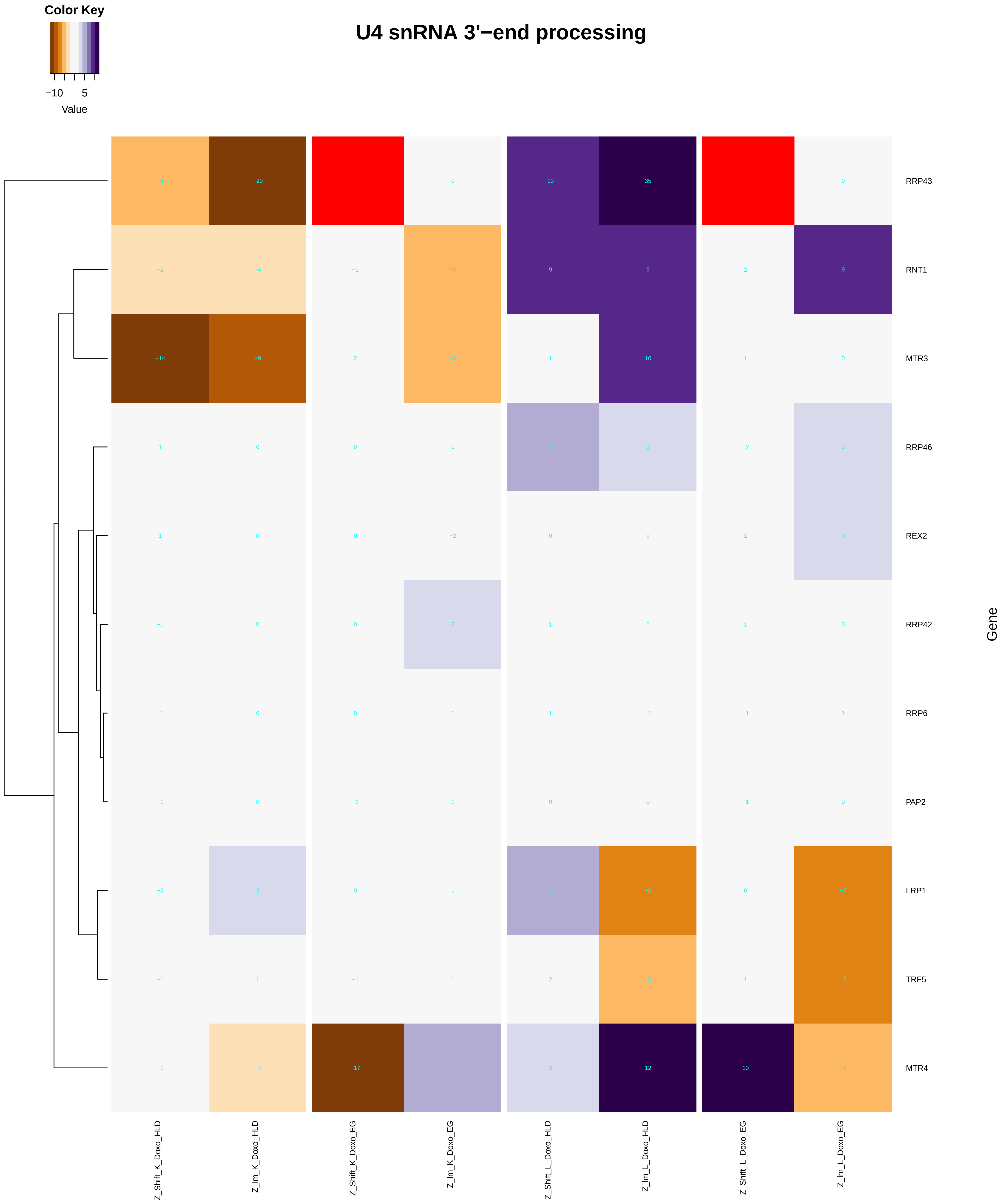
Color Key
−10
5
Value
U4 snRNA 3'−end processing
RRP43
RNT1
MTR3
RRP46
REX2
RRP42
RRP6
PAP2
LRP1
TRF5
MTR4
Gene
Z_Shift_K_Doxo_HLD
Z_lm_K_Doxo_HLD
Z_Shift_K_Doxo_EG
Z_lm_K_Doxo_EG
Z_Shift_L_Doxo_HLD
Z_lm_L_Doxo_HLD
Z_Shift_L_Doxo_EG
Z_lm_L_Doxo_EG

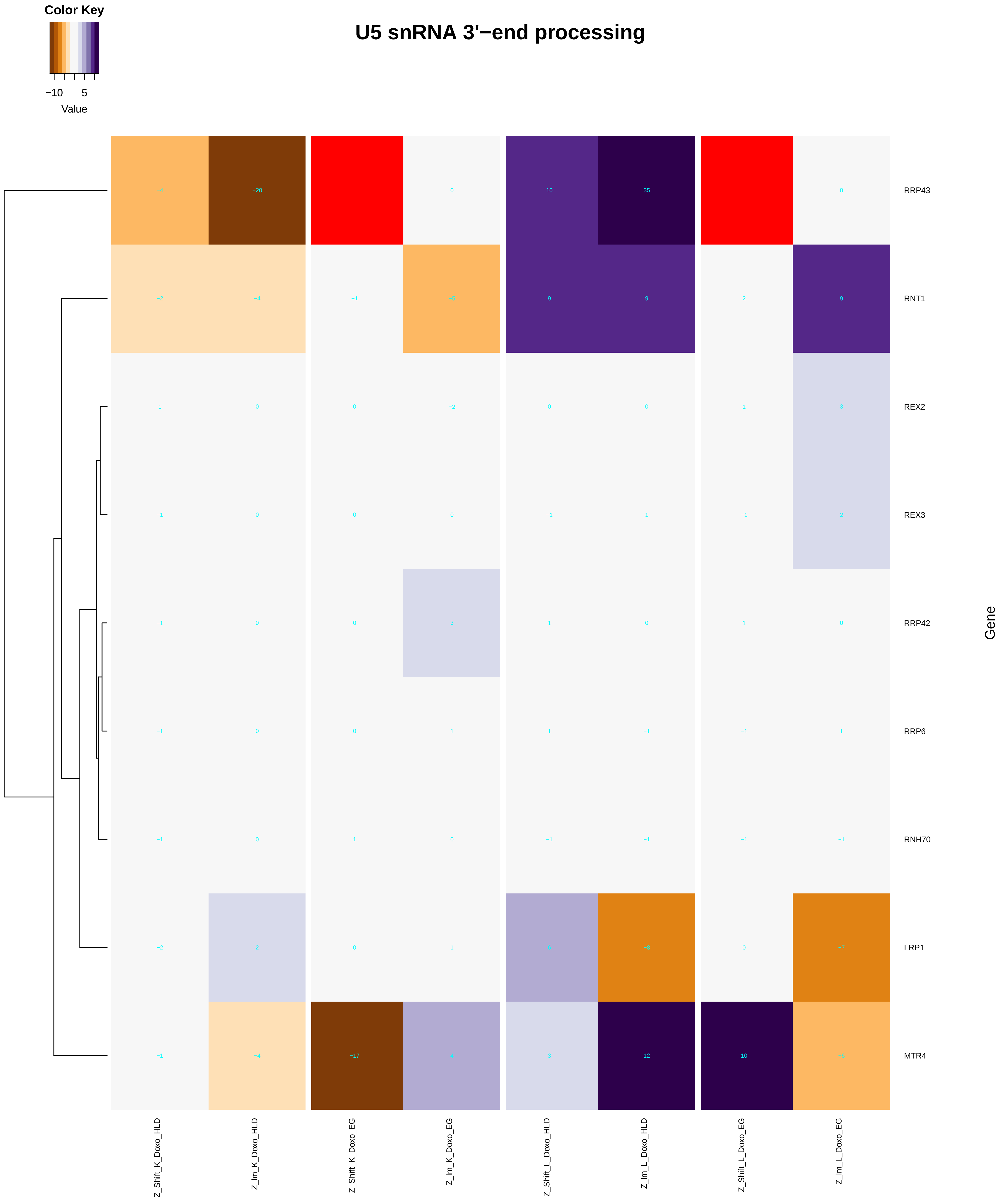

Color Key
−10
5
Value
U5 snRNA 3'−end processing
RRP43
RNT1
REX2
REX3
RRP42
RRP6
RNH70
LRP1
MTR4
Gene
Z_Shift_K_Doxo_HLD
Z_lm_K_Doxo_HLD
Z_Shift_K_Doxo_EG
Z_lm_K_Doxo_EG
Z_Shift_L_Doxo_HLD
Z_lm_L_Doxo_HLD
Z_Shift_L_Doxo_EG
Z_lm_L_Doxo_EG
