## Additional File 9 for "A yeast phenomic model for the influence of Warburg metabolism on genetic buffering of doxorubicin": negative_regulation_of_transcription_from_RNA_polymerase_II_promoter.pdf

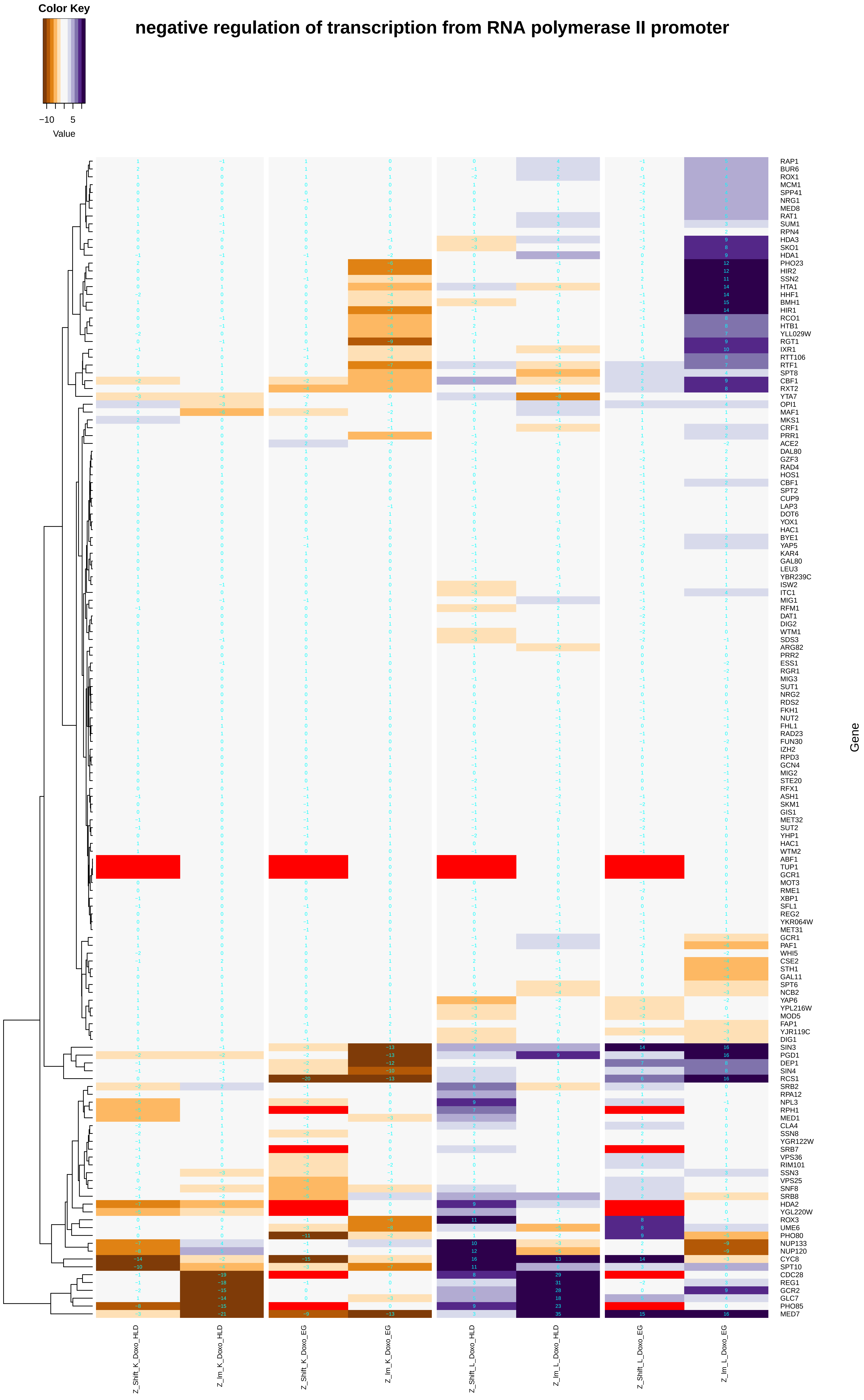

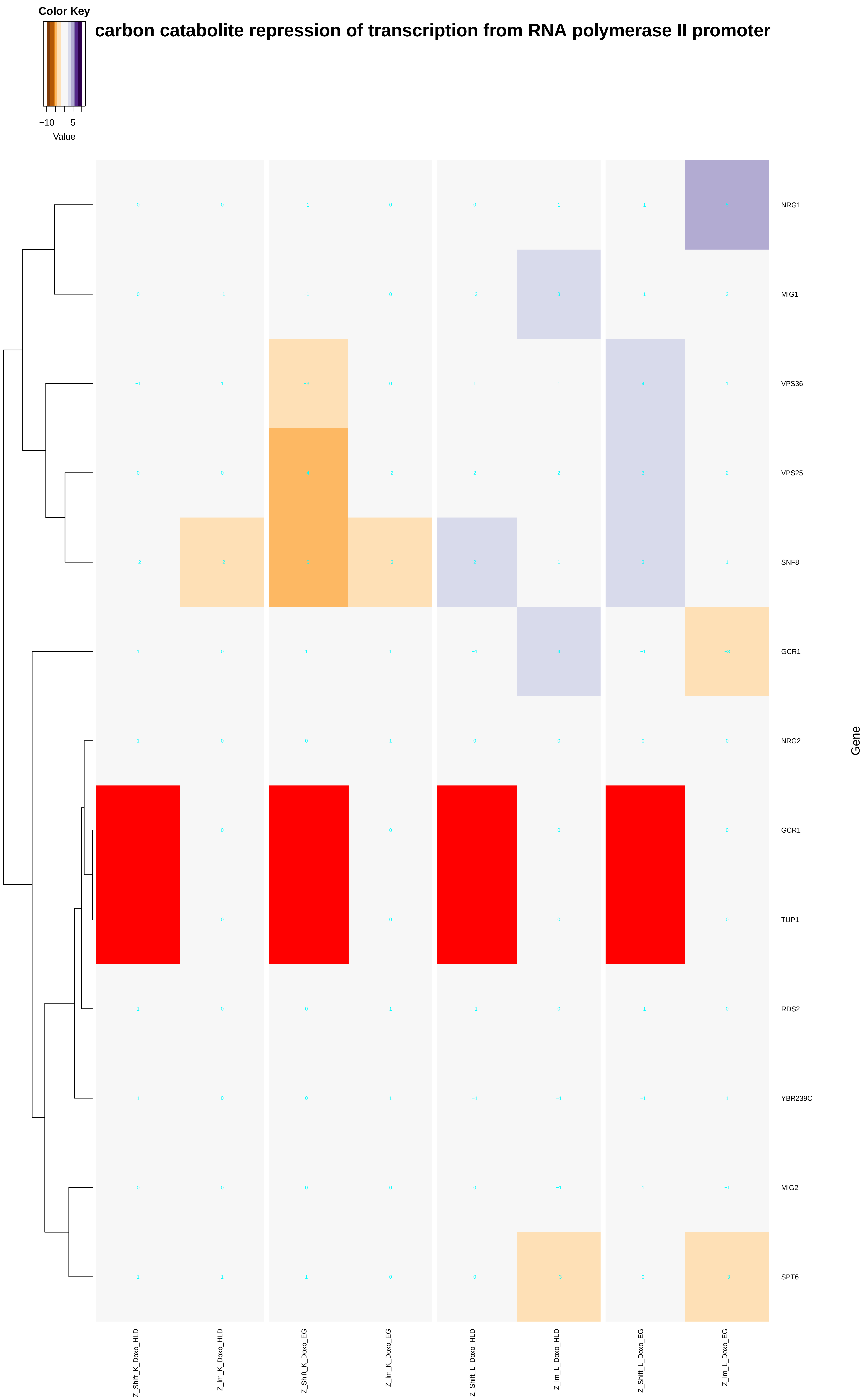

Color Key
−10
5
Value
carbon catabolite repression of transcription from RNA polymerase II promoter
NRG1
MIG1
VPS36
VPS25
SNF8
GCR1
NRG2
GCR1
TUP1
RDS2
YBR239C
MIG2
SPT6
Gene
Z_Shift_K_Doxo_HLD
Z_Im_K_Doxo_HLD
Z_Shift_K_Doxo_EG
Z_Im_K_Doxo_EG
Z_Shift_L_Doxo_HLD
Z_Im_L_Doxo_HLD
Z_Shift_L_Doxo_EG
Z_Im_L_Doxo_EG

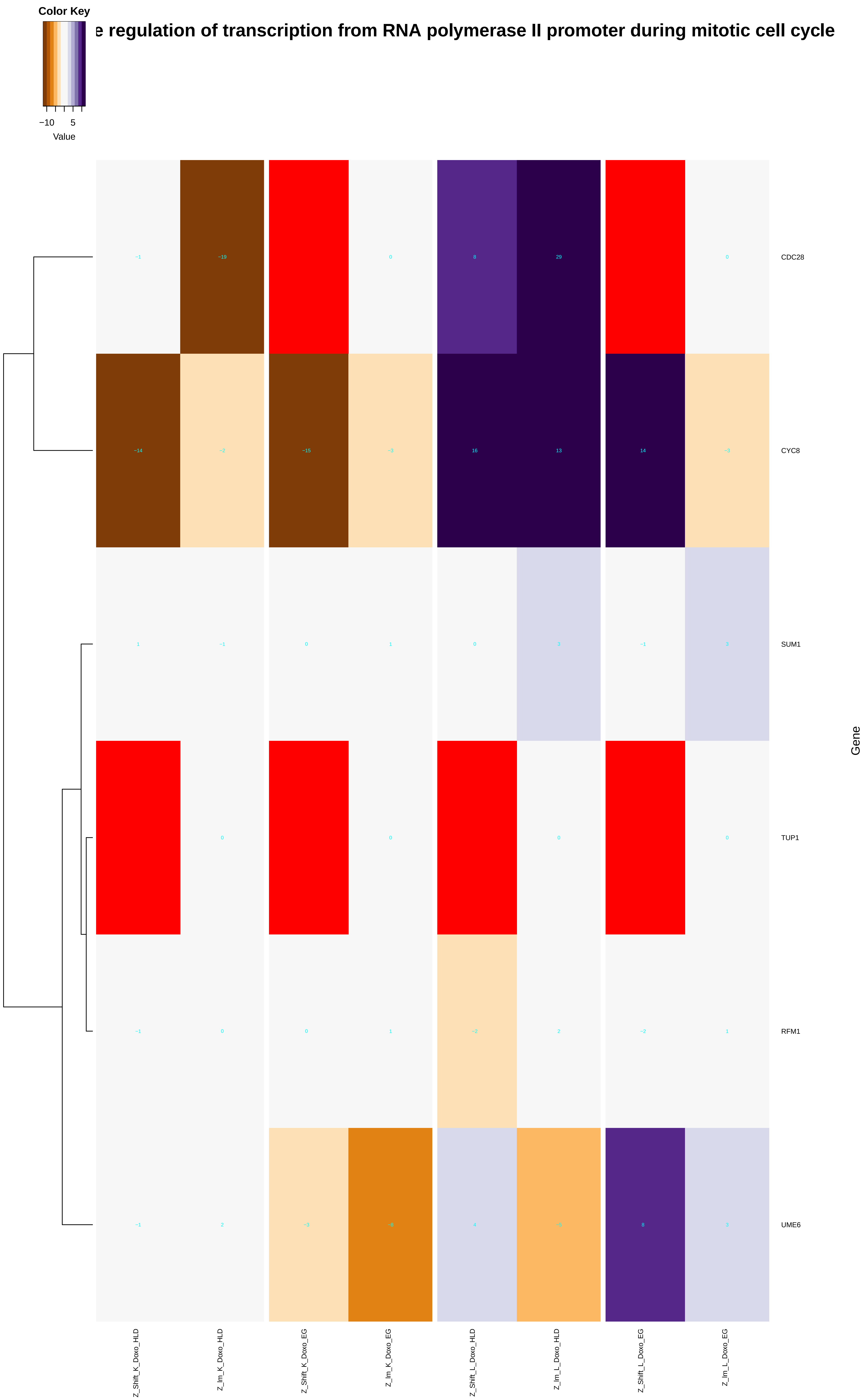

Color Key
−10
5
Value
e regulation of transcription from RNA polymerase II promoter during mitotic cell cycle
CDC28
CYC8
SUM1
TUP1
RFM1
UME6
Gene
Z_Shift_K_Doxo_HLD
Z_lm_K_Doxo_HLD
Z_Shift_K_Doxo_EG
Z_lm_K_Doxo_EG
Z_Shift_L_Doxo_HLD
Z_lm_L_Doxo_HLD
Z_Shift_L_Doxo_EG
Z_lm_L_Doxo_EG

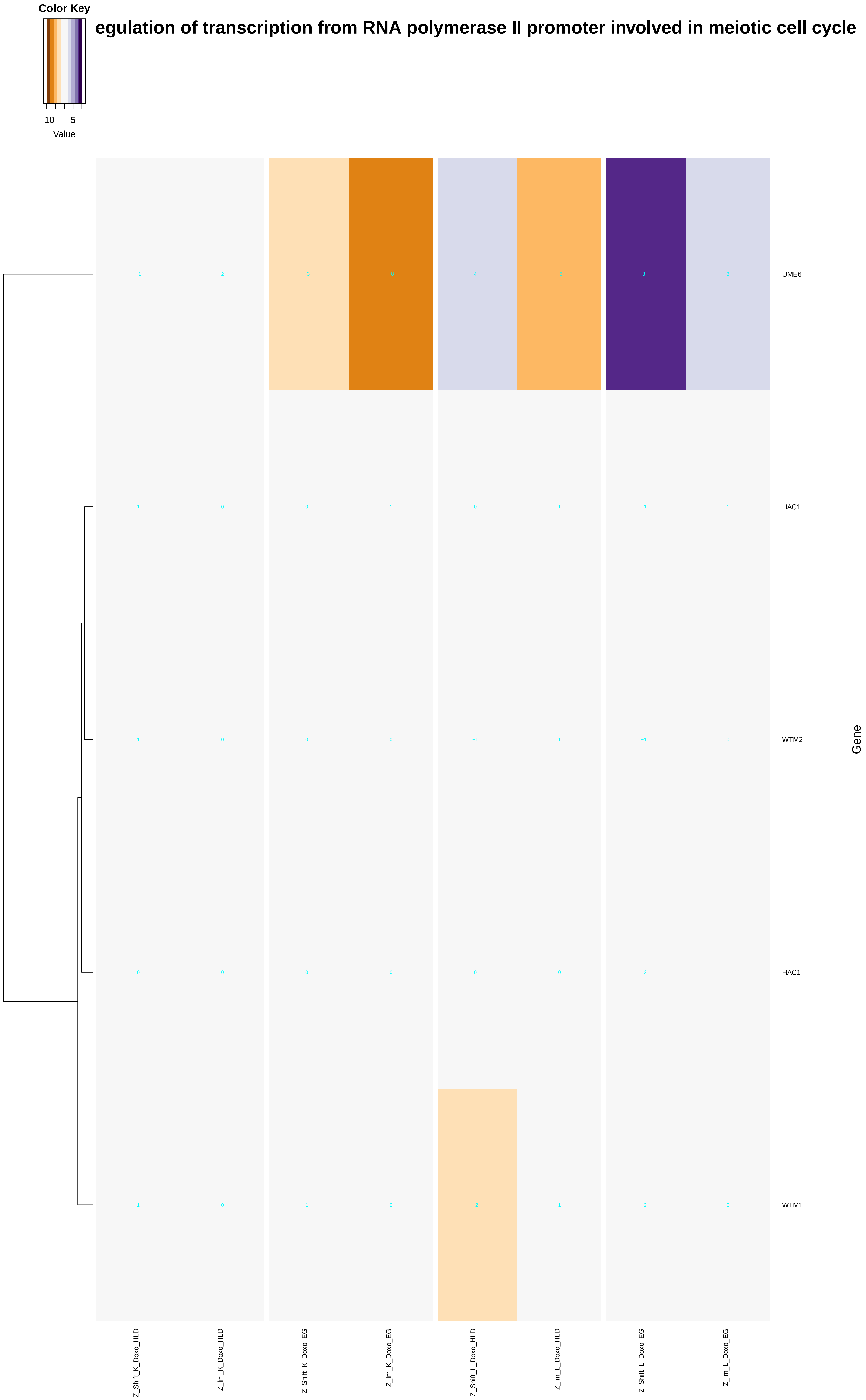
Color Key
Value
−10
5
egulation of transcription from RNA polymerase II promoter involved in meiotic cell cycle
UME6
HAC1
WTM2
HAC1
WTM1
Gene
Z_Shift_K_Doxo_HLD
Z_Im_K_Doxo_HLD
Z_Shift_K_Doxo_EG
Z_Im_K_Doxo_EG
Z_Shift_L_Doxo_HLD
Z_Im_L_Doxo_HLD
Z_Shift_L_Doxo_EG
Z_Im_L_Doxo_EG

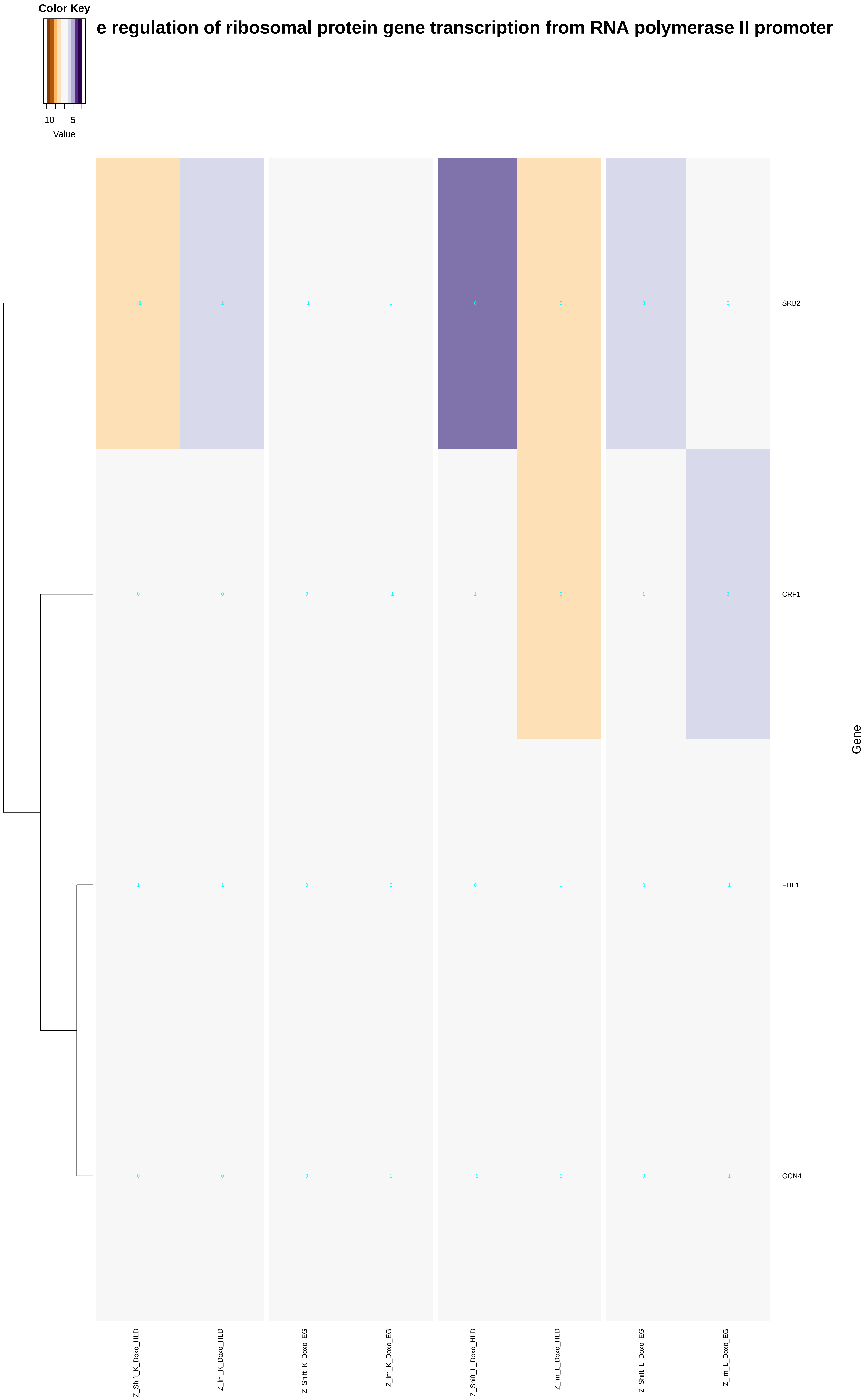

Color Key
−10
5
Value
e regulation of ribosomal protein gene transcription from RNA polymerase II promoter
SRB2
CRF1
FHL1
GCN4
Gene
Z_Shift_K_Doxo_HLD
Z_Im_K_Doxo_HLD
Z_Shift_K_Doxo_EG
Z_Im_K_Doxo_EG
Z_Shift_L_Doxo_HLD
Z_Im_L_Doxo_HLD
Z_Shift_L_Doxo_EG
Z_Im_L_Doxo_EG

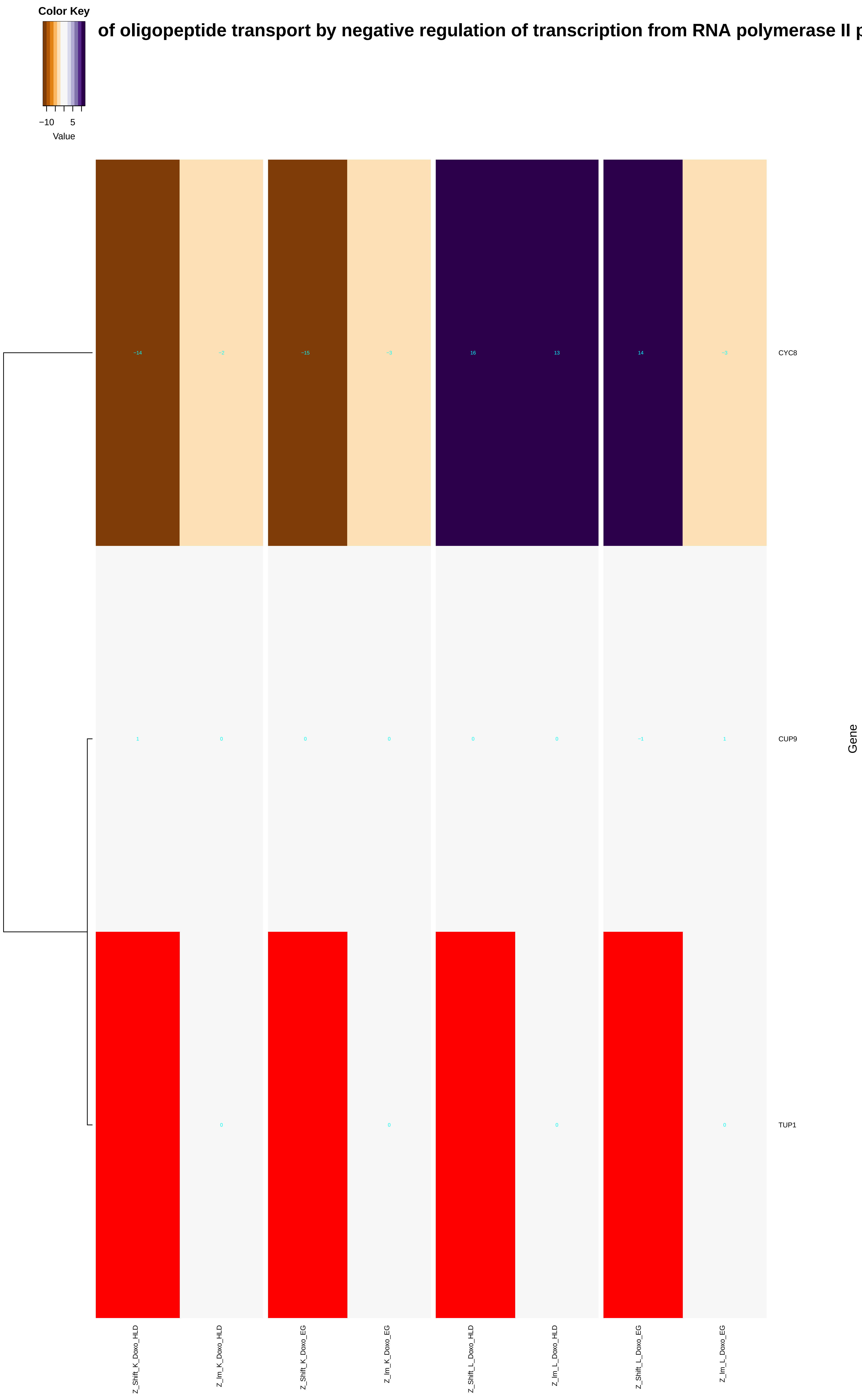
Color Key
−10
5
Value
of oligopeptide transport by negative regulation of transcription from RNA polymerase II p
−14
−2
−15
−3
16
13
14
−3
CYC8
1
0
0
0
0
0
−1
1
CUP9
0
0
0
0
TUP1
Gene
Z_Shift_K_Doxo_HLD
Z_Im_K_Doxo_HLD
Z_Shift_K_Doxo_EG
Z_Im_K_Doxo_EG
Z_Shift_L_Doxo_HLD
Z_Im_L_Doxo_HLD
Z_Shift_L_Doxo_EG
Z_Im_L_Doxo_EG

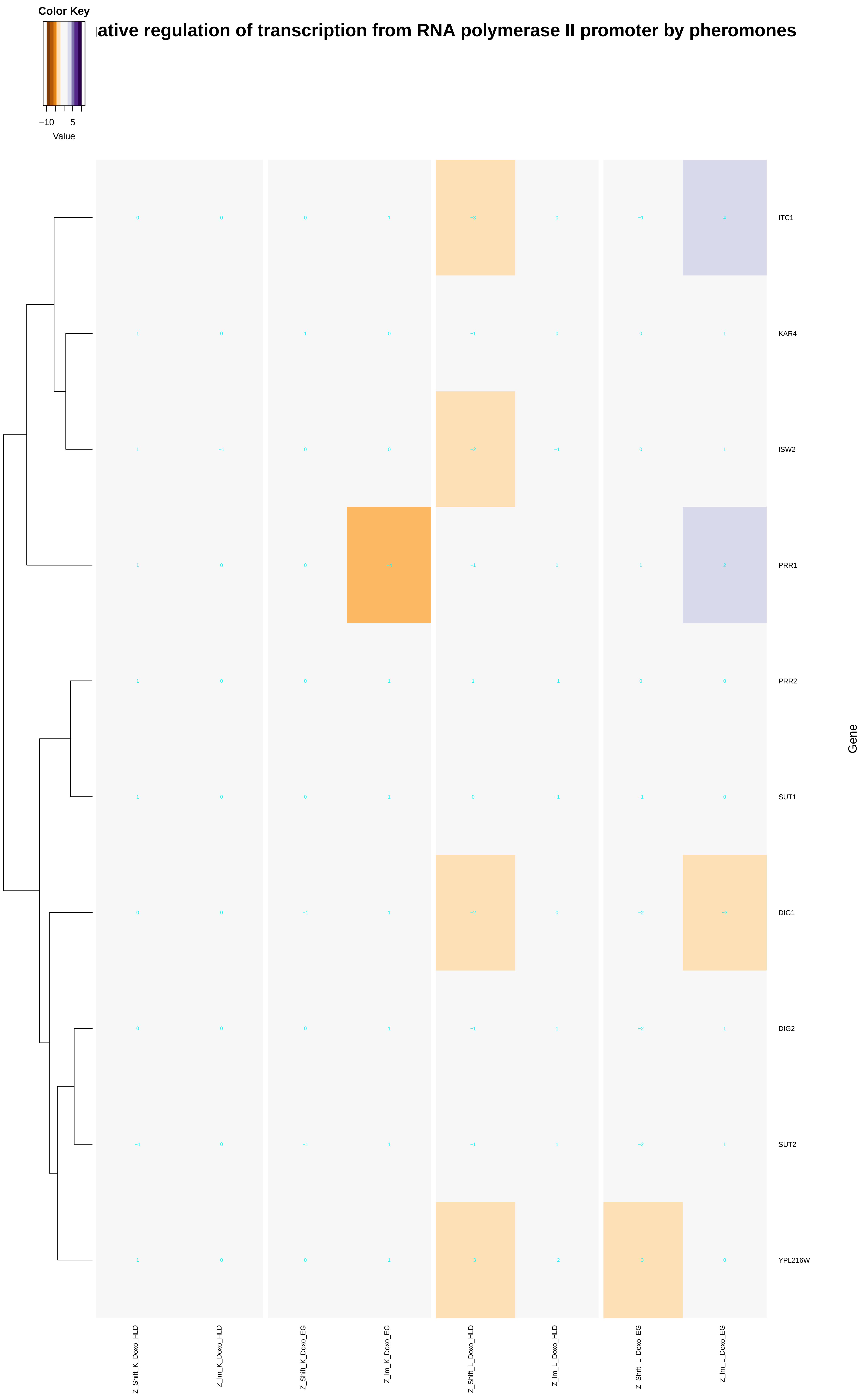

Color Key
−10
5
Value
ıative regulation of transcription from RNA polymerase II promoter by pheromones
ITC1
KAR4
ISW2
PRR1
PRR2
SUT1
DIG1
DIG2
SUT2
YPL216W
Gene
Z_Shift_K_Doxo_HLD
Z_Im_K_Doxo_HLD
Z_Shift_K_Doxo_EG
Z_Im_K_Doxo_EG
Z_Shift_L_Doxo_HLD
Z_Im_L_Doxo_HLD
Z_Shift_L_Doxo_EG
Z_Im_L_Doxo_EG

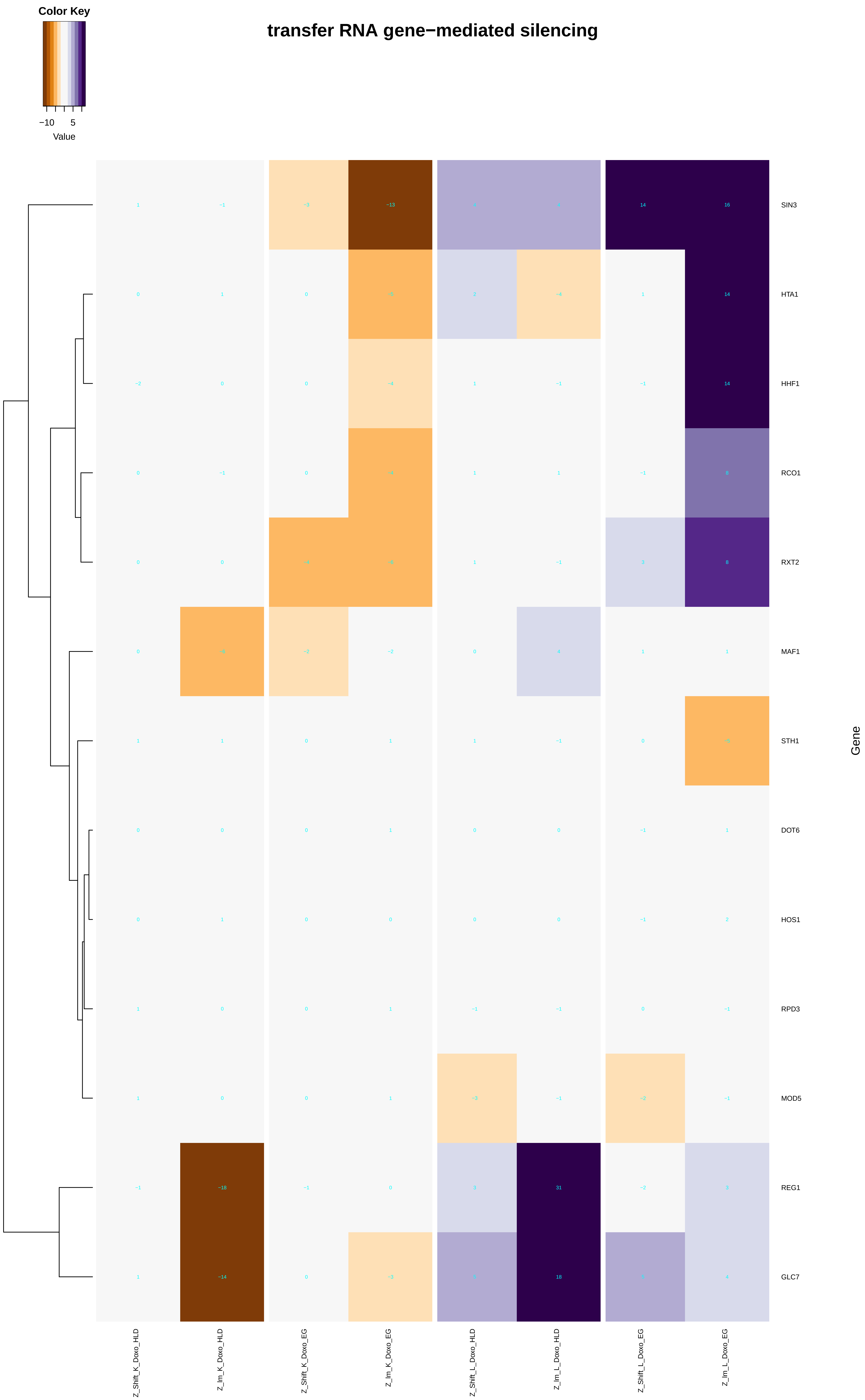

Color Key
−10
5
Value
transfer RNA gene−mediated silencing
SIN3
HTA1
HHF1
RCO1
RXT2
MAF1
STH1
DOT6
HOS1
RPD3
MOD5
REG1
GLC7
Gene
Z_Shift_K_Doxo_HLD
Z_Im_K_Doxo_HLD
Z_Shift_K_Doxo_EG
Z_Im_K_Doxo_EG
Z_Shift_L_Doxo_HLD
Z_Im_L_Doxo_HLD
Z_Shift_L_Doxo_EG
Z_Im_L_Doxo_EG

### of lipid transport by negative regulation of transcription from RNA polymerase II promoter

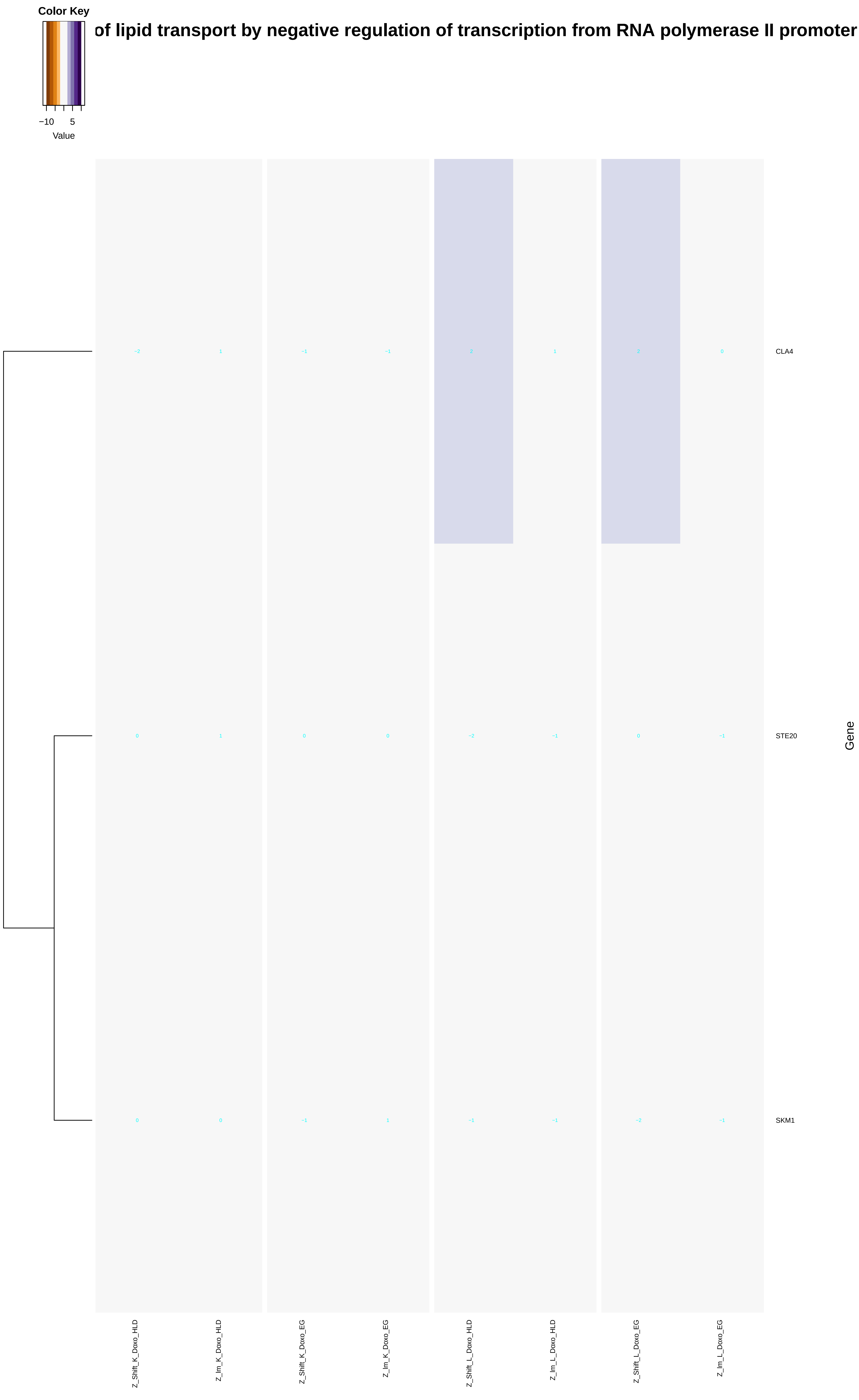

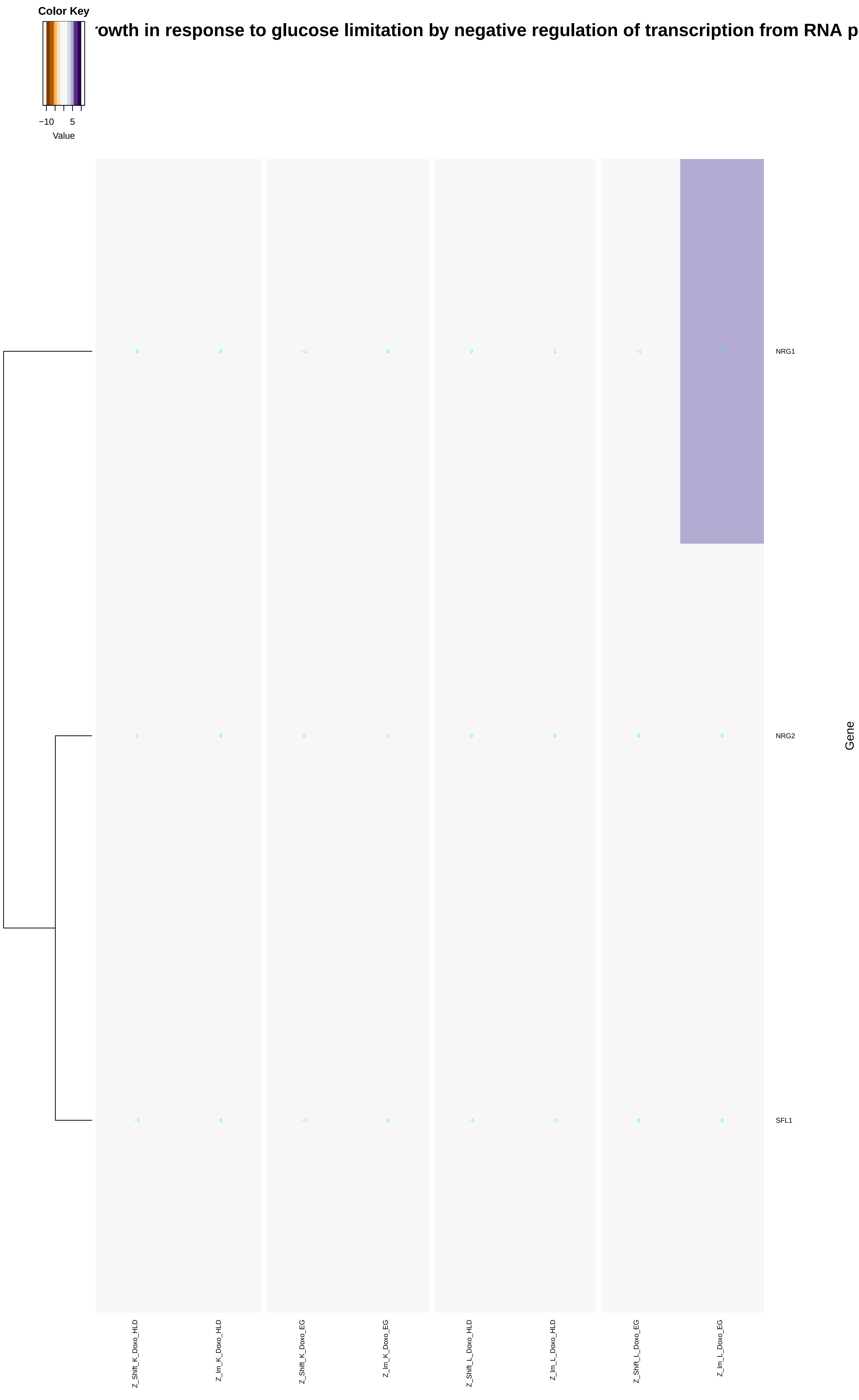

Color Key
−10
5
Value
rowth in response to glucose limitation by negative regulation of transcription from RNA p
NRG1
NRG2
SFL1
Gene
Z_Shift_K_Doxo_HLD
Z_Im_K_Doxo_HLD
Z_Shift_K_Doxo_EG
Z_Im_K_Doxo_EG
Z_Shift_L_Doxo_HLD
Z_Im_L_Doxo_HLD
Z_Shift_L_Doxo_EG
Z_Im_L_Doxo_EG

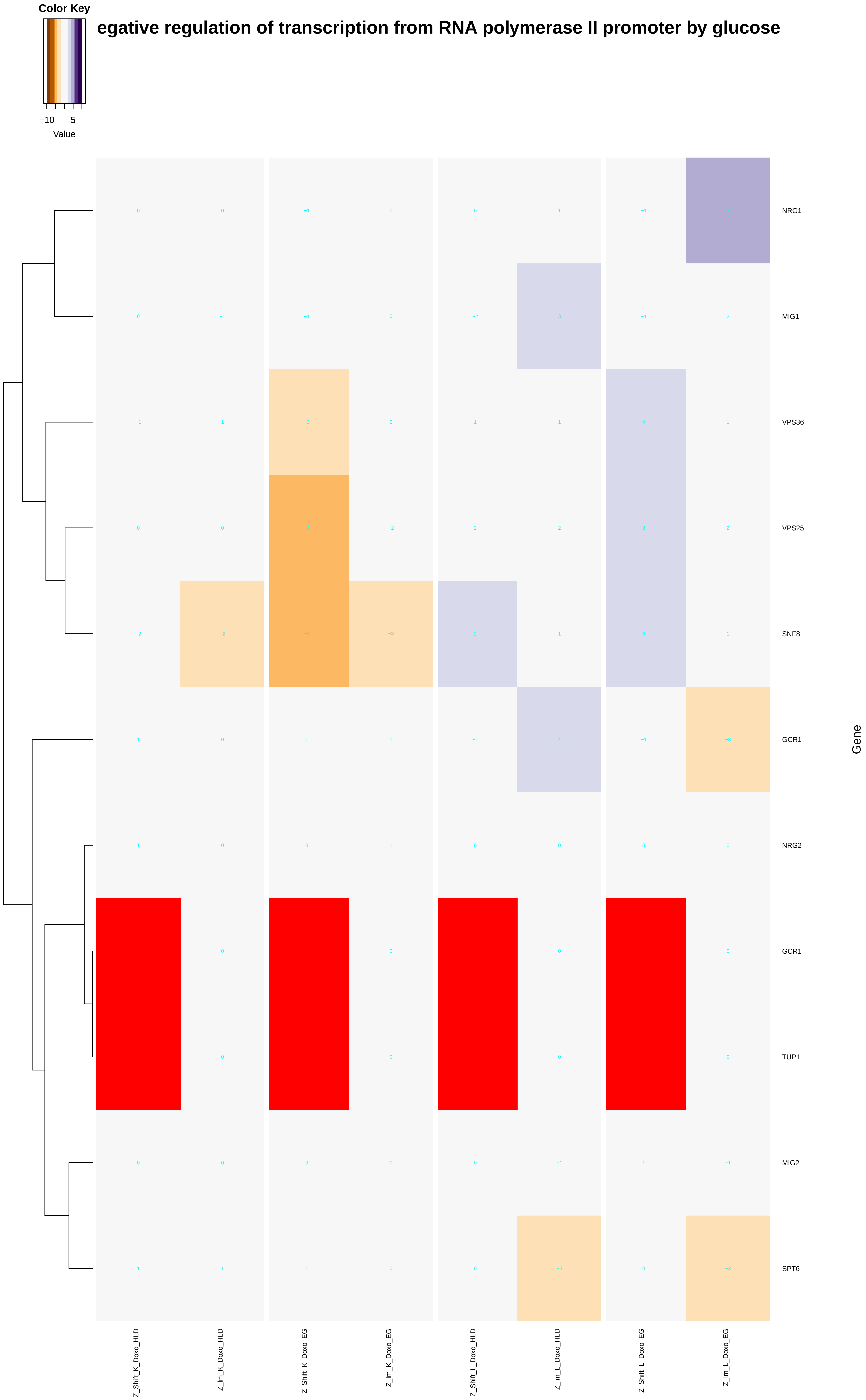
Color Key
−10
5
Value
egative regulation of transcription from RNA polymerase II promoter by glucose
NRG1
MIG1
VPS36
VPS25
SNF8
GCR1
NRG2
GCR1
TUP1
MIG2
SPT6
Gene
Z_Shift_K_Doxo_HLD
Z_lm_K_Doxo_HLD
Z_Shift_K_Doxo_EG
Z_lm_K_Doxo_EG
Z_Shift_L_Doxo_HLD
Z_lm_L_Doxo_HLD
Z_Shift_L_Doxo_EG
Z_lm_L_Doxo_EG

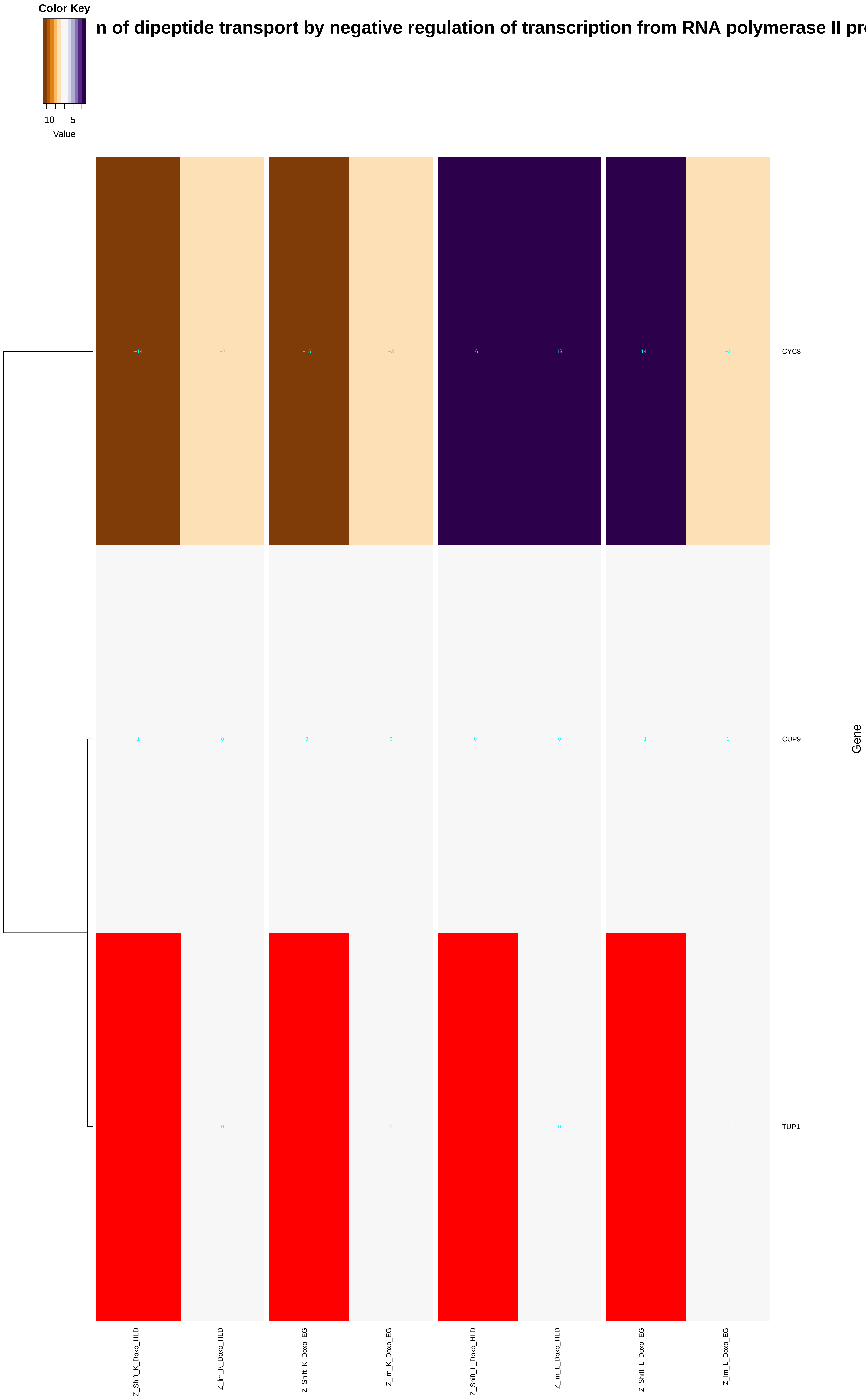
Color Key
−10
5
Value
n of dipeptide transport by negative regulation of transcription from RNA polymerase II pro
−14
−2
−15
−3
16
13
14
−3
CYC8
1
0
0
0
0
0
−1
1
CUP9
0
0
0
0
TUP1
Gene
Z_Shift_K_Doxo_HLD
Z_lm_K_Doxo_HLD
Z_Shift_K_Doxo_EG
Z_lm_K_Doxo_EG
Z_Shift_L_Doxo_HLD
Z_lm_L_Doxo_HLD
Z_Shift_L_Doxo_EG
Z_lm_L_Doxo_EG

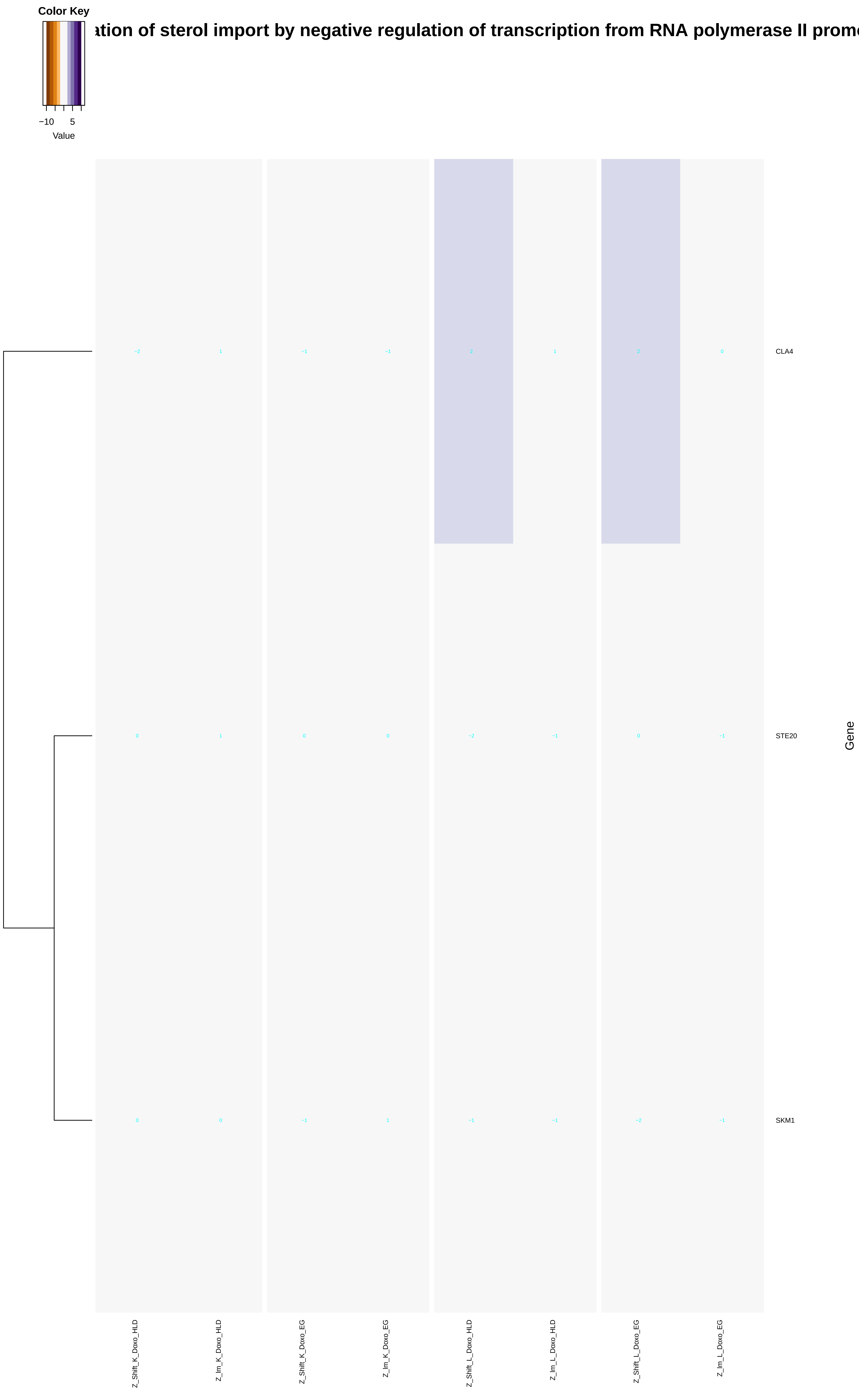
Color Key
−10
5
Value
ation of sterol import by negative regulation of transcription from RNA polymerase II prom
CLA4
STE20
SKM1
−2
1
−1
−1
2
1
2
0
0
1
0
0
−2
−1
0
−1
0
0
−1
1
−1
−1
−2
−1
Gene
Z_Shift_K_Doxo_HLD
Z_lm_K_Doxo_HLD
Z_Shift_K_Doxo_EG
Z_lm_K_Doxo_EG
Z_Shift_L_Doxo_HLD
Z_lm_L_Doxo_HLD
Z_Shift_L_Doxo_EG
Z_lm_L_Doxo_EG
