## Additional File 9 for "A yeast phenomic model for the influence of Warburg metabolism on genetic buffering of doxorubicin": nuclear-transcribed_mRNA_catabolic_process.pdf

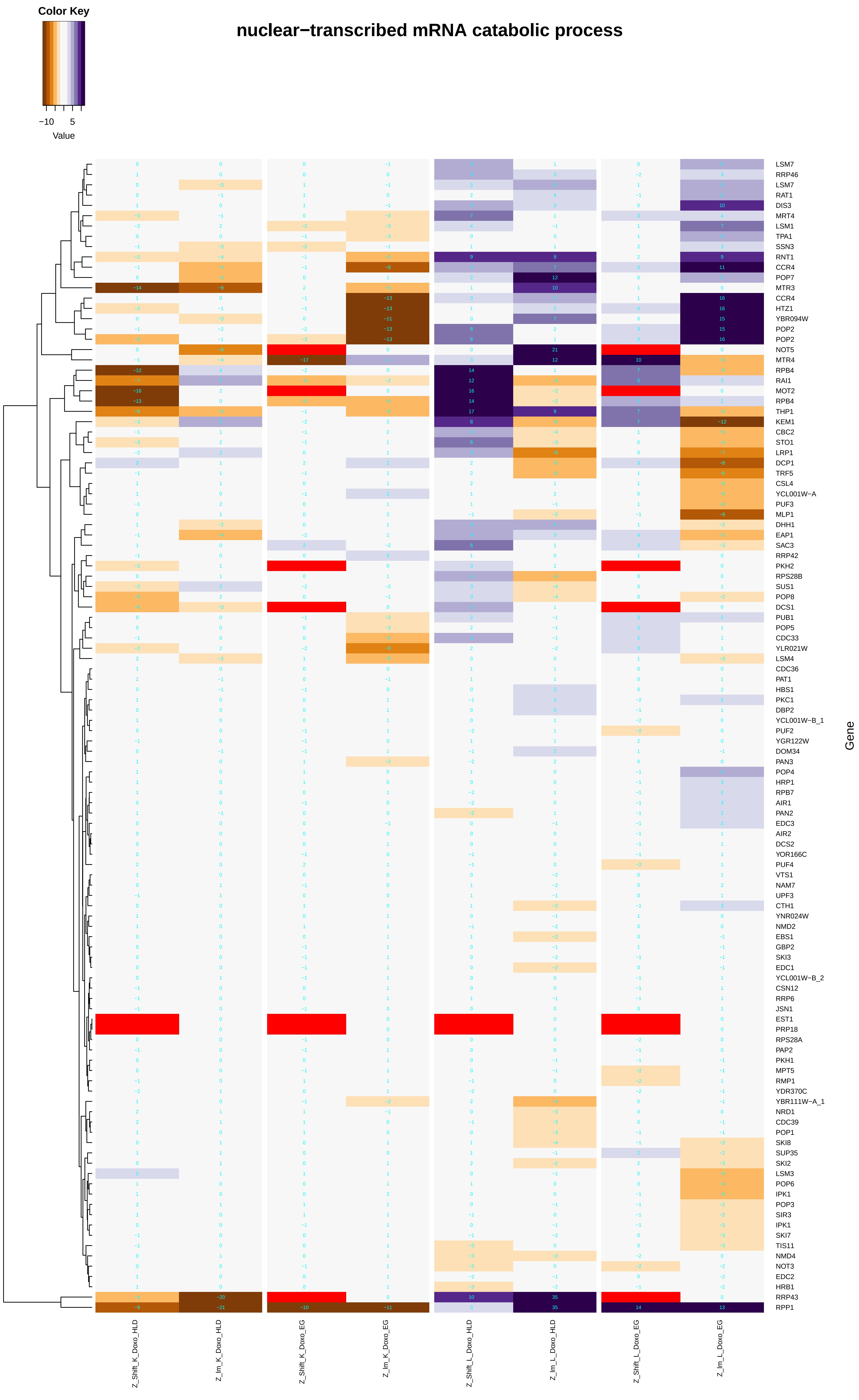

nuclear−transcribed mRNA catabolic process
Color Key
−10 5
Value
LSM7
RRP46
LSM7
RAT1
DIS3
MRT4
LSM1
TPA1
SSN3
RNT1
CCR4
POP7
MTR3
CCR4
HTZ1
YBR094W
POP2
POP2
NOT5
MTR4
RPB4
RAI1
MOT2
RPB4
THP1
KEM1
CBC2
STO1
LRP1
DCP1
TRF5
CSL4
YCL001W−A
PUF3
MLP1
DHH1
EAP1
SAC3
RRP42
PKH2
RPS28B
SUS1
POP8
DCS1
PUB1
POP5
CDC33
YLR021W
LSM4
CDC36
PAT1
HBS1
PKC1
DBP2
YCL001W−B_1
PUF2
YGR122W
DOM34
PAN3
POP4
HRP1
RPB7
AIR1
PAN2
EDC3
AIR2
DCS2
YOR166C
PUF4
VTS1
NAM7
UPF3
CTH1
YNR024W
NMD2
EBS1
GBP2
SKI3
EDC1
YCL001W−B_2
CSN12
RRP6
JSN1
EST1
PRP18
RPS28A
PAP2
PKH1
MPT5
RMP1
YDR370C
YBR111W−A_1
NRD1
CDC39
POP1
SKI8
SUP35
SKI2
LSM3
POP6
IPK1
POP3
SIR3
IPK1
SKI7
TIS11
NMD4
NOT3
EDC2
HRB1
RRP43
RPP1
Gene
Z_Shift_K_Doxo_HLD
Z_lm_K_Doxo_HLD
Z_Shift_K_Doxo_EG
Z_lm_K_Doxo_EG
Z_Shift_L_Doxo_HLD
Z_lm_L_Doxo_HLD
Z_Shift_L_Doxo_EG
Z_lm_L_Doxo_EG

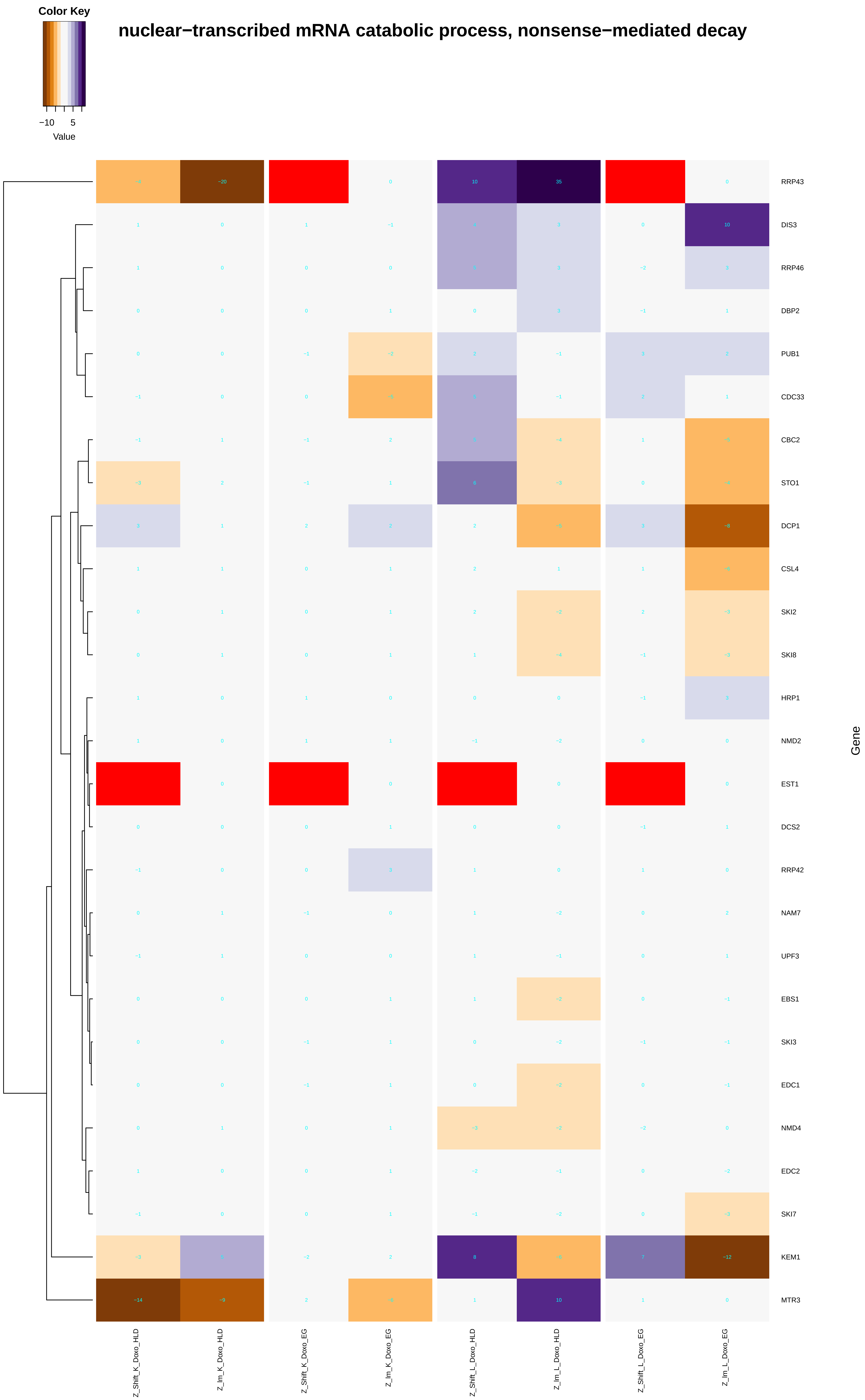
Color Key
−10
5
Value
nuclear−transcribed mRNA catabolic process, nonsense−mediated decay
RRP43
DIS3
RRP46
DBP2
PUB1
CDC33
CBC2
STO1
DCP1
CSL4
SKI2
SKI8
HRP1
NMD2
EST1
DCS2
RRP42
NAM7
UPF3
EBS1
SKI3
EDC1
NMD4
EDC2
SKI7
KEM1
MTR3
Gene
Z_Shift_K_Doxo_HLD
Z_Im_K_Doxo_HLD
Z_Shift_K_Doxo_EG
Z_Im_K_Doxo_EG
Z_Shift_L_Doxo_HLD
Z_Im_L_Doxo_HLD
Z_Shift_L_Doxo_EG
Z_Im_L_Doxo_EG

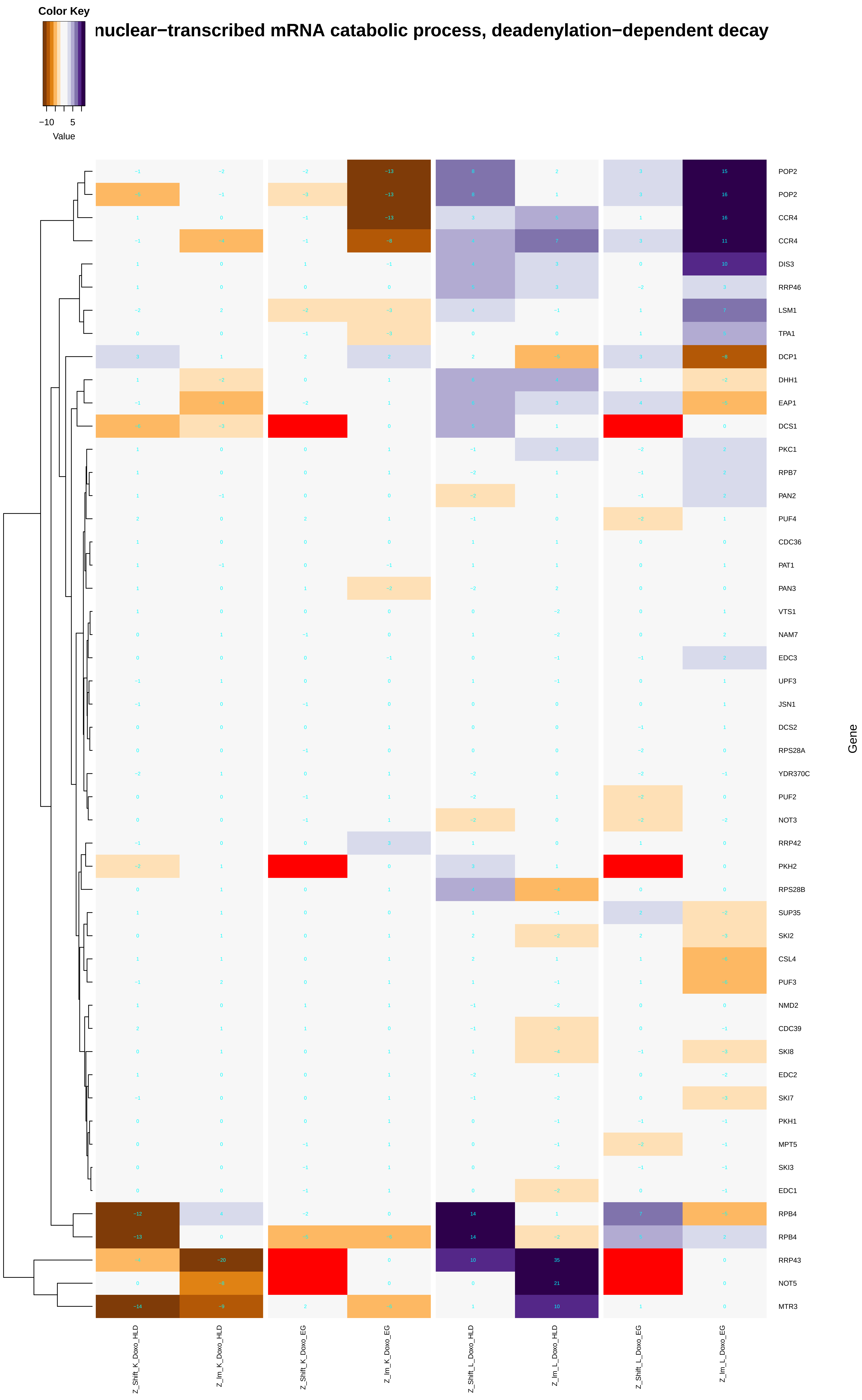

Color Key
−10
5
Value
nuclear−transcribed mRNA catabolic process, deadenylation−dependent decay
POP2
POP2
CCR4
CCR4
DIS3
RRP46
LSM1
TPA1
DCP1
DHH1
EAP1
DCS1
PKC1
RPB7
PAN2
PUF4
CDC36
PAT1
PAN3
VTS1
NAM7
EDC3
UPF3
JSN1
DCS2
RPS28A
YDR370C
PUF2
NOT3
RRP42
PKH2
RPS28B
SUP35
SKI2
CSL4
PUF3
NMD2
CDC39
SKI8
EDC2
SKI7
PKH1
MPT5
SKI3
EDC1
RPB4
RPB4
RRP43
NOT5
MTR3
Gene
Z_Shift_K_Doxo_HLD
Z_lm_K_Doxo_HLD
Z_Shift_K_Doxo_EG
Z_lm_K_Doxo_EG
Z_Shift_L_Doxo_HLD
Z_lm_L_Doxo_HLD
Z_Shift_L_Doxo_EG
Z_lm_L_Doxo_EG

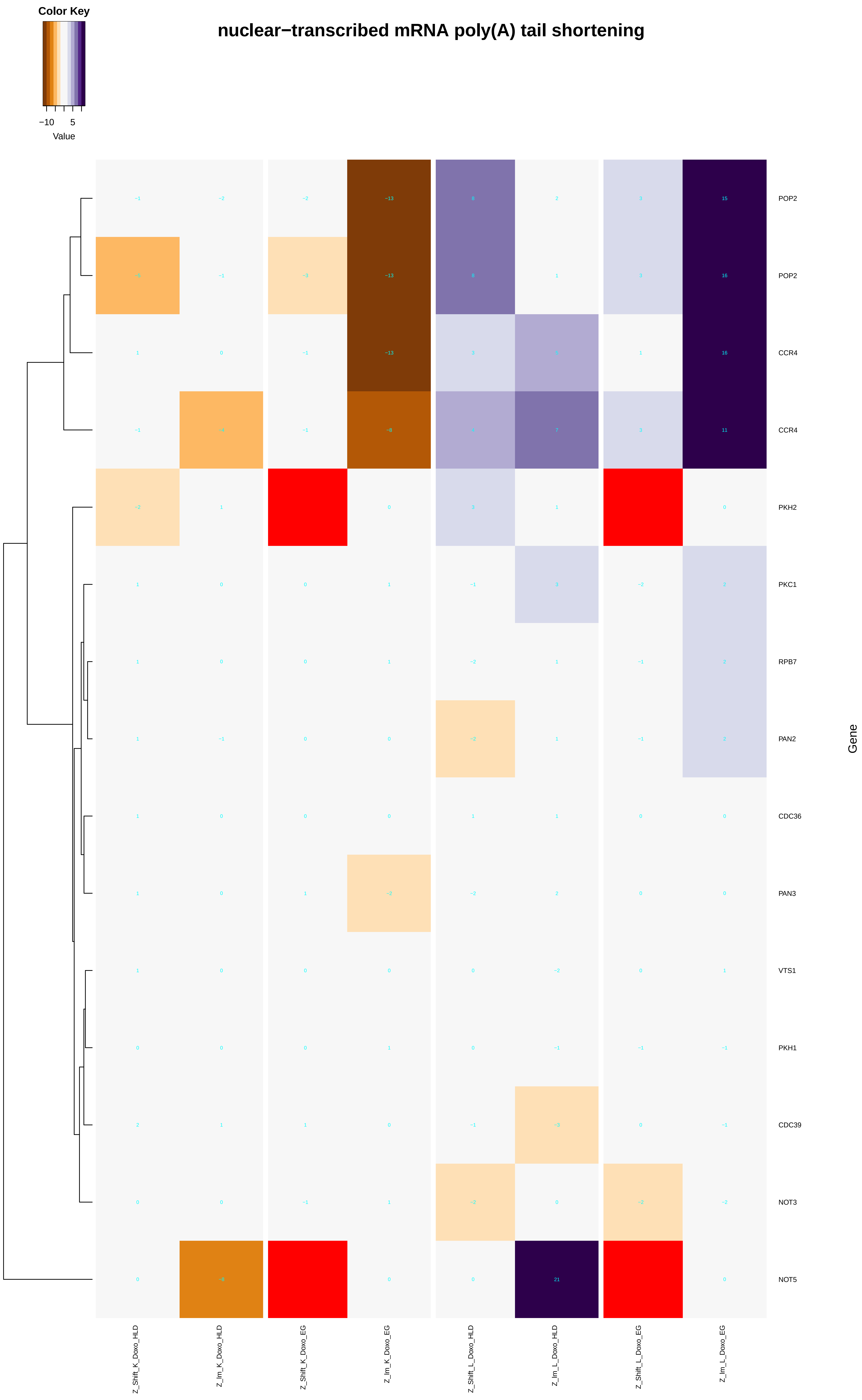

nuclear−transcribed mRNA poly(A) tail shortening
Color Key
−10
5
Value
POP2
POP2
CCR4
CCR4
PKH2
PKC1
RPB7
PAN2
CDC36
PAN3
VTS1
PKH1
CDC39
NOT3
NOT5
Gene
Z_Shift_K_Doxo_HLD
Z_Im_K_Doxo_HLD
Z_Shift_K_Doxo_EG
Z_Im_K_Doxo_EG
Z_Shift_L_Doxo_HLD
Z_Im_L_Doxo_HLD
Z_Shift_L_Doxo_EG
Z_Im_L_Doxo_EG

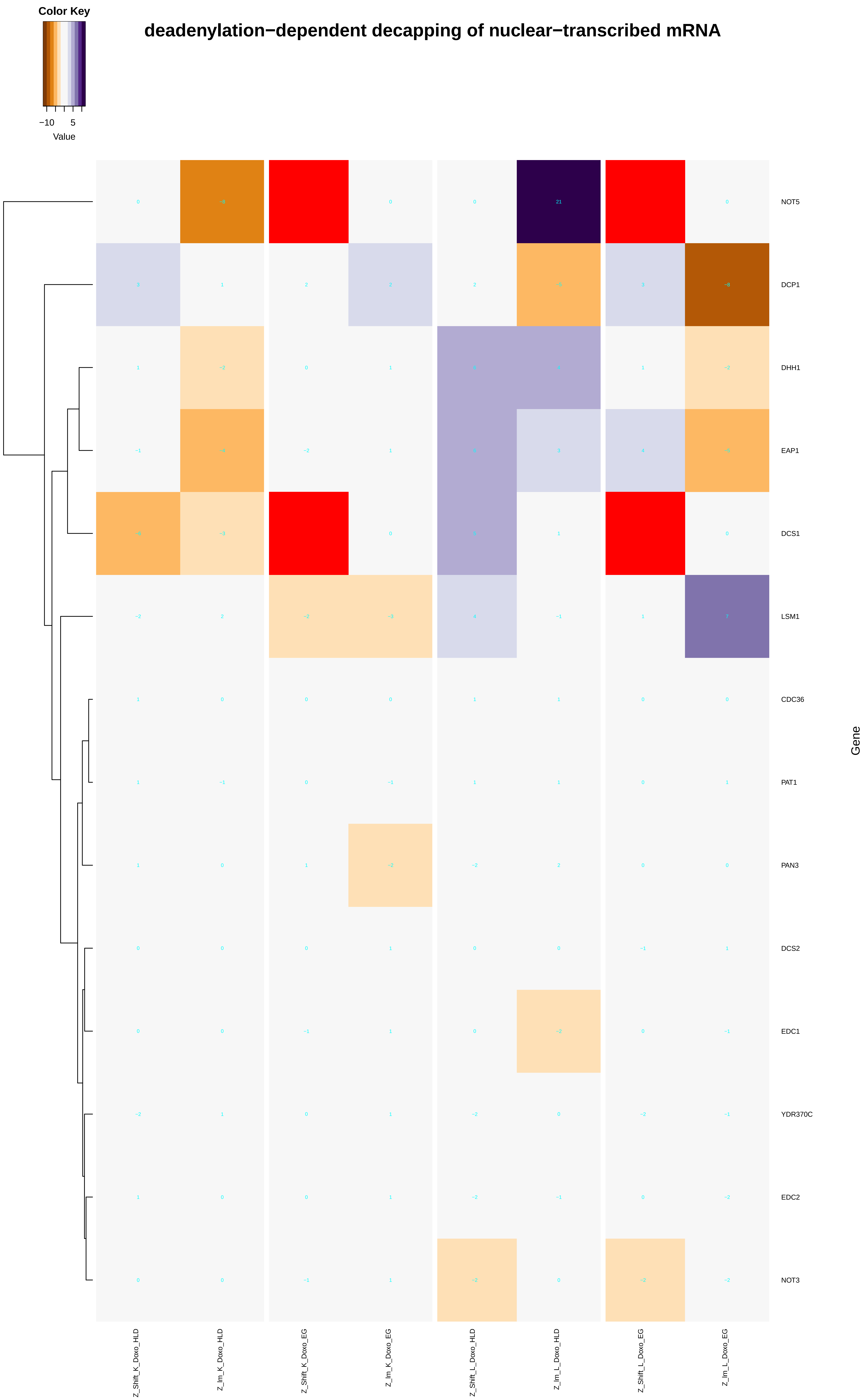

deadenylation−dependent decapping of nuclear−transcribed mRNA
Color Key
−10
5
Value
NOT5
DCP1
DHH1
EAP1
DCS1
LSM1
CDC36
PAT1
PAN3
DCS2
EDC1
YDR370C
EDC2
NOT3
Gene
Z_Shift_K_Doxo_HLD
Z_Im_K_Doxo_HLD
Z_Shift_K_Doxo_EG
Z_Im_K_Doxo_EG
Z_Shift_L_Doxo_HLD
Z_Im_L_Doxo_HLD
Z_Shift_L_Doxo_EG
Z_Im_L_Doxo_EG

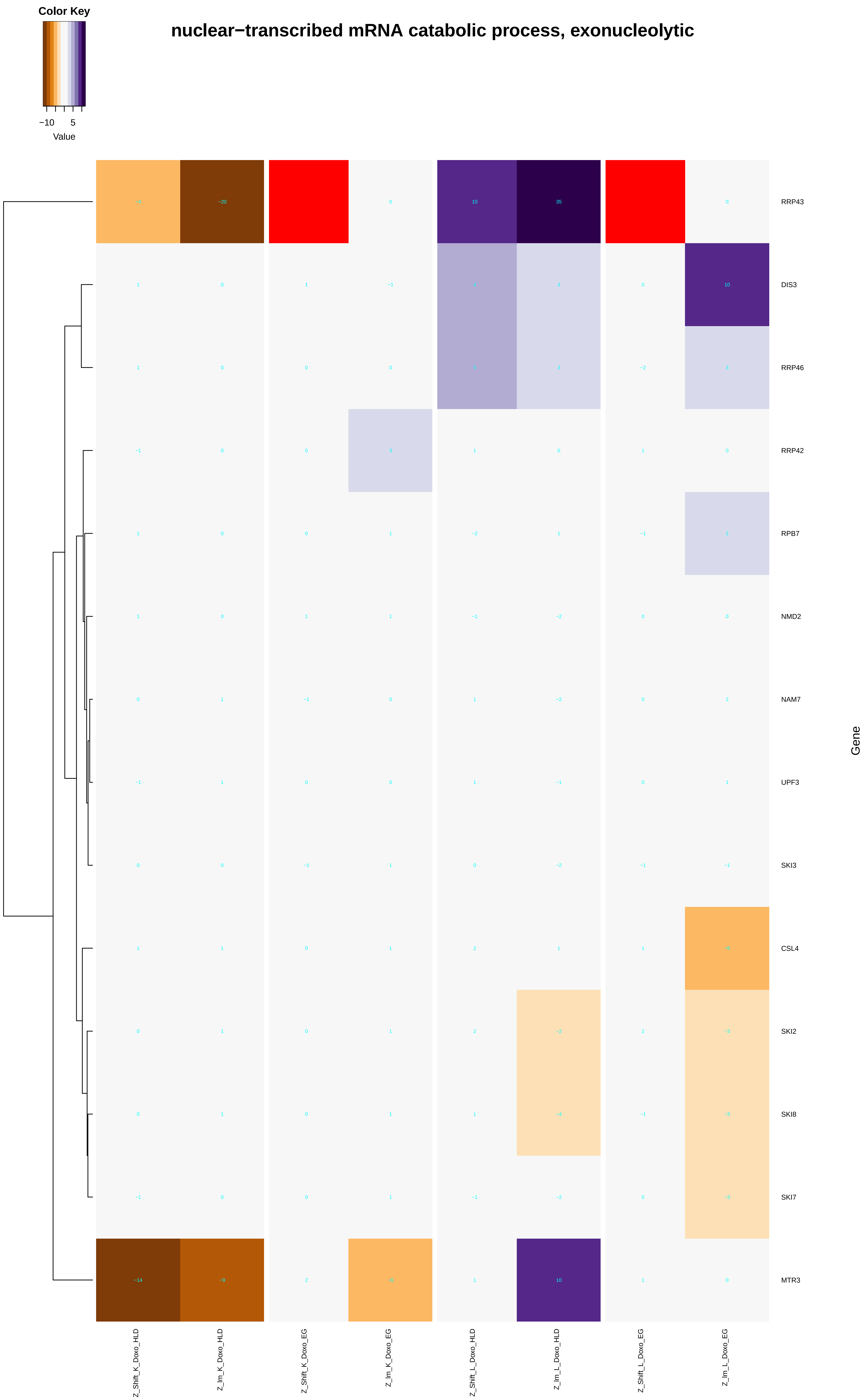

Color Key
−10
5
Value
nuclear−transcribed mRNA catabolic process, exonucleolytic
RRP43
DIS3
RRP46
RRP42
RPB7
NMD2
NAM7
UPF3
SKI3
CSL4
SKI2
SKI8
SKI7
MTR3
Gene
Z_Shift_K_Doxo_HLD
Z_Im_K_Doxo_HLD
Z_Shift_K_Doxo_EG
Z_Im_K_Doxo_EG
Z_Shift_L_Doxo_HLD
Z_Im_L_Doxo_HLD
Z_Shift_L_Doxo_EG
Z_Im_L_Doxo_EG

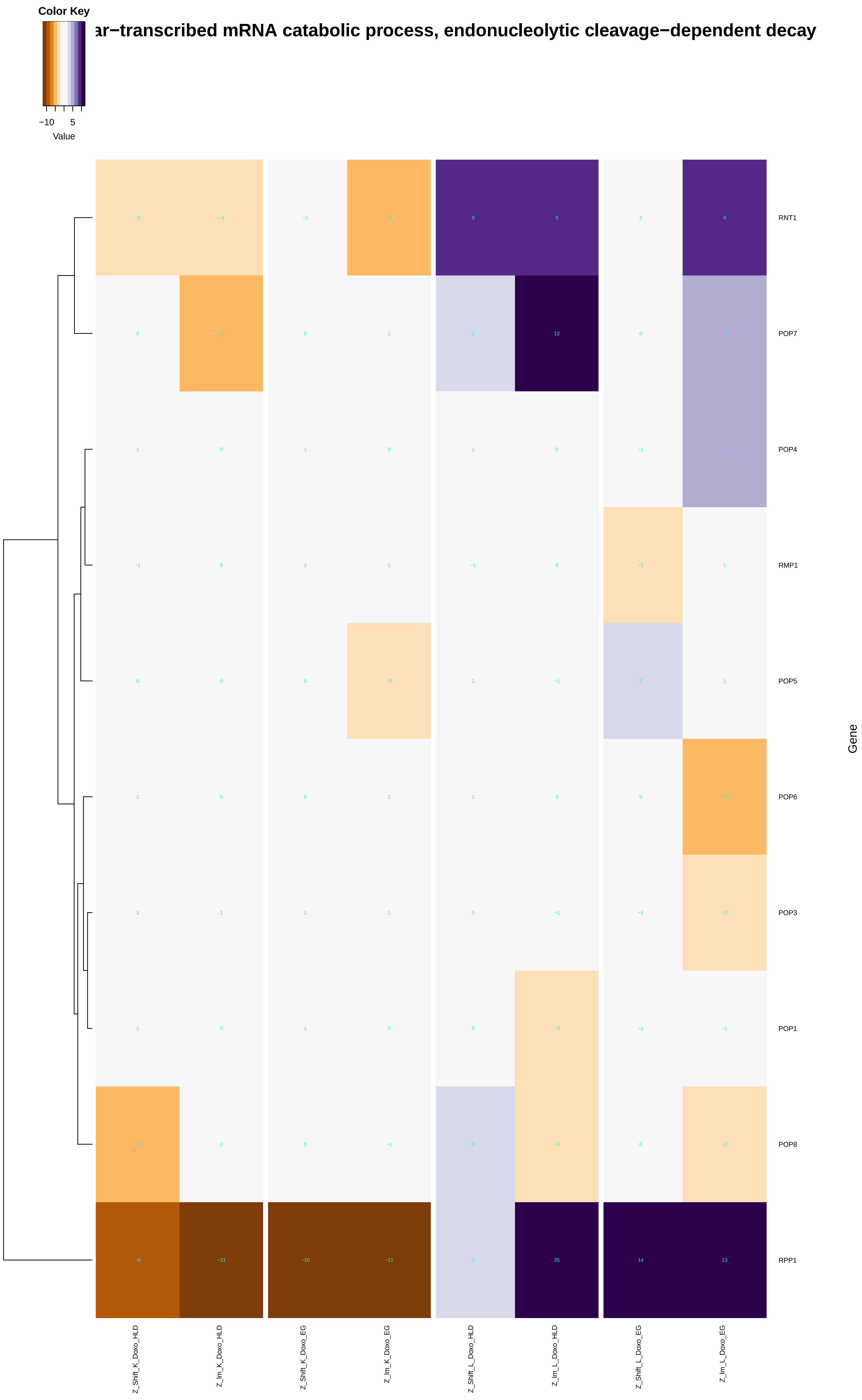

Color Key
ar−transcribed mRNA catabolic process, endonucleolytic cleavage−dependent decay
−10
5
Value
RNT1
POP7
POP4
RMP1
POP5
POP6
POP3
POP1
POP8
RPP1
Gene
Z_Shift_K_Doxo_HLD
Z_Im_K_Doxo_HLD
Z_Shift_K_Doxo_EG
Z_Im_K_Doxo_EG
Z_Shift_L_Doxo_HLD
Z_Im_L_Doxo_HLD
Z_Shift_L_Doxo_EG
Z_Im_L_Doxo_EG

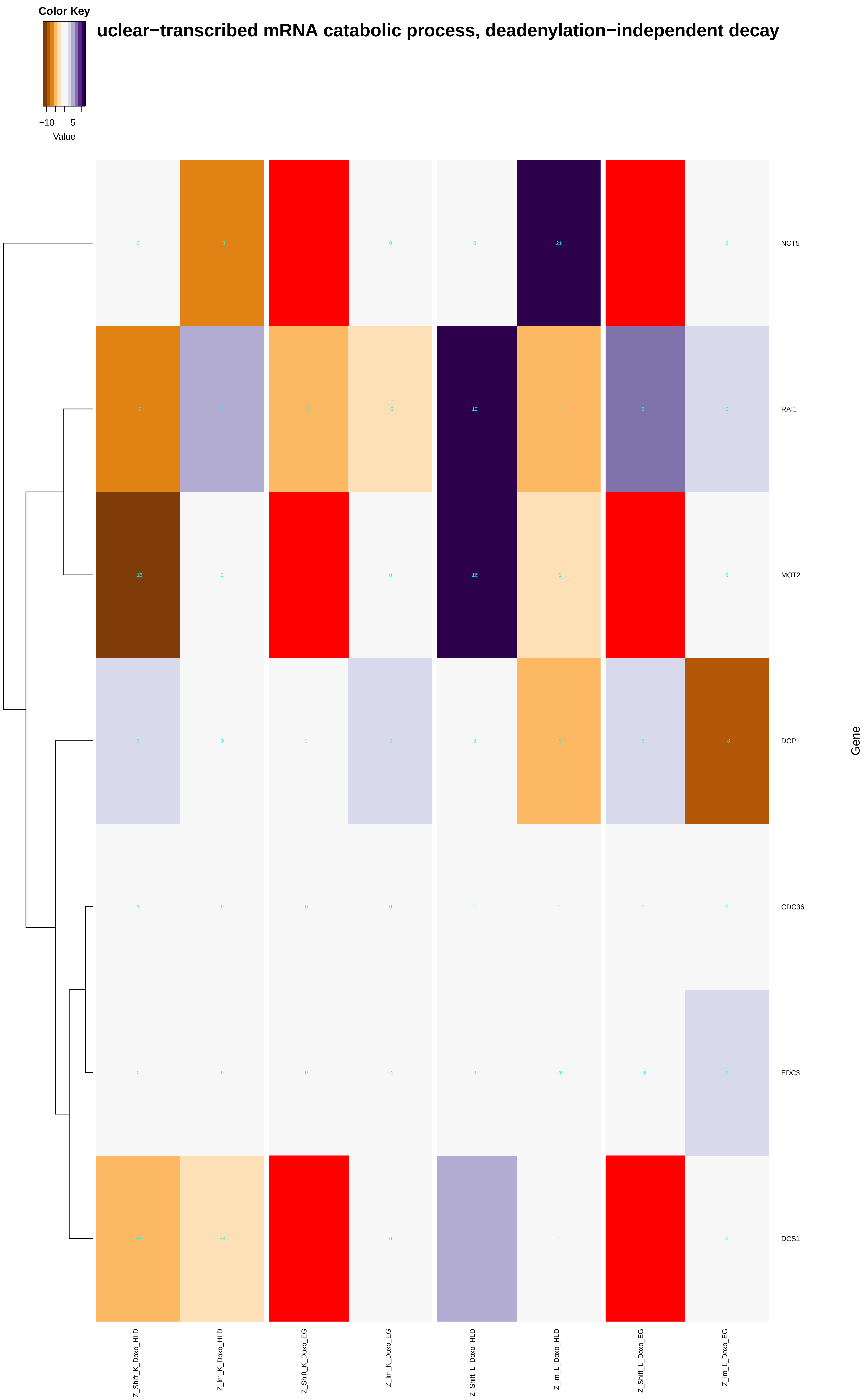

Color Key
−10
5
Value
uclear−transcribed mRNA catabolic process, deadenylation−independent decay
NOT5
RAI1
MOT2
DCP1
CDC36
EDC3
DCS1
Gene
Z_Shift_K_Doxo_HLD
Z_Im_K_Doxo_HLD
Z_Shift_K_Doxo_EG
Z_Im_K_Doxo_EG
Z_Shift_L_Doxo_HLD
Z_Im_L_Doxo_HLD
Z_Shift_L_Doxo_EG
Z_Im_L_Doxo_EG

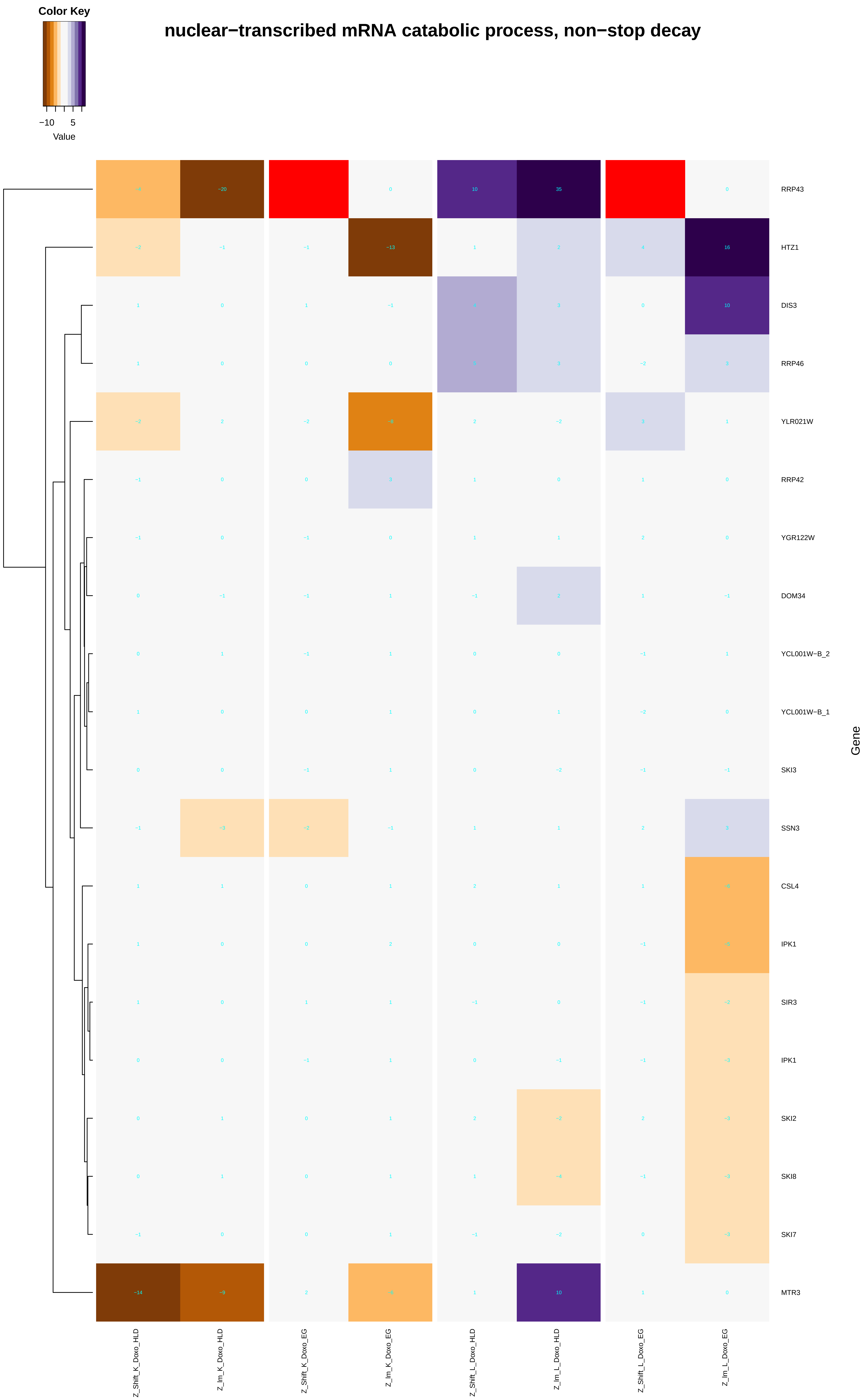

Color Key
−10
5
Value
nuclear−transcribed mRNA catabolic process, non−stop decay
RRP43
HTZ1
DIS3
RRP46
YLR021W
RRP42
YGR122W
DOM34
YCL001W−B_2
YCL001W−B_1
SKI3
SSN3
CSL4
IPK1
SIR3
IPK1
SKI2
SKI8
SKI7
MTR3
Gene
Z_Shift_K_Doxo_HLD
Z_lm_K_Doxo_HLD
Z_Shift_K_Doxo_EG
Z_lm_K_Doxo_EG
Z_Shift_L_Doxo_HLD
Z_lm_L_Doxo_HLD
Z_Shift_L_Doxo_EG
Z_lm_L_Doxo_EG

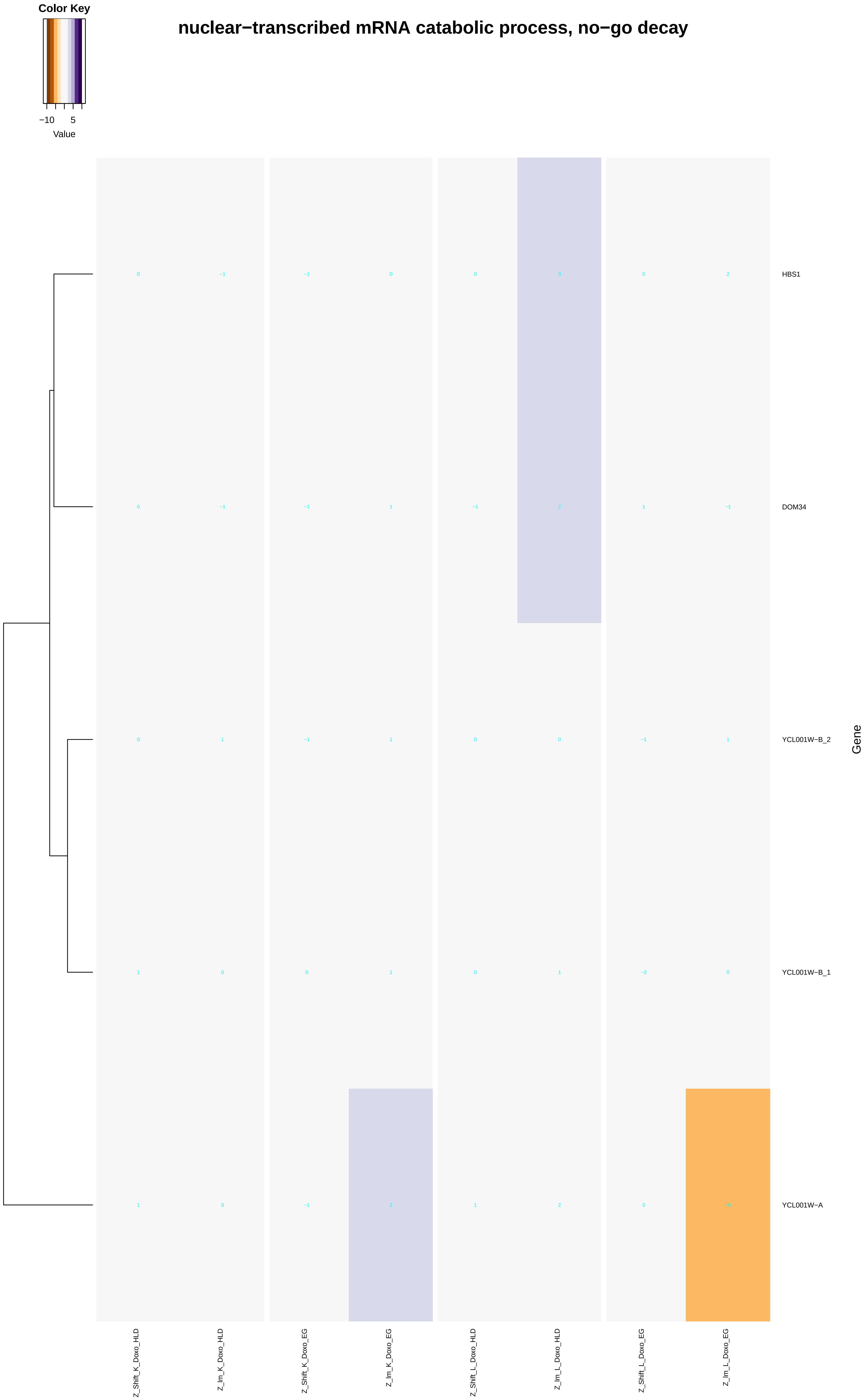
nuclear−transcribed mRNA catabolic process, no−go decay
Color Key
−10
5
Value
HBS1
DOM34
YCL001W−B_2
YCL001W−B_1
YCL001W−A
Gene
Z_Shift_K_Doxo_HLD
Z_Im_K_Doxo_HLD
Z_Shift_K_Doxo_EG
Z_Im_K_Doxo_EG
Z_Shift_L_Doxo_HLD
Z_Im_L_Doxo_HLD
Z_Shift_L_Doxo_EG
Z_Im_L_Doxo_EG

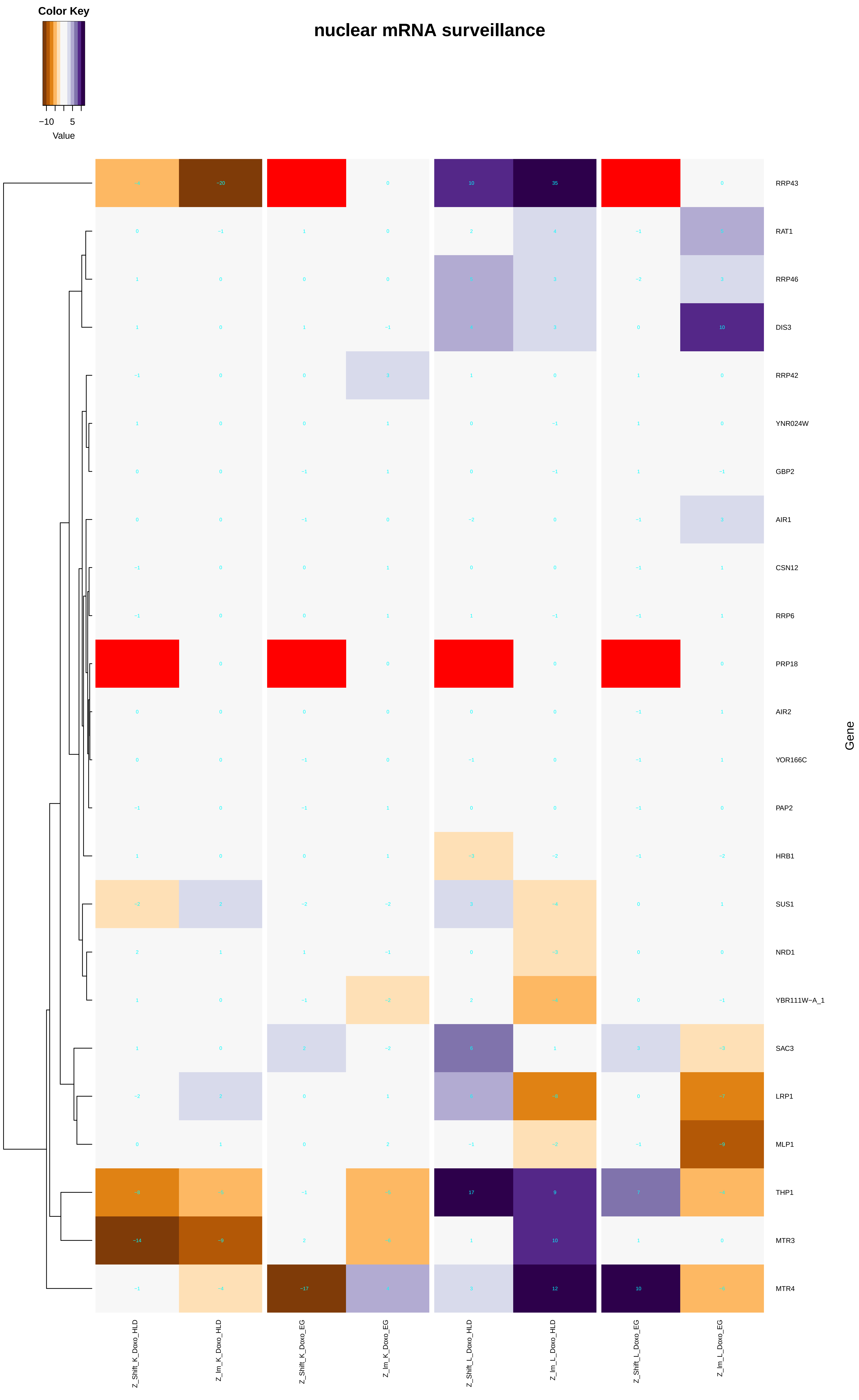
nuclear mRNA surveillance
Color Key
−10
5
Value
Gene
RRP43
RAT1
RRP46
DIS3
RRP42
YNR024W
GBP2
AIR1
CSN12
RRP6
PRP18
AIR2
YOR166C
PAP2
HRB1
SUS1
NRD1
YBR111W−A_1
SAC3
LRP1
MLP1
THP1
MTR3
MTR4
Z_Shift_K_Doxo_HLD
Z_Im_K_Doxo_HLD
Z_Shift_K_Doxo_EG
Z_Im_K_Doxo_EG
Z_Shift_L_Doxo_HLD
Z_Im_L_Doxo_HLD
Z_Shift_L_Doxo_EG
Z_Im_L_Doxo_EG

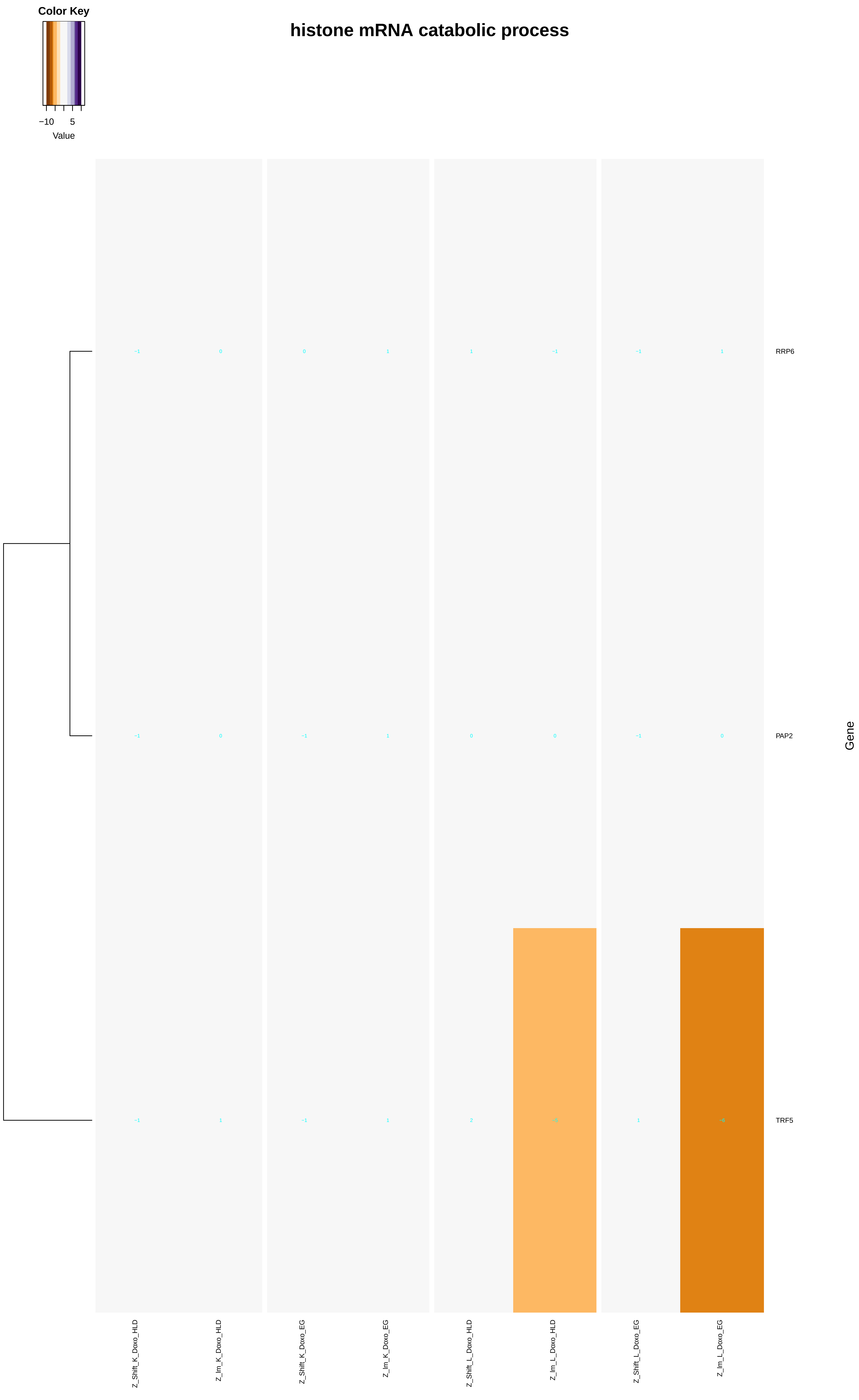

Color Key
−10
5
Value
histone mRNA catabolic process
RRP6
PAP2
TRF5
Gene
Z_Shift_K_Doxo_HLD
Z_lm_K_Doxo_HLD
Z_Shift_K_Doxo_EG
Z_lm_K_Doxo_EG
Z_Shift_L_Doxo_HLD
Z_lm_L_Doxo_HLD
Z_Shift_L_Doxo_EG
Z_lm_L_Doxo_EG

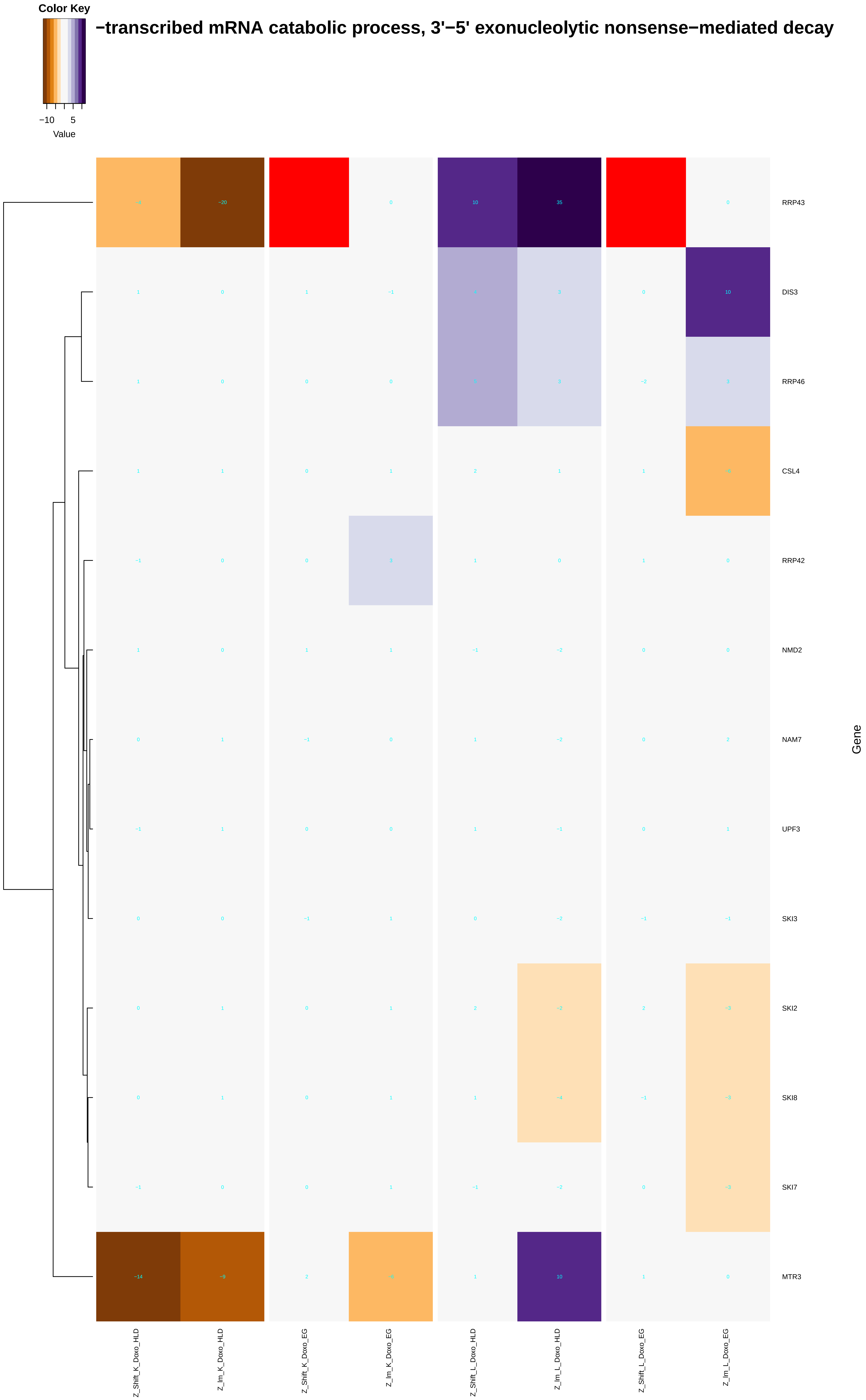
Color Key
−10
5
Value
−transcribed mRNA catabolic process, 3'−5' exonucleolytic nonsense−mediated decay
RRP43
DIS3
RRP46
CSL4
RRP42
NMD2
NAM7
UPF3
SKI3
SKI2
SKI8
SKI7
MTR3
Gene
Z_Shift_K_Doxo_HLD
Z_lm_K_Doxo_HLD
Z_Shift_K_Doxo_EG
Z_lm_K_Doxo_EG
Z_Shift_L_Doxo_HLD
Z_lm_L_Doxo_HLD
Z_Shift_L_Doxo_EG
Z_lm_L_Doxo_EG

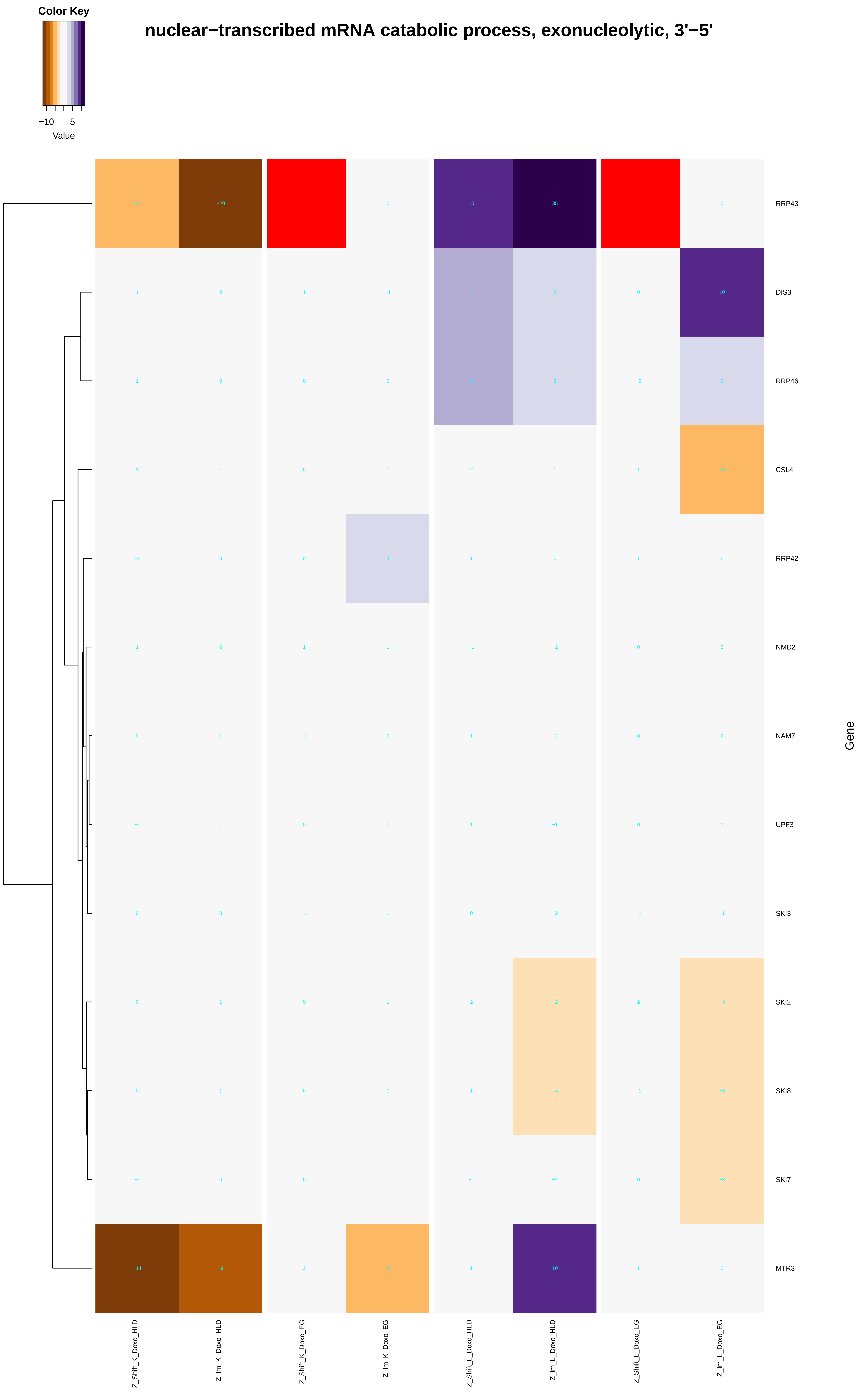
nuclear−transcribed mRNA catabolic process, exonucleolytic, 3'−5'
Color Key
−10
5
Value
RRP43
DIS3
RRP46
CSL4
RRP42
NMD2
NAM7
UPF3
SKI3
SKI2
SKI8
SKI7
MTR3
Gene
Z_Shift_K_Doxo_HLD
Z_Im_K_Doxo_HLD
Z_Shift_K_Doxo_EG
Z_Im_K_Doxo_EG
Z_Shift_L_Doxo_HLD
Z_Im_L_Doxo_HLD
Z_Shift_L_Doxo_EG
Z_Im_L_Doxo_EG

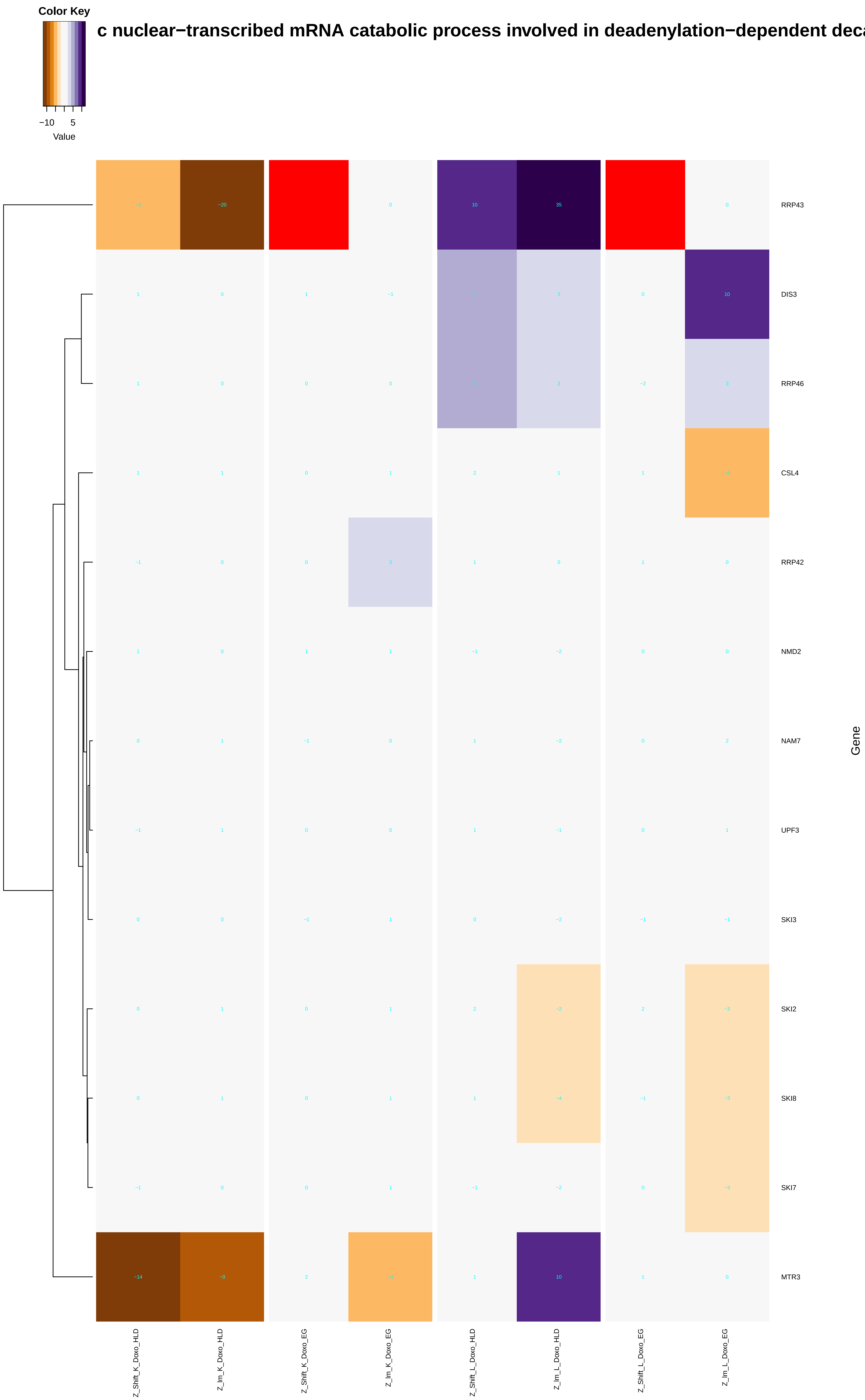
c nuclear−transcribed mRNA catabolic process involved in deadenylation−dependent deca
Color Key
−10
5
Value
RRP43
DIS3
RRP46
CSL4
RRP42
NMD2
NAM7
UPF3
SKI3
SKI2
SKI8
SKI7
MTR3
Gene
Z_Shift_K_Doxo_HLD
Z_Im_K_Doxo_HLD
Z_Shift_K_Doxo_EG
Z_Im_K_Doxo_EG
Z_Shift_L_Doxo_HLD
Z_Im_L_Doxo_HLD
Z_Shift_L_Doxo_EG
Z_Im_L_Doxo_EG

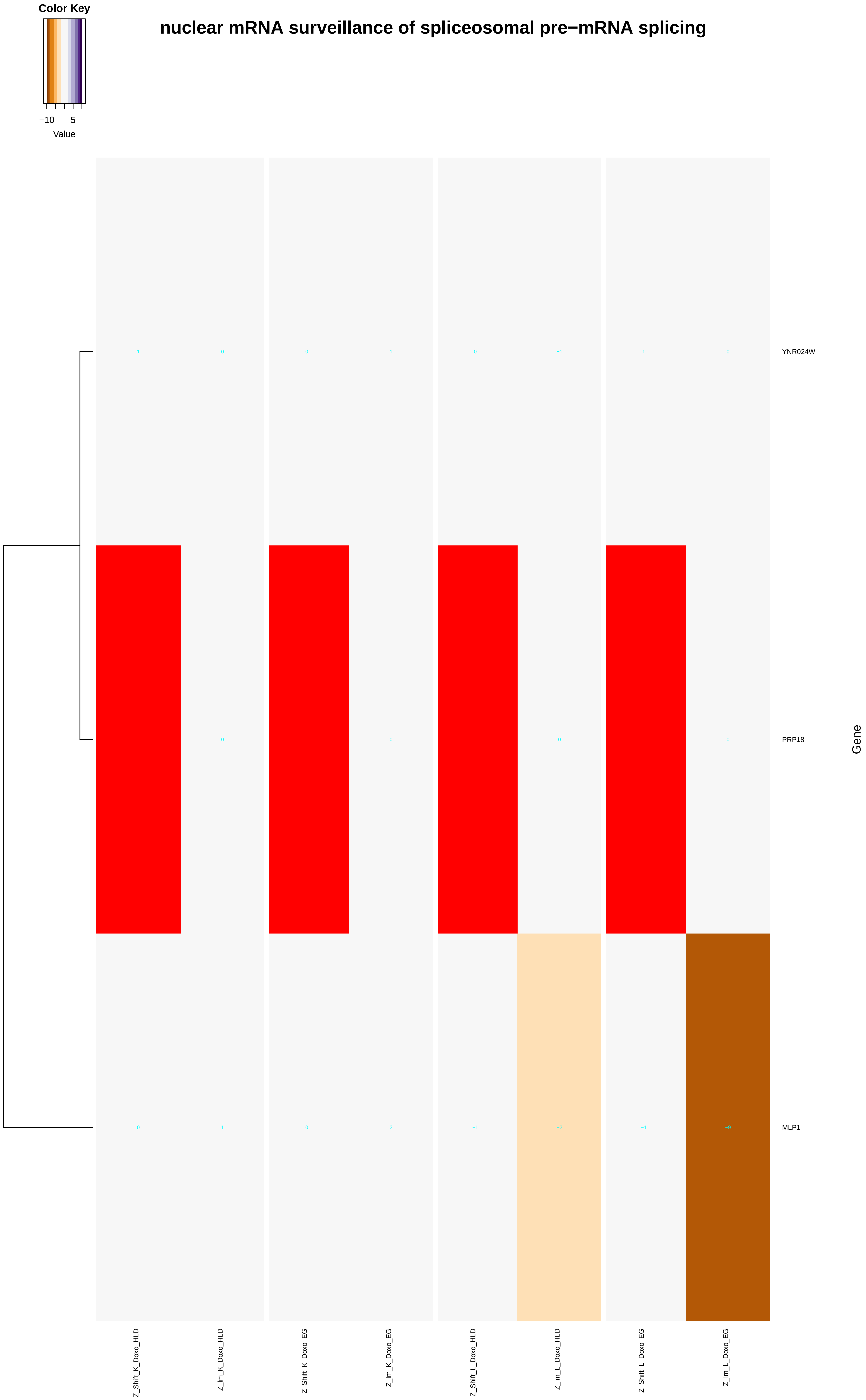

nuclear mRNA surveillance of spliceosomal pre−mRNA splicing
Color Key
−10
5
Value
YNR024W
PRP18
MLP1
Gene
Z_Shift_K_Doxo_HLD
Z_Im_K_Doxo_HLD
Z_Shift_K_Doxo_EG
Z_Im_K_Doxo_EG
Z_Shift_L_Doxo_HLD
Z_Im_L_Doxo_HLD
Z_Shift_L_Doxo_EG
Z_Im_L_Doxo_EG
1 0 0 1 0 −1 1 0
0 0 0 0
0 1 0 2 −1 −2 −1 −9

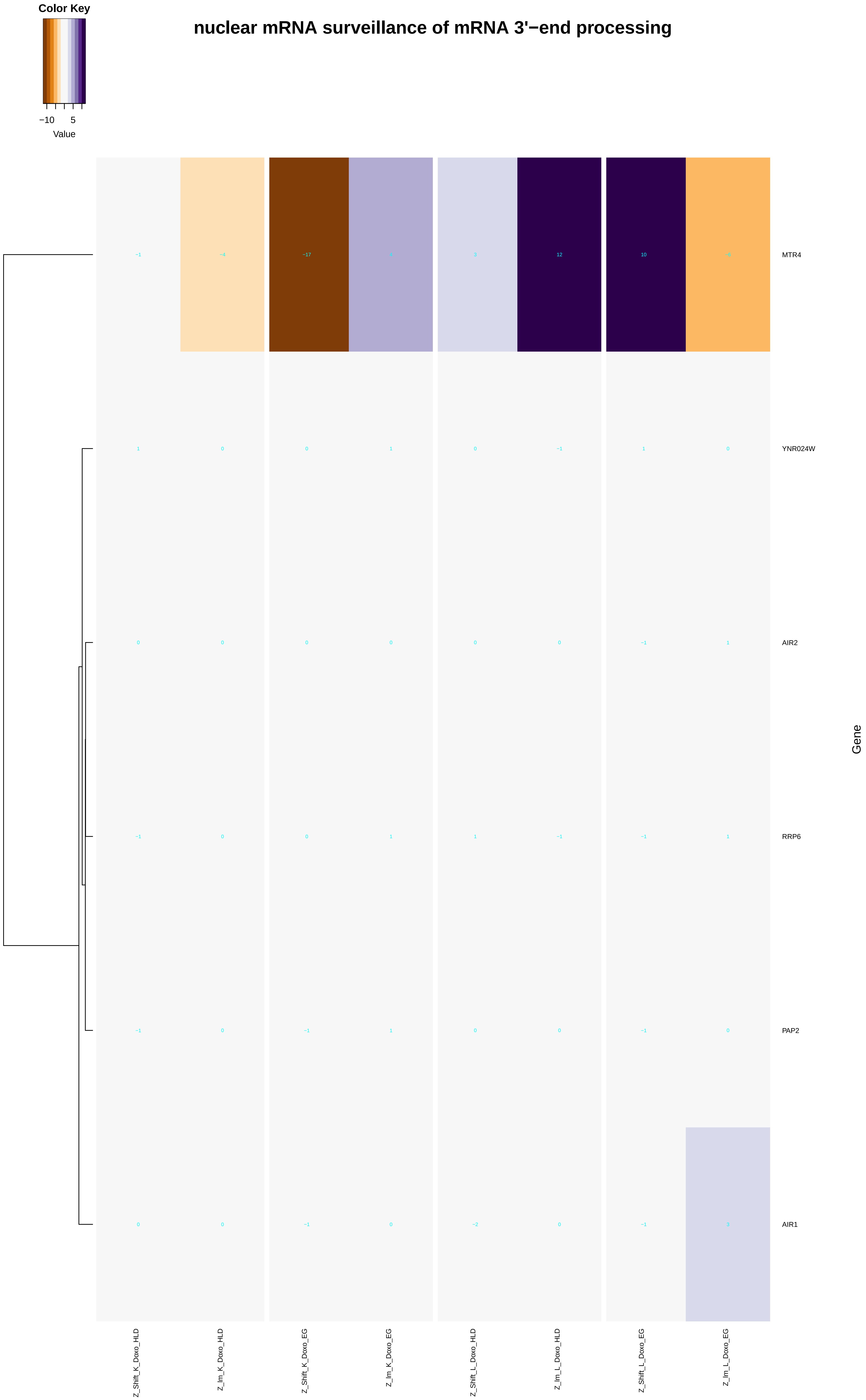

nuclear mRNA surveillance of mRNA 3'−end processing
Color Key
−10
5
Value
MTR4
YNR024W
AIR2
RRP6
PAP2
AIR1
Gene
Z_Shift_K_Doxo_HLD
Z_Im_K_Doxo_HLD
Z_Shift_K_Doxo_EG
Z_Im_K_Doxo_EG
Z_Shift_L_Doxo_HLD
Z_Im_L_Doxo_HLD
Z_Shift_L_Doxo_EG
Z_Im_L_Doxo_EG
