## Additional File 9 for "A yeast phenomic model for the influence of Warburg metabolism on genetic buffering of doxorubicin": nuclear_transport.pdf

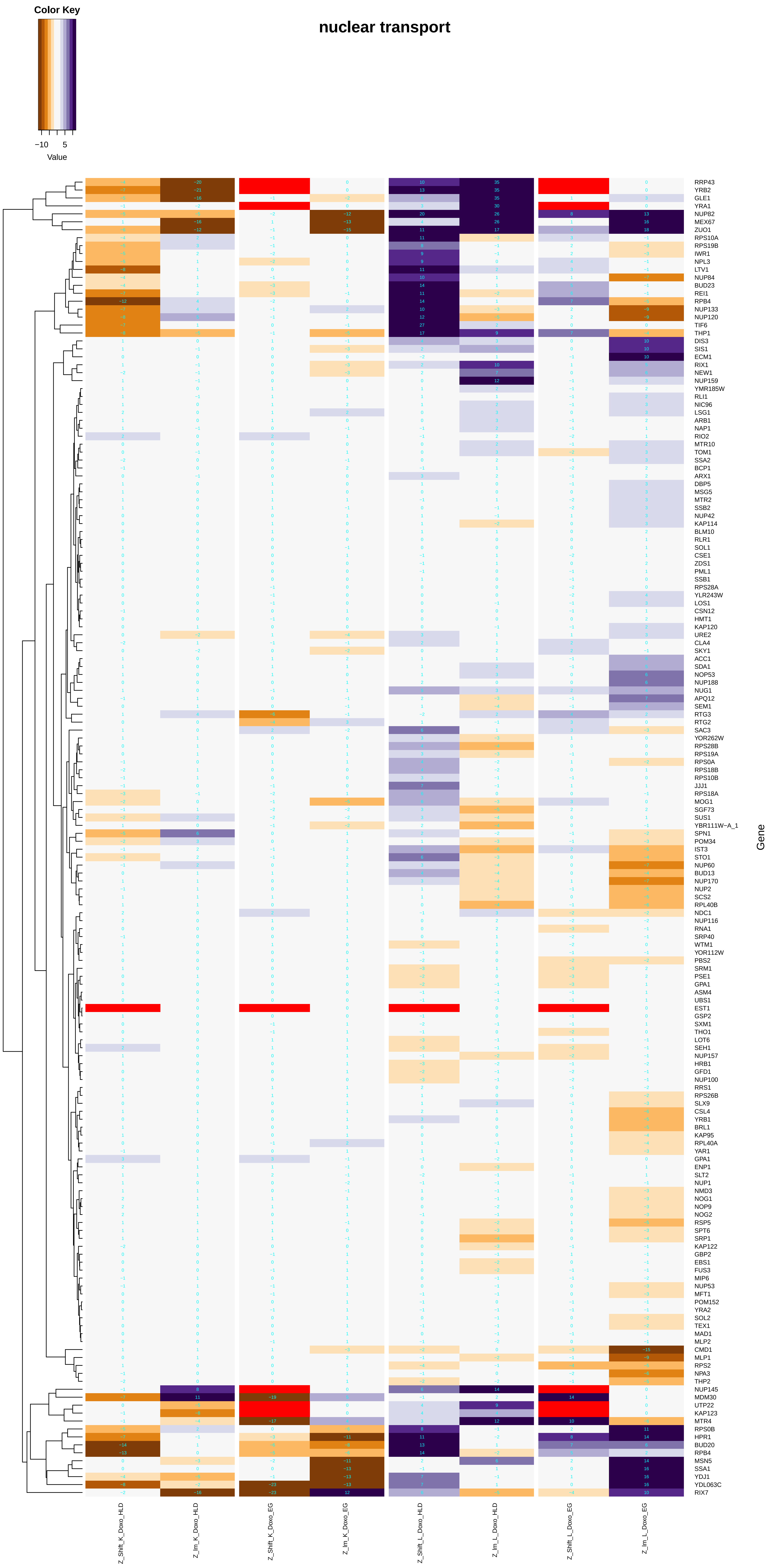
nuclear transport
Color Key
−10
5
Value
RRP43
YRB2
GLE1
YRA1
NUP82
MEX67
ZUO1
RPS10A
RPS19B
IWR1
NPL3
LTV1
NUP84
BUD23
REI1
RPB4
NUP133
NUP120
TIF6
THP1
DIS3
SIS1
ECM1
RIX1
NEW1
NUP159
YMR185W
RLI1
NIC96
LSG1
ARB1
NAP1
RIO2
MTR10
TOM1
SSA2
BCP1
ARX1
DBP5
MSG5
MTR2
SSB2
NUP42
KAP114
BLM10
RLR1
SOL1
CSE1
ZDS1
PML1
SSB1
RPS28A
YLR243W
LOS1
CSN12
HMT1
KAP120
URE2
CLA4
SKY1
ACC1
SDA1
NOP53
NUP188
NUG1
APQ12
SEM1
RTG3
RTG2
SAC3
YOR262W
RPS28B
RPS19A
RPS0A
RPS18B
RPS10B
JJJ1
RPS18A
MOG1
SGF73
SUS1
YBR111W−A_1
SPN1
POM34
IST3
STO1
NUP60
BUD13
NUP170
NUP2
SCS2
RPL40B
NDC1
NUP116
RNA1
SRP40
WTM1
YOR112W
PBS2
SRM1
PSE1
GPA1
ASM4
UBS1
EST1
GSP2
SXM1
THO1
LOT6
SEH1
NUP157
HRB1
GFD1
NUP100
RRS1
RPS26B
SLX9
CSL4
YRB1
BRL1
KAP95
RPL40A
YAR1
GPA1
ENP1
SLT2
NUP1
NMD3
NOG1
NOP9
NOG2
RSP5
SPT6
SRP1
KAP122
GBP2
EBS1
FUS3
MIP6
NUP53
MFT1
POM152
YRA2
SOL2
TEX1
MAD1
MLP2
CMD1
MLP1
RPS2
NPA3
THP2
NUP145
MDM30
UTP22
KAP123
MTR4
RPS0B
HPR1
BUD20
RPB4
MSN5
SSA1
YDJ1
YDL063C
RIX7
Gene
Z_Shift_K_Doxo_HLD
Z_Im_K_Doxo_HLD
Z_Shift_K_Doxo_EG
Z_Im_K_Doxo_EG
Z_Shift_L_Doxo_HLD
Z_Im_L_Doxo_HLD
Z_Shift_L_Doxo_EG
Z_Im_L_Doxo_EG

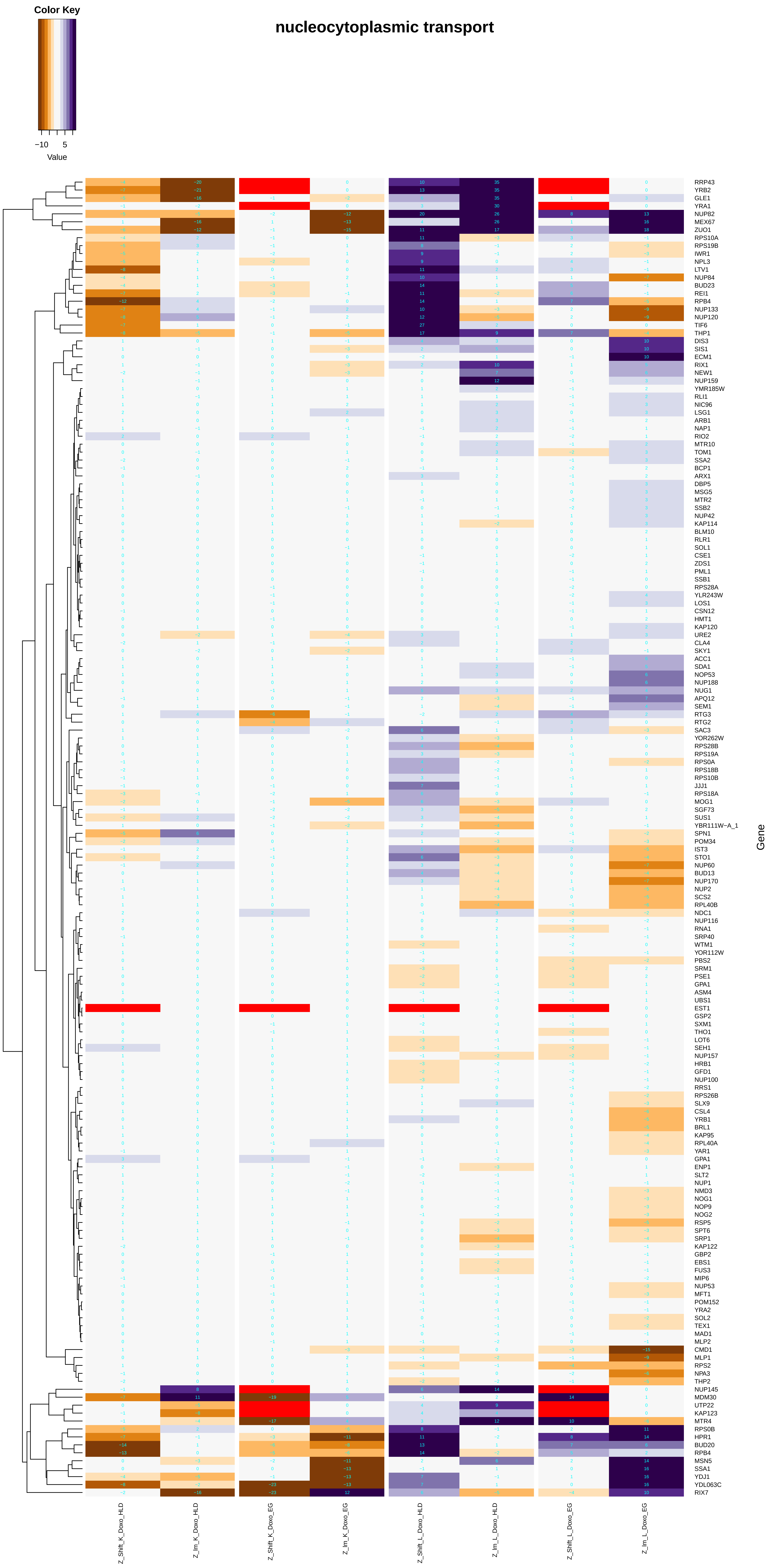

nucleocytoplasmic transport
Color Key
−10
5
Value
Gene
Z_Shift_K_Doxo_HLD
Z_lm_K_Doxo_HLD
Z_Shift_K_Doxo_EG
Z_lm_K_Doxo_EG
Z_Shift_L_Doxo_HLD
Z_lm_L_Doxo_HLD
Z_Shift_L_Doxo_EG
Z_lm_L_Doxo_EG
RRP43
YRB2
GLE1
YRA1
NUP82
MEX67
ZUO1
RPS10A
RPS19B
IWR1
NPL3
LTV1
NUP84
BUD23
REI1
RPB4
NUP133
NUP120
TIF6
THP1
DIS3
SIS1
ECM1
RIX1
NEW1
NUP159
YMR185W
RLI1
NIC96
LSG1
ARB1
NAP1
RIO2
MTR10
TOM1
SSA2
BCP1
ARX1
DBP5
MSG5
MTR2
SSB2
NUP42
KAP114
BLM10
RLR1
SOL1
CSE1
ZDS1
PML1
SSB1
RPS28A
YLR243W
LOS1
CSN12
HMT1
KAP120
URE2
CLA4
SKY1
ACC1
SDA1
NOP53
NUP188
NUG1
APQ12
SEM1
RTG3
RTG2
SAC3
YOR262W
RPS28B
RPS19A
RPS0A
RPS18B
RPS10B
JJJ1
RPS18A
MOG1
SGF73
SUS1
YBR111W-A_1
SPN1
POM34
IST3
STO1
NUP60
BUD13
NUP170
NUP2
SCS2
RPL40B
NDC1
NUP116
RNA1
SRP40
WTM1
YOR112W
PBS2
SRM1
PSE1
GPA1
ASM4
UBS1
EST1
GSP2
SXM1
THO1
LOT6
SEH1
NUP157
HRB1
GFD1
NUP100
RRS1
RPS26B
SLX9
CSL4
YRB1
BRL1
KAP95
RPL40A
YAR1
GPA1
ENP1
SLT2
NUP1
NMD3
NOG1
NOP9
NOG2
RSP5
SPT6
SRP1
KAP122
GBP2
EBS1
FUS3
MIP6
NUP53
MFT1
POM152
YRA2
SOL2
TEX1
MAD1
MLP2
CMD1
MLP1
RPS2
NPA3
THP2
NUP145
MDM30
UTP22
KAP123
MTR4
RPS0B
HPR1
BUD20
RPB4
MSN5
SSA1
YDJ1
YDL063C
RIX7

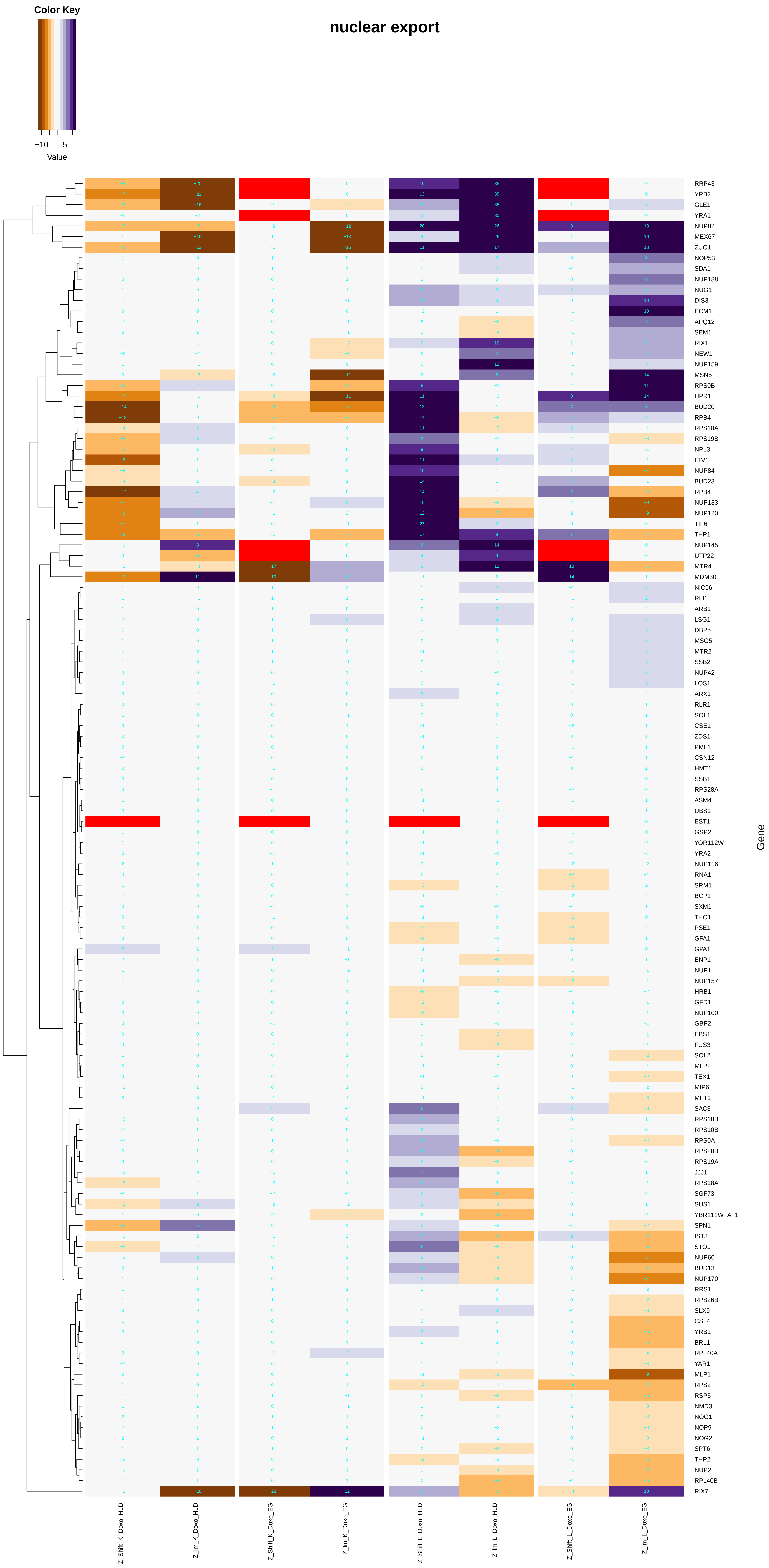

nuclear export
Color Key
−10
5
Value
RRP43
YRB2
GLE1
YRA1
NUP82
MEX67
ZUO1
NOP53
SDA1
NUP188
NUG1
DIS3
ECM1
APQ12
SEM1
RIX1
NEW1
NUP159
MSN5
RPS0B
HPR1
BUD20
RPB4
RPS10A
RPS19B
NPL3
LTV1
NUP84
BUD23
RPB4
NUP133
NUP120
TIF6
THP1
NUP145
UTP22
MTR4
MDM30
NIC96
RLI1
ARB1
LSG1
DBP5
MSG5
MTR2
SSB2
NUP42
LOS1
ARX1
RLR1
SOL1
CSE1
ZDS1
PML1
CSN12
HMT1
SSB1
RPS28A
ASM4
UBS1
EST1
GSP2
YOR112W
YRA2
NUP116
RNA1
SRM1
BCP1
SXM1
THO1
PSE1
GPA1
GPA1
ENP1
NUP1
NUP157
HRB1
GFD1
NUP100
GBP2
EBS1
FUS3
SOL2
MLP2
TEX1
MIP6
MFT1
SAC3
RPS18B
RPS10B
RPS0A
RPS28B
RPS19A
JJJ1
RPS18A
SGF73
SUS1
YBR111W−A_1
SPN1
IST3
STO1
NUP60
BUD13
NUP170
RRS1
RPS26B
SLX9
CSL4
YRB1
BRL1
RPL40A
YAR1
MLP1
RPS2
RSP5
NMD3
NOG1
NOP9
NOG2
SPT6
THP2
NUP2
RPL40B
RIX7
Gene
Z_Shift_K_Doxo_HLD
Z_Im_K_Doxo_HLD
Z_Shift_K_Doxo_EG
Z_Im_K_Doxo_EG
Z_Shift_L_Doxo_HLD
Z_Im_L_Doxo_HLD
Z_Shift_L_Doxo_EG
Z_Im_L_Doxo_EG

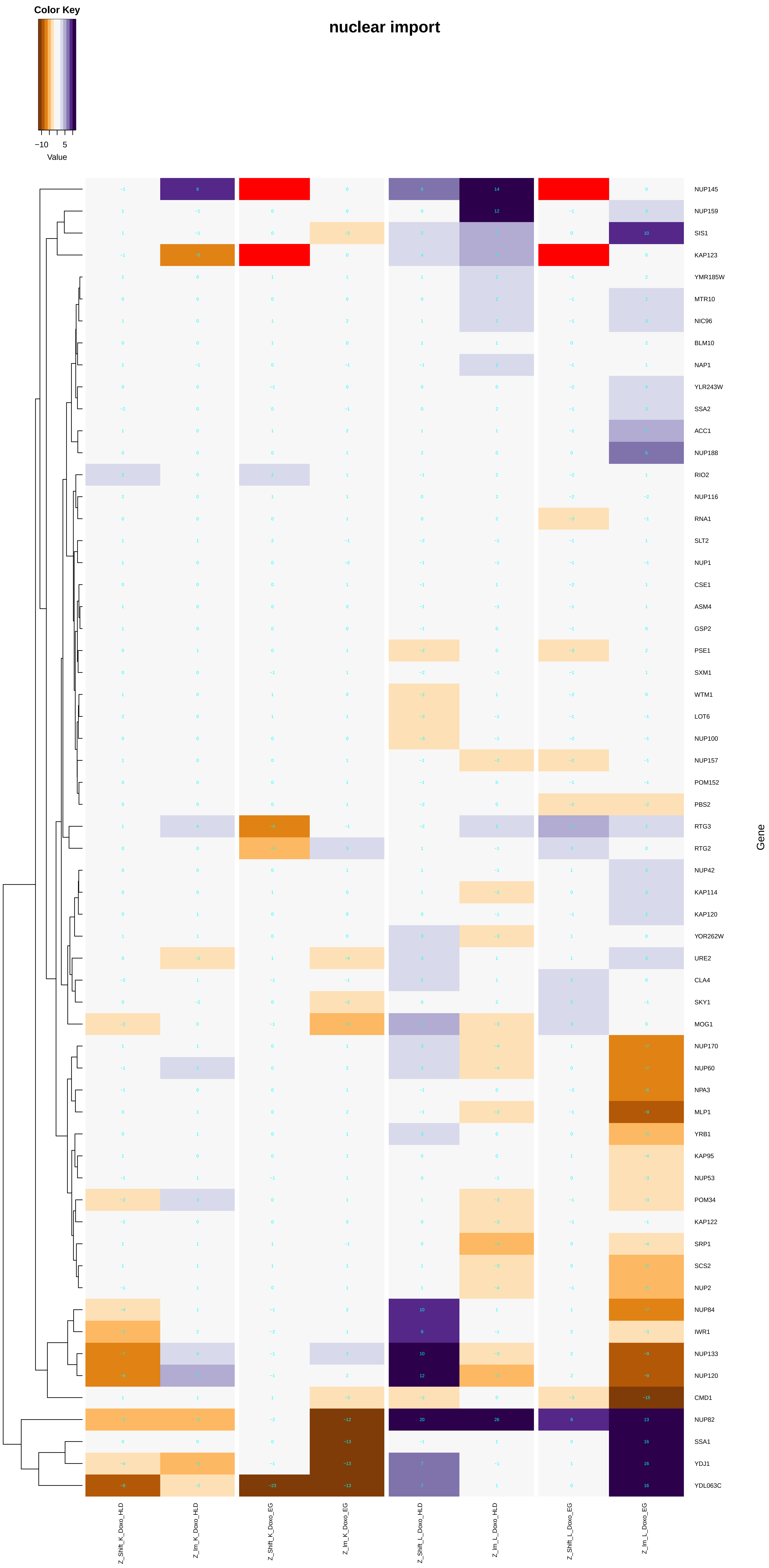

Color Key
−10
5
Value
nuclear import
Gene
NUP145
NUP159
SIS1
KAP123
YMR185W
MTR10
NIC96
BLM10
NAP1
YLR243W
SSA2
ACC1
NUP188
RIO2
NUP116
RNA1
SLT2
NUP1
CSE1
ASM4
GSP2
PSE1
SXM1
WTM1
LOT6
NUP100
NUP157
POM152
PBS2
RTG3
RTG2
NUP42
KAP114
KAP120
YOR262W
URE2
CLA4
SKY1
MOG1
NUP170
NUP60
NPA3
MLP1
YRB1
KAP95
NUP53
POM34
KAP122
SRP1
SCS2
NUP2
NUP84
IWR1
NUP133
NUP120
CMD1
NUP82
SSA1
YDJ1
YDL063C
Z_Shift_K_Doxo_HLD
Z_Im_K_Doxo_HLD
Z_Shift_K_Doxo_EG
Z_Im_K_Doxo_EG
Z_Shift_L_Doxo_HLD
Z_Im_L_Doxo_HLD
Z_Shift_L_Doxo_EG
Z_Im_L_Doxo_EG

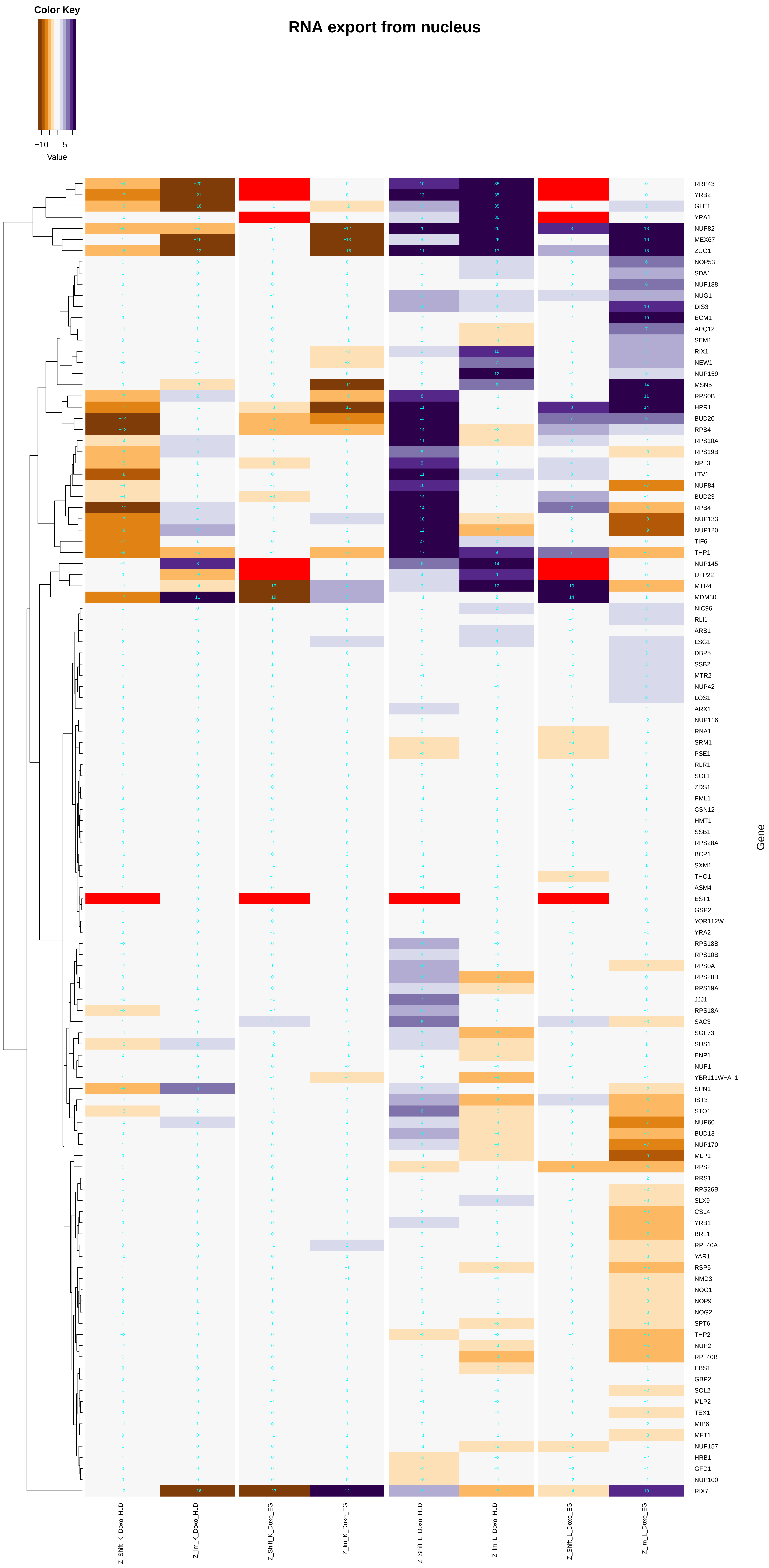
RNA export from nucleus
Color Key
−10
5
Value
RRP43
YRB2
GLE1
YRA1
NUP82
MEX67
ZUO1
NOP53
SDA1
NUP188
NUG1
DIS3
ECM1
APQ12
SEM1
RIX1
NEW1
NUP159
MSN5
RPS0B
HPR1
BUD20
RPB4
RPS10A
RPS19B
NPL3
LTV1
NUP84
BUD23
RPB4
NUP133
NUP120
TIF6
THP1
NUP145
UTP22
MTR4
MDM30
NIC96
RLI1
ARB1
LSG1
DBP5
SSB2
MTR2
NUP42
LOS1
ARX1
NUP116
RNA1
SRM1
PSE1
RLR1
SOL1
ZDS1
PML1
CSN12
HMT1
SSB1
RPS28A
BCP1
SXM1
THO1
ASM4
EST1
GSP2
YOR112W
YRA2
RPS18B
RPS10B
RPS0A
RPS28B
RPS19A
JJJ1
RPS18A
SAC3
SGF73
SUS1
ENP1
NUP1
YBR111W−A_1
SPN1
IST3
STO1
NUP60
BUD13
NUP170
MLP1
RPS2
RRS1
RPS26B
SLX9
CSL4
YRB1
BRL1
RPL40A
YAR1
RSP5
NMD3
NOG1
NOP9
NOG2
SPT6
THP2
NUP2
RPL40B
EBS1
GBP2
SOL2
MLP2
TEX1
MIP6
MFT1
NUP157
HRB1
GFD1
NUP100
RIX7
Gene
Z_Shift_K_Doxo_HLD
Z_Im_K_Doxo_HLD
Z_Shift_K_Doxo_EG
Z_Im_K_Doxo_EG
Z_Shift_L_Doxo_HLD
Z_Im_L_Doxo_HLD
Z_Shift_L_Doxo_EG
Z_Im_L_Doxo_EG

### protein export from nucleus

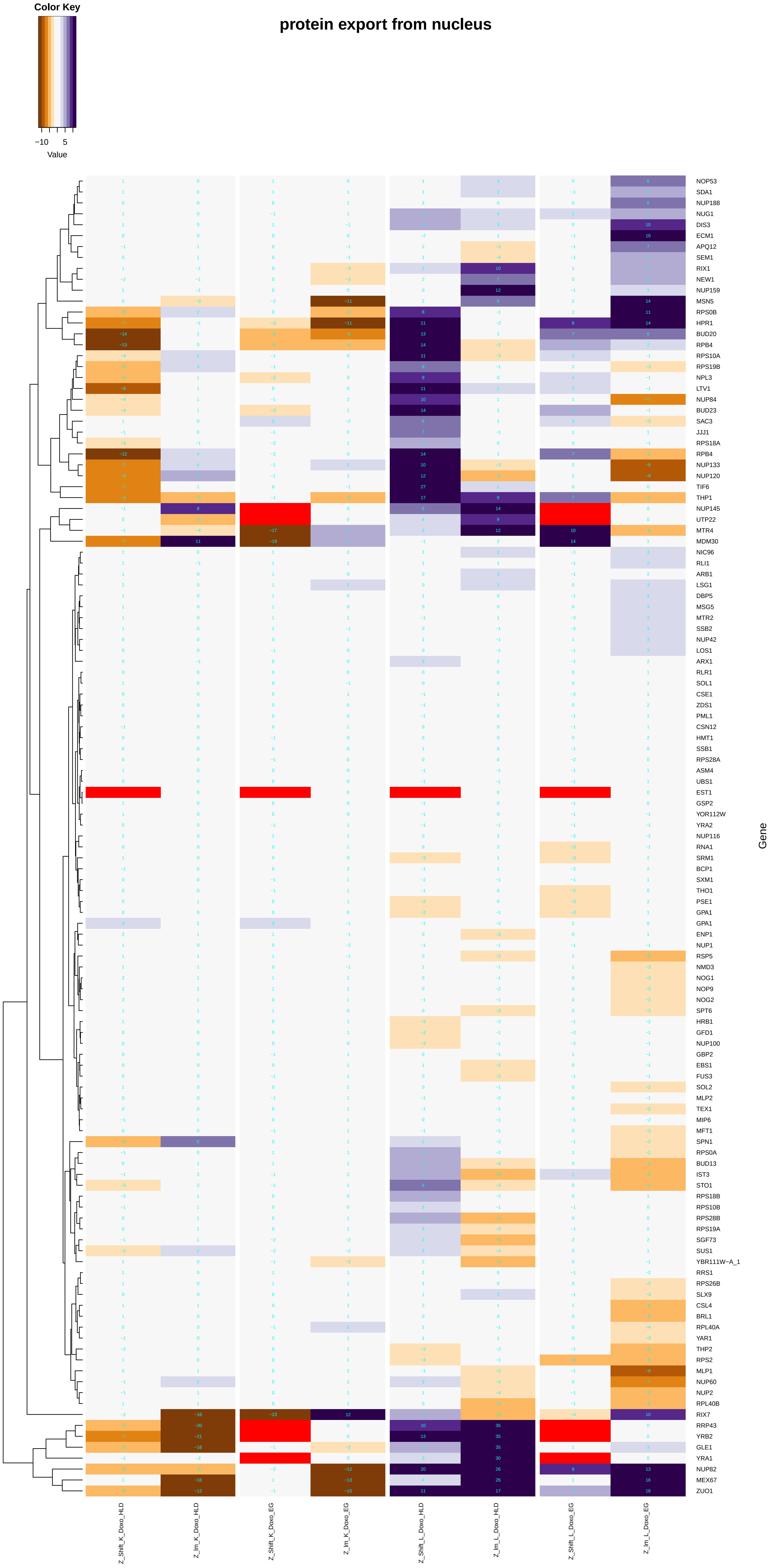

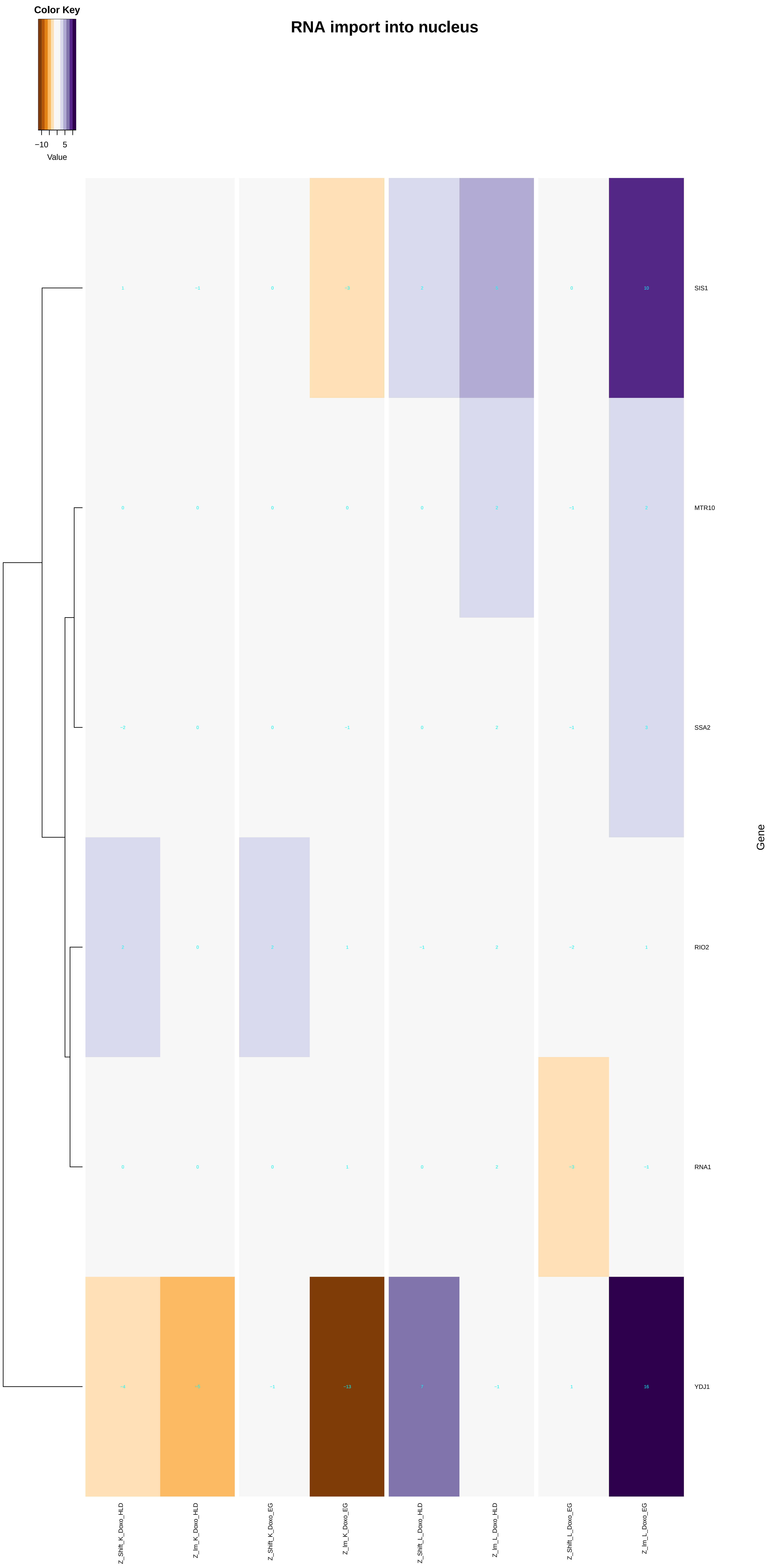
RNA import into nucleus
Color Key
−10
5
Value
SIS1
MTR10
SSA2
RIO2
RNA1
YDJ1
Gene
Z_Shift_K_Doxo_HLD
Z_Im_K_Doxo_HLD
Z_Shift_K_Doxo_EG
Z_Im_K_Doxo_EG
Z_Shift_L_Doxo_HLD
Z_Im_L_Doxo_HLD
Z_Shift_L_Doxo_EG
Z_Im_L_Doxo_EG

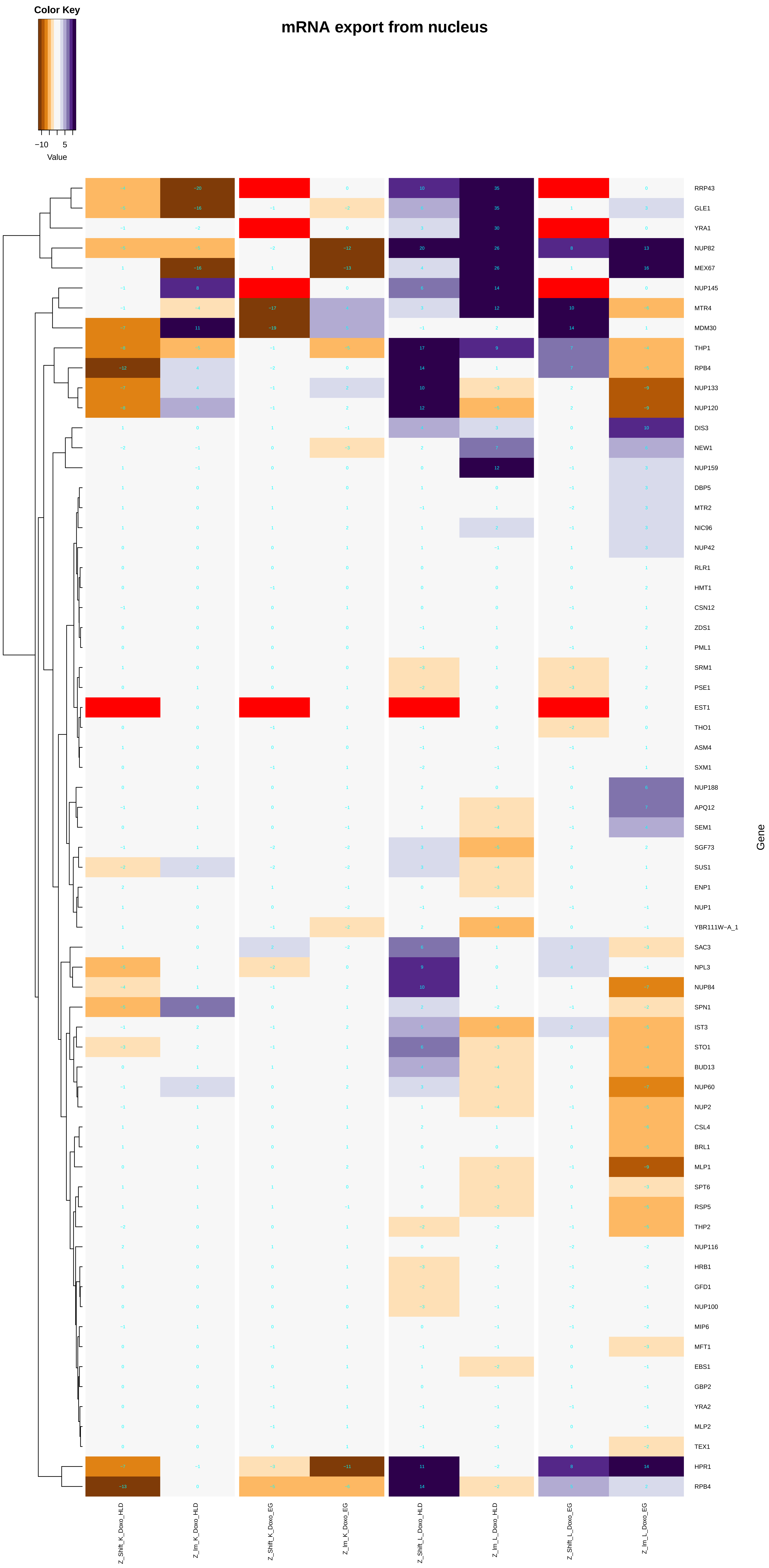

Color Key
−10
5
Value
mRNA export from nucleus
RRP43
GLE1
YRA1
NUP82
MEX67
NUP145
MTR4
MDM30
THP1
RPB4
NUP133
NUP120
DIS3
NEW1
NUP159
DBP5
MTR2
NIC96
NUP42
RLR1
HMT1
CSN12
ZDS1
PML1
SRM1
PSE1
EST1
THO1
ASM4
SXM1
NUP188
APQ12
SEM1
SGF73
SUS1
ENP1
NUP1
YBR111W−A_1
SAC3
NPL3
NUP84
SPN1
IST3
STO1
BUD13
NUP60
NUP2
CSL4
BRL1
MLP1
SPT6
RSP5
THP2
NUP116
HRB1
GFD1
NUP100
MIP6
MFT1
EBS1
GBP2
YRA2
MLP2
TEX1
HPR1
RPB4
Gene
Z_Shift_K_Doxo_HLD
Z_lm_K_Doxo_HLD
Z_Shift_K_Doxo_EG
Z_lm_K_Doxo_EG
Z_Shift_L_Doxo_HLD
Z_lm_L_Doxo_HLD
Z_Shift_L_Doxo_EG
Z_lm_L_Doxo_EG

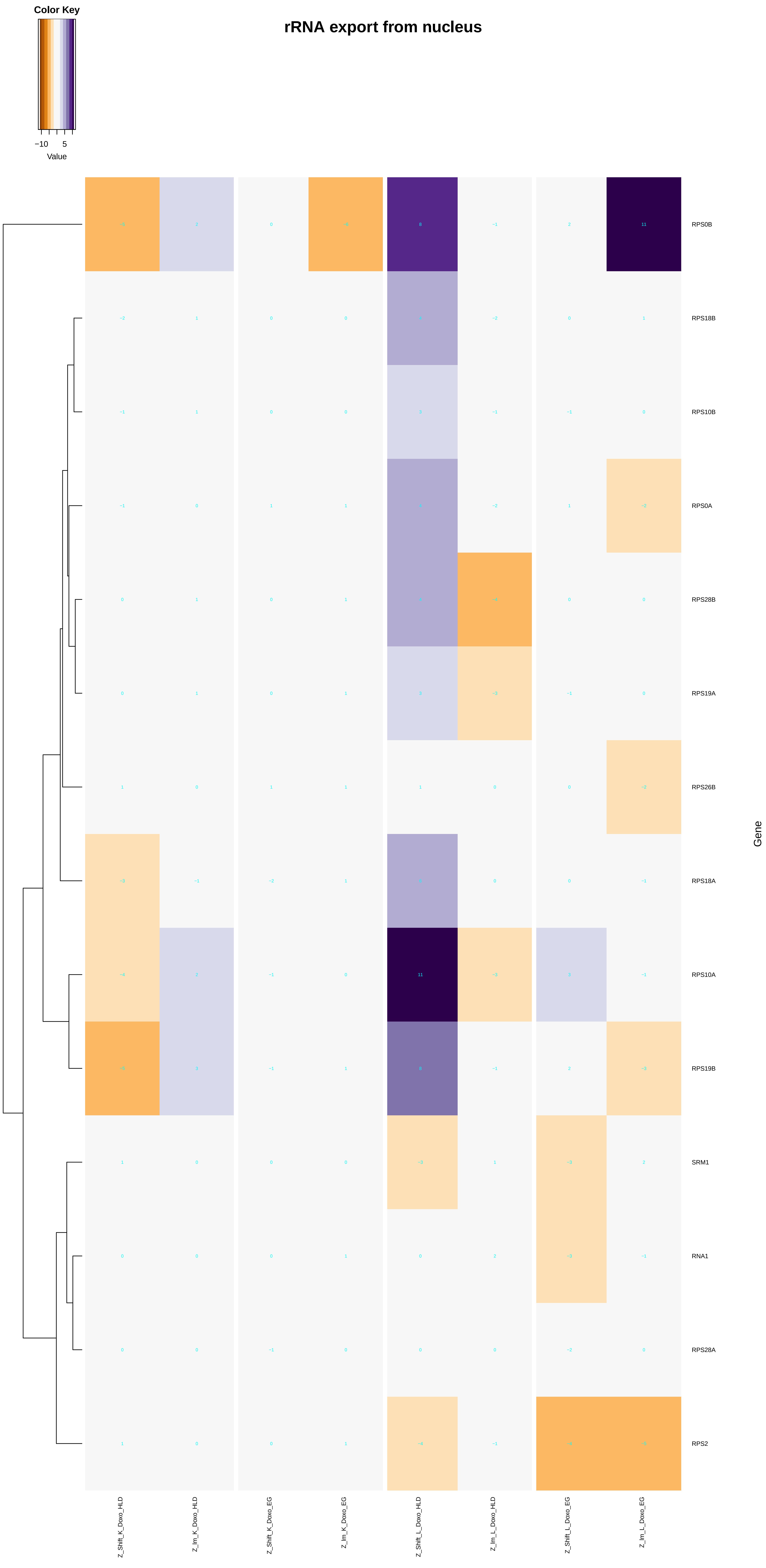

Color Key
−10
5
Value
rRNA export from nucleus
RPS0B
RPS18B
RPS10B
RPS0A
RPS28B
RPS19A
RPS26B
RPS18A
RPS10A
RPS19B
SRM1
RNA1
RPS28A
RPS2
Gene
Z_Shift_K_Doxo_HLD
Z_lm_K_Doxo_HLD
Z_Shift_K_Doxo_EG
Z_lm_K_Doxo_EG
Z_Shift_L_Doxo_HLD
Z_lm_L_Doxo_HLD
Z_Shift_L_Doxo_EG
Z_lm_L_Doxo_EG

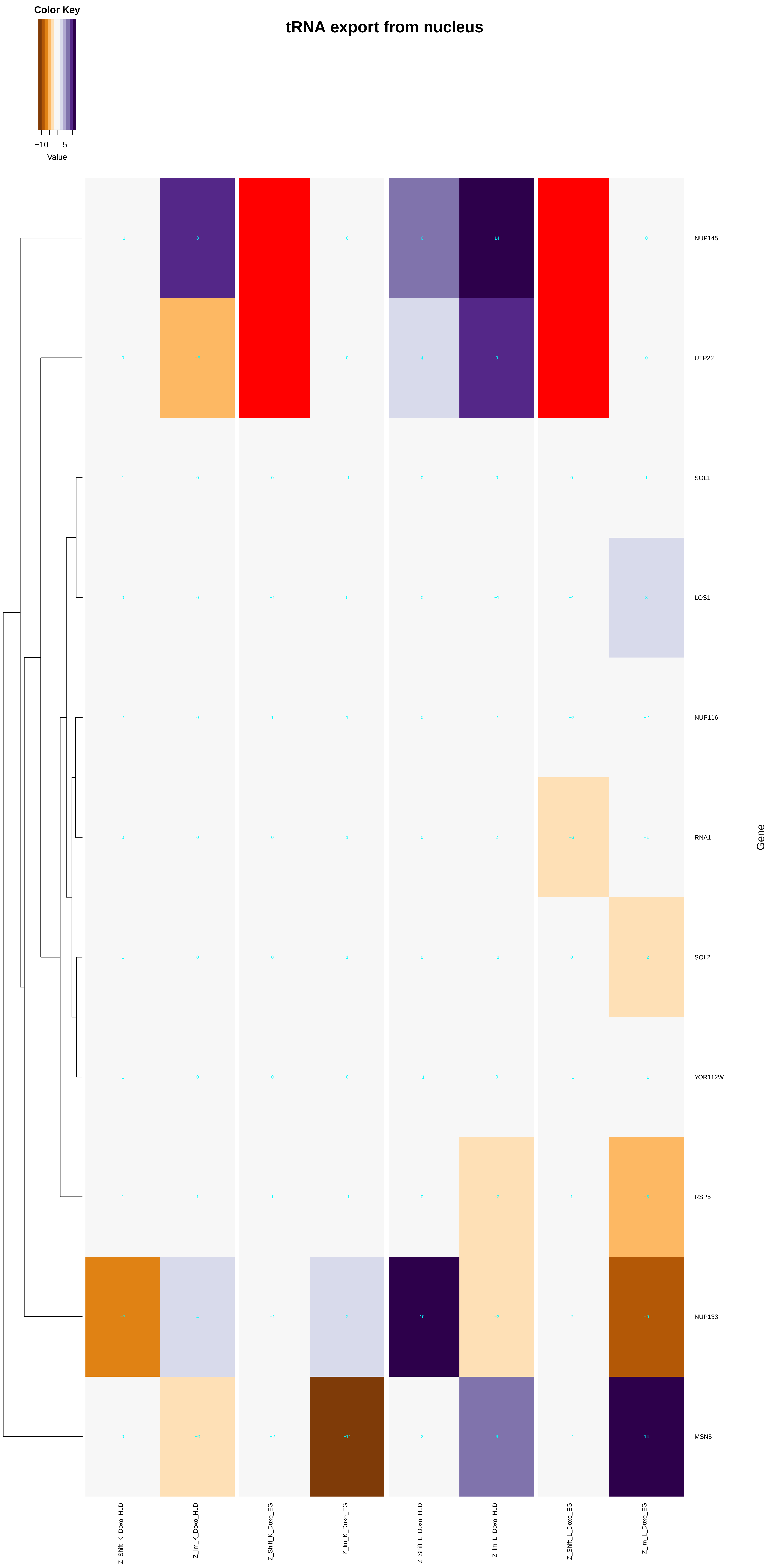

Color Key
−10
5
Value
tRNA export from nucleus
NUP145
UTP22
SOL1
LOS1
NUP116
RNA1
SOL2
YOR112W
RSP5
NUP133
MSN5
Gene
Z_Shift_K_Doxo_HLD
Z_lm_K_Doxo_HLD
Z_Shift_K_Doxo_EG
Z_lm_K_Doxo_EG
Z_Shift_L_Doxo_HLD
Z_lm_L_Doxo_HLD
Z_Shift_L_Doxo_EG
Z_lm_L_Doxo_EG

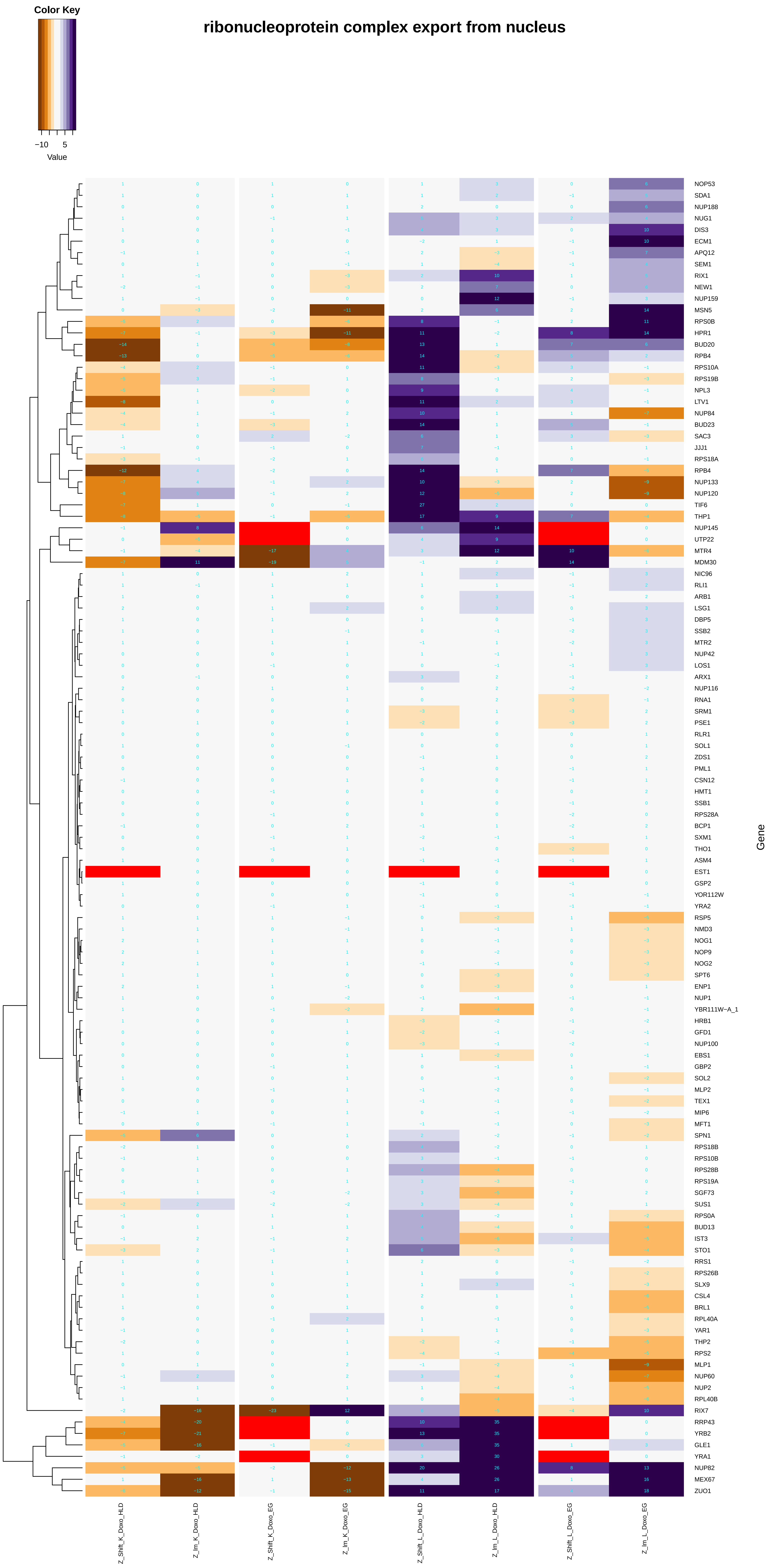
Color Key
−10
5
Value
ribonucleoprotein complex export from nucleus
NOP53
SDA1
NUP188
NUG1
DIS3
ECM1
APQ12
SEM1
RIX1
NEW1
NUP159
MSN5
RPS0B
HPR1
BUD20
RPB4
RPS10A
RPS19B
NPL3
LTV1
NUP84
BUD23
SAC3
JJJ1
RPS18A
RPB4
NUP133
NUP120
TIF6
THP1
NUP145
UTP22
MTR4
MDM30
NIC96
RLI1
ARB1
LSG1
DBP5
SSB2
MTR2
NUP42
LOS1
ARX1
NUP116
RNA1
SRM1
PSE1
RLR1
SOL1
ZDS1
PML1
CSN12
HMT1
SSB1
RPS28A
BCP1
SXM1
THO1
ASM4
EST1
GSP2
YOR112W
YRA2
RSP5
NMD3
NOG1
NOP9
NOG2
SPT6
ENP1
NUP1
YBR111W−A_1
HRB1
GFD1
NUP100
EBS1
GBP2
SOL2
MLP2
TEX1
MIP6
MFT1
SPN1
RPS18B
RPS10B
RPS28B
RPS19A
SGF73
SUS1
RPS0A
BUD13
IST3
STO1
RRS1
RPS26B
SLX9
CSL4
BRL1
RPL40A
YAR1
THP2
RPS2
MLP1
NUP60
NUP2
RPL40B
RIX7
RRP43
YRB2
GLE1
YRA1
NUP82
MEX67
ZUO1
Gene
Z_Shift_K_Doxo_HLD
Z_lm_K_Doxo_HLD
Z_Shift_K_Doxo_EG
Z_lm_K_Doxo_EG
Z_Shift_L_Doxo_HLD
Z_lm_L_Doxo_HLD
Z_Shift_L_Doxo_EG
Z_lm_L_Doxo_EG

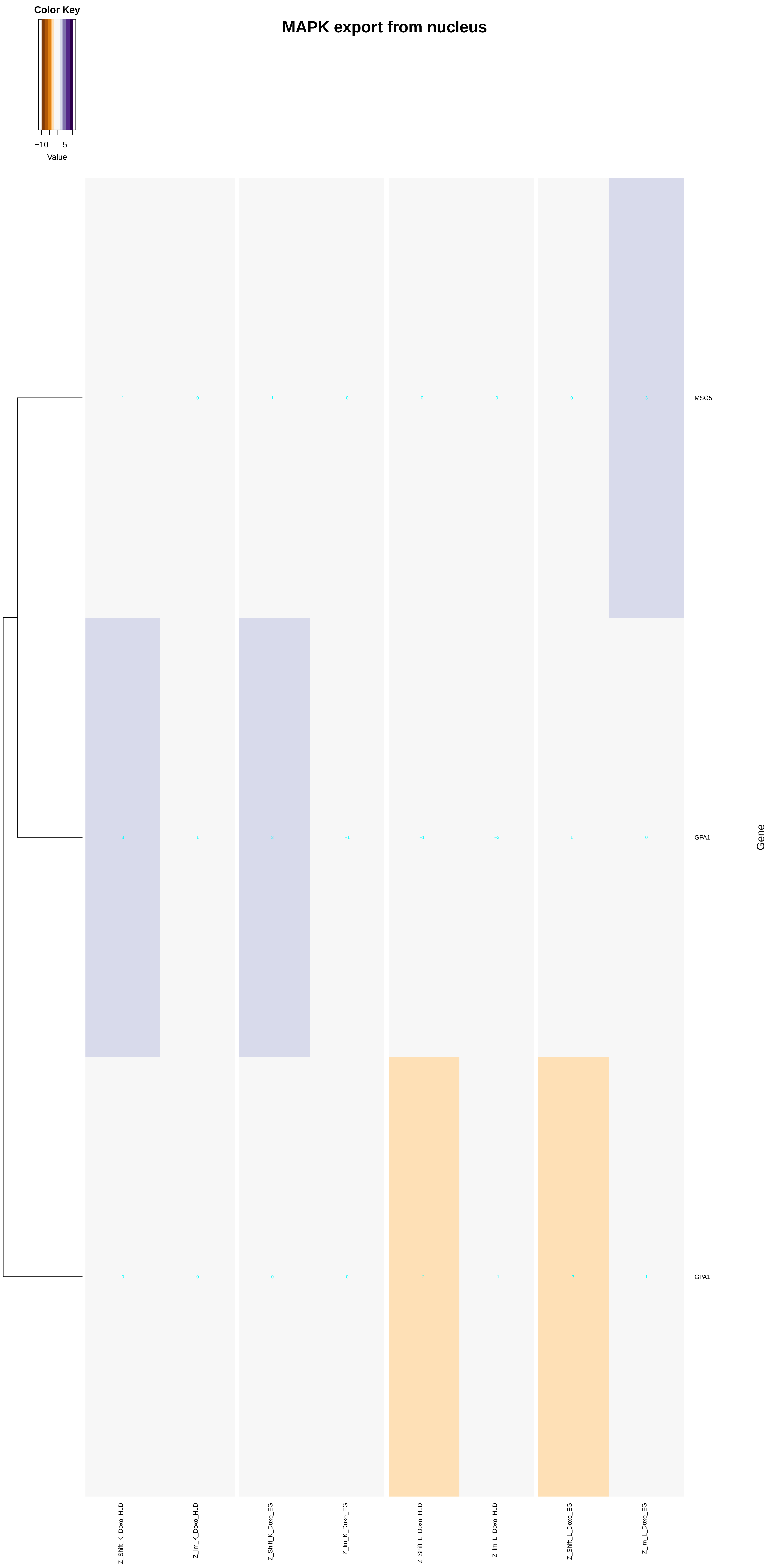
Color Key
−10
5
Value
MAPK export from nucleus
1 0 1 0 0 0 0 3
MSG5
3 1 3 −1 −1 −2 1 0
GPA1
0 0 0 0 −2 −1 −3 1
GPA1
Gene
Z_Shift_K_Doxo_HLD
Z_lm_K_Doxo_HLD
Z_Shift_K_Doxo_EG
Z_lm_K_Doxo_EG
Z_Shift_L_Doxo_HLD
Z_lm_L_Doxo_HLD
Z_Shift_L_Doxo_EG
Z_lm_L_Doxo_EG

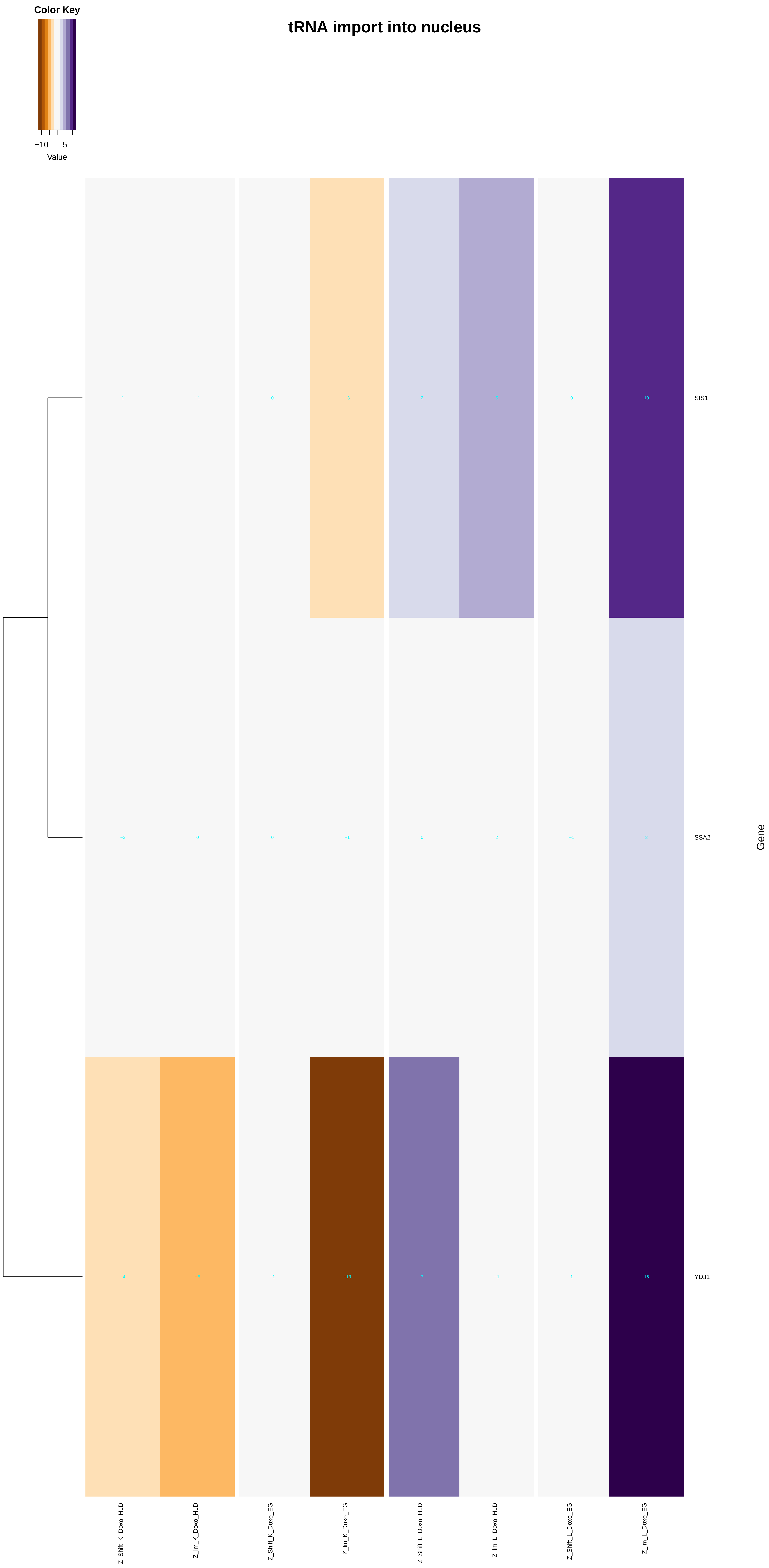
tRNA import into nucleus
Color Key
−10
5
Value
SIS1
SSA2
YDJ1
Gene
Z_Shift_K_Doxo_HLD
Z_Im_K_Doxo_HLD
Z_Shift_K_Doxo_EG
Z_Im_K_Doxo_EG
Z_Shift_L_Doxo_HLD
Z_Im_L_Doxo_HLD
Z_Shift_L_Doxo_EG
Z_Im_L_Doxo_EG

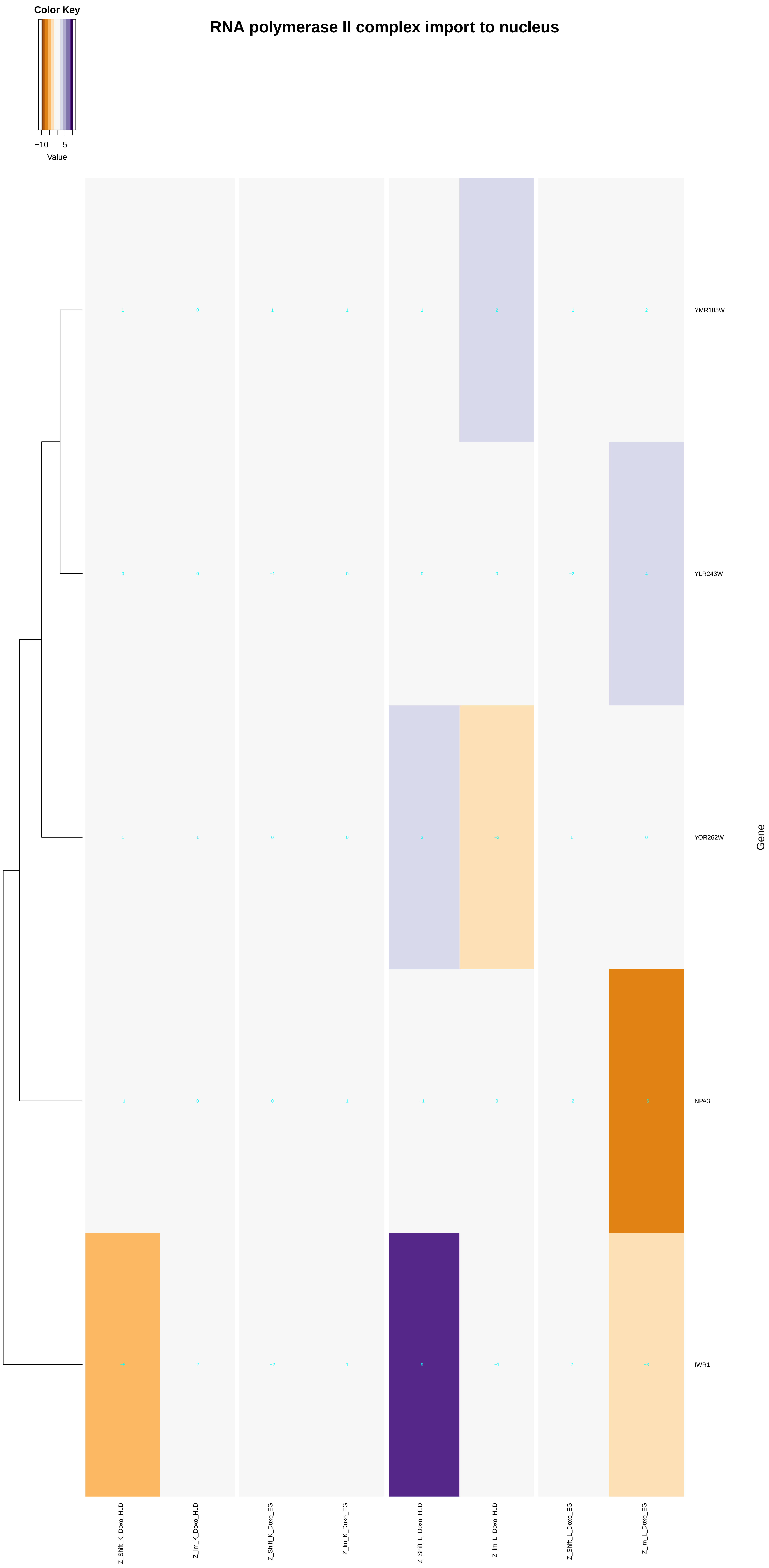

Color Key
−10
5
Value
RNA polymerase II complex import to nucleus
YMR185W
YLR243W
YOR262W
NPA3
IWR1
Gene
Z_Shift_K_Doxo_HLD
Z_Im_K_Doxo_HLD
Z_Shift_K_Doxo_EG
Z_Im_K_Doxo_EG
Z_Shift_L_Doxo_HLD
Z_Im_L_Doxo_HLD
Z_Shift_L_Doxo_EG
Z_Im_L_Doxo_EG

poly(A)+ mRNA export from nucleus

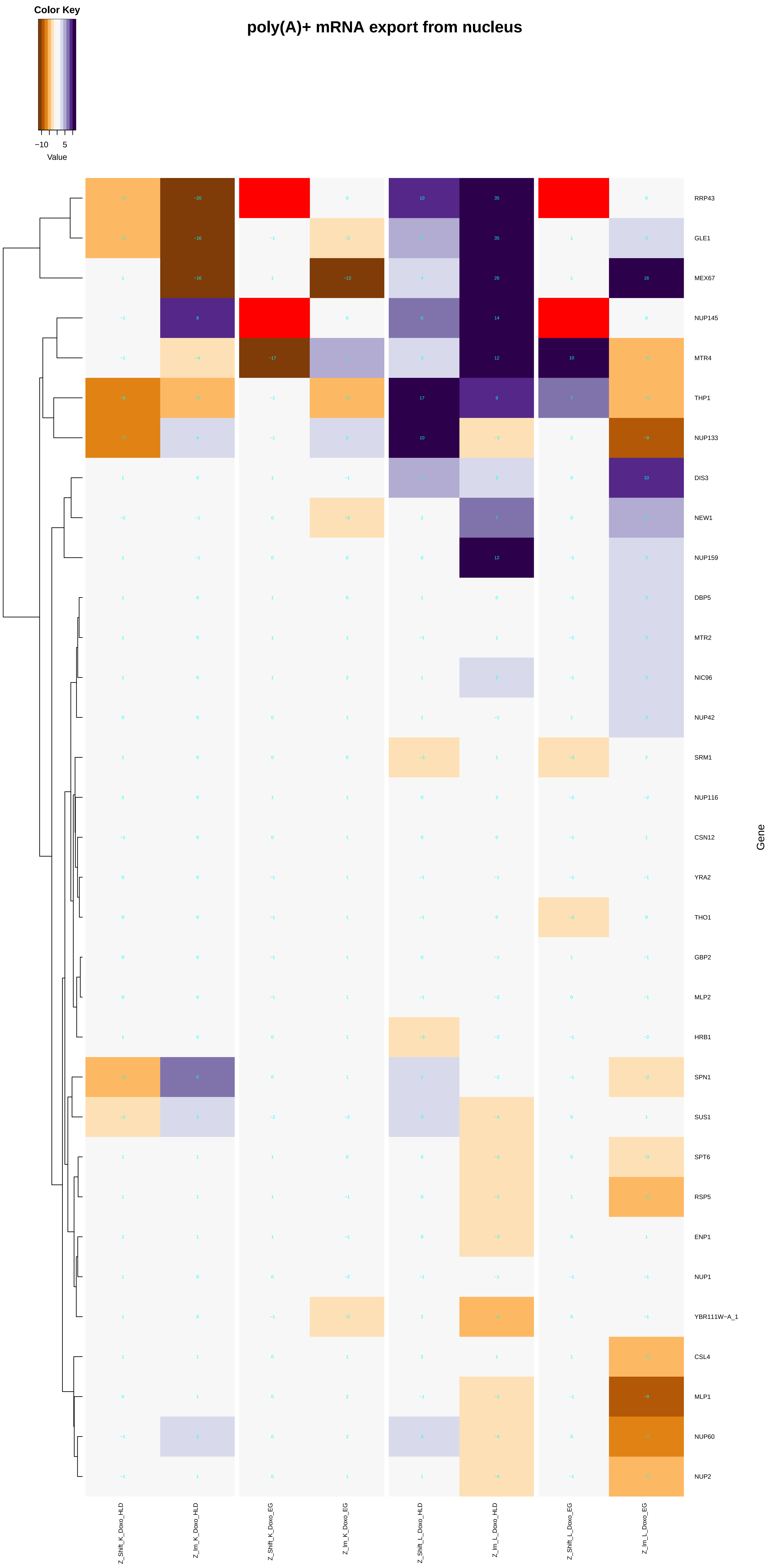

### mRNA export from nucleus in response to heat stress

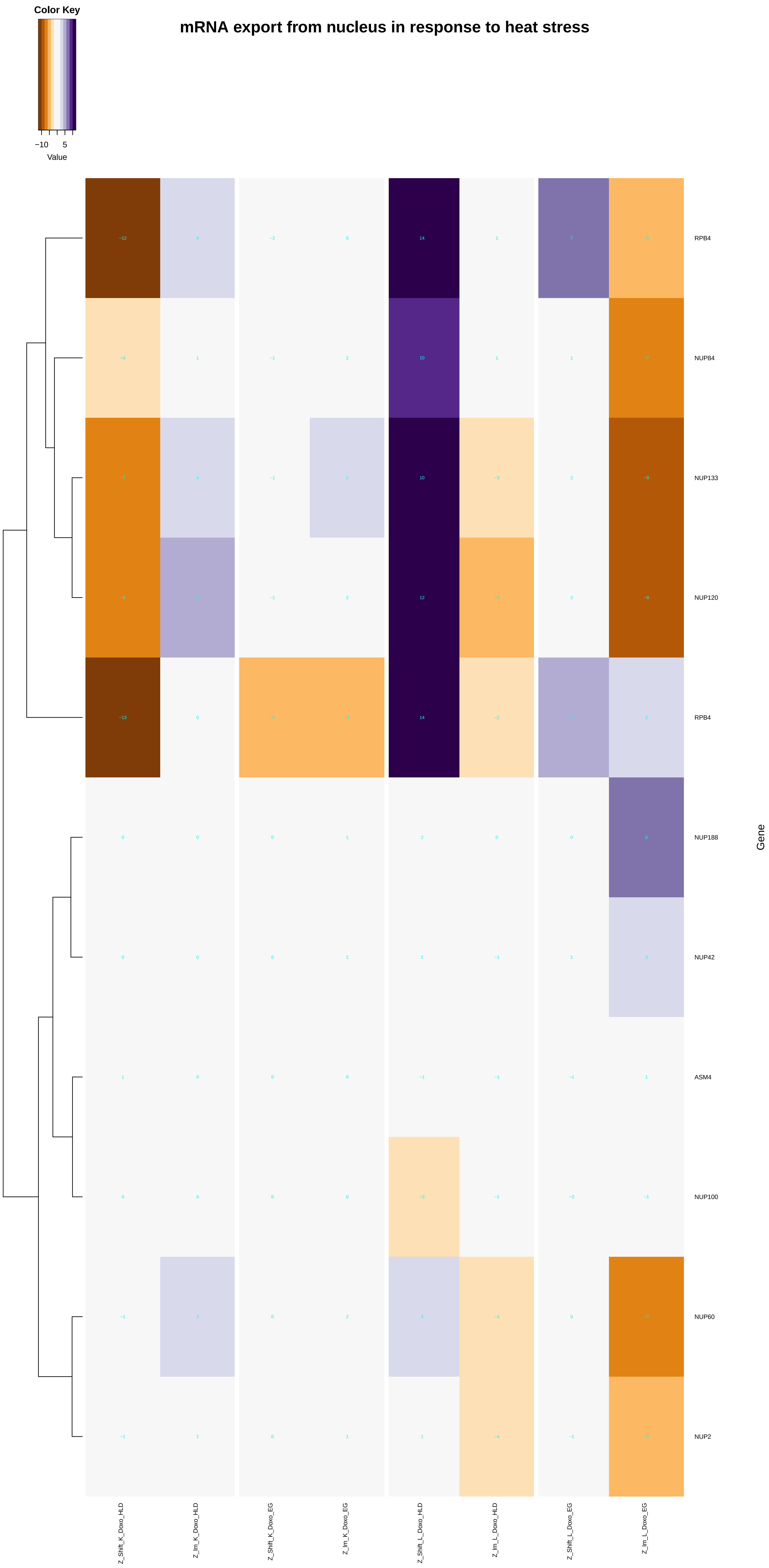

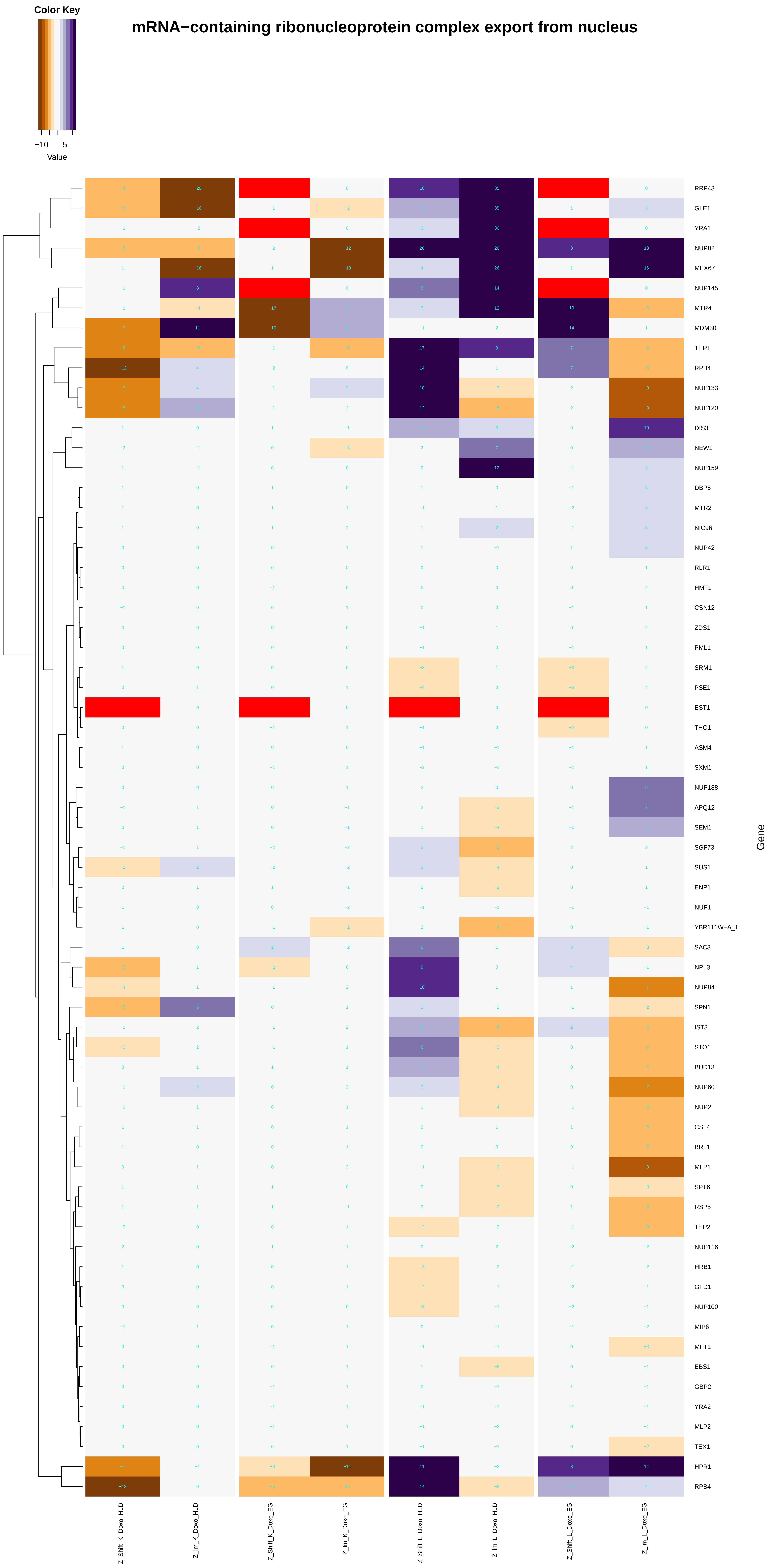

mRNA−containing ribonucleoprotein complex export from nucleus
Color Key
−10
5
Value
RRP43
GLE1
YRA1
NUP82
MEX67
NUP145
MTR4
MDM30
THP1
RPB4
NUP133
NUP120
DIS3
NEW1
NUP159
DBP5
MTR2
NIC96
NUP42
RLR1
HMT1
CSN12
ZDS1
PML1
SRM1
PSE1
EST1
THO1
ASM4
SXM1
NUP188
APQ12
SEM1
SGF73
SUS1
ENP1
NUP1
YBR111W−A_1
SAC3
NPL3
NUP84
SPN1
IST3
STO1
BUD13
NUP60
NUP2
CSL4
BRL1
MLP1
SPT6
RSP5
THP2
NUP116
HRB1
GFD1
NUP100
MIP6
MFT1
EBS1
GBP2
YRA2
MLP2
TEX1
HPR1
RPB4
Gene
Z_Shift_K_Doxo_HLD
Z_lm_K_Doxo_HLD
Z_Shift_K_Doxo_EG
Z_lm_K_Doxo_EG
Z_Shift_L_Doxo_HLD
Z_lm_L_Doxo_HLD
Z_Shift_L_Doxo_EG
Z_lm_L_Doxo_EG

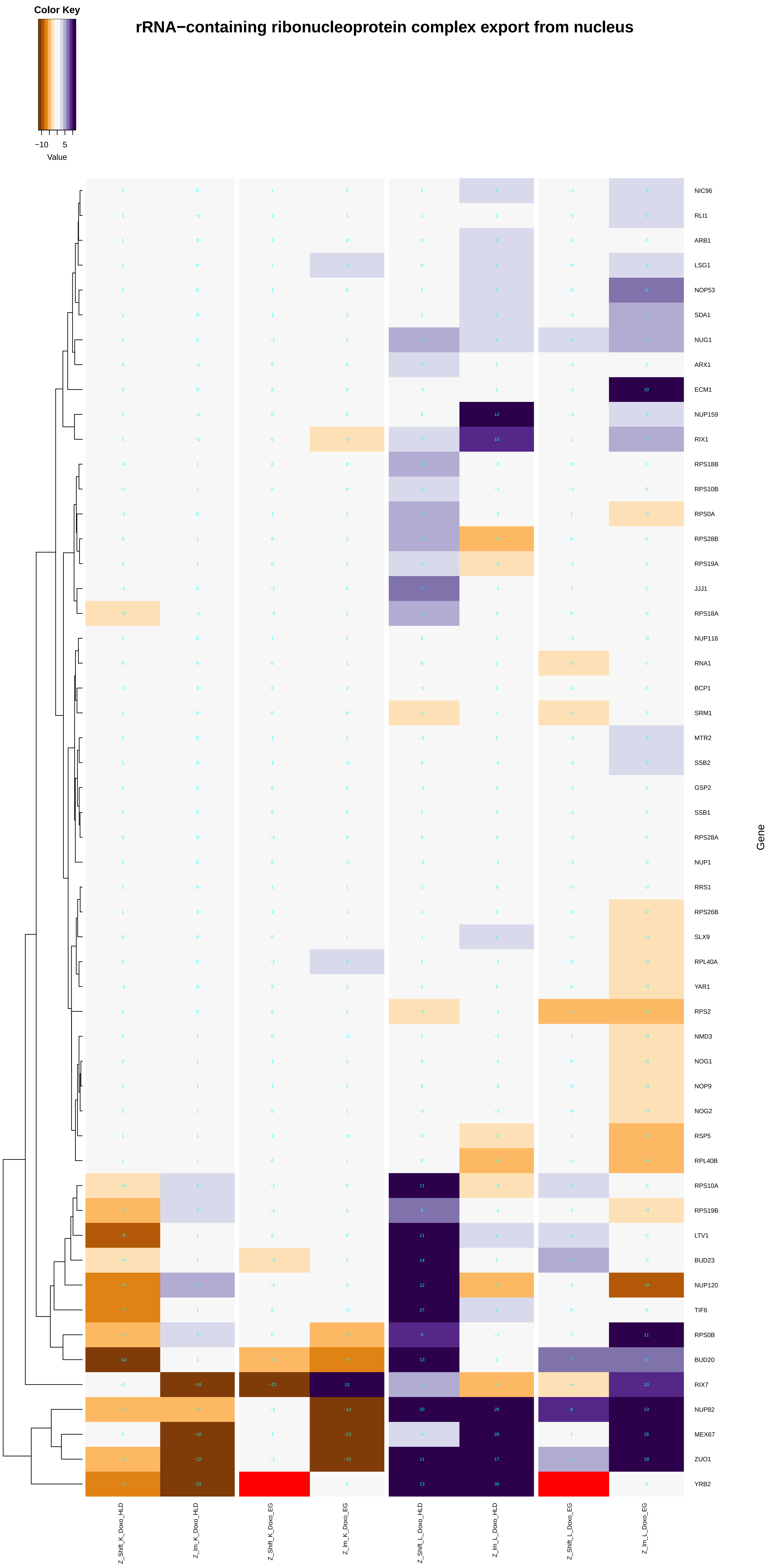

rRNA−containing ribonucleoprotein complex export from nucleus
Color Key
−10
5
Value
Gene
NIC96
RLI1
ARB1
LSG1
NOP53
SDA1
NUG1
ARX1
ECM1
NUP159
RIX1
RPS18B
RPS10B
RPS0A
RPS28B
RPS19A
JJJ1
RPS18A
NUP116
RNA1
BCP1
SRM1
MTR2
SSB2
GSP2
SSB1
RPS28A
NUP1
RRS1
RPS26B
SLX9
RPL40A
YAR1
RPS2
NMD3
NOG1
NOP9
NOG2
RSP5
RPL40B
RPS10A
RPS19B
LTV1
BUD23
NUP120
TIF6
RPS0B
BUD20
RIX7
NUP82
MEX67
ZUO1
YRB2
Z_Shift_K_Doxo_HLD
Z_Im_K_Doxo_HLD
Z_Shift_K_Doxo_EG
Z_Im_K_Doxo_EG
Z_Shift_L_Doxo_HLD
Z_Im_L_Doxo_HLD
Z_Shift_L_Doxo_EG
Z_Im_L_Doxo_EG

### tRNA−containing ribonucleoprotein complex export from nucleus

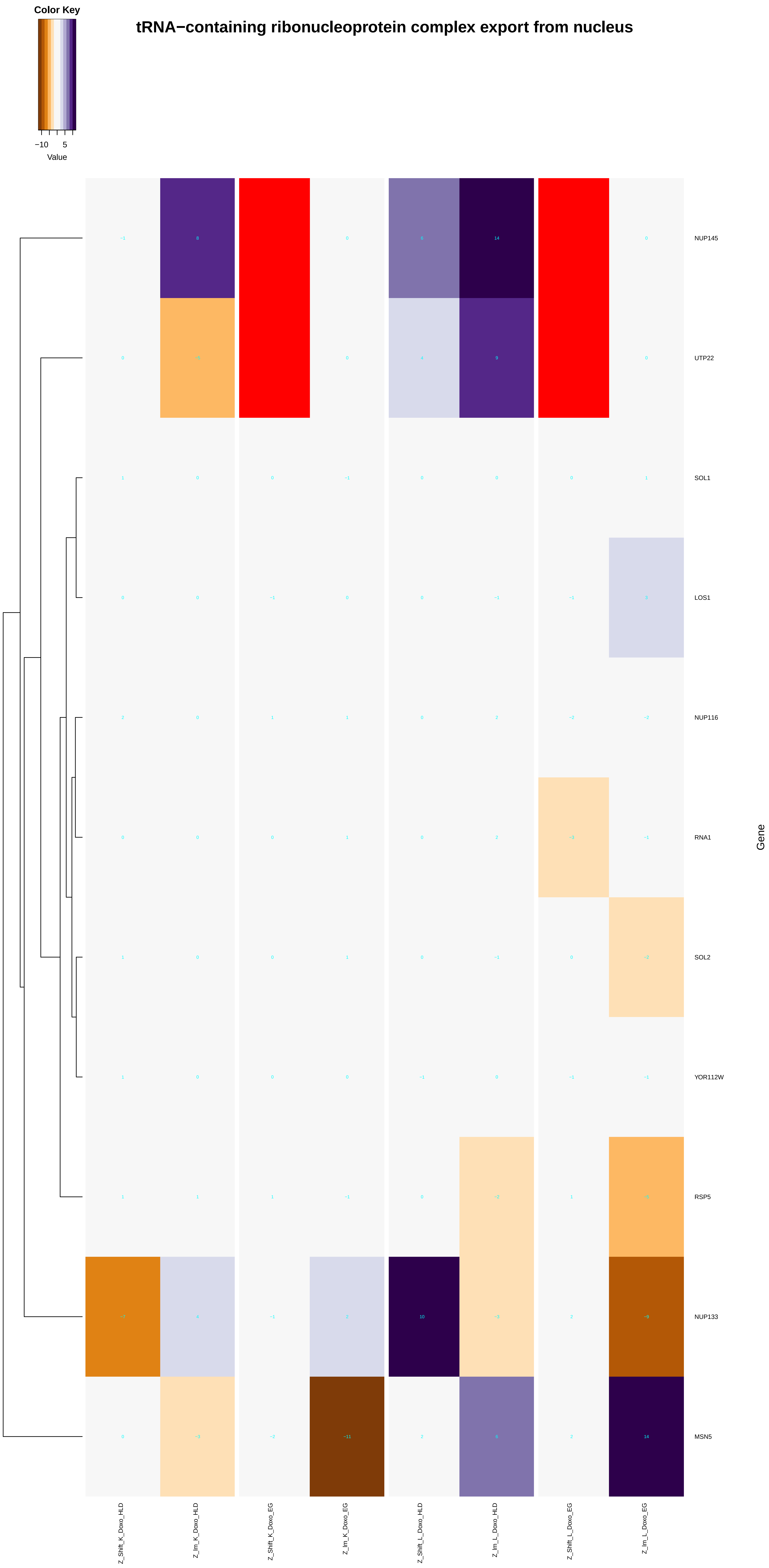

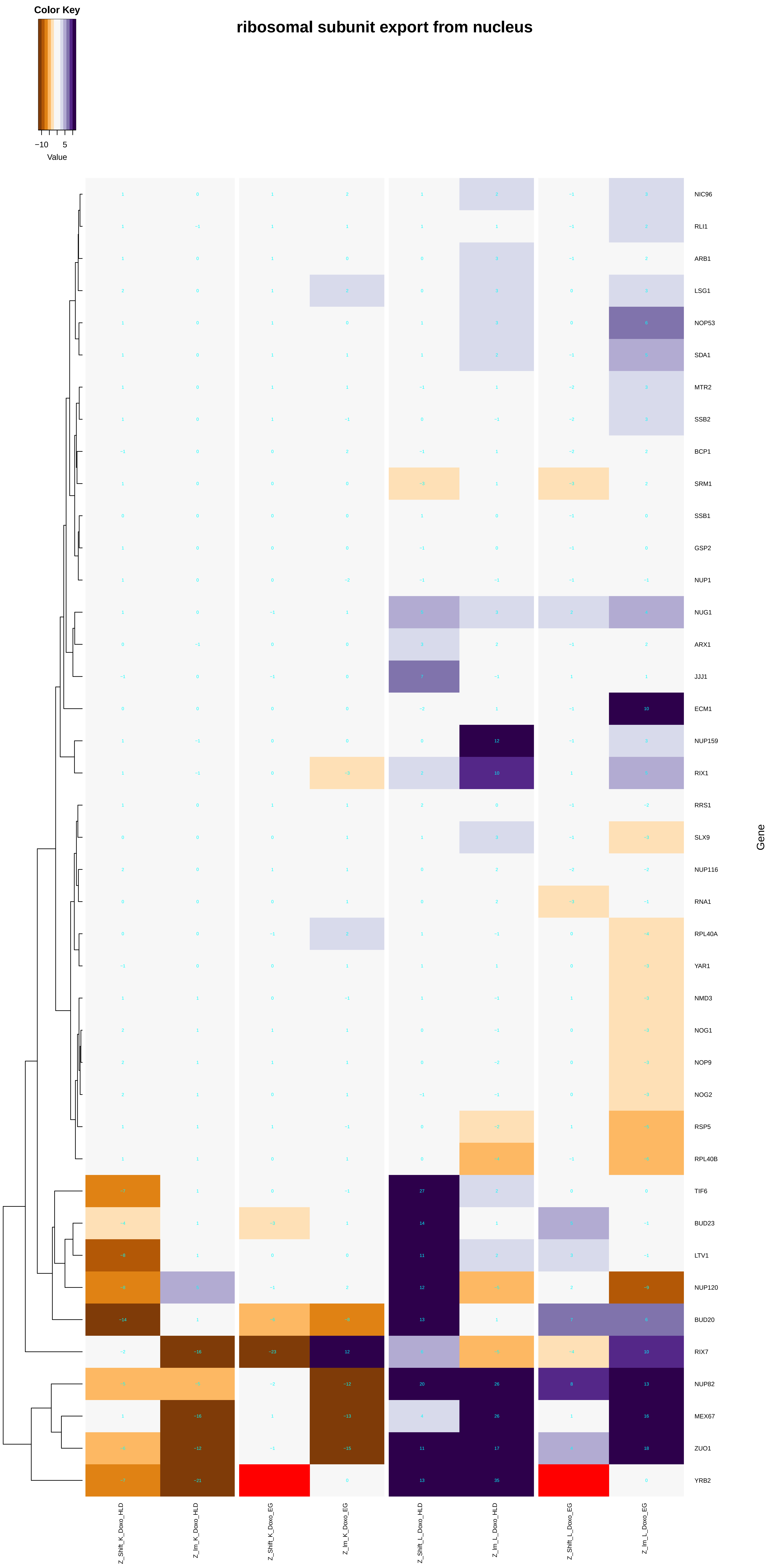

Color Key
−10
5
Value
ribosomal subunit export from nucleus
NIC96
RLI1
ARB1
LSG1
NOP53
SDA1
MTR2
SSB2
BCP1
SRM1
SSB1
GSP2
NUP1
NUG1
ARX1
JJJ1
ECM1
NUP159
RIX1
RRS1
SLX9
NUP116
RNA1
RPL40A
YAR1
NMD3
NOG1
NOP9
NOG2
RSP5
RPL40B
TIF6
BUD23
LTV1
NUP120
BUD20
RIX7
NUP82
MEX67
ZUO1
YRB2
Gene
Z_Shift_K_Doxo_HLD
Z_lm_K_Doxo_HLD
Z_Shift_K_Doxo_EG
Z_lm_K_Doxo_EG
Z_Shift_L_Doxo_HLD
Z_lm_L_Doxo_HLD
Z_Shift_L_Doxo_EG
Z_lm_L_Doxo_EG

### ribosomal large subunit export from nucleus

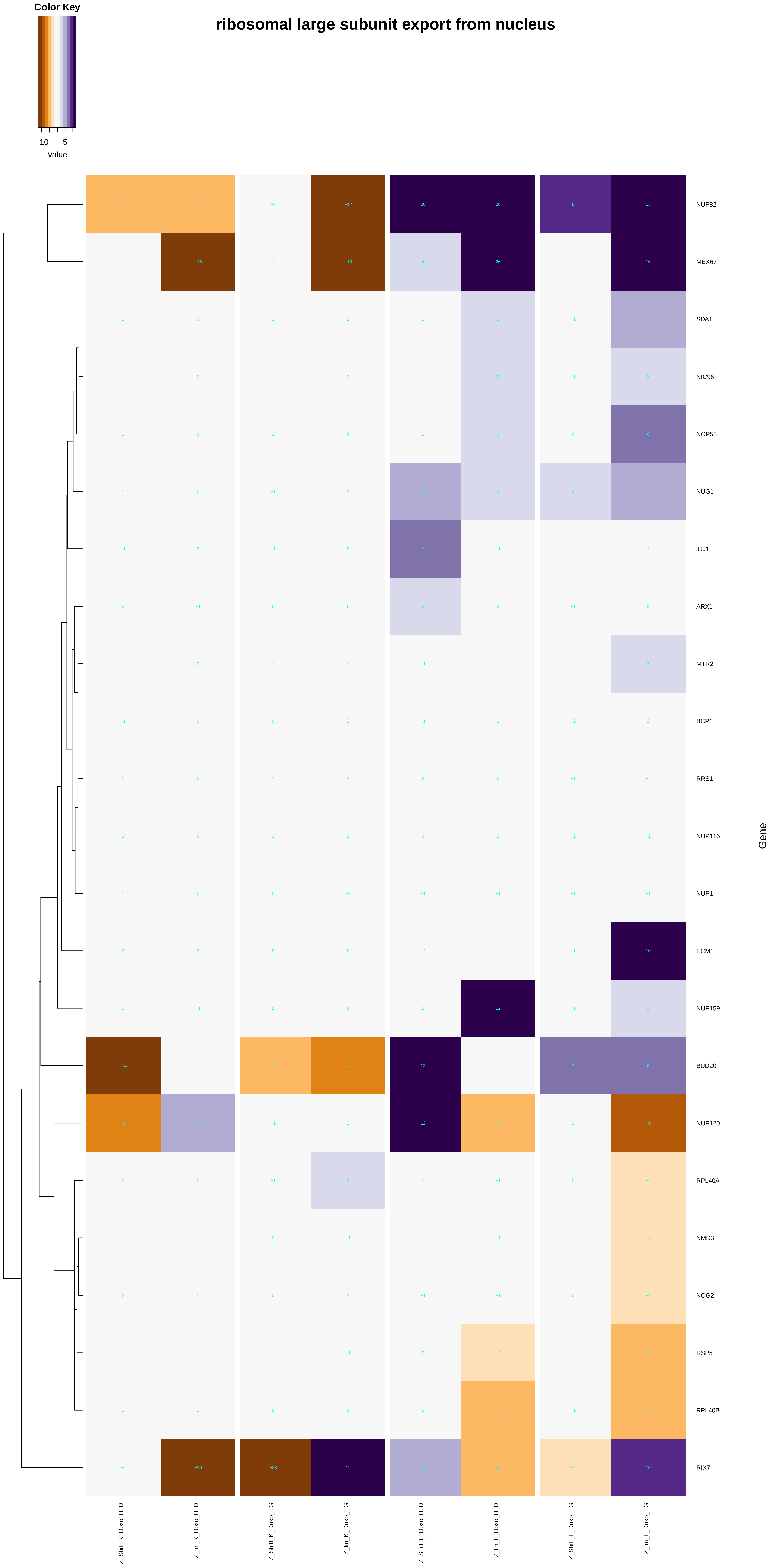

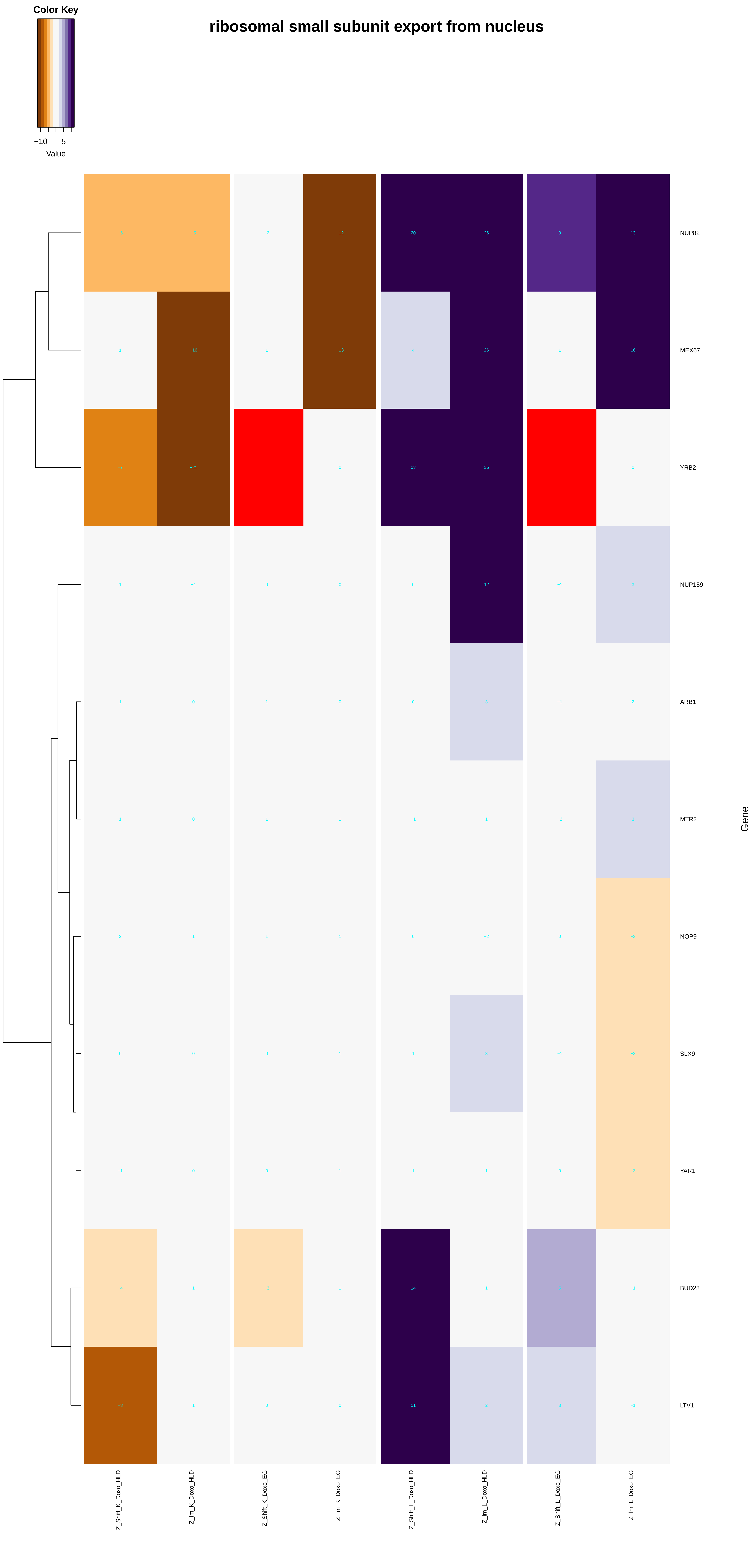
ribosomal small subunit export from nucleus
Color Key
−10
5
Value
NUP82
MEX67
YRB2
NUP159
ARB1
MTR2
NOP9
SLX9
YAR1
BUD23
LTV1
Gene
Z_Shift_K_Doxo_HLD
Z_Im_K_Doxo_HLD
Z_Shift_K_Doxo_EG
Z_Im_K_Doxo_EG
Z_Shift_L_Doxo_HLD
Z_Im_L_Doxo_HLD
Z_Shift_L_Doxo_EG
Z_Im_L_Doxo_EG
