## Additional File 9 for "A yeast phenomic model for the influence of Warburg metabolism on genetic buffering of doxorubicin": nucleic_acid-templated_transcription.pdf

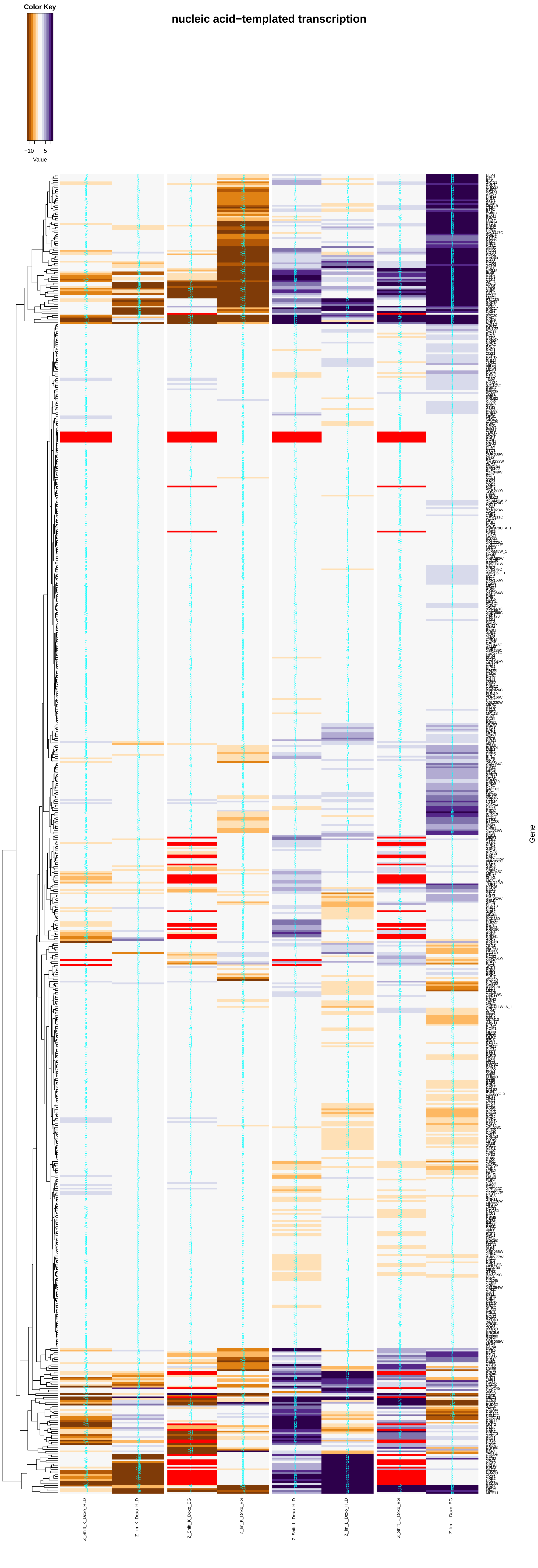

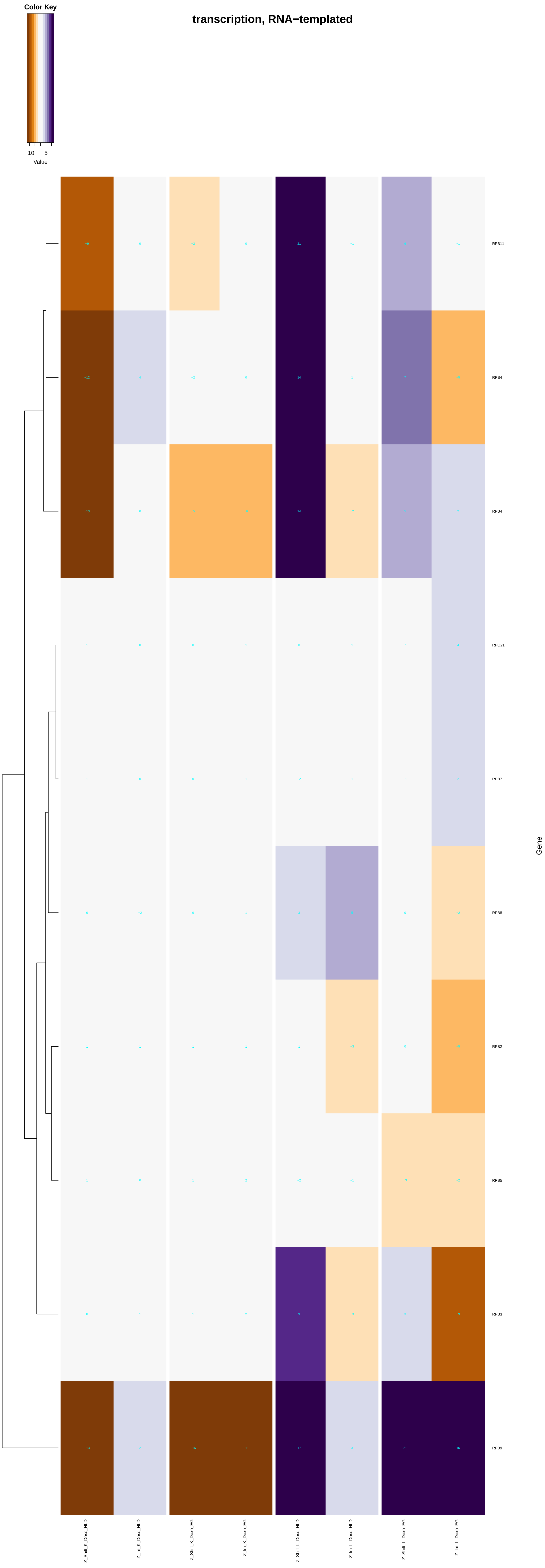

transcription, RNA−templated
Color Key
−10
5
Value
RPB11
RPB4
RPB4
RPO21
RPB7
RPB8
RPB2
RPB5
RPB3
RPB9
Gene
Z_Shift_K_Doxo_HLD
Z_Im_K_Doxo_HLD
Z_Shift_K_Doxo_EG
Z_Im_K_Doxo_EG
Z_Shift_L_Doxo_HLD
Z_Im_L_Doxo_HLD
Z_Shift_L_Doxo_EG
Z_Im_L_Doxo_EG

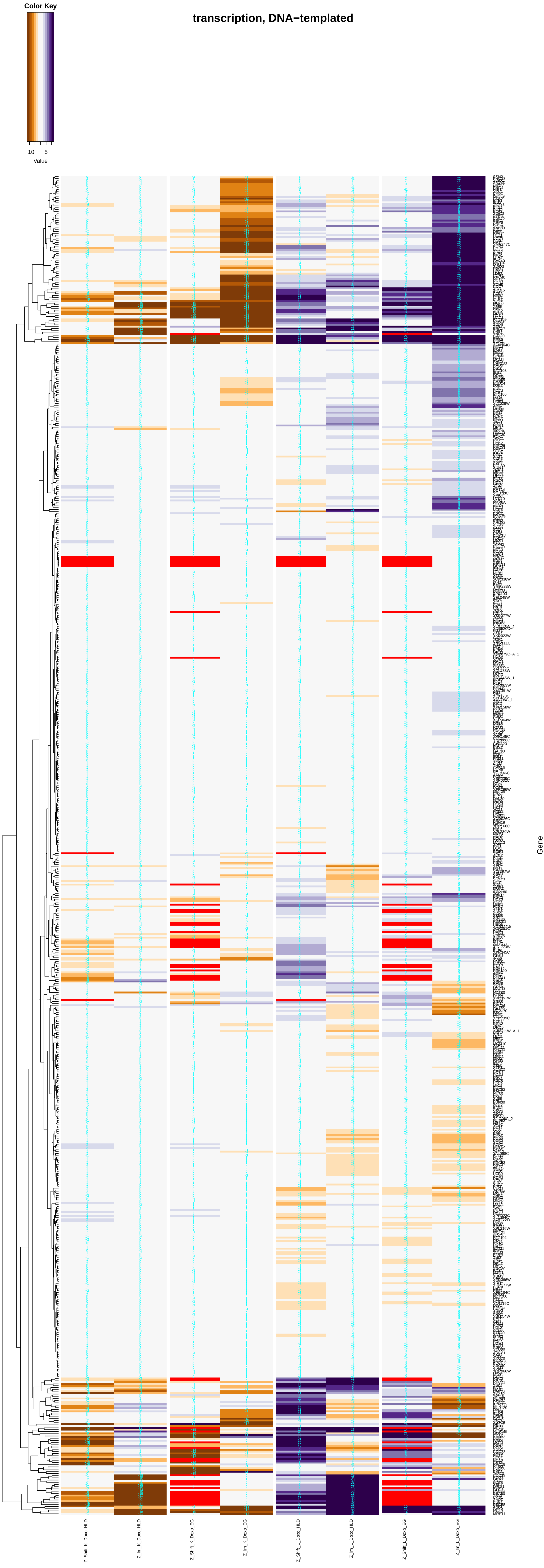
transcription, DNA−templated
Color Key
−10
5
Value
Gene
Z_Shift_K_Doxo_HLD
Z_Im_K_Doxo_HLD
Z_Shift_K_Doxo_EG
Z_Im_K_Doxo_EG
Z_Shift_L_Doxo_HLD
Z_Im_L_Doxo_HLD
Z_Shift_L_Doxo_EG
Z_Im_L_Doxo_EG

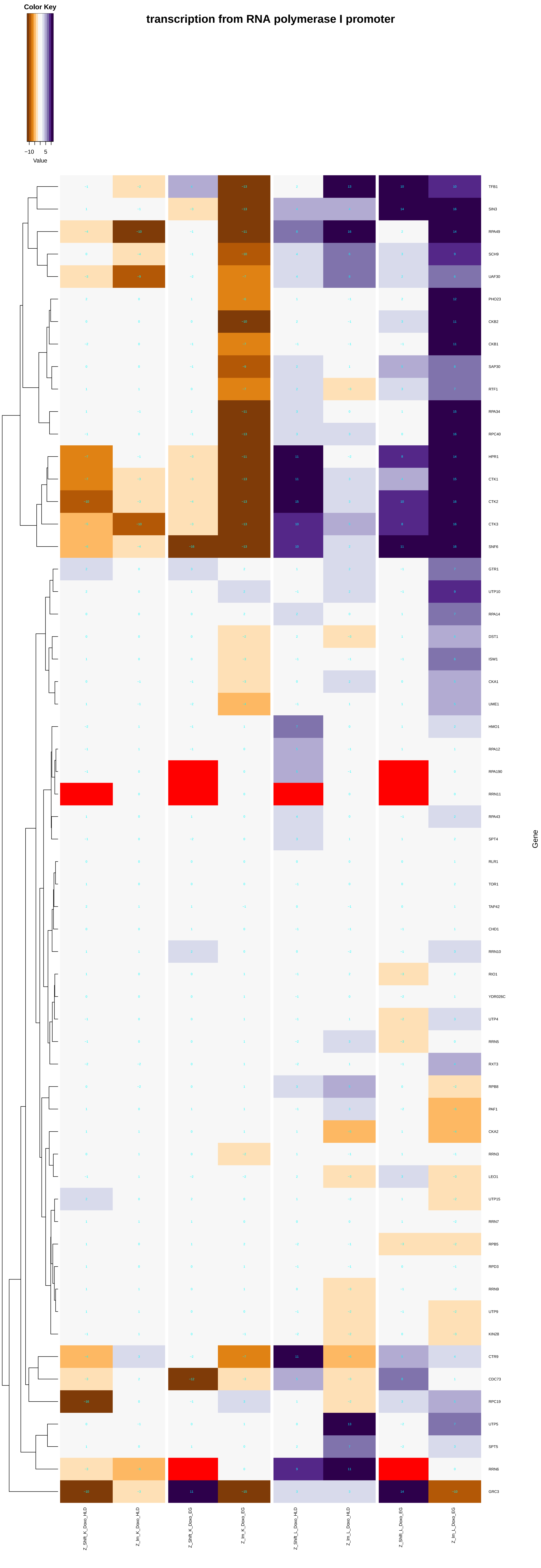

transcription from RNA polymerase I promoter
Color Key
−10
5
Value
TFB1
SIN3
RPA49
SCH9
UAF30
PHO23
CKB2
CKB1
SAP30
RTF1
RPA34
RPC40
HPR1
CTK1
CTK2
CTK3
SNF6
GTR1
UTP10
RPA14
DST1
ISW1
CKA1
UME1
HMO1
RPA12
RPA190
RRN11
RPA43
SPT4
RLR1
TOR1
TAP42
CHD1
RRN10
RIO1
YDR026C
UTP4
RRN5
RXT3
RPB8
PAF1
CKA2
RRN3
LEO1
UTP15
RRN7
RPB5
RPD3
RRN9
UTP9
KIN28
CTR9
CDC73
RPC19
UTP5
SPT5
RRN6
GRC3
Gene
Z_Shift_K_Doxo_HLD
Z_lm_K_Doxo_HLD
Z_Shift_K_Doxo_EG
Z_lm_K_Doxo_EG
Z_Shift_L_Doxo_HLD
Z_lm_L_Doxo_HLD
Z_Shift_L_Doxo_EG
Z_lm_L_Doxo_EG

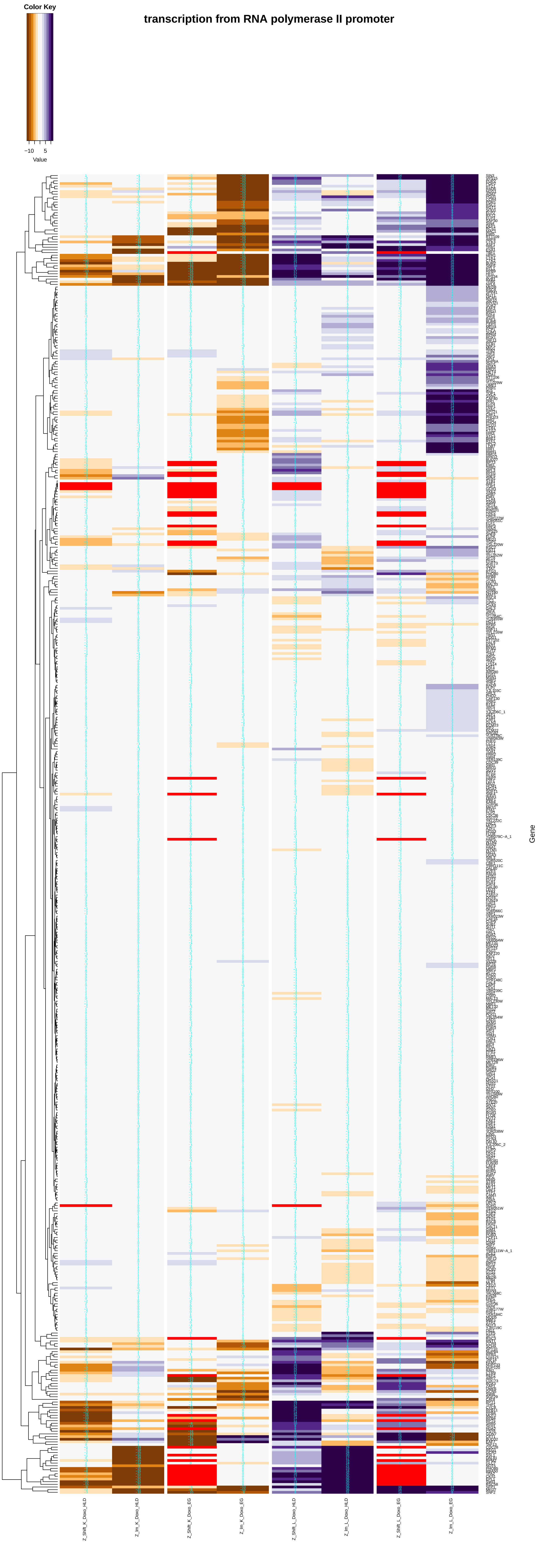
transcription from RNA polymerase II promoter
Color Key
−10
5
Value
Z_Shift_K_Doxo_HLD
Z_Im_K_Doxo_HLD
Z_Shift_K_Doxo_EG
Z_Im_K_Doxo_EG
Z_Shift_L_Doxo_HLD
Z_Im_L_Doxo_HLD
Z_Shift_L_Doxo_EG
Z_Im_L_Doxo_EG
Gene

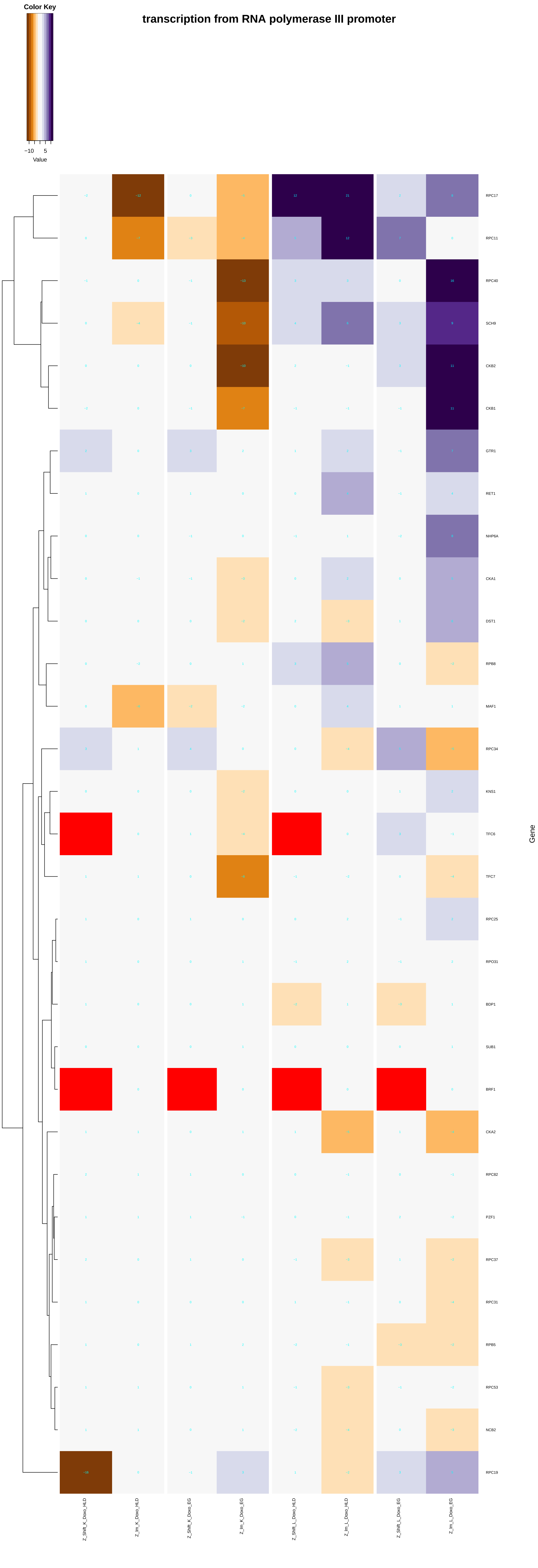

Color Key
−10
5
Value
transcription from RNA polymerase III promoter
RPC17
RPC11
RPC40
SCH9
CKB2
CKB1
GTR1
RET1
NHP6A
CKA1
DST1
RPB8
MAF1
RPC34
KNS1
TFC6
TFC7
RPC25
RPO31
BDP1
SUB1
BRF1
CKA2
RPC82
PZF1
RPC37
RPC31
RPB5
RPC53
NCB2
RPC19
Gene
Z_Shift_K_Doxo_HLD
Z_Im_K_Doxo_HLD
Z_Shift_K_Doxo_EG
Z_Im_K_Doxo_EG
Z_Shift_L_Doxo_HLD
Z_Im_L_Doxo_HLD
Z_Shift_L_Doxo_EG
Z_Im_L_Doxo_EG

### transcription from mitochondrial promoter

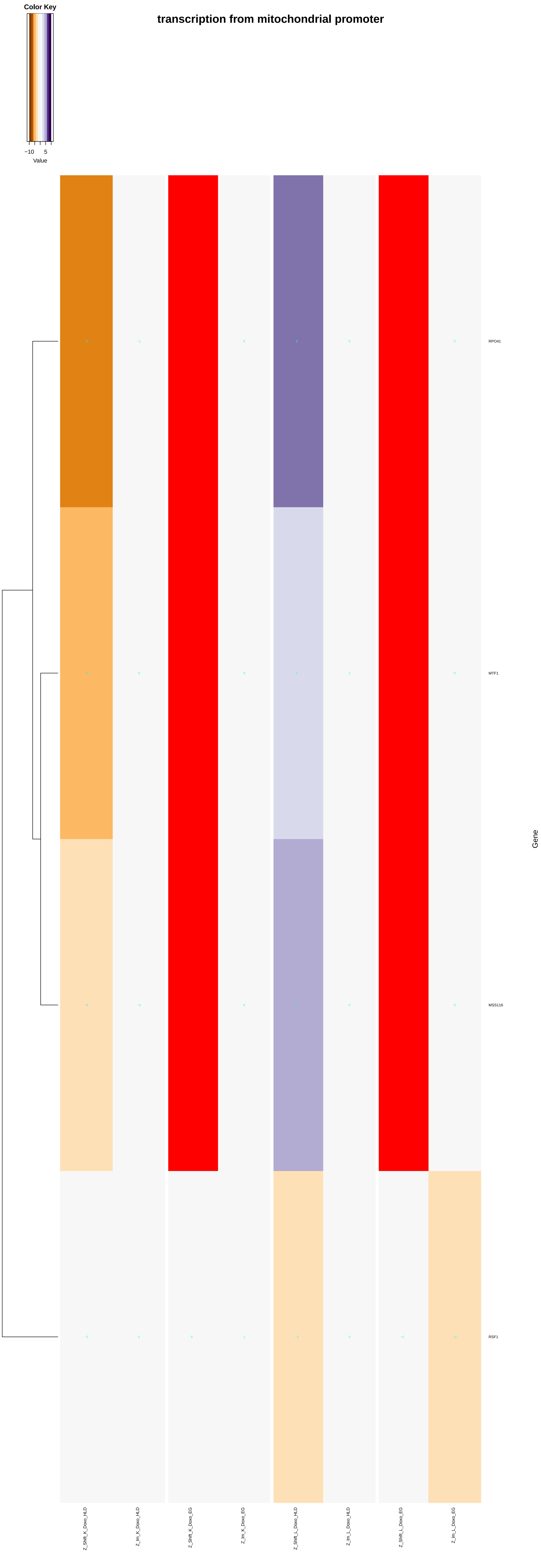

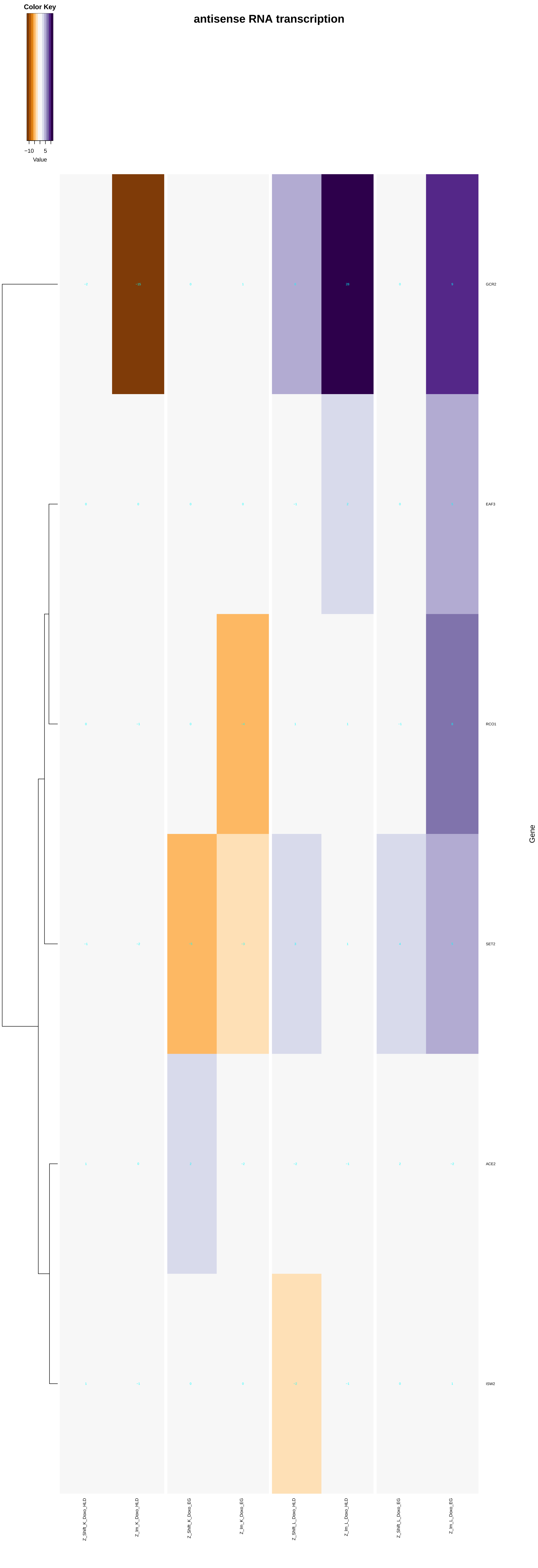
antisense RNA transcription
Color Key
−10
5
Value
GCR2
EAF3
RCO1
SET2
ACE2
ISW2
Gene
Z_Shift_K_Doxo_HLD
Z_Im_K_Doxo_HLD
Z_Shift_K_Doxo_EG
Z_Im_K_Doxo_EG
Z_Shift_L_Doxo_HLD
Z_Im_L_Doxo_HLD
Z_Shift_L_Doxo_EG
Z_Im_L_Doxo_EG

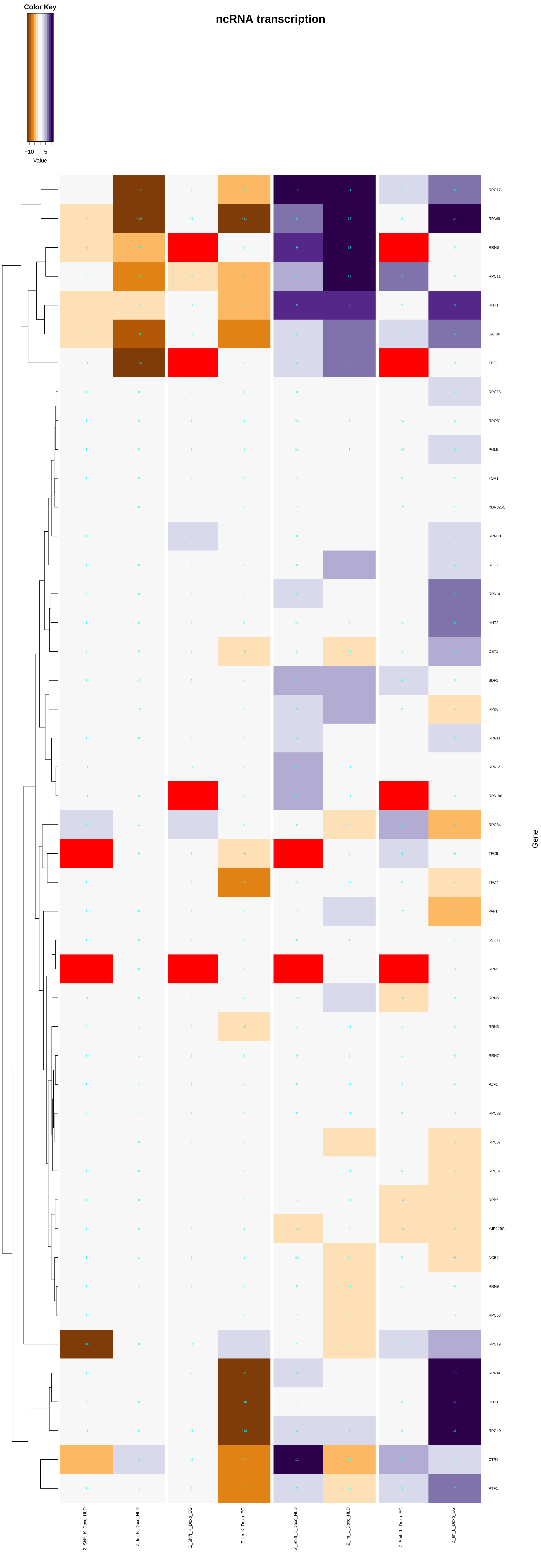
ncRNA transcription
Color Key
−10
5
Value
RPC17
RPA49
RRN6
RPC11
RNT1
UAF30
TBF1
RPC25
RPO31
POL5
TOR1
YDR026C
RRN10
RET1
RPA14
HHT2
DST1
BDF1
RPB8
RPA43
RPA12
RPA190
RPC34
TFC6
TFC7
PAF1
SSU72
RRN11
RRN5
RRN3
RRN7
PZF1
RPC82
RPC37
RPC31
RPB5
YJR119C
NCB2
RRN9
RPC53
RPC19
RPA34
HHT1
RPC40
CTR9
RTF1
Gene
Z_Shift_K_Doxo_HLD
Z_Im_K_Doxo_HLD
Z_Shift_K_Doxo_EG
Z_Im_K_Doxo_EG
Z_Shift_L_Doxo_HLD
Z_Im_L_Doxo_HLD
Z_Shift_L_Doxo_EG
Z_Im_L_Doxo_EG

### transcription of nuclear large rRNA transcript from RNA polymerase I promoter

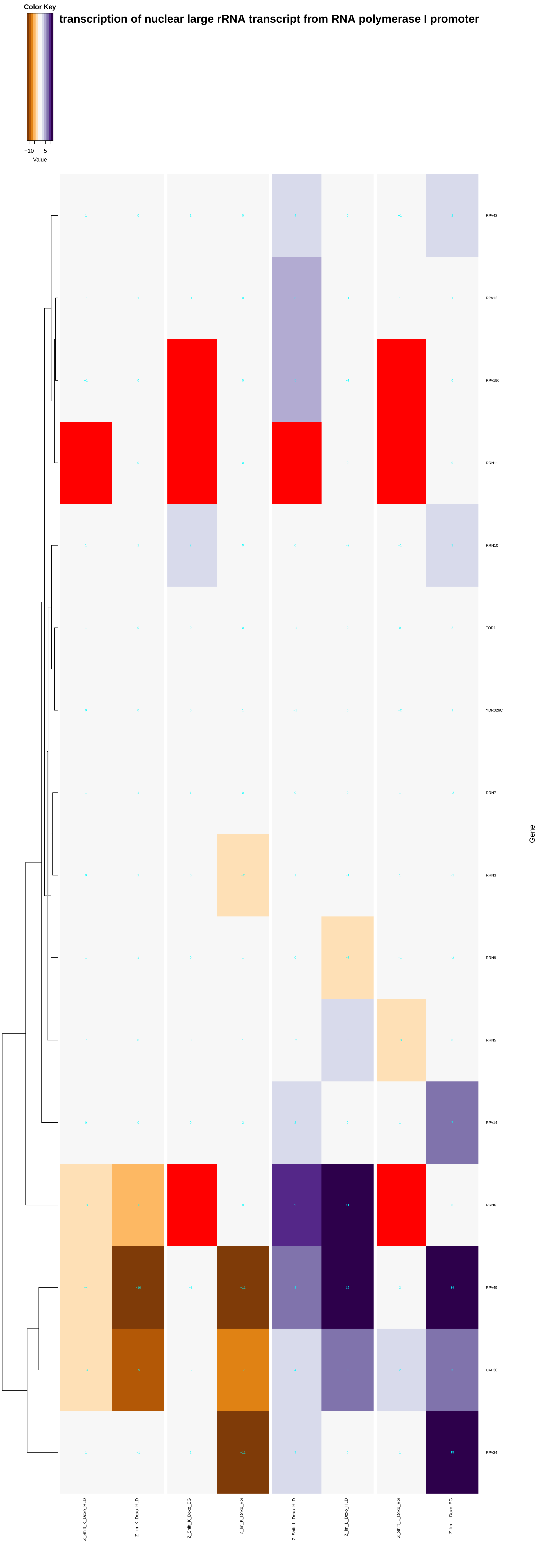

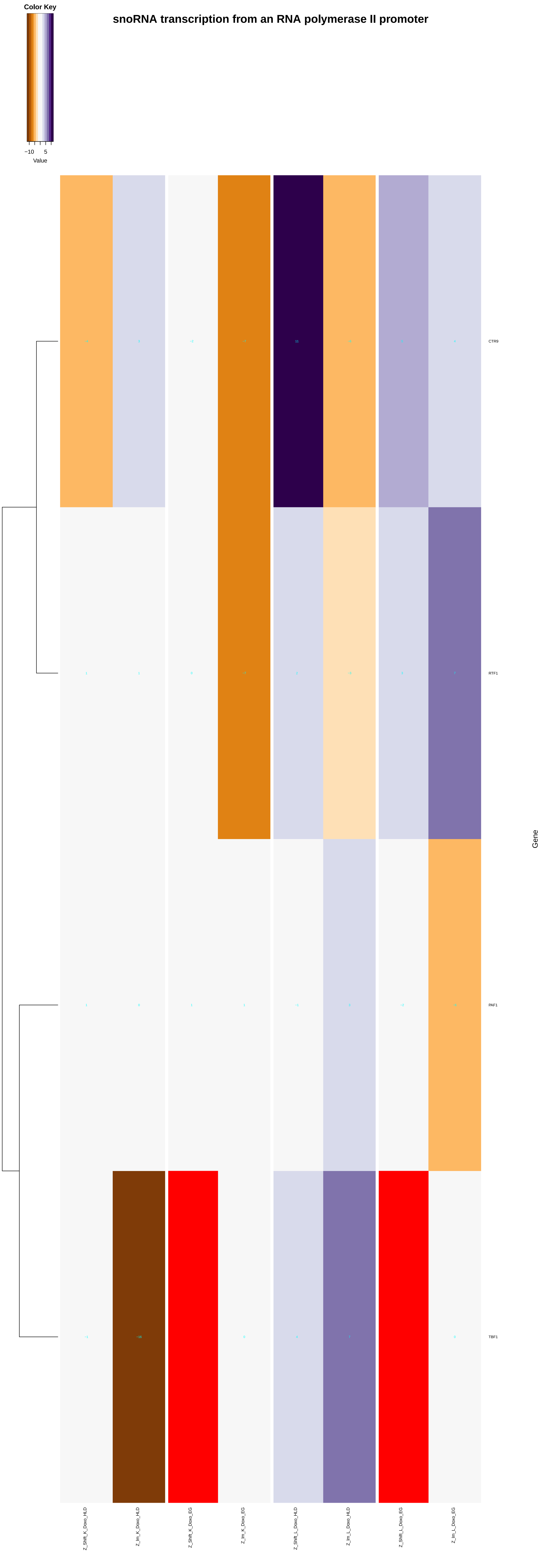
Color Key
−10
5
Value
snoRNA transcription from an RNA polymerase II promoter
CTR9
RTF1
PAF1
TBF1
Gene
Z_Shift_K_Doxo_HLD
Z_Im_K_Doxo_HLD
Z_Shift_K_Doxo_EG
Z_Im_K_Doxo_EG
Z_Shift_L_Doxo_HLD
Z_Im_L_Doxo_HLD
Z_Shift_L_Doxo_EG
Z_Im_L_Doxo_EG

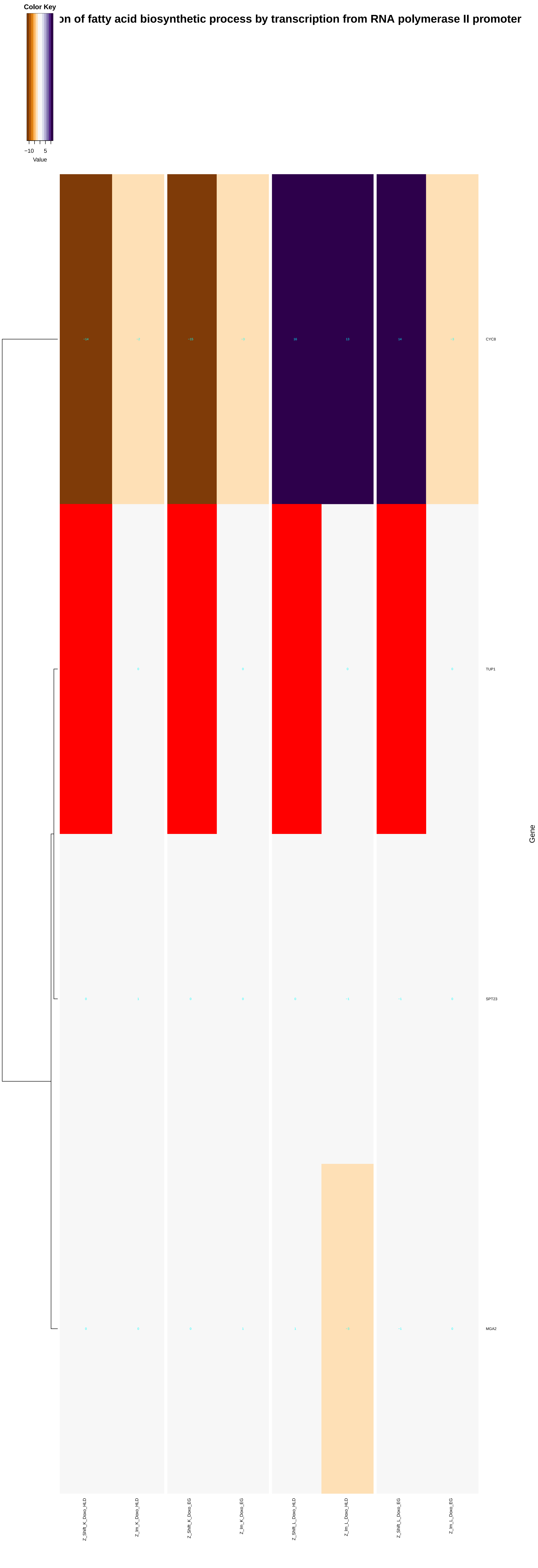

Color Key
−10
5
Value
on of fatty acid biosynthetic process by transcription from RNA polymerase II promoter
CYC8
TUP1
SPT23
MGA2
Gene
Z_Shift_K_Doxo_HLD
Z_Im_K_Doxo_HLD
Z_Shift_K_Doxo_EG
Z_Im_K_Doxo_EG
Z_Shift_L_Doxo_HLD
Z_Im_L_Doxo_HLD
Z_Shift_L_Doxo_EG
Z_Im_L_Doxo_EG

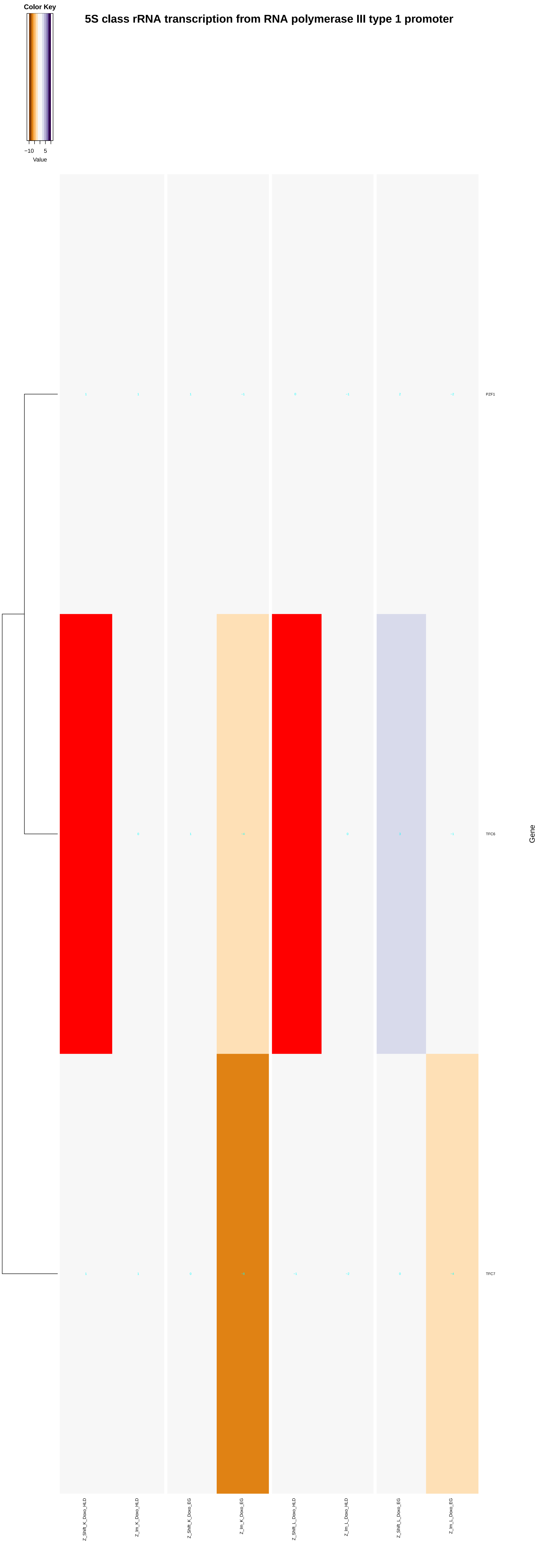

Color Key
−10
5
Value
5S class rRNA transcription from RNA polymerase III type 1 promoter
PZF1
TFC6
TFC7
Gene
Z_Shift_K_Doxo_HLD
Z_Im_K_Doxo_HLD
Z_Shift_K_Doxo_EG
Z_Im_K_Doxo_EG
Z_Shift_L_Doxo_HLD
Z_Im_L_Doxo_HLD
Z_Shift_L_Doxo_EG
Z_Im_L_Doxo_EG

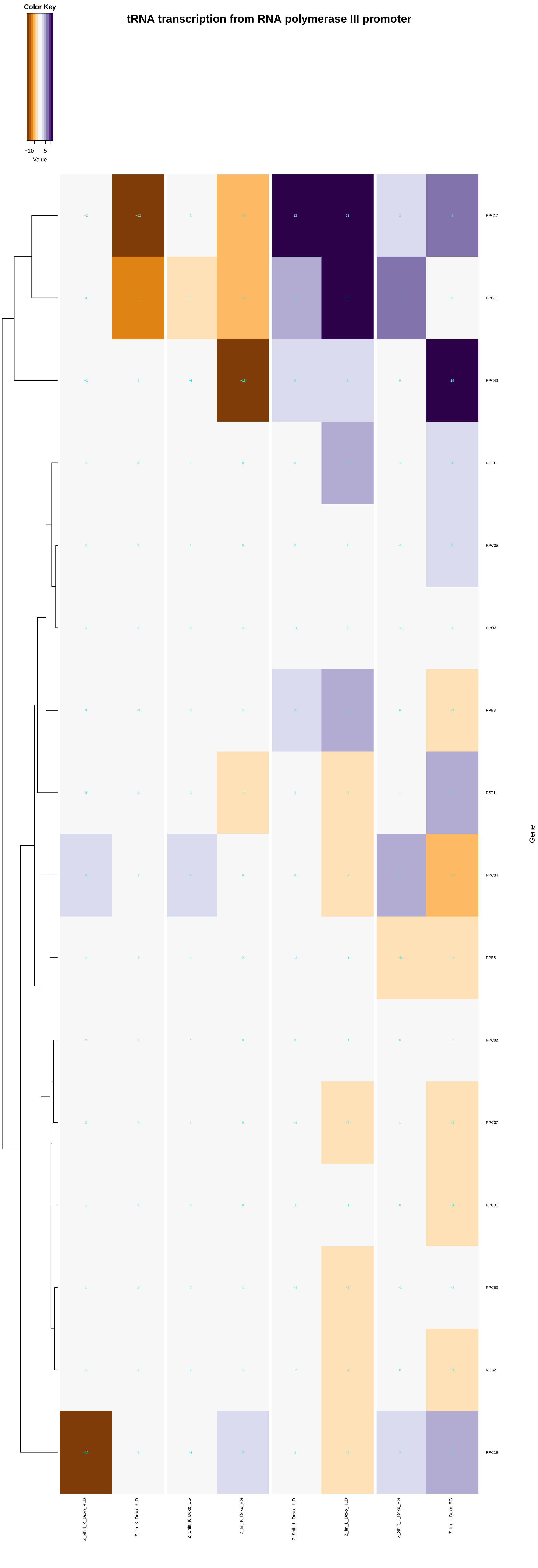
tRNA transcription from RNA polymerase III promoter
Color Key
−10
5
Value
RPC17
RPC11
RPC40
RET1
RPC25
RPO31
RPB8
DST1
RPC34
RPB5
RPC82
RPC37
RPC31
RPC53
NCB2
RPC19
Gene
Z_Shift_K_Doxo_HLD
Z_Im_K_Doxo_HLD
Z_Shift_K_Doxo_EG
Z_Im_K_Doxo_EG
Z_Shift_L_Doxo_HLD
Z_Im_L_Doxo_HLD
Z_Shift_L_Doxo_EG
Z_Im_L_Doxo_EG

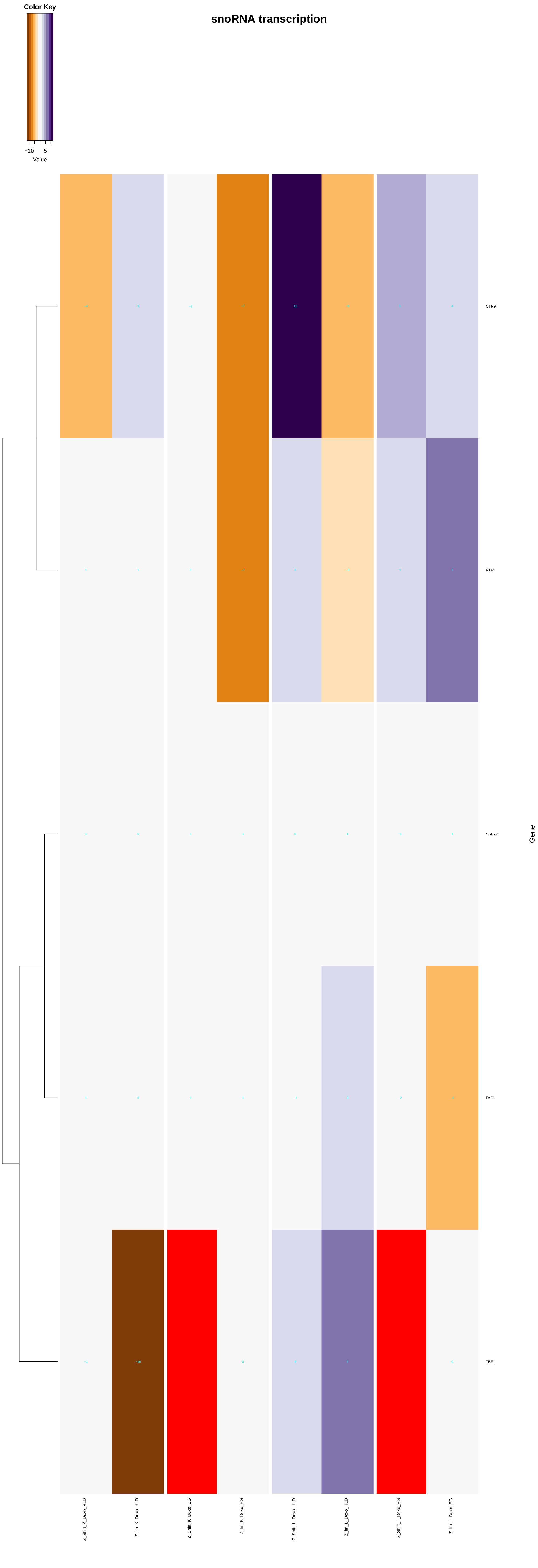
snoRNA transcription
Color Key
−10
5
Value
CTR9
RTF1
SSU72
PAF1
TBF1
Gene
Z_Shift_K_Doxo_HLD
Z_Im_K_Doxo_HLD
Z_Shift_K_Doxo_EG
Z_Im_K_Doxo_EG
Z_Shift_L_Doxo_HLD
Z_Im_L_Doxo_HLD
Z_Shift_L_Doxo_EG
Z_Im_L_Doxo_EG

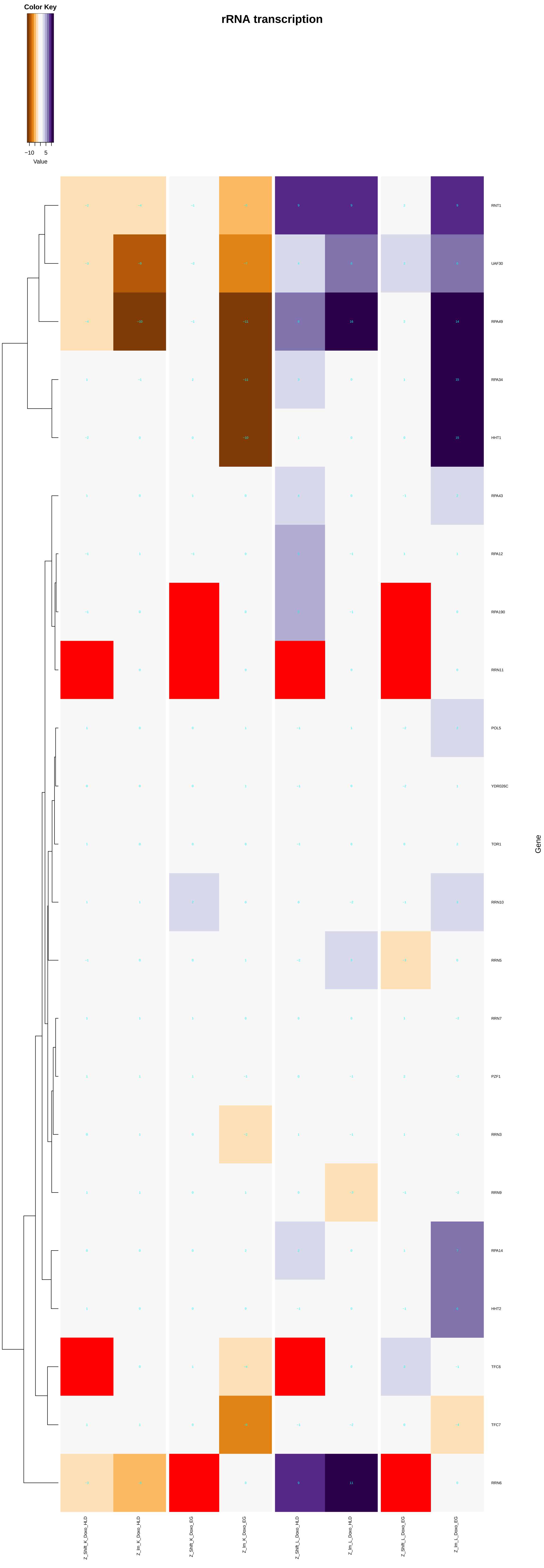
rRNA transcription
Color Key
−10
5
Value
RNT1
UAF30
RPA49
RPA34
HHT1
RPA43
RPA12
RPA190
RRN11
POL5
YDR026C
TOR1
RRN10
RRN5
RRN7
PZF1
RRN3
RRN9
RPA14
HHT2
TFC6
TFC7
RRN6
Gene
Z_Shift_K_Doxo_HLD
Z_Im_K_Doxo_HLD
Z_Shift_K_Doxo_EG
Z_Im_K_Doxo_EG
Z_Shift_L_Doxo_HLD
Z_Im_L_Doxo_HLD
Z_Shift_L_Doxo_EG
Z_Im_L_Doxo_EG

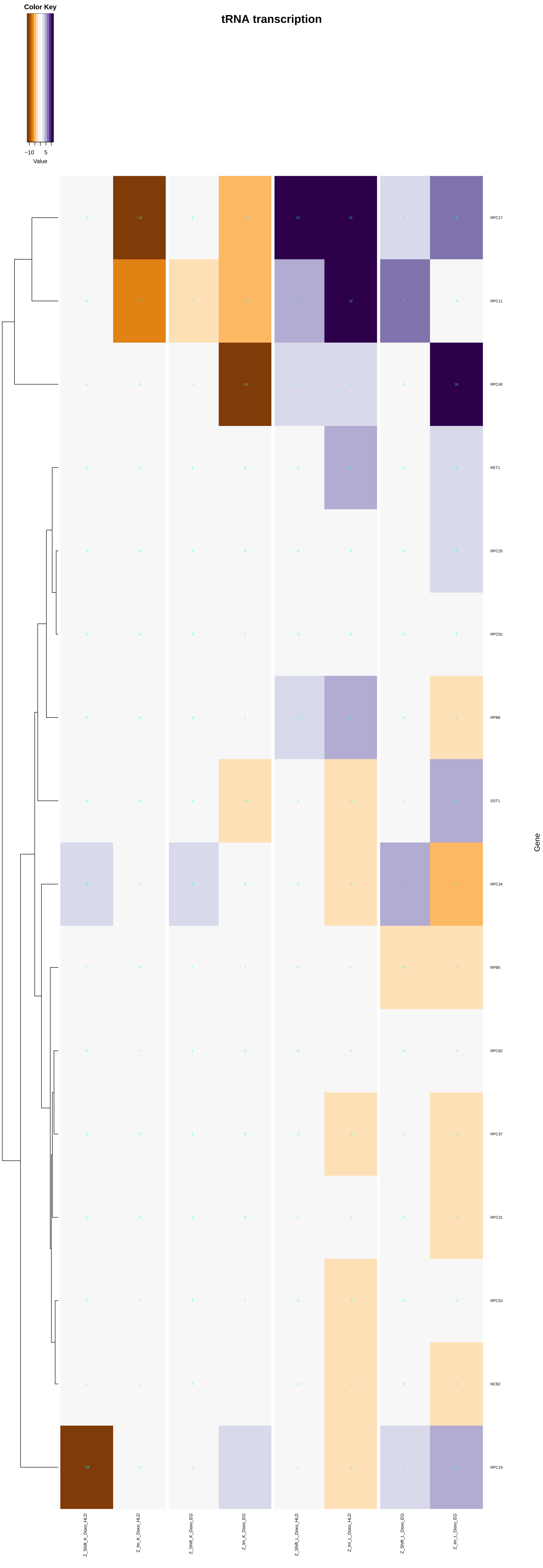
tRNA transcription
Color Key
−10
5
Value
RPC17
RPC11
RPC40
RET1
RPC25
RPO31
RPB8
DST1
RPC34
RPB5
RPC82
RPC37
RPC31
RPC53
NCB2
RPC19
Gene
Z_Shift_K_Doxo_HLD
Z_Im_K_Doxo_HLD
Z_Shift_K_Doxo_EG
Z_Im_K_Doxo_EG
Z_Shift_L_Doxo_HLD
Z_Im_L_Doxo_HLD
Z_Shift_L_Doxo_EG
Z_Im_L_Doxo_EG
