## Additional File 9 for "A yeast phenomic model for the influence of Warburg metabolism on genetic buffering of doxorubicin": nucleic_acid_phosphodiester_bond_hydrolysis.pdf

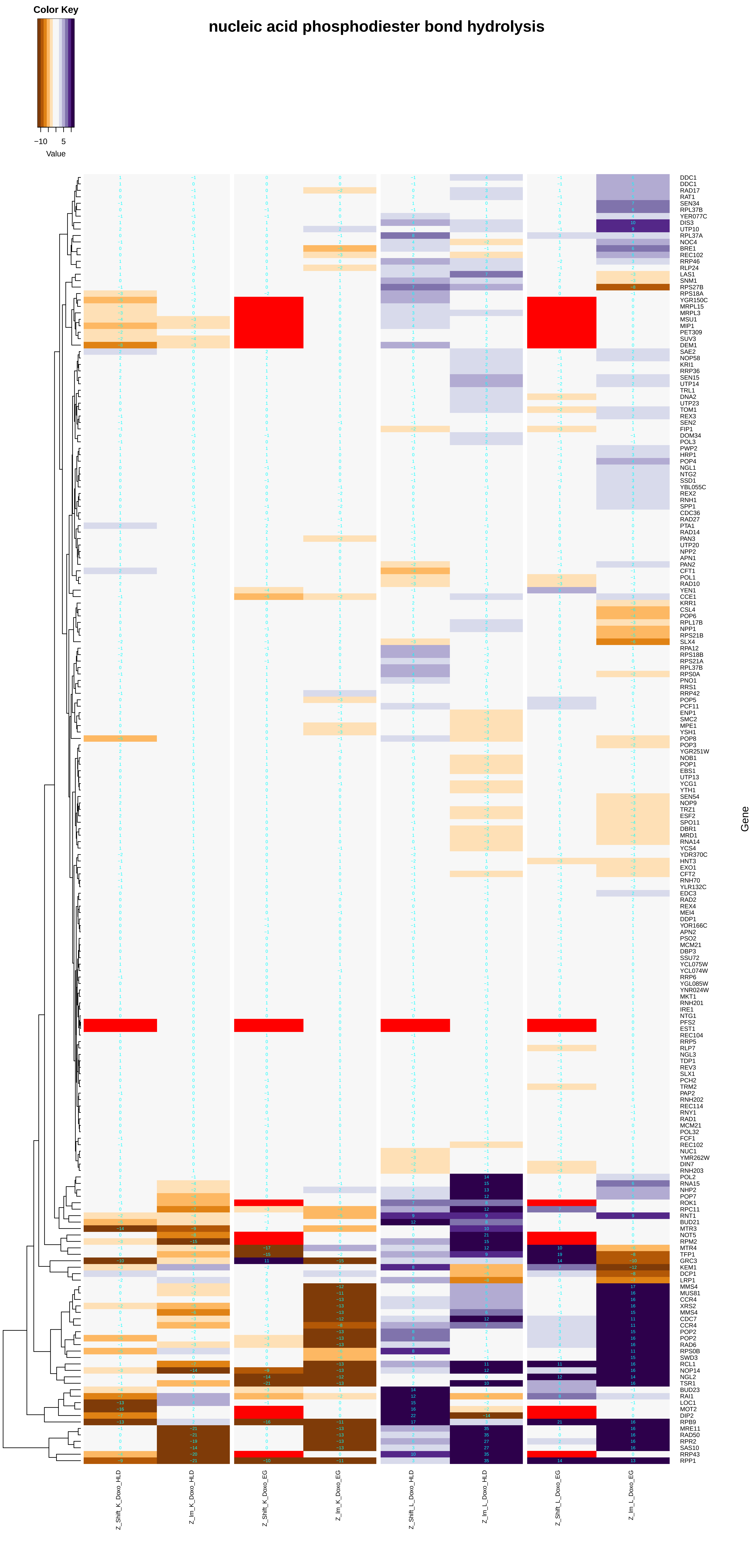

nucleic acid phosphodiester bond hydrolysis
Color Key
−10 5
Value
Gene
DDC1
DDC1
RAD17
RAT1
SEN34
RPL37B
YER077C
DIS3
UTP10
RPL37A
NOC4
BRE1
REC102
RRP46
RLP24
LAS1
SNM1
RPS27B
RPS18A
YGR150C
MRPL15
MRPL3
MSU1
MIP1
PET309
SUV3
DEM1
SAE2
NOP58
KRI1
RRP36
SEN15
UTP14
TRL1
DNA2
UTP23
TOM1
REX3
SEN2
FIP1
DOM34
POL3
PWP2
HRP1
POP4
NGL1
NTG2
SSD1
YBL055C
REX2
RNH1
SPP1
CDC36
RAD27
PTA1
RAD14
PAN3
UTP20
NPP2
APN1
PAN2
CFT1
POL1
RAD10
YEN1
CCE1
KRR1
CSL4
POP6
RPL17B
NPP1
RPS21B
SLX4
RPA12
RPS18B
RPS21A
RPL37B
RPS0A
PNO1
RRS1
RRP42
POP5
PCF11
ENP1
SMC2
MPE1
YSH1
POP8
POP3
YGR251W
NOB1
POP1
EBS1
UTP13
YCG1
YTH1
SEN54
NOP9
TRZ1
ESF2
SPO11
DBR1
MRD1
RNA14
YCS4
YDR370C
HNT3
EXO1
CFT2
RNH70
YLR132C
EDC3
RAD2
REX4
MEI4
DDP1
YOR166C
APN2
PSO2
MCM21
DBP3
SSU72
YCL075W
YCL074W
RRP6
YGL085W
YNR024W
MKT1
RNH201
IRE1
NTG1
PFS2
EST1
REC104
RRP5
RLP7
NGL3
TDP1
REV3
SLX1
PCH2
TRM2
PAP2
RNH202
REC114
RNY1
RAD1
MCM21
POL32
FCF1
REC102
NUC1
YMR262W
DIN7
RNH203
POL2
RNA15
NHP2
POP7
ROK1
RPC11
RNT1
BUD21
MTR3
NOT5
RPM2
MTR4
TFP1
GRC3
KEM1
DCP1
LRP1
MMS4
MUS81
CCR4
XRS2
MMS4
CDC7
CCR4
POP2
POP2
RAD6
RPS0B
SWD3
RCL1
NOP14
NGL2
TSR1
BUD23
RAI1
LOC1
MOT2
DIP2
RPB9
MRE11
RAD50
RPR2
SAS10
RRP43
RPP1
Z_Shift_K_Doxo_HLD
Z_lm_K_Doxo_HLD
Z_Shift_K_Doxo_EG
Z_lm_K_Doxo_EG
Z_Shift_L_Doxo_HLD
Z_lm_L_Doxo_HLD
Z_Shift_L_Doxo_EG
Z_lm_L_Doxo_EG

### DNA catabolic process, endonucleolytic

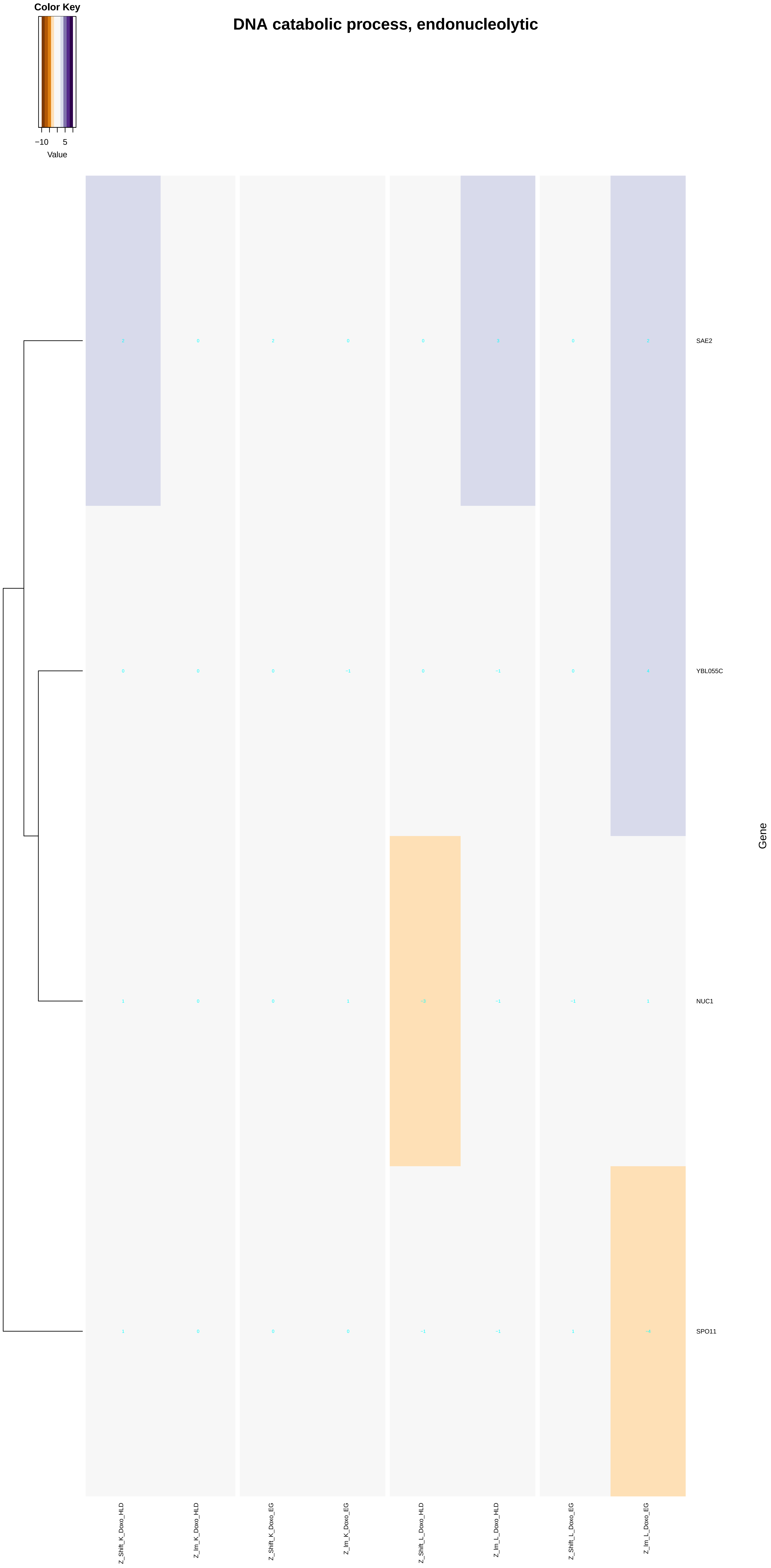

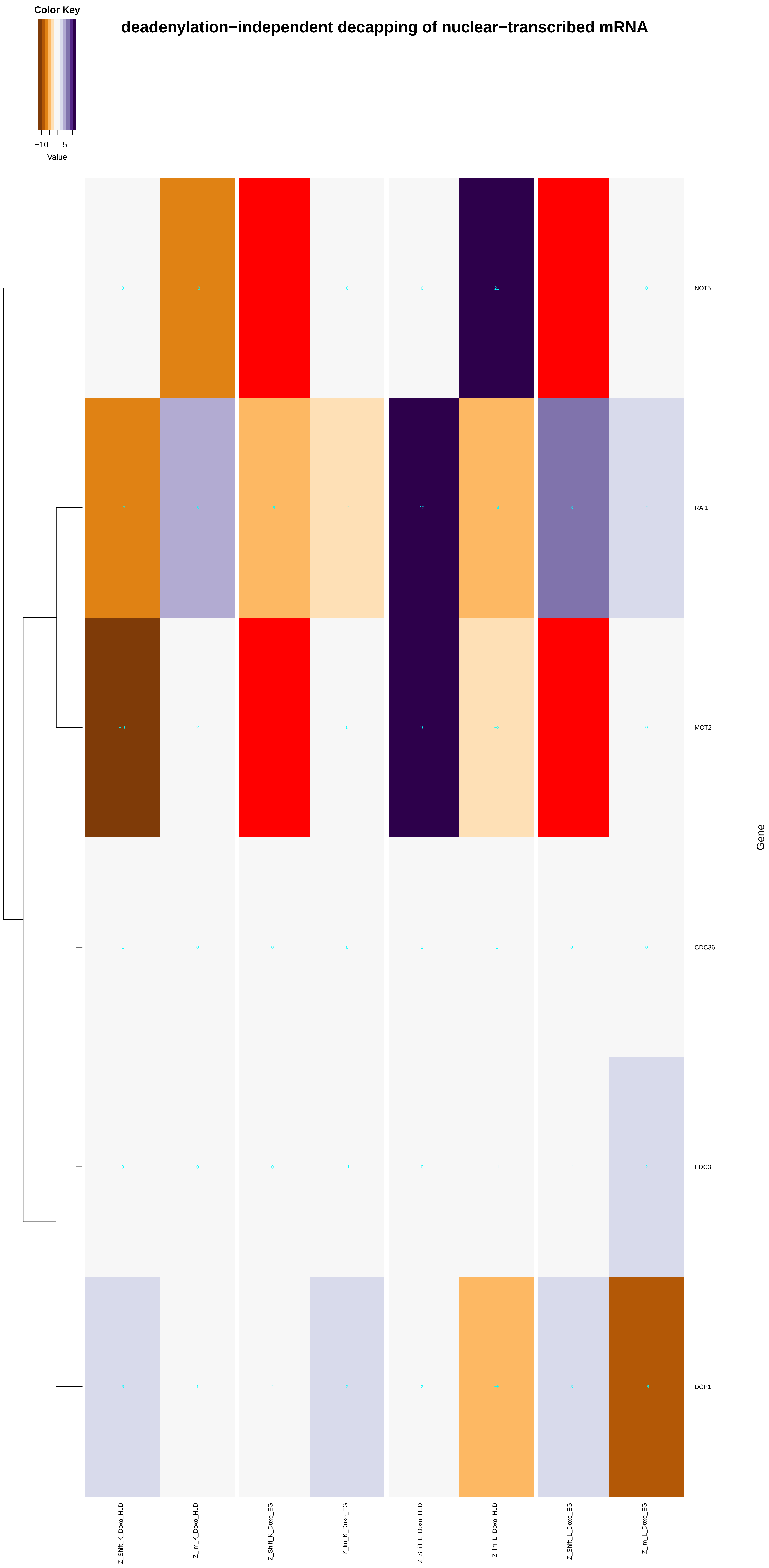
Color Key
−10
5
Value
deadenylation−independent decapping of nuclear−transcribed mRNA
NOT5
RAI1
MOT2
CDC36
EDC3
DCP1
Gene
Z_Shift_K_Doxo_HLD
Z_Im_K_Doxo_HLD
Z_Shift_K_Doxo_EG
Z_Im_K_Doxo_EG
Z_Shift_L_Doxo_HLD
Z_Im_L_Doxo_HLD
Z_Shift_L_Doxo_EG
Z_Im_L_Doxo_EG

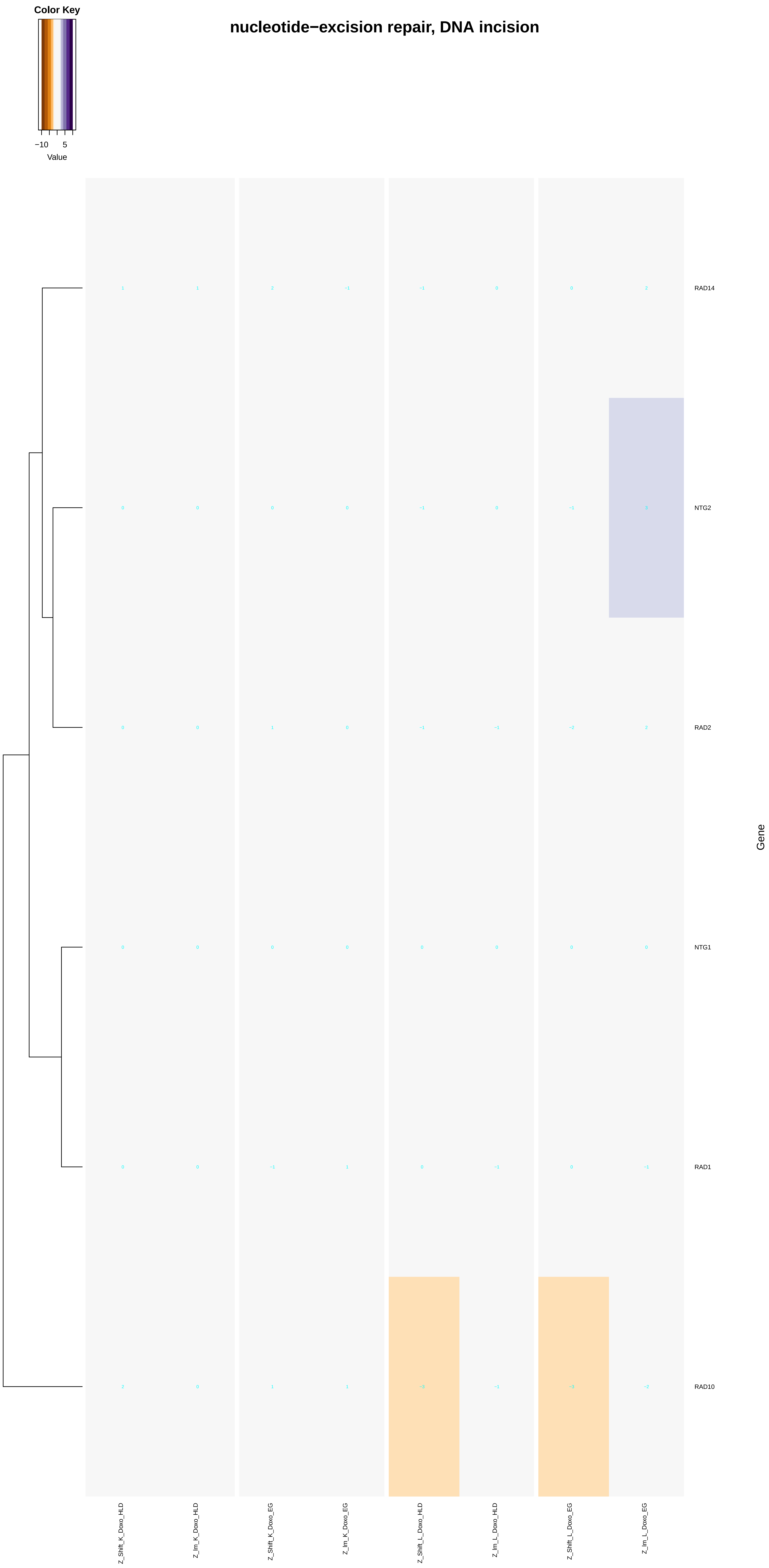
nucleotide−excision repair, DNA incision
Color Key
−10
5
Value
RAD14
NTG2
RAD2
NTG1
RAD1
RAD10
Gene
Z_Shift_K_Doxo_HLD
Z_Im_K_Doxo_HLD
Z_Shift_K_Doxo_EG
Z_Im_K_Doxo_EG
Z_Shift_L_Doxo_HLD
Z_Im_L_Doxo_HLD
Z_Shift_L_Doxo_EG
Z_Im_L_Doxo_EG

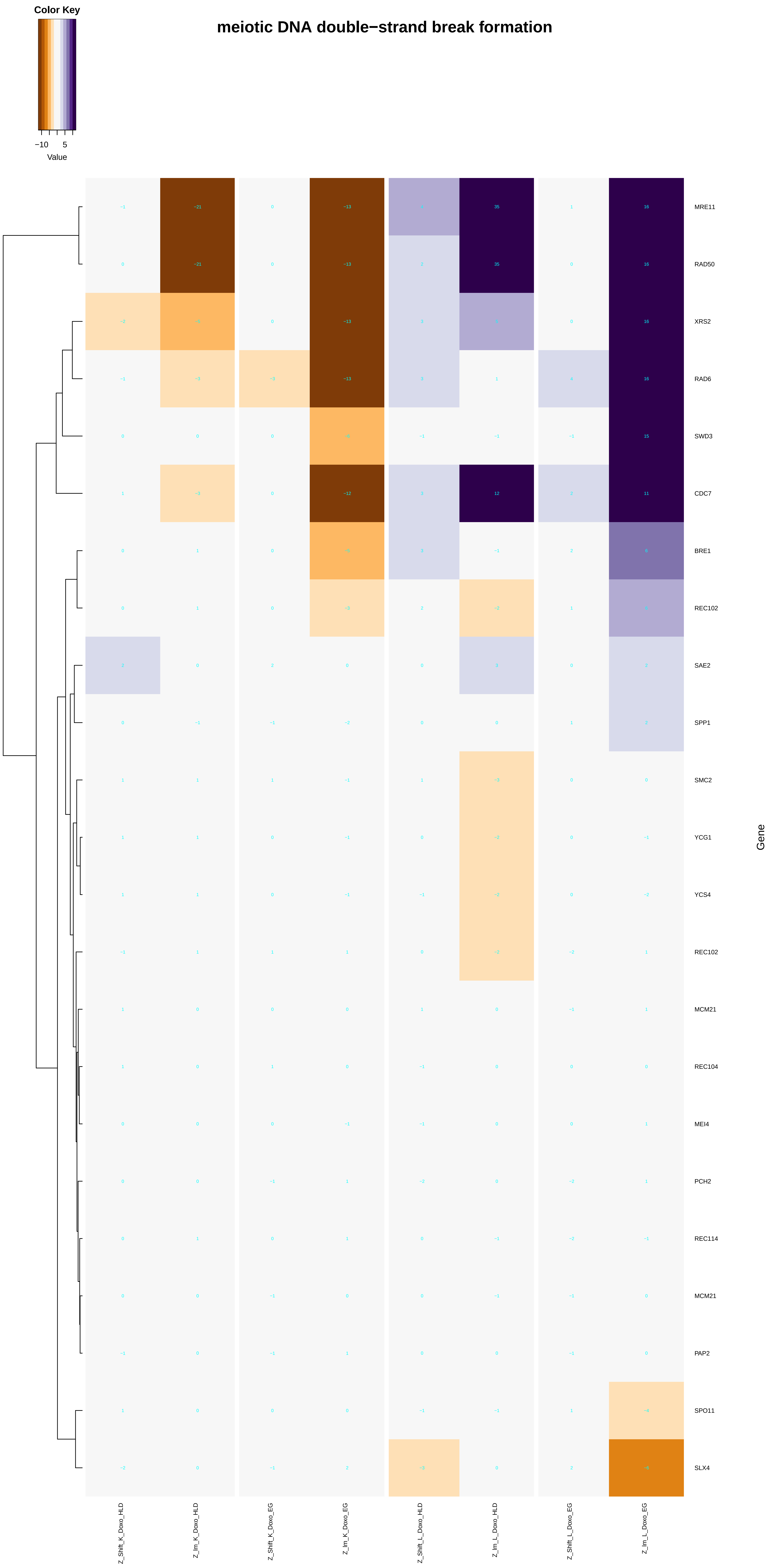
meiotic DNA double−strand break formation
Color Key
−10
5
Value
MRE11
RAD50
XRS2
RAD6
SWD3
CDC7
BRE1
REC102
SAE2
SPP1
SMC2
YCG1
YCS4
REC102
MCM21
REC104
MEI4
PCH2
REC114
MCM21
PAP2
SPO11
SLX4
Gene
Z_Shift_K_Doxo_HLD
Z_Im_K_Doxo_HLD
Z_Shift_K_Doxo_EG
Z_Im_K_Doxo_EG
Z_Shift_L_Doxo_HLD
Z_Im_L_Doxo_HLD
Z_Shift_L_Doxo_EG
Z_Im_L_Doxo_EG

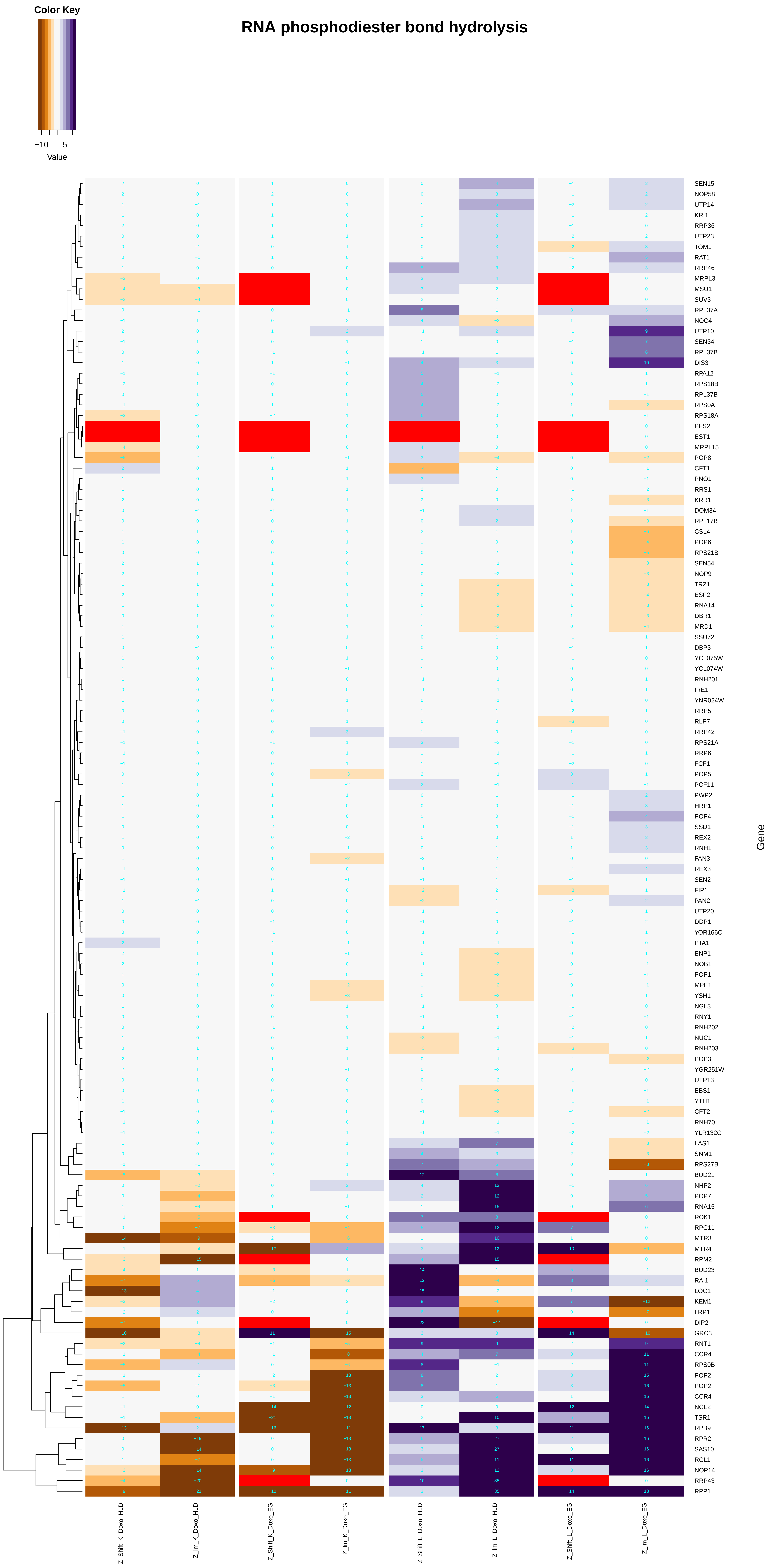
RNA phosphodiester bond hydrolysis
Color Key
−10
5
Value
SEN15
NOP58
UTP14
KRI1
RRP36
UTP23
TOM1
RAT1
RRP46
MRPL3
MSU1
SUV3
RPL37A
NOC4
UTP10
SEN34
RPL37B
DIS3
RPA12
RPS18B
RPL37B
RPS0A
RPS18A
PFS2
EST1
MRPL15
POP8
CFT1
PNO1
RRS1
KRR1
DOM34
RPL17B
CSL4
POP6
RPS21B
SEN54
NOP9
TRZ1
ESF2
RNA14
DBR1
MRD1
SSU72
DBP3
YCL075W
YCL074W
RNH201
IRE1
YNR024W
RRP5
RLP7
RRP42
RPS21A
RRP6
FCF1
POP5
PCF11
PWP2
HRP1
POP4
SSD1
REX2
RNH1
PAN3
REX3
SEN2
FIP1
PAN2
UTP20
DDP1
YOR166C
PTA1
ENP1
NOB1
POP1
MPE1
YSH1
NGL3
RNY1
RNH202
NUC1
RNH203
POP3
YGR251W
UTP13
EBS1
YTH1
CFT2
RNH70
YLR132C
LAS1
SNM1
RPS27B
BUD21
NHP2
POP7
RNA15
ROK1
RPC11
MTR3
MTR4
RPM2
BUD23
RAI1
LOC1
KEM1
LRP1
DIP2
GRC3
RNT1
CCR4
RPS0B
POP2
POP2
CCR4
NGL2
TSR1
RPB9
RPR2
SAS10
RCL1
NOP14
RRP43
RPP1
Gene
Z_Shift_K_Doxo_HLD
Z_Im_K_Doxo_HLD
Z_Shift_K_Doxo_EG
Z_Im_K_Doxo_EG
Z_Shift_L_Doxo_HLD
Z_Im_L_Doxo_HLD
Z_Shift_L_Doxo_EG
Z_Im_L_Doxo_EG

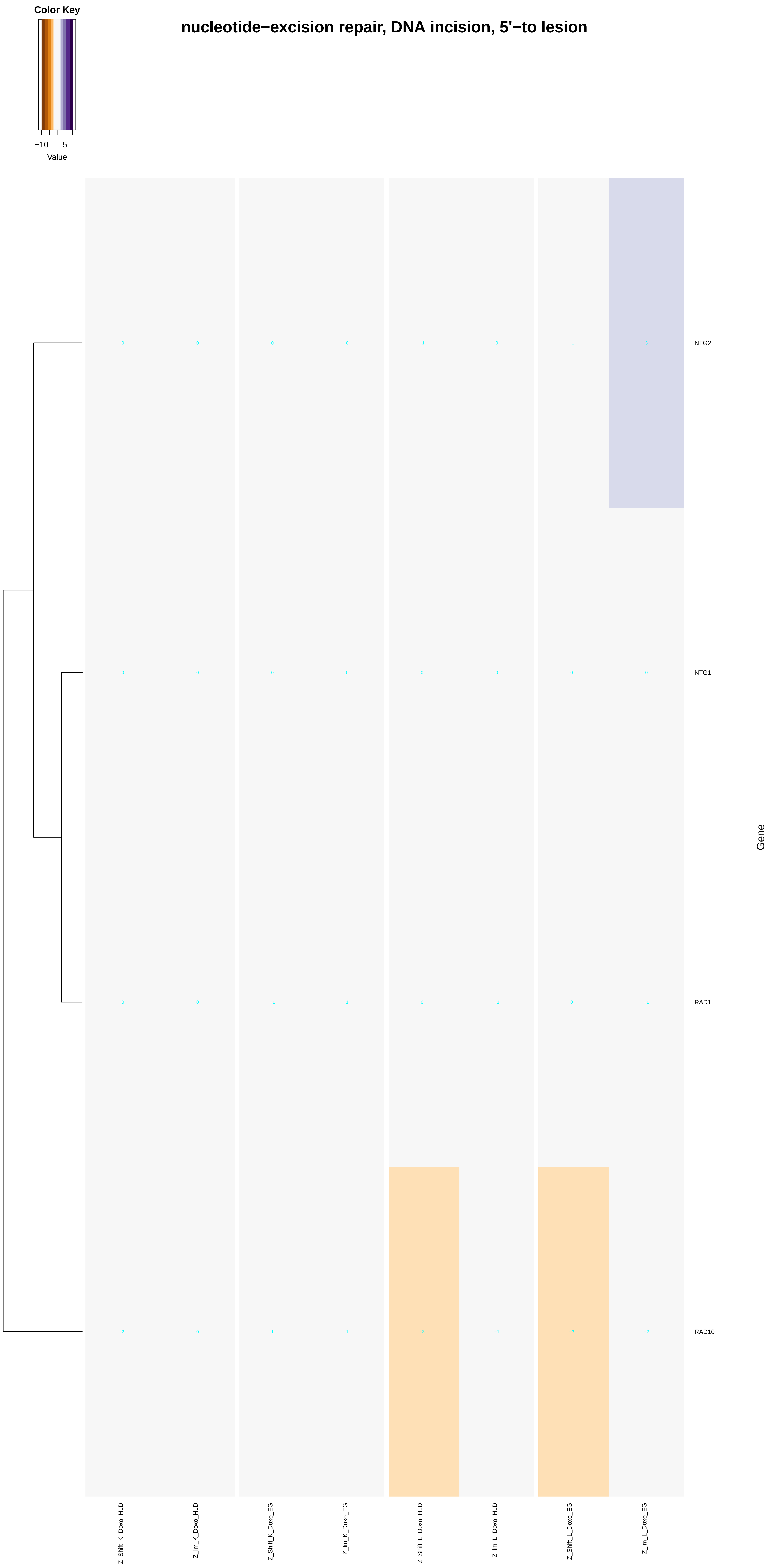
Color Key
−10
5
Value
nucleotide−excision repair, DNA incision, 5'−to lesion
NTG2
NTG1
RAD1
RAD10
Gene
Z_Shift_K_Doxo_HLD
Z_lm_K_Doxo_HLD
Z_Shift_K_Doxo_EG
Z_lm_K_Doxo_EG
Z_Shift_L_Doxo_HLD
Z_lm_L_Doxo_HLD
Z_Shift_L_Doxo_EG
Z_lm_L_Doxo_EG

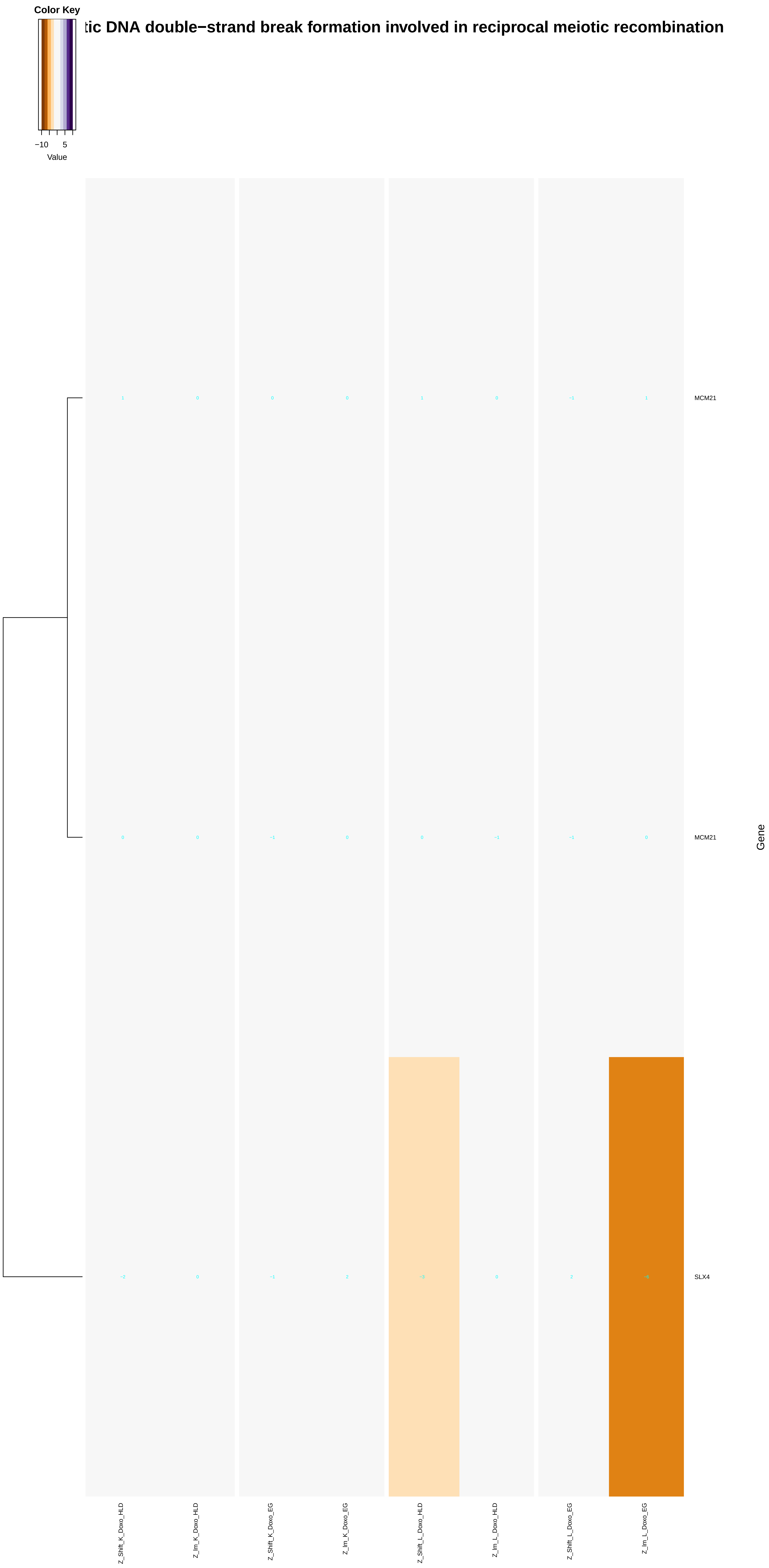

Color Key
−10
5
Value
tic DNA double−strand break formation involved in reciprocal meiotic recombination
MCM21
MCM21
SLX4
Gene
Z_Shift_K_Doxo_HLD
Z_Im_K_Doxo_HLD
Z_Shift_K_Doxo_EG
Z_Im_K_Doxo_EG
Z_Shift_L_Doxo_HLD
Z_Im_L_Doxo_HLD
Z_Shift_L_Doxo_EG
Z_Im_L_Doxo_EG

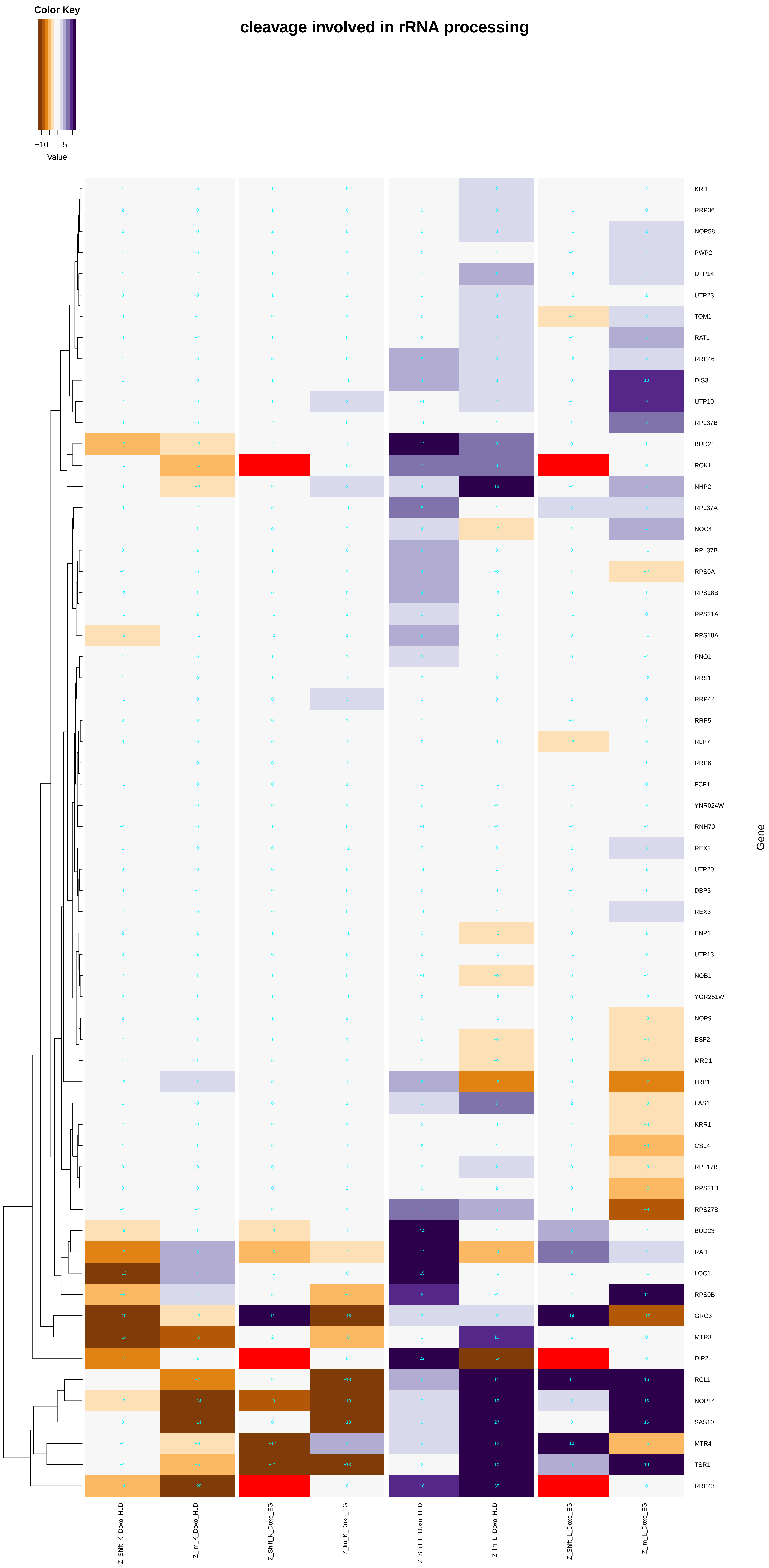
cleavage involved in rRNA processing
Color Key
−10
5
Value
Gene
KRI1
RRP36
NOP58
PWP2
UTP14
UTP23
TOM1
RAT1
RRP46
DIS3
UTP10
RPL37B
BUD21
ROK1
NHP2
RPL37A
NOC4
RPL37B
RPS0A
RPS18B
RPS21A
RPS18A
PNO1
RRS1
RRP42
RRP5
RLP7
RRP6
FCF1
YNR024W
RNH70
REX2
UTP20
DBP3
REX3
ENP1
UTP13
NOB1
YGR251W
NOP9
ESF2
MRD1
LRP1
LAS1
KRR1
CSL4
RPL17B
RPS21B
RPS27B
BUD23
RAI1
LOC1
RPS0B
GRC3
MTR3
DIP2
RCL1
NOP14
SAS10
MTR4
TSR1
RRP43
Z_Shift_K_Doxo_HLD
Z_Im_K_Doxo_HLD
Z_Shift_K_Doxo_EG
Z_Im_K_Doxo_EG
Z_Shift_L_Doxo_HLD
Z_Im_L_Doxo_HLD
Z_Shift_L_Doxo_EG
Z_Im_L_Doxo_EG

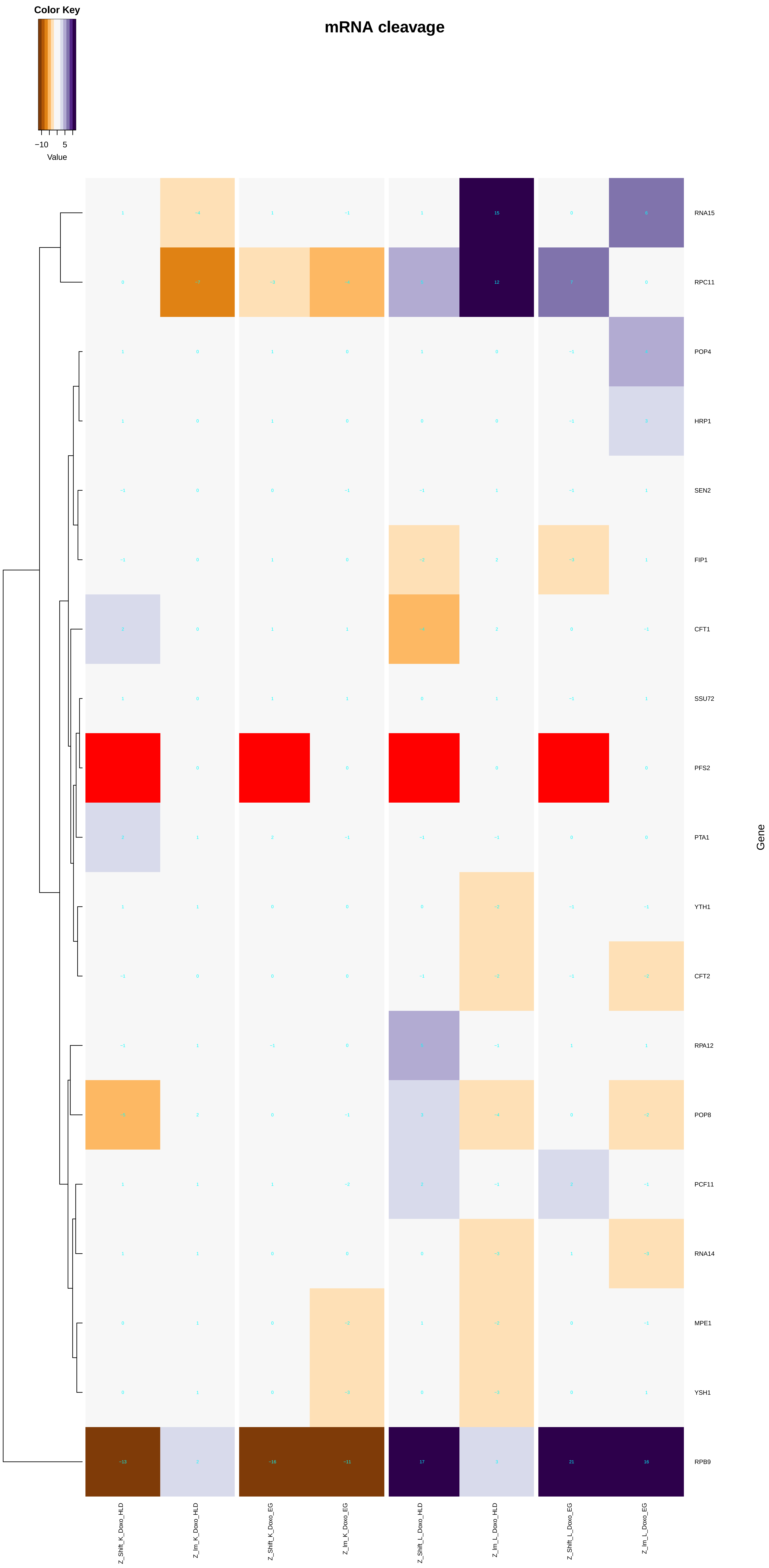

mRNA cleavage
Color Key
−10
5
Value
RNA15
RPC11
POP4
HRP1
SEN2
FIP1
CFT1
SSU72
PFS2
PTA1
YTH1
CFT2
RPA12
POP8
PCF11
RNA14
MPE1
YSH1
RPB9
Gene
Z_Shift_K_Doxo_HLD
Z_lm_K_Doxo_HLD
Z_Shift_K_Doxo_EG
Z_lm_K_Doxo_EG
Z_Shift_L_Doxo_HLD
Z_lm_L_Doxo_HLD
Z_Shift_L_Doxo_EG
Z_lm_L_Doxo_EG

### RNA phosphodiester bond hydrolysis, endonucleolytic

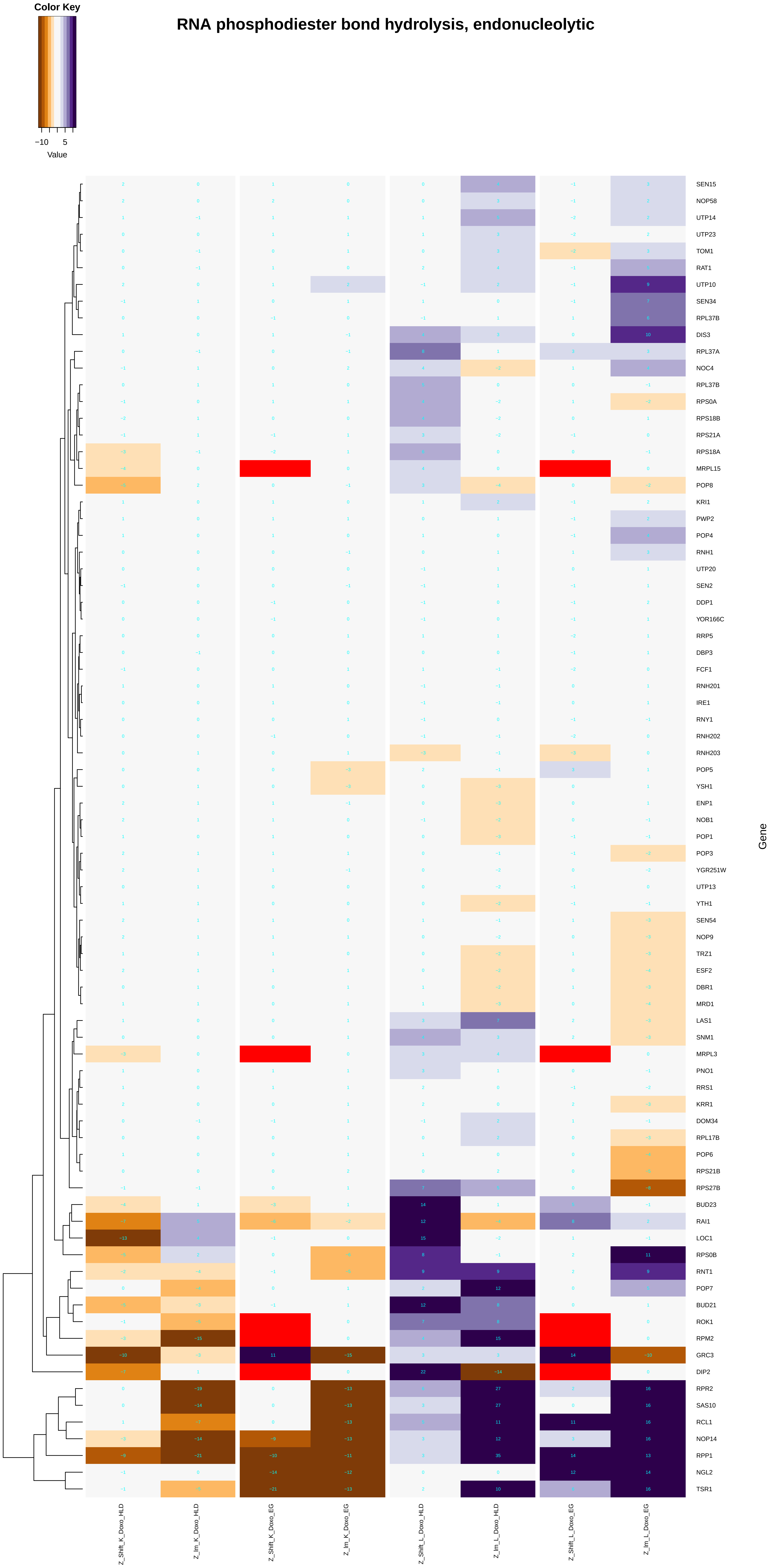

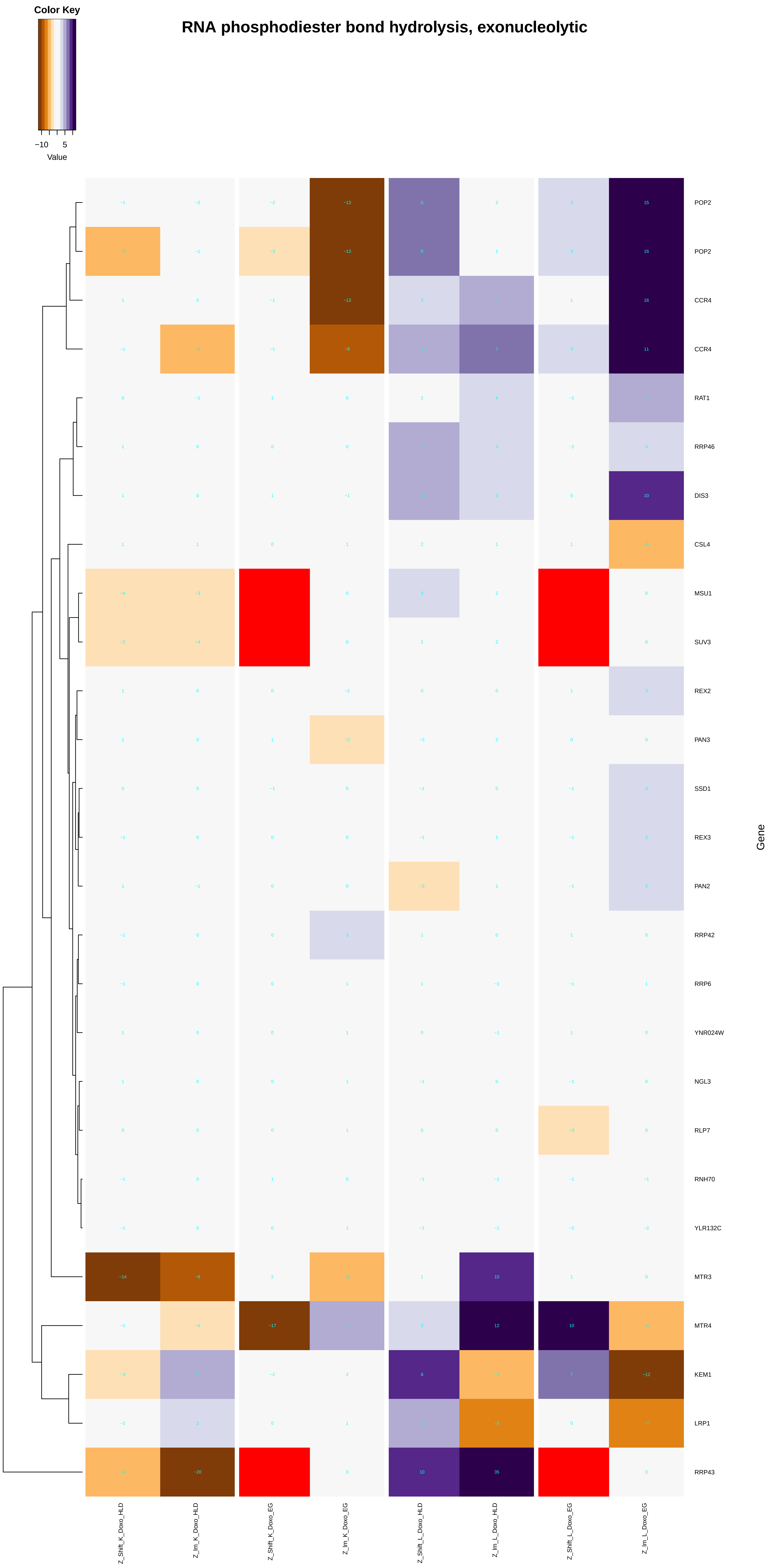
Color Key
−10
5
Value
RNA phosphodiester bond hydrolysis, exonucleolytic
POP2
POP2
CCR4
CCR4
RAT1
RRP46
DIS3
CSL4
MSU1
SUV3
REX2
PAN3
SSD1
REX3
PAN2
RRP42
RRP6
YNR024W
NGL3
RLP7
RNH70
YLR132C
MTR3
MTR4
KEM1
LRP1
RRP43
Gene
Z_Shift_K_Doxo_HLD
Z_lm_K_Doxo_HLD
Z_Shift_K_Doxo_EG
Z_lm_K_Doxo_EG
Z_Shift_L_Doxo_HLD
Z_lm_L_Doxo_HLD
Z_Shift_L_Doxo_EG
Z_lm_L_Doxo_EG

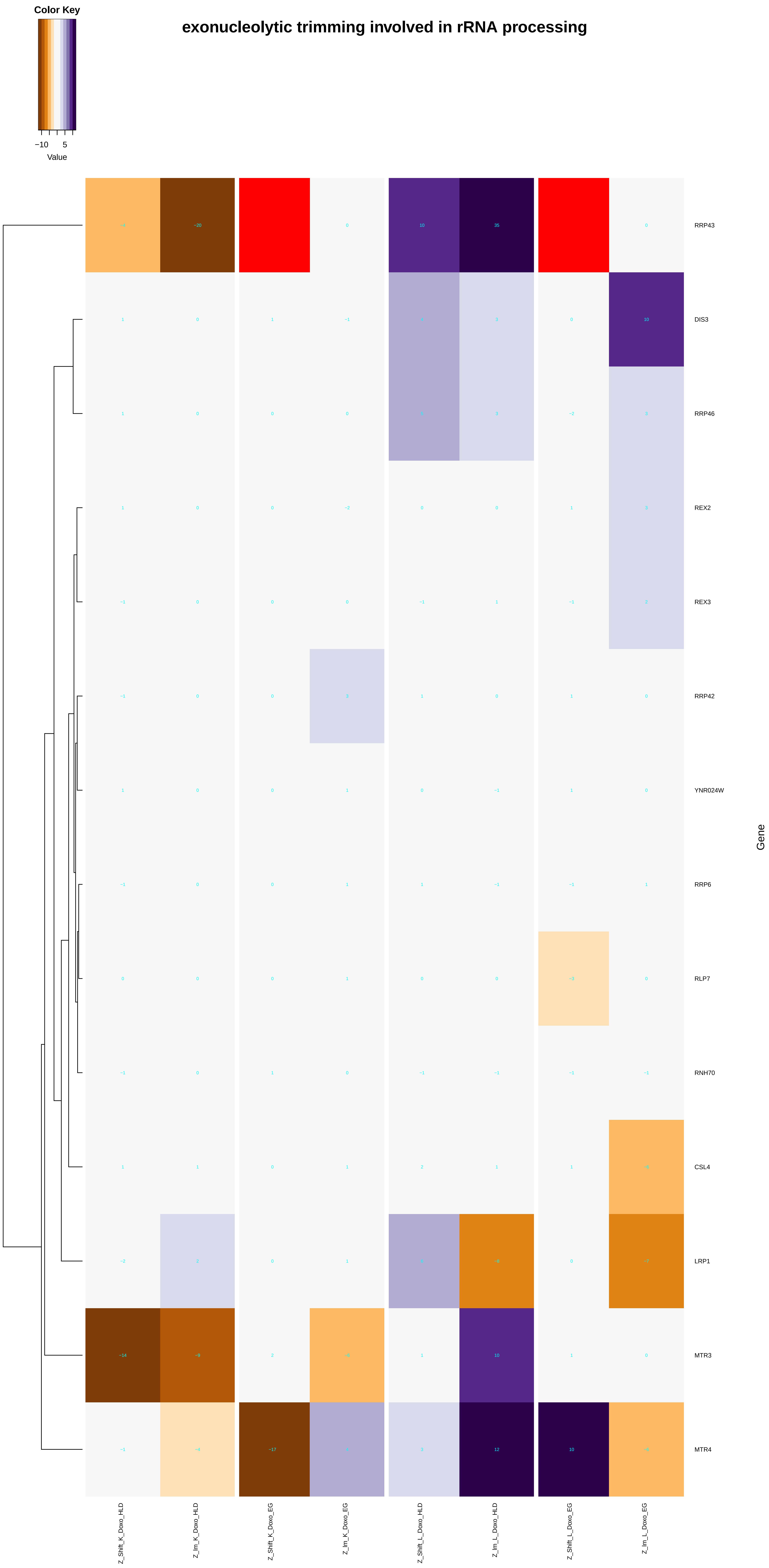
Color Key
−10
5
Value
exonucleolytic trimming involved in rRNA processing
RRP43
DIS3
RRP46
REX2
REX3
RRP42
YNR024W
RRP6
RLP7
RNH70
CSL4
LRP1
MTR3
MTR4
Gene
Z_Shift_K_Doxo_HLD
Z_lm_K_Doxo_HLD
Z_Shift_K_Doxo_EG
Z_lm_K_Doxo_EG
Z_Shift_L_Doxo_HLD
Z_lm_L_Doxo_HLD
Z_Shift_L_Doxo_EG
Z_lm_L_Doxo_EG

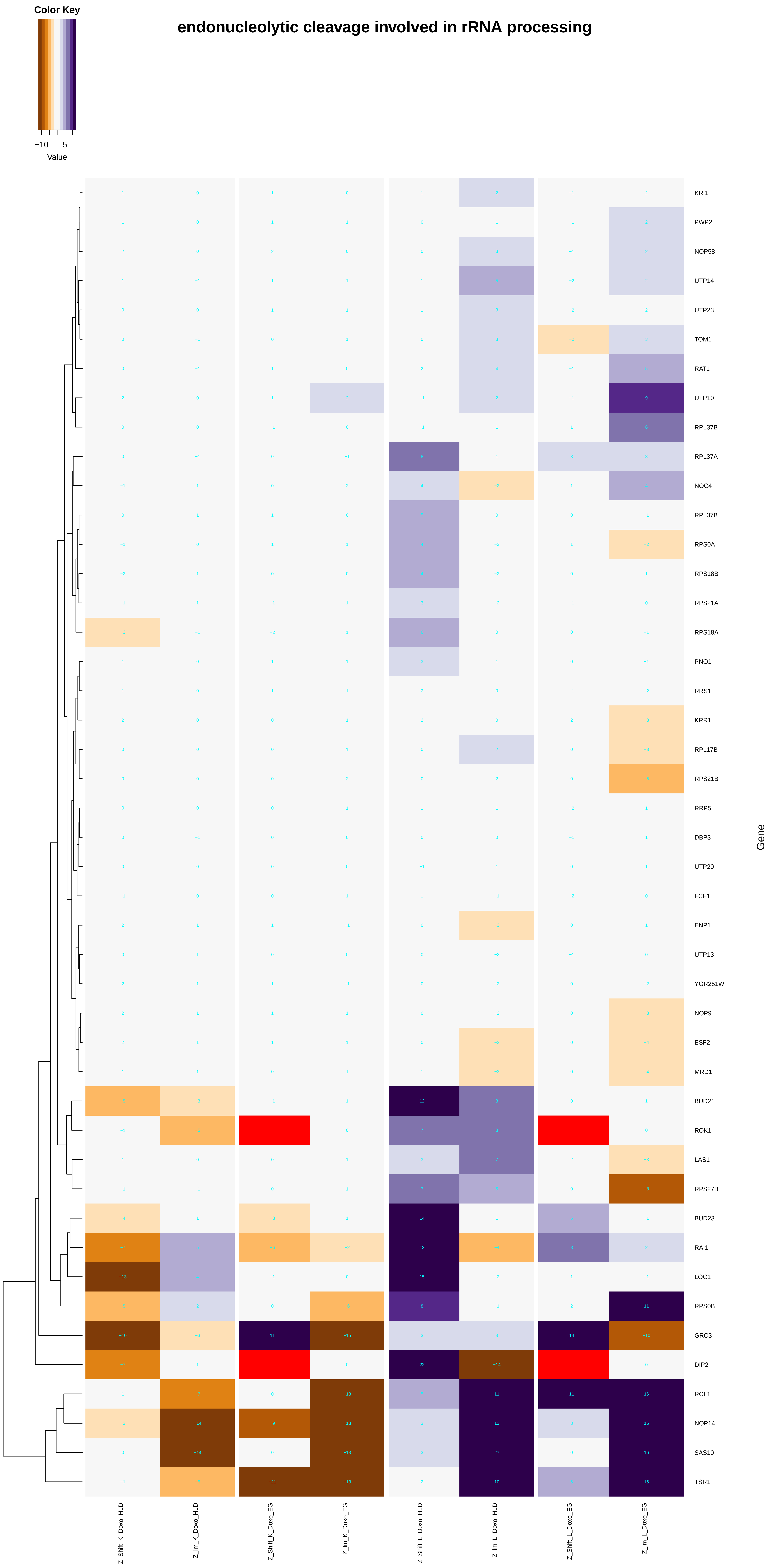

Color Key
−10
5
Value
endonucleolytic cleavage involved in rRNA processing
KRI1
PWP2
NOP58
UTP14
UTP23
TOM1
RAT1
UTP10
RPL37B
RPL37A
NOC4
RPL37B
RPS0A
RPS18B
RPS21A
RPS18A
PNO1
RRS1
KRR1
RPL17B
RPS21B
RRP5
DBP3
UTP20
FCF1
ENP1
UTP13
YGR251W
NOP9
ESF2
MRD1
BUD21
ROK1
LAS1
RPS27B
BUD23
RAI1
LOC1
RPS0B
GRC3
DIP2
RCL1
NOP14
SAS10
TSR1
Gene
Z_Shift_K_Doxo_HLD
Z_Im_K_Doxo_HLD
Z_Shift_K_Doxo_EG
Z_Im_K_Doxo_EG
Z_Shift_L_Doxo_HLD
Z_Im_L_Doxo_HLD
Z_Shift_L_Doxo_EG
Z_Im_L_Doxo_EG

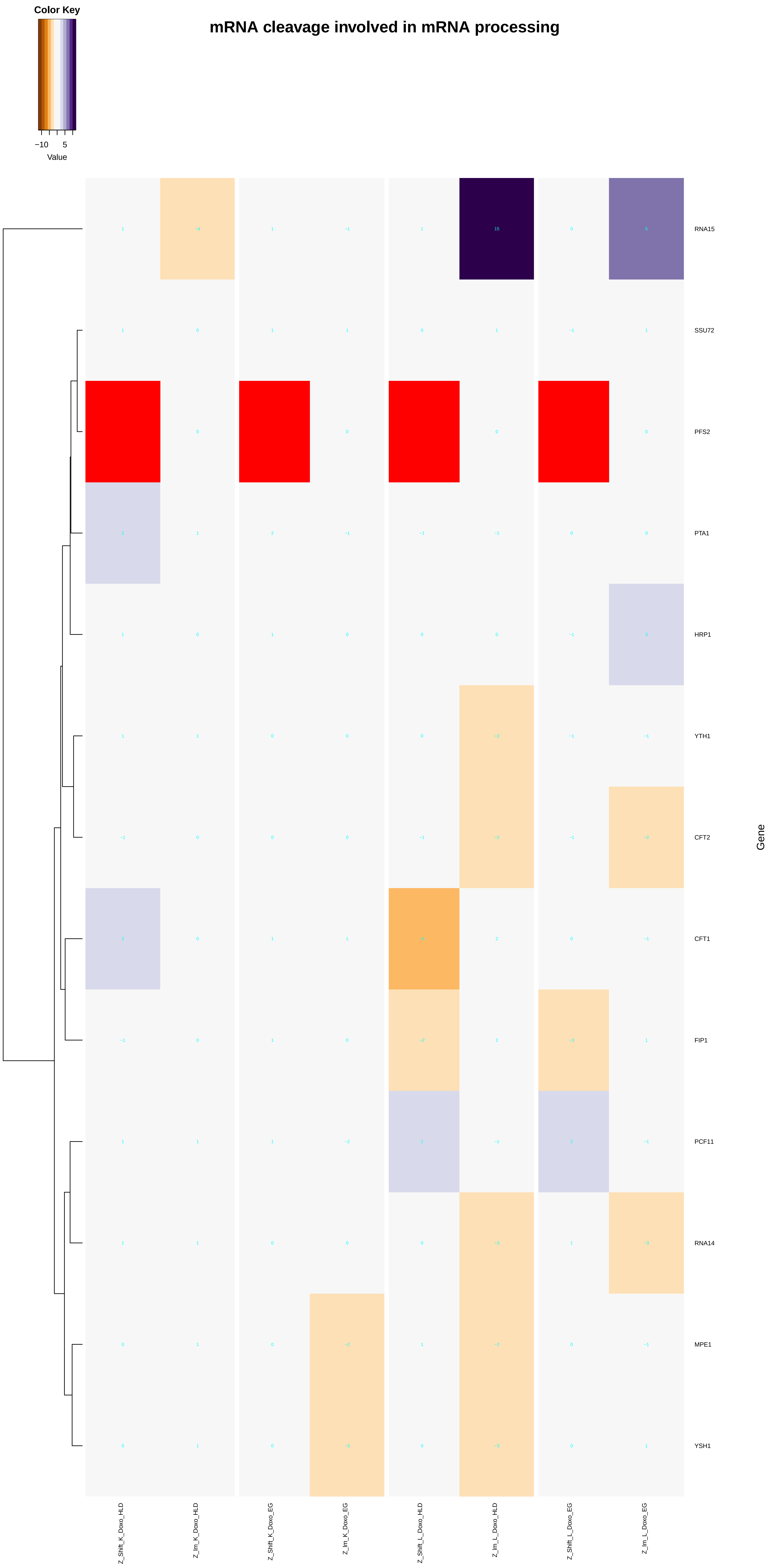

Color Key
−10
5
Value
mRNA cleavage involved in mRNA processing
RNA15
SSU72
PFS2
PTA1
HRP1
YTH1
CFT2
CFT1
FIP1
PCF11
RNA14
MPE1
YSH1
Gene
Z_Shift_K_Doxo_HLD
Z_Im_K_Doxo_HLD
Z_Shift_K_Doxo_EG
Z_Im_K_Doxo_EG
Z_Shift_L_Doxo_HLD
Z_Im_L_Doxo_HLD
Z_Shift_L_Doxo_EG
Z_Im_L_Doxo_EG

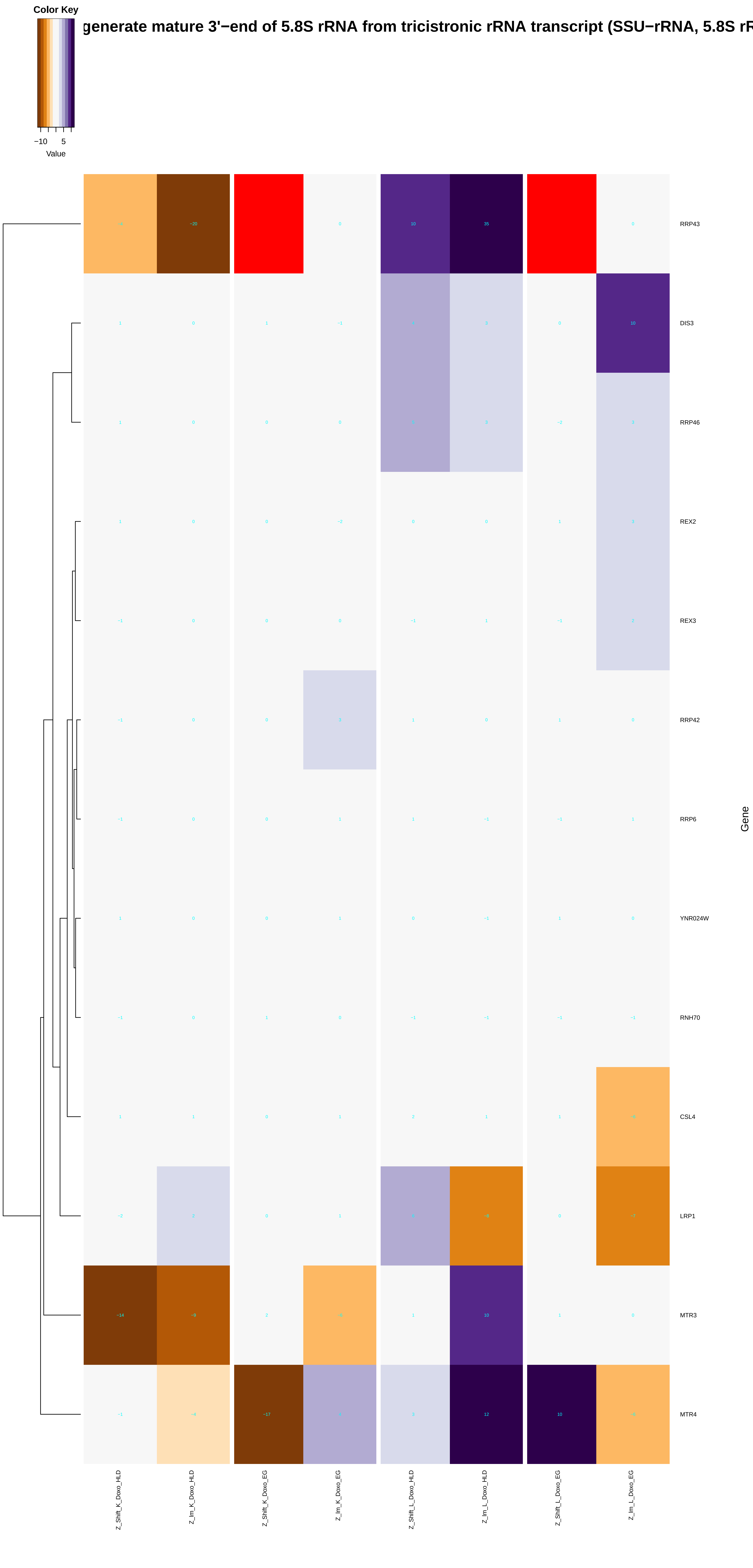

Color Key
−10
5
Value
generate mature 3'−end of 5.8S rRNA from tricistronic rRNA transcript (SSU−rRNA, 5.8S rR
RRP43
DIS3
RRP46
REX2
REX3
RRP42
RRP6
YNR024W
RNH70
CSL4
LRP1
MTR3
MTR4
Gene
Z_Shift_K_Doxo_HLD
Z_lm_K_Doxo_HLD
Z_Shift_K_Doxo_EG
Z_lm_K_Doxo_EG
Z_Shift_L_Doxo_HLD
Z_lm_L_Doxo_HLD
Z_Shift_L_Doxo_EG
Z_lm_L_Doxo_EG

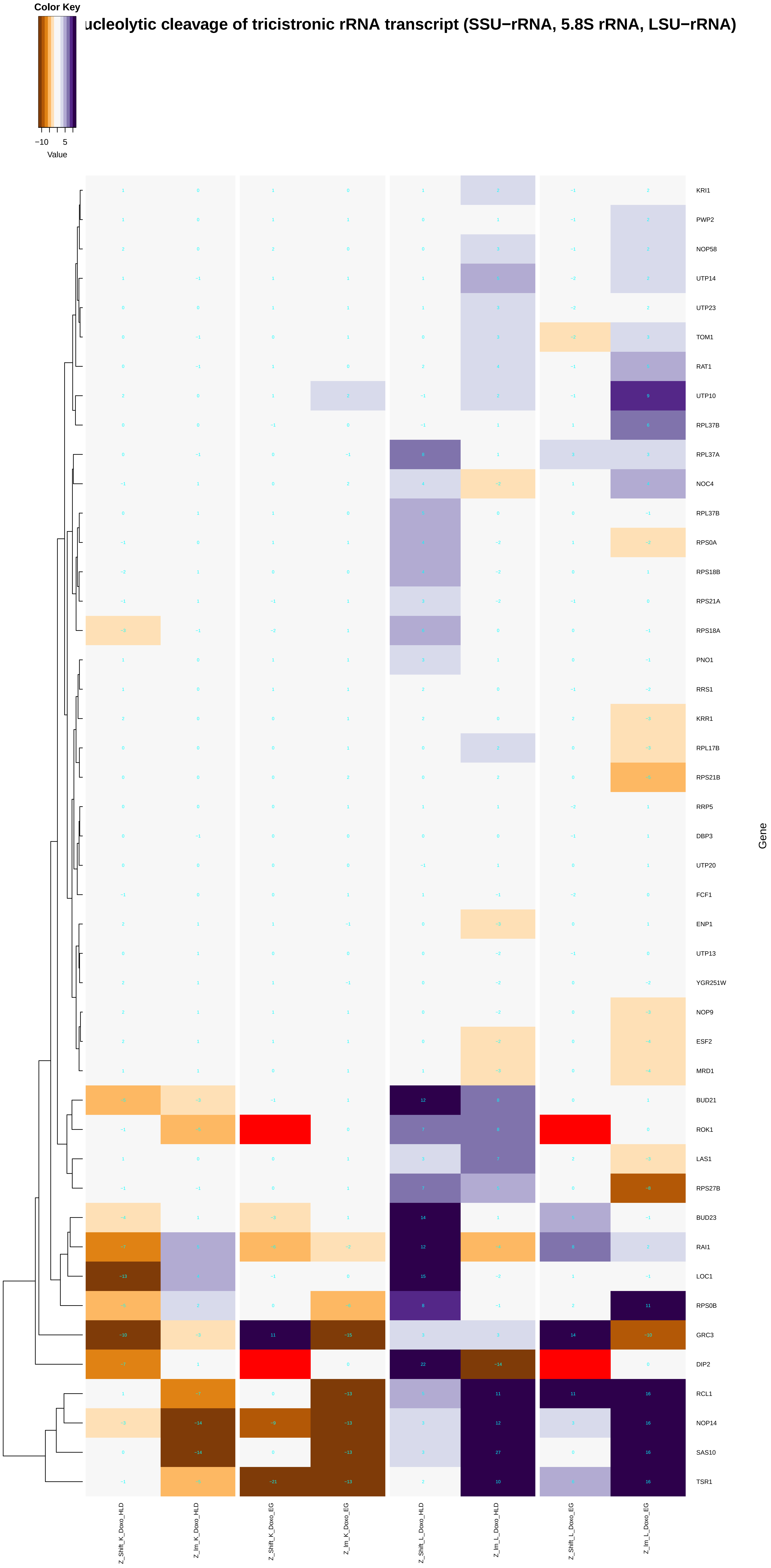
Color Key
−10
5
Value
ucleolytic cleavage of tricistronic rRNA transcript (SSU−rRNA, 5.8S rRNA, LSU−rRNA)
KRI1
PWP2
NOP58
UTP14
UTP23
TOM1
RAT1
UTP10
RPL37B
RPL37A
NOC4
RPL37B
RPS0A
RPS18B
RPS21A
RPS18A
PNO1
RRS1
KRR1
RPL17B
RPS21B
RRP5
DBP3
UTP20
FCF1
ENP1
UTP13
YGR251W
NOP9
ESF2
MRD1
BUD21
ROK1
LAS1
RPS27B
BUD23
RAI1
LOC1
RPS0B
GRC3
DIP2
RCL1
NOP14
SAS10
TSR1
Gene
Z_Shift_K_Doxo_HLD
Z_Im_K_Doxo_HLD
Z_Shift_K_Doxo_EG
Z_Im_K_Doxo_EG
Z_Shift_L_Doxo_HLD
Z_Im_L_Doxo_HLD
Z_Shift_L_Doxo_EG
Z_Im_L_Doxo_EG

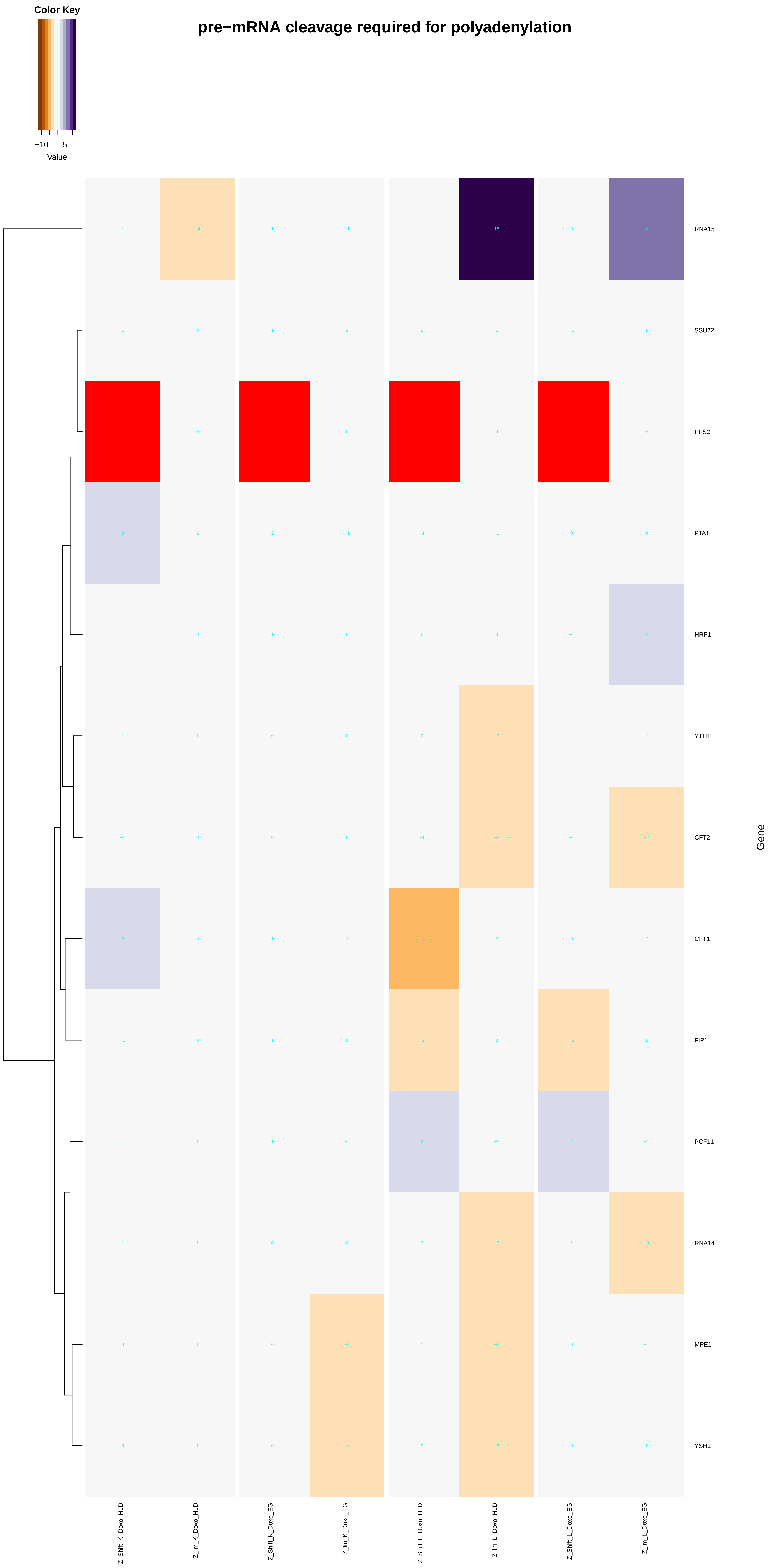

Color Key
−10
5
Value
pre−mRNA cleavage required for polyadenylation
RNA15
SSU72
PFS2
PTA1
HRP1
YTH1
CFT2
CFT1
FIP1
PCF11
RNA14
MPE1
YSH1
Gene
Z_Shift_K_Doxo_HLD
Z_lm_K_Doxo_HLD
Z_Shift_K_Doxo_EG
Z_lm_K_Doxo_EG
Z_Shift_L_Doxo_HLD
Z_lm_L_Doxo_HLD
Z_Shift_L_Doxo_EG
Z_lm_L_Doxo_EG

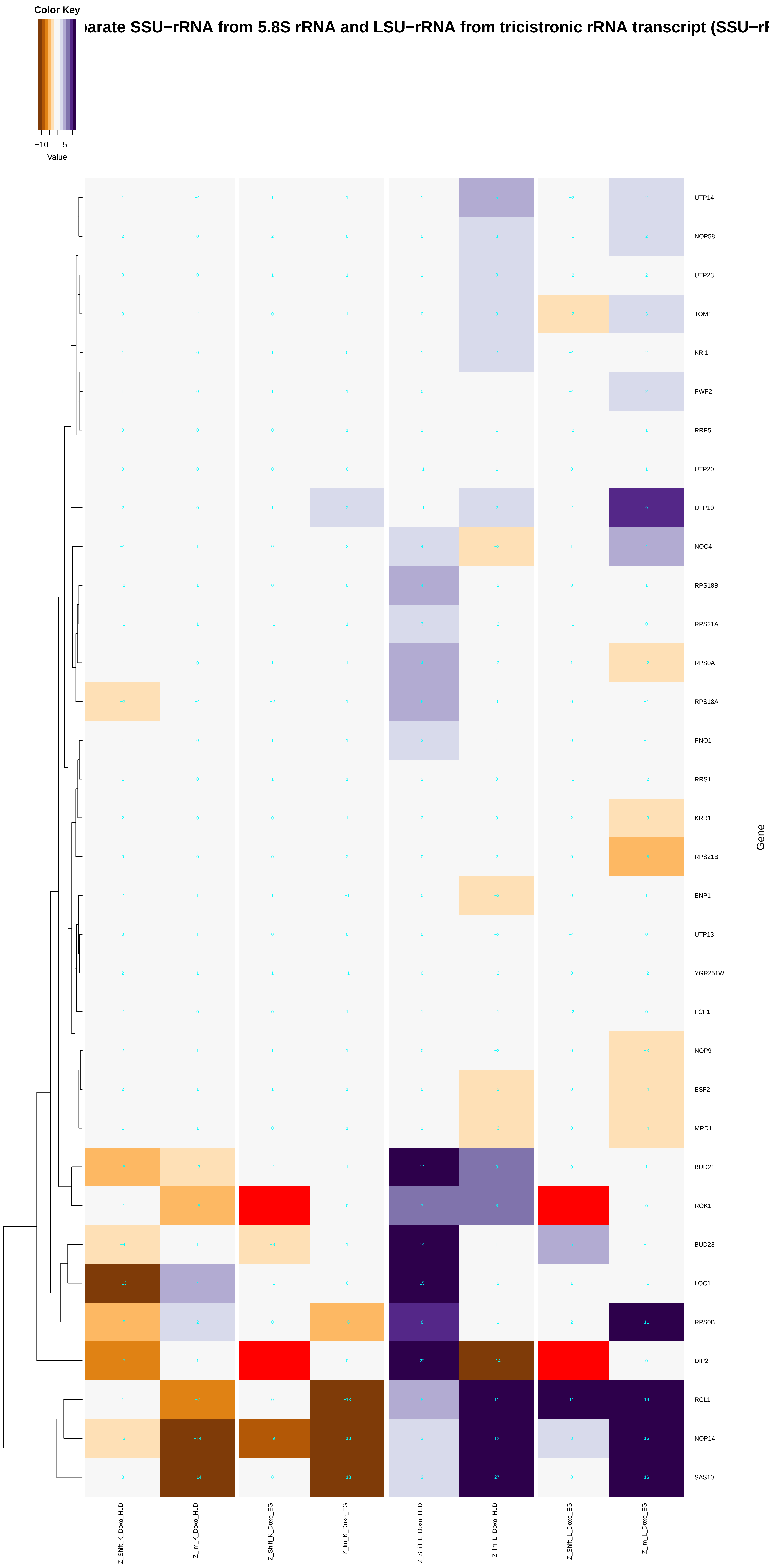

Color Key
−10
5
Value
arate SSU−rRNA from 5.8S rRNA and LSU−rRNA from tricistronic rRNA transcript (SSU−rF
UTP14
NOP58
UTP23
TOM1
KRI1
PWP2
RRP5
UTP20
UTP10
NOC4
RPS18B
RPS21A
RPS0A
RPS18A
PNO1
RRS1
KRR1
RPS21B
ENP1
UTP13
YGR251W
FCF1
NOP9
ESF2
MRD1
BUD21
ROK1
BUD23
LOC1
RPS0B
DIP2
RCL1
NOP14
SAS10
Gene
Z_Shift_K_Doxo_HLD
Z_lm_K_Doxo_HLD
Z_Shift_K_Doxo_EG
Z_lm_K_Doxo_EG
Z_Shift_L_Doxo_HLD
Z_lm_L_Doxo_HLD
Z_Shift_L_Doxo_EG
Z_lm_L_Doxo_EG

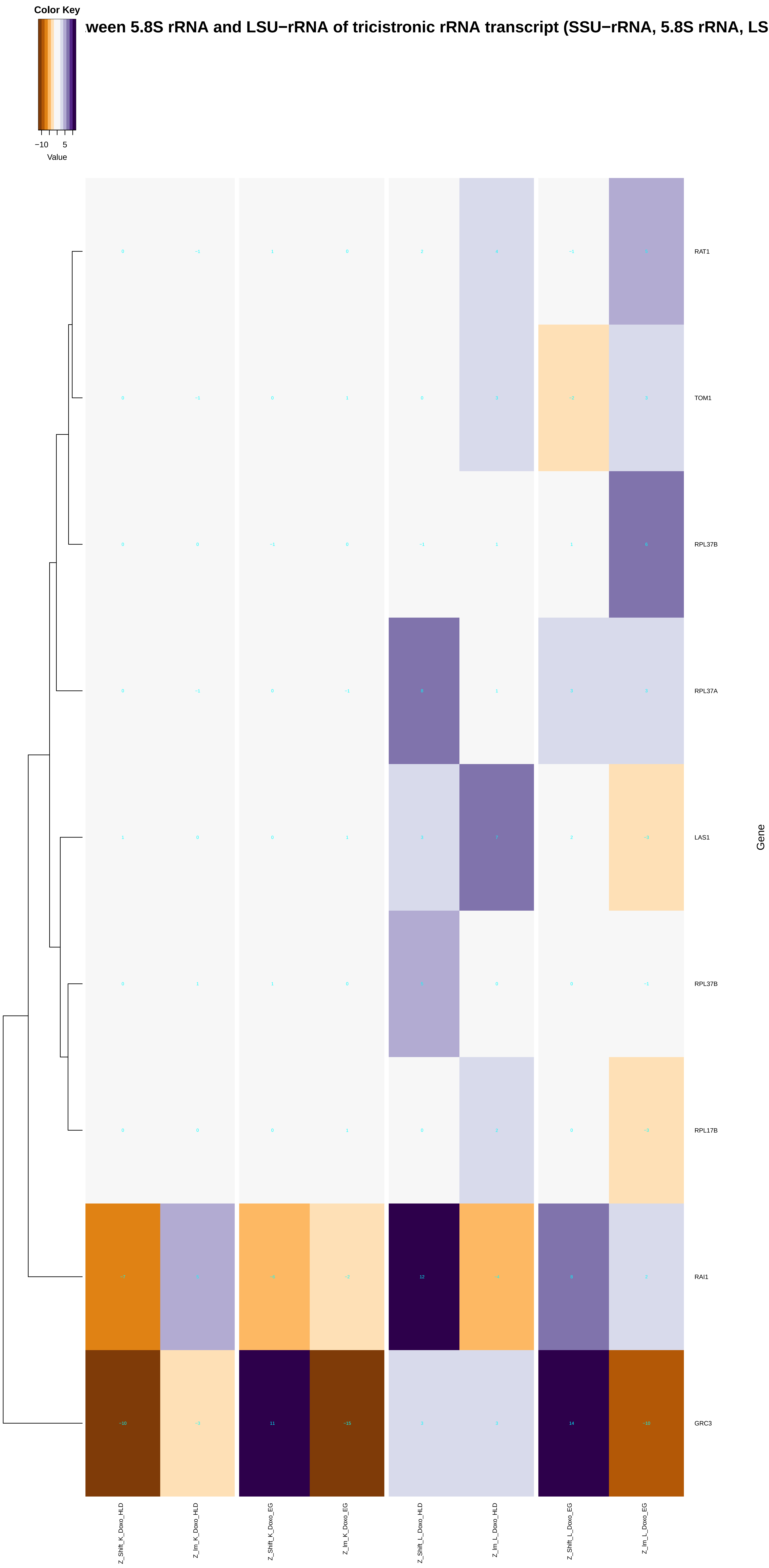

Color Key
−10
5
Value
ween 5.8S rRNA and LSU−rRNA of tricistronic rRNA transcript (SSU−rRNA, 5.8S rRNA, LS
RAT1
TOM1
RPL37B
RPL37A
LAS1
RPL37B
RPL17B
RAI1
GRC3
Gene
Z_Shift_K_Doxo_HLD
Z_Im_K_Doxo_HLD
Z_Shift_K_Doxo_EG
Z_Im_K_Doxo_EG
Z_Shift_L_Doxo_HLD
Z_Im_L_Doxo_HLD
Z_Shift_L_Doxo_EG
Z_Im_L_Doxo_EG

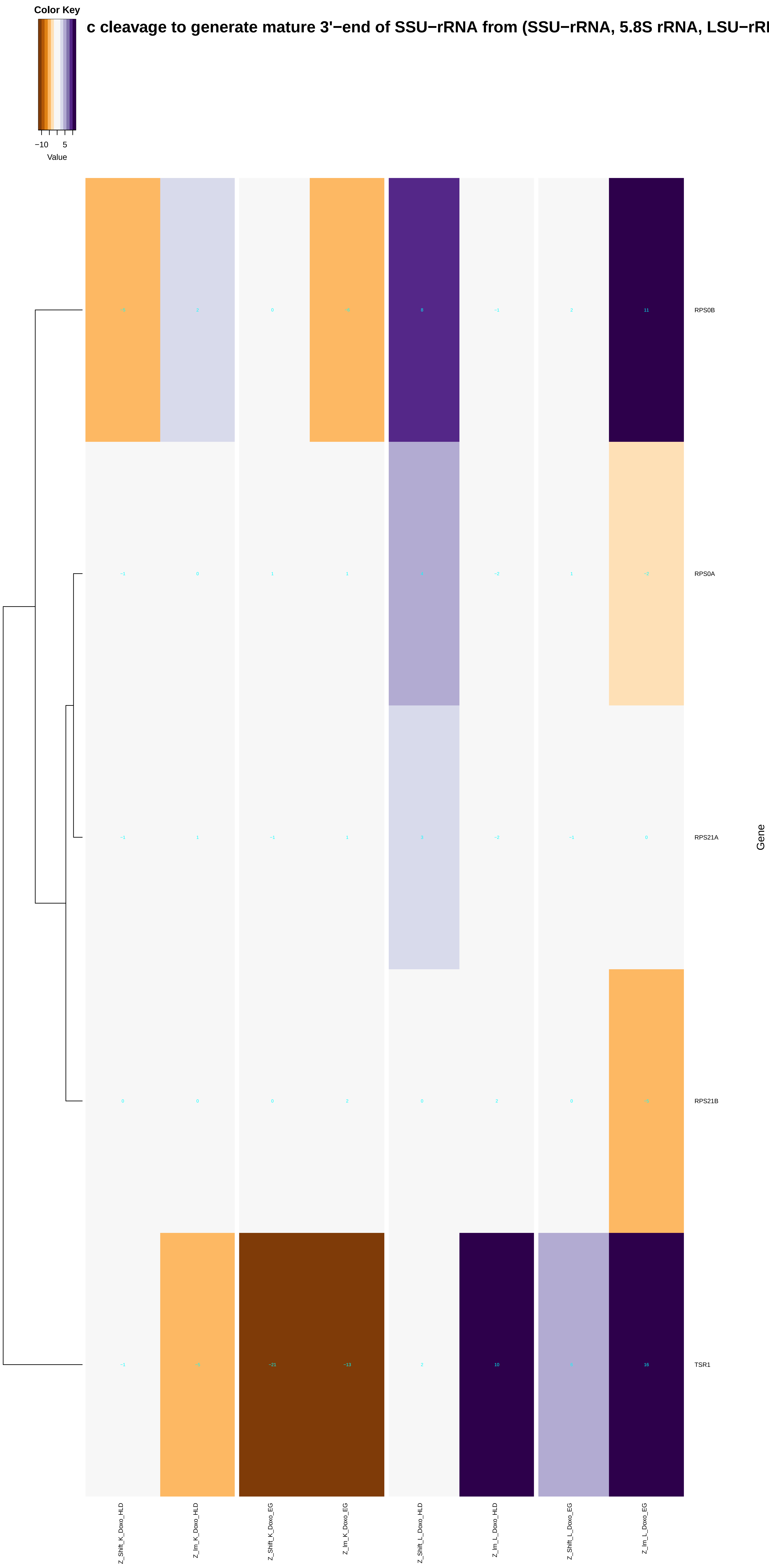

Color Key
−10
5
Value
c cleavage to generate mature 3'−end of SSU−rRNA from (SSU−rRNA, 5.8S rRNA, LSU−rRI
RPS0B
RPS0A
RPS21A
RPS21B
TSR1
Gene
Z_Shift_K_Doxo_HLD
Z_Im_K_Doxo_HLD
Z_Shift_K_Doxo_EG
Z_Im_K_Doxo_EG
Z_Shift_L_Doxo_HLD
Z_Im_L_Doxo_HLD
Z_Shift_L_Doxo_EG
Z_Im_L_Doxo_EG

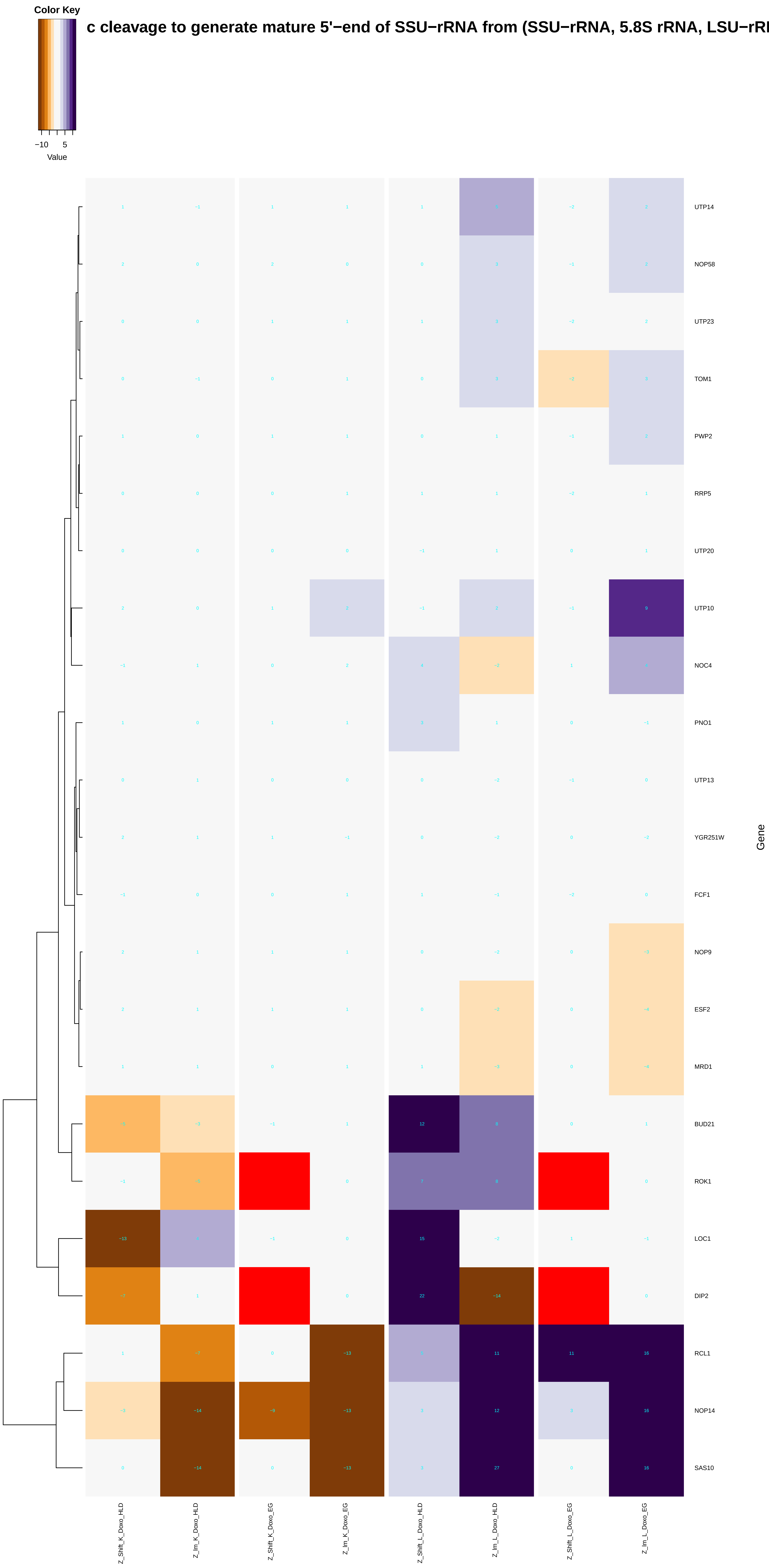

Color Key
−10 5
Value
c cleavage to generate mature 5'−end of SSU−rRNA from (SSU−rRNA, 5.8S rRNA, LSU−rR
UTP14
NOP58
UTP23
TOM1
PWP2
RRP5
UTP20
UTP10
NOC4
PNO1
UTP13
YGR251W
FCF1
NOP9
ESF2
MRD1
BUD21
ROK1
LOC1
DIP2
RCL1
NOP14
SAS10
Gene
Z_Shift_K_Doxo_HLD
Z_Im_K_Doxo_HLD
Z_Shift_K_Doxo_EG
Z_Im_K_Doxo_EG
Z_Shift_L_Doxo_HLD
Z_Im_L_Doxo_HLD
Z_Shift_L_Doxo_EG
Z_Im_L_Doxo_EG

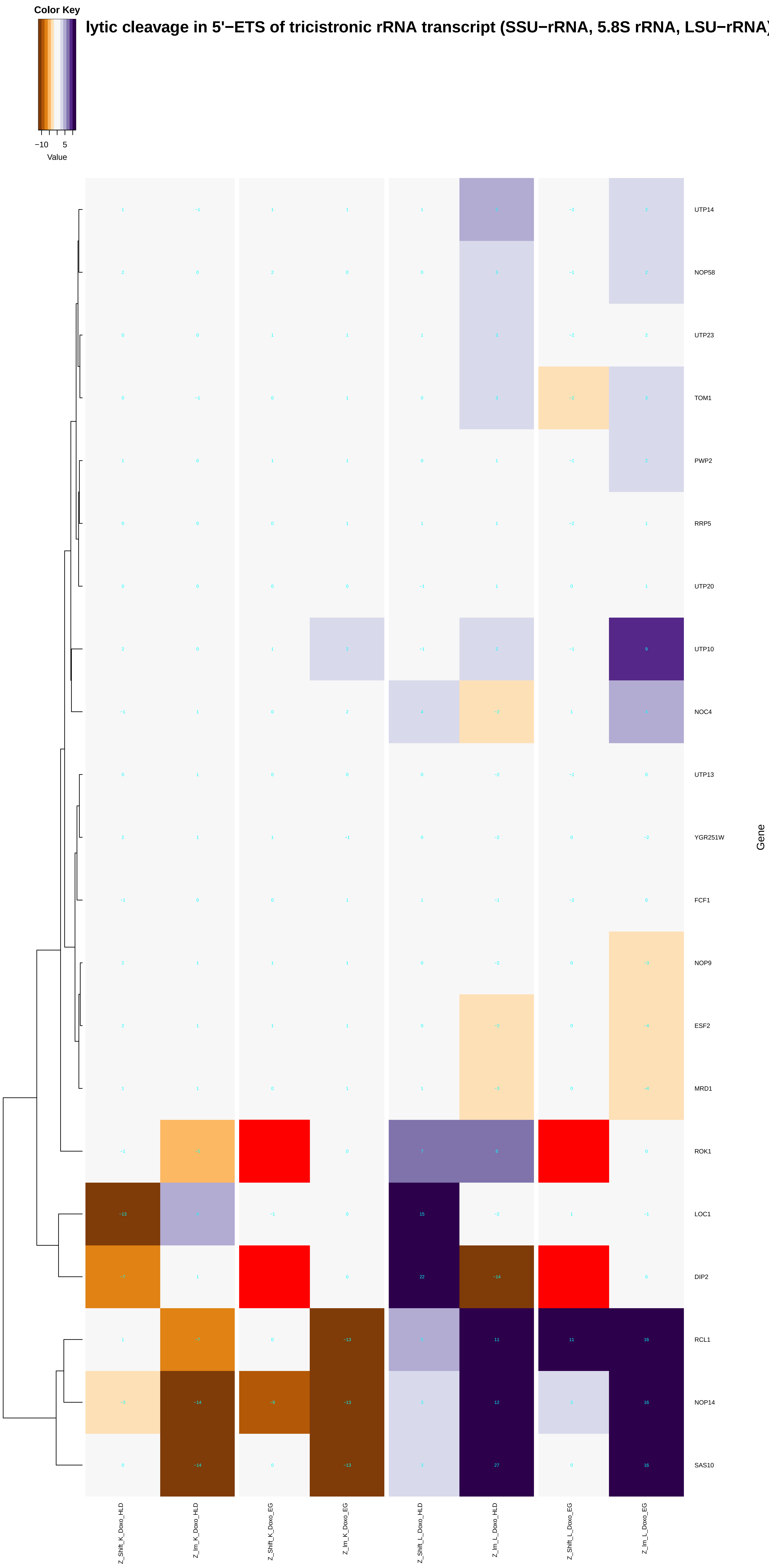

Color Key
−10
5
Value
lytic cleavage in 5'−ETS of tricistronic rRNA transcript (SSU−rRNA, 5.8S rRNA, LSU−rRNA
UTP14
NOP58
UTP23
TOM1
PWP2
RRP5
UTP20
UTP10
NOC4
UTP13
YGR251W
FCF1
NOP9
ESF2
MRD1
ROK1
LOC1
DIP2
RCL1
NOP14
SAS10
Gene
Z_Shift_K_Doxo_HLD
Z_Im_K_Doxo_HLD
Z_Shift_K_Doxo_EG
Z_Im_K_Doxo_EG
Z_Shift_L_Doxo_HLD
Z_Im_L_Doxo_HLD
Z_Shift_L_Doxo_EG
Z_Im_L_Doxo_EG
