## Additional File 9 for "A yeast phenomic model for the influence of Warburg metabolism on genetic buffering of doxorubicin": nucleic_acid_transport.pdf

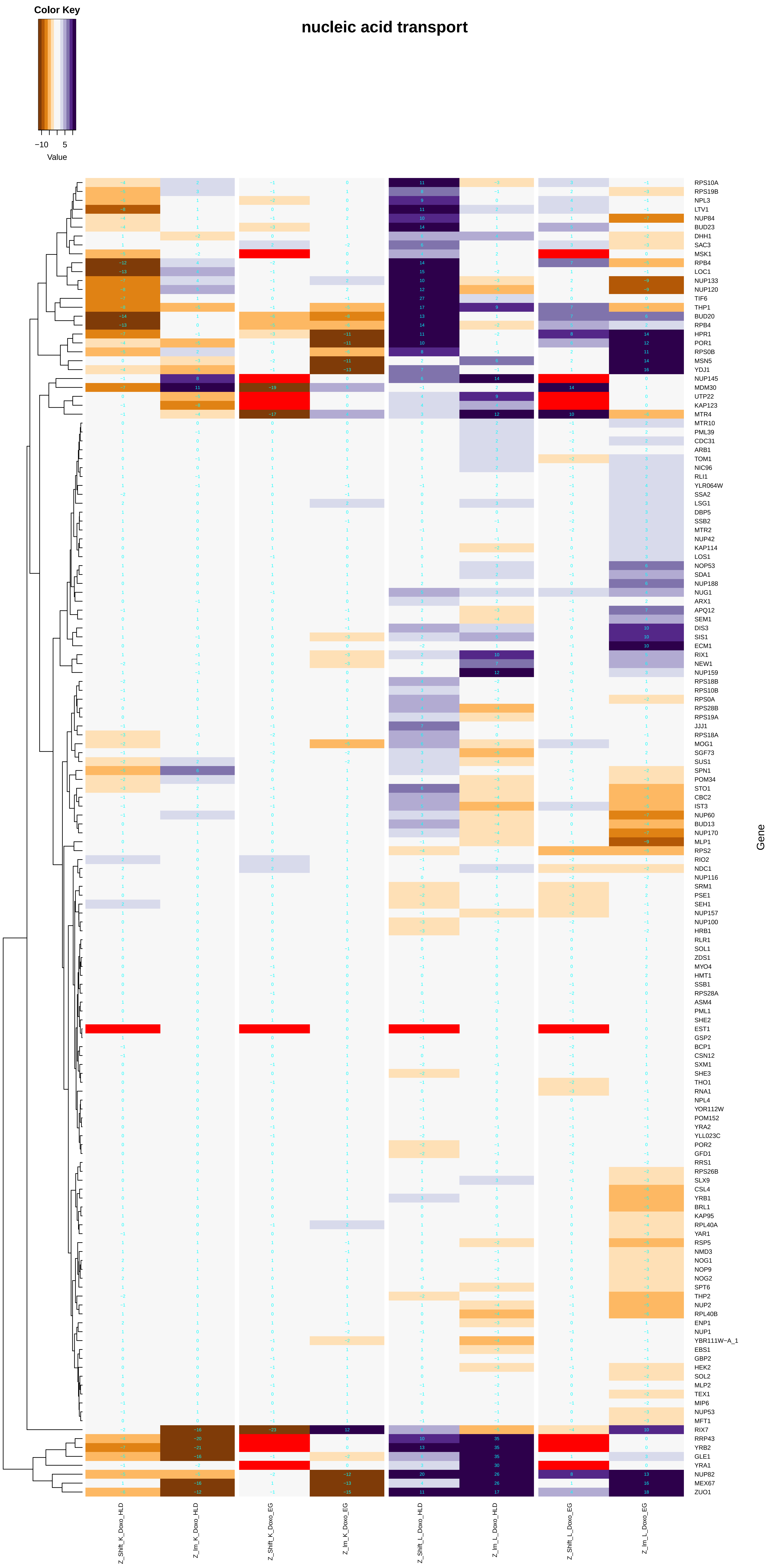

Color Key
−10
5
Value
nucleic acid transport
RPS10A
RPS19B
NPL3
LTV1
NUP84
BUD23
DHH1
SAC3
MSK1
RPB4
LOC1
NUP133
NUP120
TIF6
THP1
BUD20
RPB4
HPR1
POR1
RPS0B
MSN5
YDJ1
NUP145
MDM30
UTP22
KAP123
MTR4
MTR10
PML39
CDC31
ARB1
TOM1
NIC96
RLI1
YLR064W
SSA2
LSG1
DBP5
SSB2
MTR2
NUP42
KAP114
LOS1
NOP53
SDA1
NUP188
NUG1
ARX1
APQ12
SEM1
DIS3
SIS1
ECM1
RIX1
NEW1
NUP159
RPS18B
RPS10B
RPS0A
RPS28B
RPS19A
JJJ1
RPS18A
MOG1
SGF73
SUS1
SPN1
POM34
STO1
CBC2
IST3
NUP60
BUD13
NUP170
MLP1
RPS2
RIO2
NDC1
NUP116
SRM1
PSE1
SEH1
NUP157
NUP100
HRB1
RLR1
SOL1
ZDS1
MYO4
HMT1
SSB1
RPS28A
ASM4
PML1
SHE2
EST1
GSP2
BCP1
CSN12
SXM1
SHE3
THO1
RNA1
NPL4
YOR112W
POM152
YRA2
YLL023C
POR2
GFD1
RRS1
RPS26B
SLX9
CSL4
YRB1
BRL1
KAP95
RPL40A
YAR1
RSP5
NMD3
NOG1
NOP9
NOG2
SPT6
THP2
NUP2
RPL40B
ENP1
NUP1
YBR111W−A_1
EBS1
GBP2
HEK2
SOL2
MLP2
TEX1
MIP6
NUP53
MFT1
RIX7
RRP43
YRB2
GLE1
YRA1
NUP82
MEX67
ZUO1
Gene
Z_Shift_K_Doxo_HLD
Z_Im_K_Doxo_HLD
Z_Shift_K_Doxo_EG
Z_Im_K_Doxo_EG
Z_Shift_L_Doxo_HLD
Z_Im_L_Doxo_HLD
Z_Shift_L_Doxo_EG
Z_Im_L_Doxo_EG

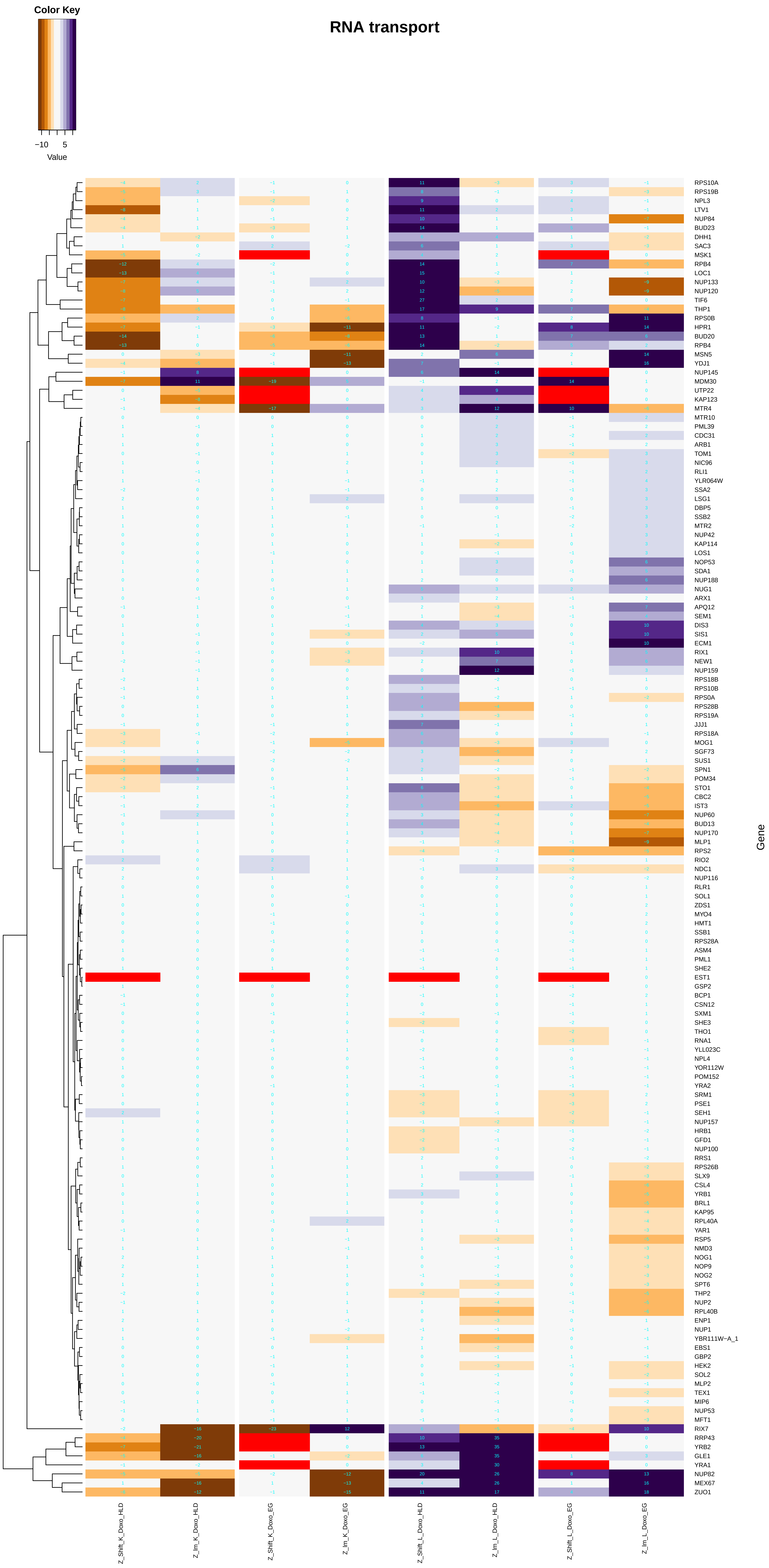

Color Key
−10
5
Value
RNA transport
Gene
Z_Shift_K_Doxo_HLD
Z_Im_K_Doxo_HLD
Z_Shift_K_Doxo_EG
Z_Im_K_Doxo_EG
Z_Shift_L_Doxo_HLD
Z_Im_L_Doxo_HLD
Z_Shift_L_Doxo_EG
Z_Im_L_Doxo_EG

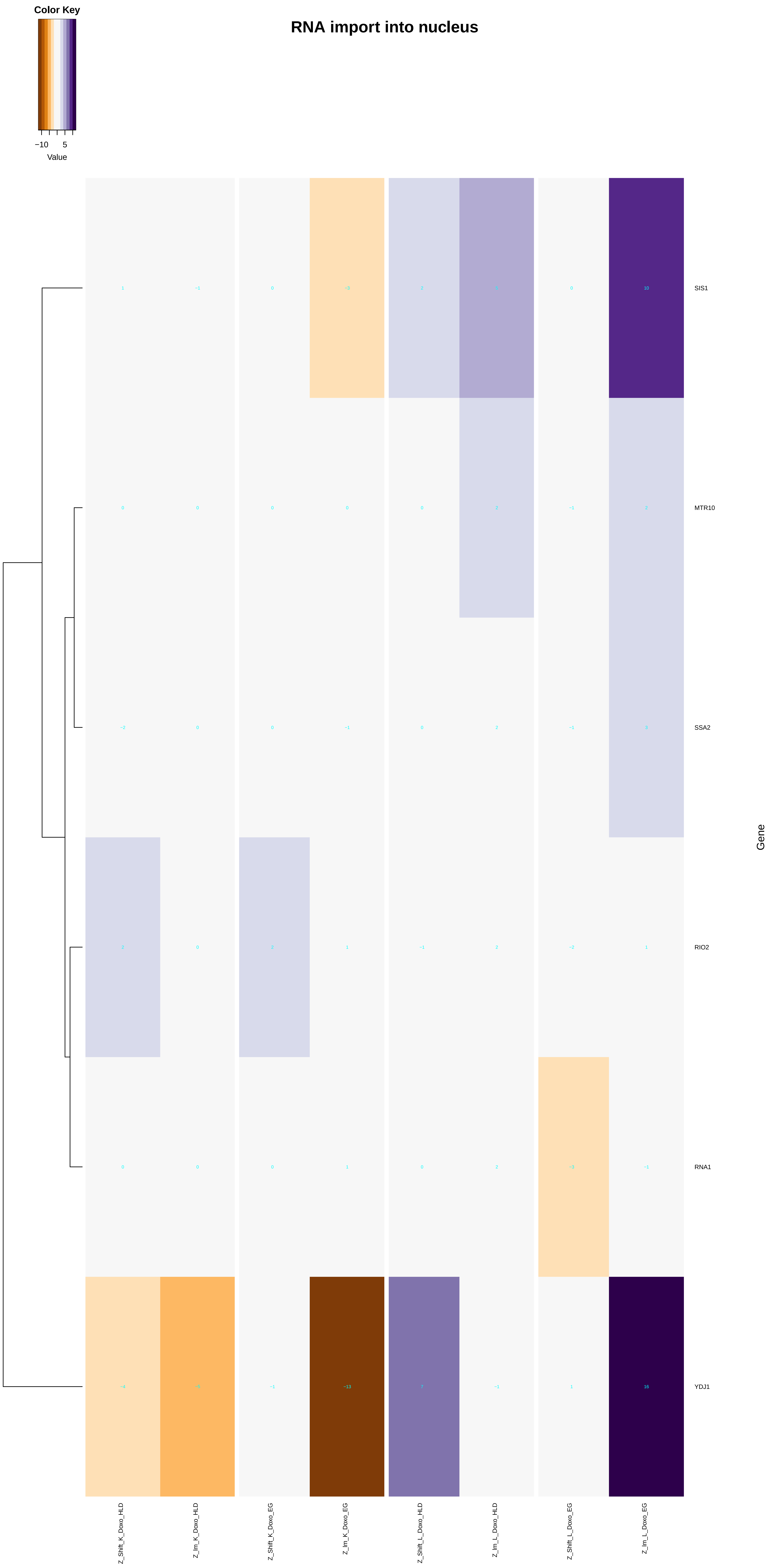

Color Key
−10
5
Value
RNA import into nucleus
SIS1
MTR10
SSA2
RIO2
RNA1
YDJ1
Gene
Z_Shift_K_Doxo_HLD
Z_Im_K_Doxo_HLD
Z_Shift_K_Doxo_EG
Z_Im_K_Doxo_EG
Z_Shift_L_Doxo_HLD
Z_Im_L_Doxo_HLD
Z_Shift_L_Doxo_EG
Z_Im_L_Doxo_EG

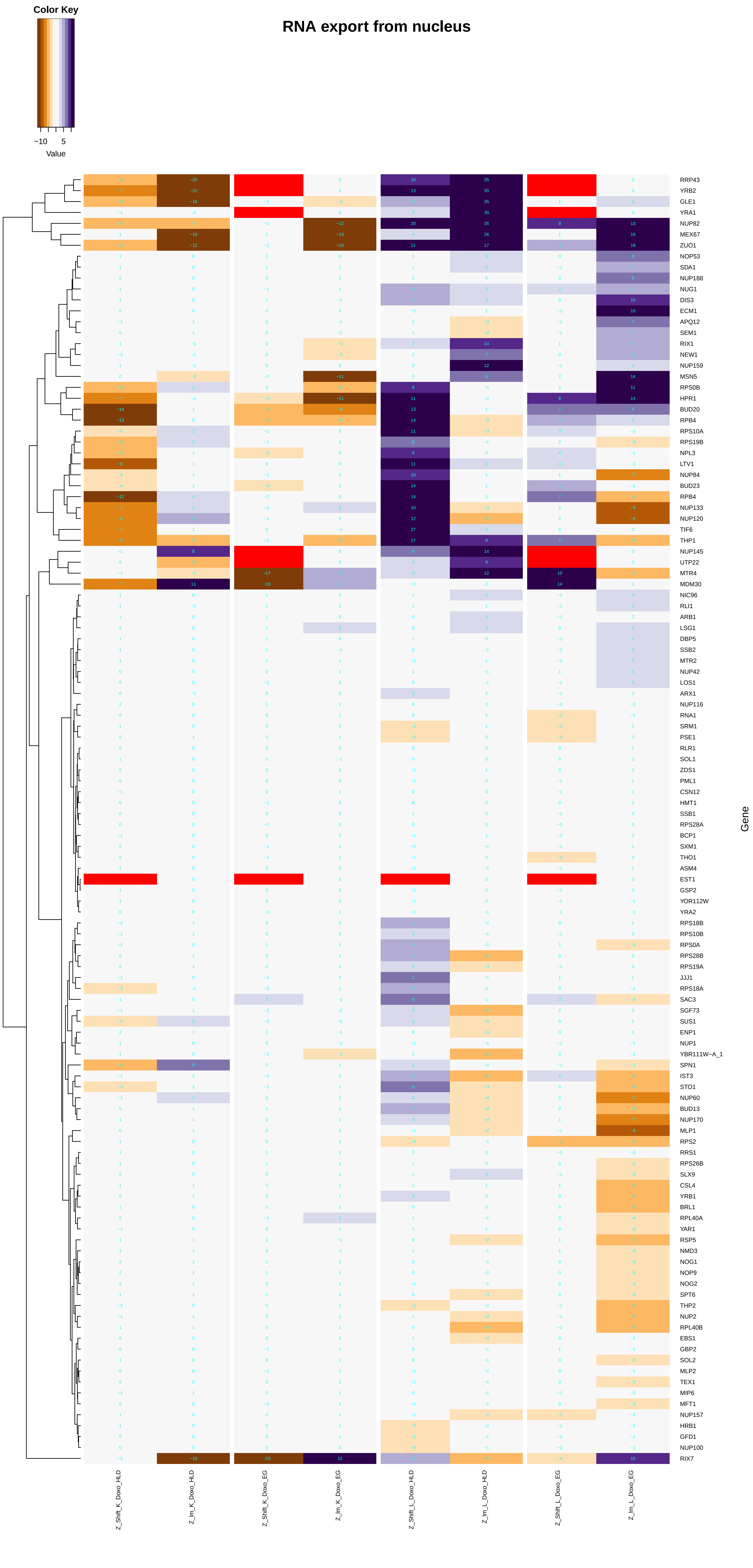

Color Key
−10
5
Value
RNA export from nucleus
Gene
RRP43
YRB2
GLE1
YRA1
NUP82
MEX67
ZUO1
NOP53
SDA1
NUP188
NUG1
DIS3
ECM1
APQ12
SEM1
RIX1
NEW1
NUP159
MSN5
RPS0B
HPR1
BUD20
RPB4
RPS10A
RPS19B
NPL3
LTV1
NUP84
BUD23
RPB4
NUP133
NUP120
TIF6
THP1
NUP145
UTP22
MTR4
MDM30
NIC96
RLI1
ARB1
LSG1
DBP5
SSB2
MTR2
NUP42
LOS1
ARX1
NUP116
RNA1
SRM1
PSE1
RLR1
SOL1
ZDS1
PML1
CSN12
HMT1
SSB1
RPS28A
BCP1
SXM1
THO1
ASM4
EST1
GSP2
YOR112W
YRA2
RPS18B
RPS10B
RPS0A
RPS28B
RPS19A
JJJ1
RPS18A
SAC3
SGF73
SUS1
ENP1
NUP1
YBR111W−A_1
SPN1
IST3
STO1
NUP60
BUD13
NUP170
MLP1
RPS2
RRS1
RPS26B
SLX9
CSL4
YRB1
BRL1
RPL40A
YAR1
RSP5
NMD3
NOG1
NOP9
NOG2
SPT6
THP2
NUP2
RPL40B
EBS1
GBP2
SOL2
MLP2
TEX1
MIP6
MFT1
NUP157
HRB1
GFD1
NUP100
RIX7
Z_Shift_K_Doxo_HLD
Z_Im_K_Doxo_HLD
Z_Shift_K_Doxo_EG
Z_Im_K_Doxo_EG
Z_Shift_L_Doxo_HLD
Z_Im_L_Doxo_HLD
Z_Shift_L_Doxo_EG
Z_Im_L_Doxo_EG

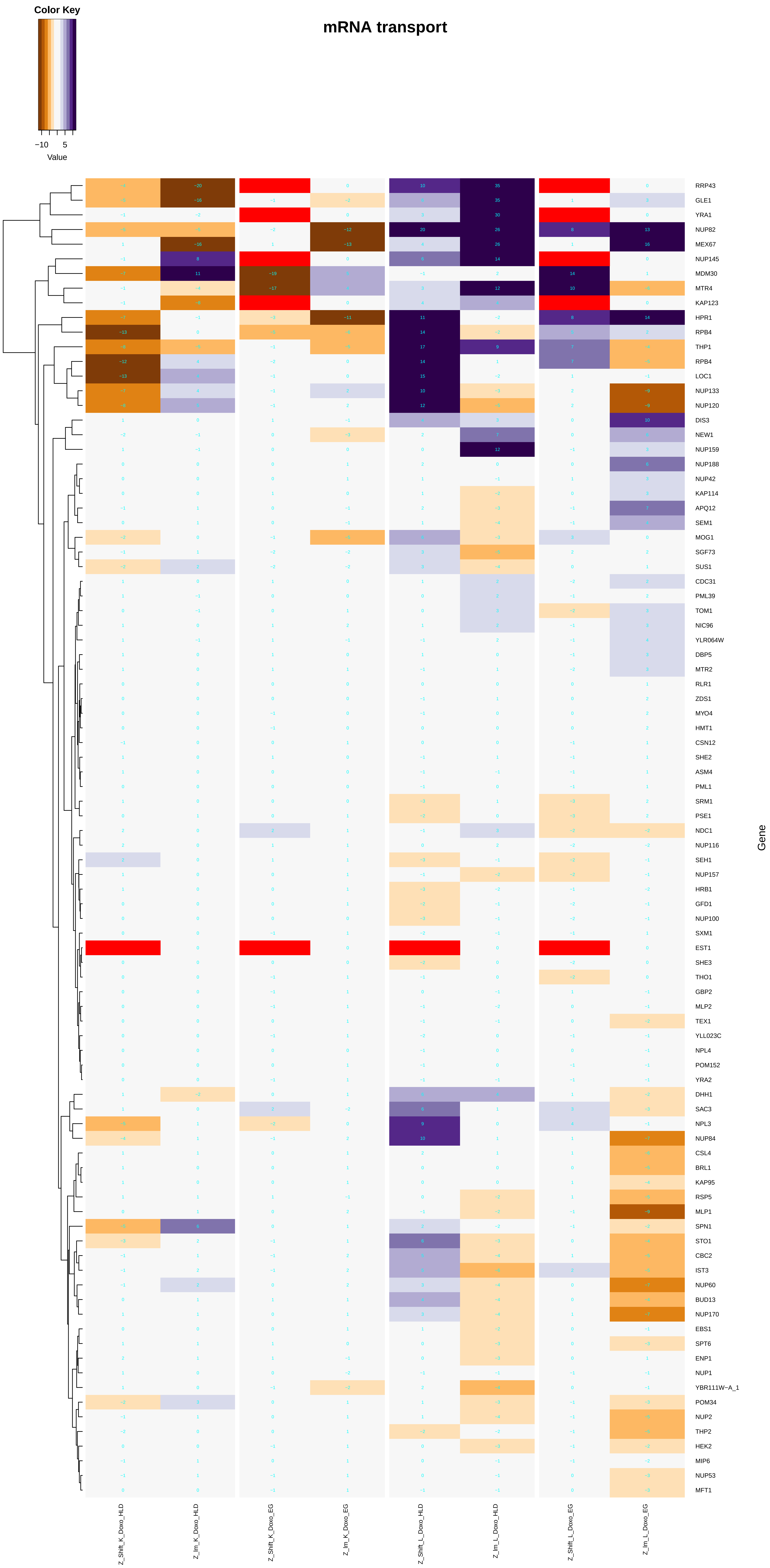

Color Key
−10
5
Value
mRNA transport
RRP43
GLE1
YRA1
NUP82
MEX67
NUP145
MDM30
MTR4
KAP123
HPR1
RPB4
THP1
RPB4
LOC1
NUP133
NUP120
DIS3
NEW1
NUP159
NUP188
NUP42
KAP114
APQ12
SEM1
MOG1
SGF73
SUS1
CDC31
PML39
TOM1
NIC96
YLR064W
DBP5
MTR2
RLR1
ZDS1
MYO4
HMT1
CSN12
SHE2
ASM4
PML1
SRM1
PSE1
NDC1
NUP116
SEH1
NUP157
HRB1
GFD1
NUP100
SXM1
EST1
SHE3
THO1
GBP2
MLP2
TEX1
YLL023C
NPL4
POM152
YRA2
DHH1
SAC3
NPL3
NUP84
CSL4
BRL1
KAP95
RSP5
MLP1
SPN1
STO1
CBC2
IST3
NUP60
BUD13
NUP170
EBS1
SPT6
ENP1
NUP1
YBR111W−A_1
POM34
NUP2
THP2
HEK2
MIP6
NUP53
MFT1
Gene
Z_Shift_K_Doxo_HLD
Z_lm_K_Doxo_HLD
Z_Shift_K_Doxo_EG
Z_lm_K_Doxo_EG
Z_Shift_L_Doxo_HLD
Z_lm_L_Doxo_HLD
Z_Shift_L_Doxo_EG
Z_lm_L_Doxo_EG

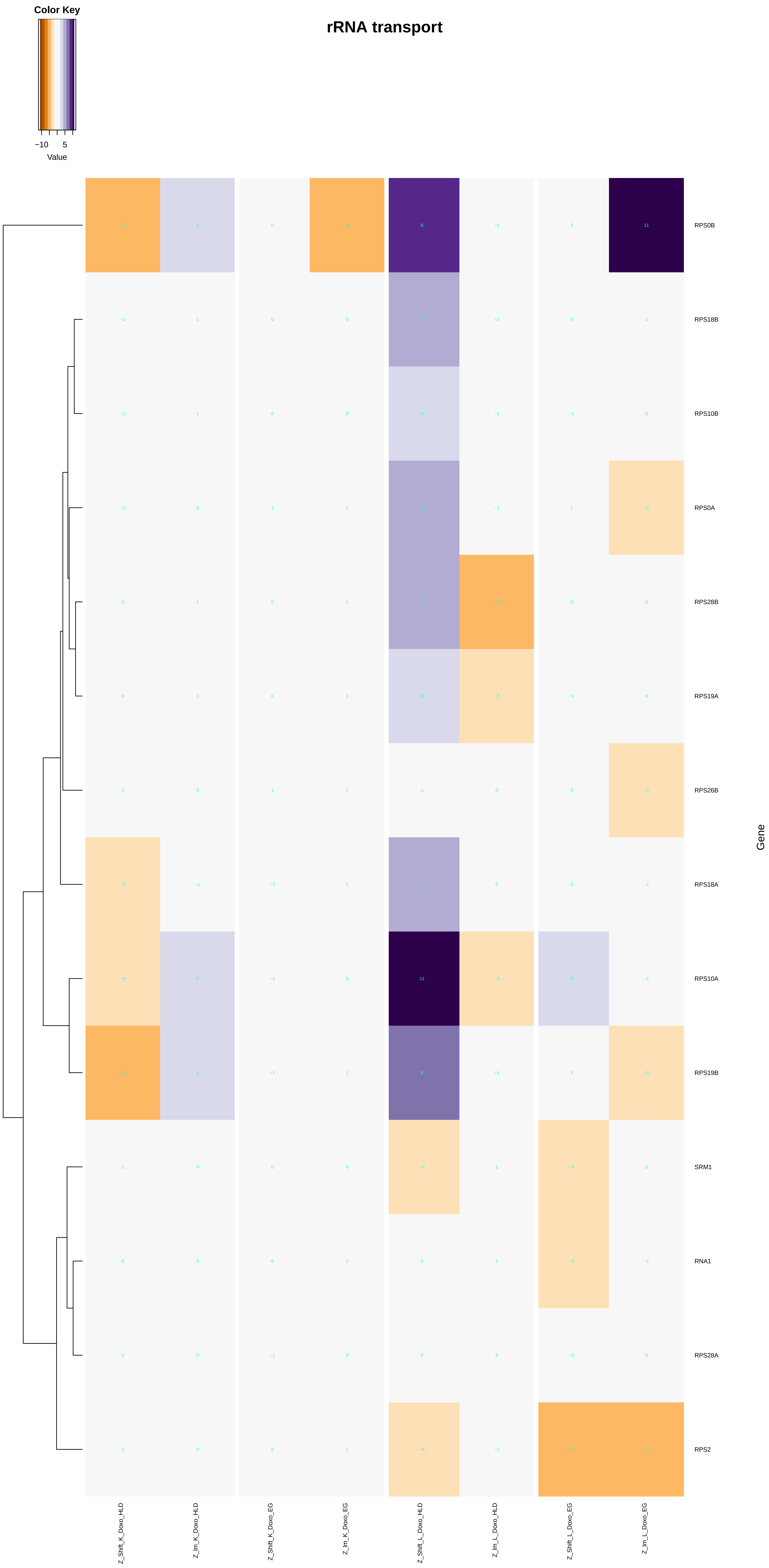
rRNA transport
Color Key
−10
5
Value
Gene
RPS0B
RPS18B
RPS10B
RPS0A
RPS28B
RPS19A
RPS26B
RPS18A
RPS10A
RPS19B
SRM1
RNA1
RPS28A
RPS2
Z_Shift_K_Doxo_HLD
Z_lm_K_Doxo_HLD
Z_Shift_K_Doxo_EG
Z_lm_K_Doxo_EG
Z_Shift_L_Doxo_HLD
Z_lm_L_Doxo_HLD
Z_Shift_L_Doxo_EG
Z_lm_L_Doxo_EG

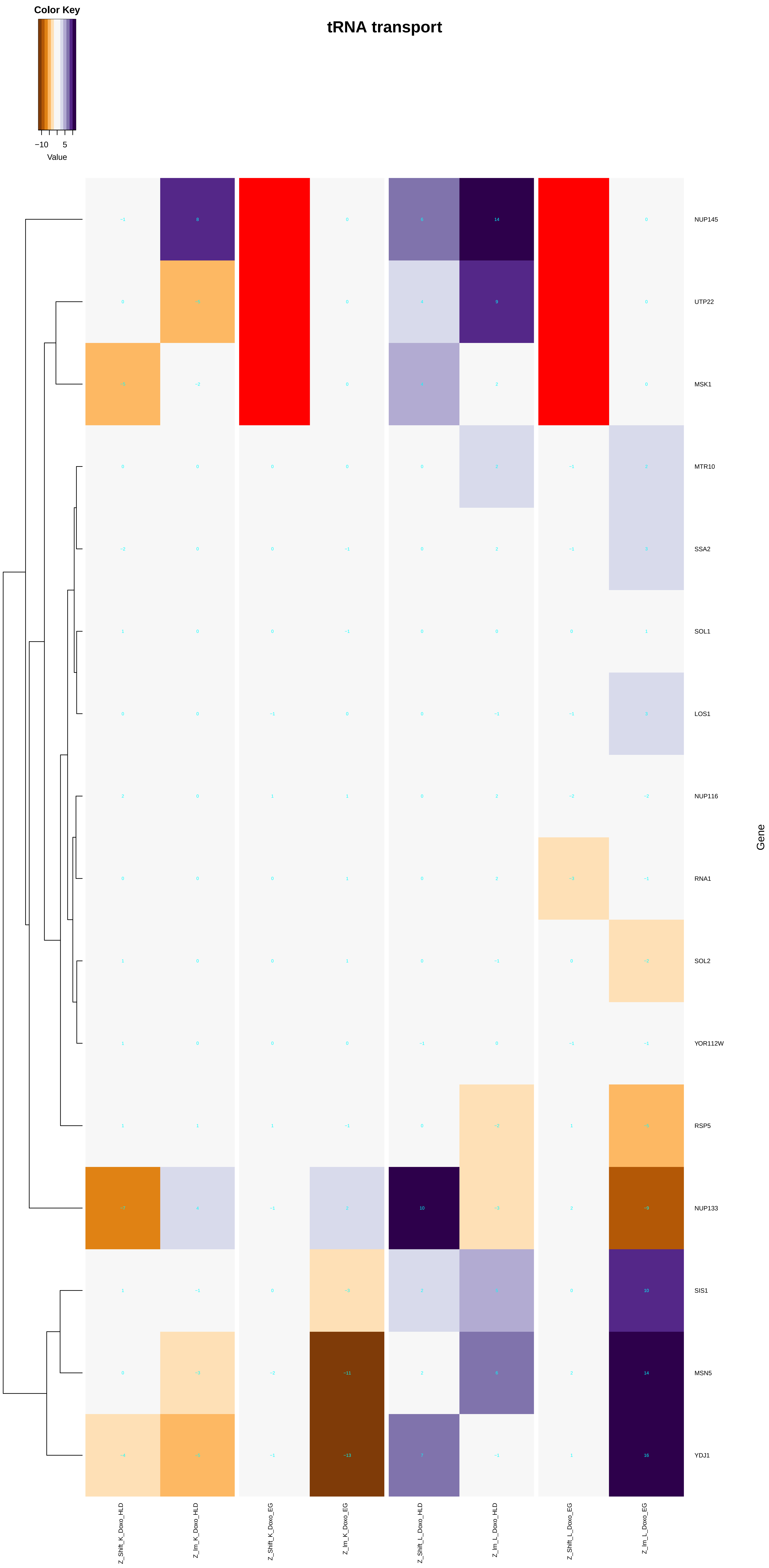

Color Key
−10
5
Value
tRNA transport
NUP145
UTP22
MSK1
MTR10
SSA2
SOL1
LOS1
NUP116
RNA1
SOL2
YOR112W
RSP5
NUP133
SIS1
MSN5
YDJ1
Gene
Z_Shift_K_Doxo_HLD
Z_Im_K_Doxo_HLD
Z_Shift_K_Doxo_EG
Z_Im_K_Doxo_EG
Z_Shift_L_Doxo_HLD
Z_Im_L_Doxo_HLD
Z_Shift_L_Doxo_EG
Z_Im_L_Doxo_EG

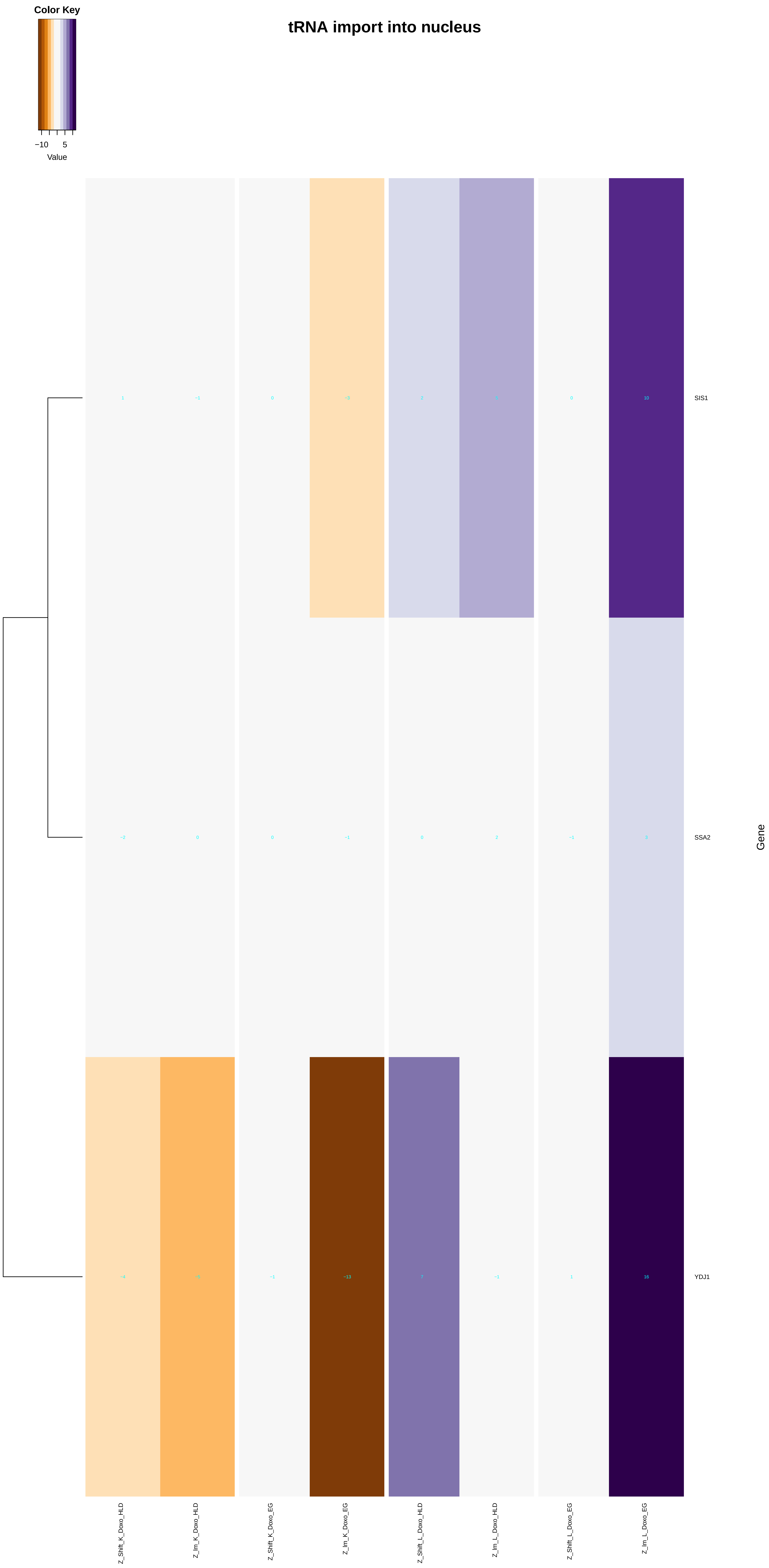

Color Key
−10
5
Value
tRNA import into nucleus
1
−1
0
−3
2
5
0
10
SIS1
−2
0
0
−1
0
2
−1
3
SSA2
−4
−5
−1
−13
7
−1
1
16
YDJ1
Gene
Z_Shift_K_Doxo_HLD
Z_Im_K_Doxo_HLD
Z_Shift_K_Doxo_EG
Z_Im_K_Doxo_EG
Z_Shift_L_Doxo_HLD
Z_Im_L_Doxo_HLD
Z_Shift_L_Doxo_EG
Z_Im_L_Doxo_EG

### mRNA export from nucleus

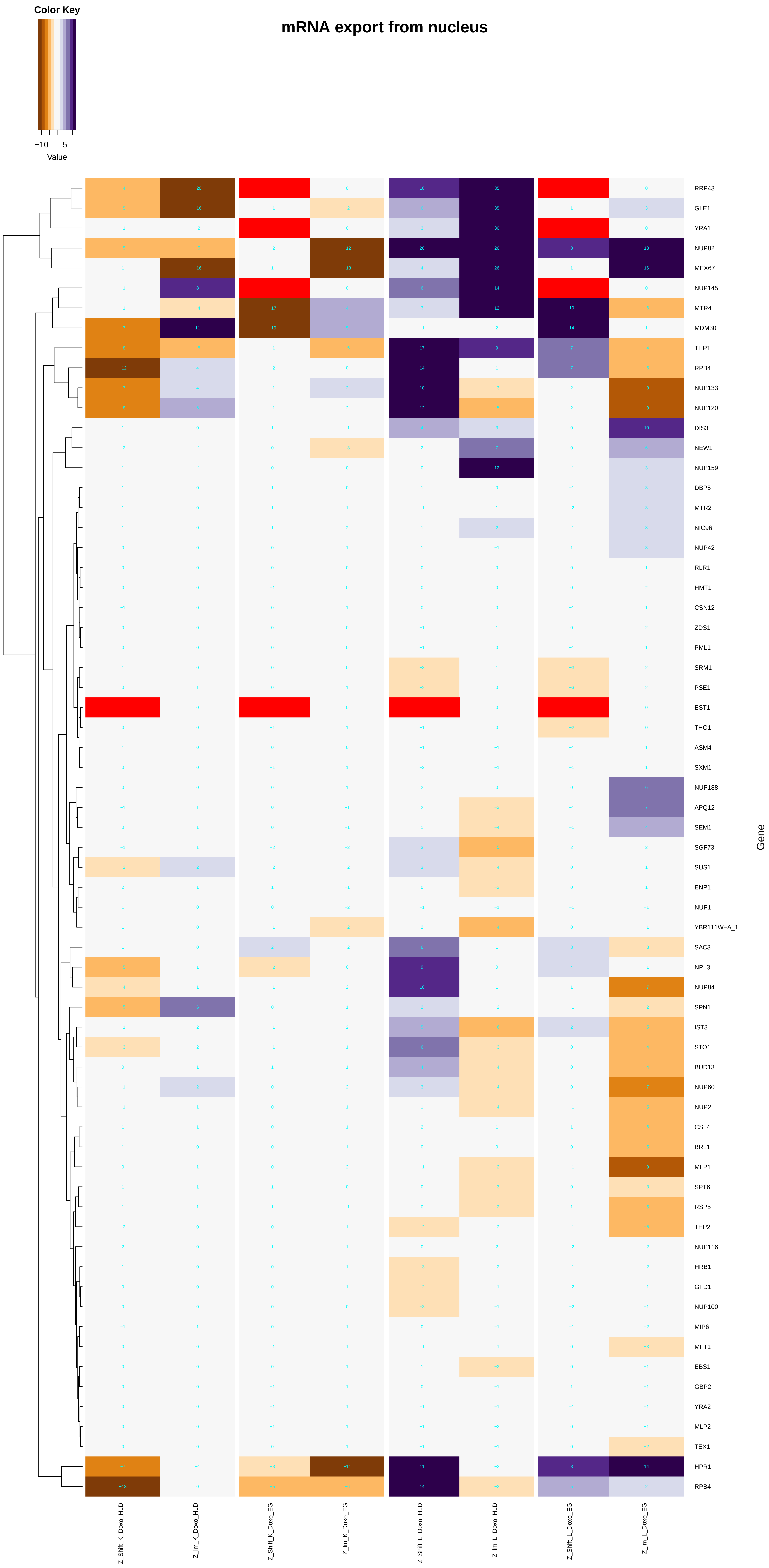

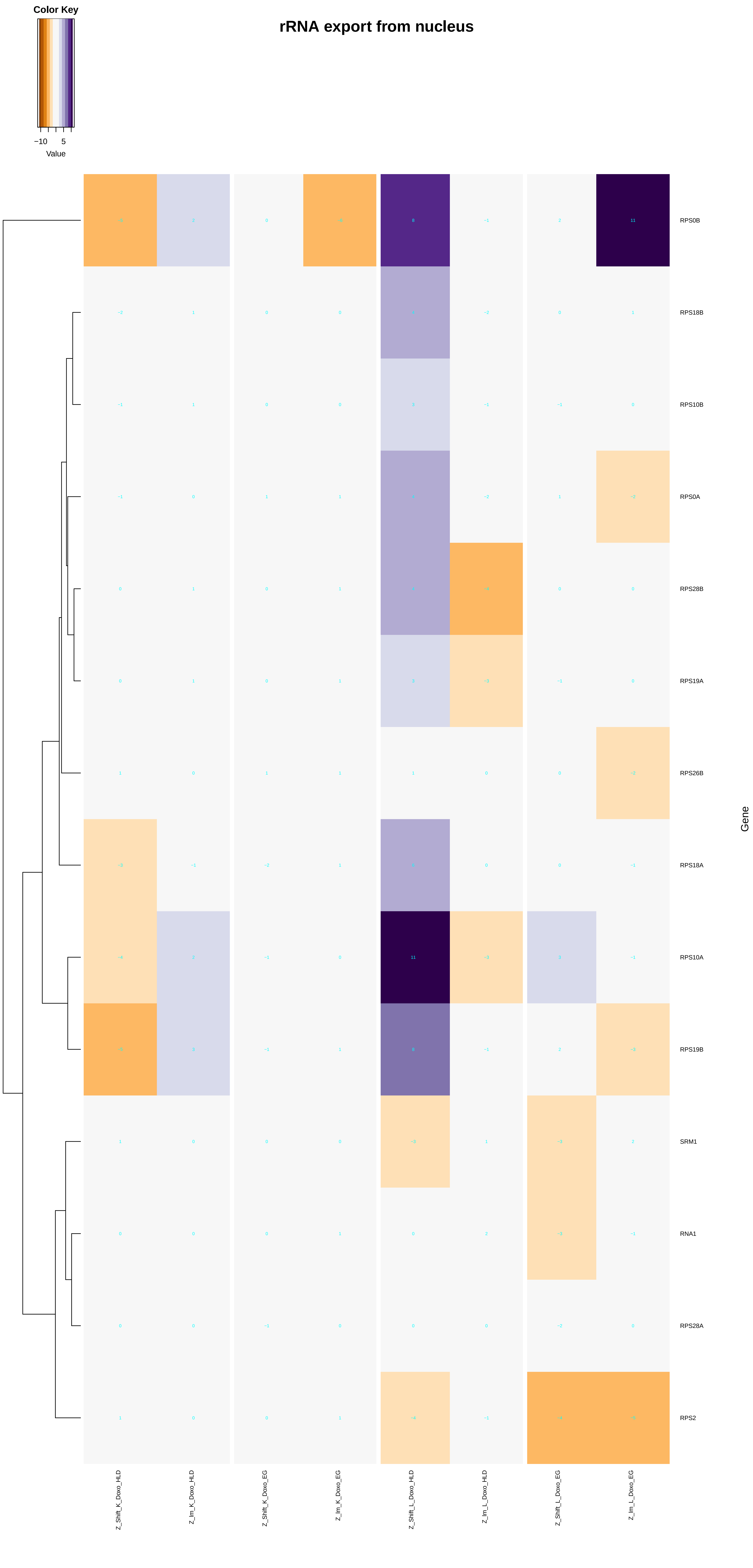

rRNA export from nucleus
Color Key
−10
5
Value
RPS0B
RPS18B
RPS10B
RPS0A
RPS28B
RPS19A
RPS26B
RPS18A
RPS10A
RPS19B
SRM1
RNA1
RPS28A
RPS2
Gene
Z_Shift_K_Doxo_HLD
Z_lm_K_Doxo_HLD
Z_Shift_K_Doxo_EG
Z_lm_K_Doxo_EG
Z_Shift_L_Doxo_HLD
Z_lm_L_Doxo_HLD
Z_Shift_L_Doxo_EG
Z_lm_L_Doxo_EG

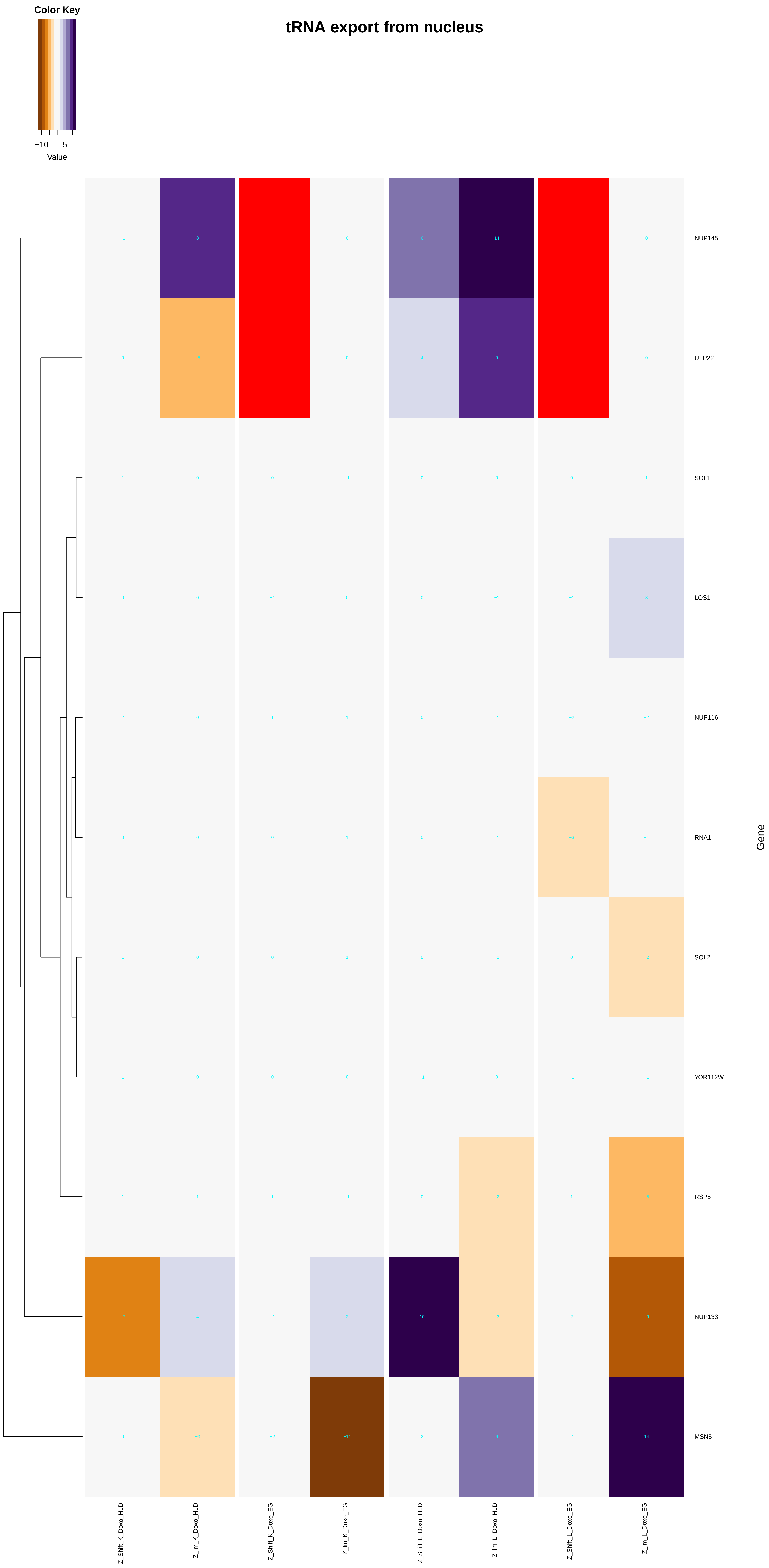

Color Key
−10
5
Value
tRNA export from nucleus
NUP145
UTP22
SOL1
LOS1
NUP116
RNA1
SOL2
YOR112W
RSP5
NUP133
MSN5
Gene
Z_Shift_K_Doxo_HLD
Z_lm_K_Doxo_HLD
Z_Shift_K_Doxo_EG
Z_lm_K_Doxo_EG
Z_Shift_L_Doxo_HLD
Z_lm_L_Doxo_HLD
Z_Shift_L_Doxo_EG
Z_lm_L_Doxo_EG

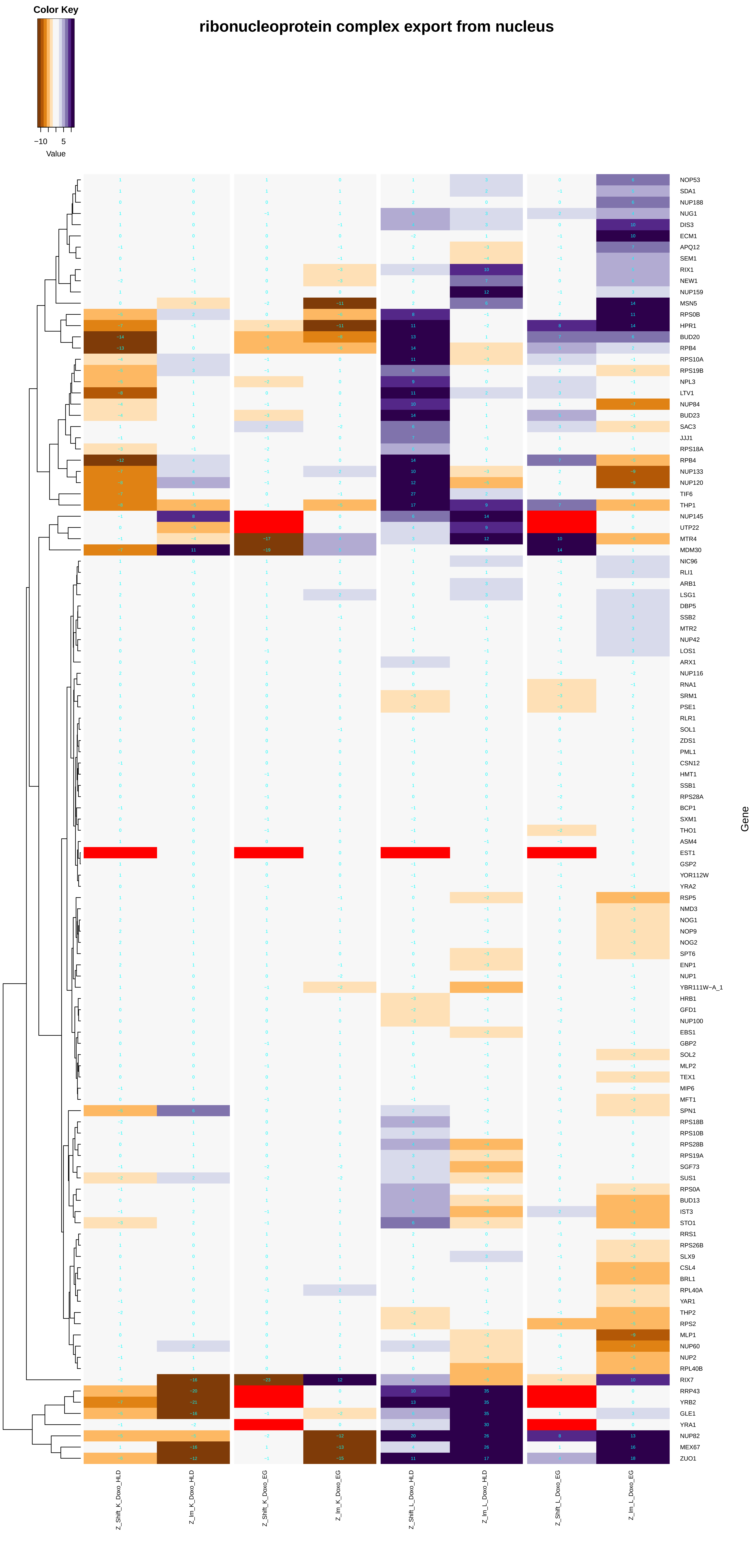

Color Key
−10
5
Value
ribonucleoprotein complex export from nucleus
NOP53
SDA1
NUP188
NUG1
DIS3
ECM1
APQ12
SEM1
RIX1
NEW1
NUP159
MSN5
RPS0B
HPR1
BUD20
RPB4
RPS10A
RPS19B
NPL3
LTV1
NUP84
BUD23
SAC3
JJJ1
RPS18A
RPB4
NUP133
NUP120
TIF6
THP1
NUP145
UTP22
MTR4
MDM30
NIC96
RLI1
ARB1
LSG1
DBP5
SSB2
MTR2
NUP42
LOS1
ARX1
NUP116
RNA1
SRM1
PSE1
RLR1
SOL1
ZDS1
PML1
CSN12
HMT1
SSB1
RPS28A
BCP1
SXM1
THO1
ASM4
EST1
GSP2
YOR112W
YRA2
RSP5
NMD3
NOG1
NOP9
NOG2
SPT6
ENP1
NUP1
YBR111W−A_1
HRB1
GFD1
NUP100
EBS1
GBP2
SOL2
MLP2
TEX1
MIP6
MFT1
SPN1
RPS18B
RPS10B
RPS28B
RPS19A
SGF73
SUS1
RPS0A
BUD13
IST3
STO1
RRS1
RPS26B
SLX9
CSL4
BRL1
RPL40A
YAR1
THP2
RPS2
MLP1
NUP60
NUP2
RPL40B
RIX7
RRP43
YRB2
GLE1
YRA1
NUP82
MEX67
ZUO1
Gene
Z_Shift_K_Doxo_HLD
Z_lm_K_Doxo_HLD
Z_Shift_K_Doxo_EG
Z_lm_K_Doxo_EG
Z_Shift_L_Doxo_HLD
Z_lm_L_Doxo_HLD
Z_Shift_L_Doxo_EG
Z_lm_L_Doxo_EG

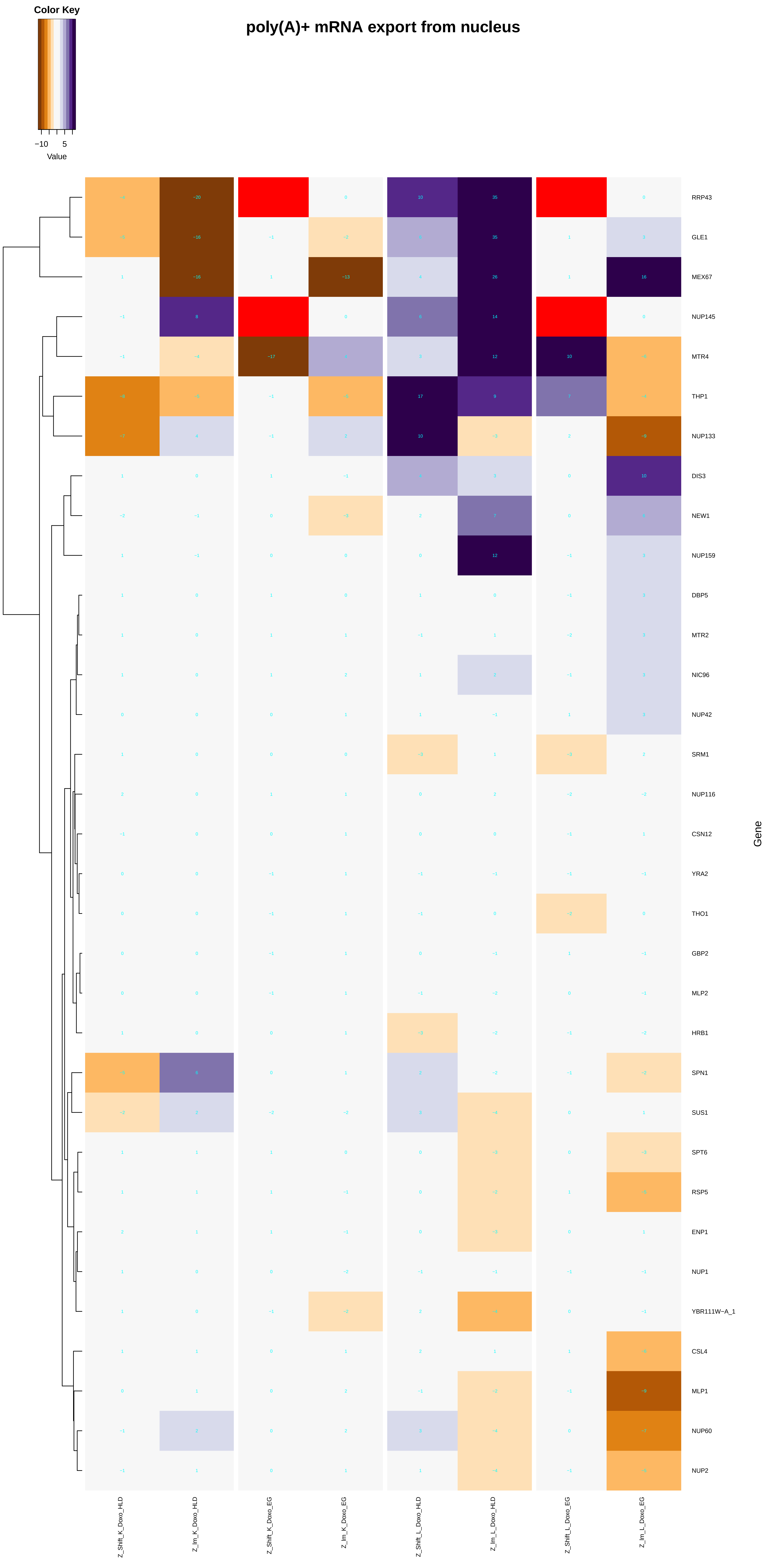
Color Key
−10
5
Value
poly(A)+ mRNA export from nucleus
RRP43
GLE1
MEX67
NUP145
MTR4
THP1
NUP133
DIS3
NEW1
NUP159
DBP5
MTR2
NIC96
NUP42
SRM1
NUP116
CSN12
YRA2
THO1
GBP2
MLP2
HRB1
SPN1
SUS1
SPT6
RSP5
ENP1
NUP1
YBR111W−A_1
CSL4
MLP1
NUP60
NUP2
Gene
Z_Shift_K_Doxo_HLD
Z_lm_K_Doxo_HLD
Z_Shift_K_Doxo_EG
Z_lm_K_Doxo_EG
Z_Shift_L_Doxo_HLD
Z_lm_L_Doxo_HLD
Z_Shift_L_Doxo_EG
Z_lm_L_Doxo_EG

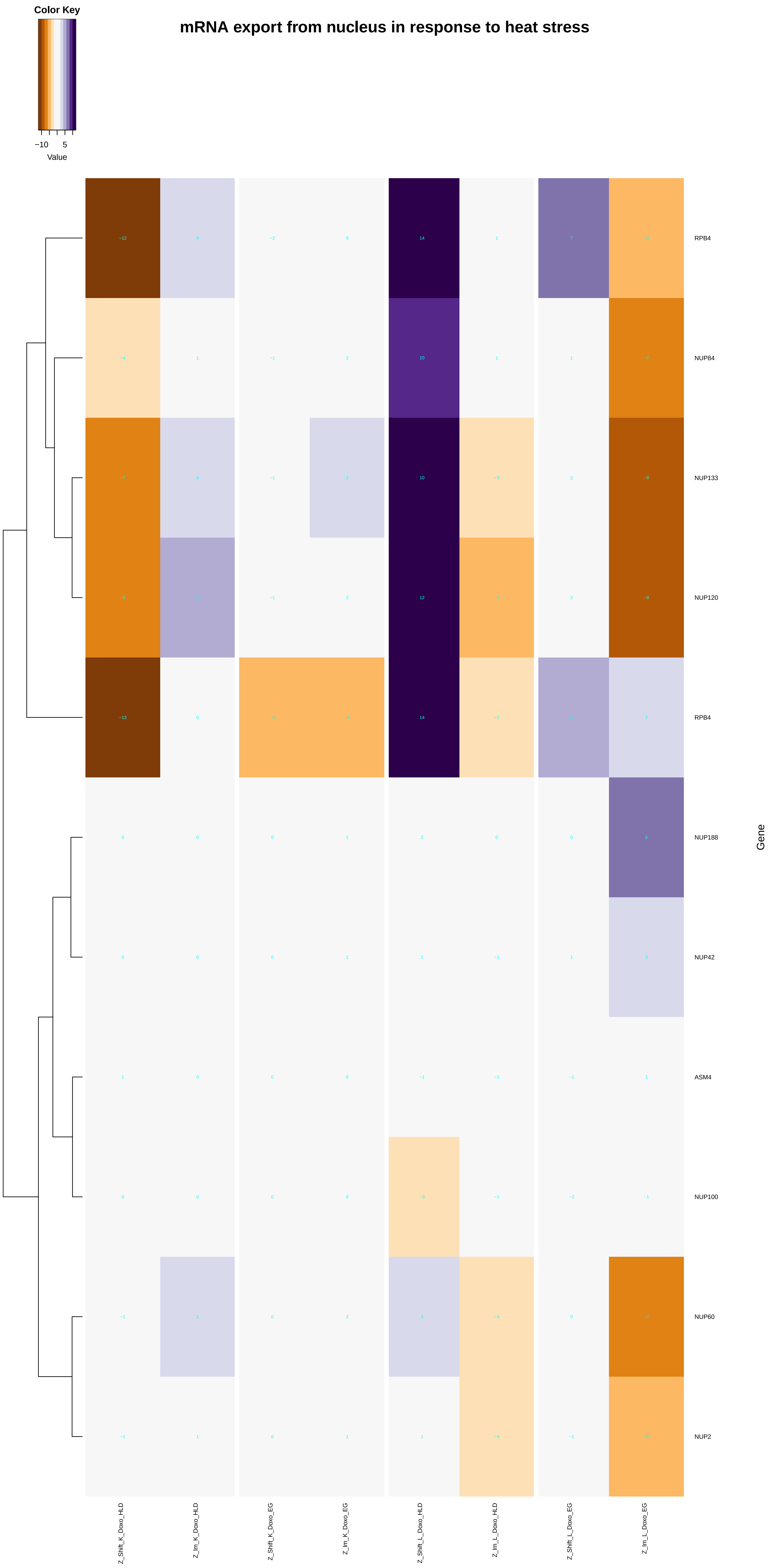
Color Key
−10
5
Value
mRNA export from nucleus in response to heat stress
RPB4
NUP84
NUP133
NUP120
RPB4
NUP188
NUP42
ASM4
NUP100
NUP60
NUP2
Gene
Z_Shift_K_Doxo_HLD
Z_Im_K_Doxo_HLD
Z_Shift_K_Doxo_EG
Z_Im_K_Doxo_EG
Z_Shift_L_Doxo_HLD
Z_Im_L_Doxo_HLD
Z_Shift_L_Doxo_EG
Z_Im_L_Doxo_EG

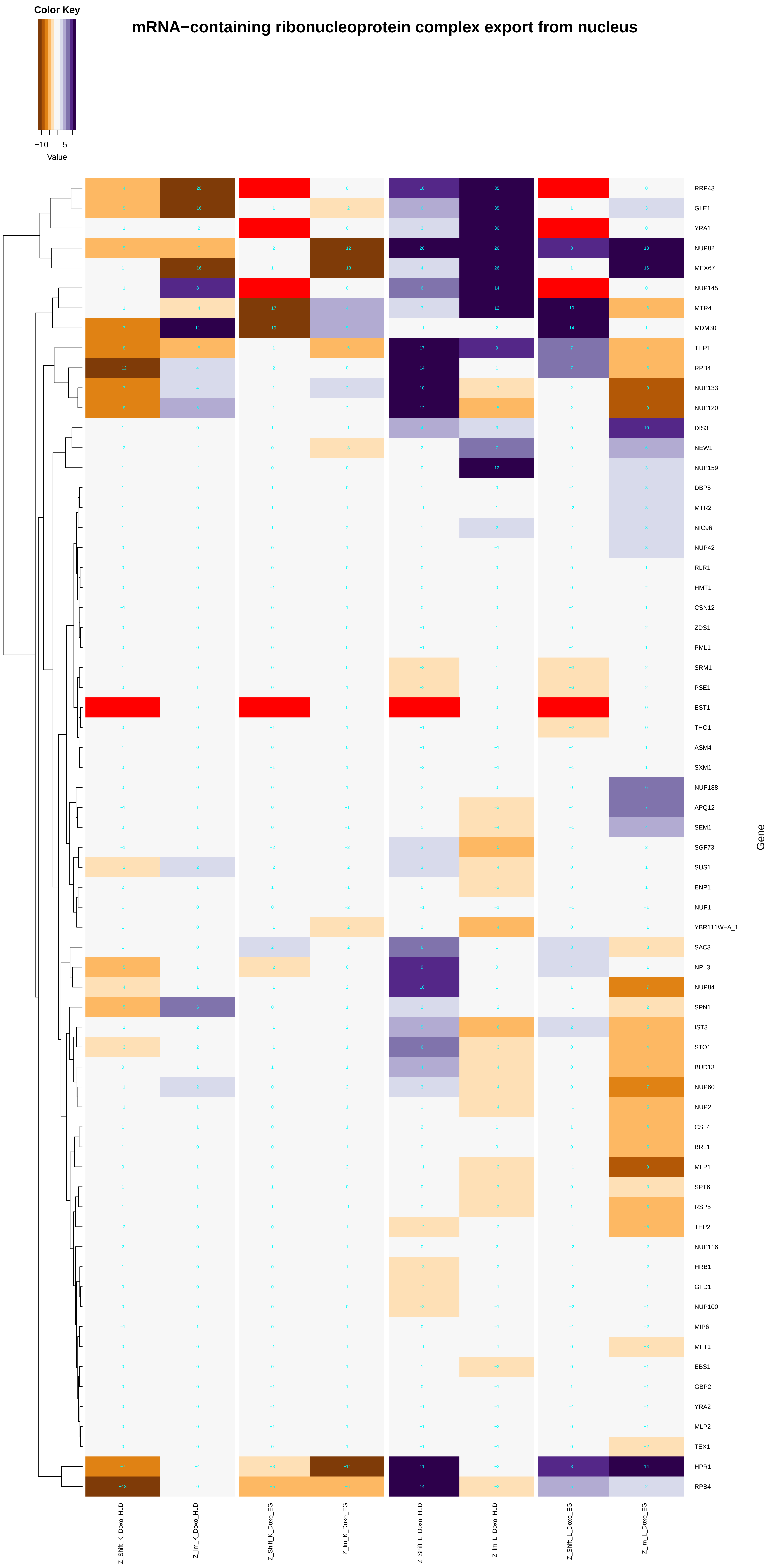
mRNA−containing ribonucleoprotein complex export from nucleus
Color Key
−10
5
Value
Gene
RRP43
GLE1
YRA1
NUP82
MEX67
NUP145
MTR4
MDM30
THP1
RPB4
NUP133
NUP120
DIS3
NEW1
NUP159
DBP5
MTR2
NIC96
NUP42
RLR1
HMT1
CSN12
ZDS1
PML1
SRM1
PSE1
EST1
THO1
ASM4
SXM1
NUP188
APQ12
SEM1
SGF73
SUS1
ENP1
NUP1
YBR111W−A_1
SAC3
NPL3
NUP84
SPN1
IST3
STO1
BUD13
NUP60
NUP2
CSL4
BRL1
MLP1
SPT6
RSP5
THP2
NUP116
HRB1
GFD1
NUP100
MIP6
MFT1
EBS1
GBP2
YRA2
MLP2
TEX1
HPR1
RPB4
Z_Shift_K_Doxo_HLD
Z_lm_K_Doxo_HLD
Z_Shift_K_Doxo_EG
Z_lm_K_Doxo_EG
Z_Shift_L_Doxo_HLD
Z_lm_L_Doxo_HLD
Z_Shift_L_Doxo_EG
Z_lm_L_Doxo_EG

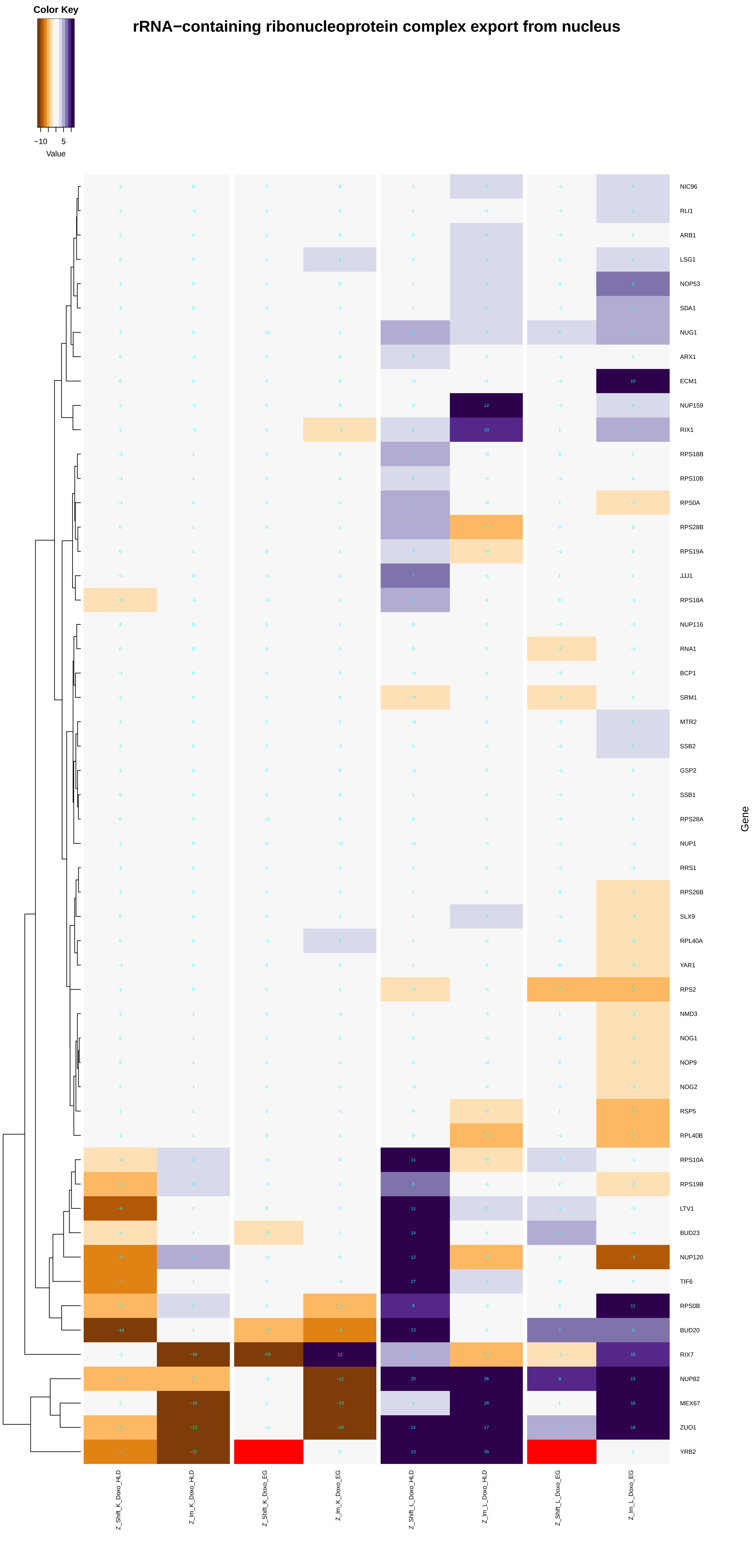

Color Key
−10 5
Value
rRNA−containing ribonucleoprotein complex export from nucleus
Gene
NIC96
RLI1
ARB1
LSG1
NOP53
SDA1
NUG1
ARX1
ECM1
NUP159
RIX1
RPS18B
RPS10B
RPS0A
RPS28B
RPS19A
JJJ1
RPS18A
NUP116
RNA1
BCP1
SRM1
MTR2
SSB2
GSP2
SSB1
RPS28A
NUP1
RRS1
RPS26B
SLX9
RPL40A
YAR1
RPS2
NMD3
NOG1
NOP9
NOG2
RSP5
RPL40B
RPS10A
RPS19B
LTV1
BUD23
NUP120
TIF6
RPS0B
BUD20
RIX7
NUP82
MEX67
ZUO1
YRB2
Z_Shift_K_Doxo_HLD
Z_lm_K_Doxo_HLD
Z_Shift_K_Doxo_EG
Z_lm_K_Doxo_EG
Z_Shift_L_Doxo_HLD
Z_lm_L_Doxo_HLD
Z_Shift_L_Doxo_EG
Z_lm_L_Doxo_EG

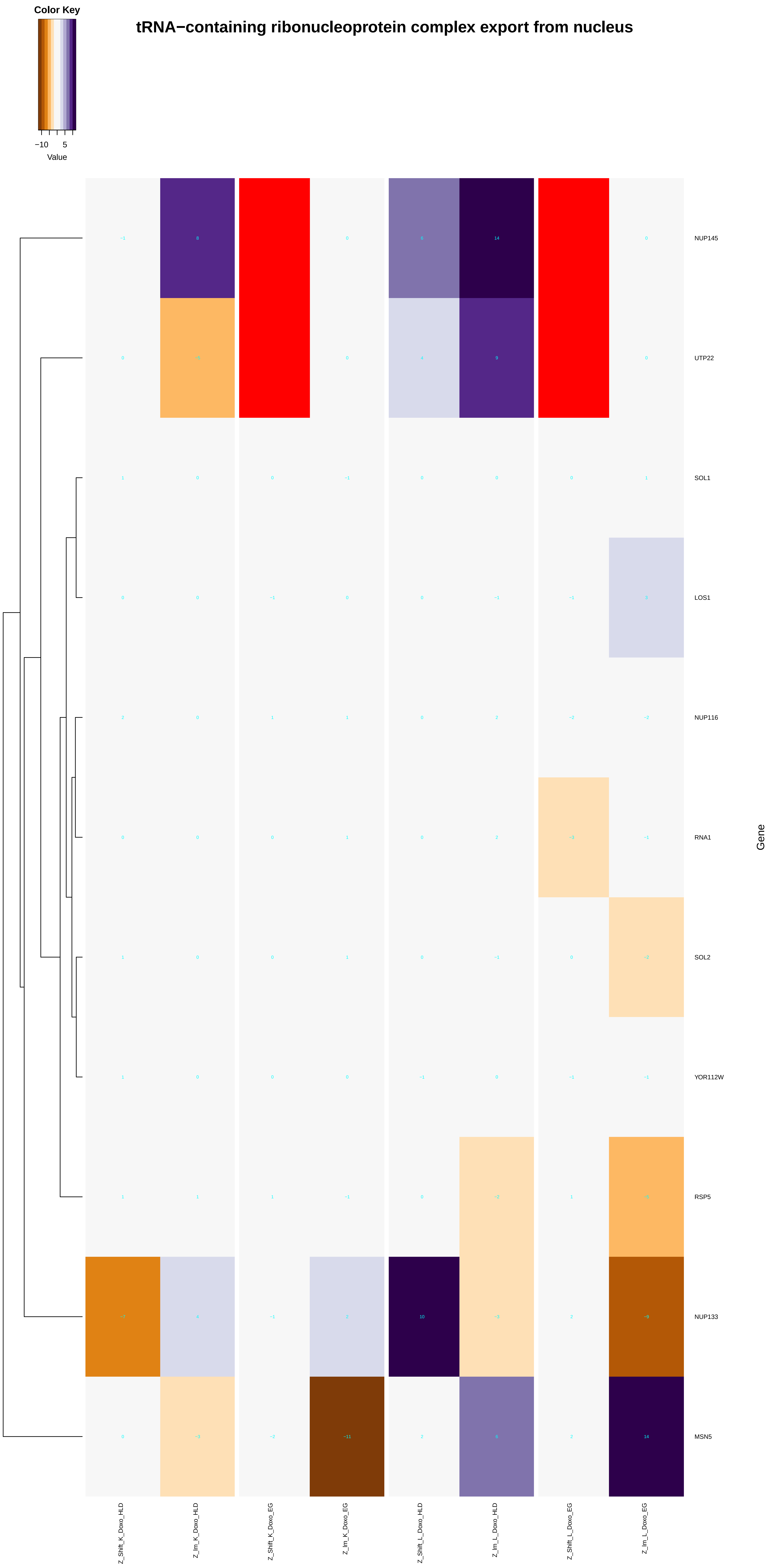
Color Key
−10
5
Value
tRNA−containing ribonucleoprotein complex export from nucleus
NUP145
UTP22
SOL1
LOS1
NUP116
RNA1
SOL2
YOR112W
RSP5
NUP133
MSN5
Gene
Z_Shift_K_Doxo_HLD
Z_Im_K_Doxo_HLD
Z_Shift_K_Doxo_EG
Z_Im_K_Doxo_EG
Z_Shift_L_Doxo_HLD
Z_Im_L_Doxo_HLD
Z_Shift_L_Doxo_EG
Z_Im_L_Doxo_EG

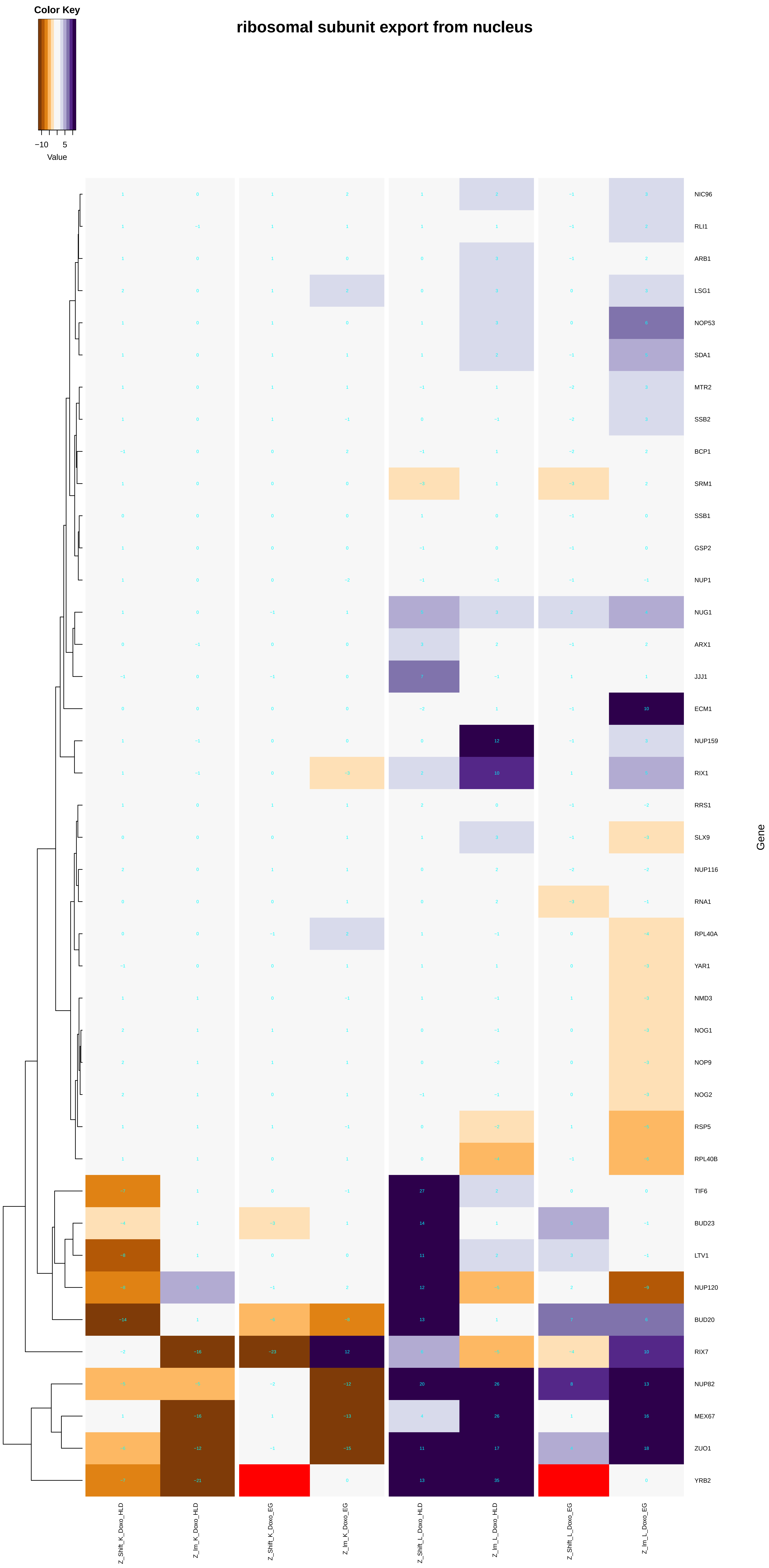

Color Key
−10
5
Value
ribosomal subunit export from nucleus
NIC96
RLI1
ARB1
LSG1
NOP53
SDA1
MTR2
SSB2
BCP1
SRM1
SSB1
GSP2
NUP1
NUG1
ARX1
JJJ1
ECM1
NUP159
RIX1
RRS1
SLX9
NUP116
RNA1
RPL40A
YAR1
NMD3
NOG1
NOP9
NOG2
RSP5
RPL40B
TIF6
BUD23
LTV1
NUP120
BUD20
RIX7
NUP82
MEX67
ZUO1
YRB2
Gene
Z_Shift_K_Doxo_HLD
Z_lm_K_Doxo_HLD
Z_Shift_K_Doxo_EG
Z_lm_K_Doxo_EG
Z_Shift_L_Doxo_HLD
Z_lm_L_Doxo_HLD
Z_Shift_L_Doxo_EG
Z_lm_L_Doxo_EG

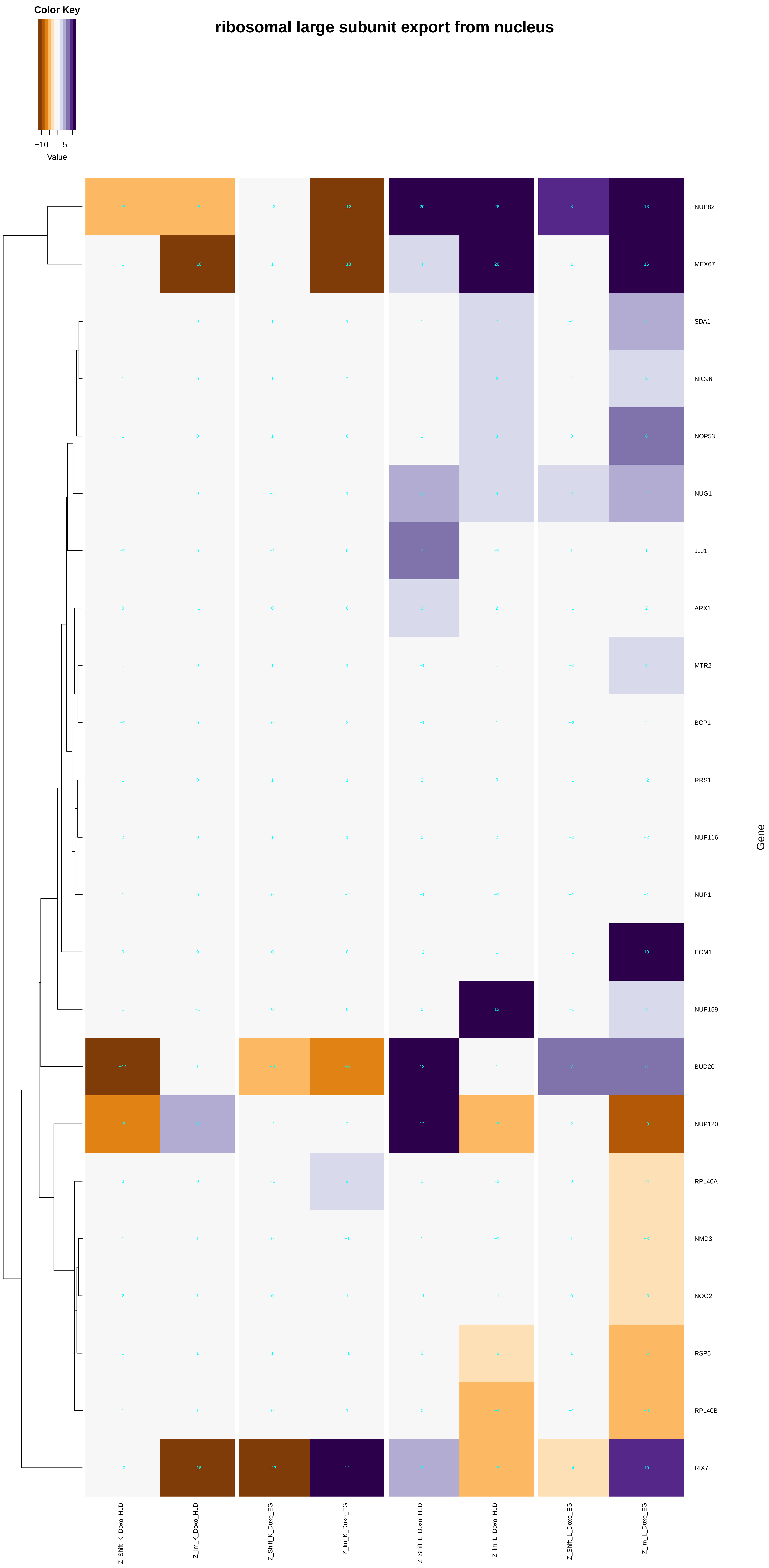

Color Key
−10
5
Value
ribosomal large subunit export from nucleus
NUP82
MEX67
SDA1
NIC96
NOP53
NUG1
JJJ1
ARX1
MTR2
BCP1
RRS1
NUP116
NUP1
ECM1
NUP159
BUD20
NUP120
RPL40A
NMD3
NOG2
RSP5
RPL40B
RIX7
Gene
Z_Shift_K_Doxo_HLD
Z_Im_K_Doxo_HLD
Z_Shift_K_Doxo_EG
Z_Im_K_Doxo_EG
Z_Shift_L_Doxo_HLD
Z_Im_L_Doxo_HLD
Z_Shift_L_Doxo_EG
Z_Im_L_Doxo_EG

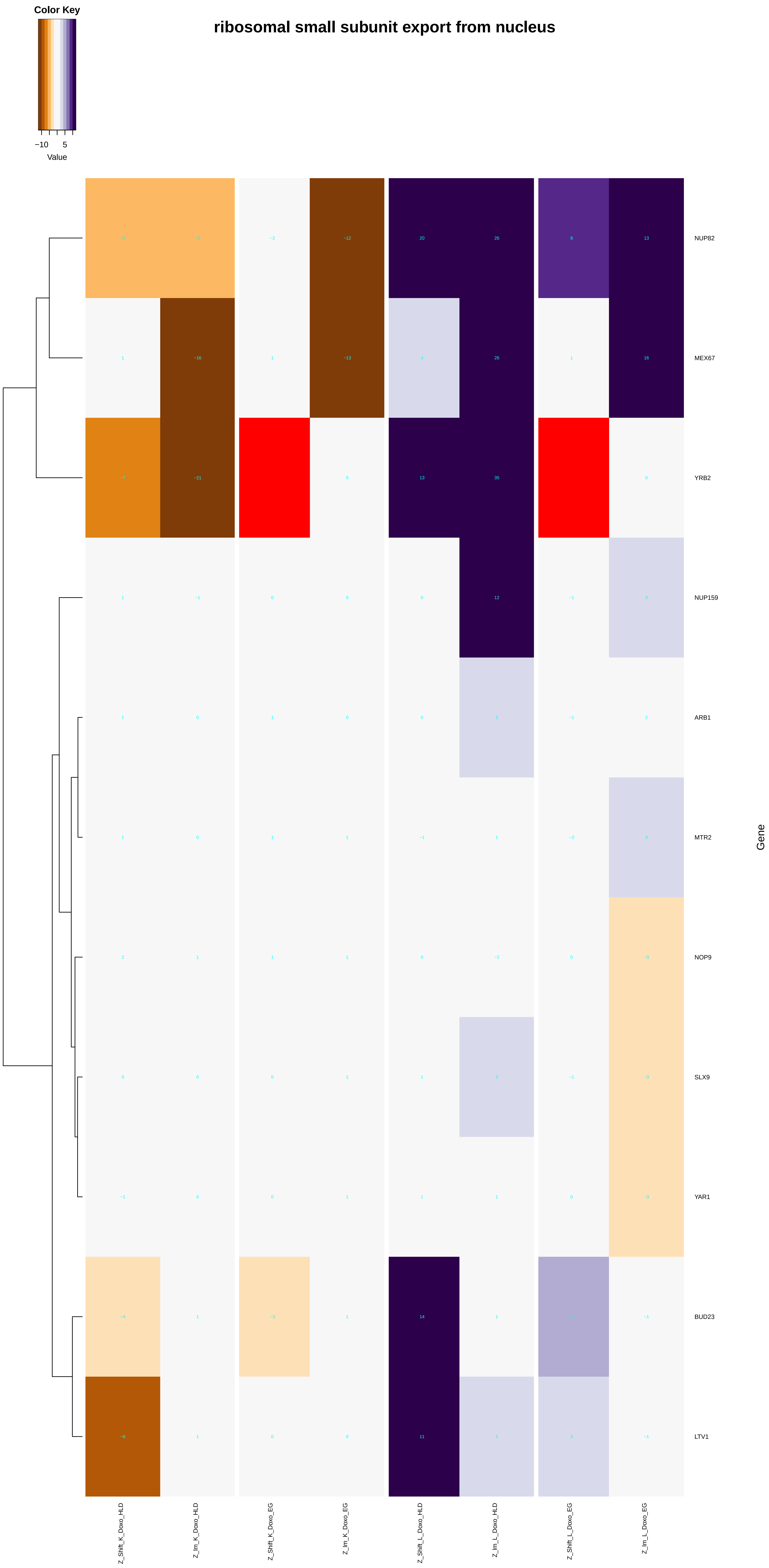

Color Key
−10
5
Value
ribosomal small subunit export from nucleus
NUP82
MEX67
YRB2
NUP159
ARB1
MTR2
NOP9
SLX9
YAR1
BUD23
LTV1
Gene
Z_Shift_K_Doxo_HLD
Z_Im_K_Doxo_HLD
Z_Shift_K_Doxo_EG
Z_Im_K_Doxo_EG
Z_Shift_L_Doxo_HLD
Z_Im_L_Doxo_HLD
Z_Shift_L_Doxo_EG
Z_Im_L_Doxo_EG
