## Additional File 9 for "A yeast phenomic model for the influence of Warburg metabolism on genetic buffering of doxorubicin": nucleoside_monophosphate_biosynthetic_process.pdf

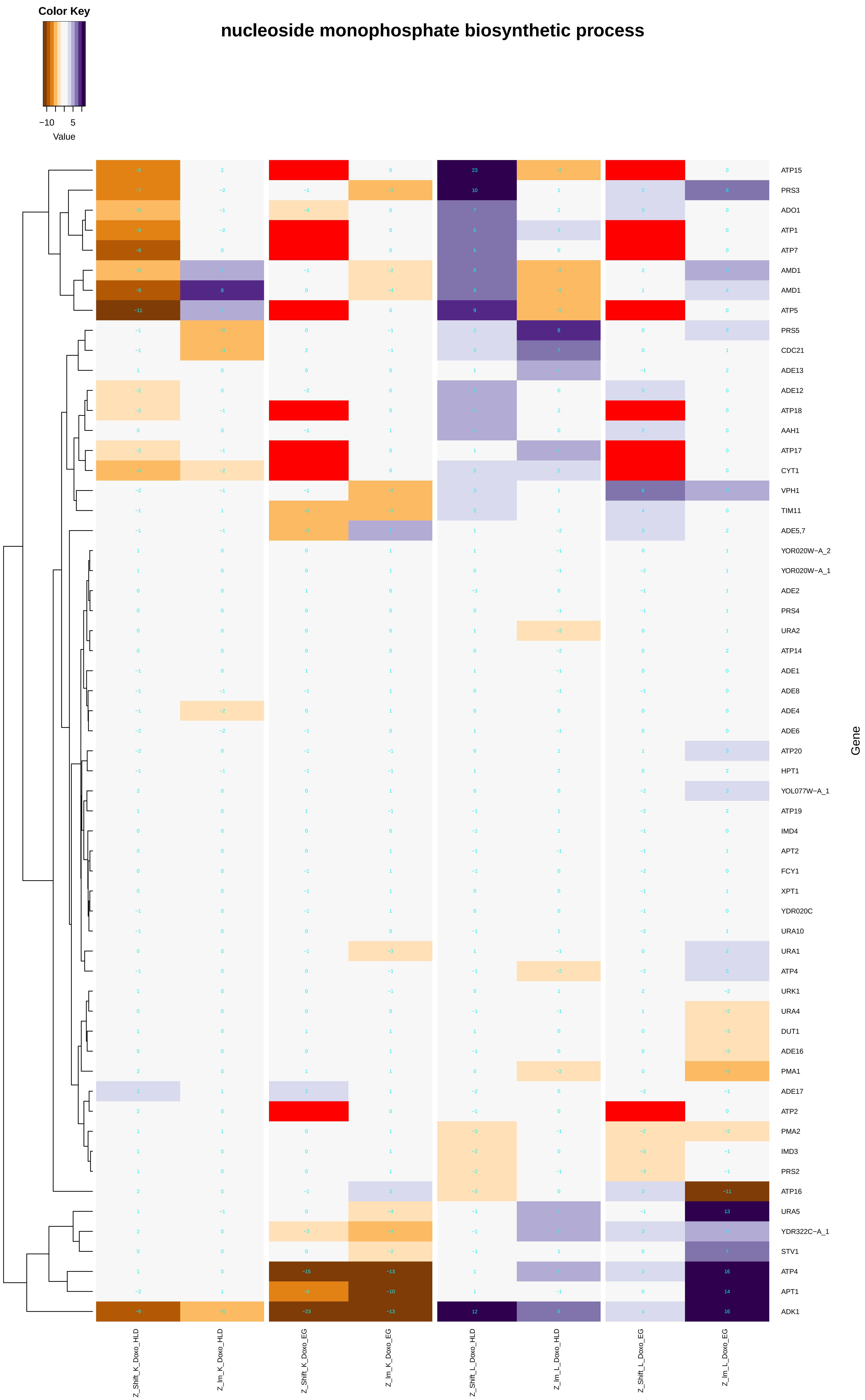

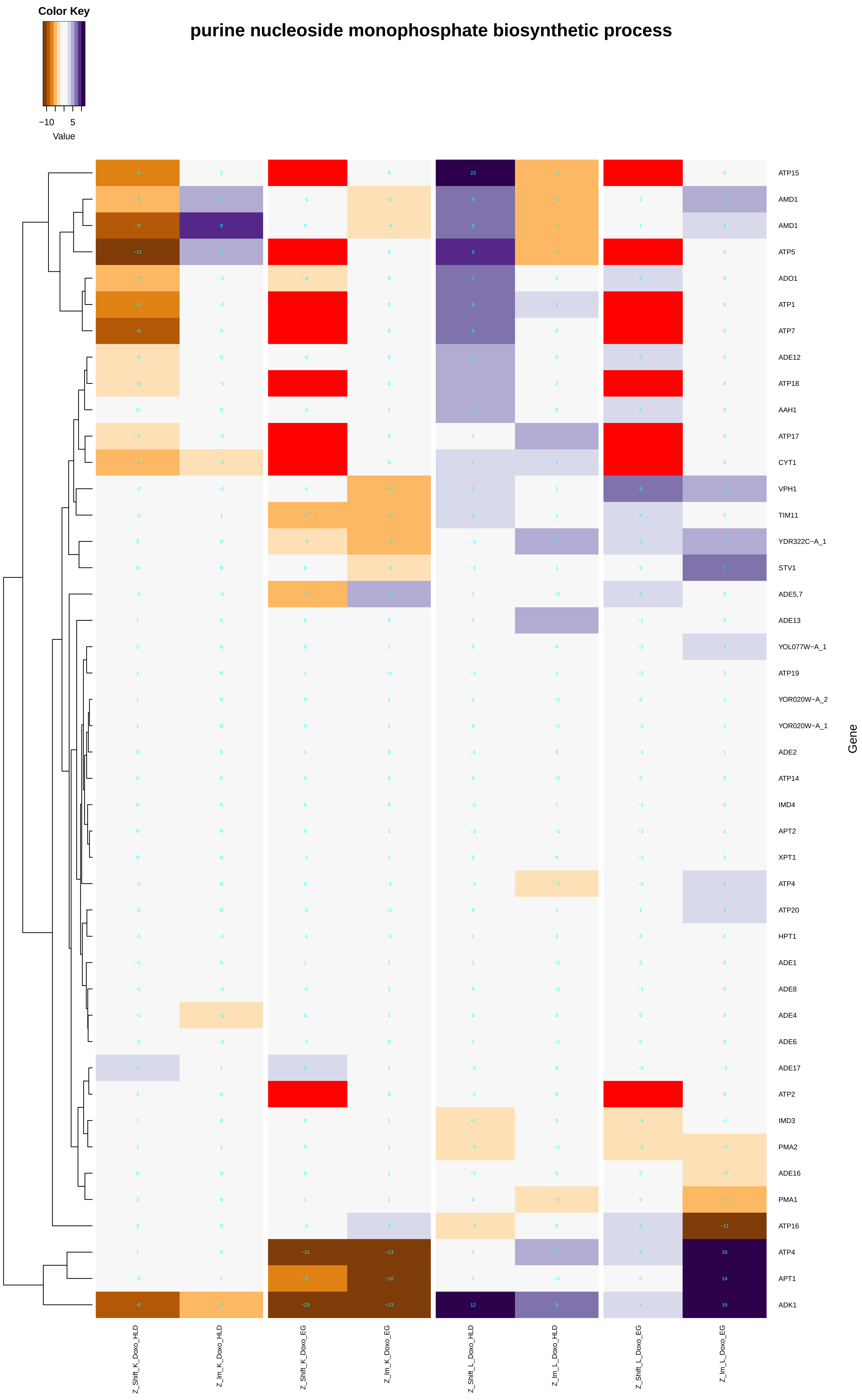

purine nucleoside monophosphate biosynthetic process
Color Key
−10
5
Value
ATP15
AMD1
AMD1
ATP5
ADO1
ATP1
ATP7
ADE12
ATP18
AAH1
ATP17
CYT1
VPH1
TIM11
YDR322C−A_1
STV1
ADE5,7
ADE13
YOL077W−A_1
ATP19
YOR020W−A_2
YOR020W−A_1
ADE2
ATP14
IMD4
APT2
XPT1
ATP4
ATP20
HPT1
ADE1
ADE8
ADE4
ADE6
ADE17
ATP2
IMD3
PMA2
ADE16
PMA1
ATP16
ATP4
APT1
ADK1
Gene
Z_Shift_K_Doxo_HLD
Z_lm_K_Doxo_HLD
Z_Shift_K_Doxo_EG
Z_lm_K_Doxo_EG
Z_Shift_L_Doxo_HLD
Z_lm_L_Doxo_HLD
Z_Shift_L_Doxo_EG
Z_lm_L_Doxo_EG

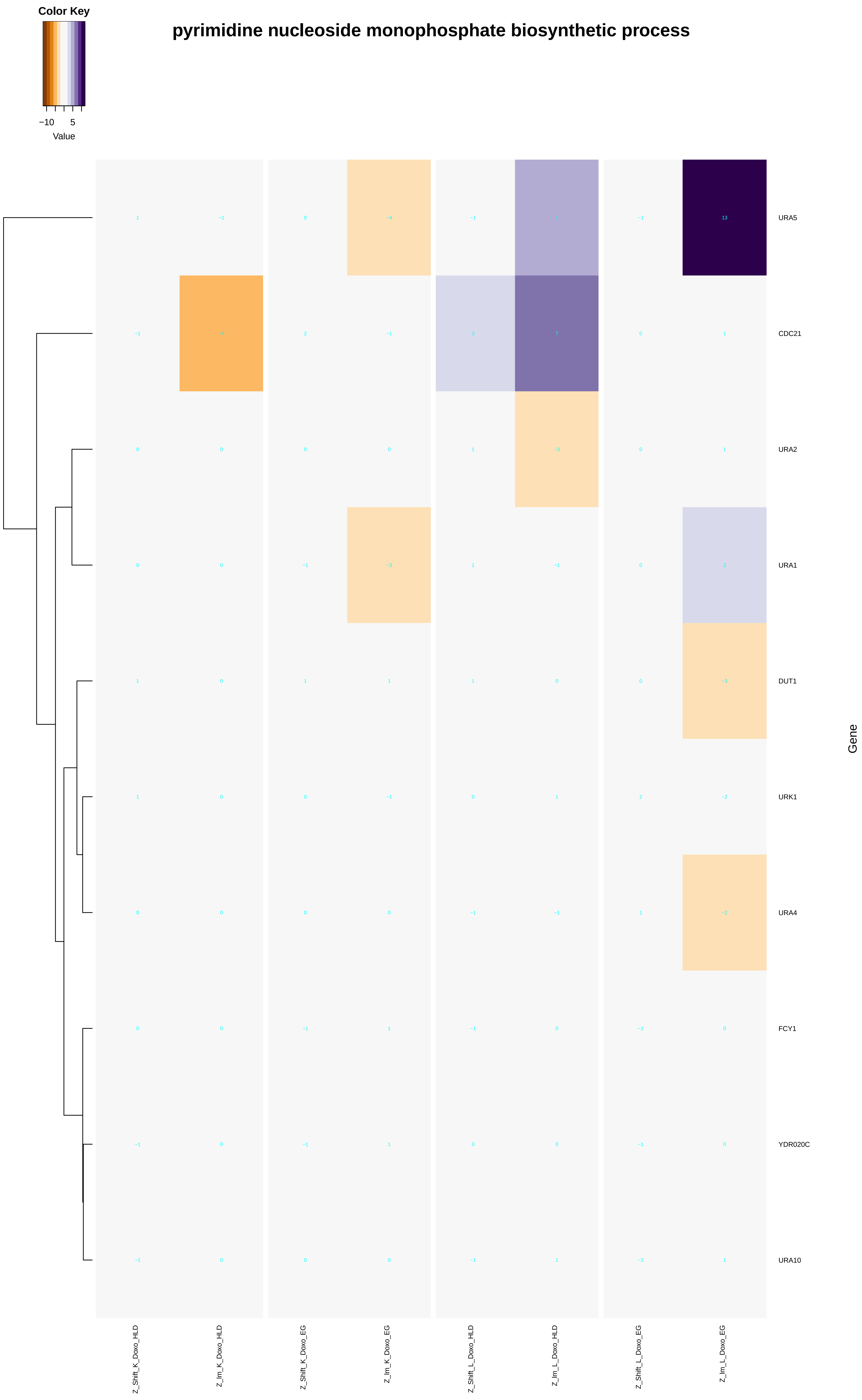

Color Key
−10
5
Value
pyrimidine nucleoside monophosphate biosynthetic process
URA5
CDC21
URA2
URA1
DUT1
URK1
URA4
FCY1
YDR020C
URA10
Gene
Z_Shift_K_Doxo_HLD
Z_lm_K_Doxo_HLD
Z_Shift_K_Doxo_EG
Z_lm_K_Doxo_EG
Z_Shift_L_Doxo_HLD
Z_lm_L_Doxo_HLD
Z_Shift_L_Doxo_EG
Z_lm_L_Doxo_EG

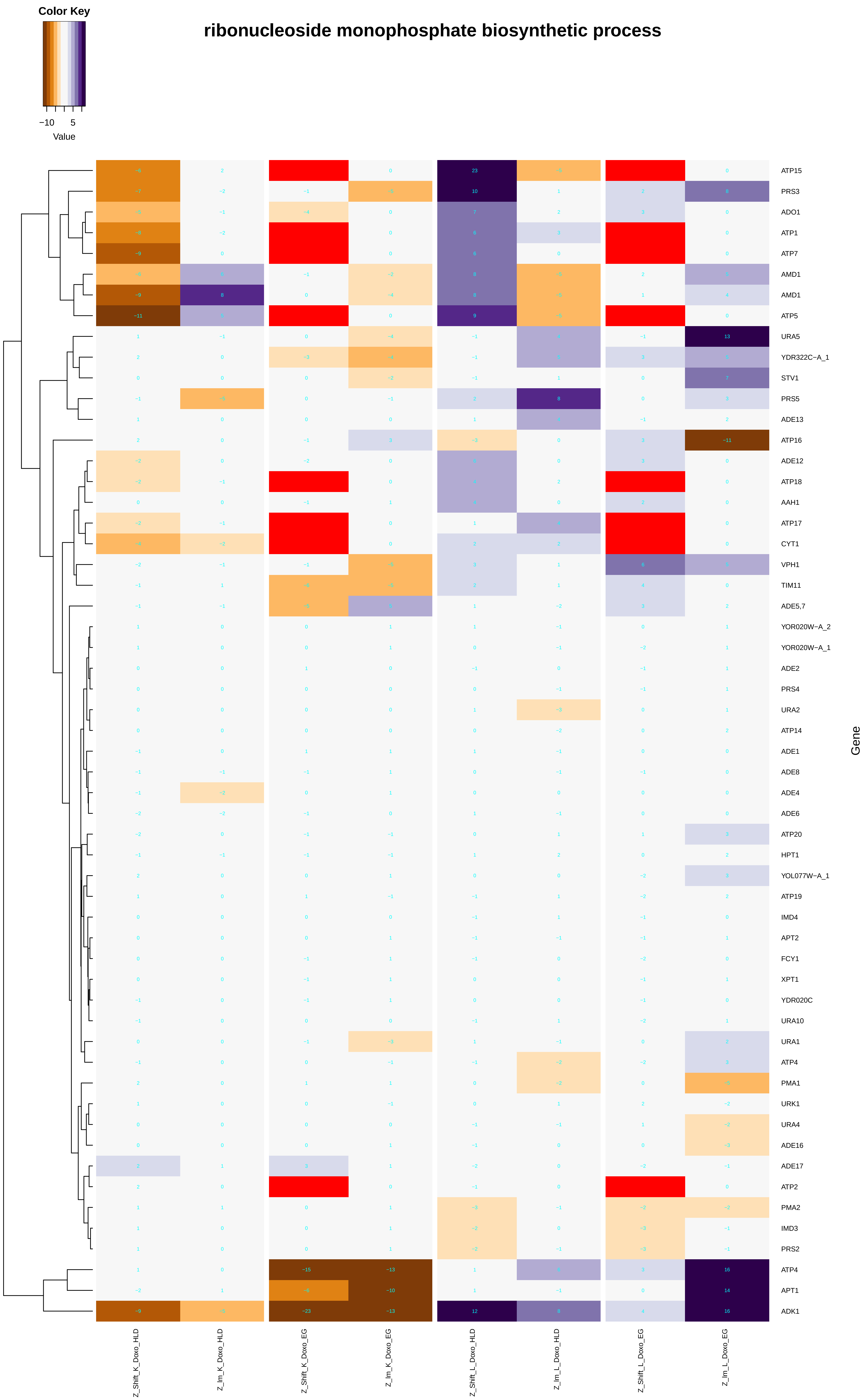

ribonucleoside monophosphate biosynthetic process
Color Key
−10
5
Value
ATP15
PRS3
ADO1
ATP1
ATP7
AMD1
AMD1
ATP5
URA5
YDR322C−A_1
STV1
PRS5
ADE13
ATP16
ADE12
ATP18
AAH1
ATP17
CYT1
VPH1
TIM11
ADE5,7
YOR020W−A_2
YOR020W−A_1
ADE2
PRS4
URA2
ATP14
ADE1
ADE8
ADE4
ADE6
ATP20
HPT1
YOL077W−A_1
ATP19
IMD4
APT2
FCY1
XPT1
YDR020C
URA10
URA1
ATP4
PMA1
URK1
URA4
ADE16
ADE17
ATP2
PMA2
IMD3
PRS2
ATP4
APT1
ADK1
Gene
Z_Shift_K_Doxo_HLD
Z_Im_K_Doxo_HLD
Z_Shift_K_Doxo_EG
Z_Im_K_Doxo_EG
Z_Shift_L_Doxo_HLD
Z_Im_L_Doxo_HLD
Z_Shift_L_Doxo_EG
Z_Im_L_Doxo_EG

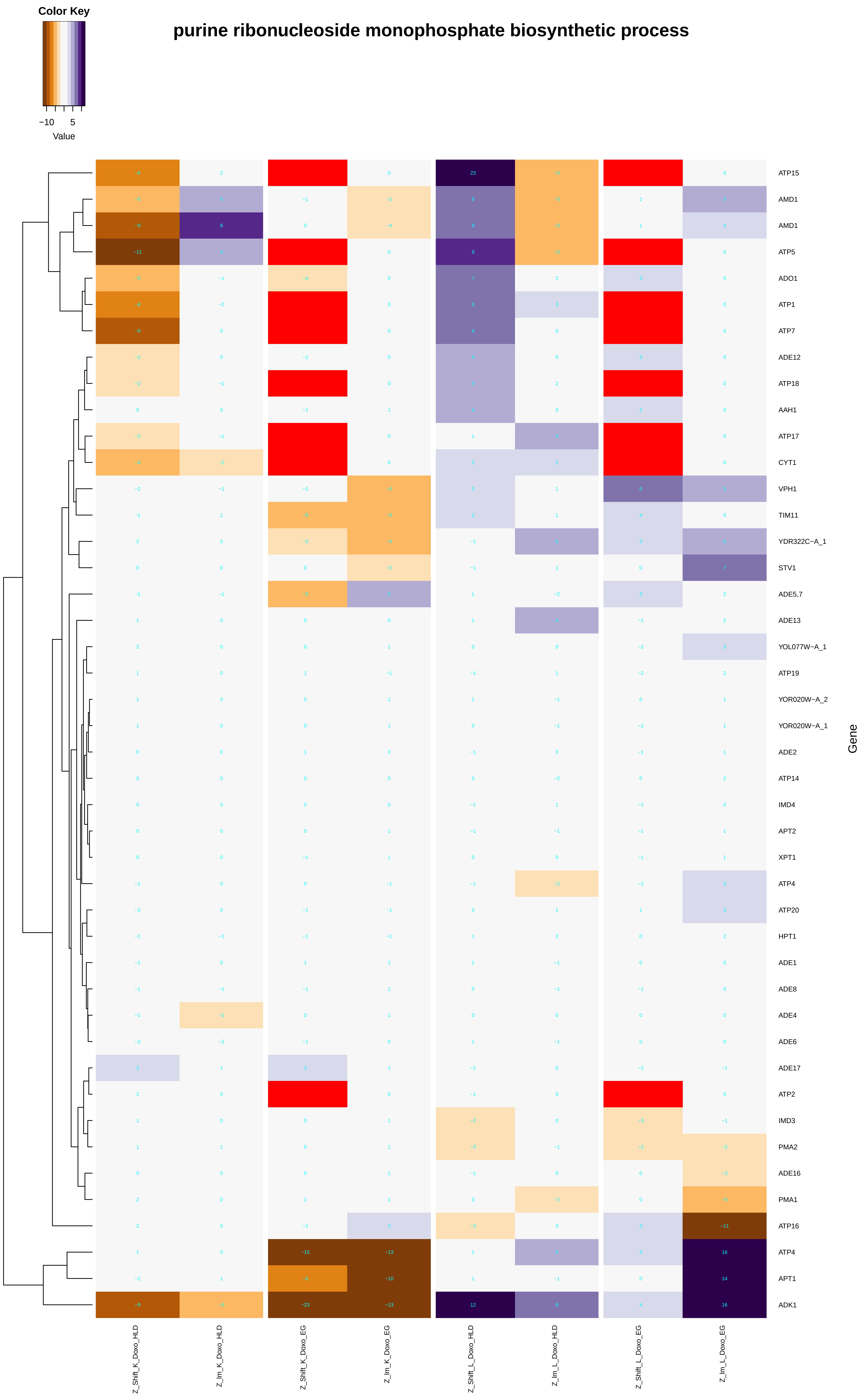
Color Key
−10
5
Value
purine ribonucleoside monophosphate biosynthetic process
ATP15
AMD1
AMD1
ATP5
ADO1
ATP1
ATP7
ADE12
ATP18
AAH1
ATP17
CYT1
VPH1
TIM11
YDR322C−A_1
STV1
ADE5,7
ADE13
YOL077W−A_1
ATP19
YOR020W−A_2
YOR020W−A_1
ADE2
ATP14
IMD4
APT2
XPT1
ATP4
ATP20
HPT1
ADE1
ADE8
ADE4
ADE6
ADE17
ATP2
IMD3
PMA2
ADE16
PMA1
ATP16
ATP4
APT1
ADK1
Gene
Z_Shift_K_Doxo_HLD
Z_Im_K_Doxo_HLD
Z_Shift_K_Doxo_EG
Z_Im_K_Doxo_EG
Z_Shift_L_Doxo_HLD
Z_Im_L_Doxo_HLD
Z_Shift_L_Doxo_EG
Z_Im_L_Doxo_EG

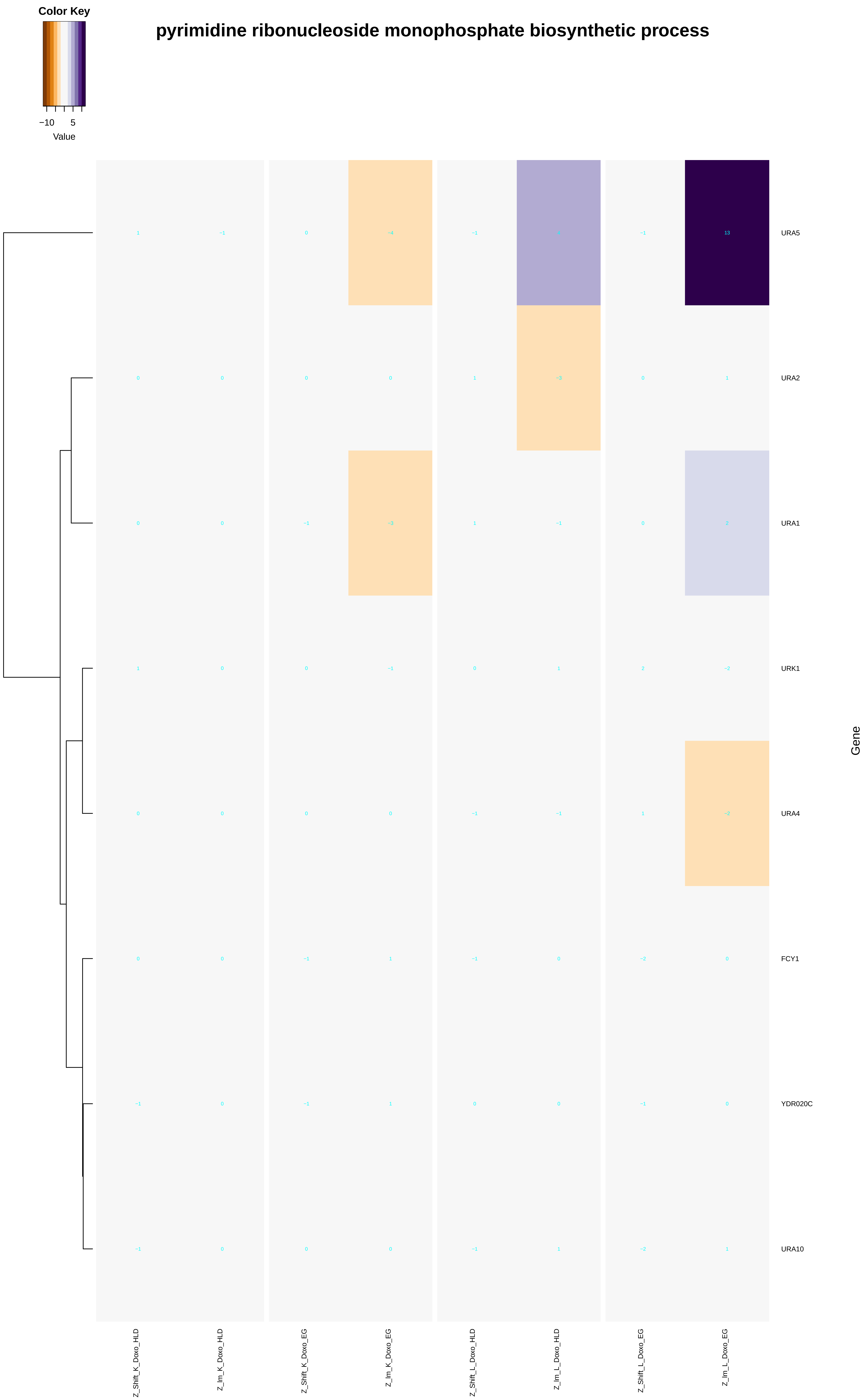

Color Key
−10
5
Value
pyrimidine ribonucleoside monophosphate biosynthetic process
URA5
URA2
URA1
URK1
URA4
FCY1
YDR020C
URA10
Gene
Z_Shift_K_Doxo_HLD
Z_Im_K_Doxo_HLD
Z_Shift_K_Doxo_EG
Z_Im_K_Doxo_EG
Z_Shift_L_Doxo_HLD
Z_Im_L_Doxo_HLD
Z_Shift_L_Doxo_EG
Z_Im_L_Doxo_EG

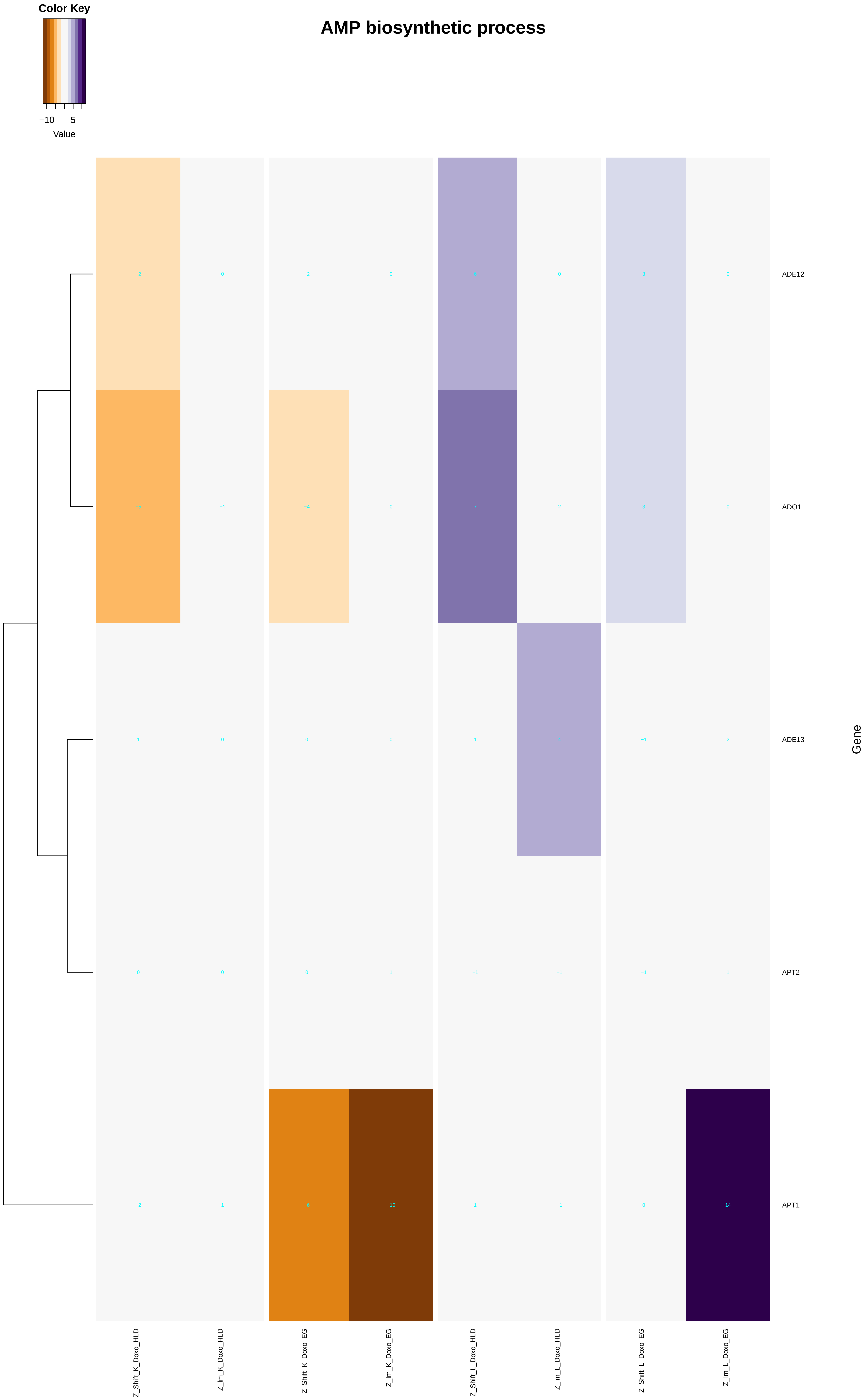
AMP biosynthetic process
Color Key
−10
5
Value
ADE12
ADO1
ADE13
APT2
APT1
Gene
Z_Shift_K_Doxo_HLD
Z_Im_K_Doxo_HLD
Z_Shift_K_Doxo_EG
Z_Im_K_Doxo_EG
Z_Shift_L_Doxo_HLD
Z_Im_L_Doxo_HLD
Z_Shift_L_Doxo_EG
Z_Im_L_Doxo_EG

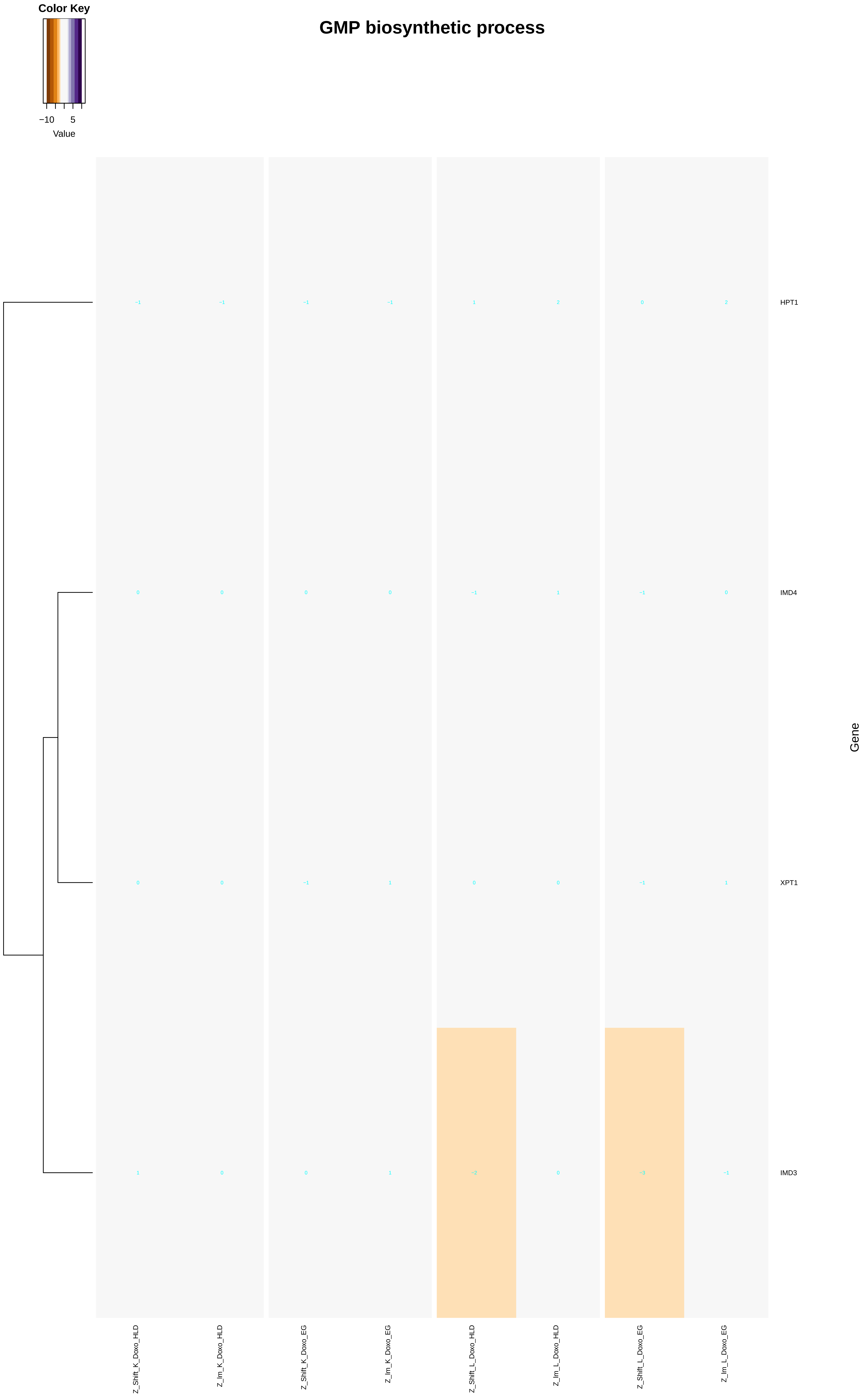

Color Key
−10
5
Value
GMP biosynthetic process
HPT1
IMD4
XPT1
IMD3
Gene
Z_Shift_K_Doxo_HLD
Z_Im_K_Doxo_HLD
Z_Shift_K_Doxo_EG
Z_Im_K_Doxo_EG
Z_Shift_L_Doxo_HLD
Z_Im_L_Doxo_HLD
Z_Shift_L_Doxo_EG
Z_Im_L_Doxo_EG

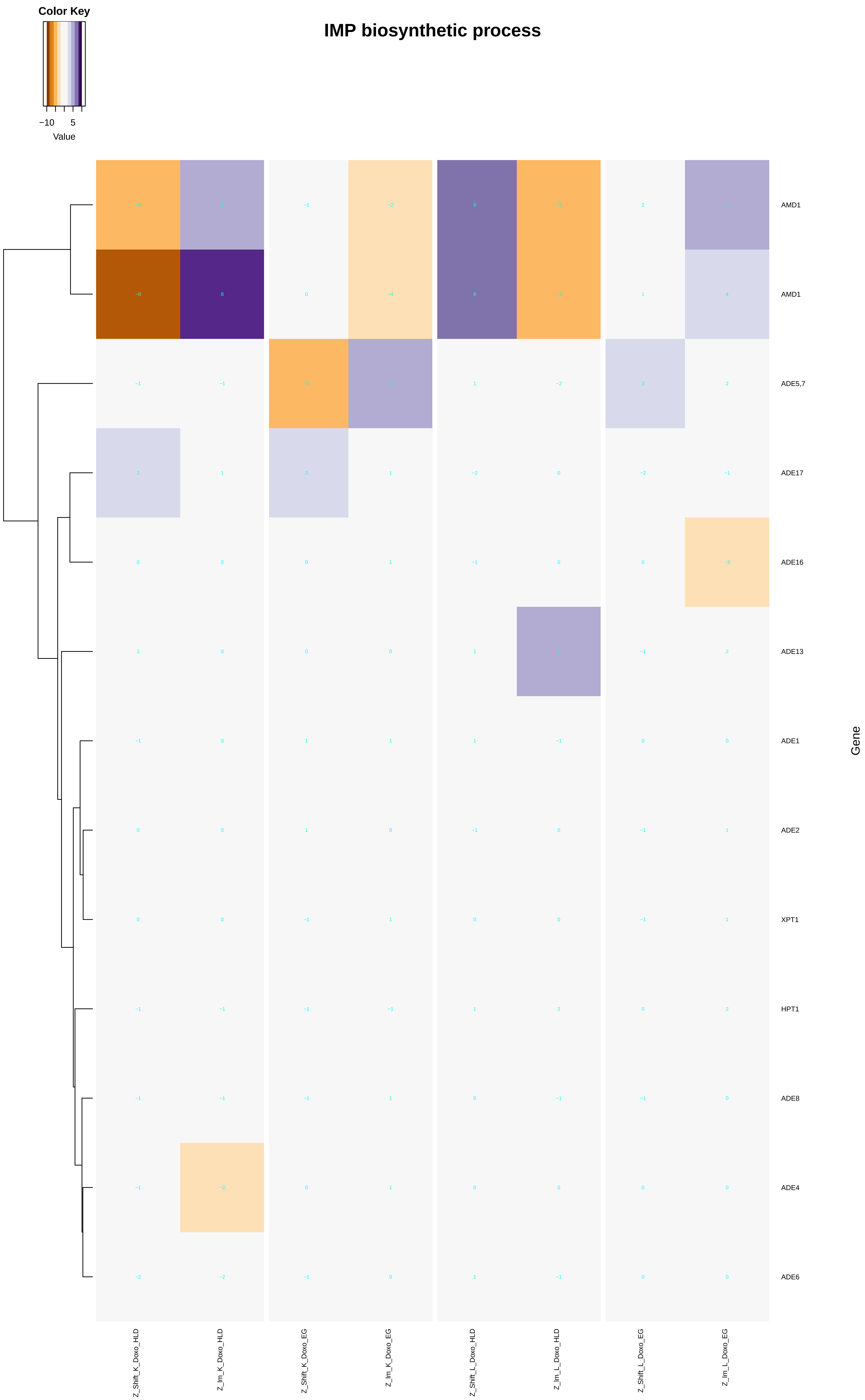

IMP biosynthetic process
Color Key
−10
5
Value
AMD1
AMD1
ADE5,7
ADE17
ADE16
ADE13
ADE1
ADE2
XPT1
HPT1
ADE8
ADE4
ADE6
Gene
Z_Shift_K_Doxo_HLD
Z_Im_K_Doxo_HLD
Z_Shift_K_Doxo_EG
Z_Im_K_Doxo_EG
Z_Shift_L_Doxo_HLD
Z_Im_L_Doxo_HLD
Z_Shift_L_Doxo_EG
Z_Im_L_Doxo_EG

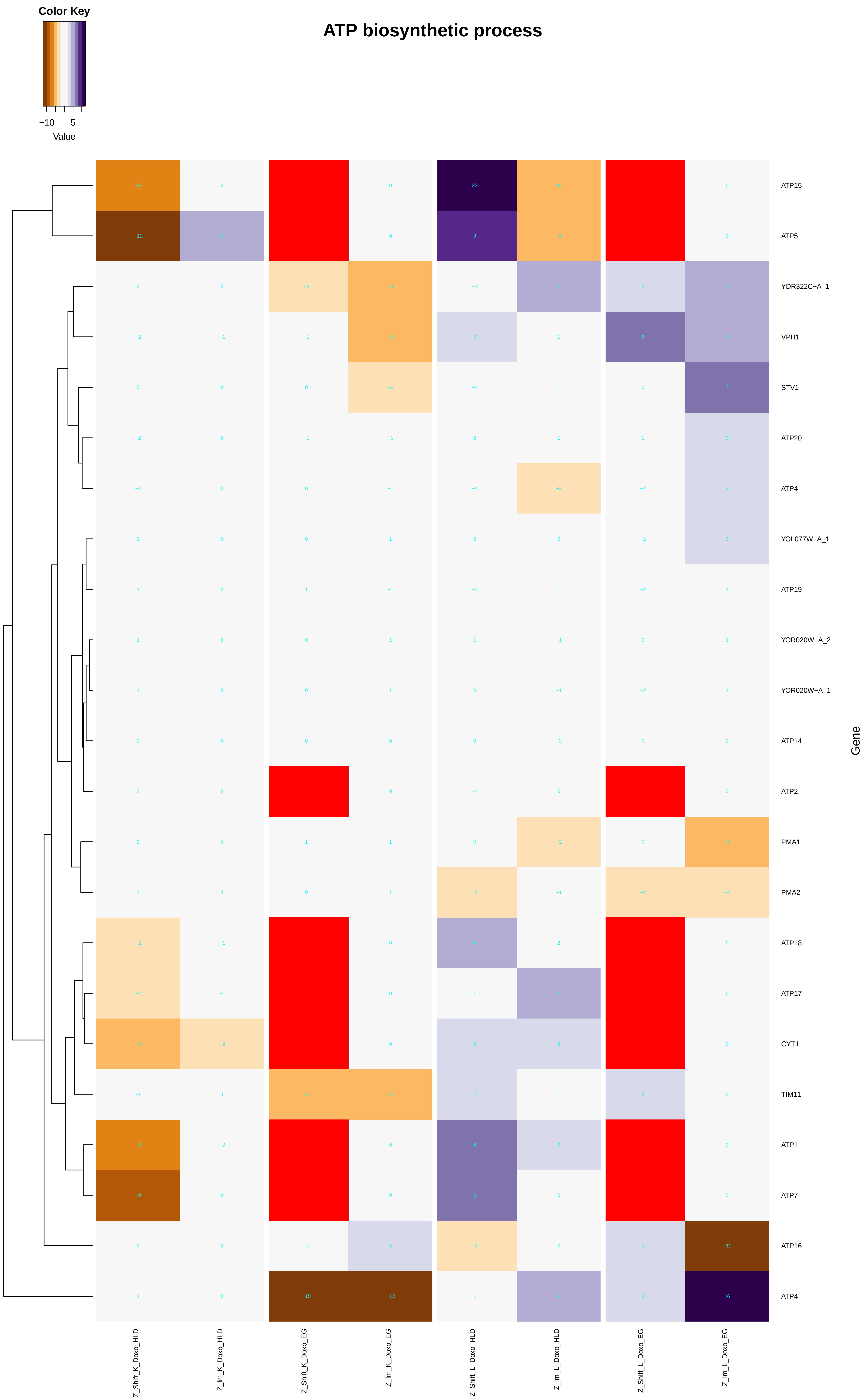
ATP biosynthetic process
Color Key
−10
5
Value
ATP15
ATP5
YDR322C−A_1
VPH1
STV1
ATP20
ATP4
YOL077W−A_1
ATP19
YOR020W−A_2
YOR020W−A_1
ATP14
ATP2
PMA1
PMA2
ATP18
ATP17
CYT1
TIM11
ATP1
ATP7
ATP16
ATP4
Gene
Z_Shift_K_Doxo_HLD
Z_Im_K_Doxo_HLD
Z_Shift_K_Doxo_EG
Z_Im_K_Doxo_EG
Z_Shift_L_Doxo_HLD
Z_Im_L_Doxo_HLD
Z_Shift_L_Doxo_EG
Z_Im_L_Doxo_EG

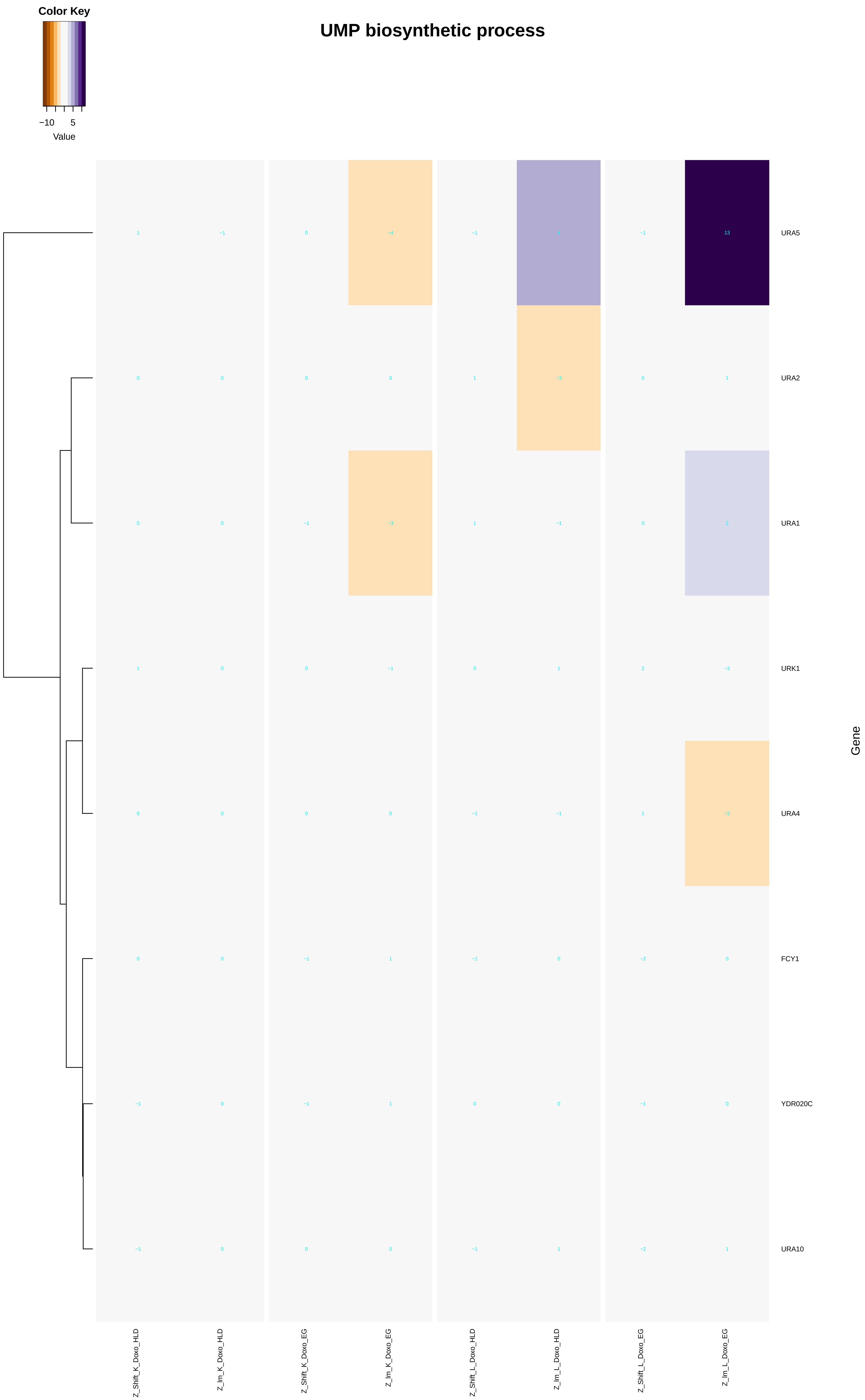

UMP biosynthetic process
Color Key
−10
5
Value
URA5
URA2
URA1
URK1
URA4
FCY1
YDR020C
URA10
Gene
Z_Shift_K_Doxo_HLD
Z_Im_K_Doxo_HLD
Z_Shift_K_Doxo_EG
Z_Im_K_Doxo_EG
Z_Shift_L_Doxo_HLD
Z_Im_L_Doxo_HLD
Z_Shift_L_Doxo_EG
Z_Im_L_Doxo_EG

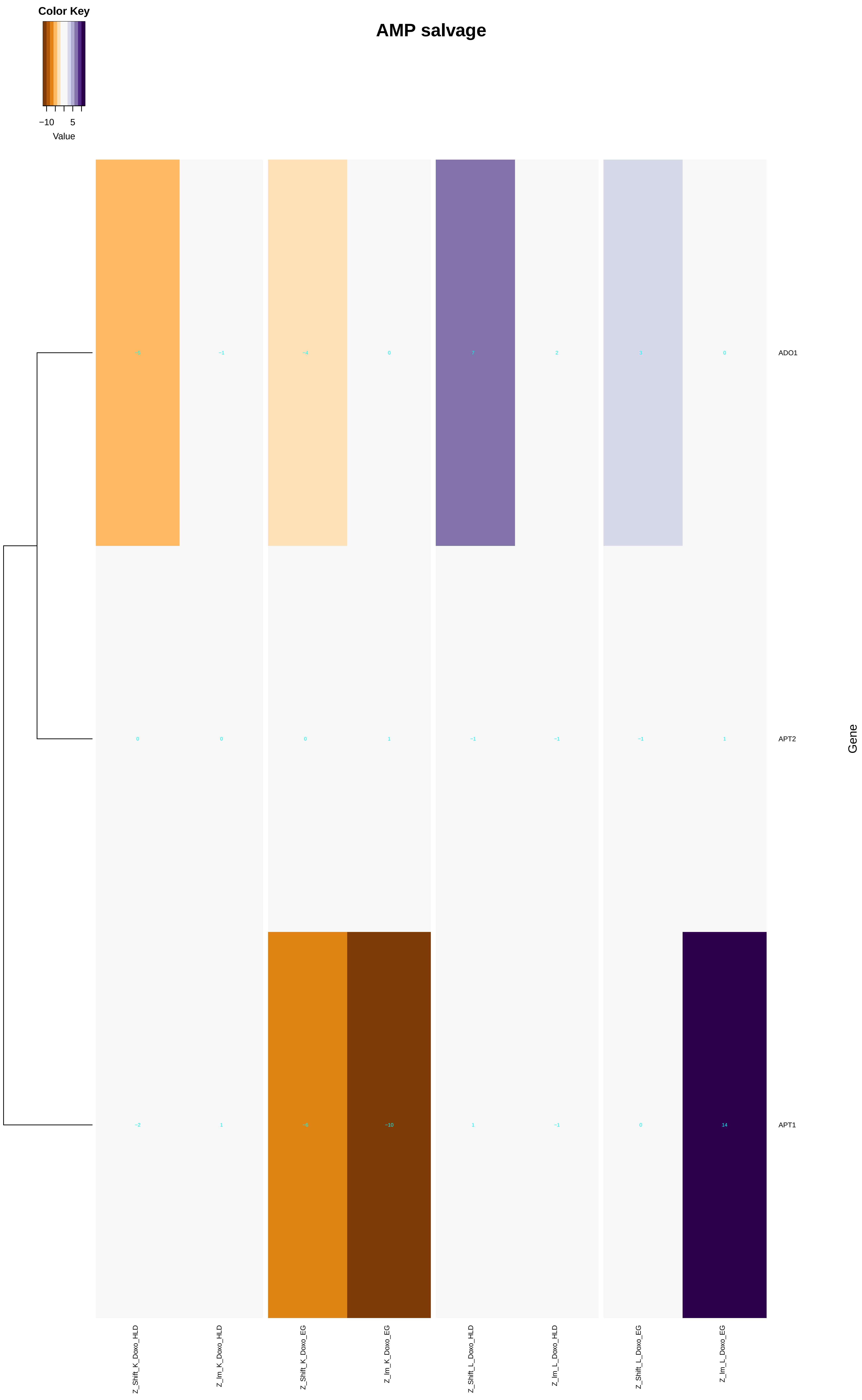
AMP salvage
Color Key
−10 5
Value
ADO1: −5 −1 −4 0 7 2 3 0
APT2: 0 0 0 1 −1 −1 −1 1
APT1: −2 1 −6 −10 1 −1 0 14
Gene
Z_Shift_K_Doxo_HLD
Z_Im_K_Doxo_HLD
Z_Shift_K_Doxo_EG
Z_Im_K_Doxo_EG
Z_Shift_L_Doxo_HLD
Z_Im_L_Doxo_HLD
Z_Shift_L_Doxo_EG
Z_Im_L_Doxo_EG

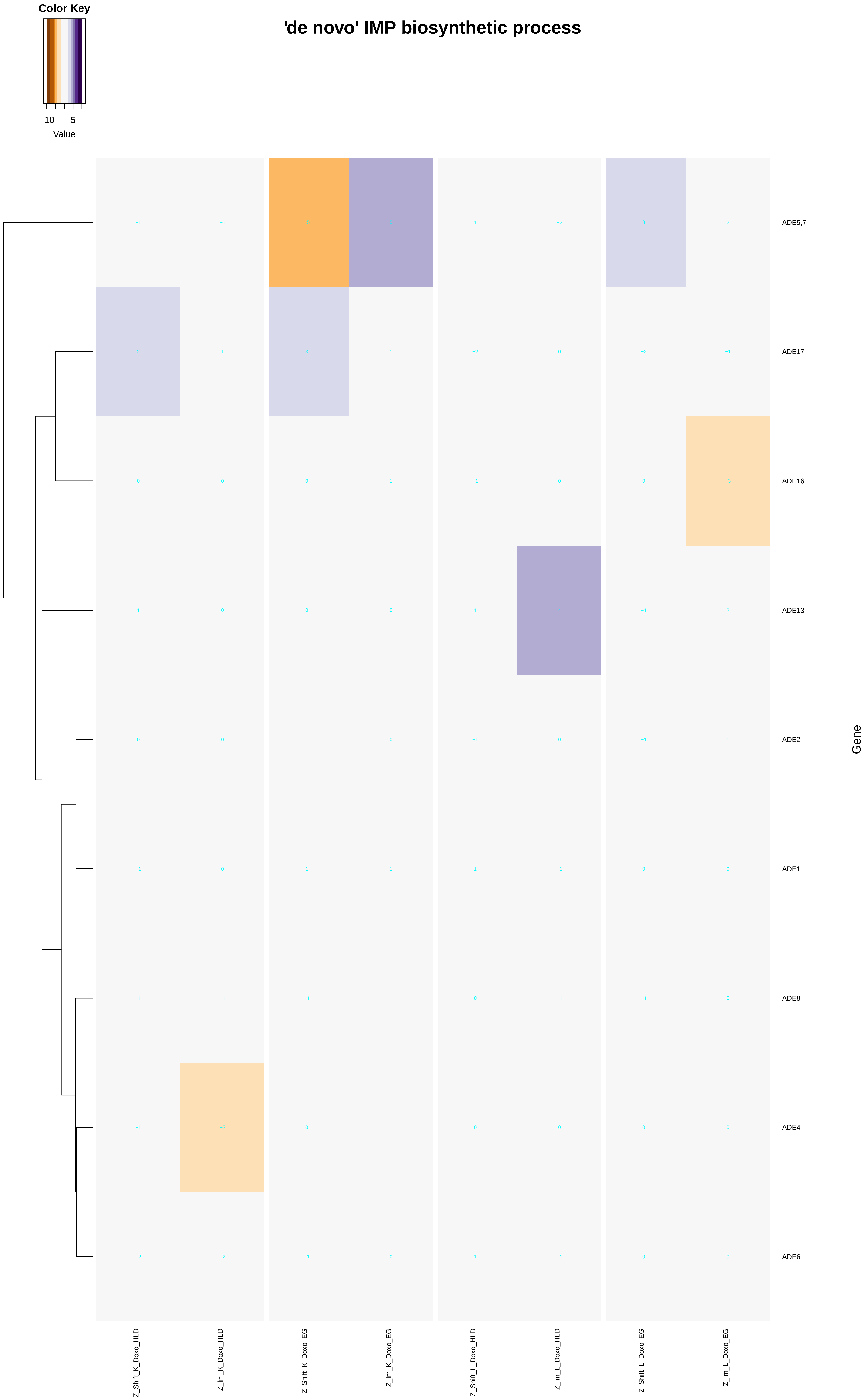

Color Key
−10
5
Value
'de novo' IMP biosynthetic process
ADE5,7
ADE17
ADE16
ADE13
ADE2
ADE1
ADE8
ADE4
ADE6
Gene
Z_Shift_K_Doxo_HLD
Z_Im_K_Doxo_HLD
Z_Shift_K_Doxo_EG
Z_Im_K_Doxo_EG
Z_Shift_L_Doxo_HLD
Z_Im_L_Doxo_HLD
Z_Shift_L_Doxo_EG
Z_Im_L_Doxo_EG

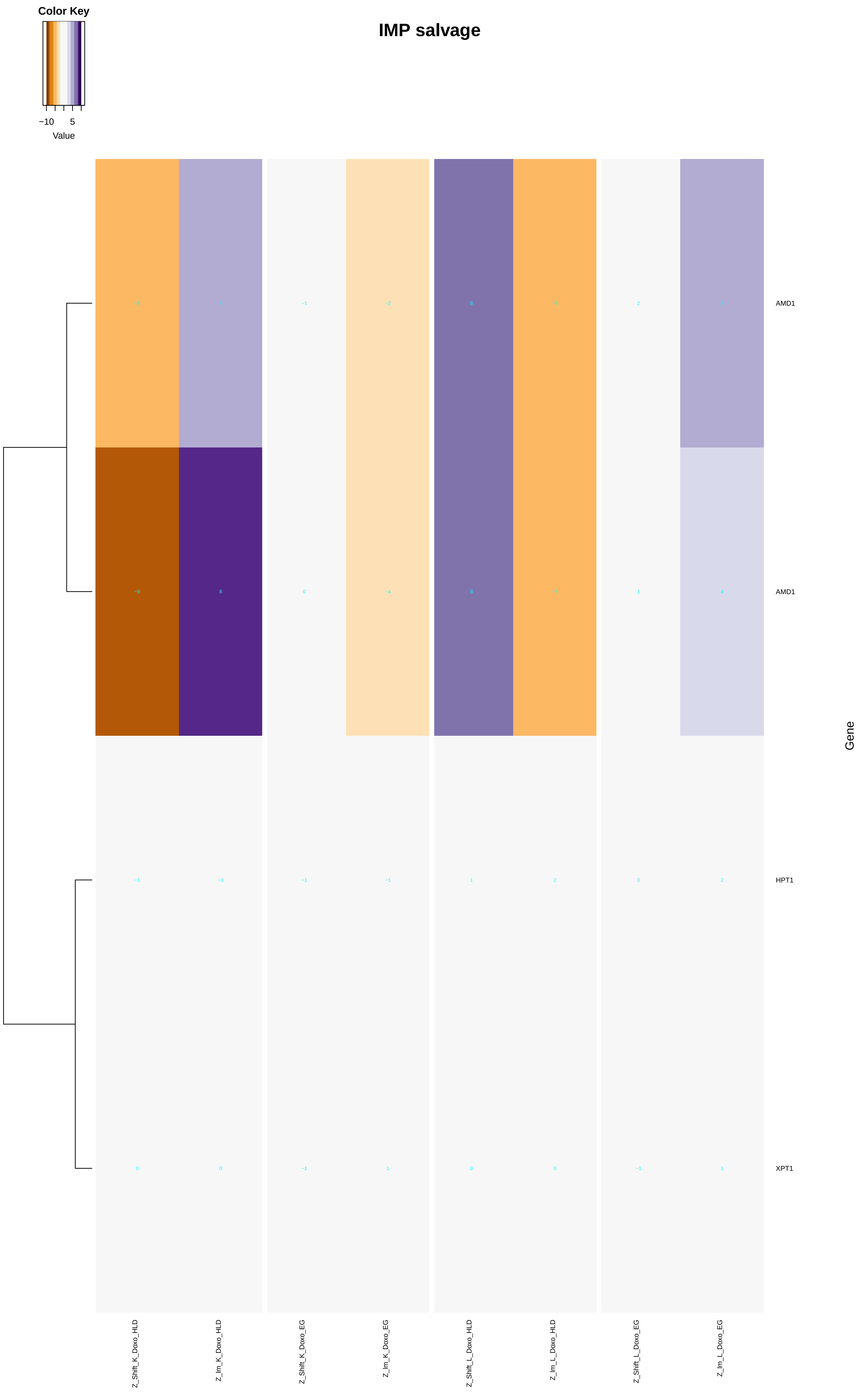
Color Key
−10
5
Value
IMP salvage
AMD1
AMD1
HPT1
XPT1
Gene
Z_Shift_K_Doxo_HLD
Z_Im_K_Doxo_HLD
Z_Shift_K_Doxo_EG
Z_Im_K_Doxo_EG
Z_Shift_L_Doxo_HLD
Z_Im_L_Doxo_HLD
Z_Shift_L_Doxo_EG
Z_Im_L_Doxo_EG

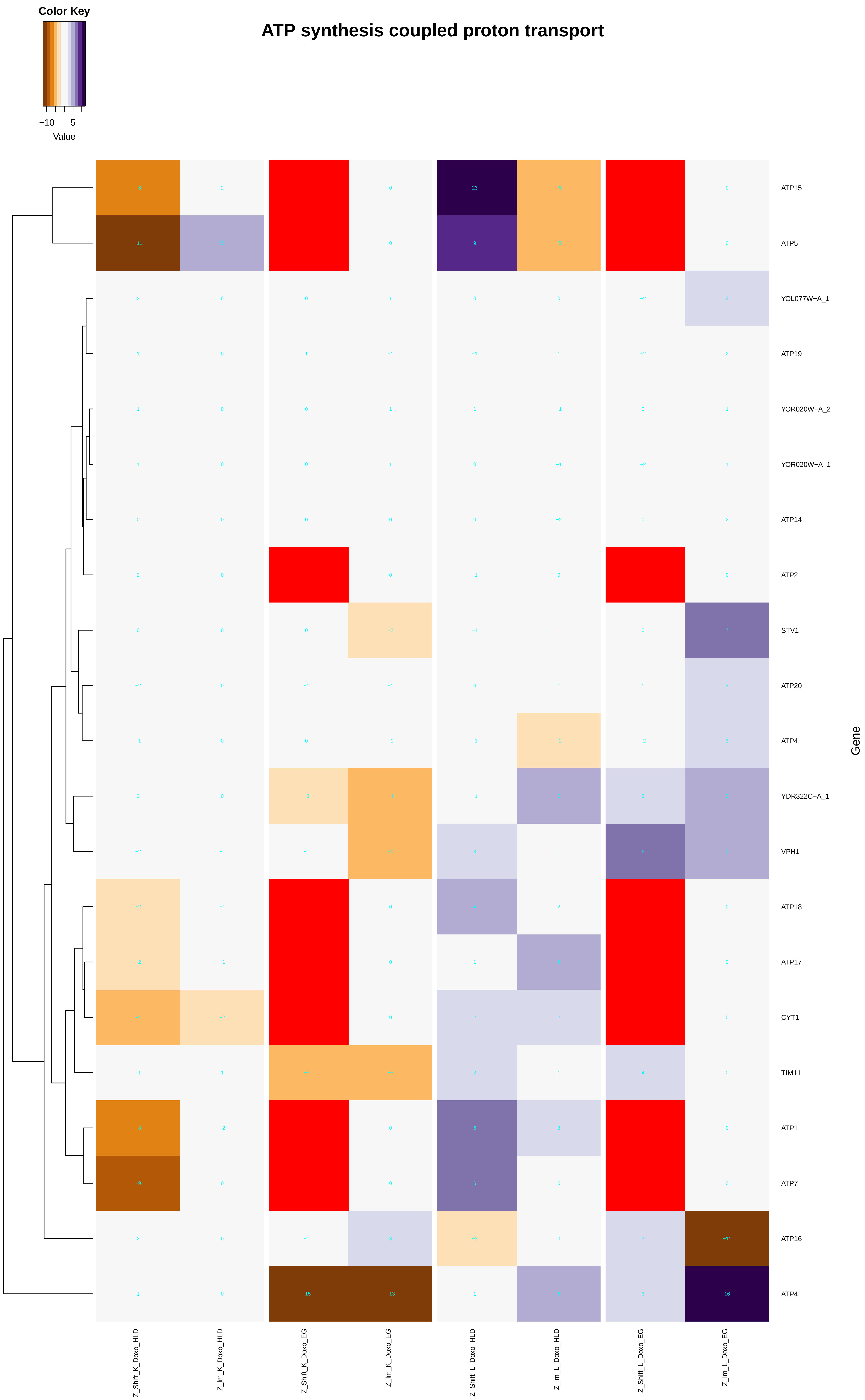

Color Key
−10
5
Value
ATP synthesis coupled proton transport
ATP15
ATP5
YOL077W−A_1
ATP19
YOR020W−A_2
YOR020W−A_1
ATP14
ATP2
STV1
ATP20
ATP4
YDR322C−A_1
VPH1
ATP18
ATP17
CYT1
TIM11
ATP1
ATP7
ATP16
ATP4
Gene
Z_Shift_K_Doxo_HLD
Z_Im_K_Doxo_HLD
Z_Shift_K_Doxo_EG
Z_Im_K_Doxo_EG
Z_Shift_L_Doxo_HLD
Z_Im_L_Doxo_HLD
Z_Shift_L_Doxo_EG
Z_Im_L_Doxo_EG

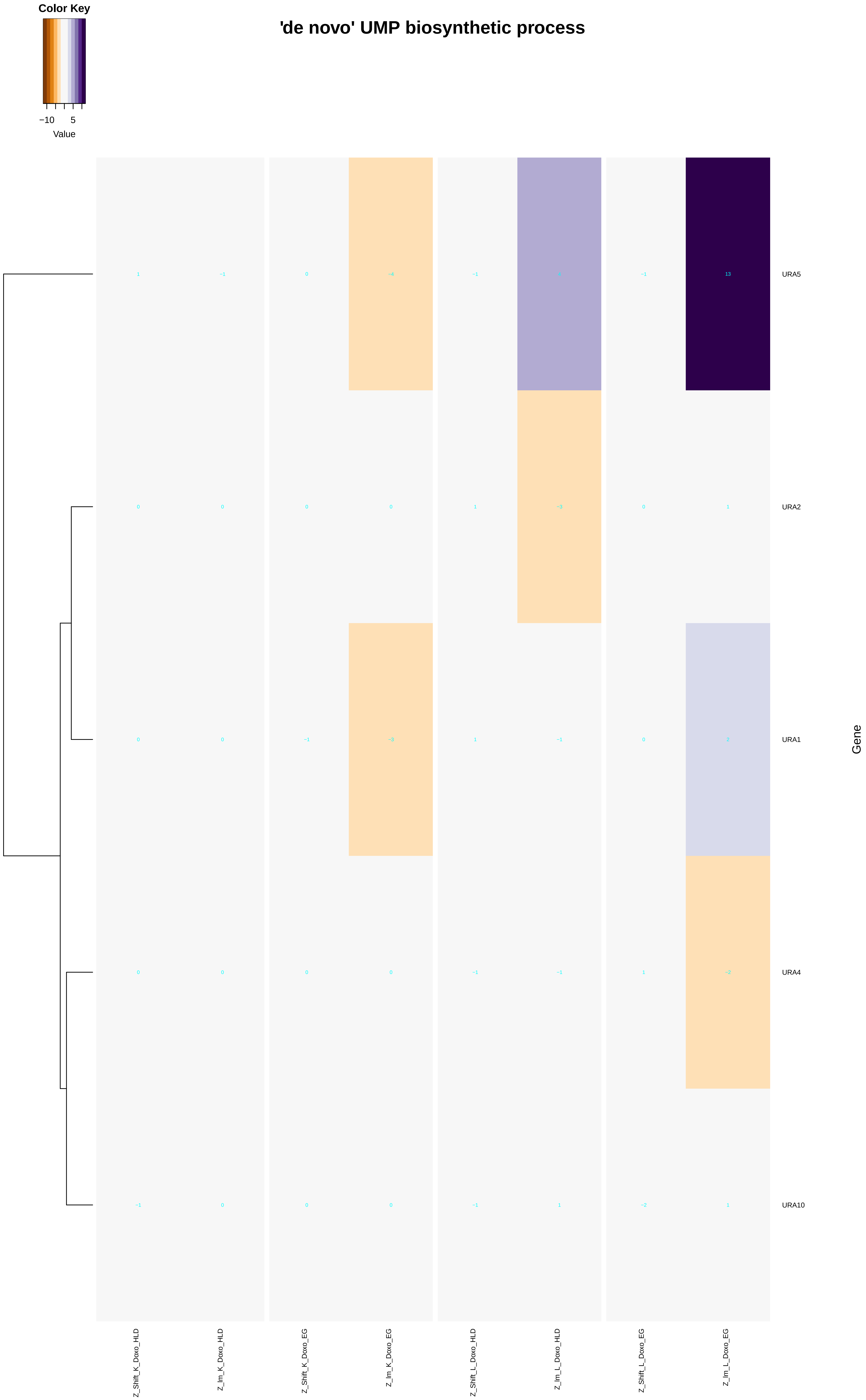

'de novo' UMP biosynthetic process
Color Key
−10
5
Value
URA5
URA2
URA1
URA4
URA10
Gene
Z_Shift_K_Doxo_HLD
Z_Im_K_Doxo_HLD
Z_Shift_K_Doxo_EG
Z_Im_K_Doxo_EG
Z_Shift_L_Doxo_HLD
Z_Im_L_Doxo_HLD
Z_Shift_L_Doxo_EG
Z_Im_L_Doxo_EG

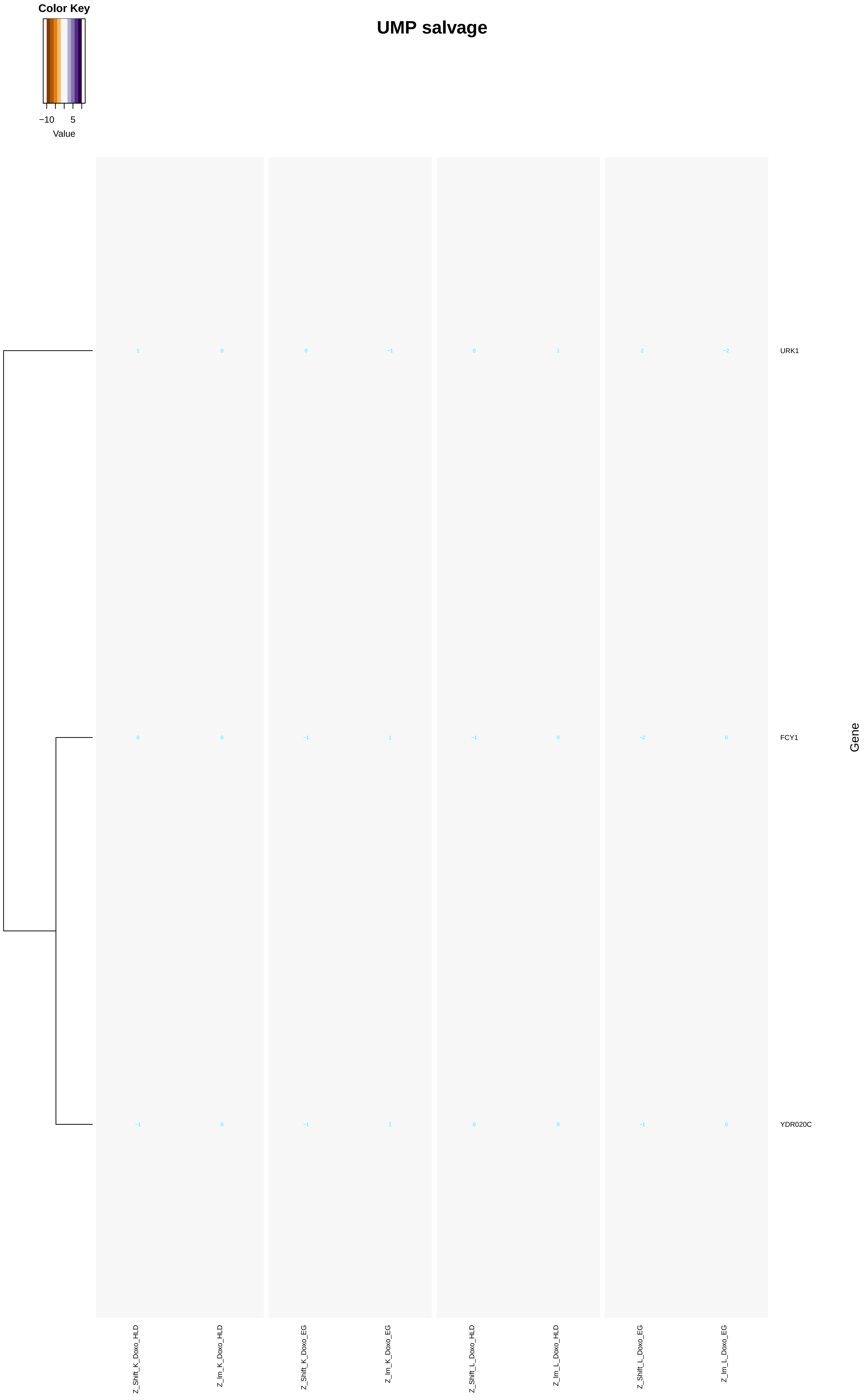
UMP salvage
Color Key
−10
5
Value
URK1
FCY1
YDR020C
Gene
1
0
0
−1
0
1
2
−2
0
0
−1
1
−1
0
−2
0
−1
0
−1
1
0
0
−1
0
Z_Shift_K_Doxo_HLD
Z_lm_K_Doxo_HLD
Z_Shift_K_Doxo_EG
Z_lm_K_Doxo_EG
Z_Shift_L_Doxo_HLD
Z_lm_L_Doxo_HLD
Z_Shift_L_Doxo_EG
Z_lm_L_Doxo_EG
