## Additional File 9 for "A yeast phenomic model for the influence of Warburg metabolism on genetic buffering of doxorubicin": nucleoside_triphosphate_biosynthetic_process.pdf

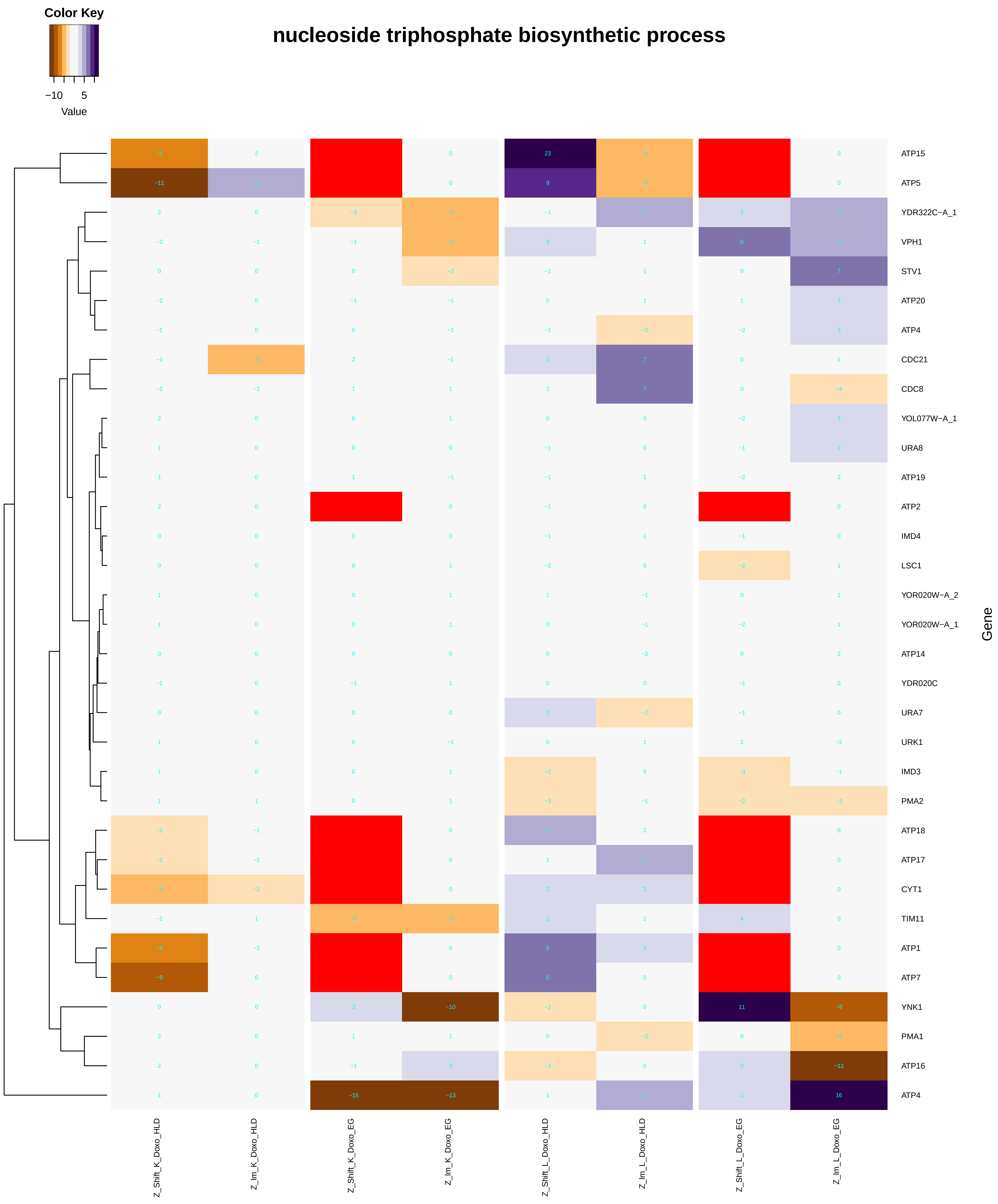

### purine nucleoside triphosphate biosynthetic process

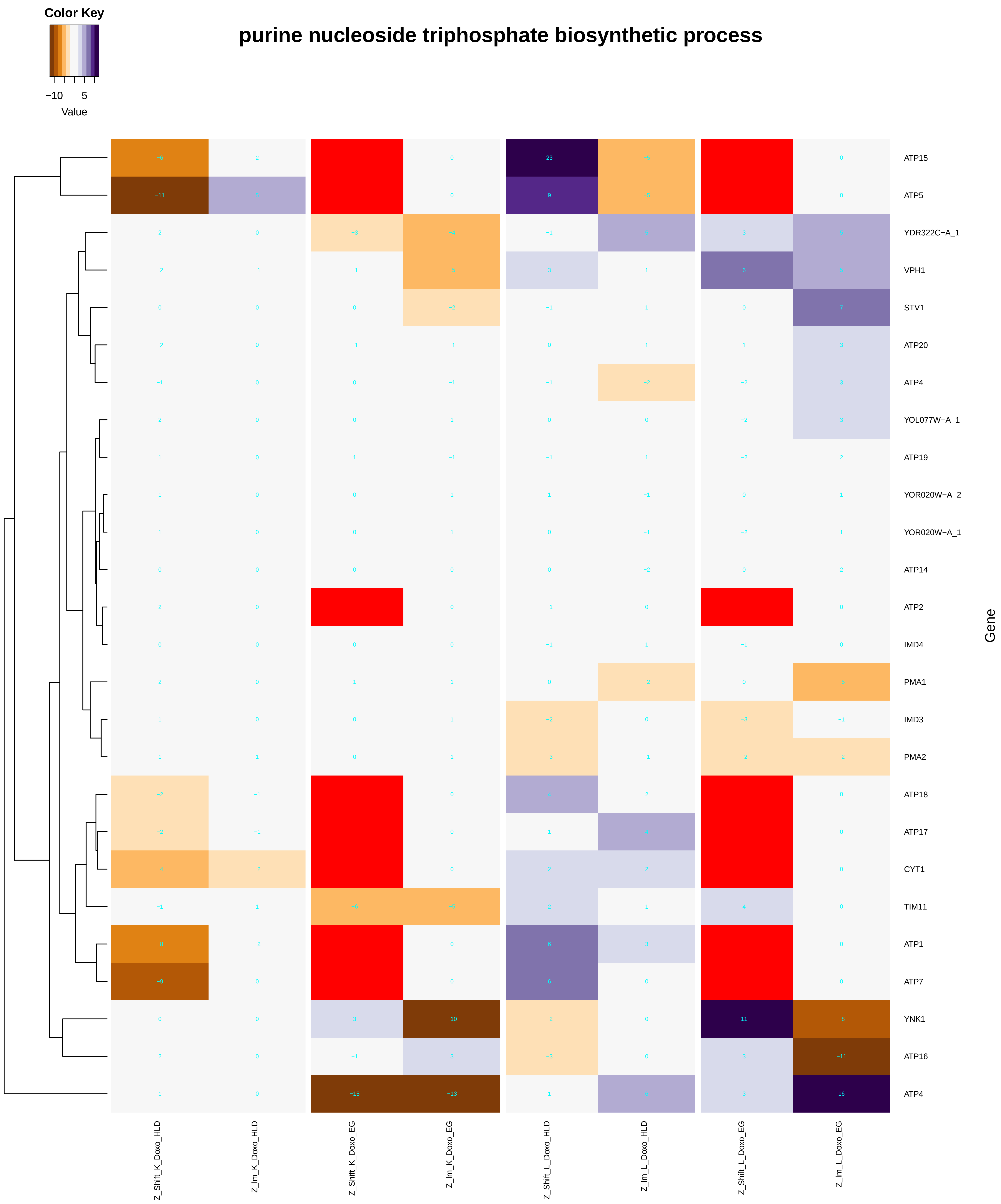

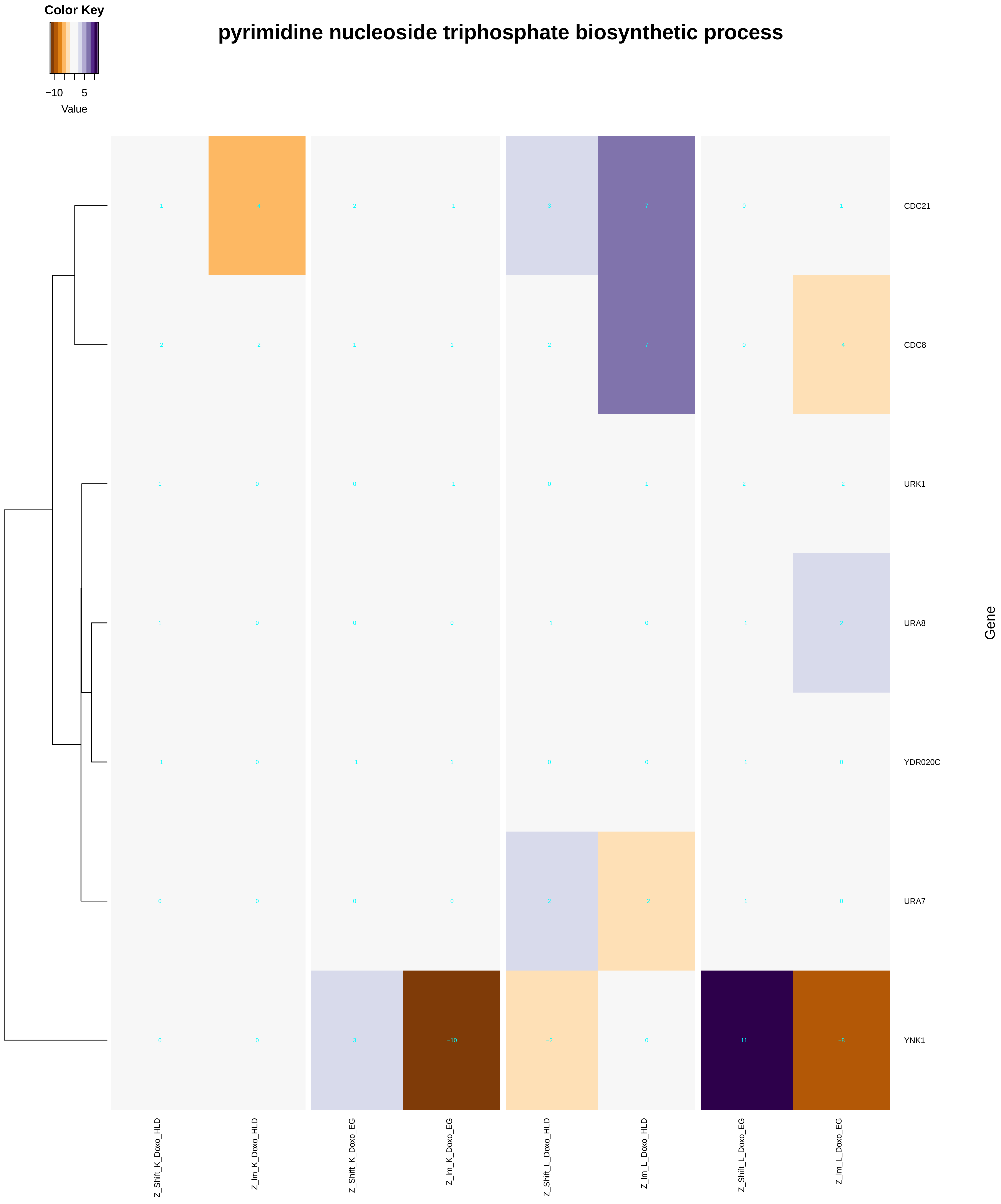
Color Key
−10
5
Value
pyrimidine nucleoside triphosphate biosynthetic process
CDC21
CDC8
URK1
URA8
YDR020C
URA7
YNK1
Gene
Z_Shift_K_Doxo_HLD
Z_lm_K_Doxo_HLD
Z_Shift_K_Doxo_EG
Z_lm_K_Doxo_EG
Z_Shift_L_Doxo_HLD
Z_lm_L_Doxo_HLD
Z_Shift_L_Doxo_EG
Z_lm_L_Doxo_EG

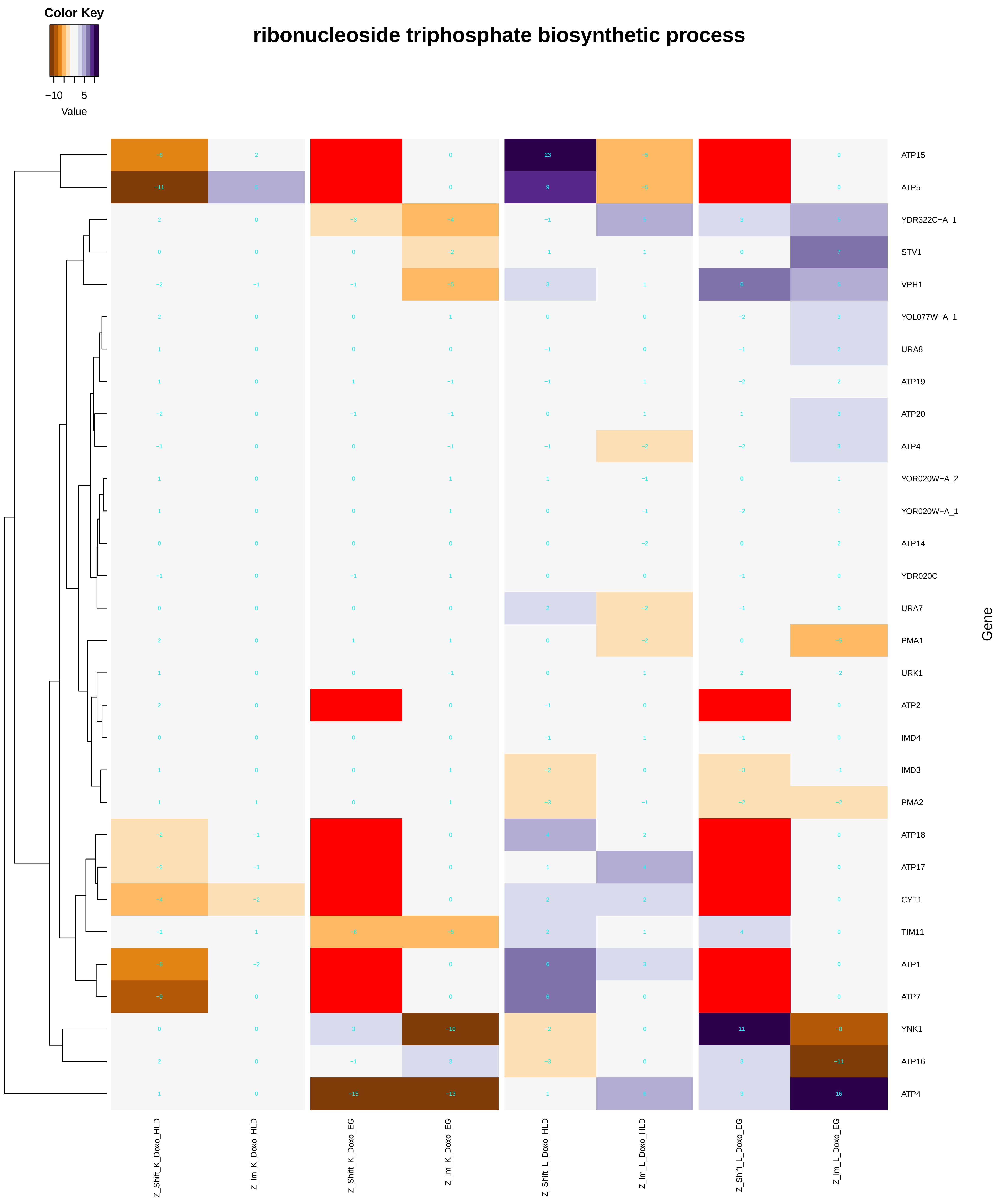
Color Key
−10
5
Value
ribonucleoside triphosphate biosynthetic process
ATP15
ATP5
YDR322C−A_1
STV1
VPH1
YOL077W−A_1
URA8
ATP19
ATP20
ATP4
YOR020W−A_2
YOR020W−A_1
ATP14
YDR020C
URA7
PMA1
URK1
ATP2
IMD4
IMD3
PMA2
ATP18
ATP17
CYT1
TIM11
ATP1
ATP7
YNK1
ATP16
ATP4
Gene
Z_Shift_K_Doxo_HLD
Z_Im_K_Doxo_HLD
Z_Shift_K_Doxo_EG
Z_Im_K_Doxo_EG
Z_Shift_L_Doxo_HLD
Z_Im_L_Doxo_HLD
Z_Shift_L_Doxo_EG
Z_Im_L_Doxo_EG

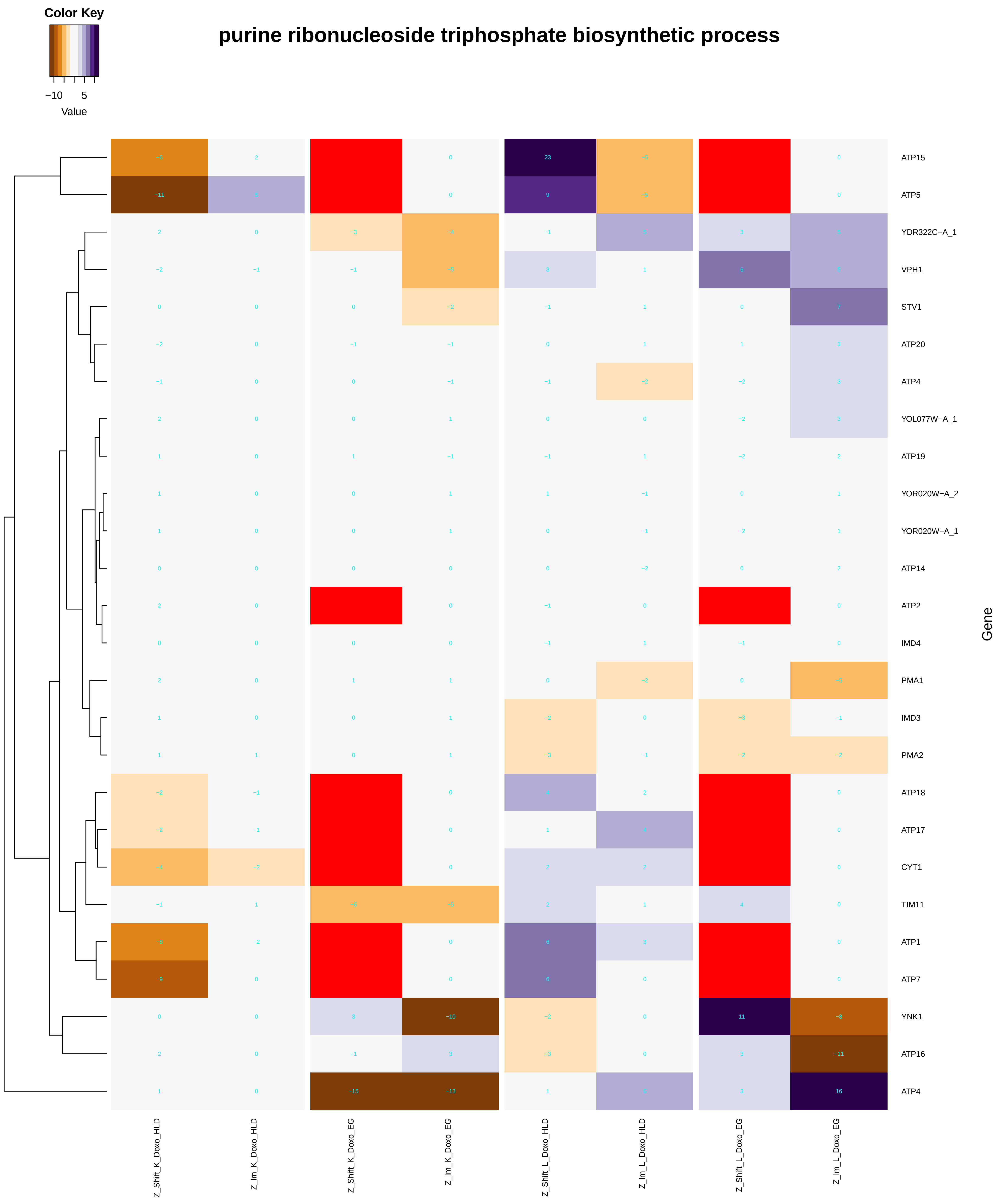
purine ribonucleoside triphosphate biosynthetic process
Color Key
−10
5
Value
ATP15
ATP5
YDR322C−A_1
VPH1
STV1
ATP20
ATP4
YOL077W−A_1
ATP19
YOR020W−A_2
YOR020W−A_1
ATP14
ATP2
IMD4
PMA1
IMD3
PMA2
ATP18
ATP17
CYT1
TIM11
ATP1
ATP7
YNK1
ATP16
ATP4
Gene
Z_Shift_K_Doxo_HLD
Z_Im_K_Doxo_HLD
Z_Shift_K_Doxo_EG
Z_Im_K_Doxo_EG
Z_Shift_L_Doxo_HLD
Z_Im_L_Doxo_HLD
Z_Shift_L_Doxo_EG
Z_Im_L_Doxo_EG

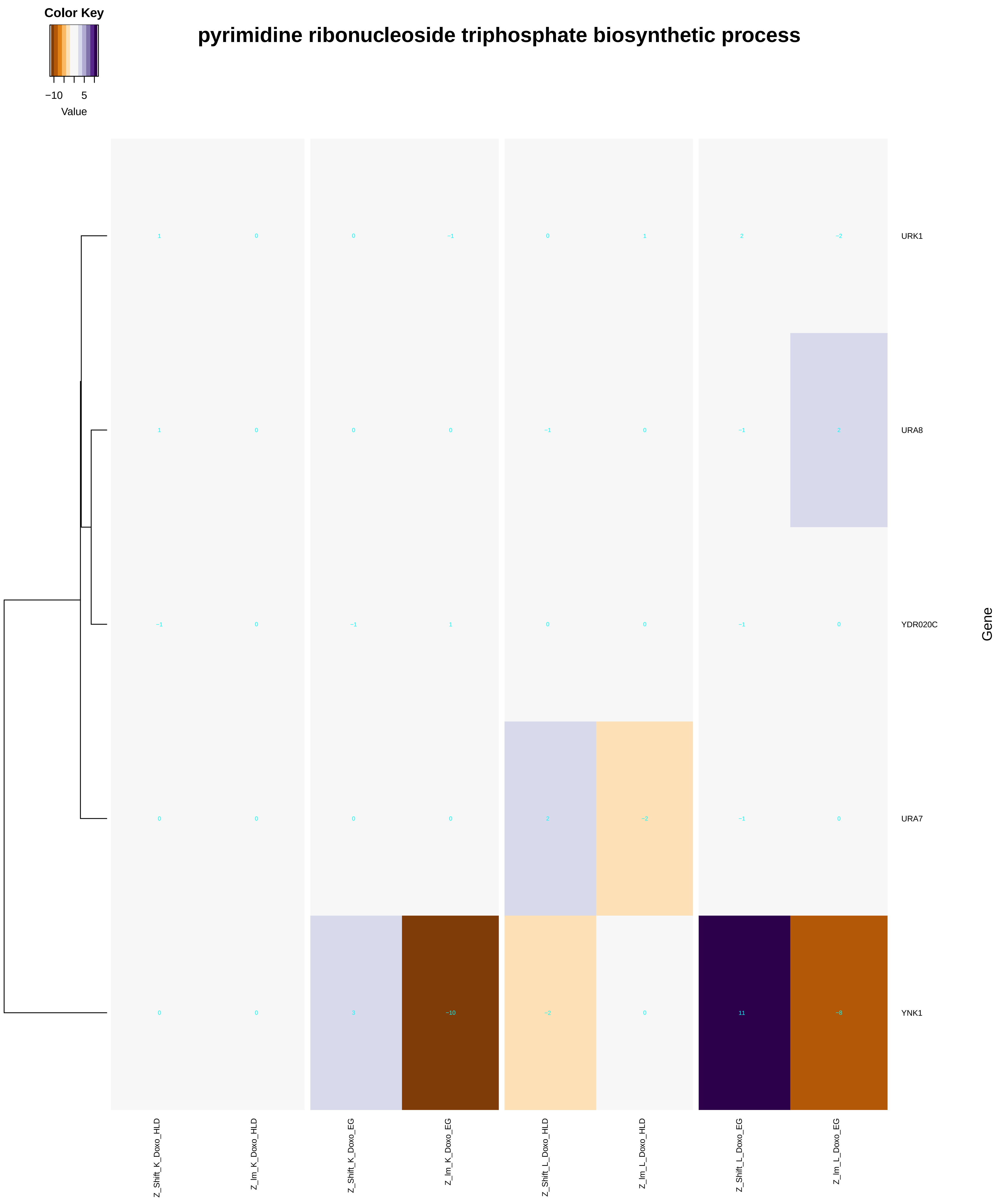
Color Key
−10
5
Value
pyrimidine ribonucleoside triphosphate biosynthetic process
URK1
URA8
YDR020C
URA7
YNK1
Gene
Z_Shift_K_Doxo_HLD
Z_Im_K_Doxo_HLD
Z_Shift_K_Doxo_EG
Z_Im_K_Doxo_EG
Z_Shift_L_Doxo_HLD
Z_Im_L_Doxo_HLD
Z_Shift_L_Doxo_EG
Z_Im_L_Doxo_EG

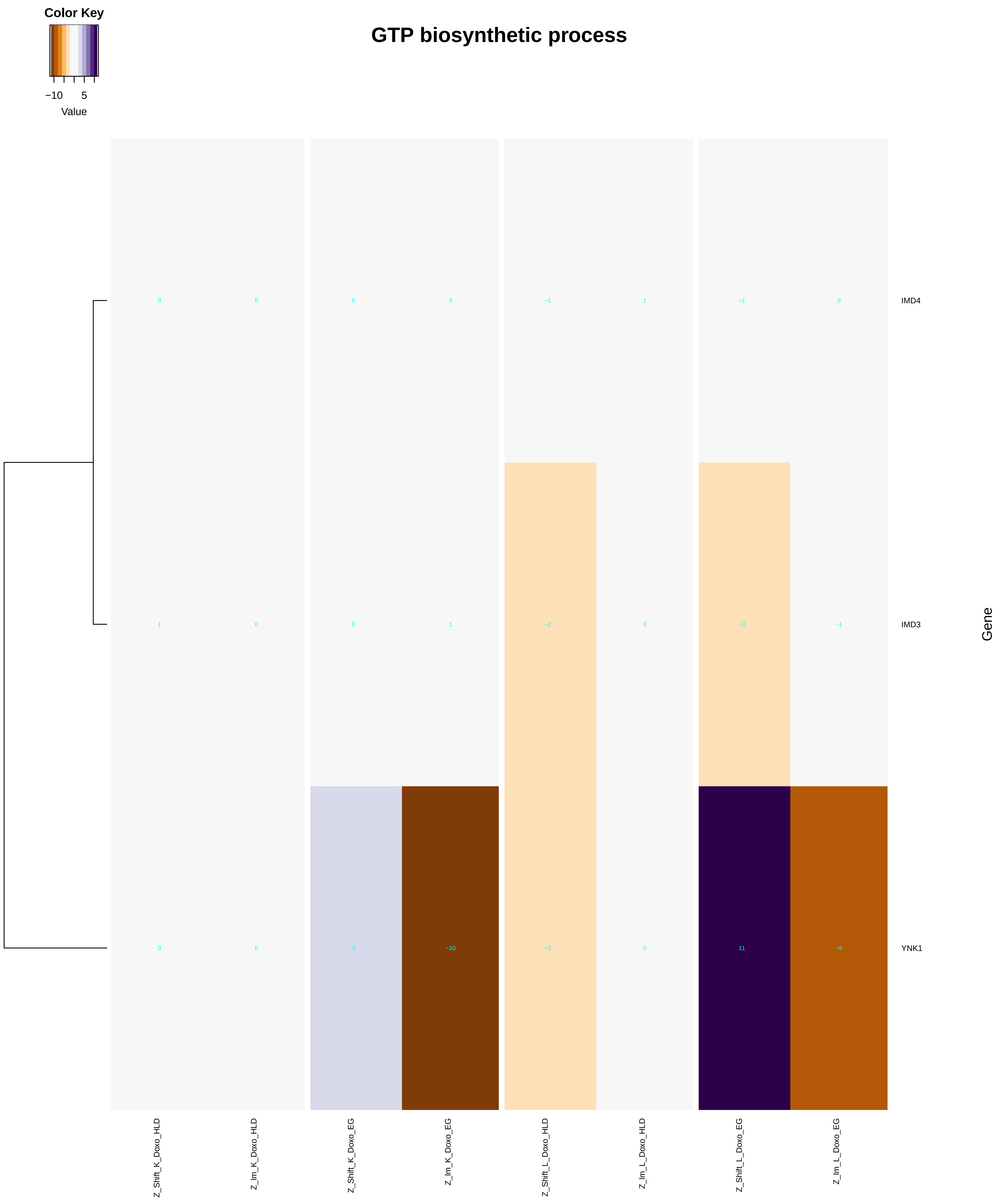
Color Key
−10
5
Value
GTP biosynthetic process
IMD4
IMD3
YNK1
Gene
Z_Shift_K_Doxo_HLD
Z_Im_K_Doxo_HLD
Z_Shift_K_Doxo_EG
Z_Im_K_Doxo_EG
Z_Shift_L_Doxo_HLD
Z_Im_L_Doxo_HLD
Z_Shift_L_Doxo_EG
Z_Im_L_Doxo_EG

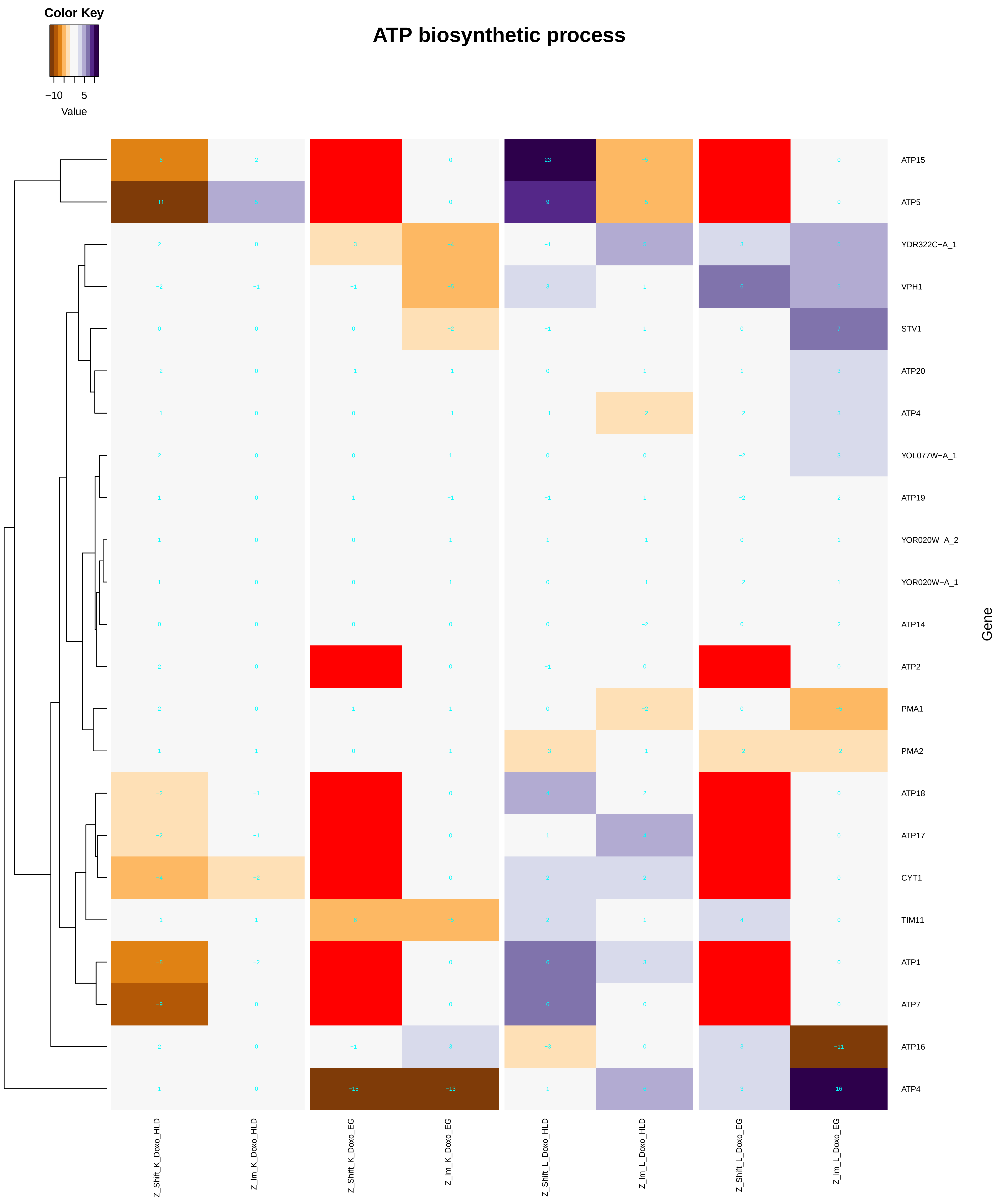
Color Key
−10
5
Value
ATP biosynthetic process
ATP15
ATP5
YDR322C−A_1
VPH1
STV1
ATP20
ATP4
YOL077W−A_1
ATP19
YOR020W−A_2
YOR020W−A_1
ATP14
ATP2
PMA1
PMA2
ATP18
ATP17
CYT1
TIM11
ATP1
ATP7
ATP16
ATP4
Gene
Z_Shift_K_Doxo_HLD
Z_Im_K_Doxo_HLD
Z_Shift_K_Doxo_EG
Z_Im_K_Doxo_EG
Z_Shift_L_Doxo_HLD
Z_Im_L_Doxo_HLD
Z_Shift_L_Doxo_EG
Z_Im_L_Doxo_EG

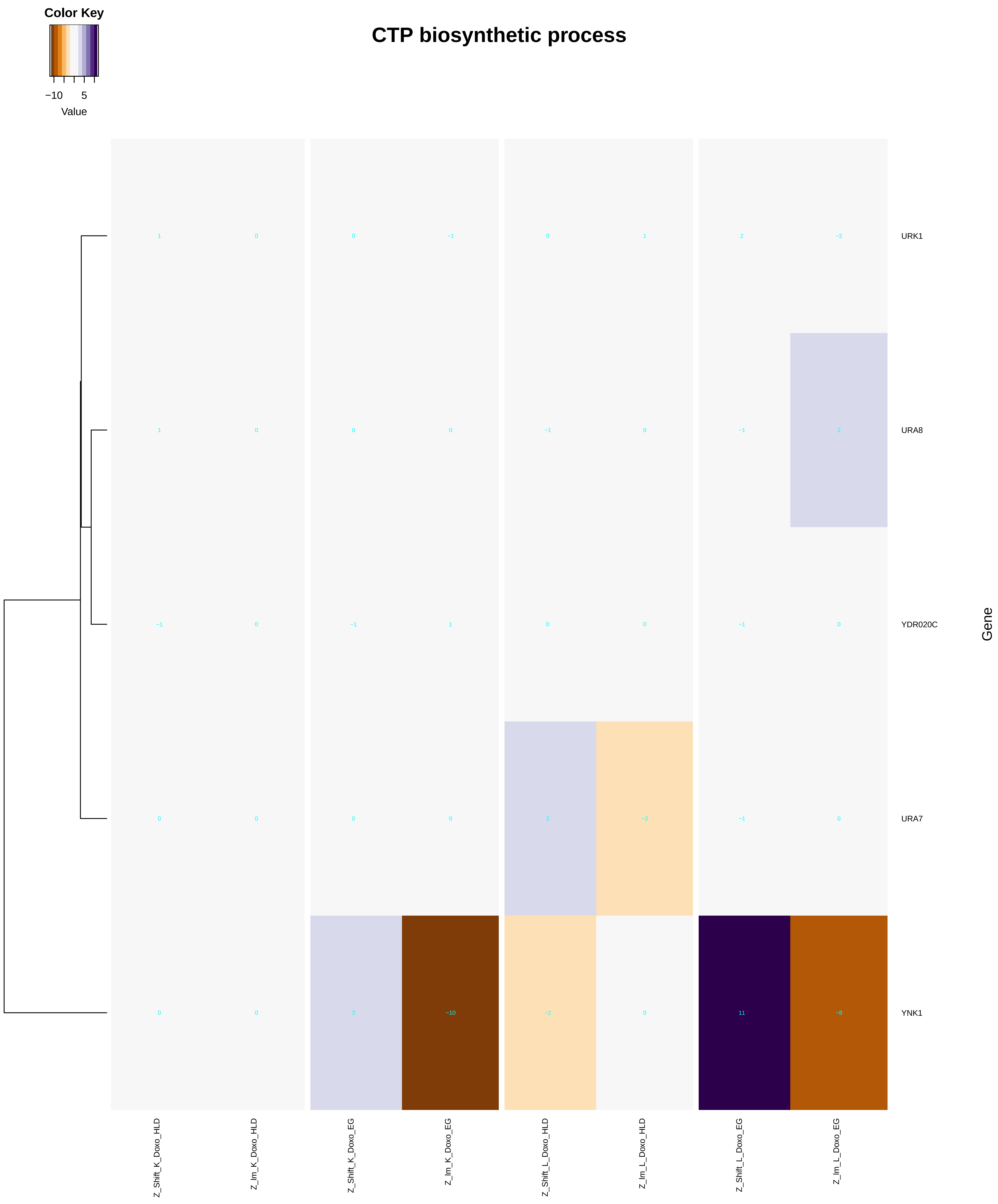
Color Key
−10
5
Value
CTP biosynthetic process
URK1
URA8
YDR020C
URA7
YNK1
Gene
Z_Shift_K_Doxo_HLD
Z_lm_K_Doxo_HLD
Z_Shift_K_Doxo_EG
Z_lm_K_Doxo_EG
Z_Shift_L_Doxo_HLD
Z_lm_L_Doxo_HLD
Z_Shift_L_Doxo_EG
Z_lm_L_Doxo_EG

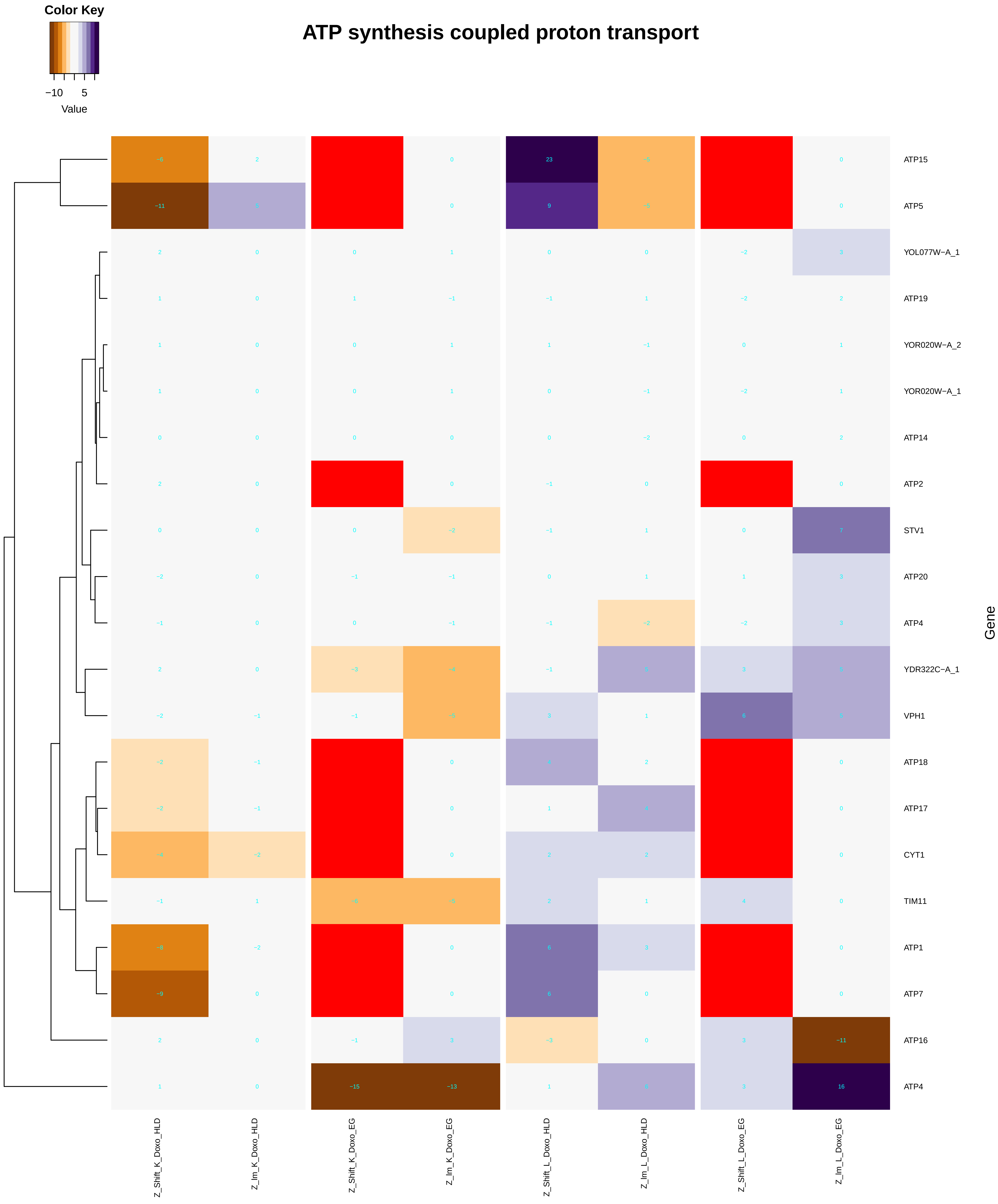

Color Key
−10 5
Value
ATP synthesis coupled proton transport
ATP15
ATP5
YOL077W−A_1
ATP19
YOR020W−A_2
YOR020W−A_1
ATP14
ATP2
STV1
ATP20
ATP4
YDR322C−A_1
VPH1
ATP18
ATP17
CYT1
TIM11
ATP1
ATP7
ATP16
ATP4
Gene
Z_Shift_K_Doxo_HLD
Z_Im_K_Doxo_HLD
Z_Shift_K_Doxo_EG
Z_Im_K_Doxo_EG
Z_Shift_L_Doxo_HLD
Z_Im_L_Doxo_HLD
Z_Shift_L_Doxo_EG
Z_Im_L_Doxo_EG
