## Additional File 9 for "A yeast phenomic model for the influence of Warburg metabolism on genetic buffering of doxorubicin": nucleosome_organization.pdf

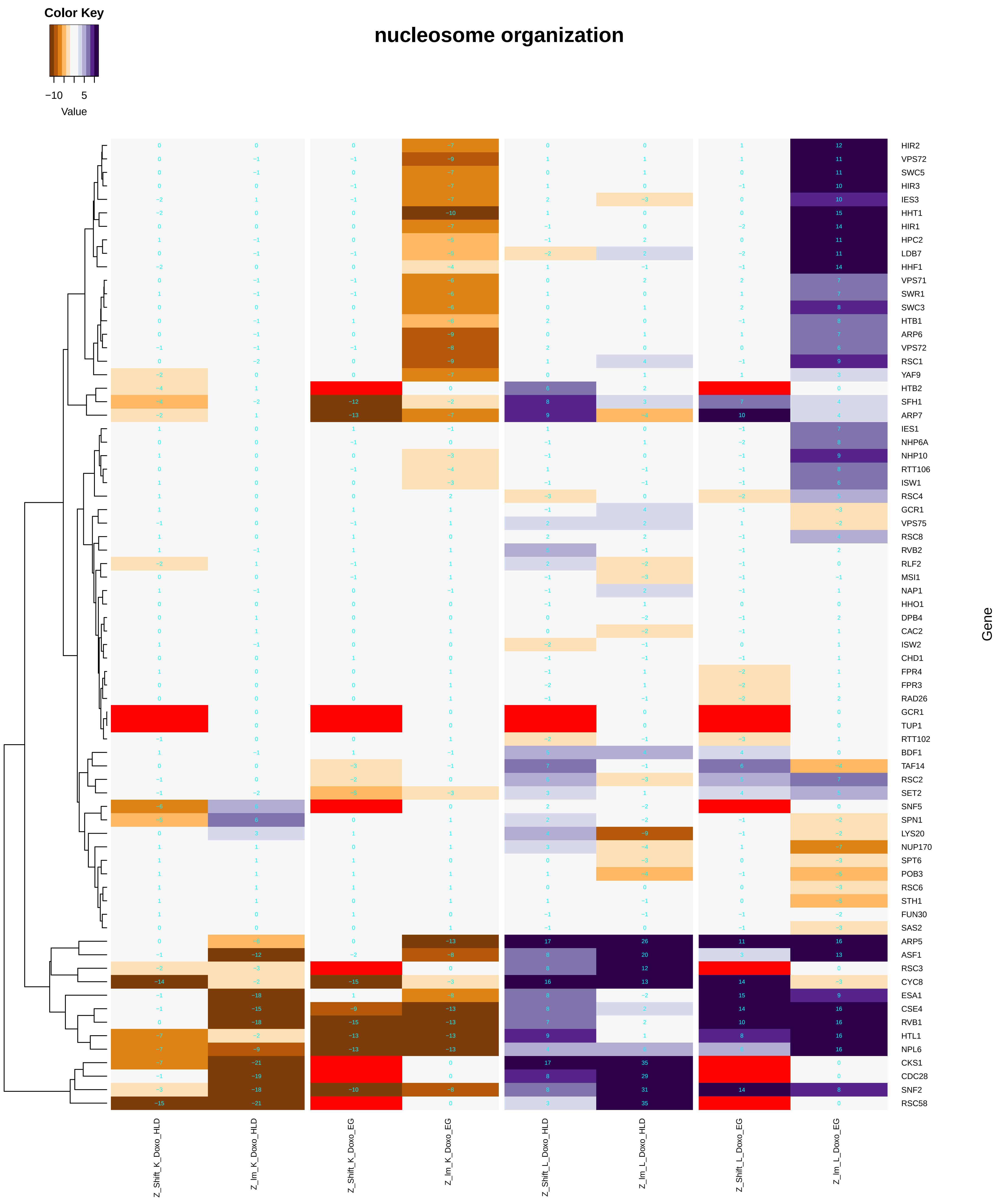
Color Key
−10 5
Value
nucleosome organization
HIR2
VPS72
SWC5
HIR3
IES3
HHT1
HIR1
HPC2
LDB7
HHF1
VPS71
SWR1
SWC3
HTB1
ARP6
VPS72
RSC1
YAF9
HTB2
SFH1
ARP7
IES1
NHP6A
NHP10
RTT106
ISW1
RSC4
GCR1
VPS75
RSC8
RVB2
RLF2
MSI1
NAP1
HHO1
DPB4
CAC2
ISW2
CHD1
FPR4
FPR3
RAD26
GCR1
TUP1
RTT102
BDF1
TAF14
RSC2
SET2
SNF5
SPN1
LYS20
NUP170
SPT6
POB3
RSC6
STH1
FUN30
SAS2
ARP5
ASF1
RSC3
CYC8
ESA1
CSE4
RVB1
HTL1
NPL6
CKS1
CDC28
SNF2
RSC58
Gene
Z_Shift_K_Doxo_HLD
Z_Im_K_Doxo_HLD
Z_Shift_K_Doxo_EG
Z_Im_K_Doxo_EG
Z_Shift_L_Doxo_HLD
Z_Im_L_Doxo_HLD
Z_Shift_L_Doxo_EG
Z_Im_L_Doxo_EG

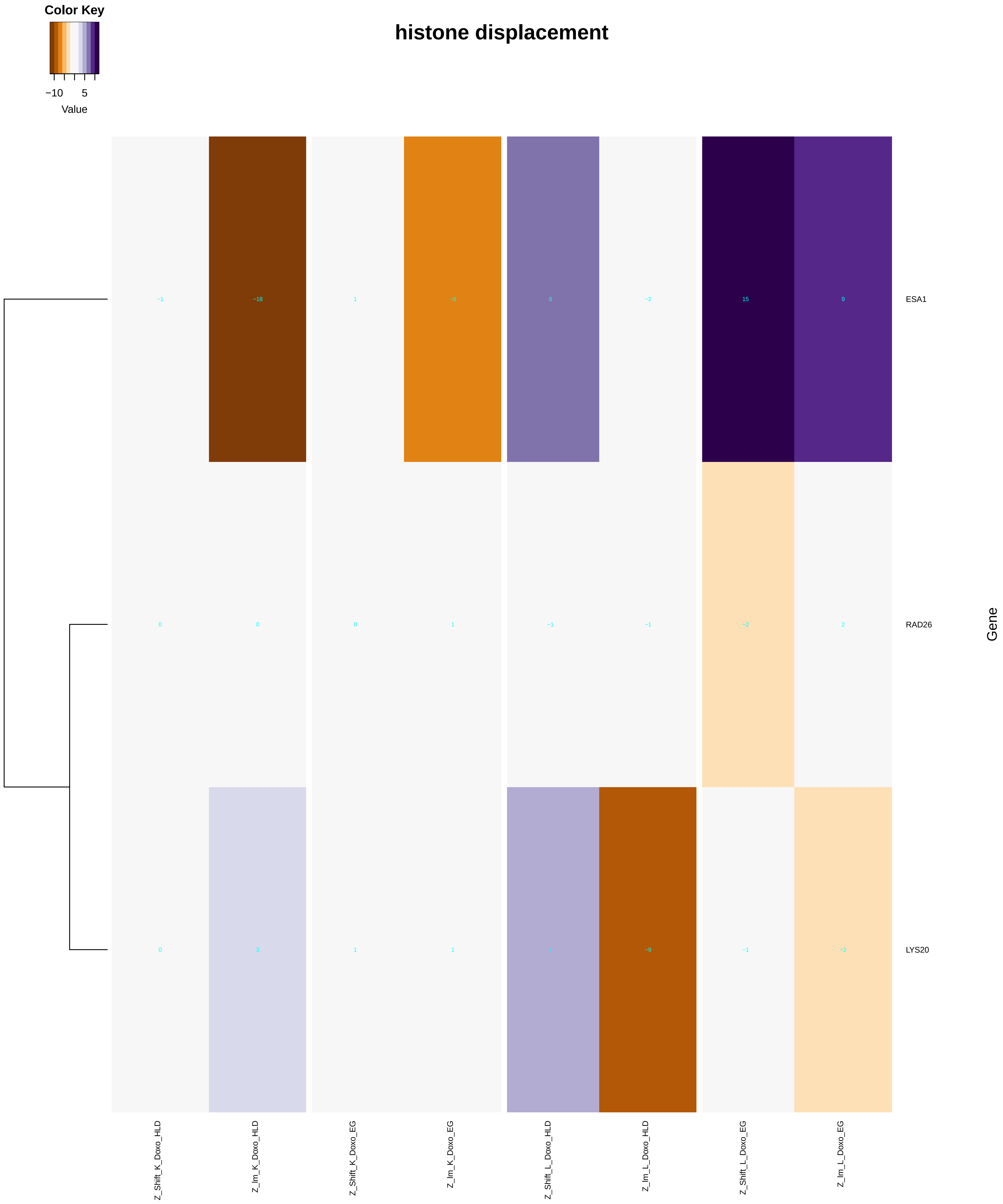
Color Key
−10
5
Value
histone displacement
−1
−18
1
−8
8
−2
15
9
ESA1
0
0
0
1
−1
−1
−2
2
RAD26
0
3
1
1
4
−9
−1
−2
LYS20
Gene
Z_Shift_K_Doxo_HLD
Z_Im_K_Doxo_HLD
Z_Shift_K_Doxo_EG
Z_Im_K_Doxo_EG
Z_Shift_L_Doxo_HLD
Z_Im_L_Doxo_HLD
Z_Shift_L_Doxo_EG
Z_Im_L_Doxo_EG

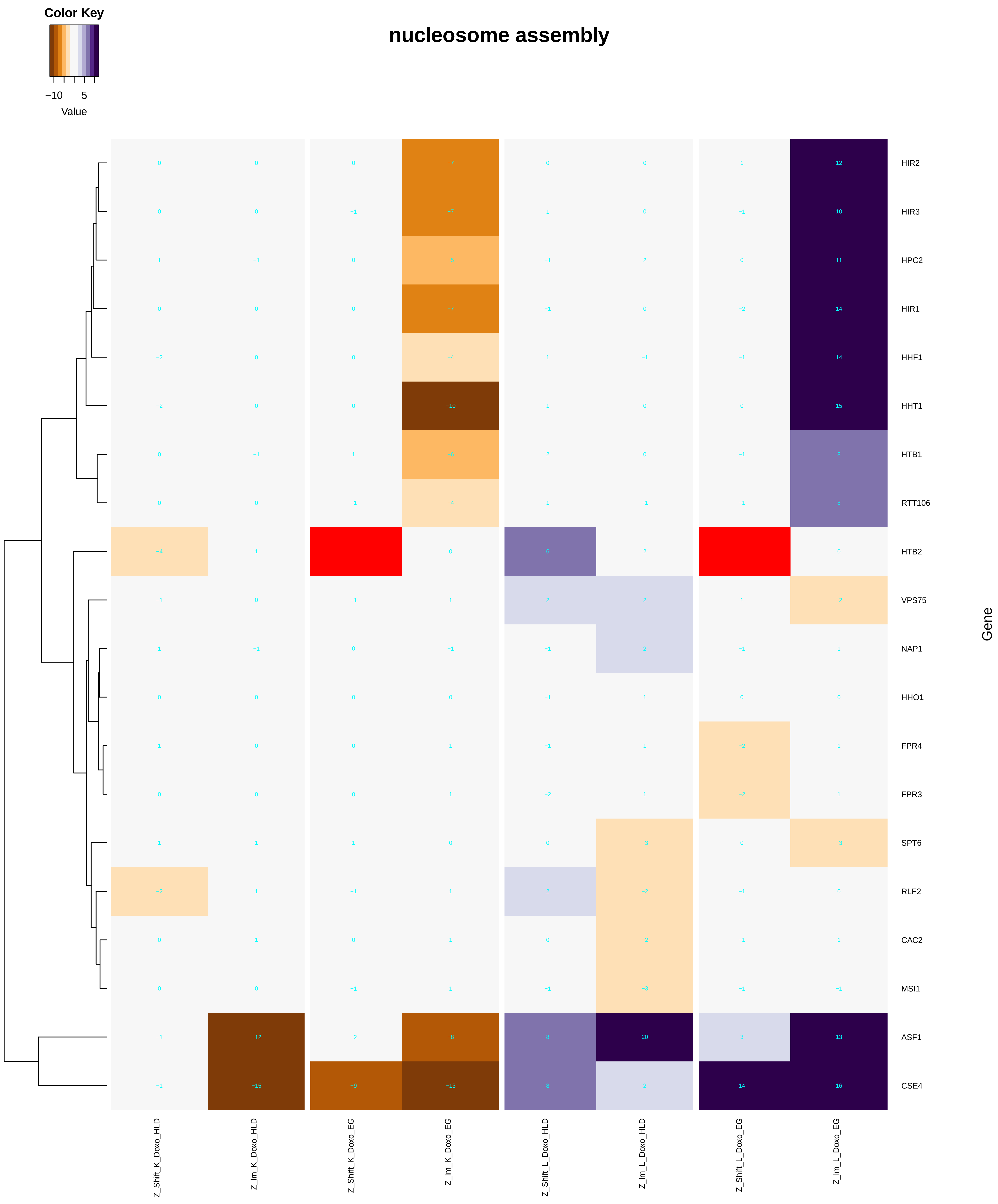

Color Key
−10
5
Value
nucleosome assembly
HIR2
HIR3
HPC2
HIR1
HHF1
HHT1
HTB1
RTT106
HTB2
VPS75
NAP1
HHO1
FPR4
FPR3
SPT6
RLF2
CAC2
MSI1
ASF1
CSE4
Gene
Z_Shift_K_Doxo_HLD
Z_Im_K_Doxo_HLD
Z_Shift_K_Doxo_EG
Z_Im_K_Doxo_EG
Z_Shift_L_Doxo_HLD
Z_Im_L_Doxo_HLD
Z_Shift_L_Doxo_EG
Z_Im_L_Doxo_EG

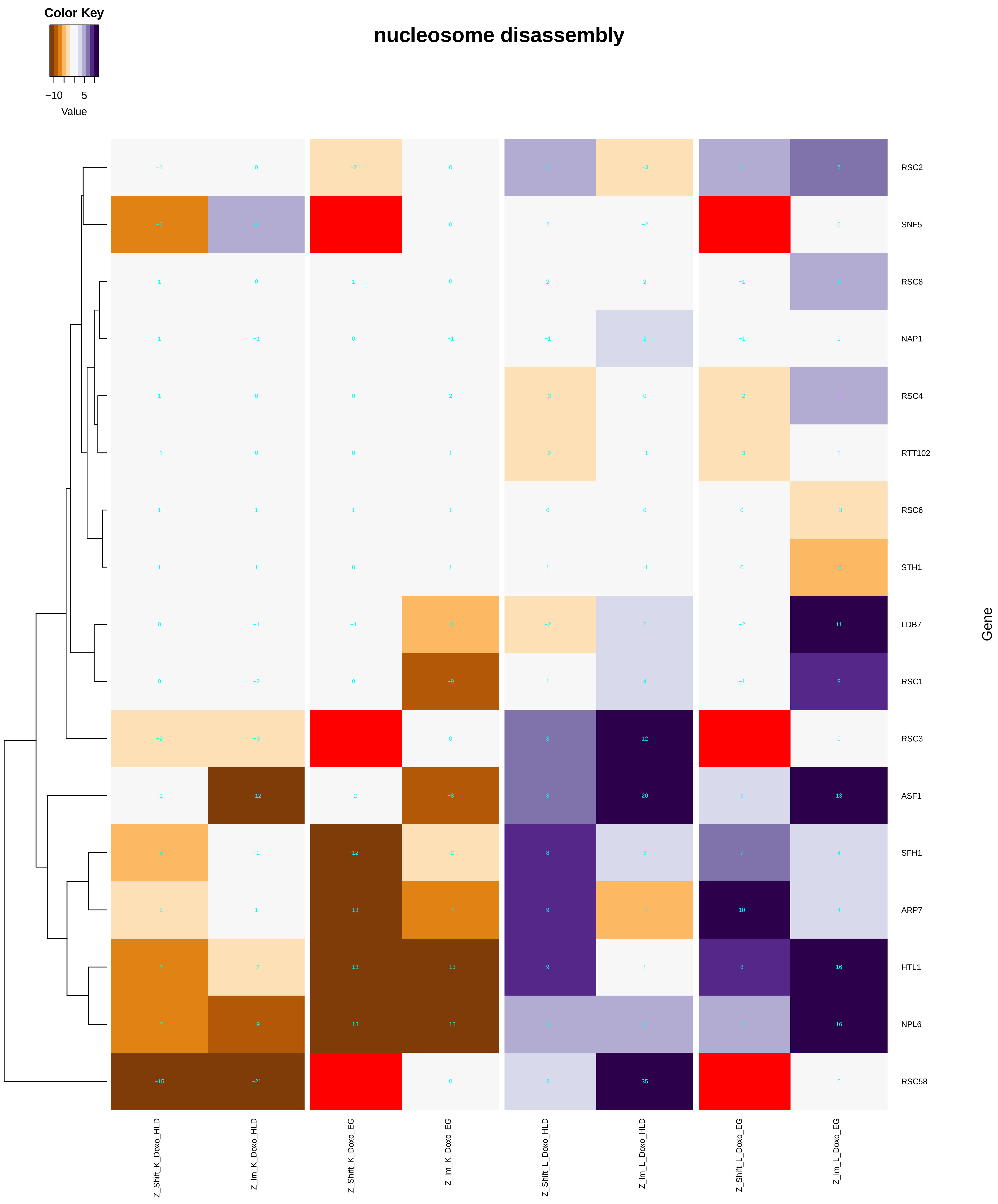
Color Key
−10 5
Value
nucleosome disassembly
RSC2
SNF5
RSC8
NAP1
RSC4
RTT102
RSC6
STH1
LDB7
RSC1
RSC3
ASF1
SFH1
ARP7
HTL1
NPL6
RSC58
Gene
Z_Shift_K_Doxo_HLD
Z_lm_K_Doxo_HLD
Z_Shift_K_Doxo_EG
Z_lm_K_Doxo_EG
Z_Shift_L_Doxo_HLD
Z_lm_L_Doxo_HLD
Z_Shift_L_Doxo_EG
Z_lm_L_Doxo_EG

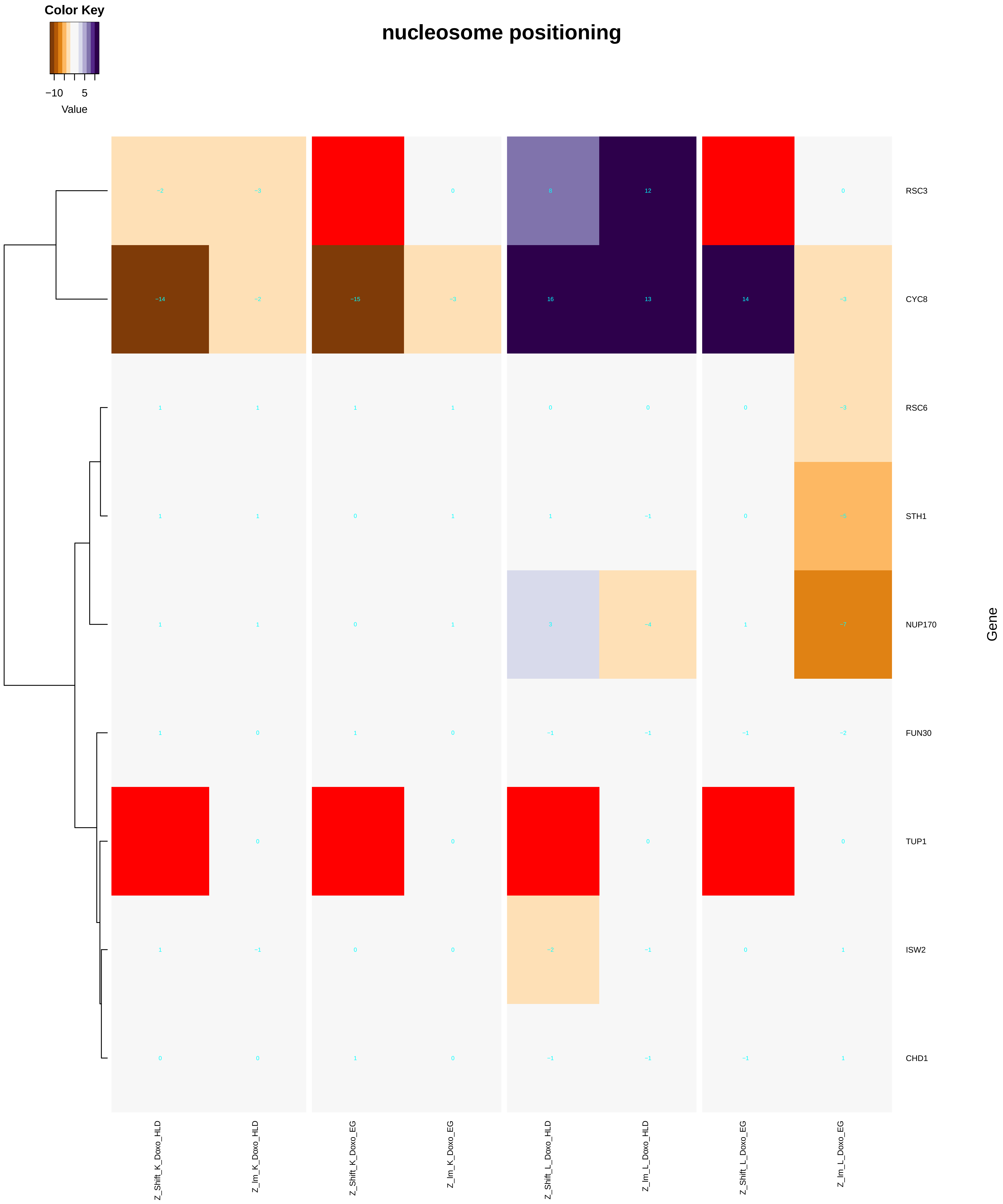

Color Key
−10
5
Value
nucleosome positioning
RSC3
CYC8
RSC6
STH1
NUP170
FUN30
TUP1
ISW2
CHD1
Gene
Z_Shift_K_Doxo_HLD
Z_Im_K_Doxo_HLD
Z_Shift_K_Doxo_EG
Z_Im_K_Doxo_EG
Z_Shift_L_Doxo_HLD
Z_Im_L_Doxo_HLD
Z_Shift_L_Doxo_EG
Z_Im_L_Doxo_EG

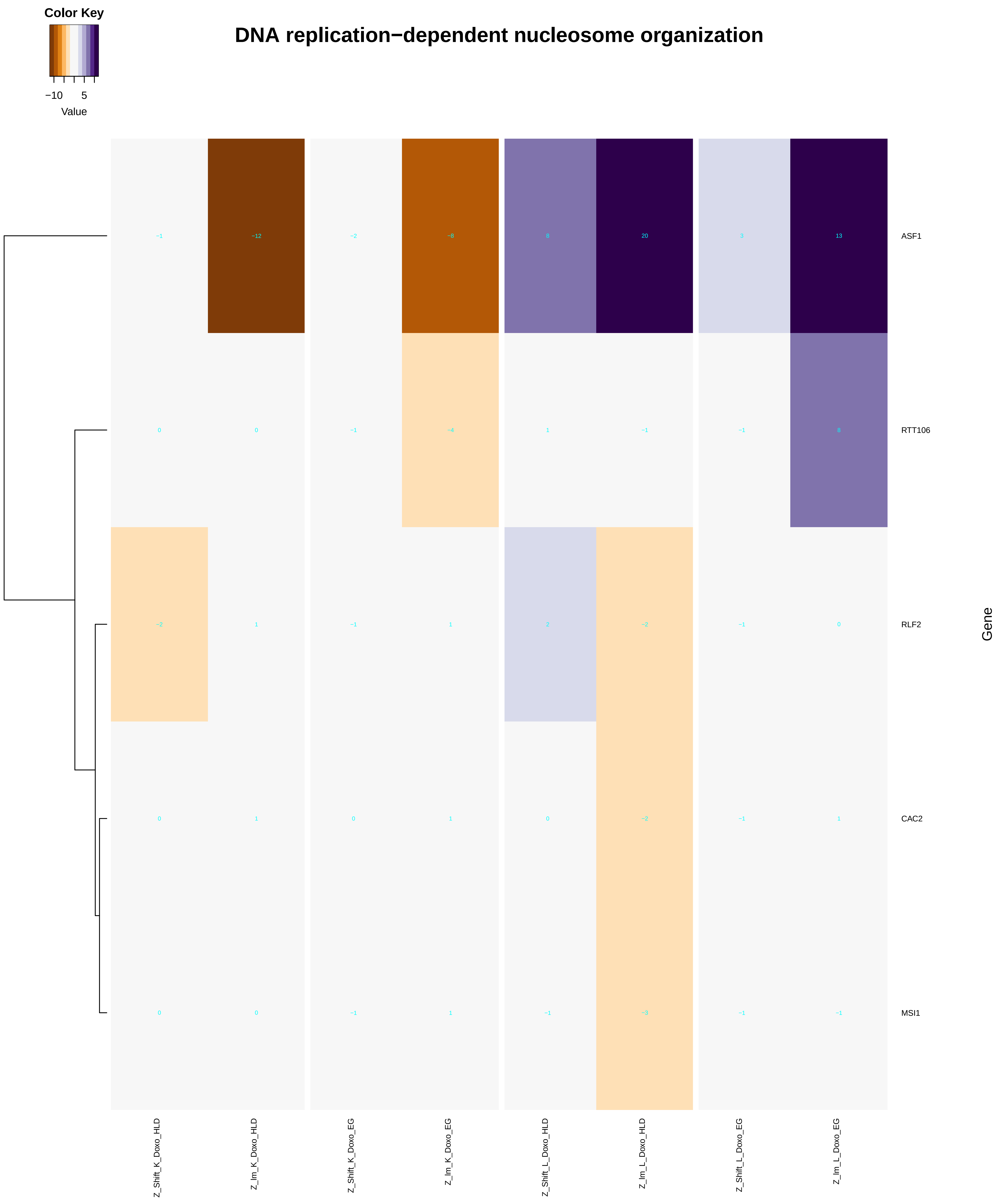

Color Key
−10
5
Value
DNA replication−dependent nucleosome organization
ASF1
RTT106
RLF2
CAC2
MSI1
Gene
Z_Shift_K_Doxo_HLD
Z_Im_K_Doxo_HLD
Z_Shift_K_Doxo_EG
Z_Im_K_Doxo_EG
Z_Shift_L_Doxo_HLD
Z_Im_L_Doxo_HLD
Z_Shift_L_Doxo_EG
Z_Im_L_Doxo_EG

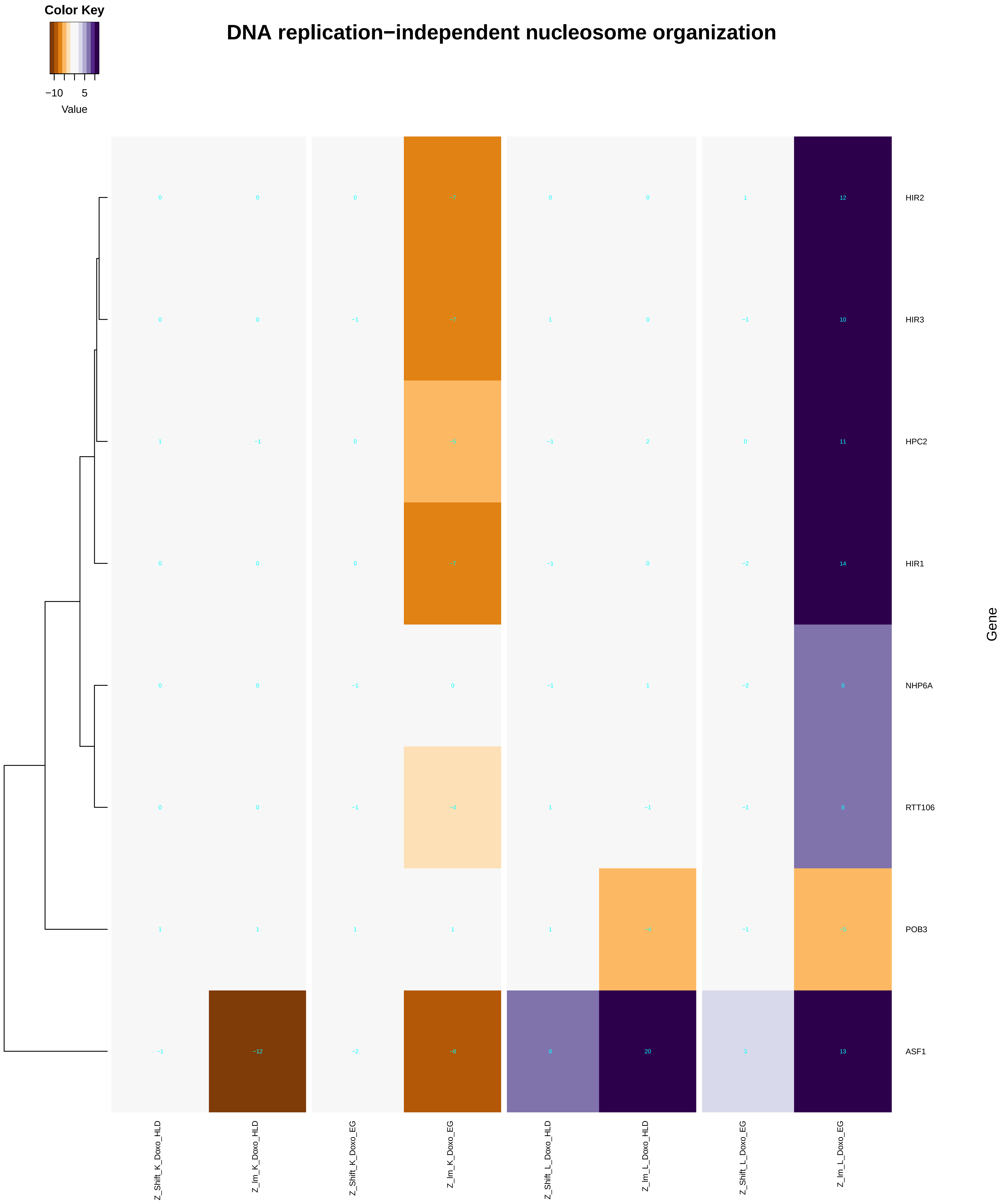

Color Key
−10
5
Value
DNA replication−independent nucleosome organization
HIR2
HIR3
HPC2
HIR1
NHP6A
RTT106
POB3
ASF1
Gene
Z_Shift_K_Doxo_HLD
Z_Im_K_Doxo_HLD
Z_Shift_K_Doxo_EG
Z_Im_K_Doxo_EG
Z_Shift_L_Doxo_HLD
Z_Im_L_Doxo_HLD
Z_Shift_L_Doxo_EG
Z_Im_L_Doxo_EG

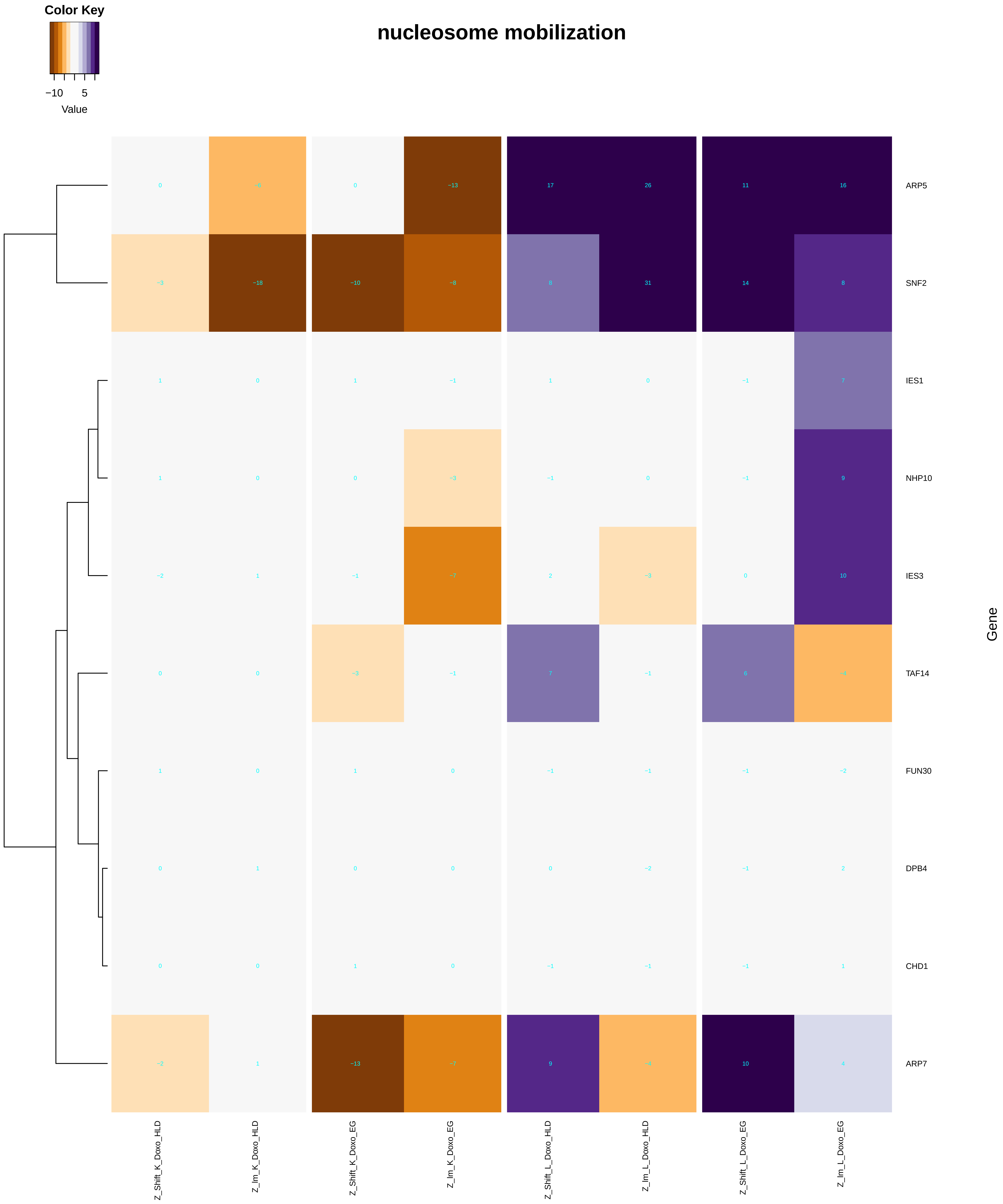

Color Key
−10
5
Value
nucleosome mobilization
ARP5
SNF2
IES1
NHP10
IES3
TAF14
FUN30
DPB4
CHD1
ARP7
Gene
Z_Shift_K_Doxo_HLD
Z_Im_K_Doxo_HLD
Z_Shift_K_Doxo_EG
Z_Im_K_Doxo_EG
Z_Shift_L_Doxo_HLD
Z_Im_L_Doxo_HLD
Z_Shift_L_Doxo_EG
Z_Im_L_Doxo_EG

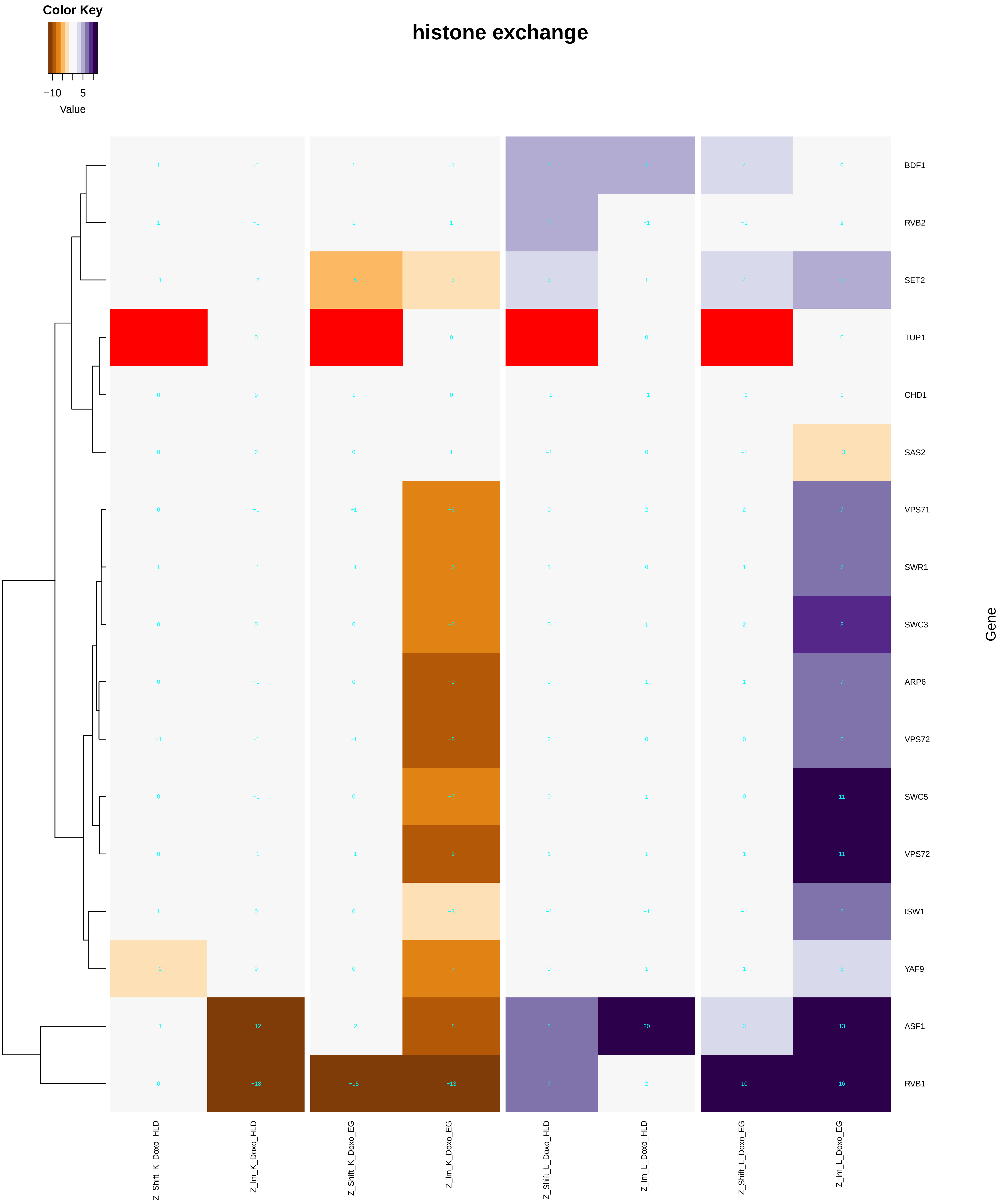
Color Key
−10
5
Value
histone exchange
BDF1
RVB2
SET2
TUP1
CHD1
SAS2
VPS71
SWR1
SWC3
ARP6
VPS72
SWC5
VPS72
ISW1
YAF9
ASF1
RVB1
Gene
Z_Shift_K_Doxo_HLD
Z_Im_K_Doxo_HLD
Z_Shift_K_Doxo_EG
Z_Im_K_Doxo_EG
Z_Shift_L_Doxo_HLD
Z_Im_L_Doxo_HLD
Z_Shift_L_Doxo_EG
Z_Im_L_Doxo_EG

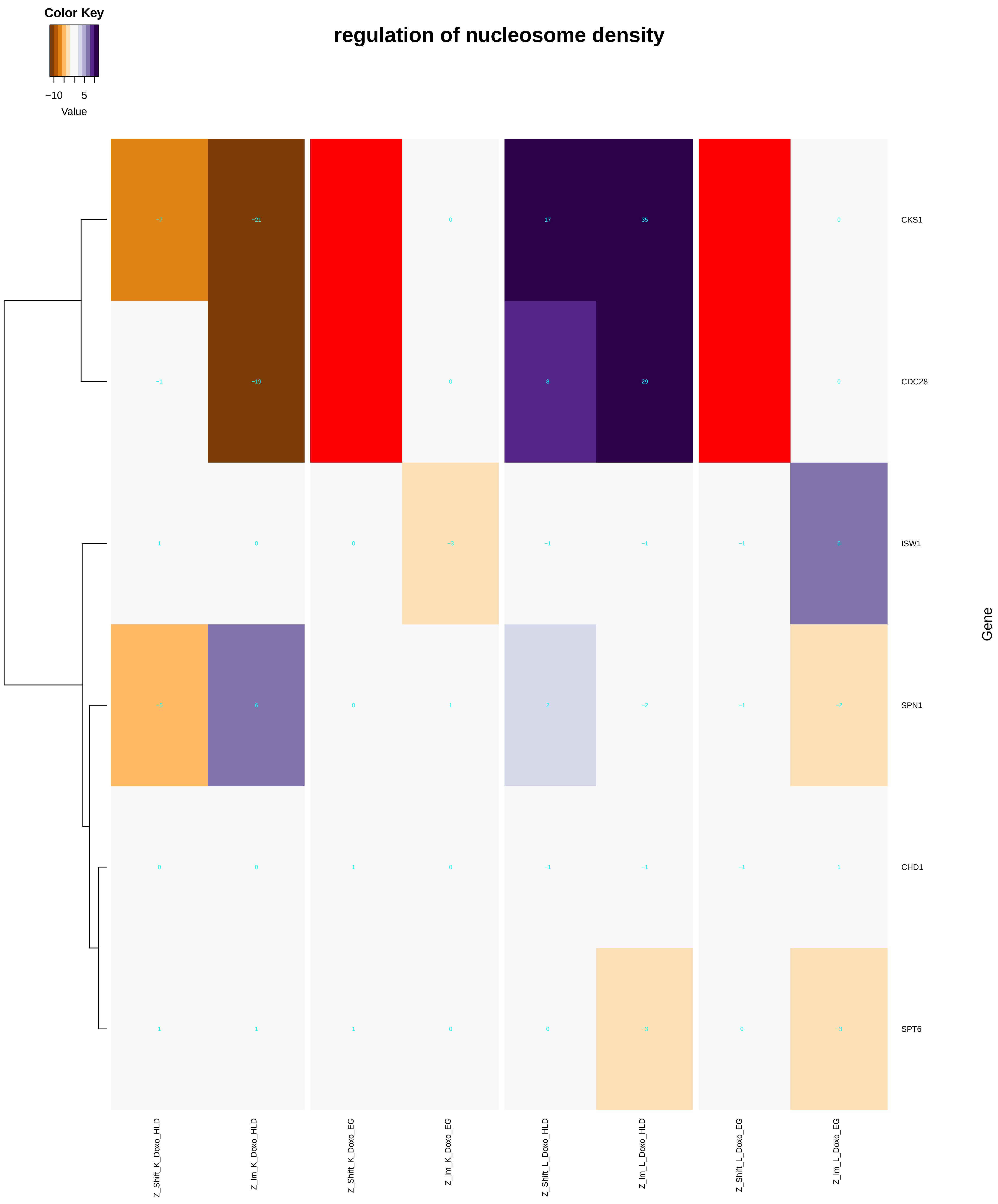
Color Key
−10
5
Value
regulation of nucleosome density
CKS1
CDC28
ISW1
SPN1
CHD1
SPT6
Gene
Z_Shift_K_Doxo_HLD
Z_lm_K_Doxo_HLD
Z_Shift_K_Doxo_EG
Z_lm_K_Doxo_EG
Z_Shift_L_Doxo_HLD
Z_lm_L_Doxo_HLD
Z_Shift_L_Doxo_EG
Z_lm_L_Doxo_EG

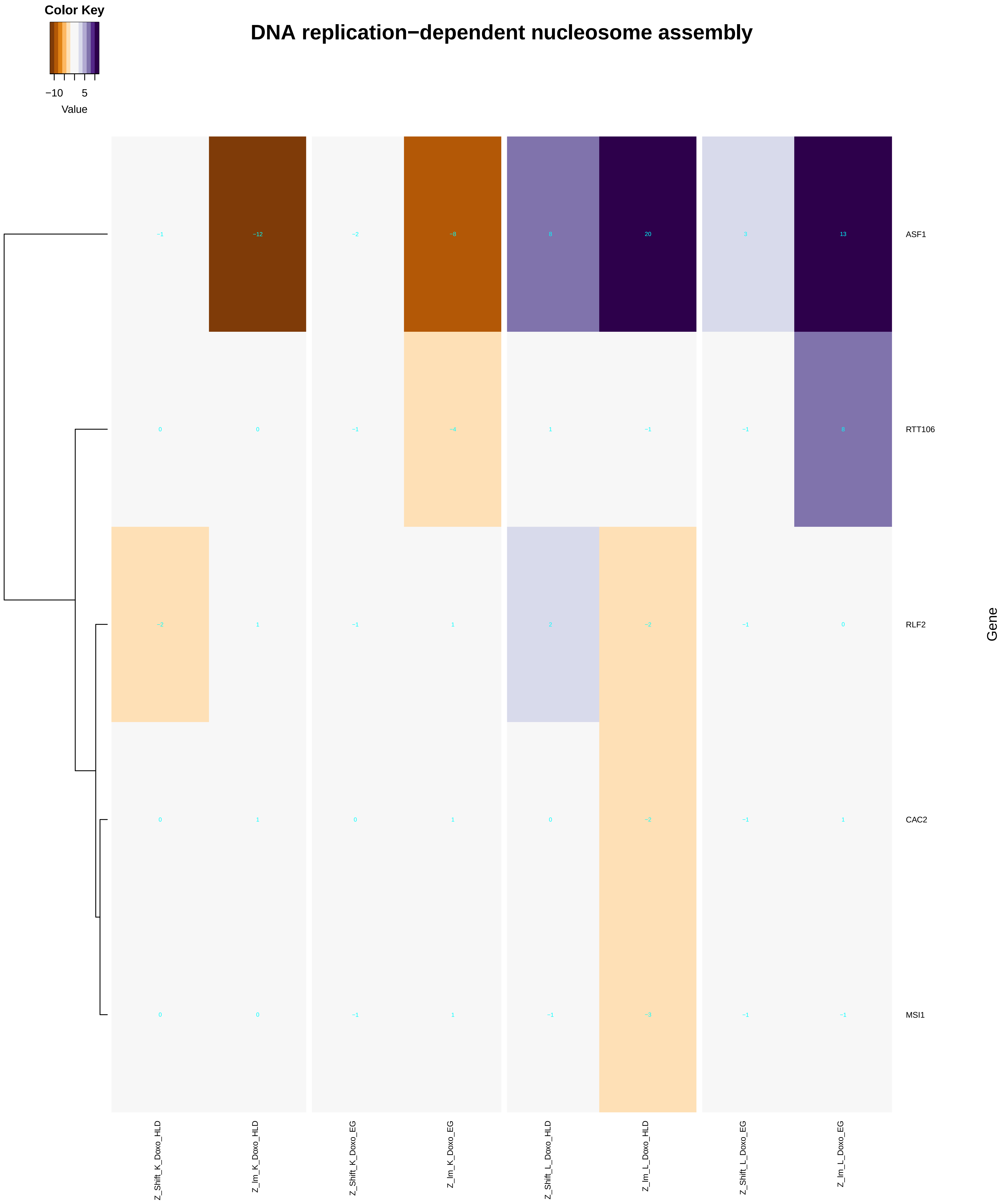
Color Key
−10
5
Value
DNA replication−dependent nucleosome assembly
ASF1
RTT106
RLF2
CAC2
MSI1
Gene
−1 −12 −2 −8 8 20 3 13
0 0 −1 −4 1 −1 −1 8
−2 1 −1 1 2 −2 −1 0
0 1 0 1 0 −2 −1 1
0 0 −1 1 −1 −3 −1 −1
Z_Shift_K_Doxo_HLD
Z_lm_K_Doxo_HLD
Z_Shift_K_Doxo_EG
Z_lm_K_Doxo_EG
Z_Shift_L_Doxo_HLD
Z_lm_L_Doxo_HLD
Z_Shift_L_Doxo_EG
Z_lm_L_Doxo_EG

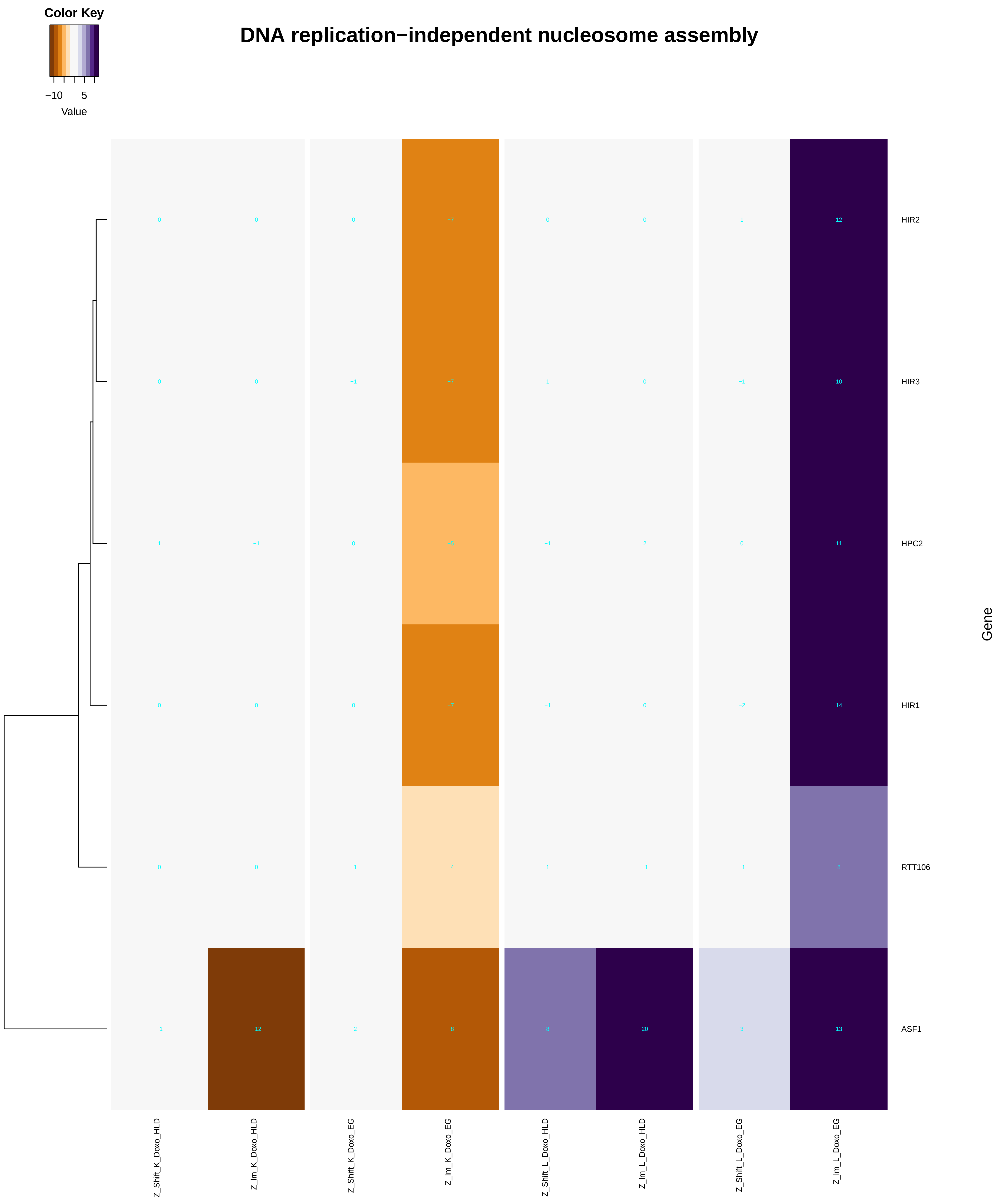

Color Key
−10 5
Value
DNA replication−independent nucleosome assembly
HIR2
HIR3
HPC2
HIR1
RTT106
ASF1
Gene
Z_Shift_K_Doxo_HLD
Z_Im_K_Doxo_HLD
Z_Shift_K_Doxo_EG
Z_Im_K_Doxo_EG
Z_Shift_L_Doxo_HLD
Z_Im_L_Doxo_HLD
Z_Shift_L_Doxo_EG
Z_Im_L_Doxo_EG
