## Additional File 9 for "A yeast phenomic model for the influence of Warburg metabolism on genetic buffering of doxorubicin": phospholipid_biosynthetic_process.pdf

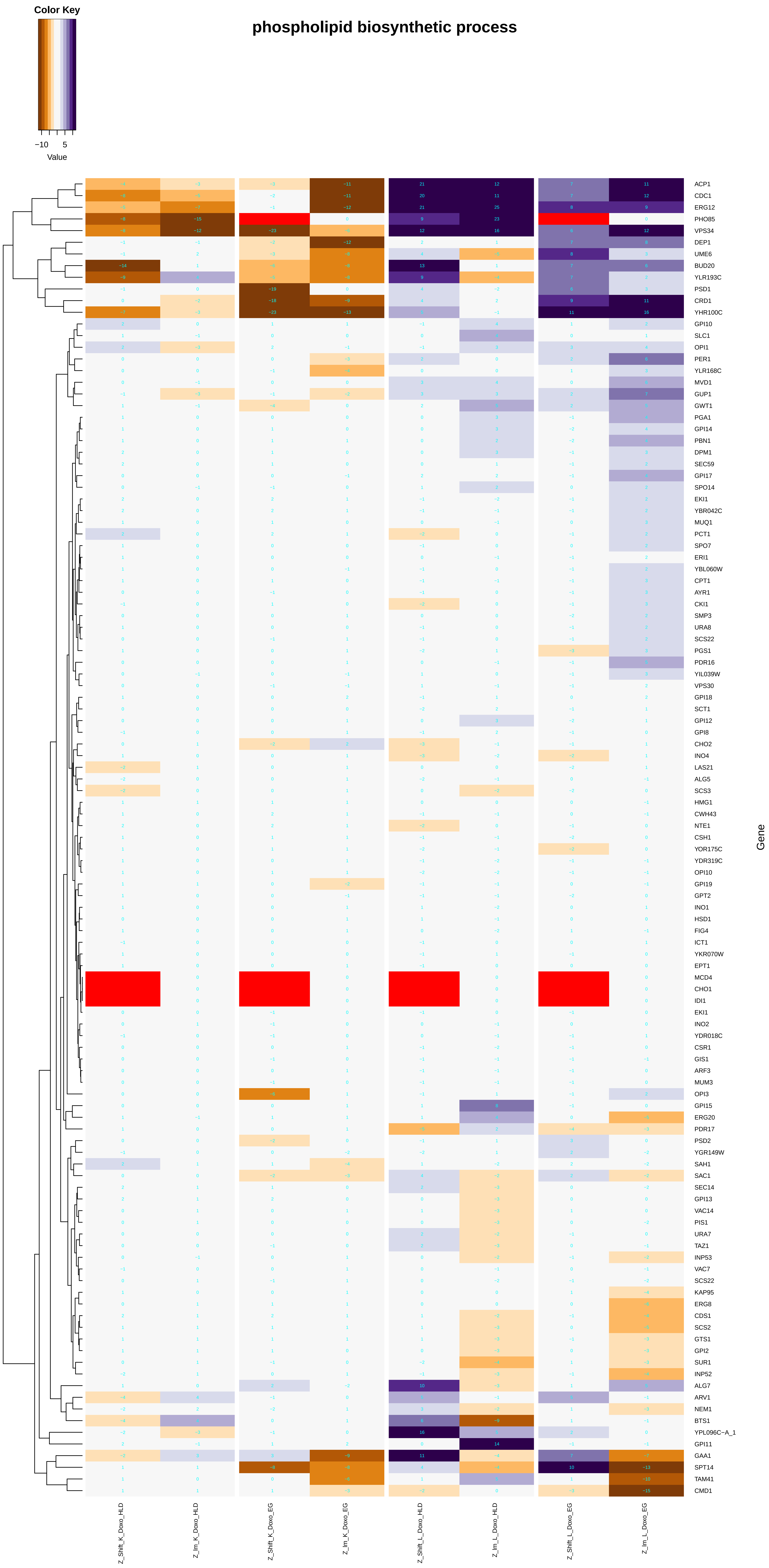
phospholipid biosynthetic process
Color Key
−10 5
Value
Gene
ACP1
CDC1
ERG12
PHO85
VPS34
DEP1
UME6
BUD20
YLR193C
PSD1
CRD1
YHR100C
GPI10
SLC1
OPI1
PER1
YLR168C
MVD1
GUP1
GWT1
PGA1
GPI14
PBN1
DPM1
SEC59
GPI17
SPO14
EKI1
YBR042C
MUQ1
PCT1
SPO7
ERI1
YBL060W
CPT1
AYR1
CKI1
SMP3
URA8
SCS22
PGS1
PDR16
YIL039W
VPS30
GPI18
SCT1
GPI12
GPI8
CHO2
INO4
LAS21
ALG5
SCS3
HMG1
CWH43
NTE1
CSH1
YOR175C
YDR319C
OPI10
GPI19
GPT2
INO1
HSD1
FIG4
ICT1
YKR070W
EPT1
MCD4
CHO1
IDI1
EKI1
INO2
YDR018C
CSR1
GIS1
ARF3
MUM3
OPI3
GPI15
ERG20
PDR17
PSD2
YGR149W
SAH1
SAC1
SEC14
GPI13
VAC14
PIS1
URA7
TAZ1
INP53
VAC7
SCS22
KAP95
ERG8
CDS1
SCS2
GTS1
GPI2
SUR1
INP52
ALG7
ARV1
NEM1
BTS1
YPL096C−A_1
GPI11
GAA1
SPT14
TAM41
CMD1
Z_Shift_K_Doxo_HLD
Z_lm_K_Doxo_HLD
Z_Shift_K_Doxo_EG
Z_lm_K_Doxo_EG
Z_Shift_L_Doxo_HLD
Z_lm_L_Doxo_HLD
Z_Shift_L_Doxo_EG
Z_lm_L_Doxo_EG

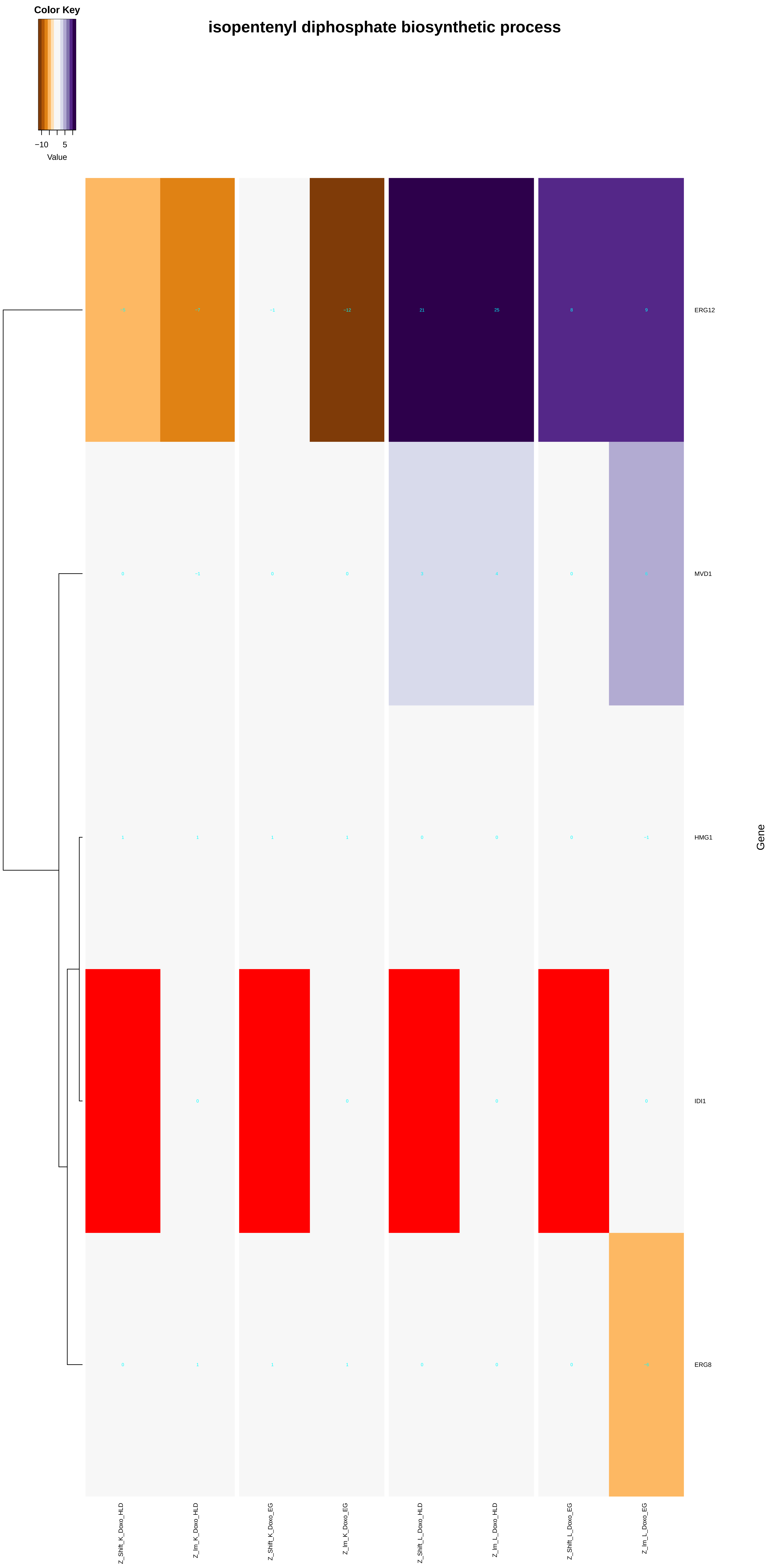

Color Key
−10
5
Value
isopentenyl diphosphate biosynthetic process
ERG12
MVD1
HMG1
IDI1
ERG8
Gene
Z_Shift_K_Doxo_HLD
Z_Im_K_Doxo_HLD
Z_Shift_K_Doxo_EG
Z_Im_K_Doxo_EG
Z_Shift_L_Doxo_HLD
Z_Im_L_Doxo_HLD
Z_Shift_L_Doxo_EG
Z_Im_L_Doxo_EG

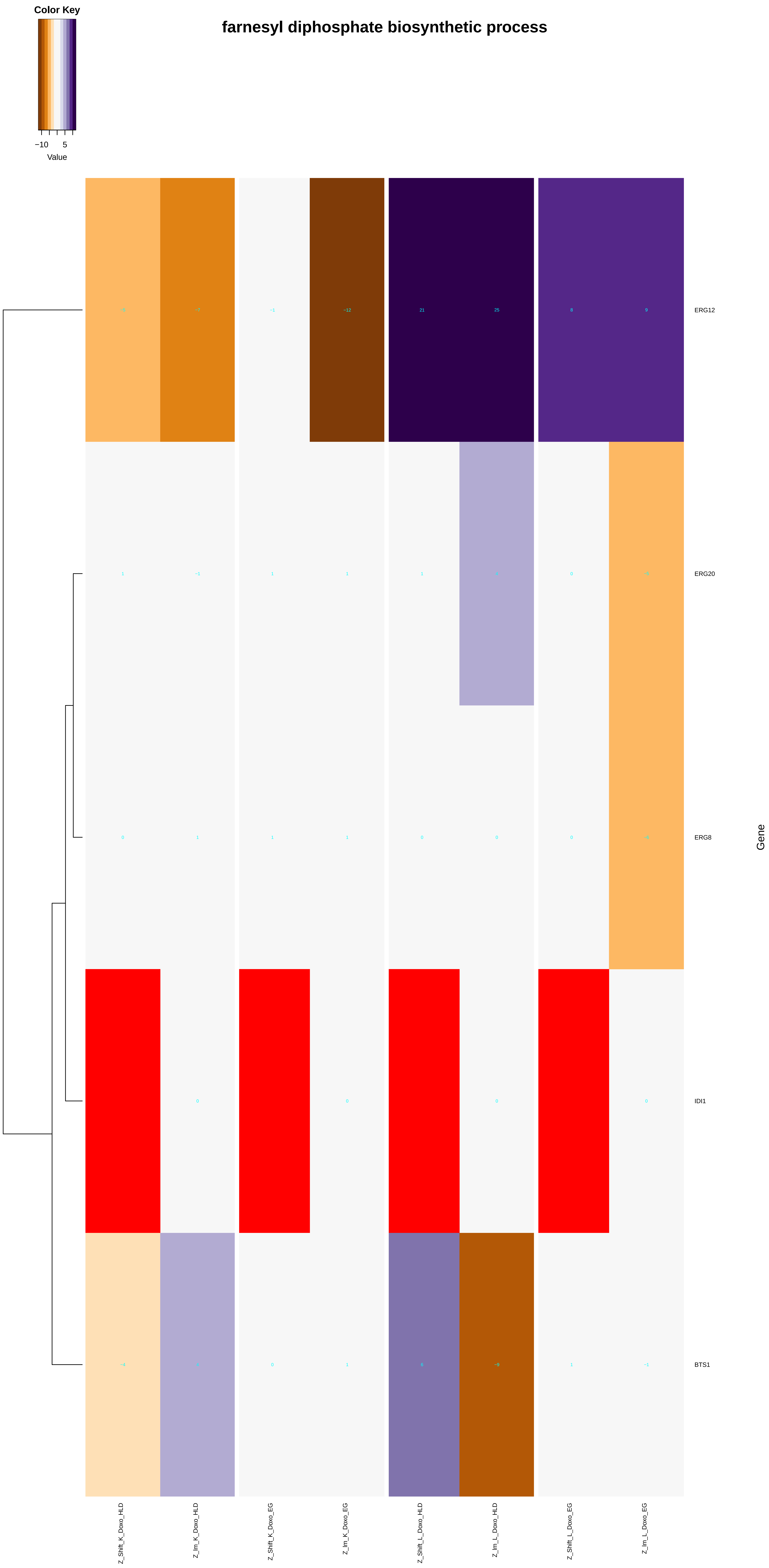

Color Key
−10
5
Value
farnesyl diphosphate biosynthetic process
ERG12
ERG20
ERG8
IDI1
BTS1
Gene
Z_Shift_K_Doxo_HLD
Z_Im_K_Doxo_HLD
Z_Shift_K_Doxo_EG
Z_Im_K_Doxo_EG
Z_Shift_L_Doxo_HLD
Z_Im_L_Doxo_HLD
Z_Shift_L_Doxo_EG
Z_Im_L_Doxo_EG

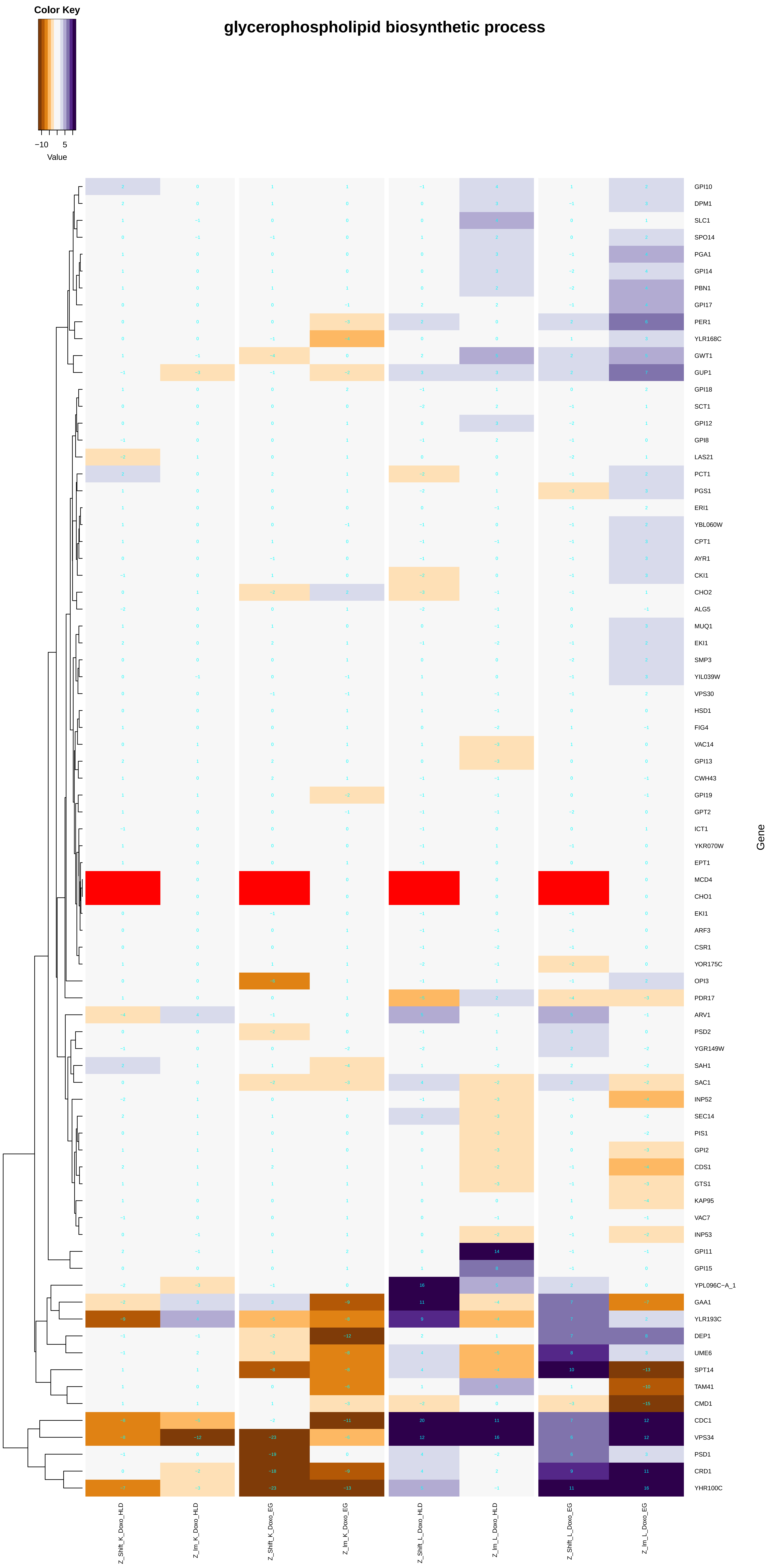
glycerophospholipid biosynthetic process
Color Key
−10
5
Value
GPI10
DPM1
SLC1
SPO14
PGA1
GPI14
PBN1
GPI17
PER1
YLR168C
GWT1
GUP1
GPI18
SCT1
GPI12
GPI8
LAS21
PCT1
PGS1
ERI1
YBL060W
CPT1
AYR1
CKI1
CHO2
ALG5
MUQ1
EKI1
SMP3
YIL039W
VPS30
HSD1
FIG4
VAC14
GPI13
CWH43
GPI19
GPT2
ICT1
YKR070W
EPT1
MCD4
CHO1
EKI1
ARF3
CSR1
YOR175C
OPI3
PDR17
ARV1
PSD2
YGR149W
SAH1
SAC1
INP52
SEC14
PIS1
GPI2
CDS1
GTS1
KAP95
VAC7
INP53
GPI11
GPI15
YPL096C−A_1
GAA1
YLR193C
DEP1
UME6
SPT14
TAM41
CMD1
CDC1
VPS34
PSD1
CRD1
YHR100C
Gene
Z_Shift_K_Doxo_HLD
Z_lm_K_Doxo_HLD
Z_Shift_K_Doxo_EG
Z_lm_K_Doxo_EG
Z_Shift_L_Doxo_HLD
Z_lm_L_Doxo_HLD
Z_Shift_L_Doxo_EG
Z_lm_L_Doxo_EG

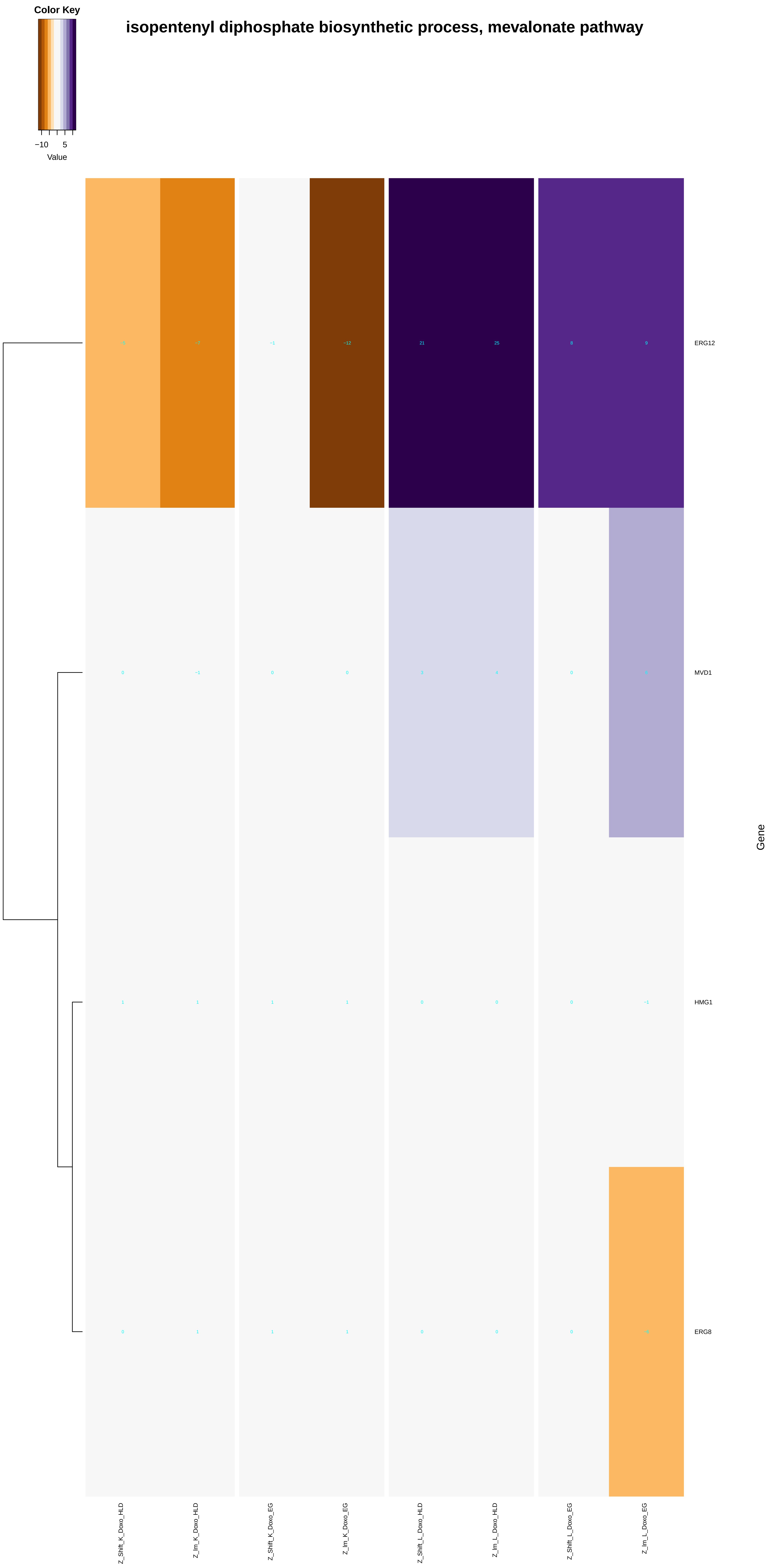
isopentenyl diphosphate biosynthetic process, mevalonate pathway
Color Key
−10
5
Value
ERG12
MVD1
HMG1
ERG8
Gene
Z_Shift_K_Doxo_HLD
Z_Im_K_Doxo_HLD
Z_Shift_K_Doxo_EG
Z_Im_K_Doxo_EG
Z_Shift_L_Doxo_HLD
Z_Im_L_Doxo_HLD
Z_Shift_L_Doxo_EG
Z_Im_L_Doxo_EG

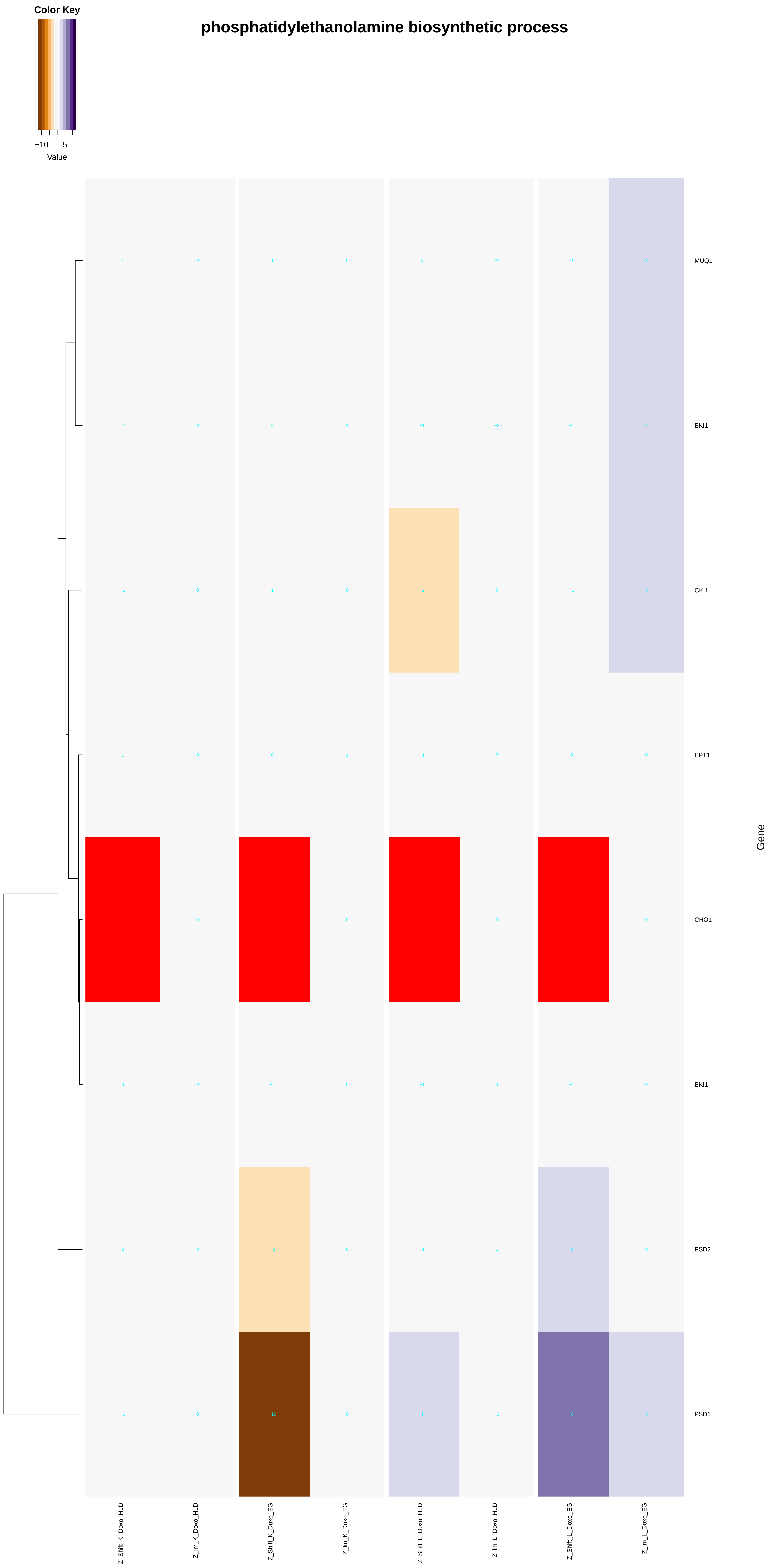
phosphatidylethanolamine biosynthetic process
Color Key
−10
5
Value
MUQ1
EKI1
CKI1
EPT1
CHO1
EKI1
PSD2
PSD1
Gene
Z_Shift_K_Doxo_HLD
Z_Im_K_Doxo_HLD
Z_Shift_K_Doxo_EG
Z_Im_K_Doxo_EG
Z_Shift_L_Doxo_HLD
Z_Im_L_Doxo_HLD
Z_Shift_L_Doxo_EG
Z_Im_L_Doxo_EG

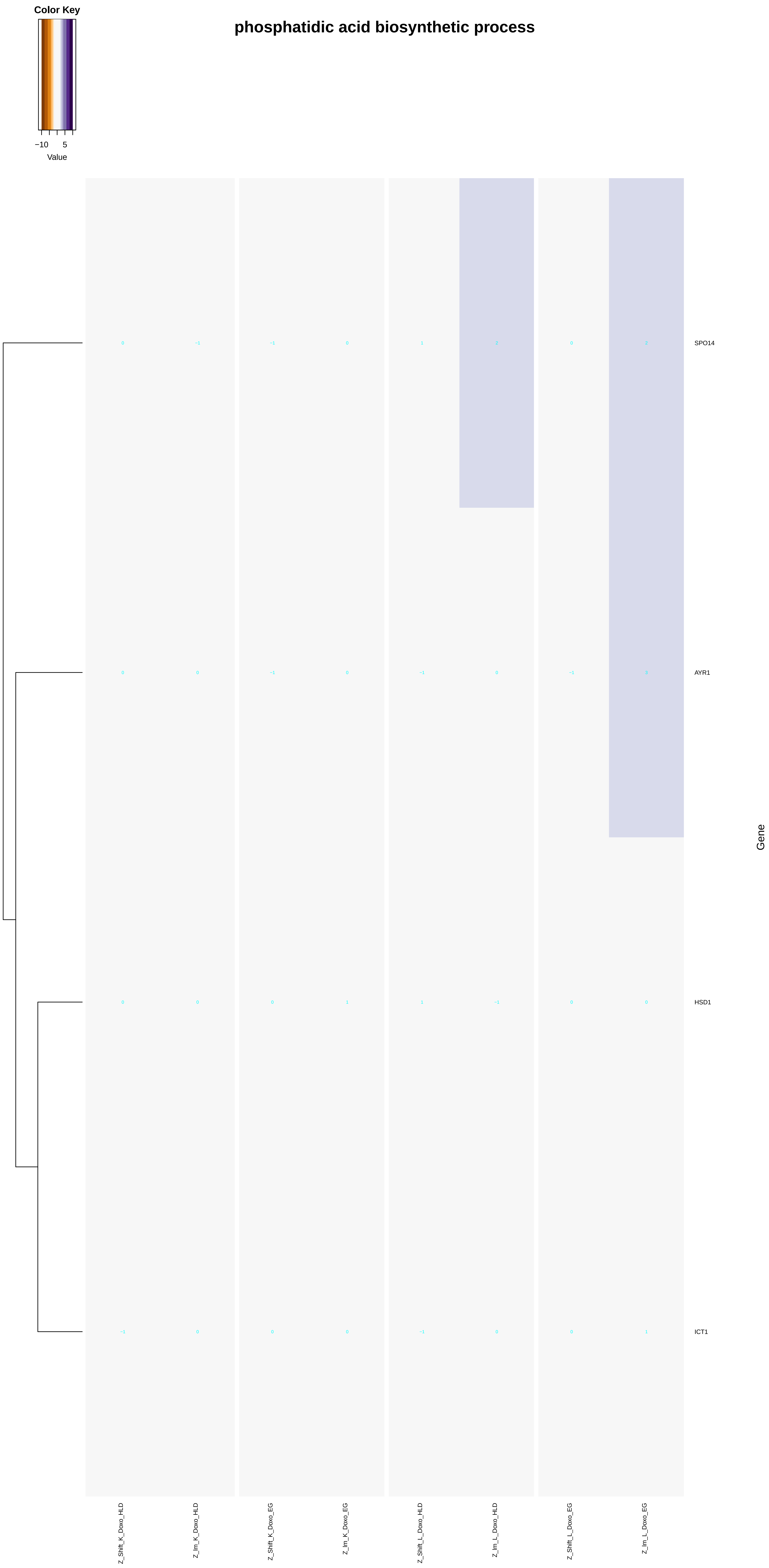
Color Key
−10
5
Value
phosphatidic acid biosynthetic process
SPO14
AYR1
HSD1
ICT1
Gene
Z_Shift_K_Doxo_HLD
Z_lm_K_Doxo_HLD
Z_Shift_K_Doxo_EG
Z_lm_K_Doxo_EG
Z_Shift_L_Doxo_HLD
Z_lm_L_Doxo_HLD
Z_Shift_L_Doxo_EG
Z_lm_L_Doxo_EG

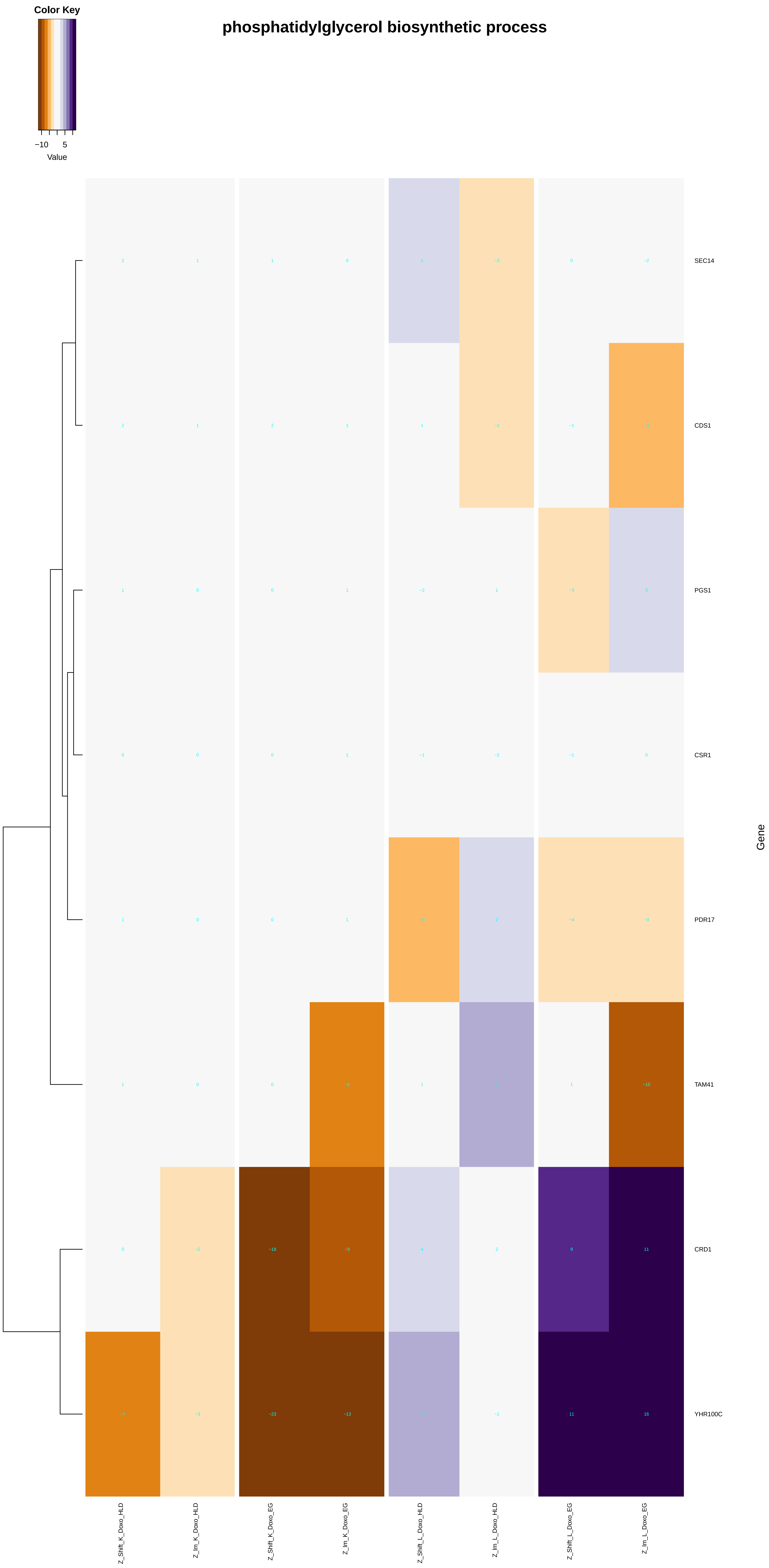
Color Key
−10
5
Value
phosphatidylglycerol biosynthetic process
SEC14
CDS1
PGS1
CSR1
PDR17
TAM41
CRD1
YHR100C
Gene
Z_Shift_K_Doxo_HLD
Z_Im_K_Doxo_HLD
Z_Shift_K_Doxo_EG
Z_Im_K_Doxo_EG
Z_Shift_L_Doxo_HLD
Z_Im_L_Doxo_HLD
Z_Shift_L_Doxo_EG
Z_Im_L_Doxo_EG

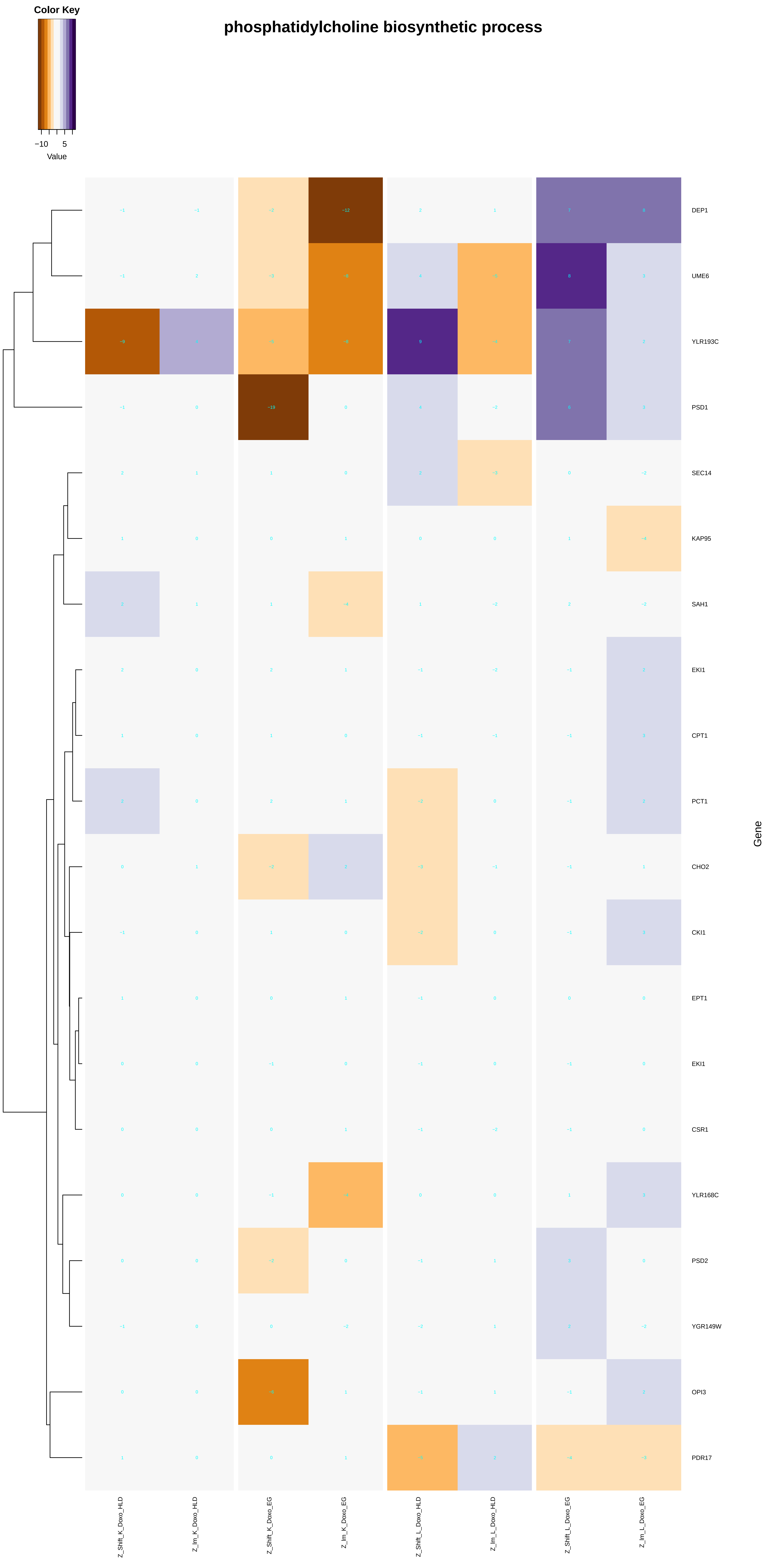

Color Key
−10
5
Value
phosphatidylcholine biosynthetic process
DEP1
UME6
YLR193C
PSD1
SEC14
KAP95
SAH1
EKI1
CPT1
PCT1
CHO2
CKI1
EPT1
EKI1
CSR1
YLR168C
PSD2
YGR149W
OPI3
PDR17
Gene
Z_Shift_K_Doxo_HLD
Z_lm_K_Doxo_HLD
Z_Shift_K_Doxo_EG
Z_lm_K_Doxo_EG
Z_Shift_L_Doxo_HLD
Z_lm_L_Doxo_HLD
Z_Shift_L_Doxo_EG
Z_lm_L_Doxo_EG

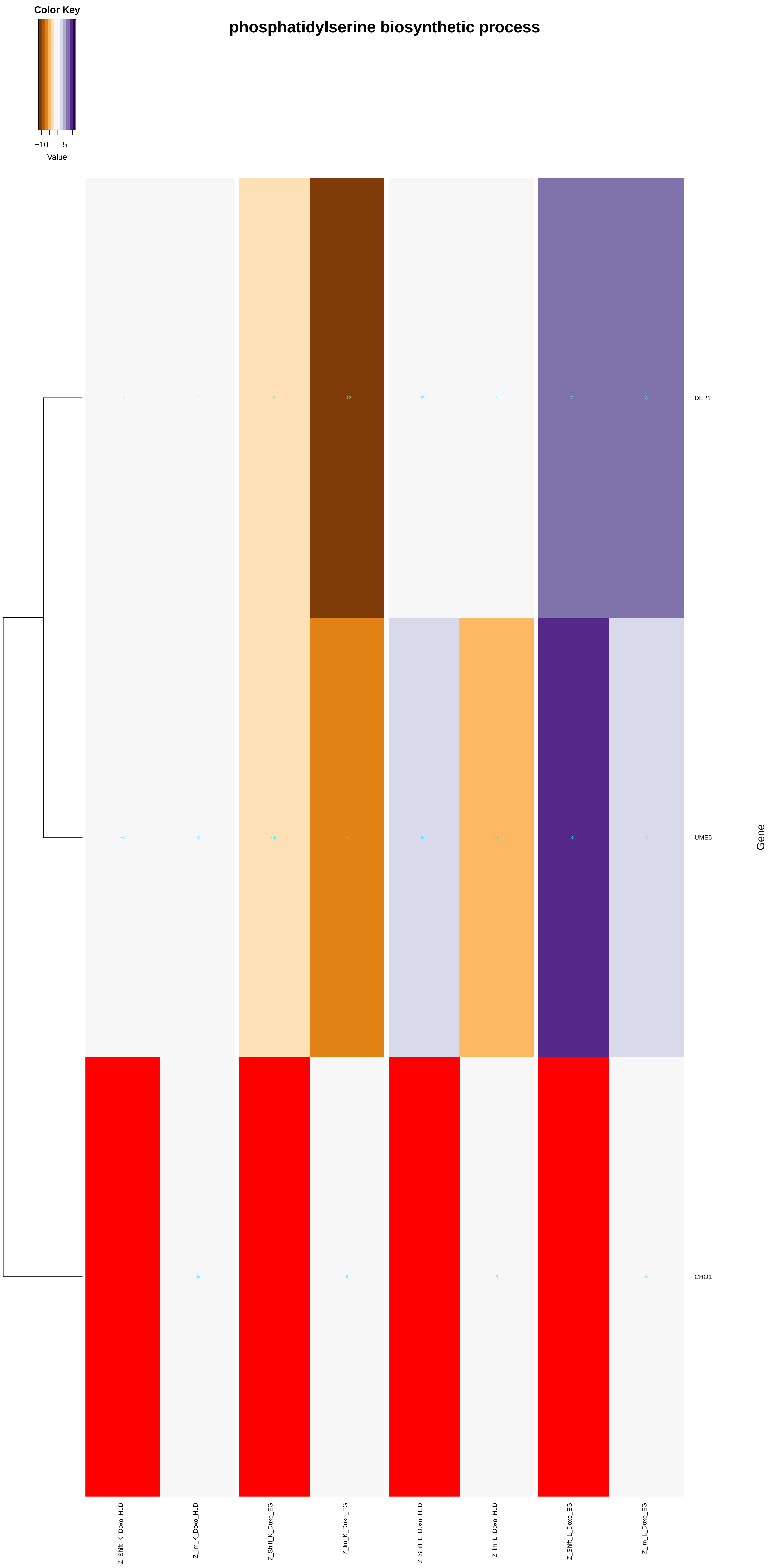
Color Key
−10
5
Value
phosphatidylserine biosynthetic process
−1
−1
−2
−12
2
1
7
8
DEP1
−1
2
−3
−8
4
−5
8
3
UME6
0
0
0
0
CHO1
Gene
Z_Shift_K_Doxo_HLD
Z_lm_K_Doxo_HLD
Z_Shift_K_Doxo_EG
Z_lm_K_Doxo_EG
Z_Shift_L_Doxo_HLD
Z_lm_L_Doxo_HLD
Z_Shift_L_Doxo_EG
Z_lm_L_Doxo_EG

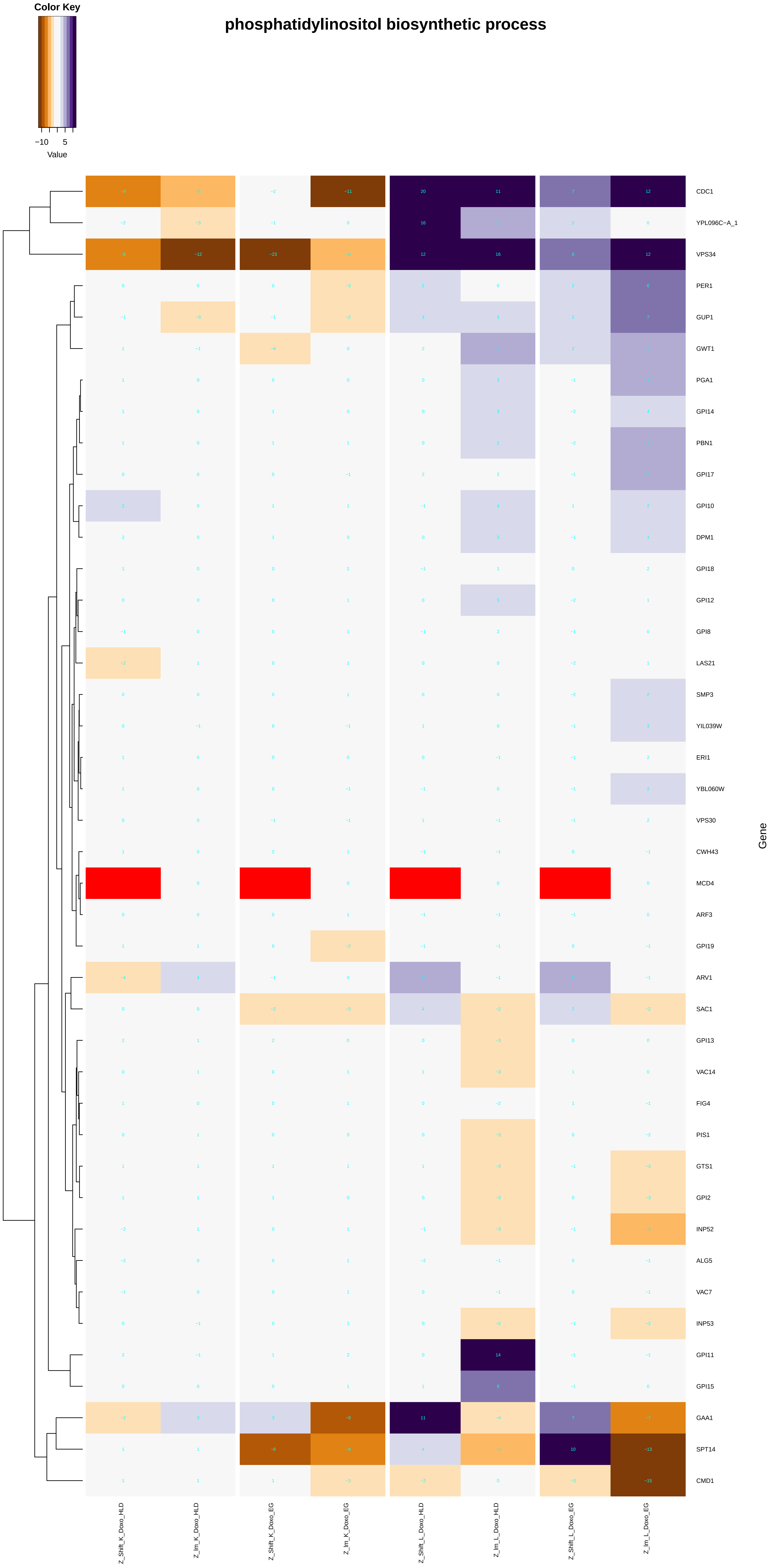
phosphatidylinositol biosynthetic process
Color Key
−10 5
Value
CDC1
YPL096C−A_1
VPS34
PER1
GUP1
GWT1
PGA1
GPI14
PBN1
GPI17
GPI10
DPM1
GPI18
GPI12
GPI8
LAS21
SMP3
YIL039W
ERI1
YBL060W
VPS30
CWH43
MCD4
ARF3
GPI19
ARV1
SAC1
GPI13
VAC14
FIG4
PIS1
GTS1
GPI2
INP52
ALG5
VAC7
INP53
GPI11
GPI15
GAA1
SPT14
CMD1
Gene
Z_Shift_K_Doxo_HLD
Z_lm_K_Doxo_HLD
Z_Shift_K_Doxo_EG
Z_lm_K_Doxo_EG
Z_Shift_L_Doxo_HLD
Z_lm_L_Doxo_HLD
Z_Shift_L_Doxo_EG
Z_lm_L_Doxo_EG

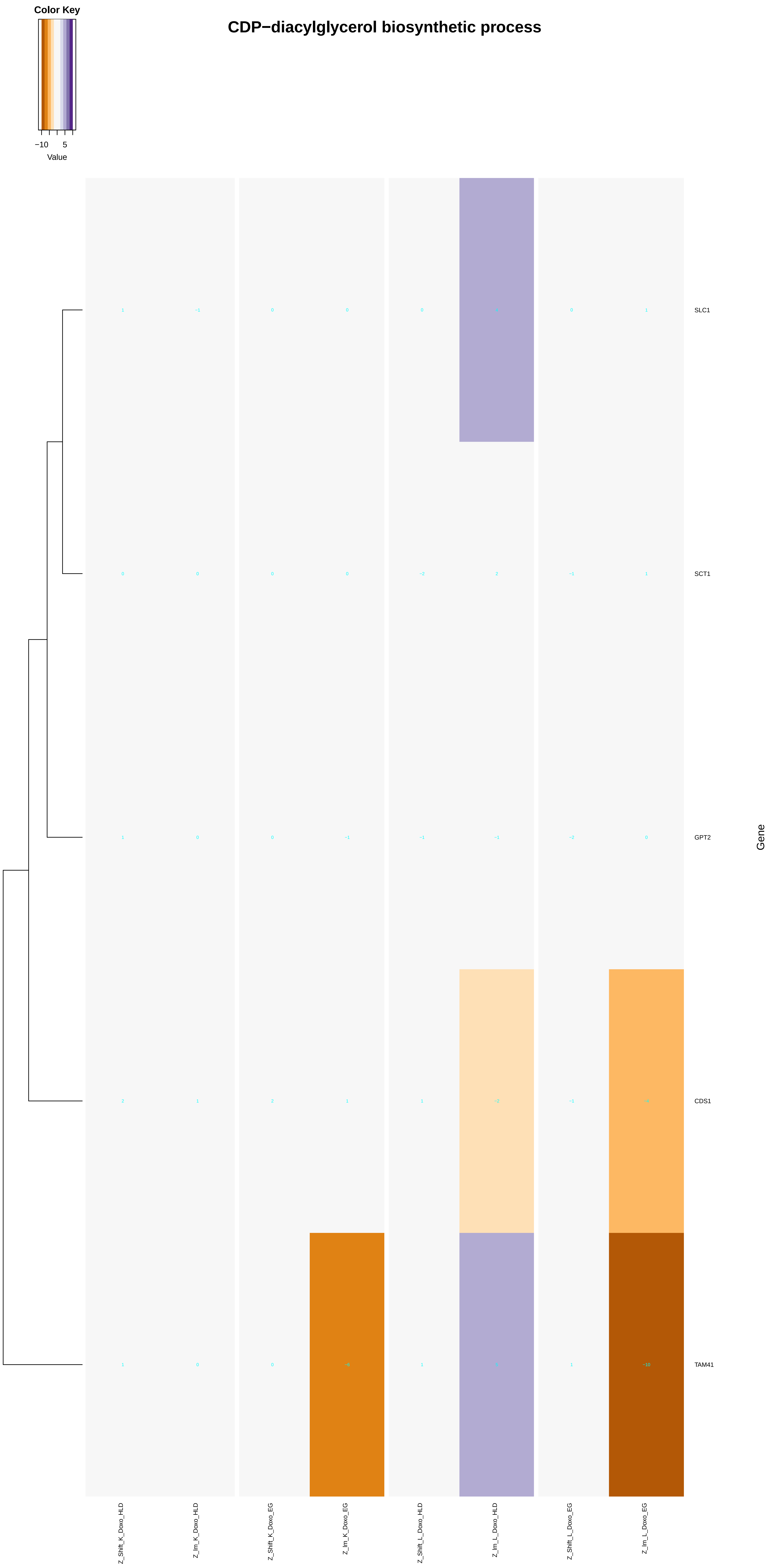
Color Key
−10
5
Value
CDP−diacylglycerol biosynthetic process
SLC1
SCT1
GPT2
CDS1
TAM41
Gene
Z_Shift_K_Doxo_HLD
Z_Im_K_Doxo_HLD
Z_Shift_K_Doxo_EG
Z_Im_K_Doxo_EG
Z_Shift_L_Doxo_HLD
Z_Im_L_Doxo_HLD
Z_Shift_L_Doxo_EG
Z_Im_L_Doxo_EG

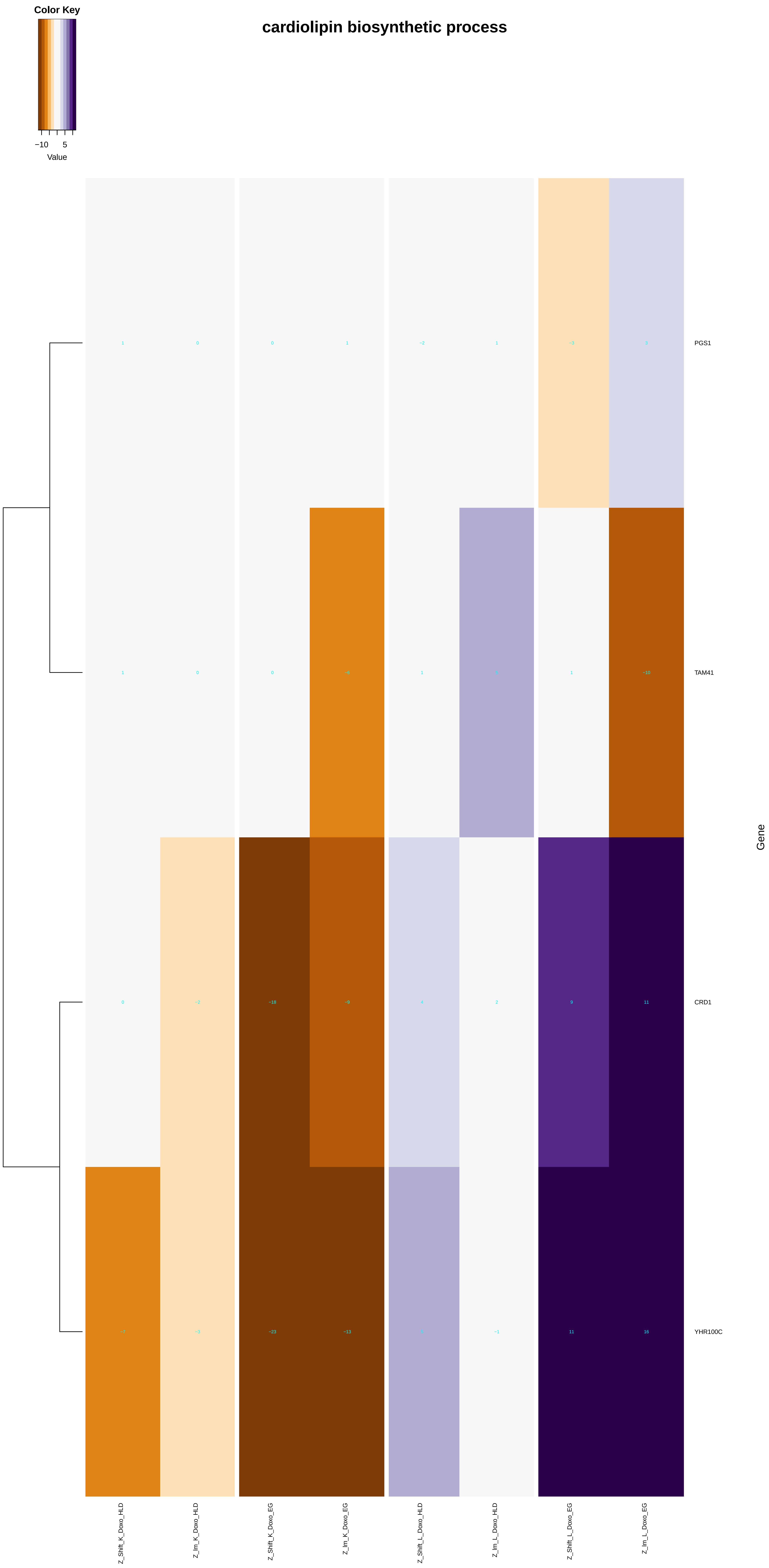

Color Key
−10
5
Value
cardiolipin biosynthetic process
PGS1
TAM41
CRD1
YHR100C
Gene
Z_Shift_K_Doxo_HLD
Z_lm_K_Doxo_HLD
Z_Shift_K_Doxo_EG
Z_lm_K_Doxo_EG
Z_Shift_L_Doxo_HLD
Z_lm_L_Doxo_HLD
Z_Shift_L_Doxo_EG
Z_lm_L_Doxo_EG

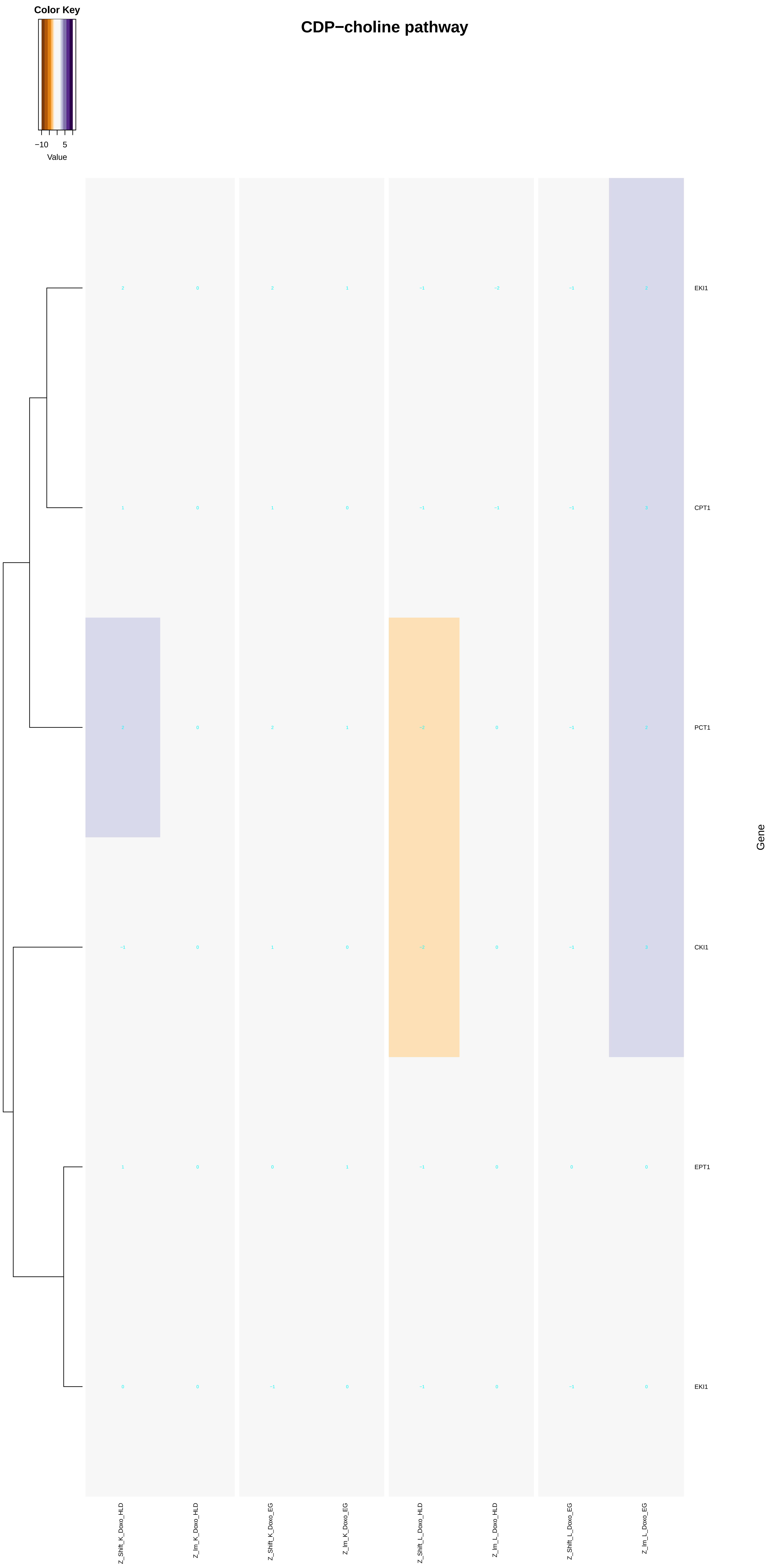
Color Key
−10
5
Value
CDP−choline pathway
EKI1
CPT1
PCT1
CKI1
EPT1
EKI1
Gene
Z_Shift_K_Doxo_HLD
Z_lm_K_Doxo_HLD
Z_Shift_K_Doxo_EG
Z_lm_K_Doxo_EG
Z_Shift_L_Doxo_HLD
Z_lm_L_Doxo_HLD
Z_Shift_L_Doxo_EG
Z_lm_L_Doxo_EG

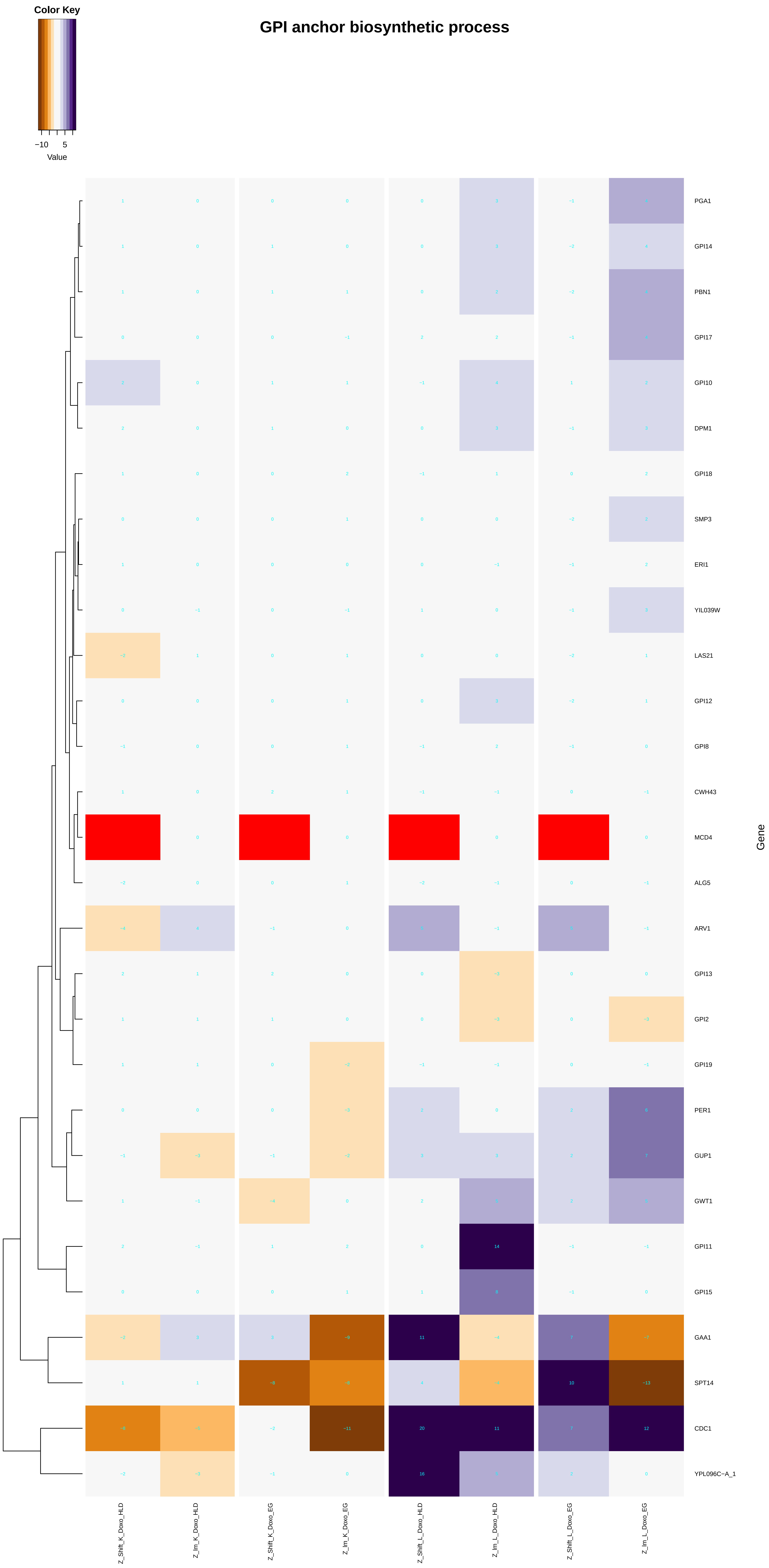

Color Key
−10
5
Value
GPI anchor biosynthetic process
PGA1
GPI14
PBN1
GPI17
GPI10
DPM1
GPI18
SMP3
ERI1
YIL039W
LAS21
GPI12
GPI8
CWH43
MCD4
ALG5
ARV1
GPI13
GPI2
GPI19
PER1
GUP1
GWT1
GPI11
GPI15
GAA1
SPT14
CDC1
YPL096C−A_1
Gene
Z_Shift_K_Doxo_HLD
Z_Im_K_Doxo_HLD
Z_Shift_K_Doxo_EG
Z_Im_K_Doxo_EG
Z_Shift_L_Doxo_HLD
Z_Im_L_Doxo_HLD
Z_Shift_L_Doxo_EG
Z_Im_L_Doxo_EG

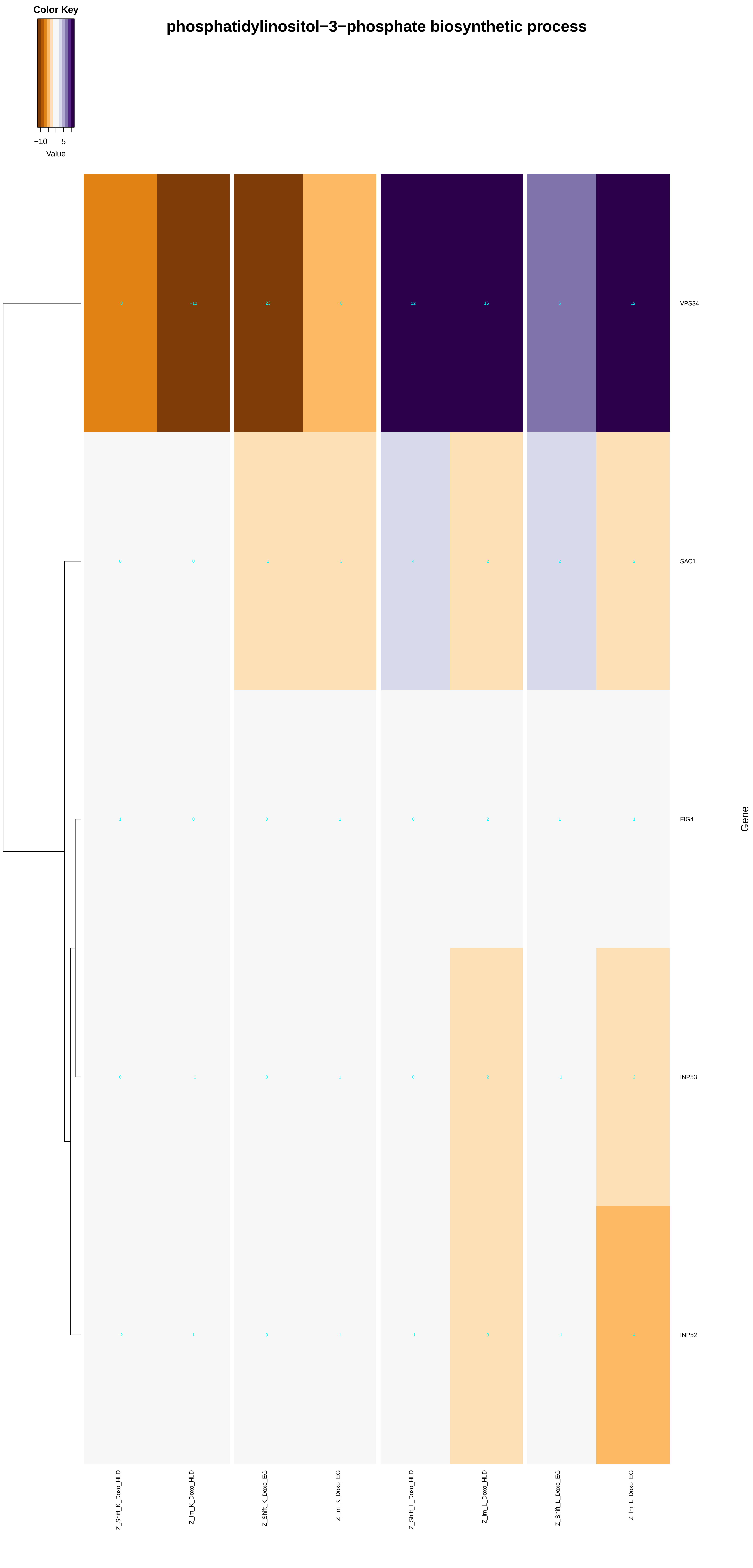

Color Key
−10
5
Value
phosphatidylinositol−3−phosphate biosynthetic process
VPS34
SAC1
FIG4
INP53
INP52
Gene
Z_Shift_K_Doxo_HLD
Z_Im_K_Doxo_HLD
Z_Shift_K_Doxo_EG
Z_Im_K_Doxo_EG
Z_Shift_L_Doxo_HLD
Z_Im_L_Doxo_HLD
Z_Shift_L_Doxo_EG
Z_Im_L_Doxo_EG
