## Supplementary figures and images for "A yeast phenomic model for the influence of Warburg metabolism on genetic buffering of doxorubicin"

### B - AcrossTissue.pdf

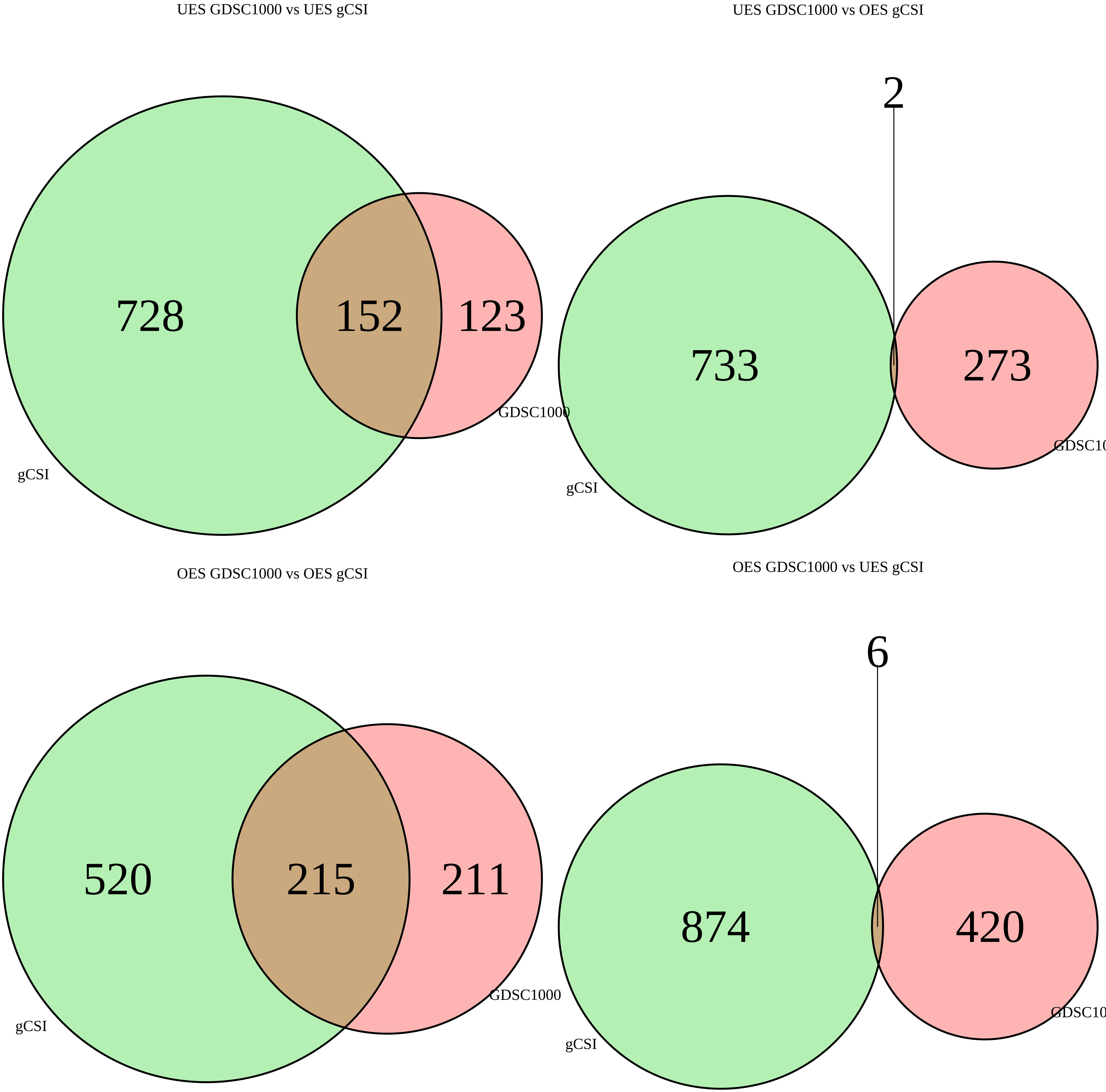

### D - gCSI_Heatmaps.pdf

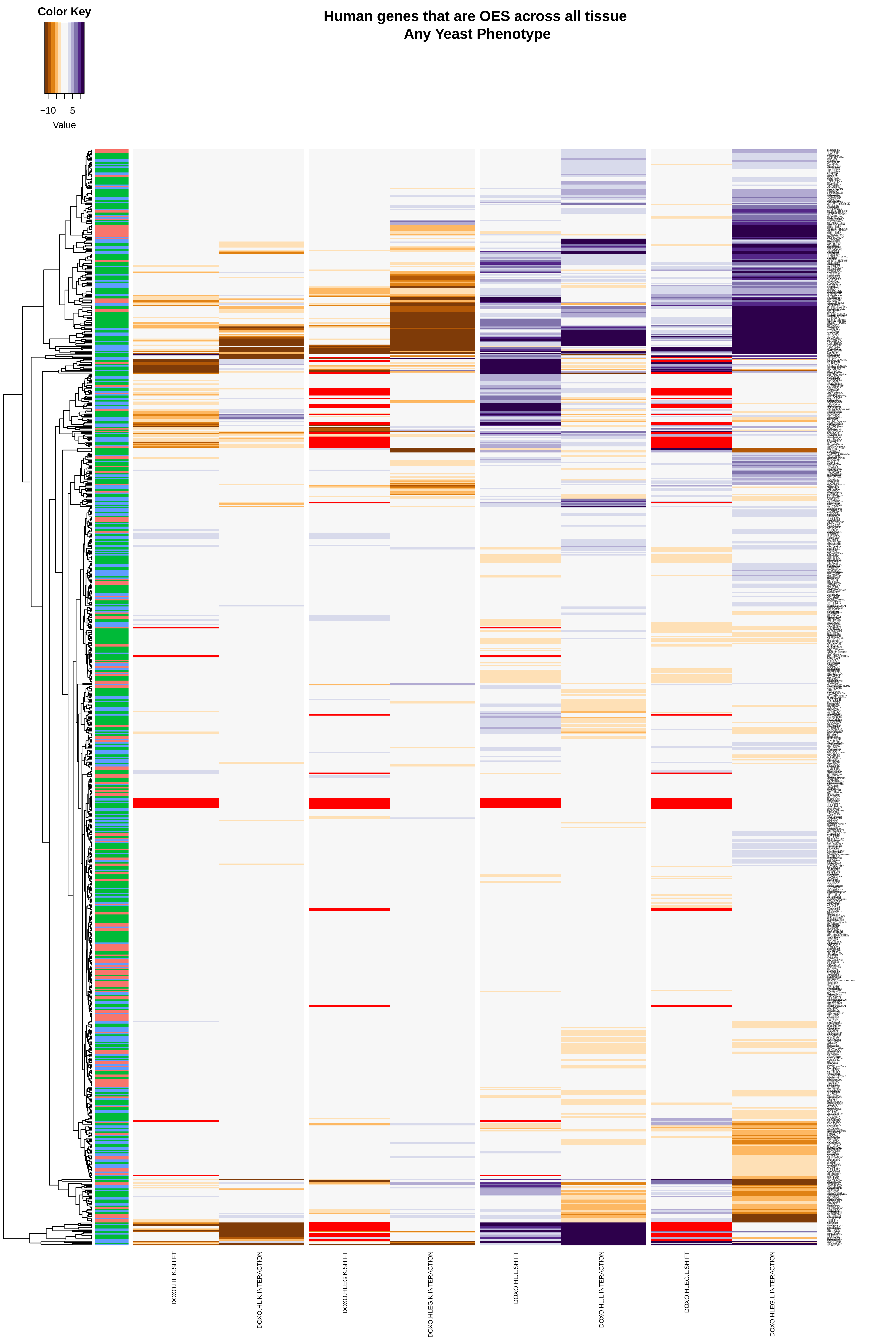

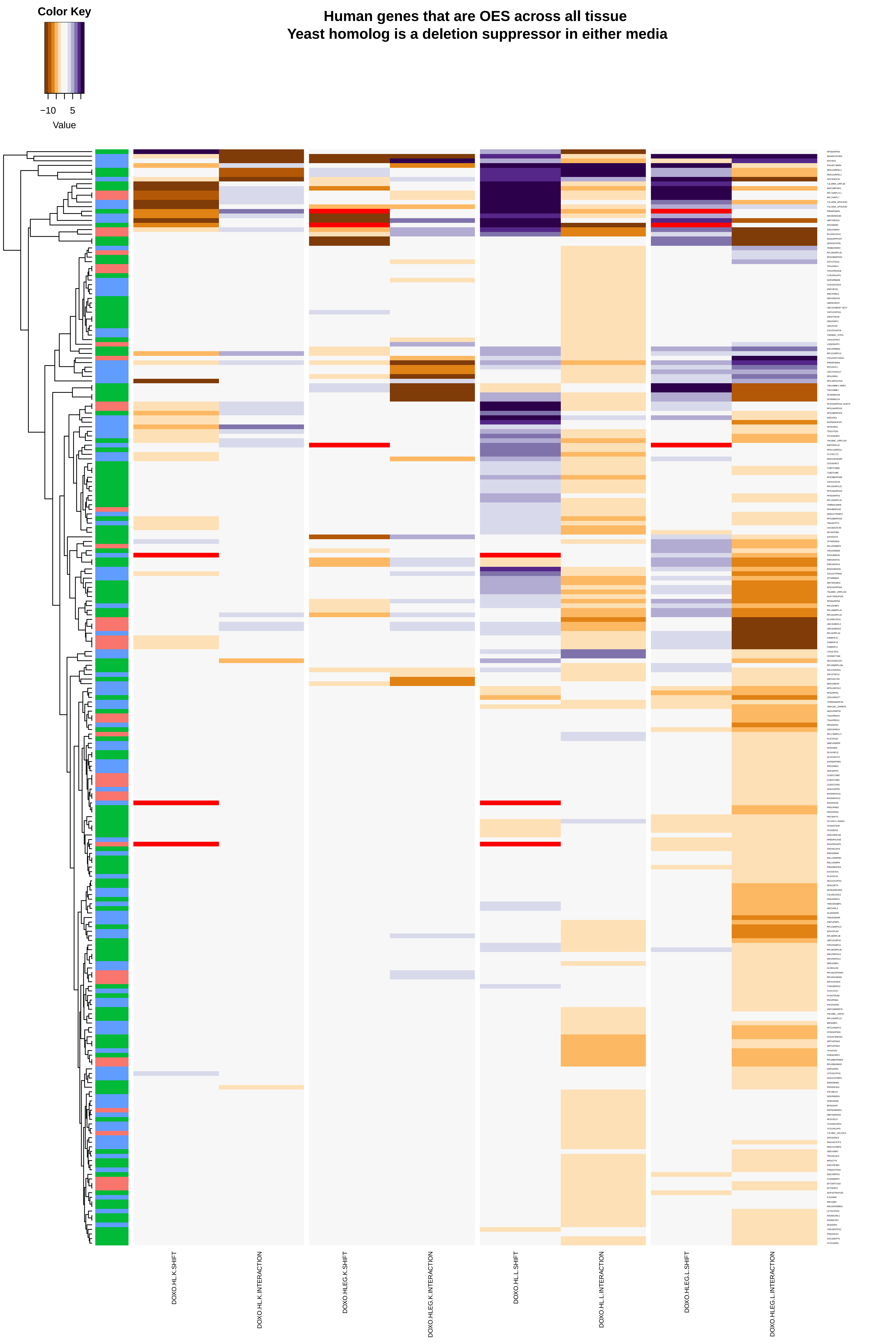

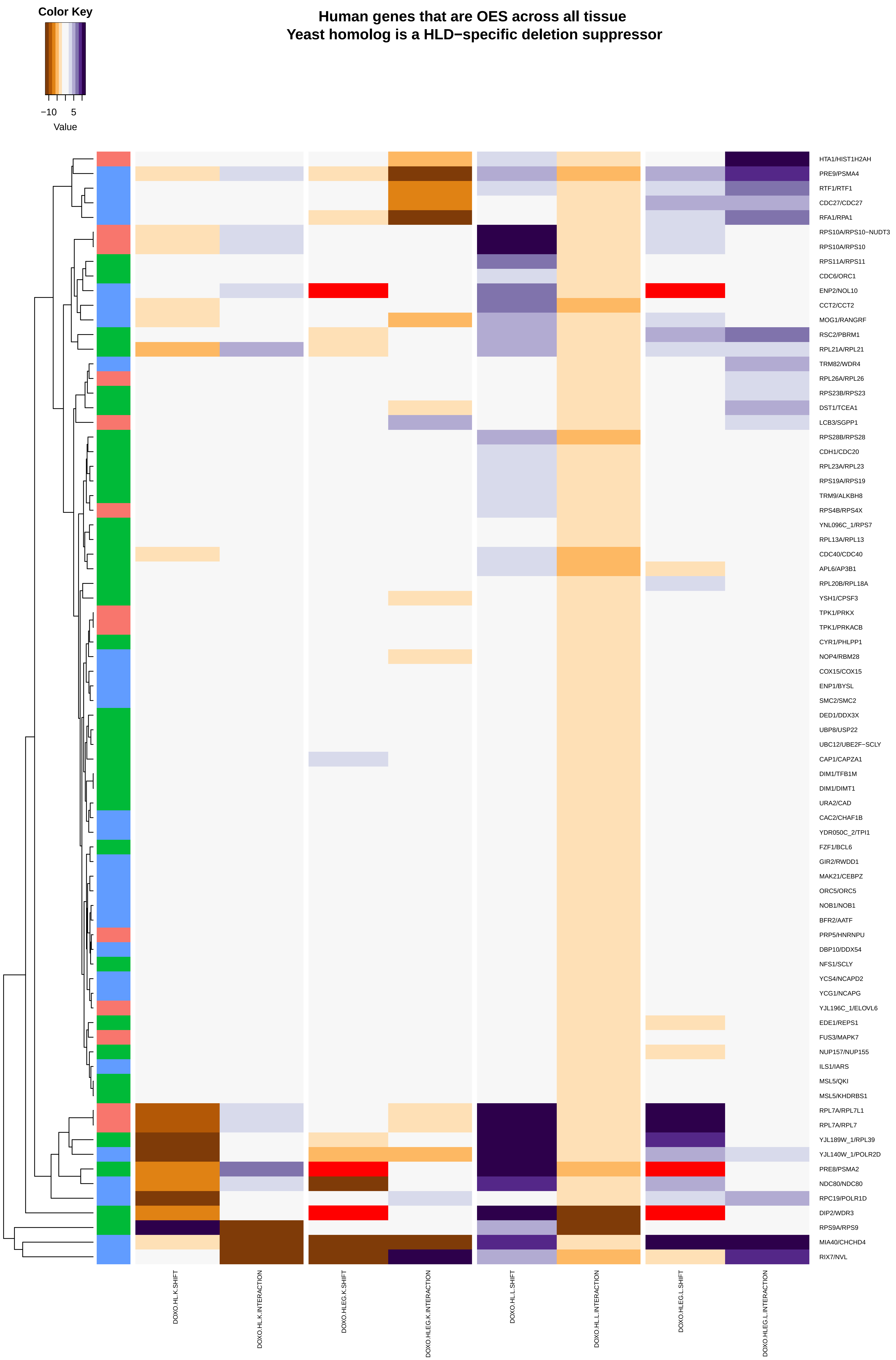

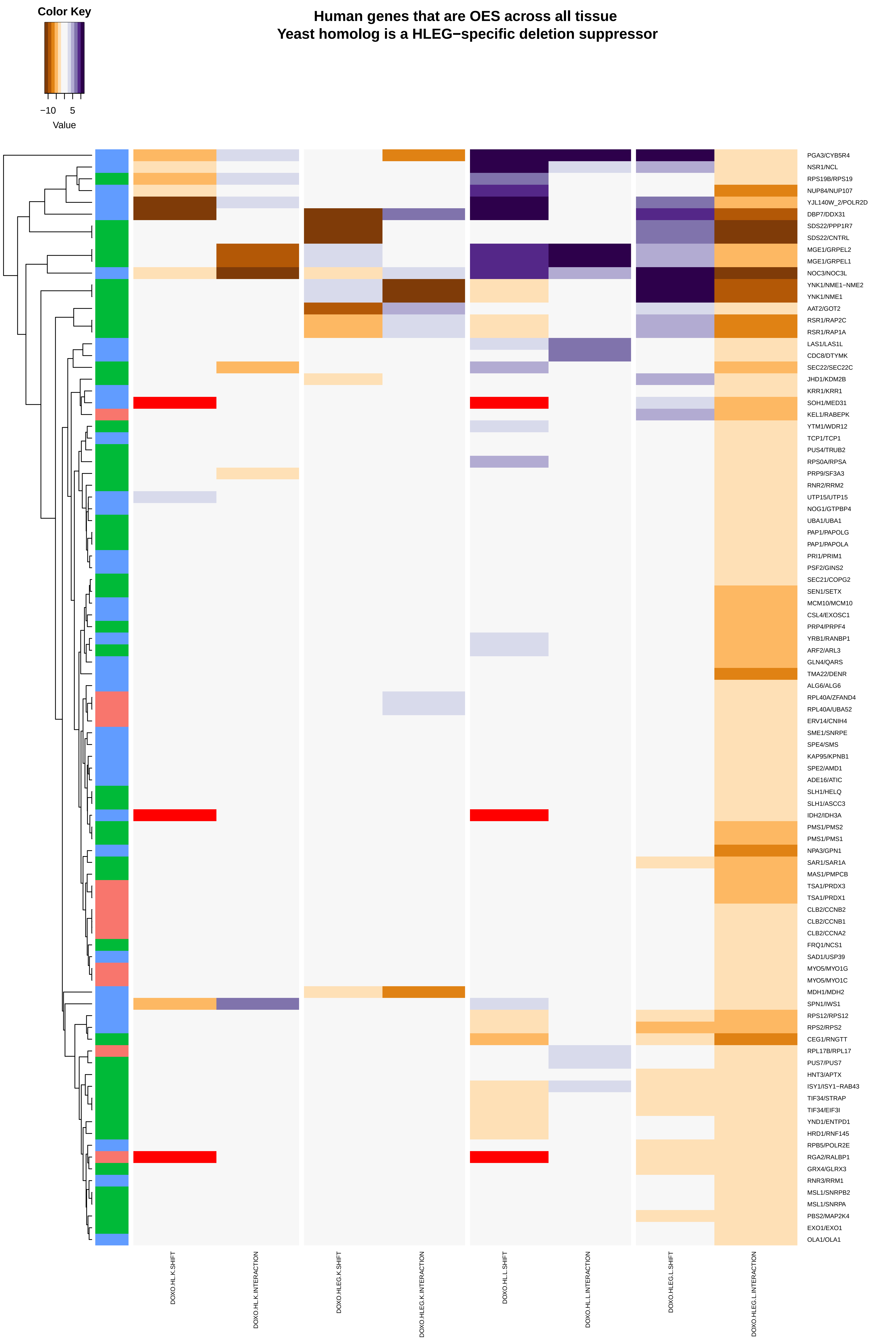

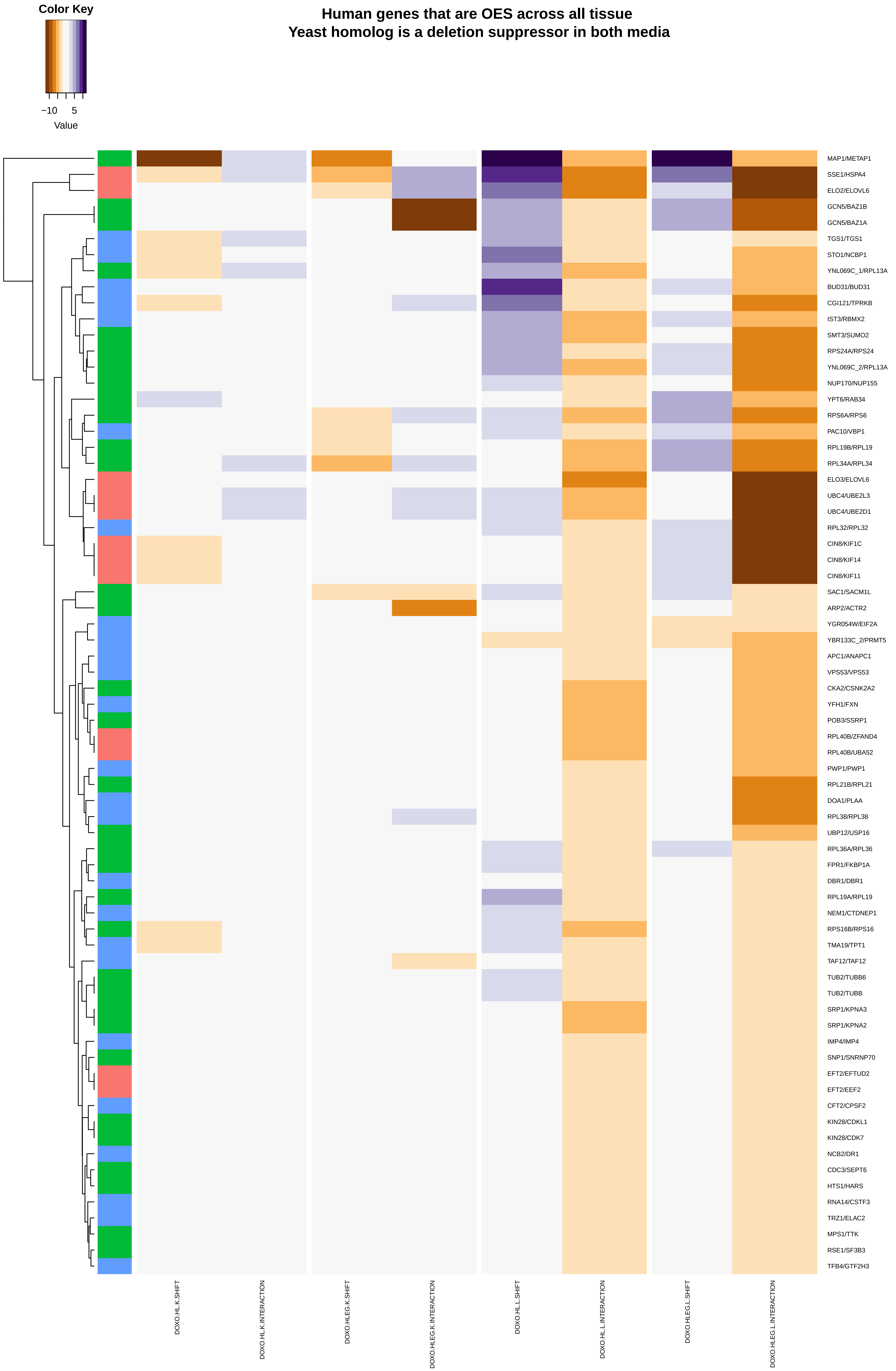

##### Human genes that are UES across all tissue
##### Any Yeast Phenotype

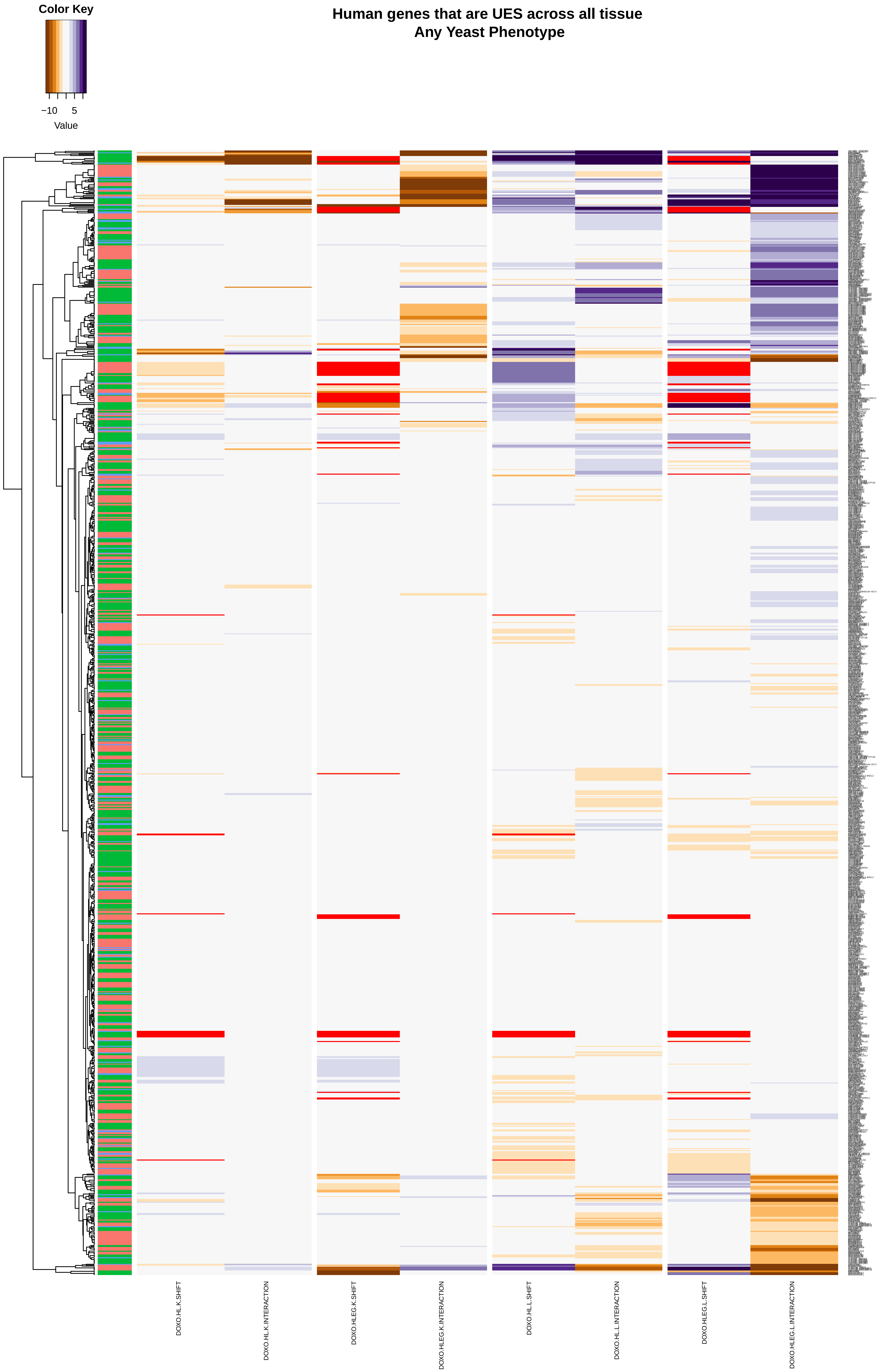

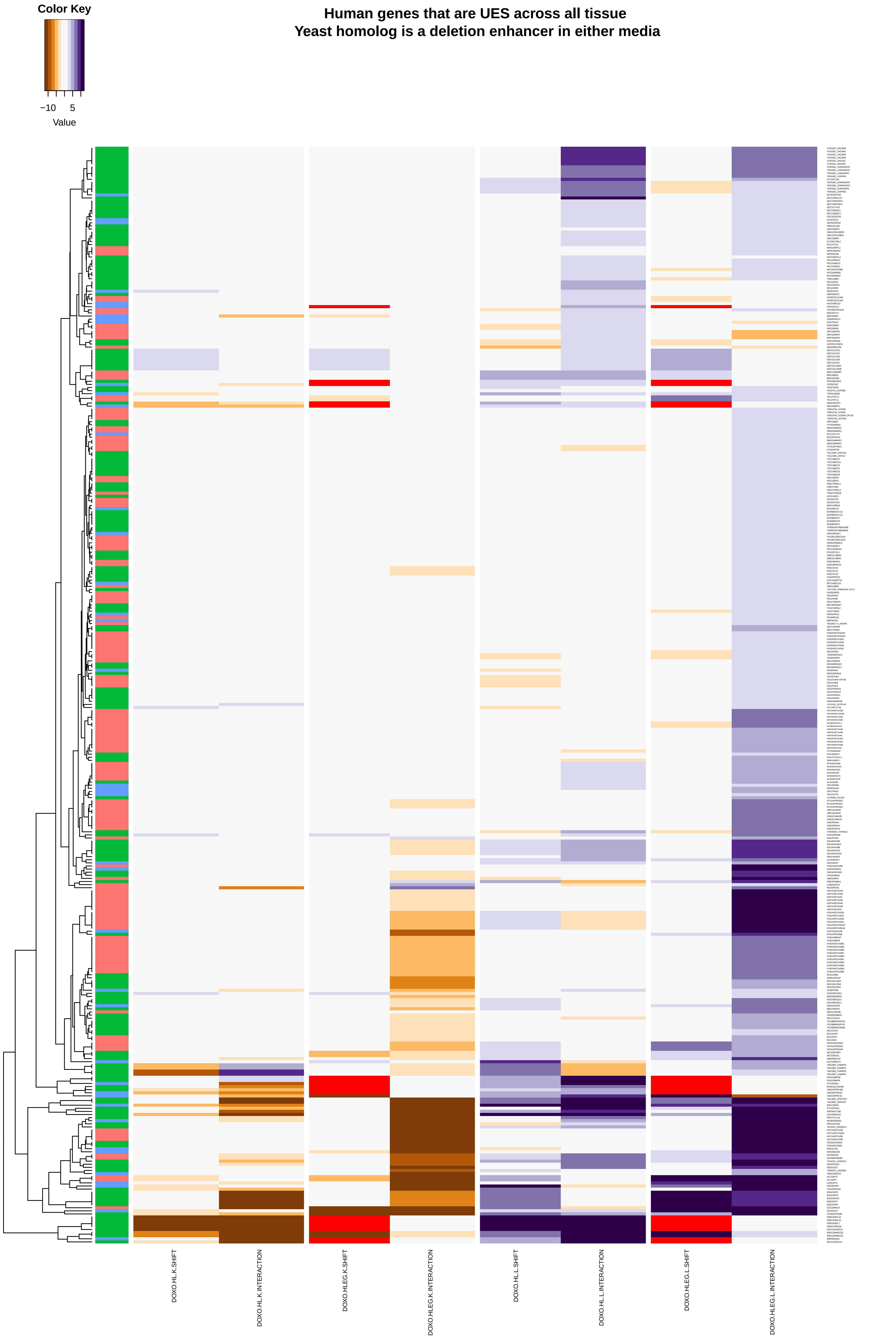

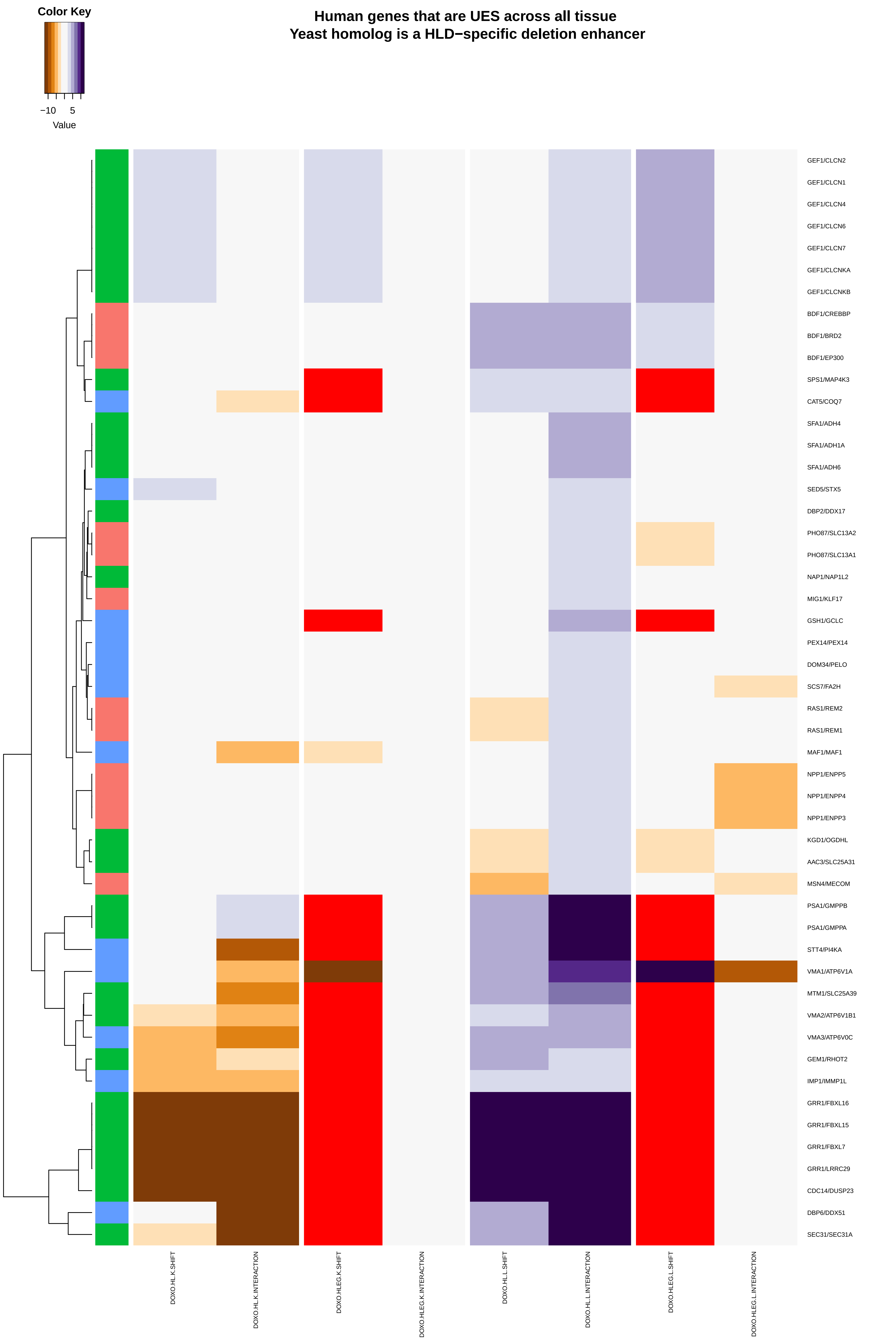

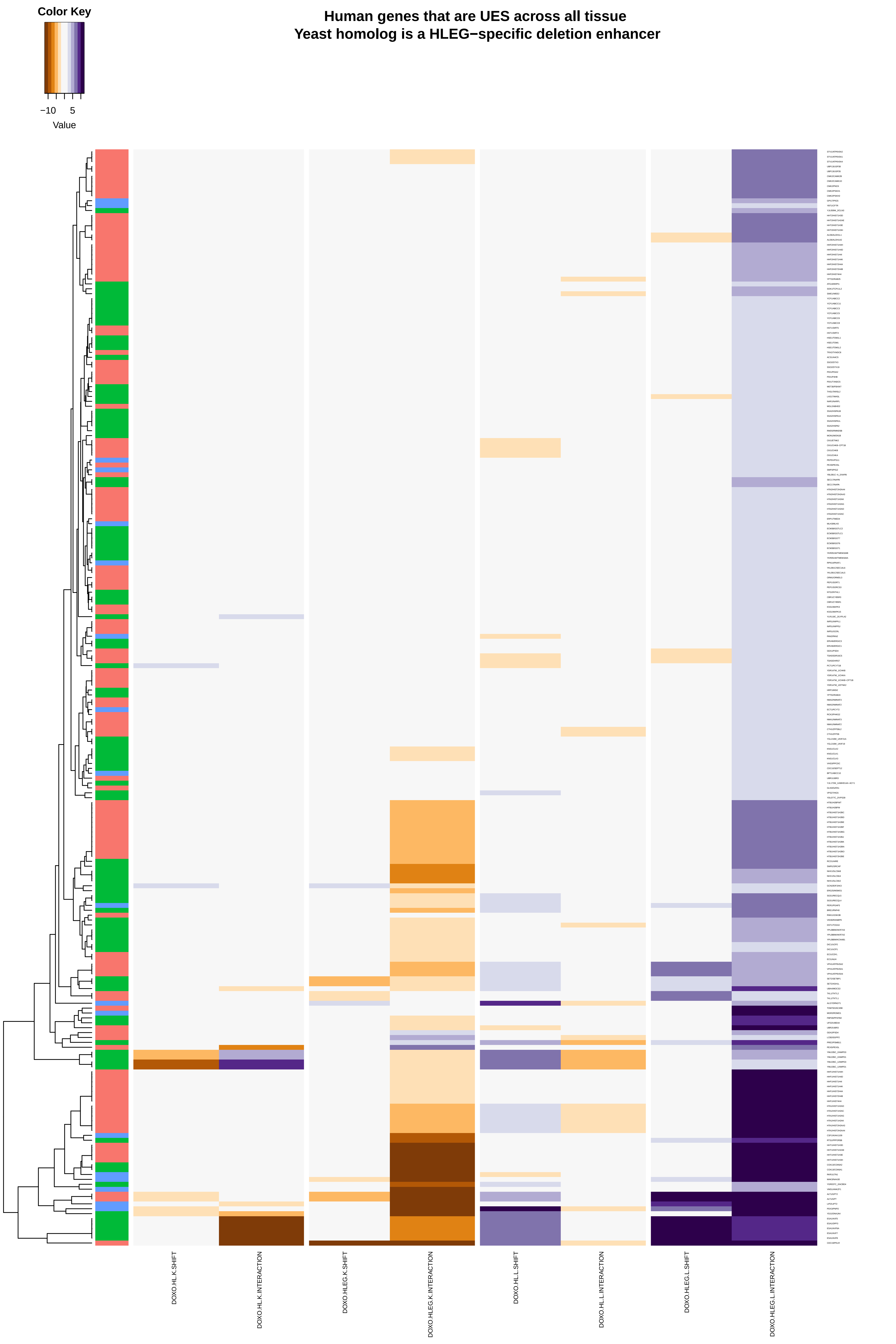

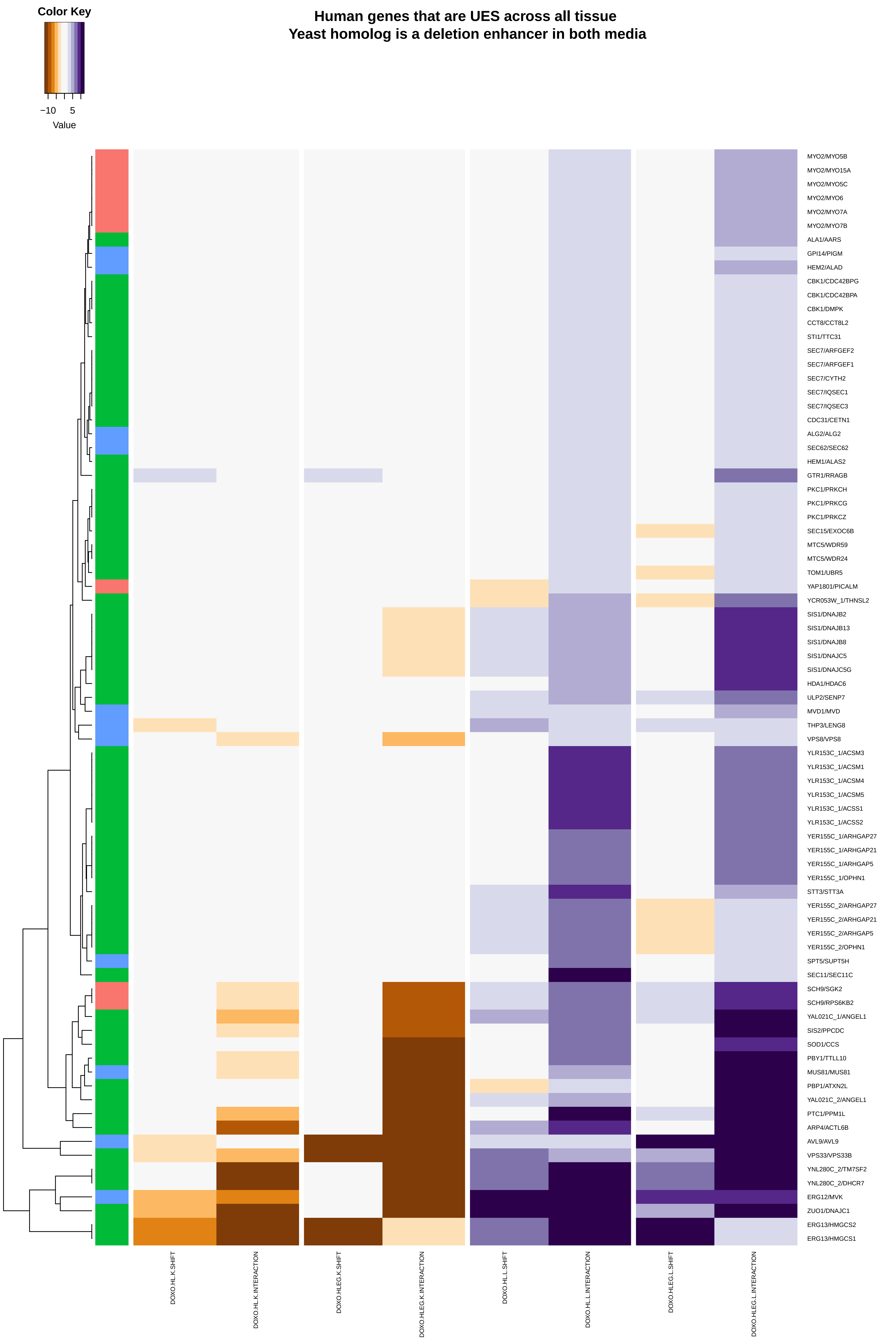

### mitochondrial_double-strand_break_repair.pdf

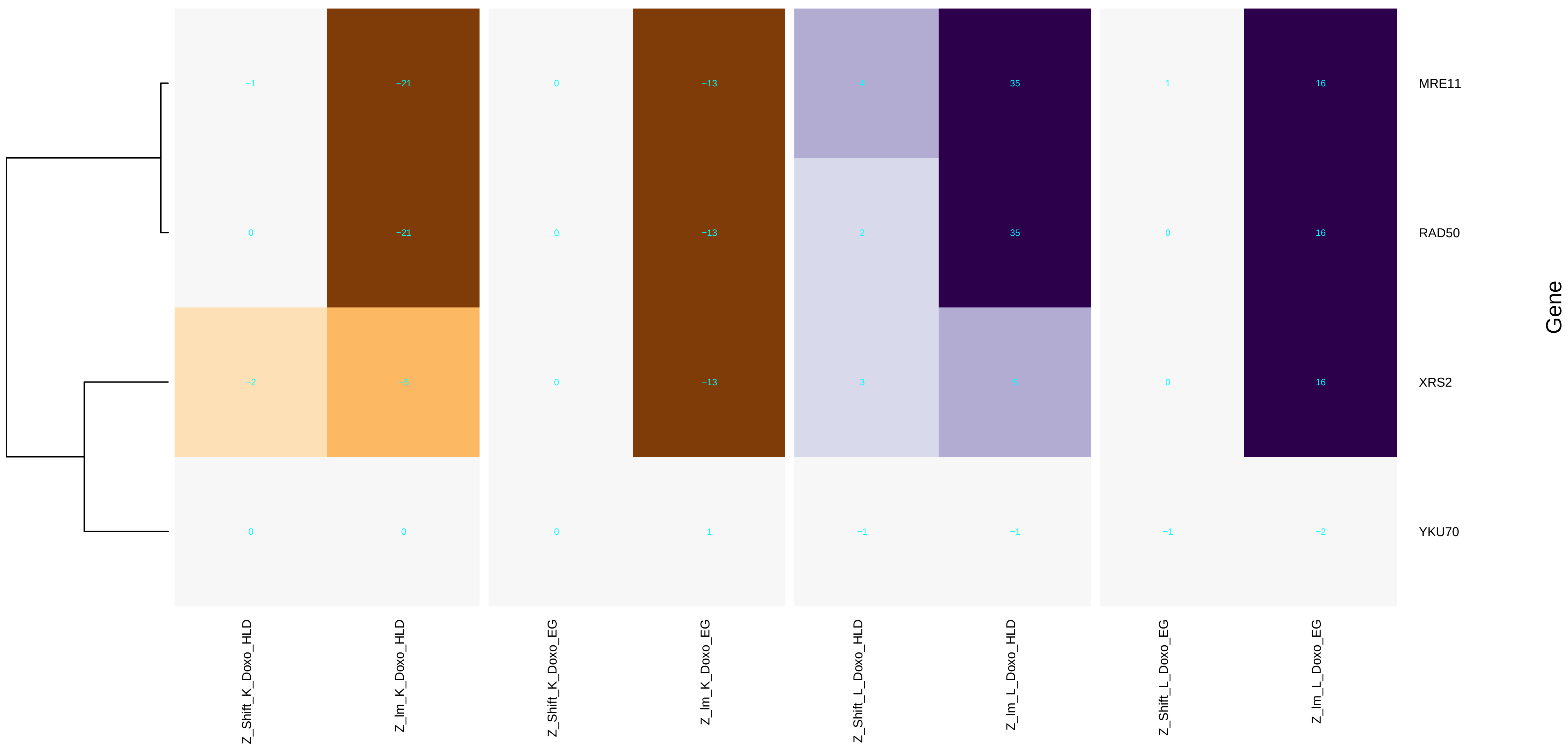

### mitochondrial_double-strand_break_repair_via_homologous_recombination.pdf

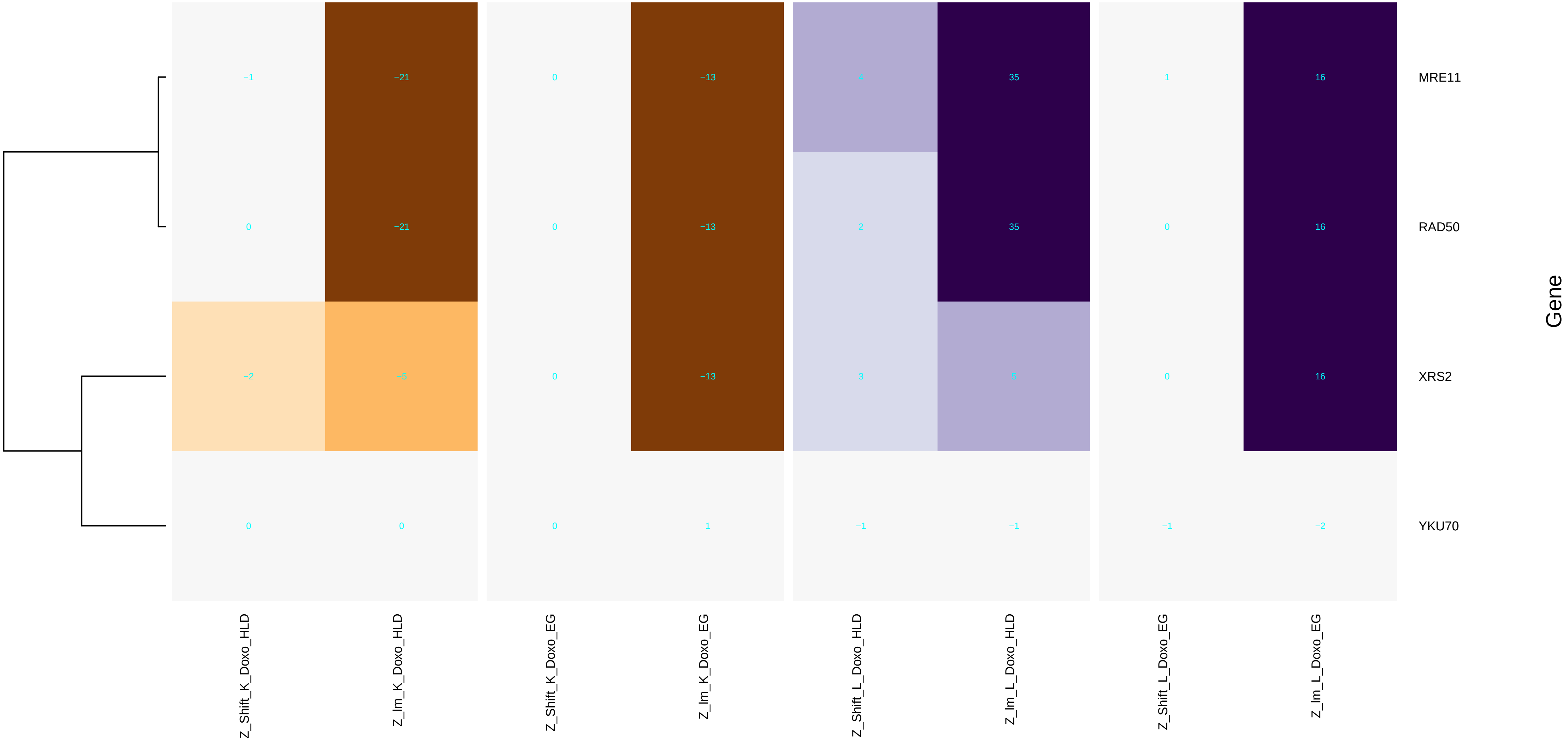

### mitochondrial_genome_maintenance.pdf

#### mitochondrial genome maintenance

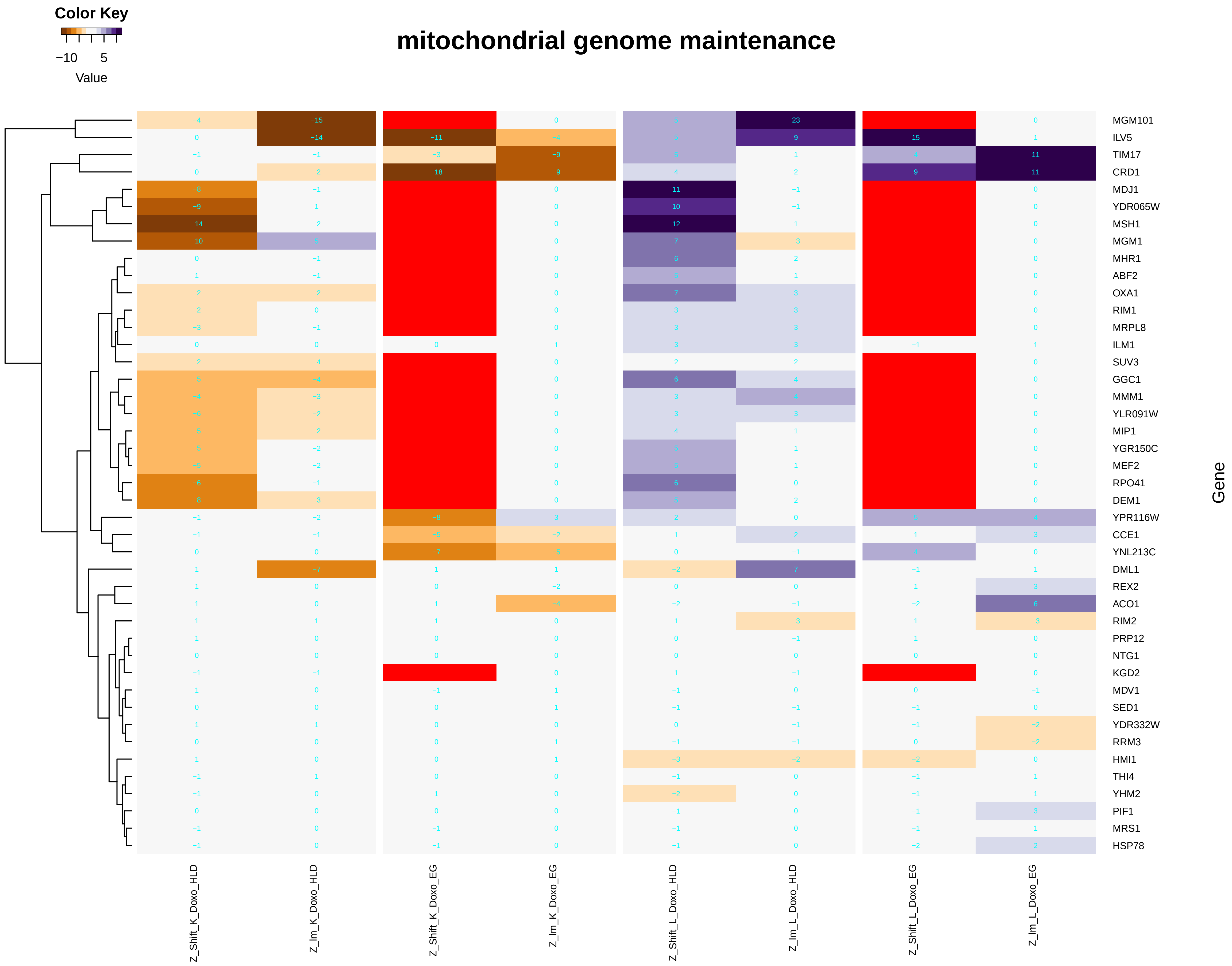

### mitochondrial_membrane_organization.pdf

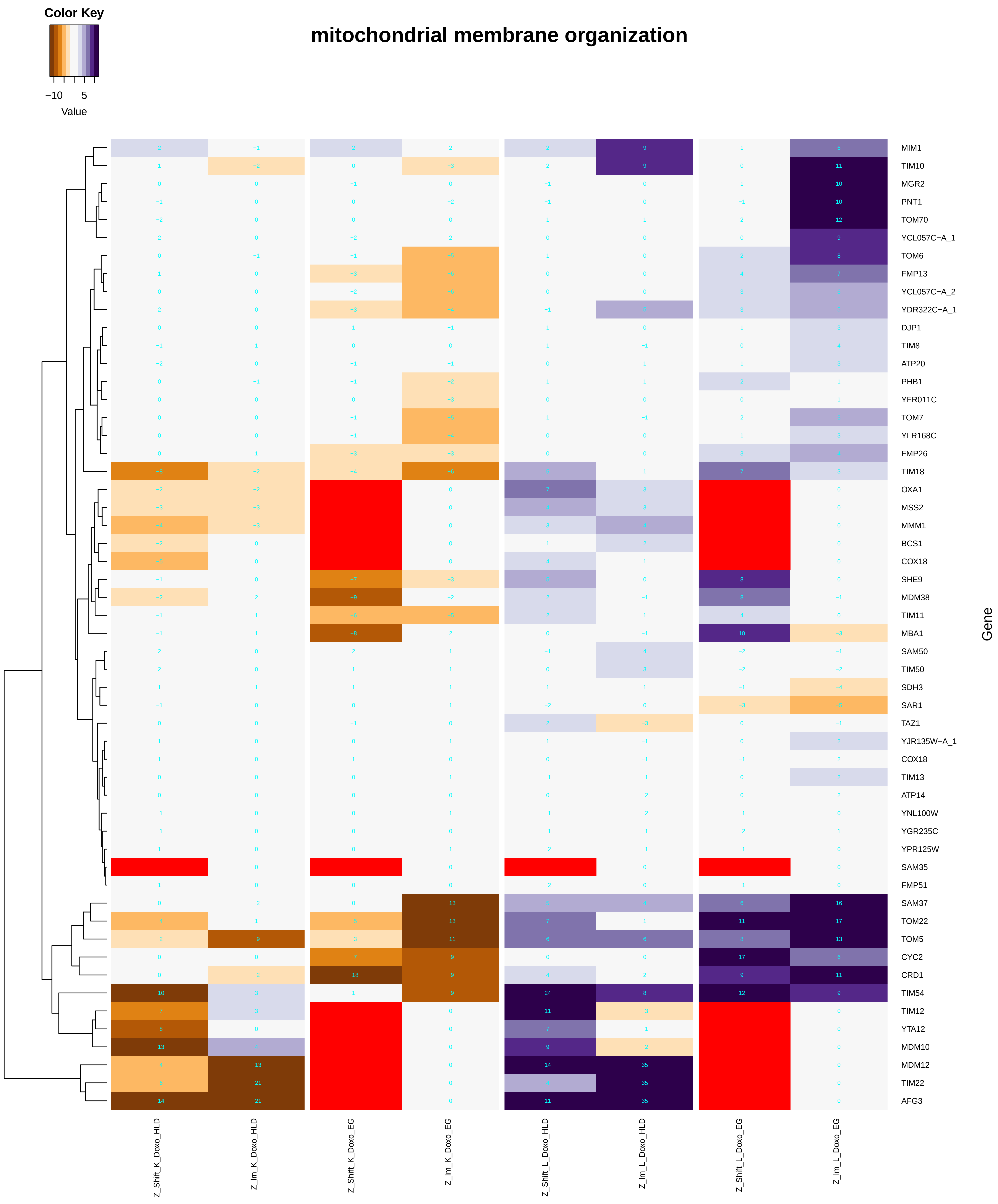

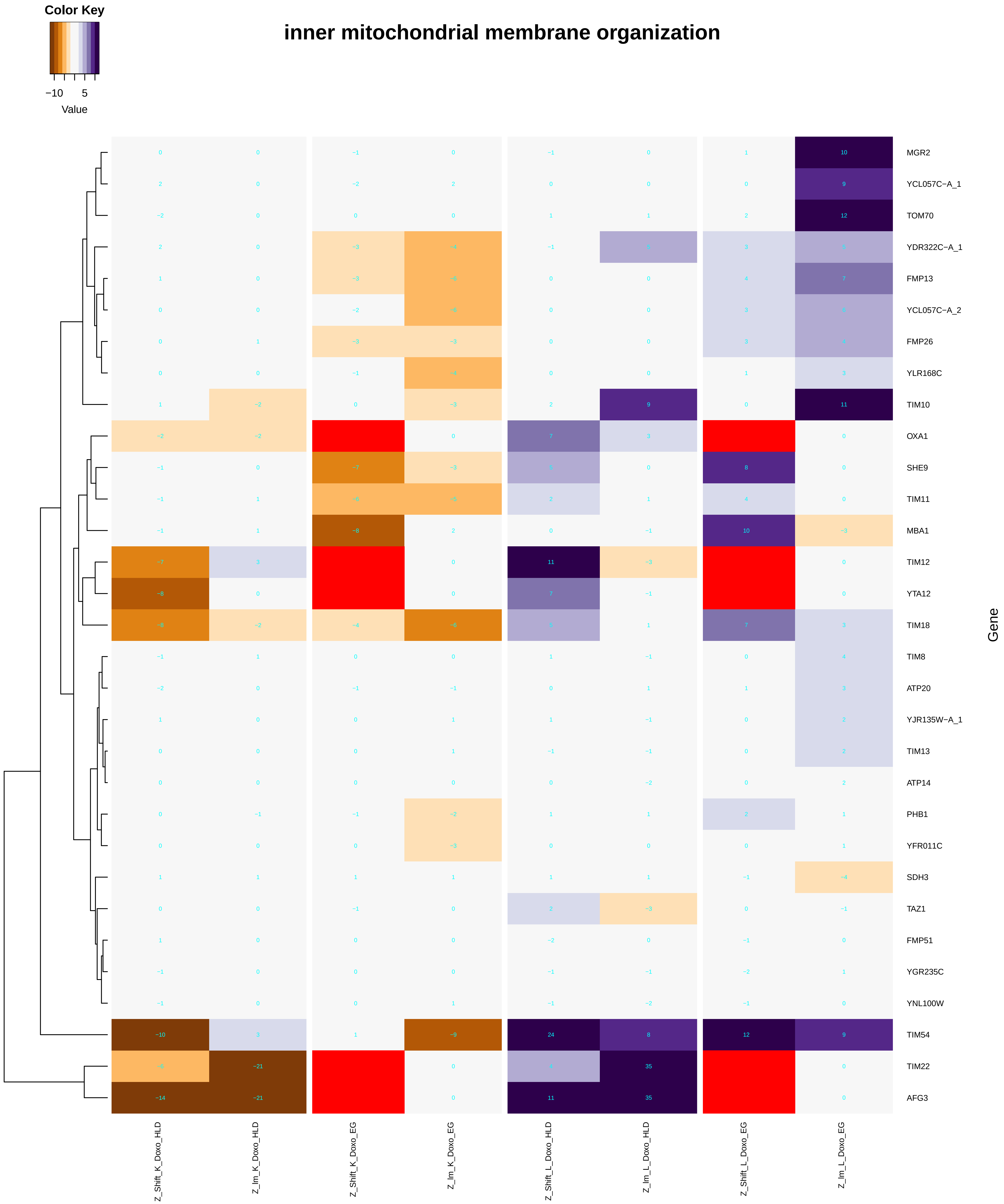

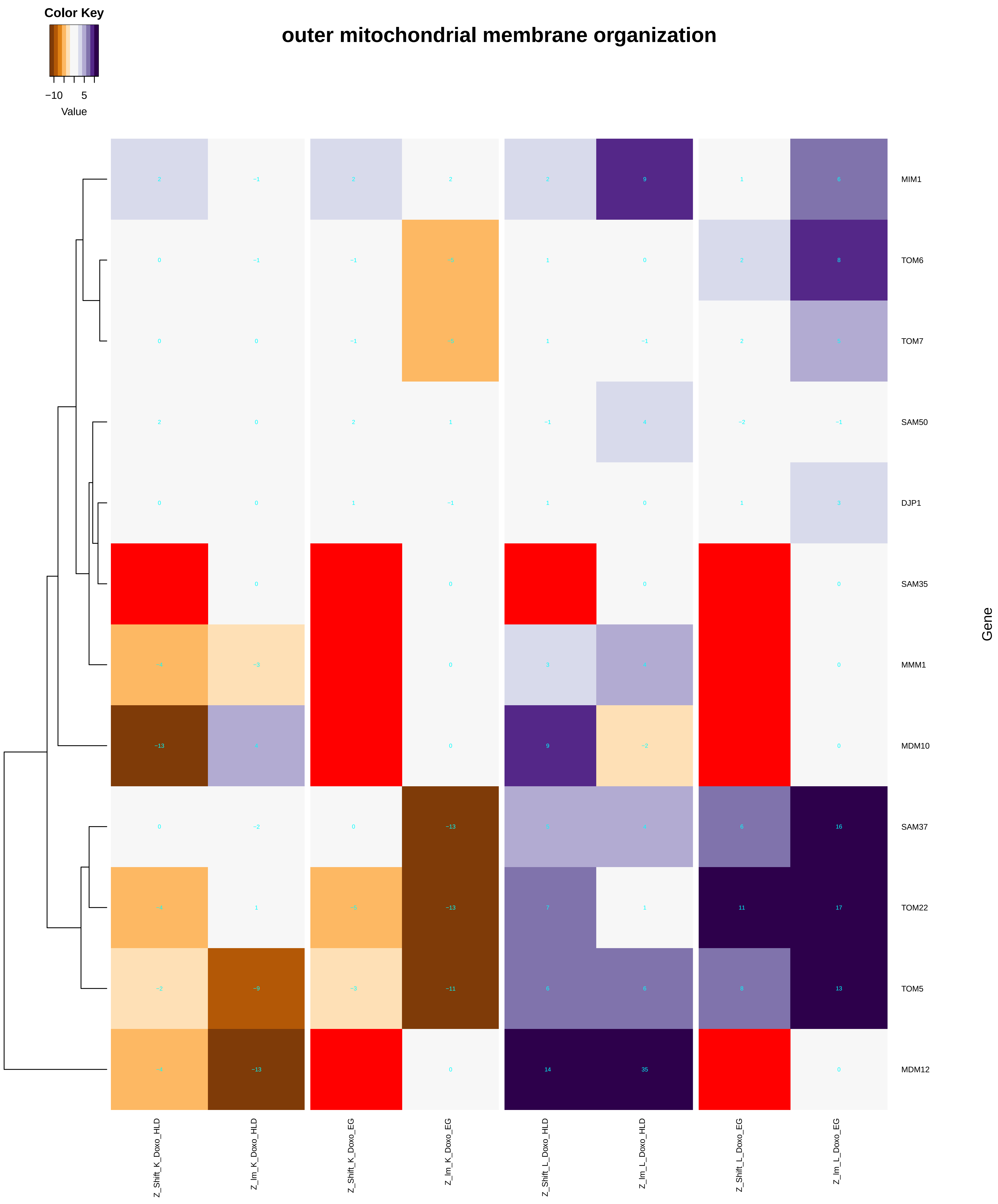

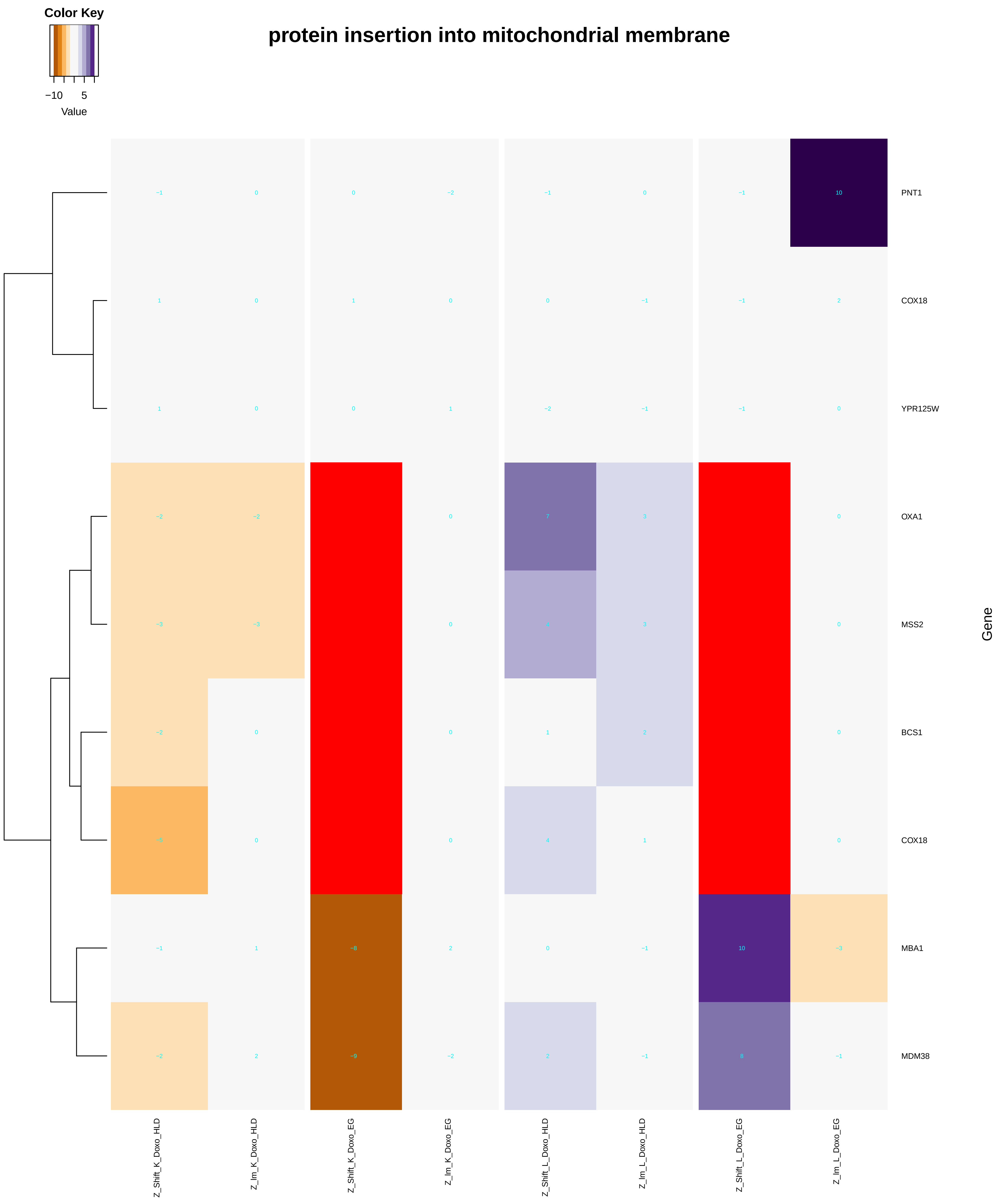

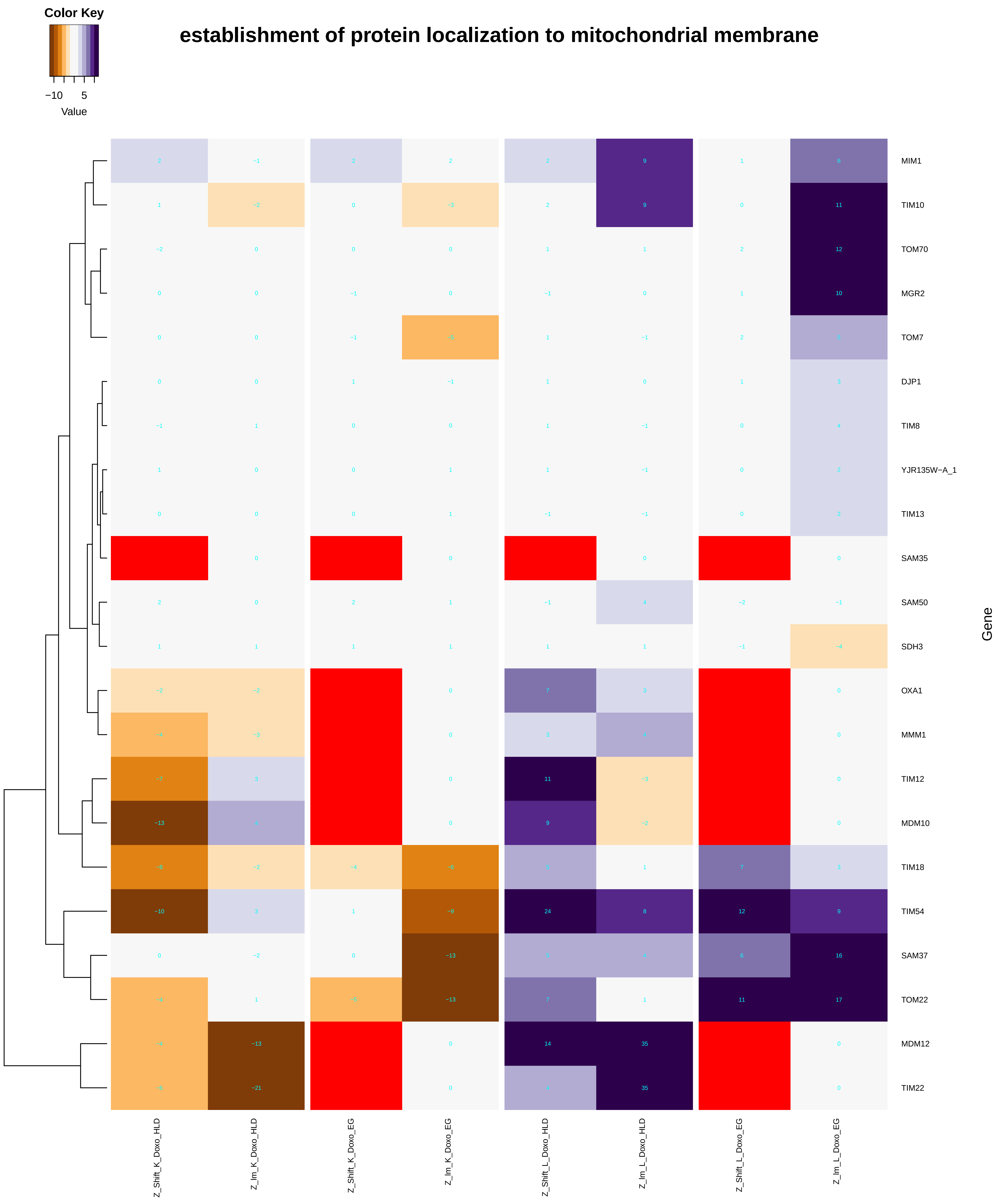

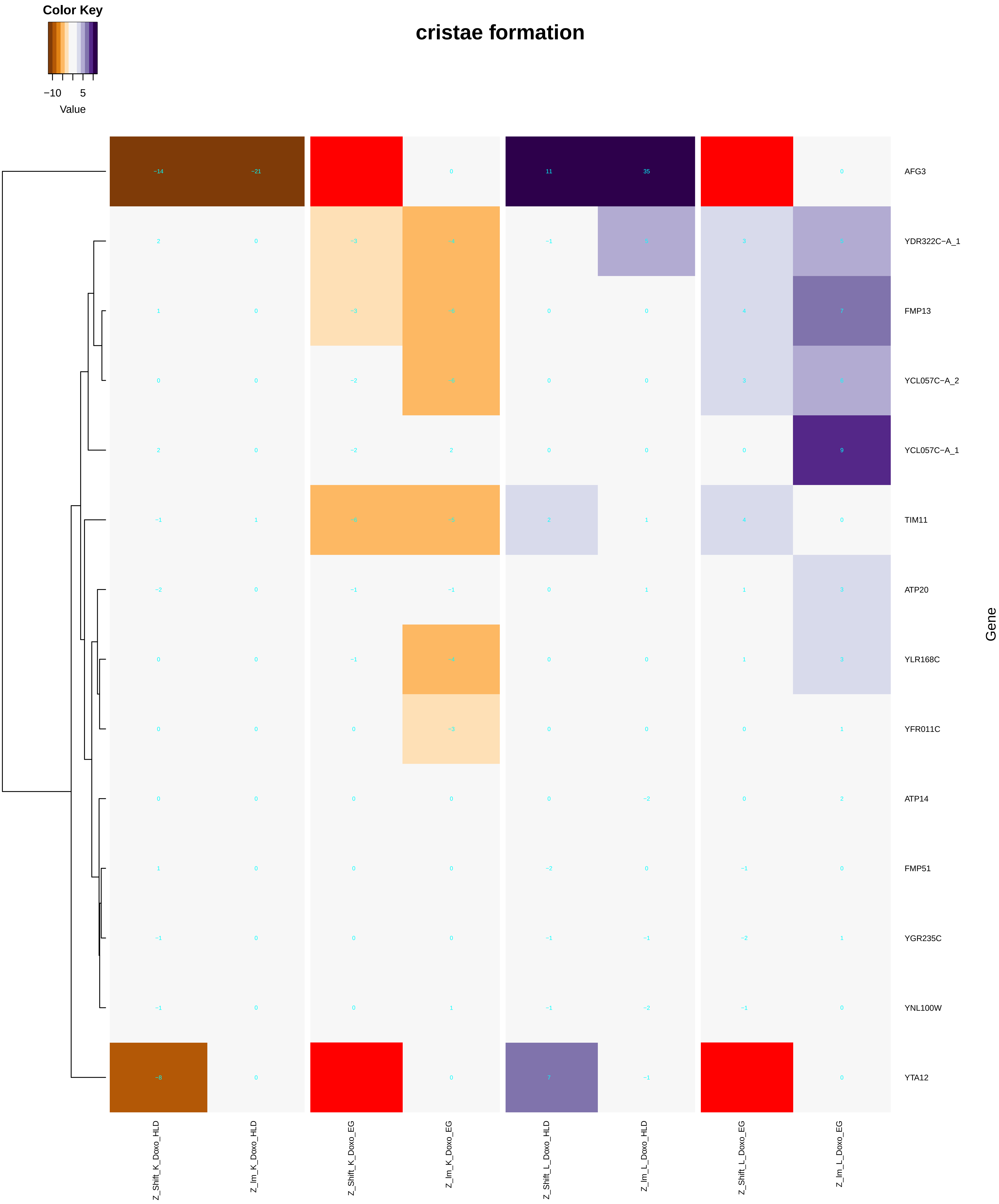

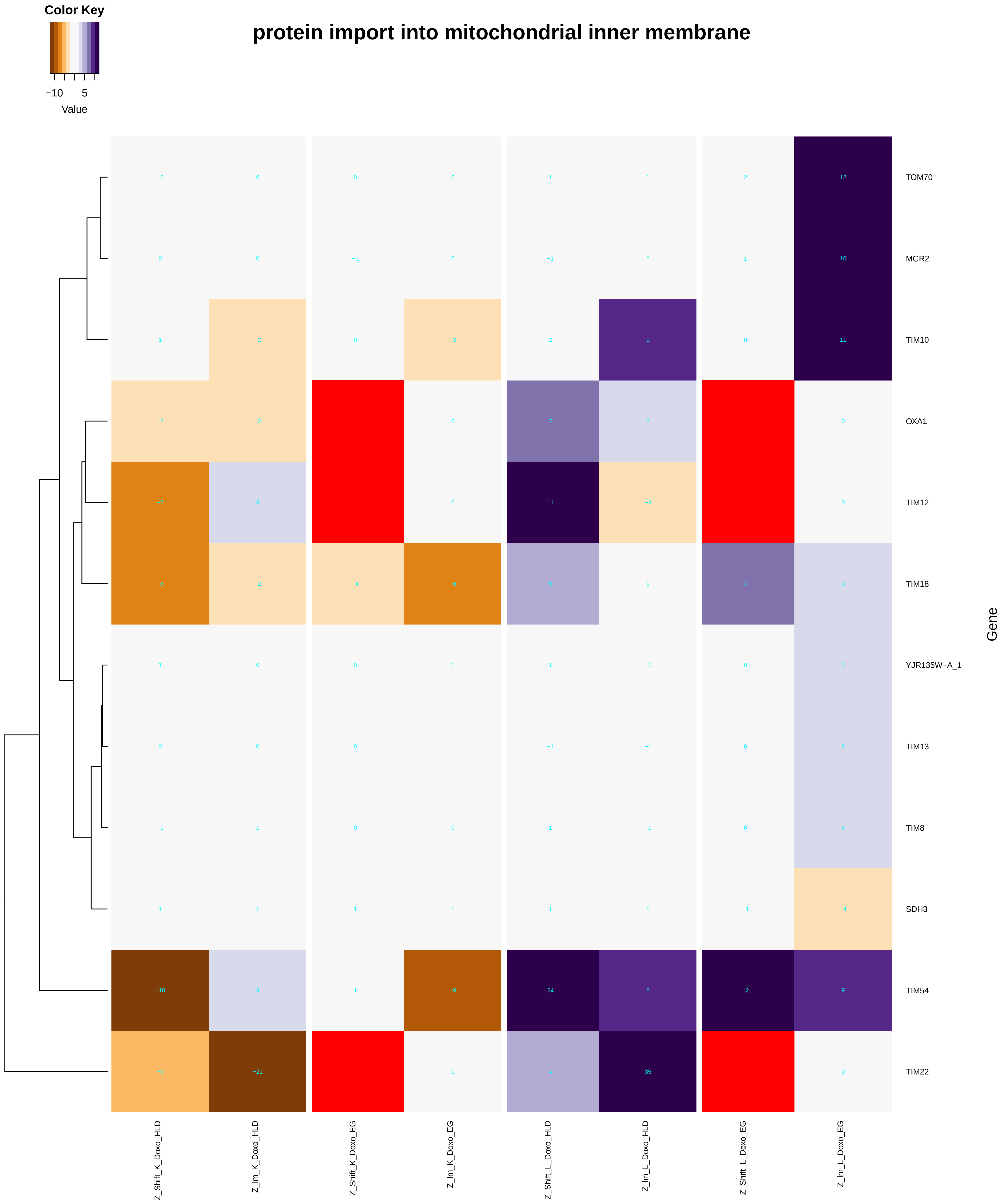

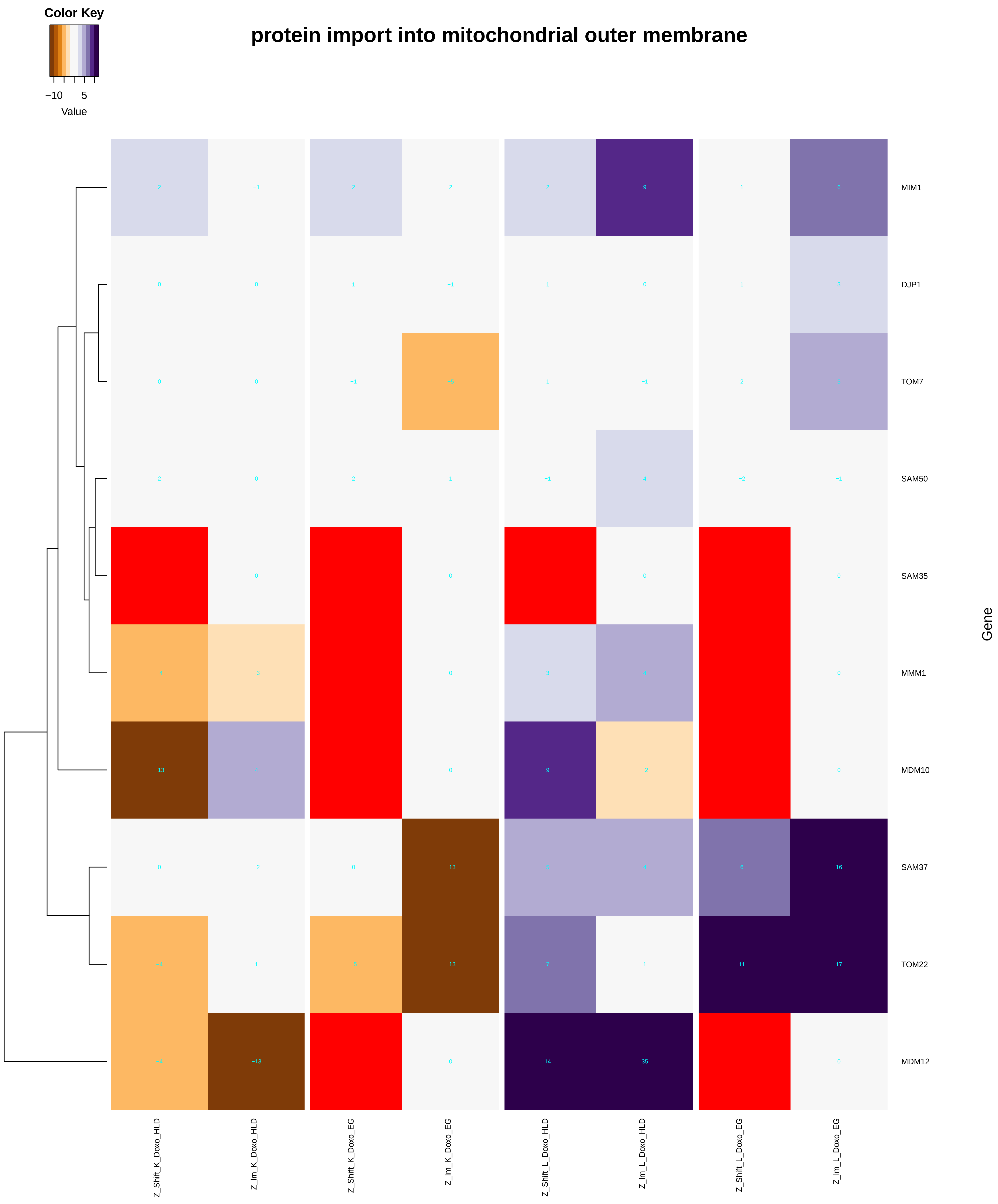

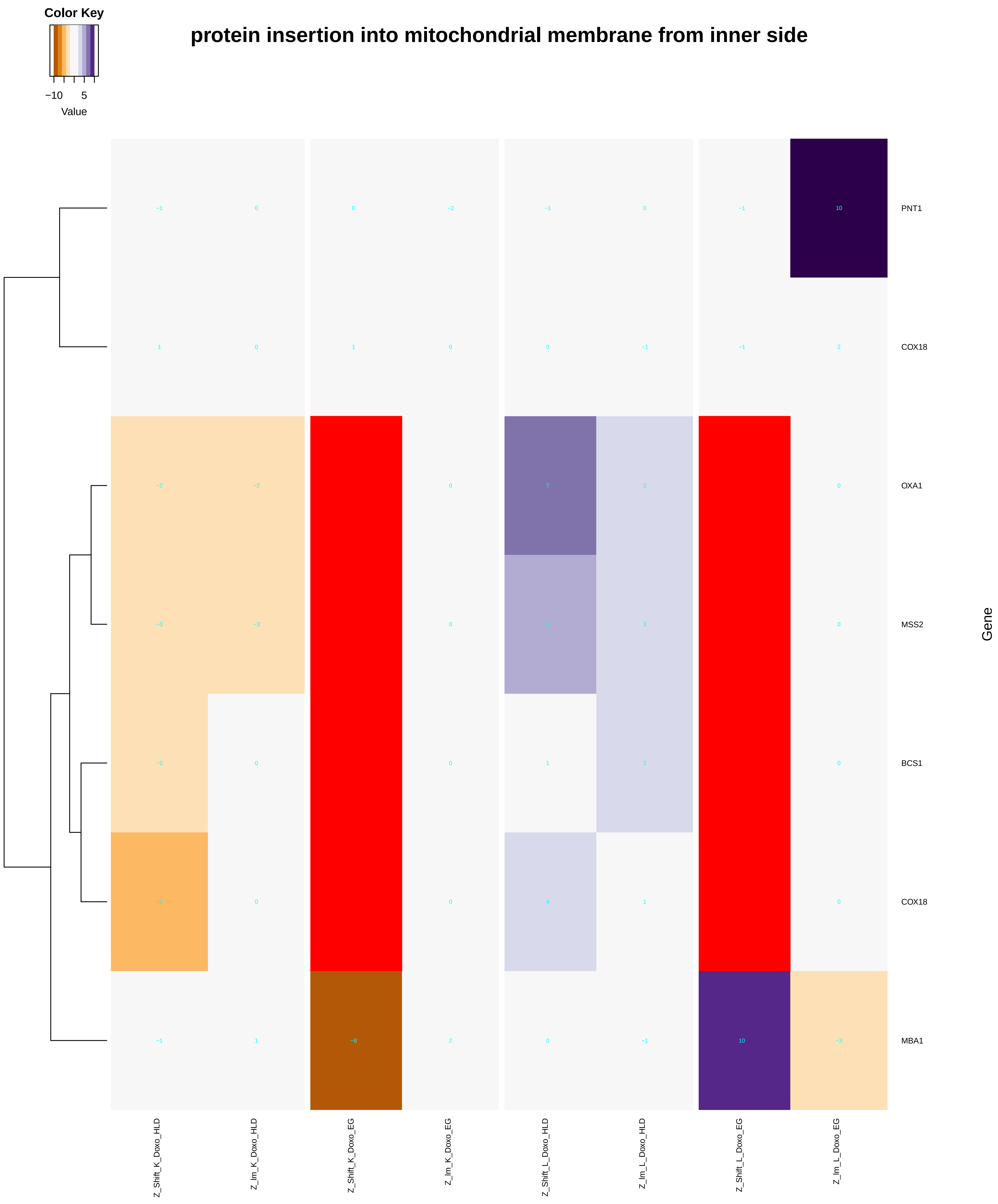

### mitochondrial_protein_processing.pdf

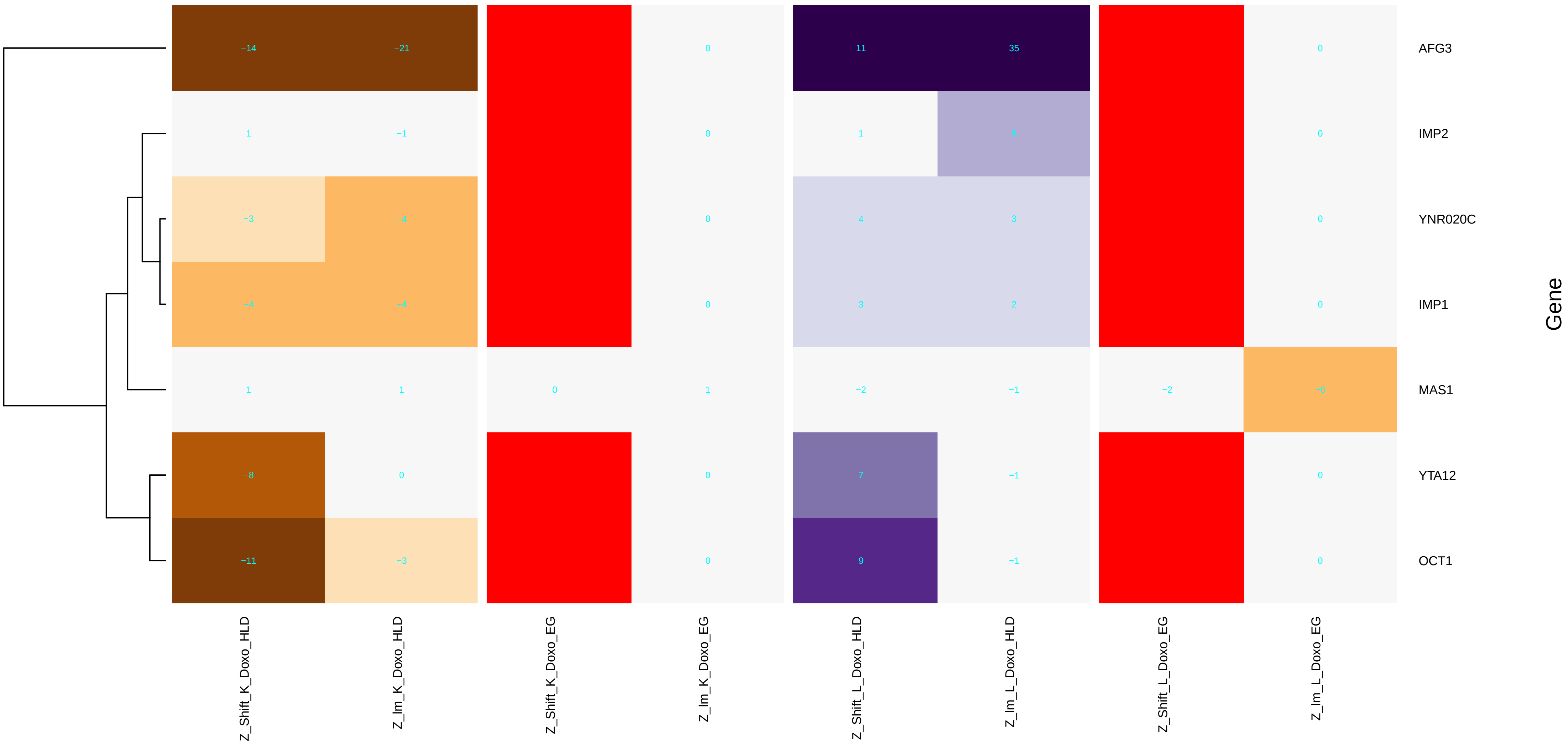

### mitochondrial_respiratory_chain_complex_assembly.pdf

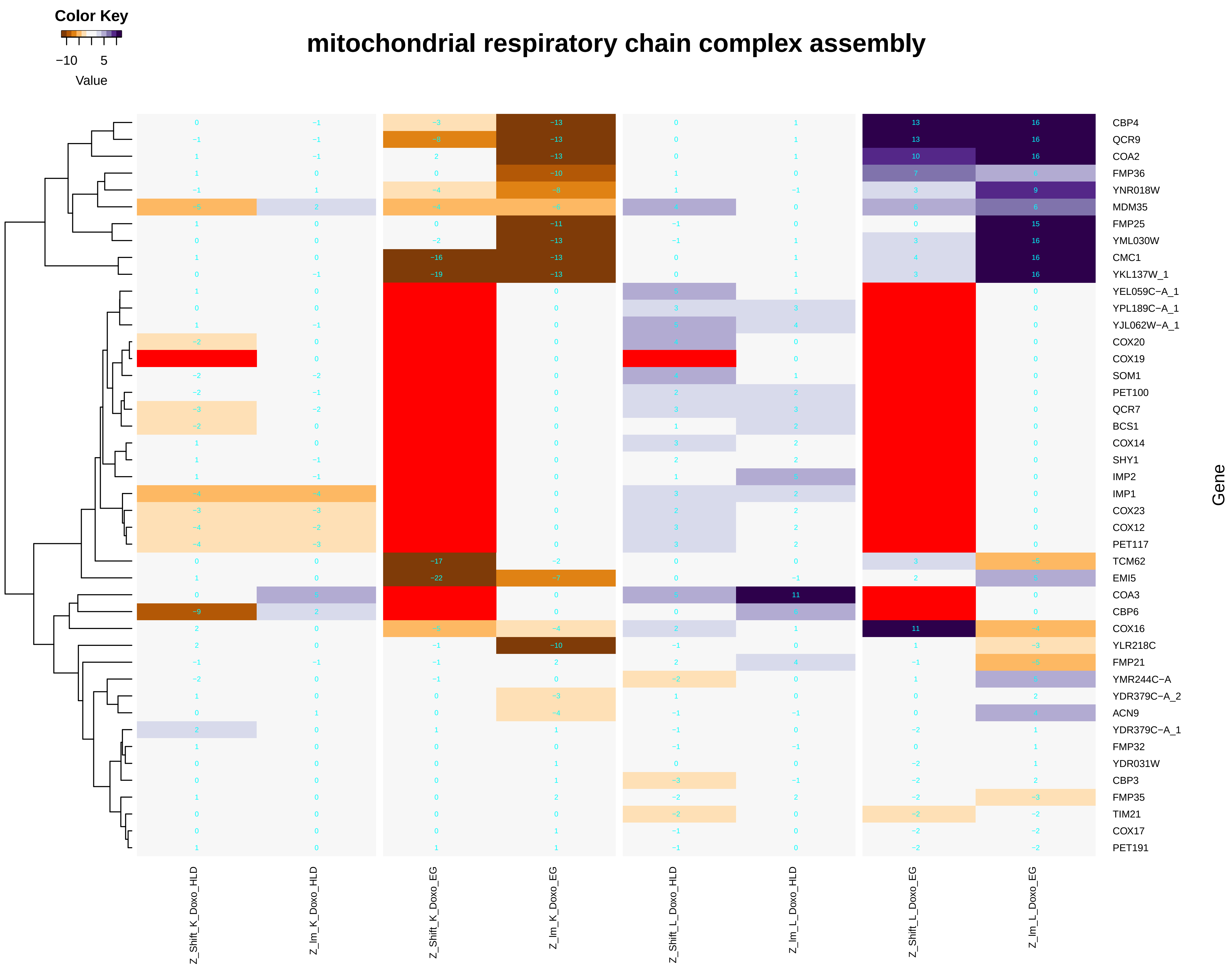

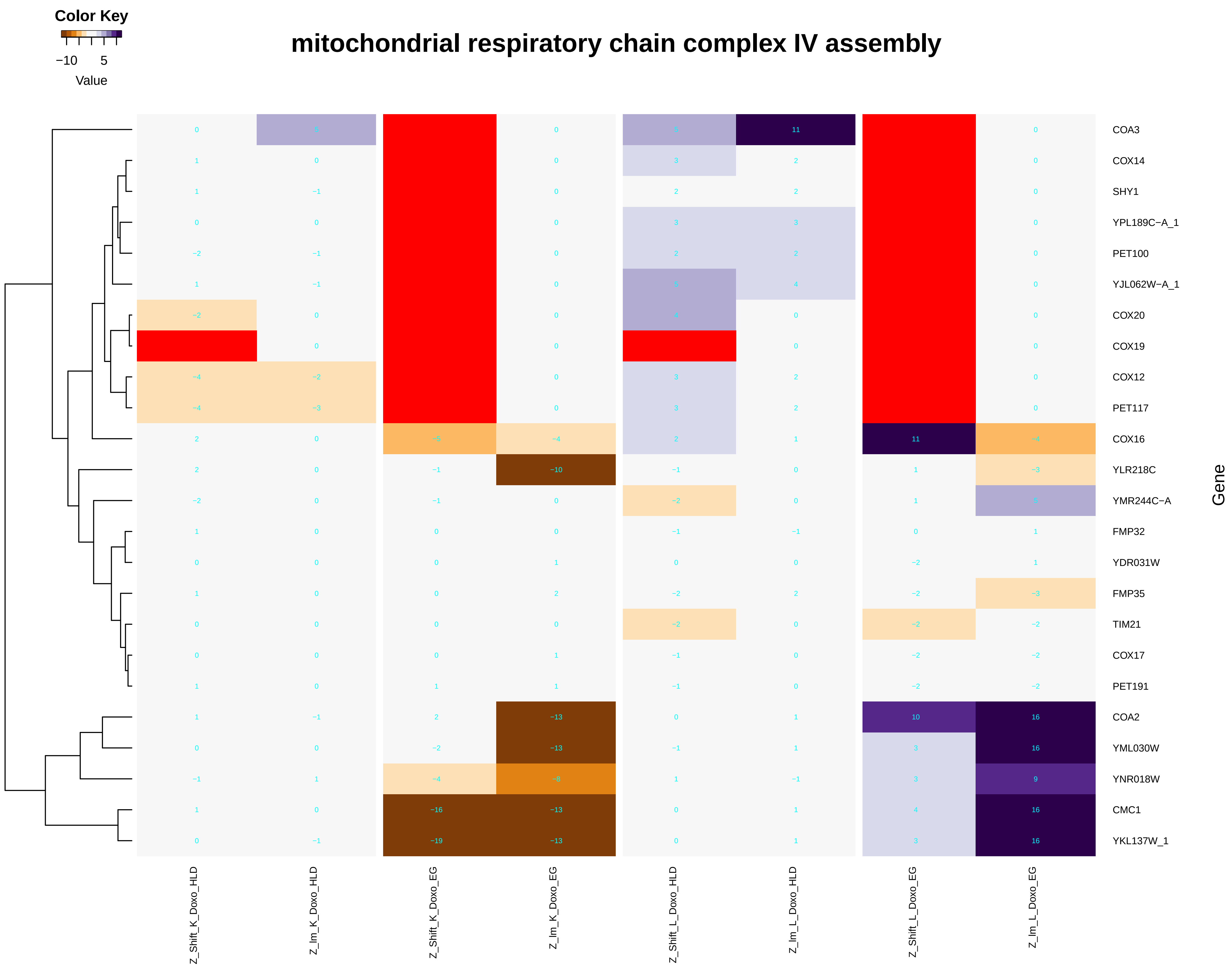

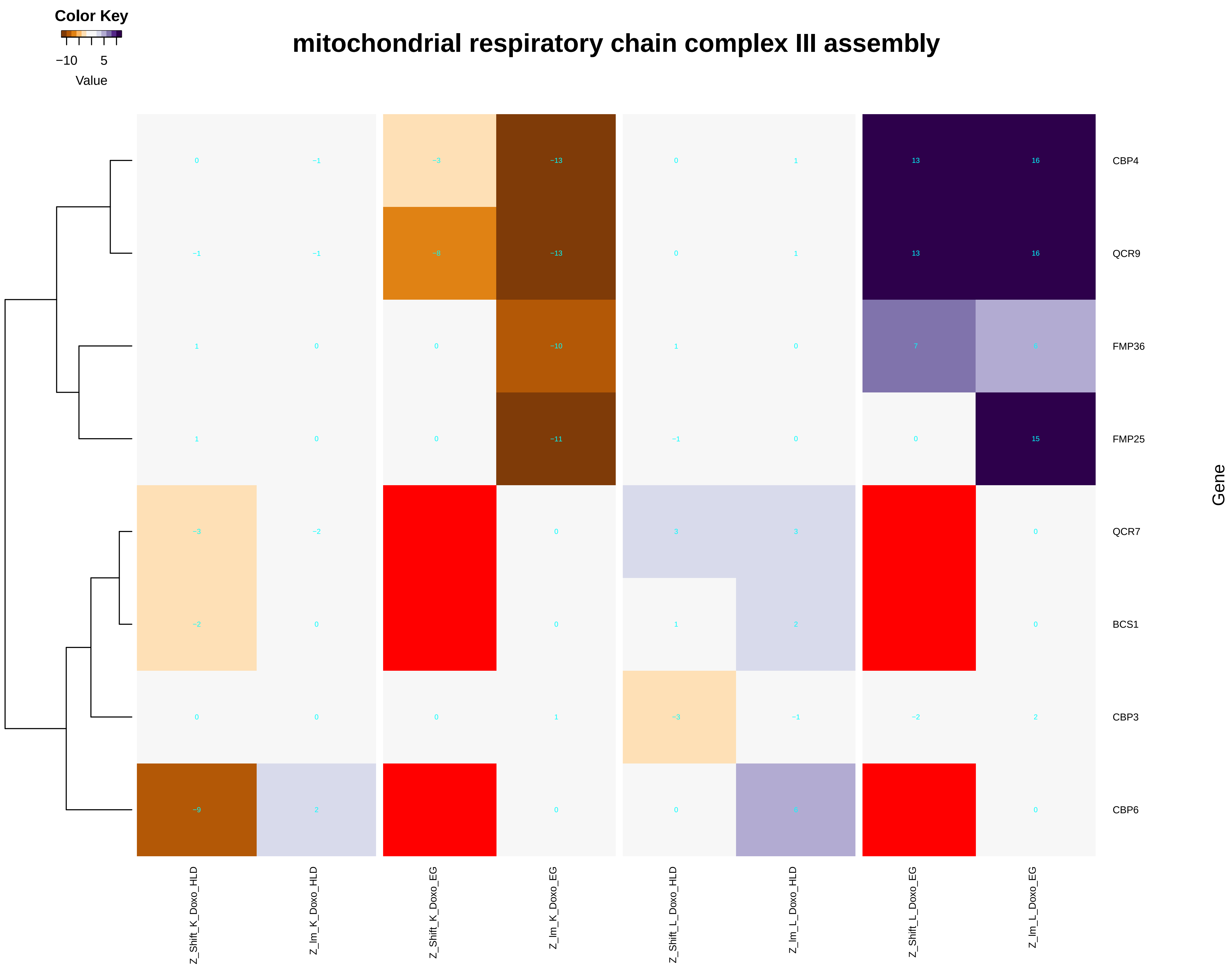

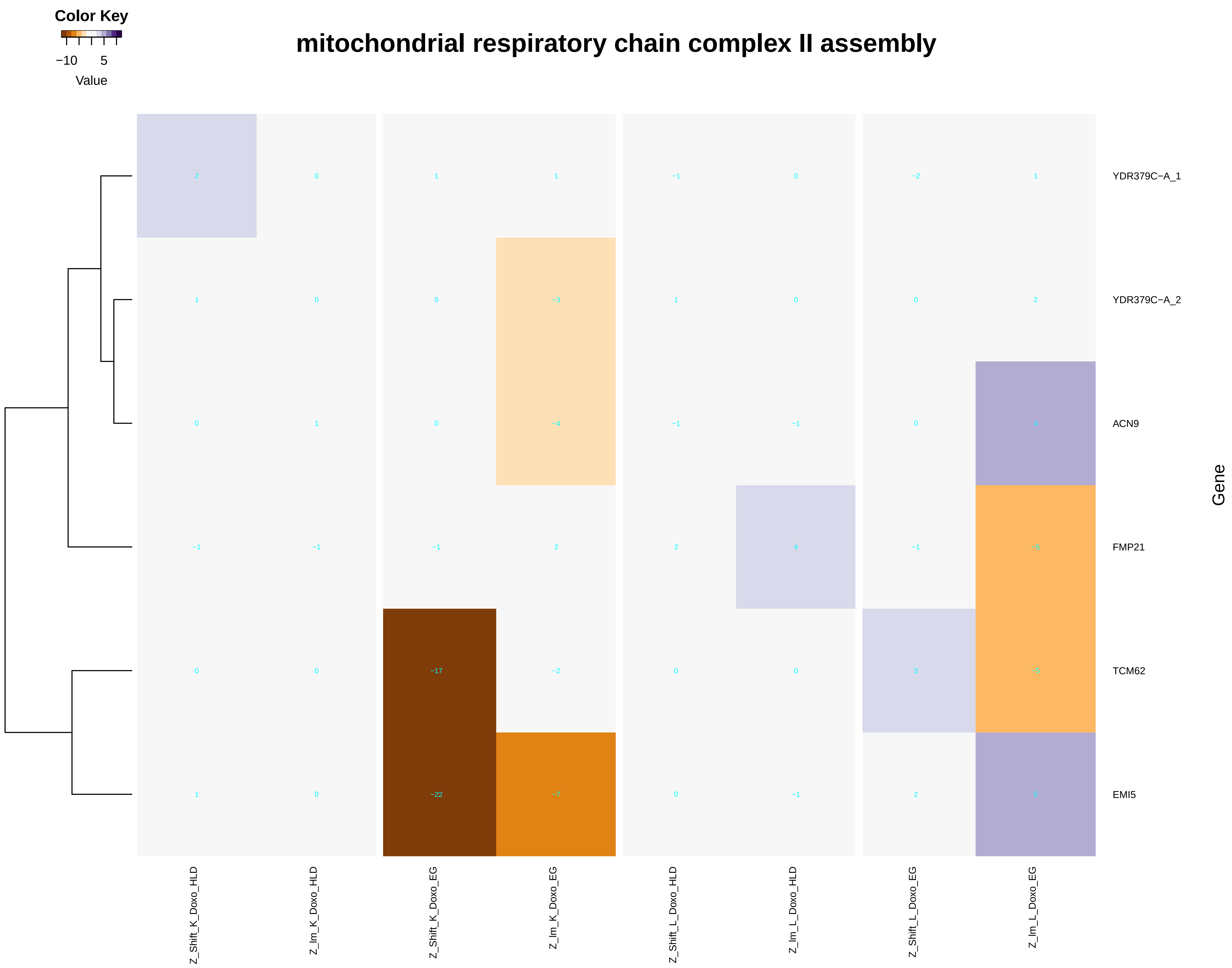

### mitochondrial_respiratory_chain_complex_III_assembly.pdf

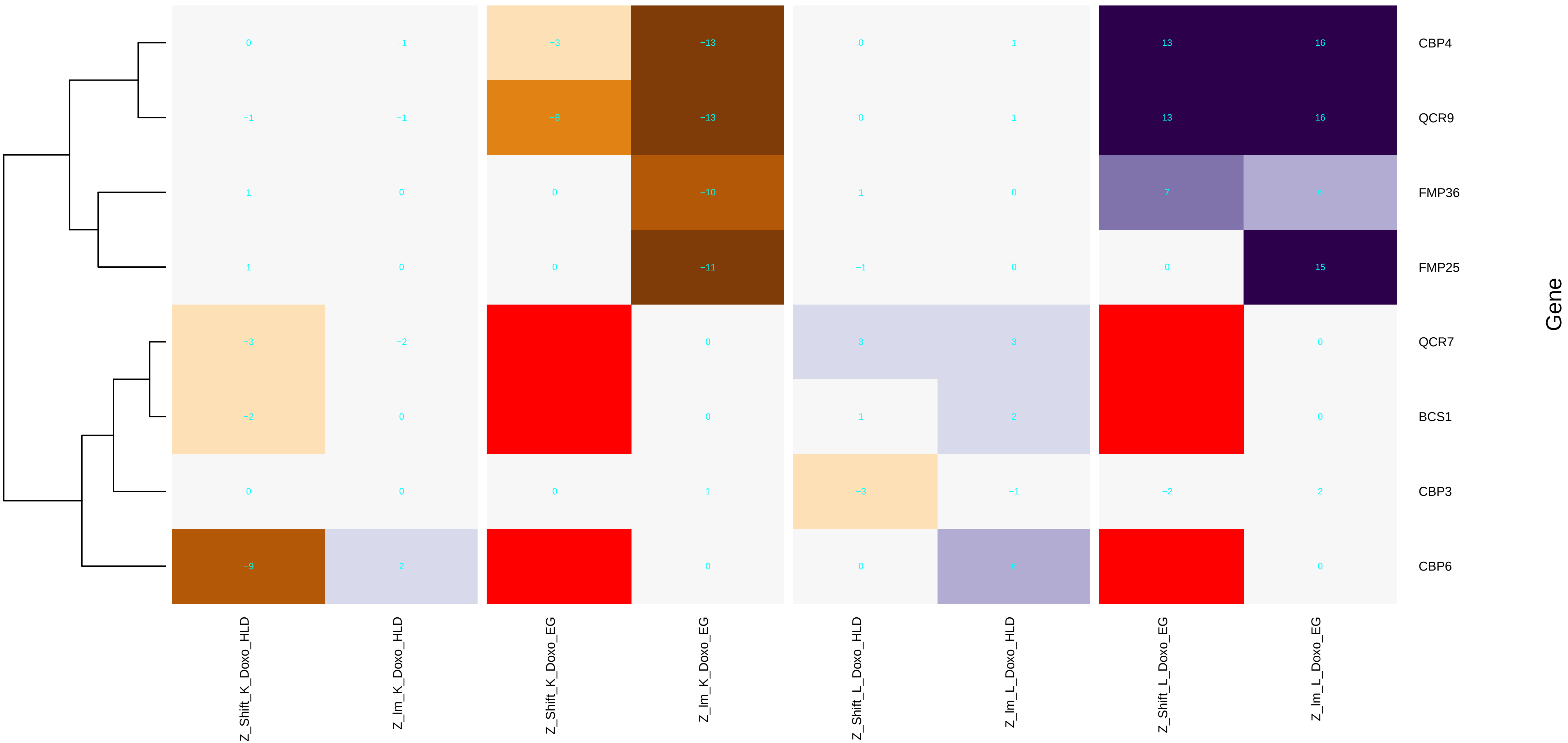

### mitochondrial_respiratory_chain_complex_IV_biogenesis.pdf

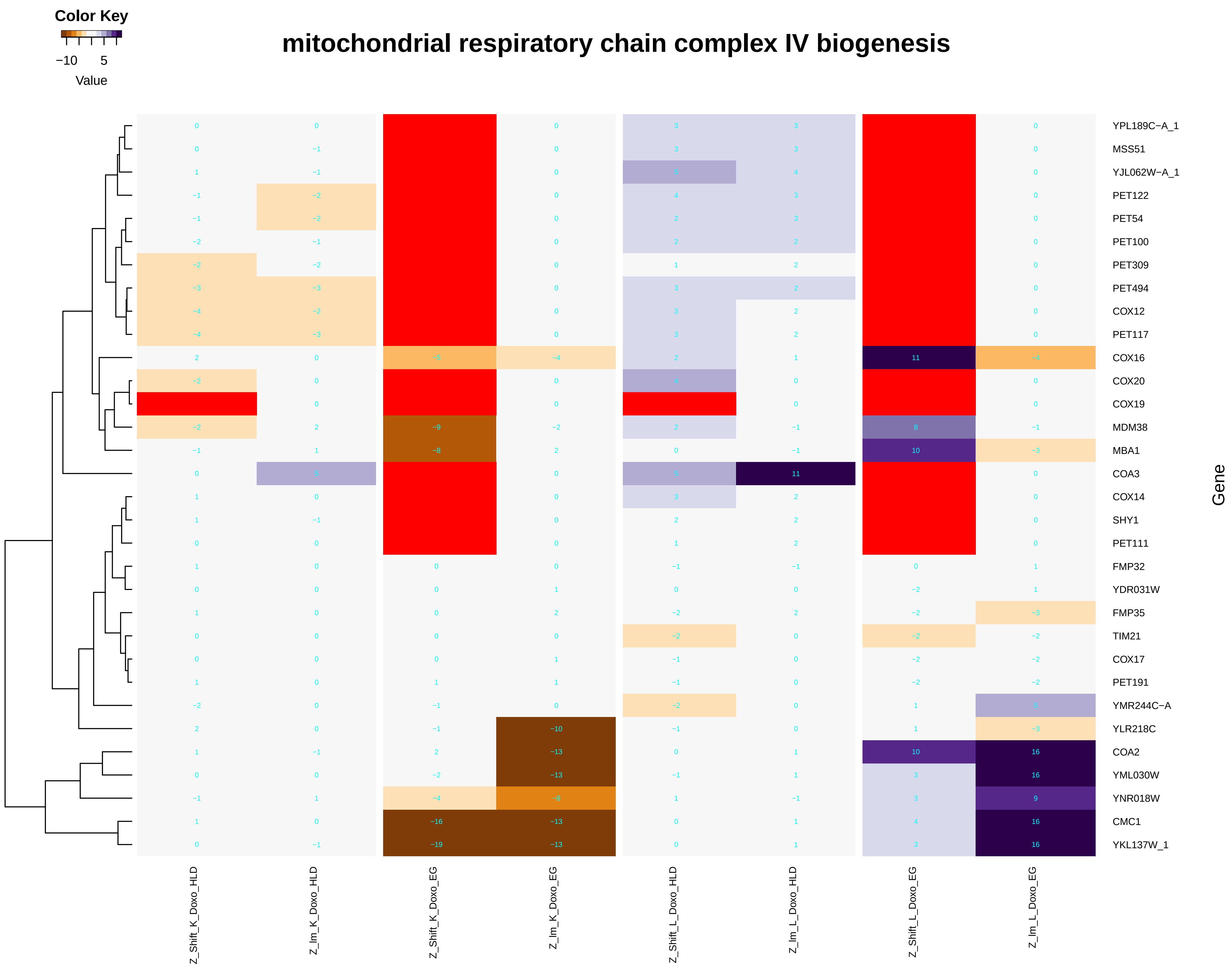

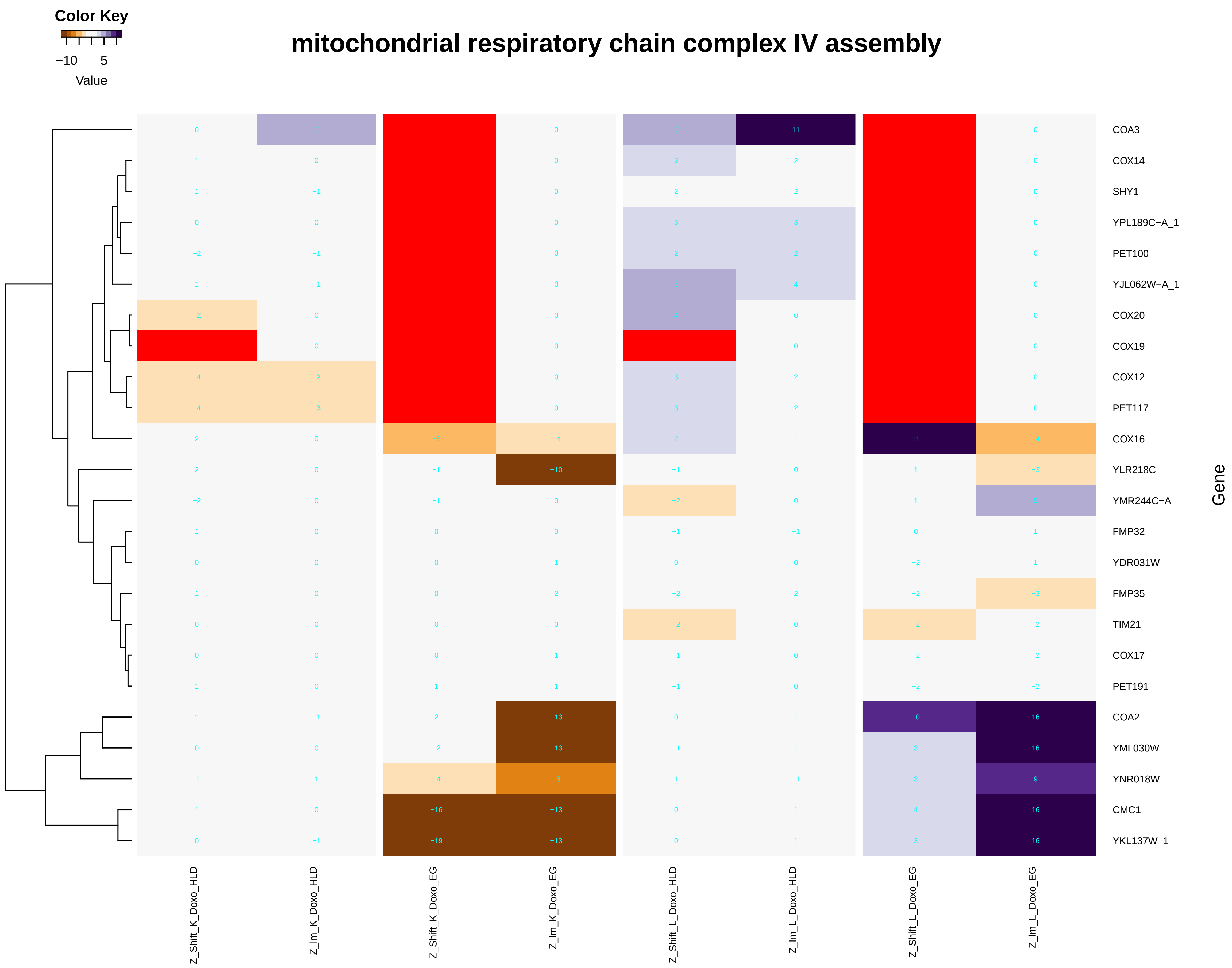

### mitochondrial_translation.pdf

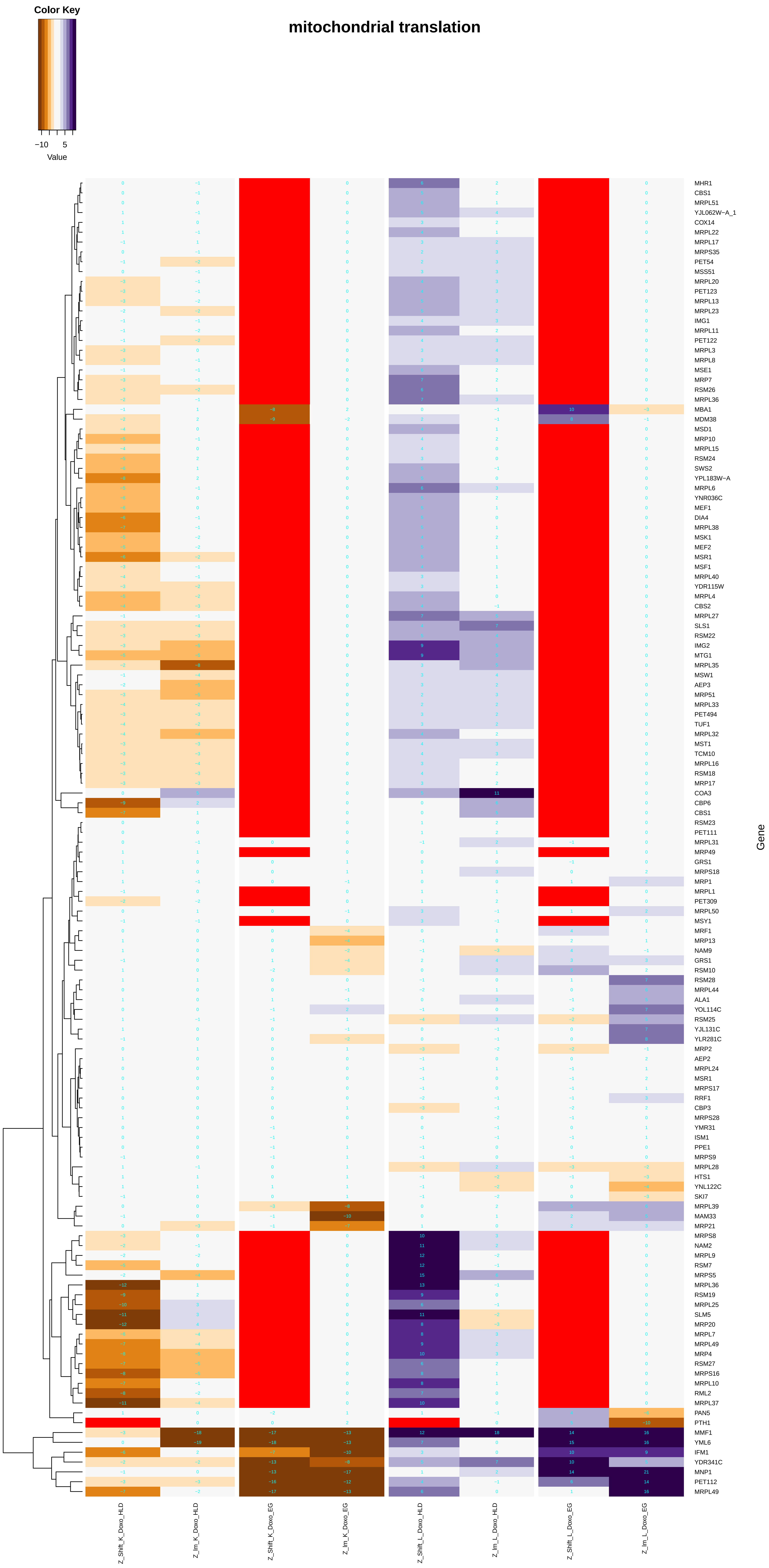

### mitochondrial_transmembrane_transport.pdf

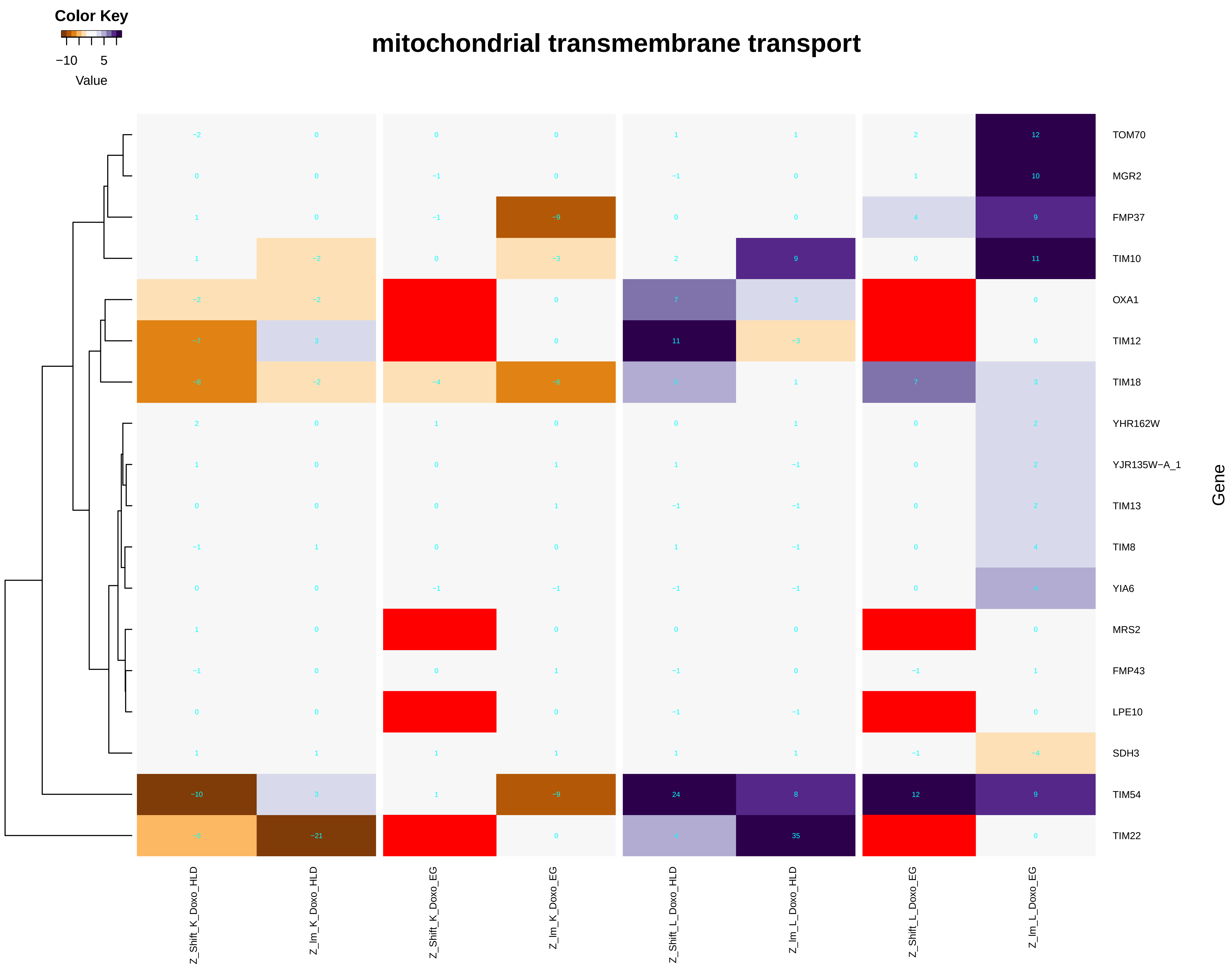

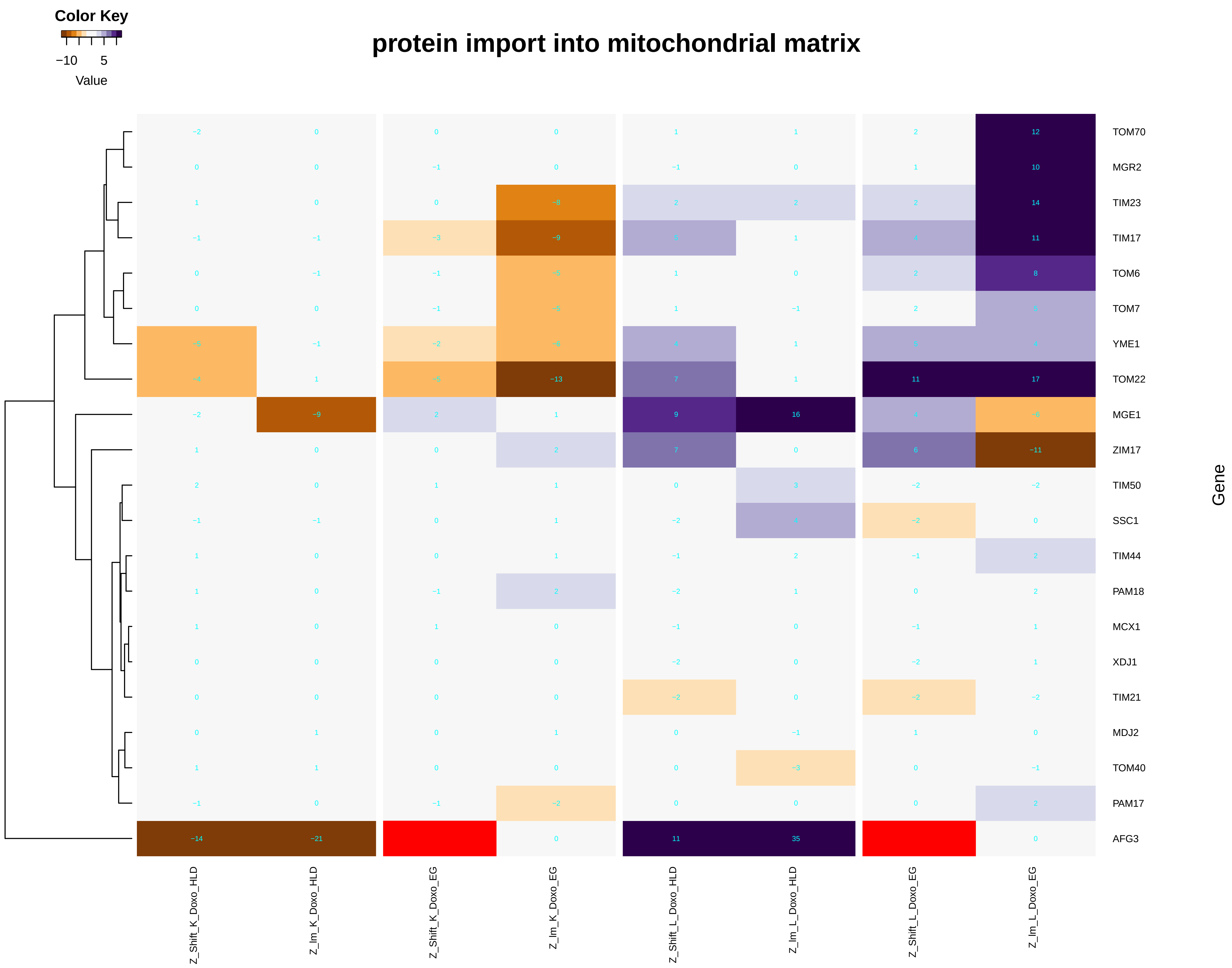

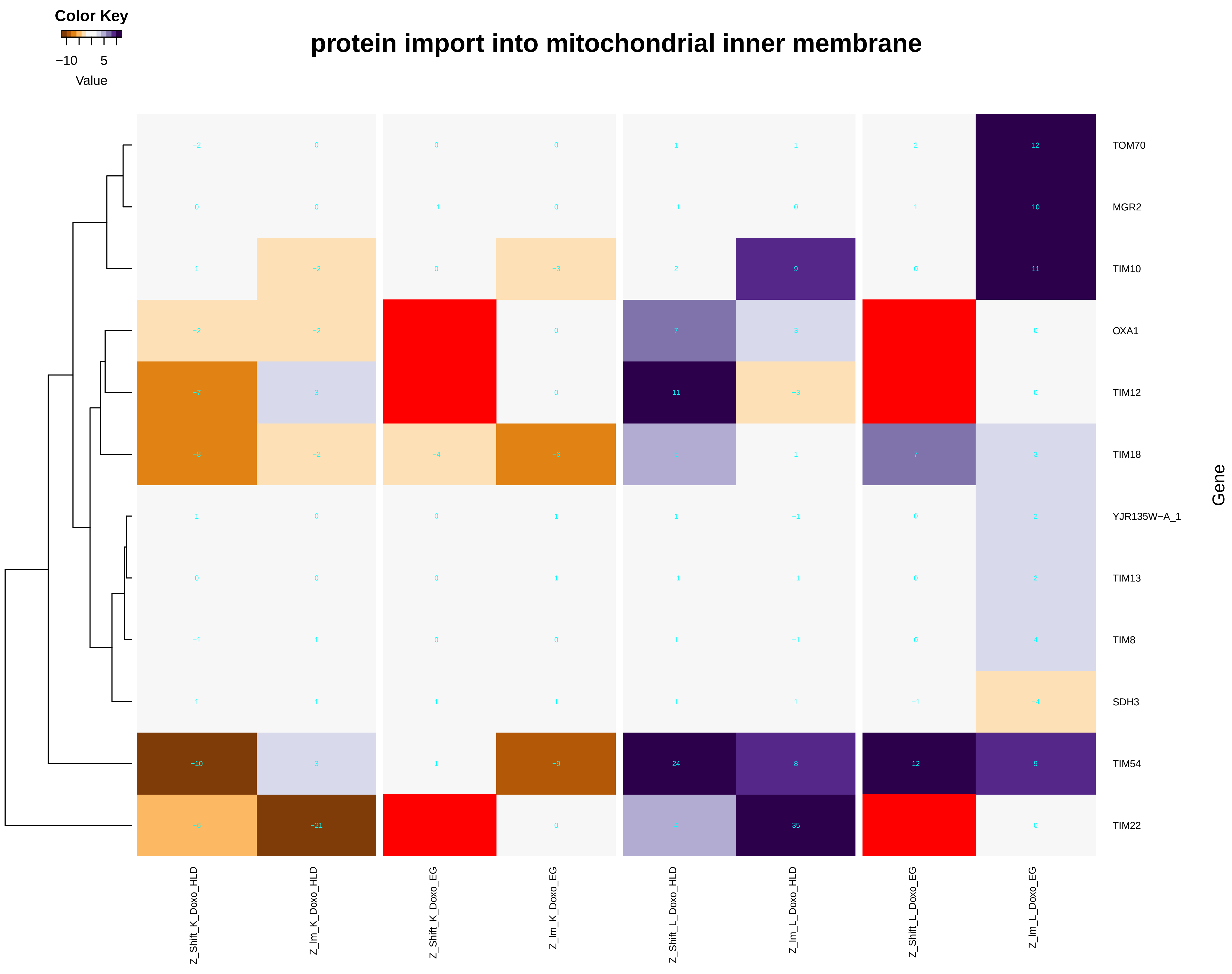

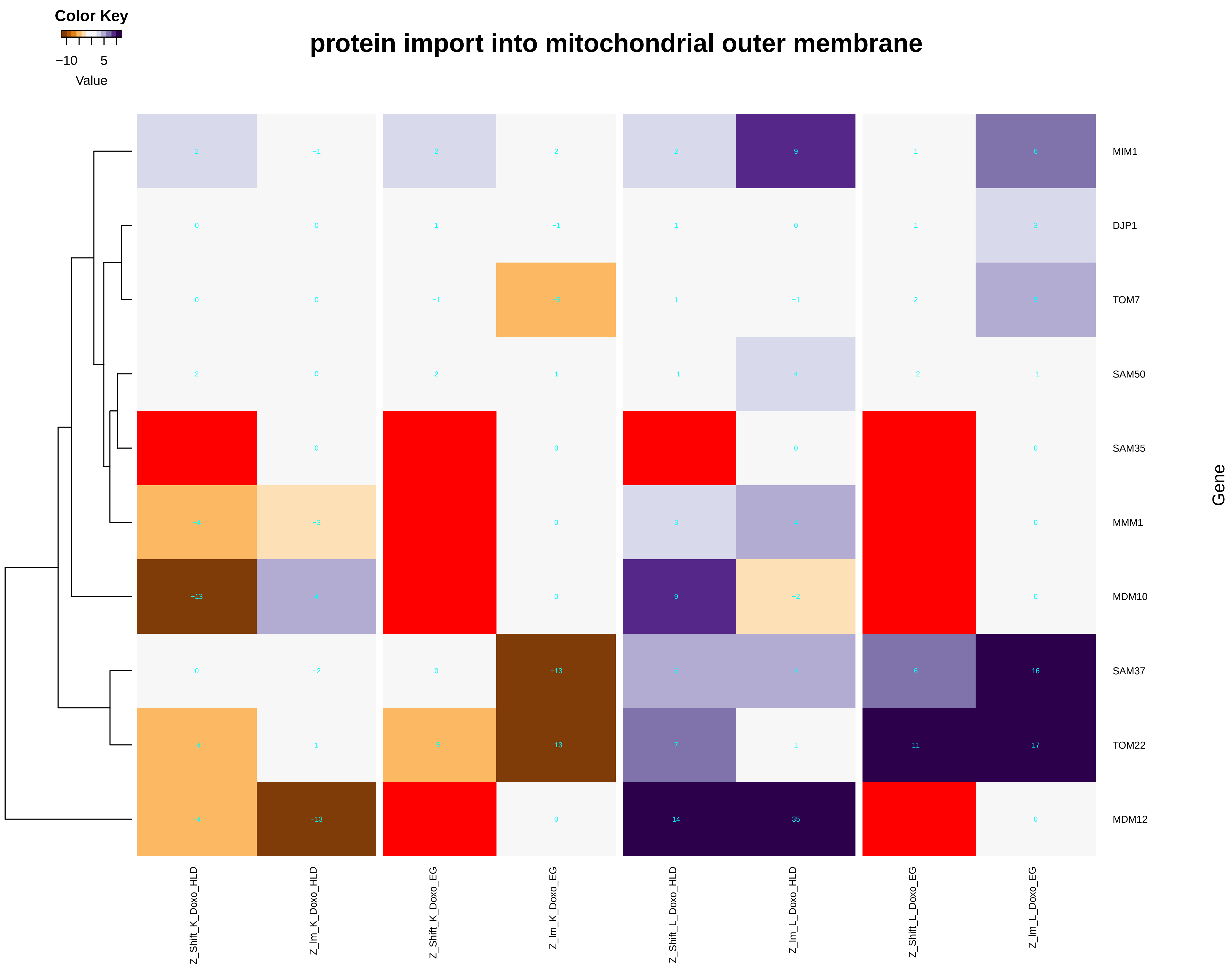

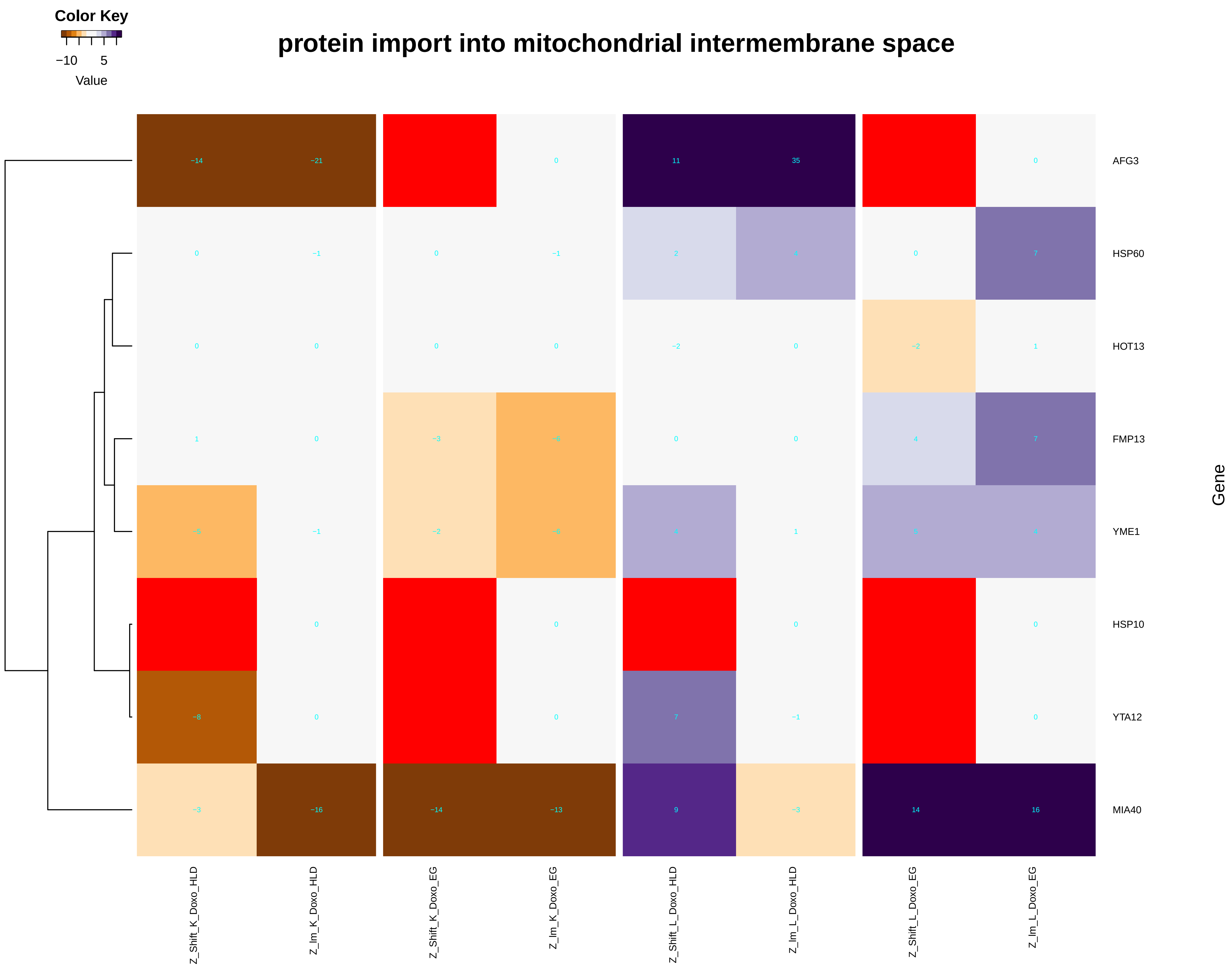

### mitochondrial_transport.pdf

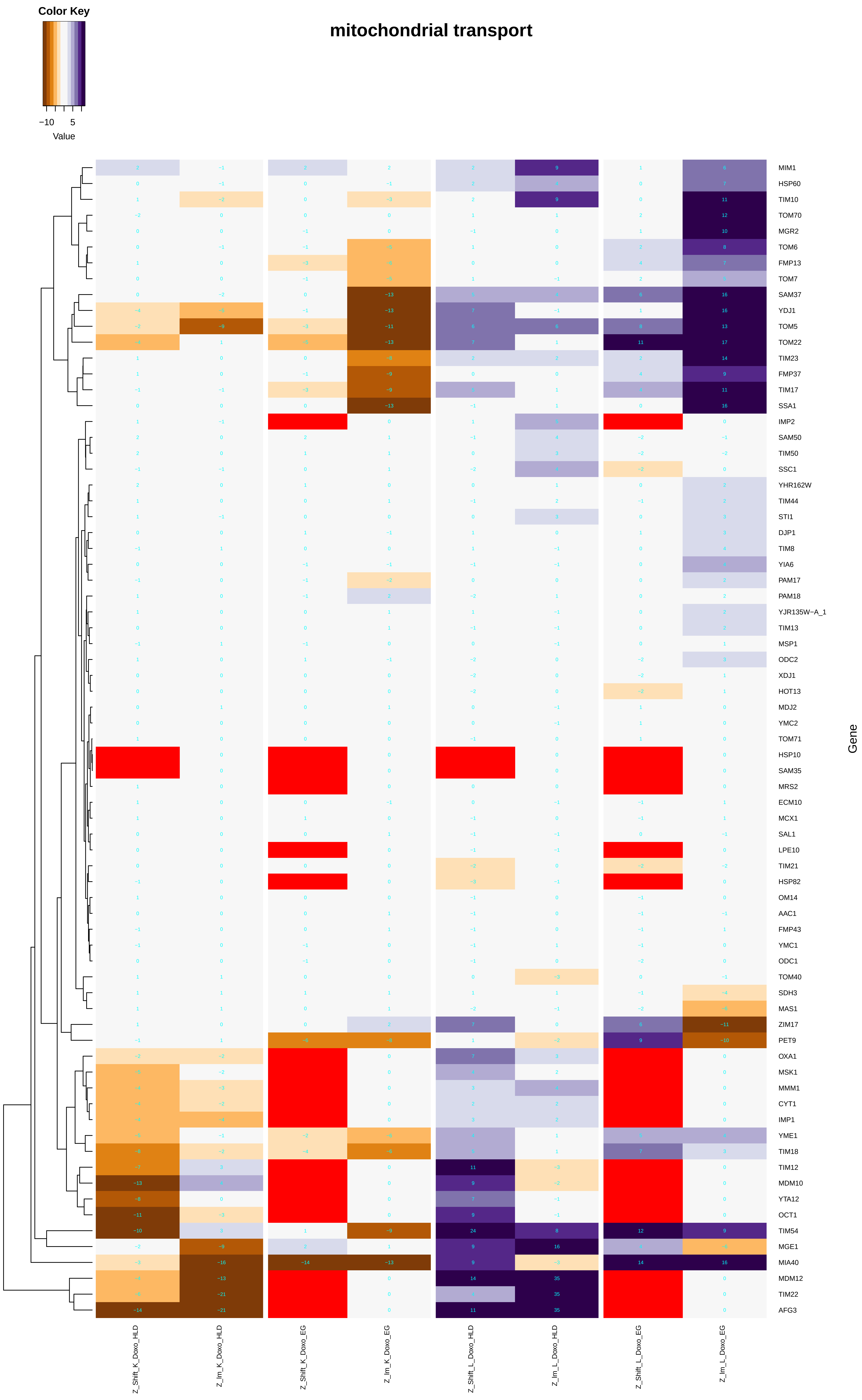

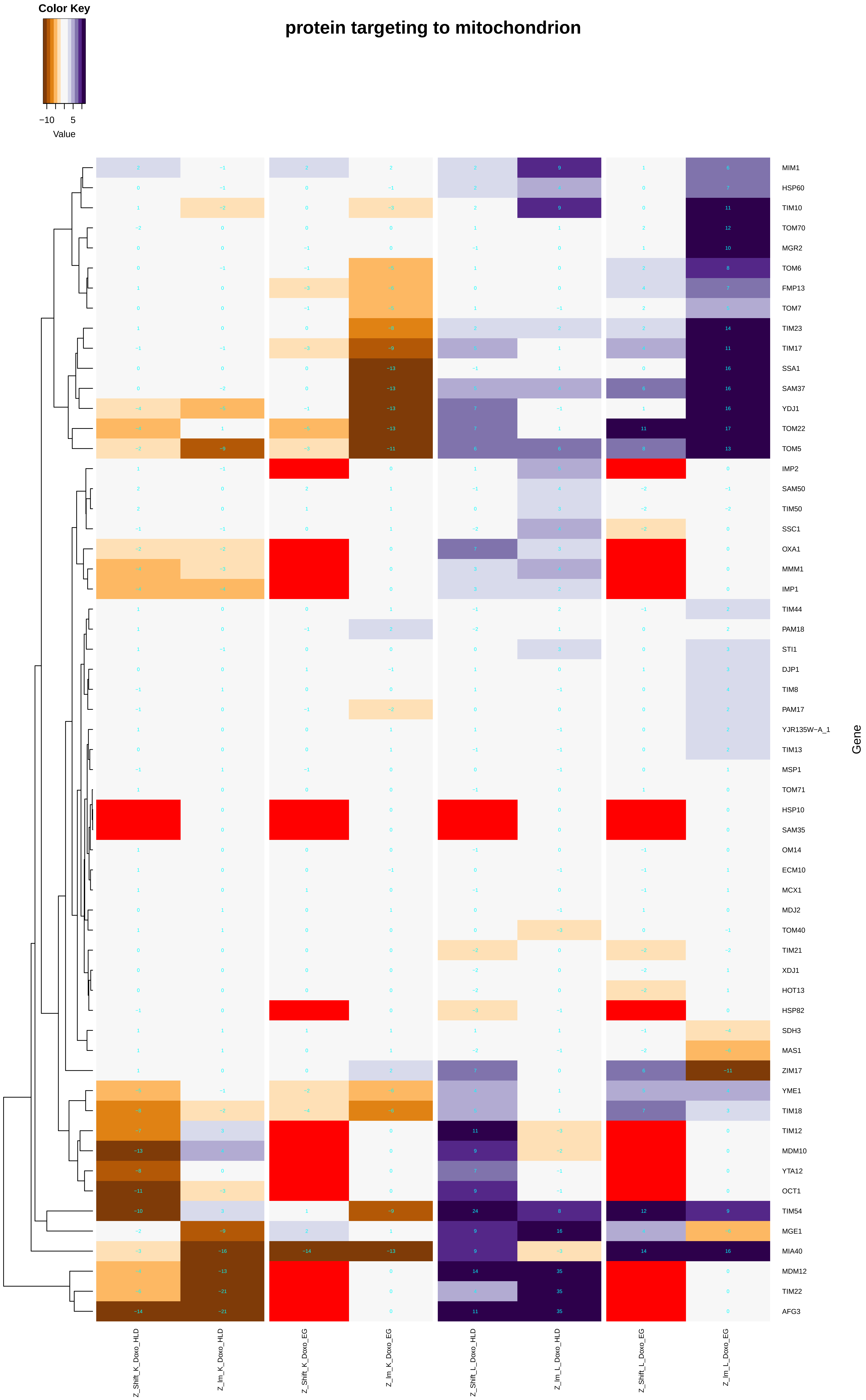

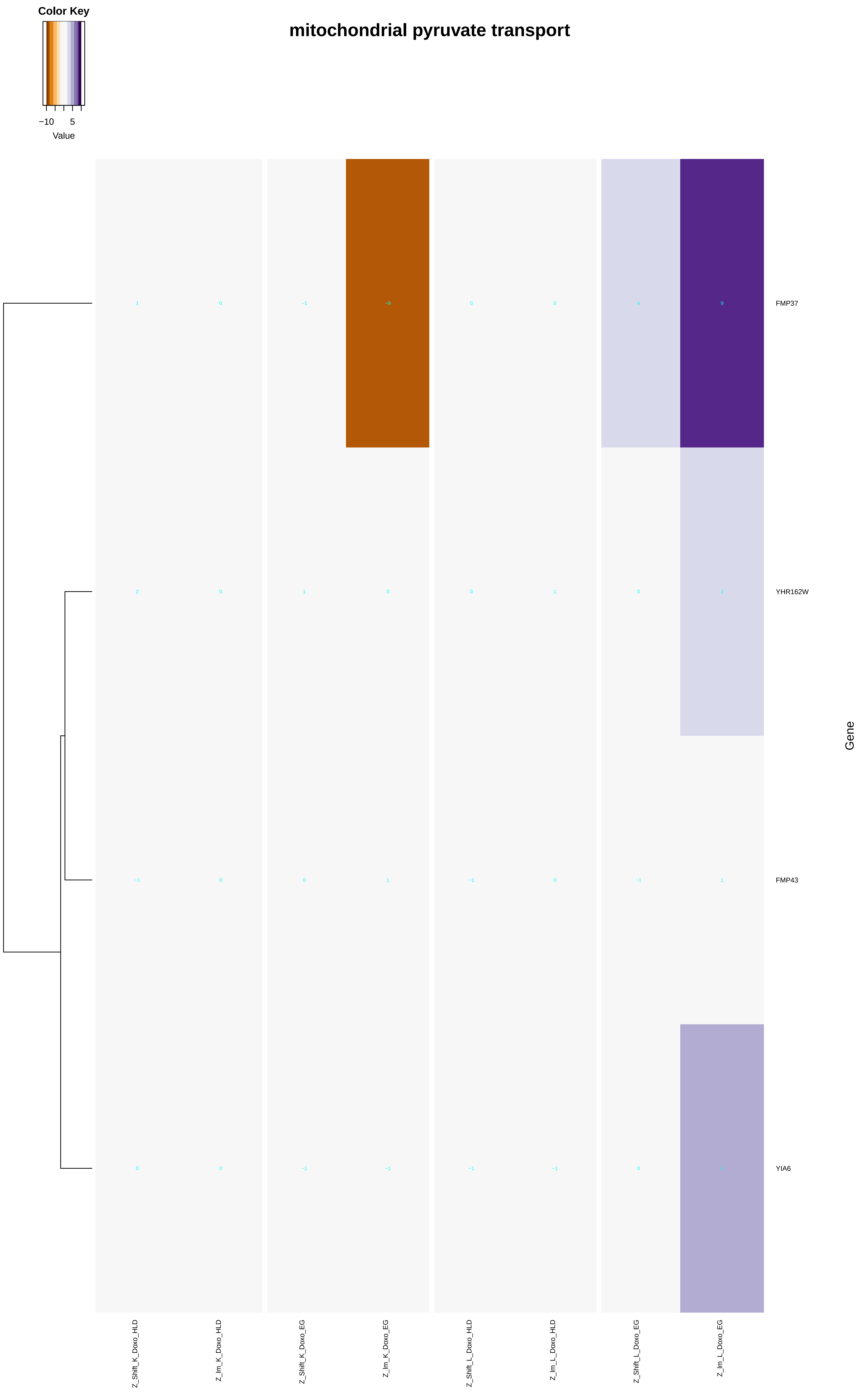

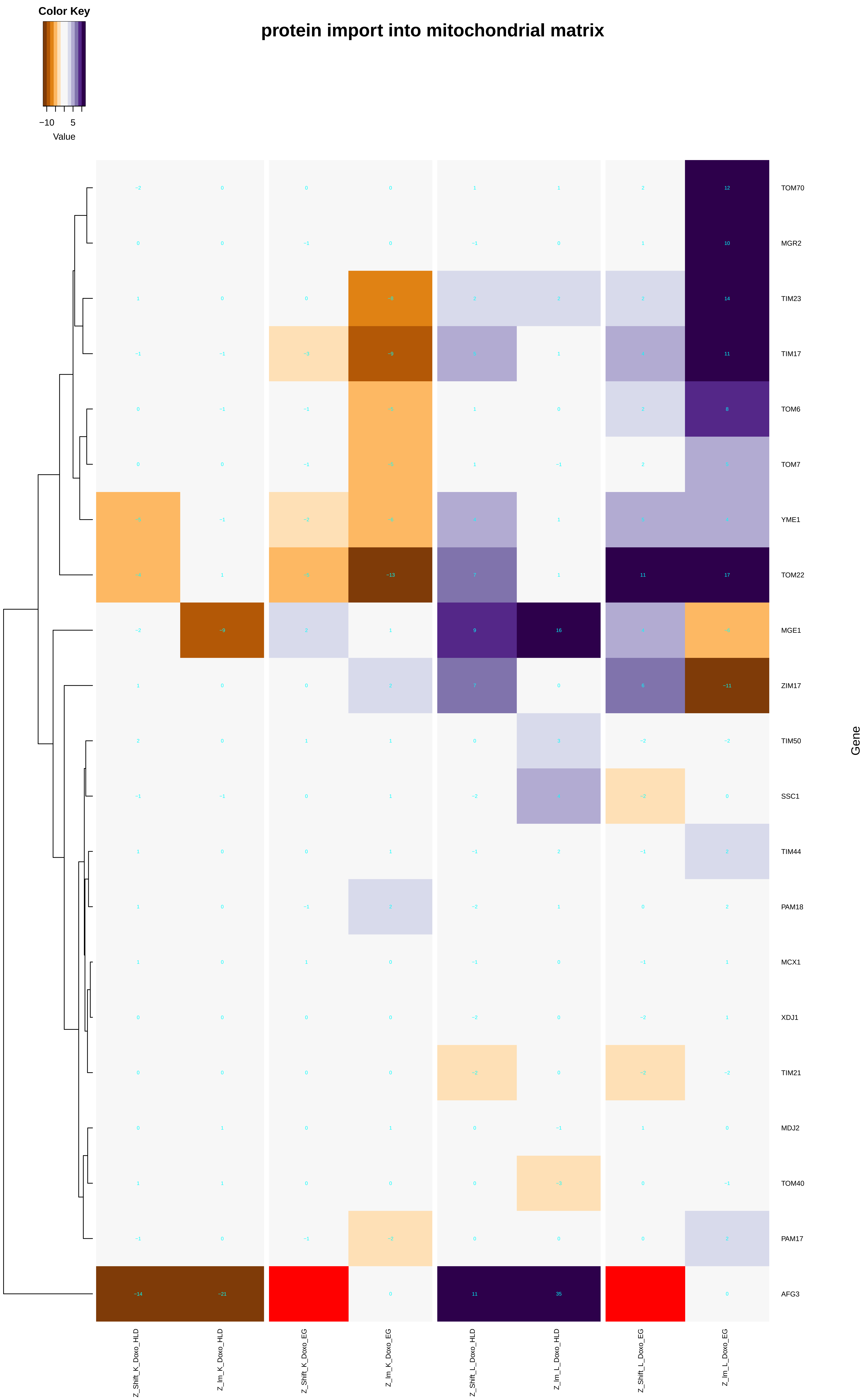

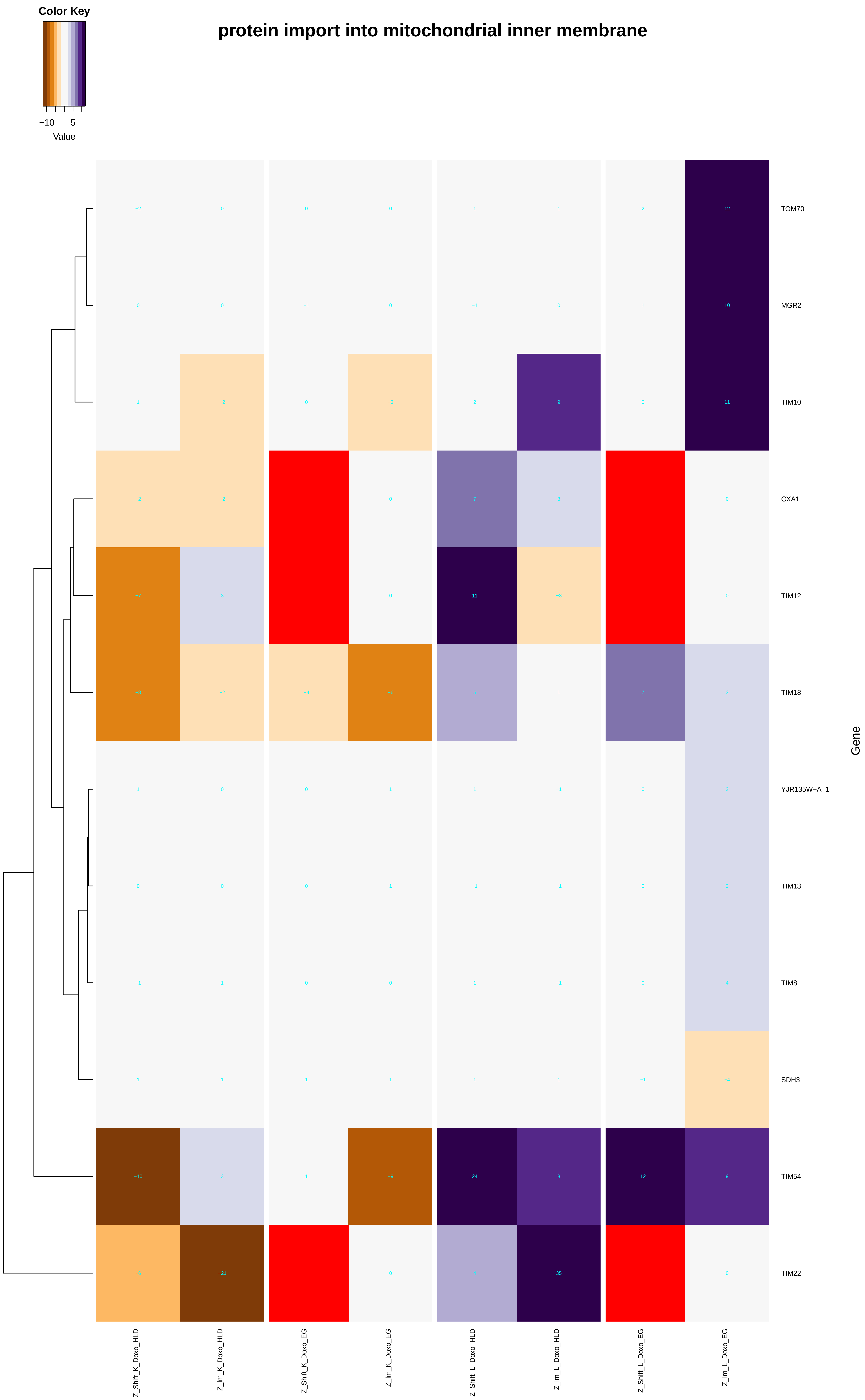

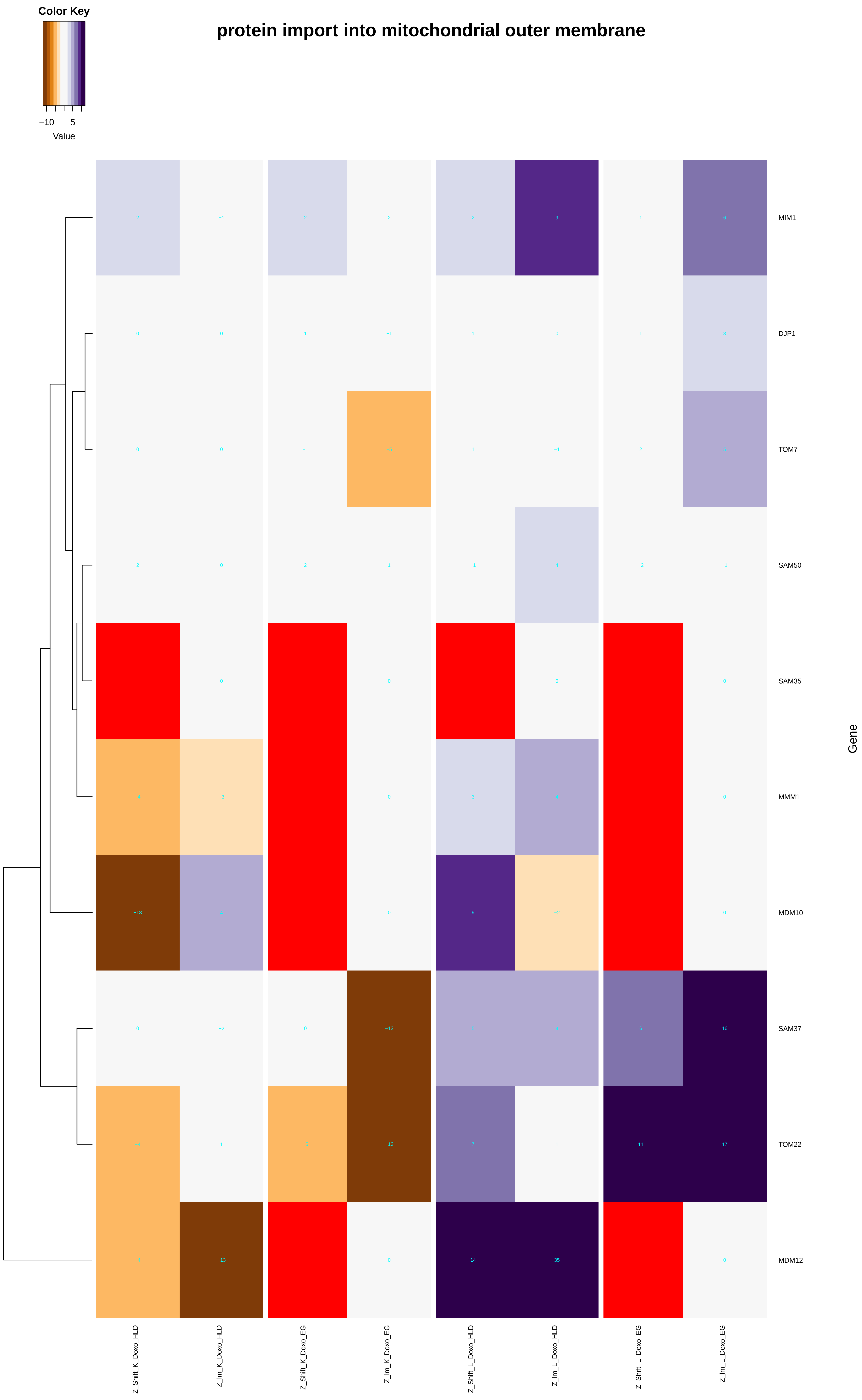

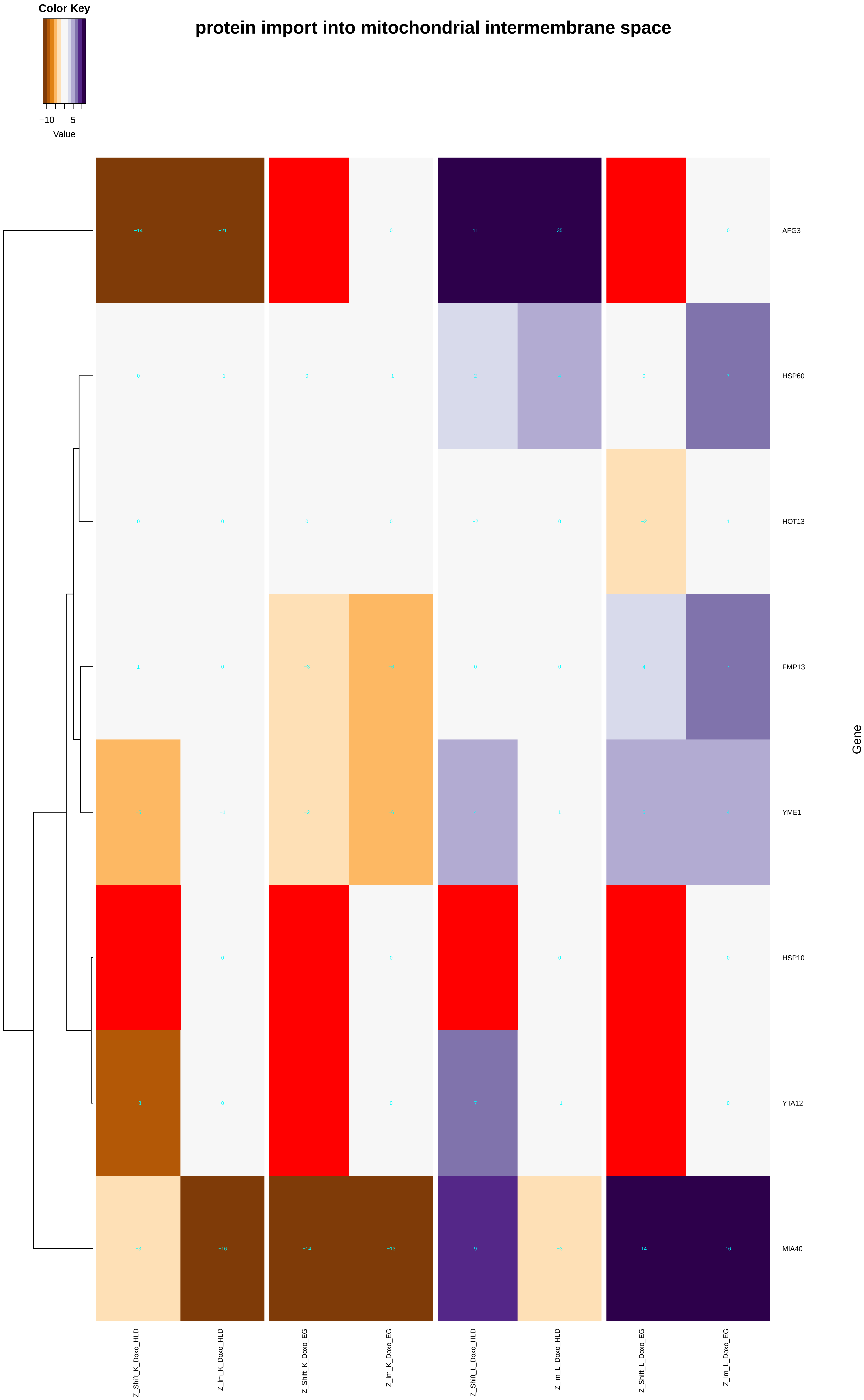

### mitochondrion-ER_tethering.pdf

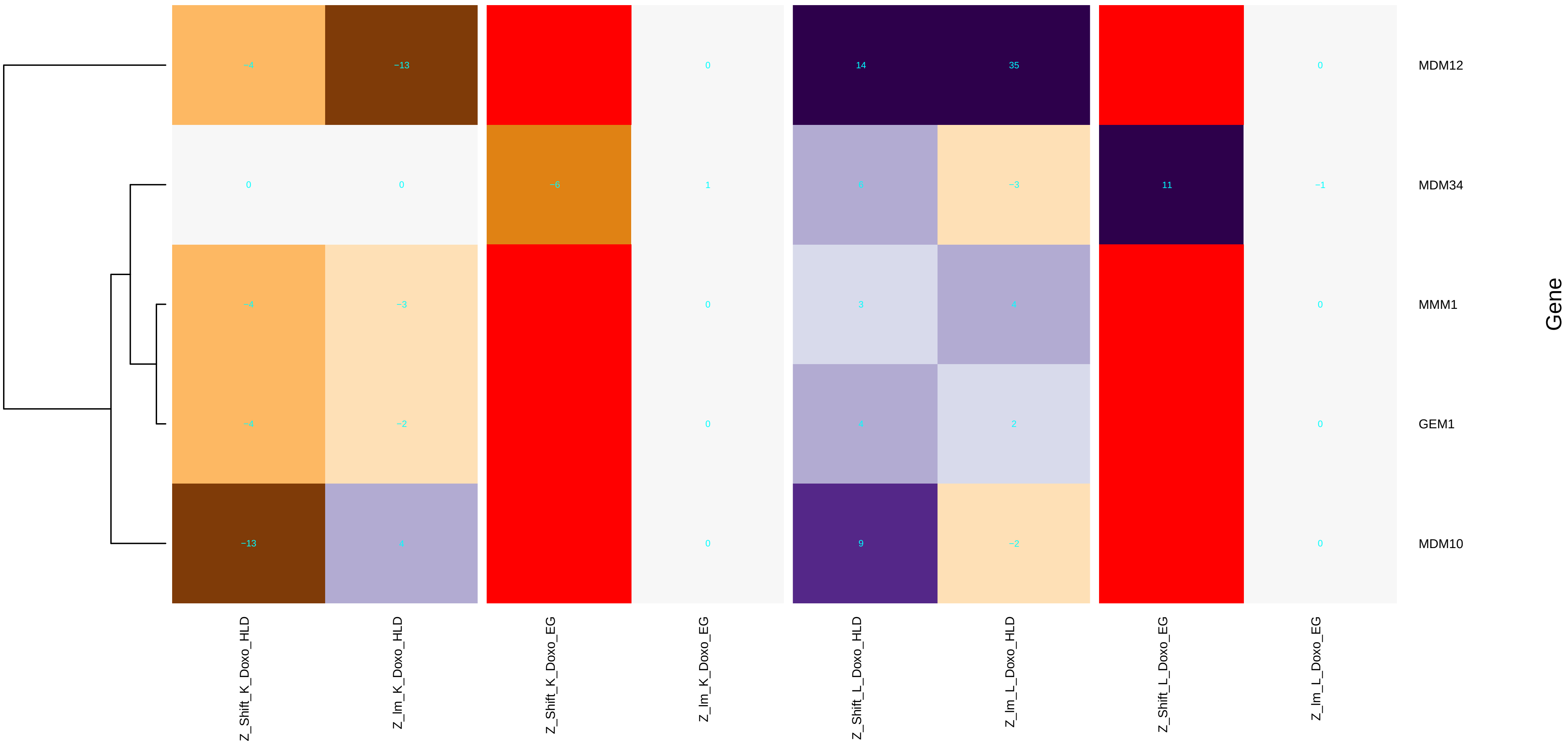

### mitotic_cell_cycle.pdf

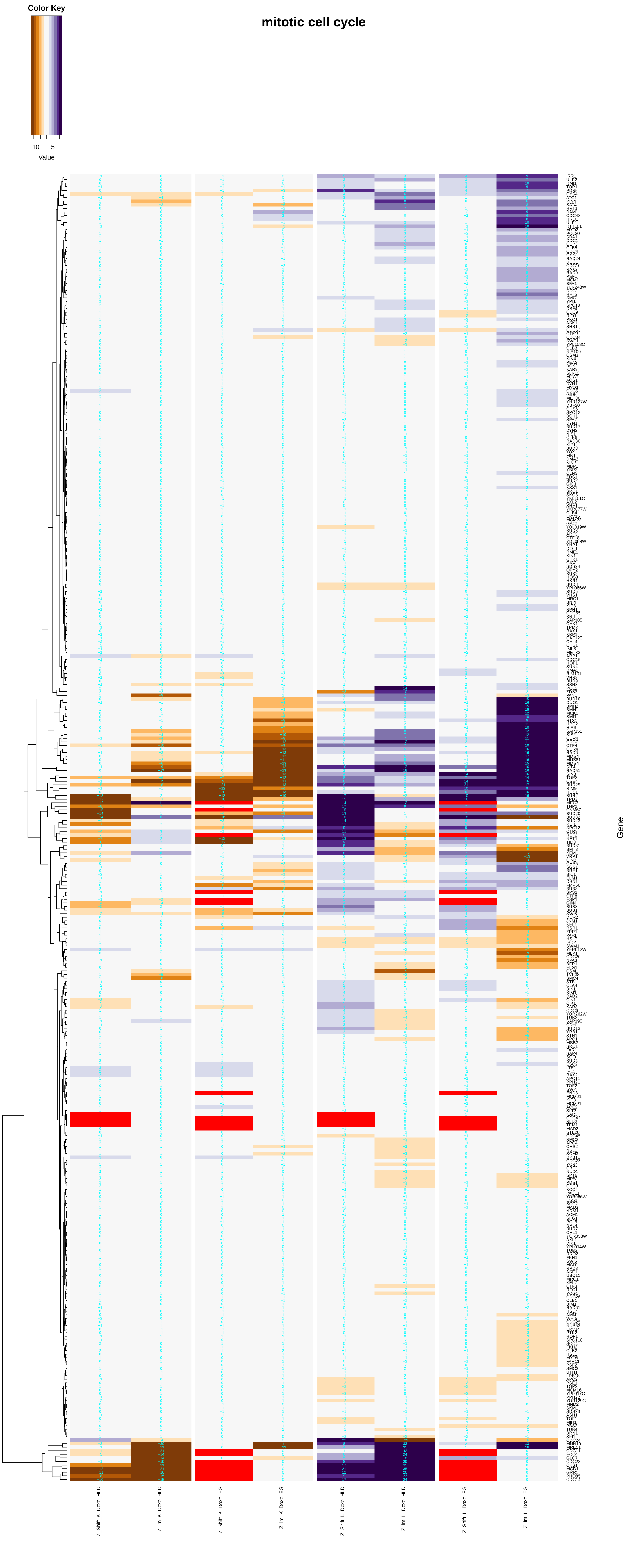

### mitotic_cell_cycle_arrest.pdf

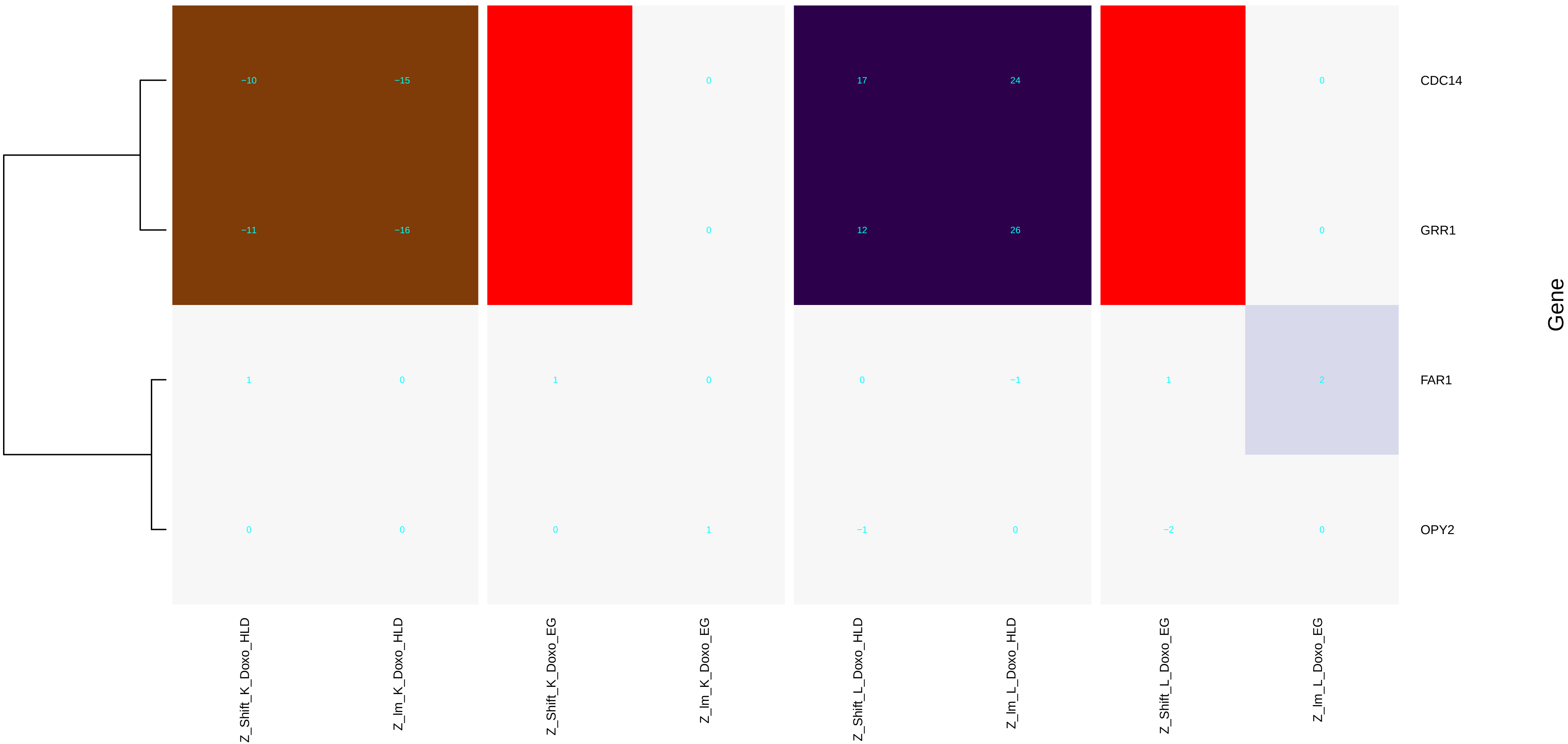

### mitotic_nuclear_division.pdf

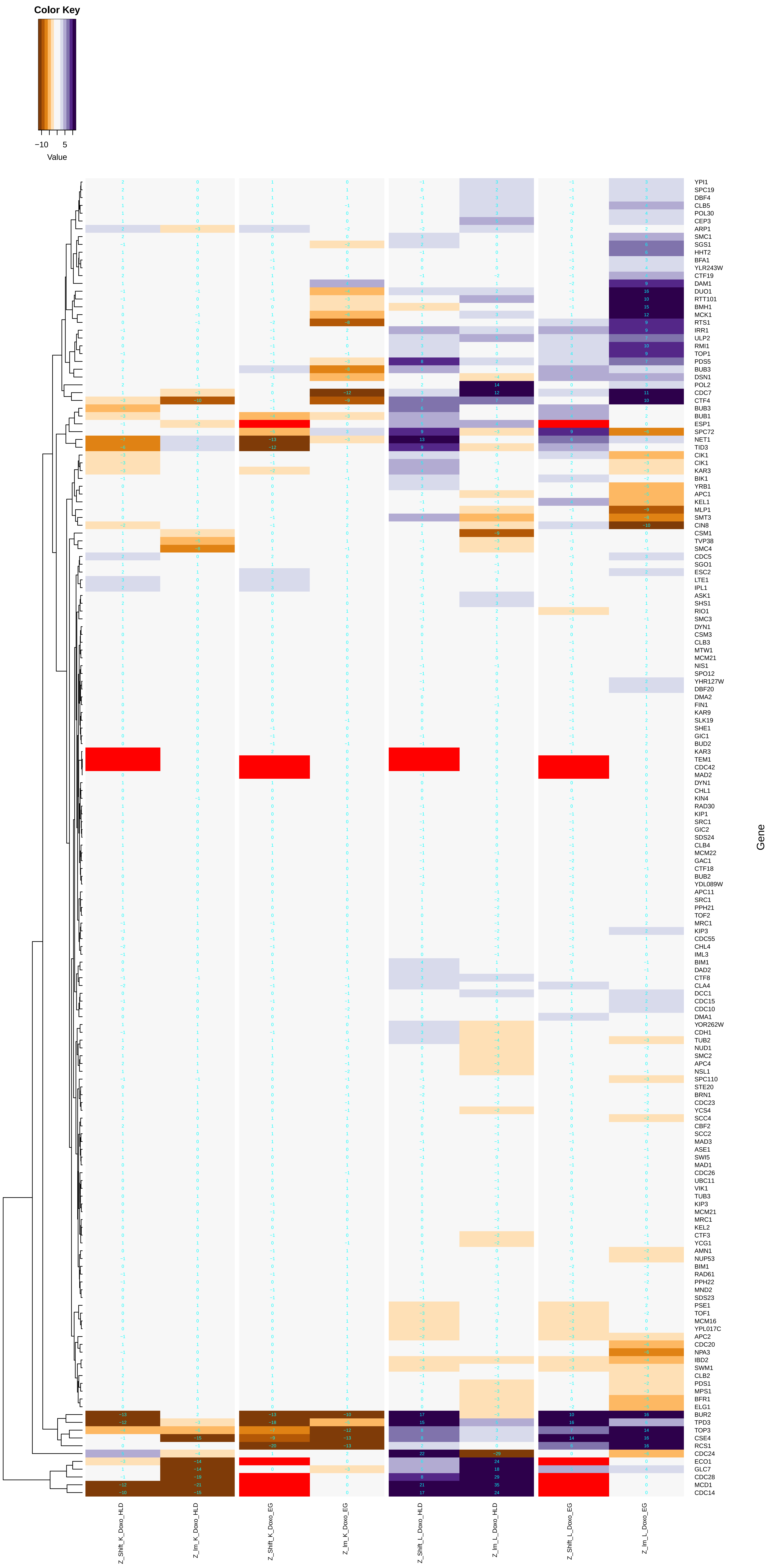

### mitotic_recombination.pdf

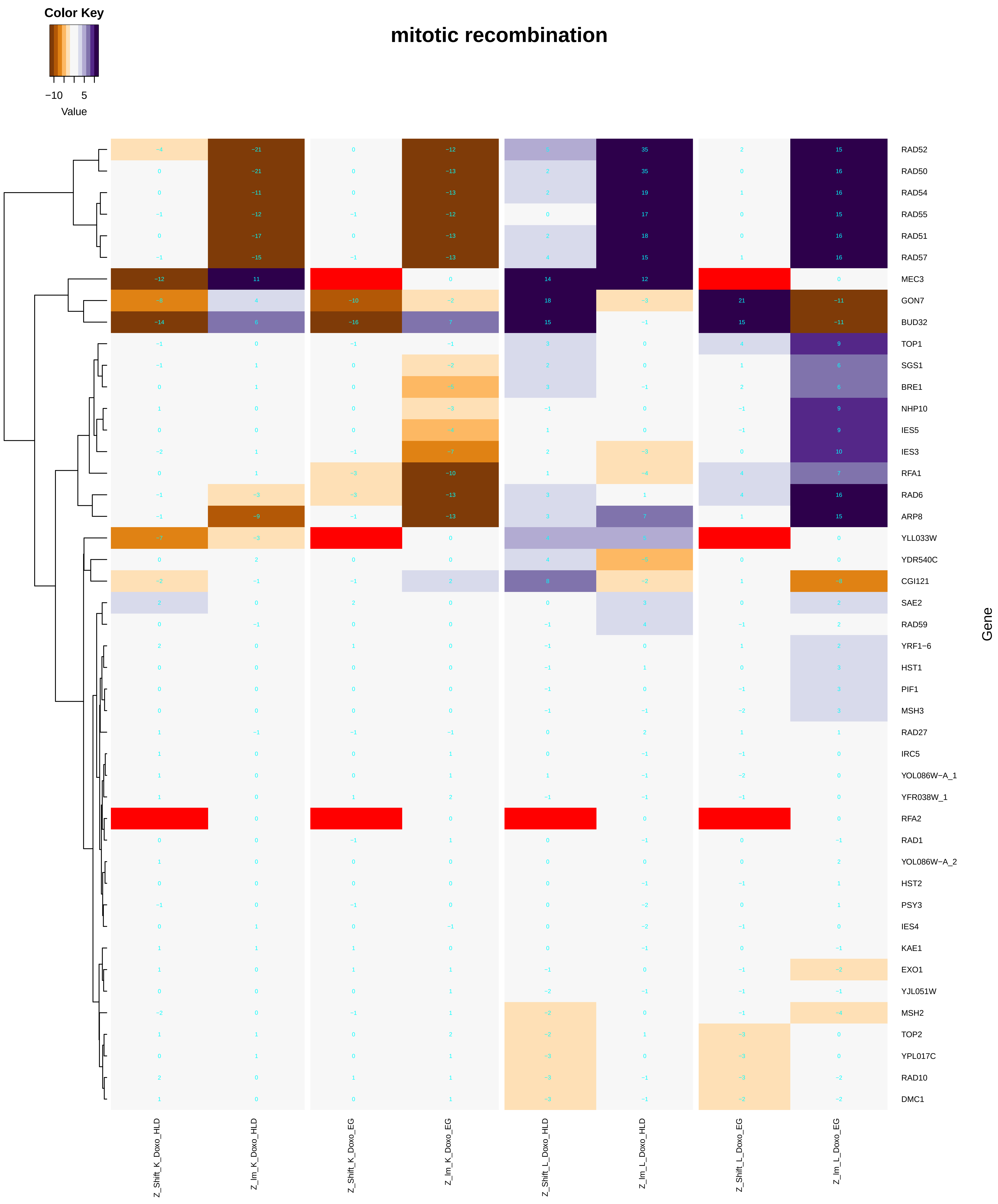

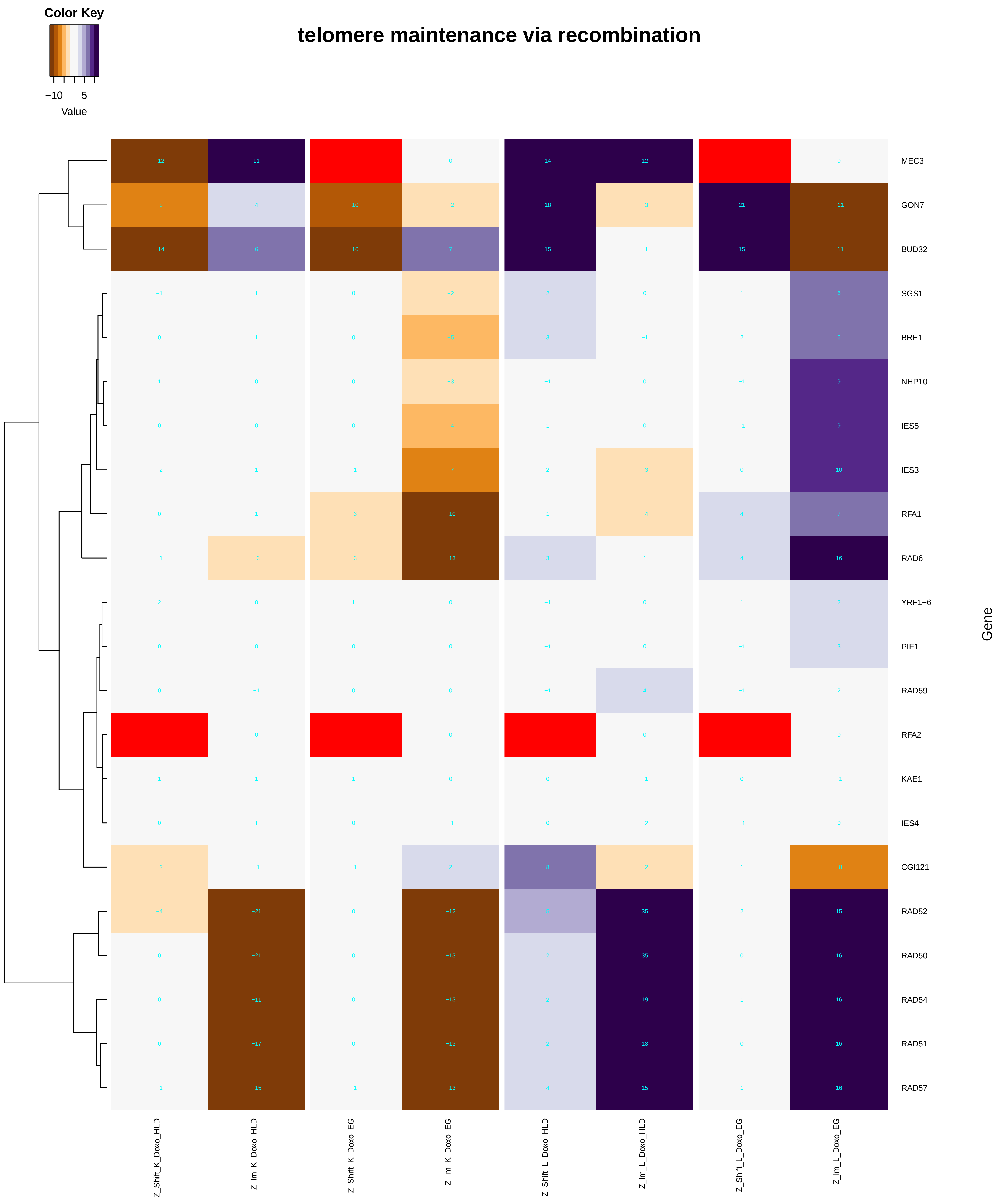

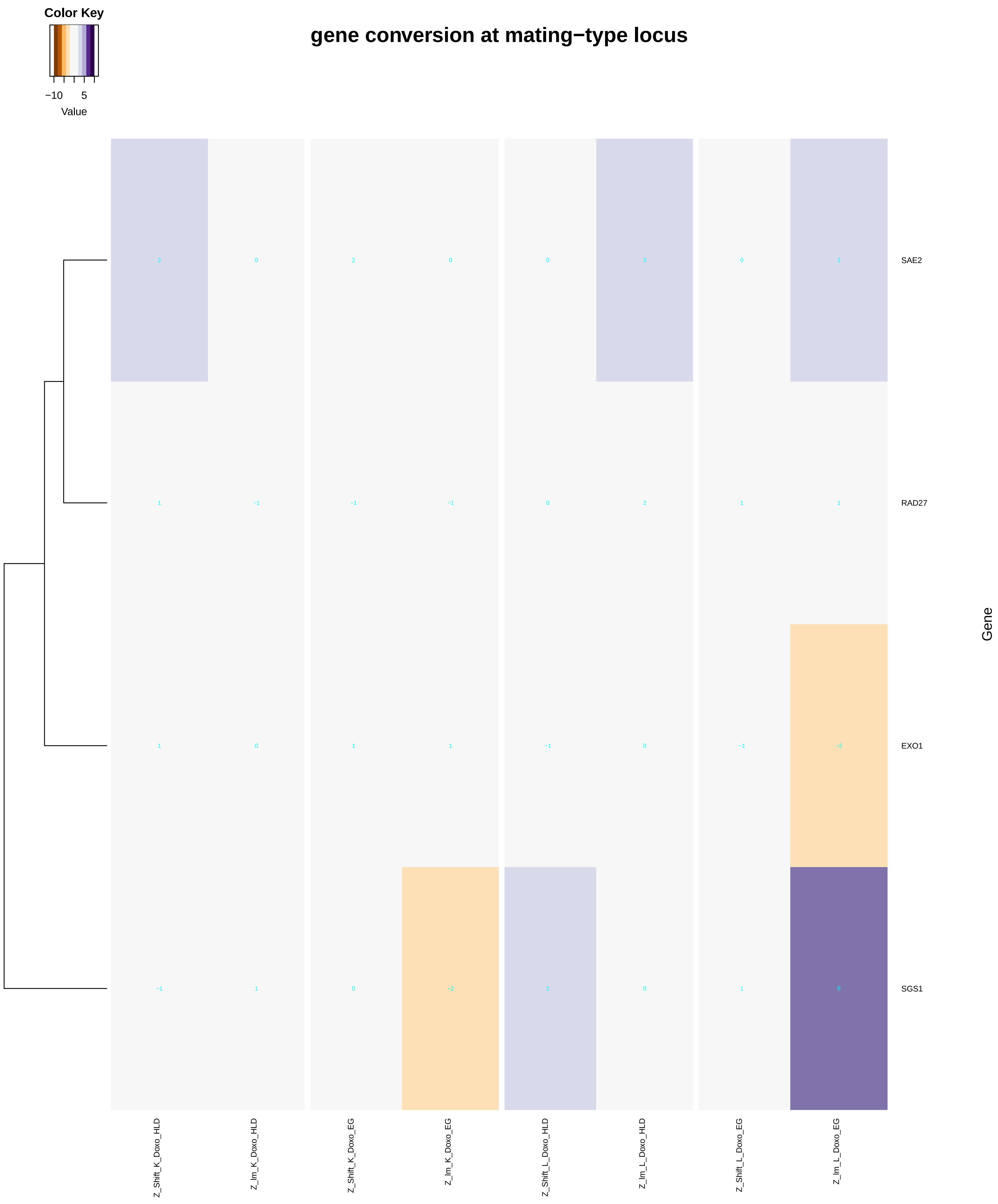

### mitotic_sister_chromatid_segregation.pdf

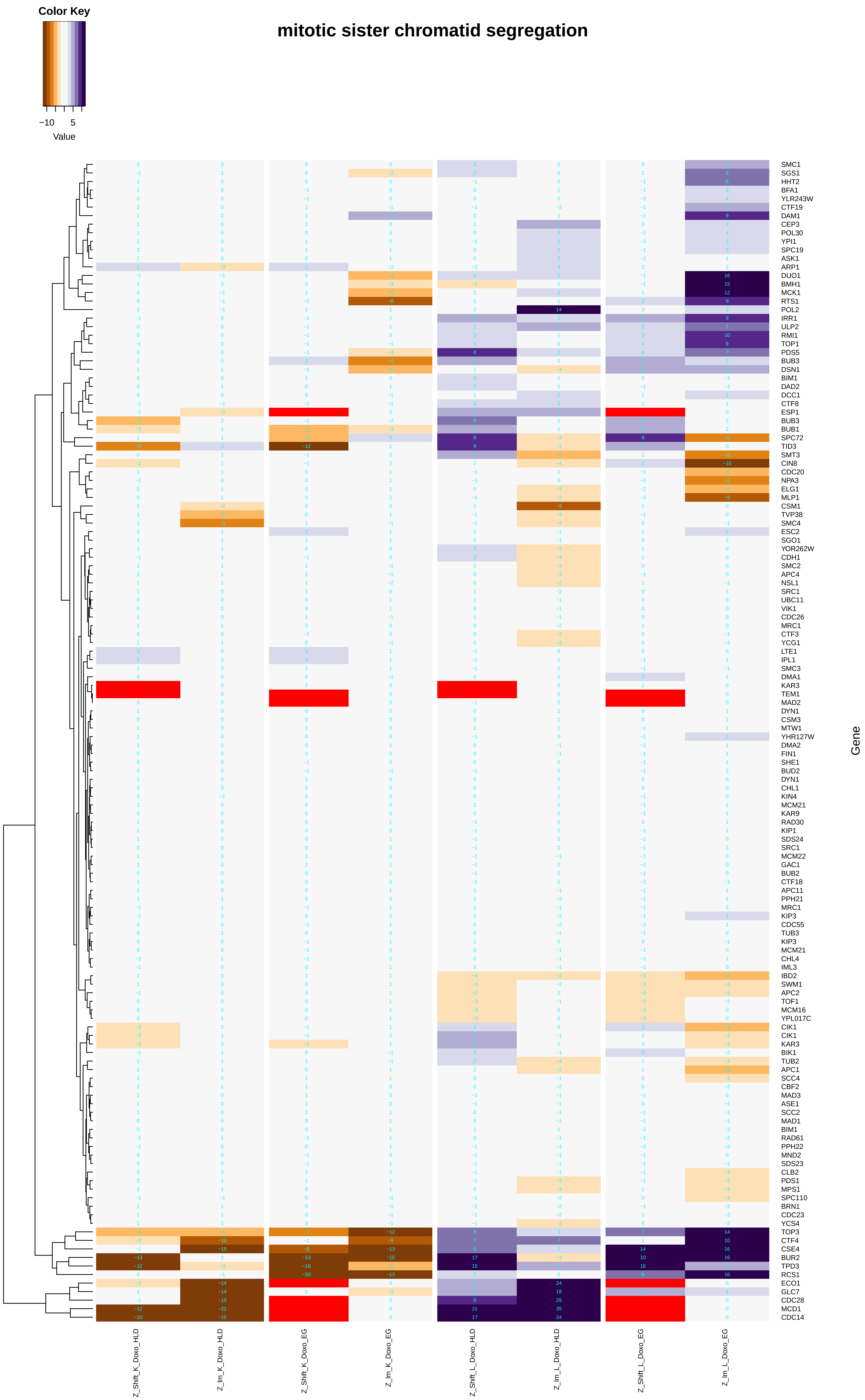

### mitotic_spindle_assembly.pdf

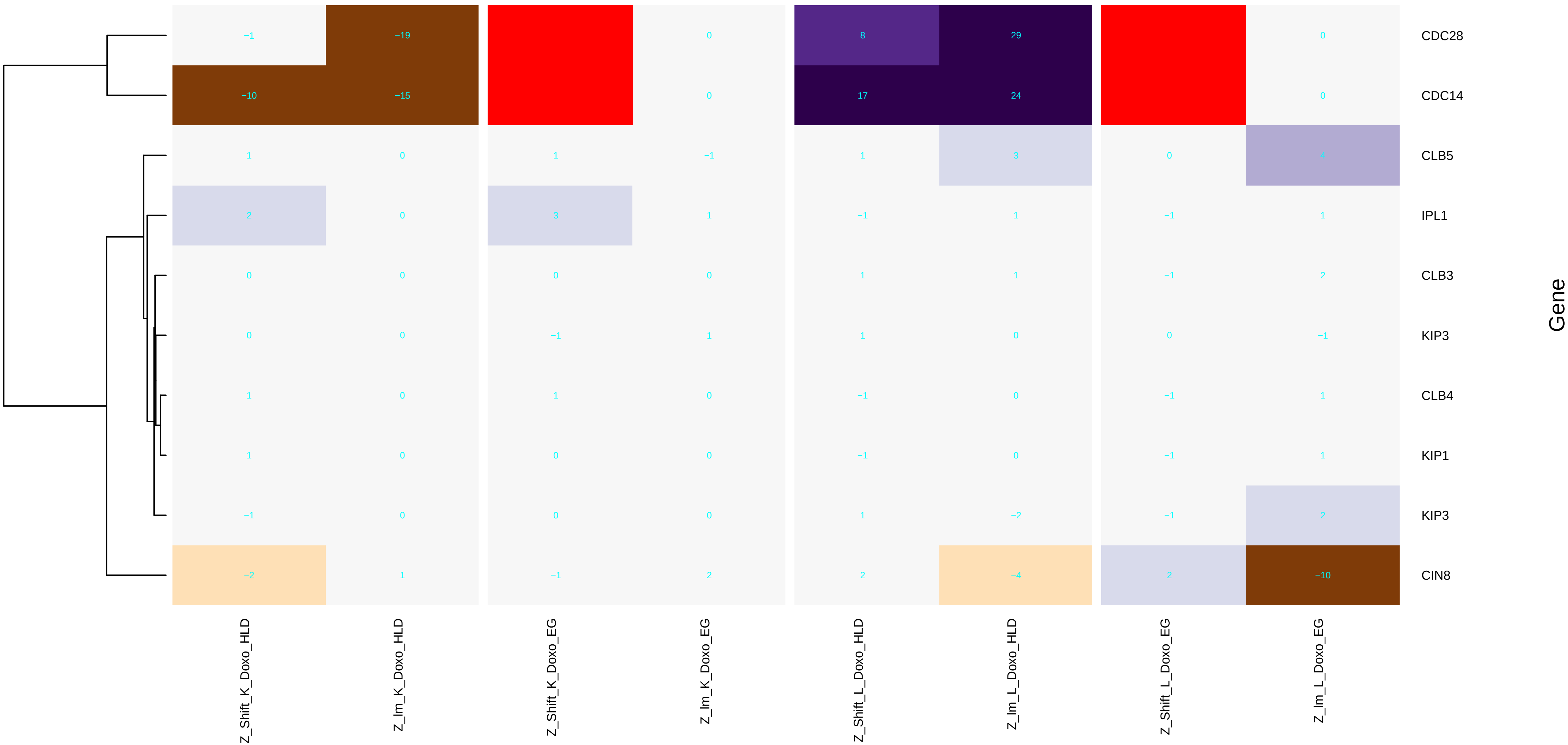

### monovalent_inorganic_cation_homeostasis.pdf

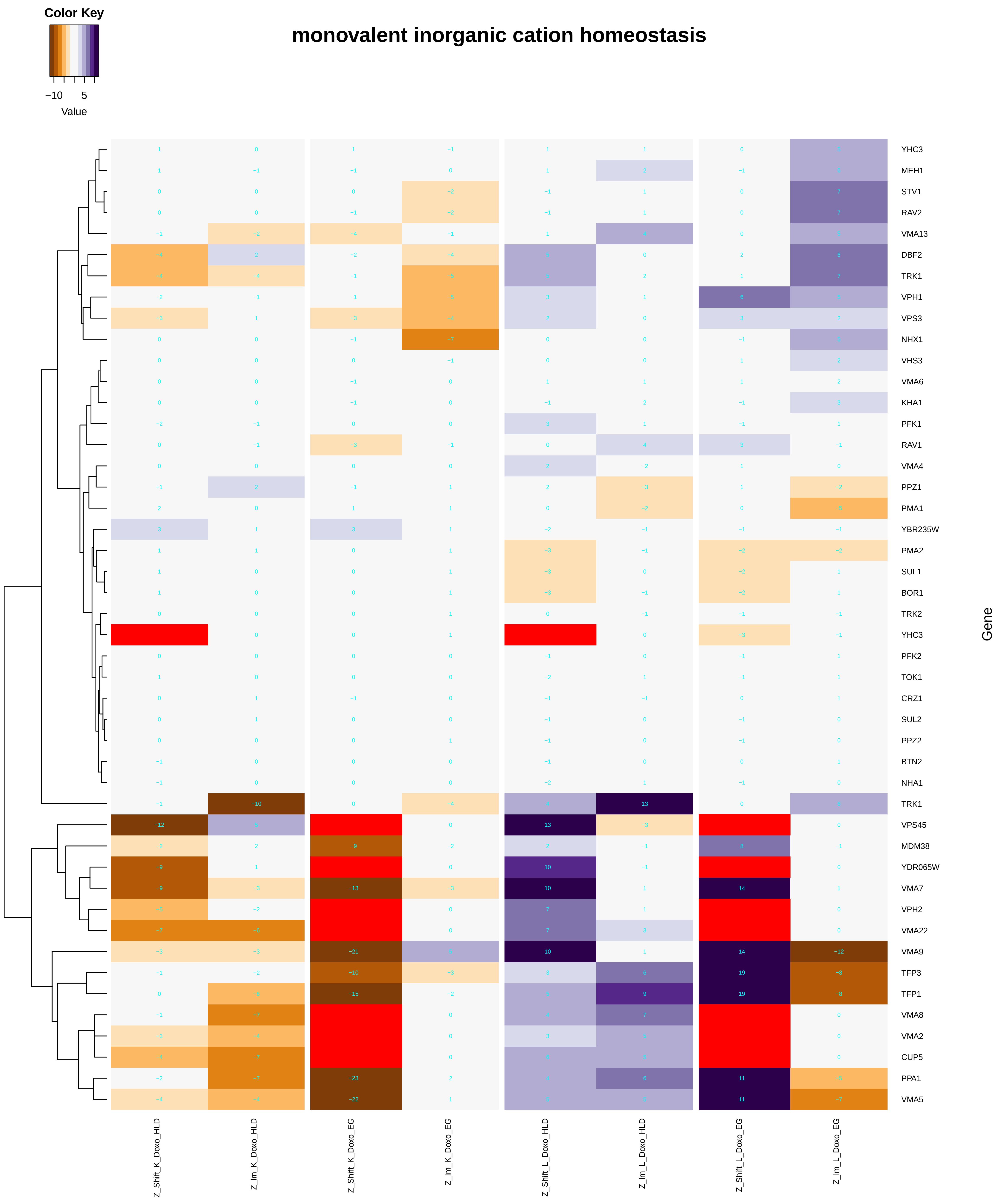

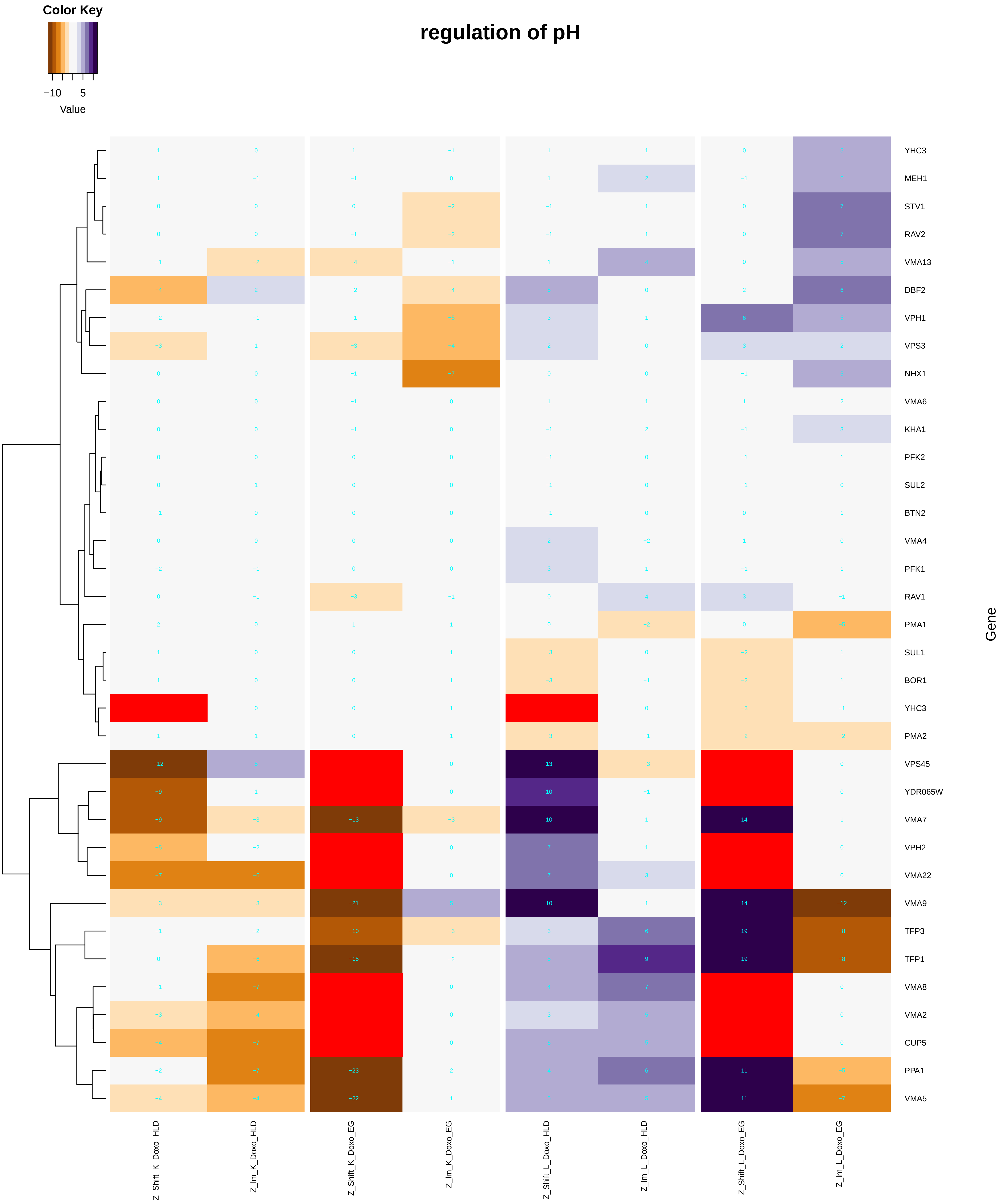

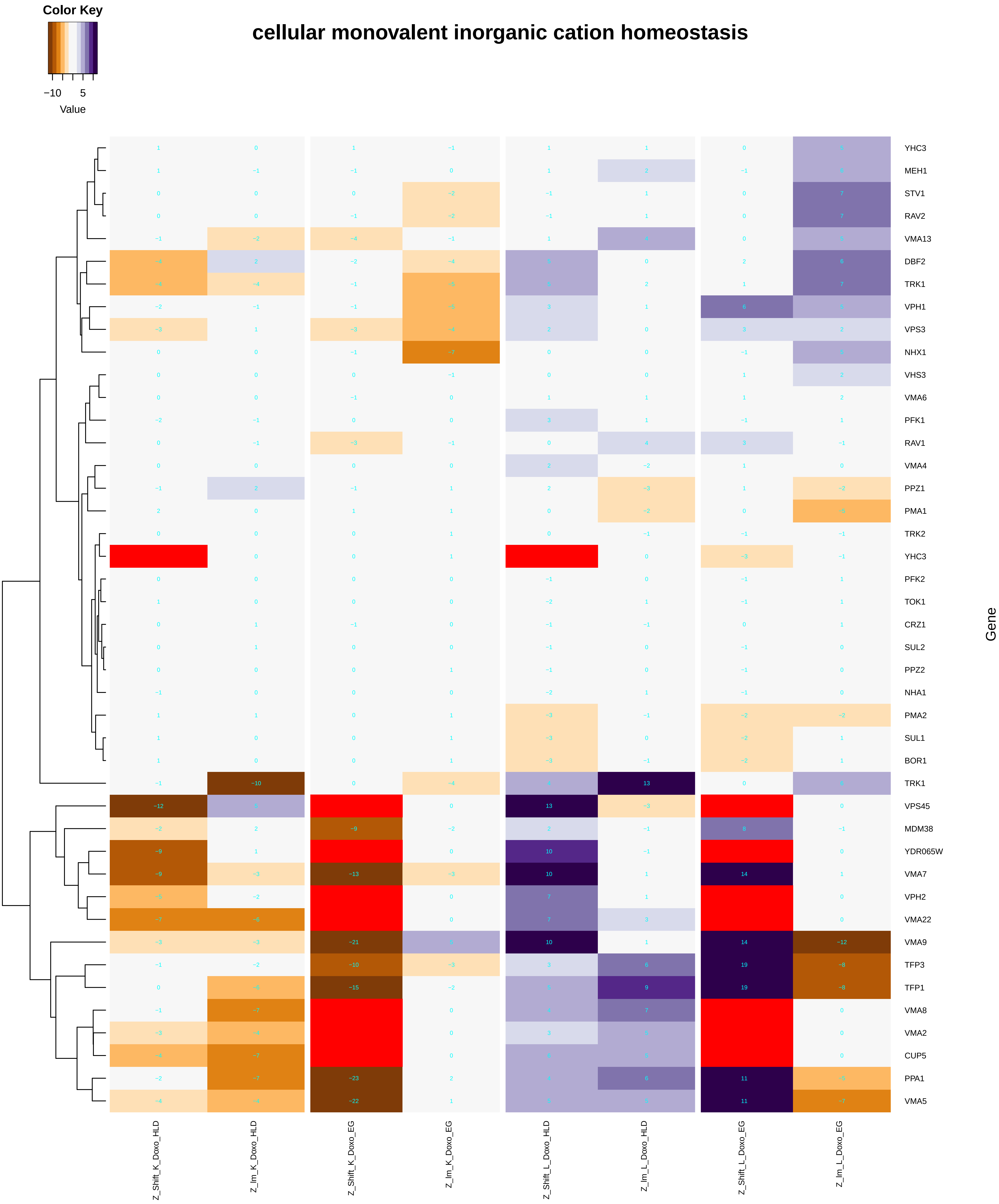

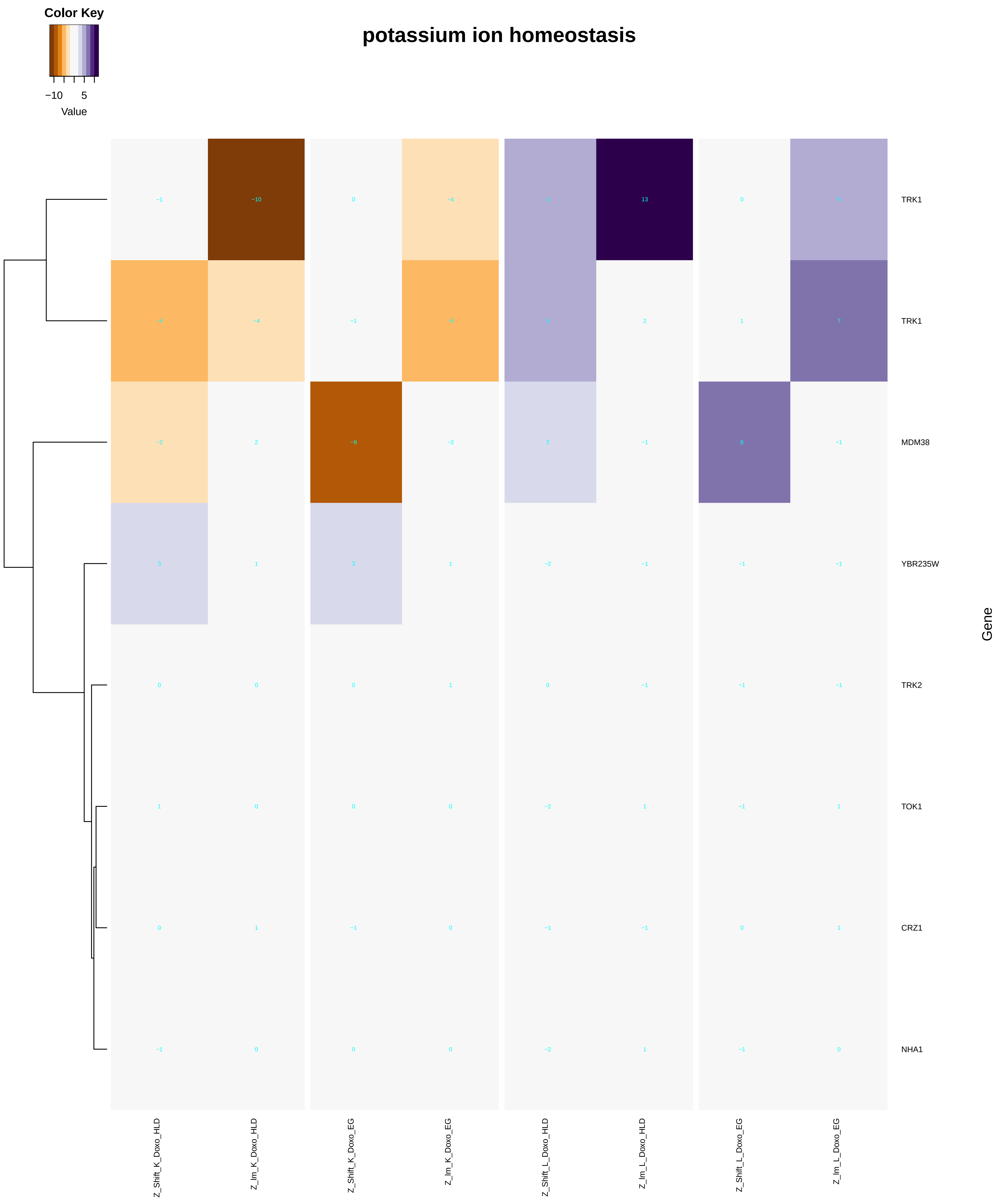

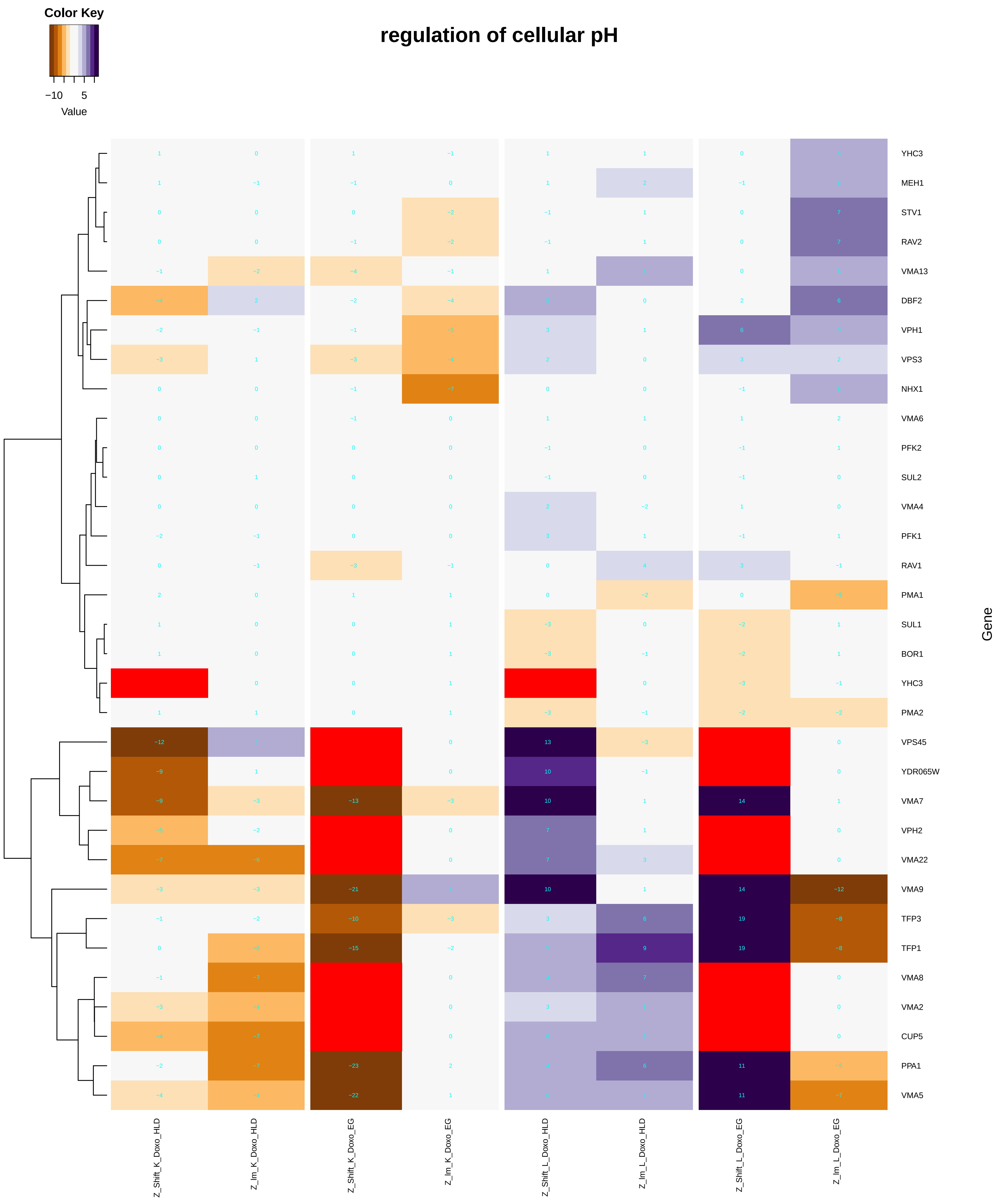

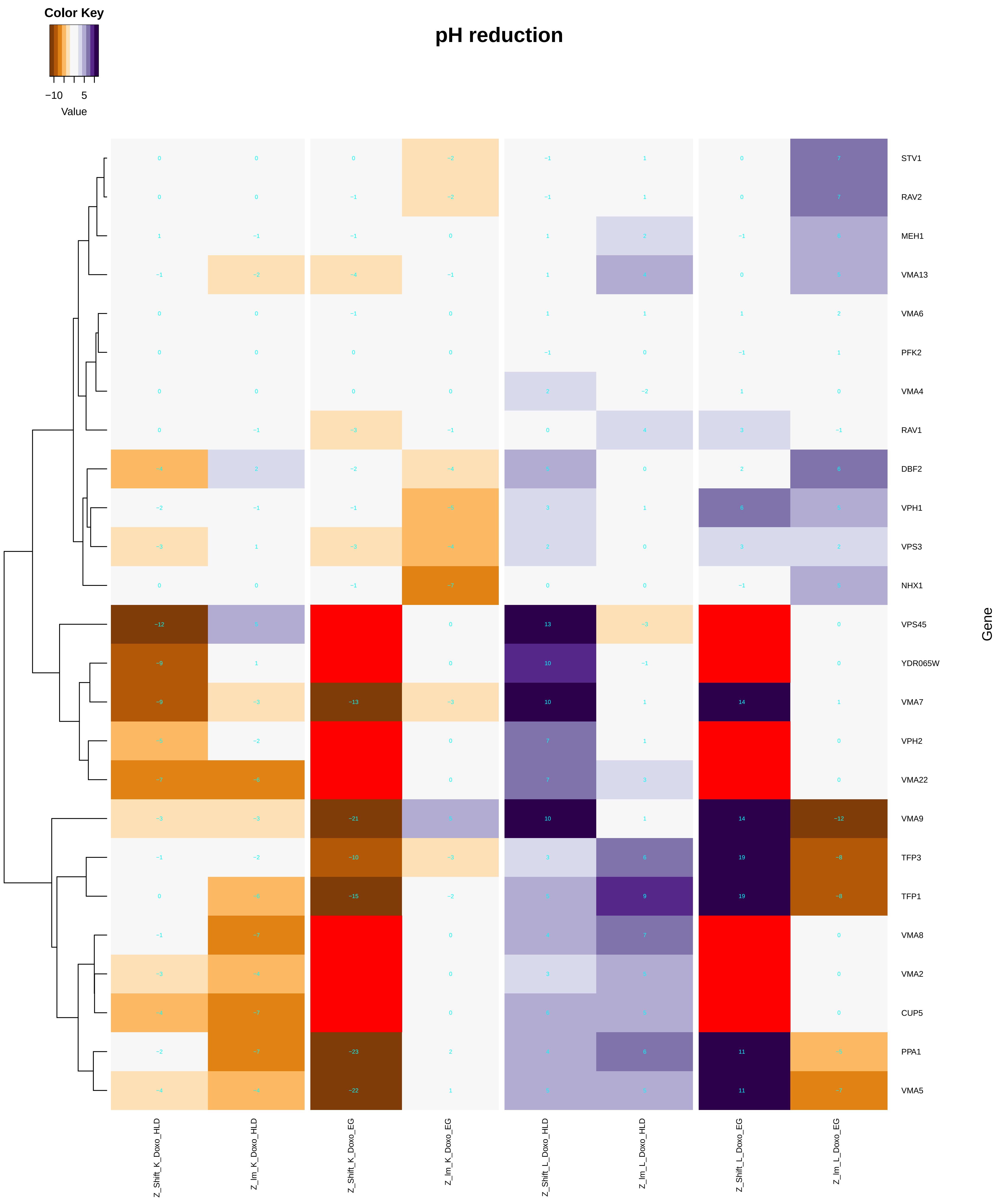

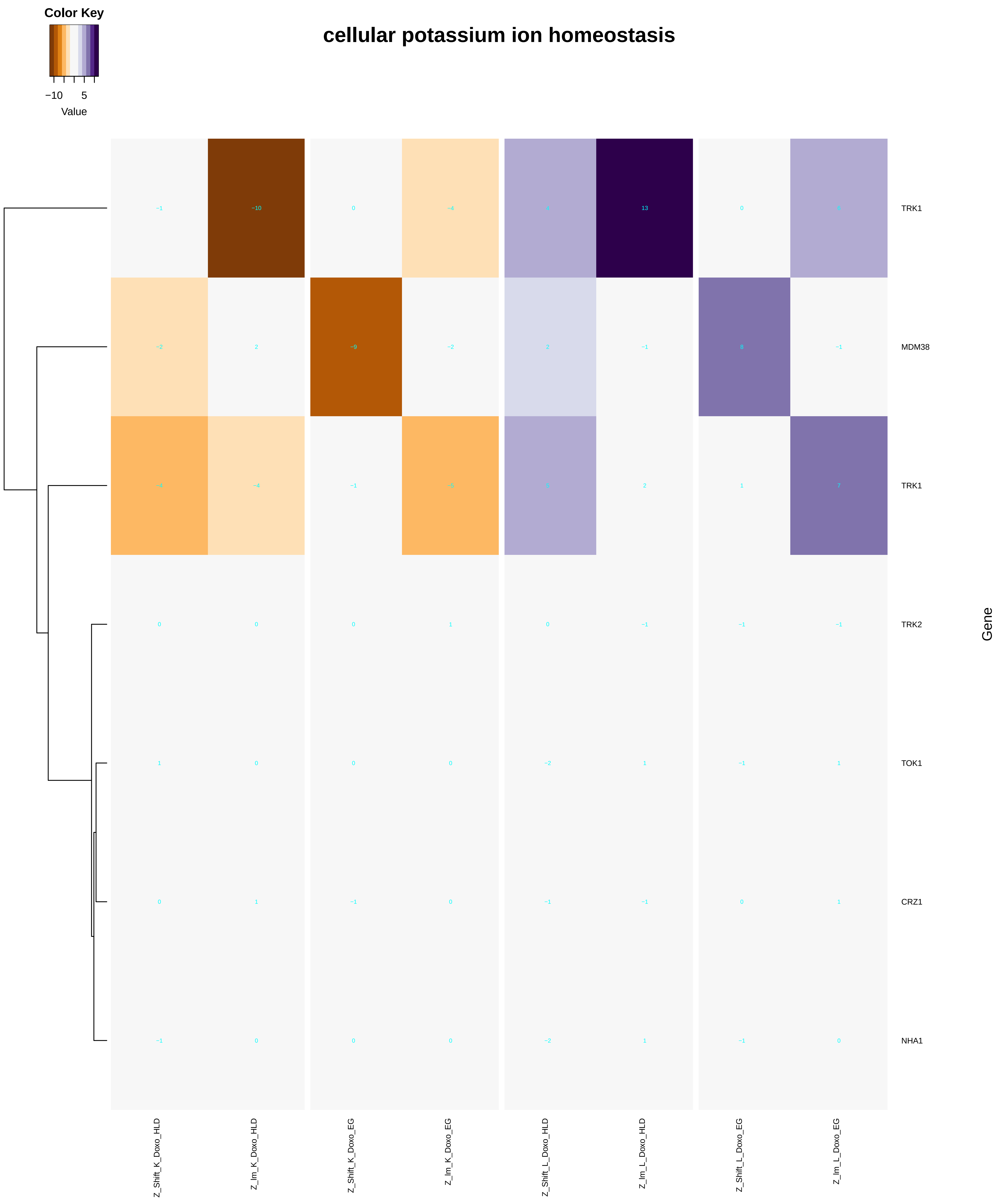

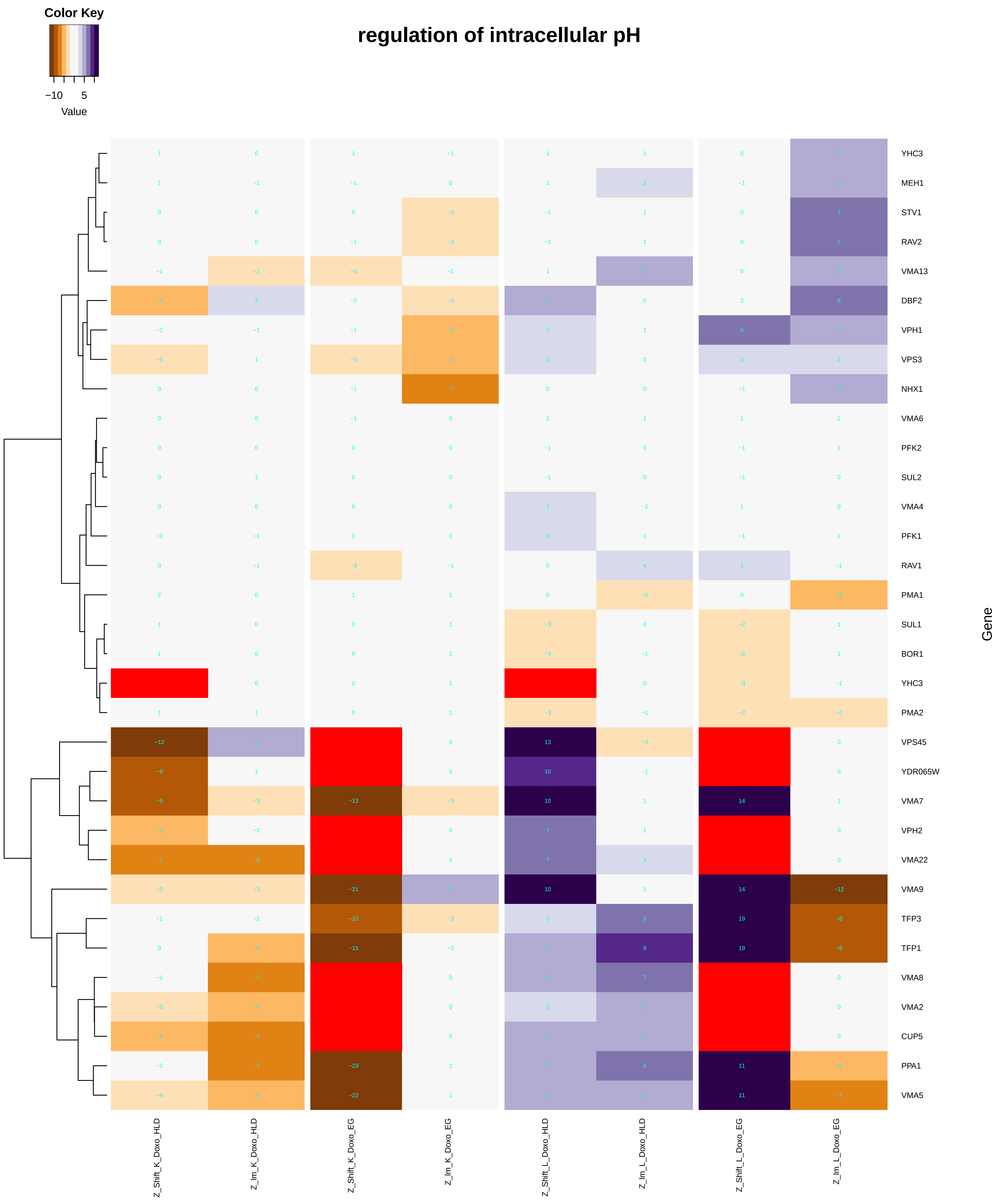

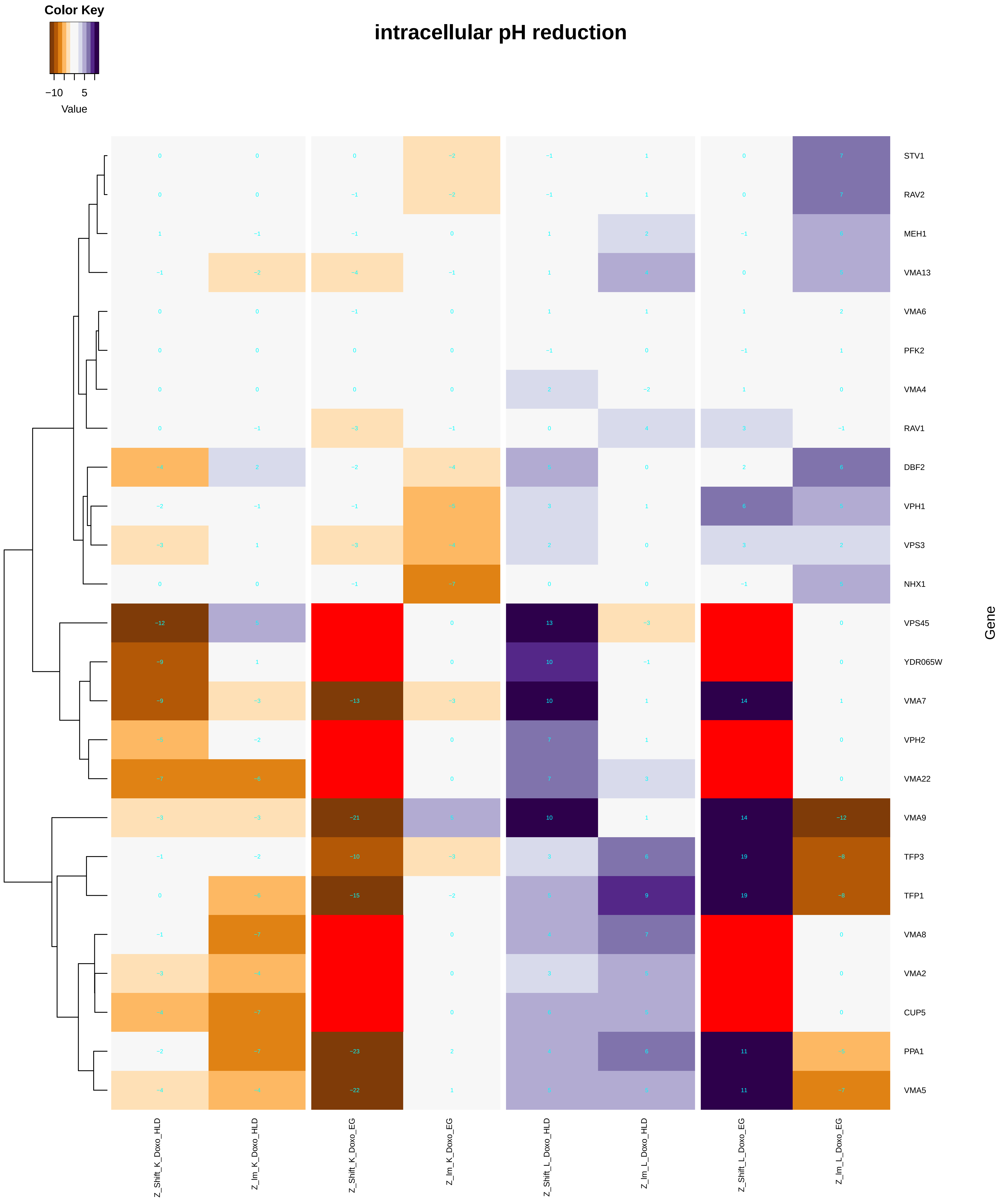

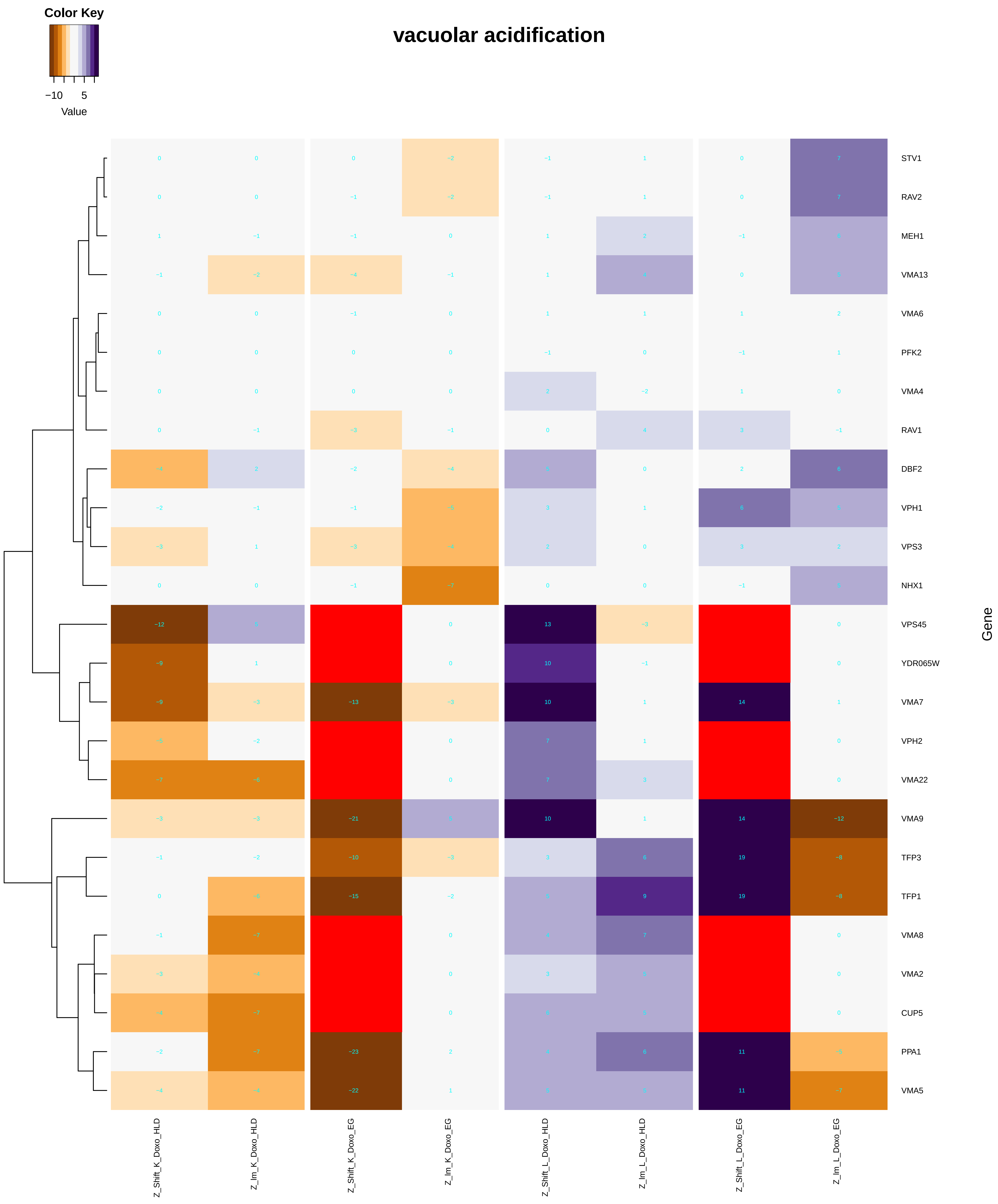

### mRNA-containing_ribonucleoprotein_complex_export_from_nucleus.pdf

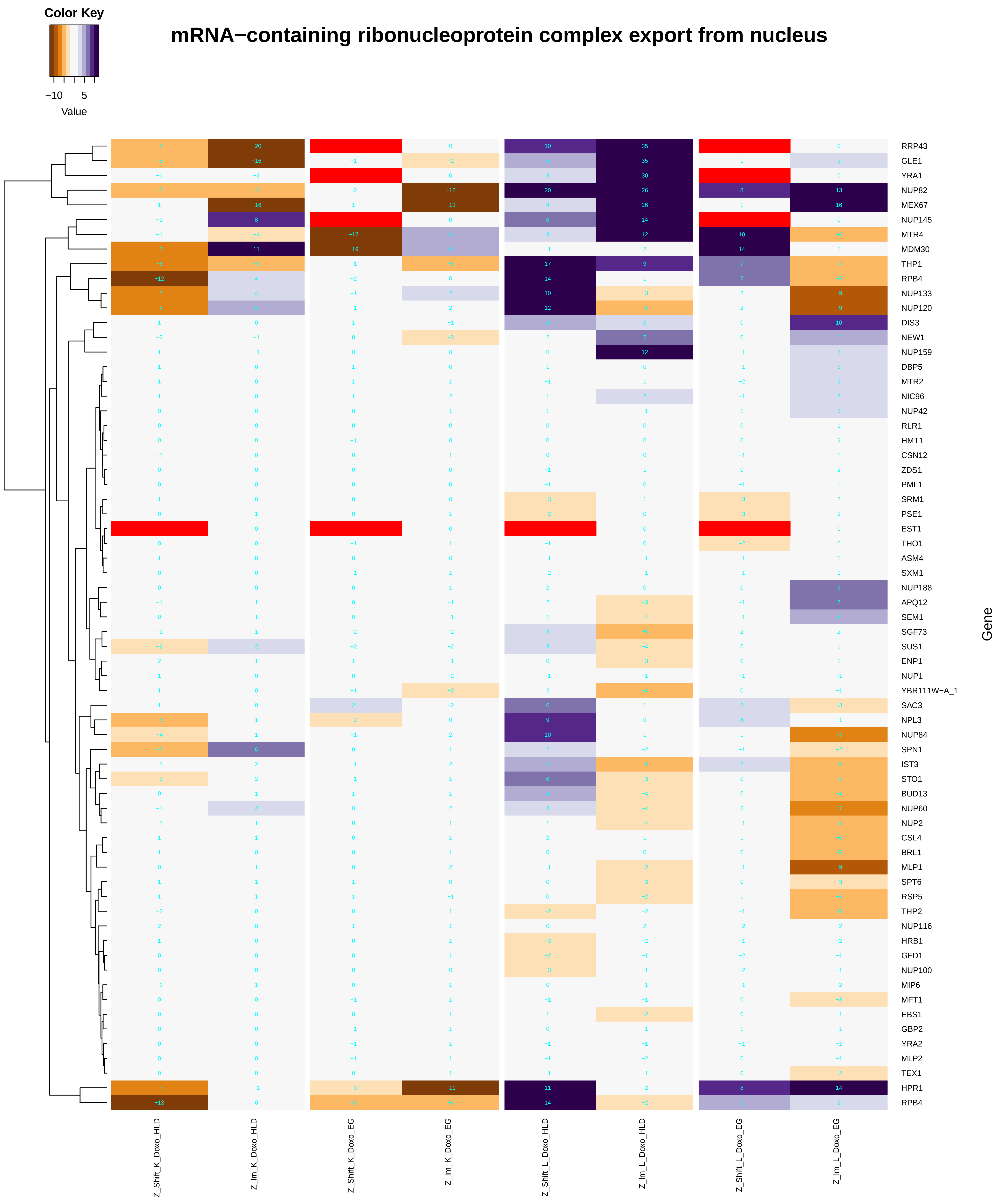

### mRNA_3'-end_processing.pdf

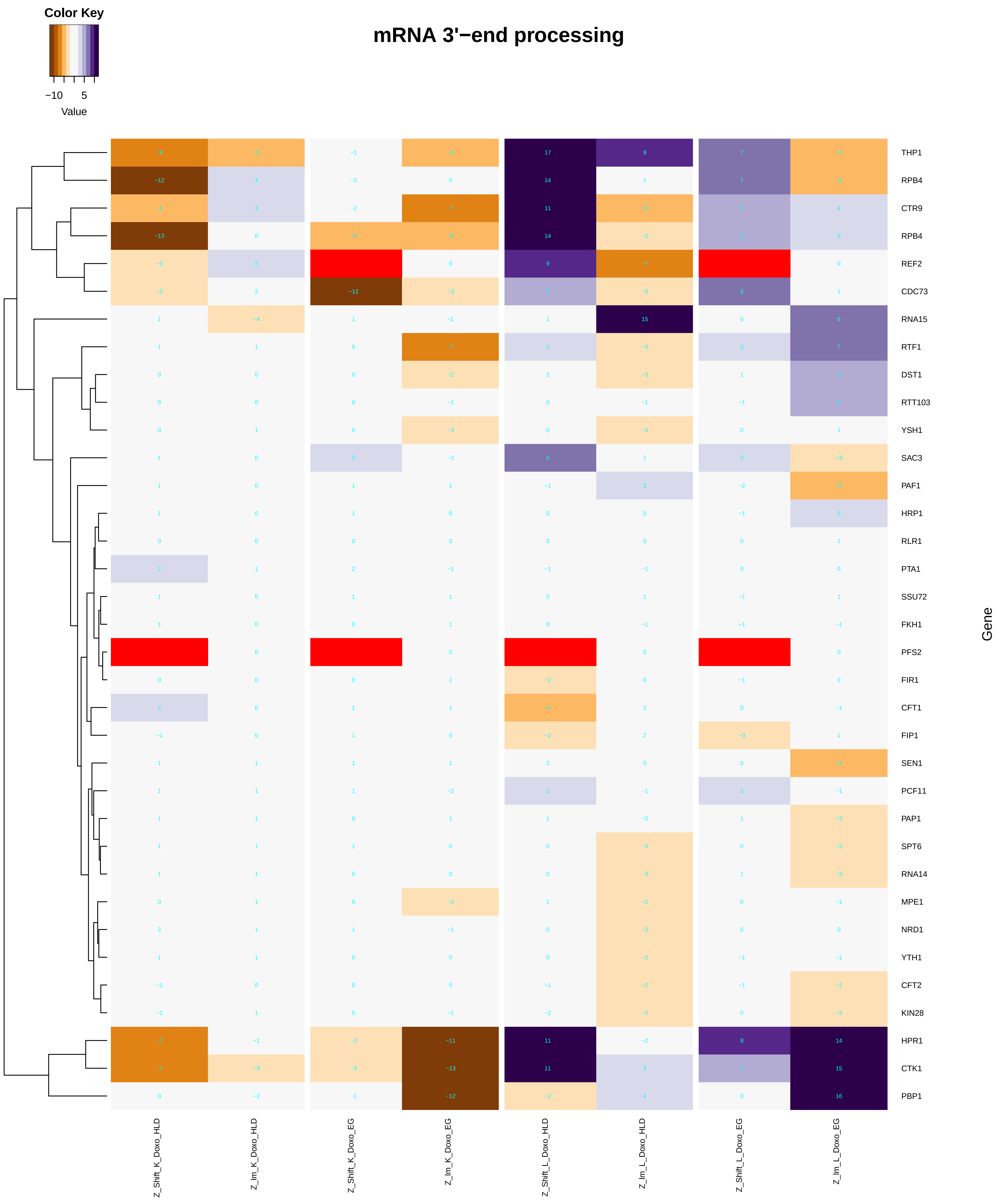

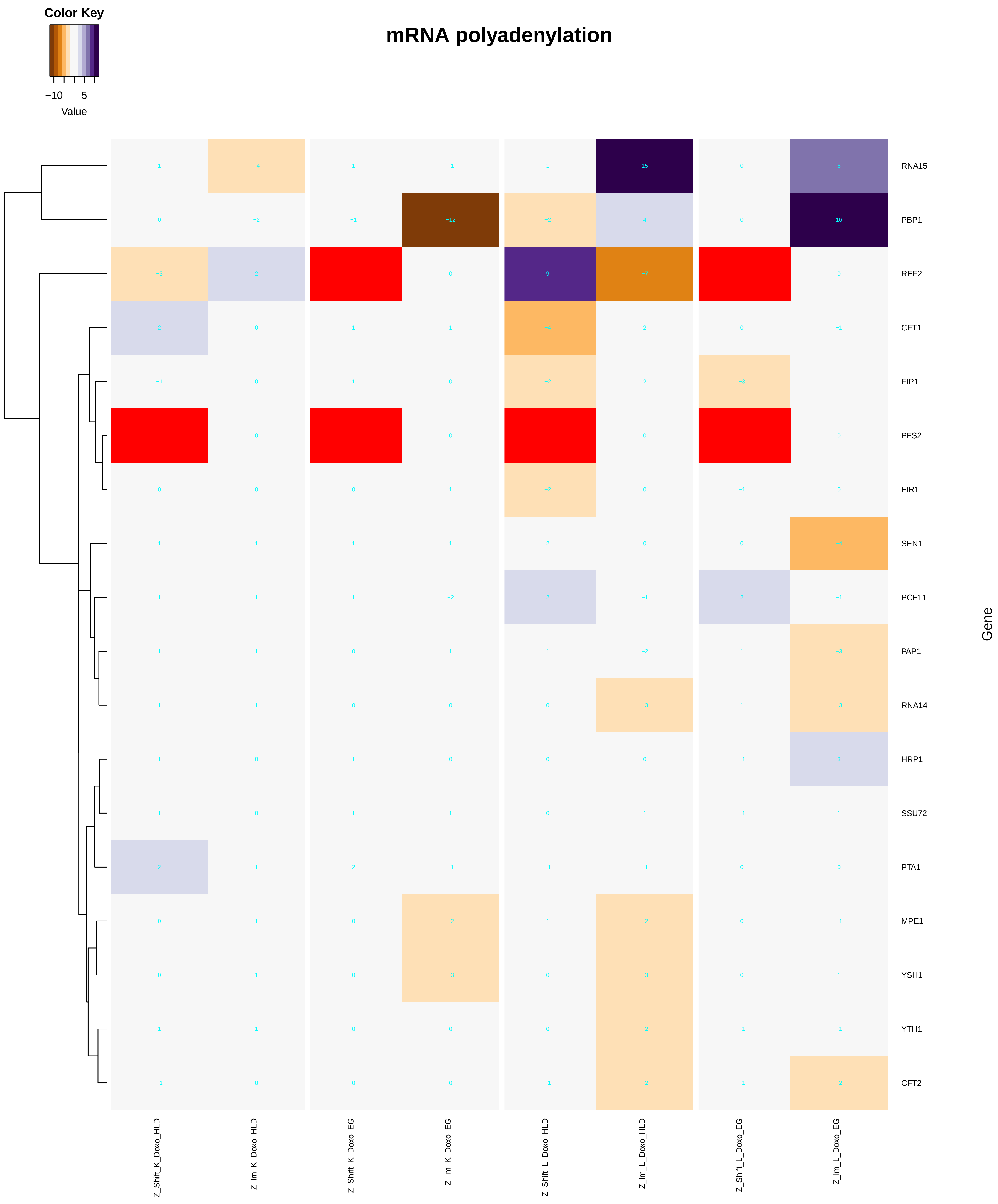

### mRNA_cleavage.pdf

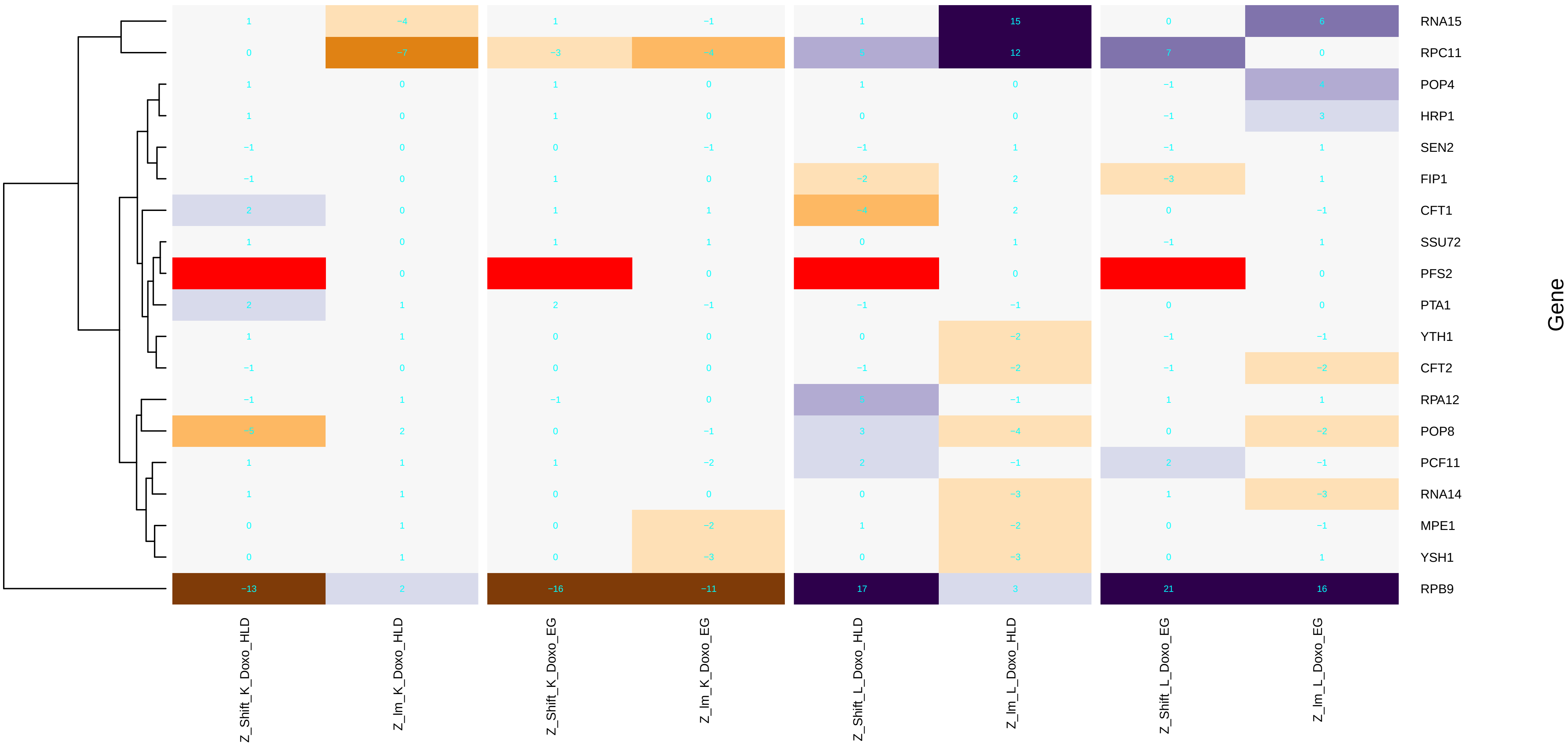

### mRNA_export_from_nucleus.pdf

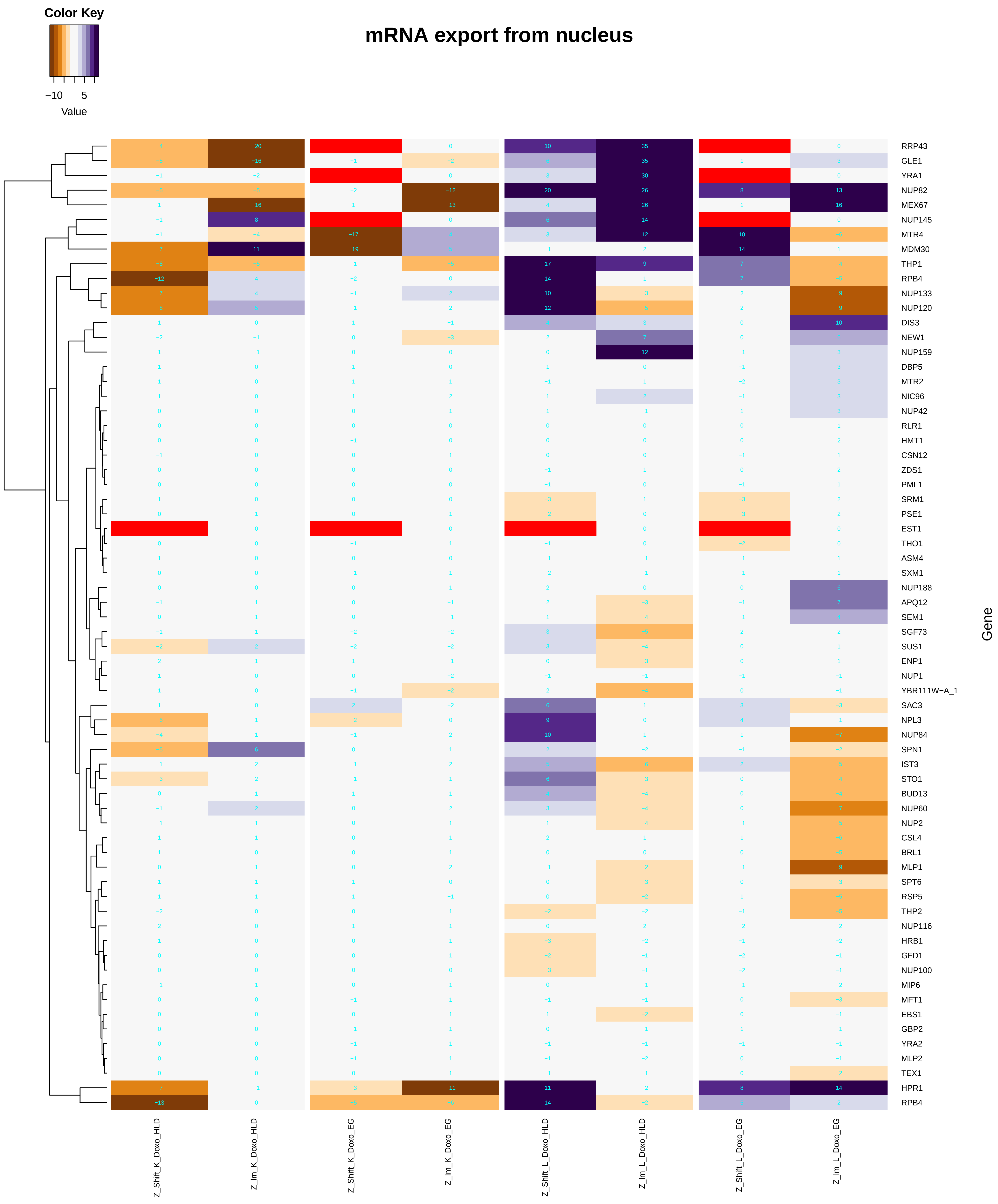

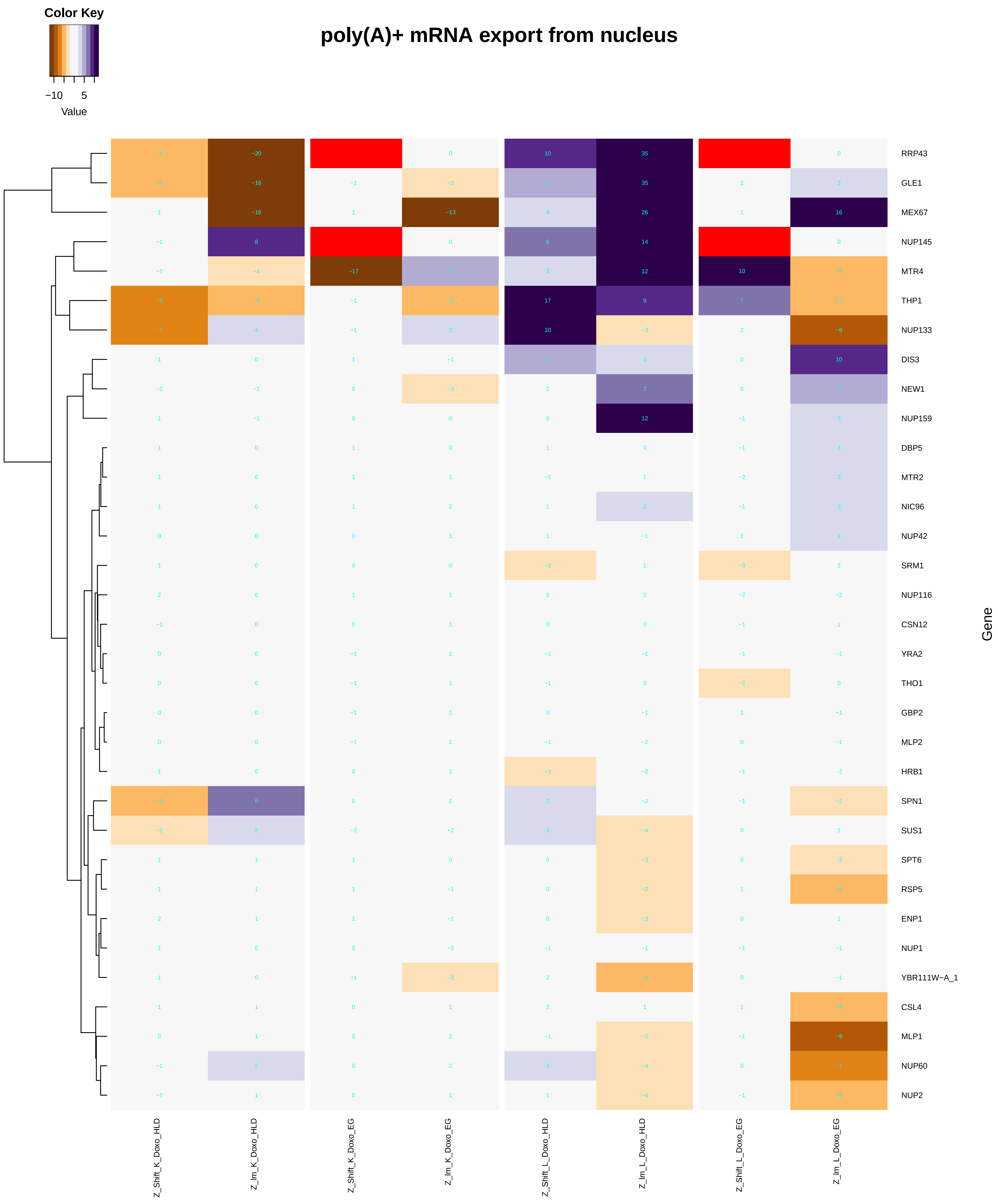

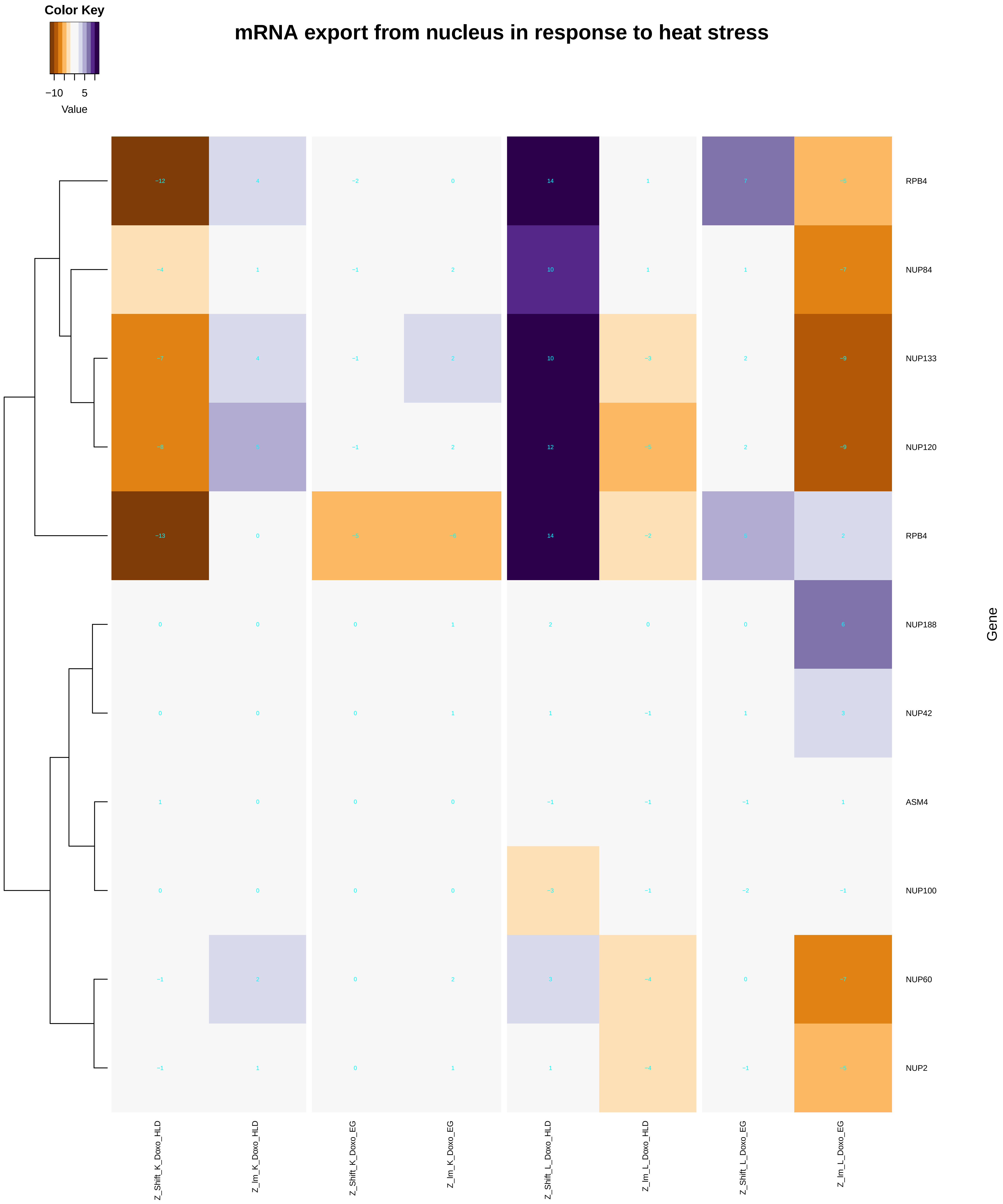

### mRNA_export_from_nucleus_in_response_to_heat_stress.pdf

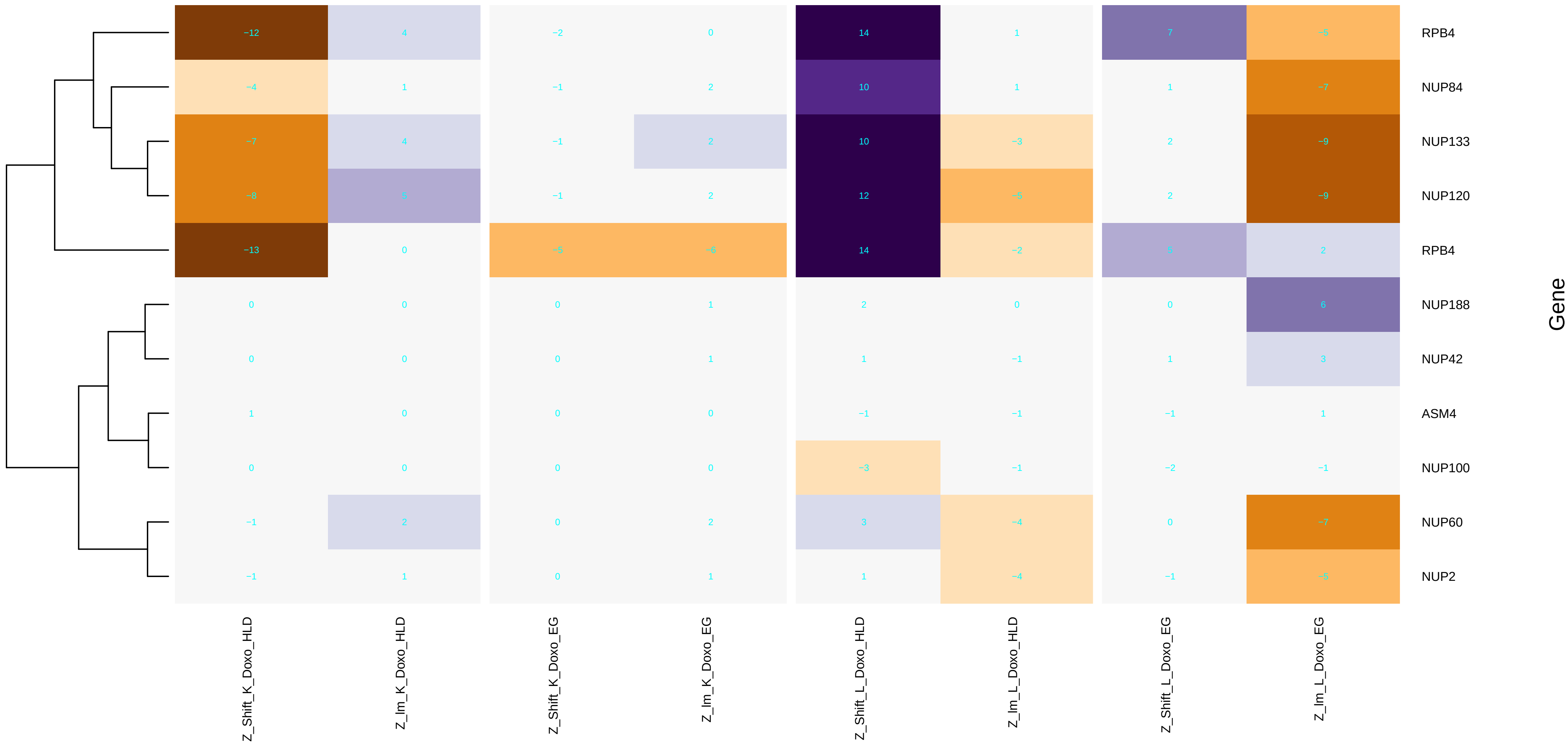

### mRNA_processing.pdf

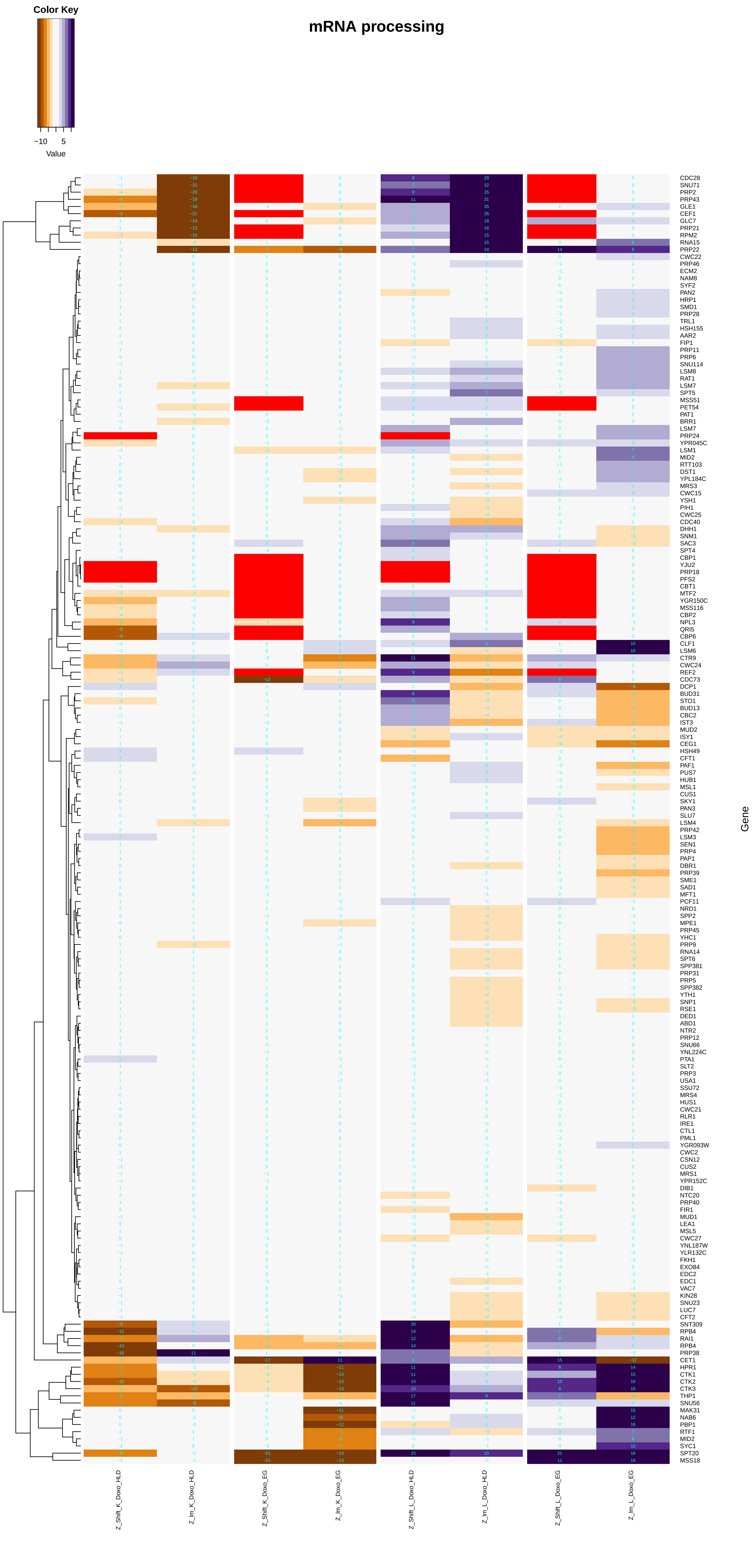

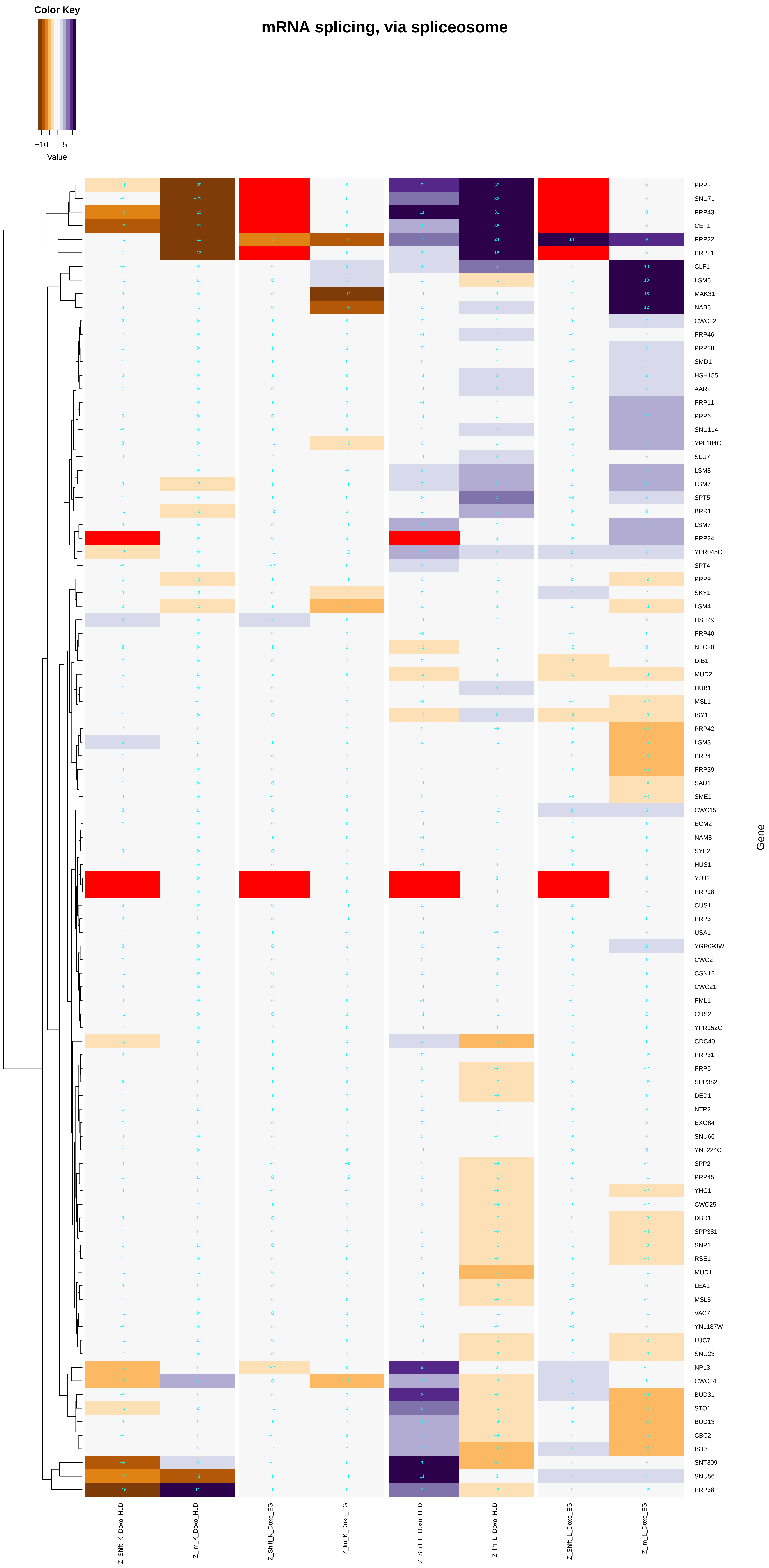

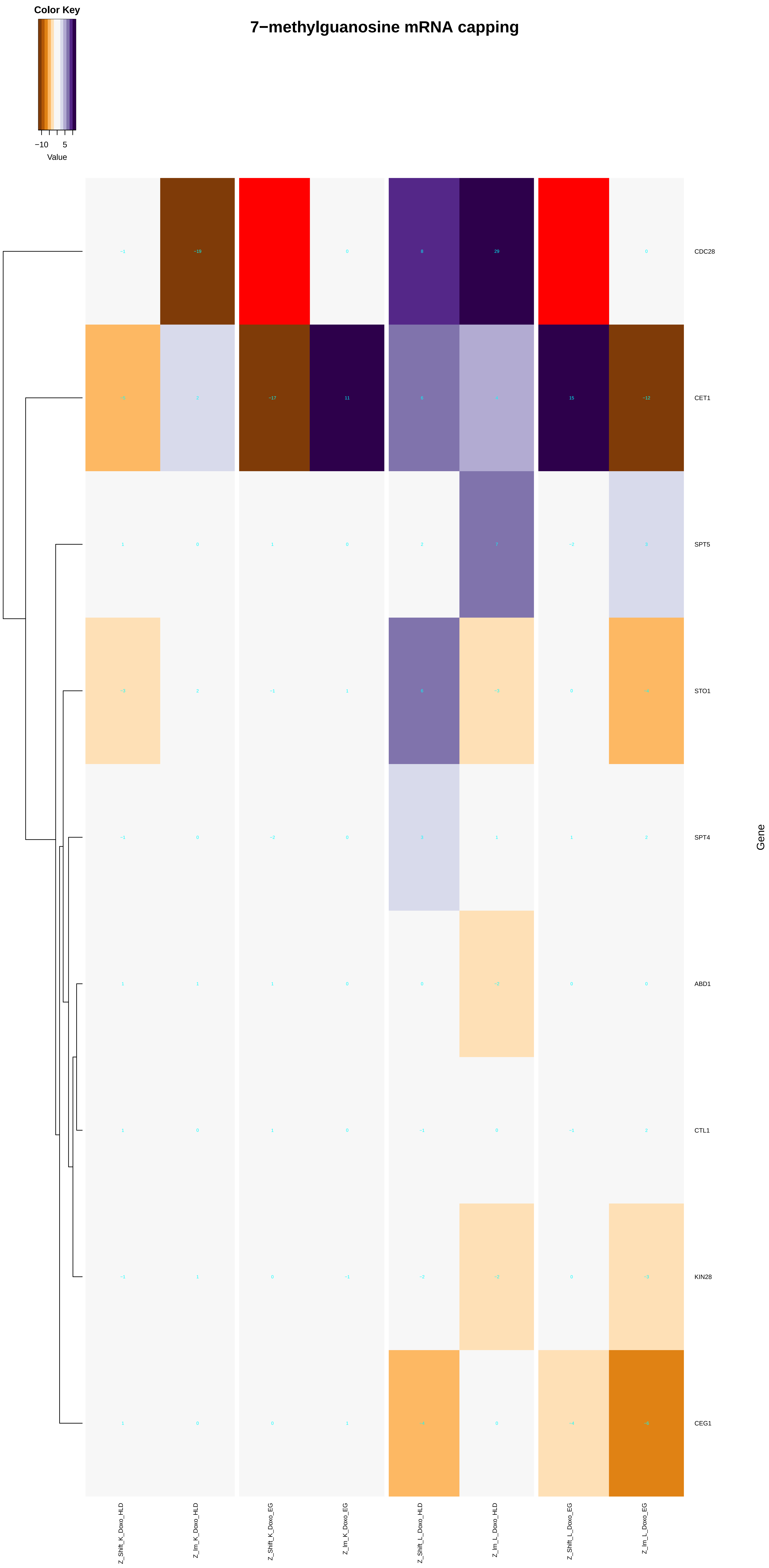

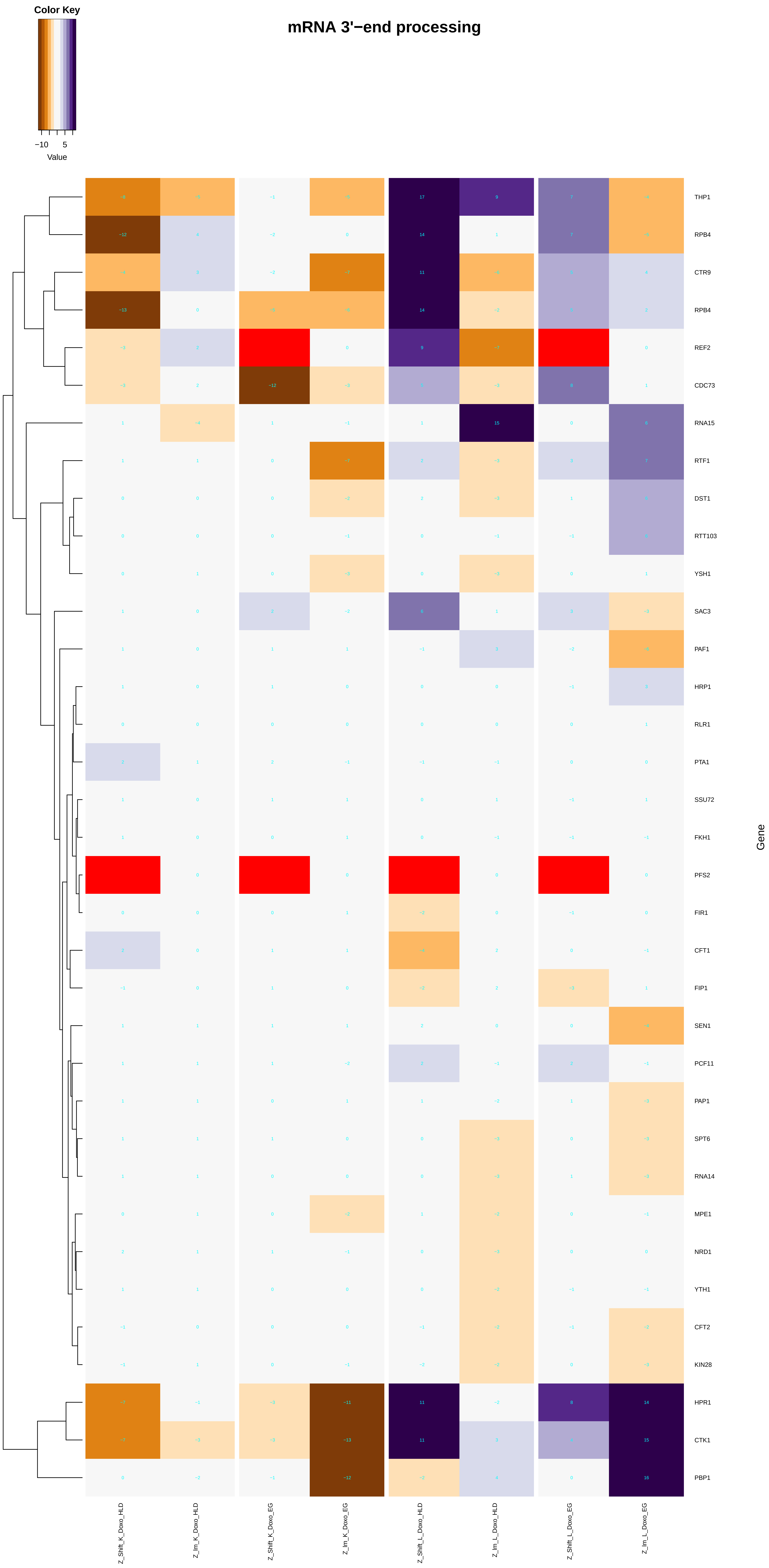

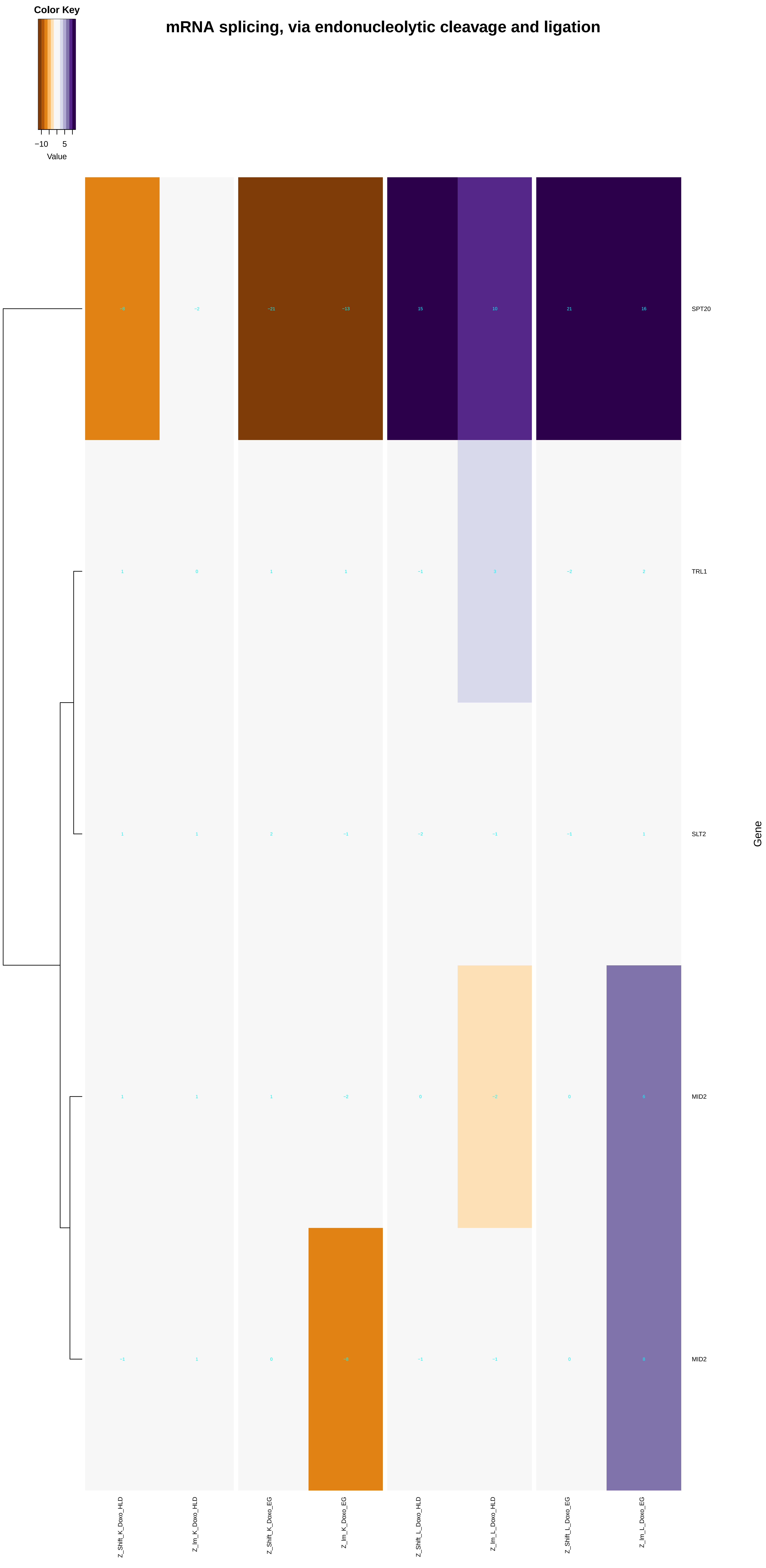

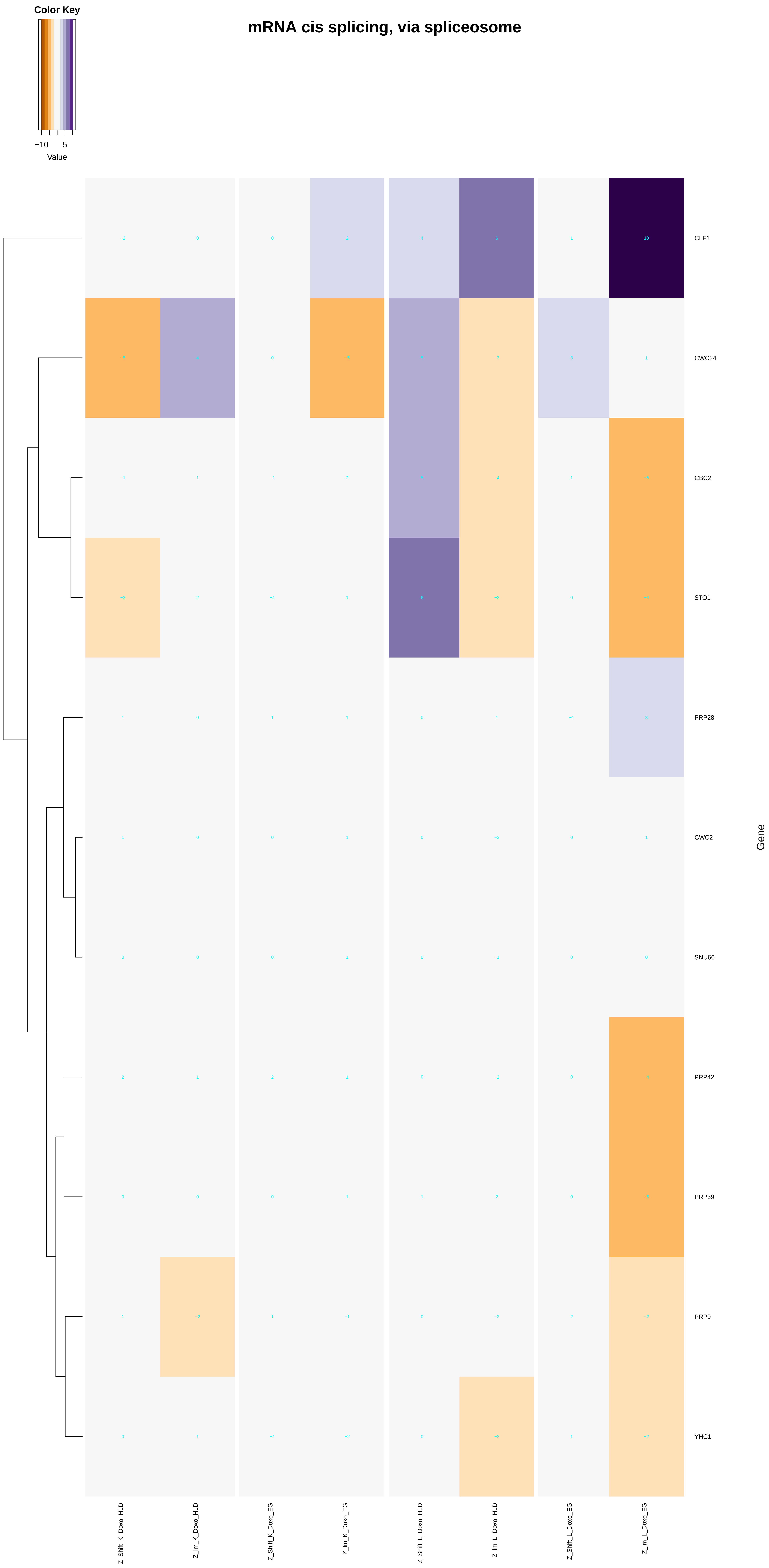

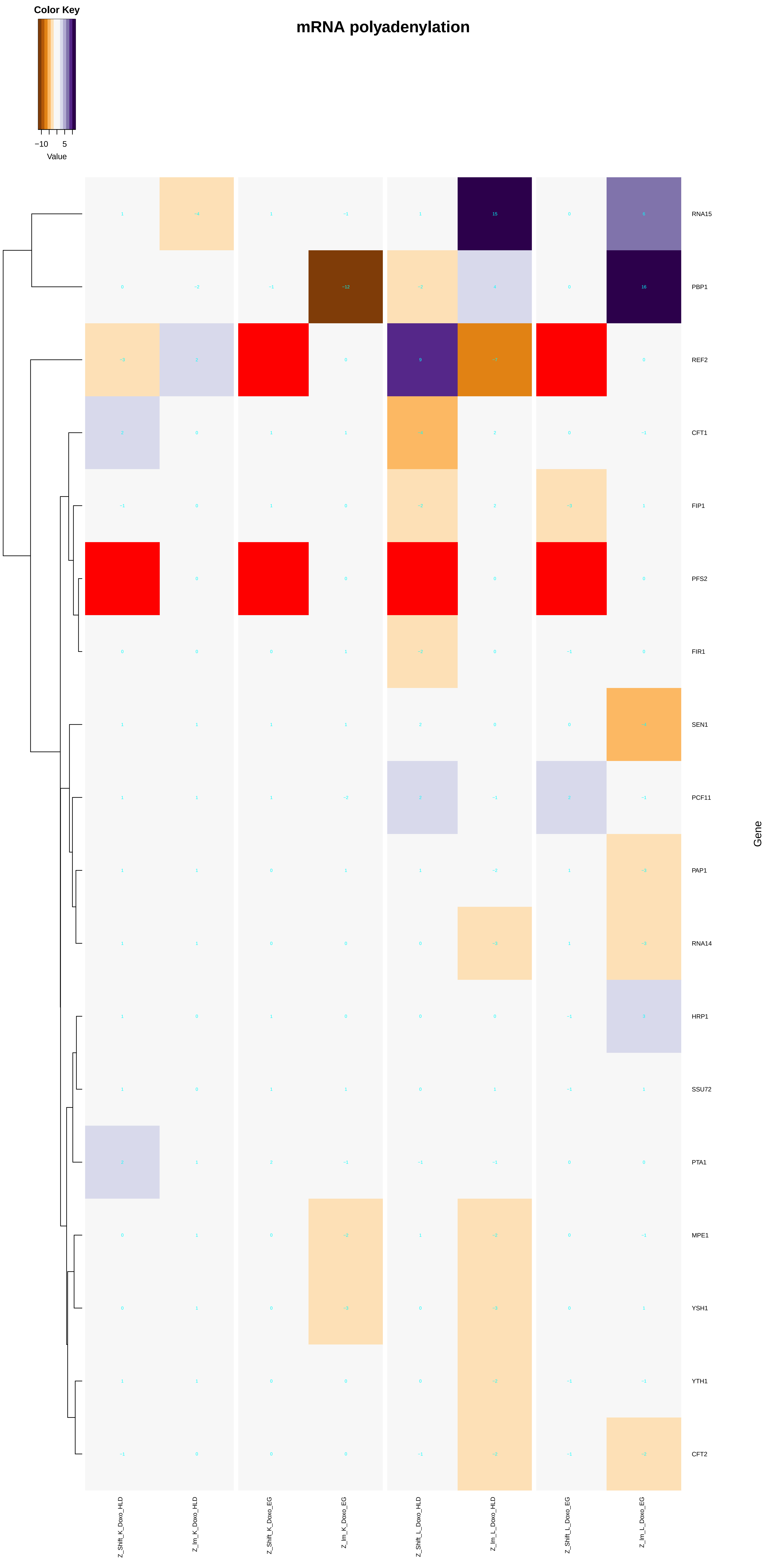

##### plicing via endonucleolytic cleavage and ligation involved in unfolded protein response

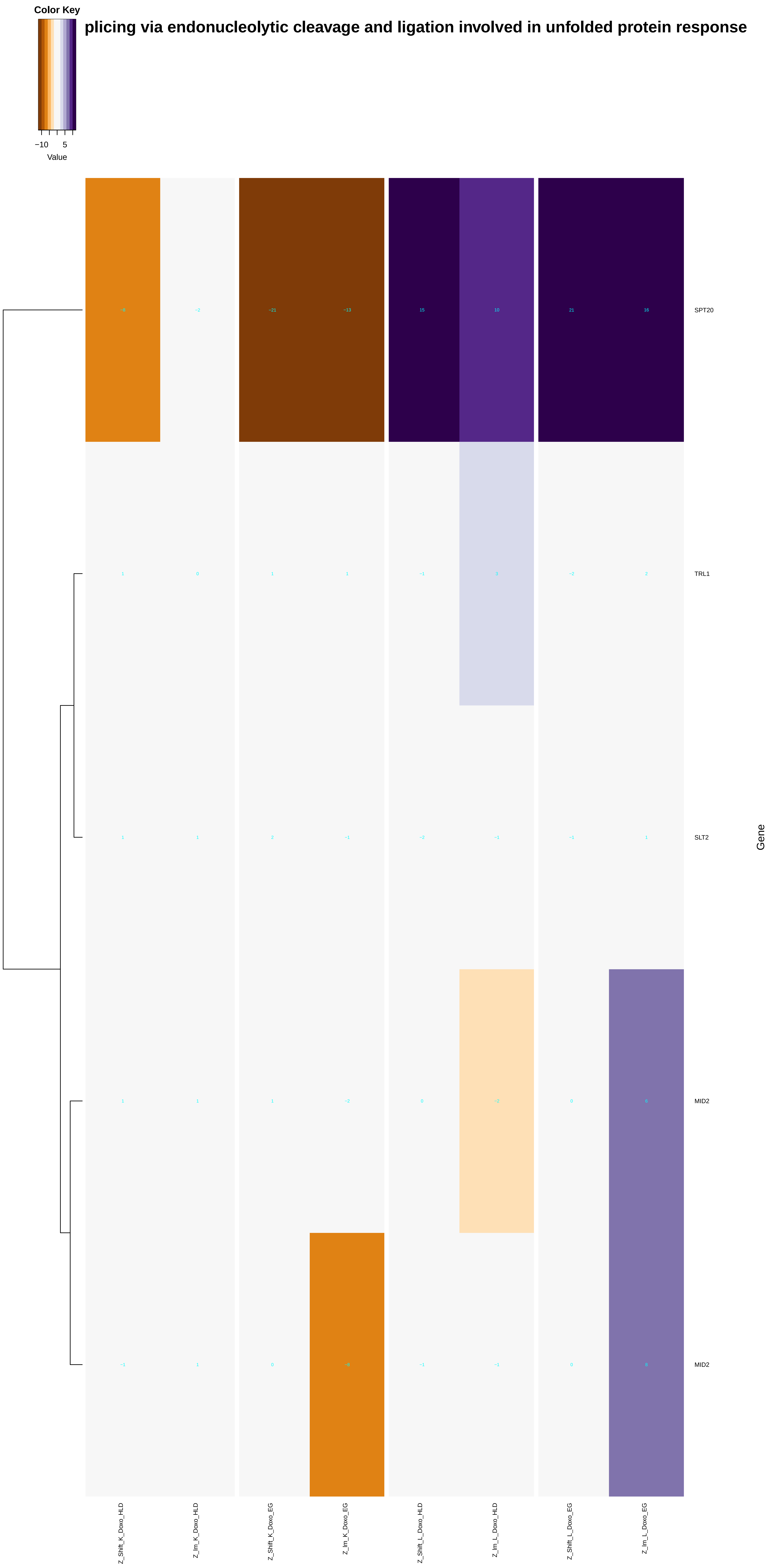

### mRNA_splicing,_via_spliceosome.pdf

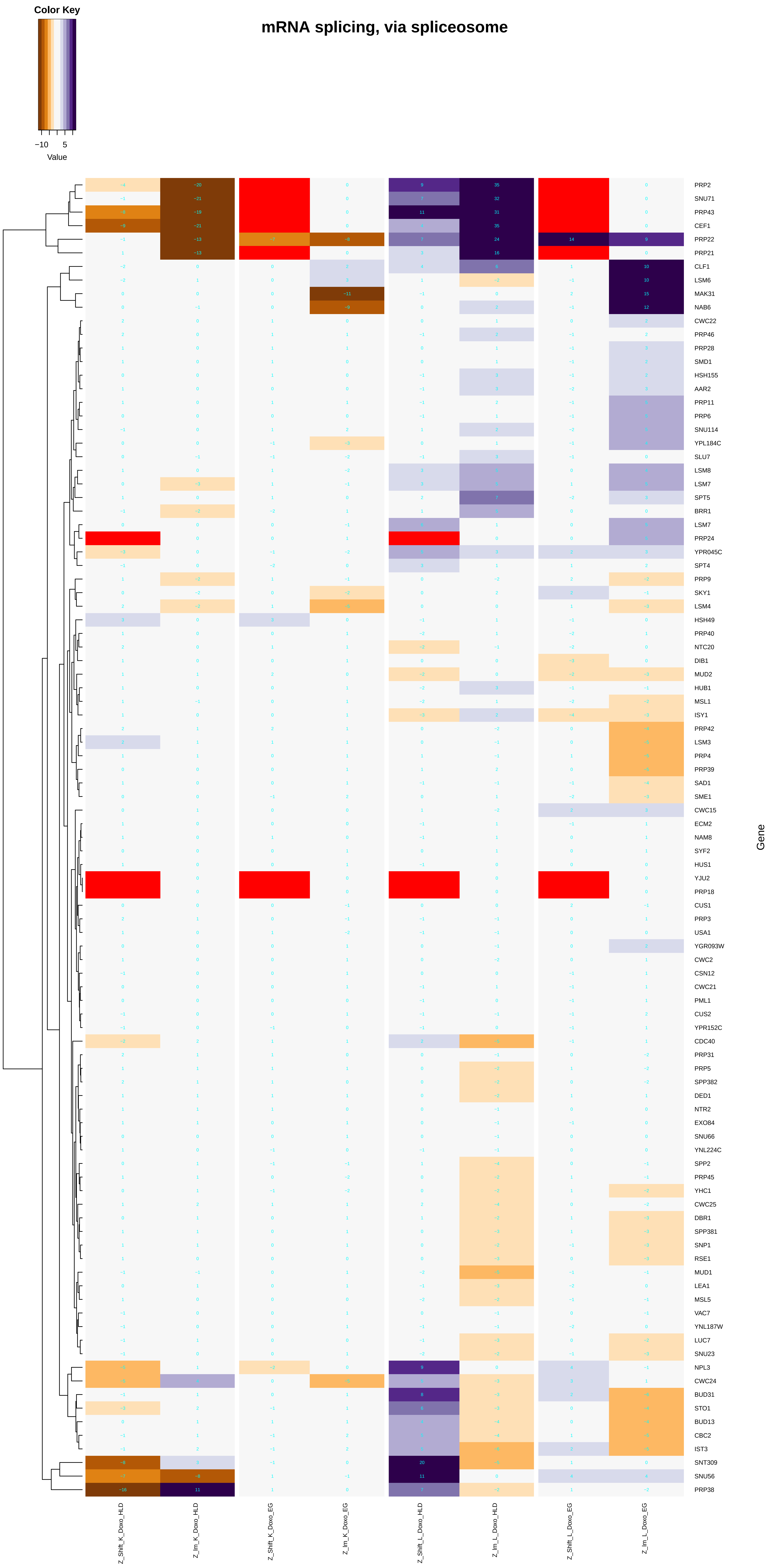

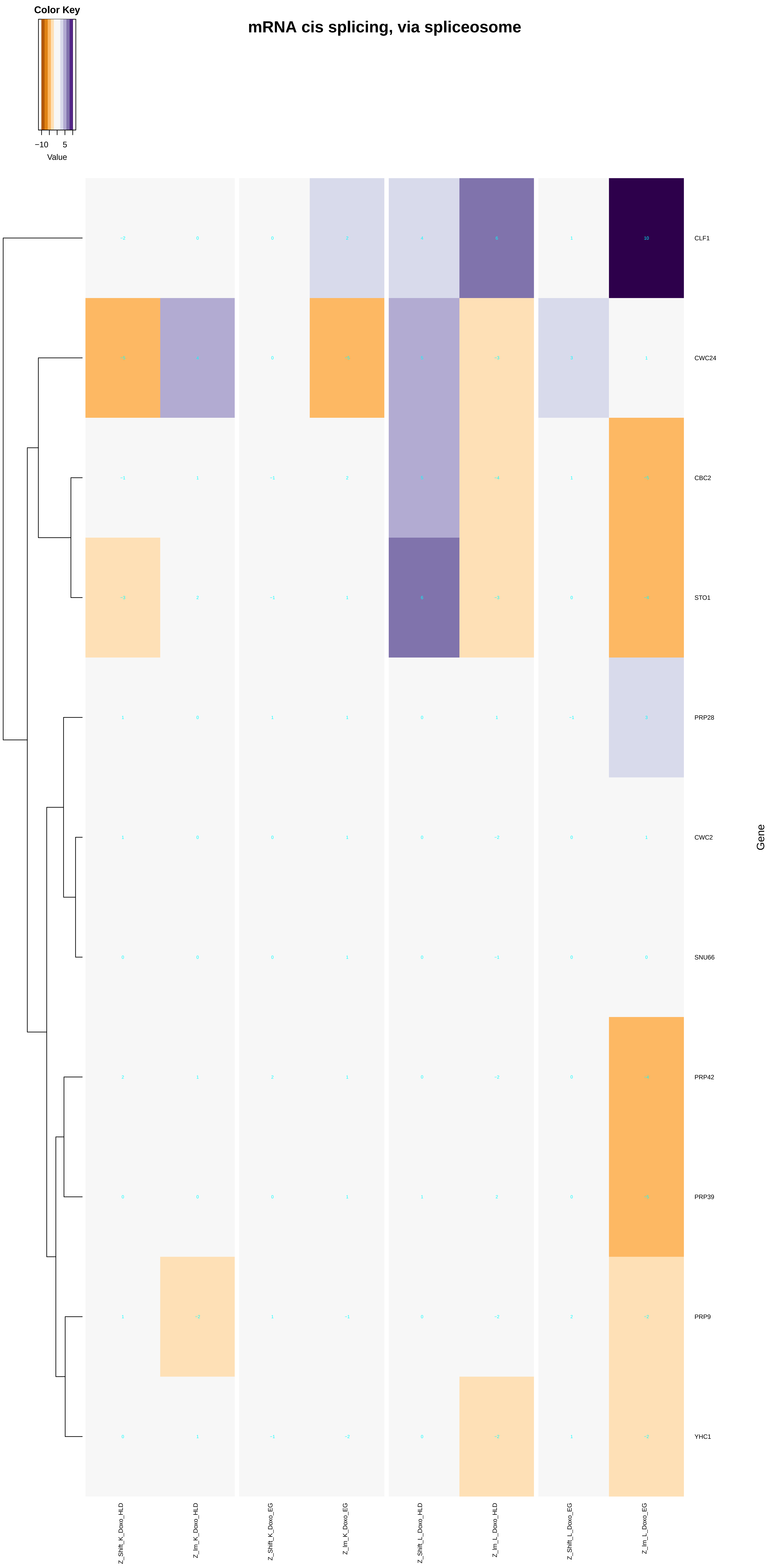

### mRNA_transport.pdf

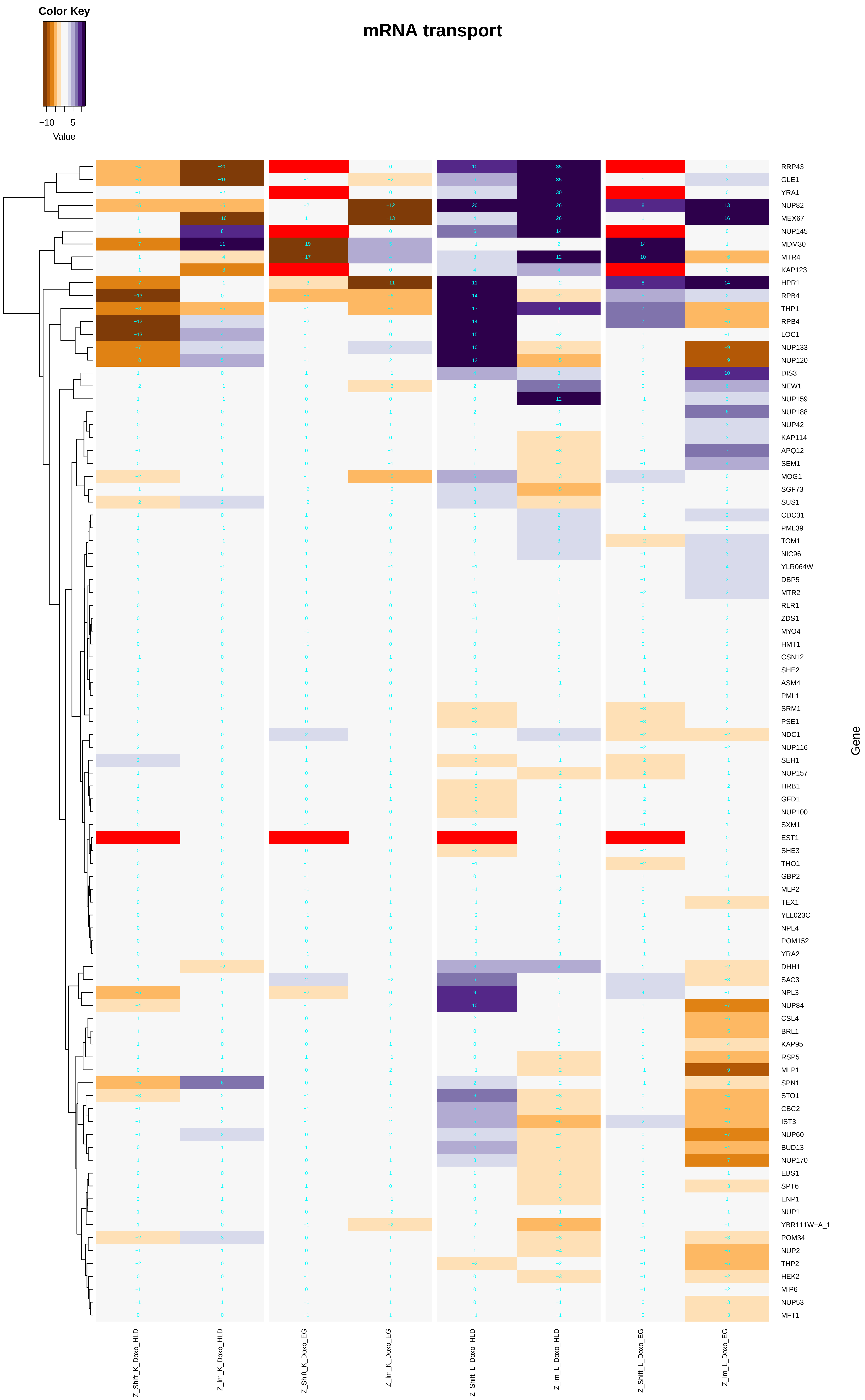

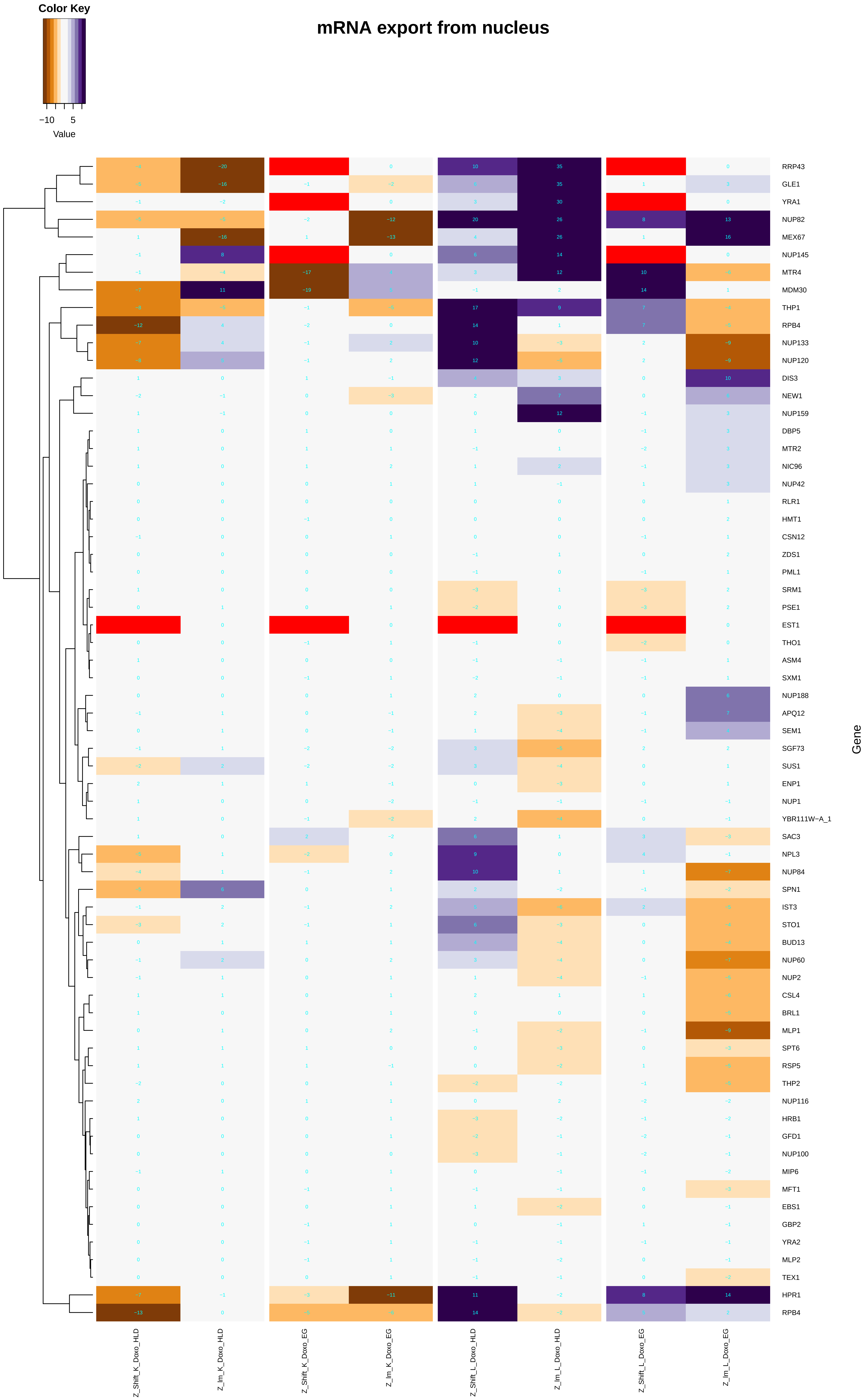

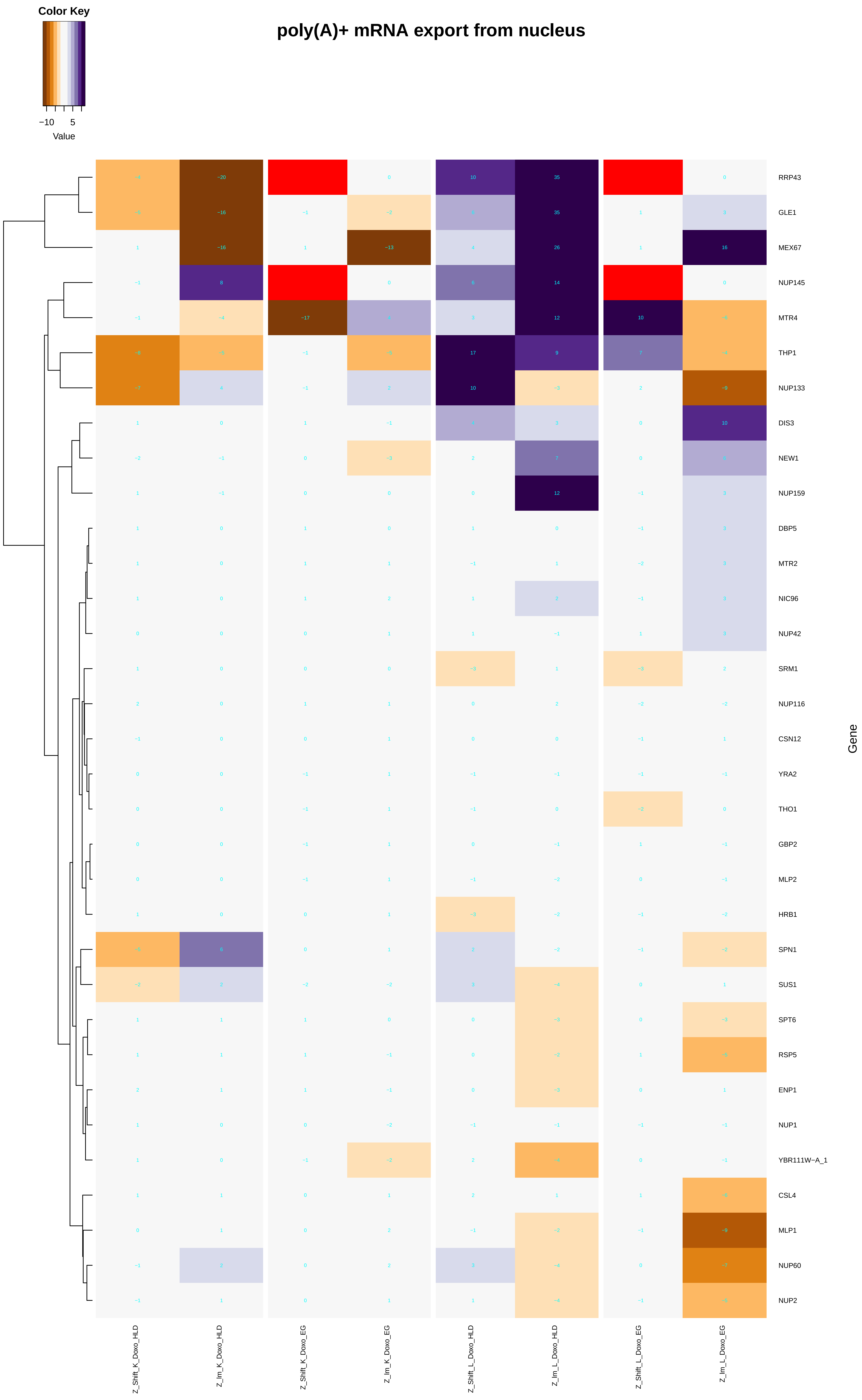

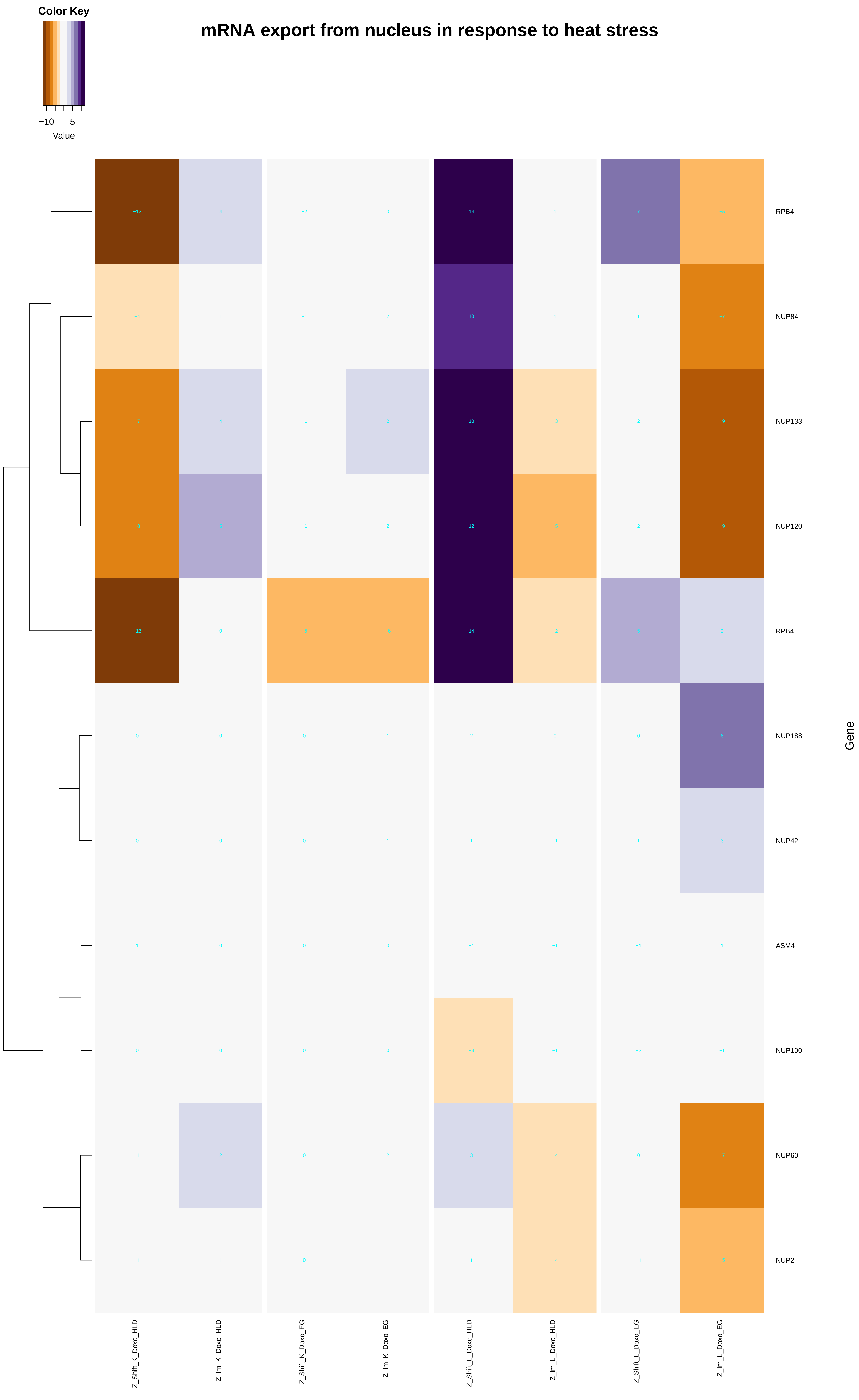

### ncRNA_transcription.pdf

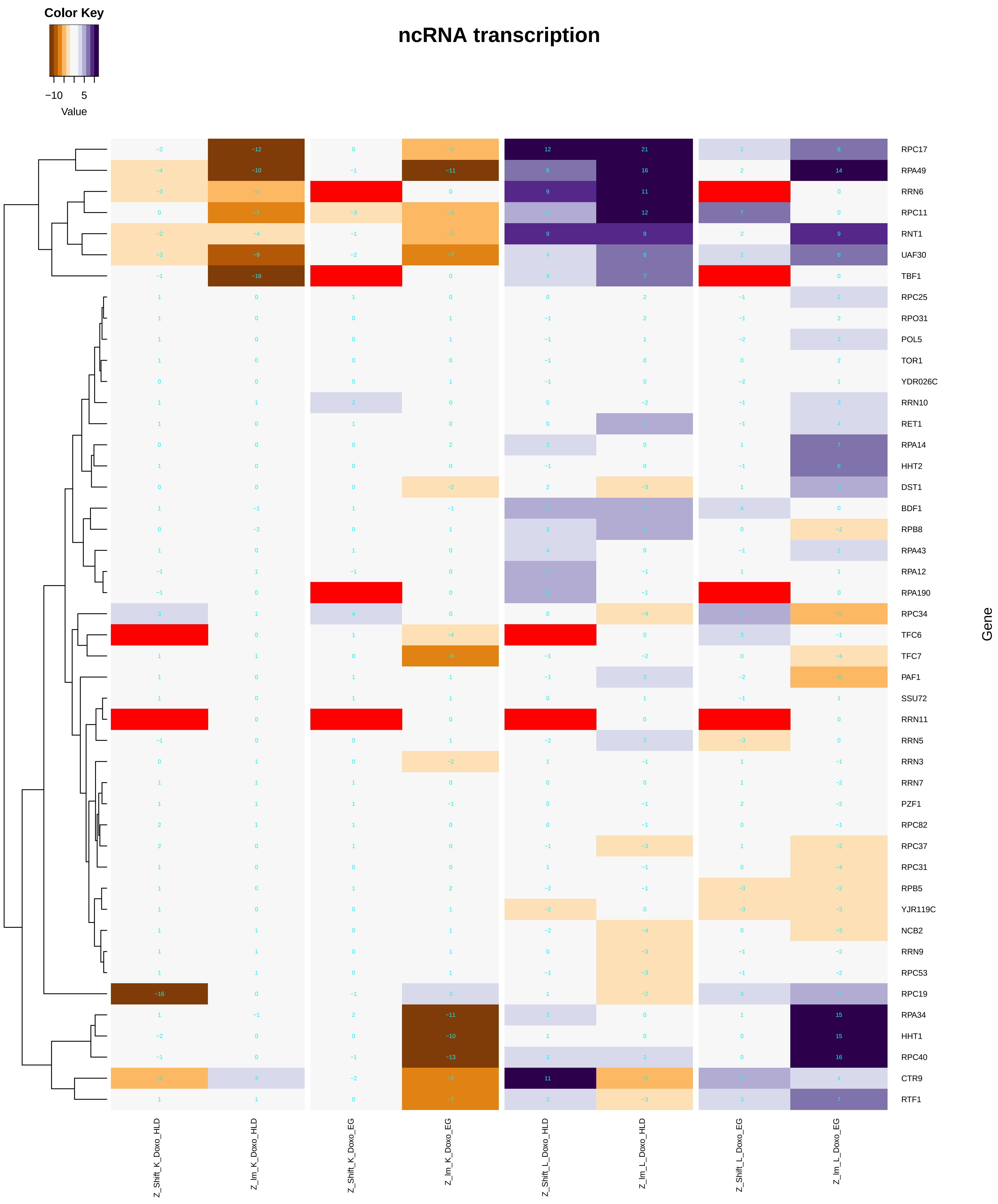

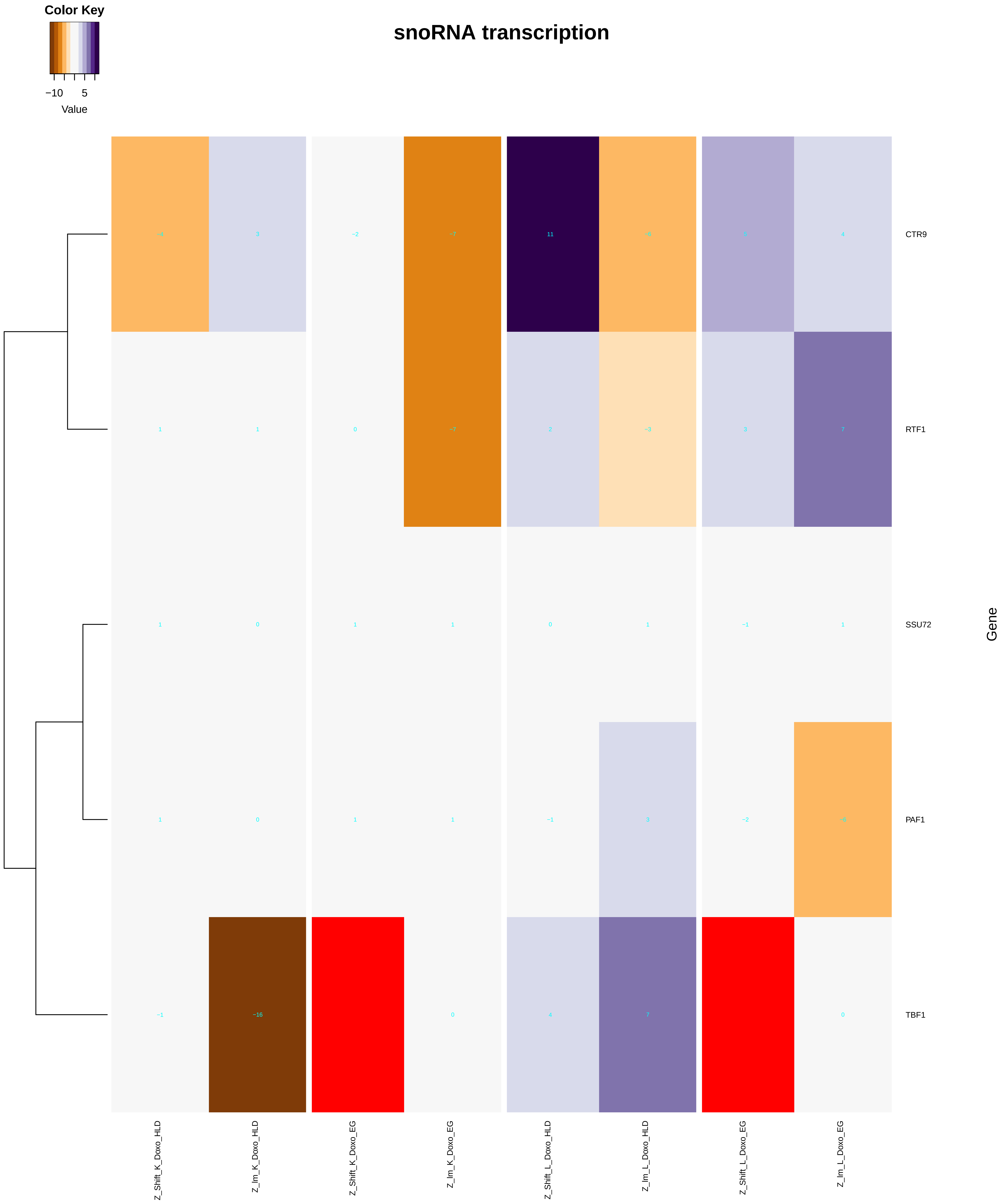

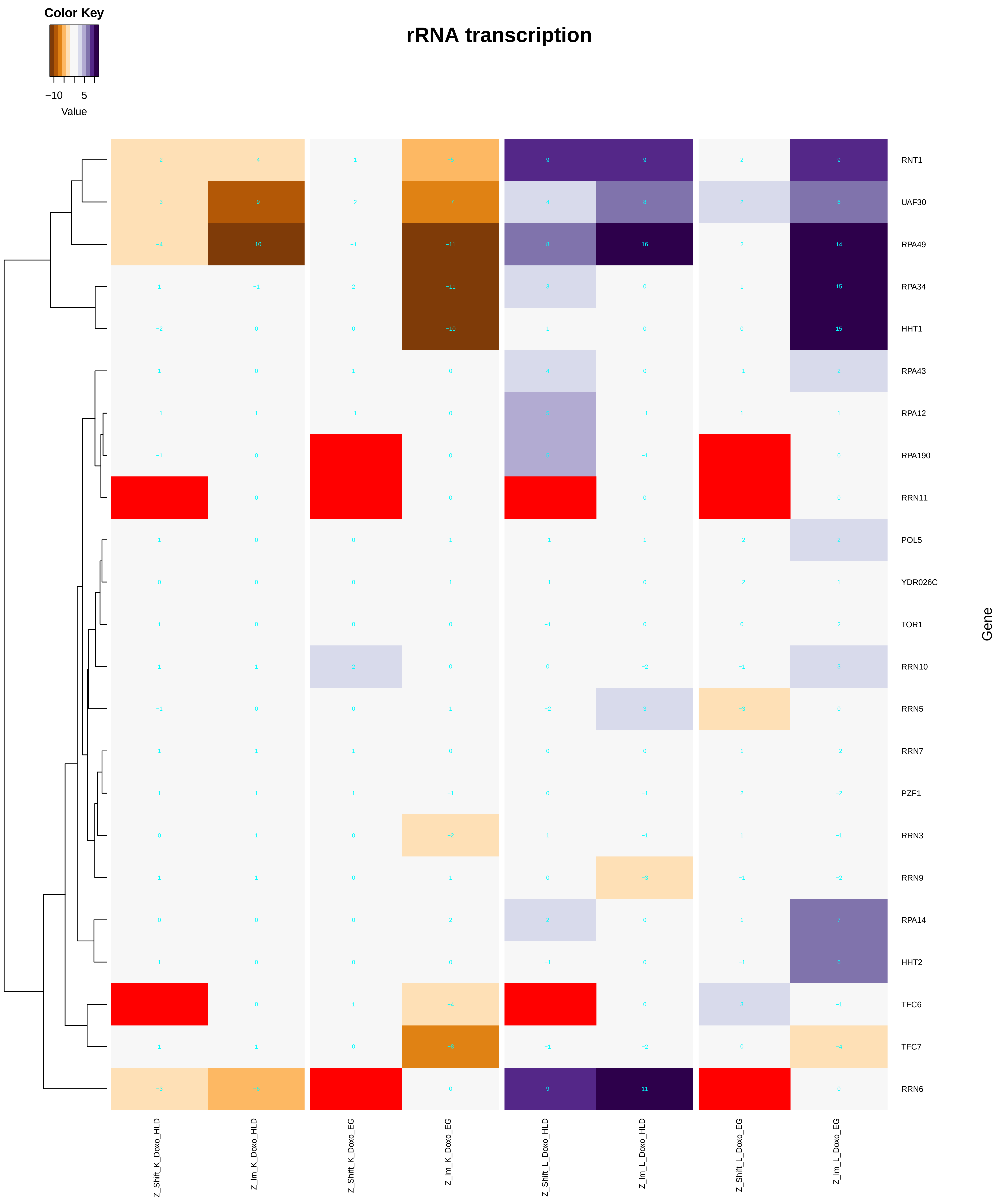

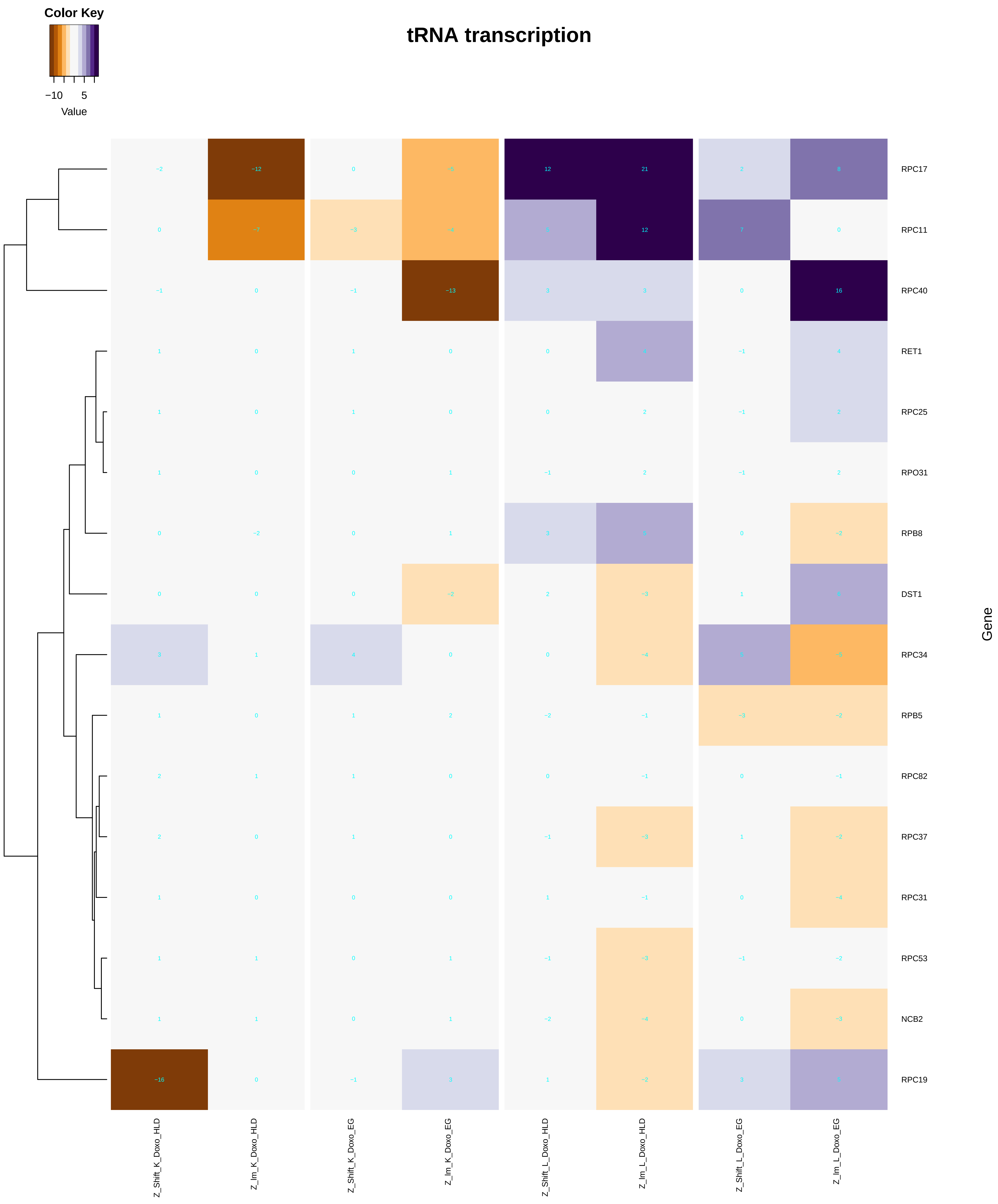

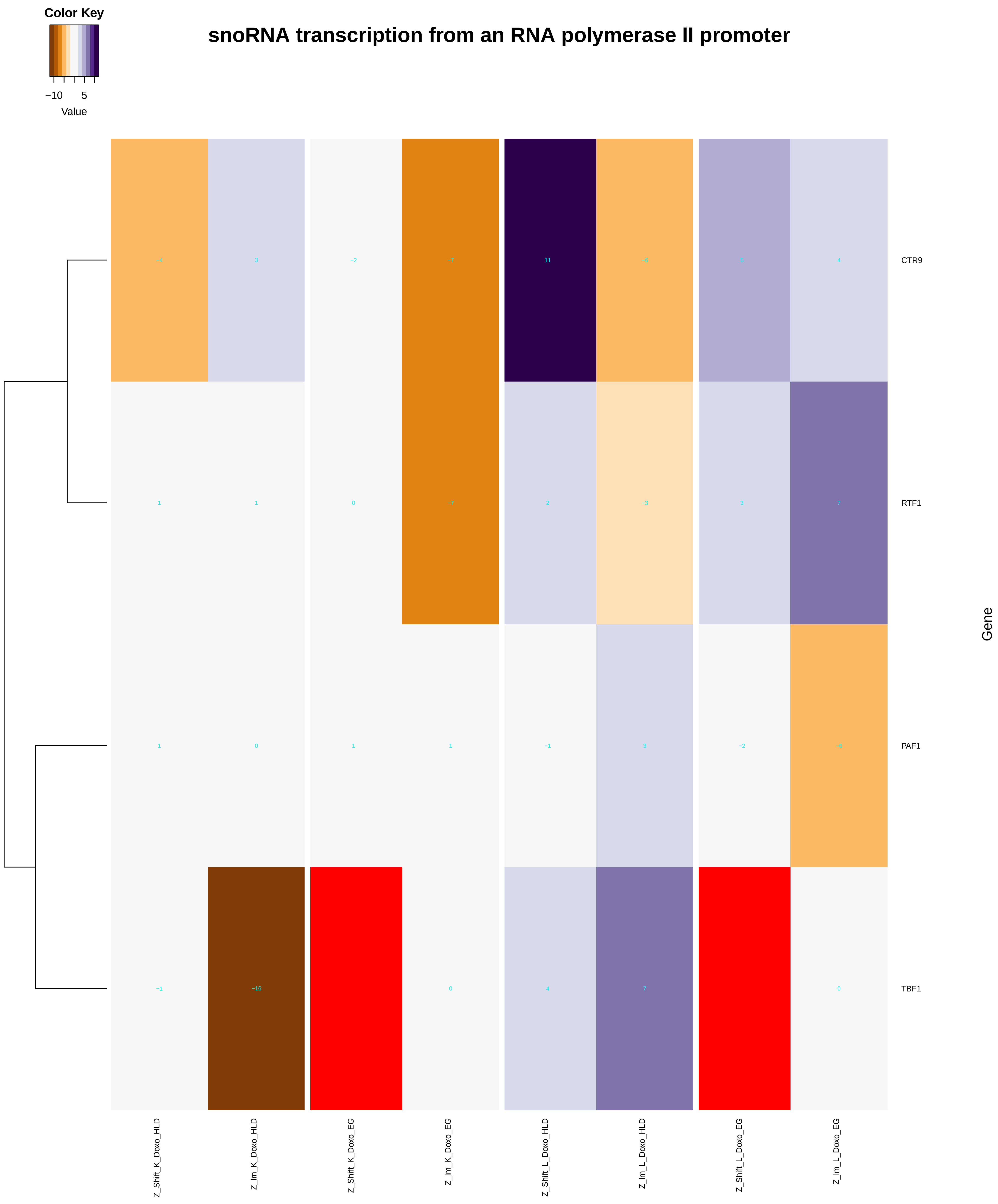

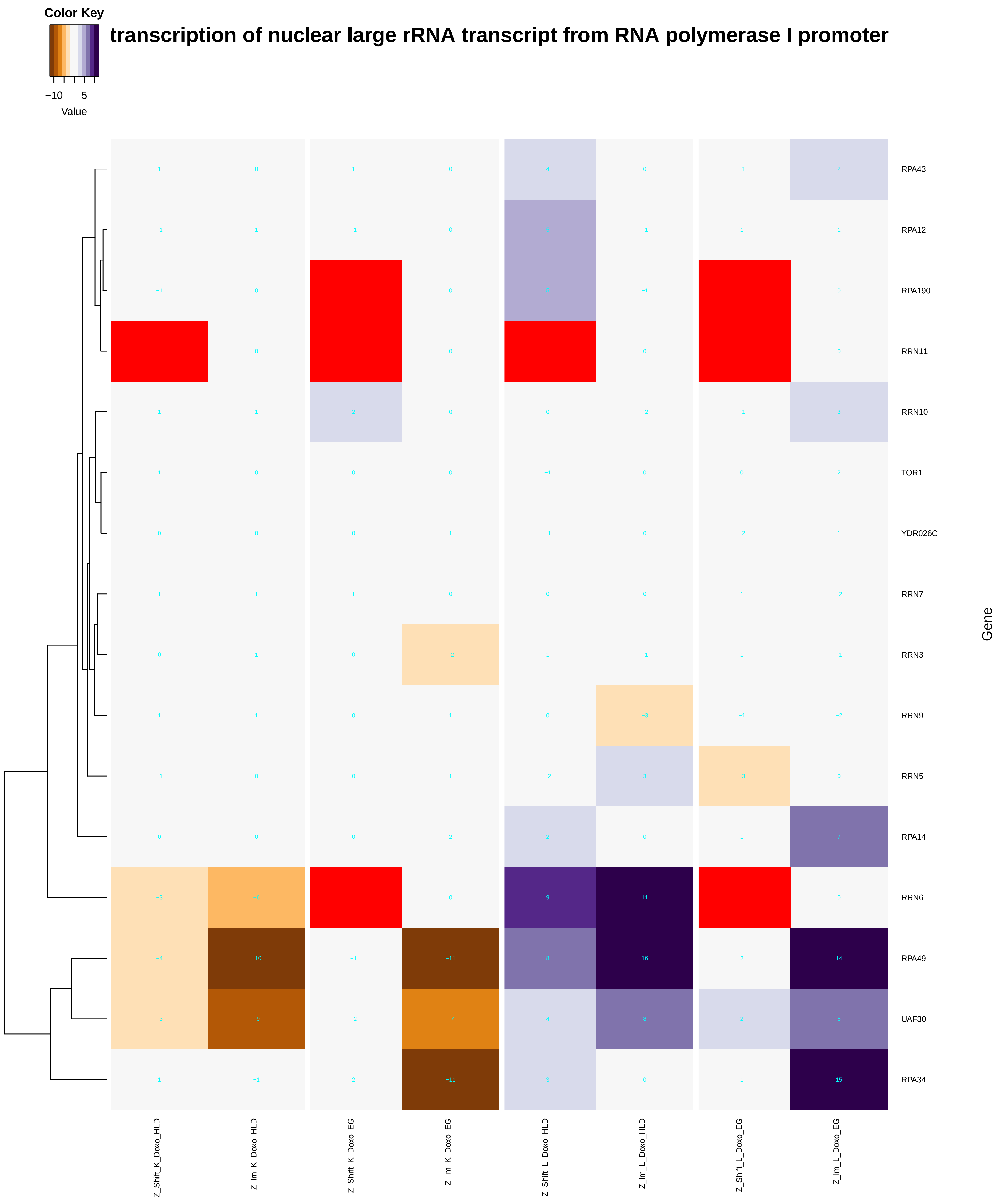

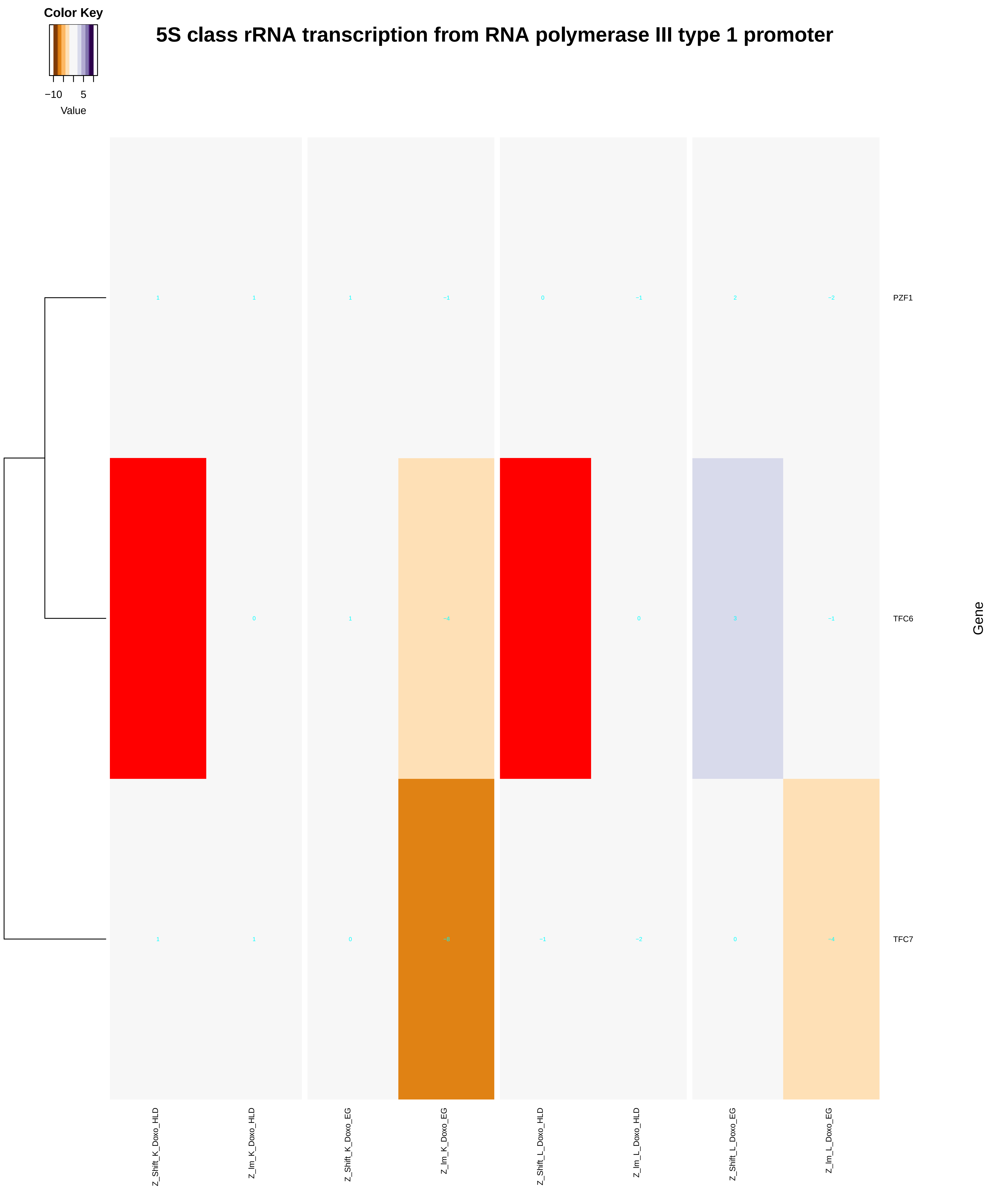

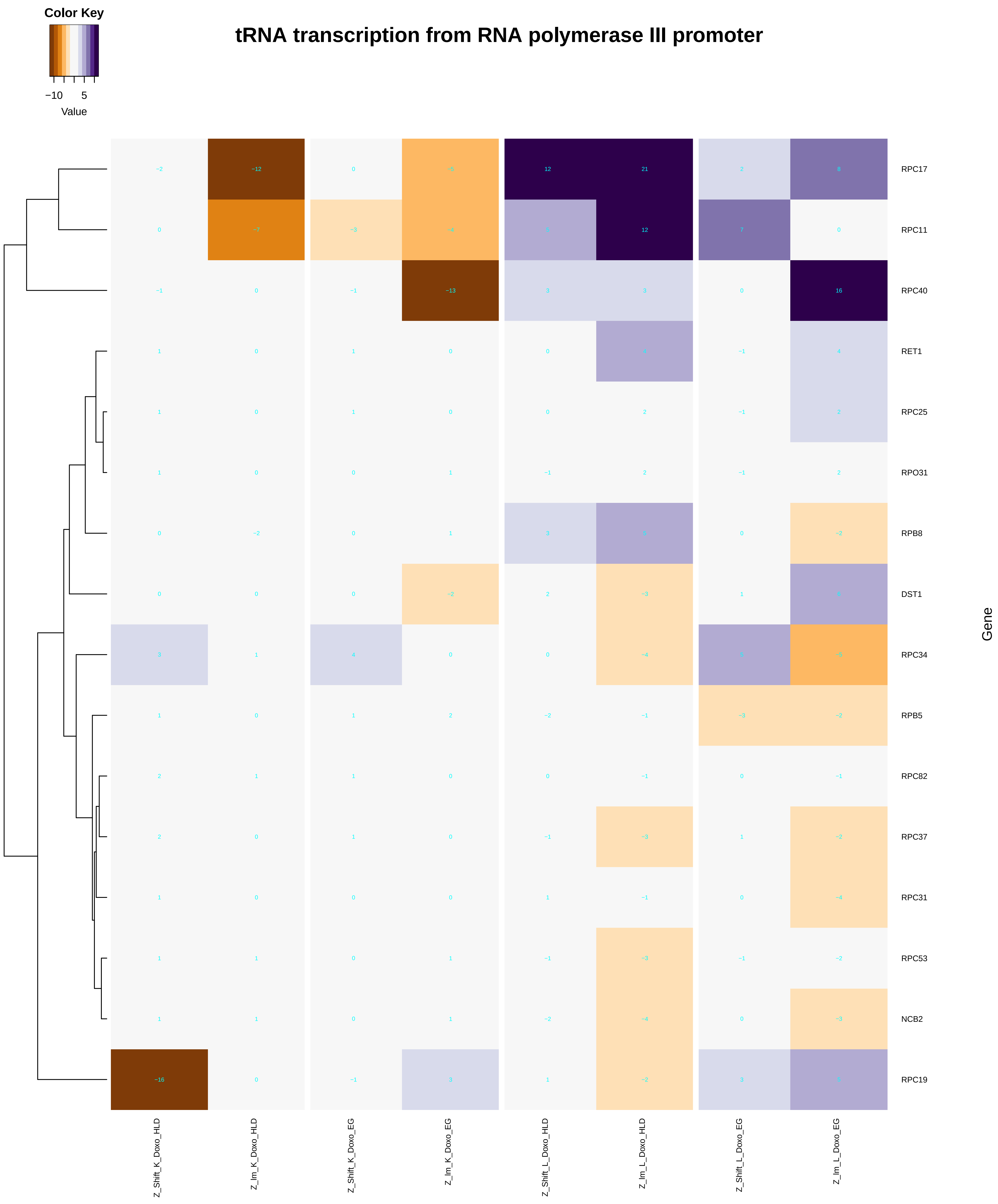

### negative_regulation_of_cellular_catabolic_process.pdf

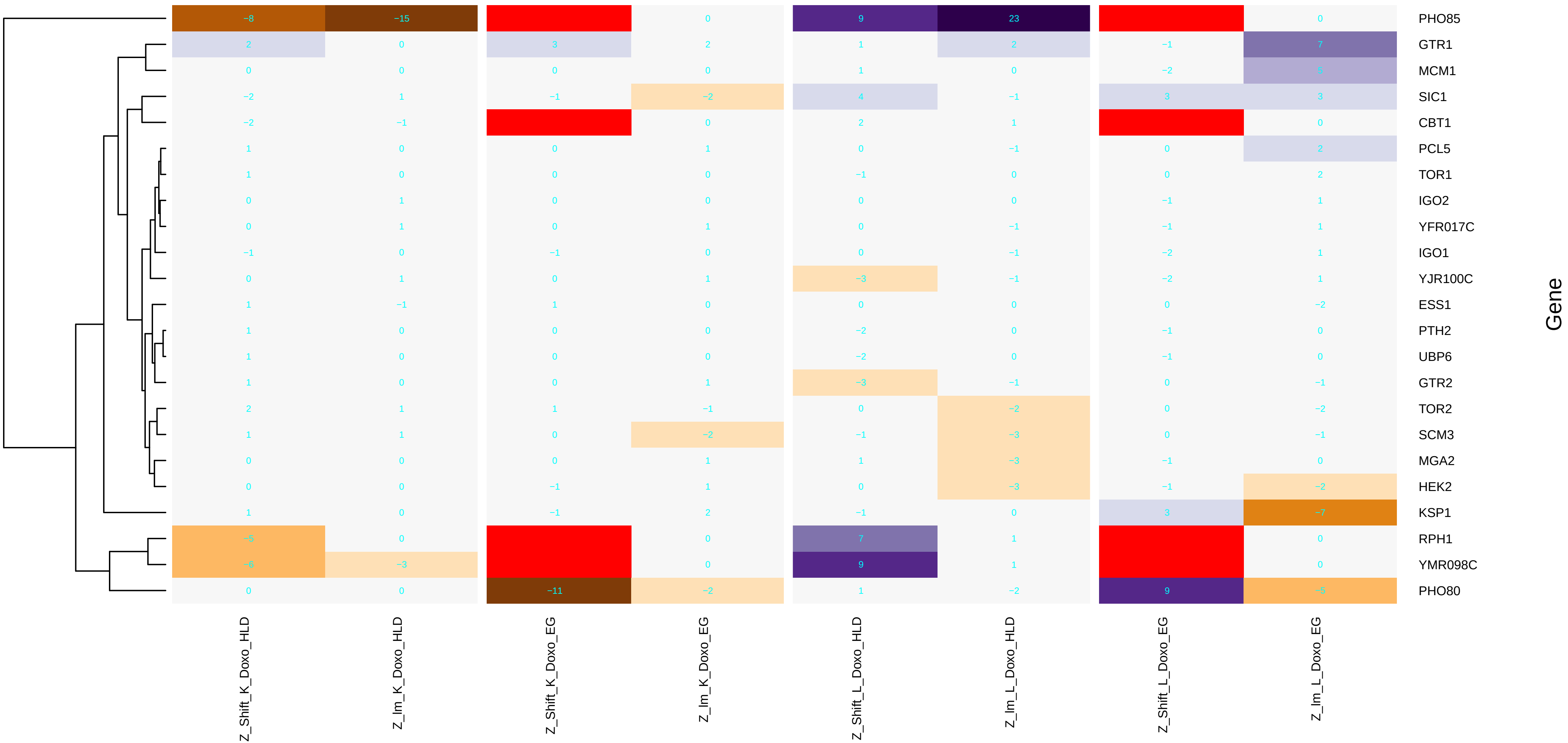

### negative_regulation_of_chromatin_silencing.pdf

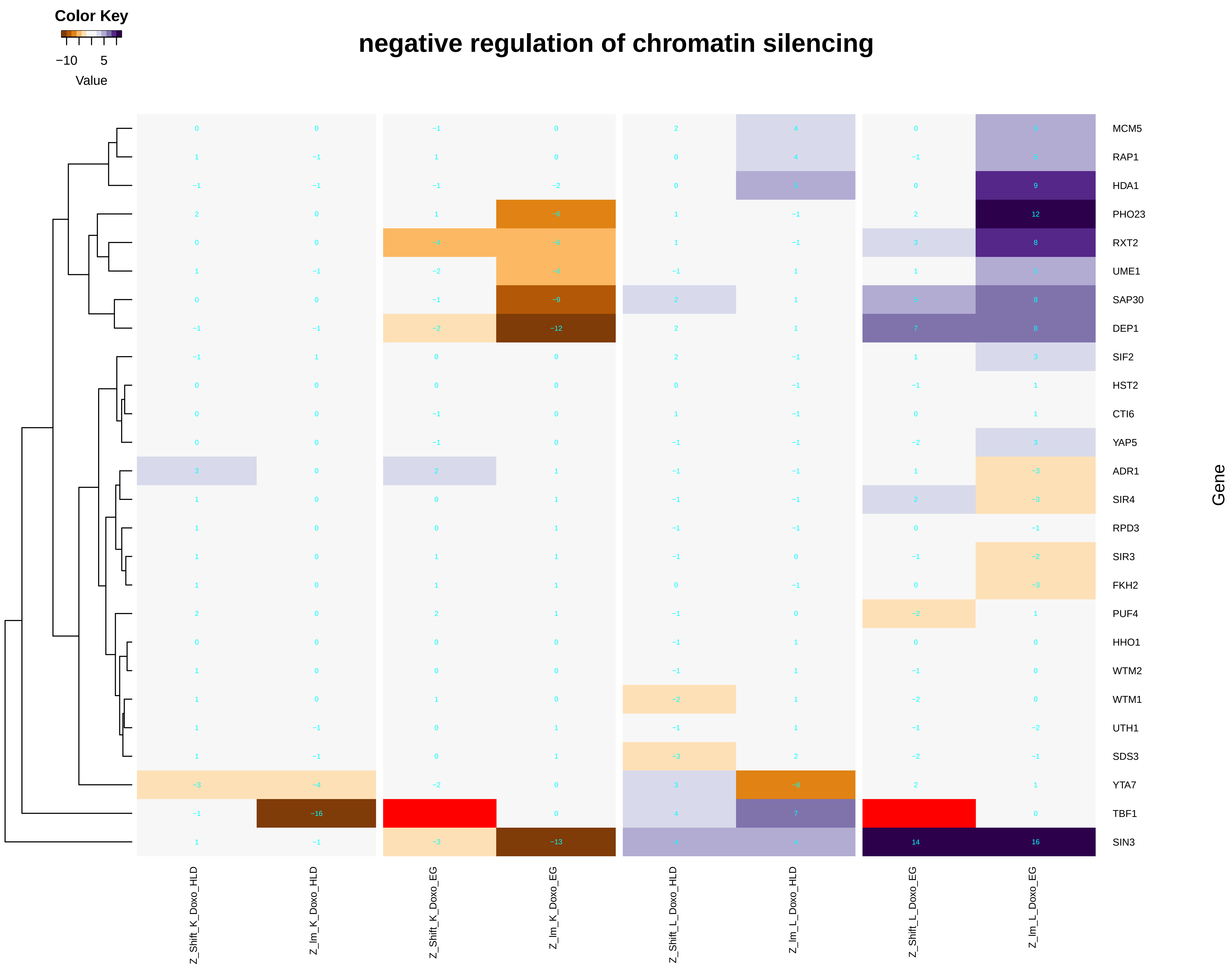

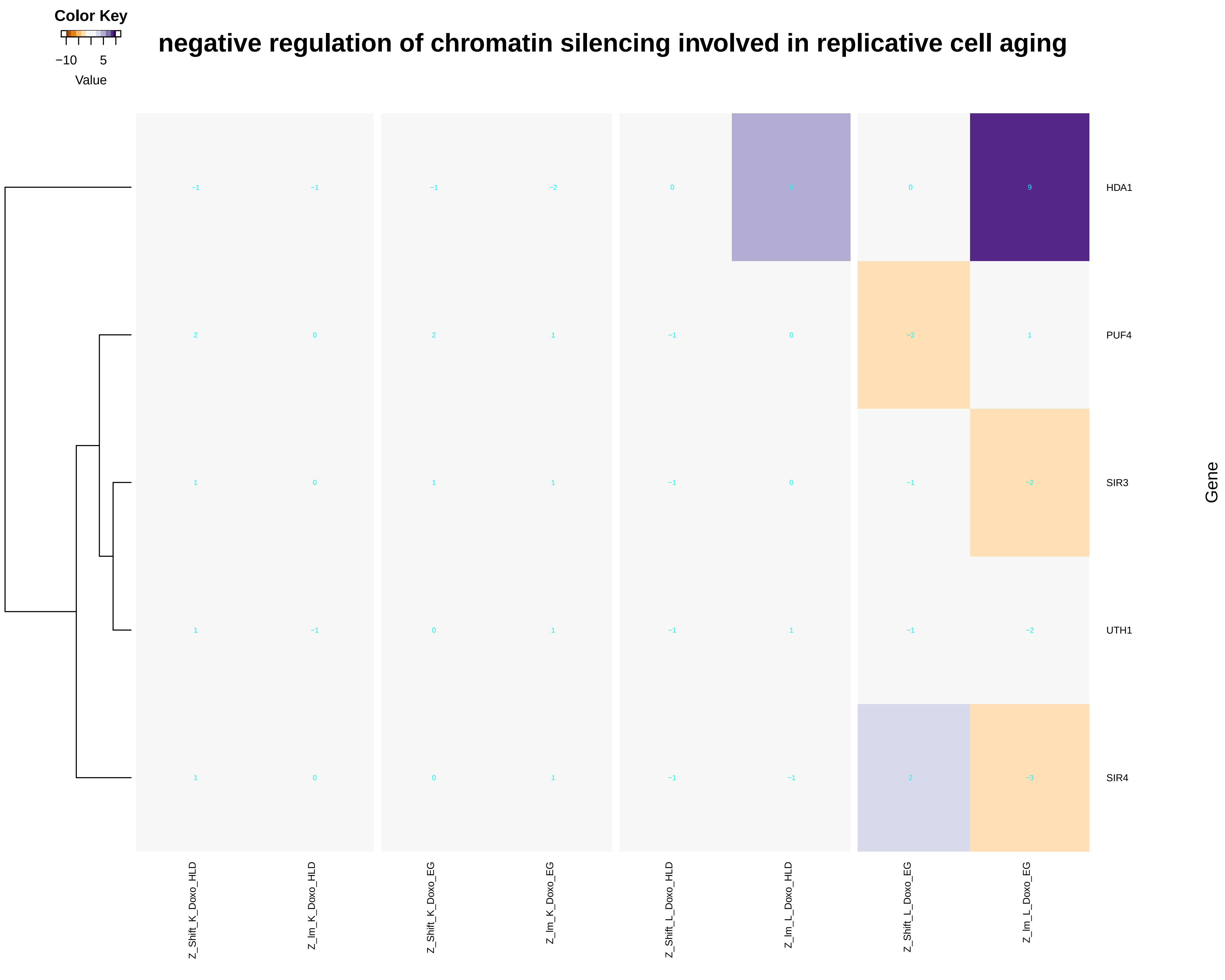

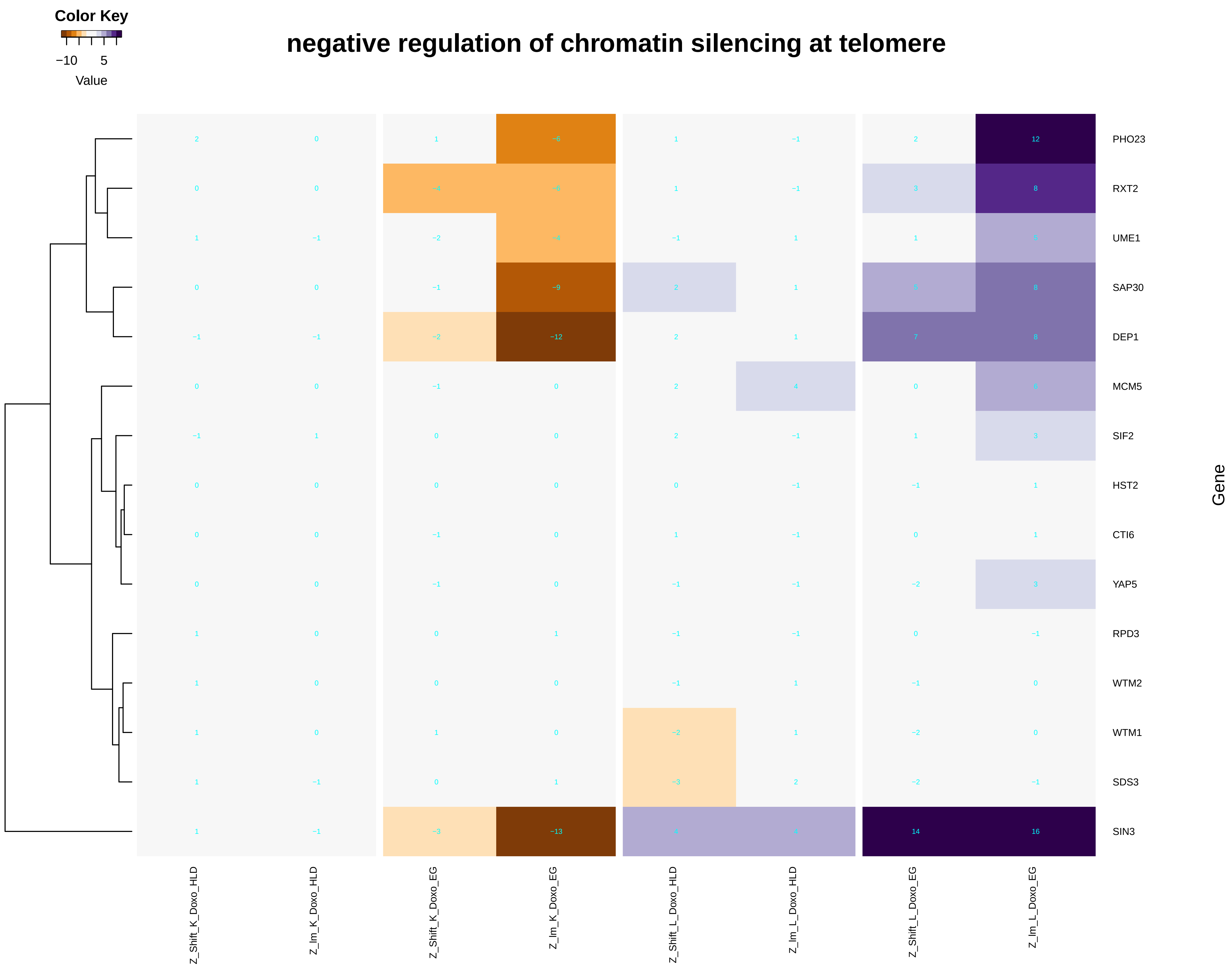

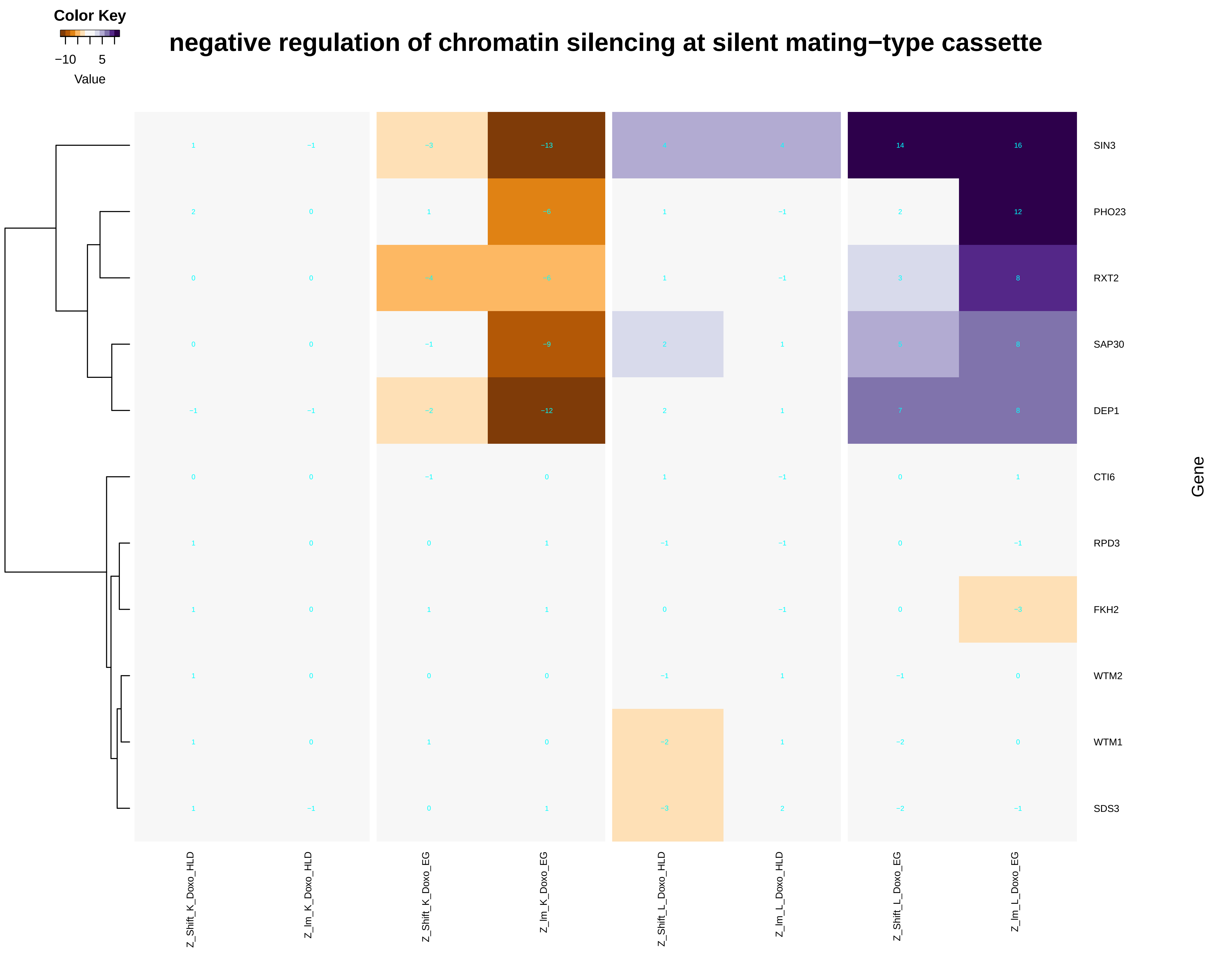

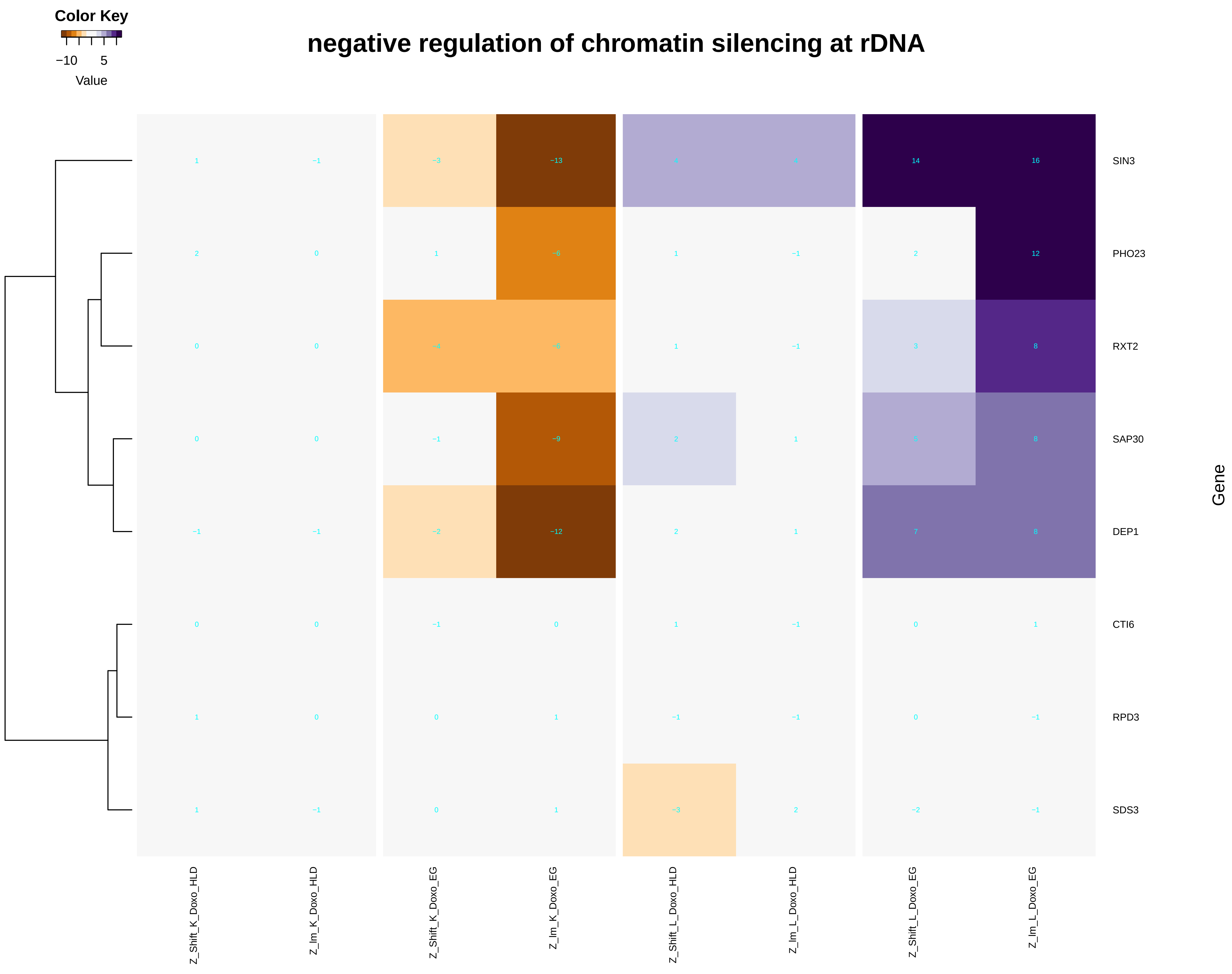

### negative_regulation_of_chromatin_silencing_at_rDNA.pdf

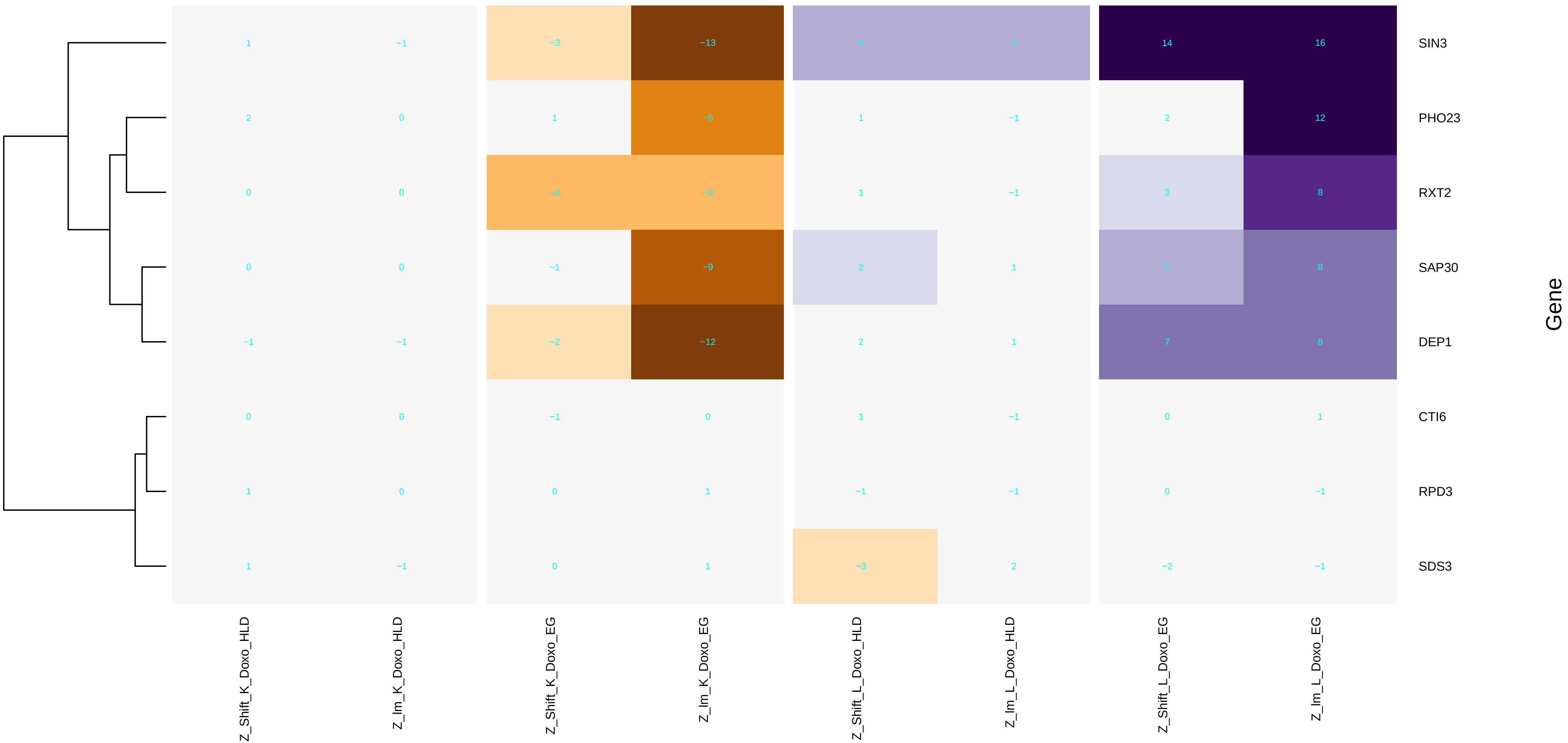

### negative_regulation_of_chromatin_silencing_at_silent_mating-type_cassette.pdf

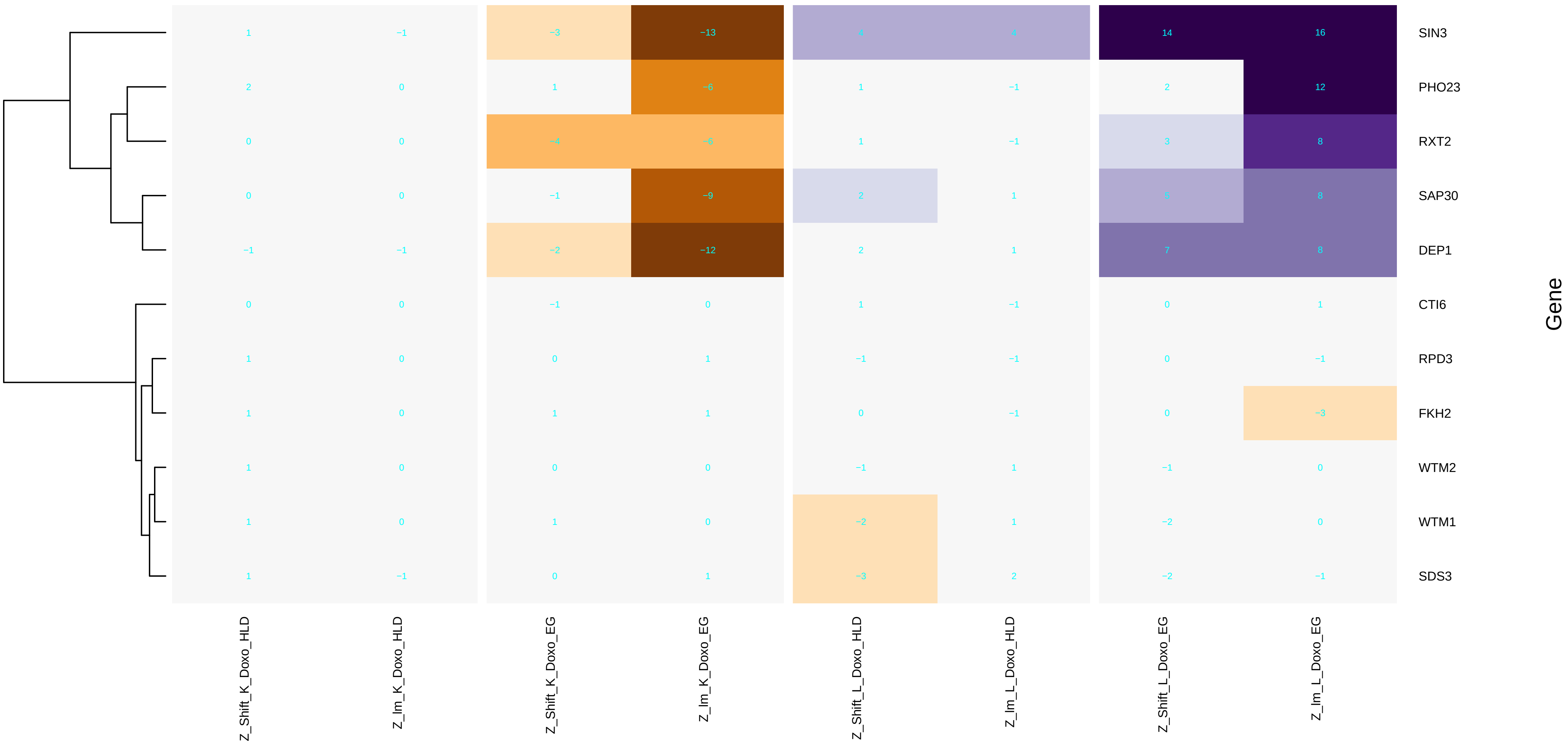

### negative_regulation_of_chromatin_silencing_at_telomere.pdf

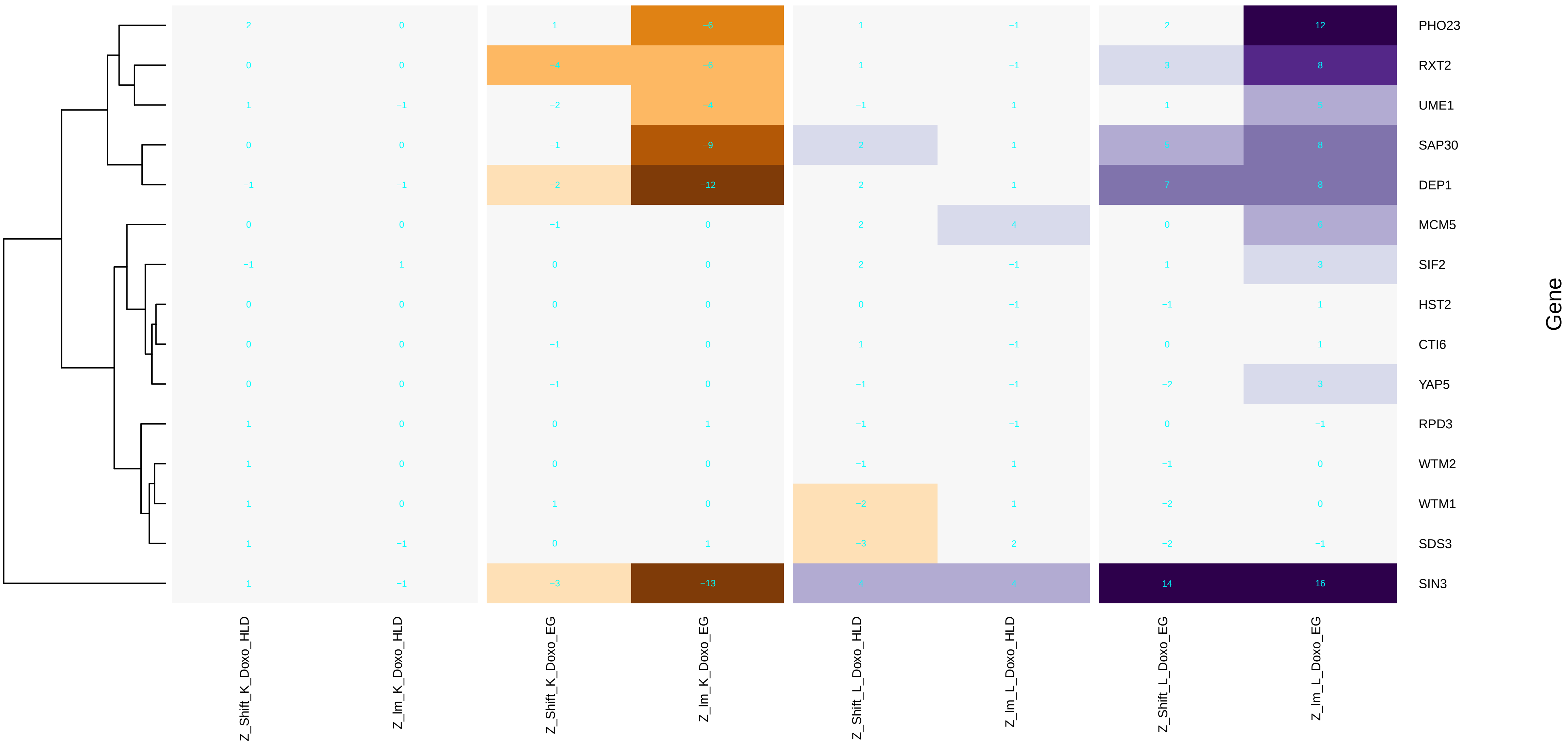

### negative_regulation_of_DNA_damage_checkpoint.pdf

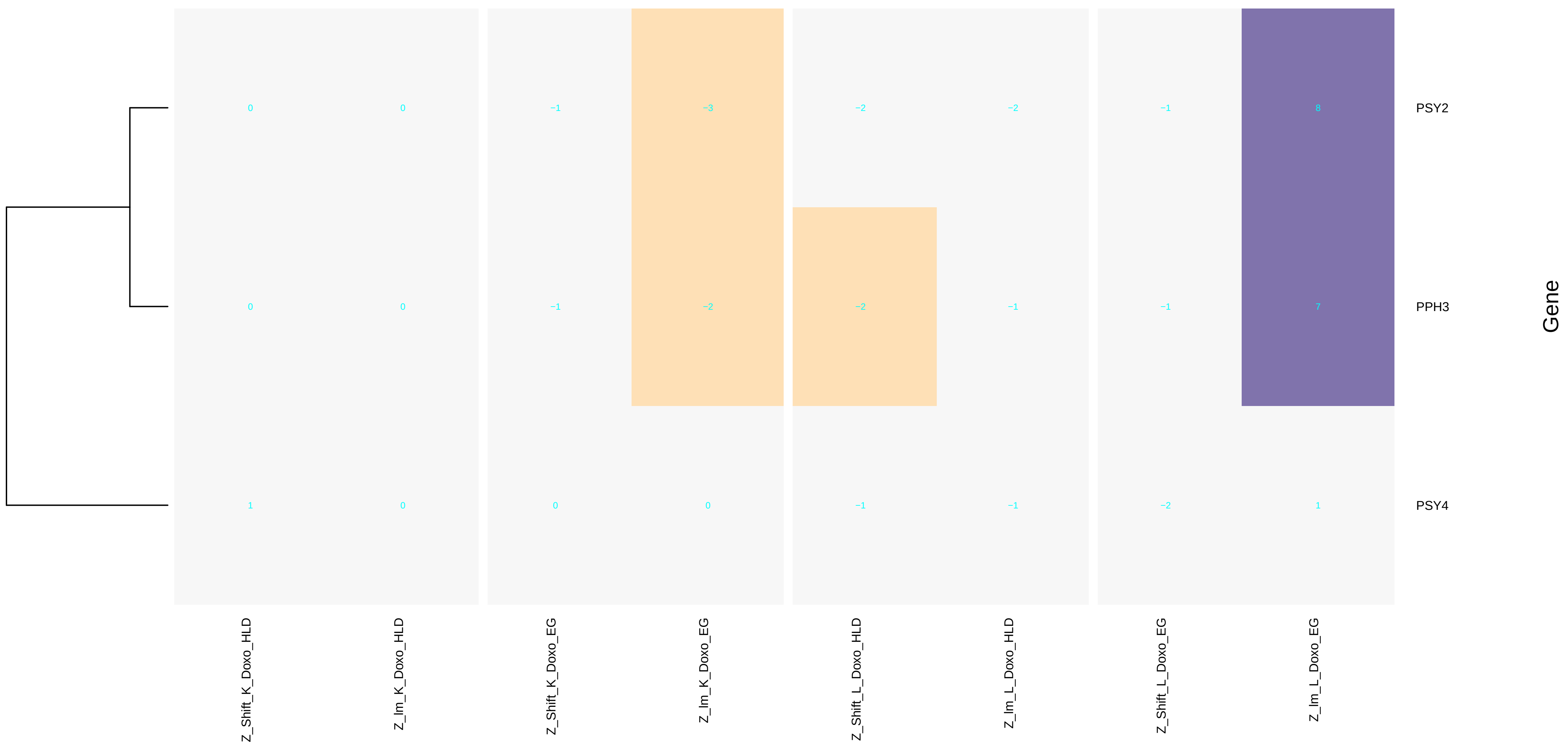

### negative_regulation_of_gene_expression,_epigenetic.pdf

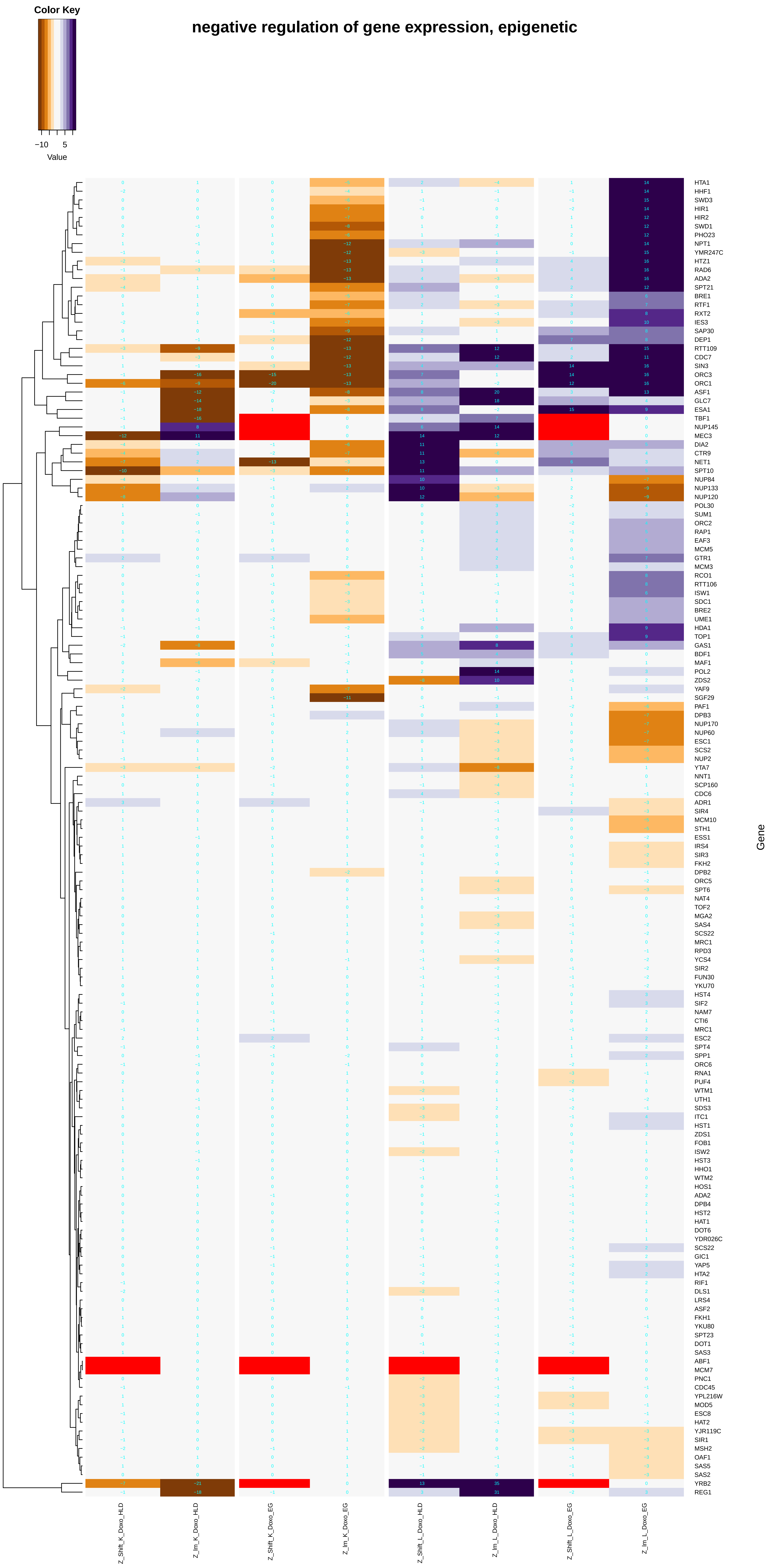

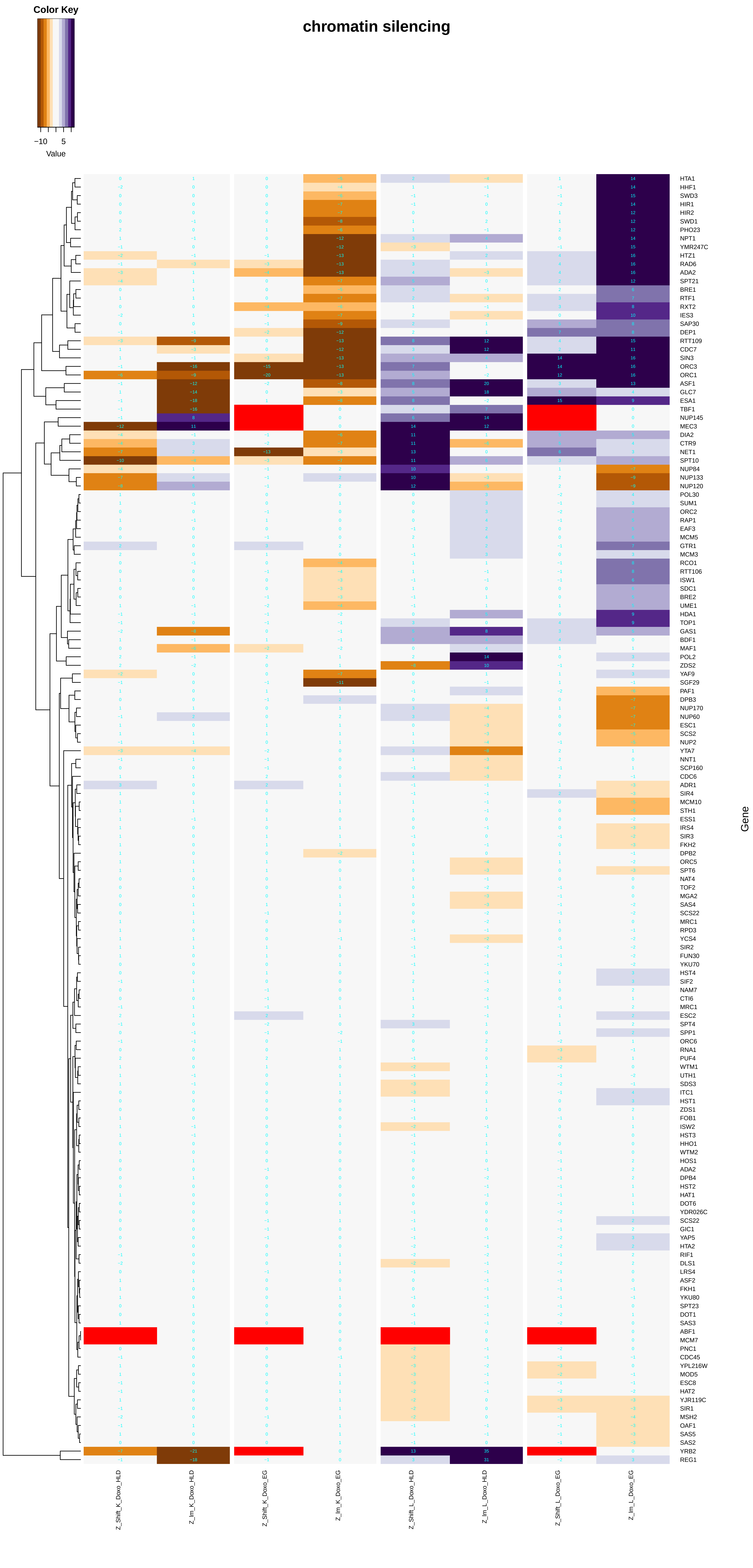

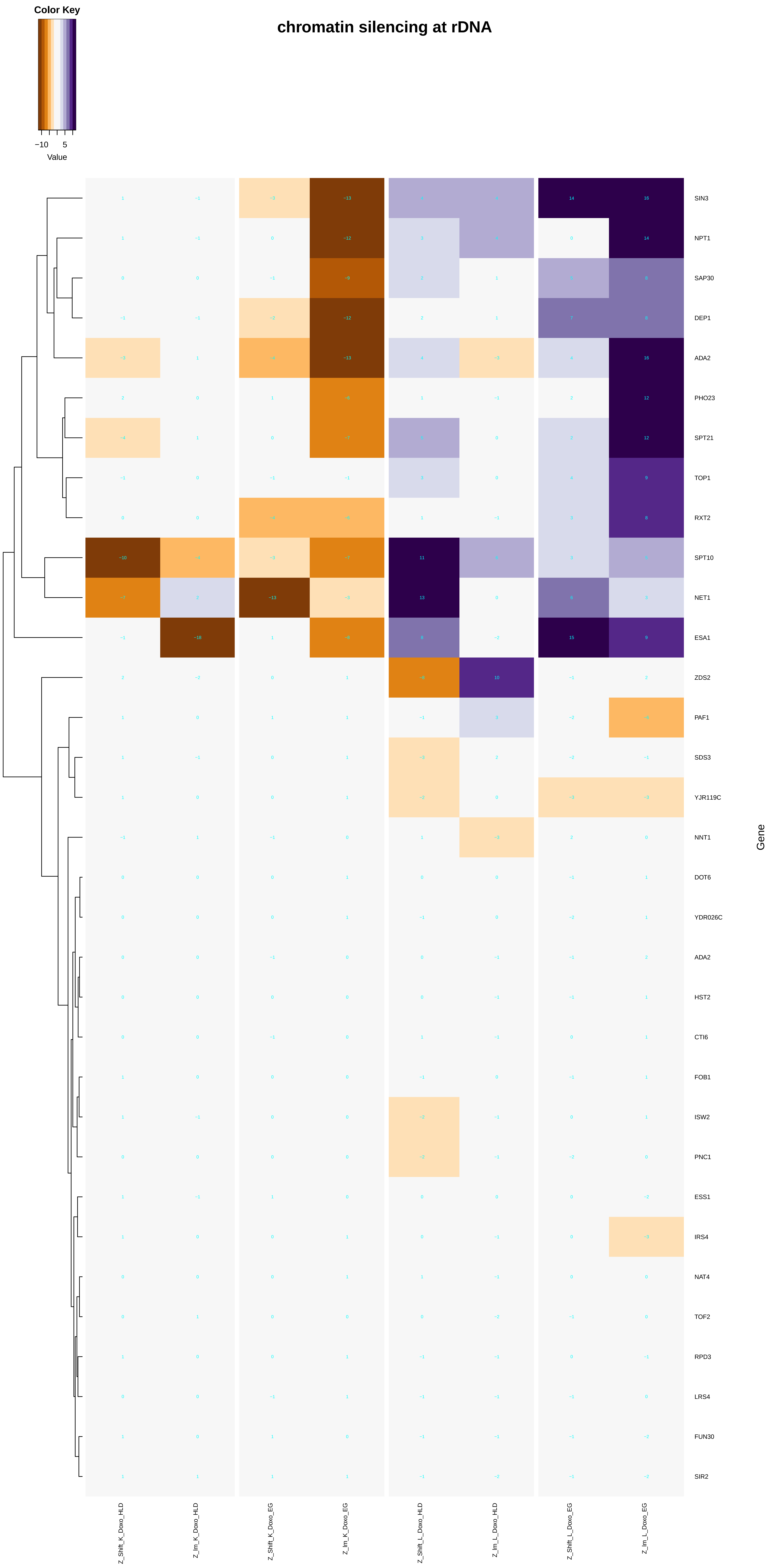

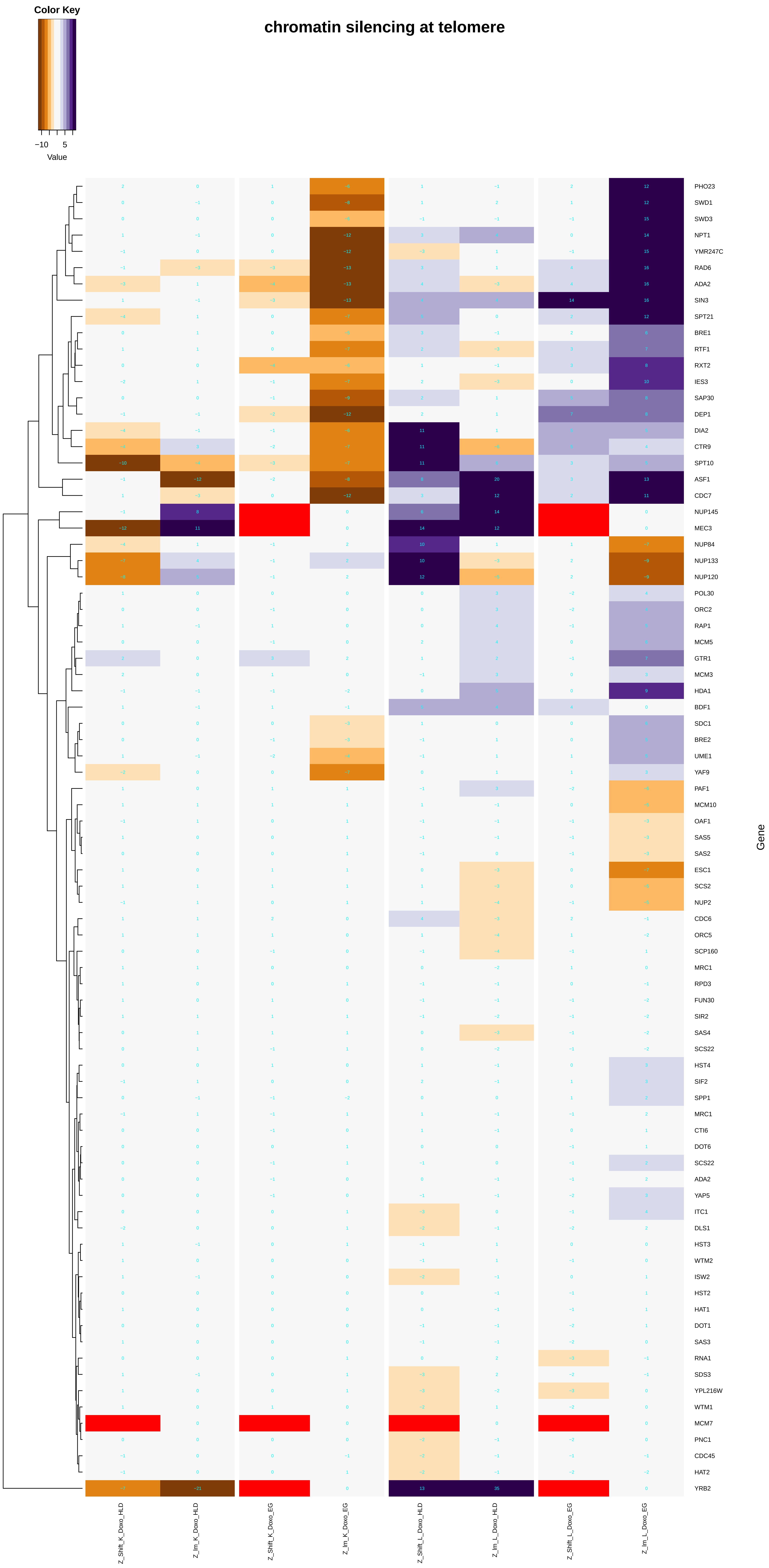

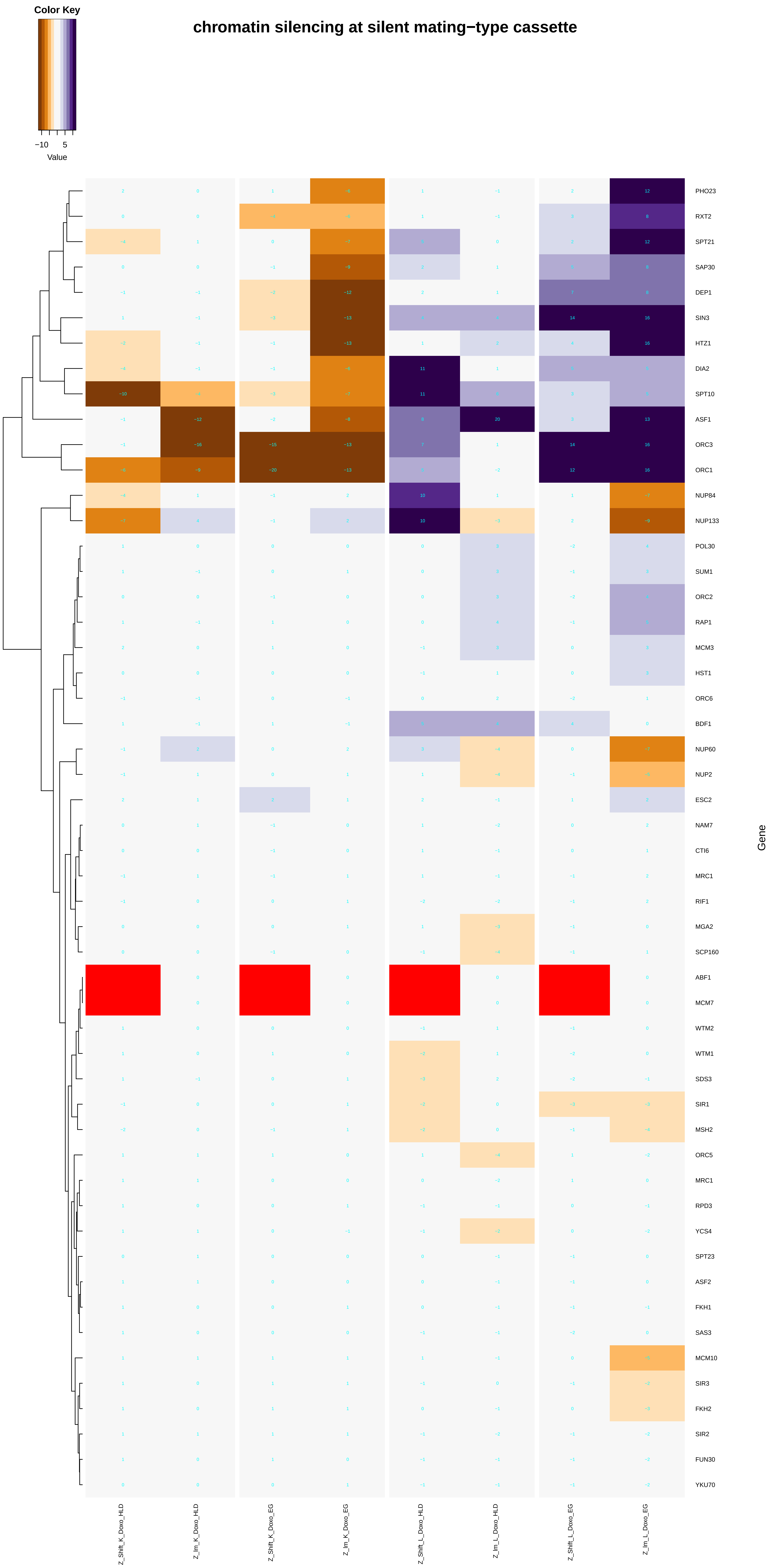

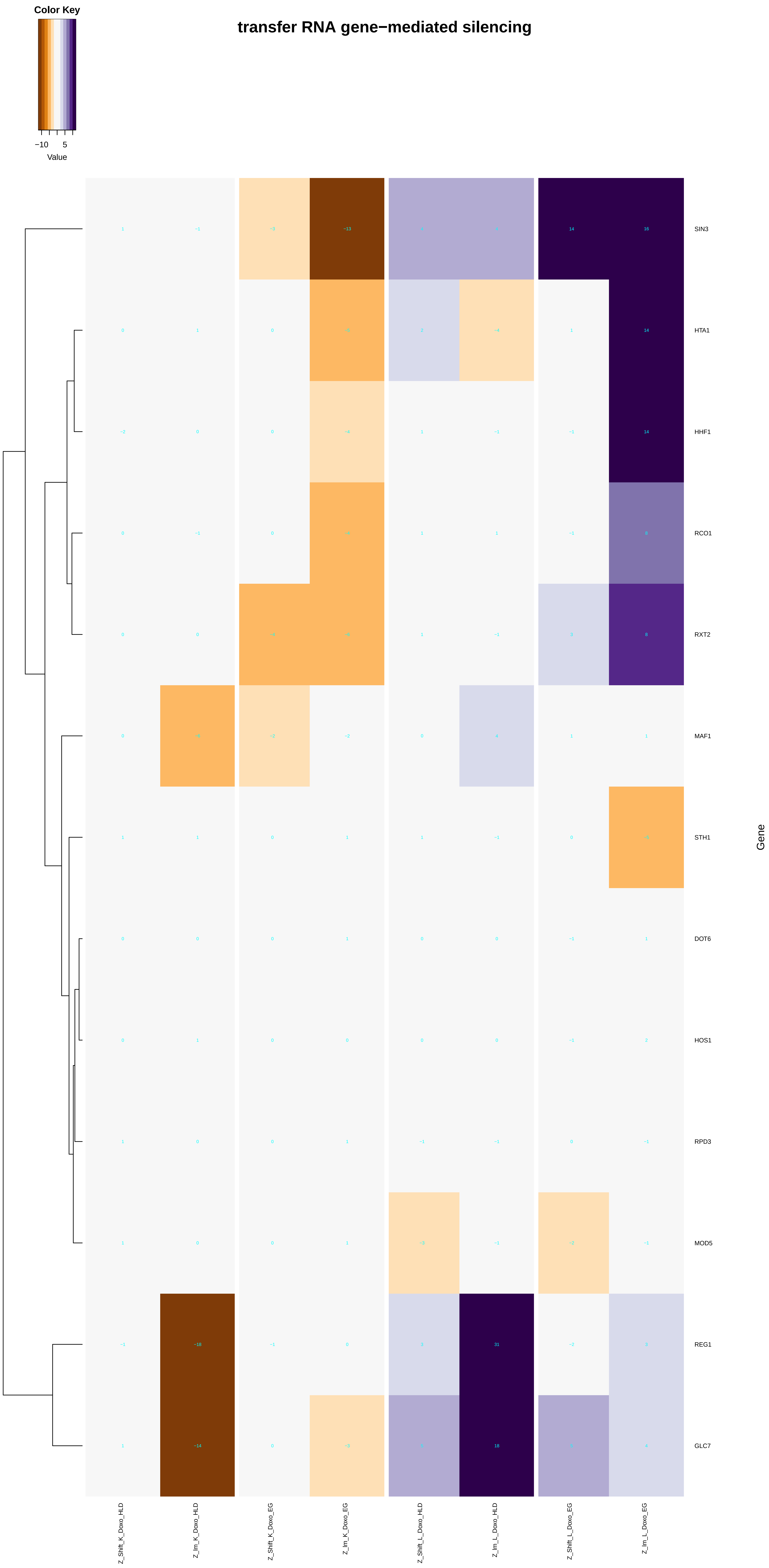

### negative_regulation_of_gene_silencing.pdf

#### negative regulation of gene silencing

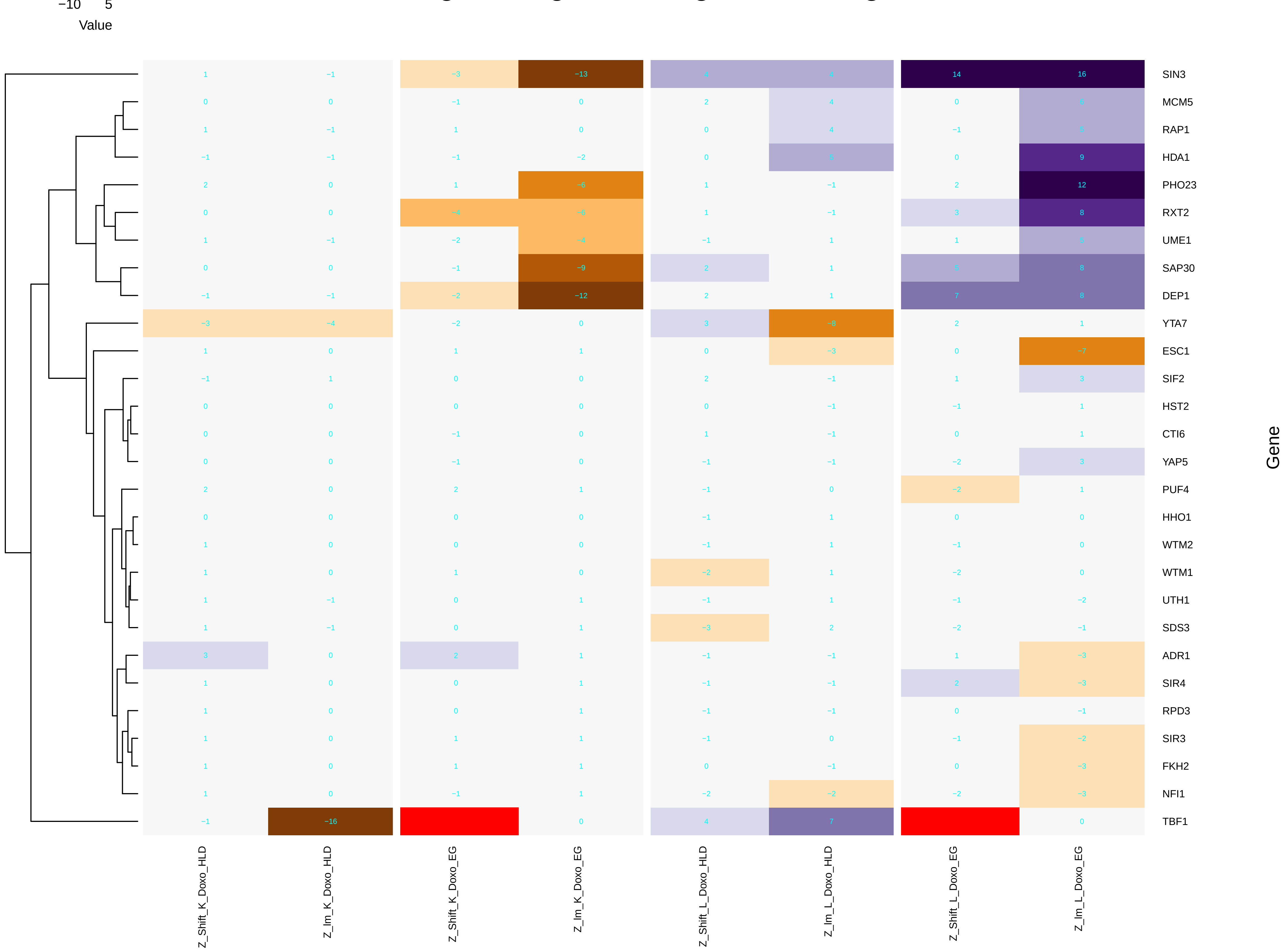

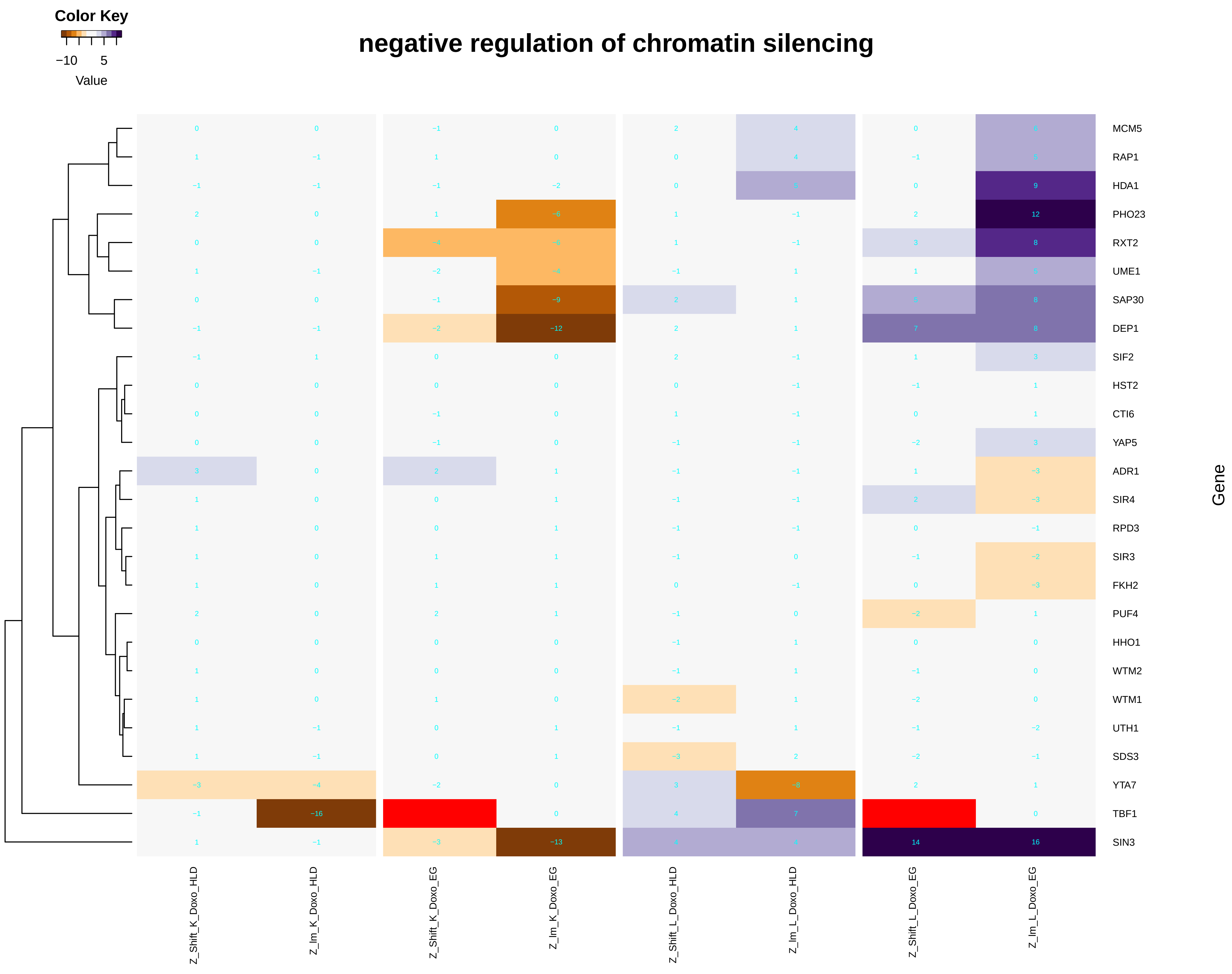

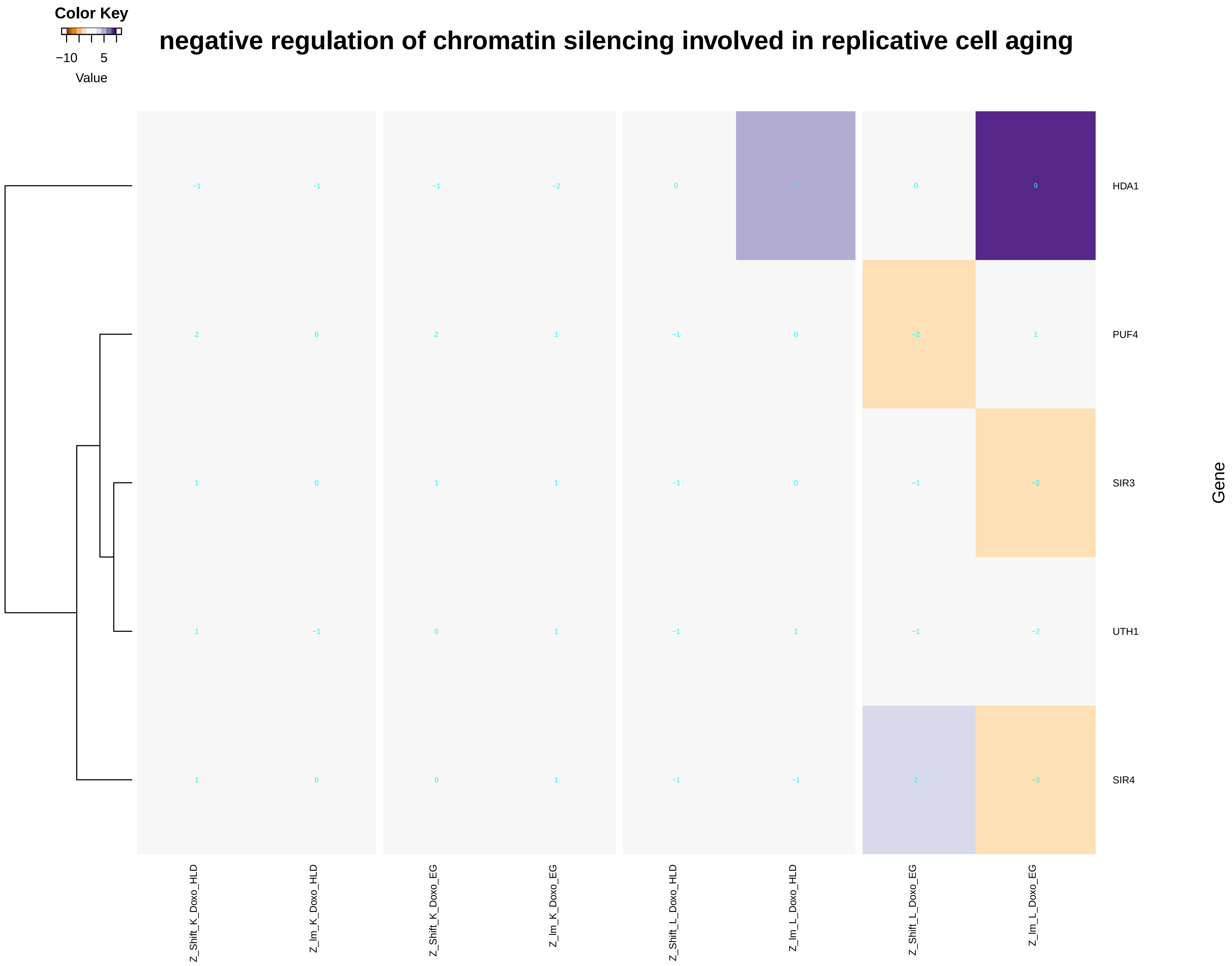

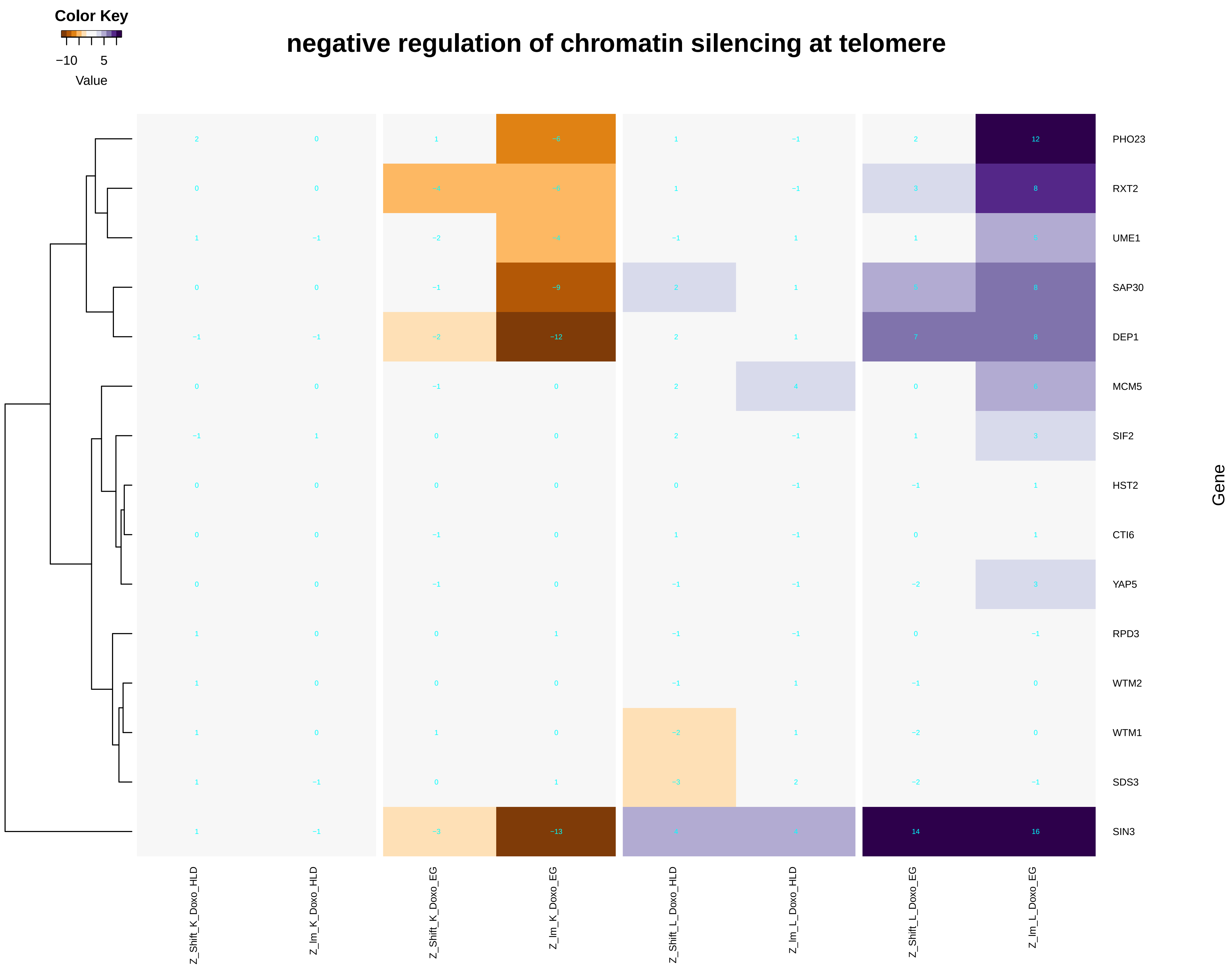

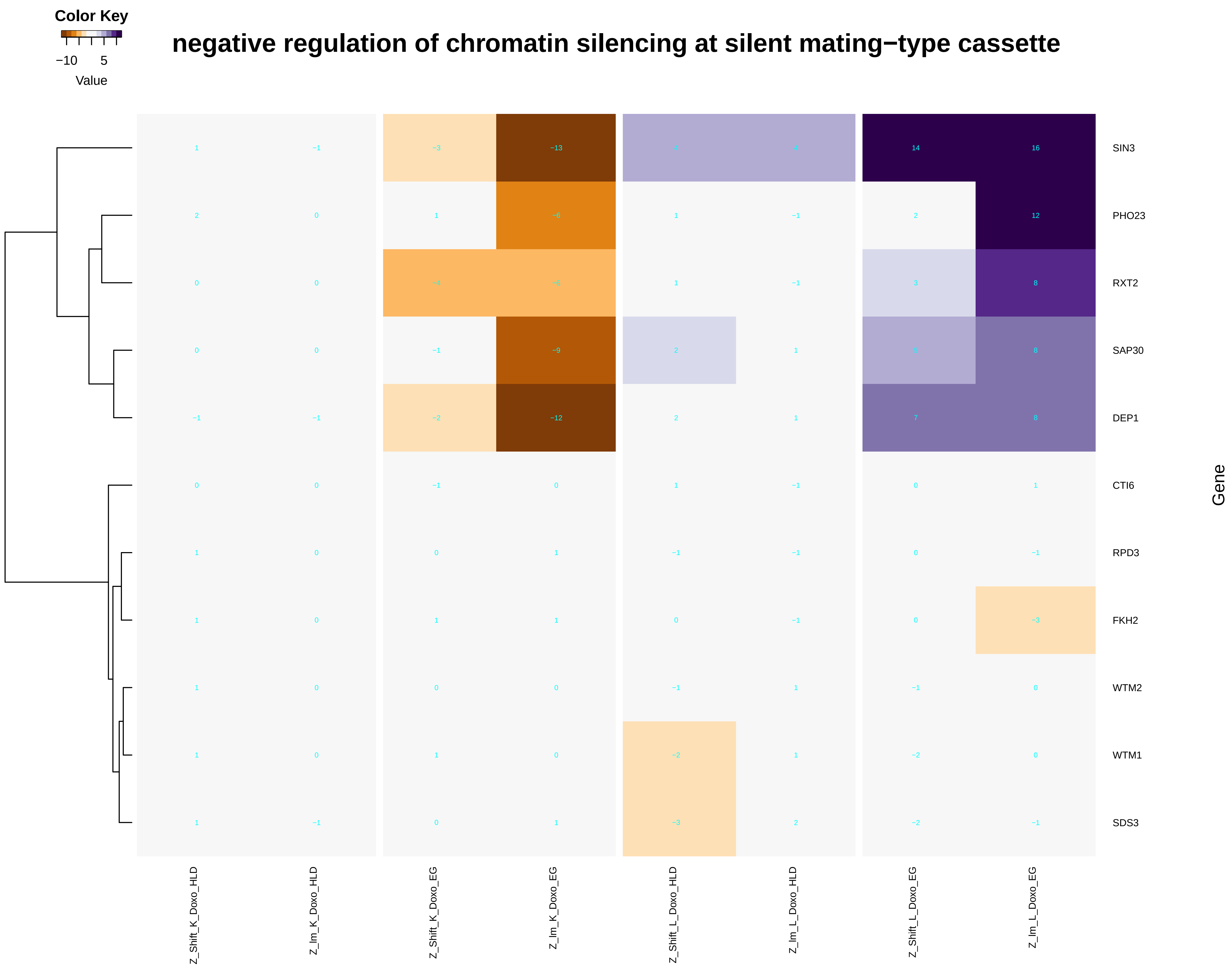

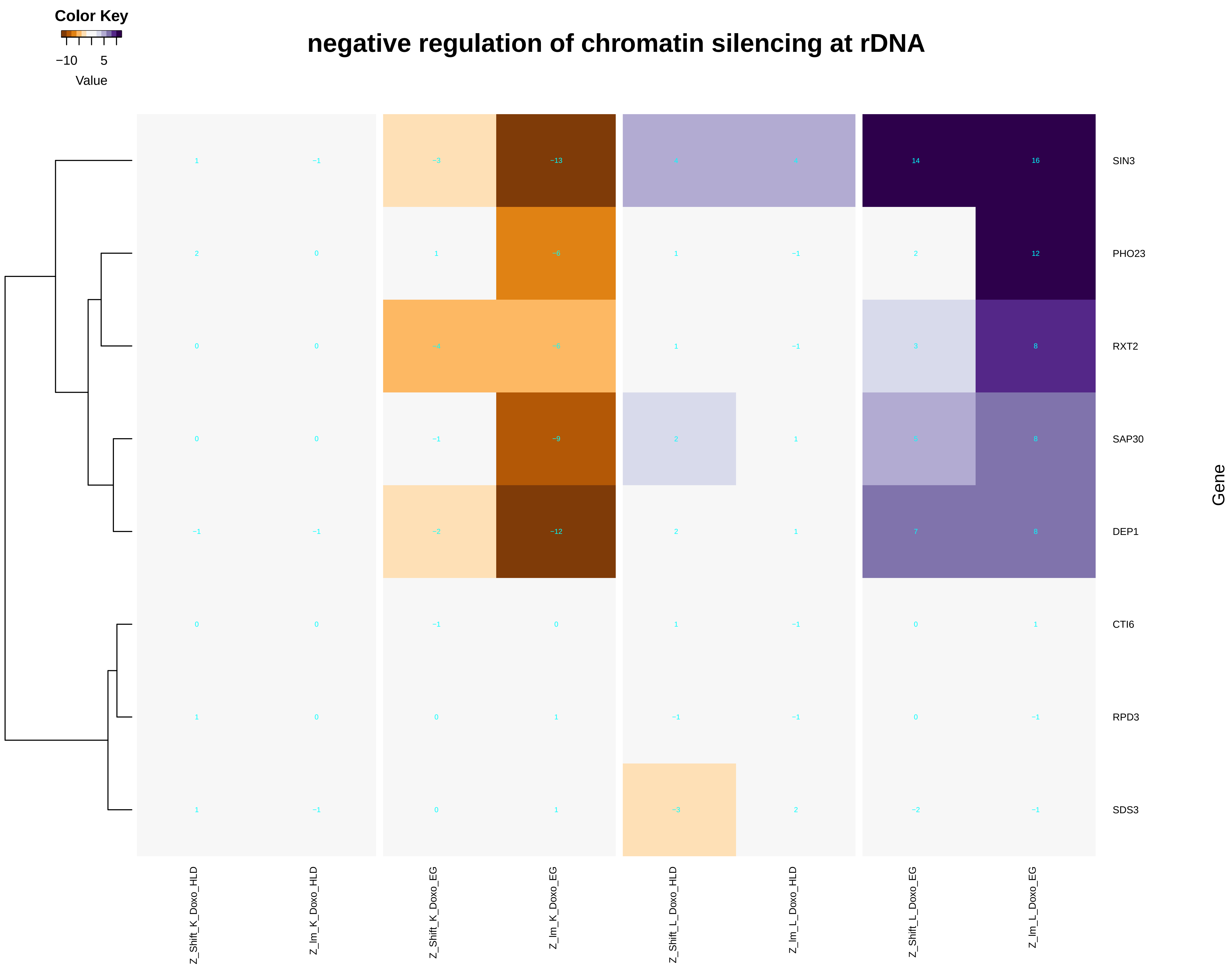

### negative_regulation_of_glucose_mediated_signaling_pathway.pdf

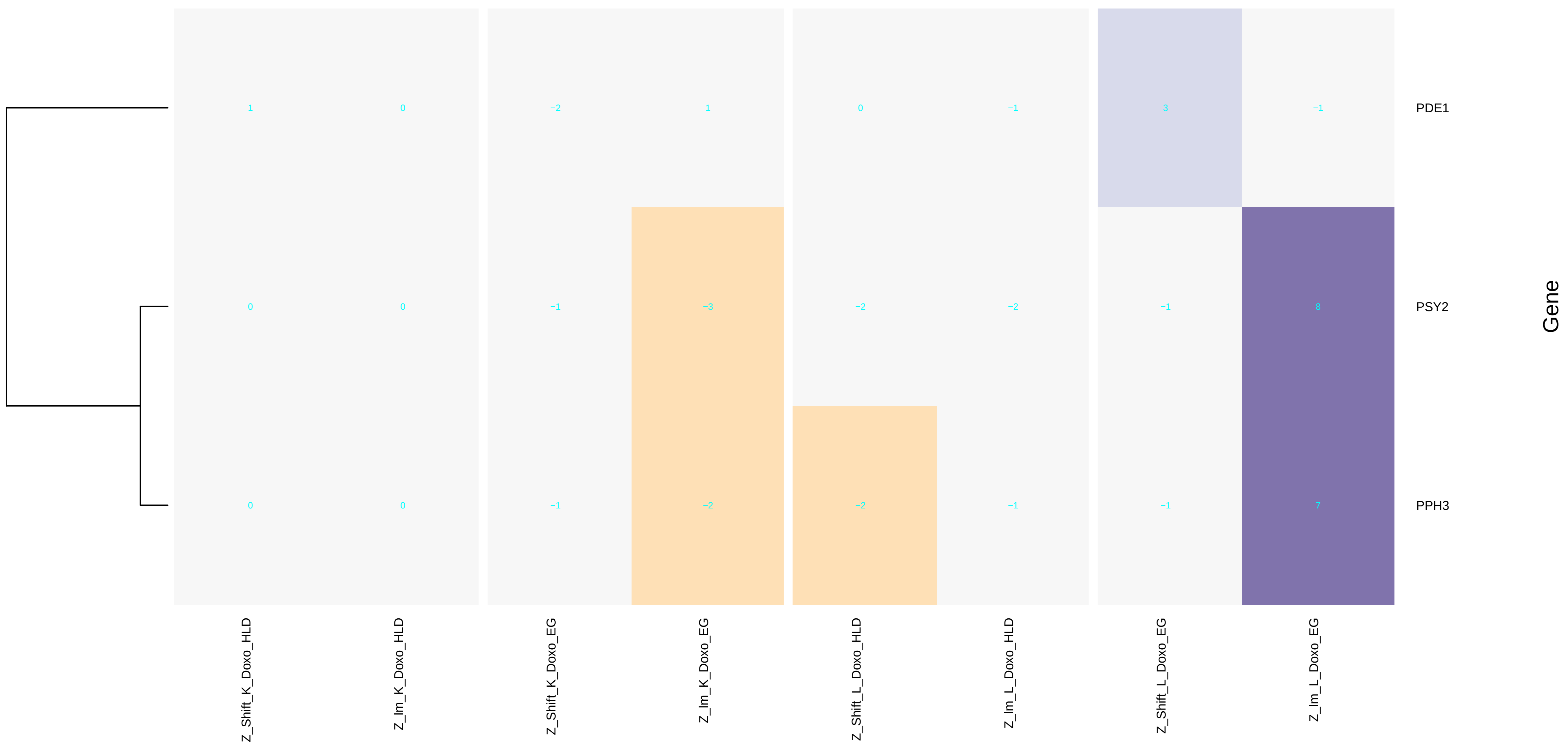

### negative_regulation_of_meiotic_DNA_double-strand_break_formation.pdf

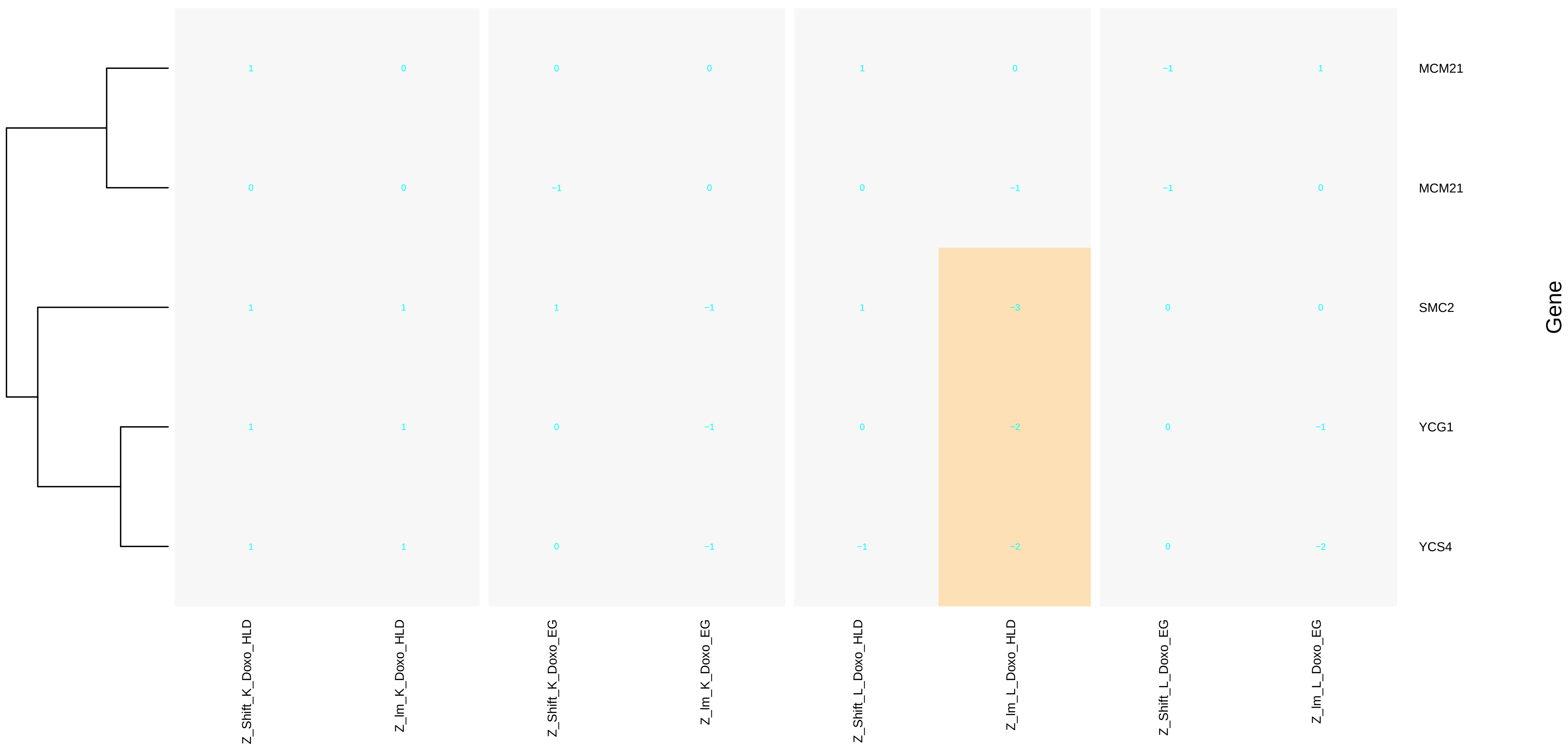

### negative_regulation_of_response_to_DNA_damage_stimulus.pdf

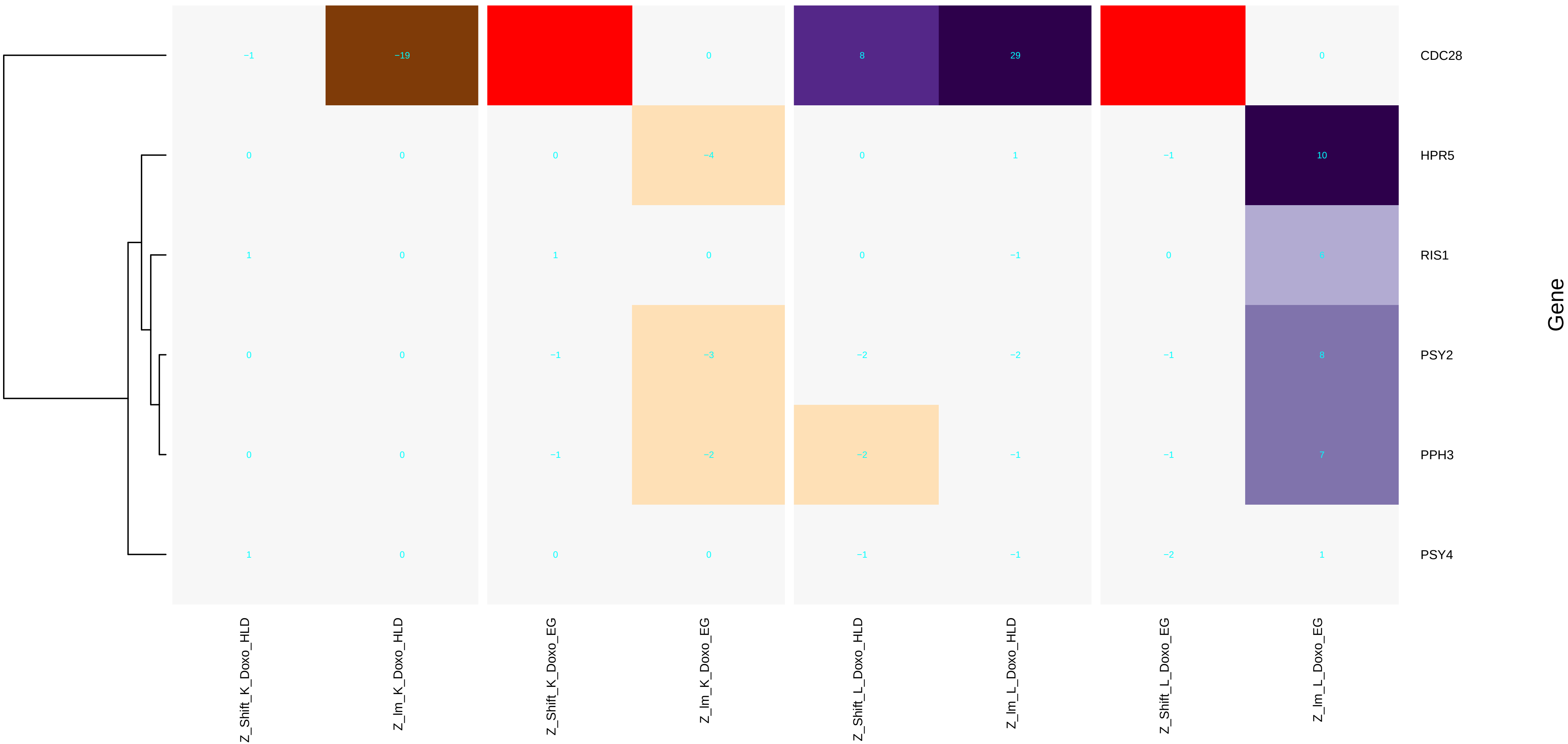

### negative_regulation_of_sequence-specific_DNA_binding_transcription_factor_activity.pdf

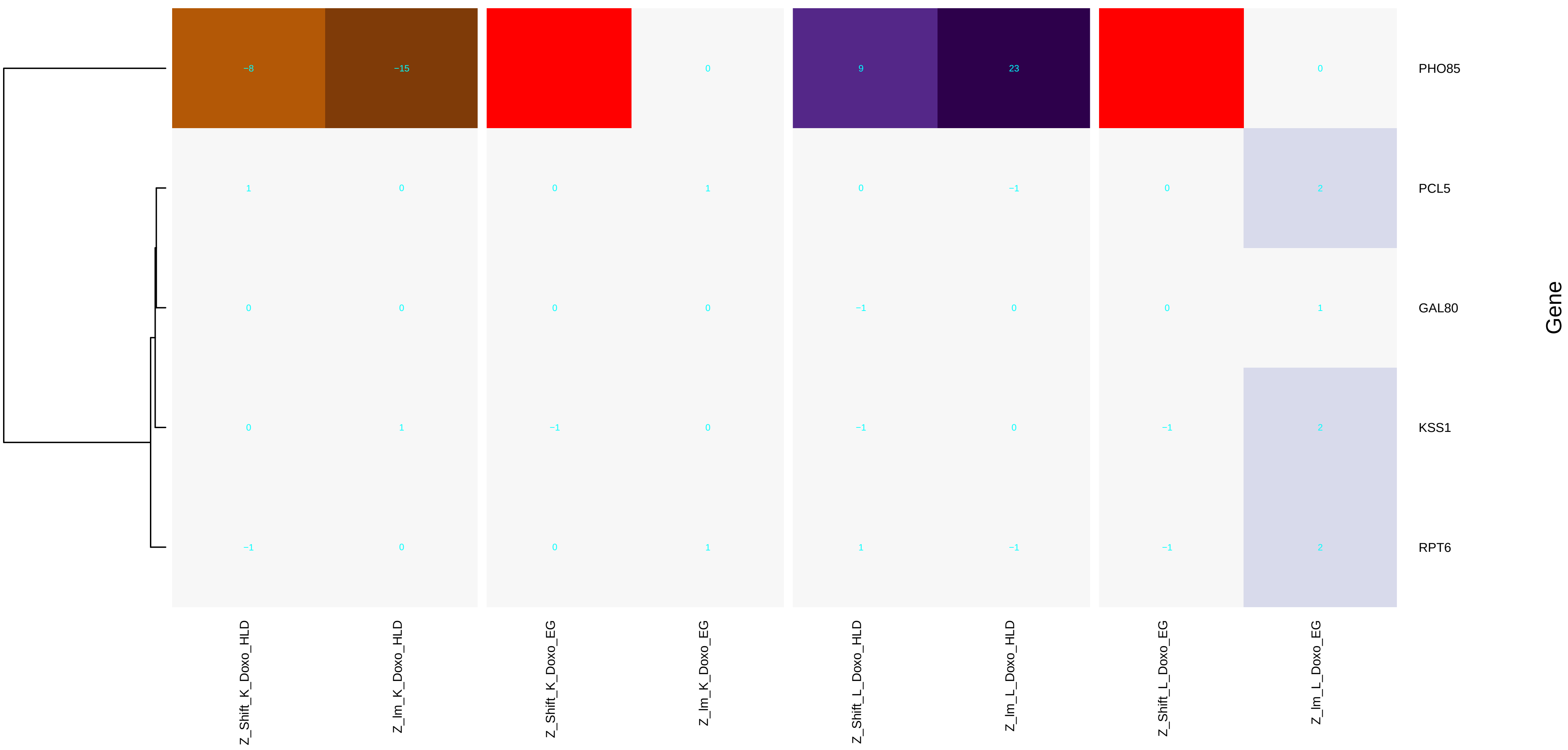

### non-recombinational_repair.pdf

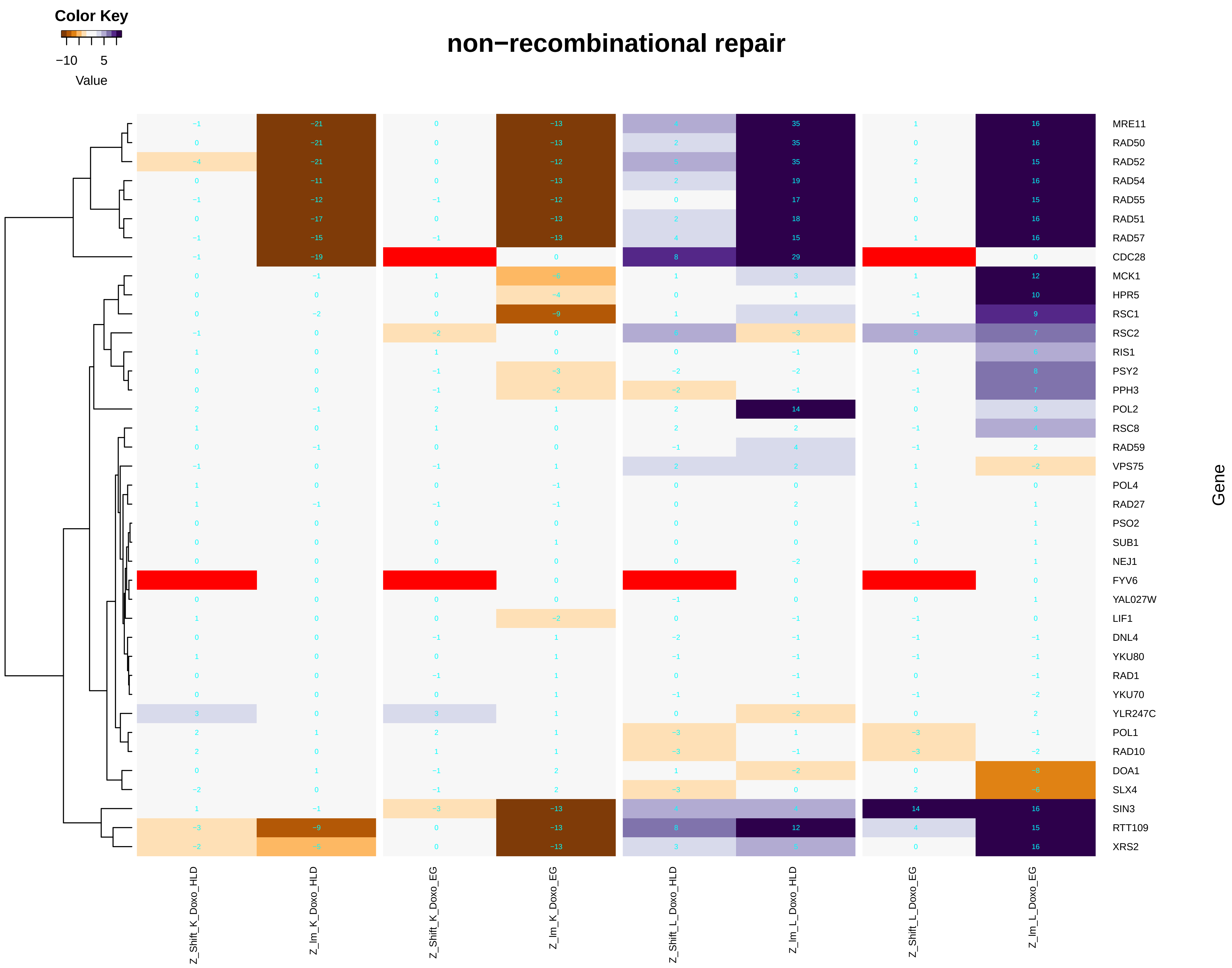

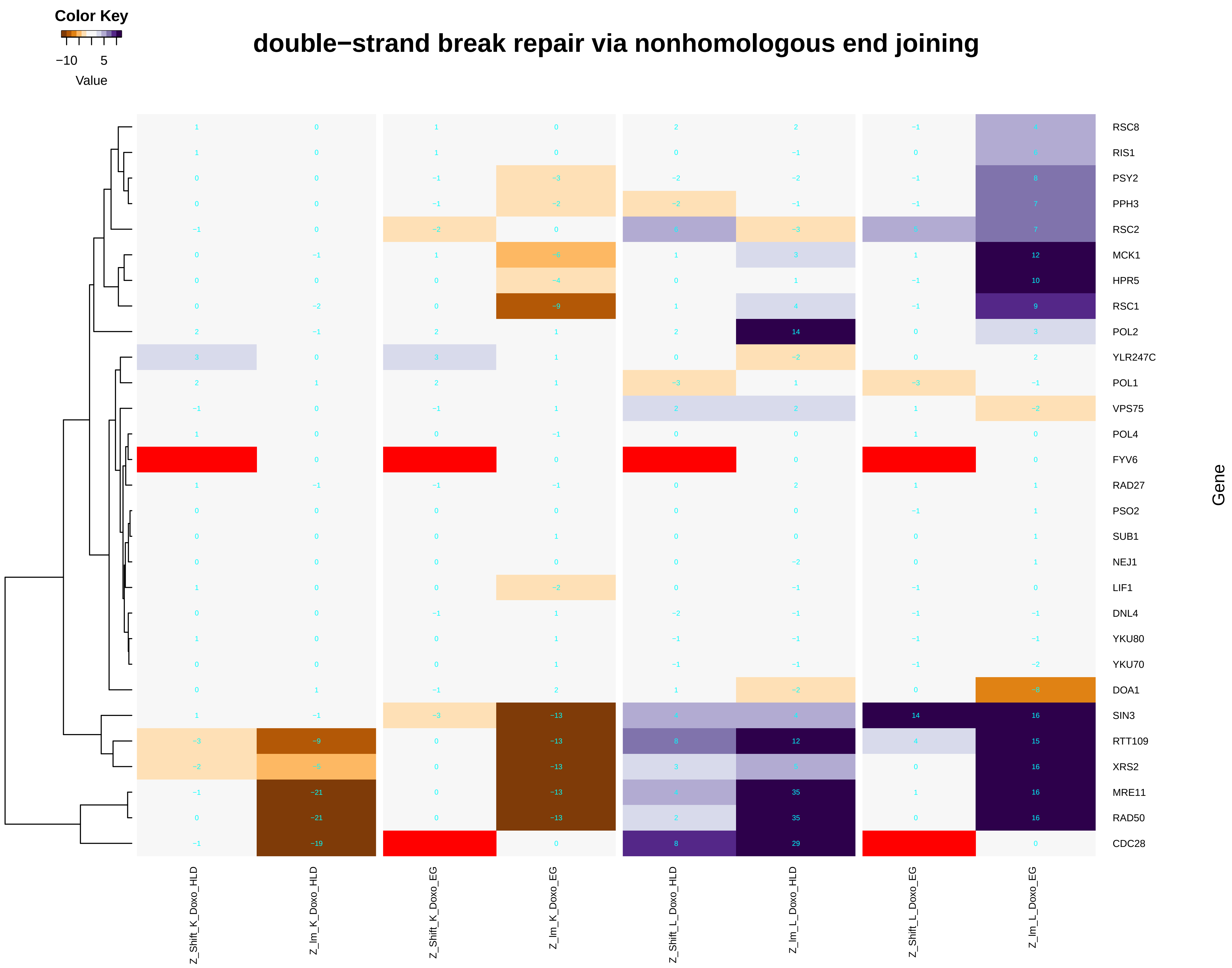

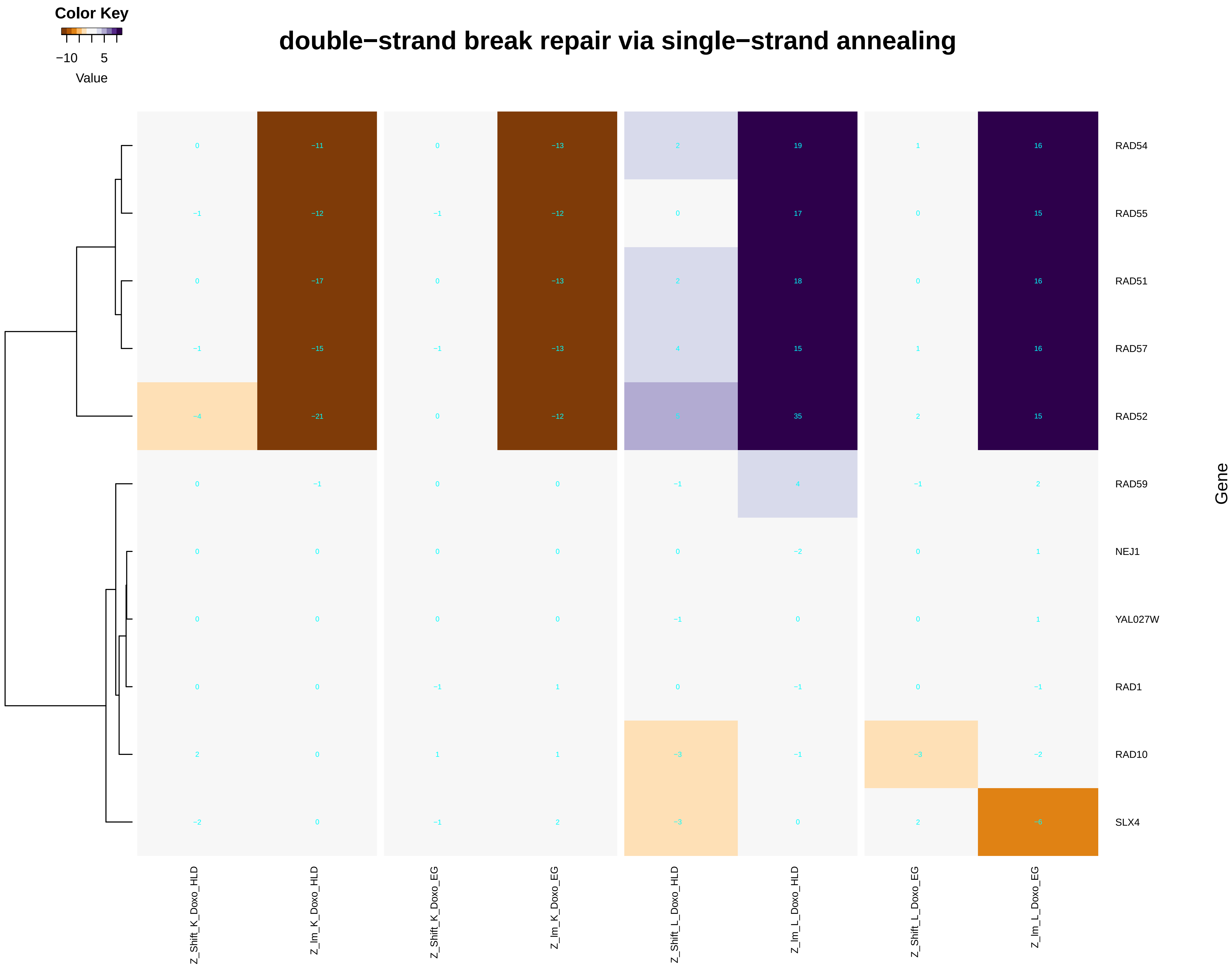

### nuclear-transcribed_mRNA_catabolic_process,_nonsense-mediated_decay.pdf

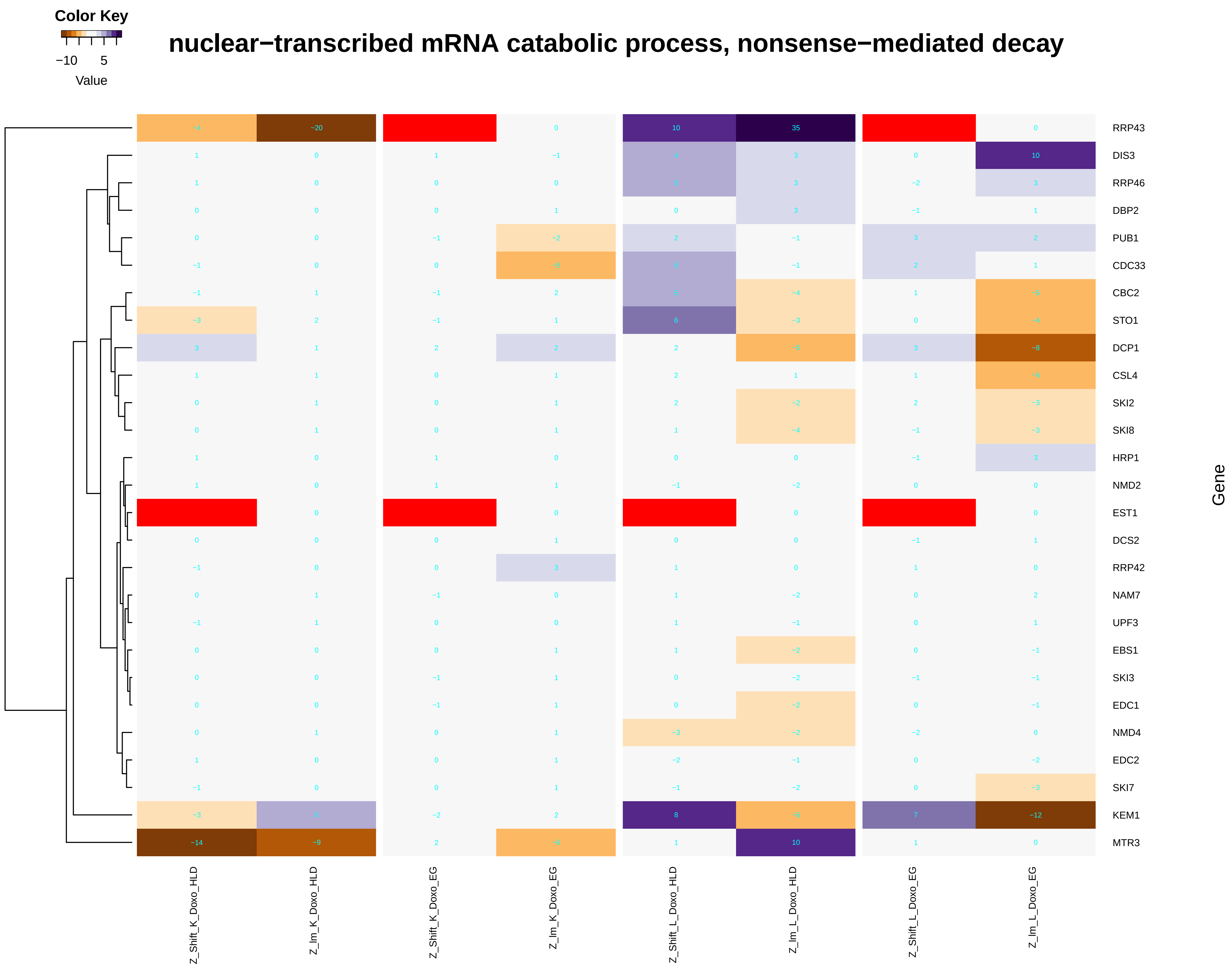

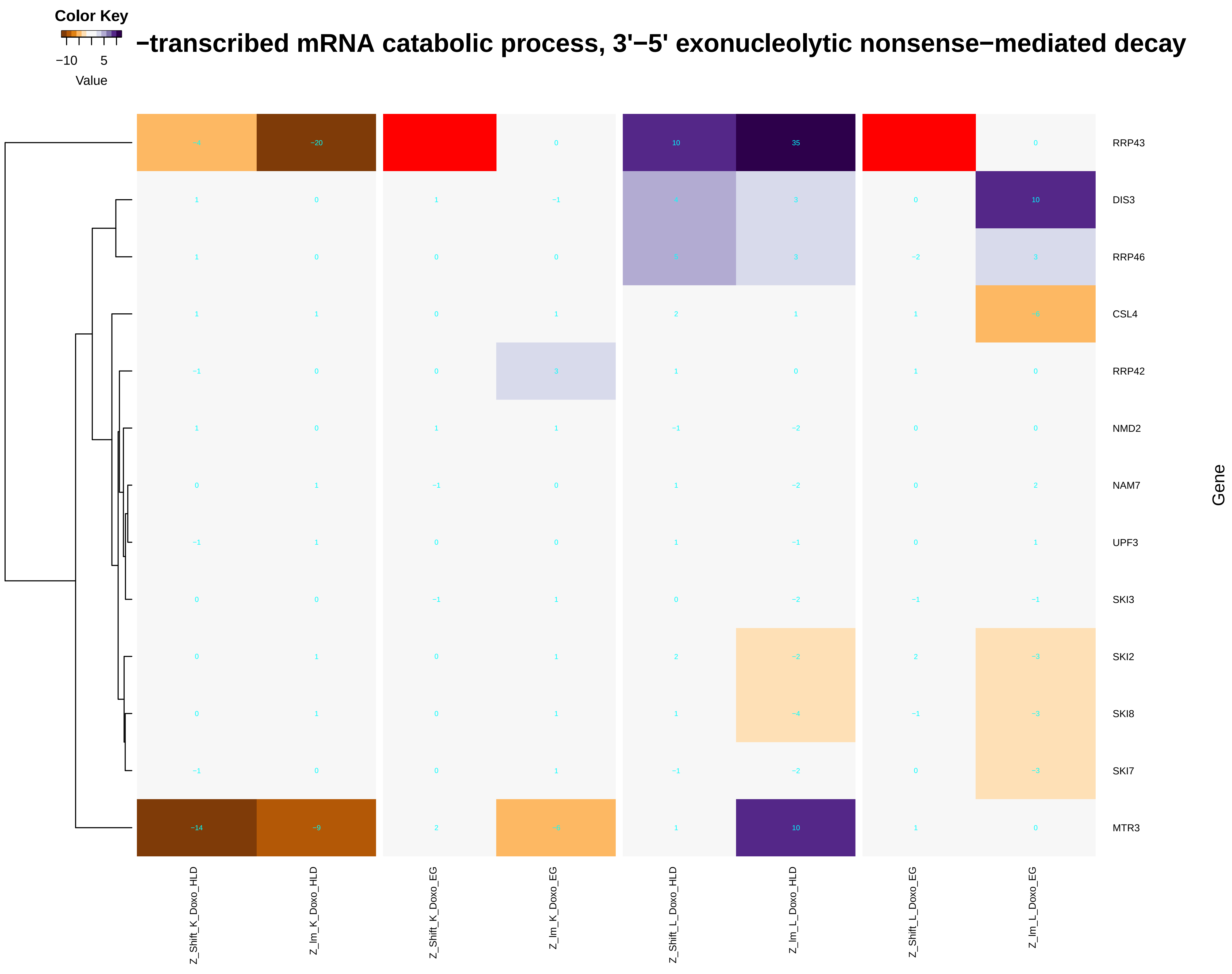

### nuclear_DNA_replication.pdf

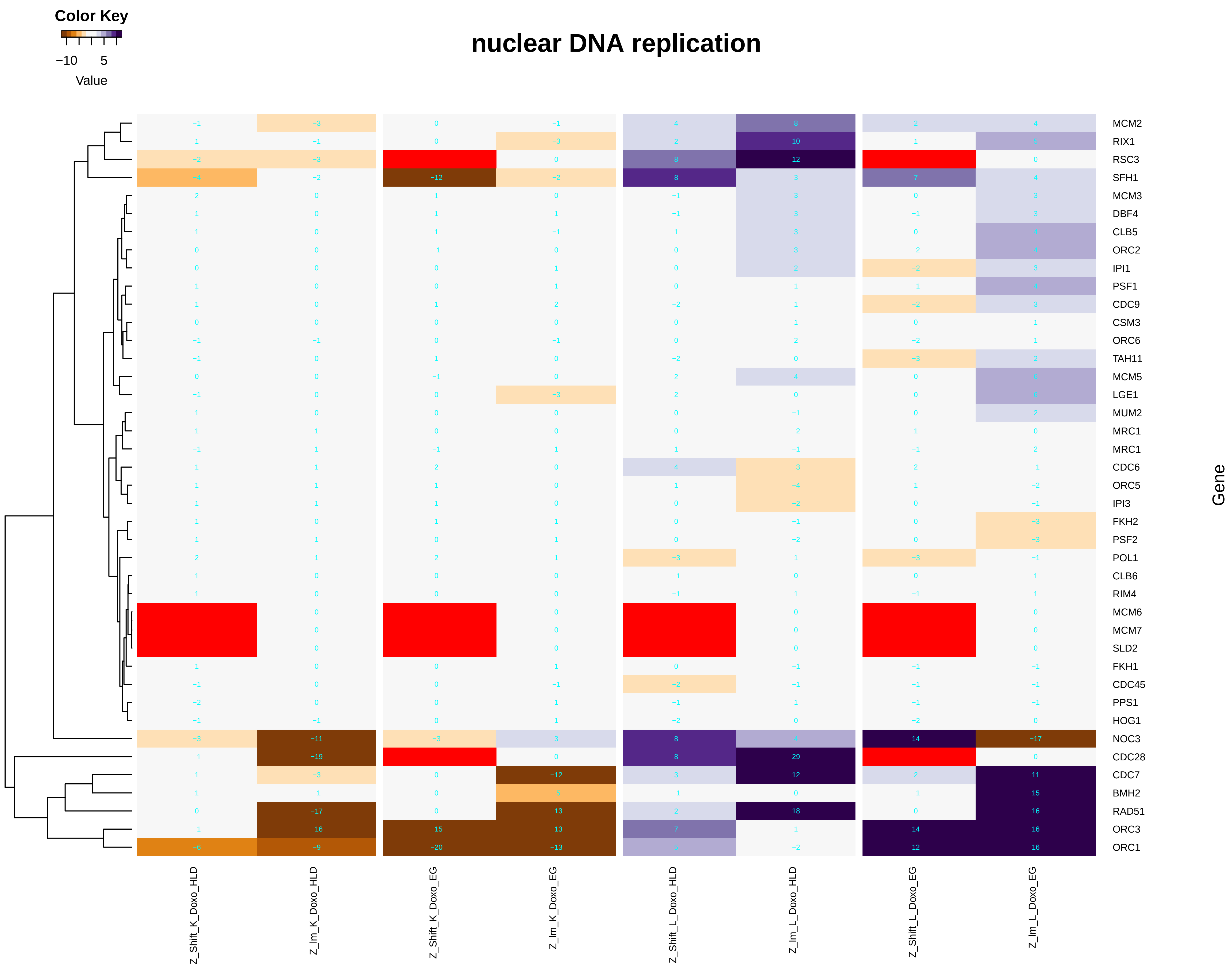

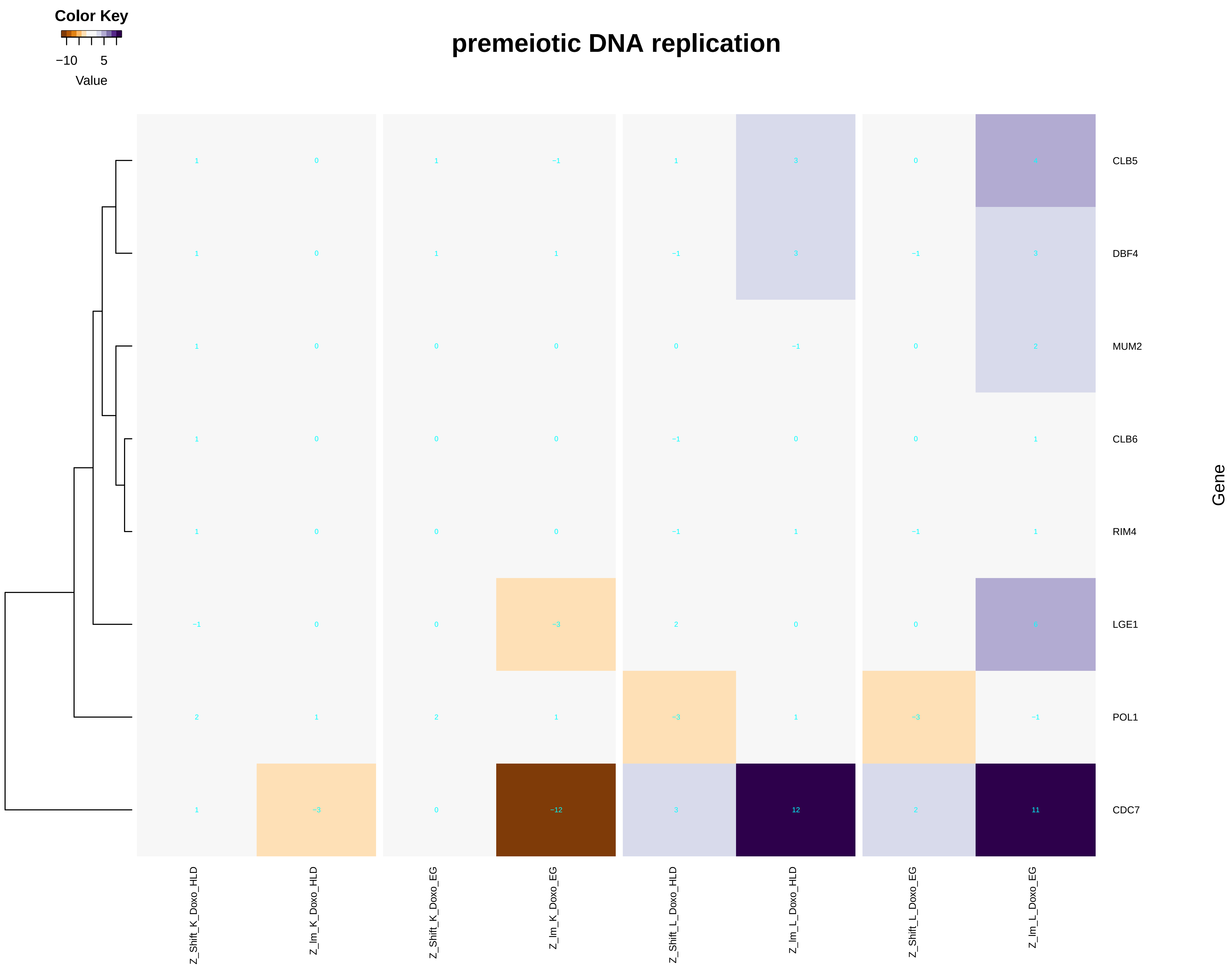

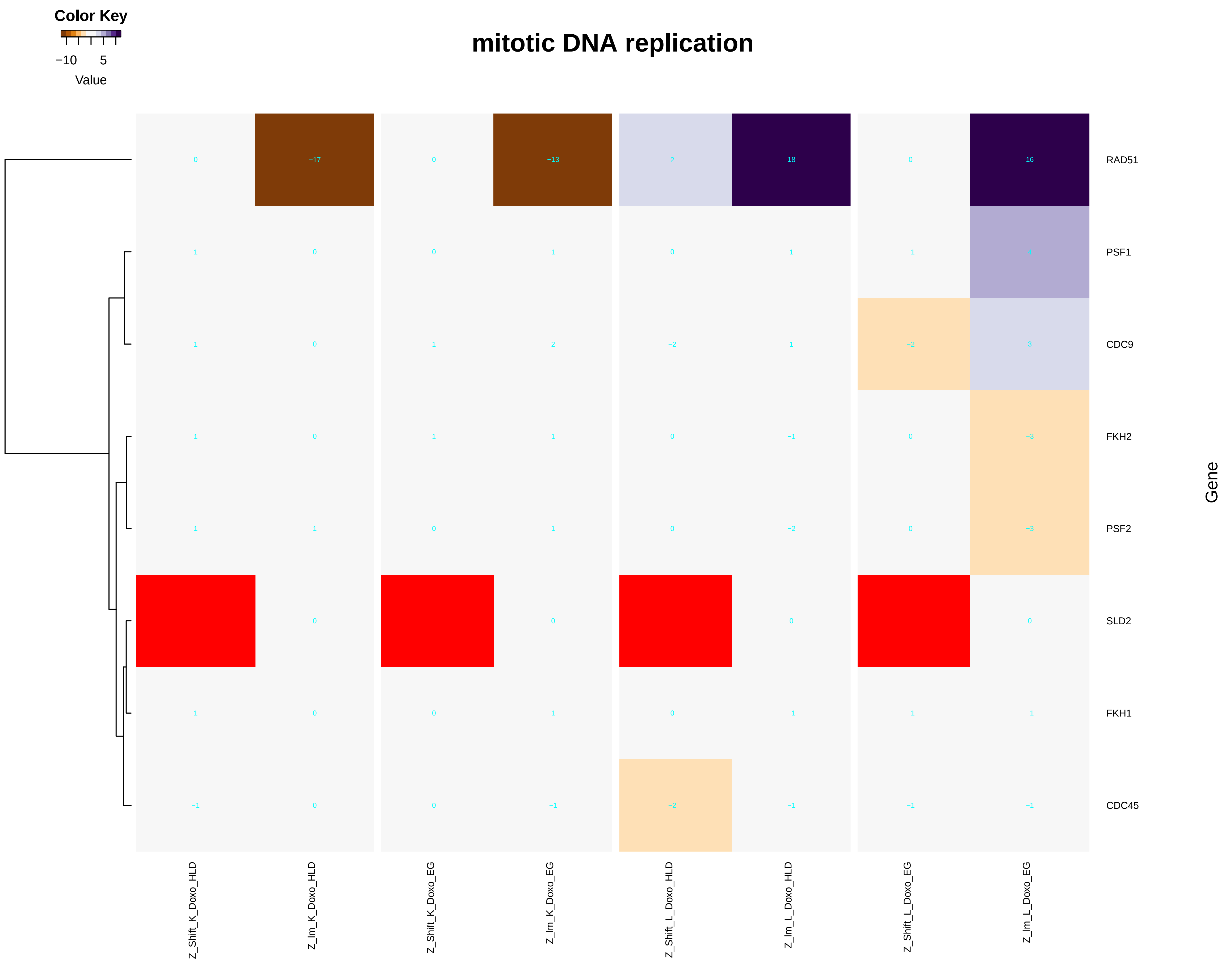

### nucleosome_assembly.pdf

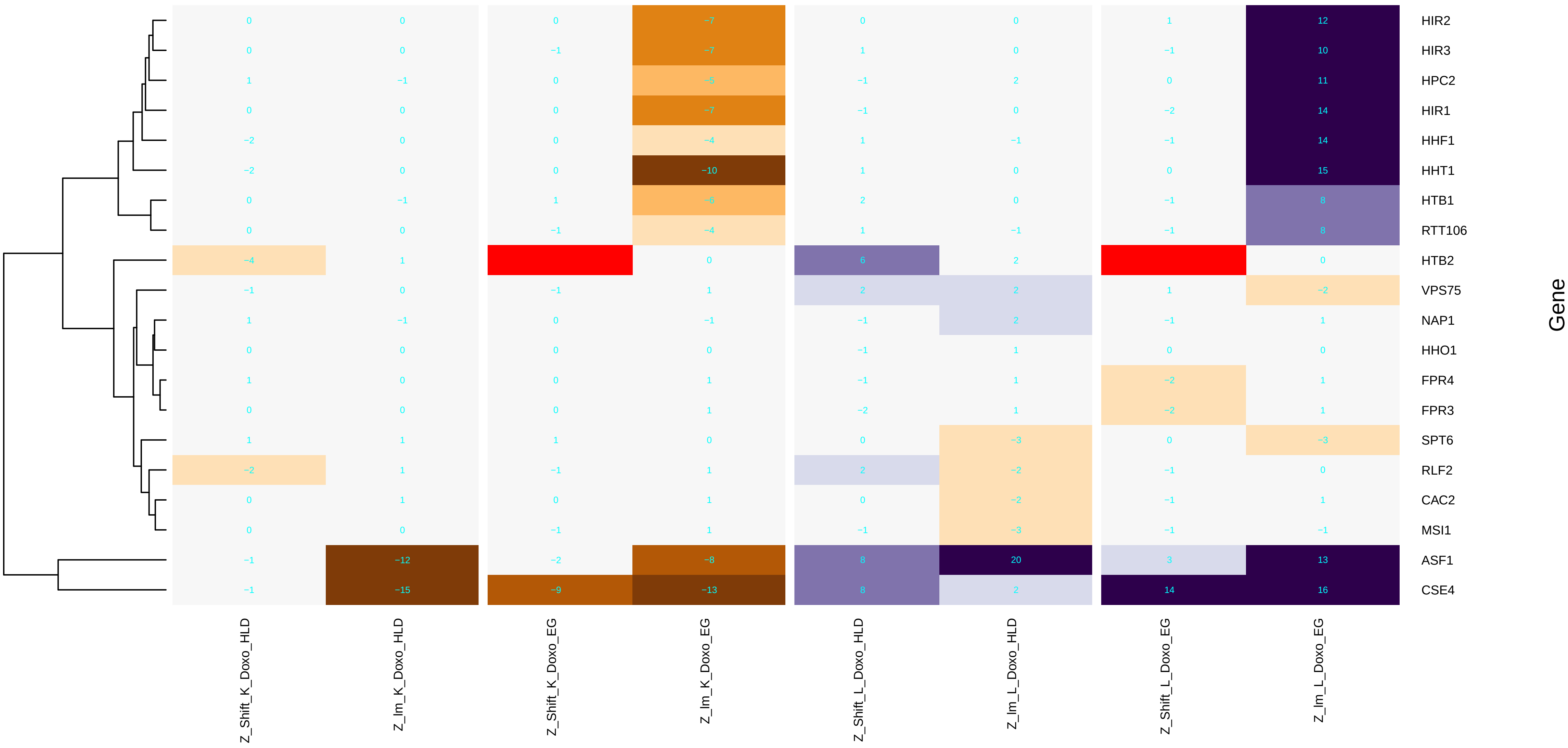

### nucleotide_phosphorylation.pdf

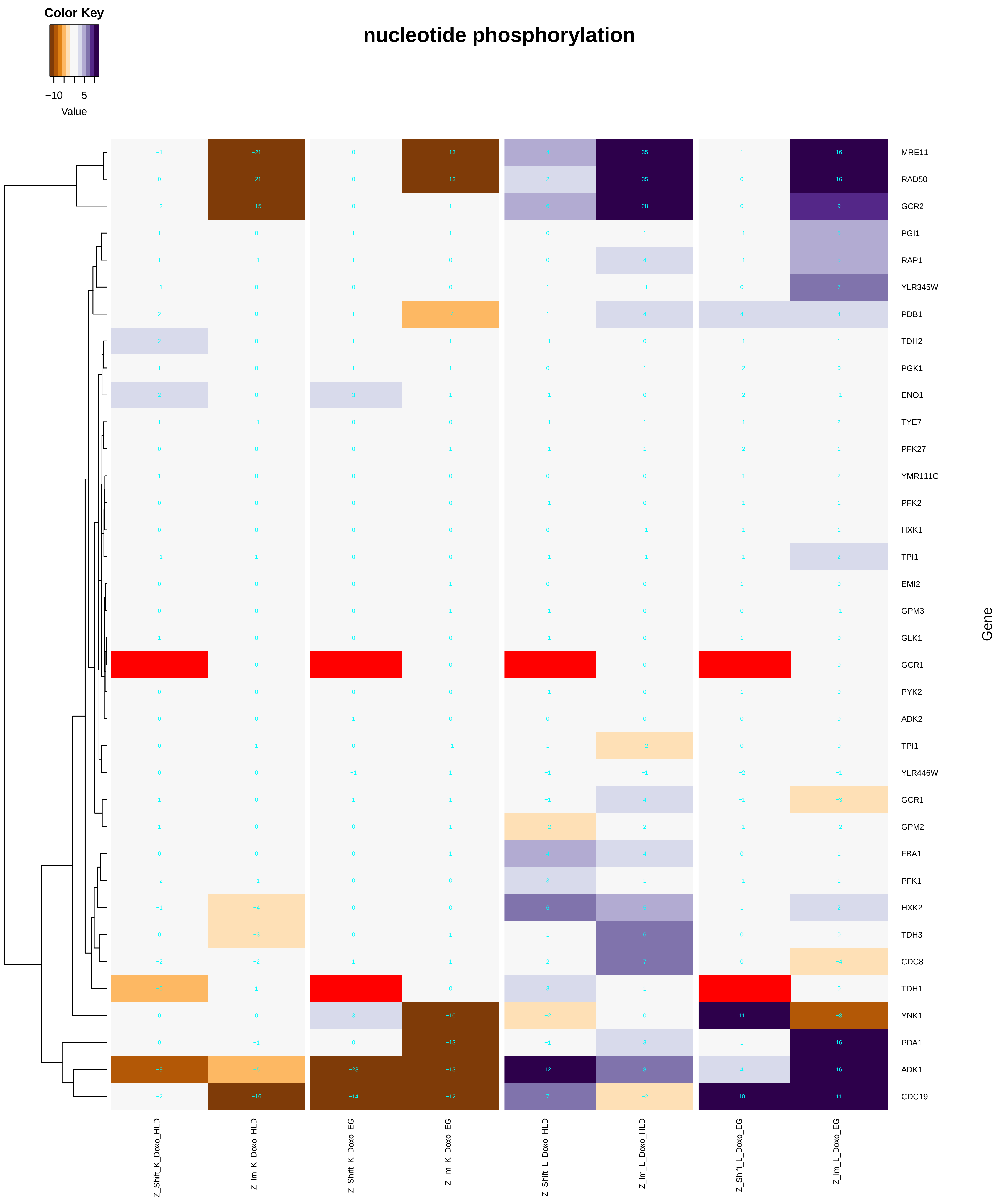

#### nucleoside diphosphate phosphorylation

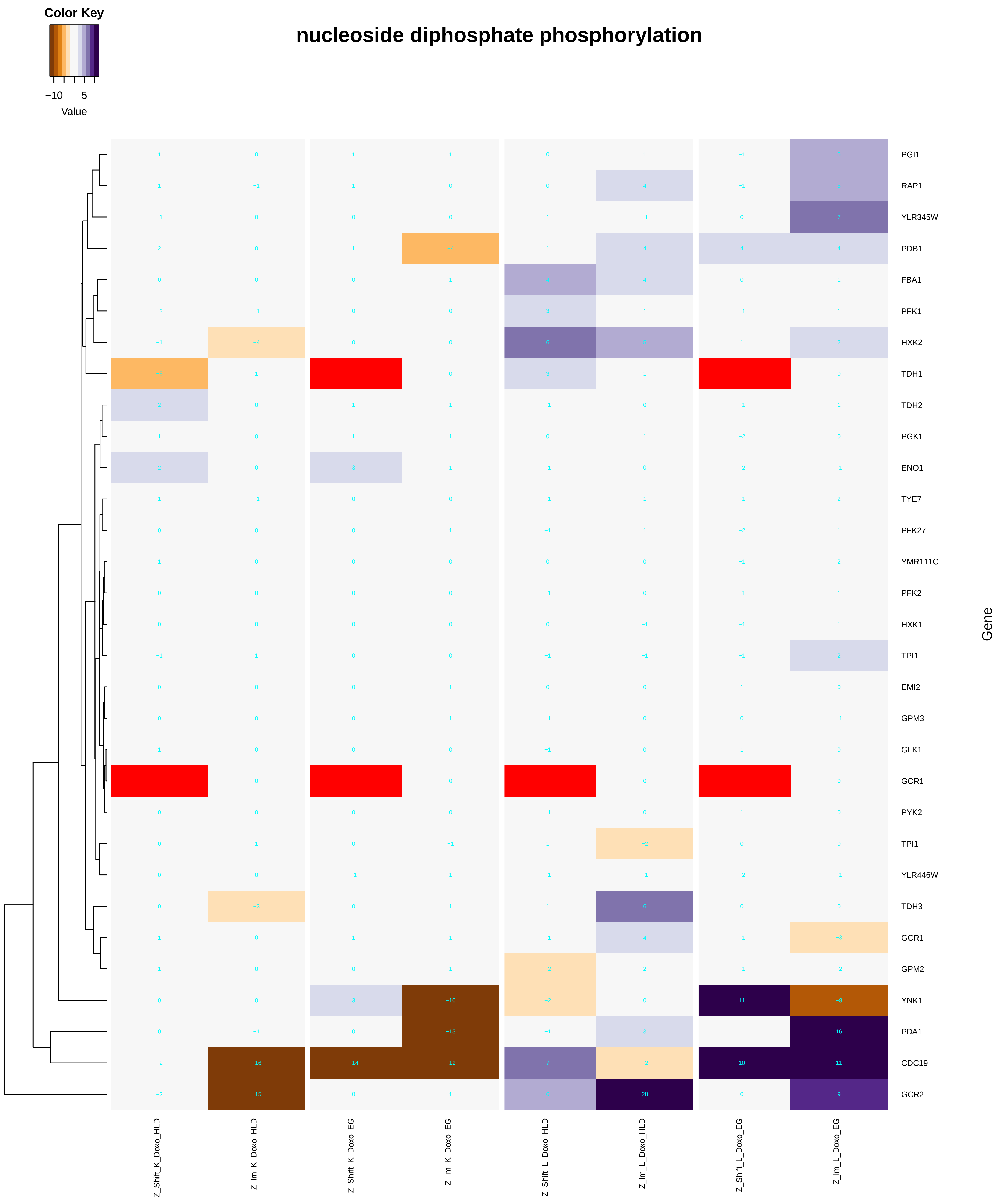

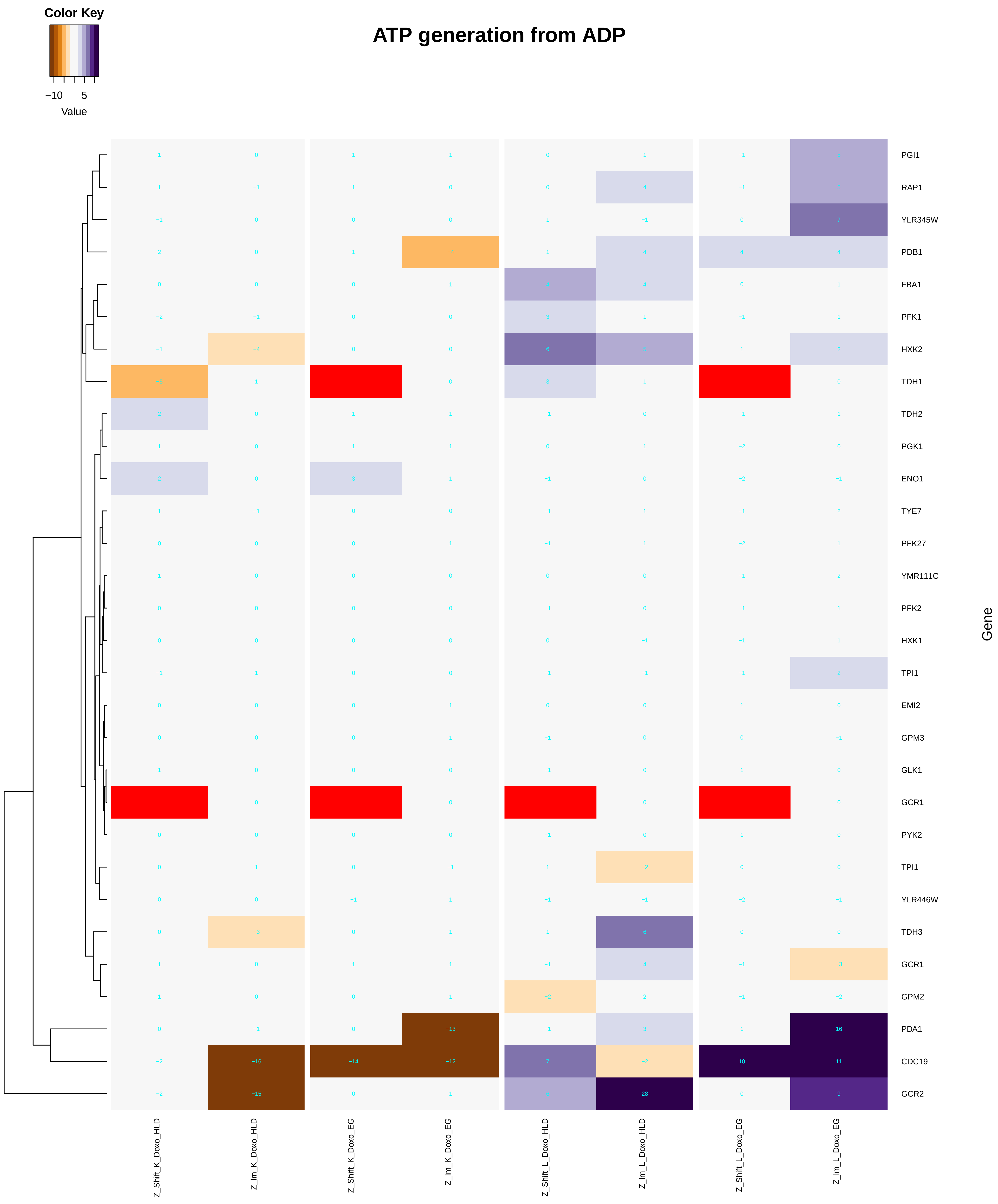

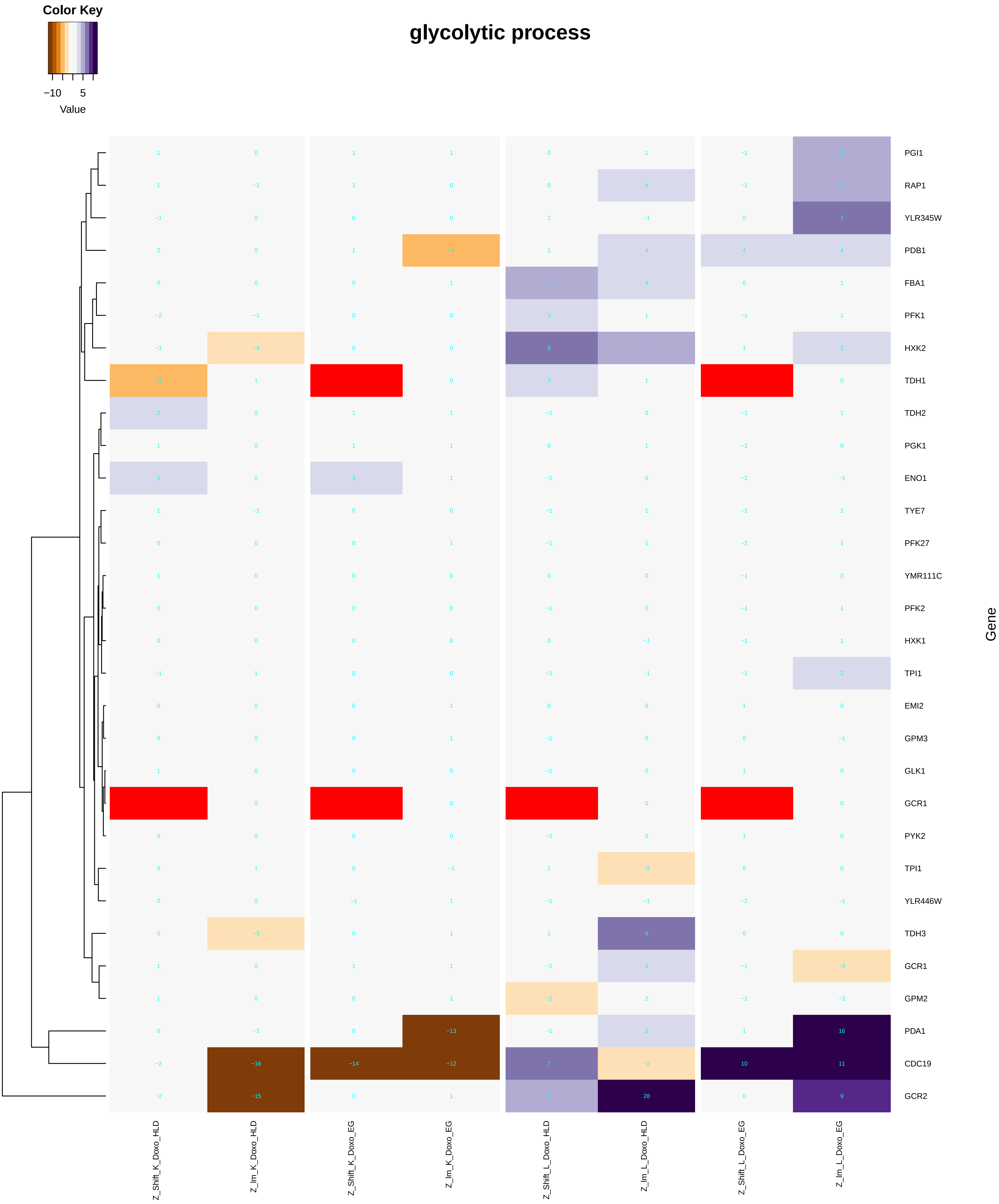

### nucleus_organization.pdf

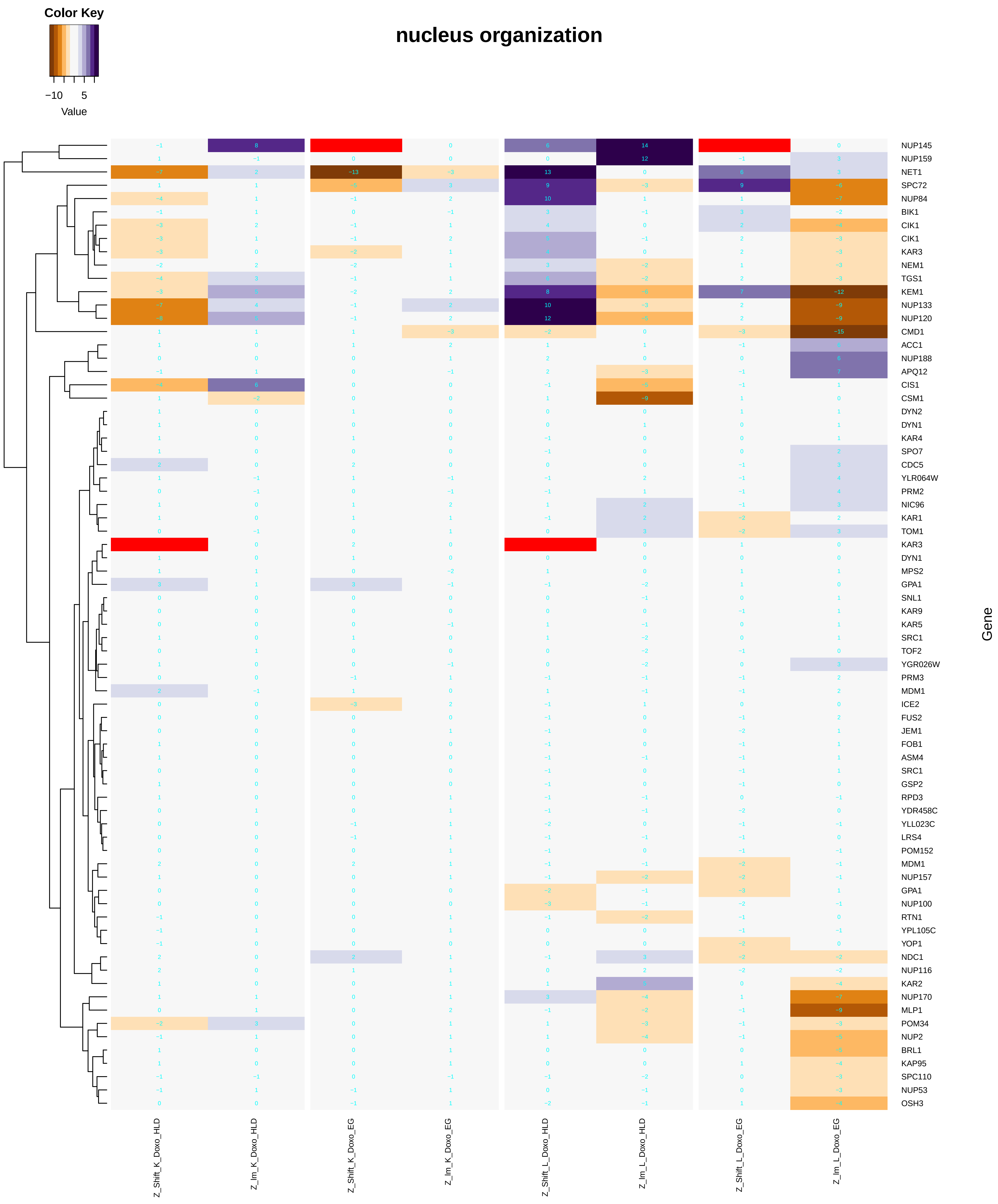

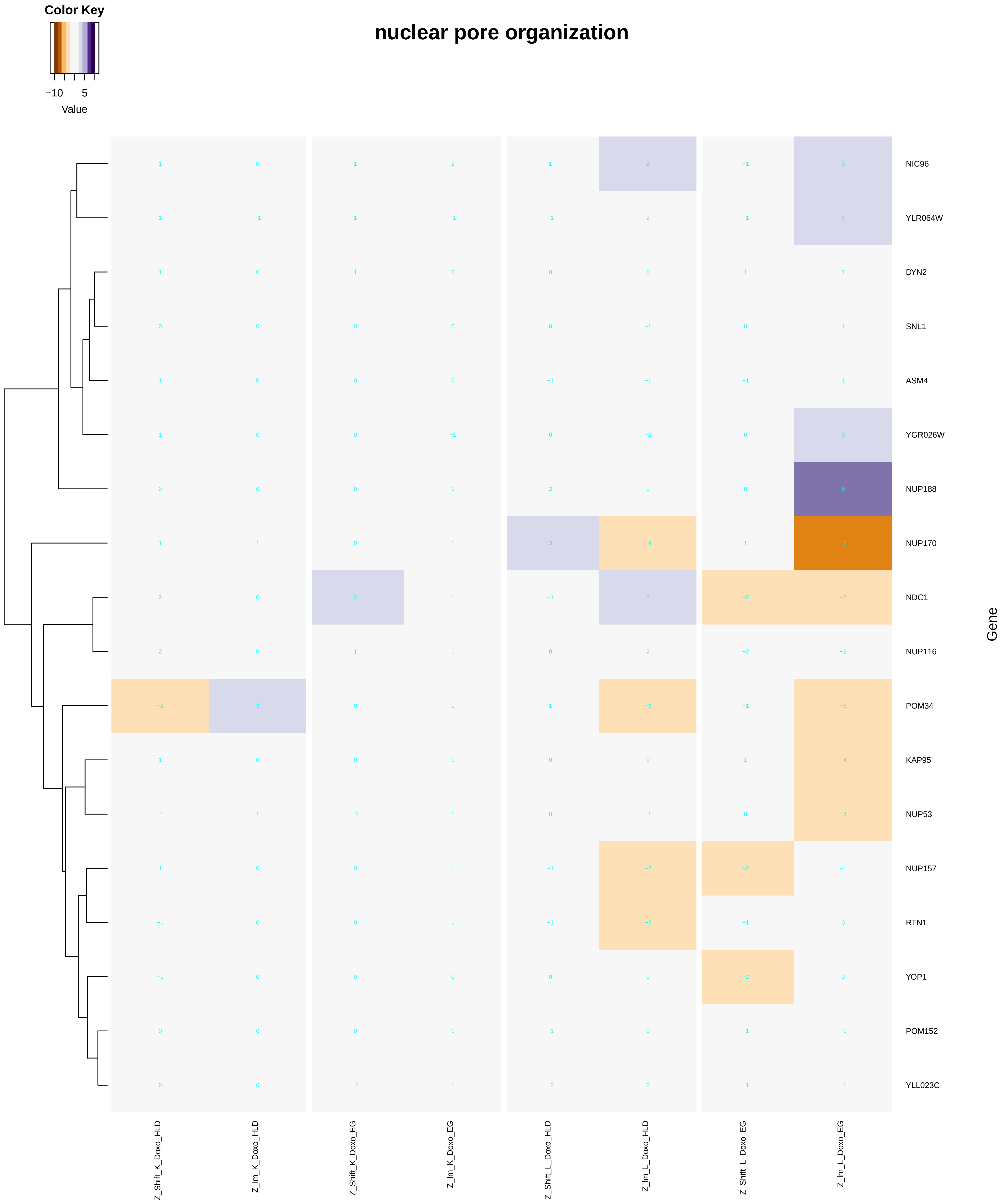

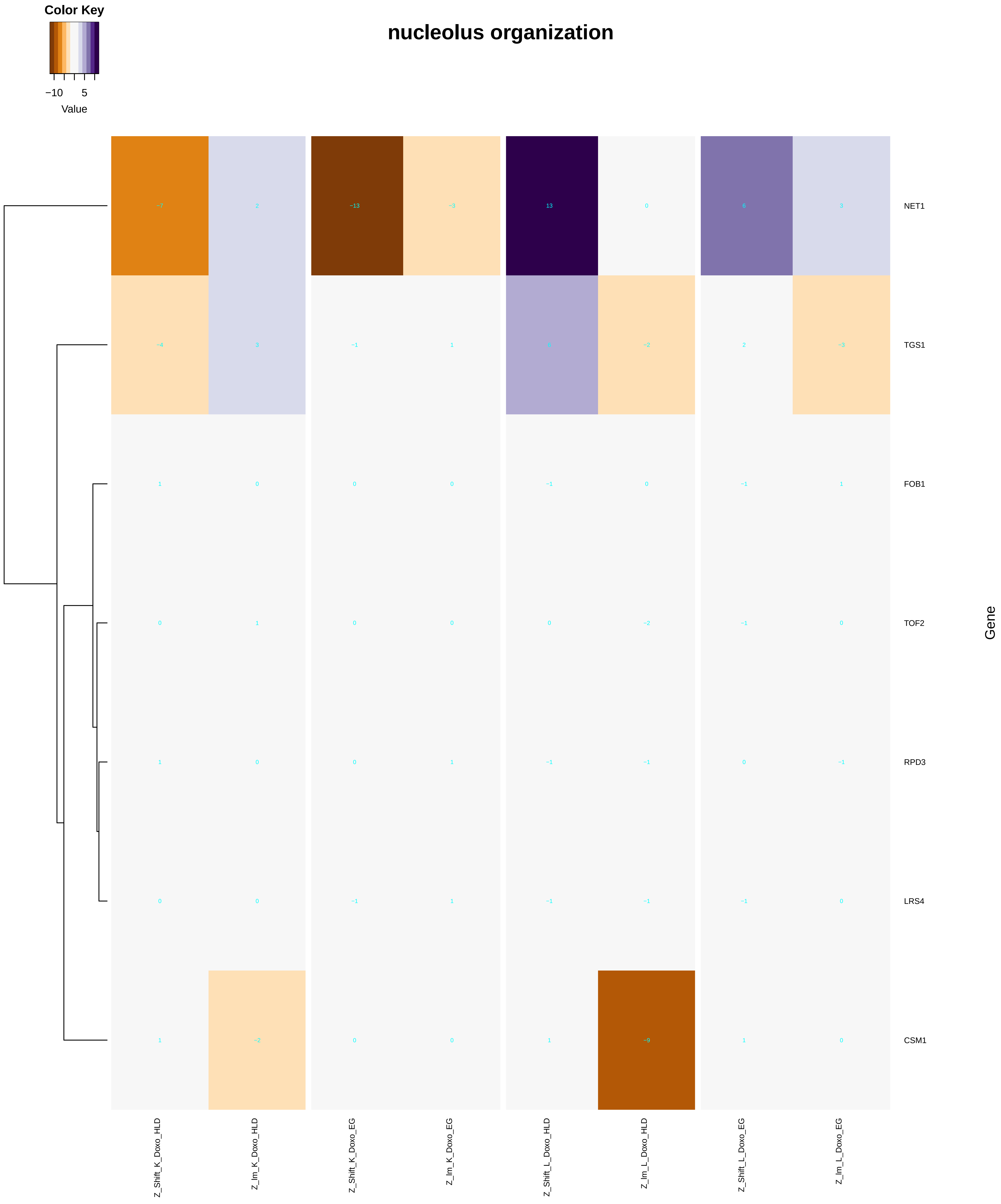

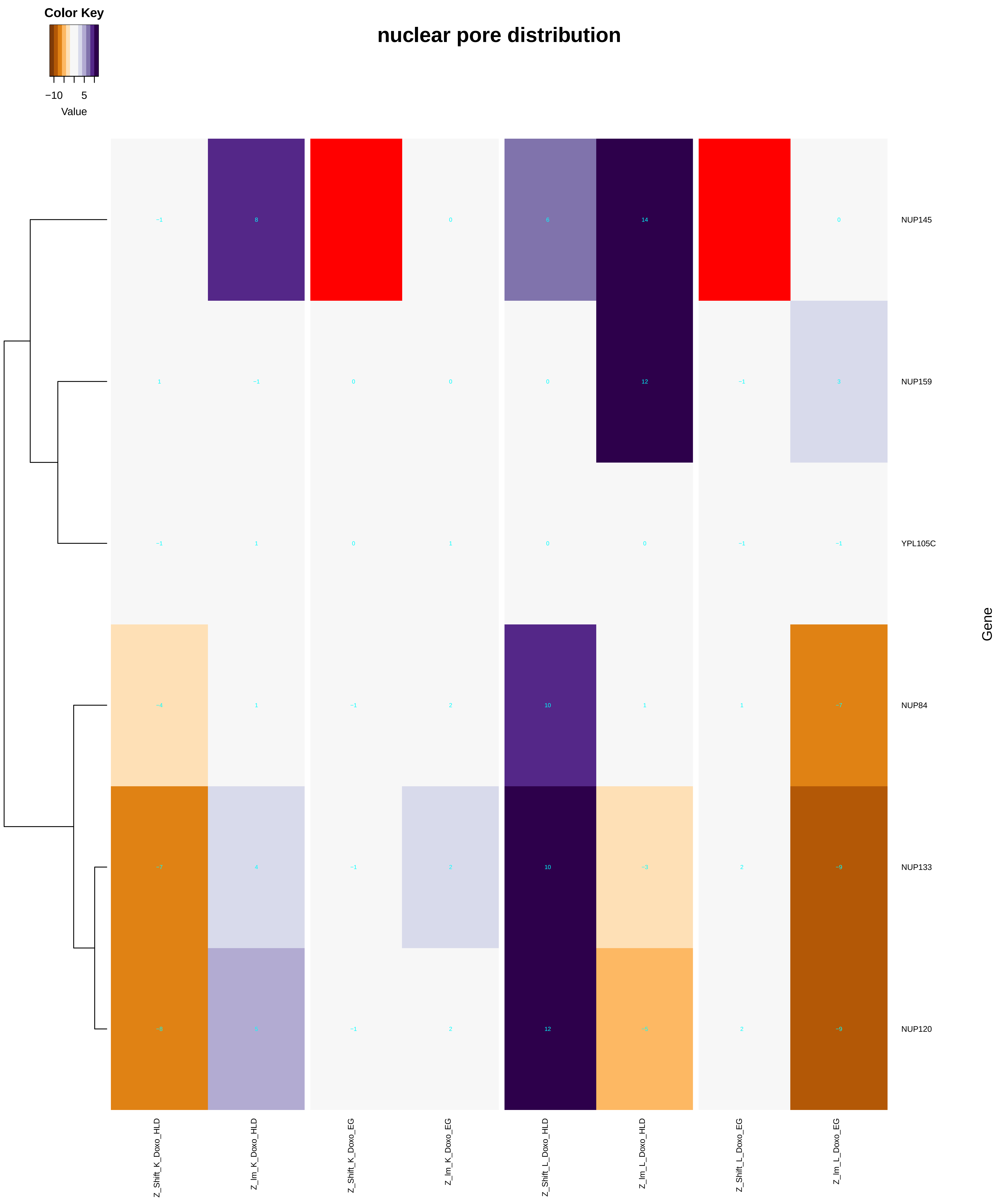

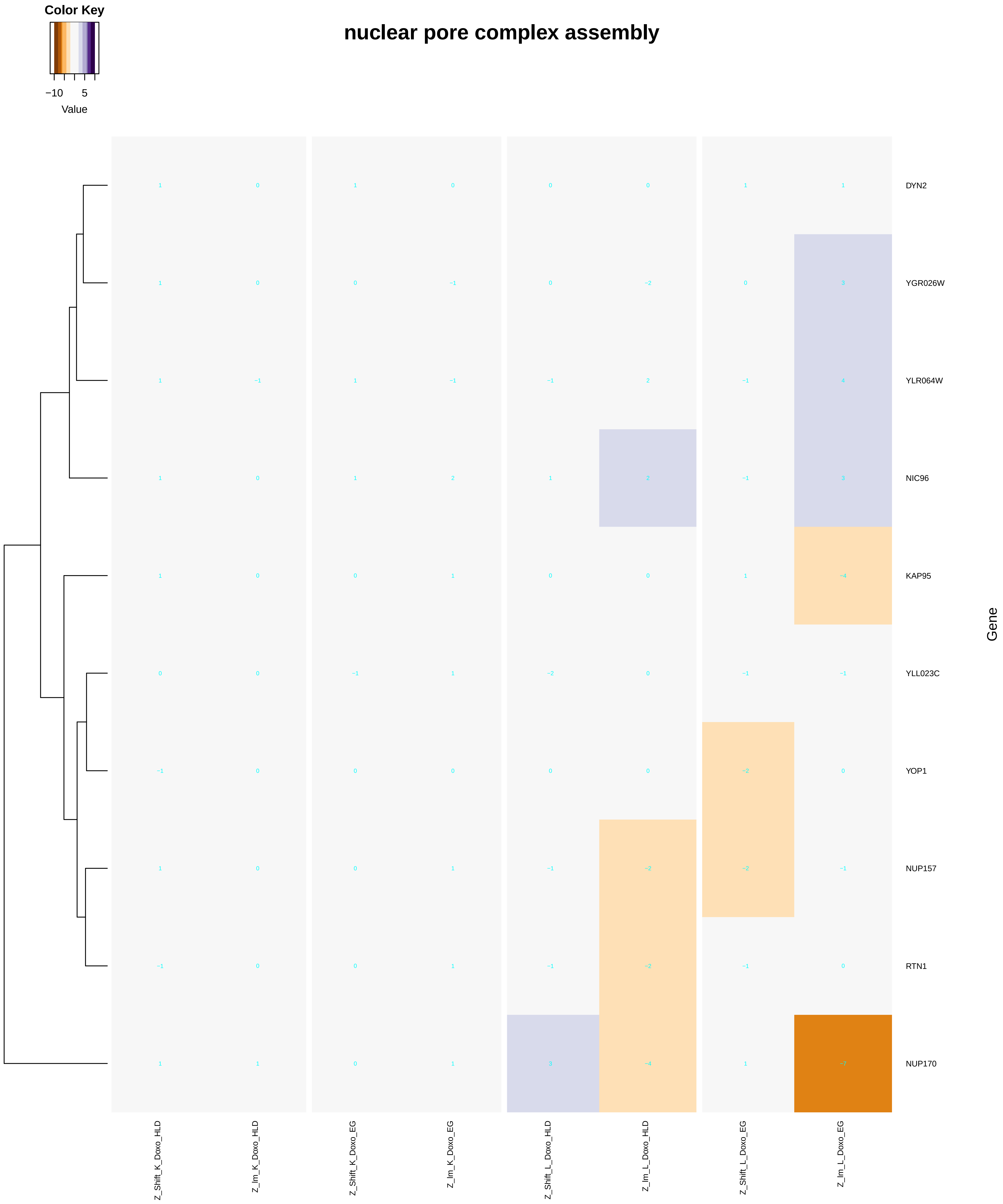

### organelle_assembly.pdf

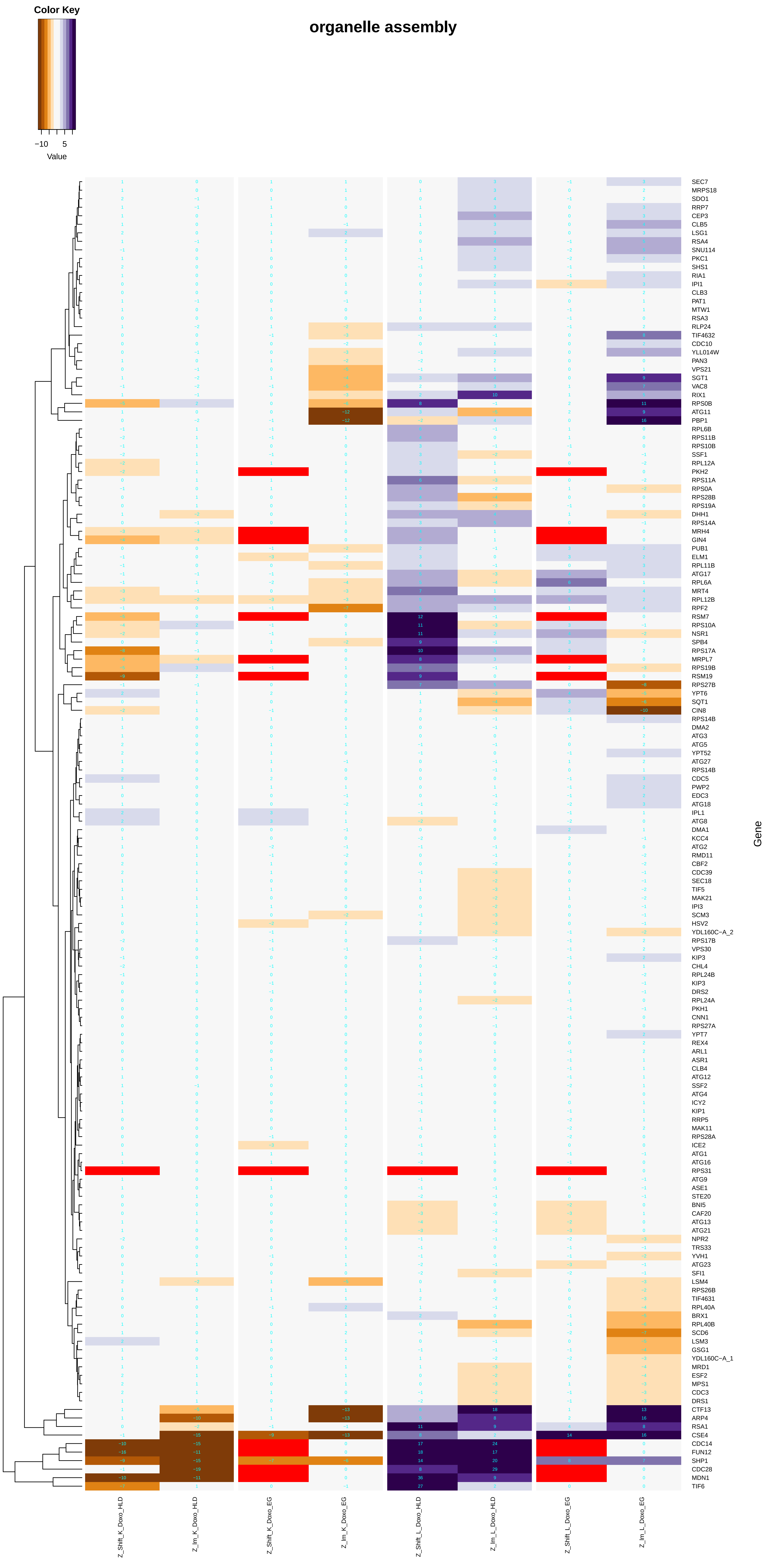

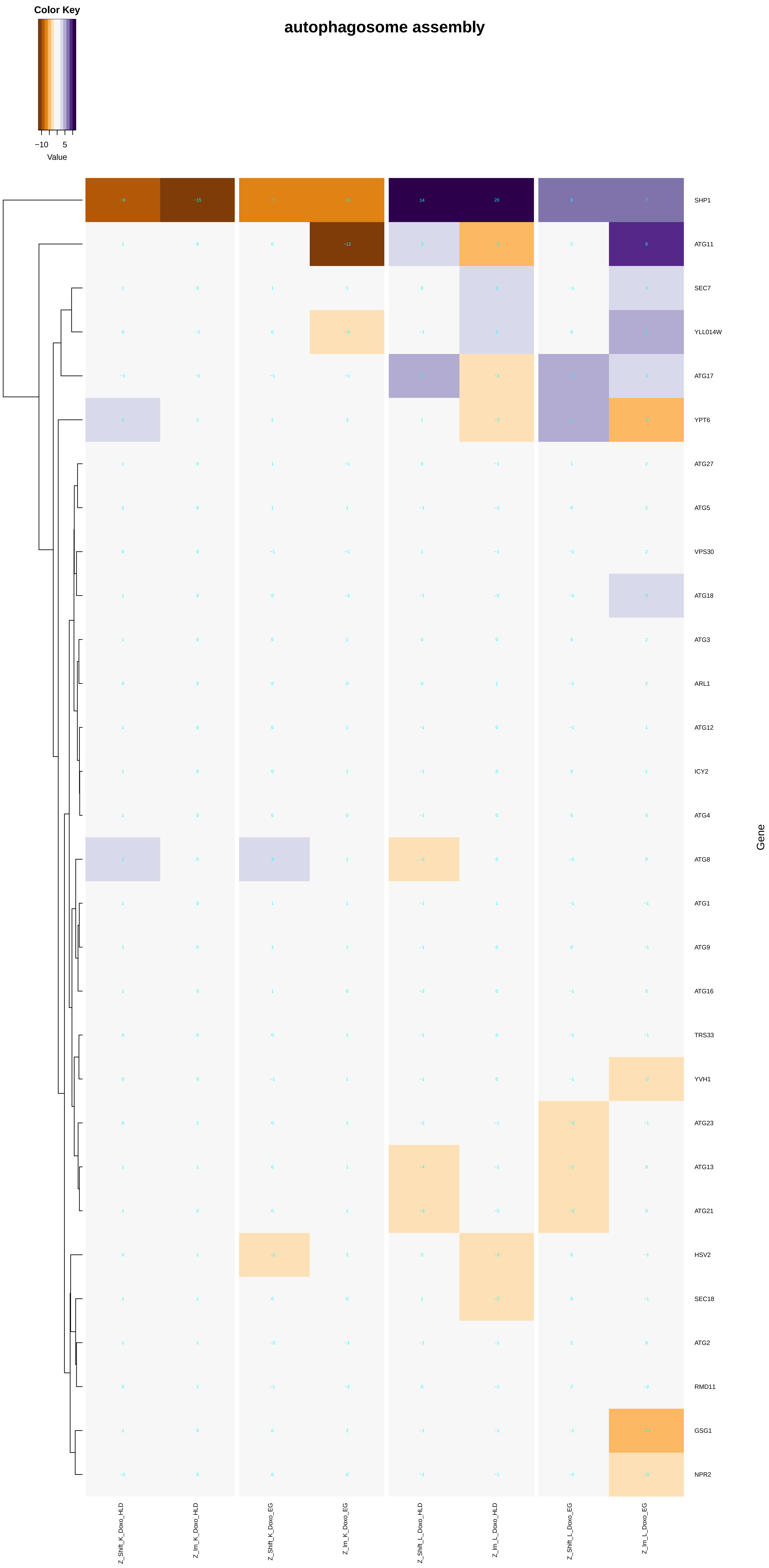

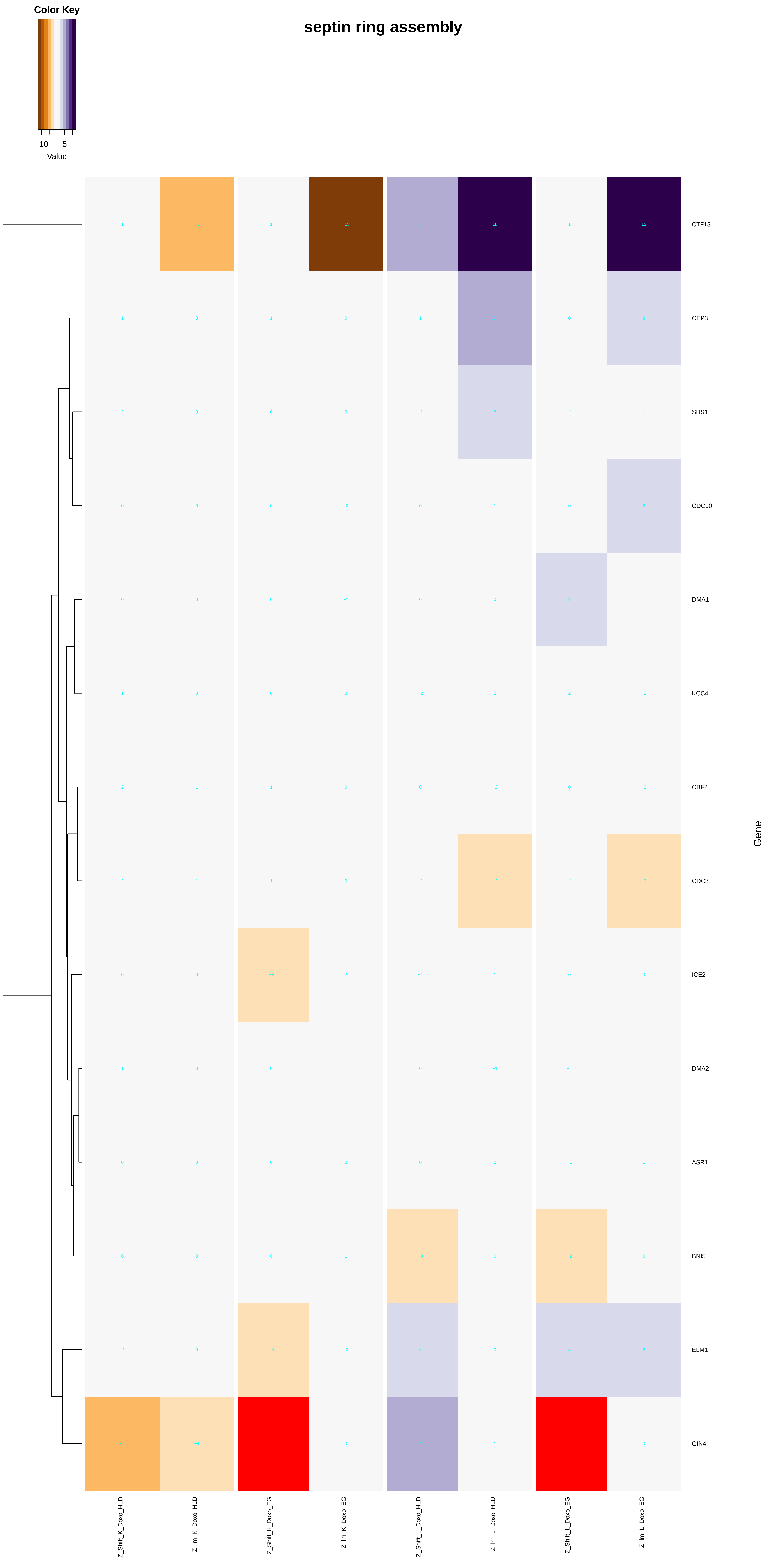

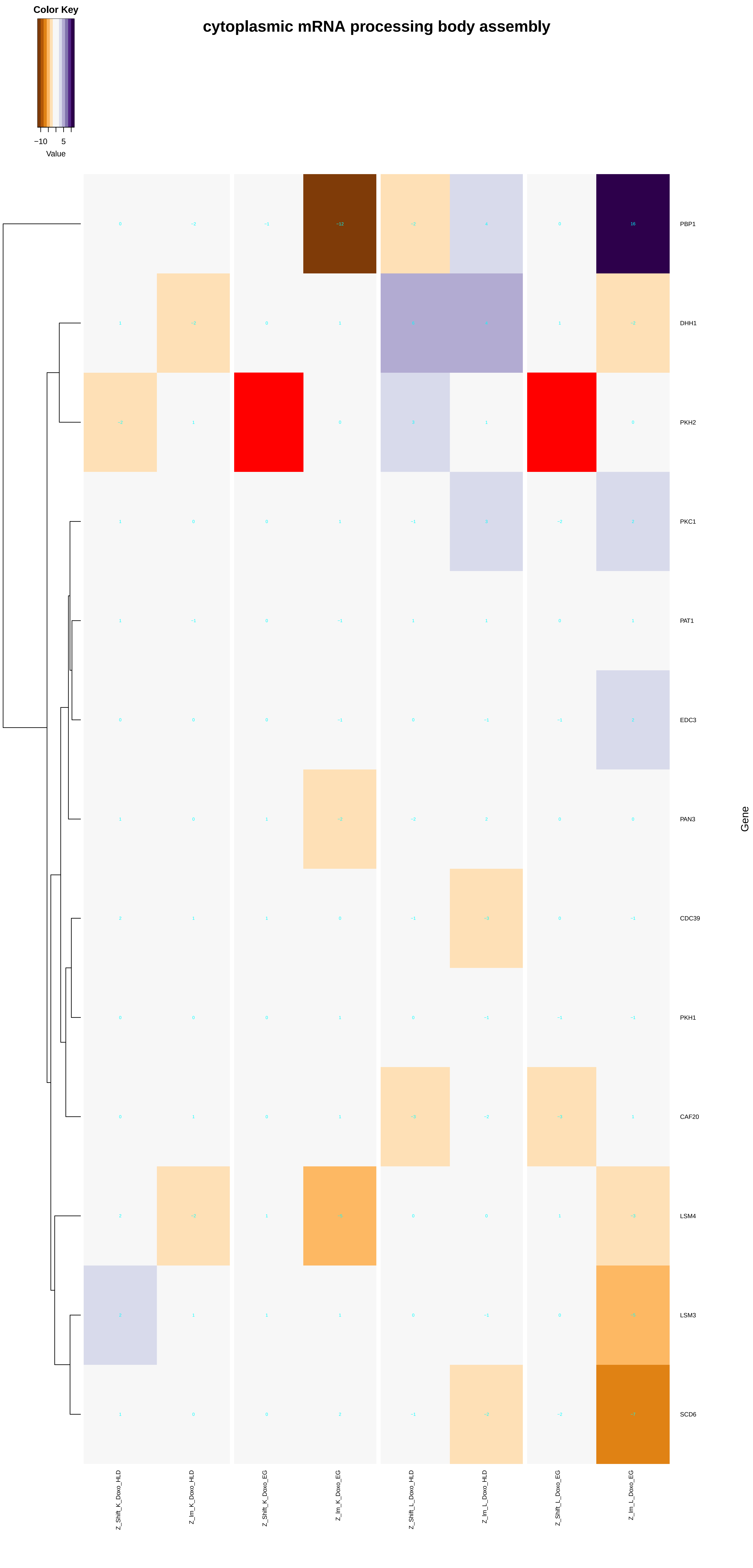

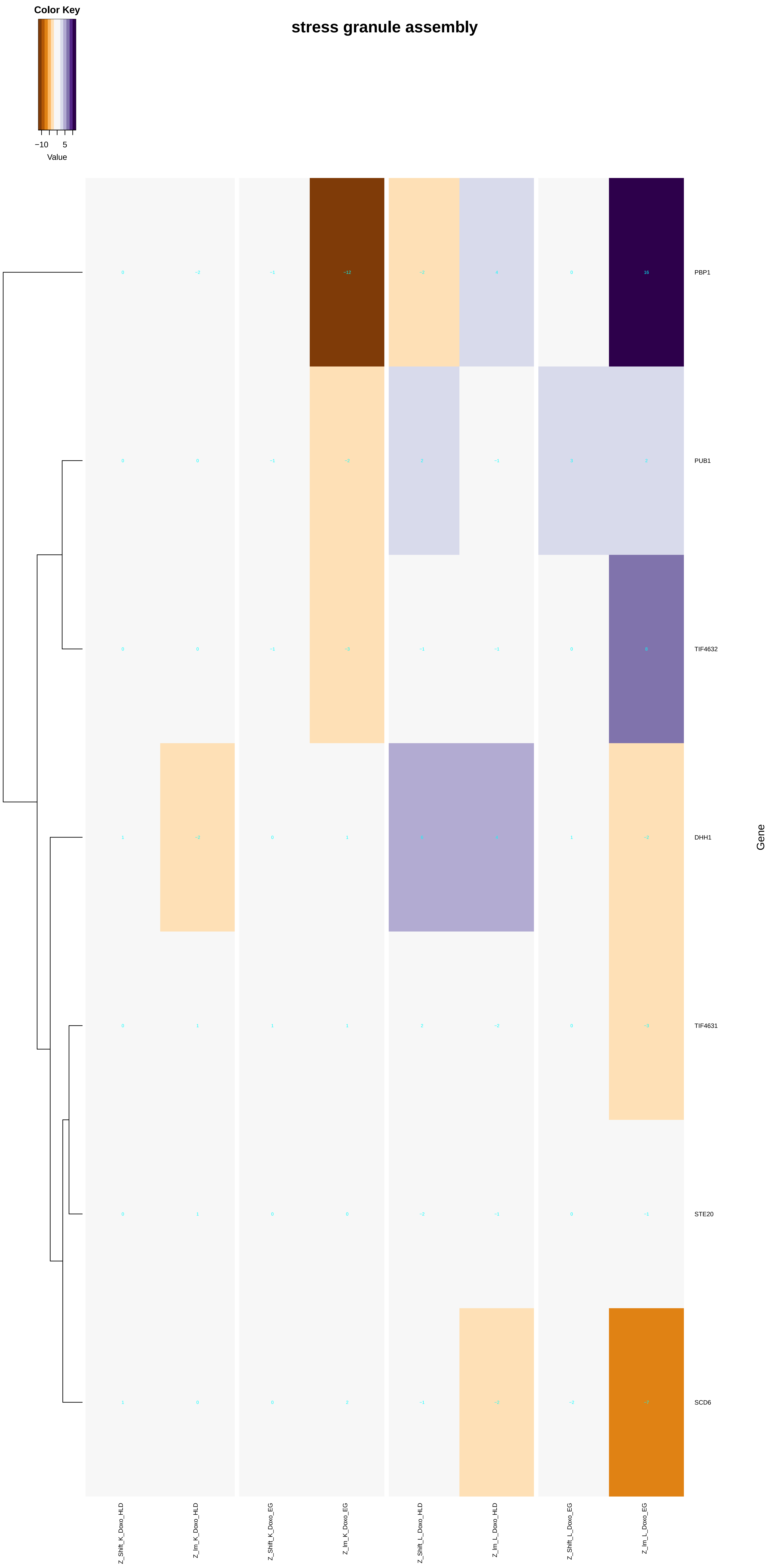

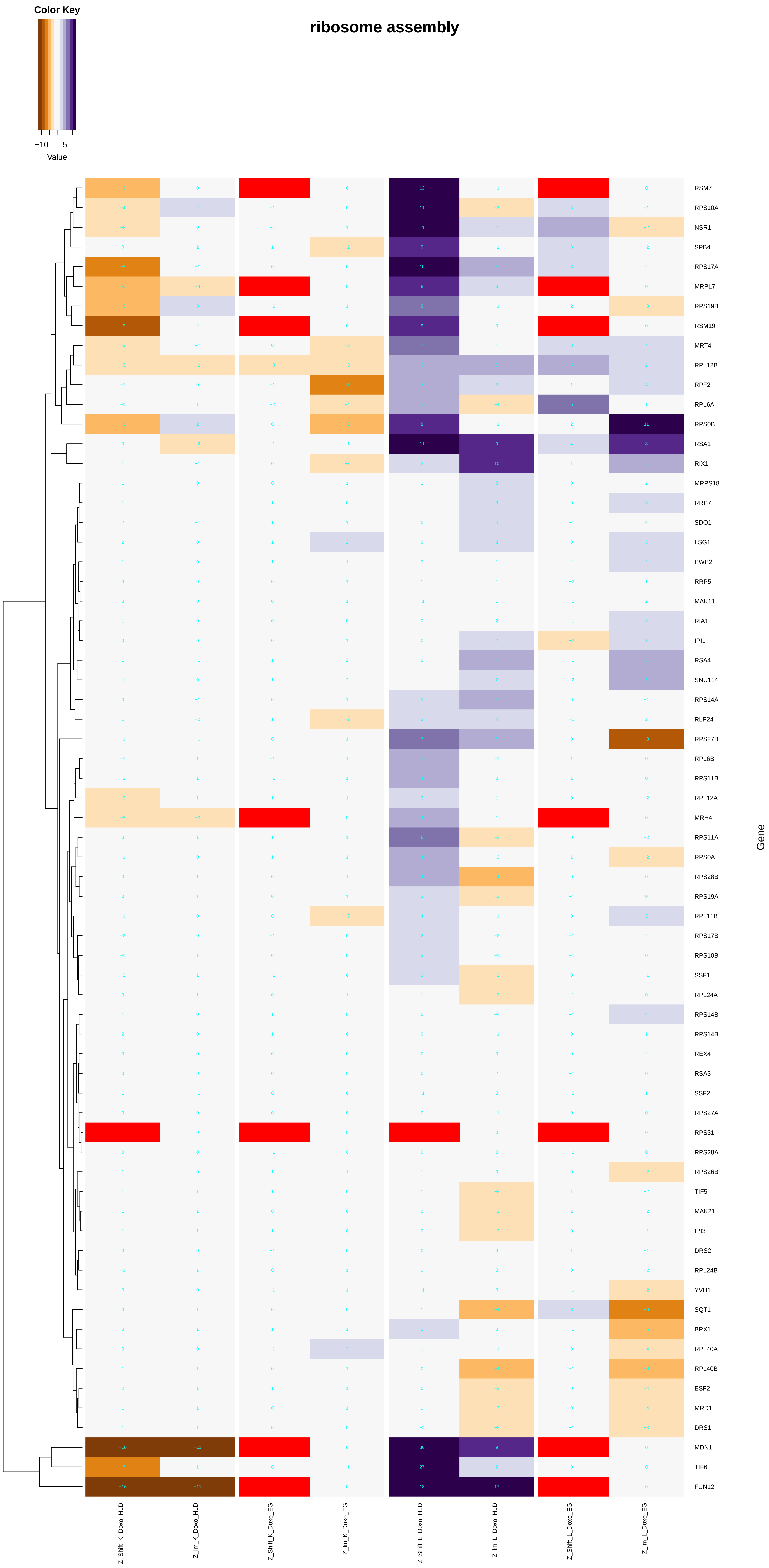

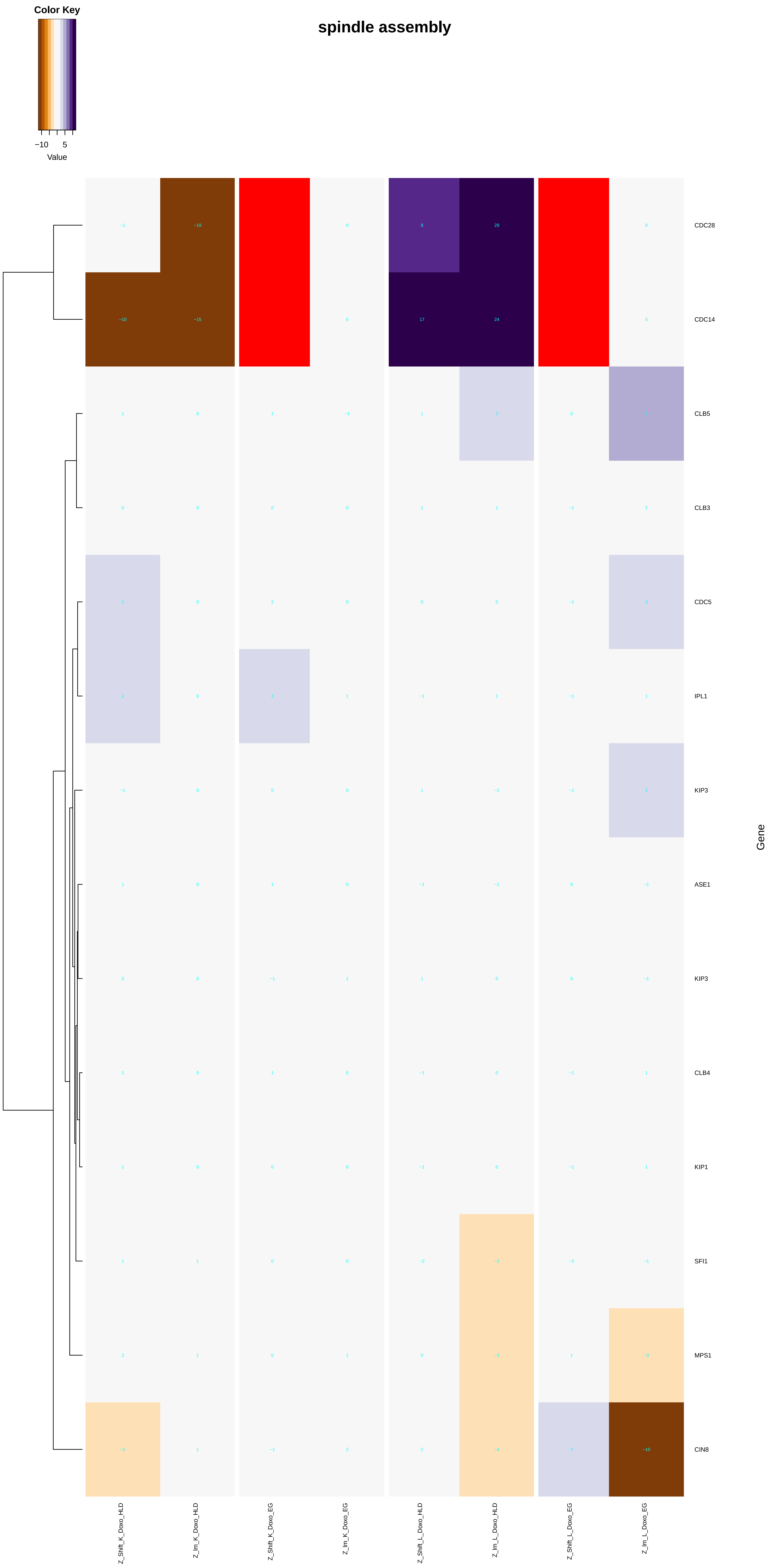

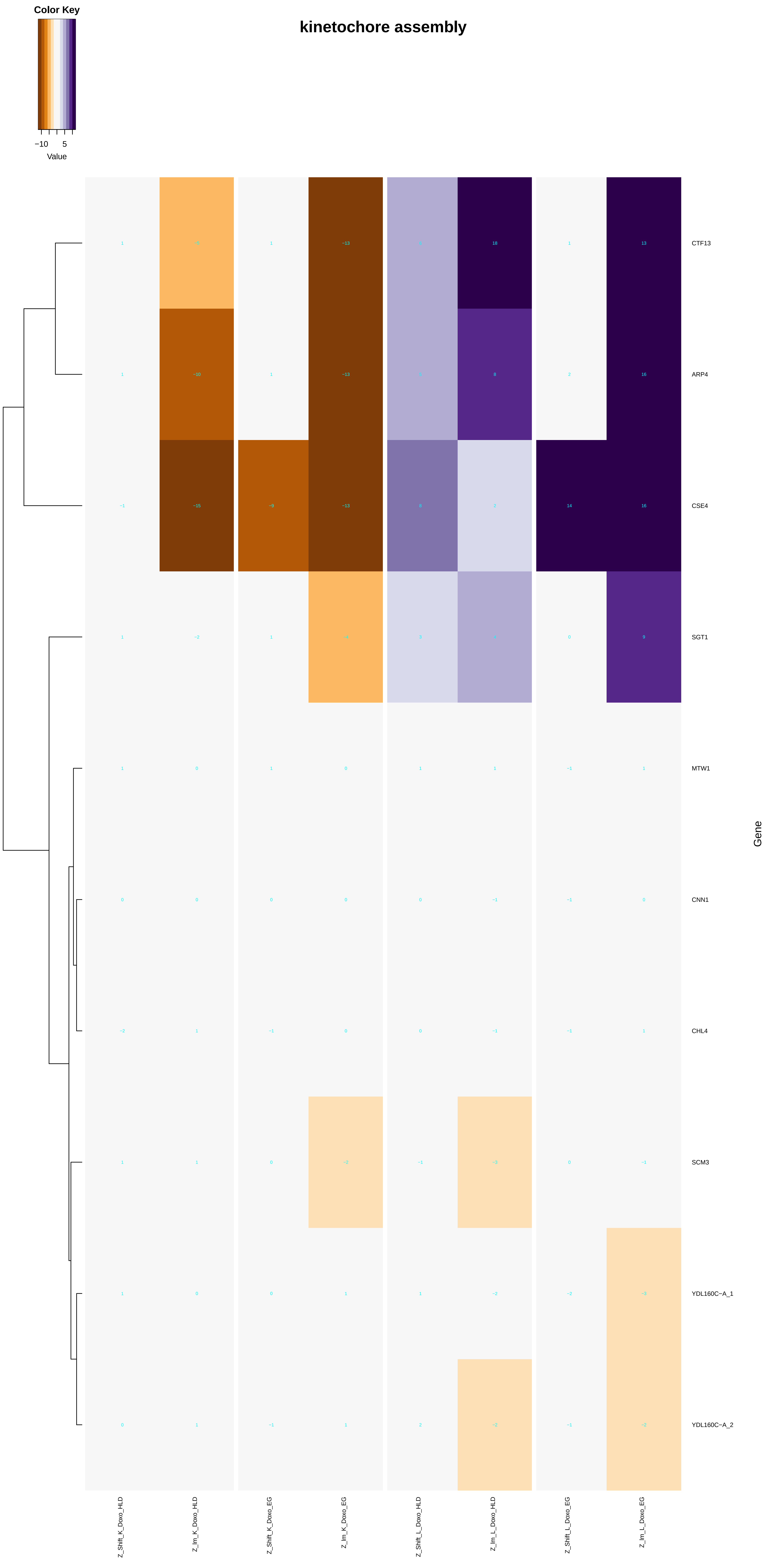

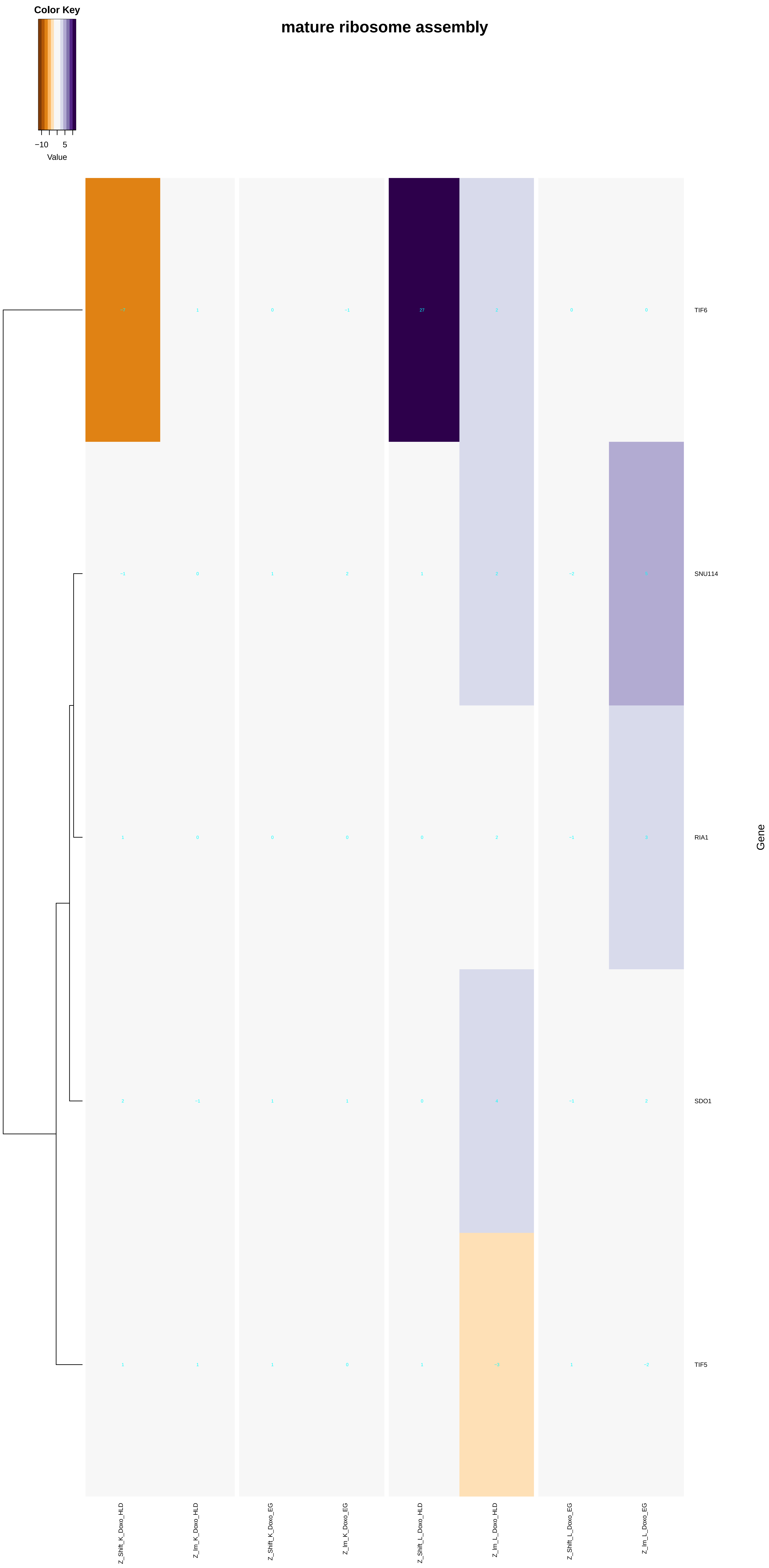

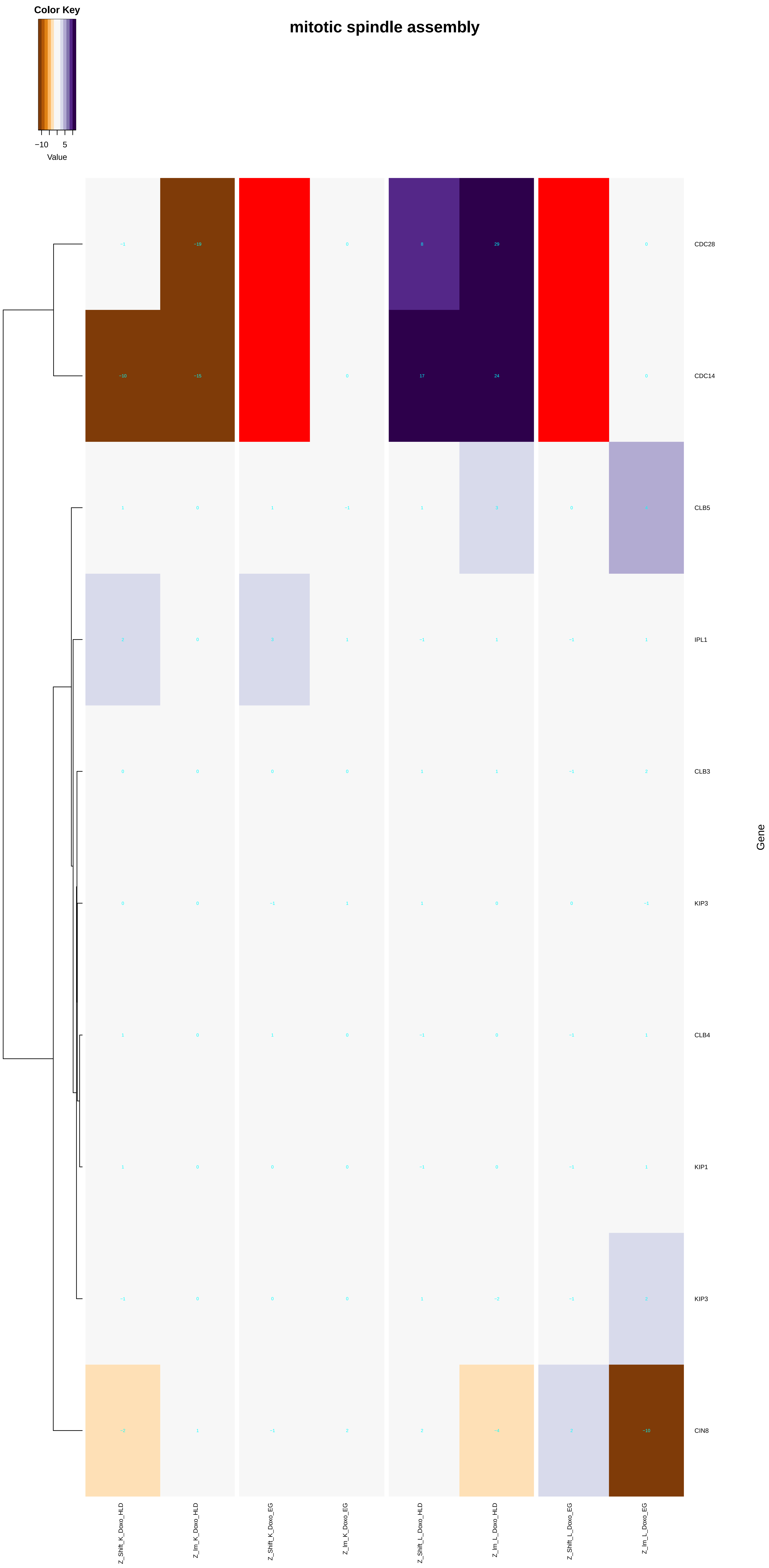

### peptide_biosynthetic_process.pdf

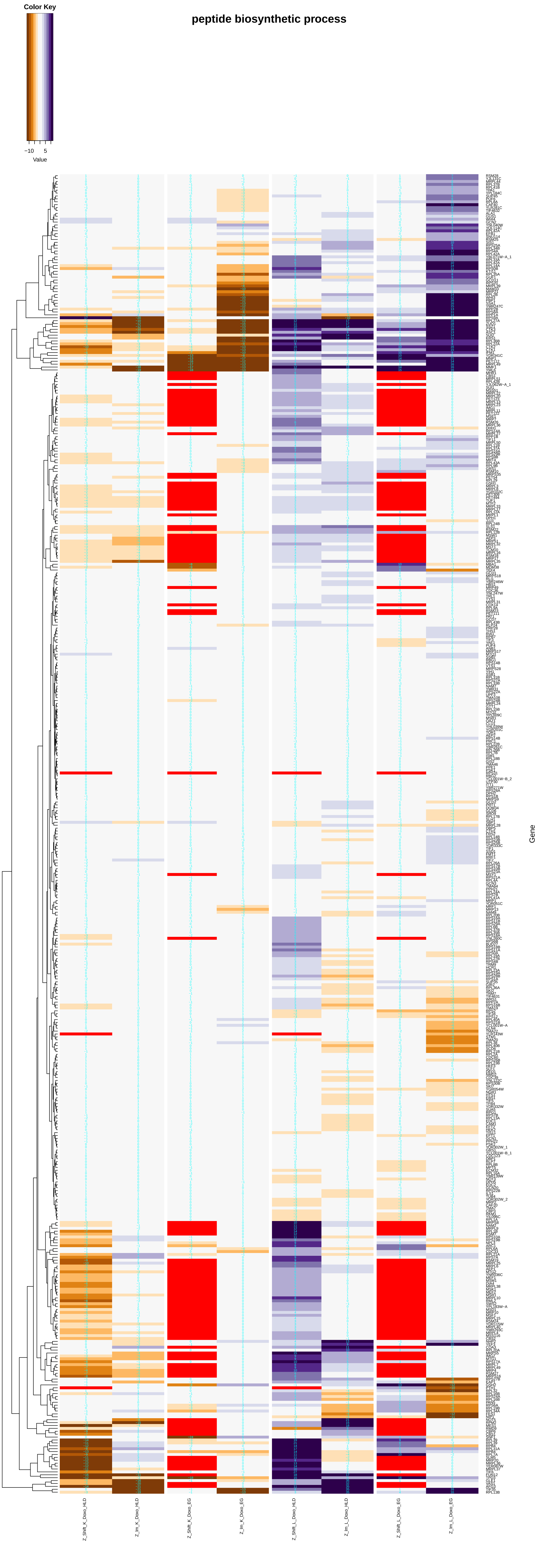

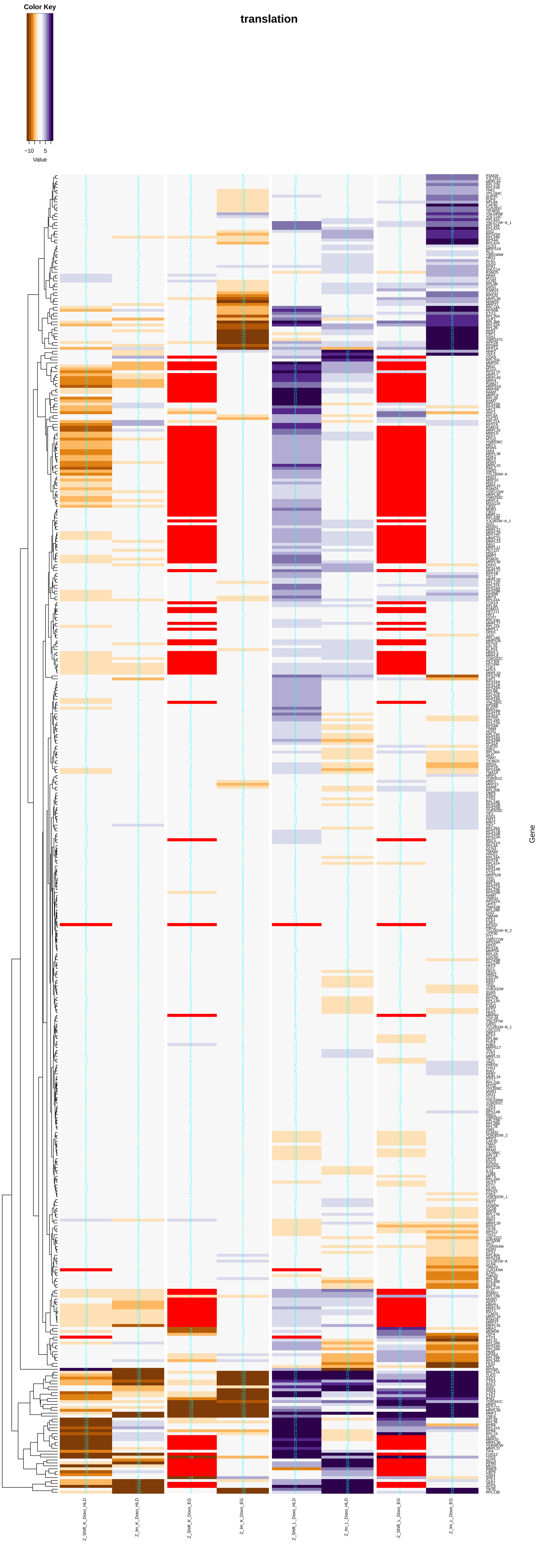

#### cytoplasmic translation

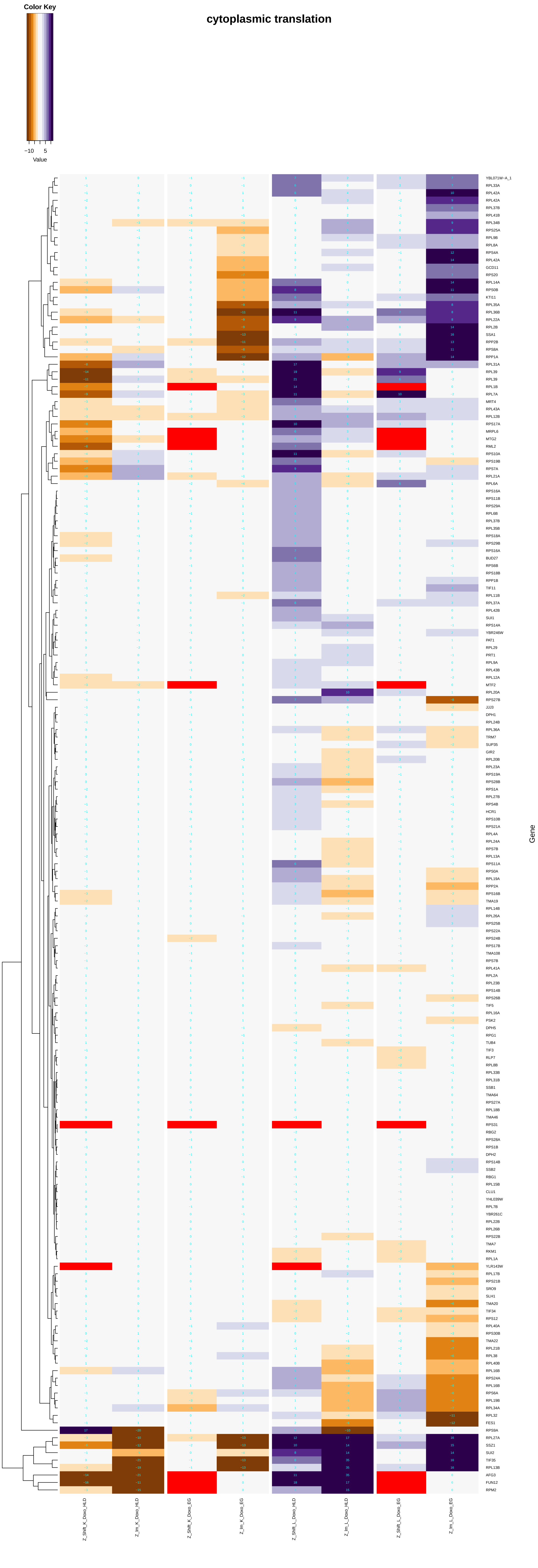

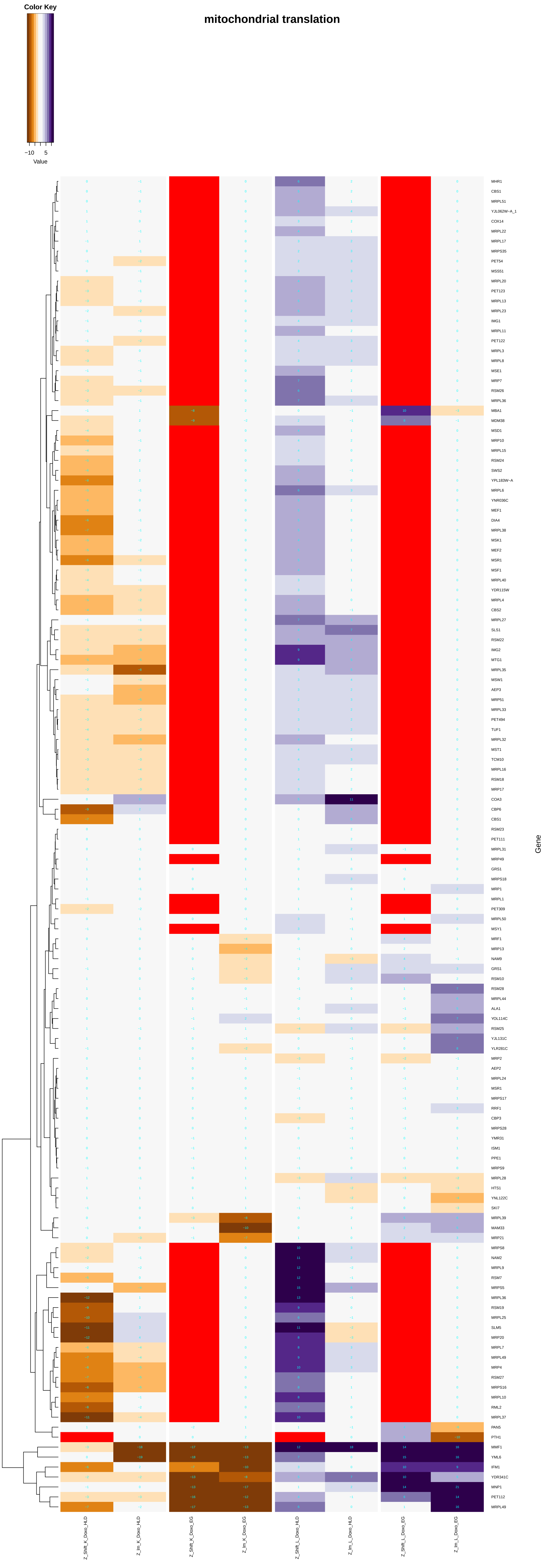

### peptide_metabolic_process.pdf

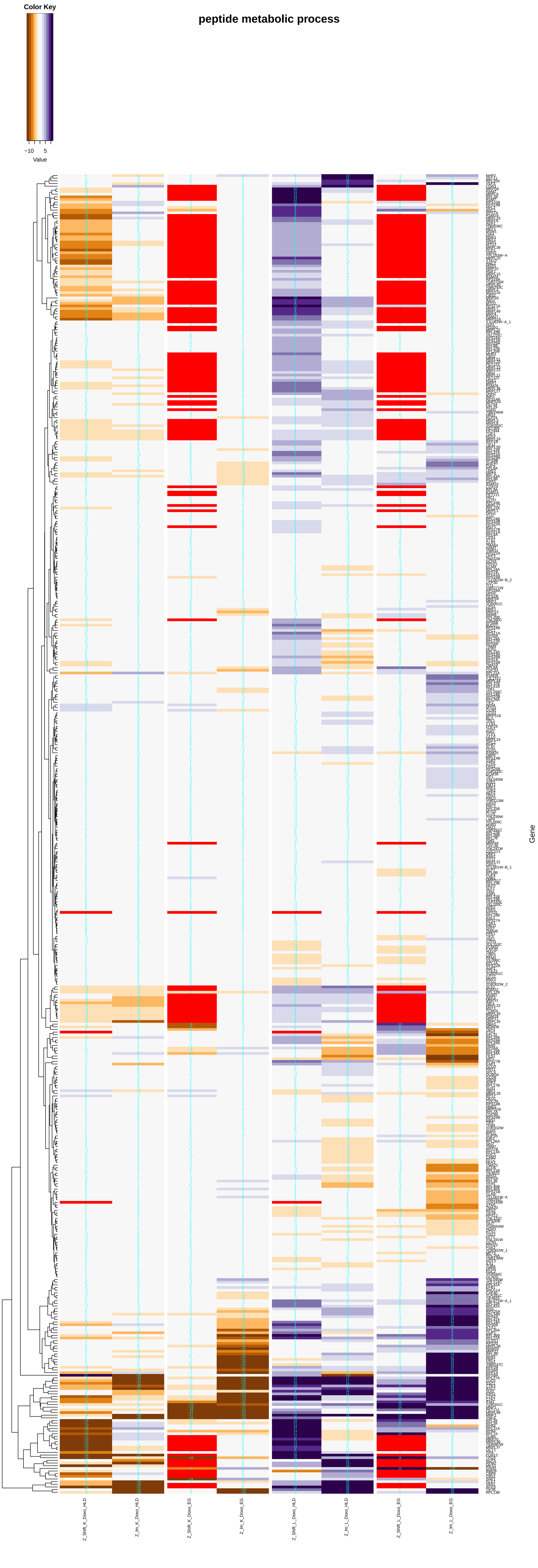

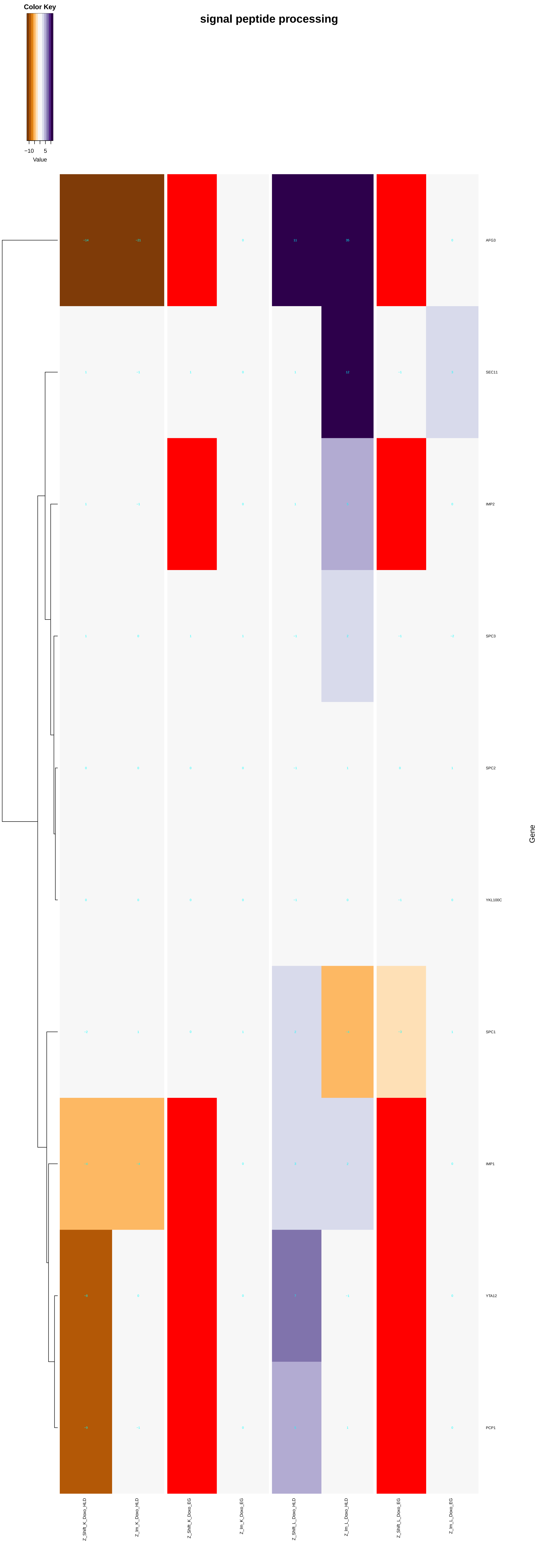

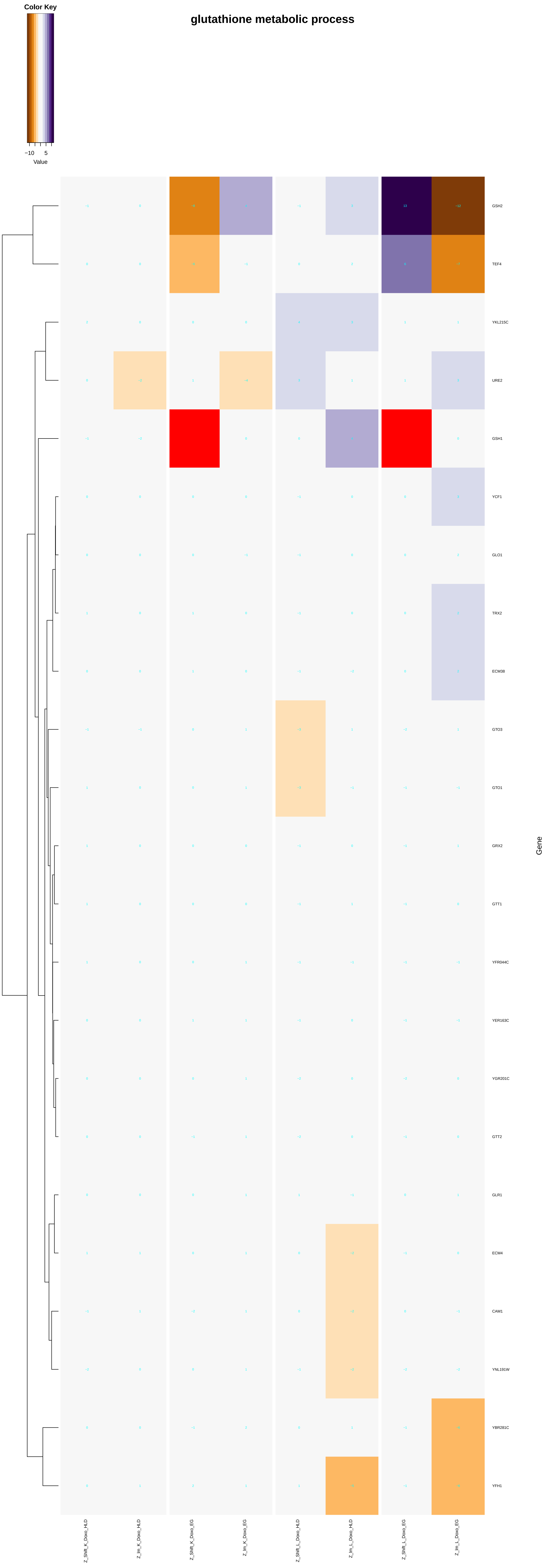

##### peptide biosynthetic process

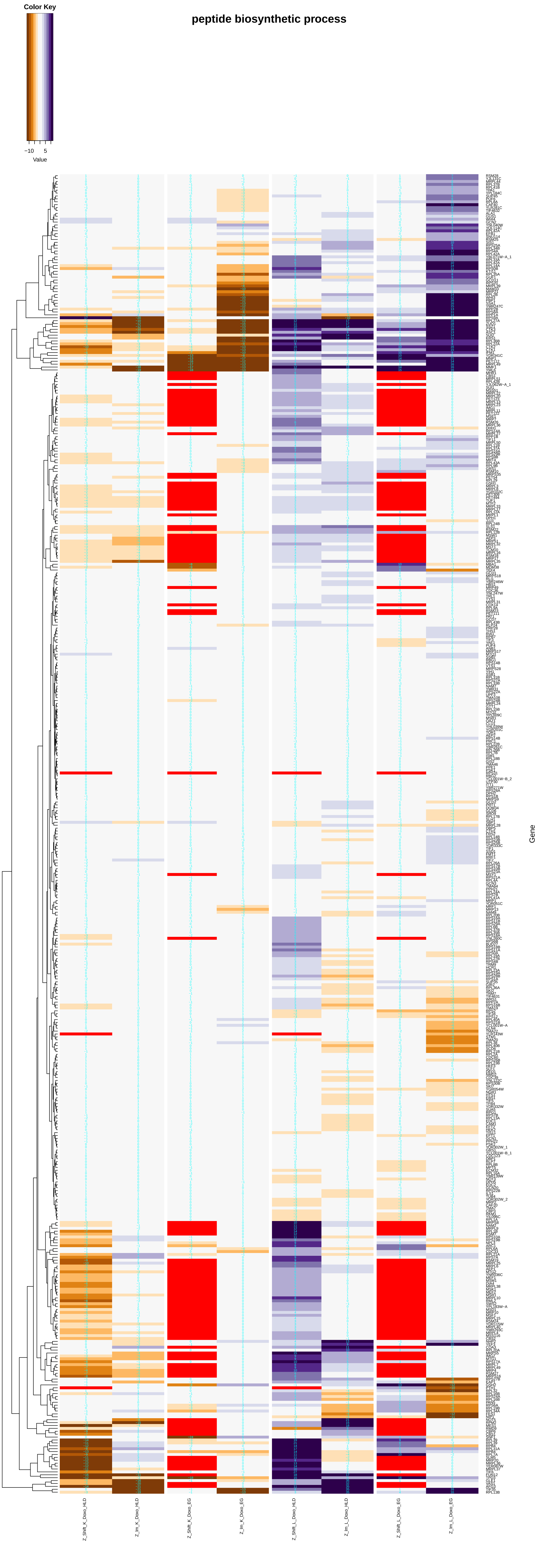

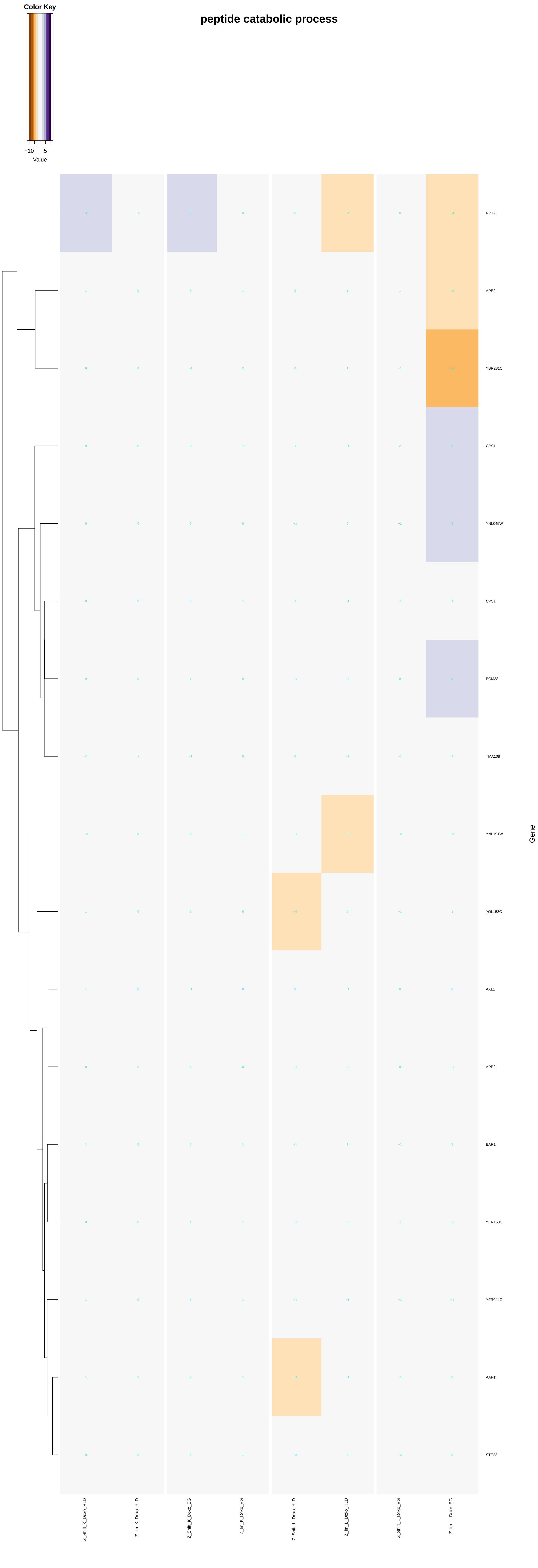

#### glutathione catabolic process

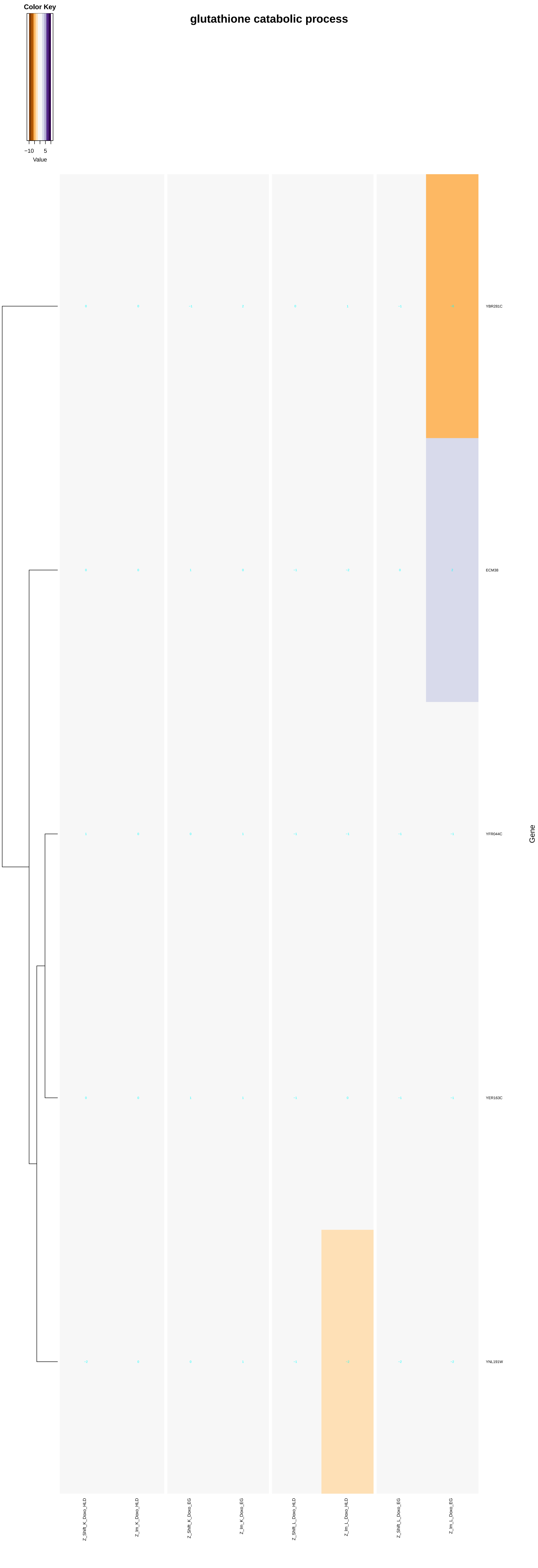

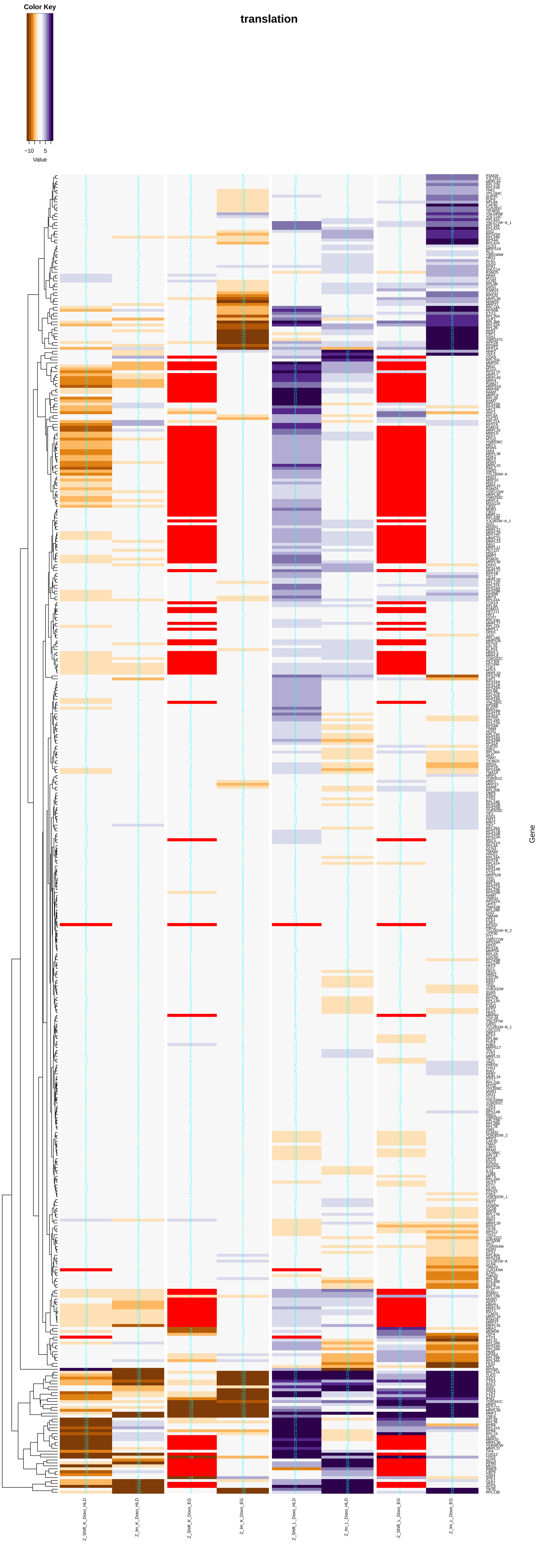

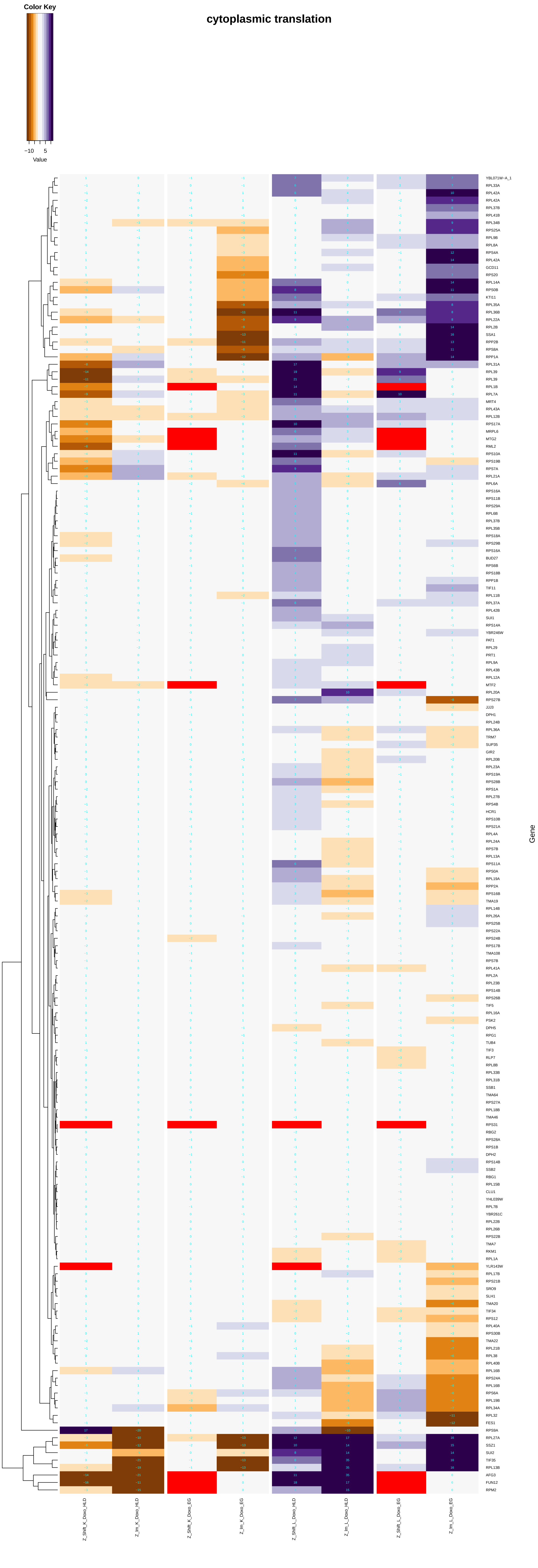

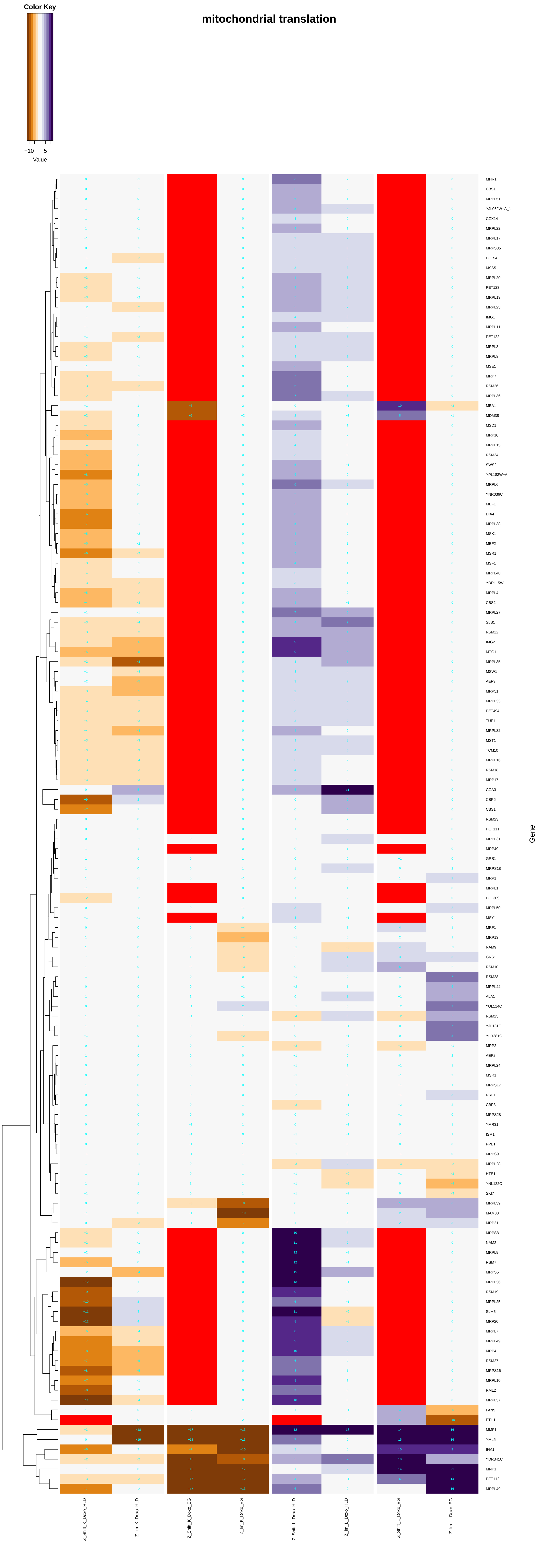

### peptidyl-lysine_acetylation.pdf

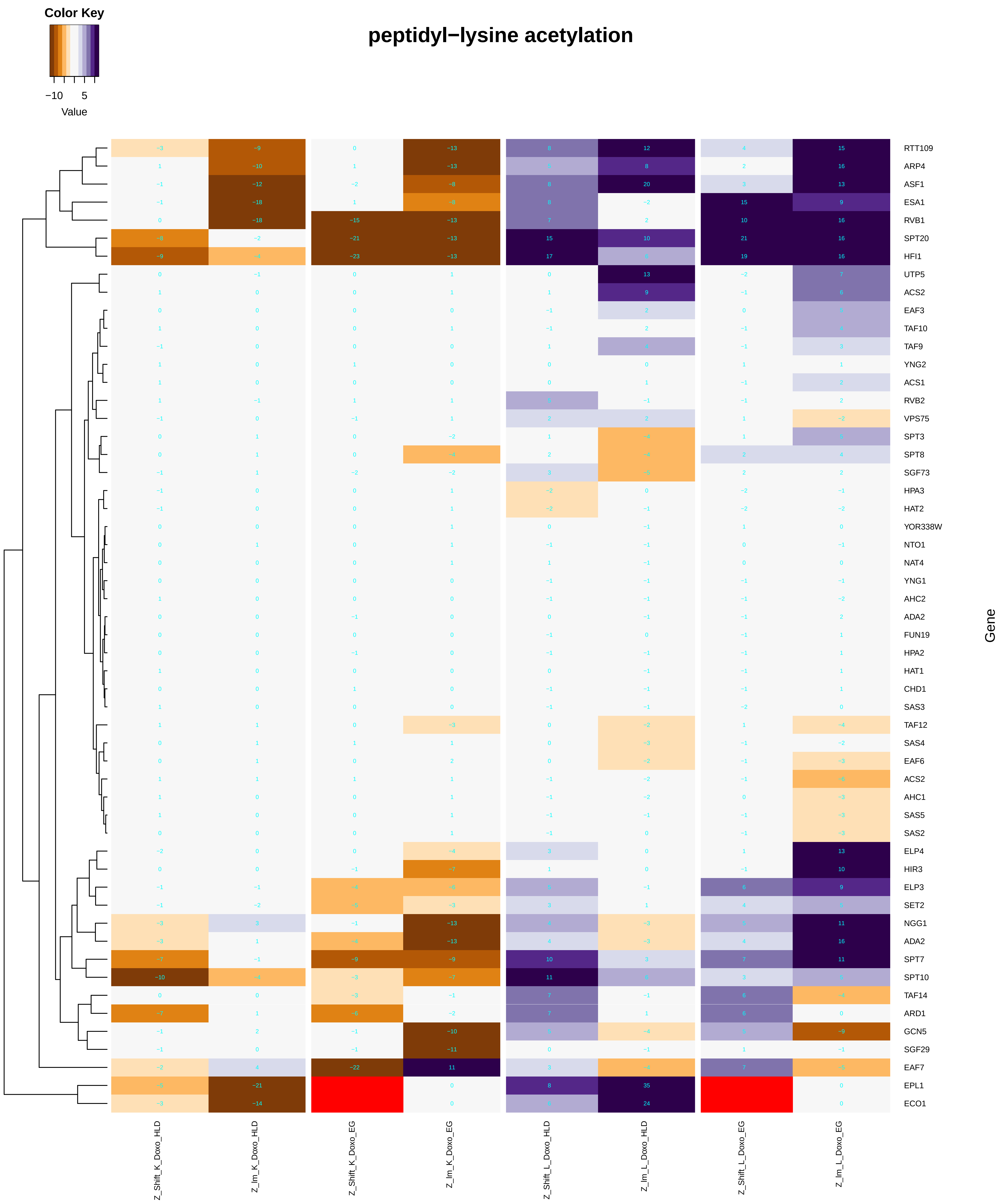

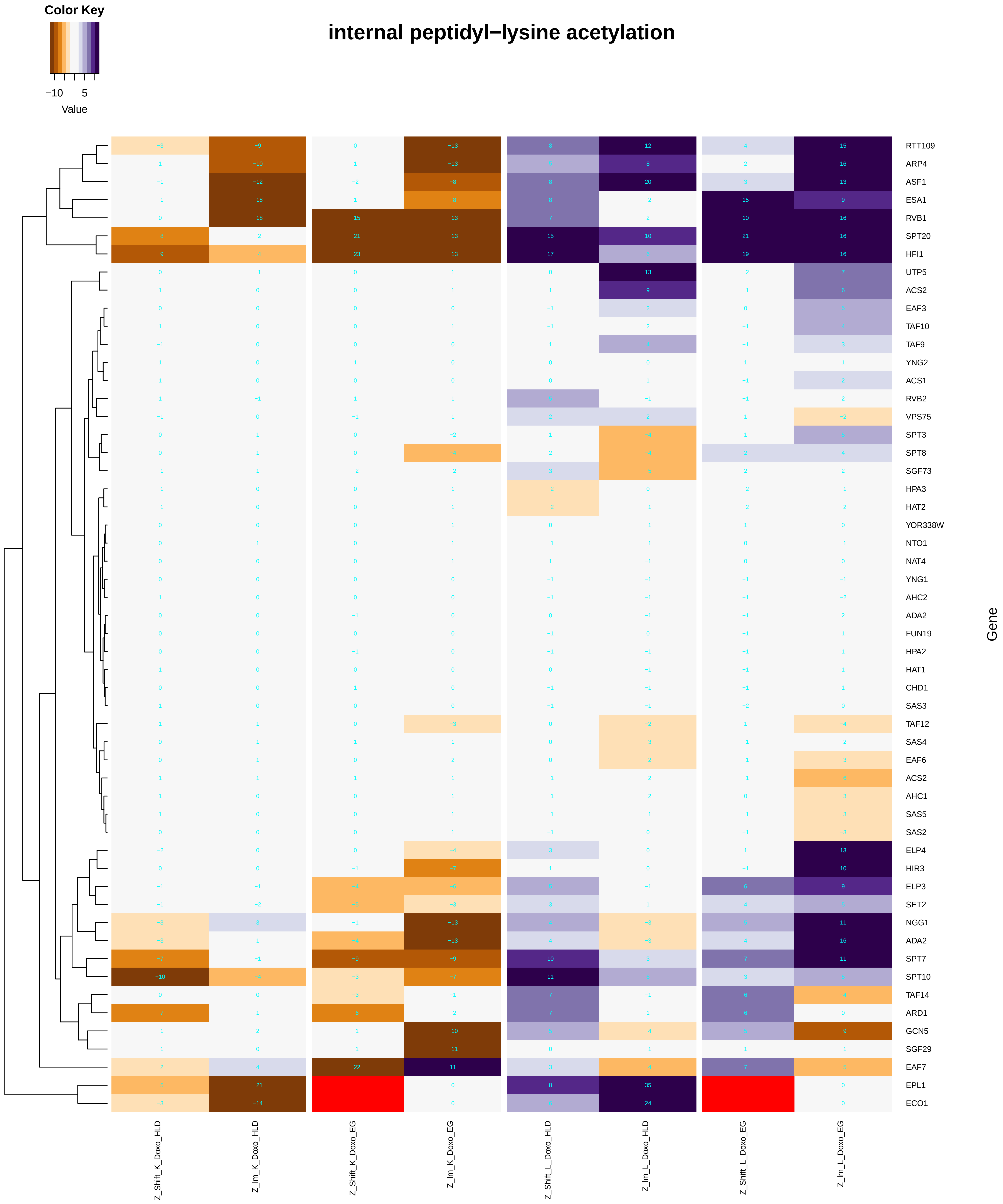

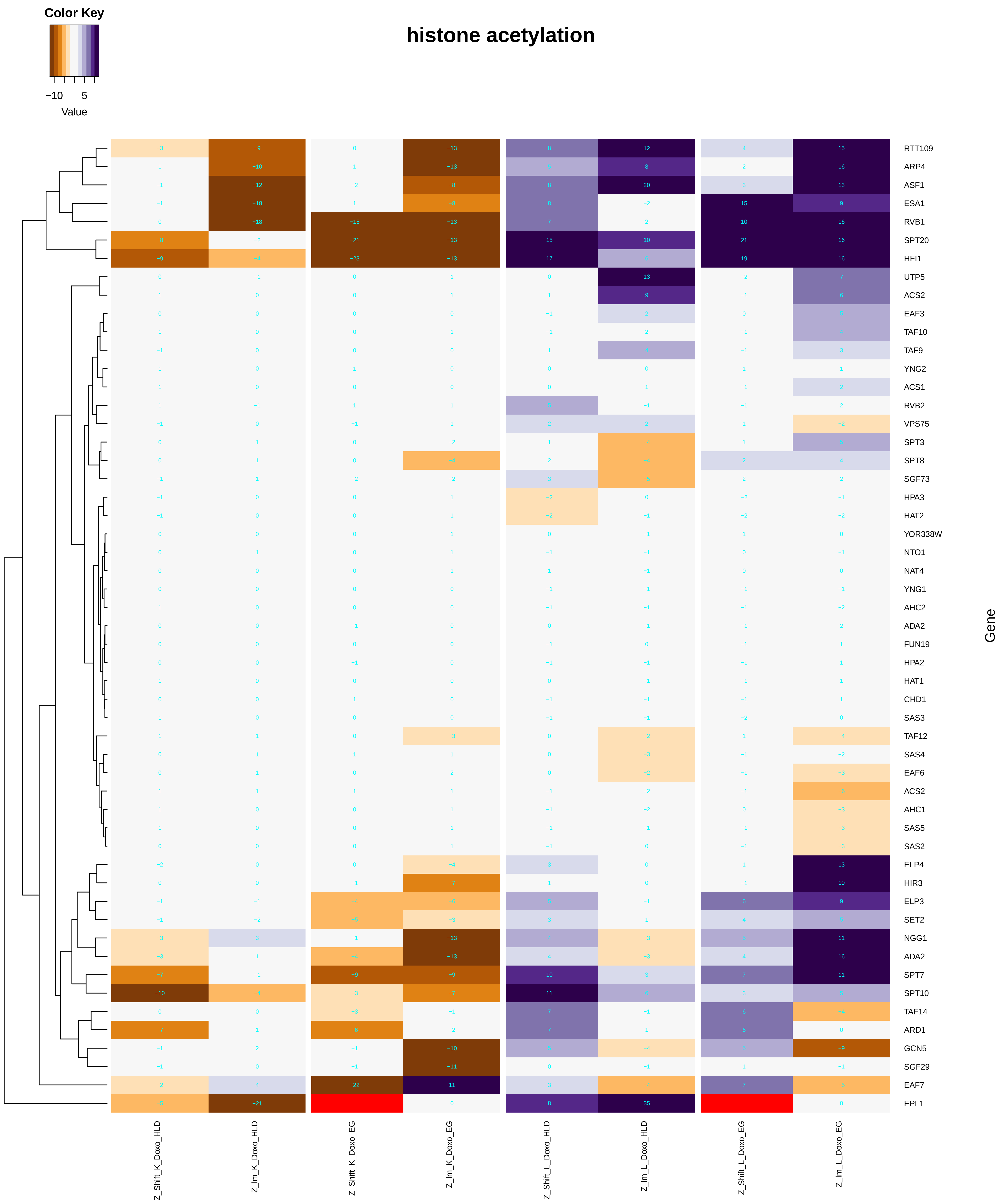

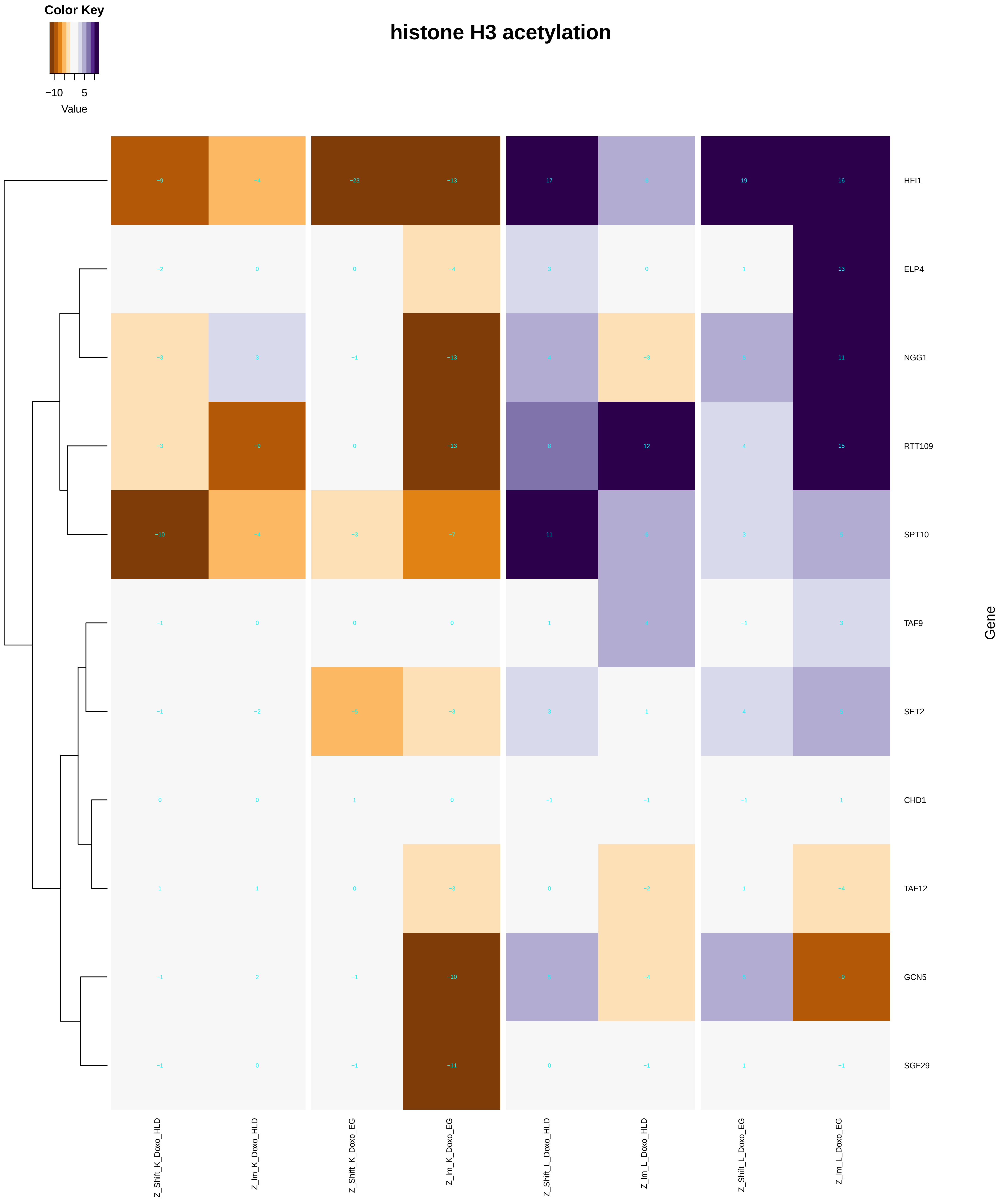

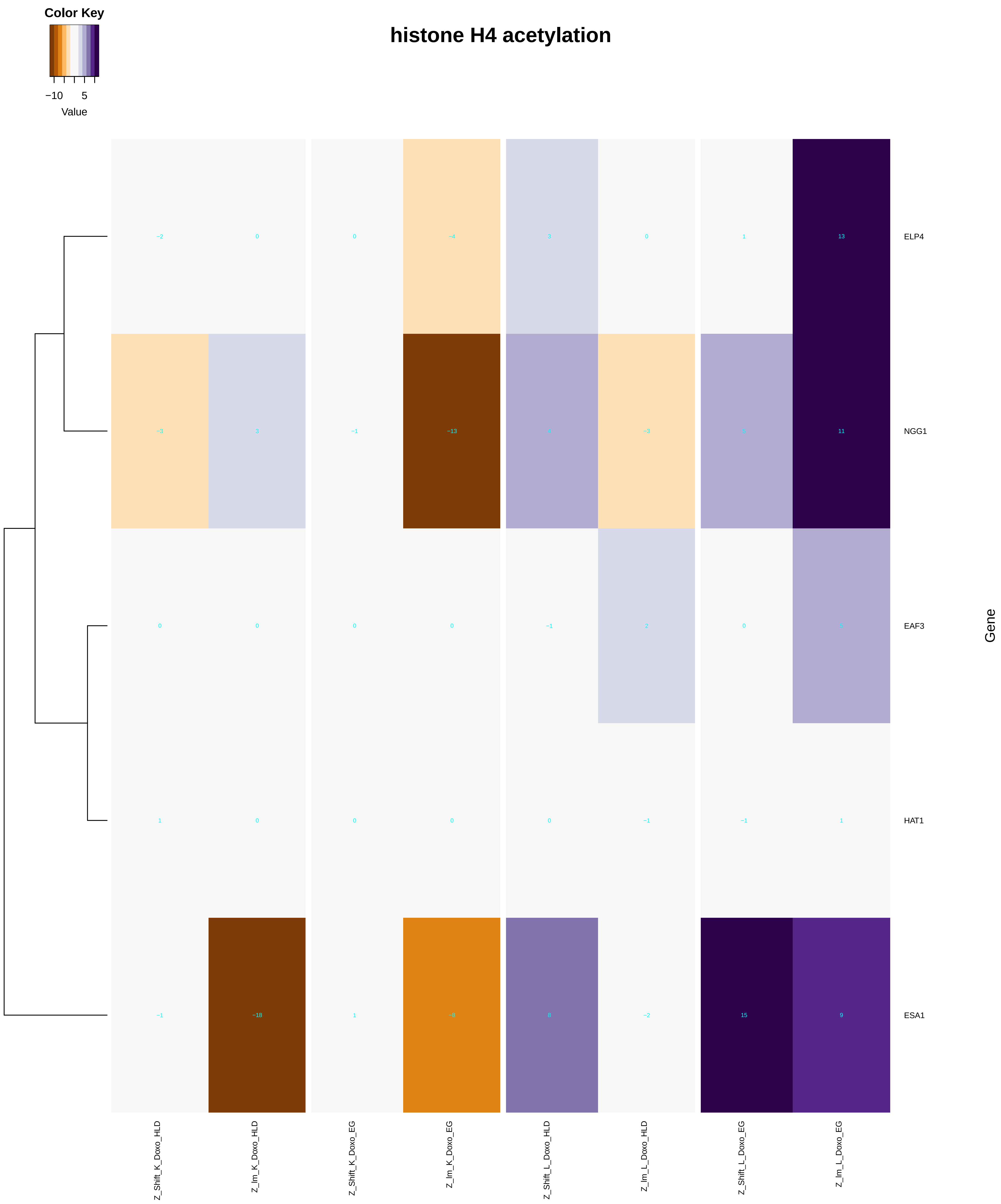

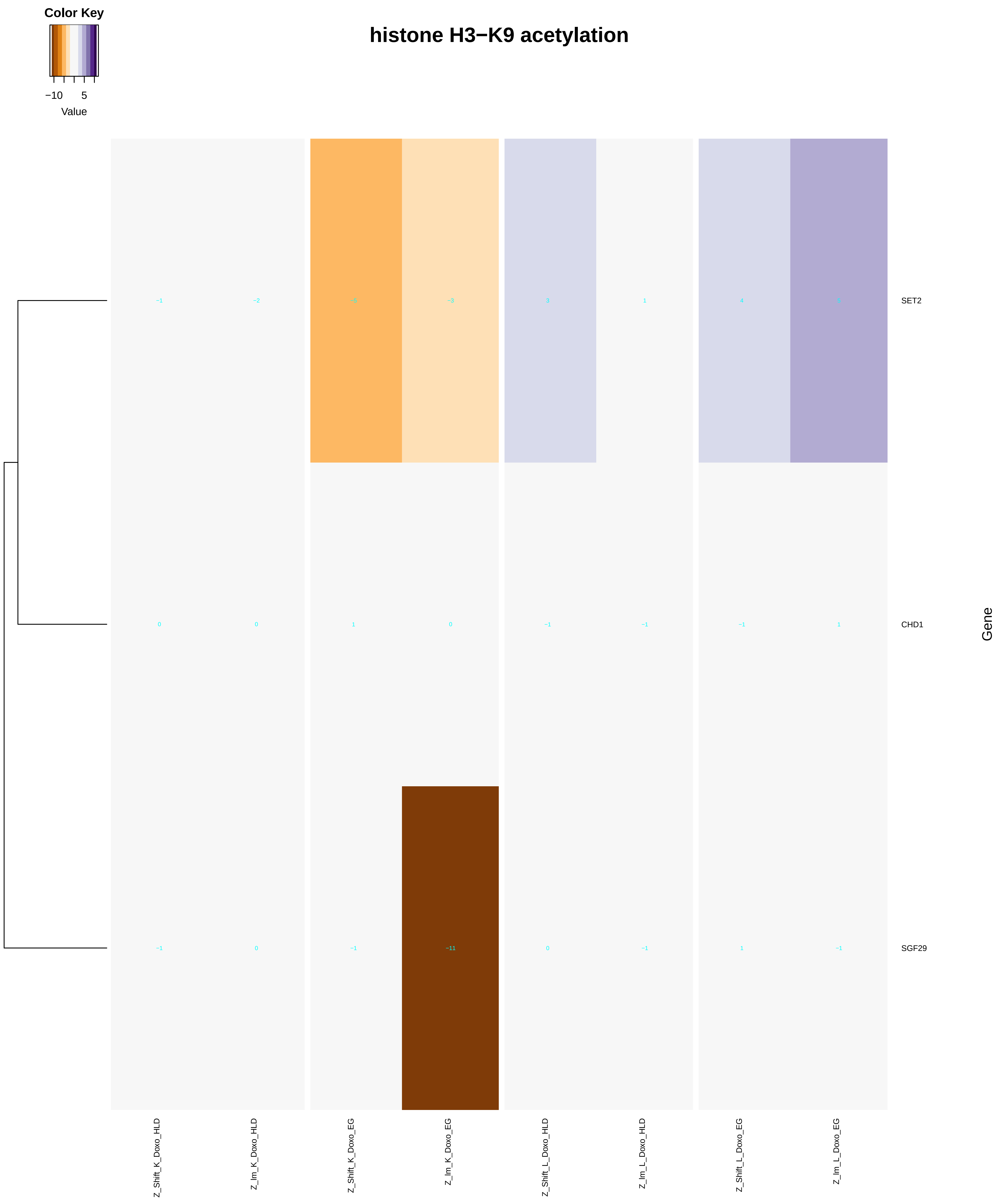

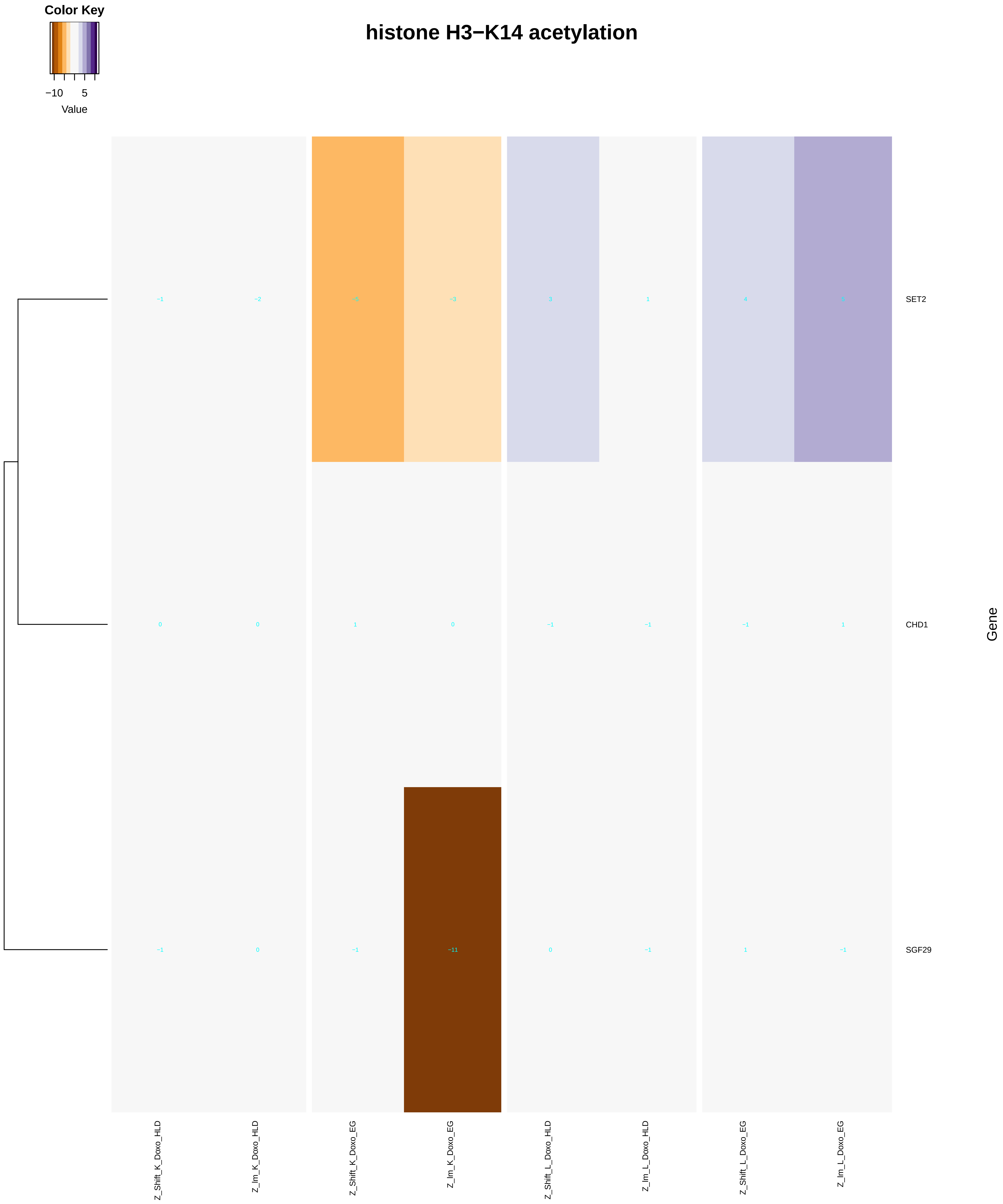

### peptidyl-serine_modification.pdf

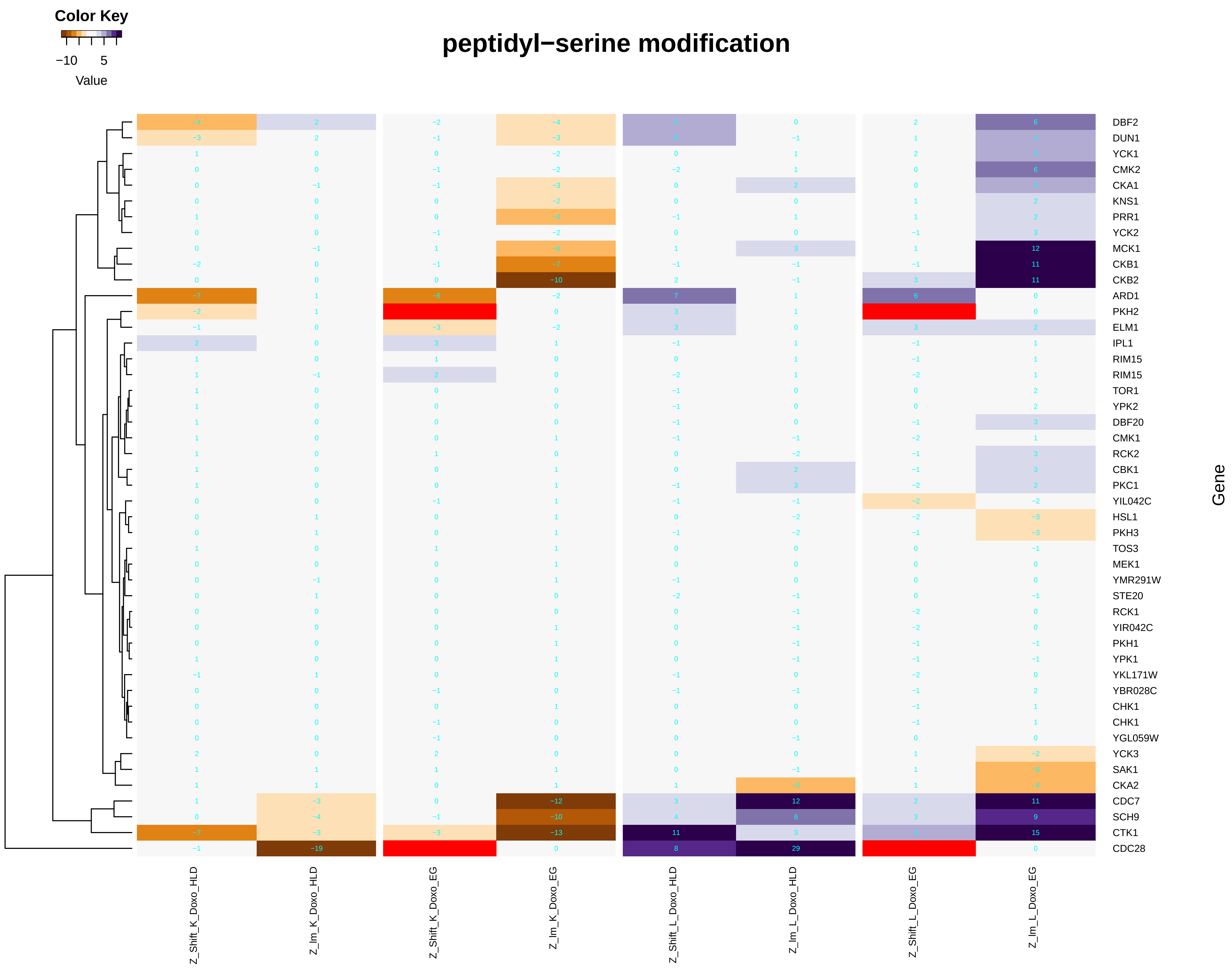

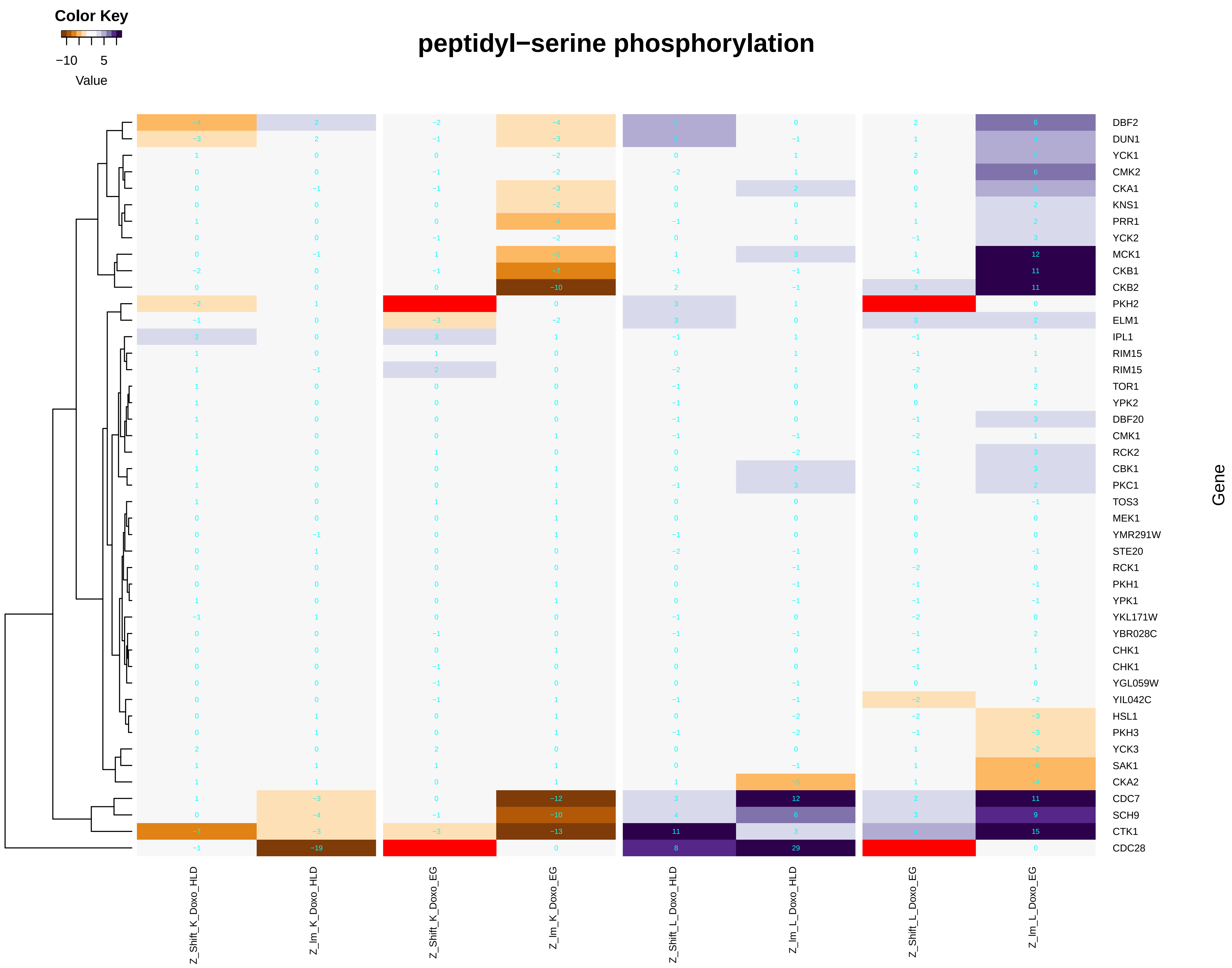

### peptidyl-serine_phosphorylation.pdf

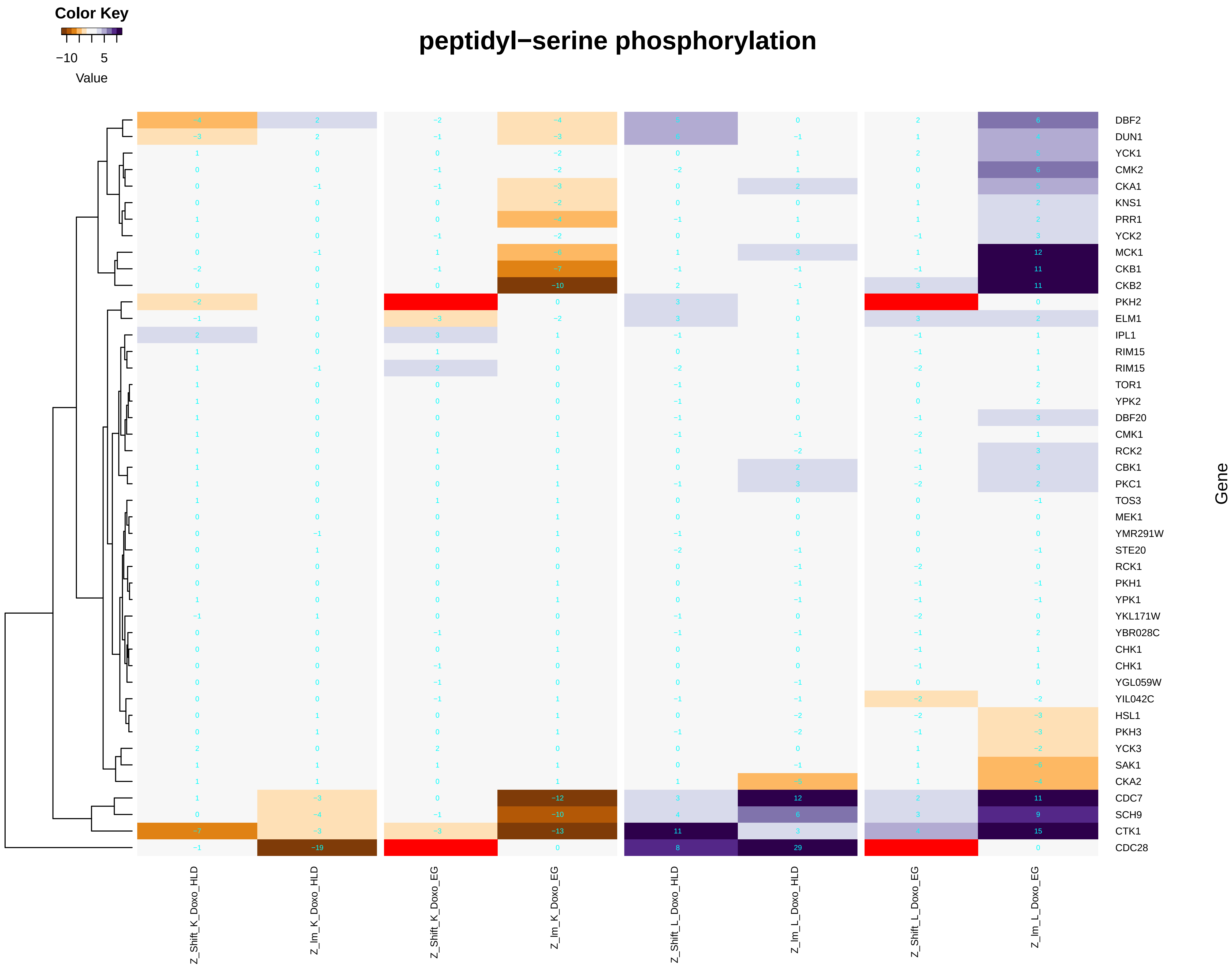

### pH_reduction.pdf

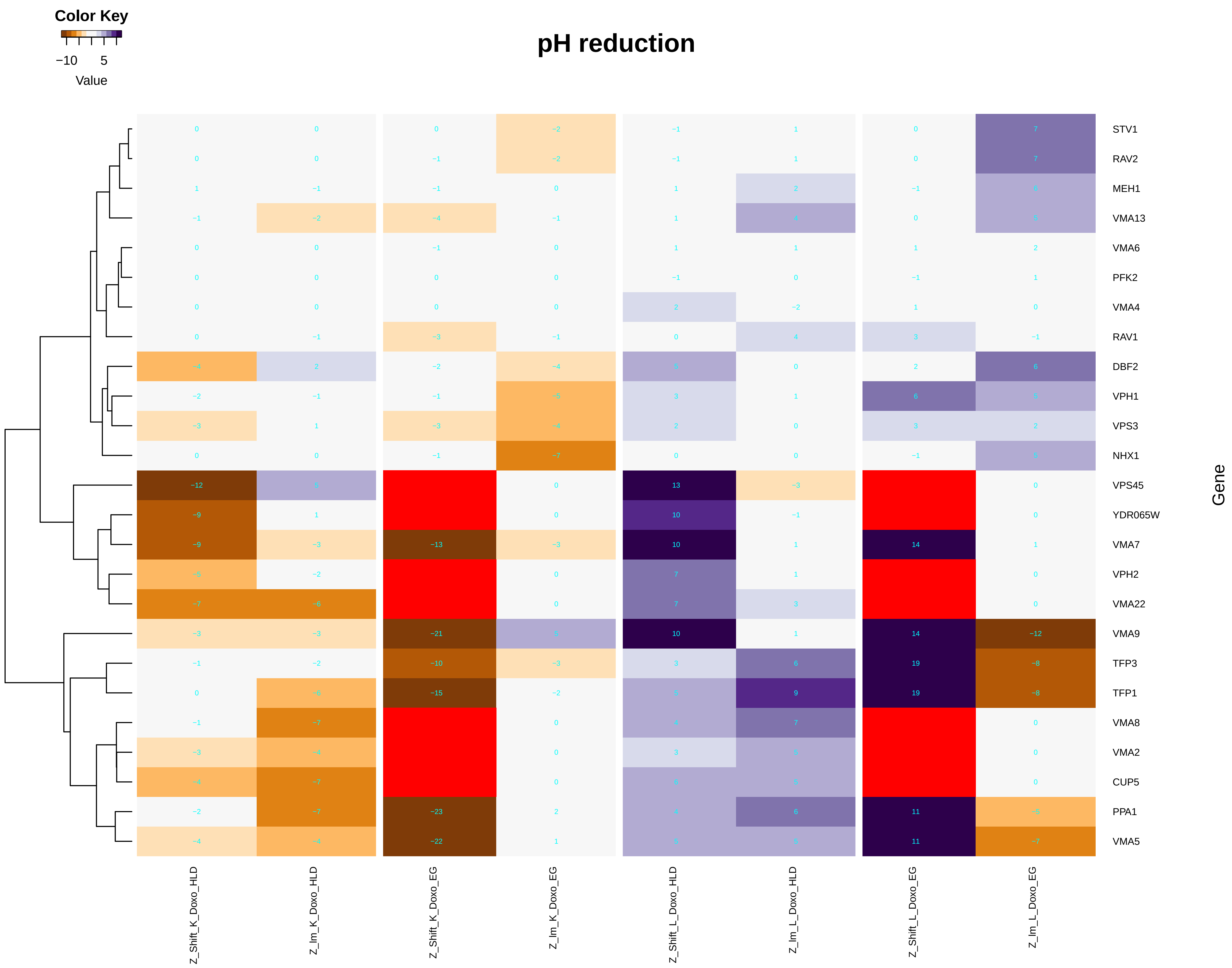

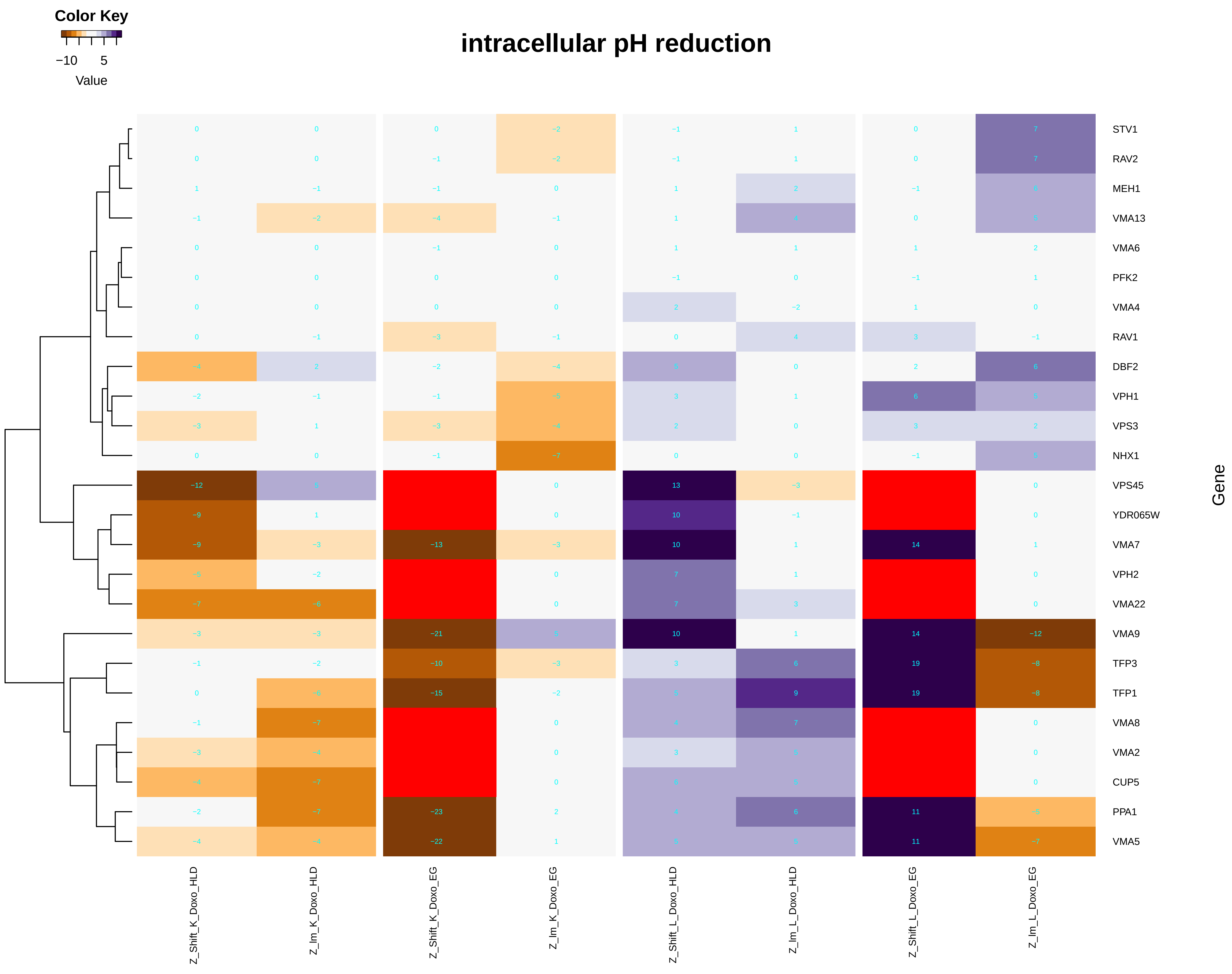

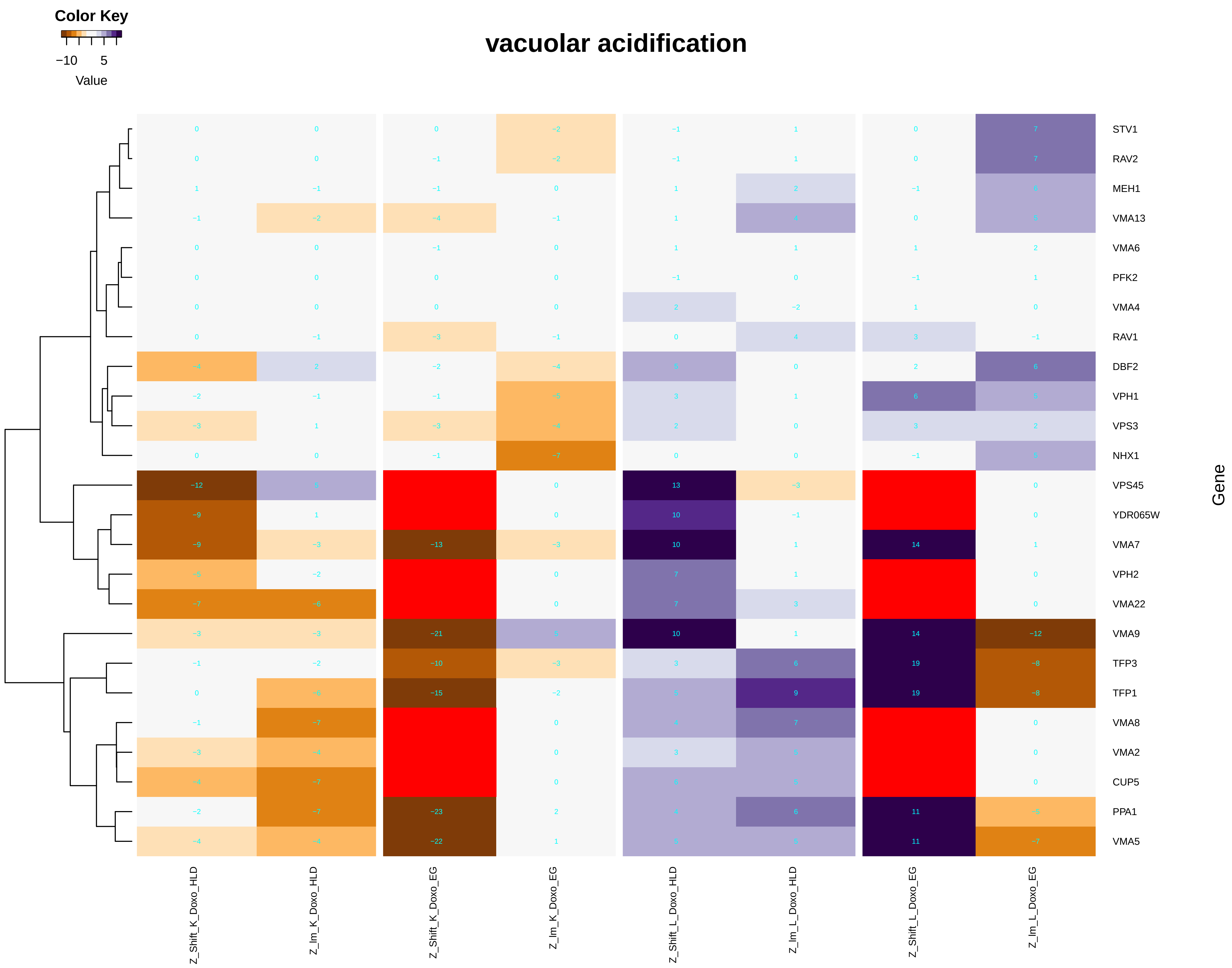

### positive_regulation_of_cellular_amide_metabolic_process.pdf

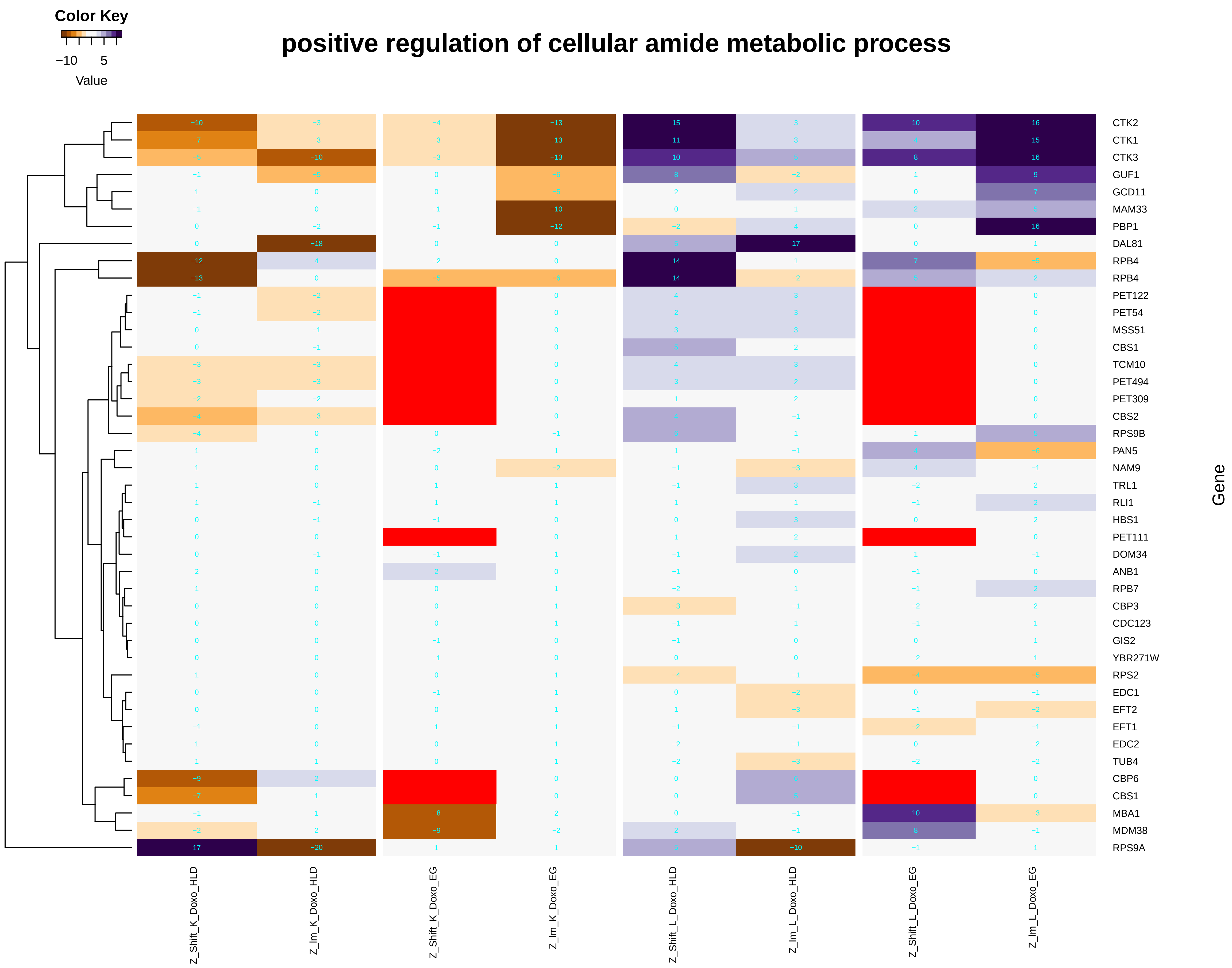

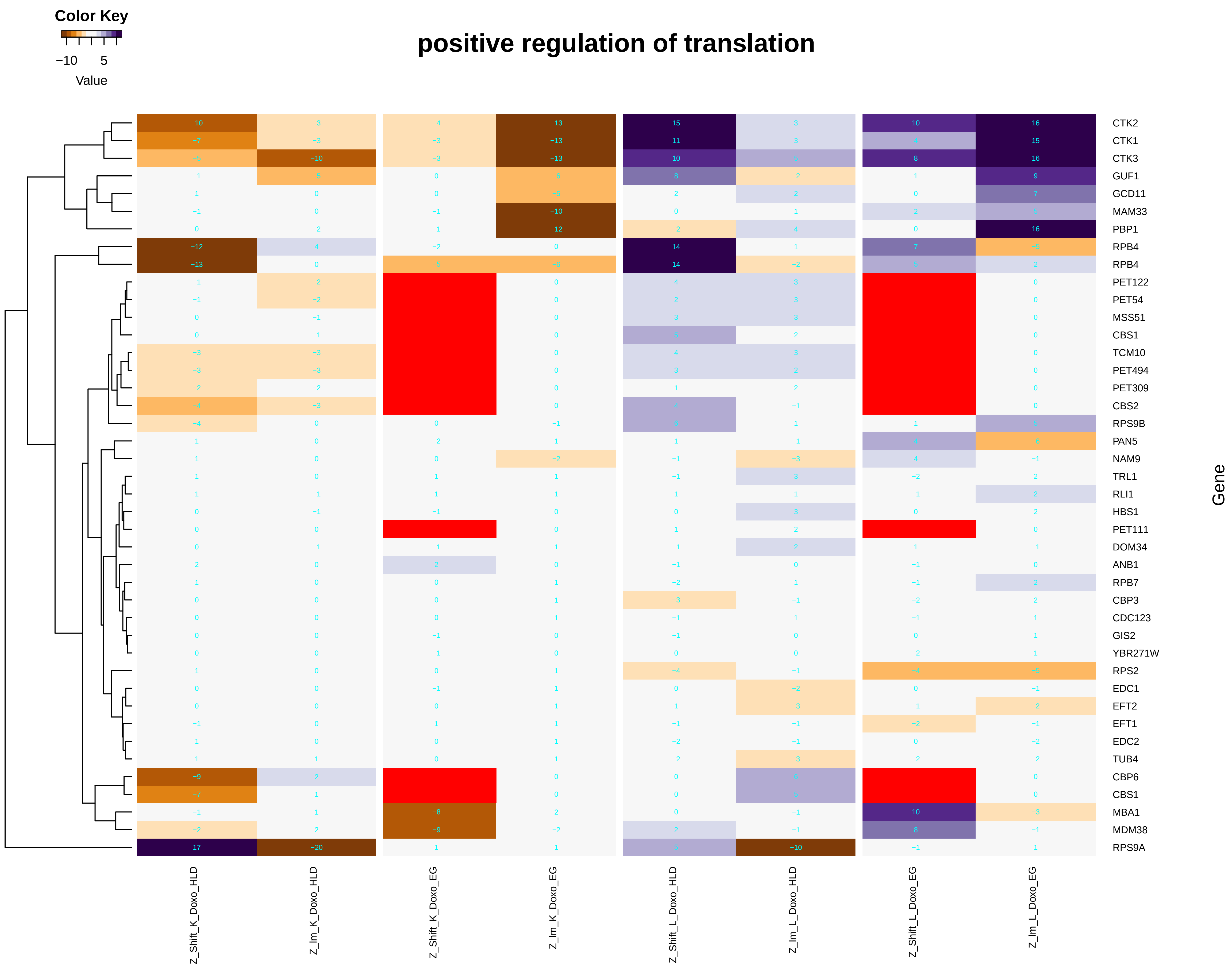

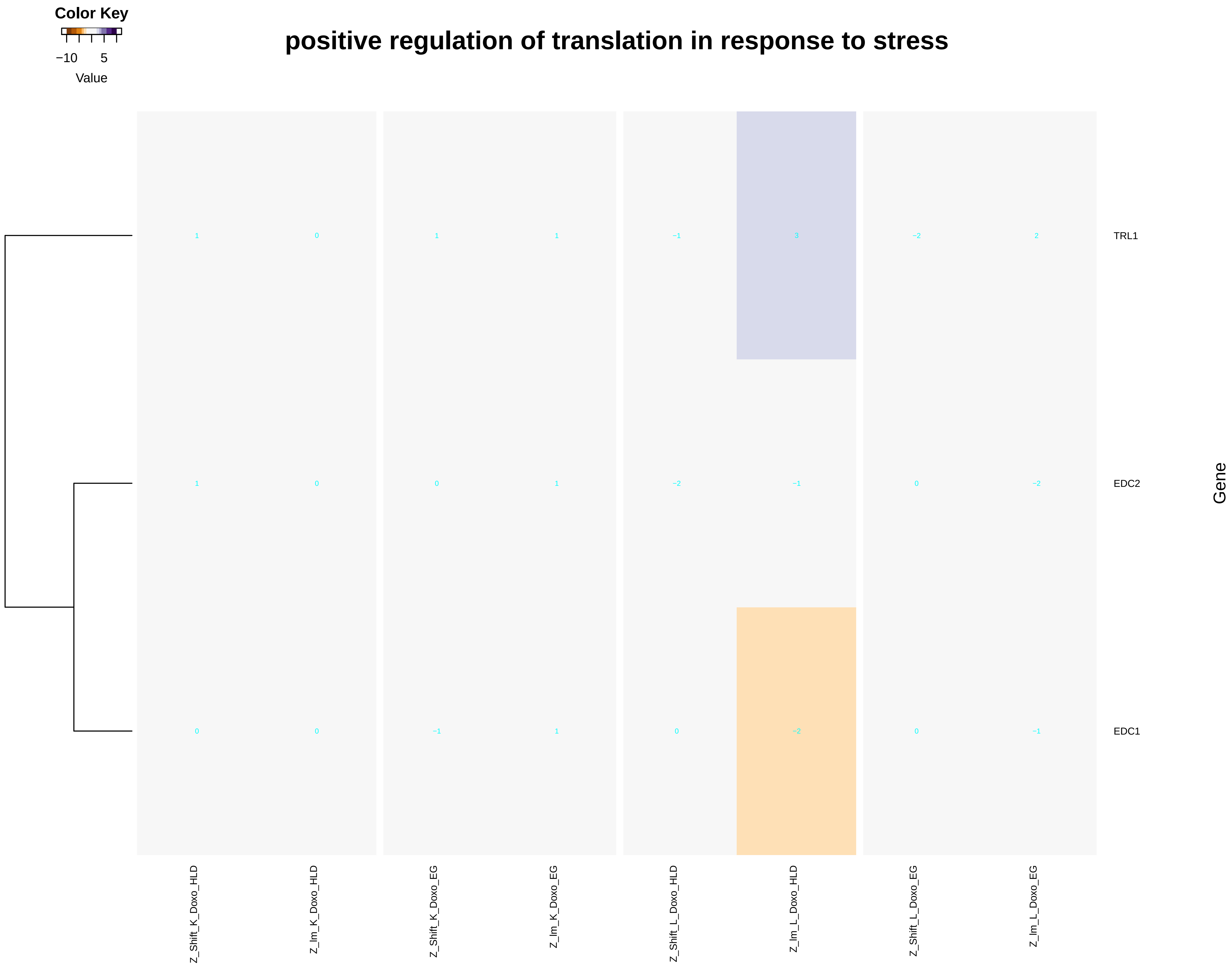

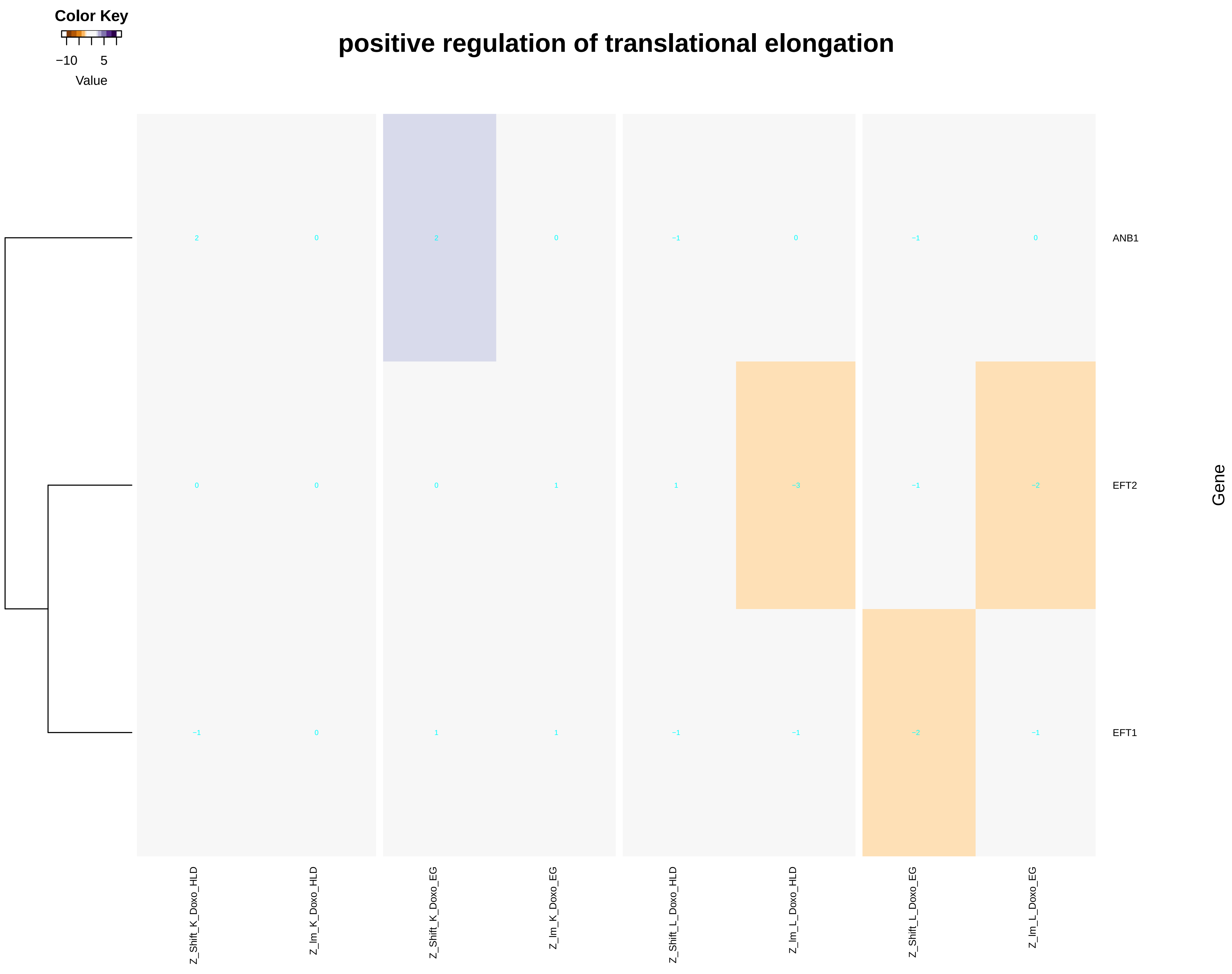

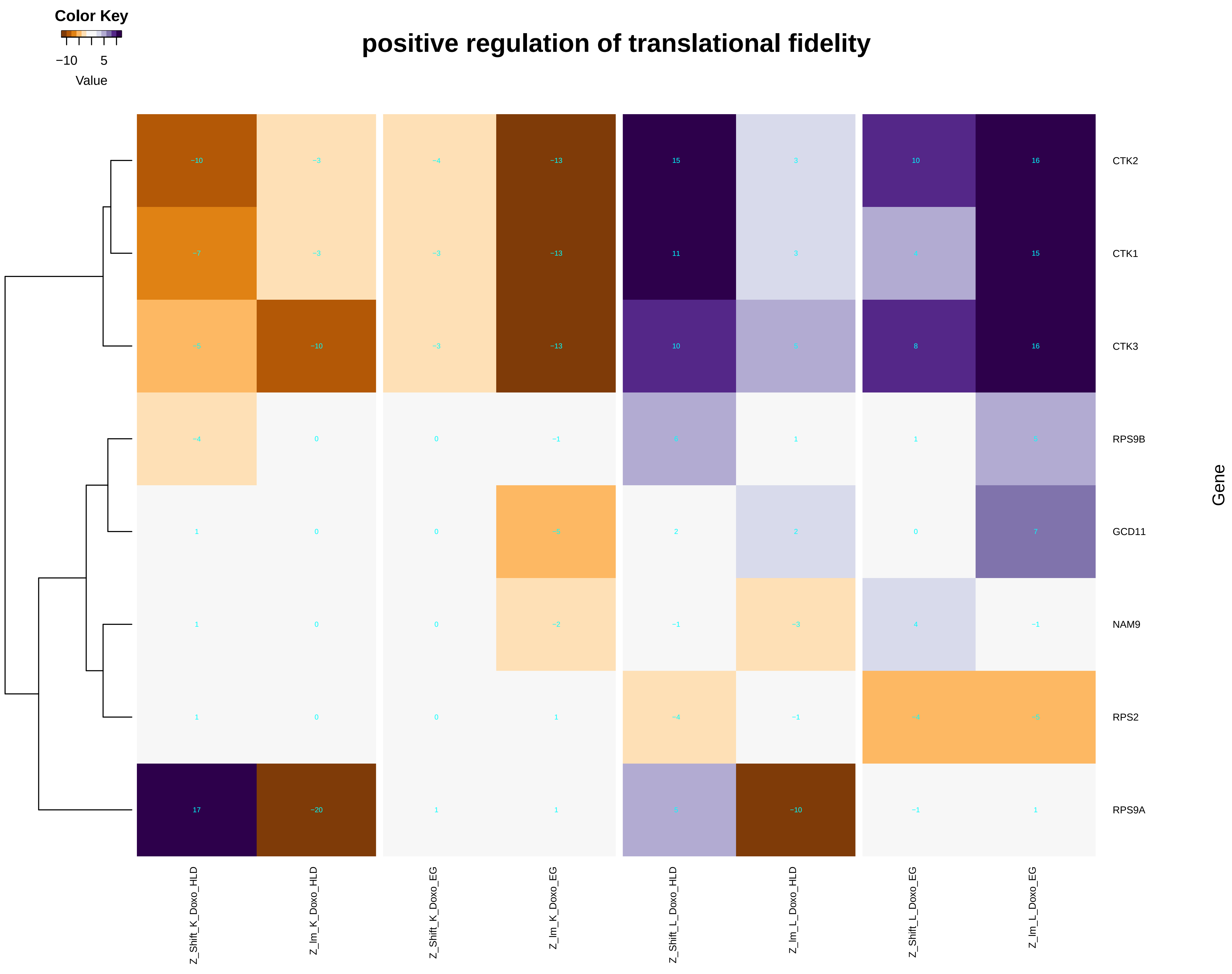

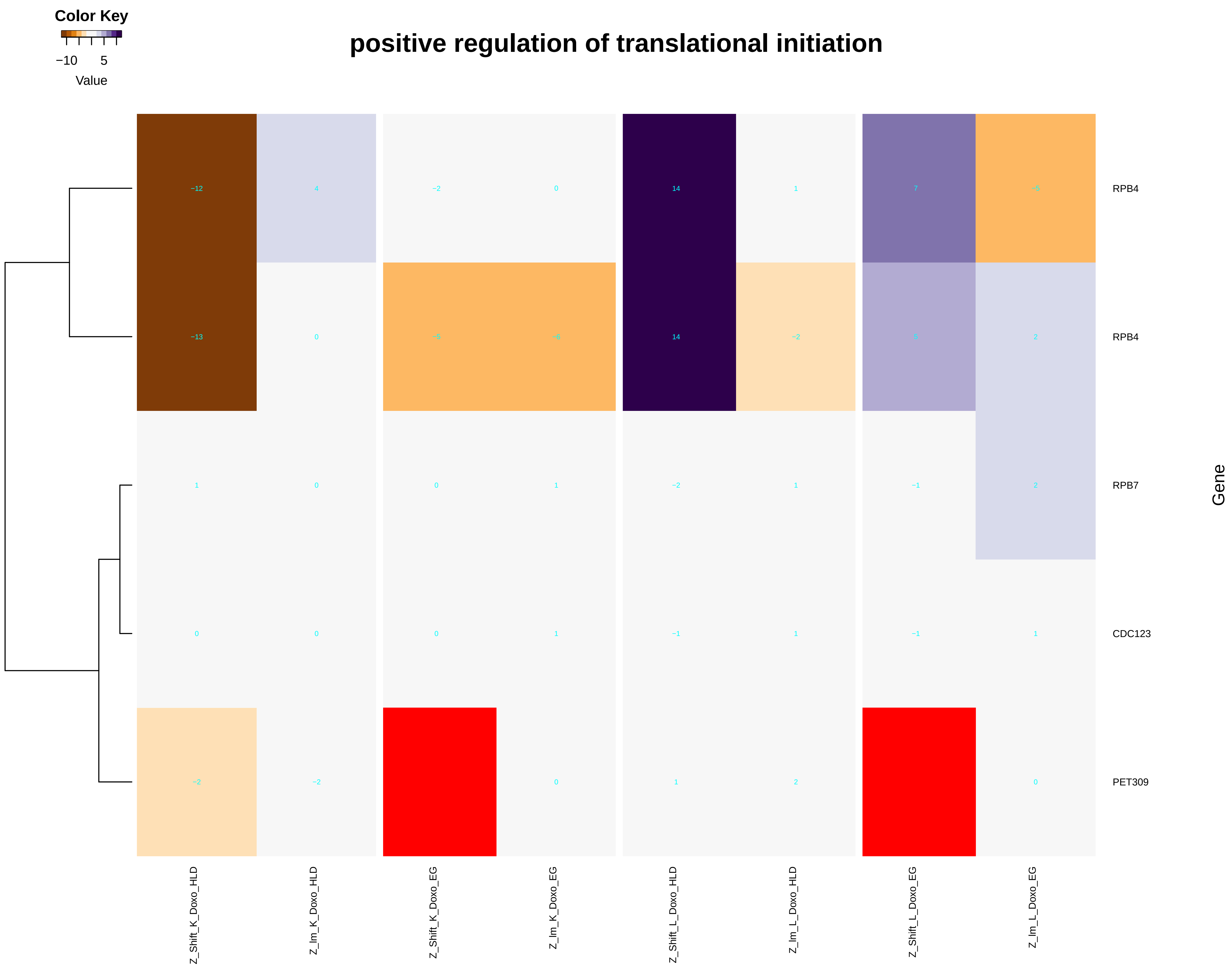

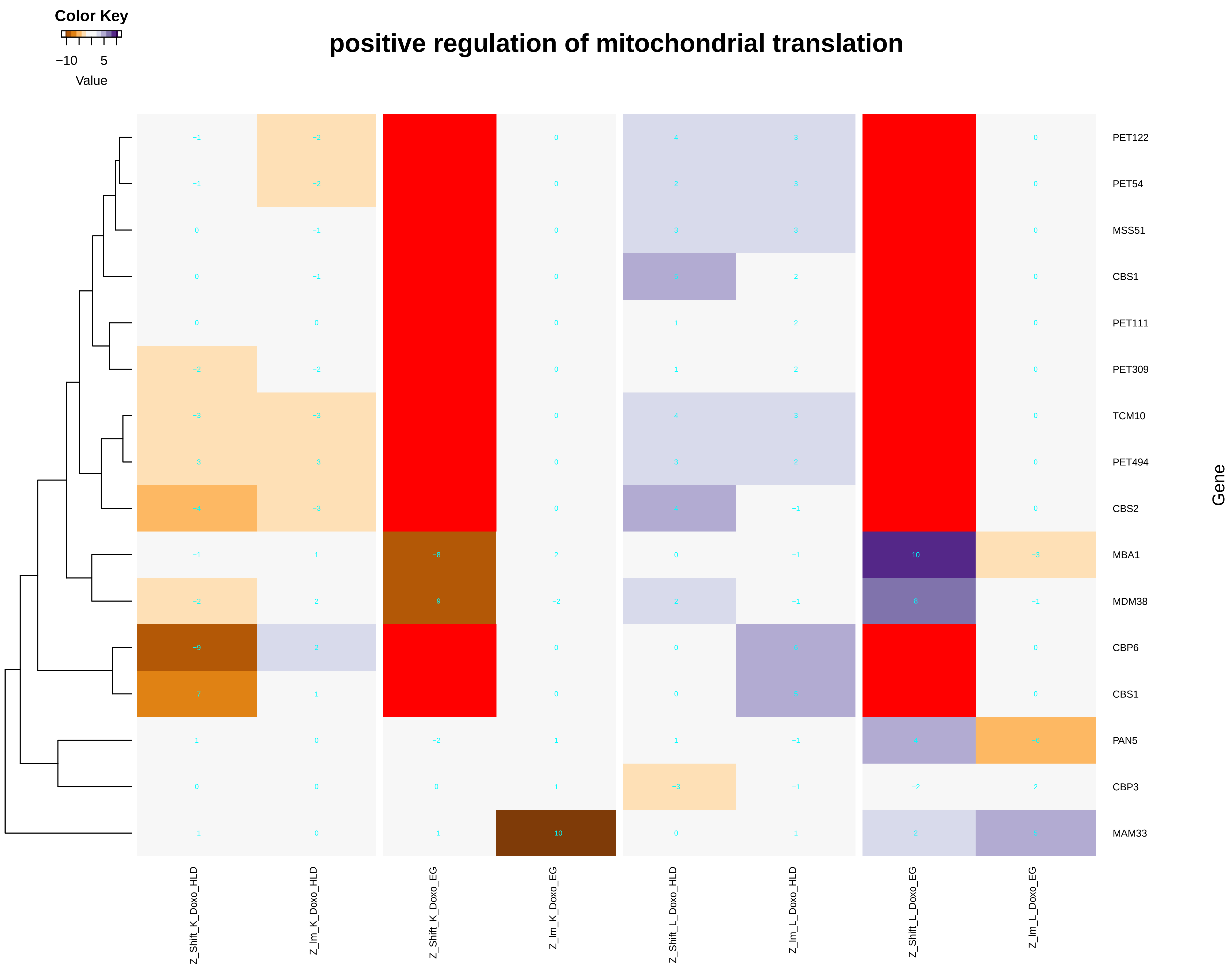

### positive_regulation_of_DNA-templated_transcription,_elongation.pdf

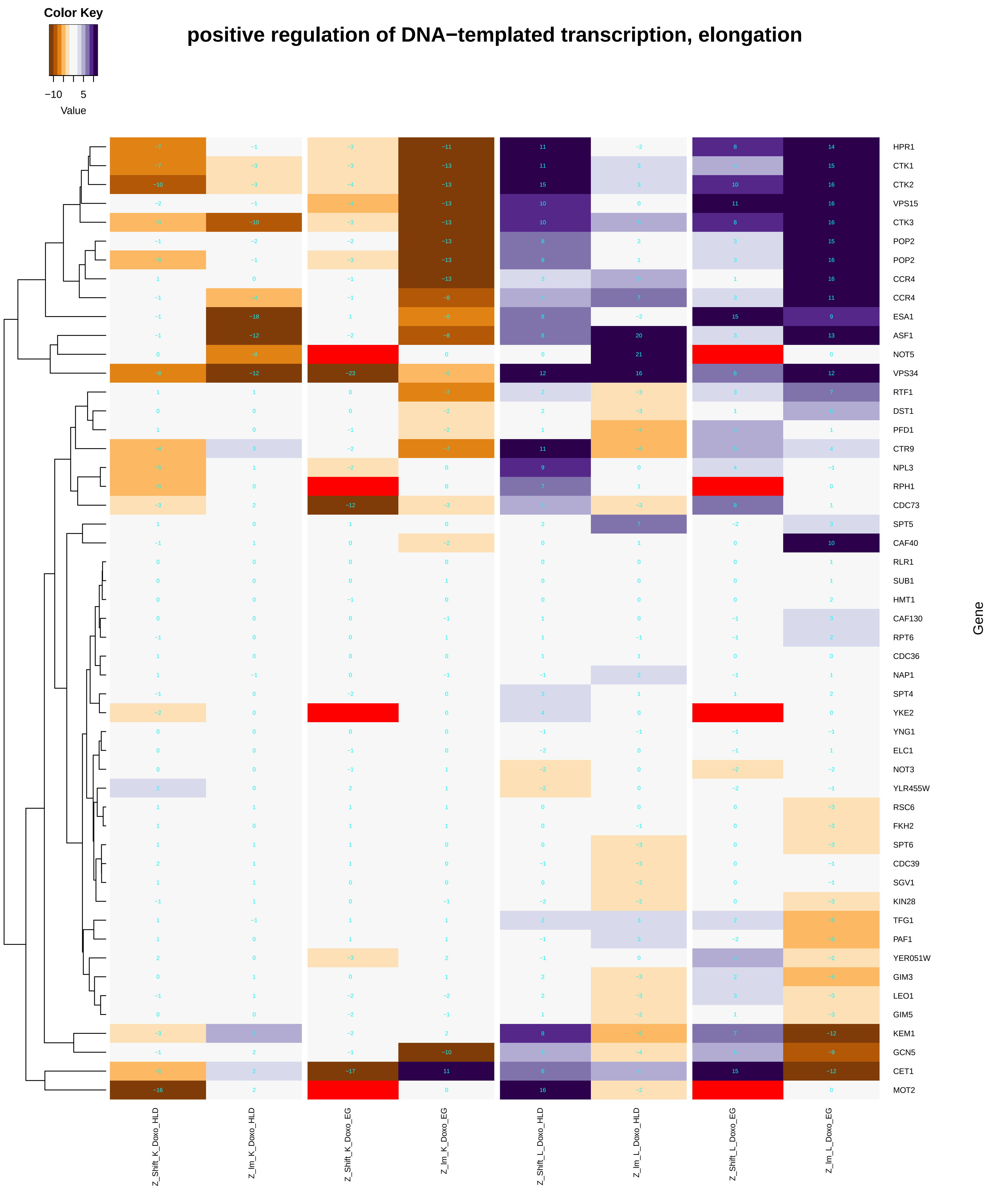

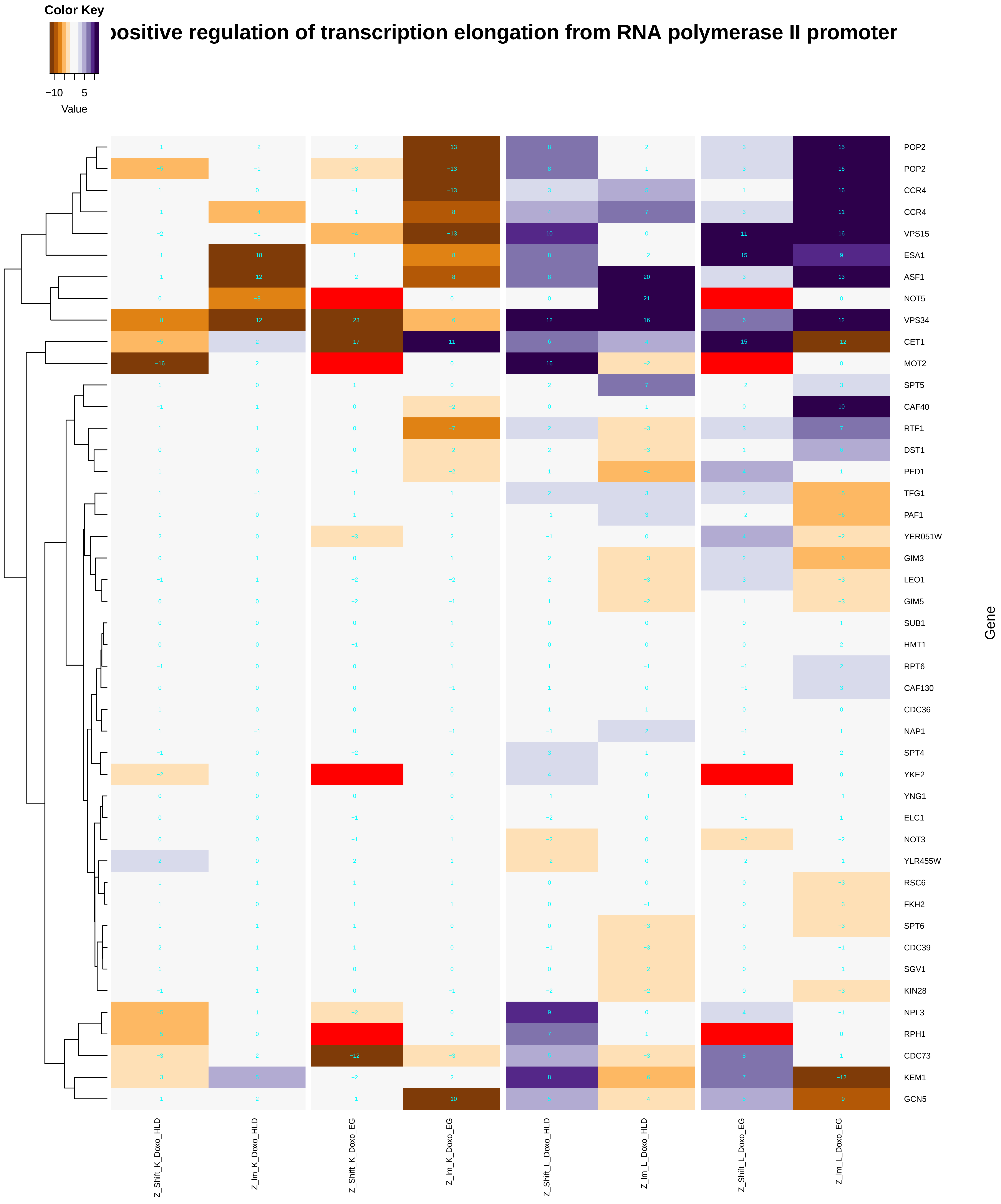

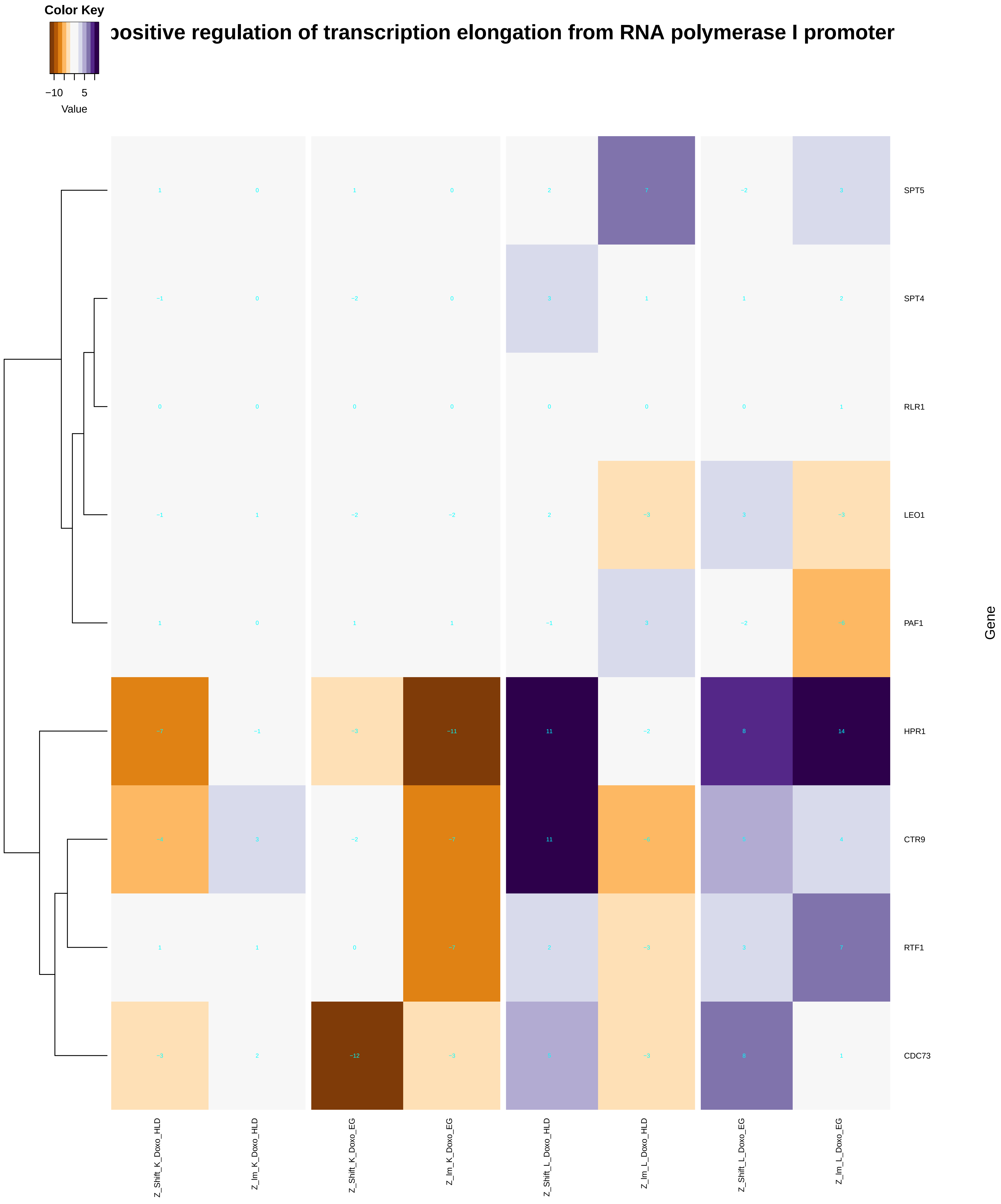

### positive_regulation_of_fatty_acid_beta-oxidation.pdf

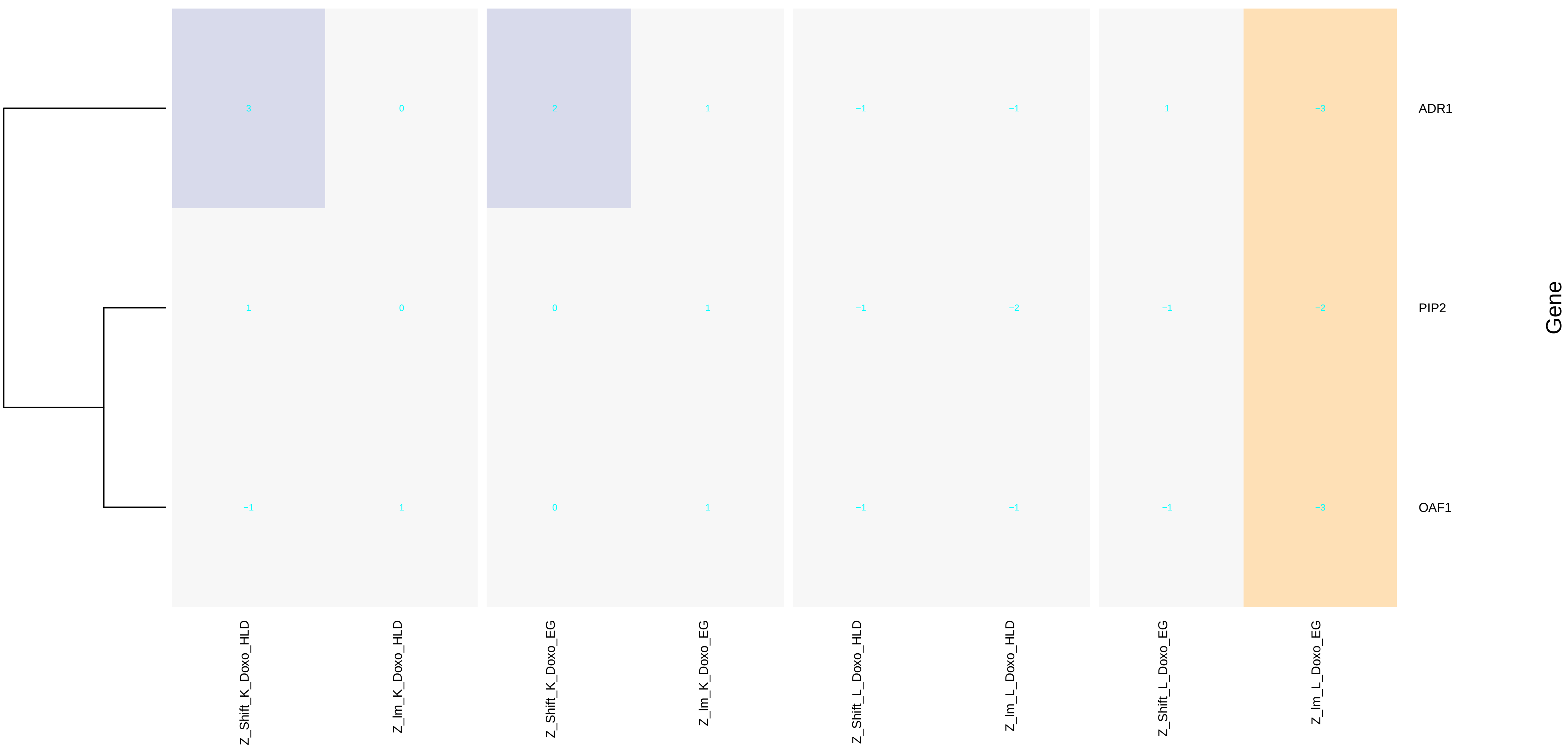

### positive_regulation_of_fatty_acid_oxidation.pdf

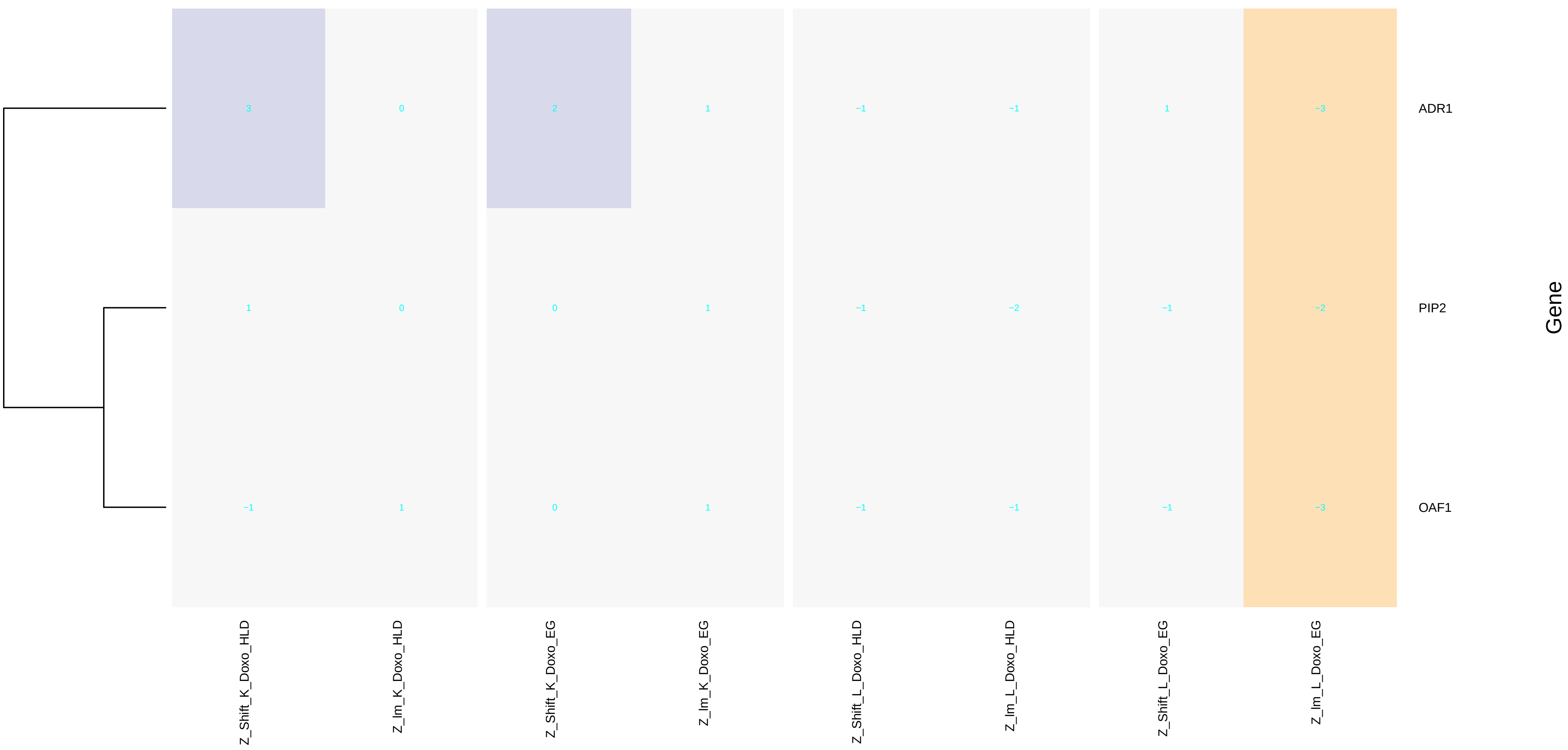

### positive_regulation_of_gene_expression,_epigenetic.pdf

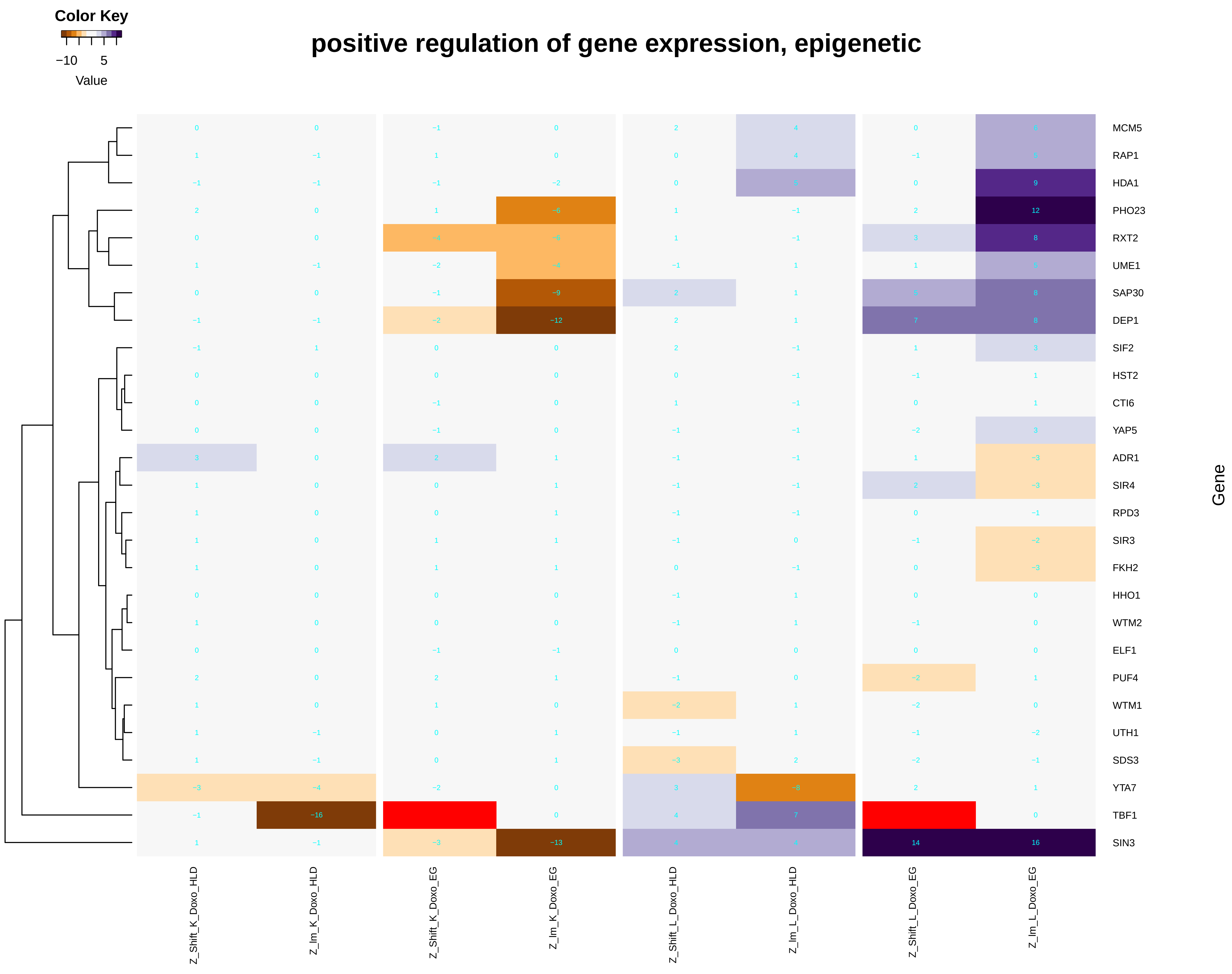

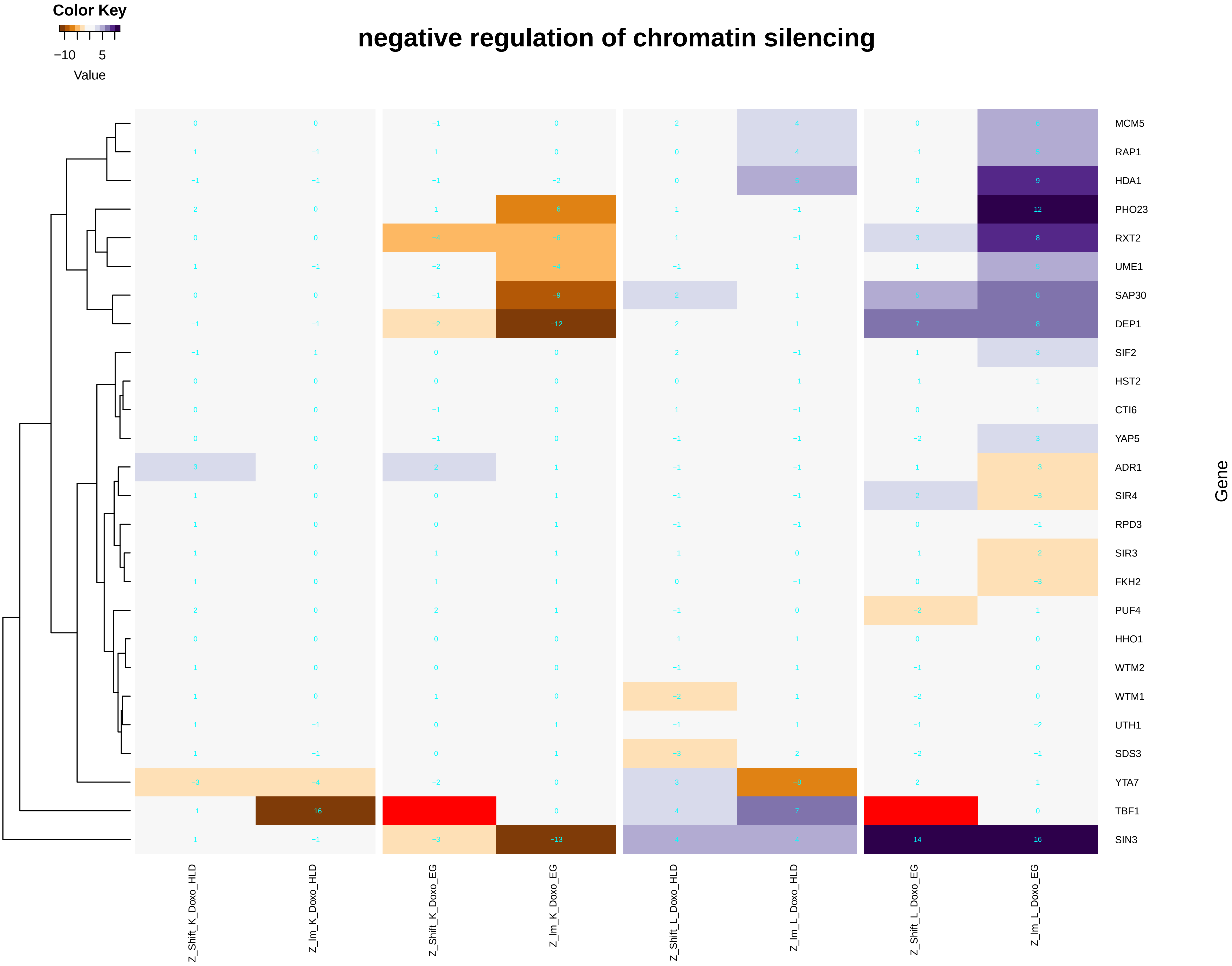

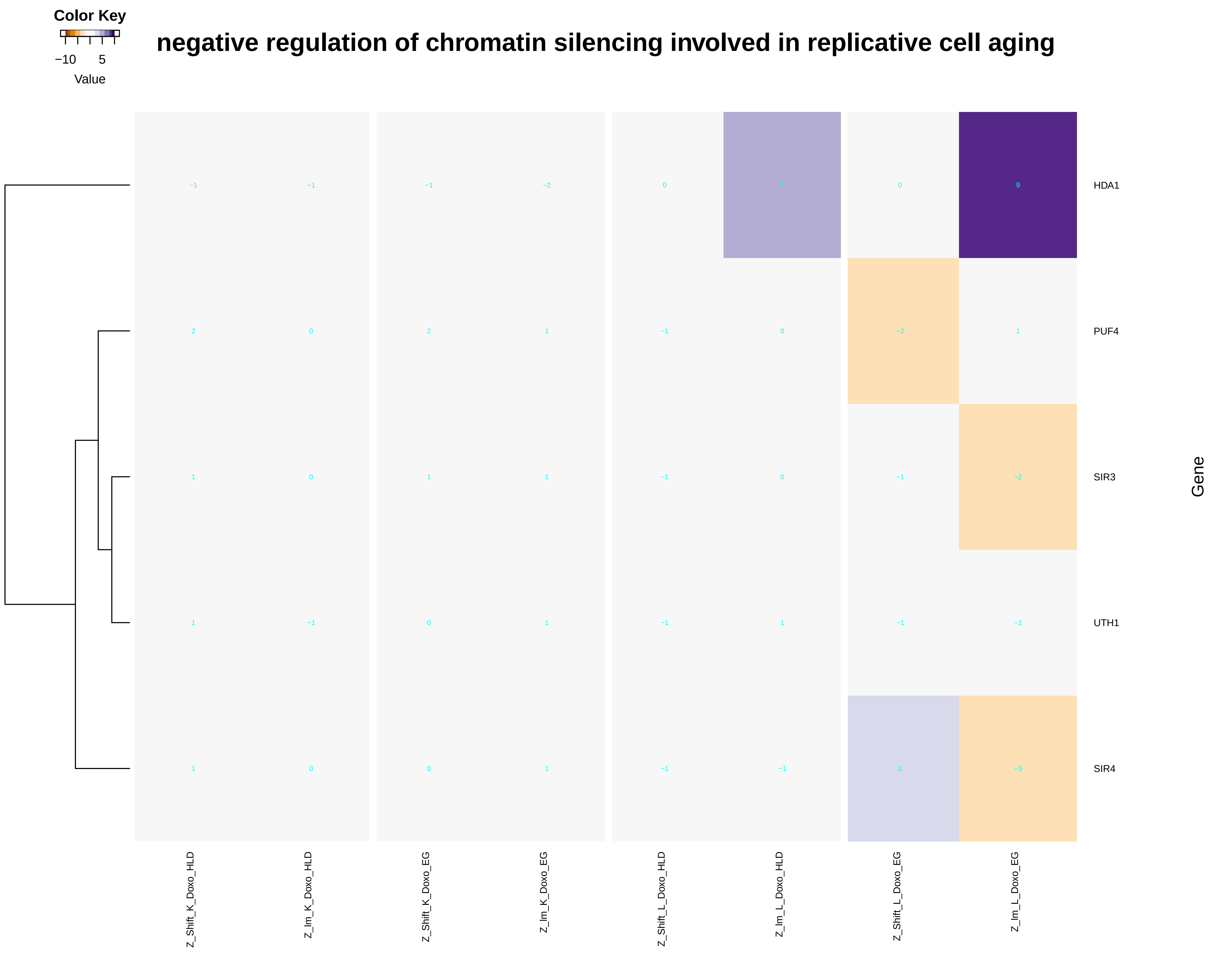

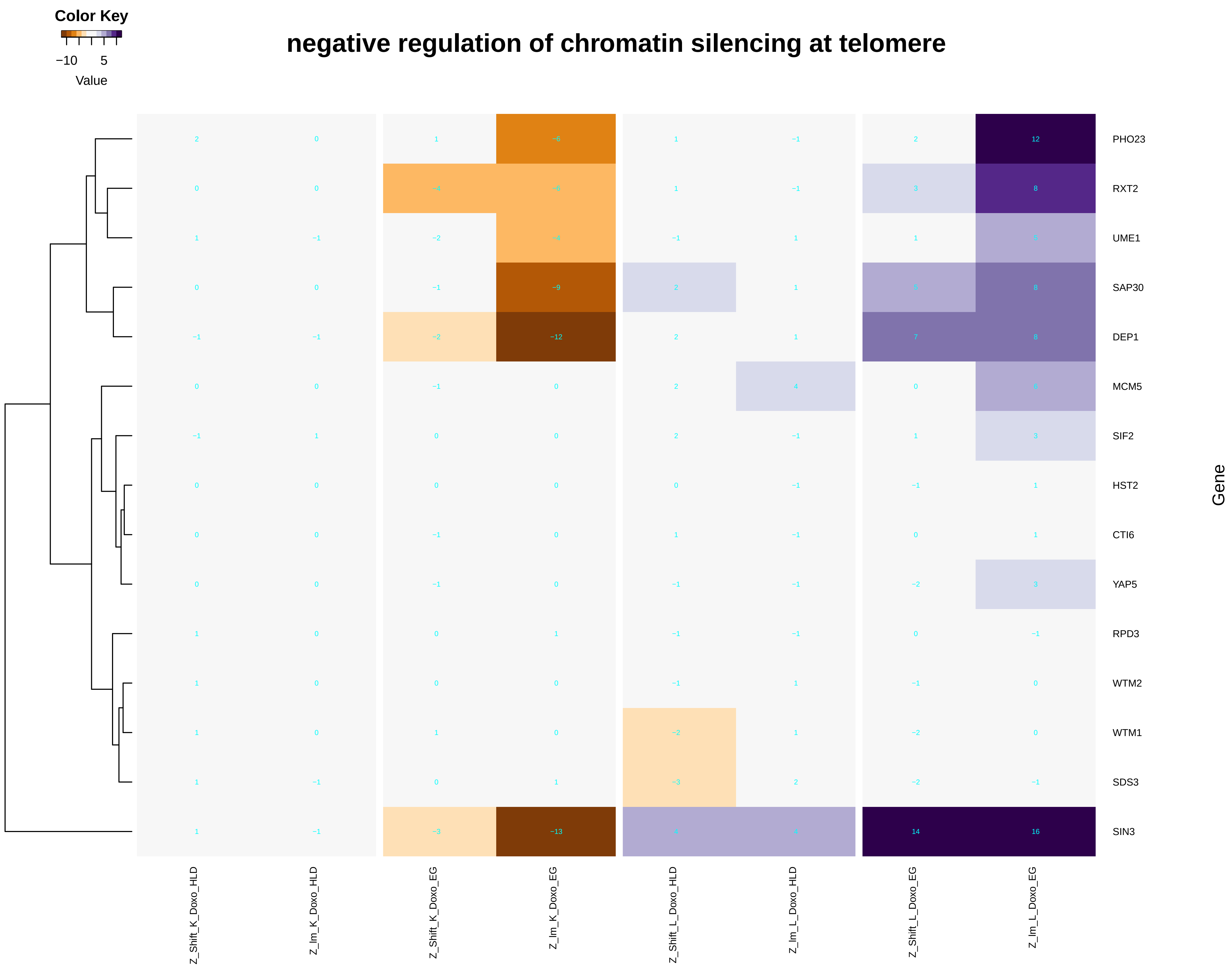

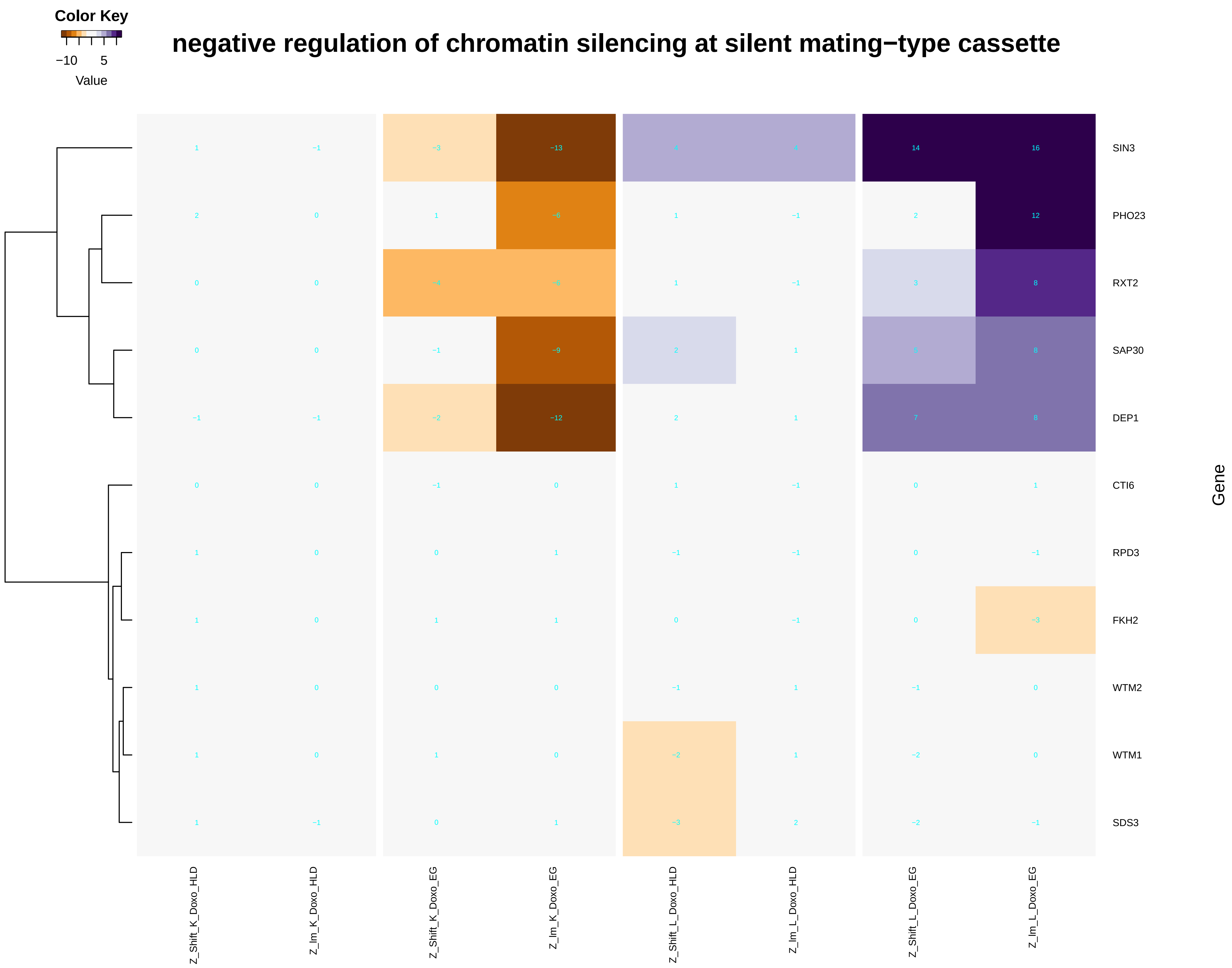

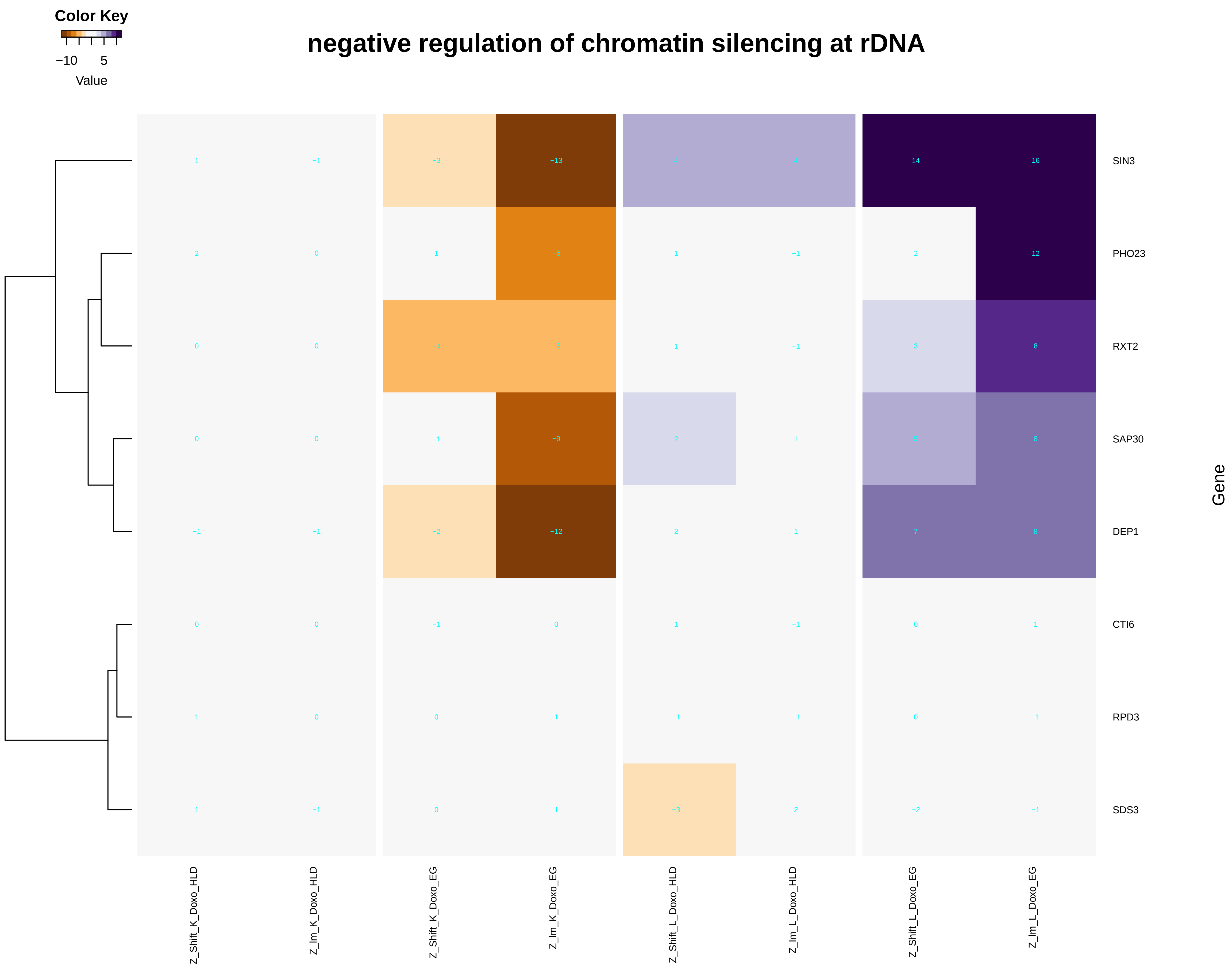

### positive_regulation_of_lipid_catabolic_process.pdf

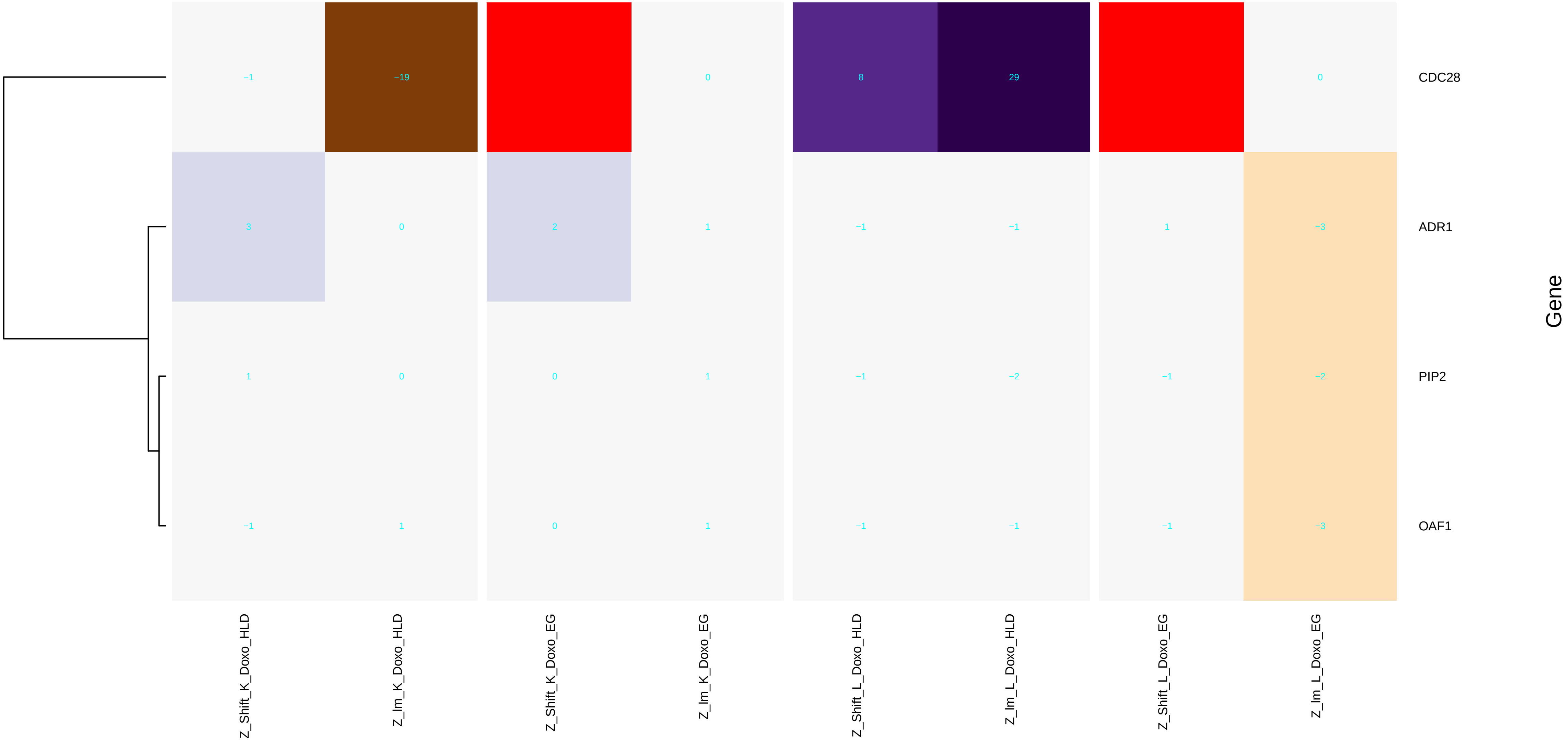

### positive_regulation_of_mitochondrial_translation.pdf

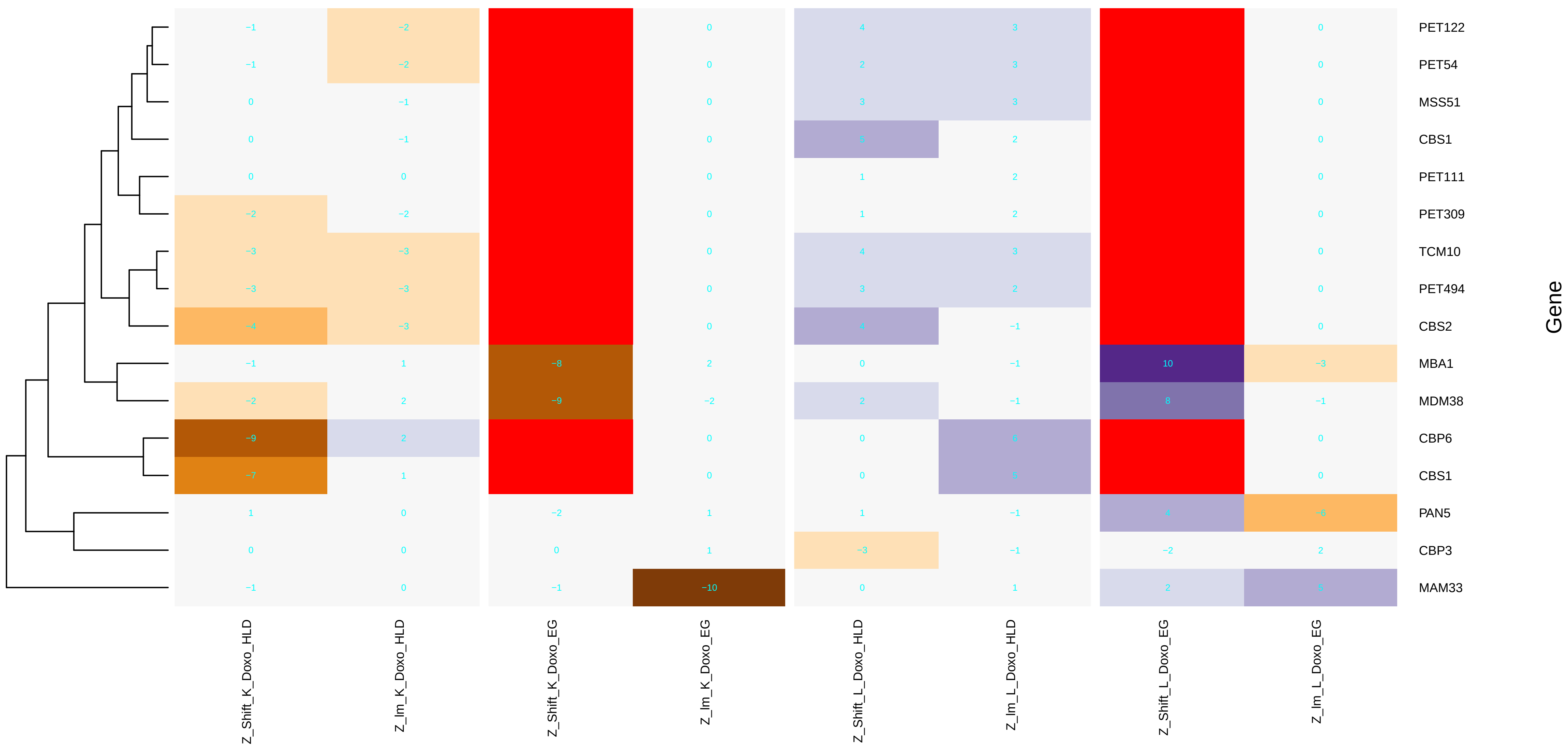

### positive_regulation_of_phosphatidylcholine_biosynthetic_process.pdf

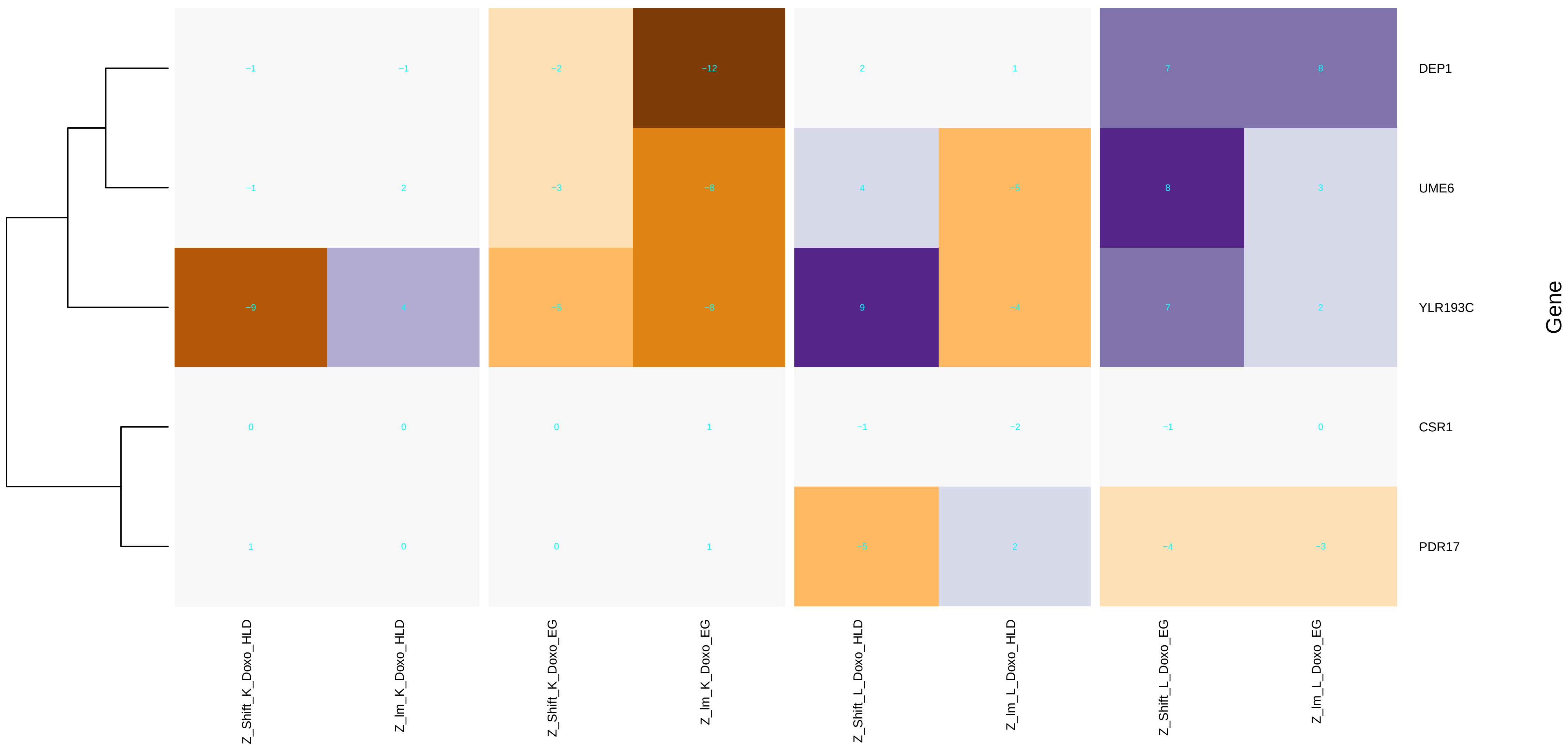

### positive_regulation_of_phospholipid_biosynthetic_process.pdf

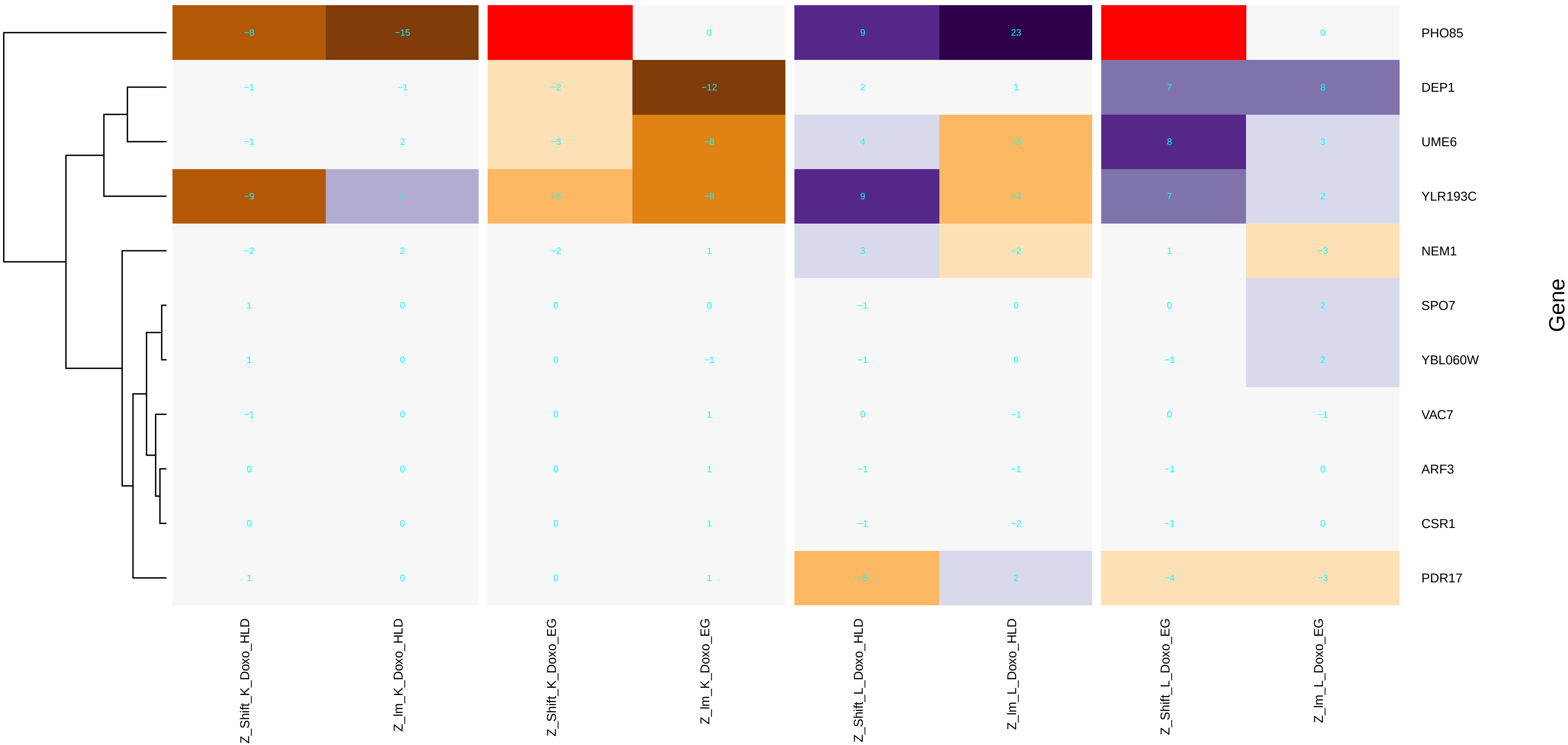
