## Additional File 11 for "A yeast phenomic model for the influence of Warburg metabolism on genetic buffering of doxorubicin": C - ByTissue.pdf

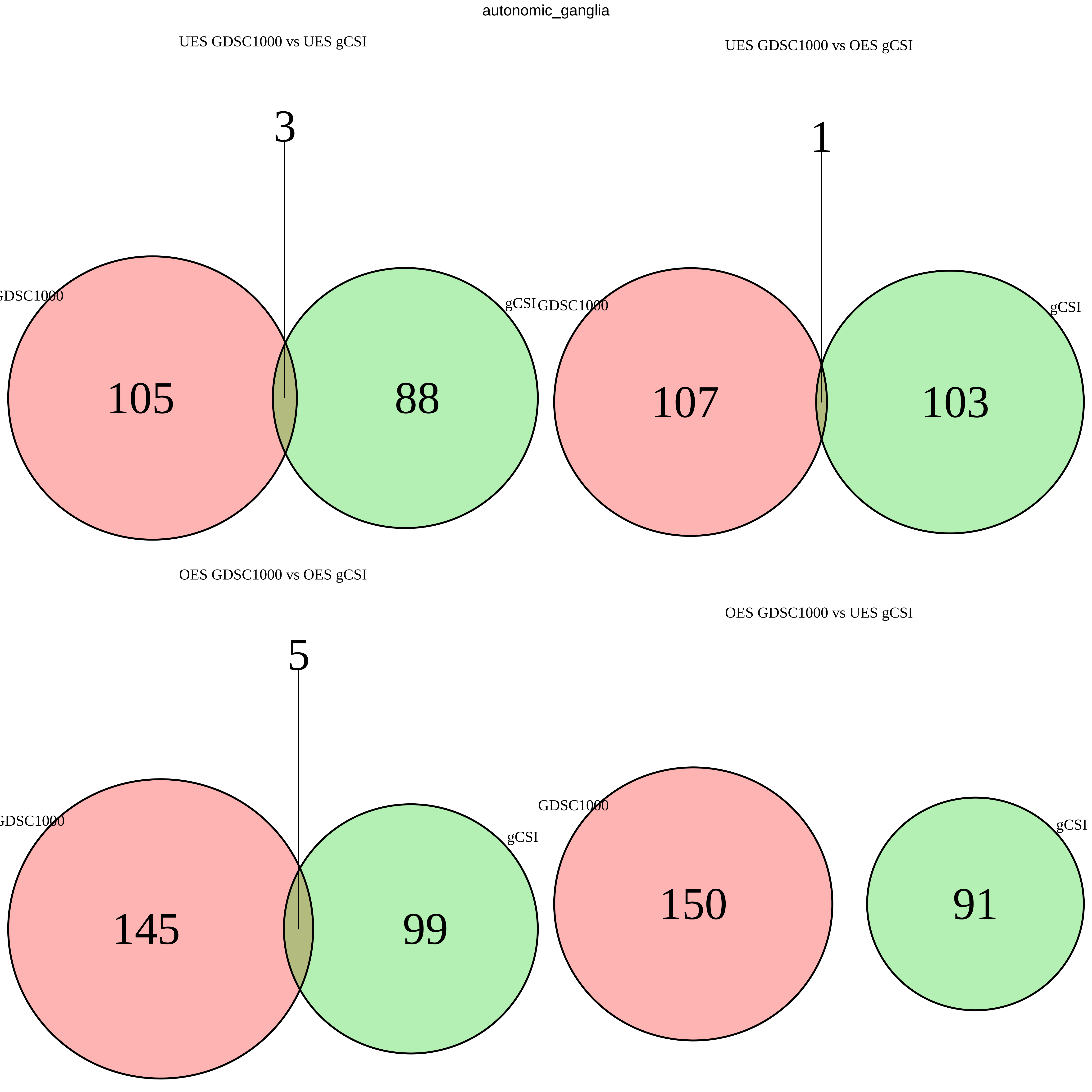
autonomic_ganglia
UES GDSC1000 vs UES gCSI
3
GDSC1000
105
88
gCSI
UES GDSC1000 vs OES gCSI
1
GDSC1000
107
103
gCSI
OES GDSC1000 vs OES gCSI
5
GDSC1000
145
99
gCSI
OES GDSC1000 vs UES gCSI
GDSC1000
150
91
gCSI

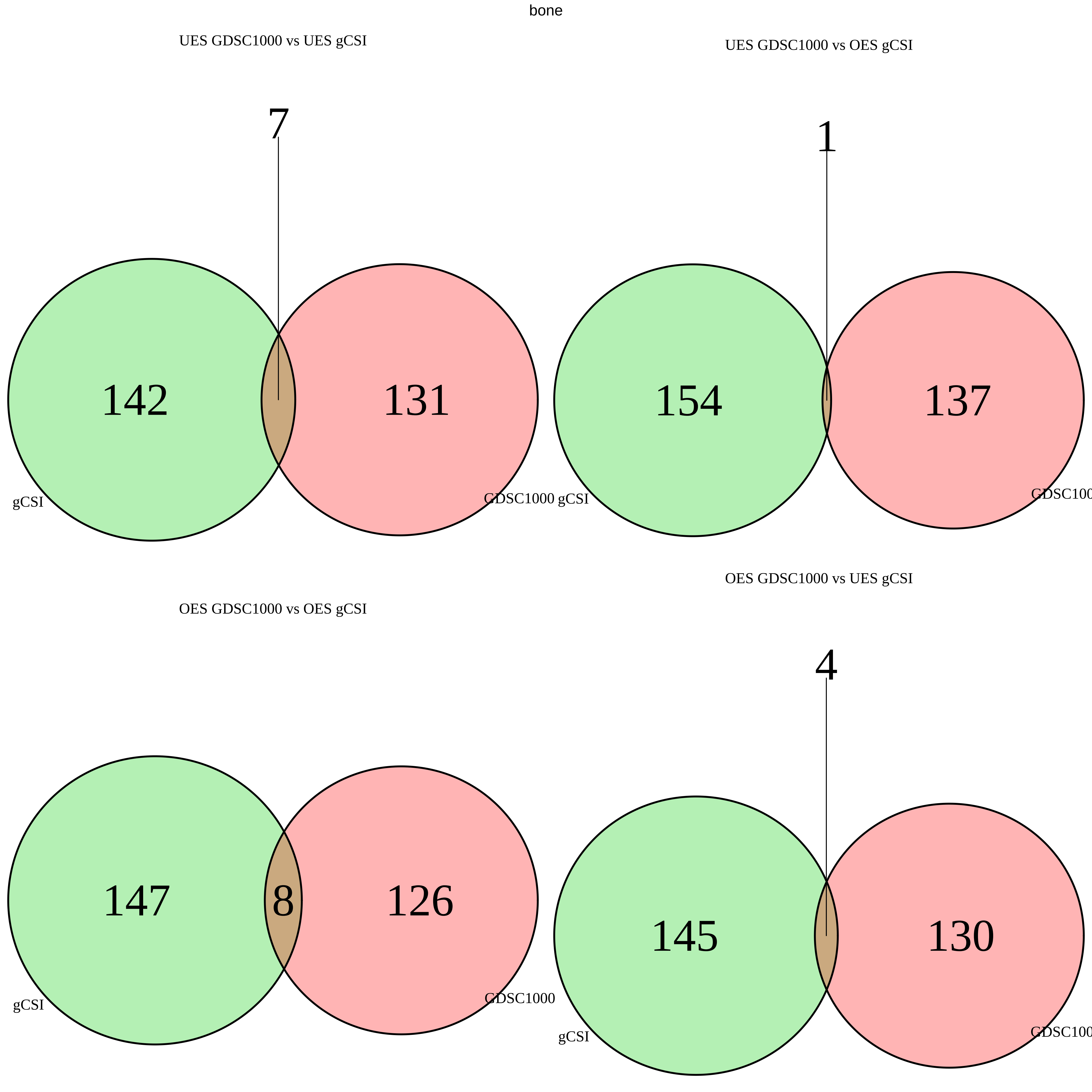
bone
UES GDSC1000 vs UES gCSI
7
142
131
gCSI
GDSC1000 gCSI
UES GDSC1000 vs OES gCSI
1
154
137
GDSC100
OES GDSC1000 vs UES gCSI
OES GDSC1000 vs OES gCSI
4
147
8
126
gCSI
GDSC1000
145
130
gCSI
GDSC100

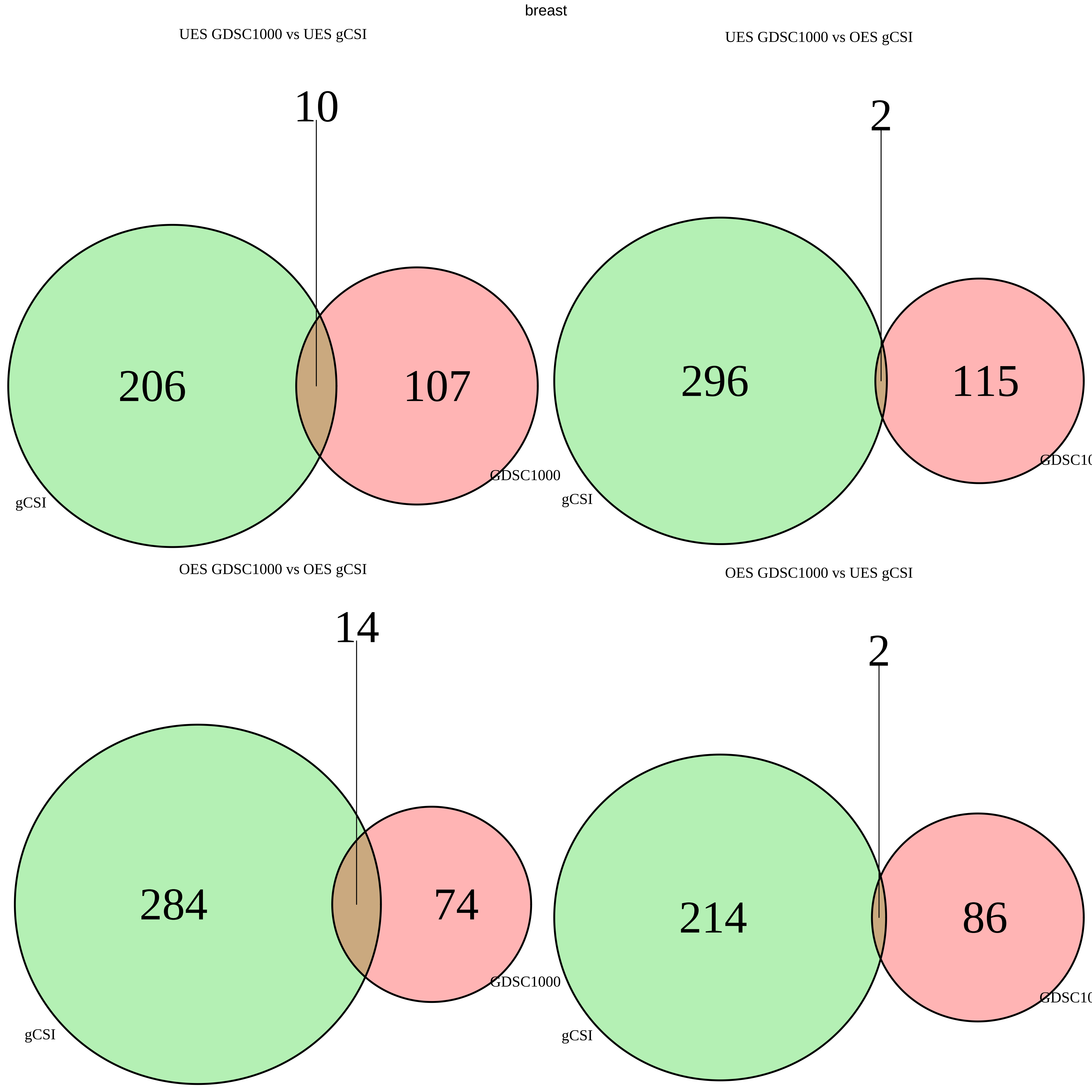
breast
UES GDSC1000 vs UES gCSI
10
206
107
gCSI
GDSC1000
UES GDSC1000 vs OES gCSI
2
296
115
gCSI
GDSC10
OES GDSC1000 vs OES gCSI
14
284
74
gCSI
GDSC1000
OES GDSC1000 vs UES gCSI
2
214
86
gCSI
GDSC10

central_nervous_system

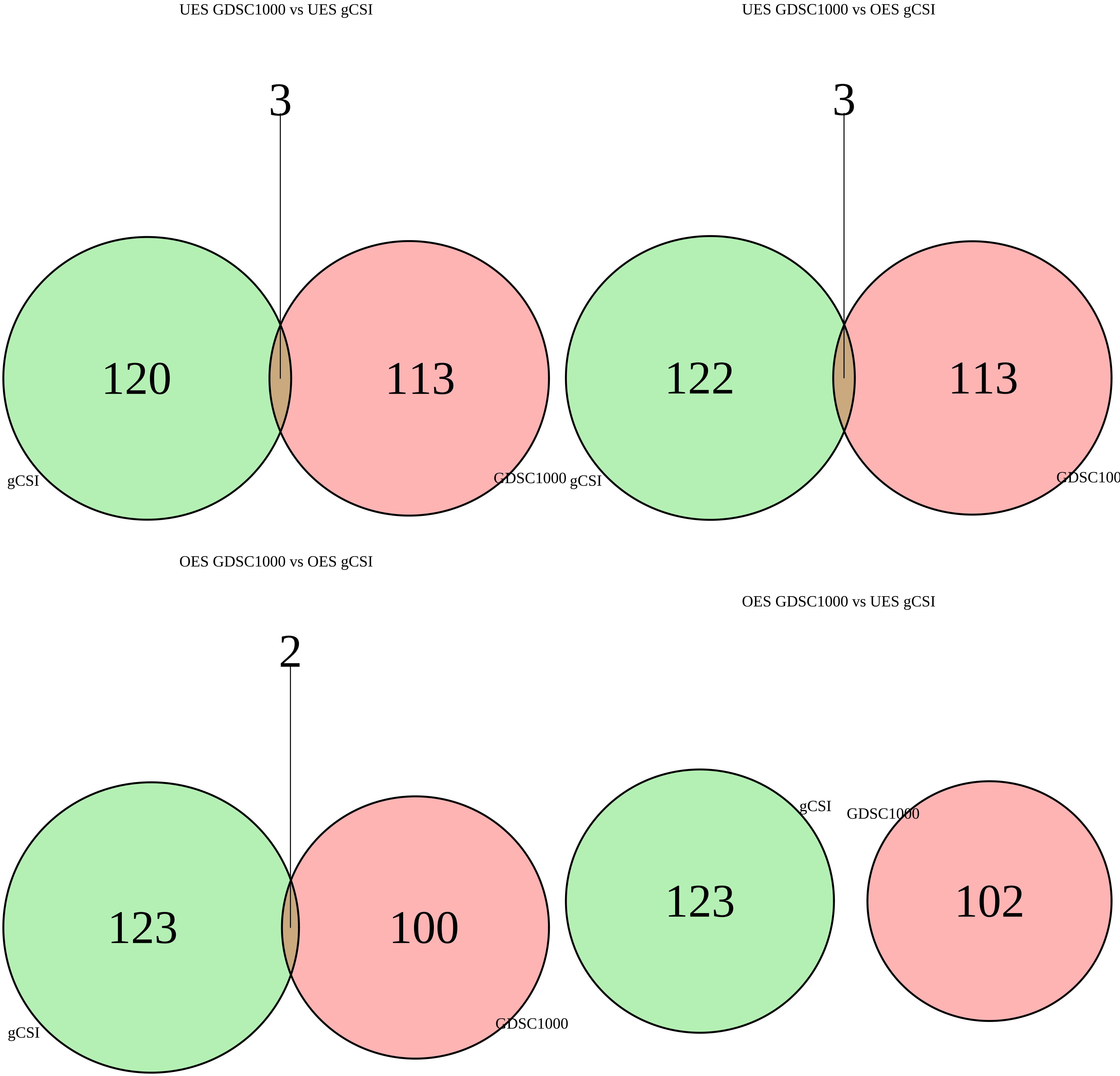

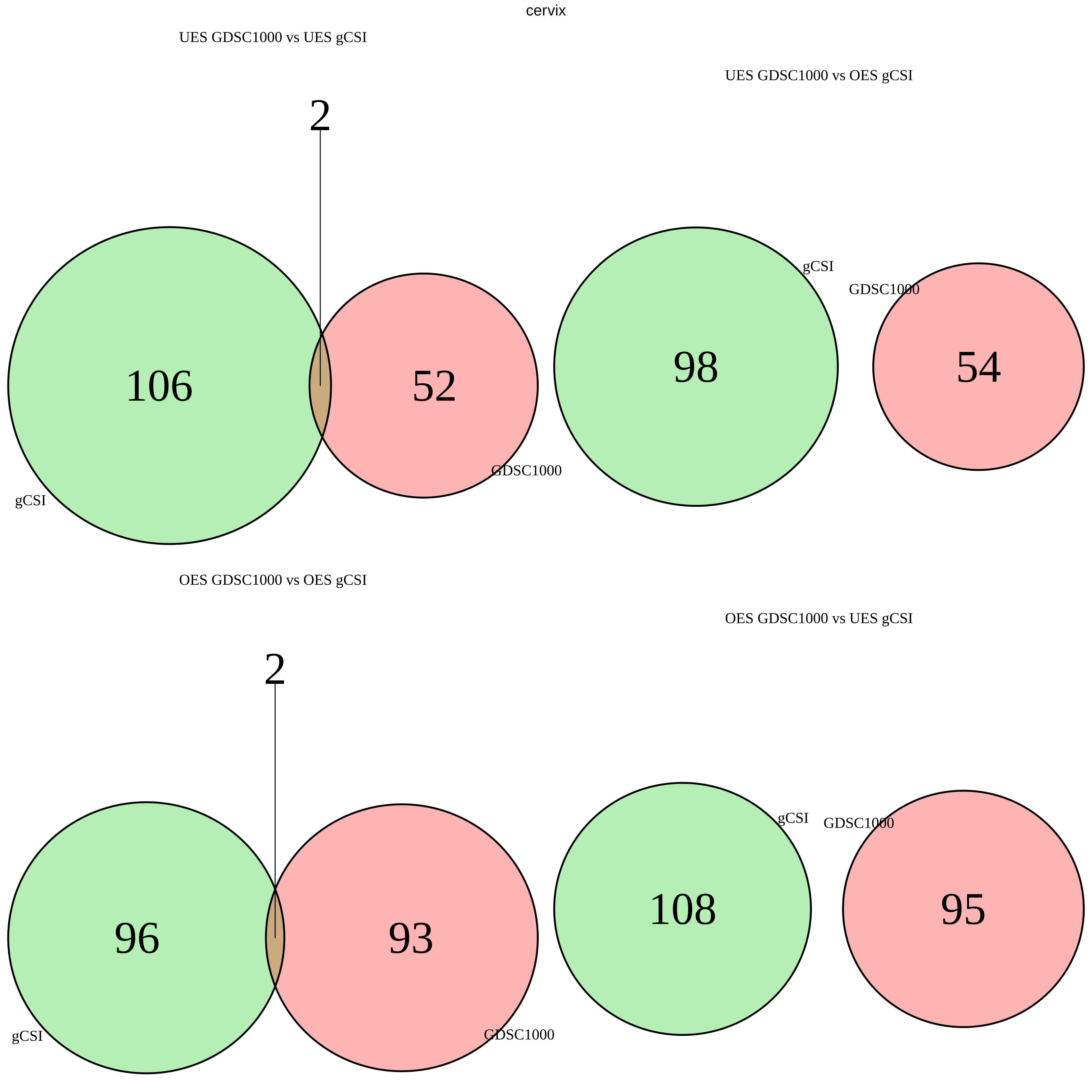
cervix
UES GDSC1000 vs UES gCSI
2
106
52
gCSI
GDSC1000
UES GDSC1000 vs OES gCSI
gCSI
GDSC1000
98
54
OES GDSC1000 vs OES gCSI
2
96
93
gCSI
GDSC1000
OES GDSC1000 vs UES gCSI
gCSI
GDSC1000
108
95

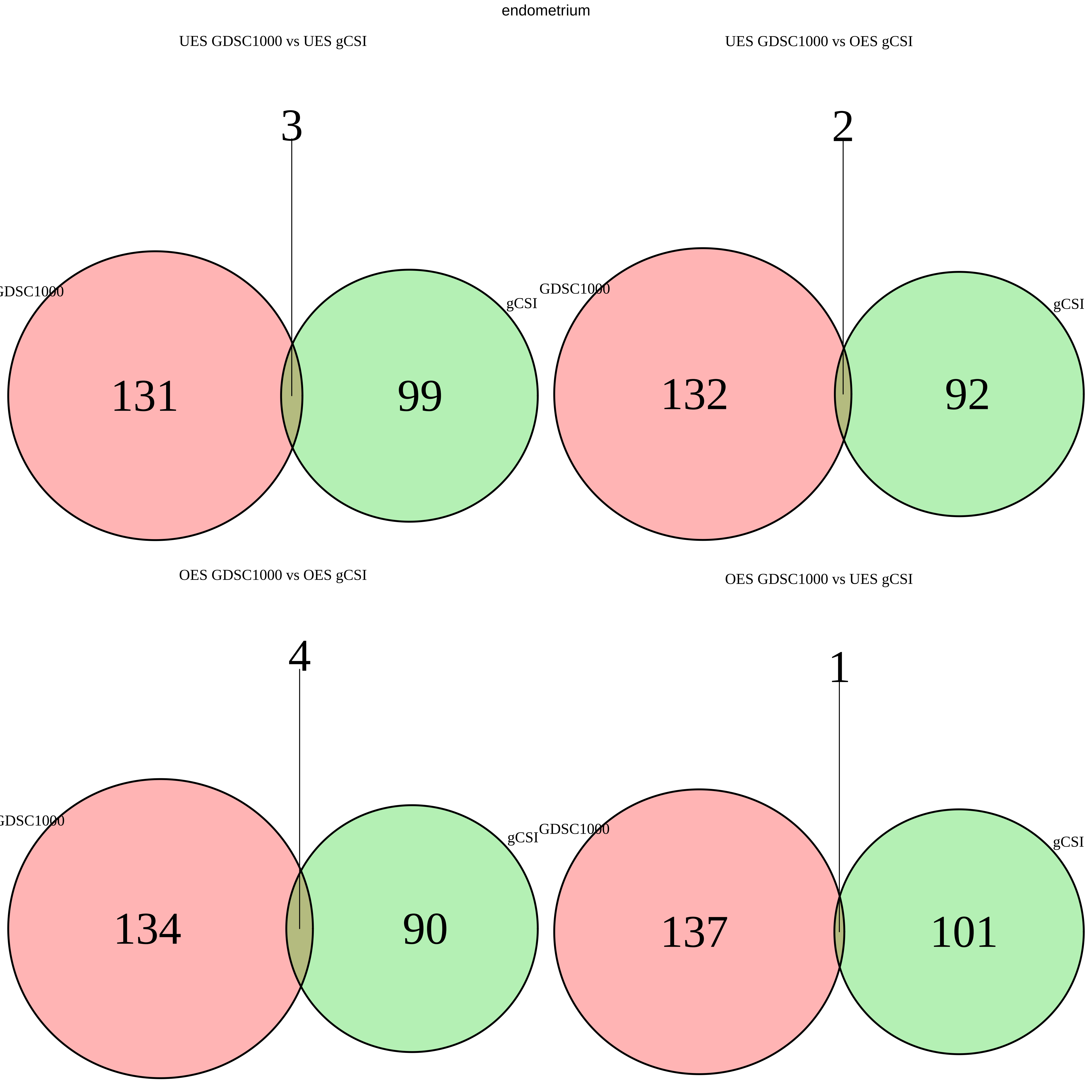
endometrium
UES GDSC1000 vs UES gCSI
3
GDSC1000
gCSI
131
99
UES GDSC1000 vs OES gCSI
2
GDSC1000
gCSI
132
92
OES GDSC1000 vs OES gCSI
4
GDSC1000
gCSI
134
90
OES GDSC1000 vs UES gCSI
1
GDSC1000
gCSI
137
101

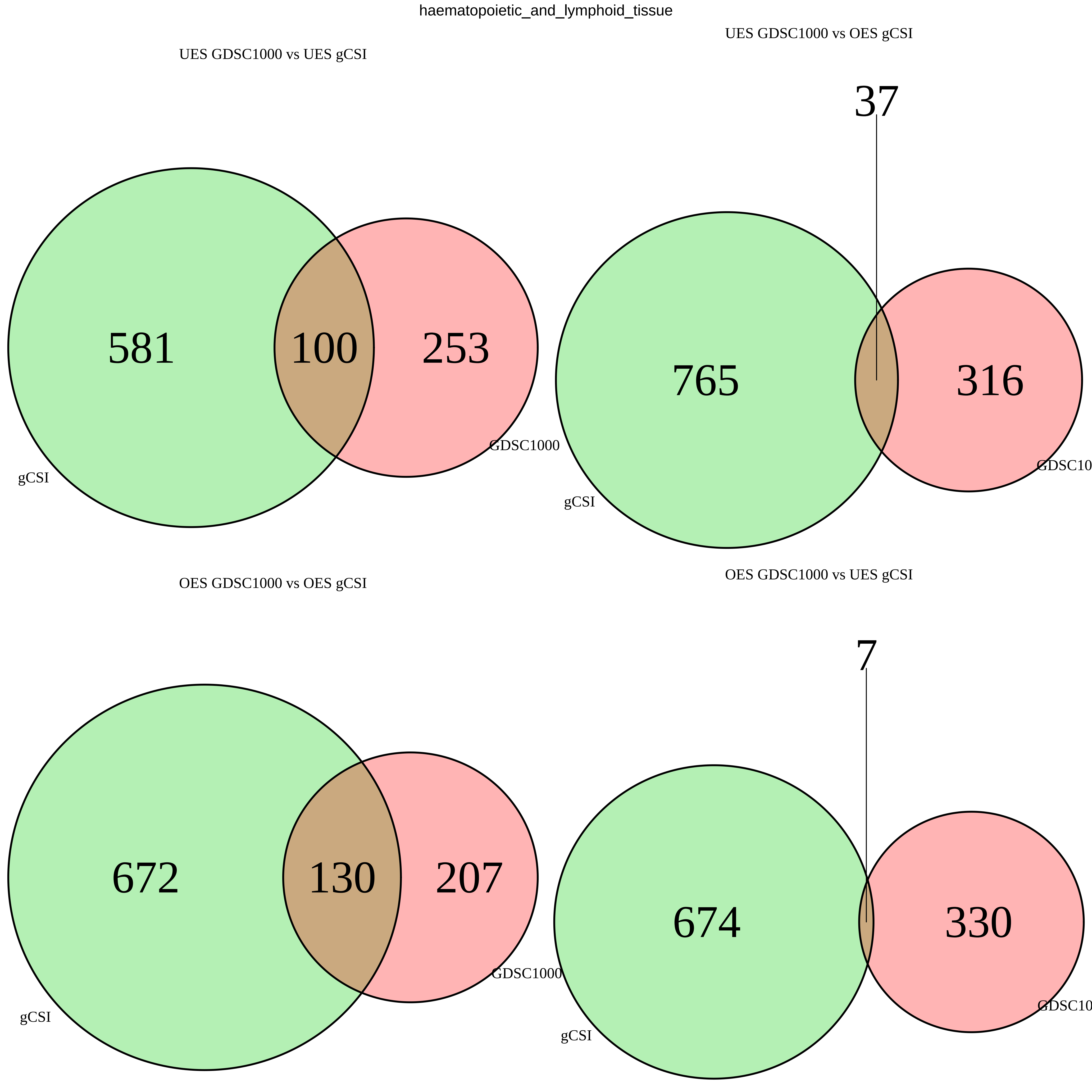
haematopoietic_and_lymphoid_tissue
UES GDSC1000 vs UES gCSI
581
100
253
GDSC1000
gCSI
UES GDSC1000 vs OES gCSI
37
765
316
GDSC10
gCSI
OES GDSC1000 vs OES gCSI
672
130
207
GDSC1000
gCSI
OES GDSC1000 vs UES gCSI
7
674
330
GDSC10
gCSI

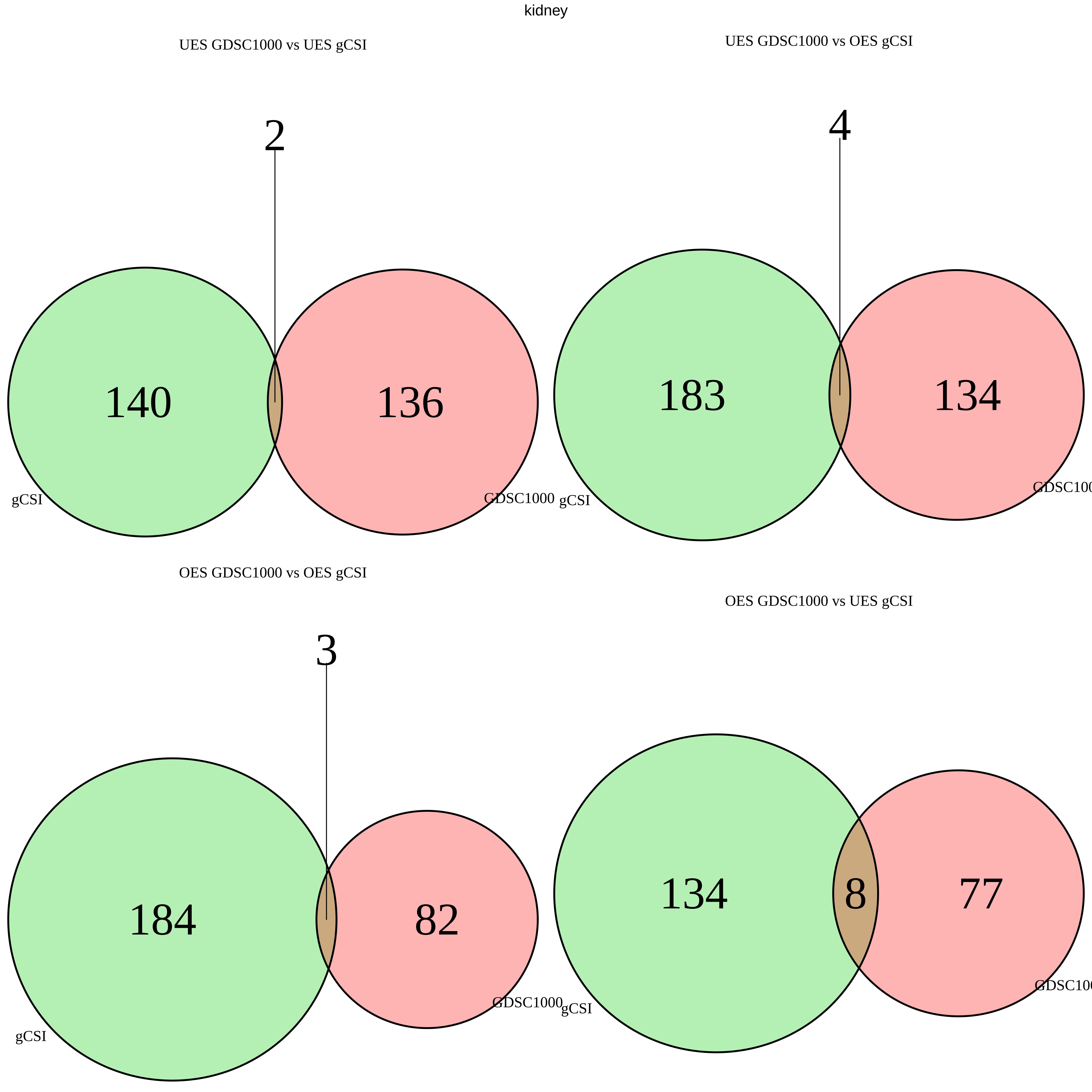

kidney
UES GDSC1000 vs UES gCSI
2
140
136
gCSI
GDSC1000
UES GDSC1000 vs OES gCSI
4
183
134
gCSI
GDSC100
OES GDSC1000 vs OES gCSI
3
184
82
gCSI
GDSC1000
OES GDSC1000 vs UES gCSI
134
8
77
gCSI
GDSC100

large_intestine

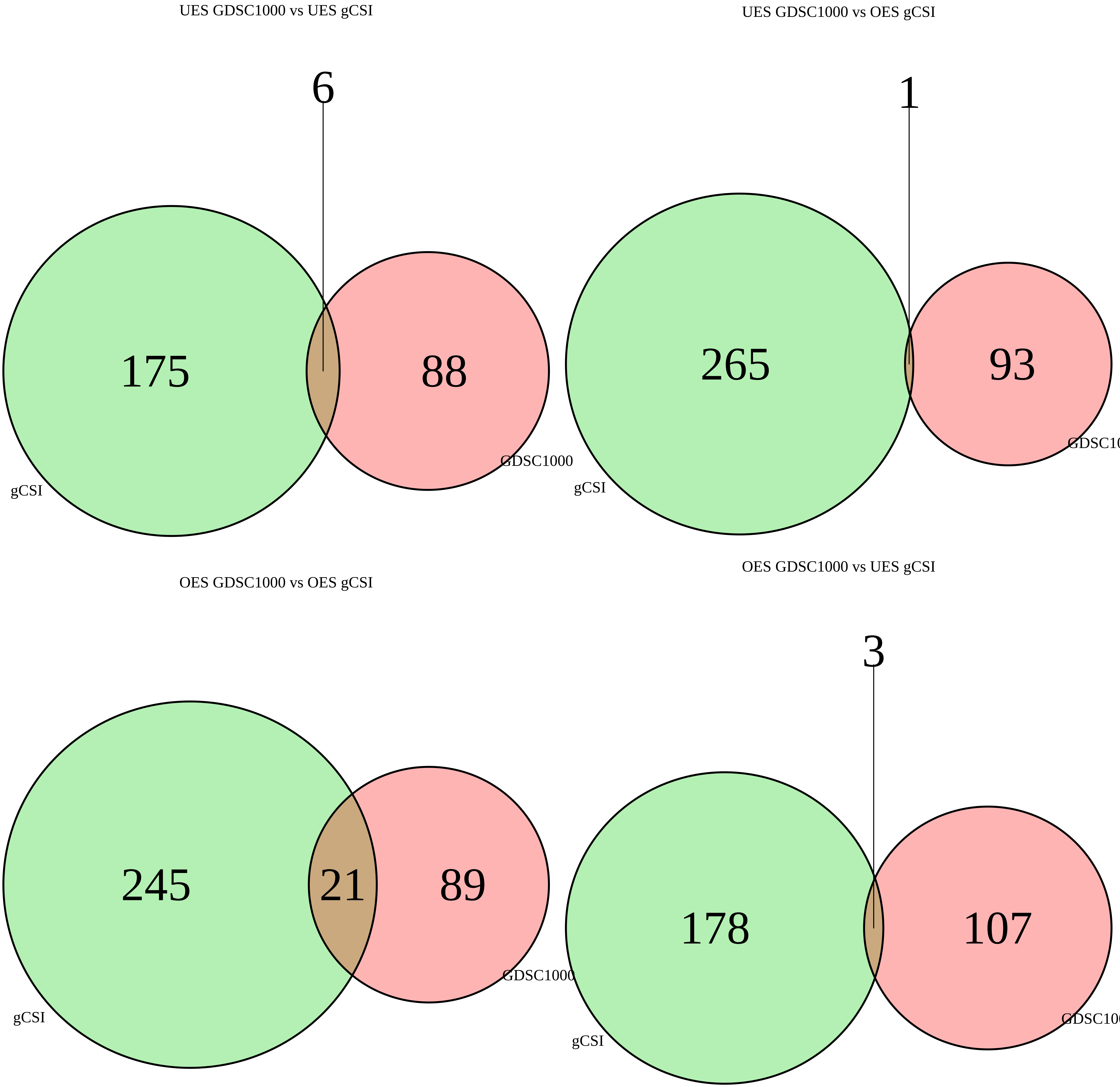

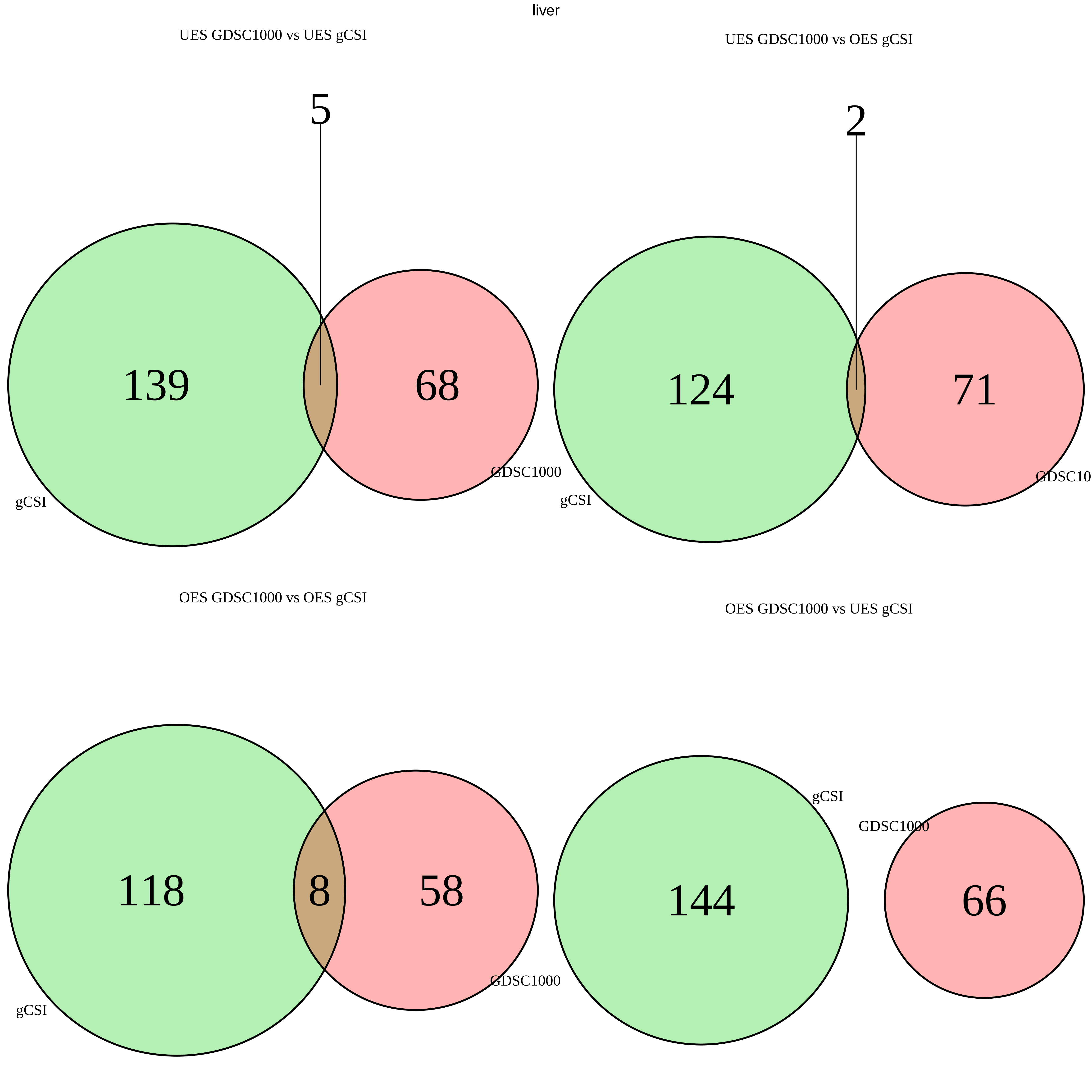
liver
UES GDSC1000 vs UES gCSI
5
139
68
GDSC1000
gCSI
UES GDSC1000 vs OES gCSI
2
124
71
gCSI
OES GDSC1000 vs OES gCSI
118
8
58
GDSC1000
gCSI
OES GDSC1000 vs UES gCSI
gCSI
GDSC1000
144
66

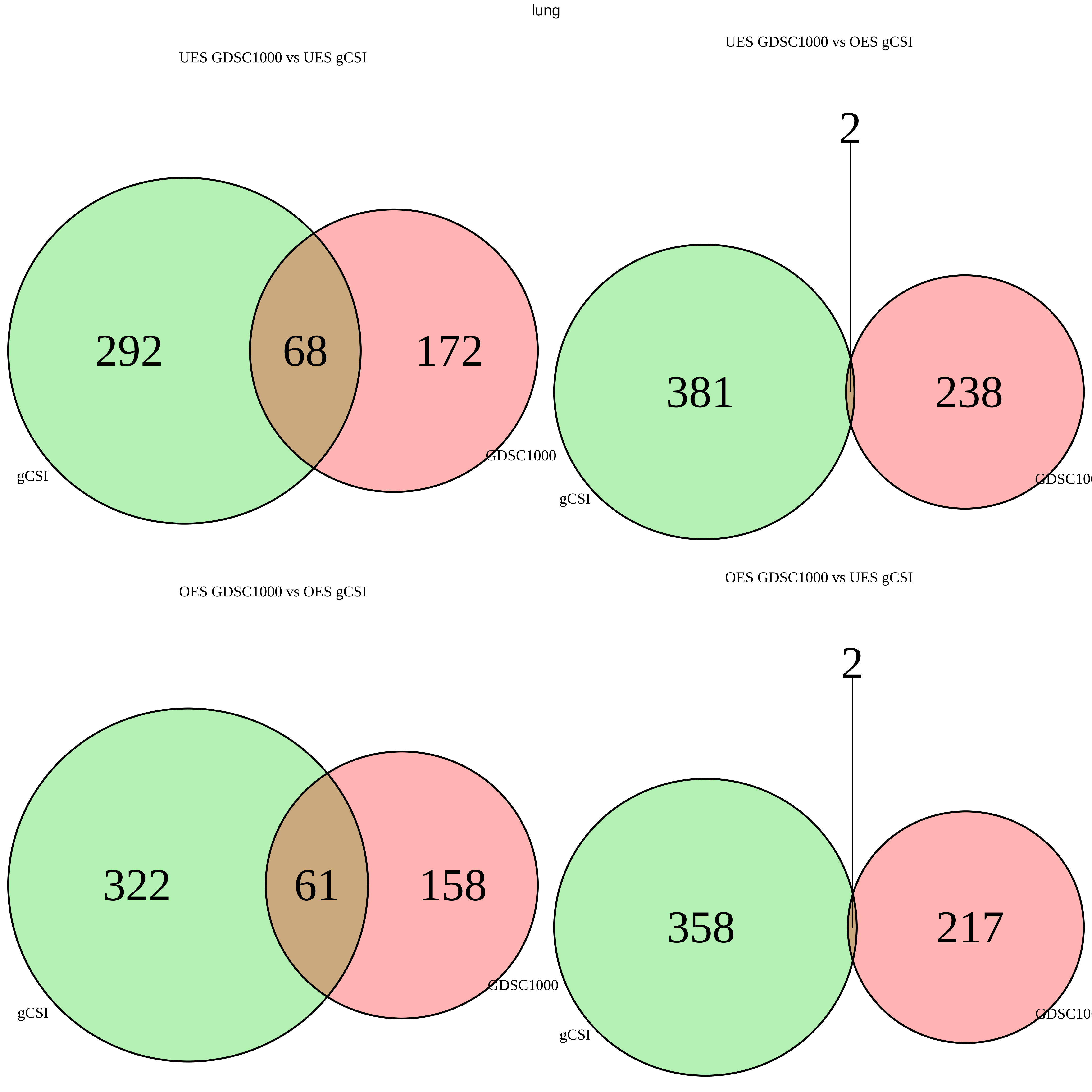

lung
UES GDSC1000 vs UES gCSI
292
68
172
gCSI
GDSC1000
UES GDSC1000 vs OES gCSI
2
381
238
gCSI
OES GDSC1000 vs OES gCSI
322
61
158
gCSI
GDSC1000
OES GDSC1000 vs UES gCSI
2
358
217
gCSI

oesophagus

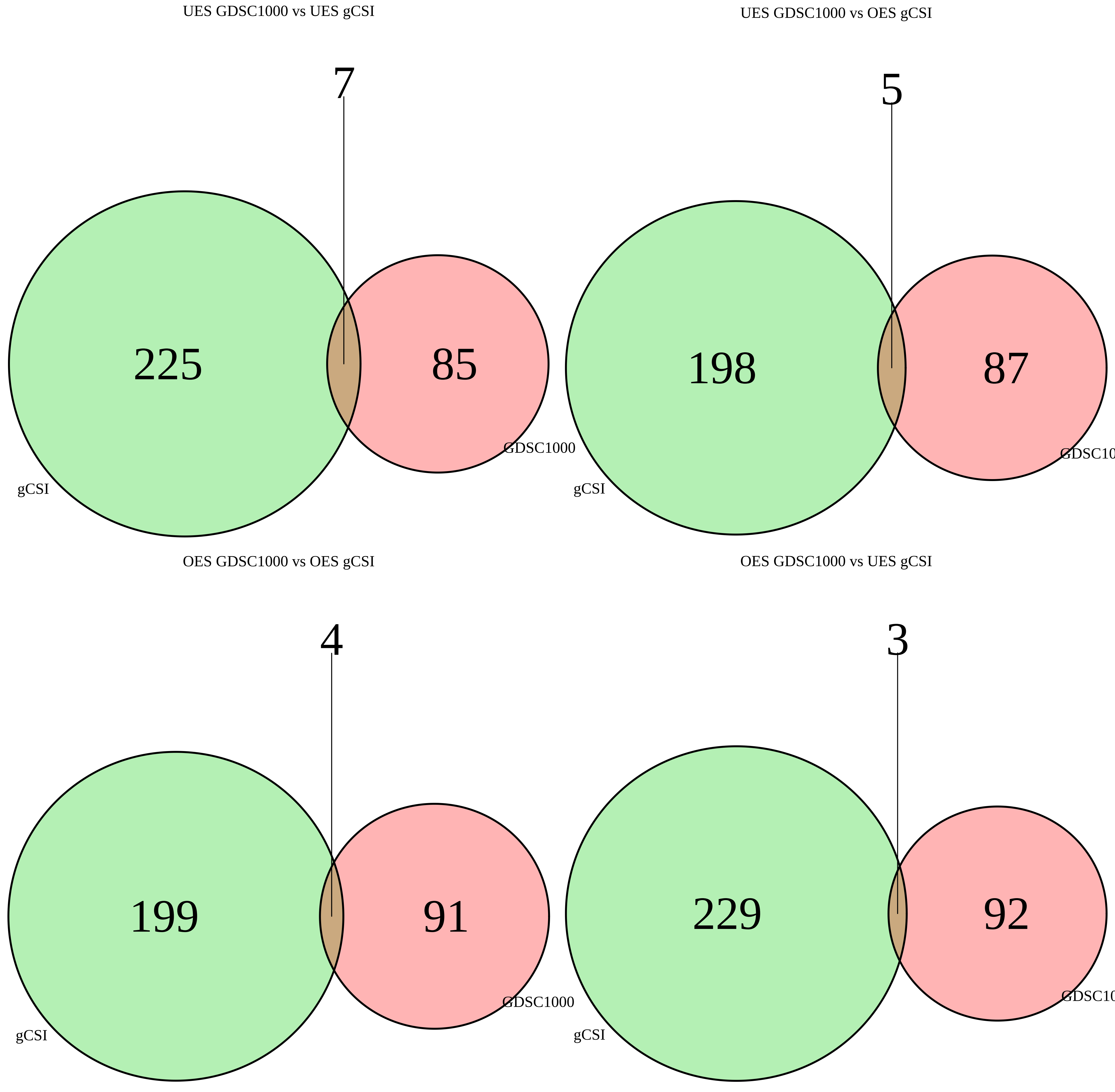
UES GDSC1000 vs UES gCSI
7
225
85
GDSC1000
gCSI
UES GDSC1000 vs OES gCSI
5
198
87
GDSC10
gCSI
OES GDSC1000 vs OES gCSI
4
199
91
GDSC1000
gCSI
OES GDSC1000 vs UES gCSI
3
229
92
GDSC10
gCSI
