## Additional File 11 for "A yeast phenomic model for the influence of Warburg metabolism on genetic buffering of doxorubicin": E - GDSC1000_Heatmaps.pdf

### Human genes that are OES across all tissue
#### Any Yeast Phenotype

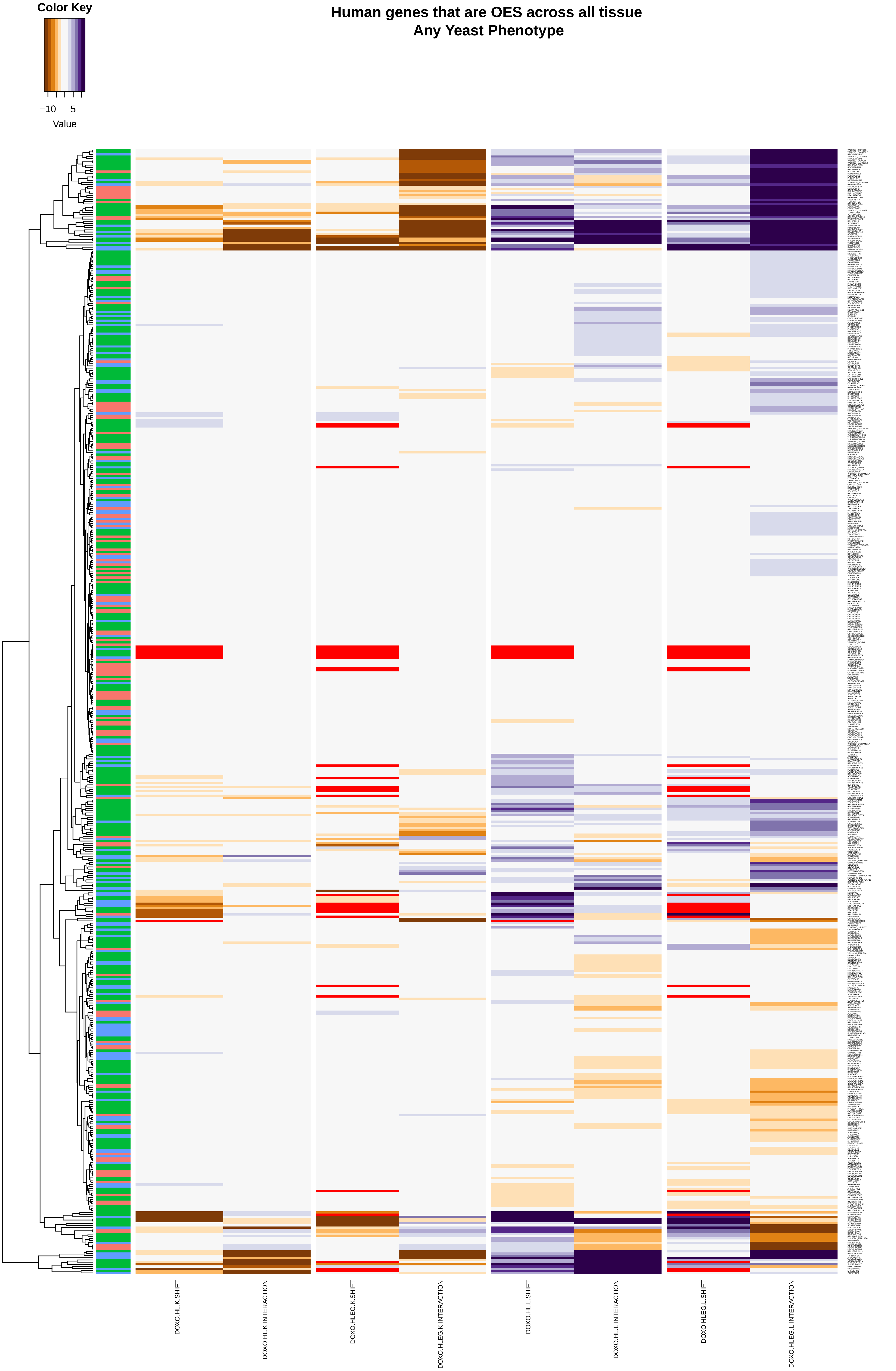

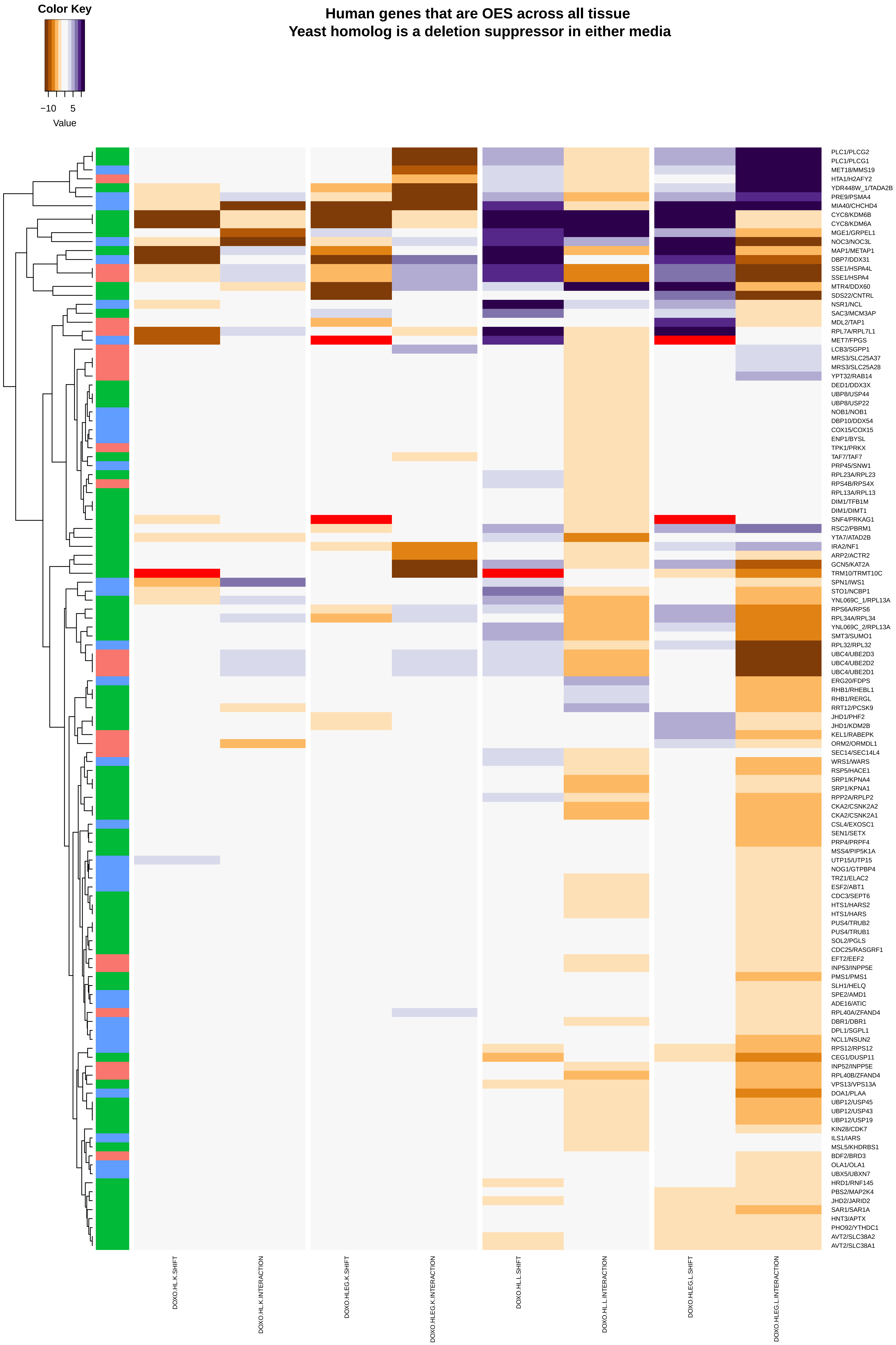
Color Key
−10
5
Value
Human genes that are OES across all tissue
Yeast homolog is a deletion suppressor in either media
PLC1/PLCG2
PLC1/PLCG1
MET18/MMS19
HTA1/H2AFY2
YDR448W_1/TADA2B
PRE9/PSMA4
MIA40/CHCHD4
CYC8/KDM6B
CYC8/KDM6A
MGE1/GRPEL1
NOC3/NOC3L
MAP1/METAP1
DBP7/DDX31
SSE1/HSPA4L
SSE1/HSPA4
MTR4/DDX60
SDS22/CNTRL
NSR1/NCL
SAC3/MCM3AP
MDL2/TAP1
RPL7A/RPL7L1
MET7/FPGS
LCB3/SGPP1
MRS3/SLC25A37
MRS3/SLC25A28
YPT32/RAB14
DED1/DDX3X
UBP8/USP44
UBP8/USP22
NOB1/NOB1
DBP10/DDX54
COX15/COX15
ENP1/BYSL
TPK1/PRKX
TAF7/TAF7
PRP45/SNW1
RPL23A/RPL23
RPS4B/RPS4X
RPL13A/RPL13
DIM1/TFB1M
DIM1/DIMT1
SNF4/PRKAG1
RSC2/PBRM1
YTA7/ATAD2B
IRA2/NF1
ARP2/ACTR2
GCN5/KAT2A
TRM10/TRMT10C
SPN1/IWS1
STO1/NCBP1
YNL069C_1/RPL13A
RPS6A/RPS6
RPL34A/RPL34
YNL069C_2/RPL13A
SMT3/SUMO1
RPL32/RPL32
UBC4/UBE2D3
UBC4/UBE2D2
UBC4/UBE2D1
ERG20/FDPS
RHB1/RHEBL1
RHB1/RERGL
RRT12/PCSK9
JHD1/PHF2
JHD1/KDM2B
KEL1/RABEPK
ORM2/ORMDL1
SEC14/SEC14L4
WRS1/WARS
RSP5/HACE1
SRP1/KPNA4
SRP1/KPNA1
RPP2A/RPLP2
CKA2/CSNK2A2
CKA2/CSNK2A1
CSL4/EXOSC1
SEN1/SETX
PRP4/PRPF4
MSS4/PIP5K1A
UTP15/UTP15
NOG1/GTPBP4
TRZ1/ELAC2
ESF2/ABT1
CDC3/SEPT6
HTS1/HARS2
HTS1/HARS
PUS4/TRUB2
PUS4/TRUB1
SOL2/PGLS
CDC25/RASGRF1
EFT2/EEF2
INP53/INPP5E
PMS1/PMS1
SLH1/HELQ
SPE2/AMD1
ADE16/ATIC
RPL40A/ZFAND4
DBR1/DBR1
DPL1/SGPL1
NCL1/NSUN2
RPS12/RPS12
CEG1/DUSP11
INP52/INPP5E
RPL40B/ZFAND4
VPS13/VPS13A
DOA1/PLAA
UBP12/USP45
UBP12/USP43
UBP12/USP19
KIN28/CDK7
ILS1/IARS
MSL5/KHDRBS1
BDF2/BRD3
OLA1/OLA1
UBX5/UBXN7
HRD1/RNF145
PBS2/MAP2K4
JHD2/JARID2
SAR1/SAR1A
HNT3/APTX
PHO92/YTHDC1
AVT2/SLC38A2
AVT2/SLC38A1
DOXO.HL.K.SHIFT
DOXO.HL.K.INTERACTION
DOXO.HLEG.K.SHIFT
DOXO.HLEG.K.INTERACTION
DOXO.HL.L.SHIFT
DOXO.HL.L.INTERACTION
DOXO.HLEG.L.SHIFT
DOXO.HLEG.L.INTERACTION

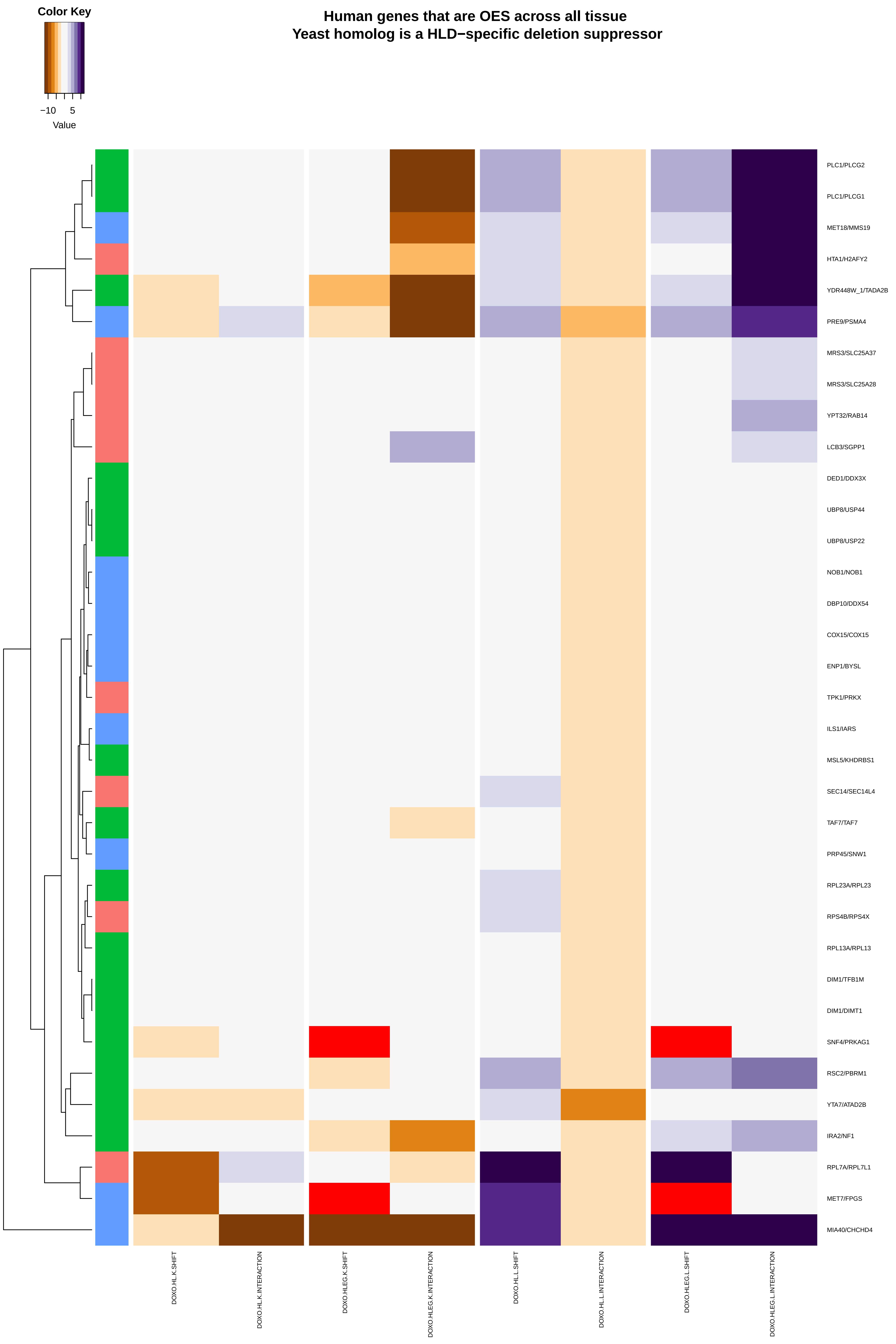
Color Key
−10
5
Value
Human genes that are OES across all tissue
Yeast homolog is a HLD−specific deletion suppressor
PLC1/PLCG2
PLC1/PLCG1
MET18/MMS19
HTA1/H2AFY2
YDR448W_1/TADA2B
PRE9/PSMA4
MRS3/SLC25A37
MRS3/SLC25A28
YPT32/RAB14
LCB3/SGPP1
DED1/DDX3X
UBP8/USP44
UBP8/USP22
NOB1/NOB1
DBP10/DDX54
COX15/COX15
ENP1/BYSL
TPK1/PRKX
ILS1/IARS
MSL5/KHDRBS1
SEC14/SEC14L4
TAF7/TAF7
PRP45/SNW1
RPL23A/RPL23
RPS4B/RPS4X
RPL13A/RPL13
DIM1/TFB1M
DIM1/DIMT1
SNF4/PRKAG1
RSC2/PBRM1
YTA7/ATAD2B
IRA2/NF1
RPL7A/RPL7L1
MET7/FPGS
MIA40/CHCHD4
DOXO.HL.K.SHIFT
DOXO.HL.K.INTERACTION
DOXO.HLEG.K.SHIFT
DOXO.HLEG.K.INTERACTION
DOXO.HL.L.SHIFT
DOXO.HL.L.INTERACTION
DOXO.HLEG.L.SHIFT
DOXO.HLEG.L.INTERACTION

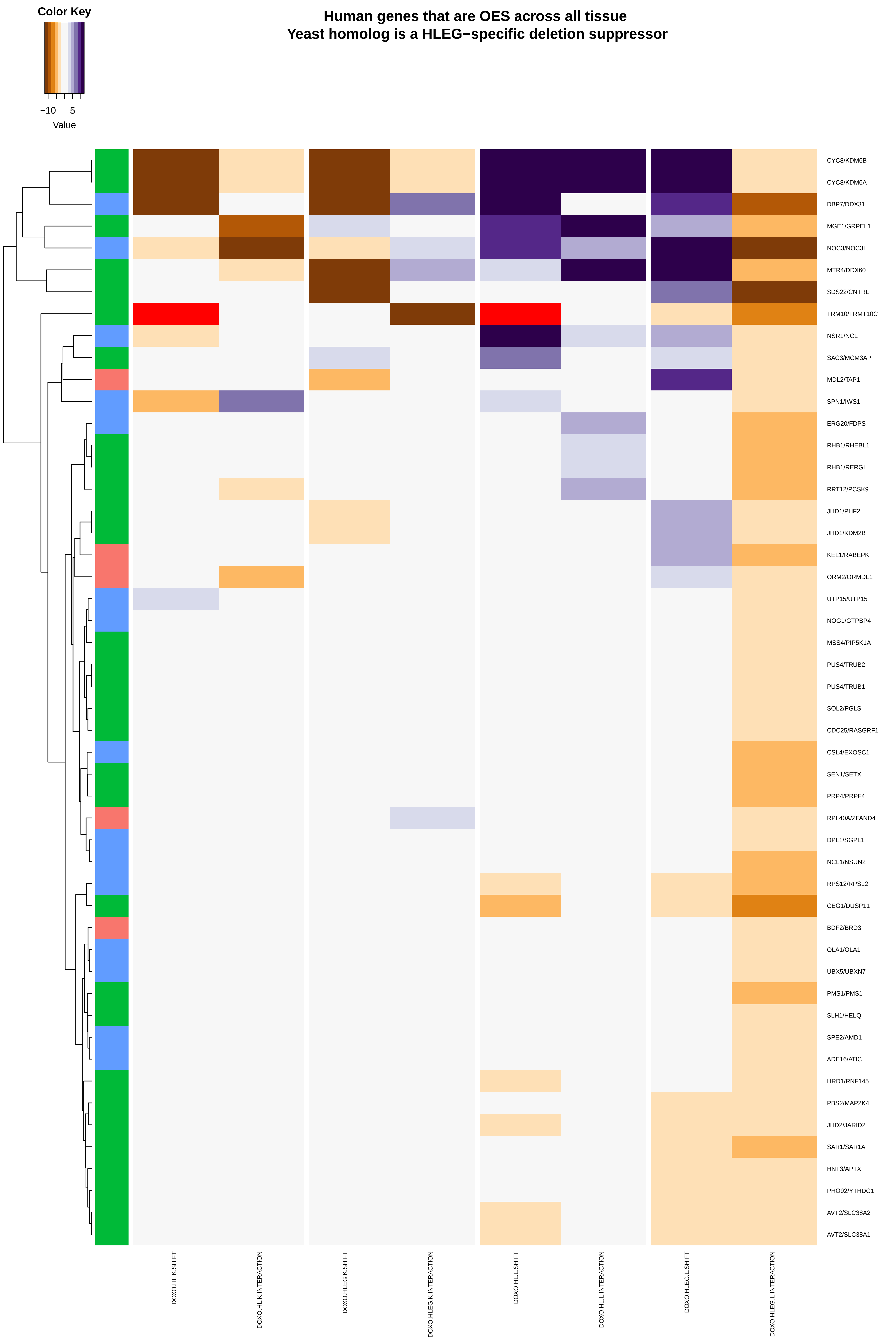

Human genes that are OES across all tissue
Yeast homolog is a HLEG−specific deletion suppressor
Color Key
−10
5
Value
CYC8/KDM6B
CYC8/KDM6A
DBP7/DDX31
MGE1/GRPEL1
NOC3/NOC3L
MTR4/DDX60
SDS22/CNTRL
TRM10/TRMT10C
NSR1/NCL
SAC3/MCM3AP
MDL2/TAP1
SPN1/IWS1
ERG20/FDPS
RHB1/RHEBL1
RHB1/RERGL
RRT12/PCSK9
JHD1/PHF2
JHD1/KDM2B
KEL1/RABEPK
ORM2/ORMDL1
UTP15/UTP15
NOG1/GTPBP4
MSS4/PIP5K1A
PUS4/TRUB2
PUS4/TRUB1
SOL2/PGLS
CDC25/RASGRF1
CSL4/EXOSC1
SEN1/SETX
PRP4/PRPF4
RPL40A/ZFAND4
DPL1/SGPL1
NCL1/NSUN2
RPS12/RPS12
CEG1/DUSP11
BDF2/BRD3
OLA1/OLA1
UBX5/UBXN7
PMS1/PMS1
SLH1/HELQ
SPE2/AMD1
ADE16/ATIC
HRD1/RNF145
PBS2/MAP2K4
JHD2/JARID2
SAR1/SAR1A
HNT3/APTX
PHO92/YTHDC1
AVT2/SLC38A2
AVT2/SLC38A1
DOXO.HL.K.SHIFT
DOXO.HL.K.INTERACTION
DOXO.HLEG.K.SHIFT
DOXO.HLEG.K.INTERACTION
DOXO.HL.L.SHIFT
DOXO.HL.L.INTERACTION
DOXO.HLEG.L.SHIFT
DOXO.HLEG.L.INTERACTION

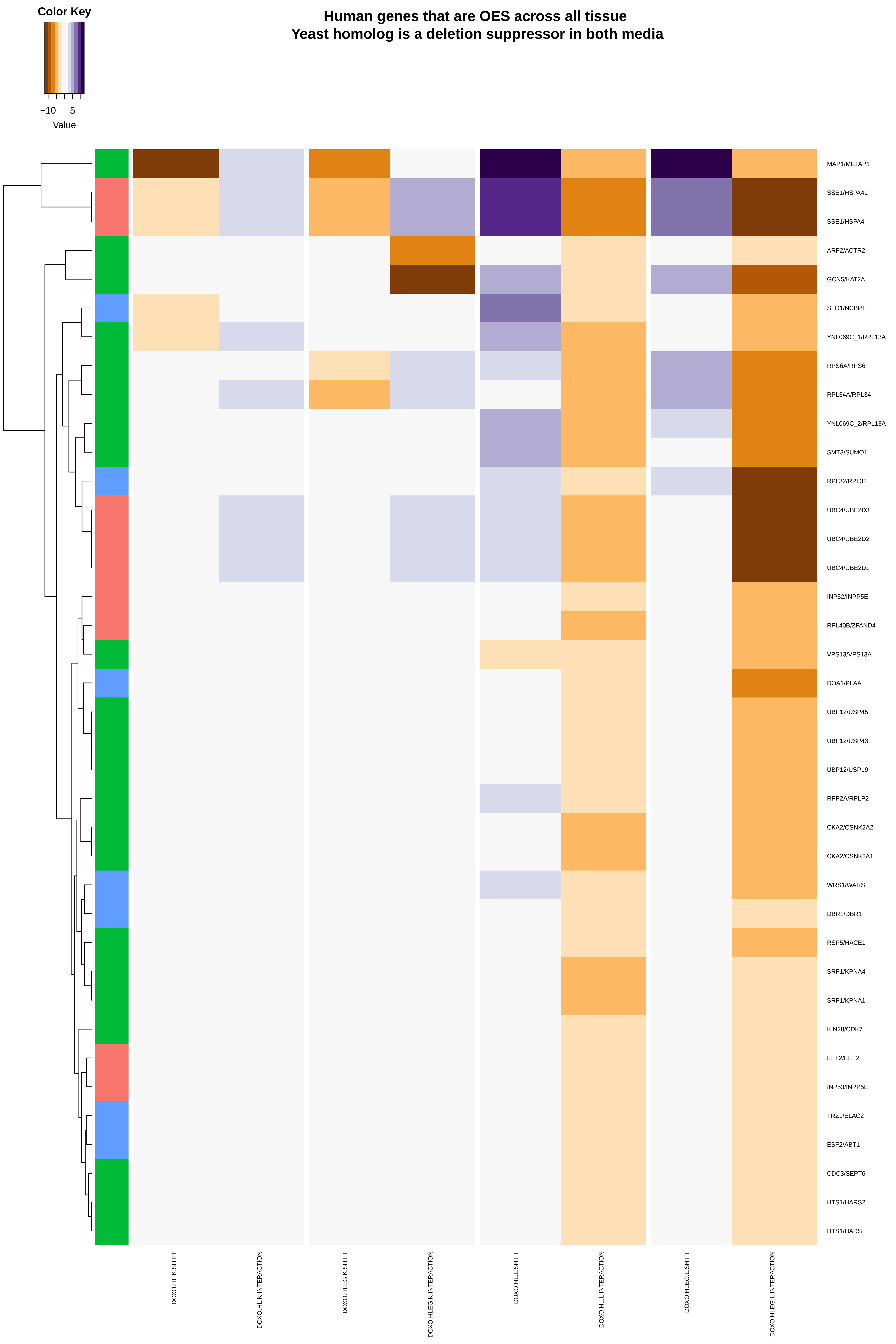

Color Key
−10
5
Value
Human genes that are OES across all tissue
Yeast homolog is a deletion suppressor in both media
MAP1/METAP1
SSE1/HSPA4L
SSE1/HSPA4
ARP2/ACTR2
GCN5/KAT2A
STO1/NCBP1
YNL069C_1/RPL13A
RPS6A/RPS6
RPL34A/RPL34
YNL069C_2/RPL13A
SMT3/SUMO1
RPL32/RPL32
UBC4/UBE2D3
UBC4/UBE2D2
UBC4/UBE2D1
INP52/INPP5E
RPL40B/ZFAND4
VPS13/VPS13A
DOA1/PLAA
UBP12/USP45
UBP12/USP43
UBP12/USP19
RPP2A/RPLP2
CKA2/CSNK2A2
CKA2/CSNK2A1
WRS1/WARS
DBR1/DBR1
RSP5/HACE1
SRP1/KPNA4
SRP1/KPNA1
KIN28/CDK7
EFT2/EEF2
INP53/INPP5E
TRZ1/ELAC2
ESF2/ABT1
CDC3/SEPT6
HTS1/HARS2
HTS1/HARS
DOXO.HL.K.SHIFT
DOXO.HL.K.INTERACTION
DOXO.HLEG.K.SHIFT
DOXO.HLEG.K.INTERACTION
DOXO.HL.L.SHIFT
DOXO.HL.L.INTERACTION
DOXO.HLEG.L.SHIFT
DOXO.HLEG.L.INTERACTION

### Human genes that are UES across all tissue
### Any Yeast Phenotype

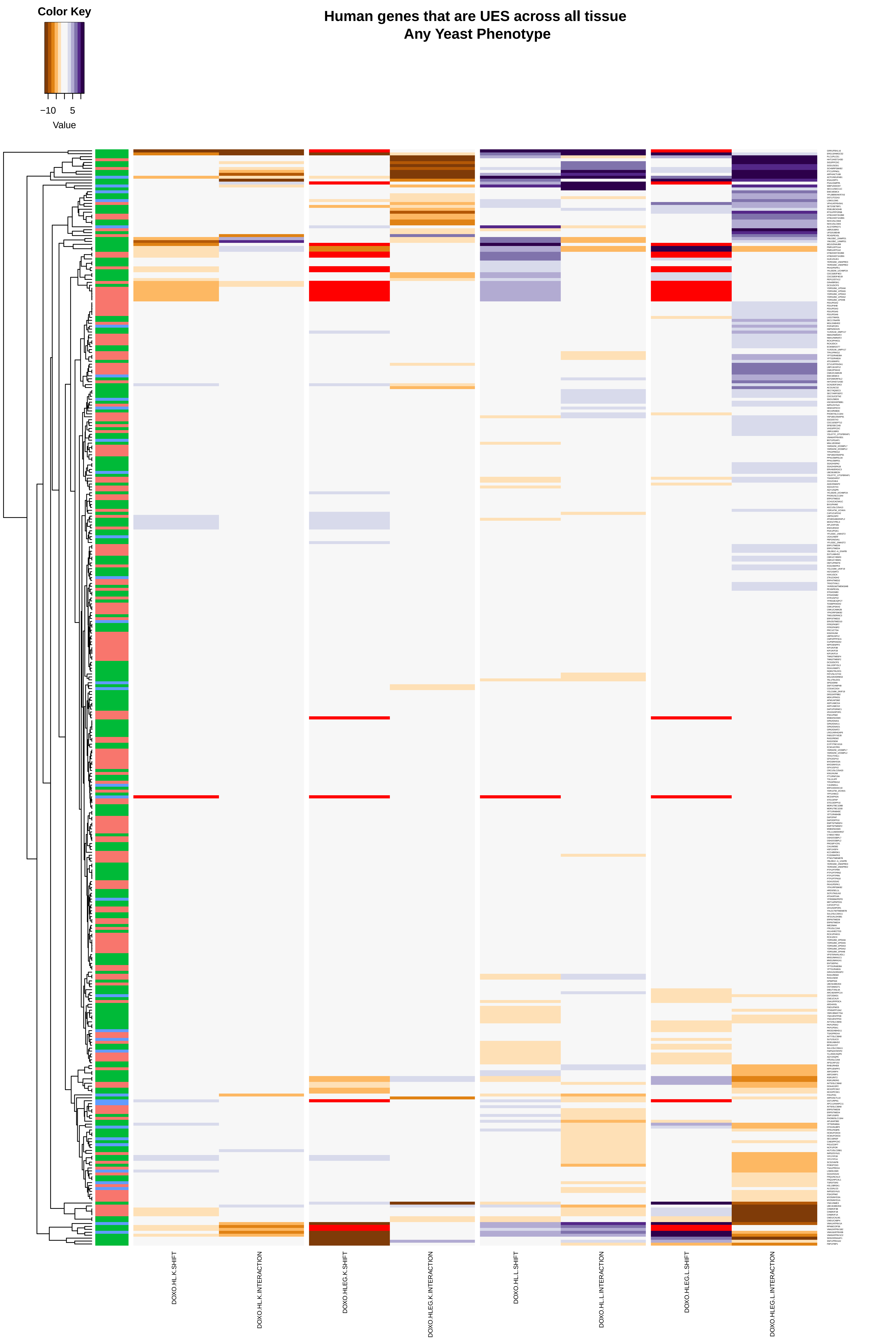

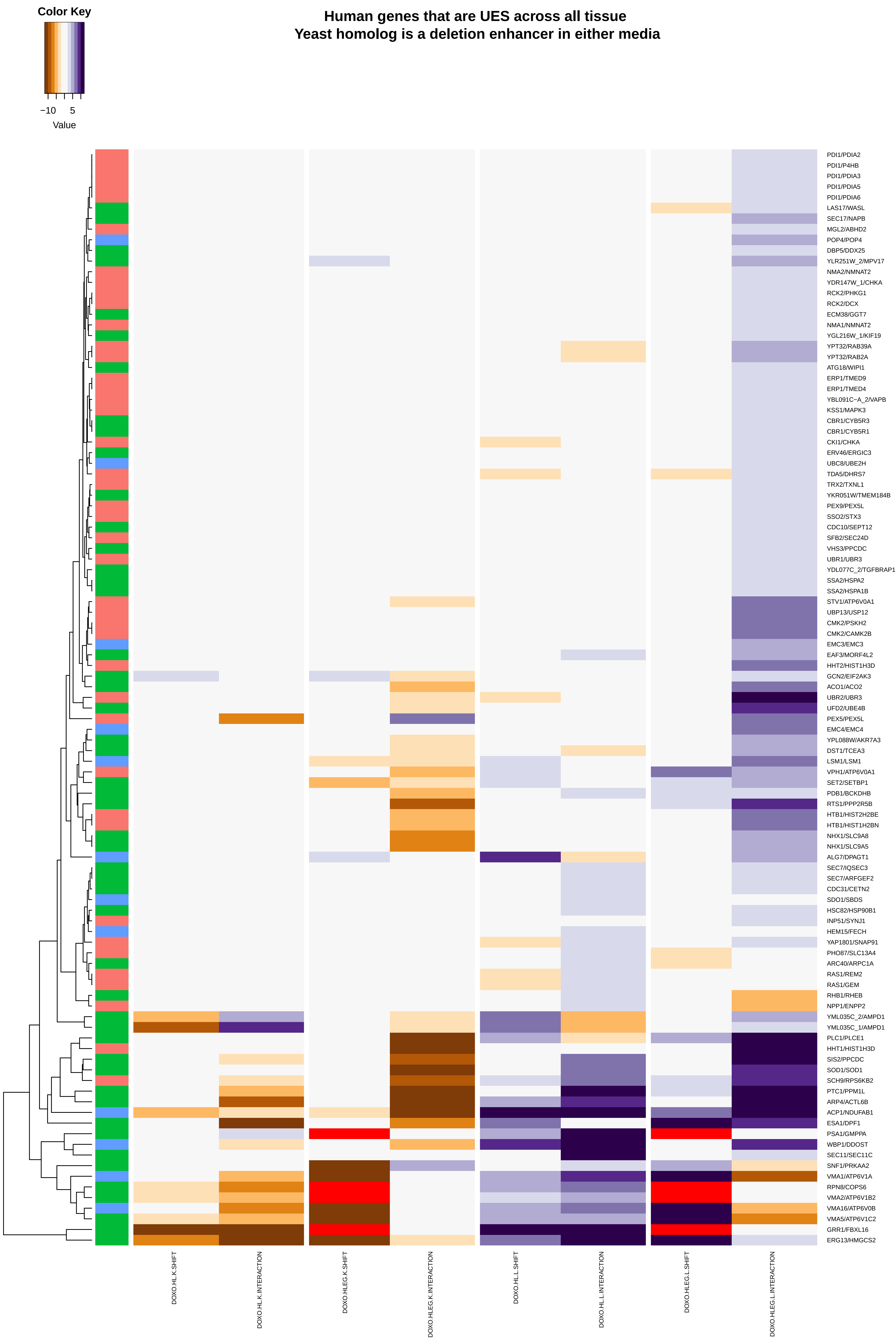
Human genes that are UES across all tissue
Yeast homolog is a deletion enhancer in either media
Color Key
−10
5
Value
PDI1/PDIA2
PDI1/P4HB
PDI1/PDIA3
PDI1/PDIA5
PDI1/PDIA6
LAS17/WASL
SEC17/NAPB
MGL2/ABHD2
POP4/POP4
DBP5/DDX25
YLR251W_2/MPV17
NMA2/NMNAT2
YDR147W_1/CHKA
RCK2/PHKG1
RCK2/DCX
ECM38/GGT7
NMA1/NMNAT2
YGL216W_1/KIF19
YPT32/RAB39A
YPT32/RAB2A
ATG18/WIPI1
ERP1/TMED9
ERP1/TMED4
YBL091C−A_2/VAPB
KSS1/MAPK3
CBR1/CYB5R3
CBR1/CYB5R1
CKI1/CHKA
ERV46/ERGIC3
UBC8/UBE2H
TDA5/DHRS7
TRX2/TXNL1
YKR051W/TMEM184B
PEX9/PEX5L
SSO2/STX3
CDC10/SEPT12
SFB2/SEC24D
VHS3/PPCDC
UBR1/UBR3
YDL077C_2/TGFBRAP1
SSA2/HSPA2
SSA2/HSPA1B
STV1/ATP6V0A1
UBP13/USP12
CMK2/PSKH2
CMK2/CAMK2B
EMC3/EMC3
EAF3/MORF4L2
HHT2/HIST1H3D
GCN2/EIF2AK3
ACO1/ACO2
UBR2/UBR3
UFD2/UBE4B
PEX5/PEX5L
EMC4/EMC4
YPL088W/AKR7A3
DST1/TCEA3
LSM1/LSM1
VPH1/ATP6V0A1
SET2/SETBP1
PDB1/BCKDHB
RTS1/PPP2R5B
HTB1/HIST2H2BE
HTB1/HIST1H2BN
NHX1/SLC9A8
NHX1/SLC9A5
ALG7/DPAGT1
SEC7/IQSEC3
SEC7/ARFGEF2
CDC31/CETN2
SDO1/SBDS
HSC82/HSP90B1
INP51/SYNJ1
HEM15/FECH
YAP1801/SNAP91
PHO87/SLC13A4
ARC40/ARPC1A
RAS1/REM2
RAS1/GEM
RHB1/RHEB
NPP1/ENPP2
YML035C_2/AMPD1
YML035C_1/AMPD1
PLC1/PLCE1
HHT1/HIST1H3D
SIS2/PPCDC
SOD1/SOD1
SCH9/RPS6KB2
PTC1/PPM1L
ARP4/ACTL6B
ACP1/NDUFAB1
ESA1/DPF1
PSA1/GMPPA
WBP1/DDOST
SEC11/SEC11C
SNF1/PRKAA2
VMA1/ATP6V1A
RPN8/COPS6
VMA2/ATP6V1B2
VMA16/ATP6V0B
VMA5/ATP6V1C2
GRR1/FBXL16
ERG13/HMGCS2
DOXO.HL.K.SHIFT
DOXO.HL.K.INTERACTION
DOXO.HLEG.K.SHIFT
DOXO.HLEG.K.INTERACTION
DOXO.HL.L.SHIFT
DOXO.HL.L.INTERACTION
DOXO.HLEG.L.SHIFT
DOXO.HLEG.L.INTERACTION

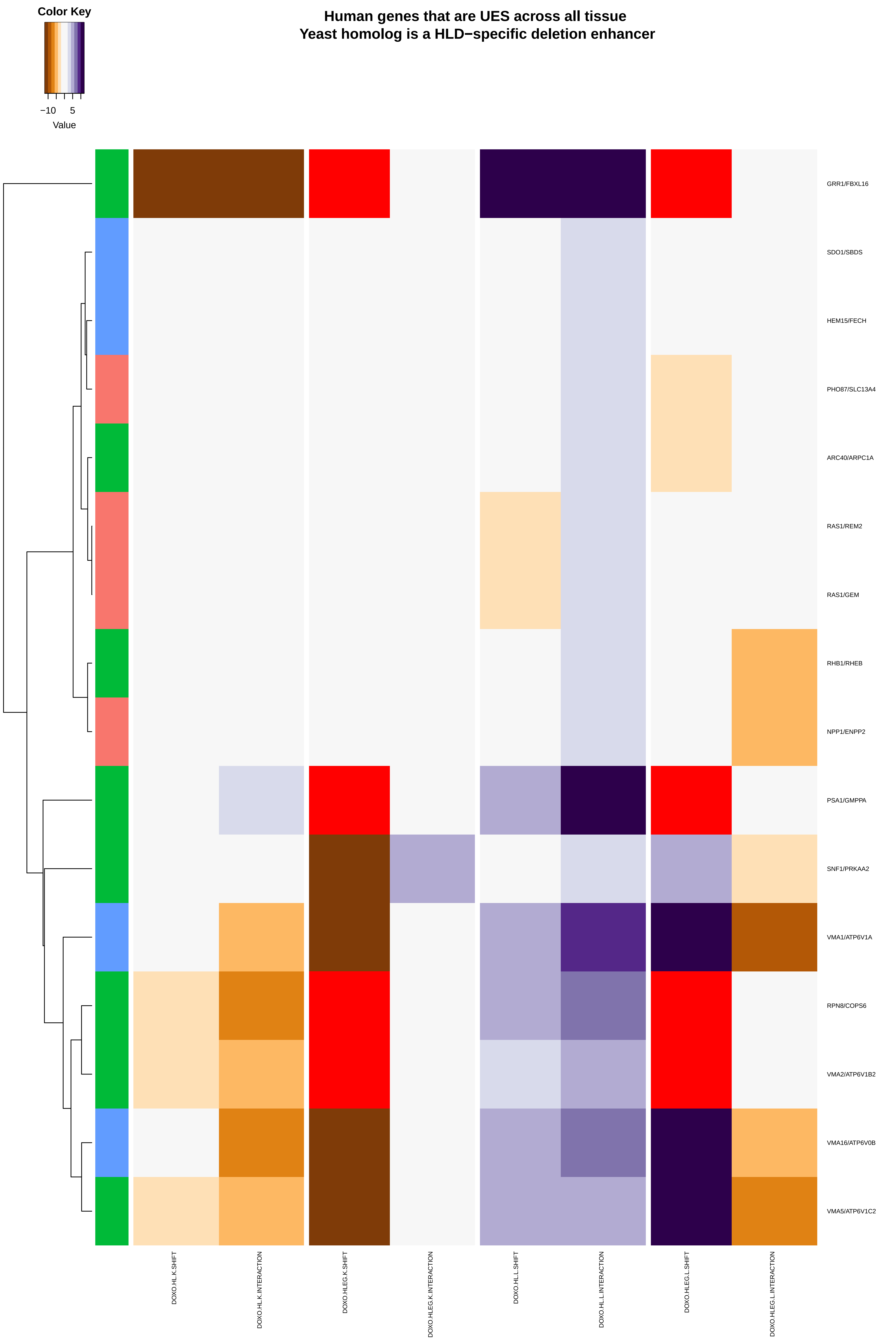

Color Key
−10
5
Value
Human genes that are UES across all tissue
Yeast homolog is a HLD−specific deletion enhancer
GRR1/FBXL16
SDO1/SBDS
HEM15/FECH
PHO87/SLC13A4
ARC40/ARPC1A
RAS1/REM2
RAS1/GEM
RHB1/RHEB
NPP1/ENPP2
PSA1/GMPPA
SNF1/PRKAA2
VMA1/ATP6V1A
RPN8/COPS6
VMA2/ATP6V1B2
VMA16/ATP6V0B
VMA5/ATP6V1C2
DOXO.HL.K.SHIFT
DOXO.HL.K.INTERACTION
DOXO.HLEG.K.SHIFT
DOXO.HLEG.K.INTERACTION
DOXO.HL.L.SHIFT
DOXO.HL.L.INTERACTION
DOXO.HLEG.L.SHIFT
DOXO.HLEG.L.INTERACTION

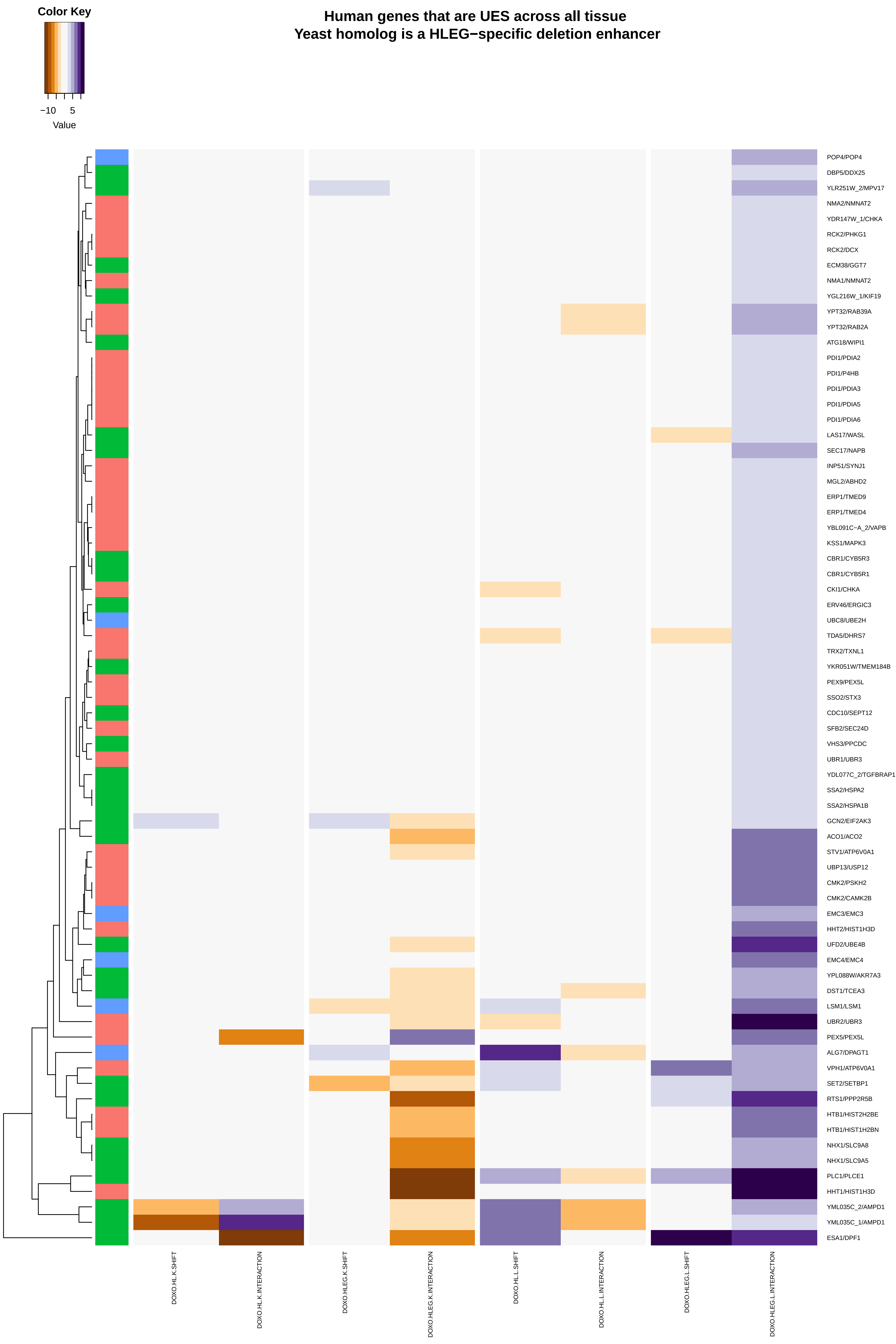
Human genes that are UES across all tissue
Yeast homolog is a HLEG−specific deletion enhancer
Color Key
−10
5
Value
POP4/POP4
DBP5/DDX25
YLR251W_2/MPV17
NMA2/NMNAT2
YDR147W_1/CHKA
RCK2/PHKG1
RCK2/DCX
ECM38/GGT7
NMA1/NMNAT2
YGL216W_1/KIF19
YPT32/RAB39A
YPT32/RAB2A
ATG18/WIPI1
PDI1/PDIA2
PDI1/P4HB
PDI1/PDIA3
PDI1/PDIA5
PDI1/PDIA6
LAS17/WASL
SEC17/NAPB
INP51/SYNJ1
MGL2/ABHD2
ERP1/TMED9
ERP1/TMED4
YBL091C−A_2/VAPB
KSS1/MAPK3
CBR1/CYB5R3
CBR1/CYB5R1
CKI1/CHKA
ERV46/ERGIC3
UBC8/UBE2H
TDA5/DHRS7
TRX2/TXNL1
YKR051W/TMEM184B
PEX9/PEX5L
SSO2/STX3
CDC10/SEPT12
SFB2/SEC24D
VHS3/PPCDC
UBR1/UBR3
YDL077C_2/TGFBRAP1
SSA2/HSPA2
SSA2/HSPA1B
GCN2/EIF2AK3
ACO1/ACO2
STV1/ATP6V0A1
UBP13/USP12
CMK2/PSKH2
CMK2/CAMK2B
EMC3/EMC3
HHT2/HIST1H3D
UFD2/UBE4B
EMC4/EMC4
YPL088W/AKR7A3
DST1/TCEA3
LSM1/LSM1
UBR2/UBR3
PEX5/PEX5L
ALG7/DPAGT1
VPH1/ATP6V0A1
SET2/SETBP1
RTS1/PPP2R5B
HTB1/HIST2H2BE
HTB1/HIST1H2BN
NHX1/SLC9A8
NHX1/SLC9A5
PLC1/PLCE1
HHT1/HIST1H3D
YML035C_2/AMPD1
YML035C_1/AMPD1
ESA1/DPF1
DOXO.HL.K.SHIFT
DOXO.HL.K.INTERACTION
DOXO.HLEG.K.SHIFT
DOXO.HLEG.K.INTERACTION
DOXO.HL.L.SHIFT
DOXO.HL.L.INTERACTION
DOXO.HLEG.L.SHIFT
DOXO.HLEG.L.INTERACTION

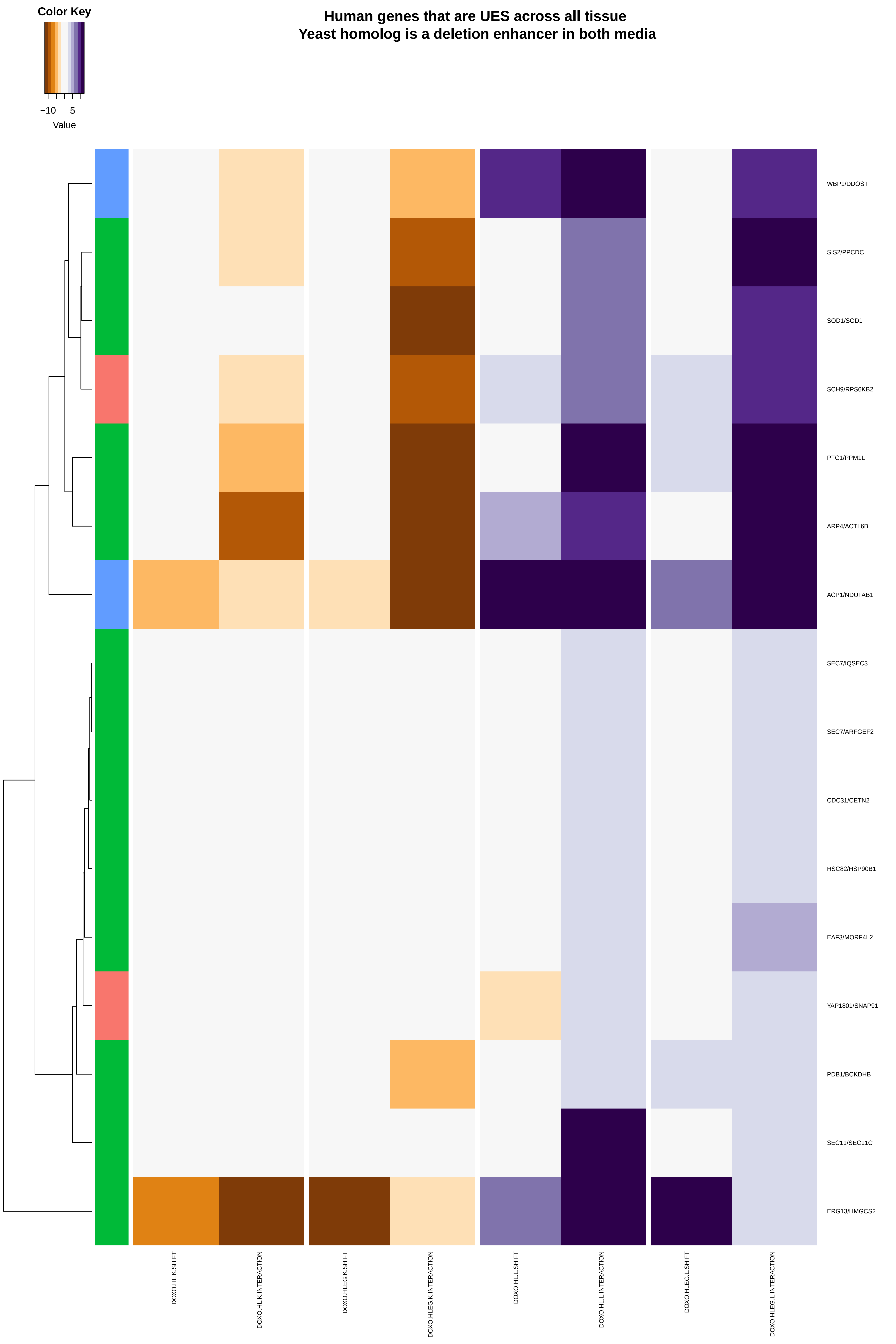

Color Key
−10
5
Value
Human genes that are UES across all tissue
Yeast homolog is a deletion enhancer in both media
WBP1/DDOST
SIS2/PPCDC
SOD1/SOD1
SCH9/RPS6KB2
PTC1/PPM1L
ARP4/ACTL6B
ACP1/NDUFAB1
SEC7/IQSEC3
SEC7/ARFGEF2
CDC31/CETN2
HSC82/HSP90B1
EAF3/MORF4L2
YAP1801/SNAP91
PDB1/BCKDHB
SEC11/SEC11C
ERG13/HMGCS2
DOXO.HL.K.SHIFT
DOXO.HL.K.INTERACTION
DOXO.HLEG.K.SHIFT
DOXO.HLEG.K.INTERACTION
DOXO.HL.L.SHIFT
DOXO.HL.L.INTERACTION
DOXO.HLEG.L.SHIFT
DOXO.HLEG.L.INTERACTION
